# Functional diversification of Ser-Arg rich protein kinases to control ubiquitin-dependent neurodevelopmental signalling

**DOI:** 10.1101/2020.04.02.005041

**Authors:** Francisco Bustos, Anna Segarra-Fas, Gino Nardocci, Andrew Cassidy, Odetta Antico, Lennart Brandenburg, Thomas Macartney, Rachel Toth, C. James Hastie, Robert Gourlay, Joby Vargese, Renata Soares, Martin Montecino, Greg M. Findlay

## Abstract

Conserved protein kinases with core cellular functions have been frequently redeployed during metazoan evolution to regulate specialized developmental processes. Ser-Arg Repeat Protein Kinase (SRPK) is one such conserved eukaryotic kinase, which controls mRNA splicing. Surprisingly, we show that SRPK has acquired a novel function in regulating a neurodevelopmental ubiquitin signalling pathway. In mammalian embryonic stem cells, SRPK phosphorylates Ser-Arg motifs in RNF12/RLIM, a key developmental E3 ubiquitin ligase that is mutated in an intellectual disability syndrome. Processive phosphorylation by SRPK stimulates RNF12-dependent ubiquitylation of transcription factor substrates, thereby acting to restrain a neural gene expression programme that is aberrantly expressed in intellectual disability. SRPK family genes are also mutated in intellectual disability disorders, and patient-derived SRPK point mutations impair RNF12 phosphorylation. Our data reveal unappreciated functional diversification of SRPK to regulate ubiquitin signalling that ensures correct regulation of neurodevelopmental gene expression.

## INTRODUCTION

Signal transduction by protein kinases controls all aspects of eukaryotic biology (Cohen, 2002), from gene expression and metabolism to complex developmental programs. As such, protein kinases that control core eukaryotic processes have been frequently redeployed during metazoan evolution to regulate specialized processes required for multicellular life. This is illustrated by increasingly complex roles of the Mitogen Activated Protein Kinase (MAPK) signalling pathway in yeast and metazoans. In yeast, MAPK signalling pathways control simple unicellular functions such as sensing mating pheromone and environmental stress (Chen and Thorner, 2007), whilst metazoan MAPK signalling has acquired the ability to regulate complex multicellular processes including lineage-specific differentiation (Cowley et al., 1994; Traverse et al., 1992). This prompts the hypothesis that other highly conserved protein kinases have undergone similar “functional diversification” to acquire new functions, thereby facilitating metazoan evolution.

In principle, functional diversification of protein kinases can be achieved via several non-mutually exclusive mechanisms; 1) evolutionary wiring of protein kinase pathways to newly-evolved cell-cell communication systems that control metazoan biology, such as receptor tyrosine kinases (Lim and Pawson, 2010); 2) evolution of new kinase-substrate relationships and 3) evolution of specific kinase activity or expression profiles that differ according to developmental time and tissue context. These mechanisms individually or in combination have the capacity to drive functional diversification, enabling highly conserved eukaryotic protein kinases to evolve novel functions in control of key metazoan processes.

The Ser-Arg Rich Protein Kinase (SRPK) family, which performs core functions in RNA splicing regulation that are conserved throughout eukaryotes (Dagher and Fu, 2001; Gui et al., 1994b; Siebel et al., 1999; Yeakley et al., 1999), represent a prominent but as yet unexplored case study for functional diversification. SRPKs phosphorylate Ser-Arg rich splicing factors, thereby modulating sub-cellular localization, spliceosome assembly and mRNA splicing (Cao et al., 1997; Koizumi et al., 1999; Mathew et al., 2008; Xiao and Manley, 1997). Few non-splicing functions of SRPKs have been reported (Hong et al., 2012; Wang et al., 2017), and it remains unclear whether SRPKs have evolved further complex regulatory roles in metazoans. However, SRPK family members exhibit highly tissue-specific expression profiles (Nakagawa et al., 2005; Wang et al., 1998), suggesting that these protein kinases may indeed perform specialized functions required for multicellular development.

Here, we show that SRPKs have undergone functional diversification to acquire a critical new role during mammalian development. Surprisingly, SRPK activity is largely dispensable for phosphorylation of Ser-Arg rich splicing factors (SRSFs) in mammalian embryo-derived stem cells. Instead, SRPK controls activity of a ubiquitin signalling pathway to regulate expression of neurodevelopmental genes. In this pathway, SRPK phosphorylates a regulatory motif on the key developmental E3 ubiquitin ligase RNF12/RLIM (Barakat et al., 2011; Shin et al., 2010; Shin et al., 2014; Zhang et al., 2012), which is mutated in the X-linked intellectual disability disorder Tonne-Kalscheuer Syndrome (TOKAS) (Frints et al., 2018; Hu et al., 2016; Tonne et al., 2015). Processive SRPK phosphorylation stimulates RNF12-dependent ubiquitylation of transcription factor substrates to modulate expression of neural genes. Data mining indicates that SRPK family genes are also mutated in neurodevelopmental disorders, and several SRPK3 point mutations identified from patients impair RNF12 phosphorylation. Thus, we uncover a previously unappreciated function for SRPK in neurodevelopmental signalling, indicating that functional diversification during eukaryotic evolution has enabled this highly conserved kinase family to govern complex metazoan processes beyond splicing regulation.

## RESULTS

### SRPK activity is largely dispensable for Ser-Arg Rich Splicing Factor phosphorylation in embryonic cells

SRPKs are key players in mRNA splicing regulation, where they control spliceosome assembly and activity (Dagher and Fu, 2001; Yeakley et al., 1999) primarily via phosphorylation of Ser-Arg Rich Splicing Factors (SRSFs) (Long and Caceres, 2009; Roscigno and Garcia-Blanco, 1995; Wu and Maniatis, 1993). Although alternative splicing plays a key role in stem cell regulation (Gabut et al., 2011; Salomonis et al., 2010), functions of SRPK during mammalian development have not yet been studied. This prompted us to examine the role of SRPK in mouse embryonic stem cells (mESCs). We first sought to confirm that SRPK activity is required for phosphorylation of the key SRSF splicing regulators. Surprisingly, and in contrast to somatic cell types (Hatcher et al., 2018), SRSF phosphorylation in mESCs is largely unaffected by the selective pan-SRPK inhibitor SRPKIN-1 (Hatcher et al., 2018) (Figure 1A). In contrast, treatment of mESCs with T3, a selective inhibitor of the closely related CLK kinases (Funnell et al., 2017), which have also been shown to phosphorylate splicing factors (Colwill et al., 1996), leads to near complete inhibition of SR-protein phosphorylation (Figure 1A). Our results suggest that SRPKs are not the major SRSF protein kinases in mESCs, and may have acquired other developmental function(s) during metazoan evolution.

**Figure 1.**
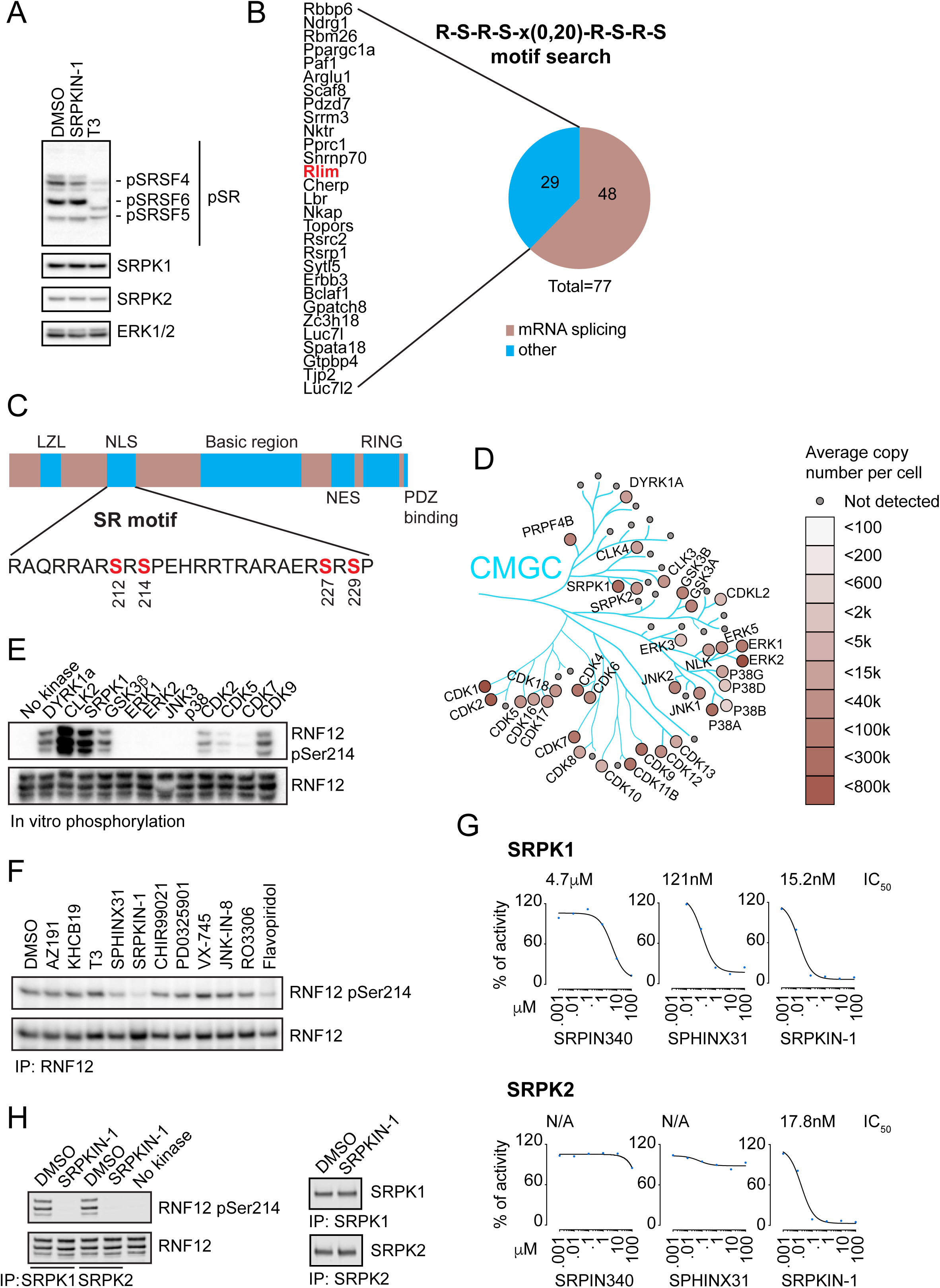
Functional diversification of SRPK to control developmental ubiquitin signalling. (A) WT mESCs were treated with either SRPKIN-1 or T3 kinase inhibitors and phosphorylation of Ser-Arg rich splicing factors (SRSF) assessed by immunoblotting. SRPK1 and SRPK2 expression is determined by immunoblotting and ERK1/2 levels are shown as a loading control. (B) Putative SRPK substrates were predicted using the ScanProsite tool and identified proteins grouped according to their UniProt functional description into mRNA splicing-related (tan) and other (cyan). (C) RNF12 domain structure showing phosphorylation sites detected via immunoprecipitation mass-spectrometry. LZL = Leucine-Zipper Like, NLS = Nuclear Localisation Signal, NES = Nuclear Export Signal, RING = RING-type E3 ubiquitin ligase catalytic domain. (D) CMGC family kinase expression in mESCs determined via quantitative total proteomics. Protein copy numbers are represented according to intensity (see right panel) and distributed in the kinase phylogenetic tree using Kinoviewer. (E) Ability of CMGC kinases (200 mU each) to phosphorylate the RNF12 SR-motif *in vitro* was determined by immunoblotting for RNF12 phospho-Ser214. RNF12 levels are shown as a loading control. (F) mESCs were treated with 10 µM of the following CMGC kinase inhibitors: AZ-191 (DYRK1B inhibitor), KH-CB19 (CLK-DYRK inhibitor), T3 (CLK inhibitor), SPHINX31 (SRPK1 inhibitor), CHIR-99021 (GSK3 inhibitor), PD-0325901 (MEK inhibitor), VX-745 (p38 inhibitor), JNK-IN-8 (JNK inhibitor), RO-3306 (CDK1 inhibitor) and Flavopiridol (CDK7/9 inhibitor) for 4 h and RNF12 SR-motif phosphorylation determined by immunoblotting for RNF12 phospho-Ser214. RNF12 levels are shown as a loading control. (G) SRPK inhibitor dose-response curves for inhibition of RNF12 phosphorylation by SRPK1 and SRPK2 *in vitro* was determined by immunoblotting for RNF12 phospho-Ser214. (H) SRPKIN-1 inhibition of SRPKs *in vivo* was determined by pre-treatment of mESCs with 10 µM SRPKIN-1 for 4 h followed by SRPK1 or SRPK2 immunoprecipitation kinase assay using recombinant RNF12 as a substrate. RNF12 SR-motif phosphorylation and SRPK1 or SRPK2 levels analysed by immunoblotting for RNF12 phospho-Ser214. RNF12 levels are shown as a loading control.

### Identification of SRPK substrates and functions in embryonic stem cells

In order to shed light on developmental functions of SRPKs, we sought to identify further SRPK substrates. Previous studies have demonstrated that SRPKs directly phosphorylate Ser-Arg repeat (SR) motifs (Gui et al., 1994a; Gui et al., 1994b; Wang et al., 1998). Therefore, we interrogated the mouse proteome database for characteristic SRPK consensus motifs of RSRS repeats separated by a linker of 0-20 residues, using the ScanProsite motif analysis tool (https://prosite.expasy.org/scanprosite). A similar approach has been employed to identify a neural-specific splicing factor (Calarco et al., 2009). Our analysis uncovered 77 predicted SRPK substrates, of which 48 have annotated splicing functions. A smaller cohort of 29 predicted substrates are not known to participate in splicing regulation (Figure 1B, Table S1). Interestingly, several have annotated roles in development, including PAF1, a key component of the PAF1 complex that controls RNA PolII and stem cell pluripotency (Ding et al., 2009; Ponnusamy et al., 2009), and TJP2/ZO-2, a component of tight junctions. Also in this dataset is RNF12/RLIM, a RING-type E3 ubiquitin ligase, which ubiquitylates substrates for proteasomal degradation to control key developmental processes including imprinted X-chromosome inactivation (Shin et al., 2014), and stem cell maintenance and differentiation (Bustos et al., 2018; Zhang et al., 2012). RNF12 mutations cause an X-linked neurodevelopmental disorder termed Tonne-Kalscheuer Syndrome (TOKAS) (Frints et al., 2018; Hu et al., 2016; Tonne et al., 2015), which is underpinned by impaired RNF12 E3 ubiquitin ligase activity resulting in deregulated neuronal differentiation (Bustos et al., 2018). Thus, we hypothesised that SRPK phosphorylates and regulates RNF12, which may represent functional diversification of SRPKs in developmental signalling.

### The RNF12 SR-motifs are phosphorylated by SRPK and other CMGC family kinases

Previous work has shown that RNF12 is phosphorylated at the SR-rich motifs identified in our screen (Jiao et al., 2013), although the kinases are not known. In order to confirm phosphorylation of the RNF12 SR-motifs *in vivo*, we performed immunoprecipitation mass-spectrometry in mESCs. Using this approach, RNF12 phosphorylation is robustly detected at two conserved sites; the SR-motifs encompassing Ser212/214/227/229 and an unstudied Ser163 site (Figure 1C, Table S2), confirming that the RNF12 SR-motifs are phosphorylated in mESCs.

The RNF12 SR-motifs consist of tandem RpSRpSP sequences (Figure 1C) flanking a central nuclear localisation signal (NLS), which resemble sequences phosphorylated by SRPKs and several other CMGC kinase sub-families, such as ERK, CDK, CLK and DYRK. Absolute quantitative proteomics shows that many CMGC family kinases are expressed in mESCs (Figure 1D). Thus, we employed a representative CMGC kinase panel to identify kinases that directly phosphorylate RNF12 *in vitro*. GSK3β, and CDK2&9, readily phosphorylate RNF12 at Ser214 within the SR-motifs (Figure 1E). However, incubation of recombinant RNF12 with active SRPK1, or related kinases CLK2 and DYRK1A, leads to a higher level of RNF12 Ser214 phosphorylation (Figure 1E). The ERK subfamily of CMGC kinases, including ERK2, JNK and p38, do not appreciably phosphorylate RNF12 at Ser214 (Figure 1E). These data identify SRPK and related kinases as strong candidates for RNF12 SR-motif phosphorylation.

### A covalent SRPK inhibitor ablates RNF12 SR-motif phosphorylation

In order to identify the kinase that phosphorylates the RNF12 SR-motifs *in vivo*, we assembled a collection of kinase inhibitors that selectively inhibit CMGC family members. Strikingly, of 12 CMGC family kinase inhibitors, only a potent and selective covalent inhibitor of SRPKs, SRPKIN-1(Hatcher et al., 2018), significantly inhibits RNF12 Ser214 phosphorylation in mESCs (Figure 1F). In contrast, the pan-CLK inhibitor T3 has little effect (Figure 1F).

Interestingly, the structurally-unrelated SRPK1 inhibitor SPHINX31(Batson et al., 2017) has a minor effect on RNF12 Ser214 phosphorylation (Figure 1F). However, we find that SRPKIN-1 is ∼10-fold and ∼300-fold more potent toward SRPK1 than SPHINX31 and another commonly used SRPK inhibitor SRPIN-340 (Fukuhara et al., 2006), respectively (Figure 1G, Figure S1A). Furthermore, although SRPKIN-1, SPHINX31 and SRPIN-340 inhibit SRPK1 phosphorylation of RNF12 *in vitro*, only SRPKIN-1 significantly inhibits SRPK2 (Figure 1G, Figure S1A), the other major SRPK isoform expressed in mESCs (Figure 1D, Figure S1B). We also confirm that SRPK1 and SRPK2 are potently inhibited by SRPKIN-1 in mESCs, as measured by RNF12 Ser214 phosphorylation by SRPK1 or SRPK2 immunoprecipitates (Figure 1H). Taken together, our data indicate that SRPK1/2 may mediate RNF12 SR-motif phosphorylation in mESCs.

### Widespread, selective RNF12 SR-motif phosphorylation by SRPK

Phosphoproteomic analysis suggests that RNF12 is phosphorylated at Ser212, Ser 214, Ser227 and Ser229 within the SR-motif (Jiao et al., 2013) (Figure 1C). In order to assess phosphorylation of these additional sites, we devised a phos-tag approach, which retards the mobility of phosphorylated proteins upon SDS-PAGE (Kinoshita et al., 2006). This analysis confirms that RNF12 is phosphorylated to high stoichiometry at all Ser residues within the SR-motif, as mutation of each increases RNF12 mobility (Figure 2A). Interestingly, mutation of Ser214 in combination with Ser229 disrupts RNF12 phosphorylation to a similar extent as when all four sites are mutated (4xSA; Figure 2A), suggesting that RNF12 SR-motifs undergo hierarchical phosphorylation with C- to N-terminal processivity characteristic of SRPK (Ma et al., 2008; Ngo et al., 2008). Importantly, an RNF12 4xSA mutant displays phos-tag mobility similar to that of dephosphorylated RNF12 (Figure 2B).

**Figure 2.**
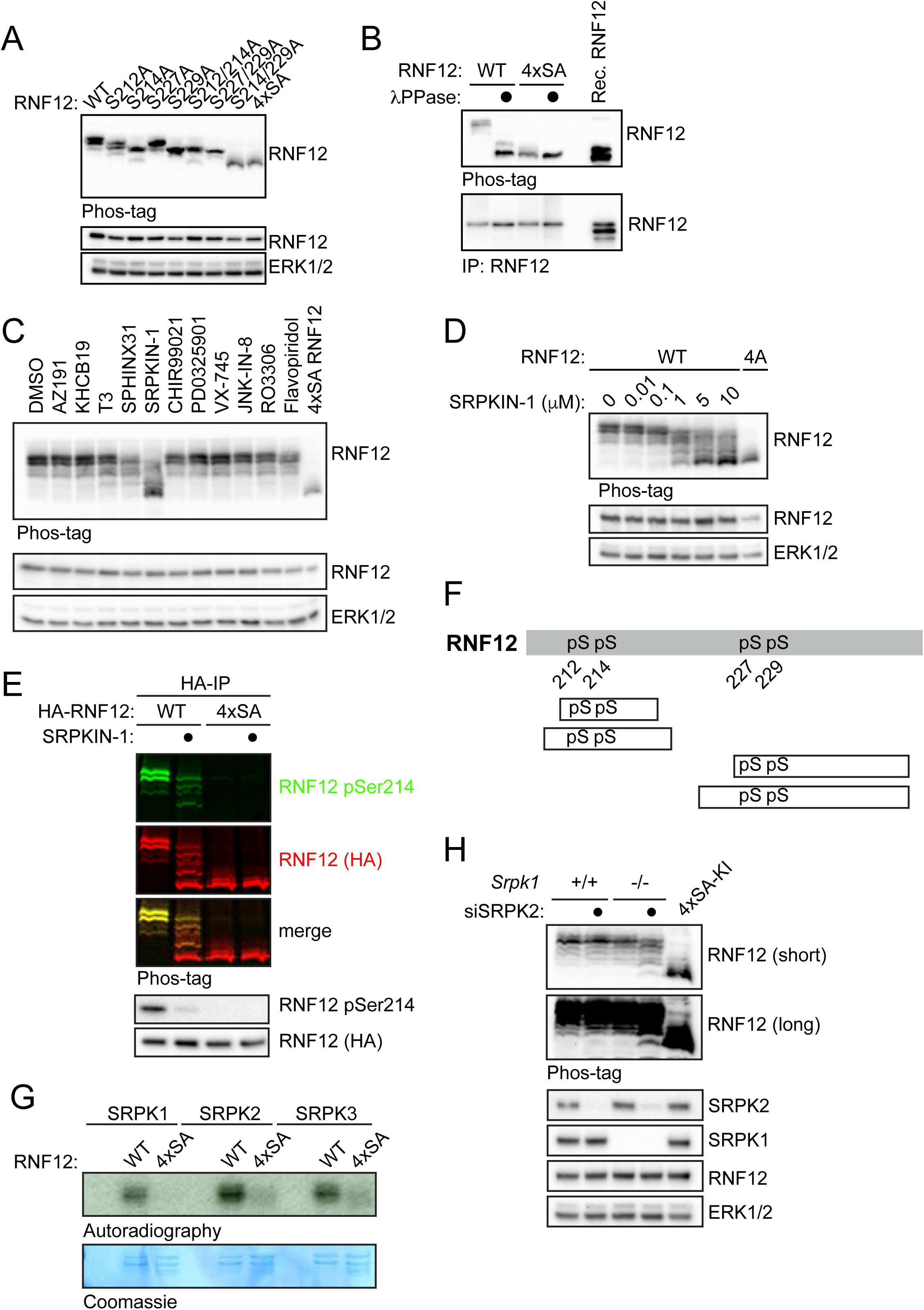
RNF12/RLIM E3 ubiquitin ligase is selectively phosphorylated by SRPKs at a SR-rich motif. (A) RNF12 knockout (*Rlim*^-/y^) mESCs were transfected with WT RNF12 or the indicated point mutants and SR-motif phosphorylation analysed by Phos-tag immunoblotting for RNF12. RNF12 4xSA = S212A/S214A/S227A/S229A. RNF12 and ERK1/2 levels are shown as a loading control. (B) RNF12 knockout (*Rlim*^-/y^) mESCs were transfected with the indicated RNF12 constructs and lysates treated with λ-phosphatase and analysed by Phos-tag immunoblotting for RNF12. Recombinant RNF12 is included as an un-phosphorylated control. (C) mESCs were treated with 10 µM of the of the following CMGC kinase inhibitors: AZ-191 (DYRK1B inhibitor), KH-CB19 (CLK-DYRK inhibitor), T3 (CLK inhibitor), SPHINX31 (SRPK1 inhibitor), CHIR-99021 (GSK3 inhibitor), PD-0325901 (MEK inhibitor), VX-745 (p38 inhibitor), JNK-IN-8 (JNK inhibitor), RO-3306 (CDK1 inhibitor) and Flavopiridol (CDK7/9 inhibitor) for 4 h and RNF12 phosphorylation analysed via Phos-tag immunoblotting for RNF12. RNF12 4xSA is included as an unphosphorylated control. RNF12 and ERK1/2 levels are shown as a control. (D) ESCs were treated with the indicated concentrations of SRPKIN-1 for 4 h and RNF12 phosphorylation analysed via Phos-tag immunoblotting for RNF12. (E) mESCs were treated with 10 µM SRPKIN-1 for 4 h and RNF12 phosphorylation analysed from HA-RNF12 immunoprecipitates via RNF12 phos-tag and phospho-Ser214 immunoblotting using multiplex infrared immunoblot. (F) Phosphorylated peptides detected via mass spectrometry analysis of RNF12 following *in vitro* phosphorylation by SRPK1. pS=phospho-Serine. (G) Autoradiography of RNF12 wild-type (WT) or S212A/S214A/S227A/S229A (4xSA) following a radioactive kinase reaction with SRPK1, SRPK2 or SRPK3. Coomassie staining of RNF12 protein is shown as a loading control. (H) *Srpk1*^-/-^ mESCs were transfected with control or SRPK2 siRNA and RNF12 phosphorylation was analysed via phos-tag immunoblotting. ERK1/2 levels are shown as a loading control.

In order to determine whether SRPKs and/or other kinases can phosphorylate further sites within the RNF12 SR-motifs, we again screened our CMGC kinase inhibitor panel using RNF12 phos-tag analysis. Of these, only SRPKIN-1 drives a major dephosphorylation of the RNF12 SR-motif (Figure 2C). In contrast, the SRPK1 selective inhibitor SPHINX31 and pan-CLK inhibitor T3 show a minor effect on RNF12 phosphorylation (Figure 2C). SRPKIN-1 treatment leads to RNF12 SR-motif de-phosphorylation at concentrations down to 1 µM (Figure 2D) and within 1-2 h (Figure S2A). Furthermore, the high-mobility form of RNF12 visualised by phos-tag SDS-PAGE is completely dephosphorylated at the Ser214 site (Figure 2E), indicating that SRPKs mediate widespread RNF12 SR-motif phosphorylation. In support of this notion, we show by mass-spectrometry that SRPKs directly phosphorylate all 4 Ser residues within the RNF12 SR-motifs (Figure 2F, Table S3). Furthermore, SRPK is highly selective for the RNF12 SR-motif, phosphorylating wild-type RNF12 but not a mutant in which SR-motif Ser residues are mutated (4xSA; Figure 2G). In summary, our data uncover a major role for SRPKs in phosphorylating the RNF12 SR-motif.

### Further evidence that SRPK1/2 are RNF12 SR-motif kinases

In order to confirm that SRPKs phosphorylate the RNF12 SR-motif kinases, we first determined SRPKIN-1 kinase inhibition specificity against a panel of representative kinases (International centre for kinase profiling). Consistent with previous kinase interaction data(Hatcher et al., 2018), SRPKIN-1 is highly specific for SRPK1 inhibition compared to 49 other kinases (Figure S2B). Inhibition of major off-target kinases, including CHK2, PLK1 and DYRK1A, does not impact on RNF12 SR-motif phosphorylation *in vivo* (Figure S2C). RNF12 SR-motif phosphorylation is also inhibited by SRPKIN-1 in wash-out assays, where SRPKIN-1 remains covalently bound to SRPKs but off-target kinases are removed (Hatcher et al., 2018) (Figure S2D).

To further substantiate the role of SRPK1/2 in RNF12 SR-motif phosphorylation, we sought to generate *Srpk1*^-/-^:*Srpk2*^-/-^ mESC lines using CRISPR/Cas9. Although we were able to obtain *Srpk1*^-/-^ and *Srpk2*^-/-^ mESC lines (Figure S2E; Appendix 1), no *Srpk1*^-/-^:*Srpk2*^-/-^ mESC lines were recovered, suggesting that SRPK1/2 perform redundant but essential functions in mESCs. RNF12 SR-motif phosphorylation is unaffected in *Srpk1*^-/-^ and *Srpk2*^-/-^ mESCs (Figure S2E), consistent with SRPK1/2 redundancy with respect to RNF12 phosphorylation. To tackle this, we sought to deplete SRPK2 in *Srpk1*^-/-^ mESCs using siRNA. Partial depletion of SRPK2 expression in the absence of SRPK1 leads to the appearance of completely dephosphorylated RNF12 (Figure 2H), providing further evidence that SRPK1/2 phosphorylate the RNF12 SR-motif in mESCs.

### RNF12 SR-motif phosphorylation drives nuclear anchoring

We next set out to explore functions of the SRPK1/2-RNF12 pathway using RNF12 SR-motif knock-in (KI) mutant mESCs. Using CRISPR/Cas9, we engineered RNF12 4xSA KI mESCs, which cannot be phosphorylated on the SR-motif, and RNF12 ΔSR-KI, in which residues 206-229 of the SR-motif are deleted entirely (Appendix 2). We also engineered control RNF12 wild-type (WT)-KI mESCs and catalytically inactive RNF12 W576Y-KI mESCs (Appendix 2). All mutants are expressed at similar levels and with a similar half-life (Figure S3A), but RNF12 4xSA is poorly phosphorylated compared to wild-type RNF12 (Figure S3B).

As the RNF12 SR-motifs flank a nuclear localisation signal(Jiao et al., 2013), we used KI mutant mESC lines to investigate the role of RNF12 SR-motif phosphorylation in nuclear localisation. Wild-type RNF12 (RNF12 WT-KI) is localised entirely in the nucleus (Figure 3A). However, RNF12 4xSA-KI and RNF12 ΔSR-KI show significant staining in both nucleus and cytosol (Figure 3A), indicating that the RNF12 SR-motif phosphorylation is not absolutely essential for nuclear localisation. In support of this, RNF12 is nuclear in 4xSA-KI ESCs treated with the CRM nuclear export inhibitor Leptomycin B (LMB) (Figure 3B). Furthermore, SRPK1 and SRPK2 are mostly located in the cytosol, with some nuclear staining particularly for SRPK2 (Figure 3C), suggesting that these kinases primarily function in the cytosol (Ding et al., 2006; Jang et al., 2009). Taken together, our data indicate that SRPK phosphorylation of the RNF12 SR-motif promotes RNF12 nuclear anchoring, but is not essential for RNF12 nuclear targeting.

**Figure 3.**
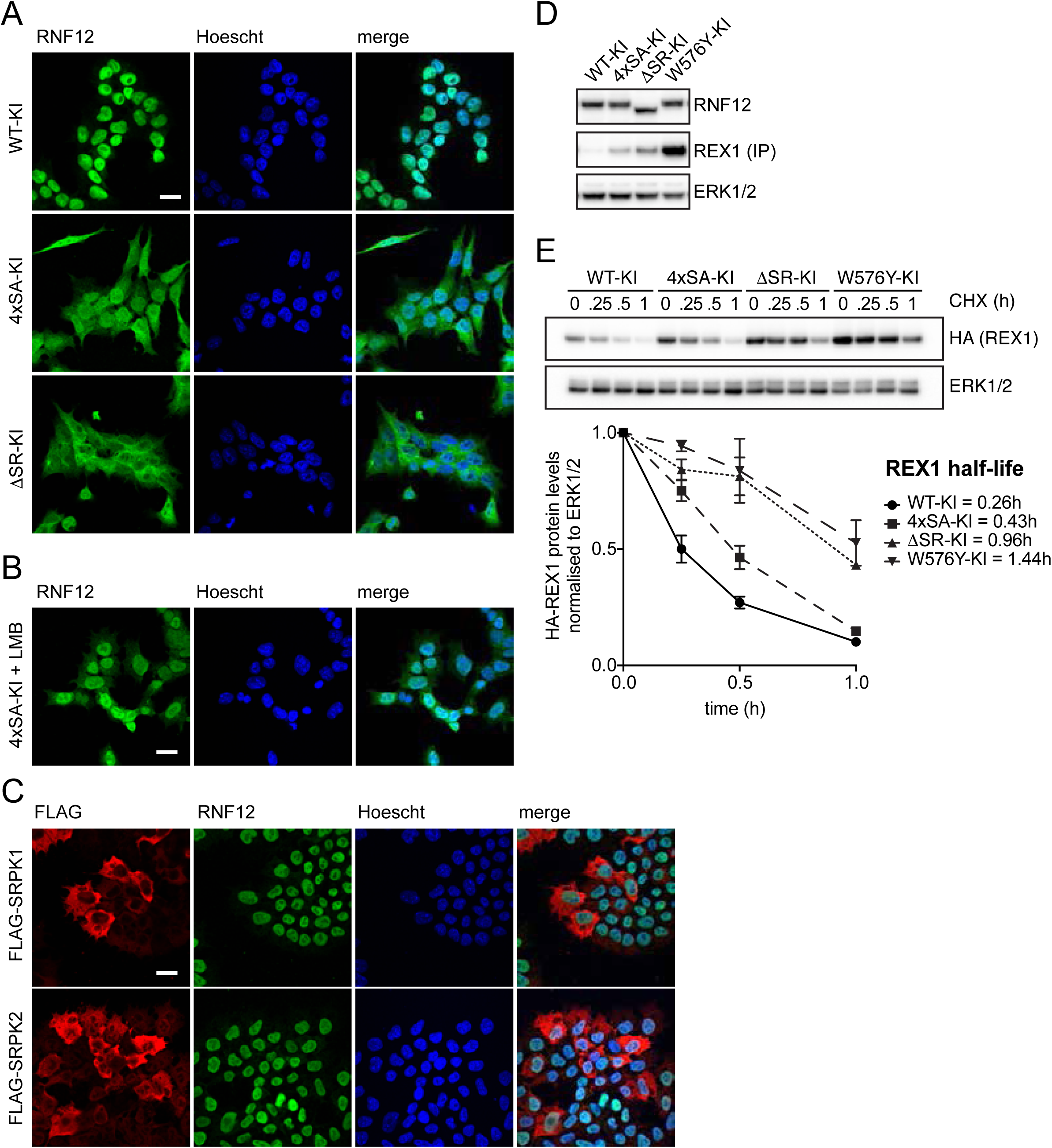
SRPK phosphorylation of RNF12 regulates nuclear anchoring and E3 ubiquitin ligase activity. (A) Subcellular localisation of RNF12 wild-type knock-in (WT-KI), SR-motif phosphorylation site knock-in (4xSA-KI) or SR-motif deletion (ΔSR-KI) in mESCs was determined by immunofluorescence. Scalebar: 20 µm. (B) RNF12 WT-KI or 4xSA-KI mESCs were treated with 30 nM Leptomycin B for 6 h and RNF12 localisation analysed via immunofluorescence. Scalebar: 20 µm. (C) FLAG-tagged SRPK1 and SRPK2 were transfected in mESCs and localisation of SRPKs and RNF12 analysed via immunofluorescence. Scalebar: 20 µm. (D) REX1 steady-state levels were analysed in RNF12 WT-KI, 4xSA-KI, ΔSR-KI and W576Y-KI mESCs by immunoprecipitation followed by immunoblotting. RNF12 and ERK1/2 levels are shown as a control. (E) REX1 half-life was determined in RNF12 WT-KI, 4xSA-KI, ΔSR-KI and W576Y-KI mESCs by immunoblotting. ERK1/2 levels are shown as a loading control.

In light of these results, we tested whether the RNF12 SR-motif is required for efficient degradation of nuclear substrates. A major substrate of RNF12 is the REX1/ZFP42 transcription factor, which mediates RNF12 function in X-chromosome inactivation (Gontan et al., 2012; Gontan et al., 2018). Increased REX1 protein levels are observed in RNF12 4xSA KI and RNF12 ΔSR-KI mESCs (Figure 3D), as well as RNF12 W576Y-KI mESCs harbouring a catalytic mutant. REX1 stability is also increased in RNF12 4xSA KI, RNF12 ΔSR-KI and RNF12 W576Y-KI mESCs, when compared to RNF12 WT-KI controls (Figure 3E). These data demonstrate that SRPK phosphorylation of RNF12 promotes degradation of REX1, and potentially other nuclear transcription factor substrates in mESCs.

### RNF12 SR-motif phosphorylation by SRPK stimulates E3 ubiquitin ligase activity

As RNF12 SR-motif phosphorylation impacts on substrate ubiquitylation and degradation, we investigated whether SRPK phosphorylation controls RNF12 catalytic activity. In order to examine the impact of RNF12 SR-motif phosphorylation on E3 ubiquitin ligase activity, we used SRPK to phosphorylate the RNF12 SR-motif to high stoichiometry in vitro (Figure S4A,B), and compared the E3 ubiquitin ligase activity of SRPK phosphorylated RNF12 and non-phosphorylated RNF12 towards REX1 substrate. Strikingly, RNF12 ubiquitylation of REX1 is enhanced by prior RNF12 phosphorylation by SRPK2 (Figure 4A), which is not observed by pre-incubation with SRPKIN-1 (Figure 4A), or by using a catalytically-inactive mutant of SRPK2 (Figure 4B). Similar results were obtained with SRPK1 (Figure S4C,D), and measuring RNF12 ubiquitylation of SMAD7, another reported substrate (Zhang et al., 2012) (Figure 4C). These results indicate that SRPK phosphorylation stimulates RNF12 substrate ubiquitylation.

**Figure 4.**
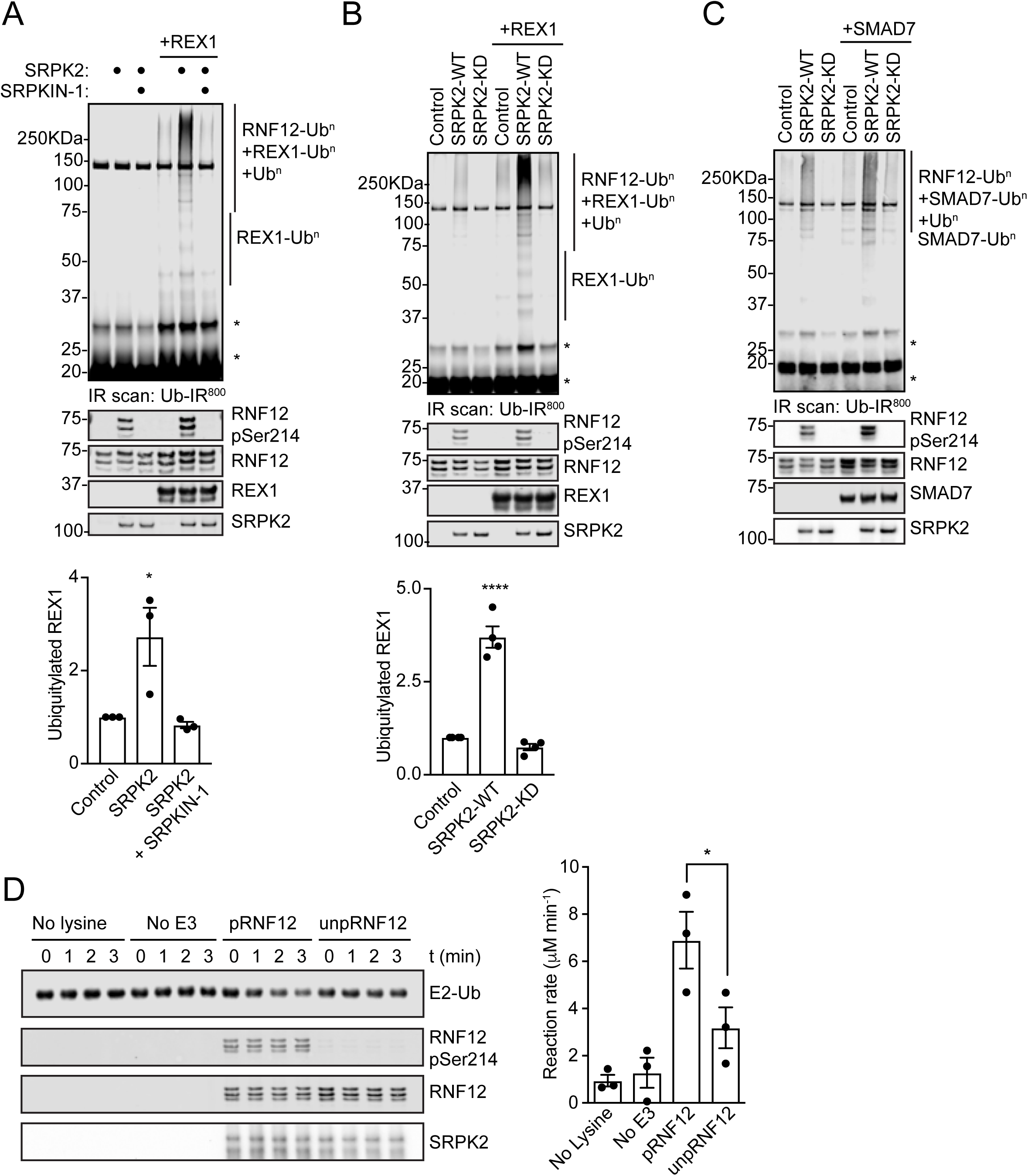
SRPK phosphorylation directly stimulates RNF12 E3 ubiquitin ligase activity. (A) Recombinant RNF12 was incubated with SRPK2 in the absence or presence of 10 µM SRPKIN-1 and subjected to REX1 fluorescent ubiquitylation assays. Infrared scans of ubiquitylated substrate signal and quantifications are shown. Data are represented as mean ± S.E.M. (n=3). One-way ANOVA followed by Tukey’s multiple comparisons test; confidence level 95%. (*) P=0.0350, (****) P<0.0001. Phosphorylated and total RNF12, REX1 and SRPK2 infrared immunoblots are showed as controls. * = non-specific fluorescent signal. (B) Recombinant RNF12 was incubated with wild-type (WT) or kinase dead (KD) SRPK2 and subjected to REX1 fluorescent ubiquitylation assays. Infrared scans of ubiquitylated substrate signal and quantifications are shown. Data are represented as mean ± S.E.M. (n=3). One-way ANOVA followed by Tukey’s multiple comparisons test; confidence level 95%. (*) P=0.0350, (****) P<0.0001. Phosphorylated and total RNF12, REX1 and SRPK2 infrared immunoblots are showed as controls. * = non-specific fluorescent signal. (C) Recombinant RNF12 was incubated with wild-type (WT) or kinase dead (KD) SRPK2 and subjected to SMAD7 fluorescent ubiquitylation assays. Infrared scans of ubiquitylated substrate signal is shown. Phosphorylated and total RNF12, REX1 and SRPK2 infrared immunoblots are showed as controls. * = non-specific fluorescent signal. (D) Recombinant RNF12 was incubated with WT or KD SRPK2 and subjected to E2 ubiquitin discharge assay for the indicated times. Infrared immunoblot scans (top) and reaction rate determinations are shown (bottom). Data are represented as mean ± S.E.M. (n=3). One-way ANOVA followed by Tukey’s multiple comparisons test; confidence level 95%. (*) P=0.0490. Phosphorylated and total RNF12, REX1 and SRPK2 infrared immunoblots are showed as controls.

We then sought to determine the mechanism by which RNF12 SR-motif phosphorylation stimulates catalytic activity. The SR-motif resides in a region proximal to a basic region implicated in the RNF12 catalytic mechanism (Bustos et al., 2018). Therefore, we investigated the impact of SR-motif phosphorylation on RNF12-dependent discharge of ubiquitin from a loaded E2 conjugating enzyme onto free lysine. At concentrations where unphosphorylated RNF12 has relatively weak capacity to discharge ubiquitin from UBE2D1, RNF12 phosphorylation by SRPK2 strongly augments E2 discharge activity (Figure 4D). Therefore, RNF12 SR-motif phosphorylation enhances substrate-independent discharge of ubiquitin from a cognate E2 conjugating enzyme. We conclude that RNF12 phosphorylation by SRPK increases its intrinsic E3 ubiquitin ligase activity.

### RNF12 E3 ubiquitin ligase activity controls a neurodevelopmental gene expression programme

As SRPK-dependent phosphorylation of the RNF12 SR-motif activates and anchors RNF12 in the nucleus to promote degradation of transcription factor substrates such as REX1, we sought to identify the gene expression network that is regulated by this emergent signalling pathway. To this end, we employed RNF12-deficient mESCs (*Rlim*^-/y^) (Bustos et al., 2018) reconstituted with either wild-type RNF12 or an E3 ubiquitin ligase catalytic mutant (W576Y), and performed RNA sequencing (RNA-SEQ) to identify genes that are specifically regulated by RNF12. As an initial validation of this experimental system, we show that degradation of REX1 substrate is restored by wild-type RNF12, but not RNF12 W576Y (Figure 5A).

**Figure 5.**
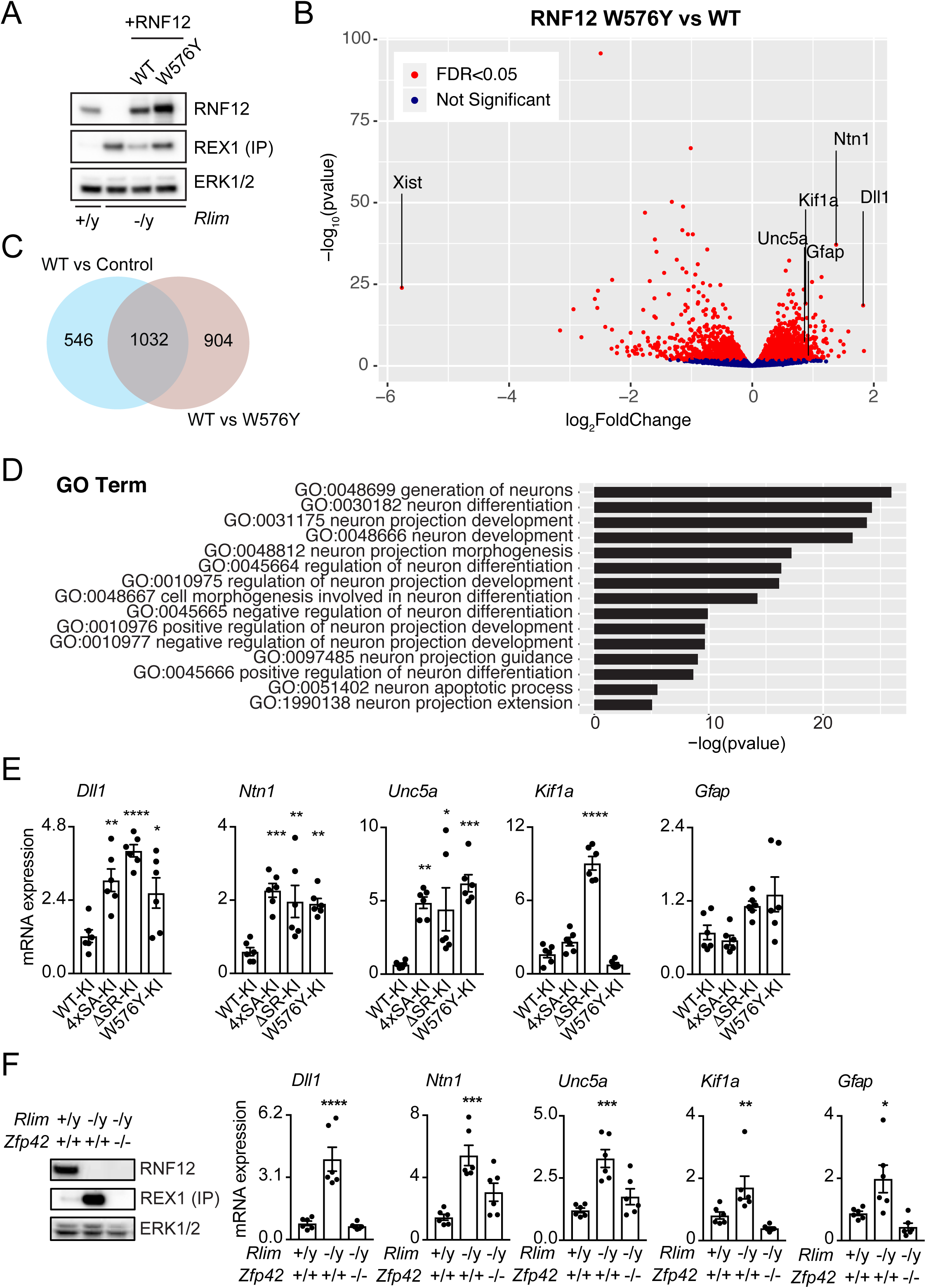
RNF12-REX1 signalling controls a neurodevelopmental gene expression programme. (A) *Rlim*^-/y^ mESCs were transfected with wild-type (WT) or W576Y catalytically inactive RNF12, and REX1 steady-state levels were analysed by immunoprecipitation followed by immunoblotting. RNF12 protein expression and ERK1/2 levels are shown as controls. (B) Volcano plot of RNA-sequencing analysis comparing mRNA expression of *Rlim*^-/y^ mESC transfected with either wild-type (WT) or W576Y catalytically inactive RNF12. mRNAs that are significantly altered by RNF12 E3 ubiquitin ligase activity are displayed in red. Key neurodevelopmental mRNAs that are inhibited by RNF12 E3 ubiquitin ligase activity are labelled (*Dll1, Ntn1, Unc5a, Kif1a, Gfap*). *Xist* is a positive control mRNA that is induced by RNF12 E3 ubiquitin ligase activity. FDR = False discovery rate. (C) Venn diagram displaying total number of mRNAs whose expression is significantly altered when comparing control vs wild-type (WT) RNF12, and wild-type (WT) RNF12 vs W576Y catalytic mutant. (D) Gene Ontology (GO) category enrichment analysis of genes/mRNAs whose expression is inhibited by RNF12. (E) RNF12 WT-KI, 4xSA-KI, ΔSR-KI and W576Y-KI mESC lines were subjected to quantitative RT-PCR analysis for relative mRNA expression of the indicated neurodevelopmental genes. Data are represented as mean ± S.E.M. (n=3). One-way ANOVA followed by Tukey’s multiple comparisons test; confidence level 95%. *Dll1* (**) P=0.0058, (****) P<0.0001, (*) P=0.0377; *Ntn1* (***) P=0.0008, (**) P=0.0057, (**) P=0.0082; *Unc5a* (**) P= 0.0079, (*) P=0.0188, (***) P=0.0006. *Kif1a* (****) P<0.0001; *Gapdh* was used as housekeeping control. (F) RNF12 and REX1 protein levels in wild-type (WT), *Rlim*^-/y^ and *Rlim*^-^/y:*Zfp42*^-/-^ mESCs were determined by immunoblotting (RNF12) and immunoprecipitation followed by immunoblotting (REX1). ERK1/2 levels are shown as loading control (Left). Wild-type (WT), *Rlim*^-/y^ and *Rlim*^-/y^:*Zfp42*^-/-^ mESCs were analysed for relative mRNA expression of indicated genes via quantitative RT-PCR (Right). Data represented as mean ± S.E.M. (n=3). One-way ANOVA followed by Tukey’s multiple comparisons test; confidence level 95%. *Dll1* (****) P<0.0001; *Ntn1* (***) P=0.0002; *Unc5a* (***) P=0.0003; *Kif1a* (*) P=0.0316; *Gfap* (*) P=0.0261. *Gapdh* was used as housekeeping control.

RNA-SEQ analysis of cells re-expressing wild-type RNF12 or the W576Y catalytic mutant reveals that RNF12 E3 ubiquitin ligase activity modulates expression of a significant cohort of mRNAs, whilst most mRNAs are not significantly altered (Figure 5B, Table S4). Additional comparison of control RNF12-deficient mESCs with those re-expressing wild-type RNF12 (Figure S5A) confirms that many RNF12 target genes are regulated in an E3 ligase activity dependent manner (Figure 5C). As proof of principle, the *Xist* long non-coding RNA, a known RNF12 target gene with a key function in X-chromosome inactivation (Barakat et al., 2011), is regulated by RNF12 E3 ubiquitin ligase activity in the expected fashion (Figure 5B).

In order to pinpoint functional clusters of genes that are regulated by RNF12 E3 ubiquitin ligase activity, we employed Gene Ontology (GO) term analysis. Enriched within the cohort of mRNAs suppressed by RNF12 re-expression are those with GO terms associated with neuronal (Figure 5D) and neural (Figure S5B) development, differentiation and function. This is consistent with a key function of RNF12 in restricting mESC differentiation to neurons (Bustos et al., 2018). The positions of genes assigned to neuronal/neural GO terms are highlighted on a further volcano plot of mRNAs that are regulated by re-expression of wild-type RNF12 (Figure S5C, Table S5). In summary, this analysis uncovers a neural gene expression programme that is suppressed by RNF12, revealing a molecular framework for RNF12-dependent regulation of neuronal differentiation (Bustos et al., 2018).

### SRPK signalling to RNF12 regulates neurodevelopmental genes

These results prompted us to investigate the function of the SRPK-RNF12 pathway in regulating expression of the neural gene network identified by RNA-SEQ. We used quantitative RT-PCR to examine expression of RNF12 responsive genes that have key functions in neural development. These are Delta-like 1 (*Dll1*), a regulator of Notch signalling in neural stem cells (Grandbarbe et al., 2003), Netrin-1 (*Ntn1*) and *Unc5a*, an axon guidance system essential for coordination of neuronal connections (Ackerman et al., 1997; Leonardo et al., 1997; Serafini et al., 1996), *Kif1a*, a motor protein for axonal transport (Okada and Hirokawa, 1999) and *Gfap*, an marker of astrocytes and radial glial cells (Middeldorp and Hol, 2011). Accordingly, each of these genes, with the exception of *Unc5a*, increases in expression during *in vitro* neural differentiation (Figure S5D). Consistent with our RNA-SEQ data, *Dll1, Ntn1, Unc5a, Kif1a* and *Gfap* are expressed at low levels in control RNF12 WT-KI mESCs, and this is augmented in catalytically inactive RNF12 W576Y-KI mESCs, with the exception of *Kif1a* (Figure 5E). These data confirm that expression of the majority of RNF12 regulated genes identified by RNA-SEQ are controlled by endogenous RNF12 E3 ubiquitin ligase activity in mESCs.

We next sought to determine the importance of the SRPK signalling to RNF12 in regulation of neurodevelopmental genes. We again employed RNF12 ΔSR-KI and RNF12 4xSA KI mESC lines (Appendix 2). When compared to RNF12 WT-KI mESCs, neural gene expression is generally augmented by mutation of the SR-motif phosphorylation sites (RNF12 4xSA KI) or deletion of the entire motif (RNF12 ΔSR-KI; Figure 5E). As further evidence of the importance of the SR-motif for RNF12-dependent transcriptional regulation, we show that induction of the known RNF12 target gene *Xist* is similarly disrupted by SR-motif mutation or deletion, or by an catalytically inactive mutant (Figure S5E). Therefore, SRPK phosphorylation of RNF12 plays a critical role in regulation of the expression of key neural genes, implicating the SRPK-RNF12 pathway in regulation of neurodevelopmental processes.

### The SRPK-RNF12 pathway regulates gene expression by promoting REX1 degradation

As RNF12 SR-motif phosphorylation is required for efficient substrate ubiquitylation and target gene regulation, we sought to define the downstream molecular pathway. The REX1 transcription factor substrate plays a critical role in RNF12-dependent regulation of *Xist* gene expression and X-chromosome activation (Gontan et al., 2012; Gontan et al., 2018). Thus, we hypothesised that REX1 ubiquitylation and degradation regulates additional genes, providing a mechanism by which RNF12 modulates neural gene expression. We therefore generated RNF12/REX1 double knock-out mESCs (*Rlim*^-/y^:*Zfp42*^-/-^; Figure 5F & Appendix 3) to investigate whether REX1 knockout reverses gene expression changes observed in RNF12-deficient mESCs (*Rlim*^-/y^). As expected, neural gene expression is augmented in RNF12-deficient mESCs (Figure 5F). Additional knockout of the REX1 substrate in an RNF12-deficient background (*Rlim*^-/y^:*Zfp42*^-/-^) reverses this gene expression profile (Figure 5F), suggesting that expression of neural RNF12 target genes is largely regulated via REX1 ubiquitylation and degradation. Our data therefore illuminate REX1 as a key substrate that controls neurodevelopmental gene expression downstream of the SRPK-RNF12 pathway.

### Intellectual disability mutations in the SRPK-RNF12 pathway lead to a deregulated neurodevelopmental gene expression programme

Mutations in RNF12 cause a neurodevelopmental disorder termed Tonne-Kalscheuer Syndrome (TOKAS), which is a syndromic form of X-linked intellectual disability (Frints et al., 2018; Hu et al., 2016; Tonne et al., 2015). We showed previously that TOKAS mutations specifically impair RNF12 E3 ubiquitin ligase activity leading to deregulated neuronal development (Bustos et al., 2018). In order to determine whether aberrant regulation of the SRPK-RNF12 dependent neurodevelopmental gene expression programme might be relevant for TOKAS pathology, we examined expression of neural genes in mESCs harbouring a TOKAS patient mutation (mouse R575C – equivalent to human R599C) (Bustos et al., 2018). Indeed, expression of *Dll1* and *Kif1a* are significantly increased in TOKAS mutant mESCs, with *Ntn1, Unc5a*, and *Gfap* showing a tendency towards increased expression. Strikingly, TOKAS mutation promotes a similar phenotype to mutating the SRPK phosphorylation sites in RNF12 (Figure 6A). Therefore, disrupting SRPK-mediated phosphorylation phenocopies RNF12 intellectual disability mutation with respect to regulation of neurodevelopmental genes.

**Figure 6.**
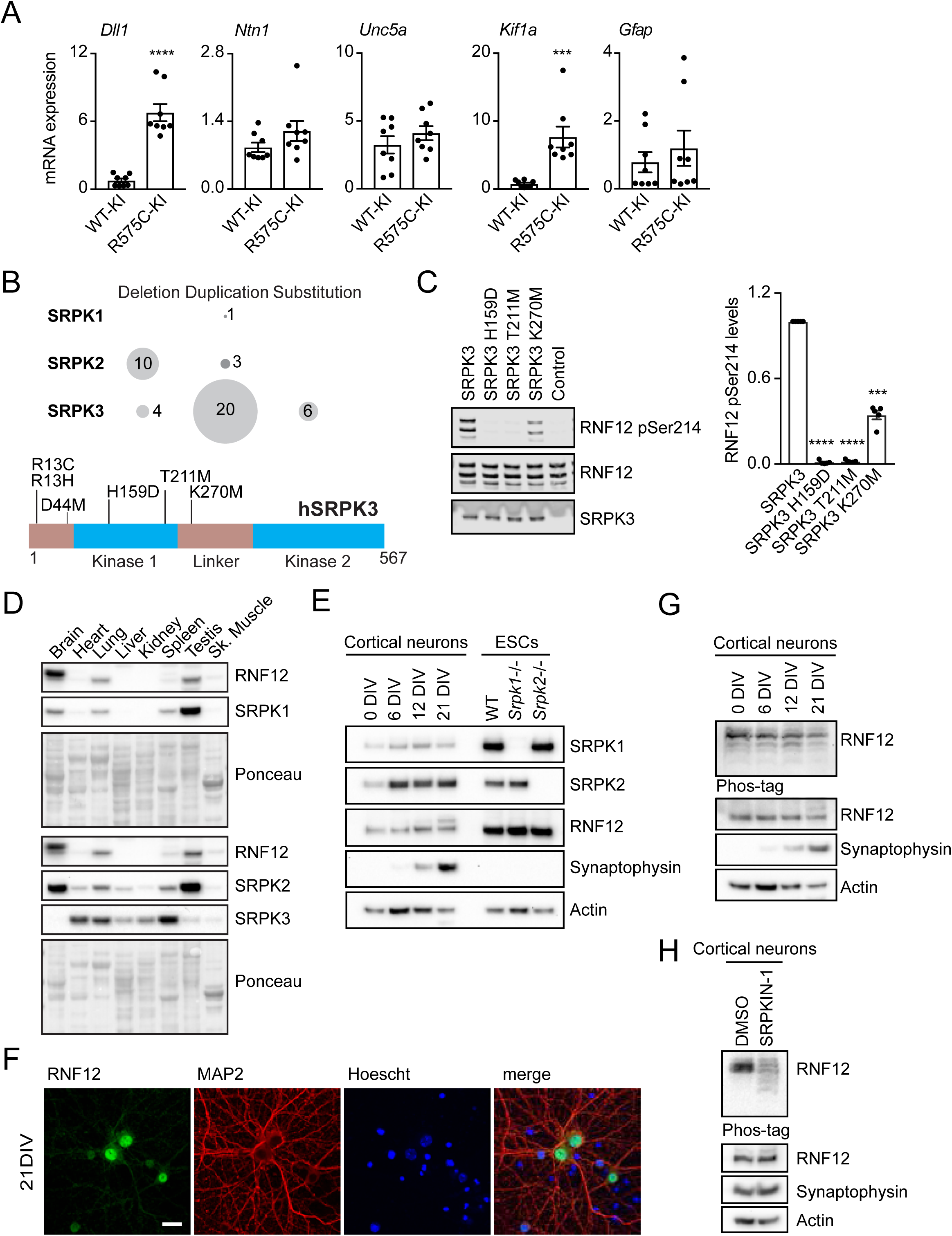
The SRPK-RNF12 signalling pathway operates in neurons and is deregulated in intellectual disability syndromes. (A) RNF12 WT-KI or R575C-KI mESC lines were analysed for relative mRNA expression of the indicated genes via quantitative RT-PCR. Data are represented as mean ± S.E.M. (n=3). Unpaired Student’s t test, two-sided, confidence level 95%. *Dll1* (****) P<0.0001; *Kif1a* (***) P=0.0005. *Gapdh* was used as housekeeping control. (B) Graphical representation of the number of SRPK mutations reported in literature grouped by type of chromosomal mutation (Top). Domain structure of SRPK3 indicating localisation of intellectual disability mutations (Bottom). (C) RNF12 phosphorylation *in vitro* by wild-type (WT) SRPK3 or the indicated mutants was analysed by immunoblotting for RNF12 phospho-Ser214. Total RNF12 and SRPK3 levels are shown as control (Left). Quantification of infrared RNF12 phospho-Ser214 immunoblotting blotting signal normalised to total RNF12 (Right). Data are represented as mean ± S.E.M. (n=3). One-way ANOVA followed by Tukey’s multiple comparisons test; confidence level 95%. (****) P>0.0001, (***) P=0.0001. (D) Expression of RNF12, SRPK1, SRPK2 and SRPK3 in adult mouse tissues analysed via immunoblotting. Ponceau S staining is shown as a loading control. (E) Primary cortical neurons isolated from E16.5 C57BL6 mice were cultured *in vitro* for the indicated number of days and RNF12, SRPK1 and SRPK2 protein expression analysed via immunoblotting alongside indicated mESC lines. Synaptophysin and Actin expression shown as neuronal maturation marker and loading control respectively. (F) Primary cortical neurons isolated from E16.5 C57BL6 mice were cultured *in vitro* for the indicated number of days and analysed by immunofluorescence. MAP2 staining is shown as neuron specific marker and Hoescht used as DNA marker. Scalebar: 20 µm. (G) RNF12 phosphorylation during in vitro mouse cortical neuron maturation was analysed via phos-tag immunoblotting. Synaptophysin and Actin expression are shown as neuron maturation marker and loading control respectively. (H) Cortical neurons were cultured for 21 days and treated with 10 µM SRPKIN-1 for 4 h whereupon RNF12 phosphorylation was analysed via phos-tag immunoblotting. Synaptophysin and Actin expression shown as neuronal maturation marker and loading control respectively.

As SRPK and RNF12 function in a pathway that is disrupted in intellectual disability, we hypothesised that mutations in SRPK family members might cause related intellectual disability syndromes. Thus, we mined molecular genetic datasets and databases of intellectual disability mutations (Deciphering Developmental Disorders, 2015, 2017; Hu et al., 2016; Niranjan et al., 2015) to determine whether SRPKs are mutated in neurodevelopmental disorders. A number of SRPK mutations have been identified in patients with intellectual disabilities or similar developmental abnormalities (Figure 6B). Of those, SRPK1 and SRPK2 are mainly deleted, suggesting that SRPK1 and SRPK2 expression is lost in these disorders. Interestingly, several point mutations in the kinase domain of the X-linked SRPK3 gene (Figure 6B) have been reported in unresolved cases of X-linked intellectual disability (Hu et al., 2016). We tested the effect of these mutations on the ability of SRPK3 to phosphorylate RNF12. SRPK3 H159D and T211M mutations strongly impair SRPK3 phosphorylation of RNF12, whilst K270M disrupts RNF12 phosphorylation to a lesser extent (Figure 6C). Therefore, SRPK mutations found in intellectual disability patients impair the ability of SRPK to phosphorylate RNF12, suggesting that the SRPK-RNF12 signalling pathway is disrupted in intellectual disability disorders.

### SRPK phosphorylates the RNF12 SR-motif in neurons

As SRPK-RNF12-REX1 signalling controls neurodevelopmental gene expression in mESCs and is disrupted in intellectual disabilities, we investigated the function of this pathway in neurons. RNF12 and SRPK1 and SRPK2, but not SRPK3, are robustly expressed in the adult mouse brain (Figure 6D). RNF12 and SRPK1, SRPK2 are also expressed during a time course of neuronal maturation of isolated mouse foetal cortical neural progenitors *in vitro* (Figure 6E). RNF12 is predominantly localised to the nucleus of cultured cortical neurons, suggesting that the RNF12 SR-motif may be phosphorylated (Figure 6F). Indeed, we show by phos-tag analysis that RNF12 SR-motif is heavily phosphorylated throughout a time-course of neuronal maturation (Figure 6G). Furthermore, treatment of mature neurons with the selective SRPK inhibitor SRPKIN-1 suppresses RNF12 phosphorylation, as measured by phos-tag (Figure 6H). Thus, our data confirm that SRPKs phosphorylate the RNF12 SR-motif during neuronal maturation *in vitro*, suggesting that SRPK activity may also regulate RNF12 function in the nervous system.

## DISCUSSION

Functional diversification of protein kinases is an important evolutionary tool, which employs pre-existing signalling cassettes for regulation of increasingly complex cellular processes. However, beyond several well-studied examples such as the MAPK signalling pathway, this concept has not been widely explored, and the importance of functional diversification for regulation of multi-cellularity remains unclear. Here, we show that Ser-Arg rich protein kinase (SRPK), a highly conserved kinase family that regulates mRNA splicing, has undergone functional diversification to control developmental ubiquitin signalling. In mammalian embryonic cells, we find that SRPK activity is largely dispensable for splicing factor phosphorylation, and instead SRPK phosphorylates a key developmental E3 ubiquitin ligase RNF12/RLIM. RNF12 phosphorylation by SRPK drives catalytic activation and nuclear anchoring, enabling RNF12 to ubiquitylate and degrade developmental transcription factors such as REX1. This pathway in turn controls expression of genes involved in control of neural development and function, such that the SRPK-RNF12 axis is mutated in patients with intellectual disability disorders (Figure 7).

**Figure 7.**
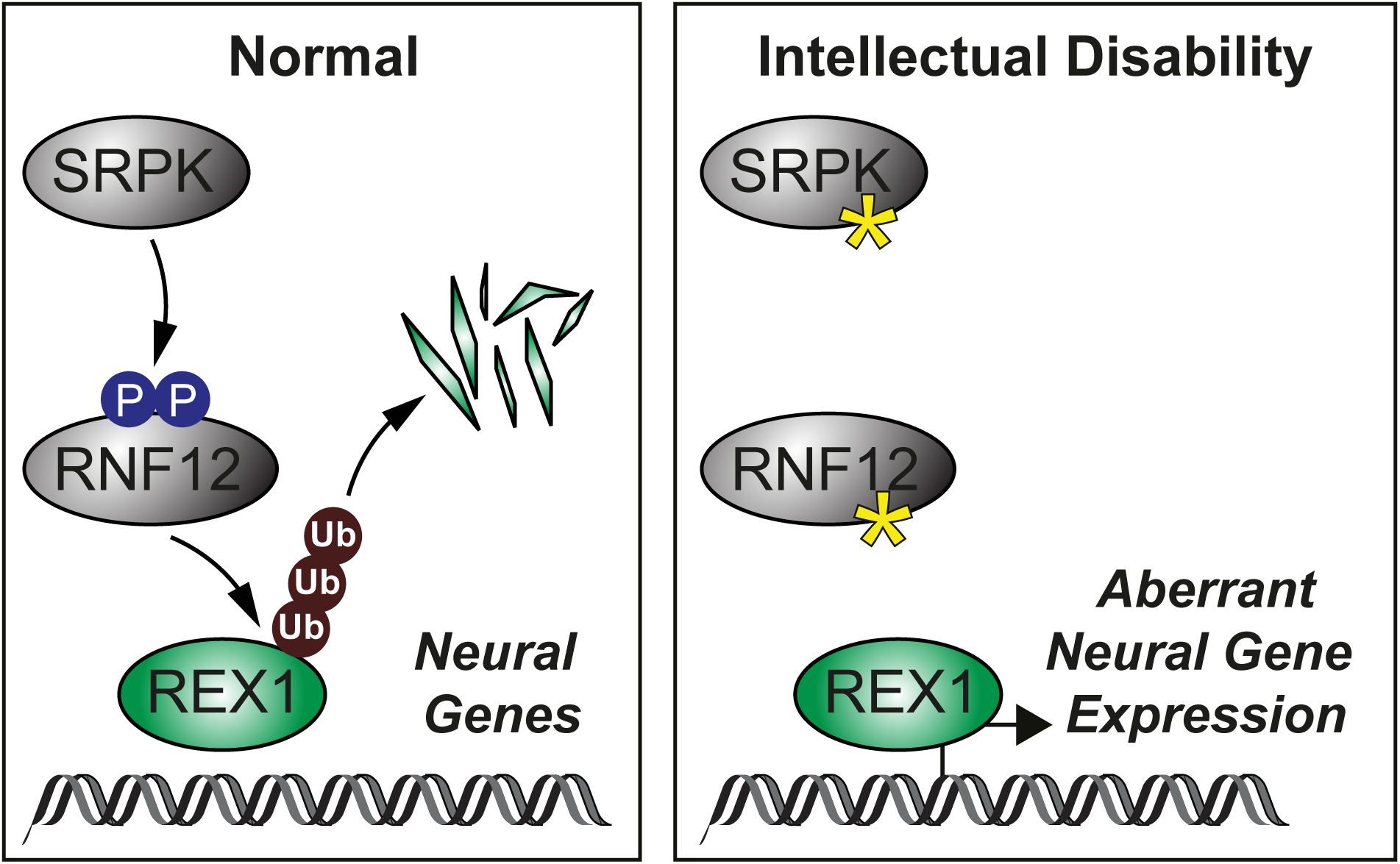
The SRPK-RNF12-REX1 signalling pathway regulates neural gene expression and is disrupted in intellectual disability disorders.

Our studies uncover the molecular mechanism by which SRPK phosphorylation controls RNF12. Although distal to the catalytic RING domain, RNF12 SR-motif phosphorylation increases substrate-independent ubiquitin discharge by a cognate E2 conjugating enzyme, indicating that phosphorylation of these motifs drives maximal catalytic activity. Previous work confirms that distal unstructured regulatory elements play important roles in RNF12 catalysis (Bustos et al., 2018; Frints et al., 2018). Furthermore, phosphorylation of distal elements in the RING E3 c-CBL mediates enzymatic activation (Dou et al., 2013). Structural investigations of full-length RNF12 in complex with cognate E2, ubiquitin and substrate will be required to determine exactly how RNF12 phosphorylation drives enzymatic activation.

Our findings propose a key role for SRPK in regulating developmental processes, although functional redundancy within the mammalian SRPK family has precluded genetic interrogation of SRPK functions during development; therefore conditional and tissue-specific mouse models of SRPK deletion and inactivation will shed light on the developmental events that are controlled by the SRPK. Nevertheless, SRPK2 is highly expressed in brain (Wang et al., 1998) and regulates processes relevant to neurodegeneration (Hong et al., 2012; Wang et al., 2017), suggesting a role for SRPK in development and maintenance of the nervous system. Furthermore, an siRNA screen indicates SRPK2 is required for efficient X chromosome inactivation (Chan et al., 2011), which is a key developmental function of RNF12. Therefore, available evidence provides support for the notion that SRPKs perform key developmental functions.

Regulation of SRPK signalling and activity in a developmental context also remains unexplored. Previous work suggests that SRPKs are constitutively auto-phosphorylated and activated (Ngo et al., 2007), with additional regulatory inputs from the AKT-mTOR pathway (Jang et al., 2009; Lee et al., 2017). Diverse temporal and tissue-specific SRPK expression patterns also suggest that transcriptional regulation may be a key mechanism to ensure that SRPK phosphorylates substrates such as RNF12 within the correct developmental time and space.

A key question relates to the function of RNF12 substrates in neuronal development. Our data indicate that RNF12 controls neural gene expression largely by ubiquitylating and degrading the REX1 transcription factor. RNF12 therefore appears to function as a ‘break’ to prevent aberrant REX1 accumulation and neural gene transcription. This could influence neuronal development in several ways; 1) transcriptional suppression of neural genes in non-neural cells, 2) by modulating the timing and levels of neural gene expression in the developing neuroepithelium or 3) by acting to regulate a specific gene at the top of the neurogenesis signalling cascade? Expression patterns in the developing brain and analysis of the affected pathways will be required to address this.

REX1 has not previously been implicated in regulation of neuronal development, which prompts the hypothesis that pathological REX1 accumulation upon RNF12 axis mutation may unleash neomorphic functions that are detrimental to neuronal development and function. Current focus is identification of REX1 transcriptional and genome occupancy profiles, to define normal and pathological REX1 functions that may be contribute to neurodevelopmental disorders. These findings suggest that approaches to activate SRPKs or normalise expression of key RNF12 substrates such as REX1, for example using protein degradation technologies such as proteolysis targeting chimeras (PROTACs), might provide therapeutic benefit in patients with neurodevelopmental disorders underpinned by deregulated SRPK-RNF12 signalling.

## MATERIALS AND METHODS

### Serine-Arginine motif search

Proteins containing tandem Serine-Arginine motifs were identified by searching the ScanProsite tool (Hulo et al., 2006) (ExPASy, Swiss Institute of Bioinformatics) for a R-S-R-S-x(0,20)-R-S-R-S motif where x is any aminoacid and (0,20) indicate the range of distance between motifs. The motif was searched in the UniProtKB database for *Mus musculus* proteome (TaxID: 10090). Resulted protein list was then categorised according to UniProt functional description. Identified proteins are listed in Table S1.

### Cell culture

Wild-type and CRISPR Cas9 edited male mouse mESCs (CCE line) were cultured in 0.1% gelatin [w/v] coated plates in DMEM containing 10% foetal calf serum [v/v], 5% Knock-Out serum replacement [v/v], 2 mM glutamine, 0.1 mM MEM non-essential amino acids, 1mM sodium pyruvate, and penicillin/streptomycin (all from Thermo Fisher Scientific), 0.1 mM beta-mercaptoethanol (Sigma Aldrich), and 100 ng/ml GST-tagged Leukaemia inhibitory factor (LIF) at 5% CO_2_ and 37°C. For 2i culture, mESCs were converted from LIF/FBS to 2i culture media composed of N2B27: 1% B27 supplement [v/v], 0.5% N2 supplement [v/v], 2 mM glutamine (all from Thermo Fisher Scientific), 0.1 mM beta-mercaptoethanol (Sigma Aldrich), and penicillin/streptomycin in 1:1 DMEM/F12:Neurobasal medium (both from Thermo Fisher Scientific with 1 µM PD0325901 and 1 µM CHIR99021) and neural differentiation induced by culturing cells in N2B27.

For primary cortical neuron cultures E16.5 C57BL6 mice brains were placed in ice cold HBSS, meninges removed, and cortex dissected. Cortex tissue was incubated with 0.125% trypsin containing DNAse at 37°C for 30 minutes. Samples were centrifuged at 1,200 rpm for 5 minutes and resuspended in complete Neurobasal media (Neurobasal containing 2 mM Glutamax, 2% B27 supplement [v/v], 10% foetal calf serum [v/v] and penicillin/streptomycin) and filtered through a 40 µm pore filter. Cells were then centrifuged for 7 minutes at 700 rpm, resuspended in complete Neurobasal media and plated at 0.5 × 10^6^ cells/well on 6-well plates coated with 0.1 mg/ml poly-L-lysine (PLL; Sigma Aldrich). Neurons were cultured at 37°C in a humidified incubator with 5% CO_2_ and medium replaced every 5 days with fresh medium containing B27.

### Mouse organ protein extraction

19-week-old C57BL/6J mice were dissected, organs collected and wrapped in tinfoil and snap frozen in liquid nitrogen. Organs were then resuspended in lysis buffer and lysed using a Polytrone PT 1200 E homogeniser (Kinematica) on ice. Samples were then clarified for 20 min at 14,000 rpm at 4°C and subjected to immunoblot analysis.

### Plasmid and siRNA transfection

mESCs were transfected with Lipofectamine LTX (Thermo Fisher Scientific) according to manufacturer instructions. Plasmids used in this study generated by MRC-PPU Reagents and Services, University of Dundee in pCAGGS puro backbone and summarised in Table S6. All cDNA clones can be found at MRC-PPU Reagents and services website http://mrcppureagents.dundee.ac.uk/. ESCs were transfected with siRNA using Lipofectamine RNAiMAX reagent (Thermo Fisher Scientific). siRNA oligos used are listed in Table S6.

### CRISPR Cas9 gene editing

*Rlim*^-/y^ mESCs were described previously (Bustos et al., 2018). To generate CRISPR Cas9 knockout mESC lines we transfected wild-type (for Srpk1 and Srpk2) or *Rlim*^-/y^ (for Zfp42) mESCs with pX335 and pKN7 vectors (Addgene) containing gRNA sequences targeting: *Srpk1* exon 3, *Srpk2* exon 5, *Zfp42* exon 4 (detailed in Table S6). *Rlim* WT-IRES-GFP (RNF12 WT-KI) and R575C-IRES-GFP (RNF12 R575C-KI) knock-in mESCs were described previously(Bustos et al., 2018). To generate *Rlim* S212A S214A S227A S229A-IRES-GFP (RNF12 4xSA-KI), *Rlim* with amino acids 206-229 deleted IRES-GFP (RNF12 ΔSR-KI) and *Rlim* W576Y-IRES-GFP (RNF W576Y-KI) knock-in mESC lines wild-type mESCs were transfected with pBABED Puro U6 and pX335 vectors encoding guide RNAs targeting *Rlim* gene as detailed in Table S6. This together with donor pMa vectors containing DNA sequence encoding RNF12 amino acids 84 to 600 harbouring the desired mutations followed by an IRES (internal ribosome entry site) and EGFP. Transfected cells were selected with 3 µg/ml puromycin for 48 h and subjected to single cell sorting. Expanded single knock-out clones were screened via immunoblot. Single EGFP positive knock-in cells were expanded and screened for EGFP expression and RNF12 size or phosphorylation via immunoblot and mutations confirmed by genomic DNA sequencing (See Appendices).

### Pharmacological inhibition

Kinase inhibitors used are listed in Table S6. All compounds were diluted in DMSO and cells were treated with 10 µM inhibitor for 4 h prior lysis unless indicated otherwise. For protein stability assays, protein synthesis was inhibited by treating cells with 350 µM cycloheximide (Sigma Aldrich).

### Kinase inhibitor profiling

SRPKIN-1 inhibition activity was analysed on in vitro kinase assays for 50 representative kinases (MRC-PPU International Centre for Kinase Profiling). Kinase activity towards specific peptides was assessed in comparison to DMSO control.

### Immunoblotting and Phos-tag

SDS-PAGE electrophoresis and immunoblotting was performed as described. Cells were lysed in lysis buffer (20 mM Tris [pH 7.4], 150 mM NaCl, 1 mM EDTA, 1% NP-40 [v/v], 0.5% sodium deoxycholate [w/v], 10 mM β-glycerophosphate, 10 mM sodium pyrophosphate, 1 mM NaF, 2 mM Na_3_VO_4_, and Roche Complete Protease Inhibitor Cocktail Tablets). Primary antibodies used are listed in Table S6. Phospho-specific antibodies were used at 1 µg/ml with 10 µg/ml Non-phosphopeptide. After secondary antibody incubation, membranes were subjected to chemiluminescence detection with Immobilon Western Chemiluminescent HRP Substrate (Millipore) using a Gel-Doc XR+ System (Bio-Rad) or Infrared detection using a LI-COR Odyssey Clx system. REX1 protein levels were determined by immunoblotting REX1 immunoprecipitates using Clean-Blot IP Detection Reagent (Thermo Fisher Scientific).

Phos-tag analyses were performed by loading protein samples containing 10 mM MnCl_2_ in 8% polyacrylamide gels containing 50 µM Phos-tag reagent (MRC-PPU reagents and services) and 0.1 mM MnCl_2_. After electrophoresis, gels were washed three times for 10 mins in Transfer buffer (48 mM Tris, 39 mM Glycine, 20% Methanol) supplemented with 20 mM EDTA. Proteins were then transferred to Nitrocellulose membranes, blocked and probed with the indicated antibodies. All protein signals were quantified using Image Studio (LI-COR Biosciences) or Image Lab software (Bio-Rad).

### Mass spectrometry

For phospho-site identification samples were separated via SDS-PAGE electrophoresis, stained with Coomassie blue and gel pieces subjected to an in-gel digestion. First, gel pieces were washed in water, 50% acetonitrile (ACN)/water, 0.1 M NH_4_HCO_3_ and 50% ACN/50 mM NH_4_HCO_3_ and then with 10 mM DTT/0.1 M NH_4_HCO_3_ (All from Sigma-Aldrich). Proteins were alkylated with 50 mM iodoacetamide/0.1 M NH_4_HCO_3_ and then washed as above. Gel pieces were then shrunk in ACN and dried using Speed-Vac. Proteins were then trypsinised by incubating with 5 µg/ml trypsin in 25 mM triethylammonium bicarbonate (Sigma-Aldrich) overnight. Supernatants were separated and gel pieces resuspended in 50% ACN/2.5% formic acid and supernatants combined. Samples were then dried via Speed-Vac and then resuspended in 30 µl 0.1 % formic acid and subjected to liquid chromatography–mass spectrometry (LC-MS) analysis using an Ultimate 3000 RSLCnano system coupled to LTQ-Orbitrap VelosPro mass spectrometer (ThermoFisher Scientific) 10 µl samples were injected and peptides were loaded onto a nanoViper C18 Trap column (5 µm particle size, 100 µm × 2 cm) and separated in a C18 reversed phase Easy-spray column (2 µm particle size, 75 µm × 50 cm) (ThermoFisher Scientific) at a flow rate of 300 nl/min. A linear gradient was used, starting at 3% B and maintained for 5 min, from 3-35% B in 40 min, 35-99% B for 2 min, maintained at 99% B for 5 min, 99-3% B in 3 min and maintained at 3% B for 5 min. Solvents used were A: 0.1% formic acid and B: 80% acetonitrile (ACN) with 0.08% formic acid.

Mass Spectrometry data was acquired in data-dependent mode using the following parameters: MS1 spectra were acquired in the Orbitrap at a resolution of 60,000 (at 400 m/z) for a mass range of 375-1600 m/z with a FTMS full AGC target of 1e6. The top 20 most intense ions (with a minimal signal threshold of 2000) were selected for MS2 analysis on the linear ion trap (with a full AGC target of 5,000) and were fragmented (using CID with a collision energy of 35%), multistage activation, and neutral loss masses of 24.4942, 32.6590, 48.9885.

Data was analysed using Proteome Discoverer v.2.0 and Mascot using MRC_Database_1 (1,950 sequences). Parameters used were the following: Variable modifications: Oxidation (M), Dioxidation (M), Phospho (STY); Fixed modifications: Carbamidomethyl (C), Enzyme: Trypsin/P, Maximum missed cleavages: 3, Precursor tolerance: 10ppm, MS2 tolerance: 0.6Da, Minimum score peptides: 18. Phospho-site assignment probability was estimated via Mascot and PhosphoRS3.1 (Proteome Discoverer v.1.4-SP1) or ptmRS (Proteome Discoverer v.2.0).

Quantitative total ESC proteomics data covering around 10000 proteins was previously described (Fernandez-Alonso et al., 2017). CMGC kinase expression representation from that dataset was generated using Kinoviewer https://peptracker.com (Brenes and Lamond, 2019).

### Protein expression and purification

All recombinant proteins were produced in *E. coli* or SF21 insect cells expression systems by MRC-PPU reagents and services and purified via standard protocols. Proteins used in this study are listed in Table S6. All proteins can be found at MRC-PPU Reagents and services website http://mrcppureagents.dundee.ac.uk/.

### *In vitro* kinase assays

For SRPK Immunoprecipitation kinase assays, mESCs were treated with 10 µM SRPKIN-1 for 4h. Cells were lysed, and 1.5 mg of protein was immunoprecipitated with 2 µg of SRPK1 or SRPK2 antibodies (BD Biosciences). Immunoprecipitated containing beads were then washed with lysis buffer supplemented with 500 mM NaCl and half sample was resuspended in loading buffer. The rest was subjected to *in vitro* phosphorylation assay containing 0.5 µg RNF12 and 2 mM ATP in kinase buffer (50 mM Tris-HCl [pH 7.5], 0.1 mM EGTA, 10 mM MgCl2, 2 mM DTT) and incubated at 30°C for 30 min. SRPK in vitro kinase assays were performed by incubating 200 mU kinase or equivalent µg of inactive kinase with 0.5 µg RNF12 and 2 mM ATP in kinase buffer. For radioactive *in vitro* kinase assays, reactions were supplemented with 1 µCi γ-^32^P ATP. Reactions were incubated at 30°C for 30 min in presence or absence of inhibitor as indicated and samples subjected to polyacrylamide electrophoresis and immunoblot or Coomassie blue staining and signal detected via ECL, infrared detection or autoradiography.

### Immunofluorescence

Immunofluorescence and confocal analysis were performed as described. mESCs were plated in 0.1% gelatin [v/v] coated coverslips. Cortical neurons were plated at a density of 1.5 × 10^5^ cells/well on poly-L-lysine German Glass Coverslips 18mm #1½ (EMSdiasum). Primary antibodies used are listed in Table S6. Cells were mounted using Fluorsave reagent (Millipore) Images were acquired in a Zeiss 710 confocal microscope and images were processed using Image J (NIH) and Photoshop CS5.1 software (Adobe).

### *In vitro* phospho-RNF12 activity assays

For substrate ubiquitylation assays, 0.5 µg RNF12 protein was subjected to a phosphorylation reaction containing 200 mU SRPK or equivalent µg of catalytically inactive kinase and 2 mM ATP in kinase buffer for 1 h at 37°C. 200 nM phosphorylated RNF12 was then incubated with a ubiquitylation mix containing 1.5 µg of REX1 or SMAD7, 0.1 µM UBE1, 0.05 µM UBE2D1, 2 µM Ub-IR^800^, 0.5 mM TCEP [pH 7.5], 5 mM ATP (both from Sigma Aldrich), 50 mM Tris-HCl [pH 7.5], 5 mM MgCl_2_ for 30 min at 30°C. Reactions were stopped with SDS sample buffer and boiled for 5 min. Samples were loaded in 4-12% Bis-Tris gradient gels (Thermo Fisher Scientific). Gels were then scanned using an Odyssey CLx Infrared Imaging System (LICOR Biosciences) for detection of fluorescently labelled ubiquitylated proteins. After scanning proteins were transferred to PVDF or nitrocellulose membranes and analysed via immunoblot and signal detected trough ECL or infrared detection.

For UBE2D1 ubiquitin discharge assays 5 µg RNF12 protein was phosphorylated as above with 2U SRPK or equivalent µg of catalytically inactive kinase and 2 mM ATP in kinase buffer for 1 h at 37°C. ATP was depleted with 4.5 U/ml apyrase (New England Biolabs) for 10 min at room temperature. UBE2D1-ubiquitin thioester was prepared by incubating 100 µM UBE2D1 with 0.2 µM UBE1, 100 µM FLAG-ubiquitin, 3 mM ATP, 0.5 mM TCEP [pH 7.5] (both from Sigma Aldrich), 5 mM MgCl2, 50 mM Tris (pH 7.5), 150 mM NaCl for 20 min at 37 °C. Reaction was stopped by depleting ATP with 4.5 U/ml apyrase (New England Biolabs) for 10 min at room temperature. Then, 40 µM UBE2D1-ubiquitin were incubated with 1 µM phosphorylated RNF12 and 150 mM L-lysine in a buffer containing 50 mM Tris [pH 7.5], 150 mM NaCl, 0.5 mM TCEP, 0.1% [v/v] NP40 at room temperature. Reactions were stopped with non-reducing SDS loading buffer and analysed via immunoblotting and membranes scanned in an Odyssey CLx Infrared Imaging System (LI-COR Biosciences). Protein signals were quantified using Image Studio software (LI-COR Biosciences). Reaction rates were determined by extrapolating protein signals in a standard curve of known concentrations of UBE2D1-ubiquitin conjugate and plotting concentration over time.

### Binding assays

For protein immunoprecipitation, protein A or G beads were incubated with 2 µg antibody and 0.5-2 µg/µl protein sample in lysis buffer overnight at 4°C. Immunoprecipitated-containing beads were then washed three times with lysis buffer supplemented with 500 µM NaCl, resuspended in 50% [v/v] loading buffer and boiled at 95°C for 5 minutes prior to immunoblotting analysis. For HA tagged protein immunoprecipitation, Anti-HA agarose conjugate (Sigma Aldrich) was used.

### RNA-Sequencing and Gene Ontology analysis

Total RNA was extracted using RNeasy Mini Kit (QIAGEN) and DNA libraries prepared using TruSeq Stranded Total RNA Sample Preparation kits (Illumina) according to manufacturer’s instructions. Sequencing was performed on Illumina NextSeq platform. Briefly, raw sequencing reads were trimmed by removing Illumina adapters sequences and low-quality bases. Trimmed reads were mapped using to mouse reference genome (mm10) using STAR software (v2.7.1a). The number of reads per transcript was counted using HTSeq (v0.11.2). The differentially expressed genes (DEGs) were estimated using SARTools (v1.6.9) and DESeq2 (v1.24) R packages. Gene Ontology (GO) analysis was carried out using the GOstats (v2.50.0) R package.

### RNA extraction and quantitative RT-PCR

Total RNA extraction and reverse transcription was performed as described. Quantitative PCR reactions using SsoFast EvaGreen Supermix (Bio-Rad) were performed in a CFX384 real time PCR system (Bio-Rad). Relative RNA expression was calculated through the ΔΔCt method and normalised to *Gapdh* expression. Data was analysed in Excel (Microsoft) and statistical analysis performed in GraphPad Prism v7.0c software (GraphPad Software Inc.). Primers used are listed in Table S6.

### Data analysis

Data is presented as mean ± standard error of the mean of at least three biological replicates. Statistical significance was estimated using ANOVA followed by Tukey’s post hoc test or t-student’s test and differences considered significant when p<0.05.

### Animal studies ethics

Mouse studies were approved by the University of Dundee ethical review committee, and further subjected to approved study plans by the Named Veterinary Surgeon and Compliance Officer (Dr. Ngaire Dennison) and performed under a UK Home Office project licence in accordance with the Animal Scientific Procedures Act (ASPA, 1986). C57BL/6J mice were housed in a SPF facility in temperature-controlled rooms at 21°C, with 45-65% relative humidity and 12-hour light/dark cycles. Mice had *ad libitum* access to food and water and regularly monitored by the School of Life Science Animal Unit Staff.

## ACKNOWLEDGEMENTS

We would like to thank Prof. Nathanael Gray and Dr. Tinghu Zhang (Dana-Farber Cancer Institute, Harvard Medical School) for providing SRPKIN-1, Prof. Helen Walden and Dr. Viduth Chaugule (University of Glasgow) for fluorescently-labelled ubiquitin, Prof. Ron Hay and Dr. Emma Branigan for recombinant UBE2D1, Prof. Miratul Muqit (MRC-PPU, School of Life Sciences, University of Dundee) for expertise in mouse neuronal culture and Prof. Vicky Cowling and Dr. Joana Silva (School of Life Sciences, University of Dundee) for mouse tissue extracts. We acknowledge the Findlay lab for comments and troubleshooting. G.M.F was supported by a Wellcome Trust/Royal Society Sir Henry Dale Fellowship (211209/Z/18/Z) and a Medical Research Council New Investigator Award (MR/N000609/1). A.S.F. was supported by an MRC-PPU prize studentship. G.N. and M.M. were supported by research grants ANID/FONDAP/15090007 and FONDECYT/11190998.

## AUTHOR CONTRIBUTIONS

F.B. and G.M.F. conceived the study and designed the experiments. F.B., A.S.-F., A.C., O.A., L.B., R.G., J.V. & G.M.F. performed experiments. G.N. and M.M. analysed data, provided expertise and prepared figures. T.M. and R.T. performed DNA cloning and CRISPR/Cas9 design. C.J.H. generated reagents. R.S. analysed mass-spectrometry data and provided expertise. F.B, and G.M.F. wrote the paper.

## FIGURE LEGENDS

**Figure S1.**
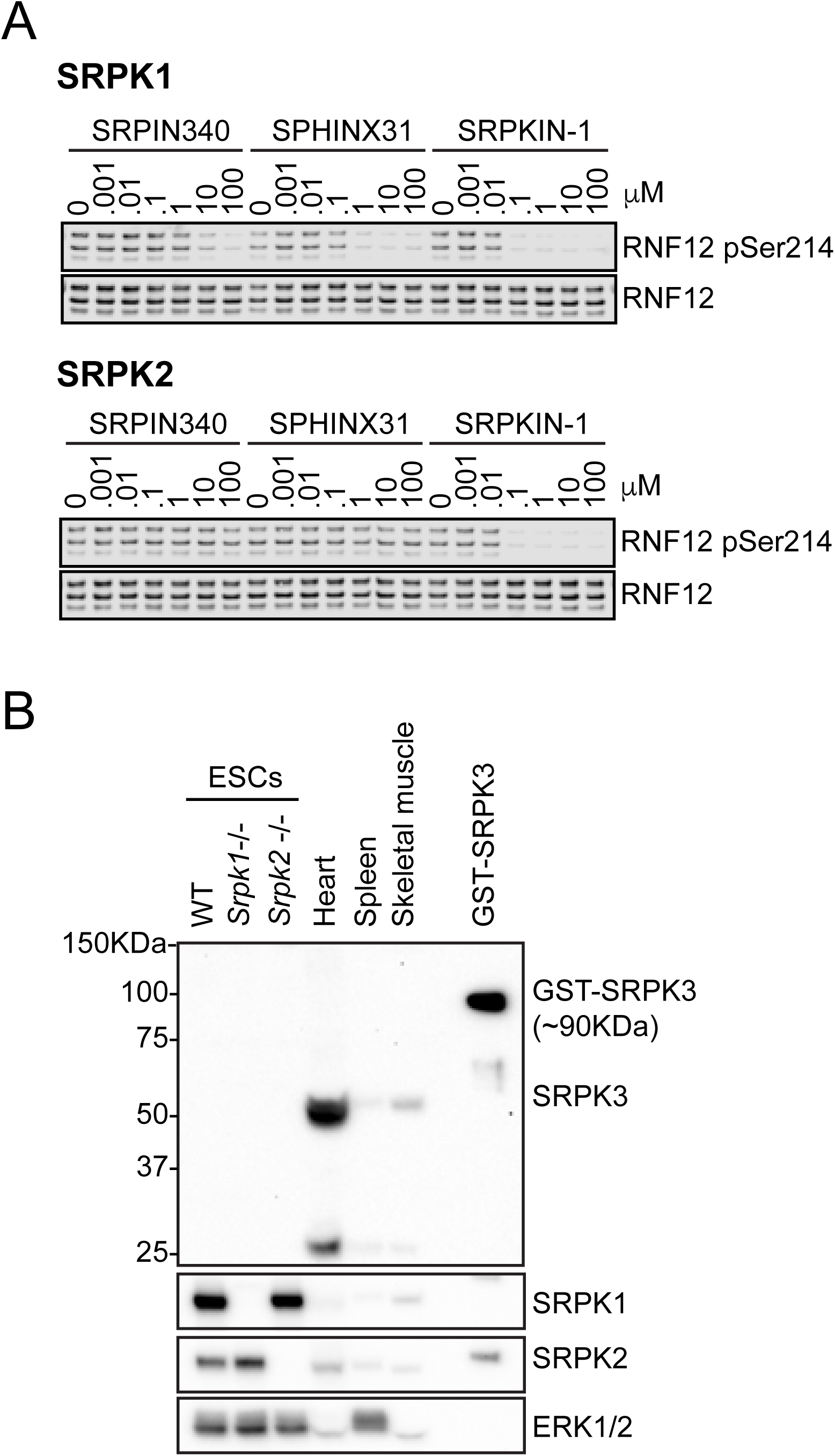
Differential inhibitor sensitivities and expression profiles of SRPK kinases (related to Figure 1). (A) Wild-type (WT), *Srpk1*^-/-^ and *Srpk2*^-/-^ mESCs were cultured and SRPK protein expression was analysed via immunoblotting. Heart, spleen and skeletal muscle tissue lysates and SRPK3 recombinant protein are shown as positive controls for SRPK3 expression. (B) Inhibition of RNF12 phosphorylation *in vitro* by SRPK1 and SRPK2 in the presence of varying concentrations of the indicated SRPK inhibitors was determined by immunoblotting for RNF12 phospho-Ser214. RNF12 levels are shown as a control

**Figure S2.**
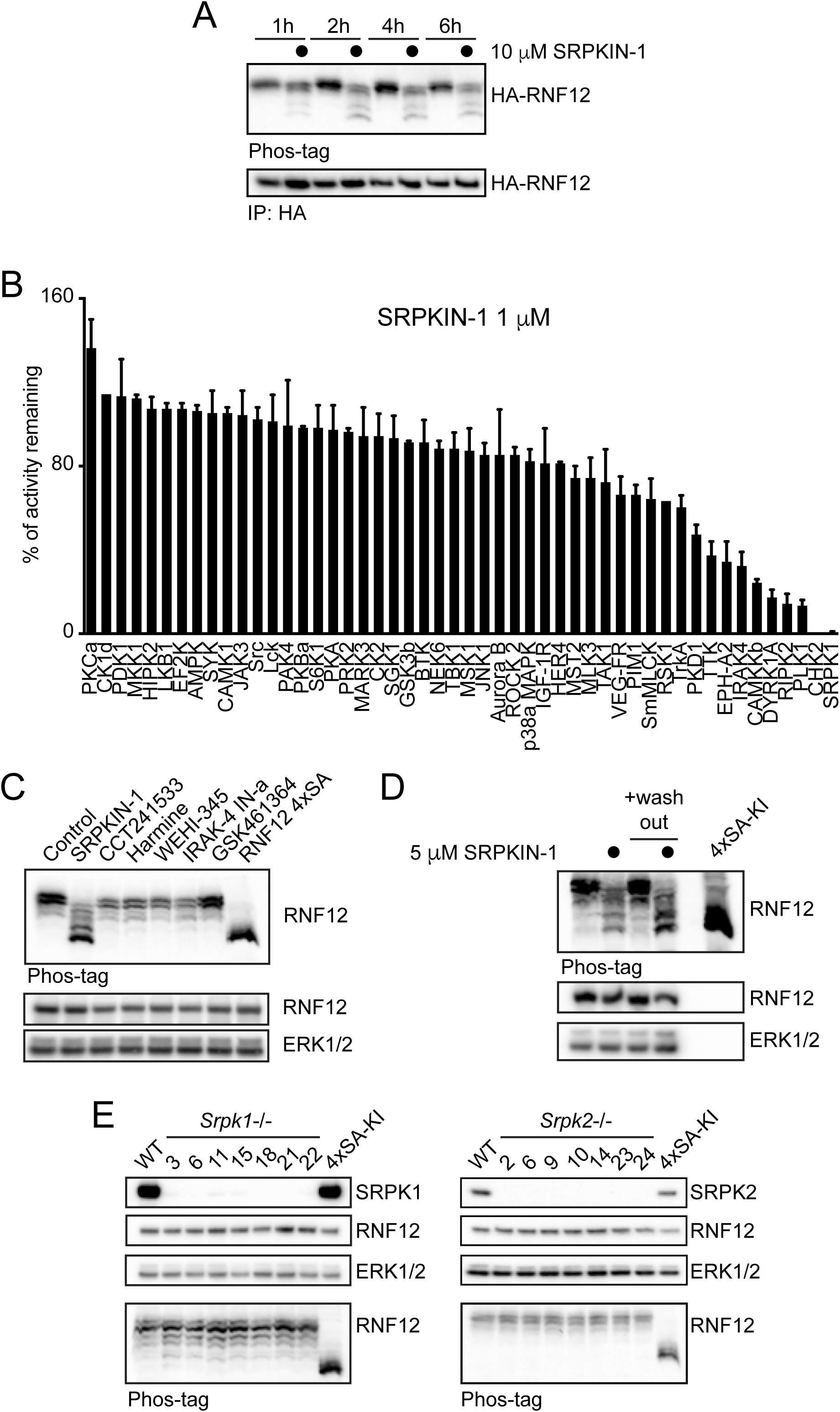
SRPK1/2 phosphorylate RNF12 SR-motif in mESCs (related to Figure 2). (A) HA-RNF12 expressing *Rlim*^-/y^ mESCs were treated with 10 µM SRPKIN-1 for the indicated times and pSR-motif phosphorylation of HA-immunoprecipitated RNF12 analysed via phos-tag immunoblotting. HA-RNF12 levels are shown as a control (B) SRPKIN-1 activity was profiled *in vitro* using 50 kinases (MRC-PPU International Centre for Kinase Profiling). Data are represented as mean ± S.D. (n=3). (C) RNF12 expressing mESCs were treated with 10 µM of the following inhibitors: SRPKIN-1 (SRPK inhibitor), CCT-241533 (CHK2 inhibitor), Harmine (DYRK1A inhibitor), WEHI-345 (RIPK2 inhibitor), IRAK-4-Inhibitor-a (IRAK4 inhibitor) and GSK-461364 (PLK1/2 inhibitor) for 4 h, and RNF12 phosphorylation analysed via phos-tag immunoblotting. HA-RNF12 and ERK1/2 levels are shown as a control. (D) RNF12 expressing mESCs were pre-treated with 5 µM SRPKIN-1 for 3 h, media changed and cells cultured for further 5 h (+ wash-out). RNF12 phosphorylation was analysed via Phos-tag gels. ERK1/2 levels are shown as a loading control. (E) Multiple *Srpk1*^-/-^ and *Srpk2*^-/-^ mESC clones were analysed for RNF12 phosphorylation via phos-tag immunoblotting. SRPK, RNF12 and ERK1/2 levels are shown as controls.

**Figure S3.**
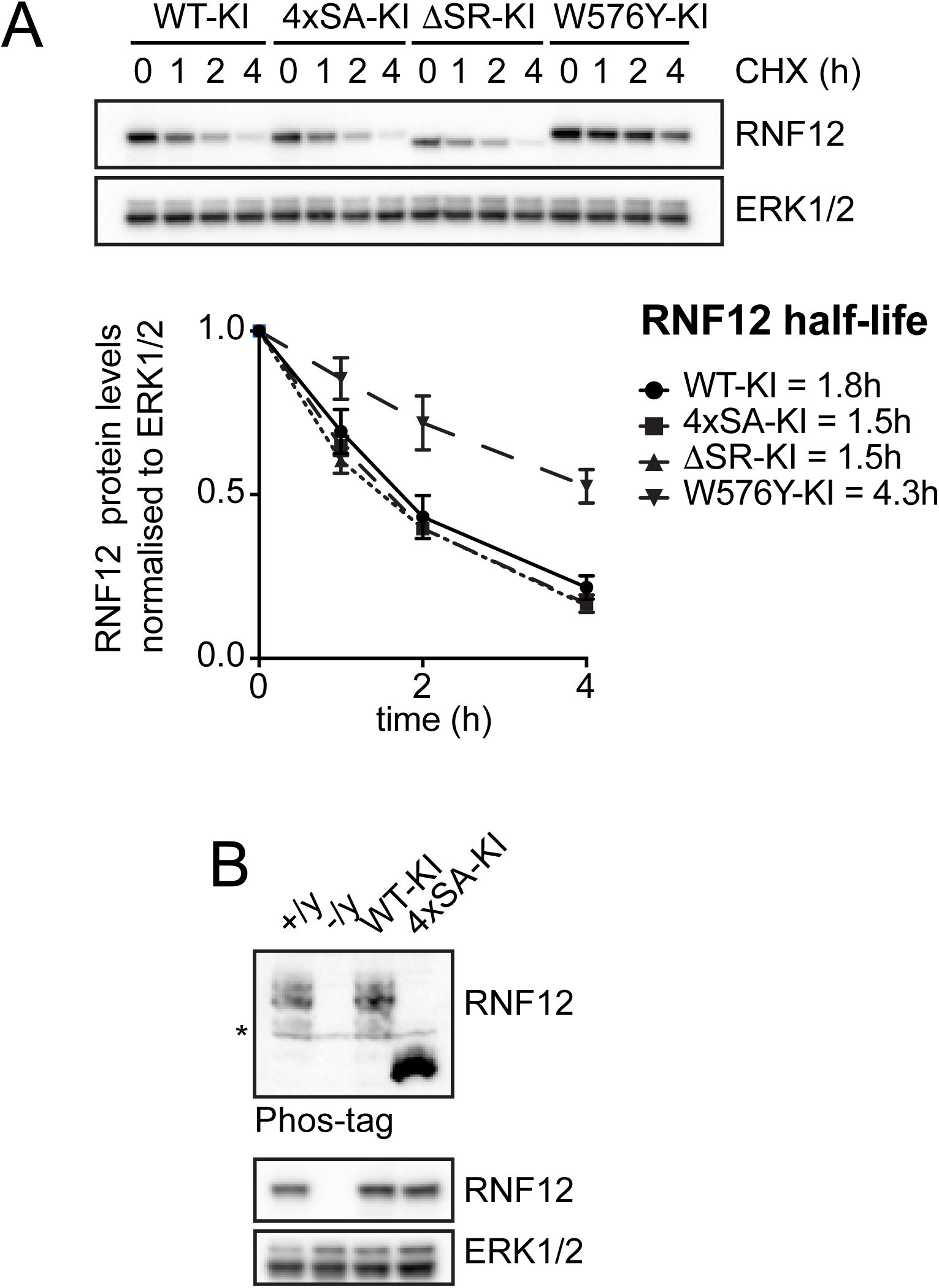
RNF12 protein stability is unaffected by SR-motif phosphorylation (related to Figure 3). (A) RNF12 knock-in mESC lines were treated with 350 µM cycloheximide for the indicated times and analysed for RNF12 levels via immunoblotting. ERK1/2 levels are shown as loading control (Top). Quantification of RNF12 signal intensity and determination of protein half-life via immunoblotting and non-linear curve fitting (Bottom). (B) Phos-tag immunoblot analysis of RNF12 SR-motif phosphorylation in the indicated mESC lines. RNF12 and ERK1/2 levels are shown as controls. (*) Indicates a non-specific band.

**Figure S4.**
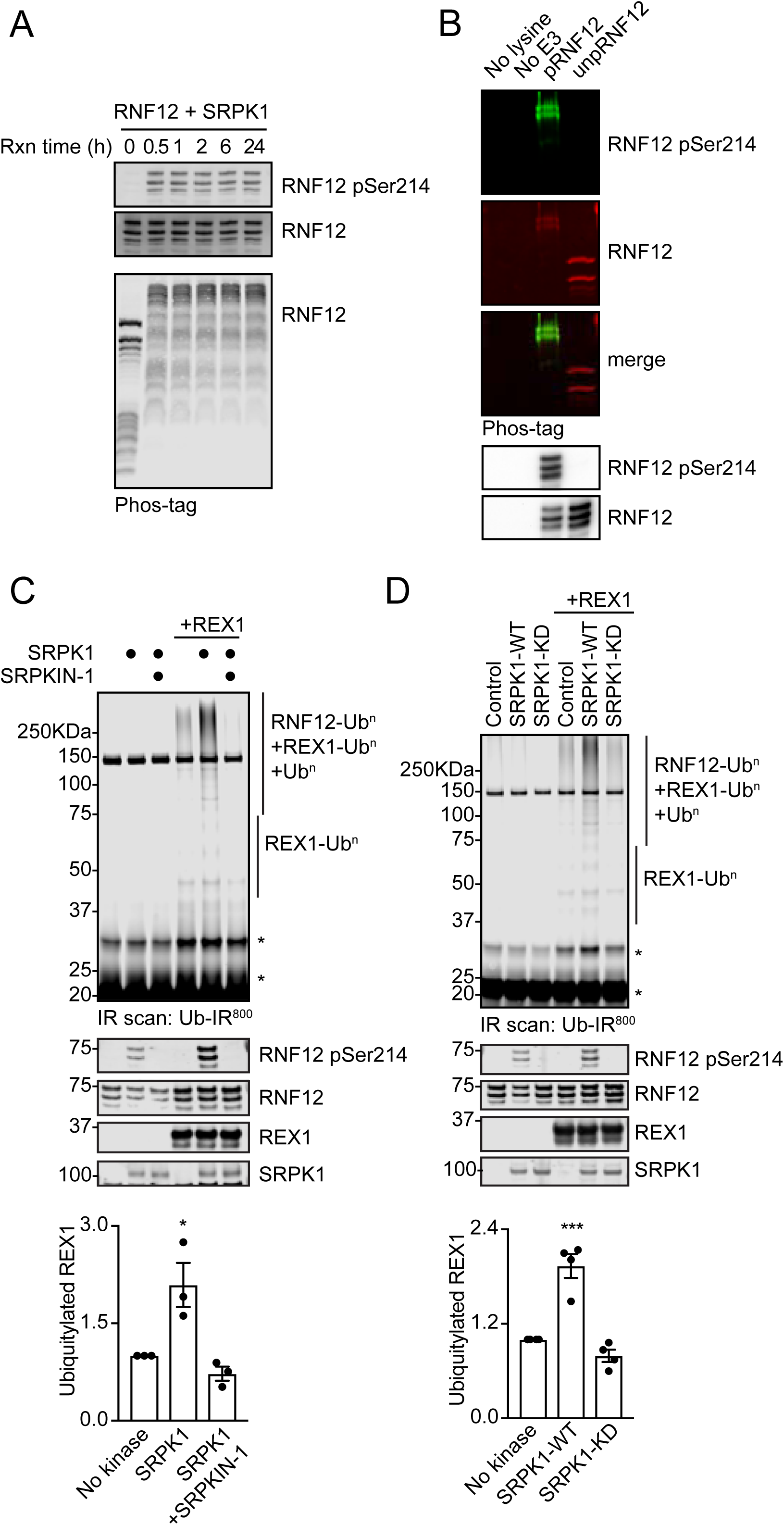
SRPK-mediated RNF12 SR-motif phosphorylation stimulates RNF12 E3 ubiquitin ligase activity (related to Figure 4). (A) Time course of RNF12 *in vitro* phosphorylation by SRPK1 analysed for SR-motif phosphorylation via pSer214 infrared and Phos-tag immunoblotting. RNF12 levels are shown as a control. (B) RNF12 *in vitro* phosphorylation by SRPK2 for 1 h was analysed via multiplex infrared Phos-tag and regular immunoblotting. This material is representative of samples used in E2 ubiquitin discharge assays displayed in Figure 4D. (C) Recombinant RNF12 was incubated with SRPK1 in absence or presence of SRPKIN-1 and subjected to REX1 fluorescent ubiquitylation assays. Infrared scans of ubiquitylated substrate signal and graphical quantifications are shown. One-way ANOVA followed by Tukey’s multiple comparisons test; confidence level 95%. (*) P=0.0221 (n=3). Phosphorylated and total RNF12, REX1 and SRPK2 infrared immunoblots are showed as controls. (D) Recombinant RNF12 was incubated with wild-type (WT) or kinase dead (KD) SRPK1 and subjected to REX1 fluorescent ubiquitylation assays. Infrared scans of ubiquitylated substrate signal and quantifications (right panel) are shown. One-way ANOVA followed by Tukey’s multiple comparisons test; confidence level 95%. (***) P=0.0002 (n=3). Phosphorylated and total RNF12, REX1 and SRPK1 infrared immunoblots are showed as controls.

**Figure S5.**
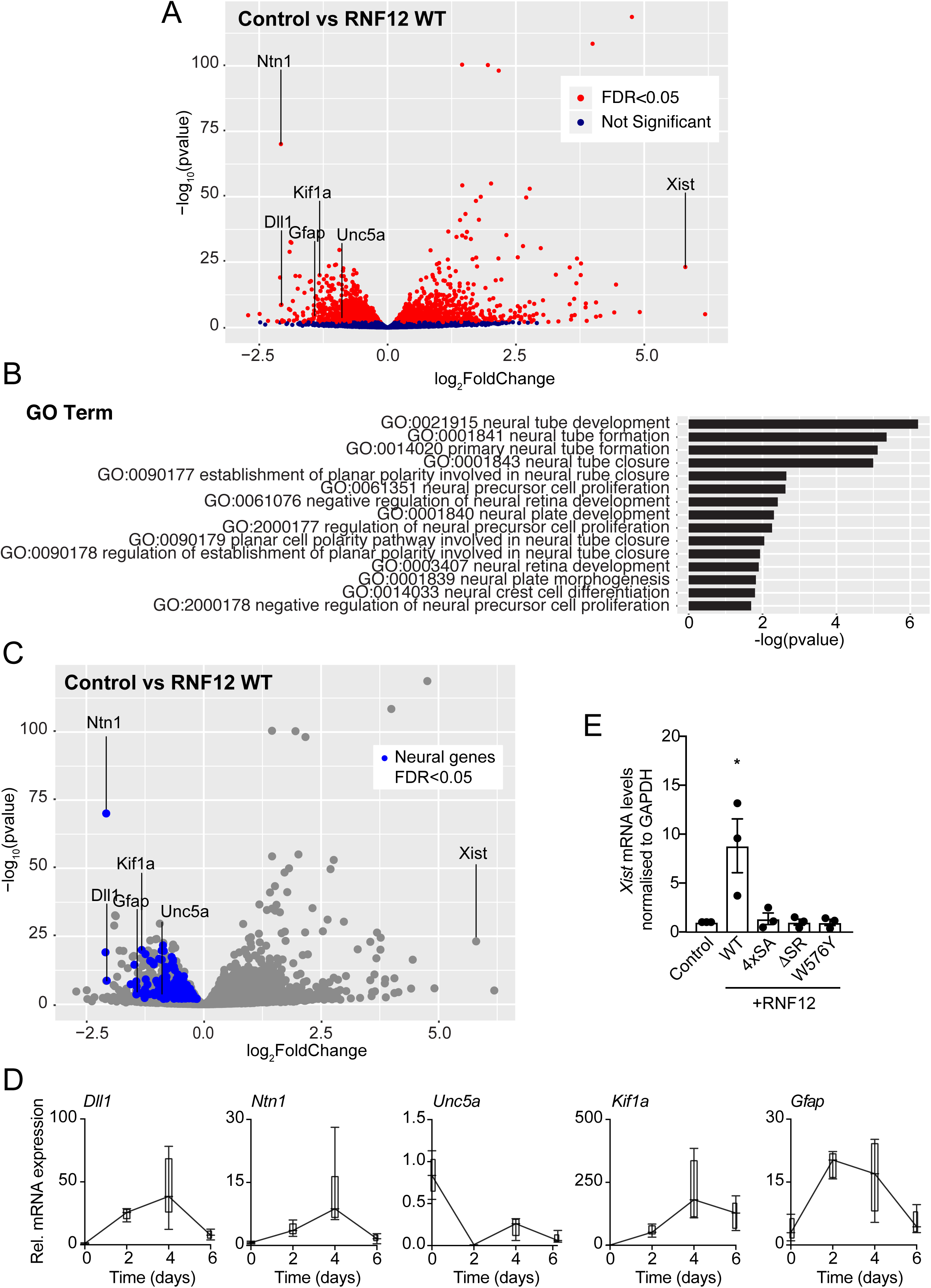
RNF12 negatively regulates neurodevelopmental gene expression in mESCs (related to Figure 5). (A) Volcano plot of an RNA-sequencing experiment comparing mRNA expression of *Rlim*^-/y^ mESCs transfected with control or wild-type (WT) RNF12. mRNAs whose expression is significantly altered by RNF12 are displayed in red. Key neurodevelopmental mRNAs that are inhibited by RNF12 E3 ubiquitin ligase activity are labelled (*Dll1, Ntn1, Gfap, Kif1a, Unc5a*). *Xist* is a known target of RNF12 activity. FDR = False discovery rate. (B) Gene Ontology analysis of RNF12 responsive genes identifies significant enrichment of genes related to neural development. (C) Volcano plot of an RNA-sequencing experiment comparing mRNA expression of *Rlim*^-/y^ mESC transfected with control or wild-type (WT) RNF12. Neurodevelopmental genes negatively regulated by RNF12 identified via Gene Ontology are indicated in blue. (D) Selected neurodevelopmental mRNA expression was analysed by quantitative RT-PCR following mESC neural differentiation using N2B27 media for the indicated times. *Gapdh* was used as housekeeping control. Data is represented as mean ± S.E.M. (n=3) (E) mESCs were transfected with the indicated vectors and cultured for 72 h prior analysis of *Xist* mRNA expression via quantitative RT-PCR. Data is represented as mean ± S.E.M. (n=3). One-way ANOVA followed by Tukey’s multiple comparisons test; confidence level 95%. (*) P=0.0292; (**) P=0.0041. *Gapdh* was used as a housekeeping control.

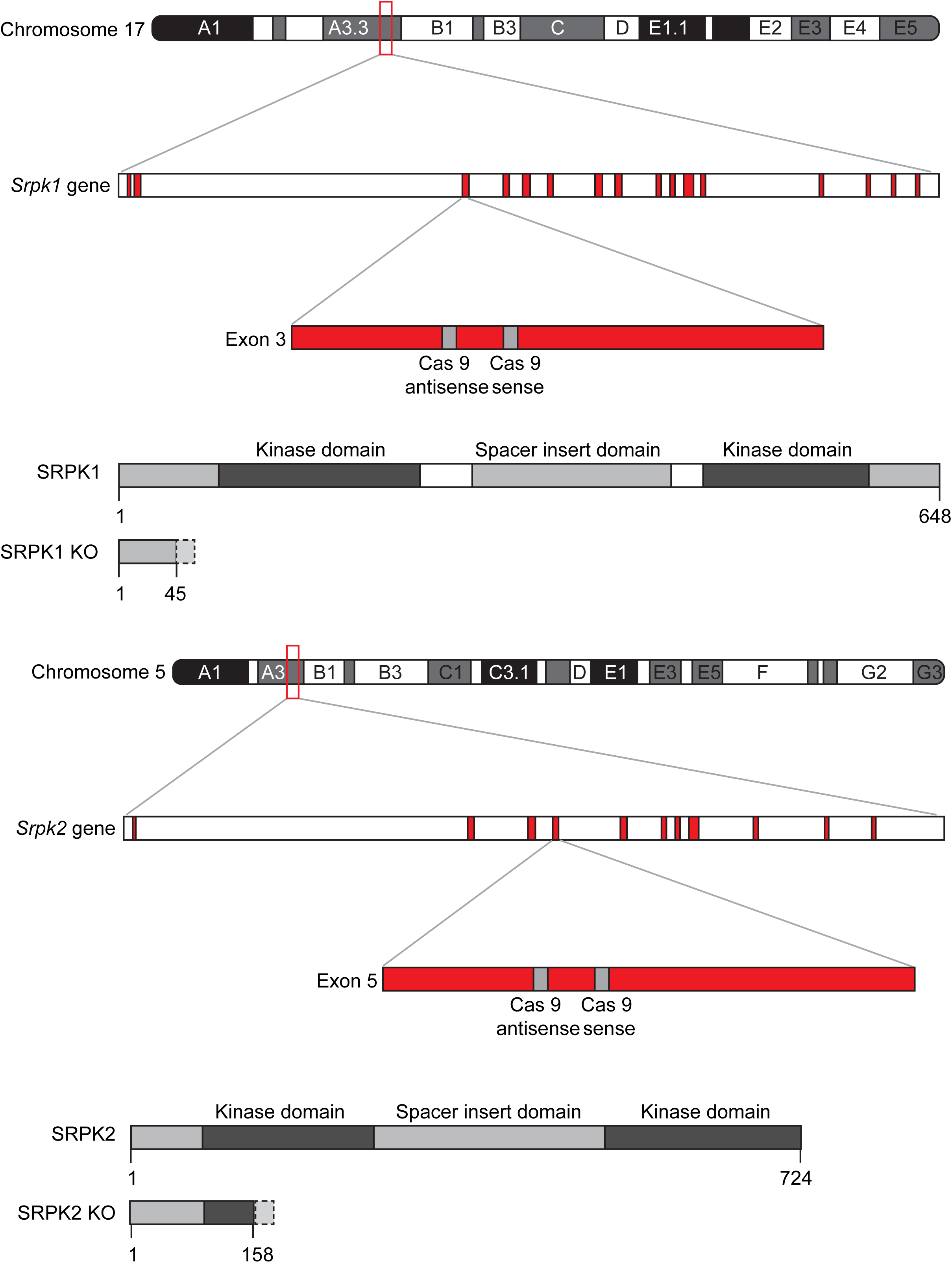

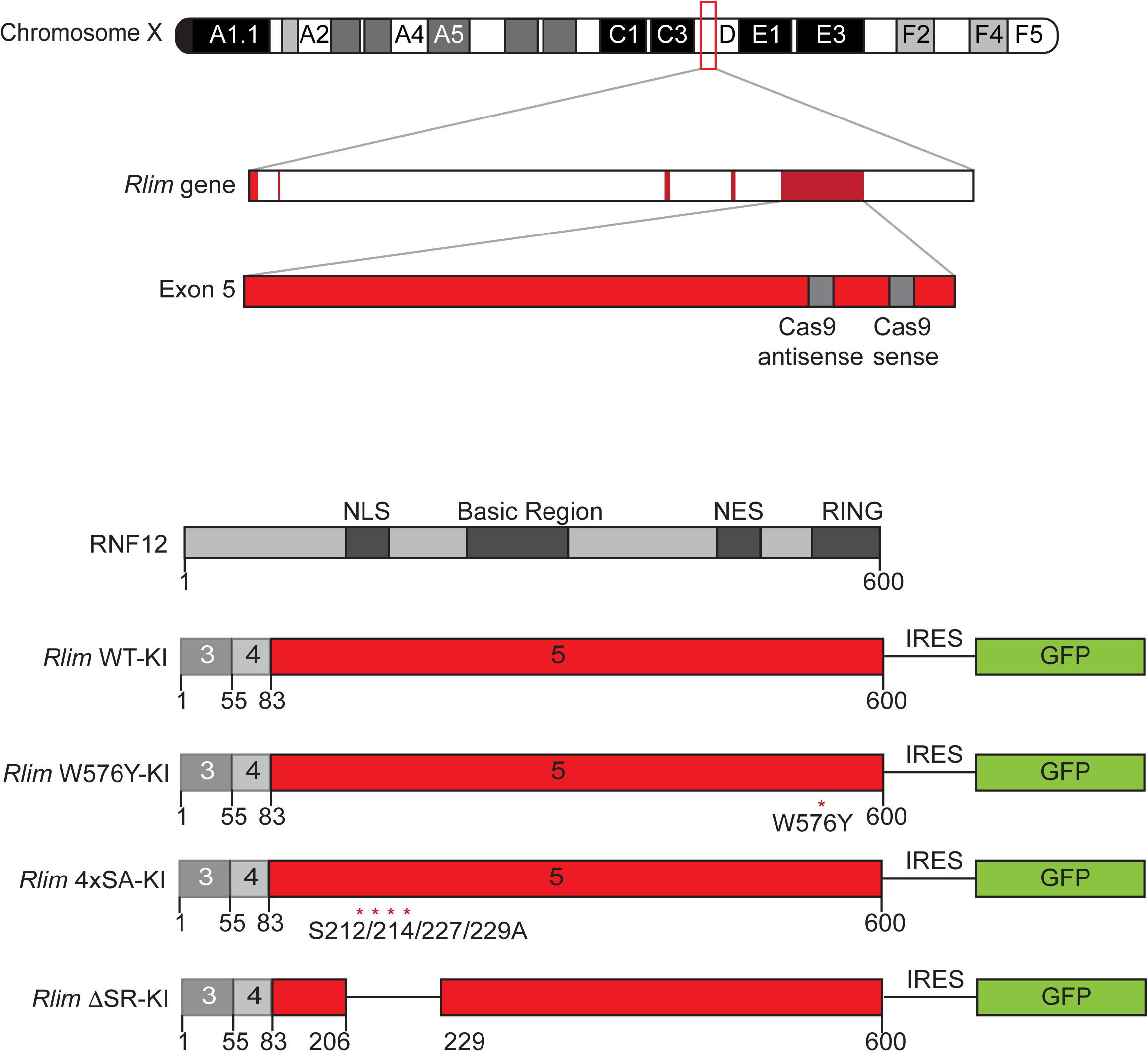

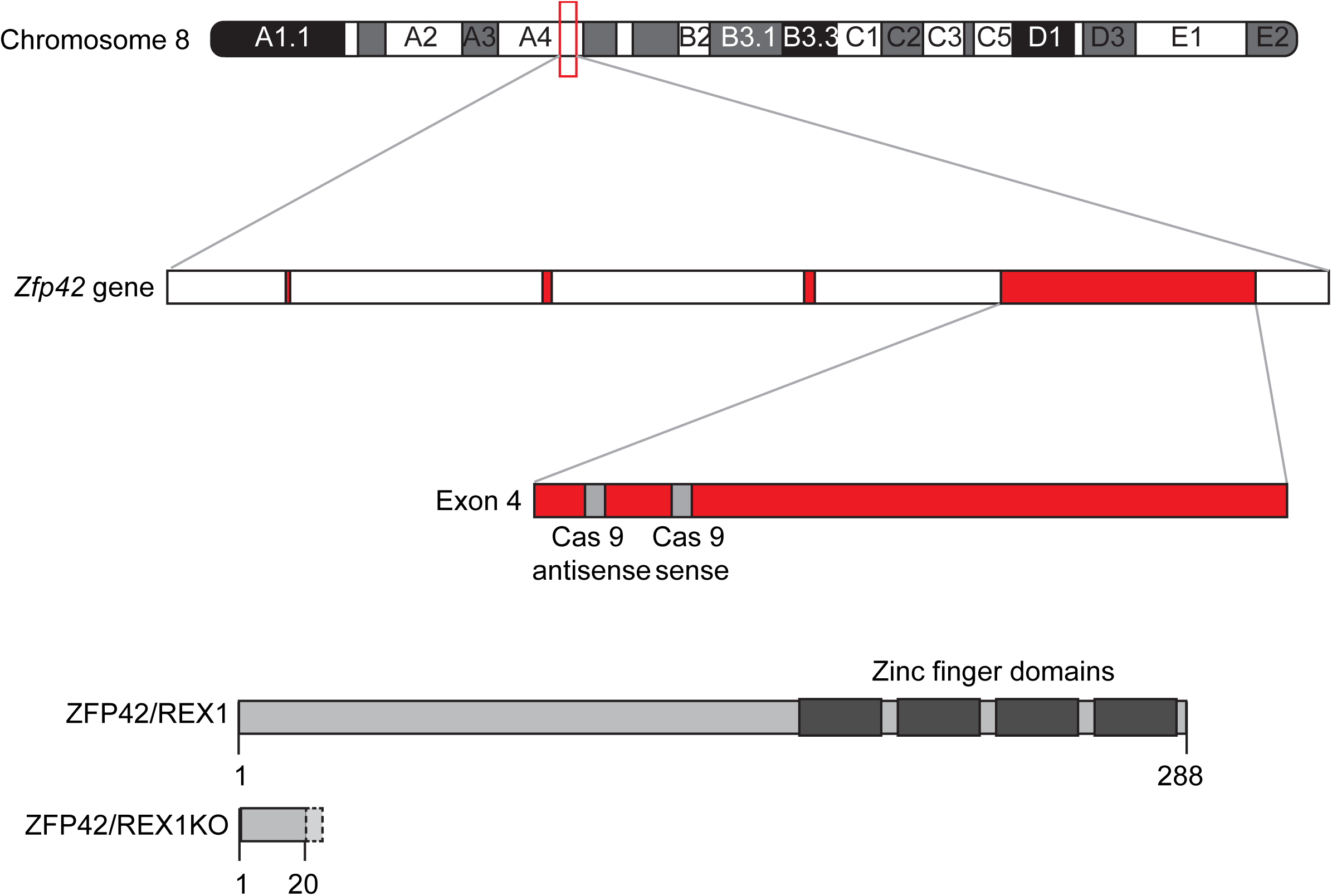

