## Supplemental Tables for "Functional diversification of Ser-Arg rich protein kinases to control ubiquitin-dependent neurodevelopmental signalling"

**Table S1. RSRS repeat-containing proteins functionally grouped**

| mRNA splicing | Other |
|---|---|
| Rbmx2 | Rbbp6 |
| Ccnl1 | Ndrg1 |
| Srsf2 | Rbm26 |
| Luc7l3 | Ppargc1a |
| Clk2 | Paf1 |
| Prpf38a | Arglu1 |
| Arl6ip4 | Scaf8 |
| Sfswap | Pdzd7 |
| Rsrc1 | Srrm3 |
| Cwc25 | Nktr |
| Rbm39 | Pprc1 |
| Srsf12 | Snrnp70 |
| Scaf4 | Rlim |
| Srek1 | Cherp |
| Pnn | Lbr |
| Clasrp | Nkap |
| Tra2b | Topors |
| Srsf5 | Rsrc2 |
| Thrap3 | Rsrp1 |
| Cactin | Sytl5 |
| Srrm1 | Erbb3 |
| Scaf1 | Bclaf1 |
| Srsf7 | Gpatch8 |
| Acin1 | Zc3h18 |
| Srsf6 | Luc7l |
| Znf638 | Spata18 |
| Prpf38b | Gtpbp4 |
| Ddx46 | Tjp2 |
| Srrm2 | Luc7l2 |
| Srsf4 | |
| Son | |
| Srsf1 | |
| U2af2 | |
| U2af1 | |
| Ppig | |
| Tra2a | |
| Ccnl2 | |
| Setd2 | |
| Cir1 | |
| Srsf3 | |
| Dhx8 | |
| Rnps1 | |
| Pnisr | |
| Cdk13 | |
| Snrnp27 | |
| Srsf10 | |
| Zranb2 | |
| Prpf4b | |

**Table S1. RSRS repeat-containing proteins identified by ScanProsite**

| Entry | Gene names (primary) | Protein names | Function [CC] |
|---|---|---|---|
| Q8R0F5 | Rbmx2 | RNA-binding motif protein, X-linked 2 | FUNCTION: Involved in pre-mRNA splicing as component of the activated spliceosome. {ECO:0000250\|UniProtKB:Q9Y388}. |
| Q52KE7 | Ccnl1 | Cyclin-L1 (Cyclin-L) (Cyclin Ania-6a) | FUNCTION: Involved in pre-mRNA splicing. Functions in association with cyclin-dependent kinases (CDKs). May play a role in the regulation of RNA polymerase II (pol II). Inhibited by the CDK-specific inhibitor CDKN1A/p21. {ECO:0000250\|UniProtKB:Q9UK58}. |
| P97868 | Rbbp6 | E3 ubiquitin-protein ligase RBBP6 (EC 2.3.2.27) (Proliferation potential-related protein) (Protein P2P-R) (RING-type E3 ubiquitin transferase RBBP6) (Retinoblastoma-binding protein 6) (p53-associated cellular protein of testis) | FUNCTION: E3 ubiquitin-protein ligase which promotes ubiquitination of YBX1, leading to its degradation by the proteasome (By similarity). May play a role as a scaffold protein to promote the assembly of the p53/TP53-MDM2 complex, resulting in increase of MDM2-mediated ubiquitination and degradation of p53/TP53; may function as negative regulator of p53/TP53, leading to both apoptosis and cell growth retardation (PubMed:17470788). Regulates DNA-replication and common fragile sites (CFS) stability in a ZBTB38- and MCM10-dependent manner. Controls ZBTB38 protein stability and abundance via ubiquitination and proteasomal degradation, and ZBTB38 in turn negatively regulates the expression of MCM10 which plays an important role in DNA-replication (PubMed:24726359). {ECO:0000250\|UniProtKB:Q7Z6E9, ECO:0000269\|PubMed:17470788, ECO:0000269\|PubMed:24726359}. |
| Q62433 | Ndrg1 | Protein NDRG1 (N-myc downstream-regulated gene 1 protein) (Protein Ndr1) | FUNCTION: Stress-responsive protein involved in hormone responses, cell growth, and differentiation. Acts as a tumor suppressor in many cell types. Necessary but not sufficient for p53/TP53-mediated caspase activation and apoptosis. Required for vesicular recycling of CDH1 and TF. May also function in lipid trafficking. Protects cells from spindle disruption damage. Functions in p53/TP53-dependent mitotic spindle checkpoint. Regulates microtubule dynamics and maintains euploidy (By similarity). Has a role in cell trafficking notably of the Schwann cell and is necessary for the maintenance and development of the peripheral nerve myelin sheath. {ECO:0000250, ECO:0000269\|PubMed:15082788, ECO:0000269\|PubMed:21303696}. |
| Q6NZN0 | Rbm26 | RNA-binding protein 26 (Protein expressed in male leptotene and zygotene spermatocytes 393) (MLZ-393) (RNA-binding motif protein 26) | |
| O70343 | Ppargc1a | Peroxisome proliferator-activated receptor gamma coactivator 1-alpha (PGC-1-alpha) (PPAR-gamma coactivator 1-alpha) (PPARGC-1-alpha) | FUNCTION: Transcriptional coactivator for steroid receptors and nuclear receptors. Greatly increases the transcriptional activity of PPARG and thyroid hormone receptor on the uncoupling protein promoter. Can regulate key mitochondrial genes that contribute to the program of adaptive thermogenesis. Plays an essential role in metabolic reprogramming in response to dietary availability through coordination of the expression of a wide array of genes involved in glucose and fatty acid metabolism. Induces the expression of PERM1 in the skeletal muscle in an ESRRA-dependent manner. Also involved in the integration of the circadian rhythms and energy metabolism. Required for oscillatory expression of clock genes, such as ARNTL/BMAL1 and NR1D1, through the coactivation of RORA and RORC, and metabolic genes, such as PDK4 and PEPCK. Isoform 4 specifically activates the expression of IGF1 and suppresses myostatin expression in skeletal muscle leading to muscle fiber hypertrophy. {ECO:0000269\|PubMed:15744310, ECO:0000269\|PubMed:17476214, ECO:0000269\|PubMed:23217713, ECO:0000269\|PubMed:9529258}. |
| Q8K2T8 | Paf1 | RNA polymerase II-associated factor 1 homolog | FUNCTION: Component of the PAF1 complex (PAF1C) which has multiple functions during transcription by RNA polymerase II and is implicated in regulation of development and maintenance of embryonic stem cell pluripotency. PAF1C associates with RNA polymerase II through interaction with POLR2A CTD non-phosphorylated and 'Ser-2'- and 'Ser-5'-phosphorylated forms and is involved in transcriptional elongation, acting both indepentently and synergistically with TCEA1 and in cooperation with the DSIF complex and HTATSF1. PAF1C is required for transcription of Hox and Wnt target genes. PAF1C is involved in hematopoiesis and stimulates transcriptional activity of KMT2A/MLL1. PAF1C is involved in histone modifications such as ubiquitination of histone H2B and methylation on histone H3 'Lys-4' (H3K4me3). PAF1C recruits the RNF20/40 E3 ubiquitin-protein ligase complex and the E2 enzyme UBE2A or UBE2B to chromatin which mediate monoubiquitination of 'Lys-120' of histone H2B (H2BK120ub1); UB2A/B-mediated H2B ubiquitination is proposed to be coupled to transcription. PAF1C is involved in mRNA 3' end formation probably through association with cleavage and poly(A) factors. Connects PAF1C with the RNF20/40 E3 ubiquitin-protein ligase complex. Involved in polyadenylation of mRNA precursors (By similarity). {ECO:0000250, ECO:0000269\|PubMed:19345177}. |
| Q62093 | Srsf2 | Serine/arginine-rich splicing factor 2 (Protein PR264) (Putative myelin regulatory factor 1) (MRF-1) (Splicing component, 35 kDa) (Splicing factor SC35) (SC-35) (Splicing factor, arginine/serine-rich 2) | FUNCTION: Necessary for the splicing of pre-mRNA. It is required for formation of the earliest ATP-dependent splicing complex and interacts with spliceosomal components bound to both the 5'- and 3'-splice sites during spliceosome assembly. It also is required for ATP-dependent interactions of both U1 and U2 snRNPs with pre-mRNA (By similarity). Can bind to the myelin basic protein (MBP) gene MB3 regulatory region and increase transcription of the mbp promoter in cells derived from the CNS. The phosphorylated form (by SRPK2) is required for cellular apoptosis in response to cisplatin treatment (By similarity). {ECO:0000250, ECO:0000269\|PubMed:7527040}. |
| Q5SUF2 | Luc7l3 | Luc7-like protein 3 (Cisplatin resistance-associated-overexpressed protein) | FUNCTION: Binds cAMP regulatory element DNA sequence. May play a role in RNA splicing (By similarity). {ECO:0000250}. |
| Q3UL36 | Arglu1 | Arginine and glutamate-rich protein 1 | |

| | | | |
|---|---|---|---|
| O35491 | Clk2 | Dual specificity protein kinase CLK2 (EC 2.7.12.1) (CDC-like kinase 2) | FUNCTION: Dual specificity kinase acting on both serine/threonine and tyrosine-containing substrates. Phosphorylates serine- and arginine-rich (SR) proteins of the spliceosomal complex. May be a constituent of a network of regulatory mechanisms that enable SR proteins to control RNA splicing and can cause redistribution of SR proteins from speckles to a diffuse nucleoplasmic distribution. Acts as a suppressor of hepatic gluconeogenesis and glucose output by repressing PPARGC1A transcriptional activity on gluconeogenic genes via its phosphorylation. Phosphorylates PPP2R5B thereby stimulating the assembly of PP2A phosphatase with the PPP2R5B-AKT1 complex leading to dephosphorylation of AKT1. Phosphorylates: PTPN1, SRSF1 and SRSF3. Regulates the alternative splicing of tissue factor (F3) pre-mRNA in endothelial cells. Phosphorylates PAGE4 at several serine and threonine residues and this phosphorylation attenuates the ability of PAGE4 to potentiate the transcriptional activator activity of JUN (By similarity). {ECO:0000250\|UniProtKB:P49760, ECO:0000269\|PubMed:20074525, ECO:0000269\|PubMed:21329884, ECO:0000269\|PubMed:9307018}. |
| Q6DID3 | Scaf8 | SR-related and CTD-associated factor 8 (RNA-binding motif protein 16) | FUNCTION: Anti-terminator protein required to prevent early mRNA termination during transcription. Together with SCAF4, acts by suppressing the use of early, alternative poly(A) sites, thereby preventing the accumulation of non-functional truncated proteins. Mechanistically, associates with the phosphorylated C-terminal heptapeptide repeat domain (CTD) of the largest RNA polymerase II subunit (POLR2A), and subsequently binds nascent RNA upstream of early polyadenylation sites to prevent premature mRNA transcript cleavage and polyadenylation. Independently of SCAF4, also acts as a positive regulator of transcript elongation. {ECO:0000250\|UniProtKB:Q9UPN6}. |
| E9Q9W7 | Pdzd7 | PDZ domain-containing protein 7 | FUNCTION: In cochlear developing hair cells, essential in organizing the USH2 complex at stereocilia ankle links (PubMed:24334608). Blocks inhibition of adenylate cyclase activity mediated by ADGRV1 (PubMed:24962568). {ECO:0000269\|PubMed:24334608, ECO:0000269\|PubMed:24962568}. |
| Q80WV7 | Srrm3 | Serine/arginine repetitive matrix protein 3 | FUNCTION: May play a role in regulating breast cancer cell invasiveness. May be involved in RYBP-mediated breast cancer progression. {ECO:0000250\|UniProtKB:A6NNA2}. |
| P30415 | Nktr | NK-tumor recognition protein (NK-TR protein) (Natural-killer cells cyclophilin-related protein) (Peptidyl-prolyl cis-trans isomerase NKTR) (PPlase) (EC 5.2.1.8) | FUNCTION: PPIase that catalyzes the cis-trans isomerization of proline imidic peptide bonds in oligopeptides and may therefore assist protein folding. Component of a putative tumor-recognition complex involved in the function of NK cells. {ECO:0000250\|UniProtKB:P30414}. |
| Q4FK66 | Prpf38a | Pre-mRNA-splicing factor 38A | FUNCTION: Involved in pre-mRNA splicing as a component of the spliceosome. {ECO:0000250\|UniProtKB:Q8NAV1}. |
| Q9JM93 | Arl6ip4 | ADP-ribosylation factor-like protein 6-interacting protein 4 (ARL-6-interacting protein 4) (Aip-4) (Splicing factor SRrp37) | FUNCTION: Involved in modulating alternative pre-mRNA splicing with either 5' distal site activation or preferential use of 3' proximal site. {ECO:0000250}. |
| Q3USH5 | Sfswap | Splicing factor, suppressor of white-apricot homolog (Splicing factor, arginine/serine-rich 8) (Suppressor of white apricot protein homolog) | FUNCTION: Plays a role as an alternative splicing regulator. Regulates its own expression at the level of RNA processing. Also regulates the splicing of fibronectin and CD45 genes. May act, at least in part, by interaction with other R/S-containing splicing factors. Represses the splicing of MAPT/Tau exon 10 (By similarity). {ECO:0000250}. |
| Q6NZN1 | Pprc1 | Peroxisome proliferator-activated receptor gamma coactivator-related protein 1 (PGC-1-related coactivator) (PRC) | FUNCTION: Acts as a coactivator during transcriptional activation of nuclear genes related to mitochondrial biogenesis and cell growth. Involved in the transcription coactivation of CREB and NRF1 target genes (By similarity). {ECO:0000250}. |
| Q62376 | Snrnp70 | U1 small nuclear ribonucleoprotein 70 kDa (U1 snRNP 70 kDa) (U1-70K) (snRNP70) | FUNCTION: Component of the spliceosomal U1 snRNP, which is essential for recognition of the pre-mRNA 5' splice-site and the subsequent assembly of the spliceosome. SNRNP70 binds to the loop I region of U1-snRNA. {ECO:0000250\|UniProtKB:P08621}.; FUNCTION: [Isoform 2]: Truncated isoforms that lack the RRM domain cannot bind U1-snRNA. {ECO:0000250\|UniProtKB:P08621}. |
| Q9DBU6 | Rsrc1 | Serine/Arginine-related protein 53 (SRrp53) (Arginine/serine-rich coiled-coil protein 1) | FUNCTION: Plays a role in pre-mRNA splicing. Involved in both constitutive and alternative pre-mRNA splicing. May have a role in the recognition of the 3' splice site during the second step of splicing (By similarity). {ECO:0000250}. |
| Q9DBF7 | Cwc25 | Pre-mRNA-splicing factor CWC25 homolog (Coiled-coil domain-containing protein 49) (Spliceosome-associated protein homolog CWC25) | FUNCTION: Involved in pre-mRNA splicing as component of the spliceosome. {ECO:0000250\|UniProtKB:Q9NXE8}. |
| Q8VH51 | Rbm39 | RNA-binding protein 39 (Coactivator of activating protein 1 and estrogen receptors) (Coactivator of AP-1 and ERs) (RNA-binding motif protein 39) (RNA-binding region-containing protein 2) (Transcription coactivator CAPER) | FUNCTION: Transcriptional coactivator for steroid nuclear receptors ESR1/ER-alpha and ESR2/ER-beta, and JUN/AP-1. May be involved in pre-mRNA splicing process. {ECO:0000269\|PubMed:11704680}. |
| Q8C8K3 | Srsf12 | Serine/arginine-rich splicing factor 12 (Splicing factor, arginine/serine-rich 13B) | FUNCTION: Splicing factor that seems to antagonize SR proteins in pre-mRNA splicing regulation. {ECO:0000250}. |

| | | | |
|---|---|---|---|
| Q7TSH6 | Scaf4 | SR-related and CTD-associated factor 4 (CTD-binding SR-like protein RA4) (Splicing factor, arginine/serine-rich 15) | FUNCTION: Anti-terminator protein required to prevent early mRNA termination during transcription. Together with SCAF8, acts by suppressing the use of early, alternative poly(A) sites, thereby preventing the accumulation of non-functional truncated proteins. Mechanistically, associates with the phosphorylated C-terminal heptapeptide repeat domain (CTD) of the largest RNA polymerase II subunit (POLR2A), and subsequently binds nascent RNA upstream of early polyadenylation sites to prevent premature mRNA transcript cleavage and polyadenylation. Independently of SCAF8, also acts as a suppressor of transcriptional readthrough. {ECO:0000250\|UniProtKB:O95104}. |
| Q9WTV7 | Rlim | E3 ubiquitin-protein ligase RLIM (EC 2.3.2.27) (LIM domain-interacting RING finger protein) (RING finger LIM domain-binding protein) (R-LIM) (RING finger protein 12) (RING-type E3 ubiquitin transferase RLIM) | FUNCTION: E3 ubiquitin-protein ligase that acts as a negative coregulator for LIM homeodomain transcription factors by mediating the ubiquitination and subsequent degradation of LIM cofactors LDB1 and LDB2 and by mediating the recruitment the SIN3a/histone deacetylase corepressor complex. Ubiquitination and degradation of LIM cofactors LDB1 and LDB2 allows DNA-bound LIM homeodomain transcription factors to interact with other protein partners such as RLIM. Plays a role in telomere length-mediated growth suppression by mediating the ubiquitination and degradation of TERF1. By targeting ZFP42 for degradation, acts as an activator of random inactivation of X chromosome in the embryo, a stochastic process in which one X chromosome is inactivated to minimize sex-related dosage differences of X-encoded genes in somatic cells of female placental mammals. {ECO:0000269\|PubMed:10431247, ECO:0000269\|PubMed:11882901, ECO:0000269\|PubMed:19945382, ECO:0000269\|PubMed:22596162}. |
| Q8BZX4 | Srek1 | Splicing regulatory glutamine/lysine-rich protein 1 (Serine/arginine-rich-splicing regulatory protein 86) (SRrp86) (Splicing factor, arginine/serine-rich 12) | FUNCTION: Participates in the regulation of alternative splicing by modulating the activity of other splice facors. Inhibits the splicing activity of SFRS1, SFRS2 and SFRS6. Augments the splicing activity of SFRS3 (By similarity). {ECO:0000250}. |
| O35691 | Pnn | Pinin | FUNCTION: Transcriptional activator binding to the E-box 1 core sequence of the E-cadherin promoter gene; the core-binding sequence is 5'CAGGTG-3'. Capable of reversing CTBP1-mediated transcription repression. Auxiliary component of the splicing-dependent multiprotein exon junction complex (EJC) deposited at splice junction on mRNAs. The EJC is a dynamic structure consisting of core proteins and several peripheral nuclear and cytoplasmic associated factors that join the complex only transiently either during EJC assembly or during subsequent mRNA metabolism. Participates in the regulation of alternative pre-mRNA splicing. Associates to spliced mRNA within 60 nt upstream of the 5'-splice sites. Component of the PSAP complex which binds RNA in a sequence-independent manner and is proposed to be recruited to the EJC prior to or during the splicing process and to regulate specific excision of introns in specific transcription subsets. Involved in the establishment and maintenance of epithelia cell-cell adhesion (By similarity). {ECO:0000250}. |
| Q8CFC7 | Clasrp | CLK4-associating serine/arginine rich protein (Clk4-associating SR-related protein) (Serine/arginine-rich splicing factor 16) (Splicing factor, arginine/serine-rich 16) (Suppressor of white-apricot homolog 2) | FUNCTION: Probably functions as an alternative splicing regulator. May regulate the mRNA splicing of genes such as CLK1. May act by regulating members of the CLK kinase family. |
| Q8CGZ0 | Cherp | Calcium homeostasis endoplasmic reticulum protein (SR-related CTD-associated factor 6) | FUNCTION: Involved in calcium homeostasis, growth and proliferation. {ECO:0000250\|UniProtKB:Q8IWX8}. |
| Q3U9G9 | Lbr | Delta(14)-sterol reductase LBR (Delta-14-SR) (EC 1.3.1.70) (3-beta-hydroxysterol Delta (14)-reductase) (C-14 sterol reductase) (C14SR) (Integral nuclear envelope inner membrane protein) (Lamin-B receptor) (Sterol C14-reductase) | FUNCTION: Catalyzes the reduction of the C14-unsaturated bond of lanosterol, as part of the metabolic pathway leading to cholesterol biosynthesis (PubMed:18785926). Plays a critical role in myeloid cell cholesterol biosynthesis which is essential to both myeloid cell growth and functional maturation (PubMed:22140257). Mediates the activation of NADPH oxidases, perhaps by maintaining critical levels of cholesterol required for membrane lipid raft formation during neutrophil differentiation (PubMed:22140257). Anchors the lamina and the heterochromatin to the inner nuclear membrane (By similarity). {ECO:0000250\|UniProtKB:Q14739, ECO:0000269\|PubMed:18785926, ECO:0000269\|PubMed:22140257}. |
| P62996 | Tra2b | Transformer-2 protein homolog beta (TRA-2 beta) (TRA2-beta) (Silica-induced gene 41 protein) (SIG-41) (Splicing factor, arginine/serine-rich 10) (Transformer-2 protein homolog B) | FUNCTION: Sequence-specific RNA-binding protein which participates in the control of pre-mRNA splicing. Can either activate or suppress exon inclusion. Acts additively with RBMX to promote exon 7 inclusion of the survival motor neuron SMN2. Activates the splicing of MAPT/Tau exon 10. Alters pre-mRNA splicing patterns by antagonizing the effects of splicing regulators, like RBMX. Binds to the AG-rich SE2 domain in the SMN exon 7 RNA. Binds to pre-mRNA (By similarity). {ECO:0000250}. |
| O35326 | Srsf5 | Serine/arginine-rich splicing factor 5 (Delayed-early protein HRS) (Pre-mRNA-splicing factor SRP40) (Splicing factor, arginine/serine-rich 5) | FUNCTION: May be required for progression through G1 and entry into S phase of cell growth. May play a regulatory role in pre-mRNA splicing. Autoregulates its own expression. Plays a role in constitutive splicing and can modulate the selection of alternative splice sites (By similarity). {ECO:0000250}. |

| | | | |
|---|---|---|---|
| Q569Z6 | Thrap3 | Thyroid hormone receptor-associated protein 3 (Thyroid hormone receptor-associated protein complex 150 kDa component) (Trap150) | FUNCTION: Involved in pre-mRNA splicing. Remains associated with spliced mRNA after splicing which probably involves interactions with the exon junction complex (EJC). Can trigger mRNA decay which seems to be independent of nonsense-mediated decay involving premature stop codons (PTC) recognition. May be involved in nuclear mRNA decay. Involved in regulation of signal-induced alternative splicing. During splicing of PTPRC/CD45 is proposed to sequester phosphorylated SFPQ from PTPRC/CD45 pre-mRNA in resting T-cells. Involved in cyclin-D1/CCND1 mRNA stability probably by acting as component of the SNARP complex which associates with both the 3'end of the CCND1 gene and its mRNA. Involved in response to DNA damage. Is excluced from DNA damage sites in a manner that parallels transcription inhibition; the function may involve the SNARP complex. Initially thought to play a role in transcriptional coactivation through its association with the TRAP complex; however, it is not regarded as a stable Mediator complex subunit. Cooperatively with HELZ2, enhances the transcriptional activation mediated by PPARG, maybe through the stabilization of the PPARG binding to DNA in presence of ligand. May play a role in the terminal stage of adipocyte differentiation. Plays a role in the positive regulation of the circadian clock. Acts as a coactivator of the CLOCK-ARNTL/BMAL1 heterodimer and promotes its transcriptional activator activity and binding to circadian target genes (PubMed:24043798). {ECO:0000269\|PubMed:23525231, ECO:0000269\|PubMed:24043798}. |
| Q9D0F4 | Nkap | NF-kappa-B-activating protein | FUNCTION: Acts as a transcriptional repressor. Plays a role as a transcriptional corepressor of the Notch-mediated signaling required for T-cell development. Also involved in the TNF and IL-1 induced NF-kappa-B activation. Associates with chromatin at the Notch-regulated SKP2 promoter (By similarity). {ECO:0000250}. |
| Q80Z37 | Topors | E3 ubiquitin-protein ligase Topors (EC 2.3.2.27) (RING-type E3 ubiquitin transferase Topors) (SUMO1-protein E3 ligase Topors) (Topoisomerase I-binding RING finger protein) (Topoisomerase I-binding arginine/serine-rich protein) (Tumor suppressor p53-binding protein 3) (p53-binding protein 3) (p53BP3) | FUNCTION: Functions as an E3 ubiquitin-protein ligase and as a E3 SUMO1-protein ligase. Probable tumor suppressor involved in cell growth, cell proliferation and apoptosis that regulates p53/TP53 stability through ubiquitin-dependent degradation. May regulate chromatin modification through sumoylation of several chromatin modification-associated proteins. May be involved in DNA-damage-induced cell death through IKBKE sumoylation. {ECO:0000269\|PubMed:15703819, ECO:0000269\|PubMed:15735665}. |
| A2RTL5 | Rsrc2 | Arginine/serine-rich coiled-coil protein 2 | |
| Q3UC65 | Rsrp1 | Arginine/serine-rich protein 1 | |
| Q80T23 | Sytl5 | Synaptotagmin-like protein 5 | FUNCTION: May act as Rab effector protein and play a role in vesicle trafficking. Binds phospholipids (By similarity). {ECO:0000250}. |
| Q61526 | Erbb3 | Receptor tyrosine-protein kinase erbB-3 (EC 2.7.10.1) (Glial growth factor receptor) (Proto-oncogene-like protein c-ErbB-3) | FUNCTION: Tyrosine-protein kinase that plays an essential role as cell surface receptor for neuregulins. Binds to neuregulin-1 (NRG1) and is activated by it; ligand-binding increases phosphorylation on tyrosine residues and promotes its association with the p85 subunit of phosphatidylinositol 3-kinase. May also be activated by CSPG5. Involved in the regulation of myeloid cell differentiation. {ECO:0000250\|UniProtKB:P21860}. |
| Q9CS00 | Cactin | Cactin | FUNCTION: Involved in the regulation of innate immune response. Acts as negative regulator of Toll-like receptor and interferon-regulatory factor (IRF) signaling pathways. Contributes to the regulation of transcriptional activation of NF-kappa-B target genes in response to endogenous proinflammatory stimuli. May play a role during early embryonic development. Probably involved in pre-mRNA splicing (By similarity). {ECO:0000250}. |
| Q52KI8 | Srrm1 | Serine/arginine repetitive matrix protein 1 (Plenty-of-prolines 101) | FUNCTION: Part of pre- and post-splicing multiprotein mRNP complexes. Involved in numerous pre-mRNA processing events. Promotes constitutive and exonic splicing enhancer (ESE)-dependent splicing activation by bridging together sequence-specific (SR family proteins, SFRS4, SFRS5 and TRA2B/SFRS10) and basal snRNP (SNRP70 and SNRPA1) factors of the spliceosome. Stimulates mRNA 3'-end cleavage independently of the formation of an exon junction complex. Binds both pre-mRNA and spliced mRNA 20-25 nt upstream of exon-exon junctions. Binds RNA and DNA with low sequence specificity and has similar preference for either double- or single-stranded nucleic acid substrates. {ECO:0000250\|UniProtKB:Q8IYB3}. |
| Q8K019 | Bclaf1 | Bcl-2-associated transcription factor 1 (Btf) | FUNCTION: Death-promoting transcriptional repressor. May be involved in cyclin-D1/CCND1 mRNA stability through the SNARP complex which associates with both the 3'end of the CCND1 gene and its mRNA (By similarity). {ECO:0000250}. |
| Q5U4C3 | Scaf1 | Splicing factor, arginine/serine-rich 19 (SR-related and CTD-associated factor 1) | FUNCTION: May function in pre-mRNA splicing. {ECO:0000250}. |
| Q8BL97 | Srsf7 | Serine/arginine-rich splicing factor 7 (Splicing factor, arginine/serine-rich 7) | FUNCTION: Required for pre-mRNA splicing. Represses the splicing of MAPT/Tau exon 10. May function as export adapter involved in mRNA nuclear export such as of histone H2A. Binds mRNA which is thought to be transferred to the NXF1-NXT1 heterodimer for export (TAP/NXF1 pathway); enhances NXF1-NXT1 RNA-binding activity. RNA-binding is semi-sequence specific (By similarity). {ECO:0000250}. |
| A2A6A1 | Gpatch8 | G patch domain-containing protein 8 | |
| Q0P678 | Zc3h18 | Zinc finger CCCH domain-containing protein 18 (Nuclear protein NHN1) | |

| | | | |
|---|---|---|---|
| Q9JIX8 | Acin1 | Apoptotic chromatin condensation inducer in the nucleus (Acinus) | FUNCTION: Auxiliary component of the splicing-dependent multiprotein exon junction complex (EJC) deposited at splice junction on mRNAs. The EJC is a dynamic structure consisting of core proteins and several peripheral nuclear and cytoplasmic associated factors that join the complex only transiently either during EJC assembly or during subsequent mRNA metabolism. Component of the ASAP complexes which bind RNA in a sequence-independent manner and are proposed to be recruited to the EJC prior to or during the splicing process and to regulate specific excision of introns in specific transcription subsets; ACIN1 confers RNA-binding to the complex. The ASAP complex can inhibit RNA processing during in vitro splicing reactions. The ASAP complex promotes apoptosis and is disassembled after induction of apoptosis. Involved in the splicing modulation of BCL2L1/Bcl-X (and probably other apoptotic genes); specifically inhibits formation of proapoptotic isoforms such as Bcl-X(S); the activity is different from the established EJC assembly and function. Induces apoptotic chromatin condensation after activation by CASP3. Regulates cyclin A1, but not cyclin A2, expression in leukemia cells (By similarity). {ECO:0000250}. |
| Q3TWW8 | Srsf6 | Serine/arginine-rich splicing factor 6 (Pre-mRNA-splicing factor SRP55) (Splicing factor, arginine/serine-rich 6) | FUNCTION: Plays a role in constitutive splicing and modulates the selection of alternative splice sites. Plays a role in the alternative splicing of MAPT/Tau exon 10. Binds to alternative exons of TNC pre-mRNA and promotes the expression of alternatively spliced TNC. Plays a role in wound healing and in the regulation of keratinocyte differentiation and proliferation via its role in alternative splicing (By similarity). {ECO:0000250}. |
| Q61464 | Znf638 | Zinc finger protein 638 (Nuclear protein 220) (Zinc finger matrin-like protein) | FUNCTION: Transcription factor that binds to cytidine clusters in double-stranded DNA (By similarity). Plays a key role in the silencing of unintegrated retroviral DNA: some part of the retroviral DNA formed immediately after infection remains unintegrated in the host genome and is transcriptionally repressed (PubMed:30487602). Mediates transcriptional repression of unintegrated viral DNA by specifically binding to the cytidine clusters of retroviral DNA and mediating the recruitment of chromatin silencers, such as the HUSH complex, SETDB1 and the histone deacetylases HDAC1 and HDAC4 (PubMed:30487602). Acts as an early regulator of adipogenesis by acting as a transcription cofactor of CEBPs (CEBPA, CEBPD and/or CEBPG), controlling the expression of PPARG and probably of other proadipogenic genes, such as SREBF1 (PubMed:21602272). May also regulate alternative splicing of target genes during adipogenesis (PubMed:25024404). {ECO:0000250\|UniProtKB:Q14966, ECO:0000269\|PubMed:21602272, ECO:0000269\|PubMed:25024404, ECO:0000269\|PubMed:30487602}. |
| Q80SY5 | Prpf38b | Pre-mRNA-splicing factor 38B | FUNCTION: May be required for pre-mRNA splicing. {ECO:0000305}. |
| Q569Z5 | Ddx46 | Probable ATP-dependent RNA helicase DDX46 (EC 3.6.4.13) (DEAD box protein 46) | FUNCTION: Plays an essential role in splicing, either prior to, or during A complex formation. {ECO:0000250}. |
| Q8BTI8 | Srrm2 | Serine/arginine repetitive matrix protein 2 | FUNCTION: Required for pre-mRNA splicing as component of the spliceosome. {ECO:0000250\|UniProtKB:Q9UQ35}. |
| Q8VE97 | Srsf4 | Serine/arginine-rich splicing factor 4 (Splicing factor, arginine/serine-rich 4) | FUNCTION: Plays a role in alternative splice site selection during pre-mRNA splicing. Represses the splicing of MAPT/Tau exon 10 (By similarity). {ECO:0000250}. |
| Q9QX47 | Son | Protein SON (Negative regulatory element-binding protein) (NRE-binding protein) | FUNCTION: RNA-binding protein that acts as a mRNA splicing cofactor by promoting efficient splicing of transcripts that possess weak splice sites. Specifically promotes splicing of many cell-cycle and DNA-repair transcripts that possess weak splice sites, such as TUBG1, KATNB1, TUBGCP2, AURKB, PCNT, AKT1, RAD23A, and FANCG. Probably acts by facilitating the interaction between Serine/arginine-rich proteins such as SRSF2 and the RNA polymerase II. Also binds to DNA; binds to the consensus DNA sequence: 5'-GA[GT]AN[CG][AG]CC-3' (By similarity). Essential for correct RNA splicing of multiple genes critical for brain development, neuronal migration and metabolism, including TUBG1, FLNA, PNKP, WDR62, PSMD3, PCK2, PFKL, IDH2, and ACY1 (By similarity). May also regulate the ghrelin signaling in hypothalamic neuron by acting as a negative regulator of GHSR expression (PubMed:20876580). {ECO:0000250\|UniProtKB:P18583, ECO:0000269\|PubMed:20876580}. |
| Q6PDM2 | Srsf1 | Serine/arginine-rich splicing factor 1 (ASF/SF2) (Pre-mRNA-splicing factor SRp30a) (Splicing factor, arginine/serine-rich 1) | FUNCTION: Plays a role in preventing exon skipping, ensuring the accuracy of splicing and regulating alternative splicing. Interacts with other spliceosomal components, via the RS domains, to form a bridge between the 5'- and 3'-splice site binding components, U1 snRNP and U2AF. Can stimulate binding of U1 snRNP to a 5'-splice site-containing pre-mRNA. Binds to purine-rich RNA sequences, either the octamer, 5'-RGAAGAAC-3' (r=A or G) or the decamers, AGGACAGAGC/AGGACGAAGC. Binds preferentially to the 5'-CGAGGCG-3' motif in vitro. Three copies of the octamer constitute a powerful splicing enhancer in vitro, the ASF/SF2 splicing enhancer (ASE) which can specifically activate ASE-dependent splicing (By similarity). Specifically regulates alternative splicing of cardiac isoforms of CAMK2D, LDB3/CYPHER and TNNT2/CTNT during heart remodeling at the juvenile to adult transition. The inappropriate accumulation of a neonatal and neuronal isoform of CAMKD2 in the adult heart results in aberrant calcium handling and defective excitation-contraction coupling in cardiomyocytes. May function as export adapter involved in mRNA nuclear export through the TAP/NXF1 pathway (PubMed:15652482). {ECO:0000250\|UniProtKB:Q07955, ECO:0000269\|PubMed:15652482}. |
| P26369 | U2af2 | Splicing factor U2AF 65 kDa subunit (U2 auxiliary factor 65 kDa subunit) (U2 snRNP auxiliary factor large subunit) | FUNCTION: Plays a role in pre-mRNA splicing and 3'-end processing. By recruiting PRPF19 and the PRP19C/Prp19 complex/NTC/Nineteen complex to the RNA polymerase II C-terminal domain (CTD), and thereby pre-mRNA, may couple transcription to splicing. Required for the export of mRNA out of the nucleus, even if the mRNA is encoded by an intron-less gene. Positively regulates pre-mRNA 3'-end processing by recruiting the CFIm complex to cleavage and polyadenylation signals. {ECO:0000250\|UniProtKB:P26368}. |

| | | | |
|---|---|---|---|
| Q9D883 | U2af1 | Splicing factor U2AF 35 kDa subunit (U2 auxiliary factor 35 kDa subunit) (U2 snRNP auxiliary factor small subunit) | FUNCTION: Plays a critical role in both constitutive and enhancer-dependent splicing by mediating protein-protein interactions and protein-RNA interactions required for accurate 3'-splice site selection. Recruits U2 snRNP to the branch point. Directly mediates interactions between U2AF2 and proteins bound to the enhancers and thus may function as a bridge between U2AF2 and the enhancer complex to recruit it to the adjacent intron (By similarity). {ECO:0000250}. |
| A2AR02 | Ppig | Peptidyl-prolyl cis-trans isomerase G (PPIase G) (Peptidyl-prolyl isomerase G) (EC 5.2.1.8) (Cyclophilin G) (Rotamase G) | FUNCTION: PPIase that catalyzes the cis-trans isomerization of proline imidic peptide bonds in oligopeptides and may therefore assist protein folding. May be implicated in the folding, transport, and assembly of proteins. May play an important role in the regulation of pre-mRNA splicing. {ECO:0000250\|UniProtKB:Q13427}. |
| Q6PFR5 | Tra2a | Transformer-2 protein homolog alpha (TRA-2 alpha) (TRA2-alpha) (Transformer-2 protein homolog A) | FUNCTION: Sequence-specific RNA-binding protein which participates in the control of pre-mRNA splicing. {ECO:0000250}. |
| Q9JJA7 | Ccnl2 | Cyclin-L2 (Cyclin Ania-6b) (Paneth cell-enhanced expression protein) (PCEE) | FUNCTION: Involved in pre-mRNA splicing. May induce cell death, possibly by acting on the transcription and RNA processing of apoptosis-related factors. {ECO:0000250\|UniProtKB:Q96S94}. |
| E9Q5F9 | Setd2 | Histone-lysine N-methyltransferase SETD2 (EC 2.1.1.-) (Lysine N-methyltransferase 3A) (Protein-lysine N-methyltransferase SETD2) (EC 2.1.1.-) (SET domain-containing protein 2) | FUNCTION: Histone methyltransferase that specifically trimethylates 'Lys-36' of histone H3 (H3K36me3) using dimethylated 'Lys-36' (H3K36me2) as substrate (PubMed:18157086, PubMed:20133625). Represents the main enzyme generating H3K36me3, a specific tag for epigenetic transcriptional activation (PubMed:18157086, PubMed:20133625). Plays a role in chromatin structure modulation during elongation by coordinating recruitment of the FACT complex and by interacting with hyperphosphorylated POLR2A (By similarity). Acts as a key regulator of DNA mismatch repair in G1 and early S phase by generating H3K36me3, a mark required to recruit MSH6 subunit of the MutS alpha complex: early recruitment of the MutS alpha complex to chromatin to be replicated allows a quick identification of mismatch DNA to initiate the mismatch repair reaction (By similarity). Required for DNA double-strand break repair in response to DNA damage: acts by mediating formation of H3K36me3, promoting recruitment of RAD51 and DNA repair via homologous recombination (HR) (By similarity). Acts as a tumor suppressor (By similarity). H3K36me3 also plays an essential role in the maintenance of a heterochromatic state, by recruiting DNA methyltransferase DNMT3A (By similarity). H3K36me3 is also enhanced in intron-containing genes, suggesting that SETD2 recruitment is enhanced by splicing and that splicing is coupled to recruitment of elongating RNA polymerase (By similarity). Required during angiogenesis (PubMed:20133625). Required for endoderm development by promoting embryonic stem cell differentiation toward endoderm: acts by mediating formation of H3K36me3 in distal promoter regions of FGFR3, leading to regulate transcription initiation of FGFR3 (PubMed:25242323). In addition to histones, also mediates methylation of other proteins, such as tubulins and STAT1 (PubMed:27518565). Trimethylates 'Lys-40' of alpha-tubulins such as TUBA1B (alpha-TubK40me3); alpha-TubK40me3 is required for normal mitosis and cytokinesis and may be a specific tag in cytoskeletal remodeling (PubMed:27518565). Involved in interferon-alpha-induced antiviral defense by mediating both monomethylation of STAT1 at 'Lys-525' and catalyzing H3K36me3 on promoters of some interferon-stimulated genes (ISGs) to activate gene transcription (By similarity). {ECO:0000250\|UniProtKB:Q9BYW2, ECO:0000269\|PubMed:18157086, ECO:0000269\|PubMed:20133625, ECO:0000269\|PubMed:25242323, ECO:0000269\|PubMed:27518565}. |
| Q9CYI4 | Luc7l | Putative RNA-binding protein Luc7-like 1 | FUNCTION: May bind to RNA via its Arg/Ser-rich domain. |
| Q0P557 | Spata18 | Mitochondria-eating protein (Spermatogenesis-associated protein 18) | FUNCTION: Key regulator of mitochondrial quality that mediates the repairing or degradation of unhealthy mitochondria in response to mitochondrial damage. Mediator of mitochondrial protein catabolic process (also named MALM) by mediating the degradation of damaged proteins inside mitochondria by promoting the accumulation in the mitochondrial matrix of hydrolases that are characteristic of the lysosomal lumen. Also involved in mitochondrion degradation of damaged mitochondria by promoting the formation of vacuole-like structures (named MIV), which engulf and degrade unhealthy mitochondria by accumulating lysosomes. May have a role in spermatogenesis, especially in cell differentiation from late elongate spermatids to mature spermatozoa (By similarity). The physical interaction of SPATA18/MIEAP, BNIP3 and BNIP3L/NIX at the mitochondrial outer membrane regulates the opening of a pore in the mitochondrial double membrane in order to mediate the translocation of lysosomal proteins from the cytoplasm to the mitochondrial matrix (By similarity). {ECO:0000250}. |
| Q9DA19 | Cir1 | Corepressor interacting with RBPJ 1 (CBF1-interacting corepressor) | FUNCTION: Regulates transcription and acts as corepressor for RBPJ. Recruits RBPJ to the Sin3-histone deacetylase complex (HDAC). Required for RBPJ-mediated repression of transcription (By similarity). May modulate splice site selection during alternative splicing of pre-mRNAs. {ECO:0000250, ECO:0000269\|PubMed:15652350}. |
| P84104 | Srsf3 | Serine/arginine-rich splicing factor 3 (Pre-mRNA-splicing factor SRP20) (Protein X16) (Splicing factor, arginine/serine-rich 3) | FUNCTION: Splicing factor that specifically promotes exon-inclusion during alternative splicing. Interaction with YTHDC1, a RNA-binding protein that recognizes and binds N6-methyladenosine (m6A)-containing RNAs, promotes recruitment of SRSF3 to its mRNA-binding elements adjacent to m6A sites, leading to exon-inclusion during alternative splicing. Also functions as export adapter involved in mRNA nuclear export. Binds mRNA which is thought to be transferred to the NXF1-NXT1 heterodimer for export (TAP/NXF1 pathway); enhances NXF1-NXT1 RNA-binding activity. Involved in nuclear export of m6A-containing mRNAs via interaction with YTHDC1: interaction with YTHDC1 facilitates m6A-containing mRNA-binding to both SRSF3 and NXF1, promoting mRNA nuclear export. RNA-binding is semi-sequence specific. {ECO:0000250\|UniProtKB:P84103}. |
| A2A4P0 | Dhx8 | ATP-dependent RNA helicase DHX8 (EC 3.6.4.13) (DEAH box protein 8) | FUNCTION: Involved in pre-mRNA splicing as component of the spliceosome. Facilitates nuclear export of spliced mRNA by releasing the RNA from the spliceosome. {ECO:0000250\|UniProtKB:Q14562}. |

| | | | |
|---|---|---|---|
| Q99M28 | Rnps1 | RNA-binding protein with serine-rich domain 1 | FUNCTION: Part of pre- and post-splicing multiprotein mRNP complexes. Auxiliary component of the splicing-dependent multiprotein exon junction complex (EJC) deposited at splice junction on mRNAs. The EJC is a dynamic structure consisting of core proteins and several peripheral nuclear and cytoplasmic associated factors that join the complex only transiently either during EJC assembly or during subsequent mRNA metabolism. Component of the ASAP and PSAP complexes which bind RNA in a sequence-independent manner and are proposed to be recruited to the EJC prior to or during the splicing process and to regulate specific excision of introns in specific transcription subsets. The ASAP complex can inhibit RNA processing during in vitro splicing reactions. The ASAP complex promotes apoptosis and is disassembled after induction of apoptosis. Enhances the formation of the ATP-dependent A complex of the spliceosome. Involved in both constitutive splicing and, in association with SRP54 and TRA2B/SFRS10, in distinctive modulation of alternative splicing in a substrate-dependent manner. Involved in the splicing modulation of BCL2L1/Bcl-X (and probably other apoptotic genes); specifically inhibits formation of proapoptotic isoforms such as Bcl-X(S); the activity is different from the established EJC assembly and function. Participates in mRNA 3'-end cleavage. Involved in UPF2-dependent nonsense-mediated decay (NMD) of mRNAs containing premature stop codons. Also mediates increase of mRNA abundance and translational efficiency. Binds spliced mRNA 20-25 nt upstream of exon-exon junctions (By similarity). {ECO:0000250}. |
| A2AJT4 | Pnisr | Arginine/serine-rich protein PNISR (Serine/arginine-rich-splicing regulatory protein 130) (SRrp130) (Splicing factor, arginine/serine-rich 130) (Splicing factor, arginine/serine-rich 18) | |
| Q69ZA1 | Cdk13 | Cyclin-dependent kinase 13 (EC 2.7.11.22) (EC 2.7.11.23) (CDC2-related protein kinase 5) (Cell division cycle 2-like protein kinase 5) (Cell division protein kinase 13) | FUNCTION: Cyclin-dependent kinase which displays CTD kinase activity and is required for RNA splicing. Has CTD kinase activity by hyperphosphorylating the C-terminal heptapeptide repeat domain (CTD) of the largest RNA polymerase II subunit RPB1, thereby acting as a key regulator of transcription elongation. Required for RNA splicing, probably by phosphorylating SRSF1/SF2. Required during hematopoiesis. {ECO:0000269\|PubMed:17261272}. |
| Q99ME9 | Gtpbp4 | Nucleolar GTP-binding protein 1 (Chronic renal failure gene protein) (GTP-binding protein NGB) | FUNCTION: Involved in the biogenesis of the 60S ribosomal subunit. {ECO:0000250}. |
| Q8K194 | Snrnp27 | U4/U6.U5 small nuclear ribonucleoprotein 27 kDa protein (U4/U6.U5 snRNP 27 kDa protein) (U4/U6.U5-27K) (U4/U6.U5 tri-snRNP-associated protein 3) | FUNCTION: May play a role in mRNA splicing. |
| Q9R0U0 | Srsf10 | Serine/arginine-rich splicing factor 10 (FUS-interacting serine-arginine-rich protein 1) (Neural-salient serine/arginine-rich protein) (Neural-specific SR protein) (Splicing factor, arginine/serine-rich 13A) (TLS-associated protein with Ser-Arg repeats) (TASR) (TLS-associated protein with SR repeats) (TLS-associated serine-arginine protein) (TLS-associated SR protein) | FUNCTION: Splicing factor that in its dephosphorylated form acts as a general repressor of pre-mRNA splicing. Seems to interfere with the U1 snRNP 5'-splice recognition of SNRNP70. Required for splicing repression in M-phase cells and after heat shock. Also acts as a splicing factor that specifically promotes exon skipping during alternative splicing. Interaction with YTHDC1, a RNA-binding protein that recognizes and binds N6-methyladenosine (m6A)-containing RNAs, prevents SRSF10 from binding to its mRNA-binding sites close to m6A-containing regions, leading to inhibit exon skipping during alternative splicing (By similarity). May be involved in regulation of alternative splicing in neurons (PubMed:10583508). {ECO:0000250\|UniProtKB:O75494, ECO:0000269\|PubMed:10583508}. |
| Q9R020 | Zranb2 | Zinc finger Ran-binding domain-containing protein 2 (Zinc finger protein 265) (Zinc finger, splicing) | FUNCTION: Splice factor required for alternative splicing of TRA2B/SFRS10 transcripts. May interfere with constitutive 5'-splice site selection (By similarity). {ECO:0000250}. |
| Q61136 | Prpf4b | Serine/threonine-protein kinase PRP4 homolog (EC 2.7.11.1) (PRP4 pre-mRNA-processing factor 4 homolog) (Pre-mRNA protein kinase) | FUNCTION: Has a role in pre-mRNA splicing. Phosphorylates SF2/ASF. |
| Q9Z0U1 | Tjp2 | Tight junction protein ZO-2 (Tight junction protein 2) (Zona occludens protein 2) (Zonula occludens protein 2) | FUNCTION: Plays a role in tight junctions and adherens junctions. |
| Q7TNC4 | Luc7l2 | Putative RNA-binding protein Luc7-like 2 (CGI-74 homolog) | FUNCTION: May bind to RNA via its Arg/Ser-rich domain. |

**Table S2. RNF12 phosphorylation sites identified by IP-MS**

Experiment 1

| pep_exp_mz | pep_exp_mr | pep_score | pep_seq | pep_var_mod | residue |
|---|---|---|---|---|---|
| 516.5920 | 1546.7504 | 23 | R.SRSPLQPTSEIPR.R | P (ST) | S227, S229 |
| 801.8524 | 1601.6868 | (26) | R.RL**S**VENMESSSQR.Q | P (ST) | S163 |
| 809.8495 | 1617.6818 | (27) | R.RL**S**VENMESSSQR.Q | O (M); P (ST) | S163 |

Experiment 2

| pep_exp_mz | pep_exp_mr | pep_score | pep_seq | pep_var_mod | residue |
|---|---|---|---|---|---|
| 701.786 | 1401.5562 | 30 | EGPPPQ**S**PDENR | P (ST) | S78 |
| 774.3826 | 1546.7504 | 40 | SR**S**PLQPTSEIPR | P (ST) | S229 |
| 801.8502 | 1601.6868 | 46 | RL**S**VENMESSSQR | P (ST) | S163 |
| 809.8483 | 1617.6818 | 62 | RL**S**VENMESSSQR | O (M); P (ST) | S163 |
| 1230.5322 | 2459.0489 | 78 | AGESSDDVTNSDSIIDWLNSVR | P (ST) | S88, S89 |
| 1255.2161 | 3762.6282 | 32 | EGPPPQSPDENRAGESSDDVTNSDSIIDWLNSVR | P (ST) | S78, S88, S89 |

Experiment 3

| pep_exp_mz | pep_exp_mr | pep_score | pep_seq | pep_var_mod | residue |
|---|---|---|---|---|---|
| 701.7868 | 1401.559 | 48 | EGPPPQ**S**PDENR | P (ST) | S78 |
| 774.3839 | 1546.7531 | 44 | SRSPLQPTSEIPR | P (ST) | S227 |
| 516.5918 | 1546.7536 | 27 | SRSPLQPTSEIPR | P (ST)?? | S227, S229, T234 |
| 809.8502 | 1617.6858 | 68 | RL**S**VENMESSSQR | O (M); P (ST) | S163 |
| 814.3668 | 1626.719 | 37 | **S**R**S**PLQPTSEIPR | 2 P (ST) | S227, S229 |
| 661.9747 | 1982.9024 | 19 | AERSRSPLQPTSEIPR | 2 P (ST)?? | S227, S229, T234, S235 |

Experiment 4

| pep_exp_mz | pep_exp_mr | pep_score | pep_seq | pep_var_mod | residue |
|---|---|---|---|---|---|
| 474.7062 | 947.3979 | 29 | SRSPEHR | P (ST)?? | S212, S214 |
| 701.7858 | 1401.5571 | 30 | EGPPPQ**S**PDENR | P (ST) | S78 |
| 774.3807 | 1546.7469 | 43 | SR**S**PLQPTSEIPR | P (ST) | S229 |
| 809.8483 | 1617.6821 | 50 | RLSVENMESSSQR | O (M); P (ST)?? | S163 |
| 814.3668 | 1626.719 | 22 | SRSPLQPTSEIPR | 2 P (ST)?? | S227, S229, T234, S235 |
| 905.9172 | 1809.8198 | 21 | AERNSAEAVTEVPTTR | P (ST)?? | S194, T199 |
| 1230.5315 | 2459.0484 | 73 | AGESSDDVTNSDSIIDWLNSVR | P (ST)?? | S88, S89, T93, |
| 1255.2163 | 3762.6271 | 39 | EGPPPQSPDENRAGESSDDVTNSDSIIDWLNSVR | P (ST) | S78 |

**Phosphosite Localisation**

Proteome Discoverer 1.4-SP1 –PhosphoRS3.1 info OR

Proteome Discoverer 2.0- –ptmRS info

1 Underlined S T is our interpretation of Mascot and MS2 data

Bold S T, is a very good assignment, S T is used where ID is not certain

?? means P is just about anywhere.

| Header | Description |
|---|---|
| pep_exp_mz | Observed or experimental m/z value |
| pep_exp_mr | Molecular mass calculated from experimental m/z value |
| pep_score | Mascot score for PSM (Peptide sequence match) |
| pep_seq | Peptide sequence in 1 letter code |
| pep_var_mod | Variable modifications from all sources as list of names |

**Table S3. RNF12 phosphorylation sites identified via SRPK in vitro phosphorylation and MS**

**SRPK1 5 min**

| pep_exp_mz | pep_exp_mr | pep_score | pep_seq | pep_var_mod |
|---|---|---|---|---|
| 460.2178 | 918.4212 | (24) | R.TYVSTIR.I | P (ST) |
| 639.3107 | 1276.6176 | (20) | R.QQI**S**GPELLGR.G | P (ST) |
| 652.8156 | 1303.6173 | (31) | R.**S**PLQPTSEIPR.R | P (ST) |
| 516.5908 | 1546.7504 | (22) | R.SRSPLQPTSEIPR.R | P (ST) |
| 774.3820 | 1546.7504 | (47) | R.SRSPLQPTSEIPR.R | P (ST) |
| 543.2460 | 1626.7168 | (26) | R.**S**R**S**PLQPTSEIPR.R | 2 P (ST) |
| 814.3652 | 1626.7168 | (41) | R.**S**R**S**PLQPTSEIPR.R | 2 P (ST) |
| 838.3497 | 1674.6886 | (46) | R.**S**QAPNNTVTYESER.G | P (ST) |
| 959.9175 | 1917.8218 | 85 | R.SR**S**QAPNNTVTYESER.G | P (ST) |
| 640.2810 | 1917.8218 | (48) | R.SRSQAPNNTVTYESER.G | P (ST) |
| 992.4553 | 1982.8976 | (28) | R.AERSRSPLQPTSEIPR.R | 2 P (ST) |
| 661.9727 | 1982.8976 | 40 | R.AERSRSPLQPTSEIPR.R | 2 P (ST) |
| 1027.4431 | 2052.8749 | (29) | R.RAPTLEQSSENEPEGSSR.T | P (ST) |
| 685.2983 | 2052.8749 | (60) | R.RAPTLEQSSENEPEGSSR.T | P (ST) |
| 689.6444 | 2065.9205 | (21) | R.DNNLLGTPGESTEEELLR.R | P (ST) |
| 771.0147 | 2310.0237 | 38 | R.RAPTLEQSSENEPEGSSRTR.H | P (ST) |

**SRPK1 60 min**

| pep_exp_mz | pep_exp_mr | pep_score | pep_seq | pep_var_mod |
|---|---|---|---|---|
| 460.2179 | 918.4212 | (19) | R.TYVSTIR.I | P (ST) |
| 514.6891 | 1027.3637 | 19 | R.**S**R**S**PEHR.R | 2 P (ST) |
| 628.2581 | 1254.5020 | 21 | R.AR**S**R**S**PEHR.R | 2 P (ST) |
| 516.5908 | 1546.7504 | (27) | R.SRSPLQPTSEIPR.R | P (ST) |
| 774.3821 | 1546.7504 | 50 | R.SRSPLQPTSEIPR.R | P (ST) |
| 814.3652 | 1626.7168 | (41) | R.**S**R**S**PLQPTSEIPR.R | 2 P (ST) |
| 543.2462 | 1626.7168 | (21) | R.**S**R**S**PLQPTSEIPR.R | 2 P (ST) |
| 838.3506 | 1674.6886 | (69) | R.**S**QAPNNTVTYESER.G | P (ST) |
| 949.3937 | 1896.7738 | (50) | R.APTLEQSSENEPEGSSR.T | P (ST) |
| 959.9170 | 1917.8218 | 82 | R.SR**S**QAPNNTVTYESER.G | P (ST) |
| 640.2809 | 1917.8218 | (29) | R.SRSQAPNNTVTYESER.G | P (ST) |
| 992.4556 | 1982.8976 | 39 | R.AER**S**R**S**PLQPTSEIPR.R | 2 P (ST) |
| 1027.4439 | 2052.8749 | (48) | R.RAPTLEQSSENEPEGSSR.T | P (ST) |
| 685.2986 | 2052.8749 | (50) | R.RAPTLEQSSENEPEGSSR.T | P (ST) |
| 752.6523 | 2254.9369 | 35 | R.TRSRSQAPNNTVTYESER.G | 2 P (ST) |
| 771.0147 | 2310.0237 | 43 | R.RAPTLEQSSENEPEGSSRTR.H | P (ST) |
| 578.5131 | 2310.0237 | (19) | R.RAPTLEQSSENEPEGSSRTR.H | P (ST) |

**SRPK2 5 min**

| pep_exp_mz | pep_exp_mr | pep_score | pep_seq | pep_var_mod |
|---|---|---|---|---|
| 473.2131 | 944.4117 | 35 | GLFAA**S**GSR | P (ST) |
| 516.5906 | 1546.7499 | 33 | SRSPLQPTSEIPR | P (ST) |
| 543.2464 | 1626.7174 | 18 | **S**R**S**PLQPTSEIPR | 2 P (ST) |
| 553.516 | 2210.0349 | 24 | ARAERSRSPLQPTSEIPR | 2 P (ST) |
| 578.5132 | 2310.0236 | 38 | RAPTLEQSSENEPEGSSRTR | P (ST) |
| 635.3174 | 1902.9303 | 38 | AERSRSPLQPTSEIPR | P (ST) |
| 640.2807 | 1917.8203 | 38 | SRSQAPNNTVTYESER | P (ST) |
| 661.9725 | 1982.8958 | 29 | AERSRSPLQPTSEIPR | 2 P (ST) |
| 685.2986 | 2052.8741 | 51 | RAPTLEQSSENEPEGSSR | P (ST) |
| 718.9813 | 2153.922 | 31 | APTLEQSSENEPEGSSRTR | P (ST) |
| 750.8328 | 1499.651 | 20 | AVSRTNPNSGDFR | P (ST) |
| 752.6522 | 2254.9348 | 42 | TRSRSQAPNNTVTYESER | 2 P (ST) |
| 771.0148 | 2310.0225 | 51 | RAPTLEQSSENEPEGSSRTR | P (ST) |
| 774.382 | 1546.7495 | 47 | SRSPLQPTSEIPR | P (ST) |
| 814.3648 | 1626.7151 | 25 | **S**R**S**PLQPTSEIPR | 2 P (ST) |
| 814.3654 | 1626.7162 | 21 | SRSPLQPTSEIPR | 2 P (ST) |
| 838.3527 | 1674.6909 | 57 | **S**QAPNNTVTYESER | P (ST) |
| 959.9179 | 1917.8213 | 69 | SRSQAPNNTVTYESER | P (ST) |

| | | | | |
|---|---|---|---|---|
| 992.455 | 1982.8955 | 27 | AERSRSPLQPTSEIPR | 2 P (ST) |
| 1027.4439 | 2052.8731 | 36 | RAPTLEQSSENEPEGSSR | P (ST) |
| 1106.0253 | 2210.036 | 18 | ARAERSRSPLQPTSEIPR | 2 P (ST) |

**SRPK2 60 min**

| pep_exp_mz | pep_exp_mr | pep_score | pep_seq | pep_var_mod |
|---|---|---|---|---|
| 460.2179 | 918.4212 | 31 | TYVSTIR | P (ST) |
| 473.213 | 944.4115 | 55 | GLFAA**S**GSR | P (ST) |
| 493.2267 | 984.4388 | 20 | DSIASRTR | P (ST) |
| 500.8911 | 1499.6514 | 25 | AVSRTNPNSGDFR | P (ST) |
| 516.5907 | 1546.7503 | 36 | SRSPLQPTSEIPR | P (ST) |
| 534.7452 | 1067.4759 | 21 | NVERVE**S**R | P (ST) |
| 534.9023 | 1601.6852 | 21 | RLSVENMESSSQR | P (ST) |
| 540.2346 | 1617.6819 | 35 | RL**S**VENMESSSQR | O (M); P (ST) |
| 543.2468 | 1626.7187 | 28 | **S**R**S**PLQPTSEIPR | 2 P (ST) |
| 578.5133 | 2310.0241 | 50 | RAPTLEQSSENEPEGSSRTR | P (ST) |
| 604.2824 | 1809.8254 | 29 | AERN**S**AEAVTEVPTTR | P (ST) |
| 633.2654 | 1896.7743 | 42 | APTLEQSSENEPEGSSR | P (ST) |
| 640.2813 | 1917.8221 | 45 | SRSQAPNNTVTYESER | P (ST) |
| 652.8159 | 1303.6172 | 25 | SPLQPTSEIPR | P (ST) |
| 661.9733 | 1982.8982 | 30 | AERSRSPLQPTSEIPR | 2 P (ST) |
| 685.2987 | 2052.8743 | 55 | RAPTLEQSSENEPEGSSR | P (ST) |
| 711.9543 | 2132.8412 | 33 | RAPTLEQSSENEPEGSSR | 2 P (ST) |
| 718.9818 | 2153.9236 | 27 | APTLEQSSENEPEGSSRTR | P (ST) |
| 725.9974 | 2174.9703 | 23 | TRSRSQAPNNTVTYESER | P (ST) |
| 750.8325 | 1499.6505 | 32 | AVSRTNPNSGDFR | P (ST) |
| 752.6528 | 2254.9365 | 49 | TRSRSQAPNNTVTYESER | 2 P (ST) |
| 771.0143 | 2310.021 | 31 | RAPTLEQSSENEPEGSSRTR | P (ST) |
| 771.0155 | 2310.0247 | 26 | RAPTLEQSSENEPEGSSRTR | P (ST) |
| 774.382 | 1546.7495 | 47 | SRSPLQPTSEIPR | P (ST) |
| 801.8499 | 1601.6853 | 70 | RL**S**VENMESSSQR | P (ST) |
| 809.8482 | 1617.6819 | 40 | RL**S**VENMESSSQR | O (M); P (ST) |
| 814.3657 | 1626.7169 | 45 | **S**R**S**PLQPTSEIPR | 2 P (ST) |
| 838.3528 | 1674.6911 | 58 | **S**QAPNNTVTYESER | P (ST) |
| 854.3486 | 1706.6827 | 19 | SRSPLQPTSEIPR | 3 P (ST) |
| 949.3945 | 1896.7744 | 58 | APTLEQSSENEPEGSSR | P (ST) |
| 959.9175 | 1917.8204 | 79 | SR**S**QAPNNTVTYESER | P (ST) |
| 992.4556 | 1982.8966 | 24 | AERSRSPLQPTSEIPR | 2 P (ST) |
| 1027.4443 | 2052.8741 | 61 | RAPTLEQSSENEPEGSSR | P (ST) |

**Phosphosite Localisation**

Proteome Discoverer 1.4-SP1 –PhosphoRS3.1 info OR

Proteome Discoverer 2.0- –ptmRS info

1 Underlined S T is our interpretation of Mascot and MS2 data

Bold S T, is a very good assignment, S T is used where ID is not certain

?? means P is just about anywhere.

| Header | Description |
|---|---|
| pep_exp_mz | Observed or experimental m/z value |
| pep_exp_mr | Molecular mass calculated from experimental m/z value |
| pep_score | Mascot score for PSM (Peptide sequence match) |
| pep_seq | Peptide sequence in 1 letter code |
| pep_var_mod | Variable modifications from all sources as list of names |

**Table S4 RNF12 W576Y vs RNF12 WT RNA-Seq analysis**

| Id | RNF12WT1 | RNF12WT2 | RNF12WT3 | RNF12W576Y1 | RNF12W576Y2 | RNF12W576Y3 | norm.RNF12WT1 | norm.RNF12WT2 | norm.RNF12WT3 | norm.RNF12W576Y1 | norm.RNF12W576Y2 | norm.RNF12W576Y3 | baseMean | Rnf12WT | Rnf12W576Y | FoldChange | log2FoldChange | pvalue | padj | dispGeneEst | dispFit | dispMAP | dispersion | betaConv | maxCooks |
|---|---|---|---|---|---|---|---|---|---|---|---|---|---|---|---|---|---|---|---|---|---|---|---|---|---|
| 0610005C13I | 26 | 18 | 2 | 5 | 7 | 6 | 20 | 19 | 2 | 5 | 8 | 6 | 10.05 | 14 | 6 | 0.447 | -1.162 | 0.12133902 | NA | 0.4188 | 0.2383 | 0.2882 | 0.2882 | TRUE | 2.1899 |
| 0610007N19 | 56 | 47 | 55 | 67 | 76 | 79 | 43 | 50 | 59 | 67 | 86 | 78 | 63.87 | 51 | 77 | 1.515 | 0.599 | 0.01793281 | 0.06372792 | 0 | 0.0395 | 0.0299 | 0.0299 | TRUE | 0.1659 |
| 0610007P14I | 2426 | 1873 | 1923 | 2265 | 2000 | 2159 | 1884 | 1992 | 2075 | 2255 | 2256 | 2131 | 2098.9 | 1984 | 2214 | 1.117 | 0.159 | 0.01578123 | 0.05798534 | 0.0013 | 0.0036 | 0.0027 | 0.0027 | TRUE | 0.0242 |
| 0610008F07I | 0 | 0 | 1 | 0 | 0 | 0 | 0 | 0 | 1 | 0 | 0 | 0 | 0.18 | 0 | 0 | 0.552 | -0.857 | 0.8336548 | NA | 0 | 13.1817 | 10 | 10 | TRUE | 0.5365 |
| 0610009B14I | 23 | 16 | 20 | 24 | 28 | 30 | 18 | 17 | 22 | 24 | 32 | 30 | 23.59 | 19 | 29 | 1.508 | 0.593 | 0.13447375 | 0.27321014 | 0 | 0.1029 | 0.0692 | 0.0692 | TRUE | 0.1254 |
| 0610009B22I | 1812 | 1293 | 1261 | 1433 | 1284 | 1388 | 1407 | 1375 | 1361 | 1426 | 1448 | 1370 | 1398.05 | 1381 | 1415 | 1.024 | 0.034 | 0.6122979 | 0.75477135 | 0 | 0.0041 | 0.0025 | 0.0025 | TRUE | 0.0093 |
| 0610009D07I | 3618 | 2593 | 2496 | 2655 | 2490 | 2768 | 2810 | 2758 | 2693 | 2643 | 2809 | 2732 | 2740.91 | 2754 | 2728 | 0.99 | -0.014 | 0.80671543 | 0.88831923 | 3.00E-04 | 0.0033 | 0.0021 | 0.0021 | TRUE | 0.0091 |
| 0610009L18F | 29 | 19 | 11 | 20 | 14 | 24 | 23 | 20 | 12 | 20 | 16 | 24 | 19 | 18 | 20 | 1.081 | 0.113 | 0.80463505 | 0.88691572 | 0 | 0.1272 | 0.0973 | 0.0973 | TRUE | 0.4391 |
| 0610009O20 | 2613 | 2007 | 2432 | 2395 | 2201 | 2624 | 2029 | 2135 | 2624 | 2384 | 2483 | 2590 | 2374.26 | 2263 | 2486 | 1.099 | 0.137 | 0.13526451 | 0.27453579 | 0.0101 | 0.0034 | 0.0056 | 0.0056 | TRUE | 0.2352 |
| 0610010B08I | 0 | 0 | 0 | 0 | 0 | 0 | 0 | 0 | 0 | 0 | 0 | 0 | 0 | 0 | 0 | NA | NA | NA | NA | NA | NA | NA | NA | NA | NA |
| 0610010F05I | 1486 | 1046 | 1062 | 1131 | 996 | 1038 | 1154 | 1113 | 1146 | 1126 | 1124 | 1025 | 1114.45 | 1138 | 1092 | 0.959 | -0.061 | 0.42008876 | 0.59229576 | 7.00E-04 | 0.0046 | 0.0032 | 0.0032 | TRUE | 0.0346 |
| 0610010K14I | 3359 | 2392 | 2683 | 2649 | 2167 | 2706 | 2609 | 2544 | 2895 | 2637 | 2445 | 2671 | 2633.45 | 2683 | 2584 | 0.964 | -0.053 | 0.45345042 | 0.62319659 | 0.0029 | 0.0033 | 0.0032 | 0.0032 | TRUE | 0.0573 |
| 0610011F06I | 824 | 636 | 694 | 715 | 646 | 810 | 640 | 677 | 749 | 712 | 729 | 800 | 717.56 | 689 | 747 | 1.087 | 0.12 | 0.20601935 | 0.36784941 | 0.0038 | 0.0057 | 0.0051 | 0.0051 | TRUE | 0.0689 |
| 0610012G03 | 955 | 752 | 793 | 861 | 703 | 880 | 742 | 800 | 856 | 857 | 793 | 869 | 819.34 | 799 | 840 | 1.053 | 0.075 | 0.39865753 | 0.57203213 | 0.0026 | 0.0053 | 0.0044 | 0.0044 | TRUE | 0.0494 |
| 0610012H03I | 17 | 10 | 6 | 17 | 15 | 12 | 13 | 11 | 6 | 17 | 17 | 12 | 12.67 | 10 | 15 | 1.484 | 0.57 | 0.30050132 | NA | 0 | 0.1895 | 0.1371 | 0.1371 | TRUE | 0.3209 |
| 0610030E20I | 1064 | 848 | 838 | 872 | 765 | 879 | 826 | 902 | 904 | 868 | 863 | 868 | 871.88 | 877 | 866 | 0.989 | -0.017 | 0.83661774 | 0.90685404 | 0 | 0.0051 | 0.0035 | 0.0035 | TRUE | 0.0328 |
| 0610031J06F | 3083 | 2316 | 2458 | 2571 | 2305 | 2639 | 2394 | 2463 | 2652 | 2559 | 2600 | 2605 | 2545.77 | 2503 | 2588 | 1.035 | 0.049 | 0.43566309 | 0.60705138 | 0.0011 | 0.0034 | 0.0024 | 0.0024 | TRUE | 0.0325 |
| 0610031O16 | 0 | 0 | 0 | 0 | 0 | 0 | 0 | 0 | 0 | 0 | 0 | 0 | 0 | 0 | 0 | NA | NA | NA | NA | NA | NA | NA | NA | NA | NA |
| 0610037L13F | 1143 | 806 | 876 | 931 | 801 | 936 | 888 | 857 | 945 | 927 | 904 | 924 | 907.43 | 897 | 918 | 1.024 | 0.035 | 0.6597805 | 0.78953806 | 0 | 0.005 | 0.0034 | 0.0034 | TRUE | 0.0255 |
| 0610038B21I | 17 | 12 | 12 | 12 | 7 | 6 | 13 | 13 | 13 | 12 | 8 | 6 | 10.78 | 13 | 9 | 0.662 | -0.594 | 0.31248265 | NA | 0 | 0.2223 | 0.152 | 0.152 | TRUE | 0.3776 |
| 0610038L08F | 0 | 0 | 0 | 0 | 0 | 0 | 0 | 0 | 0 | 0 | 0 | 0 | 0 | 0 | 0 | NA | NA | NA | NA | NA | NA | NA | NA | NA | NA |
| 0610039K10I | 2 | 1 | 0 | 1 | 0 | 6 | 2 | 1 | 0 | 1 | 0 | 6 | 1.59 | 1 | 2 | 2.579 | 1.367 | 0.43492293 | NA | 1.3161 | 1.4938 | 1.4525 | 1.4525 | TRUE | 1.531 |
| 0610040B10I | 11 | 6 | 5 | 1 | 10 | 10 | 9 | 6 | 5 | 1 | 11 | 10 | 7.08 | 7 | 7 | 1.074 | 0.103 | 0.89882495 | NA | 0.3226 | 0.3373 | 0.3338 | 0.3338 | TRUE | 1.6322 |
| 0610040F04I | 79 | 77 | 79 | 112 | 97 | 151 | 61 | 82 | 85 | 111 | 109 | 149 | 99.75 | 76 | 123 | 1.633 | 0.707 | 0.00144627 | 0.0093876 | 0.0219 | 0.0262 | 0.0249 | 0.0249 | TRUE | 0.3432 |
| 0610040J01F | 169 | 130 | 106 | 108 | 88 | 89 | 131 | 138 | 114 | 108 | 99 | 88 | 113.09 | 128 | 98 | 0.767 | -0.384 | 0.04399839 | 0.12263435 | 0 | 0.0234 | 0.0171 | 0.0171 | TRUE | 0.0859 |
| 0610043K17I | 6 | 6 | 4 | 8 | 4 | 1 | 5 | 6 | 4 | 8 | 5 | 1 | 4.8 | 5 | 5 | 0.878 | -0.187 | 0.83858125 | NA | 0.1179 | 0.4958 | 0.3996 | 0.3996 | TRUE | 0.2217 |
| 1100001G20 | 0 | 0 | 1 | 1 | 0 | 1 | 0 | 0 | 1 | 1 | 0 | 1 | 0.51 | 0 | 1 | 1.941 | 0.957 | 0.72680542 | NA | 0 | 4.6473 | 3.6356 | 3.6356 | TRUE | 0.5365 |
| 1110001A16I | 1541 | 1051 | 1222 | 1471 | 1265 | 1457 | 1197 | 1118 | 1319 | 1464 | 1427 | 1438 | 1327.14 | 1211 | 1443 | 1.192 | 0.253 | 0.00117101 | 0.00812808 | 0.0024 | 0.0042 | 0.0036 | 0.0036 | TRUE | 0.0698 |
| 1110001J03F | 586 | 408 | 456 | 539 | 470 | 574 | 455 | 434 | 492 | 537 | 530 | 567 | 502.41 | 460 | 545 | 1.184 | 0.243 | 0.01395566 | 0.05292096 | 0 | 0.0071 | 0.005 | 0.005 | TRUE | 0.0397 |
| 1110002L01F | 1224 | 824 | 724 | 802 | 691 | 753 | 951 | 876 | 781 | 798 | 780 | 743 | 821.58 | 869 | 774 | 0.888 | -0.171 | 0.0664386 | 0.16500555 | 0.0045 | 0.0053 | 0.005 | 0.005 | TRUE | 0.0929 |
| 1110004E09I | 1956 | 1504 | 1459 | 1619 | 1491 | 1642 | 1519 | 1600 | 1574 | 1612 | 1682 | 1621 | 1601.27 | 1564 | 1638 | 1.048 | 0.067 | 0.3034203 | 0.47599454 | 0 | 0.0039 | 0.0024 | 0.0024 | TRUE | 0.0081 |
| 1110004F10I | 5058 | 3354 | 3212 | 3386 | 3042 | 3283 | 3928 | 3568 | 3466 | 3370 | 3432 | 3241 | 3500.77 | 3654 | 3348 | 0.915 | -0.127 | 0.05210758 | 0.13887257 | 0.0022 | 0.0031 | 0.0028 | 0.0028 | TRUE | 0.0535 |
| 1110006O24 | 14 | 11 | 10 | 15 | 12 | 2 | 11 | 12 | 11 | 15 | 14 | 2 | 10.63 | 11 | 10 | 0.908 | -0.139 | 0.8367861 | NA | 0.2508 | 0.2253 | 0.2313 | 0.2313 | TRUE | 1.8136 |
| 1110007C09I | 844 | 584 | 659 | 815 | 738 | 962 | 655 | 621 | 711 | 811 | 833 | 950 | 763.53 | 662 | 865 | 1.306 | 0.385 | 5.57E-05 | 0.00076562 | 0.0046 | 0.0055 | 0.0052 | 0.0052 | TRUE | 0.0894 |
| 1110008F13I | 5360 | 3645 | 3352 | 3510 | 3085 | 3629 | 4163 | 3877 | 3617 | 3494 | 3480 | 3582 | 3702.21 | 3886 | 3519 | 0.905 | -0.144 | 0.02736877 | 0.08654467 | 0.0022 | 0.0031 | 0.0028 | 0.0028 | TRUE | 0.0482 |
| 1110008J03F | 1187 | 847 | 917 | 989 | 850 | 1078 | 922 | 901 | 989 | 984 | 959 | 1064 | 969.96 | 937 | 1002 | 1.07 | 0.098 | 0.22834083 | 0.39368823 | 0.0016 | 0.0049 | 0.0037 | 0.0037 | TRUE | 0.0348 |
| 1110008L16F | 1989 | 1474 | 1437 | 1502 | 1299 | 1541 | 1545 | 1568 | 1551 | 1495 | 1465 | 1521 | 1524.14 | 1555 | 1494 | 0.961 | -0.057 | 0.37584543 | 0.55040561 | 0 | 0.004 | 0.0023 | 0.0023 | TRUE | 0.0033 |
| 1110008P14I | 911 | 642 | 645 | 829 | 632 | 730 | 708 | 683 | 696 | 825 | 713 | 721 | 724.19 | 696 | 753 | 1.083 | 0.115 | 0.20588926 | 0.36775497 | 0.0022 | 0.0057 | 0.0045 | 0.0045 | TRUE | 0.0842 |
| 1110012L19F | 694 | 516 | 504 | 574 | 478 | 596 | 539 | 549 | 544 | 571 | 539 | 588 | 555.1 | 544 | 566 | 1.042 | 0.06 | 0.51715762 | 0.67926244 | 0 | 0.0067 | 0.0043 | 0.0043 | TRUE | 0.0199 |
| 1110015O18 | 0 | 0 | 0 | 0 | 0 | 0 | 0 | 0 | 0 | 0 | 0 | 0 | 0 | 0 | 0 | NA | NA | NA | NA | NA | NA | NA | NA | NA | NA |
| 1110017D15I | 5 | 3 | 2 | 2 | 4 | 1 | 4 | 3 | 2 | 2 | 5 | 1 | 2.79 | 3 | 3 | 0.791 | -0.337 | 0.76860183 | NA | 0 | 0.8528 | 0.5822 | 0.5822 | TRUE | 0.487 |
| 1110018G07 | 2751 | 1998 | 1899 | 1994 | 1847 | 2035 | 2137 | 2125 | 2049 | 1985 | 2084 | 2009 | 2064.69 | 2104 | 2026 | 0.962 | -0.055 | 0.3661298 | 0.5407764 | 0 | 0.0036 | 0.0022 | 0.0022 | TRUE | 0.0074 |
| 1110019D14I | 346 | 228 | 209 | 301 | 220 | 263 | 269 | 243 | 226 | 300 | 248 | 260 | 257.36 | 246 | 269 | 1.094 | 0.129 | 0.34640735 | 0.52074046 | 0.0052 | 0.0116 | 0.0097 | 0.0097 | TRUE | 0.1105 |
| 1110020A21I | 9 | 9 | 3 | 11 | 5 | 10 | 7 | 10 | 3 | 11 | 6 | 10 | 7.71 | 7 | 9 | 1.339 | 0.421 | 0.55431682 | NA | 0 | 0.3098 | 0.2342 | 0.2342 | TRUE | 0.3425 |
| 1110025L11F | 0 | 0 | 0 | 0 | 0 | 0 | 0 | 0 | 0 | 0 | 0 | 0 | 0 | 0 | 0 | NA | NA | NA | NA | NA | NA | NA | NA | NA | NA |
| 1110028F11I | 0 | 0 | 0 | 0 | 0 | 0 | 0 | 0 | 0 | 0 | 0 | 0 | 0 | 0 | 0 | NA | NA | NA | NA | NA | NA | NA | NA | NA | NA |
| 1110028F18I | 6 | 1 | 4 | 1 | 1 | 2 | 5 | 1 | 4 | 1 | 1 | 2 | 2.36 | 3 | 1 | 0.405 | -1.303 | 0.31608524 | NA | 0 | 1.0083 | 0.6965 | 0.6965 | TRUE | 0.4702 |
| 1110032A03I | 244 | 155 | 113 | 156 | 160 | 146 | 190 | 165 | 122 | 155 | 180 | 144 | 159.37 | 159 | 160 | 1.001 | 0.002 | 0.99280887 | 0.99592003 | 0.0247 | 0.0173 | 0.0194 | 0.0194 | TRUE | 0.4017 |
| 1110032F04I | 143 | 121 | 106 | 118 | 118 | 88 | 111 | 129 | 114 | 117 | 133 | 87 | 115.26 | 118 | 112 | 0.952 | -0.071 | 0.72742127 | 0.83661541 | 0.0164 | 0.023 | 0.0212 | 0.0212 | TRUE | 0.403 |
| 1110034G24 | 304 | 192 | 208 | 242 | 202 | 260 | 236 | 204 | 224 | 241 | 228 | 257 | 231.7 | 221 | 242 | 1.09 | 0.124 | 0.36168646 | 0.53654225 | 0 | 0.0127 | 0.009 | 0.009 | TRUE | 0.0507 |
| 1110035M17 | 4 | 1 | 1 | 3 | 2 | 1 | 3 | 1 | 1 | 3 | 2 | 1 | 1.91 | 2 | 2 | 1.147 | 0.198 | 0.88418977 | NA | 0 | 1.2413 | 0.8188 | 0.8188 | TRUE | 0.4878 |
| 1110036E04I | 0 | 3 | 1 | 2 | 0 | 0 | 0 | 3 | 1 | 2 | 0 | 0 | 1.04 | 1 | 1 | 0.485 | -1.044 | 0.6306568 | NA | 2.3232 | 2.2737 | 2.2831 | 2.2831 | TRUE | 0.547 |
| 1110037F02I | 7537 | 5165 | 4544 | 5397 | 4519 | 5358 | 5854 | 5494 | 4903 | 5372 | 5098 | 5289 | 5334.97 | 5417 | 5253 | 0.969 | -0.045 | 0.52399823 | 0.68541289 | 0.0036 | 0.0029 | 0.0034 | 0.0034 | TRUE | 0.0837 |
| 1110038B12I | 2250 | 1706 | 2594 | 2650 | 2238 | 3040 | 1747 | 1815 | 2799 | 2638 | 2525 | 3001 | 2420.75 | 2120 | 2721 | 1.284 | 0.36 | 0.13795093 | 0.27853264 | 0.0421 | 0.0034 | 0.0134 | 0.0421 | TRUE | 0.9475 |
| 1110038F14I | 1693 | 1165 | 982 | 993 | 893 | 975 | 1315 | 1239 | 1060 | 988 | 1007 | 962 | 1095.32 | 1205 | 986 | 0.817 | -0.291 | 0.00128023 | 0.00861502 | 0.0056 | 0.0046 | 0.005 | 0.005 | TRUE | 0.1314 |
| 1110046J04F | 8 | 9 | 9 | 7 | 7 | 8 | 6 | 10 | 10 | 7 | 8 | 8 | 8.04 | 9 | 8 | 0.902 | -0.149 | 0.82228791 | NA | 0 | 0.2971 | 0.1929 | 0.1929 | TRUE | 0.2258 |
| 1110051M20 | 634 | 455 | 490 | 552 | 482 | 506 | 492 | 484 | 529 | 549 | 544 | 500 | 516.3 | 502 | 531 | 1.059 | 0.082 | 0.40129923 | 0.57443865 | 0 | 0.007 | 0.0049 | 0.0049 | TRUE | 0.0319 |
| 1110054M08 | 60 | 53 | 48 | 79 | 75 | 85 | 47 | 56 | 52 | 79 | 85 | 84 | 66.99 | 52 | 83 | 1.603 | 0.681 | 0.00417351 | 0.02133476 | 0 | 0.0378 | 0.0251 | 0.0251 | TRUE | 0.0702 |
| 1110057K04I | 2433 | 1792 | 1514 | 1379 | 1260 | 1516 | 1890 | 1906 | 1634 | 1373 | 1421 | 1497 | 1619.99 | 1810 | 1430 | 0.79 | -0.34 | 1.96E-05 | 0.00032664 | 0.0041 | 0.0039 | 0.0039 | 0.0039 | TRUE | 0.0862 |
| 1110058L19F | 877 | 667 | 675 | 741 | 681 | 725 | 681 | 709 | 728 | 738 | 768 | 716 | 723.41 | 706 | 741 | 1.049 | 0.069 | 0.41344589 | 0.5860164 | 0 | 0.0057 | 0.0037 | 0.0037 | TRUE | 0.0124 |
| 1110059E24I | 2200 | 1407 | 1490 | 1541 | 1438 | 1626 | 1709 | 1497 | 1608 | 1534 | 1622 | 1605 | 1595.7 | 1605 | 1587 | 0.988 | -0.017 | 0.81324591 | 0.89214244 | 0.0021 | 0.0039 | 0.0032 | 0.0032 | TRUE | 0.0415 |
| 1110059G10 | 525 | 363 | 332 | 419 | 296 | 366 | 408 | 386 | 358 | 417 | 334 | 361 | 377.4 | 384 | 371 | 0.965 | -0.051 | 0.66917405 | 0.79664374 | 0.0058 | 0.0087 | 0.0078 | 0.0078 | TRUE | 0.1372 |
| 1110059M19 | 0 | 0 | 0 | 0 | 0 | 0 | 0 | 0 | 0 | 0 | 0 | 0 | 0 | 0 | 0 | NA | NA | NA | NA | NA | NA | NA | NA | NA | NA |
| 1110065P20I | 510 | 449 | 394 | 470 | 382 | 427 | 396 | 478 | 425 | 468 | 431 | 422 | 436.52 | 433 | 440 | 1.019 | 0.027 | 0.80769366 | 0.88878016 | 0.004 | 0.0079 | 0.0066 | 0.0066 | TRUE | 0.0937 |
| 1190002F15I | 89 | 73 | 92 | 109 | 93 | 111 | 69 | 78 | 99 | 108 | 105 | 110 | 94.84 | 82 | 108 | 1.321 | 0.402 | 0.05749915 | 0.14912554 | 0.0055 | 0.0274 | 0.0215 | 0.0215 | TRUE | 0.3084 |
| 1190002N15 | 2413 | 1933 | 1800 | 1757 | 1622 | 1817 | 1874 | 2056 | 1942 | 1749 | 1830 | 1794 | 1874.12 | 1957 | 1791 | 0.915 | -0.128 | 0.05319114 | 0.14089342 | 9.00E-04 | 0.0037 | 0.0026 | 0.0026 | TRUE | 0.0234 |

| | | | | | | | | | | | | | | | | | | | | | | | | | |
|---|---|---|---|---|---|---|---|---|---|---|---|---|---|---|---|---|---|---|---|---|---|---|---|---|---|
| 1190003J15F | 508 | 378 | 382 | 389 | 399 | 474 | 395 | 402 | 412 | 387 | 450 | 468 | 419 | 403 | 435 | 1.08 | 0.111 | 0.31571395 | 0.48947819 | 0.0029 | 0.0081 | 0.0064 | 0.0064 | TRUE | 0.109 |
| 1190003K10I | 0 | 0 | 0 | 0 | 1 | 0 | 0 | 0 | 0 | 0 | 1 | 0 | 0.19 | 0 | 0 | 2.094 | 1.067 | 0.79379329 | NA | 0 | 12.608 | 10 | 10 | TRUE | 0.5075 |
| 1190005I06R | 460 | 336 | 340 | 498 | 442 | 512 | 357 | 357 | 367 | 496 | 499 | 505 | 430.21 | 360 | 500 | 1.387 | 0.472 | 2.35E-06 | 5.73E-05 | 0 | 0.0079 | 0.0048 | 0.0048 | TRUE | 0.0026 |
| 1190007I07R | 462 | 378 | 450 | 494 | 420 | 448 | 359 | 402 | 486 | 492 | 474 | 442 | 442.37 | 416 | 469 | 1.133 | 0.18 | 0.14654021 | 0.29090266 | 0.011 | 0.0078 | 0.0088 | 0.0088 | TRUE | 0.2492 |
| 1200011I18R | 3679 | 2636 | 2692 | 2753 | 2445 | 2744 | 2857 | 2804 | 2905 | 2740 | 2758 | 2709 | 2795.52 | 2855 | 2736 | 0.958 | -0.062 | 0.26379456 | 0.43407614 | 0 | 0.0033 | 0.0018 | 0.0018 | TRUE | 0.0029 |
| 1200014J11F | 5894 | 4169 | 4071 | 4177 | 3626 | 4224 | 4578 | 4435 | 4393 | 4158 | 4091 | 4170 | 4303.79 | 4469 | 4140 | 0.926 | -0.111 | 0.03266862 | 0.09859219 | 0 | 0.003 | 0.0017 | 0.0017 | TRUE | 0.0057 |
| 1300002E11I | 459 | 388 | 259 | 307 | 268 | 292 | 356 | 413 | 279 | 306 | 302 | 288 | 324.14 | 349 | 299 | 0.854 | -0.228 | 0.11236854 | 0.24075603 | 0.0163 | 0.0097 | 0.0117 | 0.0117 | TRUE | 0.2365 |
| 1300002K09I | 21 | 15 | 12 | 9 | 16 | 10 | 16 | 16 | 13 | 9 | 18 | 10 | 13.68 | 15 | 12 | 0.807 | -0.31 | 0.55968188 | NA | 0 | 0.1756 | 0.1289 | 0.1289 | TRUE | 0.6088 |
| 1300015D01I | 4 | 2 | 3 | 4 | 1 | 2 | 3 | 2 | 3 | 4 | 1 | 2 | 2.59 | 3 | 2 | 0.841 | -0.25 | 0.83047845 | NA | 0 | 0.9166 | 0.5953 | 0.5953 | TRUE | 0.3813 |
| 1300017J02F | 84 | 70 | 50 | 98 | 71 | 62 | 65 | 74 | 54 | 98 | 80 | 61 | 72.08 | 64 | 80 | 1.232 | 0.301 | 0.23617119 | 0.40257542 | 0.026 | 0.0353 | 0.0327 | 0.0327 | TRUE | 0.1741 |
| 1300018J18F | 351 | 247 | 343 | 401 | 377 | 436 | 273 | 263 | 370 | 399 | 425 | 430 | 360.05 | 302 | 418 | 1.389 | 0.474 | 0.00057285 | 0.00479097 | 0.0159 | 0.009 | 0.0108 | 0.0108 | TRUE | 0.4299 |
| 1500002O10 | 5 | 0 | 1 | 2 | 2 | 2 | 4 | 0 | 1 | 2 | 2 | 2 | 1.86 | 2 | 2 | 1.191 | 0.253 | 0.86109851 | NA | 0 | 1.274 | 0.9752 | 0.9752 | TRUE | 1.1187 |
| 1500004A13I | 9 | 7 | 5 | 4 | 0 | 8 | 7 | 7 | 5 | 4 | 0 | 8 | 5.28 | 6 | 4 | 0.608 | -0.718 | 0.44014755 | NA | 0.2618 | 0.4509 | 0.4212 | 0.4212 | TRUE | 0.3055 |
| 1500009C09I | 15 | 7 | 15 | 28 | 12 | 23 | 12 | 7 | 16 | 28 | 14 | 23 | 16.57 | 12 | 22 | 1.827 | 0.87 | 0.08882508 | NA | 0.0593 | 0.1455 | 0.1231 | 0.1231 | TRUE | 0.2555 |
| 1500009L16F | 2197 | 1681 | 1809 | 2151 | 1767 | 2024 | 1706 | 1788 | 1952 | 2141 | 1993 | 1998 | 1929.79 | 1815 | 2044 | 1.127 | 0.173 | 0.01750537 | 0.06256671 | 0.0024 | 0.0037 | 0.0033 | 0.0033 | TRUE | 0.0519 |
| 1500011B03I | 865 | 687 | 739 | 760 | 696 | 796 | 672 | 731 | 797 | 757 | 785 | 786 | 754.56 | 733 | 776 | 1.06 | 0.084 | 0.3544753 | 0.52886383 | 0.0027 | 0.0056 | 0.0046 | 0.0046 | TRUE | 0.069 |
| 1500011K16I | 604 | 445 | 484 | 637 | 553 | 626 | 469 | 473 | 522 | 634 | 624 | 618 | 556.76 | 488 | 625 | 1.282 | 0.359 | 0.00013157 | 0.00153142 | 0 | 0.0067 | 0.0045 | 0.0045 | TRUE | 0.0419 |
| 1500012F01I | 3413 | 2743 | 4293 | 4938 | 4288 | 5515 | 2651 | 2918 | 4632 | 4915 | 4837 | 5444 | 4232.88 | 3400 | 5065 | 1.49 | 0.575 | 0.03220754 | 0.09753299 | 0.0518 | 0.003 | 0.015 | 0.0518 | TRUE | 1.2207 |
| 1500012K07I | 0 | 0 | 0 | 0 | 0 | 0 | 0 | 0 | 0 | 0 | 0 | 0 | 0 | 0 | 0 | NA | NA | NA | NA | NA | NA | NA | NA | NA | NA |
| 1500015A07I | 287 | 240 | 183 | 215 | 163 | 189 | 223 | 255 | 197 | 214 | 184 | 187 | 210.02 | 225 | 195 | 0.866 | -0.208 | 0.16971627 | 0.32133456 | 0.0068 | 0.0137 | 0.0117 | 0.0117 | TRUE | 0.1422 |
| 1500015L24F | 0 | 0 | 0 | 0 | 0 | 0 | 0 | 0 | 0 | 0 | 0 | 0 | 0 | 0 | 0 | NA | NA | NA | NA | NA | NA | NA | NA | NA | NA |
| 1500015O10 | 24 | 11 | 11 | 17 | 10 | 13 | 19 | 12 | 12 | 17 | 11 | 13 | 13.87 | 14 | 14 | 0.963 | -0.054 | 0.91729195 | NA | 0 | 0.1733 | 0.1238 | 0.1238 | TRUE | 0.439 |
| 1500016L03F | 1 | 0 | 0 | 1 | 0 | 0 | 1 | 0 | 0 | 1 | 0 | 0 | 0.3 | 0 | 0 | 1.075 | 0.105 | 0.97698525 | NA | 0 | 8.0272 | 7.5141 | 7.5141 | TRUE | 0.3906 |
| 1500017E21I | 2 | 2 | 0 | 0 | 2 | 3 | 2 | 2 | 0 | 0 | 2 | 3 | 1.48 | 1 | 2 | 1.397 | 0.482 | 0.77483248 | NA | 0 | 1.6005 | 1.3576 | 1.3576 | TRUE | 0.6254 |
| 1500032L24F | 2604 | 1882 | 1778 | 2014 | 1729 | 2111 | 2022 | 2002 | 1918 | 2005 | 1950 | 2084 | 1996.97 | 1981 | 2013 | 1.016 | 0.023 | 0.71571051 | 0.82931903 | 4.00E-04 | 0.0036 | 0.0024 | 0.0024 | TRUE | 0.0115 |
| 1600002D24I | 0 | 0 | 0 | 0 | 0 | 1 | 0 | 0 | 0 | 0 | 0 | 1 | 0.16 | 0 | 0 | 2.094 | 1.067 | 0.79379329 | NA | 0 | 14.4078 | 10 | 10 | TRUE | 0.3835 |
| 1600002H07I | 1538 | 1154 | 1117 | 1246 | 1088 | 1304 | 1194 | 1227 | 1205 | 1240 | 1227 | 1287 | 1230.35 | 1209 | 1251 | 1.036 | 0.051 | 0.46366275 | 0.63284298 | 0 | 0.0044 | 0.0026 | 0.0026 | TRUE | 0.0075 |
| 1600002K03I | 71 | 64 | 63 | 50 | 39 | 53 | 55 | 68 | 68 | 50 | 44 | 52 | 56.21 | 64 | 49 | 0.769 | -0.379 | 0.14688951 | 0.29127243 | 0 | 0.0446 | 0.031 | 0.031 | TRUE | 0.1407 |
| 1600010M07 | 1 | 0 | 0 | 0 | 1 | 0 | 1 | 0 | 0 | 0 | 1 | 0 | 0.32 | 0 | 0 | 1.075 | 0.105 | 0.97638847 | NA | 0 | 7.4681 | 7.0392 | 7.0392 | TRUE | 0.5075 |
| 1600012H06I | 1274 | 896 | 861 | 1014 | 827 | 982 | 989 | 953 | 929 | 1009 | 933 | 969 | 963.87 | 957 | 970 | 1.014 | 0.019 | 0.80206128 | 0.88530866 | 0 | 0.0049 | 0.0033 | 0.0033 | TRUE | 0.0146 |
| 1600014C10I | 477 | 321 | 328 | 390 | 314 | 346 | 370 | 341 | 354 | 388 | 354 | 342 | 358.3 | 355 | 361 | 1.016 | 0.023 | 0.83992311 | 0.90890647 | 0 | 0.009 | 0.0064 | 0.0064 | TRUE | 0.0494 |
| 1600014C23I | 21 | 22 | 12 | 14 | 13 | 18 | 16 | 23 | 13 | 14 | 15 | 18 | 16.5 | 17 | 16 | 0.883 | -0.179 | 0.70877039 | NA | 0 | 0.146 | 0.1046 | 0.1046 | TRUE | 0.3926 |
| 1600014K23I | 27 | 15 | 17 | 12 | 9 | 12 | 21 | 16 | 18 | 12 | 10 | 12 | 14.87 | 18 | 11 | 0.613 | -0.707 | 0.15382085 | NA | 0 | 0.1618 | 0.1054 | 0.1054 | TRUE | 0.0928 |
| 1600015I10R | 1 | 1 | 0 | 1 | 1 | 2 | 1 | 1 | 0 | 1 | 1 | 2 | 0.99 | 1 | 1 | 2.197 | 1.135 | 0.55299242 | NA | 0 | 2.3972 | 1.4894 | 1.4894 | TRUE | 0.1884 |
| 1600016N20 | 245 | 243 | 277 | 272 | 202 | 287 | 190 | 258 | 299 | 271 | 228 | 283 | 254.93 | 249 | 261 | 1.052 | 0.073 | 0.66631997 | 0.7948039 | 0.0285 | 0.0117 | 0.0167 | 0.0167 | TRUE | 0.4357 |
| 1600019K03I | 0 | 0 | 0 | 0 | 0 | 0 | 0 | 0 | 0 | 0 | 0 | 0 | 0 | 0 | 0 | NA | NA | NA | NA | NA | NA | NA | NA | NA | NA |
| 1600020E01I | 133 | 134 | 122 | 114 | 88 | 95 | 103 | 143 | 132 | 113 | 99 | 94 | 114 | 126 | 102 | 0.817 | -0.292 | 0.14337553 | 0.28629036 | 0.012 | 0.0232 | 0.0198 | 0.0198 | TRUE | 0.2784 |
| 1600023N17 | 5 | 14 | 8 | 9 | 6 | 9 | 4 | 15 | 9 | 9 | 7 | 9 | 8.67 | 9 | 8 | 0.918 | -0.124 | 0.85799371 | NA | 0.1214 | 0.2758 | 0.2299 | 0.2299 | TRUE | 0.2502 |
| 1600025M17 | 1 | 0 | 1 | 0 | 1 | 0 | 1 | 0 | 1 | 0 | 1 | 0 | 0.5 | 1 | 0 | 0.596 | -0.748 | 0.78656967 | NA | 0 | 4.7684 | 3.7205 | 3.7205 | TRUE | 0.5075 |
| 1600029D21I | 312 | 218 | 245 | 216 | 213 | 270 | 242 | 232 | 264 | 215 | 240 | 267 | 243.4 | 246 | 241 | 0.978 | -0.032 | 0.81678901 | 0.89487832 | 0.0037 | 0.0122 | 0.0097 | 0.0097 | TRUE | 0.1014 |
| 1600029I14R | 0 | 0 | 0 | 0 | 0 | 0 | 0 | 0 | 0 | 0 | 0 | 0 | 0 | 0 | 0 | NA | NA | NA | NA | NA | NA | NA | NA | NA | NA |
| 1600029O15 | 11 | 9 | 6 | 8 | 6 | 7 | 9 | 10 | 6 | 8 | 7 | 7 | 7.71 | 8 | 7 | 0.879 | -0.186 | 0.78319412 | NA | 0 | 0.31 | 0.1977 | 0.1977 | TRUE | 0.0902 |
| 1700001C02I | 1 | 2 | 5 | 2 | 2 | 1 | 1 | 2 | 5 | 2 | 2 | 1 | 2.26 | 3 | 2 | 0.643 | -0.637 | 0.62769968 | NA | 0 | 1.0532 | 0.766 | 0.766 | TRUE | 0.5072 |
| 1700001C19I | 42 | 24 | 19 | 35 | 29 | 21 | 33 | 26 | 21 | 35 | 33 | 21 | 27.82 | 27 | 30 | 1.111 | 0.151 | 0.69492478 | 0.81401081 | 0.0295 | 0.0876 | 0.0715 | 0.0715 | TRUE | 0.4552 |
| 1700001D01I | 0 | 0 | 0 | 0 | 0 | 0 | 0 | 0 | 0 | 0 | 0 | 0 | 0 | 0 | 0 | NA | NA | NA | NA | NA | NA | NA | NA | NA | NA |
| 1700001F09I | 2 | 0 | 0 | 0 | 0 | 1 | 2 | 0 | 0 | 0 | 0 | 1 | 0.42 | 1 | 0 | 0.682 | -0.553 | 0.86290097 | NA | 0 | 5.6 | 5.5066 | 5.5066 | TRUE | 0.9727 |
| 1700001G11 | 45 | 35 | 31 | 45 | 41 | 48 | 35 | 37 | 33 | 45 | 46 | 47 | 40.68 | 35 | 46 | 1.311 | 0.39 | 0.18854672 | 0.34603213 | 0 | 0.0607 | 0.0386 | 0.0386 | TRUE | 0.0163 |
| 1700001G17 | 7 | 7 | 6 | 6 | 2 | 5 | 5 | 7 | 6 | 6 | 2 | 5 | 5.42 | 6 | 4 | 0.69 | -0.535 | 0.51516028 | NA | 0 | 0.4397 | 0.2951 | 0.2951 | TRUE | 0.2835 |
| 1700001J03F | 5 | 5 | 3 | 5 | 4 | 2 | 4 | 5 | 3 | 5 | 5 | 2 | 3.98 | 4 | 4 | 0.92 | -0.12 | 0.89921223 | NA | 0 | 0.5974 | 0.3951 | 0.3951 | TRUE | 0.3019 |
| 1700001J11F | 0 | 0 | 0 | 1 | 0 | 1 | 0 | 0 | 0 | 1 | 0 | 1 | 0.33 | 0 | 1 | 3.763 | 1.912 | 0.57887588 | NA | 0 | 7.1752 | 6.7756 | 6.7756 | TRUE | 0.2094 |
| 1700001K19I | 1 | 0 | 0 | 0 | 0 | 0 | 1 | 0 | 0 | 0 | 0 | 0 | 0.13 | 0 | 0 | 0.552 | -0.857 | 0.8336548 | NA | 0 | 18.3121 | 10 | 10 | TRUE | 0.2566 |
| 1700001K23I | 1 | 1 | 0 | 1 | 1 | 0 | 1 | 1 | 0 | 1 | 1 | 0 | 0.66 | 1 | 1 | 1.128 | 0.173 | 0.94106052 | NA | 0 | 3.5899 | 2.4471 | 2.4471 | TRUE | 0.2681 |
| 1700001L05F | 3132 | 2716 | 2572 | 2948 | 2314 | 2709 | 2432 | 2889 | 2775 | 2934 | 2610 | 2674 | 2719.28 | 2699 | 2739 | 1.016 | 0.023 | 0.77074818 | 0.86436278 | 0.0056 | 0.0033 | 0.0041 | 0.0041 | TRUE | 0.0923 |
| 1700001L19F | 32 | 30 | 12 | 12 | 10 | 10 | 25 | 32 | 13 | 12 | 11 | 10 | 17.13 | 23 | 11 | 0.473 | -1.079 | 0.03354076 | 0.10058872 | 0 | 0.1407 | 0.118 | 0.118 | TRUE | 0.2089 |
| 1700001O22 | 43 | 38 | 29 | 48 | 38 | 29 | 33 | 40 | 31 | 48 | 43 | 29 | 37.4 | 35 | 40 | 1.135 | 0.183 | 0.58163826 | 0.73222699 | 0.0176 | 0.0658 | 0.0523 | 0.0523 | TRUE | 0.4644 |
| 1700001P01I | 8 | 11 | 10 | 11 | 11 | 8 | 14 | 6 | 12 | 11 | 9 | 14 | 10.42 | 10 | 11 | 1.198 | 0.261 | 0.66369582 | NA | 0 | 0.23 | 0.1624 | 0.1624 | TRUE | 0.4196 |
| 1700003C15I | 0 | 0 | 0 | 0 | 0 | 0 | 0 | 0 | 0 | 0 | 0 | 0 | 0 | 0 | 0 | NA | NA | NA | NA | NA | NA | NA | NA | NA | NA |
| 1700003D09I | 0 | 2 | 0 | 2 | 0 | 0 | 0 | 2 | 0 | 2 | 0 | 0 | 0.69 | 1 | 1 | 0.971 | -0.042 | 0.98762268 | NA | 8.094 | 3.4555 | 3.8966 | 3.8966 | TRUE | 0.9747 |
| 1700003E16I | 12 | 9 | 4 | 10 | 1 | 2 | 9 | 10 | 4 | 10 | 1 | 2 | 6.04 | 8 | 4 | 0.566 | -0.822 | 0.35846161 | NA | 0.4066 | 0.3945 | 0.3976 | 0.3976 | TRUE | 2.3106 |
| 1700003F12I | 15 | 3 | 11 | 21 | 13 | 19 | 12 | 3 | 12 | 21 | 15 | 19 | 13.51 | 9 | 18 | 2.013 | 1.009 | 0.07634544 | NA | 0.042 | 0.1779 | 0.1518 | 0.1518 | TRUE | 0.9126 |
| 1700003F17I | 0 | 0 | 0 | 0 | 0 | 0 | 0 | 0 | 0 | 0 | 0 | 0 | 0 | 0 | 0 | NA | NA | NA | NA | NA | NA | NA | NA | NA | NA |
| 1700003G13 | 0 | 0 | 0 | 0 | 1 | 0 | 0 | 0 | 0 | 0 | 1 | 0 | 0.19 | 0 | 0 | 2.094 | 1.067 | 0.79379329 | NA | 0 | 12.608 | 10 | 10 | TRUE | 0.5075 |
| 1700003G18 | 2 | 5 | 1 | 4 | 2 | 0 | 2 | 5 | 1 | 4 | 2 | 0 | 2.36 | 3 | 2 | 0.792 | -0.337 | 0.80399175 | NA | 0.5599 | 1.0047 | 0.8918 | 0.8918 | TRUE | 0.3516 |
| 1700003H04I | 0 | 0 | 0 | 0 | 0 | 0 | 0 | 0 | 0 | 0 | 0 | 0 | 0 | 0 | 0 | NA | NA | NA | NA | NA | NA | NA | NA | NA | NA |
| 1700003L19F | 0 | 0 | 0 | 0 | 0 | 0 | 0 | 0 | 0 | 0 | 0 | 0 | 0 | 0 | 0 | NA | NA | NA | NA | NA | NA | NA | NA | NA | NA |
| 1700003M02 | 26 | 17 | 13 | 15 | 11 | 10 | 20 | 18 | 14 | 15 | 12 | 10 | 14.92 | 17 | 12 | 0.707 | -0.5 | 0.31697306 | NA | 0 | 0.1613 | 0.1104 | 0.1104 | TRUE | 0.1371 |
| 1700003M07 | 67 | 34 | 30 | 25 | 34 | 45 | 52 | 36 | 32 | 25 | 38 | 44 | 38.04 | 40 | 36 | 0.885 | -0.177 | 0.60998999 | 0.75322701 | 0.0496 | 0.0647 | 0.0604 | 0.0604 | TRUE | 0.5729 |
| 1700003P14I | 0 | 0 | 0 | 0 | 0 | 0 | 0 | 0 | 0 | 0 | 0 | 0 | 0 | 0 | 0 | NA | NA | NA | NA | NA | NA | NA | NA | NA | NA |
| 1700006A11I | 0 | 0 | 0 | 0 | 0 | 0 | 0 | 0 | 0 | 0 | 0 | 0 | 0 | 0 | 0 | NA | NA | NA | NA | NA | NA | NA | NA | NA | NA |

| | | | | | | | | | | | | | | | | | | | | | | | | | |
|---|---|---|---|---|---|---|---|---|---|---|---|---|---|---|---|---|---|---|---|---|---|---|---|---|---|
| 1700006E09 | 1 | 0 | 1 | 1 | 0 | 1 | 1 | 0 | 1 | 1 | 0 | 1 | 0.64 | 1 | 1 | 1.077 | 0.107 | 0.96403683 | NA | 0 | 3.7074 | 2.4851 | 2.4851 | TRUE | 0.2041 |
| 1700006F04 | 0 | 0 | 0 | 0 | 0 | 0 | 0 | 0 | 0 | 0 | 0 | 0 | 0 | 0 | 0 | NA | NA | NA | NA | NA | NA | NA | NA | NA | NA |
| 1700006H21 | 0 | 2 | 0 | 0 | 1 | 1 | 0 | 2 | 0 | 0 | 1 | 1 | 0.71 | 1 | 1 | 1.018 | 0.026 | 0.99165522 | NA | 0 | 3.3542 | 2.9835 | 2.9835 | TRUE | 0.9748 |
| 1700007B14 | 2 | 0 | 2 | 1 | 0 | 0 | 2 | 0 | 2 | 1 | 0 | 0 | 0.78 | 1 | 0 | 0.298 | -1.746 | 0.45605179 | NA | 0 | 3.0237 | 2.564 | 2.564 | TRUE | 0.4004 |
| 1700007F19 | 0 | 0 | 0 | 0 | 0 | 0 | 0 | 0 | 0 | 0 | 0 | 0 | 0 | 0 | 0 | NA | NA | NA | NA | NA | NA | NA | NA | NA | NA |
| 1700007G11 | 6 | 8 | 4 | 8 | 1 | 2 | 5 | 9 | 4 | 8 | 1 | 2 | 4.76 | 6 | 4 | 0.643 | -0.637 | 0.50438872 | NA | 0.2487 | 0.5005 | 0.4334 | 0.4334 | TRUE | 1.6246 |
| 1700007J10 | 0 | 0 | 0 | 1 | 0 | 1 | 0 | 0 | 0 | 1 | 0 | 1 | 0.33 | 0 | 1 | 3.763 | 1.912 | 0.57887588 | NA | 0 | 7.1752 | 6.7756 | 6.7756 | TRUE | 0.2094 |
| 1700007K09 | 0 | 0 | 1 | 2 | 0 | 3 | 0 | 0 | 1 | 2 | 0 | 3 | 1.01 | 0 | 2 | 4.836 | 2.274 | 0.29627751 | NA | 0 | 2.3602 | 2.1051 | 2.1051 | TRUE | 0.4212 |
| 1700007K13 | 354 | 282 | 325 | 298 | 236 | 312 | 275 | 300 | 351 | 297 | 266 | 308 | 299.4 | 309 | 290 | 0.945 | -0.082 | 0.5372441 | 0.69599128 | 0.0074 | 0.0103 | 0.0094 | 0.0094 | TRUE | 0.1597 |
| 1700007L15 | 141 | 93 | 71 | 107 | 74 | 85 | 110 | 99 | 77 | 107 | 83 | 84 | 93.16 | 95 | 91 | 0.957 | -0.063 | 0.77579432 | 0.86757979 | 0.0159 | 0.0279 | 0.0245 | 0.0245 | TRUE | 0.2672 |
| 1700007P06 | 1 | 0 | 0 | 0 | 0 | 0 | 1 | 0 | 0 | 0 | 0 | 0 | 0.13 | 0 | 0 | 0.552 | -0.857 | 0.8336548 | NA | 0 | 18.3121 | 10 | 10 | TRUE | 0.2566 |
| 1700008A04 | 0 | 0 | 0 | 0 | 0 | 0 | 0 | 0 | 0 | 0 | 0 | 0 | 0 | 0 | 0 | NA | NA | NA | NA | NA | NA | NA | NA | NA | NA |
| 1700008F21 | 0 | 0 | 0 | 0 | 1 | 0 | 0 | 0 | 0 | 0 | 1 | 0 | 0.19 | 0 | 0 | 2.094 | 1.067 | 0.79379329 | NA | 0 | 12.608 | 10 | 10 | TRUE | 0.5075 |
| 1700008I05R | 96 | 76 | 50 | 76 | 54 | 67 | 75 | 81 | 54 | 76 | 61 | 66 | 68.68 | 70 | 68 | 0.967 | -0.048 | 0.84637159 | 0.91278874 | 0.0111 | 0.0369 | 0.0301 | 0.0301 | TRUE | 0.3301 |
| 1700008J07 | 67 | 54 | 56 | 45 | 47 | 48 | 52 | 57 | 60 | 45 | 53 | 47 | 52.52 | 56 | 48 | 0.856 | -0.225 | 0.3974043 | 0.57074956 | 0 | 0.0476 | 0.0315 | 0.0315 | TRUE | 0.0522 |
| 1700008K24 | 1 | 1 | 0 | 0 | 0 | 0 | 1 | 1 | 0 | 0 | 0 | 0 | 0.31 | 1 | 0 | 0.309 | -1.696 | 0.63183309 | NA | 0 | 7.7294 | 7.2472 | 7.2472 | TRUE | 0.2015 |
| 1700008O03 | 118 | 75 | 55 | 72 | 84 | 80 | 92 | 80 | 59 | 72 | 95 | 79 | 79.36 | 77 | 82 | 1.055 | 0.077 | 0.74944702 | 0.85149811 | 0.0209 | 0.0323 | 0.0291 | 0.0291 | TRUE | 0.3536 |
| 1700008P02 | 0 | 0 | 0 | 0 | 0 | 0 | 0 | 0 | 0 | 0 | 0 | 0 | 0 | 0 | 0 | NA | NA | NA | NA | NA | NA | NA | NA | NA | NA |
| 1700009C05 | 0 | 3 | 2 | 4 | 2 | 3 | 0 | 3 | 2 | 4 | 2 | 3 | 2.42 | 2 | 3 | 1.806 | 0.853 | 0.51164564 | NA | 0 | 0.9799 | 0.7654 | 0.7654 | TRUE | 0.9041 |
| 1700009J07 | 0 | 0 | 0 | 0 | 0 | 0 | 0 | 0 | 0 | 0 | 0 | 0 | 0 | 0 | 0 | NA | NA | NA | NA | NA | NA | NA | NA | NA | NA |
| 1700009N14 | 0 | 0 | 0 | 0 | 0 | 0 | 0 | 0 | 0 | 0 | 0 | 0 | 0 | 0 | 0 | NA | NA | NA | NA | NA | NA | NA | NA | NA | NA |
| 1700009P17 | 37 | 34 | 36 | 44 | 29 | 42 | 29 | 36 | 39 | 44 | 33 | 41 | 36.95 | 35 | 39 | 1.148 | 0.199 | 0.53760664 | 0.69620473 | 0 | 0.0666 | 0.0477 | 0.0477 | TRUE | 0.1868 |
| 1700010B08 | 1 | 0 | 0 | 0 | 0 | 0 | 1 | 0 | 0 | 0 | 0 | 0 | 0.13 | 0 | 0 | 0.552 | -0.857 | 0.8336548 | NA | 0 | 18.3121 | 10 | 10 | TRUE | 0.2566 |
| 1700010D01 | 2 | 6 | 1 | 0 | 0 | 0 | 2 | 6 | 1 | 0 | 0 | 0 | 1.5 | 3 | 0 | 0.064 | -3.969 | 0.03894114 | NA | 1.0986 | 1.5799 | 1.4998 | 1.4998 | TRUE | 1.1894 |
| 1700010I02R | 1 | 0 | 0 | 0 | 0 | 0 | 1 | 0 | 0 | 0 | 0 | 0 | 0.13 | 0 | 0 | 0.552 | -0.857 | 0.8336548 | NA | 0 | 18.3121 | 10 | 10 | TRUE | 0.2566 |
| 1700010I14R | 26 | 22 | 28 | 24 | 25 | 27 | 20 | 23 | 30 | 24 | 28 | 27 | 25.43 | 24 | 26 | 1.074 | 0.102 | 0.78921001 | 0.87619843 | 0 | 0.0956 | 0.0662 | 0.0662 | TRUE | 0.23 |
| 1700010J16 | 0 | 0 | 0 | 0 | 0 | 0 | 0 | 0 | 0 | 0 | 0 | 0 | 0 | 0 | 0 | NA | NA | NA | NA | NA | NA | NA | NA | NA | NA |
| 1700010K23 | 0 | 0 | 0 | 0 | 0 | 0 | 0 | 0 | 0 | 0 | 0 | 0 | 0 | 0 | 0 | NA | NA | NA | NA | NA | NA | NA | NA | NA | NA |
| 1700011A15 | 0 | 0 | 1 | 1 | 2 | 0 | 0 | 0 | 1 | 1 | 2 | 0 | 0.72 | 0 | 1 | 3.105 | 1.634 | 0.50324076 | NA | 0 | 3.2861 | 2.8125 | 2.8125 | TRUE | 0.3023 |
| 1700011B04 | 0 | 0 | 0 | 0 | 0 | 0 | 0 | 0 | 0 | 0 | 0 | 0 | 0 | 0 | 0 | NA | NA | NA | NA | NA | NA | NA | NA | NA | NA |
| 1700011E24 | 0 | 4 | 4 | 5 | 0 | 5 | 0 | 4 | 4 | 5 | 0 | 5 | 3.08 | 3 | 3 | 1.204 | 0.267 | 0.8394653 | NA | 1.6701 | 0.7718 | 0.9255 | 0.9255 | TRUE | 1.3132 |
| 1700011H14 | 0 | 0 | 0 | 0 | 0 | 0 | 0 | 0 | 0 | 0 | 0 | 0 | 0 | 0 | 0 | NA | NA | NA | NA | NA | NA | NA | NA | NA | NA |
| 1700011I03R | 0 | 0 | 0 | 0 | 0 | 0 | 0 | 0 | 0 | 0 | 0 | 0 | 0 | 0 | 0 | NA | NA | NA | NA | NA | NA | NA | NA | NA | NA |
| 1700011J10 | 330 | 185 | 209 | 242 | 198 | 180 | 256 | 197 | 226 | 241 | 223 | 178 | 220.09 | 226 | 214 | 0.942 | -0.085 | 0.59608814 | 0.74235543 | 0.0165 | 0.0132 | 0.0142 | 0.0142 | TRUE | 0.2463 |
| 1700011L22 | 1 | 0 | 0 | 0 | 0 | 0 | 1 | 0 | 0 | 0 | 0 | 0 | 0.13 | 0 | 0 | 0.552 | -0.857 | 0.8336548 | NA | 0 | 18.3121 | 10 | 10 | TRUE | 0.2566 |
| 1700011M02 | 1 | 0 | 0 | 1 | 0 | 0 | 1 | 0 | 0 | 1 | 0 | 0 | 0.3 | 0 | 0 | 1.075 | 0.105 | 0.97698525 | NA | 0 | 8.0272 | 7.5141 | 7.5141 | TRUE | 0.3906 |
| 1700012A03 | 0 | 0 | 0 | 0 | 0 | 0 | 0 | 0 | 0 | 0 | 0 | 0 | 0 | 0 | 0 | NA | NA | NA | NA | NA | NA | NA | NA | NA | NA |
| 1700012B07 | 32 | 21 | 12 | 7 | 9 | 13 | 25 | 22 | 13 | 7 | 10 | 13 | 15.02 | 20 | 10 | 0.495 | -1.016 | 0.05498907 | NA | 0 | 0.1603 | 0.1262 | 0.1262 | TRUE | 0.4855 |
| 1700012B09 | 0 | 1 | 0 | 1 | 1 | 0 | 0 | 1 | 0 | 1 | 1 | 0 | 0.53 | 0 | 1 | 2.036 | 1.026 | 0.70492654 | NA | 0 | 4.4642 | 3.5518 | 3.5518 | TRUE | 0.5217 |
| 1700012B15 | 338 | 239 | 186 | 181 | 175 | 173 | 263 | 254 | 201 | 180 | 197 | 171 | 210.96 | 239 | 183 | 0.761 | -0.393 | 0.01060235 | 0.04310301 | 0.0088 | 0.0137 | 0.0122 | 0.0122 | TRUE | 0.2118 |
| 1700012D01 | 7 | 7 | 15 | 11 | 10 | 17 | 5 | 7 | 16 | 11 | 11 | 17 | 11.35 | 9 | 13 | 1.369 | 0.453 | 0.45945168 | NA | 0.0973 | 0.2113 | 0.1795 | 0.1795 | TRUE | 1.1047 |
| 1700012D14 | 18 | 24 | 33 | 26 | 27 | 28 | 14 | 26 | 36 | 26 | 30 | 28 | 26.52 | 25 | 28 | 1.134 | 0.181 | 0.66262478 | 0.79181473 | 0.0703 | 0.0918 | 0.0857 | 0.0857 | TRUE | 0.278 |
| 1700012I11R | 0 | 0 | 0 | 0 | 0 | 0 | 0 | 0 | 0 | 0 | 0 | 0 | 0 | 0 | 0 | NA | NA | NA | NA | NA | NA | NA | NA | NA | NA |
| 1700012L04 | 0 | 0 | 0 | 0 | 0 | 0 | 0 | 0 | 0 | 0 | 0 | 0 | 0 | 0 | 0 | NA | NA | NA | NA | NA | NA | NA | NA | NA | NA |
| 1700012P22 | 0 | 0 | 0 | 0 | 0 | 0 | 0 | 0 | 0 | 0 | 0 | 0 | 0 | 0 | 0 | NA | NA | NA | NA | NA | NA | NA | NA | NA | NA |
| 1700013D24 | 0 | 0 | 0 | 0 | 0 | 0 | 0 | 0 | 0 | 0 | 0 | 0 | 0 | 0 | 0 | NA | NA | NA | NA | NA | NA | NA | NA | NA | NA |
| 1700013F07 | 10 | 3 | 4 | 8 | 3 | 3 | 8 | 3 | 4 | 8 | 3 | 3 | 4.93 | 5 | 5 | 0.924 | -0.115 | 0.89701542 | NA | 0 | 0.4831 | 0.3637 | 0.3637 | TRUE | 0.6841 |
| 1700013G23 | 13 | 9 | 19 | 9 | 8 | 11 | 10 | 10 | 21 | 9 | 9 | 11 | 11.5 | 14 | 10 | 0.725 | -0.464 | 0.42994794 | NA | 0 | 0.2085 | 0.1584 | 0.1584 | TRUE | 0.854 |
| 1700013G24 | 3 | 3 | 2 | 1 | 4 | 2 | 2 | 3 | 2 | 1 | 5 | 2 | 2.53 | 2 | 3 | 0.964 | -0.053 | 0.96462048 | NA | 0 | 0.9404 | 0.6279 | 0.6279 | TRUE | 0.4886 |
| 1700013H16 | 1723 | 1288 | 943 | 404 | 317 | 342 | 1338 | 1370 | 1017 | 402 | 358 | 338 | 803.84 | 1242 | 366 | 0.294 | -1.764 | 1.16E-47 | 3.33E-44 | 0.0164 | 0.0054 | 0.0089 | 0.0089 | TRUE | 0.2962 |
| 1700015E13 | 0 | 0 | 0 | 0 | 0 | 0 | 0 | 0 | 0 | 0 | 0 | 0 | 0 | 0 | 0 | NA | NA | NA | NA | NA | NA | NA | NA | NA | NA |
| 1700015F17 | 0 | 0 | 1 | 1 | 0 | 0 | 0 | 0 | 1 | 1 | 0 | 0 | 0.35 | 0 | 0 | 1.075 | 0.105 | 0.97567165 | NA | 0 | 6.8576 | 6.5143 | 6.5143 | TRUE | 0.5365 |
| 1700015G11 | 0 | 0 | 0 | 0 | 0 | 0 | 0 | 0 | 0 | 0 | 0 | 0 | 0 | 0 | 0 | NA | NA | NA | NA | NA | NA | NA | NA | NA | NA |
| 1700016C15 | 0 | 0 | 0 | 1 | 0 | 0 | 0 | 0 | 0 | 1 | 0 | 0 | 0.17 | 0 | 0 | 2.094 | 1.067 | 0.79379329 | NA | 0 | 14.2885 | 10 | 10 | TRUE | 0.3906 |
| 1700016D06 | 1 | 0 | 2 | 5 | 0 | 1 | 1 | 0 | 2 | 5 | 0 | 1 | 1.48 | 1 | 2 | 2.076 | 1.054 | 0.55332228 | NA | 1.2479 | 1.6004 | 1.5201 | 1.5201 | TRUE | 1.1524 |
| 1700016G22 | 3 | 1 | 2 | 1 | 1 | 1 | 2 | 1 | 2 | 1 | 1 | 1 | 1.44 | 2 | 1 | 0.554 | -0.851 | 0.58248919 | NA | 0 | 1.6441 | 0.9697 | 0.9697 | TRUE | 0.1064 |
| 1700016H13 | 30 | 17 | 20 | 11 | 13 | 18 | 23 | 18 | 22 | 11 | 15 | 18 | 17.72 | 21 | 15 | 0.687 | -0.543 | 0.23848623 | 0.40536136 | 0 | 0.1361 | 0.094 | 0.094 | TRUE | 0.2102 |
| 1700016K19 | 129 | 117 | 111 | 100 | 78 | 99 | 100 | 124 | 120 | 100 | 88 | 98 | 104.94 | 115 | 95 | 0.833 | -0.264 | 0.17835276 | 0.33291716 | 0 | 0.025 | 0.0181 | 0.0181 | TRUE | 0.1398 |
| 1700016L04 | 1 | 0 | 1 | 2 | 1 | 2 | 1 | 0 | 1 | 2 | 1 | 2 | 1.16 | 1 | 2 | 2.723 | 1.445 | 0.42461796 | NA | 0 | 2.0489 | 1.2898 | 1.2898 | TRUE | 0.1933 |
| 1700016L21 | 0 | 0 | 0 | 0 | 0 | 0 | 0 | 0 | 0 | 0 | 0 | 0 | 0 | 0 | 0 | NA | NA | NA | NA | NA | NA | NA | NA | NA | NA |
| 1700016P04 | 0 | 0 | 0 | 0 | 0 | 0 | 0 | 0 | 0 | 0 | 0 | 0 | 0 | 0 | 0 | NA | NA | NA | NA | NA | NA | NA | NA | NA | NA |
| 1700017B05 | 2633 | 1872 | 1655 | 1799 | 1581 | 1756 | 2045 | 1991 | 1786 | 1791 | 1784 | 1733 | 1854.93 | 1941 | 1769 | 0.911 | -0.135 | 0.05966265 | 0.15268725 | 0.0022 | 0.0037 | 0.0031 | 0.0031 | TRUE | 0.0582 |
| 1700017D01 | 0 | 0 | 0 | 0 | 0 | 0 | 0 | 0 | 0 | 0 | 0 | 0 | 0 | 0 | 0 | NA | NA | NA | NA | NA | NA | NA | NA | NA | NA |
| 1700017G19 | 0 | 0 | 0 | 0 | 0 | 0 | 0 | 0 | 0 | 0 | 0 | 0 | 0 | 0 | 0 | NA | NA | NA | NA | NA | NA | NA | NA | NA | NA |
| 1700017J07 | 1 | 2 | 4 | 4 | 1 | 1 | 1 | 2 | 4 | 4 | 1 | 1 | 2.22 | 2 | 2 | 0.872 | -0.198 | 0.88104639 | NA | 0 | 1.0703 | 0.8034 | 0.8034 | TRUE | 0.4372 |
| 1700017N19 | 0 | 0 | 0 | 0 | 0 | 0 | 0 | 0 | 0 | 0 | 0 | 0 | 0 | 0 | 0 | NA | NA | NA | NA | NA | NA | NA | NA | NA | NA |
| 1700018A04 | 0 | 0 | 0 | 0 | 0 | 0 | 0 | 0 | 0 | 0 | 0 | 0 | 0 | 0 | 0 | NA | NA | NA | NA | NA | NA | NA | NA | NA | NA |
| 1700018B08 | 172 | 126 | 107 | 135 | 87 | 119 | 134 | 134 | 115 | 134 | 98 | 117 | 122.18 | 128 | 116 | 0.914 | -0.129 | 0.49433054 | 0.65952267 | 0.0061 | 0.0218 | 0.0176 | 0.0176 | TRUE | 0.1838 |
| 1700018B24 | 754 | 526 | 495 | 421 | 318 | 311 | 586 | 559 | 534 | 419 | 359 | 307 | 460.67 | 560 | 362 | 0.645 | -0.632 | 1.91E-07 | 6.96E-06 | 0.0097 | 0.0076 | 0.0083 | 0.0083 | TRUE | 0.2261 |

| | | | | | | | | | | | | | | | | | | | | | | | | | |
|---|---|---|---|---|---|---|---|---|---|---|---|---|---|---|---|---|---|---|---|---|---|---|---|---|---|
| 1700018C11I | 0 | 0 | 0 | 0 | 0 | 0 | 0 | 0 | 0 | 0 | 0 | 0 | 0 | 0 | 0 | NA | NA | NA | NA | NA | NA | NA | NA | NA | NA |
| 1700018F24I | 0 | 0 | 0 | 0 | 0 | 0 | 0 | 0 | 0 | 0 | 0 | 0 | 0 | 0 | 0 | NA | NA | NA | NA | NA | NA | NA | NA | NA | NA |
| 1700018G05 | 1 | 0 | 0 | 0 | 0 | 1 | 1 | 0 | 0 | 0 | 0 | 1 | 0.29 | 0 | 0 | 1.075 | 0.105 | 0.97702352 | NA | 0 | 8.0647 | 7.5458 | 7.5458 | TRUE | 0.3835 |
| 1700018L02F | 6 | 1 | 1 | 3 | 2 | 0 | 5 | 1 | 1 | 3 | 2 | 0 | 2.01 | 2 | 2 | 0.742 | -0.43 | 0.76604242 | NA | 0.4852 | 1.183 | 0.9987 | 0.9987 | TRUE | 1.1181 |
| 1700019A02I | 114 | 105 | 96 | 86 | 55 | 50 | 89 | 112 | 104 | 86 | 62 | 49 | 83.47 | 102 | 66 | 0.651 | -0.62 | 0.01140377 | 0.04556725 | 0.0298 | 0.0308 | 0.0306 | 0.0306 | TRUE | 0.6405 |
| 1700019B03I | 2 | 2 | 5 | 1 | 4 | 3 | 2 | 2 | 5 | 1 | 5 | 3 | 2.92 | 3 | 3 | 0.942 | -0.086 | 0.94028597 | NA | 0 | 0.8129 | 0.6033 | 0.6033 | TRUE | 0.3889 |
| 1700019B21I | 202 | 144 | 153 | 123 | 111 | 94 | 157 | 153 | 165 | 122 | 125 | 93 | 135.93 | 158 | 113 | 0.715 | -0.483 | 0.00721525 | 0.03247092 | 0.003 | 0.0199 | 0.0157 | 0.0157 | TRUE | 0.2607 |
| 1700019D03I | 641 | 473 | 535 | 693 | 603 | 698 | 498 | 503 | 577 | 690 | 680 | 689 | 606.22 | 526 | 686 | 1.307 | 0.386 | 4.65E-05 | 0.00065753 | 0.0014 | 0.0063 | 0.0048 | 0.0048 | TRUE | 0.0823 |
| 1700019E08I | 34 | 24 | 32 | 16 | 11 | 11 | 26 | 26 | 35 | 16 | 12 | 11 | 20.94 | 29 | 13 | 0.455 | -1.136 | 0.00928243 | 0.03920621 | 0 | 0.1156 | 0.0811 | 0.0811 | TRUE | 0.1836 |
| 1700019G17 | 453 | 345 | 330 | 333 | 324 | 369 | 352 | 367 | 356 | 331 | 366 | 364 | 356.02 | 358 | 354 | 0.987 | -0.018 | 0.86918566 | 0.92778752 | 0 | 0.0091 | 0.006 | 0.006 | TRUE | 0.0352 |
| 1700019G24 | 7 | 2 | 5 | 1 | 2 | 4 | 5 | 2 | 5 | 1 | 2 | 4 | 3.36 | 4 | 2 | 0.552 | -0.856 | 0.42563024 | NA | 0 | 0.7078 | 0.5058 | 0.5058 | TRUE | 0.3583 |
| 1700019L03F | 3 | 1 | 2 | 3 | 0 | 3 | 2 | 1 | 2 | 3 | 0 | 3 | 1.92 | 2 | 2 | 1.078 | 0.108 | 0.93874831 | NA | 0 | 1.2391 | 0.9165 | 0.9165 | TRUE | 0.5868 |
| 1700019M22 | 0 | 0 | 0 | 0 | 0 | 0 | 0 | 0 | 0 | 0 | 0 | 0 | 0 | 0 | 0 | NA | NA | NA | NA | NA | NA | NA | NA | NA | NA |
| 1700019N19 | 41 | 23 | 23 | 35 | 39 | 28 | 32 | 24 | 25 | 35 | 44 | 28 | 31.27 | 27 | 36 | 1.299 | 0.377 | 0.28987509 | 0.4618935 | 0 | 0.0782 | 0.0591 | 0.0591 | TRUE | 0.1951 |
| 1700019O17 | 1 | 1 | 2 | 0 | 1 | 0 | 1 | 1 | 2 | 0 | 1 | 0 | 0.85 | 1 | 0 | 0.282 | -1.828 | 0.39462251 | NA | 0 | 2.7764 | 1.9548 | 1.9548 | TRUE | 0.5075 |
| 1700020A23I | 0 | 0 | 0 | 0 | 0 | 0 | 0 | 0 | 0 | 0 | 0 | 0 | 0 | 0 | 0 | NA | NA | NA | NA | NA | NA | NA | NA | NA | NA |
| 1700020D05I | 15 | 10 | 10 | 14 | 12 | 16 | 12 | 11 | 11 | 14 | 14 | 16 | 12.72 | 11 | 15 | 1.306 | 0.385 | 0.46307196 | NA | 0 | 0.1887 | 0.1193 | 0.1193 | TRUE | 0.0317 |
| 1700020G17 | 0 | 0 | 0 | 0 | 0 | 0 | 0 | 0 | 0 | 0 | 0 | 0 | 0 | 0 | 0 | NA | NA | NA | NA | NA | NA | NA | NA | NA | NA |
| 1700020I14R | 1426 | 1134 | 996 | 1185 | 1056 | 1119 | 1108 | 1206 | 1075 | 1180 | 1191 | 1105 | 1143.98 | 1130 | 1159 | 1.026 | 0.037 | 0.638421 | 0.77423614 | 0.0017 | 0.0045 | 0.0035 | 0.0035 | TRUE | 0.0412 |
| 1700020L24F | 6 | 4 | 2 | 8 | 2 | 4 | 5 | 4 | 2 | 8 | 2 | 4 | 4.21 | 4 | 5 | 1.275 | 0.35 | 0.71240963 | NA | 0 | 0.5658 | 0.4124 | 0.4124 | TRUE | 0.5797 |
| 1700020M21 | 0 | 1 | 1 | 1 | 0 | 0 | 0 | 1 | 1 | 1 | 0 | 0 | 0.52 | 1 | 0 | 0.535 | -0.904 | 0.74017109 | NA | 0 | 4.534 | 3.6534 | 3.6534 | TRUE | 0.3906 |
| 1700020N01 | 97 | 78 | 59 | 37 | 26 | 38 | 75 | 83 | 64 | 37 | 29 | 38 | 54.27 | 74 | 35 | 0.468 | -1.095 | 7.23E-05 | 0.00093916 | 0 | 0.0461 | 0.0334 | 0.0334 | TRUE | 0.1268 |
| 1700020N15 | 5 | 7 | 3 | 5 | 4 | 3 | 4 | 7 | 3 | 5 | 5 | 3 | 4.5 | 5 | 4 | 0.859 | -0.219 | 0.80728783 | NA | 0 | 0.5288 | 0.3566 | 0.3566 | TRUE | 0.3969 |
| 1700020N18 | 1 | 1 | 0 | 0 | 0 | 0 | 1 | 1 | 0 | 0 | 0 | 0 | 0.31 | 1 | 0 | 0.309 | -1.696 | 0.63183309 | NA | 0 | 7.7294 | 7.2472 | 7.2472 | TRUE | 0.2015 |
| 1700021F05I | 1459 | 990 | 1112 | 1130 | 954 | 1182 | 1133 | 1053 | 1200 | 1125 | 1076 | 1167 | 1125.64 | 1129 | 1123 | 0.995 | -0.007 | 0.92702513 | 0.95969289 | 0.0019 | 0.0045 | 0.0036 | 0.0036 | TRUE | 0.0397 |
| 1700021F07I | 5 | 1 | 3 | 7 | 5 | 3 | 4 | 1 | 3 | 7 | 6 | 3 | 3.96 | 3 | 5 | 1.864 | 0.898 | 0.36247638 | NA | 0 | 0.6011 | 0.4277 | 0.4277 | TRUE | 0.3095 |
| 1700021K19I | 2496 | 1747 | 1824 | 2027 | 1910 | 2179 | 1939 | 1858 | 1968 | 2018 | 2155 | 2151 | 2014.71 | 1922 | 2108 | 1.097 | 0.133 | 0.03753511 | 0.10938338 | 6.00E-04 | 0.0036 | 0.0024 | 0.0024 | TRUE | 0.017 |
| 1700021N21 | 0 | 0 | 0 | 0 | 0 | 1 | 0 | 0 | 0 | 0 | 0 | 1 | 0.16 | 0 | 0 | 2.094 | 1.067 | 0.79379329 | NA | 0 | 14.4078 | 10 | 10 | TRUE | 0.3835 |
| 1700022A21I | 0 | 2 | 2 | 1 | 2 | 1 | 0 | 2 | 2 | 1 | 2 | 1 | 1.42 | 1 | 1 | 1.019 | 0.027 | 0.98701488 | NA | 0 | 1.6707 | 1.2215 | 1.2215 | TRUE | 0.7215 |
| 1700022A22I | 0 | 0 | 0 | 0 | 0 | 0 | 0 | 0 | 0 | 0 | 0 | 0 | 0 | 0 | 0 | NA | NA | NA | NA | NA | NA | NA | NA | NA | NA |
| 1700022E09I | 0 | 0 | 0 | 0 | 0 | 0 | 0 | 0 | 0 | 0 | 0 | 0 | 0 | 0 | 0 | NA | NA | NA | NA | NA | NA | NA | NA | NA | NA |
| 1700022H16I | 0 | 0 | 0 | 0 | 0 | 0 | 0 | 0 | 0 | 0 | 0 | 0 | 0 | 0 | 0 | NA | NA | NA | NA | NA | NA | NA | NA | NA | NA |
| 1700022I11R | 0 | 0 | 0 | 0 | 0 | 0 | 0 | 0 | 0 | 0 | 0 | 0 | 0 | 0 | 0 | NA | NA | NA | NA | NA | NA | NA | NA | NA | NA |
| 1700022P22I | 9 | 6 | 3 | 0 | 0 | 0 | 7 | 6 | 3 | 0 | 0 | 0 | 2.77 | 5 | 0 | 0.034 | -4.87 | 0.002057 | NA | 0 | 0.8586 | 0.7127 | 0.7127 | TRUE | 0.269 |
| 1700023A16I | 2 | 1 | 1 | 0 | 0 | 0 | 2 | 1 | 1 | 0 | 0 | 0 | 0.62 | 1 | 0 | 0.154 | -2.702 | 0.27412557 | NA | 0 | 3.8499 | 3.0088 | 3.0088 | TRUE | 0.0718 |
| 1700023C21I | 0 | 0 | 0 | 1 | 0 | 0 | 0 | 0 | 0 | 1 | 0 | 0 | 0.17 | 0 | 0 | 2.094 | 1.067 | 0.79379329 | NA | 0 | 14.2885 | 10 | 10 | TRUE | 0.3906 |
| 1700023E05I | 3 | 4 | 2 | 2 | 2 | 1 | 2 | 4 | 2 | 2 | 2 | 1 | 2.33 | 3 | 2 | 0.599 | -0.74 | 0.55084868 | NA | 0 | 1.0198 | 0.6436 | 0.6436 | TRUE | 0.1899 |
| 1700023F02I | 0 | 0 | 0 | 0 | 0 | 0 | 0 | 0 | 0 | 0 | 0 | 0 | 0 | 0 | 0 | NA | NA | NA | NA | NA | NA | NA | NA | NA | NA |
| 1700023F06I | 112 | 88 | 82 | 106 | 87 | 110 | 87 | 94 | 88 | 106 | 98 | 109 | 96.89 | 90 | 104 | 1.163 | 0.218 | 0.26678915 | 0.43719529 | 0 | 0.0269 | 0.0173 | 0.0173 | TRUE | 0.0232 |
| 1700023L04F | 13 | 9 | 11 | 18 | 11 | 16 | 10 | 10 | 12 | 18 | 12 | 16 | 12.94 | 11 | 15 | 1.47 | 0.555 | 0.29351036 | NA | 0 | 0.1855 | 0.1219 | 0.1219 | TRUE | 0.1186 |
| 1700024B18I | 0 | 0 | 0 | 0 | 0 | 0 | 0 | 0 | 0 | 0 | 0 | 0 | 0 | 0 | 0 | NA | NA | NA | NA | NA | NA | NA | NA | NA | NA |
| 1700024F13I | 2 | 3 | 0 | 3 | 0 | 1 | 2 | 3 | 0 | 3 | 0 | 1 | 1.45 | 2 | 1 | 0.849 | -0.237 | 0.8905531 | NA | 0.6566 | 1.6336 | 1.4395 | 1.4395 | TRUE | 0.2416 |
| 1700024G13 | 0 | 0 | 0 | 0 | 0 | 0 | 0 | 0 | 0 | 0 | 0 | 0 | 0 | 0 | 0 | NA | NA | NA | NA | NA | NA | NA | NA | NA | NA |
| 1700024P03I | 0 | 0 | 0 | 0 | 0 | 0 | 0 | 0 | 0 | 0 | 0 | 0 | 0 | 0 | 0 | NA | NA | NA | NA | NA | NA | NA | NA | NA | NA |
| 1700024P04I | 0 | 0 | 0 | 0 | 0 | 0 | 0 | 0 | 0 | 0 | 0 | 0 | 0 | 0 | 0 | NA | NA | NA | NA | NA | NA | NA | NA | NA | NA |
| 1700024P16I | 10 | 10 | 8 | 6 | 10 | 10 | 8 | 11 | 9 | 6 | 11 | 10 | 9.03 | 9 | 9 | 1.005 | 0.007 | 0.99163408 | NA | 0 | 0.265 | 0.1798 | 0.1798 | TRUE | 0.2914 |
| 1700025B11I | 1 | 1 | 1 | 0 | 0 | 0 | 1 | 1 | 1 | 0 | 0 | 0 | 0.49 | 1 | 0 | 0.196 | -2.347 | 0.38405676 | NA | 0 | 4.8735 | 3.7415 | 3.7415 | TRUE | 0.0718 |
| 1700025C18I | 0 | 0 | 0 | 0 | 0 | 0 | 0 | 0 | 0 | 0 | 0 | 0 | 0 | 0 | 0 | NA | NA | NA | NA | NA | NA | NA | NA | NA | NA |
| 1700025E21I | 5 | 2 | 5 | 5 | 3 | 1 | 4 | 2 | 5 | 5 | 3 | 1 | 3.46 | 4 | 3 | 0.818 | -0.29 | 0.78231833 | NA | 0 | 0.6876 | 0.5013 | 0.5013 | TRUE | 0.3936 |
| 1700025F22I | 0 | 0 | 0 | 0 | 0 | 0 | 0 | 0 | 0 | 0 | 0 | 0 | 0 | 0 | 0 | NA | NA | NA | NA | NA | NA | NA | NA | NA | NA |
| 1700025F24I | 11 | 12 | 5 | 3 | 2 | 1 | 9 | 13 | 5 | 3 | 2 | 1 | 5.49 | 9 | 2 | 0.233 | -2.103 | 0.02404812 | NA | 0 | 0.4342 | 0.322 | 0.322 | TRUE | 0.1125 |
| 1700025G04 | 1000 | 758 | 707 | 709 | 636 | 696 | 777 | 806 | 763 | 706 | 717 | 687 | 742.68 | 782 | 703 | 0.899 | -0.153 | 0.0626205 | 0.158288 | 0 | 0.0056 | 0.0035 | 0.0035 | TRUE | 0.0084 |
| 1700025K24I | 0 | 0 | 0 | 0 | 0 | 0 | 0 | 0 | 0 | 0 | 0 | 0 | 0 | 0 | 0 | NA | NA | NA | NA | NA | NA | NA | NA | NA | NA |
| 1700025M24 | 0 | 0 | 0 | 0 | 0 | 0 | 0 | 0 | 0 | 0 | 0 | 0 | 0 | 0 | 0 | NA | NA | NA | NA | NA | NA | NA | NA | NA | NA |
| 1700025N23 | 0 | 0 | 0 | 0 | 0 | 0 | 0 | 0 | 0 | 0 | 0 | 0 | 0 | 0 | 0 | NA | NA | NA | NA | NA | NA | NA | NA | NA | NA |
| 1700026D08I | 8 | 4 | 8 | 8 | 4 | 11 | 6 | 4 | 9 | 8 | 5 | 11 | 7.07 | 6 | 8 | 1.232 | 0.301 | 0.68141933 | NA | 0 | 0.3375 | 0.2445 | 0.2445 | TRUE | 0.1837 |
| 1700026D11I | 0 | 0 | 0 | 0 | 0 | 0 | 0 | 0 | 0 | 0 | 0 | 0 | 0 | 0 | 0 | NA | NA | NA | NA | NA | NA | NA | NA | NA | NA |
| 1700026F02I | 0 | 0 | 0 | 0 | 0 | 0 | 0 | 0 | 0 | 0 | 0 | 0 | 0 | 0 | 0 | NA | NA | NA | NA | NA | NA | NA | NA | NA | NA |
| 1700026L06F | 18 | 29 | 20 | 18 | 17 | 8 | 14 | 31 | 22 | 18 | 19 | 8 | 18.57 | 22 | 15 | 0.681 | -0.554 | 0.2722058 | 0.44317548 | 0.1188 | 0.1301 | 0.1269 | 0.1269 | TRUE | 0.2429 |
| 1700027A15I | 0 | 0 | 0 | 0 | 0 | 0 | 0 | 0 | 0 | 0 | 0 | 0 | 0 | 0 | 0 | NA | NA | NA | NA | NA | NA | NA | NA | NA | NA |
| 1700027F09I | 0 | 0 | 0 | 0 | 0 | 0 | 0 | 0 | 0 | 0 | 0 | 0 | 0 | 0 | 0 | NA | NA | NA | NA | NA | NA | NA | NA | NA | NA |
| 1700027H10I | 0 | 0 | 0 | 0 | 0 | 0 | 0 | 0 | 0 | 0 | 0 | 0 | 0 | 0 | 0 | NA | NA | NA | NA | NA | NA | NA | NA | NA | NA |
| 1700027I24R | 1 | 1 | 0 | 0 | 0 | 1 | 1 | 1 | 0 | 0 | 0 | 1 | 0.47 | 1 | 0 | 0.599 | -0.74 | 0.79209856 | NA | 0 | 5.0317 | 3.8925 | 3.8925 | TRUE | 0.3835 |
| 1700027J07F | 0 | 0 | 0 | 0 | 0 | 0 | 0 | 0 | 0 | 0 | 0 | 0 | 0 | 0 | 0 | NA | NA | NA | NA | NA | NA | NA | NA | NA | NA |
| 1700028B04I | 50 | 48 | 42 | 33 | 37 | 48 | 39 | 51 | 45 | 33 | 42 | 47 | 42.86 | 45 | 41 | 0.906 | -0.142 | 0.63964296 | 0.77477341 | 0 | 0.0577 | 0.0429 | 0.0429 | TRUE | 0.2204 |
| 1700028D13I | 0 | 1 | 1 | 0 | 0 | 1 | 0 | 1 | 1 | 0 | 0 | 1 | 0.52 | 1 | 0 | 0.535 | -0.904 | 0.7403423 | NA | 0 | 4.5459 | 3.6616 | 3.6616 | TRUE | 0.3835 |
| 1700028E10I | 22 | 14 | 13 | 20 | 12 | 13 | 17 | 15 | 14 | 20 | 14 | 13 | 15.38 | 15 | 16 | 1.003 | 0.005 | 0.9923834 | NA | 0 | 0.1565 | 0.1068 | 0.1068 | TRUE | 0.3075 |
| 1700028I16R | 119 | 85 | 58 | 69 | 68 | 66 | 92 | 90 | 63 | 69 | 77 | 65 | 75.99 | 82 | 70 | 0.853 | -0.229 | 0.3373647 | 0.51150458 | 0.013 | 0.0336 | 0.0279 | 0.0279 | TRUE | 0.3748 |
| 1700028J19F | 36 | 31 | 25 | 24 | 29 | 30 | 28 | 33 | 27 | 24 | 33 | 30 | 29.02 | 29 | 29 | 0.98 | -0.029 | 0.93542314 | 0.96438292 | 0 | 0.0841 | 0.0572 | 0.0572 | TRUE | 0.1441 |

| | | | | | | | | | | | | | | | | | | | | | | | | | |
|---|---|---|---|---|---|---|---|---|---|---|---|---|---|---|---|---|---|---|---|---|---|---|---|---|---|
| 1700028K03I | 33 | 41 | 26 | 26 | 23 | 28 | 26 | 44 | 28 | 26 | 26 | 28 | 29.46 | 33 | 27 | 0.823 | -0.282 | 0.44575835 | 0.61648773 | 0 | 0.0829 | 0.0639 | 0.0639 | TRUE | 0.6036 |
| 1700028M03 | 0 | 0 | 0 | 0 | 0 | 0 | 0 | 0 | 0 | 0 | 0 | 0 | 0 | 0 | 0 | NA | NA | NA | NA | NA | NA | NA | NA | NA | NA |
| 1700028P14I | 2 | 1 | 1 | 2 | 1 | 2 | 2 | 1 | 1 | 2 | 1 | 2 | 1.46 | 1 | 2 | 1.369 | 0.453 | 0.76363599 | NA | 0 | 1.6204 | 0.9453 | 0.9453 | TRUE | 0.0576 |
| 1700028P15I | 1 | 3 | 2 | 0 | 3 | 0 | 1 | 3 | 2 | 0 | 3 | 0 | 1.59 | 2 | 1 | 0.542 | -0.884 | 0.60433826 | NA | 0.9218 | 1.4977 | 1.3811 | 1.3811 | TRUE | 1.2221 |
| 1700029B22I | 0 | 0 | 0 | 0 | 0 | 0 | 0 | 0 | 0 | 0 | 0 | 0 | 0 | 0 | 0 | NA | NA | NA | NA | NA | NA | NA | NA | NA | NA |
| 1700029F12I | 0 | 0 | 0 | 0 | 0 | 1 | 0 | 0 | 0 | 0 | 0 | 1 | 0.16 | 0 | 0 | 2.094 | 1.067 | 0.79379329 | NA | 0 | 14.4078 | 10 | 10 | TRUE | 0.3835 |
| 1700029H14I | 0 | 0 | 1 | 1 | 1 | 2 | 0 | 0 | 1 | 1 | 1 | 2 | 0.86 | 0 | 1 | 3.971 | 1.99 | 0.35178206 | NA | 0 | 2.7494 | 1.9121 | 1.9121 | TRUE | 0.5365 |
| 1700029I15R | 6 | 3 | 1 | 3 | 2 | 2 | 5 | 3 | 1 | 3 | 2 | 2 | 2.69 | 3 | 2 | 0.792 | -0.337 | 0.77126773 | NA | 0 | 0.8831 | 0.592 | 0.592 | TRUE | 0.2504 |
| 1700029J03F | 0 | 0 | 0 | 0 | 0 | 1 | 0 | 0 | 0 | 0 | 0 | 1 | 0.16 | 0 | 0 | 2.094 | 1.067 | 0.79379329 | NA | 0 | 14.4078 | 10 | 10 | TRUE | 0.3835 |
| 1700029J07F | 46 | 32 | 56 | 87 | 60 | 70 | 36 | 34 | 60 | 87 | 68 | 69 | 58.93 | 43 | 75 | 1.73 | 0.79 | 0.00602672 | 0.02830757 | 0.0384 | 0.0426 | 0.0415 | 0.0415 | TRUE | 0.8618 |
| 1700029M2C | 45 | 37 | 35 | 73 | 63 | 77 | 35 | 39 | 38 | 73 | 71 | 76 | 55.3 | 37 | 73 | 1.968 | 0.977 | 0.00018727 | 0.00203546 | 0 | 0.0453 | 0.0293 | 0.0293 | TRUE | 0.0284 |
| 1700029N11 | 0 | 0 | 0 | 0 | 0 | 0 | 0 | 0 | 0 | 0 | 0 | 0 | 0 | 0 | 0 | NA | NA | NA | NA | NA | NA | NA | NA | NA | NA |
| 1700029P11I | 478 | 401 | 362 | 256 | 216 | 296 | 371 | 427 | 391 | 255 | 244 | 292 | 329.84 | 396 | 264 | 0.667 | -0.584 | 2.37E-06 | 5.77E-05 | 0.0037 | 0.0096 | 0.0079 | 0.0079 | TRUE | 0.102 |
| 1700030A11I | 1 | 0 | 0 | 0 | 0 | 1 | 1 | 0 | 0 | 0 | 0 | 1 | 0.29 | 0 | 0 | 1.075 | 0.105 | 0.97702352 | NA | 0 | 8.0647 | 7.5458 | 7.5458 | TRUE | 0.3835 |
| 1700030C10I | 254 | 220 | 168 | 153 | 179 | 190 | 197 | 234 | 181 | 152 | 202 | 188 | 192.39 | 204 | 181 | 0.884 | -0.178 | 0.27943161 | 0.45103512 | 0.0132 | 0.0147 | 0.0143 | 0.0143 | TRUE | 0.2038 |
| 1700030F04I | 0 | 0 | 0 | 0 | 0 | 0 | 0 | 0 | 0 | 0 | 0 | 0 | 0 | 0 | 0 | NA | NA | NA | NA | NA | NA | NA | NA | NA | NA |
| 1700030F18I | 0 | 1 | 0 | 0 | 0 | 0 | 0 | 1 | 0 | 0 | 0 | 0 | 0.18 | 0 | 0 | 0.552 | -0.857 | 0.8336548 | NA | 0 | 13.3714 | 10 | 10 | TRUE | 0.5216 |
| 1700030J22F | 509 | 442 | 364 | 364 | 373 | 422 | 395 | 470 | 393 | 362 | 421 | 417 | 409.65 | 419 | 400 | 0.954 | -0.068 | 0.56690627 | 0.7208889 | 0.0063 | 0.0082 | 0.0076 | 0.0076 | TRUE | 0.1263 |
| 1700030K09I | 892 | 596 | 666 | 735 | 632 | 724 | 693 | 634 | 719 | 732 | 713 | 715 | 700.77 | 682 | 720 | 1.055 | 0.078 | 0.37391466 | 0.54847723 | 0 | 0.0058 | 0.0041 | 0.0041 | TRUE | 0.0426 |
| 1700030L20F | 11 | 6 | 3 | 3 | 2 | 0 | 9 | 6 | 3 | 3 | 2 | 0 | 3.9 | 6 | 2 | 0.283 | -1.82 | 0.09885333 | NA | 0 | 0.61 | 0.4984 | 0.4984 | TRUE | 0.4202 |
| 1700030M09 | 8 | 7 | 5 | 5 | 6 | 2 | 6 | 7 | 5 | 5 | 7 | 2 | 5.46 | 6 | 5 | 0.714 | -0.487 | 0.5576565 | NA | 0 | 0.4363 | 0.3059 | 0.3059 | TRUE | 0.4821 |
| 1700030N03 | 0 | 0 | 0 | 0 | 0 | 0 | 0 | 0 | 0 | 0 | 0 | 0 | 0 | 0 | 0 | NA | NA | NA | NA | NA | NA | NA | NA | NA | NA |
| 1700030O20 | 4 | 1 | 3 | 8 | 7 | 10 | 3 | 1 | 3 | 8 | 8 | 10 | 5.52 | 2 | 9 | 3.428 | 1.777 | 0.04093489 | NA | 0 | 0.4315 | 0.2926 | 0.2926 | TRUE | 0.2382 |
| 1700031A10I | 2 | 1 | 0 | 1 | 0 | 1 | 2 | 1 | 0 | 1 | 0 | 1 | 0.77 | 1 | 1 | 0.75 | -0.415 | 0.8501676 | NA | 0 | 3.0941 | 2.17 | 2.17 | TRUE | 0.2799 |
| 1700031F05I | 1 | 0 | 2 | 2 | 3 | 0 | 1 | 0 | 2 | 2 | 3 | 0 | 1.38 | 1 | 2 | 1.831 | 0.872 | 0.62309697 | NA | 0 | 1.7137 | 1.4802 | 1.4802 | TRUE | 0.2107 |
| 1700031M16 | 0 | 0 | 0 | 0 | 0 | 0 | 0 | 0 | 0 | 0 | 0 | 0 | 0 | 0 | 0 | NA | NA | NA | NA | NA | NA | NA | NA | NA | NA |
| 1700031P21I | 0 | 0 | 0 | 0 | 0 | 0 | 0 | 0 | 0 | 0 | 0 | 0 | 0 | 0 | 0 | NA | NA | NA | NA | NA | NA | NA | NA | NA | NA |
| 1700034E13I | 0 | 0 | 0 | 0 | 0 | 0 | 0 | 0 | 0 | 0 | 0 | 0 | 0 | 0 | 0 | NA | NA | NA | NA | NA | NA | NA | NA | NA | NA |
| 1700034F02I | 30 | 18 | 16 | 30 | 22 | 17 | 23 | 19 | 17 | 30 | 25 | 17 | 21.86 | 20 | 24 | 1.188 | 0.248 | 0.55744314 | 0.71234106 | 0 | 0.1108 | 0.0833 | 0.0833 | TRUE | 0.3229 |
| 1700034G24 | 0 | 0 | 0 | 0 | 0 | 0 | 0 | 0 | 0 | 0 | 0 | 0 | 0 | 0 | 0 | NA | NA | NA | NA | NA | NA | NA | NA | NA | NA |
| 1700034H15I | 31 | 25 | 21 | 17 | 19 | 14 | 24 | 27 | 23 | 17 | 21 | 14 | 20.92 | 25 | 17 | 0.709 | -0.497 | 0.23882338 | 0.40578965 | 0 | 0.1157 | 0.0785 | 0.0785 | TRUE | 0.1873 |
| 1700034I23R | 0 | 0 | 0 | 0 | 0 | 0 | 0 | 0 | 0 | 0 | 0 | 0 | 0 | 0 | 0 | NA | NA | NA | NA | NA | NA | NA | NA | NA | NA |
| 1700034J05F | 0 | 0 | 0 | 0 | 0 | 0 | 0 | 0 | 0 | 0 | 0 | 0 | 0 | 0 | 0 | NA | NA | NA | NA | NA | NA | NA | NA | NA | NA |
| 1700034K08I | 0 | 0 | 0 | 0 | 0 | 1 | 0 | 0 | 0 | 0 | 0 | 1 | 0.16 | 0 | 0 | 2.094 | 1.067 | 0.79379329 | NA | 0 | 14.4078 | 10 | 10 | TRUE | 0.3835 |
| 1700034O15 | 0 | 0 | 0 | 0 | 0 | 0 | 0 | 0 | 0 | 0 | 0 | 0 | 0 | 0 | 0 | NA | NA | NA | NA | NA | NA | NA | NA | NA | NA |
| 1700034P13I | 19 | 12 | 5 | 9 | 9 | 2 | 15 | 13 | 5 | 9 | 10 | 2 | 9 | 11 | 7 | 0.63 | -0.667 | 0.35661284 | NA | 0.2398 | 0.2658 | 0.2593 | 0.2593 | TRUE | 1.0993 |
| 1700036G14 | 0 | 0 | 0 | 0 | 0 | 0 | 0 | 0 | 0 | 0 | 0 | 0 | 0 | 0 | 0 | NA | NA | NA | NA | NA | NA | NA | NA | NA | NA |
| 1700037C18I | 55 | 41 | 27 | 51 | 40 | 34 | 43 | 44 | 29 | 51 | 45 | 34 | 40.82 | 39 | 43 | 1.116 | 0.158 | 0.62220095 | 0.7614623 | 0.0191 | 0.0605 | 0.0494 | 0.0494 | TRUE | 0.3102 |
| 1700037H04I | 6182 | 4633 | 4672 | 4987 | 4577 | 5447 | 4801 | 4928 | 5041 | 4964 | 5163 | 5377 | 5045.8 | 4923 | 5168 | 1.05 | 0.07 | 0.2074851 | 0.36911083 | 8.00E-04 | 0.0029 | 0.002 | 0.002 | TRUE | 0.0154 |
| 1700039E15I | 0 | 0 | 0 | 1 | 0 | 1 | 0 | 0 | 0 | 1 | 0 | 1 | 0.33 | 0 | 1 | 3.763 | 1.912 | 0.57887588 | NA | 0 | 7.1752 | 6.7756 | 6.7756 | TRUE | 0.2094 |
| 1700039E22I | 0 | 0 | 0 | 1 | 0 | 0 | 0 | 0 | 0 | 1 | 0 | 0 | 0.17 | 0 | 0 | 2.094 | 1.067 | 0.79379329 | NA | 0 | 14.2885 | 10 | 10 | TRUE | 0.3906 |
| 1700040L02F | 1 | 2 | 6 | 6 | 5 | 6 | 1 | 2 | 6 | 6 | 6 | 6 | 4.49 | 3 | 6 | 1.94 | 0.956 | 0.32304669 | NA | 0.0728 | 0.5308 | 0.4259 | 0.4259 | TRUE | 1.1253 |
| 1700040N02 | 2 | 1 | 2 | 4 | 0 | 2 | 2 | 1 | 2 | 4 | 0 | 2 | 1.79 | 2 | 2 | 1.267 | 0.342 | 0.81619575 | NA | 0 | 1.3276 | 0.9958 | 0.9958 | TRUE | 0.1519 |
| 1700041C23I | 0 | 0 | 0 | 0 | 0 | 0 | 0 | 0 | 0 | 0 | 0 | 0 | 0 | 0 | 0 | NA | NA | NA | NA | NA | NA | NA | NA | NA | NA |
| 1700041M19 | 0 | 0 | 0 | 0 | 0 | 0 | 0 | 0 | 0 | 0 | 0 | 0 | 0 | 0 | 0 | NA | NA | NA | NA | NA | NA | NA | NA | NA | NA |
| 1700042B14I | 0 | 0 | 0 | 0 | 0 | 0 | 0 | 0 | 0 | 0 | 0 | 0 | 0 | 0 | 0 | NA | NA | NA | NA | NA | NA | NA | NA | NA | NA |
| 1700042G07 | 0 | 1 | 1 | 0 | 0 | 0 | 0 | 1 | 1 | 0 | 0 | 0 | 0.36 | 1 | 0 | 0.271 | -1.881 | 0.5755355 | NA | 0 | 6.6391 | 6.4428 | 6.4428 | TRUE | 0.3513 |
| 1700042G15 | 7 | 7 | 4 | 5 | 3 | 5 | 5 | 7 | 4 | 5 | 3 | 5 | 5.08 | 5 | 4 | 0.778 | -0.363 | 0.66399279 | NA | 0 | 0.4687 | 0.3016 | 0.3016 | TRUE | 0.1399 |
| 1700042O10 | 58 | 43 | 35 | 19 | 25 | 30 | 45 | 46 | 38 | 19 | 28 | 30 | 34.21 | 43 | 26 | 0.595 | -0.749 | 0.0280332 | 0.0881271 | 0 | 0.0717 | 0.0523 | 0.0523 | TRUE | 0.3303 |
| 1700044C05I | 3 | 3 | 3 | 1 | 1 | 0 | 2 | 3 | 3 | 1 | 1 | 0 | 1.81 | 3 | 1 | 0.24 | -2.061 | 0.18888065 | NA | 0 | 1.3093 | 0.8628 | 0.8628 | TRUE | 0.2727 |
| 1700044K03I | 0 | 0 | 0 | 0 | 0 | 0 | 0 | 0 | 0 | 0 | 0 | 0 | 0 | 0 | 0 | NA | NA | NA | NA | NA | NA | NA | NA | NA | NA |
| 1700045H11I | 0 | 0 | 0 | 0 | 0 | 0 | 0 | 0 | 0 | 0 | 0 | 0 | 0 | 0 | 0 | NA | NA | NA | NA | NA | NA | NA | NA | NA | NA |
| 1700045I19R | 1 | 1 | 2 | 0 | 1 | 1 | 1 | 1 | 2 | 0 | 1 | 1 | 1.02 | 1 | 1 | 0.531 | -0.914 | 0.6308001 | NA | 0 | 2.3285 | 1.4951 | 1.4951 | TRUE | 0.2654 |
| 1700046C09I | 0 | 0 | 0 | 0 | 0 | 0 | 0 | 0 | 0 | 0 | 0 | 0 | 0 | 0 | 0 | NA | NA | NA | NA | NA | NA | NA | NA | NA | NA |
| 1700047E10I | 0 | 0 | 0 | 0 | 0 | 0 | 0 | 0 | 0 | 0 | 0 | 0 | 0 | 0 | 0 | NA | NA | NA | NA | NA | NA | NA | NA | NA | NA |
| 1700047G03 | 0 | 1 | 0 | 1 | 0 | 0 | 0 | 1 | 0 | 1 | 0 | 0 | 0.34 | 0 | 0 | 1.075 | 0.105 | 0.97573409 | NA | 0 | 6.9086 | 6.5582 | 6.5582 | TRUE | 0.5216 |
| 1700047I17R | 0 | 0 | 0 | 0 | 0 | 0 | 0 | 0 | 0 | 0 | 0 | 0 | 0 | 0 | 0 | NA | NA | NA | NA | NA | NA | NA | NA | NA | NA |
| 1700047L14F | 0 | 0 | 0 | 0 | 0 | 0 | 0 | 0 | 0 | 0 | 0 | 0 | 0 | 0 | 0 | NA | NA | NA | NA | NA | NA | NA | NA | NA | NA |
| 1700047M11 | 1 | 1 | 2 | 0 | 0 | 0 | 1 | 1 | 2 | 0 | 0 | 0 | 0.67 | 1 | 0 | 0.144 | -2.794 | 0.24876697 | NA | 0 | 3.559 | 2.8631 | 2.8631 | TRUE | 0.1682 |
| 1700048M11 | 0 | 0 | 0 | 0 | 0 | 0 | 0 | 0 | 0 | 0 | 0 | 0 | 0 | 0 | 0 | NA | NA | NA | NA | NA | NA | NA | NA | NA | NA |
| 1700048O20 | 125 | 100 | 93 | 117 | 73 | 101 | 97 | 106 | 100 | 116 | 82 | 100 | 100.39 | 101 | 99 | 0.986 | -0.02 | 0.92287181 | 0.95719768 | 0.0049 | 0.026 | 0.0202 | 0.0202 | TRUE | 0.1901 |
| 1700049E15I | 0 | 0 | 0 | 0 | 0 | 0 | 0 | 0 | 0 | 0 | 0 | 0 | 0 | 0 | 0 | NA | NA | NA | NA | NA | NA | NA | NA | NA | NA |
| 1700049E22I | 60 | 44 | 39 | 60 | 56 | 67 | 47 | 47 | 42 | 60 | 63 | 66 | 54.09 | 45 | 63 | 1.393 | 0.478 | 0.06580454 | 0.16391499 | 0 | 0.0462 | 0.0299 | 0.0299 | TRUE | 0.0259 |
| 1700049G17 | 255 | 171 | 182 | 247 | 175 | 222 | 198 | 182 | 196 | 246 | 197 | 219 | 206.46 | 192 | 221 | 1.15 | 0.201 | 0.16802594 | 0.31953096 | 0 | 0.0139 | 0.0105 | 0.0105 | TRUE | 0.1082 |
| 1700049L16F | 0 | 0 | 0 | 0 | 0 | 0 | 0 | 0 | 0 | 0 | 0 | 0 | 0 | 0 | 0 | NA | NA | NA | NA | NA | NA | NA | NA | NA | NA |
| 1700051A21I | 3 | 0 | 1 | 3 | 1 | 0 | 2 | 0 | 1 | 3 | 1 | 0 | 1.25 | 1 | 1 | 1.166 | 0.222 | 0.90331949 | NA | 0 | 1.8926 | 1.6217 | 1.6217 | TRUE | 0.3358 |
| 1700052I22R | 0 | 0 | 0 | 0 | 0 | 0 | 0 | 0 | 0 | 0 | 0 | 0 | 0 | 0 | 0 | NA | NA | NA | NA | NA | NA | NA | NA | NA | NA |
| 1700052K11I | 166 | 114 | 51 | 78 | 69 | 52 | 129 | 121 | 55 | 78 | 78 | 51 | 85.34 | 102 | 69 | 0.674 | -0.57 | 0.05933661 | 0.15215252 | 0.1129 | 0.0302 | 0.0535 | 0.0535 | TRUE | 1.5277 |
| 1700052N19 | 2846 | 2126 | 2378 | 2691 | 2358 | 2820 | 2210 | 2261 | 2566 | 2679 | 2660 | 2784 | 2526.68 | 2346 | 2708 | 1.155 | 0.208 | 0.00360162 | 0.01901659 | 0.0031 | 0.0034 | 0.0033 | 0.0033 | TRUE | 0.0808 |

| | | | | | | | | | | | | | | | | | | | | | | | | | |
|---|---|---|---|---|---|---|---|---|---|---|---|---|---|---|---|---|---|---|---|---|---|---|---|---|---|
| 1700054A03 | 1 | 0 | 0 | 0 | 0 | 0 | 1 | 0 | 0 | 0 | 0 | 0 | 0.13 | 0 | 0 | 0.552 | -0.857 | 0.8336548 | NA | 0 | 18.3121 | 10 | 10 | TRUE | 0.2566 |
| 1700054K19 | 1 | 0 | 0 | 0 | 0 | 0 | 1 | 0 | 0 | 0 | 0 | 0 | 0.13 | 0 | 0 | 0.552 | -0.857 | 0.8336548 | NA | 0 | 18.3121 | 10 | 10 | TRUE | 0.2566 |
| 1700054M17 | 0 | 0 | 0 | 0 | 0 | 0 | 0 | 0 | 0 | 0 | 0 | 0 | 0 | 0 | 0 | NA | NA | NA | NA | NA | NA | NA | NA | NA | NA |
| 1700054O13 | 1 | 0 | 0 | 0 | 0 | 0 | 1 | 0 | 0 | 0 | 0 | 0 | 0.13 | 0 | 0 | 0.552 | -0.857 | 0.8336548 | NA | 0 | 18.3121 | 10 | 10 | TRUE | 0.2566 |
| 1700055C04 | 1 | 2 | 0 | 0 | 0 | 0 | 1 | 2 | 0 | 0 | 0 | 0 | 0.48 | 1 | 0 | 0.197 | -2.343 | 0.43249452 | NA | 0 | 4.8992 | 4.9172 | 4.9172 | TRUE | 0.1226 |
| 1700055N04 | 0 | 2 | 2 | 1 | 3 | 1 | 0 | 2 | 2 | 1 | 3 | 1 | 1.61 | 1 | 2 | 1.288 | 0.365 | 0.81699365 | NA | 0 | 1.4757 | 1.1457 | 1.1457 | TRUE | 0.7251 |
| 1700056E22 | 38 | 34 | 21 | 35 | 23 | 28 | 30 | 36 | 23 | 35 | 26 | 28 | 29.46 | 30 | 30 | 1.002 | 0.003 | 0.99275594 | 0.99592003 | 0 | 0.0829 | 0.0614 | 0.0614 | TRUE | 0.1731 |
| 1700057G04 | 1 | 0 | 0 | 0 | 2 | 0 | 1 | 0 | 0 | 0 | 2 | 0 | 0.51 | 0 | 1 | 2.139 | 1.097 | 0.71688146 | NA | 0 | 4.6912 | 4.8901 | 4.8901 | TRUE | 0.9745 |
| 1700057H15 | 0 | 0 | 1 | 0 | 0 | 0 | 0 | 0 | 1 | 0 | 0 | 0 | 0.18 | 0 | 0 | 0.552 | -0.857 | 0.8336548 | NA | 0 | 13.1817 | 10 | 10 | TRUE | 0.5365 |
| 1700058G18 | 0 | 0 | 0 | 0 | 1 | 0 | 0 | 0 | 0 | 0 | 1 | 0 | 0.19 | 0 | 0 | 2.094 | 1.067 | 0.79379329 | NA | 0 | 12.608 | 10 | 10 | TRUE | 0.5075 |
| 1700060C16 | 0 | 0 | 0 | 0 | 0 | 0 | 0 | 0 | 0 | 0 | 0 | 0 | 0 | 0 | 0 | NA | NA | NA | NA | NA | NA | NA | NA | NA | NA |
| 1700060C20 | 0 | 1 | 0 | 0 | 0 | 0 | 0 | 1 | 0 | 0 | 0 | 0 | 0.18 | 0 | 0 | 0.552 | -0.857 | 0.8336548 | NA | 0 | 13.3714 | 10 | 10 | TRUE | 0.5216 |
| 1700061F12 | 0 | 0 | 0 | 0 | 0 | 0 | 0 | 0 | 0 | 0 | 0 | 0 | 0 | 0 | 0 | NA | NA | NA | NA | NA | NA | NA | NA | NA | NA |
| 1700061G19 | 494 | 389 | 369 | 303 | 287 | 363 | 384 | 414 | 398 | 302 | 324 | 358 | 363.22 | 399 | 328 | 0.824 | -0.28 | 0.0149324 | 0.05569644 | 0.0014 | 0.009 | 0.0067 | 0.0067 | TRUE | 0.077 |
| 1700061I17 | 0 | 1 | 0 | 0 | 0 | 0 | 0 | 1 | 0 | 0 | 0 | 0 | 0.18 | 0 | 0 | 0.552 | -0.857 | 0.8336548 | NA | 0 | 13.3714 | 10 | 10 | TRUE | 0.5216 |
| 1700063A18 | 0 | 0 | 0 | 0 | 0 | 0 | 0 | 0 | 0 | 0 | 0 | 0 | 0 | 0 | 0 | NA | NA | NA | NA | NA | NA | NA | NA | NA | NA |
| 1700063D05 | 22 | 12 | 16 | 13 | 12 | 17 | 17 | 13 | 17 | 13 | 14 | 17 | 15.06 | 16 | 15 | 0.916 | -0.126 | 0.79671903 | NA | 0 | 0.1598 | 0.1058 | 0.1058 | TRUE | 0.1169 |
| 1700063O14 | 0 | 0 | 0 | 0 | 0 | 0 | 0 | 0 | 0 | 0 | 0 | 0 | 0 | 0 | 0 | NA | NA | NA | NA | NA | NA | NA | NA | NA | NA |
| 1700064J06 | 0 | 0 | 0 | 0 | 0 | 0 | 0 | 0 | 0 | 0 | 0 | 0 | 0 | 0 | 0 | NA | NA | NA | NA | NA | NA | NA | NA | NA | NA |
| 1700064M15 | 0 | 0 | 2 | 1 | 1 | 0 | 0 | 0 | 2 | 1 | 1 | 0 | 0.71 | 1 | 1 | 1.011 | 0.016 | 0.99481353 | NA | 0 | 3.3238 | 2.9707 | 2.9707 | TRUE | 0.9749 |
| 1700065D16 | 1 | 2 | 1 | 0 | 0 | 1 | 1 | 2 | 1 | 0 | 0 | 1 | 0.83 | 1 | 0 | 0.282 | -1.824 | 0.39899776 | NA | 0 | 2.8636 | 2.0027 | 2.0027 | TRUE | 0.3835 |
| 1700065I16 | 0 | 0 | 0 | 0 | 0 | 0 | 0 | 0 | 0 | 0 | 0 | 0 | 0 | 0 | 0 | NA | NA | NA | NA | NA | NA | NA | NA | NA | NA |
| 1700065I17 | 0 | 0 | 1 | 0 | 0 | 0 | 0 | 0 | 1 | 0 | 0 | 0 | 0.18 | 0 | 0 | 0.552 | -0.857 | 0.8336548 | NA | 0 | 13.1817 | 10 | 10 | TRUE | 0.5365 |
| 1700065J11 | 0 | 0 | 0 | 0 | 0 | 0 | 0 | 0 | 0 | 0 | 0 | 0 | 0 | 0 | 0 | NA | NA | NA | NA | NA | NA | NA | NA | NA | NA |
| 1700065J18 | 0 | 0 | 0 | 0 | 0 | 0 | 0 | 0 | 0 | 0 | 0 | 0 | 0 | 0 | 0 | NA | NA | NA | NA | NA | NA | NA | NA | NA | NA |
| 1700065O20 | 0 | 0 | 0 | 0 | 0 | 0 | 0 | 0 | 0 | 0 | 0 | 0 | 0 | 0 | 0 | NA | NA | NA | NA | NA | NA | NA | NA | NA | NA |
| 1700066B17 | 9 | 2 | 2 | 5 | 0 | 2 | 7 | 2 | 2 | 5 | 0 | 2 | 3.04 | 4 | 2 | 0.612 | -0.708 | 0.56049331 | NA | 0.5334 | 0.7826 | 0.7212 | 0.7212 | TRUE | 0.5023 |
| 1700066B19 | 12 | 4 | 4 | 8 | 9 | 9 | 9 | 4 | 4 | 8 | 10 | 9 | 7.48 | 6 | 9 | 1.47 | 0.556 | 0.43269028 | NA | 0 | 0.3192 | 0.2256 | 0.2256 | TRUE | 0.7362 |
| 1700066J24 | 0 | 0 | 0 | 0 | 0 | 0 | 0 | 0 | 0 | 0 | 0 | 0 | 0 | 0 | 0 | NA | NA | NA | NA | NA | NA | NA | NA | NA | NA |
| 1700066M21 | 625 | 499 | 528 | 511 | 424 | 498 | 485 | 531 | 570 | 509 | 478 | 492 | 510.74 | 529 | 493 | 0.935 | -0.097 | 0.33631002 | 0.51055541 | 0.0021 | 0.0071 | 0.0054 | 0.0054 | TRUE | 0.0638 |
| 1700066N21 | 0 | 0 | 0 | 0 | 0 | 0 | 0 | 0 | 0 | 0 | 0 | 0 | 0 | 0 | 0 | NA | NA | NA | NA | NA | NA | NA | NA | NA | NA |
| 1700066O22 | 0 | 0 | 0 | 0 | 0 | 0 | 0 | 0 | 0 | 0 | 0 | 0 | 0 | 0 | 0 | NA | NA | NA | NA | NA | NA | NA | NA | NA | NA |
| 1700067G17 | 0 | 2 | 0 | 2 | 1 | 0 | 0 | 2 | 0 | 2 | 1 | 0 | 0.87 | 1 | 1 | 1.516 | 0.601 | 0.79248878 | NA | 0 | 2.713 | 2.5514 | 2.5514 | TRUE | 0.6486 |
| 1700067K01 | 78 | 63 | 106 | 100 | 77 | 115 | 61 | 67 | 114 | 100 | 87 | 114 | 90.32 | 81 | 100 | 1.25 | 0.321 | 0.21296347 | 0.3752326 | 0.0573 | 0.0287 | 0.0368 | 0.0368 | TRUE | 1.214 |
| 1700067P10 | 0 | 0 | 0 | 0 | 0 | 0 | 0 | 0 | 0 | 0 | 0 | 0 | 0 | 0 | 0 | NA | NA | NA | NA | NA | NA | NA | NA | NA | NA |
| 1700069L16 | 2 | 3 | 2 | 2 | 5 | 3 | 2 | 3 | 2 | 2 | 6 | 3 | 2.92 | 2 | 4 | 1.544 | 0.626 | 0.57578857 | NA | 0 | 0.8152 | 0.5421 | 0.5421 | TRUE | 0.3644 |
| 1700069P05 | 0 | 0 | 0 | 0 | 0 | 0 | 0 | 0 | 0 | 0 | 0 | 0 | 0 | 0 | 0 | NA | NA | NA | NA | NA | NA | NA | NA | NA | NA |
| 1700071K01 | 0 | 0 | 0 | 0 | 0 | 0 | 0 | 0 | 0 | 0 | 0 | 0 | 0 | 0 | 0 | NA | NA | NA | NA | NA | NA | NA | NA | NA | NA |
| 1700071M16 | 0 | 0 | 0 | 0 | 0 | 0 | 0 | 0 | 0 | 0 | 0 | 0 | 0 | 0 | 0 | NA | NA | NA | NA | NA | NA | NA | NA | NA | NA |
| 1700072B07 | 0 | 0 | 0 | 0 | 0 | 0 | 0 | 0 | 0 | 0 | 0 | 0 | 0 | 0 | 0 | NA | NA | NA | NA | NA | NA | NA | NA | NA | NA |
| 1700072O05 | 0 | 0 | 0 | 0 | 0 | 0 | 0 | 0 | 0 | 0 | 0 | 0 | 0 | 0 | 0 | NA | NA | NA | NA | NA | NA | NA | NA | NA | NA |
| 1700073E17 | 0 | 0 | 0 | 0 | 0 | 0 | 0 | 0 | 0 | 0 | 0 | 0 | 0 | 0 | 0 | NA | NA | NA | NA | NA | NA | NA | NA | NA | NA |
| 1700074H08 | 0 | 0 | 0 | 0 | 0 | 0 | 0 | 0 | 0 | 0 | 0 | 0 | 0 | 0 | 0 | NA | NA | NA | NA | NA | NA | NA | NA | NA | NA |
| 1700074P13 | 0 | 0 | 0 | 0 | 0 | 0 | 0 | 0 | 0 | 0 | 0 | 0 | 0 | 0 | 0 | NA | NA | NA | NA | NA | NA | NA | NA | NA | NA |
| 1700080E11 | 0 | 1 | 0 | 0 | 0 | 0 | 0 | 1 | 0 | 0 | 0 | 0 | 0.18 | 0 | 0 | 0.552 | -0.857 | 0.8336548 | NA | 0 | 13.3714 | 10 | 10 | TRUE | 0.5216 |
| 1700080N15 | 0 | 0 | 0 | 0 | 0 | 0 | 0 | 0 | 0 | 0 | 0 | 0 | 0 | 0 | 0 | NA | NA | NA | NA | NA | NA | NA | NA | NA | NA |
| 1700080O16 | 62 | 52 | 29 | 15 | 8 | 14 | 48 | 55 | 31 | 15 | 9 | 14 | 28.76 | 45 | 13 | 0.282 | -1.829 | 1.03E-05 | 0.00019288 | 0 | 0.0848 | 0.0723 | 0.0723 | TRUE | 0.2862 |
| 1700081H04 | 0 | 0 | 0 | 0 | 0 | 0 | 0 | 0 | 0 | 0 | 0 | 0 | 0 | 0 | 0 | NA | NA | NA | NA | NA | NA | NA | NA | NA | NA |
| 1700081H22 | 0 | 0 | 0 | 0 | 0 | 0 | 0 | 0 | 0 | 0 | 0 | 0 | 0 | 0 | 0 | NA | NA | NA | NA | NA | NA | NA | NA | NA | NA |
| 1700084C01 | 7 | 6 | 2 | 4 | 3 | 6 | 5 | 6 | 2 | 4 | 3 | 6 | 4.54 | 4 | 4 | 0.948 | -0.077 | 0.93168495 | NA | 0 | 0.524 | 0.3625 | 0.3625 | TRUE | 0.3733 |
| 1700084E18 | 10 | 10 | 10 | 9 | 8 | 3 | 8 | 11 | 11 | 9 | 9 | 3 | 8.36 | 10 | 7 | 0.718 | -0.478 | 0.48688961 | NA | 0 | 0.286 | 0.2154 | 0.2154 | TRUE | 0.7034 |
| 1700084F23 | 0 | 0 | 0 | 0 | 0 | 0 | 0 | 0 | 0 | 0 | 0 | 0 | 0 | 0 | 0 | NA | NA | NA | NA | NA | NA | NA | NA | NA | NA |
| 1700084J12 | 3 | 0 | 0 | 0 | 0 | 0 | 2 | 0 | 0 | 0 | 0 | 0 | 0.39 | 1 | 0 | 0.239 | -2.067 | 0.57190649 | NA | 10 | 6.1057 | 8.0283 | 8.0283 | TRUE | 1.4408 |
| 1700085B03 | 0 | 0 | 0 | 0 | 0 | 0 | 0 | 0 | 0 | 0 | 0 | 0 | 0 | 0 | 0 | NA | NA | NA | NA | NA | NA | NA | NA | NA | NA |
| 1700085C21 | 0 | 0 | 0 | 0 | 0 | 0 | 0 | 0 | 0 | 0 | 0 | 0 | 0 | 0 | 0 | NA | NA | NA | NA | NA | NA | NA | NA | NA | NA |
| 1700086L19 | 3 | 7 | 6 | 4 | 0 | 3 | 2 | 7 | 6 | 4 | 0 | 3 | 3.87 | 5 | 2 | 0.442 | -1.178 | 0.27640125 | NA | 0.2385 | 0.6156 | 0.5335 | 0.5335 | TRUE | 0.7123 |
| 1700086O06 | 30 | 24 | 24 | 27 | 16 | 23 | 23 | 26 | 26 | 27 | 18 | 23 | 23.73 | 25 | 23 | 0.91 | -0.136 | 0.73021588 | 0.8388371 | 0 | 0.1023 | 0.069 | 0.069 | TRUE | 0.1631 |
| 1700088E04 | 15 | 11 | 8 | 12 | 9 | 14 | 12 | 12 | 9 | 12 | 10 | 14 | 11.32 | 11 | 12 | 1.121 | 0.165 | 0.76820503 | NA | 0 | 0.2119 | 0.1383 | 0.1383 | TRUE | 0.0913 |
| 1700091E21 | 0 | 0 | 1 | 0 | 0 | 1 | 0 | 0 | 0 | 1 | 0 | 1 | 0.37 | 0 | 0 | 1.075 | 0.105 | 0.97514588 | NA | 0 | 6.4454 | 6.1578 | 6.1578 | TRUE | 0.5365 |
| 1700091H14 | 1 | 0 | 1 | 0 | 0 | 0 | 1 | 0 | 1 | 0 | 0 | 0 | 0.31 | 1 | 0 | 0.307 | -1.706 | 0.62894649 | NA | 0 | 7.6657 | 7.2 | 7.2 | TRUE | 0.2067 |
| 1700092C02 | 0 | 0 | 0 | 0 | 0 | 0 | 0 | 0 | 0 | 0 | 0 | 0 | 0 | 0 | 0 | NA | NA | NA | NA | NA | NA | NA | NA | NA | NA |
| 1700092C10 | 0 | 1 | 0 | 0 | 0 | 0 | 0 | 1 | 0 | 0 | 0 | 0 | 0.18 | 0 | 0 | 0.552 | -0.857 | 0.8336548 | NA | 0 | 13.3714 | 10 | 10 | TRUE | 0.5216 |
| 1700092E19 | 0 | 0 | 0 | 0 | 0 | 0 | 0 | 0 | 0 | 0 | 0 | 0 | 0 | 0 | 0 | NA | NA | NA | NA | NA | NA | NA | NA | NA | NA |
| 1700092K14 | 0 | 0 | 0 | 0 | 0 | 0 | 0 | 0 | 0 | 0 | 0 | 0 | 0 | 0 | 0 | NA | NA | NA | NA | NA | NA | NA | NA | NA | NA |
| 1700092M07 | 12 | 3 | 7 | 5 | 3 | 8 | 9 | 3 | 8 | 5 | 3 | 8 | 6.05 | 7 | 5 | 0.805 | -0.312 | 0.69724654 | NA | 0 | 0.3939 | 0.2982 | 0.2982 | TRUE | 0.4998 |
| 1700093K21 | 3 | 1 | 1 | 1 | 1 | 1 | 2 | 1 | 1 | 1 | 1 | 1 | 1.26 | 1 | 1 | 0.679 | -0.557 | 0.73233539 | NA | 0 | 1.8776 | 1.0978 | 1.0978 | TRUE | 0.2229 |
| 1700094D03 | 160 | 135 | 128 | 130 | 103 | 162 | 124 | 144 | 138 | 129 | 116 | 160 | 135.25 | 135 | 135 | 1.003 | 0.005 | 0.97857893 | 0.98853832 | 0.0088 | 0.02 | 0.0168 | 0.0168 | TRUE | 0.2668 |
| 1700094J05 | 0 | 0 | 0 | 1 | 0 | 0 | 0 | 0 | 0 | 1 | 0 | 0 | 0.17 | 0 | 0 | 2.094 | 1.067 | 0.79379329 | NA | 0 | 14.2885 | 10 | 10 | TRUE | 0.3906 |
| 1700094M24 | 0 | 0 | 0 | 0 | 0 | 0 | 0 | 0 | 0 | 0 | 0 | 0 | 0 | 0 | 0 | NA | NA | NA | NA | NA | NA | NA | NA | NA | NA |

| | | | | | | | | | | | | | | | | | | | | | | | | | |
|---|---|---|---|---|---|---|---|---|---|---|---|---|---|---|---|---|---|---|---|---|---|---|---|---|---|
| 1700095A21 | 0 | 0 | 0 | 0 | 0 | 0 | 0 | 0 | 0 | 0 | 0 | 0 | 0 | 0 | 0 | NA | NA | NA | NA | NA | NA | NA | NA | NA | NA |
| 1700095B10 | 3 | 2 | 1 | 3 | 3 | 1 | 2 | 2 | 1 | 3 | 3 | 1 | 2.15 | 2 | 2 | 1.304 | 0.383 | 0.76510113 | NA | 0 | 1.1053 | 0.7153 | 0.7153 | TRUE | 0.3088 |
| 1700095B22 | 1 | 0 | 0 | 1 | 0 | 1 | 1 | 0 | 0 | 1 | 0 | 1 | 0.46 | 0 | 1 | 1.94 | 0.956 | 0.73503911 | NA | 0 | 5.1562 | 3.9771 | 3.9771 | TRUE | 0.2566 |
| 1700096J18F | 0 | 0 | 0 | 0 | 0 | 0 | 0 | 0 | 0 | 0 | 0 | 0 | 0 | 0 | 0 | NA | NA | NA | NA | NA | NA | NA | NA | NA | NA |
| 1700096K18I | 70 | 37 | 43 | 49 | 43 | 40 | 54 | 39 | 46 | 49 | 49 | 39 | 46.15 | 46 | 46 | 0.969 | -0.045 | 0.87694484 | 0.93135478 | 0 | 0.0538 | 0.039 | 0.039 | TRUE | 0.1895 |
| 1700097N02 | 139 | 86 | 121 | 266 | 227 | 236 | 108 | 91 | 131 | 265 | 256 | 233 | 180.64 | 110 | 251 | 2.286 | 1.193 | 7.01E-13 | 9.13E-11 | 0.0066 | 0.0155 | 0.0135 | 0.0135 | TRUE | 0.2438 |
| 1700100L14F | 0 | 0 | 0 | 0 | 0 | 0 | 0 | 0 | 0 | 0 | 0 | 0 | 0 | 0 | 0 | NA | NA | NA | NA | NA | NA | NA | NA | NA | NA |
| 1700101E01I | 79 | 61 | 52 | 54 | 45 | 57 | 61 | 65 | 56 | 54 | 51 | 56 | 57.19 | 61 | 54 | 0.882 | -0.181 | 0.47446724 | 0.64280192 | 0 | 0.0439 | 0.0285 | 0.0285 | TRUE | 0.0358 |
| 1700101I11R | 0 | 1 | 0 | 2 | 3 | 1 | 0 | 1 | 0 | 2 | 3 | 1 | 1.24 | 0 | 2 | 6.109 | 2.611 | 0.17470329 | NA | 0 | 1.9174 | 1.421 | 1.421 | TRUE | 0.4505 |
| 1700101O22 | 0 | 0 | 0 | 0 | 0 | 0 | 0 | 0 | 0 | 0 | 0 | 0 | 0 | 0 | 0 | NA | NA | NA | NA | NA | NA | NA | NA | NA | NA |
| 1700102H20 | 3 | 2 | 4 | 10 | 7 | 0 | 2 | 2 | 4 | 10 | 8 | 0 | 4.44 | 3 | 6 | 2.044 | 1.031 | 0.3425036 | NA | 0.8023 | 0.5365 | 0.597 | 0.597 | TRUE | 1.0092 |
| 1700102P08I | 65 | 38 | 43 | 61 | 48 | 76 | 50 | 40 | 46 | 61 | 54 | 75 | 54.53 | 45 | 63 | 1.38 | 0.465 | 0.08567801 | 0.19785729 | 0 | 0.0459 | 0.0341 | 0.0341 | TRUE | 0.2324 |
| 1700104A03 | 0 | 0 | 0 | 0 | 0 | 0 | 0 | 0 | 0 | 0 | 0 | 0 | 0 | 0 | 0 | NA | NA | NA | NA | NA | NA | NA | NA | NA | NA |
| 1700105P06I | 1 | 7 | 2 | 3 | 4 | 2 | 1 | 7 | 2 | 3 | 5 | 2 | 3.31 | 3 | 3 | 0.93 | -0.105 | 0.92548823 | NA | 0.3612 | 0.7187 | 0.6067 | 0.6067 | TRUE | 1.3916 |
| 1700106J16F | 13 | 10 | 5 | 1 | 5 | 4 | 10 | 11 | 5 | 1 | 6 | 4 | 6.12 | 9 | 4 | 0.398 | -1.328 | 0.1145982 | NA | 0 | 0.3898 | 0.3069 | 0.3069 | TRUE | 0.6107 |
| 1700108F19I | 31 | 20 | 21 | 11 | 12 | 8 | 24 | 21 | 23 | 11 | 14 | 8 | 16.73 | 23 | 11 | 0.473 | -1.081 | 0.02478125 | NA | 0 | 0.1441 | 0.0977 | 0.0977 | TRUE | 0.208 |
| 1700108J01F | 0 | 0 | 0 | 0 | 0 | 0 | 0 | 0 | 0 | 0 | 0 | 0 | 0 | 0 | 0 | NA | NA | NA | NA | NA | NA | NA | NA | NA | NA |
| 1700109F18I | 0 | 0 | 0 | 0 | 0 | 0 | 0 | 0 | 0 | 0 | 0 | 0 | 0 | 0 | 0 | NA | NA | NA | NA | NA | NA | NA | NA | NA | NA |
| 1700109G14 | 1 | 1 | 1 | 0 | 0 | 0 | 1 | 1 | 1 | 0 | 0 | 0 | 0.49 | 1 | 0 | 0.196 | -2.347 | 0.38405676 | NA | 0 | 4.8735 | 3.7415 | 3.7415 | TRUE | 0.0718 |
| 1700109G15 | 1 | 1 | 1 | 1 | 1 | 1 | 1 | 1 | 1 | 1 | 1 | 1 | 1 | 1 | 1 | 1.074 | 0.103 | 0.95418264 | NA | 0 | 2.3607 | 1.3027 | 1.3027 | TRUE | 0.0201 |
| 1700109H08 | 16 | 11 | 20 | 16 | 15 | 15 | 12 | 12 | 22 | 16 | 17 | 15 | 15.56 | 15 | 16 | 1.05 | 0.07 | 0.88799828 | NA | 0 | 0.1547 | 0.112 | 0.112 | TRUE | 0.5663 |
| 1700109I08R | 0 | 0 | 0 | 0 | 0 | 0 | 0 | 0 | 0 | 0 | 0 | 0 | 0 | 0 | 0 | NA | NA | NA | NA | NA | NA | NA | NA | NA | NA |
| 1700110C19I | 4 | 0 | 1 | 1 | 2 | 4 | 3 | 0 | 1 | 1 | 2 | 4 | 1.9 | 1 | 2 | 1.639 | 0.713 | 0.62529112 | NA | 0 | 1.2514 | 0.9903 | 0.9903 | TRUE | 0.614 |
| 1700110I01R | 0 | 0 | 0 | 0 | 0 | 0 | 0 | 0 | 0 | 0 | 0 | 0 | 0 | 0 | 0 | NA | NA | NA | NA | NA | NA | NA | NA | NA | NA |
| 1700110K17I | 0 | 0 | 0 | 2 | 1 | 1 | 0 | 0 | 0 | 2 | 1 | 1 | 0.68 | 0 | 1 | 7.743 | 2.953 | 0.21812153 | NA | 0 | 3.4657 | 2.7652 | 2.7652 | TRUE | 0.1019 |
| 1700110M21 | 0 | 0 | 0 | 0 | 0 | 0 | 0 | 0 | 0 | 0 | 0 | 0 | 0 | 0 | 0 | NA | NA | NA | NA | NA | NA | NA | NA | NA | NA |
| 1700111N16 | 0 | 0 | 0 | 0 | 0 | 0 | 0 | 0 | 0 | 0 | 0 | 0 | 0 | 0 | 0 | NA | NA | NA | NA | NA | NA | NA | NA | NA | NA |
| 1700112E06I | 4 | 1 | 3 | 1 | 1 | 2 | 3 | 1 | 3 | 1 | 1 | 2 | 1.92 | 2 | 1 | 0.551 | -0.86 | 0.5313817 | NA | 0 | 1.2384 | 0.789 | 0.789 | TRUE | 0.2468 |
| 1700112H15 | 4 | 2 | 1 | 0 | 0 | 2 | 3 | 2 | 1 | 0 | 0 | 2 | 1.38 | 2 | 1 | 0.317 | -1.66 | 0.3567823 | NA | 0 | 1.7183 | 1.3433 | 1.3433 | TRUE | 0.8292 |
| 1700112J05F | 0 | 0 | 0 | 0 | 0 | 0 | 0 | 0 | 0 | 0 | 0 | 0 | 0 | 0 | 0 | NA | NA | NA | NA | NA | NA | NA | NA | NA | NA |
| 1700113A16 | 53 | 28 | 27 | 31 | 30 | 35 | 41 | 30 | 29 | 31 | 34 | 35 | 33.22 | 33 | 33 | 0.983 | -0.025 | 0.94095141 | 0.96681409 | 0 | 0.0738 | 0.0522 | 0.0522 | TRUE | 0.3435 |
| 1700113H08 | 0 | 0 | 0 | 0 | 0 | 0 | 0 | 0 | 0 | 0 | 0 | 0 | 0 | 0 | 0 | NA | NA | NA | NA | NA | NA | NA | NA | NA | NA |
| 1700119H24 | 0 | 0 | 0 | 0 | 0 | 2 | 0 | 0 | 0 | 0 | 0 | 2 | 0.33 | 0 | 1 | 3.743 | 1.904 | 0.62640105 | NA | 0 | 7.2051 | 9.2465 | 9.2465 | TRUE | 0.9739 |
| 1700120C14I | 1 | 0 | 0 | 0 | 0 | 0 | 1 | 0 | 0 | 0 | 0 | 0 | 0.13 | 0 | 0 | 0.552 | -0.857 | 0.8336548 | NA | 0 | 18.3121 | 10 | 10 | TRUE | 0.2566 |
| 1700120E14I | 0 | 0 | 0 | 0 | 0 | 0 | 0 | 0 | 0 | 0 | 0 | 0 | 0 | 0 | 0 | NA | NA | NA | NA | NA | NA | NA | NA | NA | NA |
| 1700120G07 | 0 | 0 | 0 | 0 | 0 | 0 | 0 | 0 | 0 | 0 | 0 | 0 | 0 | 0 | 0 | NA | NA | NA | NA | NA | NA | NA | NA | NA | NA |
| 1700120K04I | 0 | 0 | 0 | 2 | 0 | 0 | 0 | 0 | 0 | 2 | 0 | 0 | 0.33 | 0 | 1 | 3.77 | 1.915 | 0.62344813 | NA | 0 | 7.1455 | 9.1834 | 9.1834 | TRUE | 0.9739 |
| 1700121L16F | 0 | 0 | 0 | 0 | 0 | 0 | 0 | 0 | 0 | 0 | 0 | 0 | 0 | 0 | 0 | NA | NA | NA | NA | NA | NA | NA | NA | NA | NA |
| 1700121N20 | 2 | 0 | 1 | 1 | 4 | 2 | 2 | 0 | 1 | 1 | 5 | 2 | 1.69 | 1 | 3 | 2.705 | 1.436 | 0.3588779 | NA | 0 | 1.4084 | 1.0274 | 1.0274 | TRUE | 0.4427 |
| 1700122O11 | 1 | 2 | 0 | 0 | 0 | 0 | 1 | 2 | 0 | 0 | 0 | 0 | 0.48 | 1 | 0 | 0.197 | -2.343 | 0.43249452 | NA | 0 | 4.8992 | 4.9172 | 4.9172 | TRUE | 0.1226 |
| 1700123I01R | 55 | 38 | 43 | 32 | 33 | 36 | 43 | 40 | 46 | 32 | 37 | 36 | 39.03 | 43 | 35 | 0.807 | -0.309 | 0.31097567 | 0.48391673 | 0 | 0.0632 | 0.041 | 0.041 | TRUE | 0.0413 |
| 1700123K08I | 0 | 0 | 0 | 0 | 0 | 0 | 0 | 0 | 0 | 0 | 0 | 0 | 0 | 0 | 0 | NA | NA | NA | NA | NA | NA | NA | NA | NA | NA |
| 1700123L14F | 0 | 0 | 0 | 0 | 0 | 0 | 0 | 0 | 0 | 0 | 0 | 0 | 0 | 0 | 0 | NA | NA | NA | NA | NA | NA | NA | NA | NA | NA |
| 1700123M08 | 24 | 28 | 23 | 28 | 28 | 26 | 19 | 30 | 25 | 28 | 32 | 26 | 26.39 | 25 | 29 | 1.171 | 0.228 | 0.54959673 | 0.70603866 | 0 | 0.0922 | 0.0661 | 0.0661 | TRUE | 0.3203 |
| 1700123O12 | 0 | 0 | 0 | 0 | 0 | 0 | 0 | 0 | 0 | 0 | 0 | 0 | 0 | 0 | 0 | NA | NA | NA | NA | NA | NA | NA | NA | NA | NA |
| 1700123O20 | 1395 | 946 | 794 | 844 | 707 | 812 | 1083 | 1006 | 857 | 840 | 798 | 802 | 897.61 | 982 | 813 | 0.827 | -0.275 | 0.00444368 | 0.02244296 | 0.0066 | 0.0051 | 0.0056 | 0.0056 | TRUE | 0.1472 |
| 1700123O21 | 0 | 0 | 0 | 0 | 0 | 0 | 0 | 0 | 0 | 0 | 0 | 0 | 0 | 0 | 0 | NA | NA | NA | NA | NA | NA | NA | NA | NA | NA |
| 1700125G02 | 0 | 0 | 0 | 0 | 0 | 0 | 0 | 0 | 0 | 0 | 0 | 0 | 0 | 0 | 0 | NA | NA | NA | NA | NA | NA | NA | NA | NA | NA |
| 1700125G22 | 3 | 4 | 1 | 2 | 0 | 2 | 2 | 4 | 1 | 2 | 0 | 2 | 1.94 | 2 | 1 | 0.529 | -0.92 | 0.52047548 | NA | 0 | 1.2253 | 0.9027 | 0.9027 | TRUE | 0.2957 |
| 1700125H03 | 2 | 0 | 1 | 0 | 2 | 2 | 2 | 0 | 1 | 0 | 2 | 2 | 1.14 | 1 | 1 | 1.547 | 0.63 | 0.73713722 | NA | 0 | 2.0745 | 1.6321 | 1.6321 | TRUE | 0.5116 |
| 1700125H20 | 21 | 8 | 17 | 26 | 19 | 15 | 16 | 9 | 18 | 26 | 21 | 15 | 17.55 | 14 | 21 | 1.432 | 0.518 | 0.28777664 | 0.45998562 | 0.0401 | 0.1375 | 0.1125 | 0.1125 | TRUE | 0.3993 |
| 1700126H18 | 0 | 0 | 0 | 0 | 0 | 0 | 0 | 0 | 0 | 0 | 0 | 0 | 0 | 0 | 0 | NA | NA | NA | NA | NA | NA | NA | NA | NA | NA |
| 1700128A07 | 2 | 0 | 0 | 3 | 0 | 0 | 2 | 0 | 0 | 3 | 0 | 0 | 0.76 | 1 | 1 | 1.815 | 0.86 | 0.74367993 | NA | 8.431 | 3.135 | 3.5951 | 3.5951 | TRUE | 1.4418 |
| 1700128F08I | 0 | 0 | 1 | 0 | 0 | 0 | 0 | 0 | 1 | 0 | 0 | 0 | 0.18 | 0 | 0 | 0.552 | -0.857 | 0.8336548 | NA | 0 | 13.1817 | 10 | 10 | TRUE | 0.5365 |
| 1700129C05I | 0 | 0 | 0 | 0 | 0 | 0 | 0 | 0 | 0 | 0 | 0 | 0 | 0 | 0 | 0 | NA | NA | NA | NA | NA | NA | NA | NA | NA | NA |
| 1810006J02F | 0 | 0 | 0 | 0 | 0 | 0 | 0 | 0 | 0 | 0 | 0 | 0 | 0 | 0 | 0 | NA | NA | NA | NA | NA | NA | NA | NA | NA | NA |
| 1810007C17I | 0 | 0 | 0 | 0 | 0 | 0 | 0 | 0 | 0 | 0 | 0 | 0 | 0 | 0 | 0 | NA | NA | NA | NA | NA | NA | NA | NA | NA | NA |
| 1810007D17 | 0 | 1 | 1 | 3 | 0 | 1 | 0 | 1 | 1 | 3 | 0 | 1 | 1.02 | 1 | 1 | 1.967 | 0.976 | 0.63038931 | NA | 0 | 2.3275 | 1.8702 | 1.8702 | TRUE | 0.3453 |
| 1810007I06R | 1 | 1 | 2 | 2 | 1 | 1 | 1 | 1 | 2 | 2 | 1 | 1 | 1.35 | 1 | 1 | 1.05 | 0.07 | 0.96467497 | NA | 0 | 1.7571 | 1.0543 | 1.0543 | TRUE | 0.167 |
| 1810008I18R | 0 | 0 | 0 | 0 | 0 | 0 | 0 | 0 | 0 | 0 | 0 | 0 | 0 | 0 | 0 | NA | NA | NA | NA | NA | NA | NA | NA | NA | NA |
| 1810009A15 | 2978 | 1959 | 1695 | 1719 | 1475 | 1699 | 2313 | 2084 | 1829 | 1711 | 1664 | 1677 | 1879.63 | 2075 | 1684 | 0.811 | -0.303 | 0.00039758 | 0.00363335 | 0.0061 | 0.0037 | 0.0047 | 0.0047 | TRUE | 0.1299 |
| 1810009J06F | 0 | 0 | 0 | 0 | 1 | 0 | 0 | 0 | 0 | 0 | 1 | 0 | 0.19 | 0 | 0 | 2.094 | 1.067 | 0.79379329 | NA | 0 | 12.608 | 10 | 10 | TRUE | 0.5075 |
| 1810010D01 | 1 | 1 | 1 | 2 | 0 | 3 | 1 | 1 | 1 | 2 | 0 | 3 | 1.31 | 1 | 2 | 1.738 | 0.797 | 0.63822397 | NA | 0 | 1.809 | 1.2617 | 1.2617 | TRUE | 0.3444 |
| 1810010H24 | 125 | 101 | 115 | 158 | 128 | 149 | 97 | 107 | 124 | 157 | 144 | 147 | 129.56 | 109 | 149 | 1.373 | 0.457 | 0.01052668 | 0.0428655 | 0 | 0.0207 | 0.0151 | 0.0151 | TRUE | 0.1317 |
| 1810011H11 | 2 | 0 | 0 | 0 | 0 | 0 | 2 | 0 | 0 | 0 | 0 | 0 | 0.26 | 1 | 0 | 0.356 | -1.491 | 0.71362796 | NA | 0 | 9.1573 | 10 | 10 | TRUE | 0.968 |
| 1810011O10 | 46 | 33 | 22 | 5 | 9 | 8 | 36 | 35 | 24 | 5 | 10 | 8 | 19.6 | 32 | 8 | 0.241 | -2.054 | 3.64E-05 | 0.00054577 | 0 | 0.1234 | 0.0952 | 0.0952 | TRUE | 0.2986 |
| 1810012K16I | 9 | 4 | 7 | 10 | 8 | 10 | 7 | 4 | 8 | 10 | 9 | 10 | 7.94 | 6 | 10 | 1.529 | 0.612 | 0.36266519 | NA | 0 | 0.3009 | 0.1959 | 0.1959 | TRUE | 0.1729 |
| 1810013A23 | 0 | 0 | 0 | 1 | 0 | 0 | 0 | 0 | 0 | 1 | 0 | 0 | 0.17 | 0 | 0 | 2.094 | 1.067 | 0.79379329 | NA | 0 | 14.2885 | 10 | 10 | TRUE | 0.3906 |
| 1810013D10 | 991 | 632 | 603 | 656 | 606 | 714 | 770 | 672 | 651 | 653 | 684 | 705 | 689 | 698 | 681 | 0.973 | -0.039 | 0.68264463 | 0.80543407 | 0.0037 | 0.0059 | 0.0051 | 0.0051 | TRUE | 0.1014 |

| | | | | | | | | | | | | | | | | | | | | | | | | | |
|---|---|---|---|---|---|---|---|---|---|---|---|---|---|---|---|---|---|---|---|---|---|---|---|---|---|
| 1810013L24F | 3848 | 2862 | 2599 | 3118 | 2537 | 3014 | 2989 | 3044 | 2804 | 3104 | 2862 | 2975 | 2963.01 | 2946 | 2980 | 1.012 | 0.017 | 0.78783839 | 0.87528754 | 0.0014 | 0.0032 | 0.0025 | 0.0025 | TRUE | 0.021 |
| 1810014B01 | 218 | 152 | 145 | 157 | 133 | 171 | 169 | 162 | 156 | 156 | 150 | 169 | 160.43 | 162 | 158 | 0.974 | -0.038 | 0.80821777 | 0.8891493 | 0 | 0.0172 | 0.0114 | 0.0114 | TRUE | 0.0354 |
| 1810018F18I | 0 | 0 | 0 | 1 | 0 | 0 | 0 | 0 | 0 | 1 | 0 | 0 | 0.17 | 0 | 0 | 2.094 | 1.067 | 0.79379329 | NA | 0 | 14.2885 | 10 | 10 | TRUE | 0.3906 |
| 1810019D21 | 63 | 47 | 42 | 45 | 23 | 51 | 49 | 50 | 45 | 45 | 26 | 50 | 44.22 | 48 | 40 | 0.844 | -0.246 | 0.43391627 | 0.60572701 | 0.0236 | 0.056 | 0.0481 | 0.0481 | TRUE | 0.6772 |
| 1810019J16F | 370 | 311 | 326 | 350 | 352 | 373 | 287 | 331 | 352 | 348 | 397 | 368 | 347.27 | 323 | 371 | 1.151 | 0.203 | 0.09674139 | 0.21611827 | 0.0048 | 0.0093 | 0.0079 | 0.0079 | TRUE | 0.1162 |
| 1810020O05 | 5 | 2 | 7 | 11 | 6 | 9 | 4 | 2 | 8 | 11 | 7 | 9 | 6.69 | 5 | 9 | 1.983 | 0.988 | 0.20043278 | NA | 0 | 0.3565 | 0.2634 | 0.2634 | TRUE | 0.555 |
| 1810021B22 | 4 | 1 | 2 | 3 | 1 | 2 | 3 | 1 | 2 | 3 | 1 | 2 | 2.07 | 2 | 2 | 0.952 | -0.071 | 0.95622218 | NA | 0 | 1.1477 | 0.7349 | 0.7349 | TRUE | 0.2178 |
| 1810022K09I | 1644 | 1193 | 1157 | 1201 | 1053 | 1198 | 1277 | 1269 | 1248 | 1195 | 1188 | 1183 | 1226.7 | 1265 | 1189 | 0.94 | -0.09 | 0.18591826 | 0.34226297 | 0 | 0.0044 | 0.0025 | 0.0025 | TRUE | 0.0015 |
| 1810024B03 | 6 | 4 | 0 | 0 | 2 | 1 | 5 | 4 | 0 | 0 | 2 | 1 | 2.03 | 3 | 1 | 0.351 | -1.509 | 0.33724371 | NA | 0 | 1.1721 | 1.133 | 1.133 | TRUE | 0.8965 |
| 1810026B05 | 325 | 273 | 256 | 264 | 193 | 288 | 252 | 290 | 276 | 263 | 218 | 284 | 263.97 | 273 | 255 | 0.937 | -0.093 | 0.50395123 | 0.66805555 | 0.0074 | 0.0114 | 0.0102 | 0.0102 | TRUE | 0.1749 |
| 1810026J23F | 2104 | 1503 | 1359 | 1658 | 1410 | 1604 | 1634 | 1599 | 1466 | 1650 | 1591 | 1583 | 1587.26 | 1566 | 1608 | 1.026 | 0.037 | 0.5996892 | 0.74540027 | 0.0012 | 0.0039 | 0.0029 | 0.0029 | TRUE | 0.037 |
| 1810030O07 | 2639 | 1895 | 1821 | 1905 | 1741 | 1989 | 2050 | 2016 | 1965 | 1896 | 1964 | 1963 | 1975.64 | 2010 | 1941 | 0.965 | -0.051 | 0.40371954 | 0.57657779 | 0 | 0.0036 | 0.0022 | 0.0022 | TRUE | 0.005 |
| 1810032O08 | 1819 | 1379 | 1614 | 1899 | 1594 | 1973 | 1413 | 1467 | 1741 | 1890 | 1798 | 1948 | 1709.53 | 1540 | 1879 | 1.221 | 0.288 | 0.00079821 | 0.00613381 | 0.0065 | 0.0038 | 0.0047 | 0.0047 | TRUE | 0.1559 |
| 1810033B17 | 4 | 2 | 3 | 3 | 2 | 6 | 3 | 2 | 3 | 3 | 2 | 6 | 3.27 | 3 | 4 | 1.322 | 0.402 | 0.70011636 | NA | 0 | 0.7266 | 0.4776 | 0.4776 | TRUE | 0.4317 |
| 1810034E14I | 5 | 5 | 2 | 3 | 2 | 4 | 4 | 5 | 2 | 3 | 2 | 4 | 3.43 | 4 | 3 | 0.812 | -0.301 | 0.76759188 | NA | 0 | 0.6944 | 0.4514 | 0.4514 | TRUE | 0.1831 |
| 1810035I16R | 0 | 0 | 0 | 0 | 0 | 0 | 0 | 0 | 0 | 0 | 0 | 0 | 0 | 0 | 0 | NA | NA | NA | NA | NA | NA | NA | NA | NA | NA |
| 1810037I17R | 1843 | 1283 | 1375 | 1452 | 1179 | 1409 | 1431 | 1365 | 1484 | 1445 | 1330 | 1391 | 1407.66 | 1427 | 1389 | 0.974 | -0.038 | 0.59203887 | 0.73988035 | 9.00E-04 | 0.0041 | 0.003 | 0.003 | TRUE | 0.0167 |
| 1810041L15F | 75 | 37 | 68 | 67 | 40 | 47 | 58 | 39 | 73 | 67 | 45 | 46 | 54.86 | 57 | 53 | 0.927 | -0.11 | 0.71591083 | 0.82941668 | 0.0518 | 0.0456 | 0.0473 | 0.0473 | TRUE | 0.2197 |
| 1810043G02 | 350 | 282 | 276 | 301 | 267 | 287 | 272 | 300 | 298 | 300 | 301 | 283 | 292.29 | 290 | 295 | 1.019 | 0.026 | 0.82634767 | 0.90029631 | 0 | 0.0105 | 0.007 | 0.007 | TRUE | 0.0365 |
| 1810043H04 | 983 | 783 | 726 | 819 | 753 | 753 | 763 | 833 | 783 | 815 | 849 | 743 | 797.95 | 793 | 802 | 1.012 | 0.017 | 0.84255383 | 0.91045754 | 0.002 | 0.0054 | 0.0043 | 0.0043 | TRUE | 0.05 |
| 1810044D09 | 77 | 39 | 46 | 51 | 33 | 52 | 60 | 41 | 50 | 51 | 37 | 51 | 48.37 | 50 | 46 | 0.92 | -0.12 | 0.68089136 | 0.80416297 | 0.0115 | 0.0514 | 0.0406 | 0.0406 | TRUE | 0.2548 |
| 1810046K07I | 0 | 0 | 0 | 0 | 0 | 2 | 0 | 0 | 0 | 0 | 0 | 2 | 0.33 | 0 | 1 | 3.743 | 1.904 | 0.62640105 | NA | 0 | 7.2051 | 9.2465 | 9.2465 | TRUE | 0.9739 |
| 1810053B23 | 0 | 0 | 0 | 0 | 0 | 0 | 0 | 0 | 0 | 0 | 0 | 0 | 0 | 0 | 0 | NA | NA | NA | NA | NA | NA | NA | NA | NA | NA |
| 1810055G02 | 1761 | 1228 | 1106 | 1117 | 1069 | 1089 | 1368 | 1306 | 1193 | 1112 | 1206 | 1075 | 1210.01 | 1289 | 1131 | 0.876 | -0.191 | 0.01978338 | 0.06859871 | 0.0033 | 0.0044 | 0.004 | 0.004 | TRUE | 0.0502 |
| 1810058I24R | 2636 | 1818 | 1985 | 1953 | 1816 | 2190 | 2047 | 1934 | 2142 | 1944 | 2049 | 2162 | 2046.23 | 2041 | 2052 | 1.005 | 0.007 | 0.91527154 | 0.95330234 | 0.0022 | 0.0036 | 0.0031 | 0.0031 | TRUE | 0.0271 |
| 1810062G17 | 9 | 13 | 9 | 8 | 6 | 8 | 7 | 14 | 10 | 8 | 7 | 8 | 8.86 | 10 | 8 | 0.751 | -0.413 | 0.52273956 | NA | 0 | 0.2699 | 0.1852 | 0.1852 | TRUE | 0.2646 |
| 1810062O18 | 1 | 0 | 2 | 1 | 0 | 0 | 1 | 0 | 2 | 1 | 0 | 0 | 0.66 | 1 | 0 | 0.382 | -1.39 | 0.57951664 | NA | 0 | 3.6208 | 3.037 | 3.037 | TRUE | 0.2643 |
| 1810064F22I | 0 | 0 | 0 | 1 | 1 | 1 | 0 | 0 | 0 | 1 | 1 | 1 | 0.52 | 0 | 1 | 5.852 | 2.549 | 0.3348364 | NA | 0 | 4.5739 | 3.5345 | 3.5345 | TRUE | 0.0847 |
| 1810065E05I | 1 | 0 | 0 | 0 | 0 | 0 | 1 | 0 | 0 | 0 | 0 | 0 | 0.13 | 0 | 0 | 0.552 | -0.857 | 0.8336548 | NA | 0 | 18.3121 | 10 | 10 | TRUE | 0.2566 |
| 2010001E11I | 1 | 1 | 1 | 0 | 1 | 1 | 1 | 1 | 1 | 0 | 1 | 1 | 0.84 | 1 | 1 | 0.722 | -0.47 | 0.81690682 | NA | 0 | 2.827 | 1.7221 | 1.7221 | TRUE | 0.2649 |
| 2010001M06 | 99 | 81 | 85 | 80 | 80 | 72 | 77 | 86 | 92 | 80 | 90 | 71 | 82.62 | 85 | 80 | 0.947 | -0.078 | 0.72008212 | 0.83202375 | 0 | 0.0311 | 0.0223 | 0.0223 | TRUE | 0.1031 |
| 2010002M12 | 3 | 0 | 2 | 0 | 0 | 0 | 2 | 0 | 2 | 0 | 0 | 0 | 0.75 | 1 | 0 | 0.126 | -2.987 | 0.23840739 | NA | 0 | 3.171 | 3.3041 | 3.3041 | TRUE | 0.488 |
| 2010003K11I | 1 | 0 | 0 | 0 | 1 | 2 | 1 | 0 | 0 | 0 | 1 | 2 | 0.65 | 0 | 1 | 2.998 | 1.584 | 0.52889475 | NA | 0 | 3.6683 | 3.06 | 3.06 | TRUE | 0.14 |
| 2010003O02 | 62 | 41 | 24 | 53 | 31 | 28 | 48 | 44 | 26 | 53 | 35 | 28 | 38.84 | 39 | 39 | 0.975 | -0.037 | 0.91887153 | 0.95538599 | 0.0781 | 0.0635 | 0.0676 | 0.0676 | TRUE | 0.7238 |
| 2010005H15 | 0 | 0 | 0 | 1 | 0 | 0 | 0 | 0 | 0 | 1 | 0 | 0 | 0.17 | 0 | 0 | 2.094 | 1.067 | 0.79379329 | NA | 0 | 14.2885 | 10 | 10 | TRUE | 0.3906 |
| 2010009K17I | 0 | 0 | 0 | 0 | 0 | 0 | 0 | 0 | 0 | 0 | 0 | 0 | 0 | 0 | 0 | NA | NA | NA | NA | NA | NA | NA | NA | NA | NA |
| 2010010A06 | 1 | 8 | 3 | 6 | 6 | 4 | 1 | 9 | 3 | 6 | 7 | 4 | 4.87 | 4 | 6 | 1.368 | 0.453 | 0.63416435 | NA | 0.3064 | 0.4892 | 0.4393 | 0.4393 | TRUE | 0.7319 |
| 2010012O05 | 301 | 200 | 220 | 237 | 209 | 221 | 234 | 213 | 237 | 236 | 236 | 218 | 228.96 | 228 | 230 | 1.007 | 0.011 | 0.93603633 | 0.96464483 | 0 | 0.0128 | 0.0086 | 0.0086 | TRUE | 0.0359 |
| 2010015L04F | 96 | 98 | 69 | 92 | 81 | 76 | 75 | 104 | 74 | 92 | 91 | 75 | 85.2 | 84 | 86 | 1.021 | 0.03 | 0.89410994 | 0.94184685 | 0.0142 | 0.0302 | 0.0257 | 0.0257 | TRUE | 0.3837 |
| 2010016I18R | 0 | 0 | 1 | 0 | 0 | 0 | 0 | 0 | 1 | 0 | 0 | 0 | 0.18 | 0 | 0 | 0.552 | -0.857 | 0.8336548 | NA | 0 | 13.1817 | 10 | 10 | TRUE | 0.5365 |
| 2010106C02I | 0 | 0 | 0 | 0 | 0 | 0 | 0 | 0 | 0 | 0 | 0 | 0 | 0 | 0 | 0 | NA | NA | NA | NA | NA | NA | NA | NA | NA | NA |
| 2010106E10I | 0 | 0 | 0 | 0 | 0 | 0 | 0 | 0 | 0 | 0 | 0 | 0 | 0 | 0 | 0 | NA | NA | NA | NA | NA | NA | NA | NA | NA | NA |
| 2010107E04I | 3788 | 2626 | 2695 | 3002 | 2637 | 3018 | 2942 | 2793 | 2908 | 2988 | 2975 | 2979 | 2930.89 | 2881 | 2981 | 1.034 | 0.049 | 0.38306075 | 0.55694744 | 0 | 0.0032 | 0.0019 | 0.0019 | TRUE | 0.0085 |
| 2010107G12 | 35 | 28 | 16 | 15 | 12 | 12 | 27 | 30 | 17 | 15 | 14 | 12 | 19.09 | 25 | 14 | 0.541 | -0.886 | 0.05238868 | 0.13938991 | 0 | 0.1266 | 0.0924 | 0.0924 | TRUE | 0.3902 |
| 2010107G23 | 826 | 585 | 752 | 766 | 743 | 848 | 642 | 622 | 811 | 762 | 838 | 837 | 752.16 | 692 | 812 | 1.176 | 0.234 | 0.03159518 | 0.09616738 | 0.0107 | 0.0056 | 0.0072 | 0.0072 | TRUE | 0.2652 |
| 2010109A12 | 2 | 0 | 2 | 1 | 0 | 1 | 2 | 0 | 2 | 1 | 0 | 1 | 0.95 | 1 | 1 | 0.541 | -0.886 | 0.6640791 | NA | 0 | 2.4999 | 1.8438 | 1.8438 | TRUE | 0.3958 |
| 2010109I03R | 0 | 0 | 0 | 0 | 0 | 0 | 0 | 0 | 0 | 0 | 0 | 0 | 0 | 0 | 0 | NA | NA | NA | NA | NA | NA | NA | NA | NA | NA |
| 2010111I01R | 280 | 182 | 155 | 160 | 141 | 143 | 217 | 194 | 167 | 159 | 159 | 141 | 172.96 | 193 | 153 | 0.791 | -0.338 | 0.03768326 | 0.10965124 | 0.0064 | 0.0161 | 0.0132 | 0.0132 | TRUE | 0.1466 |
| 2010204K13I | 556 | 434 | 426 | 471 | 447 | 514 | 432 | 462 | 460 | 469 | 504 | 507 | 472.27 | 451 | 493 | 1.095 | 0.131 | 0.18580424 | 0.34218529 | 0 | 0.0074 | 0.005 | 0.005 | TRUE | 0.0226 |
| 2010300C02I | 408 | 331 | 284 | 384 | 414 | 339 | 317 | 352 | 306 | 382 | 467 | 335 | 359.88 | 325 | 395 | 1.212 | 0.278 | 0.0413859 | 0.11713439 | 0.0141 | 0.009 | 0.0106 | 0.0106 | TRUE | 0.2904 |
| 2010308F09I | 0 | 4 | 0 | 3 | 2 | 5 | 0 | 4 | 0 | 3 | 2 | 5 | 2.41 | 1 | 3 | 2.504 | 1.324 | 0.35840714 | NA | 1.0086 | 0.9877 | 0.9914 | 0.9914 | TRUE | 1.9027 |
| 2010310C07I | 0 | 1 | 0 | 1 | 1 | 0 | 0 | 1 | 0 | 1 | 1 | 0 | 0.53 | 0 | 1 | 2.036 | 1.026 | 0.70492654 | NA | 0 | 4.4642 | 3.5518 | 3.5518 | TRUE | 0.5217 |
| 2010315B03 | 28 | 22 | 20 | 16 | 25 | 12 | 22 | 23 | 22 | 16 | 28 | 12 | 20.45 | 22 | 19 | 0.833 | -0.264 | 0.55834912 | 0.71311815 | 0.0371 | 0.1183 | 0.0974 | 0.0974 | TRUE | 0.9572 |
| 2010320M18 | 281 | 252 | 221 | 271 | 204 | 252 | 218 | 268 | 238 | 270 | 230 | 249 | 245.57 | 241 | 250 | 1.037 | 0.052 | 0.70685856 | 0.82293145 | 0.0047 | 0.0121 | 0.0099 | 0.0099 | TRUE | 0.1014 |
| 2200002D01 | 158 | 84 | 84 | 83 | 76 | 72 | 123 | 89 | 91 | 83 | 86 | 71 | 90.35 | 101 | 80 | 0.785 | -0.349 | 0.11540633 | 0.24530321 | 0.0139 | 0.0287 | 0.0242 | 0.0242 | TRUE | 0.3846 |
| 2200002J24F | 2 | 4 | 1 | 6 | 4 | 12 | 2 | 4 | 1 | 6 | 5 | 12 | 4.87 | 2 | 8 | 3.312 | 1.728 | 0.07289303 | NA | 0 | 0.4891 | 0.3867 | 0.3867 | TRUE | 0.762 |
| 2210010C04I | 0 | 0 | 0 | 0 | 0 | 0 | 0 | 0 | 0 | 0 | 0 | 0 | 0 | 0 | 0 | NA | NA | NA | NA | NA | NA | NA | NA | NA | NA |
| 2210013O21 | 463 | 326 | 331 | 333 | 273 | 348 | 360 | 347 | 357 | 331 | 308 | 344 | 341.08 | 355 | 328 | 0.925 | -0.113 | 0.3150247 | 0.48870278 | 0 | 0.0094 | 0.0062 | 0.0062 | TRUE | 0.0302 |
| 2210015D19 | 515 | 342 | 332 | 359 | 339 | 359 | 400 | 364 | 358 | 357 | 382 | 354 | 369.36 | 374 | 364 | 0.972 | -0.04 | 0.71684388 | 0.82975813 | 0 | 0.0088 | 0.0062 | 0.0062 | TRUE | 0.0455 |
| 2210016F16I | 1654 | 1183 | 1010 | 972 | 824 | 856 | 1285 | 1258 | 1090 | 968 | 930 | 845 | 1062.47 | 1211 | 914 | 0.754 | -0.407 | 6.56E-06 | 0.00013308 | 0.0054 | 0.0047 | 0.0049 | 0.0049 | TRUE | 0.0904 |
| 2210016L21F | 1853 | 1384 | 1206 | 1326 | 1197 | 1381 | 1439 | 1472 | 1301 | 1320 | 1350 | 1363 | 1374.34 | 1404 | 1344 | 0.957 | -0.063 | 0.38980482 | 0.56374383 | 0.0015 | 0.0042 | 0.0032 | 0.0032 | TRUE | 0.0481 |
| 2210018M11 | 4180 | 3082 | 2843 | 3180 | 2689 | 3236 | 3246 | 3278 | 3068 | 3165 | 3033 | 3194 | 3164.26 | 3197 | 3131 | 0.979 | -0.03 | 0.60474007 | 0.74959239 | 7.00E-04 | 0.0032 | 0.0022 | 0.0022 | TRUE | 0.0151 |
| 2210019I11R | 0 | 1 | 0 | 0 | 0 | 0 | 0 | 1 | 0 | 0 | 0 | 0 | 0.18 | 0 | 0 | 0.552 | -0.857 | 0.8336548 | NA | 0 | 13.3714 | 10 | 10 | TRUE | 0.5216 |
| 2210039B01 | 4 | 5 | 9 | 12 | 9 | 11 | 3 | 5 | 10 | 12 | 10 | 11 | 8.52 | 6 | 11 | 1.863 | 0.897 | 0.18969272 | NA | 0 | 0.2808 | 0.2086 | 0.2086 | TRUE | 0.6333 |
| 2210404J11F | 362 | 227 | 185 | 253 | 183 | 221 | 281 | 241 | 200 | 252 | 206 | 218 | 233.11 | 241 | 225 | 0.934 | -0.099 | 0.52979009 | 0.6898952 | 0.016 | 0.0126 | 0.0136 | 0.0136 | TRUE | 0.1888 |
| 2210404O07 | 36 | 26 | 35 | 51 | 34 | 38 | 28 | 28 | 38 | 51 | 38 | 38 | 36.67 | 31 | 42 | 1.364 | 0.448 | 0.1728684 | 0.32523471 | 0 | 0.0671 | 0.0499 | 0.0499 | TRUE | 0.2453 |
| 2210404O09 | 211 | 124 | 107 | 102 | 98 | 106 | 164 | 132 | 115 | 102 | 111 | 105 | 121.32 | 137 | 106 | 0.765 | -0.386 | 0.04694617 | 0.12865444 | 0.0116 | 0.022 | 0.0188 | 0.0188 | TRUE | 0.328 |
| 2210407C18I | 0 | 0 | 1 | 0 | 0 | 0 | 0 | 0 | 1 | 0 | 0 | 0 | 0.18 | 0 | 0 | 0.552 | -0.857 | 0.8336548 | NA | 0 | 13.1817 | 10 | 10 | TRUE | 0.5365 |
| 2210408F21I | 2 | 1 | 2 | 1 | 3 | 2 | 2 | 1 | 2 | 1 | 3 | 2 | 1.85 | 2 | 2 | 1.32 | 0.401 | 0.77003314 | NA | 0 | 1.2802 | 0.805 | 0.805 | TRUE | 0.2286 |

| | | | | | | | | | | | | | | | | | | | | | | | | | |
|---|---|---|---|---|---|---|---|---|---|---|---|---|---|---|---|---|---|---|---|---|---|---|---|---|---|
| 2210408I21R | 148 | 108 | 77 | 89 | 58 | 87 | 115 | 115 | 83 | 89 | 65 | 86 | 92.13 | 104 | 80 | 0.766 | -0.384 | 0.08760876 | 0.20096165 | 0.0177 | 0.0282 | 0.0253 | 0.0253 | TRUE | 0.2975 |
| 2210409D07 | 0 | 0 | 0 | 0 | 0 | 0 | 0 | 0 | 0 | 0 | 0 | 0 | 0 | 0 | 0 | NA | NA | NA | NA | NA | NA | NA | NA | NA | NA |
| 2210409E12 | 119 | 61 | 32 | 40 | 30 | 26 | 92 | 65 | 35 | 40 | 34 | 26 | 48.53 | 64 | 33 | 0.513 | -0.962 | 0.00823102 | 0.03579129 | 0.1224 | 0.0513 | 0.0723 | 0.0723 | TRUE | 0.1192 |
| 2210414B05 | 0 | 0 | 0 | 0 | 0 | 0 | 0 | 0 | 0 | 0 | 0 | 0 | 0 | 0 | 0 | NA | NA | NA | NA | NA | NA | NA | NA | NA | NA |
| 2210416O15 | 12 | 6 | 8 | 6 | 5 | 5 | 9 | 6 | 9 | 6 | 6 | 5 | 6.81 | 8 | 6 | 0.676 | -0.565 | 0.43477122 | NA | 0 | 0.3503 | 0.2234 | 0.2234 | TRUE | 0.097 |
| 2210417A02 | 27 | 19 | 17 | 9 | 12 | 10 | 21 | 20 | 18 | 9 | 14 | 10 | 15.32 | 20 | 11 | 0.54 | -0.889 | 0.07249319 | NA | 0 | 0.1572 | 0.1042 | 0.1042 | TRUE | 0.1623 |
| 2210417K05 | 6 | 12 | 16 | 18 | 13 | 15 | 5 | 13 | 17 | 18 | 15 | 15 | 13.68 | 12 | 16 | 1.399 | 0.484 | 0.39515698 | NA | 0.1085 | 0.1757 | 0.158 | 0.158 | TRUE | 0.6933 |
| 2210420H20 | 6 | 10 | 5 | 10 | 6 | 6 | 5 | 11 | 5 | 10 | 7 | 6 | 7.22 | 7 | 8 | 1.109 | 0.15 | 0.83662391 | NA | 0 | 0.3306 | 0.2398 | 0.2398 | TRUE | 0.5592 |
| 2300002M23 | 7 | 11 | 12 | 12 | 10 | 12 | 5 | 12 | 13 | 12 | 11 | 12 | 10.86 | 10 | 12 | 1.188 | 0.249 | 0.67592888 | NA | 0 | 0.2207 | 0.1619 | 0.1619 | TRUE | 0.7404 |
| 2300003K06 | 0 | 0 | 0 | 0 | 0 | 0 | 0 | 0 | 0 | 0 | 0 | 0 | 0 | 0 | 0 | NA | NA | NA | NA | NA | NA | NA | NA | NA | NA |
| 2300005B03 | 0 | 0 | 0 | 0 | 1 | 0 | 0 | 0 | 0 | 0 | 1 | 0 | 0.19 | 0 | 0 | 2.094 | 1.067 | 0.79379329 | NA | 0 | 12.608 | 10 | 10 | TRUE | 0.5075 |
| 2300009A05 | 117 | 103 | 105 | 152 | 137 | 124 | 91 | 110 | 113 | 151 | 155 | 122 | 123.66 | 105 | 143 | 1.37 | 0.454 | 0.01642514 | 0.05973653 | 0.0076 | 0.0216 | 0.0176 | 0.0176 | TRUE | 0.1646 |
| 2310001H17 | 4 | 7 | 12 | 7 | 6 | 11 | 3 | 7 | 13 | 7 | 7 | 11 | 8.02 | 8 | 8 | 1.073 | 0.101 | 0.88915528 | NA | 0.1445 | 0.2981 | 0.2535 | 0.2535 | TRUE | 0.2694 |
| 2310001K24 | 1 | 0 | 0 | 1 | 0 | 0 | 1 | 0 | 0 | 1 | 0 | 0 | 0.3 | 0 | 0 | 1.075 | 0.105 | 0.97698525 | NA | 0 | 8.0272 | 7.5141 | 7.5141 | TRUE | 0.3906 |
| 2310002D06 | 0 | 0 | 0 | 0 | 0 | 0 | 0 | 0 | 0 | 0 | 0 | 0 | 0 | 0 | 0 | NA | NA | NA | NA | NA | NA | NA | NA | NA | NA |
| 2310002F09 | 5 | 6 | 8 | 11 | 6 | 12 | 4 | 6 | 9 | 11 | 7 | 12 | 8.08 | 6 | 10 | 1.6 | 0.678 | 0.32542111 | NA | 0 | 0.2959 | 0.2121 | 0.2121 | TRUE | 0.3756 |
| 2310002J15 | 8 | 5 | 9 | 14 | 8 | 13 | 6 | 5 | 10 | 14 | 9 | 13 | 9.51 | 7 | 12 | 1.702 | 0.767 | 0.22313052 | NA | 0 | 0.2517 | 0.1739 | 0.1739 | TRUE | 0.2536 |
| 2310002L09 | 0 | 0 | 0 | 0 | 0 | 0 | 0 | 0 | 0 | 0 | 0 | 0 | 0 | 0 | 0 | NA | NA | NA | NA | NA | NA | NA | NA | NA | NA |
| 2310003H01 | 2445 | 1834 | 1552 | 1836 | 1628 | 1784 | 1899 | 1951 | 1675 | 1828 | 1837 | 1761 | 1824.92 | 1842 | 1809 | 0.981 | -0.027 | 0.71684415 | 0.82975813 | 0.0029 | 0.0037 | 0.0034 | 0.0034 | TRUE | 0.0744 |
| 2310005A03 | 0 | 0 | 0 | 0 | 0 | 0 | 0 | 0 | 0 | 0 | 0 | 0 | 0 | 0 | 0 | NA | NA | NA | NA | NA | NA | NA | NA | NA | NA |
| 2310005E17 | 0 | 0 | 2 | 3 | 2 | 2 | 0 | 0 | 2 | 3 | 2 | 2 | 1.56 | 1 | 2 | 3.568 | 1.835 | 0.28504941 | NA | 0 | 1.5193 | 1.1958 | 1.1958 | TRUE | 0.9756 |
| 2310005G13 | 48 | 31 | 11 | 38 | 22 | 16 | 37 | 33 | 12 | 38 | 25 | 16 | 26.76 | 27 | 26 | 0.947 | -0.078 | 0.86801829 | 0.92704068 | 0.2066 | 0.091 | 0.1231 | 0.1231 | TRUE | 0.4965 |
| 2310007B03 | 35 | 23 | 39 | 35 | 27 | 31 | 27 | 24 | 42 | 35 | 30 | 31 | 31.6 | 31 | 32 | 1.029 | 0.041 | 0.90763215 | 0.94923397 | 0.0142 | 0.0774 | 0.06 | 0.06 | TRUE | 0.5818 |
| 2310007L24 | 1 | 0 | 2 | 0 | 4 | 0 | 1 | 0 | 2 | 0 | 5 | 0 | 1.24 | 1 | 2 | 1.518 | 0.602 | 0.76847514 | NA | 3.4394 | 1.912 | 2.1628 | 2.1628 | TRUE | 1.9008 |
| 2310008H04 | 2826 | 2025 | 2393 | 2458 | 2143 | 2553 | 2195 | 2154 | 2582 | 2447 | 2418 | 2520 | 2385.87 | 2310 | 2462 | 1.066 | 0.092 | 0.23448108 | 0.40061384 | 0.0048 | 0.0034 | 0.0039 | 0.0039 | TRUE | 0.1268 |
| 2310008H09 | 12599 | 8807 | 7914 | 8260 | 7453 | 8425 | 9785 | 9368 | 8539 | 8222 | 8408 | 8317 | 8773.11 | 9231 | 8316 | 0.901 | -0.151 | 0.01314619 | 0.05059194 | 0.0023 | 0.0027 | 0.0026 | 0.0026 | TRUE | 0.0523 |
| 2310008N11 | 4 | 1 | 1 | 0 | 0 | 0 | 3 | 1 | 1 | 0 | 0 | 0 | 0.87 | 2 | 0 | 0.107 | -3.222 | 0.14630442 | NA | 0 | 2.7114 | 2.2736 | 2.2736 | TRUE | 0.4767 |
| 2310009A05 | 173 | 133 | 118 | 124 | 123 | 123 | 134 | 141 | 127 | 123 | 139 | 121 | 131.13 | 134 | 128 | 0.95 | -0.074 | 0.66850595 | 0.79617284 | 0 | 0.0205 | 0.0138 | 0.0138 | TRUE | 0.054 |
| 2310009B15 | 142 | 117 | 86 | 83 | 75 | 81 | 110 | 124 | 93 | 83 | 85 | 80 | 95.79 | 109 | 83 | 0.754 | -0.407 | 0.04832447 | 0.13145735 | 0 | 0.0272 | 0.02 | 0.02 | TRUE | 0.159 |
| 2310010J17 | 24 | 20 | 16 | 33 | 21 | 33 | 19 | 21 | 17 | 33 | 24 | 33 | 24.38 | 19 | 30 | 1.564 | 0.646 | 0.09984706 | 0.22082633 | 0 | 0.0996 | 0.0682 | 0.0682 | TRUE | 0.1905 |
| 2310010M2 | 0 | 2 | 0 | 0 | 2 | 0 | 0 | 2 | 0 | 0 | 2 | 0 | 0.73 | 1 | 1 | 1.063 | 0.089 | 0.97363104 | NA | 9.1948 | 3.2464 | 3.7678 | 3.7678 | TRUE | 0.9747 |
| 2310011J03 | 757 | 553 | 413 | 407 | 385 | 409 | 588 | 588 | 446 | 405 | 434 | 404 | 477.49 | 541 | 414 | 0.765 | -0.387 | 0.00157598 | 0.01000689 | 0.0113 | 0.0074 | 0.0086 | 0.0086 | TRUE | 0.2694 |
| 2310014F07 | 3 | 0 | 1 | 9 | 0 | 2 | 2 | 0 | 1 | 9 | 0 | 2 | 2.39 | 1 | 4 | 3.093 | 1.629 | 0.28705748 | NA | 1.6891 | 0.994 | 1.1404 | 1.1404 | TRUE | 0.6334 |
| 2310014L17 | 72 | 52 | 54 | 74 | 29 | 53 | 56 | 55 | 58 | 74 | 33 | 52 | 54.7 | 56 | 53 | 0.941 | -0.088 | 0.77247001 | 0.86543807 | 0.0554 | 0.0458 | 0.0483 | 0.0483 | TRUE | 0.2178 |
| 2310015A10 | 252 | 203 | 211 | 202 | 188 | 222 | 196 | 216 | 228 | 201 | 212 | 219 | 211.94 | 213 | 211 | 0.992 | -0.012 | 0.93239939 | 0.96249104 | 0 | 0.0136 | 0.0095 | 0.0095 | TRUE | 0.0613 |
| 2310015B20 | 0 | 0 | 2 | 1 | 1 | 3 | 0 | 0 | 2 | 1 | 1 | 3 | 1.21 | 1 | 2 | 2.5 | 1.322 | 0.48832544 | NA | 0 | 1.9658 | 1.6105 | 1.6105 | TRUE | 0.9754 |
| 2310015D24 | 0 | 0 | 0 | 0 | 0 | 0 | 0 | 0 | 0 | 0 | 0 | 0 | 0 | 0 | 0 | NA | NA | NA | NA | NA | NA | NA | NA | NA | NA |
| 2310016D03 | 0 | 0 | 0 | 0 | 0 | 0 | 0 | 0 | 0 | 0 | 0 | 0 | 0 | 0 | 0 | NA | NA | NA | NA | NA | NA | NA | NA | NA | NA |
| 2310020H05 | 0 | 0 | 0 | 0 | 0 | 0 | 0 | 0 | 0 | 0 | 0 | 0 | 0 | 0 | 0 | NA | NA | NA | NA | NA | NA | NA | NA | NA | NA |
| 2310022A10 | 2916 | 2141 | 2178 | 2394 | 2138 | 2393 | 2265 | 2277 | 2350 | 2383 | 2412 | 2362 | 2341.54 | 2297 | 2386 | 1.039 | 0.055 | 0.34362888 | 0.51790179 | 0 | 0.0034 | 0.002 | 0.002 | TRUE | 0.0048 |
| 2310022B05 | 3924 | 3040 | 3289 | 3277 | 2885 | 3552 | 3048 | 3234 | 3549 | 3262 | 3255 | 3506 | 3308.81 | 3277 | 3341 | 1.02 | 0.029 | 0.68168213 | 0.80469794 | 0.0036 | 0.0031 | 0.0033 | 0.0033 | TRUE | 0.064 |
| 2310030A07 | 2 | 1 | 2 | 4 | 2 | 2 | 2 | 1 | 2 | 4 | 2 | 2 | 2.16 | 2 | 3 | 1.727 | 0.789 | 0.53671095 | NA | 0 | 1.0974 | 0.6821 | 0.6821 | TRUE | 0.1948 |
| 2310030G06 | 162 | 101 | 109 | 114 | 105 | 120 | 126 | 107 | 118 | 113 | 118 | 118 | 116.87 | 117 | 116 | 0.996 | -0.006 | 0.97267148 | 0.98514693 | 0 | 0.0227 | 0.0151 | 0.0151 | TRUE | 0.0502 |
| 2310033P09 | 1204 | 914 | 795 | 733 | 689 | 767 | 935 | 972 | 858 | 730 | 777 | 757 | 838.19 | 922 | 755 | 0.818 | -0.289 | 0.00060886 | 0.00502449 | 0.0013 | 0.0053 | 0.0039 | 0.0039 | TRUE | 0.0422 |
| 2310034C09 | 0 | 0 | 0 | 0 | 0 | 0 | 0 | 0 | 0 | 0 | 0 | 0 | 0 | 0 | 0 | NA | NA | NA | NA | NA | NA | NA | NA | NA | NA |
| 2310034G01 | 16 | 16 | 15 | 20 | 10 | 17 | 12 | 17 | 16 | 20 | 11 | 17 | 15.6 | 15 | 16 | 1.064 | 0.089 | 0.85606458 | NA | 0 | 0.1544 | 0.1095 | 0.1095 | TRUE | 0.2828 |
| 2310034O05 | 1 | 0 | 0 | 0 | 0 | 0 | 1 | 0 | 0 | 0 | 0 | 0 | 0.13 | 0 | 0 | 0.552 | -0.857 | 0.8336548 | NA | 0 | 18.3121 | 10 | 10 | TRUE | 0.2566 |
| 2310035C23 | 2973 | 2227 | 1912 | 1884 | 1709 | 2003 | 2309 | 2369 | 2063 | 1875 | 1928 | 1977 | 2086.9 | 2247 | 1927 | 0.857 | -0.222 | 0.00182697 | 0.01127011 | 0.0025 | 0.0036 | 0.0032 | 0.0032 | TRUE | 0.061 |
| 2310036O22 | 4520 | 3345 | 2597 | 2330 | 2181 | 2428 | 3510 | 3558 | 2802 | 2319 | 2460 | 2397 | 2841.19 | 3290 | 2392 | 0.727 | -0.461 | 1.65E-07 | 6.06E-06 | 0.0082 | 0.0033 | 0.0052 | 0.0052 | TRUE | 0.2032 |
| 2310039H08 | 228 | 212 | 215 | 210 | 197 | 212 | 177 | 226 | 232 | 209 | 222 | 209 | 212.52 | 212 | 213 | 1.014 | 0.02 | 0.89604611 | 0.94272381 | 0.0075 | 0.0136 | 0.0117 | 0.0117 | TRUE | 0.2432 |
| 2310039L15 | 44 | 27 | 31 | 34 | 26 | 20 | 34 | 29 | 33 | 34 | 29 | 20 | 29.88 | 32 | 28 | 0.858 | -0.221 | 0.54168597 | 0.69973309 | 0 | 0.0818 | 0.061 | 0.061 | TRUE | 0.4161 |
| 2310040G24 | 56 | 47 | 38 | 36 | 33 | 29 | 43 | 50 | 41 | 36 | 37 | 29 | 39.36 | 45 | 34 | 0.755 | -0.405 | 0.1907206 | 0.34885269 | 0 | 0.0626 | 0.0429 | 0.0429 | TRUE | 0.1334 |
| 2310042D19 | 0 | 0 | 0 | 0 | 0 | 1 | 0 | 0 | 0 | 0 | 0 | 1 | 0.16 | 0 | 0 | 2.094 | 1.067 | 0.79379329 | NA | 0 | 14.4078 | 10 | 10 | TRUE | 0.3835 |
| 2310042E22 | 9 | 5 | 9 | 11 | 10 | 11 | 7 | 5 | 10 | 11 | 11 | 11 | 9.18 | 7 | 11 | 1.506 | 0.591 | 0.34634615 | NA | 0 | 0.2605 | 0.1712 | 0.1712 | TRUE | 0.1935 |
| 2310043L19 | 1 | 0 | 0 | 0 | 0 | 0 | 1 | 0 | 0 | 0 | 0 | 0 | 0.13 | 0 | 0 | 0.552 | -0.857 | 0.8336548 | NA | 0 | 18.3121 | 10 | 10 | TRUE | 0.2566 |
| 2310043O21 | 0 | 0 | 0 | 0 | 0 | 0 | 0 | 0 | 0 | 0 | 0 | 0 | 0 | 0 | 0 | NA | NA | NA | NA | NA | NA | NA | NA | NA | NA |
| 2310044G17 | 1620 | 1194 | 1190 | 1252 | 1086 | 1311 | 1258 | 1270 | 1284 | 1246 | 1225 | 1294 | 1262.96 | 1271 | 1255 | 0.988 | -0.017 | 0.8037569 | 0.88642727 | 0 | 0.0043 | 0.0026 | 0.0026 | TRUE | 0.0089 |
| 2310045N01 | 350 | 268 | 251 | 245 | 225 | 291 | 272 | 285 | 271 | 244 | 254 | 287 | 268.78 | 276 | 262 | 0.949 | -0.075 | 0.55613665 | 0.71124384 | 0 | 0.0112 | 0.008 | 0.008 | TRUE | 0.0837 |
| 2310047M1 | 662 | 543 | 483 | 472 | 480 | 564 | 514 | 578 | 521 | 470 | 541 | 557 | 530.16 | 538 | 523 | 0.973 | -0.04 | 0.70502837 | 0.82140287 | 0.0042 | 0.0069 | 0.006 | 0.006 | TRUE | 0.0928 |
| 2310050B05 | 0 | 0 | 0 | 0 | 1 | 0 | 0 | 0 | 0 | 0 | 1 | 0 | 0.19 | 0 | 0 | 2.094 | 1.067 | 0.79379329 | NA | 0 | 12.608 | 10 | 10 | TRUE | 0.5075 |
| 2310050C09 | 0 | 0 | 0 | 0 | 0 | 0 | 0 | 0 | 0 | 0 | 0 | 0 | 0 | 0 | 0 | NA | NA | NA | NA | NA | NA | NA | NA | NA | NA |
| 2310057J18 | 0 | 0 | 0 | 0 | 0 | 0 | 0 | 0 | 0 | 0 | 0 | 0 | 0 | 0 | 0 | NA | NA | NA | NA | NA | NA | NA | NA | NA | NA |
| 2310057M21 | 836 | 585 | 612 | 588 | 575 | 587 | 649 | 622 | 660 | 585 | 649 | 579 | 624.22 | 644 | 604 | 0.938 | -0.093 | 0.31052612 | 0.48348038 | 6.00E-04 | 0.0062 | 0.0044 | 0.0044 | TRUE | 0.0468 |
| 2310057N15 | 0 | 0 | 0 | 0 | 0 | 0 | 0 | 0 | 0 | 0 | 0 | 0 | 0 | 0 | 0 | NA | NA | NA | NA | NA | NA | NA | NA | NA | NA |
| 2310061I04R | 2321 | 1926 | 1951 | 2171 | 2003 | 2330 | 1803 | 2049 | 2105 | 2161 | 2260 | 2300 | 2112.84 | 1986 | 2240 | 1.13 | 0.176 | 0.01824041 | 0.06451626 | 0.0031 | 0.0035 | 0.0035 | 0.0035 | TRUE | 0.0809 |
| 2310061J03 | 78 | 60 | 53 | 63 | 41 | 41 | 61 | 64 | 57 | 63 | 46 | 40 | 55.17 | 61 | 50 | 0.824 | -0.28 | 0.3039842 | 0.47651306 | 0.0061 | 0.0454 | 0.0351 | 0.0351 | TRUE | 0.4278 |
| 2310061N02 | 0 | 0 | 0 | 0 | 0 | 0 | 0 | 0 | 0 | 0 | 0 | 0 | 0 | 0 | 0 | NA | NA | NA | NA | NA | NA | NA | NA | NA | NA |
| 2310065F04 | 0 | 0 | 0 | 0 | 0 | 0 | 0 | 0 | 0 | 0 | 0 | 0 | 0 | 0 | 0 | NA | NA | NA | NA | NA | NA | NA | NA | NA | NA |
| 2310067B10 | 5147 | 3792 | 4370 | 4622 | 4355 | 5034 | 3997 | 4034 | 4715 | 4601 | 4913 | 4969 | 4538.19 | 4249 | 4828 | 1.137 | 0.185 | 0.0124122 | 0.0485343 | 0.005 | 0.0029 | 0.0037 | 0.0037 | TRUE | 0.1117 |

| | | | | | | | | | | | | | | | | | | | | | | | | | |
|---|---|---|---|---|---|---|---|---|---|---|---|---|---|---|---|---|---|---|---|---|---|---|---|---|---|
| 2310068J16F | 27 | 23 | 18 | 25 | 24 | 32 | 21 | 24 | 19 | 25 | 27 | 32 | 24.73 | 21 | 28 | 1.29 | 0.368 | 0.33801406 | 0.51223769 | 0 | 0.0982 | 0.0652 | 0.0652 | TRUE | 0.0919 |
| 2310069B03I | 23 | 15 | 11 | 26 | 17 | 17 | 18 | 16 | 12 | 26 | 19 | 17 | 17.92 | 15 | 21 | 1.345 | 0.427 | 0.35345824 | 0.52780934 | 0 | 0.1347 | 0.0962 | 0.0962 | TRUE | 0.2827 |
| 2310069G16 | 5 | 3 | 2 | 1 | 0 | 1 | 4 | 3 | 2 | 1 | 0 | 1 | 1.87 | 3 | 1 | 0.219 | -2.191 | 0.16077546 | NA | 0 | 1.2704 | 0.8413 | 0.8413 | TRUE | 0.1972 |
| 2310079G19 | 0 | 0 | 0 | 0 | 0 | 0 | 0 | 0 | 0 | 0 | 0 | 0 | 0 | 0 | 0 | NA | NA | NA | NA | NA | NA | NA | NA | NA | NA |
| 2310081J21F | 0 | 0 | 0 | 0 | 0 | 0 | 0 | 0 | 0 | 0 | 0 | 0 | 0 | 0 | 0 | NA | NA | NA | NA | NA | NA | NA | NA | NA | NA |
| 2410002F23I | 3472 | 2706 | 2538 | 2248 | 2234 | 2377 | 2697 | 2878 | 2738 | 2238 | 2520 | 2346 | 2569.61 | 2771 | 2368 | 0.854 | -0.227 | 0.00065976 | 0.00532508 | 0.0018 | 0.0033 | 0.0028 | 0.0028 | TRUE | 0.0378 |
| 2410003L11F | 1011 | 673 | 564 | 571 | 526 | 622 | 785 | 716 | 609 | 568 | 593 | 614 | 647.56 | 703 | 592 | 0.84 | -0.252 | 0.01821607 | 0.06444612 | 0.0077 | 0.0061 | 0.0066 | 0.0066 | TRUE | 0.1618 |
| 2410004B18I | 798 | 542 | 494 | 394 | 408 | 420 | 620 | 577 | 533 | 392 | 460 | 415 | 499.4 | 577 | 422 | 0.731 | -0.453 | 2.38E-05 | 0.00038379 | 0.0041 | 0.0072 | 0.0062 | 0.0062 | TRUE | 0.0692 |
| 2410004I01R | 60 | 42 | 48 | 39 | 41 | 56 | 47 | 45 | 52 | 39 | 46 | 55 | 47.24 | 48 | 47 | 0.982 | -0.026 | 0.92832808 | 0.96054762 | 0 | 0.0526 | 0.0376 | 0.0376 | TRUE | 0.2087 |
| 2410004N09 | 907 | 801 | 1147 | 1239 | 1028 | 1404 | 704 | 852 | 1238 | 1233 | 1160 | 1386 | 1095.5 | 931 | 1260 | 1.353 | 0.437 | 0.08878187 | 0.20283283 | 0.0465 | 0.0046 | 0.0158 | 0.0465 | TRUE | 0.9849 |
| 2410004P03I | 177 | 109 | 71 | 132 | 124 | 108 | 137 | 116 | 77 | 131 | 140 | 107 | 117.98 | 110 | 126 | 1.137 | 0.185 | 0.41331055 | 0.5860164 | 0.0428 | 0.0225 | 0.0282 | 0.0282 | TRUE | 0.4877 |
| 2410006H16I | 7009 | 5225 | 7883 | 9635 | 8237 | 9537 | 5444 | 5558 | 8506 | 9591 | 9292 | 9414 | 7967.4 | 6503 | 9432 | 1.451 | 0.537 | 0.01586972 | 0.05823516 | 0.0356 | 0.0027 | 0.0112 | 0.0356 | TRUE | 0.886 |
| 2410007B07I | 1461 | 1115 | 1176 | 1213 | 1071 | 1225 | 1135 | 1186 | 1269 | 1207 | 1208 | 1209 | 1202.41 | 1197 | 1208 | 1.011 | 0.016 | 0.83119463 | 0.9033735 | 9.00E-04 | 0.0044 | 0.0031 | 0.0031 | TRUE | 0.0333 |
| 2410012E07I | 1 | 0 | 0 | 0 | 1 | 0 | 1 | 0 | 0 | 0 | 1 | 0 | 0.32 | 0 | 0 | 1.075 | 0.105 | 0.97638847 | NA | 0 | 7.4681 | 7.0392 | 7.0392 | TRUE | 0.5075 |
| 2410012M07 | 720 | 525 | 454 | 502 | 417 | 487 | 559 | 558 | 490 | 500 | 470 | 481 | 509.72 | 536 | 484 | 0.902 | -0.149 | 0.13635871 | 0.27621529 | 0.0014 | 0.0071 | 0.0053 | 0.0053 | TRUE | 0.0637 |
| 2410015M20 | 2024 | 1419 | 1313 | 1513 | 1298 | 1517 | 1572 | 1509 | 1417 | 1506 | 1464 | 1498 | 1494.31 | 1499 | 1489 | 0.992 | -0.011 | 0.87584746 | 0.93088377 | 9.00E-04 | 0.004 | 0.0028 | 0.0028 | TRUE | 0.0277 |
| 2410016O06 | 2038 | 1425 | 1211 | 1283 | 1210 | 1372 | 1583 | 1516 | 1307 | 1277 | 1365 | 1354 | 1400.29 | 1469 | 1332 | 0.906 | -0.142 | 0.09186357 | 0.20788568 | 0.005 | 0.0041 | 0.0044 | 0.0044 | TRUE | 0.1105 |
| 2410017I17R | 0 | 1 | 0 | 0 | 1 | 0 | 0 | 1 | 0 | 0 | 1 | 0 | 0.37 | 0 | 0 | 1.075 | 0.105 | 0.97520514 | NA | 0 | 6.4904 | 6.1969 | 6.1969 | TRUE | 0.5216 |
| 2410018L13F | 19 | 10 | 28 | 21 | 12 | 11 | 15 | 11 | 30 | 21 | 14 | 11 | 16.82 | 19 | 15 | 0.821 | -0.285 | 0.59248019 | NA | 0.1439 | 0.1434 | 0.1435 | 0.1435 | TRUE | 1.1919 |
| 2410018M08 | 645 | 581 | 562 | 596 | 543 | 677 | 501 | 618 | 606 | 593 | 613 | 668 | 599.91 | 575 | 625 | 1.089 | 0.124 | 0.24909108 | 0.41806702 | 0.0071 | 0.0064 | 0.0066 | 0.0066 | TRUE | 0.1581 |
| 2410021H03I | 30 | 31 | 32 | 33 | 33 | 41 | 23 | 33 | 35 | 33 | 37 | 40 | 33.56 | 30 | 37 | 1.229 | 0.297 | 0.38121033 | 0.55575676 | 0 | 0.0731 | 0.0529 | 0.0529 | TRUE | 0.3316 |
| 2410057H14I | 2 | 0 | 3 | 1 | 2 | 0 | 2 | 0 | 3 | 1 | 2 | 0 | 1.34 | 2 | 1 | 0.668 | -0.583 | 0.74407913 | NA | 0 | 1.7707 | 1.5093 | 1.5093 | TRUE | 0.1443 |
| 2410066E13I | 187 | 177 | 210 | 256 | 200 | 284 | 145 | 188 | 227 | 255 | 226 | 280 | 220.15 | 187 | 254 | 1.367 | 0.451 | 0.00881859 | 0.037783 | 0.0254 | 0.0132 | 0.0168 | 0.0168 | TRUE | 0.3191 |
| 2410076I21R | 1934 | 1411 | 1568 | 1717 | 1454 | 1699 | 1502 | 1501 | 1692 | 1709 | 1640 | 1677 | 1620.22 | 1565 | 1675 | 1.072 | 0.1 | 0.16936567 | 0.32083515 | 0.0018 | 0.0039 | 0.0032 | 0.0032 | TRUE | 0.0596 |
| 2410088K16I | 99 | 81 | 71 | 58 | 42 | 74 | 77 | 86 | 77 | 58 | 47 | 73 | 69.64 | 80 | 59 | 0.747 | -0.421 | 0.08640587 | 0.19903289 | 0.0057 | 0.0365 | 0.0288 | 0.0288 | TRUE | 0.3504 |
| 2410089E03I | 3558 | 2610 | 2275 | 2551 | 2307 | 2767 | 2763 | 2776 | 2455 | 2539 | 2603 | 2731 | 2644.59 | 2665 | 2624 | 0.984 | -0.023 | 0.74483764 | 0.84828402 | 0.0027 | 0.0033 | 0.0031 | 0.0031 | TRUE | 0.0569 |
| 2410114N07 | 9 | 16 | 16 | 11 | 14 | 17 | 7 | 17 | 17 | 11 | 16 | 17 | 14.13 | 14 | 15 | 1.072 | 0.1 | 0.85419386 | NA | 0.0726 | 0.1701 | 0.1426 | 0.1426 | TRUE | 1.0318 |
| 2410124H12I | 34 | 26 | 25 | 33 | 37 | 31 | 26 | 28 | 27 | 33 | 42 | 31 | 31.04 | 27 | 35 | 1.295 | 0.373 | 0.27920395 | 0.4507316 | 0 | 0.0788 | 0.0531 | 0.0531 | TRUE | 0.1835 |
| 2410127L17F | 1044 | 893 | 912 | 936 | 794 | 897 | 811 | 950 | 984 | 932 | 896 | 885 | 909.6 | 915 | 904 | 0.991 | -0.014 | 0.88073936 | 0.93349722 | 0.0047 | 0.005 | 0.0049 | 0.0049 | TRUE | 0.1236 |
| 2410131K14I | 667 | 492 | 424 | 428 | 428 | 493 | 518 | 523 | 457 | 426 | 483 | 487 | 482.4 | 499 | 465 | 0.93 | -0.105 | 0.32333036 | 0.49734835 | 0.0033 | 0.0073 | 0.0061 | 0.0061 | TRUE | 0.0638 |
| 2410137F16I | 132 | 120 | 112 | 112 | 119 | 117 | 103 | 128 | 121 | 111 | 134 | 115 | 118.71 | 117 | 120 | 1.032 | 0.045 | 0.81045345 | 0.89071981 | 0 | 0.0224 | 0.017 | 0.017 | TRUE | 0.1328 |
| 2410137M14 | 138 | 72 | 52 | 84 | 72 | 52 | 107 | 77 | 56 | 84 | 81 | 51 | 76.01 | 80 | 72 | 0.893 | -0.163 | 0.56634852 | 0.72050017 | 0.0735 | 0.0336 | 0.0451 | 0.0451 | TRUE | 0.3088 |
| 2410141K09I | 372 | 267 | 245 | 316 | 229 | 230 | 289 | 284 | 264 | 315 | 258 | 227 | 272.87 | 279 | 267 | 0.955 | -0.067 | 0.6404414 | 0.77535397 | 0.011 | 0.0111 | 0.0111 | 0.0111 | TRUE | 0.2816 |
| 2500004C02I | 52 | 59 | 24 | 32 | 33 | 43 | 40 | 63 | 26 | 32 | 37 | 42 | 40.09 | 43 | 37 | 0.865 | -0.209 | 0.55807564 | 0.71302182 | 0.0786 | 0.0615 | 0.0663 | 0.0663 | TRUE | 0.2845 |
| 2510002D24I | 335 | 272 | 296 | 295 | 271 | 337 | 260 | 289 | 319 | 294 | 306 | 333 | 300.15 | 289 | 311 | 1.076 | 0.106 | 0.40769593 | 0.58045602 | 0.0043 | 0.0103 | 0.0085 | 0.0085 | TRUE | 0.0966 |
| 2510003D18I | 0 | 0 | 0 | 0 | 0 | 0 | 0 | 0 | 0 | 0 | 0 | 0 | 0 | 0 | 0 | NA | NA | NA | NA | NA | NA | NA | NA | NA | NA |
| 2510003E04I | 2033 | 1519 | 1581 | 1540 | 1414 | 1636 | 1579 | 1616 | 1706 | 1533 | 1595 | 1615 | 1607.27 | 1634 | 1581 | 0.968 | -0.046 | 0.49184075 | 0.65736783 | 6.00E-04 | 0.0039 | 0.0026 | 0.0026 | TRUE | 0.0179 |
| 2510009E07I | 141 | 75 | 72 | 135 | 105 | 76 | 110 | 80 | 78 | 134 | 118 | 75 | 99.14 | 89 | 109 | 1.22 | 0.287 | 0.24803371 | 0.41692883 | 0.0535 | 0.0263 | 0.0342 | 0.0342 | TRUE | 0.7259 |
| 2510039O18 | 3534 | 2702 | 2542 | 2772 | 2456 | 2691 | 2745 | 2874 | 2743 | 2759 | 2771 | 2656 | 2757.98 | 2787 | 2729 | 0.979 | -0.03 | 0.59885822 | 0.74449687 | 3.00E-04 | 0.0033 | 0.002 | 0.002 | TRUE | 0.0089 |
| 2510049J12F | 1 | 0 | 1 | 4 | 0 | 0 | 1 | 0 | 1 | 4 | 0 | 0 | 0.97 | 1 | 1 | 2.147 | 1.102 | 0.62274614 | NA | 2.5483 | 2.4386 | 2.4586 | 2.4586 | TRUE | 1.8978 |
| 2600006L11F | 0 | 1 | 0 | 0 | 0 | 0 | 0 | 1 | 0 | 0 | 0 | 0 | 0.18 | 0 | 0 | 0.552 | -0.857 | 0.8336548 | NA | 0 | 13.3714 | 10 | 10 | TRUE | 0.5216 |
| 2610001J05F | 1608 | 1115 | 1159 | 1303 | 1040 | 1169 | 1249 | 1186 | 1251 | 1297 | 1173 | 1154 | 1218.27 | 1229 | 1208 | 0.983 | -0.024 | 0.75116794 | 0.85220887 | 0.0016 | 0.0044 | 0.0034 | 0.0034 | TRUE | 0.0503 |
| 2610002J02F | 988 | 686 | 647 | 739 | 620 | 762 | 767 | 730 | 698 | 736 | 699 | 752 | 730.4 | 732 | 729 | 0.996 | -0.006 | 0.94068774 | 0.96674968 | 0 | 0.0057 | 0.0039 | 0.0039 | TRUE | 0.0227 |
| 2610002M06 | 965 | 737 | 745 | 687 | 656 | 747 | 749 | 784 | 804 | 684 | 740 | 737 | 749.76 | 779 | 720 | 0.925 | -0.112 | 0.18571348 | 0.3420622 | 0 | 0.0056 | 0.0038 | 0.0038 | TRUE | 0.0236 |
| 2610005L07F | 834 | 605 | 670 | 680 | 614 | 716 | 648 | 644 | 723 | 677 | 693 | 707 | 681.75 | 672 | 692 | 1.032 | 0.045 | 0.6111903 | 0.75399128 | 9.00E-04 | 0.0059 | 0.0043 | 0.0043 | TRUE | 0.0516 |
| 2610008E11I | 187 | 122 | 96 | 109 | 85 | 95 | 145 | 130 | 104 | 108 | 96 | 94 | 112.79 | 126 | 99 | 0.784 | -0.35 | 0.07868782 | 0.18691718 | 0.0104 | 0.0234 | 0.0196 | 0.0196 | TRUE | 0.2429 |
| 2610015P09I | 1249 | 1045 | 982 | 1152 | 927 | 1106 | 970 | 1112 | 1060 | 1147 | 1046 | 1092 | 1070.9 | 1047 | 1095 | 1.047 | 0.067 | 0.415292 | 0.58799094 | 0.0026 | 0.0046 | 0.0039 | 0.0039 | TRUE | 0.0519 |
| 2610016A17I | 2 | 5 | 3 | 1 | 3 | 1 | 2 | 5 | 3 | 1 | 3 | 1 | 2.58 | 3 | 2 | 0.53 | -0.915 | 0.45912759 | NA | 0 | 0.9213 | 0.6614 | 0.6614 | TRUE | 0.2641 |
| 2610017I09R | 3 | 2 | 2 | 0 | 0 | 2 | 2 | 2 | 2 | 0 | 0 | 2 | 1.43 | 2 | 1 | 0.306 | -1.709 | 0.33012839 | NA | 0 | 1.658 | 1.2355 | 1.2355 | TRUE | 0.9734 |
| 2610018G03 | 761 | 522 | 518 | 518 | 492 | 511 | 591 | 555 | 559 | 516 | 555 | 504 | 546.71 | 568 | 525 | 0.922 | -0.117 | 0.21504213 | 0.37765193 | 0 | 0.0068 | 0.0046 | 0.0046 | TRUE | 0.0283 |
| 2610019E17I | 212 | 177 | 143 | 136 | 122 | 169 | 165 | 188 | 154 | 135 | 138 | 167 | 157.84 | 169 | 147 | 0.869 | -0.203 | 0.22737268 | 0.39249249 | 0.0054 | 0.0174 | 0.0141 | 0.0141 | TRUE | 0.155 |
| 2610019F03I | 2286 | 1661 | 1534 | 1737 | 1569 | 1687 | 1775 | 1767 | 1655 | 1729 | 1770 | 1665 | 1726.95 | 1732 | 1721 | 0.993 | -0.01 | 0.87739004 | 0.93155072 | 6.00E-04 | 0.0038 | 0.0026 | 0.0026 | TRUE | 0.018 |
| 2610020C07I | 43 | 37 | 19 | 33 | 18 | 32 | 33 | 39 | 21 | 33 | 20 | 32 | 29.67 | 31 | 28 | 0.91 | -0.136 | 0.72288233 | 0.83413959 | 0.045 | 0.0823 | 0.0726 | 0.0726 | TRUE | 0.4934 |
| 2610020H08I | 395 | 277 | 351 | 327 | 267 | 311 | 307 | 295 | 379 | 325 | 301 | 307 | 318.98 | 327 | 311 | 0.955 | -0.067 | 0.60545772 | 0.75008085 | 0.007 | 0.0099 | 0.009 | 0.009 | TRUE | 0.2129 |
| 2610027K06I | 3 | 1 | 1 | 1 | 1 | 3 | 2 | 1 | 1 | 1 | 1 | 3 | 1.59 | 1 | 2 | 1.117 | 0.16 | 0.91339971 | NA | 0 | 1.4903 | 0.9426 | 0.9426 | TRUE | 0.3409 |
| 2610028E06I | 11 | 9 | 5 | 9 | 6 | 13 | 9 | 10 | 5 | 9 | 7 | 13 | 8.68 | 8 | 10 | 1.215 | 0.281 | 0.66729929 | NA | 0 | 0.2755 | 0.1933 | 0.1933 | TRUE | 0.3163 |
| 2610028H24I | 5 | 1 | 2 | 4 | 1 | 1 | 4 | 1 | 2 | 4 | 1 | 1 | 2.2 | 2 | 2 | 0.845 | -0.243 | 0.85210051 | NA | 0 | 1.0796 | 0.7724 | 0.7724 | TRUE | 0.6449 |
| 2610034B18I | 333 | 226 | 250 | 375 | 297 | 317 | 259 | 240 | 270 | 373 | 335 | 313 | 298.34 | 256 | 340 | 1.328 | 0.41 | 0.00113426 | 0.00794703 | 0.0026 | 0.0104 | 0.008 | 0.008 | TRUE | 0.0829 |
| 2610034M16 | 17 | 6 | 7 | 2 | 5 | 3 | 13 | 6 | 8 | 2 | 6 | 3 | 6.29 | 9 | 4 | 0.38 | -1.394 | 0.08983228 | NA | 0 | 0.3793 | 0.2857 | 0.2857 | TRUE | 0.6538 |
| 2610035D17I | 96 | 97 | 94 | 100 | 85 | 110 | 75 | 103 | 101 | 100 | 96 | 109 | 97.2 | 93 | 102 | 1.097 | 0.134 | 0.52524001 | 0.68634104 | 0.0092 | 0.0268 | 0.0217 | 0.0217 | TRUE | 0.3314 |
| 2610035F20I | 4 | 3 | 1 | 2 | 2 | 3 | 3 | 3 | 1 | 2 | 2 | 3 | 2.43 | 2 | 2 | 0.967 | -0.048 | 0.96801733 | NA | 0 | 0.9774 | 0.6197 | 0.6197 | TRUE | 0.2325 |
| 2610037D02I | 4 | 11 | 3 | 3 | 2 | 2 | 3 | 12 | 3 | 3 | 2 | 2 | 4.21 | 6 | 2 | 0.405 | -1.303 | 0.20120727 | NA | 0 | 0.5654 | 0.4522 | 0.4522 | TRUE | 1.5292 |
| 2610044O15 | 455 | 352 | 326 | 348 | 310 | 345 | 353 | 374 | 352 | 346 | 350 | 341 | 352.7 | 360 | 346 | 0.961 | -0.058 | 0.59575556 | 0.74231839 | 0 | 0.0091 | 0.0058 | 0.0058 | TRUE | 0.0138 |
| 2610100L16F | 0 | 0 | 0 | 0 | 0 | 0 | 0 | 0 | 0 | 0 | 0 | 0 | 0 | 0 | 0 | NA | NA | NA | NA | NA | NA | NA | NA | NA | NA |
| 2610203C20I | 8 | 9 | 4 | 10 | 6 | 8 | 6 | 10 | 4 | 10 | 7 | 8 | 7.45 | 7 | 8 | 1.231 | 0.3 | 0.66962622 | NA | 0 | 0.3204 | 0.2202 | 0.2202 | TRUE | 0.3249 |
| 2610203C22I | 0 | 0 | 0 | 0 | 0 | 0 | 0 | 0 | 0 | 0 | 0 | 0 | 0 | 0 | 0 | NA | NA | NA | NA | NA | NA | NA | NA | NA | NA |
| 2610204G22 | 19 | 10 | 12 | 19 | 10 | 13 | 15 | 11 | 13 | 19 | 11 | 13 | 13.56 | 13 | 14 | 1.119 | 0.162 | 0.75627629 | NA | 0 | 0.1772 | 0.1235 | 0.1235 | TRUE | 0.3496 |
| 2610206C17I | 7 | 4 | 3 | 3 | 3 | 5 | 5 | 4 | 3 | 3 | 3 | 5 | 4.04 | 4 | 4 | 0.867 | -0.205 | 0.82528856 | NA | 0 | 0.5892 | 0.3754 | 0.3754 | TRUE | 0.1239 |
| 2610301B20I | 1146 | 934 | 908 | 1040 | 920 | 1082 | 890 | 993 | 980 | 1035 | 1038 | 1068 | 1000.74 | 954 | 1047 | 1.099 | 0.136 | 0.08410297 | 0.19555616 | 0.0011 | 0.0048 | 0.0035 | 0.0035 | TRUE | 0.0439 |

| | | | | | | | | | | | | | | | | | | | | | | | | | |
|---|---|---|---|---|---|---|---|---|---|---|---|---|---|---|---|---|---|---|---|---|---|---|---|---|---|
| 2610301G19 | 3540 | 2826 | 2573 | 2364 | 2341 | 2623 | 2749 | 3006 | 2776 | 2353 | 2641 | 2589 | 2685.82 | 2844 | 2528 | 0.889 | -0.17 | 0.01409518 | 0.05332515 | 0.0025 | 0.0033 | 0.0031 | 0.0031 | TRUE | 0.0445 |
| 2610305D13 | 8606 | 6234 | 5666 | 6408 | 5296 | 6170 | 6684 | 6631 | 6114 | 6379 | 5974 | 6091 | 6312.03 | 6476 | 6148 | 0.949 | -0.075 | 0.19778951 | 0.35762245 | 0.0015 | 0.0028 | 0.0023 | 0.0023 | TRUE | 0.0291 |
| 2610306M01 | 76 | 67 | 40 | 58 | 34 | 44 | 59 | 71 | 43 | 58 | 38 | 43 | 52.16 | 58 | 46 | 0.806 | -0.312 | 0.29196504 | 0.46413706 | 0.0324 | 0.0479 | 0.0437 | 0.0437 | TRUE | 0.2037 |
| 2610307P16 | 23 | 9 | 15 | 28 | 20 | 27 | 18 | 10 | 16 | 28 | 23 | 27 | 20.12 | 15 | 26 | 1.753 | 0.81 | 0.06633806 | 0.16478449 | 0 | 0.1202 | 0.0871 | 0.0871 | TRUE | 0.3711 |
| 2610316D01 | 6 | 3 | 2 | 6 | 4 | 10 | 5 | 3 | 2 | 6 | 5 | 10 | 5.06 | 3 | 7 | 2.004 | 1.003 | 0.24958514 | NA | 0 | 0.4707 | 0.3291 | 0.3291 | TRUE | 0.4193 |
| 2610318N02 | 778 | 638 | 574 | 616 | 617 | 708 | 604 | 679 | 619 | 613 | 696 | 699 | 651.72 | 634 | 669 | 1.056 | 0.079 | 0.40901996 | 0.5818195 | 0.0031 | 0.0061 | 0.0051 | 0.0051 | TRUE | 0.0644 |
| 2610507B11 | 54177 | 38516 | 37911 | 42079 | 36458 | 41357 | 42077 | 40969 | 40906 | 41885 | 41128 | 40826 | 41298.52 | 41317 | 41280 | 0.999 | -0.001 | 0.97521113 | 0.98653073 | 1.00E-04 | 0.0025 | 0.0013 | 0.0013 | TRUE | 0.0032 |
| 2610507I01R | 10 | 12 | 12 | 20 | 14 | 15 | 8 | 13 | 13 | 20 | 16 | 15 | 14 | 11 | 17 | 1.531 | 0.614 | 0.23716819 | NA | 0 | 0.1717 | 0.12 | 0.12 | TRUE | 0.3511 |
| 2610524H06 | 783 | 544 | 696 | 657 | 602 | 672 | 608 | 579 | 751 | 654 | 679 | 663 | 655.7 | 646 | 665 | 1.031 | 0.045 | 0.67760992 | 0.80217909 | 0.0083 | 0.006 | 0.0068 | 0.0068 | TRUE | 0.2327 |
| 2610528A11 | 15 | 9 | 15 | 10 | 11 | 12 | 12 | 10 | 16 | 10 | 12 | 12 | 11.94 | 13 | 11 | 0.916 | -0.127 | 0.81853108 | NA | 0 | 0.201 | 0.1356 | 0.1356 | TRUE | 0.2482 |
| 2610528J11F | 22 | 15 | 31 | 24 | 35 | 40 | 17 | 16 | 33 | 24 | 39 | 39 | 28.22 | 22 | 34 | 1.558 | 0.64 | 0.1186537 | 0.25009841 | 0.0838 | 0.0864 | 0.0841 | 0.0841 | TRUE | 1.0641 |
| 2700029M09 | 5777 | 4237 | 4649 | 4944 | 4067 | 4750 | 4487 | 4507 | 5016 | 4921 | 4588 | 4689 | 4701.34 | 4670 | 4733 | 1.014 | 0.02 | 0.75636302 | 0.85520897 | 0.0024 | 0.0029 | 0.0027 | 0.0027 | TRUE | 0.0509 |
| 2700038G22 | 489 | 413 | 380 | 388 | 296 | 402 | 380 | 439 | 410 | 386 | 334 | 397 | 391.01 | 410 | 372 | 0.912 | -0.134 | 0.25067222 | 0.41953841 | 0.0042 | 0.0085 | 0.0072 | 0.0072 | TRUE | 0.0903 |
| 2700046A07 | 6 | 6 | 9 | 2 | 2 | 0 | 5 | 6 | 10 | 2 | 2 | 0 | 4.17 | 7 | 1 | 0.203 | -2.301 | 0.0375263 | NA | 0 | 0.5712 | 0.44 | 0.44 | TRUE | 0.4862 |
| 2700046G09 | 28 | 18 | 15 | 18 | 20 | 17 | 22 | 19 | 16 | 18 | 23 | 17 | 19.06 | 19 | 19 | 0.994 | -0.008 | 0.98515731 | 0.99244808 | 0 | 0.1268 | 0.0852 | 0.0852 | TRUE | 0.1218 |
| 2700049A03 | 3557 | 2588 | 2746 | 2906 | 2662 | 2990 | 2763 | 2753 | 2963 | 2893 | 3003 | 2952 | 2887.59 | 2826 | 2949 | 1.044 | 0.062 | 0.29910346 | 0.47138862 | 7.00E-04 | 0.0032 | 0.0022 | 0.0022 | TRUE | 0.0214 |
| 2700050L05 | 5443 | 3987 | 4137 | 4177 | 3764 | 4362 | 4227 | 4241 | 4464 | 4158 | 4246 | 4306 | 4273.66 | 4311 | 4237 | 0.983 | -0.025 | 0.64748829 | 0.78051363 | 4.00E-04 | 0.003 | 0.0019 | 0.0019 | TRUE | 0.0116 |
| 2700054A10 | 37 | 27 | 30 | 21 | 39 | 47 | 29 | 29 | 32 | 21 | 44 | 46 | 33.52 | 30 | 37 | 1.238 | 0.308 | 0.40701172 | 0.57994402 | 0.06 | 0.0731 | 0.0695 | 0.0695 | TRUE | 1.0871 |
| 2700060E02 | 7876 | 5718 | 5627 | 6751 | 5653 | 6526 | 6117 | 6082 | 6071 | 6720 | 6377 | 6442 | 6301.65 | 6090 | 6513 | 1.069 | 0.097 | 0.05247648 | 0.13956948 | 2.00E-04 | 0.0028 | 0.0016 | 0.0016 | TRUE | 0.0094 |
| 2700062C07 | 798 | 612 | 608 | 650 | 517 | 618 | 620 | 651 | 656 | 647 | 583 | 610 | 627.85 | 642 | 613 | 0.957 | -0.064 | 0.47730153 | 0.64486915 | 0 | 0.0062 | 0.0043 | 0.0043 | TRUE | 0.027 |
| 2700069I18R | 0 | 0 | 0 | 0 | 0 | 0 | 0 | 0 | 0 | 0 | 0 | 0 | 0 | 0 | 0 | NA | NA | NA | NA | NA | NA | NA | NA | NA | NA |
| 2700070H01 | 13 | 8 | 14 | 12 | 8 | 6 | 10 | 9 | 15 | 12 | 9 | 6 | 10.1 | 11 | 9 | 0.801 | -0.321 | 0.60249164 | NA | 0 | 0.2371 | 0.1727 | 0.1727 | TRUE | 0.2534 |
| 2700081L22 | 2 | 2 | 1 | 0 | 0 | 2 | 2 | 2 | 1 | 0 | 0 | 2 | 1.12 | 2 | 1 | 0.425 | -1.236 | 0.51965438 | NA | 0 | 2.1142 | 1.5828 | 1.5828 | TRUE | 0.9734 |
| 2700081O15 | 896 | 774 | 534 | 615 | 511 | 572 | 696 | 823 | 576 | 612 | 576 | 565 | 641.44 | 698 | 584 | 0.837 | -0.257 | 0.0334065 | 0.10031075 | 0.0142 | 0.0061 | 0.0089 | 0.0089 | TRUE | 0.1643 |
| 2700086A05 | 0 | 0 | 0 | 0 | 0 | 0 | 0 | 0 | 0 | 0 | 0 | 0 | 0 | 0 | 0 | NA | NA | NA | NA | NA | NA | NA | NA | NA | NA |
| 2700089E24 | 722 | 530 | 436 | 462 | 407 | 452 | 561 | 564 | 470 | 460 | 459 | 446 | 493.36 | 532 | 455 | 0.855 | -0.227 | 0.03212628 | 0.09738724 | 0.0034 | 0.0072 | 0.006 | 0.006 | TRUE | 0.1148 |
| 2700089I24R | 0 | 0 | 0 | 0 | 0 | 0 | 0 | 0 | 0 | 0 | 0 | 0 | 0 | 0 | 0 | NA | NA | NA | NA | NA | NA | NA | NA | NA | NA |
| 2700094K13 | 7230 | 4896 | 4888 | 5174 | 4556 | 5041 | 5615 | 5208 | 5274 | 5150 | 5140 | 4976 | 5227.21 | 5366 | 5089 | 0.948 | -0.077 | 0.16197887 | 0.31205405 | 7.00E-04 | 0.0029 | 0.002 | 0.002 | TRUE | 0.0205 |
| 2700097O09 | 1068 | 837 | 794 | 807 | 676 | 803 | 829 | 890 | 857 | 803 | 763 | 793 | 822.51 | 859 | 786 | 0.917 | -0.126 | 0.11774487 | 0.24899059 | 0 | 0.0053 | 0.0034 | 0.0034 | TRUE | 0.0122 |
| 2700099C18 | 442 | 389 | 399 | 358 | 318 | 352 | 343 | 414 | 431 | 356 | 359 | 347 | 375.02 | 396 | 354 | 0.898 | -0.156 | 0.19372037 | 0.35249463 | 0.0054 | 0.0087 | 0.0077 | 0.0077 | TRUE | 0.1666 |
| 2810001G20 | 178 | 116 | 101 | 91 | 91 | 130 | 138 | 123 | 109 | 91 | 103 | 128 | 115.36 | 123 | 107 | 0.865 | -0.209 | 0.29991351 | 0.47214993 | 0.0143 | 0.023 | 0.0205 | 0.0205 | TRUE | 0.3029 |
| 2810002D19 | 112 | 79 | 81 | 57 | 54 | 65 | 87 | 84 | 87 | 57 | 61 | 64 | 73.37 | 86 | 61 | 0.703 | -0.508 | 0.02401449 | 0.07897816 | 0 | 0.0347 | 0.0223 | 0.0223 | TRUE | 0.028 |
| 2810004N23 | 7721 | 5529 | 4844 | 5355 | 4788 | 5344 | 5997 | 5881 | 5227 | 5330 | 5401 | 5275 | 5518.57 | 5702 | 5335 | 0.936 | -0.096 | 0.13237278 | 0.27036327 | 0.0024 | 0.0029 | 0.0028 | 0.0028 | TRUE | 0.0643 |
| 2810006K23 | 720 | 507 | 485 | 648 | 537 | 556 | 559 | 539 | 523 | 645 | 606 | 549 | 570.24 | 540 | 600 | 1.108 | 0.149 | 0.1297002 | 0.26638397 | 0.0023 | 0.0066 | 0.0052 | 0.0052 | TRUE | 0.0665 |
| 2810007J24 | 0 | 0 | 0 | 0 | 0 | 0 | 0 | 0 | 0 | 0 | 0 | 0 | 0 | 0 | 0 | NA | NA | NA | NA | NA | NA | NA | NA | NA | NA |
| 2810008D09 | 467 | 478 | 527 | 437 | 312 | 455 | 363 | 508 | 569 | 435 | 352 | 449 | 445.98 | 480 | 412 | 0.862 | -0.214 | 0.1690033 | 0.32056364 | 0.0327 | 0.0077 | 0.0152 | 0.0152 | TRUE | 0.5536 |
| 2810011L19 | 298 | 227 | 246 | 327 | 311 | 310 | 231 | 241 | 265 | 325 | 351 | 306 | 286.78 | 246 | 327 | 1.332 | 0.414 | 0.00103169 | 0.00739827 | 0 | 0.0107 | 0.0079 | 0.0079 | TRUE | 0.0512 |
| 2810013P06 | 396 | 399 | 271 | 281 | 257 | 349 | 308 | 424 | 292 | 280 | 290 | 345 | 323.09 | 341 | 305 | 0.894 | -0.161 | 0.29760389 | 0.46975731 | 0.0243 | 0.0098 | 0.0142 | 0.0142 | TRUE | 0.505 |
| 2810021J22 | 896 | 693 | 651 | 642 | 612 | 698 | 696 | 737 | 702 | 639 | 690 | 689 | 692.32 | 712 | 673 | 0.945 | -0.081 | 0.34836119 | 0.52303033 | 0 | 0.0058 | 0.0039 | 0.0039 | TRUE | 0.023 |
| 2810025M15 | 4399 | 3187 | 3281 | 3334 | 2913 | 3384 | 3417 | 3390 | 3540 | 3319 | 3286 | 3341 | 3382 | 3449 | 3315 | 0.961 | -0.057 | 0.29221035 | 0.46437209 | 0 | 0.0031 | 0.0018 | 0.0018 | TRUE | 0.0064 |
| 2810029C07 | 19 | 24 | 22 | 24 | 23 | 29 | 15 | 26 | 24 | 24 | 26 | 29 | 23.75 | 22 | 26 | 1.241 | 0.311 | 0.44182522 | 0.61286955 | 0 | 0.1022 | 0.0755 | 0.0755 | TRUE | 0.5143 |
| 2810032G03 | 29 | 21 | 17 | 21 | 17 | 13 | 23 | 22 | 18 | 21 | 19 | 13 | 19.35 | 21 | 18 | 0.834 | -0.263 | 0.55022291 | 0.70639864 | 0 | 0.1249 | 0.0871 | 0.0871 | TRUE | 0.2981 |
| 2810047C21 | 53 | 26 | 29 | 20 | 30 | 21 | 41 | 28 | 31 | 20 | 34 | 21 | 29.1 | 33 | 25 | 0.734 | -0.445 | 0.24262775 | 0.41025416 | 0.0309 | 0.0839 | 0.0693 | 0.0693 | TRUE | 0.5648 |
| 2810049E08 | 1 | 1 | 2 | 1 | 0 | 1 | 1 | 1 | 2 | 1 | 0 | 1 | 1 | 1 | 1 | 0.514 | -0.959 | 0.61613737 | NA | 0 | 2.3801 | 1.5034 | 1.5034 | TRUE | 0.2008 |
| 2810055G20 | 0 | 0 | 0 | 0 | 0 | 0 | 0 | 0 | 0 | 0 | 0 | 0 | 0 | 0 | 0 | NA | NA | NA | NA | NA | NA | NA | NA | NA | NA |
| 2810403A07 | 4159 | 3298 | 3816 | 3891 | 3510 | 4316 | 3230 | 3508 | 4117 | 3873 | 3960 | 4261 | 3824.81 | 3618 | 4031 | 1.115 | 0.157 | 0.0671551 | 0.16625802 | 0.0087 | 0.003 | 0.005 | 0.005 | TRUE | 0.1766 |
| 2810403D21 | 12 | 4 | 5 | 8 | 7 | 5 | 9 | 4 | 5 | 8 | 8 | 5 | 6.63 | 6 | 7 | 1.074 | 0.103 | 0.89072683 | NA | 0 | 0.36 | 0.2527 | 0.2527 | TRUE | 0.5457 |
| 2810404M03 | 0 | 0 | 0 | 0 | 0 | 0 | 0 | 0 | 0 | 0 | 0 | 0 | 0 | 0 | 0 | NA | NA | NA | NA | NA | NA | NA | NA | NA | NA |
| 2810405F15 | 0 | 0 | 0 | 0 | 0 | 0 | 0 | 0 | 0 | 0 | 0 | 0 | 0 | 0 | 0 | NA | NA | NA | NA | NA | NA | NA | NA | NA | NA |
| 2810407C02 | 8156 | 5671 | 5654 | 6403 | 5517 | 6179 | 6334 | 6032 | 6101 | 6374 | 6224 | 6100 | 6194.02 | 6156 | 6233 | 1.012 | 0.017 | 0.73361121 | 0.84079179 | 4.00E-04 | 0.0028 | 0.0017 | 0.0017 | TRUE | 0.008 |
| 2810408A11 | 441 | 407 | 455 | 575 | 504 | 569 | 343 | 433 | 491 | 572 | 569 | 562 | 494.83 | 422 | 568 | 1.35 | 0.433 | 0.00058049 | 0.0048407 | 0.0141 | 0.0072 | 0.0093 | 0.0093 | TRUE | 0.3342 |
| 2810408I11R | 592 | 426 | 396 | 457 | 386 | 438 | 460 | 453 | 427 | 455 | 435 | 432 | 443.82 | 447 | 441 | 0.986 | -0.02 | 0.84126296 | 0.90968161 | 0 | 0.0078 | 0.005 | 0.005 | TRUE | 0.0163 |
| 2810408M09 | 936 | 667 | 585 | 600 | 635 | 685 | 727 | 709 | 631 | 597 | 716 | 676 | 676.24 | 689 | 663 | 0.961 | -0.058 | 0.56316787 | 0.71753952 | 0.0055 | 0.0059 | 0.0058 | 0.0058 | TRUE | 0.0899 |
| 2810410L24 | 97 | 63 | 117 | 137 | 86 | 132 | 75 | 67 | 126 | 136 | 97 | 130 | 105.38 | 89 | 121 | 1.362 | 0.446 | 0.07486905 | 0.18004295 | 0.0648 | 0.0249 | 0.0355 | 0.0355 | TRUE | 1.1823 |
| 2810417H13 | 2910 | 1887 | 1895 | 2230 | 1825 | 2309 | 2260 | 2007 | 2045 | 2220 | 2059 | 2279 | 2144.97 | 2104 | 2186 | 1.038 | 0.054 | 0.45503576 | 0.62476716 | 0.0027 | 0.0035 | 0.0033 | 0.0033 | TRUE | 0.0523 |
| 2810428I15R | 2613 | 1994 | 1965 | 2294 | 2011 | 2385 | 2029 | 2121 | 2120 | 2283 | 2269 | 2354 | 2196.17 | 2090 | 2302 | 1.102 | 0.14 | 0.01942027 | 0.06766257 | 0 | 0.0035 | 0.0021 | 0.0021 | TRUE | 0.0082 |
| 2810429I04R | 72 | 51 | 34 | 23 | 16 | 31 | 56 | 54 | 37 | 23 | 18 | 31 | 36.4 | 49 | 24 | 0.488 | -1.036 | 0.00310183 | 0.01696107 | 0.0275 | 0.0675 | 0.0571 | 0.0571 | TRUE | 0.3695 |
| 2810433D01 | 5 | 3 | 2 | 2 | 2 | 4 | 4 | 3 | 2 | 2 | 2 | 4 | 2.9 | 3 | 3 | 0.882 | -0.182 | 0.86785412 | NA | 0 | 0.8184 | 0.5179 | 0.5179 | TRUE | 0.189 |
| 2810442I21R | 21 | 11 | 9 | 14 | 9 | 9 | 16 | 12 | 10 | 14 | 10 | 9 | 11.78 | 13 | 11 | 0.866 | -0.208 | 0.71142494 | NA | 0 | 0.2036 | 0.1434 | 0.1434 | TRUE | 0.3434 |
| 2810442N19 | 0 | 0 | 0 | 0 | 0 | 0 | 0 | 0 | 0 | 0 | 0 | 0 | 0 | 0 | 0 | NA | NA | NA | NA | NA | NA | NA | NA | NA | NA |
| 2810454H06 | 30 | 17 | 14 | 20 | 19 | 13 | 23 | 18 | 15 | 20 | 21 | 13 | 18.44 | 19 | 18 | 0.948 | -0.077 | 0.86560814 | 0.92572508 | 0 | 0.1309 | 0.0969 | 0.0969 | TRUE | 0.3375 |
| 2810459M11 | 93 | 56 | 36 | 68 | 47 | 39 | 72 | 60 | 39 | 68 | 53 | 38 | 54.97 | 57 | 53 | 0.926 | -0.111 | 0.72290541 | 0.83413959 | 0.0679 | 0.0455 | 0.0519 | 0.0519 | TRUE | 0.2572 |
| 2810468N07 | 70 | 55 | 72 | 99 | 76 | 89 | 54 | 59 | 78 | 99 | 86 | 88 | 77.12 | 64 | 91 | 1.438 | 0.524 | 0.02480657 | 0.08094983 | 0.0061 | 0.0332 | 0.0259 | 0.0259 | TRUE | 0.3204 |
| 2810471M01 | 0 | 0 | 0 | 0 | 0 | 0 | 0 | 0 | 0 | 0 | 0 | 0 | 0 | 0 | 0 | NA | NA | NA | NA | NA | NA | NA | NA | NA | NA |
| 2810474O19 | 3399 | 2415 | 2117 | 2240 | 1904 | 2085 | 2640 | 2569 | 2284 | 2230 | 2148 | 2058 | 2321.45 | 2498 | 2145 | 0.858 | -0.22 | 0.00252199 | 0.01445935 | 0.003 | 0.0034 | 0.0034 | 0.0034 | TRUE | 0.0671 |
| 2900002K06 | 22 | 9 | 10 | 12 | 9 | 10 | 17 | 10 | 11 | 12 | 10 | 10 | 11.57 | 13 | 11 | 0.843 | -0.246 | 0.66375846 | NA | 0 | 0.2073 | 0.1446 | 0.1446 | TRUE | 0.5256 |
| 2900005J15 | 12 | 30 | 12 | 26 | 19 | 31 | 9 | 32 | 13 | 26 | 21 | 31 | 22.02 | 18 | 26 | 1.462 | 0.548 | 0.2631359 | 0.43335254 | 0.1668 | 0.1101 | 0.1258 | 0.1258 | TRUE | 2.2283 |
| 2900008C10 | 5 | 12 | 9 | 8 | 4 | 8 | 4 | 13 | 10 | 8 | 5 | 8 | 7.79 | 9 | 7 | 0.791 | -0.338 | 0.64152434 | NA | 0 | 0.3067 | 0.2469 | 0.2469 | TRUE | 0.4693 |
| 2900009J06 | 53 | 34 | 26 | 40 | 31 | 48 | 41 | 36 | 28 | 40 | 35 | 47 | 37.93 | 35 | 41 | 1.154 | 0.207 | 0.52051642 | 0.68185957 | 0 | 0.0649 | 0.0483 | 0.0483 | TRUE | 0.2094 |

| | | | | | | | | | | | | | | | | | | | | | | | | | |
|---|---|---|---|---|---|---|---|---|---|---|---|---|---|---|---|---|---|---|---|---|---|---|---|---|---|
| 2900011O08 | 652 | 384 | 341 | 392 | 338 | 404 | 506 | 408 | 368 | 390 | 381 | 399 | 408.85 | 427 | 390 | 0.909 | -0.137 | 0.28439403 | 0.45640291 | 0.0119 | 0.0082 | 0.0094 | 0.0094 | TRUE | 0.3142 |
| 2900026A02 | 1196 | 855 | 820 | 880 | 711 | 886 | 929 | 909 | 885 | 876 | 802 | 875 | 879.29 | 908 | 851 | 0.937 | -0.093 | 0.24508301 | 0.41332996 | 0 | 0.0051 | 0.0035 | 0.0035 | TRUE | 0.0291 |
| 2900041M22 | 148 | 102 | 110 | 152 | 152 | 170 | 115 | 108 | 119 | 151 | 171 | 168 | 138.79 | 114 | 163 | 1.433 | 0.519 | 0.00205399 | 0.01230457 | 0 | 0.0195 | 0.013 | 0.013 | TRUE | 0.0459 |
| 2900052N01 | 10 | 3 | 2 | 1 | 2 | 4 | 8 | 3 | 2 | 1 | 2 | 4 | 3.39 | 4 | 2 | 0.537 | -0.896 | 0.41312712 | NA | 0 | 0.7024 | 0.54 | 0.54 | TRUE | 1.0707 |
| 2900055J20 | 0 | 0 | 0 | 0 | 0 | 0 | 0 | 0 | 0 | 0 | 0 | 0 | 0 | 0 | 0 | NA | NA | NA | NA | NA | NA | NA | NA | NA | NA |
| 2900056M20 | 311 | 274 | 252 | 365 | 263 | 352 | 242 | 291 | 272 | 363 | 297 | 347 | 302.06 | 268 | 336 | 1.257 | 0.33 | 0.01247884 | 0.04871896 | 0.0068 | 0.0103 | 0.0092 | 0.0092 | TRUE | 0.1146 |
| 2900057B20 | 0 | 0 | 1 | 0 | 0 | 0 | 0 | 0 | 1 | 0 | 0 | 0 | 0.18 | 0 | 0 | 0.552 | -0.857 | 0.8336548 | NA | 0 | 13.1817 | 10 | 10 | TRUE | 0.5365 |
| 2900060B14 | 2 | 4 | 1 | 0 | 1 | 2 | 2 | 4 | 1 | 0 | 1 | 2 | 1.66 | 2 | 1 | 0.454 | -1.139 | 0.46744741 | NA | 0 | 1.426 | 1.0568 | 1.0568 | TRUE | 0.5286 |
| 2900076A07 | 90 | 63 | 54 | 69 | 52 | 62 | 70 | 67 | 58 | 69 | 59 | 61 | 63.95 | 65 | 63 | 0.964 | -0.053 | 0.82839995 | 0.90163878 | 0 | 0.0395 | 0.0267 | 0.0267 | TRUE | 0.0701 |
| 2900079G21 | 0 | 0 | 0 | 0 | 0 | 0 | 0 | 0 | 0 | 0 | 0 | 0 | 0 | 0 | 0 | NA | NA | NA | NA | NA | NA | NA | NA | NA | NA |
| 2900092C05 | 0 | 0 | 0 | 1 | 0 | 0 | 0 | 0 | 0 | 1 | 0 | 0 | 0.17 | 0 | 0 | 2.094 | 1.067 | 0.79379329 | NA | 0 | 14.2885 | 10 | 10 | TRUE | 0.3906 |
| 2900092D14 | 2 | 5 | 2 | 3 | 3 | 1 | 2 | 5 | 2 | 3 | 3 | 1 | 2.73 | 3 | 2 | 0.823 | -0.282 | 0.81066839 | NA | 0 | 0.8702 | 0.6196 | 0.6196 | TRUE | 0.5572 |
| 2900093L17 | 0 | 0 | 1 | 0 | 0 | 0 | 0 | 0 | 1 | 0 | 0 | 0 | 0.18 | 0 | 0 | 0.552 | -0.857 | 0.8336548 | NA | 0 | 13.1817 | 10 | 10 | TRUE | 0.5365 |
| 2900097C17 | 2930 | 2184 | 2271 | 2605 | 2129 | 2592 | 2276 | 2323 | 2450 | 2593 | 2402 | 2559 | 2433.76 | 2350 | 2518 | 1.072 | 0.101 | 0.11353345 | 0.24248857 | 0.0011 | 0.0034 | 0.0025 | 0.0025 | TRUE | 0.0195 |
| 3000002C10 | 41 | 26 | 19 | 24 | 18 | 26 | 32 | 28 | 21 | 24 | 20 | 26 | 24.98 | 27 | 23 | 0.869 | -0.203 | 0.60269399 | 0.74763976 | 0 | 0.0973 | 0.0691 | 0.0691 | TRUE | 0.2426 |
| 3010001F23 | 1 | 2 | 0 | 3 | 1 | 1 | 1 | 2 | 0 | 3 | 1 | 1 | 1.33 | 1 | 2 | 1.777 | 0.829 | 0.62394313 | NA | 0 | 1.7788 | 1.2562 | 1.2562 | TRUE | 0.4406 |
| 3010026O09 | 1457 | 1005 | 966 | 918 | 821 | 935 | 1132 | 1069 | 1042 | 914 | 926 | 923 | 1000.97 | 1081 | 921 | 0.851 | -0.233 | 0.0020963 | 0.01251564 | 0 | 0.0048 | 0.0031 | 0.0031 | TRUE | 0.0211 |
| 3100003L05 | 1 | 0 | 1 | 1 | 0 | 0 | 1 | 0 | 1 | 1 | 0 | 0 | 0.48 | 1 | 0 | 0.596 | -0.748 | 0.78941713 | NA | 0 | 4.9902 | 3.8695 | 3.8695 | TRUE | 0.3906 |
| 3110001D03 | 1530 | 1091 | 1027 | 1028 | 864 | 1061 | 1188 | 1160 | 1108 | 1023 | 975 | 1047 | 1083.7 | 1152 | 1015 | 0.881 | -0.183 | 0.01435777 | 0.05399997 | 0 | 0.0046 | 0.0031 | 0.0031 | TRUE | 0.0141 |
| 3110001I22 | 278 | 197 | 139 | 177 | 152 | 175 | 216 | 210 | 150 | 176 | 171 | 173 | 182.64 | 192 | 173 | 0.901 | -0.15 | 0.37465774 | 0.54920038 | 0.0141 | 0.0154 | 0.015 | 0.015 | TRUE | 0.379 |
| 3110002H16 | 1969 | 1474 | 1493 | 1550 | 1298 | 1559 | 1529 | 1568 | 1611 | 1543 | 1464 | 1539 | 1542.36 | 1569 | 1515 | 0.966 | -0.049 | 0.45741345 | 0.62724844 | 0 | 0.004 | 0.0025 | 0.0025 | TRUE | 0.0103 |
| 3110007F17 | 11 | 4 | 1 | 6 | 2 | 2 | 9 | 4 | 1 | 6 | 2 | 2 | 4.01 | 5 | 3 | 0.72 | -0.473 | 0.64833797 | NA | 0.3526 | 0.593 | 0.5215 | 0.5215 | TRUE | 0.227 |
| 3110009E18 | 82 | 49 | 52 | 57 | 45 | 62 | 64 | 52 | 56 | 57 | 51 | 61 | 56.77 | 57 | 56 | 0.979 | -0.031 | 0.90489322 | 0.94786653 | 0 | 0.0442 | 0.0302 | 0.0302 | TRUE | 0.0928 |
| 3110009F21 | 0 | 0 | 0 | 0 | 0 | 0 | 0 | 0 | 0 | 0 | 0 | 0 | 0 | 0 | 0 | NA | NA | NA | NA | NA | NA | NA | NA | NA | NA |
| 3110015C05 | 0 | 0 | 0 | 0 | 0 | 0 | 0 | 0 | 0 | 0 | 0 | 0 | 0 | 0 | 0 | NA | NA | NA | NA | NA | NA | NA | NA | NA | NA |
| 3110021A11 | 14 | 20 | 11 | 9 | 15 | 12 | 11 | 21 | 12 | 9 | 17 | 12 | 13.62 | 15 | 13 | 0.859 | -0.218 | 0.68965399 | NA | 0.0523 | 0.1764 | 0.1411 | 0.1411 | TRUE | 0.6645 |
| 3110021N24 | 1 | 1 | 1 | 0 | 3 | 2 | 1 | 1 | 1 | 0 | 3 | 2 | 1.38 | 1 | 2 | 1.831 | 0.873 | 0.60397437 | NA | 0 | 1.7203 | 1.2469 | 1.2469 | TRUE | 0.1952 |
| 3110035E14 | 1 | 0 | 1 | 2 | 1 | 1 | 1 | 0 | 1 | 2 | 1 | 1 | 0.99 | 1 | 1 | 2.192 | 1.132 | 0.55372876 | NA | 0 | 2.3877 | 1.4877 | 1.4877 | TRUE | 0.1946 |
| 3110039I08 | 0 | 0 | 0 | 0 | 0 | 0 | 0 | 0 | 0 | 0 | 0 | 0 | 0 | 0 | 0 | NA | NA | NA | NA | NA | NA | NA | NA | NA | NA |
| 3110039M20 | 0 | 0 | 0 | 0 | 0 | 0 | 0 | 0 | 0 | 0 | 0 | 0 | 0 | 0 | 0 | NA | NA | NA | NA | NA | NA | NA | NA | NA | NA |
| 3110040N11 | 811 | 588 | 636 | 790 | 628 | 763 | 630 | 625 | 686 | 786 | 708 | 753 | 698.26 | 647 | 749 | 1.159 | 0.213 | 0.01679137 | 0.06083625 | 0.0012 | 0.0058 | 0.0043 | 0.0043 | TRUE | 0.0318 |
| 3110043O21 | 1535 | 1188 | 1181 | 1338 | 1102 | 1301 | 1192 | 1264 | 1274 | 1332 | 1243 | 1284 | 1264.9 | 1243 | 1286 | 1.036 | 0.051 | 0.47814827 | 0.645525 | 5.00E-04 | 0.0043 | 0.0029 | 0.0029 | TRUE | 0.0164 |
| 3110045C21 | 9 | 7 | 11 | 11 | 9 | 8 | 7 | 7 | 12 | 11 | 10 | 8 | 9.22 | 9 | 10 | 1.111 | 0.152 | 0.80883987 | NA | 0 | 0.2596 | 0.1754 | 0.1754 | TRUE | 0.2796 |
| 3110047P20 | 7 | 2 | 4 | 0 | 0 | 0 | 5 | 2 | 4 | 0 | 0 | 0 | 1.98 | 4 | 0 | 0.048 | -4.389 | 0.01059143 | NA | 0 | 1.1994 | 1.0037 | 1.0037 | TRUE | 0.1895 |
| 3110052M02 | 102 | 107 | 103 | 107 | 79 | 95 | 79 | 114 | 111 | 107 | 89 | 94 | 98.93 | 101 | 97 | 0.959 | -0.061 | 0.77796113 | 0.86890666 | 0.0158 | 0.0264 | 0.0232 | 0.0232 | TRUE | 0.4042 |
| 3110056K07 | 81 | 92 | 77 | 93 | 70 | 93 | 63 | 98 | 83 | 93 | 79 | 92 | 84.53 | 81 | 88 | 1.089 | 0.123 | 0.59443121 | 0.7409266 | 0.0176 | 0.0305 | 0.0267 | 0.0267 | TRUE | 0.3447 |
| 3110057O12 | 338 | 233 | 275 | 324 | 268 | 325 | 263 | 248 | 297 | 323 | 302 | 321 | 292.12 | 269 | 315 | 1.173 | 0.231 | 0.06390545 | 0.16059394 | 0 | 0.0105 | 0.0077 | 0.0077 | TRUE | 0.0858 |
| 3110062M04 | 305 | 237 | 188 | 212 | 174 | 199 | 237 | 252 | 203 | 211 | 196 | 196 | 215.93 | 231 | 201 | 0.872 | -0.197 | 0.16913272 | 0.32065446 | 0 | 0.0134 | 0.0102 | 0.0102 | TRUE | 0.1155 |
| 3110070M22 | 2 | 2 | 4 | 0 | 1 | 4 | 2 | 2 | 4 | 0 | 1 | 4 | 2.18 | 3 | 2 | 0.648 | -0.626 | 0.65133397 | NA | 0.3066 | 1.0902 | 0.8951 | 0.8951 | TRUE | 1.0128 |
| 3110079O15 | 6 | 5 | 9 | 13 | 4 | 10 | 5 | 5 | 10 | 13 | 5 | 10 | 7.84 | 7 | 9 | 1.415 | 0.5 | 0.48594041 | NA | 6.00E-04 | 0.3049 | 0.2409 | 0.2409 | TRUE | 0.2969 |
| 3110082I17 | 1955 | 1475 | 1509 | 1398 | 1160 | 1414 | 1518 | 1569 | 1628 | 1392 | 1309 | 1396 | 1468.58 | 1572 | 1366 | 0.87 | -0.202 | 0.00354013 | 0.01874677 | 6.00E-04 | 0.004 | 0.0028 | 0.0028 | TRUE | 0.0156 |
| 3110082J24 | 5 | 4 | 9 | 6 | 6 | 7 | 4 | 4 | 10 | 6 | 7 | 7 | 6.25 | 6 | 7 | 1.117 | 0.16 | 0.83650947 | NA | 0 | 0.3816 | 0.2694 | 0.2694 | TRUE | 0.662 |
| 3110099E03 | 0 | 0 | 0 | 0 | 0 | 2 | 0 | 0 | 0 | 0 | 0 | 2 | 0.33 | 0 | 1 | 3.743 | 1.904 | 0.62640105 | NA | 0 | 7.2051 | 9.2465 | 9.2465 | TRUE | 0.9739 |
| 3200001D21 | 0 | 0 | 0 | 0 | 0 | 0 | 0 | 0 | 0 | 0 | 0 | 0 | 0 | 0 | 0 | NA | NA | NA | NA | NA | NA | NA | NA | NA | NA |
| 3230401D17 | 783 | 516 | 480 | 530 | 468 | 587 | 608 | 549 | 518 | 528 | 528 | 579 | 551.65 | 558 | 545 | 0.974 | -0.037 | 0.71162191 | 0.82672616 | 0.0033 | 0.0067 | 0.0056 | 0.0056 | TRUE | 0.0753 |
| 3300002I08 | 27 | 19 | 18 | 17 | 10 | 16 | 21 | 20 | 19 | 17 | 11 | 16 | 17.43 | 20 | 15 | 0.728 | -0.458 | 0.3164173 | 0.49040907 | 0 | 0.1384 | 0.0915 | 0.0915 | TRUE | 0.1634 |
| 3300005D01 | 0 | 0 | 1 | 0 | 0 | 1 | 0 | 0 | 1 | 0 | 0 | 1 | 0.34 | 0 | 0 | 1.075 | 0.105 | 0.97570524 | NA | 0 | 6.885 | 6.5379 | 6.5379 | TRUE | 0.5365 |
| 3425401B19 | 18 | 14 | 9 | 10 | 15 | 10 | 14 | 15 | 10 | 10 | 17 | 10 | 12.55 | 13 | 12 | 0.943 | -0.085 | 0.87748749 | NA | 0 | 0.1912 | 0.1372 | 0.1372 | TRUE | 0.4027 |
| 3632451O06 | 1693 | 1406 | 1864 | 2077 | 1756 | 2134 | 1315 | 1496 | 2011 | 2067 | 1981 | 2107 | 1829.44 | 1607 | 2052 | 1.278 | 0.354 | 0.00321678 | 0.0174223 | 0.0249 | 0.0037 | 0.0098 | 0.0098 | TRUE | 0.5817 |
| 3632454L22 | 7 | 9 | 10 | 6 | 5 | 7 | 5 | 10 | 11 | 6 | 6 | 7 | 7.39 | 9 | 6 | 0.729 | -0.456 | 0.51847185 | NA | 0 | 0.3233 | 0.2191 | 0.2191 | TRUE | 0.4244 |
| 3830403N18 | 108 | 61 | 49 | 26 | 21 | 20 | 84 | 65 | 53 | 26 | 24 | 20 | 45.16 | 67 | 23 | 0.341 | -1.551 | 1.66E-06 | 4.39E-05 | 0 | 0.0549 | 0.0459 | 0.0459 | TRUE | 0.1786 |
| 3830406C13 | 728 | 510 | 533 | 560 | 466 | 536 | 565 | 542 | 575 | 557 | 526 | 529 | 549.2 | 561 | 537 | 0.958 | -0.062 | 0.50257956 | 0.66680359 | 0 | 0.0067 | 0.0043 | 0.0043 | TRUE | 0.0125 |
| 3830408C21 | 51 | 52 | 60 | 64 | 58 | 60 | 40 | 55 | 65 | 64 | 65 | 59 | 58 | 53 | 63 | 1.19 | 0.251 | 0.35353151 | 0.52784142 | 0.0151 | 0.0433 | 0.0354 | 0.0354 | TRUE | 0.4887 |
| 3830417A13 | 77 | 57 | 41 | 10 | 15 | 11 | 60 | 61 | 44 | 10 | 17 | 11 | 33.73 | 55 | 13 | 0.227 | -2.142 | 2.91E-08 | 1.29E-06 | 0 | 0.0727 | 0.0571 | 0.0571 | TRUE | 0.3309 |
| 3930402G23 | 1 | 2 | 0 | 3 | 1 | 1 | 1 | 2 | 0 | 3 | 1 | 1 | 1.33 | 1 | 2 | 1.777 | 0.829 | 0.62394313 | NA | 0 | 1.7788 | 1.2562 | 1.2562 | TRUE | 0.4406 |
| 4430402I18 | 16 | 8 | 20 | 16 | 18 | 26 | 12 | 9 | 22 | 16 | 20 | 26 | 17.4 | 14 | 21 | 1.465 | 0.551 | 0.26754833 | 0.43808778 | 0.0603 | 0.1386 | 0.1184 | 0.1184 | TRUE | 0.6568 |
| 4632415K11 | 998 | 685 | 851 | 872 | 816 | 908 | 775 | 729 | 918 | 868 | 921 | 896 | 851.13 | 807 | 895 | 1.109 | 0.15 | 0.12214803 | 0.25535496 | 0.0062 | 0.0052 | 0.0056 | 0.0056 | TRUE | 0.1669 |
| 4632415L05 | 372 | 299 | 290 | 351 | 291 | 352 | 289 | 318 | 313 | 349 | 328 | 347 | 324.17 | 307 | 341 | 1.117 | 0.16 | 0.16531092 | 0.31621908 | 0 | 0.0097 | 0.0065 | 0.0065 | TRUE | 0.0316 |
| 4632427E13 | 15 | 7 | 5 | 11 | 7 | 11 | 12 | 7 | 5 | 11 | 8 | 11 | 9.03 | 8 | 10 | 1.196 | 0.258 | 0.68800792 | NA | 0 | 0.2648 | 0.1872 | 0.1872 | TRUE | 0.5342 |
| 4632428C04 | 11 | 9 | 6 | 9 | 7 | 12 | 9 | 10 | 6 | 9 | 8 | 12 | 8.88 | 8 | 10 | 1.168 | 0.224 | 0.72356506 | NA | 0 | 0.2693 | 0.1771 | 0.1771 | TRUE | 0.1496 |
| 4632428N05 | 319 | 252 | 246 | 198 | 195 | 234 | 248 | 268 | 265 | 197 | 220 | 231 | 238.22 | 260 | 216 | 0.831 | -0.268 | 0.04621676 | 0.1272051 | 0 | 0.0124 | 0.0087 | 0.0087 | TRUE | 0.0661 |
| 4632434I11 | 3138 | 2415 | 2486 | 2768 | 2422 | 2735 | 2437 | 2569 | 2682 | 2755 | 2732 | 2700 | 2645.95 | 2563 | 2729 | 1.066 | 0.092 | 0.1344449 | 0.27321014 | 9.00E-04 | 0.0033 | 0.0023 | 0.0023 | TRUE | 0.023 |
| 4732415M23 | 23 | 14 | 16 | 26 | 17 | 27 | 18 | 15 | 17 | 26 | 19 | 27 | 20.29 | 17 | 24 | 1.434 | 0.52 | 0.22186351 | 0.38591884 | 0 | 0.1192 | 0.08 | 0.08 | TRUE | 0.1623 |
| 4732416N19 | 3 | 5 | 2 | 2 | 0 | 0 | 2 | 5 | 2 | 2 | 0 | 0 | 1.97 | 3 | 1 | 0.209 | -2.257 | 0.1626319 | NA | 0 | 1.2078 | 0.9732 | 0.9732 | TRUE | 0.9734 |
| 4732456N10 | 1 | 0 | 1 | 0 | 0 | 0 | 1 | 0 | 1 | 0 | 0 | 0 | 0.31 | 1 | 0 | 0.307 | -1.706 | 0.62894649 | NA | 0 | 7.6657 | 7.2 | 7.2 | TRUE | 0.2067 |
| 4732471J01 | 38 | 20 | 29 | 24 | 24 | 30 | 30 | 21 | 31 | 24 | 27 | 30 | 27.11 | 27 | 27 | 0.979 | -0.03 | 0.93470557 | 0.96390669 | 0 | 0.0899 | 0.0627 | 0.0627 | TRUE | 0.2235 |
| 4732490B19 | 3 | 0 | 0 | 1 | 0 | 1 | 2 | 0 | 0 | 1 | 0 | 1 | 0.72 | 1 | 1 | 0.815 | -0.296 | 0.90325409 | NA | 0 | 3.2999 | 2.9046 | 2.9046 | TRUE | 1.439 |
| 4732491K20 | 30 | 20 | 6 | 13 | 16 | 22 | 23 | 21 | 6 | 13 | 18 | 22 | 17.29 | 17 | 18 | 1.021 | 0.029 | 0.95543891 | 0.97553943 | 0.1537 | 0.1395 | 0.1432 | 0.1432 | TRUE | 1.334 |
| 4831440E17 | 71 | 77 | 84 | 77 | 70 | 88 | 55 | 82 | 91 | 77 | 79 | 87 | 78.36 | 76 | 81 | 1.075 | 0.104 | 0.67021566 | 0.79708707 | 0.0237 | 0.0327 | 0.0301 | 0.0301 | TRUE | 0.602 |

| | | | | | | | | | | | | | | | | | | | | | | | | | |
|---|---|---|---|---|---|---|---|---|---|---|---|---|---|---|---|---|---|---|---|---|---|---|---|---|---|
| 4833403I15R | 38 | 12 | 20 | 12 | 11 | 13 | 30 | 13 | 22 | 12 | 12 | 13 | 16.84 | 22 | 12 | 0.576 | -0.796 | 0.11065674 | NA | 0 | 0.1432 | 0.1151 | 0.1151 | TRUE | 0.1353 |
| 4833412C05I | 1 | 1 | 0 | 2 | 0 | 1 | 1 | 1 | 0 | 2 | 0 | 1 | 0.8 | 1 | 1 | 1.618 | 0.694 | 0.74938576 | NA | 0 | 2.9538 | 2.1156 | 2.1156 | TRUE | 0.1303 |
| 4833417C18I | 118 | 119 | 96 | 130 | 107 | 128 | 92 | 127 | 104 | 129 | 121 | 126 | 116.38 | 108 | 125 | 1.176 | 0.234 | 0.22256541 | 0.38687503 | 0.0055 | 0.0228 | 0.0179 | 0.0179 | TRUE | 0.2428 |
| 4833418N02 | 8 | 4 | 5 | 7 | 3 | 3 | 6 | 4 | 5 | 7 | 3 | 3 | 4.86 | 5 | 4 | 0.837 | -0.257 | 0.76657266 | NA | 0 | 0.4898 | 0.3311 | 0.3311 | TRUE | 0.468 |
| 4833419F23I | 4 | 3 | 3 | 7 | 1 | 4 | 3 | 3 | 3 | 7 | 1 | 4 | 3.6 | 3 | 4 | 1.278 | 0.353 | 0.73055698 | NA | 0 | 0.6614 | 0.4797 | 0.4797 | TRUE | 0.1624 |
| 4833420G17 | 1447 | 1122 | 1091 | 1114 | 1016 | 1237 | 1124 | 1193 | 1177 | 1109 | 1146 | 1221 | 1161.77 | 1165 | 1159 | 0.996 | -0.006 | 0.93283909 | 0.96267647 | 9.00E-04 | 0.0045 | 0.0032 | 0.0032 | TRUE | 0.027 |
| 4833422C13I | 0 | 0 | 1 | 0 | 0 | 0 | 0 | 0 | 1 | 0 | 0 | 0 | 0.18 | 0 | 0 | 0.552 | -0.857 | 0.8336548 | NA | 0 | 13.1817 | 10 | 10 | TRUE | 0.5365 |
| 4833423E24I | 4 | 1 | 4 | 0 | 1 | 1 | 3 | 1 | 4 | 0 | 1 | 1 | 1.77 | 3 | 1 | 0.245 | -2.028 | 0.20852887 | NA | 0 | 1.3438 | 0.9591 | 0.9591 | TRUE | 0.2157 |
| 4833424O15 | 0 | 1 | 0 | 1 | 0 | 0 | 0 | 1 | 0 | 1 | 0 | 0 | 0.34 | 0 | 0 | 1.075 | 0.105 | 0.97573409 | NA | 0 | 6.9086 | 6.5582 | 6.5582 | TRUE | 0.5216 |
| 4833427F10I | 0 | 0 | 0 | 0 | 0 | 0 | 0 | 0 | 0 | 0 | 0 | 0 | 0 | 0 | 0 | NA | NA | NA | NA | NA | NA | NA | NA | NA | NA |
| 4833427G06 | 5 | 7 | 3 | 4 | 0 | 1 | 4 | 7 | 3 | 4 | 0 | 1 | 3.26 | 5 | 2 | 0.35 | -1.516 | 0.20264222 | NA | 0.1611 | 0.7304 | 0.6141 | 0.6141 | TRUE | 1.1399 |
| 4833428L15F | 2 | 0 | 5 | 0 | 1 | 2 | 2 | 0 | 5 | 0 | 1 | 2 | 1.68 | 2 | 1 | 0.451 | -1.15 | 0.49642791 | NA | 1.1024 | 1.4173 | 1.3495 | 1.3495 | TRUE | 0.3992 |
| 4833439L19F | 9161 | 6290 | 6045 | 6645 | 5814 | 6810 | 7115 | 6691 | 6523 | 6614 | 6559 | 6723 | 6703.97 | 6776 | 6632 | 0.978 | -0.031 | 0.56290474 | 0.71739612 | 9.00E-04 | 0.0028 | 0.002 | 0.002 | TRUE | 0.0237 |
| 4921501E09I | 0 | 0 | 0 | 0 | 0 | 0 | 0 | 0 | 0 | 0 | 0 | 0 | 0 | 0 | 0 | NA | NA | NA | NA | NA | NA | NA | NA | NA | NA |
| 4921504E06I | 1 | 1 | 1 | 0 | 0 | 0 | 1 | 1 | 1 | 0 | 0 | 0 | 0.49 | 1 | 0 | 0.196 | -2.347 | 0.38405676 | NA | 0 | 4.8735 | 3.7415 | 3.7415 | TRUE | 0.0718 |
| 4921506M07 | 25 | 12 | 10 | 7 | 9 | 18 | 19 | 13 | 11 | 7 | 10 | 18 | 12.98 | 14 | 12 | 0.805 | -0.313 | 0.58251714 | NA | 0.0822 | 0.1851 | 0.1561 | 0.1561 | TRUE | 0.8169 |
| 4921507L20F | 0 | 0 | 0 | 0 | 0 | 0 | 0 | 0 | 0 | 0 | 0 | 0 | 0 | 0 | 0 | NA | NA | NA | NA | NA | NA | NA | NA | NA | NA |
| 4921507P07I | 67 | 45 | 34 | 54 | 58 | 52 | 52 | 48 | 37 | 54 | 65 | 51 | 51.18 | 46 | 57 | 1.239 | 0.309 | 0.26927342 | 0.43990456 | 0 | 0.0487 | 0.0366 | 0.0366 | TRUE | 0.2147 |
| 4921508A21 | 0 | 0 | 0 | 0 | 0 | 0 | 0 | 0 | 0 | 0 | 0 | 0 | 0 | 0 | 0 | NA | NA | NA | NA | NA | NA | NA | NA | NA | NA |
| 4921508D12 | 0 | 0 | 0 | 0 | 0 | 0 | 0 | 0 | 0 | 0 | 0 | 0 | 0 | 0 | 0 | NA | NA | NA | NA | NA | NA | NA | NA | NA | NA |
| 4921509C19I | 14 | 7 | 9 | 8 | 7 | 8 | 11 | 7 | 10 | 8 | 8 | 8 | 8.63 | 9 | 8 | 0.842 | -0.247 | 0.6984085 | NA | 0 | 0.277 | 0.177 | 0.177 | TRUE | 0.0985 |
| 4921509O07 | 0 | 1 | 0 | 1 | 0 | 0 | 0 | 1 | 0 | 1 | 0 | 0 | 0.34 | 0 | 0 | 1.075 | 0.105 | 0.97573409 | NA | 0 | 6.9086 | 6.5582 | 6.5582 | TRUE | 0.5216 |
| 4921511C10I | 0 | 0 | 0 | 0 | 0 | 0 | 0 | 0 | 0 | 0 | 0 | 0 | 0 | 0 | 0 | NA | NA | NA | NA | NA | NA | NA | NA | NA | NA |
| 4921511C20I | 0 | 0 | 0 | 0 | 0 | 0 | 0 | 0 | 0 | 0 | 0 | 0 | 0 | 0 | 0 | NA | NA | NA | NA | NA | NA | NA | NA | NA | NA |
| 4921511H03 | 0 | 0 | 0 | 0 | 0 | 0 | 0 | 0 | 0 | 0 | 0 | 0 | 0 | 0 | 0 | NA | NA | NA | NA | NA | NA | NA | NA | NA | NA |
| 4921511I17R | 1 | 0 | 0 | 0 | 0 | 0 | 1 | 0 | 0 | 0 | 0 | 0 | 0.13 | 0 | 0 | 0.552 | -0.857 | 0.8336548 | NA | 0 | 18.3121 | 10 | 10 | TRUE | 0.2566 |
| 4921511M17 | 0 | 0 | 0 | 0 | 0 | 0 | 0 | 0 | 0 | 0 | 0 | 0 | 0 | 0 | 0 | NA | NA | NA | NA | NA | NA | NA | NA | NA | NA |
| 4921513I03R | 2 | 0 | 0 | 0 | 0 | 0 | 2 | 0 | 0 | 0 | 0 | 0 | 0.26 | 1 | 0 | 0.356 | -1.491 | 0.71362796 | NA | 0 | 9.1573 | 10 | 10 | TRUE | 0.968 |
| 4921515E04I | 0 | 0 | 0 | 0 | 0 | 0 | 0 | 0 | 0 | 0 | 0 | 0 | 0 | 0 | 0 | NA | NA | NA | NA | NA | NA | NA | NA | NA | NA |
| 4921517D22 | 0 | 0 | 2 | 2 | 1 | 0 | 0 | 0 | 2 | 2 | 1 | 0 | 0.88 | 1 | 1 | 1.503 | 0.588 | 0.79658779 | NA | 0 | 2.6973 | 2.546 | 2.546 | TRUE | 0.6542 |
| 4921524J17F | 2050 | 1668 | 1672 | 1893 | 1767 | 2043 | 1592 | 1774 | 1804 | 1884 | 1993 | 2017 | 1844.14 | 1723 | 1965 | 1.141 | 0.191 | 0.008508 | 0.03670536 | 0.0024 | 0.0037 | 0.0032 | 0.0032 | TRUE | 0.0556 |
| 4921524L21F | 0 | 0 | 0 | 0 | 0 | 0 | 0 | 0 | 0 | 0 | 0 | 0 | 0 | 0 | 0 | NA | NA | NA | NA | NA | NA | NA | NA | NA | NA |
| 4921525B02 | 0 | 1 | 0 | 0 | 0 | 0 | 0 | 1 | 0 | 0 | 0 | 0 | 0.18 | 0 | 0 | 0.552 | -0.857 | 0.8336548 | NA | 0 | 13.3714 | 10 | 10 | TRUE | 0.5216 |
| 4921525O09 | 238 | 199 | 207 | 227 | 171 | 152 | 185 | 212 | 223 | 226 | 193 | 150 | 198.13 | 207 | 190 | 0.92 | -0.121 | 0.4817569 | 0.64849796 | 0.0203 | 0.0144 | 0.0161 | 0.0161 | TRUE | 0.2934 |
| 4921530D09 | 0 | 0 | 0 | 1 | 0 | 0 | 0 | 0 | 0 | 1 | 0 | 0 | 0.17 | 0 | 0 | 2.094 | 1.067 | 0.79379329 | NA | 0 | 14.2885 | 10 | 10 | TRUE | 0.3906 |
| 4921530L18F | 16 | 13 | 8 | 6 | 14 | 8 | 12 | 14 | 9 | 6 | 16 | 8 | 10.76 | 12 | 10 | 0.84 | -0.252 | 0.68102452 | NA | 0.0449 | 0.2227 | 0.176 | 0.176 | TRUE | 0.85 |
| 4921530L21F | 0 | 0 | 0 | 0 | 0 | 0 | 0 | 0 | 0 | 0 | 0 | 0 | 0 | 0 | 0 | NA | NA | NA | NA | NA | NA | NA | NA | NA | NA |
| 4921531C22I | 22 | 22 | 12 | 15 | 15 | 17 | 17 | 23 | 13 | 15 | 17 | 17 | 17.01 | 18 | 16 | 0.91 | -0.135 | 0.77318586 | NA | 0 | 0.1417 | 0.0999 | 0.0999 | TRUE | 0.2854 |
| 4921531P14I | 0 | 0 | 0 | 0 | 0 | 0 | 0 | 0 | 0 | 0 | 0 | 0 | 0 | 0 | 0 | NA | NA | NA | NA | NA | NA | NA | NA | NA | NA |
| 4921533I20R | 2 | 0 | 0 | 0 | 0 | 0 | 2 | 0 | 0 | 0 | 0 | 0 | 0.26 | 1 | 0 | 0.356 | -1.491 | 0.71362796 | NA | 0 | 9.1573 | 10 | 10 | TRUE | 0.968 |
| 4921536K21I | 58 | 46 | 50 | 47 | 44 | 78 | 45 | 49 | 54 | 47 | 50 | 77 | 53.56 | 49 | 58 | 1.178 | 0.237 | 0.41118447 | 0.58402659 | 0.0272 | 0.0467 | 0.041 | 0.041 | TRUE | 0.7335 |
| 4921539E11I | 2 | 2 | 1 | 1 | 1 | 2 | 2 | 2 | 1 | 1 | 1 | 2 | 1.48 | 2 | 1 | 0.864 | -0.21 | 0.88889434 | NA | 0 | 1.6079 | 0.9505 | 0.9505 | TRUE | 0.0996 |
| 4922501C03I | 1489 | 1218 | 979 | 976 | 867 | 1025 | 1156 | 1296 | 1056 | 972 | 978 | 1012 | 1078.29 | 1169 | 987 | 0.844 | -0.244 | 0.00538275 | 0.02594792 | 0.0046 | 0.0046 | 0.0046 | 0.0046 | TRUE | 0.1038 |
| 4922501L14F | 4 | 4 | 10 | 2 | 1 | 3 | 3 | 4 | 11 | 2 | 1 | 3 | 4.04 | 6 | 2 | 0.343 | -1.543 | 0.14050049 | NA | 0 | 0.5893 | 0.4556 | 0.4556 | TRUE | 1.0469 |
| 4922502D21 | 1 | 0 | 0 | 0 | 0 | 0 | 1 | 0 | 0 | 0 | 0 | 0 | 0.13 | 0 | 0 | 0.552 | -0.857 | 0.8336548 | NA | 0 | 18.3121 | 10 | 10 | TRUE | 0.2566 |
| 4922502H24 | 0 | 0 | 0 | 0 | 0 | 0 | 0 | 0 | 0 | 0 | 0 | 0 | 0 | 0 | 0 | NA | NA | NA | NA | NA | NA | NA | NA | NA | NA |
| 4922502N22 | 2 | 1 | 1 | 0 | 0 | 0 | 2 | 1 | 1 | 0 | 0 | 0 | 0.62 | 1 | 0 | 0.154 | -2.702 | 0.27412557 | NA | 0 | 3.8499 | 3.0088 | 3.0088 | TRUE | 0.0718 |
| 4922505E12I | 3 | 1 | 1 | 0 | 1 | 0 | 2 | 1 | 1 | 0 | 1 | 0 | 0.93 | 1 | 0 | 0.245 | -2.03 | 0.32913391 | NA | 0 | 2.5414 | 1.7975 | 1.7975 | TRUE | 0.5075 |
| 4930401C15I | 0 | 0 | 0 | 0 | 0 | 0 | 0 | 0 | 0 | 0 | 0 | 0 | 0 | 0 | 0 | NA | NA | NA | NA | NA | NA | NA | NA | NA | NA |
| 4930401O10 | 0 | 0 | 0 | 0 | 0 | 0 | 0 | 0 | 0 | 0 | 0 | 0 | 0 | 0 | 0 | NA | NA | NA | NA | NA | NA | NA | NA | NA | NA |
| 4930401O12 | 0 | 0 | 0 | 0 | 0 | 0 | 0 | 0 | 0 | 0 | 0 | 0 | 0 | 0 | 0 | NA | NA | NA | NA | NA | NA | NA | NA | NA | NA |
| 4930402F06I | 0 | 0 | 0 | 0 | 0 | 0 | 0 | 0 | 0 | 0 | 0 | 0 | 0 | 0 | 0 | NA | NA | NA | NA | NA | NA | NA | NA | NA | NA |
| 4930402F11I | 0 | 0 | 0 | 0 | 0 | 0 | 0 | 0 | 0 | 0 | 0 | 0 | 0 | 0 | 0 | NA | NA | NA | NA | NA | NA | NA | NA | NA | NA |
| 4930402H24 | 2479 | 1884 | 2004 | 2565 | 2330 | 2591 | 1925 | 2004 | 2162 | 2553 | 2628 | 2558 | 2305.17 | 2030 | 2580 | 1.271 | 0.346 | 1.29E-07 | 4.90E-06 | 0.0012 | 0.0035 | 0.0027 | 0.0027 | TRUE | 0.0387 |
| 4930402K13I | 5 | 0 | 0 | 0 | 0 | 0 | 4 | 0 | 0 | 0 | 0 | 0 | 0.65 | 1 | 0 | 0.143 | -2.801 | 0.35655793 | NA | 10 | 3.6644 | 5.2862 | 5.2862 | TRUE | 2.3401 |
| 4930404A05 | 0 | 0 | 0 | 0 | 0 | 0 | 0 | 0 | 0 | 0 | 0 | 0 | 0 | 0 | 0 | NA | NA | NA | NA | NA | NA | NA | NA | NA | NA |
| 4930404A10 | 15 | 19 | 13 | 5 | 9 | 15 | 12 | 20 | 14 | 5 | 10 | 15 | 12.64 | 15 | 10 | 0.657 | -0.606 | 0.29691692 | NA | 0.0738 | 0.19 | 0.1595 | 0.1595 | TRUE | 0.327 |
| 4930404H11 | 0 | 0 | 0 | 0 | 0 | 0 | 0 | 0 | 0 | 0 | 0 | 0 | 0 | 0 | 0 | NA | NA | NA | NA | NA | NA | NA | NA | NA | NA |
| 4930404I05R | 1 | 0 | 0 | 0 | 0 | 0 | 1 | 0 | 0 | 0 | 0 | 0 | 0.13 | 0 | 0 | 0.552 | -0.857 | 0.8336548 | NA | 0 | 18.3121 | 10 | 10 | TRUE | 0.2566 |
| 4930404N11 | 0 | 3 | 1 | 0 | 1 | 0 | 0 | 3 | 1 | 0 | 1 | 0 | 0.9 | 1 | 0 | 0.266 | -1.909 | 0.40640649 | NA | 0 | 2.6367 | 2.4605 | 2.4605 | TRUE | 0.3216 |
| 4930405A10 | 0 | 0 | 0 | 0 | 0 | 0 | 0 | 0 | 0 | 0 | 0 | 0 | 0 | 0 | 0 | NA | NA | NA | NA | NA | NA | NA | NA | NA | NA |
| 4930405A21 | 11 | 15 | 11 | 17 | 18 | 14 | 9 | 16 | 12 | 17 | 20 | 14 | 14.57 | 12 | 17 | 1.417 | 0.503 | 0.32775923 | NA | 0 | 0.1651 | 0.1194 | 0.1194 | TRUE | 0.3476 |
| 4930405D11 | 0 | 0 | 0 | 0 | 0 | 0 | 0 | 0 | 0 | 0 | 0 | 0 | 0 | 0 | 0 | NA | NA | NA | NA | NA | NA | NA | NA | NA | NA |
| 4930405J17F | 0 | 0 | 0 | 0 | 0 | 0 | 0 | 0 | 0 | 0 | 0 | 0 | 0 | 0 | 0 | NA | NA | NA | NA | NA | NA | NA | NA | NA | NA |
| 4930405P13I | 0 | 0 | 1 | 0 | 0 | 0 | 0 | 0 | 1 | 0 | 0 | 0 | 0.18 | 0 | 0 | 0.552 | -0.857 | 0.8336548 | NA | 0 | 13.1817 | 10 | 10 | TRUE | 0.5365 |
| 4930406D18 | 0 | 0 | 0 | 0 | 0 | 0 | 0 | 0 | 0 | 0 | 0 | 0 | 0 | 0 | 0 | NA | NA | NA | NA | NA | NA | NA | NA | NA | NA |
| 4930407I10R | 1 | 0 | 0 | 1 | 0 | 0 | 1 | 0 | 0 | 1 | 0 | 0 | 0.3 | 0 | 0 | 1.075 | 0.105 | 0.97698525 | NA | 0 | 8.0272 | 7.5141 | 7.5141 | TRUE | 0.3906 |
| 4930408F14I | 3 | 5 | 2 | 1 | 3 | 8 | 2 | 5 | 2 | 1 | 3 | 8 | 3.68 | 3 | 4 | 1.268 | 0.343 | 0.74680766 | NA | 0.2857 | 0.6464 | 0.5384 | 0.5384 | TRUE | 0.3982 |
| 4930412B13 | 0 | 0 | 0 | 0 | 0 | 0 | 0 | 0 | 0 | 0 | 0 | 0 | 0 | 0 | 0 | NA | NA | NA | NA | NA | NA | NA | NA | NA | NA |

| | | | | | | | | | | | | | | | | | | | | | | | | | |
|---|---|---|---|---|---|---|---|---|---|---|---|---|---|---|---|---|---|---|---|---|---|---|---|---|---|
| 4930412C18I | 32 | 31 | 14 | 18 | 15 | 27 | 25 | 33 | 15 | 18 | 17 | 27 | 22.4 | 24 | 21 | 0.844 | -0.244 | 0.5790359 | 0.73001324 | 0.0565 | 0.1082 | 0.0942 | 0.0942 | TRUE | 0.1886 |
| 4930412O13 | 4 | 4 | 0 | 1 | 0 | 1 | 3 | 4 | 0 | 1 | 0 | 1 | 1.56 | 2 | 1 | 0.274 | -1.87 | 0.2915884 | NA | 0 | 1.5243 | 1.305 | 1.305 | TRUE | 0.3943 |
| 4930413E15I | 0 | 0 | 0 | 0 | 0 | 0 | 0 | 0 | 0 | 0 | 0 | 0 | 0 | 0 | 0 | NA | NA | NA | NA | NA | NA | NA | NA | NA | NA |
| 4930413F20I | 1 | 0 | 0 | 0 | 0 | 0 | 1 | 0 | 0 | 0 | 0 | 0 | 0.13 | 0 | 0 | 0.552 | -0.857 | 0.8336548 | NA | 0 | 18.3121 | 10 | 10 | TRUE | 0.2566 |
| 4930413G21 | 4 | 6 | 6 | 3 | 4 | 3 | 3 | 6 | 6 | 3 | 5 | 3 | 4.4 | 5 | 4 | 0.661 | -0.597 | 0.51353629 | NA | 0 | 0.5406 | 0.3595 | 0.3595 | TRUE | 0.3809 |
| 4930413M19 | 0 | 0 | 0 | 0 | 0 | 0 | 0 | 0 | 0 | 0 | 0 | 0 | 0 | 0 | 0 | NA | NA | NA | NA | NA | NA | NA | NA | NA | NA |
| 4930414L22F | 48 | 40 | 31 | 57 | 49 | 52 | 37 | 43 | 33 | 57 | 55 | 51 | 46.1 | 38 | 54 | 1.442 | 0.528 | 0.06269689 | 0.15838299 | 0 | 0.0538 | 0.0357 | 0.0357 | TRUE | 0.0819 |
| 4930414N06 | 0 | 0 | 0 | 0 | 0 | 0 | 0 | 0 | 0 | 0 | 0 | 0 | 0 | 0 | 0 | NA | NA | NA | NA | NA | NA | NA | NA | NA | NA |
| 4930415F15I | 4 | 3 | 1 | 1 | 0 | 0 | 3 | 3 | 1 | 1 | 0 | 0 | 1.4 | 2 | 0 | 0.15 | -2.739 | 0.13813771 | NA | 0 | 1.701 | 1.2643 | 1.2643 | TRUE | 0.3906 |
| 4930415L06F | 0 | 0 | 0 | 0 | 0 | 0 | 0 | 0 | 0 | 0 | 0 | 0 | 0 | 0 | 0 | NA | NA | NA | NA | NA | NA | NA | NA | NA | NA |
| 4930415O20 | 3 | 1 | 1 | 3 | 3 | 2 | 2 | 1 | 1 | 3 | 3 | 2 | 2.14 | 1 | 3 | 1.808 | 0.854 | 0.50692327 | NA | 0 | 1.1119 | 0.6931 | 0.6931 | TRUE | 0.2263 |
| 4930417O13 | 3 | 1 | 3 | 0 | 0 | 3 | 2 | 1 | 3 | 0 | 0 | 3 | 1.6 | 2 | 1 | 0.456 | -1.134 | 0.49752644 | NA | 0.3393 | 1.4849 | 1.2813 | 1.2813 | TRUE | 1.4411 |
| 4930417O22 | 41 | 49 | 18 | 23 | 17 | 17 | 32 | 52 | 19 | 23 | 19 | 17 | 27.04 | 34 | 20 | 0.57 | -0.811 | 0.05986435 | 0.15298429 | 0.101 | 0.0901 | 0.0933 | 0.0933 | TRUE | 0.2235 |
| 4930419G24 | 0 | 0 | 0 | 0 | 0 | 0 | 0 | 0 | 0 | 0 | 0 | 0 | 0 | 0 | 0 | NA | NA | NA | NA | NA | NA | NA | NA | NA | NA |
| 4930422G04 | 1894 | 1416 | 1261 | 1587 | 1337 | 1539 | 1471 | 1506 | 1361 | 1580 | 1508 | 1519 | 1490.83 | 1446 | 1536 | 1.062 | 0.087 | 0.21872315 | 0.38185049 | 9.00E-04 | 0.004 | 0.0029 | 0.0029 | TRUE | 0.0311 |
| 4930423M02 | 0 | 0 | 0 | 0 | 0 | 0 | 0 | 0 | 0 | 0 | 0 | 0 | 0 | 0 | 0 | NA | NA | NA | NA | NA | NA | NA | NA | NA | NA |
| 4930425K10I | 0 | 0 | 0 | 0 | 0 | 0 | 0 | 0 | 0 | 0 | 0 | 0 | 0 | 0 | 0 | NA | NA | NA | NA | NA | NA | NA | NA | NA | NA |
| 4930425O10 | 0 | 1 | 1 | 0 | 1 | 1 | 0 | 1 | 1 | 0 | 1 | 1 | 0.71 | 1 | 1 | 1.018 | 0.026 | 0.99107709 | NA | 0 | 3.3422 | 2.3761 | 2.3761 | TRUE | 0.3738 |
| 4930426D05I | 2 | 3 | 0 | 1 | 2 | 3 | 2 | 3 | 0 | 1 | 2 | 3 | 1.83 | 2 | 2 | 1.307 | 0.386 | 0.79066754 | NA | 0 | 1.3002 | 0.97 | 0.97 | TRUE | 0.2239 |
| 4930426L09F | 1 | 1 | 1 | 0 | 0 | 1 | 1 | 1 | 1 | 0 | 0 | 1 | 0.65 | 1 | 0 | 0.383 | -1.383 | 0.55329037 | NA | 0 | 3.6426 | 2.4227 | 2.4227 | TRUE | 0.3835 |
| 4930427A07I | 2755 | 2124 | 2159 | 2408 | 2224 | 2530 | 2140 | 2259 | 2330 | 2397 | 2509 | 2498 | 2355.3 | 2243 | 2468 | 1.101 | 0.139 | 0.02680055 | 0.0854473 | 9.00E-04 | 0.0034 | 0.0024 | 0.0024 | TRUE | 0.0203 |
| 4930428D18I | 0 | 0 | 0 | 0 | 0 | 0 | 0 | 0 | 0 | 0 | 0 | 0 | 0 | 0 | 0 | NA | NA | NA | NA | NA | NA | NA | NA | NA | NA |
| 4930428E07I | 0 | 0 | 0 | 0 | 0 | 0 | 0 | 0 | 0 | 0 | 0 | 0 | 0 | 0 | 0 | NA | NA | NA | NA | NA | NA | NA | NA | NA | NA |
| 4930428G15 | 0 | 0 | 0 | 0 | 0 | 0 | 0 | 0 | 0 | 0 | 0 | 0 | 0 | 0 | 0 | NA | NA | NA | NA | NA | NA | NA | NA | NA | NA |
| 4930428O21 | 0 | 0 | 0 | 0 | 0 | 0 | 0 | 0 | 0 | 0 | 0 | 0 | 0 | 0 | 0 | NA | NA | NA | NA | NA | NA | NA | NA | NA | NA |
| 4930429B21I | 44 | 38 | 16 | 28 | 17 | 17 | 34 | 40 | 17 | 28 | 19 | 17 | 25.95 | 30 | 21 | 0.693 | -0.529 | 0.21615927 | 0.37885523 | 0.0857 | 0.0938 | 0.0915 | 0.0915 | TRUE | 0.5921 |
| 4930429D17I | 0 | 1 | 1 | 0 | 0 | 0 | 0 | 1 | 1 | 0 | 0 | 0 | 0.36 | 1 | 0 | 0.271 | -1.881 | 0.5755355 | NA | 0 | 6.6391 | 6.4428 | 6.4428 | TRUE | 0.3513 |
| 4930429F11I | 0 | 0 | 0 | 0 | 0 | 0 | 0 | 0 | 0 | 0 | 0 | 0 | 0 | 0 | 0 | NA | NA | NA | NA | NA | NA | NA | NA | NA | NA |
| 4930429F24I | 0 | 1 | 1 | 0 | 0 | 1 | 0 | 1 | 1 | 0 | 0 | 1 | 0.52 | 1 | 0 | 0.535 | -0.904 | 0.7403423 | NA | 0 | 4.5459 | 3.6616 | 3.6616 | TRUE | 0.3835 |
| 4930430A15I | 0 | 0 | 0 | 0 | 0 | 0 | 0 | 0 | 0 | 0 | 0 | 0 | 0 | 0 | 0 | NA | NA | NA | NA | NA | NA | NA | NA | NA | NA |
| 4930430D24I | 0 | 0 | 0 | 0 | 0 | 0 | 0 | 0 | 0 | 0 | 0 | 0 | 0 | 0 | 0 | NA | NA | NA | NA | NA | NA | NA | NA | NA | NA |
| 4930430F08I | 1707 | 1196 | 1364 | 1425 | 1216 | 1377 | 1326 | 1272 | 1472 | 1418 | 1372 | 1359 | 1369.87 | 1357 | 1383 | 1.02 | 0.029 | 0.70700249 | 0.82296495 | 0.0023 | 0.0042 | 0.0035 | 0.0035 | TRUE | 0.0646 |
| 4930430F21I | 0 | 0 | 0 | 0 | 0 | 0 | 0 | 0 | 0 | 0 | 0 | 0 | 0 | 0 | 0 | NA | NA | NA | NA | NA | NA | NA | NA | NA | NA |
| 4930430J02F | 0 | 0 | 0 | 0 | 0 | 0 | 0 | 0 | 0 | 0 | 0 | 0 | 0 | 0 | 0 | NA | NA | NA | NA | NA | NA | NA | NA | NA | NA |
| 4930431F12I | 1 | 0 | 0 | 0 | 1 | 2 | 1 | 0 | 0 | 0 | 1 | 2 | 0.65 | 0 | 1 | 2.998 | 1.584 | 0.52889475 | NA | 0 | 3.6683 | 3.06 | 3.06 | TRUE | 0.14 |
| 4930431P03I | 0 | 0 | 0 | 0 | 1 | 0 | 0 | 0 | 0 | 0 | 1 | 0 | 0.19 | 0 | 0 | 2.094 | 1.067 | 0.79379329 | NA | 0 | 12.608 | 10 | 10 | TRUE | 0.5075 |
| 4930432J09F | 0 | 0 | 0 | 0 | 0 | 0 | 0 | 0 | 0 | 0 | 0 | 0 | 0 | 0 | 0 | NA | NA | NA | NA | NA | NA | NA | NA | NA | NA |
| 4930432K09I | 0 | 0 | 0 | 0 | 0 | 0 | 0 | 0 | 0 | 0 | 0 | 0 | 0 | 0 | 0 | NA | NA | NA | NA | NA | NA | NA | NA | NA | NA |
| 4930432K21I | 198 | 151 | 121 | 126 | 129 | 121 | 154 | 161 | 131 | 125 | 146 | 119 | 139.22 | 149 | 130 | 0.875 | -0.193 | 0.27099463 | 0.44196113 | 0.0032 | 0.0194 | 0.015 | 0.015 | TRUE | 0.1083 |
| 4930432M17 | 0 | 0 | 0 | 0 | 0 | 0 | 0 | 0 | 0 | 0 | 0 | 0 | 0 | 0 | 0 | NA | NA | NA | NA | NA | NA | NA | NA | NA | NA |
| 4930433B08I | 0 | 0 | 0 | 0 | 0 | 0 | 0 | 0 | 0 | 0 | 0 | 0 | 0 | 0 | 0 | NA | NA | NA | NA | NA | NA | NA | NA | NA | NA |
| 4930433I11R | 0 | 0 | 0 | 0 | 0 | 0 | 0 | 0 | 0 | 0 | 0 | 0 | 0 | 0 | 0 | NA | NA | NA | NA | NA | NA | NA | NA | NA | NA |
| 4930433N12 | 95 | 67 | 84 | 103 | 79 | 86 | 74 | 71 | 91 | 103 | 89 | 85 | 85.37 | 79 | 92 | 1.177 | 0.235 | 0.2801192 | 0.45153307 | 0 | 0.0302 | 0.0223 | 0.0223 | TRUE | 0.1577 |
| 4930434J06F | 0 | 0 | 0 | 0 | 0 | 0 | 0 | 0 | 0 | 0 | 0 | 0 | 0 | 0 | 0 | NA | NA | NA | NA | NA | NA | NA | NA | NA | NA |
| 4930435E12I | 0 | 0 | 0 | 0 | 1 | 1 | 0 | 0 | 0 | 0 | 1 | 1 | 0.35 | 0 | 1 | 3.965 | 1.987 | 0.55529285 | NA | 0 | 6.7252 | 6.4426 | 6.4426 | TRUE | 0.261 |
| 4930438E09I | 0 | 0 | 0 | 0 | 0 | 0 | 0 | 0 | 0 | 0 | 0 | 0 | 0 | 0 | 0 | NA | NA | NA | NA | NA | NA | NA | NA | NA | NA |
| 4930440C22I | 0 | 0 | 0 | 0 | 0 | 0 | 0 | 0 | 0 | 0 | 0 | 0 | 0 | 0 | 0 | NA | NA | NA | NA | NA | NA | NA | NA | NA | NA |
| 4930441J16F | 0 | 0 | 0 | 0 | 0 | 0 | 0 | 0 | 0 | 0 | 0 | 0 | 0 | 0 | 0 | NA | NA | NA | NA | NA | NA | NA | NA | NA | NA |
| 4930441O14 | 9 | 7 | 4 | 7 | 6 | 6 | 7 | 7 | 4 | 7 | 7 | 6 | 6.4 | 6 | 7 | 1.042 | 0.06 | 0.93593803 | NA | 0 | 0.3726 | 0.2398 | 0.2398 | TRUE | 0.159 |
| 4930442J19F | 2 | 3 | 1 | 1 | 2 | 0 | 2 | 3 | 1 | 1 | 2 | 0 | 1.51 | 2 | 1 | 0.549 | -0.865 | 0.59002952 | NA | 0 | 1.5694 | 1.1193 | 1.1193 | TRUE | 0.3427 |
| 4930442L01F | 0 | 0 | 1 | 0 | 0 | 0 | 0 | 0 | 1 | 0 | 0 | 0 | 0.18 | 0 | 0 | 0.552 | -0.857 | 0.8336548 | NA | 0 | 13.1817 | 10 | 10 | TRUE | 0.5365 |
| 4930443O20 | 0 | 0 | 0 | 0 | 0 | 1 | 0 | 0 | 0 | 0 | 0 | 1 | 0.16 | 0 | 0 | 2.094 | 1.067 | 0.79379329 | NA | 0 | 14.4078 | 10 | 10 | TRUE | 0.3835 |
| 4930444A02I | 1369 | 1019 | 1043 | 1128 | 1079 | 1179 | 1063 | 1084 | 1125 | 1123 | 1217 | 1164 | 1129.4 | 1091 | 1168 | 1.071 | 0.099 | 0.18020008 | 0.33508136 | 0 | 0.0045 | 0.003 | 0.003 | TRUE | 0.016 |
| 4930444F02I | 1 | 0 | 1 | 4 | 0 | 1 | 1 | 0 | 1 | 4 | 0 | 1 | 1.14 | 1 | 2 | 2.677 | 1.421 | 0.47190338 | NA | 0 | 2.0862 | 1.7319 | 1.7319 | TRUE | 0.5809 |
| 4930444G20 | 0 | 0 | 0 | 0 | 0 | 0 | 0 | 0 | 0 | 0 | 0 | 0 | 0 | 0 | 0 | NA | NA | NA | NA | NA | NA | NA | NA | NA | NA |
| 4930444M15 | 123 | 91 | 82 | 135 | 97 | 143 | 96 | 97 | 88 | 134 | 109 | 141 | 110.96 | 94 | 128 | 1.373 | 0.457 | 0.01762643 | 0.06293645 | 0 | 0.0238 | 0.0175 | 0.0175 | TRUE | 0.1616 |
| 4930444P10I | 1 | 1 | 3 | 2 | 1 | 1 | 1 | 1 | 3 | 2 | 1 | 1 | 1.53 | 2 | 1 | 0.829 | -0.27 | 0.85906504 | NA | 0 | 1.5509 | 0.9947 | 0.9947 | TRUE | 0.4555 |
| 4930447A16I | 0 | 0 | 0 | 0 | 0 | 0 | 0 | 0 | 0 | 0 | 0 | 0 | 0 | 0 | 0 | NA | NA | NA | NA | NA | NA | NA | NA | NA | NA |
| 4930447C04I | 1274 | 1016 | 1030 | 1161 | 894 | 1065 | 989 | 1081 | 1111 | 1156 | 1009 | 1051 | 1066.17 | 1060 | 1072 | 1.012 | 0.018 | 0.83344724 | 0.90506468 | 0.0034 | 0.0046 | 0.0042 | 0.0042 | TRUE | 0.0562 |
| 4930447J18F | 0 | 0 | 0 | 0 | 0 | 0 | 0 | 0 | 0 | 0 | 0 | 0 | 0 | 0 | 0 | NA | NA | NA | NA | NA | NA | NA | NA | NA | NA |
| 4930447K03I | 2 | 0 | 2 | 9 | 7 | 4 | 2 | 0 | 2 | 9 | 8 | 4 | 4.09 | 1 | 7 | 5.513 | 2.463 | 0.02778516 | NA | 0 | 0.5825 | 0.446 | 0.446 | TRUE | 0.3882 |
| 4930447N08 | 0 | 1 | 0 | 0 | 1 | 1 | 0 | 1 | 0 | 0 | 1 | 1 | 0.53 | 0 | 1 | 2.03 | 1.022 | 0.70625413 | NA | 0 | 4.4757 | 3.5573 | 3.5573 | TRUE | 0.5217 |
| 4930448C13I | 0 | 0 | 0 | 0 | 0 | 0 | 0 | 0 | 0 | 0 | 0 | 0 | 0 | 0 | 0 | NA | NA | NA | NA | NA | NA | NA | NA | NA | NA |
| 4930448F12I | 0 | 0 | 0 | 0 | 0 | 0 | 0 | 0 | 0 | 0 | 0 | 0 | 0 | 0 | 0 | NA | NA | NA | NA | NA | NA | NA | NA | NA | NA |
| 4930448H16I | 0 | 0 | 0 | 0 | 0 | 0 | 0 | 0 | 0 | 0 | 0 | 0 | 0 | 0 | 0 | NA | NA | NA | NA | NA | NA | NA | NA | NA | NA |
| 4930448I06R | 0 | 0 | 0 | 0 | 0 | 0 | 0 | 0 | 0 | 0 | 0 | 0 | 0 | 0 | 0 | NA | NA | NA | NA | NA | NA | NA | NA | NA | NA |
| 4930448I18R | 0 | 0 | 0 | 0 | 0 | 0 | 0 | 0 | 0 | 0 | 0 | 0 | 0 | 0 | 0 | NA | NA | NA | NA | NA | NA | NA | NA | NA | NA |
| 4930448K20I | 0 | 0 | 2 | 0 | 0 | 0 | 0 | 0 | 2 | 0 | 0 | 0 | 0.36 | 1 | 0 | 0.269 | -1.896 | 0.61680529 | NA | 10 | 6.5921 | 8.6623 | 8.6623 | TRUE | 0.9744 |
| 4930449E01I | 2 | 0 | 1 | 0 | 0 | 2 | 2 | 0 | 1 | 0 | 0 | 2 | 0.77 | 1 | 1 | 0.747 | -0.422 | 0.85833462 | NA | 0 | 3.0894 | 2.7108 | 2.7108 | TRUE | 0.9736 |

| | | | | | | | | | | | | | | | | | | | | | | | | |
|---|---|---|---|---|---|---|---|---|---|---|---|---|---|---|---|---|---|---|---|---|---|---|---|---|
| 4930449E18I | 0 | 0 | 0 | 0 | 0 | 0 | 0 | 0 | 0 | 0 | 0 | 0 | 0 | 0 | 0 | NA | NA | NA | NA | NA | NA | NA | NA | NA | NA |
| 4930449I24R | 0 | 0 | 0 | 0 | 0 | 0 | 0 | 0 | 0 | 0 | 0 | 0 | 0 | 0 | 0 | NA | NA | NA | NA | NA | NA | NA | NA | NA | NA |
| 4930451C15I | 19 | 14 | 13 | 27 | 28 | 35 | 15 | 15 | 14 | 27 | 32 | 35 | 22.78 | 15 | 31 | 2.128 | 1.089 | 0.00757523 | 0.03364767 | 0 | 0.1065 | 0.0702 | 0.0702 | TRUE | 0.0944 |
| 4930451G09 | 52 | 38 | 28 | 37 | 24 | 49 | 40 | 40 | 30 | 37 | 27 | 48 | 37.22 | 37 | 37 | 1.012 | 0.017 | 0.9586541 | 0.9773909 | 0.0244 | 0.0661 | 0.055 | 0.055 | TRUE | 0.2286 |
| 4930451I11R | 2 | 0 | 1 | 0 | 0 | 0 | 2 | 0 | 1 | 0 | 0 | 0 | 0.44 | 1 | 0 | 0.214 | -2.223 | 0.47051571 | NA | 0 | 5.4047 | 5.2946 | 5.2946 | TRUE | 0.2788 |
| 4930452A19I | 0 | 0 | 0 | 0 | 0 | 0 | 0 | 0 | 0 | 0 | 0 | 0 | 0 | 0 | 0 | NA | NA | NA | NA | NA | NA | NA | NA | NA | NA |
| 4930452B06I | 1842 | 1351 | 1445 | 1238 | 1175 | 1376 | 1431 | 1437 | 1559 | 1232 | 1326 | 1358 | 1390.49 | 1476 | 1305 | 0.885 | -0.176 | 0.01783666 | 0.06346511 | 0.0017 | 0.0041 | 0.0033 | 0.0033 | TRUE | 0.0292 |
| 4930452G13 | 1 | 1 | 0 | 0 | 0 | 0 | 1 | 1 | 0 | 0 | 0 | 0 | 0.31 | 1 | 0 | 0.309 | -1.696 | 0.63183309 | NA | 0 | 7.7294 | 7.2472 | 7.2472 | TRUE | 0.2015 |
| 4930452N14 | 0 | 0 | 0 | 0 | 0 | 0 | 0 | 0 | 0 | 0 | 0 | 0 | 0 | 0 | 0 | NA | NA | NA | NA | NA | NA | NA | NA | NA | NA |
| 4930453H23I | 2 | 4 | 0 | 1 | 2 | 2 | 2 | 4 | 0 | 1 | 2 | 2 | 1.84 | 2 | 2 | 0.9 | -0.151 | 0.91774588 | NA | 0 | 1.2912 | 1.0006 | 1.0006 | TRUE | 0.3507 |
| 4930453L07F | 0 | 0 | 0 | 1 | 0 | 0 | 0 | 0 | 0 | 1 | 0 | 0 | 0.17 | 0 | 0 | 2.094 | 1.067 | 0.79379329 | NA | 0 | 14.2885 | 10 | 10 | TRUE | 0.3906 |
| 4930453N24 | 2147 | 1415 | 1210 | 1228 | 1127 | 1231 | 1667 | 1505 | 1306 | 1222 | 1271 | 1215 | 1364.52 | 1493 | 1236 | 0.827 | -0.274 | 0.00260184 | 0.01480007 | 0.0066 | 0.0042 | 0.0052 | 0.0052 | TRUE | 0.144 |
| 4930455B14I | 5 | 2 | 9 | 5 | 7 | 3 | 4 | 2 | 10 | 5 | 8 | 3 | 5.26 | 5 | 5 | 1.01 | 0.014 | 0.98727611 | NA | 0.1848 | 0.4531 | 0.3759 | 0.3759 | TRUE | 0.8337 |
| 4930455C13I | 0 | 0 | 0 | 0 | 0 | 0 | 0 | 0 | 0 | 0 | 0 | 0 | 0 | 0 | 0 | NA | NA | NA | NA | NA | NA | NA | NA | NA | NA |
| 4930455D15I | 0 | 0 | 0 | 0 | 0 | 0 | 0 | 0 | 0 | 0 | 0 | 0 | 0 | 0 | 0 | NA | NA | NA | NA | NA | NA | NA | NA | NA | NA |
| 4930455F16I | 0 | 0 | 0 | 0 | 0 | 0 | 0 | 0 | 0 | 0 | 0 | 0 | 0 | 0 | 0 | NA | NA | NA | NA | NA | NA | NA | NA | NA | NA |
| 4930455F23I | 812 | 584 | 545 | 632 | 497 | 611 | 631 | 621 | 588 | 629 | 561 | 603 | 605.47 | 613 | 598 | 0.974 | -0.037 | 0.68535558 | 0.80709763 | 0 | 0.0063 | 0.0045 | 0.0045 | TRUE | 0.0334 |
| 4930455H04I | 0 | 0 | 0 | 0 | 0 | 0 | 0 | 0 | 0 | 0 | 0 | 0 | 0 | 0 | 0 | NA | NA | NA | NA | NA | NA | NA | NA | NA | NA |
| 4930455J16F | 0 | 0 | 0 | 0 | 0 | 0 | 0 | 0 | 0 | 0 | 0 | 0 | 0 | 0 | 0 | NA | NA | NA | NA | NA | NA | NA | NA | NA | NA |
| 4930456L15F | 1 | 0 | 0 | 0 | 0 | 0 | 1 | 0 | 0 | 0 | 0 | 0 | 0.13 | 0 | 0 | 0.552 | -0.857 | 0.8336548 | NA | 0 | 18.3121 | 10 | 10 | TRUE | 0.2566 |
| 4930459L07F | 0 | 0 | 0 | 0 | 0 | 0 | 0 | 0 | 0 | 0 | 0 | 0 | 0 | 0 | 0 | NA | NA | NA | NA | NA | NA | NA | NA | NA | NA |
| 4930461G14 | 608 | 416 | 457 | 495 | 428 | 506 | 472 | 442 | 493 | 493 | 483 | 500 | 480.48 | 469 | 492 | 1.048 | 0.068 | 0.49217252 | 0.6574413 | 0 | 0.0074 | 0.0049 | 0.0049 | TRUE | 0.0274 |
| 4930465A12I | 0 | 0 | 0 | 0 | 0 | 0 | 0 | 0 | 0 | 0 | 0 | 0 | 0 | 0 | 0 | NA | NA | NA | NA | NA | NA | NA | NA | NA | NA |
| 4930465K10I | 1 | 3 | 2 | 1 | 1 | 2 | 1 | 3 | 2 | 1 | 1 | 2 | 1.7 | 2 | 1 | 0.687 | -0.542 | 0.70832811 | NA | 0 | 1.3934 | 0.891 | 0.891 | TRUE | 0.2936 |
| 4930465M2C | 4 | 1 | 0 | 1 | 0 | 0 | 3 | 1 | 0 | 1 | 0 | 0 | 0.86 | 1 | 0 | 0.259 | -1.946 | 0.39766866 | NA | 0 | 2.7553 | 2.477 | 2.477 | TRUE | 0.2295 |
| 4930467D21I | 20 | 12 | 14 | 15 | 12 | 11 | 16 | 13 | 15 | 15 | 14 | 11 | 13.79 | 15 | 13 | 0.903 | -0.147 | 0.77197478 | NA | 0 | 0.1743 | 0.1134 | 0.1134 | TRUE | 0.0989 |
| 4930467E23I | 0 | 0 | 0 | 0 | 0 | 0 | 0 | 0 | 0 | 0 | 0 | 0 | 0 | 0 | 0 | NA | NA | NA | NA | NA | NA | NA | NA | NA | NA |
| 4930467K11I | 1 | 2 | 0 | 0 | 1 | 0 | 1 | 2 | 0 | 0 | 1 | 0 | 0.67 | 1 | 0 | 0.385 | -1.379 | 0.57963618 | NA | 0 | 3.5292 | 2.9645 | 2.9645 | TRUE | 0.3656 |
| 4930468A15I | 0 | 0 | 0 | 0 | 0 | 0 | 0 | 0 | 0 | 0 | 0 | 0 | 0 | 0 | 0 | NA | NA | NA | NA | NA | NA | NA | NA | NA | NA |
| 4930469G21 | 0 | 2 | 0 | 1 | 1 | 0 | 0 | 2 | 0 | 1 | 1 | 0 | 0.71 | 1 | 1 | 1.021 | 0.03 | 0.99038186 | NA | 0 | 3.3477 | 2.9808 | 2.9808 | TRUE | 0.9748 |
| 4930470H14I | 0 | 0 | 0 | 0 | 0 | 0 | 0 | 0 | 0 | 0 | 0 | 0 | 0 | 0 | 0 | NA | NA | NA | NA | NA | NA | NA | NA | NA | NA |
| 4930470P17I | 4 | 3 | 4 | 1 | 2 | 2 | 3 | 3 | 4 | 1 | 2 | 2 | 2.64 | 3 | 2 | 0.492 | -1.025 | 0.38362975 | NA | 0 | 0.9002 | 0.5608 | 0.5608 | TRUE | 0.1147 |
| 4930471C04I | 0 | 0 | 0 | 0 | 0 | 0 | 0 | 0 | 0 | 0 | 0 | 0 | 0 | 0 | 0 | NA | NA | NA | NA | NA | NA | NA | NA | NA | NA |
| 4930471G03 | 0 | 0 | 0 | 0 | 0 | 0 | 0 | 0 | 0 | 0 | 0 | 0 | 0 | 0 | 0 | NA | NA | NA | NA | NA | NA | NA | NA | NA | NA |
| 4930471I20R | 0 | 0 | 0 | 0 | 0 | 0 | 0 | 0 | 0 | 0 | 0 | 0 | 0 | 0 | 0 | NA | NA | NA | NA | NA | NA | NA | NA | NA | NA |
| 4930471M09 | 1 | 0 | 0 | 0 | 0 | 0 | 1 | 0 | 0 | 0 | 0 | 0 | 0.13 | 0 | 0 | 0.552 | -0.857 | 0.8336548 | NA | 0 | 18.3121 | 10 | 10 | TRUE | 0.2566 |
| 4930473A02I | 1 | 0 | 1 | 2 | 1 | 2 | 1 | 0 | 1 | 2 | 1 | 2 | 1.16 | 1 | 2 | 2.723 | 1.445 | 0.42461796 | NA | 0 | 2.0489 | 1.2898 | 1.2898 | TRUE | 0.1933 |
| 4930473O22 | 0 | 0 | 0 | 0 | 0 | 0 | 0 | 0 | 0 | 0 | 0 | 0 | 0 | 0 | 0 | NA | NA | NA | NA | NA | NA | NA | NA | NA | NA |
| 4930474G06 | 0 | 0 | 0 | 0 | 0 | 0 | 0 | 0 | 0 | 0 | 0 | 0 | 0 | 0 | 0 | NA | NA | NA | NA | NA | NA | NA | NA | NA | NA |
| 4930474H20I | 0 | 1 | 0 | 0 | 0 | 0 | 0 | 1 | 0 | 0 | 0 | 0 | 0.18 | 0 | 0 | 0.552 | -0.857 | 0.8336548 | NA | 0 | 13.3714 | 10 | 10 | TRUE | 0.5216 |
| 4930474M22 | 0 | 0 | 0 | 0 | 0 | 0 | 0 | 0 | 0 | 0 | 0 | 0 | 0 | 0 | 0 | NA | NA | NA | NA | NA | NA | NA | NA | NA | NA |
| 4930474N05 | 0 | 0 | 0 | 0 | 0 | 0 | 0 | 0 | 0 | 0 | 0 | 0 | 0 | 0 | 0 | NA | NA | NA | NA | NA | NA | NA | NA | NA | NA |
| 4930474N09 | 0 | 0 | 1 | 0 | 0 | 0 | 0 | 0 | 1 | 0 | 0 | 0 | 0.18 | 0 | 0 | 0.552 | -0.857 | 0.8336548 | NA | 0 | 13.1817 | 10 | 10 | TRUE | 0.5365 |
| 4930478L05F | 0 | 0 | 0 | 0 | 0 | 0 | 0 | 0 | 0 | 0 | 0 | 0 | 0 | 0 | 0 | NA | NA | NA | NA | NA | NA | NA | NA | NA | NA |
| 4930478P22I | 0 | 0 | 0 | 0 | 0 | 0 | 0 | 0 | 0 | 0 | 0 | 0 | 0 | 0 | 0 | NA | NA | NA | NA | NA | NA | NA | NA | NA | NA |
| 4930479D17I | 2 | 1 | 1 | 0 | 0 | 1 | 2 | 1 | 1 | 0 | 0 | 1 | 0.78 | 1 | 0 | 0.299 | -1.743 | 0.4255496 | NA | 0 | 3.0389 | 2.0589 | 2.0589 | TRUE | 0.3835 |
| 4930479M11 | 13 | 6 | 10 | 12 | 8 | 10 | 10 | 6 | 11 | 12 | 9 | 10 | 9.68 | 9 | 10 | 1.129 | 0.174 | 0.77474778 | NA | 0 | 0.2471 | 0.1646 | 0.1646 | TRUE | 0.2005 |
| 4930480E11I | 0 | 0 | 0 | 0 | 0 | 0 | 0 | 0 | 0 | 0 | 0 | 0 | 0 | 0 | 0 | NA | NA | NA | NA | NA | NA | NA | NA | NA | NA |
| 4930480G23 | 0 | 1 | 0 | 0 | 2 | 0 | 0 | 1 | 0 | 0 | 2 | 0 | 0.55 | 0 | 1 | 2.136 | 1.095 | 0.70925113 | NA | 0 | 4.2858 | 4.5115 | 4.5115 | TRUE | 0.9746 |
| 4930480K15I | 0 | 0 | 0 | 0 | 0 | 0 | 0 | 0 | 0 | 0 | 0 | 0 | 0 | 0 | 0 | NA | NA | NA | NA | NA | NA | NA | NA | NA | NA |
| 4930480M12 | 0 | 0 | 0 | 0 | 0 | 0 | 0 | 0 | 0 | 0 | 0 | 0 | 0 | 0 | 0 | NA | NA | NA | NA | NA | NA | NA | NA | NA | NA |
| 4930481A15I | 8 | 9 | 4 | 5 | 3 | 3 | 6 | 10 | 4 | 5 | 3 | 3 | 5.24 | 7 | 4 | 0.565 | -0.823 | 0.33234824 | NA | 0 | 0.4549 | 0.3103 | 0.3103 | TRUE | 0.278 |
| 4930483J18F | 1 | 2 | 1 | 3 | 0 | 2 | 1 | 2 | 1 | 3 | 0 | 2 | 1.49 | 1 | 2 | 1.285 | 0.362 | 0.82041438 | NA | 0 | 1.5924 | 1.1516 | 1.1516 | TRUE | 0.3748 |
| 4930483K19I | 0 | 0 | 0 | 1 | 0 | 0 | 0 | 0 | 0 | 1 | 0 | 0 | 0.17 | 0 | 0 | 2.094 | 1.067 | 0.79379329 | NA | 0 | 14.2885 | 10 | 10 | TRUE | 0.3906 |
| 4930483O08 | 0 | 0 | 1 | 0 | 0 | 0 | 0 | 0 | 1 | 0 | 0 | 0 | 0.18 | 0 | 0 | 0.552 | -0.857 | 0.8336548 | NA | 0 | 13.1817 | 10 | 10 | TRUE | 0.5365 |
| 4930486F22I | 0 | 0 | 0 | 0 | 0 | 0 | 0 | 0 | 0 | 0 | 0 | 0 | 0 | 0 | 0 | NA | NA | NA | NA | NA | NA | NA | NA | NA | NA |
| 4930486I03R | 0 | 0 | 0 | 0 | 0 | 0 | 0 | 0 | 0 | 0 | 0 | 0 | 0 | 0 | 0 | NA | NA | NA | NA | NA | NA | NA | NA | NA | NA |
| 4930486L24F | 119 | 87 | 90 | 30 | 31 | 33 | 92 | 93 | 97 | 30 | 35 | 33 | 63.25 | 94 | 33 | 0.345 | -1.536 | 2.06E-09 | 1.27E-07 | 0 | 0.0399 | 0.0262 | 0.0262 | TRUE | 0.0343 |
| 4930487D11I | 0 | 0 | 0 | 0 | 0 | 0 | 0 | 0 | 0 | 0 | 0 | 0 | 0 | 0 | 0 | NA | NA | NA | NA | NA | NA | NA | NA | NA | NA |
| 4930487H11I | 0 | 0 | 1 | 0 | 0 | 0 | 0 | 0 | 1 | 0 | 0 | 0 | 0.18 | 0 | 0 | 0.552 | -0.857 | 0.8336548 | NA | 0 | 13.1817 | 10 | 10 | TRUE | 0.5365 |
| 4930488B22I | 0 | 0 | 0 | 0 | 0 | 0 | 0 | 0 | 0 | 0 | 0 | 0 | 0 | 0 | 0 | NA | NA | NA | NA | NA | NA | NA | NA | NA | NA |
| 4930488L21F | 0 | 0 | 0 | 0 | 0 | 0 | 0 | 0 | 0 | 0 | 0 | 0 | 0 | 0 | 0 | NA | NA | NA | NA | NA | NA | NA | NA | NA | NA |
| 4930500F04I | 0 | 0 | 0 | 0 | 0 | 0 | 0 | 0 | 0 | 0 | 0 | 0 | 0 | 0 | 0 | NA | NA | NA | NA | NA | NA | NA | NA | NA | NA |
| 4930500J02F | 1157 | 912 | 960 | 770 | 665 | 636 | 899 | 970 | 1036 | 766 | 750 | 628 | 841.5 | 968 | 715 | 0.739 | -0.437 | 1.02E-05 | 0.00019123 | 0.007 | 0.0052 | 0.0058 | 0.0058 | TRUE | 0.1358 |
| 4930500L23F | 0 | 0 | 0 | 0 | 0 | 0 | 0 | 0 | 0 | 0 | 0 | 0 | 0 | 0 | 0 | NA | NA | NA | NA | NA | NA | NA | NA | NA | NA |
| 4930502A04I | 0 | 0 | 0 | 0 | 0 | 0 | 0 | 0 | 0 | 0 | 0 | 0 | 0 | 0 | 0 | NA | NA | NA | NA | NA | NA | NA | NA | NA | NA |
| 4930502E09I | 3 | 6 | 8 | 7 | 3 | 4 | 2 | 6 | 9 | 7 | 3 | 4 | 5.27 | 6 | 5 | 0.846 | -0.241 | 0.78075864 | NA | 0 | 0.4518 | 0.3489 | 0.3489 | TRUE | 0.4669 |
| 4930502E18I | 378 | 294 | 230 | 178 | 174 | 182 | 294 | 313 | 248 | 177 | 196 | 180 | 234.6 | 285 | 184 | 0.646 | -0.63 | 1.04E-05 | 0.00019326 | 0.0048 | 0.0125 | 0.0102 | 0.0102 | TRUE | 0.1359 |
| 4930503B20I | 5 | 6 | 1 | 6 | 5 | 6 | 4 | 6 | 1 | 6 | 6 | 6 | 4.81 | 4 | 6 | 1.543 | 0.625 | 0.48420795 | NA | 0 | 0.4948 | 0.3601 | 0.3601 | TRUE | 0.2787 |

| | | | | | | | | | | | | | | | | | | | | | | | | | |
|---|---|---|---|---|---|---|---|---|---|---|---|---|---|---|---|---|---|---|---|---|---|---|---|---|---|
| 4930503E14I | 3 | 4 | 0 | 1 | 3 | 1 | 2 | 4 | 0 | 1 | 3 | 1 | 1.99 | 2 | 2 | 0.801 | -0.321 | 0.82377183 | NA | 0.3393 | 1.1923 | 0.9843 | 0.9843 | TRUE | 0.1848 |
| 4930503E24I | 0 | 0 | 0 | 0 | 1 | 1 | 0 | 0 | 0 | 0 | 1 | 1 | 0.35 | 0 | 1 | 3.965 | 1.987 | 0.55529285 | NA | 0 | 6.7252 | 6.4426 | 6.4426 | TRUE | 0.261 |
| 4930503H13 | 0 | 0 | 0 | 0 | 0 | 0 | 0 | 0 | 0 | 0 | 0 | 0 | 0 | 0 | 0 | NA | NA | NA | NA | NA | NA | NA | NA | NA | NA |
| 4930503L19F | 1299 | 906 | 868 | 951 | 801 | 929 | 1009 | 964 | 937 | 947 | 904 | 917 | 946.07 | 970 | 923 | 0.95 | -0.073 | 0.34117568 | 0.51526036 | 0 | 0.0049 | 0.0032 | 0.0032 | TRUE | 0.0157 |
| 4930503O07 | 0 | 0 | 0 | 0 | 0 | 0 | 0 | 0 | 0 | 0 | 0 | 0 | 0 | 0 | 0 | NA | NA | NA | NA | NA | NA | NA | NA | NA | NA |
| 4930504O13 | 16 | 10 | 3 | 8 | 6 | 8 | 12 | 11 | 3 | 8 | 7 | 8 | 8.15 | 9 | 8 | 0.849 | -0.235 | 0.73768082 | NA | 0 | 0.2931 | 0.2332 | 0.2332 | TRUE | 0.8755 |
| 4930505A04 | 11 | 15 | 5 | 5 | 7 | 2 | 9 | 16 | 5 | 5 | 8 | 2 | 7.46 | 10 | 5 | 0.493 | -1.022 | 0.18886382 | NA | 0.175 | 0.3203 | 0.2811 | 0.2811 | TRUE | 0.3896 |
| 4930505G20 | 0 | 0 | 0 | 0 | 0 | 0 | 0 | 0 | 0 | 0 | 0 | 0 | 0 | 0 | 0 | NA | NA | NA | NA | NA | NA | NA | NA | NA | NA |
| 4930506C21I | 0 | 0 | 0 | 2 | 0 | 0 | 0 | 0 | 0 | 2 | 0 | 0 | 0.33 | 0 | 1 | 3.77 | 1.915 | 0.62344813 | NA | 0 | 7.1455 | 9.1834 | 9.1834 | TRUE | 0.9739 |
| 4930506M07 | 857 | 677 | 629 | 764 | 615 | 679 | 666 | 720 | 679 | 760 | 694 | 670 | 698.16 | 688 | 708 | 1.03 | 0.043 | 0.63482955 | 0.77160293 | 0.0016 | 0.0058 | 0.0044 | 0.0044 | TRUE | 0.0501 |
| 4930507D05 | 4 | 5 | 1 | 2 | 5 | 4 | 3 | 5 | 1 | 2 | 6 | 4 | 3.51 | 3 | 4 | 1.21 | 0.275 | 0.7926126 | NA | 0 | 0.6769 | 0.4984 | 0.4984 | TRUE | 0.219 |
| 4930509E16I | 23 | 12 | 13 | 34 | 29 | 34 | 18 | 13 | 14 | 34 | 33 | 34 | 24.13 | 15 | 34 | 2.22 | 1.15 | 0.00390312 | 0.02023994 | 0 | 0.1006 | 0.067 | 0.067 | TRUE | 0.1774 |
| 4930509J09F | 0 | 0 | 0 | 0 | 0 | 0 | 0 | 0 | 0 | 0 | 0 | 0 | 0 | 0 | 0 | NA | NA | NA | NA | NA | NA | NA | NA | NA | NA |
| 4930509K18I | 0 | 0 | 0 | 1 | 0 | 0 | 0 | 0 | 0 | 1 | 0 | 0 | 0.17 | 0 | 0 | 2.094 | 1.067 | 0.79379329 | NA | 0 | 14.2885 | 10 | 10 | TRUE | 0.3906 |
| 4930511A02 | 1 | 1 | 1 | 2 | 1 | 0 | 1 | 1 | 1 | 2 | 1 | 0 | 1.01 | 1 | 1 | 1.075 | 0.104 | 0.95576718 | NA | 0 | 2.3575 | 1.5354 | 1.5354 | TRUE | 0.2671 |
| 4930511E03I | 1 | 0 | 0 | 0 | 0 | 1 | 1 | 0 | 0 | 0 | 0 | 1 | 0.29 | 0 | 0 | 1.075 | 0.105 | 0.97702352 | NA | 0 | 8.0647 | 7.5458 | 7.5458 | TRUE | 0.3835 |
| 4930511M06 | 24 | 9 | 15 | 10 | 9 | 12 | 19 | 10 | 16 | 10 | 10 | 12 | 12.73 | 15 | 11 | 0.714 | -0.485 | 0.37454116 | NA | 0 | 0.1887 | 0.1344 | 0.1344 | TRUE | 0.4066 |
| 4930511M11 | 0 | 0 | 0 | 0 | 0 | 0 | 0 | 0 | 0 | 0 | 0 | 0 | 0 | 0 | 0 | NA | NA | NA | NA | NA | NA | NA | NA | NA | NA |
| 4930512B01 | 0 | 0 | 0 | 0 | 0 | 0 | 0 | 0 | 0 | 0 | 0 | 0 | 0 | 0 | 0 | NA | NA | NA | NA | NA | NA | NA | NA | NA | NA |
| 4930513D17 | 3 | 0 | 0 | 0 | 0 | 0 | 2 | 0 | 0 | 0 | 0 | 0 | 0.39 | 1 | 0 | 0.239 | -2.067 | 0.57190649 | NA | 10 | 6.1057 | 8.0283 | 8.0283 | TRUE | 1.4408 |
| 4930513N10 | 221 | 146 | 108 | 114 | 84 | 97 | 172 | 155 | 117 | 113 | 95 | 96 | 124.58 | 148 | 101 | 0.683 | -0.55 | 0.00600404 | 0.02821959 | 0.0181 | 0.0215 | 0.0205 | 0.0205 | TRUE | 0.3474 |
| 4930513O06 | 1 | 0 | 0 | 0 | 0 | 0 | 1 | 0 | 0 | 0 | 0 | 0 | 0.13 | 0 | 0 | 0.552 | -0.857 | 0.8336548 | NA | 0 | 18.3121 | 10 | 10 | TRUE | 0.2566 |
| 4930515B02 | 0 | 0 | 0 | 0 | 0 | 0 | 0 | 0 | 0 | 0 | 0 | 0 | 0 | 0 | 0 | NA | NA | NA | NA | NA | NA | NA | NA | NA | NA |
| 4930515G01 | 0 | 1 | 0 | 0 | 1 | 0 | 0 | 1 | 0 | 0 | 1 | 0 | 0.37 | 0 | 0 | 1.075 | 0.105 | 0.97520514 | NA | 0 | 6.4904 | 6.1969 | 6.1969 | TRUE | 0.5216 |
| 4930515G16 | 0 | 0 | 0 | 0 | 0 | 0 | 0 | 0 | 0 | 0 | 0 | 0 | 0 | 0 | 0 | NA | NA | NA | NA | NA | NA | NA | NA | NA | NA |
| 4930515L03F | 0 | 0 | 0 | 0 | 0 | 0 | 0 | 0 | 0 | 0 | 0 | 0 | 0 | 0 | 0 | NA | NA | NA | NA | NA | NA | NA | NA | NA | NA |
| 4930515L19F | 0 | 0 | 0 | 0 | 0 | 0 | 0 | 0 | 0 | 0 | 0 | 0 | 0 | 0 | 0 | NA | NA | NA | NA | NA | NA | NA | NA | NA | NA |
| 4930517E11I | 0 | 0 | 0 | 0 | 0 | 0 | 0 | 0 | 0 | 0 | 0 | 0 | 0 | 0 | 0 | NA | NA | NA | NA | NA | NA | NA | NA | NA | NA |
| 4930518P08I | 9 | 4 | 5 | 3 | 6 | 4 | 7 | 4 | 5 | 3 | 7 | 4 | 5.06 | 5 | 5 | 0.809 | -0.306 | 0.71786872 | NA | 0 | 0.4711 | 0.3169 | 0.3169 | TRUE | 0.2951 |
| 4930519D14 | 0 | 0 | 0 | 0 | 0 | 0 | 0 | 0 | 0 | 0 | 0 | 0 | 0 | 0 | 0 | NA | NA | NA | NA | NA | NA | NA | NA | NA | NA |
| 4930519F09I | 4 | 1 | 1 | 1 | 0 | 1 | 3 | 1 | 1 | 1 | 0 | 1 | 1.21 | 2 | 1 | 0.378 | -1.405 | 0.43600426 | NA | 0 | 1.9688 | 1.3109 | 1.3109 | TRUE | 0.4821 |
| 4930519F16I | 13 | 18 | 10 | 8 | 3 | 6 | 10 | 19 | 11 | 8 | 3 | 6 | 9.55 | 13 | 6 | 0.438 | -1.191 | 0.07281323 | NA | 0 | 0.2506 | 0.1922 | 0.1922 | TRUE | 0.5835 |
| 4930519F24I | 0 | 0 | 0 | 0 | 0 | 0 | 0 | 0 | 0 | 0 | 0 | 0 | 0 | 0 | 0 | NA | NA | NA | NA | NA | NA | NA | NA | NA | NA |
| 4930519G04 | 0 | 0 | 0 | 0 | 1 | 1 | 0 | 0 | 0 | 0 | 1 | 1 | 0.35 | 0 | 1 | 3.965 | 1.987 | 0.55529285 | NA | 0 | 6.7252 | 6.4426 | 6.4426 | TRUE | 0.261 |
| 4930519H02 | 0 | 0 | 0 | 0 | 0 | 0 | 0 | 0 | 0 | 0 | 0 | 0 | 0 | 0 | 0 | NA | NA | NA | NA | NA | NA | NA | NA | NA | NA |
| 4930520O04 | 28 | 20 | 16 | 27 | 13 | 17 | 22 | 21 | 17 | 27 | 15 | 17 | 19.77 | 20 | 20 | 0.968 | -0.047 | 0.91571024 | 0.95343864 | 0 | 0.1223 | 0.0915 | 0.0915 | TRUE | 0.604 |
| 4930520P13I | 0 | 0 | 0 | 0 | 0 | 0 | 0 | 0 | 0 | 0 | 0 | 0 | 0 | 0 | 0 | NA | NA | NA | NA | NA | NA | NA | NA | NA | NA |
| 4930521E06I | 0 | 0 | 0 | 0 | 0 | 0 | 0 | 0 | 0 | 0 | 0 | 0 | 0 | 0 | 0 | NA | NA | NA | NA | NA | NA | NA | NA | NA | NA |
| 4930522H14 | 0 | 0 | 0 | 0 | 0 | 0 | 0 | 0 | 0 | 0 | 0 | 0 | 0 | 0 | 0 | NA | NA | NA | NA | NA | NA | NA | NA | NA | NA |
| 4930522O17 | 0 | 0 | 0 | 0 | 0 | 0 | 0 | 0 | 0 | 0 | 0 | 0 | 0 | 0 | 0 | NA | NA | NA | NA | NA | NA | NA | NA | NA | NA |
| 4930523C07I | 29 | 21 | 12 | 13 | 19 | 14 | 23 | 22 | 13 | 13 | 21 | 14 | 17.67 | 19 | 16 | 0.824 | -0.279 | 0.55812873 | 0.71302591 | 0 | 0.1366 | 0.1062 | 0.1062 | TRUE | 0.4023 |
| 4930523O13 | 0 | 0 | 0 | 0 | 0 | 0 | 0 | 0 | 0 | 0 | 0 | 0 | 0 | 0 | 0 | NA | NA | NA | NA | NA | NA | NA | NA | NA | NA |
| 4930524B15 | 230 | 156 | 122 | 108 | 100 | 79 | 179 | 166 | 132 | 108 | 113 | 78 | 129.08 | 159 | 100 | 0.623 | -0.682 | 0.00073691 | 0.00578963 | 0.0222 | 0.0208 | 0.0212 | 0.0212 | TRUE | 0.355 |
| 4930524C18I | 1 | 0 | 0 | 0 | 0 | 0 | 1 | 0 | 0 | 0 | 0 | 0 | 0.13 | 0 | 0 | 0.552 | -0.857 | 0.8336548 | NA | 0 | 18.3121 | 10 | 10 | TRUE | 0.2566 |
| 4930524L23F | 16 | 7 | 4 | 7 | 2 | 4 | 12 | 7 | 4 | 7 | 2 | 4 | 6.23 | 8 | 4 | 0.541 | -0.887 | 0.27874891 | NA | 0.1191 | 0.383 | 0.3075 | 0.3075 | TRUE | 0.2502 |
| 4930524N10 | 0 | 0 | 0 | 0 | 0 | 0 | 0 | 0 | 0 | 0 | 0 | 0 | 0 | 0 | 0 | NA | NA | NA | NA | NA | NA | NA | NA | NA | NA |
| 4930524O05 | 0 | 0 | 0 | 0 | 0 | 0 | 0 | 0 | 0 | 0 | 0 | 0 | 0 | 0 | 0 | NA | NA | NA | NA | NA | NA | NA | NA | NA | NA |
| 4930524O08 | 0 | 0 | 0 | 0 | 0 | 0 | 0 | 0 | 0 | 0 | 0 | 0 | 0 | 0 | 0 | NA | NA | NA | NA | NA | NA | NA | NA | NA | NA |
| 4930525D18 | 0 | 0 | 1 | 0 | 1 | 0 | 0 | 0 | 1 | 0 | 1 | 0 | 0.37 | 0 | 0 | 1.075 | 0.105 | 0.97514588 | NA | 0 | 6.4454 | 6.1578 | 6.1578 | TRUE | 0.5365 |
| 4930525G20 | 5 | 3 | 3 | 6 | 3 | 5 | 4 | 3 | 3 | 6 | 3 | 5 | 4.1 | 3 | 5 | 1.384 | 0.469 | 0.6109546 | NA | 0 | 0.5804 | 0.365 | 0.365 | TRUE | 0.1093 |
| 4930525M21 | 0 | 0 | 0 | 0 | 0 | 0 | 0 | 0 | 0 | 0 | 0 | 0 | 0 | 0 | 0 | NA | NA | NA | NA | NA | NA | NA | NA | NA | NA |
| 4930526D03 | 2 | 2 | 1 | 3 | 0 | 1 | 2 | 2 | 1 | 3 | 0 | 1 | 1.46 | 2 | 1 | 0.849 | -0.237 | 0.88245499 | NA | 0 | 1.6308 | 1.1578 | 1.1578 | TRUE | 0.5795 |
| 4930526I15R | 106 | 97 | 72 | 90 | 88 | 94 | 82 | 103 | 78 | 90 | 99 | 93 | 90.81 | 88 | 94 | 1.071 | 0.099 | 0.63729072 | 0.77337377 | 0 | 0.0285 | 0.021 | 0.021 | TRUE | 0.2139 |
| 4930526L06F | 30 | 16 | 22 | 31 | 19 | 26 | 23 | 17 | 24 | 31 | 21 | 26 | 23.67 | 21 | 26 | 1.216 | 0.282 | 0.47924792 | 0.64621609 | 0 | 0.1026 | 0.0722 | 0.0722 | TRUE | 0.1676 |
| 4930527F14I | 0 | 1 | 0 | 0 | 0 | 0 | 0 | 1 | 0 | 0 | 0 | 0 | 0.18 | 0 | 0 | 0.552 | -0.857 | 0.8336548 | NA | 0 | 13.3714 | 10 | 10 | TRUE | 0.5216 |
| 4930527G23 | 0 | 0 | 0 | 0 | 0 | 0 | 0 | 0 | 0 | 0 | 0 | 0 | 0 | 0 | 0 | NA | NA | NA | NA | NA | NA | NA | NA | NA | NA |
| 4930528A17 | 41 | 18 | 14 | 34 | 17 | 15 | 32 | 19 | 15 | 34 | 19 | 15 | 22.32 | 22 | 23 | 1.015 | 0.021 | 0.9641839 | 0.98029759 | 0.1253 | 0.1086 | 0.1135 | 0.1135 | TRUE | 1.1173 |
| 4930528D03 | 6 | 9 | 5 | 6 | 2 | 6 | 5 | 10 | 5 | 6 | 2 | 6 | 5.63 | 7 | 5 | 0.735 | -0.445 | 0.58954185 | NA | 0 | 0.4234 | 0.307 | 0.307 | TRUE | 0.3921 |
| 4930528P14I | 0 | 0 | 0 | 0 | 0 | 0 | 0 | 0 | 0 | 0 | 0 | 0 | 0 | 0 | 0 | NA | NA | NA | NA | NA | NA | NA | NA | NA | NA |
| 4930529C04I | 0 | 1 | 0 | 0 | 1 | 0 | 0 | 1 | 0 | 0 | 1 | 0 | 0.37 | 0 | 0 | 1.075 | 0.105 | 0.97520514 | NA | 0 | 6.4904 | 6.1969 | 6.1969 | TRUE | 0.5216 |
| 4930529F24I | 0 | 0 | 0 | 0 | 0 | 0 | 0 | 0 | 0 | 0 | 0 | 0 | 0 | 0 | 0 | NA | NA | NA | NA | NA | NA | NA | NA | NA | NA |
| 4930529K09I | 0 | 0 | 0 | 0 | 0 | 0 | 0 | 0 | 0 | 0 | 0 | 0 | 0 | 0 | 0 | NA | NA | NA | NA | NA | NA | NA | NA | NA | NA |
| 4930529L06F | 0 | 0 | 0 | 0 | 0 | 0 | 0 | 0 | 0 | 0 | 0 | 0 | 0 | 0 | 0 | NA | NA | NA | NA | NA | NA | NA | NA | NA | NA |
| 4930529M08 | 12 | 15 | 17 | 24 | 21 | 17 | 9 | 16 | 18 | 24 | 24 | 17 | 18 | 14 | 22 | 1.496 | 0.581 | 0.21885563 | 0.38194182 | 0 | 0.1341 | 0.103 | 0.103 | TRUE | 0.5724 |
| 4930533B01 | 0 | 0 | 0 | 0 | 0 | 0 | 0 | 0 | 0 | 0 | 0 | 0 | 0 | 0 | 0 | NA | NA | NA | NA | NA | NA | NA | NA | NA | NA |
| 4930533P14I | 0 | 0 | 0 | 1 | 0 | 0 | 0 | 0 | 0 | 1 | 0 | 0 | 0.17 | 0 | 0 | 2.094 | 1.067 | 0.79379329 | NA | 0 | 14.2885 | 10 | 10 | TRUE | 0.3906 |
| 4930538K18I | 10 | 11 | 18 | 21 | 19 | 16 | 8 | 12 | 19 | 21 | 21 | 16 | 16.17 | 13 | 19 | 1.518 | 0.602 | 0.2357378 | NA | 0.044 | 0.149 | 0.1213 | 0.1213 | TRUE | 0.657 |
| 4930539C22I | 0 | 0 | 0 | 0 | 0 | 0 | 0 | 0 | 0 | 0 | 0 | 0 | 0 | 0 | 0 | NA | NA | NA | NA | NA | NA | NA | NA | NA | NA |
| 4930539E08I | 474 | 373 | 393 | 404 | 350 | 420 | 368 | 397 | 424 | 402 | 395 | 415 | 400.09 | 396 | 404 | 1.022 | 0.031 | 0.77505326 | 0.86721285 | 0 | 0.0084 | 0.0059 | 0.0059 | TRUE | 0.0481 |

| | | | | | | | | | | | | | | | | | | | | | | | | | |
|---|---|---|---|---|---|---|---|---|---|---|---|---|---|---|---|---|---|---|---|---|---|---|---|---|---|
| 4930539J05F | 4 | 1 | 2 | 3 | 4 | 3 | 3 | 1 | 2 | 3 | 5 | 3 | 2.8 | 2 | 4 | 1.61 | 0.687 | 0.54268952 | NA | 0 | 0.8495 | 0.5451 | 0.5451 | TRUE | | 0.2202 |
| 4930539M17 | 7 | 5 | 5 | 2 | 0 | 2 | 5 | 5 | 5 | 2 | 0 | 2 | 3.35 | 5 | 1 | 0.252 | -1.989 | 0.08578827 | NA | 0 | 0.7094 | 0.4951 | 0.4951 | TRUE | | 0.3873 |
| 4930539N22 | 0 | 0 | 0 | 0 | 0 | 0 | 0 | 0 | 0 | 0 | 0 | 0 | 0 | 0 | 0 | NA | NA | NA | NA | NA | NA | NA | NA | NA | NA | |
| 4930540M03 | 0 | 0 | 0 | 0 | 0 | 0 | 0 | 0 | 0 | 0 | 0 | 0 | 0 | 0 | 0 | NA | NA | NA | NA | NA | NA | NA | NA | NA | NA | |
| 4930542C21I | 0 | 0 | 0 | 0 | 0 | 0 | 0 | 0 | 0 | 0 | 0 | 0 | 0 | 0 | 0 | NA | NA | NA | NA | NA | NA | NA | NA | NA | NA | |
| 4930542D17I | 0 | 1 | 0 | 1 | 0 | 0 | 0 | 1 | 0 | 1 | 0 | 0 | 0.34 | 0 | 0 | 1.075 | 0.105 | 0.97573409 | NA | 0 | 6.9086 | 6.5582 | 6.5582 | TRUE | | 0.5216 |
| 4930542H20I | 0 | 0 | 0 | 0 | 0 | 0 | 0 | 0 | 0 | 0 | 0 | 0 | 0 | 0 | 0 | NA | NA | NA | NA | NA | NA | NA | NA | NA | NA | |
| 4930543E12I | 6 | 3 | 1 | 4 | 0 | 3 | 5 | 3 | 1 | 4 | 0 | 3 | 2.65 | 3 | 2 | 0.776 | -0.366 | 0.7702915 | NA | 0.2223 | 0.8983 | 0.7551 | 0.7551 | TRUE | | 0.4813 |
| 4930544D05I | 1 | 0 | 0 | 0 | 0 | 0 | 1 | 0 | 0 | 0 | 0 | 0 | 0.13 | 0 | 0 | 0.552 | -0.857 | 0.8336548 | NA | 0 | 18.3121 | 10 | 10 | TRUE | | 0.2566 |
| 4930544G11 | 0 | 0 | 0 | 0 | 0 | 0 | 0 | 0 | 0 | 0 | 0 | 0 | 0 | 0 | 0 | NA | NA | NA | NA | NA | NA | NA | NA | NA | NA | |
| 4930544M13 | 0 | 0 | 0 | 0 | 0 | 0 | 0 | 0 | 0 | 0 | 0 | 0 | 0 | 0 | 0 | NA | NA | NA | NA | NA | NA | NA | NA | NA | NA | |
| 4930545E07I | 0 | 0 | 0 | 0 | 0 | 0 | 0 | 0 | 0 | 0 | 0 | 0 | 0 | 0 | 0 | NA | NA | NA | NA | NA | NA | NA | NA | NA | NA | |
| 4930545H06I | 0 | 0 | 1 | 0 | 0 | 0 | 0 | 0 | 1 | 0 | 0 | 0 | 0.18 | 0 | 0 | 0.552 | -0.857 | 0.8336548 | NA | 0 | 13.1817 | 10 | 10 | TRUE | | 0.5365 |
| 4930545L23F | 0 | 0 | 0 | 0 | 0 | 0 | 0 | 0 | 0 | 0 | 0 | 0 | 0 | 0 | 0 | NA | NA | NA | NA | NA | NA | NA | NA | NA | NA | |
| 4930546C10I | 0 | 0 | 0 | 0 | 0 | 0 | 0 | 0 | 0 | 0 | 0 | 0 | 0 | 0 | 0 | NA | NA | NA | NA | NA | NA | NA | NA | NA | NA | |
| 4930546K05I | 3 | 2 | 2 | 1 | 0 | 0 | 2 | 2 | 2 | 1 | 0 | 0 | 1.27 | 2 | 0 | 0.168 | -2.574 | 0.16863336 | NA | 0 | 1.8709 | 1.3012 | 1.3012 | TRUE | | 0.3906 |
| 4930547C10I | 8 | 1 | 0 | 5 | 4 | 4 | 6 | 1 | 0 | 5 | 5 | 4 | 3.45 | 2 | 5 | 1.741 | 0.8 | 0.48640016 | NA | 0.4992 | 0.6889 | 0.6478 | 0.6478 | TRUE | | 2.5595 |
| 4930547E08I | 2 | 3 | 5 | 7 | 5 | 4 | 2 | 3 | 5 | 7 | 6 | 4 | 4.45 | 3 | 6 | 1.675 | 0.744 | 0.42140138 | NA | 0 | 0.5351 | 0.3774 | 0.3774 | TRUE | | 0.4374 |
| 4930547E14I | 0 | 0 | 0 | 0 | 0 | 0 | 0 | 0 | 0 | 0 | 0 | 0 | 0 | 0 | 0 | NA | NA | NA | NA | NA | NA | NA | NA | NA | NA | |
| 4930548G14 | 0 | 0 | 0 | 0 | 0 | 0 | 0 | 0 | 0 | 0 | 0 | 0 | 0 | 0 | 0 | NA | NA | NA | NA | NA | NA | NA | NA | NA | NA | |
| 4930548H24I | 0 | 0 | 1 | 3 | 0 | 0 | 0 | 0 | 1 | 3 | 0 | 0 | 0.68 | 0 | 1 | 2.911 | 1.541 | 0.57330413 | NA | 7.7556 | 3.5005 | 3.8827 | 3.8827 | TRUE | | 1.4418 |
| 4930548J01F | 0 | 0 | 0 | 0 | 0 | 0 | 0 | 0 | 0 | 0 | 0 | 0 | 0 | 0 | 0 | NA | NA | NA | NA | NA | NA | NA | NA | NA | NA | |
| 4930548K13I | 0 | 0 | 0 | 0 | 0 | 0 | 0 | 0 | 0 | 0 | 0 | 0 | 0 | 0 | 0 | NA | NA | NA | NA | NA | NA | NA | NA | NA | NA | |
| 4930549C01I | 1 | 0 | 0 | 0 | 0 | 0 | 1 | 0 | 0 | 0 | 0 | 0 | 0.13 | 0 | 0 | 0.552 | -0.857 | 0.8336548 | NA | 0 | 18.3121 | 10 | 10 | TRUE | | 0.2566 |
| 4930549G23 | 1 | 1 | 2 | 0 | 0 | 0 | 1 | 1 | 2 | 0 | 0 | 0 | 0.67 | 1 | 0 | 0.144 | -2.794 | 0.24876697 | NA | 0 | 3.559 | 2.8631 | 2.8631 | TRUE | | 0.1682 |
| 4930550C14I | 24 | 16 | 13 | 19 | 17 | 16 | 19 | 17 | 14 | 19 | 19 | 16 | 17.26 | 17 | 18 | 1.077 | 0.108 | 0.81305073 | 0.89199678 | 0 | 0.1397 | 0.0915 | 0.0915 | TRUE | | 0.0813 |
| 4930550L24F | 161 | 134 | 83 | 23 | 14 | 8 | 125 | 143 | 90 | 23 | 16 | 8 | 67.29 | 119 | 16 | 0.13 | -2.942 | 4.43E-18 | 1.04E-15 | 0.0851 | 0.0377 | 0.0456 | 0.0456 | TRUE | | 0.5117 |
| 4930552N02 | 0 | 0 | 0 | 0 | 0 | 0 | 0 | 0 | 0 | 0 | 0 | 0 | 0 | 0 | 0 | NA | NA | NA | NA | NA | NA | NA | NA | NA | NA | |
| 4930552P12I | 23 | 15 | 28 | 12 | 15 | 17 | 18 | 16 | 30 | 12 | 17 | 17 | 18.28 | 21 | 15 | 0.716 | -0.482 | 0.30681052 | 0.47941868 | 0 | 0.1321 | 0.1038 | 0.1038 | TRUE | | 0.6917 |
| 4930553E22I | 0 | 0 | 0 | 0 | 0 | 0 | 0 | 0 | 0 | 0 | 0 | 0 | 0 | 0 | 0 | NA | NA | NA | NA | NA | NA | NA | NA | NA | NA | |
| 4930554C24I | 0 | 0 | 0 | 0 | 0 | 0 | 0 | 0 | 0 | 0 | 0 | 0 | 0 | 0 | 0 | NA | NA | NA | NA | NA | NA | NA | NA | NA | NA | |
| 4930555B11I | 0 | 0 | 0 | 0 | 0 | 0 | 0 | 0 | 0 | 0 | 0 | 0 | 0 | 0 | 0 | NA | NA | NA | NA | NA | NA | NA | NA | NA | NA | |
| 4930555G01 | 1 | 2 | 3 | 1 | 4 | 2 | 1 | 2 | 3 | 1 | 5 | 2 | 2.27 | 2 | 3 | 1.236 | 0.305 | 0.81280371 | NA | 0 | 1.0463 | 0.7463 | 0.7463 | TRUE | | 0.4831 |
| 4930556C24I | 0 | 0 | 0 | 0 | 0 | 0 | 0 | 0 | 0 | 0 | 0 | 0 | 0 | 0 | 0 | NA | NA | NA | NA | NA | NA | NA | NA | NA | NA | |
| 4930556G01 | 2 | 0 | 1 | 0 | 0 | 0 | 2 | 0 | 1 | 0 | 0 | 0 | 0.44 | 1 | 0 | 0.214 | -2.223 | 0.47051571 | NA | 0 | 5.4047 | 5.2946 | 5.2946 | TRUE | | 0.2788 |
| 4930556J02F | 0 | 0 | 0 | 0 | 0 | 0 | 0 | 0 | 0 | 0 | 0 | 0 | 0 | 0 | 0 | NA | NA | NA | NA | NA | NA | NA | NA | NA | NA | |
| 4930556M19 | 18 | 17 | 19 | 32 | 22 | 22 | 14 | 18 | 21 | 32 | 25 | 22 | 21.83 | 18 | 26 | 1.506 | 0.59 | 0.1597751 | 0.30927214 | 0 | 0.111 | 0.0795 | 0.0795 | TRUE | | 0.2351 |
| 4930556N09 | 0 | 0 | 0 | 0 | 0 | 0 | 0 | 0 | 0 | 0 | 0 | 0 | 0 | 0 | 0 | NA | NA | NA | NA | NA | NA | NA | NA | NA | NA | |
| 4930557A04I | 0 | 0 | 0 | 0 | 0 | 0 | 0 | 0 | 0 | 0 | 0 | 0 | 0 | 0 | 0 | NA | NA | NA | NA | NA | NA | NA | NA | NA | NA | |
| 4930557J02F | 0 | 0 | 0 | 0 | 0 | 0 | 0 | 0 | 0 | 0 | 0 | 0 | 0 | 0 | 0 | NA | NA | NA | NA | NA | NA | NA | NA | NA | NA | |
| 4930558C23I | 8 | 4 | 7 | 2 | 0 | 1 | 6 | 4 | 8 | 2 | 0 | 1 | 3.5 | 6 | 1 | 0.17 | -2.558 | 0.03653739 | NA | 0 | 0.6796 | 0.4916 | 0.4916 | TRUE | | 0.3518 |
| 4930558G05 | 0 | 0 | 0 | 0 | 0 | 0 | 0 | 0 | 0 | 0 | 0 | 0 | 0 | 0 | 0 | NA | NA | NA | NA | NA | NA | NA | NA | NA | NA | |
| 4930558J18F | 204 | 148 | 167 | 194 | 144 | 163 | 158 | 157 | 180 | 193 | 162 | 161 | 168.75 | 165 | 172 | 1.043 | 0.061 | 0.70085227 | 0.81860599 | 0 | 0.0165 | 0.0125 | 0.0125 | TRUE | | 0.1229 |
| 4930558K02I | 1 | 2 | 1 | 3 | 5 | 2 | 1 | 2 | 1 | 3 | 6 | 2 | 2.43 | 1 | 4 | 2.701 | 1.433 | 0.26202577 | NA | 0 | 0.9775 | 0.6587 | 0.6587 | TRUE | | 0.3645 |
| 4930562C15I | 11 | 3 | 5 | 7 | 6 | 7 | 9 | 3 | 5 | 7 | 7 | 7 | 6.3 | 6 | 7 | 1.182 | 0.241 | 0.75176873 | NA | 0 | 0.3789 | 0.2612 | 0.2612 | TRUE | | 0.4285 |
| 4930562F07I | 2 | 0 | 1 | 0 | 0 | 0 | 2 | 0 | 1 | 0 | 0 | 0 | 0.44 | 1 | 0 | 0.214 | -2.223 | 0.47051571 | NA | 0 | 5.4047 | 5.2946 | 5.2946 | TRUE | | 0.2788 |
| 4930563D23I | 0 | 1 | 0 | 1 | 1 | 0 | 0 | 1 | 0 | 1 | 1 | 0 | 0.53 | 0 | 1 | 2.036 | 1.026 | 0.70492654 | NA | 0 | 4.4642 | 3.5518 | 3.5518 | TRUE | | 0.5217 |
| 4930563E18I | 3 | 0 | 1 | 2 | 2 | 0 | 2 | 0 | 1 | 2 | 2 | 0 | 1.28 | 1 | 1 | 1.19 | 0.251 | 0.88877104 | NA | 0 | 1.8598 | 1.5429 | 1.5429 | TRUE | | 0.3097 |
| 4930563E22I | 101 | 108 | 87 | 92 | 81 | 96 | 78 | 115 | 94 | 92 | 91 | 95 | 94.15 | 96 | 93 | 0.973 | -0.04 | 0.85260076 | 0.9167369 | 0.01 | 0.0276 | 0.0224 | 0.0224 | TRUE | | 0.273 |
| 4930563F08I | 0 | 0 | 0 | 0 | 0 | 0 | 0 | 0 | 0 | 0 | 0 | 0 | 0 | 0 | 0 | NA | NA | NA | NA | NA | NA | NA | NA | NA | NA | |
| 4930563M20 | 1 | 1 | 0 | 2 | 2 | 1 | 1 | 1 | 0 | 2 | 2 | 1 | 1.18 | 1 | 2 | 2.782 | 1.476 | 0.4136547 | NA | 0 | 2.0125 | 1.2823 | 1.2823 | TRUE | | 0.187 |
| 4930564B18I | 0 | 0 | 0 | 0 | 0 | 0 | 0 | 0 | 0 | 0 | 0 | 0 | 0 | 0 | 0 | NA | NA | NA | NA | NA | NA | NA | NA | NA | NA | |
| 4930564C03I | 0 | 0 | 0 | 0 | 0 | 0 | 0 | 0 | 0 | 0 | 0 | 0 | 0 | 0 | 0 | NA | NA | NA | NA | NA | NA | NA | NA | NA | NA | |
| 4930564D02I | 0 | 0 | 0 | 0 | 0 | 0 | 0 | 0 | 0 | 0 | 0 | 0 | 0 | 0 | 0 | NA | NA | NA | NA | NA | NA | NA | NA | NA | NA | |
| 4930564G21 | 0 | 1 | 2 | 0 | 2 | 1 | 0 | 1 | 2 | 0 | 2 | 1 | 1.08 | 1 | 1 | 1.028 | 0.04 | 0.98369621 | NA | 0 | 2.202 | 1.8158 | 1.8158 | TRUE | | 0.2386 |
| 4930564K09I | 0 | 0 | 0 | 1 | 0 | 0 | 0 | 0 | 0 | 1 | 0 | 0 | 0.17 | 0 | 0 | 2.094 | 1.067 | 0.79379329 | NA | 0 | 14.2885 | 10 | 10 | TRUE | | 0.3906 |
| 4930565D16I | 0 | 0 | 0 | 0 | 0 | 0 | 0 | 0 | 0 | 0 | 0 | 0 | 0 | 0 | 0 | NA | NA | NA | NA | NA | NA | NA | NA | NA | NA | |
| 4930565N06 | 0 | 0 | 0 | 0 | 0 | 0 | 0 | 0 | 0 | 0 | 0 | 0 | 0 | 0 | 0 | NA | NA | NA | NA | NA | NA | NA | NA | NA | NA | |
| 4930567H12I | 0 | 0 | 0 | 0 | 0 | 0 | 0 | 0 | 0 | 0 | 0 | 0 | 0 | 0 | 0 | NA | NA | NA | NA | NA | NA | NA | NA | NA | NA | |
| 4930567H17I | 0 | 0 | 0 | 0 | 0 | 0 | 0 | 0 | 0 | 0 | 0 | 0 | 0 | 0 | 0 | NA | NA | NA | NA | NA | NA | NA | NA | NA | NA | |
| 4930567J20F | 0 | 0 | 0 | 0 | 0 | 0 | 0 | 0 | 0 | 0 | 0 | 0 | 0 | 0 | 0 | NA | NA | NA | NA | NA | NA | NA | NA | NA | NA | |
| 4930567K20I | 0 | 0 | 0 | 0 | 0 | 0 | 0 | 0 | 0 | 0 | 0 | 0 | 0 | 0 | 0 | NA | NA | NA | NA | NA | NA | NA | NA | NA | NA | |
| 4930568D16I | 0 | 0 | 0 | 0 | 0 | 0 | 0 | 0 | 0 | 0 | 0 | 0 | 0 | 0 | 0 | NA | NA | NA | NA | NA | NA | NA | NA | NA | NA | |
| 4930568E12I | 23 | 13 | 8 | 1 | 4 | 0 | 18 | 14 | 9 | 1 | 5 | 0 | 7.64 | 14 | 2 | 0.13 | -2.944 | 0.00151498 | NA | 0.1437 | 0.3127 | 0.2911 | 0.2911 | TRUE | | 0.759 |
| 4930568G15 | 0 | 0 | 0 | 0 | 0 | 0 | 0 | 0 | 0 | 0 | 0 | 0 | 0 | 0 | 0 | NA | NA | NA | NA | NA | NA | NA | NA | NA | NA | |
| 4930568K20I | 0 | 0 | 0 | 0 | 0 | 0 | 0 | 0 | 0 | 0 | 0 | 0 | 0 | 0 | 0 | NA | NA | NA | NA | NA | NA | NA | NA | NA | NA | |
| 4930570G19 | 0 | 0 | 0 | 0 | 0 | 0 | 0 | 0 | 0 | 0 | 0 | 0 | 0 | 0 | 0 | NA | NA | NA | NA | NA | NA | NA | NA | NA | NA | |
| 4930571K23I | 0 | 0 | 0 | 0 | 0 | 0 | 0 | 0 | 0 | 0 | 0 | 0 | 0 | 0 | 0 | NA | NA | NA | NA | NA | NA | NA | NA | NA | NA | |
| 4930572K03I | 0 | 0 | 0 | 0 | 0 | 0 | 0 | 0 | 0 | 0 | 0 | 0 | 0 | 0 | 0 | NA | NA | NA | NA | NA | NA | NA | NA | NA | NA | |

| | | | | | | | | | | | | | | | | | | | | | | | | | |
|---|---|---|---|---|---|---|---|---|---|---|---|---|---|---|---|---|---|---|---|---|---|---|---|---|---|
| 4930572O03 | 1 | 0 | 0 | 0 | 0 | 0 | 1 | 0 | 0 | 0 | 0 | 0 | 0.13 | 0 | 0 | 0.552 | -0.857 | 0.8336548 | NA | 0 | 18.3121 | 10 | 10 | TRUE | 0.2566 |
| 4930572O13 | 0 | 0 | 0 | 0 | 0 | 0 | 0 | 0 | 0 | 0 | 0 | 0 | 0 | 0 | 0 | NA | NA | NA | NA | NA | NA | NA | NA | NA | NA |
| 4930573O16 | 0 | 0 | 0 | 0 | 0 | 0 | 0 | 0 | 0 | 0 | 0 | 0 | 0 | 0 | 0 | NA | NA | NA | NA | NA | NA | NA | NA | NA | NA |
| 4930577N17 | 11 | 3 | 1 | 5 | 2 | 4 | 9 | 3 | 1 | 5 | 2 | 4 | 4 | 4 | 4 | 0.85 | -0.234 | 0.81832671 | NA | 0.2757 | 0.5951 | 0.499 | 0.499 | TRUE | 0.7559 |
| 4930578C19I | 0 | 0 | 0 | 0 | 0 | 0 | 0 | 0 | 0 | 0 | 0 | 0 | 0 | 0 | 0 | NA | NA | NA | NA | NA | NA | NA | NA | NA | NA |
| 4930578E11I | 0 | 0 | 0 | 0 | 0 | 0 | 0 | 0 | 0 | 0 | 0 | 0 | 0 | 0 | 0 | NA | NA | NA | NA | NA | NA | NA | NA | NA | NA |
| 4930578I06R | 0 | 0 | 0 | 0 | 0 | 0 | 0 | 0 | 0 | 0 | 0 | 0 | 0 | 0 | 0 | NA | NA | NA | NA | NA | NA | NA | NA | NA | NA |
| 4930578M01 | 3 | 1 | 0 | 0 | 1 | 2 | 2 | 1 | 0 | 0 | 1 | 2 | 1.08 | 1 | 1 | 0.882 | -0.182 | 0.92471943 | NA | 0 | 2.1915 | 1.7495 | 1.7495 | TRUE | 0.2446 |
| 4930578N18 | 0 | 0 | 0 | 0 | 0 | 0 | 0 | 0 | 0 | 0 | 0 | 0 | 0 | 0 | 0 | NA | NA | NA | NA | NA | NA | NA | NA | NA | NA |
| 4930579F01I | 0 | 0 | 1 | 0 | 0 | 0 | 0 | 0 | 1 | 0 | 0 | 0 | 0.18 | 0 | 0 | 0.552 | -0.857 | 0.8336548 | NA | 0 | 13.1817 | 10 | 10 | TRUE | 0.5365 |
| 4930579G18 | 5 | 4 | 5 | 6 | 6 | 4 | 4 | 4 | 5 | 6 | 7 | 4 | 5.04 | 4 | 6 | 1.236 | 0.306 | 0.7140117 | NA | 0 | 0.4729 | 0.3023 | 0.3023 | TRUE | 0.1474 |
| 4930579G24 | 1172 | 855 | 796 | 789 | 670 | 799 | 910 | 909 | 859 | 785 | 756 | 789 | 834.75 | 893 | 777 | 0.87 | -0.201 | 0.01124847 | 0.04504745 | 0 | 0.0053 | 0.0033 | 0.0033 | TRUE | 0.0128 |
| 4930579K19I | 18 | 13 | 13 | 11 | 12 | 15 | 14 | 14 | 14 | 11 | 14 | 15 | 13.52 | 14 | 13 | 0.939 | -0.091 | 0.85837964 | NA | 0 | 0.1777 | 0.1138 | 0.1138 | TRUE | 0.0894 |
| 4930581F22I | 11 | 13 | 11 | 10 | 5 | 7 | 9 | 14 | 12 | 10 | 6 | 7 | 9.46 | 12 | 8 | 0.666 | -0.587 | 0.35199763 | NA | 0 | 0.253 | 0.175 | 0.175 | TRUE | 0.2376 |
| 4930583K01I | 2 | 0 | 1 | 0 | 2 | 1 | 2 | 0 | 1 | 0 | 2 | 1 | 0.98 | 1 | 1 | 1.179 | 0.238 | 0.90541994 | NA | 0 | 2.4226 | 1.8634 | 1.8634 | TRUE | 0.2099 |
| 4930583P06I | 0 | 0 | 0 | 0 | 0 | 0 | 0 | 0 | 0 | 0 | 0 | 0 | 0 | 0 | 0 | NA | NA | NA | NA | NA | NA | NA | NA | NA | NA |
| 4930584F24I | 0 | 0 | 0 | 0 | 0 | 0 | 0 | 0 | 0 | 0 | 0 | 0 | 0 | 0 | 0 | NA | NA | NA | NA | NA | NA | NA | NA | NA | NA |
| 4930590J08F | 21 | 19 | 14 | 16 | 10 | 19 | 16 | 20 | 15 | 16 | 11 | 19 | 16.26 | 17 | 15 | 0.896 | -0.159 | 0.73996199 | NA | 0 | 0.1481 | 0.1027 | 0.1027 | TRUE | 0.2204 |
| 4930590L20F | 4 | 3 | 3 | 0 | 2 | 1 | 3 | 3 | 3 | 0 | 2 | 1 | 2.13 | 3 | 1 | 0.332 | -1.592 | 0.25629765 | NA | 0 | 1.1153 | 0.7739 | 0.7739 | TRUE | 0.3992 |
| 4930591A17I | 239 | 160 | 130 | 110 | 86 | 102 | 186 | 170 | 140 | 109 | 97 | 101 | 133.88 | 165 | 102 | 0.618 | -0.695 | 0.00014242 | 0.00163372 | 0 | 0.0201 | 0.0161 | 0.0161 | TRUE | 0.1818 |
| 4930592A05I | 0 | 0 | 0 | 0 | 0 | 0 | 0 | 0 | 0 | 0 | 0 | 0 | 0 | 0 | 0 | NA | NA | NA | NA | NA | NA | NA | NA | NA | NA |
| 4930592I03R | 0 | 0 | 0 | 0 | 0 | 0 | 0 | 0 | 0 | 0 | 0 | 0 | 0 | 0 | 0 | NA | NA | NA | NA | NA | NA | NA | NA | NA | NA |
| 4930593A02I | 0 | 0 | 0 | 0 | 1 | 0 | 0 | 0 | 0 | 0 | 1 | 0 | 0.19 | 0 | 0 | 2.094 | 1.067 | 0.79379329 | NA | 0 | 12.608 | 10 | 10 | TRUE | 0.5075 |
| 4930593C16I | 0 | 0 | 0 | 0 | 0 | 0 | 0 | 0 | 0 | 0 | 0 | 0 | 0 | 0 | 0 | NA | NA | NA | NA | NA | NA | NA | NA | NA | NA |
| 4930594C11I | 4 | 0 | 3 | 4 | 1 | 3 | 3 | 0 | 3 | 4 | 1 | 3 | 2.4 | 2 | 3 | 1.262 | 0.335 | 0.79551919 | NA | 0 | 0.9889 | 0.7913 | 0.7913 | TRUE | 0.6529 |
| 4930595M18 | 0 | 0 | 0 | 0 | 0 | 0 | 0 | 0 | 0 | 0 | 0 | 0 | 0 | 0 | 0 | NA | NA | NA | NA | NA | NA | NA | NA | NA | NA |
| 4930596D02I | 0 | 0 | 0 | 0 | 0 | 0 | 0 | 0 | 0 | 0 | 0 | 0 | 0 | 0 | 0 | NA | NA | NA | NA | NA | NA | NA | NA | NA | NA |
| 4930596M17 | 0 | 0 | 0 | 0 | 0 | 0 | 0 | 0 | 0 | 0 | 0 | 0 | 0 | 0 | 0 | NA | NA | NA | NA | NA | NA | NA | NA | NA | NA |
| 4930597G03 | 0 | 0 | 0 | 0 | 0 | 0 | 0 | 0 | 0 | 0 | 0 | 0 | 0 | 0 | 0 | NA | NA | NA | NA | NA | NA | NA | NA | NA | NA |
| 4930598F16I | 0 | 0 | 0 | 0 | 0 | 0 | 0 | 0 | 0 | 0 | 0 | 0 | 0 | 0 | 0 | NA | NA | NA | NA | NA | NA | NA | NA | NA | NA |
| 4930599N23 | 4 | 4 | 2 | 1 | 2 | 2 | 3 | 4 | 2 | 1 | 2 | 2 | 2.46 | 3 | 2 | 0.546 | -0.873 | 0.47136235 | NA | 0 | 0.9668 | 0.6088 | 0.6088 | TRUE | 0.1146 |
| 4931402G19 | 0 | 0 | 0 | 0 | 0 | 0 | 0 | 0 | 0 | 0 | 0 | 0 | 0 | 0 | 0 | NA | NA | NA | NA | NA | NA | NA | NA | NA | NA |
| 4931403E22I | 0 | 0 | 0 | 1 | 0 | 0 | 0 | 0 | 0 | 1 | 0 | 0 | 0.17 | 0 | 0 | 2.094 | 1.067 | 0.79379329 | NA | 0 | 14.2885 | 10 | 10 | TRUE | 0.3906 |
| 4931403G20 | 0 | 0 | 0 | 0 | 0 | 0 | 0 | 0 | 0 | 0 | 0 | 0 | 0 | 0 | 0 | NA | NA | NA | NA | NA | NA | NA | NA | NA | NA |
| 4931406B18I | 2 | 1 | 3 | 2 | 2 | 3 | 2 | 1 | 3 | 2 | 2 | 3 | 2.18 | 2 | 2 | 1.246 | 0.317 | 0.8018456 | NA | 0 | 1.0911 | 0.6845 | 0.6845 | TRUE | 0.26 |
| 4931406C07I | 1006 | 686 | 753 | 981 | 848 | 884 | 781 | 730 | 812 | 976 | 957 | 873 | 854.88 | 774 | 935 | 1.207 | 0.272 | 0.00154879 | 0.00988252 | 0.002 | 0.0052 | 0.0041 | 0.0041 | TRUE | 0.0415 |
| 4931406H21I | 3 | 1 | 0 | 1 | 1 | 0 | 2 | 1 | 0 | 1 | 1 | 0 | 0.92 | 1 | 1 | 0.598 | -0.741 | 0.72094137 | NA | 0 | 2.5799 | 1.9516 | 1.9516 | TRUE | 0.1813 |
| 4931406P16I | 5795 | 4281 | 3923 | 4304 | 3779 | 4116 | 4501 | 4554 | 4233 | 4284 | 4263 | 4063 | 4316.28 | 4429 | 4203 | 0.949 | -0.076 | 0.18505181 | 0.34128329 | 8.00E-04 | 0.003 | 0.0021 | 0.0021 | TRUE | 0.018 |
| 4931408C20I | 0 | 0 | 0 | 0 | 0 | 0 | 0 | 0 | 0 | 0 | 0 | 0 | 0 | 0 | 0 | NA | NA | NA | NA | NA | NA | NA | NA | NA | NA |
| 4931408D14I | 7 | 6 | 4 | 8 | 6 | 2 | 5 | 6 | 4 | 8 | 7 | 2 | 5.47 | 5 | 6 | 1.031 | 0.043 | 0.95868631 | NA | 0 | 0.4354 | 0.3233 | 0.3233 | TRUE | 0.7388 |
| 4931409K22I | 14 | 9 | 13 | 13 | 9 | 10 | 11 | 10 | 14 | 13 | 10 | 10 | 11.24 | 12 | 11 | 0.96 | -0.059 | 0.91631692 | NA | 0 | 0.2133 | 0.1403 | 0.1403 | TRUE | 0.1285 |
| 4931414P19I | 442 | 343 | 377 | 359 | 297 | 352 | 343 | 365 | 407 | 357 | 335 | 347 | 359.13 | 372 | 346 | 0.935 | -0.097 | 0.40201104 | 0.5750704 | 0.0017 | 0.009 | 0.0068 | 0.0068 | TRUE | 0.0774 |
| 4931417E11I | 0 | 0 | 0 | 0 | 0 | 0 | 0 | 0 | 0 | 0 | 0 | 0 | 0 | 0 | 0 | NA | NA | NA | NA | NA | NA | NA | NA | NA | NA |
| 4931417G12 | 36 | 26 | 16 | 15 | 18 | 22 | 28 | 28 | 17 | 15 | 20 | 22 | 21.64 | 24 | 19 | 0.776 | -0.365 | 0.39147762 | 0.56513379 | 0 | 0.112 | 0.0836 | 0.0836 | TRUE | 0.3562 |
| 4931419H13I | 0 | 0 | 0 | 0 | 0 | 0 | 0 | 0 | 0 | 0 | 0 | 0 | 0 | 0 | 0 | NA | NA | NA | NA | NA | NA | NA | NA | NA | NA |
| 4931420L22F | 0 | 1 | 1 | 0 | 0 | 0 | 0 | 1 | 1 | 0 | 0 | 0 | 0.36 | 1 | 0 | 0.271 | -1.881 | 0.5755355 | NA | 0 | 6.6391 | 6.4428 | 6.4428 | TRUE | 0.3513 |
| 4931423N10 | 3 | 2 | 3 | 0 | 0 | 1 | 2 | 2 | 3 | 0 | 0 | 1 | 1.45 | 2 | 0 | 0.145 | -2.786 | 0.12277827 | NA | 0 | 1.6404 | 1.1603 | 1.1603 | TRUE | 0.3835 |
| 4931428F04I | 942 | 780 | 744 | 823 | 748 | 844 | 732 | 830 | 803 | 819 | 844 | 833 | 810.04 | 788 | 832 | 1.058 | 0.081 | 0.33893173 | 0.51300976 | 0.0012 | 0.0054 | 0.0039 | 0.0039 | TRUE | 0.0494 |
| 4931428L18F | 3 | 1 | 3 | 2 | 1 | 1 | 2 | 1 | 3 | 2 | 1 | 1 | 1.79 | 2 | 1 | 0.62 | -0.689 | 0.62437276 | NA | 0 | 1.3269 | 0.8297 | 0.8297 | TRUE | 0.1815 |
| 4931429I11R | 9 | 3 | 8 | 6 | 10 | 14 | 7 | 3 | 9 | 6 | 11 | 14 | 8.31 | 6 | 10 | 1.644 | 0.717 | 0.30502473 | NA | 0 | 0.2875 | 0.2254 | 0.2254 | TRUE | 0.4135 |
| 4931429L15F | 1 | 3 | 0 | 5 | 0 | 3 | 1 | 3 | 0 | 5 | 0 | 3 | 1.98 | 1 | 3 | 2.051 | 1.036 | 0.50721916 | NA | 1.2017 | 1.1968 | 1.1962 | 1.1962 | TRUE | 0.3017 |
| 4931429P17I | 0 | 0 | 0 | 0 | 0 | 0 | 0 | 0 | 0 | 0 | 0 | 0 | 0 | 0 | 0 | NA | NA | NA | NA | NA | NA | NA | NA | NA | NA |
| 4931430N09 | 0 | 0 | 1 | 0 | 0 | 0 | 0 | 0 | 1 | 0 | 0 | 0 | 0.18 | 0 | 0 | 0.552 | -0.857 | 0.8336548 | NA | 0 | 13.1817 | 10 | 10 | TRUE | 0.5365 |
| 4931431B13I | 0 | 0 | 0 | 0 | 0 | 0 | 0 | 0 | 0 | 0 | 0 | 0 | 0 | 0 | 0 | NA | NA | NA | NA | NA | NA | NA | NA | NA | NA |
| 4931431C16I | 0 | 0 | 0 | 1 | 1 | 1 | 0 | 0 | 0 | 1 | 1 | 1 | 0.52 | 0 | 1 | 5.852 | 2.549 | 0.3348364 | NA | 0 | 4.5739 | 3.5345 | 3.5345 | TRUE | 0.0847 |
| 4931431F19I | 0 | 0 | 0 | 0 | 0 | 0 | 0 | 0 | 0 | 0 | 0 | 0 | 0 | 0 | 0 | NA | NA | NA | NA | NA | NA | NA | NA | NA | NA |
| 4931440F15I | 12 | 2 | 4 | 8 | 3 | 6 | 9 | 2 | 4 | 8 | 3 | 6 | 5.51 | 5 | 6 | 1.073 | 0.102 | 0.90573393 | NA | 0.1547 | 0.4329 | 0.3526 | 0.3526 | TRUE | 0.8013 |
| 4931440J10F | 0 | 0 | 0 | 0 | 0 | 0 | 0 | 0 | 0 | 0 | 0 | 0 | 0 | 0 | 0 | NA | NA | NA | NA | NA | NA | NA | NA | NA | NA |
| 4931440L10F | 27 | 27 | 21 | 34 | 24 | 28 | 21 | 29 | 23 | 34 | 27 | 28 | 26.82 | 24 | 30 | 1.232 | 0.301 | 0.41922205 | 0.59168956 | 0 | 0.0908 | 0.0623 | 0.0623 | TRUE | 0.1598 |
| 4931440P22I | 25 | 13 | 23 | 23 | 17 | 18 | 19 | 14 | 25 | 23 | 19 | 18 | 19.65 | 19 | 20 | 1.032 | 0.045 | 0.91897997 | 0.95541029 | 0 | 0.123 | 0.0889 | 0.0889 | TRUE | 0.1797 |
| 4932411E22I | 0 | 1 | 0 | 0 | 0 | 0 | 0 | 1 | 0 | 0 | 0 | 0 | 0.18 | 0 | 0 | 0.552 | -0.857 | 0.8336548 | NA | 0 | 13.3714 | 10 | 10 | TRUE | 0.5216 |
| 4932411N23 | 6 | 3 | 0 | 1 | 4 | 1 | 5 | 3 | 0 | 1 | 5 | 1 | 2.39 | 3 | 2 | 0.795 | -0.332 | 0.80749493 | NA | 0.657 | 0.9937 | 0.9108 | 0.9108 | TRUE | 0.5013 |
| 4932412D23I | 51 | 54 | 34 | 20 | 29 | 24 | 40 | 57 | 37 | 20 | 33 | 24 | 35.01 | 45 | 26 | 0.569 | -0.812 | 0.0221539 | 0.07432997 | 0.0315 | 0.0701 | 0.0596 | 0.0596 | TRUE | 0.4752 |
| 4932413F04I | 1 | 1 | 2 | 2 | 1 | 2 | 1 | 1 | 2 | 2 | 1 | 2 | 1.52 | 1 | 2 | 1.308 | 0.387 | 0.79622987 | NA | 0 | 1.5666 | 0.9456 | 0.9456 | TRUE | 0.1668 |
| 4932414J04F | 0 | 0 | 0 | 0 | 0 | 0 | 0 | 0 | 0 | 0 | 0 | 0 | 0 | 0 | 0 | NA | NA | NA | NA | NA | NA | NA | NA | NA | NA |
| 4932414N04 | 1 | 0 | 0 | 0 | 0 | 0 | 1 | 0 | 0 | 0 | 0 | 0 | 0.13 | 0 | 0 | 0.552 | -0.857 | 0.8336548 | NA | 0 | 18.3121 | 10 | 10 | TRUE | 0.2566 |
| 4932415G12 | 86 | 75 | 42 | 67 | 52 | 57 | 67 | 80 | 45 | 67 | 59 | 56 | 62.25 | 64 | 61 | 0.945 | -0.081 | 0.76316769 | 0.85943643 | 0.0253 | 0.0405 | 0.0364 | 0.0364 | TRUE | 0.3361 |
| 4932415M13 | 9 | 4 | 2 | 2 | 0 | 0 | 7 | 4 | 2 | 2 | 0 | 0 | 2.57 | 4 | 1 | 0.15 | -2.733 | 0.07059841 | NA | 0 | 0.9262 | 0.7799 | 0.7799 | TRUE | 0.6006 |
| 4932416H05I | 79 | 55 | 52 | 75 | 69 | 63 | 61 | 59 | 56 | 75 | 78 | 62 | 65.11 | 59 | 72 | 1.216 | 0.282 | 0.24191663 | 0.40948773 | 0 | 0.0388 | 0.0264 | 0.0264 | TRUE | 0.1212 |

| | | | | | | | | | | | | | | | | | | | | | | | | | |
|---|---|---|---|---|---|---|---|---|---|---|---|---|---|---|---|---|---|---|---|---|---|---|---|---|---|
| 4932416K20I | 0 | 0 | 0 | 0 | 0 | 0 | 0 | 0 | 0 | 0 | 0 | 0 | 0 | 0 | 0 | NA | NA | NA | NA | NA | NA | NA | NA | NA | NA |
| 4932418E24I | 0 | 0 | 0 | 0 | 2 | 0 | 0 | 0 | 0 | 0 | 2 | 0 | 0.38 | 0 | 1 | 4.195 | 2.069 | 0.57837236 | NA | 10 | 6.3052 | 8.2866 | 8.2866 | TRUE | 0.9743 |
| 4932429P05I | 0 | 0 | 0 | 0 | 0 | 0 | 0 | 0 | 0 | 0 | 0 | 0 | 0 | 0 | 0 | NA | NA | NA | NA | NA | NA | NA | NA | NA | NA |
| 4932435O22 | 1 | 0 | 0 | 2 | 0 | 0 | 1 | 0 | 0 | 2 | 0 | 0 | 0.46 | 0 | 1 | 1.942 | 0.958 | 0.7580687 | NA | 0 | 5.1409 | 5.1921 | 5.1921 | TRUE | 0.9738 |
| 4932438A13 | 5083 | 3567 | 3520 | 4188 | 3632 | 4134 | 3948 | 3794 | 3798 | 4169 | 4097 | 4081 | 3981.15 | 3847 | 4116 | 1.07 | 0.097 | 0.065177 | 0.16297672 | 0 | 0.003 | 0.0017 | 0.0017 | TRUE | 0.0066 |
| 4932438H23 | 0 | 1 | 1 | 0 | 4 | 1 | 0 | 1 | 1 | 0 | 5 | 1 | 1.27 | 1 | 2 | 2.655 | 1.409 | 0.46508953 | NA | 0 | 1.8632 | 1.6836 | 1.6836 | TRUE | 0.7589 |
| 4932441J04F | 0 | 0 | 0 | 0 | 0 | 0 | 0 | 0 | 0 | 0 | 0 | 0 | 0 | 0 | 0 | NA | NA | NA | NA | NA | NA | NA | NA | NA | NA |
| 4932443I19R | 0 | 0 | 1 | 0 | 0 | 1 | 0 | 0 | 1 | 0 | 0 | 1 | 0.34 | 0 | 0 | 1.075 | 0.105 | 0.97570524 | NA | 0 | 6.885 | 6.5379 | 6.5379 | TRUE | 0.5365 |
| 4932702P03I | 0 | 0 | 0 | 0 | 0 | 1 | 0 | 0 | 0 | 0 | 0 | 1 | 0.16 | 0 | 0 | 2.094 | 1.067 | 0.79379329 | NA | 0 | 14.4078 | 10 | 10 | TRUE | 0.3835 |
| 4933400A11 | 2 | 0 | 0 | 1 | 1 | 0 | 2 | 0 | 0 | 1 | 1 | 0 | 0.61 | 1 | 1 | 1.262 | 0.336 | 0.89669251 | NA | 0 | 3.87 | 3.2065 | 3.2065 | TRUE | 0.9721 |
| 4933400B14 | 0 | 0 | 0 | 0 | 0 | 0 | 0 | 0 | 0 | 0 | 0 | 0 | 0 | 0 | 0 | NA | NA | NA | NA | NA | NA | NA | NA | NA | NA |
| 4933400C23I | 0 | 0 | 0 | 0 | 0 | 0 | 0 | 0 | 0 | 0 | 0 | 0 | 0 | 0 | 0 | NA | NA | NA | NA | NA | NA | NA | NA | NA | NA |
| 4933400F21I | 0 | 0 | 0 | 0 | 0 | 0 | 0 | 0 | 0 | 0 | 0 | 0 | 0 | 0 | 0 | NA | NA | NA | NA | NA | NA | NA | NA | NA | NA |
| 4933400L20F | 0 | 0 | 0 | 0 | 0 | 0 | 0 | 0 | 0 | 0 | 0 | 0 | 0 | 0 | 0 | NA | NA | NA | NA | NA | NA | NA | NA | NA | NA |
| 4933401B06 | 0 | 0 | 0 | 0 | 0 | 0 | 0 | 0 | 0 | 0 | 0 | 0 | 0 | 0 | 0 | NA | NA | NA | NA | NA | NA | NA | NA | NA | NA |
| 4933401D09 | 2 | 3 | 0 | 1 | 1 | 1 | 2 | 3 | 0 | 1 | 1 | 1 | 1.31 | 2 | 1 | 0.654 | -0.613 | 0.718172 | NA | 0 | 1.8128 | 1.273 | 1.273 | TRUE | 0.1306 |
| 4933401H06 | 0 | 0 | 0 | 0 | 0 | 0 | 0 | 0 | 0 | 0 | 0 | 0 | 0 | 0 | 0 | NA | NA | NA | NA | NA | NA | NA | NA | NA | NA |
| 4933401P06I | 0 | 0 | 0 | 0 | 0 | 0 | 0 | 0 | 0 | 0 | 0 | 0 | 0 | 0 | 0 | NA | NA | NA | NA | NA | NA | NA | NA | NA | NA |
| 4933402C06I | 0 | 0 | 0 | 0 | 0 | 0 | 0 | 0 | 0 | 0 | 0 | 0 | 0 | 0 | 0 | NA | NA | NA | NA | NA | NA | NA | NA | NA | NA |
| 4933402D24 | 0 | 0 | 0 | 0 | 0 | 0 | 0 | 0 | 0 | 0 | 0 | 0 | 0 | 0 | 0 | NA | NA | NA | NA | NA | NA | NA | NA | NA | NA |
| 4933402E13I | 140 | 118 | 91 | 79 | 47 | 50 | 109 | 126 | 98 | 79 | 53 | 49 | 85.58 | 111 | 60 | 0.546 | -0.874 | 0.00026211 | 0.00262533 | 0.0235 | 0.0301 | 0.0284 | 0.0284 | TRUE | 0.6197 |
| 4933402J07F | 2 | 0 | 0 | 4 | 2 | 2 | 2 | 0 | 0 | 4 | 2 | 2 | 1.63 | 1 | 3 | 4.689 | 2.229 | 0.19281605 | NA | 0 | 1.4586 | 1.1019 | 1.1019 | TRUE | 0.9707 |
| 4933402J10F | 0 | 0 | 0 | 0 | 0 | 0 | 0 | 0 | 0 | 0 | 0 | 0 | 0 | 0 | 0 | NA | NA | NA | NA | NA | NA | NA | NA | NA | NA |
| 4933402J15F | 1 | 0 | 0 | 0 | 0 | 0 | 1 | 0 | 0 | 0 | 0 | 0 | 0.13 | 0 | 0 | 0.552 | -0.857 | 0.8336548 | NA | 0 | 18.3121 | 10 | 10 | TRUE | 0.2566 |
| 4933402N03 | 0 | 0 | 0 | 0 | 0 | 0 | 0 | 0 | 0 | 0 | 0 | 0 | 0 | 0 | 0 | NA | NA | NA | NA | NA | NA | NA | NA | NA | NA |
| 4933402N22 | 1 | 0 | 0 | 0 | 0 | 0 | 1 | 0 | 0 | 0 | 0 | 0 | 0.13 | 0 | 0 | 0.552 | -0.857 | 0.8336548 | NA | 0 | 18.3121 | 10 | 10 | TRUE | 0.2566 |
| 4933402P03I | 0 | 0 | 0 | 0 | 0 | 0 | 0 | 0 | 0 | 0 | 0 | 0 | 0 | 0 | 0 | NA | NA | NA | NA | NA | NA | NA | NA | NA | NA |
| 4933403G14 | 823 | 676 | 548 | 517 | 512 | 536 | 639 | 719 | 591 | 515 | 578 | 529 | 595.14 | 650 | 541 | 0.831 | -0.267 | 0.00964221 | 0.04032067 | 0.005 | 0.0064 | 0.0059 | 0.0059 | TRUE | 0.0988 |
| 4933403O08 | 9 | 4 | 2 | 7 | 0 | 1 | 7 | 4 | 2 | 7 | 0 | 1 | 3.56 | 4 | 3 | 0.594 | -0.752 | 0.52436344 | NA | 0.8345 | 0.6682 | 0.7065 | 0.7065 | TRUE | 1.2242 |
| 4933404G15 | 0 | 0 | 0 | 0 | 0 | 0 | 0 | 0 | 0 | 0 | 0 | 0 | 0 | 0 | 0 | NA | NA | NA | NA | NA | NA | NA | NA | NA | NA |
| 4933404K08I | 0 | 0 | 0 | 0 | 0 | 0 | 0 | 0 | 0 | 0 | 0 | 0 | 0 | 0 | 0 | NA | NA | NA | NA | NA | NA | NA | NA | NA | NA |
| 4933404O12 | 2187 | 1631 | 1868 | 1900 | 1647 | 2065 | 1699 | 1735 | 2016 | 1891 | 1858 | 2038 | 1872.78 | 1817 | 1929 | 1.063 | 0.088 | 0.27484804 | 0.44590037 | 0.005 | 0.0037 | 0.0042 | 0.0042 | TRUE | 0.1102 |
| 4933405D12 | 3 | 4 | 4 | 8 | 9 | 7 | 2 | 4 | 4 | 8 | 10 | 7 | 5.99 | 3 | 8 | 2.334 | 1.223 | 0.12655295 | NA | 0 | 0.3982 | 0.2644 | 0.2644 | TRUE | 0.2156 |
| 4933405E24I | 0 | 0 | 0 | 0 | 0 | 0 | 0 | 0 | 0 | 0 | 0 | 0 | 0 | 0 | 0 | NA | NA | NA | NA | NA | NA | NA | NA | NA | NA |
| 4933405L10F | 26 | 12 | 19 | 28 | 18 | 14 | 20 | 13 | 21 | 28 | 20 | 14 | 19.24 | 18 | 21 | 1.155 | 0.207 | 0.65418737 | 0.78521163 | 0.0404 | 0.1256 | 0.1023 | 0.1023 | TRUE | 0.2053 |
| 4933405O20 | 0 | 0 | 0 | 0 | 0 | 0 | 0 | 0 | 0 | 0 | 0 | 0 | 0 | 0 | 0 | NA | NA | NA | NA | NA | NA | NA | NA | NA | NA |
| 4933406C10I | 1 | 3 | 5 | 2 | 4 | 2 | 1 | 3 | 5 | 2 | 5 | 2 | 2.97 | 3 | 3 | 0.922 | -0.117 | 0.91901449 | NA | 0 | 0.7995 | 0.6091 | 0.6091 | TRUE | 0.2303 |
| 4933406D12 | 0 | 0 | 0 | 0 | 0 | 0 | 0 | 0 | 0 | 0 | 0 | 0 | 0 | 0 | 0 | NA | NA | NA | NA | NA | NA | NA | NA | NA | NA |
| 4933406F09I | 0 | 0 | 0 | 0 | 0 | 0 | 0 | 0 | 0 | 0 | 0 | 0 | 0 | 0 | 0 | NA | NA | NA | NA | NA | NA | NA | NA | NA | NA |
| 4933406G16 | 0 | 0 | 0 | 0 | 0 | 0 | 0 | 0 | 0 | 0 | 0 | 0 | 0 | 0 | 0 | NA | NA | NA | NA | NA | NA | NA | NA | NA | NA |
| 4933406I18R | 0 | 0 | 0 | 1 | 1 | 0 | 0 | 0 | 0 | 1 | 1 | 0 | 0.35 | 0 | 1 | 3.978 | 1.992 | 0.55384016 | NA | 0 | 6.6991 | 6.4234 | 6.4234 | TRUE | 0.2647 |
| 4933406J08F | 181 | 138 | 141 | 157 | 112 | 118 | 141 | 147 | 152 | 156 | 126 | 116 | 139.77 | 147 | 133 | 0.91 | -0.136 | 0.44080381 | 0.61198726 | 0.005 | 0.0194 | 0.0154 | 0.0154 | TRUE | 0.2455 |
| 4933406J10F | 0 | 0 | 1 | 0 | 1 | 0 | 0 | 0 | 1 | 0 | 1 | 0 | 0.37 | 0 | 0 | 1.075 | 0.105 | 0.97514588 | NA | 0 | 6.4454 | 6.1578 | 6.1578 | TRUE | 0.5365 |
| 4933406K04I | 0 | 0 | 0 | 0 | 0 | 0 | 0 | 0 | 0 | 0 | 0 | 0 | 0 | 0 | 0 | NA | NA | NA | NA | NA | NA | NA | NA | NA | NA |
| 4933406M09 | 1 | 1 | 2 | 3 | 3 | 2 | 1 | 1 | 2 | 3 | 3 | 2 | 2.06 | 1 | 3 | 2.13 | 1.091 | 0.4128957 | NA | 0 | 1.1545 | 0.7229 | 0.7229 | TRUE | 0.1664 |
| 4933407E24I | 0 | 0 | 0 | 0 | 0 | 0 | 0 | 0 | 0 | 0 | 0 | 0 | 0 | 0 | 0 | NA | NA | NA | NA | NA | NA | NA | NA | NA | NA |
| 4933407G14 | 0 | 0 | 0 | 0 | 0 | 0 | 0 | 0 | 0 | 0 | 0 | 0 | 0 | 0 | 0 | NA | NA | NA | NA | NA | NA | NA | NA | NA | NA |
| 4933407I05R | 0 | 0 | 0 | 0 | 0 | 0 | 0 | 0 | 0 | 0 | 0 | 0 | 0 | 0 | 0 | NA | NA | NA | NA | NA | NA | NA | NA | NA | NA |
| 4933407K13I | 60 | 38 | 51 | 59 | 32 | 44 | 47 | 40 | 55 | 59 | 36 | 43 | 46.72 | 47 | 46 | 0.977 | -0.034 | 0.91138988 | 0.95132517 | 0.0188 | 0.0532 | 0.0439 | 0.0439 | TRUE | 0.4577 |
| 4933407L21F | 0 | 0 | 0 | 0 | 0 | 0 | 0 | 0 | 0 | 0 | 0 | 0 | 0 | 0 | 0 | NA | NA | NA | NA | NA | NA | NA | NA | NA | NA |
| 4933408B17 | 4 | 7 | 4 | 6 | 9 | 11 | 3 | 7 | 4 | 6 | 10 | 11 | 6.98 | 5 | 9 | 1.843 | 0.882 | 0.23853928 | NA | 0 | 0.3422 | 0.2473 | 0.2473 | TRUE | 0.3707 |
| 4933408J17F | 1 | 0 | 0 | 2 | 0 | 2 | 1 | 0 | 0 | 2 | 0 | 2 | 0.79 | 0 | 1 | 3.872 | 1.953 | 0.40305747 | NA | 0 | 3.0014 | 2.5443 | 2.5443 | TRUE | 0.4068 |
| 4933408N05 | 22 | 11 | 11 | 8 | 5 | 4 | 17 | 12 | 12 | 8 | 6 | 4 | 9.7 | 14 | 6 | 0.428 | -1.223 | 0.05717279 | NA | 0 | 0.2467 | 0.174 | 0.174 | TRUE | 0.2749 |
| 4933409G03 | 0 | 0 | 0 | 0 | 0 | 0 | 0 | 0 | 0 | 0 | 0 | 0 | 0 | 0 | 0 | NA | NA | NA | NA | NA | NA | NA | NA | NA | NA |
| 4933409K07I | 0 | 0 | 0 | 0 | 0 | 0 | 0 | 0 | 0 | 0 | 0 | 0 | 0 | 0 | 0 | NA | NA | NA | NA | NA | NA | NA | NA | NA | NA |
| 4933411E08I | 1 | 2 | 2 | 2 | 1 | 0 | 1 | 2 | 2 | 2 | 1 | 0 | 1.36 | 2 | 1 | 0.626 | -0.676 | 0.68475361 | NA | 0 | 1.7407 | 1.2066 | 1.2066 | TRUE | 0.378 |
| 4933411G06 | 0 | 0 | 0 | 0 | 0 | 0 | 0 | 0 | 0 | 0 | 0 | 0 | 0 | 0 | 0 | NA | NA | NA | NA | NA | NA | NA | NA | NA | NA |
| 4933411G11 | 7 | 3 | 3 | 3 | 4 | 3 | 5 | 3 | 3 | 3 | 5 | 3 | 3.72 | 4 | 4 | 0.864 | -0.211 | 0.82811808 | NA | 0 | 0.6394 | 0.408 | 0.408 | TRUE | 0.2361 |
| 4933411K16I | 8 | 8 | 3 | 5 | 3 | 6 | 6 | 9 | 3 | 5 | 3 | 6 | 5.37 | 6 | 5 | 0.798 | -0.326 | 0.69460184 | NA | 0 | 0.4435 | 0.3076 | 0.3076 | TRUE | 0.2009 |
| 4933411K20I | 2770 | 2006 | 1868 | 2028 | 1741 | 1982 | 2151 | 2134 | 2016 | 2019 | 1964 | 1957 | 2039.98 | 2100 | 1980 | 0.942 | -0.086 | 0.16772468 | 0.3191704 | 3.00E-04 | 0.0036 | 0.0023 | 0.0023 | TRUE | 0.0148 |
| 4933412E12I | 69 | 28 | 34 | 53 | 30 | 46 | 54 | 30 | 37 | 53 | 34 | 45 | 42.01 | 40 | 44 | 1.091 | 0.126 | 0.70401896 | 0.82094194 | 0.0472 | 0.0588 | 0.0555 | 0.0555 | TRUE | 0.7586 |
| 4933412E24I | 0 | 2 | 0 | 0 | 0 | 1 | 0 | 2 | 0 | 0 | 0 | 1 | 0.52 | 1 | 0 | 0.534 | -0.904 | 0.76321451 | NA | 0 | 4.5682 | 4.7886 | 4.7886 | TRUE | 0.9746 |
| 4933412O06 | 0 | 0 | 0 | 0 | 0 | 0 | 0 | 0 | 0 | 0 | 0 | 0 | 0 | 0 | 0 | NA | NA | NA | NA | NA | NA | NA | NA | NA | NA |
| 4933413G19 | 2 | 1 | 0 | 1 | 0 | 0 | 2 | 1 | 0 | 1 | 0 | 0 | 0.6 | 1 | 0 | 0.416 | -1.265 | 0.62159733 | NA | 0 | 3.939 | 3.1896 | 3.1896 | TRUE | 0.3906 |
| 4933413J09F | 17 | 14 | 8 | 4 | 4 | 2 | 13 | 15 | 9 | 4 | 5 | 2 | 7.87 | 12 | 4 | 0.282 | -1.826 | 0.01491378 | NA | 0 | 0.3037 | 0.2165 | 0.2165 | TRUE | 0.243 |
| 4933413L06F | 0 | 0 | 0 | 0 | 0 | 0 | 0 | 0 | 0 | 0 | 0 | 0 | 0 | 0 | 0 | NA | NA | NA | NA | NA | NA | NA | NA | NA | NA |
| 4933415F23I | 1 | 1 | 1 | 1 | 1 | 1 | 1 | 1 | 1 | 1 | 1 | 1 | 1 | 1 | 1 | 1.074 | 0.103 | 0.95418264 | NA | 0 | 2.3607 | 1.3027 | 1.3027 | TRUE | 0.0201 |
| 4933416C03I | 169 | 140 | 121 | 61 | 61 | 59 | 131 | 149 | 131 | 61 | 69 | 58 | 99.75 | 137 | 63 | 0.457 | -1.131 | 2.91E-08 | 1.29E-06 | 0 | 0.0262 | 0.0181 | 0.0181 | TRUE | 0.0615 |
| 4933416E03I | 0 | 0 | 0 | 0 | 0 | 0 | 0 | 0 | 0 | 0 | 0 | 0 | 0 | 0 | 0 | NA | NA | NA | NA | NA | NA | NA | NA | NA | NA |

| | | | | | | | | | | | | | | | | | | | | | | | | | |
|---|---|---|---|---|---|---|---|---|---|---|---|---|---|---|---|---|---|---|---|---|---|---|---|---|---|
| 4933416I08R | 4 | 2 | 3 | 0 | 0 | 0 | 3 | 2 | 3 | 0 | 0 | 0 | 1.41 | 3 | 0 | 0.067 | -3.893 | 0.03710637 | NA | 0 | 1.6811 | 1.3428 | 1.3428 | TRUE | 0.0718 |
| 4933416M06 | 1 | 0 | 0 | 0 | 0 | 0 | 1 | 0 | 0 | 0 | 0 | 0 | 0.13 | 0 | 0 | 0.552 | -0.857 | 0.8336548 | NA | 0 | 18.3121 | 10 | 10 | TRUE | 0.2566 |
| 4933416M07 | 49 | 24 | 23 | 19 | 23 | 20 | 38 | 26 | 25 | 19 | 26 | 20 | 25.5 | 30 | 22 | 0.721 | -0.471 | 0.23333133 | 0.39940175 | 0 | 0.0954 | 0.0723 | 0.0723 | TRUE | 0.5036 |
| 4933417A18 | 0 | 0 | 0 | 0 | 0 | 0 | 0 | 0 | 0 | 0 | 0 | 0 | 0 | 0 | 0 | NA | NA | NA | NA | NA | NA | NA | NA | NA | NA |
| 4933417D19 | 5 | 2 | 0 | 1 | 1 | 1 | 4 | 2 | 0 | 1 | 1 | 1 | 1.52 | 2 | 1 | 0.502 | -0.994 | 0.54036614 | NA | 0 | 1.5615 | 1.1647 | 1.1647 | TRUE | 0.1359 |
| 4933417E11I | 1 | 0 | 0 | 0 | 0 | 1 | 1 | 0 | 0 | 0 | 0 | 1 | 0.29 | 0 | 0 | 1.075 | 0.105 | 0.97702352 | NA | 0 | 8.0647 | 7.5458 | 7.5458 | TRUE | 0.3835 |
| 4933417G07 | 0 | 0 | 0 | 0 | 0 | 0 | 0 | 0 | 0 | 0 | 0 | 0 | 0 | 0 | 0 | NA | NA | NA | NA | NA | NA | NA | NA | NA | NA |
| 4933417O13 | 0 | 0 | 0 | 0 | 0 | 1 | 0 | 0 | 0 | 0 | 0 | 1 | 0.16 | 0 | 0 | 2.094 | 1.067 | 0.79379329 | NA | 0 | 14.4078 | 10 | 10 | TRUE | 0.3835 |
| 4933421I07R | 1 | 1 | 0 | 1 | 0 | 0 | 1 | 1 | 0 | 1 | 0 | 0 | 0.47 | 1 | 0 | 0.599 | -0.74 | 0.79191656 | NA | 0 | 5.0171 | 3.8827 | 3.8827 | TRUE | 0.3906 |
| 4933421O10 | 15 | 11 | 9 | 5 | 12 | 10 | 12 | 12 | 10 | 5 | 14 | 10 | 10.24 | 11 | 10 | 0.851 | -0.233 | 0.70390687 | NA | 0 | 0.2339 | 0.1725 | 0.1725 | TRUE | 0.3171 |
| 4933422A05 | 2 | 0 | 0 | 0 | 1 | 0 | 2 | 0 | 0 | 0 | 1 | 0 | 0.45 | 1 | 0 | 0.68 | -0.556 | 0.85993441 | NA | 0 | 5.3057 | 5.2576 | 5.2576 | TRUE | 0.9726 |
| 4933422H20 | 23 | 17 | 14 | 20 | 23 | 25 | 18 | 18 | 15 | 20 | 26 | 25 | 20.26 | 17 | 24 | 1.376 | 0.461 | 0.27597811 | 0.4470764 | 0 | 0.1194 | 0.0786 | 0.0786 | TRUE | 0.1081 |
| 4933424G05 | 0 | 0 | 2 | 0 | 0 | 1 | 0 | 0 | 2 | 0 | 0 | 1 | 0.52 | 1 | 0 | 0.529 | -0.92 | 0.75853134 | NA | 0 | 4.5238 | 4.7579 | 4.7579 | TRUE | 0.9746 |
| 4933424G06 | 1 | 0 | 2 | 4 | 1 | 1 | 1 | 0 | 2 | 4 | 1 | 1 | 1.51 | 1 | 2 | 2.115 | 1.08 | 0.50975914 | NA | 0 | 1.577 | 1.1837 | 1.1837 | TRUE | 0.6559 |
| 4933425B07 | 1 | 0 | 0 | 0 | 0 | 1 | 1 | 0 | 0 | 0 | 0 | 1 | 0.29 | 0 | 0 | 1.075 | 0.105 | 0.97702352 | NA | 0 | 8.0647 | 7.5458 | 7.5458 | TRUE | 0.3835 |
| 4933425H06 | 0 | 0 | 0 | 0 | 0 | 0 | 0 | 0 | 0 | 0 | 0 | 0 | 0 | 0 | 0 | NA | NA | NA | NA | NA | NA | NA | NA | NA | NA |
| 4933425L06F | 26 | 16 | 15 | 4 | 8 | 2 | 20 | 17 | 16 | 4 | 9 | 2 | 11.4 | 18 | 5 | 0.274 | -1.867 | 0.00398077 | NA | 0 | 0.2104 | 0.1702 | 0.1702 | TRUE | 0.896 |
| 4933426D04 | 0 | 0 | 0 | 0 | 0 | 0 | 0 | 0 | 0 | 0 | 0 | 0 | 0 | 0 | 0 | NA | NA | NA | NA | NA | NA | NA | NA | NA | NA |
| 4933426M11 | 4112 | 2964 | 3229 | 3646 | 3209 | 3587 | 3194 | 3153 | 3484 | 3629 | 3620 | 3541 | 3436.78 | 3277 | 3597 | 1.098 | 0.135 | 0.02634492 | 0.08444673 | 0.0012 | 0.0031 | 0.0024 | 0.0024 | TRUE | 0.0367 |
| 4933427D06 | 689 | 483 | 375 | 318 | 274 | 282 | 535 | 514 | 405 | 317 | 309 | 278 | 392.92 | 485 | 301 | 0.621 | -0.688 | 1.07E-07 | 4.18E-06 | 0.0114 | 0.0085 | 0.0094 | 0.0094 | TRUE | 0.2369 |
| 4933427D14 | 969 | 888 | 815 | 832 | 691 | 894 | 753 | 945 | 879 | 828 | 780 | 883 | 844.45 | 859 | 830 | 0.969 | -0.046 | 0.64831817 | 0.78131632 | 0.0076 | 0.0052 | 0.006 | 0.006 | TRUE | 0.1455 |
| 4933427E11I | 3 | 1 | 0 | 4 | 1 | 0 | 2 | 1 | 0 | 4 | 1 | 0 | 1.42 | 1 | 2 | 1.453 | 0.539 | 0.76012243 | NA | 1.0365 | 1.6747 | 1.5318 | 1.5318 | TRUE | 0.5801 |
| 4933427E13I | 0 | 0 | 0 | 0 | 0 | 0 | 0 | 0 | 0 | 0 | 0 | 0 | 0 | 0 | 0 | NA | NA | NA | NA | NA | NA | NA | NA | NA | NA |
| 4933427G17 | 8 | 5 | 0 | 11 | 9 | 3 | 6 | 5 | 0 | 11 | 10 | 3 | 5.93 | 4 | 8 | 2.031 | 1.022 | 0.27230306 | NA | 0.589 | 0.4019 | 0.4373 | 0.4373 | TRUE | 1.101 |
| 4933427I22R | 0 | 0 | 0 | 0 | 0 | 0 | 0 | 0 | 0 | 0 | 0 | 0 | 0 | 0 | 0 | NA | NA | NA | NA | NA | NA | NA | NA | NA | NA |
| 4933428C19I | 2 | 3 | 2 | 0 | 0 | 0 | 2 | 3 | 2 | 0 | 0 | 0 | 1.15 | 2 | 0 | 0.083 | -3.586 | 0.07265248 | NA | 0 | 2.0626 | 1.6639 | 1.6639 | TRUE | 0.112 |
| 4933429K18I | 0 | 0 | 0 | 0 | 0 | 0 | 0 | 0 | 0 | 0 | 0 | 0 | 0 | 0 | 0 | NA | NA | NA | NA | NA | NA | NA | NA | NA | NA |
| 4933429O19 | 1 | 0 | 0 | 0 | 0 | 0 | 1 | 0 | 0 | 0 | 0 | 0 | 0.13 | 0 | 0 | 0.552 | -0.857 | 0.8336548 | NA | 0 | 18.3121 | 10 | 10 | TRUE | 0.2566 |
| 4933430H16 | 0 | 1 | 0 | 0 | 0 | 0 | 0 | 1 | 0 | 0 | 0 | 0 | 0.18 | 0 | 0 | 0.552 | -0.857 | 0.8336548 | NA | 0 | 13.3714 | 10 | 10 | TRUE | 0.5216 |
| 4933430I17R | 38 | 29 | 27 | 30 | 37 | 39 | 30 | 31 | 29 | 30 | 42 | 38 | 33.27 | 30 | 37 | 1.228 | 0.297 | 0.37473566 | 0.54920038 | 0 | 0.0737 | 0.0502 | 0.0502 | TRUE | 0.1958 |
| 4933430M04 | 0 | 0 | 0 | 0 | 0 | 0 | 0 | 0 | 0 | 0 | 0 | 0 | 0 | 0 | 0 | NA | NA | NA | NA | NA | NA | NA | NA | NA | NA |
| 4933430N04 | 0 | 0 | 0 | 0 | 0 | 0 | 0 | 0 | 0 | 0 | 0 | 0 | 0 | 0 | 0 | NA | NA | NA | NA | NA | NA | NA | NA | NA | NA |
| 4933431E20I | 434 | 273 | 317 | 418 | 303 | 406 | 337 | 290 | 342 | 416 | 342 | 401 | 354.7 | 323 | 386 | 1.195 | 0.257 | 0.03784737 | 0.10997204 | 0.0062 | 0.0091 | 0.0082 | 0.0082 | TRUE | 0.1128 |
| 4933431G14 | 0 | 0 | 0 | 0 | 0 | 0 | 0 | 0 | 0 | 0 | 0 | 0 | 0 | 0 | 0 | NA | NA | NA | NA | NA | NA | NA | NA | NA | NA |
| 4933432G23 | 3 | 2 | 9 | 5 | 5 | 3 | 2 | 2 | 10 | 5 | 6 | 3 | 4.62 | 5 | 5 | 0.973 | -0.04 | 0.96654099 | NA | 0.2332 | 0.5149 | 0.4287 | 0.4287 | TRUE | 1.5318 |
| 4933432I03R | 0 | 1 | 1 | 1 | 0 | 0 | 0 | 1 | 1 | 1 | 0 | 0 | 0.52 | 1 | 0 | 0.535 | -0.904 | 0.74017109 | NA | 0 | 4.534 | 3.6534 | 3.6534 | TRUE | 0.3906 |
| 4933432I09R | 0 | 0 | 0 | 0 | 0 | 1 | 0 | 0 | 0 | 0 | 0 | 1 | 0.16 | 0 | 0 | 2.094 | 1.067 | 0.79379329 | NA | 0 | 14.4078 | 10 | 10 | TRUE | 0.3835 |
| 4933433C11I | 25 | 21 | 19 | 22 | 13 | 19 | 19 | 22 | 21 | 22 | 15 | 19 | 19.6 | 21 | 19 | 0.894 | -0.162 | 0.70694269 | 0.82296237 | 0 | 0.1234 | 0.0824 | 0.0824 | TRUE | 0.149 |
| 4933433F19I | 0 | 0 | 0 | 0 | 0 | 0 | 0 | 0 | 0 | 0 | 0 | 0 | 0 | 0 | 0 | NA | NA | NA | NA | NA | NA | NA | NA | NA | NA |
| 4933433G15 | 5 | 10 | 6 | 21 | 12 | 16 | 4 | 11 | 6 | 21 | 14 | 16 | 11.87 | 7 | 17 | 2.451 | 1.294 | 0.03090973 | NA | 0 | 0.2021 | 0.1577 | 0.1577 | TRUE | 0.5585 |
| 4933433G19 | 9 | 10 | 2 | 5 | 5 | 6 | 7 | 11 | 2 | 5 | 6 | 6 | 6.05 | 7 | 6 | 0.833 | -0.263 | 0.74537375 | NA | 0 | 0.3939 | 0.3064 | 0.3064 | TRUE | 0.2144 |
| 4933433H22 | 0 | 0 | 0 | 0 | 0 | 0 | 0 | 0 | 0 | 0 | 0 | 0 | 0 | 0 | 0 | NA | NA | NA | NA | NA | NA | NA | NA | NA | NA |
| 4933434E20I | 1242 | 997 | 1031 | 1012 | 877 | 1141 | 965 | 1060 | 1112 | 1007 | 989 | 1126 | 1043.43 | 1046 | 1041 | 0.997 | -0.004 | 0.96262616 | 0.97941456 | 0.0042 | 0.0047 | 0.0045 | 0.0045 | TRUE | 0.0619 |
| 4933434I20R | 10 | 6 | 4 | 9 | 8 | 11 | 8 | 6 | 4 | 9 | 9 | 11 | 7.88 | 6 | 10 | 1.544 | 0.627 | 0.35426161 | NA | 0 | 0.303 | 0.199 | 0.199 | TRUE | 0.166 |
| 4933436C20I | 101 | 84 | 126 | 115 | 85 | 100 | 78 | 89 | 136 | 114 | 96 | 99 | 102.14 | 101 | 103 | 1.025 | 0.035 | 0.88085615 | 0.93349722 | 0.0381 | 0.0256 | 0.0293 | 0.0293 | TRUE | 0.8617 |
| 4933436E23I | 0 | 0 | 0 | 0 | 0 | 0 | 0 | 0 | 0 | 0 | 0 | 0 | 0 | 0 | 0 | NA | NA | NA | NA | NA | NA | NA | NA | NA | NA |
| 4933436H12 | 0 | 0 | 0 | 0 | 0 | 0 | 0 | 0 | 0 | 0 | 0 | 0 | 0 | 0 | 0 | NA | NA | NA | NA | NA | NA | NA | NA | NA | NA |
| 4933436I01R | 0 | 0 | 0 | 0 | 0 | 0 | 0 | 0 | 0 | 0 | 0 | 0 | 0 | 0 | 0 | NA | NA | NA | NA | NA | NA | NA | NA | NA | NA |
| 4933438B17 | 0 | 0 | 0 | 0 | 0 | 0 | 0 | 0 | 0 | 0 | 0 | 0 | 0 | 0 | 0 | NA | NA | NA | NA | NA | NA | NA | NA | NA | NA |
| 4933438K21I | 4 | 2 | 1 | 1 | 0 | 1 | 3 | 2 | 1 | 1 | 0 | 1 | 1.38 | 2 | 1 | 0.317 | -1.657 | 0.33459241 | NA | 0 | 1.7166 | 1.134 | 1.134 | TRUE | 0.1727 |
| 4933439C10I | 95 | 88 | 130 | 121 | 96 | 113 | 74 | 94 | 140 | 120 | 108 | 112 | 107.99 | 103 | 113 | 1.115 | 0.156 | 0.50640177 | 0.6698751 | 0.0459 | 0.0244 | 0.0306 | 0.0306 | TRUE | 0.724 |
| 4933439K11I | 1 | 1 | 1 | 0 | 0 | 1 | 1 | 1 | 1 | 0 | 0 | 1 | 0.65 | 1 | 0 | 0.383 | -1.383 | 0.55329037 | NA | 0 | 3.6426 | 2.4227 | 2.4227 | TRUE | 0.3835 |
| 4933440J02F | 0 | 1 | 0 | 0 | 0 | 0 | 0 | 1 | 0 | 0 | 0 | 0 | 0.18 | 0 | 0 | 0.552 | -0.857 | 0.8336548 | NA | 0 | 13.3714 | 10 | 10 | TRUE | 0.5216 |
| 4933440M02 | 7 | 3 | 11 | 20 | 12 | 13 | 5 | 3 | 12 | 20 | 14 | 13 | 11.13 | 7 | 16 | 2.286 | 1.193 | 0.05854312 | NA | 0.0685 | 0.2154 | 0.1816 | 0.1816 | TRUE | 0.9717 |
| 5031410I06R | 1 | 2 | 0 | 2 | 0 | 0 | 1 | 2 | 0 | 2 | 0 | 0 | 0.82 | 1 | 1 | 0.698 | -0.518 | 0.82391604 | NA | 0 | 2.9076 | 2.6343 | 2.6343 | TRUE | 0.7035 |
| 5031414D18 | 26 | 13 | 14 | 22 | 17 | 18 | 20 | 14 | 15 | 22 | 19 | 18 | 18 | 16 | 20 | 1.187 | 0.247 | 0.58352321 | 0.73325466 | 0 | 0.1341 | 0.0908 | 0.0908 | TRUE | 0.2491 |
| 5031425E22I | 164 | 134 | 148 | 152 | 129 | 199 | 127 | 143 | 160 | 151 | 146 | 196 | 153.81 | 143 | 164 | 1.154 | 0.206 | 0.25087347 | 0.41967867 | 0.0142 | 0.0178 | 0.0168 | 0.0168 | TRUE | 0.3117 |
| 5031425F14I | 0 | 0 | 0 | 0 | 0 | 0 | 0 | 0 | 0 | 0 | 0 | 0 | 0 | 0 | 0 | NA | NA | NA | NA | NA | NA | NA | NA | NA | NA |
| 5031426D15 | 0 | 1 | 0 | 0 | 0 | 0 | 0 | 1 | 0 | 0 | 0 | 0 | 0.18 | 0 | 0 | 0.552 | -0.857 | 0.8336548 | NA | 0 | 13.3714 | 10 | 10 | TRUE | 0.5216 |
| 5031434C07I | 8 | 11 | 12 | 12 | 6 | 10 | 6 | 12 | 13 | 12 | 7 | 10 | 9.91 | 10 | 10 | 0.944 | -0.083 | 0.89448617 | NA | 0 | 0.2416 | 0.1785 | 0.1785 | TRUE | 0.5571 |
| 5031434O11 | 10 | 9 | 6 | 8 | 7 | 8 | 8 | 10 | 6 | 8 | 8 | 8 | 7.93 | 8 | 8 | 0.998 | -0.003 | 0.99685158 | NA | 0 | 0.3014 | 0.1911 | 0.1911 | TRUE | 0.0852 |
| 5031439G07 | 4090 | 2955 | 3039 | 3211 | 2930 | 3263 | 3177 | 3143 | 3279 | 3196 | 3305 | 3221 | 3220.24 | 3200 | 3241 | 1.013 | 0.018 | 0.7364236 | 0.84273038 | 0 | 0.0032 | 0.0019 | 0.0019 | TRUE | 0.0056 |
| 5033403H07 | 0 | 0 | 0 | 0 | 0 | 0 | 0 | 0 | 0 | 0 | 0 | 0 | 0 | 0 | 0 | NA | NA | NA | NA | NA | NA | NA | NA | NA | NA |
| 5033404E19I | 8 | 12 | 10 | 21 | 11 | 15 | 6 | 13 | 11 | 21 | 12 | 15 | 12.98 | 10 | 16 | 1.648 | 0.721 | 0.19430986 | NA | 0 | 0.185 | 0.1405 | 0.1405 | TRUE | 0.4929 |
| 5033406O09 | 48 | 36 | 29 | 34 | 41 | 38 | 37 | 38 | 31 | 34 | 46 | 38 | 37.41 | 35 | 39 | 1.095 | 0.131 | 0.67913461 | 0.8028453 | 0 | 0.0658 | 0.0458 | 0.0458 | TRUE | 0.1714 |
| 5033411D12 | 15 | 5 | 6 | 16 | 14 | 10 | 12 | 5 | 6 | 16 | 16 | 10 | 10.84 | 8 | 14 | 1.733 | 0.793 | 0.19119377 | NA | 0 | 0.2211 | 0.1677 | 0.1677 | TRUE | 0.687 |
| 5330411J11F | 74 | 46 | 42 | 42 | 41 | 42 | 57 | 49 | 45 | 42 | 46 | 41 | 46.87 | 50 | 43 | 0.848 | -0.237 | 0.40032047 | 0.57345805 | 0 | 0.053 | 0.0359 | 0.0359 | TRUE | 0.134 |
| 5330413P13I | 7 | 1 | 11 | 3 | 2 | 7 | 5 | 1 | 12 | 3 | 2 | 7 | 5.09 | 6 | 4 | 0.669 | -0.58 | 0.54843067 | NA | 0.4612 | 0.4683 | 0.4663 | 0.4663 | TRUE | 0.2132 |
| 5330417C22I | 48 | 40 | 48 | 46 | 42 | 52 | 37 | 43 | 52 | 46 | 47 | 51 | 46.02 | 44 | 48 | 1.105 | 0.144 | 0.61687141 | 0.75818583 | 0 | 0.0539 | 0.0378 | 0.0378 | TRUE | 0.1876 |

| | | | | | | | | | | | | | | | | | | | | | | | | | |
|---|---|---|---|---|---|---|---|---|---|---|---|---|---|---|---|---|---|---|---|---|---|---|---|---|---|
| 5330426P16I | 107 | 86 | 72 | 81 | 68 | 69 | 83 | 91 | 78 | 81 | 77 | 68 | 79.62 | 84 | 75 | 0.894 | -0.162 | 0.45840366 | 0.6281242 | 0 | 0.0322 | 0.0219 | 0.0219 | TRUE | 0.0636 |
| 5330434G04 | 0 | 0 | 0 | 0 | 0 | 0 | 0 | 0 | 0 | 0 | 0 | 0 | 0 | 0 | 0 | NA | NA | NA | NA | NA | NA | NA | NA | NA | NA |
| 5330439B14I | 0 | 0 | 0 | 0 | 0 | 0 | 0 | 0 | 0 | 0 | 0 | 0 | 0 | 0 | 0 | NA | NA | NA | NA | NA | NA | NA | NA | NA | NA |
| 5430401F13I | 0 | 1 | 1 | 0 | 0 | 0 | 0 | 1 | 1 | 0 | 0 | 0 | 0.36 | 1 | 0 | 0.271 | -1.881 | 0.5755355 | NA | 0 | 6.6391 | 6.4428 | 6.4428 | TRUE | 0.3513 |
| 5430402E10I | 0 | 0 | 0 | 0 | 0 | 0 | 0 | 0 | 0 | 0 | 0 | 0 | 0 | 0 | 0 | NA | NA | NA | NA | NA | NA | NA | NA | NA | NA |
| 5430402O13 | 36 | 27 | 7 | 8 | 3 | 5 | 28 | 29 | 8 | 8 | 3 | 5 | 13.42 | 22 | 5 | 0.254 | -1.978 | 0.00280775 | NA | 0.2491 | 0.179 | 0.1987 | 0.1987 | TRUE | 1.6853 |
| 5430403N17 | 0 | 0 | 0 | 0 | 0 | 0 | 0 | 0 | 0 | 0 | 0 | 0 | 0 | 0 | 0 | NA | NA | NA | NA | NA | NA | NA | NA | NA | NA |
| 5430405H02I | 30 | 11 | 19 | 11 | 8 | 12 | 23 | 12 | 21 | 11 | 9 | 12 | 14.55 | 19 | 11 | 0.571 | -0.809 | 0.12241924 | NA | 0 | 0.1653 | 0.1233 | 0.1233 | TRUE | 0.5057 |
| 5430416N02 | 424 | 343 | 367 | 481 | 440 | 474 | 329 | 365 | 396 | 479 | 496 | 468 | 422.2 | 363 | 481 | 1.327 | 0.408 | 0.00019329 | 0.00207725 | 0.0022 | 0.008 | 0.0062 | 0.0062 | TRUE | 0.0835 |
| 5430416O09 | 3 | 4 | 1 | 1 | 2 | 3 | 2 | 4 | 1 | 1 | 2 | 3 | 2.31 | 2 | 2 | 0.812 | -0.301 | 0.81023721 | NA | 0 | 1.0272 | 0.6937 | 0.6937 | TRUE | 0.2693 |
| 5430417L22F | 402 | 351 | 386 | 369 | 372 | 368 | 312 | 373 | 416 | 367 | 420 | 363 | 375.38 | 367 | 383 | 1.046 | 0.065 | 0.61547697 | 0.7572023 | 0.0113 | 0.0087 | 0.0095 | 0.0095 | TRUE | 0.2109 |
| 5430419D17I | 2 | 1 | 2 | 2 | 2 | 1 | 2 | 1 | 2 | 2 | 2 | 1 | 1.67 | 2 | 2 | 1.092 | 0.127 | 0.92887276 | NA | 0 | 1.4232 | 0.8567 | 0.8567 | TRUE | 0.119 |
| 5430421F17I | 18 | 10 | 8 | 4 | 6 | 6 | 14 | 11 | 9 | 4 | 7 | 6 | 8.32 | 11 | 6 | 0.495 | -1.014 | 0.13526002 | NA | 0 | 0.2873 | 0.1956 | 0.1956 | TRUE | 0.2446 |
| 5430421N21 | 0 | 1 | 0 | 0 | 1 | 0 | 0 | 1 | 0 | 0 | 1 | 0 | 0.37 | 0 | 0 | 1.075 | 0.105 | 0.97520514 | NA | 0 | 6.4904 | 6.1969 | 6.1969 | TRUE | 0.5216 |
| 5430425J12F | 0 | 0 | 0 | 0 | 0 | 0 | 0 | 0 | 0 | 0 | 0 | 0 | 0 | 0 | 0 | NA | NA | NA | NA | NA | NA | NA | NA | NA | NA |
| 5430427M07 | 0 | 0 | 0 | 0 | 0 | 0 | 0 | 0 | 0 | 0 | 0 | 0 | 0 | 0 | 0 | NA | NA | NA | NA | NA | NA | NA | NA | NA | NA |
| 5430427O19 | 0 | 0 | 0 | 0 | 0 | 0 | 0 | 0 | 0 | 0 | 0 | 0 | 0 | 0 | 0 | NA | NA | NA | NA | NA | NA | NA | NA | NA | NA |
| 5430428K19I | 0 | 0 | 0 | 0 | 0 | 0 | 0 | 0 | 0 | 0 | 0 | 0 | 0 | 0 | 0 | NA | NA | NA | NA | NA | NA | NA | NA | NA | NA |
| 5430434I15R | 0 | 0 | 0 | 0 | 0 | 0 | 0 | 0 | 0 | 0 | 0 | 0 | 0 | 0 | 0 | NA | NA | NA | NA | NA | NA | NA | NA | NA | NA |
| 5430435G22 | 5 | 1 | 1 | 3 | 5 | 2 | 4 | 1 | 1 | 3 | 6 | 2 | 2.77 | 2 | 4 | 1.674 | 0.743 | 0.52857751 | NA | 0 | 0.8577 | 0.6246 | 0.6246 | TRUE | 0.7977 |
| 5430437J10F | 0 | 0 | 0 | 0 | 0 | 0 | 0 | 0 | 0 | 0 | 0 | 0 | 0 | 0 | 0 | NA | NA | NA | NA | NA | NA | NA | NA | NA | NA |
| 5430440P10I | 7 | 3 | 3 | 2 | 2 | 1 | 5 | 3 | 3 | 2 | 2 | 1 | 2.85 | 4 | 2 | 0.433 | -1.208 | 0.2942044 | NA | 0 | 0.8341 | 0.5369 | 0.5369 | TRUE | 0.2338 |
| 5530400C23I | 0 | 0 | 0 | 0 | 0 | 0 | 0 | 0 | 0 | 0 | 0 | 0 | 0 | 0 | 0 | NA | NA | NA | NA | NA | NA | NA | NA | NA | NA |
| 5530401A14I | 0 | 0 | 0 | 0 | 0 | 0 | 0 | 0 | 0 | 0 | 0 | 0 | 0 | 0 | 0 | NA | NA | NA | NA | NA | NA | NA | NA | NA | NA |
| 5530601H04I | 184 | 124 | 101 | 117 | 113 | 128 | 143 | 132 | 109 | 116 | 127 | 126 | 125.68 | 128 | 123 | 0.961 | -0.058 | 0.75076058 | 0.85197467 | 0 | 0.0213 | 0.0161 | 0.0161 | TRUE | 0.1656 |
| 5730403I07R | 3 | 6 | 3 | 3 | 8 | 3 | 2 | 6 | 3 | 3 | 9 | 3 | 4.49 | 4 | 5 | 1.258 | 0.331 | 0.72646316 | NA | 0.157 | 0.5306 | 0.4169 | 0.4169 | TRUE | 0.9064 |
| 5730405O15 | 57 | 47 | 42 | 57 | 50 | 61 | 44 | 50 | 45 | 57 | 56 | 60 | 52.16 | 46 | 58 | 1.245 | 0.316 | 0.23059143 | 0.39635909 | 0 | 0.0479 | 0.0308 | 0.0308 | TRUE | 0.0323 |
| 5730408K05I | 140 | 127 | 102 | 112 | 89 | 113 | 109 | 135 | 110 | 111 | 100 | 112 | 112.89 | 118 | 108 | 0.917 | -0.125 | 0.51179344 | 0.67469688 | 0 | 0.0234 | 0.0171 | 0.0171 | TRUE | 0.1573 |
| 5730409E04I | 223 | 211 | 176 | 211 | 189 | 233 | 173 | 224 | 190 | 210 | 213 | 230 | 206.8 | 196 | 218 | 1.116 | 0.159 | 0.29070275 | 0.46274801 | 0.0054 | 0.0139 | 0.0114 | 0.0114 | TRUE | 0.1751 |
| 5730412P04I | 0 | 0 | 0 | 0 | 0 | 0 | 0 | 0 | 0 | 0 | 0 | 0 | 0 | 0 | 0 | NA | NA | NA | NA | NA | NA | NA | NA | NA | NA |
| 5730416F02I | 2 | 0 | 0 | 2 | 0 | 1 | 2 | 0 | 0 | 2 | 0 | 1 | 0.76 | 1 | 1 | 1.792 | 0.842 | 0.72383071 | NA | 0 | 3.1407 | 2.7141 | 2.7141 | TRUE | 0.3907 |
| 5730420D15I | 3 | 4 | 4 | 3 | 5 | 5 | 2 | 4 | 4 | 3 | 6 | 5 | 4.08 | 3 | 5 | 1.259 | 0.333 | 0.72223153 | NA | 0 | 0.5837 | 0.3811 | 0.3811 | TRUE | 0.2107 |
| 5730422E09I | 1 | 2 | 1 | 3 | 0 | 2 | 1 | 2 | 1 | 3 | 0 | 2 | 1.49 | 1 | 2 | 1.285 | 0.362 | 0.82041438 | NA | 0 | 1.5924 | 1.1516 | 1.1516 | TRUE | 0.3748 |
| 5730435O14 | 1 | 0 | 1 | 0 | 0 | 0 | 1 | 0 | 1 | 0 | 0 | 0 | 0.31 | 1 | 0 | 0.307 | -1.706 | 0.62894649 | NA | 0 | 7.6657 | 7.2 | 7.2 | TRUE | 0.2067 |
| 5730455P16I | 2212 | 1544 | 1608 | 2196 | 1822 | 1956 | 1718 | 1642 | 1735 | 2186 | 2055 | 1931 | 1877.92 | 1698 | 2057 | 1.211 | 0.276 | 7.99E-05 | 0.00100913 | 0.0019 | 0.0037 | 0.003 | 0.003 | TRUE | 0.0366 |
| 5730457N03 | 3 | 5 | 1 | 0 | 0 | 2 | 2 | 5 | 1 | 0 | 0 | 2 | 1.78 | 3 | 1 | 0.233 | -2.1 | 0.21486876 | NA | 0.3393 | 1.3312 | 1.1337 | 1.1337 | TRUE | 0.7673 |
| 5730460C07I | 0 | 0 | 0 | 1 | 0 | 0 | 0 | 0 | 0 | 1 | 0 | 0 | 0.17 | 0 | 0 | 2.094 | 1.067 | 0.79379329 | NA | 0 | 14.2885 | 10 | 10 | TRUE | 0.3906 |
| 5730480H06I | 62 | 38 | 35 | 40 | 26 | 48 | 48 | 40 | 38 | 40 | 29 | 47 | 40.48 | 42 | 39 | 0.921 | -0.118 | 0.7084446 | 0.82410273 | 0 | 0.061 | 0.0472 | 0.0472 | TRUE | 0.2865 |
| 5730488B01I | 0 | 0 | 2 | 0 | 1 | 0 | 0 | 0 | 2 | 0 | 1 | 0 | 0.55 | 1 | 0 | 0.529 | -0.918 | 0.75547616 | NA | 0 | 4.3299 | 4.5761 | 4.5761 | TRUE | 0.9747 |
| 5730507C01I | 115 | 64 | 73 | 61 | 51 | 67 | 89 | 68 | 79 | 61 | 58 | 66 | 70.09 | 79 | 62 | 0.778 | -0.362 | 0.12701174 | 0.26250766 | 0 | 0.0362 | 0.026 | 0.026 | TRUE | 0.1445 |
| 5730508B09I | 178 | 119 | 110 | 134 | 97 | 127 | 138 | 127 | 119 | 133 | 109 | 125 | 125.28 | 128 | 122 | 0.959 | -0.061 | 0.7361645 | 0.8425688 | 0 | 0.0213 | 0.0154 | 0.0154 | TRUE | 0.0865 |
| 5730522E02I | 6 | 1 | 4 | 0 | 3 | 0 | 5 | 1 | 4 | 0 | 3 | 0 | 2.24 | 3 | 1 | 0.32 | -1.642 | 0.27920954 | NA | 0.9029 | 1.0618 | 1.0367 | 1.0367 | TRUE | 1.4446 |
| 5730559C18I | 717 | 525 | 575 | 601 | 609 | 668 | 557 | 558 | 620 | 598 | 687 | 659 | 613.4 | 578 | 648 | 1.121 | 0.165 | 0.0895429 | 0.20402307 | 0.0027 | 0.0063 | 0.0051 | 0.0051 | TRUE | 0.0539 |
| 5730577I03R | 9 | 3 | 4 | 2 | 4 | 4 | 7 | 3 | 4 | 2 | 5 | 4 | 4.16 | 5 | 4 | 0.706 | -0.502 | 0.59376126 | NA | 0 | 0.5724 | 0.3899 | 0.3899 | TRUE | 0.3917 |
| 5830403L16F | 1 | 1 | 1 | 4 | 1 | 0 | 1 | 1 | 1 | 4 | 1 | 0 | 1.34 | 1 | 2 | 1.773 | 0.826 | 0.63206573 | NA | 0 | 1.7735 | 1.3418 | 1.3418 | TRUE | 0.5285 |
| 5830403M04 | 1 | 2 | 1 | 0 | 2 | 0 | 1 | 2 | 1 | 0 | 2 | 0 | 1.04 | 1 | 1 | 0.551 | -0.859 | 0.66722185 | NA | 0 | 2.2816 | 1.7914 | 1.7914 | TRUE | 0.975 |
| 5830411N06 | 0 | 0 | 0 | 2 | 0 | 0 | 0 | 0 | 0 | 2 | 0 | 0 | 0.33 | 0 | 1 | 3.77 | 1.915 | 0.62344813 | NA | 0 | 7.1455 | 9.1834 | 9.1834 | TRUE | 0.9739 |
| 5830415F09I | 739 | 563 | 506 | 518 | 417 | 513 | 574 | 599 | 546 | 516 | 470 | 506 | 535.2 | 573 | 497 | 0.869 | -0.203 | 0.03459073 | 0.10297964 | 0 | 0.0069 | 0.0048 | 0.0048 | TRUE | 0.0255 |
| 5830416I19R | 2 | 1 | 0 | 2 | 2 | 0 | 2 | 1 | 0 | 2 | 2 | 0 | 1.14 | 1 | 1 | 1.556 | 0.638 | 0.73376063 | NA | 0 | 2.0742 | 1.6319 | 1.6319 | TRUE | 0.5196 |
| 5830416P10I | 15 | 23 | 11 | 19 | 33 | 25 | 12 | 24 | 12 | 19 | 37 | 25 | 21.47 | 16 | 27 | 1.693 | 0.76 | 0.10579776 | 0.23061878 | 0.0999 | 0.1128 | 0.1093 | 0.1093 | TRUE | 0.6173 |
| 5830417I10R | 1997 | 1446 | 1426 | 1379 | 1304 | 1470 | 1551 | 1538 | 1539 | 1373 | 1471 | 1451 | 1487.09 | 1543 | 1432 | 0.928 | -0.108 | 0.10361345 | 0.22703466 | 0 | 0.004 | 0.0025 | 0.0025 | TRUE | 0.0157 |
| 5830418K08I | 4690 | 3479 | 3545 | 3925 | 3626 | 4080 | 3643 | 3701 | 3825 | 3907 | 4091 | 4028 | 3865.53 | 3723 | 4009 | 1.077 | 0.107 | 0.04971375 | 0.13427033 | 3.00E-04 | 0.003 | 0.0019 | 0.0019 | TRUE | 0.007 |
| 5830418P13I | 85 | 45 | 34 | 44 | 53 | 30 | 66 | 48 | 37 | 44 | 60 | 30 | 47.29 | 50 | 45 | 0.874 | -0.195 | 0.56342358 | 0.71780135 | 0.0805 | 0.0525 | 0.0604 | 0.0604 | TRUE | 0.2362 |
| 5830428M24 | 3 | 1 | 2 | 2 | 1 | 0 | 2 | 1 | 2 | 2 | 1 | 0 | 1.45 | 2 | 1 | 0.556 | -0.848 | 0.59994311 | NA | 0 | 1.6425 | 1.1248 | 1.1248 | TRUE | 0.395 |
| 5830432E09I | 0 | 0 | 0 | 0 | 0 | 0 | 0 | 0 | 0 | 0 | 0 | 0 | 0 | 0 | 0 | NA | NA | NA | NA | NA | NA | NA | NA | NA | NA |
| 5830454E08I | 15 | 12 | 16 | 14 | 18 | 16 | 12 | 13 | 17 | 14 | 20 | 16 | 15.29 | 14 | 17 | 1.205 | 0.269 | 0.58474262 | NA | 0 | 0.1575 | 0.1084 | 0.1084 | TRUE | 0.1834 |
| 5830473C10I | 3 | 4 | 2 | 0 | 0 | 1 | 2 | 4 | 2 | 0 | 0 | 1 | 1.62 | 3 | 0 | 0.128 | -2.968 | 0.09173646 | NA | 0 | 1.4639 | 1.0655 | 1.0655 | TRUE | 0.3835 |
| 5930403L14F | 52 | 33 | 26 | 39 | 33 | 35 | 40 | 35 | 28 | 39 | 37 | 35 | 35.69 | 34 | 37 | 1.061 | 0.085 | 0.79301892 | 0.87869058 | 0 | 0.0688 | 0.0478 | 0.0478 | TRUE | 0.1847 |
| 5930412G12 | 62 | 52 | 63 | 87 | 85 | 86 | 48 | 55 | 68 | 87 | 96 | 85 | 73.14 | 57 | 89 | 1.569 | 0.65 | 0.00599507 | 0.02820837 | 0 | 0.0348 | 0.0259 | 0.0259 | TRUE | 0.2269 |
| 5930438M14 | 5 | 1 | 1 | 3 | 0 | 1 | 4 | 1 | 1 | 3 | 0 | 1 | 1.67 | 2 | 1 | 0.652 | -0.618 | 0.69115956 | NA | 0.3393 | 1.4246 | 1.1237 | 1.1237 | TRUE | 0.6713 |
| 6030407O03 | 6 | 8 | 3 | 2 | 1 | 1 | 5 | 9 | 3 | 2 | 1 | 1 | 3.42 | 6 | 1 | 0.252 | -1.988 | 0.08352202 | NA | 0 | 0.6957 | 0.4855 | 0.4855 | TRUE | 0.4382 |
| 6030408B16I | 3 | 5 | 3 | 2 | 4 | 4 | 2 | 5 | 3 | 2 | 5 | 4 | 3.56 | 3 | 4 | 0.969 | -0.046 | 0.96379366 | NA | 0 | 0.6689 | 0.4464 | 0.4464 | TRUE | 0.2439 |
| 6030419C18I | 272 | 199 | 141 | 214 | 149 | 162 | 211 | 212 | 152 | 213 | 168 | 160 | 186.01 | 192 | 180 | 0.939 | -0.091 | 0.6113819 | 0.75404219 | 0.0236 | 0.0152 | 0.0176 | 0.0176 | TRUE | 0.3395 |
| 6030440G07 | 1 | 2 | 0 | 0 | 3 | 0 | 1 | 2 | 0 | 0 | 3 | 0 | 1.05 | 1 | 1 | 1.142 | 0.191 | 0.92902053 | NA | 2.7233 | 2.2638 | 2.3395 | 2.3395 | TRUE | 1.2909 |
| 6030458C11I | 3948 | 2834 | 2756 | 2744 | 2603 | 2813 | 3066 | 3014 | 2974 | 2731 | 2936 | 2777 | 2916.52 | 3018 | 2815 | 0.932 | -0.101 | 0.08252849 | 0.19342383 | 5.00E-04 | 0.0032 | 0.0021 | 0.0021 | TRUE | 0.0171 |
| 6030466F02I | 3 | 0 | 2 | 1 | 3 | 0 | 2 | 0 | 2 | 1 | 3 | 0 | 1.48 | 1 | 1 | 0.941 | -0.088 | 0.95918289 | NA | 0.7906 | 1.606 | 1.4349 | 1.4349 | TRUE | 0.635 |
| 6030468B19I | 1 | 1 | 0 | 0 | 0 | 0 | 1 | 1 | 0 | 0 | 0 | 0 | 0.31 | 1 | 0 | 0.309 | -1.696 | 0.63183309 | NA | 0 | 7.7294 | 7.2472 | 7.2472 | TRUE | 0.2015 |
| 6030498E09I | 0 | 0 | 0 | 0 | 0 | 0 | 0 | 0 | 0 | 0 | 0 | 0 | 0 | 0 | 0 | NA | NA | NA | NA | NA | NA | NA | NA | NA | NA |
| 6230400D17I | 32 | 23 | 28 | 29 | 26 | 19 | 25 | 24 | 30 | 29 | 29 | 19 | 26.08 | 26 | 26 | 0.968 | -0.048 | 0.90124743 | 0.94568228 | 0 | 0.0933 | 0.0674 | 0.0674 | TRUE | 0.3524 |

| | | | | | | | | | | | | | | | | | | | | | | | | | |
|---|---|---|---|---|---|---|---|---|---|---|---|---|---|---|---|---|---|---|---|---|---|---|---|---|---|
| 6330403A02 | 3 | 3 | 2 | 2 | 1 | 3 | 2 | 3 | 2 | 2 | 1 | 3 | 2.29 | 2 | 2 | 0.801 | -0.32 | 0.79429444 | NA | 0 | 1.0359 | 0.6409 | 0.6409 | TRUE | 0.1504 |
| 6330403K07 | 234 | 156 | 157 | 164 | 160 | 166 | 182 | 166 | 169 | 163 | 180 | 164 | 170.78 | 172 | 169 | 0.979 | -0.031 | 0.8389939 | 0.90825872 | 0 | 0.0163 | 0.0108 | 0.0108 | TRUE | 0.0347 |
| 6330403M23 | 131 | 108 | 104 | 144 | 102 | 134 | 102 | 115 | 112 | 143 | 115 | 132 | 119.92 | 110 | 130 | 1.193 | 0.255 | 0.16577579 | 0.31676916 | 0 | 0.0222 | 0.016 | 0.016 | TRUE | 0.1011 |
| 6330407A03 | 0 | 0 | 0 | 0 | 1 | 0 | 0 | 0 | 0 | 0 | 1 | 0 | 0.19 | 0 | 0 | 2.094 | 1.067 | 0.79379329 | NA | 0 | 12.608 | 10 | 10 | TRUE | 0.5075 |
| 6330408A02 | 298 | 187 | 191 | 200 | 189 | 224 | 231 | 199 | 206 | 199 | 213 | 221 | 211.64 | 212 | 211 | 0.992 | -0.011 | 0.93616677 | 0.96464794 | 0 | 0.0136 | 0.0097 | 0.0097 | TRUE | 0.0754 |
| 6330409D20 | 1 | 0 | 0 | 0 | 0 | 2 | 1 | 0 | 0 | 0 | 0 | 2 | 0.46 | 0 | 1 | 1.93 | 0.948 | 0.7608301 | NA | 0 | 5.1717 | 5.216 | 5.216 | TRUE | 0.9737 |
| 6330410L21 | 160 | 115 | 60 | 62 | 48 | 38 | 124 | 122 | 65 | 62 | 54 | 38 | 77.45 | 104 | 51 | 0.49 | -1.028 | 0.000428 | 0.00382814 | 0.081 | 0.033 | 0.0466 | 0.0466 | TRUE | 1.0276 |
| 6330415B21 | 0 | 0 | 0 | 0 | 0 | 0 | 0 | 0 | 0 | 0 | 0 | 0 | 0 | 0 | 0 | NA | NA | NA | NA | NA | NA | NA | NA | NA | NA |
| 6330416G13 | 705 | 577 | 582 | 628 | 549 | 583 | 548 | 614 | 628 | 625 | 619 | 576 | 601.54 | 597 | 607 | 1.019 | 0.027 | 0.77743183 | 0.8684512 | 0.0023 | 0.0064 | 0.005 | 0.005 | TRUE | 0.0643 |
| 6330418K02 | 36 | 20 | 14 | 19 | 18 | 14 | 28 | 21 | 15 | 19 | 20 | 14 | 19.56 | 21 | 18 | 0.814 | -0.297 | 0.50943422 | 0.67264141 | 0 | 0.1236 | 0.0948 | 0.0948 | TRUE | 0.1925 |
| 6330549D23 | 15 | 7 | 7 | 5 | 9 | 6 | 12 | 7 | 8 | 5 | 10 | 6 | 7.95 | 9 | 7 | 0.775 | -0.367 | 0.59315645 | NA | 0 | 0.3005 | 0.2123 | 0.2123 | TRUE | 0.3629 |
| 6430411K18 | 31 | 29 | 18 | 17 | 20 | 38 | 24 | 31 | 19 | 17 | 23 | 38 | 25.22 | 25 | 26 | 1.038 | 0.054 | 0.89785698 | 0.94349536 | 0.0695 | 0.0964 | 0.0888 | 0.0888 | TRUE | 0.8067 |
| 6430531B16 | 62 | 55 | 53 | 53 | 53 | 77 | 48 | 59 | 57 | 53 | 60 | 76 | 58.73 | 55 | 63 | 1.157 | 0.21 | 0.42546572 | 0.59812939 | 0 | 0.0428 | 0.033 | 0.033 | TRUE | 0.3015 |
| 6430548M08 | 220 | 159 | 166 | 184 | 178 | 212 | 171 | 169 | 179 | 183 | 201 | 209 | 185.39 | 173 | 198 | 1.143 | 0.193 | 0.18908847 | 0.34662582 | 0 | 0.0152 | 0.0102 | 0.0102 | TRUE | 0.0464 |
| 6430550D23 | 14 | 6 | 16 | 11 | 7 | 10 | 11 | 6 | 17 | 11 | 8 | 10 | 10.54 | 11 | 10 | 0.837 | -0.258 | 0.67391585 | NA | 0 | 0.2273 | 0.174 | 0.174 | TRUE | 0.2398 |
| 6430562O15 | 1 | 0 | 2 | 0 | 0 | 1 | 1 | 0 | 2 | 0 | 0 | 1 | 0.65 | 1 | 0 | 0.382 | -1.39 | 0.57974155 | NA | 0 | 3.6284 | 3.0424 | 3.0424 | TRUE | 0.2583 |
| 6430571L13 | 21 | 16 | 17 | 18 | 17 | 8 | 16 | 17 | 18 | 18 | 19 | 8 | 16.11 | 17 | 15 | 0.868 | -0.204 | 0.68323225 | NA | 0 | 0.1495 | 0.1173 | 0.1173 | TRUE | 0.8114 |
| 6430573F11 | 54 | 49 | 36 | 50 | 53 | 57 | 42 | 52 | 39 | 50 | 60 | 56 | 49.79 | 44 | 55 | 1.249 | 0.32 | 0.2483722 | 0.41735055 | 0 | 0.05 | 0.0352 | 0.0352 | TRUE | 0.1734 |
| 6430584L05 | 1 | 0 | 0 | 0 | 0 | 0 | 1 | 0 | 0 | 0 | 0 | 0 | 0.13 | 0 | 0 | 0.552 | -0.857 | 0.8336548 | NA | 0 | 18.3121 | 10 | 10 | TRUE | 0.2566 |
| 6430706D22 | 0 | 0 | 0 | 0 | 0 | 0 | 0 | 0 | 0 | 0 | 0 | 0 | 0 | 0 | 0 | NA | NA | NA | NA | NA | NA | NA | NA | NA | NA |
| 6530402F18 | 14 | 10 | 13 | 15 | 23 | 15 | 11 | 11 | 14 | 15 | 26 | 15 | 15.2 | 12 | 19 | 1.564 | 0.645 | 0.20383851 | NA | 0 | 0.1583 | 0.117 | 0.117 | TRUE | 0.577 |
| 6530411M01 | 1 | 1 | 0 | 0 | 2 | 1 | 1 | 1 | 0 | 0 | 2 | 1 | 0.85 | 1 | 1 | 1.713 | 0.776 | 0.71789357 | NA | 0 | 2.7997 | 2.0659 | 2.0659 | TRUE | 0.1674 |
| 6720401G13 | 2987 | 2822 | 2501 | 2521 | 2447 | 2649 | 2320 | 3002 | 2699 | 2509 | 2760 | 2615 | 2650.83 | 2674 | 2628 | 0.984 | -0.024 | 0.79134644 | 0.87744725 | 0.0091 | 0.0033 | 0.0053 | 0.0053 | TRUE | 0.1652 |
| 6720456H20 | 142 | 104 | 67 | 66 | 43 | 70 | 110 | 111 | 72 | 66 | 49 | 69 | 79.42 | 98 | 61 | 0.625 | -0.678 | 0.00700525 | 0.03175581 | 0.0319 | 0.0323 | 0.0322 | 0.0322 | TRUE | 0.4818 |
| 6720468P15 | 29 | 25 | 34 | 32 | 31 | 28 | 23 | 27 | 37 | 32 | 35 | 28 | 30.04 | 29 | 32 | 1.108 | 0.148 | 0.68163104 | 0.80469794 | 0 | 0.0813 | 0.0607 | 0.0607 | TRUE | 0.3816 |
| 6720483E21 | 3 | 5 | 8 | 11 | 10 | 7 | 2 | 5 | 9 | 11 | 11 | 7 | 7.57 | 5 | 10 | 1.834 | 0.875 | 0.23504366 | NA | 0 | 0.3155 | 0.2467 | 0.2467 | TRUE | 0.421 |
| 6720489N17 | 262 | 184 | 150 | 178 | 156 | 157 | 203 | 196 | 162 | 177 | 176 | 155 | 178.2 | 187 | 169 | 0.903 | -0.148 | 0.3515911 | 0.52609837 | 0.0046 | 0.0157 | 0.0125 | 0.0125 | TRUE | 0.1441 |
| 6820408C15 | 12 | 7 | 13 | 5 | 7 | 7 | 9 | 7 | 14 | 5 | 8 | 7 | 8.43 | 10 | 7 | 0.643 | -0.638 | 0.33989876 | NA | 0 | 0.2836 | 0.1963 | 0.1963 | TRUE | 0.3307 |
| 6820431F20 | 218 | 185 | 200 | 176 | 167 | 192 | 169 | 197 | 216 | 175 | 188 | 190 | 189.17 | 194 | 184 | 0.954 | -0.068 | 0.65940866 | 0.78925812 | 0.004 | 0.015 | 0.0117 | 0.0117 | TRUE | 0.1473 |
| 7420426K07 | 3 | 3 | 1 | 1 | 2 | 3 | 2 | 3 | 1 | 1 | 2 | 3 | 2.14 | 2 | 2 | 0.937 | -0.093 | 0.94205064 | NA | 0 | 1.1123 | 0.7196 | 0.7196 | TRUE | 0.2 |
| 7420461P10 | 0 | 0 | 1 | 0 | 0 | 0 | 0 | 0 | 1 | 0 | 0 | 0 | 0.18 | 0 | 0 | 0.552 | -0.857 | 0.8336548 | NA | 0 | 13.1817 | 10 | 10 | TRUE | 0.5365 |
| 7420700N18 | 0 | 0 | 0 | 0 | 0 | 0 | 0 | 0 | 0 | 0 | 0 | 0 | 0 | 0 | 0 | NA | NA | NA | NA | NA | NA | NA | NA | NA | NA |
| 7420701I03R | 0 | 0 | 0 | 0 | 0 | 0 | 0 | 0 | 0 | 0 | 0 | 0 | 0 | 0 | 0 | NA | NA | NA | NA | NA | NA | NA | NA | NA | NA |
| 7530416G11 | 0 | 0 | 0 | 0 | 0 | 0 | 0 | 0 | 0 | 0 | 0 | 0 | 0 | 0 | 0 | NA | NA | NA | NA | NA | NA | NA | NA | NA | NA |
| 7530420F21 | 0 | 0 | 0 | 0 | 0 | 0 | 0 | 0 | 0 | 0 | 0 | 0 | 0 | 0 | 0 | NA | NA | NA | NA | NA | NA | NA | NA | NA | NA |
| 7630403G23 | 0 | 0 | 0 | 0 | 0 | 0 | 0 | 0 | 0 | 0 | 0 | 0 | 0 | 0 | 0 | NA | NA | NA | NA | NA | NA | NA | NA | NA | NA |
| 8030411F24 | 1 | 0 | 0 | 1 | 0 | 0 | 1 | 0 | 0 | 1 | 0 | 0 | 0.3 | 0 | 0 | 1.075 | 0.105 | 0.97698525 | NA | 0 | 8.0272 | 7.5141 | 7.5141 | TRUE | 0.3906 |
| 8030423F21 | 1 | 1 | 0 | 1 | 3 | 1 | 1 | 1 | 0 | 1 | 3 | 1 | 1.2 | 1 | 2 | 2.832 | 1.502 | 0.41030437 | NA | 0 | 1.9755 | 1.3347 | 1.3347 | TRUE | 0.4384 |
| 8030423J24 | 0 | 0 | 0 | 0 | 0 | 0 | 0 | 0 | 0 | 0 | 0 | 0 | 0 | 0 | 0 | NA | NA | NA | NA | NA | NA | NA | NA | NA | NA |
| 8030442B05 | 0 | 0 | 0 | 0 | 0 | 0 | 0 | 0 | 0 | 0 | 0 | 0 | 0 | 0 | 0 | NA | NA | NA | NA | NA | NA | NA | NA | NA | NA |
| 8030443G20 | 0 | 0 | 0 | 0 | 0 | 0 | 0 | 0 | 0 | 0 | 0 | 0 | 0 | 0 | 0 | NA | NA | NA | NA | NA | NA | NA | NA | NA | NA |
| 8030462N17 | 3450 | 2493 | 2456 | 2486 | 2292 | 2640 | 2679 | 2652 | 2650 | 2475 | 2586 | 2606 | 2607.92 | 2660 | 2556 | 0.96 | -0.058 | 0.30803831 | 0.48060171 | 0 | 0.0033 | 0.002 | 0.002 | TRUE | 0.0094 |
| 8430403D17 | 0 | 0 | 0 | 0 | 0 | 0 | 0 | 0 | 0 | 0 | 0 | 0 | 0 | 0 | 0 | NA | NA | NA | NA | NA | NA | NA | NA | NA | NA |
| 8430408G22 | 1 | 0 | 0 | 0 | 1 | 0 | 1 | 0 | 0 | 0 | 1 | 0 | 0.32 | 0 | 0 | 1.075 | 0.105 | 0.97638847 | NA | 0 | 7.4681 | 7.0392 | 7.0392 | TRUE | 0.5075 |
| 8430410A17 | 2174 | 1686 | 1705 | 1588 | 1485 | 1700 | 1688 | 1793 | 1840 | 1581 | 1675 | 1678 | 1709.27 | 1774 | 1645 | 0.928 | -0.108 | 0.11243499 | 0.24079013 | 0.001 | 0.0038 | 0.0028 | 0.0028 | TRUE | 0.0222 |
| 8430419L09 | 4170 | 3372 | 3749 | 4823 | 4337 | 4963 | 3239 | 3587 | 4045 | 4801 | 4893 | 4899 | 4243.87 | 3624 | 4864 | 1.343 | 0.426 | 3.83E-08 | 1.67E-06 | 0.0059 | 0.003 | 0.0041 | 0.0041 | TRUE | 0.1259 |
| 8430422H06 | 0 | 0 | 0 | 0 | 0 | 0 | 0 | 0 | 0 | 0 | 0 | 0 | 0 | 0 | 0 | NA | NA | NA | NA | NA | NA | NA | NA | NA | NA |
| 8430423G03 | 0 | 1 | 0 | 0 | 0 | 1 | 0 | 1 | 0 | 0 | 0 | 1 | 0.34 | 0 | 0 | 1.075 | 0.105 | 0.97576789 | NA | 0 | 6.9363 | 6.5821 | 6.5821 | TRUE | 0.5216 |
| 8430427H17 | 1166 | 886 | 1034 | 1121 | 934 | 1136 | 906 | 942 | 1116 | 1116 | 1054 | 1121 | 1042.43 | 988 | 1097 | 1.112 | 0.154 | 0.09350793 | 0.21037444 | 0.0056 | 0.0047 | 0.0051 | 0.0051 | TRUE | 0.1511 |
| 8430429K09 | 66 | 38 | 17 | 30 | 40 | 33 | 51 | 40 | 18 | 30 | 45 | 33 | 36.26 | 36 | 36 | 0.965 | -0.052 | 0.89416399 | 0.94184685 | 0.1176 | 0.0678 | 0.0815 | 0.0815 | TRUE | 0.4928 |
| 8430431K14 | 0 | 0 | 0 | 0 | 0 | 0 | 0 | 0 | 0 | 0 | 0 | 0 | 0 | 0 | 0 | NA | NA | NA | NA | NA | NA | NA | NA | NA | NA |
| 8430436N08 | 0 | 0 | 0 | 0 | 0 | 0 | 0 | 0 | 0 | 0 | 0 | 0 | 0 | 0 | 0 | NA | NA | NA | NA | NA | NA | NA | NA | NA | NA |
| 8430437L04 | 0 | 0 | 0 | 0 | 0 | 0 | 0 | 0 | 0 | 0 | 0 | 0 | 0 | 0 | 0 | NA | NA | NA | NA | NA | NA | NA | NA | NA | NA |
| 9030025P20 | 82 | 74 | 48 | 68 | 55 | 72 | 64 | 79 | 52 | 68 | 62 | 71 | 65.83 | 65 | 67 | 1.035 | 0.05 | 0.84082459 | 0.90934518 | 0.0061 | 0.0384 | 0.0299 | 0.0299 | TRUE | 0.2423 |
| 9030204H09 | 0 | 0 | 0 | 1 | 0 | 1 | 0 | 0 | 0 | 1 | 0 | 1 | 0.33 | 0 | 1 | 3.763 | 1.912 | 0.57887588 | NA | 0 | 7.1752 | 6.7756 | 6.7756 | TRUE | 0.2094 |
| 9030404E10 | 11 | 1 | 3 | 4 | 3 | 3 | 9 | 1 | 3 | 4 | 3 | 3 | 3.86 | 4 | 3 | 0.78 | -0.359 | 0.72638824 | NA | 0.2365 | 0.6161 | 0.4988 | 0.4988 | TRUE | 0.6606 |
| 9030612E09 | 5 | 5 | 7 | 13 | 7 | 6 | 4 | 5 | 8 | 13 | 8 | 6 | 7.25 | 6 | 9 | 1.622 | 0.698 | 0.33962678 | NA | 0 | 0.3292 | 0.2393 | 0.2393 | TRUE | 0.5118 |
| 9030617O03 | 385 | 255 | 306 | 385 | 308 | 343 | 299 | 271 | 330 | 383 | 347 | 339 | 328.28 | 300 | 356 | 1.188 | 0.249 | 0.04303552 | 0.12079863 | 0.0034 | 0.0096 | 0.0078 | 0.0078 | TRUE | 0.0811 |
| 9030619P08 | 2 | 2 | 1 | 0 | 0 | 0 | 2 | 2 | 1 | 0 | 0 | 0 | 0.79 | 2 | 0 | 0.12 | -3.059 | 0.17609562 | NA | 0 | 2.9901 | 2.379 | 2.379 | TRUE | 0.0718 |
| 9030624G23 | 506 | 399 | 413 | 477 | 433 | 504 | 393 | 424 | 446 | 475 | 488 | 498 | 453.97 | 421 | 487 | 1.159 | 0.213 | 0.03700326 | 0.1083564 | 0 | 0.0076 | 0.0053 | 0.0053 | TRUE | 0.0423 |
| 9030624J02 | 3873 | 2939 | 3723 | 3337 | 3383 | 3919 | 3008 | 3126 | 4017 | 3322 | 3816 | 3869 | 3526.32 | 3384 | 3669 | 1.085 | 0.117 | 0.24920639 | 0.41810537 | 0.0144 | 0.0031 | 0.0072 | 0.0072 | TRUE | 0.325 |
| 9130008F23 | 11 | 7 | 9 | 4 | 4 | 4 | 9 | 7 | 10 | 4 | 5 | 4 | 6.36 | 9 | 4 | 0.483 | -1.049 | 0.16872992 | NA | 0 | 0.3752 | 0.2371 | 0.2371 | TRUE | 0.0377 |
| 9130011E15 | 1611 | 1155 | 1161 | 1244 | 1140 | 1288 | 1251 | 1229 | 1253 | 1238 | 1286 | 1271 | 1254.71 | 1244 | 1265 | 1.017 | 0.024 | 0.72599453 | 0.83588205 | 0 | 0.0043 | 0.0025 | 0.0025 | TRUE | 0.0042 |
| 9130015A21 | 0 | 1 | 0 | 3 | 1 | 2 | 0 | 1 | 0 | 3 | 1 | 2 | 1.19 | 0 | 2 | 5.915 | 2.564 | 0.18484089 | NA | 0 | 1.9906 | 1.4414 | 1.4414 | TRUE | 0.4979 |
| 9130015L21 | 0 | 0 | 0 | 0 | 0 | 0 | 0 | 0 | 0 | 0 | 0 | 0 | 0 | 0 | 0 | NA | NA | NA | NA | NA | NA | NA | NA | NA | NA |
| 9130017N09 | 137 | 138 | 190 | 277 | 358 | 354 | 106 | 147 | 205 | 276 | 404 | 349 | 247.87 | 153 | 343 | 2.258 | 1.175 | 1.44E-08 | 7.02E-07 | 0.0631 | 0.012 | 0.0263 | 0.0263 | TRUE | 0.4359 |
| 9130019O22 | 170 | 118 | 76 | 111 | 94 | 97 | 132 | 126 | 82 | 110 | 106 | 96 | 108.64 | 113 | 104 | 0.915 | -0.128 | 0.5541898 | 0.70983375 | 0.0252 | 0.0242 | 0.0245 | 0.0245 | TRUE | 0.5563 |
| 9130019P16 | 0 | 0 | 0 | 1 | 0 | 0 | 0 | 0 | 0 | 1 | 0 | 0 | 0.17 | 0 | 0 | 2.094 | 1.067 | 0.79379329 | NA | 0 | 14.2885 | 10 | 10 | TRUE | 0.3906 |
| 9130023H24 | 105 | 54 | 30 | 57 | 41 | 34 | 82 | 57 | 32 | 57 | 46 | 34 | 51.32 | 57 | 46 | 0.79 | -0.341 | 0.32853468 | 0.50307973 | 0.1156 | 0.0486 | 0.068 | 0.068 | TRUE | 0.1577 |

| | | | | | | | | | | | | | | | | | | | | | | | | | |
|---|---|---|---|---|---|---|---|---|---|---|---|---|---|---|---|---|---|---|---|---|---|---|---|---|---|
| 9130024F11I | 6 | 7 | 3 | 3 | 2 | 2 | 5 | 7 | 3 | 3 | 2 | 2 | 3.76 | 5 | 2 | 0.472 | -1.083 | 0.28079789 | NA | 0 | 0.6328 | 0.4175 | 0.4175 | TRUE | 0.2994 |
| 9130204L05F | 0 | 1 | 0 | 0 | 0 | 2 | 0 | 1 | 0 | 0 | 0 | 2 | 0.51 | 0 | 1 | 1.931 | 0.949 | 0.75306031 | NA | 0 | 4.6833 | 4.7842 | 4.7842 | TRUE | 0.9737 |
| 9130206I24R | 23 | 12 | 11 | 6 | 6 | 9 | 18 | 13 | 12 | 6 | 7 | 9 | 10.69 | 14 | 7 | 0.505 | -0.984 | 0.100289 | NA | 0 | 0.2242 | 0.1531 | 0.1531 | TRUE | 0.2832 |
| 9130209A04 | 0 | 0 | 0 | 0 | 0 | 0 | 0 | 0 | 0 | 0 | 0 | 0 | 0 | 0 | 0 | NA | NA | NA | NA | NA | NA | NA | NA | NA | NA |
| 9130221F21I | 0 | 0 | 0 | 0 | 0 | 0 | 0 | 0 | 0 | 0 | 0 | 0 | 0 | 0 | 0 | NA | NA | NA | NA | NA | NA | NA | NA | NA | NA |
| 9130221H12 | 231 | 178 | 178 | 166 | 150 | 182 | 179 | 189 | 192 | 165 | 169 | 180 | 179.15 | 187 | 171 | 0.918 | -0.123 | 0.40524813 | 0.57800733 | 0 | 0.0157 | 0.0101 | 0.0101 | TRUE | 0.0197 |
| 9130227L01F | 11 | 7 | 6 | 0 | 1 | 2 | 9 | 7 | 6 | 0 | 1 | 2 | 4.26 | 7 | 1 | 0.137 | -2.863 | 0.01351796 | NA | 0 | 0.5587 | 0.4033 | 0.4033 | TRUE | 0.3947 |
| 9130230L23F | 12 | 5 | 8 | 3 | 2 | 5 | 9 | 5 | 9 | 3 | 2 | 5 | 5.57 | 8 | 3 | 0.438 | -1.19 | 0.15544825 | NA | 0 | 0.4276 | 0.2928 | 0.2928 | TRUE | 0.2355 |
| 9130401M01 | 4610 | 3369 | 3287 | 3536 | 3023 | 3525 | 3580 | 3584 | 3547 | 3520 | 3410 | 3480 | 3520.05 | 3570 | 3470 | 0.972 | -0.041 | 0.43482503 | 0.6065737 | 0 | 0.0031 | 0.0017 | 0.0017 | TRUE | 0.0027 |
| 9130409I23R | 1 | 2 | 1 | 1 | 0 | 1 | 1 | 2 | 1 | 1 | 0 | 1 | 0.99 | 1 | 1 | 0.516 | -0.956 | 0.61758228 | NA | 0 | 2.3862 | 1.5048 | 1.5048 | TRUE | 0.2008 |
| 9230009I02R | 0 | 0 | 0 | 0 | 0 | 2 | 0 | 0 | 0 | 0 | 0 | 2 | 0.33 | 0 | 1 | 3.743 | 1.904 | 0.62640105 | NA | 0 | 7.2051 | 9.2465 | 9.2465 | TRUE | 0.9739 |
| 9230102K24I | 1 | 0 | 0 | 0 | 0 | 0 | 1 | 0 | 0 | 0 | 0 | 0 | 0.13 | 0 | 0 | 0.552 | -0.857 | 0.8336548 | NA | 0 | 18.3121 | 10 | 10 | TRUE | 0.2566 |
| 9230102O04 | 3 | 0 | 1 | 0 | 0 | 0 | 2 | 0 | 1 | 0 | 0 | 0 | 0.57 | 1 | 0 | 0.165 | -2.602 | 0.35448349 | NA | 0 | 4.1739 | 4.2733 | 4.2733 | TRUE | 0.066 |
| 9230104L09F | 0 | 1 | 0 | 0 | 1 | 0 | 0 | 1 | 0 | 0 | 1 | 0 | 0.37 | 0 | 0 | 1.075 | 0.105 | 0.97520514 | NA | 0 | 6.4904 | 6.1969 | 6.1969 | TRUE | 0.5216 |
| 9230105E05I | 7 | 7 | 8 | 9 | 6 | 15 | 5 | 7 | 9 | 9 | 7 | 15 | 8.67 | 7 | 10 | 1.442 | 0.528 | 0.42676123 | NA | 0 | 0.2756 | 0.1993 | 0.1993 | TRUE | 0.5676 |
| 9230110C19I | 436 | 313 | 322 | 389 | 320 | 356 | 339 | 333 | 347 | 387 | 361 | 351 | 353.1 | 340 | 366 | 1.079 | 0.11 | 0.31861058 | 0.4925266 | 0 | 0.0091 | 0.006 | 0.006 | TRUE | 0.0284 |
| 9230110F15I | 0 | 0 | 0 | 0 | 0 | 0 | 0 | 0 | 0 | 0 | 0 | 0 | 0 | 0 | 0 | NA | NA | NA | NA | NA | NA | NA | NA | NA | NA |
| 9230112D13 | 1 | 0 | 0 | 1 | 0 | 1 | 1 | 0 | 0 | 1 | 0 | 1 | 0.46 | 0 | 1 | 1.94 | 0.956 | 0.73503911 | NA | 0 | 5.1562 | 3.9771 | 3.9771 | TRUE | 0.2566 |
| 9230112J17F | 2 | 1 | 2 | 0 | 2 | 1 | 2 | 1 | 2 | 0 | 2 | 1 | 1.34 | 2 | 1 | 0.666 | -0.586 | 0.72539072 | NA | 0 | 1.7759 | 1.211 | 1.211 | TRUE | 0.3101 |
| 9230114K14I | 99 | 82 | 65 | 89 | 64 | 64 | 77 | 87 | 70 | 89 | 72 | 63 | 76.37 | 78 | 75 | 0.957 | -0.064 | 0.78494305 | 0.87364767 | 0.0075 | 0.0335 | 0.0262 | 0.0262 | TRUE | 0.2507 |
| 9230115E21I | 3 | 0 | 0 | 0 | 0 | 1 | 2 | 0 | 0 | 0 | 0 | 1 | 0.55 | 1 | 0 | 0.458 | -1.128 | 0.69846567 | NA | 0 | 4.2894 | 4.5119 | 4.5119 | TRUE | 1.4399 |
| 9230116N13 | 46 | 33 | 46 | 59 | 36 | 41 | 36 | 35 | 50 | 59 | 41 | 40 | 43.38 | 40 | 47 | 1.167 | 0.223 | 0.47435157 | 0.64280192 | 0.0214 | 0.0571 | 0.0471 | 0.0471 | TRUE | 0.4178 |
| 9330020H09 | 26 | 36 | 22 | 14 | 20 | 15 | 20 | 38 | 24 | 14 | 23 | 15 | 22.25 | 27 | 17 | 0.625 | -0.678 | 0.12619147 | 0.26136135 | 0.0556 | 0.1089 | 0.0934 | 0.0934 | TRUE | 0.7455 |
| 9330111N05 | 0 | 0 | 0 | 0 | 0 | 0 | 0 | 0 | 0 | 0 | 0 | 0 | 0 | 0 | 0 | NA | NA | NA | NA | NA | NA | NA | NA | NA | NA |
| 9330117O12 | 0 | 0 | 0 | 0 | 0 | 0 | 0 | 0 | 0 | 0 | 0 | 0 | 0 | 0 | 0 | NA | NA | NA | NA | NA | NA | NA | NA | NA | NA |
| 9330133O14 | 102 | 89 | 50 | 67 | 69 | 44 | 79 | 95 | 54 | 67 | 78 | 43 | 69.3 | 76 | 63 | 0.821 | -0.284 | 0.32144588 | 0.49524911 | 0.0648 | 0.0366 | 0.0444 | 0.0444 | TRUE | 0.3476 |
| 9330151L19F | 128 | 137 | 157 | 150 | 111 | 157 | 99 | 146 | 169 | 149 | 125 | 155 | 140.68 | 138 | 143 | 1.044 | 0.063 | 0.76355101 | 0.85969854 | 0.0358 | 0.0193 | 0.0242 | 0.0242 | TRUE | 0.5555 |
| 9330158H04 | 0 | 0 | 0 | 0 | 0 | 0 | 0 | 0 | 0 | 0 | 0 | 0 | 0 | 0 | 0 | NA | NA | NA | NA | NA | NA | NA | NA | NA | NA |
| 9330159F19I | 36 | 31 | 34 | 54 | 43 | 57 | 28 | 33 | 37 | 54 | 49 | 56 | 42.69 | 33 | 53 | 1.637 | 0.711 | 0.01715006 | 0.06177565 | 0 | 0.0579 | 0.0394 | 0.0394 | TRUE | 0.1281 |
| 9330159M07 | 69 | 58 | 43 | 54 | 45 | 45 | 54 | 62 | 46 | 54 | 51 | 44 | 51.77 | 54 | 50 | 0.921 | -0.119 | 0.66118944 | 0.79076066 | 0 | 0.0482 | 0.0339 | 0.0339 | TRUE | 0.1228 |
| 9330162B11 | 0 | 0 | 0 | 0 | 0 | 0 | 0 | 0 | 0 | 0 | 0 | 0 | 0 | 0 | 0 | NA | NA | NA | NA | NA | NA | NA | NA | NA | NA |
| 9330175E14I | 1 | 1 | 0 | 1 | 0 | 0 | 1 | 1 | 0 | 1 | 0 | 0 | 0.47 | 1 | 0 | 0.599 | -0.74 | 0.79191656 | NA | 0 | 5.0171 | 3.8827 | 3.8827 | TRUE | 0.3906 |
| 9330175M20 | 0 | 0 | 0 | 0 | 0 | 0 | 0 | 0 | 0 | 0 | 0 | 0 | 0 | 0 | 0 | NA | NA | NA | NA | NA | NA | NA | NA | NA | NA |
| 9330178D15 | 0 | 0 | 0 | 0 | 0 | 0 | 0 | 0 | 0 | 0 | 0 | 0 | 0 | 0 | 0 | NA | NA | NA | NA | NA | NA | NA | NA | NA | NA |
| 9330179D12 | 1 | 0 | 0 | 0 | 0 | 0 | 1 | 0 | 0 | 0 | 0 | 0 | 0.13 | 0 | 0 | 0.552 | -0.857 | 0.8336548 | NA | 0 | 18.3121 | 10 | 10 | TRUE | 0.2566 |
| 9330182L06F | 437 | 311 | 308 | 468 | 395 | 405 | 339 | 331 | 332 | 466 | 446 | 400 | 385.63 | 334 | 437 | 1.307 | 0.386 | 0.0004831 | 0.00421031 | 0 | 0.0086 | 0.0062 | 0.0062 | TRUE | 0.0659 |
| 9330182O14 | 3 | 0 | 2 | 5 | 0 | 0 | 2 | 0 | 2 | 5 | 0 | 0 | 1.58 | 1 | 2 | 1.104 | 0.143 | 0.93777213 | NA | 3.5114 | 1.5048 | 1.7999 | 1.7999 | TRUE | 2.3423 |
| 9430007A20 | 0 | 0 | 0 | 0 | 0 | 0 | 0 | 0 | 0 | 0 | 0 | 0 | 0 | 0 | 0 | NA | NA | NA | NA | NA | NA | NA | NA | NA | NA |
| 9430008C03I | 1194 | 1040 | 932 | 912 | 815 | 922 | 927 | 1106 | 1006 | 908 | 919 | 910 | 962.76 | 1013 | 912 | 0.902 | -0.149 | 0.08339817 | 0.19442342 | 0.0032 | 0.0049 | 0.0043 | 0.0043 | TRUE | 0.0774 |
| 9430014N10 | 0 | 0 | 0 | 0 | 0 | 0 | 0 | 0 | 0 | 0 | 0 | 0 | 0 | 0 | 0 | NA | NA | NA | NA | NA | NA | NA | NA | NA | NA |
| 9430015G10 | 1705 | 1485 | 1303 | 1566 | 1362 | 1489 | 1324 | 1580 | 1406 | 1559 | 1536 | 1470 | 1479.14 | 1437 | 1522 | 1.06 | 0.085 | 0.29270693 | 0.46492953 | 0.0036 | 0.004 | 0.004 | 0.004 | TRUE | 0.0909 |
| 9430016H08 | 1051 | 758 | 722 | 743 | 595 | 695 | 816 | 806 | 779 | 740 | 671 | 686 | 749.74 | 800 | 699 | 0.873 | -0.196 | 0.02033935 | 0.0700311 | 0 | 0.0056 | 0.0038 | 0.0038 | TRUE | 0.0307 |
| 9430018G01 | 76 | 62 | 40 | 29 | 22 | 27 | 59 | 66 | 43 | 29 | 25 | 27 | 41.41 | 56 | 27 | 0.478 | -1.066 | 0.000817 | 0.00624456 | 0 | 0.0597 | 0.0452 | 0.0452 | TRUE | 0.3184 |
| 9430019J16F | 0 | 0 | 0 | 0 | 0 | 0 | 0 | 0 | 0 | 0 | 0 | 0 | 0 | 0 | 0 | NA | NA | NA | NA | NA | NA | NA | NA | NA | NA |
| 9430020K01I | 2257 | 1699 | 1665 | 1846 | 1594 | 1644 | 1753 | 1807 | 1797 | 1838 | 1798 | 1623 | 1769.2 | 1786 | 1753 | 0.982 | -0.027 | 0.7060871 | 0.82216725 | 0.0017 | 0.0038 | 0.003 | 0.003 | TRUE | 0.0514 |
| 9430021M05 | 0 | 0 | 1 | 0 | 0 | 3 | 0 | 0 | 1 | 0 | 0 | 3 | 0.67 | 0 | 1 | 2.892 | 1.532 | 0.57632544 | NA | 7.6973 | 3.5219 | 3.8977 | 3.8977 | TRUE | 1.4418 |
| 9430023L20F | 3141 | 2206 | 1946 | 2139 | 1865 | 2106 | 2439 | 2346 | 2100 | 2129 | 2104 | 2079 | 2199.62 | 2295 | 2104 | 0.916 | -0.127 | 0.07474966 | 0.17993429 | 0.0027 | 0.0035 | 0.0032 | 0.0032 | TRUE | 0.0666 |
| 9430037G07 | 38 | 29 | 12 | 28 | 29 | 23 | 30 | 31 | 13 | 28 | 33 | 23 | 26.1 | 25 | 28 | 1.125 | 0.17 | 0.68248168 | 0.80530922 | 0.0658 | 0.0932 | 0.0863 | 0.0863 | TRUE | 0.9367 |
| 9430038I01R | 676 | 483 | 318 | 419 | 403 | 419 | 525 | 514 | 343 | 417 | 455 | 414 | 444.54 | 461 | 429 | 0.928 | -0.108 | 0.46009193 | 0.62988358 | 0.026 | 0.0078 | 0.0132 | 0.0132 | TRUE | 0.5661 |
| 9430041J12F | 7 | 12 | 5 | 6 | 3 | 1 | 5 | 13 | 5 | 6 | 3 | 1 | 5.66 | 8 | 3 | 0.442 | -1.179 | 0.18517082 | NA | 0.1857 | 0.4214 | 0.3589 | 0.3589 | TRUE | 0.442 |
| 9430060I03R | 10 | 9 | 5 | 1 | 8 | 4 | 8 | 10 | 5 | 1 | 9 | 4 | 6.12 | 8 | 5 | 0.604 | -0.727 | 0.3908192 | NA | 0.1735 | 0.3899 | 0.3409 | 0.3409 | TRUE | 0.5891 |
| 9430069I07R | 3 | 4 | 6 | 6 | 5 | 14 | 2 | 4 | 6 | 6 | 6 | 14 | 6.42 | 4 | 9 | 2 | 1 | 0.21633523 | NA | 0.1179 | 0.3719 | 0.2976 | 0.2976 | TRUE | 0.9588 |
| 9430076C15I | 4 | 1 | 3 | 0 | 0 | 1 | 3 | 1 | 3 | 0 | 0 | 1 | 1.4 | 2 | 0 | 0.149 | -2.744 | 0.13743044 | NA | 0 | 1.6964 | 1.2638 | 1.2638 | TRUE | 0.3835 |
| 9430076G02 | 4 | 0 | 1 | 0 | 0 | 0 | 3 | 0 | 1 | 0 | 0 | 0 | 0.7 | 1 | 0 | 0.134 | -2.902 | 0.26925366 | NA | 6.5513 | 3.3999 | 3.6307 | 3.6307 | TRUE | 0.1404 |
| 9430083A17 | 9 | 21 | 5 | 4 | 12 | 10 | 7 | 22 | 5 | 4 | 14 | 10 | 10.35 | 11 | 9 | 0.791 | -0.338 | 0.63623242 | NA | 0.3555 | 0.2314 | 0.2685 | 0.2685 | TRUE | 1.2757 |
| 9430091E24I | 17 | 11 | 5 | 11 | 14 | 13 | 13 | 12 | 5 | 11 | 16 | 13 | 11.65 | 10 | 13 | 1.286 | 0.363 | 0.53034958 | NA | 0 | 0.2059 | 0.1544 | 0.1544 | TRUE | 0.5283 |
| 9530002B09 | 0 | 0 | 0 | 0 | 0 | 0 | 0 | 0 | 0 | 0 | 0 | 0 | 0 | 0 | 0 | NA | NA | NA | NA | NA | NA | NA | NA | NA | NA |
| 9530003J23F | 0 | 0 | 0 | 0 | 0 | 0 | 0 | 0 | 0 | 0 | 0 | 0 | 0 | 0 | 0 | NA | NA | NA | NA | NA | NA | NA | NA | NA | NA |
| 9530026F06I | 0 | 0 | 0 | 0 | 0 | 0 | 0 | 0 | 0 | 0 | 0 | 0 | 0 | 0 | 0 | NA | NA | NA | NA | NA | NA | NA | NA | NA | NA |
| 9530026P05I | 8 | 8 | 9 | 10 | 4 | 9 | 6 | 9 | 10 | 10 | 5 | 9 | 7.96 | 8 | 8 | 0.971 | -0.042 | 0.95085441 | NA | 0 | 0.3 | 0.2101 | 0.2101 | TRUE | 0.3432 |
| 9530027J09F | 5 | 4 | 1 | 9 | 5 | 7 | 4 | 4 | 1 | 9 | 6 | 7 | 5.12 | 3 | 7 | 2.306 | 1.205 | 0.16941625 | NA | 0 | 0.4652 | 0.328 | 0.328 | TRUE | 0.3717 |
| 9530036O11 | 0 | 0 | 1 | 1 | 1 | 0 | 0 | 0 | 1 | 1 | 1 | 0 | 0.53 | 0 | 1 | 2.036 | 1.026 | 0.70454668 | NA | 0 | 4.4428 | 3.5371 | 3.5371 | TRUE | 0.5365 |
| 9530048J24F | 0 | 0 | 0 | 0 | 0 | 0 | 0 | 0 | 0 | 0 | 0 | 0 | 0 | 0 | 0 | NA | NA | NA | NA | NA | NA | NA | NA | NA | NA |
| 9530051G07 | 9 | 11 | 12 | 23 | 10 | 15 | 7 | 12 | 13 | 23 | 11 | 15 | 13.44 | 11 | 16 | 1.577 | 0.657 | 0.23391191 | NA | 0 | 0.1788 | 0.1415 | 0.1415 | TRUE | 0.4768 |
| 9530052E02I | 0 | 0 | 1 | 0 | 0 | 0 | 0 | 0 | 1 | 0 | 0 | 0 | 0.18 | 0 | 0 | 0.552 | -0.857 | 0.8336548 | NA | 0 | 13.1817 | 10 | 10 | TRUE | 0.5365 |
| 9530053A07 | 51 | 31 | 31 | 44 | 29 | 36 | 40 | 33 | 33 | 44 | 33 | 36 | 36.35 | 35 | 38 | 1.054 | 0.076 | 0.81364867 | 0.89251577 | 0 | 0.0676 | 0.0468 | 0.0468 | TRUE | 0.1684 |
| 9530059O14 | 2 | 0 | 0 | 1 | 1 | 0 | 2 | 0 | 0 | 1 | 1 | 0 | 0.61 | 1 | 1 | 1.262 | 0.336 | 0.89669251 | NA | 0 | 3.87 | 3.2065 | 3.2065 | TRUE | 0.9721 |
| 9530068E07I | 2524 | 1941 | 1992 | 2281 | 1981 | 2132 | 1960 | 2065 | 2149 | 2271 | 2235 | 2105 | 2130.69 | 2058 | 2204 | 1.071 | 0.099 | 0.13576446 | 0.27520656 | 0.0014 | 0.0035 | 0.0027 | 0.0027 | TRUE | 0.0216 |
| 9530077C05I | 242 | 165 | 143 | 167 | 135 | 169 | 188 | 176 | 154 | 166 | 152 | 167 | 167.18 | 173 | 162 | 0.935 | -0.096 | 0.54203131 | 0.6999739 | 0 | 0.0166 | 0.012 | 0.012 | TRUE | 0.0892 |

| | | | | | | | | | | | | | | | | | | | | | | | | | |
|---|---|---|---|---|---|---|---|---|---|---|---|---|---|---|---|---|---|---|---|---|---|---|---|---|---|
| 9530080O11 | 9 | 5 | 5 | 8 | 13 | 7 | 7 | 5 | 5 | 8 | 15 | 7 | 7.87 | 6 | 10 | 1.643 | 0.716 | 0.30352312 | NA | 0 | 0.3034 | 0.2153 | 0.2153 | TRUE | 0.5697 |
| 9530082P21 | 40 | 32 | 24 | 39 | 32 | 41 | 31 | 34 | 26 | 39 | 36 | 40 | 34.4 | 30 | 38 | 1.268 | 0.342 | 0.29393531 | 0.46590435 | 0 | 0.0713 | 0.0473 | 0.0473 | TRUE | 0.0984 |
| 9530091C08 | 0 | 0 | 1 | 4 | 3 | 2 | 0 | 0 | 1 | 4 | 3 | 2 | 1.74 | 0 | 3 | 9.019 | 3.173 | 0.06730035 | NA | 0 | 1.3672 | 1.0009 | 1.0009 | TRUE | 0.5365 |
| 9630013A20 | 0 | 0 | 0 | 0 | 0 | 0 | 0 | 0 | 0 | 0 | 0 | 0 | 0 | 0 | 0 | NA | NA | NA | NA | NA | NA | NA | NA | NA | NA |
| 9630028B13 | 0 | 1 | 1 | 0 | 0 | 0 | 0 | 1 | 1 | 0 | 0 | 0 | 0.36 | 1 | 0 | 0.271 | -1.881 | 0.5755355 | NA | 0 | 6.6391 | 6.4428 | 6.4428 | TRUE | 0.3513 |
| 9630028H03 | 0 | 0 | 0 | 0 | 0 | 0 | 0 | 0 | 0 | 0 | 0 | 0 | 0 | 0 | 0 | NA | NA | NA | NA | NA | NA | NA | NA | NA | NA |
| 9630033F20 | 3696 | 2632 | 2108 | 2444 | 2179 | 2271 | 2871 | 2800 | 2275 | 2433 | 2458 | 2242 | 2512.9 | 2649 | 2378 | 0.897 | -0.157 | 0.07696853 | 0.18362731 | 0.0082 | 0.0034 | 0.0052 | 0.0052 | TRUE | 0.1835 |
| 9830001H06 | 1118 | 760 | 706 | 766 | 618 | 710 | 868 | 808 | 762 | 762 | 697 | 701 | 766.5 | 813 | 720 | 0.885 | -0.176 | 0.04800719 | 0.13077845 | 0.0023 | 0.0055 | 0.0044 | 0.0044 | TRUE | 0.0444 |
| 9830107B12 | 0 | 0 | 0 | 0 | 0 | 0 | 0 | 0 | 0 | 0 | 0 | 0 | 0 | 0 | 0 | NA | NA | NA | NA | NA | NA | NA | NA | NA | NA |
| 9830132P13 | 0 | 0 | 0 | 0 | 0 | 0 | 0 | 0 | 0 | 0 | 0 | 0 | 0 | 0 | 0 | NA | NA | NA | NA | NA | NA | NA | NA | NA | NA |
| 9830147E19 | 1223 | 923 | 848 | 984 | 824 | 959 | 950 | 982 | 915 | 979 | 930 | 947 | 950.39 | 949 | 952 | 1.003 | 0.005 | 0.94939913 | 0.97225491 | 0 | 0.0049 | 0.0032 | 0.0032 | TRUE | 0.0111 |
| 9830166K06 | 2 | 1 | 2 | 0 | 0 | 0 | 2 | 1 | 2 | 0 | 0 | 0 | 0.8 | 2 | 0 | 0.12 | -3.062 | 0.17528122 | NA | 0 | 2.9805 | 2.3743 | 2.3743 | TRUE | 0.0718 |
| 9930012K11 | 262 | 199 | 158 | 220 | 184 | 229 | 203 | 212 | 170 | 219 | 208 | 226 | 206.38 | 195 | 218 | 1.113 | 0.155 | 0.28836329 | 0.46051134 | 0 | 0.0139 | 0.0105 | 0.0105 | TRUE | 0.1257 |
| 9930013L23 | 37 | 19 | 16 | 26 | 12 | 15 | 29 | 20 | 17 | 26 | 14 | 15 | 20.07 | 22 | 18 | 0.814 | -0.297 | 0.51772471 | 0.67975742 | 0.0524 | 0.1205 | 0.1013 | 0.1013 | TRUE | 0.7402 |
| 9930014A18 | 14 | 6 | 5 | 2 | 10 | 8 | 11 | 6 | 5 | 2 | 11 | 8 | 7.3 | 7 | 7 | 0.914 | -0.13 | 0.86652074 | NA | 0.1954 | 0.3269 | 0.2921 | 0.2921 | TRUE | 0.4447 |
| 9930021J03 | 3159 | 2327 | 2389 | 2600 | 2156 | 2595 | 2453 | 2475 | 2578 | 2588 | 2432 | 2562 | 2514.71 | 2502 | 2527 | 1.011 | 0.015 | 0.79980549 | 0.88333244 | 5.00E-04 | 0.0034 | 0.0022 | 0.0022 | TRUE | 0.0129 |
| 9930104L06 | 1064 | 789 | 639 | 643 | 631 | 688 | 826 | 839 | 689 | 640 | 712 | 679 | 731.02 | 785 | 677 | 0.861 | -0.216 | 0.02986775 | 0.09240255 | 0.0059 | 0.0057 | 0.0057 | 0.0057 | TRUE | 0.1311 |
| 9930111J21 | 3 | 0 | 1 | 1 | 0 | 0 | 2 | 0 | 1 | 1 | 0 | 0 | 0.73 | 1 | 0 | 0.318 | -1.651 | 0.49291572 | NA | 0 | 3.2311 | 2.7619 | 2.7619 | TRUE | 0.1857 |
| 9930111J21 | 0 | 0 | 0 | 0 | 0 | 0 | 0 | 0 | 0 | 0 | 0 | 0 | 0 | 0 | 0 | NA | NA | NA | NA | NA | NA | NA | NA | NA | NA |
| a | 2 | 4 | 8 | 4 | 2 | 4 | 2 | 4 | 9 | 4 | 2 | 4 | 4.1 | 5 | 3 | 0.726 | -0.462 | 0.64154562 | NA | 0 | 0.5799 | 0.455 | 0.455 | TRUE | 0.2804 |
| A030009H04 | 402 | 276 | 230 | 271 | 217 | 242 | 312 | 294 | 248 | 270 | 245 | 239 | 267.9 | 285 | 251 | 0.88 | -0.184 | 0.17530454 | 0.32873626 | 0.0056 | 0.0113 | 0.0096 | 0.0096 | TRUE | 0.1377 |
| A130010J15 | 925 | 699 | 675 | 847 | 696 | 749 | 718 | 744 | 728 | 843 | 785 | 739 | 759.65 | 730 | 789 | 1.081 | 0.113 | 0.19027493 | 0.34831415 | 0.0011 | 0.0055 | 0.004 | 0.004 | TRUE | 0.0432 |
| A130049A11 | 29 | 40 | 48 | 25 | 35 | 41 | 23 | 43 | 52 | 25 | 39 | 40 | 36.95 | 39 | 35 | 0.906 | -0.142 | 0.70555193 | 0.82174868 | 0.0958 | 0.0666 | 0.0751 | 0.0751 | TRUE | 0.594 |
| A130077B15 | 0 | 0 | 0 | 0 | 0 | 0 | 0 | 0 | 0 | 0 | 0 | 0 | 0 | 0 | 0 | NA | NA | NA | NA | NA | NA | NA | NA | NA | NA |
| A1bg | 0 | 0 | 1 | 1 | 0 | 0 | 0 | 0 | 1 | 1 | 0 | 0 | 0.35 | 0 | 0 | 1.075 | 0.105 | 0.97567165 | NA | 0 | 6.8576 | 6.5143 | 6.5143 | TRUE | 0.5365 |
| A1cf | 0 | 1 | 0 | 0 | 0 | 0 | 0 | 1 | 0 | 0 | 0 | 0 | 0.18 | 0 | 0 | 0.552 | -0.857 | 0.8336548 | NA | 0 | 13.3714 | 10 | 10 | TRUE | 0.5216 |
| A230001M1 | 0 | 1 | 0 | 0 | 0 | 0 | 0 | 1 | 0 | 0 | 0 | 0 | 0.18 | 0 | 0 | 0.552 | -0.857 | 0.8336548 | NA | 0 | 13.3714 | 10 | 10 | TRUE | 0.5216 |
| A230020J21 | 6 | 1 | 0 | 2 | 0 | 0 | 5 | 1 | 0 | 2 | 0 | 0 | 1.29 | 2 | 1 | 0.343 | -1.542 | 0.44850371 | NA | 2.6237 | 1.8458 | 1.9696 | 1.9696 | TRUE | 0.7551 |
| A230028O05 | 10 | 1 | 3 | 4 | 2 | 2 | 8 | 1 | 3 | 4 | 2 | 2 | 3.38 | 4 | 3 | 0.664 | -0.592 | 0.58921099 | NA | 0 | 0.7037 | 0.5597 | 0.5597 | TRUE | 0.444 |
| A230046K03 | 9262 | 6722 | 7062 | 7829 | 6692 | 7621 | 7193 | 7150 | 7620 | 7793 | 7549 | 7523 | 7471.45 | 7321 | 7622 | 1.041 | 0.058 | 0.26271903 | 0.43318169 | 6.00E-04 | 0.0027 | 0.0018 | 0.0018 | TRUE | 0.0154 |
| A230050P20 | 660 | 589 | 601 | 516 | 528 | 634 | 513 | 627 | 648 | 514 | 596 | 626 | 587.12 | 596 | 579 | 0.973 | -0.04 | 0.73344333 | 0.84073346 | 0.0116 | 0.0065 | 0.0081 | 0.0081 | TRUE | 0.1843 |
| A230056J06 | 2 | 5 | 1 | 4 | 4 | 2 | 2 | 5 | 1 | 4 | 5 | 2 | 3.07 | 3 | 4 | 1.331 | 0.413 | 0.71264762 | NA | 0 | 0.7745 | 0.5719 | 0.5719 | TRUE | 0.8361 |
| A230056P14 | 19 | 13 | 9 | 13 | 8 | 14 | 15 | 14 | 10 | 13 | 9 | 14 | 12.35 | 13 | 12 | 0.933 | -0.101 | 0.85320874 | NA | 0 | 0.1944 | 0.1327 | 0.1327 | TRUE | 0.1658 |
| A230057D06 | 2 | 1 | 1 | 0 | 3 | 0 | 2 | 1 | 1 | 0 | 3 | 0 | 1.18 | 1 | 1 | 0.88 | -0.185 | 0.92210897 | NA | 0.6585 | 2.0109 | 1.7084 | 1.7084 | TRUE | 1.444 |
| A230065H16 | 13 | 2 | 2 | 4 | 4 | 3 | 10 | 2 | 2 | 4 | 5 | 3 | 4.31 | 5 | 4 | 0.768 | -0.381 | 0.6987602 | NA | 0.284 | 0.5528 | 0.4652 | 0.4652 | TRUE | 2.3161 |
| A230070E04 | 39 | 21 | 19 | 36 | 19 | 31 | 30 | 22 | 21 | 36 | 21 | 31 | 26.83 | 24 | 29 | 1.195 | 0.257 | 0.50669906 | 0.67002033 | 0 | 0.0908 | 0.0709 | 0.0709 | TRUE | 0.3303 |
| A230072C01 | 105 | 95 | 89 | 106 | 93 | 124 | 82 | 101 | 96 | 106 | 105 | 122 | 101.91 | 93 | 111 | 1.201 | 0.264 | 0.18485804 | 0.34114602 | 0 | 0.0257 | 0.0187 | 0.0187 | TRUE | 0.1252 |
| A230072E10 | 0 | 0 | 1 | 0 | 0 | 0 | 0 | 0 | 1 | 0 | 0 | 0 | 0.18 | 0 | 0 | 0.552 | -0.857 | 0.8336548 | NA | 0 | 13.1817 | 10 | 10 | TRUE | 0.5365 |
| A230073K19 | 9 | 1 | 9 | 4 | 7 | 6 | 7 | 1 | 10 | 4 | 8 | 6 | 5.93 | 6 | 6 | 0.993 | -0.011 | 0.99018918 | NA | 0.2179 | 0.4023 | 0.3576 | 0.3576 | TRUE | 0.5408 |
| A230077H06 | 6 | 6 | 3 | 2 | 2 | 3 | 5 | 6 | 3 | 2 | 2 | 3 | 3.58 | 5 | 2 | 0.506 | -0.984 | 0.33236891 | NA | 0 | 0.6642 | 0.4254 | 0.4254 | TRUE | 0.1538 |
| A230108P19 | 0 | 0 | 0 | 0 | 0 | 1 | 0 | 0 | 0 | 0 | 0 | 1 | 0.16 | 0 | 0 | 2.094 | 1.067 | 0.79379329 | NA | 0 | 14.4078 | 10 | 10 | TRUE | 0.3835 |
| A2m | 11379 | 10307 | 14205 | 10480 | 11233 | 12242 | 8838 | 10963 | 15327 | 10432 | 12672 | 12085 | 11719.43 | 11709 | 11730 | 1.002 | 0.003 | 0.99085759 | 0.99545425 | 0.0357 | 0.0026 | 0.0137 | 0.0357 | TRUE | 0.5332 |
| A330009N23 | 46 | 26 | 17 | 28 | 33 | 30 | 36 | 28 | 18 | 28 | 37 | 30 | 29.41 | 27 | 32 | 1.143 | 0.193 | 0.60898396 | 0.75256855 | 0.0297 | 0.083 | 0.0685 | 0.0685 | TRUE | 0.2572 |
| A330021E22 | 136 | 127 | 107 | 111 | 105 | 115 | 106 | 135 | 115 | 110 | 118 | 114 | 116.44 | 119 | 114 | 0.965 | -0.052 | 0.78040906 | 0.87053897 | 0 | 0.0228 | 0.0165 | 0.0165 | TRUE | 0.1448 |
| A330023F24 | 13 | 17 | 9 | 8 | 9 | 7 | 10 | 18 | 10 | 8 | 10 | 7 | 10.49 | 13 | 8 | 0.662 | -0.595 | 0.32671382 | NA | 0 | 0.2284 | 0.1648 | 0.1648 | TRUE | 0.5484 |
| A330032B11 | 0 | 0 | 0 | 0 | 1 | 0 | 0 | 0 | 0 | 0 | 1 | 0 | 0.19 | 0 | 0 | 2.094 | 1.067 | 0.79379329 | NA | 0 | 12.608 | 10 | 10 | TRUE | 0.5075 |
| A330035P11 | 18 | 14 | 15 | 16 | 11 | 11 | 14 | 15 | 16 | 16 | 12 | 11 | 14.04 | 15 | 13 | 0.873 | -0.196 | 0.69793203 | NA | 0 | 0.1712 | 0.1123 | 0.1123 | TRUE | 0.1588 |
| A330040F15 | 2 | 0 | 0 | 0 | 0 | 0 | 2 | 0 | 0 | 0 | 0 | 0 | 0.26 | 1 | 0 | 0.356 | -1.491 | 0.71362796 | NA | 0 | 9.1573 | 10 | 10 | TRUE | 0.968 |
| A330041J22 | 1 | 2 | 1 | 5 | 3 | 1 | 1 | 2 | 1 | 5 | 3 | 1 | 2.22 | 1 | 3 | 2.394 | 1.259 | 0.34618011 | NA | 0 | 1.0691 | 0.7509 | 0.7509 | TRUE | 0.1966 |
| A330048O09 | 17 | 14 | 9 | 11 | 16 | 20 | 13 | 15 | 10 | 11 | 18 | 20 | 14.42 | 13 | 16 | 1.284 | 0.361 | 0.48344382 | NA | 0 | 0.1667 | 0.1213 | 0.1213 | TRUE | 0.4019 |
| A330049N07 | 0 | 0 | 0 | 0 | 0 | 0 | 0 | 0 | 0 | 0 | 0 | 0 | 0 | 0 | 0 | NA | NA | NA | NA | NA | NA | NA | NA | NA | NA |
| A330050B17 | 6 | 1 | 1 | 2 | 2 | 3 | 5 | 1 | 1 | 2 | 2 | 3 | 2.34 | 2 | 2 | 1.02 | 0.028 | 0.9821844 | NA | 0 | 1.0174 | 0.7122 | 0.7122 | TRUE | 1.1246 |
| A330050F15 | 7 | 7 | 6 | 24 | 19 | 23 | 5 | 7 | 6 | 24 | 21 | 23 | 14.56 | 6 | 23 | 3.551 | 1.828 | 0.00064392 | NA | 0 | 0.1652 | 0.1089 | 0.1089 | TRUE | 0.0661 |
| A330069E16 | 54 | 51 | 54 | 71 | 38 | 64 | 42 | 54 | 58 | 71 | 43 | 63 | 55.2 | 51 | 59 | 1.155 | 0.208 | 0.46562013 | 0.63463117 | 0.028 | 0.0454 | 0.0406 | 0.0406 | TRUE | 0.4548 |
| A330070K13 | 0 | 0 | 0 | 0 | 0 | 0 | 0 | 0 | 0 | 0 | 0 | 0 | 0 | 0 | 0 | NA | NA | NA | NA | NA | NA | NA | NA | NA | NA |
| A330076C08 | 0 | 0 | 0 | 0 | 0 | 0 | 0 | 0 | 0 | 0 | 0 | 0 | 0 | 0 | 0 | NA | NA | NA | NA | NA | NA | NA | NA | NA | NA |
| A330076H08 | 3 | 3 | 5 | 3 | 2 | 3 | 2 | 3 | 5 | 3 | 2 | 3 | 3.19 | 3 | 3 | 0.765 | -0.387 | 0.71368077 | NA | 0 | 0.7462 | 0.477 | 0.477 | TRUE | 0.2576 |
| A330093E20 | 1 | 0 | 0 | 0 | 0 | 0 | 1 | 0 | 0 | 0 | 0 | 0 | 0.13 | 0 | 0 | 0.552 | -0.857 | 0.8336548 | NA | 0 | 18.3121 | 10 | 10 | TRUE | 0.2566 |
| A330102I10 | 26 | 23 | 18 | 22 | 14 | 21 | 20 | 24 | 19 | 22 | 16 | 21 | 20.42 | 21 | 20 | 0.916 | -0.126 | 0.76556192 | 0.86097856 | 0 | 0.1185 | 0.0795 | 0.0795 | TRUE | 0.1314 |
| A3galt2 | 0 | 0 | 0 | 0 | 0 | 0 | 0 | 0 | 0 | 0 | 0 | 0 | 0 | 0 | 0 | NA | NA | NA | NA | NA | NA | NA | NA | NA | NA |
| A430005L14 | 2629 | 1940 | 1792 | 1622 | 1509 | 1744 | 2042 | 2064 | 1934 | 1615 | 1702 | 1722 | 1846.23 | 2013 | 1680 | 0.834 | -0.262 | 6.23E-05 | 0.00083881 | 6.00E-04 | 0.0037 | 0.0025 | 0.0025 | TRUE | 0.0141 |
| A430033K04 | 292 | 205 | 219 | 277 | 236 | 280 | 227 | 218 | 236 | 276 | 266 | 276 | 249.92 | 227 | 273 | 1.202 | 0.265 | 0.03638809 | 0.10703744 | 0 | 0.0119 | 0.0076 | 0.0076 | TRUE | 0.0129 |
| A430035B10 | 1 | 1 | 3 | 1 | 1 | 0 | 1 | 1 | 3 | 1 | 1 | 0 | 1.2 | 2 | 1 | 0.42 | -1.252 | 0.49196762 | NA | 0 | 1.9773 | 1.3614 | 1.3614 | TRUE | 0.4566 |
| A430071A18 | 1 | 0 | 0 | 0 | 0 | 0 | 1 | 0 | 0 | 0 | 0 | 0 | 0.13 | 0 | 0 | 0.552 | -0.857 | 0.8336548 | NA | 0 | 18.3121 | 10 | 10 | TRUE | 0.2566 |
| A430078G23 | 26 | 21 | 9 | 30 | 29 | 19 | 20 | 22 | 10 | 30 | 33 | 19 | 22.26 | 17 | 27 | 1.542 | 0.625 | 0.16311301 | 0.31365186 | 0.066 | 0.1089 | 0.0981 | 0.0981 | TRUE | 0.6772 |
| A430088P11 | 0 | 0 | 0 | 0 | 0 | 0 | 0 | 0 | 0 | 0 | 0 | 0 | 0 | 0 | 0 | NA | NA | NA | NA | NA | NA | NA | NA | NA | NA |
| A430089I19 | 0 | 0 | 1 | 0 | 0 | 0 | 0 | 0 | 1 | 0 | 0 | 0 | 0.18 | 0 | 0 | 0.552 | -0.857 | 0.8336548 | NA | 0 | 13.1817 | 10 | 10 | TRUE | 0.5365 |
| A430090L17 | 0 | 1 | 0 | 0 | 0 | 0 | 0 | 1 | 0 | 0 | 0 | 0 | 0.18 | 0 | 0 | 0.552 | -0.857 | 0.8336548 | NA | 0 | 13.3714 | 10 | 10 | TRUE | 0.5216 |
| A430092G05 | 0 | 0 | 0 | 0 | 0 | 0 | 0 | 0 | 0 | 0 | 0 | 0 | 0 | 0 | 0 | NA | NA | NA | NA | NA | NA | NA | NA | NA | NA |

| | | | | | | | | | | | | | | | | | | | | | | | | | |
|---|---|---|---|---|---|---|---|---|---|---|---|---|---|---|---|---|---|---|---|---|---|---|---|---|---|
| A430093F15 | 2 | 0 | 1 | 3 | 1 | 1 | 2 | 0 | 1 | 3 | 1 | 1 | 1.29 | 1 | 2 | 1.878 | 0.909 | 0.59350075 | NA | 0 | 1.8412 | 1.2591 | 1.2591 | TRUE | 0.3462 |
| A430105I19F | 61 | 57 | 73 | 78 | 65 | 77 | 47 | 61 | 79 | 78 | 73 | 76 | 68.96 | 62 | 76 | 1.226 | 0.294 | 0.24504746 | 0.41331878 | 0.0178 | 0.0368 | 0.0315 | 0.0315 | TRUE | 0.3386 |
| A430107P09 | 3 | 5 | 6 | 4 | 0 | 5 | 2 | 5 | 6 | 4 | 0 | 5 | 3.84 | 4 | 3 | 0.652 | -0.618 | 0.5627337 | NA | 0.2527 | 0.6196 | 0.5454 | 0.5454 | TRUE | 0.8486 |
| A4galt | 566 | 366 | 407 | 415 | 359 | 452 | 440 | 389 | 439 | 413 | 405 | 446 | 422.05 | 423 | 421 | 0.996 | -0.005 | 0.96220892 | 0.97919915 | 0.0014 | 0.008 | 0.006 | 0.006 | TRUE | 0.0534 |
| A4gnt | 0 | 0 | 0 | 1 | 0 | 0 | 0 | 0 | 0 | 1 | 0 | 0 | 0.17 | 0 | 0 | 2.094 | 1.067 | 0.79379329 | NA | 0 | 14.2885 | 10 | 10 | TRUE | 0.3906 |
| A530006G24 | 0 | 0 | 0 | 0 | 0 | 0 | 0 | 0 | 0 | 0 | 0 | 0 | 0 | 0 | 0 | NA | NA | NA | NA | NA | NA | NA | NA | NA | NA |
| A530013C23 | 1 | 0 | 0 | 0 | 0 | 0 | 1 | 0 | 0 | 0 | 0 | 0 | 0.13 | 0 | 0 | 0.552 | -0.857 | 0.8336548 | NA | 0 | 18.3121 | 10 | 10 | TRUE | 0.2566 |
| A530016L24I | 1 | 1 | 0 | 0 | 0 | 0 | 1 | 1 | 0 | 0 | 0 | 0 | 0.31 | 1 | 0 | 0.309 | -1.696 | 0.63183309 | NA | 0 | 7.7294 | 7.2472 | 7.2472 | TRUE | 0.2015 |
| A530032D15 | 4 | 2 | 4 | 3 | 0 | 0 | 3 | 2 | 4 | 3 | 0 | 0 | 2.09 | 3 | 1 | 0.319 | -1.647 | 0.27365753 | NA | 0 | 1.1368 | 0.97 | 0.97 | TRUE | 1.4412 |
| A530046M15 | 1 | 1 | 0 | 0 | 0 | 0 | 1 | 1 | 0 | 0 | 0 | 0 | 0.31 | 1 | 0 | 0.309 | -1.696 | 0.63183309 | NA | 0 | 7.7294 | 7.2472 | 7.2472 | TRUE | 0.2015 |
| A530050N04 | 0 | 0 | 0 | 0 | 0 | 0 | 0 | 0 | 0 | 0 | 0 | 0 | 0 | 0 | 0 | NA | NA | NA | NA | NA | NA | NA | NA | NA | NA |
| A530053G22 | 0 | 0 | 0 | 0 | 0 | 0 | 0 | 0 | 0 | 0 | 0 | 0 | 0 | 0 | 0 | NA | NA | NA | NA | NA | NA | NA | NA | NA | NA |
| A530054K11 | 966 | 791 | 817 | 996 | 812 | 961 | 750 | 841 | 882 | 991 | 916 | 949 | 888.21 | 824 | 952 | 1.157 | 0.211 | 0.01629013 | 0.05939566 | 0.0032 | 0.0051 | 0.0044 | 0.0044 | TRUE | 0.0776 |
| A530058N18 | 0 | 0 | 0 | 0 | 0 | 0 | 0 | 0 | 0 | 0 | 0 | 0 | 0 | 0 | 0 | NA | NA | NA | NA | NA | NA | NA | NA | NA | NA |
| A530064D06 | 3 | 3 | 1 | 1 | 0 | 1 | 2 | 3 | 1 | 1 | 0 | 1 | 1.43 | 2 | 1 | 0.307 | -1.703 | 0.31641515 | NA | 0 | 1.6593 | 1.0996 | 1.0996 | TRUE | 0.1988 |
| A530065N20 | 0 | 0 | 0 | 0 | 0 | 0 | 0 | 0 | 0 | 0 | 0 | 0 | 0 | 0 | 0 | NA | NA | NA | NA | NA | NA | NA | NA | NA | NA |
| A530072M11 | 18 | 13 | 9 | 11 | 7 | 10 | 14 | 14 | 10 | 11 | 8 | 10 | 11.04 | 13 | 10 | 0.764 | -0.388 | 0.49803426 | NA | 0 | 0.2171 | 0.1436 | 0.1436 | TRUE | 0.142 |
| A530088E08 | 0 | 2 | 1 | 0 | 2 | 1 | 0 | 2 | 1 | 0 | 2 | 1 | 1.07 | 1 | 1 | 1.031 | 0.045 | 0.98188495 | NA | 0 | 2.2072 | 1.8171 | 1.8171 | TRUE | 0.2451 |
| A530099J19F | 0 | 0 | 0 | 0 | 0 | 0 | 0 | 0 | 0 | 0 | 0 | 0 | 0 | 0 | 0 | NA | NA | NA | NA | NA | NA | NA | NA | NA | NA |
| A630001G21 | 36 | 18 | 19 | 23 | 26 | 20 | 28 | 19 | 21 | 23 | 29 | 20 | 23.26 | 23 | 24 | 1.051 | 0.072 | 0.85921014 | 0.92142036 | 0 | 0.1043 | 0.0761 | 0.0761 | TRUE | 0.3096 |
| A630007B06 | 1240 | 864 | 909 | 869 | 847 | 963 | 963 | 919 | 981 | 865 | 956 | 951 | 939 | 954 | 924 | 0.967 | -0.048 | 0.55274631 | 0.70844002 | 9.00E-04 | 0.005 | 0.0036 | 0.0036 | TRUE | 0.0372 |
| A630010A05 | 1 | 0 | 0 | 0 | 0 | 2 | 1 | 0 | 0 | 0 | 0 | 2 | 0.46 | 0 | 1 | 1.93 | 0.948 | 0.7608301 | NA | 0 | 5.1717 | 5.216 | 5.216 | TRUE | 0.9737 |
| A630012P03 | 0 | 0 | 0 | 0 | 0 | 0 | 0 | 0 | 0 | 0 | 0 | 0 | 0 | 0 | 0 | NA | NA | NA | NA | NA | NA | NA | NA | NA | NA |
| A630019I02F | 10 | 19 | 7 | 11 | 7 | 6 | 8 | 20 | 8 | 11 | 8 | 6 | 10.05 | 12 | 8 | 0.704 | -0.506 | 0.44146939 | NA | 0.1382 | 0.2383 | 0.2078 | 0.2078 | TRUE | 1.4185 |
| A630020A06 | 91 | 44 | 41 | 61 | 52 | 54 | 71 | 47 | 44 | 61 | 59 | 53 | 55.73 | 54 | 58 | 1.057 | 0.079 | 0.77621473 | 0.86766476 | 0.0223 | 0.045 | 0.0383 | 0.0383 | TRUE | 0.6986 |
| A630023A22 | 0 | 0 | 0 | 0 | 0 | 0 | 0 | 0 | 0 | 0 | 0 | 0 | 0 | 0 | 0 | NA | NA | NA | NA | NA | NA | NA | NA | NA | NA |
| A630033H20 | 0 | 0 | 0 | 0 | 0 | 0 | 0 | 0 | 0 | 0 | 0 | 0 | 0 | 0 | 0 | NA | NA | NA | NA | NA | NA | NA | NA | NA | NA |
| A630066F11 | 21 | 18 | 8 | 8 | 12 | 5 | 16 | 19 | 9 | 8 | 14 | 5 | 11.75 | 15 | 9 | 0.592 | -0.757 | 0.21351714 | NA | 0.0909 | 0.2041 | 0.1747 | 0.1747 | TRUE | 0.5913 |
| A630072M18 | 164 | 149 | 76 | 99 | 91 | 91 | 127 | 158 | 82 | 99 | 103 | 90 | 109.82 | 122 | 97 | 0.79 | -0.341 | 0.14659723 | 0.29093505 | 0.0463 | 0.024 | 0.0304 | 0.0304 | TRUE | 0.2415 |
| A630073D07 | 0 | 0 | 0 | 0 | 0 | 0 | 0 | 0 | 0 | 0 | 0 | 0 | 0 | 0 | 0 | NA | NA | NA | NA | NA | NA | NA | NA | NA | NA |
| A630075F10 | 2 | 0 | 1 | 2 | 2 | 4 | 2 | 0 | 1 | 2 | 2 | 4 | 1.8 | 1 | 3 | 2.993 | 1.581 | 0.2903249 | NA | 0 | 1.3157 | 0.8974 | 0.8974 | TRUE | 0.2797 |
| A630076J17F | 0 | 0 | 0 | 0 | 0 | 0 | 0 | 0 | 0 | 0 | 0 | 0 | 0 | 0 | 0 | NA | NA | NA | NA | NA | NA | NA | NA | NA | NA |
| A630077J23F | 1 | 2 | 1 | 0 | 0 | 0 | 1 | 2 | 1 | 0 | 0 | 0 | 0.66 | 1 | 0 | 0.145 | -2.79 | 0.24993894 | NA | 0 | 3.5727 | 2.8701 | 2.8701 | TRUE | 0.1613 |
| A630089N07 | 279 | 192 | 132 | 130 | 110 | 139 | 217 | 204 | 142 | 129 | 124 | 137 | 159.01 | 188 | 130 | 0.691 | -0.532 | 0.00420401 | 0.02144464 | 0.0208 | 0.0173 | 0.0184 | 0.0184 | TRUE | 0.4662 |
| A630095E13 | 0 | 0 | 0 | 0 | 0 | 0 | 0 | 0 | 0 | 0 | 0 | 0 | 0 | 0 | 0 | NA | NA | NA | NA | NA | NA | NA | NA | NA | NA |
| A630095N17 | 2 | 0 | 0 | 3 | 4 | 3 | 2 | 0 | 0 | 3 | 5 | 3 | 2 | 1 | 4 | 5.886 | 2.557 | 0.11048162 | NA | 0 | 1.1861 | 0.8889 | 0.8889 | TRUE | 0.9705 |
| A730008H23 | 0 | 0 | 0 | 0 | 0 | 0 | 0 | 0 | 0 | 0 | 0 | 0 | 0 | 0 | 0 | NA | NA | NA | NA | NA | NA | NA | NA | NA | NA |
| A730017C20 | 45 | 30 | 18 | 10 | 9 | 5 | 35 | 32 | 19 | 10 | 10 | 5 | 18.55 | 29 | 8 | 0.287 | -1.8 | 0.00041524 | 0.00373742 | 0 | 0.1302 | 0.1084 | 0.1084 | TRUE | 0.4963 |
| A730017L22I | 25 | 14 | 11 | 20 | 15 | 14 | 19 | 15 | 12 | 20 | 17 | 14 | 16.14 | 15 | 17 | 1.085 | 0.118 | 0.80617466 | NA | 0 | 0.1493 | 0.1054 | 0.1054 | TRUE | 0.3005 |
| A730018C14 | 0 | 0 | 0 | 0 | 0 | 1 | 0 | 0 | 0 | 0 | 0 | 1 | 0.16 | 0 | 0 | 2.094 | 1.067 | 0.79379329 | NA | 0 | 14.4078 | 10 | 10 | TRUE | 0.3835 |
| A730020E08 | 1 | 1 | 0 | 0 | 1 | 0 | 1 | 1 | 0 | 0 | 1 | 0 | 0.49 | 1 | 0 | 0.599 | -0.74 | 0.78905321 | NA | 0 | 4.7929 | 3.7323 | 3.7323 | TRUE | 0.5075 |
| A730020M07 | 1 | 1 | 0 | 2 | 0 | 2 | 1 | 1 | 0 | 2 | 0 | 2 | 0.97 | 1 | 1 | 2.15 | 1.105 | 0.58648156 | NA | 0 | 2.4519 | 1.8115 | 1.8115 | TRUE | 0.4033 |
| A730036I17F | 43 | 30 | 20 | 25 | 28 | 28 | 33 | 32 | 22 | 25 | 32 | 28 | 28.5 | 29 | 28 | 0.96 | -0.058 | 0.87399786 | 0.93005063 | 0 | 0.0856 | 0.062 | 0.062 | TRUE | 0.3024 |
| A730043L09I | 0 | 0 | 0 | 1 | 2 | 0 | 0 | 0 | 0 | 1 | 2 | 0 | 0.54 | 0 | 1 | 6.07 | 2.602 | 0.36554818 | NA | 0 | 4.3758 | 4.4761 | 4.4761 | TRUE | 0.0722 |
| A730046J19F | 9 | 7 | 1 | 3 | 2 | 0 | 7 | 7 | 1 | 3 | 2 | 0 | 3.46 | 5 | 2 | 0.333 | -1.588 | 0.18619131 | NA | 0.5067 | 0.6875 | 0.6493 | 0.6493 | TRUE | 0.9096 |
| A730056A06 | 0 | 1 | 0 | 0 | 2 | 0 | 0 | 1 | 0 | 0 | 2 | 0 | 0.55 | 0 | 1 | 2.136 | 1.095 | 0.70925113 | NA | 0 | 4.2858 | 4.5115 | 4.5115 | TRUE | 0.9746 |
| A730082K24 | 3 | 5 | 4 | 2 | 1 | 2 | 2 | 5 | 4 | 2 | 1 | 2 | 2.84 | 4 | 2 | 0.435 | -1.202 | 0.29949591 | NA | 0 | 0.8361 | 0.5415 | 0.5415 | TRUE | 0.2992 |
| A730085A09 | 225 | 174 | 156 | 95 | 75 | 80 | 175 | 185 | 168 | 95 | 85 | 79 | 131.05 | 176 | 86 | 0.489 | -1.032 | 6.60E-09 | 3.51E-07 | 0 | 0.0205 | 0.0141 | 0.0141 | TRUE | 0.0735 |
| A730085K08 | 0 | 0 | 0 | 0 | 0 | 0 | 0 | 0 | 0 | 0 | 0 | 0 | 0 | 0 | 0 | NA | NA | NA | NA | NA | NA | NA | NA | NA | NA |
| A730090H04 | 4 | 4 | 3 | 3 | 5 | 5 | 3 | 4 | 3 | 3 | 6 | 5 | 4.03 | 3 | 5 | 1.281 | 0.358 | 0.70155654 | NA | 0 | 0.591 | 0.3754 | 0.3754 | TRUE | 0.168 |
| A730090N16 | 5 | 5 | 4 | 0 | 1 | 2 | 4 | 5 | 4 | 0 | 1 | 2 | 2.77 | 4 | 1 | 0.23 | -2.119 | 0.1021581 | NA | 0 | 0.858 | 0.6032 | 0.6032 | TRUE | 0.3922 |
| A730098P11 | 330 | 249 | 245 | 210 | 254 | 244 | 256 | 265 | 264 | 209 | 287 | 241 | 253.66 | 262 | 246 | 0.936 | -0.095 | 0.5045614 | 0.66826706 | 0.008 | 0.0118 | 0.0107 | 0.0107 | TRUE | 0.2302 |
| A830009L08I | 0 | 0 | 0 | 0 | 0 | 0 | 0 | 0 | 0 | 0 | 0 | 0 | 0 | 0 | 0 | NA | NA | NA | NA | NA | NA | NA | NA | NA | NA |
| A830010M20 | 616 | 475 | 450 | 471 | 440 | 432 | 478 | 505 | 486 | 469 | 496 | 426 | 476.81 | 490 | 464 | 0.947 | -0.079 | 0.44122955 | 0.61245438 | 9.00E-04 | 0.0074 | 0.0054 | 0.0054 | TRUE | 0.0583 |
| A830018L16I | 156 | 109 | 127 | 132 | 114 | 131 | 121 | 116 | 137 | 131 | 129 | 129 | 127.24 | 125 | 130 | 1.042 | 0.059 | 0.73378214 | 0.84079179 | 0 | 0.0211 | 0.0141 | 0.0141 | TRUE | 0.0713 |
| A830019L24I | 0 | 0 | 0 | 0 | 0 | 0 | 0 | 0 | 0 | 0 | 0 | 0 | 0 | 0 | 0 | NA | NA | NA | NA | NA | NA | NA | NA | NA | NA |
| A830052D11 | 1 | 0 | 0 | 0 | 1 | 1 | 1 | 0 | 0 | 0 | 1 | 1 | 0.48 | 0 | 1 | 2.031 | 1.022 | 0.71376068 | NA | 0 | 4.9197 | 3.859 | 3.859 | TRUE | 0.2623 |
| A830080D01 | 2774 | 2063 | 2048 | 2579 | 2032 | 2289 | 2154 | 2194 | 2210 | 2567 | 2292 | 2260 | 2279.61 | 2186 | 2373 | 1.086 | 0.119 | 0.08381939 | 0.19518255 | 0.002 | 0.0035 | 0.003 | 0.003 | TRUE | 0.0624 |
| A830082K12 | 4 | 1 | 1 | 2 | 6 | 2 | 3 | 1 | 1 | 2 | 7 | 2 | 2.66 | 2 | 4 | 1.949 | 0.963 | 0.43094962 | NA | 0 | 0.8921 | 0.6652 | 0.6652 | TRUE | 0.828 |
| A830082N09 | 51 | 32 | 22 | 32 | 29 | 33 | 40 | 34 | 24 | 32 | 33 | 33 | 32.42 | 33 | 33 | 0.989 | -0.016 | 0.9638433 | 0.98009479 | 0 | 0.0755 | 0.0557 | 0.0557 | TRUE | 0.3588 |
| A930001A20 | 0 | 0 | 0 | 0 | 0 | 0 | 0 | 0 | 0 | 0 | 0 | 0 | 0 | 0 | 0 | NA | NA | NA | NA | NA | NA | NA | NA | NA | NA |
| A930001C03 | 35 | 27 | 31 | 45 | 39 | 47 | 27 | 29 | 33 | 45 | 44 | 46 | 37.42 | 30 | 45 | 1.52 | 0.604 | 0.05360738 | 0.14156329 | 0 | 0.0658 | 0.0428 | 0.0428 | TRUE | 0.0726 |
| A930003A15 | 0 | 0 | 0 | 0 | 0 | 0 | 0 | 0 | 0 | 0 | 0 | 0 | 0 | 0 | 0 | NA | NA | NA | NA | NA | NA | NA | NA | NA | NA |
| A930003O13 | 0 | 0 | 0 | 0 | 0 | 0 | 0 | 0 | 0 | 0 | 0 | 0 | 0 | 0 | 0 | NA | NA | NA | NA | NA | NA | NA | NA | NA | NA |
| A930004D18 | 330 | 271 | 183 | 146 | 156 | 187 | 256 | 288 | 197 | 145 | 176 | 185 | 207.99 | 247 | 169 | 0.681 | -0.555 | 0.00116479 | 0.00809747 | 0.0212 | 0.0138 | 0.016 | 0.016 | TRUE | 0.3081 |
| A930005H10 | 256 | 157 | 148 | 164 | 142 | 169 | 199 | 167 | 160 | 163 | 160 | 167 | 169.3 | 175 | 163 | 0.929 | -0.106 | 0.50445088 | 0.66826706 | 0 | 0.0164 | 0.0123 | 0.0123 | TRUE | 0.1624 |
| A930006I01F | 0 | 0 | 0 | 0 | 0 | 0 | 0 | 0 | 0 | 0 | 0 | 0 | 0 | 0 | 0 | NA | NA | NA | NA | NA | NA | NA | NA | NA | NA |
| A930006K02 | 9 | 1 | 9 | 5 | 2 | 8 | 7 | 1 | 10 | 5 | 2 | 8 | 5.48 | 6 | 5 | 0.855 | -0.226 | 0.80389676 | NA | 0.3526 | 0.4347 | 0.4137 | 0.4137 | TRUE | 0.4797 |
| A930007I19F | 4 | 4 | 8 | 10 | 5 | 6 | 3 | 4 | 9 | 10 | 6 | 6 | 6.25 | 5 | 7 | 1.374 | 0.459 | 0.56043526 | NA | 0 | 0.3815 | 0.2831 | 0.2831 | TRUE | 0.5869 |

| | | | | | | | | | | | | | | | | | | | | | | | | | |
|---|---|---|---|---|---|---|---|---|---|---|---|---|---|---|---|---|---|---|---|---|---|---|---|---|---|
| A930009A15 | 0 | 0 | 0 | 0 | 0 | 0 | 0 | 0 | 0 | 0 | 0 | 0 | 0 | 0 | 0 | NA | NA | NA | NA | NA | NA | NA | NA | NA | NA |
| A930011G23 | 0 | 0 | 0 | 0 | 1 | 0 | 0 | 0 | 0 | 0 | 1 | 0 | 0.19 | 0 | 0 | 2.094 | 1.067 | 0.79379329 | NA | 0 | 12.608 | 10 | 10 | TRUE | 0.5075 |
| A930011O12 | 99 | 94 | 74 | 140 | 124 | 134 | 77 | 100 | 80 | 139 | 140 | 132 | 111.37 | 86 | 137 | 1.609 | 0.686 | 0.00036617 | 0.00339407 | 0 | 0.0237 | 0.0173 | 0.0173 | TRUE | 0.1981 |
| A930012L18 | 2 | 4 | 1 | 0 | 1 | 2 | 2 | 4 | 1 | 0 | 1 | 2 | 1.66 | 2 | 1 | 0.454 | -1.139 | 0.46744741 | NA | 0 | 1.426 | 1.0568 | 1.0568 | TRUE | 0.5286 |
| A930013F10 | 172 | 168 | 147 | 189 | 122 | 232 | 134 | 179 | 159 | 188 | 138 | 229 | 170.95 | 157 | 185 | 1.185 | 0.245 | 0.21505754 | 0.37765193 | 0.0365 | 0.0163 | 0.0222 | 0.0222 | TRUE | 0.2052 |
| A930015D03 | 127 | 99 | 57 | 64 | 59 | 54 | 99 | 105 | 62 | 64 | 67 | 53 | 74.84 | 89 | 61 | 0.688 | -0.539 | 0.0370198 | 0.10838262 | 0.0342 | 0.0341 | 0.0341 | 0.0341 | TRUE | 0.6424 |
| A930016O22 | 19 | 20 | 33 | 20 | 24 | 22 | 15 | 21 | 36 | 20 | 27 | 22 | 23.39 | 24 | 23 | 0.969 | -0.046 | 0.91651552 | 0.95388891 | 0.0739 | 0.1038 | 0.095 | 0.095 | TRUE | 0.5647 |
| A930017M01 | 7 | 1 | 7 | 1 | 1 | 3 | 5 | 1 | 8 | 1 | 1 | 3 | 3.19 | 5 | 2 | 0.364 | -1.46 | 0.21731795 | NA | 0 | 0.7455 | 0.6034 | 0.6034 | TRUE | 0.2635 |
| A930018P22 | 1 | 1 | 1 | 1 | 1 | 5 | 1 | 1 | 1 | 1 | 1 | 5 | 1.66 | 1 | 2 | 2.464 | 1.301 | 0.39682188 | NA | 0 | 1.4275 | 0.98 | 0.98 | TRUE | 0.9899 |
| A930019D19 | 0 | 0 | 0 | 0 | 0 | 0 | 0 | 0 | 0 | 0 | 0 | 0 | 0 | 0 | 0 | NA | NA | NA | NA | NA | NA | NA | NA | NA | NA |
| A930024E05 | 66 | 32 | 50 | 57 | 55 | 75 | 51 | 34 | 54 | 57 | 62 | 74 | 55.34 | 46 | 64 | 1.381 | 0.465 | 0.09501666 | 0.21299766 | 0.0154 | 0.0453 | 0.0376 | 0.0376 | TRUE | 0.4034 |
| A930041C12 | 0 | 0 | 0 | 0 | 0 | 0 | 0 | 0 | 0 | 0 | 0 | 0 | 0 | 0 | 0 | NA | NA | NA | NA | NA | NA | NA | NA | NA | NA |
| AA387883 | 0 | 0 | 0 | 0 | 0 | 0 | 0 | 0 | 0 | 0 | 0 | 0 | 0 | 0 | 0 | NA | NA | NA | NA | NA | NA | NA | NA | NA | NA |
| AA388235 | 45 | 27 | 16 | 21 | 26 | 28 | 35 | 29 | 17 | 21 | 29 | 28 | 26.47 | 27 | 26 | 0.951 | -0.073 | 0.85564945 | 0.91884279 | 0.0381 | 0.092 | 0.0772 | 0.0772 | TRUE | 0.4085 |
| AA414768 | 330 | 160 | 101 | 103 | 99 | 97 | 256 | 170 | 109 | 103 | 112 | 96 | 140.9 | 178 | 104 | 0.575 | -0.798 | 0.0017893 | 0.01109301 | 0.0869 | 0.0192 | 0.0393 | 0.0393 | TRUE | 0.1889 |
| AA415398 | 29 | 19 | 10 | 25 | 14 | 16 | 23 | 20 | 11 | 25 | 16 | 16 | 18.33 | 18 | 19 | 1.047 | 0.067 | 0.88835442 | 0.93803559 | 0.0469 | 0.1317 | 0.1088 | 0.1088 | TRUE | 0.56 |
| AA465934 | 600 | 358 | 308 | 311 | 266 | 337 | 466 | 381 | 332 | 310 | 300 | 333 | 353.57 | 393 | 314 | 0.797 | -0.328 | 0.01684696 | 0.06097947 | 0.014 | 0.0091 | 0.0107 | 0.0107 | TRUE | 0.3172 |
| AA467197 | 87 | 86 | 67 | 51 | 41 | 59 | 68 | 91 | 72 | 51 | 46 | 58 | 64.43 | 77 | 52 | 0.675 | -0.567 | 0.02508571 | 0.08150719 | 0 | 0.0392 | 0.0299 | 0.0299 | TRUE | 0.2389 |
| AA474331 | 84 | 68 | 67 | 69 | 54 | 80 | 65 | 72 | 72 | 69 | 61 | 79 | 69.74 | 70 | 70 | 0.998 | -0.002 | 0.99213486 | 0.99575757 | 0 | 0.0364 | 0.0253 | 0.0253 | TRUE | 0.1279 |
| AA536875 | 0 | 0 | 0 | 0 | 0 | 0 | 0 | 0 | 0 | 0 | 0 | 0 | 0 | 0 | 0 | NA | NA | NA | NA | NA | NA | NA | NA | NA | NA |
| AA543186 | 6 | 11 | 6 | 9 | 5 | 6 | 5 | 12 | 6 | 9 | 6 | 6 | 7.23 | 8 | 7 | 0.914 | -0.13 | 0.85779214 | NA | 0 | 0.3304 | 0.2424 | 0.2424 | TRUE | 0.6011 |
| AA545190 | 0 | 0 | 0 | 0 | 0 | 0 | 0 | 0 | 0 | 0 | 0 | 0 | 0 | 0 | 0 | NA | NA | NA | NA | NA | NA | NA | NA | NA | NA |
| AA619741 | 4 | 2 | 0 | 0 | 1 | 0 | 3 | 2 | 0 | 0 | 1 | 0 | 1.06 | 2 | 0 | 0.209 | -2.26 | 0.29098819 | NA | 0 | 2.2376 | 2.0276 | 2.0276 | TRUE | 0.4075 |
| AA792892 | 0 | 0 | 0 | 0 | 0 | 0 | 0 | 0 | 0 | 0 | 0 | 0 | 0 | 0 | 0 | NA | NA | NA | NA | NA | NA | NA | NA | NA | NA |
| AA986860 | 62 | 40 | 32 | 44 | 33 | 51 | 48 | 43 | 35 | 44 | 37 | 50 | 42.77 | 42 | 44 | 1.045 | 0.064 | 0.8320618 | 0.90401574 | 0 | 0.0578 | 0.0426 | 0.0426 | TRUE | 0.1664 |
| AA987161 | 2096 | 1658 | 1835 | 1992 | 1741 | 1974 | 1628 | 1764 | 1980 | 1983 | 1964 | 1949 | 1877.82 | 1791 | 1965 | 1.099 | 0.136 | 0.08457104 | 0.19613401 | 0.0042 | 0.0037 | 0.0039 | 0.0039 | TRUE | 0.1032 |
| Aaas | 7334 | 5391 | 5583 | 5757 | 5024 | 5749 | 5696 | 5734 | 6024 | 5731 | 5668 | 5675 | 5754.6 | 5818 | 5691 | 0.978 | -0.032 | 0.53759289 | 0.69620473 | 3.00E-04 | 0.0028 | 0.0017 | 0.0017 | TRUE | 0.0116 |
| Aacs | 3177 | 2492 | 2330 | 2669 | 2399 | 2843 | 2467 | 2651 | 2514 | 2657 | 2706 | 2806 | 2633.62 | 2544 | 2723 | 1.071 | 0.099 | 0.1034957 | 0.22688099 | 7.00E-04 | 0.0033 | 0.0023 | 0.0023 | TRUE | 0.0162 |
| Aadac | 1 | 1 | 1 | 1 | 1 | 1 | 0 | 1 | 1 | 1 | 1 | 0 | 0.84 | 1 | 1 | 0.724 | -0.466 | 0.8180547 | NA | 0 | 2.8223 | 1.7212 | 1.7212 | TRUE | 0.2697 |
| Aadacl2 | 6 | 2 | 0 | 0 | 0 | 2 | 5 | 2 | 0 | 0 | 0 | 2 | 1.46 | 2 | 1 | 0.291 | -1.781 | 0.35523316 | NA | 2.1331 | 1.6255 | 1.6962 | 1.6962 | TRUE | 0.287 |
| Aadacl3 | 2 | 1 | 1 | 0 | 1 | 1 | 2 | 1 | 1 | 0 | 1 | 1 | 0.97 | 1 | 1 | 0.56 | -0.836 | 0.66301683 | NA | 0 | 2.4495 | 1.5188 | 1.5188 | TRUE | 0.2653 |
| Aadat | 27 | 12 | 10 | 12 | 13 | 12 | 21 | 13 | 11 | 12 | 15 | 12 | 13.83 | 15 | 13 | 0.849 | -0.236 | 0.6549506 | NA | 0 | 0.1738 | 0.128 | 0.128 | TRUE | 0.7254 |
| Aaed1 | 728 | 568 | 571 | 566 | 565 | 599 | 565 | 604 | 616 | 563 | 637 | 591 | 596.29 | 595 | 597 | 1.004 | 0.006 | 0.94934036 | 0.97225491 | 0.0013 | 0.0064 | 0.0047 | 0.0047 | TRUE | 0.0393 |
| Aagab | 4236 | 3232 | 3160 | 3395 | 3110 | 3595 | 3290 | 3438 | 3410 | 3379 | 3508 | 3549 | 3429 | 3379 | 3479 | 1.03 | 0.042 | 0.44577329 | 0.61648773 | 3.00E-04 | 0.0031 | 0.0019 | 0.0019 | TRUE | 0.0077 |
| Aak1 | 3444 | 2141 | 1964 | 1977 | 1728 | 1965 | 2675 | 2277 | 2119 | 1968 | 1949 | 1940 | 2154.72 | 2357 | 1952 | 0.827 | -0.273 | 0.00131235 | 0.00876107 | 0.0067 | 0.0035 | 0.0047 | 0.0047 | TRUE | 0.1713 |
| Aamdc | 360 | 283 | 257 | 330 | 295 | 342 | 280 | 301 | 277 | 328 | 333 | 338 | 309.47 | 286 | 333 | 1.165 | 0.221 | 0.05724902 | 0.14866207 | 0 | 0.0101 | 0.0065 | 0.0065 | TRUE | 0.0229 |
| Aamp | 8580 | 6274 | 5995 | 6023 | 5470 | 6612 | 6664 | 6674 | 6469 | 5995 | 6171 | 6527 | 6416.49 | 6602 | 6231 | 0.944 | -0.083 | 0.12573581 | 0.26082056 | 9.00E-04 | 0.0028 | 0.002 | 0.002 | TRUE | 0.0212 |
| Aanat | 4 | 3 | 1 | 2 | 4 | 2 | 3 | 3 | 1 | 2 | 5 | 2 | 2.64 | 2 | 3 | 1.125 | 0.17 | 0.88438569 | NA | 0 | 0.8994 | 0.6008 | 0.6008 | TRUE | 0.2973 |
| Aar2 | 2720 | 1776 | 1505 | 1658 | 1389 | 1599 | 2113 | 1889 | 1624 | 1650 | 1567 | 1578 | 1736.88 | 1875 | 1598 | 0.852 | -0.232 | 0.01126958 | 0.0450941 | 0.008 | 0.0038 | 0.0054 | 0.0054 | TRUE | 0.1653 |
| Aard | 3109 | 1754 | 1135 | 1361 | 1107 | 1141 | 2415 | 1866 | 1225 | 1355 | 1249 | 1126 | 1539.15 | 1835 | 1243 | 0.677 | -0.562 | 0.04483719 | 0.12441428 | 0.0559 | 0.004 | 0.0189 | 0.0559 | TRUE | 0.1598 |
| Aars | 21778 | 15294 | 13210 | 12823 | 10739 | 12697 | 16914 | 16268 | 14253 | 12764 | 12115 | 12534 | 14141.36 | 15812 | 12471 | 0.789 | -0.343 | 4.46E-07 | 1.48E-05 | 0.0041 | 0.0026 | 0.0032 | 0.0032 | TRUE | 0.0908 |
| Aars2 | 3607 | 2728 | 2432 | 2736 | 2701 | 2854 | 2801 | 2902 | 2624 | 2723 | 3047 | 2817 | 2819.17 | 2776 | 2862 | 1.031 | 0.044 | 0.52049891 | 0.68185957 | 0.0023 | 0.0033 | 0.003 | 0.003 | TRUE | 0.0382 |
| Aarsd1 | 5520 | 4072 | 4272 | 4614 | 3949 | 4587 | 4287 | 4331 | 4609 | 4593 | 4455 | 4528 | 4467.28 | 4409 | 4525 | 1.027 | 0.038 | 0.49201596 | 0.65741649 | 6.00E-04 | 0.003 | 0.002 | 0.002 | TRUE | 0.019 |
| Aasdh | 1534 | 1250 | 1337 | 1487 | 1330 | 1411 | 1191 | 1330 | 1443 | 1480 | 1500 | 1393 | 1389.5 | 1321 | 1458 | 1.105 | 0.144 | 0.08595327 | 0.19818198 | 0.0047 | 0.0041 | 0.0043 | 0.0043 | TRUE | 0.0921 |
| Aasdhppt | 3846 | 2536 | 2848 | 2881 | 2498 | 2890 | 2987 | 2698 | 3073 | 2868 | 2818 | 2853 | 2882.69 | 2919 | 2846 | 0.975 | -0.037 | 0.57535201 | 0.72717057 | 0.0017 | 0.0032 | 0.0028 | 0.0028 | TRUE | 0.0528 |
| Aass | 1027 | 788 | 1149 | 933 | 815 | 1044 | 798 | 838 | 1240 | 929 | 919 | 1031 | 959.05 | 959 | 960 | 1.003 | 0.004 | 0.97838672 | 0.98841401 | 0.0311 | 0.0049 | 0.0127 | 0.0127 | TRUE | 0.7815 |
| Aatf | 4442 | 3185 | 2744 | 2869 | 2453 | 2915 | 3450 | 3388 | 2961 | 2856 | 2767 | 2878 | 3049.85 | 3266 | 2834 | 0.867 | -0.206 | 0.00356949 | 0.01887485 | 0.003 | 0.0032 | 0.0033 | 0.0033 | TRUE | 0.0806 |
| Aatk | 145 | 116 | 101 | 154 | 125 | 161 | 113 | 123 | 109 | 153 | 141 | 159 | 133.04 | 115 | 151 | 1.316 | 0.396 | 0.02092872 | 0.07159746 | 0 | 0.0202 | 0.0136 | 0.0136 | TRUE | 0.0384 |
| AB099516 | 0 | 0 | 1 | 0 | 0 | 0 | 0 | 0 | 1 | 0 | 0 | 0 | 0.18 | 0 | 0 | 0.552 | -0.857 | 0.8336548 | NA | 0 | 13.1817 | 10 | 10 | TRUE | 0.5365 |
| AB124611 | 2 | 1 | 2 | 2 | 5 | 1 | 2 | 1 | 2 | 2 | 6 | 1 | 2.23 | 2 | 3 | 1.783 | 0.835 | 0.52405754 | NA | 0 | 1.0642 | 0.7505 | 0.7505 | TRUE | 0.7938 |
| Abat | 505 | 302 | 254 | 281 | 270 | 272 | 392 | 321 | 274 | 280 | 305 | 269 | 306.72 | 329 | 285 | 0.86 | -0.218 | 0.13239156 | 0.27036327 | 0.0157 | 0.0102 | 0.0119 | 0.0119 | TRUE | 0.2998 |
| Abca1 | 1180 | 794 | 1006 | 1345 | 1313 | 1379 | 916 | 845 | 1085 | 1339 | 1481 | 1361 | 1171.3 | 949 | 1394 | 1.469 | 0.555 | 7.43E-09 | 3.88E-07 | 0.0087 | 0.0044 | 0.0058 | 0.0058 | TRUE | 0.1847 |
| Abca12 | 12 | 4 | 9 | 4 | 9 | 1 | 9 | 4 | 10 | 4 | 10 | 1 | 6.4 | 8 | 5 | 0.636 | -0.654 | 0.44528117 | NA | 0.3293 | 0.3727 | 0.362 | 0.362 | TRUE | 0.7461 |
| Abca13 | 141 | 95 | 87 | 82 | 81 | 62 | 110 | 101 | 94 | 82 | 91 | 61 | 89.77 | 102 | 78 | 0.765 | -0.386 | 0.08030945 | 0.18966672 | 0.009 | 0.0288 | 0.0237 | 0.0237 | TRUE | 0.3367 |
| Abca14 | 5 | 2 | 0 | 3 | 0 | 1 | 4 | 2 | 0 | 3 | 0 | 1 | 1.66 | 2 | 1 | 0.654 | -0.613 | 0.70898259 | NA | 0.9106 | 1.4267 | 1.3233 | 1.3233 | TRUE | 0.2386 |
| Abca15 | 1 | 0 | 0 | 0 | 0 | 0 | 1 | 0 | 0 | 0 | 0 | 0 | 0.13 | 0 | 0 | 0.552 | -0.857 | 0.8336548 | NA | 0 | 18.3121 | 10 | 10 | TRUE | 0.2566 |
| Abca16 | 45 | 48 | 32 | 29 | 14 | 18 | 35 | 51 | 35 | 29 | 16 | 18 | 30.49 | 40 | 21 | 0.522 | -0.939 | 0.01437629 | 0.0540027 | 0.0383 | 0.0801 | 0.0691 | 0.0691 | TRUE | 0.6437 |
| Abca17 | 5 | 6 | 5 | 1 | 1 | 5 | 4 | 6 | 5 | 1 | 1 | 5 | 3.79 | 5 | 2 | 0.46 | -1.122 | 0.28147866 | NA | 0 | 0.6283 | 0.471 | 0.471 | TRUE | 0.9936 |
| Abca2 | 2808 | 2109 | 2153 | 2540 | 2176 | 2368 | 2181 | 2243 | 2323 | 2528 | 2455 | 2338 | 2344.65 | 2249 | 2440 | 1.085 | 0.118 | 0.06011808 | 0.15349547 | 9.00E-04 | 0.0034 | 0.0024 | 0.0024 | TRUE | 0.0166 |
| Abca3 | 3045 | 2161 | 1971 | 2400 | 2079 | 2184 | 2365 | 2299 | 2127 | 2389 | 2345 | 2156 | 2280.08 | 2264 | 2297 | 1.014 | 0.02 | 0.77523247 | 0.86721285 | 0.0025 | 0.0035 | 0.0031 | 0.0031 | TRUE | 0.0351 |
| Abca4 | 230 | 169 | 168 | 155 | 153 | 157 | 179 | 180 | 181 | 154 | 173 | 155 | 170.26 | 180 | 161 | 0.892 | -0.165 | 0.27671102 | 0.44785761 | 0 | 0.0163 | 0.0106 | 0.0106 | TRUE | 0.043 |
| Abca5 | 1953 | 1480 | 1350 | 1682 | 1505 | 1601 | 1517 | 1574 | 1457 | 1674 | 1698 | 1580 | 1583.37 | 1516 | 1651 | 1.089 | 0.123 | 0.0739871 | 0.17857102 | 8.00E-04 | 0.0039 | 0.0028 | 0.0028 | TRUE | 0.0169 |
| Abca6 | 22 | 18 | 5 | 13 | 8 | 9 | 17 | 19 | 5 | 13 | 9 | 9 | 12.08 | 14 | 10 | 0.737 | -0.441 | 0.46616812 | NA | 0.1236 | 0.1986 | 0.1785 | 0.1785 | TRUE | 1.1333 |
| Abca7 | 710 | 682 | 767 | 772 | 678 | 870 | 551 | 725 | 828 | 768 | 765 | 859 | 749.43 | 701 | 797 | 1.14 | 0.189 | 0.14165601 | 0.28396765 | 0.0217 | 0.0056 | 0.0106 | 0.0106 | TRUE | 0.4294 |
| Abca8a | 12 | 18 | 9 | 14 | 4 | 9 | 9 | 19 | 10 | 14 | 5 | 9 | 10.92 | 13 | 9 | 0.727 | -0.46 | 0.46628172 | NA | 0.1224 | 0.2195 | 0.1928 | 0.1928 | TRUE | 0.3009 |
| Abca8b | 39 | 24 | 19 | 10 | 17 | 16 | 30 | 26 | 21 | 10 | 19 | 16 | 20.21 | 26 | 15 | 0.582 | -0.78 | 0.08176454 | 0.1921823 | 0 | 0.1197 | 0.0912 | 0.0912 | TRUE | 0.3274 |
| Abca9 | 7 | 1 | 3 | 0 | 2 | 2 | 5 | 1 | 3 | 0 | 2 | 2 | 2.33 | 3 | 1 | 0.422 | -1.245 | 0.36087811 | NA | 0 | 1.0205 | 0.8204 | 0.8204 | TRUE | 0.1887 |
| Abcb10 | 3363 | 2438 | 2401 | 2616 | 2331 | 2629 | 2612 | 2593 | 2591 | 2604 | 2630 | 2595 | 2604.1 | 2599 | 2610 | 1.004 | 0.006 | 0.91528751 | 0.95330234 | 0 | 0.0033 | 0.0018 | 0.0018 | TRUE | 5.00E-04 |
| Abcb11 | 11 | 10 | 5 | 2 | 2 | 0 | 9 | 11 | 5 | 2 | 2 | 0 | 4.8 | 8 | 1 | 0.169 | -2.561 | 0.01589964 | NA | 0 | 0.4958 | 0.382 | 0.382 | TRUE | 0.4485 |

| | | | | | | | | | | | | | | | | | | | | | | | | | |
|---|---|---|---|---|---|---|---|---|---|---|---|---|---|---|---|---|---|---|---|---|---|---|---|---|---|
| Abcb1a | 1385 | 1099 | 948 | 730 | 656 | 838 | 1076 | 1169 | 1023 | 727 | 740 | 827 | 926.91 | 1089 | 765 | 0.702 | -0.51 | 8.55E-09 | 4.35E-07 | 0.0036 | 0.005 | 0.0045 | 0.0045 | TRUE | 0.0616 |
| Abcb1b | 2046 | 1216 | 954 | 1128 | 868 | 957 | 1589 | 1293 | 1029 | 1123 | 979 | 945 | 1159.76 | 1304 | 1016 | 0.778 | -0.362 | 0.00517247 | 0.02518381 | 0.0261 | 0.0045 | 0.0112 | 0.0112 | TRUE | 0.1715 |
| Abcb4 | 52 | 32 | 54 | 61 | 38 | 49 | 40 | 34 | 58 | 61 | 43 | 48 | 47.44 | 44 | 51 | 1.151 | 0.203 | 0.50699711 | 0.67035243 | 0.0305 | 0.0524 | 0.0465 | 0.0465 | TRUE | 0.5602 |
| Abcb5 | 133 | 85 | 28 | 24 | 20 | 14 | 103 | 90 | 30 | 24 | 23 | 14 | 47.37 | 74 | 20 | 0.267 | -1.903 | 7.61E-06 | 0.00015018 | 0.2115 | 0.0525 | 0.098 | 0.098 | TRUE | 0.9041 |
| Abcb6 | 1551 | 1175 | 1245 | 1681 | 1566 | 1725 | 1205 | 1250 | 1343 | 1673 | 1767 | 1703 | 1490.08 | 1266 | 1714 | 1.355 | 0.439 | 7.92E-10 | 5.47E-08 | 0.0012 | 0.004 | 0.003 | 0.003 | TRUE | 0.034 |
| Abcb7 | 5139 | 3621 | 3907 | 4246 | 3607 | 4139 | 3991 | 3852 | 4216 | 4226 | 4069 | 4086 | 4073.31 | 4020 | 4127 | 1.027 | 0.038 | 0.50740989 | 0.67077409 | 9.00E-04 | 0.003 | 0.0022 | 0.0022 | TRUE | 0.0218 |
| Abcb8 | 1564 | 1130 | 1215 | 1402 | 1274 | 1352 | 1215 | 1202 | 1311 | 1396 | 1437 | 1335 | 1315.84 | 1243 | 1389 | 1.118 | 0.161 | 0.02701596 | 0.08586594 | 0.001 | 0.0042 | 0.0031 | 0.0031 | TRUE | 0.0271 |
| Abcb9 | 35 | 30 | 18 | 34 | 32 | 23 | 27 | 32 | 19 | 34 | 36 | 23 | 28.53 | 26 | 31 | 1.175 | 0.232 | 0.53954136 | 0.69782505 | 0.0217 | 0.0855 | 0.0682 | 0.0682 | TRUE | 0.3712 |
| Abcc1 | 6027 | 4824 | 6024 | 6623 | 6215 | 6864 | 4681 | 5131 | 6500 | 6593 | 7011 | 6776 | 6115.25 | 5437 | 6793 | 1.25 | 0.322 | 0.00099967 | 0.00723822 | 0.0154 | 0.0028 | 0.0067 | 0.0067 | TRUE | 0.3559 |
| Abcc10 | 1274 | 960 | 971 | 1241 | 1081 | 1212 | 989 | 1021 | 1048 | 1235 | 1219 | 1196 | 1118.25 | 1019 | 1217 | 1.195 | 0.257 | 0.00033793 | 0.00318185 | 0 | 0.0045 | 0.0028 | 0.0028 | TRUE | 0.0084 |
| Abcc12 | 4 | 2 | 5 | 6 | 3 | 5 | 3 | 2 | 5 | 6 | 3 | 5 | 4.15 | 3 | 5 | 1.359 | 0.443 | 0.63503295 | NA | 0 | 0.573 | 0.3822 | 0.3822 | TRUE | 0.279 |
| Abcc2 | 283 | 291 | 251 | 262 | 306 | 330 | 220 | 310 | 271 | 261 | 345 | 326 | 288.65 | 267 | 311 | 1.168 | 0.224 | 0.15142992 | 0.29763309 | 0.0219 | 0.0106 | 0.014 | 0.014 | TRUE | 0.2843 |
| Abcc3 | 108 | 85 | 71 | 98 | 82 | 82 | 84 | 90 | 77 | 98 | 93 | 81 | 86.98 | 84 | 91 | 1.079 | 0.11 | 0.60152692 | 0.74667517 | 0 | 0.0297 | 0.0206 | 0.0206 | TRUE | 0.0816 |
| Abcc4 | 6823 | 5149 | 5793 | 5989 | 6030 | 6891 | 5299 | 5477 | 6251 | 5961 | 6802 | 6802 | 6098.84 | 5676 | 6522 | 1.149 | 0.201 | 0.00968406 | 0.04044352 | 0.0065 | 0.0028 | 0.0042 | 0.0042 | TRUE | 0.0955 |
| Abcc5 | 4964 | 3553 | 3509 | 4051 | 3555 | 4146 | 3855 | 3779 | 3786 | 4032 | 4010 | 4093 | 3926.05 | 3807 | 4045 | 1.062 | 0.087 | 0.08872359 | 0.20280286 | 0 | 0.003 | 0.0016 | 0.0016 | TRUE | 0.0015 |
| Abcc6 | 42 | 34 | 39 | 40 | 33 | 40 | 33 | 36 | 42 | 40 | 37 | 39 | 37.9 | 37 | 39 | 1.057 | 0.08 | 0.79707409 | 0.88160891 | 0 | 0.065 | 0.0431 | 0.0431 | TRUE | 0.1032 |
| Abcc8 | 493 | 459 | 491 | 534 | 543 | 622 | 383 | 488 | 530 | 532 | 613 | 614 | 526.5 | 467 | 586 | 1.259 | 0.332 | 0.00846115 | 0.03654737 | 0.0153 | 0.0069 | 0.0095 | 0.0095 | TRUE | 0.3044 |
| Abcc9 | 7 | 1 | 0 | 0 | 1 | 1 | 5 | 1 | 0 | 0 | 1 | 1 | 1.44 | 2 | 1 | 0.311 | -1.685 | 0.36640519 | NA | 1.2368 | 1.653 | 1.5505 | 1.5505 | TRUE | 1.0837 |
| Abcd1 | 780 | 549 | 598 | 706 | 557 | 622 | 606 | 584 | 645 | 703 | 628 | 614 | 630.02 | 612 | 648 | 1.061 | 0.085 | 0.37180263 | 0.54649756 | 0.0023 | 0.0062 | 0.0049 | 0.0049 | TRUE | 0.0643 |
| Abcd2 | 34 | 13 | 13 | 6 | 13 | 15 | 26 | 14 | 14 | 6 | 15 | 15 | 14.95 | 18 | 12 | 0.643 | -0.638 | 0.24616123 | NA | 0.1121 | 0.1609 | 0.1474 | 0.1474 | TRUE | 0.9905 |
| Abcd3 | 3935 | 2761 | 2404 | 2738 | 2326 | 2639 | 3056 | 2937 | 2594 | 2725 | 2624 | 2605 | 2756.89 | 2862 | 2651 | 0.926 | -0.111 | 0.1233304 | 0.25718725 | 0.0033 | 0.0033 | 0.0034 | 0.0034 | TRUE | 0.0811 |
| Abcd4 | 1653 | 1296 | 1369 | 1391 | 1256 | 1499 | 1284 | 1379 | 1477 | 1385 | 1417 | 1480 | 1403.46 | 1380 | 1427 | 1.036 | 0.05 | 0.50875936 | 0.67201243 | 0.0024 | 0.0041 | 0.0035 | 0.0035 | TRUE | 0.0464 |
| Abce1 | 38769 | 26324 | 26580 | 27116 | 22938 | 26906 | 30110 | 28000 | 28680 | 26991 | 25876 | 26560 | 27703.06 | 28930 | 26476 | 0.915 | -0.128 | 0.01022897 | 0.04198815 | 9.00E-04 | 0.0025 | 0.0018 | 0.0018 | TRUE | 0.0156 |
| Abcf1 | 16698 | 12748 | 11843 | 12917 | 11579 | 13981 | 12969 | 13560 | 12778 | 12858 | 13062 | 13801 | 13171.38 | 13102 | 13240 | 1.011 | 0.015 | 0.7730908 | 0.86583504 | 9.00E-04 | 0.0026 | 0.0019 | 0.0019 | TRUE | 0.0168 |
| Abcf2 | 12876 | 9098 | 8042 | 8569 | 7602 | 8937 | 10000 | 9677 | 8677 | 8530 | 8576 | 8822 | 9047.09 | 9451 | 8643 | 0.914 | -0.129 | 0.0392743 | 0.11301542 | 0.0027 | 0.0027 | 0.0027 | 0.0027 | TRUE | 0.0626 |
| Abcf3 | 2909 | 2124 | 1931 | 2110 | 1847 | 2124 | 2259 | 2259 | 2084 | 2100 | 2084 | 2097 | 2147.12 | 2201 | 2094 | 0.951 | -0.073 | 0.24888646 | 0.41805458 | 6.00E-04 | 0.0035 | 0.0024 | 0.0024 | TRUE | 0.0258 |
| Abcg1 | 1014 | 970 | 1387 | 1879 | 1907 | 2185 | 788 | 1032 | 1497 | 1870 | 2151 | 2157 | 1582.41 | 1106 | 2059 | 1.864 | 0.899 | 0.00125537 | 0.00850382 | 0.0552 | 0.0039 | 0.0173 | 0.0552 | TRUE | 0.9603 |
| Abcg2 | 3150 | 2050 | 2047 | 2183 | 1829 | 2134 | 2446 | 2181 | 2209 | 2173 | 2063 | 2107 | 2196.43 | 2279 | 2114 | 0.927 | -0.109 | 0.11111344 | 0.23885463 | 0.002 | 0.0035 | 0.0029 | 0.0029 | TRUE | 0.0517 |
| Abcg3 | 42 | 35 | 31 | 39 | 32 | 36 | 33 | 37 | 33 | 39 | 36 | 36 | 35.63 | 34 | 37 | 1.072 | 0.1 | 0.75213405 | 0.85292383 | 0 | 0.069 | 0.0442 | 0.0442 | TRUE | 0.0341 |
| Abcg4 | 34 | 26 | 38 | 46 | 24 | 27 | 26 | 28 | 41 | 46 | 27 | 27 | 32.43 | 32 | 33 | 1.055 | 0.078 | 0.83469977 | 0.90601463 | 0.0512 | 0.0755 | 0.0686 | 0.0686 | TRUE | 0.782 |
| Abcg5 | 16 | 13 | 11 | 5 | 10 | 2 | 12 | 14 | 12 | 5 | 11 | 2 | 9.39 | 13 | 6 | 0.471 | -1.088 | 0.11464419 | NA | 0.0593 | 0.2548 | 0.2179 | 0.2179 | TRUE | 0.941 |
| Abcg8 | 24 | 11 | 14 | 25 | 20 | 9 | 19 | 12 | 15 | 25 | 23 | 9 | 16.96 | 15 | 19 | 1.226 | 0.294 | 0.56653056 | NA | 0.1013 | 0.1421 | 0.1309 | 0.1309 | TRUE | 1.1485 |
| Abhd1 | 425 | 265 | 280 | 351 | 314 | 320 | 330 | 282 | 302 | 349 | 354 | 316 | 322.26 | 305 | 340 | 1.112 | 0.153 | 0.20651459 | 0.36831958 | 0.0022 | 0.0098 | 0.0075 | 0.0075 | TRUE | 0.0648 |
| Abhd10 | 1642 | 1084 | 1316 | 1389 | 1209 | 1482 | 1275 | 1153 | 1420 | 1383 | 1364 | 1463 | 1342.95 | 1283 | 1403 | 1.094 | 0.13 | 0.12936464 | 0.26608271 | 0.005 | 0.0042 | 0.0045 | 0.0045 | TRUE | 0.1021 |
| Abhd11 | 1910 | 1442 | 1464 | 1559 | 1412 | 1568 | 1483 | 1534 | 1580 | 1552 | 1593 | 1548 | 1548.24 | 1532 | 1564 | 1.021 | 0.03 | 0.64270592 | 0.7773516 | 0 | 0.004 | 0.0025 | 0.0025 | TRUE | 0.0098 |
| Abhd12 | 1895 | 1495 | 1535 | 1751 | 1427 | 1594 | 1472 | 1590 | 1656 | 1743 | 1610 | 1574 | 1607.42 | 1573 | 1642 | 1.045 | 0.064 | 0.39582755 | 0.56928589 | 0.0027 | 0.0039 | 0.0035 | 0.0035 | TRUE | 0.0394 |
| Abhd12b | 1 | 1 | 3 | 1 | 2 | 1 | 1 | 1 | 3 | 1 | 2 | 1 | 1.55 | 2 | 1 | 0.844 | -0.244 | 0.87215008 | NA | 0 | 1.5289 | 0.9939 | 0.9939 | TRUE | 0.4555 |
| Abhd13 | 3896 | 2954 | 2906 | 3141 | 2614 | 3048 | 3026 | 3142 | 3136 | 3127 | 2949 | 3009 | 3064.63 | 3101 | 3028 | 0.977 | -0.034 | 0.55443309 | 0.70999991 | 4.00E-04 | 0.0032 | 0.002 | 0.002 | TRUE | 0.0098 |
| Abhd14a | 1784 | 1321 | 1320 | 1486 | 1256 | 1599 | 1386 | 1405 | 1424 | 1479 | 1417 | 1578 | 1448.25 | 1405 | 1491 | 1.062 | 0.087 | 0.21583229 | 0.37851422 | 9.00E-04 | 0.0041 | 0.0029 | 0.0029 | TRUE | 0.0314 |
| Abhd14b | 432 | 339 | 468 | 589 | 473 | 707 | 336 | 361 | 505 | 586 | 534 | 698 | 503.15 | 401 | 606 | 1.518 | 0.602 | 6.18E-05 | 0.00083283 | 0.0326 | 0.0071 | 0.0142 | 0.0142 | TRUE | 0.595 |
| Abhd15 | 60 | 30 | 34 | 45 | 56 | 59 | 47 | 32 | 37 | 45 | 63 | 58 | 46.9 | 39 | 55 | 1.428 | 0.514 | 0.08417378 | 0.19565717 | 0.0142 | 0.053 | 0.0421 | 0.0421 | TRUE | 0.3024 |
| Abhd16a | 704 | 572 | 585 | 629 | 633 | 730 | 547 | 608 | 631 | 626 | 714 | 721 | 641.2 | 595 | 687 | 1.155 | 0.208 | 0.03586461 | 0.10578153 | 0.0044 | 0.0061 | 0.0055 | 0.0055 | TRUE | 0.0717 |
| Abhd16b | 13 | 9 | 3 | 8 | 8 | 12 | 10 | 10 | 3 | 8 | 9 | 12 | 8.62 | 8 | 10 | 1.244 | 0.314 | 0.64127694 | NA | 0 | 0.2773 | 0.212 | 0.212 | TRUE | 0.65 |
| Abhd2 | 1647 | 1221 | 1324 | 1362 | 1292 | 1536 | 1279 | 1299 | 1429 | 1356 | 1458 | 1516 | 1389.34 | 1336 | 1443 | 1.081 | 0.113 | 0.14569409 | 0.28966556 | 0.0027 | 0.0041 | 0.0036 | 0.0036 | TRUE | 0.044 |
| Abhd3 | 343 | 238 | 244 | 230 | 187 | 226 | 266 | 253 | 263 | 229 | 211 | 223 | 240.97 | 261 | 221 | 0.847 | -0.24 | 0.06385796 | 0.16057739 | 0 | 0.0123 | 0.0079 | 0.0079 | TRUE | 0.0166 |
| Abhd4 | 2440 | 1729 | 2126 | 2183 | 1966 | 2305 | 1895 | 1839 | 2294 | 2173 | 2218 | 2275 | 2115.72 | 2009 | 2222 | 1.107 | 0.146 | 0.08682094 | 0.19976363 | 0.0072 | 0.0035 | 0.0048 | 0.0048 | TRUE | 0.1836 |
| Abhd5 | 842 | 510 | 531 | 476 | 412 | 433 | 654 | 542 | 573 | 474 | 465 | 427 | 522.57 | 590 | 455 | 0.77 | -0.377 | 0.00040776 | 0.00369807 | 0.0048 | 0.007 | 0.0062 | 0.0062 | TRUE | 0.1116 |
| Abhd6 | 431 | 337 | 324 | 365 | 342 | 398 | 335 | 358 | 350 | 363 | 386 | 393 | 364.14 | 348 | 381 | 1.096 | 0.133 | 0.22440253 | 0.38882112 | 0 | 0.0089 | 0.0058 | 0.0058 | TRUE | 0.0186 |
| Abhd8 | 1055 | 738 | 910 | 1049 | 882 | 978 | 819 | 785 | 982 | 1044 | 995 | 965 | 931.81 | 862 | 1001 | 1.163 | 0.218 | 0.02304452 | 0.07661367 | 0.0066 | 0.005 | 0.0055 | 0.0055 | TRUE | 0.1718 |
| Abi1 | 4257 | 3096 | 3109 | 3460 | 3006 | 3338 | 3306 | 3293 | 3355 | 3444 | 3391 | 3295 | 3347.38 | 3318 | 3377 | 1.018 | 0.025 | 0.63858476 | 0.77423614 | 0 | 0.0031 | 0.0018 | 0.0018 | TRUE | 0.0055 |
| Abi2 | 5051 | 3830 | 3802 | 4511 | 3908 | 4483 | 3923 | 4074 | 4102 | 4490 | 4409 | 4425 | 4237.24 | 4033 | 4441 | 1.102 | 0.14 | 0.00745304 | 0.03327908 | 0 | 0.003 | 0.0017 | 0.0017 | TRUE | 0.0071 |
| Abi3 | 83 | 73 | 75 | 107 | 103 | 146 | 64 | 78 | 81 | 107 | 116 | 144 | 98.31 | 74 | 122 | 1.655 | 0.726 | 0.00067227 | 0.00539515 | 0.0115 | 0.0265 | 0.0221 | 0.0221 | TRUE | 0.2533 |
| Abi3bp | 44 | 38 | 25 | 17 | 21 | 23 | 34 | 40 | 27 | 17 | 24 | 23 | 27.48 | 34 | 21 | 0.622 | -0.686 | 0.07032609 | 0.17206121 | 0 | 0.0887 | 0.0646 | 0.0646 | TRUE | 0.144 |
| Abl1 | 6205 | 4819 | 4534 | 4678 | 4248 | 4916 | 4819 | 5126 | 4892 | 4656 | 4792 | 4853 | 4856.45 | 4946 | 4767 | 0.964 | -0.053 | 0.32766134 | 0.5019575 | 4.00E-04 | 0.0029 | 0.0019 | 0.0019 | TRUE | 0.0123 |
| Abl2 | 8812 | 6426 | 6658 | 7416 | 6836 | 7909 | 6844 | 6835 | 7184 | 7382 | 7712 | 7807 | 7294.01 | 6954 | 7634 | 1.098 | 0.135 | 0.01012063 | 0.04167506 | 6.00E-04 | 0.0028 | 0.0018 | 0.0018 | TRUE | 0.0102 |
| Ablim1 | 666 | 579 | 863 | 690 | 797 | 866 | 517 | 616 | 931 | 687 | 899 | 855 | 750.85 | 688 | 814 | 1.183 | 0.243 | 0.3848073 | 0.55883466 | 0.0548 | 0.0056 | 0.0192 | 0.0548 | TRUE | 1.1252 |
| Ablim2 | 264 | 237 | 299 | 317 | 339 | 338 | 205 | 252 | 323 | 316 | 382 | 334 | 301.9 | 260 | 344 | 1.328 | 0.409 | 0.01084454 | 0.04383511 | 0.0259 | 0.0103 | 0.0152 | 0.0152 | TRUE | 0.4335 |
| Ablim3 | 193 | 168 | 164 | 224 | 208 | 258 | 150 | 179 | 177 | 223 | 235 | 255 | 202.97 | 169 | 238 | 1.415 | 0.501 | 0.00070108 | 0.00557603 | 0 | 0.0141 | 0.0107 | 0.0107 | TRUE | 0.1111 |
| Abo | 25 | 18 | 12 | 19 | 28 | 13 | 19 | 19 | 13 | 19 | 32 | 13 | 19.14 | 17 | 21 | 1.215 | 0.281 | 0.55584842 | 0.71100245 | 0.0742 | 0.1263 | 0.1117 | 0.1117 | TRUE | 0.4396 |
| Abp1 | 34 | 19 | 21 | 13 | 11 | 17 | 26 | 20 | 23 | 13 | 12 | 17 | 18.57 | 23 | 14 | 0.607 | -0.721 | 0.10819983 | 0.23424723 | 0 | 0.1301 | 0.0875 | 0.0875 | TRUE | 0.1311 |
| Abr | 5570 | 4106 | 3855 | 4401 | 4033 | 4453 | 4326 | 4367 | 4160 | 4381 | 4550 | 4396 | 4363.2 | 4284 | 4442 | 1.037 | 0.052 | 0.33110349 | 0.50549692 | 3.00E-04 | 0.003 | 0.0018 | 0.0018 | TRUE | 0.0078 |
| Abra | 6 | 1 | 1 | 1 | 2 | 2 | 5 | 1 | 1 | 1 | 2 | 2 | 2 | 2 | 2 | 0.737 | -0.44 | 0.74681281 | NA | 0 | 1.1846 | 0.8317 | 0.8317 | TRUE | 1.1215 |
| Abracl | 1346 | 831 | 905 | 1154 | 879 | 1139 | 1045 | 884 | 976 | 1149 | 992 | 1124 | 1028.41 | 968 | 1088 | 1.123 | 0.167 | 0.06752932 | 0.16698904 | 0.0056 | 0.0047 | 0.005 | 0.005 | TRUE | 0.0709 |
| Abt1 | 1897 | 1360 | 1132 | 1202 | 1089 | 1255 | 1473 | 1447 | 1221 | 1196 | 1229 | 1239 | 1300.87 | 1380 | 1221 | 0.884 | -0.178 | 0.03536183 | 0.10468653 | 0.0047 | 0.0042 | 0.0044 | 0.0044 | TRUE | 0.1202 |
| Abtb1 | 1455 | 1093 | 1076 | 946 | 929 | 1063 | 1130 | 1163 | 1161 | 942 | 1048 | 1049 | 1082.11 | 1151 | 1013 | 0.88 | -0.184 | 0.01684288 | 0.06097947 | 0.001 | 0.0046 | 0.0034 | 0.0034 | TRUE | 0.0459 |
| Abtb2 | 624 | 458 | 457 | 607 | 512 | 593 | 485 | 487 | 493 | 604 | 578 | 585 | 538.68 | 488 | 589 | 1.207 | 0.271 | 0.00302914 | 0.01665854 | 0 | 0.0068 | 0.0042 | 0.0042 | TRUE | 0.006 |
| Acaa1a | 2059 | 1660 | 1397 | 1528 | 1338 | 1543 | 1599 | 1766 | 1507 | 1521 | 1509 | 1523 | 1570.96 | 1624 | 1518 | 0.935 | -0.097 | 0.19586572 | 0.35547397 | 0.0023 | 0.0039 | 0.0034 | 0.0034 | TRUE | 0.0685 |
| Acaa1b | 243 | 191 | 188 | 193 | 184 | 218 | 189 | 203 | 203 | 192 | 208 | 215 | 201.6 | 198 | 205 | 1.035 | 0.05 | 0.72210592 | 0.83368819 | 0 | 0.0142 | 0.0095 | 0.0095 | TRUE | 0.0336 |
| Acaa2 | 2153 | 1549 | 1627 | 1778 | 1550 | 1760 | 1672 | 1648 | 1756 | 1770 | 1749 | 1737 | 1721.85 | 1692 | 1752 | 1.036 | 0.051 | 0.42536686 | 0.59804923 | 0 | 0.0038 | 0.0023 | 0.0023 | TRUE | 0.0127 |

| | | | | | | | | | | | | | | | | | | | | | | | | | |
|---|---|---|---|---|---|---|---|---|---|---|---|---|---|---|---|---|---|---|---|---|---|---|---|---|---|
| Acaca | 10467 | 7409 | 7077 | 8842 | 7574 | 8711 | 8129 | 7881 | 7636 | 8801 | 8544 | 8599 | 8265.14 | 7882 | 8648 | 1.097 | 0.134 | 0.00793502 | 0.03485564 | 4.00E-04 | 0.0027 | 0.0017 | 0.0017 | TRUE | 0.0093 |
| Acacb | 2111 | 1541 | 1601 | 1864 | 1811 | 2103 | 1640 | 1639 | 1727 | 1855 | 2043 | 2076 | 1830.09 | 1669 | 1991 | 1.194 | 0.255 | 0.00026722 | 0.00266156 | 0.0018 | 0.0037 | 0.003 | 0.003 | TRUE | 0.0435 |
| Acad10 | 1120 | 739 | 846 | 840 | 747 | 878 | 870 | 786 | 913 | 836 | 843 | 867 | 852.38 | 856 | 849 | 0.991 | -0.013 | 0.8752389 | 0.93071867 | 0.0017 | 0.0052 | 0.0041 | 0.0041 | TRUE | 0.0589 |
| Acad11 | 2527 | 1726 | 1938 | 2054 | 1955 | 2144 | 1963 | 1836 | 2091 | 2045 | 2205 | 2116 | 2042.68 | 1963 | 2122 | 1.081 | 0.112 | 0.11208268 | 0.24035975 | 0.0021 | 0.0036 | 0.0031 | 0.0031 | TRUE | 0.0381 |
| Acad12 | 471 | 352 | 247 | 385 | 308 | 344 | 366 | 374 | 267 | 383 | 347 | 340 | 346.17 | 336 | 357 | 1.061 | 0.086 | 0.54087026 | 0.6991642 | 0.0159 | 0.0093 | 0.0112 | 0.0112 | TRUE | 0.3569 |
| Acad8 | 1672 | 1225 | 1214 | 1323 | 1107 | 1282 | 1299 | 1303 | 1310 | 1317 | 1249 | 1266 | 1290.46 | 1304 | 1277 | 0.98 | -0.029 | 0.66513311 | 0.7937289 | 0 | 0.0043 | 0.0026 | 0.0026 | TRUE | 0.009 |
| Acad9 | 4843 | 3398 | 3002 | 3056 | 2784 | 3095 | 3761 | 3614 | 3239 | 3042 | 3141 | 3055 | 3308.79 | 3538 | 3079 | 0.87 | -0.201 | 0.00306179 | 0.01679284 | 0.0026 | 0.0031 | 0.003 | 0.003 | TRUE | 0.0661 |
| Acadl | 1540 | 1148 | 1326 | 1525 | 1309 | 1492 | 1196 | 1221 | 1431 | 1518 | 1477 | 1473 | 1385.9 | 1283 | 1489 | 1.162 | 0.217 | 0.00840974 | 0.03640229 | 0.0041 | 0.0041 | 0.0042 | 0.0042 | TRUE | 0.1209 |
| Acadm | 6986 | 5140 | 4983 | 4894 | 4548 | 5379 | 5426 | 5467 | 5377 | 4871 | 5131 | 5310 | 5263.62 | 5423 | 5104 | 0.941 | -0.088 | 0.10987378 | 0.23713917 | 7.00E-04 | 0.0029 | 0.002 | 0.002 | TRUE | 0.0195 |
| Acads | 5778 | 4223 | 4546 | 5288 | 4813 | 5432 | 4488 | 4492 | 4905 | 5264 | 5430 | 5362 | 4990 | 4628 | 5352 | 1.157 | 0.21 | 0.00027015 | 0.00268144 | 0.0012 | 0.0029 | 0.0022 | 0.0022 | TRUE | 0.0331 |
| Acadsb | 2237 | 1602 | 1755 | 1943 | 1649 | 1912 | 1737 | 1704 | 1894 | 1934 | 1860 | 1887 | 1836.13 | 1778 | 1894 | 1.066 | 0.092 | 0.17628483 | 0.32996846 | 0.0012 | 0.0037 | 0.0028 | 0.0028 | TRUE | 0.038 |
| Acadvl | 4313 | 2861 | 2761 | 3347 | 2872 | 3071 | 3350 | 3043 | 2979 | 3332 | 3240 | 3032 | 3162.52 | 3124 | 3201 | 1.024 | 0.034 | 0.61868197 | 0.75917079 | 0.0029 | 0.0032 | 0.0031 | 0.0031 | TRUE | 0.0495 |
| Acan | 2172 | 1471 | 1207 | 1504 | 1333 | 1378 | 1687 | 1565 | 1302 | 1497 | 1504 | 1360 | 1485.84 | 1518 | 1454 | 0.956 | -0.064 | 0.50073499 | 0.66509628 | 0.0096 | 0.004 | 0.0059 | 0.0059 | TRUE | 0.1845 |
| Acap1 | 711 | 587 | 571 | 938 | 795 | 1005 | 552 | 624 | 616 | 934 | 897 | 992 | 769.22 | 597 | 941 | 1.579 | 0.659 | 1.92E-13 | 2.72E-11 | 0.0022 | 0.0055 | 0.0044 | 0.0044 | TRUE | 0.0555 |
| Acap2 | 5231 | 3855 | 4280 | 4801 | 4218 | 4731 | 4063 | 4101 | 4618 | 4779 | 4758 | 4670 | 4498.13 | 4261 | 4736 | 1.112 | 0.153 | 0.01689491 | 0.06111811 | 0.0024 | 0.003 | 0.0027 | 0.0027 | TRUE | 0.0652 |
| Acap3 | 579 | 413 | 299 | 356 | 385 | 422 | 450 | 439 | 323 | 354 | 434 | 417 | 402.81 | 404 | 402 | 0.992 | -0.012 | 0.93446115 | 0.96372416 | 0.0194 | 0.0083 | 0.0116 | 0.0116 | TRUE | 0.3483 |
| Acat1 | 2148 | 1610 | 1757 | 2321 | 2019 | 2354 | 1668 | 1713 | 1896 | 2310 | 2278 | 2324 | 2031.38 | 1759 | 2304 | 1.311 | 0.391 | 1.33E-08 | 6.48E-07 | 0.0017 | 0.0036 | 0.0029 | 0.0029 | TRUE | 0.0554 |
| Acat2 | 6444 | 4643 | 4412 | 5115 | 4411 | 5199 | 5005 | 4939 | 4761 | 5091 | 4976 | 5132 | 4983.96 | 4902 | 5066 | 1.034 | 0.048 | 0.3582681 | 0.53274302 | 3.00E-04 | 0.0029 | 0.0017 | 0.0017 | TRUE | 0.0076 |
| Acat3 | 17 | 13 | 17 | 13 | 13 | 27 | 13 | 14 | 18 | 13 | 15 | 27 | 16.61 | 15 | 18 | 1.206 | 0.27 | 0.58605978 | NA | 0.0445 | 0.1452 | 0.1161 | 0.1161 | TRUE | 0.9021 |
| Acbd3 | 3401 | 2627 | 2723 | 2879 | 2593 | 2891 | 2641 | 2794 | 2938 | 2866 | 2925 | 2854 | 2836.43 | 2791 | 2882 | 1.033 | 0.047 | 0.45072901 | 0.62076217 | 0.0012 | 0.0033 | 0.0024 | 0.0024 | TRUE | 0.0275 |
| Acbd4 | 1279 | 939 | 1058 | 1049 | 909 | 1063 | 993 | 999 | 1142 | 1044 | 1025 | 1049 | 1042.11 | 1045 | 1039 | 0.996 | -0.005 | 0.95012146 | 0.97257652 | 0.0023 | 0.0047 | 0.0039 | 0.0039 | TRUE | 0.0774 |
| Acbd5 | 1919 | 1418 | 1553 | 1795 | 1431 | 1809 | 1490 | 1508 | 1676 | 1787 | 1614 | 1786 | 1643.53 | 1558 | 1729 | 1.111 | 0.152 | 0.04726359 | 0.1293144 | 0.0029 | 0.0039 | 0.0036 | 0.0036 | TRUE | 0.0518 |
| Acbd6 | 3104 | 2325 | 2335 | 2526 | 2330 | 2674 | 2411 | 2473 | 2519 | 2514 | 2628 | 2640 | 2530.96 | 2468 | 2594 | 1.052 | 0.073 | 0.216774 | 0.37953812 | 3.00E-04 | 0.0034 | 0.0021 | 0.0021 | TRUE | 0.0089 |
| Acbd7 | 153 | 91 | 127 | 145 | 132 | 148 | 119 | 97 | 137 | 144 | 149 | 146 | 132 | 118 | 146 | 1.245 | 0.317 | 0.08204941 | 0.19260401 | 0.0041 | 0.0204 | 0.0162 | 0.0162 | TRUE | 0.2234 |
| Accs | 115 | 102 | 78 | 91 | 88 | 77 | 89 | 108 | 84 | 91 | 99 | 76 | 91.31 | 94 | 89 | 0.942 | -0.085 | 0.69099901 | 0.81113382 | 0.006 | 0.0284 | 0.0223 | 0.0223 | TRUE | 0.1671 |
| Accsl | 222 | 143 | 94 | 76 | 87 | 101 | 172 | 152 | 101 | 76 | 98 | 100 | 116.57 | 142 | 91 | 0.639 | -0.647 | 0.00430697 | 0.02186838 | 0.0404 | 0.0228 | 0.0279 | 0.0279 | TRUE | 0.4918 |
| Acd | 2657 | 1923 | 1879 | 1865 | 1727 | 1978 | 2064 | 2045 | 2027 | 1856 | 1948 | 1953 | 1982.29 | 2045 | 1919 | 0.938 | -0.092 | 0.12954458 | 0.26633784 | 0 | 0.0036 | 0.0022 | 0.0022 | TRUE | 0.01 |
| Ace | 859 | 796 | 811 | 864 | 762 | 807 | 667 | 847 | 875 | 860 | 860 | 797 | 817.53 | 796 | 839 | 1.056 | 0.078 | 0.46512729 | 0.63411526 | 0.0105 | 0.0053 | 0.007 | 0.007 | TRUE | 0.2492 |
| Ace2 | 0 | 0 | 0 | 0 | 0 | 2 | 0 | 0 | 0 | 0 | 0 | 2 | 0.33 | 0 | 1 | 3.743 | 1.904 | 0.62640105 | NA | 0 | 7.2051 | 9.2465 | 9.2465 | TRUE | 0.9739 |
| Ace3 | 4 | 1 | 2 | 3 | 2 | 5 | 3 | 1 | 2 | 3 | 2 | 5 | 2.75 | 2 | 3 | 1.583 | 0.662 | 0.56271046 | NA | 0 | 0.8639 | 0.5686 | 0.5686 | TRUE | 0.2352 |
| Acer1 | 22 | 13 | 19 | 46 | 34 | 30 | 17 | 14 | 21 | 46 | 38 | 30 | 27.53 | 17 | 38 | 2.213 | 1.146 | 0.00312901 | 0.01705684 | 0 | 0.0885 | 0.0666 | 0.0666 | TRUE | 0.1333 |
| Acer2 | 714 | 557 | 492 | 590 | 492 | 609 | 555 | 592 | 531 | 587 | 555 | 601 | 570.23 | 559 | 581 | 1.04 | 0.056 | 0.54926475 | 0.7056756 | 0 | 0.0066 | 0.0046 | 0.0046 | TRUE | 0.0301 |
| Acer3 | 1091 | 784 | 819 | 973 | 808 | 972 | 847 | 834 | 884 | 969 | 912 | 960 | 900.75 | 855 | 947 | 1.108 | 0.148 | 0.05788859 | 0.14986445 | 0 | 0.0051 | 0.0033 | 0.0033 | TRUE | 0.0122 |
| Ache | 2 | 1 | 1 | 3 | 4 | 3 | 2 | 1 | 1 | 3 | 5 | 3 | 2.36 | 1 | 4 | 2.775 | 1.473 | 0.24478154 | NA | 0 | 1.007 | 0.622 | 0.622 | TRUE | 0.0872 |
| Acin1 | 17640 | 13485 | 12964 | 12445 | 11524 | 13780 | 13700 | 14344 | 13988 | 12388 | 13000 | 13603 | 13503.85 | 14011 | 12997 | 0.928 | -0.108 | 0.04451398 | 0.12378178 | 0.0012 | 0.0026 | 0.002 | 0.002 | TRUE | 0.0206 |
| Acly | 41911 | 30725 | 32008 | 33931 | 30061 | 35783 | 32550 | 32682 | 34536 | 33775 | 33912 | 35323 | 33796.47 | 33256 | 34337 | 1.033 | 0.046 | 0.34774237 | 0.52236276 | 8.00E-04 | 0.0025 | 0.0017 | 0.0017 | TRUE | 0.0139 |
| Acmsd | 1 | 0 | 0 | 1 | 0 | 0 | 1 | 0 | 0 | 1 | 0 | 0 | 0.3 | 0 | 0 | 1.075 | 0.105 | 0.97698525 | NA | 0 | 8.0272 | 7.5141 | 7.5141 | TRUE | 0.3906 |
| Acn9 | 423 | 300 | 300 | 374 | 301 | 349 | 329 | 319 | 324 | 372 | 340 | 345 | 337.95 | 324 | 352 | 1.088 | 0.121 | 0.28119518 | 0.45275857 | 0 | 0.0094 | 0.0061 | 0.0061 | TRUE | 0.0289 |
| Acnat1 | 0 | 3 | 1 | 3 | 0 | 1 | 0 | 3 | 1 | 3 | 0 | 1 | 1.37 | 1 | 1 | 0.973 | -0.039 | 0.98249369 | NA | 1.1511 | 1.7275 | 1.5972 | 1.5972 | TRUE | 0.4682 |
| Acnat2 | 0 | 0 | 0 | 0 | 0 | 0 | 0 | 0 | 0 | 0 | 0 | 0 | 0 | 0 | 0 | NA | NA | NA | NA | NA | NA | NA | NA | NA | NA |
| Aco1 | 4794 | 3530 | 3859 | 4354 | 4031 | 4665 | 3723 | 3755 | 4164 | 4334 | 4547 | 4605 | 4188.06 | 3881 | 4495 | 1.159 | 0.213 | 0.0008334 | 0.00633834 | 0.0021 | 0.003 | 0.0027 | 0.0027 | TRUE | 0.0492 |
| Aco2 | 22995 | 17017 | 18160 | 20987 | 18814 | 21002 | 17859 | 18101 | 19594 | 20890 | 21224 | 20732 | 19733.55 | 18518 | 20949 | 1.131 | 0.178 | 0.00079522 | 0.00612399 | 0.0013 | 0.0025 | 0.002 | 0.002 | TRUE | 0.0316 |
| Acot1 | 1533 | 1405 | 1787 | 1671 | 1612 | 2024 | 1191 | 1494 | 1928 | 1663 | 1819 | 1998 | 1682.18 | 1538 | 1827 | 1.188 | 0.249 | 0.23902166 | 0.40603002 | 0.0316 | 0.0038 | 0.0122 | 0.0316 | TRUE | 0.3626 |
| Acot10 | 4 | 5 | 8 | 4 | 8 | 4 | 3 | 5 | 9 | 4 | 9 | 4 | 5.67 | 6 | 6 | 1.002 | 0.003 | 0.99744988 | NA | 0 | 0.4205 | 0.3212 | 0.3212 | TRUE | 0.3927 |
| Acot11 | 522 | 315 | 298 | 317 | 284 | 294 | 405 | 335 | 322 | 316 | 320 | 290 | 331.36 | 354 | 309 | 0.869 | -0.203 | 0.11219671 | 0.2405321 | 0.0071 | 0.0096 | 0.0088 | 0.0088 | TRUE | 0.1952 |
| Acot12 | 5 | 8 | 1 | 0 | 1 | 1 | 4 | 9 | 1 | 0 | 1 | 1 | 2.6 | 5 | 1 | 0.155 | -2.691 | 0.07364591 | NA | 0 | 0.9147 | 0.7773 | 0.7773 | TRUE | 0.1304 |
| Acot13 | 1121 | 799 | 999 | 978 | 889 | 1179 | 871 | 850 | 1078 | 974 | 1003 | 1164 | 989.78 | 933 | 1047 | 1.123 | 0.168 | 0.11645337 | 0.24699035 | 0.0124 | 0.0048 | 0.0072 | 0.0072 | TRUE | 0.2174 |
| Acot2 | 357 | 262 | 235 | 232 | 232 | 255 | 277 | 279 | 254 | 231 | 262 | 252 | 258.98 | 270 | 248 | 0.918 | -0.124 | 0.33639472 | 0.51061908 | 0 | 0.0116 | 0.008 | 0.008 | TRUE | 0.0411 |
| Acot3 | 45 | 25 | 23 | 21 | 27 | 21 | 35 | 27 | 25 | 21 | 30 | 21 | 26.41 | 29 | 24 | 0.825 | -0.278 | 0.47081984 | 0.63967946 | 0 | 0.0922 | 0.0685 | 0.0685 | TRUE | 0.3002 |
| Acot4 | 10 | 10 | 9 | 10 | 7 | 8 | 8 | 11 | 10 | 10 | 8 | 8 | 8.98 | 10 | 9 | 0.923 | -0.115 | 0.85412348 | NA | 0 | 0.2664 | 0.1711 | 0.1711 | TRUE | 0.0976 |
| Acot5 | 7 | 6 | 3 | 5 | 2 | 4 | 5 | 6 | 3 | 5 | 2 | 4 | 4.37 | 5 | 4 | 0.745 | -0.424 | 0.64108687 | NA | 0 | 0.5444 | 0.3622 | 0.3622 | TRUE | 0.1741 |
| Acot6 | 13 | 16 | 10 | 14 | 11 | 21 | 10 | 17 | 11 | 14 | 12 | 21 | 14.16 | 13 | 16 | 1.256 | 0.328 | 0.53065559 | NA | 0 | 0.1698 | 0.1262 | 0.1262 | TRUE | 0.3784 |
| Acot7 | 3442 | 2566 | 2509 | 2994 | 2688 | 3139 | 2673 | 2729 | 2707 | 2980 | 3032 | 3099 | 2870.19 | 2703 | 3037 | 1.124 | 0.168 | 0.00234891 | 0.01369855 | 0 | 0.0033 | 0.0019 | 0.0019 | TRUE | 0.0039 |
| Acot8 | 685 | 465 | 441 | 537 | 447 | 483 | 532 | 495 | 476 | 535 | 504 | 477 | 503.01 | 501 | 505 | 1.007 | 0.01 | 0.9209134 | 0.95651589 | 0.0015 | 0.0071 | 0.0053 | 0.0053 | TRUE | 0.0368 |
| Acot9 | 4052 | 2953 | 3320 | 3437 | 3012 | 3546 | 3147 | 3141 | 3582 | 3421 | 3398 | 3500 | 3364.97 | 3290 | 3440 | 1.046 | 0.065 | 0.33414455 | 0.50818557 | 0.0026 | 0.0031 | 0.003 | 0.003 | TRUE | 0.0726 |
| Acox1 | 1782 | 1260 | 1303 | 1462 | 1237 | 1469 | 1384 | 1340 | 1406 | 1455 | 1395 | 1450 | 1405.17 | 1377 | 1433 | 1.041 | 0.059 | 0.38150731 | 0.55579288 | 0 | 0.0041 | 0.0025 | 0.0025 | TRUE | 0.0064 |
| Acox2 | 2 | 0 | 1 | 3 | 1 | 1 | 2 | 0 | 1 | 3 | 1 | 1 | 1.29 | 1 | 2 | 1.878 | 0.909 | 0.59350075 | NA | 0 | 1.8412 | 1.2591 | 1.2591 | TRUE | 0.3462 |
| Acox3 | 319 | 235 | 265 | 286 | 208 | 280 | 248 | 250 | 286 | 285 | 235 | 276 | 263.23 | 261 | 265 | 1.019 | 0.027 | 0.8445448 | 0.91195135 | 0.0044 | 0.0114 | 0.0094 | 0.0094 | TRUE | 0.1096 |
| Acoxl | 25 | 20 | 16 | 21 | 28 | 27 | 19 | 21 | 17 | 21 | 32 | 27 | 22.85 | 19 | 27 | 1.361 | 0.445 | 0.27165311 | 0.44262858 | 0 | 0.1062 | 0.0733 | 0.0733 | TRUE | 0.1822 |
| Acp1 | 2686 | 1884 | 1856 | 1971 | 1627 | 1920 | 2086 | 2004 | 2003 | 1962 | 1835 | 1895 | 1964.23 | 2031 | 1897 | 0.934 | -0.098 | 0.11842706 | 0.24987864 | 3.00E-04 | 0.0036 | 0.0023 | 0.0023 | TRUE | 0.0106 |
| Acp2 | 838 | 598 | 467 | 574 | 471 | 519 | 651 | 636 | 504 | 571 | 531 | 512 | 567.64 | 597 | 538 | 0.9 | -0.151 | 0.18252199 | 0.33810094 | 0.0095 | 0.0066 | 0.0075 | 0.0075 | TRUE | 0.2135 |
| Acp5 | 858 | 631 | 612 | 802 | 739 | 804 | 666 | 671 | 660 | 798 | 834 | 794 | 737.26 | 666 | 809 | 1.214 | 0.279 | 0.00062617 | 0.00513384 | 0 | 0.0056 | 0.0034 | 0.0034 | TRUE | 0.0085 |
| Acp6 | 11431 | 8739 | 8924 | 9477 | 8714 | 9866 | 8878 | 9296 | 9629 | 9433 | 9830 | 9739 | 9467.57 | 9268 | 9667 | 1.043 | 0.061 | 0.24695383 | 0.41568896 | 9.00E-04 | 0.0027 | 0.0019 | 0.0019 | TRUE | 0.0167 |
| Acpl2 | 910 | 641 | 750 | 838 | 713 | 851 | 707 | 682 | 809 | 834 | 804 | 840 | 779.4 | 733 | 826 | 1.129 | 0.175 | 0.053247 | 0.14096303 | 0.0029 | 0.0055 | 0.0046 | 0.0046 | TRUE | 0.096 |
| Acpp | 384 | 319 | 363 | 264 | 249 | 299 | 298 | 339 | 392 | 263 | 281 | 295 | 311.34 | 343 | 280 | 0.818 | -0.291 | 0.02958905 | 0.0917199 | 0.0086 | 0.01 | 0.0096 | 0.0096 | TRUE | 0.1743 |
| Acpt | 0 | 0 | 0 | 0 | 0 | 0 | 0 | 0 | 0 | 0 | 0 | 0 | 0 | 0 | 0 | NA | NA | NA | NA | NA | NA | NA | NA | NA | NA |
| Acr | 2 | 3 | 2 | 1 | 1 | 0 | 2 | 3 | 2 | 1 | 1 | 0 | 1.5 | 2 | 1 | 0.306 | -1.71 | 0.30391297 | NA | 0 | 1.5779 | 1.0396 | 1.0396 | TRUE | 0.2719 |
| Acrbp | 532 | 433 | 559 | 566 | 531 | 651 | 413 | 461 | 603 | 563 | 599 | 643 | 547 | 492 | 602 | 1.226 | 0.293 | 0.02592635 | 0.08349825 | 0.0196 | 0.0068 | 0.0107 | 0.0107 | TRUE | 0.4481 |
| Acrv1 | 0 | 0 | 0 | 0 | 0 | 0 | 0 | 0 | 0 | 0 | 0 | 0 | 0 | 0 | 0 | NA | NA | NA | NA | NA | NA | NA | NA | NA | NA |

| | | | | | | | | | | | | | | | | | | | | | | | | | |
|---|---|---|---|---|---|---|---|---|---|---|---|---|---|---|---|---|---|---|---|---|---|---|---|---|---|
| Acsbg1 | 111 | 71 | 59 | 73 | 66 | 77 | 86 | 76 | 64 | 73 | 74 | 76 | 74.75 | 75 | 74 | 0.985 | -0.022 | 0.9222146 | 0.95711845 | 0 | 0.0341 | 0.0245 | 0.0245 | TRUE | 0.1717 |
| Acsbg2 | 30 | 21 | 24 | 24 | 17 | 16 | 23 | 22 | 26 | 24 | 19 | 16 | 21.73 | 24 | 20 | 0.824 | -0.28 | 0.49719461 | 0.66159903 | 0 | 0.1115 | 0.0756 | 0.0756 | TRUE | 0.2017 |
| Acsf2 | 2601 | 2084 | 2307 | 3016 | 2645 | 2984 | 2020 | 2217 | 2489 | 3002 | 2984 | 2946 | 2609.61 | 2242 | 2977 | 1.329 | 0.411 | 1.31E-07 | 4.95E-06 | 0.005 | 0.0033 | 0.004 | 0.004 | TRUE | 0.1125 |
| Acsf3 | 1587 | 1133 | 1089 | 1032 | 1027 | 1090 | 1233 | 1205 | 1175 | 1027 | 1159 | 1076 | 1145.76 | 1204 | 1087 | 0.902 | -0.149 | 0.05091507 | 0.13663708 | 0.0011 | 0.0045 | 0.0033 | 0.0033 | TRUE | 0.0386 |
| Acsl1 | 5776 | 4618 | 5251 | 5838 | 5482 | 6490 | 4486 | 4912 | 5666 | 5811 | 6184 | 6407 | 5577.65 | 5021 | 6134 | 1.222 | 0.289 | 0.00042444 | 0.00380825 | 0.008 | 0.0029 | 0.0047 | 0.0047 | TRUE | 0.1535 |
| Acsl3 | 7573 | 5629 | 6121 | 6052 | 5232 | 6129 | 5882 | 5987 | 6605 | 6024 | 5902 | 6050 | 6075.05 | 6158 | 5992 | 0.973 | -0.039 | 0.51763504 | 0.67970212 | 0.0018 | 0.0028 | 0.0024 | 0.0024 | TRUE | 0.0489 |
| Acsl4 | 5090 | 3474 | 3797 | 4300 | 3412 | 4172 | 3953 | 3695 | 4097 | 4280 | 3849 | 4118 | 3998.85 | 3915 | 4082 | 1.043 | 0.061 | 0.35302699 | 0.52775572 | 0.0024 | 0.003 | 0.0028 | 0.0028 | TRUE | 0.0302 |
| Acsl5 | 679 | 504 | 594 | 577 | 547 | 691 | 527 | 536 | 641 | 574 | 617 | 682 | 596.32 | 568 | 624 | 1.101 | 0.139 | 0.20336828 | 0.36472149 | 0.008 | 0.0064 | 0.0069 | 0.0069 | TRUE | 0.1445 |
| Acsl6 | 910 | 650 | 628 | 865 | 776 | 803 | 707 | 691 | 678 | 861 | 875 | 793 | 767.48 | 692 | 843 | 1.217 | 0.283 | 0.00075197 | 0.00587317 | 0 | 0.0055 | 0.0038 | 0.0038 | TRUE | 0.0328 |
| Acsm1 | 0 | 0 | 0 | 0 | 0 | 0 | 0 | 0 | 0 | 0 | 0 | 0 | 0 | 0 | 0 | NA | NA | NA | NA | NA | NA | NA | NA | NA | NA |
| Acsm2 | 0 | 0 | 0 | 0 | 0 | 0 | 0 | 0 | 0 | 0 | 0 | 0 | 0 | 0 | 0 | NA | NA | NA | NA | NA | NA | NA | NA | NA | NA |
| Acsm3 | 4 | 3 | 3 | 5 | 5 | 3 | 3 | 3 | 3 | 5 | 6 | 3 | 3.85 | 3 | 5 | 1.42 | 0.506 | 0.5960575 | NA | 0 | 0.6177 | 0.389 | 0.389 | TRUE | 0.1764 |
| Acsm4 | 3 | 4 | 1 | 0 | 0 | 1 | 2 | 4 | 1 | 0 | 0 | 1 | 1.44 | 2 | 0 | 0.145 | -2.782 | 0.13070355 | NA | 0 | 1.6462 | 1.2529 | 1.2529 | TRUE | 0.2957 |
| Acsm5 | 0 | 0 | 0 | 0 | 0 | 0 | 0 | 0 | 0 | 0 | 0 | 0 | 0 | 0 | 0 | NA | NA | NA | NA | NA | NA | NA | NA | NA | NA |
| Acss1 | 699 | 616 | 986 | 1122 | 1338 | 1475 | 543 | 655 | 1064 | 1117 | 1509 | 1456 | 1057.38 | 754 | 1361 | 1.806 | 0.852 | 0.00949362 | 0.03986261 | 0.0768 | 0.0047 | 0.0224 | 0.0768 | TRUE | 1.5285 |
| Acss2 | 2438 | 1848 | 1909 | 2403 | 2162 | 2624 | 1893 | 1966 | 2060 | 2392 | 2439 | 2590 | 2223.36 | 1973 | 2474 | 1.255 | 0.327 | 6.18E-07 | 1.96E-05 | 0.0013 | 0.0035 | 0.0027 | 0.0027 | TRUE | 0.0208 |
| Acss3 | 72 | 44 | 23 | 49 | 31 | 36 | 56 | 47 | 25 | 49 | 35 | 36 | 41.14 | 43 | 40 | 0.928 | -0.107 | 0.75910872 | 0.8571376 | 0.0742 | 0.06 | 0.064 | 0.064 | TRUE | 0.6216 |
| Acta1 | 13 | 10 | 11 | 11 | 10 | 21 | 10 | 11 | 12 | 11 | 11 | 21 | 12.59 | 11 | 14 | 1.326 | 0.407 | 0.46018524 | NA | 0 | 0.1906 | 0.1388 | 0.1388 | TRUE | 0.6981 |
| Acta2 | 62 | 31 | 27 | 28 | 26 | 20 | 48 | 33 | 29 | 28 | 29 | 20 | 31.2 | 37 | 26 | 0.69 | -0.536 | 0.14637534 | 0.2907369 | 0.0323 | 0.0784 | 0.0649 | 0.0649 | TRUE | 0.6156 |
| Actb | 80203 | 54840 | 42302 | 40924 | 35443 | 41534 | 62290 | 58333 | 45644 | 40736 | 39983 | 41000 | 47997.65 | 55422 | 40573 | 0.732 | -0.45 | 8.60E-07 | 2.57E-05 | 0.0124 | 0.0025 | 0.006 | 0.006 | TRUE | 0.2916 |
| Actbl2 | 0 | 0 | 0 | 0 | 0 | 0 | 0 | 0 | 0 | 0 | 0 | 0 | 0 | 0 | 0 | NA | NA | NA | NA | NA | NA | NA | NA | NA | NA |
| Actc1 | 10 | 3 | 2 | 2 | 3 | 4 | 8 | 3 | 2 | 2 | 3 | 4 | 3.74 | 4 | 3 | 0.692 | -0.531 | 0.60007147 | NA | 0 | 0.6362 | 0.4649 | 0.4649 | TRUE | 1.0741 |
| Actg1 | 67970 | 45735 | 44380 | 39610 | 36959 | 44363 | 52789 | 48648 | 47886 | 39428 | 41694 | 43793 | 45706.21 | 49774 | 41638 | 0.837 | -0.258 | 1.86E-05 | 0.00031463 | 0.0027 | 0.0025 | 0.0026 | 0.0026 | TRUE | 0.0344 |
| Actg2 | 3 | 1 | 0 | 2 | 1 | 0 | 2 | 1 | 0 | 2 | 1 | 0 | 1.09 | 1 | 1 | 0.885 | -0.177 | 0.92674766 | NA | 0 | 2.186 | 1.7483 | 1.7483 | TRUE | 0.2457 |
| Actl10 | 7 | 3 | 7 | 16 | 6 | 10 | 5 | 3 | 8 | 16 | 7 | 10 | 8.12 | 5 | 11 | 2.023 | 1.016 | 0.15191403 | NA | 0 | 0.2941 | 0.2258 | 0.2258 | TRUE | 0.3061 |
| Actl11 | 0 | 0 | 0 | 0 | 0 | 0 | 0 | 0 | 0 | 0 | 0 | 0 | 0 | 0 | 0 | NA | NA | NA | NA | NA | NA | NA | NA | NA | NA |
| Actl6a | 9147 | 6640 | 7084 | 6309 | 5808 | 6784 | 7104 | 7063 | 7644 | 6280 | 6552 | 6697 | 6889.9 | 7270 | 6510 | 0.896 | -0.159 | 0.00496149 | 0.02436988 | 0.0013 | 0.0028 | 0.0022 | 0.0022 | TRUE | 0.0245 |
| Actl6b | 0 | 0 | 0 | 1 | 2 | 0 | 0 | 0 | 0 | 1 | 2 | 0 | 0.54 | 0 | 1 | 6.07 | 2.602 | 0.36554818 | NA | 0 | 4.3758 | 4.4761 | 4.4761 | TRUE | 0.0722 |
| Actl7a | 0 | 1 | 0 | 0 | 0 | 0 | 0 | 1 | 0 | 0 | 0 | 0 | 0.18 | 0 | 0 | 0.552 | -0.857 | 0.8336548 | NA | 0 | 13.3714 | 10 | 10 | TRUE | 0.5216 |
| Actl7b | 67 | 42 | 34 | 54 | 40 | 49 | 52 | 45 | 37 | 54 | 45 | 48 | 46.77 | 45 | 49 | 1.098 | 0.135 | 0.6392091 | 0.77458378 | 0 | 0.0531 | 0.0382 | 0.0382 | TRUE | 0.2009 |
| Actl9 | 0 | 0 | 0 | 0 | 0 | 0 | 0 | 0 | 0 | 0 | 0 | 0 | 0 | 0 | 0 | NA | NA | NA | NA | NA | NA | NA | NA | NA | NA |
| Actn1 | 13359 | 10435 | 10749 | 10257 | 9735 | 11681 | 10375 | 11100 | 11598 | 10210 | 10982 | 11531 | 10965.98 | 11024 | 10908 | 0.99 | -0.015 | 0.81545832 | 0.89395197 | 0.0027 | 0.0026 | 0.0029 | 0.0029 | TRUE | 0.0384 |
| Actn2 | 523 | 423 | 294 | 295 | 262 | 293 | 406 | 450 | 317 | 294 | 296 | 289 | 341.97 | 391 | 293 | 0.748 | -0.419 | 0.00222007 | 0.01308687 | 0.0131 | 0.0094 | 0.0105 | 0.0105 | TRUE | 0.3025 |
| Actn3 | 611 | 456 | 410 | 889 | 792 | 1008 | 475 | 485 | 442 | 885 | 893 | 995 | 695.9 | 467 | 924 | 1.978 | 0.984 | 1.87E-26 | 1.12E-23 | 0.0021 | 0.0058 | 0.0046 | 0.0046 | TRUE | 0.0537 |
| Actn4 | 33580 | 23110 | 20685 | 21260 | 19144 | 22357 | 26080 | 24582 | 22319 | 21162 | 21596 | 22070 | 22968.22 | 24327 | 21609 | 0.888 | -0.171 | 0.006714 | 0.03076728 | 0.0032 | 0.0025 | 0.0028 | 0.0028 | TRUE | 0.0638 |
| Actr10 | 3559 | 2460 | 2276 | 2281 | 1959 | 2257 | 2764 | 2617 | 2456 | 2271 | 2210 | 2228 | 2424.17 | 2612 | 2236 | 0.855 | -0.225 | 0.00056752 | 0.00475195 | 0.0015 | 0.0034 | 0.0027 | 0.0027 | TRUE | 0.0331 |
| Actr1a | 6598 | 4410 | 4508 | 5188 | 4414 | 5356 | 5124 | 4691 | 4864 | 5164 | 4979 | 5287 | 5018.35 | 4893 | 5143 | 1.051 | 0.072 | 0.21509248 | 0.37765193 | 0.0011 | 0.0029 | 0.0022 | 0.0022 | TRUE | 0.0212 |
| Actr1b | 4741 | 3465 | 3323 | 3257 | 2845 | 3303 | 3682 | 3686 | 3585 | 3242 | 3209 | 3261 | 3444.22 | 3651 | 3237 | 0.887 | -0.174 | 0.0010175 | 0.00732866 | 0 | 0.0031 | 0.0017 | 0.0017 | TRUE | 0.003 |
| Actr2 | 23691 | 16974 | 17659 | 20358 | 17363 | 19613 | 18400 | 18055 | 19054 | 20264 | 19587 | 19361 | 19120.23 | 18503 | 19737 | 1.067 | 0.093 | 0.05382068 | 0.14182504 | 6.00E-04 | 0.0026 | 0.0016 | 0.0016 | TRUE | 0.0083 |
| Actr3 | 16933 | 12282 | 13686 | 13414 | 11644 | 13655 | 13151 | 13064 | 14767 | 13352 | 13136 | 13480 | 13491.66 | 13661 | 13323 | 0.975 | -0.036 | 0.54895915 | 0.70553666 | 0.0022 | 0.0026 | 0.0025 | 0.0025 | TRUE | 0.0612 |
| Actr3b | 1176 | 866 | 793 | 1002 | 826 | 971 | 913 | 921 | 856 | 997 | 932 | 959 | 929.65 | 897 | 963 | 1.073 | 0.102 | 0.19330503 | 0.35205259 | 0 | 0.005 | 0.0034 | 0.0034 | TRUE | 0.0185 |
| Actr5 | 1362 | 1044 | 974 | 1065 | 941 | 1114 | 1058 | 1110 | 1051 | 1060 | 1062 | 1100 | 1073.43 | 1073 | 1074 | 1.001 | 0.002 | 0.98252075 | 0.99089635 | 0 | 0.0046 | 0.0029 | 0.0029 | TRUE | 0.0108 |
| Actr6 | 2054 | 1477 | 1447 | 1485 | 1299 | 1434 | 1595 | 1571 | 1561 | 1478 | 1465 | 1416 | 1514.46 | 1576 | 1453 | 0.922 | -0.118 | 0.06946097 | 0.17071391 | 0 | 0.004 | 0.0024 | 0.0024 | TRUE | 0.0062 |
| Actr8 | 2814 | 2154 | 2324 | 2441 | 2192 | 2680 | 2186 | 2291 | 2508 | 2430 | 2473 | 2646 | 2422.07 | 2328 | 2516 | 1.082 | 0.113 | 0.11385096 | 0.24287638 | 0.0029 | 0.0034 | 0.0033 | 0.0033 | TRUE | 0.0548 |
| Actrt1 | 0 | 0 | 0 | 0 | 0 | 0 | 0 | 0 | 0 | 0 | 0 | 0 | 0 | 0 | 0 | NA | NA | NA | NA | NA | NA | NA | NA | NA | NA |
| Actrt2 | 0 | 0 | 0 | 0 | 2 | 1 | 0 | 0 | 0 | 0 | 2 | 1 | 0.54 | 0 | 1 | 6.057 | 2.599 | 0.3664873 | NA | 0 | 4.3869 | 4.4843 | 4.4843 | TRUE | 0.0738 |
| Actrt3 | 8 | 3 | 2 | 6 | 4 | 3 | 6 | 3 | 2 | 6 | 5 | 3 | 4.17 | 4 | 5 | 1.134 | 0.182 | 0.84668225 | NA | 0 | 0.571 | 0.3998 | 0.3998 | TRUE | 0.6102 |
| Acvr1 | 578 | 444 | 437 | 494 | 422 | 510 | 449 | 472 | 472 | 492 | 476 | 503 | 477.32 | 464 | 490 | 1.058 | 0.081 | 0.40210285 | 0.57507729 | 0 | 0.0074 | 0.0047 | 0.0047 | TRUE | 0.0105 |
| Acvr1b | 3813 | 2957 | 3549 | 3564 | 3446 | 3932 | 2961 | 3145 | 3829 | 3548 | 3887 | 3881 | 3542.1 | 3312 | 3772 | 1.14 | 0.188 | 0.03598984 | 0.1060869 | 0.0104 | 0.0031 | 0.0055 | 0.0055 | TRUE | 0.2263 |
| Acvr1c | 7 | 9 | 8 | 4 | 3 | 1 | 5 | 10 | 9 | 4 | 3 | 1 | 5.33 | 8 | 3 | 0.354 | -1.496 | 0.09302512 | NA | 0 | 0.4469 | 0.3232 | 0.3232 | TRUE | 0.4117 |
| Acvr2a | 1271 | 911 | 897 | 980 | 858 | 946 | 987 | 969 | 968 | 975 | 968 | 934 | 966.88 | 975 | 959 | 0.984 | -0.024 | 0.74712251 | 0.84993112 | 0 | 0.0049 | 0.0029 | 0.0029 | TRUE | 0.0064 |
| Acvr2b | 1360 | 1022 | 1049 | 1134 | 1075 | 1222 | 1056 | 1087 | 1132 | 1129 | 1213 | 1206 | 1137.17 | 1092 | 1183 | 1.084 | 0.116 | 0.11776845 | 0.24900362 | 6.00E-04 | 0.0045 | 0.0031 | 0.0031 | TRUE | 0.0192 |
| Acvrl1 | 131 | 131 | 151 | 196 | 237 | 234 | 102 | 139 | 163 | 195 | 267 | 231 | 182.91 | 135 | 231 | 1.726 | 0.787 | 3.94E-05 | 0.00058163 | 0.0337 | 0.0154 | 0.0206 | 0.0206 | TRUE | 0.3033 |
| Acy1 | 2363 | 1958 | 2092 | 2243 | 2171 | 2606 | 1835 | 2083 | 2257 | 2233 | 2449 | 2573 | 2238.25 | 2058 | 2418 | 1.176 | 0.234 | 0.0067064 | 0.03074338 | 0.0072 | 0.0035 | 0.0049 | 0.0049 | TRUE | 0.1113 |
| Acy3 | 350 | 244 | 269 | 277 | 246 | 263 | 272 | 260 | 290 | 276 | 278 | 260 | 272.41 | 274 | 271 | 0.989 | -0.016 | 0.90043416 | 0.94534343 | 0 | 0.0111 | 0.0074 | 0.0074 | TRUE | 0.0287 |
| Acyp1 | 578 | 439 | 545 | 637 | 474 | 577 | 449 | 467 | 588 | 634 | 535 | 570 | 540.38 | 501 | 580 | 1.159 | 0.213 | 0.07776744 | 0.18524843 | 0.0124 | 0.0068 | 0.0086 | 0.0086 | TRUE | 0.263 |
| Acyp2 | 83 | 53 | 33 | 37 | 31 | 43 | 64 | 56 | 36 | 37 | 35 | 42 | 45.12 | 52 | 38 | 0.726 | -0.462 | 0.14160183 | 0.28393869 | 0.0318 | 0.055 | 0.0483 | 0.0483 | TRUE | 0.5716 |
| Ada | 1299 | 968 | 846 | 949 | 828 | 972 | 1009 | 1030 | 913 | 945 | 934 | 960 | 964.93 | 984 | 946 | 0.961 | -0.057 | 0.47468918 | 0.64293333 | 0.001 | 0.0049 | 0.0036 | 0.0036 | TRUE | 0.046 |
| Adad1 | 68 | 70 | 64 | 60 | 51 | 62 | 53 | 74 | 69 | 60 | 58 | 61 | 62.46 | 65 | 60 | 0.916 | -0.127 | 0.61458684 | 0.75648379 | 0 | 0.0404 | 0.0299 | 0.0299 | TRUE | 0.2917 |
| Adad2 | 168 | 99 | 92 | 140 | 126 | 144 | 130 | 105 | 99 | 139 | 142 | 142 | 126.45 | 111 | 141 | 1.257 | 0.329 | 0.07111723 | 0.17351974 | 0 | 0.0212 | 0.016 | 0.016 | TRUE | 0.2399 |
| Adal | 1056 | 831 | 775 | 823 | 753 | 809 | 820 | 884 | 836 | 819 | 849 | 799 | 834.6 | 847 | 822 | 0.972 | -0.042 | 0.60683478 | 0.75114657 | 0 | 0.0053 | 0.0035 | 0.0035 | TRUE | 0.0173 |
| Adam10 | 4653 | 3254 | 3272 | 3795 | 3095 | 3486 | 3614 | 3461 | 3530 | 3778 | 3491 | 3441 | 3552.62 | 3535 | 3570 | 1.01 | 0.014 | 0.81644585 | 0.89469144 | 0.0012 | 0.0031 | 0.0023 | 0.0023 | TRUE | 0.0316 |
| Adam11 | 235 | 196 | 193 | 254 | 244 | 252 | 183 | 208 | 208 | 253 | 275 | 249 | 229.35 | 200 | 259 | 1.299 | 0.378 | 0.00577323 | 0.027396 | 0 | 0.0128 | 0.0091 | 0.0091 | TRUE | 0.0687 |
| Adam12 | 4 | 5 | 2 | 2 | 2 | 2 | 3 | 5 | 2 | 2 | 2 | 2 | 2.8 | 3 | 2 | 0.589 | -0.764 | 0.50037839 | NA | 0 | 0.8487 | 0.539 | 0.539 | TRUE | 0.2695 |
| Adam15 | 1964 | 1494 | 1405 | 1578 | 1416 | 1609 | 1525 | 1589 | 1516 | 1571 | 1597 | 1588 | 1564.49 | 1543 | 1585 | 1.027 | 0.039 | 0.54477933 | 0.70214926 | 0 | 0.0039 | 0.0023 | 0.0023 | TRUE | 0.0079 |
| Adam17 | 4007 | 2834 | 2898 | 3165 | 2919 | 3292 | 3112 | 3014 | 3127 | 3150 | 3293 | 3250 | 3157.76 | 3084 | 3231 | 1.047 | 0.067 | 0.22966277 | 0.39520467 | 0 | 0.0032 | 0.0019 | 0.0019 | TRUE | 0.0058 |
| Adam18 | 5 | 3 | 1 | 7 | 0 | 1 | 4 | 3 | 1 | 7 | 0 | 1 | 2.68 | 3 | 3 | 0.975 | -0.037 | 0.97770915 | NA | 0.9135 | 0.8852 | 0.8924 | 0.8924 | TRUE | 2.3809 |
| Adam19 | 7656 | 6319 | 9163 | 9346 | 9311 | 10272 | 5946 | 6721 | 9887 | 9303 | 10504 | 10140 | 8750.19 | 7518 | 9982 | 1.328 | 0.409 | 0.08401087 | 0.19540558 | 0.0403 | 0.0027 | 0.0124 | 0.0403 | TRUE | 0.9272 |
| Adam1a | 12 | 7 | 6 | 10 | 4 | 10 | 9 | 7 | 6 | 10 | 5 | 10 | 7.93 | 7 | 8 | 1.046 | 0.065 | 0.92366867 | NA | 0 | 0.3013 | 0.2112 | 0.2112 | TRUE | 0.3956 |

| | | | | | | | | | | | | | | | | | | | | | | | | | |
|---|---|---|---|---|---|---|---|---|---|---|---|---|---|---|---|---|---|---|---|---|---|---|---|---|---|
| Adam1b | 1 | 1 | 0 | 0 | 0 | 0 | 1 | 1 | 0 | 0 | 0 | 0 | 0.31 | 1 | 0 | 0.309 | -1.696 | 0.63183309 | NA | 0 | 7.7294 | 7.2472 | 7.2472 | TRUE | 0.2015 |
| Adam2 | 9 | 3 | 7 | 12 | 10 | 6 | 7 | 3 | 8 | 12 | 11 | 6 | 7.81 | 6 | 10 | 1.628 | 0.703 | 0.32111363 | NA | 0 | 0.3057 | 0.2272 | 0.2272 | TRUE | 0.4121 |
| Adam20 | 0 | 1 | 0 | 0 | 0 | 0 | 0 | 1 | 0 | 0 | 0 | 0 | 0.18 | 0 | 0 | 0.552 | -0.857 | 0.8336548 | NA | 0 | 13.3714 | 10 | 10 | TRUE | 0.5216 |
| Adam21 | 0 | 0 | 0 | 1 | 0 | 0 | 0 | 0 | 0 | 1 | 0 | 0 | 0.17 | 0 | 0 | 2.094 | 1.067 | 0.79379329 | NA | 0 | 14.2885 | 10 | 10 | TRUE | 0.3906 |
| Adam22 | 750 | 570 | 602 | 694 | 626 | 669 | 582 | 606 | 650 | 691 | 706 | 660 | 649.29 | 613 | 686 | 1.12 | 0.164 | 0.06800121 | 0.16786551 | 0 | 0.0061 | 0.0043 | 0.0043 | TRUE | 0.032 |
| Adam23 | 5022 | 3937 | 4438 | 4804 | 4548 | 5055 | 3900 | 4188 | 4789 | 4782 | 5131 | 4990 | 4629.88 | 4292 | 4968 | 1.158 | 0.211 | 0.00581842 | 0.02757381 | 0.0059 | 0.0029 | 0.004 | 0.004 | TRUE | 0.1244 |
| Adam24 | 0 | 0 | 0 | 0 | 0 | 0 | 0 | 0 | 0 | 0 | 0 | 0 | 0 | 0 | 0 | NA | NA | NA | NA | NA | NA | NA | NA | NA | NA |
| Adam25 | 2 | 2 | 1 | 1 | 1 | 1 | 2 | 2 | 1 | 1 | 1 | 1 | 1.31 | 2 | 1 | 0.652 | -0.616 | 0.70036224 | NA | 0 | 1.8093 | 1.0423 | 1.0423 | TRUE | 0.0571 |
| Adam26a | 0 | 0 | 0 | 0 | 0 | 0 | 0 | 0 | 0 | 0 | 0 | 0 | 0 | 0 | 0 | NA | NA | NA | NA | NA | NA | NA | NA | NA | NA |
| Adam26b | 0 | 0 | 0 | 0 | 0 | 0 | 0 | 0 | 0 | 0 | 0 | 0 | 0 | 0 | 0 | NA | NA | NA | NA | NA | NA | NA | NA | NA | NA |
| Adam28 | 1 | 0 | 0 | 0 | 0 | 0 | 1 | 0 | 0 | 0 | 0 | 0 | 0.13 | 0 | 0 | 0.552 | -0.857 | 0.8336548 | NA | 0 | 18.3121 | 10 | 10 | TRUE | 0.2566 |
| Adam29 | 0 | 0 | 0 | 0 | 0 | 0 | 0 | 0 | 0 | 0 | 0 | 0 | 0 | 0 | 0 | NA | NA | NA | NA | NA | NA | NA | NA | NA | NA |
| Adam3 | 8 | 5 | 1 | 6 | 5 | 4 | 6 | 5 | 1 | 6 | 6 | 4 | 4.7 | 4 | 5 | 1.208 | 0.273 | 0.76343151 | NA | 0 | 0.5072 | 0.3809 | 0.3809 | TRUE | 0.6615 |
| Adam30 | 10 | 5 | 3 | 5 | 4 | 3 | 8 | 5 | 3 | 5 | 5 | 3 | 4.8 | 5 | 4 | 0.75 | -0.416 | 0.63406165 | NA | 0 | 0.4967 | 0.3373 | 0.3373 | TRUE | 0.2247 |
| Adam32 | 24 | 23 | 17 | 15 | 15 | 17 | 19 | 24 | 18 | 15 | 17 | 17 | 18.35 | 20 | 16 | 0.793 | -0.334 | 0.45224106 | 0.6220654 | 0 | 0.1316 | 0.0867 | 0.0867 | TRUE | 0.1491 |
| Adam33 | 31 | 20 | 22 | 32 | 23 | 34 | 24 | 21 | 24 | 32 | 26 | 34 | 26.74 | 23 | 31 | 1.323 | 0.404 | 0.27323273 | 0.44421357 | 0 | 0.0911 | 0.06 | 0.06 | TRUE | 0.103 |
| Adam34 | 0 | 0 | 0 | 0 | 0 | 0 | 0 | 0 | 0 | 0 | 0 | 0 | 0 | 0 | 0 | NA | NA | NA | NA | NA | NA | NA | NA | NA | NA |
| Adam39 | 0 | 0 | 0 | 0 | 0 | 0 | 0 | 0 | 0 | 0 | 0 | 0 | 0 | 0 | 0 | NA | NA | NA | NA | NA | NA | NA | NA | NA | NA |
| Adam4 | 7 | 1 | 3 | 5 | 3 | 4 | 5 | 1 | 3 | 5 | 3 | 4 | 3.67 | 3 | 4 | 1.234 | 0.304 | 0.76285058 | NA | 0 | 0.6474 | 0.4589 | 0.4589 | TRUE | 0.2603 |
| Adam5 | 2 | 1 | 0 | 0 | 3 | 1 | 2 | 1 | 0 | 0 | 3 | 1 | 1.16 | 1 | 1 | 1.59 | 0.669 | 0.72443691 | NA | 0 | 2.0373 | 1.696 | 1.696 | TRUE | 0.4739 |
| Adam6a | 0 | 0 | 0 | 0 | 0 | 0 | 0 | 0 | 0 | 0 | 0 | 0 | 0 | 0 | 0 | NA | NA | NA | NA | NA | NA | NA | NA | NA | NA |
| Adam6b | 0 | 0 | 0 | 0 | 0 | 0 | 0 | 0 | 0 | 0 | 0 | 0 | 0 | 0 | 0 | NA | NA | NA | NA | NA | NA | NA | NA | NA | NA |
| Adam7 | 1 | 0 | 0 | 1 | 0 | 1 | 1 | 0 | 0 | 1 | 0 | 1 | 0.46 | 0 | 1 | 1.94 | 0.956 | 0.73503911 | NA | 0 | 5.1562 | 3.9771 | 3.9771 | TRUE | 0.2566 |
| Adam8 | 78 | 54 | 62 | 71 | 66 | 53 | 61 | 57 | 67 | 71 | 74 | 52 | 63.73 | 62 | 66 | 1.066 | 0.093 | 0.71278129 | 0.82740055 | 0 | 0.0396 | 0.0299 | 0.0299 | TRUE | 0.2913 |
| Adam9 | 4162 | 2958 | 2976 | 3265 | 2762 | 3001 | 3232 | 3146 | 3211 | 3250 | 3116 | 2962 | 3153.03 | 3196 | 3109 | 0.973 | -0.04 | 0.49948024 | 0.66397611 | 9.00E-04 | 0.0032 | 0.0022 | 0.0022 | TRUE | 0.021 |
| Adamdec1 | 0 | 0 | 0 | 0 | 0 | 0 | 0 | 0 | 0 | 0 | 0 | 0 | 0 | 0 | 0 | NA | NA | NA | NA | NA | NA | NA | NA | NA | NA |
| Adamts1 | 115 | 101 | 78 | 83 | 64 | 87 | 89 | 107 | 84 | 83 | 72 | 86 | 86.93 | 93 | 80 | 0.859 | -0.219 | 0.30932441 | 0.48202942 | 0 | 0.0297 | 0.0217 | 0.0217 | TRUE | 0.1519 |
| Adamts10 | 929 | 832 | 806 | 895 | 767 | 939 | 722 | 885 | 870 | 891 | 865 | 927 | 859.87 | 826 | 894 | 1.086 | 0.119 | 0.21115286 | 0.37314636 | 0.0059 | 0.0052 | 0.0054 | 0.0054 | TRUE | 0.1511 |
| Adamts12 | 2 | 3 | 1 | 3 | 3 | 5 | 2 | 3 | 1 | 3 | 3 | 5 | 2.85 | 2 | 4 | 1.963 | 0.973 | 0.39153508 | NA | 0 | 0.8326 | 0.5413 | 0.5413 | TRUE | 0.2493 |
| Adamts13 | 0 | 1 | 1 | 0 | 1 | 2 | 0 | 1 | 1 | 0 | 1 | 2 | 0.87 | 1 | 1 | 1.512 | 0.597 | 0.77900931 | NA | 0 | 2.7136 | 2.0546 | 2.0546 | TRUE | 0.2336 |
| Adamts14 | 1042 | 968 | 1056 | 1208 | 1180 | 1236 | 809 | 1030 | 1139 | 1202 | 1331 | 1220 | 1122.01 | 993 | 1251 | 1.263 | 0.336 | 0.00256015 | 0.01464736 | 0.0145 | 0.0045 | 0.0081 | 0.0081 | TRUE | 0.3223 |
| Adamts15 | 264 | 222 | 204 | 336 | 282 | 327 | 205 | 236 | 220 | 334 | 318 | 323 | 272.78 | 220 | 325 | 1.479 | 0.565 | 6.82E-06 | 0.00013754 | 0 | 0.0111 | 0.0076 | 0.0076 | TRUE | 0.0456 |
| Adamts16 | 1 | 4 | 1 | 2 | 0 | 0 | 1 | 4 | 1 | 2 | 0 | 0 | 1.35 | 2 | 1 | 0.337 | -1.569 | 0.4012003 | NA | 0.7058 | 1.7578 | 1.5106 | 1.5106 | TRUE | 0.9735 |
| Adamts17 | 106 | 66 | 70 | 97 | 107 | 92 | 82 | 70 | 76 | 97 | 121 | 91 | 89.36 | 76 | 103 | 1.343 | 0.426 | 0.04809984 | 0.13092572 | 0 | 0.029 | 0.0221 | 0.0221 | TRUE | 0.2212 |
| Adamts18 | 2 | 5 | 0 | 3 | 2 | 0 | 2 | 5 | 0 | 3 | 2 | 0 | 2.02 | 2 | 2 | 0.765 | -0.387 | 0.80126912 | NA | 1.2645 | 1.1763 | 1.194 | 1.194 | TRUE | 0.5414 |
| Adamts19 | 104 | 61 | 51 | 67 | 64 | 89 | 81 | 65 | 55 | 67 | 72 | 88 | 71.24 | 67 | 76 | 1.123 | 0.167 | 0.50011125 | 0.66457109 | 0.0163 | 0.0357 | 0.0302 | 0.0302 | TRUE | 0.3292 |
| Adamts2 | 140 | 107 | 80 | 127 | 95 | 113 | 109 | 114 | 86 | 126 | 107 | 112 | 109 | 103 | 115 | 1.116 | 0.158 | 0.42038989 | 0.59260736 | 0.0032 | 0.0242 | 0.0186 | 0.0186 | TRUE | 0.184 |
| Adamts20 | 167 | 152 | 127 | 114 | 114 | 126 | 130 | 162 | 137 | 113 | 129 | 124 | 132.48 | 143 | 122 | 0.857 | -0.222 | 0.2102769 | 0.37224273 | 0 | 0.0203 | 0.0151 | 0.0151 | TRUE | 0.1363 |
| Adamts3 | 110 | 77 | 84 | 112 | 93 | 106 | 85 | 82 | 91 | 111 | 105 | 105 | 96.5 | 86 | 107 | 1.245 | 0.316 | 0.10715622 | 0.23265586 | 0 | 0.027 | 0.0174 | 0.0174 | TRUE | 0.0193 |
| Adamts4 | 88 | 79 | 70 | 68 | 67 | 74 | 68 | 84 | 76 | 68 | 76 | 73 | 74.04 | 76 | 72 | 0.952 | -0.071 | 0.75351246 | 0.85367419 | 0 | 0.0344 | 0.0234 | 0.0234 | TRUE | 0.0815 |
| Adamts5 | 25 | 30 | 9 | 14 | 18 | 14 | 19 | 32 | 10 | 14 | 20 | 14 | 18.18 | 20 | 16 | 0.785 | -0.35 | 0.49345176 | 0.65871915 | 0.1284 | 0.1328 | 0.1316 | 0.1316 | TRUE | 0.1995 |
| Adamts6 | 3 | 1 | 2 | 2 | 2 | 1 | 2 | 1 | 2 | 2 | 2 | 1 | 1.8 | 2 | 2 | 0.928 | -0.108 | 0.93758572 | NA | 0 | 1.3209 | 0.8079 | 0.8079 | TRUE | 0.119 |
| Adamts7 | 557 | 448 | 424 | 445 | 416 | 428 | 433 | 477 | 457 | 443 | 469 | 423 | 450.23 | 456 | 445 | 0.977 | -0.033 | 0.74836572 | 0.85076667 | 0 | 0.0077 | 0.0054 | 0.0054 | TRUE | 0.0258 |
| Adamts8 | 210 | 144 | 153 | 115 | 106 | 106 | 163 | 153 | 165 | 114 | 120 | 105 | 136.67 | 160 | 113 | 0.702 | -0.509 | 0.00262395 | 0.01488051 | 0 | 0.0198 | 0.0131 | 0.0131 | TRUE | 0.0416 |
| Adamts9 | 806 | 617 | 586 | 701 | 573 | 641 | 626 | 656 | 632 | 698 | 646 | 633 | 648.59 | 638 | 659 | 1.033 | 0.047 | 0.59417448 | 0.74089299 | 0 | 0.0061 | 0.0041 | 0.0041 | TRUE | 0.0318 |
| Adamtsl1 | 265 | 173 | 215 | 193 | 222 | 246 | 206 | 184 | 232 | 192 | 250 | 243 | 217.87 | 207 | 228 | 1.102 | 0.139 | 0.37057182 | 0.54525253 | 0.0119 | 0.0133 | 0.0129 | 0.0129 | TRUE | 0.2161 |
| Adamtsl2 | 0 | 0 | 0 | 0 | 0 | 0 | 0 | 0 | 0 | 0 | 0 | 0 | 0 | 0 | 0 | NA | NA | NA | NA | NA | NA | NA | NA | NA | NA |
| Adamtsl3 | 31 | 12 | 19 | 18 | 11 | 20 | 24 | 13 | 21 | 18 | 12 | 20 | 17.9 | 19 | 17 | 0.869 | -0.202 | 0.66823387 | 0.79609449 | 0 | 0.1348 | 0.1039 | 0.1039 | TRUE | 0.422 |
| Adamtsl4 | 351 | 297 | 340 | 418 | 367 | 413 | 273 | 316 | 367 | 416 | 414 | 408 | 365.53 | 319 | 413 | 1.301 | 0.379 | 0.00260074 | 0.01480007 | 0.0081 | 0.0089 | 0.0087 | 0.0087 | TRUE | 0.1999 |
| Adamtsl5 | 132 | 103 | 122 | 176 | 142 | 163 | 103 | 110 | 132 | 175 | 160 | 161 | 140 | 115 | 165 | 1.451 | 0.537 | 0.00205406 | 0.01230457 | 0 | 0.0194 | 0.0145 | 0.0145 | TRUE | 0.1651 |
| Adap1 | 1381 | 979 | 954 | 1191 | 1039 | 1213 | 1073 | 1041 | 1029 | 1186 | 1172 | 1197 | 1116.38 | 1048 | 1185 | 1.13 | 0.177 | 0.01238344 | 0.04849221 | 0 | 0.0045 | 0.0027 | 0.0027 | TRUE | 0.0054 |
| Adap2 | 388 | 283 | 264 | 307 | 268 | 311 | 301 | 301 | 285 | 306 | 302 | 307 | 300.36 | 296 | 305 | 1.031 | 0.044 | 0.70867875 | 0.82415787 | 0 | 0.0103 | 0.0064 | 0.0064 | TRUE | 0.0112 |
| Adar | 4540 | 3309 | 3099 | 3428 | 2951 | 3496 | 3526 | 3520 | 3344 | 3412 | 3329 | 3451 | 3430.32 | 3463 | 3397 | 0.981 | -0.028 | 0.61709492 | 0.75833012 | 3.00E-04 | 0.0031 | 0.0019 | 0.0019 | TRUE | 0.0109 |
| Adarb1 | 1188 | 952 | 936 | 1122 | 974 | 1154 | 923 | 1013 | 1010 | 1117 | 1099 | 1139 | 1050 | 982 | 1118 | 1.141 | 0.19 | 0.01292708 | 0.04996189 | 7.00E-04 | 0.0047 | 0.0033 | 0.0033 | TRUE | 0.035 |
| Adarb2 | 5 | 1 | 2 | 4 | 1 | 3 | 4 | 1 | 2 | 4 | 1 | 3 | 2.53 | 2 | 3 | 1.118 | 0.161 | 0.89329991 | NA | 0 | 0.9394 | 0.6511 | 0.6511 | TRUE | 0.4388 |
| Adat1 | 1665 | 1200 | 917 | 1007 | 925 | 1113 | 1293 | 1276 | 989 | 1002 | 1043 | 1099 | 1117.26 | 1186 | 1048 | 0.883 | -0.18 | 0.07996986 | 0.18919567 | 0.0112 | 0.0045 | 0.0067 | 0.0067 | TRUE | 0.2492 |
| Adat2 | 678 | 537 | 538 | 605 | 522 | 575 | 527 | 571 | 580 | 602 | 589 | 568 | 572.83 | 559 | 586 | 1.05 | 0.07 | 0.45268083 | 0.62249059 | 0 | 0.0066 | 0.0045 | 0.0045 | TRUE | 0.0332 |
| Adat3 | 342 | 275 | 280 | 263 | 226 | 278 | 266 | 293 | 302 | 262 | 255 | 274 | 275.24 | 287 | 264 | 0.922 | -0.117 | 0.35148874 | 0.52609837 | 0 | 0.011 | 0.0076 | 0.0076 | TRUE | 0.051 |
| Adc | 992 | 813 | 922 | 991 | 835 | 945 | 770 | 865 | 995 | 986 | 942 | 933 | 915.22 | 877 | 954 | 1.09 | 0.125 | 0.20570457 | 0.36751692 | 0.0071 | 0.005 | 0.0059 | 0.0059 | TRUE | 0.165 |
| Adck1 | 1218 | 838 | 905 | 906 | 807 | 937 | 946 | 891 | 976 | 902 | 910 | 925 | 925.17 | 938 | 912 | 0.973 | -0.04 | 0.60682805 | 0.75114657 | 0 | 0.005 | 0.0033 | 0.0033 | TRUE | 0.0216 |
| Adck2 | 315 | 235 | 250 | 275 | 233 | 277 | 245 | 250 | 270 | 274 | 263 | 273 | 262.4 | 255 | 270 | 1.062 | 0.086 | 0.48975296 | 0.65555713 | 0 | 0.0115 | 0.0075 | 0.0075 | TRUE | 0.0285 |
| Adck3 | 1310 | 1211 | 1440 | 1777 | 1752 | 1875 | 1017 | 1288 | 1554 | 1769 | 1976 | 1851 | 1575.91 | 1286 | 1865 | 1.452 | 0.538 | 7.02E-06 | 0.00014017 | 0.0223 | 0.0039 | 0.0097 | 0.0097 | TRUE | 0.2881 |
| Adck4 | 612 | 440 | 424 | 427 | 373 | 412 | 475 | 468 | 457 | 425 | 421 | 407 | 442.23 | 467 | 418 | 0.893 | -0.162 | 0.10154901 | 0.22365982 | 0 | 0.0078 | 0.0048 | 0.0048 | TRUE | 0.0061 |
| Adck5 | 878 | 604 | 505 | 597 | 538 | 593 | 682 | 642 | 545 | 594 | 607 | 585 | 609.3 | 623 | 595 | 0.954 | -0.069 | 0.50454016 | 0.66826706 | 0.0052 | 0.0063 | 0.006 | 0.006 | TRUE | 0.1392 |
| Adcy1 | 1464 | 1108 | 1171 | 1330 | 1324 | 1491 | 1137 | 1179 | 1263 | 1324 | 1494 | 1472 | 1311.4 | 1193 | 1430 | 1.199 | 0.262 | 0.0009212 | 0.00683273 | 0.0028 | 0.0042 | 0.0037 | 0.0037 | TRUE | 0.0508 |
| Adcy10 | 14 | 13 | 5 | 7 | 6 | 7 | 11 | 14 | 5 | 7 | 7 | 7 | 8.46 | 10 | 7 | 0.684 | -0.548 | 0.41507367 | NA | 0 | 0.2827 | 0.203 | 0.203 | TRUE | 0.3157 |
| Adcy2 | 186 | 104 | 91 | 87 | 85 | 81 | 144 | 111 | 98 | 87 | 96 | 80 | 102.62 | 118 | 88 | 0.737 | -0.44 | 0.0401969 | 0.11483794 | 0.0176 | 0.0255 | 0.0231 | 0.0231 | TRUE | 0.431 |
| Adcy3 | 1774 | 1310 | 1392 | 1626 | 1460 | 1594 | 1378 | 1393 | 1502 | 1619 | 1647 | 1574 | 1518.71 | 1424 | 1613 | 1.133 | 0.18 | 0.00894163 | 0.03817284 | 7.00E-04 | 0.004 | 0.0028 | 0.0028 | TRUE | 0.0268 |
| Adcy4 | 6 | 19 | 11 | 19 | 16 | 14 | 5 | 20 | 12 | 19 | 18 | 14 | 14.59 | 12 | 17 | 1.41 | 0.496 | 0.3856461 | NA | 0.1642 | 0.1649 | 0.1647 | 0.1647 | TRUE | 0.3251 |

| | | | | | | | | | | | | | | | | | | | | | | | | |
|---|---|---|---|---|---|---|---|---|---|---|---|---|---|---|---|---|---|---|---|---|---|---|---|---|
| Adcy5 | 149 | 123 | 134 | 155 | 154 | 169 | 116 | 131 | 145 | 154 | 174 | 167 | 147.66 | 131 | 165 | 1.27 | 0.344 | 0.04093366 | 0.11638447 | 0 | 0.0185 | 0.0136 | 0.0136 | TRUE | 0.1115 |
| Adcy6 | 467 | 359 | 414 | 585 | 481 | 524 | 363 | 382 | 447 | 582 | 543 | 517 | 472.24 | 397 | 547 | 1.382 | 0.467 | 2.89E-05 | 0.0004496 | 0.0054 | 0.0074 | 0.0068 | 0.0068 | TRUE | 0.135 |
| Adcy7 | 775 | 494 | 388 | 487 | 389 | 464 | 602 | 525 | 419 | 485 | 439 | 458 | 487.94 | 515 | 461 | 0.891 | -0.166 | 0.20011624 | 0.3607342 | 0.0159 | 0.0073 | 0.01 | 0.01 | TRUE | 0.2606 |
| Adcy8 | 3 | 1 | 1 | 0 | 0 | 0 | 2 | 1 | 1 | 0 | 0 | 0 | 0.75 | 1 | 0 | 0.126 | -2.986 | 0.19880652 | NA | 0 | 3.1818 | 2.5705 | 2.5705 | TRUE | 0.2178 |
| Adcy9 | 453 | 351 | 294 | 368 | 301 | 369 | 352 | 373 | 317 | 366 | 340 | 364 | 352.09 | 347 | 357 | 1.027 | 0.038 | 0.74143126 | 0.8460229 | 0 | 0.0092 | 0.0067 | 0.0067 | TRUE | 0.0624 |
| Adcyap1 | 27 | 26 | 20 | 24 | 23 | 29 | 21 | 28 | 22 | 24 | 26 | 29 | 24.78 | 24 | 26 | 1.122 | 0.167 | 0.66404913 | 0.7929864 | 0 | 0.0981 | 0.0655 | 0.0655 | TRUE | 0.1411 |
| Adcyap1r1 | 203 | 127 | 105 | 113 | 121 | 129 | 158 | 135 | 113 | 112 | 137 | 127 | 130.39 | 135 | 125 | 0.921 | -0.119 | 0.52963297 | 0.68981629 | 0.0115 | 0.0206 | 0.0179 | 0.0179 | TRUE | 0.194 |
| Add1 | 7421 | 5043 | 4298 | 4073 | 3601 | 3968 | 5764 | 5364 | 4638 | 4054 | 4062 | 3917 | 4633.14 | 5255 | 4011 | 0.763 | -0.39 | 4.71E-07 | 1.54E-05 | 0.0058 | 0.0029 | 0.0041 | 0.0041 | TRUE | 0.1285 |
| Add2 | 80 | 59 | 56 | 66 | 67 | 73 | 62 | 63 | 60 | 66 | 76 | 72 | 66.44 | 62 | 71 | 1.15 | 0.201 | 0.39075178 | 0.5645332 | 0 | 0.0381 | 0.0246 | 0.0246 | TRUE | 0.0405 |
| Add3 | 4235 | 3196 | 3140 | 2922 | 2748 | 3145 | 3289 | 3400 | 3388 | 2909 | 3100 | 3105 | 3198.32 | 3359 | 3038 | 0.905 | -0.145 | 0.01197149 | 0.04726755 | 5.00E-04 | 0.0032 | 0.0021 | 0.0021 | TRUE | 0.0169 |
| Adgb | 2 | 5 | 4 | 3 | 1 | 2 | 2 | 5 | 4 | 3 | 1 | 2 | 2.88 | 4 | 2 | 0.56 | -0.838 | 0.46761231 | NA | 0 | 0.8255 | 0.5732 | 0.5732 | TRUE | 0.561 |
| Adh1 | 16 | 15 | 13 | 9 | 9 | 7 | 12 | 16 | 14 | 9 | 10 | 7 | 11.41 | 14 | 9 | 0.614 | -0.704 | 0.21459433 | NA | 0 | 0.2102 | 0.1376 | 0.1376 | TRUE | 0.1021 |
| Adh4 | 110 | 83 | 61 | 40 | 38 | 40 | 85 | 88 | 66 | 40 | 43 | 39 | 60.28 | 80 | 41 | 0.508 | -0.976 | 0.00018325 | 0.00199783 | 0 | 0.0417 | 0.0303 | 0.0303 | TRUE | 0.2073 |
| Adh5 | 17431 | 12778 | 13864 | 14837 | 13191 | 15926 | 13538 | 13592 | 14959 | 14769 | 14881 | 15721 | 14576.64 | 14030 | 15124 | 1.078 | 0.109 | 0.06329852 | 0.15945113 | 0.0021 | 0.0026 | 0.0024 | 0.0024 | TRUE | 0.041 |
| Adh6-ps1 | 0 | 0 | 0 | 0 | 0 | 0 | 0 | 0 | 0 | 0 | 0 | 0 | 0 | 0 | 0 | NA | NA | NA | NA | NA | NA | NA | NA | NA | NA |
| Adh6a | 0 | 0 | 0 | 0 | 1 | 1 | 0 | 0 | 0 | 0 | 1 | 1 | 0.35 | 0 | 1 | 3.965 | 1.987 | 0.55529285 | NA | 0 | 6.7252 | 6.4426 | 6.4426 | TRUE | 0.261 |
| Adh7 | 54 | 46 | 21 | 21 | 20 | 17 | 42 | 49 | 23 | 21 | 23 | 17 | 28.96 | 38 | 20 | 0.528 | -0.922 | 0.02123665 | 0.07225315 | 0.0582 | 0.0843 | 0.0769 | 0.0769 | TRUE | 0.5114 |
| Adhfe1 | 90 | 100 | 132 | 150 | 148 | 168 | 70 | 106 | 142 | 149 | 167 | 166 | 133.47 | 106 | 161 | 1.526 | 0.609 | 0.00739132 | 0.03308625 | 0.0537 | 0.0202 | 0.0295 | 0.0295 | TRUE | 0.3144 |
| Adi1 | 2925 | 2051 | 2308 | 2239 | 1919 | 2306 | 2272 | 2182 | 2490 | 2229 | 2165 | 2276 | 2268.93 | 2315 | 2223 | 0.961 | -0.057 | 0.40448247 | 0.57722147 | 0.002 | 0.0035 | 0.003 | 0.003 | TRUE | 0.0524 |
| Adig | 0 | 0 | 0 | 1 | 0 | 1 | 0 | 0 | 0 | 1 | 0 | 1 | 0.33 | 0 | 1 | 3.763 | 1.912 | 0.57887588 | NA | 0 | 7.1752 | 6.7756 | 6.7756 | TRUE | 0.2094 |
| Adipoq | 5 | 4 | 1 | 2 | 0 | 2 | 4 | 4 | 1 | 2 | 0 | 2 | 2.2 | 3 | 1 | 0.436 | -1.199 | 0.3818782 | NA | 0 | 1.0812 | 0.8155 | 0.8155 | TRUE | 0.3935 |
| Adipor1 | 6998 | 5054 | 5854 | 6160 | 5500 | 6121 | 5435 | 5376 | 6316 | 6132 | 6205 | 6042 | 5917.66 | 5709 | 6126 | 1.073 | 0.102 | 0.14069318 | 0.28258882 | 0.0041 | 0.0028 | 0.0033 | 0.0033 | TRUE | 0.105 |
| Adipor2 | 14787 | 10780 | 11588 | 13129 | 11803 | 13929 | 11484 | 11467 | 12503 | 13069 | 13315 | 13750 | 12598 | 11818 | 13378 | 1.132 | 0.179 | 0.00119168 | 0.00821245 | 0.0015 | 0.0026 | 0.0021 | 0.0021 | TRUE | 0.0313 |
| Adk | 3350 | 2312 | 2206 | 2561 | 2243 | 2832 | 2602 | 2459 | 2380 | 2549 | 2530 | 2796 | 2552.75 | 2480 | 2625 | 1.058 | 0.081 | 0.23070775 | 0.3964778 | 0.002 | 0.0034 | 0.0029 | 0.0029 | TRUE | 0.0394 |
| Adm | 133 | 111 | 82 | 149 | 129 | 145 | 103 | 118 | 88 | 148 | 146 | 143 | 124.47 | 103 | 146 | 1.41 | 0.496 | 0.00651725 | 0.03004687 | 0 | 0.0215 | 0.0157 | 0.0157 | TRUE | 0.1414 |
| Adm2 | 29 | 25 | 34 | 41 | 40 | 41 | 23 | 27 | 37 | 41 | 45 | 40 | 35.37 | 29 | 42 | 1.486 | 0.571 | 0.08778518 | 0.20125342 | 0 | 0.0694 | 0.0513 | 0.0513 | TRUE | 0.3811 |
| Adnp | 10241 | 7244 | 7078 | 7071 | 6583 | 7272 | 7954 | 7705 | 7637 | 7038 | 7426 | 7179 | 7489.93 | 7765 | 7214 | 0.929 | -0.107 | 0.03522075 | 0.10437698 | 4.00E-04 | 0.0027 | 0.0017 | 0.0017 | TRUE | 0.008 |
| Adnp2 | 4302 | 3264 | 3376 | 3130 | 2831 | 3188 | 3341 | 3472 | 3643 | 3116 | 3194 | 3147 | 3318.67 | 3485 | 3152 | 0.905 | -0.144 | 0.01335428 | 0.05126852 | 8.00E-04 | 0.0031 | 0.0021 | 0.0021 | TRUE | 0.0189 |
| Ado | 3258 | 2554 | 2533 | 2685 | 2455 | 2864 | 2530 | 2717 | 2733 | 2673 | 2769 | 2827 | 2708.24 | 2660 | 2756 | 1.037 | 0.052 | 0.39623315 | 0.56958266 | 0.001 | 0.0033 | 0.0024 | 0.0024 | TRUE | 0.0228 |
| Adora1 | 22 | 23 | 18 | 37 | 51 | 41 | 17 | 24 | 19 | 37 | 58 | 40 | 32.63 | 20 | 45 | 2.22 | 1.151 | 0.00152741 | 0.00978545 | 0 | 0.075 | 0.0595 | 0.0595 | TRUE | 0.4432 |
| Adora2a | 71 | 61 | 42 | 50 | 31 | 58 | 55 | 65 | 45 | 50 | 35 | 57 | 51.22 | 55 | 47 | 0.862 | -0.214 | 0.46289611 | 0.63217944 | 0.0221 | 0.0487 | 0.0418 | 0.0418 | TRUE | 0.3158 |
| Adora2b | 404 | 279 | 246 | 335 | 293 | 289 | 314 | 297 | 265 | 333 | 331 | 285 | 304.21 | 292 | 316 | 1.081 | 0.112 | 0.37965748 | 0.5542278 | 0.0041 | 0.0102 | 0.0084 | 0.0084 | TRUE | 0.0855 |
| Adora3 | 0 | 0 | 0 | 0 | 0 | 0 | 0 | 0 | 0 | 0 | 0 | 0 | 0 | 0 | 0 | NA | NA | NA | NA | NA | NA | NA | NA | NA | NA |
| Adpgk | 5246 | 3788 | 3509 | 3906 | 3458 | 3873 | 4074 | 4029 | 3786 | 3888 | 3901 | 3823 | 3917.01 | 3963 | 3871 | 0.976 | -0.035 | 0.53556231 | 0.69450118 | 6.00E-04 | 0.003 | 0.002 | 0.002 | TRUE | 0.0184 |
| Adprh | 9752 | 7120 | 7395 | 7776 | 6916 | 7824 | 7574 | 7573 | 7979 | 7740 | 7802 | 7723 | 7732.04 | 7709 | 7755 | 1.006 | 0.009 | 0.8585842 | 0.92109464 | 3.00E-04 | 0.0027 | 0.0016 | 0.0016 | TRUE | 0.0114 |
| Adprhl1 | 37 | 26 | 12 | 42 | 39 | 30 | 29 | 28 | 13 | 42 | 44 | 30 | 30.79 | 23 | 39 | 1.646 | 0.719 | 0.06362577 | 0.16013438 | 0.057 | 0.0794 | 0.074 | 0.074 | TRUE | 0.7948 |
| Adprhl2 | 2158 | 1527 | 1657 | 1703 | 1524 | 1708 | 1676 | 1624 | 1788 | 1695 | 1719 | 1686 | 1698.1 | 1696 | 1700 | 1.003 | 0.004 | 0.95549676 | 0.97553943 | 6.00E-04 | 0.0038 | 0.0026 | 0.0026 | TRUE | 0.0263 |
| Adprm | 1503 | 1036 | 1008 | 1179 | 1069 | 1150 | 1167 | 1102 | 1088 | 1174 | 1206 | 1135 | 1145.28 | 1119 | 1172 | 1.046 | 0.064 | 0.38013783 | 0.5547024 | 0 | 0.0045 | 0.003 | 0.003 | TRUE | 0.018 |
| Adra1a | 1 | 0 | 0 | 0 | 0 | 1 | 1 | 0 | 0 | 0 | 0 | 1 | 0.29 | 0 | 0 | 1.075 | 0.105 | 0.97702352 | NA | 0 | 8.0647 | 7.5458 | 7.5458 | TRUE | 0.3835 |
| Adra1b | 0 | 1 | 0 | 1 | 0 | 0 | 0 | 1 | 0 | 1 | 0 | 0 | 0.34 | 0 | 0 | 1.075 | 0.105 | 0.97573409 | NA | 0 | 6.9086 | 6.5582 | 6.5582 | TRUE | 0.5216 |
| Adra1d | 3 | 2 | 2 | 4 | 4 | 3 | 2 | 2 | 2 | 4 | 5 | 3 | 3.01 | 2 | 4 | 1.722 | 0.784 | 0.46675529 | NA | 0 | 0.7894 | 0.4821 | 0.4821 | TRUE | 0.064 |
| Adra2a | 43 | 36 | 36 | 48 | 46 | 46 | 33 | 38 | 39 | 48 | 52 | 45 | 42.6 | 37 | 48 | 1.316 | 0.397 | 0.17570684 | 0.32921205 | 0 | 0.0581 | 0.038 | 0.038 | TRUE | 0.0588 |
| Adra2b | 73 | 55 | 45 | 62 | 34 | 47 | 57 | 59 | 49 | 62 | 38 | 46 | 51.7 | 55 | 49 | 0.896 | -0.158 | 0.57599085 | 0.72769249 | 0.0109 | 0.0483 | 0.0385 | 0.0385 | TRUE | 0.4324 |
| Adra2c | 122 | 87 | 73 | 61 | 54 | 67 | 95 | 93 | 79 | 61 | 61 | 66 | 75.64 | 89 | 63 | 0.704 | -0.505 | 0.02481626 | 0.08096297 | 0 | 0.0338 | 0.0229 | 0.0229 | TRUE | 0.0869 |
| Adrb1 | 102 | 99 | 101 | 145 | 145 | 138 | 79 | 105 | 109 | 144 | 164 | 136 | 122.94 | 98 | 148 | 1.522 | 0.606 | 0.00175331 | 0.01092419 | 0.0104 | 0.0217 | 0.0186 | 0.0186 | TRUE | 0.3078 |
| Adrb2 | 269 | 218 | 198 | 191 | 182 | 175 | 209 | 232 | 214 | 190 | 205 | 173 | 203.77 | 218 | 189 | 0.868 | -0.204 | 0.15755485 | 0.30642692 | 0 | 0.0141 | 0.0101 | 0.0101 | TRUE | 0.0644 |
| Adrb3 | 1335 | 994 | 857 | 881 | 782 | 963 | 1037 | 1057 | 925 | 877 | 882 | 951 | 954.77 | 1006 | 903 | 0.897 | -0.156 | 0.06417015 | 0.16107015 | 0.0025 | 0.0049 | 0.0041 | 0.0041 | TRUE | 0.0584 |
| Adrbk1 | 4433 | 3189 | 3238 | 3507 | 3096 | 3475 | 3443 | 3392 | 3494 | 3491 | 3493 | 3430 | 3457.11 | 3443 | 3471 | 1.008 | 0.012 | 0.82343714 | 0.89863229 | 0 | 0.0031 | 0.0017 | 0.0017 | TRUE | 0.002 |
| Adrbk2 | 735 | 602 | 594 | 737 | 729 | 754 | 571 | 640 | 641 | 734 | 822 | 744 | 692.07 | 617 | 767 | 1.244 | 0.314 | 0.00076542 | 0.00595219 | 0.0028 | 0.0059 | 0.0048 | 0.0048 | TRUE | 0.0545 |
| Adrm1 | 6312 | 4523 | 4129 | 4237 | 3896 | 4585 | 4902 | 4811 | 4455 | 4218 | 4395 | 4526 | 4551.2 | 4723 | 4380 | 0.927 | -0.109 | 0.07136785 | 0.17395306 | 0.0015 | 0.0029 | 0.0024 | 0.0024 | TRUE | 0.0297 |
| Adsl | 8061 | 5800 | 6085 | 6473 | 5879 | 6608 | 6261 | 6169 | 6566 | 6443 | 6632 | 6523 | 6432.36 | 6332 | 6533 | 1.032 | 0.045 | 0.3810662 | 0.55560331 | 4.00E-04 | 0.0028 | 0.0018 | 0.0018 | TRUE | 0.0126 |
| Adss | 10308 | 7303 | 7447 | 7997 | 6916 | 7900 | 8006 | 7768 | 8035 | 7960 | 7802 | 7799 | 7894.97 | 7936 | 7854 | 0.99 | -0.015 | 0.74974734 | 0.85153057 | 1.00E-04 | 0.0027 | 0.0015 | 0.0015 | TRUE | 0.0042 |
| Adssl1 | 212 | 209 | 261 | 405 | 394 | 400 | 165 | 222 | 282 | 403 | 444 | 395 | 318.51 | 223 | 414 | 1.869 | 0.902 | 4.71E-08 | 2.01E-06 | 0.0313 | 0.0099 | 0.0162 | 0.0162 | TRUE | 0.4016 |
| Adtrp | 58 | 35 | 40 | 30 | 22 | 21 | 45 | 37 | 43 | 30 | 25 | 21 | 33.47 | 42 | 25 | 0.6 | -0.738 | 0.02990456 | 0.09247641 | 0 | 0.0732 | 0.051 | 0.051 | TRUE | 0.1723 |
| Aebp1 | 212 | 168 | 118 | 142 | 138 | 155 | 165 | 179 | 127 | 141 | 156 | 153 | 153.45 | 157 | 150 | 0.954 | -0.068 | 0.69753916 | 0.81631335 | 0.0091 | 0.0179 | 0.0154 | 0.0154 | TRUE | 0.2705 |
| Aebp2 | 10778 | 7915 | 7448 | 7029 | 6115 | 7071 | 8371 | 8419 | 8036 | 6997 | 6898 | 6980 | 7616.9 | 8275 | 6958 | 0.841 | -0.25 | 2.80E-07 | 9.76E-06 | 2.00E-04 | 0.0027 | 0.0016 | 0.0016 | TRUE | 0.0077 |
| Aen | 4837 | 3564 | 3734 | 3683 | 3382 | 3955 | 3757 | 3791 | 4029 | 3666 | 3815 | 3904 | 3827.02 | 3859 | 3795 | 0.984 | -0.024 | 0.68525525 | 0.80704955 | 8.00E-04 | 0.003 | 0.0022 | 0.0022 | TRUE | 0.0179 |
| Aes | 16672 | 11884 | 11863 | 14274 | 11890 | 13506 | 12948 | 12641 | 12800 | 14208 | 13413 | 13333 | 13223.89 | 12796 | 13651 | 1.067 | 0.093 | 0.05917068 | 0.15186326 | 6.00E-04 | 0.0026 | 0.0017 | 0.0017 | TRUE | 0.0156 |
| AF067061 | 370 | 272 | 110 | 150 | 136 | 113 | 287 | 289 | 119 | 149 | 153 | 112 | 184.94 | 232 | 138 | 0.595 | -0.749 | 0.0760002 | 0.18186353 | 0.1225 | 0.0152 | 0.0447 | 0.1225 | TRUE | 1.9888 |
| AF067063 | 29 | 25 | 17 | 7 | 8 | 10 | 23 | 27 | 18 | 7 | 9 | 10 | 15.55 | 23 | 9 | 0.383 | -1.385 | 0.00669578 | NA | 0 | 0.1548 | 0.1064 | 0.1064 | TRUE | 0.0971 |
| AF251705 | 0 | 1 | 0 | 0 | 0 | 0 | 0 | 1 | 0 | 0 | 0 | 0 | 0.18 | 0 | 0 | 0.552 | -0.857 | 0.8336548 | NA | 0 | 13.3714 | 10 | 10 | TRUE | 0.5216 |
| AF357355 | 1 | 1 | 0 | 2 | 1 | 1 | 1 | 1 | 0 | 2 | 1 | 1 | 0.99 | 1 | 1 | 2.2 | 1.137 | 0.55221173 | NA | 0 | 2.3939 | 1.4887 | 1.4887 | TRUE | 0.1884 |
| AF357359 | 70 | 47 | 50 | 85 | 56 | 62 | 54 | 50 | 54 | 85 | 63 | 61 | 61.22 | 53 | 70 | 1.32 | 0.401 | 0.11755947 | 0.24874574 | 0 | 0.0411 | 0.0307 | 0.0307 | TRUE | 0.322 |
| AF357399 | 5 | 3 | 5 | 3 | 4 | 4 | 4 | 3 | 5 | 3 | 5 | 4 | 3.99 | 4 | 4 | 0.918 | -0.123 | 0.89483471 | NA | 0 | 0.597 | 0.3756 | 0.3756 | TRUE | 0.1044 |
| AF357425 | 2 | 3 | 2 | 2 | 1 | 1 | 2 | 3 | 2 | 2 | 1 | 1 | 1.83 | 2 | 1 | 0.604 | -0.728 | 0.59891961 | NA | 0 | 1.2942 | 0.7914 | 0.7914 | TRUE | 0.1143 |
| AF357426 | 4 | 1 | 1 | 4 | 3 | 3 | 3 | 1 | 1 | 4 | 3 | 3 | 2.6 | 2 | 3 | 1.894 | 0.921 | 0.43673779 | NA | 0 | 0.9154 | 0.5963 | 0.5963 | TRUE | 0.492 |
| AF366264 | 0 | 0 | 0 | 0 | 0 | 0 | 0 | 0 | 0 | 0 | 0 | 0 | 0 | 0 | 0 | NA | NA | NA | NA | NA | NA | NA | NA | NA | NA |
| AF529169 | 103 | 73 | 69 | 95 | 77 | 111 | 80 | 78 | 74 | 95 | 87 | 110 | 87.18 | 77 | 97 | 1.254 | 0.327 | 0.12187281 | 0.25492874 | 0 | 0.0296 | 0.0206 | 0.0206 | TRUE | 0.1261 |
| Afap1 | 1369 | 959 | 1013 | 1125 | 965 | 1037 | 1063 | 1020 | 1093 | 1120 | 1089 | 1024 | 1068.08 | 1059 | 1078 | 1.017 | 0.025 | 0.74458396 | 0.84821935 | 7.00E-04 | 0.0046 | 0.0033 | 0.0033 | TRUE | 0.0231 |

| | | | | | | | | | | | | | | | | | | | | | | | | | |
|---|---|---|---|---|---|---|---|---|---|---|---|---|---|---|---|---|---|---|---|---|---|---|---|---|---|
| Afap1l1 | 786 | 632 | 668 | 1034 | 1004 | 1030 | 610 | 672 | 721 | 1029 | 1133 | 1017 | 863.68 | 668 | 1060 | 1.589 | 0.668 | 2.27E-13 | 3.15E-11 | 0.004 | 0.0052 | 0.0048 | 0.0048 | TRUE | 0.0707 |
| Afap1l2 | 271 | 202 | 179 | 142 | 175 | 154 | 210 | 215 | 193 | 141 | 197 | 152 | 184.88 | 206 | 163 | 0.791 | -0.339 | 0.03908216 | 0.11264394 | 0.0099 | 0.0152 | 0.0139 | 0.0139 | TRUE | 0.3341 |
| Aff1 | 4134 | 3179 | 3490 | 3977 | 3720 | 4396 | 3211 | 3381 | 3766 | 3959 | 4197 | 4340 | 3808.77 | 3453 | 4165 | 1.207 | 0.271 | 0.00016071 | 0.0018002 | 0.0041 | 0.003 | 0.0035 | 0.0035 | TRUE | 0.0762 |
| Aff2 | 14 | 4 | 6 | 5 | 3 | 3 | 11 | 4 | 6 | 5 | 3 | 3 | 5.49 | 7 | 4 | 0.517 | -0.952 | 0.259547 | NA | 0 | 0.4343 | 0.3107 | 0.3107 | TRUE | 0.3903 |
| Aff3 | 724 | 520 | 531 | 613 | 560 | 686 | 562 | 553 | 573 | 610 | 632 | 677 | 601.24 | 563 | 640 | 1.137 | 0.185 | 0.04172941 | 0.11791136 | 0 | 0.0064 | 0.0043 | 0.0043 | TRUE | 0.0316 |
| Aff4 | 22205 | 15020 | 14934 | 16601 | 14533 | 16826 | 17246 | 15977 | 16114 | 16525 | 16395 | 16610 | 16477.52 | 16446 | 16510 | 1.004 | 0.005 | 0.91447429 | 0.9529197 | 7.00E-04 | 0.0026 | 0.0018 | 0.0018 | TRUE | 0.0223 |
| Afg3l1 | 13484 | 9802 | 9148 | 10708 | 9187 | 10629 | 10472 | 10426 | 9871 | 10659 | 10364 | 10492 | 10380.73 | 10256 | 10505 | 1.024 | 0.034 | 0.48832813 | 0.65426265 | 5.00E-04 | 0.0027 | 0.0017 | 0.0017 | TRUE | 0.0132 |
| Afg3l2 | 10650 | 8059 | 8493 | 8990 | 8133 | 9263 | 8271 | 8572 | 9164 | 8949 | 9175 | 9144 | 8879.17 | 8669 | 9089 | 1.049 | 0.069 | 0.21642662 | 0.37913784 | 0.0013 | 0.0027 | 0.0021 | 0.0021 | TRUE | 0.0304 |
| Afm | 0 | 0 | 0 | 0 | 0 | 0 | 0 | 0 | 0 | 0 | 0 | 0 | 0 | 0 | 0 | NA | NA | NA | NA | NA | NA | NA | NA | NA | NA |
| Afmid | 284 | 205 | 196 | 252 | 228 | 248 | 221 | 218 | 211 | 251 | 257 | 245 | 233.83 | 217 | 251 | 1.157 | 0.21 | 0.10583417 | 0.23064549 | 0 | 0.0126 | 0.0079 | 0.0079 | TRUE | 0.0051 |
| Afp | 11 | 4 | 1 | 3 | 1 | 1 | 9 | 4 | 1 | 3 | 1 | 1 | 3.16 | 5 | 2 | 0.362 | -1.466 | 0.22090623 | NA | 0 | 0.7517 | 0.6289 | 0.6289 | TRUE | 0.142 |
| Aftph | 2927 | 2134 | 2244 | 2528 | 2155 | 2571 | 2273 | 2270 | 2421 | 2516 | 2431 | 2538 | 2408.31 | 2321 | 2495 | 1.075 | 0.105 | 0.08495523 | 0.19667397 | 5.00E-04 | 0.0034 | 0.0023 | 0.0023 | TRUE | 0.0168 |
| Aga | 1616 | 1218 | 1356 | 1279 | 1137 | 1193 | 1255 | 1296 | 1463 | 1273 | 1283 | 1178 | 1291.2 | 1338 | 1245 | 0.931 | -0.103 | 0.20738475 | 0.36908856 | 0.0037 | 0.0043 | 0.0041 | 0.0041 | TRUE | 0.0797 |
| Agap1 | 3355 | 2549 | 2558 | 2742 | 2596 | 2972 | 2606 | 2711 | 2760 | 2729 | 2929 | 2934 | 2778.14 | 2692 | 2864 | 1.064 | 0.09 | 0.14279204 | 0.28548421 | 9.00E-04 | 0.0033 | 0.0023 | 0.0023 | TRUE | 0.0206 |
| Agap2 | 76 | 48 | 39 | 65 | 73 | 66 | 59 | 51 | 42 | 65 | 82 | 65 | 60.73 | 51 | 71 | 1.382 | 0.467 | 0.07273344 | 0.17646904 | 0 | 0.0415 | 0.0321 | 0.0321 | TRUE | 0.1934 |
| Agap3 | 2843 | 2149 | 2145 | 2205 | 2209 | 2362 | 2208 | 2286 | 2314 | 2195 | 2492 | 2332 | 2304.47 | 2269 | 2340 | 1.031 | 0.044 | 0.51556636 | 0.67798234 | 0.0017 | 0.0035 | 0.0028 | 0.0028 | TRUE | 0.0389 |
| Agbl1 | 0 | 0 | 0 | 0 | 0 | 1 | 0 | 0 | 0 | 0 | 0 | 1 | 0.16 | 0 | 0 | 2.094 | 1.067 | 0.79379329 | NA | 0 | 14.4078 | 10 | 10 | TRUE | 0.3835 |
| Agbl2 | 40 | 37 | 59 | 78 | 62 | 63 | 31 | 39 | 64 | 78 | 70 | 62 | 57.31 | 45 | 70 | 1.581 | 0.661 | 0.02684272 | 0.08552453 | 0.0519 | 0.0438 | 0.0459 | 0.0459 | TRUE | 1.0473 |
| Agbl3 | 73 | 56 | 45 | 59 | 49 | 59 | 57 | 60 | 49 | 59 | 55 | 58 | 56.18 | 55 | 57 | 1.044 | 0.062 | 0.80733809 | 0.8885257 | 0 | 0.0446 | 0.0294 | 0.0294 | TRUE | 0.0798 |
| Agbl4 | 0 | 0 | 0 | 0 | 1 | 0 | 0 | 0 | 0 | 0 | 1 | 0 | 0.19 | 0 | 0 | 2.094 | 1.067 | 0.79379329 | NA | 0 | 12.608 | 10 | 10 | TRUE | 0.5075 |
| Agbl5 | 1359 | 1053 | 1082 | 1198 | 1050 | 1233 | 1055 | 1120 | 1167 | 1192 | 1185 | 1217 | 1156.19 | 1114 | 1198 | 1.077 | 0.106 | 0.15000624 | 0.29565051 | 6.00E-04 | 0.0045 | 0.0031 | 0.0031 | TRUE | 0.0269 |
| Ager | 10 | 10 | 10 | 11 | 9 | 7 | 8 | 11 | 11 | 11 | 10 | 7 | 9.53 | 10 | 9 | 0.966 | -0.051 | 0.93452418 | NA | 0 | 0.251 | 0.1671 | 0.1671 | TRUE | 0.1786 |
| Agfg1 | 10103 | 7538 | 7814 | 8412 | 7597 | 8572 | 7847 | 8018 | 8431 | 8373 | 8570 | 8462 | 8283.55 | 8099 | 8468 | 1.046 | 0.065 | 0.20712129 | 0.36896996 | 6.00E-04 | 0.0027 | 0.0018 | 0.0018 | TRUE | 0.0157 |
| Agfg2 | 1416 | 1060 | 1200 | 1522 | 1257 | 1435 | 1100 | 1128 | 1295 | 1515 | 1418 | 1417 | 1311.94 | 1174 | 1450 | 1.237 | 0.306 | 0.00019086 | 0.00206661 | 0.0036 | 0.0042 | 0.0041 | 0.0041 | TRUE | 0.0958 |
| Aggf1 | 7192 | 5061 | 4488 | 4823 | 4162 | 4756 | 5586 | 5383 | 4843 | 4801 | 4695 | 4695 | 5000.41 | 5271 | 4730 | 0.897 | -0.157 | 0.01508556 | 0.05606288 | 0.0024 | 0.0029 | 0.0028 | 0.0028 | TRUE | 0.0612 |
| Agk | 2025 | 1424 | 1432 | 1317 | 1197 | 1427 | 1573 | 1515 | 1545 | 1311 | 1350 | 1409 | 1450.41 | 1544 | 1357 | 0.878 | -0.187 | 0.00564486 | 0.02693891 | 0 | 0.0041 | 0.0026 | 0.0026 | TRUE | 0.0138 |
| Agl | 2272 | 1600 | 1683 | 1822 | 1623 | 1808 | 1765 | 1702 | 1816 | 1814 | 1831 | 1785 | 1785.29 | 1761 | 1810 | 1.028 | 0.039 | 0.5331051 | 0.69263607 | 0 | 0.0038 | 0.0023 | 0.0023 | TRUE | 0.01 |
| Agmat | 62 | 41 | 44 | 62 | 43 | 59 | 48 | 44 | 47 | 62 | 49 | 58 | 51.28 | 46 | 56 | 1.211 | 0.276 | 0.30636545 | 0.47877557 | 0 | 0.0486 | 0.0328 | 0.0328 | TRUE | 0.1117 |
| Agmo | 0 | 0 | 0 | 0 | 0 | 0 | 0 | 0 | 0 | 0 | 0 | 0 | 0 | 0 | 0 | NA | NA | NA | NA | NA | NA | NA | NA | NA | NA |
| Agpat1 | 4845 | 3743 | 3624 | 3460 | 3249 | 3797 | 3763 | 3981 | 3910 | 3444 | 3665 | 3748 | 3752.01 | 3885 | 3619 | 0.932 | -0.102 | 0.08512641 | 0.19700643 | 0.001 | 0.0031 | 0.0023 | 0.0023 | TRUE | 0.0219 |
| Agpat2 | 2323 | 1770 | 1669 | 1811 | 1670 | 1870 | 1804 | 1883 | 1801 | 1803 | 1884 | 1846 | 1836.72 | 1829 | 1844 | 1.008 | 0.012 | 0.85009104 | 0.91534965 | 0 | 0.0037 | 0.0023 | 0.0023 | TRUE | 0.0078 |
| Agpat3 | 1791 | 1269 | 1634 | 2546 | 2172 | 2436 | 1391 | 1350 | 1763 | 2534 | 2450 | 2405 | 1982.19 | 1501 | 2463 | 1.642 | 0.715 | 3.11E-14 | 4.89E-12 | 0.011 | 0.0036 | 0.0059 | 0.0059 | TRUE | 0.277 |
| Agpat4 | 912 | 746 | 950 | 817 | 713 | 876 | 708 | 794 | 1025 | 813 | 804 | 865 | 834.87 | 842 | 827 | 0.984 | -0.023 | 0.85045085 | 0.91553016 | 0.0179 | 0.0053 | 0.0093 | 0.0093 | TRUE | 0.4262 |
| Agpat5 | 8246 | 6174 | 6419 | 7386 | 6249 | 7026 | 6404 | 6567 | 6926 | 7352 | 7050 | 6936 | 6872.48 | 6632 | 7113 | 1.073 | 0.101 | 0.06704228 | 0.16607199 | 0.001 | 0.0028 | 0.0021 | 0.0021 | TRUE | 0.0182 |
| Agpat6 | 4804 | 3478 | 3163 | 3280 | 2818 | 3205 | 3731 | 3700 | 3413 | 3265 | 3179 | 3164 | 3408.53 | 3615 | 3203 | 0.886 | -0.175 | 0.0033947 | 0.01824077 | 0.0011 | 0.0031 | 0.0023 | 0.0023 | TRUE | 0.0286 |
| Agpat9 | 256 | 175 | 145 | 50 | 54 | 47 | 199 | 186 | 156 | 50 | 61 | 46 | 116.42 | 180 | 52 | 0.288 | -1.794 | 7.03E-18 | 1.62E-15 | 0 | 0.0228 | 0.0187 | 0.0187 | TRUE | 0.1565 |
| Agphd1 | 544 | 403 | 392 | 520 | 398 | 466 | 423 | 429 | 423 | 518 | 449 | 460 | 450.12 | 425 | 476 | 1.121 | 0.164 | 0.11374553 | 0.24283269 | 0 | 0.0077 | 0.0055 | 0.0055 | TRUE | 0.0702 |
| Agps | 14159 | 10342 | 10669 | 12042 | 10585 | 11638 | 10997 | 11001 | 11512 | 11987 | 11941 | 11489 | 11487.53 | 11170 | 11806 | 1.057 | 0.08 | 0.10459781 | 0.22856085 | 5.00E-04 | 0.0026 | 0.0017 | 0.0017 | TRUE | 0.0087 |
| Agr2 | 0 | 0 | 0 | 0 | 0 | 1 | 0 | 0 | 0 | 0 | 0 | 1 | 0.16 | 0 | 0 | 2.094 | 1.067 | 0.79379329 | NA | 0 | 14.4078 | 10 | 10 | TRUE | 0.3835 |
| Agr3 | 0 | 0 | 0 | 0 | 0 | 0 | 0 | 0 | 0 | 0 | 0 | 0 | 0 | 0 | 0 | NA | NA | NA | NA | NA | NA | NA | NA | NA | NA |
| Agrn | 12261 | 9609 | 9103 | 10962 | 10077 | 11160 | 9523 | 10221 | 9822 | 10912 | 11368 | 11017 | 10476.96 | 9855 | 11099 | 1.126 | 0.172 | 0.00081255 | 0.00622392 | 7.00E-04 | 0.0027 | 0.0018 | 0.0018 | TRUE | 0.0129 |
| Agrp | 11 | 7 | 4 | 7 | 9 | 6 | 9 | 7 | 4 | 7 | 10 | 6 | 7.22 | 7 | 8 | 1.117 | 0.159 | 0.8233394 | NA | 0 | 0.3305 | 0.2278 | 0.2278 | TRUE | 0.2372 |
| Agt | 4 | 4 | 4 | 2 | 3 | 2 | 3 | 4 | 4 | 2 | 3 | 2 | 3.17 | 4 | 2 | 0.629 | -0.669 | 0.52694389 | NA | 0 | 0.7498 | 0.467 | 0.467 | TRUE | 0.1054 |
| Agtpbp1 | 1317 | 1052 | 1035 | 1233 | 1101 | 1267 | 1023 | 1119 | 1117 | 1227 | 1242 | 1251 | 1163.12 | 1086 | 1240 | 1.143 | 0.193 | 0.00885698 | 0.03792473 | 6.00E-04 | 0.0045 | 0.0031 | 0.0031 | TRUE | 0.0329 |
| Agtr1a | 117 | 75 | 64 | 85 | 78 | 79 | 91 | 80 | 69 | 85 | 88 | 78 | 81.71 | 80 | 84 | 1.039 | 0.055 | 0.80088317 | 0.88434975 | 0 | 0.0314 | 0.0226 | 0.0226 | TRUE | 0.1508 |
| Agtr1b | 3 | 2 | 3 | 4 | 6 | 4 | 2 | 2 | 3 | 4 | 7 | 4 | 3.73 | 2 | 5 | 1.909 | 0.933 | 0.34476604 | NA | 0 | 0.6375 | 0.408 | 0.408 | TRUE | 0.1972 |
| Agtr2 | 0 | 0 | 0 | 0 | 0 | 0 | 0 | 0 | 0 | 0 | 0 | 0 | 0 | 0 | 0 | NA | NA | NA | NA | NA | NA | NA | NA | NA | NA |
| Agtrap | 4431 | 3326 | 3123 | 3492 | 3248 | 3585 | 3441 | 3538 | 3370 | 3476 | 3664 | 3539 | 3504.64 | 3450 | 3560 | 1.032 | 0.045 | 0.41801986 | 0.59074133 | 3.00E-04 | 0.0031 | 0.0019 | 0.0019 | TRUE | 0.0079 |
| Agxt | 7 | 1 | 0 | 7 | 2 | 0 | 5 | 1 | 0 | 7 | 2 | 0 | 2.62 | 2 | 3 | 1.364 | 0.448 | 0.76034446 | NA | 2.1757 | 0.9068 | 1.1695 | 1.1695 | TRUE | 0.5625 |
| Agxt2 | 0 | 0 | 0 | 0 | 0 | 0 | 0 | 0 | 0 | 0 | 0 | 0 | 0 | 0 | 0 | NA | NA | NA | NA | NA | NA | NA | NA | NA | NA |
| Agxt2l1 | 0 | 0 | 1 | 0 | 0 | 0 | 0 | 0 | 1 | 0 | 0 | 0 | 0.18 | 0 | 0 | 0.552 | -0.857 | 0.8336548 | NA | 0 | 13.1817 | 10 | 10 | TRUE | 0.5365 |
| Agxt2l2 | 468 | 349 | 352 | 402 | 377 | 454 | 363 | 371 | 380 | 400 | 425 | 448 | 398.02 | 371 | 424 | 1.144 | 0.194 | 0.06867037 | 0.16907935 | 0 | 0.0084 | 0.0056 | 0.0056 | TRUE | 0.0299 |
| Ahctf1 | 19839 | 13688 | 12755 | 13909 | 12186 | 13897 | 15408 | 14560 | 13763 | 13845 | 13747 | 13718 | 14173.49 | 14577 | 13770 | 0.944 | -0.082 | 0.1363784 | 0.27621602 | 0.0015 | 0.0026 | 0.0021 | 0.0021 | TRUE | 0.0306 |
| Ahcy | 6064 | 4231 | 4437 | 4685 | 4175 | 4771 | 4710 | 4500 | 4787 | 4663 | 4710 | 4710 | 4680.1 | 4666 | 4694 | 1.006 | 0.009 | 0.8697745 | 0.92801725 | 3.00E-04 | 0.0029 | 0.0018 | 0.0018 | TRUE | 0.0115 |
| Ahcyl1 | 10682 | 7950 | 8796 | 9116 | 8057 | 9079 | 8296 | 8456 | 9491 | 9074 | 9089 | 8962 | 8894.82 | 8748 | 9042 | 1.034 | 0.048 | 0.4384982 | 0.60989659 | 0.0024 | 0.0027 | 0.0027 | 0.0027 | TRUE | 0.0672 |
| Ahcyl2 | 2642 | 1875 | 2026 | 2409 | 2072 | 2402 | 2052 | 1994 | 2186 | 2398 | 2337 | 2371 | 2223.15 | 2077 | 2369 | 1.141 | 0.19 | 0.00251895 | 0.01445915 | 5.00E-04 | 0.0035 | 0.0024 | 0.0024 | TRUE | 0.0246 |
| Ahdc1 | 5462 | 4177 | 3655 | 4622 | 3950 | 4465 | 4242 | 4443 | 3944 | 4601 | 4456 | 4408 | 4348.87 | 4210 | 4488 | 1.066 | 0.092 | 0.13273833 | 0.27080058 | 0.0017 | 0.003 | 0.0025 | 0.0025 | TRUE | 0.0366 |
| Ahi1 | 914 | 705 | 724 | 859 | 684 | 854 | 710 | 750 | 781 | 855 | 772 | 843 | 785.11 | 747 | 823 | 1.104 | 0.143 | 0.09751141 | 0.21736995 | 0.0014 | 0.0054 | 0.0041 | 0.0041 | TRUE | 0.0348 |
| Ahnak | 3433 | 2345 | 2180 | 3049 | 2778 | 3244 | 2666 | 2494 | 2352 | 3035 | 3134 | 3202 | 2814 | 2504 | 3124 | 1.246 | 0.318 | 1.39E-06 | 3.80E-05 | 0.0018 | 0.0033 | 0.0028 | 0.0028 | TRUE | 0.0398 |
| Ahr | 598 | 460 | 410 | 448 | 425 | 481 | 464 | 489 | 442 | 446 | 479 | 475 | 466.05 | 465 | 467 | 1.003 | 0.004 | 0.97047929 | 0.98397988 | 0 | 0.0075 | 0.0051 | 0.0051 | TRUE | 0.0221 |
| Ahrr | 182 | 141 | 132 | 182 | 154 | 149 | 141 | 150 | 142 | 181 | 174 | 147 | 155.96 | 144 | 167 | 1.157 | 0.211 | 0.19573652 | 0.35534892 | 0 | 0.0176 | 0.0127 | 0.0127 | TRUE | 0.1222 |
| Ahsa1 | 11198 | 8132 | 7542 | 7193 | 6565 | 7661 | 8697 | 8650 | 8138 | 7160 | 7406 | 7563 | 7935.53 | 8495 | 7376 | 0.868 | -0.204 | 0.00013659 | 0.00157335 | 9.00E-04 | 0.0027 | 0.0019 | 0.0019 | TRUE | 0.0164 |
| Ahsa2 | 1938 | 1406 | 1286 | 1308 | 1110 | 1376 | 1505 | 1496 | 1388 | 1302 | 1252 | 1358 | 1383.47 | 1463 | 1304 | 0.891 | -0.166 | 0.02162588 | 0.07312679 | 0.0011 | 0.0041 | 0.003 | 0.003 | TRUE | 0.0239 |
| Ahsg | 120 | 101 | 70 | 56 | 59 | 54 | 93 | 107 | 76 | 56 | 67 | 53 | 75.29 | 92 | 59 | 0.634 | -0.657 | 0.00599573 | 0.02820837 | 0.0093 | 0.0339 | 0.0271 | 0.0271 | TRUE | 0.16 |
| AI115009 | 0 | 0 | 0 | 0 | 1 | 0 | 0 | 0 | 0 | 0 | 1 | 0 | 0.19 | 0 | 0 | 2.094 | 1.067 | 0.79379329 | NA | 0 | 12.608 | 10 | 10 | TRUE | 0.5075 |
| AI118078 | 102 | 77 | 49 | 66 | 59 | 51 | 79 | 82 | 53 | 66 | 67 | 50 | 66.1 | 71 | 61 | 0.849 | -0.236 | 0.36773154 | 0.54235754 | 0.0235 | 0.0383 | 0.0342 | 0.0342 | TRUE | 0.4367 |
| AI182371 | 15 | 10 | 8 | 5 | 6 | 12 | 12 | 11 | 9 | 5 | 7 | 12 | 9.09 | 11 | 8 | 0.762 | -0.393 | 0.54380946 | NA | 0 | 0.2633 | 0.1893 | 0.1893 | TRUE | 0.5946 |
| AI197445 | 11 | 6 | 3 | 4 | 7 | 7 | 9 | 6 | 3 | 4 | 8 | 7 | 6.16 | 6 | 6 | 1.014 | 0.02 | 0.97936999 | NA | 0 | 0.3873 | 0.2798 | 0.2798 | TRUE | 0.3064 |
| AI314180 | 5305 | 3533 | 3490 | 3599 | 3014 | 3455 | 4120 | 3758 | 3766 | 3582 | 3400 | 3411 | 3672.84 | 3881 | 3464 | 0.892 | -0.165 | 0.00768143 | 0.03395939 | 0.0014 | 0.0031 | 0.0025 | 0.0025 | TRUE | 0.036 |

| | | | | | | | | | | | | | | | | | | | | | | | | | |
|---|---|---|---|---|---|---|---|---|---|---|---|---|---|---|---|---|---|---|---|---|---|---|---|---|---|
| AI314831 | 3 | 1 | 0 | 2 | 1 | 1 | 2 | 1 | 0 | 2 | 1 | 1 | 1.25 | 1 | 1 | 1.162 | 0.216 | 0.89888599 | NA | 0 | 1.8985 | 1.3002 | 1.3002 | TRUE | 0.3124 |
| AI316807 | 654 | 491 | 435 | 496 | 426 | 477 | 508 | 522 | 469 | 494 | 481 | 471 | 490.79 | 500 | 482 | 0.963 | -0.054 | 0.58103941 | 0.73170012 | 0 | 0.0073 | 0.0049 | 0.0049 | TRUE | 0.0316 |
| AI317395 | 67 | 63 | 49 | 23 | 29 | 36 | 52 | 67 | 53 | 23 | 33 | 36 | 43.84 | 57 | 31 | 0.531 | -0.914 | 0.00321236 | 0.01741311 | 0 | 0.0565 | 0.0438 | 0.0438 | TRUE | 0.3214 |
| AI413582 | 492 | 371 | 443 | 541 | 505 | 585 | 382 | 395 | 478 | 539 | 570 | 577 | 490.07 | 418 | 562 | 1.347 | 0.429 | 0.00010772 | 0.00128728 | 0.0053 | 0.0073 | 0.0068 | 0.0068 | TRUE | 0.1764 |
| AI414108 | 1011 | 799 | 892 | 951 | 805 | 962 | 785 | 850 | 962 | 947 | 908 | 950 | 900.32 | 866 | 935 | 1.082 | 0.114 | 0.21338483 | 0.37565086 | 0.0046 | 0.0051 | 0.0049 | 0.0049 | TRUE | 0.1126 |
| AI427809 | 163 | 109 | 88 | 99 | 97 | 91 | 127 | 116 | 95 | 99 | 109 | 90 | 105.88 | 113 | 99 | 0.877 | -0.189 | 0.34790653 | 0.52249953 | 0.0065 | 0.0248 | 0.0196 | 0.0196 | TRUE | 0.1796 |
| AI429214 | 47 | 40 | 33 | 46 | 28 | 27 | 37 | 43 | 36 | 46 | 32 | 27 | 36.45 | 39 | 35 | 0.91 | -0.137 | 0.68312774 | 0.80580546 | 0.0149 | 0.0675 | 0.0533 | 0.0533 | TRUE | 0.5715 |
| AI450353 | 115 | 95 | 109 | 83 | 64 | 79 | 89 | 101 | 118 | 83 | 72 | 78 | 90.13 | 103 | 78 | 0.76 | -0.396 | 0.06312294 | 0.15913899 | 0 | 0.0287 | 0.0213 | 0.0213 | TRUE | 0.157 |
| AI462493 | 710 | 439 | 455 | 450 | 409 | 438 | 551 | 467 | 491 | 448 | 461 | 432 | 475.17 | 503 | 447 | 0.886 | -0.174 | 0.09797609 | 0.2181605 | 0.0027 | 0.0074 | 0.0058 | 0.0058 | TRUE | 0.0872 |
| AI463170 | 2 | 2 | 2 | 0 | 0 | 0 | 2 | 2 | 2 | 0 | 0 | 0 | 0.97 | 2 | 0 | 0.098 | -3.348 | 0.1108432 | NA | 0 | 2.438 | 1.9269 | 1.9269 | TRUE | 0.0718 |
| AI464131 | 1326 | 1059 | 1055 | 1342 | 1203 | 1361 | 1030 | 1126 | 1138 | 1336 | 1357 | 1344 | 1221.85 | 1098 | 1346 | 1.227 | 0.295 | 5.89E-05 | 0.00079896 | 8.00E-04 | 0.0044 | 0.0031 | 0.0031 | TRUE | 0.0375 |
| AI467606 | 298 | 205 | 187 | 256 | 219 | 263 | 231 | 218 | 202 | 255 | 247 | 260 | 235.46 | 217 | 254 | 1.167 | 0.223 | 0.09320011 | 0.2100007 | 0 | 0.0125 | 0.0084 | 0.0084 | TRUE | 0.0409 |
| AI480653 | 2209 | 1755 | 1824 | 1918 | 1801 | 2011 | 1716 | 1867 | 1968 | 1909 | 2032 | 1985 | 1912.76 | 1850 | 1975 | 1.069 | 0.096 | 0.1823759 | 0.33796172 | 0.0024 | 0.0037 | 0.0032 | 0.0032 | TRUE | 0.0506 |
| AI504432 | 84 | 49 | 24 | 44 | 42 | 33 | 65 | 52 | 26 | 44 | 47 | 33 | 44.5 | 48 | 41 | 0.855 | -0.227 | 0.5216713 | 0.68299422 | 0.0981 | 0.0557 | 0.0676 | 0.0676 | TRUE | 0.4302 |
| AI506816 | 37 | 32 | 22 | 35 | 28 | 26 | 29 | 34 | 24 | 35 | 32 | 26 | 29.77 | 29 | 31 | 1.064 | 0.089 | 0.80303024 | 0.88603607 | 0 | 0.082 | 0.0582 | 0.0582 | TRUE | 0.1473 |
| AI507597 | 1 | 0 | 0 | 1 | 0 | 0 | 1 | 0 | 0 | 1 | 0 | 0 | 0.3 | 0 | 0 | 1.075 | 0.105 | 0.97698525 | NA | 0 | 8.0272 | 7.5141 | 7.5141 | TRUE | 0.3906 |
| AI593442 | 46 | 36 | 33 | 17 | 11 | 5 | 36 | 38 | 36 | 17 | 12 | 5 | 23.98 | 37 | 11 | 0.312 | -1.681 | 0.00019597 | 0.00209952 | 0 | 0.1013 | 0.0881 | 0.0881 | TRUE | 0.9976 |
| AI597468 | 2447 | 1842 | 1846 | 2041 | 1686 | 1857 | 1900 | 1959 | 1992 | 2032 | 1902 | 1833 | 1936.39 | 1950 | 1922 | 0.986 | -0.02 | 0.75984061 | 0.8573099 | 0.0012 | 0.0037 | 0.0027 | 0.0027 | TRUE | 0.0303 |
| AI597479 | 3501 | 2587 | 2552 | 2587 | 2172 | 2557 | 2719 | 2752 | 2754 | 2575 | 2450 | 2524 | 2628.99 | 2742 | 2516 | 0.918 | -0.123 | 0.03012818 | 0.09296091 | 0 | 0.0033 | 0.0019 | 0.0019 | TRUE | 0.0063 |
| AI606473 | 0 | 0 | 0 | 0 | 0 | 0 | 0 | 0 | 0 | 0 | 0 | 0 | 0 | 0 | 0 | NA | NA | NA | NA | NA | NA | NA | NA | NA | NA |
| AI607873 | 0 | 2 | 0 | 0 | 1 | 0 | 0 | 2 | 0 | 0 | 1 | 0 | 0.54 | 1 | 0 | 0.535 | -0.903 | 0.76014553 | NA | 0 | 4.3706 | 4.6038 | 4.6038 | TRUE | 0.9746 |
| AI646519 | 0 | 1 | 0 | 1 | 0 | 2 | 0 | 1 | 0 | 1 | 0 | 2 | 0.67 | 0 | 1 | 2.905 | 1.538 | 0.53584396 | NA | 0 | 3.5281 | 2.9323 | 2.9323 | TRUE | 0.2752 |
| AI661453 | 398 | 327 | 325 | 257 | 258 | 286 | 309 | 348 | 351 | 256 | 291 | 282 | 306.13 | 336 | 276 | 0.824 | -0.279 | 0.0234497 | 0.0776208 | 0 | 0.0102 | 0.0076 | 0.0076 | TRUE | 0.0597 |
| AI662270 | 1099 | 863 | 720 | 162 | 122 | 155 | 854 | 918 | 777 | 161 | 138 | 153 | 500.05 | 850 | 151 | 0.178 | -2.493 | 1.83E-96 | 2.62E-92 | 0.0043 | 0.0072 | 0.0063 | 0.0063 | TRUE | 0.0643 |
| AI747448 | 12 | 10 | 15 | 21 | 16 | 12 | 9 | 11 | 16 | 21 | 18 | 12 | 14.49 | 12 | 17 | 1.418 | 0.504 | 0.33404352 | NA | 0 | 0.166 | 0.1249 | 0.1249 | TRUE | 0.353 |
| AI836003 | 69 | 49 | 54 | 18 | 31 | 28 | 54 | 52 | 58 | 18 | 35 | 28 | 40.75 | 55 | 27 | 0.489 | -1.033 | 0.00154568 | 0.00986774 | 0 | 0.0606 | 0.0485 | 0.0485 | TRUE | 0.5327 |
| AI837181 | 1270 | 903 | 945 | 963 | 870 | 1041 | 986 | 961 | 1020 | 959 | 981 | 1028 | 989.03 | 989 | 989 | 1.001 | 0.001 | 0.99109056 | 0.99553503 | 0 | 0.0048 | 0.0032 | 0.0032 | TRUE | 0.014 |
| AI846148 | 2778 | 1967 | 1778 | 1796 | 1550 | 1712 | 2158 | 2092 | 1918 | 1788 | 1749 | 1690 | 1899.1 | 2056 | 1742 | 0.847 | -0.24 | 0.00055209 | 0.00464725 | 0.0018 | 0.0037 | 0.003 | 0.003 | TRUE | 0.041 |
| AI847159 | 2 | 2 | 2 | 1 | 1 | 1 | 2 | 2 | 2 | 1 | 1 | 1 | 1.49 | 2 | 1 | 0.536 | -0.9 | 0.55440082 | NA | 0 | 1.5914 | 0.9219 | 0.9219 | TRUE | 0.0377 |
| AI848285 | 382 | 282 | 290 | 386 | 332 | 401 | 297 | 300 | 313 | 384 | 375 | 396 | 344.03 | 303 | 385 | 1.271 | 0.346 | 0.00177298 | 0.01103231 | 0 | 0.0093 | 0.0059 | 0.0059 | TRUE | 0.0086 |
| AI854517 | 108 | 69 | 65 | 55 | 50 | 82 | 84 | 73 | 70 | 55 | 56 | 81 | 69.92 | 76 | 64 | 0.844 | -0.245 | 0.32419971 | 0.49809594 | 0.0141 | 0.0363 | 0.0302 | 0.0302 | TRUE | 0.479 |
| AI854703 | 149 | 118 | 131 | 143 | 128 | 133 | 116 | 126 | 141 | 142 | 144 | 131 | 133.44 | 128 | 139 | 1.096 | 0.132 | 0.44754237 | 0.61788255 | 0 | 0.0202 | 0.0143 | 0.0143 | TRUE | 0.0901 |
| AI987944 | 846 | 635 | 718 | 849 | 711 | 821 | 657 | 675 | 775 | 845 | 802 | 810 | 760.81 | 702 | 819 | 1.168 | 0.225 | 0.01383441 | 0.05264516 | 0.0029 | 0.0055 | 0.0047 | 0.0047 | TRUE | 0.0941 |
| Aicda | 45 | 27 | 16 | 15 | 11 | 13 | 35 | 29 | 17 | 15 | 12 | 13 | 20.18 | 27 | 13 | 0.493 | -1.021 | 0.02604343 | 0.08378093 | 0 | 0.1198 | 0.0953 | 0.0953 | TRUE | 0.2556 |
| Aida | 1772 | 1263 | 1087 | 1084 | 941 | 1055 | 1376 | 1343 | 1173 | 1079 | 1062 | 1041 | 1179.09 | 1297 | 1061 | 0.817 | -0.292 | 0.0003509 | 0.00327809 | 0.0031 | 0.0044 | 0.004 | 0.004 | TRUE | 0.0834 |
| Aif1 | 0 | 0 | 0 | 0 | 0 | 2 | 0 | 0 | 0 | 0 | 0 | 2 | 0.33 | 0 | 1 | 3.743 | 1.904 | 0.62640105 | NA | 0 | 7.2051 | 9.2465 | 9.2465 | TRUE | 0.9739 |
| Aif1l | 1598 | 1087 | 1148 | 1216 | 1122 | 1262 | 1241 | 1156 | 1239 | 1210 | 1266 | 1246 | 1226.32 | 1212 | 1241 | 1.023 | 0.033 | 0.64748437 | 0.78051363 | 0 | 0.0044 | 0.0029 | 0.0029 | TRUE | 0.019 |
| Aifm1 | 5052 | 3433 | 3564 | 3846 | 3225 | 3961 | 3924 | 3652 | 3846 | 3828 | 3638 | 3910 | 3799.57 | 3807 | 3792 | 0.996 | -0.006 | 0.92214673 | 0.95711845 | 0.0011 | 0.0031 | 0.0022 | 0.0022 | TRUE | 0.0154 |
| Aifm2 | 1453 | 1052 | 1141 | 1294 | 1129 | 1367 | 1128 | 1119 | 1231 | 1288 | 1274 | 1349 | 1231.62 | 1159 | 1304 | 1.125 | 0.17 | 0.0218054 | 0.07356156 | 0.001 | 0.0044 | 0.0032 | 0.0032 | TRUE | 0.0341 |
| Aifm3 | 655 | 525 | 564 | 728 | 606 | 736 | 509 | 558 | 609 | 725 | 684 | 727 | 635.09 | 559 | 712 | 1.278 | 0.353 | 0.00025159 | 0.00254128 | 0.003 | 0.0062 | 0.0051 | 0.0051 | TRUE | 0.0762 |
| Aig1 | 182 | 154 | 142 | 119 | 159 | 155 | 141 | 164 | 153 | 118 | 179 | 153 | 151.53 | 153 | 150 | 0.983 | -0.024 | 0.8946647 | 0.94219294 | 0.0168 | 0.0181 | 0.0177 | 0.0177 | TRUE | 0.2736 |
| Aim1 | 334 | 228 | 198 | 155 | 173 | 187 | 259 | 243 | 214 | 154 | 195 | 185 | 208.27 | 239 | 178 | 0.744 | -0.427 | 0.0050758 | 0.02479438 | 0.0069 | 0.0138 | 0.0118 | 0.0118 | TRUE | 0.1474 |
| Aim1l | 437 | 344 | 343 | 420 | 413 | 451 | 339 | 366 | 370 | 418 | 466 | 445 | 400.76 | 358 | 443 | 1.237 | 0.307 | 0.00427702 | 0.02177823 | 0 | 0.0083 | 0.0058 | 0.0058 | TRUE | 0.0283 |
| Aim2 | 949 | 711 | 760 | 374 | 301 | 336 | 737 | 756 | 820 | 372 | 340 | 332 | 559.48 | 771 | 348 | 0.452 | -1.147 | 1.46E-30 | 1.30E-27 | 0.0016 | 0.0067 | 0.0051 | 0.0051 | TRUE | 0.044 |
| Aimp1 | 6205 | 4515 | 4725 | 5049 | 4667 | 5256 | 4819 | 4803 | 5098 | 5026 | 5265 | 5188 | 5033.18 | 4907 | 5160 | 1.052 | 0.073 | 0.18009678 | 0.33499782 | 6.00E-04 | 0.0029 | 0.0019 | 0.0019 | TRUE | 0.0141 |
| Aimp2 | 3728 | 2583 | 2382 | 2323 | 2088 | 2437 | 2895 | 2748 | 2570 | 2312 | 2355 | 2406 | 2547.75 | 2738 | 2358 | 0.861 | -0.216 | 0.00093132 | 0.0068899 | 0.0017 | 0.0034 | 0.0027 | 0.0027 | TRUE | 0.0346 |
| Aip | 1735 | 1197 | 1312 | 1620 | 1396 | 1569 | 1348 | 1273 | 1416 | 1613 | 1575 | 1549 | 1462.1 | 1346 | 1579 | 1.173 | 0.231 | 0.00101981 | 0.00733942 | 7.00E-04 | 0.004 | 0.0029 | 0.0029 | TRUE | 0.0255 |
| Aipl1 | 48 | 43 | 51 | 75 | 73 | 60 | 37 | 46 | 55 | 75 | 82 | 59 | 59.05 | 46 | 72 | 1.576 | 0.656 | 0.01549183 | 0.05723049 | 0.0156 | 0.0426 | 0.0351 | 0.0351 | TRUE | 0.2593 |
| Aire | 851 | 841 | 964 | 974 | 859 | 946 | 661 | 895 | 1040 | 970 | 969 | 934 | 911.34 | 865 | 958 | 1.109 | 0.149 | 0.25111007 | 0.41997616 | 0.025 | 0.005 | 0.0111 | 0.0111 | TRUE | 0.4124 |
| Airn | 3 | 0 | 3 | 1 | 1 | 0 | 2 | 0 | 3 | 1 | 1 | 0 | 1.28 | 2 | 1 | 0.374 | -1.418 | 0.44694565 | NA | 0 | 1.8515 | 1.5066 | 1.5066 | TRUE | 0.4398 |
| Ajap1 | 20 | 15 | 11 | 16 | 15 | 20 | 16 | 16 | 12 | 16 | 17 | 20 | 15.99 | 15 | 18 | 1.21 | 0.275 | 0.56202736 | NA | 0 | 0.1506 | 0.0988 | 0.0988 | TRUE | 0.0986 |
| Ajuba | 5827 | 4304 | 4009 | 4230 | 3638 | 4108 | 4526 | 4578 | 4326 | 4211 | 4104 | 4055 | 4299.86 | 4477 | 4123 | 0.921 | -0.119 | 0.02778651 | 0.08755567 | 4.00E-04 | 0.003 | 0.0019 | 0.0019 | TRUE | 0.0104 |
| AK010878 | 386 | 266 | 242 | 277 | 201 | 280 | 300 | 283 | 261 | 276 | 227 | 276 | 270.45 | 281 | 260 | 0.923 | -0.116 | 0.38697105 | 0.56071448 | 0.0044 | 0.0112 | 0.0092 | 0.0092 | TRUE | 0.1316 |
| Ak1 | 2424 | 1883 | 2080 | 2303 | 2194 | 2475 | 1883 | 2003 | 2244 | 2292 | 2475 | 2443 | 2223.42 | 2043 | 2403 | 1.177 | 0.235 | 0.00222036 | 0.01308687 | 0.0042 | 0.0035 | 0.0038 | 0.0038 | TRUE | 0.0892 |
| AK129341 | 741 | 581 | 545 | 614 | 520 | 635 | 576 | 618 | 588 | 611 | 587 | 627 | 601.03 | 594 | 608 | 1.026 | 0.036 | 0.68581919 | 0.80738108 | 0 | 0.0064 | 0.0042 | 0.0042 | TRUE | 0.0147 |
| Ak2 | 14575 | 10477 | 10257 | 11161 | 9745 | 11463 | 11320 | 11144 | 11067 | 11110 | 10993 | 11316 | 11158.34 | 11177 | 11140 | 0.997 | -0.005 | 0.91443794 | 0.9529197 | 1.00E-04 | 0.0026 | 0.0014 | 0.0014 | TRUE | 0.0023 |
| Ak3 | 908 | 616 | 526 | 591 | 555 | 613 | 705 | 655 | 568 | 588 | 626 | 605 | 624.58 | 643 | 606 | 0.941 | -0.087 | 0.39161813 | 0.56522246 | 0.0051 | 0.0062 | 0.0058 | 0.0058 | TRUE | 0.1212 |
| Ak4 | 18891 | 14328 | 14733 | 10676 | 10005 | 11000 | 14672 | 15241 | 15897 | 10627 | 11287 | 10859 | 13096.9 | 15270 | 10924 | 0.715 | -0.483 | 1.62E-19 | 4.78E-17 | 0.0012 | 0.0026 | 0.002 | 0.002 | TRUE | 0.0158 |
| Ak5 | 30 | 21 | 16 | 24 | 14 | 16 | 23 | 22 | 17 | 24 | 16 | 16 | 19.73 | 21 | 19 | 0.88 | -0.184 | 0.67409973 | 0.79984289 | 0 | 0.1226 | 0.0872 | 0.0872 | TRUE | 0.3445 |
| Ak7 | 629 | 400 | 328 | 257 | 205 | 242 | 489 | 425 | 354 | 256 | 231 | 239 | 332.31 | 423 | 242 | 0.571 | -0.808 | 5.38E-09 | 2.98E-07 | 0.0126 | 0.0096 | 0.0105 | 0.0105 | TRUE | 0.1346 |
| Ak8 | 38 | 30 | 48 | 46 | 39 | 55 | 30 | 32 | 52 | 46 | 44 | 54 | 42.88 | 38 | 48 | 1.285 | 0.361 | 0.25811815 | 0.42814004 | 0.0291 | 0.0577 | 0.0499 | 0.0499 | TRUE | 0.743 |
| Akap1 | 18861 | 13987 | 13654 | 13199 | 12629 | 13966 | 14649 | 14878 | 14733 | 13138 | 14247 | 13787 | 14238.43 | 14753 | 13724 | 0.93 | -0.104 | 0.03843529 | 0.11125063 | 7.00E-04 | 0.0026 | 0.0018 | 0.0018 | TRUE | 0.017 |
| Akap10 | 4226 | 3005 | 3058 | 3336 | 2996 | 3480 | 3282 | 3196 | 3300 | 3321 | 3380 | 3435 | 3318.97 | 3259 | 3379 | 1.036 | 0.052 | 0.33836443 | 0.51249222 | 0 | 0.0031 | 0.0018 | 0.0018 | TRUE | 0.0034 |
| Akap11 | 11186 | 7642 | 6857 | 7872 | 6811 | 7553 | 8688 | 8129 | 7399 | 7836 | 7684 | 7456 | 7865.05 | 8072 | 7659 | 0.949 | -0.076 | 0.24781093 | 0.41665236 | 0.0033 | 0.0027 | 0.003 | 0.003 | TRUE | 0.0649 |
| Akap12 | 35783 | 27082 | 26890 | 29155 | 26683 | 31087 | 27791 | 28807 | 29014 | 29021 | 30101 | 30688 | 29236.95 | 28537 | 29937 | 1.049 | 0.069 | 0.14860431 | 0.29381677 | 6.00E-04 | 0.0025 | 0.0016 | 0.0016 | TRUE | 0.0088 |
| Akap13 | 3945 | 2707 | 2512 | 2998 | 2704 | 3095 | 3064 | 2879 | 2710 | 2984 | 3050 | 3055 | 2957.26 | 2884 | 3030 | 1.05 | 0.07 | 0.27445907 | 0.44542335 | 0.0015 | 0.0032 | 0.0026 | 0.0026 | TRUE | 0.0368 |
| Akap14 | 3 | 2 | 1 | 0 | 1 | 2 | 2 | 2 | 1 | 0 | 1 | 2 | 1.44 | 2 | 1 | 0.555 | -0.85 | 0.59944244 | NA | 0 | 1.6485 | 1.1253 | 1.1253 | TRUE | 0.3878 |
| Akap17b | 1310 | 878 | 753 | 927 | 748 | 861 | 1017 | 934 | 812 | 923 | 844 | 850 | 896.72 | 921 | 872 | 0.945 | -0.081 | 0.39984976 | 0.57316745 | 0.0066 | 0.0051 | 0.0056 | 0.0056 | TRUE | 0.1259 |
| Akap2 | 3527 | 2710 | 3663 | 3522 | 3535 | 4430 | 2739 | 2883 | 3952 | 3506 | 3988 | 4373 | 3573.49 | 3191 | 3956 | 1.24 | 0.31 | 0.11235884 | 0.24075603 | 0.0272 | 0.0031 | 0.0099 | 0.0272 | TRUE | 0.5282 |

| | | | | | | | | | | | | | | | | | | | | | | | | | |
|---|---|---|---|---|---|---|---|---|---|---|---|---|---|---|---|---|---|---|---|---|---|---|---|---|---|
| Akap3 | 7 | 4 | 2 | 5 | 4 | 3 | 5 | 4 | 2 | 5 | 5 | 3 | 4.05 | 4 | 4 | 1.033 | 0.047 | 0.96047063 | NA | 0 | 0.5876 | 0.3899 | 0.3899 | TRUE | 0.2376 |
| Akap4 | 2 | 1 | 1 | 0 | 0 | 0 | 2 | 1 | 1 | 0 | 0 | 0 | 0.62 | 1 | 0 | 0.154 | -2.702 | 0.27412557 | NA | 0 | 3.8499 | 3.0088 | 3.0088 | TRUE | 0.0718 |
| Akap5 | 319 | 287 | 299 | 326 | 286 | 352 | 248 | 305 | 323 | 325 | 323 | 347 | 311.71 | 292 | 332 | 1.141 | 0.19 | 0.14764913 | 0.2923496 | 0.0074 | 0.01 | 0.0092 | 0.0092 | TRUE | 0.2137 |
| Akap6 | 22 | 14 | 10 | 5 | 8 | 7 | 17 | 15 | 11 | 5 | 9 | 7 | 10.61 | 14 | 7 | 0.483 | -1.05 | 0.08390922 | NA | 0 | 0.2257 | 0.1576 | 0.1576 | TRUE | 0.1871 |
| Akap7 | 1202 | 911 | 1027 | 1124 | 979 | 985 | 934 | 969 | 1108 | 1119 | 1104 | 972 | 1034.38 | 1004 | 1065 | 1.062 | 0.087 | 0.34523136 | 0.51936368 | 0.0056 | 0.0047 | 0.0052 | 0.0052 | TRUE | 0.0981 |
| Akap8 | 8701 | 6416 | 5841 | 5648 | 5216 | 6062 | 6758 | 6825 | 6302 | 5622 | 5884 | 5984 | 6229.17 | 6628 | 5830 | 0.879 | -0.185 | 0.00109025 | 0.00769531 | 0.0012 | 0.0028 | 0.0022 | 0.0022 | TRUE | 0.0224 |
| Akap8l | 3237 | 2696 | 2739 | 3117 | 2712 | 3179 | 2514 | 2868 | 2955 | 3103 | 3059 | 3138 | 2939.56 | 2779 | 3100 | 1.117 | 0.159 | 0.02527277 | 0.08191015 | 0.0031 | 0.0032 | 0.0033 | 0.0033 | TRUE | 0.0865 |
| Akap9 | 2421 | 1783 | 1712 | 2302 | 2001 | 2440 | 1880 | 1897 | 1847 | 2291 | 2257 | 2409 | 2096.91 | 1875 | 2319 | 1.237 | 0.307 | 5.70E-07 | 1.82E-05 | 0 | 0.0036 | 0.0022 | 0.0022 | TRUE | 0.0139 |
| Akip1 | 718 | 470 | 511 | 556 | 472 | 499 | 558 | 500 | 551 | 553 | 532 | 493 | 531.24 | 536 | 526 | 0.98 | -0.029 | 0.76809304 | 0.86284342 | 0.0016 | 0.0069 | 0.0052 | 0.0052 | TRUE | 0.0402 |
| Akirin1 | 4431 | 2998 | 2880 | 3350 | 2835 | 3200 | 3441 | 3189 | 3107 | 3335 | 3198 | 3159 | 3238.24 | 3246 | 3231 | 0.995 | -0.008 | 0.90149494 | 0.94568228 | 0.0016 | 0.0032 | 0.0025 | 0.0025 | TRUE | 0.0345 |
| Akirin2 | 6338 | 4643 | 4529 | 5101 | 4416 | 5151 | 4922 | 4939 | 4887 | 5078 | 4982 | 5085 | 4982 | 4916 | 5048 | 1.027 | 0.038 | 0.43532985 | 0.60698451 | 0 | 0.0029 | 0.0015 | 0.0015 | TRUE | 0.0016 |
| Akna | 1240 | 970 | 1036 | 999 | 879 | 985 | 963 | 1032 | 1118 | 994 | 992 | 972 | 1011.84 | 1038 | 986 | 0.952 | -0.071 | 0.3823132 | 0.55640702 | 0.0021 | 0.0048 | 0.0038 | 0.0038 | TRUE | 0.0546 |
| Aknad1 | 1 | 4 | 2 | 1 | 1 | 1 | 1 | 4 | 2 | 1 | 1 | 1 | 1.72 | 2 | 1 | 0.441 | -1.18 | 0.43138537 | NA | 0 | 1.383 | 0.9161 | 0.9161 | TRUE | 0.1962 |
| Akp3 | 3 | 2 | 4 | 1 | 1 | 1 | 2 | 2 | 4 | 1 | 1 | 1 | 1.98 | 3 | 1 | 0.356 | -1.489 | 0.28709552 | NA | 0 | 1.199 | 0.7429 | 0.7429 | TRUE | 0.2011 |
| Akr1a1 | 14920 | 10158 | 9744 | 11721 | 10270 | 11861 | 11588 | 10805 | 10514 | 11667 | 11586 | 11709 | 11311.28 | 10969 | 11654 | 1.062 | 0.087 | 0.10589927 | 0.2307345 | 0.0012 | 0.0026 | 0.002 | 0.002 | TRUE | 0.03 |
| Akr1b10 | 459 | 366 | 280 | 306 | 252 | 264 | 356 | 389 | 302 | 305 | 284 | 261 | 316.23 | 349 | 283 | 0.81 | -0.304 | 0.021276 | 0.07231489 | 0.008 | 0.0099 | 0.0093 | 0.0093 | TRUE | 0.1521 |
| Akr1b3 | 0 | 0 | 0 | 0 | 0 | 0 | 0 | 0 | 0 | 0 | 0 | 0 | 0 | 0 | 0 | NA | NA | NA | NA | NA | NA | NA | NA | NA | NA |
| Akr1b7 | 2 | 12 | 16 | 21 | 20 | 28 | 2 | 13 | 17 | 21 | 23 | 28 | 17.11 | 11 | 24 | 2.322 | 1.215 | 0.04373529 | 0.12213922 | 0.3868 | 0.1409 | 0.1925 | 0.1925 | TRUE | 1.9496 |
| Akr1b8 | 1518 | 941 | 929 | 1196 | 1126 | 1272 | 1179 | 1001 | 1002 | 1190 | 1270 | 1256 | 1149.78 | 1061 | 1239 | 1.165 | 0.221 | 0.01010664 | 0.04165893 | 0.0043 | 0.0045 | 0.0044 | 0.0044 | TRUE | 0.1179 |
| Akr1c12 | 12 | 8 | 3 | 3 | 2 | 2 | 9 | 9 | 3 | 3 | 2 | 2 | 4.71 | 7 | 2 | 0.34 | -1.558 | 0.1000212 | NA | 0 | 0.5052 | 0.3636 | 0.3636 | TRUE | 0.5423 |
| Akr1c13 | 68 | 43 | 37 | 25 | 13 | 11 | 53 | 46 | 40 | 25 | 15 | 11 | 31.48 | 46 | 17 | 0.363 | -1.46 | 0.0001734 | 0.00191087 | 0.0308 | 0.0777 | 0.0678 | 0.0678 | TRUE | 0.8968 |
| Akr1c14 | 14 | 12 | 7 | 9 | 4 | 5 | 11 | 13 | 8 | 9 | 5 | 5 | 8.27 | 11 | 6 | 0.592 | -0.757 | 0.26689253 | NA | 0 | 0.2891 | 0.2045 | 0.2045 | TRUE | 0.396 |
| Akr1c18 | 6 | 9 | 14 | 2 | 3 | 0 | 5 | 10 | 15 | 2 | 3 | 0 | 5.79 | 10 | 2 | 0.181 | -2.467 | 0.01403353 | NA | 0.2676 | 0.4121 | 0.3809 | 0.3809 | TRUE | 0.2131 |
| Akr1c19 | 2 | 0 | 0 | 1 | 0 | 0 | 2 | 0 | 0 | 1 | 0 | 0 | 0.42 | 1 | 0 | 0.681 | -0.553 | 0.8627148 | NA | 0 | 5.5819 | 5.4906 | 5.4906 | TRUE | 0.9727 |
| Akr1c20 | 0 | 0 | 0 | 0 | 0 | 0 | 0 | 0 | 0 | 0 | 0 | 0 | 0 | 0 | 0 | NA | NA | NA | NA | NA | NA | NA | NA | NA | NA |
| Akr1c21 | 0 | 1 | 1 | 0 | 0 | 0 | 0 | 1 | 1 | 0 | 0 | 0 | 0.36 | 1 | 0 | 0.271 | -1.881 | 0.5755355 | NA | 0 | 6.6391 | 6.4428 | 6.4428 | TRUE | 0.3513 |
| Akr1c6 | 0 | 0 | 0 | 0 | 0 | 0 | 0 | 0 | 0 | 0 | 0 | 0 | 0 | 0 | 0 | NA | NA | NA | NA | NA | NA | NA | NA | NA | NA |
| Akr1cl | 93 | 42 | 20 | 37 | 26 | 14 | 72 | 45 | 22 | 37 | 29 | 14 | 36.41 | 46 | 27 | 0.572 | -0.807 | 0.07560802 | 0.18125887 | 0.2472 | 0.0675 | 0.1187 | 0.1187 | TRUE | 0.1494 |
| Akr1d1 | 0 | 0 | 0 | 0 | 0 | 0 | 0 | 0 | 0 | 0 | 0 | 0 | 0 | 0 | 0 | NA | NA | NA | NA | NA | NA | NA | NA | NA | NA |
| Akr1e1 | 721 | 473 | 584 | 610 | 526 | 547 | 560 | 503 | 630 | 607 | 593 | 540 | 572.3 | 564 | 580 | 1.028 | 0.04 | 0.70933099 | 0.82473504 | 0.0063 | 0.0066 | 0.0065 | 0.0065 | TRUE | 0.1164 |
| Akr7a5 | 1533 | 1239 | 1323 | 1448 | 1343 | 1435 | 1191 | 1318 | 1428 | 1441 | 1515 | 1417 | 1384.83 | 1312 | 1458 | 1.112 | 0.154 | 0.06063637 | 0.15437755 | 0.0041 | 0.0041 | 0.0041 | 0.0041 | TRUE | 0.0818 |
| Akt1 | 3264 | 2517 | 2546 | 2459 | 2338 | 2716 | 2535 | 2677 | 2747 | 2448 | 2638 | 2681 | 2620.95 | 2653 | 2589 | 0.976 | -0.035 | 0.59321593 | 0.74070371 | 0.0017 | 0.0033 | 0.0027 | 0.0027 | TRUE | 0.0278 |
| Akt1s1 | 2684 | 1878 | 1525 | 1603 | 1596 | 1784 | 2085 | 1998 | 1645 | 1596 | 1800 | 1761 | 1814.13 | 1909 | 1719 | 0.899 | -0.153 | 0.09842931 | 0.21859076 | 0.0086 | 0.0037 | 0.0056 | 0.0056 | TRUE | 0.175 |
| Akt2 | 3971 | 2953 | 2944 | 2845 | 2494 | 2892 | 3084 | 3141 | 3177 | 2832 | 2813 | 2855 | 2983.66 | 3134 | 2833 | 0.904 | -0.145 | 0.00753322 | 0.03355324 | 0 | 0.0032 | 0.0018 | 0.0018 | TRUE | 0.0024 |
| Akt3 | 2915 | 2085 | 2313 | 2135 | 1843 | 2048 | 2264 | 2218 | 2496 | 2125 | 2079 | 2022 | 2200.57 | 2326 | 2075 | 0.893 | -0.164 | 0.01604455 | 0.05869586 | 0.0017 | 0.0035 | 0.0029 | 0.0029 | TRUE | 0.0486 |
| Aktip | 4662 | 3576 | 3906 | 4402 | 3823 | 4476 | 3621 | 3804 | 4215 | 4382 | 4313 | 4419 | 4125.34 | 3880 | 4371 | 1.127 | 0.173 | 0.00931072 | 0.03931405 | 0.0028 | 0.003 | 0.0029 | 0.0029 | TRUE | 0.0691 |
| Alad | 2917 | 2037 | 1841 | 1916 | 1667 | 1880 | 2266 | 2167 | 1986 | 1907 | 1881 | 1856 | 2010.37 | 2140 | 1881 | 0.879 | -0.187 | 0.00692768 | 0.03148598 | 0.0019 | 0.0036 | 0.0029 | 0.0029 | TRUE | 0.0471 |
| Alas1 | 2841 | 2071 | 1856 | 2029 | 1766 | 2091 | 2206 | 2203 | 2003 | 2020 | 1992 | 2064 | 2081.34 | 2137 | 2025 | 0.947 | -0.078 | 0.23563849 | 0.40209897 | 0.0012 | 0.0036 | 0.0027 | 0.0027 | TRUE | 0.0362 |
| Alas2 | 6 | 2 | 0 | 1 | 2 | 1 | 5 | 2 | 0 | 1 | 2 | 1 | 1.84 | 2 | 1 | 0.599 | -0.74 | 0.62136133 | NA | 0 | 1.2921 | 1.0419 | 1.0419 | TRUE | 0.1658 |
| Alb | 0 | 0 | 0 | 0 | 0 | 0 | 0 | 0 | 0 | 0 | 0 | 0 | 0 | 0 | 0 | NA | NA | NA | NA | NA | NA | NA | NA | NA | NA |
| Alcam | 74 | 62 | 61 | 57 | 31 | 41 | 57 | 66 | 66 | 57 | 35 | 40 | 53.57 | 63 | 44 | 0.703 | -0.509 | 0.06935526 | 0.17056 | 0.0095 | 0.0467 | 0.0372 | 0.0372 | TRUE | 0.497 |
| Aldh16a1 | 2607 | 1949 | 1766 | 1814 | 1672 | 1782 | 2025 | 2073 | 1905 | 1806 | 1886 | 1759 | 1909.05 | 2001 | 1817 | 0.908 | -0.14 | 0.03502199 | 0.10389666 | 0.001 | 0.0037 | 0.0027 | 0.0027 | TRUE | 0.0206 |
| Aldh18a1 | 3107 | 2307 | 2139 | 2067 | 1821 | 2082 | 2413 | 2454 | 2308 | 2057 | 2054 | 2055 | 2223.66 | 2392 | 2055 | 0.859 | -0.219 | 0.00025439 | 0.00256418 | 0 | 0.0035 | 0.0021 | 0.0021 | TRUE | 0.0111 |
| Aldh1a1 | 15 | 7 | 10 | 2 | 2 | 2 | 12 | 7 | 11 | 2 | 2 | 2 | 6.02 | 10 | 2 | 0.207 | -2.274 | 0.01017257 | NA | 0 | 0.3963 | 0.266 | 0.266 | TRUE | 0.1594 |
| Aldh1a2 | 319 | 217 | 189 | 177 | 168 | 196 | 248 | 231 | 204 | 176 | 190 | 193 | 206.95 | 228 | 186 | 0.817 | -0.292 | 0.04455018 | 0.1238294 | 0 | 0.0139 | 0.0103 | 0.0103 | TRUE | 0.0884 |
| Aldh1a3 | 31 | 39 | 35 | 40 | 28 | 35 | 24 | 41 | 38 | 40 | 32 | 35 | 34.88 | 34 | 36 | 1.038 | 0.054 | 0.87630587 | 0.93096567 | 0 | 0.0704 | 0.0565 | 0.0565 | TRUE | 0.5915 |
| Aldh1a7 | 28 | 20 | 15 | 5 | 12 | 5 | 22 | 21 | 16 | 5 | 14 | 5 | 13.78 | 20 | 8 | 0.389 | -1.362 | 0.01731333 | NA | 0.0217 | 0.1745 | 0.1445 | 0.1445 | TRUE | 1.064 |
| Aldh1b1 | 883 | 648 | 685 | 699 | 608 | 672 | 686 | 689 | 739 | 696 | 686 | 663 | 693.2 | 705 | 682 | 0.968 | -0.047 | 0.58296894 | 0.73290307 | 0 | 0.0058 | 0.0038 | 0.0038 | TRUE | 0.021 |
| Aldh1l1 | 802 | 668 | 940 | 1283 | 1157 | 1356 | 623 | 711 | 1014 | 1277 | 1305 | 1339 | 1044.76 | 783 | 1307 | 1.673 | 0.743 | 5.08E-08 | 2.13E-06 | 0.0332 | 0.0047 | 0.0124 | 0.0124 | TRUE | 0.7922 |
| Aldh1l2 | 128 | 108 | 87 | 75 | 68 | 84 | 99 | 115 | 94 | 75 | 77 | 83 | 90.41 | 103 | 78 | 0.761 | -0.394 | 0.05819905 | 0.15039577 | 0 | 0.0286 | 0.0198 | 0.0198 | TRUE | 0.0996 |
| Aldh2 | 7866 | 6064 | 7109 | 9008 | 8509 | 9700 | 6109 | 6450 | 7671 | 8967 | 9599 | 9575 | 8061.82 | 6743 | 9380 | 1.391 | 0.477 | 2.95E-09 | 1.71E-07 | 0.0079 | 0.0027 | 0.0045 | 0.0045 | TRUE | 0.1763 |
| Aldh3a1 | 60 | 41 | 52 | 49 | 55 | 52 | 47 | 44 | 56 | 49 | 62 | 51 | 51.41 | 49 | 54 | 1.108 | 0.148 | 0.58948761 | 0.7382741 | 0 | 0.0485 | 0.0345 | 0.0345 | TRUE | 0.1311 |
| Aldh3a2 | 3200 | 2288 | 2262 | 2199 | 1886 | 2301 | 2485 | 2434 | 2441 | 2189 | 2128 | 2271 | 2324.6 | 2453 | 2196 | 0.895 | -0.16 | 0.00737386 | 0.03305908 | 0 | 0.0034 | 0.0021 | 0.0021 | TRUE | 0.011 |
| Aldh3b1 | 80 | 56 | 71 | 115 | 75 | 93 | 62 | 60 | 77 | 114 | 85 | 92 | 81.53 | 66 | 97 | 1.473 | 0.559 | 0.01506635 | 0.05600605 | 0.0097 | 0.0315 | 0.0254 | 0.0254 | TRUE | 0.2441 |
| Aldh3b2 | 25 | 16 | 21 | 19 | 16 | 22 | 19 | 17 | 23 | 19 | 18 | 22 | 19.63 | 20 | 20 | 0.995 | -0.007 | 0.98705338 | 0.99373352 | 0 | 0.1232 | 0.0808 | 0.0808 | TRUE | 0.0831 |
| Aldh4a1 | 4887 | 3683 | 3909 | 4337 | 3858 | 4479 | 3796 | 3918 | 4218 | 4317 | 4352 | 4421 | 4170.27 | 3977 | 4363 | 1.098 | 0.135 | 0.02335286 | 0.07740779 | 0.0012 | 0.003 | 0.0023 | 0.0023 | TRUE | 0.034 |
| Aldh5a1 | 2304 | 1708 | 1705 | 2001 | 1672 | 1957 | 1789 | 1817 | 1840 | 1992 | 1886 | 1932 | 1875.95 | 1815 | 1937 | 1.067 | 0.094 | 0.12871071 | 0.26519564 | 0 | 0.0037 | 0.0022 | 0.0022 | TRUE | 0.0075 |
| Aldh6a1 | 3389 | 2569 | 3153 | 3829 | 3461 | 3980 | 2632 | 2733 | 3402 | 3811 | 3904 | 3929 | 3401.9 | 2922 | 3881 | 1.329 | 0.41 | 3.37E-06 | 7.68E-05 | 0.0099 | 0.0031 | 0.0053 | 0.0053 | TRUE | 0.2494 |
| Aldh7a1 | 5716 | 4138 | 4392 | 4744 | 4092 | 5056 | 4439 | 4402 | 4739 | 4722 | 4616 | 4991 | 4651.54 | 4527 | 4776 | 1.056 | 0.078 | 0.18691197 | 0.34364985 | 0.0013 | 0.0029 | 0.0023 | 0.0023 | TRUE | 0.0203 |
| Aldh8a1 | 1 | 2 | 3 | 0 | 3 | 1 | 1 | 2 | 3 | 0 | 3 | 1 | 1.75 | 2 | 1 | 0.71 | -0.494 | 0.7466262 | NA | 0.2746 | 1.3551 | 1.0773 | 1.0773 | TRUE | 0.6372 |
| Aldh9a1 | 6406 | 4833 | 5307 | 5526 | 4903 | 5797 | 4975 | 5141 | 5726 | 5501 | 5531 | 5723 | 5432.75 | 5281 | 5585 | 1.058 | 0.081 | 0.20863265 | 0.3702951 | 0.0027 | 0.0029 | 0.0028 | 0.0028 | TRUE | 0.0661 |
| Aldoa | 104176 | 83317 | 109329 | 114124 | 105438 | 120526 | 80909 | 88623 | 117965 | 113599 | 118945 | 118978 | 106503.17 | 95832 | 117174 | 1.223 | 0.29 | 0.09045016 | 0.20537001 | 0.0211 | 0.0024 | 0.0079 | 0.0211 | TRUE | 0.4999 |
| Aldoart1 | 21 | 16 | 22 | 28 | 20 | 28 | 16 | 17 | 24 | 28 | 23 | 28 | 22.52 | 19 | 26 | 1.38 | 0.464 | 0.25435523 | 0.42366849 | 0 | 0.1077 | 0.0743 | 0.0743 | TRUE | 0.2308 |
| Aldoart2 | 15 | 2 | 6 | 8 | 4 | 7 | 12 | 2 | 6 | 8 | 5 | 7 | 6.61 | 7 | 7 | 0.94 | -0.09 | 0.91027078 | NA | 0.1725 | 0.3612 | 0.3082 | 0.3082 | TRUE | 0.2384 |
| Aldob | 0 | 0 | 0 | 2 | 0 | 0 | 0 | 0 | 0 | 2 | 0 | 0 | 0.33 | 0 | 1 | 3.77 | 1.915 | 0.62344813 | NA | 0 | 7.1455 | 9.1834 | 9.1834 | TRUE | 0.9739 |
| Aldoc | 2983 | 2611 | 4103 | 4520 | 4470 | 5020 | 2317 | 2777 | 4427 | 4499 | 5043 | 4956 | 4003.08 | 3174 | 4833 | 1.523 | 0.607 | 0.0398501 | 0.11423534 | 0.0626 | 0.003 | 0.0174 | 0.0626 | TRUE | 1.4498 |
| Alg1 | 1176 | 881 | 951 | 965 | 852 | 974 | 913 | 937 | 1026 | 961 | 961 | 961 | 959.96 | 959 | 961 | 1.003 | 0.005 | 0.94999263 | 0.97253743 | 9.00E-04 | 0.0049 | 0.0035 | 0.0035 | TRUE | 0.0442 |
| Alg10b | 4013 | 2799 | 2893 | 2983 | 2534 | 2869 | 3117 | 2977 | 3122 | 2969 | 2859 | 2832 | 2979.26 | 3072 | 2887 | 0.94 | -0.09 | 0.11682922 | 0.24767726 | 3.00E-04 | 0.0032 | 0.002 | 0.002 | TRUE | 0.0087 |
| Alg11 | 4130 | 2994 | 2845 | 3057 | 2912 | 2958 | 3208 | 3185 | 3070 | 3043 | 3285 | 2920 | 3118.33 | 3154 | 3083 | 0.977 | -0.034 | 0.5961167 | 0.74235543 | 0.0015 | 0.0032 | 0.0026 | 0.0026 | TRUE | 0.0397 |

| | | | | | | | | | | | | | | | | | | | | | | | | | |
|---|---|---|---|---|---|---|---|---|---|---|---|---|---|---|---|---|---|---|---|---|---|---|---|---|---|
| Alg12 | 662 | 436 | 501 | 590 | 438 | 554 | 514 | 464 | 541 | 587 | 494 | 547 | 524.46 | 506 | 543 | 1.073 | 0.102 | 0.33876295 | 0.51295317 | 0.0046 | 0.0069 | 0.0062 | 0.0062 | TRUE | 0.07 |
| Alg13 | 13373 | 10811 | 12187 | 10672 | 10902 | 12624 | 10386 | 11500 | 13150 | 10623 | 12299 | 12462 | 11736.46 | 11679 | 11795 | 1.01 | 0.014 | 0.86855552 | 0.92747584 | 0.0086 | 0.0026 | 0.0054 | 0.0054 | TRUE | 0.1484 |
| Alg14 | 835 | 532 | 474 | 546 | 506 | 655 | 649 | 566 | 511 | 543 | 571 | 647 | 581.12 | 575 | 587 | 1.018 | 0.026 | 0.81920341 | 0.89620164 | 0.0097 | 0.0065 | 0.0075 | 0.0075 | TRUE | 0.1517 |
| Alg2 | 1817 | 1372 | 1164 | 1202 | 1205 | 1307 | 1411 | 1459 | 1256 | 1196 | 1359 | 1290 | 1328.76 | 1375 | 1282 | 0.931 | -0.103 | 0.21684194 | 0.37958627 | 0.0039 | 0.0042 | 0.0042 | 0.0042 | TRUE | 0.0679 |
| Alg3 | 1520 | 1129 | 1205 | 1330 | 1206 | 1411 | 1181 | 1201 | 1300 | 1324 | 1360 | 1393 | 1293.14 | 1227 | 1359 | 1.108 | 0.149 | 0.04111125 | 0.1166342 | 9.00E-04 | 0.0043 | 0.003 | 0.003 | TRUE | 0.0319 |
| Alg5 | 1009 | 747 | 741 | 821 | 678 | 734 | 784 | 795 | 800 | 817 | 765 | 725 | 780.73 | 793 | 769 | 0.97 | -0.043 | 0.6064334 | 0.7508545 | 0 | 0.0055 | 0.0038 | 0.0038 | TRUE | 0.0366 |
| Alg6 | 1932 | 1338 | 1360 | 1579 | 1288 | 1462 | 1501 | 1423 | 1467 | 1572 | 1453 | 1443 | 1476.52 | 1464 | 1489 | 1.017 | 0.025 | 0.72493283 | 0.83533255 | 8.00E-04 | 0.004 | 0.0028 | 0.0028 | TRUE | 0.0285 |
| Alg8 | 721 | 509 | 579 | 599 | 556 | 611 | 560 | 541 | 625 | 596 | 627 | 603 | 592.12 | 575 | 609 | 1.059 | 0.082 | 0.38555791 | 0.55936317 | 0.0013 | 0.0064 | 0.0048 | 0.0048 | TRUE | 0.0635 |
| Alg9 | 2019 | 1538 | 1559 | 1723 | 1418 | 1608 | 1568 | 1636 | 1682 | 1715 | 1600 | 1587 | 1631.37 | 1629 | 1634 | 1.004 | 0.006 | 0.93150698 | 0.96216772 | 0.001 | 0.0039 | 0.0028 | 0.0028 | TRUE | 0.0229 |
| Alk | 99 | 75 | 62 | 67 | 69 | 63 | 77 | 80 | 67 | 67 | 78 | 62 | 71.71 | 75 | 69 | 0.921 | -0.118 | 0.61145941 | 0.75404219 | 0 | 0.0355 | 0.0249 | 0.0249 | TRUE | 0.1086 |
| Alkbh1 | 1181 | 859 | 1002 | 1098 | 1032 | 1192 | 917 | 914 | 1081 | 1093 | 1164 | 1177 | 1057.65 | 971 | 1145 | 1.18 | 0.239 | 0.00633297 | 0.02934103 | 0.0042 | 0.0047 | 0.0046 | 0.0046 | TRUE | 0.1159 |
| Alkbh2 | 875 | 671 | 719 | 725 | 673 | 794 | 680 | 714 | 776 | 722 | 759 | 784 | 738.96 | 723 | 755 | 1.046 | 0.064 | 0.47003558 | 0.63909994 | 0.0019 | 0.0056 | 0.0044 | 0.0044 | TRUE | 0.0477 |
| Alkbh3 | 504 | 402 | 320 | 362 | 318 | 363 | 391 | 428 | 345 | 360 | 359 | 358 | 373.62 | 388 | 359 | 0.925 | -0.112 | 0.33175563 | 0.50595211 | 0.0028 | 0.0088 | 0.007 | 0.007 | TRUE | 0.1012 |
| Alkbh4 | 606 | 465 | 433 | 492 | 446 | 503 | 471 | 495 | 467 | 490 | 503 | 497 | 486.98 | 478 | 497 | 1.04 | 0.057 | 0.5541408 | 0.70983375 | 0 | 0.0073 | 0.0045 | 0.0045 | TRUE | 0.011 |
| Alkbh5 | 16020 | 12987 | 14507 | 14118 | 12558 | 14603 | 12442 | 13814 | 15653 | 14053 | 14167 | 14415 | 14090.71 | 13970 | 14212 | 1.018 | 0.025 | 0.74149086 | 0.8460229 | 0.0053 | 0.0026 | 0.0041 | 0.0041 | TRUE | 0.1358 |
| Alkbh6 | 383 | 269 | 268 | 268 | 248 | 292 | 297 | 286 | 289 | 267 | 280 | 288 | 284.59 | 291 | 278 | 0.956 | -0.065 | 0.58591386 | 0.73524731 | 0 | 0.0108 | 0.0068 | 0.0068 | TRUE | 0.0151 |
| Alkbh7 | 1323 | 1011 | 947 | 1191 | 1046 | 1178 | 1028 | 1075 | 1022 | 1186 | 1180 | 1163 | 1108.85 | 1042 | 1176 | 1.13 | 0.176 | 0.01382639 | 0.05264267 | 0 | 0.0046 | 0.0028 | 0.0028 | TRUE | 0.0094 |
| Alkbh8 | 1058 | 811 | 806 | 824 | 722 | 864 | 822 | 863 | 870 | 820 | 814 | 853 | 840.27 | 852 | 829 | 0.975 | -0.036 | 0.64625341 | 0.78007769 | 0 | 0.0052 | 0.0033 | 0.0033 | TRUE | 0.0117 |
| Allc | 2 | 1 | 0 | 2 | 1 | 0 | 2 | 1 | 0 | 2 | 1 | 0 | 0.96 | 1 | 1 | 1.153 | 0.206 | 0.91844446 | NA | 0 | 2.4816 | 1.8799 | 1.8799 | TRUE | 0.2473 |
| Alms1 | 4519 | 3248 | 3052 | 3538 | 3172 | 3771 | 3510 | 3455 | 3293 | 3522 | 3578 | 3723 | 3513.38 | 3419 | 3608 | 1.055 | 0.077 | 0.18021202 | 0.33508136 | 7.00E-04 | 0.0031 | 0.0021 | 0.0021 | TRUE | 0.0125 |
| Alms1-ps2 | 3 | 0 | 1 | 1 | 1 | 2 | 2 | 0 | 1 | 1 | 1 | 2 | 1.25 | 1 | 1 | 1.157 | 0.211 | 0.90134218 | NA | 0 | 1.8968 | 1.2998 | 1.2998 | TRUE | 0.3005 |
| Alox12 | 136 | 95 | 71 | 128 | 90 | 120 | 106 | 101 | 77 | 127 | 102 | 118 | 105.11 | 95 | 116 | 1.223 | 0.29 | 0.15786514 | 0.30682166 | 0.01 | 0.025 | 0.0209 | 0.0209 | TRUE | 0.2503 |
| Alox12b | 105 | 80 | 106 | 143 | 136 | 146 | 82 | 85 | 114 | 142 | 153 | 144 | 120.15 | 94 | 146 | 1.573 | 0.653 | 0.00071451 | 0.00565165 | 0.0062 | 0.0222 | 0.0181 | 0.0181 | TRUE | 0.3463 |
| Alox12e | 19 | 13 | 12 | 9 | 5 | 5 | 15 | 14 | 13 | 9 | 6 | 5 | 10.18 | 14 | 7 | 0.47 | -1.088 | 0.07665046 | NA | 0 | 0.2353 | 0.1579 | 0.1579 | TRUE | 0.2809 |
| Alox15 | 200 | 133 | 90 | 144 | 120 | 135 | 155 | 141 | 97 | 143 | 135 | 133 | 134.31 | 131 | 137 | 1.041 | 0.057 | 0.77094444 | 0.86441946 | 0.0217 | 0.0201 | 0.0205 | 0.0205 | TRUE | 0.5125 |
| Alox5 | 193 | 105 | 122 | 129 | 131 | 135 | 150 | 112 | 132 | 128 | 148 | 133 | 133.78 | 131 | 136 | 1.035 | 0.05 | 0.78198522 | 0.87149433 | 0.0065 | 0.0201 | 0.0162 | 0.0162 | TRUE | 0.1781 |
| Alox5ap | 72 | 34 | 36 | 74 | 44 | 71 | 56 | 36 | 39 | 74 | 50 | 70 | 54.05 | 44 | 65 | 1.466 | 0.552 | 0.05821244 | 0.15040317 | 0.0319 | 0.0463 | 0.0423 | 0.0423 | TRUE | 0.5412 |
| Alox8 | 54 | 39 | 29 | 42 | 41 | 39 | 42 | 41 | 31 | 42 | 46 | 38 | 40.21 | 38 | 42 | 1.097 | 0.134 | 0.66224458 | 0.79149275 | 0 | 0.0614 | 0.0427 | 0.0427 | TRUE | 0.1732 |
| Aloxe3 | 97 | 67 | 113 | 103 | 101 | 134 | 75 | 71 | 122 | 103 | 114 | 132 | 102.88 | 89 | 116 | 1.305 | 0.384 | 0.1026555 | 0.22548871 | 0.0429 | 0.0255 | 0.0301 | 0.0301 | TRUE | 0.9215 |
| Alpi | 0 | 0 | 0 | 0 | 0 | 0 | 0 | 0 | 0 | 0 | 0 | 0 | 0 | 0 | 0 | NA | NA | NA | NA | NA | NA | NA | NA | NA | NA |
| Alpk1 | 57 | 44 | 29 | 36 | 31 | 32 | 44 | 47 | 31 | 36 | 35 | 32 | 37.46 | 41 | 34 | 0.834 | -0.262 | 0.41415916 | 0.58679114 | 0 | 0.0657 | 0.0475 | 0.0475 | TRUE | 0.2904 |
| Alpk2 | 4 | 3 | 1 | 3 | 0 | 1 | 3 | 3 | 1 | 3 | 0 | 1 | 1.89 | 2 | 1 | 0.544 | -0.879 | 0.54623101 | NA | 0 | 1.2553 | 0.9497 | 0.9497 | TRUE | 0.7356 |
| Alpk3 | 764 | 658 | 670 | 698 | 706 | 736 | 593 | 700 | 723 | 695 | 796 | 727 | 705.66 | 672 | 739 | 1.102 | 0.14 | 0.16982971 | 0.32137906 | 0.0066 | 0.0058 | 0.0061 | 0.0061 | TRUE | 0.1305 |
| Alpl | 6768 | 4767 | 5010 | 4786 | 4434 | 4961 | 5256 | 5071 | 5406 | 4764 | 5002 | 4897 | 5066 | 5244 | 4888 | 0.932 | -0.102 | 0.05949788 | 0.15237471 | 5.00E-04 | 0.0029 | 0.0019 | 0.0019 | TRUE | 0.0101 |
| Alppl2 | 154 | 111 | 71 | 57 | 53 | 53 | 120 | 118 | 77 | 57 | 60 | 52 | 80.52 | 105 | 56 | 0.535 | -0.903 | 0.00023085 | 0.00238752 | 0.024 | 0.0319 | 0.0295 | 0.0295 | TRUE | 0.5213 |
| Als2 | 8676 | 5889 | 5372 | 5200 | 4690 | 5363 | 6738 | 6264 | 5796 | 5176 | 5291 | 5294 | 5759.94 | 6266 | 5254 | 0.838 | -0.255 | 7.64E-05 | 0.00098107 | 0.0026 | 0.0028 | 0.0028 | 0.0028 | TRUE | 0.0535 |
| Als2cl | 1564 | 1228 | 1122 | 1336 | 1247 | 1361 | 1215 | 1306 | 1211 | 1330 | 1407 | 1344 | 1301.94 | 1244 | 1360 | 1.094 | 0.129 | 0.07102022 | 0.17334221 | 5.00E-04 | 0.0042 | 0.0029 | 0.0029 | TRUE | 0.0226 |
| Als2cr11 | 35 | 21 | 26 | 15 | 12 | 17 | 27 | 22 | 28 | 15 | 14 | 17 | 20.47 | 26 | 15 | 0.584 | -0.777 | 0.06813273 | 0.16796046 | 0 | 0.1182 | 0.0778 | 0.0778 | TRUE | 0.0821 |
| Als2cr12 | 8 | 6 | 4 | 3 | 3 | 4 | 6 | 6 | 4 | 3 | 3 | 4 | 4.54 | 5 | 3 | 0.608 | -0.718 | 0.41794659 | NA | 0 | 0.5246 | 0.33 | 0.33 | TRUE | 0.0848 |
| Alx1 | 14 | 16 | 18 | 16 | 13 | 16 | 11 | 17 | 19 | 16 | 15 | 16 | 15.62 | 16 | 16 | 0.993 | -0.01 | 0.98393188 | NA | 0 | 0.1542 | 0.1081 | 0.1081 | TRUE | 0.4437 |
| Alx3 | 11 | 4 | 1 | 8 | 5 | 7 | 9 | 4 | 1 | 8 | 6 | 7 | 5.73 | 5 | 7 | 1.433 | 0.52 | 0.5414126 | NA | 0.13 | 0.4159 | 0.345 | 0.345 | TRUE | 0.4412 |
| Alx4 | 10 | 4 | 5 | 9 | 4 | 7 | 8 | 4 | 5 | 9 | 5 | 7 | 6.3 | 6 | 7 | 1.161 | 0.215 | 0.77769119 | NA | 0 | 0.3787 | 0.2592 | 0.2592 | TRUE | 0.2387 |
| Alyref | 8053 | 6215 | 5171 | 5314 | 4916 | 5544 | 6254 | 6611 | 5579 | 5290 | 5546 | 5473 | 5792.13 | 6148 | 5436 | 0.884 | -0.178 | 0.00953638 | 0.03997169 | 0.0037 | 0.0028 | 0.0032 | 0.0032 | TRUE | 0.0792 |
| Alyref2 | 143 | 101 | 90 | 102 | 72 | 97 | 111 | 107 | 97 | 102 | 81 | 96 | 99.02 | 105 | 93 | 0.882 | -0.18 | 0.36671977 | 0.5414766 | 0 | 0.0264 | 0.0186 | 0.0186 | TRUE | 0.1084 |
| Amacr | 939 | 663 | 633 | 749 | 740 | 709 | 729 | 705 | 683 | 746 | 835 | 700 | 732.96 | 706 | 760 | 1.075 | 0.104 | 0.2607933 | 0.43080723 | 0.0032 | 0.0057 | 0.0048 | 0.0048 | TRUE | 0.0859 |
| Ambn | 0 | 1 | 0 | 0 | 0 | 0 | 0 | 1 | 0 | 0 | 0 | 0 | 0.18 | 0 | 0 | 0.552 | -0.857 | 0.8336548 | NA | 0 | 13.3714 | 10 | 10 | TRUE | 0.5216 |
| Ambp | 0 | 0 | 1 | 0 | 1 | 0 | 0 | 0 | 1 | 0 | 1 | 0 | 0.37 | 0 | 0 | 1.075 | 0.105 | 0.97514588 | NA | 0 | 6.4454 | 6.1578 | 6.1578 | TRUE | 0.5365 |
| Ambra1 | 4241 | 3001 | 2691 | 2855 | 2658 | 2848 | 3294 | 3192 | 2904 | 2842 | 2998 | 2811 | 3006.88 | 3130 | 2884 | 0.921 | -0.119 | 0.07542693 | 0.18094614 | 0.0022 | 0.0032 | 0.0029 | 0.0029 | TRUE | 0.0482 |
| Amd1 | 0 | 0 | 0 | 0 | 0 | 0 | 0 | 0 | 0 | 0 | 0 | 0 | 0 | 0 | 0 | NA | NA | NA | NA | NA | NA | NA | NA | NA | NA |
| Amd2 | 0 | 0 | 0 | 0 | 0 | 0 | 0 | 0 | 0 | 0 | 0 | 0 | 0 | 0 | 0 | NA | NA | NA | NA | NA | NA | NA | NA | NA | NA |
| Amdhd1 | 43 | 40 | 48 | 54 | 35 | 39 | 33 | 43 | 52 | 54 | 39 | 38 | 43.24 | 43 | 44 | 1.041 | 0.057 | 0.85427117 | 0.91805247 | 0.0212 | 0.0572 | 0.047 | 0.047 | TRUE | 0.2181 |
| Amdhd2 | 1071 | 805 | 770 | 833 | 702 | 806 | 832 | 856 | 831 | 829 | 792 | 796 | 822.61 | 840 | 806 | 0.96 | -0.059 | 0.45277689 | 0.62256278 | 0 | 0.0053 | 0.0032 | 0.0032 | TRUE | 0.0079 |
| Amelx | 0 | 0 | 0 | 0 | 0 | 0 | 0 | 0 | 0 | 0 | 0 | 0 | 0 | 0 | 0 | NA | NA | NA | NA | NA | NA | NA | NA | NA | NA |
| Amer1 | 1444 | 1160 | 968 | 975 | 929 | 958 | 1121 | 1234 | 1044 | 971 | 1048 | 946 | 1060.68 | 1133 | 988 | 0.872 | -0.198 | 0.02149291 | 0.07284496 | 0.0036 | 0.0047 | 0.0044 | 0.0044 | TRUE | 0.0702 |
| Amer2 | 24 | 19 | 13 | 17 | 8 | 19 | 19 | 20 | 14 | 17 | 9 | 19 | 16.26 | 18 | 15 | 0.849 | -0.236 | 0.63066929 | NA | 0 | 0.1482 | 0.1119 | 0.1119 | TRUE | 0.4832 |
| Amer3 | 5 | 1 | 3 | 2 | 1 | 4 | 4 | 1 | 3 | 2 | 1 | 4 | 2.54 | 3 | 2 | 0.859 | -0.219 | 0.8553518 | NA | 0 | 0.9348 | 0.6503 | 0.6503 | TRUE | 0.3732 |
| Amfr | 5558 | 3957 | 4170 | 5449 | 4595 | 5112 | 4317 | 4209 | 4499 | 5424 | 5184 | 5046 | 4779.83 | 4342 | 5218 | 1.202 | 0.265 | 3.03E-06 | 6.98E-05 | 9.00E-04 | 0.0029 | 0.0021 | 0.0021 | TRUE | 0.0146 |
| Amh | 14 | 7 | 16 | 16 | 11 | 10 | 11 | 7 | 17 | 16 | 12 | 10 | 12.3 | 12 | 13 | 1.078 | 0.108 | 0.84815287 | NA | 0 | 0.1951 | 0.1495 | 0.1495 | TRUE | 0.4369 |
| Amhr2 | 387 | 310 | 328 | 382 | 358 | 400 | 301 | 330 | 354 | 380 | 404 | 395 | 360.53 | 328 | 393 | 1.201 | 0.264 | 0.02064831 | 0.07082464 | 0 | 0.009 | 0.0066 | 0.0066 | TRUE | 0.0665 |
| Amica1 | 8 | 7 | 5 | 5 | 2 | 1 | 6 | 7 | 5 | 5 | 2 | 1 | 4.55 | 6 | 3 | 0.433 | -1.208 | 0.20020942 | NA | 0 | 0.5238 | 0.3766 | 0.3766 | TRUE | 0.6379 |
| Amigo1 | 486 | 395 | 445 | 502 | 485 | 511 | 377 | 420 | 480 | 500 | 547 | 504 | 471.5 | 426 | 517 | 1.217 | 0.283 | 0.01241817 | 0.0485351 | 0.0064 | 0.0075 | 0.0071 | 0.0071 | TRUE | 0.1426 |
| Amigo2 | 342 | 237 | 227 | 272 | 233 | 263 | 266 | 252 | 245 | 271 | 263 | 260 | 259.31 | 254 | 265 | 1.038 | 0.054 | 0.66301068 | 0.7920772 | 0 | 0.0116 | 0.0074 | 0.0074 | TRUE | 0.0187 |
| Amigo3 | 18 | 12 | 7 | 8 | 10 | 9 | 14 | 13 | 8 | 8 | 11 | 9 | 10.4 | 12 | 9 | 0.81 | -0.303 | 0.61165586 | NA | 0 | 0.2302 | 0.1599 | 0.1599 | TRUE | 0.3089 |
| Ammecr1 | 1375 | 960 | 916 | 964 | 788 | 868 | 1068 | 1021 | 988 | 960 | 889 | 857 | 963.79 | 1026 | 902 | 0.878 | -0.187 | 0.02098233 | 0.0716608 | 0.0014 | 0.0049 | 0.0037 | 0.0037 | TRUE | 0.038 |
| Ammecr1l | 2270 | 1589 | 1377 | 1371 | 1184 | 1387 | 1763 | 1690 | 1486 | 1365 | 1336 | 1369 | 1501.42 | 1646 | 1357 | 0.823 | -0.281 | 0.00037351 | 0.00345311 | 0.0035 | 0.004 | 0.0038 | 0.0038 | TRUE | 0.0868 |
| Amn | 320 | 260 | 267 | 348 | 341 | 380 | 249 | 277 | 288 | 346 | 385 | 375 | 319.9 | 271 | 369 | 1.364 | 0.447 | 0.00020341 | 0.00215841 | 0 | 0.0098 | 0.0073 | 0.0073 | TRUE | 0.0649 |
| Amn1 | 171 | 112 | 165 | 184 | 155 | 156 | 133 | 119 | 178 | 183 | 175 | 154 | 157 | 143 | 171 | 1.193 | 0.255 | 0.16420508 | 0.31511589 | 0.0184 | 0.0175 | 0.0178 | 0.0178 | TRUE | 0.4455 |
| Amot | 966 | 635 | 537 | 600 | 518 | 427 | 750 | 675 | 579 | 597 | 584 | 422 | 601.37 | 668 | 534 | 0.798 | -0.326 | 0.01740146 | 0.06230756 | 0.0246 | 0.0064 | 0.0118 | 0.0118 | TRUE | 0.4032 |
| Amotl1 | 1939 | 1412 | 1287 | 1269 | 1071 | 1142 | 1506 | 1502 | 1389 | 1263 | 1208 | 1127 | 1332.54 | 1466 | 1199 | 0.818 | -0.29 | 0.00013199 | 0.00153504 | 0.0019 | 0.0042 | 0.0034 | 0.0034 | TRUE | 0.0339 |

| | | | | | | | | | | | | | | | | | | | | | | | | | |
|---|---|---|---|---|---|---|---|---|---|---|---|---|---|---|---|---|---|---|---|---|---|---|---|---|---|
| Amotl2 | 2378 | 1520 | 1269 | 1281 | 1121 | 1153 | 1847 | 1617 | 1369 | 1275 | 1265 | 1138 | 1418.47 | 1611 | 1226 | 0.76 | -0.396 | 0.0001181 | 0.00139624 | 0.0118 | 0.0041 | 0.0069 | 0.0069 | TRUE | 0.1568 |
| Ampd1 | 20 | 14 | 17 | 14 | 10 | 17 | 16 | 15 | 18 | 14 | 11 | 17 | 15.13 | 16 | 14 | 0.866 | -0.208 | 0.67066737 | NA | 0 | 0.1591 | 0.1054 | 0.1054 | TRUE | 0.1337 |
| Ampd2 | 3743 | 2835 | 2290 | 2183 | 1939 | 2434 | 2907 | 3016 | 2471 | 2173 | 2187 | 2403 | 2526.09 | 2798 | 2254 | 0.806 | -0.312 | 0.00016976 | 0.00188033 | 0.0063 | 0.0034 | 0.0046 | 0.0046 | TRUE | 0.1254 |
| Ampd3 | 847 | 735 | 867 | 954 | 919 | 1093 | 658 | 782 | 935 | 950 | 1037 | 1079 | 906.74 | 792 | 1022 | 1.293 | 0.371 | 0.00139318 | 0.00913848 | 0.0158 | 0.005 | 0.0086 | 0.0086 | TRUE | 0.2994 |
| Amph | 1752 | 1265 | 1207 | 1401 | 1370 | 1432 | 1361 | 1346 | 1302 | 1395 | 1546 | 1414 | 1393.71 | 1336 | 1452 | 1.085 | 0.118 | 0.10195528 | 0.22420607 | 0.0011 | 0.0041 | 0.003 | 0.003 | TRUE | 0.0382 |
| Amt | 2550 | 1874 | 1851 | 1932 | 1634 | 1845 | 1980 | 1993 | 1997 | 1923 | 1843 | 1821 | 1926.46 | 1990 | 1862 | 0.936 | -0.095 | 0.11929991 | 0.25101629 | 0 | 0.0037 | 0.0022 | 0.0022 | TRUE | 0.0099 |
| Amtn | 0 | 0 | 0 | 0 | 0 | 0 | 0 | 0 | 0 | 0 | 0 | 0 | 0 | 0 | 0 | NA | NA | NA | NA | NA | NA | NA | NA | NA | NA |
| Amy1 | 6 | 3 | 1 | 4 | 1 | 2 | 5 | 3 | 1 | 4 | 1 | 2 | 2.67 | 3 | 2 | 0.784 | -0.351 | 0.76752347 | NA | 0 | 0.8904 | 0.6377 | 0.6377 | TRUE | 0.2641 |
| Amy2a2 | 0 | 0 | 0 | 0 | 0 | 0 | 0 | 0 | 0 | 0 | 0 | 0 | 0 | 0 | 0 | NA | NA | NA | NA | NA | NA | NA | NA | NA | NA |
| Amy2a3 | 0 | 0 | 0 | 0 | 0 | 0 | 0 | 0 | 0 | 0 | 0 | 0 | 0 | 0 | 0 | NA | NA | NA | NA | NA | NA | NA | NA | NA | NA |
| Amy2a4 | 0 | 0 | 0 | 0 | 0 | 0 | 0 | 0 | 0 | 0 | 0 | 0 | 0 | 0 | 0 | NA | NA | NA | NA | NA | NA | NA | NA | NA | NA |
| Amy2a5 | 0 | 0 | 0 | 0 | 0 | 0 | 0 | 0 | 0 | 0 | 0 | 0 | 0 | 0 | 0 | NA | NA | NA | NA | NA | NA | NA | NA | NA | NA |
| Amy2b | 0 | 0 | 0 | 0 | 0 | 0 | 0 | 0 | 0 | 0 | 0 | 0 | 0 | 0 | 0 | NA | NA | NA | NA | NA | NA | NA | NA | NA | NA |
| Amz1 | 105 | 92 | 66 | 125 | 109 | 117 | 82 | 98 | 71 | 124 | 123 | 115 | 102.25 | 84 | 121 | 1.449 | 0.535 | 0.00776682 | 0.03425 | 0 | 0.0256 | 0.019 | 0.019 | TRUE | 0.1956 |
| Amz2 | 3633 | 2713 | 3157 | 3230 | 2884 | 3358 | 2822 | 2886 | 3406 | 3215 | 3253 | 3315 | 3149.54 | 3038 | 3261 | 1.074 | 0.103 | 0.17813719 | 0.33260808 | 0.0052 | 0.0032 | 0.0039 | 0.0039 | TRUE | 0.1359 |
| Anapc1 | 15244 | 10727 | 10988 | 11606 | 10332 | 12171 | 11839 | 11410 | 11856 | 11553 | 11656 | 12015 | 11721.39 | 11702 | 11741 | 1.003 | 0.005 | 0.92067731 | 0.95643782 | 3.00E-04 | 0.0026 | 0.0016 | 0.0016 | TRUE | 0.0058 |
| Anapc10 | 1861 | 1299 | 1094 | 1023 | 805 | 934 | 1445 | 1382 | 1180 | 1018 | 908 | 922 | 1142.65 | 1336 | 949 | 0.71 | -0.494 | 9.63E-08 | 3.80E-06 | 0.0067 | 0.0045 | 0.0053 | 0.0053 | TRUE | 0.1231 |
| Anapc11 | 4624 | 3204 | 3092 | 3601 | 3103 | 3558 | 3591 | 3408 | 3336 | 3584 | 3501 | 3512 | 3488.8 | 3445 | 3532 | 1.025 | 0.035 | 0.53286983 | 0.69245464 | 6.00E-04 | 0.0031 | 0.002 | 0.002 | TRUE | 0.0172 |
| Anapc13 | 1261 | 919 | 929 | 1008 | 937 | 1054 | 979 | 978 | 1002 | 1003 | 1057 | 1040 | 1010.02 | 986 | 1033 | 1.048 | 0.067 | 0.35950465 | 0.53419275 | 0 | 0.0048 | 0.0029 | 0.0029 | TRUE | 0.0079 |
| Anapc15 | 2985 | 2158 | 2036 | 2266 | 1894 | 2202 | 2318 | 2295 | 2197 | 2256 | 2137 | 2174 | 2229.42 | 2270 | 2189 | 0.964 | -0.053 | 0.38515081 | 0.55910628 | 3.00E-04 | 0.0035 | 0.0022 | 0.0022 | TRUE | 0.0095 |
| Anapc16 | 1088 | 794 | 746 | 842 | 720 | 843 | 845 | 845 | 805 | 838 | 812 | 832 | 829.51 | 832 | 827 | 0.995 | -0.007 | 0.9259618 | 0.95917949 | 0 | 0.0053 | 0.0032 | 0.0032 | TRUE | 0.009 |
| Anapc2 | 6735 | 4987 | 4742 | 4629 | 4270 | 4897 | 5231 | 5305 | 5117 | 4608 | 4817 | 4834 | 4985.13 | 5218 | 4753 | 0.911 | -0.135 | 0.00993957 | 0.04117865 | 3.00E-04 | 0.0029 | 0.0018 | 0.0018 | TRUE | 0.0088 |
| Anapc4 | 4314 | 3072 | 3057 | 3152 | 2699 | 3167 | 3351 | 3268 | 3298 | 3137 | 3045 | 3126 | 3204.2 | 3306 | 3103 | 0.939 | -0.091 | 0.09008514 | 0.20486665 | 0 | 0.0032 | 0.0018 | 0.0018 | TRUE | 0.0032 |
| Anapc5 | 22986 | 16651 | 17260 | 18772 | 16535 | 18975 | 17852 | 17711 | 18623 | 18686 | 18653 | 18731 | 18376.2 | 18062 | 18690 | 1.035 | 0.049 | 0.28359125 | 0.45548598 | 3.00E-04 | 0.0026 | 0.0015 | 0.0015 | TRUE | 0.009 |
| Anapc7 | 4676 | 3517 | 3623 | 3749 | 3354 | 4005 | 3632 | 3741 | 3909 | 3732 | 3784 | 3954 | 3791.8 | 3761 | 3823 | 1.017 | 0.024 | 0.673628 | 0.79961507 | 9.00E-04 | 0.0031 | 0.0022 | 0.0022 | TRUE | 0.0145 |
| Ang | 11 | 6 | 8 | 7 | 16 | 9 | 9 | 6 | 9 | 7 | 18 | 9 | 9.58 | 8 | 11 | 1.422 | 0.508 | 0.4314072 | NA | 0 | 0.2499 | 0.1934 | 0.1934 | TRUE | 0.9395 |
| Ang2 | 0 | 0 | 0 | 0 | 0 | 0 | 0 | 0 | 0 | 0 | 0 | 0 | 0 | 0 | 0 | NA | NA | NA | NA | NA | NA | NA | NA | NA | NA |
| Ang3 | 0 | 0 | 0 | 0 | 0 | 0 | 0 | 0 | 0 | 0 | 0 | 0 | 0 | 0 | 0 | NA | NA | NA | NA | NA | NA | NA | NA | NA | NA |
| Ang4 | 0 | 0 | 0 | 0 | 1 | 0 | 0 | 0 | 0 | 0 | 1 | 0 | 0.19 | 0 | 0 | 2.094 | 1.067 | 0.79379329 | NA | 0 | 12.608 | 10 | 10 | TRUE | 0.5075 |
| Ang5 | 0 | 0 | 0 | 0 | 0 | 0 | 0 | 0 | 0 | 0 | 0 | 0 | 0 | 0 | 0 | NA | NA | NA | NA | NA | NA | NA | NA | NA | NA |
| Ang6 | 0 | 0 | 0 | 0 | 0 | 0 | 0 | 0 | 0 | 0 | 0 | 0 | 0 | 0 | 0 | NA | NA | NA | NA | NA | NA | NA | NA | NA | NA |
| Angel1 | 2596 | 1935 | 1885 | 2235 | 1921 | 2205 | 2016 | 2058 | 2034 | 2225 | 2167 | 2177 | 2112.8 | 2036 | 2190 | 1.076 | 0.105 | 0.07295624 | 0.17685949 | 0 | 0.0035 | 0.002 | 0.002 | TRUE | 0.0024 |
| Angel2 | 2371 | 1647 | 1501 | 1445 | 1311 | 1360 | 1841 | 1752 | 1620 | 1438 | 1479 | 1343 | 1578.79 | 1738 | 1420 | 0.816 | -0.293 | 0.00010913 | 0.00130097 | 0.0027 | 0.0039 | 0.0035 | 0.0035 | TRUE | 0.0423 |
| Angpt1 | 0 | 0 | 0 | 1 | 0 | 0 | 0 | 0 | 0 | 1 | 0 | 0 | 0.17 | 0 | 0 | 2.094 | 1.067 | 0.79379329 | NA | 0 | 14.2885 | 10 | 10 | TRUE | 0.3906 |
| Angpt2 | 12 | 8 | 6 | 6 | 8 | 9 | 9 | 9 | 6 | 6 | 9 | 9 | 8.03 | 8 | 8 | 0.974 | -0.037 | 0.95532108 | NA | 0 | 0.2975 | 0.1947 | 0.1947 | TRUE | 0.1441 |
| Angpt4 | 2 | 0 | 0 | 3 | 4 | 1 | 2 | 0 | 0 | 3 | 5 | 1 | 1.67 | 1 | 3 | 4.816 | 2.268 | 0.19020573 | NA | 0 | 1.4189 | 1.1525 | 1.1525 | TRUE | 0.5476 |
| Angptl1 | 1 | 0 | 0 | 0 | 0 | 0 | 1 | 0 | 0 | 0 | 0 | 0 | 0.13 | 0 | 0 | 0.552 | -0.857 | 0.8336548 | NA | 0 | 18.3121 | 10 | 10 | TRUE | 0.2566 |
| Angptl2 | 39 | 24 | 25 | 28 | 24 | 25 | 30 | 26 | 27 | 28 | 27 | 25 | 27.07 | 28 | 27 | 0.958 | -0.062 | 0.86359571 | 0.92466653 | 0 | 0.09 | 0.0581 | 0.0581 | TRUE | 0.0562 |
| Angptl3 | 0 | 0 | 0 | 0 | 0 | 0 | 0 | 0 | 0 | 0 | 0 | 0 | 0 | 0 | 0 | NA | NA | NA | NA | NA | NA | NA | NA | NA | NA |
| Angptl4 | 1212 | 851 | 720 | 766 | 654 | 828 | 941 | 905 | 777 | 762 | 738 | 817 | 823.5 | 874 | 772 | 0.883 | -0.18 | 0.05845155 | 0.15071711 | 0.0053 | 0.0053 | 0.0053 | 0.0053 | TRUE | 0.1115 |
| Angptl6 | 317 | 231 | 275 | 310 | 272 | 307 | 246 | 246 | 297 | 309 | 307 | 303 | 284.52 | 263 | 306 | 1.167 | 0.223 | 0.08076278 | 0.19036098 | 0.002 | 0.0108 | 0.0082 | 0.0082 | TRUE | 0.1366 |
| Angptl7 | 0 | 0 | 0 | 0 | 0 | 0 | 0 | 0 | 0 | 0 | 0 | 0 | 0 | 0 | 0 | NA | NA | NA | NA | NA | NA | NA | NA | NA | NA |
| Ank | 1054 | 734 | 808 | 756 | 700 | 723 | 819 | 781 | 872 | 753 | 790 | 714 | 787.85 | 824 | 752 | 0.913 | -0.132 | 0.12652266 | 0.26185756 | 0.0014 | 0.0054 | 0.0041 | 0.0041 | TRUE | 0.0297 |
| Ank1 | 61 | 32 | 32 | 38 | 27 | 36 | 47 | 34 | 35 | 38 | 30 | 36 | 36.63 | 39 | 35 | 0.89 | -0.168 | 0.60520583 | 0.74984457 | 0 | 0.0671 | 0.0488 | 0.0488 | TRUE | 0.3364 |
| Ank2 | 613 | 435 | 433 | 498 | 439 | 557 | 476 | 463 | 467 | 496 | 495 | 550 | 491.13 | 469 | 514 | 1.096 | 0.132 | 0.17980946 | 0.33464802 | 0 | 0.0073 | 0.005 | 0.005 | TRUE | 0.0451 |
| Ank3 | 872 | 642 | 831 | 846 | 879 | 1029 | 677 | 683 | 897 | 842 | 992 | 1016 | 851.04 | 752 | 950 | 1.264 | 0.338 | 0.00413908 | 0.02120423 | 0.0169 | 0.0052 | 0.0088 | 0.0088 | TRUE | 0.3298 |
| Ankar | 2 | 0 | 1 | 0 | 0 | 1 | 2 | 0 | 1 | 0 | 0 | 1 | 0.6 | 1 | 0 | 0.414 | -1.271 | 0.61985863 | NA | 0 | 3.9313 | 3.1876 | 3.1876 | TRUE | 0.3835 |
| Ankdd1b | 5 | 4 | 5 | 3 | 5 | 3 | 4 | 4 | 5 | 3 | 6 | 3 | 4.19 | 4 | 4 | 0.854 | -0.228 | 0.80414499 | NA | 0 | 0.5685 | 0.3658 | 0.3658 | TRUE | 0.2326 |
| Ankef1 | 41 | 32 | 28 | 29 | 21 | 26 | 32 | 34 | 30 | 29 | 24 | 26 | 29.05 | 32 | 26 | 0.815 | -0.294 | 0.4029296 | 0.57589577 | 0 | 0.084 | 0.0544 | 0.0544 | TRUE | 0.055 |
| Ankfn1 | 0 | 0 | 0 | 0 | 0 | 0 | 0 | 0 | 0 | 0 | 0 | 0 | 0 | 0 | 0 | NA | NA | NA | NA | NA | NA | NA | NA | NA | NA |
| Ankfy1 | 16716 | 12052 | 10668 | 11735 | 10576 | 11886 | 12983 | 12820 | 11511 | 11681 | 11931 | 11733 | 12109.67 | 12438 | 11782 | 0.947 | -0.078 | 0.18323549 | 0.33907105 | 0.002 | 0.0026 | 0.0024 | 0.0024 | TRUE | 0.0518 |
| Ankhd1 | 7177 | 5203 | 5079 | 5604 | 5049 | 5847 | 5574 | 5534 | 5480 | 5578 | 5696 | 5772 | 5605.76 | 5529 | 5682 | 1.027 | 0.039 | 0.42587718 | 0.59841355 | 0 | 0.0028 | 0.0016 | 0.0016 | TRUE | 0.0031 |
| Ankib1 | 2857 | 1989 | 1968 | 2003 | 1764 | 2018 | 2219 | 2116 | 2123 | 1994 | 1990 | 1992 | 2072.31 | 2153 | 1992 | 0.925 | -0.113 | 0.06049698 | 0.15413251 | 0 | 0.0036 | 0.0021 | 0.0021 | TRUE | 0.0091 |
| Ankk1 | 12 | 8 | 2 | 5 | 9 | 9 | 9 | 9 | 2 | 5 | 10 | 9 | 7.33 | 7 | 8 | 1.179 | 0.238 | 0.75166857 | NA | 0.1026 | 0.3256 | 0.2706 | 0.2706 | TRUE | 0.8061 |
| Ankle1 | 543 | 507 | 410 | 534 | 455 | 503 | 422 | 539 | 442 | 532 | 513 | 497 | 490.79 | 468 | 514 | 1.101 | 0.138 | 0.22281624 | 0.38710497 | 0.0072 | 0.0073 | 0.0072 | 0.0072 | TRUE | 0.205 |
| Ankle2 | 4506 | 3329 | 3466 | 3709 | 3138 | 3610 | 3500 | 3541 | 3740 | 3692 | 3540 | 3564 | 3595.99 | 3594 | 3599 | 1.002 | 0.003 | 0.96329282 | 0.97974413 | 6.00E-04 | 0.0031 | 0.002 | 0.002 | TRUE | 0.0153 |
| Ankmy1 | 84 | 52 | 58 | 63 | 52 | 71 | 65 | 55 | 63 | 63 | 59 | 70 | 62.43 | 61 | 64 | 1.044 | 0.062 | 0.7987846 | 0.88244151 | 0 | 0.0404 | 0.0273 | 0.0273 | TRUE | 0.0655 |
| Ankmy2 | 1967 | 1635 | 1804 | 2012 | 1780 | 2058 | 1528 | 1739 | 1947 | 2003 | 2008 | 2032 | 1875.94 | 1738 | 2014 | 1.161 | 0.215 | 0.01243762 | 0.04859784 | 0.0065 | 0.0037 | 0.0048 | 0.0048 | TRUE | 0.1387 |
| Ankra2 | 1314 | 960 | 1064 | 1235 | 914 | 1203 | 1021 | 1021 | 1148 | 1229 | 1031 | 1188 | 1106.28 | 1063 | 1149 | 1.082 | 0.114 | 0.20339991 | 0.36472149 | 0.0052 | 0.0046 | 0.0049 | 0.0049 | TRUE | 0.0954 |
| Ankrd1 | 11 | 3 | 0 | 6 | 3 | 1 | 9 | 3 | 0 | 6 | 3 | 1 | 3.68 | 4 | 3 | 0.855 | -0.227 | 0.84734152 | NA | 0.9812 | 0.6465 | 0.7278 | 0.7278 | TRUE | 0.3572 |
| Ankrd10 | 6532 | 5534 | 5657 | 5486 | 4951 | 6381 | 5073 | 5886 | 6104 | 5461 | 5585 | 6299 | 5734.74 | 5688 | 5782 | 1.017 | 0.024 | 0.76376212 | 0.85969854 | 0.0075 | 0.0028 | 0.0045 | 0.0045 | TRUE | 0.11 |
| Ankrd11 | 18918 | 14101 | 12788 | 12901 | 11619 | 13386 | 14693 | 14999 | 13798 | 12842 | 13107 | 13214 | 13775.52 | 14497 | 13054 | 0.9 | -0.151 | 0.00351326 | 0.0186618 | 9.00E-04 | 0.0026 | 0.0019 | 0.0019 | TRUE | 0.0216 |
| Ankrd12 | 888 | 684 | 723 | 851 | 752 | 853 | 690 | 728 | 780 | 847 | 848 | 842 | 789.13 | 733 | 846 | 1.157 | 0.21 | 0.01254384 | 0.04893262 | 7.00E-04 | 0.0054 | 0.0038 | 0.0038 | TRUE | 0.0381 |
| Ankrd13a | 5749 | 4275 | 4281 | 3983 | 3649 | 4336 | 4465 | 4547 | 4619 | 3965 | 4116 | 4280 | 4332.14 | 4544 | 4120 | 0.907 | -0.141 | 0.01103878 | 0.04441929 | 5.00E-04 | 0.003 | 0.002 | 0.002 | TRUE | 0.0142 |
| Ankrd13b | 1267 | 1015 | 946 | 1232 | 1011 | 1294 | 984 | 1080 | 1021 | 1226 | 1141 | 1277 | 1121.44 | 1028 | 1215 | 1.183 | 0.243 | 0.00210275 | 0.01254367 | 0.0018 | 0.0045 | 0.0036 | 0.0036 | TRUE | 0.0336 |
| Ankrd13c | 3144 | 2160 | 2023 | 1995 | 1807 | 1928 | 2442 | 2298 | 2183 | 1986 | 2038 | 1903 | 2141.62 | 2308 | 1976 | 0.855 | -0.225 | 0.00091588 | 0.00680386 | 0.0018 | 0.0035 | 0.0029 | 0.0029 | TRUE | 0.0323 |
| Ankrd13d | 472 | 393 | 385 | 443 | 462 | 502 | 367 | 418 | 415 | 441 | 521 | 496 | 442.95 | 400 | 486 | 1.217 | 0.283 | 0.0110802 | 0.04449825 | 0.0043 | 0.0078 | 0.0067 | 0.0067 | TRUE | 0.0766 |
| Ankrd16 | 389 | 321 | 305 | 286 | 254 | 276 | 302 | 341 | 329 | 285 | 287 | 272 | 302.72 | 324 | 281 | 0.869 | -0.203 | 0.09034227 | 0.20525546 | 0 | 0.0103 | 0.007 | 0.007 | TRUE | 0.0439 |
| Ankrd17 | 14889 | 10698 | 9893 | 9893 | 8812 | 10168 | 11564 | 11379 | 10674 | 9847 | 9941 | 10037 | 10573.86 | 11206 | 9942 | 0.887 | -0.173 | 0.00084012 | 0.00636848 | 8.00E-04 | 0.0027 | 0.0018 | 0.0018 | TRUE | 0.021 |

| | | | | | | | | | | | | | | | | | | | | | | | | | |
|---|---|---|---|---|---|---|---|---|---|---|---|---|---|---|---|---|---|---|---|---|---|---|---|---|---|
| Ankrd2 | 2 | 1 | 1 | 3 | 1 | 0 | 2 | 1 | 1 | 3 | 1 | 0 | 1.3 | 1 | 1 | 1.104 | 0.142 | 0.93281931 | NA | 0 | 1.8231 | 1.2916 | 1.2916 | TRUE | 0.3572 |
| Ankrd22 | 37 | 27 | 2 | 5 | 6 | 5 | 29 | 29 | 2 | 5 | 7 | 5 | 12.72 | 20 | 6 | 0.277 | -1.853 | 0.0147804 | NA | 0.5796 | 0.1888 | 0.2985 | 0.2985 | TRUE | 3.081 |
| Ankrd23 | 99 | 100 | 89 | 87 | 84 | 109 | 77 | 106 | 96 | 87 | 95 | 108 | 94.71 | 93 | 97 | 1.041 | 0.058 | 0.78733224 | 0.87519506 | 0.0104 | 0.0275 | 0.0225 | 0.0225 | TRUE | 0.2536 |
| Ankrd24 | 462 | 440 | 441 | 524 | 434 | 527 | 359 | 468 | 476 | 522 | 490 | 520 | 472.35 | 434 | 511 | 1.18 | 0.239 | 0.04876042 | 0.13233939 | 0.0108 | 0.0074 | 0.0085 | 0.0085 | TRUE | 0.2844 |
| Ankrd26 | 3559 | 2611 | 2514 | 2763 | 2378 | 2697 | 2764 | 2777 | 2713 | 2750 | 2683 | 2662 | 2724.88 | 2751 | 2698 | 0.981 | -0.028 | 0.61427876 | 0.75623483 | 0 | 0.0033 | 0.0019 | 0.0019 | TRUE | 0.0035 |
| Ankrd27 | 5643 | 4345 | 4247 | 4910 | 4216 | 4918 | 4383 | 4622 | 4582 | 4887 | 4756 | 4855 | 4680.87 | 4529 | 4833 | 1.068 | 0.094 | 0.07309045 | 0.17700121 | 3.00E-04 | 0.0029 | 0.0018 | 0.0018 | TRUE | 0.01 |
| Ankrd28 | 5652 | 4063 | 3897 | 3937 | 3522 | 4024 | 4390 | 4322 | 4205 | 3919 | 3973 | 3972 | 4130.11 | 4306 | 3955 | 0.918 | -0.123 | 0.017997 | 0.06387663 | 0 | 0.003 | 0.0017 | 0.0017 | TRUE | 0.0051 |
| Ankrd29 | 38 | 17 | 24 | 24 | 20 | 39 | 30 | 18 | 26 | 24 | 23 | 38 | 26.41 | 25 | 28 | 1.151 | 0.203 | 0.61100955 | 0.75392458 | 0.0396 | 0.0922 | 0.0775 | 0.0775 | TRUE | 0.6533 |
| Ankrd32 | 1241 | 896 | 806 | 860 | 779 | 711 | 964 | 953 | 870 | 856 | 879 | 702 | 870.54 | 929 | 812 | 0.873 | -0.195 | 0.04903792 | 0.13279633 | 0.0076 | 0.0051 | 0.0059 | 0.0059 | TRUE | 0.1694 |
| Ankrd33 | 0 | 0 | 0 | 0 | 0 | 0 | 0 | 0 | 0 | 0 | 0 | 0 | 0 | 0 | 0 | NA | NA | NA | NA | NA | NA | NA | NA | NA | NA |
| Ankrd33b | 873 | 617 | 583 | 1028 | 864 | 977 | 678 | 656 | 629 | 1023 | 975 | 964 | 820.96 | 654 | 987 | 1.507 | 0.592 | 3.55E-13 | 4.84E-11 | 0 | 0.0053 | 0.0035 | 0.0035 | TRUE | 0.0134 |
| Ankrd34a | 30 | 14 | 24 | 29 | 22 | 28 | 23 | 15 | 26 | 29 | 25 | 28 | 24.24 | 21 | 27 | 1.267 | 0.341 | 0.39085684 | 0.56457977 | 0 | 0.1002 | 0.0721 | 0.0721 | TRUE | 0.361 |
| Ankrd34b | 116 | 100 | 106 | 119 | 117 | 117 | 90 | 106 | 114 | 118 | 132 | 115 | 112.8 | 103 | 122 | 1.182 | 0.241 | 0.20641051 | 0.3682718 | 0 | 0.0234 | 0.0172 | 0.0172 | TRUE | 0.1455 |
| Ankrd34c | 0 | 0 | 1 | 0 | 0 | 0 | 0 | 0 | 1 | 0 | 0 | 0 | 0.18 | 0 | 0 | 0.552 | -0.857 | 0.8336548 | NA | 0 | 13.1817 | 10 | 10 | TRUE | 0.5365 |
| Ankrd35 | 393 | 293 | 335 | 310 | 318 | 334 | 305 | 312 | 361 | 309 | 359 | 330 | 329.23 | 326 | 333 | 1.02 | 0.029 | 0.81635571 | 0.89466126 | 0.0041 | 0.0096 | 0.0079 | 0.0079 | TRUE | 0.0995 |
| Ankrd36 | 35 | 14 | 22 | 21 | 11 | 15 | 27 | 15 | 24 | 21 | 12 | 15 | 18.99 | 22 | 16 | 0.729 | -0.457 | 0.32370081 | 0.49775747 | 0 | 0.1272 | 0.1005 | 0.1005 | TRUE | 0.4264 |
| Ankrd37 | 1098 | 848 | 1081 | 1066 | 912 | 1113 | 853 | 902 | 1166 | 1061 | 1029 | 1099 | 1018.3 | 974 | 1063 | 1.093 | 0.129 | 0.23932307 | 0.40630058 | 0.0128 | 0.0048 | 0.0077 | 0.0077 | TRUE | 0.3547 |
| Ankrd39 | 516 | 355 | 375 | 427 | 375 | 426 | 401 | 378 | 405 | 425 | 423 | 421 | 408.6 | 395 | 423 | 1.072 | 0.1 | 0.33021639 | 0.50457375 | 0 | 0.0082 | 0.0052 | 0.0052 | TRUE | 0.0152 |
| Ankrd40 | 4978 | 3566 | 3437 | 3836 | 3420 | 3873 | 3866 | 3793 | 3708 | 3818 | 3858 | 3823 | 3811.25 | 3789 | 3833 | 1.011 | 0.016 | 0.75736919 | 0.85587269 | 0 | 0.003 | 0.0017 | 0.0017 | TRUE | 0.0042 |
| Ankrd42 | 106 | 64 | 51 | 70 | 71 | 63 | 82 | 68 | 55 | 70 | 80 | 62 | 69.57 | 68 | 71 | 1.023 | 0.033 | 0.89606338 | 0.94272381 | 0.0147 | 0.0365 | 0.0303 | 0.0303 | TRUE | 0.2346 |
| Ankrd44 | 838 | 523 | 458 | 722 | 487 | 574 | 651 | 556 | 494 | 719 | 549 | 567 | 589.34 | 567 | 612 | 1.077 | 0.107 | 0.4099952 | 0.58285871 | 0.019 | 0.0064 | 0.0104 | 0.0104 | TRUE | 0.2773 |
| Ankrd45 | 198 | 139 | 113 | 119 | 84 | 70 | 154 | 148 | 122 | 118 | 95 | 69 | 117.65 | 141 | 94 | 0.665 | -0.589 | 0.00670434 | 0.03074338 | 0.0318 | 0.0226 | 0.0251 | 0.0251 | TRUE | 0.3211 |
| Ankrd46 | 4590 | 3358 | 3529 | 3676 | 3192 | 3607 | 3565 | 3572 | 3808 | 3659 | 3601 | 3561 | 3627.52 | 3648 | 3607 | 0.989 | -0.016 | 0.77634692 | 0.86766476 | 5.00E-04 | 0.0031 | 0.002 | 0.002 | TRUE | 0.0176 |
| Ankrd49 | 1311 | 899 | 724 | 781 | 771 | 824 | 1018 | 956 | 781 | 777 | 870 | 813 | 869.37 | 918 | 820 | 0.891 | -0.166 | 0.11119361 | 0.23898904 | 0.0099 | 0.0052 | 0.0067 | 0.0067 | TRUE | 0.2012 |
| Ankrd50 | 1222 | 833 | 917 | 962 | 854 | 1021 | 949 | 886 | 989 | 958 | 963 | 1008 | 958.9 | 941 | 976 | 1.037 | 0.052 | 0.50930176 | 0.67252864 | 8.00E-04 | 0.0049 | 0.0035 | 0.0035 | TRUE | 0.0304 |
| Ankrd52 | 2017 | 1619 | 1655 | 1603 | 1484 | 1751 | 1567 | 1722 | 1786 | 1596 | 1674 | 1729 | 1678.77 | 1692 | 1666 | 0.986 | -0.02 | 0.78525905 | 0.87383344 | 0.0026 | 0.0038 | 0.0034 | 0.0034 | TRUE | 0.0522 |
| Ankrd53 | 0 | 0 | 0 | 0 | 0 | 1 | 0 | 0 | 0 | 0 | 0 | 1 | 0.16 | 0 | 0 | 2.094 | 1.067 | 0.79379329 | NA | 0 | 14.4078 | 10 | 10 | TRUE | 0.3835 |
| Ankrd54 | 984 | 692 | 689 | 723 | 675 | 855 | 764 | 736 | 743 | 720 | 761 | 844 | 761.48 | 748 | 775 | 1.036 | 0.051 | 0.56764666 | 0.72138115 | 0.0023 | 0.0055 | 0.0044 | 0.0044 | TRUE | 0.0731 |
| Ankrd55 | 10 | 10 | 8 | 14 | 10 | 5 | 8 | 11 | 9 | 14 | 11 | 5 | 9.53 | 9 | 10 | 1.118 | 0.161 | 0.80127289 | NA | 0 | 0.2511 | 0.1909 | 0.1909 | TRUE | 0.6587 |
| Ankrd6 | 1814 | 1314 | 1282 | 1348 | 1298 | 1371 | 1409 | 1398 | 1383 | 1342 | 1464 | 1353 | 1391.55 | 1397 | 1386 | 0.992 | -0.012 | 0.86888559 | 0.92762038 | 4.00E-04 | 0.0041 | 0.0028 | 0.0028 | TRUE | 0.0284 |
| Ankrd60 | 0 | 0 | 0 | 0 | 0 | 0 | 0 | 0 | 0 | 0 | 0 | 0 | 0 | 0 | 0 | NA | NA | NA | NA | NA | NA | NA | NA | NA | NA |
| Ankrd61 | 3 | 0 | 0 | 0 | 1 | 0 | 2 | 0 | 0 | 0 | 1 | 0 | 0.58 | 1 | 0 | 0.457 | -1.13 | 0.69406848 | NA | 0 | 4.1146 | 4.3478 | 4.3478 | TRUE | 1.4398 |
| Ankrd63 | 11 | 13 | 8 | 7 | 12 | 17 | 9 | 14 | 9 | 7 | 14 | 17 | 11.38 | 11 | 13 | 1.209 | 0.274 | 0.64434462 | NA | 0 | 0.2107 | 0.1649 | 0.1649 | TRUE | 0.4691 |
| Ankrd66 | 2 | 2 | 2 | 2 | 4 | 4 | 2 | 2 | 2 | 2 | 5 | 4 | 2.72 | 2 | 4 | 1.801 | 0.849 | 0.46005815 | NA | 0 | 0.8754 | 0.5521 | 0.5521 | TRUE | 0.2141 |
| Ankrd7 | 0 | 0 | 0 | 0 | 0 | 0 | 0 | 0 | 0 | 0 | 0 | 0 | 0 | 0 | 0 | NA | NA | NA | NA | NA | NA | NA | NA | NA | NA |
| Ankrd9 | 692 | 502 | 414 | 507 | 456 | 529 | 537 | 534 | 447 | 505 | 514 | 522 | 509.9 | 506 | 514 | 1.014 | 0.019 | 0.85266269 | 0.9167369 | 0.0034 | 0.0071 | 0.0059 | 0.0059 | TRUE | 0.1188 |
| Anks1 | 3125 | 2293 | 2356 | 2232 | 2068 | 2303 | 2427 | 2439 | 2542 | 2222 | 2333 | 2273 | 2372.71 | 2469 | 2276 | 0.922 | -0.117 | 0.04834039 | 0.13145735 | 0 | 0.0034 | 0.0021 | 0.0021 | TRUE | 0.0079 |
| Anks1b | 49 | 22 | 37 | 20 | 34 | 23 | 38 | 23 | 40 | 20 | 38 | 23 | 30.39 | 34 | 27 | 0.791 | -0.338 | 0.38649005 | 0.56036744 | 0.0645 | 0.0804 | 0.0761 | 0.0761 | TRUE | 0.7964 |
| Anks3 | 808 | 661 | 622 | 670 | 601 | 706 | 628 | 703 | 671 | 667 | 678 | 697 | 673.93 | 667 | 681 | 1.022 | 0.031 | 0.72609291 | 0.83589639 | 0 | 0.0059 | 0.0041 | 0.0041 | TRUE | 0.0342 |
| Anks4b | 1 | 0 | 0 | 0 | 0 | 0 | 1 | 0 | 0 | 0 | 0 | 0 | 0.13 | 0 | 0 | 0.552 | -0.857 | 0.8336548 | NA | 0 | 18.3121 | 10 | 10 | TRUE | 0.2566 |
| Anks6 | 92 | 92 | 71 | 77 | 60 | 90 | 71 | 98 | 77 | 77 | 68 | 89 | 79.85 | 82 | 78 | 0.954 | -0.068 | 0.76882975 | 0.86325859 | 0.0113 | 0.0321 | 0.0262 | 0.0262 | TRUE | 0.2624 |
| Ankzf1 | 1801 | 1674 | 1628 | 1534 | 1400 | 1740 | 1399 | 1781 | 1757 | 1527 | 1579 | 1718 | 1626.65 | 1646 | 1608 | 0.979 | -0.031 | 0.74700589 | 0.84993112 | 0.0103 | 0.0039 | 0.0061 | 0.0061 | TRUE | 0.2124 |
| Anln | 19947 | 14661 | 16993 | 16091 | 14586 | 16406 | 15492 | 15595 | 18335 | 16017 | 16455 | 16195 | 16348.12 | 16474 | 16222 | 0.985 | -0.022 | 0.74951646 | 0.8514989 | 0.004 | 0.0026 | 0.0034 | 0.0034 | TRUE | 0.1193 |
| Ano1 | 63 | 45 | 55 | 57 | 55 | 76 | 49 | 48 | 59 | 57 | 62 | 75 | 58.32 | 52 | 65 | 1.245 | 0.316 | 0.22400757 | 0.38840902 | 0 | 0.0431 | 0.0315 | 0.0315 | TRUE | 0.1804 |
| Ano10 | 499 | 342 | 416 | 566 | 544 | 590 | 388 | 364 | 449 | 563 | 614 | 582 | 493.28 | 400 | 586 | 1.467 | 0.553 | 2.78E-07 | 9.71E-06 | 0.0038 | 0.0072 | 0.0062 | 0.0062 | TRUE | 0.1251 |
| Ano2 | 14 | 14 | 3 | 12 | 13 | 10 | 11 | 15 | 3 | 12 | 15 | 10 | 10.91 | 10 | 12 | 1.248 | 0.319 | 0.61228969 | NA | 0.1072 | 0.2196 | 0.1933 | 0.1933 | TRUE | 0.5221 |
| Ano3 | 454 | 329 | 408 | 410 | 328 | 386 | 353 | 350 | 440 | 408 | 370 | 381 | 383.66 | 381 | 386 | 1.017 | 0.024 | 0.8446507 | 0.91195135 | 0.0073 | 0.0086 | 0.0082 | 0.0082 | TRUE | 0.2071 |
| Ano4 | 25 | 16 | 14 | 22 | 22 | 25 | 19 | 17 | 15 | 22 | 25 | 25 | 20.49 | 17 | 24 | 1.376 | 0.461 | 0.27130187 | 0.44220751 | 0 | 0.1181 | 0.0767 | 0.0767 | TRUE | 0.0804 |
| Ano5 | 14 | 8 | 7 | 9 | 0 | 6 | 11 | 9 | 8 | 9 | 0 | 6 | 6.97 | 9 | 5 | 0.558 | -0.841 | 0.31259228 | NA | 0.3476 | 0.3425 | 0.3433 | 0.3433 | TRUE | 0.7517 |
| Ano6 | 3090 | 2424 | 2364 | 2619 | 2399 | 2919 | 2400 | 2578 | 2551 | 2607 | 2706 | 2882 | 2620.63 | 2510 | 2732 | 1.089 | 0.123 | 0.05902574 | 0.15156286 | 0.0017 | 0.0033 | 0.0027 | 0.0027 | TRUE | 0.0282 |
| Ano7 | 39 | 34 | 30 | 36 | 46 | 41 | 30 | 36 | 32 | 36 | 52 | 40 | 37.84 | 33 | 43 | 1.297 | 0.376 | 0.2394918 | 0.40642666 | 0 | 0.0651 | 0.0466 | 0.0466 | TRUE | 0.2529 |
| Ano8 | 773 | 683 | 654 | 815 | 701 | 805 | 600 | 726 | 706 | 811 | 791 | 795 | 738.2 | 677 | 799 | 1.182 | 0.242 | 0.01072296 | 0.04347895 | 0.004 | 0.0056 | 0.0051 | 0.0051 | TRUE | 0.1242 |
| Ano9 | 1149 | 1066 | 1184 | 1225 | 1050 | 1259 | 892 | 1134 | 1278 | 1219 | 1185 | 1243 | 1158.42 | 1101 | 1216 | 1.106 | 0.145 | 0.19212113 | 0.35037387 | 0.0158 | 0.0045 | 0.0081 | 0.0081 | TRUE | 0.3393 |
| Anp32a | 16040 | 13140 | 14058 | 15279 | 14148 | 16590 | 12458 | 13977 | 15168 | 15209 | 15960 | 16377 | 14858.15 | 13868 | 15849 | 1.143 | 0.193 | 0.00755538 | 0.03361879 | 0.0054 | 0.0026 | 0.0037 | 0.0037 | TRUE | 0.0972 |
| Anp32b | 22662 | 16568 | 14758 | 13949 | 12478 | 14829 | 17601 | 17623 | 15924 | 13885 | 14076 | 14639 | 15624.56 | 17049 | 14200 | 0.833 | -0.264 | 4.60E-06 | 9.91E-05 | 0.0019 | 0.0026 | 0.0023 | 0.0023 | TRUE | 0.0407 |
| Anp32e | 15690 | 11097 | 10327 | 10946 | 9427 | 10602 | 12186 | 11804 | 11143 | 10896 | 10635 | 10466 | 11188.05 | 11711 | 10666 | 0.911 | -0.135 | 0.01160304 | 0.04615704 | 0.0011 | 0.0026 | 0.002 | 0.002 | TRUE | 0.022 |
| Anpep | 3116 | 2140 | 2121 | 3216 | 2681 | 3107 | 2420 | 2276 | 2289 | 3201 | 3024 | 3067 | 2712.94 | 2328 | 3097 | 1.33 | 0.411 | 8.50E-12 | 8.93E-10 | 7.00E-04 | 0.0033 | 0.0022 | 0.0022 | TRUE | 0.0149 |
| Antxr1 | 299 | 199 | 152 | 154 | 125 | 166 | 232 | 212 | 164 | 153 | 141 | 164 | 177.68 | 203 | 153 | 0.752 | -0.412 | 0.01571327 | 0.05781029 | 0.0138 | 0.0158 | 0.0152 | 0.0152 | TRUE | 0.2943 |
| Antxr2 | 36 | 17 | 23 | 23 | 19 | 12 | 28 | 18 | 25 | 23 | 21 | 12 | 21.17 | 24 | 19 | 0.786 | -0.347 | 0.4305591 | 0.60268154 | 0.0268 | 0.1144 | 0.0917 | 0.0917 | TRUE | 0.5596 |
| Antxrl | 0 | 0 | 0 | 0 | 0 | 0 | 0 | 0 | 0 | 0 | 0 | 0 | 0 | 0 | 0 | NA | NA | NA | NA | NA | NA | NA | NA | NA | NA |
| Anxa1 | 275 | 199 | 158 | 114 | 74 | 70 | 214 | 212 | 170 | 113 | 83 | 69 | 143.63 | 199 | 88 | 0.446 | -1.164 | 8.78E-09 | 4.43E-07 | 0.0279 | 0.0189 | 0.0212 | 0.0212 | TRUE | 0.5847 |
| Anxa10 | 5 | 1 | 1 | 3 | 3 | 6 | 4 | 1 | 1 | 3 | 3 | 6 | 3.05 | 2 | 4 | 1.958 | 0.969 | 0.38904478 | NA | 0 | 0.7787 | 0.5568 | 0.5568 | TRUE | 0.7998 |
| Anxa11 | 3554 | 2510 | 2796 | 3204 | 2887 | 3534 | 2760 | 2670 | 3017 | 3189 | 3257 | 3489 | 3063.61 | 2816 | 3312 | 1.177 | 0.235 | 0.00059122 | 0.00490441 | 0.0027 | 0.0032 | 0.003 | 0.003 | TRUE | 0.0467 |
| Anxa13 | 1 | 1 | 0 | 0 | 0 | 1 | 1 | 1 | 0 | 0 | 0 | 1 | 0.47 | 1 | 0 | 0.599 | -0.74 | 0.79209856 | NA | 0 | 5.0317 | 3.8925 | 3.8925 | TRUE | 0.3835 |
| Anxa2 | 4399 | 3428 | 4910 | 5580 | 5093 | 6046 | 3417 | 3646 | 5298 | 5554 | 5745 | 5968 | 4938.13 | 4120 | 5756 | 1.397 | 0.483 | 0.02112798 | 0.07203775 | 0.0314 | 0.0029 | 0.0105 | 0.0314 | TRUE | 0.7606 |
| Anxa3 | 407 | 252 | 200 | 177 | 164 | 118 | 316 | 268 | 216 | 176 | 185 | 116 | 212.94 | 267 | 159 | 0.594 | -0.75 | 9.75E-05 | 0.00118574 | 0.0424 | 0.0136 | 0.0217 | 0.0217 | TRUE | 0.2596 |
| Anxa4 | 1757 | 1283 | 1376 | 1818 | 1463 | 1616 | 1365 | 1365 | 1485 | 1810 | 1650 | 1595 | 1544.88 | 1405 | 1685 | 1.2 | 0.264 | 0.0005262 | 0.00447749 | 0.0027 | 0.004 | 0.0035 | 0.0035 | TRUE | 0.0509 |
| Anxa5 | 836 | 688 | 870 | 1076 | 1004 | 1234 | 649 | 732 | 939 | 1071 | 1133 | 1218 | 956.94 | 773 | 1141 | 1.478 | 0.564 | 2.21E-06 | 5.46E-05 | 0.0191 | 0.0049 | 0.0091 | 0.0091 | TRUE | 0.4133 |
| Anxa6 | 6939 | 5032 | 4851 | 5977 | 5365 | 6226 | 5389 | 5352 | 5234 | 5949 | 6052 | 6146 | 5687.28 | 5325 | 6049 | 1.136 | 0.184 | 0.00020592 | 0.00217956 | 0 | 0.0028 | 0.0016 | 0.0016 | TRUE | 0.0027 |

| | | | | | | | | | | | | | | | | | | | | | | | | | |
|---|---|---|---|---|---|---|---|---|---|---|---|---|---|---|---|---|---|---|---|---|---|---|---|---|---|
| Anxa7 | 6115 | 4525 | 5015 | 5908 | 5638 | 6460 | 4749 | 4813 | 5411 | 5881 | 6360 | 6377 | 5598.61 | 4991 | 6206 | 1.244 | 0.315 | 2.96E-06 | 6.84E-05 | 0.0034 | 0.0028 | 0.0031 | 0.0031 | TRUE | 0.0657 |
| Anxa8 | 316 | 272 | 351 | 365 | 348 | 325 | 245 | 289 | 379 | 363 | 393 | 321 | 331.7 | 304 | 359 | 1.182 | 0.242 | 0.12200116 | 0.25508522 | 0.0254 | 0.0096 | 0.0145 | 0.0145 | TRUE | 0.5107 |
| Anxa9 | 28 | 18 | 10 | 10 | 11 | 10 | 22 | 19 | 11 | 10 | 12 | 10 | 13.99 | 17 | 11 | 0.617 | -0.698 | 0.18899081 | NA | 0 | 0.1719 | 0.1272 | 0.1272 | TRUE | 0.4929 |
| Aoah | 113 | 79 | 54 | 22 | 19 | 33 | 88 | 84 | 58 | 22 | 21 | 33 | 50.99 | 77 | 25 | 0.33 | -1.6 | 3.14E-07 | 1.08E-05 | 0.0297 | 0.0489 | 0.0438 | 0.0438 | TRUE | 0.3985 |
| Aoc2 | 228 | 192 | 187 | 219 | 178 | 215 | 177 | 204 | 202 | 218 | 201 | 212 | 202.35 | 194 | 210 | 1.086 | 0.119 | 0.40435251 | 0.5771907 | 0 | 0.0141 | 0.0098 | 0.0098 | TRUE | 0.0726 |
| Aoc3 | 388 | 371 | 518 | 374 | 404 | 550 | 301 | 395 | 559 | 372 | 456 | 543 | 437.64 | 418 | 457 | 1.094 | 0.129 | 0.6673101 | 0.7954185 | 0.0628 | 0.0078 | 0.0233 | 0.0628 | TRUE | 0.456 |
| Aox1 | 93 | 62 | 71 | 106 | 88 | 109 | 72 | 66 | 77 | 106 | 99 | 108 | 87.86 | 72 | 104 | 1.455 | 0.541 | 0.00905143 | 0.0384746 | 0 | 0.0294 | 0.0193 | 0.0193 | TRUE | 0.0394 |
| Aox3 | 48 | 32 | 36 | 22 | 17 | 16 | 37 | 34 | 39 | 22 | 19 | 16 | 27.84 | 37 | 19 | 0.516 | -0.955 | 0.0099933 | 0.04135328 | 0 | 0.0876 | 0.0586 | 0.0586 | TRUE | 0.1181 |
| Aox3l1 | 2 | 3 | 1 | 2 | 5 | 5 | 2 | 3 | 1 | 2 | 6 | 5 | 3.07 | 2 | 4 | 2.168 | 1.116 | 0.32113126 | NA | 0 | 0.7757 | 0.538 | 0.538 | TRUE | 0.3768 |
| Aox4 | 112 | 81 | 76 | 65 | 46 | 51 | 87 | 86 | 82 | 65 | 52 | 50 | 70.35 | 85 | 56 | 0.654 | -0.612 | 0.00934203 | 0.03939973 | 0 | 0.0361 | 0.0249 | 0.0249 | TRUE | 0.1746 |
| Ap1ar | 3186 | 2318 | 2175 | 2084 | 1900 | 2199 | 2474 | 2466 | 2347 | 2074 | 2143 | 2171 | 2279.24 | 2429 | 2129 | 0.876 | -0.19 | 0.00165096 | 0.0103967 | 3.00E-04 | 0.0035 | 0.0022 | 0.0022 | TRUE | 0.0104 |
| Ap1b1 | 7577 | 5455 | 5129 | 6025 | 5377 | 6488 | 5885 | 5802 | 5534 | 5997 | 6066 | 6405 | 5948.17 | 5740 | 6156 | 1.072 | 0.101 | 0.06810821 | 0.16796046 | 9.00E-04 | 0.0028 | 0.002 | 0.002 | TRUE | 0.0153 |
| Ap1g1 | 14978 | 10494 | 10327 | 11377 | 9709 | 11168 | 11633 | 11162 | 11143 | 11325 | 10953 | 11025 | 11206.63 | 11313 | 11101 | 0.981 | -0.027 | 0.56776833 | 0.72147162 | 3.00E-04 | 0.0026 | 0.0016 | 0.0016 | TRUE | 0.0076 |
| Ap1g2 | 795 | 577 | 597 | 567 | 586 | 655 | 617 | 614 | 644 | 564 | 661 | 647 | 624.57 | 625 | 624 | 0.998 | -0.003 | 0.97541649 | 0.98666864 | 0.0022 | 0.0062 | 0.0049 | 0.0049 | TRUE | 0.083 |
| Ap1m1 | 3456 | 2490 | 2260 | 2311 | 2067 | 2343 | 2684 | 2649 | 2439 | 2300 | 2332 | 2313 | 2452.72 | 2591 | 2315 | 0.893 | -0.163 | 0.00949723 | 0.03986267 | 0.001 | 0.0034 | 0.0024 | 0.0024 | TRUE | 0.0315 |
| Ap1m2 | 309 | 269 | 343 | 346 | 369 | 410 | 240 | 286 | 370 | 344 | 416 | 405 | 343.6 | 299 | 388 | 1.305 | 0.384 | 0.01290084 | 0.04991653 | 0.0249 | 0.0093 | 0.0142 | 0.0142 | TRUE | 0.4899 |
| Ap1s1 | 2774 | 1874 | 1911 | 1974 | 1701 | 1922 | 2154 | 1993 | 2062 | 1965 | 1919 | 1897 | 1998.48 | 2070 | 1927 | 0.93 | -0.104 | 0.09991666 | 0.22083019 | 5.00E-04 | 0.0036 | 0.0024 | 0.0024 | TRUE | 0.016 |
| Ap1s2 | 1304 | 804 | 764 | 789 | 585 | 703 | 1013 | 855 | 824 | 785 | 660 | 694 | 805.27 | 897 | 713 | 0.794 | -0.334 | 0.00142167 | 0.00929123 | 0.0086 | 0.0054 | 0.0066 | 0.0066 | TRUE | 0.1553 |
| Ap1s3 | 1075 | 792 | 811 | 911 | 747 | 849 | 835 | 842 | 875 | 907 | 843 | 838 | 856.67 | 851 | 863 | 1.015 | 0.021 | 0.7951299 | 0.87996838 | 0 | 0.0052 | 0.0035 | 0.0035 | TRUE | 0.0243 |
| Ap2a1 | 14493 | 10402 | 10286 | 11097 | 10152 | 11721 | 11256 | 11064 | 11099 | 11046 | 11453 | 11570 | 11247.99 | 11140 | 11356 | 1.019 | 0.028 | 0.55642054 | 0.71154323 | 2.00E-04 | 0.0026 | 0.0015 | 0.0015 | TRUE | 0.007 |
| Ap2a2 | 15361 | 10959 | 11530 | 11763 | 10539 | 12346 | 11930 | 11657 | 12441 | 11709 | 11889 | 12187 | 11968.89 | 12009 | 11928 | 0.993 | -0.01 | 0.84671531 | 0.91302165 | 5.00E-04 | 0.0026 | 0.0017 | 0.0017 | TRUE | 0.012 |
| Ap2b1 | 20438 | 14405 | 15342 | 16442 | 14574 | 16811 | 15873 | 15322 | 16554 | 16366 | 16441 | 16595 | 16192 | 15916 | 16467 | 1.035 | 0.049 | 0.32148062 | 0.49524925 | 7.00E-04 | 0.0026 | 0.0017 | 0.0017 | TRUE | 0.0149 |
| Ap2m1 | 7192 | 5020 | 4651 | 5013 | 4223 | 5143 | 5586 | 5340 | 5018 | 4990 | 4764 | 5077 | 5129.11 | 5315 | 4944 | 0.93 | -0.105 | 0.08307853 | 0.19410862 | 0.0017 | 0.0029 | 0.0024 | 0.0024 | TRUE | 0.0289 |
| Ap2s1 | 2938 | 2097 | 2099 | 2217 | 1784 | 2331 | 2282 | 2231 | 2265 | 2207 | 2013 | 2301 | 2216.26 | 2259 | 2174 | 0.962 | -0.055 | 0.41653634 | 0.58934408 | 0.0017 | 0.0035 | 0.0029 | 0.0029 | TRUE | 0.0501 |
| Ap3b1 | 8217 | 6118 | 6157 | 5665 | 5225 | 6076 | 6382 | 6508 | 6643 | 5639 | 5894 | 5998 | 6177.34 | 6511 | 5844 | 0.898 | -0.156 | 0.0028734 | 0.0159928 | 5.00E-04 | 0.0028 | 0.0018 | 0.0018 | TRUE | 0.0115 |
| Ap3b2 | 176 | 137 | 153 | 196 | 204 | 214 | 137 | 146 | 165 | 195 | 230 | 211 | 180.66 | 149 | 212 | 1.425 | 0.511 | 0.00102463 | 0.00735933 | 0 | 0.0155 | 0.0118 | 0.0118 | TRUE | 0.0891 |
| Ap3d1 | 10098 | 7640 | 6049 | 5935 | 5691 | 6203 | 7843 | 8127 | 6527 | 5908 | 6420 | 6123 | 6824.52 | 7499 | 6150 | 0.82 | -0.286 | 0.00034905 | 0.00326289 | 0.0071 | 0.0028 | 0.0045 | 0.0045 | TRUE | 0.1563 |
| Ap3m1 | 4842 | 3546 | 3600 | 4023 | 3437 | 3792 | 3761 | 3772 | 3884 | 4004 | 3877 | 3743 | 3840.31 | 3806 | 3875 | 1.018 | 0.026 | 0.63534922 | 0.77174822 | 5.00E-04 | 0.003 | 0.002 | 0.002 | TRUE | 0.0109 |
| Ap3m2 | 930 | 842 | 1112 | 727 | 717 | 846 | 722 | 896 | 1200 | 724 | 809 | 835 | 864.23 | 939 | 789 | 0.842 | -0.249 | 0.08453386 | 0.19609882 | 0.0341 | 0.0052 | 0.0138 | 0.0138 | TRUE | 0.3519 |
| Ap3s1 | 1908 | 1346 | 1422 | 1567 | 1443 | 1481 | 1482 | 1432 | 1534 | 1560 | 1628 | 1462 | 1516.26 | 1483 | 1550 | 1.045 | 0.063 | 0.37476691 | 0.54920038 | 0.0013 | 0.004 | 0.003 | 0.003 | TRUE | 0.0298 |
| Ap3s2 | 5579 | 3863 | 3736 | 4067 | 3746 | 4297 | 4333 | 4109 | 4031 | 4048 | 4226 | 4242 | 4164.85 | 4158 | 4172 | 1.003 | 0.004 | 0.94043808 | 0.96671811 | 8.00E-04 | 0.003 | 0.0021 | 0.0021 | TRUE | 0.0169 |
| Ap4b1 | 1188 | 805 | 797 | 913 | 719 | 854 | 923 | 856 | 860 | 909 | 811 | 843 | 866.97 | 880 | 854 | 0.971 | -0.043 | 0.60717437 | 0.75124163 | 0.0015 | 0.0052 | 0.0039 | 0.0039 | TRUE | 0.0371 |
| Ap4e1 | 2898 | 2085 | 2081 | 2202 | 1804 | 2103 | 2251 | 2218 | 2245 | 2192 | 2035 | 2076 | 2169.48 | 2238 | 2101 | 0.939 | -0.091 | 0.13799794 | 0.27858822 | 2.00E-04 | 0.0035 | 0.0022 | 0.0022 | TRUE | 0.0174 |
| Ap4m1 | 756 | 523 | 488 | 509 | 490 | 524 | 587 | 556 | 527 | 507 | 553 | 517 | 541.12 | 557 | 526 | 0.942 | -0.086 | 0.37236864 | 0.54688296 | 0 | 0.0068 | 0.0049 | 0.0049 | TRUE | 0.0285 |
| Ap4s1 | 343 | 192 | 170 | 189 | 159 | 202 | 266 | 204 | 183 | 188 | 179 | 199 | 203.49 | 218 | 189 | 0.863 | -0.213 | 0.19952886 | 0.36017505 | 0.0168 | 0.0141 | 0.0149 | 0.0149 | TRUE | 0.4392 |
| Ap5b1 | 330 | 184 | 149 | 166 | 167 | 124 | 256 | 196 | 161 | 165 | 188 | 122 | 181.47 | 204 | 158 | 0.772 | -0.373 | 0.06677065 | 0.16557129 | 0.0455 | 0.0155 | 0.0242 | 0.0242 | TRUE | 0.3371 |
| Ap5m1 | 1533 | 1092 | 1120 | 1190 | 1045 | 1160 | 1191 | 1162 | 1208 | 1185 | 1179 | 1145 | 1178.19 | 1187 | 1170 | 0.985 | -0.022 | 0.75799781 | 0.85637995 | 0 | 0.0044 | 0.0027 | 0.0027 | TRUE | 0.0041 |
| Ap5s1 | 400 | 277 | 235 | 247 | 270 | 266 | 311 | 295 | 254 | 246 | 305 | 263 | 278.65 | 287 | 271 | 0.943 | -0.085 | 0.53794595 | 0.69645484 | 0.0078 | 0.0109 | 0.01 | 0.01 | TRUE | 0.1275 |
| Ap5z1 | 691 | 587 | 430 | 400 | 401 | 423 | 537 | 624 | 464 | 398 | 452 | 418 | 482.19 | 542 | 423 | 0.78 | -0.359 | 0.00313836 | 0.01709474 | 0.0111 | 0.0073 | 0.0085 | 0.0085 | TRUE | 0.1829 |
| Apaf1 | 5247 | 3655 | 4059 | 4277 | 3569 | 4115 | 4075 | 3888 | 4380 | 4257 | 4026 | 4062 | 4114.7 | 4114 | 4115 | 1 | 0.001 | 0.99194974 | 0.99565418 | 0.002 | 0.003 | 0.0026 | 0.0026 | TRUE | 0.0382 |
| Apba1 | 47 | 18 | 20 | 25 | 20 | 29 | 37 | 19 | 22 | 25 | 23 | 29 | 25.55 | 26 | 26 | 0.973 | -0.039 | 0.92265546 | 0.95719768 | 0.0396 | 0.0952 | 0.0788 | 0.0788 | TRUE | 0.9751 |
| Apba2 | 362 | 327 | 325 | 232 | 248 | 278 | 281 | 348 | 351 | 231 | 280 | 274 | 294.13 | 327 | 262 | 0.803 | -0.316 | 0.0227001 | 0.07568289 | 0.0101 | 0.0105 | 0.0104 | 0.0104 | TRUE | 0.18 |
| Apba3 | 2166 | 1592 | 1463 | 1385 | 1189 | 1432 | 1682 | 1693 | 1579 | 1379 | 1341 | 1414 | 1514.62 | 1651 | 1378 | 0.834 | -0.261 | 0.00012007 | 0.00141475 | 4.00E-04 | 0.004 | 0.0027 | 0.0027 | TRUE | 0.0175 |
| Apbb1 | 1235 | 932 | 944 | 889 | 784 | 897 | 959 | 991 | 1019 | 885 | 884 | 885 | 937.32 | 990 | 885 | 0.895 | -0.16 | 0.03380705 | 0.10127944 | 0 | 0.005 | 0.003 | 0.003 | TRUE | 0.0092 |
| Apbb1ip | 40 | 31 | 21 | 27 | 27 | 27 | 31 | 33 | 23 | 27 | 30 | 27 | 28.45 | 29 | 28 | 0.964 | -0.053 | 0.88450869 | 0.93598006 | 0 | 0.0857 | 0.0591 | 0.0591 | TRUE | 0.2135 |
| Apbb2 | 1702 | 1224 | 1223 | 1383 | 1318 | 1422 | 1322 | 1302 | 1320 | 1377 | 1487 | 1404 | 1368.44 | 1315 | 1423 | 1.082 | 0.113 | 0.09947168 | 0.22032368 | 0 | 0.0042 | 0.0027 | 0.0027 | TRUE | 0.0185 |
| Apbb3 | 469 | 364 | 311 | 297 | 214 | 279 | 364 | 387 | 336 | 296 | 241 | 275 | 316.58 | 362 | 271 | 0.748 | -0.418 | 0.00088321 | 0.00663014 | 0.0037 | 0.0099 | 0.0082 | 0.0082 | TRUE | 0.097 |
| Apc | 9660 | 6867 | 6691 | 7506 | 6734 | 7940 | 7503 | 7304 | 7220 | 7471 | 7597 | 7838 | 7488.75 | 7342 | 7635 | 1.04 | 0.056 | 0.25887519 | 0.42878846 | 3.00E-04 | 0.0027 | 0.0017 | 0.0017 | TRUE | 0.0066 |
| Apc2 | 155 | 142 | 150 | 212 | 181 | 186 | 120 | 151 | 162 | 211 | 204 | 184 | 172.02 | 144 | 200 | 1.39 | 0.475 | 0.0042817 | 0.0217943 | 0.0083 | 0.0162 | 0.0139 | 0.0139 | TRUE | 0.2471 |
| Apcdd1 | 529 | 407 | 411 | 405 | 343 | 375 | 411 | 433 | 443 | 403 | 387 | 370 | 407.92 | 429 | 387 | 0.902 | -0.148 | 0.15871437 | 0.30792785 | 0 | 0.0082 | 0.0055 | 0.0055 | TRUE | 0.0172 |
| Apcs | 0 | 0 | 0 | 0 | 0 | 0 | 0 | 0 | 0 | 0 | 0 | 0 | 0 | 0 | 0 | NA | NA | NA | NA | NA | NA | NA | NA | NA | NA |
| Apeh | 2385 | 1893 | 1765 | 1843 | 1717 | 1861 | 1852 | 2014 | 1904 | 1835 | 1937 | 1837 | 1896.48 | 1923 | 1870 | 0.972 | -0.041 | 0.53827307 | 0.69678096 | 9.00E-04 | 0.0037 | 0.0026 | 0.0026 | TRUE | 0.0202 |
| Apex1 | 9544 | 6388 | 5302 | 5475 | 5153 | 5890 | 7412 | 6795 | 5721 | 5450 | 5813 | 5814 | 6167.56 | 6643 | 5692 | 0.857 | -0.223 | 0.00844209 | 0.03648709 | 0.0087 | 0.0028 | 0.005 | 0.005 | TRUE | 0.1795 |
| Apex2 | 603 | 420 | 351 | 327 | 325 | 368 | 468 | 447 | 379 | 325 | 367 | 363 | 391.53 | 431 | 352 | 0.814 | -0.298 | 0.01267032 | 0.04924675 | 0.0059 | 0.0085 | 0.0077 | 0.0077 | TRUE | 0.1283 |
| Aph1a | 1753 | 1244 | 1376 | 1477 | 1359 | 1489 | 1361 | 1323 | 1485 | 1470 | 1533 | 1470 | 1440.43 | 1390 | 1491 | 1.073 | 0.102 | 0.15955287 | 0.30888381 | 0.0013 | 0.0041 | 0.0031 | 0.0031 | TRUE | 0.0417 |
| Aph1b | 216 | 152 | 134 | 163 | 104 | 157 | 168 | 162 | 145 | 162 | 117 | 155 | 151.43 | 158 | 145 | 0.917 | -0.124 | 0.47987434 | 0.64663372 | 0.0095 | 0.0181 | 0.0157 | 0.0157 | TRUE | 0.2744 |
| Aph1c | 163 | 125 | 120 | 160 | 139 | 125 | 127 | 133 | 129 | 159 | 157 | 123 | 138.08 | 130 | 146 | 1.129 | 0.175 | 0.31764642 | 0.4915678 | 0 | 0.0196 | 0.0149 | 0.0149 | TRUE | 0.2032 |
| Api5 | 15126 | 10950 | 10423 | 10457 | 9306 | 10415 | 11748 | 11647 | 11246 | 10409 | 10498 | 10281 | 10971.61 | 11547 | 10396 | 0.9 | -0.152 | 0.00125144 | 0.00849327 | 2.00E-04 | 0.0026 | 0.0015 | 0.0015 | TRUE | 0.0063 |
| Apip | 1126 | 799 | 729 | 841 | 775 | 870 | 875 | 850 | 787 | 837 | 874 | 859 | 846.87 | 837 | 857 | 1.022 | 0.032 | 0.69857929 | 0.81695252 | 0 | 0.0052 | 0.0036 | 0.0036 | TRUE | 0.0324 |
| Apitd1 | 1083 | 776 | 776 | 691 | 623 | 697 | 841 | 825 | 837 | 688 | 703 | 688 | 763.75 | 834 | 693 | 0.83 | -0.269 | 0.00076328 | 0.00594206 | 0 | 0.0055 | 0.0033 | 0.0033 | TRUE | 0.0018 |
| Aplf | 868 | 606 | 570 | 590 | 583 | 631 | 674 | 645 | 615 | 587 | 658 | 623 | 633.6 | 645 | 623 | 0.964 | -0.053 | 0.5662493 | 0.72043806 | 0.0011 | 0.0062 | 0.0045 | 0.0045 | TRUE | 0.029 |
| Apln | 150 | 131 | 144 | 251 | 189 | 198 | 116 | 139 | 155 | 250 | 213 | 195 | 178.29 | 137 | 219 | 1.61 | 0.687 | 5.21E-05 | 0.00072392 | 0.0129 | 0.0157 | 0.0149 | 0.0149 | TRUE | 0.1992 |
| Aplnr | 0 | 0 | 0 | 0 | 0 | 0 | 0 | 0 | 0 | 0 | 0 | 0 | 0 | 0 | 0 | NA | NA | NA | NA | NA | NA | NA | NA | NA | NA |
| Aplp1 | 3018 | 2132 | 2674 | 4065 | 3330 | 3930 | 2344 | 2268 | 2885 | 4046 | 3757 | 3880 | 3196.56 | 2499 | 3894 | 1.559 | 0.641 | 1.74E-13 | 2.48E-11 | 0.0093 | 0.0032 | 0.0051 | 0.0051 | TRUE | 0.2198 |
| Aplp2 | 21023 | 15631 | 15769 | 17510 | 15534 | 17535 | 16328 | 16626 | 17015 | 17429 | 17524 | 17310 | 17038.66 | 16656 | 17421 | 1.046 | 0.065 | 0.14911467 | 0.29437485 | 1.00E-04 | 0.0026 | 0.0014 | 0.0014 | TRUE | 0.0043 |
| Apmap | 1059 | 678 | 572 | 679 | 564 | 631 | 822 | 721 | 617 | 676 | 636 | 623 | 682.64 | 720 | 645 | 0.893 | -0.162 | 0.13927692 | 0.28048626 | 0.01 | 0.0059 | 0.0072 | 0.0072 | TRUE | 0.174 |
| Apoa1 | 9 | 4 | 4 | 4 | 5 | 7 | 7 | 4 | 4 | 4 | 6 | 7 | 5.35 | 5 | 6 | 1.048 | 0.067 | 0.93451918 | NA | 0 | 0.4455 | 0.294 | 0.294 | TRUE | 0.2539 |
| Apoa1bp | 2653 | 2006 | 2010 | 2415 | 2067 | 2396 | 2060 | 2134 | 2169 | 2404 | 2332 | 2365 | 2243.98 | 2121 | 2367 | 1.117 | 0.159 | 0.0073673 | 0.03305113 | 0 | 0.0035 | 0.0021 | 0.0021 | TRUE | 0.0078 |
| Apoa2 | 6 | 7 | 8 | 3 | 6 | 12 | 5 | 7 | 9 | 3 | 7 | 12 | 7.06 | 7 | 7 | 1.056 | 0.078 | 0.9183047 | NA | 0.1143 | 0.3383 | 0.2746 | 0.2746 | TRUE | 0.2753 |

| | | | | | | | | | | | | | | | | | | | | | | | | | |
|---|---|---|---|---|---|---|---|---|---|---|---|---|---|---|---|---|---|---|---|---|---|---|---|---|---|
| Apoa4 | 1 | 0 | 0 | 1 | 0 | 0 | 1 | 0 | 0 | 1 | 0 | 0 | 0.3 | 0 | 0 | 1.075 | 0.105 | 0.97698525 | NA | 0 | 8.0272 | 7.5141 | 7.5141 | TRUE | 0.3906 |
| Apoa5 | 4 | 0 | 3 | 2 | 0 | 1 | 3 | 0 | 3 | 2 | 0 | 1 | 1.55 | 2 | 1 | 0.473 | -1.081 | 0.52212622 | NA | 0 | 1.528 | 1.3186 | 1.3186 | TRUE | 0.527 |
| Apob | 142 | 75 | 45 | 53 | 49 | 44 | 110 | 80 | 49 | 53 | 55 | 43 | 65.01 | 80 | 50 | 0.629 | -0.669 | 0.02792911 | 0.08790811 | 0.0776 | 0.0389 | 0.0504 | 0.0504 | TRUE | 0.1292 |
| Apobec1 | 383 | 288 | 352 | 484 | 395 | 461 | 297 | 306 | 380 | 482 | 446 | 455 | 394.34 | 328 | 461 | 1.41 | 0.496 | 4.12E-05 | 0.00059971 | 0.0064 | 0.0084 | 0.0079 | 0.0079 | TRUE | 0.2129 |
| Apobec2 | 137 | 100 | 101 | 130 | 150 | 151 | 106 | 106 | 109 | 129 | 169 | 149 | 128.24 | 107 | 149 | 1.389 | 0.474 | 0.00868022 | 0.03725718 | 0 | 0.0209 | 0.0156 | 0.0156 | TRUE | 0.1415 |
| Apobec3 | 7321 | 5817 | 6799 | 7190 | 6761 | 7722 | 5686 | 6187 | 7336 | 7157 | 7627 | 7623 | 6936.05 | 6403 | 7469 | 1.167 | 0.223 | 0.00817321 | 0.03561576 | 0.0092 | 0.0028 | 0.005 | 0.005 | TRUE | 0.198 |
| Apobec4 | 0 | 0 | 0 | 0 | 0 | 0 | 0 | 0 | 0 | 0 | 0 | 0 | 0 | 0 | 0 | NA | NA | NA | NA | NA | NA | NA | NA | NA | NA |
| Apobr | 102 | 66 | 51 | 94 | 69 | 87 | 79 | 70 | 55 | 94 | 78 | 86 | 76.96 | 68 | 86 | 1.252 | 0.325 | 0.16352783 | 0.31417234 | 0.0061 | 0.0332 | 0.026 | 0.026 | TRUE | 0.2424 |
| Apoc1 | 377 | 283 | 347 | 407 | 322 | 373 | 293 | 301 | 374 | 405 | 363 | 368 | 350.8 | 323 | 379 | 1.177 | 0.236 | 0.06435189 | 0.16139351 | 0.008 | 0.0092 | 0.0088 | 0.0088 | TRUE | 0.2174 |
| Apoc2 | 6 | 4 | 8 | 9 | 7 | 3 | 5 | 4 | 9 | 9 | 8 | 3 | 6.23 | 6 | 7 | 1.135 | 0.183 | 0.81777763 | NA | 0 | 0.383 | 0.2919 | 0.2919 | TRUE | 0.6177 |
| Apoc3 | 1 | 1 | 0 | 0 | 0 | 0 | 1 | 1 | 0 | 0 | 0 | 0 | 0.31 | 1 | 0 | 0.309 | -1.696 | 0.63183309 | NA | 0 | 7.7294 | 7.2472 | 7.2472 | TRUE | 0.2015 |
| Apoc4 | 1 | 4 | 2 | 7 | 4 | 3 | 1 | 4 | 2 | 7 | 5 | 3 | 3.61 | 2 | 5 | 2.076 | 1.054 | 0.31926619 | NA | 0 | 0.6598 | 0.4917 | 0.4917 | TRUE | 0.4627 |
| Apod | 211 | 150 | 162 | 178 | 160 | 187 | 164 | 160 | 175 | 177 | 180 | 185 | 173.42 | 166 | 181 | 1.089 | 0.123 | 0.40964992 | 0.58242577 | 0 | 0.0161 | 0.0103 | 0.0103 | TRUE | 0.021 |
| Apoe | 4131 | 3542 | 4833 | 6092 | 5924 | 6839 | 3208 | 3768 | 5215 | 6064 | 6683 | 6751 | 5281.45 | 4064 | 6499 | 1.6 | 0.678 | 0.00182292 | 0.01125971 | 0.0339 | 0.0029 | 0.0113 | 0.0339 | TRUE | 0.7474 |
| Apof | 47 | 43 | 25 | 28 | 31 | 34 | 37 | 46 | 27 | 28 | 35 | 34 | 34.27 | 37 | 32 | 0.881 | -0.183 | 0.59381353 | 0.74083813 | 0 | 0.0716 | 0.0551 | 0.0551 | TRUE | 0.1578 |
| Apoh | 2 | 0 | 0 | 0 | 0 | 0 | 2 | 0 | 0 | 0 | 0 | 0 | 0.26 | 1 | 0 | 0.356 | -1.491 | 0.71362796 | NA | 0 | 9.1573 | 10 | 10 | TRUE | 0.968 |
| Apol10a | 0 | 1 | 0 | 0 | 0 | 0 | 0 | 1 | 0 | 0 | 0 | 0 | 0.18 | 0 | 0 | 0.552 | -0.857 | 0.8336548 | NA | 0 | 13.3714 | 10 | 10 | TRUE | 0.5216 |
| Apol10b | 11 | 3 | 6 | 6 | 3 | 4 | 9 | 3 | 6 | 6 | 3 | 4 | 5.25 | 6 | 4 | 0.724 | -0.466 | 0.5809559 | NA | 0 | 0.4537 | 0.3196 | 0.3196 | TRUE | 0.2723 |
| Apol11a | 0 | 0 | 0 | 0 | 0 | 0 | 0 | 0 | 0 | 0 | 0 | 0 | 0 | 0 | 0 | NA | NA | NA | NA | NA | NA | NA | NA | NA | NA |
| Apol11b | 0 | 0 | 0 | 0 | 0 | 0 | 0 | 0 | 0 | 0 | 0 | 0 | 0 | 0 | 0 | NA | NA | NA | NA | NA | NA | NA | NA | NA | NA |
| Apol6 | 12 | 7 | 2 | 1 | 4 | 10 | 9 | 7 | 2 | 1 | 5 | 10 | 5.72 | 6 | 5 | 0.804 | -0.315 | 0.73196388 | NA | 0.476 | 0.417 | 0.4329 | 0.4329 | TRUE | 0.4154 |
| Apol7a | 31 | 23 | 21 | 13 | 28 | 24 | 24 | 24 | 23 | 13 | 32 | 24 | 23.24 | 24 | 23 | 0.952 | -0.071 | 0.86879525 | 0.92759322 | 0.0411 | 0.1044 | 0.0881 | 0.0881 | TRUE | 0.298 |
| Apol7b | 49 | 40 | 14 | 22 | 16 | 24 | 38 | 43 | 15 | 22 | 18 | 24 | 26.56 | 32 | 21 | 0.663 | -0.593 | 0.17177949 | 0.32382541 | 0.1084 | 0.0917 | 0.0963 | 0.0963 | TRUE | 1.351 |
| Apol7c | 1 | 0 | 0 | 1 | 1 | 1 | 1 | 0 | 0 | 1 | 1 | 1 | 0.65 | 0 | 1 | 3.004 | 1.587 | 0.49664754 | NA | 0 | 3.6606 | 2.4271 | 2.4271 | TRUE | 0.2566 |
| Apol7d | 23 | 24 | 16 | 24 | 31 | 39 | 18 | 26 | 17 | 24 | 35 | 38 | 26.34 | 20 | 32 | 1.611 | 0.688 | 0.08064689 | 0.19021222 | 0 | 0.0924 | 0.0718 | 0.0718 | TRUE | 0.3766 |
| Apol7e | 14 | 11 | 2 | 11 | 13 | 9 | 11 | 12 | 2 | 11 | 15 | 9 | 9.87 | 8 | 12 | 1.373 | 0.457 | 0.49507268 | NA | 0.1437 | 0.2425 | 0.2205 | 0.2205 | TRUE | 1.1268 |
| Apol8 | 29 | 18 | 30 | 40 | 21 | 25 | 23 | 19 | 32 | 40 | 24 | 25 | 27.04 | 25 | 30 | 1.198 | 0.261 | 0.51258588 | 0.67504942 | 0.0436 | 0.0901 | 0.0771 | 0.0771 | TRUE | 0.6427 |
| Apol9a | 0 | 0 | 0 | 0 | 1 | 0 | 0 | 0 | 0 | 0 | 1 | 0 | 0.19 | 0 | 0 | 2.094 | 1.067 | 0.79379329 | NA | 0 | 12.608 | 10 | 10 | TRUE | 0.5075 |
| Apol9b | 0 | 0 | 0 | 0 | 0 | 0 | 0 | 0 | 0 | 0 | 0 | 0 | 0 | 0 | 0 | NA | NA | NA | NA | NA | NA | NA | NA | NA | NA |
| Apold1 | 84 | 67 | 58 | 48 | 36 | 38 | 65 | 71 | 63 | 48 | 41 | 38 | 54.17 | 66 | 42 | 0.633 | -0.66 | 0.01309546 | 0.05046465 | 0 | 0.0462 | 0.0313 | 0.0313 | TRUE | 0.1159 |
| Apom | 26 | 24 | 20 | 20 | 14 | 14 | 20 | 26 | 22 | 20 | 16 | 14 | 19.47 | 23 | 17 | 0.739 | -0.437 | 0.31605549 | 0.48990144 | 0 | 0.1242 | 0.0838 | 0.0838 | TRUE | 0.163 |
| Apon | 3 | 0 | 1 | 1 | 0 | 2 | 2 | 0 | 1 | 1 | 0 | 2 | 1.06 | 1 | 1 | 0.856 | -0.224 | 0.90756607 | NA | 0 | 2.2318 | 1.7592 | 1.7592 | TRUE | 0.2268 |
| Apoo | 239 | 142 | 150 | 167 | 130 | 159 | 186 | 151 | 162 | 166 | 147 | 157 | 161.39 | 166 | 157 | 0.94 | -0.09 | 0.58069207 | 0.73139159 | 0 | 0.0171 | 0.0128 | 0.0128 | TRUE | 0.1218 |
| Apoo-ps | 0 | 0 | 0 | 0 | 0 | 0 | 0 | 0 | 0 | 0 | 0 | 0 | 0 | 0 | 0 | NA | NA | NA | NA | NA | NA | NA | NA | NA | NA |
| Apool | 1214 | 904 | 875 | 1098 | 877 | 977 | 943 | 962 | 944 | 1093 | 989 | 964 | 982.55 | 950 | 1015 | 1.07 | 0.098 | 0.22331247 | 0.38773143 | 0.0013 | 0.0048 | 0.0036 | 0.0036 | TRUE | 0.0537 |
| Apopt1 | 787 | 622 | 588 | 658 | 621 | 645 | 611 | 662 | 634 | 655 | 701 | 637 | 649.92 | 636 | 664 | 1.045 | 0.063 | 0.47862713 | 0.64580449 | 0 | 0.0061 | 0.0042 | 0.0042 | TRUE | 0.0265 |
| App | 1562 | 1137 | 1316 | 1216 | 1169 | 1287 | 1213 | 1209 | 1420 | 1210 | 1319 | 1270 | 1273.69 | 1281 | 1266 | 0.989 | -0.015 | 0.85561369 | 0.91884279 | 0.0044 | 0.0043 | 0.0043 | 0.0043 | TRUE | 0.1067 |
| Appbp2 | 7455 | 5543 | 5119 | 5614 | 4852 | 5438 | 5790 | 5896 | 5523 | 5588 | 5474 | 5368 | 5606.54 | 5736 | 5477 | 0.955 | -0.067 | 0.2072462 | 0.36907176 | 5.00E-04 | 0.0028 | 0.0019 | 0.0019 | TRUE | 0.0127 |
| Appl1 | 4768 | 3461 | 3914 | 3881 | 3507 | 3821 | 3703 | 3681 | 4223 | 3863 | 3956 | 3772 | 3866.5 | 3869 | 3864 | 0.999 | -0.002 | 0.98065939 | 0.9899403 | 0.0031 | 0.003 | 0.0031 | 0.0031 | TRUE | 0.0773 |
| Appl2 | 1908 | 1497 | 1719 | 1798 | 1660 | 1768 | 1482 | 1592 | 1855 | 1790 | 1873 | 1745 | 1722.78 | 1643 | 1803 | 1.098 | 0.135 | 0.11849915 | 0.24997369 | 0.0066 | 0.0038 | 0.0048 | 0.0048 | TRUE | 0.1526 |
| Aprt | 7824 | 5587 | 5645 | 5836 | 5258 | 6294 | 6077 | 5943 | 6091 | 5809 | 5932 | 6213 | 6010.69 | 6037 | 5985 | 0.991 | -0.013 | 0.81056277 | 0.89074498 | 5.00E-04 | 0.0028 | 0.0018 | 0.0018 | TRUE | 0.0137 |
| Aptx | 976 | 655 | 668 | 809 | 673 | 775 | 758 | 697 | 721 | 805 | 759 | 765 | 750.84 | 725 | 776 | 1.07 | 0.097 | 0.24700527 | 0.41568896 | 0 | 0.0056 | 0.0038 | 0.0038 | TRUE | 0.0197 |
| Aqp1 | 7 | 2 | 1 | 2 | 2 | 0 | 5 | 2 | 1 | 2 | 2 | 0 | 2.15 | 3 | 1 | 0.473 | -1.079 | 0.44343921 | NA | 0 | 1.1056 | 0.8941 | 0.8941 | TRUE | 0.9781 |
| Aqp11 | 378 | 274 | 280 | 327 | 321 | 354 | 294 | 291 | 302 | 325 | 362 | 349 | 320.7 | 296 | 345 | 1.169 | 0.225 | 0.05127098 | 0.13733434 | 0 | 0.0098 | 0.0065 | 0.0065 | TRUE | 0.03 |
| Aqp12 | 0 | 0 | 0 | 0 | 0 | 0 | 0 | 0 | 0 | 0 | 0 | 0 | 0 | 0 | 0 | NA | NA | NA | NA | NA | NA | NA | NA | NA | NA |
| Aqp2 | 5 | 4 | 3 | 12 | 9 | 9 | 4 | 4 | 3 | 12 | 10 | 9 | 7.06 | 4 | 10 | 2.719 | 1.443 | 0.05125027 | NA | 0 | 0.3382 | 0.2183 | 0.2183 | TRUE | 0.069 |
| Aqp3 | 7408 | 4546 | 3410 | 3510 | 3076 | 3190 | 5753 | 4836 | 3679 | 3494 | 3470 | 3149 | 4063.55 | 4756 | 3371 | 0.709 | -0.497 | 0.0081063 | 0.0354194 | 0.0251 | 0.003 | 0.0099 | 0.0251 | TRUE | 0.1846 |
| Aqp4 | 3 | 2 | 0 | 1 | 0 | 1 | 2 | 2 | 0 | 1 | 0 | 1 | 1.07 | 1 | 1 | 0.446 | -1.164 | 0.55264457 | NA | 0 | 2.2106 | 1.683 | 1.683 | TRUE | 0.4676 |
| Aqp5 | 200 | 131 | 111 | 233 | 203 | 226 | 155 | 139 | 120 | 232 | 229 | 223 | 183.08 | 138 | 228 | 1.642 | 0.716 | 3.97E-06 | 8.78E-05 | 0 | 0.0154 | 0.0116 | 0.0116 | TRUE | 0.1361 |
| Aqp6 | 57 | 54 | 57 | 66 | 65 | 67 | 44 | 57 | 62 | 66 | 73 | 66 | 61.4 | 54 | 68 | 1.265 | 0.339 | 0.18140822 | 0.33695486 | 0 | 0.041 | 0.0299 | 0.0299 | TRUE | 0.2618 |
| Aqp7 | 5 | 5 | 2 | 1 | 1 | 5 | 4 | 5 | 2 | 1 | 1 | 5 | 3.07 | 4 | 2 | 0.627 | -0.673 | 0.55347318 | NA | 0.0625 | 0.7745 | 0.5851 | 0.5851 | TRUE | 0.8439 |
| Aqp8 | 10 | 8 | 3 | 7 | 3 | 3 | 8 | 9 | 3 | 7 | 3 | 3 | 5.47 | 7 | 4 | 0.68 | -0.556 | 0.51046749 | NA | 0 | 0.4356 | 0.3247 | 0.3247 | TRUE | 0.4681 |
| Aqp9 | 32 | 21 | 17 | 4 | 6 | 9 | 25 | 22 | 18 | 4 | 7 | 9 | 14.19 | 22 | 7 | 0.298 | -1.745 | 0.00173966 | NA | 0 | 0.1694 | 0.1226 | 0.1226 | TRUE | 0.3125 |
| Aqr | 10460 | 7468 | 7510 | 7578 | 6529 | 7927 | 8124 | 7944 | 8103 | 7543 | 7365 | 7825 | 7817.4 | 8057 | 7578 | 0.941 | -0.088 | 0.077074 | 0.1837869 | 4.00E-04 | 0.0027 | 0.0017 | 0.0017 | TRUE | 0.01 |
| Ar | 20 | 12 | 14 | 20 | 6 | 9 | 16 | 13 | 15 | 20 | 7 | 9 | 13.16 | 15 | 12 | 0.823 | -0.282 | 0.61858478 | NA | 0.0752 | 0.1825 | 0.154 | 0.154 | TRUE | 1.3987 |
| Araf | 3395 | 2659 | 2698 | 3085 | 2650 | 3259 | 2637 | 2828 | 2911 | 3071 | 2989 | 3217 | 2942.27 | 2792 | 3092 | 1.109 | 0.149 | 0.02015049 | 0.06956788 | 0.0015 | 0.0032 | 0.0026 | 0.0026 | TRUE | 0.0295 |
| Arap1 | 2915 | 2219 | 2386 | 2632 | 2482 | 2954 | 2264 | 2360 | 2574 | 2620 | 2800 | 2916 | 2589.11 | 2399 | 2779 | 1.159 | 0.213 | 0.00299033 | 0.01650223 | 0.0031 | 0.0033 | 0.0033 | 0.0033 | TRUE | 0.049 |
| Arap2 | 770 | 537 | 527 | 601 | 480 | 543 | 598 | 571 | 569 | 598 | 541 | 536 | 568.93 | 579 | 558 | 0.964 | -0.053 | 0.5715447 | 0.72478826 | 0 | 0.0066 | 0.0046 | 0.0046 | TRUE | 0.0458 |
| Arap3 | 932 | 625 | 616 | 672 | 618 | 657 | 724 | 665 | 665 | 669 | 697 | 649 | 677.99 | 685 | 672 | 0.979 | -0.03 | 0.73117334 | 0.83927567 | 0 | 0.0059 | 0.0041 | 0.0041 | TRUE | 0.0317 |
| Arc | 705 | 629 | 666 | 667 | 608 | 753 | 548 | 669 | 719 | 664 | 686 | 743 | 671.39 | 645 | 698 | 1.085 | 0.117 | 0.28857115 | 0.46071562 | 0.0101 | 0.006 | 0.0073 | 0.0073 | TRUE | 0.2164 |
| Arcn1 | 17551 | 12310 | 12986 | 14177 | 12601 | 14373 | 13631 | 13094 | 14012 | 14112 | 14215 | 14188 | 13875.38 | 13579 | 14172 | 1.044 | 0.062 | 0.20332038 | 0.36472149 | 5.00E-04 | 0.0026 | 0.0016 | 0.0016 | TRUE | 0.0118 |
| Areg | 23 | 17 | 12 | 5 | 7 | 6 | 18 | 18 | 13 | 5 | 8 | 6 | 11.28 | 16 | 6 | 0.381 | -1.392 | 0.01974161 | NA | 0 | 0.2125 | 0.1444 | 0.1444 | TRUE | 0.1434 |
| Arf1 | 25051 | 17709 | 18326 | 18550 | 16470 | 18750 | 19456 | 18837 | 19774 | 18465 | 18580 | 18509 | 18936.69 | 19356 | 18518 | 0.957 | -0.064 | 0.15996039 | 0.30945258 | 2.00E-04 | 0.0026 | 0.0014 | 0.0014 | TRUE | 0.0067 |
| Arf2 | 2627 | 1735 | 1575 | 1816 | 1593 | 1660 | 2040 | 1845 | 1699 | 1808 | 1797 | 1639 | 1804.76 | 1861 | 1748 | 0.938 | -0.093 | 0.25621041 | 0.42612273 | 0.0049 | 0.0037 | 0.0043 | 0.0043 | TRUE | 0.087 |
| Arf3 | 9526 | 6529 | 5693 | 5790 | 5112 | 5925 | 7398 | 6945 | 6143 | 5763 | 5767 | 5849 | 6310.84 | 6829 | 5793 | 0.848 | -0.238 | 0.00070991 | 0.00562462 | 0.0041 | 0.0028 | 0.0034 | 0.0034 | TRUE | 0.094 |
| Arf4 | 8914 | 6029 | 6464 | 7341 | 6031 | 6980 | 6923 | 6413 | 6975 | 7307 | 6804 | 6890 | 6885.31 | 6770 | 7000 | 1.034 | 0.048 | 0.40804018 | 0.58080247 | 0.0016 | 0.0028 | 0.0023 | 0.0023 | TRUE | 0.0258 |
| Arf5 | 4193 | 3175 | 3539 | 3495 | 3237 | 3762 | 3257 | 3377 | 3819 | 3479 | 3652 | 3714 | 3549.42 | 3484 | 3615 | 1.038 | 0.054 | 0.44948157 | 0.61958145 | 0.0034 | 0.0031 | 0.0034 | 0.0034 | TRUE | 0.0853 |
| Arf6 | 8037 | 5889 | 5777 | 6125 | 5204 | 5804 | 6242 | 6264 | 6233 | 6097 | 5871 | 5729 | 6072.72 | 6246 | 5899 | 0.944 | -0.083 | 0.10471038 | 0.22870193 | 2.00E-04 | 0.0028 | 0.0017 | 0.0017 | TRUE | 0.0106 |
| Arfgap1 | 2164 | 1435 | 1234 | 1348 | 1165 | 1279 | 1681 | 1526 | 1331 | 1342 | 1314 | 1263 | 1409.53 | 1513 | 1306 | 0.862 | -0.214 | 0.01646494 | 0.05985082 | 0.0061 | 0.0041 | 0.005 | 0.005 | TRUE | 0.1316 |

| | | | | | | | | | | | | | | | | | | | | | | | | | |
|---|---|---|---|---|---|---|---|---|---|---|---|---|---|---|---|---|---|---|---|---|---|---|---|---|---|
| Arfgap2 | 2484 | 1773 | 1500 | 1528 | 1280 | 1518 | 1929 | 1886 | 1618 | 1521 | 1444 | 1499 | 1649.51 | 1811 | 1488 | 0.821 | -0.285 | 0.00038474 | 0.00352959 | 0.0043 | 0.0039 | 0.004 | 0.004 | TRUE | 0.1033 |
| Arfgap3 | 874 | 732 | 824 | 860 | 756 | 917 | 679 | 779 | 889 | 856 | 853 | 905 | 826.77 | 782 | 871 | 1.117 | 0.159 | 0.11987775 | 0.25155727 | 0.0085 | 0.0053 | 0.0063 | 0.0063 | TRUE | 0.1696 |
| Arfgef1 | 7510 | 5572 | 5912 | 6343 | 5466 | 5895 | 5833 | 5927 | 6379 | 6314 | 6166 | 5819 | 6072.98 | 6046 | 6100 | 1.009 | 0.013 | 0.8279514 | 0.90144951 | 0.0019 | 0.0028 | 0.0024 | 0.0024 | TRUE | 0.0281 |
| Arfgef2 | 7527 | 5533 | 5365 | 5689 | 5209 | 5982 | 5846 | 5885 | 5789 | 5663 | 5876 | 5905 | 5827.39 | 5840 | 5815 | 0.996 | -0.006 | 0.89854603 | 0.94384643 | 1.00E-04 | 0.0028 | 0.0016 | 0.0016 | TRUE | 0.0064 |
| Arfip1 | 2435 | 1831 | 1707 | 1959 | 1664 | 1897 | 1891 | 1948 | 1842 | 1950 | 1877 | 1873 | 1896.73 | 1894 | 1900 | 1.003 | 0.005 | 0.9364332 | 0.96471383 | 0 | 0.0037 | 0.0023 | 0.0023 | TRUE | 0.0073 |
| Arfip2 | 1427 | 968 | 941 | 971 | 885 | 940 | 1108 | 1030 | 1015 | 967 | 998 | 928 | 1007.68 | 1051 | 964 | 0.916 | -0.127 | 0.10529684 | 0.22977218 | 0.001 | 0.0048 | 0.0034 | 0.0034 | TRUE | 0.0284 |
| Arfrp1 | 1560 | 1071 | 1062 | 1029 | 838 | 983 | 1212 | 1139 | 1146 | 1024 | 945 | 970 | 1072.78 | 1166 | 980 | 0.84 | -0.251 | 0.0009065 | 0.00674471 | 6.00E-04 | 0.0046 | 0.0032 | 0.0032 | TRUE | 0.0187 |
| Arg1 | 14 | 8 | 6 | 5 | 6 | 8 | 11 | 9 | 6 | 5 | 7 | 8 | 7.58 | 9 | 7 | 0.752 | -0.411 | 0.55256555 | NA | 0 | 0.315 | 0.2105 | 0.2105 | TRUE | 0.2085 |
| Arg2 | 114 | 76 | 41 | 28 | 38 | 43 | 89 | 81 | 44 | 28 | 43 | 42 | 54.47 | 71 | 38 | 0.526 | -0.927 | 0.00401054 | 0.02066426 | 0.0751 | 0.0459 | 0.0542 | 0.0542 | TRUE | 0.9503 |
| Arglu1 | 4110 | 3471 | 2881 | 2371 | 1941 | 2462 | 3192 | 3692 | 3109 | 2360 | 2190 | 2430 | 2828.8 | 3331 | 2327 | 0.699 | -0.517 | 4.45E-11 | 3.87E-09 | 0.0054 | 0.0033 | 0.0041 | 0.0041 | TRUE | 0.1085 |
| Arhgap1 | 2583 | 1850 | 1779 | 1803 | 1611 | 1852 | 2006 | 1968 | 1920 | 1795 | 1817 | 1828 | 1888.96 | 1965 | 1813 | 0.923 | -0.116 | 0.05722357 | 0.14865441 | 0 | 0.0037 | 0.0022 | 0.0022 | TRUE | 0.0048 |
| Arhgap10 | 2359 | 1821 | 1847 | 1657 | 1479 | 1655 | 1832 | 1937 | 1993 | 1649 | 1668 | 1634 | 1785.6 | 1921 | 1650 | 0.86 | -0.218 | 0.00084056 | 0.00636848 | 6.00E-04 | 0.0038 | 0.0025 | 0.0025 | TRUE | 0.0204 |
| Arhgap11a | 6991 | 5132 | 5179 | 5135 | 4853 | 5462 | 5430 | 5459 | 5588 | 5111 | 5475 | 5392 | 5409.08 | 5492 | 5326 | 0.97 | -0.044 | 0.40544843 | 0.57815607 | 5.00E-04 | 0.0029 | 0.0019 | 0.0019 | TRUE | 0.0152 |
| Arhgap12 | 5507 | 4266 | 4749 | 5262 | 4857 | 5590 | 4277 | 4538 | 5124 | 5238 | 5479 | 5518 | 5029.01 | 4646 | 5412 | 1.165 | 0.221 | 0.00206257 | 0.01235035 | 0.0045 | 0.0029 | 0.0035 | 0.0035 | TRUE | 0.0984 |
| Arhgap15 | 0 | 0 | 0 | 0 | 0 | 0 | 0 | 0 | 0 | 0 | 0 | 0 | 0 | 0 | 0 | NA | NA | NA | NA | NA | NA | NA | NA | NA | NA |
| Arhgap17 | 2578 | 1887 | 1863 | 1822 | 1625 | 1848 | 2002 | 2007 | 2010 | 1814 | 1833 | 1824 | 1915.1 | 2006 | 1824 | 0.909 | -0.138 | 0.02043592 | 0.07024786 | 0 | 0.0037 | 0.002 | 0.002 | TRUE | 3.00E-04 |
| Arhgap18 | 514 | 357 | 408 | 473 | 374 | 466 | 399 | 380 | 440 | 471 | 422 | 460 | 428.65 | 406 | 451 | 1.111 | 0.152 | 0.15999545 | 0.30945258 | 0.0018 | 0.008 | 0.0061 | 0.0061 | TRUE | 0.0582 |
| Arhgap19 | 9397 | 7035 | 7998 | 8975 | 7601 | 8918 | 7298 | 7483 | 8630 | 8934 | 8575 | 8803 | 8287.15 | 7804 | 8771 | 1.124 | 0.169 | 0.01424098 | 0.05367784 | 0.0043 | 0.0027 | 0.0033 | 0.0033 | TRUE | 0.1044 |
| Arhgap20 | 89 | 68 | 79 | 62 | 45 | 56 | 69 | 72 | 85 | 62 | 51 | 55 | 65.74 | 75 | 56 | 0.743 | -0.428 | 0.07859533 | 0.18679133 | 0 | 0.0385 | 0.027 | 0.027 | TRUE | 0.1109 |
| Arhgap21 | 4574 | 3177 | 2742 | 2903 | 2559 | 3022 | 3552 | 3379 | 2959 | 2890 | 2887 | 2983 | 3108.33 | 3297 | 2920 | 0.885 | -0.176 | 0.01716323 | 0.06180752 | 0.004 | 0.0032 | 0.0036 | 0.0036 | TRUE | 0.0973 |
| Arhgap22 | 370 | 346 | 376 | 405 | 385 | 421 | 287 | 368 | 406 | 403 | 434 | 416 | 385.69 | 354 | 418 | 1.185 | 0.245 | 0.06476391 | 0.16225603 | 0.0136 | 0.0086 | 0.0101 | 0.0101 | TRUE | 0.3289 |
| Arhgap23 | 4648 | 3825 | 3741 | 4358 | 4178 | 4874 | 3610 | 4069 | 4037 | 4338 | 4713 | 4811 | 4262.93 | 3905 | 4621 | 1.184 | 0.244 | 0.00040305 | 0.00366927 | 0.0033 | 0.003 | 0.0032 | 0.0032 | TRUE | 0.0541 |
| Arhgap24 | 35 | 30 | 26 | 18 | 29 | 22 | 27 | 32 | 28 | 18 | 33 | 22 | 26.58 | 29 | 24 | 0.827 | -0.274 | 0.47858833 | 0.64580449 | 0 | 0.0916 | 0.0694 | 0.0694 | TRUE | 0.5354 |
| Arhgap25 | 50 | 25 | 22 | 35 | 24 | 24 | 39 | 27 | 24 | 35 | 27 | 24 | 29.13 | 30 | 29 | 0.951 | -0.073 | 0.84583775 | 0.91235068 | 0.0264 | 0.0838 | 0.0672 | 0.0672 | TRUE | 0.5594 |
| Arhgap26 | 617 | 422 | 429 | 547 | 498 | 549 | 479 | 449 | 463 | 544 | 562 | 542 | 506.53 | 464 | 549 | 1.183 | 0.243 | 0.01030809 | 0.04222802 | 0 | 0.0071 | 0.0045 | 0.0045 | TRUE | 0.0108 |
| Arhgap27 | 1174 | 846 | 810 | 911 | 802 | 906 | 912 | 900 | 874 | 907 | 905 | 894 | 898.59 | 895 | 902 | 1.007 | 0.01 | 0.89484087 | 0.94223961 | 0 | 0.0051 | 0.003 | 0.003 | TRUE | 0.005 |
| Arhgap28 | 1932 | 1385 | 1472 | 1679 | 1493 | 1769 | 1501 | 1473 | 1588 | 1671 | 1684 | 1746 | 1610.63 | 1521 | 1700 | 1.119 | 0.162 | 0.0151451 | 0.0562257 | 3.00E-04 | 0.0039 | 0.0026 | 0.0026 | TRUE | 0.0177 |
| Arhgap29 | 3196 | 2509 | 2509 | 2512 | 2268 | 2633 | 2482 | 2669 | 2707 | 2500 | 2559 | 2599 | 2586.06 | 2619 | 2553 | 0.975 | -0.036 | 0.5592982 | 0.71420076 | 0.001 | 0.0033 | 0.0024 | 0.0024 | TRUE | 0.0263 |
| Arhgap30 | 991 | 760 | 747 | 817 | 668 | 808 | 770 | 808 | 806 | 813 | 754 | 798 | 791.42 | 795 | 788 | 0.993 | -0.01 | 0.90268319 | 0.94610624 | 0 | 0.0054 | 0.0036 | 0.0036 | TRUE | 0.017 |
| Arhgap31 | 90 | 56 | 50 | 50 | 38 | 30 | 70 | 60 | 54 | 50 | 43 | 30 | 50.94 | 61 | 41 | 0.663 | -0.593 | 0.04233551 | 0.1192735 | 0.018 | 0.0489 | 0.0409 | 0.0409 | TRUE | 0.4493 |
| Arhgap32 | 4604 | 3552 | 3666 | 3879 | 3846 | 4182 | 3576 | 3778 | 3956 | 3861 | 4339 | 4128 | 3939.61 | 3770 | 4109 | 1.09 | 0.125 | 0.05976919 | 0.15285042 | 0.0026 | 0.003 | 0.0029 | 0.0029 | TRUE | 0.034 |
| Arhgap33 | 416 | 386 | 398 | 579 | 505 | 592 | 323 | 411 | 429 | 576 | 570 | 584 | 482.26 | 388 | 577 | 1.494 | 0.579 | 8.59E-07 | 2.57E-05 | 0.009 | 0.0073 | 0.0078 | 0.0078 | TRUE | 0.2625 |
| Arhgap36 | 28 | 24 | 11 | 15 | 18 | 20 | 22 | 26 | 12 | 15 | 20 | 20 | 19.02 | 20 | 18 | 0.924 | -0.113 | 0.80552834 | 0.88769596 | 0 | 0.127 | 0.1004 | 0.1004 | TRUE | 0.5909 |
| Arhgap39 | 1666 | 1380 | 1439 | 1455 | 1421 | 1714 | 1294 | 1468 | 1553 | 1448 | 1603 | 1692 | 1509.63 | 1438 | 1581 | 1.101 | 0.138 | 0.11750509 | 0.24870412 | 0.0063 | 0.004 | 0.005 | 0.005 | TRUE | 0.0955 |
| Arhgap4 | 328 | 279 | 298 | 327 | 335 | 352 | 255 | 297 | 322 | 325 | 378 | 347 | 320.66 | 291 | 350 | 1.207 | 0.271 | 0.03552179 | 0.10498615 | 0.0067 | 0.0098 | 0.0089 | 0.0089 | TRUE | 0.1452 |
| Arhgap40 | 29 | 19 | 13 | 19 | 25 | 22 | 23 | 20 | 14 | 19 | 28 | 22 | 20.93 | 19 | 23 | 1.199 | 0.262 | 0.54114076 | 0.69945062 | 0 | 0.1157 | 0.0846 | 0.0846 | TRUE | 0.2493 |
| Arhgap42 | 1176 | 800 | 720 | 588 | 518 | 551 | 913 | 851 | 777 | 585 | 584 | 544 | 709.12 | 847 | 571 | 0.673 | -0.571 | 1.13E-09 | 7.43E-08 | 0.0031 | 0.0058 | 0.0049 | 0.0049 | TRUE | 0.0619 |
| Arhgap44 | 2720 | 2075 | 2120 | 1980 | 1841 | 2099 | 2113 | 2207 | 2287 | 1971 | 2077 | 2072 | 2121.15 | 2202 | 2040 | 0.927 | -0.11 | 0.08566311 | 0.19785729 | 8.00E-04 | 0.0035 | 0.0025 | 0.0025 | TRUE | 0.016 |
| Arhgap5 | 6826 | 5028 | 4886 | 5652 | 4734 | 5394 | 5301 | 5348 | 5272 | 5626 | 5340 | 5325 | 5368.8 | 5307 | 5430 | 1.023 | 0.033 | 0.52069303 | 0.68196329 | 3.00E-04 | 0.0029 | 0.0017 | 0.0017 | TRUE | 0.0122 |
| Arhgap6 | 59 | 32 | 23 | 25 | 22 | 18 | 46 | 34 | 25 | 25 | 25 | 18 | 28.69 | 35 | 23 | 0.637 | -0.65 | 0.09324949 | 0.2100007 | 0.0377 | 0.085 | 0.0712 | 0.0712 | TRUE | 0.1929 |
| Arhgap8 | 626 | 521 | 637 | 669 | 648 | 775 | 486 | 554 | 687 | 666 | 731 | 765 | 648.28 | 576 | 721 | 1.255 | 0.327 | 0.00755493 | 0.03361879 | 0.0159 | 0.0061 | 0.0092 | 0.0092 | TRUE | 0.3343 |
| Arhgap9 | 251 | 165 | 193 | 226 | 168 | 192 | 195 | 176 | 208 | 225 | 190 | 190 | 197.12 | 193 | 202 | 1.044 | 0.063 | 0.67803815 | 0.80231161 | 0.0033 | 0.0145 | 0.0113 | 0.0113 | TRUE | 0.1161 |
| Arhgdia | 23829 | 16686 | 15255 | 14523 | 13257 | 15828 | 18507 | 17749 | 16460 | 14456 | 14955 | 15625 | 16291.97 | 17572 | 15012 | 0.854 | -0.227 | 0.00014771 | 0.00168497 | 0.0024 | 0.0026 | 0.0025 | 0.0025 | TRUE | 0.0374 |
| Arhgdib | 325 | 172 | 131 | 140 | 127 | 123 | 252 | 183 | 141 | 139 | 143 | 121 | 163.46 | 192 | 134 | 0.696 | -0.522 | 0.01164522 | 0.04626172 | 0.0419 | 0.0169 | 0.0245 | 0.0245 | TRUE | 0.289 |
| Arhgdig | 660 | 467 | 425 | 528 | 515 | 558 | 513 | 497 | 459 | 526 | 581 | 551 | 520.88 | 490 | 553 | 1.127 | 0.172 | 0.08022304 | 0.18952526 | 0 | 0.007 | 0.0051 | 0.0051 | TRUE | 0.0343 |
| Arhgef1 | 4397 | 3271 | 3494 | 3340 | 3341 | 3815 | 3415 | 3479 | 3770 | 3325 | 3769 | 3766 | 3587.32 | 3555 | 3620 | 1.019 | 0.026 | 0.70772326 | 0.82353566 | 0.0037 | 0.0031 | 0.0033 | 0.0033 | TRUE | 0.0621 |
| Arhgef10 | 2723 | 2015 | 2186 | 2424 | 2222 | 2480 | 2115 | 2143 | 2359 | 2413 | 2507 | 2448 | 2330.75 | 2206 | 2456 | 1.114 | 0.156 | 0.01805999 | 0.06406836 | 0.0014 | 0.0034 | 0.0027 | 0.0027 | TRUE | 0.0441 |
| Arhgef10l | 2317 | 1934 | 2089 | 2051 | 1992 | 2393 | 1800 | 2057 | 2254 | 2042 | 2247 | 2362 | 2126.95 | 2037 | 2217 | 1.089 | 0.124 | 0.16739756 | 0.31884506 | 0.0082 | 0.0035 | 0.0053 | 0.0053 | TRUE | 0.1285 |
| Arhgef11 | 2552 | 1846 | 1868 | 1835 | 1672 | 1847 | 1982 | 1964 | 2016 | 1827 | 1886 | 1823 | 1916.2 | 1987 | 1845 | 0.929 | -0.107 | 0.07777764 | 0.18524843 | 0 | 0.0037 | 0.0021 | 0.0021 | TRUE | 0.0044 |
| Arhgef12 | 4564 | 3355 | 3183 | 3581 | 3423 | 3659 | 3545 | 3569 | 3434 | 3565 | 3861 | 3612 | 3597.63 | 3516 | 3679 | 1.046 | 0.065 | 0.26305283 | 0.43335254 | 7.00E-04 | 0.0031 | 0.0022 | 0.0022 | TRUE | 0.0226 |
| Arhgef15 | 440 | 301 | 351 | 421 | 313 | 396 | 342 | 320 | 379 | 419 | 353 | 391 | 367.28 | 347 | 388 | 1.119 | 0.163 | 0.1716378 | 0.32364368 | 0.0042 | 0.0089 | 0.0075 | 0.0075 | TRUE | 0.0699 |
| Arhgef16 | 654 | 503 | 589 | 531 | 515 | 646 | 508 | 535 | 636 | 529 | 581 | 638 | 570.95 | 560 | 583 | 1.043 | 0.061 | 0.59390061 | 0.74083813 | 0.0097 | 0.0066 | 0.0076 | 0.0076 | TRUE | 0.1634 |
| Arhgef17 | 2249 | 1764 | 1776 | 1813 | 1735 | 1903 | 1747 | 1876 | 1916 | 1805 | 1957 | 1879 | 1863.3 | 1846 | 1880 | 1.019 | 0.027 | 0.69470773 | 0.81382326 | 0.0015 | 0.0037 | 0.0029 | 0.0029 | TRUE | 0.028 |
| Arhgef18 | 6096 | 4959 | 5417 | 5332 | 5169 | 5920 | 4735 | 5275 | 5845 | 5307 | 5831 | 5844 | 5472.8 | 5285 | 5661 | 1.072 | 0.1 | 0.20659059 | 0.36835958 | 0.0066 | 0.0029 | 0.0043 | 0.0043 | TRUE | 0.1048 |
| Arhgef19 | 338 | 299 | 289 | 358 | 313 | 366 | 263 | 318 | 312 | 356 | 353 | 361 | 327.19 | 298 | 357 | 1.204 | 0.268 | 0.02694767 | 0.08576331 | 0.0027 | 0.0097 | 0.0075 | 0.0075 | TRUE | 0.1301 |
| Arhgef2 | 4129 | 3291 | 3534 | 3663 | 3570 | 4160 | 3207 | 3501 | 3813 | 3646 | 4027 | 4107 | 3716.77 | 3507 | 3927 | 1.12 | 0.164 | 0.03248561 | 0.09816424 | 0.0053 | 0.0031 | 0.004 | 0.004 | TRUE | 0.071 |
| Arhgef25 | 1822 | 1519 | 1588 | 1735 | 1762 | 2132 | 1415 | 1616 | 1713 | 1727 | 1988 | 2105 | 1760.6 | 1581 | 1940 | 1.228 | 0.296 | 0.00145654 | 0.00944142 | 0.0087 | 0.0038 | 0.0057 | 0.0057 | TRUE | 0.112 |
| Arhgef26 | 1017 | 634 | 346 | 308 | 297 | 236 | 790 | 674 | 373 | 307 | 335 | 233 | 452.03 | 612 | 292 | 0.476 | -1.072 | 0.00224311 | 0.0132046 | 0.0862 | 0.0077 | 0.0289 | 0.0862 | TRUE | 0.43 |
| Arhgef28 | 524 | 382 | 495 | 548 | 523 | 598 | 407 | 406 | 534 | 545 | 590 | 590 | 512.2 | 449 | 575 | 1.283 | 0.36 | 0.00266729 | 0.0150522 | 0.0115 | 0.0071 | 0.0084 | 0.0084 | TRUE | 0.3106 |
| Arhgef3 | 675 | 618 | 629 | 829 | 728 | 913 | 524 | 657 | 679 | 825 | 821 | 901 | 734.67 | 620 | 849 | 1.374 | 0.458 | 1.97E-05 | 0.0003271 | 0.0096 | 0.0057 | 0.0069 | 0.0069 | TRUE | 0.2268 |
| Arhgef33 | 84 | 60 | 60 | 63 | 54 | 52 | 65 | 64 | 65 | 63 | 61 | 51 | 61.46 | 65 | 58 | 0.902 | -0.149 | 0.54311748 | 0.7009121 | 0 | 0.041 | 0.0271 | 0.0271 | TRUE | 0.0969 |
| Arhgef37 | 49 | 34 | 32 | 40 | 43 | 34 | 38 | 36 | 35 | 40 | 49 | 34 | 38.44 | 36 | 41 | 1.115 | 0.157 | 0.61623738 | 0.75773235 | 0 | 0.0641 | 0.0448 | 0.0448 | TRUE | 0.2027 |
| Arhgef38 | 12 | 9 | 17 | 4 | 6 | 6 | 9 | 10 | 18 | 4 | 7 | 6 | 8.98 | 12 | 6 | 0.449 | -1.154 | 0.088421 | NA | 0 | 0.2662 | 0.1978 | 0.1978 | TRUE | 0.6608 |
| Arhgef39 | 1168 | 881 | 914 | 991 | 892 | 988 | 907 | 937 | 986 | 986 | 1006 | 975 | 966.41 | 943 | 989 | 1.049 | 0.07 | 0.36230224 | 0.53696431 | 0 | 0.0049 | 0.0032 | 0.0032 | TRUE | 0.0185 |
| Arhgef4 | 596 | 474 | 546 | 540 | 519 | 594 | 463 | 504 | 589 | 538 | 585 | 586 | 544.26 | 519 | 570 | 1.101 | 0.139 | 0.20674453 | 0.36845382 | 0.007 | 0.0068 | 0.0069 | 0.0069 | TRUE | 0.163 |
| Arhgef40 | 689 | 493 | 440 | 567 | 472 | 574 | 535 | 524 | 475 | 564 | 532 | 567 | 532.96 | 511 | 554 | 1.083 | 0.115 | 0.23448602 | 0.40061384 | 0 | 0.0069 | 0.0049 | 0.0049 | TRUE | 0.0446 |
| Arhgef5 | 367 | 310 | 333 | 409 | 358 | 393 | 285 | 330 | 359 | 407 | 404 | 388 | 362.17 | 325 | 400 | 1.235 | 0.305 | 0.01099715 | 0.04427671 | 0.0044 | 0.009 | 0.0076 | 0.0076 | TRUE | 0.1406 |
| Arhgef6 | 108 | 63 | 51 | 54 | 39 | 51 | 84 | 67 | 55 | 54 | 44 | 50 | 59 | 69 | 49 | 0.716 | -0.483 | 0.07219825 | 0.17558781 | 0.0153 | 0.0426 | 0.0345 | 0.0345 | TRUE | 0.2462 |
| Arhgef7 | 5363 | 3993 | 3810 | 4028 | 3558 | 4055 | 4165 | 4247 | 4111 | 4009 | 4014 | 4003 | 4091.61 | 4174 | 4009 | 0.96 | -0.058 | 0.25274485 | 0.4216249 | 0 | 0.003 | 0.0016 | 0.0016 | TRUE | 0.0028 |

| | | | | | | | | | | | | | | | | | | | | | | | | | |
|---|---|---|---|---|---|---|---|---|---|---|---|---|---|---|---|---|---|---|---|---|---|---|---|---|---|
| Arhgef9 | 233 | 162 | 150 | 244 | 214 | 225 | 181 | 172 | 162 | 243 | 241 | 222 | 203.59 | 172 | 235 | 1.368 | 0.452 | 0.00142276 | 0.00929408 | 0 | 0.0141 | 0.0095 | 0.0095 | TRUE | 0.0278 |
| Arid1a | 18615 | 12677 | 10579 | 11658 | 10656 | 11984 | 14457 | 13484 | 11415 | 11604 | 12021 | 11830 | 12468.66 | 13119 | 11818 | 0.901 | -0.151 | 0.0538026 | 0.14180352 | 0.0071 | 0.0026 | 0.0043 | 0.0043 | TRUE | 0.1577 |
| Arid1b | 1688 | 1253 | 1113 | 1086 | 1015 | 1125 | 1311 | 1333 | 1201 | 1081 | 1145 | 1111 | 1196.88 | 1282 | 1112 | 0.867 | -0.206 | 0.00624585 | 0.02906383 | 0.0011 | 0.0044 | 0.0032 | 0.0032 | TRUE | 0.0354 |
| Arid2 | 5941 | 4167 | 4227 | 4548 | 4062 | 4505 | 4614 | 4432 | 4561 | 4527 | 4582 | 4447 | 4527.33 | 4536 | 4519 | 0.996 | -0.006 | 0.91076719 | 0.95109194 | 0 | 0.0029 | 0.0017 | 0.0017 | TRUE | 0.0048 |
| Arid3a | 1073 | 958 | 1022 | 1091 | 1123 | 1280 | 833 | 1019 | 1103 | 1086 | 1267 | 1264 | 1095.25 | 985 | 1206 | 1.226 | 0.294 | 0.00612492 | 0.02866518 | 0.0121 | 0.0046 | 0.0074 | 0.0074 | TRUE | 0.224 |
| Arid3b | 4381 | 3009 | 3102 | 3481 | 3155 | 3602 | 3403 | 3201 | 3347 | 3465 | 3559 | 3556 | 3421.68 | 3317 | 3527 | 1.063 | 0.088 | 0.11413817 | 0.24326351 | 3.00E-04 | 0.0031 | 0.0019 | 0.0019 | TRUE | 0.0113 |
| Arid3c | 44 | 23 | 30 | 45 | 40 | 55 | 34 | 24 | 32 | 45 | 45 | 54 | 39.2 | 30 | 48 | 1.578 | 0.658 | 0.03624937 | 0.10669527 | 0 | 0.0629 | 0.0446 | 0.0446 | TRUE | 0.1804 |
| Arid4a | 2262 | 1703 | 1559 | 1560 | 1412 | 1595 | 1757 | 1811 | 1682 | 1553 | 1593 | 1575 | 1661.77 | 1750 | 1574 | 0.899 | -0.154 | 0.01812022 | 0.0642342 | 0 | 0.0039 | 0.0024 | 0.0024 | TRUE | 0.0134 |
| Arid4b | 2774 | 2118 | 2119 | 2128 | 1918 | 2240 | 2154 | 2253 | 2286 | 2118 | 2164 | 2211 | 2197.81 | 2231 | 2164 | 0.971 | -0.043 | 0.48046853 | 0.64736224 | 3.00E-04 | 0.0035 | 0.0022 | 0.0022 | TRUE | 0.0114 |
| Arid5a | 318 | 215 | 170 | 199 | 187 | 219 | 247 | 229 | 183 | 198 | 211 | 216 | 214.05 | 220 | 208 | 0.945 | -0.082 | 0.59127576 | 0.73937922 | 0.0082 | 0.0135 | 0.0119 | 0.0119 | TRUE | 0.2222 |
| Arid5b | 5641 | 4250 | 4001 | 3790 | 3837 | 4188 | 4381 | 4521 | 4317 | 3773 | 4329 | 4134 | 4242.36 | 4406 | 4079 | 0.925 | -0.112 | 0.08323999 | 0.19418135 | 0.0023 | 0.003 | 0.0028 | 0.0028 | TRUE | 0.0525 |
| Arih1 | 10952 | 8223 | 8868 | 9169 | 8437 | 9522 | 8506 | 8747 | 9568 | 9127 | 9518 | 9400 | 9144.24 | 8940 | 9348 | 1.046 | 0.065 | 0.2744181 | 0.44540743 | 0.002 | 0.0027 | 0.0024 | 0.0024 | TRUE | 0.046 |
| Arih2 | 12398 | 8759 | 8279 | 7502 | 6706 | 7233 | 9629 | 9317 | 8933 | 7467 | 7565 | 7140 | 8341.91 | 9293 | 7391 | 0.795 | -0.331 | 9.09E-10 | 6.19E-08 | 0.001 | 0.0027 | 0.002 | 0.002 | TRUE | 0.014 |
| Arl1 | 2699 | 1903 | 1925 | 2155 | 1898 | 1965 | 2096 | 2024 | 2077 | 2145 | 2141 | 1940 | 2070.57 | 2066 | 2075 | 1.004 | 0.006 | 0.92839732 | 0.96054762 | 0.0013 | 0.0036 | 0.0027 | 0.0027 | TRUE | 0.0399 |
| Arl10 | 406 | 286 | 348 | 407 | 330 | 401 | 315 | 304 | 375 | 405 | 372 | 396 | 361.38 | 331 | 391 | 1.182 | 0.241 | 0.04427214 | 0.12322915 | 0.0041 | 0.009 | 0.0075 | 0.0075 | TRUE | 0.146 |
| Arl11 | 2 | 0 | 1 | 6 | 2 | 6 | 2 | 0 | 1 | 6 | 2 | 6 | 2.8 | 1 | 5 | 5.149 | 2.364 | 0.07376358 | NA | 0 | 0.8497 | 0.63 | 0.63 | TRUE | 0.3551 |
| Arl13a | 0 | 0 | 0 | 0 | 0 | 0 | 0 | 0 | 0 | 0 | 0 | 0 | 0 | 0 | 0 | NA | NA | NA | NA | NA | NA | NA | NA | NA | NA |
| Arl13b | 1573 | 1107 | 1194 | 1374 | 1103 | 1292 | 1222 | 1178 | 1288 | 1368 | 1244 | 1275 | 1262.48 | 1229 | 1296 | 1.055 | 0.077 | 0.30553191 | 0.47820495 | 0.0014 | 0.0043 | 0.0033 | 0.0033 | TRUE | 0.0284 |
| Arl14 | 1 | 0 | 0 | 0 | 0 | 1 | 1 | 0 | 0 | 0 | 0 | 1 | 0.29 | 0 | 0 | 1.075 | 0.105 | 0.97702352 | NA | 0 | 8.0647 | 7.5458 | 7.5458 | TRUE | 0.3835 |
| Arl14ep | 2134 | 1484 | 1401 | 1344 | 1237 | 1418 | 1657 | 1579 | 1512 | 1338 | 1395 | 1400 | 1480.11 | 1583 | 1378 | 0.87 | -0.201 | 0.00384313 | 0.01998176 | 8.00E-04 | 0.004 | 0.0028 | 0.0028 | TRUE | 0.0214 |
| Arl14epl | 4071 | 2783 | 2494 | 1954 | 1659 | 1916 | 3162 | 2960 | 2691 | 1945 | 1872 | 1891 | 2420.16 | 2938 | 1903 | 0.647 | -0.628 | 4.20E-18 | 1.02E-15 | 0.0032 | 0.0034 | 0.0033 | 0.0033 | TRUE | 0.0652 |
| Arl15 | 2533 | 1730 | 1754 | 1865 | 1624 | 1822 | 1967 | 1840 | 1893 | 1856 | 1832 | 1799 | 1864.51 | 1900 | 1829 | 0.962 | -0.056 | 0.37608657 | 0.5506459 | 0 | 0.0037 | 0.0023 | 0.0023 | TRUE | 0.0121 |
| Arl16 | 470 | 347 | 246 | 255 | 239 | 259 | 365 | 369 | 265 | 254 | 270 | 256 | 296.45 | 333 | 260 | 0.777 | -0.363 | 0.01192283 | 0.04712354 | 0.0143 | 0.0104 | 0.0116 | 0.0116 | TRUE | 0.3488 |
| Arl2 | 1528 | 1068 | 1099 | 1386 | 1169 | 1379 | 1187 | 1136 | 1186 | 1380 | 1319 | 1361 | 1261.37 | 1170 | 1353 | 1.157 | 0.21 | 0.00236302 | 0.01375748 | 0 | 0.0043 | 0.0027 | 0.0027 | TRUE | 0.0074 |
| Arl2bp | 3098 | 1986 | 1963 | 2028 | 1708 | 1859 | 2406 | 2112 | 2118 | 2019 | 1927 | 1835 | 2069.54 | 2212 | 1927 | 0.87 | -0.201 | 0.00750722 | 0.03345826 | 0.0033 | 0.0036 | 0.0036 | 0.0036 | TRUE | 0.0728 |
| Arl3 | 1547 | 1075 | 1163 | 1337 | 1128 | 1410 | 1201 | 1143 | 1255 | 1331 | 1273 | 1392 | 1265.84 | 1200 | 1332 | 1.11 | 0.151 | 0.04248972 | 0.11961899 | 0.0012 | 0.0043 | 0.0032 | 0.0032 | TRUE | 0.0195 |
| Arl4a | 1108 | 783 | 732 | 795 | 714 | 729 | 861 | 833 | 790 | 791 | 805 | 720 | 799.94 | 828 | 772 | 0.931 | -0.102 | 0.23371942 | 0.39987452 | 0.0015 | 0.0054 | 0.0041 | 0.0041 | TRUE | 0.0424 |
| Arl4c | 326 | 259 | 294 | 279 | 260 | 304 | 253 | 275 | 317 | 278 | 293 | 300 | 286.17 | 282 | 290 | 1.033 | 0.047 | 0.71963464 | 0.83170841 | 0.0042 | 0.0107 | 0.0087 | 0.0087 | TRUE | 0.1326 |
| Arl4d | 1178 | 862 | 637 | 597 | 519 | 554 | 915 | 917 | 687 | 594 | 585 | 547 | 707.62 | 840 | 575 | 0.684 | -0.547 | 2.15E-06 | 5.33E-05 | 0.0131 | 0.0058 | 0.0081 | 0.0081 | TRUE | 0.2943 |
| Arl5a | 4296 | 3083 | 3047 | 3097 | 2779 | 3164 | 3337 | 3279 | 3288 | 3083 | 3135 | 3123 | 3207.44 | 3301 | 3114 | 0.943 | -0.085 | 0.1109151 | 0.23852327 | 0 | 0.0032 | 0.0017 | 0.0017 | TRUE | 0.0011 |
| Arl5b | 1486 | 1095 | 1134 | 1148 | 970 | 1134 | 1154 | 1165 | 1224 | 1143 | 1094 | 1119 | 1149.81 | 1181 | 1119 | 0.948 | -0.077 | 0.28453943 | 0.45649591 | 0 | 0.0045 | 0.0028 | 0.0028 | TRUE | 0.0118 |
| Arl5c | 664 | 485 | 418 | 534 | 489 | 575 | 516 | 516 | 451 | 532 | 552 | 568 | 522.23 | 494 | 551 | 1.112 | 0.153 | 0.12369115 | 0.2577515 | 0.0013 | 0.007 | 0.0052 | 0.0052 | TRUE | 0.0656 |
| Arl6 | 1254 | 864 | 862 | 910 | 748 | 929 | 974 | 919 | 930 | 906 | 844 | 917 | 914.96 | 941 | 889 | 0.944 | -0.083 | 0.29638973 | 0.46830626 | 0 | 0.005 | 0.0034 | 0.0034 | TRUE | 0.0228 |
| Arl6ip1 | 23877 | 17041 | 17511 | 19758 | 17166 | 18378 | 18544 | 18126 | 18894 | 19667 | 19365 | 18142 | 18789.8 | 18521 | 19058 | 1.029 | 0.041 | 0.42688561 | 0.59901775 | 0.001 | 0.0026 | 0.0019 | 0.0019 | TRUE | 0.0217 |
| Arl6ip4 | 2833 | 1979 | 1852 | 2009 | 1816 | 2107 | 2200 | 2105 | 1998 | 2000 | 2049 | 2080 | 2071.99 | 2101 | 2043 | 0.971 | -0.042 | 0.51846995 | 0.68017357 | 9.00E-04 | 0.0036 | 0.0025 | 0.0025 | TRUE | 0.0221 |
| Arl6ip5 | 4650 | 3346 | 3751 | 3983 | 3750 | 4385 | 3611 | 3559 | 4047 | 3965 | 4230 | 4329 | 3956.93 | 3739 | 4175 | 1.117 | 0.159 | 0.01987156 | 0.06880432 | 0.0032 | 0.003 | 0.0031 | 0.0031 | TRUE | 0.0626 |
| Arl6ip6 | 3137 | 2219 | 2385 | 2200 | 1967 | 2454 | 2436 | 2360 | 2573 | 2190 | 2219 | 2422 | 2366.9 | 2456 | 2277 | 0.927 | -0.109 | 0.10796765 | 0.23381539 | 0.0019 | 0.0034 | 0.0029 | 0.0029 | TRUE | 0.0381 |
| Arl8a | 2018 | 1530 | 1741 | 1898 | 1563 | 1728 | 1567 | 1627 | 1879 | 1889 | 1763 | 1706 | 1738.59 | 1691 | 1786 | 1.057 | 0.08 | 0.33277553 | 0.50702065 | 0.0051 | 0.0038 | 0.0044 | 0.0044 | TRUE | 0.1125 |
| Arl8b | 9223 | 6781 | 7222 | 7747 | 6700 | 7632 | 7163 | 7213 | 7792 | 7711 | 7558 | 7534 | 7495.34 | 7389 | 7601 | 1.029 | 0.041 | 0.45044624 | 0.62049229 | 0.001 | 0.0027 | 0.002 | 0.002 | TRUE | 0.0276 |
| Arl9 | 0 | 0 | 0 | 0 | 1 | 0 | 0 | 0 | 0 | 0 | 1 | 0 | 0.19 | 0 | 0 | 2.094 | 1.067 | 0.79379329 | NA | 0 | 12.608 | 10 | 10 | TRUE | 0.5075 |
| Armc1 | 6647 | 4859 | 4692 | 5008 | 4430 | 4970 | 5162 | 5168 | 5063 | 4985 | 4997 | 4906 | 5047.02 | 5131 | 4963 | 0.967 | -0.048 | 0.32754902 | 0.50194683 | 0 | 0.0029 | 0.0016 | 0.0016 | TRUE | 0.0016 |
| Armc10 | 4767 | 3539 | 3712 | 3849 | 3387 | 4029 | 3702 | 3764 | 4005 | 3831 | 3821 | 3977 | 3850.23 | 3824 | 3876 | 1.014 | 0.02 | 0.7243941 | 0.83484639 | 9.00E-04 | 0.003 | 0.0021 | 0.0021 | TRUE | 0.0208 |
| Armc12 | 6 | 6 | 8 | 1 | 3 | 4 | 5 | 6 | 9 | 1 | 3 | 4 | 4.67 | 7 | 3 | 0.425 | -1.233 | 0.1855755 | NA | 0 | 0.5103 | 0.3636 | 0.3636 | TRUE | 0.3772 |
| Armc2 | 251 | 177 | 151 | 223 | 195 | 198 | 195 | 188 | 163 | 222 | 220 | 195 | 197.26 | 182 | 212 | 1.164 | 0.218 | 0.14081792 | 0.28268406 | 0 | 0.0144 | 0.0108 | 0.0108 | TRUE | 0.0873 |
| Armc3 | 19 | 5 | 12 | 13 | 8 | 7 | 15 | 5 | 13 | 13 | 9 | 7 | 10.32 | 11 | 10 | 0.866 | -0.208 | 0.74138917 | NA | 0.0772 | 0.2322 | 0.1891 | 0.1891 | TRUE | 0.7084 |
| Armc4 | 13 | 4 | 13 | 11 | 9 | 5 | 10 | 4 | 14 | 11 | 10 | 5 | 9.07 | 9 | 9 | 0.914 | -0.13 | 0.84801799 | NA | 0.1085 | 0.2638 | 0.2228 | 0.2228 | TRUE | 0.2778 |
| Armc5 | 785 | 561 | 461 | 532 | 493 | 548 | 610 | 597 | 497 | 530 | 556 | 541 | 555.08 | 568 | 542 | 0.953 | -0.07 | 0.50302223 | 0.66713341 | 0.0046 | 0.0067 | 0.006 | 0.006 | TRUE | 0.1347 |
| Armc6 | 2302 | 1629 | 1404 | 1546 | 1323 | 1452 | 1788 | 1733 | 1515 | 1539 | 1492 | 1433 | 1583.37 | 1679 | 1488 | 0.886 | -0.175 | 0.02748636 | 0.08682044 | 0.0039 | 0.0039 | 0.0039 | 0.0039 | TRUE | 0.0866 |
| Armc7 | 153 | 135 | 121 | 132 | 105 | 122 | 119 | 144 | 131 | 131 | 118 | 120 | 127.21 | 131 | 123 | 0.946 | -0.08 | 0.65014488 | 0.7827712 | 0 | 0.0211 | 0.0148 | 0.0148 | TRUE | 0.0759 |
| Armc8 | 4616 | 3133 | 2969 | 3337 | 2874 | 3280 | 3585 | 3333 | 3204 | 3322 | 3242 | 3238 | 3320.47 | 3374 | 3267 | 0.968 | -0.047 | 0.44687793 | 0.61748127 | 0.0013 | 0.0031 | 0.0025 | 0.0025 | TRUE | 0.0373 |
| Armc9 | 1178 | 866 | 960 | 886 | 864 | 929 | 915 | 921 | 1036 | 882 | 975 | 917 | 940.93 | 957 | 925 | 0.966 | -0.05 | 0.56084314 | 0.71547046 | 0.0027 | 0.0049 | 0.0042 | 0.0042 | TRUE | 0.0602 |
| Armcx1 | 1781 | 1308 | 1530 | 1710 | 1459 | 1820 | 1383 | 1391 | 1651 | 1702 | 1646 | 1797 | 1595.01 | 1475 | 1715 | 1.164 | 0.219 | 0.00904965 | 0.0384746 | 0.0054 | 0.0039 | 0.0044 | 0.0044 | TRUE | 0.1291 |
| Armcx2 | 3113 | 2226 | 2426 | 2557 | 2438 | 2802 | 2418 | 2368 | 2618 | 2545 | 2750 | 2766 | 2577.45 | 2468 | 2687 | 1.089 | 0.123 | 0.06628902 | 0.16471991 | 0.0019 | 0.0033 | 0.0029 | 0.0029 | TRUE | 0.0337 |
| Armcx3 | 218 | 164 | 175 | 237 | 187 | 199 | 169 | 174 | 189 | 236 | 211 | 196 | 195.98 | 177 | 214 | 1.21 | 0.275 | 0.06287214 | 0.15868471 | 0 | 0.0145 | 0.0106 | 0.0106 | TRUE | 0.0861 |
| Armcx4 | 2669 | 1757 | 1419 | 1807 | 1610 | 1937 | 2073 | 1869 | 1531 | 1799 | 1816 | 1912 | 1833.32 | 1824 | 1842 | 1.009 | 0.013 | 0.89675946 | 0.94321753 | 0.0116 | 0.0037 | 0.0063 | 0.0063 | TRUE | 0.2374 |
| Armcx5 | 907 | 628 | 650 | 679 | 622 | 725 | 704 | 668 | 701 | 676 | 702 | 716 | 694.5 | 691 | 698 | 1.009 | 0.013 | 0.88082223 | 0.93349722 | 0 | 0.0058 | 0.0037 | 0.0037 | TRUE | 0.01 |
| Armcx6 | 204 | 157 | 147 | 209 | 195 | 224 | 158 | 167 | 159 | 208 | 220 | 221 | 188.86 | 161 | 216 | 1.342 | 0.424 | 0.0032214 | 0.01744071 | 0 | 0.015 | 0.0095 | 0.0095 | TRUE | 0.0127 |
| Arnt | 6409 | 4706 | 5404 | 5716 | 5092 | 6281 | 4978 | 5006 | 5831 | 5690 | 5744 | 6200 | 5574.75 | 5272 | 5878 | 1.115 | 0.158 | 0.03095301 | 0.09481598 | 0.0051 | 0.0029 | 0.0037 | 0.0037 | TRUE | 0.1046 |
| Arnt2 | 994 | 679 | 671 | 890 | 773 | 936 | 772 | 722 | 724 | 886 | 872 | 924 | 816.69 | 739 | 894 | 1.208 | 0.272 | 0.00078864 | 0.00608167 | 0 | 0.0053 | 0.0035 | 0.0035 | TRUE | 0.0187 |
| Arntl | 1292 | 889 | 814 | 951 | 848 | 1034 | 1003 | 946 | 878 | 947 | 957 | 1021 | 958.56 | 942 | 975 | 1.033 | 0.047 | 0.57383055 | 0.72601222 | 0.0021 | 0.0049 | 0.0039 | 0.0039 | TRUE | 0.0419 |
| Arntl2 | 202 | 177 | 147 | 108 | 95 | 104 | 157 | 188 | 159 | 108 | 107 | 103 | 136.85 | 168 | 106 | 0.631 | -0.664 | 0.00013387 | 0.00155121 | 0 | 0.0197 | 0.014 | 0.014 | TRUE | 0.1162 |
| Arpc1a | 5539 | 3919 | 4070 | 4408 | 3832 | 4340 | 4302 | 4169 | 4391 | 4388 | 4323 | 4284 | 4309.48 | 4287 | 4332 | 1.01 | 0.015 | 0.77812689 | 0.86902388 | 2.00E-04 | 0.003 | 0.0018 | 0.0018 | TRUE | 0.007 |
| Arpc1b | 6845 | 5091 | 5053 | 5216 | 4519 | 5166 | 5316 | 5415 | 5452 | 5192 | 5098 | 5100 | 5262.19 | 5394 | 5130 | 0.951 | -0.072 | 0.14163954 | 0.28396765 | 0 | 0.0029 | 0.0016 | 0.0016 | TRUE | 0.002 |
| Arpc2 | 7610 | 5559 | 5262 | 5222 | 4747 | 5336 | 5910 | 5913 | 5678 | 5198 | 5355 | 5267 | 5553.6 | 5834 | 5273 | 0.904 | -0.146 | 0.00393262 | 0.02033732 | 2.00E-04 | 0.0029 | 0.0017 | 0.0017 | TRUE | 0.0066 |
| Arpc3 | 5259 | 3766 | 3928 | 3875 | 3455 | 4180 | 4084 | 4006 | 4238 | 3857 | 3898 | 4126 | 4034.94 | 4109 | 3960 | 0.964 | -0.053 | 0.35143687 | 0.52609837 | 7.00E-04 | 0.003 | 0.0021 | 0.0021 | TRUE | 0.0165 |
| Arpc4 | 7835 | 5506 | 4783 | 4834 | 4099 | 4864 | 6085 | 5857 | 5161 | 4812 | 4624 | 4802 | 5223.33 | 5701 | 4746 | 0.832 | -0.265 | 0.00011828 | 0.00139718 | 0.0035 | 0.0029 | 0.0032 | 0.0032 | TRUE | 0.0834 |
| Arpc5 | 4618 | 3136 | 2792 | 2664 | 2355 | 2558 | 3587 | 3336 | 3013 | 2652 | 2657 | 2525 | 2961.41 | 3312 | 2611 | 0.788 | -0.344 | 2.59E-06 | 6.19E-05 | 0.0037 | 0.0032 | 0.0035 | 0.0035 | TRUE | 0.0756 |
| Arpc5l | 2321 | 1779 | 1574 | 1747 | 1382 | 1702 | 1803 | 1892 | 1698 | 1739 | 1559 | 1680 | 1728.57 | 1798 | 1659 | 0.923 | -0.115 | 0.11542527 | 0.24530321 | 0.0021 | 0.0038 | 0.0033 | 0.0033 | TRUE | 0.0332 |
| Arpp19 | 5309 | 3581 | 3483 | 3757 | 3064 | 3664 | 4123 | 3809 | 3758 | 3740 | 3457 | 3617 | 3750.6 | 3897 | 3605 | 0.925 | -0.113 | 0.07070741 | 0.17281482 | 0.0017 | 0.0031 | 0.0026 | 0.0026 | TRUE | 0.032 |

| | | | | | | | | | | | | | | | | | | | | | | | | | |
|---|---|---|---|---|---|---|---|---|---|---|---|---|---|---|---|---|---|---|---|---|---|---|---|---|---|
| Arpp21 | 12 | 11 | 10 | 5 | 3 | 3 | 9 | 12 | 11 | 5 | 3 | 3 | 7.19 | 11 | 4 | 0.358 | -1.482 | 0.04676497 | NA | 0 | 0.3321 | 0.2178 | 0.2178 | TRUE | 0.1289 |
| Arr3 | 0 | 0 | 0 | 0 | 0 | 0 | 0 | 0 | 0 | 0 | 0 | 0 | 0 | 0 | 0 | NA | NA | NA | NA | NA | NA | NA | NA | NA | NA |
| Arrb1 | 2247 | 1842 | 2098 | 2419 | 2460 | 2681 | 1745 | 1959 | 2264 | 2408 | 2775 | 2647 | 2299.63 | 1989 | 2610 | 1.313 | 0.393 | 2.69E-05 | 0.00042457 | 0.0105 | 0.0035 | 0.0059 | 0.0059 | TRUE | 0.1758 |
| Arrb2 | 3751 | 2762 | 2707 | 2455 | 2287 | 2903 | 2913 | 2938 | 2921 | 2444 | 2580 | 2866 | 2776.89 | 2924 | 2630 | 0.9 | -0.153 | 0.02852933 | 0.08926657 | 0.0028 | 0.0033 | 0.0031 | 0.0031 | TRUE | 0.0753 |
| Arrdc1 | 724 | 543 | 613 | 680 | 579 | 733 | 562 | 578 | 661 | 677 | 653 | 724 | 642.49 | 600 | 685 | 1.142 | 0.192 | 0.04882483 | 0.13241999 | 0.0036 | 0.0061 | 0.0053 | 0.0053 | TRUE | 0.091 |
| Arrdc2 | 1226 | 1096 | 1204 | 1395 | 1274 | 1530 | 952 | 1166 | 1299 | 1389 | 1437 | 1510 | 1292.2 | 1139 | 1445 | 1.271 | 0.346 | 0.00078279 | 0.00606152 | 0.0115 | 0.0043 | 0.0069 | 0.0069 | TRUE | 0.2545 |
| Arrdc3 | 4351 | 3090 | 2795 | 3248 | 2727 | 2829 | 3379 | 3287 | 3016 | 3233 | 3076 | 2793 | 3130.64 | 3227 | 3034 | 0.94 | -0.09 | 0.22086066 | 0.384502 | 0.0039 | 0.0032 | 0.0036 | 0.0036 | TRUE | 0.0592 |
| Arrdc4 | 3731 | 2796 | 3222 | 3692 | 3203 | 3691 | 2898 | 2974 | 3477 | 3675 | 3613 | 3644 | 3380.03 | 3116 | 3644 | 1.17 | 0.227 | 0.00229989 | 0.01346123 | 0.0047 | 0.0031 | 0.0037 | 0.0037 | TRUE | 0.1236 |
| Arrdc5 | 5 | 2 | 0 | 2 | 1 | 2 | 4 | 2 | 0 | 2 | 1 | 2 | 1.85 | 2 | 2 | 0.825 | -0.277 | 0.84789776 | NA | 0 | 1.2831 | 0.9715 | 0.9715 | TRUE | 0.1974 |
| Arsa | 807 | 583 | 744 | 815 | 763 | 908 | 627 | 620 | 803 | 811 | 861 | 896 | 769.67 | 683 | 856 | 1.255 | 0.328 | 0.00246522 | 0.01421923 | 0.0108 | 0.0055 | 0.0071 | 0.0071 | TRUE | 0.2716 |
| Arsb | 192 | 116 | 92 | 93 | 89 | 78 | 149 | 123 | 99 | 93 | 100 | 77 | 106.96 | 124 | 90 | 0.721 | -0.472 | 0.02837031 | 0.08888575 | 0.0217 | 0.0246 | 0.0237 | 0.0237 | TRUE | 0.16 |
| Arsg | 126 | 109 | 106 | 124 | 85 | 100 | 98 | 116 | 114 | 123 | 96 | 99 | 107.7 | 109 | 106 | 0.974 | -0.038 | 0.84663464 | 0.91300354 | 0.0061 | 0.0244 | 0.019 | 0.019 | TRUE | 0.2067 |
| Arsi | 9 | 6 | 9 | 14 | 7 | 19 | 7 | 6 | 10 | 14 | 8 | 19 | 10.61 | 8 | 14 | 1.777 | 0.829 | 0.17693029 | NA | 0 | 0.2258 | 0.1707 | 0.1707 | TRUE | 0.1528 |
| Arsj | 3 | 2 | 5 | 0 | 1 | 5 | 2 | 2 | 5 | 0 | 1 | 5 | 2.65 | 3 | 2 | 0.626 | -0.675 | 0.60146662 | NA | 0.4965 | 0.8959 | 0.8006 | 0.8006 | TRUE | 1.4982 |
| Arsk | 107 | 82 | 78 | 84 | 76 | 73 | 83 | 87 | 84 | 84 | 86 | 72 | 82.65 | 85 | 81 | 0.948 | -0.077 | 0.71886657 | 0.83129126 | 0 | 0.0311 | 0.0206 | 0.0206 | TRUE | 0.0796 |
| Art1 | 60 | 47 | 61 | 40 | 37 | 55 | 47 | 50 | 66 | 40 | 42 | 54 | 49.71 | 54 | 45 | 0.842 | -0.249 | 0.38900698 | 0.56310277 | 0.0131 | 0.0501 | 0.0397 | 0.0397 | TRUE | 0.286 |
| Art2a-ps | 0 | 0 | 0 | 0 | 0 | 0 | 0 | 0 | 0 | 0 | 0 | 0 | 0 | 0 | 0 | NA | NA | NA | NA | NA | NA | NA | NA | NA | NA |
| Art2b | 0 | 0 | 0 | 0 | 0 | 0 | 0 | 0 | 0 | 0 | 0 | 0 | 0 | 0 | 0 | NA | NA | NA | NA | NA | NA | NA | NA | NA | NA |
| Art3 | 32 | 18 | 28 | 21 | 19 | 13 | 25 | 19 | 30 | 21 | 21 | 13 | 21.56 | 25 | 18 | 0.741 | -0.432 | 0.31520456 | 0.48874146 | 0 | 0.1123 | 0.0855 | 0.0855 | TRUE | 0.2626 |
| Art4 | 12 | 8 | 7 | 5 | 2 | 2 | 9 | 9 | 8 | 5 | 2 | 2 | 5.76 | 9 | 3 | 0.364 | -1.46 | 0.08167629 | NA | 0 | 0.4135 | 0.2819 | 0.2819 | TRUE | 0.4051 |
| Art5 | 91 | 67 | 71 | 109 | 75 | 100 | 71 | 71 | 77 | 108 | 85 | 99 | 85.06 | 73 | 97 | 1.339 | 0.422 | 0.04955639 | 0.13392125 | 0 | 0.0303 | 0.0213 | 0.0213 | TRUE | 0.1192 |
| Artn | 51 | 42 | 66 | 100 | 90 | 94 | 40 | 45 | 71 | 100 | 102 | 93 | 74.89 | 52 | 98 | 1.907 | 0.931 | 0.00028674 | 0.0028071 | 0.029 | 0.0341 | 0.0328 | 0.0328 | TRUE | 0.848 |
| Arv1 | 693 | 594 | 590 | 707 | 580 | 724 | 538 | 632 | 637 | 704 | 654 | 715 | 646.57 | 602 | 691 | 1.151 | 0.203 | 0.03963748 | 0.11380845 | 0.0041 | 0.0061 | 0.0054 | 0.0054 | TRUE | 0.108 |
| Arvcf | 4614 | 3386 | 3428 | 4007 | 3307 | 3772 | 3583 | 3602 | 3699 | 3989 | 3731 | 3724 | 3721.11 | 3628 | 3815 | 1.052 | 0.073 | 0.20071748 | 0.36140779 | 7.00E-04 | 0.0031 | 0.0021 | 0.0021 | TRUE | 0.0195 |
| Arx | 9 | 12 | 5 | 5 | 4 | 7 | 7 | 13 | 5 | 5 | 5 | 7 | 6.92 | 8 | 6 | 0.658 | -0.605 | 0.41645842 | NA | 0 | 0.3447 | 0.247 | 0.247 | TRUE | 0.591 |
| Arxes1 | 397 | 271 | 188 | 205 | 183 | 194 | 308 | 288 | 203 | 204 | 206 | 192 | 233.58 | 266 | 201 | 0.75 | -0.414 | 0.01199101 | 0.04733152 | 0.0213 | 0.0126 | 0.0152 | 0.0152 | TRUE | 0.4749 |
| Arxes2 | 580 | 381 | 320 | 346 | 281 | 301 | 450 | 405 | 345 | 344 | 317 | 297 | 359.92 | 400 | 319 | 0.796 | -0.329 | 0.01049883 | 0.04278865 | 0.0093 | 0.009 | 0.0091 | 0.0091 | TRUE | 0.1642 |
| As3mt | 1023 | 706 | 654 | 662 | 594 | 686 | 795 | 751 | 706 | 659 | 670 | 677 | 709.56 | 751 | 669 | 0.89 | -0.168 | 0.05302586 | 0.14069029 | 0 | 0.0058 | 0.004 | 0.004 | TRUE | 0.0331 |
| Asah1 | 8635 | 6308 | 7497 | 9080 | 7905 | 8879 | 6706 | 6710 | 8089 | 9038 | 8918 | 8765 | 8037.7 | 7168 | 8907 | 1.243 | 0.314 | 2.77E-05 | 0.00043384 | 0.0061 | 0.0027 | 0.0039 | 0.0039 | TRUE | 0.1536 |
| Asah2 | 100 | 52 | 79 | 81 | 74 | 70 | 78 | 55 | 85 | 81 | 83 | 69 | 75.24 | 73 | 78 | 1.065 | 0.091 | 0.70404481 | 0.82094194 | 0.0134 | 0.0339 | 0.0285 | 0.0285 | TRUE | 0.3751 |
| Asap1 | 3830 | 2803 | 2818 | 2772 | 2583 | 3055 | 2975 | 2982 | 3041 | 2759 | 2914 | 3016 | 2947.6 | 2999 | 2896 | 0.966 | -0.05 | 0.40000216 | 0.57332841 | 7.00E-04 | 0.0032 | 0.0022 | 0.0022 | TRUE | 0.021 |
| Asap2 | 2239 | 1765 | 1953 | 2185 | 2039 | 2386 | 1739 | 1877 | 2107 | 2175 | 2300 | 2355 | 2092.35 | 1908 | 2277 | 1.195 | 0.257 | 0.00128747 | 0.00865561 | 0.0049 | 0.0036 | 0.0041 | 0.0041 | TRUE | 0.1007 |
| Asap3 | 1003 | 782 | 770 | 816 | 716 | 797 | 779 | 832 | 831 | 812 | 808 | 787 | 808.06 | 814 | 802 | 0.987 | -0.019 | 0.81108397 | 0.89107019 | 0 | 0.0054 | 0.0034 | 0.0034 | TRUE | 0.0178 |
| Asb1 | 691 | 439 | 470 | 598 | 471 | 573 | 537 | 467 | 507 | 595 | 531 | 566 | 533.83 | 504 | 564 | 1.119 | 0.162 | 0.1061726 | 0.23118906 | 0.0022 | 0.0069 | 0.0054 | 0.0054 | TRUE | 0.0468 |
| Asb10 | 38 | 19 | 38 | 43 | 32 | 42 | 30 | 20 | 41 | 43 | 36 | 41 | 35.18 | 30 | 40 | 1.33 | 0.412 | 0.23595365 | 0.40240602 | 0.0221 | 0.0698 | 0.0582 | 0.0582 | TRUE | 0.2955 |
| Asb11 | 2 | 3 | 2 | 1 | 2 | 2 | 2 | 3 | 2 | 1 | 2 | 2 | 2.02 | 2 | 2 | 0.762 | -0.391 | 0.76499342 | NA | 0 | 1.1749 | 0.7247 | 0.7247 | TRUE | 0.1148 |
| Asb12 | 3 | 1 | 3 | 1 | 5 | 1 | 2 | 1 | 3 | 1 | 6 | 1 | 2.38 | 2 | 3 | 1.127 | 0.172 | 0.89339354 | NA | 0.2223 | 1.0001 | 0.7679 | 0.7679 | TRUE | 1.1407 |
| Asb13 | 220 | 187 | 193 | 200 | 201 | 231 | 171 | 199 | 208 | 199 | 227 | 228 | 205.31 | 193 | 218 | 1.135 | 0.183 | 0.21796968 | 0.38095385 | 0.0038 | 0.014 | 0.0109 | 0.0109 | TRUE | 0.1171 |
| Asb14 | 12 | 10 | 7 | 3 | 7 | 9 | 9 | 11 | 8 | 3 | 8 | 9 | 7.88 | 9 | 7 | 0.716 | -0.482 | 0.49065044 | NA | 0 | 0.3032 | 0.2216 | 0.2216 | TRUE | 0.6016 |
| Asb15 | 1 | 0 | 0 | 0 | 0 | 0 | 1 | 0 | 0 | 0 | 0 | 0 | 0.13 | 0 | 0 | 0.552 | -0.857 | 0.8336548 | NA | 0 | 18.3121 | 10 | 10 | TRUE | 0.2566 |
| Asb16 | 102 | 79 | 90 | 90 | 81 | 80 | 79 | 84 | 97 | 90 | 91 | 79 | 86.72 | 87 | 87 | 1.001 | 0.001 | 0.99579502 | 0.99753911 | 0 | 0.0298 | 0.0208 | 0.0208 | TRUE | 0.0994 |
| Asb17 | 5 | 2 | 0 | 2 | 0 | 1 | 4 | 2 | 0 | 2 | 0 | 1 | 1.5 | 2 | 1 | 0.492 | -1.024 | 0.55096757 | NA | 0 | 1.5845 | 1.3777 | 1.3777 | TRUE | 0.1332 |
| Asb18 | 0 | 1 | 0 | 0 | 1 | 0 | 0 | 1 | 0 | 0 | 1 | 0 | 0.37 | 0 | 0 | 1.075 | 0.105 | 0.97520514 | NA | 0 | 6.4904 | 6.1969 | 6.1969 | TRUE | 0.5216 |
| Asb2 | 17 | 8 | 16 | 24 | 9 | 9 | 13 | 9 | 17 | 24 | 10 | 9 | 13.65 | 13 | 14 | 1.105 | 0.144 | 0.80230405 | NA | 0.14 | 0.1761 | 0.1656 | 0.1656 | TRUE | 0.9586 |
| Asb3 | 1314 | 981 | 933 | 947 | 853 | 963 | 1021 | 1043 | 1007 | 943 | 962 | 951 | 987.71 | 1024 | 952 | 0.93 | -0.105 | 0.15233023 | 0.2990186 | 0 | 0.0048 | 0.0029 | 0.0029 | TRUE | 0.0033 |
| Asb4 | 11 | 20 | 9 | 6 | 5 | 8 | 9 | 21 | 10 | 6 | 6 | 8 | 9.84 | 13 | 7 | 0.5 | -1.001 | 0.12872483 | NA | 0 | 0.2433 | 0.1965 | 0.1965 | TRUE | 1.1615 |
| Asb5 | 2 | 0 | 1 | 2 | 0 | 1 | 2 | 0 | 1 | 2 | 0 | 1 | 0.94 | 1 | 1 | 1.117 | 0.159 | 0.93729373 | NA | 0 | 2.5371 | 1.8957 | 1.8957 | TRUE | 0.1676 |
| Asb6 | 1650 | 1131 | 879 | 902 | 814 | 911 | 1281 | 1203 | 948 | 898 | 918 | 899 | 1024.73 | 1144 | 905 | 0.79 | -0.34 | 0.00120291 | 0.00826939 | 0.0115 | 0.0047 | 0.007 | 0.007 | TRUE | 0.2656 |
| Asb7 | 1291 | 939 | 821 | 866 | 747 | 844 | 1003 | 999 | 886 | 862 | 843 | 833 | 904.2 | 963 | 846 | 0.878 | -0.187 | 0.02377021 | 0.07842742 | 0.0016 | 0.005 | 0.0038 | 0.0038 | TRUE | 0.0564 |
| Asb8 | 1022 | 660 | 724 | 678 | 592 | 682 | 794 | 702 | 781 | 675 | 668 | 673 | 715.49 | 759 | 672 | 0.884 | -0.177 | 0.04330425 | 0.12124421 | 9.00E-04 | 0.0057 | 0.0041 | 0.0041 | TRUE | 0.05 |
| Asb9 | 6 | 4 | 5 | 2 | 2 | 2 | 5 | 4 | 5 | 2 | 2 | 2 | 3.42 | 5 | 2 | 0.434 | -1.203 | 0.24727444 | NA | 0 | 0.6951 | 0.4281 | 0.4281 | TRUE | 0.0229 |
| Ascc1 | 2179 | 1737 | 2036 | 2243 | 2127 | 2343 | 1692 | 1848 | 2197 | 2233 | 2399 | 2313 | 2113.64 | 1912 | 2315 | 1.212 | 0.277 | 0.0021979 | 0.01297588 | 0.0091 | 0.0035 | 0.0054 | 0.0054 | TRUE | 0.2042 |
| Ascc2 | 2683 | 1779 | 1821 | 1875 | 1626 | 1913 | 2084 | 1892 | 1965 | 1866 | 1834 | 1888 | 1921.67 | 1980 | 1863 | 0.94 | -0.089 | 0.17369457 | 0.3264453 | 9.00E-04 | 0.0037 | 0.0026 | 0.0026 | TRUE | 0.0261 |
| Ascc3 | 5994 | 4137 | 3692 | 3880 | 3269 | 3842 | 4655 | 4400 | 3984 | 3862 | 3688 | 3793 | 4063.66 | 4346 | 3781 | 0.87 | -0.202 | 0.00292877 | 0.01624402 | 0.0029 | 0.003 | 0.0031 | 0.0031 | TRUE | 0.0647 |
| Ascl1 | 16 | 15 | 16 | 15 | 12 | 17 | 12 | 16 | 17 | 15 | 14 | 17 | 15.15 | 15 | 15 | 1.001 | 0.001 | 0.99868429 | NA | 0 | 0.1589 | 0.1045 | 0.1045 | TRUE | 0.1509 |
| Ascl2 | 222 | 199 | 200 | 265 | 224 | 287 | 172 | 212 | 216 | 264 | 253 | 283 | 233.28 | 200 | 267 | 1.34 | 0.423 | 0.00309893 | 0.01695748 | 0.005 | 0.0126 | 0.0103 | 0.0103 | TRUE | 0.1745 |
| Ascl3 | 0 | 0 | 0 | 0 | 0 | 0 | 0 | 0 | 0 | 0 | 0 | 0 | 0 | 0 | 0 | NA | NA | NA | NA | NA | NA | NA | NA | NA | NA |
| Ascl4 | 9 | 6 | 6 | 9 | 4 | 4 | 7 | 6 | 6 | 9 | 5 | 4 | 6.21 | 6 | 6 | 0.879 | -0.186 | 0.80791496 | NA | 0 | 0.384 | 0.2613 | 0.2613 | TRUE | 0.5305 |
| Ascl5 | 0 | 1 | 0 | 4 | 3 | 1 | 0 | 1 | 0 | 4 | 3 | 1 | 1.57 | 0 | 3 | 8.054 | 3.01 | 0.09635669 | NA | 0 | 1.5125 | 1.1747 | 1.1747 | TRUE | 0.5217 |
| Asf1a | 2276 | 1588 | 1287 | 1327 | 1096 | 1335 | 1768 | 1689 | 1389 | 1321 | 1236 | 1318 | 1453.44 | 1615 | 1292 | 0.799 | -0.323 | 0.00046364 | 0.00408309 | 0.0081 | 0.0041 | 0.0055 | 0.0055 | TRUE | 0.1796 |
| Asf1b | 3584 | 2554 | 2496 | 2483 | 2369 | 2617 | 2784 | 2717 | 2693 | 2472 | 2672 | 2583 | 2653.47 | 2731 | 2576 | 0.943 | -0.085 | 0.15337722 | 0.30026306 | 5.00E-04 | 0.0033 | 0.0022 | 0.0022 | TRUE | 0.0152 |
| Asgr1 | 18 | 20 | 16 | 17 | 13 | 11 | 14 | 21 | 17 | 17 | 15 | 11 | 15.83 | 17 | 14 | 0.813 | -0.298 | 0.54080199 | NA | 0 | 0.1522 | 0.107 | 0.107 | TRUE | 0.1957 |
| Asgr2 | 72 | 56 | 62 | 63 | 69 | 67 | 56 | 60 | 67 | 63 | 78 | 66 | 64.85 | 61 | 69 | 1.135 | 0.182 | 0.45337915 | 0.62319659 | 0 | 0.039 | 0.0271 | 0.0271 | TRUE | 0.1089 |
| Ash1l | 1810 | 1375 | 1406 | 1590 | 1386 | 1573 | 1406 | 1463 | 1517 | 1583 | 1564 | 1553 | 1514.07 | 1462 | 1567 | 1.072 | 0.101 | 0.13039552 | 0.26731829 | 0 | 0.004 | 0.0025 | 0.0025 | TRUE | 0.0142 |
| Ash2l | 33783 | 24428 | 25609 | 27331 | 25355 | 28672 | 26238 | 25984 | 27632 | 27205 | 28603 | 28304 | 27327.58 | 26618 | 28037 | 1.053 | 0.075 | 0.1312737 | 0.2686689 | 8.00E-04 | 0.0025 | 0.0017 | 0.0017 | TRUE | 0.0136 |
| Asic1 | 1598 | 1176 | 1130 | 1559 | 1332 | 1563 | 1241 | 1251 | 1219 | 1552 | 1503 | 1543 | 1384.77 | 1237 | 1533 | 1.239 | 0.309 | 2.93E-06 | 6.78E-05 | 0 | 0.0041 | 0.0024 | 0.0024 | TRUE | 0.0034 |
| Asic2 | 82 | 70 | 53 | 48 | 58 | 58 | 64 | 74 | 57 | 48 | 65 | 57 | 60.97 | 65 | 57 | 0.871 | -0.199 | 0.43852986 | 0.60989659 | 0 | 0.0413 | 0.031 | 0.031 | TRUE | 0.1656 |
| Asic3 | 4 | 9 | 10 | 13 | 13 | 10 | 3 | 10 | 11 | 13 | 15 | 10 | 10.16 | 8 | 13 | 1.634 | 0.709 | 0.27090709 | NA | 0.0625 | 0.2358 | 0.1939 | 0.1939 | TRUE | 1.0953 |
| Asic4 | 1 | 1 | 0 | 2 | 1 | 3 | 1 | 1 | 0 | 2 | 1 | 3 | 1.32 | 1 | 2 | 3.263 | 1.706 | 0.32825016 | NA | 0 | 1.7978 | 1.1723 | 1.1723 | TRUE | 0.1787 |

| | | | | | | | | | | | | | | | | | | | | | | | | | |
|---|---|---|---|---|---|---|---|---|---|---|---|---|---|---|---|---|---|---|---|---|---|---|---|---|---|
| Asic5 | 0 | 0 | 0 | 0 | 0 | 0 | 0 | 0 | 0 | 0 | 0 | 0 | 0 | 0 | 0 | NA | NA | NA | NA | NA | NA | NA | NA | NA | NA |
| Asl | 2347 | 1729 | 1912 | 2267 | 2149 | 2307 | 1823 | 1839 | 2063 | 2257 | 2424 | 2277 | 2113.87 | 1908 | 2319 | 1.216 | 0.282 | 8.05E-05 | 0.001015 | 0.0025 | 0.0035 | 0.0032 | 0.0032 | TRUE | 0.06 |
| Asmt | 0 | 0 | 0 | 0 | 0 | 0 | 0 | 0 | 0 | 0 | 0 | 0 | 0 | 0 | 0 | NA | NA | NA | NA | NA | NA | NA | NA | NA | NA |
| Asna1 | 12276 | 8992 | 8632 | 9182 | 8436 | 9992 | 9534 | 9565 | 9314 | 9140 | 9517 | 9864 | 9488.81 | 9471 | 9507 | 1.004 | 0.005 | 0.91648878 | 0.95388891 | 6.00E-04 | 0.0027 | 0.0018 | 0.0018 | TRUE | 0.014 |
| Asns | 11318 | 8338 | 8310 | 8746 | 7642 | 8784 | 8790 | 8869 | 8966 | 8706 | 8621 | 8671 | 8770.59 | 8875 | 8666 | 0.977 | -0.034 | 0.45209287 | 0.62194332 | 0 | 0.0027 | 0.0014 | 0.0014 | TRUE | 0.001 |
| Asnsd1 | 10110 | 7351 | 7345 | 7954 | 6670 | 7554 | 7852 | 7819 | 7925 | 7917 | 7524 | 7457 | 7749.2 | 7865 | 7633 | 0.97 | -0.043 | 0.38833861 | 0.56236309 | 4.00E-04 | 0.0027 | 0.0017 | 0.0017 | TRUE | 0.013 |
| Aspa | 153 | 123 | 77 | 31 | 33 | 29 | 119 | 131 | 83 | 31 | 37 | 29 | 71.58 | 111 | 32 | 0.289 | -1.79 | 5.23E-11 | 4.51E-09 | 0.0272 | 0.0355 | 0.0331 | 0.0331 | TRUE | 0.4288 |
| Aspdh | 0 | 0 | 1 | 1 | 0 | 0 | 0 | 0 | 1 | 1 | 0 | 0 | 0.35 | 0 | 0 | 1.075 | 0.105 | 0.97567165 | NA | 0 | 6.8576 | 6.5143 | 6.5143 | TRUE | 0.5365 |
| Aspg | 553 | 564 | 601 | 680 | 638 | 734 | 429 | 600 | 648 | 677 | 720 | 725 | 633.18 | 559 | 707 | 1.268 | 0.343 | 0.00948674 | 0.03984544 | 0.022 | 0.0062 | 0.011 | 0.011 | TRUE | 0.5048 |
| Asph | 4585 | 3161 | 3228 | 3451 | 3031 | 3476 | 3561 | 3362 | 3483 | 3435 | 3419 | 3431 | 3448.67 | 3469 | 3428 | 0.988 | -0.017 | 0.75119517 | 0.85220887 | 0 | 0.0031 | 0.0019 | 0.0019 | TRUE | 0.0087 |
| Asphd1 | 9 | 3 | 7 | 8 | 5 | 15 | 7 | 3 | 8 | 8 | 6 | 15 | 7.69 | 6 | 10 | 1.6 | 0.678 | 0.35151686 | NA | 0.0772 | 0.3106 | 0.2464 | 0.2464 | TRUE | 0.7439 |
| Asphd2 | 293 | 212 | 208 | 228 | 187 | 234 | 228 | 226 | 224 | 227 | 211 | 231 | 224.4 | 226 | 223 | 0.988 | -0.018 | 0.89479146 | 0.94223961 | 0 | 0.013 | 0.0083 | 0.0083 | TRUE | 0.0234 |
| Aspm | 6243 | 4363 | 4353 | 4865 | 4257 | 4591 | 4849 | 4641 | 4697 | 4843 | 4802 | 4532 | 4727.23 | 4729 | 4726 | 0.999 | -0.001 | 0.97939628 | 0.98908436 | 7.00E-04 | 0.0029 | 0.002 | 0.002 | TRUE | 0.0157 |
| Aspn | 0 | 0 | 0 | 0 | 0 | 0 | 0 | 0 | 0 | 0 | 0 | 0 | 0 | 0 | 0 | NA | NA | NA | NA | NA | NA | NA | NA | NA | NA |
| Asprv1 | 40 | 22 | 21 | 30 | 23 | 21 | 31 | 23 | 23 | 30 | 26 | 21 | 25.61 | 26 | 26 | 0.984 | -0.023 | 0.9526306 | 0.97381985 | 0 | 0.095 | 0.068 | 0.068 | TRUE | 0.2462 |
| Aspscr1 | 5919 | 4393 | 4022 | 4338 | 4068 | 4829 | 4597 | 4673 | 4340 | 4318 | 4589 | 4767 | 4547.28 | 4537 | 4558 | 1.005 | 0.007 | 0.91341139 | 0.95246365 | 0.0018 | 0.0029 | 0.0025 | 0.0025 | TRUE | 0.026 |
| Asrgl1 | 2384 | 1795 | 1803 | 2085 | 1867 | 2110 | 1852 | 1909 | 1945 | 2075 | 2106 | 2083 | 1995.13 | 1902 | 2088 | 1.098 | 0.135 | 0.02516397 | 0.08165009 | 0 | 0.0036 | 0.0021 | 0.0021 | TRUE | 0.0068 |
| Ass1 | 10 | 8 | 5 | 1 | 4 | 2 | 8 | 9 | 5 | 1 | 5 | 2 | 4.86 | 7 | 3 | 0.338 | -1.565 | 0.09436844 | NA | 0 | 0.4902 | 0.3526 | 0.3526 | TRUE | 0.4849 |
| Aste1 | 516 | 394 | 400 | 350 | 321 | 385 | 401 | 419 | 432 | 348 | 362 | 380 | 390.33 | 417 | 363 | 0.873 | -0.197 | 0.06558969 | 0.16366505 | 0 | 0.0085 | 0.0056 | 0.0056 | TRUE | 0.0187 |
| Astl | 2 | 0 | 1 | 1 | 1 | 1 | 2 | 0 | 1 | 1 | 1 | 1 | 0.96 | 1 | 1 | 1.143 | 0.193 | 0.91871967 | NA | 0 | 2.4786 | 1.5559 | 1.5559 | TRUE | 0.2848 |
| Astn1 | 12 | 15 | 9 | 5 | 5 | 3 | 9 | 16 | 10 | 5 | 6 | 3 | 8.09 | 12 | 5 | 0.388 | -1.366 | 0.05533155 | NA | 0 | 0.2952 | 0.2094 | 0.2094 | TRUE | 0.3804 |
| Astn2 | 13 | 5 | 4 | 1 | 4 | 3 | 10 | 5 | 4 | 1 | 5 | 3 | 4.7 | 6 | 3 | 0.418 | -1.26 | 0.18285306 | NA | 0 | 0.5067 | 0.3866 | 0.3866 | TRUE | 0.7041 |
| Asun | 2006 | 1386 | 1309 | 1462 | 1302 | 1477 | 1558 | 1474 | 1412 | 1455 | 1469 | 1458 | 1471.12 | 1481 | 1461 | 0.985 | -0.022 | 0.75000118 | 0.85158618 | 7.00E-04 | 0.004 | 0.0028 | 0.0028 | TRUE | 0.0255 |
| Asxl1 | 13263 | 9442 | 9789 | 10418 | 9373 | 11127 | 10301 | 10043 | 10562 | 10370 | 10574 | 10984 | 10472.37 | 10302 | 10643 | 1.033 | 0.047 | 0.3518568 | 0.52644082 | 6.00E-04 | 0.0027 | 0.0017 | 0.0017 | TRUE | 0.0097 |
| Asxl2 | 2740 | 1967 | 1726 | 1745 | 1635 | 2005 | 2128 | 2092 | 1862 | 1737 | 1844 | 1979 | 1940.55 | 2027 | 1853 | 0.914 | -0.13 | 0.09461822 | 0.21230427 | 0.0039 | 0.0036 | 0.0039 | 0.0039 | TRUE | 0.0608 |
| Asxl3 | 323 | 220 | 153 | 175 | 192 | 162 | 251 | 234 | 165 | 174 | 217 | 160 | 200.11 | 217 | 184 | 0.843 | -0.246 | 0.17904737 | 0.33361023 | 0.0311 | 0.0143 | 0.0192 | 0.0192 | TRUE | 0.4624 |
| Asz1 | 686 | 492 | 422 | 374 | 311 | 261 | 533 | 523 | 455 | 372 | 351 | 258 | 415.37 | 504 | 327 | 0.648 | -0.626 | 5.91E-06 | 0.00012146 | 0.0189 | 0.0081 | 0.0112 | 0.0112 | TRUE | 0.3973 |
| Atad1 | 5006 | 3709 | 4088 | 4488 | 3769 | 4536 | 3888 | 3945 | 4411 | 4467 | 4252 | 4478 | 4240.16 | 4081 | 4399 | 1.078 | 0.109 | 0.09499088 | 0.21299223 | 0.0025 | 0.003 | 0.0028 | 0.0028 | TRUE | 0.0603 |
| Atad2 | 11571 | 8772 | 8740 | 9242 | 8224 | 9541 | 8987 | 9331 | 9430 | 9199 | 9278 | 9418 | 9273.87 | 9249 | 9298 | 1.006 | 0.008 | 0.86923688 | 0.92778752 | 3.00E-04 | 0.0027 | 0.0016 | 0.0016 | TRUE | 0.0076 |
| Atad2b | 5893 | 4038 | 3986 | 5001 | 4393 | 4999 | 4577 | 4295 | 4301 | 4978 | 4956 | 4935 | 4673.57 | 4391 | 4956 | 1.128 | 0.174 | 0.00119822 | 0.00824565 | 5.00E-04 | 0.0029 | 0.0019 | 0.0019 | TRUE | 0.0171 |
| Atad3a | 8574 | 6492 | 5969 | 6765 | 5744 | 6811 | 6659 | 6905 | 6441 | 6734 | 6480 | 6724 | 6657.04 | 6668 | 6646 | 0.997 | -0.005 | 0.92777671 | 0.960172 | 5.00E-04 | 0.0028 | 0.0019 | 0.0019 | TRUE | 0.0116 |
| Atad5 | 5020 | 3697 | 3712 | 4137 | 3476 | 3938 | 3899 | 3932 | 4005 | 4118 | 3921 | 3887 | 3960.53 | 3945 | 3975 | 1.008 | 0.011 | 0.83573396 | 0.90644601 | 3.00E-04 | 0.003 | 0.0019 | 0.0019 | TRUE | 0.012 |
| Atat1 | 471 | 354 | 393 | 377 | 378 | 444 | 366 | 377 | 424 | 375 | 426 | 438 | 401.06 | 389 | 413 | 1.065 | 0.09 | 0.43167495 | 0.60365229 | 0.004 | 0.0083 | 0.007 | 0.007 | TRUE | 0.0761 |
| Atcay | 40 | 28 | 15 | 39 | 43 | 43 | 31 | 30 | 16 | 39 | 49 | 42 | 34.47 | 26 | 43 | 1.667 | 0.737 | 0.03586492 | 0.10578153 | 0.0149 | 0.0712 | 0.0583 | 0.0583 | TRUE | 0.5925 |
| Ate1 | 4276 | 3023 | 3044 | 3208 | 2902 | 3351 | 3321 | 3216 | 3284 | 3193 | 3274 | 3308 | 3265.99 | 3274 | 3258 | 0.995 | -0.007 | 0.89587102 | 0.94272381 | 0 | 0.0032 | 0.0018 | 0.0018 | TRUE | 0.0037 |
| Atf1 | 4765 | 3492 | 3427 | 3611 | 3151 | 3578 | 3701 | 3714 | 3698 | 3594 | 3555 | 3532 | 3632.33 | 3704 | 3560 | 0.961 | -0.057 | 0.26770808 | 0.43829916 | 0 | 0.0031 | 0.0016 | 0.0016 | TRUE | 9.00E-04 |
| Atf2 | 6782 | 4893 | 4541 | 4439 | 3988 | 4709 | 5267 | 5205 | 4900 | 4419 | 4499 | 4649 | 4822.93 | 5124 | 4522 | 0.882 | -0.181 | 0.0012459 | 0.00846776 | 8.00E-04 | 0.0029 | 0.002 | 0.002 | TRUE | 0.0177 |
| Atf3 | 95 | 43 | 45 | 62 | 48 | 54 | 74 | 46 | 49 | 62 | 54 | 53 | 56.21 | 56 | 56 | 0.997 | -0.004 | 0.98746799 | 0.99373352 | 0.0254 | 0.0446 | 0.0389 | 0.0389 | TRUE | 0.7313 |
| Atf4 | 2309 | 1594 | 1377 | 788 | 746 | 940 | 1793 | 1696 | 1486 | 784 | 842 | 928 | 1254.74 | 1658 | 851 | 0.513 | -0.963 | 5.11E-25 | 2.61E-22 | 0.0068 | 0.0043 | 0.0054 | 0.0054 | TRUE | 0.099 |
| Atf5 | 612 | 458 | 346 | 315 | 269 | 303 | 475 | 487 | 373 | 314 | 303 | 299 | 375.32 | 445 | 305 | 0.685 | -0.546 | 1.53E-05 | 0.00026983 | 0.0087 | 0.0087 | 0.0087 | 0.0087 | TRUE | 0.2242 |
| Atf6 | 2861 | 1964 | 2070 | 2305 | 2042 | 2265 | 2222 | 2089 | 2234 | 2294 | 2304 | 2236 | 2229.75 | 2182 | 2278 | 1.044 | 0.062 | 0.31189787 | 0.48519332 | 3.00E-04 | 0.0035 | 0.0022 | 0.0022 | TRUE | 0.0163 |
| Atf6b | 2841 | 2157 | 2102 | 2117 | 1993 | 2336 | 2206 | 2294 | 2268 | 2107 | 2248 | 2306 | 2238.41 | 2256 | 2220 | 0.984 | -0.023 | 0.7203955 | 0.83218402 | 8.00E-04 | 0.0035 | 0.0024 | 0.0024 | TRUE | 0.0244 |
| Atf7 | 396 | 268 | 311 | 286 | 261 | 309 | 308 | 285 | 336 | 285 | 294 | 305 | 302.06 | 310 | 295 | 0.953 | -0.07 | 0.56740618 | 0.72126792 | 0 | 0.0103 | 0.0074 | 0.0074 | TRUE | 0.0581 |
| Atf7ip | 10062 | 7429 | 7691 | 7053 | 6732 | 8225 | 7815 | 7902 | 8299 | 7021 | 7594 | 8119 | 7791.61 | 8005 | 7578 | 0.947 | -0.079 | 0.21949724 | 0.3825944 | 0.003 | 0.0027 | 0.0028 | 0.0028 | TRUE | 0.0508 |
| Atf7ip2 | 983 | 671 | 636 | 786 | 650 | 728 | 763 | 714 | 686 | 782 | 733 | 719 | 732.95 | 721 | 745 | 1.031 | 0.045 | 0.61118584 | 0.75399128 | 0.0013 | 0.0057 | 0.0042 | 0.0042 | TRUE | 0.0329 |
| Atg10 | 403 | 270 | 312 | 334 | 308 | 311 | 313 | 287 | 337 | 332 | 347 | 307 | 320.63 | 312 | 329 | 1.053 | 0.074 | 0.53963281 | 0.69788018 | 0.0017 | 0.0098 | 0.0074 | 0.0074 | TRUE | 0.0526 |
| Atg12 | 3795 | 2791 | 3115 | 3782 | 3192 | 3820 | 2947 | 2969 | 3361 | 3765 | 3601 | 3771 | 3402.28 | 3092 | 3712 | 1.201 | 0.264 | 8.86E-05 | 0.0010933 | 0.0027 | 0.0031 | 0.003 | 0.003 | TRUE | 0.0695 |
| Atg13 | 3441 | 2403 | 2328 | 2378 | 2160 | 2358 | 2672 | 2556 | 2512 | 2367 | 2437 | 2328 | 2478.65 | 2580 | 2377 | 0.921 | -0.119 | 0.04652561 | 0.12783363 | 4.00E-04 | 0.0034 | 0.0022 | 0.0022 | TRUE | 0.0123 |
| Atg14 | 3231 | 2402 | 2394 | 2468 | 2191 | 2528 | 2509 | 2555 | 2583 | 2457 | 2472 | 2496 | 2511.88 | 2549 | 2475 | 0.971 | -0.042 | 0.45115079 | 0.62104384 | 0 | 0.0034 | 0.0019 | 0.0019 | TRUE | 0.0023 |
| Atg16l1 | 2644 | 1890 | 1640 | 1830 | 1536 | 1749 | 2053 | 2010 | 1770 | 1822 | 1733 | 1727 | 1852.38 | 1944 | 1761 | 0.905 | -0.145 | 0.05344319 | 0.14127307 | 0.0031 | 0.0037 | 0.0035 | 0.0035 | TRUE | 0.0739 |
| Atg16l2 | 376 | 254 | 216 | 244 | 169 | 235 | 292 | 270 | 233 | 243 | 191 | 232 | 243.46 | 265 | 222 | 0.836 | -0.258 | 0.08001627 | 0.18922445 | 0.01 | 0.0122 | 0.0115 | 0.0115 | TRUE | 0.1597 |
| Atg2a | 4675 | 3421 | 3015 | 2878 | 2819 | 3342 | 3631 | 3639 | 3253 | 2865 | 3180 | 3299 | 3311.14 | 3508 | 3115 | 0.888 | -0.172 | 0.01944869 | 0.06768447 | 0.0041 | 0.0031 | 0.0036 | 0.0036 | TRUE | 0.0602 |
| Atg2b | 7371 | 5411 | 5499 | 6318 | 6000 | 7162 | 5725 | 5756 | 5933 | 6289 | 6769 | 7070 | 6256.88 | 5805 | 6709 | 1.156 | 0.209 | 0.00042648 | 0.00382179 | 0.0017 | 0.0028 | 0.0024 | 0.0024 | TRUE | 0.0368 |
| Atg3 | 1611 | 1148 | 1091 | 1166 | 935 | 1157 | 1251 | 1221 | 1177 | 1161 | 1055 | 1142 | 1167.84 | 1216 | 1119 | 0.92 | -0.12 | 0.10766077 | 0.23350461 | 8.00E-04 | 0.0045 | 0.0032 | 0.0032 | TRUE | 0.0297 |
| Atg4a | 215 | 145 | 135 | 159 | 127 | 153 | 167 | 154 | 146 | 158 | 143 | 151 | 153.24 | 156 | 151 | 0.968 | -0.048 | 0.76708456 | 0.86221607 | 0 | 0.0179 | 0.0121 | 0.0121 | TRUE | 0.0472 |
| Atg4b | 2543 | 1982 | 2016 | 1969 | 1884 | 2126 | 1975 | 2108 | 2175 | 1960 | 2125 | 2099 | 2073.75 | 2086 | 2061 | 0.989 | -0.016 | 0.80916315 | 0.88984911 | 0.0017 | 0.0036 | 0.0029 | 0.0029 | TRUE | 0.0271 |
| Atg4c | 845 | 630 | 658 | 861 | 710 | 864 | 656 | 670 | 710 | 857 | 801 | 853 | 757.88 | 679 | 837 | 1.235 | 0.304 | 0.00029424 | 0.00286874 | 0 | 0.0056 | 0.0038 | 0.0038 | TRUE | 0.0187 |
| Atg4d | 2237 | 1666 | 1640 | 1601 | 1554 | 1707 | 1737 | 1772 | 1770 | 1594 | 1753 | 1685 | 1718.47 | 1760 | 1677 | 0.953 | -0.069 | 0.2954028 | 0.46710834 | 5.00E-04 | 0.0038 | 0.0026 | 0.0026 | TRUE | 0.023 |
| Atg5 | 3020 | 2219 | 2192 | 2174 | 1916 | 2044 | 2346 | 2360 | 2365 | 2164 | 2161 | 2018 | 2235.69 | 2357 | 2114 | 0.897 | -0.157 | 0.01022747 | 0.04198815 | 3.00E-04 | 0.0035 | 0.0022 | 0.0022 | TRUE | 0.0195 |
| Atg7 | 1473 | 1141 | 1060 | 1236 | 1121 | 1217 | 1144 | 1214 | 1144 | 1230 | 1265 | 1201 | 1199.62 | 1167 | 1232 | 1.056 | 0.078 | 0.27240046 | 0.44329042 | 0 | 0.0044 | 0.0028 | 0.0028 | TRUE | 0.0143 |
| Atg9a | 1629 | 1219 | 1292 | 1473 | 1336 | 1513 | 1265 | 1297 | 1394 | 1466 | 1507 | 1494 | 1403.8 | 1319 | 1489 | 1.13 | 0.176 | 0.01201977 | 0.04741888 | 6.00E-04 | 0.0041 | 0.0028 | 0.0028 | TRUE | 0.0297 |
| Atg9b | 324 | 296 | 313 | 327 | 292 | 379 | 252 | 315 | 338 | 325 | 329 | 374 | 322.21 | 302 | 343 | 1.143 | 0.193 | 0.15820674 | 0.3072349 | 0.0117 | 0.0098 | 0.0104 | 0.0104 | TRUE | 0.2538 |
| Athl1 | 1016 | 766 | 703 | 721 | 658 | 772 | 789 | 815 | 759 | 718 | 742 | 762 | 764.08 | 788 | 741 | 0.94 | -0.089 | 0.28342727 | 0.45527376 | 0 | 0.0055 | 0.0036 | 0.0036 | TRUE | 0.0116 |
| Atic | 14347 | 10256 | 9806 | 9107 | 8316 | 9840 | 11143 | 10909 | 10581 | 9065 | 9381 | 9714 | 10132.08 | 10878 | 9387 | 0.863 | -0.213 | 4.07E-05 | 0.00059527 | 8.00E-04 | 0.0027 | 0.0018 | 0.0018 | TRUE | 0.0114 |
| Atl1 | 294 | 230 | 264 | 273 | 261 | 310 | 228 | 245 | 285 | 272 | 294 | 306 | 271.67 | 253 | 291 | 1.154 | 0.207 | 0.12192957 | 0.25501014 | 0.0047 | 0.0112 | 0.0092 | 0.0092 | TRUE | 0.1362 |
| Atl2 | 5539 | 4149 | 4112 | 4649 | 4092 | 4726 | 4302 | 4413 | 4437 | 4628 | 4616 | 4665 | 4510.18 | 4384 | 4636 | 1.058 | 0.081 | 0.10770478 | 0.23352795 | 0 | 0.003 | 0.0016 | 0.0016 | TRUE | 0.0034 |
| Atl3 | 3252 | 2450 | 2780 | 3321 | 2841 | 3112 | 2526 | 2606 | 3000 | 3306 | 3205 | 3072 | 2952.33 | 2711 | 3194 | 1.179 | 0.238 | 0.00153821 | 0.00983036 | 0.0045 | 0.0032 | 0.0037 | 0.0037 | TRUE | 0.105 |
| Atm | 5516 | 3696 | 3139 | 3463 | 3177 | 3571 | 4284 | 3931 | 3387 | 3447 | 3584 | 3525 | 3693.09 | 3867 | 3519 | 0.909 | -0.137 | 0.09033814 | 0.20525546 | 0.0066 | 0.0031 | 0.0045 | 0.0045 | TRUE | 0.1433 |
| Atmin | 9313 | 6924 | 7091 | 7158 | 6683 | 7739 | 7233 | 7365 | 7651 | 7125 | 7539 | 7640 | 7425.48 | 7416 | 7435 | 1.003 | 0.004 | 0.94492793 | 0.96969091 | 0.001 | 0.0027 | 0.002 | 0.002 | TRUE | 0.0163 |

| | | | | | | | | | | | | | | | | | | | | | | | | | |
|---|---|---|---|---|---|---|---|---|---|---|---|---|---|---|---|---|---|---|---|---|---|---|---|---|---|
| Atn1 | 4305 | 3217 | 2567 | 2558 | 2464 | 2813 | 3344 | 3422 | 2770 | 2546 | 2780 | 2777 | 2939.65 | 3179 | 2701 | 0.849 | -0.235 | 0.0050106 | 0.02456827 | 0.007 | 0.0032 | 0.0047 | 0.0047 | TRUE | 0.1523 |
| Atoh1 | 16 | 17 | 5 | 7 | 15 | 4 | 12 | 18 | 5 | 7 | 17 | 4 | 10.62 | 12 | 9 | 0.765 | -0.386 | 0.57622003 | NA | 0.3017 | 0.2255 | 0.2468 | 0.2468 | TRUE | 0.4171 |
| Atoh7 | 1 | 6 | 0 | 0 | 0 | 0 | 1 | 6 | 0 | 0 | 0 | 0 | 1.19 | 2 | 0 | 0.08 | -3.635 | 0.11010291 | NA | 8.8819 | 1.9888 | 2.5258 | 2.5258 | TRUE | 2.1502 |
| Atoh8 | 13 | 11 | 9 | 13 | 12 | 8 | 10 | 12 | 10 | 13 | 14 | 8 | 10.98 | 11 | 12 | 1.089 | 0.123 | 0.83067633 | NA | 0 | 0.2183 | 0.147 | 0.147 | TRUE | 0.2928 |
| Atox1 | 5144 | 3674 | 3818 | 4473 | 3817 | 4637 | 3995 | 3908 | 4120 | 4452 | 4306 | 4577 | 4226.42 | 4008 | 4445 | 1.109 | 0.15 | 0.00661031 | 0.03038959 | 5.00E-04 | 0.003 | 0.002 | 0.002 | TRUE | 0.0091 |
| Atp10a | 764 | 583 | 756 | 693 | 658 | 698 | 593 | 620 | 816 | 690 | 742 | 689 | 691.73 | 676 | 707 | 1.047 | 0.066 | 0.57607504 | 0.72769249 | 0.0147 | 0.0059 | 0.0087 | 0.0087 | TRUE | 0.3789 |
| Atp10b | 3 | 0 | 0 | 1 | 0 | 0 | 2 | 0 | 0 | 1 | 0 | 0 | 0.55 | 1 | 0 | 0.458 | -1.128 | 0.69820437 | NA | 0 | 4.2787 | 4.502 | 4.502 | TRUE | 1.4399 |
| Atp10d | 4067 | 3015 | 3281 | 4508 | 4011 | 4853 | 3159 | 3207 | 3540 | 4487 | 4525 | 4791 | 3951.43 | 3302 | 4601 | 1.394 | 0.479 | 9.87E-14 | 1.49E-11 | 0.0022 | 0.003 | 0.0027 | 0.0027 | TRUE | 0.048 |
| Atp11a | 4199 | 3030 | 2908 | 2934 | 2663 | 2962 | 3261 | 3223 | 3138 | 2921 | 3004 | 2924 | 3078.41 | 3207 | 2950 | 0.919 | -0.121 | 0.02725072 | 0.08632413 | 0 | 0.0032 | 0.0019 | 0.0019 | TRUE | 0.0043 |
| Atp11b | 10555 | 7789 | 7570 | 8721 | 7920 | 8924 | 8198 | 8285 | 8168 | 8681 | 8935 | 8809 | 8512.58 | 8217 | 8808 | 1.072 | 0.1 | 0.03060146 | 0.0939001 | 0 | 0.0027 | 0.0014 | 0.0014 | TRUE | 0.002 |
| Atp11c | 1930 | 1333 | 1450 | 1636 | 1309 | 1499 | 1499 | 1418 | 1565 | 1628 | 1477 | 1480 | 1511.05 | 1494 | 1528 | 1.023 | 0.033 | 0.65399528 | 0.78517888 | 0.0021 | 0.004 | 0.0033 | 0.0033 | TRUE | 0.0399 |
| Atp12a | 207 | 159 | 212 | 352 | 277 | 281 | 161 | 169 | 229 | 350 | 312 | 277 | 249.82 | 186 | 313 | 1.69 | 0.757 | 2.50E-06 | 6.00E-05 | 0.0202 | 0.0119 | 0.0144 | 0.0144 | TRUE | 0.4198 |
| Atp13a1 | 5645 | 4265 | 3705 | 3769 | 3367 | 4020 | 4384 | 4537 | 3998 | 3752 | 3798 | 3968 | 4072.81 | 4306 | 3839 | 0.892 | -0.166 | 0.00991082 | 0.04110189 | 0.0022 | 0.003 | 0.0027 | 0.0027 | TRUE | 0.0472 |
| Atp13a2 | 2622 | 2012 | 1848 | 2143 | 1856 | 2062 | 2036 | 2140 | 1994 | 2133 | 2094 | 2036 | 2072.15 | 2057 | 2088 | 1.015 | 0.021 | 0.7317785 | 0.83929626 | 4.00E-04 | 0.0036 | 0.0023 | 0.0023 | TRUE | 0.0148 |
| Atp13a3 | 14294 | 10034 | 9894 | 11234 | 9353 | 10697 | 11102 | 10673 | 10676 | 11182 | 10551 | 10560 | 10790.53 | 10817 | 10764 | 0.995 | -0.007 | 0.88857399 | 0.93805954 | 6.00E-04 | 0.0026 | 0.0018 | 0.0018 | TRUE | 0.0141 |
| Atp13a4 | 32 | 23 | 19 | 11 | 11 | 10 | 25 | 24 | 21 | 11 | 12 | 10 | 17.17 | 23 | 11 | 0.474 | -1.078 | 0.02185972 | 0.0736368 | 0 | 0.1404 | 0.0919 | 0.0919 | TRUE | 0.0595 |
| Atp13a5 | 0 | 0 | 0 | 0 | 0 | 0 | 0 | 0 | 0 | 0 | 0 | 0 | 0 | 0 | 0 | NA | NA | NA | NA | NA | NA | NA | NA | NA | NA |
| Atp1a1 | 39704 | 29276 | 31523 | 31594 | 30336 | 33536 | 30836 | 31141 | 34013 | 31449 | 34222 | 33105 | 32460.99 | 31997 | 32925 | 1.029 | 0.041 | 0.47977408 | 0.64660659 | 0.0021 | 0.0025 | 0.0024 | 0.0024 | TRUE | 0.0372 |
| Atp1a2 | 98 | 82 | 177 | 233 | 180 | 199 | 76 | 87 | 191 | 232 | 203 | 196 | 164.29 | 118 | 210 | 1.793 | 0.843 | 0.0015915 | 0.01007854 | 0.1219 | 0.0169 | 0.0447 | 0.0447 | TRUE | 2.8942 |
| Atp1a3 | 4655 | 3719 | 4826 | 7305 | 6655 | 7960 | 3615 | 3956 | 5207 | 7271 | 7508 | 7858 | 5902.51 | 4259 | 7546 | 1.772 | 0.826 | 4.69E-15 | 7.99E-13 | 0.0199 | 0.0028 | 0.0078 | 0.0078 | TRUE | 0.4607 |
| Atp1a4 | 16 | 11 | 14 | 14 | 16 | 18 | 12 | 12 | 15 | 14 | 18 | 18 | 14.83 | 13 | 17 | 1.27 | 0.344 | 0.483855 | NA | 0 | 0.1622 | 0.1061 | 0.1061 | TRUE | 0.0971 |
| Atp1b1 | 5871 | 4733 | 5193 | 6900 | 6385 | 7122 | 4560 | 5034 | 5603 | 6868 | 7203 | 7031 | 6049.85 | 5066 | 7034 | 1.389 | 0.474 | 1.50E-10 | 1.16E-08 | 0.0053 | 0.0028 | 0.0038 | 0.0038 | TRUE | 0.1049 |
| Atp1b2 | 4332 | 3220 | 3818 | 4004 | 4005 | 4408 | 3364 | 3425 | 4120 | 3986 | 4518 | 4351 | 3960.69 | 3636 | 4285 | 1.179 | 0.237 | 0.00474341 | 0.02353263 | 0.0083 | 0.003 | 0.0048 | 0.0048 | TRUE | 0.1636 |
| Atp1b3 | 7830 | 5246 | 5030 | 4926 | 4156 | 4821 | 6081 | 5580 | 5427 | 4903 | 4688 | 4759 | 5239.91 | 5696 | 4783 | 0.839 | -0.252 | 3.29E-05 | 0.00050283 | 0.0018 | 0.0029 | 0.0025 | 0.0025 | TRUE | 0.0432 |
| Atp1b4 | 0 | 0 | 1 | 0 | 0 | 0 | 0 | 0 | 1 | 0 | 0 | 0 | 0.18 | 0 | 0 | 0.552 | -0.857 | 0.8336548 | NA | 0 | 13.1817 | 10 | 10 | TRUE | 0.5365 |
| Atp2a1 | 8 | 5 | 7 | 12 | 12 | 4 | 6 | 5 | 8 | 12 | 14 | 4 | 8.09 | 6 | 10 | 1.536 | 0.619 | 0.38410216 | NA | 0 | 0.2955 | 0.2353 | 0.2353 | TRUE | 0.9694 |
| Atp2a2 | 35590 | 25045 | 23785 | 27614 | 23995 | 27207 | 27641 | 26640 | 25664 | 27487 | 27069 | 26858 | 26893.07 | 26648 | 27138 | 1.018 | 0.026 | 0.59134722 | 0.73939153 | 7.00E-04 | 0.0025 | 0.0017 | 0.0017 | TRUE | 0.0131 |
| Atp2a3 | 2722 | 2165 | 2151 | 3512 | 3020 | 3484 | 2114 | 2303 | 2321 | 3496 | 3407 | 3439 | 2846.64 | 2246 | 3447 | 1.536 | 0.62 | 8.16E-24 | 3.64E-21 | 0.001 | 0.0033 | 0.0024 | 0.0024 | TRUE | 0.0331 |
| Atp2b1 | 6088 | 4353 | 4323 | 4758 | 4167 | 4671 | 4728 | 4630 | 4664 | 4736 | 4701 | 4611 | 4678.49 | 4674 | 4683 | 1.002 | 0.002 | 0.96291113 | 0.97947032 | 0 | 0.0029 | 0.0016 | 0.0016 | TRUE | 0.0022 |
| Atp2b2 | 177 | 119 | 132 | 211 | 195 | 199 | 137 | 127 | 142 | 210 | 220 | 196 | 172.15 | 135 | 209 | 1.539 | 0.622 | 5.00E-05 | 0.00070114 | 0 | 0.0162 | 0.0109 | 0.0109 | TRUE | 0.0318 |
| Atp2b3 | 35 | 19 | 20 | 23 | 18 | 16 | 27 | 20 | 22 | 23 | 20 | 16 | 21.33 | 23 | 20 | 0.849 | -0.236 | 0.57255305 | 0.72523048 | 0 | 0.1136 | 0.0789 | 0.0789 | TRUE | 0.1795 |
| Atp2b4 | 2137 | 1784 | 1851 | 1627 | 1531 | 1840 | 1660 | 1898 | 1997 | 1620 | 1727 | 1816 | 1786.26 | 1852 | 1721 | 0.931 | -0.104 | 0.21318474 | 0.37543735 | 0.0052 | 0.0038 | 0.0044 | 0.0044 | TRUE | 0.1019 |
| Atp2c1 | 5486 | 4223 | 3979 | 3785 | 3407 | 3927 | 4261 | 4492 | 4293 | 3768 | 3843 | 3877 | 4088.93 | 4349 | 3829 | 0.881 | -0.183 | 0.00064104 | 0.00521479 | 3.00E-04 | 0.003 | 0.0018 | 0.0018 | TRUE | 0.01 |
| Atp2c2 | 83 | 70 | 88 | 87 | 99 | 92 | 64 | 74 | 95 | 87 | 112 | 91 | 87.16 | 78 | 97 | 1.241 | 0.312 | 0.17300056 | 0.32542077 | 0.0171 | 0.0296 | 0.0261 | 0.0261 | TRUE | 0.3286 |
| Atp4a | 62 | 61 | 70 | 99 | 109 | 113 | 48 | 65 | 76 | 99 | 123 | 112 | 86.94 | 63 | 111 | 1.781 | 0.832 | 0.00033259 | 0.00314399 | 0.0171 | 0.0297 | 0.0264 | 0.0264 | TRUE | 0.4289 |
| Atp4b | 0 | 0 | 0 | 0 | 0 | 0 | 0 | 0 | 0 | 0 | 0 | 0 | 0 | 0 | 0 | NA | NA | NA | NA | NA | NA | NA | NA | NA | NA |
| Atp5a1 | 72181 | 52773 | 56516 | 60607 | 53421 | 62438 | 56060 | 56134 | 60980 | 60328 | 60264 | 61636 | 59233.78 | 57725 | 60743 | 1.052 | 0.074 | 0.15396121 | 0.30107642 | 0.0012 | 0.0025 | 0.0019 | 0.0019 | TRUE | 0.0298 |
| Atp5b | 128724 | 92155 | 98253 | 110752 | 98060 | 112425 | 99974 | 98024 | 106014 | 110242 | 110622 | 110981 | 105976.3 | 101337 | 110615 | 1.092 | 0.126 | 0.00905163 | 0.0384746 | 8.00E-04 | 0.0024 | 0.0017 | 0.0017 | TRUE | 0.0199 |
| Atp5c1 | 16036 | 11579 | 12718 | 12866 | 11259 | 12777 | 12454 | 12316 | 13723 | 12807 | 12701 | 12613 | 12769.09 | 12831 | 12707 | 0.99 | -0.014 | 0.80647285 | 0.88818896 | 0.0014 | 0.0026 | 0.0022 | 0.0022 | TRUE | 0.045 |
| Atp5d | 10241 | 7369 | 7463 | 8085 | 7180 | 8355 | 7954 | 7838 | 8053 | 8048 | 8100 | 8248 | 8039.97 | 7948 | 8132 | 1.023 | 0.033 | 0.48280332 | 0.64923397 | 0 | 0.0027 | 0.0015 | 0.0015 | TRUE | 0.0019 |
| Atp5e | 4444 | 3088 | 3338 | 3824 | 3239 | 3800 | 3451 | 3285 | 3602 | 3806 | 3654 | 3751 | 3591.55 | 3446 | 3737 | 1.085 | 0.117 | 0.0461556 | 0.12710159 | 9.00E-04 | 0.0031 | 0.0022 | 0.0022 | TRUE | 0.0199 |
| Atp5f1 | 17192 | 11686 | 11958 | 12949 | 10919 | 12897 | 13352 | 12430 | 12903 | 12889 | 12318 | 12731 | 12770.61 | 12895 | 12646 | 0.981 | -0.028 | 0.58048825 | 0.73119931 | 8.00E-04 | 0.0026 | 0.0018 | 0.0018 | TRUE | 0.0122 |
| Atp5g1 | 13036 | 8527 | 7796 | 9033 | 7386 | 9065 | 10125 | 9070 | 8412 | 8991 | 8332 | 8949 | 8979.76 | 9202 | 8757 | 0.951 | -0.072 | 0.31935514 | 0.49330412 | 0.005 | 0.0027 | 0.0036 | 0.0036 | TRUE | 0.0945 |
| Atp5g2 | 4069 | 3060 | 3265 | 3469 | 3253 | 3886 | 3160 | 3255 | 3523 | 3453 | 3670 | 3836 | 3482.81 | 3313 | 3653 | 1.103 | 0.142 | 0.03446734 | 0.10271929 | 0.0026 | 0.0031 | 0.0029 | 0.0029 | TRUE | 0.0373 |
| Atp5g3 | 18992 | 13146 | 11825 | 12766 | 10939 | 12363 | 14750 | 13983 | 12759 | 12707 | 12340 | 12204 | 13124.06 | 13831 | 12417 | 0.898 | -0.156 | 0.01203653 | 0.04744572 | 0.0027 | 0.0026 | 0.0027 | 0.0027 | TRUE | 0.0561 |
| Atp5h | 9035 | 6417 | 6636 | 7210 | 6288 | 7229 | 7017 | 6826 | 7160 | 7177 | 7094 | 7136 | 7068.24 | 7001 | 7136 | 1.019 | 0.027 | 0.57231668 | 0.72505421 | 1.00E-04 | 0.0028 | 0.0016 | 0.0016 | TRUE | 0.0058 |
| Atp5j | 6062 | 4075 | 4189 | 4660 | 3773 | 4328 | 4708 | 4335 | 4520 | 4639 | 4256 | 4272 | 4454.97 | 4521 | 4389 | 0.971 | -0.043 | 0.48433103 | 0.65044468 | 0.0017 | 0.003 | 0.0025 | 0.0025 | TRUE | 0.0302 |
| Atp5j2 | 7598 | 5579 | 5464 | 5743 | 4986 | 5798 | 5901 | 5934 | 5896 | 5717 | 5625 | 5724 | 5799.3 | 5910 | 5689 | 0.962 | -0.055 | 0.24905383 | 0.41806702 | 0 | 0.0028 | 0.0015 | 0.0015 | TRUE | 0.0012 |
| Atp5k | 2738 | 1788 | 1763 | 1800 | 1400 | 1775 | 2126 | 1902 | 1902 | 1792 | 1579 | 1752 | 1842.31 | 1977 | 1708 | 0.863 | -0.212 | 0.00609496 | 0.02855303 | 0.0035 | 0.0037 | 0.0037 | 0.0037 | TRUE | 0.0543 |
| Atp5l | 1044 | 636 | 643 | 711 | 591 | 643 | 811 | 677 | 694 | 708 | 667 | 635 | 698.38 | 727 | 670 | 0.919 | -0.122 | 0.21962981 | 0.38273212 | 0.0053 | 0.0058 | 0.0056 | 0.0056 | TRUE | 0.1255 |
| Atp5o | 12972 | 9059 | 8889 | 9943 | 8709 | 10321 | 10075 | 9636 | 9591 | 9897 | 9825 | 10188 | 9868.7 | 9767 | 9970 | 1.021 | 0.029 | 0.55007509 | 0.70633575 | 4.00E-04 | 0.0027 | 0.0016 | 0.0016 | TRUE | 0.0094 |
| Atp5s | 523 | 423 | 433 | 522 | 441 | 560 | 406 | 450 | 467 | 520 | 497 | 553 | 482.21 | 441 | 523 | 1.19 | 0.251 | 0.01560956 | 0.05754641 | 0.0021 | 0.0073 | 0.0057 | 0.0057 | TRUE | 0.0598 |
| Atp5sl | 1859 | 1416 | 1539 | 1317 | 1273 | 1589 | 1444 | 1506 | 1661 | 1311 | 1436 | 1569 | 1487.69 | 1537 | 1439 | 0.937 | -0.094 | 0.27505083 | 0.44599502 | 0.0056 | 0.004 | 0.0047 | 0.0047 | TRUE | 0.0762 |
| Atp6ap1 | 2319 | 1782 | 2023 | 2275 | 1961 | 2292 | 1801 | 1895 | 2183 | 2265 | 2212 | 2263 | 2103.11 | 1960 | 2247 | 1.147 | 0.198 | 0.01119546 | 0.04486432 | 0.0045 | 0.0036 | 0.0039 | 0.0039 | TRUE | 0.1191 |
| Atp6ap1l | 27 | 11 | 16 | 21 | 10 | 16 | 21 | 12 | 17 | 21 | 11 | 16 | 16.32 | 17 | 16 | 0.957 | -0.064 | 0.89679215 | NA | 0 | 0.1477 | 0.1142 | 0.1142 | TRUE | 0.2233 |
| Atp6ap2 | 3772 | 2704 | 2980 | 3537 | 3149 | 3663 | 2930 | 2876 | 3215 | 3521 | 3552 | 3616 | 3285.04 | 3007 | 3563 | 1.185 | 0.245 | 8.59E-05 | 0.00106707 | 0.0015 | 0.0031 | 0.0025 | 0.0025 | TRUE | 0.0439 |
| Atp6v0a1 | 3403 | 2255 | 2078 | 2627 | 2137 | 2483 | 2643 | 2399 | 2242 | 2615 | 2411 | 2451 | 2460.08 | 2428 | 2492 | 1.026 | 0.037 | 0.62424122 | 0.76326288 | 0.004 | 0.0034 | 0.0036 | 0.0036 | TRUE | 0.0743 |
| Atp6v0a2 | 916 | 614 | 544 | 663 | 591 | 682 | 711 | 653 | 587 | 660 | 667 | 673 | 658.57 | 650 | 667 | 1.023 | 0.032 | 0.73785262 | 0.84383266 | 0.0033 | 0.006 | 0.0051 | 0.0051 | TRUE | 0.0851 |
| Atp6v0a4 | 1096 | 1080 | 1650 | 1467 | 1629 | 1815 | 851 | 1149 | 1780 | 1460 | 1838 | 1792 | 1478.33 | 1260 | 1697 | 1.347 | 0.43 | 0.18340589 | 0.33925461 | 0.0744 | 0.004 | 0.0223 | 0.0744 | TRUE | 0.7717 |
| Atp6v0b | 3970 | 2798 | 3010 | 3423 | 2986 | 3479 | 3083 | 2976 | 3248 | 3407 | 3369 | 3434 | 3252.89 | 3102 | 3403 | 1.097 | 0.134 | 0.02150294 | 0.07284702 | 6.00E-04 | 0.0032 | 0.0021 | 0.0021 | TRUE | 0.0199 |
| Atp6v0c | 1 | 1 | 0 | 1 | 0 | 1 | 1 | 1 | 0 | 1 | 0 | 1 | 0.64 | 1 | 1 | 1.082 | 0.113 | 0.96183421 | NA | 0 | 3.7222 | 2.4899 | 2.4899 | TRUE | 0.2041 |
| Atp6v0c-ps2 | 0 | 0 | 0 | 0 | 0 | 0 | 0 | 0 | 0 | 0 | 0 | 0 | 0 | 0 | 0 | NA | NA | NA | NA | NA | NA | NA | NA | NA | NA |
| Atp6v0d1 | 8080 | 5729 | 5971 | 6907 | 6206 | 7158 | 6275 | 6094 | 6443 | 6875 | 7001 | 7066 | 6625.7 | 6271 | 6981 | 1.113 | 0.155 | 0.00209421 | 0.01251219 | 3.00E-04 | 0.0028 | 0.0017 | 0.0017 | TRUE | 0.0073 |
| Atp6v0d2 | 0 | 0 | 0 | 0 | 0 | 0 | 0 | 0 | 0 | 0 | 0 | 0 | 0 | 0 | 0 | NA | NA | NA | NA | NA | NA | NA | NA | NA | NA |
| Atp6v0e | 2221 | 1447 | 1464 | 1808 | 1515 | 1722 | 1725 | 1539 | 1580 | 1800 | 1709 | 1700 | 1675.4 | 1615 | 1736 | 1.074 | 0.103 | 0.14769558 | 0.29238406 | 0.0018 | 0.0038 | 0.0031 | 0.0031 | TRUE | 0.0447 |
| Atp6v0e2 | 510 | 345 | 420 | 464 | 421 | 471 | 396 | 367 | 453 | 462 | 475 | 465 | 436.33 | 405 | 467 | 1.153 | 0.206 | 0.06123291 | 0.15559147 | 0.0029 | 0.0079 | 0.0064 | 0.0064 | TRUE | 0.1165 |
| Atp6v1a | 14109 | 9955 | 9796 | 11871 | 9970 | 10980 | 10958 | 10589 | 10570 | 11816 | 11247 | 10839 | 11003.2 | 10706 | 11301 | 1.055 | 0.078 | 0.14185117 | 0.28419944 | 0.001 | 0.0026 | 0.0019 | 0.0019 | TRUE | 0.0195 |
| Atp6v1b1 | 0 | 0 | 1 | 0 | 0 | 0 | 0 | 0 | 1 | 0 | 0 | 0 | 0.18 | 0 | 0 | 0.552 | -0.857 | 0.8336548 | NA | 0 | 13.1817 | 10 | 10 | TRUE | 0.5365 |
| Atp6v1b2 | 16913 | 12172 | 12221 | 12449 | 11134 | 13077 | 13136 | 12947 | 13186 | 12392 | 12560 | 12909 | 12855.03 | 13090 | 12620 | 0.964 | -0.053 | 0.25196015 | 0.42090538 | 1.00E-04 | 0.0026 | 0.0014 | 0.0014 | TRUE | 0.0049 |

| | | | | | | | | | | | | | | | | | | | | | | | | | |
|---|---|---|---|---|---|---|---|---|---|---|---|---|---|---|---|---|---|---|---|---|---|---|---|---|---|
| Atp6v1c1 | 5649 | 4127 | 4181 | 4601 | 4219 | 4579 | 4387 | 4390 | 4511 | 4580 | 4759 | 4520 | 4524.65 | 4429 | 4620 | 1.043 | 0.061 | 0.24933504 | 0.41813278 | 2.00E-04 | 0.003 | 0.0018 | 0.0018 | TRUE | 0.0084 |
| Atp6v1c2 | 146 | 134 | 112 | 137 | 119 | 122 | 113 | 143 | 121 | 136 | 134 | 120 | 127.97 | 126 | 130 | 1.041 | 0.058 | 0.74906445 | 0.85119878 | 0 | 0.0209 | 0.0155 | 0.0155 | TRUE | 0.1392 |
| Atp6v1d | 3187 | 2444 | 2546 | 2632 | 2462 | 2938 | 2475 | 2600 | 2747 | 2620 | 2777 | 2900 | 2686.59 | 2607 | 2766 | 1.061 | 0.086 | 0.20338807 | 0.36472149 | 0.0021 | 0.0033 | 0.0029 | 0.0029 | TRUE | 0.0266 |
| Atp6v1e1 | 6715 | 4545 | 4577 | 4884 | 4435 | 4678 | 5215 | 4834 | 4939 | 4862 | 5003 | 4618 | 4911.81 | 4996 | 4828 | 0.966 | -0.05 | 0.39189009 | 0.56555786 | 0.0013 | 0.0029 | 0.0023 | 0.0023 | TRUE | 0.0183 |
| Atp6v1e2 | 3 | 2 | 2 | 2 | 0 | 0 | 2 | 2 | 2 | 2 | 0 | 0 | 1.43 | 2 | 1 | 0.307 | -1.704 | 0.33134919 | NA | 0 | 1.6548 | 1.2354 | 1.2354 | TRUE | 0.9735 |
| Atp6v1f | 4338 | 3005 | 3210 | 3724 | 3408 | 3884 | 3369 | 3196 | 3464 | 3707 | 3845 | 3834 | 3569.11 | 3343 | 3795 | 1.135 | 0.183 | 0.00149469 | 0.00963628 | 6.00E-04 | 0.0031 | 0.0021 | 0.0021 | TRUE | 0.0175 |
| Atp6v1g1 | 3382 | 2393 | 2783 | 2808 | 2477 | 2831 | 2627 | 2545 | 3003 | 2795 | 2794 | 2795 | 2759.82 | 2725 | 2795 | 1.026 | 0.037 | 0.60696766 | 0.75124163 | 0.0035 | 0.0033 | 0.0034 | 0.0034 | TRUE | 0.0953 |
| Atp6v1g2 | 22 | 21 | 24 | 26 | 19 | 20 | 17 | 22 | 26 | 26 | 21 | 20 | 22.06 | 22 | 22 | 1.036 | 0.051 | 0.90170512 | 0.94568228 | 0 | 0.1098 | 0.077 | 0.077 | TRUE | 0.2503 |
| Atp6v1g3 | 0 | 0 | 0 | 1 | 0 | 0 | 0 | 0 | 0 | 1 | 0 | 0 | 0.17 | 0 | 0 | 2.094 | 1.067 | 0.79379329 | NA | 0 | 14.2885 | 10 | 10 | TRUE | 0.3906 |
| Atp6v1h | 4043 | 3014 | 3070 | 3327 | 2871 | 3209 | 3140 | 3206 | 3313 | 3312 | 3239 | 3168 | 3229.46 | 3220 | 3240 | 1.007 | 0.009 | 0.86732305 | 0.92678292 | 3.00E-04 | 0.0032 | 0.002 | 0.002 | TRUE | 0.0077 |
| Atp7a | 2731 | 1788 | 2105 | 2567 | 2241 | 2599 | 2121 | 1902 | 2271 | 2555 | 2528 | 2566 | 2323.85 | 2098 | 2550 | 1.215 | 0.281 | 0.00012435 | 0.00145678 | 0.0033 | 0.0034 | 0.0034 | 0.0034 | TRUE | 0.0793 |
| Atp7b | 1022 | 797 | 902 | 862 | 824 | 934 | 794 | 848 | 973 | 858 | 930 | 922 | 887.39 | 872 | 903 | 1.038 | 0.054 | 0.56751475 | 0.72134178 | 0.0054 | 0.0051 | 0.0052 | 0.0052 | TRUE | 0.1229 |
| Atp8a1 | 1492 | 1116 | 1326 | 1479 | 1380 | 1589 | 1159 | 1187 | 1431 | 1472 | 1557 | 1569 | 1395.69 | 1259 | 1533 | 1.219 | 0.285 | 0.00130524 | 0.00872171 | 0.0066 | 0.0041 | 0.005 | 0.005 | TRUE | 0.1693 |
| Atp8a2 | 87 | 62 | 86 | 54 | 70 | 70 | 68 | 66 | 93 | 54 | 79 | 69 | 71.36 | 76 | 67 | 0.893 | -0.164 | 0.51750895 | 0.67959899 | 0.0225 | 0.0356 | 0.032 | 0.032 | TRUE | 0.355 |
| Atp8b1 | 23 | 18 | 19 | 28 | 16 | 13 | 18 | 19 | 21 | 28 | 18 | 13 | 19.38 | 19 | 20 | 1.025 | 0.036 | 0.93759811 | 0.96535728 | 0.0268 | 0.1247 | 0.0978 | 0.0978 | TRUE | 0.4667 |
| Atp8b2 | 3112 | 2298 | 2030 | 2117 | 1961 | 2087 | 2417 | 2444 | 2190 | 2107 | 2212 | 2060 | 2238.55 | 2350 | 2126 | 0.904 | -0.145 | 0.03333629 | 0.10020516 | 0.0018 | 0.0035 | 0.0029 | 0.0029 | TRUE | 0.0423 |
| Atp8b3 | 225 | 177 | 218 | 291 | 275 | 291 | 175 | 188 | 235 | 290 | 310 | 287 | 247.57 | 199 | 296 | 1.488 | 0.574 | 7.10E-05 | 0.00092808 | 0.0079 | 0.012 | 0.0108 | 0.0108 | TRUE | 0.2621 |
| Atp8b4 | 13 | 13 | 10 | 1 | 1 | 4 | 10 | 14 | 11 | 1 | 1 | 4 | 6.8 | 12 | 2 | 0.178 | -2.491 | 0.00450558 | NA | 0 | 0.3511 | 0.2603 | 0.2603 | TRUE | 0.6539 |
| Atp8b5 | 63 | 72 | 66 | 101 | 84 | 101 | 49 | 77 | 71 | 101 | 95 | 100 | 81.95 | 66 | 99 | 1.515 | 0.599 | 0.01025452 | 0.04206885 | 0.0132 | 0.0313 | 0.0265 | 0.0265 | TRUE | 0.5102 |
| Atp9a | 2465 | 1869 | 2047 | 2351 | 2056 | 2321 | 1914 | 1988 | 2209 | 2340 | 2319 | 2291 | 2176.99 | 2037 | 2317 | 1.138 | 0.187 | 0.00760862 | 0.03373138 | 0.0022 | 0.0035 | 0.0031 | 0.0031 | TRUE | 0.0652 |
| Atp9b | 2108 | 1539 | 1419 | 1396 | 1343 | 1497 | 1637 | 1637 | 1531 | 1390 | 1515 | 1478 | 1531.28 | 1602 | 1461 | 0.911 | -0.134 | 0.05613223 | 0.14651169 | 0.001 | 0.004 | 0.0029 | 0.0029 | TRUE | 0.022 |
| Atpaf1 | 1558 | 1064 | 1056 | 1209 | 1009 | 1114 | 1210 | 1132 | 1139 | 1203 | 1138 | 1100 | 1153.77 | 1160 | 1147 | 0.988 | -0.018 | 0.81235981 | 0.89151249 | 0.001 | 0.0045 | 0.0032 | 0.0032 | TRUE | 0.0225 |
| Atpaf2 | 2482 | 1749 | 1566 | 1662 | 1447 | 1767 | 1928 | 1860 | 1690 | 1654 | 1632 | 1744 | 1751.46 | 1826 | 1677 | 0.918 | -0.124 | 0.09007824 | 0.20486665 | 0.0024 | 0.0038 | 0.0033 | 0.0033 | TRUE | 0.0509 |
| Atpbd4 | 2476 | 1818 | 1814 | 1838 | 1597 | 1890 | 1923 | 1934 | 1957 | 1830 | 1802 | 1866 | 1885.15 | 1938 | 1833 | 0.946 | -0.08 | 0.18371068 | 0.33973046 | 0 | 0.0037 | 0.0021 | 0.0021 | TRUE | 0.0031 |
| Atpif1 | 7389 | 5410 | 5630 | 6652 | 5785 | 6951 | 5739 | 5755 | 6075 | 6621 | 6526 | 6862 | 6262.86 | 5856 | 6670 | 1.139 | 0.188 | 0.00039808 | 0.00363566 | 6.00E-04 | 0.0028 | 0.0019 | 0.0019 | TRUE | 0.0129 |
| Atr | 7133 | 5566 | 5639 | 6368 | 5601 | 6631 | 5540 | 5920 | 6084 | 6339 | 6319 | 6546 | 6124.64 | 5848 | 6401 | 1.095 | 0.131 | 0.02003701 | 0.06927648 | 0.0011 | 0.0028 | 0.0021 | 0.0021 | TRUE | 0.0264 |
| Atraid | 768 | 470 | 532 | 623 | 529 | 587 | 596 | 500 | 574 | 620 | 597 | 579 | 577.8 | 557 | 599 | 1.074 | 0.102 | 0.30361438 | 0.47614223 | 0.003 | 0.0065 | 0.0054 | 0.0054 | TRUE | 0.0918 |
| Atrip | 1284 | 1016 | 973 | 1078 | 969 | 1121 | 997 | 1081 | 1050 | 1073 | 1093 | 1107 | 1066.76 | 1043 | 1091 | 1.047 | 0.067 | 0.3669312 | 0.54162426 | 0 | 0.0046 | 0.003 | 0.003 | TRUE | 0.0183 |
| Atrn | 2426 | 1792 | 1857 | 2210 | 1879 | 1963 | 1884 | 1906 | 2004 | 2200 | 2120 | 1938 | 2008.55 | 1931 | 2086 | 1.08 | 0.111 | 0.11326845 | 0.24206728 | 0.0022 | 0.0036 | 0.0031 | 0.0031 | TRUE | 0.0471 |
| Atrnl1 | 856 | 665 | 702 | 820 | 700 | 780 | 665 | 707 | 757 | 816 | 790 | 770 | 750.92 | 710 | 792 | 1.118 | 0.16 | 0.06594411 | 0.16411964 | 0.0013 | 0.0056 | 0.0041 | 0.0041 | TRUE | 0.0405 |
| Atrx | 12109 | 8558 | 7616 | 7340 | 6550 | 7702 | 9405 | 9103 | 8218 | 7306 | 7389 | 7603 | 8170.59 | 8909 | 7433 | 0.834 | -0.262 | 2.50E-05 | 0.00039933 | 0.0024 | 0.0027 | 0.0027 | 0.0027 | TRUE | 0.056 |
| Atxn1 | 367 | 242 | 246 | 272 | 248 | 265 | 285 | 257 | 265 | 271 | 280 | 262 | 270 | 269 | 271 | 1.003 | 0.004 | 0.97451223 | 0.98624254 | 0 | 0.0112 | 0.0074 | 0.0074 | TRUE | 0.0318 |
| Atxn10 | 17201 | 12799 | 14259 | 12949 | 12347 | 13996 | 13359 | 13614 | 15385 | 12889 | 13929 | 13816 | 13832.18 | 14119 | 13545 | 0.959 | -0.06 | 0.36565675 | 0.54031585 | 0.0037 | 0.0026 | 0.0031 | 0.0031 | TRUE | 0.0751 |
| Atxn1l | 3308 | 2309 | 2266 | 2081 | 2034 | 2326 | 2569 | 2456 | 2445 | 2071 | 2295 | 2296 | 2355.39 | 2490 | 2221 | 0.891 | -0.166 | 0.01240012 | 0.04853092 | 0.0014 | 0.0034 | 0.0027 | 0.0027 | TRUE | 0.0422 |
| Atxn2 | 6251 | 4774 | 4594 | 4849 | 4162 | 4786 | 4855 | 5078 | 4957 | 4827 | 4695 | 4725 | 4856.03 | 4963 | 4749 | 0.957 | -0.063 | 0.21744255 | 0.38021854 | 2.00E-04 | 0.0029 | 0.0017 | 0.0017 | TRUE | 0.005 |
| Atxn2l | 15102 | 11787 | 12173 | 13179 | 11638 | 14044 | 11729 | 12538 | 13135 | 13118 | 13129 | 13864 | 12918.69 | 12467 | 13370 | 1.073 | 0.101 | 0.08241716 | 0.19327127 | 0.002 | 0.0026 | 0.0024 | 0.0024 | TRUE | 0.033 |
| Atxn3 | 5074 | 3869 | 3944 | 4536 | 4160 | 4826 | 3941 | 4115 | 4256 | 4515 | 4693 | 4764 | 4380.63 | 4104 | 4657 | 1.135 | 0.183 | 0.00126522 | 0.0085341 | 8.00E-04 | 0.003 | 0.0021 | 0.0021 | TRUE | 0.0151 |
| Atxn7 | 600 | 438 | 453 | 444 | 362 | 419 | 466 | 466 | 489 | 442 | 408 | 414 | 447.44 | 474 | 421 | 0.891 | -0.167 | 0.09578182 | 0.21444386 | 0 | 0.0077 | 0.005 | 0.005 | TRUE | 0.0214 |
| Atxn7l1 | 484 | 322 | 368 | 442 | 414 | 495 | 376 | 343 | 397 | 440 | 467 | 489 | 418.52 | 372 | 465 | 1.251 | 0.323 | 0.00291475 | 0.01617882 | 0.0013 | 0.0081 | 0.0061 | 0.0061 | TRUE | 0.0516 |
| Atxn7l2 | 509 | 427 | 428 | 491 | 403 | 518 | 395 | 454 | 462 | 489 | 455 | 511 | 461.01 | 437 | 485 | 1.113 | 0.155 | 0.15234405 | 0.2990186 | 0.0034 | 0.0076 | 0.0062 | 0.0062 | TRUE | 0.087 |
| Atxn7l3 | 8711 | 6220 | 5334 | 5622 | 5013 | 5699 | 6765 | 6616 | 5755 | 5596 | 5655 | 5626 | 6002.34 | 6379 | 5626 | 0.882 | -0.182 | 0.00731639 | 0.03285363 | 0.0034 | 0.0028 | 0.0031 | 0.0031 | TRUE | 0.0888 |
| Atxn7l3b | 5041 | 3576 | 3233 | 3247 | 2769 | 3059 | 3915 | 3804 | 3488 | 3232 | 3124 | 3020 | 3430.46 | 3736 | 3125 | 0.836 | -0.258 | 6.46E-05 | 0.00086276 | 0.0019 | 0.0031 | 0.0027 | 0.0027 | TRUE | 0.0405 |
| AU015228 | 62 | 41 | 9 | 8 | 11 | 5 | 48 | 44 | 10 | 8 | 12 | 5 | 21.13 | 34 | 8 | 0.246 | -2.025 | 0.00070281 | 0.00558071 | 0.3499 | 0.1146 | 0.1812 | 0.1812 | TRUE | 2.3644 |
| AU015836 | 786 | 501 | 536 | 372 | 315 | 401 | 610 | 533 | 578 | 370 | 355 | 396 | 473.86 | 574 | 374 | 0.651 | -0.619 | 3.33E-09 | 1.93E-07 | 0.002 | 0.0074 | 0.0057 | 0.0057 | TRUE | 0.0454 |
| AU016765 | 0 | 0 | 0 | 0 | 0 | 1 | 0 | 0 | 0 | 0 | 0 | 1 | 0.16 | 0 | 0 | 2.094 | 1.067 | 0.79379329 | NA | 0 | 14.4078 | 10 | 10 | TRUE | 0.3835 |
| AU018091 | 12792 | 8622 | 9727 | 9037 | 7884 | 8989 | 9935 | 9171 | 10495 | 8995 | 8894 | 8874 | 9394.06 | 9867 | 8921 | 0.904 | -0.145 | 0.01523087 | 0.05645613 | 0.0021 | 0.0027 | 0.0025 | 0.0025 | TRUE | 0.0462 |
| AU018829 | 0 | 0 | 0 | 0 | 0 | 0 | 0 | 0 | 0 | 0 | 0 | 0 | 0 | 0 | 0 | NA | NA | NA | NA | NA | NA | NA | NA | NA | NA |
| AU019823 | 2322 | 1706 | 1427 | 1565 | 1321 | 1410 | 1803 | 1815 | 1540 | 1558 | 1490 | 1392 | 1599.61 | 1719 | 1480 | 0.86 | -0.217 | 0.00916108 | 0.03878534 | 0.0047 | 0.0039 | 0.0044 | 0.0044 | TRUE | 0.0992 |
| AU019990 | 22 | 19 | 1 | 2 | 3 | 2 | 17 | 20 | 1 | 2 | 3 | 2 | 7.62 | 13 | 2 | 0.189 | -2.405 | 0.0121367 | NA | 0.6684 | 0.3134 | 0.412 | 0.412 | TRUE | 1.0298 |
| AU021063 | 1 | 0 | 2 | 1 | 0 | 0 | 1 | 0 | 2 | 1 | 0 | 0 | 0.66 | 1 | 0 | 0.382 | -1.39 | 0.57951664 | NA | 0 | 3.6208 | 3.037 | 3.037 | TRUE | 0.2643 |
| AU021092 | 7 | 2 | 1 | 1 | 2 | 1 | 5 | 2 | 1 | 1 | 2 | 1 | 2.15 | 3 | 1 | 0.471 | -1.085 | 0.42245975 | NA | 0 | 1.1063 | 0.7847 | 0.7847 | TRUE | 0.9805 |
| AU022252 | 1138 | 870 | 884 | 987 | 825 | 994 | 884 | 925 | 954 | 982 | 931 | 981 | 942.91 | 921 | 965 | 1.049 | 0.069 | 0.37404664 | 0.54861034 | 0 | 0.0049 | 0.0033 | 0.0033 | TRUE | 0.0157 |
| AU022751 | 872 | 596 | 511 | 385 | 310 | 279 | 677 | 634 | 551 | 383 | 350 | 275 | 478.49 | 621 | 336 | 0.54 | -0.888 | 1.34E-11 | 1.31E-09 | 0.0167 | 0.0074 | 0.0101 | 0.0101 | TRUE | 0.2899 |
| AU022754 | 0 | 1 | 1 | 0 | 0 | 0 | 0 | 1 | 1 | 0 | 0 | 0 | 0.36 | 1 | 0 | 0.271 | -1.881 | 0.5755355 | NA | 0 | 6.6391 | 6.4428 | 6.4428 | TRUE | 0.3513 |
| AU022793 | 0 | 0 | 0 | 0 | 0 | 0 | 0 | 0 | 0 | 0 | 0 | 0 | 0 | 0 | 0 | NA | NA | NA | NA | NA | NA | NA | NA | NA | NA |
| AU023762 | 1 | 2 | 3 | 1 | 0 | 1 | 1 | 2 | 3 | 1 | 0 | 1 | 1.35 | 2 | 1 | 0.337 | -1.569 | 0.36866479 | NA | 0 | 1.7529 | 1.1898 | 1.1898 | TRUE | 0.1561 |
| AU023871 | 43 | 50 | 72 | 55 | 49 | 73 | 33 | 53 | 78 | 55 | 55 | 72 | 57.73 | 55 | 61 | 1.121 | 0.165 | 0.60151375 | 0.74667517 | 0.0814 | 0.0435 | 0.0545 | 0.0545 | TRUE | 0.2753 |
| AU040320 | 888 | 578 | 619 | 723 | 651 | 735 | 690 | 615 | 668 | 720 | 734 | 726 | 692 | 658 | 727 | 1.103 | 0.142 | 0.10305655 | 0.22616353 | 0 | 0.0059 | 0.004 | 0.004 | TRUE | 0.0374 |
| AU040972 | 18 | 9 | 17 | 5 | 2 | 5 | 14 | 10 | 18 | 5 | 2 | 5 | 9.01 | 14 | 4 | 0.294 | -1.768 | 0.01199248 | NA | 0 | 0.2655 | 0.1959 | 0.1959 | TRUE | 0.1071 |
| AU041133 | 88 | 79 | 57 | 83 | 48 | 70 | 68 | 84 | 62 | 83 | 54 | 69 | 69.96 | 71 | 69 | 0.966 | -0.049 | 0.84462327 | 0.91195135 | 0.0184 | 0.0363 | 0.0315 | 0.0315 | TRUE | 0.2058 |
| Auh | 710 | 598 | 712 | 692 | 622 | 747 | 551 | 636 | 768 | 689 | 702 | 737 | 680.61 | 652 | 709 | 1.091 | 0.125 | 0.27921203 | 0.4507316 | 0.0131 | 0.0059 | 0.0082 | 0.0082 | TRUE | 0.2863 |
| Aup1 | 2122 | 1526 | 1623 | 1607 | 1429 | 1664 | 1648 | 1623 | 1751 | 1600 | 1612 | 1643 | 1646.12 | 1674 | 1618 | 0.967 | -0.049 | 0.46026055 | 0.62988358 | 0 | 0.0039 | 0.0025 | 0.0025 | TRUE | 0.019 |
| Aurka | 8674 | 5989 | 6276 | 6213 | 5645 | 6598 | 6737 | 6370 | 6772 | 6184 | 6368 | 6513 | 6490.78 | 6626 | 6355 | 0.959 | -0.06 | 0.25711258 | 0.42711648 | 7.00E-04 | 0.0028 | 0.0019 | 0.0019 | TRUE | 0.0138 |
| Aurkaip1 | 3595 | 2473 | 2512 | 2659 | 2441 | 2757 | 2792 | 2630 | 2710 | 2647 | 2754 | 2722 | 2709.18 | 2711 | 2708 | 0.998 | -0.003 | 0.96242793 | 0.9793523 | 3.00E-04 | 0.0033 | 0.0021 | 0.0021 | TRUE | 0.0086 |
| Aurkb | 6674 | 4925 | 5622 | 5972 | 5237 | 5947 | 5183 | 5239 | 6066 | 5945 | 5908 | 5871 | 5701.86 | 5496 | 5908 | 1.075 | 0.105 | 0.12607488 | 0.26127132 | 0.0038 | 0.0028 | 0.0032 | 0.0032 | TRUE | 0.0999 |
| Aurkc | 621 | 463 | 485 | 430 | 388 | 441 | 482 | 492 | 523 | 428 | 438 | 435 | 466.53 | 499 | 434 | 0.869 | -0.202 | 0.04001021 | 0.11444986 | 0 | 0.0075 | 0.0048 | 0.0048 | TRUE | 0.0201 |
| Auts2 | 852 | 671 | 700 | 632 | 546 | 651 | 662 | 714 | 755 | 629 | 616 | 643 | 669.74 | 710 | 629 | 0.888 | -0.172 | 0.05678272 | 0.14778944 | 0.0013 | 0.006 | 0.0044 | 0.0044 | TRUE | 0.0448 |
| AV039307 | 2 | 3 | 1 | 1 | 4 | 2 | 2 | 3 | 1 | 1 | 5 | 2 | 2.22 | 2 | 3 | 1.278 | 0.354 | 0.78377603 | NA | 0 | 1.0712 | 0.7405 | 0.7405 | TRUE | 0.4896 |
| AV051173 | 2 | 2 | 1 | 0 | 1 | 1 | 2 | 2 | 1 | 0 | 1 | 1 | 1.15 | 2 | 1 | 0.439 | -1.188 | 0.5118568 | NA | 0 | 2.0709 | 1.3166 | 1.3166 | TRUE | 0.2658 |

| | | | | | | | | | | | | | | | | | | | | | | | | | |
|---|---|---|---|---|---|---|---|---|---|---|---|---|---|---|---|---|---|---|---|---|---|---|---|---|---|
| AV320801 | 0 | 0 | 0 | 0 | 0 | 0 | 0 | 0 | 0 | 0 | 0 | 0 | 0 | 0 | 0 | NA | NA | NA | NA | NA | NA | NA | NA | NA | NA |
| Aven | 1645 | 1142 | 965 | 1087 | 1019 | 1166 | 1278 | 1215 | 1041 | 1082 | 1150 | 1151 | 1152.69 | 1178 | 1128 | 0.956 | -0.065 | 0.46244653 | 0.63177429 | 0.0054 | 0.0045 | 0.0048 | 0.0048 | TRUE | 0.1218 |
| Avil | 52 | 37 | 39 | 37 | 24 | 24 | 40 | 39 | 42 | 37 | 27 | 24 | 34.9 | 40 | 29 | 0.72 | -0.474 | 0.15438217 | 0.30177577 | 0 | 0.0703 | 0.0502 | 0.0502 | TRUE | 0.3536 |
| Avl9 | 2829 | 1962 | 2104 | 2255 | 2021 | 2215 | 2197 | 2087 | 2270 | 2245 | 2280 | 2187 | 2210.9 | 2185 | 2237 | 1.024 | 0.034 | 0.58945367 | 0.7382741 | 6.00E-04 | 0.0035 | 0.0024 | 0.0024 | TRUE | 0.0181 |
| Avp | 0 | 1 | 0 | 0 | 1 | 0 | 0 | 1 | 0 | 0 | 1 | 0 | 0.37 | 0 | 0 | 1.075 | 0.105 | 0.97520514 | NA | 0 | 6.4904 | 6.1969 | 6.1969 | TRUE | 0.5216 |
| Avpi1 | 1542 | 1101 | 1074 | 988 | 857 | 1002 | 1198 | 1171 | 1159 | 983 | 967 | 989 | 1077.82 | 1176 | 980 | 0.833 | -0.264 | 0.0002214 | 0.00230816 | 0 | 0.0046 | 0.0027 | 0.0027 | TRUE | 0.0033 |
| Avpr1a | 8 | 2 | 1 | 5 | 0 | 3 | 6 | 2 | 1 | 5 | 0 | 3 | 2.89 | 3 | 3 | 0.831 | -0.267 | 0.83050068 | NA | 0.6542 | 0.8217 | 0.7812 | 0.7812 | TRUE | 0.4058 |
| Avpr1b | 5 | 2 | 4 | 4 | 4 | 3 | 4 | 2 | 4 | 4 | 5 | 3 | 3.63 | 3 | 4 | 1.1 | 0.138 | 0.88792826 | NA | 0 | 0.6553 | 0.4158 | 0.4158 | TRUE | 0.1477 |
| Avpr2 | 1 | 0 | 1 | 2 | 0 | 1 | 1 | 0 | 1 | 2 | 0 | 1 | 0.81 | 1 | 1 | 1.611 | 0.688 | 0.75133267 | NA | 0 | 2.9444 | 2.113 | 2.113 | TRUE | 0.131 |
| AW011738 | 1 | 3 | 3 | 1 | 1 | 0 | 1 | 3 | 3 | 1 | 1 | 0 | 1.55 | 2 | 1 | 0.296 | -1.757 | 0.29557974 | NA | 0 | 1.5269 | 1.0863 | 1.0863 | TRUE | 0.5104 |
| AW046200 | 4 | 1 | 0 | 1 | 2 | 1 | 3 | 1 | 0 | 1 | 2 | 1 | 1.4 | 1 | 1 | 0.963 | -0.055 | 0.97327594 | NA | 0 | 1.6935 | 1.2463 | 1.2463 | TRUE | 0.5628 |
| AW112010 | 0 | 0 | 0 | 0 | 0 | 0 | 0 | 0 | 0 | 0 | 0 | 0 | 0 | 0 | 0 | NA | NA | NA | NA | NA | NA | NA | NA | NA | NA |
| AW146154 | 279 | 197 | 155 | 189 | 168 | 179 | 217 | 210 | 167 | 188 | 190 | 177 | 191.3 | 198 | 185 | 0.931 | -0.103 | 0.50445068 | 0.66826706 | 0.0052 | 0.0148 | 0.012 | 0.012 | TRUE | 0.191 |
| AW209491 | 1091 | 748 | 714 | 782 | 614 | 743 | 847 | 796 | 770 | 778 | 693 | 733 | 769.65 | 804 | 735 | 0.913 | -0.132 | 0.13216138 | 0.27004756 | 0.0016 | 0.0055 | 0.0042 | 0.0042 | TRUE | 0.0318 |
| AW495222 | 4 | 3 | 1 | 0 | 1 | 0 | 3 | 3 | 1 | 0 | 1 | 0 | 1.42 | 2 | 0 | 0.15 | -2.739 | 0.13678552 | NA | 0 | 1.6745 | 1.2477 | 1.2477 | TRUE | 0.5075 |
| AW549542 | 33 | 33 | 33 | 26 | 26 | 38 | 26 | 35 | 36 | 26 | 29 | 38 | 31.51 | 32 | 31 | 0.971 | -0.043 | 0.90263989 | 0.94610624 | 0 | 0.0776 | 0.0578 | 0.0578 | TRUE | 0.2584 |
| AW549877 | 5202 | 4202 | 4828 | 5261 | 4405 | 5167 | 4040 | 4470 | 5209 | 5237 | 4969 | 5101 | 4837.65 | 4573 | 5102 | 1.116 | 0.159 | 0.0578316 | 0.14974404 | 0.0084 | 0.0029 | 0.0048 | 0.0048 | TRUE | 0.1803 |
| AW551984 | 157 | 130 | 141 | 171 | 154 | 191 | 122 | 138 | 152 | 170 | 174 | 189 | 157.47 | 137 | 178 | 1.297 | 0.376 | 0.02182698 | 0.07356156 | 0 | 0.0175 | 0.0129 | 0.0129 | TRUE | 0.1132 |
| AW554918 | 585 | 400 | 456 | 444 | 376 | 445 | 454 | 425 | 492 | 442 | 424 | 439 | 446.21 | 457 | 435 | 0.952 | -0.071 | 0.49530798 | 0.65996429 | 0 | 0.0077 | 0.0055 | 0.0055 | TRUE | 0.0484 |
| Awat1 | 0 | 0 | 0 | 0 | 0 | 0 | 0 | 0 | 0 | 0 | 0 | 0 | 0 | 0 | 0 | NA | NA | NA | NA | NA | NA | NA | NA | NA | NA |
| Awat2 | 44 | 24 | 21 | 24 | 28 | 28 | 34 | 26 | 23 | 24 | 32 | 28 | 27.58 | 28 | 28 | 0.998 | -0.003 | 0.99334425 | 0.99603371 | 0 | 0.0884 | 0.064 | 0.064 | TRUE | 0.348 |
| Axin1 | 5002 | 3861 | 3854 | 3876 | 3496 | 4109 | 3885 | 4107 | 4158 | 3858 | 3944 | 4056 | 4001.4 | 4050 | 3953 | 0.976 | -0.034 | 0.54287778 | 0.70074524 | 8.00E-04 | 0.003 | 0.0021 | 0.0021 | TRUE | 0.0159 |
| Axin2 | 867 | 638 | 605 | 724 | 709 | 778 | 673 | 679 | 653 | 721 | 800 | 768 | 715.55 | 668 | 763 | 1.141 | 0.19 | 0.02677032 | 0.08539328 | 0 | 0.0057 | 0.0039 | 0.0039 | TRUE | 0.0278 |
| Axl | 253 | 198 | 260 | 278 | 277 | 295 | 196 | 211 | 281 | 277 | 312 | 291 | 261.34 | 229 | 293 | 1.285 | 0.361 | 0.01736898 | 0.0622638 | 0.0153 | 0.0115 | 0.0127 | 0.0127 | TRUE | 0.4147 |
| AY074887 | 3 | 4 | 4 | 4 | 2 | 3 | 2 | 4 | 4 | 4 | 2 | 3 | 3.35 | 3 | 3 | 0.86 | -0.218 | 0.83163986 | NA | 0 | 0.7099 | 0.4518 | 0.4518 | TRUE | 0.2086 |
| AY358078 | 3 | 7 | 2 | 3 | 0 | 1 | 2 | 7 | 2 | 3 | 0 | 1 | 2.65 | 4 | 1 | 0.343 | -1.543 | 0.24317603 | NA | 0.3423 | 0.8964 | 0.754 | 0.754 | TRUE | 0.9141 |
| AY512915 | 0 | 0 | 0 | 0 | 0 | 1 | 0 | 0 | 0 | 0 | 0 | 1 | 0.16 | 0 | 0 | 2.094 | 1.067 | 0.79379329 | NA | 0 | 14.4078 | 10 | 10 | TRUE | 0.3835 |
| AY512931 | 17 | 20 | 13 | 29 | 24 | 24 | 13 | 21 | 14 | 29 | 27 | 24 | 21.36 | 16 | 27 | 1.654 | 0.726 | 0.08704545 | 0.20002261 | 0 | 0.1134 | 0.0802 | 0.0802 | TRUE | 0.3448 |
| AY761184 | 1 | 0 | 3 | 0 | 0 | 1 | 1 | 0 | 3 | 0 | 0 | 1 | 0.83 | 1 | 0 | 0.28 | -1.835 | 0.43280827 | NA | 0 | 2.8461 | 2.5799 | 2.5799 | TRUE | 0.5383 |
| AY761185 | 0 | 0 | 0 | 0 | 0 | 0 | 0 | 0 | 0 | 0 | 0 | 0 | 0 | 0 | 0 | NA | NA | NA | NA | NA | NA | NA | NA | NA | NA |
| Aym1 | 2 | 3 | 10 | 5 | 5 | 10 | 2 | 3 | 11 | 5 | 6 | 10 | 6 | 5 | 7 | 1.357 | 0.441 | 0.61329983 | NA | 0.3284 | 0.3972 | 0.3771 | 0.3771 | TRUE | 1.7765 |
| Azgp1 | 0 | 0 | 0 | 0 | 0 | 0 | 0 | 0 | 0 | 0 | 0 | 0 | 0 | 0 | 0 | NA | NA | NA | NA | NA | NA | NA | NA | NA | NA |
| Azi1 | 3577 | 2901 | 2960 | 3218 | 2976 | 3537 | 2778 | 3086 | 3194 | 3203 | 3357 | 3492 | 3184.95 | 3019 | 3351 | 1.111 | 0.151 | 0.0303483 | 0.09338392 | 0.003 | 0.0032 | 0.0032 | 0.0032 | TRUE | 0.0606 |
| Azi2 | 3052 | 2148 | 2078 | 2034 | 1774 | 1981 | 2370 | 2285 | 2242 | 2025 | 2001 | 1956 | 2146.46 | 2299 | 1994 | 0.867 | -0.206 | 0.00066235 | 0.00533953 | 0 | 0.0035 | 0.0022 | 0.0022 | TRUE | 0.0092 |
| Azin1 | 11537 | 7979 | 6591 | 7209 | 6199 | 6913 | 8960 | 8487 | 7112 | 7176 | 6993 | 6824 | 7592.04 | 8186 | 6998 | 0.855 | -0.227 | 0.00428826 | 0.02181992 | 0.007 | 0.0027 | 0.0044 | 0.0044 | TRUE | 0.1607 |
| B020004C17 | 1 | 0 | 0 | 0 | 0 | 0 | 1 | 0 | 0 | 0 | 0 | 0 | 0.13 | 0 | 0 | 0.552 | -0.857 | 0.8336548 | NA | 0 | 18.3121 | 10 | 10 | TRUE | 0.2566 |
| B020004J07 | 88 | 67 | 26 | 30 | 29 | 12 | 68 | 71 | 28 | 30 | 33 | 12 | 40.35 | 56 | 25 | 0.44 | -1.183 | 0.00542882 | 0.02613475 | 0.2112 | 0.0612 | 0.1009 | 0.1009 | TRUE | 1.5314 |
| B020014A21 | 0 | 0 | 0 | 0 | 0 | 0 | 0 | 0 | 0 | 0 | 0 | 0 | 0 | 0 | 0 | NA | NA | NA | NA | NA | NA | NA | NA | NA | NA |
| B020018J22 | 1 | 1 | 0 | 0 | 0 | 0 | 1 | 1 | 0 | 0 | 0 | 0 | 0.31 | 1 | 0 | 0.309 | -1.696 | 0.63183309 | NA | 0 | 7.7294 | 7.2472 | 7.2472 | TRUE | 0.2015 |
| B020031M17 | 0 | 0 | 0 | 0 | 0 | 0 | 0 | 0 | 0 | 0 | 0 | 0 | 0 | 0 | 0 | NA | NA | NA | NA | NA | NA | NA | NA | NA | NA |
| B130006D01 | 4 | 4 | 9 | 32 | 15 | 16 | 3 | 4 | 10 | 32 | 17 | 16 | 13.61 | 6 | 22 | 3.89 | 1.96 | 0.0015942 | NA | 0.1384 | 0.1766 | 0.1669 | 0.1669 | TRUE | 1.0088 |
| B130024G19 | 0 | 1 | 0 | 0 | 0 | 0 | 0 | 1 | 0 | 0 | 0 | 0 | 0.18 | 0 | 0 | 0.552 | -0.857 | 0.8336548 | NA | 0 | 13.3714 | 10 | 10 | TRUE | 0.5216 |
| B130034C11 | 8 | 8 | 1 | 4 | 6 | 5 | 6 | 9 | 1 | 4 | 7 | 5 | 5.25 | 5 | 5 | 0.981 | -0.027 | 0.97527608 | NA | 0.1179 | 0.454 | 0.3734 | 0.3734 | TRUE | 0.5479 |
| B230118H07 | 833 | 559 | 512 | 626 | 523 | 604 | 647 | 595 | 552 | 623 | 590 | 596 | 600.56 | 598 | 603 | 1.007 | 0.01 | 0.92064131 | 0.95643782 | 0.0021 | 0.0064 | 0.005 | 0.005 | TRUE | 0.0636 |
| B230119M05 | 3 | 1 | 0 | 3 | 1 | 2 | 2 | 1 | 0 | 3 | 1 | 2 | 1.58 | 1 | 2 | 1.72 | 0.782 | 0.61325739 | NA | 0 | 1.5021 | 1.0658 | 1.0658 | TRUE | 0.4405 |
| B230120H23 | 1572 | 1122 | 1224 | 1410 | 1191 | 1379 | 1221 | 1193 | 1321 | 1404 | 1344 | 1361 | 1307.24 | 1245 | 1370 | 1.101 | 0.138 | 0.05607693 | 0.14639407 | 8.00E-04 | 0.0042 | 0.003 | 0.003 | TRUE | 0.0329 |
| B230206H07 | 2 | 0 | 0 | 4 | 1 | 1 | 2 | 0 | 0 | 4 | 1 | 1 | 1.28 | 1 | 2 | 3.542 | 1.825 | 0.33556405 | NA | 0 | 1.8613 | 1.4989 | 1.4989 | TRUE | 0.9711 |
| B230208H11 | 22 | 15 | 18 | 18 | 11 | 20 | 17 | 16 | 19 | 18 | 12 | 20 | 17.09 | 17 | 17 | 0.96 | -0.06 | 0.89768469 | 0.94349536 | 0 | 0.1411 | 0.0958 | 0.0958 | TRUE | 0.2203 |
| B230209E15 | 0 | 0 | 0 | 0 | 0 | 0 | 0 | 0 | 0 | 0 | 0 | 0 | 0 | 0 | 0 | NA | NA | NA | NA | NA | NA | NA | NA | NA | NA |
| B230209K01 | 8 | 3 | 5 | 2 | 8 | 10 | 6 | 3 | 5 | 2 | 9 | 10 | 5.95 | 5 | 7 | 1.393 | 0.478 | 0.5680016 | NA | 0 | 0.4009 | 0.3339 | 0.3339 | TRUE | 1.0648 |
| B230214G05 | 0 | 0 | 0 | 0 | 0 | 0 | 0 | 0 | 0 | 0 | 0 | 0 | 0 | 0 | 0 | NA | NA | NA | NA | NA | NA | NA | NA | NA | NA |
| B230216G23 | 7 | 5 | 0 | 1 | 2 | 1 | 5 | 5 | 0 | 1 | 2 | 1 | 2.5 | 3 | 1 | 0.384 | -1.38 | 0.31345358 | NA | 0 | 0.9508 | 0.8476 | 0.8476 | TRUE | 1.0552 |
| B230216N24 | 2 | 5 | 2 | 7 | 4 | 5 | 2 | 5 | 2 | 7 | 5 | 5 | 4.24 | 3 | 6 | 1.863 | 0.898 | 0.34575446 | NA | 0 | 0.5613 | 0.3956 | 0.3956 | TRUE | 0.5559 |
| B230217C12 | 463 | 327 | 315 | 355 | 297 | 381 | 360 | 348 | 340 | 353 | 335 | 376 | 351.97 | 349 | 355 | 1.016 | 0.023 | 0.83510384 | 0.90624396 | 0 | 0.0092 | 0.0062 | 0.0062 | TRUE | 0.0315 |
| B230217O12 | 15 | 18 | 12 | 9 | 3 | 8 | 12 | 19 | 13 | 9 | 3 | 8 | 10.66 | 15 | 7 | 0.47 | -1.089 | 0.08042646 | NA | 0 | 0.2247 | 0.1705 | 0.1705 | TRUE | 0.426 |
| B230219D22 | 5474 | 4378 | 4821 | 5949 | 4860 | 5729 | 4251 | 4657 | 5202 | 5922 | 5483 | 5655 | 5194.95 | 4703 | 5687 | 1.21 | 0.275 | 0.00027498 | 0.0027174 | 0.0056 | 0.0029 | 0.0039 | 0.0039 | TRUE | 0.1046 |
| B230312C02 | 1 | 1 | 1 | 0 | 1 | 0 | 1 | 1 | 1 | 0 | 1 | 0 | 0.67 | 1 | 0 | 0.383 | -1.383 | 0.54981825 | NA | 0 | 3.5158 | 2.3538 | 2.3538 | TRUE | 0.5075 |
| B230319C09 | 2 | 1 | 1 | 0 | 1 | 0 | 2 | 1 | 1 | 0 | 1 | 0 | 0.8 | 1 | 0 | 0.299 | -1.743 | 0.42212658 | NA | 0 | 2.9502 | 2.0092 | 2.0092 | TRUE | 0.5075 |
| B230323A14 | 0 | 0 | 0 | 0 | 0 | 0 | 0 | 0 | 0 | 0 | 0 | 0 | 0 | 0 | 0 | NA | NA | NA | NA | NA | NA | NA | NA | NA | NA |
| B230378P21 | 0 | 0 | 0 | 0 | 0 | 0 | 0 | 0 | 0 | 0 | 0 | 0 | 0 | 0 | 0 | NA | NA | NA | NA | NA | NA | NA | NA | NA | NA |
| B2m | 680 | 455 | 501 | 577 | 464 | 516 | 528 | 484 | 541 | 574 | 523 | 509 | 526.64 | 518 | 535 | 1.035 | 0.049 | 0.62075543 | 0.76065413 | 0.0017 | 0.0069 | 0.0053 | 0.0053 | TRUE | 0.0473 |
| B330016D10 | 32 | 52 | 49 | 58 | 50 | 61 | 25 | 55 | 53 | 58 | 56 | 60 | 51.23 | 44 | 58 | 1.329 | 0.41 | 0.19985057 | 0.36048598 | 0.0673 | 0.0487 | 0.0538 | 0.0538 | TRUE | 1.3674 |
| B3galnt1 | 922 | 629 | 703 | 808 | 673 | 835 | 716 | 669 | 759 | 804 | 759 | 824 | 755.24 | 715 | 796 | 1.114 | 0.156 | 0.07358337 | 0.17783684 | 0.0013 | 0.0056 | 0.0042 | 0.0042 | TRUE | 0.0349 |
| B3galnt2 | 2774 | 2135 | 2204 | 2354 | 2058 | 2329 | 2154 | 2271 | 2378 | 2343 | 2322 | 2299 | 2294.57 | 2268 | 2321 | 1.024 | 0.035 | 0.58216347 | 0.73253195 | 9.00E-04 | 0.0035 | 0.0024 | 0.0024 | TRUE | 0.024 |
| B3galt1 | 57 | 33 | 42 | 58 | 51 | 76 | 44 | 35 | 45 | 58 | 58 | 75 | 52.5 | 41 | 64 | 1.524 | 0.608 | 0.02782652 | 0.08764311 | 0 | 0.0476 | 0.0354 | 0.0354 | TRUE | 0.2283 |
| B3galt2 | 0 | 0 | 0 | 0 | 0 | 0 | 0 | 0 | 0 | 0 | 0 | 0 | 0 | 0 | 0 | NA | NA | NA | NA | NA | NA | NA | NA | NA | NA |
| B3galt4 | 138 | 103 | 115 | 82 | 87 | 105 | 107 | 110 | 124 | 82 | 98 | 104 | 104.04 | 114 | 95 | 0.833 | -0.264 | 0.18212936 | 0.33776393 | 0 | 0.0252 | 0.0185 | 0.0185 | TRUE | 0.1404 |
| B3galt5 | 16 | 13 | 12 | 14 | 15 | 15 | 12 | 14 | 13 | 14 | 17 | 15 | 14.14 | 13 | 15 | 1.165 | 0.22 | 0.6582689 | NA | 0 | 0.17 | 0.1079 | 0.1079 | TRUE | 0.0397 |
| B3galt6 | 422 | 258 | 224 | 235 | 239 | 242 | 328 | 274 | 242 | 234 | 270 | 239 | 264.38 | 281 | 248 | 0.875 | -0.192 | 0.18732018 | 0.3442231 | 0.0117 | 0.0114 | 0.0115 | 0.0115 | TRUE | 0.2485 |

| | | | | | | | | | | | | | | | | | | | | | | | | | |
|---|---|---|---|---|---|---|---|---|---|---|---|---|---|---|---|---|---|---|---|---|---|---|---|---|---|
| B3galtl | 1268 | 910 | 931 | 979 | 907 | 1014 | 985 | 968 | 1005 | 974 | 1023 | 1001 | 992.66 | 986 | 999 | 1.014 | 0.02 | 0.78960366 | 0.87639639 | 0 | 0.0048 | 0.0029 | 0.0029 | TRUE | 0.0058 |
| B3gat1 | 50 | 33 | 31 | 27 | 33 | 31 | 39 | 35 | 33 | 27 | 37 | 31 | 33.68 | 36 | 32 | 0.876 | -0.19 | 0.56689886 | 0.7208889 | 0 | 0.0728 | 0.0496 | 0.0496 | TRUE | 0.1542 |
| B3gat2 | 6 | 4 | 8 | 8 | 9 | 10 | 5 | 4 | 9 | 8 | 10 | 10 | 7.59 | 6 | 9 | 1.61 | 0.687 | 0.32627448 | NA | 0 | 0.3147 | 0.2138 | 0.2138 | TRUE | 0.3615 |
| B3gat3 | 1040 | 728 | 725 | 757 | 747 | 838 | 808 | 774 | 782 | 754 | 843 | 827 | 797.97 | 788 | 808 | 1.024 | 0.034 | 0.68499951 | 0.80694783 | 7.00E-04 | 0.0054 | 0.0038 | 0.0038 | TRUE | 0.0414 |
| B3gnt1 | 548 | 407 | 330 | 428 | 315 | 359 | 426 | 433 | 356 | 426 | 355 | 354 | 391.73 | 405 | 378 | 0.934 | -0.098 | 0.43096403 | 0.60313022 | 0.0087 | 0.0085 | 0.0085 | 0.0085 | TRUE | 0.1393 |
| B3gnt2 | 2314 | 1795 | 1849 | 2156 | 1919 | 2119 | 1797 | 1909 | 1995 | 2146 | 2165 | 2092 | 2017.38 | 1900 | 2134 | 1.124 | 0.169 | 0.01040273 | 0.0424939 | 0.0011 | 0.0036 | 0.0026 | 0.0026 | TRUE | 0.0284 |
| B3gnt3 | 101 | 82 | 83 | 86 | 93 | 92 | 78 | 87 | 90 | 86 | 105 | 91 | 89.43 | 85 | 94 | 1.104 | 0.142 | 0.49527856 | 0.65996429 | 0 | 0.0289 | 0.0201 | 0.0201 | TRUE | 0.0988 |
| B3gnt4 | 22 | 18 | 12 | 6 | 8 | 9 | 17 | 19 | 13 | 6 | 9 | 9 | 12.18 | 16 | 8 | 0.484 | -1.048 | 0.06288128 | NA | 0 | 0.1971 | 0.1343 | 0.1343 | TRUE | 0.1481 |
| B3gnt5 | 2068 | 1503 | 1551 | 1984 | 1541 | 1704 | 1606 | 1599 | 1674 | 1975 | 1738 | 1682 | 1712.29 | 1626 | 1798 | 1.106 | 0.146 | 0.05770788 | 0.14953204 | 0.0032 | 0.0038 | 0.0037 | 0.0037 | TRUE | 0.0896 |
| B3gnt6 | 4 | 0 | 4 | 4 | 4 | 1 | 3 | 0 | 4 | 4 | 5 | 1 | 2.82 | 2 | 3 | 1.26 | 0.333 | 0.78721459 | NA | 0.3393 | 0.8437 | 0.7389 | 0.7389 | TRUE | 0.7512 |
| B3gnt7 | 5645 | 4085 | 4127 | 5156 | 4906 | 5514 | 4384 | 4345 | 4453 | 5132 | 5534 | 5443 | 4882.05 | 4394 | 5370 | 1.222 | 0.289 | 1.16E-07 | 4.47E-06 | 5.00E-04 | 0.0029 | 0.0019 | 0.0019 | TRUE | 0.0183 |
| B3gnt8 | 24 | 12 | 9 | 17 | 15 | 19 | 19 | 13 | 10 | 17 | 17 | 19 | 15.62 | 14 | 18 | 1.261 | 0.335 | 0.49589982 | NA | 0 | 0.1542 | 0.1101 | 0.1101 | TRUE | 0.5321 |
| B3gnt9 | 76 | 65 | 87 | 107 | 80 | 112 | 59 | 69 | 94 | 107 | 90 | 111 | 88.23 | 74 | 103 | 1.396 | 0.481 | 0.03766271 | 0.10964193 | 0.0211 | 0.0293 | 0.027 | 0.027 | TRUE | 0.4907 |
| B3gntl1 | 1312 | 1076 | 995 | 1052 | 947 | 1069 | 1019 | 1145 | 1074 | 1047 | 1068 | 1055 | 1067.97 | 1079 | 1057 | 0.981 | -0.028 | 0.71261976 | 0.82728023 | 0.001 | 0.0046 | 0.0033 | 0.0033 | TRUE | 0.0337 |
| B430010I23F | 2 | 0 | 0 | 2 | 1 | 1 | 2 | 0 | 0 | 2 | 1 | 1 | 0.94 | 1 | 1 | 2.402 | 1.264 | 0.53839702 | NA | 0 | 2.5151 | 1.8224 | 1.8224 | TRUE | 0.9714 |
| B430212C06 | 2 | 6 | 1 | 5 | 1 | 2 | 2 | 6 | 1 | 5 | 1 | 2 | 2.85 | 3 | 3 | 0.917 | -0.124 | 0.91778796 | NA | 0.3781 | 0.8343 | 0.6919 | 0.6919 | TRUE | 1.1785 |
| B430306N03 | 33 | 19 | 14 | 13 | 3 | 19 | 26 | 20 | 15 | 13 | 3 | 19 | 16 | 20 | 12 | 0.578 | -0.79 | 0.15995825 | NA | 0.195 | 0.1505 | 0.1603 | 0.1603 | TRUE | 0.5943 |
| B430319G15 | 10 | 5 | 2 | 1 | 3 | 1 | 8 | 5 | 2 | 1 | 3 | 1 | 3.43 | 5 | 2 | 0.34 | -1.557 | 0.16813255 | NA | 0 | 0.6924 | 0.5303 | 0.5303 | TRUE | 0.2136 |
| B4galnt1 | 508 | 394 | 389 | 439 | 352 | 429 | 395 | 419 | 420 | 437 | 397 | 423 | 415.15 | 411 | 419 | 1.022 | 0.031 | 0.76731242 | 0.86240437 | 0 | 0.0081 | 0.0054 | 0.0054 | TRUE | 0.0237 |
| B4galnt2 | 259 | 173 | 322 | 400 | 419 | 425 | 201 | 184 | 347 | 398 | 473 | 420 | 337.16 | 244 | 430 | 1.766 | 0.82 | 1.81E-05 | 0.00030897 | 0.0604 | 0.0095 | 0.0232 | 0.0232 | TRUE | 1.4889 |
| B4galnt3 | 1750 | 1421 | 1467 | 1322 | 1268 | 1523 | 1359 | 1511 | 1583 | 1316 | 1430 | 1503 | 1450.55 | 1484 | 1416 | 0.955 | -0.066 | 0.42731962 | 0.5992621 | 0.0047 | 0.0041 | 0.0043 | 0.0043 | TRUE | 0.0679 |
| B4galnt4 | 1074 | 832 | 832 | 865 | 950 | 966 | 834 | 885 | 898 | 861 | 1072 | 954 | 917.19 | 872 | 962 | 1.103 | 0.141 | 0.1312987 | 0.2686689 | 0.0057 | 0.005 | 0.0052 | 0.0052 | TRUE | 0.1166 |
| B4galt1 | 2314 | 1688 | 1681 | 1720 | 1505 | 1705 | 1797 | 1796 | 1814 | 1712 | 1698 | 1683 | 1749.91 | 1802 | 1698 | 0.942 | -0.086 | 0.15868422 | 0.30791115 | 0 | 0.0038 | 0.0021 | 0.0021 | TRUE | 7.00E-04 |
| B4galt2 | 763 | 622 | 679 | 652 | 561 | 601 | 593 | 662 | 733 | 649 | 633 | 593 | 643.66 | 663 | 625 | 0.946 | -0.08 | 0.42824592 | 0.600425 | 0.0054 | 0.0061 | 0.0059 | 0.0059 | TRUE | 0.1054 |
| B4galt3 | 1611 | 1137 | 1128 | 1171 | 1018 | 1199 | 1251 | 1209 | 1217 | 1166 | 1148 | 1184 | 1195.89 | 1226 | 1166 | 0.951 | -0.073 | 0.29219021 | 0.46437209 | 0 | 0.0044 | 0.0026 | 0.0026 | TRUE | 0.0041 |
| B4galt4 | 526 | 343 | 430 | 402 | 341 | 381 | 409 | 365 | 464 | 400 | 385 | 376 | 399.71 | 413 | 387 | 0.939 | -0.091 | 0.43492987 | 0.60666074 | 0.005 | 0.0084 | 0.0073 | 0.0073 | TRUE | 0.131 |
| B4galt5 | 5138 | 3717 | 3588 | 3617 | 3220 | 3509 | 3990 | 3954 | 3871 | 3600 | 3632 | 3464 | 3752.06 | 3938 | 3565 | 0.905 | -0.144 | 0.00748148 | 0.03338676 | 2.00E-04 | 0.0031 | 0.0018 | 0.0018 | TRUE | 0.0076 |
| B4galt6 | 8018 | 5968 | 6313 | 7243 | 6461 | 7250 | 6227 | 6348 | 6812 | 7210 | 7289 | 7157 | 6840.37 | 6462 | 7219 | 1.117 | 0.16 | 0.00342838 | 0.01835841 | 0.001 | 0.0028 | 0.002 | 0.002 | TRUE | 0.0272 |
| B4galt7 | 586 | 432 | 433 | 497 | 458 | 467 | 455 | 460 | 467 | 495 | 517 | 461 | 475.7 | 461 | 491 | 1.065 | 0.091 | 0.3577156 | 0.53214291 | 0 | 0.0074 | 0.005 | 0.005 | TRUE | 0.0331 |
| B630005N14 | 2293 | 1683 | 1983 | 2015 | 1642 | 1947 | 1781 | 1790 | 2140 | 2006 | 1852 | 1922 | 1915.13 | 1904 | 1927 | 1.013 | 0.019 | 0.82127458 | 0.89729992 | 0.0058 | 0.0037 | 0.0044 | 0.0044 | TRUE | 0.1408 |
| B630019K06 | 85 | 56 | 49 | 57 | 47 | 55 | 66 | 60 | 53 | 57 | 53 | 54 | 57.08 | 60 | 55 | 0.916 | -0.127 | 0.61951665 | 0.75986884 | 0 | 0.0439 | 0.0294 | 0.0294 | TRUE | 0.0907 |
| B830017H08 | 4 | 6 | 3 | 5 | 1 | 0 | 3 | 6 | 3 | 5 | 1 | 0 | 3.14 | 4 | 2 | 0.486 | -1.04 | 0.38751631 | NA | 0.3742 | 0.7576 | 0.6771 | 0.6771 | TRUE | 1.506 |
| B930003M22 | 33 | 30 | 46 | 54 | 40 | 46 | 26 | 32 | 50 | 54 | 45 | 45 | 41.91 | 36 | 48 | 1.362 | 0.445 | 0.17367764 | 0.3264453 | 0.0361 | 0.059 | 0.0528 | 0.0528 | TRUE | 0.804 |
| B930018H19 | 1 | 0 | 0 | 0 | 0 | 0 | 1 | 0 | 0 | 0 | 0 | 0 | 0.13 | 0 | 0 | 0.552 | -0.857 | 0.8336548 | NA | 0 | 18.3121 | 10 | 10 | TRUE | 0.2566 |
| B930025P03 | 0 | 0 | 0 | 0 | 0 | 0 | 0 | 0 | 0 | 0 | 0 | 0 | 0 | 0 | 0 | NA | NA | NA | NA | NA | NA | NA | NA | NA | NA |
| B930041F14 | 304 | 257 | 295 | 339 | 293 | 363 | 236 | 273 | 318 | 337 | 331 | 358 | 309.01 | 276 | 342 | 1.245 | 0.317 | 0.01814877 | 0.06426088 | 0.0087 | 0.0101 | 0.0097 | 0.0097 | TRUE | 0.2017 |
| B930059L03I | 18 | 13 | 9 | 12 | 7 | 12 | 14 | 14 | 10 | 12 | 8 | 12 | 11.53 | 13 | 11 | 0.844 | -0.244 | 0.66414348 | NA | 0 | 0.2079 | 0.1403 | 0.1403 | TRUE | 0.156 |
| B930092H01 | 4 | 4 | 3 | 6 | 7 | 5 | 3 | 4 | 3 | 6 | 8 | 5 | 4.9 | 3 | 6 | 1.777 | 0.83 | 0.33329144 | NA | 0 | 0.4861 | 0.31 | 0.31 | TRUE | 0.1122 |
| B9d1 | 1654 | 1222 | 1133 | 1135 | 1004 | 1236 | 1285 | 1300 | 1222 | 1130 | 1133 | 1220 | 1214.91 | 1269 | 1161 | 0.915 | -0.129 | 0.07916572 | 0.18767882 | 6.00E-04 | 0.0044 | 0.003 | 0.003 | TRUE | 0.0242 |
| B9d2 | 195 | 155 | 157 | 164 | 147 | 184 | 151 | 165 | 169 | 163 | 166 | 182 | 166.07 | 162 | 170 | 1.054 | 0.076 | 0.62177742 | 0.76142749 | 0 | 0.0167 | 0.0113 | 0.0113 | TRUE | 0.0376 |
| Baalc | 11 | 9 | 17 | 17 | 10 | 18 | 9 | 10 | 18 | 17 | 11 | 18 | 13.74 | 12 | 15 | 1.281 | 0.358 | 0.50990586 | NA | 0.039 | 0.1749 | 0.1383 | 0.1383 | TRUE | 0.7403 |
| Baat | 8 | 4 | 3 | 4 | 3 | 5 | 6 | 4 | 3 | 4 | 3 | 5 | 4.33 | 4 | 4 | 0.888 | -0.172 | 0.84877397 | NA | 0 | 0.5492 | 0.3535 | 0.3535 | TRUE | 0.2451 |
| Baat1 | 13 | 11 | 6 | 3 | 3 | 4 | 10 | 12 | 6 | 3 | 3 | 4 | 6.43 | 9 | 3 | 0.364 | -1.457 | 0.0653131 | NA | 0 | 0.3709 | 0.2488 | 0.2488 | TRUE | 0.2257 |
| Babam1 | 2900 | 2233 | 2345 | 2157 | 2065 | 2393 | 2252 | 2375 | 2530 | 2147 | 2330 | 2362 | 2332.77 | 2386 | 2280 | 0.956 | -0.065 | 0.35556872 | 0.53010782 | 0.0024 | 0.0034 | 0.0031 | 0.0031 | TRUE | 0.0339 |
| Bace1 | 371 | 235 | 222 | 263 | 242 | 289 | 288 | 250 | 240 | 262 | 273 | 285 | 266.29 | 259 | 273 | 1.051 | 0.072 | 0.5849088 | 0.73437293 | 0.0026 | 0.0113 | 0.0086 | 0.0086 | TRUE | 0.1146 |
| Bace2 | 13 | 8 | 10 | 8 | 6 | 4 | 10 | 9 | 11 | 8 | 7 | 4 | 8.01 | 10 | 6 | 0.634 | -0.658 | 0.33215289 | NA | 0 | 0.2982 | 0.1996 | 0.1996 | TRUE | 0.2616 |
| Bach1 | 2522 | 1908 | 2264 | 2264 | 2255 | 2687 | 1959 | 2030 | 2443 | 2254 | 2544 | 2652 | 2313.5 | 2144 | 2483 | 1.159 | 0.213 | 0.02141222 | 0.07264297 | 0.0102 | 0.0035 | 0.0057 | 0.0057 | TRUE | 0.1794 |
| Bach2 | 193 | 154 | 158 | 210 | 161 | 184 | 150 | 164 | 170 | 209 | 182 | 182 | 176.08 | 161 | 191 | 1.186 | 0.246 | 0.11029008 | 0.23758494 | 0 | 0.0159 | 0.0114 | 0.0114 | TRUE | 0.0769 |
| Bad | 392 | 244 | 259 | 242 | 190 | 192 | 304 | 260 | 279 | 241 | 214 | 190 | 248.03 | 281 | 215 | 0.763 | -0.391 | 0.00572395 | 0.02721639 | 0.0062 | 0.012 | 0.0103 | 0.0103 | TRUE | 0.1259 |
| Bag1 | 6110 | 4354 | 4388 | 4719 | 4114 | 4705 | 4745 | 4631 | 4735 | 4697 | 4641 | 4645 | 4682.36 | 4704 | 4661 | 0.991 | -0.013 | 0.78958184 | 0.87639639 | 0 | 0.0029 | 0.0016 | 0.0016 | TRUE | 0.0022 |
| Bag2 | 620 | 474 | 506 | 458 | 486 | 570 | 482 | 504 | 546 | 456 | 548 | 563 | 516.42 | 511 | 522 | 1.024 | 0.034 | 0.75814363 | 0.85640931 | 0.0065 | 0.007 | 0.0069 | 0.0069 | TRUE | 0.1463 |
| Bag3 | 3153 | 2326 | 2306 | 2367 | 2219 | 2573 | 2449 | 2474 | 2488 | 2356 | 2503 | 2540 | 2468.4 | 2470 | 2466 | 0.998 | -0.002 | 0.97048656 | 0.98397988 | 4.00E-04 | 0.0034 | 0.0022 | 0.0022 | TRUE | 0.0187 |
| Bag4 | 9100 | 6916 | 7375 | 8140 | 7235 | 7911 | 7068 | 7356 | 7958 | 8103 | 8162 | 7809 | 7742.56 | 7461 | 8025 | 1.076 | 0.105 | 0.07596128 | 0.18180083 | 0.0019 | 0.0027 | 0.0024 | 0.0024 | TRUE | 0.0414 |
| Bag5 | 721 | 564 | 454 | 551 | 452 | 484 | 560 | 600 | 490 | 548 | 510 | 478 | 530.98 | 550 | 512 | 0.931 | -0.104 | 0.33630115 | 0.51055541 | 0.0056 | 0.0069 | 0.0065 | 0.0065 | TRUE | 0.1025 |
| Bag6 | 19206 | 14136 | 13271 | 13668 | 12529 | 14168 | 14916 | 15036 | 14319 | 13605 | 14134 | 13986 | 14332.86 | 14757 | 13908 | 0.942 | -0.086 | 0.07446163 | 0.17945519 | 4.00E-04 | 0.0026 | 0.0016 | 0.0016 | TRUE | 0.0082 |
| Bahcc1 | 3155 | 2600 | 3445 | 4307 | 3977 | 4744 | 2450 | 2766 | 3717 | 4287 | 4486 | 4683 | 3731.63 | 2978 | 4485 | 1.507 | 0.591 | 0.00161615 | 0.01020301 | 0.0251 | 0.0031 | 0.0093 | 0.0251 | TRUE | 0.5727 |
| Bahd1 | 1574 | 1166 | 1218 | 1244 | 1160 | 1266 | 1222 | 1240 | 1314 | 1238 | 1309 | 1250 | 1262.26 | 1259 | 1266 | 1.006 | 0.008 | 0.91090513 | 0.95115633 | 0 | 0.0043 | 0.0029 | 0.0029 | TRUE | 0.0174 |
| Bai1 | 429 | 343 | 356 | 447 | 417 | 426 | 333 | 365 | 384 | 445 | 470 | 421 | 403.01 | 361 | 445 | 1.237 | 0.306 | 0.0055006 | 0.02640031 | 0.0018 | 0.0083 | 0.0063 | 0.0063 | TRUE | 0.0553 |
| Bai2 | 1466 | 1233 | 1215 | 1664 | 1456 | 1619 | 1139 | 1312 | 1311 | 1656 | 1643 | 1598 | 1443.02 | 1254 | 1632 | 1.304 | 0.383 | 7.04E-07 | 2.18E-05 | 0.0028 | 0.0041 | 0.0036 | 0.0036 | TRUE | 0.0809 |
| Bai3 | 64 | 35 | 51 | 31 | 38 | 24 | 50 | 37 | 55 | 31 | 43 | 24 | 39.9 | 47 | 33 | 0.682 | -0.553 | 0.09755082 | 0.2173829 | 0.0313 | 0.0618 | 0.0539 | 0.0539 | TRUE | 0.4971 |
| Baiap2 | 1379 | 1123 | 1144 | 1228 | 1086 | 1264 | 1071 | 1195 | 1234 | 1222 | 1225 | 1248 | 1199.19 | 1167 | 1232 | 1.058 | 0.081 | 0.30162861 | 0.47401625 | 0.0021 | 0.0044 | 0.0036 | 0.0036 | TRUE | 0.0646 |
| Baiap2l1 | 689 | 532 | 706 | 781 | 689 | 818 | 535 | 566 | 762 | 777 | 777 | 807 | 704.15 | 621 | 787 | 1.271 | 0.346 | 0.00453068 | 0.02276971 | 0.0177 | 0.0058 | 0.0093 | 0.0093 | TRUE | 0.4583 |
| Baiap2l2 | 75 | 54 | 44 | 60 | 72 | 54 | 58 | 57 | 47 | 60 | 81 | 53 | 59.57 | 54 | 65 | 1.183 | 0.243 | 0.36328942 | 0.53786674 | 0.0142 | 0.0422 | 0.0344 | 0.0344 | TRUE | 0.4136 |
| Baiap3 | 215 | 143 | 158 | 151 | 139 | 135 | 167 | 152 | 170 | 150 | 157 | 133 | 154.99 | 163 | 147 | 0.898 | -0.155 | 0.33642341 | 0.51061908 | 0 | 0.0177 | 0.0123 | 0.0123 | TRUE | 0.069 |
| Bak1 | 3614 | 2383 | 2161 | 2288 | 2006 | 2463 | 2807 | 2535 | 2332 | 2277 | 2263 | 2431 | 2440.85 | 2558 | 2324 | 0.908 | -0.139 | 0.07182722 | 0.17492358 | 0.0045 | 0.0034 | 0.0039 | 0.0039 | TRUE | 0.0896 |
| Bambi | 709 | 452 | 506 | 615 | 549 | 582 | 551 | 481 | 546 | 612 | 619 | 575 | 563.9 | 526 | 602 | 1.143 | 0.193 | 0.04768367 | 0.13016477 | 0.0017 | 0.0066 | 0.0051 | 0.0051 | TRUE | 0.0638 |
| Bambi-ps1 | 78 | 33 | 34 | 48 | 43 | 47 | 61 | 35 | 37 | 48 | 49 | 46 | 45.84 | 44 | 48 | 1.065 | 0.09 | 0.77051884 | 0.8642217 | 0.0314 | 0.0541 | 0.0475 | 0.0475 | TRUE | 0.9456 |
| Banf1 | 9864 | 7022 | 7904 | 8799 | 7631 | 9037 | 7661 | 7469 | 8528 | 8759 | 8609 | 8921 | 8324.42 | 7886 | 8763 | 1.111 | 0.152 | 0.01374695 | 0.05241 | 0.0025 | 0.0027 | 0.0026 | 0.0026 | TRUE | 0.0616 |
| Banf2 | 2 | 0 | 0 | 2 | 0 | 0 | 2 | 0 | 0 | 2 | 0 | 0 | 0.59 | 1 | 1 | 1.22 | 0.287 | 0.92000925 | NA | 0 | 4.0148 | 4.2481 | 4.2481 | TRUE | 0.9738 |

| | | | | | | | | | | | | | | | | | | | | | | | | | |
|---|---|---|---|---|---|---|---|---|---|---|---|---|---|---|---|---|---|---|---|---|---|---|---|---|---|
| Bank1 | 4 | 4 | 1 | 0 | 0 | 2 | 3 | 4 | 1 | 0 | 0 | 2 | 1.74 | 3 | 1 | 0.24 | -2.062 | 0.22252284 | NA | 0 | 1.3678 | 1.1222 | 1.1222 | TRUE | 0.8219 |
| Banp | 2953 | 2435 | 2637 | 2699 | 2451 | 3002 | 2293 | 2590 | 2845 | 2687 | 2765 | 2963 | 2690.64 | 2576 | 2805 | 1.09 | 0.124 | 0.13282705 | 0.27090425 | 0.0065 | 0.0033 | 0.0045 | 0.0045 | TRUE | 0.114 |
| Bap1 | 7356 | 5272 | 4995 | 4922 | 4437 | 5008 | 5713 | 5608 | 5390 | 4899 | 5005 | 4944 | 5259.81 | 5570 | 4949 | 0.888 | -0.171 | 0.00097702 | 0.00712034 | 3.00E-04 | 0.0029 | 0.0017 | 0.0017 | TRUE | 0.0098 |
| Bard1 | 4245 | 3266 | 3112 | 3384 | 3043 | 3454 | 3297 | 3474 | 3358 | 3368 | 3433 | 3410 | 3389.94 | 3376 | 3404 | 1.008 | 0.012 | 0.82532403 | 0.8999218 | 0 | 0.0031 | 0.0018 | 0.0018 | TRUE | 0.0078 |
| Barhl1 | 6 | 3 | 1 | 3 | 2 | 1 | 5 | 3 | 1 | 3 | 2 | 1 | 2.53 | 3 | 2 | 0.682 | -0.551 | 0.65009986 | NA | 0 | 0.9405 | 0.6552 | 0.6552 | TRUE | 0.2237 |
| Barhl2 | 27 | 17 | 16 | 16 | 13 | 18 | 21 | 18 | 17 | 16 | 15 | 18 | 17.45 | 19 | 16 | 0.856 | -0.224 | 0.6201202 | 0.76038736 | 0 | 0.1383 | 0.0893 | 0.0893 | TRUE | 0.0675 |
| Barx1 | 149 | 114 | 160 | 116 | 125 | 128 | 116 | 121 | 173 | 115 | 141 | 126 | 132.08 | 137 | 127 | 0.937 | -0.093 | 0.64037948 | 0.77534468 | 0.0228 | 0.0204 | 0.0211 | 0.0211 | TRUE | 0.538 |
| Barx2 | 109 | 92 | 90 | 131 | 110 | 144 | 85 | 98 | 97 | 130 | 124 | 142 | 112.71 | 93 | 132 | 1.425 | 0.511 | 0.00632416 | 0.02933817 | 0 | 0.0235 | 0.0161 | 0.0161 | TRUE | 0.0712 |
| Basp1 | 7063 | 5132 | 4134 | 5030 | 4247 | 5009 | 5486 | 5459 | 4461 | 5007 | 4791 | 4945 | 5024.58 | 5135 | 4914 | 0.957 | -0.064 | 0.42078122 | 0.5929435 | 0.006 | 0.0029 | 0.0043 | 0.0043 | TRUE | 0.1601 |
| Batf | 28 | 20 | 15 | 22 | 17 | 13 | 22 | 21 | 16 | 22 | 19 | 13 | 18.85 | 20 | 18 | 0.906 | -0.143 | 0.75041529 | 0.85178572 | 0 | 0.1281 | 0.0921 | 0.0921 | TRUE | 0.3322 |
| Batf2 | 5 | 4 | 6 | 11 | 9 | 8 | 4 | 4 | 6 | 11 | 10 | 8 | 7.27 | 5 | 10 | 2.005 | 1.003 | 0.16141749 | NA | 0 | 0.3285 | 0.2165 | 0.2165 | TRUE | 0.1504 |
| Batf3 | 59 | 51 | 32 | 43 | 45 | 44 | 46 | 54 | 35 | 43 | 51 | 43 | 45.27 | 45 | 46 | 1.015 | 0.021 | 0.94247546 | 0.96793862 | 0 | 0.0548 | 0.0415 | 0.0415 | TRUE | 0.2558 |
| Bax | 8207 | 5972 | 6015 | 6410 | 5888 | 6771 | 6374 | 6352 | 6490 | 6381 | 6642 | 6684 | 6487.22 | 6405 | 6569 | 1.026 | 0.036 | 0.46456599 | 0.63371271 | 2.00E-04 | 0.0028 | 0.0016 | 0.0016 | TRUE | 0.0077 |
| Baz1a | 4071 | 2820 | 2482 | 2500 | 2143 | 2512 | 3162 | 3000 | 2678 | 2488 | 2418 | 2480 | 2704.2 | 2947 | 2462 | 0.835 | -0.26 | 0.00028905 | 0.00282781 | 0.0031 | 0.0033 | 0.0033 | 0.0033 | TRUE | 0.0766 |
| Baz1b | 9934 | 7167 | 6228 | 6037 | 5557 | 6570 | 7715 | 7623 | 6720 | 6009 | 6269 | 6486 | 6803.74 | 7353 | 6255 | 0.85 | -0.234 | 0.00045041 | 0.00398377 | 0.0033 | 0.0028 | 0.003 | 0.003 | TRUE | 0.0689 |
| Baz2a | 35225 | 26648 | 27188 | 28684 | 27058 | 30693 | 27358 | 28345 | 29336 | 28552 | 30524 | 30299 | 29068.92 | 28346 | 29792 | 1.051 | 0.072 | 0.16925836 | 0.32079683 | 0.0012 | 0.0025 | 0.0019 | 0.0019 | TRUE | 0.0162 |
| Baz2b | 2287 | 1614 | 1653 | 1653 | 1479 | 1693 | 1776 | 1717 | 1784 | 1645 | 1668 | 1671 | 1710.28 | 1759 | 1661 | 0.945 | -0.082 | 0.18715074 | 0.34395597 | 0 | 0.0038 | 0.0022 | 0.0022 | TRUE | 0.0052 |
| BB014433 | 0 | 1 | 0 | 0 | 0 | 0 | 0 | 1 | 0 | 0 | 0 | 0 | 0.18 | 0 | 0 | 0.552 | -0.857 | 0.8336548 | NA | 0 | 13.3714 | 10 | 10 | TRUE | 0.5216 |
| BB019430 | 0 | 0 | 0 | 0 | 0 | 1 | 0 | 0 | 0 | 0 | 0 | 1 | 0.16 | 0 | 0 | 2.094 | 1.067 | 0.79379329 | NA | 0 | 14.4078 | 10 | 10 | TRUE | 0.3835 |
| BB031773 | 0 | 0 | 0 | 0 | 0 | 0 | 0 | 0 | 0 | 0 | 0 | 0 | 0 | 0 | 0 | NA | NA | NA | NA | NA | NA | NA | NA | NA | NA |
| BB123696 | 1 | 0 | 1 | 0 | 0 | 0 | 1 | 0 | 1 | 0 | 0 | 0 | 0.31 | 1 | 0 | 0.307 | -1.706 | 0.62894649 | NA | 0 | 7.6657 | 7.2 | 7.2 | TRUE | 0.2067 |
| BB283400 | 12 | 13 | 22 | 18 | 23 | 20 | 9 | 14 | 24 | 18 | 26 | 20 | 18.42 | 16 | 21 | 1.374 | 0.459 | 0.34484795 | 0.51910603 | 0.0684 | 0.1311 | 0.114 | 0.114 | TRUE | 0.7389 |
| BB287469 | 92 | 85 | 39 | 42 | 26 | 38 | 71 | 90 | 42 | 42 | 29 | 38 | 52.1 | 68 | 36 | 0.533 | -0.907 | 0.00514912 | 0.02510964 | 0.0703 | 0.0479 | 0.0543 | 0.0543 | TRUE | 0.1957 |
| BB557941 | 9 | 4 | 4 | 4 | 3 | 12 | 7 | 4 | 4 | 4 | 3 | 12 | 5.8 | 5 | 6 | 1.228 | 0.296 | 0.72344235 | NA | 0.1477 | 0.4114 | 0.3314 | 0.3314 | TRUE | 1.402 |
| Bbc3 | 1617 | 1394 | 1845 | 2008 | 1793 | 2213 | 1256 | 1483 | 1991 | 1999 | 2023 | 2185 | 1822.57 | 1577 | 2069 | 1.313 | 0.393 | 0.05187396 | 0.13850823 | 0.0288 | 0.0037 | 0.0109 | 0.0288 | TRUE | 0.6364 |
| Bbip1 | 587 | 399 | 381 | 445 | 336 | 429 | 456 | 424 | 411 | 443 | 379 | 423 | 422.81 | 430 | 415 | 0.964 | -0.053 | 0.62655805 | 0.76482743 | 0.0021 | 0.008 | 0.0062 | 0.0062 | TRUE | 0.0646 |
| Bbox1 | 0 | 0 | 0 | 0 | 0 | 0 | 0 | 0 | 0 | 0 | 0 | 0 | 0 | 0 | 0 | NA | NA | NA | NA | NA | NA | NA | NA | NA | NA |
| Bbs1 | 382 | 287 | 218 | 250 | 215 | 247 | 297 | 305 | 235 | 249 | 243 | 244 | 262.07 | 279 | 245 | 0.877 | -0.19 | 0.16906922 | 0.32057659 | 0.0062 | 0.0115 | 0.0099 | 0.0099 | TRUE | 0.2031 |
| Bbs10 | 558 | 375 | 346 | 339 | 340 | 328 | 433 | 399 | 373 | 337 | 384 | 324 | 375.06 | 402 | 348 | 0.864 | -0.211 | 0.07270943 | 0.17644072 | 0.0041 | 0.0087 | 0.0073 | 0.0073 | TRUE | 0.0871 |
| Bbs12 | 231 | 164 | 120 | 145 | 130 | 142 | 179 | 174 | 129 | 144 | 147 | 140 | 152.42 | 161 | 144 | 0.889 | -0.17 | 0.33138308 | 0.50569314 | 0.0095 | 0.018 | 0.0155 | 0.0155 | TRUE | 0.2993 |
| Bbs2 | 1458 | 1093 | 1130 | 1257 | 1165 | 1344 | 1132 | 1163 | 1219 | 1251 | 1314 | 1327 | 1234.4 | 1171 | 1297 | 1.108 | 0.148 | 0.03920323 | 0.11285634 | 0 | 0.0043 | 0.0029 | 0.0029 | TRUE | 0.0151 |
| Bbs4 | 867 | 641 | 733 | 685 | 658 | 786 | 673 | 682 | 791 | 682 | 742 | 776 | 724.36 | 715 | 733 | 1.026 | 0.038 | 0.69878992 | 0.8170087 | 0.0048 | 0.0057 | 0.0054 | 0.0054 | TRUE | 0.099 |
| Bbs5 | 545 | 445 | 408 | 387 | 365 | 405 | 423 | 473 | 440 | 385 | 412 | 400 | 422.27 | 445 | 399 | 0.896 | -0.158 | 0.13074838 | 0.26784959 | 0 | 0.008 | 0.0055 | 0.0055 | TRUE | 0.034 |
| Bbs7 | 459 | 348 | 380 | 346 | 279 | 343 | 356 | 370 | 410 | 344 | 315 | 339 | 355.73 | 379 | 333 | 0.88 | -0.184 | 0.10777762 | 0.23352795 | 0 | 0.0091 | 0.0066 | 0.0066 | TRUE | 0.0581 |
| Bbs9 | 1587 | 1136 | 1197 | 1075 | 1071 | 1163 | 1233 | 1208 | 1292 | 1070 | 1208 | 1148 | 1193.13 | 1244 | 1142 | 0.918 | -0.124 | 0.10571019 | 0.23056856 | 0.0015 | 0.0044 | 0.0034 | 0.0034 | TRUE | 0.0366 |
| Bbx | 2319 | 1672 | 1655 | 1692 | 1454 | 1659 | 1801 | 1778 | 1786 | 1684 | 1640 | 1638 | 1721.24 | 1788 | 1654 | 0.925 | -0.113 | 0.06795584 | 0.16778249 | 0 | 0.0038 | 0.0022 | 0.0022 | TRUE | 0.0031 |
| BC002163 | 934 | 664 | 626 | 766 | 700 | 809 | 725 | 706 | 675 | 762 | 790 | 799 | 742.98 | 702 | 784 | 1.114 | 0.156 | 0.05935224 | 0.15216529 | 0 | 0.0056 | 0.0036 | 0.0036 | TRUE | 0.0131 |
| BC003331 | 1590 | 1179 | 1256 | 1457 | 1294 | 1443 | 1235 | 1254 | 1355 | 1450 | 1460 | 1424 | 1363.12 | 1281 | 1445 | 1.128 | 0.174 | 0.01350758 | 0.05173199 | 6.00E-04 | 0.0042 | 0.0028 | 0.0028 | TRUE | 0.03 |
| BC003965 | 1319 | 959 | 782 | 872 | 801 | 870 | 1024 | 1020 | 844 | 868 | 904 | 859 | 919.78 | 963 | 877 | 0.909 | -0.137 | 0.13925806 | 0.28048626 | 0.0053 | 0.005 | 0.0051 | 0.0051 | TRUE | 0.1365 |
| BC004004 | 2671 | 1924 | 2100 | 2319 | 2116 | 2293 | 2074 | 2047 | 2266 | 2308 | 2387 | 2264 | 2224.3 | 2129 | 2320 | 1.09 | 0.124 | 0.060153 | 0.15353327 | 0.0013 | 0.0035 | 0.0027 | 0.0027 | TRUE | 0.0376 |
| BC005537 | 8285 | 6051 | 4719 | 5106 | 4376 | 4934 | 6435 | 6436 | 5092 | 5083 | 4937 | 4871 | 5475.41 | 5988 | 4964 | 0.829 | -0.271 | 0.00138195 | 0.00908566 | 0.0084 | 0.0029 | 0.005 | 0.005 | TRUE | 0.2081 |
| BC005561 | 680 | 471 | 441 | 540 | 425 | 508 | 528 | 501 | 476 | 538 | 479 | 501 | 503.9 | 502 | 506 | 1.008 | 0.012 | 0.90785076 | 0.94923397 | 0.0011 | 0.0071 | 0.0052 | 0.0052 | TRUE | 0.0344 |
| BC005624 | 1941 | 1475 | 1469 | 1379 | 1250 | 1453 | 1507 | 1569 | 1585 | 1373 | 1410 | 1434 | 1479.76 | 1554 | 1406 | 0.905 | -0.144 | 0.03055013 | 0.09378286 | 0 | 0.004 | 0.0025 | 0.0025 | TRUE | 0.0086 |
| BC005764 | 186 | 179 | 233 | 303 | 274 | 340 | 144 | 190 | 251 | 302 | 309 | 336 | 255.43 | 195 | 316 | 1.624 | 0.7 | 6.66E-05 | 0.0008844 | 0.0331 | 0.0117 | 0.0181 | 0.0181 | TRUE | 0.4575 |
| BC006965 | 0 | 0 | 0 | 0 | 0 | 0 | 0 | 0 | 0 | 0 | 0 | 0 | 0 | 0 | 0 | NA | NA | NA | NA | NA | NA | NA | NA | NA | NA |
| BC016423 | 6002 | 3960 | 3308 | 3505 | 2904 | 3135 | 4661 | 4212 | 3569 | 3489 | 3276 | 3095 | 3717.1 | 4147 | 3287 | 0.792 | -0.336 | 0.00019822 | 0.00211901 | 0.0102 | 0.0031 | 0.0056 | 0.0056 | TRUE | 0.1807 |
| BC016579 | 4 | 1 | 0 | 1 | 1 | 1 | 3 | 1 | 0 | 1 | 1 | 1 | 1.21 | 1 | 1 | 0.714 | -0.486 | 0.78184158 | NA | 0 | 1.9555 | 1.383 | 1.383 | TRUE | 0.4387 |
| BC017158 | 389 | 306 | 226 | 297 | 250 | 251 | 302 | 325 | 244 | 296 | 282 | 248 | 282.82 | 290 | 275 | 0.946 | -0.08 | 0.57034168 | 0.72384067 | 0.0112 | 0.0108 | 0.0109 | 0.0109 | TRUE | 0.2124 |
| BC017612 | 292 | 237 | 251 | 235 | 231 | 274 | 227 | 252 | 271 | 234 | 261 | 270 | 252.45 | 250 | 255 | 1.023 | 0.033 | 0.80728976 | 0.8885257 | 0.0033 | 0.0118 | 0.0092 | 0.0092 | TRUE | 0.0794 |
| BC017643 | 1387 | 973 | 948 | 963 | 853 | 1007 | 1077 | 1035 | 1023 | 959 | 962 | 994 | 1008.33 | 1045 | 972 | 0.929 | -0.106 | 0.15260278 | 0.29927984 | 0 | 0.0048 | 0.003 | 0.003 | TRUE | 0.0092 |
| BC018242 | 348 | 302 | 309 | 409 | 374 | 447 | 270 | 321 | 333 | 407 | 422 | 441 | 365.87 | 308 | 423 | 1.379 | 0.463 | 0.0001022 | 0.00122979 | 0.0042 | 0.0089 | 0.0075 | 0.0075 | TRUE | 0.1435 |
| BC018473 | 0 | 0 | 0 | 0 | 1 | 0 | 0 | 0 | 0 | 0 | 1 | 0 | 0.19 | 0 | 0 | 2.094 | 1.067 | 0.79379329 | NA | 0 | 12.608 | 10 | 10 | TRUE | 0.5075 |
| BC018507 | 6897 | 5065 | 5302 | 5627 | 5150 | 5975 | 5357 | 5388 | 5721 | 5601 | 5810 | 5898 | 5629.01 | 5489 | 5770 | 1.051 | 0.072 | 0.18557314 | 0.34184777 | 8.00E-04 | 0.0028 | 0.002 | 0.002 | TRUE | 0.0166 |
| BC020402 | 25 | 23 | 28 | 25 | 16 | 21 | 19 | 24 | 30 | 25 | 18 | 21 | 22.96 | 24 | 21 | 0.868 | -0.204 | 0.61719111 | 0.75833714 | 0 | 0.1057 | 0.0759 | 0.0759 | TRUE | 0.2148 |
| BC021614 | 299 | 231 | 287 | 427 | 334 | 379 | 232 | 246 | 310 | 425 | 377 | 374 | 327.26 | 263 | 392 | 1.499 | 0.584 | 1.56E-05 | 0.00027449 | 0.0103 | 0.0097 | 0.01 | 0.01 | TRUE | 0.2696 |
| BC021767 | 73 | 50 | 48 | 66 | 42 | 67 | 57 | 53 | 52 | 66 | 47 | 66 | 56.81 | 54 | 60 | 1.11 | 0.15 | 0.56729842 | 0.72119508 | 0 | 0.0441 | 0.0322 | 0.0322 | TRUE | 0.2629 |
| BC021785 | 3 | 1 | 4 | 1 | 3 | 2 | 2 | 1 | 4 | 1 | 3 | 2 | 2.34 | 2 | 2 | 0.819 | -0.288 | 0.81846734 | NA | 0 | 1.0136 | 0.6959 | 0.6959 | TRUE | 0.2769 |
| BC021891 | 45 | 31 | 41 | 51 | 49 | 51 | 35 | 33 | 44 | 51 | 55 | 50 | 44.76 | 37 | 52 | 1.397 | 0.483 | 0.09631605 | 0.21537003 | 0 | 0.0554 | 0.0378 | 0.0378 | TRUE | 0.1733 |
| BC022687 | 215 | 139 | 151 | 137 | 140 | 155 | 167 | 148 | 163 | 136 | 158 | 153 | 154.18 | 159 | 149 | 0.934 | -0.099 | 0.54219975 | 0.70012296 | 0 | 0.0178 | 0.0123 | 0.0123 | TRUE | 0.0592 |
| BC023829 | 1354 | 948 | 830 | 863 | 783 | 850 | 1052 | 1008 | 896 | 859 | 883 | 839 | 922.82 | 985 | 860 | 0.872 | -0.198 | 0.02154066 | 0.07294022 | 0.0028 | 0.005 | 0.0042 | 0.0042 | TRUE | 0.0744 |
| BC024139 | 2 | 1 | 3 | 1 | 11 | 1 | 2 | 1 | 3 | 1 | 12 | 1 | 3.37 | 2 | 5 | 2.444 | 1.289 | 0.30449416 | NA | 0.9191 | 0.7048 | 0.777 | 0.777 | TRUE | 3.425 |
| BC024386 | 2 | 0 | 1 | 0 | 1 | 0 | 2 | 0 | 1 | 0 | 1 | 0 | 0.63 | 1 | 0 | 0.414 | -1.271 | 0.61575572 | NA | 0 | 3.784 | 3.0839 | 3.0839 | TRUE | 0.5075 |
| BC024582 | 2 | 0 | 0 | 0 | 1 | 0 | 2 | 0 | 0 | 0 | 1 | 0 | 0.45 | 1 | 0 | 0.68 | -0.556 | 0.85993441 | NA | 0 | 5.3057 | 5.2576 | 5.2576 | TRUE | 0.9726 |
| BC024978 | 272 | 211 | 212 | 227 | 193 | 230 | 211 | 224 | 229 | 226 | 218 | 227 | 222.53 | 221 | 224 | 1.012 | 0.017 | 0.90111244 | 0.94563878 | 0 | 0.0131 | 0.0084 | 0.0084 | TRUE | 0.0198 |
| BC025920 | 43 | 30 | 29 | 34 | 34 | 44 | 33 | 32 | 31 | 34 | 38 | 43 | 35.37 | 32 | 38 | 1.195 | 0.258 | 0.4222975 | 0.59466977 | 0 | 0.0694 | 0.0458 | 0.0458 | TRUE | 0.0949 |
| BC026585 | 753 | 525 | 593 | 615 | 577 | 574 | 585 | 558 | 640 | 612 | 651 | 567 | 602.14 | 594 | 610 | 1.026 | 0.037 | 0.70745765 | 0.82342771 | 0.003 | 0.0064 | 0.0053 | 0.0053 | TRUE | 0.0505 |
| BC026590 | 866 | 556 | 471 | 512 | 432 | 475 | 673 | 591 | 508 | 510 | 487 | 469 | 539.68 | 591 | 489 | 0.825 | -0.278 | 0.01559658 | 0.0575134 | 0.0092 | 0.0068 | 0.0076 | 0.0076 | TRUE | 0.1626 |
| BC027072 | 21 | 7 | 9 | 4 | 9 | 5 | 16 | 7 | 10 | 4 | 10 | 5 | 8.76 | 11 | 6 | 0.556 | -0.846 | 0.22227632 | NA | 0.0917 | 0.2731 | 0.2214 | 0.2214 | TRUE | 0.7594 |
| BC027231 | 2809 | 1945 | 2086 | 2221 | 2004 | 2361 | 2182 | 2069 | 2251 | 2211 | 2261 | 2331 | 2217.24 | 2167 | 2268 | 1.046 | 0.065 | 0.30257785 | 0.4751423 | 8.00E-04 | 0.0035 | 0.0024 | 0.0024 | TRUE | 0.0185 |

| | | | | | | | | | | | | | | | | | | | | | | | | | |
|---|---|---|---|---|---|---|---|---|---|---|---|---|---|---|---|---|---|---|---|---|---|---|---|---|---|
| BC028528 | 68 | 45 | 52 | 86 | 74 | 86 | 53 | 48 | 56 | 86 | 83 | 85 | 68.46 | 52 | 85 | 1.62 | 0.696 | 0.00289437 | 0.01609071 | 0 | 0.037 | 0.024 | 0.024 | TRUE | 0.0406 |
| BC029214 | 71 | 50 | 35 | 56 | 49 | 57 | 55 | 53 | 38 | 56 | 55 | 56 | 52.23 | 49 | 56 | 1.139 | 0.188 | 0.49210136 | 0.6574413 | 0 | 0.0478 | 0.0348 | 0.0348 | TRUE | 0.291 |
| BC029722 | 485 | 359 | 385 | 375 | 356 | 403 | 377 | 382 | 415 | 373 | 402 | 398 | 391.11 | 391 | 391 | 1 | 0 | 0.99811162 | 0.99895031 | 0 | 0.0085 | 0.0058 | 0.0058 | TRUE | 0.0324 |
| BC030307 | 79 | 53 | 42 | 67 | 45 | 55 | 61 | 56 | 45 | 67 | 51 | 54 | 55.8 | 54 | 57 | 1.05 | 0.07 | 0.79269623 | 0.87856756 | 0 | 0.0449 | 0.0338 | 0.0338 | TRUE | 0.1786 |
| BC030336 | 1481 | 1102 | 1086 | 1388 | 1085 | 1317 | 1150 | 1172 | 1172 | 1382 | 1224 | 1300 | 1233.31 | 1165 | 1302 | 1.119 | 0.162 | 0.02967944 | 0.09189954 | 0.0011 | 0.0043 | 0.0032 | 0.0032 | TRUE | 0.0344 |
| BC030476 | 23 | 21 | 12 | 28 | 17 | 23 | 18 | 22 | 13 | 28 | 19 | 23 | 20.48 | 18 | 23 | 1.315 | 0.395 | 0.36331675 | 0.53786674 | 0 | 0.1181 | 0.0864 | 0.0864 | TRUE | 0.2285 |
| BC030500 | 0 | 1 | 0 | 1 | 1 | 0 | 0 | 1 | 0 | 1 | 1 | 0 | 0.53 | 0 | 1 | 2.036 | 1.026 | 0.70492654 | NA | 0 | 4.4642 | 3.5518 | 3.5518 | TRUE | 0.5217 |
| BC030867 | 3254 | 2374 | 2451 | 2146 | 2054 | 2429 | 2527 | 2525 | 2645 | 2136 | 2317 | 2398 | 2424.68 | 2566 | 2284 | 0.89 | -0.168 | 0.01106089 | 0.04445772 | 0.0015 | 0.0034 | 0.0027 | 0.0027 | TRUE | 0.0391 |
| BC030870 | 1 | 8 | 0 | 0 | 2 | 0 | 1 | 9 | 0 | 0 | 2 | 0 | 1.92 | 3 | 1 | 0.236 | -2.083 | 0.26943375 | NA | 4.2026 | 1.2344 | 1.69 | 1.69 | TRUE | 2.9748 |
| BC031181 | 2559 | 1701 | 1633 | 1985 | 1730 | 1971 | 1987 | 1809 | 1762 | 1976 | 1952 | 1946 | 1905.33 | 1853 | 1958 | 1.055 | 0.078 | 0.25849496 | 0.42846671 | 0.0016 | 0.0037 | 0.0029 | 0.0029 | TRUE | 0.0503 |
| BC031361 | 22 | 12 | 12 | 21 | 12 | 14 | 17 | 13 | 13 | 21 | 14 | 14 | 15.18 | 14 | 16 | 1.121 | 0.165 | 0.73849732 | NA | 0 | 0.1586 | 0.1111 | 0.1111 | TRUE | 0.3333 |
| BC033916 | 395 | 274 | 258 | 224 | 203 | 223 | 307 | 291 | 278 | 223 | 229 | 220 | 258.12 | 292 | 224 | 0.765 | -0.386 | 0.0022576 | 0.01325169 | 0 | 0.0116 | 0.0076 | 0.0076 | TRUE | 0.0232 |
| BC035044 | 0 | 0 | 1 | 1 | 0 | 0 | 0 | 0 | 1 | 1 | 0 | 0 | 0.35 | 0 | 0 | 1.075 | 0.105 | 0.97567165 | NA | 0 | 6.8576 | 6.5143 | 6.5143 | TRUE | 0.5365 |
| BC037032 | 0 | 1 | 2 | 0 | 0 | 3 | 0 | 1 | 2 | 0 | 0 | 3 | 1.03 | 1 | 1 | 0.962 | -0.057 | 0.97905301 | NA | 2.6483 | 2.3023 | 2.36 | 2.36 | TRUE | 0.6755 |
| BC037034 | 1185 | 1006 | 934 | 958 | 870 | 1047 | 920 | 1070 | 1008 | 954 | 981 | 1034 | 994.46 | 999 | 990 | 0.992 | -0.012 | 0.88719965 | 0.93716439 | 0.0029 | 0.0048 | 0.0041 | 0.0041 | TRUE | 0.06 |
| BC037704 | 98 | 86 | 73 | 105 | 65 | 85 | 76 | 91 | 79 | 105 | 73 | 84 | 84.68 | 82 | 87 | 1.067 | 0.094 | 0.67544749 | 0.80071088 | 0.0091 | 0.0304 | 0.0245 | 0.0245 | TRUE | 0.2856 |
| BC039771 | 8 | 5 | 4 | 9 | 4 | 7 | 6 | 5 | 4 | 9 | 5 | 7 | 6.04 | 5 | 7 | 1.283 | 0.359 | 0.64112524 | NA | 0 | 0.3949 | 0.2615 | 0.2615 | TRUE | 0.1746 |
| BC039966 | 15 | 6 | 10 | 5 | 3 | 8 | 12 | 6 | 11 | 5 | 3 | 8 | 7.51 | 10 | 5 | 0.565 | -0.824 | 0.25235097 | NA | 0 | 0.3179 | 0.2288 | 0.2288 | TRUE | 0.3528 |
| BC048403 | 1590 | 1293 | 1276 | 1306 | 1255 | 1366 | 1235 | 1375 | 1377 | 1300 | 1416 | 1348 | 1341.87 | 1329 | 1355 | 1.02 | 0.029 | 0.70440709 | 0.82107628 | 0.0022 | 0.0042 | 0.0035 | 0.0035 | TRUE | 0.0481 |
| BC048502 | 1 | 2 | 3 | 1 | 1 | 0 | 1 | 2 | 3 | 1 | 1 | 0 | 1.38 | 2 | 1 | 0.347 | -1.526 | 0.37918523 | NA | 0 | 1.7231 | 1.1882 | 1.1882 | TRUE | 0.1962 |
| BC048507 | 1 | 1 | 0 | 0 | 0 | 3 | 1 | 1 | 0 | 0 | 0 | 3 | 0.8 | 1 | 1 | 1.611 | 0.688 | 0.77122419 | NA | 1.7276 | 2.9639 | 2.7421 | 2.7421 | TRUE | 1.4416 |
| BC048546 | 21 | 18 | 13 | 11 | 11 | 10 | 16 | 19 | 14 | 11 | 12 | 10 | 13.79 | 16 | 11 | 0.67 | -0.578 | 0.26053843 | NA | 0 | 0.1744 | 0.1139 | 0.1139 | TRUE | 0.087 |
| BC048562 | 0 | 0 | 0 | 0 | 0 | 0 | 0 | 0 | 0 | 0 | 0 | 0 | 0 | 0 | 0 | NA | NA | NA | NA | NA | NA | NA | NA | NA | NA |
| BC048602 | 0 | 0 | 0 | 0 | 0 | 0 | 0 | 0 | 0 | 0 | 0 | 0 | 0 | 0 | 0 | NA | NA | NA | NA | NA | NA | NA | NA | NA | NA |
| BC048609 | 1 | 0 | 2 | 1 | 2 | 3 | 1 | 0 | 2 | 1 | 2 | 3 | 1.52 | 1 | 2 | 2.143 | 1.1 | 0.49232419 | NA | 0 | 1.557 | 1.0999 | 1.0999 | TRUE | 0.4431 |
| BC048644 | 3 | 3 | 3 | 3 | 2 | 4 | 2 | 3 | 3 | 3 | 2 | 4 | 2.99 | 3 | 3 | 1.064 | 0.089 | 0.93356123 | NA | 0 | 0.7947 | 0.489 | 0.489 | TRUE | 0.0863 |
| BC048671 | 0 | 0 | 0 | 0 | 0 | 0 | 0 | 0 | 0 | 0 | 0 | 0 | 0 | 0 | 0 | NA | NA | NA | NA | NA | NA | NA | NA | NA | NA |
| BC048679 | 3 | 1 | 5 | 7 | 5 | 6 | 2 | 1 | 5 | 7 | 6 | 6 | 4.55 | 3 | 6 | 2.136 | 1.095 | 0.23928092 | NA | 0 | 0.5229 | 0.374 | 0.374 | TRUE | 0.6086 |
| BC048943 | 0 | 0 | 0 | 0 | 0 | 0 | 0 | 0 | 0 | 0 | 0 | 0 | 0 | 0 | 0 | NA | NA | NA | NA | NA | NA | NA | NA | NA | NA |
| BC049352 | 1 | 0 | 0 | 1 | 1 | 0 | 1 | 0 | 0 | 1 | 1 | 0 | 0.48 | 0 | 1 | 2.037 | 1.027 | 0.71241723 | NA | 0 | 4.9057 | 3.8521 | 3.8521 | TRUE | 0.2664 |
| BC049635 | 6 | 8 | 5 | 5 | 8 | 10 | 5 | 9 | 5 | 5 | 9 | 10 | 7.07 | 6 | 8 | 1.297 | 0.375 | 0.60617908 | NA | 0 | 0.3375 | 0.2369 | 0.2369 | TRUE | 0.3253 |
| BC049715 | 3 | 0 | 0 | 0 | 0 | 0 | 2 | 0 | 0 | 0 | 0 | 0 | 0.39 | 1 | 0 | 0.239 | -2.067 | 0.57190649 | NA | 10 | 6.1057 | 8.0283 | 8.0283 | TRUE | 1.4408 |
| BC049730 | 1 | 0 | 0 | 0 | 0 | 0 | 1 | 0 | 0 | 0 | 0 | 0 | 0.13 | 0 | 0 | 0.552 | -0.857 | 0.8336548 | NA | 0 | 18.3121 | 10 | 10 | TRUE | 0.2566 |
| BC049762 | 21 | 10 | 8 | 12 | 9 | 8 | 16 | 11 | 9 | 12 | 10 | 8 | 10.93 | 12 | 10 | 0.831 | -0.266 | 0.65046101 | NA | 0 | 0.2193 | 0.157 | 0.157 | TRUE | 0.5259 |
| BC051019 | 91 | 81 | 72 | 52 | 71 | 74 | 71 | 86 | 78 | 52 | 80 | 73 | 73.24 | 78 | 68 | 0.874 | -0.194 | 0.42711403 | 0.59915014 | 0.014 | 0.0348 | 0.0292 | 0.0292 | TRUE | 0.4074 |
| BC051070 | 0 | 0 | 0 | 0 | 0 | 0 | 0 | 0 | 0 | 0 | 0 | 0 | 0 | 0 | 0 | NA | NA | NA | NA | NA | NA | NA | NA | NA | NA |
| BC051142 | 4 | 4 | 5 | 4 | 3 | 4 | 3 | 4 | 5 | 4 | 3 | 4 | 4.01 | 4 | 4 | 0.899 | -0.154 | 0.86823294 | NA | 0 | 0.5932 | 0.3708 | 0.3708 | TRUE | 0.1336 |
| BC051226 | 186 | 157 | 163 | 174 | 172 | 210 | 144 | 167 | 176 | 173 | 194 | 207 | 176.98 | 162 | 191 | 1.183 | 0.243 | 0.12518288 | 0.26021509 | 0.0041 | 0.0158 | 0.0124 | 0.0124 | TRUE | 0.1104 |
| BC051537 | 4 | 5 | 4 | 9 | 8 | 6 | 3 | 5 | 4 | 9 | 9 | 6 | 6.11 | 4 | 8 | 1.897 | 0.924 | 0.23549423 | NA | 0 | 0.3905 | 0.2573 | 0.2573 | TRUE | 0.1536 |
| BC051628 | 12 | 4 | 5 | 4 | 4 | 2 | 9 | 4 | 5 | 4 | 5 | 2 | 4.91 | 6 | 4 | 0.54 | -0.888 | 0.31636815 | NA | 0 | 0.4855 | 0.3418 | 0.3418 | TRUE | 0.5406 |
| BC051665 | 0 | 0 | 0 | 0 | 0 | 0 | 0 | 0 | 0 | 0 | 0 | 0 | 0 | 0 | 0 | NA | NA | NA | NA | NA | NA | NA | NA | NA | NA |
| BC052040 | 3628 | 2758 | 3362 | 3046 | 2949 | 3246 | 2818 | 2934 | 3628 | 3032 | 3327 | 3204 | 3157 | 3127 | 3188 | 1.02 | 0.029 | 0.75059371 | 0.85185293 | 0.0102 | 0.0032 | 0.0056 | 0.0056 | TRUE | 0.2381 |
| BC052688 | 60 | 56 | 34 | 37 | 32 | 41 | 47 | 60 | 37 | 37 | 36 | 40 | 42.71 | 48 | 38 | 0.795 | -0.331 | 0.2814545 | 0.45301905 | 0 | 0.0579 | 0.0444 | 0.0444 | TRUE | 0.2047 |
| BC053393 | 3 | 3 | 0 | 0 | 0 | 1 | 2 | 3 | 0 | 0 | 0 | 1 | 1.08 | 2 | 0 | 0.2 | -2.321 | 0.27423319 | NA | 0 | 2.1874 | 1.9858 | 1.9858 | TRUE | 0.3784 |
| BC053749 | 607 | 514 | 532 | 520 | 466 | 599 | 471 | 547 | 574 | 518 | 526 | 591 | 537.8 | 531 | 545 | 1.03 | 0.042 | 0.69823647 | 0.81661843 | 0.0063 | 0.0068 | 0.0066 | 0.0066 | TRUE | 0.1185 |
| BC055111 | 127 | 80 | 81 | 90 | 82 | 109 | 99 | 85 | 87 | 90 | 93 | 108 | 93.47 | 90 | 97 | 1.066 | 0.092 | 0.6541073 | 0.78520683 | 0 | 0.0278 | 0.0195 | 0.0195 | TRUE | 0.0994 |
| BC055324 | 6449 | 4884 | 4812 | 5239 | 4638 | 5268 | 5009 | 5195 | 5192 | 5215 | 5232 | 5200 | 5173.86 | 5132 | 5216 | 1.017 | 0.024 | 0.63473228 | 0.77160293 | 0 | 0.0029 | 0.0016 | 0.0016 | TRUE | 0.0055 |
| BC055402 | 1 | 2 | 0 | 0 | 1 | 0 | 1 | 2 | 0 | 0 | 1 | 0 | 0.67 | 1 | 0 | 0.385 | -1.379 | 0.57963618 | NA | 0 | 3.5292 | 2.9645 | 2.9645 | TRUE | 0.3656 |
| BC056474 | 1623 | 1136 | 1191 | 1236 | 1098 | 1239 | 1261 | 1208 | 1285 | 1230 | 1239 | 1223 | 1241 | 1251 | 1231 | 0.983 | -0.024 | 0.72582542 | 0.83575467 | 0 | 0.0043 | 0.0026 | 0.0026 | TRUE | 0.0104 |
| BC061194 | 1 | 0 | 0 | 0 | 1 | 3 | 1 | 0 | 0 | 0 | 1 | 3 | 0.81 | 0 | 1 | 3.963 | 1.987 | 0.39811512 | NA | 0 | 2.9247 | 2.6042 | 2.6042 | TRUE | 0.1519 |
| BC061212 | 0 | 0 | 0 | 0 | 0 | 0 | 0 | 0 | 0 | 0 | 0 | 0 | 0 | 0 | 0 | NA | NA | NA | NA | NA | NA | NA | NA | NA | NA |
| BC061237 | 0 | 0 | 0 | 0 | 0 | 0 | 0 | 0 | 0 | 0 | 0 | 0 | 0 | 0 | 0 | NA | NA | NA | NA | NA | NA | NA | NA | NA | NA |
| BC064078 | 6 | 4 | 3 | 2 | 1 | 1 | 5 | 4 | 3 | 2 | 1 | 1 | 2.71 | 4 | 1 | 0.338 | -1.567 | 0.19385247 | NA | 0 | 0.8771 | 0.553 | 0.553 | TRUE | 0.1034 |
| BC065397 | 131 | 136 | 137 | 112 | 121 | 161 | 102 | 145 | 148 | 111 | 137 | 159 | 133.52 | 132 | 136 | 1.038 | 0.054 | 0.79285128 | 0.87867126 | 0.0298 | 0.0202 | 0.023 | 0.023 | TRUE | 0.3625 |
| BC068157 | 442 | 403 | 489 | 465 | 445 | 554 | 343 | 429 | 528 | 463 | 502 | 547 | 468.55 | 433 | 504 | 1.167 | 0.223 | 0.11859993 | 0.25005318 | 0.0229 | 0.0075 | 0.0126 | 0.0126 | TRUE | 0.2625 |
| BC068281 | 317 | 251 | 227 | 235 | 208 | 253 | 246 | 267 | 245 | 234 | 235 | 250 | 246.07 | 253 | 240 | 0.949 | -0.076 | 0.55498013 | 0.7102732 | 0 | 0.0121 | 0.0079 | 0.0079 | TRUE | 0.0262 |
| BC080695 | 50 | 53 | 42 | 37 | 32 | 26 | 39 | 56 | 45 | 37 | 36 | 26 | 39.85 | 47 | 33 | 0.704 | -0.506 | 0.11534144 | 0.24528718 | 0 | 0.0619 | 0.0484 | 0.0484 | TRUE | 0.2629 |
| BC089491 | 2 | 2 | 1 | 3 | 2 | 2 | 2 | 2 | 1 | 3 | 2 | 2 | 2 | 2 | 2 | 1.518 | 0.602 | 0.64543124 | NA | 0 | 1.1898 | 0.7151 | 0.7151 | TRUE | 0.0561 |
| BC089597 | 1 | 1 | 0 | 3 | 0 | 0 | 1 | 1 | 0 | 3 | 0 | 0 | 0.8 | 1 | 1 | 1.621 | 0.697 | 0.76811698 | NA | 1.7582 | 2.9487 | 2.7352 | 2.7352 | TRUE | 1.4417 |
| BC090627 | 1 | 0 | 0 | 0 | 0 | 0 | 1 | 0 | 0 | 0 | 0 | 0 | 0.13 | 0 | 0 | 0.552 | -0.857 | 0.8336548 | NA | 0 | 18.3121 | 10 | 10 | TRUE | 0.2566 |
| BC094916 | 0 | 0 | 0 | 0 | 0 | 0 | 0 | 0 | 0 | 0 | 0 | 0 | 0 | 0 | 0 | NA | NA | NA | NA | NA | NA | NA | NA | NA | NA |
| BC096441 | 16 | 7 | 22 | 12 | 14 | 13 | 12 | 7 | 24 | 12 | 16 | 13 | 14.03 | 14 | 14 | 0.933 | -0.1 | 0.8575935 | NA | 0.0972 | 0.1714 | 0.1508 | 0.1508 | TRUE | 0.5538 |
| Bc1 | 0 | 0 | 0 | 0 | 0 | 1 | 0 | 0 | 0 | 0 | 0 | 1 | 0.16 | 0 | 0 | 2.094 | 1.067 | 0.79379329 | NA | 0 | 14.4078 | 10 | 10 | TRUE | 0.3835 |
| BC100451 | 11 | 12 | 5 | 4 | 3 | 11 | 9 | 13 | 5 | 4 | 3 | 11 | 7.49 | 9 | 6 | 0.688 | -0.539 | 0.47438027 | NA | 0.1389 | 0.3189 | 0.2698 | 0.2698 | TRUE | 0.599 |
| BC100530 | 0 | 0 | 0 | 0 | 0 | 0 | 0 | 0 | 0 | 0 | 0 | 0 | 0 | 0 | 0 | NA | NA | NA | NA | NA | NA | NA | NA | NA | NA |
| BC107364 | 4 | 15 | 11 | 9 | 8 | 16 | 3 | 16 | 12 | 9 | 9 | 16 | 10.78 | 10 | 11 | 1.119 | 0.162 | 0.80836328 | NA | 0.2422 | 0.2222 | 0.2279 | 0.2279 | TRUE | 0.733 |
| BC117090 | 0 | 0 | 0 | 0 | 0 | 0 | 0 | 0 | 0 | 0 | 0 | 0 | 0 | 0 | 0 | NA | NA | NA | NA | NA | NA | NA | NA | NA | NA |
| BC125332 | 39 | 25 | 19 | 22 | 20 | 20 | 30 | 27 | 21 | 22 | 23 | 20 | 23.6 | 26 | 22 | 0.823 | -0.28 | 0.47935518 | 0.64629975 | 0 | 0.1029 | 0.0703 | 0.0703 | TRUE | 0.1882 |
| BC147527 | 96 | 93 | 39 | 48 | 51 | 38 | 75 | 99 | 42 | 48 | 58 | 38 | 59.73 | 72 | 48 | 0.66 | -0.599 | 0.06164879 | 0.15645352 | 0.0928 | 0.0421 | 0.0565 | 0.0565 | TRUE | 0.1833 |

| | | | | | | | | | | | | | | | | | | | | | | | | | |
|---|---|---|---|---|---|---|---|---|---|---|---|---|---|---|---|---|---|---|---|---|---|---|---|---|---|
| Bcam | 2747 | 1747 | 1661 | 2444 | 1913 | 2245 | 2133 | 1858 | 1792 | 2433 | 2158 | 2216 | 2098.49 | 1928 | 2269 | 1.176 | 0.234 | 0.0044457 | 0.02244522 | 0.0057 | 0.0036 | 0.0044 | 0.0044 | TRUE | 0.1074 |
| Bcan | 355 | 337 | 278 | 493 | 396 | 439 | 276 | 358 | 300 | 491 | 447 | 433 | 384.16 | 311 | 457 | 1.472 | 0.558 | 7.42E-06 | 0.00014708 | 0.0082 | 0.0086 | 0.0085 | 0.0085 | TRUE | 0.1963 |
| Bcap29 | 1557 | 1197 | 1276 | 1660 | 1312 | 1358 | 1209 | 1273 | 1377 | 1652 | 1480 | 1341 | 1388.71 | 1286 | 1491 | 1.16 | 0.214 | 0.01737618 | 0.0622638 | 0.0066 | 0.0041 | 0.0051 | 0.0051 | TRUE | 0.109 |
| Bcap31 | 6050 | 4609 | 4885 | 5552 | 4973 | 5520 | 4699 | 4903 | 5271 | 5526 | 5610 | 5449 | 5242.97 | 4958 | 5528 | 1.116 | 0.158 | 0.00785886 | 0.034594 | 0.0015 | 0.0029 | 0.0023 | 0.0023 | TRUE | 0.0372 |
| Bcar1 | 5169 | 3790 | 3469 | 3837 | 3506 | 3791 | 4015 | 4031 | 3743 | 3819 | 3955 | 3742 | 3884.29 | 3930 | 3839 | 0.977 | -0.034 | 0.55603919 | 0.71118283 | 9.00E-04 | 0.003 | 0.0022 | 0.0022 | TRUE | 0.0207 |
| Bcar3 | 1667 | 1177 | 1426 | 1424 | 1297 | 1486 | 1295 | 1252 | 1539 | 1417 | 1463 | 1467 | 1405.47 | 1362 | 1449 | 1.065 | 0.091 | 0.29191804 | 0.46411396 | 0.0054 | 0.0041 | 0.0046 | 0.0046 | TRUE | 0.1526 |
| Bcas1 | 812 | 670 | 742 | 781 | 689 | 850 | 631 | 713 | 801 | 777 | 777 | 839 | 756.28 | 715 | 798 | 1.119 | 0.163 | 0.10637987 | 0.23148179 | 0.0068 | 0.0056 | 0.006 | 0.006 | TRUE | 0.1314 |
| Bcas2 | 9040 | 6442 | 6730 | 6968 | 6293 | 7266 | 7021 | 6852 | 7262 | 6936 | 7099 | 7173 | 7057.1 | 7045 | 7069 | 1.003 | 0.005 | 0.92130822 | 0.95666871 | 3.00E-04 | 0.0028 | 0.0017 | 0.0017 | TRUE | 0.0087 |
| Bcas3 | 3954 | 2899 | 3153 | 2655 | 2553 | 2966 | 3071 | 3084 | 3402 | 2643 | 2880 | 2928 | 3001.22 | 3186 | 2817 | 0.885 | -0.177 | 0.0101101 | 0.04165893 | 0.0027 | 0.0032 | 0.0031 | 0.0031 | TRUE | 0.0426 |
| Bcat1 | 8529 | 6102 | 6059 | 6284 | 5669 | 6496 | 6624 | 6491 | 6538 | 6255 | 6395 | 6413 | 6452.53 | 6551 | 6354 | 0.97 | -0.044 | 0.35740143 | 0.53189698 | 0 | 0.0028 | 0.0015 | 0.0015 | TRUE | 0.0023 |
| Bcat2 | 6656 | 4962 | 5872 | 6568 | 6430 | 7957 | 5169 | 5278 | 6336 | 6538 | 7254 | 7855 | 6404.93 | 5594 | 7216 | 1.29 | 0.367 | 2.90E-05 | 0.0004509 | 0.0106 | 0.0028 | 0.0054 | 0.0054 | TRUE | 0.1635 |
| Bccip | 5573 | 4148 | 4062 | 4469 | 3869 | 4587 | 4328 | 4412 | 4383 | 4448 | 4365 | 4528 | 4410.75 | 4374 | 4447 | 1.017 | 0.024 | 0.63710377 | 0.77324184 | 0 | 0.003 | 0.0016 | 0.0016 | TRUE | 0.0032 |
| Bcdin3d | 326 | 226 | 209 | 218 | 191 | 208 | 253 | 240 | 226 | 217 | 215 | 205 | 226.15 | 240 | 212 | 0.885 | -0.176 | 0.18965552 | 0.3474425 | 0 | 0.0129 | 0.0086 | 0.0086 | TRUE | 0.0292 |
| Bche | 466 | 282 | 160 | 158 | 120 | 116 | 362 | 300 | 173 | 157 | 135 | 115 | 206.95 | 278 | 136 | 0.486 | -1.04 | 5.33E-06 | 0.00011188 | 0.077 | 0.0139 | 0.0321 | 0.0321 | TRUE | 0.2933 |
| Bckdha | 3913 | 2946 | 3178 | 3419 | 3275 | 3833 | 3039 | 3134 | 3429 | 3403 | 3695 | 3784 | 3413.88 | 3201 | 3627 | 1.134 | 0.181 | 0.00891639 | 0.03811064 | 0.0031 | 0.0031 | 0.0032 | 0.0032 | TRUE | 0.0471 |
| Bckdhb | 2547 | 1861 | 2076 | 2284 | 2028 | 2388 | 1978 | 1980 | 2240 | 2273 | 2288 | 2357 | 2186.04 | 2066 | 2306 | 1.117 | 0.16 | 0.02201077 | 0.07395391 | 0.0022 | 0.0035 | 0.003 | 0.003 | TRUE | 0.0648 |
| Bckdk | 4561 | 3310 | 3156 | 3253 | 2924 | 3399 | 3542 | 3521 | 3405 | 3238 | 3299 | 3355 | 3393.4 | 3489 | 3297 | 0.945 | -0.082 | 0.13157679 | 0.26908385 | 0 | 0.0031 | 0.0018 | 0.0018 | TRUE | 0.0054 |
| Bcl10 | 2177 | 1460 | 1524 | 1546 | 1306 | 1551 | 1691 | 1553 | 1644 | 1539 | 1473 | 1531 | 1571.9 | 1629 | 1514 | 0.929 | -0.106 | 0.11791152 | 0.24916845 | 6.00E-04 | 0.0039 | 0.0027 | 0.0027 | TRUE | 0.0201 |
| Bcl11a | 52 | 44 | 60 | 48 | 66 | 91 | 40 | 47 | 65 | 48 | 74 | 90 | 60.67 | 51 | 71 | 1.404 | 0.49 | 0.10230312 | 0.22481834 | 0.0635 | 0.0415 | 0.0478 | 0.0478 | TRUE | 0.3114 |
| Bcl11b | 289 | 226 | 250 | 310 | 291 | 367 | 224 | 240 | 270 | 309 | 328 | 362 | 288.96 | 245 | 333 | 1.364 | 0.448 | 0.00059043 | 0.00490069 | 0.0043 | 0.0106 | 0.0087 | 0.0087 | TRUE | 0.0863 |
| Bcl2 | 40 | 22 | 14 | 25 | 16 | 11 | 31 | 23 | 15 | 25 | 18 | 11 | 20.56 | 23 | 18 | 0.765 | -0.386 | 0.41352797 | 0.58607391 | 0.0935 | 0.1177 | 0.111 | 0.111 | TRUE | 0.2075 |
| Bcl2a1a | 0 | 0 | 0 | 0 | 0 | 0 | 0 | 0 | 0 | 0 | 0 | 0 | 0 | 0 | 0 | NA | NA | NA | NA | NA | NA | NA | NA | NA | NA |
| Bcl2a1b | 0 | 0 | 0 | 0 | 0 | 0 | 0 | 0 | 0 | 0 | 0 | 0 | 0 | 0 | 0 | NA | NA | NA | NA | NA | NA | NA | NA | NA | NA |
| Bcl2a1c | 0 | 0 | 0 | 0 | 0 | 0 | 0 | 0 | 0 | 0 | 0 | 0 | 0 | 0 | 0 | NA | NA | NA | NA | NA | NA | NA | NA | NA | NA |
| Bcl2a1d | 0 | 0 | 0 | 0 | 0 | 0 | 0 | 0 | 0 | 0 | 0 | 0 | 0 | 0 | 0 | NA | NA | NA | NA | NA | NA | NA | NA | NA | NA |
| Bcl2l1 | 1542 | 1020 | 955 | 1027 | 787 | 954 | 1198 | 1085 | 1030 | 1022 | 888 | 942 | 1027.47 | 1104 | 951 | 0.86 | -0.218 | 0.01407637 | 0.05328687 | 0.0046 | 0.0047 | 0.0047 | 0.0047 | TRUE | 0.0676 |
| Bcl2l10 | 14 | 11 | 4 | 9 | 5 | 11 | 11 | 12 | 4 | 9 | 6 | 11 | 8.72 | 9 | 9 | 0.944 | -0.083 | 0.90194497 | NA | 0 | 0.2741 | 0.2123 | 0.2123 | TRUE | 0.5952 |
| Bcl2l11 | 1621 | 1233 | 1280 | 1298 | 1134 | 1383 | 1259 | 1312 | 1381 | 1292 | 1279 | 1365 | 1314.69 | 1317 | 1312 | 0.997 | -0.004 | 0.95784419 | 0.97691359 | 0.001 | 0.0042 | 0.003 | 0.003 | TRUE | 0.0215 |
| Bcl2l12 | 448 | 354 | 361 | 355 | 350 | 363 | 348 | 377 | 390 | 353 | 395 | 358 | 370.09 | 372 | 369 | 0.994 | -0.008 | 0.94148852 | 0.967203 | 0 | 0.0088 | 0.0064 | 0.0064 | TRUE | 0.0422 |
| Bcl2l13 | 7249 | 5029 | 5546 | 5953 | 5194 | 6124 | 5630 | 5349 | 5984 | 5926 | 5859 | 6045 | 5798.95 | 5654 | 5943 | 1.051 | 0.072 | 0.21656854 | 0.37929348 | 0.0014 | 0.0028 | 0.0023 | 0.0023 | TRUE | 0.0313 |
| Bcl2l14 | 527 | 368 | 382 | 237 | 228 | 255 | 409 | 391 | 412 | 236 | 257 | 252 | 326.29 | 404 | 248 | 0.614 | -0.705 | 9.57E-10 | 6.42E-08 | 0 | 0.0097 | 0.0063 | 0.0063 | TRUE | 0.0213 |
| Bcl2l15 | 1 | 1 | 1 | 0 | 1 | 1 | 1 | 1 | 1 | 0 | 1 | 1 | 0.84 | 1 | 1 | 0.722 | -0.47 | 0.81690682 | NA | 0 | 2.827 | 1.7221 | 1.7221 | TRUE | 0.2649 |
| Bcl2l2 | 929 | 683 | 907 | 1087 | 963 | 1135 | 722 | 726 | 979 | 1082 | 1086 | 1120 | 952.57 | 809 | 1096 | 1.357 | 0.441 | 7.52E-05 | 0.00097057 | 0.0151 | 0.0049 | 0.0079 | 0.0079 | TRUE | 0.3951 |
| Bcl3 | 3036 | 2415 | 2480 | 2007 | 1981 | 2395 | 2358 | 2569 | 2676 | 1998 | 2235 | 2364 | 2366.57 | 2534 | 2199 | 0.868 | -0.204 | 0.0101138 | 0.04165893 | 0.005 | 0.0034 | 0.0041 | 0.0041 | TRUE | 0.0781 |
| Bcl6 | 253 | 177 | 211 | 156 | 191 | 166 | 196 | 188 | 228 | 155 | 215 | 164 | 191.18 | 204 | 178 | 0.872 | -0.198 | 0.23516874 | 0.40153707 | 0.0147 | 0.0148 | 0.0148 | 0.0148 | TRUE | 0.3468 |
| Bcl6b | 450 | 338 | 281 | 369 | 344 | 451 | 349 | 360 | 303 | 367 | 388 | 445 | 368.8 | 337 | 400 | 1.185 | 0.245 | 0.04509826 | 0.12487202 | 0.0063 | 0.0089 | 0.0081 | 0.0081 | TRUE | 0.1135 |
| Bcl7a | 1034 | 868 | 916 | 899 | 792 | 974 | 803 | 923 | 988 | 895 | 893 | 961 | 910.75 | 905 | 916 | 1.015 | 0.022 | 0.81580352 | 0.89426181 | 0.0055 | 0.005 | 0.0052 | 0.0052 | TRUE | 0.1208 |
| Bcl7b | 1063 | 765 | 780 | 804 | 726 | 905 | 826 | 814 | 842 | 800 | 819 | 893 | 832.27 | 827 | 837 | 1.013 | 0.019 | 0.819343 | 0.89628574 | 6.00E-04 | 0.0053 | 0.0037 | 0.0037 | TRUE | 0.041 |
| Bcl7c | 1271 | 1035 | 965 | 1104 | 999 | 1142 | 987 | 1101 | 1041 | 1099 | 1127 | 1127 | 1080.42 | 1043 | 1118 | 1.073 | 0.101 | 0.18244914 | 0.33805362 | 8.00E-04 | 0.0046 | 0.0032 | 0.0032 | TRUE | 0.0282 |
| Bcl9 | 2584 | 1990 | 1785 | 1920 | 1724 | 1924 | 2007 | 2117 | 1926 | 1911 | 1945 | 1899 | 1967.49 | 2017 | 1918 | 0.951 | -0.072 | 0.26394327 | 0.43407614 | 6.00E-04 | 0.0036 | 0.0025 | 0.0025 | TRUE | 0.0222 |
| Bcl9l | 553 | 444 | 428 | 421 | 374 | 422 | 429 | 472 | 462 | 419 | 422 | 417 | 436.85 | 454 | 419 | 0.924 | -0.114 | 0.25957949 | 0.42956694 | 0 | 0.0079 | 0.0051 | 0.0051 | TRUE | 0.0291 |
| Bclaf1 | 16624 | 11233 | 7988 | 7343 | 6400 | 7213 | 12911 | 11948 | 8619 | 7309 | 7220 | 7120 | 9187.99 | 11159 | 7216 | 0.647 | -0.629 | 0.00018471 | 0.00200923 | 0.0203 | 0.0027 | 0.0086 | 0.0203 | TRUE | 0.4845 |
| Bcmo1 | 13 | 6 | 6 | 3 | 1 | 14 | 10 | 6 | 6 | 3 | 1 | 14 | 6.81 | 7 | 6 | 0.784 | -0.351 | 0.68351644 | NA | 0.482 | 0.3502 | 0.3859 | 0.3859 | TRUE | 3.1383 |
| Bco2 | 15 | 9 | 6 | 3 | 4 | 5 | 12 | 10 | 6 | 3 | 5 | 5 | 6.69 | 9 | 4 | 0.444 | -1.17 | 0.12502897 | NA | 0 | 0.3568 | 0.2422 | 0.2422 | TRUE | 0.2054 |
| Bcor | 2231 | 1847 | 1781 | 1799 | 1721 | 2054 | 1733 | 1965 | 1922 | 1791 | 1941 | 2028 | 1896.47 | 1873 | 1920 | 1.026 | 0.037 | 0.62910889 | 0.76644848 | 0.0037 | 0.0037 | 0.0037 | 0.0037 | TRUE | 0.0535 |
| Bcorl1 | 2098 | 1521 | 1195 | 1289 | 1170 | 1231 | 1629 | 1618 | 1289 | 1283 | 1320 | 1215 | 1392.47 | 1512 | 1273 | 0.841 | -0.25 | 0.00806077 | 0.03524398 | 0.0088 | 0.0041 | 0.0057 | 0.0057 | TRUE | 0.1975 |
| Bcr | 2517 | 1871 | 1856 | 1932 | 1814 | 2073 | 1955 | 1990 | 2003 | 1923 | 2046 | 2046 | 1993.91 | 1983 | 2005 | 1.012 | 0.017 | 0.78938156 | 0.87632082 | 0 | 0.0036 | 0.0023 | 0.0023 | TRUE | 0.0157 |
| Bcs1l | 1506 | 1017 | 1054 | 1021 | 934 | 1081 | 1170 | 1082 | 1137 | 1016 | 1054 | 1067 | 1087.62 | 1130 | 1046 | 0.925 | -0.113 | 0.1294538 | 0.26619936 | 0 | 0.0046 | 0.003 | 0.003 | TRUE | 0.0163 |
| Bdh1 | 1474 | 1061 | 1270 | 1404 | 1255 | 1515 | 1145 | 1129 | 1370 | 1398 | 1416 | 1496 | 1325.42 | 1215 | 1437 | 1.184 | 0.243 | 0.00531104 | 0.02565964 | 0.0057 | 0.0042 | 0.0047 | 0.0047 | TRUE | 0.1485 |
| Bdh2 | 252 | 157 | 182 | 190 | 185 | 222 | 196 | 167 | 196 | 189 | 209 | 219 | 196.01 | 186 | 206 | 1.102 | 0.14 | 0.34779693 | 0.52238982 | 0 | 0.0145 | 0.0108 | 0.0108 | TRUE | 0.0854 |
| Bdkrb1 | 6 | 2 | 4 | 1 | 2 | 6 | 5 | 2 | 4 | 1 | 2 | 6 | 3.38 | 4 | 3 | 0.824 | -0.28 | 0.79429931 | NA | 0 | 0.7037 | 0.5322 | 0.5322 | TRUE | 0.925 |
| Bdkrb2 | 38 | 31 | 37 | 44 | 44 | 47 | 30 | 33 | 40 | 44 | 50 | 46 | 40.37 | 34 | 47 | 1.372 | 0.457 | 0.13431877 | 0.27320595 | 0 | 0.0611 | 0.0417 | 0.0417 | TRUE | 0.1485 |
| Bdnf | 81 | 60 | 33 | 39 | 37 | 42 | 63 | 64 | 36 | 39 | 42 | 41 | 47.39 | 54 | 41 | 0.748 | -0.42 | 0.1727441 | 0.32507828 | 0.0317 | 0.0524 | 0.0466 | 0.0466 | TRUE | 0.7022 |
| Bdp1 | 3988 | 2972 | 2892 | 2994 | 2635 | 3101 | 3097 | 3161 | 3120 | 2980 | 2973 | 3061 | 3065.5 | 3126 | 3005 | 0.961 | -0.057 | 0.29332901 | 0.46521877 | 0 | 0.0032 | 0.0018 | 0.0018 | TRUE | 0.0033 |
| Bean1 | 8 | 6 | 4 | 13 | 8 | 10 | 6 | 6 | 4 | 13 | 9 | 10 | 8.12 | 5 | 11 | 1.876 | 0.908 | 0.17761699 | NA | 0 | 0.2941 | 0.1931 | 0.1931 | TRUE | 0.1355 |
| Becn1 | 4704 | 3471 | 3086 | 3319 | 2960 | 3465 | 3653 | 3692 | 3330 | 3304 | 3339 | 3420 | 3456.44 | 3558 | 3354 | 0.942 | -0.086 | 0.16485307 | 0.31559669 | 0.0013 | 0.0031 | 0.0024 | 0.0024 | TRUE | 0.0379 |
| Begain | 118 | 115 | 123 | 158 | 136 | 152 | 92 | 122 | 133 | 157 | 153 | 150 | 134.57 | 116 | 153 | 1.339 | 0.421 | 0.02382553 | 0.07855554 | 0.0106 | 0.02 | 0.0174 | 0.0174 | TRUE | 0.372 |
| Bend3 | 2398 | 1581 | 1204 | 1150 | 1057 | 1228 | 1862 | 1682 | 1299 | 1145 | 1192 | 1212 | 1398.76 | 1614 | 1183 | 0.732 | -0.45 | 4.96E-05 | 0.00069708 | 0.0152 | 0.0041 | 0.0081 | 0.0081 | TRUE | 0.3497 |
| Bend4 | 17263 | 12531 | 12148 | 13079 | 11556 | 12886 | 13407 | 13329 | 13108 | 13019 | 13036 | 12720 | 13103.29 | 13281 | 12925 | 0.973 | -0.039 | 0.38241264 | 0.55640702 | 1.00E-04 | 0.0026 | 0.0014 | 0.0014 | TRUE | 0.0024 |
| Bend5 | 147 | 120 | 114 | 126 | 115 | 124 | 114 | 128 | 123 | 125 | 130 | 122 | 123.73 | 122 | 126 | 1.037 | 0.052 | 0.76592111 | 0.86116938 | 0 | 0.0216 | 0.014 | 0.014 | TRUE | 0.0327 |
| Bend6 | 154 | 108 | 126 | 125 | 121 | 142 | 120 | 115 | 136 | 124 | 137 | 140 | 128.59 | 124 | 134 | 1.084 | 0.116 | 0.50823867 | 0.67156838 | 0 | 0.0209 | 0.0145 | 0.0145 | TRUE | 0.0746 |
| Bend7 | 385 | 257 | 283 | 298 | 271 | 317 | 299 | 273 | 305 | 297 | 306 | 313 | 298.83 | 292 | 305 | 1.042 | 0.059 | 0.61778512 | 0.75859223 | 0 | 0.0104 | 0.0069 | 0.0069 | TRUE | 0.0354 |
| Best1 | 57 | 36 | 39 | 53 | 52 | 53 | 44 | 38 | 42 | 53 | 59 | 52 | 48.06 | 41 | 55 | 1.309 | 0.388 | 0.15940508 | 0.30872315 | 0 | 0.0517 | 0.0337 | 0.0337 | TRUE | 0.0345 |
| Best2 | 58 | 45 | 63 | 95 | 66 | 102 | 45 | 48 | 68 | 95 | 74 | 101 | 71.77 | 54 | 90 | 1.691 | 0.758 | 0.0028176 | 0.01572508 | 0.0208 | 0.0355 | 0.0316 | 0.0316 | TRUE | 0.4359 |
| Best3 | 22 | 22 | 10 | 20 | 17 | 15 | 17 | 23 | 11 | 20 | 19 | 15 | 17.53 | 17 | 18 | 1.049 | 0.069 | 0.88427971 | 0.93584543 | 0 | 0.1376 | 0.1058 | 0.1058 | TRUE | 0.177 |
| Bet1 | 1127 | 802 | 827 | 935 | 777 | 982 | 875 | 853 | 892 | 931 | 877 | 969 | 899.55 | 873 | 926 | 1.06 | 0.084 | 0.29021467 | 0.46233151 | 0 | 0.0051 | 0.0035 | 0.0035 | TRUE | 0.0249 |
| Bet1l | 583 | 414 | 409 | 381 | 339 | 415 | 453 | 440 | 441 | 379 | 382 | 410 | 417.63 | 445 | 390 | 0.878 | -0.188 | 0.06647887 | 0.1650274 | 0 | 0.0081 | 0.0052 | 0.0052 | TRUE | 0.0217 |
| Bet3l | 1 | 1 | 0 | 1 | 0 | 0 | 1 | 1 | 0 | 1 | 0 | 0 | 0.47 | 1 | 0 | 0.599 | -0.74 | 0.79191656 | NA | 0 | 5.0171 | 3.8827 | 3.8827 | TRUE | 0.3906 |
| Beta-s | 0 | 0 | 0 | 0 | 0 | 0 | 0 | 0 | 0 | 0 | 0 | 0 | 0 | 0 | 0 | NA | NA | NA | NA | NA | NA | NA | NA | NA | NA |

| | | | | | | | | | | | | | | | | | | | | | | | | | |
|---|---|---|---|---|---|---|---|---|---|---|---|---|---|---|---|---|---|---|---|---|---|---|---|---|---|
| Bex1 | 12754 | 9113 | 8861 | 9212 | 7824 | 9322 | 9905 | 9693 | 9561 | 9170 | 8826 | 9202 | 9392.99 | 9720 | 9066 | 0.933 | -0.101 | 0.03774643 | 0.10981268 | 3.00E-04 | 0.0027 | 0.0016 | 0.0016 | TRUE | 0.0065 |
| Bex2 | 2101 | 1544 | 1265 | 1391 | 1248 | 1379 | 1632 | 1642 | 1365 | 1385 | 1408 | 1361 | 1465.46 | 1546 | 1385 | 0.895 | -0.161 | 0.05411892 | 0.14242712 | 0.0049 | 0.004 | 0.0043 | 0.0043 | TRUE | 0.1248 |
| Bex4 | 4205 | 3148 | 3120 | 3353 | 2988 | 3315 | 3266 | 3348 | 3366 | 3338 | 3371 | 3272 | 3326.92 | 3327 | 3327 | 1 | 0 | 0.99624398 | 0.99770986 | 0 | 0.0031 | 0.0018 | 0.0018 | TRUE | 0.0032 |
| Bex6 | 14 | 21 | 10 | 3 | 1 | 2 | 11 | 22 | 11 | 3 | 1 | 2 | 8.35 | 15 | 2 | 0.141 | -2.827 | 0.00083283 | NA | 0 | 0.2863 | 0.2311 | 0.2311 | TRUE | 0.8985 |
| Bfar | 2012 | 1447 | 1443 | 1656 | 1469 | 1701 | 1563 | 1539 | 1557 | 1648 | 1657 | 1679 | 1607.25 | 1553 | 1661 | 1.07 | 0.097 | 0.11993845 | 0.251619 | 0 | 0.0039 | 0.0022 | 0.0022 | TRUE | 0.001 |
| Bfsp1 | 52 | 41 | 35 | 35 | 34 | 36 | 40 | 44 | 38 | 35 | 38 | 36 | 38.42 | 41 | 36 | 0.892 | -0.165 | 0.59035083 | 0.73882577 | 0 | 0.0641 | 0.0412 | 0.0412 | TRUE | 0.0293 |
| Bfsp2 | 8 | 6 | 3 | 0 | 5 | 3 | 6 | 6 | 3 | 0 | 6 | 3 | 4.07 | 5 | 3 | 0.531 | -0.912 | 0.38438665 | NA | 0.1924 | 0.5844 | 0.5174 | 0.5174 | TRUE | 0.3911 |
| Bglap | 0 | 0 | 0 | 1 | 0 | 0 | 0 | 0 | 0 | 1 | 0 | 0 | 0.17 | 0 | 0 | 2.094 | 1.067 | 0.79379329 | NA | 0 | 14.2885 | 10 | 10 | TRUE | 0.3906 |
| Bglap2 | 0 | 0 | 0 | 0 | 0 | 0 | 0 | 0 | 0 | 0 | 0 | 0 | 0 | 0 | 0 | NA | NA | NA | NA | NA | NA | NA | NA | NA | NA |
| Bglap3 | 10 | 21 | 18 | 12 | 15 | 16 | 8 | 22 | 19 | 12 | 17 | 16 | 15.7 | 16 | 15 | 0.915 | -0.128 | 0.80962543 | NA | 0.1011 | 0.1534 | 0.1382 | 0.1382 | TRUE | 1.3164 |
| Bgn | 91 | 54 | 61 | 118 | 65 | 104 | 71 | 57 | 66 | 117 | 73 | 103 | 81.23 | 65 | 98 | 1.513 | 0.597 | 0.01311092 | 0.050497 | 0.0225 | 0.0316 | 0.029 | 0.029 | TRUE | 0.3414 |
| Bhlha15 | 410 | 353 | 428 | 610 | 620 | 702 | 318 | 375 | 462 | 607 | 699 | 693 | 525.89 | 385 | 666 | 1.736 | 0.796 | 1.00E-09 | 6.68E-08 | 0.0175 | 0.0069 | 0.0102 | 0.0102 | TRUE | 0.346 |
| Bhlha9 | 5 | 5 | 1 | 2 | 5 | 1 | 4 | 5 | 1 | 2 | 6 | 1 | 3.15 | 3 | 3 | 0.821 | -0.285 | 0.80191187 | NA | 0.22 | 0.7549 | 0.6082 | 0.6082 | TRUE | 0.6887 |
| Bhlhb9 | 3659 | 2526 | 2409 | 3013 | 2478 | 2846 | 2842 | 2687 | 2599 | 2999 | 2795 | 2809 | 2788.66 | 2709 | 2868 | 1.058 | 0.081 | 0.20063463 | 0.36134964 | 0.0013 | 0.0033 | 0.0026 | 0.0026 | TRUE | 0.0228 |
| Bhlhe22 | 38 | 17 | 12 | 8 | 9 | 16 | 30 | 18 | 13 | 8 | 10 | 16 | 15.74 | 20 | 11 | 0.554 | -0.851 | 0.1109784 | NA | 0.0959 | 0.153 | 0.1362 | 0.1362 | TRUE | 0.4299 |
| Bhlhe23 | 56 | 27 | 15 | 24 | 13 | 30 | 43 | 29 | 16 | 24 | 15 | 30 | 26.09 | 29 | 23 | 0.766 | -0.385 | 0.38934554 | 0.5633074 | 0.1356 | 0.0933 | 0.1053 | 0.1053 | TRUE | 0.1897 |
| Bhlhe40 | 5256 | 5383 | 7796 | 7819 | 7726 | 9557 | 4082 | 5726 | 8412 | 7783 | 8716 | 9434 | 7358.79 | 6073 | 8644 | 1.423 | 0.509 | 0.10037842 | 0.22161088 | 0.0691 | 0.0027 | 0.0192 | 0.0691 | TRUE | 0.5461 |
| Bhlhe41 | 58 | 65 | 62 | 100 | 89 | 110 | 45 | 69 | 67 | 100 | 100 | 109 | 81.6 | 60 | 103 | 1.722 | 0.784 | 0.00081189 | 0.0062222 | 0.0116 | 0.0315 | 0.0264 | 0.0264 | TRUE | 0.5023 |
| Bhmt | 91 | 59 | 37 | 58 | 46 | 39 | 71 | 63 | 40 | 58 | 52 | 38 | 53.58 | 58 | 49 | 0.848 | -0.237 | 0.42884028 | 0.60086404 | 0.0446 | 0.0467 | 0.0461 | 0.0461 | TRUE | 0.5932 |
| Bhmt2 | 87 | 60 | 67 | 93 | 77 | 64 | 68 | 64 | 72 | 93 | 87 | 63 | 74.38 | 68 | 81 | 1.19 | 0.251 | 0.29157134 | 0.4636659 | 0.0093 | 0.0343 | 0.0273 | 0.0273 | TRUE | 0.3513 |
| Bicc1 | 363 | 278 | 277 | 285 | 262 | 247 | 282 | 296 | 299 | 284 | 296 | 244 | 283.27 | 292 | 275 | 0.939 | -0.091 | 0.4753298 | 0.64342464 | 0.0017 | 0.0108 | 0.0081 | 0.0081 | TRUE | 0.1078 |
| Bicd1 | 206 | 133 | 82 | 104 | 86 | 84 | 160 | 141 | 88 | 104 | 97 | 83 | 112.23 | 130 | 95 | 0.723 | -0.468 | 0.04368257 | 0.12204243 | 0.0442 | 0.0235 | 0.0296 | 0.0296 | TRUE | 0.6801 |
| Bicd2 | 2179 | 1602 | 1500 | 1578 | 1340 | 1453 | 1692 | 1704 | 1618 | 1571 | 1512 | 1434 | 1588.6 | 1671 | 1506 | 0.9 | -0.151 | 0.02706089 | 0.08591231 | 8.00E-04 | 0.0039 | 0.0027 | 0.0027 | TRUE | 0.021 |
| Bid | 4075 | 3116 | 3195 | 3722 | 3417 | 3884 | 3165 | 3314 | 3447 | 3705 | 3855 | 3834 | 3553.4 | 3309 | 3798 | 1.148 | 0.2 | 0.00062916 | 0.00515332 | 9.00E-04 | 0.0031 | 0.0022 | 0.0022 | TRUE | 0.0181 |
| Bik | 178 | 142 | 141 | 149 | 171 | 188 | 138 | 151 | 152 | 148 | 193 | 186 | 161.37 | 147 | 176 | 1.194 | 0.256 | 0.12349818 | 0.257462 | 0.0054 | 0.0171 | 0.0137 | 0.0137 | TRUE | 0.2003 |
| Bin1 | 437 | 328 | 297 | 399 | 370 | 365 | 339 | 349 | 320 | 397 | 417 | 360 | 363.94 | 336 | 391 | 1.163 | 0.218 | 0.05426857 | 0.14268959 | 0 | 0.0089 | 0.0065 | 0.0065 | TRUE | 0.0568 |
| Bin2 | 2428 | 1813 | 1876 | 2062 | 1965 | 2307 | 1886 | 1928 | 2024 | 2053 | 2217 | 2277 | 2064.16 | 1946 | 2182 | 1.122 | 0.166 | 0.01459737 | 0.05464647 | 0.0016 | 0.0036 | 0.0028 | 0.0028 | TRUE | 0.033 |
| Bin3 | 581 | 455 | 409 | 369 | 337 | 384 | 451 | 484 | 441 | 367 | 380 | 379 | 417.18 | 459 | 375 | 0.819 | -0.289 | 0.00531217 | 0.02565964 | 0 | 0.0081 | 0.0053 | 0.0053 | TRUE | 0.0256 |
| Birc2 | 2728 | 2025 | 2286 | 2649 | 2273 | 2416 | 2119 | 2154 | 2467 | 2637 | 2564 | 2385 | 2387.54 | 2247 | 2529 | 1.126 | 0.172 | 0.02475212 | 0.08080904 | 0.0043 | 0.0034 | 0.0038 | 0.0038 | TRUE | 0.0882 |
| Birc3 | 303 | 190 | 186 | 242 | 210 | 222 | 235 | 202 | 201 | 241 | 237 | 219 | 222.51 | 213 | 232 | 1.088 | 0.122 | 0.38493828 | 0.55896808 | 0 | 0.0131 | 0.0096 | 0.0096 | TRUE | 0.103 |
| Birc5 | 10658 | 7583 | 7289 | 7699 | 6753 | 7973 | 8278 | 8066 | 7865 | 7664 | 7618 | 7871 | 7893.43 | 8070 | 7718 | 0.956 | -0.065 | 0.19249999 | 0.35094418 | 3.00E-04 | 0.0027 | 0.0016 | 0.0016 | TRUE | 0.0063 |
| Birc6 | 10221 | 6714 | 5704 | 6882 | 6068 | 6948 | 7938 | 7142 | 6155 | 6850 | 6845 | 6859 | 6964.8 | 7078 | 6851 | 0.968 | -0.047 | 0.55944527 | 0.71432475 | 0.0068 | 0.0028 | 0.0046 | 0.0046 | TRUE | 0.1588 |
| Birc7 | 29 | 21 | 24 | 18 | 14 | 18 | 23 | 22 | 26 | 18 | 16 | 18 | 20.37 | 24 | 17 | 0.73 | -0.454 | 0.27891556 | 0.45050741 | 0 | 0.1188 | 0.0761 | 0.0761 | TRUE | 0.0403 |
| Bivm | 1663 | 1189 | 1374 | 1639 | 1313 | 1463 | 1292 | 1265 | 1483 | 1631 | 1481 | 1444 | 1432.62 | 1347 | 1519 | 1.129 | 0.175 | 0.03814382 | 0.1105637 | 0.005 | 0.0041 | 0.0044 | 0.0044 | TRUE | 0.0926 |
| Blcap | 1369 | 1021 | 1004 | 1099 | 999 | 1152 | 1063 | 1086 | 1083 | 1094 | 1127 | 1137 | 1098.45 | 1077 | 1119 | 1.039 | 0.055 | 0.43551955 | 0.60705138 | 0 | 0.0046 | 0.0027 | 0.0027 | TRUE | 0.0048 |
| Blk | 0 | 1 | 0 | 0 | 0 | 0 | 0 | 1 | 0 | 0 | 0 | 0 | 0.18 | 0 | 0 | 0.552 | -0.857 | 0.8336548 | NA | 0 | 13.3714 | 10 | 10 | TRUE | 0.5216 |
| Blm | 11850 | 9080 | 9185 | 10383 | 9008 | 10182 | 9203 | 9658 | 9911 | 10335 | 10162 | 10051 | 9886.77 | 9591 | 10183 | 1.062 | 0.087 | 0.08927455 | 0.2036997 | 6.00E-04 | 0.0027 | 0.0018 | 0.0018 | TRUE | 0.0154 |
| Blmh | 13713 | 9480 | 8690 | 9840 | 8283 | 9561 | 10650 | 10084 | 9376 | 9795 | 9344 | 9438 | 9781.25 | 10037 | 9526 | 0.949 | -0.076 | 0.20732607 | 0.36908856 | 0.0022 | 0.0027 | 0.0025 | 0.0025 | TRUE | 0.0404 |
| Blnk | 123 | 84 | 107 | 77 | 75 | 88 | 96 | 89 | 115 | 77 | 85 | 87 | 91.41 | 100 | 83 | 0.827 | -0.273 | 0.19266956 | 0.35111896 | 0 | 0.0284 | 0.0207 | 0.0207 | TRUE | 0.1656 |
| Bloc1s1 | 959 | 649 | 645 | 677 | 591 | 631 | 745 | 690 | 696 | 674 | 667 | 623 | 682.43 | 710 | 655 | 0.92 | -0.12 | 0.17015397 | 0.32177966 | 0 | 0.0059 | 0.0041 | 0.0041 | TRUE | 0.0225 |
| Bloc1s2a | 2090 | 1570 | 1497 | 1611 | 1363 | 1580 | 1623 | 1670 | 1615 | 1604 | 1538 | 1560 | 1601.56 | 1636 | 1567 | 0.958 | -0.062 | 0.33412816 | 0.50818557 | 0 | 0.0039 | 0.0023 | 0.0023 | TRUE | 0.0051 |
| Bloc1s3 | 580 | 448 | 499 | 592 | 550 | 616 | 450 | 477 | 538 | 589 | 620 | 608 | 547.21 | 488 | 606 | 1.243 | 0.314 | 0.00178628 | 0.01109093 | 0.0025 | 0.0068 | 0.0054 | 0.0054 | TRUE | 0.0918 |
| Bloc1s4 | 697 | 539 | 470 | 558 | 485 | 558 | 541 | 573 | 507 | 555 | 547 | 551 | 545.86 | 540 | 551 | 1.019 | 0.028 | 0.76862872 | 0.86320502 | 0 | 0.0068 | 0.0046 | 0.0046 | TRUE | 0.0323 |
| Bloc1s5 | 686 | 513 | 520 | 534 | 496 | 566 | 533 | 546 | 561 | 532 | 560 | 559 | 548.22 | 547 | 550 | 1.007 | 0.01 | 0.91462047 | 0.9529197 | 0 | 0.0067 | 0.0043 | 0.0043 | TRUE | 0.0102 |
| Bloc1s6 | 2279 | 1511 | 1546 | 1348 | 1240 | 1372 | 1770 | 1607 | 1668 | 1342 | 1399 | 1354 | 1523.4 | 1682 | 1365 | 0.811 | -0.303 | 1.35E-05 | 0.00024122 | 0.001 | 0.004 | 0.0028 | 0.0028 | TRUE | 0.0264 |
| Blvra | 645 | 406 | 419 | 466 | 436 | 471 | 501 | 432 | 452 | 464 | 492 | 465 | 467.59 | 462 | 474 | 1.023 | 0.033 | 0.75273614 | 0.85340348 | 0.0017 | 0.0075 | 0.0056 | 0.0056 | TRUE | 0.0687 |
| Blvrb | 1153 | 896 | 1003 | 1202 | 1140 | 1337 | 895 | 953 | 1082 | 1196 | 1286 | 1320 | 1122.19 | 977 | 1267 | 1.299 | 0.378 | 1.86E-05 | 0.00031463 | 0.0047 | 0.0045 | 0.0047 | 0.0047 | TRUE | 0.1053 |
| Blzf1 | 1777 | 1331 | 1267 | 1221 | 1125 | 1315 | 1380 | 1416 | 1367 | 1215 | 1269 | 1298 | 1324.26 | 1388 | 1261 | 0.909 | -0.138 | 0.04465209 | 0.12399694 | 0 | 0.0042 | 0.0027 | 0.0027 | TRUE | 0.0122 |
| Bmf | 1586 | 1205 | 1657 | 1899 | 1884 | 2100 | 1232 | 1282 | 1788 | 1890 | 2125 | 2073 | 1731.67 | 1434 | 2029 | 1.417 | 0.503 | 2.18E-05 | 0.00035797 | 0.024 | 0.0038 | 0.0095 | 0.0095 | TRUE | 0.5587 |
| Bmi1 | 1211 | 915 | 816 | 801 | 602 | 701 | 941 | 973 | 880 | 797 | 679 | 692 | 827.12 | 931 | 723 | 0.776 | -0.365 | 6.46E-05 | 0.00086276 | 0.0035 | 0.0053 | 0.0048 | 0.0048 | TRUE | 0.0969 |
| Bmp1 | 400 | 255 | 239 | 242 | 191 | 222 | 311 | 271 | 258 | 241 | 215 | 219 | 252.55 | 280 | 225 | 0.802 | -0.318 | 0.01917677 | 0.06698277 | 0.0036 | 0.0118 | 0.0092 | 0.0092 | TRUE | 0.1109 |
| Bmp10 | 0 | 0 | 0 | 1 | 1 | 2 | 0 | 0 | 0 | 1 | 1 | 2 | 0.68 | 0 | 1 | 7.73 | 2.951 | 0.21871019 | NA | 0 | 3.4727 | 2.7687 | 2.7687 | TRUE | 0.0985 |
| Bmp15 | 4 | 2 | 3 | 0 | 0 | 0 | 3 | 2 | 3 | 0 | 0 | 0 | 1.41 | 3 | 0 | 0.067 | -3.893 | 0.03710637 | NA | 0 | 1.6811 | 1.3428 | 1.3428 | TRUE | 0.0718 |
| Bmp2 | 27 | 10 | 22 | 20 | 13 | 16 | 21 | 11 | 24 | 20 | 15 | 16 | 17.62 | 19 | 17 | 0.908 | -0.139 | 0.77165579 | 0.86486091 | 0 | 0.1369 | 0.1088 | 0.1088 | TRUE | 0.6548 |
| Bmp2k | 625 | 528 | 475 | 526 | 430 | 533 | 485 | 562 | 513 | 524 | 485 | 526 | 515.73 | 520 | 512 | 0.986 | -0.02 | 0.84509315 | 0.91202932 | 0.002 | 0.007 | 0.0054 | 0.0054 | TRUE | 0.0573 |
| Bmp3 | 1 | 1 | 1 | 0 | 1 | 1 | 1 | 1 | 1 | 0 | 1 | 1 | 0.84 | 1 | 1 | 0.722 | -0.47 | 0.81690682 | NA | 0 | 2.827 | 1.7221 | 1.7221 | TRUE | 0.2649 |
| Bmp4 | 1983 | 1502 | 1495 | 2178 | 1914 | 2185 | 1540 | 1598 | 1613 | 2168 | 2159 | 2157 | 1872.49 | 1584 | 2161 | 1.366 | 0.45 | 2.19E-13 | 3.07E-11 | 0 | 0.0037 | 0.0022 | 0.0022 | TRUE | 0.0073 |
| Bmp5 | 0 | 0 | 0 | 0 | 0 | 0 | 0 | 0 | 0 | 0 | 0 | 0 | 0 | 0 | 0 | NA | NA | NA | NA | NA | NA | NA | NA | NA | NA |
| Bmp6 | 120 | 82 | 77 | 107 | 89 | 83 | 93 | 87 | 83 | 107 | 100 | 82 | 92.06 | 88 | 96 | 1.093 | 0.128 | 0.53917075 | 0.69753507 | 0 | 0.0282 | 0.0205 | 0.0205 | TRUE | 0.1697 |
| Bmp7 | 719 | 619 | 830 | 934 | 864 | 1039 | 558 | 658 | 896 | 930 | 975 | 1026 | 840.41 | 704 | 977 | 1.39 | 0.475 | 0.00049846 | 0.00429704 | 0.0296 | 0.0052 | 0.0122 | 0.0122 | TRUE | 0.6663 |
| Bmp8a | 98 | 91 | 39 | 102 | 99 | 72 | 76 | 97 | 42 | 102 | 112 | 71 | 83.21 | 72 | 95 | 1.318 | 0.398 | 0.16675152 | 0.31778393 | 0.093 | 0.0309 | 0.0476 | 0.0476 | TRUE | 0.4738 |
| Bmp8b | 235 | 185 | 209 | 281 | 268 | 259 | 183 | 197 | 226 | 280 | 302 | 256 | 240.42 | 202 | 279 | 1.388 | 0.473 | 0.00081531 | 0.00623837 | 0.005 | 0.0123 | 0.0101 | 0.0101 | TRUE | 0.115 |
| Bmper | 39 | 20 | 14 | 15 | 21 | 20 | 30 | 21 | 15 | 15 | 24 | 20 | 20.84 | 22 | 20 | 0.862 | -0.214 | 0.63185536 | 0.76906316 | 0.0436 | 0.1162 | 0.0954 | 0.0954 | TRUE | 0.3105 |
| Bmpr1a | 5940 | 4215 | 4364 | 4612 | 3884 | 4468 | 4613 | 4483 | 4709 | 4591 | 4382 | 4411 | 4531.41 | 4602 | 4461 | 0.969 | -0.045 | 0.40292088 | 0.57589577 | 3.00E-04 | 0.0029 | 0.0018 | 0.0018 | TRUE | 0.0079 |
| Bmpr1b | 115 | 86 | 91 | 74 | 90 | 110 | 89 | 91 | 98 | 74 | 102 | 109 | 93.79 | 93 | 95 | 1.018 | 0.025 | 0.90684226 | 0.94912712 | 0.0105 | 0.0277 | 0.023 | 0.023 | TRUE | 0.3705 |
| Bmpr2 | 956 | 715 | 628 | 731 | 565 | 653 | 742 | 761 | 678 | 728 | 637 | 645 | 698.38 | 727 | 670 | 0.922 | -0.118 | 0.20900541 | 0.37076598 | 0.003 | 0.0058 | 0.0049 | 0.0049 | TRUE | 0.0677 |
| Bms1 | 22809 | 16605 | 15396 | 17427 | 15204 | 17830 | 17715 | 17663 | 16612 | 17347 | 17152 | 17601 | 17348.15 | 17330 | 17367 | 1.002 | 0.003 | 0.95190398 | 0.97342517 | 5.00E-04 | 0.0026 | 0.0017 | 0.0017 | TRUE | 0.016 |
| Bmx | 0 | 0 | 0 | 0 | 0 | 0 | 0 | 0 | 0 | 0 | 0 | 0 | 0 | 0 | 0 | NA | NA | NA | NA | NA | NA | NA | NA | NA | NA |
| Bmyc | 790 | 633 | 638 | 697 | 614 | 643 | 614 | 673 | 688 | 694 | 693 | 635 | 666.08 | 658 | 674 | 1.025 | 0.035 | 0.70196784 | 0.81950718 | 0.0018 | 0.006 | 0.0046 | 0.0046 | TRUE | 0.0445 |

| | | | | | | | | | | | | | | | | | | | | | | | | | |
|---|---|---|---|---|---|---|---|---|---|---|---|---|---|---|---|---|---|---|---|---|---|---|---|---|---|
| Bnc1 | 35 | 29 | 24 | 22 | 33 | 29 | 27 | 31 | 26 | 22 | 37 | 29 | 28.61 | 28 | 29 | 1.042 | 0.06 | 0.87149092 | 0.92866769 | 0 | 0.0853 | 0.0629 | 0.0629 | TRUE | 0.2334 |
| Bnc2 | 1034 | 839 | 858 | 930 | 885 | 1010 | 803 | 892 | 926 | 926 | 998 | 997 | 923.73 | 874 | 974 | 1.116 | 0.159 | 0.06371365 | 0.16029909 | 0.0027 | 0.005 | 0.0042 | 0.0042 | TRUE | 0.0628 |
| Bnip1 | 945 | 714 | 763 | 821 | 775 | 834 | 734 | 759 | 823 | 817 | 874 | 823 | 805.25 | 772 | 838 | 1.086 | 0.12 | 0.15934147 | 0.30864178 | 0.0012 | 0.0054 | 0.004 | 0.004 | TRUE | 0.0391 |
| Bnip2 | 1880 | 1376 | 1189 | 1245 | 1030 | 1232 | 1460 | 1464 | 1283 | 1239 | 1162 | 1216 | 1304.01 | 1402 | 1206 | 0.86 | -0.218 | 0.00533545 | 0.02574597 | 0.0026 | 0.0042 | 0.0037 | 0.0037 | TRUE | 0.0653 |
| Bnip3 | 9654 | 7982 | 11115 | 11354 | 10516 | 12210 | 7498 | 8490 | 11993 | 11302 | 11863 | 12053 | 10533.21 | 9327 | 11739 | 1.259 | 0.332 | 0.11833917 | 0.24976694 | 0.0325 | 0.0027 | 0.0107 | 0.0325 | TRUE | 0.7636 |
| Bnip3l | 8817 | 6897 | 7855 | 8331 | 7467 | 8856 | 6848 | 7336 | 8475 | 8293 | 8424 | 8742 | 8019.66 | 7553 | 8486 | 1.124 | 0.168 | 0.02626568 | 0.08426833 | 0.0063 | 0.0027 | 0.004 | 0.004 | TRUE | 0.1391 |
| Bnipl | 34 | 20 | 28 | 39 | 36 | 47 | 26 | 21 | 30 | 39 | 41 | 46 | 33.95 | 26 | 42 | 1.616 | 0.692 | 0.03851881 | 0.1114247 | 0 | 0.0722 | 0.0497 | 0.0497 | TRUE | 0.1394 |
| Boc | 382 | 334 | 341 | 400 | 413 | 489 | 297 | 355 | 368 | 398 | 466 | 483 | 394.45 | 340 | 449 | 1.324 | 0.405 | 0.00120085 | 0.00825972 | 0.0093 | 0.0084 | 0.0087 | 0.0087 | TRUE | 0.152 |
| Bod1 | 4450 | 3257 | 3086 | 3301 | 2922 | 3404 | 3456 | 3464 | 3330 | 3286 | 3296 | 3360 | 3365.45 | 3417 | 3314 | 0.97 | -0.044 | 0.41363075 | 0.58616145 | 0 | 0.0031 | 0.0018 | 0.0018 | TRUE | 0.0059 |
| Bod1l | 6943 | 4905 | 4428 | 5588 | 4792 | 5557 | 5392 | 5217 | 4778 | 5562 | 5406 | 5486 | 5306.88 | 5129 | 5485 | 1.069 | 0.096 | 0.11196437 | 0.24021415 | 0.0017 | 0.0029 | 0.0024 | 0.0024 | TRUE | 0.0435 |
| Bok | 809 | 540 | 445 | 476 | 419 | 497 | 628 | 574 | 480 | 474 | 473 | 491 | 519.99 | 561 | 479 | 0.852 | -0.231 | 0.04095432 | 0.11642008 | 0.0079 | 0.007 | 0.0073 | 0.0073 | TRUE | 0.1831 |
| Bola1 | 975 | 695 | 606 | 691 | 631 | 644 | 757 | 739 | 654 | 688 | 712 | 636 | 697.63 | 717 | 679 | 0.945 | -0.082 | 0.3852958 | 0.55920317 | 0.0033 | 0.0058 | 0.005 | 0.005 | TRUE | 0.0682 |
| Bola2 | 2296 | 1576 | 1294 | 1455 | 1140 | 1368 | 1783 | 1676 | 1396 | 1448 | 1286 | 1350 | 1490.09 | 1618 | 1361 | 0.841 | -0.251 | 0.00792863 | 0.03484743 | 0.0084 | 0.004 | 0.0057 | 0.0057 | TRUE | 0.1727 |
| Bola3 | 706 | 467 | 437 | 497 | 393 | 453 | 548 | 497 | 472 | 495 | 443 | 447 | 483.64 | 506 | 462 | 0.912 | -0.133 | 0.20695169 | 0.36873105 | 0.0031 | 0.0073 | 0.0059 | 0.0059 | TRUE | 0.0676 |
| Boll | 42 | 53 | 45 | 41 | 31 | 31 | 33 | 56 | 49 | 41 | 35 | 31 | 40.66 | 46 | 36 | 0.78 | -0.358 | 0.27350555 | 0.44443097 | 0.0306 | 0.0607 | 0.0518 | 0.0518 | TRUE | 0.4962 |
| Bop1 | 9037 | 6572 | 5381 | 5558 | 4799 | 5829 | 7019 | 6991 | 5806 | 5532 | 5414 | 5754 | 6085.93 | 6605 | 5567 | 0.843 | -0.247 | 0.00118541 | 0.00819694 | 0.0057 | 0.0028 | 0.004 | 0.004 | TRUE | 0.1361 |
| Bora | 2704 | 1933 | 1656 | 1846 | 1609 | 1746 | 2100 | 2056 | 1787 | 1838 | 1815 | 1724 | 1886.53 | 1981 | 1792 | 0.904 | -0.146 | 0.05841356 | 0.15071711 | 0.0034 | 0.0037 | 0.0037 | 0.0037 | TRUE | 0.0878 |
| Bpgm | 1417 | 946 | 805 | 991 | 876 | 1077 | 1101 | 1006 | 869 | 986 | 988 | 1063 | 1002.2 | 992 | 1012 | 1.019 | 0.028 | 0.77104839 | 0.86442816 | 0.007 | 0.0048 | 0.0055 | 0.0055 | TRUE | 0.1396 |
| Bphl | 664 | 464 | 407 | 504 | 426 | 459 | 516 | 494 | 439 | 502 | 481 | 453 | 480.63 | 483 | 479 | 0.989 | -0.016 | 0.87995097 | 0.93298562 | 0.0027 | 0.0074 | 0.0059 | 0.0059 | TRUE | 0.0712 |
| Bpi | 10 | 6 | 12 | 13 | 8 | 11 | 8 | 6 | 13 | 13 | 9 | 11 | 9.99 | 9 | 11 | 1.222 | 0.29 | 0.63579616 | NA | 0 | 0.2397 | 0.1688 | 0.1688 | TRUE | 0.4201 |
| Bpifa1 | 0 | 0 | 0 | 0 | 0 | 0 | 0 | 0 | 0 | 0 | 0 | 0 | 0 | 0 | 0 | NA | NA | NA | NA | NA | NA | NA | NA | NA | NA |
| Bpifa2 | 1 | 0 | 1 | 0 | 0 | 0 | 1 | 0 | 1 | 0 | 0 | 0 | 0.31 | 1 | 0 | 0.307 | -1.706 | 0.62894649 | NA | 0 | 7.6657 | 7.2 | 7.2 | TRUE | 0.2067 |
| Bpifa3 | 0 | 0 | 0 | 0 | 0 | 0 | 0 | 0 | 0 | 0 | 0 | 0 | 0 | 0 | 0 | NA | NA | NA | NA | NA | NA | NA | NA | NA | NA |
| Bpifa5 | 0 | 0 | 0 | 0 | 0 | 0 | 0 | 0 | 0 | 0 | 0 | 0 | 0 | 0 | 0 | NA | NA | NA | NA | NA | NA | NA | NA | NA | NA |
| Bpifa6 | 0 | 0 | 0 | 0 | 0 | 0 | 0 | 0 | 0 | 0 | 0 | 0 | 0 | 0 | 0 | NA | NA | NA | NA | NA | NA | NA | NA | NA | NA |
| Bpifb1 | 8 | 6 | 1 | 3 | 3 | 3 | 6 | 6 | 1 | 3 | 3 | 3 | 3.83 | 4 | 3 | 0.673 | -0.572 | 0.57105398 | NA | 0 | 0.6205 | 0.4628 | 0.4628 | TRUE | 0.7498 |
| Bpifb2 | 1 | 2 | 1 | 0 | 0 | 1 | 1 | 2 | 1 | 0 | 0 | 1 | 0.83 | 1 | 0 | 0.282 | -1.824 | 0.39899776 | NA | 0 | 2.8636 | 2.0027 | 2.0027 | TRUE | 0.3835 |
| Bpifb3 | 0 | 1 | 0 | 0 | 0 | 0 | 0 | 1 | 0 | 0 | 0 | 0 | 0.18 | 0 | 0 | 0.552 | -0.857 | 0.8336548 | NA | 0 | 13.3714 | 10 | 10 | TRUE | 0.5216 |
| Bpifb4 | 110 | 89 | 65 | 70 | 62 | 87 | 85 | 95 | 70 | 70 | 70 | 86 | 79.29 | 83 | 75 | 0.901 | -0.15 | 0.51171696 | 0.67469683 | 0 | 0.0323 | 0.025 | 0.025 | TRUE | 0.169 |
| Bpifb5 | 36 | 20 | 9 | 21 | 18 | 19 | 28 | 21 | 10 | 21 | 20 | 19 | 19.82 | 20 | 20 | 1.003 | 0.004 | 0.99285416 | 0.99592003 | 0.0738 | 0.122 | 0.109 | 0.109 | TRUE | 0.3898 |
| Bpifb6 | 0 | 0 | 0 | 1 | 0 | 0 | 0 | 0 | 0 | 1 | 0 | 0 | 0.17 | 0 | 0 | 2.094 | 1.067 | 0.79379329 | NA | 0 | 14.2885 | 10 | 10 | TRUE | 0.3906 |
| Bpifb9a | 0 | 0 | 0 | 0 | 0 | 0 | 0 | 0 | 0 | 0 | 0 | 0 | 0 | 0 | 0 | NA | NA | NA | NA | NA | NA | NA | NA | NA | NA |
| Bpifb9b | 1 | 0 | 0 | 0 | 0 | 0 | 1 | 0 | 0 | 0 | 0 | 0 | 0.13 | 0 | 0 | 0.552 | -0.857 | 0.8336548 | NA | 0 | 18.3121 | 10 | 10 | TRUE | 0.2566 |
| Bpifc | 2 | 2 | 0 | 2 | 1 | 0 | 2 | 2 | 0 | 2 | 1 | 0 | 1.13 | 1 | 1 | 0.837 | -0.257 | 0.891055 | NA | 0 | 2.0938 | 1.6517 | 1.6517 | TRUE | 0.3105 |
| Bpnt1 | 2494 | 1826 | 1673 | 1673 | 1377 | 1717 | 1937 | 1942 | 1805 | 1665 | 1553 | 1695 | 1766.35 | 1895 | 1638 | 0.864 | -0.21 | 0.00224823 | 0.01322383 | 0.0013 | 0.0038 | 0.0028 | 0.0028 | TRUE | 0.0241 |
| Bptf | 17150 | 12484 | 11633 | 12776 | 11494 | 13923 | 13320 | 13279 | 12552 | 12717 | 12966 | 13744 | 13096.41 | 13050 | 13142 | 1.007 | 0.01 | 0.85174456 | 0.91640026 | 0.001 | 0.0026 | 0.002 | 0.002 | TRUE | 0.0197 |
| Braf | 4038 | 2919 | 3076 | 3831 | 3258 | 3793 | 3136 | 3105 | 3319 | 3813 | 3675 | 3744 | 3465.51 | 3187 | 3744 | 1.175 | 0.233 | 3.88E-05 | 0.00057527 | 4.00E-04 | 0.0031 | 0.002 | 0.002 | TRUE | 0.0158 |
| Brap | 3171 | 2210 | 2147 | 2114 | 1882 | 2346 | 2463 | 2351 | 2317 | 2104 | 2123 | 2316 | 2278.89 | 2377 | 2181 | 0.917 | -0.124 | 0.05978196 | 0.15285573 | 0.0013 | 0.0035 | 0.0027 | 0.0027 | TRUE | 0.0357 |
| Brat1 | 8 | 5 | 6 | 9 | 4 | 3 | 6 | 5 | 6 | 9 | 5 | 3 | 5.74 | 6 | 6 | 0.914 | -0.13 | 0.87186589 | NA | 0 | 0.4154 | 0.2943 | 0.2943 | TRUE | 0.697 |
| Brca1 | 10911 | 7917 | 8194 | 8310 | 7364 | 8491 | 8474 | 8421 | 8841 | 8272 | 8307 | 8382 | 8449.6 | 8579 | 8320 | 0.97 | -0.044 | 0.36333194 | 0.53786674 | 2.00E-04 | 0.0027 | 0.0016 | 0.0016 | TRUE | 0.0087 |
| Brca2 | 10554 | 7728 | 6795 | 7991 | 7201 | 8389 | 8197 | 8220 | 7332 | 7954 | 8123 | 8281 | 8017.95 | 7916 | 8119 | 1.026 | 0.036 | 0.54647823 | 0.70367688 | 0.0018 | 0.0027 | 0.0025 | 0.0025 | TRUE | 0.0506 |
| Brcc3 | 1484 | 1050 | 1033 | 1034 | 887 | 1051 | 1153 | 1117 | 1115 | 1029 | 1001 | 1037 | 1075.23 | 1128 | 1022 | 0.906 | -0.142 | 0.04785395 | 0.13052987 | 0 | 0.0046 | 0.0028 | 0.0028 | TRUE | 0.0046 |
| Brd1 | 8744 | 6780 | 7115 | 6902 | 6250 | 6957 | 6791 | 7212 | 7677 | 6870 | 7051 | 6868 | 7078.07 | 7227 | 6930 | 0.959 | -0.06 | 0.30991899 | 0.48274544 | 0.0018 | 0.0028 | 0.0024 | 0.0024 | TRUE | 0.0363 |
| Brd2 | 19719 | 14187 | 12825 | 12916 | 11511 | 12943 | 15315 | 15091 | 13838 | 12857 | 12986 | 12777 | 13810.4 | 14748 | 12873 | 0.873 | -0.196 | 0.00034201 | 0.00320551 | 0.0014 | 0.0026 | 0.0021 | 0.0021 | TRUE | 0.0355 |
| Brd3 | 8305 | 6361 | 6048 | 5961 | 5458 | 6233 | 6450 | 6766 | 6526 | 5934 | 6157 | 6153 | 6330.95 | 6581 | 6081 | 0.924 | -0.114 | 0.02569355 | 0.08285449 | 3.00E-04 | 0.0028 | 0.0017 | 0.0017 | TRUE | 0.0074 |
| Brd4 | 13687 | 10195 | 9374 | 9543 | 8726 | 10400 | 10630 | 10844 | 10114 | 9499 | 9844 | 10266 | 10199.7 | 10529 | 9870 | 0.937 | -0.093 | 0.08789164 | 0.20135202 | 0.0012 | 0.0027 | 0.0021 | 0.0021 | TRUE | 0.0152 |
| Brd7 | 9234 | 6863 | 6661 | 6789 | 5831 | 6925 | 7172 | 7300 | 7187 | 6758 | 6578 | 6836 | 6971.78 | 7220 | 6724 | 0.931 | -0.102 | 0.03374422 | 0.1011336 | 1.00E-04 | 0.0028 | 0.0015 | 0.0015 | TRUE | 0.0044 |
| Brd8 | 7838 | 5842 | 5534 | 5840 | 5146 | 6017 | 6087 | 6214 | 5971 | 5813 | 5805 | 5940 | 5971.78 | 6091 | 5853 | 0.961 | -0.058 | 0.24459358 | 0.41279679 | 1.00E-04 | 0.0028 | 0.0016 | 0.0016 | TRUE | 0.0038 |
| Brd9 | 1552 | 1076 | 877 | 846 | 758 | 907 | 1205 | 1145 | 946 | 842 | 855 | 895 | 981.46 | 1099 | 864 | 0.786 | -0.348 | 0.00034319 | 0.00321447 | 0.0076 | 0.0048 | 0.0058 | 0.0058 | TRUE | 0.174 |
| Brdt | 275 | 180 | 170 | 182 | 128 | 148 | 214 | 191 | 183 | 181 | 144 | 146 | 176.69 | 196 | 157 | 0.8 | -0.321 | 0.04585529 | 0.12645373 | 0.0057 | 0.0158 | 0.0129 | 0.0129 | TRUE | 0.1896 |
| Bre | 2759 | 1931 | 2046 | 2208 | 2056 | 2377 | 2143 | 2054 | 2208 | 2198 | 2319 | 2346 | 2211.35 | 2135 | 2288 | 1.072 | 0.1 | 0.11523888 | 0.24514202 | 8.00E-04 | 0.0035 | 0.0024 | 0.0024 | TRUE | 0.0145 |
| Brf1 | 1258 | 997 | 881 | 833 | 796 | 890 | 977 | 1060 | 951 | 829 | 898 | 879 | 932.3 | 996 | 869 | 0.872 | -0.198 | 0.015667 | 0.05766892 | 0.0014 | 0.005 | 0.0037 | 0.0037 | TRUE | 0.0372 |
| Brf2 | 1345 | 910 | 799 | 912 | 759 | 885 | 1045 | 968 | 862 | 908 | 856 | 874 | 918.72 | 958 | 879 | 0.916 | -0.126 | 0.15949005 | 0.30880403 | 0.0041 | 0.005 | 0.0047 | 0.0047 | TRUE | 0.0909 |
| Bri3 | 1848 | 1518 | 1583 | 1660 | 1537 | 1824 | 1435 | 1615 | 1708 | 1652 | 1734 | 1801 | 1657.47 | 1586 | 1729 | 1.092 | 0.126 | 0.11445945 | 0.2438469 | 0.0043 | 0.0039 | 0.004 | 0.004 | TRUE | 0.0861 |
| Bri3bp | 4282 | 3280 | 3493 | 3575 | 3180 | 3553 | 3326 | 3489 | 3769 | 3559 | 3587 | 3507 | 3539.46 | 3528 | 3551 | 1.007 | 0.01 | 0.87064711 | 0.92839146 | 0.0018 | 0.0031 | 0.0026 | 0.0026 | TRUE | 0.0433 |
| Bricd5 | 4 | 3 | 3 | 3 | 2 | 9 | 3 | 3 | 3 | 3 | 2 | 9 | 3.94 | 3 | 5 | 1.495 | 0.58 | 0.55754856 | NA | 0 | 0.6034 | 0.4441 | 0.4441 | TRUE | 1.1856 |
| Brip1 | 3518 | 2404 | 2294 | 2375 | 1960 | 2300 | 2732 | 2557 | 2475 | 2364 | 2211 | 2270 | 2435.03 | 2588 | 2282 | 0.881 | -0.182 | 0.00511644 | 0.02496733 | 0.0015 | 0.0034 | 0.0026 | 0.0026 | TRUE | 0.0295 |
| Brix1 | 8062 | 5641 | 5507 | 5612 | 4822 | 5817 | 6261 | 6000 | 5942 | 5586 | 5440 | 5742 | 5828.64 | 6068 | 5589 | 0.921 | -0.119 | 0.0248638 | 0.08109958 | 5.00E-04 | 0.0028 | 0.0018 | 0.0018 | TRUE | 0.0097 |
| Brk1 | 6971 | 4644 | 4822 | 5548 | 4679 | 5475 | 5414 | 4940 | 5203 | 5522 | 5278 | 5405 | 5293.71 | 5186 | 5402 | 1.041 | 0.059 | 0.30213455 | 0.47465477 | 0.001 | 0.0029 | 0.0021 | 0.0021 | TRUE | 0.0209 |
| Brms1 | 1316 | 897 | 680 | 638 | 596 | 596 | 1022 | 954 | 734 | 635 | 672 | 588 | 767.61 | 903 | 632 | 0.698 | -0.518 | 1.31E-05 | 0.00023548 | 0.0159 | 0.0055 | 0.0088 | 0.0088 | TRUE | 0.3175 |
| Brms1l | 2272 | 1570 | 1537 | 1576 | 1411 | 1602 | 1765 | 1670 | 1658 | 1569 | 1592 | 1581 | 1639.15 | 1698 | 1581 | 0.93 | -0.104 | 0.10777512 | 0.23352795 | 0 | 0.0039 | 0.0024 | 0.0024 | TRUE | 0.0149 |
| Brox | 2277 | 1620 | 1495 | 1614 | 1427 | 1597 | 1768 | 1723 | 1613 | 1607 | 1610 | 1576 | 1649.6 | 1701 | 1598 | 0.938 | -0.092 | 0.16952052 | 0.32103858 | 6.00E-04 | 0.0039 | 0.0026 | 0.0026 | TRUE | 0.0246 |
| Brpf1 | 5268 | 3703 | 3312 | 3059 | 2856 | 3211 | 4091 | 3939 | 3574 | 3045 | 3222 | 3170 | 3506.74 | 3868 | 3146 | 0.813 | -0.299 | 6.90E-06 | 0.00013845 | 0.0024 | 0.0031 | 0.0029 | 0.0029 | TRUE | 0.0536 |
| Brpf3 | 4219 | 3142 | 3276 | 2446 | 2372 | 2988 | 3277 | 3342 | 3535 | 2435 | 2676 | 2950 | 3035.64 | 3385 | 2687 | 0.794 | -0.333 | 1.30E-05 | 0.00023351 | 0.0049 | 0.0032 | 0.0039 | 0.0039 | TRUE | 0.0896 |
| Brs3 | 0 | 0 | 0 | 0 | 0 | 0 | 0 | 0 | 0 | 0 | 0 | 0 | 0 | 0 | 0 | NA | NA | NA | NA | NA | NA | NA | NA | NA | NA |
| Brsk1 | 1115 | 784 | 730 | 901 | 751 | 897 | 866 | 834 | 788 | 897 | 847 | 885 | 852.85 | 829 | 876 | 1.056 | 0.079 | 0.33076389 | 0.50524829 | 0 | 0.0052 | 0.0036 | 0.0036 | TRUE | 0.0225 |
| Brsk2 | 477 | 394 | 355 | 318 | 261 | 271 | 370 | 419 | 383 | 317 | 294 | 268 | 341.85 | 391 | 293 | 0.75 | -0.415 | 0.00052628 | 0.00447749 | 0.0026 | 0.0094 | 0.0073 | 0.0073 | TRUE | 0.0664 |
| Brwd1 | 6427 | 4854 | 4883 | 5389 | 4945 | 5434 | 4992 | 5163 | 5269 | 5364 | 5578 | 5364 | 5288.38 | 5141 | 5435 | 1.057 | 0.081 | 0.12546507 | 0.26066038 | 5.00E-04 | 0.0029 | 0.0018 | 0.0018 | TRUE | 0.0081 |
| Brwd3 | 2421 | 1618 | 1555 | 1710 | 1409 | 1668 | 1880 | 1721 | 1678 | 1702 | 1589 | 1647 | 1702.89 | 1760 | 1646 | 0.935 | -0.098 | 0.17276251 | 0.32507828 | 0.0019 | 0.0038 | 0.0031 | 0.0031 | TRUE | 0.0448 |

| | | | | | | | | | | | | | | | | | | | | | | | | |
|---|---|---|---|---|---|---|---|---|---|---|---|---|---|---|---|---|---|---|---|---|---|---|---|---|
| Bscl2 | 1095 | 742 | 679 | 668 | 664 | 710 | 850 | 789 | 733 | 665 | 749 | 701 | 747.87 | 791 | 705 | 0.889 | -0.169 | 0.06803975 | 0.16791649 | 0.0033 | 0.0056 | 0.0048 | 0.0048 | TRUE | 0.054 |
| Bsdc1 | 2723 | 1961 | 2013 | 2218 | 1925 | 2244 | 2115 | 2086 | 2172 | 2208 | 2172 | 2215 | 2161.22 | 2124 | 2198 | 1.035 | 0.05 | 0.39897928 | 0.57224403 | 0 | 0.0035 | 0.002 | 0.002 | TRUE | 0.0045 |
| Bsg | 23220 | 17605 | 20909 | 22063 | 19787 | 21850 | 18034 | 18726 | 22561 | 21961 | 22322 | 21569 | 20862.24 | 19774 | 21951 | 1.11 | 0.151 | 0.0525167 | 0.13965046 | 0.0077 | 0.0025 | 0.0043 | 0.0043 | TRUE | 0.1859 |
| Bsn | 130 | 96 | 76 | 105 | 77 | 98 | 101 | 102 | 82 | 105 | 87 | 97 | 95.53 | 95 | 96 | 1.01 | 0.014 | 0.94603372 | 0.97029019 | 0 | 0.0272 | 0.0199 | 0.0199 | TRUE | 0.131 |
| Bsnd | 0 | 1 | 1 | 3 | 0 | 1 | 0 | 1 | 1 | 3 | 0 | 1 | 1.02 | 1 | 1 | 1.967 | 0.976 | 0.63038931 | NA | 0 | 2.3275 | 1.8702 | 1.8702 | TRUE | 0.3453 |
| Bsph1 | 0 | 0 | 0 | 0 | 0 | 0 | 0 | 0 | 0 | 0 | 0 | 0 | 0 | 0 | 0 | NA | NA | NA | NA | NA | NA | NA | NA | NA | NA |
| Bsph2 | 0 | 0 | 0 | 0 | 0 | 0 | 0 | 0 | 0 | 0 | 0 | 0 | 0 | 0 | 0 | NA | NA | NA | NA | NA | NA | NA | NA | NA | NA |
| Bspry | 544 | 448 | 442 | 436 | 392 | 472 | 423 | 477 | 477 | 434 | 442 | 466 | 453.02 | 459 | 447 | 0.978 | -0.032 | 0.75579424 | 0.85511255 | 0 | 0.0077 | 0.0056 | 0.0056 | TRUE | 0.0593 |
| Bst1 | 137 | 113 | 101 | 105 | 110 | 108 | 106 | 120 | 109 | 105 | 124 | 107 | 111.8 | 112 | 112 | 0.999 | -0.002 | 0.99326503 | 0.99603371 | 0 | 0.0236 | 0.0164 | 0.0164 | TRUE | 0.0889 |
| Bst2 | 2178 | 1650 | 2261 | 2209 | 1907 | 2220 | 1692 | 1755 | 2440 | 2199 | 2151 | 2191 | 2071.31 | 1962 | 2180 | 1.112 | 0.153 | 0.17622748 | 0.3299043 | 0.019 | 0.0036 | 0.0088 | 0.0088 | TRUE | 0.5458 |
| Bsx | 1 | 0 | 0 | 1 | 0 | 1 | 1 | 0 | 0 | 1 | 0 | 1 | 0.46 | 0 | 1 | 1.94 | 0.956 | 0.73503911 | NA | 0 | 5.1562 | 3.9771 | 3.9771 | TRUE | 0.2566 |
| Btaf1 | 8629 | 6035 | 5339 | 5917 | 5395 | 6004 | 6702 | 6419 | 5761 | 5890 | 6086 | 5927 | 6130.77 | 6294 | 5968 | 0.948 | -0.077 | 0.23452824 | 0.40061384 | 0.0029 | 0.0028 | 0.0029 | 0.0029 | TRUE | 0.0668 |
| Btbd1 | 3507 | 2678 | 2921 | 3518 | 2911 | 3450 | 2724 | 2849 | 3152 | 3502 | 3284 | 3406 | 3152.57 | 2908 | 3397 | 1.169 | 0.225 | 0.00108452 | 0.00767615 | 0.0029 | 0.0032 | 0.0031 | 0.0031 | TRUE | 0.065 |
| Btbd10 | 2931 | 2040 | 1951 | 2052 | 1815 | 2083 | 2276 | 2170 | 2105 | 2043 | 2048 | 2056 | 2116.29 | 2184 | 2049 | 0.937 | -0.093 | 0.132641 | 0.27071795 | 4.00E-04 | 0.0035 | 0.0023 | 0.0023 | TRUE | 0.0172 |
| Btbd11 | 1157 | 812 | 706 | 805 | 748 | 862 | 899 | 864 | 762 | 801 | 844 | 851 | 836.69 | 842 | 832 | 0.987 | -0.018 | 0.83580642 | 0.90645327 | 0.003 | 0.0053 | 0.0045 | 0.0045 | TRUE | 0.0799 |
| Btbd16 | 3 | 1 | 1 | 1 | 1 | 1 | 2 | 1 | 1 | 1 | 1 | 1 | 1.26 | 1 | 1 | 0.679 | -0.557 | 0.73233539 | NA | 0 | 1.8776 | 1.0978 | 1.0978 | TRUE | 0.2229 |
| Btbd17 | 27 | 28 | 22 | 26 | 25 | 30 | 21 | 30 | 24 | 26 | 28 | 30 | 26.36 | 25 | 28 | 1.13 | 0.177 | 0.63618587 | 0.77255774 | 0 | 0.0923 | 0.0625 | 0.0625 | TRUE | 0.1789 |
| Btbd19 | 859 | 796 | 698 | 633 | 565 | 664 | 667 | 847 | 753 | 630 | 637 | 655 | 698.32 | 756 | 641 | 0.85 | -0.234 | 0.0210184 | 0.07174968 | 0.0063 | 0.0058 | 0.006 | 0.006 | TRUE | 0.132 |
| Btbd2 | 3430 | 2992 | 3427 | 3650 | 3360 | 3748 | 2664 | 3183 | 3698 | 3633 | 3790 | 3700 | 3444.61 | 3182 | 3708 | 1.166 | 0.222 | 0.02190591 | 0.07375763 | 0.013 | 0.0031 | 0.0065 | 0.0065 | TRUE | 0.2199 |
| Btbd3 | 1813 | 1374 | 1447 | 1503 | 1261 | 1546 | 1408 | 1462 | 1561 | 1496 | 1423 | 1526 | 1479.28 | 1477 | 1482 | 1.004 | 0.006 | 0.93060732 | 0.96190625 | 0.0014 | 0.004 | 0.003 | 0.003 | TRUE | 0.0298 |
| Btbd6 | 841 | 603 | 499 | 526 | 491 | 568 | 653 | 641 | 538 | 524 | 554 | 561 | 578.53 | 611 | 546 | 0.892 | -0.165 | 0.10961894 | 0.23667424 | 0.0046 | 0.0065 | 0.0059 | 0.0059 | TRUE | 0.1237 |
| Btbd7 | 2221 | 1604 | 1784 | 1730 | 1694 | 1912 | 1725 | 1706 | 1925 | 1722 | 1911 | 1887 | 1812.75 | 1785 | 1840 | 1.031 | 0.044 | 0.5600558 | 0.71470543 | 0.0033 | 0.0037 | 0.0036 | 0.0036 | TRUE | 0.0556 |
| Btbd8 | 149 | 117 | 114 | 139 | 116 | 115 | 116 | 124 | 123 | 138 | 131 | 114 | 124.32 | 121 | 128 | 1.055 | 0.077 | 0.6651707 | 0.7937289 | 0 | 0.0215 | 0.0149 | 0.0149 | TRUE | 0.0978 |
| Btbd9 | 2796 | 2022 | 1975 | 2140 | 1914 | 2216 | 2172 | 2151 | 2131 | 2130 | 2159 | 2188 | 2155.03 | 2151 | 2159 | 1.003 | 0.005 | 0.93164468 | 0.96216772 | 0 | 0.0035 | 0.002 | 0.002 | TRUE | 0.0017 |
| Btc | 22 | 15 | 18 | 13 | 19 | 16 | 17 | 16 | 19 | 13 | 21 | 16 | 17.11 | 17 | 17 | 0.953 | -0.07 | 0.88071203 | 0.93349722 | 0 | 0.141 | 0.097 | 0.097 | TRUE | 0.2691 |
| Btd | 533 | 375 | 311 | 389 | 343 | 353 | 414 | 399 | 336 | 387 | 387 | 348 | 378.5 | 383 | 374 | 0.975 | -0.037 | 0.75958502 | 0.8571787 | 0.0053 | 0.0087 | 0.0076 | 0.0076 | TRUE | 0.1301 |
| Btf3 | 19926 | 14228 | 14896 | 15908 | 13381 | 16148 | 15476 | 15134 | 16073 | 15835 | 15095 | 15941 | 15592.17 | 15561 | 15624 | 1.004 | 0.006 | 0.90689352 | 0.94912712 | 7.00E-04 | 0.0026 | 0.0018 | 0.0018 | TRUE | 0.0107 |
| Btf3l4 | 1669 | 1225 | 1277 | 1268 | 1074 | 1193 | 1296 | 1303 | 1378 | 1262 | 1212 | 1178 | 1271.43 | 1326 | 1217 | 0.919 | -0.123 | 0.08592128 | 0.19817217 | 4.00E-04 | 0.0043 | 0.0029 | 0.0029 | TRUE | 0.014 |
| Btg1 | 1879 | 1519 | 1851 | 1762 | 1567 | 1826 | 1459 | 1616 | 1997 | 1754 | 1768 | 1803 | 1732.74 | 1691 | 1775 | 1.051 | 0.072 | 0.47540128 | 0.64342464 | 0.0125 | 0.0038 | 0.0067 | 0.0067 | TRUE | 0.3024 |
| Btg2 | 2407 | 1795 | 1826 | 1992 | 1705 | 1815 | 1869 | 1909 | 1970 | 1983 | 1923 | 1792 | 1907.82 | 1916 | 1899 | 0.991 | -0.012 | 0.85286722 | 0.91681943 | 0.0012 | 0.0037 | 0.0027 | 0.0027 | TRUE | 0.0301 |
| Btg3 | 1 | 0 | 1 | 0 | 0 | 0 | 1 | 0 | 1 | 0 | 0 | 0 | 0.31 | 1 | 0 | 0.307 | -1.706 | 0.62894649 | NA | 0 | 7.6657 | 7.2 | 7.2 | TRUE | 0.2067 |
| Btg4 | 0 | 0 | 0 | 0 | 0 | 0 | 0 | 0 | 0 | 0 | 0 | 0 | 0 | 0 | 0 | NA | NA | NA | NA | NA | NA | NA | NA | NA | NA |
| Btk | 22 | 9 | 12 | 12 | 12 | 12 | 17 | 10 | 13 | 12 | 14 | 12 | 12.82 | 13 | 13 | 0.931 | -0.103 | 0.84681301 | NA | 0 | 0.1873 | 0.1285 | 0.1285 | TRUE | 0.2816 |
| Btla | 1 | 1 | 1 | 1 | 0 | 2 | 1 | 1 | 1 | 1 | 0 | 2 | 0.98 | 1 | 1 | 1.044 | 0.062 | 0.97363744 | NA | 0 | 2.4171 | 1.5475 | 1.5475 | TRUE | 0.1802 |
| Btn1a1 | 14 | 8 | 4 | 14 | 12 | 17 | 11 | 9 | 4 | 14 | 14 | 17 | 11.33 | 8 | 15 | 1.836 | 0.877 | 0.13538143 | NA | 0 | 0.2117 | 0.1535 | 0.1535 | TRUE | 0.4182 |
| Btn2a2 | 110 | 81 | 75 | 84 | 77 | 107 | 85 | 86 | 81 | 84 | 87 | 106 | 88.1 | 84 | 92 | 1.094 | 0.129 | 0.53911844 | 0.69753507 | 0 | 0.0293 | 0.0206 | 0.0206 | TRUE | 0.1642 |
| Btnl1 | 14 | 14 | 13 | 7 | 16 | 12 | 11 | 15 | 14 | 7 | 18 | 12 | 12.78 | 13 | 12 | 0.925 | -0.112 | 0.84146286 | NA | 0 | 0.1879 | 0.1458 | 0.1458 | TRUE | 0.2597 |
| Btnl10 | 0 | 0 | 0 | 1 | 0 | 0 | 0 | 0 | 0 | 1 | 0 | 0 | 0.17 | 0 | 0 | 2.094 | 1.067 | 0.79379329 | NA | 0 | 14.2885 | 10 | 10 | TRUE | 0.3906 |
| Btnl2 | 0 | 0 | 0 | 0 | 0 | 0 | 0 | 0 | 0 | 0 | 0 | 0 | 0 | 0 | 0 | NA | NA | NA | NA | NA | NA | NA | NA | NA | NA |
| Btnl4 | 4 | 4 | 3 | 9 | 6 | 5 | 3 | 4 | 3 | 9 | 7 | 5 | 5.21 | 3 | 7 | 1.961 | 0.972 | 0.2484955 | NA | 0 | 0.4573 | 0.2986 | 0.2986 | TRUE | 0.1698 |
| Btnl5 | 2 | 0 | 2 | 1 | 0 | 3 | 2 | 0 | 2 | 1 | 0 | 3 | 1.28 | 1 | 1 | 1.074 | 0.103 | 0.95409135 | NA | 0 | 1.8569 | 1.5459 | 1.5459 | TRUE | 0.4767 |
| Btnl6 | 1 | 1 | 0 | 1 | 0 | 4 | 1 | 1 | 0 | 1 | 0 | 4 | 1.13 | 1 | 2 | 2.678 | 1.421 | 0.4723908 | NA | 0 | 2.0985 | 1.735 | 1.735 | TRUE | 0.558 |
| Btnl9 | 18 | 10 | 10 | 12 | 5 | 12 | 14 | 11 | 11 | 12 | 6 | 12 | 10.81 | 12 | 10 | 0.832 | -0.266 | 0.65291461 | NA | 0 | 0.2217 | 0.1583 | 0.1583 | TRUE | 0.4182 |
| Btrc | 3101 | 2208 | 2293 | 2443 | 2155 | 2290 | 2408 | 2349 | 2474 | 2432 | 2431 | 2261 | 2392.43 | 2410 | 2375 | 0.985 | -0.022 | 0.72538878 | 0.83572689 | 8.00E-04 | 0.0034 | 0.0024 | 0.0024 | TRUE | 0.0215 |
| Bub1 | 14124 | 10330 | 11451 | 13379 | 11392 | 13151 | 10970 | 10988 | 12356 | 13317 | 12851 | 12982 | 12243.97 | 11438 | 13050 | 1.141 | 0.191 | 0.00158805 | 0.01006643 | 0.0025 | 0.0026 | 0.0025 | 0.0025 | TRUE | 0.06 |
| Bub1b | 18220 | 13506 | 14056 | 14771 | 13409 | 15590 | 14151 | 14366 | 15166 | 14703 | 15127 | 15390 | 14817.11 | 14561 | 15073 | 1.035 | 0.05 | 0.32577354 | 0.49986916 | 8.00E-04 | 0.0026 | 0.0018 | 0.0018 | TRUE | 0.0161 |
| Bub3 | 7989 | 5783 | 5926 | 6007 | 5414 | 6435 | 6205 | 6151 | 6394 | 5979 | 6108 | 6352 | 6198.23 | 6250 | 6146 | 0.984 | -0.024 | 0.64501733 | 0.77898037 | 4.00E-04 | 0.0028 | 0.0018 | 0.0018 | TRUE | 0.0105 |
| Bud13 | 1901 | 1398 | 1344 | 1407 | 1217 | 1512 | 1476 | 1487 | 1450 | 1401 | 1373 | 1493 | 1446.61 | 1471 | 1422 | 0.967 | -0.049 | 0.47651973 | 0.64430176 | 3.00E-04 | 0.0041 | 0.0027 | 0.0027 | TRUE | 0.0228 |
| Bud31 | 1717 | 1148 | 844 | 852 | 659 | 832 | 1334 | 1221 | 911 | 848 | 743 | 821 | 979.69 | 1155 | 804 | 0.696 | -0.524 | 1.98E-05 | 0.00032822 | 0.0207 | 0.0048 | 0.0098 | 0.0098 | TRUE | 0.4071 |
| Bves | 153 | 98 | 95 | 141 | 105 | 107 | 119 | 104 | 103 | 140 | 118 | 106 | 115 | 109 | 121 | 1.116 | 0.158 | 0.41225085 | 0.5851342 | 0.0061 | 0.023 | 0.0181 | 0.0181 | TRUE | 0.192 |
| Bysl | 6758 | 4628 | 3699 | 4318 | 3822 | 4129 | 5249 | 4923 | 3991 | 4298 | 4312 | 4076 | 4474.72 | 4721 | 4229 | 0.895 | -0.16 | 0.07265894 | 0.1763781 | 0.0097 | 0.003 | 0.0055 | 0.0055 | TRUE | 0.222 |
| Bzrap1 | 73 | 76 | 74 | 85 | 91 | 110 | 57 | 81 | 80 | 85 | 103 | 109 | 85.54 | 73 | 99 | 1.371 | 0.455 | 0.04799626 | 0.13077845 | 0.0163 | 0.0301 | 0.0262 | 0.0262 | TRUE | 0.3801 |
| Bzw1 | 20484 | 14129 | 12613 | 13202 | 11289 | 13209 | 15909 | 15029 | 13609 | 13141 | 12735 | 13039 | 13910.49 | 14849 | 12972 | 0.873 | -0.195 | 0.00210423 | 0.01254725 | 0.0031 | 0.0026 | 0.0028 | 0.0028 | TRUE | 0.0652 |
| Bzw2 | 12381 | 9421 | 10577 | 11256 | 10537 | 12495 | 9616 | 10021 | 11412 | 11204 | 11887 | 12334 | 11079.14 | 10350 | 11808 | 1.141 | 0.191 | 0.00766089 | 0.03387903 | 0.0052 | 0.0026 | 0.0036 | 0.0036 | TRUE | 0.0984 |
| C030006K11I | 279 | 159 | 143 | 197 | 168 | 219 | 217 | 169 | 154 | 196 | 190 | 216 | 190.32 | 180 | 201 | 1.109 | 0.149 | 0.3688728 | 0.54353597 | 0.0136 | 0.0149 | 0.0145 | 0.0145 | TRUE | 0.3662 |
| C030007H22 | 0 | 0 | 0 | 0 | 0 | 0 | 0 | 0 | 0 | 0 | 0 | 0 | 0 | 0 | 0 | NA | NA | NA | NA | NA | NA | NA | NA | NA | NA |
| C030013G03 | 0 | 0 | 0 | 0 | 0 | 0 | 0 | 0 | 0 | 0 | 0 | 0 | 0 | 0 | 0 | NA | NA | NA | NA | NA | NA | NA | NA | NA | NA |
| C030016D13 | 4 | 6 | 8 | 7 | 9 | 9 | 3 | 6 | 9 | 7 | 10 | 9 | 7.35 | 6 | 9 | 1.463 | 0.549 | 0.44674406 | NA | 0 | 0.3247 | 0.2338 | 0.2338 | TRUE | 0.5381 |
| C030018K13I | 1 | 1 | 2 | 0 | 0 | 2 | 1 | 1 | 2 | 0 | 0 | 2 | 1 | 1 | 1 | 0.511 | -0.968 | 0.63227045 | NA | 0 | 2.3833 | 1.811 | 1.811 | TRUE | 0.9735 |
| C030023E24I | 0 | 0 | 0 | 0 | 0 | 0 | 0 | 0 | 0 | 0 | 0 | 0 | 0 | 0 | 0 | NA | NA | NA | NA | NA | NA | NA | NA | NA | NA |
| C030030A07 | 3 | 0 | 0 | 0 | 0 | 1 | 2 | 0 | 0 | 0 | 0 | 1 | 0.55 | 1 | 0 | 0.458 | -1.128 | 0.69846567 | NA | 0 | 4.2894 | 4.5119 | 4.5119 | TRUE | 1.4399 |
| C030034I22R | 40 | 39 | 39 | 67 | 44 | 47 | 31 | 41 | 42 | 67 | 50 | 46 | 46.23 | 38 | 54 | 1.432 | 0.518 | 0.08473301 | 0.19644588 | 0 | 0.0537 | 0.0427 | 0.0427 | TRUE | 0.3434 |
| C030034L19F | 1 | 0 | 0 | 3 | 0 | 0 | 1 | 0 | 0 | 3 | 0 | 0 | 0.63 | 0 | 1 | 2.91 | 1.541 | 0.58287418 | NA | 7.7564 | 3.7816 | 4.1567 | 4.1567 | TRUE | 1.4419 |
| C030037D09 | 47 | 42 | 48 | 55 | 46 | 64 | 37 | 45 | 52 | 55 | 52 | 63 | 50.46 | 45 | 57 | 1.288 | 0.365 | 0.19143416 | 0.34953607 | 0 | 0.0494 | 0.0362 | 0.0362 | TRUE | 0.2214 |
| C030039L03F | 853 | 653 | 461 | 358 | 303 | 331 | 662 | 695 | 497 | 356 | 342 | 327 | 479.9 | 618 | 342 | 0.552 | -0.857 | 4.25E-11 | 3.78E-09 | 0.0153 | 0.0074 | 0.0099 | 0.0099 | TRUE | 0.3356 |
| C030046E11I | 3240 | 2304 | 2266 | 2569 | 2089 | 2419 | 2516 | 2451 | 2445 | 2557 | 2357 | 2388 | 2452.3 | 2471 | 2434 | 0.985 | -0.022 | 0.72341464 | 0.83439037 | 7.00E-04 | 0.0034 | 0.0023 | 0.0023 | TRUE | 0.0239 |
| C130021I20R | 37 | 26 | 16 | 11 | 19 | 12 | 29 | 28 | 17 | 11 | 21 | 12 | 19.65 | 25 | 15 | 0.593 | -0.755 | 0.10651898 | 0.23166096 | 0.0443 | 0.1231 | 0.1026 | 0.1026 | TRUE | 0.6463 |
| C130026I21R | 20 | 17 | 7 | 3 | 14 | 6 | 16 | 18 | 8 | 3 | 16 | 6 | 10.98 | 14 | 8 | 0.59 | -0.762 | 0.26230071 | NA | 0.2795 | 0.2183 | 0.2343 | 0.2343 | TRUE | 1.625 |
| C130026L21F | 2 | 2 | 3 | 4 | 10 | 7 | 2 | 2 | 3 | 4 | 11 | 7 | 4.85 | 2 | 7 | 3.249 | 1.7 | 0.07194229 | NA | 0 | 0.4913 | 0.3599 | 0.3599 | TRUE | 0.1483 |

| | | | | | | | | | | | | | | | | | | | | | | | | | |
|---|---|---|---|---|---|---|---|---|---|---|---|---|---|---|---|---|---|---|---|---|---|---|---|---|---|
| C130030K03I | 1 | 0 | 0 | 1 | 0 | 0 | 1 | 0 | 0 | 1 | 0 | 0 | 0.3 | 0 | 0 | 1.075 | 0.105 | 0.97698525 | NA | 0 | 8.0272 | 7.5141 | 7.5141 | TRUE | 0.3906 |
| C130036L24I | 27 | 20 | 12 | 15 | 20 | 16 | 21 | 21 | 13 | 15 | 23 | 16 | 18.08 | 18 | 18 | 0.956 | -0.065 | 0.88854096 | 0.93805954 | 0 | 0.1335 | 0.0986 | 0.0986 | TRUE | 0.3175 |
| C130050O18 | 0 | 0 | 1 | 1 | 1 | 0 | 0 | 0 | 1 | 1 | 1 | 0 | 0.53 | 0 | 1 | 2.036 | 1.026 | 0.70454668 | NA | 0 | 4.4428 | 3.5371 | 3.5371 | TRUE | 0.5365 |
| C130060C02I | 4 | 2 | 2 | 2 | 4 | 5 | 3 | 2 | 2 | 2 | 5 | 5 | 3.14 | 2 | 4 | 1.525 | 0.609 | 0.5682614 | NA | 0 | 0.7576 | 0.4913 | 0.4913 | TRUE | 0.2903 |
| C130060K24I | 1 | 2 | 3 | 0 | 1 | 1 | 1 | 2 | 3 | 0 | 1 | 1 | 1.38 | 2 | 1 | 0.346 | -1.529 | 0.3785048 | NA | 0 | 1.7248 | 1.1883 | 1.1883 | TRUE | 0.1928 |
| C130071C03I | 1 | 0 | 0 | 0 | 1 | 0 | 1 | 0 | 0 | 0 | 1 | 0 | 0.32 | 0 | 0 | 1.075 | 0.105 | 0.97638847 | NA | 0 | 7.4681 | 7.0392 | 7.0392 | TRUE | 0.5075 |
| C130074G19 | 598 | 418 | 458 | 467 | 357 | 468 | 464 | 445 | 494 | 465 | 403 | 462 | 455.47 | 468 | 443 | 0.949 | -0.076 | 0.47546709 | 0.64342464 | 0.002 | 0.0076 | 0.0059 | 0.0059 | TRUE | 0.0715 |
| C130079G13 | 0 | 0 | 0 | 0 | 0 | 0 | 0 | 0 | 0 | 0 | 0 | 0 | 0 | 0 | 0 | NA | NA | NA | NA | NA | NA | NA | NA | NA | NA |
| C130080G10 | 0 | 0 | 0 | 0 | 0 | 0 | 0 | 0 | 0 | 0 | 0 | 0 | 0 | 0 | 0 | NA | NA | NA | NA | NA | NA | NA | NA | NA | NA |
| C130083M11 | 3 | 2 | 1 | 2 | 1 | 0 | 2 | 2 | 1 | 2 | 1 | 0 | 1.44 | 2 | 1 | 0.557 | -0.845 | 0.6011101 | NA | 0 | 1.6454 | 1.1252 | 1.1252 | TRUE | 0.395 |
| C1d | 2157 | 1519 | 1364 | 1362 | 1185 | 1321 | 1675 | 1616 | 1472 | 1356 | 1337 | 1304 | 1459.88 | 1588 | 1332 | 0.838 | -0.254 | 0.00056252 | 0.00472113 | 0.0018 | 0.004 | 0.0032 | 0.0032 | TRUE | 0.0485 |
| C1galt1 | 1157 | 785 | 805 | 927 | 753 | 825 | 899 | 835 | 869 | 923 | 849 | 814 | 864.8 | 868 | 862 | 0.993 | -0.01 | 0.9086552 | 0.94963864 | 0.0016 | 0.0052 | 0.0039 | 0.0039 | TRUE | 0.0456 |
| C1galt1c1 | 401 | 261 | 277 | 292 | 213 | 266 | 311 | 278 | 299 | 291 | 240 | 263 | 280.24 | 296 | 265 | 0.893 | -0.163 | 0.20713741 | 0.36896996 | 0.0023 | 0.0109 | 0.0084 | 0.0084 | TRUE | 0.0836 |
| C1qa | 0 | 0 | 0 | 0 | 0 | 0 | 0 | 0 | 0 | 0 | 0 | 0 | 0 | 0 | 0 | NA | NA | NA | NA | NA | NA | NA | NA | NA | NA |
| C1qb | 0 | 0 | 0 | 0 | 0 | 0 | 0 | 0 | 0 | 0 | 0 | 0 | 0 | 0 | 0 | NA | NA | NA | NA | NA | NA | NA | NA | NA | NA |
| C1qbp | 28550 | 20071 | 20241 | 22745 | 19620 | 22719 | 22174 | 21349 | 21840 | 22640 | 22133 | 22427 | 22093.93 | 21788 | 22400 | 1.028 | 0.04 | 0.36987679 | 0.54467844 | 2.00E-04 | 0.0025 | 0.0014 | 0.0014 | TRUE | 0.0038 |
| C1qc | 0 | 0 | 0 | 0 | 0 | 0 | 0 | 0 | 0 | 0 | 0 | 0 | 0 | 0 | 0 | NA | NA | NA | NA | NA | NA | NA | NA | NA | NA |
| C1ql1 | 305 | 251 | 291 | 374 | 382 | 427 | 237 | 267 | 314 | 372 | 431 | 422 | 340.43 | 273 | 408 | 1.503 | 0.587 | 8.53E-06 | 0.00016458 | 0.0098 | 0.0094 | 0.0095 | 0.0095 | TRUE | 0.1957 |
| C1ql2 | 11 | 4 | 1 | 4 | 4 | 2 | 9 | 4 | 1 | 4 | 5 | 2 | 4.06 | 5 | 4 | 0.731 | -0.452 | 0.65510719 | NA | 0.2468 | 0.5865 | 0.4869 | 0.4869 | TRUE | 0.2319 |
| C1ql3 | 6 | 9 | 7 | 6 | 0 | 1 | 5 | 10 | 8 | 6 | 0 | 1 | 4.79 | 8 | 2 | 0.328 | -1.608 | 0.12023872 | NA | 0.4251 | 0.4971 | 0.4854 | 0.4854 | TRUE | 1.5763 |
| C1ql4 | 95 | 70 | 76 | 80 | 57 | 66 | 74 | 74 | 82 | 80 | 64 | 65 | 73.22 | 77 | 70 | 0.911 | -0.135 | 0.55465273 | 0.71014436 | 0 | 0.0348 | 0.024 | 0.024 | TRUE | 0.142 |
| C1qtnf1 | 7 | 7 | 10 | 8 | 5 | 5 | 5 | 7 | 11 | 8 | 6 | 5 | 7.04 | 8 | 6 | 0.793 | -0.334 | 0.6453151 | NA | 0 | 0.3393 | 0.2334 | 0.2334 | TRUE | 0.286 |
| C1qtnf2 | 3 | 0 | 2 | 2 | 1 | 1 | 2 | 0 | 2 | 2 | 1 | 1 | 1.43 | 1 | 1 | 0.9 | -0.152 | 0.92429029 | NA | 0 | 1.6571 | 1.1591 | 1.1591 | TRUE | 0.4744 |
| C1qtnf3 | 0 | 0 | 0 | 0 | 0 | 1 | 0 | 0 | 0 | 0 | 0 | 1 | 0.16 | 0 | 0 | 2.094 | 1.067 | 0.79379329 | NA | 0 | 14.4078 | 10 | 10 | TRUE | 0.3835 |
| C1qtnf4 | 90 | 79 | 81 | 133 | 89 | 140 | 70 | 84 | 87 | 132 | 100 | 138 | 102.05 | 80 | 123 | 1.548 | 0.631 | 0.00278135 | 0.01557139 | 0.0123 | 0.0257 | 0.0218 | 0.0218 | TRUE | 0.2626 |
| C1qtnf5 | 0 | 0 | 0 | 0 | 0 | 0 | 0 | 0 | 0 | 0 | 0 | 0 | 0 | 0 | 0 | NA | NA | NA | NA | NA | NA | NA | NA | NA | NA |
| C1qtnf6 | 16 | 20 | 22 | 20 | 20 | 28 | 12 | 21 | 24 | 20 | 23 | 28 | 21.26 | 19 | 24 | 1.238 | 0.308 | 0.47892027 | 0.64589568 | 0 | 0.1139 | 0.089 | 0.089 | TRUE | 0.6298 |
| C1qtnf7 | 0 | 0 | 0 | 0 | 0 | 0 | 0 | 0 | 0 | 0 | 0 | 0 | 0 | 0 | 0 | NA | NA | NA | NA | NA | NA | NA | NA | NA | NA |
| C1qtnf9 | 4 | 5 | 3 | 7 | 5 | 2 | 3 | 5 | 3 | 7 | 6 | 2 | 4.37 | 4 | 5 | 1.257 | 0.329 | 0.72269298 | NA | 0 | 0.5443 | 0.3892 | 0.3892 | TRUE | 0.5589 |
| C1ra | 15 | 16 | 10 | 15 | 13 | 13 | 12 | 17 | 11 | 15 | 15 | 13 | 13.65 | 13 | 14 | 1.079 | 0.11 | 0.83141042 | NA | 0 | 0.1761 | 0.1182 | 0.1182 | TRUE | 0.257 |
| C1rb | 6 | 16 | 5 | 5 | 1 | 1 | 5 | 17 | 5 | 5 | 1 | 1 | 5.69 | 9 | 2 | 0.267 | -1.904 | 0.05177119 | NA | 0.4224 | 0.4186 | 0.4197 | 0.4197 | TRUE | 1.8532 |
| C1rl | 269 | 197 | 191 | 268 | 215 | 219 | 209 | 210 | 206 | 267 | 243 | 216 | 225.01 | 208 | 242 | 1.161 | 0.216 | 0.12229993 | 0.25548566 | 0 | 0.013 | 0.0096 | 0.0096 | TRUE | 0.0983 |
| C1s | 463 | 364 | 284 | 191 | 187 | 214 | 360 | 387 | 306 | 190 | 211 | 211 | 277.59 | 351 | 204 | 0.581 | -0.784 | 7.93E-09 | 4.08E-07 | 0.0058 | 0.011 | 0.0094 | 0.0094 | TRUE | 0.1351 |
| C2 | 42 | 29 | 63 | 61 | 65 | 98 | 33 | 31 | 68 | 61 | 73 | 97 | 60.37 | 44 | 77 | 1.771 | 0.824 | 0.01165888 | 0.04628827 | 0.106 | 0.0417 | 0.0591 | 0.0591 | TRUE | 1.1529 |
| C230004F18I | 0 | 0 | 0 | 0 | 0 | 0 | 0 | 0 | 0 | 0 | 0 | 0 | 0 | 0 | 0 | NA | NA | NA | NA | NA | NA | NA | NA | NA | NA |
| C230024C17I | 0 | 0 | 1 | 0 | 0 | 0 | 0 | 0 | 1 | 0 | 0 | 0 | 0.18 | 0 | 0 | 0.552 | -0.857 | 0.8336548 | NA | 0 | 13.1817 | 10 | 10 | TRUE | 0.5365 |
| C230035I16R | 9 | 12 | 10 | 9 | 9 | 2 | 7 | 13 | 11 | 9 | 10 | 2 | 8.6 | 10 | 7 | 0.691 | -0.533 | 0.45492881 | NA | 0.1372 | 0.2779 | 0.2442 | 0.2442 | TRUE | 1.1002 |
| C230052I12R | 712 | 494 | 362 | 416 | 400 | 499 | 553 | 525 | 391 | 414 | 451 | 493 | 471.16 | 490 | 453 | 0.922 | -0.116 | 0.38521748 | 0.55914628 | 0.0185 | 0.0075 | 0.0108 | 0.0108 | TRUE | 0.3576 |
| C230079O03 | 0 | 0 | 0 | 0 | 0 | 0 | 0 | 0 | 0 | 0 | 0 | 0 | 0 | 0 | 0 | NA | NA | NA | NA | NA | NA | NA | NA | NA | NA |
| C230081A13 | 3136 | 2152 | 2078 | 2533 | 2256 | 2545 | 2436 | 2289 | 2242 | 2521 | 2545 | 2512 | 2424.24 | 2322 | 2526 | 1.087 | 0.12 | 0.04887346 | 0.13246864 | 6.00E-04 | 0.0034 | 0.0023 | 0.0023 | TRUE | 0.0228 |
| C230091D08 | 1 | 1 | 1 | 1 | 0 | 1 | 1 | 1 | 1 | 1 | 0 | 1 | 0.82 | 1 | 1 | 0.698 | -0.518 | 0.79971927 | NA | 0 | 2.9034 | 1.738 | 1.738 | TRUE | 0.2018 |
| C2cd2 | 772 | 579 | 533 | 661 | 547 | 622 | 600 | 616 | 575 | 658 | 617 | 614 | 613.27 | 597 | 630 | 1.055 | 0.077 | 0.38797024 | 0.56194351 | 0 | 0.0063 | 0.0041 | 0.0041 | TRUE | 0.0184 |
| C2cd2l | 549 | 381 | 393 | 405 | 364 | 482 | 426 | 405 | 424 | 403 | 411 | 476 | 424.21 | 418 | 430 | 1.027 | 0.039 | 0.72387631 | 0.83455196 | 0.0023 | 0.008 | 0.0062 | 0.0062 | TRUE | 0.1027 |
| C2cd3 | 2821 | 2052 | 2016 | 2033 | 1889 | 2198 | 2191 | 2183 | 2175 | 2024 | 2131 | 2170 | 2145.55 | 2183 | 2108 | 0.966 | -0.051 | 0.40709915 | 0.58001078 | 0 | 0.0035 | 0.0022 | 0.0022 | TRUE | 0.015 |
| C2cd4a | 2 | 6 | 2 | 1 | 5 | 3 | 2 | 6 | 2 | 1 | 6 | 3 | 3.28 | 3 | 3 | 0.957 | -0.064 | 0.95458938 | NA | 0.2806 | 0.7246 | 0.5904 | 0.5904 | TRUE | 0.4066 |
| C2cd4b | 78 | 70 | 42 | 37 | 51 | 41 | 61 | 74 | 45 | 37 | 58 | 40 | 52.53 | 60 | 45 | 0.744 | -0.426 | 0.15367813 | 0.30072858 | 0.0372 | 0.0475 | 0.0447 | 0.0447 | TRUE | 0.1937 |
| C2cd4c | 97 | 84 | 91 | 120 | 105 | 100 | 75 | 89 | 98 | 119 | 118 | 99 | 99.91 | 87 | 112 | 1.286 | 0.363 | 0.07679246 | 0.18333062 | 0 | 0.0261 | 0.0202 | 0.0202 | TRUE | 0.1636 |
| C2cd4d | 0 | 0 | 0 | 1 | 0 | 0 | 0 | 0 | 0 | 1 | 0 | 0 | 0.17 | 0 | 0 | 2.094 | 1.067 | 0.79379329 | NA | 0 | 14.2885 | 10 | 10 | TRUE | 0.3906 |
| C2cd5 | 3816 | 2891 | 2904 | 3322 | 3245 | 3511 | 2964 | 3075 | 3133 | 3307 | 3661 | 3466 | 3267.59 | 3057 | 3478 | 1.138 | 0.186 | 0.00260112 | 0.01480007 | 0.0012 | 0.0032 | 0.0024 | 0.0024 | TRUE | 0.0255 |
| C3 | 160 | 136 | 216 | 293 | 271 | 285 | 124 | 145 | 233 | 292 | 306 | 281 | 230.12 | 167 | 293 | 1.76 | 0.816 | 2.94E-05 | 0.0004571 | 0.0496 | 0.0127 | 0.0228 | 0.0228 | TRUE | 1.2385 |
| C330005M16 | 21 | 27 | 29 | 30 | 32 | 35 | 16 | 29 | 31 | 30 | 36 | 35 | 29.47 | 25 | 34 | 1.334 | 0.416 | 0.26923552 | 0.43989292 | 0.0268 | 0.0828 | 0.0674 | 0.0674 | TRUE | 0.7531 |
| C330006A16 | 1716 | 1291 | 1322 | 1379 | 1262 | 1463 | 1333 | 1373 | 1426 | 1373 | 1424 | 1444 | 1395.49 | 1377 | 1414 | 1.027 | 0.038 | 0.57788421 | 0.7293317 | 0 | 0.0041 | 0.0027 | 0.0027 | TRUE | 0.0116 |
| C330007P06I | 1346 | 977 | 867 | 961 | 791 | 857 | 1045 | 1039 | 935 | 957 | 892 | 846 | 952.5 | 1006 | 898 | 0.892 | -0.165 | 0.05224882 | 0.13917115 | 0.0028 | 0.0049 | 0.0042 | 0.0042 | TRUE | 0.0447 |
| C330008G21 | 24 | 19 | 12 | 5 | 3 | 3 | 19 | 20 | 13 | 5 | 3 | 3 | 10.52 | 17 | 4 | 0.219 | -2.194 | 0.00110323 | NA | 0 | 0.2277 | 0.1611 | 0.1611 | TRUE | 0.2194 |
| C330011F03I | 0 | 0 | 0 | 0 | 0 | 0 | 0 | 0 | 0 | 0 | 0 | 0 | 0 | 0 | 0 | NA | NA | NA | NA | NA | NA | NA | NA | NA | NA |
| C330013E15I | 36 | 31 | 21 | 33 | 25 | 30 | 28 | 33 | 23 | 33 | 28 | 30 | 29.04 | 28 | 30 | 1.085 | 0.118 | 0.74109913 | 0.84598937 | 0 | 0.084 | 0.0574 | 0.0574 | TRUE | 0.1533 |
| C330013F16I | 201 | 155 | 141 | 124 | 110 | 132 | 156 | 165 | 152 | 123 | 124 | 130 | 141.82 | 158 | 126 | 0.799 | -0.324 | 0.04869018 | 0.13218021 | 0 | 0.0191 | 0.0123 | 0.0123 | TRUE | 0.0157 |
| C330018D20 | 678 | 484 | 459 | 333 | 330 | 329 | 527 | 515 | 495 | 331 | 372 | 325 | 427.53 | 512 | 343 | 0.668 | -0.582 | 4.21E-08 | 1.83E-06 | 0 | 0.008 | 0.0057 | 0.0057 | TRUE | 0.0621 |
| C330021F23I | 449 | 314 | 402 | 462 | 420 | 484 | 349 | 334 | 434 | 460 | 474 | 478 | 421.32 | 372 | 471 | 1.266 | 0.341 | 0.00415832 | 0.02127991 | 0.0071 | 0.0081 | 0.0078 | 0.0078 | TRUE | 0.2322 |
| C330022C24I | 0 | 0 | 0 | 1 | 1 | 0 | 0 | 0 | 0 | 1 | 1 | 0 | 0.35 | 0 | 1 | 3.978 | 1.992 | 0.55384016 | NA | 0 | 6.6991 | 6.4234 | 6.4234 | TRUE | 0.2647 |
| C330024C12I | 1 | 4 | 1 | 2 | 0 | 0 | 1 | 4 | 1 | 2 | 0 | 0 | 1.35 | 2 | 1 | 0.337 | -1.569 | 0.4012003 | NA | 0.7058 | 1.7578 | 1.5106 | 1.5106 | TRUE | 0.9735 |
| C330024D21 | 888 | 676 | 569 | 646 | 537 | 604 | 690 | 719 | 614 | 643 | 606 | 596 | 644.62 | 674 | 615 | 0.912 | -0.133 | 0.16006912 | 0.30951354 | 0.0025 | 0.0061 | 0.0049 | 0.0049 | TRUE | 0.0694 |
| C330027C09I | 8037 | 5723 | 5882 | 6620 | 5841 | 6813 | 6242 | 6087 | 6347 | 6590 | 6589 | 6725 | 6430.06 | 6225 | 6635 | 1.066 | 0.092 | 0.06091871 | 0.15501355 | 1.00E-04 | 0.0028 | 0.0016 | 0.0016 | TRUE | 0.0045 |
| C330046G13 | 402 | 316 | 439 | 461 | 472 | 471 | 312 | 336 | 474 | 459 | 532 | 465 | 429.72 | 374 | 485 | 1.301 | 0.38 | 0.01012505 | 0.04168127 | 0.0271 | 0.0079 | 0.0133 | 0.0133 | TRUE | 0.6157 |
| C3ar1 | 0 | 0 | 0 | 0 | 0 | 0 | 0 | 0 | 0 | 0 | 0 | 0 | 0 | 0 | 0 | NA | NA | NA | NA | NA | NA | NA | NA | NA | NA |
| C430002E04I | 241 | 187 | 186 | 193 | 139 | 159 | 187 | 199 | 201 | 192 | 157 | 157 | 182.11 | 196 | 169 | 0.864 | -0.211 | 0.17166764 | 0.32365725 | 0 | 0.0154 | 0.0116 | 0.0116 | TRUE | 0.1603 |
| C430049B03 | 39 | 44 | 31 | 36 | 27 | 27 | 30 | 47 | 33 | 36 | 30 | 27 | 33.91 | 37 | 31 | 0.846 | -0.242 | 0.48260157 | 0.6491459 | 0 | 0.0723 | 0.0555 | 0.0555 | TRUE | 0.3927 |
| C4a | 56 | 70 | 41 | 62 | 44 | 44 | 43 | 74 | 44 | 62 | 50 | 43 | 52.83 | 54 | 52 | 0.96 | -0.059 | 0.84492367 | 0.91201028 | 0.0473 | 0.0473 | 0.0473 | 0.0473 | TRUE | 0.879 |
| C4b | 30 | 16 | 18 | 16 | 15 | 17 | 23 | 17 | 19 | 16 | 17 | 17 | 18.23 | 20 | 17 | 0.825 | -0.277 | 0.53267209 | 0.69226061 | 0 | 0.1324 | 0.0868 | 0.0868 | TRUE | 0.1464 |

| | | | | | | | | | | | | | | | | | | | | | | | | | |
|---|---|---|---|---|---|---|---|---|---|---|---|---|---|---|---|---|---|---|---|---|---|---|---|---|---|
| C4bp | 0 | 0 | 0 | 0 | 0 | 0 | 0 | 0 | 0 | 0 | 0 | 0 | 0 | 0 | 0 | NA | NA | NA | NA | NA | NA | NA | NA | NA | NA |
| C4bp-ps1 | 2 | 1 | 1 | 1 | 3 | 2 | 2 | 1 | 1 | 1 | 3 | 2 | 1.68 | 1 | 2 | 1.682 | 0.75 | 0.60454026 | NA | 0 | 1.4174 | 0.8785 | 0.8785 | TRUE | 0.2074 |
| C530005A16 | 13 | 17 | 12 | 18 | 8 | 19 | 10 | 18 | 13 | 18 | 9 | 19 | 14.47 | 14 | 15 | 1.128 | 0.174 | 0.74191349 | NA | 0.0409 | 0.1662 | 0.1324 | 0.1324 | TRUE | 0.5258 |
| C530008M17 | 369 | 337 | 288 | 350 | 323 | 372 | 287 | 358 | 311 | 348 | 364 | 367 | 339.3 | 319 | 360 | 1.133 | 0.18 | 0.13964067 | 0.28083355 | 0.004 | 0.0094 | 0.0078 | 0.0078 | TRUE | 0.1349 |
| C530044C16 | 1 | 0 | 0 | 0 | 0 | 0 | 1 | 0 | 0 | 0 | 0 | 0 | 0.13 | 0 | 0 | 0.552 | -0.857 | 0.8336548 | NA | 0 | 18.3121 | 10 | 10 | TRUE | 0.2566 |
| C5ar1 | 0 | 0 | 0 | 0 | 0 | 0 | 0 | 0 | 0 | 0 | 0 | 0 | 0 | 0 | 0 | NA | NA | NA | NA | NA | NA | NA | NA | NA | NA |
| C5ar2 | 2 | 2 | 0 | 0 | 0 | 0 | 2 | 2 | 0 | 0 | 0 | 0 | 0.61 | 1 | 0 | 0.154 | -2.696 | 0.32099274 | NA | 0 | 3.8659 | 3.9247 | 3.9247 | TRUE | 0.3505 |
| C6 | 2 | 0 | 2 | 0 | 1 | 1 | 2 | 0 | 2 | 0 | 1 | 1 | 0.97 | 1 | 1 | 0.56 | -0.836 | 0.67994454 | NA | 0 | 2.443 | 1.8311 | 1.8311 | TRUE | 0.3957 |
| C630020P19 | 21 | 13 | 17 | 8 | 7 | 20 | 16 | 14 | 18 | 8 | 8 | 20 | 14.01 | 16 | 12 | 0.739 | -0.437 | 0.42650337 | NA | 0.0676 | 0.1715 | 0.1443 | 0.1443 | TRUE | 1.3536 |
| C630028M04 | 3 | 1 | 0 | 2 | 0 | 0 | 2 | 1 | 0 | 2 | 0 | 0 | 0.9 | 1 | 1 | 0.579 | -0.789 | 0.7251066 | NA | 0 | 2.6434 | 2.4396 | 2.4396 | TRUE | 0.4576 |
| C630031E19 | 0 | 0 | 0 | 0 | 0 | 0 | 0 | 0 | 0 | 0 | 0 | 0 | 0 | 0 | 0 | NA | NA | NA | NA | NA | NA | NA | NA | NA | NA |
| C630043F03 | 79 | 62 | 40 | 58 | 48 | 59 | 61 | 66 | 43 | 58 | 54 | 58 | 56.76 | 57 | 57 | 0.996 | -0.006 | 0.98089602 | 0.9900737 | 0 | 0.0442 | 0.0338 | 0.0338 | TRUE | 0.3479 |
| C7 | 51 | 26 | 33 | 11 | 14 | 18 | 40 | 28 | 36 | 11 | 16 | 18 | 24.56 | 35 | 15 | 0.43 | -1.216 | 0.00312997 | 0.01705684 | 0 | 0.0989 | 0.0732 | 0.0732 | TRUE | 0.2481 |
| C730002L08 | 0 | 0 | 0 | 0 | 0 | 0 | 0 | 0 | 0 | 0 | 0 | 0 | 0 | 0 | 0 | NA | NA | NA | NA | NA | NA | NA | NA | NA | NA |
| C730027H18 | 2 | 0 | 0 | 2 | 1 | 0 | 2 | 0 | 0 | 2 | 1 | 0 | 0.78 | 1 | 1 | 1.851 | 0.888 | 0.70714822 | NA | 0 | 3.046 | 2.6704 | 2.6704 | TRUE | 0.5318 |
| C730036E19 | 4 | 1 | 2 | 2 | 2 | 1 | 3 | 1 | 2 | 2 | 2 | 1 | 1.93 | 2 | 2 | 0.808 | -0.308 | 0.81870767 | NA | 0 | 1.2323 | 0.7763 | 0.7763 | TRUE | 0.2174 |
| C77080 | 5082 | 4138 | 4868 | 5364 | 5079 | 5621 | 3947 | 4402 | 5253 | 5339 | 5730 | 5549 | 5036.47 | 4534 | 5539 | 1.222 | 0.29 | 0.00125385 | 0.00849884 | 0.011 | 0.0029 | 0.0056 | 0.0056 | TRUE | 0.2341 |
| C77370 | 1958 | 1322 | 1187 | 1217 | 1099 | 1182 | 1521 | 1406 | 1281 | 1211 | 1240 | 1167 | 1304.28 | 1403 | 1206 | 0.859 | -0.22 | 0.00680589 | 0.03110861 | 0.0035 | 0.0042 | 0.004 | 0.004 | TRUE | 0.069 |
| C78339 | 1385 | 1012 | 976 | 1124 | 988 | 1130 | 1076 | 1076 | 1053 | 1119 | 1115 | 1115 | 1092.35 | 1068 | 1116 | 1.045 | 0.063 | 0.37102076 | 0.54568838 | 0 | 0.0046 | 0.0027 | 0.0027 | TRUE | 0.0018 |
| C86187 | 1 | 0 | 0 | 0 | 0 | 0 | 1 | 0 | 0 | 0 | 0 | 0 | 0.13 | 0 | 0 | 0.552 | -0.857 | 0.8336548 | NA | 0 | 18.3121 | 10 | 10 | TRUE | 0.2566 |
| C86695 | 21 | 16 | 4 | 3 | 7 | 4 | 16 | 17 | 4 | 3 | 8 | 4 | 8.75 | 12 | 5 | 0.387 | -1.371 | 0.06819946 | NA | 0.2366 | 0.2734 | 0.2632 | 0.2632 | TRUE | 1.2342 |
| C87198 | 0 | 0 | 0 | 0 | 0 | 0 | 0 | 0 | 0 | 0 | 0 | 0 | 0 | 0 | 0 | NA | NA | NA | NA | NA | NA | NA | NA | NA | NA |
| C87414 | 1 | 0 | 0 | 0 | 0 | 0 | 1 | 0 | 0 | 0 | 0 | 0 | 0.13 | 0 | 0 | 0.552 | -0.857 | 0.8336548 | NA | 0 | 18.3121 | 10 | 10 | TRUE | 0.2566 |
| C87436 | 422 | 380 | 330 | 345 | 349 | 396 | 328 | 404 | 356 | 343 | 394 | 391 | 369.34 | 363 | 376 | 1.039 | 0.055 | 0.65173026 | 0.7837103 | 0.006 | 0.0088 | 0.008 | 0.008 | TRUE | 0.1143 |
| C87499 | 0 | 2 | 0 | 1 | 1 | 0 | 0 | 2 | 0 | 1 | 1 | 0 | 0.71 | 1 | 1 | 1.021 | 0.03 | 0.99038186 | NA | 0 | 3.3477 | 2.9808 | 2.9808 | TRUE | 0.9748 |
| C87977 | 0 | 0 | 0 | 0 | 0 | 0 | 0 | 0 | 0 | 0 | 0 | 0 | 0 | 0 | 0 | NA | NA | NA | NA | NA | NA | NA | NA | NA | NA |
| C8a | 6 | 3 | 4 | 9 | 5 | 2 | 5 | 3 | 4 | 9 | 6 | 2 | 4.79 | 4 | 6 | 1.353 | 0.436 | 0.6283364 | NA | 0 | 0.4972 | 0.3732 | 0.3732 | TRUE | 0.1683 |
| C8b | 99 | 88 | 84 | 61 | 50 | 60 | 77 | 94 | 91 | 61 | 56 | 59 | 72.91 | 87 | 59 | 0.678 | -0.56 | 0.01501142 | 0.05588909 | 0 | 0.0349 | 0.0238 | 0.0238 | TRUE | 0.1121 |
| C8g | 121 | 74 | 98 | 136 | 125 | 153 | 94 | 79 | 106 | 135 | 141 | 151 | 117.64 | 93 | 142 | 1.535 | 0.618 | 0.00105598 | 0.00752776 | 0 | 0.0226 | 0.0168 | 0.0168 | TRUE | 0.1563 |
| C9 | 1 | 1 | 1 | 1 | 0 | 0 | 1 | 1 | 1 | 1 | 0 | 0 | 0.65 | 1 | 0 | 0.383 | -1.383 | 0.55308337 | NA | 0 | 3.6349 | 2.4186 | 2.4186 | TRUE | 0.3906 |
| C920006O11 | 153 | 111 | 103 | 150 | 93 | 141 | 119 | 118 | 111 | 149 | 105 | 139 | 123.57 | 116 | 131 | 1.132 | 0.179 | 0.34589981 | 0.52014162 | 0.0085 | 0.0216 | 0.018 | 0.018 | TRUE | 0.2989 |
| C920009B18 | 3 | 8 | 5 | 1 | 1 | 4 | 2 | 9 | 5 | 1 | 1 | 4 | 3.72 | 5 | 2 | 0.384 | -1.382 | 0.20471713 | NA | 0 | 0.6399 | 0.5116 | 0.5116 | TRUE | 0.2127 |
| C920021L13 | 3 | 2 | 4 | 3 | 4 | 4 | 2 | 2 | 4 | 3 | 5 | 4 | 3.37 | 3 | 4 | 1.314 | 0.394 | 0.69943563 | NA | 0 | 0.7057 | 0.4484 | 0.4484 | TRUE | 0.1985 |
| C920025E04 | 9 | 5 | 5 | 6 | 8 | 5 | 7 | 5 | 5 | 6 | 9 | 5 | 6.27 | 6 | 7 | 1.111 | 0.151 | 0.84080676 | NA | 0 | 0.3803 | 0.2496 | 0.2496 | TRUE | 0.2235 |
| Caap1 | 1543 | 1073 | 872 | 963 | 805 | 905 | 1198 | 1141 | 941 | 959 | 908 | 893 | 1006.78 | 1093 | 920 | 0.84 | -0.251 | 0.0097452 | 0.04062063 | 0.0078 | 0.0048 | 0.0058 | 0.0058 | TRUE | 0.1761 |
| Cab39 | 7643 | 5549 | 5096 | 5127 | 4638 | 5029 | 5936 | 5902 | 5499 | 5103 | 5232 | 4964 | 5439.48 | 5779 | 5100 | 0.882 | -0.181 | 0.00130374 | 0.00871753 | 0.001 | 0.0029 | 0.0021 | 0.0021 | TRUE | 0.0218 |
| Cab39l | 3843 | 2767 | 2749 | 3245 | 2918 | 3160 | 2985 | 2943 | 2966 | 3230 | 3292 | 3119 | 3089.22 | 2965 | 3214 | 1.084 | 0.116 | 0.03617265 | 0.10653527 | 0 | 0.0032 | 0.0019 | 0.0019 | TRUE | 0.008 |
| Cabin1 | 5892 | 4490 | 4001 | 4161 | 3960 | 4407 | 4576 | 4776 | 4317 | 4142 | 4467 | 4350 | 4438.1 | 4556 | 4320 | 0.948 | -0.077 | 0.207442 | 0.36909854 | 0.0017 | 0.003 | 0.0025 | 0.0025 | TRUE | 0.0253 |
| Cables1 | 165 | 118 | 111 | 112 | 100 | 108 | 128 | 126 | 120 | 111 | 113 | 107 | 117.39 | 125 | 110 | 0.885 | -0.177 | 0.32233901 | 0.49614381 | 0 | 0.0226 | 0.0144 | 0.0144 | TRUE | 0.0108 |
| Cables2 | 1426 | 1121 | 994 | 1034 | 972 | 1159 | 1108 | 1192 | 1073 | 1029 | 1097 | 1144 | 1107.05 | 1124 | 1090 | 0.97 | -0.044 | 0.5780871 | 0.72933171 | 0.002 | 0.0046 | 0.0037 | 0.0037 | TRUE | 0.0328 |
| Cabp1 | 348 | 271 | 240 | 350 | 283 | 377 | 270 | 288 | 259 | 348 | 319 | 372 | 309.55 | 272 | 346 | 1.273 | 0.349 | 0.00414072 | 0.02120505 | 0 | 0.0101 | 0.0074 | 0.0074 | TRUE | 0.0526 |
| Cabp2 | 5 | 4 | 2 | 7 | 12 | 4 | 4 | 4 | 2 | 7 | 14 | 4 | 5.79 | 3 | 8 | 2.339 | 1.226 | 0.15111714 | NA | 0 | 0.4116 | 0.3218 | 0.3218 | TRUE | 0.3116 |
| Cabp4 | 28 | 20 | 26 | 19 | 15 | 16 | 22 | 21 | 28 | 19 | 17 | 16 | 20.45 | 24 | 17 | 0.729 | -0.456 | 0.2810343 | 0.4526508 | 0 | 0.1183 | 0.0786 | 0.0786 | TRUE | 0.1431 |
| Cabp5 | 1 | 1 | 1 | 0 | 0 | 0 | 1 | 1 | 1 | 0 | 0 | 0 | 0.49 | 1 | 0 | 0.196 | -2.347 | 0.38405676 | NA | 0 | 4.8735 | 3.7415 | 3.7415 | TRUE | 0.0718 |
| Cabp7 | 64 | 41 | 48 | 47 | 52 | 50 | 50 | 44 | 52 | 47 | 59 | 49 | 49.99 | 49 | 52 | 1.063 | 0.089 | 0.7455214 | 0.84872461 | 0 | 0.0498 | 0.0339 | 0.0339 | TRUE | 0.1102 |
| Cabs1 | 2 | 0 | 0 | 0 | 0 | 0 | 2 | 0 | 0 | 0 | 0 | 0 | 0.26 | 1 | 0 | 0.356 | -1.491 | 0.71362796 | NA | 0 | 9.1573 | 10 | 10 | TRUE | 0.968 |
| Cabyr | 31 | 27 | 13 | 14 | 14 | 14 | 24 | 29 | 14 | 14 | 16 | 14 | 18.4 | 22 | 15 | 0.649 | -0.624 | 0.18154519 | 0.33712166 | 0 | 0.1313 | 0.1001 | 0.1001 | TRUE | 0.3917 |
| Cacfd1 | 431 | 333 | 345 | 357 | 328 | 349 | 335 | 354 | 372 | 355 | 370 | 345 | 355.18 | 354 | 357 | 1.009 | 0.014 | 0.90318718 | 0.94645925 | 0 | 0.0091 | 0.0061 | 0.0061 | TRUE | 0.0273 |
| Cachd1 | 459 | 327 | 235 | 331 | 306 | 338 | 356 | 348 | 254 | 329 | 345 | 334 | 327.7 | 319 | 336 | 1.049 | 0.07 | 0.6169593 | 0.75822866 | 0.014 | 0.0097 | 0.0109 | 0.0109 | TRUE | 0.3566 |
| Cacna1a | 2389 | 1906 | 2028 | 2393 | 2269 | 2625 | 1855 | 2027 | 2188 | 2382 | 2560 | 2591 | 2267.33 | 2023 | 2511 | 1.242 | 0.313 | 3.46E-05 | 0.00052377 | 0.0037 | 0.0035 | 0.0037 | 0.0037 | TRUE | 0.0658 |
| Cacna1b | 546 | 436 | 455 | 628 | 603 | 693 | 424 | 464 | 491 | 625 | 680 | 684 | 561.37 | 460 | 663 | 1.446 | 0.532 | 7.48E-08 | 3.02E-06 | 0.0022 | 0.0066 | 0.0052 | 0.0052 | TRUE | 0.0572 |
| Cacna1c | 140 | 81 | 91 | 86 | 107 | 129 | 109 | 86 | 98 | 86 | 121 | 127 | 104.46 | 98 | 111 | 1.134 | 0.181 | 0.3989953 | 0.57224403 | 0.0196 | 0.0251 | 0.0235 | 0.0235 | TRUE | 0.4129 |
| Cacna1d | 112 | 65 | 61 | 94 | 88 | 88 | 87 | 69 | 66 | 94 | 99 | 87 | 83.61 | 74 | 93 | 1.251 | 0.323 | 0.14251181 | 0.28511991 | 0 | 0.0308 | 0.0228 | 0.0228 | TRUE | 0.2439 |
| Cacna1e | 33 | 21 | 16 | 7 | 7 | 9 | 26 | 22 | 17 | 7 | 8 | 9 | 14.83 | 22 | 8 | 0.362 | -1.465 | 0.00527059 | NA | 0 | 0.1622 | 0.1113 | 0.1113 | TRUE | 0.1742 |
| Cacna1f | 3 | 3 | 6 | 4 | 5 | 6 | 2 | 3 | 6 | 4 | 6 | 6 | 4.59 | 4 | 5 | 1.319 | 0.4 | 0.65557841 | NA | 0 | 0.5188 | 0.3559 | 0.3559 | TRUE | 0.4586 |
| Cacna1g | 143 | 123 | 94 | 239 | 188 | 215 | 111 | 131 | 101 | 238 | 212 | 212 | 167.59 | 114 | 221 | 1.932 | 0.95 | 6.46E-09 | 3.47E-07 | 0.0017 | 0.0166 | 0.0127 | 0.0127 | TRUE | 0.1465 |
| Cacna1h | 490 | 429 | 471 | 726 | 713 | 785 | 381 | 456 | 508 | 723 | 804 | 775 | 607.83 | 448 | 767 | 1.717 | 0.78 | 4.68E-12 | 5.15E-10 | 0.0099 | 0.0063 | 0.0074 | 0.0074 | TRUE | 0.2165 |
| Cacna1i | 118 | 74 | 68 | 90 | 92 | 95 | 92 | 79 | 73 | 90 | 104 | 94 | 88.48 | 81 | 96 | 1.171 | 0.228 | 0.28064027 | 0.45221999 | 0 | 0.0292 | 0.0207 | 0.0207 | TRUE | 0.1322 |
| Cacna1s | 25 | 23 | 24 | 32 | 38 | 36 | 19 | 24 | 26 | 32 | 43 | 36 | 30.01 | 23 | 37 | 1.588 | 0.667 | 0.06173278 | 0.15657854 | 0 | 0.0814 | 0.057 | 0.057 | TRUE | 0.1536 |
| Cacna2d1 | 431 | 320 | 305 | 237 | 220 | 244 | 335 | 340 | 329 | 236 | 248 | 241 | 288.19 | 335 | 242 | 0.722 | -0.471 | 7.80E-05 | 0.00098999 | 0 | 0.0107 | 0.0066 | 0.0066 | TRUE | 0.0059 |
| Cacna2d2 | 275 | 258 | 282 | 367 | 352 | 355 | 214 | 274 | 304 | 365 | 397 | 350 | 317.52 | 264 | 371 | 1.411 | 0.496 | 0.00047504 | 0.0041553 | 0.0146 | 0.0099 | 0.0113 | 0.0113 | TRUE | 0.3388 |
| Cacna2d3 | 29 | 20 | 18 | 22 | 25 | 25 | 23 | 21 | 19 | 22 | 28 | 25 | 23 | 21 | 25 | 1.177 | 0.235 | 0.55205543 | 0.7078682 | 0 | 0.1055 | 0.0689 | 0.0689 | TRUE | 0.0737 |
| Cacna2d4 | 6 | 3 | 2 | 6 | 5 | 6 | 5 | 3 | 2 | 6 | 6 | 6 | 4.59 | 3 | 6 | 1.719 | 0.782 | 0.37678616 | NA | 0 | 0.5187 | 0.3334 | 0.3334 | TRUE | 0.2318 |
| Cacnb1 | 633 | 540 | 631 | 679 | 675 | 761 | 492 | 574 | 681 | 676 | 761 | 751 | 655.9 | 582 | 729 | 1.256 | 0.329 | 0.00527145 | 0.02552332 | 0.013 | 0.006 | 0.0085 | 0.0085 | TRUE | 0.2569 |
| Cacnb2 | 78 | 50 | 57 | 53 | 38 | 43 | 61 | 53 | 62 | 53 | 43 | 42 | 52.22 | 59 | 46 | 0.788 | -0.344 | 0.20052842 | 0.36124939 | 0 | 0.0478 | 0.0326 | 0.0326 | TRUE | 0.1322 |
| Cacnb3 | 1716 | 1442 | 1576 | 1583 | 1470 | 1767 | 1333 | 1534 | 1700 | 1576 | 1658 | 1744 | 1590.9 | 1522 | 1659 | 1.092 | 0.126 | 0.1658981 | 0.31683429 | 0.0077 | 0.0039 | 0.0054 | 0.0054 | TRUE | 0.1471 |
| Cacnb4 | 17 | 16 | 20 | 19 | 17 | 18 | 13 | 17 | 22 | 19 | 19 | 18 | 17.94 | 17 | 19 | 1.088 | 0.122 | 0.78768707 | 0.87526281 | 0 | 0.1345 | 0.092 | 0.092 | TRUE | 0.2673 |
| Cacng1 | 12 | 5 | 2 | 8 | 5 | 4 | 9 | 5 | 2 | 8 | 6 | 4 | 5.72 | 5 | 6 | 1.021 | 0.029 | 0.97182114 | NA | 0.1179 | 0.4164 | 0.3294 | 0.3294 | TRUE | 0.29 |
| Cacng2 | 5 | 0 | 1 | 4 | 2 | 1 | 4 | 0 | 1 | 4 | 2 | 1 | 2.03 | 2 | 2 | 1.386 | 0.471 | 0.74179429 | NA | 0.4488 | 1.1693 | 0.9801 | 0.9801 | TRUE | 0.885 |

| | | | | | | | | | | | | | | | | | | | | | | | | | |
|---|---|---|---|---|---|---|---|---|---|---|---|---|---|---|---|---|---|---|---|---|---|---|---|---|---|
| Cacng3 | 0 | 1 | 0 | 0 | 0 | 0 | 0 | 1 | 0 | 0 | 0 | 0 | 0.18 | 0 | 0 | 0.552 | -0.857 | 0.8336548 | NA | 0 | 13.3714 | 10 | 10 | TRUE | 0.5216 |
| Cacng4 | 104 | 95 | 95 | 77 | 68 | 92 | 81 | 101 | 103 | 77 | 77 | 91 | 88.08 | 95 | 82 | 0.864 | -0.211 | 0.32803327 | 0.50241958 | 0 | 0.0293 | 0.0221 | 0.0221 | TRUE | 0.1828 |
| Cacng5 | 275 | 195 | 206 | 350 | 349 | 343 | 214 | 207 | 222 | 348 | 394 | 339 | 287.33 | 214 | 360 | 1.678 | 0.747 | 2.86E-09 | 1.68E-07 | 0 | 0.0107 | 0.0077 | 0.0077 | TRUE | 0.0735 |
| Cacng6 | 146 | 106 | 117 | 193 | 166 | 178 | 113 | 113 | 126 | 192 | 187 | 176 | 151.25 | 117 | 185 | 1.577 | 0.658 | 5.02E-05 | 0.00070171 | 0 | 0.0181 | 0.012 | 0.012 | TRUE | 0.0406 |
| Cacng7 | 689 | 478 | 481 | 495 | 388 | 502 | 535 | 508 | 519 | 493 | 438 | 496 | 498.09 | 521 | 476 | 0.913 | -0.132 | 0.18553465 | 0.34182093 | 0 | 0.0072 | 0.0051 | 0.0051 | TRUE | 0.0539 |
| Cacng8 | 92 | 61 | 60 | 92 | 54 | 67 | 71 | 65 | 65 | 92 | 61 | 66 | 69.95 | 67 | 73 | 1.087 | 0.121 | 0.624804 | 0.76346484 | 0.0124 | 0.0363 | 0.0295 | 0.0295 | TRUE | 0.4623 |
| Cactin | 3718 | 2647 | 2070 | 2079 | 1867 | 2168 | 2888 | 2816 | 2234 | 2069 | 2106 | 2140 | 2375.41 | 2646 | 2105 | 0.795 | -0.33 | 0.00027228 | 0.00270069 | 0.0087 | 0.0034 | 0.0055 | 0.0055 | TRUE | 0.2236 |
| Cacul1 | 5601 | 4031 | 3885 | 4190 | 3781 | 4287 | 4350 | 4288 | 4192 | 4171 | 4265 | 4232 | 4249.61 | 4277 | 4223 | 0.987 | -0.019 | 0.71680161 | 0.82975813 | 0 | 0.003 | 0.0017 | 0.0017 | TRUE | 0.0036 |
| Cacybp | 15218 | 10590 | 10527 | 9962 | 8190 | 9940 | 11819 | 11264 | 11359 | 9916 | 9239 | 9812 | 10568.3 | 11481 | 9656 | 0.841 | -0.25 | 1.96E-06 | 5.01E-05 | 9.00E-04 | 0.0027 | 0.0019 | 0.0019 | TRUE | 0.0173 |
| Cad | 32873 | 23549 | 23957 | 25971 | 22853 | 26838 | 25531 | 25049 | 25849 | 25852 | 25781 | 26493 | 25759.09 | 25476 | 26042 | 1.022 | 0.032 | 0.47375558 | 0.64226392 | 2.00E-04 | 0.0025 | 0.0014 | 0.0014 | TRUE | 0.0028 |
| Cadm1 | 2682 | 2239 | 2478 | 2222 | 2198 | 2445 | 2083 | 2382 | 2674 | 2212 | 2480 | 2414 | 2373.88 | 2380 | 2369 | 0.996 | -0.006 | 0.94984301 | 0.97250044 | 0.0092 | 0.0034 | 0.0054 | 0.0054 | TRUE | 0.1466 |
| Cadm2 | 8 | 6 | 6 | 10 | 5 | 11 | 6 | 6 | 6 | 10 | 6 | 11 | 7.59 | 6 | 9 | 1.396 | 0.482 | 0.48548887 | NA | 0 | 0.3148 | 0.2095 | 0.2095 | TRUE | 0.279 |
| Cadm3 | 28 | 14 | 14 | 14 | 18 | 13 | 22 | 15 | 15 | 14 | 20 | 13 | 16.47 | 17 | 16 | 0.897 | -0.157 | 0.74376945 | NA | 0 | 0.1463 | 0.1057 | 0.1057 | TRUE | 0.3173 |
| Cadm4 | 606 | 536 | 602 | 953 | 809 | 893 | 471 | 570 | 650 | 949 | 913 | 882 | 738.85 | 564 | 915 | 1.628 | 0.703 | 3.10E-10 | 2.29E-08 | 0.0112 | 0.0056 | 0.0076 | 0.0076 | TRUE | 0.2566 |
| Cadps | 35 | 28 | 28 | 23 | 16 | 23 | 27 | 30 | 30 | 23 | 18 | 23 | 25.14 | 29 | 21 | 0.734 | -0.446 | 0.24010705 | 0.40734126 | 0 | 0.0967 | 0.0631 | 0.0631 | TRUE | 0.0859 |
| Cadps2 | 69 | 54 | 43 | 66 | 61 | 47 | 54 | 57 | 46 | 66 | 69 | 46 | 56.39 | 52 | 60 | 1.146 | 0.196 | 0.468547 | 0.63749644 | 0 | 0.0445 | 0.0349 | 0.0349 | TRUE | 0.3595 |
| Cage1 | 43 | 35 | 32 | 48 | 51 | 51 | 33 | 37 | 35 | 48 | 58 | 50 | 43.47 | 35 | 52 | 1.481 | 0.566 | 0.05201533 | 0.13879494 | 0 | 0.057 | 0.0375 | 0.0375 | TRUE | 0.0699 |
| Calb1 | 19 | 19 | 17 | 9 | 12 | 8 | 15 | 20 | 18 | 9 | 14 | 8 | 13.95 | 18 | 10 | 0.569 | -0.813 | 0.12182374 | NA | 0 | 0.1723 | 0.12 | 0.12 | TRUE | 0.2713 |
| Calb2 | 324 | 270 | 217 | 247 | 229 | 235 | 252 | 287 | 234 | 246 | 258 | 232 | 251.53 | 258 | 245 | 0.952 | -0.071 | 0.601044 | 0.74643513 | 0.0026 | 0.0118 | 0.0092 | 0.0092 | TRUE | 0.1067 |
| Calca | 263 | 229 | 304 | 443 | 344 | 426 | 204 | 244 | 328 | 441 | 388 | 421 | 337.57 | 259 | 417 | 1.619 | 0.695 | 8.74E-06 | 0.00016784 | 0.0272 | 0.0094 | 0.0145 | 0.0145 | TRUE | 0.6096 |
| Calcb | 0 | 0 | 0 | 2 | 1 | 2 | 0 | 0 | 0 | 2 | 1 | 2 | 0.85 | 0 | 2 | 9.62 | 3.266 | 0.1405906 | NA | 0 | 2.7945 | 2.2362 | 2.2362 | TRUE | 0.0847 |
| Calcoco1 | 2298 | 1817 | 1833 | 1746 | 1667 | 1906 | 1785 | 1933 | 1978 | 1738 | 1881 | 1882 | 1865.88 | 1899 | 1834 | 0.966 | -0.049 | 0.48520606 | 0.65111743 | 0.002 | 0.0037 | 0.0031 | 0.0031 | TRUE | 0.0343 |
| Calcoco2 | 4991 | 2524 | 1501 | 938 | 732 | 890 | 3876 | 2685 | 1620 | 934 | 826 | 879 | 1803.11 | 2727 | 880 | 0.322 | -1.633 | 2.68E-06 | 6.34E-05 | 0.0865 | 0.0037 | 0.0258 | 0.0865 | TRUE | 0.0937 |
| Calcr | 47 | 32 | 28 | 17 | 18 | 19 | 37 | 34 | 30 | 17 | 20 | 19 | 26.12 | 34 | 19 | 0.553 | -0.855 | 0.02395386 | 0.07888767 | 0 | 0.0932 | 0.0611 | 0.0611 | TRUE | 0.0515 |
| Calcrl | 18 | 12 | 13 | 9 | 7 | 7 | 14 | 13 | 14 | 9 | 8 | 7 | 10.76 | 14 | 8 | 0.582 | -0.78 | 0.17879184 | NA | 0 | 0.2228 | 0.1418 | 0.1418 | TRUE | 0.0403 |
| Cald1 | 3066 | 2029 | 2126 | 2352 | 1863 | 2016 | 2381 | 2158 | 2294 | 2341 | 2102 | 1990 | 2211.05 | 2278 | 2144 | 0.941 | -0.088 | 0.25195907 | 0.42090538 | 0.004 | 0.0035 | 0.0038 | 0.0038 | TRUE | 0.0787 |
| Calhm1 | 1 | 6 | 0 | 1 | 2 | 4 | 1 | 6 | 0 | 1 | 2 | 4 | 2.39 | 2 | 2 | 1.026 | 0.038 | 0.97858688 | NA | 0.9837 | 0.9928 | 0.9903 | 0.9903 | TRUE | 1.7499 |
| Calhm2 | 124 | 94 | 108 | 95 | 87 | 86 | 96 | 100 | 117 | 95 | 98 | 85 | 98.4 | 104 | 93 | 0.889 | -0.17 | 0.39649465 | 0.56990124 | 0 | 0.0265 | 0.0186 | 0.0186 | TRUE | 0.1006 |
| Calm1 | 13992 | 9089 | 7090 | 7142 | 5887 | 6824 | 10867 | 9668 | 7650 | 7109 | 6641 | 6736 | 8111.92 | 9395 | 6829 | 0.727 | -0.461 | 3.82E-06 | 8.50E-05 | 0.0154 | 0.0027 | 0.007 | 0.007 | TRUE | 0.2899 |
| Calm2 | 21312 | 15393 | 15457 | 16213 | 14376 | 16333 | 16552 | 16373 | 16678 | 16138 | 16218 | 16123 | 16347.11 | 16534 | 16160 | 0.977 | -0.033 | 0.4440636 | 0.61501948 | 0 | 0.0026 | 0.0013 | 0.0013 | TRUE | 9.00E-04 |
| Calm3 | 11395 | 8239 | 8057 | 8137 | 7187 | 8020 | 8850 | 8764 | 8693 | 8100 | 8108 | 7917 | 8405.23 | 8769 | 8042 | 0.917 | -0.125 | 0.00703023 | 0.03184097 | 0 | 0.0027 | 0.0014 | 0.0014 | TRUE | 0.0022 |
| Calm4 | 0 | 0 | 0 | 0 | 0 | 0 | 0 | 0 | 0 | 0 | 0 | 0 | 0 | 0 | 0 | NA | NA | NA | NA | NA | NA | NA | NA | NA | NA |
| Calm5 | 0 | 0 | 0 | 0 | 0 | 0 | 0 | 0 | 0 | 0 | 0 | 0 | 0 | 0 | 0 | NA | NA | NA | NA | NA | NA | NA | NA | NA | NA |
| Calml3 | 0 | 0 | 1 | 0 | 0 | 0 | 0 | 0 | 1 | 0 | 0 | 0 | 0.18 | 0 | 0 | 0.552 | -0.857 | 0.8336548 | NA | 0 | 13.1817 | 10 | 10 | TRUE | 0.5365 |
| Calml4 | 189 | 110 | 152 | 198 | 165 | 196 | 147 | 117 | 164 | 197 | 186 | 193 | 167.42 | 143 | 192 | 1.348 | 0.43 | 0.00955002 | 0.04001714 | 0.0056 | 0.0166 | 0.0138 | 0.0138 | TRUE | 0.2403 |
| Caln1 | 6 | 5 | 0 | 1 | 2 | 0 | 5 | 5 | 0 | 1 | 2 | 0 | 2.2 | 3 | 1 | 0.315 | -1.665 | 0.27969148 | NA | 0.9768 | 1.0773 | 1.0763 | 1.0763 | TRUE | 0.9901 |
| Calr | 41574 | 29677 | 28175 | 33504 | 27657 | 31726 | 32289 | 31567 | 30401 | 33350 | 31200 | 31318 | 31687.44 | 31419 | 31956 | 1.017 | 0.024 | 0.63508809 | 0.77168359 | 0.001 | 0.0025 | 0.0019 | 0.0019 | TRUE | 0.0178 |
| Calr3 | 120 | 91 | 75 | 114 | 95 | 79 | 93 | 97 | 81 | 113 | 107 | 78 | 94.92 | 90 | 99 | 1.1 | 0.137 | 0.5277348 | 0.68871694 | 0.0131 | 0.0274 | 0.0234 | 0.0234 | TRUE | 0.3596 |
| Calr4 | 74 | 41 | 42 | 55 | 65 | 48 | 57 | 44 | 45 | 55 | 73 | 47 | 53.64 | 49 | 58 | 1.187 | 0.247 | 0.38257164 | 0.55654925 | 0.0198 | 0.0466 | 0.039 | 0.039 | TRUE | 0.3987 |
| Calu | 16866 | 11957 | 12604 | 14823 | 12877 | 14068 | 13099 | 12719 | 13600 | 14755 | 14527 | 13887 | 13764.32 | 13139 | 14390 | 1.095 | 0.131 | 0.0114616 | 0.04572163 | 9.00E-04 | 0.0026 | 0.0019 | 0.0019 | TRUE | 0.0114 |
| Caly | 34 | 22 | 33 | 22 | 23 | 23 | 26 | 23 | 36 | 22 | 26 | 23 | 25.99 | 28 | 24 | 0.827 | -0.274 | 0.47241996 | 0.64096316 | 0 | 0.0936 | 0.0658 | 0.0658 | TRUE | 0.2879 |
| Camk1 | 796 | 507 | 419 | 570 | 433 | 532 | 618 | 539 | 452 | 567 | 488 | 525 | 531.77 | 536 | 527 | 0.98 | -0.029 | 0.81264054 | 0.8917342 | 0.0132 | 0.0069 | 0.0089 | 0.0089 | TRUE | 0.2043 |
| Camk1d | 996 | 769 | 877 | 968 | 993 | 1073 | 774 | 818 | 946 | 964 | 1120 | 1059 | 946.8 | 846 | 1048 | 1.24 | 0.31 | 0.00136875 | 0.00903212 | 0.0069 | 0.0049 | 0.0057 | 0.0057 | TRUE | 0.1268 |
| Camk1g | 0 | 1 | 1 | 3 | 0 | 1 | 0 | 1 | 1 | 3 | 0 | 1 | 1.02 | 1 | 1 | 1.967 | 0.976 | 0.63038931 | NA | 0 | 2.3275 | 1.8702 | 1.8702 | TRUE | 0.3453 |
| Camk2a | 304 | 183 | 208 | 365 | 314 | 300 | 236 | 195 | 224 | 363 | 354 | 296 | 278.15 | 218 | 338 | 1.542 | 0.625 | 5.56E-06 | 0.00011583 | 0.0075 | 0.0109 | 0.0099 | 0.0099 | TRUE | 0.1353 |
| Camk2b | 2218 | 1685 | 1820 | 1979 | 1880 | 2227 | 1723 | 1792 | 1964 | 1970 | 2121 | 2198 | 1961.3 | 1826 | 2096 | 1.149 | 0.2 | 0.00749027 | 0.03340357 | 0.0031 | 0.0036 | 0.0035 | 0.0035 | TRUE | 0.0521 |
| Camk2d | 416 | 262 | 273 | 280 | 227 | 231 | 323 | 279 | 295 | 279 | 256 | 228 | 276.53 | 299 | 254 | 0.849 | -0.236 | 0.07486348 | 0.18004295 | 0.0044 | 0.011 | 0.009 | 0.009 | TRUE | 0.0935 |
| Camk2g | 2993 | 2131 | 2153 | 2434 | 2092 | 2415 | 2325 | 2267 | 2323 | 2423 | 2360 | 2384 | 2346.85 | 2305 | 2389 | 1.036 | 0.052 | 0.36805295 | 0.54268049 | 0 | 0.0034 | 0.0019 | 0.0019 | TRUE | 0.0025 |
| Camk2n1 | 411 | 378 | 583 | 848 | 764 | 871 | 319 | 402 | 629 | 844 | 862 | 860 | 652.69 | 450 | 855 | 1.902 | 0.928 | 0.0016637 | 0.01045928 | 0.061 | 0.0061 | 0.0199 | 0.061 | TRUE | 1.3977 |
| Camk2n2 | 117 | 86 | 97 | 105 | 73 | 125 | 91 | 91 | 105 | 105 | 82 | 123 | 99.55 | 96 | 103 | 1.086 | 0.119 | 0.57558669 | 0.72733274 | 0.0126 | 0.0262 | 0.0225 | 0.0225 | TRUE | 0.2032 |
| Camk4 | 39 | 24 | 23 | 18 | 12 | 21 | 30 | 26 | 25 | 18 | 14 | 21 | 22.14 | 27 | 18 | 0.647 | -0.628 | 0.12811643 | 0.2643494 | 0 | 0.1095 | 0.0751 | 0.0751 | TRUE | 0.1683 |
| Camkk1 | 823 | 709 | 705 | 814 | 708 | 810 | 639 | 754 | 761 | 810 | 799 | 800 | 760.43 | 718 | 803 | 1.121 | 0.165 | 0.07586208 | 0.18165592 | 0.0036 | 0.0055 | 0.0049 | 0.0049 | TRUE | 0.1155 |
| Camkk2 | 1088 | 833 | 912 | 999 | 902 | 1063 | 845 | 886 | 984 | 994 | 1018 | 1049 | 962.73 | 905 | 1020 | 1.129 | 0.175 | 0.03655992 | 0.10745444 | 0.0024 | 0.0049 | 0.004 | 0.004 | TRUE | 0.0687 |
| Camkmt | 308 | 223 | 175 | 193 | 156 | 180 | 239 | 237 | 189 | 192 | 176 | 178 | 201.84 | 222 | 182 | 0.819 | -0.288 | 0.05724383 | 0.14866207 | 0.0053 | 0.0142 | 0.0116 | 0.0116 | TRUE | 0.1781 |
| Camkv | 838 | 691 | 727 | 918 | 806 | 1030 | 651 | 735 | 784 | 914 | 909 | 1017 | 835.01 | 723 | 947 | 1.312 | 0.391 | 3.74E-05 | 0.00055539 | 0.0053 | 0.0053 | 0.0053 | 0.0053 | TRUE | 0.0961 |
| Caml | 772 | 593 | 494 | 551 | 466 | 566 | 600 | 631 | 533 | 548 | 526 | 559 | 566.04 | 588 | 544 | 0.926 | -0.111 | 0.25945856 | 0.42946626 | 0.0022 | 0.0066 | 0.0052 | 0.0052 | TRUE | 0.0746 |
| Camp | 2 | 0 | 0 | 0 | 0 | 0 | 2 | 0 | 0 | 0 | 0 | 0 | 0.26 | 1 | 0 | 0.356 | -1.491 | 0.71362796 | NA | 0 | 9.1573 | 10 | 10 | TRUE | 0.968 |
| Camsap1 | 5469 | 3989 | 4072 | 4462 | 3855 | 4358 | 4248 | 4243 | 4394 | 4441 | 4349 | 4302 | 4329.43 | 4295 | 4364 | 1.016 | 0.023 | 0.65324647 | 0.78460939 | 0 | 0.003 | 0.0017 | 0.0017 | TRUE | 0.0049 |
| Camsap2 | 3449 | 2408 | 2272 | 2340 | 2071 | 2372 | 2679 | 2561 | 2451 | 2329 | 2336 | 2342 | 2449.76 | 2564 | 2336 | 0.91 | -0.135 | 0.02680666 | 0.08544773 | 7.00E-04 | 0.0034 | 0.0023 | 0.0023 | TRUE | 0.0192 |
| Camsap3 | 4210 | 3239 | 3134 | 2908 | 2690 | 2983 | 3270 | 3445 | 3382 | 2895 | 3035 | 2945 | 3161.75 | 3366 | 2958 | 0.879 | -0.186 | 0.0010178 | 0.00732866 | 4.00E-04 | 0.0032 | 0.002 | 0.002 | TRUE | 0.0078 |
| Camta1 | 2511 | 1955 | 2318 | 2576 | 2309 | 2705 | 1950 | 2080 | 2501 | 2564 | 2605 | 2670 | 2395 | 2177 | 2613 | 1.201 | 0.265 | 0.00243721 | 0.01408179 | 0.0083 | 0.0034 | 0.0051 | 0.0051 | TRUE | 0.2045 |
| Camta2 | 1084 | 818 | 800 | 850 | 759 | 888 | 842 | 870 | 863 | 846 | 856 | 877 | 859.02 | 858 | 860 | 1.002 | 0.003 | 0.96966044 | 0.9834812 | 0 | 0.0052 | 0.0031 | 0.0031 | TRUE | 0.0036 |
| Cand1 | 24463 | 17518 | 17571 | 17544 | 15849 | 17879 | 18999 | 18634 | 18959 | 17463 | 17879 | 17649 | 18263.99 | 18864 | 17664 | 0.936 | -0.095 | 0.03013278 | 0.09296091 | 0 | 0.0026 | 0.0013 | 0.0013 | TRUE | 0.0014 |
| Cand2 | 1798 | 1324 | 1116 | 964 | 967 | 991 | 1396 | 1408 | 1204 | 960 | 1091 | 978 | 1172.94 | 1336 | 1010 | 0.755 | -0.406 | 4.21E-06 | 9.21E-05 | 0.0048 | 0.0044 | 0.0047 | 0.0047 | TRUE | 0.0883 |
| Cant1 | 2825 | 2041 | 2058 | 2225 | 1958 | 2278 | 2194 | 2171 | 2221 | 2215 | 2209 | 2249 | 2209.66 | 2195 | 2224 | 1.013 | 0.019 | 0.74239206 | 0.84665858 | 0 | 0.0035 | 0.002 | 0.002 | TRUE | 0.0012 |
| Canx | 67262 | 49614 | 52957 | 59987 | 50413 | 57087 | 52240 | 52774 | 57140 | 59711 | 56871 | 56354 | 55848.21 | 54051 | 57645 | 1.067 | 0.093 | 0.08800343 | 0.20157581 | 0.0017 | 0.0025 | 0.0021 | 0.0021 | TRUE | 0.0306 |
| Cap1 | 14767 | 10490 | 10172 | 9835 | 8905 | 10085 | 11469 | 11158 | 10976 | 9790 | 10046 | 9955 | 10565.57 | 11201 | 9930 | 0.886 | -0.174 | 0.0002325 | 0.0023994 | 2.00E-04 | 0.0027 | 0.0015 | 0.0015 | TRUE | 0.0054 |
| Cap2 | 164 | 89 | 121 | 178 | 154 | 143 | 127 | 95 | 131 | 177 | 174 | 141 | 140.78 | 118 | 164 | 1.391 | 0.476 | 0.01081444 | 0.04375059 | 0.0142 | 0.0193 | 0.0179 | 0.0179 | TRUE | 0.2779 |
| Capg | 1062 | 814 | 984 | 1130 | 1056 | 1232 | 825 | 866 | 1062 | 1125 | 1191 | 1216 | 1047.44 | 918 | 1177 | 1.285 | 0.362 | 0.00024315 | 0.00247705 | 0.0089 | 0.0047 | 0.0061 | 0.0061 | TRUE | 0.2231 |

| | | | | | | | | | | | | | | | | | | | | | | | | | |
|---|---|---|---|---|---|---|---|---|---|---|---|---|---|---|---|---|---|---|---|---|---|---|---|---|---|
| Capn1 | 1932 | 1386 | 1449 | 1378 | 1254 | 1449 | 1501 | 1474 | 1563 | 1372 | 1415 | 1430 | 1459.15 | 1513 | 1406 | 0.929 | -0.106 | 0.11388688 | 0.24288051 | 0 | 0.0041 | 0.0025 | 0.0025 | TRUE | 0.01 |
| Capn10 | 1871 | 1405 | 1244 | 1167 | 1100 | 1210 | 1453 | 1494 | 1342 | 1162 | 1241 | 1194 | 1314.48 | 1430 | 1199 | 0.838 | -0.255 | 0.00056501 | 0.00473647 | 0.0013 | 0.0042 | 0.0032 | 0.0032 | TRUE | 0.0336 |
| Capn11 | 109 | 84 | 82 | 121 | 114 | 137 | 85 | 89 | 88 | 120 | 129 | 135 | 107.79 | 87 | 128 | 1.466 | 0.552 | 0.00330401 | 0.01780033 | 0 | 0.0244 | 0.0159 | 0.0159 | TRUE | 0.0288 |
| Capn12 | 110 | 67 | 70 | 121 | 110 | 119 | 85 | 71 | 76 | 120 | 124 | 117 | 99.04 | 77 | 120 | 1.551 | 0.633 | 0.00135311 | 0.00895785 | 0 | 0.0264 | 0.0177 | 0.0177 | TRUE | 0.0869 |
| Capn13 | 1 | 0 | 0 | 0 | 0 | 1 | 1 | 0 | 0 | 0 | 0 | 1 | 0.29 | 0 | 0 | 1.075 | 0.105 | 0.97702352 | NA | 0 | 8.0647 | 7.5458 | 7.5458 | TRUE | 0.3835 |
| Capn2 | 1012 | 756 | 958 | 1022 | 924 | 1111 | 786 | 804 | 1034 | 1017 | 1042 | 1097 | 963.37 | 875 | 1052 | 1.205 | 0.269 | 0.01082865 | 0.04379566 | 0.0116 | 0.0049 | 0.007 | 0.007 | TRUE | 0.2977 |
| Capn3 | 120 | 86 | 70 | 108 | 84 | 91 | 93 | 91 | 76 | 108 | 95 | 90 | 92.05 | 87 | 98 | 1.12 | 0.163 | 0.433635 | 0.60556376 | 0 | 0.0282 | 0.0204 | 0.0204 | TRUE | 0.1147 |
| Capn5 | 2126 | 1737 | 2215 | 2407 | 2373 | 2647 | 1651 | 1848 | 2390 | 2396 | 2677 | 2613 | 2262.45 | 1963 | 2562 | 1.306 | 0.385 | 0.00051332 | 0.00439381 | 0.02 | 0.0035 | 0.0084 | 0.0084 | TRUE | 0.437 |
| Capn6 | 226 | 168 | 165 | 71 | 46 | 52 | 176 | 179 | 178 | 71 | 52 | 51 | 117.69 | 178 | 58 | 0.327 | -1.611 | 6.81E-16 | 1.30E-13 | 0 | 0.0226 | 0.017 | 0.017 | TRUE | 0.3207 |
| Capn7 | 3368 | 2380 | 2272 | 2482 | 2061 | 2460 | 2616 | 2532 | 2451 | 2471 | 2325 | 2428 | 2470.47 | 2533 | 2408 | 0.95 | -0.073 | 0.22930329 | 0.39472864 | 6.00E-04 | 0.0034 | 0.0023 | 0.0023 | TRUE | 0.0108 |
| Capn8 | 4 | 1 | 1 | 3 | 2 | 0 | 3 | 1 | 1 | 3 | 2 | 0 | 1.75 | 2 | 2 | 0.967 | -0.049 | 0.97390668 | NA | 0 | 1.3578 | 1.0492 | 1.0492 | TRUE | 0.6383 |
| Capn9 | 20 | 9 | 15 | 22 | 9 | 15 | 16 | 10 | 16 | 22 | 10 | 15 | 14.69 | 14 | 16 | 1.136 | 0.183 | 0.72615587 | NA | 0.0331 | 0.1637 | 0.1296 | 0.1296 | TRUE | 0.3005 |
| Capns1 | 12511 | 9059 | 9153 | 10984 | 9355 | 10944 | 9717 | 9636 | 9876 | 10933 | 10553 | 10803 | 10253.17 | 9743 | 10763 | 1.105 | 0.144 | 0.00199164 | 0.01207745 | 1.00E-04 | 0.0027 | 0.0015 | 0.0015 | TRUE | 0.0035 |
| Capns2 | 6 | 3 | 2 | 6 | 5 | 5 | 5 | 3 | 2 | 6 | 6 | 5 | 4.43 | 3 | 6 | 1.621 | 0.697 | 0.43770477 | NA | 0 | 0.5379 | 0.346 | 0.346 | TRUE | 0.2315 |
| Caprin1 | 43107 | 31900 | 32132 | 33084 | 29814 | 35460 | 33479 | 33932 | 34670 | 32932 | 33633 | 35004 | 33941.78 | 34027 | 33856 | 0.995 | -0.007 | 0.87949443 | 0.93288415 | 5.00E-04 | 0.0025 | 0.0016 | 0.0016 | TRUE | 0.0108 |
| Caprin2 | 1714 | 1303 | 1306 | 1436 | 1168 | 1383 | 1331 | 1386 | 1409 | 1429 | 1318 | 1365 | 1373.1 | 1375 | 1371 | 0.998 | -0.003 | 0.96062776 | 0.97828661 | 6.00E-04 | 0.0042 | 0.0028 | 0.0028 | TRUE | 0.0169 |
| Caps2 | 80 | 93 | 65 | 79 | 71 | 80 | 62 | 99 | 70 | 79 | 80 | 79 | 78.15 | 77 | 79 | 1.034 | 0.048 | 0.84003378 | 0.90890647 | 0.0185 | 0.0328 | 0.0287 | 0.0287 | TRUE | 0.5563 |
| Capsl | 442 | 322 | 295 | 349 | 240 | 307 | 343 | 343 | 318 | 347 | 271 | 303 | 320.88 | 335 | 307 | 0.918 | -0.123 | 0.32994623 | 0.50437663 | 0.0051 | 0.0098 | 0.0084 | 0.0084 | TRUE | 0.1493 |
| Capza1 | 2955 | 1993 | 2045 | 2076 | 1933 | 2136 | 2295 | 2120 | 2207 | 2066 | 2181 | 2109 | 2162.85 | 2207 | 2119 | 0.959 | -0.06 | 0.34055987 | 0.51467152 | 7.00E-04 | 0.0035 | 0.0024 | 0.0024 | TRUE | 0.0152 |
| Capza2 | 8676 | 5953 | 5870 | 6502 | 5370 | 6156 | 6738 | 6332 | 6334 | 6472 | 6058 | 6077 | 6335.17 | 6468 | 6202 | 0.959 | -0.061 | 0.27860286 | 0.4501033 | 0.0011 | 0.0028 | 0.0021 | 0.0021 | TRUE | 0.0177 |
| Capza3 | 0 | 0 | 0 | 0 | 0 | 0 | 0 | 0 | 0 | 0 | 0 | 0 | 0 | 0 | 0 | NA | NA | NA | NA | NA | NA | NA | NA | NA | NA |
| Capzb | 13196 | 9244 | 9389 | 9512 | 8674 | 9993 | 10249 | 9833 | 10131 | 9468 | 9785 | 9865 | 9888.36 | 10071 | 9706 | 0.964 | -0.053 | 0.27115761 | 0.44203202 | 3.00E-04 | 0.0027 | 0.0016 | 0.0016 | TRUE | 0.0056 |
| Car1 | 0 | 0 | 1 | 0 | 0 | 0 | 0 | 0 | 1 | 0 | 0 | 0 | 0.18 | 0 | 0 | 0.552 | -0.857 | 0.8336548 | NA | 0 | 13.1817 | 10 | 10 | TRUE | 0.5365 |
| Car10 | 8 | 6 | 4 | 6 | 4 | 2 | 6 | 6 | 4 | 6 | 5 | 2 | 4.9 | 5 | 4 | 0.732 | -0.45 | 0.60481225 | NA | 0 | 0.4866 | 0.3337 | 0.3337 | TRUE | 0.3848 |
| Car11 | 22 | 20 | 25 | 25 | 28 | 35 | 17 | 21 | 27 | 25 | 32 | 35 | 26.06 | 22 | 31 | 1.406 | 0.491 | 0.20463104 | 0.36610233 | 0 | 0.0934 | 0.0686 | 0.0686 | TRUE | 0.2522 |
| Car12 | 797 | 589 | 707 | 622 | 587 | 657 | 619 | 627 | 763 | 619 | 662 | 649 | 656.38 | 670 | 643 | 0.962 | -0.056 | 0.58873648 | 0.73762904 | 0.0062 | 0.006 | 0.0061 | 0.0061 | TRUE | 0.1725 |
| Car13 | 667 | 552 | 661 | 662 | 595 | 685 | 518 | 587 | 713 | 659 | 671 | 676 | 637.46 | 606 | 669 | 1.106 | 0.146 | 0.20406979 | 0.36544644 | 0.011 | 0.0061 | 0.0079 | 0.0079 | TRUE | 0.2793 |
| Car14 | 62 | 32 | 38 | 78 | 53 | 85 | 48 | 34 | 41 | 78 | 60 | 84 | 57.42 | 41 | 74 | 1.788 | 0.839 | 0.00215234 | 0.01275434 | 0.0115 | 0.0437 | 0.0352 | 0.0352 | TRUE | 0.2369 |
| Car15 | 5 | 3 | 5 | 1 | 3 | 1 | 4 | 3 | 5 | 1 | 3 | 1 | 2.97 | 4 | 2 | 0.423 | -1.242 | 0.28070661 | NA | 0 | 0.7997 | 0.5433 | 0.5433 | TRUE | 0.434 |
| Car2 | 2678 | 1772 | 1412 | 1110 | 969 | 1090 | 2080 | 1885 | 1524 | 1105 | 1093 | 1076 | 1460.38 | 1830 | 1091 | 0.596 | -0.747 | 2.40E-13 | 3.30E-11 | 0.0121 | 0.004 | 0.0068 | 0.0068 | TRUE | 0.2569 |
| Car3 | 21 | 14 | 9 | 20 | 7 | 11 | 16 | 15 | 10 | 20 | 8 | 11 | 13.26 | 14 | 13 | 0.944 | -0.084 | 0.88112286 | NA | 0.0657 | 0.1811 | 0.1496 | 0.1496 | TRUE | 0.9487 |
| Car4 | 709 | 349 | 304 | 244 | 164 | 199 | 551 | 371 | 328 | 243 | 185 | 196 | 312.37 | 417 | 208 | 0.498 | -1.005 | 4.82E-08 | 2.05E-06 | 0.0456 | 0.01 | 0.0208 | 0.0208 | TRUE | 0.9396 |
| Car5a | 30 | 25 | 37 | 26 | 29 | 30 | 23 | 27 | 40 | 26 | 33 | 30 | 29.67 | 30 | 30 | 0.989 | -0.017 | 0.9642485 | 0.98029759 | 0 | 0.0823 | 0.064 | 0.064 | TRUE | 0.5404 |
| Car5b | 53 | 27 | 43 | 50 | 47 | 56 | 41 | 29 | 46 | 50 | 53 | 55 | 45.73 | 39 | 53 | 1.357 | 0.44 | 0.13654583 | 0.27647677 | 0 | 0.0543 | 0.0407 | 0.0407 | TRUE | 0.3506 |
| Car6 | 137 | 69 | 64 | 81 | 81 | 97 | 106 | 73 | 69 | 81 | 91 | 96 | 86.1 | 83 | 89 | 1.066 | 0.093 | 0.69162899 | 0.81149322 | 0.0229 | 0.03 | 0.0279 | 0.0279 | TRUE | 0.6342 |
| Car7 | 100 | 70 | 56 | 68 | 53 | 58 | 78 | 74 | 60 | 68 | 60 | 57 | 66.21 | 71 | 62 | 0.866 | -0.207 | 0.39377369 | 0.56735961 | 0 | 0.0382 | 0.0272 | 0.0272 | TRUE | 0.1421 |
| Car8 | 222 | 173 | 205 | 202 | 178 | 200 | 172 | 184 | 221 | 201 | 201 | 197 | 196.16 | 192 | 200 | 1.041 | 0.058 | 0.70066966 | 0.8185838 | 0.0038 | 0.0145 | 0.0114 | 0.0114 | TRUE | 0.1792 |
| Car9 | 106 | 71 | 108 | 167 | 162 | 170 | 82 | 76 | 117 | 166 | 183 | 168 | 131.86 | 92 | 172 | 1.89 | 0.919 | 2.43E-06 | 5.87E-05 | 0.0148 | 0.0204 | 0.019 | 0.019 | TRUE | 0.5209 |
| Card10 | 1088 | 873 | 757 | 861 | 745 | 943 | 845 | 929 | 817 | 857 | 840 | 931 | 869.79 | 864 | 876 | 1.015 | 0.022 | 0.79815315 | 0.88215303 | 0.0025 | 0.0052 | 0.0043 | 0.0043 | TRUE | 0.0503 |
| Card11 | 48 | 29 | 32 | 41 | 52 | 60 | 37 | 31 | 35 | 41 | 59 | 59 | 43.56 | 34 | 53 | 1.538 | 0.621 | 0.04101417 | 0.11654644 | 0 | 0.0568 | 0.0428 | 0.0428 | TRUE | 0.3381 |
| Card14 | 17 | 11 | 13 | 9 | 14 | 17 | 13 | 12 | 14 | 9 | 16 | 17 | 13.41 | 13 | 14 | 1.064 | 0.09 | 0.86490331 | NA | 0 | 0.1792 | 0.1259 | 0.1259 | TRUE | 0.4261 |
| Card6 | 164 | 103 | 109 | 83 | 95 | 69 | 127 | 110 | 118 | 83 | 107 | 68 | 102.07 | 118 | 86 | 0.723 | -0.468 | 0.02910814 | 0.09060163 | 0.0153 | 0.0256 | 0.023 | 0.023 | TRUE | 0.4196 |
| Card9 | 30 | 33 | 32 | 28 | 33 | 29 | 23 | 35 | 35 | 28 | 37 | 29 | 31.11 | 31 | 31 | 1.015 | 0.021 | 0.95287194 | 0.9739969 | 0 | 0.0786 | 0.0594 | 0.0594 | TRUE | 0.3853 |
| Carf | 311 | 227 | 232 | 292 | 233 | 233 | 242 | 241 | 250 | 291 | 263 | 230 | 252.81 | 244 | 261 | 1.069 | 0.096 | 0.47950443 | 0.64643999 | 0.0033 | 0.0118 | 0.0093 | 0.0093 | TRUE | 0.1251 |
| Carhsp1 | 4669 | 3454 | 3496 | 4163 | 3492 | 3984 | 3626 | 3674 | 3772 | 4144 | 3939 | 3933 | 3848.06 | 3691 | 4005 | 1.086 | 0.118 | 0.03103101 | 0.09501418 | 4.00E-04 | 0.003 | 0.0019 | 0.0019 | TRUE | 0.0112 |
| Carkd | 1668 | 1231 | 1175 | 1302 | 1126 | 1269 | 1295 | 1309 | 1268 | 1296 | 1270 | 1253 | 1281.94 | 1291 | 1273 | 0.986 | -0.02 | 0.76455346 | 0.86031826 | 0 | 0.0043 | 0.0025 | 0.0025 | TRUE | 0.0031 |
| Carm1 | 9927 | 7128 | 7405 | 7827 | 6896 | 8647 | 7710 | 7582 | 7990 | 7791 | 7779 | 8536 | 7898.01 | 7761 | 8035 | 1.036 | 0.05 | 0.38243494 | 0.55640702 | 0.0016 | 0.0027 | 0.0023 | 0.0023 | TRUE | 0.0363 |
| Carns1 | 51 | 35 | 27 | 50 | 37 | 46 | 40 | 37 | 29 | 50 | 42 | 45 | 40.48 | 35 | 46 | 1.287 | 0.364 | 0.23308948 | 0.39927467 | 0 | 0.061 | 0.0422 | 0.0422 | TRUE | 0.1572 |
| Cars | 4604 | 3400 | 3147 | 3043 | 2543 | 3170 | 3576 | 3617 | 3396 | 3029 | 2869 | 3129 | 3269.15 | 3530 | 3009 | 0.853 | -0.23 | 0.00015504 | 0.00174873 | 0.001 | 0.0032 | 0.0024 | 0.0024 | TRUE | 0.0199 |
| Cars2 | 2822 | 1982 | 2105 | 2351 | 2061 | 2448 | 2192 | 2108 | 2271 | 2340 | 2325 | 2417 | 2275.5 | 2190 | 2361 | 1.078 | 0.108 | 0.07779727 | 0.18526432 | 4.00E-04 | 0.0035 | 0.0023 | 0.0023 | TRUE | 0.0126 |
| Cartpt | 0 | 0 | 1 | 0 | 0 | 0 | 0 | 0 | 1 | 0 | 0 | 0 | 0.18 | 0 | 0 | 0.552 | -0.857 | 0.8336548 | NA | 0 | 13.1817 | 10 | 10 | TRUE | 0.5365 |
| Casc1 | 2 | 2 | 2 | 4 | 3 | 3 | 2 | 2 | 2 | 4 | 3 | 3 | 2.69 | 2 | 3 | 1.788 | 0.838 | 0.46199594 | NA | 0 | 0.8821 | 0.5351 | 0.5351 | TRUE | 0.0388 |
| Casc3 | 8703 | 6943 | 6435 | 6580 | 5970 | 6844 | 6759 | 7385 | 6943 | 6550 | 6735 | 6756 | 6854.72 | 7029 | 6680 | 0.951 | -0.073 | 0.18215325 | 0.33776393 | 0.001 | 0.0028 | 0.002 | 0.002 | TRUE | 0.0239 |
| Casc4 | 3631 | 2617 | 2734 | 2915 | 2477 | 2778 | 2820 | 2784 | 2950 | 2902 | 2794 | 2742 | 2831.98 | 2851 | 2813 | 0.987 | -0.019 | 0.74297572 | 0.84697336 | 5.00E-04 | 0.0033 | 0.0021 | 0.0021 | TRUE | 0.0109 |
| Casc5 | 4587 | 3251 | 3569 | 3729 | 3372 | 3853 | 3563 | 3458 | 3851 | 3712 | 3804 | 3804 | 3698.47 | 3624 | 3773 | 1.041 | 0.058 | 0.33397175 | 0.50810999 | 0.0013 | 0.0031 | 0.0024 | 0.0024 | TRUE | 0.036 |
| Casd1 | 770 | 534 | 498 | 512 | 508 | 502 | 598 | 568 | 537 | 510 | 573 | 496 | 546.94 | 568 | 526 | 0.924 | -0.114 | 0.25763373 | 0.427535 | 0.0025 | 0.0068 | 0.0054 | 0.0054 | TRUE | 0.0695 |
| Cask | 1615 | 1182 | 1135 | 1143 | 1050 | 1105 | 1254 | 1257 | 1225 | 1138 | 1185 | 1091 | 1191.55 | 1245 | 1138 | 0.913 | -0.131 | 0.06649358 | 0.1650274 | 0 | 0.0044 | 0.0029 | 0.0029 | TRUE | 0.0156 |
| Caskin1 | 339 | 300 | 261 | 286 | 247 | 271 | 263 | 319 | 282 | 285 | 279 | 268 | 282.48 | 288 | 277 | 0.964 | -0.053 | 0.67812114 | 0.80231161 | 0.0021 | 0.0108 | 0.0082 | 0.0082 | TRUE | 0.0995 |
| Caskin2 | 1647 | 1250 | 1118 | 1049 | 975 | 1117 | 1279 | 1330 | 1206 | 1044 | 1100 | 1103 | 1176.97 | 1272 | 1082 | 0.851 | -0.233 | 0.00174775 | 0.01089904 | 8.00E-04 | 0.0044 | 0.0031 | 0.0031 | TRUE | 0.0234 |
| Casp1 | 0 | 0 | 0 | 0 | 0 | 0 | 0 | 0 | 0 | 0 | 0 | 0 | 0 | 0 | 0 | NA | NA | NA | NA | NA | NA | NA | NA | NA | NA |
| Casp12 | 0 | 0 | 0 | 0 | 0 | 0 | 0 | 0 | 0 | 0 | 0 | 0 | 0 | 0 | 0 | NA | NA | NA | NA | NA | NA | NA | NA | NA | NA |
| Casp14 | 34 | 19 | 13 | 20 | 13 | 10 | 26 | 20 | 14 | 20 | 15 | 10 | 17.51 | 20 | 15 | 0.726 | -0.462 | 0.34453415 | 0.51885225 | 0.0516 | 0.1377 | 0.1138 | 0.1138 | TRUE | 0.173 |
| Casp2 | 1075 | 818 | 785 | 741 | 595 | 760 | 835 | 870 | 847 | 738 | 671 | 750 | 785.18 | 851 | 720 | 0.847 | -0.239 | 0.00440864 | 0.02232119 | 4.00E-04 | 0.0054 | 0.0038 | 0.0038 | TRUE | 0.0399 |
| Casp3 | 2966 | 2144 | 2620 | 2998 | 2569 | 3167 | 2304 | 2281 | 2827 | 2984 | 2898 | 3126 | 2736.62 | 2471 | 3003 | 1.216 | 0.283 | 0.00090617 | 0.00674471 | 0.008 | 0.0033 | 0.0049 | 0.0049 | TRUE | 0.1918 |
| Casp4 | 0 | 0 | 0 | 0 | 0 | 0 | 0 | 0 | 0 | 0 | 0 | 0 | 0 | 0 | 0 | NA | NA | NA | NA | NA | NA | NA | NA | NA | NA |
| Casp6 | 998 | 784 | 938 | 893 | 784 | 893 | 775 | 834 | 1012 | 889 | 884 | 882 | 879.33 | 874 | 885 | 1.015 | 0.021 | 0.83545137 | 0.90637245 | 0.0086 | 0.0051 | 0.0063 | 0.0063 | TRUE | 0.2266 |
| Casp7 | 379 | 279 | 261 | 269 | 237 | 296 | 294 | 297 | 282 | 268 | 267 | 292 | 283.34 | 291 | 276 | 0.948 | -0.077 | 0.52549029 | 0.68648504 | 0 | 0.0108 | 0.0071 | 0.0071 | TRUE | 0.031 |
| Casp8 | 438 | 304 | 316 | 290 | 264 | 281 | 340 | 323 | 341 | 289 | 298 | 277 | 311.4 | 335 | 288 | 0.859 | -0.219 | 0.05845718 | 0.15071711 | 0 | 0.01 | 0.0064 | 0.0064 | TRUE | 0.0117 |
| Casp8ap2 | 5706 | 4121 | 4000 | 4327 | 3768 | 4453 | 4432 | 4383 | 4316 | 4307 | 4251 | 4396 | 4347.44 | 4377 | 4318 | 0.986 | -0.02 | 0.70042224 | 0.81850931 | 0 | 0.003 | 0.0017 | 0.0017 | TRUE | 0.003 |

| | | | | | | | | | | | | | | | | | | | | | | | | | |
|---|---|---|---|---|---|---|---|---|---|---|---|---|---|---|---|---|---|---|---|---|---|---|---|---|---|
| Casp9 | 2749 | 2071 | 2047 | 2282 | 1924 | 2297 | 2135 | 2203 | 2209 | 2272 | 2170 | 2267 | 2209.35 | 2182 | 2236 | 1.026 | 0.036 | 0.54356858 | 0.70106712 | 0 | 0.0035 | 0.0021 | 0.0021 | TRUE | 0.0079 |
| Casq1 | 73 | 47 | 49 | 78 | 47 | 76 | 57 | 50 | 53 | 78 | 53 | 75 | 60.87 | 53 | 69 | 1.29 | 0.367 | 0.15668866 | 0.30499129 | 0 | 0.0414 | 0.0319 | 0.0319 | TRUE | 0.33 |
| Casq2 | 14 | 6 | 5 | 6 | 6 | 5 | 11 | 6 | 5 | 6 | 7 | 5 | 6.72 | 7 | 6 | 0.766 | -0.385 | 0.60289341 | NA | 0 | 0.3551 | 0.2426 | 0.2426 | TRUE | 0.5338 |
| Casr | 19 | 11 | 9 | 4 | 7 | 9 | 15 | 12 | 10 | 4 | 8 | 9 | 9.49 | 12 | 7 | 0.569 | -0.815 | 0.20269328 | NA | 0 | 0.2522 | 0.1799 | 0.1799 | TRUE | 0.3781 |
| Cass4 | 185 | 148 | 110 | 221 | 201 | 227 | 144 | 157 | 119 | 220 | 227 | 224 | 181.77 | 140 | 224 | 1.596 | 0.674 | 1.59E-05 | 0.00027848 | 0 | 0.0155 | 0.0118 | 0.0118 | TRUE | 0.1691 |
| Cast | 2139 | 1472 | 1505 | 1605 | 1336 | 1537 | 1661 | 1566 | 1624 | 1598 | 1507 | 1517 | 1578.82 | 1617 | 1541 | 0.952 | -0.07 | 0.29277608 | 0.46496095 | 4.00E-04 | 0.0039 | 0.0026 | 0.0026 | TRUE | 0.0127 |
| Casz1 | 1415 | 1074 | 1134 | 1284 | 1272 | 1355 | 1099 | 1142 | 1224 | 1278 | 1435 | 1338 | 1252.6 | 1155 | 1350 | 1.17 | 0.226 | 0.00396442 | 0.02047092 | 0.0024 | 0.0043 | 0.0036 | 0.0036 | TRUE | 0.0356 |
| Cat | 5949 | 4433 | 4563 | 5045 | 4717 | 5491 | 4620 | 4715 | 4923 | 5022 | 5321 | 5420 | 5003.77 | 4753 | 5254 | 1.106 | 0.145 | 0.01106388 | 0.04445772 | 0.001 | 0.0029 | 0.0021 | 0.0021 | TRUE | 0.0184 |
| Catsper1 | 0 | 0 | 0 | 0 | 0 | 0 | 0 | 0 | 0 | 0 | 0 | 0 | 0 | 0 | 0 | NA | NA | NA | NA | NA | NA | NA | NA | NA | NA |
| Catsper2 | 120 | 83 | 57 | 82 | 57 | 57 | 93 | 88 | 62 | 82 | 64 | 56 | 74.2 | 81 | 67 | 0.829 | -0.271 | 0.28568772 | 0.45767031 | 0.0285 | 0.0344 | 0.0327 | 0.0327 | TRUE | 0.3949 |
| Catsper3 | 7 | 2 | 3 | 3 | 5 | 4 | 5 | 2 | 3 | 3 | 6 | 4 | 3.9 | 3 | 4 | 1.134 | 0.182 | 0.84997379 | NA | 0 | 0.6108 | 0.4107 | 0.4107 | TRUE | 0.4037 |
| Catsper4 | 1 | 1 | 2 | 2 | 1 | 1 | 1 | 1 | 2 | 2 | 1 | 1 | 1.35 | 1 | 1 | 1.05 | 0.07 | 0.96467497 | NA | 0 | 1.7571 | 1.0543 | 1.0543 | TRUE | 0.167 |
| Catsperb | 1 | 0 | 0 | 0 | 2 | 0 | 1 | 0 | 0 | 0 | 2 | 0 | 0.51 | 0 | 1 | 2.139 | 1.097 | 0.71688146 | NA | 0 | 4.6912 | 4.8901 | 4.8901 | TRUE | 0.9745 |
| Catsperd | 95 | 72 | 108 | 164 | 144 | 158 | 74 | 77 | 117 | 163 | 162 | 156 | 124.76 | 89 | 160 | 1.816 | 0.861 | 2.37E-05 | 0.0003829 | 0.0206 | 0.0214 | 0.0212 | 0.0212 | TRUE | 0.6742 |
| Catsperg1 | 161 | 166 | 190 | 159 | 115 | 172 | 125 | 177 | 205 | 158 | 130 | 170 | 160.74 | 169 | 153 | 0.91 | -0.136 | 0.49596722 | 0.66060083 | 0.0349 | 0.0172 | 0.0225 | 0.0225 | TRUE | 0.4274 |
| Catsperg2 | 46 | 48 | 49 | 39 | 36 | 44 | 36 | 51 | 53 | 39 | 41 | 43 | 43.75 | 47 | 41 | 0.887 | -0.173 | 0.56623511 | 0.72043806 | 0 | 0.0566 | 0.0422 | 0.0422 | TRUE | 0.3879 |
| Cav1 | 7 | 1 | 4 | 7 | 9 | 5 | 5 | 1 | 4 | 7 | 10 | 5 | 5.48 | 3 | 7 | 1.984 | 0.988 | 0.25174233 | NA | 0 | 0.435 | 0.3344 | 0.3344 | TRUE | 0.45 |
| Cav2 | 26 | 18 | 23 | 16 | 8 | 8 | 20 | 19 | 25 | 16 | 9 | 8 | 16.17 | 21 | 11 | 0.515 | -0.959 | 0.05541755 | NA | 0 | 0.149 | 0.1107 | 0.1107 | TRUE | 0.6029 |
| Cav3 | 0 | 1 | 0 | 1 | 1 | 1 | 0 | 1 | 0 | 1 | 1 | 1 | 0.7 | 0 | 1 | 3.004 | 1.587 | 0.48898579 | NA | 0 | 3.409 | 2.2901 | 2.2901 | TRUE | 0.5217 |
| Cbfa2t2 | 7408 | 5336 | 5911 | 6741 | 6044 | 7000 | 5753 | 5676 | 6378 | 6710 | 6818 | 6910 | 6374.26 | 5936 | 6813 | 1.148 | 0.199 | 0.00095259 | 0.00700018 | 0.002 | 0.0028 | 0.0025 | 0.0025 | TRUE | 0.0514 |
| Cbfa2t3 | 125 | 97 | 87 | 118 | 124 | 143 | 97 | 103 | 94 | 117 | 140 | 141 | 115.44 | 98 | 133 | 1.354 | 0.438 | 0.01852037 | 0.0652964 | 0 | 0.023 | 0.0161 | 0.0161 | TRUE | 0.1074 |
| Cbfb | 5336 | 4031 | 3547 | 3678 | 2998 | 3533 | 4144 | 4288 | 3827 | 3661 | 3382 | 3488 | 3798.32 | 4086 | 3510 | 0.859 | -0.219 | 0.0006528 | 0.00528341 | 0.0021 | 0.0031 | 0.0027 | 0.0027 | TRUE | 0.037 |
| Cbl | 2252 | 1547 | 1479 | 1832 | 1593 | 1898 | 1749 | 1646 | 1596 | 1824 | 1797 | 1874 | 1747.44 | 1664 | 1832 | 1.1 | 0.138 | 0.03939192 | 0.11327412 | 8.00E-04 | 0.0038 | 0.0026 | 0.0026 | TRUE | 0.0254 |
| Cblb | 1697 | 1242 | 1347 | 1342 | 1177 | 1342 | 1318 | 1321 | 1453 | 1336 | 1328 | 1325 | 1346.81 | 1364 | 1330 | 0.975 | -0.036 | 0.61469808 | 0.75655556 | 9.00E-04 | 0.0042 | 0.003 | 0.003 | TRUE | 0.0386 |
| Cblc | 260 | 224 | 303 | 359 | 299 | 392 | 202 | 238 | 327 | 357 | 337 | 387 | 308.12 | 256 | 360 | 1.417 | 0.503 | 0.00190539 | 0.01163833 | 0.029 | 0.0101 | 0.0156 | 0.0156 | TRUE | 0.6556 |
| Cbll1 | 2592 | 1976 | 1872 | 1967 | 1806 | 2122 | 2013 | 2102 | 2020 | 1958 | 2037 | 2095 | 2037.48 | 2045 | 2030 | 0.993 | -0.01 | 0.87035648 | 0.92822005 | 4.00E-04 | 0.0036 | 0.0023 | 0.0023 | TRUE | 0.0118 |
| Cbln1 | 1610 | 1390 | 1461 | 1460 | 1357 | 1463 | 1250 | 1479 | 1576 | 1453 | 1531 | 1444 | 1455.61 | 1435 | 1476 | 1.03 | 0.043 | 0.63351902 | 0.77062871 | 0.007 | 0.0041 | 0.0051 | 0.0051 | TRUE | 0.1573 |
| Cbln2 | 32 | 20 | 20 | 25 | 24 | 16 | 25 | 21 | 22 | 25 | 27 | 16 | 22.58 | 23 | 23 | 0.994 | -0.009 | 0.98189309 | 0.99062596 | 0 | 0.1074 | 0.0776 | 0.0776 | TRUE | 0.4121 |
| Cbln3 | 47 | 44 | 56 | 31 | 53 | 60 | 37 | 47 | 60 | 31 | 60 | 59 | 48.93 | 48 | 50 | 1.048 | 0.067 | 0.83841327 | 0.907974 | 0.0742 | 0.0509 | 0.0574 | 0.0574 | TRUE | 0.6185 |
| Cbln4 | 0 | 1 | 2 | 1 | 2 | 3 | 0 | 1 | 2 | 1 | 2 | 3 | 1.57 | 1 | 2 | 2.029 | 1.021 | 0.52141865 | NA | 0 | 1.5097 | 1.1037 | 1.1037 | TRUE | 0.4301 |
| Cbr1 | 1962 | 1370 | 1394 | 1264 | 1225 | 1483 | 1524 | 1457 | 1504 | 1258 | 1382 | 1464 | 1431.54 | 1495 | 1368 | 0.915 | -0.129 | 0.09162279 | 0.20745819 | 0.0024 | 0.0041 | 0.0035 | 0.0035 | TRUE | 0.0601 |
| Cbr2 | 0 | 2 | 2 | 0 | 0 | 1 | 0 | 2 | 2 | 0 | 0 | 1 | 0.88 | 1 | 0 | 0.266 | -1.912 | 0.40283109 | NA | 0 | 2.6995 | 2.4177 | 2.4177 | TRUE | 0.6891 |
| Cbr3 | 216 | 150 | 165 | 173 | 164 | 176 | 168 | 160 | 178 | 172 | 185 | 174 | 172.72 | 169 | 177 | 1.05 | 0.071 | 0.63913886 | 0.77456433 | 0 | 0.0161 | 0.0106 | 0.0106 | TRUE | 0.0244 |
| Cbr4 | 1422 | 953 | 1070 | 1265 | 1010 | 1169 | 1104 | 1014 | 1155 | 1259 | 1139 | 1154 | 1137.53 | 1091 | 1184 | 1.086 | 0.119 | 0.14462509 | 0.28812968 | 0.0027 | 0.0045 | 0.0039 | 0.0039 | TRUE | 0.0444 |
| Cbs | 1394 | 1058 | 1390 | 1829 | 1737 | 1978 | 1083 | 1125 | 1500 | 1821 | 1960 | 1953 | 1573.42 | 1236 | 1911 | 1.548 | 0.63 | 5.68E-09 | 3.12E-07 | 0.0171 | 0.0039 | 0.0078 | 0.0078 | TRUE | 0.4155 |
| Cbwd1 | 2684 | 1895 | 1788 | 2287 | 1856 | 2111 | 2085 | 2016 | 1929 | 2276 | 2094 | 2084 | 2080.6 | 2010 | 2151 | 1.07 | 0.098 | 0.14751087 | 0.29213979 | 0.0015 | 0.0036 | 0.0028 | 0.0028 | TRUE | 0.0315 |
| Cbx1 | 8843 | 6604 | 5867 | 6743 | 5771 | 6942 | 6868 | 7025 | 6330 | 6712 | 6510 | 6853 | 6716.35 | 6741 | 6692 | 0.993 | -0.011 | 0.85494644 | 0.91850192 | 0.0016 | 0.0028 | 0.0023 | 0.0023 | TRUE | 0.0343 |
| Cbx2 | 3137 | 2353 | 2148 | 1954 | 1884 | 2176 | 2436 | 2503 | 2318 | 1945 | 2125 | 2148 | 2245.88 | 2419 | 2073 | 0.857 | -0.223 | 0.0009134 | 0.00678896 | 0.0015 | 0.0035 | 0.0028 | 0.0028 | TRUE | 0.0355 |
| Cbx3 | 3675 | 2915 | 2948 | 2921 | 2481 | 2932 | 2854 | 3101 | 3181 | 2908 | 2799 | 2894 | 2956.08 | 3045 | 2867 | 0.942 | -0.086 | 0.1749961 | 0.32841639 | 0.0015 | 0.0032 | 0.0025 | 0.0025 | TRUE | 0.0375 |
| Cbx4 | 391 | 275 | 275 | 282 | 244 | 301 | 304 | 293 | 297 | 281 | 275 | 297 | 291 | 298 | 284 | 0.955 | -0.066 | 0.57824446 | 0.72941333 | 0 | 0.0106 | 0.0067 | 0.0067 | TRUE | 0.0176 |
| Cbx5 | 19692 | 13957 | 13954 | 16334 | 13394 | 15738 | 15294 | 14846 | 15056 | 16259 | 15110 | 15536 | 15350.1 | 15065 | 15635 | 1.038 | 0.054 | 0.28288836 | 0.45466359 | 7.00E-04 | 0.0026 | 0.0017 | 0.0017 | TRUE | 0.0149 |
| Cbx6 | 2599 | 2017 | 1900 | 1909 | 1866 | 2080 | 2019 | 2145 | 2050 | 1900 | 2105 | 2053 | 2045.44 | 2071 | 2019 | 0.975 | -0.036 | 0.58699819 | 0.73617813 | 0.0014 | 0.0036 | 0.0028 | 0.0028 | TRUE | 0.0324 |
| Cbx7 | 9037 | 6968 | 7194 | 7115 | 6805 | 7725 | 7019 | 7412 | 7762 | 7082 | 7677 | 7626 | 7429.58 | 7398 | 7462 | 1.009 | 0.013 | 0.83295994 | 0.90481048 | 0.0021 | 0.0027 | 0.0025 | 0.0025 | TRUE | 0.0248 |
| Cbx8 | 76 | 71 | 72 | 51 | 50 | 60 | 59 | 76 | 78 | 51 | 56 | 59 | 63.11 | 71 | 55 | 0.788 | -0.343 | 0.1709027 | 0.32259143 | 0 | 0.04 | 0.0291 | 0.0291 | TRUE | 0.2175 |
| Cby1 | 1000 | 737 | 674 | 789 | 697 | 795 | 777 | 784 | 727 | 785 | 786 | 785 | 774.05 | 763 | 785 | 1.029 | 0.042 | 0.6074003 | 0.75128613 | 0 | 0.0055 | 0.0035 | 0.0035 | TRUE | 0.0188 |
| Cc2d1a | 2391 | 1689 | 1680 | 1721 | 1538 | 1751 | 1857 | 1797 | 1813 | 1713 | 1735 | 1729 | 1773.81 | 1822 | 1726 | 0.947 | -0.079 | 0.19749724 | 0.35727582 | 0 | 0.0038 | 0.0022 | 0.0022 | TRUE | 0.0035 |
| Cc2d1b | 4144 | 3165 | 2903 | 2748 | 2559 | 3001 | 3218 | 3367 | 3132 | 2735 | 2887 | 2962 | 3050.33 | 3239 | 2861 | 0.883 | -0.179 | 0.00352049 | 0.01868486 | 0.001 | 0.0032 | 0.0024 | 0.0024 | TRUE | 0.0182 |
| Cc2d2a | 1691 | 1152 | 1364 | 1013 | 889 | 1033 | 1313 | 1225 | 1472 | 1008 | 1003 | 1020 | 1173.57 | 1337 | 1010 | 0.756 | -0.403 | 1.37E-06 | 3.77E-05 | 0.0036 | 0.0044 | 0.0042 | 0.0042 | TRUE | 0.0913 |
| Ccar1 | 6030 | 4366 | 3552 | 3725 | 3109 | 3741 | 4683 | 4644 | 3833 | 3708 | 3507 | 3693 | 4011.33 | 4387 | 3636 | 0.829 | -0.271 | 0.00063625 | 0.00519352 | 0.0061 | 0.003 | 0.0043 | 0.0043 | TRUE | 0.1478 |
| Ccbe1 | 109 | 92 | 90 | 148 | 127 | 127 | 85 | 98 | 97 | 147 | 143 | 125 | 115.93 | 93 | 138 | 1.492 | 0.578 | 0.0019655 | 0.01194932 | 0 | 0.0229 | 0.0162 | 0.0162 | TRUE | 0.0745 |
| Ccbl1 | 464 | 388 | 325 | 398 | 385 | 401 | 360 | 413 | 351 | 396 | 434 | 396 | 391.68 | 375 | 409 | 1.092 | 0.126 | 0.26385393 | 0.43407614 | 0.0025 | 0.0085 | 0.0067 | 0.0067 | TRUE | 0.0877 |
| Ccbl2 | 1054 | 691 | 839 | 1026 | 920 | 1065 | 819 | 735 | 905 | 1021 | 1038 | 1051 | 928.22 | 820 | 1037 | 1.265 | 0.339 | 0.0001461 | 0.00166922 | 0.0039 | 0.005 | 0.0047 | 0.0047 | TRUE | 0.095 |
| Ccbp2 | 17 | 10 | 11 | 14 | 10 | 14 | 13 | 11 | 12 | 14 | 11 | 14 | 12.46 | 12 | 13 | 1.091 | 0.125 | 0.81349951 | NA | 0 | 0.1927 | 0.1235 | 0.1235 | TRUE | 0.0505 |
| Ccdc101 | 1508 | 1039 | 1002 | 1032 | 987 | 1165 | 1171 | 1105 | 1081 | 1027 | 1113 | 1150 | 1108.04 | 1119 | 1097 | 0.979 | -0.031 | 0.69776763 | 0.81640412 | 0.0017 | 0.0046 | 0.0035 | 0.0035 | TRUE | 0.0375 |
| Ccdc102a | 628 | 449 | 393 | 607 | 514 | 585 | 488 | 478 | 424 | 604 | 580 | 577 | 525.15 | 463 | 587 | 1.266 | 0.34 | 0.0005514 | 0.00464695 | 9.00E-04 | 0.0069 | 0.0051 | 0.0051 | TRUE | 0.0613 |
| Ccdc103 | 66 | 31 | 27 | 28 | 25 | 27 | 51 | 33 | 29 | 28 | 28 | 27 | 32.68 | 38 | 28 | 0.722 | -0.471 | 0.18886222 | 0.34643323 | 0.0302 | 0.0749 | 0.0611 | 0.0611 | TRUE | 0.8197 |
| Ccdc104 | 2646 | 2014 | 2067 | 2187 | 1903 | 2247 | 2055 | 2142 | 2230 | 2177 | 2147 | 2218 | 2161.57 | 2142 | 2181 | 1.019 | 0.027 | 0.67007306 | 0.79708707 | 6.00E-04 | 0.0035 | 0.0023 | 0.0023 | TRUE | 0.016 |
| Ccdc105 | 4 | 1 | 0 | 3 | 3 | 0 | 3 | 1 | 0 | 3 | 3 | 0 | 1.76 | 1 | 2 | 1.451 | 0.537 | 0.73833922 | NA | 0.9941 | 1.3515 | 1.2818 | 1.2818 | TRUE | 0.786 |
| Ccdc106 | 381 | 284 | 199 | 290 | 274 | 280 | 296 | 302 | 215 | 289 | 309 | 276 | 281.15 | 271 | 291 | 1.072 | 0.101 | 0.49235878 | 0.65756719 | 0.0148 | 0.0109 | 0.012 | 0.012 | TRUE | 0.3549 |
| Ccdc107 | 695 | 528 | 555 | 549 | 528 | 571 | 540 | 562 | 599 | 546 | 596 | 564 | 567.67 | 567 | 569 | 1.004 | 0.006 | 0.95062759 | 0.97276276 | 0 | 0.0066 | 0.0047 | 0.0047 | TRUE | 0.0284 |
| Ccdc108 | 12 | 7 | 1 | 4 | 2 | 6 | 9 | 7 | 1 | 4 | 2 | 6 | 5 | 6 | 4 | 0.675 | -0.567 | 0.54550674 | NA | 0.289 | 0.4764 | 0.4266 | 0.4266 | TRUE | 0.8988 |
| Ccdc109b | 20 | 12 | 6 | 9 | 7 | 5 | 16 | 13 | 6 | 9 | 8 | 5 | 9.43 | 12 | 7 | 0.619 | -0.693 | 0.28819928 | NA | 0 | 0.2538 | 0.1939 | 0.1939 | TRUE | 0.4335 |
| Ccdc11 | 45 | 32 | 38 | 52 | 52 | 43 | 35 | 34 | 41 | 52 | 59 | 42 | 43.81 | 37 | 51 | 1.39 | 0.475 | 0.10988405 | 0.23713917 | 0 | 0.0565 | 0.0402 | 0.0402 | TRUE | 0.1775 |
| Ccdc110 | 3 | 1 | 2 | 3 | 2 | 0 | 2 | 1 | 2 | 3 | 2 | 0 | 1.8 | 2 | 2 | 0.931 | -0.104 | 0.94326775 | NA | 0 | 1.3199 | 0.9747 | 0.9747 | TRUE | 0.6392 |
| Ccdc111 | 1406 | 1204 | 1509 | 1593 | 1347 | 1682 | 1092 | 1281 | 1628 | 1586 | 1520 | 1660 | 1461.08 | 1334 | 1589 | 1.193 | 0.255 | 0.02947996 | 0.09148004 | 0.0207 | 0.004 | 0.0092 | 0.0092 | TRUE | 0.4481 |
| Ccdc112 | 2255 | 1684 | 1400 | 2015 | 1829 | 1974 | 1751 | 1791 | 1511 | 2006 | 2063 | 1949 | 1845.15 | 1684 | 2006 | 1.19 | 0.251 | 0.00120712 | 0.00829092 | 0.0037 | 0.0037 | 0.0038 | 0.0038 | TRUE | 0.0964 |
| Ccdc113 | 220 | 171 | 205 | 211 | 189 | 227 | 171 | 182 | 221 | 210 | 213 | 224 | 203.55 | 191 | 216 | 1.132 | 0.179 | 0.2341941 | 0.40038472 | 0.0049 | 0.0141 | 0.0114 | 0.0114 | TRUE | 0.197 |
| Ccdc114 | 706 | 472 | 392 | 511 | 437 | 484 | 548 | 502 | 423 | 509 | 493 | 478 | 492.13 | 491 | 493 | 1.001 | 0.002 | 0.98552699 | 0.99275053 | 0.0071 | 0.0072 | 0.0072 | 0.0072 | TRUE | 0.1686 |
| Ccdc115 | 1911 | 1438 | 1387 | 1455 | 1244 | 1464 | 1484 | 1530 | 1497 | 1448 | 1403 | 1445 | 1467.87 | 1504 | 1432 | 0.953 | -0.069 | 0.28746887 | 0.45954509 | 0 | 0.004 | 0.0024 | 0.0024 | TRUE | 0.0037 |

| | | | | | | | | | | | | | | | | | | | | | | | | | |
|---|---|---|---|---|---|---|---|---|---|---|---|---|---|---|---|---|---|---|---|---|---|---|---|---|---|
| Ccdc116 | 115 | 76 | 36 | 62 | 42 | 61 | 89 | 81 | 39 | 62 | 47 | 60 | 63.05 | 70 | 56 | 0.806 | -0.31 | 0.31428718 | 0.48779527 | 0.0846 | 0.04 | 0.0525 | 0.0525 | TRUE | 1.2855 |
| Ccdc117 | 6850 | 5030 | 5234 | 5215 | 4335 | 5094 | 5320 | 5350 | 5647 | 5191 | 4890 | 5029 | 5237.97 | 5439 | 5037 | 0.926 | -0.111 | 0.04415284 | 0.12299289 | 7.00E-04 | 0.0029 | 0.002 | 0.002 | TRUE | 0.0136 |
| Ccdc12 | 1394 | 1087 | 1015 | 989 | 965 | 1134 | 1083 | 1156 | 1095 | 984 | 1089 | 1119 | 1087.76 | 1111 | 1064 | 0.958 | -0.062 | 0.43634755 | 0.60763012 | 0.0019 | 0.0046 | 0.0037 | 0.0037 | TRUE | 0.052 |
| Ccdc120 | 960 | 719 | 747 | 760 | 733 | 818 | 746 | 765 | 806 | 757 | 827 | 807 | 784.55 | 772 | 797 | 1.033 | 0.046 | 0.58028692 | 0.73107457 | 0 | 0.0054 | 0.0038 | 0.0038 | TRUE | 0.0236 |
| Ccdc121 | 13 | 8 | 3 | 4 | 3 | 4 | 10 | 9 | 3 | 4 | 3 | 4 | 5.53 | 7 | 4 | 0.513 | -0.964 | 0.2539467 | NA | 0 | 0.4313 | 0.312 | 0.312 | TRUE | 0.5923 |
| Ccdc122 | 310 | 237 | 216 | 257 | 190 | 272 | 241 | 252 | 233 | 256 | 214 | 269 | 244.1 | 242 | 246 | 1.019 | 0.028 | 0.83981835 | 0.90890647 | 0.0028 | 0.0121 | 0.0095 | 0.0095 | TRUE | 0.1369 |
| Ccdc124 | 5223 | 3695 | 3074 | 3562 | 2840 | 3397 | 4056 | 3930 | 3317 | 3546 | 3204 | 3353 | 3567.74 | 3768 | 3368 | 0.893 | -0.162 | 0.0439676 | 0.12257244 | 0.0064 | 0.0031 | 0.0044 | 0.0044 | TRUE | 0.1327 |
| Ccdc125 | 355 | 262 | 293 | 362 | 295 | 378 | 276 | 279 | 316 | 360 | 333 | 373 | 322.8 | 290 | 355 | 1.228 | 0.296 | 0.01354435 | 0.05184504 | 0 | 0.0098 | 0.0073 | 0.0073 | TRUE | 0.0667 |
| Ccdc126 | 376 | 261 | 255 | 257 | 230 | 275 | 292 | 278 | 275 | 256 | 259 | 271 | 271.92 | 282 | 262 | 0.93 | -0.104 | 0.39460361 | 0.56811659 | 0 | 0.0111 | 0.0071 | 0.0071 | TRUE | 0.0128 |
| Ccdc127 | 3016 | 2167 | 1967 | 2285 | 1924 | 2065 | 2342 | 2305 | 2122 | 2274 | 2170 | 2038 | 2208.87 | 2256 | 2161 | 0.957 | -0.063 | 0.36855914 | 0.54321707 | 0.0022 | 0.0035 | 0.0031 | 0.0031 | TRUE | 0.0324 |
| Ccdc129 | 1 | 0 | 3 | 0 | 0 | 1 | 1 | 0 | 3 | 0 | 0 | 1 | 0.83 | 1 | 0 | 0.28 | -1.835 | 0.43280827 | NA | 0 | 2.8461 | 2.5799 | 2.5799 | TRUE | 0.5383 |
| Ccdc13 | 44 | 35 | 27 | 28 | 22 | 33 | 34 | 37 | 29 | 28 | 25 | 33 | 30.97 | 33 | 29 | 0.849 | -0.236 | 0.49552199 | 0.66012637 | 0 | 0.079 | 0.0536 | 0.0536 | TRUE | 0.1075 |
| Ccdc130 | 862 | 673 | 678 | 799 | 772 | 878 | 669 | 716 | 732 | 795 | 871 | 867 | 774.97 | 706 | 844 | 1.198 | 0.26 | 0.00243372 | 0.01407738 | 0.0012 | 0.0055 | 0.004 | 0.004 | TRUE | 0.0309 |
| Ccdc132 | 4180 | 3105 | 2799 | 2994 | 2744 | 3092 | 3246 | 3303 | 3020 | 2980 | 3096 | 3052 | 3116.22 | 3190 | 3043 | 0.954 | -0.069 | 0.25402206 | 0.42321218 | 8.00E-04 | 0.0032 | 0.0023 | 0.0023 | TRUE | 0.0258 |
| Ccdc134 | 483 | 368 | 402 | 402 | 339 | 386 | 375 | 391 | 434 | 400 | 382 | 381 | 393.99 | 400 | 388 | 0.971 | -0.042 | 0.70257179 | 0.81985628 | 0 | 0.0084 | 0.0061 | 0.0061 | TRUE | 0.0612 |
| Ccdc135 | 73 | 42 | 52 | 67 | 42 | 44 | 57 | 45 | 56 | 67 | 47 | 43 | 52.5 | 53 | 52 | 0.999 | -0.002 | 0.99547921 | 0.99730837 | 0.0166 | 0.0476 | 0.039 | 0.039 | TRUE | 0.4717 |
| Ccdc136 | 1822 | 1566 | 1794 | 1933 | 1968 | 2314 | 1415 | 1666 | 1936 | 1924 | 2220 | 2284 | 1907.5 | 1672 | 2143 | 1.283 | 0.359 | 0.00069043 | 0.00552535 | 0.0153 | 0.0037 | 0.0075 | 0.0075 | TRUE | 0.2294 |
| Ccdc137 | 845 | 658 | 578 | 565 | 454 | 597 | 656 | 700 | 624 | 562 | 512 | 589 | 607.29 | 660 | 554 | 0.841 | -0.249 | 0.0097625 | 0.04067618 | 0.0022 | 0.0063 | 0.005 | 0.005 | TRUE | 0.051 |
| Ccdc138 | 719 | 615 | 703 | 716 | 636 | 813 | 558 | 654 | 759 | 713 | 717 | 803 | 700.64 | 657 | 744 | 1.136 | 0.184 | 0.10778334 | 0.23352795 | 0.0118 | 0.0058 | 0.008 | 0.008 | TRUE | 0.2153 |
| Ccdc14 | 1857 | 1390 | 1387 | 1376 | 1197 | 1363 | 1442 | 1479 | 1497 | 1370 | 1350 | 1345 | 1413.81 | 1473 | 1355 | 0.921 | -0.119 | 0.06990416 | 0.17145226 | 0 | 0.0041 | 0.0024 | 0.0024 | TRUE | 0.0041 |
| Ccdc141 | 404 | 386 | 512 | 463 | 459 | 569 | 314 | 411 | 552 | 461 | 518 | 562 | 469.53 | 426 | 514 | 1.21 | 0.275 | 0.09595023 | 0.21470336 | 0.0423 | 0.0075 | 0.0176 | 0.0176 | TRUE | 0.3831 |
| Ccdc142 | 108 | 114 | 90 | 57 | 67 | 77 | 84 | 121 | 97 | 57 | 76 | 76 | 85.1 | 101 | 70 | 0.691 | -0.532 | 0.02324474 | 0.07719263 | 0.0202 | 0.0303 | 0.0274 | 0.0274 | TRUE | 0.3074 |
| Ccdc144b | 0 | 0 | 0 | 0 | 0 | 0 | 0 | 0 | 0 | 0 | 0 | 0 | 0 | 0 | 0 | NA | NA | NA | NA | NA | NA | NA | NA | NA | NA |
| Ccdc146 | 8 | 5 | 4 | 4 | 2 | 2 | 6 | 5 | 4 | 4 | 2 | 2 | 4.01 | 5 | 3 | 0.517 | -0.953 | 0.32035928 | NA | 0 | 0.5935 | 0.3828 | 0.3828 | TRUE | 0.1959 |
| Ccdc147 | 1 | 1 | 2 | 2 | 2 | 1 | 1 | 1 | 2 | 2 | 2 | 1 | 1.54 | 1 | 2 | 1.329 | 0.411 | 0.78362382 | NA | 0 | 1.5427 | 0.944 | 0.944 | TRUE | 0.1668 |
| Ccdc148 | 3 | 0 | 1 | 1 | 1 | 1 | 2 | 0 | 1 | 1 | 1 | 1 | 1.09 | 1 | 1 | 0.877 | -0.19 | 0.91667219 | NA | 0 | 2.1836 | 1.4553 | 1.4553 | TRUE | 0.2362 |
| Ccdc149 | 20 | 16 | 12 | 7 | 8 | 9 | 16 | 17 | 13 | 7 | 9 | 9 | 11.73 | 15 | 8 | 0.545 | -0.875 | 0.11935904 | NA | 0 | 0.2045 | 0.1331 | 0.1331 | TRUE | 0.068 |
| Ccdc15 | 469 | 322 | 330 | 484 | 348 | 404 | 364 | 343 | 356 | 482 | 393 | 399 | 389.33 | 354 | 425 | 1.198 | 0.26 | 0.02670949 | 0.08529911 | 0.0049 | 0.0085 | 0.0074 | 0.0074 | TRUE | 0.1631 |
| Ccdc150 | 43 | 30 | 21 | 47 | 33 | 34 | 33 | 32 | 23 | 47 | 37 | 34 | 34.26 | 29 | 39 | 1.33 | 0.411 | 0.22687114 | 0.391816 | 0 | 0.0716 | 0.0538 | 0.0538 | TRUE | 0.2413 |
| Ccdc151 | 113 | 78 | 92 | 99 | 109 | 142 | 88 | 83 | 99 | 99 | 123 | 140 | 105.28 | 90 | 121 | 1.341 | 0.423 | 0.04038332 | 0.11523233 | 0.0114 | 0.0249 | 0.021 | 0.021 | TRUE | 0.2399 |
| Ccdc152 | 235 | 165 | 152 | 177 | 149 | 150 | 183 | 176 | 164 | 176 | 168 | 148 | 169.06 | 174 | 164 | 0.941 | -0.088 | 0.57486857 | 0.72681084 | 0 | 0.0164 | 0.0116 | 0.0116 | TRUE | 0.0798 |
| Ccdc153 | 3 | 1 | 0 | 0 | 0 | 4 | 2 | 1 | 0 | 0 | 0 | 4 | 1.22 | 1 | 1 | 1.142 | 0.192 | 0.92412136 | NA | 3.0466 | 1.9392 | 2.1166 | 2.1166 | TRUE | 1.0112 |
| Ccdc154 | 1 | 1 | 0 | 0 | 2 | 1 | 1 | 1 | 0 | 0 | 2 | 1 | 0.85 | 1 | 1 | 1.713 | 0.776 | 0.71789357 | NA | 0 | 2.7997 | 2.0659 | 2.0659 | TRUE | 0.1674 |
| Ccdc155 | 102 | 88 | 82 | 137 | 130 | 160 | 79 | 94 | 88 | 136 | 147 | 158 | 117.04 | 87 | 147 | 1.694 | 0.761 | 4.31E-05 | 0.00062014 | 0 | 0.0227 | 0.0158 | 0.0158 | TRUE | 0.0688 |
| Ccdc157 | 322 | 239 | 172 | 211 | 171 | 235 | 250 | 254 | 186 | 210 | 193 | 232 | 220.8 | 230 | 212 | 0.919 | -0.122 | 0.44240851 | 0.61332152 | 0.0144 | 0.0132 | 0.0135 | 0.0135 | TRUE | 0.3022 |
| Ccdc158 | 241 | 152 | 196 | 203 | 150 | 225 | 187 | 162 | 211 | 202 | 169 | 222 | 192.29 | 187 | 198 | 1.061 | 0.085 | 0.60156152 | 0.74667517 | 0.0123 | 0.0148 | 0.0141 | 0.0141 | TRUE | 0.1668 |
| Ccdc159 | 47 | 60 | 43 | 58 | 41 | 58 | 37 | 64 | 46 | 58 | 46 | 57 | 51.33 | 49 | 54 | 1.11 | 0.151 | 0.609139 | 0.75263876 | 0.0286 | 0.0486 | 0.043 | 0.043 | TRUE | 0.4489 |
| Ccdc160 | 301 | 234 | 238 | 206 | 196 | 245 | 234 | 249 | 257 | 205 | 221 | 242 | 234.58 | 247 | 223 | 0.905 | -0.144 | 0.28858791 | 0.46071562 | 0 | 0.0125 | 0.009 | 0.009 | TRUE | 0.0644 |
| Ccdc162 | 135 | 98 | 145 | 222 | 173 | 212 | 105 | 104 | 156 | 221 | 195 | 209 | 165.16 | 122 | 208 | 1.72 | 0.783 | 2.00E-05 | 0.00033086 | 0.0207 | 0.0168 | 0.0178 | 0.0178 | TRUE | 0.6026 |
| Ccdc163 | 477 | 389 | 411 | 420 | 337 | 389 | 370 | 414 | 443 | 418 | 380 | 384 | 401.66 | 409 | 394 | 0.966 | -0.05 | 0.6582853 | 0.78834284 | 0.0034 | 0.0083 | 0.0067 | 0.0067 | TRUE | 0.085 |
| Ccdc164 | 231 | 193 | 227 | 213 | 222 | 235 | 179 | 205 | 245 | 212 | 250 | 232 | 220.68 | 210 | 231 | 1.107 | 0.146 | 0.34403224 | 0.51831483 | 0.0114 | 0.0132 | 0.0126 | 0.0126 | TRUE | 0.2311 |
| Ccdc166 | 263 | 215 | 210 | 173 | 176 | 204 | 204 | 229 | 227 | 172 | 199 | 201 | 205.28 | 220 | 191 | 0.869 | -0.203 | 0.16122923 | 0.31099182 | 0 | 0.014 | 0.0102 | 0.0102 | TRUE | 0.0794 |
| Ccdc167 | 402 | 305 | 287 | 311 | 259 | 285 | 312 | 324 | 310 | 310 | 292 | 281 | 304.9 | 315 | 294 | 0.934 | -0.099 | 0.39777888 | 0.57100066 | 0 | 0.0102 | 0.0066 | 0.0066 | TRUE | 0.0236 |
| Ccdc169 | 167 | 124 | 89 | 73 | 77 | 97 | 130 | 132 | 96 | 73 | 87 | 96 | 102.15 | 119 | 85 | 0.712 | -0.49 | 0.02147701 | 0.07282821 | 0.015 | 0.0256 | 0.0226 | 0.0226 | TRUE | 0.2785 |
| Ccdc17 | 9 | 11 | 10 | 13 | 5 | 7 | 7 | 12 | 11 | 13 | 6 | 7 | 9.16 | 10 | 9 | 0.878 | -0.188 | 0.77324932 | NA | 0 | 0.2611 | 0.1952 | 0.1952 | TRUE | 0.6548 |
| Ccdc170 | 50 | 35 | 62 | 46 | 56 | 57 | 39 | 37 | 67 | 46 | 63 | 56 | 51.36 | 48 | 55 | 1.161 | 0.215 | 0.48308125 | 0.6494855 | 0.0477 | 0.0486 | 0.0483 | 0.0483 | TRUE | 0.9511 |
| Ccdc171 | 272 | 225 | 180 | 175 | 159 | 179 | 211 | 239 | 194 | 174 | 179 | 177 | 195.84 | 215 | 177 | 0.823 | -0.282 | 0.05637355 | 0.14706099 | 0 | 0.0145 | 0.0106 | 0.0106 | TRUE | 0.1022 |
| Ccdc172 | 703 | 529 | 513 | 536 | 407 | 466 | 546 | 563 | 554 | 534 | 459 | 460 | 519.15 | 554 | 484 | 0.875 | -0.193 | 0.05502444 | 0.14422598 | 0.0017 | 0.007 | 0.0053 | 0.0053 | TRUE | 0.0934 |
| Ccdc173 | 84 | 69 | 69 | 77 | 48 | 69 | 65 | 73 | 74 | 77 | 54 | 68 | 68.67 | 71 | 66 | 0.939 | -0.091 | 0.70591336 | 0.82203191 | 0 | 0.0369 | 0.0274 | 0.0274 | TRUE | 0.2129 |
| Ccdc174 | 692 | 483 | 408 | 423 | 368 | 373 | 537 | 514 | 440 | 421 | 415 | 368 | 449.31 | 497 | 401 | 0.806 | -0.312 | 0.00638005 | 0.02952089 | 0.006 | 0.0077 | 0.0071 | 0.0071 | TRUE | 0.1144 |
| Ccdc175 | 14 | 15 | 10 | 14 | 9 | 11 | 11 | 16 | 11 | 14 | 10 | 11 | 12.09 | 13 | 12 | 0.935 | -0.096 | 0.86049393 | NA | 0 | 0.1984 | 0.1338 | 0.1338 | TRUE | 0.2161 |
| Ccdc176 | 79 | 82 | 57 | 68 | 57 | 84 | 61 | 87 | 62 | 68 | 64 | 83 | 70.83 | 70 | 72 | 1.028 | 0.04 | 0.87094556 | 0.92839601 | 0.0167 | 0.0359 | 0.0305 | 0.0305 | TRUE | 0.4 |
| Ccdc177 | 8 | 14 | 4 | 5 | 5 | 1 | 6 | 15 | 4 | 5 | 6 | 1 | 6.17 | 8 | 4 | 0.455 | -1.136 | 0.19375727 | NA | 0.2808 | 0.3865 | 0.3581 | 0.3581 | TRUE | 1.269 |
| Ccdc178 | 0 | 1 | 0 | 1 | 0 | 0 | 0 | 1 | 0 | 1 | 0 | 0 | 0.34 | 0 | 0 | 1.075 | 0.105 | 0.97573409 | NA | 0 | 6.9086 | 6.5582 | 6.5582 | TRUE | 0.5216 |
| Ccdc18 | 1420 | 961 | 923 | 1028 | 937 | 1074 | 1103 | 1022 | 996 | 1023 | 1057 | 1060 | 1043.58 | 1040 | 1047 | 1.005 | 0.007 | 0.92864532 | 0.96063744 | 9.00E-04 | 0.0047 | 0.0033 | 0.0033 | TRUE | 0.0347 |
| Ccdc19 | 53 | 47 | 47 | 46 | 51 | 70 | 41 | 50 | 51 | 46 | 58 | 69 | 52.38 | 47 | 58 | 1.221 | 0.289 | 0.30297278 | 0.47552416 | 0 | 0.0477 | 0.0373 | 0.0373 | TRUE | 0.163 |
| Ccdc22 | 1321 | 1021 | 861 | 953 | 860 | 1048 | 1026 | 1086 | 929 | 949 | 970 | 1035 | 999.05 | 1014 | 985 | 0.971 | -0.042 | 0.61735326 | 0.75838678 | 0.003 | 0.0048 | 0.0042 | 0.0042 | TRUE | 0.0615 |
| Ccdc23 | 860 | 581 | 547 | 584 | 520 | 578 | 668 | 618 | 590 | 581 | 587 | 571 | 602.44 | 625 | 580 | 0.925 | -0.113 | 0.22185447 | 0.38591884 | 0 | 0.0064 | 0.0045 | 0.0045 | TRUE | 0.0441 |
| Ccdc24 | 93 | 74 | 94 | 105 | 110 | 108 | 72 | 79 | 101 | 105 | 124 | 107 | 97.93 | 84 | 112 | 1.334 | 0.415 | 0.04962126 | 0.13406899 | 0.0092 | 0.0266 | 0.0218 | 0.0218 | TRUE | 0.2952 |
| Ccdc25 | 2690 | 1879 | 1911 | 1791 | 1531 | 1924 | 2089 | 1999 | 2062 | 1783 | 1727 | 1899 | 1926.5 | 2050 | 1803 | 0.88 | -0.185 | 0.00489164 | 0.02409243 | 9.00E-04 | 0.0037 | 0.0026 | 0.0026 | TRUE | 0.0264 |
| Ccdc27 | 0 | 1 | 0 | 0 | 0 | 0 | 0 | 1 | 0 | 0 | 0 | 0 | 0.18 | 0 | 0 | 0.552 | -0.857 | 0.8336548 | NA | 0 | 13.3714 | 10 | 10 | TRUE | 0.5216 |
| Ccdc28a | 424 | 302 | 311 | 337 | 311 | 366 | 329 | 321 | 336 | 335 | 351 | 361 | 338.95 | 329 | 349 | 1.062 | 0.087 | 0.4339638 | 0.60572701 | 0 | 0.0094 | 0.006 | 0.006 | TRUE | 0.0139 |
| Ccdc28b | 109 | 82 | 90 | 101 | 90 | 92 | 85 | 87 | 97 | 101 | 102 | 91 | 93.65 | 90 | 98 | 1.091 | 0.125 | 0.53427866 | 0.69340279 | 0 | 0.0277 | 0.0185 | 0.0185 | TRUE | 0.0485 |
| Ccdc3 | 216 | 166 | 165 | 311 | 264 | 312 | 168 | 177 | 178 | 310 | 298 | 308 | 239.62 | 174 | 305 | 1.756 | 0.812 | 4.97E-10 | 3.61E-08 | 0 | 0.0123 | 0.0078 | 0.0078 | TRUE | 0.0123 |
| Ccdc30 | 160 | 126 | 99 | 110 | 84 | 96 | 124 | 134 | 107 | 109 | 95 | 95 | 110.69 | 122 | 100 | 0.819 | -0.288 | 0.13373777 | 0.27225664 | 0 | 0.0238 | 0.0175 | 0.0175 | TRUE | 0.1083 |
| Ccdc32 | 620 | 442 | 422 | 487 | 406 | 442 | 482 | 470 | 455 | 485 | 458 | 436 | 464.35 | 469 | 460 | 0.979 | -0.03 | 0.76360492 | 0.85969854 | 0 | 0.0075 | 0.0051 | 0.0051 | TRUE | 0.0271 |
| Ccdc33 | 36 | 23 | 20 | 51 | 36 | 37 | 28 | 24 | 22 | 51 | 41 | 37 | 33.65 | 25 | 43 | 1.719 | 0.781 | 0.0218306 | 0.07356156 | 0 | 0.0729 | 0.0521 | 0.0521 | TRUE | 0.2194 |
| Ccdc34 | 2489 | 1932 | 1911 | 1878 | 1698 | 1997 | 1933 | 2055 | 2062 | 1869 | 1916 | 1971 | 1967.72 | 2017 | 1919 | 0.952 | -0.071 | 0.26786299 | 0.43840213 | 6.00E-04 | 0.0036 | 0.0024 | 0.0024 | TRUE | 0.0165 |
| Ccdc36 | 209 | 139 | 120 | 138 | 114 | 129 | 162 | 148 | 129 | 137 | 129 | 127 | 138.83 | 146 | 131 | 0.892 | -0.166 | 0.33649944 | 0.51068025 | 0 | 0.0195 | 0.0142 | 0.0142 | TRUE | 0.1057 |

| | | | | | | | | | | | | | | | | | | | | | | | | |
|---|---|---|---|---|---|---|---|---|---|---|---|---|---|---|---|---|---|---|---|---|---|---|---|---|
| Ccdc37 | 21 | 32 | 23 | 16 | 13 | 19 | 16 | 34 | 25 | 16 | 15 | 19 | 20.75 | 25 | 17 | 0.665 | -0.589 | 0.19003913 | 0.34801144 | 0 | 0.1166 | 0.0948 | 0.0948 | TRUE | 0.1521 |
| Ccdc38 | 173 | 120 | 120 | 115 | 74 | 111 | 134 | 128 | 129 | 114 | 83 | 110 | 116.5 | 130 | 102 | 0.787 | -0.345 | 0.07006049 | 0.17161517 | 5.00E-04 | 0.0228 | 0.0175 | 0.0175 | TRUE | 0.244 |
| Ccdc39 | 131 | 140 | 129 | 132 | 105 | 116 | 102 | 149 | 139 | 131 | 118 | 115 | 125.7 | 130 | 121 | 0.941 | -0.088 | 0.65308065 | 0.78454209 | 0.0163 | 0.0213 | 0.0198 | 0.0198 | TRUE | 0.4112 |
| Ccdc40 | 481 | 342 | 311 | 363 | 367 | 436 | 374 | 364 | 336 | 361 | 414 | 430 | 379.78 | 358 | 402 | 1.122 | 0.166 | 0.15051079 | 0.29623392 | 0.0032 | 0.0087 | 0.007 | 0.007 | TRUE | 0.0915 |
| Ccdc41 | 2856 | 2231 | 2216 | 2085 | 1885 | 2133 | 2218 | 2373 | 2391 | 2075 | 2126 | 2106 | 2214.96 | 2327 | 2102 | 0.904 | -0.146 | 0.01889728 | 0.06636334 | 6.00E-04 | 0.0035 | 0.0023 | 0.0023 | TRUE | 0.0212 |
| Ccdc42 | 2 | 3 | 2 | 3 | 2 | 1 | 2 | 3 | 2 | 3 | 2 | 1 | 2.19 | 2 | 2 | 0.912 | -0.133 | 0.91641615 | NA | 0 | 1.0853 | 0.6892 | 0.6892 | TRUE | 0.2071 |
| Ccdc42b | 0 | 1 | 0 | 0 | 0 | 0 | 0 | 1 | 0 | 0 | 0 | 0 | 0.18 | 0 | 0 | 0.552 | -0.857 | 0.8336548 | NA | 0 | 13.3714 | 10 | 10 | TRUE | 0.5216 |
| Ccdc43 | 7879 | 5847 | 5847 | 5977 | 5120 | 5928 | 6119 | 6219 | 6309 | 5949 | 5776 | 5852 | 6037.46 | 6216 | 5859 | 0.943 | -0.085 | 0.08266112 | 0.19363001 | 0 | 0.0028 | 0.0016 | 0.0016 | TRUE | 0.0023 |
| Ccdc47 | 8291 | 5850 | 5652 | 6097 | 5255 | 6067 | 6439 | 6223 | 6098 | 6069 | 5928 | 5989 | 6124.42 | 6253 | 5995 | 0.959 | -0.061 | 0.22691534 | 0.39184498 | 3.00E-04 | 0.0028 | 0.0017 | 0.0017 | TRUE | 0.0084 |
| Ccdc50 | 4489 | 3314 | 3448 | 3760 | 3277 | 3728 | 3486 | 3525 | 3720 | 3743 | 3697 | 3680 | 3641.91 | 3577 | 3707 | 1.036 | 0.052 | 0.34997268 | 0.52455531 | 4.00E-04 | 0.0031 | 0.0019 | 0.0019 | TRUE | 0.0147 |
| Ccdc51 | 604 | 438 | 403 | 492 | 401 | 452 | 469 | 466 | 435 | 490 | 452 | 446 | 459.69 | 457 | 463 | 1.013 | 0.018 | 0.85549043 | 0.91884279 | 0 | 0.0076 | 0.0052 | 0.0052 | TRUE | 0.0307 |
| Ccdc53 | 1461 | 1024 | 1054 | 1069 | 978 | 1121 | 1135 | 1089 | 1137 | 1064 | 1103 | 1107 | 1105.86 | 1120 | 1091 | 0.974 | -0.038 | 0.5916358 | 0.73955467 | 0 | 0.0046 | 0.0028 | 0.0028 | TRUE | 0.0069 |
| Ccdc54 | 0 | 0 | 0 | 0 | 0 | 0 | 0 | 0 | 0 | 0 | 0 | 0 | 0 | 0 | 0 | NA | NA | NA | NA | NA | NA | NA | NA | NA | NA |
| Ccdc55 | 3398 | 2561 | 2370 | 2306 | 2087 | 2277 | 2639 | 2724 | 2557 | 2295 | 2354 | 2248 | 2469.65 | 2640 | 2299 | 0.871 | -0.2 | 0.00083054 | 0.00632441 | 3.00E-04 | 0.0034 | 0.0022 | 0.0022 | TRUE | 0.0091 |
| Ccdc57 | 207 | 181 | 228 | 200 | 184 | 239 | 161 | 193 | 246 | 199 | 208 | 236 | 206.98 | 200 | 214 | 1.078 | 0.108 | 0.52957686 | 0.68980608 | 0.0224 | 0.0139 | 0.0164 | 0.0164 | TRUE | 0.413 |
| Ccdc58 | 4857 | 3451 | 4049 | 4637 | 4062 | 4827 | 3772 | 3671 | 4369 | 4616 | 4582 | 4765 | 4295.81 | 3937 | 4654 | 1.183 | 0.242 | 0.00078672 | 0.00607768 | 0.0045 | 0.003 | 0.0035 | 0.0035 | TRUE | 0.111 |
| Ccdc59 | 1323 | 964 | 897 | 836 | 744 | 888 | 1028 | 1025 | 968 | 832 | 839 | 877 | 928.14 | 1007 | 849 | 0.843 | -0.246 | 0.0014622 | 0.00946525 | 0 | 0.005 | 0.0032 | 0.0032 | TRUE | 0.0134 |
| Ccdc6 | 9732 | 7541 | 7623 | 8148 | 7578 | 8637 | 7558 | 8021 | 8225 | 8111 | 8549 | 8526 | 8165.03 | 7935 | 8395 | 1.058 | 0.082 | 0.13996239 | 0.28132225 | 0.0012 | 0.0027 | 0.0021 | 0.0021 | TRUE | 0.0213 |
| Ccdc60 | 50 | 53 | 49 | 69 | 69 | 76 | 39 | 56 | 53 | 69 | 78 | 75 | 61.6 | 49 | 74 | 1.508 | 0.593 | 0.02117941 | 0.07213125 | 0 | 0.0409 | 0.0308 | 0.0308 | TRUE | 0.3365 |
| Ccdc61 | 2174 | 1781 | 2421 | 2598 | 2390 | 2783 | 1688 | 1894 | 2612 | 2586 | 2696 | 2747 | 2370.76 | 2065 | 2676 | 1.297 | 0.375 | 0.05660856 | 0.14748564 | 0.0274 | 0.0034 | 0.0102 | 0.0274 | TRUE | 0.6494 |
| Ccdc62 | 84 | 54 | 35 | 60 | 59 | 48 | 65 | 57 | 38 | 60 | 67 | 47 | 55.68 | 53 | 58 | 1.072 | 0.101 | 0.72574283 | 0.83572689 | 0.0332 | 0.045 | 0.0418 | 0.0418 | TRUE | 0.5216 |
| Ccdc63 | 86 | 69 | 60 | 92 | 85 | 81 | 67 | 73 | 65 | 92 | 96 | 80 | 78.73 | 68 | 89 | 1.304 | 0.383 | 0.0815638 | 0.19189983 | 0 | 0.0325 | 0.022 | 0.022 | TRUE | 0.0788 |
| Ccdc64 | 1104 | 880 | 778 | 822 | 764 | 823 | 857 | 936 | 839 | 818 | 862 | 812 | 854.24 | 877 | 831 | 0.947 | -0.079 | 0.34056973 | 0.51467152 | 0.001 | 0.0052 | 0.0038 | 0.0038 | TRUE | 0.0392 |
| Ccdc64b | 48 | 30 | 40 | 45 | 43 | 44 | 37 | 32 | 43 | 45 | 49 | 43 | 41.51 | 37 | 46 | 1.216 | 0.283 | 0.34495785 | 0.51916211 | 0 | 0.0595 | 0.0403 | 0.0403 | TRUE | 0.1167 |
| Ccdc65 | 54 | 38 | 42 | 44 | 29 | 49 | 42 | 40 | 45 | 44 | 33 | 48 | 42.09 | 42 | 42 | 0.982 | -0.026 | 0.9314461 | 0.96216772 | 0 | 0.0587 | 0.0418 | 0.0418 | TRUE | 0.2467 |
| Ccdc66 | 1704 | 1252 | 1092 | 1090 | 986 | 1100 | 1323 | 1332 | 1178 | 1085 | 1112 | 1086 | 1186.1 | 1278 | 1094 | 0.856 | -0.225 | 0.00354206 | 0.01874677 | 0.0016 | 0.0044 | 0.0034 | 0.0034 | TRUE | 0.0544 |
| Ccdc67 | 10 | 7 | 3 | 6 | 5 | 5 | 8 | 7 | 3 | 6 | 6 | 5 | 5.83 | 6 | 6 | 0.887 | -0.173 | 0.82676692 | NA | 0 | 0.4087 | 0.2772 | 0.2772 | TRUE | 0.3715 |
| Ccdc68 | 294 | 238 | 291 | 279 | 237 | 286 | 228 | 253 | 314 | 278 | 267 | 282 | 270.48 | 265 | 276 | 1.044 | 0.063 | 0.66010545 | 0.78986079 | 0.0104 | 0.0112 | 0.0109 | 0.0109 | TRUE | 0.2861 |
| Ccdc69 | 73 | 39 | 67 | 63 | 52 | 59 | 57 | 41 | 72 | 63 | 59 | 58 | 58.35 | 57 | 60 | 1.054 | 0.076 | 0.77902557 | 0.86964954 | 0.0174 | 0.043 | 0.0363 | 0.0363 | TRUE | 0.1849 |
| Ccdc7 | 0 | 0 | 0 | 0 | 0 | 0 | 0 | 0 | 0 | 0 | 0 | 0 | 0 | 0 | 0 | NA | NA | NA | NA | NA | NA | NA | NA | NA | NA |
| Ccdc70 | 0 | 0 | 0 | 0 | 0 | 0 | 0 | 0 | 0 | 0 | 0 | 0 | 0 | 0 | 0 | NA | NA | NA | NA | NA | NA | NA | NA | NA | NA |
| Ccdc71 | 2473 | 1774 | 1513 | 1463 | 1433 | 1552 | 1921 | 1887 | 1633 | 1456 | 1617 | 1532 | 1674.18 | 1814 | 1535 | 0.846 | -0.242 | 0.00285896 | 0.01593106 | 0.0042 | 0.0038 | 0.0041 | 0.0041 | TRUE | 0.0908 |
| Ccdc71l | 794 | 559 | 410 | 379 | 333 | 355 | 617 | 595 | 442 | 377 | 376 | 350 | 459.5 | 551 | 368 | 0.666 | -0.587 | 6.84E-06 | 0.00013765 | 0.0151 | 0.0076 | 0.01 | 0.01 | TRUE | 0.3424 |
| Ccdc73 | 510 | 381 | 368 | 355 | 277 | 272 | 396 | 405 | 397 | 353 | 312 | 269 | 355.46 | 399 | 311 | 0.78 | -0.359 | 0.00372812 | 0.01954014 | 0.0059 | 0.0091 | 0.0082 | 0.0082 | TRUE | 0.1695 |
| Ccdc74a | 36 | 38 | 38 | 26 | 32 | 32 | 28 | 40 | 41 | 26 | 36 | 32 | 33.83 | 36 | 31 | 0.86 | -0.217 | 0.52703665 | 0.68806493 | 0 | 0.0725 | 0.055 | 0.055 | TRUE | 0.36 |
| Ccdc77 | 2941 | 2065 | 2087 | 2400 | 2144 | 2492 | 2284 | 2197 | 2252 | 2389 | 2419 | 2460 | 2333.35 | 2244 | 2423 | 1.079 | 0.11 | 0.05850381 | 0.15072007 | 0 | 0.0034 | 0.002 | 0.002 | TRUE | 0.0042 |
| Ccdc78 | 0 | 1 | 0 | 1 | 1 | 3 | 0 | 1 | 0 | 1 | 1 | 3 | 1.02 | 0 | 2 | 4.937 | 2.304 | 0.25879915 | NA | 0 | 2.3152 | 1.6948 | 1.6948 | TRUE | 0.5217 |
| Ccdc79 | 115 | 77 | 59 | 98 | 84 | 90 | 89 | 82 | 64 | 98 | 95 | 89 | 86.01 | 78 | 94 | 1.19 | 0.251 | 0.24972584 | 0.41849354 | 0 | 0.03 | 0.0227 | 0.0227 | TRUE | 0.2359 |
| Ccdc8 | 156 | 159 | 140 | 171 | 153 | 191 | 121 | 169 | 151 | 170 | 173 | 189 | 162.12 | 147 | 177 | 1.211 | 0.276 | 0.10673854 | 0.23203852 | 0.0098 | 0.017 | 0.0149 | 0.0149 | TRUE | 0.2776 |
| Ccdc80 | 1 | 0 | 0 | 0 | 1 | 3 | 1 | 0 | 0 | 0 | 1 | 3 | 0.81 | 0 | 1 | 3.963 | 1.987 | 0.39811512 | NA | 0 | 2.9247 | 2.6042 | 2.6042 | TRUE | 0.1519 |
| Ccdc81 | 5 | 4 | 8 | 3 | 6 | 1 | 4 | 4 | 9 | 3 | 7 | 1 | 4.59 | 6 | 4 | 0.637 | -0.652 | 0.49563454 | NA | 0.1657 | 0.5193 | 0.4249 | 0.4249 | TRUE | 0.6051 |
| Ccdc82 | 288 | 221 | 193 | 311 | 263 | 312 | 224 | 235 | 208 | 310 | 297 | 308 | 263.54 | 222 | 305 | 1.371 | 0.455 | 0.00029611 | 0.00287978 | 0 | 0.0114 | 0.0075 | 0.0075 | TRUE | 0.0312 |
| Ccdc83 | 1 | 2 | 0 | 1 | 1 | 0 | 1 | 2 | 0 | 1 | 1 | 0 | 0.84 | 1 | 1 | 0.726 | -0.461 | 0.83029328 | NA | 0 | 2.8309 | 2.0932 | 2.0932 | TRUE | 0.3091 |
| Ccdc84 | 198 | 140 | 145 | 112 | 95 | 154 | 154 | 149 | 156 | 111 | 107 | 152 | 138.3 | 153 | 123 | 0.809 | -0.306 | 0.09642933 | 0.21558962 | 0.0104 | 0.0196 | 0.0171 | 0.0171 | TRUE | 0.4187 |
| Ccdc85a | 260 | 180 | 195 | 160 | 143 | 174 | 202 | 191 | 210 | 159 | 161 | 172 | 182.69 | 201 | 164 | 0.816 | -0.294 | 0.04569721 | 0.12616385 | 0 | 0.0154 | 0.01 | 0.01 | TRUE | 0.0185 |
| Ccdc85b | 726 | 564 | 586 | 621 | 539 | 614 | 564 | 600 | 632 | 618 | 608 | 606 | 604.73 | 599 | 611 | 1.022 | 0.031 | 0.73096092 | 0.83916662 | 0 | 0.0063 | 0.0043 | 0.0043 | TRUE | 0.0326 |
| Ccdc85c | 1323 | 1102 | 1036 | 1061 | 955 | 1175 | 1028 | 1172 | 1118 | 1056 | 1077 | 1160 | 1101.82 | 1106 | 1098 | 0.994 | -0.008 | 0.91827877 | 0.95516763 | 0.0027 | 0.0046 | 0.0039 | 0.0039 | TRUE | 0.0481 |
| Ccdc86 | 1795 | 1258 | 958 | 879 | 870 | 873 | 1394 | 1338 | 1034 | 875 | 981 | 862 | 1080.68 | 1255 | 906 | 0.721 | -0.473 | 1.78E-05 | 0.00030491 | 0.0144 | 0.0046 | 0.0078 | 0.0078 | TRUE | 0.2833 |
| Ccdc87 | 4 | 5 | 3 | 1 | 4 | 3 | 3 | 5 | 3 | 1 | 5 | 3 | 3.36 | 4 | 3 | 0.723 | -0.467 | 0.6574095 | NA | 0 | 0.7088 | 0.49 | 0.49 | TRUE | 0.325 |
| Ccdc88a | 7625 | 5387 | 5214 | 6510 | 5388 | 6027 | 5922 | 5730 | 5626 | 6480 | 6078 | 5950 | 5964.3 | 5759 | 6169 | 1.071 | 0.099 | 0.0789128 | 0.18729269 | 0.0011 | 0.0028 | 0.0021 | 0.0021 | TRUE | 0.0238 |
| Ccdc88b | 156 | 117 | 115 | 194 | 128 | 159 | 121 | 124 | 124 | 193 | 144 | 157 | 144.03 | 123 | 165 | 1.34 | 0.423 | 0.01585576 | 0.0582138 | 0.006 | 0.0189 | 0.0151 | 0.0151 | TRUE | 0.2406 |
| Ccdc88c | 4057 | 3115 | 3101 | 3402 | 3240 | 3802 | 3151 | 3313 | 3346 | 3386 | 3655 | 3753 | 3434.13 | 3270 | 3598 | 1.101 | 0.138 | 0.02696862 | 0.08578747 | 0.0015 | 0.0031 | 0.0025 | 0.0025 | TRUE | 0.0325 |
| Ccdc89 | 56 | 41 | 24 | 21 | 19 | 20 | 43 | 44 | 26 | 21 | 21 | 20 | 29.18 | 38 | 21 | 0.546 | -0.872 | 0.02086429 | 0.07141123 | 0 | 0.0836 | 0.0649 | 0.0649 | TRUE | 0.5125 |
| Ccdc9 | 2244 | 1653 | 1574 | 1477 | 1264 | 1672 | 1743 | 1758 | 1698 | 1470 | 1426 | 1651 | 1624.35 | 1733 | 1516 | 0.875 | -0.193 | 0.009502 | 0.03986267 | 0.0023 | 0.0039 | 0.0034 | 0.0034 | TRUE | 0.0735 |
| Ccdc90a | 691 | 521 | 487 | 518 | 426 | 507 | 537 | 554 | 525 | 516 | 481 | 500 | 518.83 | 539 | 499 | 0.927 | -0.11 | 0.24257703 | 0.41021693 | 0 | 0.007 | 0.0045 | 0.0045 | TRUE | 0.0116 |
| Ccdc90b | 1749 | 1252 | 1338 | 1528 | 1368 | 1526 | 1358 | 1332 | 1444 | 1521 | 1543 | 1506 | 1450.74 | 1378 | 1523 | 1.106 | 0.145 | 0.03263474 | 0.09851075 | 0 | 0.0041 | 0.0026 | 0.0026 | TRUE | 0.0203 |
| Ccdc91 | 749 | 501 | 517 | 531 | 494 | 572 | 582 | 533 | 558 | 529 | 557 | 565 | 553.83 | 558 | 550 | 0.986 | -0.021 | 0.82191715 | 0.89779608 | 0 | 0.0067 | 0.0045 | 0.0045 | TRUE | 0.0181 |
| Ccdc92 | 1494 | 1116 | 992 | 998 | 888 | 1069 | 1160 | 1187 | 1070 | 993 | 1002 | 1055 | 1078.03 | 1139 | 1017 | 0.892 | -0.164 | 0.03337931 | 0.10028062 | 0.001 | 0.0046 | 0.0034 | 0.0034 | TRUE | 0.0325 |
| Ccdc93 | 2301 | 1745 | 1754 | 1962 | 1748 | 1979 | 1787 | 1856 | 1893 | 1953 | 1972 | 1954 | 1902.38 | 1845 | 1960 | 1.063 | 0.087 | 0.1556717 | 0.30342501 | 0 | 0.0037 | 0.0022 | 0.0022 | TRUE | 0.0096 |
| Ccdc94 | 489 | 345 | 293 | 268 | 283 | 269 | 380 | 367 | 316 | 267 | 319 | 266 | 319.08 | 354 | 284 | 0.798 | -0.325 | 0.01240902 | 0.0485343 | 0.0069 | 0.0099 | 0.009 | 0.009 | TRUE | 0.1297 |
| Ccdc96 | 72 | 47 | 34 | 30 | 26 | 17 | 56 | 50 | 37 | 30 | 29 | 17 | 36.43 | 48 | 25 | 0.529 | -0.92 | 0.00911617 | 0.03866878 | 0.0344 | 0.0675 | 0.059 | 0.059 | TRUE | 0.554 |
| Ccdc97 | 2127 | 1515 | 1432 | 1434 | 1393 | 1571 | 1652 | 1611 | 1545 | 1427 | 1571 | 1551 | 1559.7 | 1603 | 1516 | 0.945 | -0.081 | 0.25113971 | 0.41997658 | 0.0012 | 0.0039 | 0.0029 | 0.0029 | TRUE | 0.0321 |
| Ccer1 | 7 | 3 | 0 | 3 | 3 | 0 | 5 | 3 | 0 | 3 | 3 | 0 | 2.5 | 3 | 2 | 0.715 | -0.484 | 0.72992583 | NA | 1.2727 | 0.9506 | 1.0072 | 1.0072 | TRUE | 0.2255 |
| Cchcr1 | 1437 | 1121 | 1080 | 986 | 931 | 992 | 1116 | 1192 | 1165 | 981 | 1050 | 979 | 1080.79 | 1158 | 1003 | 0.867 | -0.206 | 0.00616143 | 0.02877995 | 0 | 0.0046 | 0.0031 | 0.0031 | TRUE | 0.0193 |
| Ccin | 4 | 1 | 1 | 0 | 1 | 2 | 3 | 1 | 1 | 0 | 1 | 2 | 1.39 | 2 | 1 | 0.576 | -0.795 | 0.6319054 | NA | 0 | 1.7051 | 1.2151 | 1.2151 | TRUE | 0.483 |
| Cck | 2 | 1 | 2 | 1 | 0 | 2 | 2 | 1 | 2 | 1 | 0 | 2 | 1.29 | 2 | 1 | 0.635 | -0.654 | 0.69698437 | NA | 0 | 1.8386 | 1.217 | 1.217 | TRUE | 0.2593 |
| Cckar | 1 | 0 | 2 | 0 | 2 | 1 | 1 | 0 | 2 | 0 | 2 | 1 | 1.03 | 1 | 1 | 1.098 | 0.135 | 0.94560368 | NA | 0 | 2.3042 | 1.8365 | 1.8365 | TRUE | 0.2585 |
| Cckbr | 55 | 36 | 34 | 44 | 29 | 40 | 43 | 38 | 37 | 44 | 33 | 39 | 38.95 | 39 | 39 | 0.985 | -0.022 | 0.94245641 | 0.96793862 | 0 | 0.0633 | 0.0431 | 0.0431 | TRUE | 0.1236 |
| Ccl1 | 0 | 0 | 0 | 0 | 0 | 0 | 0 | 0 | 0 | 0 | 0 | 0 | 0 | 0 | 0 | NA | NA | NA | NA | NA | NA | NA | NA | NA | NA |

| | | | | | | | | | | | | | | | | | | | | | | | | | |
|---|---|---|---|---|---|---|---|---|---|---|---|---|---|---|---|---|---|---|---|---|---|---|---|---|---|
| Ccl11 | 0 | 0 | 0 | 0 | 0 | 0 | 0 | 0 | 0 | 0 | 0 | 0 | 0 | 0 | 0 | NA | NA | NA | NA | NA | NA | NA | NA | NA | NA |
| Ccl12 | 0 | 0 | 0 | 0 | 0 | 0 | 0 | 0 | 0 | 0 | 0 | 0 | 0 | 0 | 0 | NA | NA | NA | NA | NA | NA | NA | NA | NA | NA |
| Ccl17 | 0 | 0 | 0 | 0 | 0 | 2 | 0 | 0 | 0 | 0 | 0 | 2 | 0.33 | 0 | 1 | 3.743 | 1.904 | 0.62640105 | NA | 0 | 7.2051 | 9.2465 | 9.2465 | TRUE | 0.9739 |
| Ccl19 | 0 | 0 | 0 | 0 | 0 | 0 | 0 | 0 | 0 | 0 | 0 | 0 | 0 | 0 | 0 | NA | NA | NA | NA | NA | NA | NA | NA | NA | NA |
| Ccl2 | 1 | 0 | 0 | 0 | 0 | 0 | 1 | 0 | 0 | 0 | 0 | 0 | 0.13 | 0 | 0 | 0.552 | -0.857 | 0.8336548 | NA | 0 | 18.3121 | 10 | 10 | TRUE | 0.2566 |
| Ccl20 | 6 | 4 | 0 | 0 | 0 | 0 | 5 | 4 | 0 | 0 | 0 | 0 | 1.49 | 3 | 0 | 0.063 | -3.978 | 0.05197378 | NA | 6.5736 | 1.5976 | 1.8574 | 1.8574 | TRUE | 0.9077 |
| Ccl21a | 0 | 0 | 0 | 0 | 0 | 0 | 0 | 0 | 0 | 0 | 0 | 0 | 0 | 0 | 0 | NA | NA | NA | NA | NA | NA | NA | NA | NA | NA |
| Ccl21b | 0 | 0 | 0 | 0 | 0 | 0 | 0 | 0 | 0 | 0 | 0 | 0 | 0 | 0 | 0 | NA | NA | NA | NA | NA | NA | NA | NA | NA | NA |
| Ccl21c | 0 | 0 | 0 | 0 | 0 | 0 | 0 | 0 | 0 | 0 | 0 | 0 | 0 | 0 | 0 | NA | NA | NA | NA | NA | NA | NA | NA | NA | NA |
| Ccl22 | 0 | 1 | 0 | 0 | 2 | 1 | 0 | 1 | 0 | 0 | 2 | 1 | 0.72 | 0 | 1 | 3.099 | 1.632 | 0.50458863 | NA | 0 | 3.3041 | 2.8241 | 2.8241 | TRUE | 0.2937 |
| Ccl24 | 0 | 0 | 0 | 0 | 0 | 0 | 0 | 0 | 0 | 0 | 0 | 0 | 0 | 0 | 0 | NA | NA | NA | NA | NA | NA | NA | NA | NA | NA |
| Ccl25 | 84 | 63 | 66 | 65 | 45 | 66 | 65 | 67 | 71 | 65 | 51 | 65 | 64.01 | 68 | 60 | 0.891 | -0.166 | 0.49623522 | 0.66061075 | 0 | 0.0395 | 0.0272 | 0.0272 | TRUE | 0.1511 |
| Ccl26 | 0 | 0 | 0 | 0 | 0 | 0 | 0 | 0 | 0 | 0 | 0 | 0 | 0 | 0 | 0 | NA | NA | NA | NA | NA | NA | NA | NA | NA | NA |
| Ccl27a | 5 | 3 | 5 | 5 | 3 | 5 | 4 | 3 | 5 | 5 | 3 | 5 | 4.29 | 4 | 4 | 1.074 | 0.103 | 0.90856305 | NA | 0 | 0.5543 | 0.3507 | 0.3507 | TRUE | 0.1041 |
| Ccl27b | 0 | 0 | 0 | 0 | 0 | 0 | 0 | 0 | 0 | 0 | 0 | 0 | 0 | 0 | 0 | NA | NA | NA | NA | NA | NA | NA | NA | NA | NA |
| Ccl28 | 2 | 1 | 4 | 5 | 2 | 5 | 2 | 1 | 4 | 5 | 2 | 5 | 3.18 | 2 | 4 | 1.797 | 0.845 | 0.44159607 | NA | 0 | 0.7469 | 0.5307 | 0.5307 | TRUE | 0.5327 |
| Ccl3 | 68 | 40 | 8 | 14 | 10 | 16 | 53 | 43 | 9 | 14 | 11 | 16 | 24.17 | 35 | 14 | 0.392 | -1.35 | 0.01437423 | 0.0540027 | 0.3321 | 0.1005 | 0.1677 | 0.1677 | TRUE | 0.5762 |
| Ccl4 | 0 | 0 | 0 | 0 | 0 | 0 | 0 | 0 | 0 | 0 | 0 | 0 | 0 | 0 | 0 | NA | NA | NA | NA | NA | NA | NA | NA | NA | NA |
| Ccl5 | 0 | 0 | 0 | 0 | 0 | 0 | 0 | 0 | 0 | 0 | 0 | 0 | 0 | 0 | 0 | NA | NA | NA | NA | NA | NA | NA | NA | NA | NA |
| Ccl6 | 0 | 0 | 0 | 0 | 1 | 0 | 0 | 0 | 0 | 0 | 1 | 0 | 0.19 | 0 | 0 | 2.094 | 1.067 | 0.79379329 | NA | 0 | 12.608 | 10 | 10 | TRUE | 0.5075 |
| Ccl7 | 1 | 0 | 0 | 0 | 0 | 0 | 1 | 0 | 0 | 0 | 0 | 0 | 0.13 | 0 | 0 | 0.552 | -0.857 | 0.8336548 | NA | 0 | 18.3121 | 10 | 10 | TRUE | 0.2566 |
| Ccl8 | 1 | 0 | 0 | 0 | 1 | 0 | 1 | 0 | 0 | 0 | 0 | 0 | 0.13 | 0 | 0 | 0.552 | -0.857 | 0.8336548 | NA | 0 | 18.3121 | 10 | 10 | TRUE | 0.2566 |
| Ccl9 | 0 | 3 | 1 | 0 | 1 | 0 | 0 | 3 | 1 | 0 | 1 | 0 | 0.9 | 1 | 0 | 0.266 | -1.909 | 0.40640649 | NA | 0 | 2.6367 | 2.4605 | 2.4605 | TRUE | 0.3216 |
| Ccm2 | 2305 | 1473 | 1375 | 1684 | 1489 | 1544 | 1790 | 1567 | 1484 | 1676 | 1680 | 1524 | 1620.13 | 1614 | 1627 | 1.007 | 0.01 | 0.91012233 | 0.95062689 | 0.0057 | 0.0039 | 0.0045 | 0.0045 | TRUE | 0.1134 |
| Ccm2l | 29 | 18 | 24 | 41 | 36 | 46 | 23 | 19 | 26 | 41 | 41 | 45 | 32.4 | 23 | 42 | 1.879 | 0.91 | 0.00771673 | 0.03409433 | 0 | 0.0756 | 0.0504 | 0.0504 | TRUE | 0.0878 |
| Ccna1 | 130 | 84 | 108 | 148 | 132 | 135 | 101 | 89 | 117 | 147 | 149 | 133 | 122.72 | 102 | 143 | 1.4 | 0.485 | 0.00831906 | 0.03607533 | 0 | 0.0217 | 0.016 | 0.016 | TRUE | 0.134 |
| Ccna2 | 19068 | 13936 | 15333 | 15773 | 14701 | 16346 | 14809 | 14824 | 16544 | 15700 | 16584 | 16136 | 15766.29 | 15392 | 16140 | 1.049 | 0.069 | 0.24890817 | 0.41805458 | 0.0024 | 0.0026 | 0.0025 | 0.0025 | TRUE | 0.0523 |
| Ccnb1 | 20226 | 14096 | 14383 | 15621 | 13616 | 15122 | 15709 | 14994 | 15519 | 15549 | 15360 | 14928 | 15343.11 | 15407 | 15279 | 0.992 | -0.012 | 0.79832656 | 0.88226146 | 4.00E-04 | 0.0026 | 0.0016 | 0.0016 | TRUE | 0.0067 |
| Ccnb1ip1 | 2166 | 1519 | 1419 | 1421 | 1199 | 1451 | 1682 | 1616 | 1531 | 1414 | 1353 | 1432 | 1504.75 | 1610 | 1400 | 0.869 | -0.202 | 0.003866 | 0.02008605 | 0.001 | 0.004 | 0.0029 | 0.0029 | TRUE | 0.0218 |
| Ccnb2 | 11610 | 8537 | 9612 | 9965 | 8989 | 10154 | 9017 | 9081 | 10371 | 9919 | 10141 | 10024 | 9758.7 | 9490 | 10028 | 1.057 | 0.08 | 0.21201803 | 0.37430505 | 0.0031 | 0.0027 | 0.0028 | 0.0028 | TRUE | 0.0804 |
| Ccnb3 | 8 | 8 | 3 | 3 | 2 | 2 | 6 | 9 | 3 | 3 | 2 | 2 | 4.2 | 6 | 2 | 0.402 | -1.316 | 0.1763412 | NA | 0 | 0.5673 | 0.3876 | 0.3876 | TRUE | 0.1346 |
| Ccnc | 2459 | 1794 | 1696 | 1926 | 1544 | 1799 | 1910 | 1908 | 1830 | 1917 | 1742 | 1776 | 1847.14 | 1883 | 1812 | 0.962 | -0.055 | 0.40752259 | 0.58032484 | 0.001 | 0.0037 | 0.0027 | 0.0027 | TRUE | 0.0315 |
| Ccnd1 | 3911 | 2924 | 2896 | 3378 | 3012 | 3404 | 3038 | 3110 | 3125 | 3362 | 3398 | 3360 | 3232.18 | 3091 | 3373 | 1.092 | 0.127 | 0.01765087 | 0.06296079 | 0 | 0.0032 | 0.0017 | 0.0017 | TRUE | 0.0029 |
| Ccnd2 | 131 | 106 | 78 | 133 | 106 | 142 | 102 | 113 | 84 | 132 | 120 | 140 | 115.13 | 100 | 131 | 1.313 | 0.393 | 0.04029741 | 0.1150791 | 0.0026 | 0.023 | 0.0177 | 0.0177 | TRUE | 0.1647 |
| Ccnd3 | 12624 | 8758 | 8868 | 8494 | 7126 | 8467 | 9805 | 9316 | 9568 | 8455 | 8039 | 8358 | 8923.47 | 9563 | 8284 | 0.866 | -0.207 | 4.04E-05 | 0.0005917 | 5.00E-04 | 0.0027 | 0.0017 | 0.0017 | TRUE | 0.0081 |
| Ccndbp1 | 1484 | 1089 | 1094 | 1105 | 1011 | 1150 | 1153 | 1158 | 1180 | 1100 | 1141 | 1135 | 1144.5 | 1164 | 1125 | 0.967 | -0.048 | 0.48936581 | 0.65510026 | 0 | 0.0045 | 0.0027 | 0.0027 | TRUE | 0.0047 |
| Ccne1 | 12226 | 8644 | 7764 | 8274 | 7693 | 9048 | 9495 | 9195 | 8377 | 8236 | 8679 | 8932 | 8818.9 | 9022 | 8616 | 0.955 | -0.067 | 0.28848466 | 0.4606537 | 0.0027 | 0.0027 | 0.0027 | 0.0027 | TRUE | 0.0477 |
| Ccne2 | 1156 | 813 | 650 | 640 | 467 | 559 | 898 | 865 | 701 | 637 | 527 | 552 | 696.61 | 821 | 572 | 0.696 | -0.523 | 4.35E-06 | 9.46E-05 | 0.0122 | 0.0058 | 0.0079 | 0.0079 | TRUE | 0.1911 |
| Ccnf | 13456 | 9839 | 10466 | 11952 | 10668 | 11547 | 10451 | 10466 | 11293 | 11897 | 12035 | 11399 | 11256.56 | 10737 | 11777 | 1.097 | 0.134 | 0.01423824 | 0.05367784 | 0.0013 | 0.0026 | 0.0021 | 0.0021 | TRUE | 0.025 |
| Ccng1 | 22259 | 15917 | 17667 | 21342 | 17356 | 19436 | 17288 | 16931 | 19063 | 21244 | 19579 | 19186 | 18881.73 | 17761 | 20003 | 1.126 | 0.172 | 0.00740804 | 0.03313193 | 0.0035 | 0.0026 | 0.0029 | 0.0029 | TRUE | 0.0502 |
| Ccng2 | 1488 | 1422 | 1956 | 2295 | 2038 | 2368 | 1156 | 1513 | 2111 | 2284 | 2299 | 2338 | 1949.97 | 1593 | 2307 | 1.449 | 0.535 | 0.03359743 | 0.10071477 | 0.0451 | 0.0036 | 0.0149 | 0.0451 | TRUE | 0.5781 |
| Ccnh | 3558 | 2591 | 2714 | 2669 | 2467 | 2808 | 2763 | 2756 | 2928 | 2657 | 2783 | 2772 | 2776.57 | 2816 | 2737 | 0.972 | -0.041 | 0.49332272 | 0.65860841 | 5.00E-04 | 0.0033 | 0.0022 | 0.0022 | TRUE | 0.0146 |
| Ccni | 16548 | 12303 | 13392 | 15348 | 13356 | 15035 | 12852 | 13087 | 14450 | 15277 | 15067 | 14842 | 14262.45 | 13463 | 15062 | 1.119 | 0.162 | 0.00521072 | 0.02530978 | 0.0021 | 0.0026 | 0.0024 | 0.0024 | TRUE | 0.0502 |
| Ccnj | 634 | 423 | 398 | 398 | 332 | 408 | 492 | 450 | 429 | 396 | 375 | 403 | 424.21 | 457 | 391 | 0.854 | -0.228 | 0.03313325 | 0.09970366 | 0 | 0.008 | 0.0058 | 0.0058 | TRUE | 0.0558 |
| Ccnjl | 754 | 554 | 505 | 567 | 509 | 566 | 586 | 589 | 545 | 564 | 574 | 559 | 569.52 | 573 | 566 | 0.986 | -0.02 | 0.82427781 | 0.89901206 | 0 | 0.0066 | 0.0042 | 0.0042 | TRUE | 0.0211 |
| Ccnk | 7195 | 5539 | 5561 | 5893 | 5517 | 6365 | 5588 | 5892 | 6000 | 5866 | 6224 | 6283 | 5975.49 | 5827 | 6124 | 1.051 | 0.072 | 0.20114988 | 0.36195835 | 0.0011 | 0.0028 | 0.0021 | 0.0021 | TRUE | 0.0167 |
| Ccnl1 | 1634 | 1245 | 1245 | 1033 | 951 | 1174 | 1269 | 1324 | 1343 | 1028 | 1073 | 1159 | 1199.45 | 1312 | 1087 | 0.829 | -0.271 | 0.00037772 | 0.00348529 | 0.0014 | 0.0044 | 0.0033 | 0.0033 | TRUE | 0.0412 |
| Ccnl2 | 3971 | 3310 | 2960 | 2730 | 2378 | 2749 | 3084 | 3521 | 3194 | 2717 | 2683 | 2714 | 2985.41 | 3266 | 2705 | 0.829 | -0.271 | 3.87E-05 | 0.0005731 | 0.0019 | 0.0032 | 0.0028 | 0.0028 | TRUE | 0.0563 |
| Ccno | 317 | 221 | 207 | 202 | 213 | 219 | 246 | 235 | 223 | 201 | 240 | 216 | 227.03 | 235 | 219 | 0.93 | -0.104 | 0.45108686 | 0.62102725 | 0 | 0.0129 | 0.0093 | 0.0093 | TRUE | 0.0749 |
| Ccnt1 | 1807 | 1237 | 1151 | 1253 | 1128 | 1218 | 1403 | 1316 | 1242 | 1247 | 1273 | 1202 | 1280.54 | 1320 | 1241 | 0.938 | -0.092 | 0.22259647 | 0.38687503 | 0.0016 | 0.0043 | 0.0033 | 0.0033 | TRUE | 0.0378 |
| Ccnt2 | 4878 | 3864 | 3859 | 4008 | 3718 | 4187 | 3789 | 4110 | 4164 | 3990 | 4194 | 4133 | 4063.25 | 4021 | 4106 | 1.022 | 0.031 | 0.60781294 | 0.75164132 | 0.0014 | 0.003 | 0.0024 | 0.0024 | TRUE | 0.0318 |
| Ccny | 4547 | 3276 | 3539 | 3607 | 3053 | 3426 | 3531 | 3485 | 3819 | 3590 | 3444 | 3382 | 3541.86 | 3612 | 3472 | 0.962 | -0.056 | 0.3567086 | 0.5312531 | 0.0013 | 0.0031 | 0.0024 | 0.0024 | TRUE | 0.0302 |
| Ccnyl1 | 1529 | 1129 | 868 | 991 | 801 | 931 | 1188 | 1201 | 937 | 986 | 904 | 919 | 1022.34 | 1109 | 936 | 0.844 | -0.244 | 0.01565032 | 0.05765882 | 0.0097 | 0.0047 | 0.0064 | 0.0064 | TRUE | 0.2163 |
| Ccp110 | 5036 | 3727 | 3320 | 3355 | 2930 | 3370 | 3911 | 3964 | 3582 | 3340 | 3305 | 3327 | 3571.58 | 3819 | 3324 | 0.87 | -0.201 | 0.00085879 | 0.00647743 | 0.0011 | 0.0031 | 0.0023 | 0.0023 | TRUE | 0.0354 |
| Ccpg1 | 4403 | 3268 | 3635 | 3953 | 3588 | 4251 | 3420 | 3476 | 3922 | 3935 | 4048 | 4196 | 3832.79 | 3606 | 4060 | 1.126 | 0.172 | 0.01146479 | 0.04572163 | 0.003 | 0.003 | 0.0031 | 0.0031 | TRUE | 0.0711 |
| Ccr1 | 0 | 0 | 0 | 0 | 0 | 0 | 0 | 0 | 0 | 0 | 0 | 0 | 0 | 0 | 0 | NA | NA | NA | NA | NA | NA | NA | NA | NA | NA |
| Ccr10 | 0 | 0 | 1 | 0 | 0 | 0 | 0 | 0 | 1 | 0 | 0 | 0 | 0.18 | 0 | 0 | 0.552 | -0.857 | 0.8336548 | NA | 0 | 13.1817 | 10 | 10 | TRUE | 0.5365 |
| Ccr1l1 | 1 | 0 | 1 | 0 | 0 | 0 | 1 | 0 | 1 | 0 | 0 | 0 | 0.31 | 1 | 0 | 0.307 | -1.706 | 0.62894649 | NA | 0 | 7.6657 | 7.2 | 7.2 | TRUE | 0.2067 |
| Ccr2 | 0 | 0 | 0 | 0 | 0 | 0 | 0 | 0 | 0 | 0 | 0 | 0 | 0 | 0 | 0 | NA | NA | NA | NA | NA | NA | NA | NA | NA | NA |
| Ccr3 | 0 | 0 | 0 | 0 | 0 | 0 | 0 | 0 | 0 | 0 | 0 | 0 | 0 | 0 | 0 | NA | NA | NA | NA | NA | NA | NA | NA | NA | NA |
| Ccr4 | 27 | 23 | 17 | 19 | 8 | 6 | 21 | 24 | 18 | 19 | 9 | 6 | 16.27 | 21 | 11 | 0.533 | -0.909 | 0.08425972 | NA | 0.0759 | 0.1481 | 0.1313 | 0.1313 | TRUE | 1.3618 |
| Ccr5 | 0 | 0 | 0 | 0 | 0 | 0 | 0 | 0 | 0 | 0 | 0 | 0 | 0 | 0 | 0 | NA | NA | NA | NA | NA | NA | NA | NA | NA | NA |
| Ccr6 | 0 | 0 | 0 | 0 | 0 | 0 | 0 | 0 | 0 | 0 | 0 | 0 | 0 | 0 | 0 | NA | NA | NA | NA | NA | NA | NA | NA | NA | NA |
| Ccr7 | 28 | 27 | 26 | 35 | 45 | 42 | 22 | 29 | 28 | 35 | 51 | 41 | 34.26 | 26 | 42 | 1.626 | 0.701 | 0.0393959 | 0.11327412 | 0 | 0.0716 | 0.0527 | 0.0527 | TRUE | 0.196 |
| Ccr8 | 7 | 4 | 2 | 3 | 3 | 4 | 5 | 4 | 2 | 3 | 3 | 4 | 3.69 | 4 | 3 | 0.858 | -0.22 | 0.82174705 | NA | 0 | 0.6439 | 0.418 | 0.418 | TRUE | 0.238 |
| Ccr9 | 0 | 0 | 0 | 0 | 0 | 0 | 0 | 0 | 0 | 0 | 0 | 0 | 0 | 0 | 0 | NA | NA | NA | NA | NA | NA | NA | NA | NA | NA |
| Ccrl1 | 4 | 1 | 5 | 10 | 1 | 8 | 3 | 1 | 5 | 10 | 1 | 8 | 4.76 | 3 | 6 | 2.007 | 1.005 | 0.31173729 | NA | 0.4133 | 0.5006 | 0.4782 | 0.4782 | TRUE | 0.6991 |
| Ccrl2 | 3 | 6 | 2 | 3 | 4 | 5 | 2 | 6 | 2 | 3 | 5 | 5 | 3.88 | 3 | 4 | 1.158 | 0.212 | 0.82831786 | NA | 0 | 0.6126 | 0.4278 | 0.4278 | TRUE | 0.6349 |
| Ccrn4l | 5118 | 3432 | 2889 | 3144 | 2904 | 3185 | 3975 | 3651 | 3117 | 3130 | 3276 | 3144 | 3382.06 | 3581 | 3183 | 0.888 | -0.171 | 0.04031061 | 0.1150938 | 0.0072 | 0.0031 | 0.0047 | 0.0047 | TRUE | 0.1555 |

| | | | | | | | | | | | | | | | | | | | | | | | | | |
|---|---|---|---|---|---|---|---|---|---|---|---|---|---|---|---|---|---|---|---|---|---|---|---|---|---|
| Ccs | 948 | 629 | 684 | 766 | 676 | 713 | 736 | 669 | 738 | 762 | 763 | 704 | 728.71 | 714 | 743 | 1.039 | 0.055 | 0.53140864 | 0.691373 | 0.0012 | 0.0057 | 0.0042 | 0.0042 | TRUE | 0.0355 |
| Ccsap | 1002 | 760 | 742 | 715 | 692 | 714 | 778 | 808 | 801 | 712 | 781 | 705 | 764.07 | 796 | 733 | 0.92 | -0.12 | 0.15489674 | 0.3024758 | 0 | 0.0055 | 0.0038 | 0.0038 | TRUE | 0.0383 |
| Ccser1 | 44 | 18 | 11 | 14 | 23 | 15 | 34 | 19 | 12 | 14 | 26 | 15 | 19.98 | 22 | 18 | 0.822 | -0.283 | 0.57146004 | 0.72474519 | 0.1517 | 0.1211 | 0.1301 | 0.1301 | TRUE | 0.4905 |
| Ccser2 | 3584 | 2695 | 2905 | 3343 | 2898 | 3400 | 2784 | 2867 | 3134 | 3328 | 3269 | 3356 | 3122.97 | 2928 | 3318 | 1.134 | 0.181 | 0.00435634 | 0.02210337 | 0.0016 | 0.0032 | 0.0026 | 0.0026 | TRUE | 0.0458 |
| Cct2 | 54421 | 39483 | 40418 | 43987 | 38075 | 44172 | 42266 | 41998 | 43611 | 43785 | 42953 | 43605 | 43036.08 | 42625 | 43448 | 1.019 | 0.028 | 0.52817776 | 0.68874916 | 2.00E-04 | 0.0025 | 0.0014 | 0.0014 | TRUE | 0.005 |
| Cct3 | 36410 | 25456 | 23372 | 24625 | 22074 | 25088 | 28278 | 27077 | 25218 | 24512 | 24902 | 24766 | 25792.12 | 26858 | 24727 | 0.921 | -0.119 | 0.02987865 | 0.09241626 | 0.0016 | 0.0025 | 0.0021 | 0.0021 | TRUE | 0.0349 |
| Cct4 | 38301 | 28786 | 30938 | 32932 | 29569 | 33767 | 29747 | 30619 | 33382 | 32780 | 33357 | 33333 | 32203.09 | 31249 | 33157 | 1.061 | 0.086 | 0.12464236 | 0.25931778 | 0.0018 | 0.0025 | 0.0022 | 0.0022 | TRUE | 0.0436 |
| Cct5 | 50808 | 36878 | 37171 | 38381 | 34232 | 39951 | 39460 | 39227 | 40107 | 38204 | 38617 | 39438 | 39175.62 | 39598 | 38753 | 0.979 | -0.031 | 0.47222837 | 0.64086802 | 2.00E-04 | 0.0025 | 0.0013 | 0.0013 | TRUE | 0.0029 |
| Cct6a | 28660 | 19915 | 18957 | 21764 | 18267 | 21087 | 22259 | 21183 | 20454 | 21664 | 20607 | 20816 | 21163.97 | 21299 | 21029 | 0.987 | -0.019 | 0.72420799 | 0.83476652 | 0.001 | 0.0025 | 0.0019 | 0.0019 | TRUE | 0.0191 |
| Cct6b | 115 | 71 | 64 | 72 | 55 | 57 | 89 | 76 | 69 | 72 | 62 | 56 | 70.65 | 78 | 63 | 0.808 | -0.307 | 0.19981222 | 0.36048598 | 0 | 0.036 | 0.0269 | 0.0269 | TRUE | 0.168 |
| Cct7 | 51518 | 37322 | 38162 | 41264 | 36628 | 41937 | 40012 | 39699 | 41176 | 41074 | 41320 | 41398 | 40779.98 | 40296 | 41264 | 1.024 | 0.034 | 0.42730969 | 0.5992621 | 2.00E-04 | 0.0025 | 0.0013 | 0.0013 | TRUE | 0.0045 |
| Cct8 | 45760 | 32165 | 32011 | 31748 | 27403 | 32226 | 35540 | 34213 | 34540 | 31602 | 30913 | 31812 | 33103.39 | 34764 | 31442 | 0.904 | -0.145 | 0.00114952 | 0.00802253 | 3.00E-04 | 0.0025 | 0.0014 | 0.0014 | TRUE | 0.0047 |
| Cct8l1 | 3 | 2 | 7 | 3 | 5 | 2 | 2 | 2 | 8 | 3 | 6 | 2 | 3.77 | 4 | 4 | 0.888 | -0.171 | 0.86728983 | NA | 0.1626 | 0.6313 | 0.4882 | 0.4882 | TRUE | 0.9314 |
| Ccz1 | 5171 | 3626 | 3471 | 3837 | 3125 | 3640 | 4016 | 3857 | 3745 | 3819 | 3525 | 3593 | 3759.35 | 3873 | 3646 | 0.941 | -0.087 | 0.14380773 | 0.28679278 | 0.0011 | 0.0031 | 0.0023 | 0.0023 | TRUE | 0.0211 |
| Cd101 | 1 | 4 | 4 | 2 | 0 | 3 | 1 | 4 | 4 | 2 | 0 | 3 | 2.38 | 3 | 2 | 0.552 | -0.858 | 0.52254129 | NA | 0 | 0.9969 | 0.8315 | 0.8315 | TRUE | 0.7922 |
| Cd109 | 454 | 369 | 473 | 452 | 388 | 444 | 353 | 393 | 510 | 450 | 438 | 438 | 430.23 | 419 | 442 | 1.059 | 0.083 | 0.53315484 | 0.69263607 | 0.0163 | 0.0079 | 0.0105 | 0.0105 | TRUE | 0.4218 |
| Cd14 | 13 | 12 | 6 | 10 | 4 | 5 | 10 | 13 | 6 | 10 | 5 | 5 | 8.12 | 10 | 7 | 0.664 | -0.591 | 0.39499457 | NA | 0 | 0.2942 | 0.2194 | 0.2194 | TRUE | 0.4376 |
| Cd151 | 2787 | 1976 | 2049 | 2580 | 2269 | 2561 | 2165 | 2102 | 2211 | 2568 | 2560 | 2528 | 2355.52 | 2159 | 2552 | 1.182 | 0.241 | 3.39E-05 | 0.0005151 | 0 | 0.0034 | 0.002 | 0.002 | TRUE | 0.0063 |
| Cd160 | 4 | 0 | 0 | 3 | 4 | 1 | 3 | 0 | 0 | 3 | 5 | 1 | 1.93 | 1 | 3 | 2.49 | 1.316 | 0.40098815 | NA | 0.8572 | 1.2291 | 1.1578 | 1.1578 | TRUE | 0.9533 |
| Cd163 | 9 | 7 | 1 | 1 | 2 | 4 | 7 | 7 | 1 | 1 | 2 | 4 | 3.79 | 5 | 2 | 0.46 | -1.119 | 0.30040545 | NA | 0 | 0.6285 | 0.5343 | 0.5343 | TRUE | 0.904 |
| Cd163l1 | 6 | 0 | 3 | 0 | 0 | 0 | 5 | 0 | 3 | 0 | 0 | 0 | 1.32 | 3 | 0 | 0.072 | -3.806 | 0.07300904 | NA | 6.4483 | 1.8032 | 2.0656 | 2.0656 | TRUE | 0.18 |
| Cd164 | 17417 | 12943 | 14032 | 15424 | 13181 | 15194 | 13527 | 13767 | 15140 | 15353 | 14870 | 14999 | 14609.36 | 14145 | 15074 | 1.066 | 0.092 | 0.10883015 | 0.23536228 | 0.0019 | 0.0026 | 0.0023 | 0.0023 | TRUE | 0.0462 |
| Cd164l2 | 193 | 134 | 174 | 252 | 208 | 252 | 150 | 143 | 188 | 251 | 235 | 249 | 202.4 | 160 | 245 | 1.534 | 0.617 | 4.99E-05 | 0.0006993 | 0.0044 | 0.0141 | 0.0115 | 0.0115 | TRUE | 0.2297 |
| Cd177 | 7 | 3 | 1 | 1 | 0 | 4 | 5 | 3 | 1 | 1 | 0 | 4 | 2.44 | 3 | 2 | 0.51 | -0.972 | 0.47092319 | NA | 0.4853 | 0.9731 | 0.8559 | 0.8559 | TRUE | 0.4178 |
| Cd180 | 6 | 8 | 0 | 3 | 7 | 2 | 5 | 9 | 0 | 3 | 8 | 2 | 4.34 | 5 | 4 | 0.963 | -0.054 | 0.95979228 | NA | 0.8012 | 0.5488 | 0.6052 | 0.6052 | TRUE | 0.2179 |
| Cd19 | 3 | 4 | 2 | 1 | 1 | 3 | 2 | 4 | 2 | 1 | 1 | 3 | 2.3 | 3 | 2 | 0.59 | -0.762 | 0.54550169 | NA | 0 | 1.0308 | 0.6743 | 0.6743 | TRUE | 0.3419 |
| Cd1d1 | 568 | 463 | 457 | 441 | 434 | 487 | 441 | 492 | 493 | 439 | 490 | 481 | 472.67 | 475 | 470 | 0.989 | -0.016 | 0.88109002 | 0.93360662 | 0.0018 | 0.0074 | 0.0056 | 0.0056 | TRUE | 0.05 |
| Cd1d2 | 60 | 30 | 53 | 42 | 36 | 47 | 47 | 32 | 57 | 42 | 41 | 46 | 44.09 | 45 | 43 | 0.95 | -0.075 | 0.80912842 | 0.88984911 | 0.0182 | 0.0562 | 0.0461 | 0.0461 | TRUE | 0.2588 |
| Cd2 | 59 | 46 | 47 | 57 | 47 | 59 | 46 | 49 | 51 | 57 | 53 | 58 | 52.24 | 49 | 56 | 1.159 | 0.212 | 0.41934849 | 0.59169931 | 0 | 0.0478 | 0.0307 | 0.0307 | TRUE | 0.022 |
| Cd200 | 102 | 66 | 49 | 38 | 27 | 23 | 79 | 70 | 53 | 38 | 30 | 23 | 48.88 | 67 | 30 | 0.448 | -1.16 | 0.00018067 | 0.00197881 | 0.0283 | 0.0509 | 0.0449 | 0.0449 | TRUE | 0.3022 |
| Cd200r1 | 5 | 2 | 2 | 1 | 2 | 0 | 4 | 2 | 2 | 1 | 2 | 0 | 1.9 | 3 | 1 | 0.383 | -1.384 | 0.34574121 | NA | 0 | 1.2476 | 0.8869 | 0.8869 | TRUE | 0.4044 |
| Cd200r2 | 27 | 17 | 7 | 8 | 9 | 10 | 21 | 18 | 8 | 8 | 10 | 10 | 12.43 | 16 | 9 | 0.593 | -0.753 | 0.1962186 | NA | 0 | 0.1931 | 0.1584 | 0.1584 | TRUE | 0.8723 |
| Cd200r3 | 1 | 1 | 0 | 1 | 2 | 2 | 1 | 1 | 0 | 1 | 2 | 2 | 1.18 | 1 | 2 | 2.779 | 1.475 | 0.41423757 | NA | 0 | 2.0149 | 1.2827 | 1.2827 | TRUE | 0.187 |
| Cd200r4 | 0 | 1 | 0 | 0 | 0 | 0 | 0 | 1 | 0 | 0 | 0 | 0 | 0.18 | 0 | 0 | 0.552 | -0.857 | 0.8336548 | NA | 0 | 13.3714 | 10 | 10 | TRUE | 0.5216 |
| Cd207 | 0 | 0 | 2 | 1 | 0 | 1 | 0 | 0 | 2 | 1 | 0 | 1 | 0.69 | 1 | 1 | 0.966 | -0.049 | 0.98423749 | NA | 0 | 3.4369 | 3.0198 | 3.0198 | TRUE | 0.9749 |
| Cd209a | 0 | 1 | 1 | 2 | 1 | 0 | 0 | 1 | 1 | 2 | 1 | 0 | 0.88 | 1 | 1 | 1.518 | 0.602 | 0.77693369 | NA | 0 | 2.7051 | 2.052 | 2.052 | TRUE | 0.2258 |
| Cd209b | 0 | 0 | 0 | 0 | 0 | 0 | 0 | 0 | 0 | 0 | 0 | 0 | 0 | 0 | 0 | NA | NA | NA | NA | NA | NA | NA | NA | NA | NA |
| Cd209c | 0 | 1 | 0 | 0 | 1 | 0 | 0 | 1 | 0 | 0 | 1 | 0 | 0.37 | 0 | 0 | 1.075 | 0.105 | 0.97520514 | NA | 0 | 6.4904 | 6.1969 | 6.1969 | TRUE | 0.5216 |
| Cd209d | 0 | 0 | 0 | 0 | 0 | 0 | 0 | 0 | 0 | 0 | 0 | 0 | 0 | 0 | 0 | NA | NA | NA | NA | NA | NA | NA | NA | NA | NA |
| Cd209e | 6 | 5 | 1 | 1 | 0 | 1 | 5 | 5 | 1 | 1 | 0 | 1 | 2.17 | 4 | 1 | 0.183 | -2.451 | 0.11235003 | NA | 0 | 1.093 | 0.825 | 0.825 | TRUE | 0.5409 |
| Cd209f | 3 | 3 | 2 | 0 | 0 | 3 | 2 | 3 | 2 | 0 | 0 | 3 | 1.77 | 2 | 1 | 0.395 | -1.339 | 0.39845181 | NA | 0 | 1.3389 | 1.1107 | 1.1107 | TRUE | 1.441 |
| Cd209g | 0 | 1 | 1 | 0 | 0 | 0 | 0 | 1 | 1 | 0 | 0 | 0 | 0.36 | 1 | 0 | 0.271 | -1.881 | 0.5755355 | NA | 0 | 6.6391 | 6.4428 | 6.4428 | TRUE | 0.3513 |
| Cd22 | 23 | 10 | 11 | 19 | 15 | 17 | 18 | 11 | 12 | 19 | 17 | 17 | 15.5 | 14 | 18 | 1.286 | 0.363 | 0.45756831 | NA | 0 | 0.1554 | 0.1076 | 0.1076 | TRUE | 0.4379 |
| Cd226 | 0 | 0 | 0 | 0 | 1 | 0 | 0 | 0 | 0 | 0 | 1 | 0 | 0.19 | 0 | 0 | 2.094 | 1.067 | 0.79379329 | NA | 0 | 12.608 | 10 | 10 | TRUE | 0.5075 |
| Cd244 | 1 | 1 | 1 | 2 | 2 | 0 | 1 | 1 | 1 | 2 | 2 | 0 | 1.19 | 1 | 1 | 1.454 | 0.54 | 0.7585334 | NA | 0 | 1.9868 | 1.3533 | 1.3533 | TRUE | 0.521 |
| Cd247 | 8 | 4 | 4 | 5 | 4 | 8 | 6 | 4 | 4 | 5 | 5 | 8 | 5.36 | 5 | 6 | 1.168 | 0.224 | 0.78327691 | NA | 0 | 0.4445 | 0.2912 | 0.2912 | TRUE | 0.2367 |
| Cd248 | 20 | 11 | 4 | 12 | 19 | 20 | 16 | 12 | 4 | 12 | 21 | 20 | 14.11 | 11 | 18 | 1.649 | 0.721 | 0.20287241 | NA | 0.1149 | 0.1704 | 0.1568 | 0.1568 | TRUE | 0.683 |
| Cd24a | 2076 | 1555 | 1490 | 1912 | 1661 | 1926 | 1612 | 1654 | 1608 | 1903 | 1874 | 1901 | 1758.72 | 1625 | 1893 | 1.165 | 0.221 | 0.00033243 | 0.00314399 | 0 | 0.0038 | 0.0022 | 0.0022 | TRUE | 0.0029 |
| Cd27 | 0 | 0 | 0 | 1 | 0 | 0 | 0 | 0 | 0 | 1 | 0 | 0 | 0.17 | 0 | 0 | 2.094 | 1.067 | 0.79379329 | NA | 0 | 14.2885 | 10 | 10 | TRUE | 0.3906 |
| Cd274 | 59 | 38 | 36 | 57 | 45 | 58 | 46 | 40 | 39 | 57 | 51 | 57 | 48.31 | 42 | 55 | 1.313 | 0.393 | 0.15426279 | 0.30158366 | 0 | 0.0515 | 0.0339 | 0.0339 | TRUE | 0.0675 |
| Cd276 | 3392 | 2485 | 2451 | 2597 | 2408 | 2677 | 2634 | 2643 | 2645 | 2585 | 2716 | 2643 | 2644.41 | 2641 | 2648 | 1.003 | 0.004 | 0.94695741 | 0.97100669 | 0 | 0.0033 | 0.0019 | 0.0019 | TRUE | 0.0061 |
| Cd28 | 2 | 3 | 1 | 2 | 0 | 1 | 2 | 3 | 1 | 2 | 0 | 1 | 1.47 | 2 | 1 | 0.525 | -0.931 | 0.56462803 | NA | 0 | 1.6181 | 1.1211 | 1.1211 | TRUE | 0.2966 |
| Cd2ap | 9272 | 7016 | 7162 | 7518 | 6660 | 7853 | 7201 | 7463 | 7728 | 7483 | 7513 | 7752 | 7523.41 | 7464 | 7583 | 1.016 | 0.023 | 0.65578062 | 0.78626561 | 6.00E-04 | 0.0027 | 0.0018 | 0.0018 | TRUE | 0.0117 |
| Cd2bp2 | 5821 | 4120 | 3894 | 3886 | 3341 | 3837 | 4521 | 4382 | 4202 | 3868 | 3769 | 3788 | 4088.29 | 4368 | 3808 | 0.872 | -0.198 | 0.00033315 | 0.00314722 | 6.00E-04 | 0.003 | 0.002 | 0.002 | TRUE | 0.0135 |
| Cd300a | 17 | 15 | 6 | 19 | 7 | 17 | 13 | 16 | 6 | 19 | 8 | 17 | 13.2 | 12 | 15 | 1.225 | 0.293 | 0.60841195 | NA | 0.0948 | 0.1819 | 0.1596 | 0.1596 | TRUE | 0.6413 |
| Cd300c | 0 | 0 | 0 | 0 | 1 | 0 | 0 | 0 | 0 | 0 | 1 | 0 | 0.19 | 0 | 0 | 2.094 | 1.067 | 0.79379329 | NA | 0 | 12.608 | 10 | 10 | TRUE | 0.5075 |
| Cd300e | 5 | 0 | 1 | 1 | 1 | 1 | 4 | 0 | 1 | 1 | 1 | 1 | 1.35 | 2 | 1 | 0.601 | -0.735 | 0.66734624 | NA | 0 | 1.7639 | 1.3254 | 1.3254 | TRUE | 0.6547 |
| Cd300lb | 6 | 5 | 6 | 2 | 1 | 5 | 5 | 5 | 6 | 2 | 1 | 5 | 4.08 | 5 | 3 | 0.498 | -1.005 | 0.30378471 | NA | 0 | 0.5827 | 0.4094 | 0.4094 | TRUE | 0.647 |
| Cd300ld | 1 | 1 | 1 | 0 | 0 | 0 | 1 | 1 | 1 | 0 | 0 | 0 | 0.49 | 1 | 0 | 0.196 | -2.347 | 0.38405676 | NA | 0 | 4.8735 | 3.7415 | 3.7415 | TRUE | 0.0718 |
| Cd300lf | 24 | 13 | 23 | 31 | 22 | 30 | 19 | 14 | 25 | 31 | 25 | 30 | 23.76 | 19 | 29 | 1.494 | 0.579 | 0.15212076 | 0.29872099 | 0 | 0.1022 | 0.0744 | 0.0744 | TRUE | 0.3172 |
| Cd300lg | 28 | 24 | 22 | 40 | 43 | 34 | 22 | 26 | 24 | 40 | 49 | 34 | 32.15 | 24 | 41 | 1.718 | 0.781 | 0.02447803 | 0.08009717 | 0 | 0.0761 | 0.0536 | 0.0536 | TRUE | 0.1867 |
| Cd300lh | 0 | 0 | 0 | 0 | 0 | 0 | 0 | 0 | 0 | 0 | 0 | 0 | 0 | 0 | 0 | NA | NA | NA | NA | NA | NA | NA | NA | NA | NA |
| Cd302 | 347 | 268 | 314 | 351 | 270 | 316 | 270 | 285 | 339 | 349 | 305 | 312 | 309.88 | 298 | 322 | 1.085 | 0.117 | 0.36906771 | 0.54371107 | 0.0068 | 0.0101 | 0.0091 | 0.0091 | TRUE | 0.1607 |
| Cd320 | 1893 | 1375 | 1441 | 1688 | 1433 | 1712 | 1470 | 1463 | 1555 | 1680 | 1617 | 1690 | 1579.07 | 1496 | 1662 | 1.112 | 0.153 | 0.02085428 | 0.07139406 | 0 | 0.0039 | 0.0025 | 0.0025 | TRUE | 0.014 |
| Cd33 | 0 | 1 | 0 | 3 | 1 | 1 | 0 | 1 | 0 | 3 | 1 | 1 | 1.03 | 0 | 2 | 4.949 | 2.307 | 0.2578665 | NA | 0 | 2.309 | 1.6929 | 1.6929 | TRUE | 0.5217 |
| Cd34 | 754 | 576 | 534 | 730 | 613 | 711 | 586 | 613 | 576 | 727 | 692 | 702 | 649.08 | 592 | 707 | 1.195 | 0.257 | 0.00293327 | 0.0162564 | 0 | 0.0061 | 0.0038 | 0.0038 | TRUE | 0.011 |
| Cd36 | 2 | 0 | 1 | 0 | 0 | 0 | 2 | 0 | 1 | 0 | 0 | 0 | 0.44 | 1 | 0 | 0.214 | -2.223 | 0.47051571 | NA | 0 | 5.4047 | 5.2946 | 5.2946 | TRUE | 0.2788 |
| Cd37 | 391 | 293 | 268 | 492 | 436 | 502 | 304 | 312 | 289 | 490 | 492 | 496 | 396.94 | 302 | 493 | 1.633 | 0.707 | 1.54E-11 | 1.47E-09 | 0 | 0.0084 | 0.0053 | 0.0053 | TRUE | 0.0134 |
| Cd38 | 723 | 577 | 686 | 679 | 655 | 754 | 562 | 614 | 740 | 676 | 739 | 744 | 679.09 | 639 | 720 | 1.13 | 0.176 | 0.1100092 | 0.23719438 | 0.0095 | 0.0059 | 0.0072 | 0.0072 | TRUE | 0.2268 |

| | | | | | | | | | | | | | | | | | | | | | | | | |
|---|---|---|---|---|---|---|---|---|---|---|---|---|---|---|---|---|---|---|---|---|---|---|---|---|
| Cd3d | 2 | 3 | 1 | 4 | 3 | 1 | 2 | 3 | 1 | 4 | 3 | 1 | 2.36 | 2 | 3 | 1.44 | 0.526 | 0.67392834 | NA | 0 | 1.0055 | 0.6904 | 0.6904 | TRUE | 0.4063 |
| Cd3e | 52 | 33 | 35 | 24 | 19 | 20 | 40 | 35 | 38 | 24 | 21 | 20 | 29.72 | 38 | 22 | 0.573 | -0.803 | 0.02334284 | 0.07740444 | 0 | 0.0822 | 0.0537 | 0.0537 | TRUE | 0.0465 |
| Cd3eap | 1833 | 1319 | 1093 | 1017 | 981 | 1068 | 1424 | 1403 | 1179 | 1012 | 1107 | 1054 | 1196.54 | 1335 | 1058 | 0.791 | -0.338 | 0.00013851 | 0.00159399 | 0.0056 | 0.0044 | 0.0048 | 0.0048 | TRUE | 0.1236 |
| Cd3g | 5 | 5 | 1 | 1 | 5 | 2 | 4 | 5 | 1 | 1 | 6 | 2 | 3.15 | 3 | 3 | 0.82 | -0.286 | 0.80128229 | NA | 0.2183 | 0.7552 | 0.608 | 0.608 | TRUE | 0.6897 |
| Cd4 | 5 | 5 | 2 | 3 | 4 | 3 | 4 | 5 | 2 | 3 | 5 | 3 | 3.64 | 4 | 4 | 0.914 | -0.129 | 0.89582548 | NA | 0 | 0.6541 | 0.425 | 0.425 | TRUE | 0.1957 |
| Cd40 | 112 | 78 | 77 | 134 | 93 | 132 | 87 | 83 | 83 | 133 | 105 | 130 | 103.61 | 84 | 123 | 1.458 | 0.544 | 0.00595016 | 0.02804936 | 0 | 0.0253 | 0.0182 | 0.0182 | TRUE | 0.1577 |
| Cd40lg | 0 | 0 | 0 | 0 | 0 | 0 | 0 | 0 | 0 | 0 | 0 | 0 | 0 | 0 | 0 | NA | NA | NA | NA | NA | NA | NA | NA | NA | NA |
| Cd44 | 68 | 58 | 44 | 80 | 54 | 63 | 53 | 62 | 47 | 80 | 61 | 62 | 60.79 | 54 | 68 | 1.254 | 0.327 | 0.20420332 | 0.36556589 | 0 | 0.0414 | 0.0311 | 0.0311 | TRUE | 0.2199 |
| Cd46 | 1 | 2 | 3 | 1 | 1 | 0 | 1 | 2 | 3 | 1 | 1 | 0 | 1.38 | 2 | 1 | 0.347 | -1.526 | 0.37918523 | NA | 0 | 1.7231 | 1.1882 | 1.1882 | TRUE | 0.1962 |
| Cd47 | 546 | 406 | 447 | 596 | 436 | 538 | 424 | 432 | 482 | 593 | 492 | 531 | 492.4 | 446 | 539 | 1.21 | 0.275 | 0.0114109 | 0.04558301 | 0.0049 | 0.0072 | 0.0065 | 0.0065 | TRUE | 0.0918 |
| Cd48 | 0 | 0 | 0 | 0 | 0 | 0 | 0 | 0 | 0 | 0 | 0 | 0 | 0 | 0 | 0 | NA | NA | NA | NA | NA | NA | NA | NA | NA | NA |
| Cd5 | 1 | 2 | 1 | 0 | 0 | 0 | 1 | 2 | 1 | 0 | 0 | 0 | 0.66 | 1 | 0 | 0.145 | -2.79 | 0.24993894 | NA | 0 | 3.5727 | 2.8701 | 2.8701 | TRUE | 0.1613 |
| Cd52 | 2 | 5 | 3 | 1 | 1 | 0 | 2 | 5 | 3 | 1 | 1 | 0 | 2.04 | 3 | 1 | 0.209 | -2.256 | 0.14535735 | NA | 0 | 1.165 | 0.8408 | 0.8408 | TRUE | 0.1716 |
| Cd53 | 0 | 0 | 0 | 0 | 0 | 2 | 0 | 0 | 0 | 0 | 0 | 2 | 0.33 | 0 | 1 | 3.743 | 1.904 | 0.62640105 | NA | 0 | 7.2051 | 9.2465 | 9.2465 | TRUE | 0.9739 |
| Cd55 | 759 | 516 | 651 | 762 | 626 | 661 | 589 | 549 | 702 | 758 | 706 | 653 | 659.66 | 613 | 706 | 1.151 | 0.203 | 0.06262637 | 0.158288 | 0.0089 | 0.006 | 0.007 | 0.007 | TRUE | 0.1832 |
| Cd59a | 138 | 98 | 112 | 86 | 85 | 99 | 107 | 104 | 121 | 86 | 96 | 98 | 101.91 | 111 | 93 | 0.841 | -0.25 | 0.20030284 | 0.36102503 | 0 | 0.0257 | 0.0175 | 0.0175 | TRUE | 0.0598 |
| Cd59b | 14 | 18 | 7 | 13 | 17 | 20 | 11 | 19 | 8 | 13 | 19 | 20 | 14.91 | 13 | 17 | 1.384 | 0.469 | 0.37390297 | NA | 0.0475 | 0.1614 | 0.132 | 0.132 | TRUE | 0.7975 |
| Cd5l | 0 | 1 | 1 | 0 | 0 | 0 | 0 | 1 | 1 | 0 | 0 | 0 | 0.36 | 1 | 0 | 0.271 | -1.881 | 0.5755355 | NA | 0 | 6.6391 | 6.4428 | 6.4428 | TRUE | 0.3513 |
| Cd6 | 270 | 237 | 253 | 273 | 323 | 372 | 210 | 252 | 273 | 272 | 364 | 367 | 289.69 | 245 | 334 | 1.369 | 0.453 | 0.00311725 | 0.01701864 | 0.0199 | 0.0106 | 0.0134 | 0.0134 | TRUE | 0.3095 |
| Cd63 | 3260 | 2282 | 2173 | 2180 | 1629 | 1911 | 2532 | 2427 | 2345 | 2170 | 1838 | 1886 | 2199.66 | 2435 | 1965 | 0.807 | -0.31 | 5.50E-05 | 0.00075607 | 0.0042 | 0.0035 | 0.0038 | 0.0038 | TRUE | 0.1013 |
| Cd68 | 226 | 119 | 203 | 186 | 195 | 240 | 176 | 127 | 219 | 185 | 220 | 237 | 193.86 | 174 | 214 | 1.232 | 0.301 | 0.11539835 | 0.24530321 | 0.0378 | 0.0147 | 0.0211 | 0.0211 | TRUE | 0.2556 |
| Cd69 | 1 | 0 | 1 | 0 | 0 | 1 | 1 | 0 | 1 | 0 | 0 | 1 | 0.47 | 1 | 0 | 0.596 | -0.748 | 0.78959815 | NA | 0 | 5.0047 | 3.8792 | 3.8792 | TRUE | 0.3835 |
| Cd7 | 58 | 51 | 68 | 70 | 49 | 58 | 45 | 54 | 73 | 70 | 55 | 57 | 59.15 | 57 | 61 | 1.064 | 0.09 | 0.7422942 | 0.84665858 | 0.0232 | 0.0425 | 0.037 | 0.037 | TRUE | 0.4796 |
| Cd70 | 0 | 0 | 0 | 0 | 0 | 0 | 0 | 0 | 0 | 0 | 0 | 0 | 0 | 0 | 0 | NA | NA | NA | NA | NA | NA | NA | NA | NA | NA |
| Cd72 | 2 | 0 | 0 | 0 | 1 | 1 | 2 | 0 | 0 | 0 | 1 | 1 | 0.61 | 1 | 1 | 1.258 | 0.332 | 0.89795466 | NA | 0 | 3.8787 | 3.2104 | 3.2104 | TRUE | 0.9721 |
| Cd74 | 274 | 142 | 139 | 277 | 179 | 230 | 213 | 151 | 150 | 276 | 202 | 227 | 203.09 | 171 | 235 | 1.365 | 0.449 | 0.01267776 | 0.0492534 | 0.0285 | 0.0141 | 0.0184 | 0.0184 | TRUE | 0.5133 |
| Cd79a | 64 | 41 | 32 | 56 | 30 | 63 | 50 | 44 | 35 | 56 | 34 | 62 | 46.6 | 43 | 51 | 1.186 | 0.246 | 0.43567712 | 0.60705138 | 0.0414 | 0.0533 | 0.0501 | 0.0501 | TRUE | 0.6389 |
| Cd79b | 226 | 136 | 168 | 259 | 236 | 272 | 176 | 145 | 181 | 258 | 266 | 269 | 215.67 | 167 | 264 | 1.578 | 0.658 | 4.51E-06 | 9.77E-05 | 0 | 0.0134 | 0.01 | 0.01 | TRUE | 0.1393 |
| Cd80 | 507 | 302 | 335 | 544 | 447 | 551 | 394 | 321 | 361 | 541 | 504 | 544 | 444.36 | 359 | 530 | 1.473 | 0.559 | 3.98E-07 | 1.33E-05 | 0.0034 | 0.0078 | 0.0064 | 0.0064 | TRUE | 0.095 |
| Cd81 | 6944 | 5301 | 6795 | 7973 | 6978 | 7943 | 5393 | 5639 | 7332 | 7936 | 7872 | 7841 | 7002.11 | 6121 | 7883 | 1.288 | 0.365 | 0.00012953 | 0.00151129 | 0.0147 | 0.0028 | 0.0064 | 0.0064 | TRUE | 0.3645 |
| Cd82 | 87 | 87 | 87 | 115 | 99 | 113 | 68 | 93 | 94 | 114 | 112 | 112 | 98.61 | 85 | 113 | 1.34 | 0.422 | 0.0423795 | 0.1193557 | 0 | 0.0265 | 0.0208 | 0.0208 | TRUE | 0.3387 |
| Cd83 | 23 | 10 | 13 | 20 | 15 | 12 | 18 | 11 | 14 | 20 | 17 | 12 | 15.2 | 14 | 16 | 1.132 | 0.179 | 0.72090384 | NA | 0 | 0.1583 | 0.116 | 0.116 | TRUE | 0.2849 |
| Cd84 | 0 | 1 | 0 | 0 | 0 | 0 | 0 | 1 | 0 | 0 | 0 | 0 | 0.18 | 0 | 0 | 0.552 | -0.857 | 0.8336548 | NA | 0 | 13.3714 | 10 | 10 | TRUE | 0.5216 |
| Cd86 | 3 | 4 | 0 | 1 | 4 | 3 | 2 | 4 | 0 | 1 | 5 | 3 | 2.51 | 2 | 3 | 1.271 | 0.346 | 0.7890011 | NA | 0.2746 | 0.9471 | 0.8003 | 0.8003 | TRUE | 0.268 |
| Cd8a | 53 | 27 | 26 | 51 | 25 | 30 | 41 | 29 | 28 | 51 | 28 | 30 | 34.42 | 33 | 36 | 1.102 | 0.141 | 0.69920262 | 0.8173471 | 0.0549 | 0.0713 | 0.0665 | 0.0665 | TRUE | 0.906 |
| Cd8b1 | 37 | 13 | 12 | 22 | 12 | 13 | 29 | 14 | 13 | 22 | 14 | 13 | 17.3 | 19 | 16 | 0.857 | -0.222 | 0.66246386 | 0.79168863 | 0.1053 | 0.1394 | 0.1292 | 0.1292 | TRUE | 1.4462 |
| Cd9 | 16725 | 12303 | 15097 | 17036 | 15695 | 16931 | 12990 | 13087 | 16290 | 16958 | 17706 | 16714 | 15623.73 | 14122 | 17126 | 1.213 | 0.278 | 0.00068961 | 0.00552187 | 0.0092 | 0.0026 | 0.0048 | 0.0048 | TRUE | 0.2202 |
| Cd93 | 3 | 2 | 1 | 1 | 4 | 0 | 2 | 2 | 1 | 1 | 5 | 0 | 1.84 | 2 | 2 | 0.963 | -0.054 | 0.97090814 | NA | 0.3393 | 1.29 | 1.0496 | 1.0496 | TRUE | 1.1862 |
| Cd96 | 5 | 1 | 4 | 3 | 1 | 2 | 4 | 1 | 4 | 3 | 1 | 2 | 2.56 | 3 | 2 | 0.657 | -0.605 | 0.61547536 | NA | 0 | 0.9287 | 0.6378 | 0.6378 | TRUE | 0.3992 |
| Cd97 | 7738 | 5871 | 6104 | 7781 | 7196 | 8285 | 6010 | 6245 | 6586 | 7745 | 8118 | 8179 | 7147.08 | 6280 | 8014 | 1.276 | 0.352 | 4.44E-10 | 3.26E-08 | 0.0013 | 0.0028 | 0.0022 | 0.0022 | TRUE | 0.0221 |
| Cd99l2 | 891 | 584 | 682 | 680 | 649 | 690 | 692 | 621 | 736 | 677 | 732 | 681 | 689.87 | 683 | 697 | 1.019 | 0.028 | 0.76913992 | 0.86332904 | 0.0029 | 0.0059 | 0.0049 | 0.0049 | TRUE | 0.0711 |
| Cda | 92 | 42 | 69 | 59 | 70 | 70 | 71 | 45 | 74 | 59 | 79 | 69 | 66.23 | 63 | 69 | 1.079 | 0.11 | 0.67908768 | 0.8028453 | 0.0298 | 0.0382 | 0.036 | 0.036 | TRUE | 0.559 |
| Cdadc1 | 476 | 349 | 295 | 271 | 252 | 285 | 370 | 371 | 318 | 270 | 284 | 281 | 315.77 | 353 | 278 | 0.787 | -0.345 | 0.00429817 | 0.02186252 | 0 | 0.0099 | 0.0073 | 0.0073 | TRUE | 0.0816 |
| Cdan1 | 2124 | 1572 | 1534 | 1494 | 1310 | 1543 | 1650 | 1672 | 1655 | 1487 | 1478 | 1523 | 1577.51 | 1659 | 1496 | 0.902 | -0.149 | 0.01879788 | 0.06606298 | 0 | 0.0039 | 0.0023 | 0.0023 | TRUE | 0.0031 |
| Cdc123 | 4133 | 2715 | 2604 | 2836 | 2408 | 2869 | 3210 | 2888 | 2810 | 2823 | 2716 | 2832 | 2879.85 | 2969 | 2790 | 0.939 | -0.091 | 0.18126284 | 0.33681614 | 0.0023 | 0.0032 | 0.003 | 0.003 | TRUE | 0.0624 |
| Cdc14a | 311 | 190 | 180 | 161 | 137 | 139 | 242 | 202 | 194 | 160 | 155 | 137 | 181.65 | 213 | 151 | 0.706 | -0.503 | 0.00161024 | 0.0101792 | 0.0062 | 0.0155 | 0.0126 | 0.0126 | TRUE | 0.1664 |
| Cdc14b | 2927 | 2256 | 2597 | 3220 | 2814 | 3359 | 2273 | 2400 | 2802 | 3205 | 3174 | 3316 | 2861.77 | 2492 | 3232 | 1.298 | 0.376 | 2.06E-06 | 5.18E-05 | 0.0059 | 0.0033 | 0.0042 | 0.0042 | TRUE | 0.1433 |
| Cdc16 | 6948 | 4691 | 4936 | 5211 | 4616 | 5135 | 5396 | 4990 | 5326 | 5187 | 5207 | 5069 | 5195.88 | 5237 | 5154 | 0.984 | -0.023 | 0.66959063 | 0.79697335 | 7.00E-04 | 0.0029 | 0.002 | 0.002 | TRUE | 0.0207 |
| Cdc20 | 12629 | 9395 | 9243 | 8962 | 8451 | 9446 | 9808 | 9993 | 9973 | 8921 | 9534 | 9325 | 9592.32 | 9925 | 9260 | 0.933 | -0.1 | 0.04436369 | 0.12341186 | 5.00E-04 | 0.0027 | 0.0017 | 0.0017 | TRUE | 0.0125 |
| Cdc23 | 8395 | 6100 | 5839 | 6286 | 5425 | 6207 | 6520 | 6488 | 6300 | 6257 | 6120 | 6127 | 6302.18 | 6436 | 6168 | 0.958 | -0.062 | 0.20811344 | 0.36974089 | 1.00E-04 | 0.0028 | 0.0016 | 0.0016 | TRUE | 0.0041 |
| Cdc25a | 6689 | 4828 | 4969 | 5348 | 4620 | 5281 | 5195 | 5135 | 5362 | 5323 | 5212 | 5213 | 5240.07 | 5231 | 5249 | 1.004 | 0.005 | 0.91595175 | 0.95357989 | 1.00E-04 | 0.0029 | 0.0016 | 0.0016 | TRUE | 0.0057 |
| Cdc25b | 2381 | 1755 | 1912 | 2422 | 2130 | 2433 | 1849 | 1867 | 2063 | 2411 | 2403 | 2402 | 2165.75 | 1926 | 2405 | 1.249 | 0.321 | 1.15E-06 | 3.30E-05 | 0.0012 | 0.0035 | 0.0027 | 0.0027 | TRUE | 0.0459 |
| Cdc25c | 1457 | 1080 | 1170 | 1248 | 1045 | 1221 | 1132 | 1149 | 1262 | 1242 | 1179 | 1205 | 1194.87 | 1181 | 1209 | 1.025 | 0.035 | 0.64113532 | 0.77592588 | 0.0013 | 0.0044 | 0.0033 | 0.0033 | TRUE | 0.0429 |
| Cdc26 | 1267 | 824 | 821 | 858 | 688 | 898 | 984 | 876 | 886 | 854 | 776 | 886 | 877.17 | 915 | 839 | 0.916 | -0.127 | 0.15124933 | 0.29740083 | 0.0033 | 0.0051 | 0.0045 | 0.0045 | TRUE | 0.0534 |
| Cdc27 | 7199 | 4661 | 3989 | 4634 | 3884 | 4125 | 5591 | 4958 | 4304 | 4613 | 4382 | 4072 | 4653.22 | 4951 | 4356 | 0.879 | -0.186 | 0.03650434 | 0.10735732 | 0.0101 | 0.0029 | 0.0055 | 0.0055 | TRUE | 0.159 |
| Cdc34 | 15068 | 10557 | 9346 | 9949 | 8899 | 9738 | 11703 | 11229 | 10084 | 9903 | 10039 | 9613 | 10428.57 | 11005 | 9852 | 0.895 | -0.16 | 0.01192513 | 0.04712354 | 0.003 | 0.0027 | 0.0028 | 0.0028 | TRUE | 0.0654 |
| Cdc37 | 14465 | 10846 | 10306 | 10719 | 9209 | 11051 | 11234 | 11537 | 11120 | 10670 | 10389 | 10909 | 10976.44 | 11297 | 10656 | 0.943 | -0.084 | 0.08115981 | 0.19110662 | 3.00E-04 | 0.0026 | 0.0016 | 0.0016 | TRUE | 0.0058 |
| Cdc37l1 | 3190 | 2375 | 2470 | 2397 | 2179 | 2626 | 2478 | 2526 | 2665 | 2386 | 2458 | 2592 | 2517.55 | 2556 | 2479 | 0.97 | -0.044 | 0.49216052 | 0.6574413 | 0.0012 | 0.0034 | 0.0025 | 0.0025 | TRUE | 0.0197 |
| Cdc40 | 2370 | 1733 | 1780 | 1787 | 1671 | 1895 | 1841 | 1843 | 1921 | 1779 | 1885 | 1871 | 1856.52 | 1868 | 1845 | 0.988 | -0.018 | 0.77682915 | 0.86798148 | 0 | 0.0037 | 0.0024 | 0.0024 | TRUE | 0.0119 |
| Cdc42 | 19519 | 13140 | 13502 | 14571 | 12417 | 14040 | 15160 | 13977 | 14569 | 14504 | 14008 | 13860 | 14346.04 | 14569 | 14124 | 0.969 | -0.045 | 0.38968192 | 0.56368016 | 0.001 | 0.0026 | 0.0019 | 0.0019 | TRUE | 0.0155 |
| Cdc42bpa | 2614 | 1835 | 1633 | 1702 | 1398 | 1695 | 2030 | 1952 | 1762 | 1694 | 1577 | 1673 | 1781.42 | 1915 | 1648 | 0.86 | -0.217 | 0.00353485 | 0.01873325 | 0.0029 | 0.0038 | 0.0034 | 0.0034 | TRUE | 0.0582 |
| Cdc42bpb | 6446 | 4731 | 4108 | 3860 | 3600 | 4265 | 5006 | 5032 | 4432 | 3842 | 4061 | 4210 | 4430.79 | 4823 | 4038 | 0.837 | -0.257 | 0.00016468 | 0.00183463 | 0.0032 | 0.003 | 0.0031 | 0.0031 | TRUE | 0.0609 |
| Cdc42bpg | 4991 | 4041 | 4240 | 5229 | 4603 | 5336 | 3876 | 4298 | 4575 | 5205 | 5193 | 5267 | 4735.77 | 4250 | 5222 | 1.229 | 0.298 | 1.01E-05 | 0.00018998 | 0.0031 | 0.0029 | 0.0031 | 0.0031 | TRUE | 0.0732 |
| Cdc42ep1 | 144 | 99 | 102 | 144 | 152 | 155 | 112 | 105 | 110 | 143 | 171 | 153 | 132.5 | 109 | 156 | 1.426 | 0.512 | 0.00314403 | 0.01711909 | 0 | 0.0203 | 0.0139 | 0.0139 | TRUE | 0.0767 |
| Cdc42ep2 | 198 | 159 | 232 | 223 | 204 | 257 | 154 | 169 | 250 | 222 | 230 | 254 | 213.17 | 191 | 235 | 1.237 | 0.307 | 0.08937902 | 0.20381211 | 0.0322 | 0.0135 | 0.0188 | 0.0188 | TRUE | 0.7821 |
| Cdc42ep3 | 731 | 554 | 503 | 451 | 455 | 455 | 568 | 589 | 543 | 449 | 500 | 449 | 516.26 | 567 | 466 | 0.821 | -0.284 | 0.00392213 | 0.02029651 | 0 | 0.007 | 0.005 | 0.005 | TRUE | 0.0454 |
| Cdc42ep4 | 4888 | 3732 | 3838 | 4446 | 3939 | 4376 | 3796 | 3970 | 4141 | 4426 | 4444 | 4320 | 4182.68 | 3969 | 4397 | 1.108 | 0.148 | 0.00903491 | 0.03845621 | 7.00E-04 | 0.003 | 0.0021 | 0.0021 | TRUE | 0.0181 |
| Cdc42ep5 | 431 | 281 | 357 | 362 | 370 | 443 | 335 | 299 | 385 | 360 | 417 | 437 | 372.31 | 340 | 405 | 1.193 | 0.254 | 0.04736665 | 0.12953195 | 0.01 | 0.0088 | 0.0092 | 0.0092 | TRUE | 0.1497 |
| Cdc42se1 | 1755 | 1182 | 1217 | 1303 | 1119 | 1259 | 1363 | 1257 | 1313 | 1297 | 1262 | 1243 | 1289.27 | 1311 | 1267 | 0.966 | -0.05 | 0.47825681 | 0.64561056 | 0 | 0.0043 | 0.0028 | 0.0028 | TRUE | 0.0155 |

| | | | | | | | | | | | | | | | | | | | | | | | | | |
|---|---|---|---|---|---|---|---|---|---|---|---|---|---|---|---|---|---|---|---|---|---|---|---|---|---|
| Cdc42se2 | 9957 | 7122 | 6641 | 7728 | 6522 | 7511 | 7733 | 7576 | 7166 | 7692 | 7357 | 7415 | 7489.8 | 7492 | 7488 | 0.999 | -0.001 | 0.98698572 | 0.99373352 | 9.00E-04 | 0.0027 | 0.0019 | 0.0019 | TRUE | 0.0176 |
| Cdc45 | 3386 | 2462 | 2547 | 2531 | 2160 | 2552 | 2630 | 2619 | 2748 | 2519 | 2437 | 2519 | 2578.67 | 2666 | 2492 | 0.935 | -0.097 | 0.09599445 | 0.21471894 | 0 | 0.0033 | 0.002 | 0.002 | TRUE | 0.0088 |
| Cdc5l | 20288 | 14155 | 12644 | 13749 | 11890 | 12731 | 15757 | 15056 | 13643 | 13686 | 13413 | 12567 | 14020.41 | 14819 | 13222 | 0.892 | -0.165 | 0.01166194 | 0.04628827 | 0.0035 | 0.0026 | 0.003 | 0.003 | TRUE | 0.0588 |
| Cdc6 | 5942 | 4110 | 3908 | 3708 | 3261 | 3830 | 4615 | 4372 | 4217 | 3691 | 3679 | 3781 | 4058.97 | 4401 | 3717 | 0.844 | -0.244 | 2.20E-05 | 0.00036186 | 0.001 | 0.003 | 0.0021 | 0.0021 | TRUE | 0.0224 |
| Cdc7 | 3150 | 2456 | 2632 | 2717 | 2257 | 2725 | 2446 | 2612 | 2840 | 2704 | 2546 | 2690 | 2639.9 | 2633 | 2647 | 1.006 | 0.009 | 0.89851189 | 0.94384643 | 0.003 | 0.0033 | 0.0032 | 0.0032 | TRUE | 0.0572 |
| Cdc73 | 5248 | 3698 | 3077 | 3262 | 2807 | 3169 | 4076 | 3934 | 3320 | 3247 | 3167 | 3128 | 3478.56 | 3777 | 3181 | 0.842 | -0.248 | 0.00145748 | 0.00944325 | 0.0054 | 0.0031 | 0.0041 | 0.0041 | TRUE | 0.1352 |
| Cdca2 | 6195 | 4753 | 5000 | 5415 | 4962 | 5578 | 4811 | 5056 | 5395 | 5390 | 5598 | 5506 | 5292.69 | 5087 | 5498 | 1.081 | 0.113 | 0.05869001 | 0.15103642 | 0.0016 | 0.0029 | 0.0024 | 0.0024 | TRUE | 0.034 |
| Cdca3 | 8165 | 5999 | 6336 | 6209 | 5649 | 6528 | 6341 | 6381 | 6836 | 6180 | 6373 | 6444 | 6426.03 | 6519 | 6332 | 0.971 | -0.042 | 0.44456658 | 0.61547754 | 9.00E-04 | 0.0028 | 0.002 | 0.002 | TRUE | 0.0219 |
| Cdca4 | 3490 | 2506 | 2588 | 2571 | 2383 | 2607 | 2711 | 2666 | 2792 | 2559 | 2688 | 2574 | 2664.92 | 2723 | 2607 | 0.957 | -0.063 | 0.27976026 | 0.45138501 | 2.00E-04 | 0.0033 | 0.0021 | 0.0021 | TRUE | 0.0089 |
| Cdca5 | 3649 | 2567 | 2415 | 2554 | 2136 | 2408 | 2834 | 2730 | 2606 | 2542 | 2410 | 2377 | 2583.2 | 2723 | 2443 | 0.897 | -0.157 | 0.01251517 | 0.04884745 | 0.0012 | 0.0033 | 0.0025 | 0.0025 | TRUE | 0.0173 |
| Cdca7 | 8151 | 6434 | 6977 | 6911 | 6075 | 7391 | 6331 | 6844 | 7528 | 6879 | 6853 | 7296 | 6955.15 | 6901 | 7009 | 1.016 | 0.023 | 0.73983162 | 0.84538261 | 0.0043 | 0.0028 | 0.0033 | 0.0033 | TRUE | 0.0771 |
| Cdca7l | 2487 | 1898 | 1824 | 1667 | 1450 | 1635 | 1932 | 2019 | 1968 | 1659 | 1636 | 1614 | 1804.6 | 1973 | 1636 | 0.83 | -0.269 | 1.44E-05 | 0.00025631 | 0 | 0.0037 | 0.0022 | 0.0022 | TRUE | 0.005 |
| Cdca8 | 9229 | 6674 | 6571 | 6461 | 5796 | 6548 | 7168 | 7099 | 7090 | 6431 | 6538 | 6464 | 6798.42 | 7119 | 6478 | 0.91 | -0.136 | 0.00363787 | 0.01916552 | 0 | 0.0028 | 0.0014 | 0.0014 | TRUE | 8.00E-04 |
| Cdcp1 | 229 | 219 | 299 | 279 | 267 | 286 | 178 | 233 | 323 | 278 | 301 | 282 | 265.78 | 245 | 287 | 1.18 | 0.238 | 0.19062648 | 0.3487296 | 0.0413 | 0.0113 | 0.0201 | 0.0201 | TRUE | 0.435 |
| Cdcp2 | 8 | 3 | 5 | 7 | 4 | 6 | 6 | 3 | 5 | 7 | 5 | 6 | 5.37 | 5 | 6 | 1.168 | 0.224 | 0.78317587 | NA | 0 | 0.444 | 0.2911 | 0.2911 | TRUE | 0.1771 |
| Cdh1 | 21555 | 16523 | 18049 | 19301 | 18134 | 21574 | 16741 | 17575 | 19475 | 19212 | 20457 | 21297 | 19126.16 | 17930 | 20322 | 1.134 | 0.181 | 0.00743917 | 0.0332483 | 0.0043 | 0.0026 | 0.0032 | 0.0032 | TRUE | 0.0694 |
| Cdh10 | 10 | 5 | 9 | 14 | 15 | 12 | 8 | 5 | 10 | 14 | 17 | 12 | 10.92 | 8 | 14 | 1.867 | 0.901 | 0.12590538 | NA | 0 | 0.2195 | 0.1499 | 0.1499 | TRUE | 0.1716 |
| Cdh11 | 13 | 6 | 7 | 11 | 6 | 7 | 10 | 6 | 8 | 11 | 7 | 7 | 8.11 | 8 | 8 | 1.017 | 0.025 | 0.97056076 | NA | 0 | 0.2947 | 0.1997 | 0.1997 | TRUE | 0.2619 |
| Cdh12 | 4 | 2 | 1 | 0 | 0 | 0 | 3 | 2 | 1 | 0 | 0 | 0 | 1.05 | 2 | 0 | 0.09 | -3.481 | 0.09287895 | NA | 0 | 2.255 | 1.8642 | 1.8642 | TRUE | 0.0734 |
| Cdh13 | 1 | 1 | 2 | 3 | 2 | 1 | 1 | 1 | 2 | 3 | 2 | 1 | 1.7 | 1 | 2 | 1.589 | 0.668 | 0.64355948 | NA | 0 | 1.3928 | 0.8824 | 0.8824 | TRUE | 0.2062 |
| Cdh15 | 161 | 127 | 95 | 171 | 129 | 143 | 125 | 135 | 103 | 170 | 146 | 141 | 136.59 | 121 | 152 | 1.259 | 0.332 | 0.06601587 | 0.16424106 | 0.0064 | 0.0198 | 0.0161 | 0.0161 | TRUE | 0.167 |
| Cdh16 | 1 | 0 | 0 | 0 | 2 | 2 | 1 | 0 | 0 | 0 | 2 | 2 | 0.83 | 0 | 1 | 4.067 | 2.024 | 0.3803708 | NA | 0 | 2.8424 | 2.4672 | 2.4672 | TRUE | 0.5093 |
| Cdh17 | 5 | 7 | 4 | 0 | 1 | 0 | 4 | 7 | 4 | 0 | 1 | 0 | 2.8 | 5 | 0 | 0.072 | -3.806 | 0.01420283 | NA | 0 | 0.8502 | 0.6428 | 0.6428 | TRUE | 0.5075 |
| Cdh18 | 13 | 9 | 9 | 7 | 4 | 3 | 10 | 10 | 10 | 7 | 5 | 3 | 7.3 | 10 | 5 | 0.491 | -1.025 | 0.15643141 | NA | 0 | 0.3269 | 0.2202 | 0.2202 | TRUE | 0.3148 |
| Cdh19 | 3 | 1 | 0 | 1 | 1 | 1 | 2 | 1 | 0 | 1 | 1 | 1 | 1.08 | 1 | 1 | 0.879 | -0.186 | 0.91830054 | NA | 0 | 2.1887 | 1.4564 | 1.4564 | TRUE | 0.2453 |
| Cdh2 | 293 | 184 | 197 | 161 | 122 | 127 | 228 | 196 | 213 | 160 | 138 | 125 | 176.52 | 212 | 141 | 0.665 | -0.59 | 0.0002373 | 0.00243309 | 0.0042 | 0.0159 | 0.0126 | 0.0126 | TRUE | 0.1508 |
| Cdh20 | 5 | 4 | 5 | 0 | 0 | 1 | 4 | 4 | 5 | 0 | 0 | 1 | 2.42 | 4 | 0 | 0.082 | -3.6 | 0.02346142 | NA | 0 | 0.9818 | 0.7113 | 0.7113 | TRUE | 0.3835 |
| Cdh22 | 47 | 46 | 70 | 119 | 130 | 171 | 37 | 49 | 76 | 118 | 147 | 169 | 99.15 | 54 | 145 | 2.728 | 1.448 | 2.52E-08 | 1.14E-06 | 0.0651 | 0.0263 | 0.036 | 0.036 | TRUE | 0.5585 |
| Cdh23 | 103 | 86 | 121 | 150 | 147 | 168 | 80 | 91 | 131 | 149 | 166 | 166 | 130.5 | 101 | 160 | 1.603 | 0.681 | 0.00078708 | 0.00607768 | 0.0243 | 0.0206 | 0.0216 | 0.0216 | TRUE | 0.6443 |
| Cdh24 | 279 | 232 | 176 | 166 | 136 | 183 | 217 | 247 | 190 | 165 | 153 | 181 | 192.11 | 218 | 166 | 0.765 | -0.386 | 0.01381748 | 0.05263538 | 0.0066 | 0.0148 | 0.0124 | 0.0124 | TRUE | 0.1406 |
| Cdh26 | 2 | 0 | 1 | 4 | 3 | 3 | 2 | 0 | 1 | 4 | 3 | 3 | 2.16 | 1 | 3 | 3.748 | 1.906 | 0.1737571 | NA | 0 | 1.0997 | 0.7418 | 0.7418 | TRUE | 0.2779 |
| Cdh3 | 3203 | 2977 | 3487 | 2935 | 3144 | 3913 | 2488 | 3167 | 3762 | 2921 | 3547 | 3863 | 3291.28 | 3139 | 3444 | 1.097 | 0.134 | 0.5190186 | 0.68051856 | 0.0308 | 0.0031 | 0.0112 | 0.0308 | TRUE | 0.2422 |
| Cdh4 | 145 | 104 | 105 | 108 | 88 | 98 | 113 | 111 | 113 | 108 | 99 | 97 | 106.68 | 112 | 101 | 0.902 | -0.149 | 0.42638941 | 0.5987801 | 0 | 0.0246 | 0.0158 | 0.0158 | TRUE | 0.0301 |
| Cdh5 | 39 | 21 | 17 | 17 | 17 | 25 | 30 | 22 | 18 | 17 | 19 | 25 | 21.96 | 23 | 20 | 0.849 | -0.236 | 0.57619705 | 0.72771813 | 0 | 0.1104 | 0.0829 | 0.0829 | TRUE | 0.4257 |
| Cdh6 | 27 | 22 | 19 | 19 | 6 | 15 | 21 | 23 | 21 | 19 | 7 | 15 | 17.56 | 22 | 14 | 0.63 | -0.666 | 0.17035469 | 0.32195322 | 0 | 0.1374 | 0.1099 | 0.1099 | TRUE | 0.6425 |
| Cdh7 | 1 | 4 | 1 | 4 | 1 | 1 | 1 | 4 | 1 | 4 | 1 | 1 | 2.03 | 2 | 2 | 1.026 | 0.037 | 0.97851985 | NA | 0.3023 | 1.1673 | 0.8894 | 0.8894 | TRUE | 0.7856 |
| Cdh8 | 12 | 3 | 7 | 5 | 0 | 3 | 9 | 3 | 8 | 5 | 0 | 3 | 4.67 | 7 | 3 | 0.4 | -1.324 | 0.18697842 | NA | 0.2498 | 0.5103 | 0.4616 | 0.4616 | TRUE | 0.5929 |
| Cdh9 | 23 | 27 | 18 | 14 | 7 | 18 | 18 | 29 | 19 | 14 | 8 | 18 | 17.6 | 22 | 13 | 0.608 | -0.719 | 0.14130053 | 0.28345383 | 0.0367 | 0.1371 | 0.1111 | 0.1111 | TRUE | 0.4612 |
| Cdhr1 | 352 | 304 | 447 | 461 | 471 | 549 | 273 | 323 | 482 | 459 | 531 | 542 | 435.2 | 359 | 511 | 1.424 | 0.51 | 0.00258893 | 0.0147601 | 0.0464 | 0.0079 | 0.0183 | 0.0183 | TRUE | 1.0109 |
| Cdhr2 | 12 | 4 | 7 | 11 | 5 | 3 | 9 | 4 | 8 | 11 | 6 | 3 | 6.78 | 7 | 7 | 0.917 | -0.125 | 0.87213151 | NA | 0.1173 | 0.352 | 0.2863 | 0.2863 | TRUE | 0.5472 |
| Cdhr3 | 8 | 1 | 5 | 6 | 2 | 5 | 6 | 1 | 5 | 6 | 2 | 5 | 4.31 | 4 | 4 | 1.03 | 0.043 | 0.96470219 | NA | 0 | 0.5528 | 0.4372 | 0.4372 | TRUE | 0.6894 |
| Cdhr5 | 6 | 4 | 2 | 3 | 1 | 0 | 5 | 4 | 2 | 3 | 1 | 0 | 2.53 | 4 | 1 | 0.37 | -1.436 | 0.27121067 | NA | 0 | 0.9388 | 0.7208 | 0.7208 | TRUE | 0.7017 |
| Cdip1 | 1747 | 1252 | 1291 | 1276 | 1105 | 1339 | 1357 | 1332 | 1393 | 1270 | 1247 | 1322 | 1320 | 1361 | 1280 | 0.941 | -0.088 | 0.20032879 | 0.36102628 | 0 | 0.0042 | 0.0026 | 0.0026 | TRUE | 0.0101 |
| Cdipt | 2295 | 1497 | 1423 | 1654 | 1300 | 1497 | 1782 | 1592 | 1535 | 1646 | 1467 | 1478 | 1583.48 | 1636 | 1530 | 0.934 | -0.098 | 0.22868034 | 0.39402174 | 0.0043 | 0.0039 | 0.0042 | 0.0042 | TRUE | 0.0751 |
| Cdk1 | 10575 | 7520 | 7884 | 8217 | 7256 | 8216 | 8213 | 7999 | 8507 | 8179 | 8186 | 8110 | 8199 | 8240 | 8158 | 0.99 | -0.014 | 0.77280739 | 0.86564989 | 3.00E-04 | 0.0027 | 0.0016 | 0.0016 | TRUE | 0.0097 |
| Cdk10 | 2378 | 1827 | 1619 | 1647 | 1398 | 1554 | 1847 | 1943 | 1747 | 1639 | 1577 | 1534 | 1714.61 | 1846 | 1583 | 0.858 | -0.221 | 0.00148844 | 0.00960028 | 0.0014 | 0.0038 | 0.0029 | 0.0029 | TRUE | 0.0257 |
| Cdk11b | 8140 | 6188 | 5605 | 5787 | 5208 | 6170 | 6322 | 6582 | 6048 | 5760 | 5875 | 6091 | 6113.02 | 6317 | 5909 | 0.935 | -0.096 | 0.08458958 | 0.19614518 | 0.0011 | 0.0028 | 0.0021 | 0.0021 | TRUE | 0.0168 |
| Cdk12 | 8717 | 5831 | 4864 | 5712 | 5102 | 5768 | 6770 | 6202 | 5248 | 5686 | 5756 | 5694 | 5892.65 | 6073 | 5712 | 0.94 | -0.089 | 0.27818751 | 0.44963632 | 0.0078 | 0.0028 | 0.0047 | 0.0047 | TRUE | 0.172 |
| Cdk13 | 5573 | 3985 | 3648 | 3701 | 3401 | 3832 | 4328 | 4239 | 3936 | 3684 | 3837 | 3783 | 3967.78 | 4168 | 3768 | 0.904 | -0.146 | 0.01350704 | 0.05173199 | 0.001 | 0.003 | 0.0023 | 0.0023 | TRUE | 0.0285 |
| Cdk14 | 186 | 127 | 103 | 91 | 106 | 115 | 144 | 135 | 111 | 91 | 120 | 114 | 119.06 | 130 | 108 | 0.825 | -0.278 | 0.15494477 | 0.3024758 | 0.0107 | 0.0223 | 0.0191 | 0.0191 | TRUE | 0.1982 |
| Cdk15 | 3 | 5 | 1 | 0 | 0 | 4 | 2 | 5 | 1 | 0 | 0 | 4 | 2.11 | 3 | 1 | 0.463 | -1.111 | 0.46884079 | NA | 1.2255 | 1.1243 | 1.1434 | 1.1434 | TRUE | 1.5052 |
| Cdk16 | 8116 | 5997 | 5886 | 6776 | 5887 | 6616 | 6303 | 6379 | 6351 | 6745 | 6641 | 6531 | 6491.7 | 6344 | 6639 | 1.046 | 0.066 | 0.17123663 | 0.32305771 | 0 | 0.0028 | 0.0015 | 0.0015 | TRUE | 0.0025 |
| Cdk17 | 2937 | 2082 | 1976 | 2282 | 2046 | 2294 | 2281 | 2215 | 2132 | 2272 | 2308 | 2265 | 2245.31 | 2209 | 2282 | 1.032 | 0.045 | 0.45120223 | 0.62105484 | 0 | 0.0035 | 0.0022 | 0.0022 | TRUE | 0.0112 |
| Cdk18 | 1661 | 1232 | 1187 | 1362 | 1274 | 1585 | 1290 | 1310 | 1281 | 1356 | 1437 | 1565 | 1373.14 | 1294 | 1453 | 1.123 | 0.167 | 0.02677166 | 0.08539328 | 0.002 | 0.0042 | 0.0034 | 0.0034 | TRUE | 0.0557 |
| Cdk19 | 1059 | 896 | 898 | 927 | 887 | 987 | 822 | 953 | 969 | 923 | 1001 | 974 | 940.36 | 915 | 966 | 1.058 | 0.081 | 0.36218862 | 0.53695279 | 0.0039 | 0.0049 | 0.0046 | 0.0046 | TRUE | 0.0975 |
| Cdk2 | 4903 | 3733 | 3652 | 3914 | 3492 | 4090 | 3808 | 3971 | 3940 | 3896 | 3939 | 4037 | 3931.99 | 3906 | 3957 | 1.013 | 0.019 | 0.71765689 | 0.83029545 | 2.00E-04 | 0.003 | 0.0018 | 0.0018 | TRUE | 0.0061 |
| Cdk20 | 342 | 247 | 265 | 256 | 232 | 290 | 266 | 263 | 286 | 255 | 262 | 286 | 269.52 | 272 | 268 | 0.987 | -0.019 | 0.88123311 | 0.93368902 | 0 | 0.0112 | 0.0077 | 0.0077 | TRUE | 0.0426 |
| Cdk2ap1 | 1495 | 1006 | 885 | 1107 | 941 | 1175 | 1161 | 1070 | 955 | 1102 | 1062 | 1160 | 1084.91 | 1062 | 1108 | 1.042 | 0.059 | 0.50452408 | 0.66826706 | 0.0049 | 0.0046 | 0.0047 | 0.0047 | TRUE | 0.0923 |
| Cdk2ap2 | 1160 | 839 | 962 | 914 | 893 | 1008 | 901 | 892 | 1038 | 910 | 1007 | 995 | 957.26 | 944 | 971 | 1.029 | 0.042 | 0.63932068 | 0.77465332 | 0.0041 | 0.0049 | 0.0046 | 0.0046 | TRUE | 0.089 |
| Cdk3-ps | 66 | 45 | 31 | 59 | 31 | 48 | 51 | 48 | 33 | 59 | 35 | 47 | 45.61 | 44 | 47 | 1.062 | 0.087 | 0.78149609 | 0.87115299 | 0.0333 | 0.0544 | 0.0487 | 0.0487 | TRUE | 0.1857 |
| Cdk4 | 14948 | 10610 | 9915 | 9615 | 8868 | 10286 | 11609 | 11286 | 10698 | 9571 | 10004 | 10154 | 10553.68 | 11198 | 9910 | 0.885 | -0.177 | 0.00114467 | 0.00799647 | 0.0012 | 0.0027 | 0.002 | 0.002 | TRUE | 0.0186 |
| Cdk5 | 937 | 658 | 694 | 698 | 600 | 695 | 728 | 700 | 749 | 695 | 677 | 686 | 705.7 | 726 | 686 | 0.946 | -0.081 | 0.33381371 | 0.50806085 | 0 | 0.0058 | 0.0036 | 0.0036 | TRUE | 0.0107 |
| Cdk5r1 | 146 | 146 | 111 | 108 | 84 | 79 | 113 | 155 | 120 | 108 | 95 | 78 | 111.45 | 129 | 94 | 0.723 | -0.467 | 0.0256828 | 0.08285449 | 0.0189 | 0.0237 | 0.0223 | 0.0223 | TRUE | 0.3053 |
| Cdk5r2 | 160 | 129 | 114 | 157 | 154 | 148 | 124 | 137 | 123 | 156 | 174 | 146 | 143.43 | 128 | 159 | 1.238 | 0.308 | 0.06661672 | 0.16521825 | 0 | 0.019 | 0.0133 | 0.0133 | TRUE | 0.0695 |
| Cdk5rap1 | 1650 | 1242 | 1037 | 1316 | 1128 | 1275 | 1281 | 1321 | 1119 | 1310 | 1273 | 1259 | 1260.43 | 1240 | 1281 | 1.032 | 0.045 | 0.57933461 | 0.730261 | 0.0031 | 0.0043 | 0.0039 | 0.0039 | TRUE | 0.0859 |
| Cdk5rap2 | 4279 | 3194 | 3071 | 2972 | 2861 | 3353 | 3323 | 3397 | 3314 | 2958 | 3228 | 3310 | 3255.01 | 3345 | 3165 | 0.946 | -0.08 | 0.20103452 | 0.36180686 | 0.0013 | 0.0032 | 0.0025 | 0.0025 | TRUE | 0.04 |
| Cdk5rap3 | 7787 | 6320 | 6395 | 7362 | 6946 | 7761 | 6048 | 6723 | 6900 | 7328 | 7836 | 7661 | 7082.62 | 6557 | 7608 | 1.161 | 0.215 | 0.00076023 | 0.00592476 | 0.0027 | 0.0028 | 0.0028 | 0.0028 | TRUE | 0.0569 |
| Cdk6 | 630 | 555 | 613 | 806 | 632 | 762 | 489 | 590 | 661 | 802 | 713 | 752 | 668.09 | 580 | 756 | 1.307 | 0.386 | 0.00067093 | 0.00538745 | 0.0116 | 0.006 | 0.0078 | 0.0078 | TRUE | 0.2327 |
| Cdk7 | 3795 | 2744 | 2601 | 2718 | 2410 | 2811 | 2947 | 2919 | 2806 | 2705 | 2719 | 2775 | 2811.96 | 2891 | 2733 | 0.945 | -0.081 | 0.1501155 | 0.29569004 | 0 | 0.0033 | 0.0019 | 0.0019 | TRUE | 0.0078 |

| | | | | | | | | | | | | | | | | | | | | | | | | | |
|---|---|---|---|---|---|---|---|---|---|---|---|---|---|---|---|---|---|---|---|---|---|---|---|---|---|
| Cdk8 | 4879 | 3548 | 3516 | 4130 | 3468 | 4016 | 3789 | 3774 | 3794 | 4111 | 3912 | 3964 | 3890.78 | 3786 | 3996 | 1.056 | 0.078 | 0.13940705 | 0.28067967 | 0 | 0.003 | 0.0018 | 0.0018 | TRUE | 0.0077 |
| Cdk9 | 5489 | 4040 | 3642 | 3577 | 3361 | 3784 | 4263 | 4297 | 3930 | 3561 | 3792 | 3735 | 3929.59 | 4163 | 3696 | 0.887 | -0.172 | 0.00451745 | 0.02273518 | 0.0013 | 0.003 | 0.0024 | 0.0024 | TRUE | 0.029 |
| Cdkal1 | 1465 | 1015 | 1075 | 1128 | 1073 | 1165 | 1138 | 1080 | 1160 | 1123 | 1210 | 1150 | 1143.44 | 1126 | 1161 | 1.031 | 0.044 | 0.55533319 | 0.71053418 | 5.00E-04 | 0.0045 | 0.0031 | 0.0031 | TRUE | 0.0162 |
| Cdkl1 | 18 | 16 | 13 | 7 | 2 | 5 | 14 | 17 | 14 | 7 | 2 | 5 | 9.86 | 15 | 5 | 0.319 | -1.649 | 0.01250873 | NA | 0 | 0.2427 | 0.1736 | 0.1736 | TRUE | 0.3435 |
| Cdkl2 | 1388 | 1075 | 1067 | 1248 | 1093 | 1253 | 1078 | 1143 | 1151 | 1242 | 1233 | 1237 | 1180.82 | 1124 | 1237 | 1.102 | 0.14 | 0.04861542 | 0.13202741 | 0 | 0.0044 | 0.0028 | 0.0028 | TRUE | 0.0164 |
| Cdkl3 | 220 | 136 | 113 | 163 | 141 | 137 | 171 | 145 | 122 | 162 | 159 | 135 | 149 | 146 | 152 | 1.038 | 0.053 | 0.76764462 | 0.86257577 | 0.0129 | 0.0183 | 0.0167 | 0.0167 | TRUE | 0.2513 |
| Cdkl4 | 206 | 140 | 151 | 264 | 215 | 274 | 160 | 149 | 163 | 263 | 243 | 270 | 207.94 | 157 | 259 | 1.645 | 0.718 | 3.64E-07 | 1.24E-05 | 0 | 0.0138 | 0.0093 | 0.0093 | TRUE | 0.0317 |
| Cdkl5 | 412 | 268 | 233 | 249 | 243 | 285 | 320 | 285 | 251 | 248 | 274 | 281 | 276.63 | 285 | 268 | 0.935 | -0.098 | 0.47063705 | 0.63963211 | 0.0064 | 0.011 | 0.0096 | 0.0096 | TRUE | 0.1341 |
| Cdkn1a | 3578 | 2735 | 3779 | 4384 | 3971 | 4428 | 2779 | 2909 | 4078 | 4364 | 4480 | 4371 | 3830.04 | 3255 | 4405 | 1.353 | 0.437 | 0.0169686 | 0.06131944 | 0.0238 | 0.003 | 0.0089 | 0.0238 | TRUE | 0.5929 |
| Cdkn1b | 2606 | 2171 | 2488 | 2785 | 2620 | 3121 | 2024 | 2309 | 2685 | 2772 | 2956 | 3081 | 2637.75 | 2339 | 2936 | 1.256 | 0.329 | 0.00042229 | 0.0037937 | 0.0109 | 0.0033 | 0.0059 | 0.0059 | TRUE | 0.2019 |
| Cdkn1c | 354 | 288 | 339 | 347 | 302 | 363 | 275 | 306 | 366 | 345 | 341 | 358 | 331.91 | 316 | 348 | 1.107 | 0.147 | 0.25761568 | 0.427535 | 0.0079 | 0.0096 | 0.0091 | 0.0091 | TRUE | 0.2162 |
| Cdkn2a | 919 | 621 | 532 | 599 | 472 | 470 | 714 | 661 | 574 | 596 | 532 | 464 | 590.17 | 650 | 531 | 0.816 | -0.293 | 0.01263956 | 0.04921199 | 0.0122 | 0.0064 | 0.0082 | 0.0082 | TRUE | 0.1448 |
| Cdkn2aip | 1837 | 1337 | 1292 | 1367 | 1190 | 1337 | 1427 | 1422 | 1394 | 1361 | 1342 | 1320 | 1377.65 | 1414 | 1341 | 0.948 | -0.077 | 0.24253302 | 0.41021693 | 0 | 0.0041 | 0.0024 | 0.0024 | TRUE | 0.0023 |
| Cdkn2aipnl | 6664 | 4639 | 4600 | 4269 | 3906 | 4529 | 5176 | 4934 | 4963 | 4249 | 4406 | 4471 | 4700 | 5024 | 4375 | 0.871 | -0.2 | 0.00019876 | 0.00212322 | 5.00E-04 | 0.0029 | 0.0019 | 0.0019 | TRUE | 0.0086 |
| Cdkn2b | 243 | 138 | 123 | 151 | 128 | 125 | 189 | 147 | 133 | 150 | 144 | 123 | 147.72 | 156 | 139 | 0.887 | -0.172 | 0.35296107 | 0.52775572 | 0.0169 | 0.0185 | 0.018 | 0.018 | TRUE | 0.38 |
| Cdkn2c | 54 | 32 | 31 | 38 | 30 | 42 | 42 | 34 | 33 | 38 | 34 | 41 | 37.09 | 36 | 38 | 1.03 | 0.042 | 0.89399418 | 0.94183378 | 0 | 0.0663 | 0.0451 | 0.0451 | TRUE | 0.1465 |
| Cdkn2d | 606 | 464 | 526 | 550 | 528 | 551 | 471 | 494 | 568 | 547 | 596 | 544 | 536.46 | 511 | 562 | 1.103 | 0.141 | 0.17618676 | 0.32987128 | 0.0042 | 0.0068 | 0.006 | 0.006 | TRUE | 0.1093 |
| Cdkn3 | 636 | 449 | 414 | 468 | 425 | 437 | 494 | 478 | 447 | 466 | 479 | 431 | 465.82 | 473 | 459 | 0.969 | -0.046 | 0.65412843 | 0.78520683 | 0 | 0.0075 | 0.0054 | 0.0054 | TRUE | 0.0323 |
| Cdnf | 69 | 41 | 42 | 64 | 60 | 62 | 54 | 44 | 45 | 64 | 68 | 61 | 55.85 | 48 | 64 | 1.343 | 0.425 | 0.10009707 | 0.221058 | 0 | 0.0449 | 0.0301 | 0.0301 | TRUE | 0.1174 |
| Cdo1 | 316 | 217 | 223 | 179 | 176 | 165 | 245 | 231 | 241 | 178 | 199 | 163 | 209.41 | 239 | 180 | 0.751 | -0.413 | 0.00380957 | 0.01984809 | 0 | 0.0137 | 0.0098 | 0.0098 | TRUE | 0.0849 |
| Cdon | 2583 | 1866 | 1783 | 1859 | 1565 | 1843 | 2006 | 1985 | 1924 | 1850 | 1765 | 1819 | 1891.67 | 1972 | 1811 | 0.919 | -0.122 | 0.04888889 | 0.13246864 | 0 | 0.0037 | 0.0022 | 0.0022 | TRUE | 0.0059 |
| Cdpf1 | 734 | 593 | 548 | 696 | 648 | 660 | 570 | 631 | 591 | 693 | 731 | 652 | 644.58 | 597 | 692 | 1.159 | 0.213 | 0.0212801 | 0.07231489 | 0.0015 | 0.0061 | 0.0046 | 0.0046 | TRUE | 0.0307 |
| Cdr1 | 0 | 0 | 0 | 0 | 0 | 0 | 0 | 0 | 0 | 0 | 0 | 0 | 0 | 0 | 0 | NA | NA | NA | NA | NA | NA | NA | NA | NA | NA |
| Cdr2 | 2542 | 1683 | 1649 | 1712 | 1519 | 1701 | 1974 | 1790 | 1779 | 1704 | 1714 | 1679 | 1773.43 | 1848 | 1699 | 0.918 | -0.123 | 0.07378797 | 0.17818463 | 0.0014 | 0.0038 | 0.0028 | 0.0028 | TRUE | 0.0447 |
| Cdr2l | 1983 | 1641 | 1679 | 1788 | 1541 | 1916 | 1540 | 1746 | 1812 | 1780 | 1738 | 1891 | 1751.14 | 1699 | 1803 | 1.063 | 0.088 | 0.26284855 | 0.43322121 | 0.004 | 0.0038 | 0.0039 | 0.0039 | TRUE | 0.0835 |
| Cdrt4 | 14 | 13 | 12 | 16 | 7 | 9 | 11 | 14 | 13 | 16 | 8 | 9 | 11.73 | 13 | 11 | 0.877 | -0.19 | 0.73848157 | NA | 0 | 0.2045 | 0.1471 | 0.1471 | TRUE | 0.6102 |
| Cds1 | 737 | 554 | 622 | 648 | 641 | 733 | 572 | 589 | 671 | 645 | 723 | 724 | 654.09 | 611 | 697 | 1.143 | 0.192 | 0.05083218 | 0.13649151 | 0.0042 | 0.006 | 0.0055 | 0.0055 | TRUE | 0.0858 |
| Cds2 | 6668 | 5053 | 5020 | 5520 | 4984 | 5909 | 5179 | 5375 | 5417 | 5495 | 5622 | 5833 | 5486.71 | 5324 | 5650 | 1.062 | 0.086 | 0.10446421 | 0.22844046 | 5.00E-04 | 0.0029 | 0.0019 | 0.0019 | TRUE | 0.0099 |
| Cdsn | 95 | 68 | 83 | 88 | 94 | 111 | 74 | 72 | 90 | 88 | 106 | 110 | 89.81 | 79 | 101 | 1.289 | 0.366 | 0.08665069 | 0.19946825 | 0 | 0.0288 | 0.0217 | 0.0217 | TRUE | 0.1361 |
| Cdt1 | 12673 | 10073 | 10444 | 10927 | 9975 | 11924 | 9843 | 10715 | 11269 | 10877 | 11253 | 11771 | 10954.41 | 10609 | 11300 | 1.065 | 0.091 | 0.14879155 | 0.29406494 | 0.0029 | 0.0026 | 0.0028 | 0.0028 | TRUE | 0.0491 |
| Cdv3 | 22343 | 15666 | 15177 | 16410 | 13858 | 16003 | 17353 | 16664 | 16376 | 16334 | 15633 | 15797 | 16359.6 | 16798 | 15921 | 0.948 | -0.077 | 0.11490447 | 0.2446127 | 6.00E-04 | 0.0026 | 0.0017 | 0.0017 | TRUE | 0.0103 |
| Cdx1 | 120 | 89 | 77 | 231 | 170 | 195 | 93 | 95 | 83 | 230 | 192 | 192 | 147.53 | 90 | 205 | 2.265 | 1.179 | 1.12E-11 | 1.15E-09 | 0 | 0.0185 | 0.0139 | 0.0139 | TRUE | 0.1278 |
| Cdx2 | 28 | 12 | 8 | 7 | 5 | 8 | 22 | 13 | 9 | 7 | 6 | 8 | 10.61 | 15 | 7 | 0.47 | -1.089 | 0.0847057 | NA | 0 | 0.2258 | 0.1789 | 0.1789 | TRUE | 0.3756 |
| Cdx4 | 8 | 2 | 3 | 4 | 3 | 0 | 6 | 2 | 3 | 4 | 3 | 0 | 3.16 | 4 | 2 | 0.621 | -0.688 | 0.55593556 | NA | 0.2746 | 0.7531 | 0.6477 | 0.6477 | TRUE | 0.8748 |
| Cdyl | 3141 | 2427 | 2357 | 2310 | 2107 | 2425 | 2439 | 2582 | 2543 | 2299 | 2377 | 2394 | 2439.06 | 2521 | 2357 | 0.935 | -0.097 | 0.10359365 | 0.22702607 | 3.00E-04 | 0.0034 | 0.0021 | 0.0021 | TRUE | 0.0101 |
| Cdyl2 | 17839 | 13415 | 14833 | 16637 | 15046 | 16334 | 13855 | 14269 | 16005 | 16560 | 16973 | 16124 | 15631.15 | 14710 | 16552 | 1.125 | 0.17 | 0.00714807 | 0.03224979 | 0.0032 | 0.0026 | 0.0028 | 0.0028 | TRUE | 0.0724 |
| Ceacam-ps1 | 0 | 1 | 0 | 0 | 0 | 0 | 0 | 1 | 0 | 0 | 0 | 0 | 0.18 | 0 | 0 | 0.552 | -0.857 | 0.8336548 | NA | 0 | 13.3714 | 10 | 10 | TRUE | 0.5216 |
| Ceacam1 | 933 | 631 | 638 | 614 | 549 | 630 | 725 | 671 | 688 | 611 | 619 | 622 | 656.1 | 695 | 617 | 0.888 | -0.172 | 0.04679982 | 0.12834033 | 0 | 0.006 | 0.0038 | 0.0038 | TRUE | 0.0178 |
| Ceacam10 | 45 | 25 | 37 | 50 | 25 | 46 | 35 | 27 | 40 | 50 | 28 | 45 | 37.47 | 34 | 41 | 1.22 | 0.287 | 0.39978917 | 0.57313807 | 0.0322 | 0.0657 | 0.0569 | 0.0569 | TRUE | 0.531 |
| Ceacam11 | 0 | 0 | 0 | 0 | 0 | 0 | 0 | 0 | 0 | 0 | 0 | 0 | 0 | 0 | 0 | NA | NA | NA | NA | NA | NA | NA | NA | NA | NA |
| Ceacam12 | 0 | 0 | 0 | 0 | 0 | 0 | 0 | 0 | 0 | 0 | 0 | 0 | 0 | 0 | 0 | NA | NA | NA | NA | NA | NA | NA | NA | NA | NA |
| Ceacam13 | 1 | 0 | 0 | 0 | 0 | 0 | 1 | 0 | 0 | 0 | 0 | 0 | 0.13 | 0 | 0 | 0.552 | -0.857 | 0.8336548 | NA | 0 | 18.3121 | 10 | 10 | TRUE | 0.2566 |
| Ceacam14 | 0 | 0 | 0 | 0 | 0 | 0 | 0 | 0 | 0 | 0 | 0 | 0 | 0 | 0 | 0 | NA | NA | NA | NA | NA | NA | NA | NA | NA | NA |
| Ceacam15 | 0 | 0 | 0 | 0 | 0 | 0 | 0 | 0 | 0 | 0 | 0 | 0 | 0 | 0 | 0 | NA | NA | NA | NA | NA | NA | NA | NA | NA | NA |
| Ceacam16 | 43 | 40 | 45 | 68 | 58 | 60 | 33 | 43 | 49 | 68 | 65 | 59 | 52.81 | 42 | 64 | 1.558 | 0.639 | 0.01984053 | 0.06873017 | 0 | 0.0473 | 0.0345 | 0.0345 | TRUE | 0.2678 |
| Ceacam18 | 25 | 11 | 10 | 16 | 12 | 19 | 19 | 12 | 11 | 16 | 14 | 19 | 15.02 | 14 | 16 | 1.136 | 0.184 | 0.71500502 | NA | 0 | 0.1602 | 0.1176 | 0.1176 | TRUE | 0.6289 |
| Ceacam19 | 40 | 30 | 19 | 23 | 29 | 23 | 31 | 32 | 21 | 23 | 33 | 23 | 26.97 | 28 | 26 | 0.93 | -0.104 | 0.78596783 | 0.87416062 | 0 | 0.0903 | 0.0691 | 0.0691 | TRUE | 0.3142 |
| Ceacam2 | 41 | 30 | 26 | 15 | 14 | 22 | 32 | 32 | 28 | 15 | 16 | 22 | 24.04 | 31 | 18 | 0.572 | -0.807 | 0.04254477 | 0.11971622 | 0 | 0.101 | 0.0687 | 0.0687 | TRUE | 0.2335 |
| Ceacam20 | 415 | 277 | 230 | 215 | 164 | 226 | 322 | 295 | 248 | 214 | 185 | 223 | 247.87 | 288 | 207 | 0.718 | -0.478 | 0.00110188 | 0.0077659 | 0.0096 | 0.012 | 0.0113 | 0.0113 | TRUE | 0.1641 |
| Ceacam3 | 0 | 0 | 0 | 0 | 0 | 0 | 0 | 0 | 0 | 0 | 0 | 0 | 0 | 0 | 0 | NA | NA | NA | NA | NA | NA | NA | NA | NA | NA |
| Ceacam5 | 0 | 1 | 2 | 1 | 1 | 3 | 0 | 1 | 2 | 1 | 1 | 3 | 1.38 | 1 | 2 | 1.664 | 0.735 | 0.66159436 | NA | 0 | 1.7144 | 1.2556 | 1.2556 | TRUE | 0.362 |
| Ceacam9 | 21 | 11 | 17 | 10 | 9 | 12 | 16 | 12 | 18 | 10 | 10 | 12 | 13.05 | 15 | 11 | 0.689 | -0.538 | 0.31089377 | NA | 0 | 0.184 | 0.1231 | 0.1231 | TRUE | 0.1921 |
| Cebpa | 527 | 435 | 307 | 368 | 354 | 338 | 409 | 463 | 331 | 366 | 399 | 334 | 383.76 | 401 | 366 | 0.912 | -0.132 | 0.32792508 | 0.5023077 | 0.0151 | 0.0086 | 0.0106 | 0.0106 | TRUE | 0.2569 |
| Cebpb | 347 | 304 | 229 | 240 | 221 | 236 | 270 | 323 | 247 | 239 | 249 | 233 | 260.19 | 280 | 240 | 0.859 | -0.219 | 0.11400318 | 0.24303355 | 0.0065 | 0.0115 | 0.01 | 0.01 | TRUE | 0.1979 |
| Cebpd | 93 | 71 | 53 | 74 | 59 | 66 | 72 | 76 | 57 | 74 | 67 | 65 | 68.38 | 68 | 69 | 1 | 0 | 0.99924278 | 0.99952251 | 0 | 0.0371 | 0.0264 | 0.0264 | TRUE | 0.1688 |
| Cebpe | 17 | 14 | 12 | 15 | 10 | 10 | 13 | 15 | 13 | 15 | 11 | 10 | 12.85 | 14 | 12 | 0.881 | -0.183 | 0.72934734 | NA | 0 | 0.1868 | 0.1228 | 0.1228 | TRUE | 0.1823 |
| Cebpg | 1586 | 1259 | 1291 | 1444 | 1323 | 1541 | 1232 | 1339 | 1393 | 1437 | 1492 | 1521 | 1402.5 | 1321 | 1483 | 1.124 | 0.169 | 0.02250444 | 0.0751884 | 0.0017 | 0.0041 | 0.0032 | 0.0032 | TRUE | 0.0441 |
| Cebpz | 13874 | 10311 | 10242 | 10350 | 9103 | 10652 | 10775 | 10968 | 11051 | 10302 | 10269 | 10515 | 10646.79 | 10931 | 10362 | 0.948 | -0.077 | 0.09252045 | 0.20904122 | 1.00E-04 | 0.0026 | 0.0014 | 0.0014 | TRUE | 0.002 |
| Cecr2 | 9982 | 7229 | 7266 | 7247 | 6970 | 7926 | 7753 | 7689 | 7840 | 7214 | 7863 | 7824 | 7697.11 | 7761 | 7634 | 0.984 | -0.024 | 0.66054146 | 0.79018406 | 9.00E-04 | 0.0027 | 0.002 | 0.002 | TRUE | 0.0283 |
| Cecr5 | 3183 | 2375 | 2166 | 2669 | 2240 | 2500 | 2472 | 2526 | 2337 | 2657 | 2527 | 2468 | 2497.84 | 2445 | 2551 | 1.043 | 0.061 | 0.33700354 | 0.51122815 | 0.0011 | 0.0034 | 0.0025 | 0.0025 | TRUE | 0.0177 |
| Cecr6 | 5 | 3 | 0 | 4 | 2 | 7 | 4 | 3 | 0 | 4 | 2 | 7 | 3.37 | 2 | 4 | 1.818 | 0.862 | 0.44374594 | NA | 0.1799 | 0.7056 | 0.5973 | 0.5973 | TRUE | 0.5015 |
| Cel | 0 | 0 | 1 | 0 | 3 | 0 | 0 | 0 | 1 | 0 | 3 | 0 | 0.74 | 0 | 1 | 3.214 | 1.684 | 0.52794035 | NA | 8.5559 | 3.1885 | 3.6646 | 3.6646 | TRUE | 1.4433 |
| Cela1 | 166 | 113 | 135 | 188 | 152 | 195 | 129 | 120 | 146 | 187 | 171 | 192 | 157.65 | 132 | 183 | 1.399 | 0.484 | 0.00281304 | 0.01570932 | 0 | 0.0175 | 0.0124 | 0.0124 | TRUE | 0.0834 |
| Cela2a | 193 | 192 | 158 | 118 | 92 | 109 | 150 | 204 | 170 | 117 | 104 | 108 | 142.24 | 175 | 110 | 0.63 | -0.667 | 0.00023632 | 0.00242655 | 0.0099 | 0.0191 | 0.0162 | 0.0162 | TRUE | 0.2295 |
| Cela3b | 0 | 0 | 0 | 0 | 0 | 0 | 0 | 0 | 0 | 0 | 0 | 0 | 0 | 0 | 0 | NA | NA | NA | NA | NA | NA | NA | NA | NA | NA |
| Celf1 | 23301 | 17015 | 16701 | 17076 | 14875 | 17282 | 18097 | 18099 | 18020 | 16997 | 16781 | 17060 | 17508.96 | 18072 | 16946 | 0.938 | -0.093 | 0.0308501 | 0.09454128 | 0 | 0.0026 | 0.0013 | 0.0013 | TRUE | 9.00E-04 |
| Celf2 | 774 | 593 | 640 | 674 | 586 | 616 | 601 | 631 | 691 | 671 | 661 | 608 | 643.75 | 641 | 647 | 1.01 | 0.015 | 0.87378293 | 0.9300007 | 0.0024 | 0.0061 | 0.0049 | 0.0049 | TRUE | 0.0537 |
| Celf3 | 6 | 10 | 8 | 7 | 8 | 12 | 5 | 11 | 9 | 7 | 9 | 12 | 8.63 | 8 | 9 | 1.184 | 0.243 | 0.71486922 | NA | 0 | 0.2771 | 0.2019 | 0.2019 | TRUE | 0.5212 |
| Celf4 | 602 | 497 | 489 | 773 | 579 | 748 | 468 | 529 | 528 | 769 | 653 | 738 | 614.14 | 508 | 720 | 1.422 | 0.508 | 5.05E-07 | 1.64E-05 | 0.0045 | 0.0063 | 0.0057 | 0.0057 | TRUE | 0.0769 |

| | | | | | | | | | | | | | | | | | | | | | | | | | |
|---|---|---|---|---|---|---|---|---|---|---|---|---|---|---|---|---|---|---|---|---|---|---|---|---|---|
| Celf5 | 876 | 744 | 742 | 1092 | 909 | 1091 | 680 | 791 | 801 | 1087 | 1025 | 1077 | 910.29 | 757 | 1063 | 1.407 | 0.493 | 2.23E-08 | 1.02E-06 | 0.0034 | 0.005 | 0.0045 | 0.0045 | TRUE | 0.0995 |
| Celf6 | 37 | 25 | 27 | 36 | 27 | 40 | 29 | 27 | 29 | 36 | 30 | 39 | 31.71 | 28 | 35 | 1.254 | 0.327 | 0.33480189 | 0.50886042 | 0 | 0.0772 | 0.0509 | 0.0509 | TRUE | 0.0933 |
| Celsr1 | 2344 | 1812 | 1746 | 2414 | 2236 | 2543 | 1820 | 1927 | 1884 | 2403 | 2522 | 2510 | 2177.91 | 1877 | 2478 | 1.321 | 0.402 | 6.26E-11 | 5.33E-09 | 3.00E-04 | 0.0035 | 0.0023 | 0.0023 | TRUE | 0.0088 |
| Celsr2 | 980 | 696 | 523 | 589 | 493 | 548 | 761 | 740 | 564 | 586 | 556 | 541 | 624.86 | 688 | 561 | 0.814 | -0.298 | 0.01096292 | 0.04416248 | 0.0126 | 0.0062 | 0.0082 | 0.0082 | TRUE | 0.2889 |
| Celsr3 | 672 | 534 | 473 | 596 | 543 | 714 | 522 | 568 | 510 | 593 | 613 | 705 | 585.16 | 533 | 637 | 1.195 | 0.257 | 0.0116535 | 0.0462805 | 0.0043 | 0.0065 | 0.0058 | 0.0058 | TRUE | 0.104 |
| Cend1 | 13 | 10 | 14 | 1 | 10 | 10 | 10 | 11 | 15 | 1 | 11 | 10 | 9.66 | 12 | 7 | 0.616 | -0.699 | 0.3261735 | NA | 0.281 | 0.2477 | 0.2543 | 0.2543 | TRUE | 1.6375 |
| Cenpa | 7362 | 5546 | 6241 | 6424 | 5899 | 6625 | 5718 | 5899 | 6734 | 6394 | 6655 | 6540 | 6323.33 | 6117 | 6530 | 1.068 | 0.095 | 0.16665678 | 0.31777285 | 0.0039 | 0.0028 | 0.0032 | 0.0032 | TRUE | 0.0947 |
| Cenpb | 4434 | 2897 | 2135 | 2115 | 1943 | 2034 | 3444 | 3082 | 2304 | 2105 | 2192 | 2008 | 2522.32 | 2943 | 2102 | 0.714 | -0.487 | 1.95E-05 | 0.00032626 | 0.0199 | 0.0034 | 0.009 | 0.009 | TRUE | 0.4196 |
| Cenpc1 | 3179 | 2281 | 2307 | 2247 | 1818 | 2246 | 2469 | 2426 | 2489 | 2237 | 2051 | 2217 | 2314.87 | 2461 | 2168 | 0.881 | -0.182 | 0.00347839 | 0.01853312 | 5.00E-04 | 0.0035 | 0.0024 | 0.0024 | TRUE | 0.0267 |
| Cenpe | 10380 | 7570 | 6718 | 6826 | 6170 | 7040 | 8062 | 8052 | 7249 | 6795 | 6960 | 6950 | 7344.51 | 7788 | 6902 | 0.886 | -0.175 | 0.00295502 | 0.01632631 | 0.0017 | 0.0027 | 0.0023 | 0.0023 | TRUE | 0.0445 |
| Cenpf | 14781 | 10908 | 11291 | 13217 | 12177 | 13762 | 11480 | 11603 | 12183 | 13156 | 13737 | 13585 | 12623.95 | 11755 | 13493 | 1.148 | 0.199 | 6.84E-05 | 0.00090277 | 6.00E-04 | 0.0026 | 0.0017 | 0.0017 | TRUE | 0.0123 |
| Cenph | 2719 | 1984 | 1929 | 2052 | 1742 | 1982 | 2112 | 2110 | 2081 | 2043 | 1965 | 1957 | 2044.62 | 2101 | 1988 | 0.946 | -0.08 | 0.18282583 | 0.33854092 | 0 | 0.0036 | 0.0021 | 0.0021 | TRUE | 0.007 |
| Cenpi | 2290 | 1632 | 1702 | 1628 | 1328 | 1535 | 1779 | 1736 | 1836 | 1621 | 1498 | 1515 | 1664.14 | 1784 | 1545 | 0.866 | -0.207 | 0.00207711 | 0.01243224 | 6.00E-04 | 0.0039 | 0.0027 | 0.0027 | TRUE | 0.0224 |
| Cenpj | 1641 | 1116 | 951 | 1082 | 950 | 1154 | 1274 | 1187 | 1026 | 1077 | 1072 | 1139 | 1129.26 | 1162 | 1096 | 0.941 | -0.087 | 0.33435933 | 0.50840404 | 0.0058 | 0.0045 | 0.005 | 0.005 | TRUE | 0.125 |
| Cenpk | 937 | 667 | 754 | 751 | 681 | 801 | 728 | 709 | 814 | 748 | 768 | 791 | 759.54 | 750 | 769 | 1.026 | 0.036 | 0.67842248 | 0.80244062 | 0.0017 | 0.0055 | 0.0043 | 0.0043 | TRUE | 0.0624 |
| Cenpl | 929 | 604 | 624 | 692 | 566 | 647 | 722 | 642 | 673 | 689 | 639 | 639 | 667.21 | 679 | 656 | 0.964 | -0.053 | 0.55754724 | 0.71241039 | 0.0014 | 0.006 | 0.0044 | 0.0044 | TRUE | 0.0373 |
| Cenpm | 2038 | 1281 | 1281 | 1283 | 1005 | 1159 | 1583 | 1363 | 1382 | 1277 | 1134 | 1144 | 1313.76 | 1443 | 1185 | 0.821 | -0.285 | 0.00088313 | 0.00663014 | 0.0047 | 0.0042 | 0.0045 | 0.0045 | TRUE | 0.0896 |
| Cenpn | 3778 | 2882 | 2626 | 2878 | 2536 | 2933 | 2934 | 3066 | 2833 | 2865 | 2861 | 2895 | 2909.02 | 2944 | 2874 | 0.976 | -0.035 | 0.5472338 | 0.7040944 | 4.00E-04 | 0.0032 | 0.0021 | 0.0021 | TRUE | 0.0153 |
| Cenpo | 1392 | 1009 | 949 | 1036 | 873 | 996 | 1081 | 1073 | 1024 | 1031 | 985 | 983 | 1029.6 | 1059 | 1000 | 0.943 | -0.084 | 0.25704737 | 0.42708724 | 0 | 0.0047 | 0.003 | 0.003 | TRUE | 0.01 |
| Cenpp | 1139 | 886 | 748 | 810 | 656 | 772 | 885 | 942 | 807 | 806 | 740 | 762 | 823.75 | 878 | 769 | 0.876 | -0.19 | 0.03173992 | 0.0965026 | 0.0027 | 0.0053 | 0.0044 | 0.0044 | TRUE | 0.0571 |
| Cenpq | 739 | 565 | 563 | 571 | 505 | 537 | 574 | 601 | 607 | 568 | 570 | 530 | 575.1 | 594 | 556 | 0.936 | -0.095 | 0.30136934 | 0.47371296 | 0 | 0.0065 | 0.0043 | 0.0043 | TRUE | 0.0198 |
| Cenpt | 4897 | 3690 | 3263 | 3458 | 3179 | 3596 | 3803 | 3925 | 3521 | 3442 | 3586 | 3550 | 3637.86 | 3750 | 3526 | 0.94 | -0.089 | 0.14624195 | 0.2905931 | 0.0013 | 0.0031 | 0.0024 | 0.0024 | TRUE | 0.0341 |
| Cenpv | 4152 | 2970 | 2782 | 3190 | 2871 | 3161 | 3225 | 3159 | 3002 | 3175 | 3239 | 3120 | 3153.35 | 3129 | 3178 | 1.015 | 0.022 | 0.70409836 | 0.82094194 | 5.00E-04 | 0.0032 | 0.0021 | 0.0021 | TRUE | 0.0151 |
| Cenpw | 1340 | 982 | 1029 | 1131 | 889 | 1049 | 1041 | 1045 | 1110 | 1126 | 1003 | 1036 | 1059.96 | 1065 | 1055 | 0.991 | -0.013 | 0.87086221 | 0.92839601 | 0.0015 | 0.0047 | 0.0035 | 0.0035 | TRUE | 0.0417 |
| Cep104 | 1500 | 1007 | 1021 | 1034 | 907 | 1047 | 1165 | 1071 | 1102 | 1029 | 1023 | 1034 | 1070.63 | 1113 | 1029 | 0.924 | -0.115 | 0.12197441 | 0.2550666 | 0 | 0.0046 | 0.003 | 0.003 | TRUE | 0.0214 |
| Cep110 | 4648 | 3544 | 3426 | 3974 | 3576 | 4141 | 3610 | 3770 | 3697 | 3956 | 4034 | 4088 | 3858.98 | 3692 | 4026 | 1.091 | 0.125 | 0.01852657 | 0.06530217 | 0 | 0.003 | 0.0018 | 0.0018 | TRUE | 0.0047 |
| Cep112 | 297 | 236 | 254 | 259 | 233 | 262 | 231 | 251 | 274 | 258 | 263 | 259 | 255.84 | 252 | 260 | 1.034 | 0.048 | 0.71113091 | 0.8262229 | 0 | 0.0117 | 0.0083 | 0.0083 | TRUE | 0.0663 |
| Cep120 | 4303 | 3139 | 3306 | 3344 | 2745 | 3269 | 3342 | 3339 | 3567 | 3329 | 3097 | 3227 | 3316.71 | 3416 | 3218 | 0.942 | -0.086 | 0.1531197 | 0.30004659 | 9.00E-04 | 0.0031 | 0.0023 | 0.0023 | TRUE | 0.018 |
| Cep128 | 1762 | 1276 | 1147 | 1155 | 1012 | 1158 | 1368 | 1357 | 1238 | 1150 | 1142 | 1143 | 1232.97 | 1321 | 1145 | 0.866 | -0.208 | 0.00464141 | 0.02315519 | 8.00E-04 | 0.0043 | 0.0031 | 0.0031 | TRUE | 0.036 |
| Cep135 | 1710 | 1221 | 1079 | 902 | 869 | 999 | 1328 | 1299 | 1164 | 898 | 980 | 986 | 1109.24 | 1264 | 955 | 0.755 | -0.406 | 9.10E-07 | 2.71E-05 | 0.003 | 0.0046 | 0.004 | 0.004 | TRUE | 0.056 |
| Cep152 | 968 | 805 | 782 | 704 | 583 | 771 | 752 | 856 | 844 | 701 | 658 | 761 | 761.9 | 817 | 707 | 0.866 | -0.207 | 0.02739626 | 0.08659326 | 0.004 | 0.0055 | 0.005 | 0.005 | TRUE | 0.0613 |
| Cep164 | 4322 | 3305 | 3113 | 2669 | 2498 | 2943 | 3357 | 3515 | 3359 | 2657 | 2818 | 2905 | 3101.84 | 3410 | 2793 | 0.819 | -0.288 | 2.04E-06 | 5.15E-05 | 9.00E-04 | 0.0032 | 0.0023 | 0.0023 | TRUE | 0.0224 |
| Cep170 | 3171 | 2449 | 2479 | 2939 | 2431 | 2607 | 2463 | 2605 | 2675 | 2925 | 2742 | 2574 | 2664 | 2581 | 2747 | 1.065 | 0.091 | 0.18709954 | 0.3439061 | 0.0024 | 0.0033 | 0.003 | 0.003 | TRUE | 0.0395 |
| Cep170b | 5811 | 4440 | 4946 | 5465 | 5017 | 5716 | 4513 | 4723 | 5337 | 5440 | 5660 | 5643 | 5219.12 | 4858 | 5581 | 1.149 | 0.201 | 0.00367194 | 0.01930231 | 0.0038 | 0.0029 | 0.0033 | 0.0033 | TRUE | 0.0904 |
| Cep19 | 1159 | 808 | 799 | 795 | 676 | 741 | 900 | 859 | 862 | 791 | 763 | 731 | 817.86 | 874 | 762 | 0.871 | -0.199 | 0.01382079 | 0.05263538 | 0 | 0.0053 | 0.0035 | 0.0035 | TRUE | 0.0147 |
| Cep192 | 11469 | 8299 | 8766 | 9710 | 8950 | 10357 | 8907 | 8828 | 9458 | 9665 | 10097 | 10224 | 9529.88 | 9064 | 9995 | 1.103 | 0.141 | 0.00815897 | 0.03556455 | 0.001 | 0.0027 | 0.0019 | 0.0019 | TRUE | 0.0175 |
| Cep250 | 2473 | 2137 | 1980 | 1891 | 1664 | 1984 | 1921 | 2273 | 2136 | 1882 | 1877 | 1959 | 2008.03 | 2110 | 1906 | 0.904 | -0.145 | 0.05363373 | 0.14156329 | 0.0035 | 0.0036 | 0.0036 | 0.0036 | TRUE | 0.0768 |
| Cep290 | 1636 | 1104 | 918 | 937 | 756 | 868 | 1271 | 1174 | 991 | 933 | 853 | 857 | 1012.97 | 1145 | 881 | 0.768 | -0.38 | 0.00011606 | 0.00137434 | 0.0084 | 0.0048 | 0.006 | 0.006 | TRUE | 0.1655 |
| Cep350 | 4696 | 3185 | 3134 | 3674 | 3524 | 3960 | 3647 | 3388 | 3382 | 3657 | 3975 | 3909 | 3659.71 | 3472 | 3847 | 1.107 | 0.147 | 0.01775421 | 0.06323473 | 0.0015 | 0.0031 | 0.0025 | 0.0025 | TRUE | 0.0241 |
| Cep44 | 1651 | 1111 | 1197 | 1231 | 1143 | 1250 | 1282 | 1182 | 1292 | 1225 | 1289 | 1234 | 1250.71 | 1252 | 1249 | 0.997 | -0.004 | 0.95911823 | 0.97739513 | 7.00E-04 | 0.0043 | 0.003 | 0.003 | TRUE | 0.0281 |
| Cep55 | 4895 | 3731 | 3847 | 4030 | 3581 | 3940 | 3802 | 3969 | 4151 | 4011 | 4040 | 3889 | 3976.97 | 3974 | 3980 | 1.002 | 0.003 | 0.95990756 | 0.97776217 | 9.00E-04 | 0.003 | 0.0021 | 0.0021 | TRUE | 0.0185 |
| Cep57 | 4317 | 3239 | 3559 | 3486 | 3155 | 3485 | 3353 | 3445 | 3840 | 3470 | 3559 | 3440 | 3517.93 | 3546 | 3490 | 0.985 | -0.022 | 0.73378412 | 0.84079179 | 0.0025 | 0.0031 | 0.0029 | 0.0029 | TRUE | 0.0636 |
| Cep57l1 | 2402 | 1884 | 1676 | 1749 | 1546 | 1937 | 1866 | 2004 | 1808 | 1741 | 1744 | 1912 | 1845.84 | 1893 | 1799 | 0.951 | -0.073 | 0.31247878 | 0.48572717 | 0.0021 | 0.0037 | 0.0032 | 0.0032 | TRUE | 0.0368 |
| Cep63 | 1282 | 949 | 923 | 974 | 781 | 946 | 996 | 1009 | 996 | 970 | 881 | 934 | 964.24 | 1000 | 928 | 0.928 | -0.107 | 0.1633687 | 0.31398857 | 0 | 0.0049 | 0.0032 | 0.0032 | TRUE | 0.0229 |
| Cep68 | 3764 | 2903 | 2984 | 3225 | 2876 | 3356 | 2923 | 3088 | 3220 | 3210 | 3244 | 3313 | 3166.4 | 3077 | 3256 | 1.059 | 0.082 | 0.1703381 | 0.32195322 | 0.001 | 0.0032 | 0.0023 | 0.0023 | TRUE | 0.0238 |
| Cep70 | 1117 | 814 | 885 | 870 | 759 | 897 | 868 | 866 | 955 | 866 | 856 | 885 | 882.66 | 896 | 869 | 0.971 | -0.042 | 0.59903506 | 0.74465195 | 6.00E-04 | 0.0051 | 0.0036 | 0.0036 | TRUE | 0.0382 |
| Cep72 | 2086 | 1634 | 1614 | 1794 | 1575 | 1827 | 1620 | 1738 | 1741 | 1786 | 1777 | 1804 | 1744.28 | 1700 | 1789 | 1.053 | 0.075 | 0.24901076 | 0.41806702 | 0 | 0.0038 | 0.0025 | 0.0025 | TRUE | 0.0212 |
| Cep76 | 2154 | 1435 | 1312 | 1517 | 1329 | 1489 | 1673 | 1526 | 1416 | 1510 | 1499 | 1470 | 1515.68 | 1538 | 1493 | 0.969 | -0.045 | 0.5599797 | 0.7146879 | 0.0031 | 0.004 | 0.0037 | 0.0037 | TRUE | 0.0728 |
| Cep78 | 3027 | 2228 | 2168 | 2158 | 2079 | 2551 | 2351 | 2370 | 2339 | 2148 | 2345 | 2518 | 2345.29 | 2353 | 2337 | 0.993 | -0.01 | 0.88855395 | 0.93805954 | 0.0028 | 0.0034 | 0.0032 | 0.0032 | TRUE | 0.0614 |
| Cep85 | 7294 | 5336 | 5318 | 5713 | 5137 | 5766 | 5665 | 5676 | 5738 | 5687 | 5795 | 5692 | 5708.76 | 5693 | 5725 | 1.006 | 0.008 | 0.86830846 | 0.92728129 | 0 | 0.0028 | 0.0015 | 0.0015 | TRUE | 0.0014 |
| Cep85l | 366 | 258 | 243 | 281 | 241 | 258 | 284 | 274 | 262 | 280 | 272 | 255 | 271.19 | 273 | 269 | 0.981 | -0.028 | 0.8202568 | 0.89651557 | 0 | 0.0112 | 0.0074 | 0.0074 | TRUE | 0.0241 |
| Cep89 | 1217 | 1013 | 919 | 971 | 892 | 1018 | 945 | 1078 | 992 | 967 | 1006 | 1005 | 998.67 | 1005 | 993 | 0.989 | -0.016 | 0.84236746 | 0.91032497 | 0.0016 | 0.0048 | 0.0037 | 0.0037 | TRUE | 0.0477 |
| Cep95 | 1608 | 1124 | 1135 | 1359 | 1126 | 1383 | 1249 | 1196 | 1225 | 1353 | 1270 | 1365 | 1276.22 | 1223 | 1329 | 1.087 | 0.12 | 0.08821296 | 0.20181091 | 0 | 0.0043 | 0.0028 | 0.0028 | TRUE | 0.0178 |
| Cep97 | 1132 | 819 | 844 | 1022 | 882 | 1048 | 879 | 871 | 911 | 1017 | 995 | 1035 | 951.3 | 887 | 1016 | 1.146 | 0.196 | 0.00893565 | 0.03817014 | 0 | 0.0049 | 0.003 | 0.003 | TRUE | 0.0062 |
| Cept1 | 1781 | 1273 | 1420 | 1538 | 1339 | 1405 | 1383 | 1354 | 1532 | 1531 | 1511 | 1387 | 1449.65 | 1423 | 1476 | 1.038 | 0.053 | 0.49289355 | 0.6581584 | 0.0026 | 0.0041 | 0.0036 | 0.0036 | TRUE | 0.0529 |
| Cer1 | 0 | 0 | 0 | 0 | 0 | 1 | 0 | 0 | 0 | 0 | 0 | 1 | 0.16 | 0 | 0 | 2.094 | 1.067 | 0.79379329 | NA | 0 | 14.4078 | 10 | 10 | TRUE | 0.3835 |
| Cercam | 1397 | 1071 | 1222 | 1479 | 1283 | 1411 | 1085 | 1139 | 1319 | 1472 | 1447 | 1393 | 1309.19 | 1181 | 1437 | 1.219 | 0.286 | 0.00076989 | 0.00598045 | 0.0046 | 0.0042 | 0.0044 | 0.0044 | TRUE | 0.1234 |
| Cerk | 440 | 297 | 318 | 330 | 302 | 334 | 342 | 316 | 343 | 328 | 341 | 330 | 333.27 | 334 | 333 | 0.997 | -0.004 | 0.96962654 | 0.9834812 | 0 | 0.0095 | 0.0062 | 0.0062 | TRUE | 0.0235 |
| Cerkl | 32 | 32 | 32 | 35 | 22 | 31 | 25 | 34 | 35 | 35 | 25 | 31 | 30.61 | 31 | 30 | 0.976 | -0.035 | 0.92167241 | 0.95683559 | 0 | 0.0798 | 0.058 | 0.058 | TRUE | 0.2557 |
| Cers1 | 0 | 0 | 0 | 0 | 0 | 0 | 0 | 0 | 0 | 0 | 0 | 0 | 0 | 0 | 0 | NA | NA | NA | NA | NA | NA | NA | NA | NA | NA |
| Cers2 | 2824 | 2129 | 2209 | 2158 | 1933 | 2278 | 2193 | 2265 | 2383 | 2148 | 2181 | 2249 | 2236.47 | 2280 | 2193 | 0.962 | -0.056 | 0.3749484 | 0.54931708 | 8.00E-04 | 0.0035 | 0.0024 | 0.0024 | TRUE | 0.0188 |
| Cers3 | 58 | 28 | 44 | 40 | 33 | 27 | 45 | 30 | 47 | 40 | 37 | 27 | 37.67 | 41 | 35 | 0.844 | -0.244 | 0.46570958 | 0.63463117 | 0.0232 | 0.0653 | 0.0541 | 0.0541 | TRUE | 0.395 |
| Cers4 | 3808 | 2951 | 3431 | 3869 | 3589 | 4097 | 2958 | 3139 | 3702 | 3851 | 4049 | 4044 | 3623.8 | 3266 | 3981 | 1.22 | 0.287 | 0.0004129 | 0.00372572 | 0.007 | 0.0031 | 0.0045 | 0.0045 | TRUE | 0.1651 |
| Cers5 | 2720 | 1897 | 1987 | 2042 | 1831 | 2133 | 2113 | 2018 | 2144 | 2033 | 2066 | 2106 | 2079.67 | 2092 | 2068 | 0.989 | -0.017 | 0.787787 | 0.87528754 | 0 | 0.0036 | 0.0022 | 0.0022 | TRUE | 0.0112 |
| Cers6 | 1351 | 942 | 1016 | 1259 | 1112 | 1221 | 1049 | 1002 | 1096 | 1253 | 1254 | 1205 | 1143.41 | 1049 | 1237 | 1.18 | 0.238 | 0.0011838 | 0.00818977 | 0 | 0.0045 | 0.003 | 0.003 | TRUE | 0.0176 |
| Ces1a | 2 | 0 | 0 | 1 | 1 | 1 | 2 | 0 | 0 | 1 | 1 | 1 | 0.78 | 1 | 1 | 1.828 | 0.87 | 0.69255718 | NA | 0 | 3.0514 | 2.1415 | 2.1415 | TRUE | 0.9716 |
| Ces1b | 0 | 0 | 0 | 1 | 1 | 1 | 0 | 0 | 0 | 1 | 1 | 1 | 0.52 | 0 | 1 | 5.852 | 2.549 | 0.3348364 | NA | 0 | 4.5739 | 3.5345 | 3.5345 | TRUE | 0.0847 |
| Ces1c | 7 | 1 | 0 | 1 | 2 | 2 | 5 | 1 | 0 | 1 | 2 | 2 | 1.95 | 2 | 2 | 0.765 | -0.386 | 0.7935699 | NA | 0.6515 | 1.2151 | 1.0537 | 1.0537 | TRUE | 1.9007 |
| Ces1d | 0 | 0 | 0 | 0 | 0 | 0 | 0 | 0 | 0 | 0 | 0 | 0 | 0 | 0 | 0 | NA | NA | NA | NA | NA | NA | NA | NA | NA | NA |

| | | | | | | | | | | | | | | | | | | | | | | | | | |
|---|---|---|---|---|---|---|---|---|---|---|---|---|---|---|---|---|---|---|---|---|---|---|---|---|---|
| Ces1e | 0 | 0 | 0 | 0 | 0 | 0 | 0 | 0 | 0 | 0 | 0 | 0 | 0 | 0 | 0 | NA | NA | NA | NA | NA | NA | NA | NA | NA | NA |
| Ces1f | 0 | 0 | 0 | 0 | 0 | 0 | 0 | 0 | 0 | 0 | 0 | 0 | 0 | 0 | 0 | NA | NA | NA | NA | NA | NA | NA | NA | NA | NA |
| Ces1g | 0 | 0 | 0 | 0 | 0 | 0 | 0 | 0 | 0 | 0 | 0 | 0 | 0 | 0 | 0 | NA | NA | NA | NA | NA | NA | NA | NA | NA | NA |
| Ces2a | 0 | 0 | 0 | 0 | 0 | 0 | 0 | 0 | 0 | 0 | 0 | 0 | 0 | 0 | 0 | NA | NA | NA | NA | NA | NA | NA | NA | NA | NA |
| Ces2b | 0 | 0 | 1 | 0 | 0 | 0 | 0 | 0 | 1 | 0 | 0 | 0 | 0.18 | 0 | 0 | 0.552 | -0.857 | 0.8336548 | NA | 0 | 13.1817 | 10 | 10 | TRUE | 0.5365 |
| Ces2c | 0 | 0 | 0 | 0 | 0 | 0 | 0 | 0 | 0 | 0 | 0 | 0 | 0 | 0 | 0 | NA | NA | NA | NA | NA | NA | NA | NA | NA | NA |
| Ces2d-ps | 0 | 0 | 0 | 0 | 0 | 0 | 0 | 0 | 0 | 0 | 0 | 0 | 0 | 0 | 0 | NA | NA | NA | NA | NA | NA | NA | NA | NA | NA |
| Ces2e | 31 | 20 | 15 | 18 | 6 | 13 | 24 | 21 | 16 | 18 | 7 | 13 | 16.51 | 20 | 13 | 0.611 | -0.71 | 0.15887659 | NA | 0.0296 | 0.146 | 0.1186 | 0.1186 | TRUE | 0.3367 |
| Ces2f | 3 | 0 | 1 | 2 | 1 | 0 | 2 | 0 | 1 | 2 | 1 | 0 | 1.09 | 1 | 1 | 0.882 | -0.181 | 0.92509194 | NA | 0 | 2.1808 | 1.7468 | 1.7468 | TRUE | 0.2364 |
| Ces2g | 9 | 5 | 3 | 1 | 0 | 1 | 7 | 5 | 3 | 1 | 0 | 1 | 2.92 | 5 | 1 | 0.13 | -2.94 | 0.03681251 | NA | 0 | 0.8137 | 0.5777 | 0.5777 | TRUE | 0.1671 |
| Ces2h | 0 | 0 | 0 | 0 | 0 | 0 | 0 | 0 | 0 | 0 | 0 | 0 | 0 | 0 | 0 | NA | NA | NA | NA | NA | NA | NA | NA | NA | NA |
| Ces3a | 0 | 0 | 0 | 0 | 0 | 0 | 0 | 0 | 0 | 0 | 0 | 0 | 0 | 0 | 0 | NA | NA | NA | NA | NA | NA | NA | NA | NA | NA |
| Ces3b | 0 | 0 | 0 | 0 | 0 | 0 | 0 | 0 | 0 | 0 | 0 | 0 | 0 | 0 | 0 | NA | NA | NA | NA | NA | NA | NA | NA | NA | NA |
| Ces4a | 1 | 2 | 1 | 0 | 0 | 1 | 1 | 2 | 1 | 0 | 0 | 1 | 0.83 | 1 | 0 | 0.282 | -1.824 | 0.39899776 | NA | 0 | 2.8636 | 2.0027 | 2.0027 | TRUE | 0.3835 |
| Ces5a | 2 | 3 | 0 | 0 | 0 | 0 | 2 | 3 | 0 | 0 | 0 | 0 | 0.79 | 2 | 0 | 0.12 | -3.055 | 0.22090216 | NA | 5.3026 | 2.9997 | 3.1827 | 3.1827 | TRUE | 0.0516 |
| Cetn1 | 0 | 0 | 0 | 0 | 0 | 0 | 0 | 0 | 0 | 0 | 0 | 0 | 0 | 0 | 0 | NA | NA | NA | NA | NA | NA | NA | NA | NA | NA |
| Cetn2 | 1469 | 1110 | 1039 | 1243 | 1038 | 1172 | 1141 | 1181 | 1121 | 1237 | 1171 | 1157 | 1167.98 | 1148 | 1188 | 1.036 | 0.051 | 0.48197368 | 0.64860652 | 0 | 0.0045 | 0.0029 | 0.0029 | TRUE | 0.0157 |
| Cetn3 | 3484 | 2358 | 2675 | 2675 | 2183 | 2509 | 2706 | 2508 | 2886 | 2663 | 2463 | 2477 | 2617.08 | 2700 | 2534 | 0.939 | -0.091 | 0.19559147 | 0.35527584 | 0.0028 | 0.0033 | 0.0032 | 0.0032 | TRUE | 0.046 |
| Cetn4 | 44 | 23 | 12 | 35 | 21 | 19 | 34 | 24 | 13 | 35 | 24 | 19 | 24.81 | 24 | 26 | 1.067 | 0.093 | 0.83470234 | 0.90601463 | 0.1166 | 0.0979 | 0.1032 | 0.1032 | TRUE | 0.2398 |
| Cfb | 0 | 1 | 1 | 1 | 3 | 2 | 0 | 1 | 1 | 1 | 3 | 2 | 1.42 | 1 | 2 | 3.129 | 1.646 | 0.33803814 | NA | 0 | 1.6761 | 1.1642 | 1.1642 | TRUE | 0.3327 |
| Cfc1 | 3 | 4 | 3 | 5 | 3 | 2 | 2 | 4 | 3 | 5 | 3 | 2 | 3.36 | 3 | 3 | 1.065 | 0.09 | 0.93008751 | NA | 0 | 0.7079 | 0.4628 | 0.4628 | TRUE | 0.2183 |
| Cfd | 0 | 0 | 0 | 0 | 0 | 0 | 0 | 0 | 0 | 0 | 0 | 0 | 0 | 0 | 0 | NA | NA | NA | NA | NA | NA | NA | NA | NA | NA |
| Cfdp1 | 15312 | 10854 | 10506 | 10980 | 9446 | 11315 | 11892 | 11545 | 11336 | 10929 | 10656 | 11170 | 11254.75 | 11591 | 10918 | 0.942 | -0.086 | 0.07757299 | 0.18488423 | 4.00E-04 | 0.0026 | 0.0016 | 0.0016 | TRUE | 0.0064 |
| Cfh | 6 | 5 | 0 | 0 | 0 | 3 | 5 | 5 | 0 | 0 | 0 | 3 | 2.16 | 3 | 1 | 0.3 | -1.735 | 0.29692125 | NA | 3.1156 | 1.1014 | 1.335 | 1.335 | TRUE | 1.4412 |
| Cfhr1 | 0 | 0 | 0 | 0 | 0 | 0 | 0 | 0 | 0 | 0 | 0 | 0 | 0 | 0 | 0 | NA | NA | NA | NA | NA | NA | NA | NA | NA | NA |
| Cfhr2 | 0 | 2 | 0 | 1 | 0 | 0 | 0 | 2 | 0 | 1 | 0 | 0 | 0.52 | 1 | 0 | 0.534 | -0.904 | 0.76303124 | NA | 0 | 4.5562 | 4.7774 | 4.7774 | TRUE | 0.9746 |
| Cfi | 2 | 9 | 2 | 7 | 3 | 2 | 2 | 10 | 2 | 7 | 3 | 2 | 4.27 | 5 | 4 | 0.948 | -0.078 | 0.9398792 | NA | 0.4532 | 0.5577 | 0.5245 | 0.5245 | TRUE | 1.7998 |
| Cfl1 | 35591 | 24604 | 21533 | 20453 | 17553 | 21563 | 27642 | 26171 | 23234 | 20359 | 19802 | 21286 | 23082.23 | 25682 | 20482 | 0.797 | -0.327 | 1.72E-06 | 4.50E-05 | 0.0044 | 0.0025 | 0.0033 | 0.0033 | TRUE | 0.0851 |
| Cfl2 | 3166 | 2254 | 2310 | 2505 | 2050 | 2386 | 2459 | 2398 | 2492 | 2493 | 2313 | 2355 | 2418.39 | 2450 | 2387 | 0.975 | -0.037 | 0.54339005 | 0.70096336 | 5.00E-04 | 0.0034 | 0.0023 | 0.0023 | TRUE | 0.0185 |
| Cflar | 305 | 218 | 202 | 216 | 204 | 261 | 237 | 232 | 218 | 215 | 230 | 258 | 231.58 | 229 | 234 | 1.023 | 0.032 | 0.81477647 | 0.8934786 | 0 | 0.0127 | 0.0092 | 0.0092 | TRUE | 0.0865 |
| Cfp | 72 | 58 | 56 | 49 | 40 | 46 | 56 | 62 | 60 | 49 | 45 | 45 | 52.89 | 59 | 46 | 0.784 | -0.35 | 0.18199998 | 0.33758977 | 0 | 0.0472 | 0.0303 | 0.0303 | TRUE | 0.0254 |
| Cftr | 38 | 31 | 10 | 16 | 9 | 8 | 30 | 33 | 11 | 16 | 10 | 8 | 17.88 | 25 | 11 | 0.462 | -1.114 | 0.03856384 | 0.1115098 | 0.1657 | 0.135 | 0.1437 | 0.1437 | TRUE | 1.3376 |
| Cga | 5 | 3 | 3 | 10 | 1 | 5 | 4 | 3 | 3 | 10 | 1 | 5 | 4.39 | 3 | 5 | 1.561 | 0.642 | 0.50925086 | NA | 0 | 0.5425 | 0.4465 | 0.4465 | TRUE | 0.1841 |
| Cggbp1 | 8259 | 5861 | 5905 | 6197 | 5564 | 6437 | 6414 | 6234 | 6371 | 6168 | 6277 | 6354 | 6303.28 | 6340 | 6266 | 0.988 | -0.017 | 0.72716248 | 0.83661541 | 0 | 0.0028 | 0.0015 | 0.0015 | TRUE | 0.0026 |
| Cgn | 2786 | 2075 | 2071 | 2474 | 2118 | 2509 | 2164 | 2207 | 2235 | 2463 | 2389 | 2477 | 2322.37 | 2202 | 2443 | 1.11 | 0.151 | 0.0096831 | 0.04044352 | 0 | 0.0034 | 0.002 | 0.002 | TRUE | 0.0044 |
| Cgnl1 | 1632 | 1164 | 964 | 987 | 863 | 1177 | 1268 | 1238 | 1040 | 982 | 974 | 1162 | 1110.61 | 1182 | 1039 | 0.879 | -0.186 | 0.06250561 | 0.15815058 | 0.0093 | 0.0046 | 0.0063 | 0.0063 | TRUE | 0.1303 |
| Cgref1 | 562 | 373 | 405 | 543 | 407 | 463 | 436 | 397 | 437 | 541 | 459 | 457 | 454.49 | 423 | 486 | 1.147 | 0.197 | 0.07301772 | 0.17694849 | 0.0041 | 0.0076 | 0.0065 | 0.0065 | TRUE | 0.1152 |
| Cgrrf1 | 260 | 194 | 258 | 301 | 266 | 298 | 202 | 206 | 278 | 300 | 300 | 294 | 263.42 | 229 | 298 | 1.306 | 0.385 | 0.00844566 | 0.03649147 | 0.0118 | 0.0114 | 0.0116 | 0.0116 | TRUE | 0.3838 |
| Ch25h | 3 | 1 | 3 | 1 | 1 | 5 | 2 | 1 | 3 | 1 | 1 | 5 | 2.28 | 2 | 2 | 1.073 | 0.102 | 0.93718262 | NA | 0 | 1.0412 | 0.7621 | 0.7621 | TRUE | 0.9911 |
| Chac1 | 119 | 84 | 48 | 52 | 37 | 46 | 92 | 89 | 52 | 52 | 42 | 45 | 62.08 | 78 | 46 | 0.593 | -0.754 | 0.00796808 | 0.0349456 | 0.0417 | 0.0406 | 0.0409 | 0.0409 | TRUE | 0.7559 |
| Chac2 | 372 | 258 | 247 | 299 | 264 | 293 | 289 | 274 | 267 | 298 | 298 | 289 | 285.76 | 277 | 295 | 1.064 | 0.09 | 0.4536973 | 0.62346846 | 0 | 0.0107 | 0.0068 | 0.0068 | TRUE | 0.0185 |
| Chad | 7 | 5 | 5 | 3 | 6 | 3 | 5 | 5 | 5 | 3 | 7 | 3 | 4.81 | 5 | 4 | 0.778 | -0.361 | 0.67701381 | NA | 0 | 0.4951 | 0.3294 | 0.3294 | TRUE | 0.4255 |
| Chadl | 81 | 44 | 66 | 74 | 49 | 58 | 63 | 47 | 71 | 74 | 55 | 57 | 61.19 | 60 | 62 | 1.029 | 0.042 | 0.87486983 | 0.9305339 | 0.0166 | 0.0412 | 0.0346 | 0.0346 | TRUE | 0.3092 |
| Chaf1a | 8054 | 5871 | 5086 | 4871 | 4453 | 4953 | 6255 | 6245 | 5488 | 4849 | 5023 | 4889 | 5458.21 | 5996 | 4920 | 0.82 | -0.286 | 1.00E-05 | 0.00018932 | 0.0026 | 0.0029 | 0.0028 | 0.0028 | TRUE | 0.0666 |
| Chaf1b | 5473 | 4173 | 3884 | 3900 | 3589 | 4112 | 4251 | 4439 | 4191 | 3882 | 4049 | 4059 | 4145.04 | 4294 | 3997 | 0.931 | -0.103 | 0.06044271 | 0.15407036 | 4.00E-04 | 0.003 | 0.0019 | 0.0019 | TRUE | 0.0105 |
| Chat | 4 | 3 | 2 | 2 | 2 | 1 | 3 | 3 | 2 | 2 | 2 | 1 | 2.28 | 3 | 2 | 0.613 | -0.707 | 0.56869779 | NA | 0 | 1.0412 | 0.6402 | 0.6402 | TRUE | 0.1193 |
| Chchd1 | 2444 | 1601 | 1612 | 1923 | 1640 | 1860 | 1898 | 1703 | 1739 | 1914 | 1850 | 1836 | 1823.47 | 1780 | 1867 | 1.048 | 0.067 | 0.32944182 | 0.50382109 | 0.0015 | 0.0037 | 0.0029 | 0.0029 | TRUE | 0.042 |
| Chchd10 | 17284 | 13510 | 14584 | 19792 | 17262 | 20635 | 13424 | 14370 | 15736 | 19701 | 19473 | 20370 | 17179.06 | 14510 | 19848 | 1.368 | 0.452 | 1.70E-12 | 2.02E-10 | 0.0034 | 0.0026 | 0.0029 | 0.0029 | TRUE | 0.0668 |
| Chchd2 | 10747 | 7607 | 7292 | 8491 | 7049 | 8235 | 8347 | 8091 | 7868 | 8452 | 7952 | 8129 | 8139.89 | 8102 | 8178 | 1.009 | 0.013 | 0.80071356 | 0.88423077 | 6.00E-04 | 0.0027 | 0.0019 | 0.0019 | TRUE | 0.0105 |
| Chchd3 | 6852 | 4888 | 4635 | 5499 | 4688 | 5454 | 5322 | 5199 | 5001 | 5474 | 5289 | 5384 | 5278.05 | 5174 | 5382 | 1.04 | 0.057 | 0.28297585 | 0.45474428 | 5.00E-04 | 0.0029 | 0.0018 | 0.0018 | TRUE | 0.0104 |
| Chchd4 | 3361 | 2465 | 2596 | 2821 | 2417 | 2972 | 2610 | 2622 | 2801 | 2808 | 2727 | 2934 | 2750.31 | 2678 | 2823 | 1.055 | 0.077 | 0.21521031 | 0.37769249 | 9.00E-04 | 0.0033 | 0.0024 | 0.0024 | TRUE | 0.0194 |
| Chchd5 | 1425 | 992 | 1174 | 1250 | 1101 | 1322 | 1107 | 1055 | 1267 | 1244 | 1242 | 1305 | 1203.33 | 1143 | 1264 | 1.107 | 0.146 | 0.08186339 | 0.19231975 | 0.0038 | 0.0044 | 0.0042 | 0.0042 | TRUE | 0.105 |
| Chchd6 | 1189 | 918 | 888 | 977 | 920 | 1024 | 923 | 976 | 958 | 973 | 1038 | 1011 | 979.88 | 952 | 1007 | 1.058 | 0.081 | 0.28722556 | 0.45936163 | 0 | 0.0048 | 0.0031 | 0.0031 | TRUE | 0.0108 |
| Chchd7 | 889 | 637 | 632 | 672 | 611 | 668 | 690 | 678 | 682 | 669 | 689 | 659 | 677.92 | 683 | 672 | 0.984 | -0.024 | 0.77586265 | 0.86757979 | 0 | 0.0059 | 0.0036 | 0.0036 | TRUE | 0.0054 |
| Chd1 | 10321 | 7237 | 6977 | 6729 | 5855 | 6904 | 8016 | 7698 | 7528 | 6698 | 6605 | 6815 | 7226.72 | 7747 | 6706 | 0.865 | -0.208 | 4.49E-05 | 0.00063731 | 4.00E-04 | 0.0028 | 0.0017 | 0.0017 | TRUE | 0.0114 |
| Chd1l | 4252 | 3337 | 3181 | 3378 | 3080 | 3537 | 3302 | 3550 | 3432 | 3362 | 3475 | 3492 | 3435.45 | 3428 | 3443 | 1.005 | 0.007 | 0.90531495 | 0.94816931 | 6.00E-04 | 0.0031 | 0.0021 | 0.0021 | TRUE | 0.0129 |
| Chd2 | 5883 | 4182 | 4973 | 5733 | 5271 | 6318 | 4569 | 4448 | 5366 | 5707 | 5946 | 6237 | 5378.82 | 4794 | 5963 | 1.244 | 0.315 | 3.67E-05 | 0.00054842 | 0.0061 | 0.0029 | 0.004 | 0.004 | TRUE | 0.1317 |
| Chd3 | 2859 | 2128 | 2135 | 2129 | 1829 | 2265 | 2220 | 2264 | 2304 | 2119 | 2063 | 2236 | 2201.01 | 2263 | 2139 | 0.946 | -0.08 | 0.19968101 | 0.36040418 | 5.00E-04 | 0.0035 | 0.0023 | 0.0023 | TRUE | 0.0189 |
| Chd4 | 50094 | 35623 | 32851 | 33267 | 30977 | 35960 | 38906 | 37892 | 35446 | 33114 | 34945 | 35498 | 35966.81 | 37415 | 34519 | 0.923 | -0.116 | 0.03559617 | 0.10516247 | 0.0017 | 0.0025 | 0.0022 | 0.0022 | TRUE | 0.0259 |
| Chd5 | 525 | 418 | 424 | 605 | 592 | 705 | 408 | 445 | 457 | 602 | 668 | 696 | 545.98 | 437 | 655 | 1.504 | 0.589 | 6.73E-09 | 3.55E-07 | 0.003 | 0.0068 | 0.0055 | 0.0055 | TRUE | 0.0602 |
| Chd6 | 4173 | 3099 | 3254 | 3730 | 3439 | 3804 | 3241 | 3296 | 3511 | 3713 | 3880 | 3755 | 3565.99 | 3349 | 3783 | 1.13 | 0.176 | 0.00256926 | 0.01467136 | 7.00E-04 | 0.0031 | 0.0022 | 0.0022 | TRUE | 0.0215 |
| Chd7 | 4568 | 3252 | 3003 | 3779 | 3520 | 4144 | 3548 | 3459 | 3240 | 3762 | 3971 | 4091 | 3678.4 | 3416 | 3941 | 1.153 | 0.206 | 0.00095816 | 0.00703753 | 0.0016 | 0.0031 | 0.0025 | 0.0025 | TRUE | 0.0244 |
| Chd8 | 9746 | 6665 | 5853 | 6837 | 5765 | 6748 | 7569 | 7089 | 6315 | 6806 | 6504 | 6661 | 6824.08 | 6991 | 6657 | 0.952 | -0.071 | 0.30760752 | 0.48024087 | 0.0041 | 0.0028 | 0.0034 | 0.0034 | TRUE | 0.0871 |
| Chd9 | 9642 | 6937 | 7164 | 10189 | 8962 | 10637 | 7489 | 7379 | 7730 | 10142 | 10110 | 10500 | 8891.63 | 7533 | 10251 | 1.361 | 0.445 | 1.64E-19 | 4.78E-17 | 3.00E-04 | 0.0027 | 0.0016 | 0.0016 | TRUE | 0.0063 |
| Chdh | 26 | 29 | 12 | 11 | 25 | 16 | 20 | 31 | 13 | 11 | 28 | 16 | 19.82 | 21 | 18 | 0.853 | -0.229 | 0.64672377 | 0.78033832 | 0.1447 | 0.122 | 0.1283 | 0.1283 | TRUE | 0.3796 |
| Chek1 | 4410 | 3305 | 3608 | 3720 | 3146 | 3716 | 3425 | 3515 | 3893 | 3703 | 3549 | 3668 | 3625.62 | 3611 | 3640 | 1.009 | 0.012 | 0.8493834 | 0.91479443 | 0.0023 | 0.0031 | 0.0028 | 0.0028 | TRUE | 0.0563 |
| Chek2 | 3673 | 2663 | 2822 | 2877 | 2696 | 3096 | 2853 | 2833 | 3045 | 2864 | 3041 | 3056 | 2948.59 | 2910 | 2987 | 1.027 | 0.038 | 0.53604559 | 0.69493874 | 9.00E-04 | 0.0032 | 0.0024 | 0.0024 | TRUE | 0.0197 |
| Cherp | 13752 | 10044 | 9060 | 8616 | 8104 | 9343 | 10681 | 10684 | 9776 | 8576 | 9142 | 9223 | 9680.24 | 10380 | 8980 | 0.865 | -0.209 | 0.00036388 | 0.00338159 | 0.0019 | 0.0027 | 0.0024 | 0.0024 | TRUE | 0.0316 |
| Chfr | 4628 | 3221 | 3204 | 3402 | 2900 | 3321 | 3594 | 3426 | 3457 | 3386 | 3272 | 3278 | 3402.3 | 3492 | 3312 | 0.948 | -0.077 | 0.16324291 | 0.31385942 | 3.00E-04 | 0.0031 | 0.0019 | 0.0019 | TRUE | 0.0081 |

| | | | | | | | | | | | | | | | | | | | | | | | | | |
|---|---|---|---|---|---|---|---|---|---|---|---|---|---|---|---|---|---|---|---|---|---|---|---|---|---|
| Chga | 874 | 668 | 704 | 969 | 869 | 989 | 679 | 711 | 760 | 965 | 980 | 976 | 845.02 | 717 | 974 | 1.361 | 0.445 | 4.82E-08 | 2.05E-06 | 0 | 0.0052 | 0.0036 | 0.0036 | TRUE | 0.0327 |
| Chgb | 15 | 13 | 11 | 8 | 8 | 18 | 12 | 14 | 12 | 8 | 9 | 18 | 12.02 | 13 | 12 | 0.936 | -0.095 | 0.86771327 | NA | 0 | 0.1996 | 0.15 | 0.15 | TRUE | 0.8659 |
| Chi3l1 | 41 | 36 | 29 | 60 | 46 | 68 | 32 | 38 | 31 | 60 | 52 | 67 | 46.69 | 34 | 60 | 1.77 | 0.824 | 0.00436161 | 0.02211637 | 0 | 0.0532 | 0.0372 | 0.0372 | TRUE | 0.1061 |
| Chi3l3 | 2 | 2 | 3 | 0 | 2 | 1 | 2 | 2 | 3 | 0 | 2 | 1 | 1.69 | 2 | 1 | 0.463 | -1.111 | 0.46744084 | NA | 0 | 1.4019 | 0.9816 | 0.9816 | TRUE | 0.3789 |
| Chi3l4 | 1 | 0 | 1 | 0 | 0 | 0 | 1 | 0 | 1 | 0 | 0 | 0 | 0.31 | 1 | 0 | 0.307 | -1.706 | 0.62894649 | NA | 0 | 7.6657 | 7.2 | 7.2 | TRUE | 0.2067 |
| Chia | 0 | 0 | 0 | 0 | 0 | 0 | 0 | 0 | 0 | 0 | 0 | 0 | 0 | 0 | 0 | NA | NA | NA | NA | NA | NA | NA | NA | NA | NA |
| Chic1 | 1389 | 983 | 1100 | 1145 | 997 | 1146 | 1079 | 1046 | 1187 | 1140 | 1125 | 1131 | 1117.83 | 1104 | 1132 | 1.026 | 0.037 | 0.62943173 | 0.76676619 | 0.0012 | 0.0045 | 0.0034 | 0.0034 | TRUE | 0.0504 |
| Chic2 | 1562 | 1106 | 968 | 1095 | 948 | 1129 | 1213 | 1176 | 1044 | 1090 | 1069 | 1114 | 1117.99 | 1144 | 1091 | 0.952 | -0.07 | 0.38380269 | 0.55782915 | 0.0024 | 0.0045 | 0.0038 | 0.0038 | TRUE | 0.0691 |
| Chid1 | 927 | 677 | 610 | 869 | 805 | 803 | 720 | 720 | 658 | 865 | 908 | 793 | 777.35 | 699 | 855 | 1.221 | 0.288 | 0.00123403 | 0.00841908 | 0.0024 | 0.0055 | 0.0044 | 0.0044 | TRUE | 0.049 |
| Chit1 | 24 | 24 | 23 | 22 | 29 | 32 | 19 | 26 | 25 | 22 | 33 | 32 | 25.86 | 23 | 29 | 1.257 | 0.33 | 0.39421906 | 0.56782959 | 0 | 0.0941 | 0.0687 | 0.0687 | TRUE | 0.2883 |
| Chka | 2383 | 1911 | 1960 | 1816 | 1717 | 2247 | 1851 | 2033 | 2115 | 1808 | 1937 | 2218 | 1993.51 | 2000 | 1988 | 0.995 | -0.007 | 0.93208965 | 0.96225044 | 0.0073 | 0.0036 | 0.0049 | 0.0049 | TRUE | 0.1254 |
| Chkb | 17 | 15 | 12 | 14 | 17 | 14 | 13 | 16 | 13 | 14 | 19 | 14 | 14.84 | 14 | 16 | 1.113 | 0.154 | 0.75406419 | NA | 0 | 0.1621 | 0.1074 | 0.1074 | TRUE | 0.1658 |
| Chl1 | 18 | 2 | 5 | 4 | 4 | 5 | 14 | 2 | 5 | 4 | 5 | 5 | 5.82 | 7 | 5 | 0.609 | -0.716 | 0.41241765 | NA | 0.2795 | 0.4095 | 0.3681 | 0.3681 | TRUE | 0.5895 |
| Chm | 2118 | 1552 | 1435 | 1711 | 1373 | 1608 | 1645 | 1651 | 1548 | 1703 | 1549 | 1587 | 1613.92 | 1615 | 1613 | 0.999 | -0.002 | 0.98268051 | 0.99089635 | 0.0012 | 0.0039 | 0.0029 | 0.0029 | TRUE | 0.0288 |
| Chml | 1427 | 1116 | 1152 | 1130 | 1001 | 1193 | 1108 | 1187 | 1243 | 1125 | 1129 | 1178 | 1161.68 | 1179 | 1144 | 0.971 | -0.042 | 0.57955821 | 0.73041403 | 0.0013 | 0.0045 | 0.0033 | 0.0033 | TRUE | 0.035 |
| Chmp1a | 8245 | 5903 | 5427 | 5259 | 4792 | 5438 | 6404 | 6279 | 5856 | 5235 | 5406 | 5368 | 5757.83 | 6180 | 5336 | 0.863 | -0.212 | 0.00015115 | 0.00172006 | 0.001 | 0.0028 | 0.0021 | 0.0021 | TRUE | 0.0255 |
| Chmp1b | 1506 | 1048 | 945 | 979 | 857 | 986 | 1170 | 1115 | 1020 | 974 | 967 | 973 | 1036.44 | 1102 | 971 | 0.881 | -0.183 | 0.02193679 | 0.07380025 | 0.0016 | 0.0047 | 0.0036 | 0.0036 | TRUE | 0.0498 |
| Chmp2a | 4472 | 3378 | 3886 | 4408 | 3936 | 4257 | 3473 | 3593 | 4193 | 4388 | 4440 | 4202 | 4048.26 | 3753 | 4343 | 1.158 | 0.211 | 0.00503656 | 0.02467865 | 0.0053 | 0.003 | 0.0038 | 0.0038 | TRUE | 0.1273 |
| Chmp2b | 2043 | 1395 | 1456 | 1456 | 1198 | 1389 | 1587 | 1484 | 1571 | 1449 | 1351 | 1371 | 1468.92 | 1547 | 1390 | 0.899 | -0.154 | 0.02624313 | 0.08423379 | 6.00E-04 | 0.004 | 0.0028 | 0.0028 | TRUE | 0.0165 |
| Chmp3 | 6005 | 4122 | 4104 | 4532 | 3911 | 4455 | 4664 | 4385 | 4428 | 4511 | 4412 | 4398 | 4466.24 | 4492 | 4440 | 0.988 | -0.017 | 0.74838658 | 0.85076667 | 5.00E-04 | 0.003 | 0.0019 | 0.0019 | TRUE | 0.0139 |
| Chmp4b | 6015 | 4458 | 4164 | 4068 | 3592 | 4179 | 4672 | 4742 | 4493 | 4049 | 4052 | 4125 | 4355.53 | 4636 | 4075 | 0.879 | -0.186 | 0.00042014 | 0.00377674 | 2.00E-04 | 0.003 | 0.0018 | 0.0018 | TRUE | 0.0087 |
| Chmp4c | 1371 | 1103 | 1457 | 1225 | 1157 | 1326 | 1065 | 1173 | 1572 | 1219 | 1305 | 1309 | 1273.95 | 1270 | 1278 | 1.007 | 0.011 | 0.92925603 | 0.9609898 | 0.0211 | 0.0043 | 0.0095 | 0.0095 | TRUE | 0.5177 |
| Chmp5 | 3568 | 2531 | 2616 | 2836 | 2390 | 2807 | 2771 | 2692 | 2823 | 2823 | 2696 | 2771 | 2762.67 | 2762 | 2763 | 1.001 | 0.001 | 0.98741673 | 0.99373352 | 1.00E-04 | 0.0033 | 0.002 | 0.002 | TRUE | 0.0058 |
| Chmp6 | 2883 | 2035 | 1914 | 2017 | 1760 | 1948 | 2239 | 2165 | 2065 | 2008 | 1985 | 1923 | 2064.18 | 2156 | 1972 | 0.914 | -0.13 | 0.04089647 | 0.11630189 | 7.00E-04 | 0.0036 | 0.0024 | 0.0024 | TRUE | 0.0164 |
| Chmp7 | 4234 | 3039 | 2923 | 3111 | 2536 | 2924 | 3288 | 3233 | 3154 | 3097 | 2861 | 2886 | 3086.47 | 3225 | 2948 | 0.914 | -0.13 | 0.02951988 | 0.09154428 | 7.00E-04 | 0.0032 | 0.0022 | 0.0022 | TRUE | 0.0237 |
| Chn1 | 1605 | 1272 | 1331 | 1412 | 1190 | 1320 | 1247 | 1353 | 1436 | 1406 | 1342 | 1303 | 1347.78 | 1345 | 1350 | 1.005 | 0.007 | 0.92464167 | 0.95836863 | 0.0026 | 0.0042 | 0.0036 | 0.0036 | TRUE | 0.0515 |
| Chn2 | 210 | 112 | 119 | 158 | 140 | 145 | 163 | 119 | 128 | 157 | 158 | 143 | 144.83 | 137 | 153 | 1.109 | 0.149 | 0.40291063 | 0.57589577 | 0.0095 | 0.0188 | 0.016 | 0.016 | TRUE | 0.3171 |
| Chodl | 0 | 1 | 0 | 1 | 0 | 0 | 0 | 1 | 0 | 1 | 0 | 0 | 0.34 | 0 | 1 | 1.075 | 0.105 | 0.97573409 | NA | 0 | 6.9086 | 6.5582 | 6.5582 | TRUE | 0.5216 |
| Chordc1 | 4552 | 3253 | 2791 | 2641 | 2201 | 2631 | 3535 | 3460 | 3011 | 2629 | 2483 | 2597 | 2952.67 | 3335 | 2570 | 0.77 | -0.377 | 2.29E-07 | 8.23E-06 | 0.0035 | 0.0032 | 0.0035 | 0.0035 | TRUE | 0.0872 |
| Chp1 | 4202 | 2766 | 3014 | 3450 | 2907 | 3248 | 3264 | 2942 | 3252 | 3434 | 3279 | 3206 | 3229.59 | 3153 | 3306 | 1.049 | 0.068 | 0.29040208 | 0.4624239 | 0.0018 | 0.0032 | 0.0027 | 0.0027 | TRUE | 0.041 |
| Chp2 | 6 | 6 | 3 | 3 | 3 | 1 | 5 | 6 | 3 | 3 | 3 | 1 | 3.61 | 5 | 2 | 0.511 | -0.969 | 0.3466846 | NA | 0 | 0.6597 | 0.4492 | 0.4492 | TRUE | 0.3104 |
| Chpf | 2179 | 1603 | 1510 | 1803 | 1632 | 1896 | 1692 | 1705 | 1629 | 1795 | 1841 | 1872 | 1755.69 | 1675 | 1836 | 1.095 | 0.131 | 0.03736891 | 0.10904734 | 0 | 0.0038 | 0.0023 | 0.0023 | TRUE | 0.0068 |
| Chpf2 | 272 | 214 | 169 | 222 | 194 | 195 | 211 | 228 | 182 | 221 | 219 | 192 | 208.93 | 207 | 211 | 1.016 | 0.024 | 0.8732565 | 0.9297124 | 0.0038 | 0.0138 | 0.0109 | 0.0109 | TRUE | 0.1119 |
| Chpt1 | 1040 | 766 | 661 | 807 | 672 | 730 | 808 | 815 | 713 | 803 | 758 | 721 | 769.62 | 779 | 761 | 0.976 | -0.035 | 0.70073494 | 0.8185838 | 0.0029 | 0.0055 | 0.0046 | 0.0046 | TRUE | 0.0621 |
| Chrac1 | 1173 | 869 | 891 | 862 | 745 | 918 | 911 | 924 | 961 | 858 | 840 | 906 | 900.24 | 932 | 868 | 0.932 | -0.101 | 0.19723354 | 0.35697847 | 0 | 0.0051 | 0.0033 | 0.0033 | TRUE | 0.0176 |
| Chrd | 162 | 136 | 134 | 168 | 137 | 160 | 126 | 145 | 145 | 167 | 155 | 158 | 149.13 | 139 | 160 | 1.16 | 0.214 | 0.18893299 | 0.34647412 | 0 | 0.0183 | 0.0125 | 0.0125 | TRUE | 0.0731 |
| Chrdl1 | 62 | 54 | 30 | 40 | 30 | 34 | 48 | 57 | 32 | 40 | 34 | 34 | 40.86 | 46 | 36 | 0.776 | -0.365 | 0.25656559 | 0.42653739 | 0.0213 | 0.0604 | 0.0497 | 0.0497 | TRUE | 0.2975 |
| Chrdl2 | 17 | 15 | 11 | 11 | 16 | 10 | 13 | 16 | 12 | 11 | 18 | 10 | 13.32 | 14 | 13 | 0.942 | -0.086 | 0.87156089 | NA | 0 | 0.1804 | 0.1282 | 0.1282 | TRUE | 0.4447 |
| Chrm1 | 0 | 0 | 0 | 0 | 0 | 1 | 0 | 0 | 0 | 0 | 0 | 1 | 0.16 | 0 | 0 | 2.094 | 1.067 | 0.79379329 | NA | 0 | 14.4078 | 10 | 10 | TRUE | 0.3835 |
| Chrm2 | 0 | 0 | 0 | 0 | 0 | 0 | 0 | 0 | 0 | 0 | 0 | 0 | 0 | 0 | 0 | NA | NA | NA | NA | NA | NA | NA | NA | NA | NA |
| Chrm3 | 103 | 82 | 40 | 38 | 37 | 37 | 80 | 87 | 43 | 38 | 42 | 37 | 54.41 | 70 | 39 | 0.549 | -0.865 | 0.00519997 | 0.02529719 | 0.0554 | 0.046 | 0.0487 | 0.0487 | TRUE | 0.965 |
| Chrm4 | 508 | 392 | 382 | 431 | 432 | 486 | 395 | 417 | 412 | 429 | 487 | 480 | 436.63 | 408 | 465 | 1.141 | 0.191 | 0.06811418 | 0.16796046 | 0 | 0.0079 | 0.0056 | 0.0056 | TRUE | 0.0548 |
| Chrm5 | 0 | 1 | 0 | 0 | 0 | 1 | 0 | 1 | 0 | 0 | 0 | 1 | 0.34 | 0 | 0 | 1.075 | 0.105 | 0.97576789 | NA | 0 | 6.9363 | 6.5821 | 6.5821 | TRUE | 0.5216 |
| Chrna1 | 2 | 3 | 2 | 2 | 1 | 5 | 2 | 3 | 2 | 2 | 1 | 5 | 2.49 | 2 | 3 | 1.194 | 0.256 | 0.83259114 | NA | 0 | 0.9532 | 0.6527 | 0.6527 | TRUE | 0.6469 |
| Chrna10 | 13 | 10 | 7 | 11 | 9 | 9 | 10 | 11 | 8 | 11 | 10 | 9 | 9.71 | 10 | 10 | 1.056 | 0.079 | 0.89560043 | NA | 0 | 0.2465 | 0.1584 | 0.1584 | TRUE | 0.0904 |
| Chrna2 | 1 | 0 | 3 | 1 | 2 | 0 | 1 | 0 | 3 | 1 | 2 | 0 | 1.21 | 1 | 1 | 0.808 | -0.307 | 0.87018566 | NA | 0.7906 | 1.9597 | 1.6967 | 1.6967 | TRUE | 0.5419 |
| Chrna3 | 2 | 0 | 0 | 0 | 1 | 2 | 2 | 0 | 0 | 0 | 1 | 2 | 0.78 | 1 | 1 | 1.844 | 0.883 | 0.7091433 | NA | 0 | 3.0568 | 2.6753 | 2.6753 | TRUE | 0.5491 |
| Chrna4 | 193 | 152 | 165 | 253 | 191 | 220 | 150 | 162 | 178 | 252 | 215 | 217 | 195.68 | 163 | 228 | 1.403 | 0.489 | 0.00118286 | 0.00818977 | 0 | 0.0145 | 0.0111 | 0.0111 | TRUE | 0.0919 |
| Chrna5 | 139 | 110 | 93 | 141 | 121 | 106 | 108 | 117 | 100 | 140 | 137 | 105 | 117.8 | 108 | 127 | 1.171 | 0.228 | 0.23613786 | 0.40256661 | 0.0073 | 0.0225 | 0.0182 | 0.0182 | TRUE | 0.251 |
| Chrna6 | 1 | 0 | 0 | 1 | 0 | 0 | 1 | 0 | 0 | 1 | 0 | 0 | 0.3 | 0 | 0 | 1.075 | 0.105 | 0.97698525 | NA | 0 | 8.0272 | 7.5141 | 7.5141 | TRUE | 0.3906 |
| Chrna7 | 328 | 268 | 259 | 235 | 208 | 225 | 255 | 285 | 279 | 234 | 235 | 222 | 251.66 | 273 | 230 | 0.845 | -0.244 | 0.05872214 | 0.15109191 | 0 | 0.0118 | 0.008 | 0.008 | TRUE | 0.0423 |
| Chrna9 | 214 | 144 | 128 | 146 | 113 | 145 | 166 | 153 | 138 | 145 | 127 | 143 | 145.57 | 152 | 138 | 0.908 | -0.14 | 0.40467924 | 0.57736878 | 0 | 0.0187 | 0.0135 | 0.0135 | TRUE | 0.0712 |
| Chrnb1 | 137 | 135 | 121 | 210 | 148 | 191 | 106 | 144 | 131 | 209 | 167 | 189 | 157.52 | 127 | 188 | 1.493 | 0.579 | 0.0009973 | 0.00723207 | 0.0113 | 0.0175 | 0.0157 | 0.0157 | TRUE | 0.2284 |
| Chrnb2 | 40 | 34 | 26 | 58 | 37 | 41 | 31 | 36 | 28 | 58 | 42 | 40 | 39.21 | 32 | 47 | 1.472 | 0.557 | 0.08050419 | 0.18996968 | 0 | 0.0629 | 0.047 | 0.047 | TRUE | 0.3494 |
| Chrnb3 | 5 | 3 | 4 | 6 | 4 | 3 | 4 | 3 | 4 | 6 | 5 | 3 | 4.14 | 4 | 5 | 1.179 | 0.238 | 0.7960636 | NA | 0 | 0.575 | 0.3673 | 0.3673 | TRUE | 0.1716 |
| Chrnb4 | 332 | 227 | 265 | 347 | 281 | 322 | 258 | 241 | 286 | 345 | 317 | 318 | 294.25 | 262 | 327 | 1.25 | 0.321 | 0.00968856 | 0.04044352 | 0 | 0.0105 | 0.0077 | 0.0077 | TRUE | 0.0685 |
| Chrnd | 16 | 10 | 8 | 16 | 15 | 20 | 12 | 11 | 9 | 16 | 17 | 20 | 14.05 | 11 | 18 | 1.646 | 0.719 | 0.15858011 | NA | 0 | 0.1711 | 0.1128 | 0.1128 | TRUE | 0.1107 |
| Chrne | 10 | 8 | 11 | 11 | 16 | 8 | 8 | 9 | 12 | 11 | 18 | 8 | 10.84 | 10 | 12 | 1.311 | 0.391 | 0.51569265 | NA | 0 | 0.2211 | 0.1667 | 0.1667 | TRUE | 0.499 |
| Chrng | 15 | 7 | 9 | 4 | 12 | 7 | 12 | 7 | 10 | 4 | 14 | 7 | 8.87 | 10 | 8 | 0.833 | -0.263 | 0.6974549 | NA | 0.0654 | 0.2695 | 0.2156 | 0.2156 | TRUE | 0.6514 |
| Chst1 | 49 | 38 | 60 | 50 | 43 | 39 | 38 | 40 | 65 | 50 | 49 | 38 | 46.67 | 48 | 46 | 0.961 | -0.058 | 0.85150374 | 0.91638754 | 0.0344 | 0.0532 | 0.0478 | 0.0478 | TRUE | 0.7455 |
| Chst10 | 480 | 366 | 359 | 416 | 357 | 390 | 373 | 389 | 387 | 414 | 403 | 385 | 391.88 | 383 | 401 | 1.046 | 0.065 | 0.53395913 | 0.69315568 | 0 | 0.0085 | 0.0054 | 0.0054 | TRUE | 0.0138 |
| Chst11 | 1058 | 709 | 862 | 961 | 905 | 950 | 822 | 754 | 930 | 957 | 1021 | 938 | 903.54 | 835 | 972 | 1.163 | 0.218 | 0.01866491 | 0.06569256 | 0.005 | 0.005 | 0.0051 | 0.0051 | TRUE | 0.1131 |
| Chst12 | 790 | 552 | 646 | 639 | 631 | 720 | 614 | 587 | 697 | 636 | 712 | 711 | 659.4 | 633 | 686 | 1.085 | 0.118 | 0.23152229 | 0.39754301 | 0.0044 | 0.006 | 0.0055 | 0.0055 | TRUE | 0.0904 |
| Chst13 | 58 | 54 | 65 | 47 | 49 | 57 | 45 | 57 | 70 | 47 | 55 | 56 | 55.16 | 57 | 53 | 0.923 | -0.115 | 0.67684891 | 0.80190662 | 0.0147 | 0.0454 | 0.0365 | 0.0365 | TRUE | 0.194 |
| Chst14 | 236 | 174 | 206 | 209 | 163 | 188 | 183 | 185 | 222 | 208 | 184 | 186 | 194.69 | 197 | 193 | 0.981 | -0.028 | 0.85174024 | 0.91640026 | 0.0034 | 0.0146 | 0.0113 | 0.0113 | TRUE | 0.1333 |
| Chst15 | 120 | 87 | 71 | 81 | 64 | 71 | 93 | 93 | 77 | 81 | 72 | 70 | 80.88 | 88 | 74 | 0.848 | -0.238 | 0.27748 | 0.44874651 | 0 | 0.0317 | 0.0221 | 0.0221 | TRUE | 0.1058 |
| Chst2 | 294 | 192 | 173 | 210 | 186 | 165 | 228 | 204 | 187 | 209 | 210 | 163 | 200.16 | 206 | 194 | 0.936 | -0.096 | 0.54480063 | 0.70214926 | 0.0103 | 0.0143 | 0.0131 | 0.0131 | TRUE | 0.2163 |
| Chst3 | 147 | 118 | 117 | 121 | 108 | 139 | 114 | 126 | 126 | 120 | 122 | 137 | 124.24 | 122 | 126 | 1.04 | 0.057 | 0.74795476 | 0.85041106 | 0 | 0.0215 | 0.0145 | 0.0145 | TRUE | 0.0573 |
| Chst4 | 1 | 0 | 0 | 2 | 0 | 0 | 1 | 0 | 0 | 2 | 0 | 0 | 0.46 | 0 | 1 | 1.942 | 0.958 | 0.7580687 | NA | 0 | 5.1409 | 5.1921 | 5.1921 | TRUE | 0.9738 |

| | | | | | | | | | | | | | | | | | | | | | | | | | |
|---|---|---|---|---|---|---|---|---|---|---|---|---|---|---|---|---|---|---|---|---|---|---|---|---|---|
| Chst5 | 106 | 101 | 66 | 121 | 104 | 101 | 82 | 107 | 71 | 120 | 117 | 100 | 99.74 | 87 | 112 | 1.294 | 0.372 | 0.08311375 | 0.19410862 | 0.0145 | 0.0262 | 0.0231 | 0.0231 | TRUE | 0.3778 |
| Chst7 | 144 | 89 | 113 | 133 | 111 | 121 | 112 | 95 | 122 | 132 | 125 | 119 | 117.58 | 110 | 125 | 1.147 | 0.198 | 0.28579627 | 0.4577416 | 0 | 0.0226 | 0.0164 | 0.0164 | TRUE | 0.1298 |
| Chst8 | 87 | 52 | 56 | 56 | 53 | 57 | 68 | 55 | 60 | 56 | 60 | 56 | 59.18 | 61 | 57 | 0.933 | -0.1 | 0.68880725 | 0.80956516 | 0 | 0.0425 | 0.0282 | 0.0282 | TRUE | 0.0853 |
| Chst9 | 1 | 1 | 6 | 2 | 0 | 0 | 1 | 1 | 6 | 2 | 0 | 0 | 1.72 | 3 | 1 | 0.248 | -2.009 | 0.25989347 | NA | 1.2414 | 1.3824 | 1.3492 | 1.3492 | TRUE | 1.5534 |
| Chsy1 | 3428 | 2588 | 2670 | 3115 | 2723 | 3142 | 2662 | 2753 | 2881 | 3101 | 3072 | 3102 | 2928.37 | 2765 | 3092 | 1.119 | 0.162 | 0.00546545 | 0.02627572 | 5.00E-04 | 0.0032 | 0.0021 | 0.0021 | TRUE | 0.0161 |
| Chsy3 | 66 | 44 | 28 | 43 | 43 | 55 | 51 | 47 | 30 | 43 | 49 | 54 | 45.65 | 43 | 49 | 1.127 | 0.173 | 0.57097293 | 0.72425596 | 0.019 | 0.0543 | 0.0449 | 0.0449 | TRUE | 0.4765 |
| Chtf18 | 4766 | 3522 | 3133 | 3198 | 2925 | 3238 | 3702 | 3746 | 3380 | 3183 | 3300 | 3196 | 3417.96 | 3609 | 3226 | 0.894 | -0.162 | 0.00847858 | 0.03660754 | 0.0013 | 0.0031 | 0.0024 | 0.0024 | TRUE | 0.0369 |
| Chtf8 | 10410 | 7600 | 7546 | 7435 | 6819 | 7953 | 8085 | 8084 | 8142 | 7401 | 7693 | 7851 | 7875.88 | 8104 | 7648 | 0.944 | -0.083 | 0.09060445 | 0.20562233 | 3.00E-04 | 0.0027 | 0.0016 | 0.0016 | TRUE | 0.0098 |
| Chtop | 10420 | 7632 | 7152 | 7221 | 6640 | 7393 | 8093 | 8118 | 7717 | 7188 | 7491 | 7298 | 7650.7 | 7976 | 7326 | 0.918 | -0.123 | 0.01516161 | 0.05624316 | 4.00E-04 | 0.0027 | 0.0017 | 0.0017 | TRUE | 0.0098 |
| Chuk | 5048 | 3493 | 3031 | 3113 | 2678 | 2989 | 3921 | 3715 | 3270 | 3099 | 3021 | 2951 | 3329.47 | 3635 | 3024 | 0.831 | -0.267 | 0.00028492 | 0.00279307 | 0.004 | 0.0031 | 0.0036 | 0.0036 | TRUE | 0.0934 |
| Churc1 | 1749 | 1214 | 1203 | 1452 | 1189 | 1395 | 1358 | 1291 | 1298 | 1445 | 1341 | 1377 | 1351.91 | 1316 | 1388 | 1.054 | 0.076 | 0.27564766 | 0.44669297 | 4.00E-04 | 0.0042 | 0.0028 | 0.0028 | TRUE | 0.0158 |
| Ciao1 | 2895 | 2009 | 1954 | 2023 | 1740 | 2053 | 2248 | 2137 | 2108 | 2014 | 1963 | 2027 | 2082.82 | 2164 | 2001 | 0.924 | -0.114 | 0.06496836 | 0.16259723 | 4.00E-04 | 0.0036 | 0.0023 | 0.0023 | TRUE | 0.0144 |
| Ciapin1 | 12304 | 8626 | 7543 | 7868 | 7289 | 8393 | 9556 | 9175 | 8139 | 7832 | 8223 | 8285 | 8534.99 | 8957 | 8113 | 0.906 | -0.143 | 0.03319273 | 0.09981562 | 0.0036 | 0.0027 | 0.0031 | 0.0031 | TRUE | 0.0777 |
| Cib1 | 1945 | 1384 | 1236 | 1443 | 1204 | 1363 | 1511 | 1472 | 1334 | 1436 | 1358 | 1345 | 1409.41 | 1439 | 1380 | 0.958 | -0.061 | 0.41540608 | 0.5880942 | 0.0021 | 0.0041 | 0.0034 | 0.0034 | TRUE | 0.0485 |
| Cib2 | 182 | 130 | 147 | 192 | 179 | 235 | 141 | 138 | 159 | 191 | 202 | 232 | 177.21 | 146 | 208 | 1.429 | 0.515 | 0.0010413 | 0.00745281 | 0 | 0.0158 | 0.012 | 0.012 | TRUE | 0.1099 |
| Cib3 | 2 | 1 | 0 | 0 | 3 | 0 | 2 | 1 | 0 | 0 | 3 | 0 | 1 | 1 | 1 | 1.229 | 0.297 | 0.89095244 | NA | 2.3937 | 2.372 | 2.3763 | 2.3763 | TRUE | 1.4437 |
| Cib4 | 0 | 0 | 0 | 0 | 0 | 2 | 0 | 0 | 0 | 0 | 0 | 2 | 0.33 | 0 | 1 | 3.743 | 1.904 | 0.62640105 | NA | 0 | 7.2051 | 9.2465 | 9.2465 | TRUE | 0.9739 |
| Cic | 9547 | 7287 | 7046 | 6994 | 6341 | 7391 | 7415 | 7751 | 7603 | 6962 | 7153 | 7296 | 7363.27 | 7590 | 7137 | 0.941 | -0.088 | 0.07797019 | 0.18558628 | 3.00E-04 | 0.0027 | 0.0017 | 0.0017 | TRUE | 0.0057 |
| Cidea | 206 | 89 | 77 | 97 | 78 | 78 | 160 | 95 | 83 | 97 | 88 | 77 | 99.88 | 113 | 87 | 0.768 | -0.381 | 0.13743133 | 0.27771893 | 0.0623 | 0.0262 | 0.0372 | 0.0372 | TRUE | 1.4504 |
| Cideb | 22 | 16 | 29 | 29 | 33 | 39 | 17 | 17 | 31 | 29 | 37 | 38 | 28.33 | 22 | 35 | 1.613 | 0.69 | 0.07603815 | 0.18189343 | 0.0303 | 0.0861 | 0.0718 | 0.0718 | TRUE | 0.7703 |
| Cidec | 40 | 25 | 18 | 26 | 7 | 15 | 31 | 27 | 19 | 26 | 8 | 15 | 20.94 | 26 | 16 | 0.632 | -0.663 | 0.16673081 | 0.31778393 | 0.114 | 0.1156 | 0.1152 | 0.1152 | TRUE | 0.5628 |
| Ciita | 5 | 2 | 6 | 11 | 3 | 5 | 4 | 2 | 6 | 11 | 3 | 5 | 5.29 | 4 | 6 | 1.559 | 0.64 | 0.46563698 | NA | 0.1326 | 0.4503 | 0.3584 | 0.3584 | TRUE | 0.9552 |
| Cilp | 23 | 17 | 8 | 12 | 13 | 4 | 18 | 18 | 9 | 12 | 15 | 4 | 12.52 | 15 | 10 | 0.677 | -0.564 | 0.35364191 | NA | 0.1555 | 0.1917 | 0.1826 | 0.1826 | TRUE | 1.0474 |
| Cilp2 | 238 | 147 | 129 | 174 | 141 | 164 | 185 | 156 | 139 | 173 | 159 | 162 | 162.43 | 160 | 165 | 1.024 | 0.034 | 0.83699969 | 0.90706168 | 0.0061 | 0.017 | 0.0138 | 0.0138 | TRUE | 0.2096 |
| Cinp | 3676 | 2608 | 2436 | 2728 | 2407 | 2802 | 2855 | 2774 | 2628 | 2715 | 2715 | 2766 | 2742.39 | 2752 | 2732 | 0.992 | -0.011 | 0.84857404 | 0.91423444 | 6.00E-04 | 0.0033 | 0.0022 | 0.0022 | TRUE | 0.0187 |
| Cir1 | 720 | 594 | 576 | 529 | 484 | 559 | 559 | 632 | 621 | 527 | 546 | 552 | 572.82 | 604 | 542 | 0.898 | -0.155 | 0.10190291 | 0.22414563 | 0 | 0.0066 | 0.0047 | 0.0047 | TRUE | 0.0532 |
| Cirbp | 2876 | 2304 | 1953 | 2286 | 1993 | 2096 | 2234 | 2451 | 2107 | 2275 | 2248 | 2069 | 2230.76 | 2264 | 2197 | 0.971 | -0.043 | 0.56651512 | 0.72064797 | 0.0038 | 0.0035 | 0.0036 | 0.0036 | TRUE | 0.062 |
| Cirh1a | 16758 | 11802 | 10287 | 10248 | 9175 | 10586 | 13015 | 12554 | 11100 | 10201 | 10350 | 10450 | 11278.28 | 12223 | 10334 | 0.845 | -0.242 | 0.00019236 | 0.00207325 | 0.0033 | 0.0026 | 0.003 | 0.003 | TRUE | 0.0788 |
| Cisd1 | 3058 | 2103 | 2292 | 2589 | 2173 | 2595 | 2375 | 2237 | 2473 | 2577 | 2451 | 2562 | 2445.85 | 2362 | 2530 | 1.071 | 0.1 | 0.11890029 | 0.25043274 | 0.001 | 0.0034 | 0.0025 | 0.0025 | TRUE | 0.0252 |
| Cisd2 | 1137 | 735 | 689 | 719 | 649 | 702 | 883 | 782 | 743 | 716 | 732 | 693 | 758.18 | 803 | 714 | 0.887 | -0.173 | 0.06032838 | 0.1538675 | 0.0034 | 0.0056 | 0.0048 | 0.0048 | TRUE | 0.0949 |
| Cisd3 | 1831 | 1263 | 1283 | 1343 | 1247 | 1626 | 1422 | 1343 | 1384 | 1337 | 1407 | 1605 | 1416.42 | 1383 | 1450 | 1.048 | 0.067 | 0.41294498 | 0.58582854 | 0.004 | 0.0041 | 0.0042 | 0.0042 | TRUE | 0.1071 |
| Cish | 211 | 131 | 182 | 213 | 163 | 226 | 164 | 139 | 196 | 212 | 184 | 223 | 186.43 | 166 | 206 | 1.241 | 0.312 | 0.06046622 | 0.15408161 | 0.0128 | 0.0151 | 0.0145 | 0.0145 | TRUE | 0.2453 |
| Cit | 2974 | 2238 | 2228 | 2236 | 2192 | 2479 | 2310 | 2381 | 2404 | 2226 | 2473 | 2447 | 2373.33 | 2365 | 2382 | 1.007 | 0.011 | 0.87164977 | 0.92876771 | 0.0014 | 0.0034 | 0.0027 | 0.0027 | TRUE | 0.0401 |
| Cited1 | 130 | 98 | 95 | 156 | 118 | 124 | 101 | 104 | 103 | 155 | 133 | 122 | 119.75 | 103 | 137 | 1.336 | 0.418 | 0.02334643 | 0.07740444 | 0 | 0.0222 | 0.016 | 0.016 | TRUE | 0.1454 |
| Cited2 | 1426 | 1108 | 843 | 729 | 690 | 794 | 1108 | 1179 | 910 | 726 | 778 | 784 | 913.92 | 1066 | 763 | 0.715 | -0.483 | 1.95E-06 | 5.01E-05 | 0.0089 | 0.005 | 0.0063 | 0.0063 | TRUE | 0.1911 |
| Cited4 | 126 | 106 | 129 | 106 | 116 | 163 | 98 | 113 | 139 | 106 | 131 | 161 | 124.51 | 117 | 133 | 1.141 | 0.19 | 0.36732996 | 0.54210089 | 0.0304 | 0.0215 | 0.0241 | 0.0241 | TRUE | 0.2108 |
| Ciz1 | 1002 | 842 | 884 | 758 | 733 | 843 | 778 | 896 | 954 | 755 | 827 | 832 | 840.21 | 876 | 805 | 0.92 | -0.12 | 0.21172225 | 0.37392144 | 0.0059 | 0.0052 | 0.0055 | 0.0055 | TRUE | 0.1184 |
| CK137956 | 1 | 0 | 0 | 1 | 3 | 1 | 1 | 0 | 0 | 1 | 3 | 1 | 1.02 | 0 | 2 | 5.137 | 2.361 | 0.24864342 | NA | 0 | 2.3171 | 1.7262 | 1.7262 | TRUE | 0.4395 |
| Ckap2 | 9572 | 6968 | 7126 | 7479 | 6453 | 7433 | 7434 | 7412 | 7689 | 7445 | 7280 | 7338 | 7432.77 | 7512 | 7354 | 0.979 | -0.03 | 0.52761264 | 0.6886911 | 1.00E-04 | 0.0027 | 0.0015 | 0.0015 | TRUE | 0.0052 |
| Ckap2l | 4398 | 3161 | 3369 | 3592 | 3163 | 3604 | 3416 | 3362 | 3635 | 3575 | 3568 | 3558 | 3519.09 | 3471 | 3567 | 1.028 | 0.04 | 0.48438629 | 0.65044468 | 4.00E-04 | 0.0031 | 0.002 | 0.002 | TRUE | 0.0204 |
| Ckap4 | 5553 | 4202 | 4023 | 4972 | 4469 | 4769 | 4313 | 4470 | 4341 | 4949 | 5041 | 4708 | 4636.92 | 4375 | 4899 | 1.12 | 0.164 | 0.00271047 | 0.01522827 | 5.00E-04 | 0.0029 | 0.0019 | 0.0019 | TRUE | 0.0143 |
| Ckap5 | 17984 | 13377 | 14019 | 15751 | 13718 | 15862 | 13967 | 14229 | 15126 | 15679 | 15475 | 15658 | 15022.47 | 14441 | 15604 | 1.081 | 0.112 | 0.02703537 | 0.08588944 | 8.00E-04 | 0.0026 | 0.0018 | 0.0018 | TRUE | 0.021 |
| Ckb | 8363 | 6445 | 5864 | 5357 | 5030 | 5791 | 6495 | 6855 | 6327 | 5332 | 5674 | 5717 | 6066.86 | 6559 | 5574 | 0.85 | -0.235 | 4.41E-05 | 0.0006306 | 0.0013 | 0.0028 | 0.0022 | 0.0022 | TRUE | 0.0189 |
| Cklf | 471 | 293 | 246 | 320 | 273 | 302 | 366 | 312 | 265 | 319 | 308 | 298 | 311.25 | 314 | 308 | 0.977 | -0.034 | 0.80318026 | 0.88613128 | 0.0106 | 0.01 | 0.0102 | 0.0102 | TRUE | 0.2363 |
| Ckm | 8 | 4 | 6 | 16 | 8 | 8 | 6 | 4 | 6 | 16 | 9 | 8 | 8.3 | 5 | 11 | 1.934 | 0.952 | 0.16501068 | NA | 0 | 0.288 | 0.2066 | 0.2066 | TRUE | 0.5994 |
| Ckmt1 | 505 | 417 | 535 | 693 | 597 | 752 | 392 | 444 | 577 | 690 | 673 | 742 | 586.44 | 471 | 702 | 1.495 | 0.58 | 6.76E-06 | 0.00013649 | 0.0192 | 0.0065 | 0.0102 | 0.0102 | TRUE | 0.449 |
| Ckmt2 | 0 | 0 | 2 | 1 | 1 | 0 | 0 | 0 | 2 | 1 | 1 | 0 | 0.71 | 1 | 1 | 1.011 | 0.016 | 0.99481353 | NA | 0 | 3.3238 | 2.9707 | 2.9707 | TRUE | 0.9749 |
| Cks1b | 4511 | 3067 | 3108 | 3239 | 2814 | 3308 | 3504 | 3262 | 3354 | 3224 | 3174 | 3266 | 3297.24 | 3373 | 3221 | 0.955 | -0.067 | 0.23913925 | 0.40618151 | 5.00E-04 | 0.0031 | 0.002 | 0.002 | TRUE | 0.0143 |
| Cks2 | 4247 | 3016 | 3331 | 3144 | 2785 | 3250 | 3298 | 3208 | 3594 | 3130 | 3142 | 3208 | 3263.37 | 3367 | 3160 | 0.939 | -0.091 | 0.14468421 | 0.28817885 | 0.0014 | 0.0032 | 0.0025 | 0.0025 | TRUE | 0.0417 |
| Clasp1 | 4078 | 2757 | 2400 | 2354 | 2092 | 2393 | 3167 | 2933 | 2590 | 2343 | 2360 | 2362 | 2625.8 | 2897 | 2355 | 0.812 | -0.3 | 0.00010101 | 0.00122245 | 0.0045 | 0.0033 | 0.0039 | 0.0039 | TRUE | 0.1038 |
| Clasp2 | 7481 | 5579 | 5739 | 6479 | 5761 | 6674 | 5810 | 5934 | 6192 | 6449 | 6499 | 6588 | 6245.55 | 5979 | 6512 | 1.089 | 0.124 | 0.01614739 | 0.05902551 | 3.00E-04 | 0.0028 | 0.0017 | 0.0017 | TRUE | 0.0118 |
| Clasrp | 2155 | 1904 | 1753 | 1636 | 1552 | 1866 | 1674 | 2025 | 1891 | 1628 | 1751 | 1842 | 1801.96 | 1863 | 1740 | 0.935 | -0.097 | 0.24997214 | 0.41880809 | 0.0056 | 0.0037 | 0.0046 | 0.0046 | TRUE | 0.0985 |
| Clca1 | 6 | 8 | 3 | 3 | 1 | 6 | 5 | 9 | 3 | 3 | 1 | 6 | 4.41 | 6 | 3 | 0.622 | -0.685 | 0.47488997 | NA | 0.0954 | 0.5402 | 0.4214 | 0.4214 | TRUE | 0.5116 |
| Clca2 | 3 | 0 | 3 | 2 | 0 | 3 | 2 | 0 | 3 | 2 | 0 | 3 | 1.75 | 2 | 2 | 0.897 | -0.157 | 0.92023231 | NA | 0 | 1.3543 | 1.2042 | 1.2042 | TRUE | 0.5664 |
| Clca3 | 53 | 50 | 54 | 38 | 31 | 36 | 41 | 53 | 58 | 38 | 35 | 36 | 43.49 | 51 | 36 | 0.715 | -0.484 | 0.10576819 | 0.23060527 | 0 | 0.0569 | 0.0407 | 0.0407 | TRUE | 0.2664 |
| Clca4 | 282 | 262 | 228 | 108 | 122 | 137 | 219 | 279 | 246 | 108 | 138 | 135 | 187.35 | 248 | 127 | 0.512 | -0.965 | 6.05E-09 | 3.30E-07 | 0.0107 | 0.0151 | 0.0138 | 0.0138 | TRUE | 0.184 |
| Clca5 | 1 | 1 | 1 | 0 | 2 | 1 | 1 | 1 | 1 | 0 | 2 | 1 | 1.03 | 1 | 1 | 1.1 | 0.138 | 0.94104339 | NA | 0 | 2.3099 | 1.5261 | 1.5261 | TRUE | 0.2247 |
| Clca6 | 0 | 0 | 1 | 0 | 0 | 0 | 0 | 0 | 1 | 0 | 0 | 0 | 0.18 | 0 | 0 | 0.552 | -0.857 | 0.8336548 | NA | 0 | 13.1817 | 10 | 10 | TRUE | 0.5365 |
| Clcc1 | 1292 | 939 | 1164 | 1229 | 1110 | 1304 | 1003 | 999 | 1256 | 1223 | 1252 | 1287 | 1170.16 | 1086 | 1254 | 1.156 | 0.209 | 0.02779405 | 0.08756014 | 0.0082 | 0.0045 | 0.0057 | 0.0057 | TRUE | 0.2209 |
| Clcf1 | 5 | 7 | 9 | 12 | 5 | 12 | 4 | 7 | 10 | 12 | 6 | 12 | 8.41 | 7 | 10 | 1.434 | 0.52 | 0.45131817 | NA | 0 | 0.2842 | 0.2207 | 0.2207 | TRUE | 0.5483 |
| Clcn1 | 71 | 77 | 66 | 81 | 57 | 67 | 55 | 82 | 71 | 81 | 64 | 66 | 69.89 | 69 | 70 | 1.021 | 0.031 | 0.90188971 | 0.94570564 | 0.0151 | 0.0363 | 0.0302 | 0.0302 | TRUE | 0.3348 |
| Clcn2 | 1171 | 1048 | 1014 | 1022 | 962 | 1127 | 909 | 1115 | 1094 | 1017 | 1085 | 1113 | 1055.56 | 1039 | 1072 | 1.033 | 0.047 | 0.61521099 | 0.75699128 | 0.0065 | 0.0047 | 0.0053 | 0.0053 | TRUE | 0.1489 |
| Clcn3 | 12179 | 9085 | 9234 | 10204 | 8551 | 9864 | 9459 | 9664 | 9963 | 10157 | 9646 | 9737 | 9771.11 | 9695 | 9847 | 1.016 | 0.023 | 0.65356577 | 0.78486099 | 5.00E-04 | 0.0027 | 0.0017 | 0.0017 | TRUE | 0.0093 |
| Clcn4-2 | 1025 | 686 | 573 | 595 | 505 | 641 | 796 | 730 | 618 | 592 | 570 | 633 | 656.46 | 715 | 598 | 0.836 | -0.259 | 0.01587767 | 0.05824938 | 0.0082 | 0.006 | 0.0068 | 0.0068 | TRUE | 0.1626 |
| Clcn5 | 1257 | 863 | 738 | 701 | 586 | 654 | 976 | 918 | 796 | 698 | 661 | 646 | 782.49 | 897 | 668 | 0.744 | -0.427 | 9.29E-06 | 0.0001772 | 0.0052 | 0.0055 | 0.0054 | 0.0054 | TRUE | 0.113 |
| Clcn6 | 2145 | 1847 | 1782 | 1942 | 1854 | 2056 | 1666 | 1965 | 1923 | 1933 | 2092 | 2030 | 1934.58 | 1851 | 2018 | 1.091 | 0.126 | 0.10677411 | 0.23207457 | 0.0038 | 0.0037 | 0.0039 | 0.0039 | TRUE | 0.0954 |
| Clcn7 | 1692 | 1350 | 1244 | 1250 | 1142 | 1361 | 1314 | 1436 | 1342 | 1244 | 1288 | 1344 | 1328.07 | 1364 | 1292 | 0.948 | -0.077 | 0.28925176 | 0.46130681 | 0.0011 | 0.0042 | 0.0031 | 0.0031 | TRUE | 0.0253 |
| Clcnka | 257 | 207 | 172 | 178 | 196 | 204 | 200 | 220 | 186 | 177 | 221 | 201 | 200.84 | 202 | 200 | 0.989 | -0.015 | 0.91888975 | 0.95538599 | 0.0042 | 0.0142 | 0.0114 | 0.0114 | TRUE | 0.1079 |
| Clcnkb | 161 | 143 | 143 | 134 | 101 | 125 | 125 | 152 | 154 | 133 | 114 | 123 | 133.69 | 144 | 123 | 0.864 | -0.211 | 0.23591687 | 0.40240602 | 0 | 0.0202 | 0.0153 | 0.0153 | TRUE | 0.1509 |

| | | | | | | | | | | | | | | | | | | | | | | | | | |
|---|---|---|---|---|---|---|---|---|---|---|---|---|---|---|---|---|---|---|---|---|---|---|---|---|---|
| Cldn1 | 37 | 18 | 17 | 11 | 6 | 12 | 29 | 19 | 18 | 11 | 7 | 12 | 15.96 | 22 | 10 | 0.445 | -1.167 | 0.02196043 | NA | 0 | 0.1509 | 0.113 | 0.113 | TRUE | 0.474 |
| Cldn10 | 27 | 20 | 25 | 11 | 14 | 14 | 21 | 21 | 27 | 11 | 16 | 14 | 18.3 | 23 | 14 | 0.586 | -0.771 | 0.08963845 | 0.20415483 | 0 | 0.132 | 0.0893 | 0.0893 | TRUE | 0.1201 |
| Cldn11 | 29 | 28 | 32 | 28 | 41 | 33 | 23 | 30 | 35 | 28 | 46 | 33 | 32.26 | 29 | 36 | 1.234 | 0.303 | 0.40096775 | 0.57407915 | 0.0284 | 0.0759 | 0.0626 | 0.0626 | TRUE | 0.4617 |
| Cldn12 | 2675 | 1848 | 1791 | 1998 | 1620 | 1850 | 2078 | 1966 | 1932 | 1989 | 1828 | 1826 | 1936.38 | 1992 | 1881 | 0.944 | -0.083 | 0.21945672 | 0.3825944 | 0.0015 | 0.0037 | 0.0028 | 0.0028 | TRUE | 0.0307 |
| Cldn13 | 0 | 0 | 0 | 1 | 0 | 0 | 0 | 0 | 0 | 1 | 0 | 0 | 0.17 | 0 | 0 | 2.094 | 1.067 | 0.79379329 | NA | 0 | 14.2885 | 10 | 10 | TRUE | 0.3906 |
| Cldn14 | 4 | 3 | 5 | 3 | 5 | 4 | 3 | 3 | 5 | 3 | 6 | 4 | 4.04 | 4 | 4 | 1.079 | 0.11 | 0.90664001 | NA | 0 | 0.5884 | 0.3817 | 0.3817 | TRUE | 0.169 |
| Cldn15 | 9 | 7 | 5 | 18 | 14 | 14 | 7 | 7 | 5 | 18 | 16 | 14 | 11.23 | 6 | 16 | 2.388 | 1.256 | 0.03129512 | NA | 0 | 0.2135 | 0.14 | 0.14 | TRUE | 0.0641 |
| Cldn16 | 3 | 3 | 3 | 2 | 1 | 0 | 2 | 3 | 3 | 2 | 1 | 0 | 1.98 | 3 | 1 | 0.357 | -1.486 | 0.30148917 | NA | 0 | 1.1997 | 0.8241 | 0.8241 | TRUE | 0.3977 |
| Cldn17 | 0 | 0 | 0 | 0 | 0 | 0 | 0 | 0 | 0 | 0 | 0 | 0 | 0 | 0 | 0 | NA | NA | NA | NA | NA | NA | NA | NA | NA | NA |
| Cldn18 | 25 | 20 | 15 | 27 | 16 | 10 | 19 | 21 | 16 | 27 | 18 | 10 | 18.61 | 19 | 18 | 0.962 | -0.055 | 0.90809084 | 0.94933751 | 0.0666 | 0.1298 | 0.1127 | 0.1127 | TRUE | 0.2171 |
| Cldn19 | 31 | 21 | 24 | 48 | 40 | 46 | 24 | 22 | 26 | 48 | 45 | 45 | 35.1 | 24 | 46 | 1.914 | 0.936 | 0.00397951 | 0.02051919 | 0 | 0.0699 | 0.045 | 0.045 | TRUE | 0.0221 |
| Cldn2 | 23 | 12 | 9 | 9 | 9 | 12 | 18 | 13 | 10 | 9 | 10 | 12 | 11.88 | 14 | 10 | 0.759 | -0.398 | 0.47852855 | NA | 0 | 0.2019 | 0.1417 | 0.1417 | TRUE | 0.3926 |
| Cldn20 | 0 | 0 | 1 | 0 | 0 | 1 | 0 | 0 | 1 | 0 | 0 | 1 | 0.34 | 0 | 0 | 1.075 | 0.105 | 0.97570524 | NA | 0 | 6.885 | 6.5379 | 6.5379 | TRUE | 0.5365 |
| Cldn22 | 0 | 0 | 0 | 0 | 1 | 0 | 0 | 0 | 0 | 0 | 1 | 0 | 0.19 | 0 | 0 | 2.094 | 1.067 | 0.79379329 | NA | 0 | 12.608 | 10 | 10 | TRUE | 0.5075 |
| Cldn23 | 20 | 17 | 19 | 11 | 11 | 7 | 16 | 18 | 21 | 11 | 12 | 7 | 14.06 | 18 | 10 | 0.559 | -0.839 | 0.10862939 | NA | 0 | 0.1709 | 0.1184 | 0.1184 | TRUE | 0.2754 |
| Cldn24 | 0 | 0 | 0 | 0 | 0 | 0 | 0 | 0 | 0 | 0 | 0 | 0 | 0 | 0 | 0 | NA | NA | NA | NA | NA | NA | NA | NA | NA | NA |
| Cldn25 | 2114 | 1456 | 1616 | 1782 | 1489 | 1684 | 1642 | 1549 | 1744 | 1774 | 1680 | 1662 | 1675.02 | 1645 | 1705 | 1.037 | 0.052 | 0.46157035 | 0.63095143 | 0.0015 | 0.0038 | 0.0031 | 0.0031 | TRUE | 0.0323 |
| Cldn26 | 10 | 7 | 5 | 9 | 3 | 5 | 8 | 7 | 5 | 9 | 3 | 5 | 6.31 | 7 | 6 | 0.84 | -0.252 | 0.74298151 | NA | 0 | 0.3778 | 0.265 | 0.265 | TRUE | 0.5429 |
| Cldn3 | 106 | 80 | 115 | 138 | 116 | 122 | 82 | 85 | 124 | 137 | 131 | 120 | 113.36 | 97 | 129 | 1.34 | 0.422 | 0.04173467 | 0.11791136 | 0.0185 | 0.0233 | 0.022 | 0.022 | TRUE | 0.5503 |
| Cldn4 | 869 | 729 | 1140 | 1089 | 1158 | 1255 | 675 | 775 | 1230 | 1084 | 1306 | 1239 | 1051.6 | 893 | 1210 | 1.355 | 0.438 | 0.12517354 | 0.26021509 | 0.0578 | 0.0047 | 0.0181 | 0.0578 | TRUE | 1.2908 |
| Cldn5 | 76 | 65 | 73 | 125 | 110 | 159 | 59 | 69 | 79 | 124 | 124 | 157 | 102.07 | 69 | 135 | 1.974 | 0.981 | 3.49E-06 | 7.87E-05 | 0.0105 | 0.0256 | 0.0213 | 0.0213 | TRUE | 0.2084 |
| Cldn6 | 900 | 798 | 1137 | 1152 | 1036 | 1397 | 699 | 849 | 1227 | 1147 | 1169 | 1379 | 1078.18 | 925 | 1232 | 1.332 | 0.414 | 0.10945309 | 0.23635187 | 0.0473 | 0.0046 | 0.016 | 0.0473 | TRUE | 0.9707 |
| Cldn7 | 632 | 603 | 1002 | 1150 | 1053 | 1306 | 491 | 641 | 1081 | 1145 | 1188 | 1289 | 972.54 | 738 | 1207 | 1.637 | 0.711 | 0.04153287 | 0.11748057 | 0.0867 | 0.0049 | 0.0246 | 0.0867 | TRUE | 1.6466 |
| Cldn8 | 0 | 0 | 1 | 0 | 0 | 0 | 0 | 0 | 1 | 0 | 0 | 0 | 0.18 | 0 | 0 | 0.552 | -0.857 | 0.8336548 | NA | 0 | 13.1817 | 10 | 10 | TRUE | 0.5365 |
| Cldn9 | 54 | 33 | 47 | 60 | 72 | 68 | 42 | 35 | 51 | 60 | 81 | 67 | 55.97 | 43 | 69 | 1.627 | 0.702 | 0.01022524 | 0.04198815 | 0.0075 | 0.0448 | 0.035 | 0.035 | TRUE | 0.1935 |
| Cldnd2 | 2 | 4 | 3 | 2 | 0 | 5 | 2 | 4 | 3 | 2 | 0 | 5 | 2.66 | 3 | 2 | 0.791 | -0.339 | 0.78658425 | NA | 0.2753 | 0.8928 | 0.7466 | 0.7466 | TRUE | 0.3982 |
| Clec10a | 71 | 44 | 19 | 13 | 10 | 8 | 55 | 47 | 21 | 13 | 11 | 8 | 25.76 | 41 | 11 | 0.26 | -1.943 | 4.94E-05 | 0.00069418 | 0.1304 | 0.0944 | 0.1049 | 0.1049 | TRUE | 0.4493 |
| Clec11a | 61 | 43 | 55 | 61 | 56 | 37 | 47 | 46 | 59 | 61 | 63 | 37 | 52.15 | 51 | 54 | 1.052 | 0.073 | 0.80469622 | 0.88691572 | 0.0323 | 0.0479 | 0.0436 | 0.0436 | TRUE | 0.6555 |
| Clec12a | 0 | 3 | 0 | 4 | 2 | 1 | 0 | 3 | 0 | 4 | 2 | 1 | 1.74 | 1 | 2 | 2.364 | 1.241 | 0.45331009 | NA | 0.9412 | 1.3677 | 1.2857 | 1.2857 | TRUE | 1.005 |
| Clec12b | 0 | 0 | 0 | 0 | 0 | 0 | 0 | 0 | 0 | 0 | 0 | 0 | 0 | 0 | 0 | NA | NA | NA | NA | NA | NA | NA | NA | NA | NA |
| Clec14a | 1 | 2 | 1 | 0 | 2 | 1 | 1 | 2 | 1 | 0 | 2 | 1 | 1.2 | 1 | 1 | 0.811 | -0.303 | 0.86289797 | NA | 0 | 1.9703 | 1.3597 | 1.3597 | TRUE | 0.2764 |
| Clec16a | 1825 | 1203 | 928 | 1008 | 906 | 1088 | 1417 | 1280 | 1001 | 1003 | 1022 | 1074 | 1132.96 | 1233 | 1033 | 0.837 | -0.257 | 0.02083873 | 0.07135794 | 0.0154 | 0.0045 | 0.008 | 0.008 | TRUE | 0.321 |
| Clec18a | 1 | 0 | 0 | 1 | 2 | 3 | 1 | 0 | 0 | 1 | 2 | 3 | 1.16 | 0 | 2 | 6.003 | 2.586 | 0.18409684 | NA | 0 | 2.0369 | 1.4806 | 1.4806 | TRUE | 0.2566 |
| Clec1a | 1 | 3 | 0 | 1 | 0 | 0 | 1 | 3 | 0 | 1 | 0 | 0 | 0.83 | 1 | 0 | 0.283 | -1.823 | 0.43666934 | NA | 0 | 2.8676 | 2.5891 | 2.5891 | TRUE | 0.5266 |
| Clec1b | 1 | 1 | 0 | 0 | 0 | 0 | 1 | 1 | 0 | 0 | 0 | 0 | 0.31 | 1 | 0 | 0.309 | -1.696 | 0.63183309 | NA | 0 | 7.7294 | 7.2472 | 7.2472 | TRUE | 0.2015 |
| Clec2d | 8 | 6 | 2 | 7 | 4 | 5 | 6 | 6 | 2 | 7 | 5 | 5 | 5.19 | 5 | 6 | 1.103 | 0.142 | 0.86658622 | NA | 0 | 0.4587 | 0.3213 | 0.3213 | TRUE | 0.4293 |
| Clec2e | 0 | 0 | 0 | 0 | 0 | 0 | 0 | 0 | 0 | 0 | 0 | 0 | 0 | 0 | 0 | NA | NA | NA | NA | NA | NA | NA | NA | NA | NA |
| Clec2g | 3 | 7 | 2 | 1 | 0 | 0 | 2 | 7 | 2 | 1 | 0 | 0 | 2.15 | 4 | 0 | 0.094 | -3.411 | 0.04201177 | NA | 0 | 1.1023 | 0.9053 | 0.9053 | TRUE | 0.9159 |
| Clec2h | 0 | 0 | 0 | 0 | 0 | 0 | 0 | 0 | 0 | 0 | 0 | 0 | 0 | 0 | 0 | NA | NA | NA | NA | NA | NA | NA | NA | NA | NA |
| Clec2i | 54 | 27 | 28 | 16 | 11 | 16 | 42 | 29 | 30 | 16 | 12 | 16 | 24.17 | 34 | 15 | 0.436 | -1.198 | 0.00362195 | 0.01910277 | 0 | 0.1005 | 0.0729 | 0.0729 | TRUE | 0.3805 |
| Clec2l | 196 | 135 | 152 | 203 | 152 | 165 | 152 | 144 | 164 | 202 | 171 | 163 | 166.04 | 153 | 179 | 1.168 | 0.223 | 0.1663385 | 0.31738434 | 0 | 0.0167 | 0.0127 | 0.0127 | TRUE | 0.1416 |
| Clec3a | 1 | 2 | 0 | 0 | 0 | 0 | 1 | 2 | 0 | 0 | 0 | 0 | 0.48 | 1 | 0 | 0.197 | -2.343 | 0.43249452 | NA | 0 | 4.8992 | 4.9172 | 4.9172 | TRUE | 0.1226 |
| Clec3b | 2 | 2 | 2 | 0 | 2 | 2 | 2 | 2 | 2 | 0 | 2 | 2 | 1.68 | 2 | 1 | 0.722 | -0.469 | 0.75249826 | NA | 0 | 1.4147 | 0.976 | 0.976 | TRUE | 0.5149 |
| Clec4a1 | 14 | 12 | 9 | 1 | 2 | 3 | 11 | 13 | 10 | 1 | 2 | 3 | 6.59 | 11 | 2 | 0.186 | -2.426 | 0.00519326 | NA | 0 | 0.3619 | 0.2494 | 0.2494 | TRUE | 0.2036 |
| Clec4a2 | 3 | 1 | 2 | 1 | 1 | 0 | 2 | 1 | 2 | 1 | 1 | 0 | 1.28 | 2 | 1 | 0.374 | -1.42 | 0.41745247 | NA | 0 | 1.8552 | 1.2075 | 1.2075 | TRUE | 0.2712 |
| Clec4a3 | 4 | 2 | 2 | 0 | 0 | 0 | 3 | 2 | 2 | 0 | 0 | 0 | 1.23 | 2 | 0 | 0.077 | -3.702 | 0.05731828 | NA | 0 | 1.9262 | 1.5379 | 1.5379 | TRUE | 0.0744 |
| Clec4a4 | 0 | 1 | 0 | 0 | 0 | 0 | 0 | 1 | 0 | 0 | 0 | 0 | 0.18 | 0 | 0 | 0.552 | -0.857 | 0.8336548 | NA | 0 | 13.3714 | 10 | 10 | TRUE | 0.5216 |
| Clec4b1 | 0 | 0 | 0 | 0 | 0 | 0 | 0 | 0 | 0 | 0 | 0 | 0 | 0 | 0 | 0 | NA | NA | NA | NA | NA | NA | NA | NA | NA | NA |
| Clec4b2 | 0 | 0 | 0 | 0 | 0 | 0 | 0 | 0 | 0 | 0 | 0 | 0 | 0 | 0 | 0 | NA | NA | NA | NA | NA | NA | NA | NA | NA | NA |
| Clec4d | 0 | 0 | 0 | 0 | 0 | 0 | 0 | 0 | 0 | 0 | 0 | 0 | 0 | 0 | 0 | NA | NA | NA | NA | NA | NA | NA | NA | NA | NA |
| Clec4e | 5 | 7 | 7 | 1 | 1 | 2 | 4 | 7 | 8 | 1 | 1 | 2 | 3.83 | 6 | 1 | 0.221 | -2.181 | 0.04834858 | NA | 0 | 0.6212 | 0.425 | 0.425 | TRUE | 0.3602 |
| Clec4f | 8 | 2 | 5 | 1 | 5 | 1 | 6 | 2 | 5 | 1 | 6 | 1 | 3.56 | 4 | 3 | 0.537 | -0.896 | 0.41444646 | NA | 0.24 | 0.6682 | 0.5563 | 0.5563 | TRUE | 1.138 |
| Clec4g | 1 | 2 | 2 | 4 | 4 | 3 | 1 | 2 | 2 | 4 | 5 | 3 | 2.75 | 2 | 4 | 2.326 | 1.218 | 0.29737611 | NA | 0 | 0.8634 | 0.5546 | 0.5546 | TRUE | 0.2504 |
| Clec4n | 0 | 0 | 0 | 0 | 0 | 0 | 0 | 0 | 0 | 0 | 0 | 0 | 0 | 0 | 0 | NA | NA | NA | NA | NA | NA | NA | NA | NA | NA |
| Clec5a | 0 | 0 | 0 | 0 | 0 | 0 | 0 | 0 | 0 | 0 | 0 | 0 | 0 | 0 | 0 | NA | NA | NA | NA | NA | NA | NA | NA | NA | NA |
| Clec7a | 0 | 0 | 0 | 0 | 0 | 0 | 0 | 0 | 0 | 0 | 0 | 0 | 0 | 0 | 0 | NA | NA | NA | NA | NA | NA | NA | NA | NA | NA |
| Clec9a | 0 | 0 | 0 | 0 | 0 | 0 | 0 | 0 | 0 | 0 | 0 | 0 | 0 | 0 | 0 | NA | NA | NA | NA | NA | NA | NA | NA | NA | NA |
| Clgn | 2790 | 1910 | 1975 | 2454 | 2205 | 2506 | 2167 | 2032 | 2131 | 2443 | 2487 | 2474 | 2288.92 | 2110 | 2468 | 1.169 | 0.225 | 0.00016389 | 0.00182994 | 0 | 0.0035 | 0.0021 | 0.0021 | TRUE | 0.0126 |
| Clhc1 | 1476 | 1047 | 1062 | 1147 | 1020 | 1060 | 1146 | 1114 | 1146 | 1142 | 1151 | 1046 | 1124.11 | 1135 | 1113 | 0.98 | -0.03 | 0.69274398 | 0.81225511 | 6.00E-04 | 0.0045 | 0.0031 | 0.0031 | TRUE | 0.0331 |
| Clic1 | 2040 | 1641 | 2017 | 2027 | 1833 | 2120 | 1584 | 1746 | 2176 | 2018 | 2068 | 2093 | 1947.41 | 1835 | 2060 | 1.123 | 0.168 | 0.09466108 | 0.2123671 | 0.0133 | 0.0036 | 0.0067 | 0.0067 | TRUE | 0.318 |
| Clic3 | 16 | 14 | 12 | 13 | 12 | 11 | 12 | 15 | 13 | 13 | 14 | 11 | 12.93 | 13 | 13 | 0.928 | -0.107 | 0.83713688 | NA | 0 | 0.1857 | 0.1187 | 0.1187 | TRUE | 0.0519 |
| Clic4 | 12827 | 9238 | 9271 | 9171 | 8015 | 9201 | 9962 | 9826 | 10003 | 9129 | 9042 | 9083 | 9507.54 | 9930 | 9085 | 0.915 | -0.128 | 0.00445978 | 0.02249664 | 0 | 0.0027 | 0.0014 | 0.0014 | TRUE | 0.001 |
| Clic5 | 11 | 11 | 7 | 15 | 7 | 7 | 9 | 12 | 8 | 15 | 8 | 7 | 9.59 | 10 | 10 | 1.076 | 0.105 | 0.86812289 | NA | 0 | 0.2496 | 0.1844 | 0.1844 | TRUE | 0.6939 |
| Clic6 | 143 | 105 | 97 | 123 | 121 | 112 | 111 | 112 | 105 | 122 | 137 | 111 | 116.15 | 109 | 123 | 1.126 | 0.171 | 0.3533737 | 0.52780934 | 0 | 0.0228 | 0.0158 | 0.0158 | TRUE | 0.0871 |
| Clint1 | 10694 | 7668 | 7134 | 7738 | 6799 | 7837 | 8306 | 8156 | 7698 | 7702 | 7670 | 7736 | 7878.03 | 8053 | 7703 | 0.956 | -0.064 | 0.21330423 | 0.37556494 | 6.00E-04 | 0.0027 | 0.0018 | 0.0018 | TRUE | 0.0181 |
| Clip1 | 1190 | 912 | 912 | 1136 | 983 | 1134 | 924 | 970 | 984 | 1131 | 1109 | 1119 | 1039.58 | 959 | 1120 | 1.168 | 0.224 | 0.00223452 | 0.01315944 | 0 | 0.0047 | 0.0029 | 0.0029 | TRUE | 0.0131 |
| Clip2 | 729 | 585 | 636 | 705 | 659 | 804 | 566 | 622 | 686 | 702 | 743 | 794 | 685.59 | 625 | 746 | 1.197 | 0.259 | 0.00901351 | 0.03838799 | 0.0051 | 0.0059 | 0.0056 | 0.0056 | TRUE | 0.0866 |
| Clip3 | 851 | 585 | 610 | 582 | 570 | 716 | 661 | 622 | 658 | 579 | 643 | 707 | 645.09 | 647 | 643 | 0.993 | -0.01 | 0.92151333 | 0.95679138 | 0.004 | 0.0061 | 0.0054 | 0.0054 | TRUE | 0.0906 |
| Clip4 | 1533 | 1160 | 1222 | 1535 | 1339 | 1483 | 1191 | 1234 | 1319 | 1528 | 1511 | 1464 | 1374.24 | 1248 | 1501 | 1.204 | 0.268 | 0.00017709 | 0.00194405 | 8.00E-04 | 0.0042 | 0.0029 | 0.0029 | TRUE | 0.0293 |

| | | | | | | | | | | | | | | | | | | | | | | | | | |
|---|---|---|---|---|---|---|---|---|---|---|---|---|---|---|---|---|---|---|---|---|---|---|---|---|---|
| Clk1 | 4311 | 3755 | 4688 | 5213 | 4377 | 6100 | 3348 | 3994 | 5058 | 5189 | 4938 | 6022 | 4758.17 | 4133 | 5383 | 1.302 | 0.381 | 0.04999917 | 0.13477101 | 0.0271 | 0.0029 | 0.0099 | 0.0271 | TRUE | 0.4662 |
| Clk2 | 3115 | 2582 | 2427 | 2369 | 2142 | 2507 | 2419 | 2746 | 2619 | 2358 | 2416 | 2475 | 2505.62 | 2595 | 2416 | 0.932 | -0.102 | 0.12994161 | 0.26669377 | 0.002 | 0.0034 | 0.0028 | 0.0028 | TRUE | 0.0437 |
| Clk3 | 2387 | 1719 | 2056 | 2162 | 1906 | 2138 | 1854 | 1828 | 2218 | 2152 | 2150 | 2111 | 2052.25 | 1967 | 2138 | 1.087 | 0.121 | 0.1355184 | 0.2748637 | 0.0055 | 0.0036 | 0.0042 | 0.0042 | TRUE | 0.1495 |
| Clk4 | 1402 | 1081 | 1247 | 1207 | 924 | 1264 | 1089 | 1150 | 1346 | 1201 | 1042 | 1248 | 1179.3 | 1195 | 1164 | 0.976 | -0.035 | 0.72024049 | 0.83208603 | 0.0097 | 0.0044 | 0.0062 | 0.0062 | TRUE | 0.1451 |
| Clmn | 1829 | 1462 | 1649 | 2146 | 1892 | 2201 | 1421 | 1555 | 1779 | 2136 | 2134 | 2173 | 1866.35 | 1585 | 2148 | 1.357 | 0.44 | 1.26E-07 | 4.83E-06 | 0.0058 | 0.0037 | 0.0045 | 0.0045 | TRUE | 0.1382 |
| Clmp | 116 | 79 | 92 | 72 | 64 | 68 | 90 | 84 | 99 | 72 | 72 | 67 | 80.73 | 91 | 70 | 0.772 | -0.374 | 0.08397694 | 0.19539026 | 0 | 0.0318 | 0.021 | 0.021 | TRUE | 0.0542 |
| Cln3 | 1009 | 762 | 823 | 817 | 751 | 860 | 784 | 811 | 888 | 813 | 847 | 849 | 831.93 | 828 | 836 | 1.012 | 0.017 | 0.83570899 | 0.90644601 | 0.0013 | 0.0053 | 0.0039 | 0.0039 | TRUE | 0.0481 |
| Cln5 | 2121 | 1512 | 1553 | 1538 | 1452 | 1651 | 1647 | 1608 | 1676 | 1531 | 1638 | 1630 | 1621.66 | 1644 | 1600 | 0.973 | -0.04 | 0.54857658 | 0.70517178 | 0 | 0.0039 | 0.0025 | 0.0025 | TRUE | 0.0171 |
| Cln6 | 1992 | 1564 | 1864 | 1895 | 1783 | 1891 | 1547 | 1664 | 2011 | 1886 | 2011 | 1867 | 1831.06 | 1741 | 1921 | 1.105 | 0.144 | 0.12253376 | 0.25586196 | 0.0096 | 0.0037 | 0.0057 | 0.0057 | TRUE | 0.2222 |
| Cln8 | 3421 | 2539 | 2444 | 2463 | 2216 | 2436 | 2657 | 2701 | 2637 | 2452 | 2500 | 2405 | 2558.49 | 2665 | 2452 | 0.92 | -0.12 | 0.0339019 | 0.10152103 | 0 | 0.0034 | 0.0019 | 0.0019 | TRUE | 0.0035 |
| Clnk | 10 | 8 | 4 | 3 | 7 | 1 | 8 | 9 | 4 | 3 | 8 | 1 | 5.41 | 7 | 4 | 0.564 | -0.827 | 0.35591644 | NA | 0.1622 | 0.4405 | 0.3752 | 0.3752 | TRUE | 0.9689 |
| Clns1a | 8362 | 5805 | 5470 | 5407 | 4685 | 5169 | 6494 | 6175 | 5902 | 5382 | 5285 | 5103 | 5723.51 | 6190 | 5257 | 0.849 | -0.236 | 4.04E-05 | 0.0005917 | 0.0012 | 0.0028 | 0.0022 | 0.0022 | TRUE | 0.0228 |
| Clock | 1488 | 1104 | 1064 | 1152 | 912 | 1059 | 1156 | 1174 | 1148 | 1147 | 1029 | 1045 | 1116.49 | 1159 | 1074 | 0.926 | -0.11 | 0.1462095 | 0.29056901 | 8.00E-04 | 0.0045 | 0.0032 | 0.0032 | TRUE | 0.0427 |
| Clp1 | 3073 | 2129 | 1521 | 1685 | 1473 | 1639 | 2387 | 2265 | 1641 | 1677 | 1662 | 1618 | 1874.88 | 2098 | 1652 | 0.787 | -0.345 | 0.00217444 | 0.01286393 | 0.0175 | 0.0037 | 0.0086 | 0.0086 | TRUE | 0.4356 |
| Clpb | 6848 | 5079 | 5144 | 5888 | 5309 | 5991 | 5319 | 5402 | 5550 | 5861 | 5989 | 5914 | 5672.57 | 5424 | 5921 | 1.092 | 0.127 | 0.01089502 | 0.0439894 | 1.00E-04 | 0.0028 | 0.0016 | 0.0016 | TRUE | 0.0051 |
| Clpp | 3876 | 2827 | 2541 | 2779 | 2407 | 2908 | 3010 | 3007 | 2742 | 2766 | 2715 | 2871 | 2851.88 | 2920 | 2784 | 0.953 | -0.069 | 0.27666929 | 0.4478408 | 0.0013 | 0.0033 | 0.0025 | 0.0025 | TRUE | 0.0341 |
| Clps | 4 | 7 | 5 | 4 | 3 | 6 | 3 | 7 | 5 | 4 | 3 | 6 | 4.87 | 5 | 4 | 0.85 | -0.235 | 0.78688112 | NA | 0 | 0.4888 | 0.334 | 0.334 | TRUE | 0.2361 |
| Clpsl2 | 0 | 4 | 2 | 1 | 0 | 1 | 0 | 4 | 2 | 1 | 0 | 1 | 1.4 | 2 | 1 | 0.324 | -1.627 | 0.38165852 | NA | 0 | 1.6963 | 1.4956 | 1.4956 | TRUE | 0.0976 |
| Clptm1 | 7849 | 5642 | 5541 | 5617 | 5186 | 6151 | 6096 | 6001 | 5979 | 5591 | 5850 | 6072 | 5931.58 | 6025 | 5838 | 0.969 | -0.046 | 0.39469071 | 0.56811659 | 7.00E-04 | 0.0028 | 0.0019 | 0.0019 | TRUE | 0.0168 |
| Clptm1l | 3634 | 2637 | 2673 | 2793 | 2438 | 2751 | 2822 | 2805 | 2884 | 2780 | 2750 | 2716 | 2792.93 | 2837 | 2749 | 0.969 | -0.046 | 0.40888974 | 0.58169215 | 0 | 0.0033 | 0.0018 | 0.0018 | TRUE | 0.0025 |
| Clpx | 3355 | 2462 | 2568 | 2804 | 2369 | 2803 | 2606 | 2619 | 2771 | 2791 | 2672 | 2767 | 2704.32 | 2665 | 2743 | 1.03 | 0.043 | 0.47176971 | 0.64057116 | 5.00E-04 | 0.0033 | 0.0022 | 0.0022 | TRUE | 0.0144 |
| Clrn1 | 2 | 1 | 1 | 1 | 0 | 0 | 2 | 1 | 1 | 1 | 0 | 0 | 0.78 | 1 | 0 | 0.299 | -1.743 | 0.425346 | NA | 0 | 3.0336 | 2.0559 | 2.0559 | TRUE | 0.3906 |
| Clrn2 | 0 | 0 | 0 | 0 | 0 | 0 | 0 | 0 | 0 | 0 | 0 | 0 | 0 | 0 | 0 | NA | NA | NA | NA | NA | NA | NA | NA | NA | NA |
| Clrn3 | 0 | 0 | 0 | 0 | 1 | 0 | 0 | 0 | 0 | 0 | 1 | 0 | 0.19 | 0 | 0 | 2.094 | 1.067 | 0.79379329 | NA | 0 | 12.608 | 10 | 10 | TRUE | 0.5075 |
| Clspn | 4375 | 3038 | 2736 | 2859 | 2508 | 2950 | 3398 | 3231 | 2952 | 2846 | 2829 | 2912 | 3028.12 | 3194 | 2862 | 0.896 | -0.159 | 0.01722062 | 0.061905 | 0.0021 | 0.0032 | 0.0029 | 0.0029 | TRUE | 0.053 |
| Clstn1 | 8722 | 6614 | 7253 | 7986 | 7443 | 8062 | 6774 | 7035 | 7826 | 7949 | 8396 | 7958 | 7656.55 | 7212 | 8101 | 1.124 | 0.168 | 0.00968383 | 0.04044352 | 0.0032 | 0.0027 | 0.0029 | 0.0029 | TRUE | 0.0676 |
| Clstn2 | 33 | 26 | 22 | 18 | 19 | 21 | 26 | 28 | 24 | 18 | 21 | 21 | 22.85 | 26 | 20 | 0.779 | -0.36 | 0.36319047 | 0.53786674 | 0 | 0.1061 | 0.0684 | 0.0684 | TRUE | 0.047 |
| Clstn3 | 1950 | 1389 | 1357 | 1584 | 1331 | 1653 | 1514 | 1477 | 1464 | 1577 | 1502 | 1632 | 1527.69 | 1485 | 1570 | 1.057 | 0.08 | 0.23634142 | 0.40276952 | 4.00E-04 | 0.004 | 0.0026 | 0.0026 | TRUE | 0.0172 |
| Clta | 10661 | 8187 | 8013 | 8503 | 7569 | 9007 | 8280 | 8708 | 8646 | 8464 | 8539 | 8891 | 8588.02 | 8545 | 8631 | 1.01 | 0.015 | 0.7705619 | 0.8642217 | 5.00E-04 | 0.0027 | 0.0018 | 0.0018 | TRUE | 0.0091 |
| Cltb | 1382 | 997 | 1070 | 1364 | 1249 | 1455 | 1073 | 1060 | 1155 | 1358 | 1409 | 1436 | 1248.57 | 1096 | 1401 | 1.279 | 0.355 | 1.02E-06 | 3.00E-05 | 5.00E-04 | 0.0043 | 0.003 | 0.003 | TRUE | 0.0252 |
| Cltc | 50456 | 34169 | 32936 | 39129 | 35187 | 40410 | 39187 | 36345 | 35538 | 38949 | 39695 | 39891 | 38267.38 | 37023 | 39512 | 1.067 | 0.094 | 0.07611686 | 0.18200333 | 0.0014 | 0.0025 | 0.002 | 0.002 | TRUE | 0.0321 |
| Clu | 1837 | 1367 | 1288 | 1263 | 1099 | 1165 | 1427 | 1454 | 1390 | 1257 | 1240 | 1150 | 1319.59 | 1424 | 1216 | 0.854 | -0.228 | 0.00138615 | 0.00910488 | 5.00E-04 | 0.0042 | 0.0029 | 0.0029 | TRUE | 0.0272 |
| Cluap1 | 689 | 546 | 592 | 590 | 535 | 542 | 535 | 581 | 639 | 587 | 604 | 535 | 580.09 | 585 | 575 | 0.985 | -0.022 | 0.83161739 | 0.90369553 | 0.0043 | 0.0065 | 0.0058 | 0.0058 | TRUE | 0.0759 |
| Cluh | 17523 | 13242 | 11883 | 12586 | 11628 | 13309 | 13609 | 14085 | 12822 | 12528 | 13118 | 13138 | 13216.68 | 13505 | 12928 | 0.957 | -0.063 | 0.24819277 | 0.41714714 | 0.0014 | 0.0026 | 0.0021 | 0.0021 | TRUE | 0.0238 |
| Clvs1 | 51 | 27 | 25 | 14 | 7 | 4 | 40 | 29 | 27 | 14 | 8 | 4 | 20.18 | 32 | 9 | 0.269 | -1.893 | 0.0001826 | 0.0019938 | 0.0647 | 0.1199 | 0.1091 | 0.1091 | TRUE | 0.9642 |
| Clvs2 | 67 | 41 | 30 | 31 | 40 | 45 | 52 | 44 | 32 | 31 | 45 | 44 | 41.4 | 43 | 40 | 0.932 | -0.101 | 0.75244463 | 0.85314065 | 0.0257 | 0.0597 | 0.0503 | 0.0503 | TRUE | 0.2497 |
| Clybl | 1777 | 1311 | 1688 | 1871 | 1764 | 1901 | 1380 | 1394 | 1821 | 1862 | 1990 | 1877 | 1720.82 | 1532 | 1910 | 1.248 | 0.319 | 0.00148169 | 0.00956538 | 0.0132 | 0.0038 | 0.0067 | 0.0067 | TRUE | 0.3263 |
| Cma1 | 0 | 0 | 1 | 0 | 0 | 0 | 0 | 0 | 1 | 0 | 0 | 0 | 0.18 | 0 | 0 | 0.552 | -0.857 | 0.8336548 | NA | 0 | 13.1817 | 10 | 10 | TRUE | 0.5365 |
| Cma2 | 0 | 0 | 0 | 0 | 0 | 0 | 0 | 0 | 0 | 0 | 0 | 0 | 0 | 0 | 0 | NA | NA | NA | NA | NA | NA | NA | NA | NA | NA |
| Cmah | 19 | 10 | 13 | 12 | 8 | 7 | 15 | 11 | 14 | 12 | 9 | 7 | 11.22 | 13 | 9 | 0.705 | -0.505 | 0.37914457 | NA | 0 | 0.2137 | 0.1456 | 0.1456 | TRUE | 0.2143 |
| Cmas | 1156 | 803 | 684 | 794 | 658 | 715 | 898 | 854 | 738 | 790 | 742 | 706 | 788.07 | 830 | 746 | 0.898 | -0.156 | 0.10717178 | 0.23265586 | 0.0055 | 0.0054 | 0.0055 | 0.0055 | TRUE | 0.1099 |
| Cmbl | 47 | 21 | 31 | 36 | 35 | 44 | 37 | 22 | 33 | 36 | 39 | 43 | 35.17 | 31 | 39 | 1.278 | 0.354 | 0.29131492 | 0.4634644 | 0 | 0.0698 | 0.0524 | 0.0524 | TRUE | 0.3647 |
| Cmc1 | 1069 | 730 | 786 | 970 | 852 | 943 | 830 | 776 | 848 | 966 | 961 | 931 | 885.4 | 818 | 953 | 1.164 | 0.219 | 0.00570589 | 0.02714857 | 0 | 0.0051 | 0.0034 | 0.0034 | TRUE | 0.0228 |
| Cmc2 | 1204 | 816 | 704 | 796 | 679 | 764 | 935 | 868 | 760 | 792 | 766 | 754 | 812.53 | 854 | 771 | 0.901 | -0.151 | 0.10919125 | 0.23596469 | 0.0048 | 0.0053 | 0.0051 | 0.0051 | TRUE | 0.1103 |
| Cmip | 11125 | 8367 | 8293 | 8536 | 7782 | 9050 | 8640 | 8900 | 8948 | 8497 | 8779 | 8934 | 8782.94 | 8829 | 8737 | 0.99 | -0.015 | 0.75939066 | 0.8571376 | 3.00E-04 | 0.0027 | 0.0016 | 0.0016 | TRUE | 0.0071 |
| Cmklr1 | 1261 | 983 | 1031 | 1074 | 1027 | 1184 | 979 | 1046 | 1112 | 1069 | 1159 | 1169 | 1088.97 | 1046 | 1132 | 1.084 | 0.116 | 0.15218839 | 0.29879514 | 0.0024 | 0.0046 | 0.0038 | 0.0038 | TRUE | 0.0387 |
| Cml1 | 48 | 35 | 45 | 47 | 46 | 51 | 37 | 37 | 49 | 47 | 52 | 50 | 45.35 | 41 | 50 | 1.215 | 0.281 | 0.32924355 | 0.50362565 | 0 | 0.0547 | 0.0375 | 0.0375 | TRUE | 0.1836 |
| Cml2 | 138 | 141 | 69 | 66 | 45 | 52 | 107 | 150 | 74 | 66 | 51 | 51 | 83.23 | 110 | 56 | 0.507 | -0.981 | 0.00032356 | 0.00307895 | 0.0617 | 0.0309 | 0.04 | 0.04 | TRUE | 0.1477 |
| Cml3 | 0 | 0 | 0 | 1 | 1 | 0 | 0 | 0 | 0 | 1 | 1 | 0 | 0.35 | 0 | 1 | 3.978 | 1.992 | 0.55384016 | NA | 0 | 6.6991 | 6.4234 | 6.4234 | TRUE | 0.2647 |
| Cml5 | 0 | 1 | 0 | 0 | 0 | 0 | 0 | 1 | 0 | 0 | 0 | 0 | 0.18 | 0 | 0 | 0.552 | -0.857 | 0.8336548 | NA | 0 | 13.3714 | 10 | 10 | TRUE | 0.5216 |
| Cmpk1 | 4521 | 3088 | 3024 | 3221 | 2736 | 3039 | 3511 | 3285 | 3263 | 3206 | 3086 | 3000 | 3225.24 | 3353 | 3097 | 0.923 | -0.115 | 0.05716372 | 0.1485259 | 0.0011 | 0.0032 | 0.0023 | 0.0023 | TRUE | 0.0212 |
| Cmpk2 | 156 | 93 | 77 | 85 | 71 | 78 | 121 | 99 | 83 | 85 | 80 | 77 | 90.81 | 101 | 81 | 0.792 | -0.336 | 0.12550652 | 0.26066038 | 0.0115 | 0.0285 | 0.0234 | 0.0234 | TRUE | 0.2646 |
| Cmss1 | 1135 | 807 | 676 | 631 | 578 | 684 | 882 | 858 | 729 | 628 | 652 | 675 | 737.44 | 823 | 652 | 0.791 | -0.339 | 0.0004388 | 0.00390039 | 0.0046 | 0.0056 | 0.0053 | 0.0053 | TRUE | 0.1155 |
| Cmtm1 | 0 | 0 | 0 | 0 | 1 | 0 | 0 | 0 | 0 | 0 | 1 | 0 | 0.19 | 0 | 0 | 2.094 | 1.067 | 0.79379329 | NA | 0 | 12.608 | 10 | 10 | TRUE | 0.5075 |
| Cmtm2a | 0 | 0 | 1 | 0 | 1 | 0 | 0 | 0 | 1 | 0 | 1 | 0 | 0.37 | 0 | 0 | 1.075 | 0.105 | 0.97514588 | NA | 0 | 6.4454 | 6.1578 | 6.1578 | TRUE | 0.5365 |
| Cmtm2b | 1 | 2 | 1 | 1 | 2 | 6 | 1 | 2 | 1 | 1 | 2 | 6 | 2.19 | 1 | 3 | 2.361 | 1.24 | 0.3614695 | NA | 0 | 1.0832 | 0.7907 | 0.7907 | TRUE | 0.6167 |
| Cmtm3 | 334 | 269 | 250 | 252 | 225 | 264 | 259 | 286 | 270 | 251 | 254 | 261 | 263.43 | 272 | 255 | 0.94 | -0.089 | 0.47564942 | 0.64354953 | 0 | 0.0114 | 0.0074 | 0.0074 | TRUE | 0.0238 |
| Cmtm4 | 13007 | 9661 | 10690 | 11970 | 10566 | 11961 | 10102 | 10276 | 11534 | 11915 | 11920 | 11807 | 11259.08 | 10637 | 11881 | 1.117 | 0.16 | 0.00895753 | 0.0382179 | 0.0026 | 0.0026 | 0.0026 | 0.0026 | TRUE | 0.0663 |
| Cmtm5 | 0 | 1 | 0 | 1 | 0 | 1 | 0 | 1 | 0 | 1 | 0 | 1 | 0.51 | 0 | 1 | 1.941 | 0.957 | 0.72720315 | NA | 0 | 4.6706 | 3.6514 | 3.6514 | TRUE | 0.5217 |
| Cmtm6 | 7681 | 5770 | 6545 | 7320 | 6632 | 7505 | 5966 | 6137 | 7062 | 7286 | 7482 | 7409 | 6890.25 | 6388 | 7392 | 1.157 | 0.211 | 0.00219162 | 0.01294415 | 0.0041 | 0.0028 | 0.0033 | 0.0033 | TRUE | 0.1035 |
| Cmtm7 | 1443 | 989 | 903 | 939 | 863 | 917 | 1121 | 1052 | 974 | 935 | 974 | 905 | 993.41 | 1049 | 938 | 0.892 | -0.164 | 0.04695037 | 0.12865444 | 0.0023 | 0.0048 | 0.0039 | 0.0039 | TRUE | 0.046 |
| Cmtm8 | 103 | 66 | 116 | 98 | 100 | 147 | 80 | 70 | 125 | 98 | 113 | 145 | 105.14 | 92 | 119 | 1.296 | 0.374 | 0.12798159 | 0.26411217 | 0.0575 | 0.025 | 0.0339 | 0.0339 | TRUE | 0.932 |
| Cmya5 | 51 | 32 | 20 | 15 | 16 | 13 | 40 | 34 | 22 | 15 | 18 | 13 | 23.51 | 32 | 15 | 0.476 | -1.071 | 0.0121099 | 0.04769545 | 0 | 0.1033 | 0.0822 | 0.0822 | TRUE | 0.3525 |
| Cnbd2 | 43 | 44 | 32 | 48 | 42 | 43 | 33 | 47 | 35 | 48 | 47 | 42 | 42.06 | 38 | 46 | 1.204 | 0.268 | 0.37408216 | 0.54861034 | 0 | 0.0588 | 0.0418 | 0.0418 | TRUE | 0.2702 |
| Cnbp | 27872 | 19464 | 20289 | 22292 | 19221 | 21369 | 21647 | 20704 | 21892 | 22189 | 21683 | 21094 | 21534.91 | 21414 | 21655 | 1.011 | 0.016 | 0.74137079 | 0.8460229 | 6.00E-04 | 0.0025 | 0.0017 | 0.0017 | TRUE | 0.0103 |
| Cndp1 | 3 | 4 | 2 | 4 | 0 | 5 | 2 | 4 | 2 | 4 | 0 | 5 | 2.94 | 3 | 3 | 1.042 | 0.059 | 0.96011989 | NA | 0 | 0.8077 | 0.6687 | 0.6687 | TRUE | 0.8539 |
| Cndp2 | 5327 | 3755 | 3687 | 3899 | 3281 | 3626 | 4137 | 3994 | 3978 | 3881 | 3701 | 3579 | 3878.57 | 4036 | 3720 | 0.921 | -0.118 | 0.03938229 | 0.11327412 | 7.00E-04 | 0.003 | 0.0021 | 0.0021 | TRUE | 0.0175 |
| Cnep1r1 | 3261 | 2209 | 2296 | 2329 | 2091 | 2383 | 2533 | 2350 | 2477 | 2318 | 2359 | 2352 | 2398.21 | 2453 | 2343 | 0.955 | -0.067 | 0.26635635 | 0.43674458 | 4.00E-04 | 0.0034 | 0.0022 | 0.0022 | TRUE | 0.0164 |
| Cnfn | 1 | 1 | 0 | 1 | 0 | 0 | 1 | 1 | 0 | 1 | 0 | 0 | 0.47 | 1 | 0 | 0.599 | -0.74 | 0.79191656 | NA | 0 | 5.0171 | 3.8827 | 3.8827 | TRUE | 0.3906 |

| | | | | | | | | | | | | | | | | | | | | | | | | | |
|---|---|---|---|---|---|---|---|---|---|---|---|---|---|---|---|---|---|---|---|---|---|---|---|---|---|
| Cnga1 | 0 | 0 | 0 | 0 | 0 | 0 | 0 | 0 | 0 | 0 | 0 | 0 | 0 | 0 | 0 | NA | NA | NA | NA | NA | NA | NA | NA | NA | NA |
| Cnga2 | 0 | 0 | 0 | 0 | 0 | 0 | 0 | 0 | 0 | 0 | 0 | 0 | 0 | 0 | 0 | NA | NA | NA | NA | NA | NA | NA | NA | NA | NA |
| Cnga3 | 594 | 436 | 362 | 372 | 325 | 349 | 461 | 464 | 391 | 370 | 367 | 345 | 399.52 | 439 | 361 | 0.821 | -0.285 | 0.01178333 | 0.0466923 | 0.0031 | 0.0084 | 0.0067 | 0.0067 | TRUE | 0.1023 |
| Cnga4 | 12 | 2 | 7 | 5 | 3 | 6 | 9 | 2 | 8 | 5 | 3 | 6 | 5.55 | 6 | 5 | 0.745 | -0.425 | 0.61573575 | NA | 0 | 0.4297 | 0.3342 | 0.3342 | TRUE | 0.7685 |
| Cngb1 | 330 | 244 | 249 | 270 | 273 | 340 | 256 | 260 | 269 | 269 | 308 | 336 | 282.81 | 262 | 304 | 1.164 | 0.219 | 0.09151583 | 0.20726283 | 0.0034 | 0.0108 | 0.0085 | 0.0085 | TRUE | 0.1197 |
| Cngb3 | 0 | 0 | 0 | 0 | 0 | 0 | 0 | 0 | 0 | 0 | 0 | 0 | 0 | 0 | 0 | NA | NA | NA | NA | NA | NA | NA | NA | NA | NA |
| Cnih | 4114 | 2978 | 3420 | 3783 | 3219 | 3720 | 3195 | 3168 | 3690 | 3766 | 3631 | 3672 | 3520.36 | 3351 | 3690 | 1.102 | 0.14 | 0.04737029 | 0.12953195 | 0.0036 | 0.0031 | 0.0033 | 0.0033 | TRUE | 0.0944 |
| Cnih2 | 205 | 166 | 130 | 133 | 107 | 111 | 159 | 177 | 140 | 132 | 121 | 110 | 139.79 | 159 | 121 | 0.762 | -0.393 | 0.02553479 | 0.08253476 | 0.004 | 0.0194 | 0.015 | 0.015 | TRUE | 0.1015 |
| Cnih3 | 6 | 3 | 6 | 9 | 2 | 4 | 5 | 3 | 6 | 9 | 2 | 4 | 4.91 | 5 | 5 | 1.069 | 0.096 | 0.91436082 | NA | 0 | 0.4847 | 0.3688 | 0.3688 | TRUE | 0.9419 |
| Cnih4 | 2930 | 1977 | 2406 | 2802 | 2479 | 2709 | 2276 | 2103 | 2596 | 2789 | 2797 | 2674 | 2539.07 | 2325 | 2753 | 1.185 | 0.244 | 0.0020443 | 0.01227055 | 0.0055 | 0.0034 | 0.0041 | 0.0041 | TRUE | 0.1242 |
| Cnksr1 | 159 | 121 | 146 | 174 | 151 | 182 | 123 | 129 | 158 | 173 | 170 | 180 | 155.49 | 137 | 174 | 1.282 | 0.359 | 0.03043957 | 0.09356404 | 0 | 0.0177 | 0.0133 | 0.0133 | TRUE | 0.1799 |
| Cnksr2 | 431 | 315 | 263 | 359 | 282 | 307 | 335 | 335 | 284 | 357 | 318 | 303 | 322.02 | 318 | 326 | 1.025 | 0.035 | 0.77904781 | 0.86964954 | 0.0049 | 0.0098 | 0.0083 | 0.0083 | TRUE | 0.0961 |
| Cnksr3 | 1565 | 1064 | 941 | 1009 | 931 | 1113 | 1215 | 1132 | 1015 | 1004 | 1050 | 1099 | 1085.98 | 1121 | 1051 | 0.936 | -0.095 | 0.27440649 | 0.44540743 | 0.0043 | 0.0046 | 0.0045 | 0.0045 | TRUE | 0.0805 |
| Cnn1 | 66 | 53 | 52 | 126 | 104 | 126 | 51 | 56 | 56 | 125 | 117 | 124 | 88.48 | 54 | 122 | 2.25 | 1.17 | 2.94E-08 | 1.30E-06 | 0 | 0.0292 | 0.019 | 0.019 | TRUE | 0.0284 |
| Cnn2 | 2129 | 1533 | 1537 | 1550 | 1490 | 1787 | 1654 | 1631 | 1658 | 1543 | 1681 | 1764 | 1655.06 | 1648 | 1663 | 1.009 | 0.013 | 0.85555022 | 0.91884279 | 0.0017 | 0.0039 | 0.0031 | 0.0031 | TRUE | 0.0485 |
| Cnn3 | 17730 | 12870 | 13418 | 14815 | 12584 | 14657 | 13770 | 13690 | 14478 | 14747 | 14196 | 14469 | 14224.89 | 13979 | 14471 | 1.035 | 0.05 | 0.30726501 | 0.47991901 | 6.00E-04 | 0.0026 | 0.0017 | 0.0017 | TRUE | 0.0118 |
| Cnnm1 | 229 | 198 | 134 | 166 | 123 | 170 | 178 | 211 | 145 | 165 | 139 | 168 | 167.48 | 178 | 157 | 0.886 | -0.174 | 0.3241695 | 0.49809594 | 0.0164 | 0.0166 | 0.0165 | 0.0165 | TRUE | 0.1579 |
| Cnnm2 | 2309 | 1659 | 1838 | 2157 | 1992 | 2271 | 1793 | 1765 | 1983 | 2147 | 2247 | 2242 | 2029.54 | 1847 | 2212 | 1.198 | 0.261 | 0.00014884 | 0.00169648 | 0.0016 | 0.0036 | 0.0029 | 0.0029 | TRUE | 0.0492 |
| Cnnm3 | 2113 | 1728 | 1462 | 1579 | 1564 | 1692 | 1641 | 1838 | 1577 | 1572 | 1764 | 1670 | 1677.16 | 1685 | 1669 | 0.99 | -0.014 | 0.85580508 | 0.91887185 | 0.0042 | 0.0038 | 0.004 | 0.004 | TRUE | 0.0743 |
| Cnnm4 | 1327 | 987 | 898 | 859 | 819 | 830 | 1031 | 1050 | 969 | 855 | 924 | 819 | 941.29 | 1017 | 866 | 0.851 | -0.232 | 0.00462908 | 0.02311791 | 0.0015 | 0.0049 | 0.0038 | 0.0038 | TRUE | 0.0397 |
| Cnot1 | 59189 | 42857 | 46396 | 50289 | 46039 | 53238 | 45970 | 45586 | 50061 | 50058 | 51937 | 52554 | 49360.9 | 47206 | 51516 | 1.091 | 0.126 | 0.02036372 | 0.07008396 | 0.0017 | 0.0025 | 0.0021 | 0.0021 | TRUE | 0.0342 |
| Cnot10 | 5431 | 4205 | 3991 | 4091 | 3850 | 4402 | 4218 | 4473 | 4306 | 4072 | 4343 | 4345 | 4292.99 | 4332 | 4253 | 0.982 | -0.026 | 0.64509919 | 0.77901341 | 8.00E-04 | 0.003 | 0.0021 | 0.0021 | TRUE | 0.017 |
| Cnot11 | 3531 | 2367 | 2184 | 2002 | 1953 | 2114 | 2742 | 2518 | 2357 | 1993 | 2203 | 2087 | 2316.58 | 2539 | 2094 | 0.824 | -0.279 | 0.00018287 | 0.0019952 | 0.0035 | 0.0034 | 0.0036 | 0.0036 | TRUE | 0.0608 |
| Cnot2 | 5846 | 4361 | 4230 | 4462 | 4042 | 4732 | 4540 | 4639 | 4564 | 4441 | 4560 | 4671 | 4569.28 | 4581 | 4557 | 0.995 | -0.007 | 0.88815748 | 0.93792008 | 2.00E-04 | 0.0029 | 0.0017 | 0.0017 | TRUE | 0.0061 |
| Cnot3 | 8247 | 6163 | 5910 | 5654 | 5459 | 6308 | 6405 | 6556 | 6377 | 5628 | 6158 | 6227 | 6225.12 | 6446 | 6004 | 0.931 | -0.102 | 0.0759062 | 0.18169943 | 0.0014 | 0.0028 | 0.0022 | 0.0022 | TRUE | 0.0368 |
| Cnot4 | 2783 | 2064 | 1893 | 2056 | 1756 | 2182 | 2161 | 2195 | 2043 | 2047 | 1981 | 2154 | 2096.81 | 2133 | 2061 | 0.966 | -0.05 | 0.44581384 | 0.61648773 | 0.0011 | 0.0036 | 0.0026 | 0.0026 | TRUE | 0.0191 |
| Cnot6 | 5826 | 4023 | 3600 | 3678 | 3217 | 3536 | 4525 | 4279 | 3884 | 3661 | 3629 | 3491 | 3911.53 | 4229 | 3594 | 0.849 | -0.236 | 0.00049713 | 0.00428926 | 0.0028 | 0.003 | 0.003 | 0.003 | TRUE | 0.0618 |
| Cnot6l | 1329 | 1079 | 1137 | 1182 | 935 | 1139 | 1032 | 1148 | 1227 | 1177 | 1055 | 1124 | 1127.07 | 1136 | 1119 | 0.987 | -0.019 | 0.82655801 | 0.90038821 | 0.0045 | 0.0045 | 0.0045 | 0.0045 | TRUE | 0.0791 |
| Cnot7 | 7959 | 5758 | 5879 | 6108 | 5280 | 5983 | 6181 | 6125 | 6343 | 6080 | 5956 | 5906 | 6098.66 | 6216 | 5981 | 0.962 | -0.056 | 0.25860771 | 0.42850468 | 1.00E-04 | 0.0028 | 0.0016 | 0.0016 | TRUE | 0.0038 |
| Cnot8 | 5180 | 3792 | 4043 | 4493 | 3925 | 4635 | 4023 | 4034 | 4362 | 4472 | 4428 | 4575 | 4315.76 | 4140 | 4492 | 1.085 | 0.118 | 0.03876057 | 0.11194279 | 9.00E-04 | 0.003 | 0.0021 | 0.0021 | TRUE | 0.0267 |
| Cnp | 3192 | 2170 | 2113 | 2510 | 2213 | 2581 | 2479 | 2308 | 2280 | 2498 | 2496 | 2548 | 2435 | 2356 | 2514 | 1.066 | 0.093 | 0.13242845 | 0.27039998 | 8.00E-04 | 0.0034 | 0.0023 | 0.0023 | TRUE | 0.0262 |
| Cnppd1 | 1337 | 976 | 965 | 1138 | 998 | 998 | 1038 | 1038 | 1041 | 1133 | 1126 | 985 | 1060.26 | 1039 | 1081 | 1.04 | 0.057 | 0.48417715 | 0.65044468 | 0.0021 | 0.0047 | 0.0038 | 0.0038 | TRUE | 0.0732 |
| Cnpy1 | 449 | 467 | 497 | 552 | 458 | 614 | 349 | 497 | 536 | 549 | 517 | 606 | 508.99 | 461 | 557 | 1.215 | 0.281 | 0.05055275 | 0.13591994 | 0.0264 | 0.0071 | 0.0129 | 0.0129 | TRUE | 0.5496 |
| Cnpy2 | 2215 | 1610 | 1406 | 1459 | 1355 | 1447 | 1720 | 1713 | 1517 | 1452 | 1529 | 1428 | 1559.86 | 1650 | 1470 | 0.89 | -0.168 | 0.02496888 | 0.08133095 | 0.0025 | 0.0039 | 0.0034 | 0.0034 | TRUE | 0.0588 |
| Cnpy3 | 2284 | 1753 | 1877 | 2107 | 1895 | 2228 | 1774 | 1865 | 2025 | 2097 | 2138 | 2199 | 2016.37 | 1888 | 2145 | 1.137 | 0.185 | 0.00789217 | 0.03471924 | 0.0018 | 0.0036 | 0.003 | 0.003 | TRUE | 0.0487 |
| Cnpy4 | 929 | 694 | 641 | 654 | 536 | 630 | 722 | 738 | 692 | 651 | 605 | 622 | 671.48 | 717 | 626 | 0.873 | -0.196 | 0.02352708 | 0.07776887 | 0 | 0.006 | 0.0039 | 0.0039 | TRUE | 0.0146 |
| Cnr1 | 143 | 90 | 81 | 123 | 100 | 116 | 111 | 96 | 87 | 122 | 113 | 115 | 107.32 | 98 | 117 | 1.183 | 0.243 | 0.20819726 | 0.36975192 | 0 | 0.0245 | 0.0175 | 0.0175 | TRUE | 0.1463 |
| Cnr2 | 3 | 3 | 0 | 3 | 1 | 1 | 2 | 3 | 0 | 3 | 1 | 1 | 1.77 | 2 | 2 | 0.919 | -0.123 | 0.9341462 | NA | 0 | 1.3411 | 1.0247 | 1.0247 | TRUE | 0.5606 |
| Cnrip1 | 6 | 9 | 2 | 6 | 5 | 2 | 5 | 10 | 2 | 6 | 6 | 2 | 5 | 6 | 5 | 0.828 | -0.272 | 0.76519315 | NA | 0.1731 | 0.4768 | 0.3922 | 0.3922 | TRUE | 0.4008 |
| Cnst | 1400 | 998 | 1005 | 1162 | 978 | 1125 | 1087 | 1062 | 1084 | 1157 | 1103 | 1111 | 1100.63 | 1078 | 1124 | 1.042 | 0.06 | 0.40170235 | 0.5748316 | 0 | 0.0046 | 0.0028 | 0.0028 | TRUE | 0.008 |
| Cntd1 | 397 | 317 | 455 | 542 | 483 | 516 | 308 | 337 | 491 | 540 | 545 | 509 | 455.04 | 379 | 531 | 1.407 | 0.492 | 0.00102232 | 0.00735014 | 0.0308 | 0.0076 | 0.0139 | 0.0139 | TRUE | 0.7611 |
| Cntf | 8 | 5 | 6 | 4 | 5 | 2 | 6 | 5 | 6 | 4 | 6 | 2 | 4.93 | 6 | 4 | 0.637 | -0.65 | 0.45185359 | NA | 0 | 0.4828 | 0.3222 | 0.3222 | TRUE | 0.3085 |
| Cntfr | 513 | 382 | 376 | 684 | 556 | 667 | 398 | 406 | 406 | 681 | 627 | 658 | 529.49 | 403 | 655 | 1.626 | 0.701 | 1.11E-13 | 1.61E-11 | 0 | 0.0069 | 0.0044 | 0.0044 | TRUE | 0.0163 |
| Cntln | 327 | 213 | 185 | 143 | 141 | 123 | 254 | 227 | 200 | 142 | 159 | 121 | 183.83 | 227 | 141 | 0.619 | -0.693 | 2.85E-05 | 0.00044393 | 0.0106 | 0.0153 | 0.0139 | 0.0139 | TRUE | 0.1542 |
| Cntn1 | 21 | 11 | 9 | 12 | 12 | 4 | 16 | 12 | 10 | 12 | 14 | 4 | 11.19 | 13 | 10 | 0.768 | -0.381 | 0.53711439 | NA | 0.0905 | 0.2142 | 0.183 | 0.183 | TRUE | 0.9726 |
| Cntn2 | 906 | 715 | 819 | 1198 | 1182 | 1228 | 704 | 761 | 884 | 1192 | 1333 | 1212 | 1014.34 | 783 | 1246 | 1.595 | 0.673 | 2.41E-12 | 2.78E-10 | 0.0072 | 0.0048 | 0.0056 | 0.0056 | TRUE | 0.1502 |
| Cntn3 | 60 | 28 | 29 | 29 | 40 | 37 | 47 | 30 | 31 | 29 | 45 | 37 | 36.36 | 36 | 37 | 1.013 | 0.019 | 0.95609768 | 0.9757358 | 0.0319 | 0.0676 | 0.0574 | 0.0574 | TRUE | 0.4137 |
| Cntn4 | 5 | 4 | 4 | 3 | 3 | 6 | 4 | 4 | 4 | 3 | 3 | 6 | 4.12 | 4 | 4 | 0.993 | -0.01 | 0.99141744 | NA | 0 | 0.577 | 0.3683 | 0.3683 | TRUE | 0.2699 |
| Cntn5 | 3 | 1 | 3 | 5 | 2 | 0 | 2 | 1 | 3 | 5 | 2 | 0 | 2.31 | 2 | 2 | 1.09 | 0.124 | 0.92655853 | NA | 0.3393 | 1.0281 | 0.869 | 0.869 | TRUE | 0.2264 |
| Cntn6 | 12 | 12 | 12 | 3 | 8 | 5 | 9 | 13 | 13 | 3 | 9 | 5 | 8.66 | 12 | 6 | 0.481 | -1.056 | 0.12303749 | NA | 0 | 0.276 | 0.2037 | 0.2037 | TRUE | 0.543 |
| Cntnap1 | 44 | 42 | 45 | 62 | 62 | 60 | 34 | 45 | 49 | 62 | 70 | 59 | 53.05 | 43 | 64 | 1.509 | 0.593 | 0.03033573 | 0.09336534 | 0 | 0.0471 | 0.0345 | 0.0345 | TRUE | 0.2697 |
| Cntnap2 | 898 | 603 | 483 | 643 | 496 | 533 | 697 | 641 | 521 | 640 | 560 | 526 | 597.62 | 620 | 575 | 0.926 | -0.111 | 0.36137912 | 0.53636467 | 0.0144 | 0.0064 | 0.0089 | 0.0089 | TRUE | 0.2256 |
| Cntnap3 | 71 | 43 | 20 | 31 | 30 | 19 | 55 | 46 | 22 | 31 | 34 | 19 | 34.32 | 41 | 28 | 0.674 | -0.569 | 0.15707088 | 0.30565201 | 0.1237 | 0.0715 | 0.0862 | 0.0862 | TRUE | 0.5382 |
| Cntnap4 | 8 | 3 | 4 | 3 | 1 | 3 | 6 | 3 | 4 | 3 | 1 | 3 | 3.47 | 4 | 2 | 0.515 | -0.956 | 0.35742773 | NA | 0 | 0.6862 | 0.4567 | 0.4567 | TRUE | 0.2418 |
| Cntnap5a | 1 | 0 | 0 | 0 | 0 | 0 | 1 | 0 | 0 | 0 | 0 | 0 | 0.13 | 0 | 0 | 0.552 | -0.857 | 0.8336548 | NA | 0 | 18.3121 | 10 | 10 | TRUE | 0.2566 |
| Cntnap5b | 0 | 0 | 0 | 0 | 0 | 0 | 0 | 0 | 0 | 0 | 0 | 0 | 0 | 0 | 0 | NA | NA | NA | NA | NA | NA | NA | NA | NA | NA |
| Cntnap5c | 0 | 0 | 0 | 0 | 0 | 0 | 0 | 0 | 0 | 0 | 0 | 0 | 0 | 0 | 0 | NA | NA | NA | NA | NA | NA | NA | NA | NA | NA |
| Cntrob | 3272 | 2448 | 2418 | 2544 | 2336 | 2683 | 2541 | 2604 | 2609 | 2532 | 2635 | 2649 | 2595.04 | 2585 | 2605 | 1.008 | 0.012 | 0.83736064 | 0.9072782 | 0 | 0.0033 | 0.002 | 0.002 | TRUE | 0.0074 |
| Coa3 | 2089 | 1354 | 1353 | 1418 | 1126 | 1471 | 1622 | 1440 | 1460 | 1411 | 1270 | 1452 | 1442.73 | 1507 | 1378 | 0.914 | -0.13 | 0.10696449 | 0.23241766 | 0.0036 | 0.0041 | 0.004 | 0.004 | TRUE | 0.0553 |
| Coa4 | 509 | 343 | 322 | 442 | 389 | 420 | 395 | 365 | 347 | 440 | 439 | 415 | 400.17 | 369 | 431 | 1.165 | 0.22 | 0.04118158 | 0.11674104 | 0 | 0.0083 | 0.0059 | 0.0059 | TRUE | 0.0472 |
| Coa5 | 7726 | 5477 | 5580 | 6209 | 5254 | 6253 | 6000 | 5826 | 6021 | 6180 | 5927 | 6173 | 6021.2 | 5949 | 6093 | 1.024 | 0.035 | 0.49302211 | 0.65826857 | 2.00E-04 | 0.0028 | 0.0017 | 0.0017 | TRUE | 0.0069 |
| Coa6 | 1151 | 796 | 692 | 781 | 675 | 790 | 894 | 847 | 747 | 777 | 761 | 780 | 801 | 829 | 773 | 0.931 | -0.103 | 0.25233612 | 0.42118886 | 0.0032 | 0.0054 | 0.0046 | 0.0046 | TRUE | 0.0885 |
| Coasy | 2418 | 1820 | 1996 | 2201 | 1929 | 2246 | 1878 | 1936 | 2154 | 2191 | 2176 | 2217 | 2091.94 | 1989 | 2195 | 1.104 | 0.143 | 0.04035671 | 0.11517939 | 0.002 | 0.0036 | 0.003 | 0.003 | TRUE | 0.0627 |
| Cobl | 7403 | 5295 | 5228 | 8530 | 7227 | 8423 | 5750 | 5632 | 5641 | 8491 | 8153 | 8315 | 6996.86 | 5674 | 8320 | 1.466 | 0.552 | 6.87E-30 | 5.77E-27 | 2.00E-04 | 0.0028 | 0.0016 | 0.0016 | TRUE | 0.004 |
| Cobll1 | 459 | 319 | 275 | 259 | 224 | 244 | 356 | 339 | 297 | 258 | 253 | 241 | 290.65 | 331 | 251 | 0.755 | -0.405 | 0.00137331 | 0.00904552 | 0 | 0.0106 | 0.008 | 0.008 | TRUE | 0.0903 |
| Coch | 1277 | 848 | 790 | 1217 | 952 | 1101 | 992 | 902 | 852 | 1211 | 1074 | 1087 | 1019.74 | 915 | 1124 | 1.226 | 0.294 | 0.00081848 | 0.00624654 | 0.0043 | 0.0048 | 0.0046 | 0.0046 | TRUE | 0.0661 |
| Cog1 | 3602 | 2630 | 2946 | 2889 | 2579 | 3020 | 2798 | 2797 | 3179 | 2876 | 2909 | 2981 | 2923.34 | 2925 | 2922 | 1 | 0 | 0.99414771 | 0.99663461 | 0.0026 | 0.0032 | 0.003 | 0.003 | TRUE | 0.0694 |
| Cog2 | 3279 | 2413 | 2285 | 2355 | 2156 | 2508 | 2547 | 2567 | 2465 | 2344 | 2432 | 2476 | 2471.83 | 2526 | 2417 | 0.957 | -0.064 | 0.27910599 | 0.45068084 | 0 | 0.0034 | 0.0021 | 0.0021 | TRUE | 0.0086 |
| Cog3 | 3159 | 2298 | 2292 | 2464 | 2195 | 2404 | 2453 | 2444 | 2473 | 2453 | 2476 | 2373 | 2445.47 | 2457 | 2434 | 0.991 | -0.014 | 0.81073271 | 0.8908004 | 0 | 0.0034 | 0.002 | 0.002 | TRUE | 0.0058 |

| | | | | | | | | | | | | | | | | | | | | | | | | | |
|---|---|---|---|---|---|---|---|---|---|---|---|---|---|---|---|---|---|---|---|---|---|---|---|---|---|
| Cog4 | 9855 | 7423 | 7750 | 7942 | 7029 | 8272 | 7654 | 7896 | 8362 | 7905 | 7929 | 8166 | 7985.42 | 7971 | 8000 | 1.004 | 0.006 | 0.91574936 | 0.95343864 | 0.0011 | 0.0027 | 0.002 | 0.002 | TRUE | 0.0225 |
| Cog5 | 2212 | 1619 | 1787 | 1964 | 1698 | 1973 | 1718 | 1722 | 1928 | 1955 | 1916 | 1948 | 1864.4 | 1789 | 1940 | 1.085 | 0.117 | 0.09163112 | 0.20745819 | 0.0015 | 0.0037 | 0.0029 | 0.0029 | TRUE | 0.0547 |
| Cog6 | 3975 | 2759 | 2947 | 3292 | 2900 | 3451 | 3087 | 2935 | 3180 | 3277 | 3272 | 3407 | 3192.79 | 3067 | 3319 | 1.082 | 0.114 | 0.05330513 | 0.14101245 | 6.00E-04 | 0.0032 | 0.0022 | 0.0022 | TRUE | 0.017 |
| Cog7 | 2544 | 1879 | 1858 | 1727 | 1565 | 1823 | 1976 | 1999 | 2005 | 1719 | 1765 | 1800 | 1877.23 | 1993 | 1761 | 0.884 | -0.178 | 0.0035667 | 0.01886709 | 0 | 0.0037 | 0.0022 | 0.0022 | TRUE | 0.0054 |
| Cog8 | 3154 | 2086 | 1701 | 1836 | 1710 | 1799 | 2450 | 2219 | 1835 | 1828 | 1929 | 1776 | 2006.05 | 2168 | 1844 | 0.85 | -0.235 | 0.0141272 | 0.05339076 | 0.0104 | 0.0036 | 0.0061 | 0.0061 | TRUE | 0.2169 |
| Coil | 5473 | 3915 | 4102 | 4451 | 4019 | 4620 | 4251 | 4164 | 4426 | 4431 | 4534 | 4561 | 4394.34 | 4280 | 4509 | 1.053 | 0.075 | 0.16144399 | 0.31132203 | 3.00E-04 | 0.003 | 0.0018 | 0.0018 | TRUE | 0.0106 |
| Col10a1 | 1 | 0 | 0 | 0 | 0 | 0 | 1 | 0 | 0 | 0 | 0 | 0 | 0.13 | 0 | 0 | 0.552 | -0.857 | 0.8336548 | NA | 0 | 18.3121 | 10 | 10 | TRUE | 0.2566 |
| Col11a1 | 551 | 343 | 294 | 273 | 253 | 268 | 428 | 365 | 317 | 272 | 285 | 265 | 321.95 | 370 | 274 | 0.737 | -0.44 | 0.00108486 | 0.00767615 | 0.01 | 0.0098 | 0.0098 | 0.0098 | TRUE | 0.2182 |
| Col11a2 | 18 | 25 | 19 | 29 | 17 | 28 | 14 | 27 | 21 | 29 | 19 | 28 | 22.79 | 21 | 25 | 1.258 | 0.331 | 0.43532783 | 0.60698451 | 0.0268 | 0.1064 | 0.0854 | 0.0854 | TRUE | 0.2891 |
| Col12a1 | 85 | 52 | 48 | 42 | 34 | 25 | 66 | 55 | 52 | 42 | 38 | 25 | 46.33 | 58 | 35 | 0.602 | -0.732 | 0.01642293 | 0.05973653 | 0.0154 | 0.0536 | 0.044 | 0.044 | TRUE | 0.4877 |
| Col13a1 | 214 | 133 | 152 | 239 | 183 | 178 | 166 | 141 | 164 | 238 | 206 | 176 | 181.96 | 157 | 207 | 1.312 | 0.392 | 0.01733081 | 0.06217602 | 0.0105 | 0.0155 | 0.014 | 0.014 | TRUE | 0.1521 |
| Col14a1 | 15 | 12 | 13 | 6 | 13 | 6 | 12 | 13 | 14 | 6 | 15 | 6 | 10.83 | 13 | 9 | 0.685 | -0.546 | 0.37007576 | NA | 0.0179 | 0.2212 | 0.1703 | 0.1703 | TRUE | 0.9446 |
| Col15a1 | 46 | 37 | 35 | 19 | 23 | 19 | 36 | 39 | 38 | 19 | 26 | 19 | 29.41 | 38 | 21 | 0.562 | -0.831 | 0.02103199 | 0.07177894 | 0 | 0.083 | 0.0561 | 0.0561 | TRUE | 0.1943 |
| Col16a1 | 50 | 43 | 37 | 40 | 26 | 37 | 39 | 46 | 40 | 40 | 29 | 37 | 38.36 | 42 | 35 | 0.853 | -0.229 | 0.46312595 | 0.63236939 | 0 | 0.0642 | 0.044 | 0.044 | TRUE | 0.14 |
| Col17a1 | 176 | 149 | 163 | 160 | 145 | 182 | 137 | 158 | 176 | 159 | 164 | 180 | 162.26 | 157 | 168 | 1.072 | 0.1 | 0.54332565 | 0.70095114 | 0.0046 | 0.017 | 0.0134 | 0.0134 | TRUE | 0.1501 |
| Col18a1 | 11973 | 9100 | 9379 | 10417 | 10132 | 11113 | 9299 | 9680 | 10120 | 10369 | 11430 | 10970 | 10311.27 | 9700 | 10923 | 1.126 | 0.172 | 0.00340453 | 0.0182763 | 0.0019 | 0.0027 | 0.0024 | 0.0024 | TRUE | 0.0241 |
| Col19a1 | 4 | 1 | 1 | 2 | 3 | 0 | 3 | 1 | 1 | 2 | 3 | 0 | 1.77 | 2 | 2 | 0.982 | -0.026 | 0.98615169 | NA | 0 | 1.3409 | 1.0489 | 1.0489 | TRUE | 0.3085 |
| Col1a1 | 182 | 122 | 98 | 184 | 153 | 163 | 141 | 130 | 106 | 183 | 173 | 161 | 148.92 | 126 | 172 | 1.364 | 0.448 | 0.00947302 | 0.03982292 | 0.005 | 0.0183 | 0.0147 | 0.0147 | TRUE | 0.188 |
| Col1a2 | 81 | 64 | 42 | 58 | 51 | 69 | 63 | 68 | 45 | 58 | 58 | 68 | 59.95 | 59 | 61 | 1.038 | 0.054 | 0.83633835 | 0.90661995 | 0.0075 | 0.042 | 0.0328 | 0.0328 | TRUE | 0.3183 |
| Col20a1 | 59 | 59 | 55 | 64 | 57 | 52 | 46 | 63 | 59 | 64 | 64 | 51 | 57.88 | 56 | 60 | 1.074 | 0.103 | 0.69763294 | 0.81631335 | 0 | 0.0434 | 0.0329 | 0.0329 | TRUE | 0.2486 |
| Col22a1 | 585 | 450 | 403 | 417 | 409 | 340 | 454 | 479 | 435 | 415 | 461 | 336 | 429.99 | 456 | 404 | 0.885 | -0.176 | 0.15790688 | 0.30686105 | 0.0112 | 0.0079 | 0.0089 | 0.0089 | TRUE | 0.255 |
| Col23a1 | 118 | 80 | 77 | 98 | 81 | 64 | 92 | 85 | 83 | 98 | 91 | 63 | 85.32 | 87 | 84 | 0.967 | -0.048 | 0.83295347 | 0.90481048 | 0.0144 | 0.0302 | 0.0259 | 0.0259 | TRUE | 0.4562 |
| Col24a1 | 23 | 12 | 15 | 10 | 2 | 4 | 18 | 13 | 16 | 10 | 2 | 4 | 10.5 | 16 | 5 | 0.348 | -1.523 | 0.02141852 | NA | 0.0142 | 0.2283 | 0.1899 | 0.1899 | TRUE | 1.1963 |
| Col25a1 | 31 | 24 | 24 | 30 | 25 | 16 | 24 | 26 | 26 | 30 | 28 | 16 | 24.89 | 25 | 25 | 0.977 | -0.033 | 0.93388975 | 0.96320438 | 0 | 0.0976 | 0.0748 | 0.0748 | TRUE | 0.616 |
| Col26a1 | 62 | 48 | 72 | 88 | 70 | 87 | 48 | 51 | 78 | 88 | 79 | 86 | 71.56 | 59 | 84 | 1.438 | 0.524 | 0.03711042 | 0.10853677 | 0.019 | 0.0355 | 0.0312 | 0.0312 | TRUE | 0.6341 |
| Col27a1 | 254 | 181 | 170 | 174 | 143 | 156 | 197 | 193 | 183 | 173 | 161 | 154 | 176.96 | 191 | 163 | 0.851 | -0.232 | 0.12078889 | 0.25306883 | 0 | 0.0158 | 0.0104 | 0.0104 | TRUE | 0.0338 |
| Col28a1 | 8 | 9 | 6 | 8 | 4 | 8 | 6 | 10 | 6 | 8 | 5 | 8 | 7.11 | 7 | 7 | 0.926 | -0.111 | 0.87648752 | NA | 0 | 0.336 | 0.2256 | 0.2256 | TRUE | 0.1963 |
| Col2a1 | 297 | 246 | 207 | 168 | 159 | 148 | 231 | 262 | 223 | 167 | 179 | 146 | 201.4 | 239 | 164 | 0.688 | -0.54 | 0.00033183 | 0.00314095 | 0.0032 | 0.0142 | 0.0111 | 0.0111 | TRUE | 0.1003 |
| Col3a1 | 1 | 2 | 0 | 0 | 0 | 0 | 1 | 2 | 0 | 0 | 0 | 0 | 0.48 | 1 | 0 | 0.197 | -2.343 | 0.43249452 | NA | 0 | 4.8992 | 4.9172 | 4.9172 | TRUE | 0.1226 |
| Col4a1 | 8951 | 5991 | 5910 | 8458 | 6797 | 6842 | 6952 | 6373 | 6377 | 8419 | 7668 | 6754 | 7090.36 | 6567 | 7614 | 1.159 | 0.213 | 0.00712783 | 0.03218897 | 0.0071 | 0.0028 | 0.0044 | 0.0044 | TRUE | 0.1194 |
| Col4a2 | 3201 | 2066 | 2034 | 2820 | 2322 | 2158 | 2486 | 2198 | 2195 | 2807 | 2619 | 2130 | 2405.85 | 2293 | 2519 | 1.098 | 0.135 | 0.16444109 | 0.31523023 | 0.0117 | 0.0034 | 0.0064 | 0.0064 | TRUE | 0.222 |
| Col4a3 | 2 | 0 | 0 | 0 | 2 | 2 | 2 | 0 | 0 | 0 | 2 | 2 | 0.96 | 1 | 1 | 2.479 | 1.31 | 0.54958602 | NA | 0 | 2.4611 | 2.2311 | 2.2311 | TRUE | 0.9717 |
| Col4a3bp | 4031 | 2850 | 2639 | 2728 | 2271 | 2644 | 3131 | 3032 | 2847 | 2715 | 2562 | 2610 | 2816.18 | 3003 | 2629 | 0.875 | -0.192 | 0.0021177 | 0.01259604 | 0.0013 | 0.0033 | 0.0025 | 0.0025 | TRUE | 0.0248 |
| Col4a4 | 6 | 6 | 3 | 2 | 6 | 6 | 5 | 6 | 3 | 2 | 7 | 6 | 4.83 | 5 | 5 | 1.023 | 0.032 | 0.97104377 | NA | 0 | 0.4934 | 0.3562 | 0.3562 | TRUE | 0.5512 |
| Col4a5 | 109 | 67 | 61 | 53 | 52 | 52 | 85 | 71 | 66 | 53 | 59 | 51 | 64.08 | 74 | 54 | 0.729 | -0.455 | 0.06503966 | 0.16271869 | 0 | 0.0394 | 0.0278 | 0.0278 | TRUE | 0.1657 |
| Col4a6 | 34 | 28 | 15 | 24 | 14 | 17 | 26 | 30 | 16 | 24 | 16 | 17 | 21.47 | 24 | 19 | 0.779 | -0.36 | 0.40859481 | 0.58139857 | 0 | 0.1128 | 0.0891 | 0.0891 | TRUE | 0.4544 |
| Col5a1 | 1908 | 1306 | 1265 | 1392 | 1168 | 1181 | 1482 | 1389 | 1365 | 1386 | 1318 | 1166 | 1350.84 | 1412 | 1290 | 0.913 | -0.132 | 0.10906195 | 0.23575657 | 0.0041 | 0.0042 | 0.0041 | 0.0041 | TRUE | 0.0859 |
| Col5a2 | 172 | 103 | 110 | 50 | 42 | 47 | 134 | 110 | 119 | 50 | 47 | 46 | 84.23 | 121 | 48 | 0.396 | -1.338 | 2.32E-09 | 1.40E-07 | 0 | 0.0306 | 0.0213 | 0.0213 | TRUE | 0.0983 |
| Col5a3 | 618 | 460 | 541 | 926 | 773 | 873 | 480 | 489 | 584 | 922 | 872 | 862 | 701.43 | 518 | 885 | 1.714 | 0.777 | 2.11E-15 | 3.67E-13 | 0.0045 | 0.0058 | 0.0054 | 0.0054 | TRUE | 0.142 |
| Col6a1 | 33 | 36 | 23 | 35 | 37 | 43 | 26 | 38 | 25 | 35 | 42 | 42 | 34.63 | 30 | 40 | 1.347 | 0.43 | 0.20387402 | 0.36525999 | 0 | 0.0709 | 0.053 | 0.053 | TRUE | 0.4138 |
| Col6a2 | 208 | 135 | 142 | 160 | 132 | 158 | 162 | 144 | 153 | 159 | 149 | 156 | 153.75 | 153 | 155 | 1.011 | 0.016 | 0.91998073 | 0.956103 | 0 | 0.0178 | 0.0117 | 0.0117 | TRUE | 0.0293 |
| Col6a3 | 34 | 47 | 50 | 27 | 35 | 31 | 26 | 50 | 54 | 27 | 39 | 31 | 37.89 | 43 | 32 | 0.749 | -0.416 | 0.2474791 | 0.41629031 | 0.0685 | 0.065 | 0.066 | 0.066 | TRUE | 1.0681 |
| Col6a4 | 300 | 260 | 243 | 221 | 222 | 239 | 233 | 277 | 262 | 220 | 250 | 236 | 246.35 | 257 | 235 | 0.917 | -0.125 | 0.35618069 | 0.53063584 | 0 | 0.012 | 0.0091 | 0.0091 | TRUE | 0.0827 |
| Col6a5 | 210 | 138 | 111 | 149 | 150 | 172 | 163 | 147 | 120 | 148 | 169 | 170 | 152.83 | 143 | 162 | 1.128 | 0.174 | 0.31513482 | 0.48870278 | 0.0082 | 0.0179 | 0.0152 | 0.0152 | TRUE | 0.2069 |
| Col6a6 | 187 | 182 | 143 | 142 | 132 | 174 | 145 | 194 | 154 | 141 | 149 | 172 | 159.19 | 164 | 154 | 0.94 | -0.089 | 0.61059134 | 0.75370775 | 0.0113 | 0.0173 | 0.0155 | 0.0155 | TRUE | 0.2519 |
| Col7a1 | 188 | 159 | 120 | 144 | 124 | 151 | 146 | 169 | 129 | 143 | 140 | 149 | 146.15 | 148 | 144 | 0.973 | -0.04 | 0.81632855 | 0.89466126 | 0 | 0.0186 | 0.014 | 0.014 | TRUE | 0.1495 |
| Col8a1 | 0 | 1 | 0 | 1 | 0 | 1 | 0 | 1 | 0 | 1 | 0 | 1 | 0.51 | 0 | 1 | 1.941 | 0.957 | 0.72720315 | NA | 0 | 4.6706 | 3.6514 | 3.6514 | TRUE | 0.5217 |
| Col8a2 | 111 | 91 | 82 | 152 | 117 | 118 | 86 | 97 | 88 | 151 | 132 | 116 | 111.88 | 90 | 133 | 1.475 | 0.561 | 0.0037224 | 0.01952447 | 0 | 0.0236 | 0.0177 | 0.0177 | TRUE | 0.1483 |
| Col9a1 | 43 | 28 | 18 | 9 | 13 | 13 | 33 | 30 | 19 | 9 | 15 | 13 | 19.84 | 27 | 12 | 0.437 | -1.195 | 0.00986068 | 0.04095282 | 0 | 0.1219 | 0.0943 | 0.0943 | TRUE | 0.3997 |
| Col9a2 | 893 | 639 | 508 | 589 | 482 | 540 | 694 | 680 | 548 | 586 | 544 | 533 | 597.41 | 641 | 554 | 0.864 | -0.21 | 0.05380179 | 0.14180352 | 0.0079 | 0.0064 | 0.0069 | 0.0069 | TRUE | 0.1832 |
| Col9a3 | 399 | 317 | 253 | 441 | 357 | 408 | 310 | 337 | 273 | 439 | 403 | 403 | 360.75 | 307 | 415 | 1.352 | 0.435 | 0.00023983 | 0.00245022 | 0.0031 | 0.009 | 0.0073 | 0.0073 | TRUE | 0.098 |
| Colec10 | 0 | 0 | 0 | 0 | 0 | 0 | 0 | 0 | 0 | 0 | 0 | 0 | 0 | 0 | 0 | NA | NA | NA | NA | NA | NA | NA | NA | NA | NA |
| Colec11 | 1 | 0 | 1 | 1 | 0 | 2 | 1 | 0 | 1 | 1 | 0 | 2 | 0.8 | 1 | 1 | 1.608 | 0.686 | 0.7523688 | NA | 0 | 2.9494 | 2.1146 | 2.1146 | TRUE | 0.1306 |
| Colec12 | 566 | 387 | 334 | 381 | 315 | 330 | 440 | 412 | 360 | 379 | 355 | 326 | 378.66 | 404 | 353 | 0.873 | -0.196 | 0.10246431 | 0.22510335 | 0.0055 | 0.0087 | 0.0077 | 0.0077 | TRUE | 0.1003 |
| Colq | 16 | 13 | 12 | 15 | 16 | 12 | 12 | 14 | 13 | 15 | 18 | 12 | 14 | 13 | 15 | 1.142 | 0.191 | 0.70580298 | NA | 0 | 0.1717 | 0.1131 | 0.1131 | TRUE | 0.1528 |
| Commd1 | 2644 | 1982 | 2110 | 2354 | 2019 | 2246 | 2053 | 2108 | 2277 | 2343 | 2278 | 2217 | 2212.72 | 2146 | 2279 | 1.063 | 0.088 | 0.18283592 | 0.33854092 | 0.0014 | 0.0035 | 0.0027 | 0.0027 | TRUE | 0.034 |
| Commd10 | 3747 | 2667 | 2933 | 2866 | 2574 | 2935 | 2910 | 2837 | 3165 | 2853 | 2904 | 2897 | 2927.59 | 2971 | 2885 | 0.971 | -0.042 | 0.50204191 | 0.66632789 | 0.0012 | 0.0032 | 0.0025 | 0.0025 | TRUE | 0.039 |
| Commd2 | 1975 | 1381 | 1307 | 1332 | 1115 | 1358 | 1534 | 1469 | 1410 | 1326 | 1258 | 1341 | 1389.56 | 1471 | 1308 | 0.889 | -0.17 | 0.01642005 | 0.05973653 | 8.00E-04 | 0.0041 | 0.0029 | 0.0029 | TRUE | 0.0175 |
| Commd3 | 1268 | 874 | 941 | 934 | 796 | 864 | 985 | 930 | 1015 | 930 | 898 | 853 | 935.06 | 977 | 894 | 0.915 | -0.129 | 0.10802213 | 0.23389794 | 8.00E-04 | 0.005 | 0.0035 | 0.0035 | TRUE | 0.0203 |
| Commd4 | 1718 | 1340 | 1364 | 1244 | 1172 | 1357 | 1334 | 1425 | 1472 | 1238 | 1322 | 1340 | 1355.23 | 1410 | 1300 | 0.922 | -0.116 | 0.1147114 | 0.24431083 | 0.0015 | 0.0042 | 0.0032 | 0.0032 | TRUE | 0.028 |
| Commd5 | 1103 | 688 | 601 | 556 | 525 | 552 | 857 | 732 | 648 | 553 | 592 | 545 | 654.59 | 746 | 563 | 0.754 | -0.408 | 0.00023006 | 0.0023845 | 0.0097 | 0.006 | 0.0073 | 0.0073 | TRUE | 0.2077 |
| Commd6 | 1048 | 717 | 655 | 726 | 572 | 671 | 814 | 763 | 707 | 723 | 645 | 662 | 718.94 | 761 | 677 | 0.888 | -0.171 | 0.065378 | 0.1633079 | 0.003 | 0.0057 | 0.0048 | 0.0048 | TRUE | 0.0459 |
| Commd7 | 677 | 509 | 484 | 412 | 414 | 476 | 526 | 541 | 522 | 410 | 467 | 470 | 489.41 | 530 | 449 | 0.847 | -0.239 | 0.01763364 | 0.06294645 | 0 | 0.0073 | 0.0053 | 0.0053 | TRUE | 0.0676 |
| Commd8 | 2902 | 2232 | 2169 | 2351 | 1913 | 2300 | 2254 | 2374 | 2340 | 2340 | 2158 | 2270 | 2289.51 | 2323 | 2256 | 0.972 | -0.041 | 0.51444681 | 0.67707074 | 7.00E-04 | 0.0035 | 0.0024 | 0.0024 | TRUE | 0.0173 |
| Commd9 | 1106 | 783 | 726 | 695 | 650 | 817 | 859 | 833 | 783 | 692 | 733 | 807 | 784.46 | 825 | 744 | 0.901 | -0.15 | 0.09496216 | 0.2129757 | 0.0028 | 0.0054 | 0.0046 | 0.0046 | TRUE | 0.0655 |
| Comp | 68 | 62 | 74 | 96 | 102 | 89 | 53 | 66 | 80 | 96 | 115 | 88 | 82.85 | 66 | 100 | 1.512 | 0.596 | 0.01138779 | 0.04551612 | 0.0179 | 0.031 | 0.0274 | 0.0274 | TRUE | 0.322 |
| Comt | 1580 | 1060 | 1135 | 1257 | 1174 | 1208 | 1227 | 1128 | 1225 | 1251 | 1324 | 1192 | 1224.56 | 1193 | 1256 | 1.052 | 0.073 | 0.34219072 | 0.51631201 | 0.0017 | 0.0044 | 0.0034 | 0.0034 | TRUE | 0.0272 |
| Comtd1 | 720 | 605 | 616 | 676 | 643 | 718 | 559 | 644 | 665 | 673 | 725 | 709 | 662.4 | 623 | 702 | 1.131 | 0.177 | 0.06591481 | 0.16411964 | 0.0035 | 0.006 | 0.0052 | 0.0052 | TRUE | 0.0989 |
| Copa | 17368 | 12328 | 12858 | 14003 | 12726 | 14272 | 13489 | 13113 | 13874 | 13939 | 14356 | 14089 | 13809.88 | 13492 | 14128 | 1.047 | 0.066 | 0.1651813 | 0.31601343 | 4.00E-04 | 0.0026 | 0.0016 | 0.0016 | TRUE | 0.0074 |

| | | | | | | | | | | | | | | | | | | | | | | | | | |
|---|---|---|---|---|---|---|---|---|---|---|---|---|---|---|---|---|---|---|---|---|---|---|---|---|---|
| Copb1 | 9975 | 6909 | 7798 | 8531 | 7340 | 8212 | 7747 | 7349 | 8414 | 8492 | 8280 | 8107 | 8064.78 | 7837 | 8293 | 1.058 | 0.082 | 0.18493523 | 0.34120038 | 0.0024 | 0.0027 | 0.0026 | 0.0026 | TRUE | 0.0503 |
| Copb2 | 16305 | 11556 | 11429 | 12688 | 11454 | 12782 | 12663 | 12292 | 12332 | 12630 | 12921 | 12618 | 12575.98 | 12429 | 12723 | 1.024 | 0.034 | 0.46346495 | 0.63270253 | 1.00E-04 | 0.0026 | 0.0014 | 0.0014 | TRUE | 0.0034 |
| Cope | 5137 | 3948 | 4332 | 4656 | 4380 | 4959 | 3990 | 4199 | 4674 | 4635 | 4941 | 4895 | 4555.71 | 4288 | 4824 | 1.125 | 0.171 | 0.01362163 | 0.05200158 | 0.0036 | 0.0029 | 0.0032 | 0.0032 | TRUE | 0.0755 |
| Copg1 | 10499 | 7429 | 7114 | 7338 | 6387 | 7547 | 8154 | 7902 | 7676 | 7304 | 7205 | 7450 | 7615.28 | 7911 | 7320 | 0.925 | -0.112 | 0.02649289 | 0.08474997 | 4.00E-04 | 0.0027 | 0.0017 | 0.0017 | TRUE | 0.009 |
| Copg2 | 5309 | 4039 | 4426 | 5210 | 4459 | 5085 | 4123 | 4296 | 4776 | 5186 | 5030 | 5020 | 4738.51 | 4398 | 5079 | 1.155 | 0.208 | 0.00151758 | 0.00974866 | 0.0028 | 0.0029 | 0.0029 | 0.0029 | TRUE | 0.0683 |
| Coprs | 2985 | 2287 | 2100 | 2226 | 2036 | 2279 | 2318 | 2433 | 2266 | 2216 | 2297 | 2250 | 2296.53 | 2339 | 2254 | 0.964 | -0.053 | 0.38064993 | 0.55516627 | 3.00E-04 | 0.0035 | 0.0022 | 0.0022 | TRUE | 0.0145 |
| Cops2 | 14192 | 9811 | 9688 | 10931 | 9423 | 10630 | 11022 | 10436 | 10453 | 10881 | 10630 | 10493 | 10652.62 | 10637 | 10668 | 1.003 | 0.004 | 0.93709679 | 0.96508948 | 4.00E-04 | 0.0026 | 0.0017 | 0.0017 | TRUE | 0.0124 |
| Cops3 | 10109 | 6849 | 6048 | 6318 | 5509 | 6180 | 7851 | 7285 | 6526 | 6289 | 6215 | 6101 | 6711.07 | 7221 | 6202 | 0.859 | -0.22 | 0.0016362 | 0.01031594 | 0.0041 | 0.0028 | 0.0034 | 0.0034 | TRUE | 0.0864 |
| Cops4 | 5471 | 3948 | 3817 | 4133 | 3472 | 4111 | 4249 | 4199 | 4119 | 4114 | 3917 | 4058 | 4109.33 | 4189 | 4030 | 0.962 | -0.056 | 0.29160394 | 0.46366616 | 2.00E-04 | 0.003 | 0.0018 | 0.0018 | TRUE | 0.0072 |
| Cops5 | 6319 | 4365 | 4358 | 4587 | 3980 | 4708 | 4908 | 4643 | 4702 | 4566 | 4490 | 4648 | 4659.37 | 4751 | 4568 | 0.961 | -0.057 | 0.28227551 | 0.45400943 | 4.00E-04 | 0.0029 | 0.0018 | 0.0018 | TRUE | 0.0104 |
| Cops6 | 7331 | 5304 | 5182 | 5308 | 4640 | 5476 | 5694 | 5642 | 5591 | 5284 | 5234 | 5406 | 5475.07 | 5642 | 5308 | 0.941 | -0.088 | 0.07313862 | 0.17706136 | 0 | 0.0029 | 0.0016 | 0.0016 | TRUE | 0.0032 |
| Cops7a | 8985 | 6780 | 6532 | 5574 | 5350 | 6117 | 6978 | 7212 | 7048 | 5548 | 6035 | 6038 | 6476.7 | 7079 | 5874 | 0.83 | -0.269 | 1.37E-06 | 3.76E-05 | 0.0011 | 0.0028 | 0.0021 | 0.0021 | TRUE | 0.0288 |
| Cops7b | 2658 | 1893 | 1920 | 1857 | 1620 | 1950 | 2064 | 2014 | 2072 | 1848 | 1828 | 1925 | 1958.42 | 2050 | 1867 | 0.911 | -0.135 | 0.0282155 | 0.08853919 | 0 | 0.0036 | 0.0022 | 0.0022 | TRUE | 0.009 |
| Cops8 | 4191 | 2932 | 2738 | 2797 | 2448 | 2816 | 3255 | 3119 | 2954 | 2784 | 2762 | 2780 | 2942.26 | 3109 | 2775 | 0.892 | -0.165 | 0.00628179 | 0.02921756 | 9.00E-04 | 0.0032 | 0.0023 | 0.0023 | TRUE | 0.023 |
| Copz1 | 8837 | 6106 | 6011 | 6802 | 6027 | 7029 | 6863 | 6495 | 6486 | 6771 | 6799 | 6939 | 6725.42 | 6615 | 6836 | 1.033 | 0.047 | 0.3578638 | 0.53225257 | 4.00E-04 | 0.0028 | 0.0017 | 0.0017 | TRUE | 0.0134 |
| Copz2 | 257 | 172 | 342 | 373 | 357 | 423 | 200 | 183 | 369 | 371 | 403 | 418 | 323.86 | 251 | 397 | 1.59 | 0.669 | 0.0010483 | 0.0074917 | 0.073 | 0.0097 | 0.0268 | 0.0268 | TRUE | 1.8739 |
| Coq10a | 559 | 437 | 588 | 702 | 620 | 727 | 434 | 465 | 634 | 699 | 699 | 718 | 608.21 | 511 | 705 | 1.383 | 0.468 | 0.00025788 | 0.00259391 | 0.0199 | 0.0063 | 0.0101 | 0.0101 | TRUE | 0.5137 |
| Coq10b | 1150 | 782 | 820 | 863 | 725 | 788 | 893 | 832 | 885 | 859 | 818 | 778 | 844.09 | 870 | 818 | 0.94 | -0.089 | 0.27974952 | 0.45138501 | 8.00E-04 | 0.0052 | 0.0037 | 0.0037 | TRUE | 0.0231 |
| Coq2 | 2163 | 1617 | 1681 | 1883 | 1632 | 1803 | 1680 | 1720 | 1814 | 1874 | 1841 | 1780 | 1784.82 | 1738 | 1832 | 1.055 | 0.077 | 0.24191038 | 0.40948773 | 6.00E-04 | 0.0038 | 0.0025 | 0.0025 | TRUE | 0.0174 |
| Coq3 | 2061 | 1586 | 1874 | 2104 | 1892 | 1969 | 1601 | 1687 | 2022 | 2094 | 2134 | 1944 | 1913.69 | 1770 | 2057 | 1.163 | 0.218 | 0.01473681 | 0.05508191 | 0.0083 | 0.0037 | 0.0053 | 0.0053 | TRUE | 0.1864 |
| Coq4 | 875 | 649 | 560 | 571 | 508 | 611 | 680 | 690 | 604 | 568 | 573 | 603 | 619.79 | 658 | 581 | 0.883 | -0.179 | 0.05580958 | 0.14581649 | 0.0015 | 0.0063 | 0.0047 | 0.0047 | TRUE | 0.0583 |
| Coq5 | 3527 | 2551 | 2578 | 3050 | 2743 | 3287 | 2739 | 2713 | 2782 | 3036 | 3094 | 3245 | 2934.92 | 2745 | 3125 | 1.139 | 0.187 | 0.00111823 | 0.00786948 | 4.00E-04 | 0.0032 | 0.002 | 0.002 | TRUE | 0.0138 |
| Coq6 | 2185 | 1603 | 1658 | 1872 | 1601 | 1867 | 1697 | 1705 | 1789 | 1863 | 1806 | 1843 | 1783.93 | 1730 | 1837 | 1.063 | 0.087 | 0.16513275 | 0.31596284 | 0 | 0.0038 | 0.0023 | 0.0023 | TRUE | 0.0104 |
| Coq7 | 1915 | 1391 | 1410 | 1605 | 1273 | 1498 | 1487 | 1480 | 1521 | 1598 | 1436 | 1479 | 1500.12 | 1496 | 1504 | 1.006 | 0.008 | 0.90405375 | 0.9470566 | 9.00E-04 | 0.004 | 0.0029 | 0.0029 | TRUE | 0.0357 |
| Coq9 | 3240 | 2323 | 2392 | 2731 | 2543 | 2828 | 2516 | 2471 | 2581 | 2718 | 2869 | 2792 | 2657.86 | 2523 | 2793 | 1.107 | 0.147 | 0.01155082 | 0.04600054 | 2.00E-04 | 0.0033 | 0.0021 | 0.0021 | TRUE | 0.0067 |
| Corin | 24 | 16 | 11 | 11 | 12 | 10 | 19 | 17 | 12 | 11 | 14 | 10 | 13.65 | 16 | 12 | 0.716 | -0.482 | 0.35492883 | NA | 0 | 0.1761 | 0.1203 | 0.1203 | TRUE | 0.211 |
| Coro1a | 382 | 248 | 263 | 341 | 292 | 363 | 297 | 264 | 284 | 339 | 329 | 358 | 311.9 | 282 | 342 | 1.215 | 0.281 | 0.01757246 | 0.06277508 | 0 | 0.01 | 0.0069 | 0.0069 | TRUE | 0.0332 |
| Coro1b | 5402 | 3804 | 4053 | 4072 | 3684 | 4346 | 4196 | 4046 | 4373 | 4053 | 4156 | 4290 | 4185.72 | 4205 | 4166 | 0.991 | -0.013 | 0.81767493 | 0.89531267 | 9.00E-04 | 0.003 | 0.0021 | 0.0021 | TRUE | 0.0146 |
| Coro1c | 15518 | 11391 | 9577 | 10403 | 9437 | 10890 | 12052 | 12116 | 10334 | 10355 | 10646 | 10750 | 11042.22 | 11501 | 10584 | 0.92 | -0.12 | 0.07629415 | 0.18235321 | 0.0039 | 0.0026 | 0.0032 | 0.0032 | TRUE | 0.096 |
| Coro2a | 637 | 439 | 550 | 672 | 614 | 681 | 495 | 467 | 593 | 669 | 693 | 672 | 598.16 | 518 | 678 | 1.309 | 0.389 | 0.00020906 | 0.00220038 | 0.0058 | 0.0064 | 0.0062 | 0.0062 | TRUE | 0.1809 |
| Coro2b | 2289 | 1888 | 2363 | 2885 | 2828 | 3297 | 1778 | 2008 | 2550 | 2872 | 3190 | 3255 | 2608.72 | 2112 | 3106 | 1.472 | 0.558 | 3.01E-07 | 1.04E-05 | 0.0192 | 0.0033 | 0.0081 | 0.0081 | TRUE | 0.3972 |
| Coro6 | 158 | 120 | 101 | 118 | 108 | 155 | 123 | 128 | 109 | 117 | 122 | 153 | 125.27 | 120 | 131 | 1.092 | 0.127 | 0.4961046 | 0.66061075 | 0.0062 | 0.0213 | 0.017 | 0.017 | TRUE | 0.2318 |
| Coro7 | 1068 | 796 | 754 | 738 | 747 | 803 | 829 | 847 | 814 | 735 | 843 | 793 | 809.95 | 830 | 790 | 0.951 | -0.072 | 0.3957076 | 0.56928522 | 0.0011 | 0.0054 | 0.004 | 0.004 | TRUE | 0.0447 |
| Cort | 20 | 20 | 32 | 18 | 20 | 20 | 16 | 21 | 35 | 18 | 23 | 20 | 21.93 | 24 | 20 | 0.852 | -0.23 | 0.60154416 | 0.74667517 | 0.0543 | 0.1105 | 0.0938 | 0.0938 | TRUE | 0.5494 |
| Cotl1 | 7365 | 5253 | 6441 | 7446 | 6694 | 8282 | 5720 | 5588 | 6950 | 7412 | 7552 | 8176 | 6899.38 | 6086 | 7713 | 1.268 | 0.342 | 3.55E-05 | 0.00053401 | 0.0088 | 0.0028 | 0.0048 | 0.0048 | TRUE | 0.1874 |
| Cox10 | 3529 | 2523 | 2327 | 2459 | 2337 | 2588 | 2741 | 2684 | 2511 | 2448 | 2636 | 2555 | 2595.69 | 2645 | 2546 | 0.962 | -0.056 | 0.38102251 | 0.55559628 | 0.0013 | 0.0033 | 0.0026 | 0.0026 | TRUE | 0.0237 |
| Cox11 | 3264 | 2538 | 2198 | 2200 | 2019 | 2367 | 2535 | 2700 | 2372 | 2190 | 2278 | 2337 | 2401.73 | 2536 | 2268 | 0.894 | -0.161 | 0.01862175 | 0.06558913 | 0.002 | 0.0034 | 0.0029 | 0.0029 | TRUE | 0.038 |
| Cox14 | 1344 | 995 | 1059 | 1193 | 1037 | 1248 | 1044 | 1058 | 1143 | 1188 | 1170 | 1232 | 1139.03 | 1082 | 1197 | 1.107 | 0.147 | 0.04879249 | 0.13235738 | 6.00E-04 | 0.0045 | 0.0031 | 0.0031 | TRUE | 0.0286 |
| Cox15 | 3305 | 2359 | 2317 | 2471 | 2021 | 2446 | 2567 | 2509 | 2500 | 2460 | 2280 | 2415 | 2455.04 | 2525 | 2385 | 0.944 | -0.083 | 0.17087992 | 0.32259143 | 4.00E-04 | 0.0034 | 0.0022 | 0.0022 | TRUE | 0.0176 |
| Cox16 | 805 | 529 | 579 | 638 | 537 | 589 | 625 | 563 | 625 | 635 | 606 | 581 | 605.82 | 604 | 607 | 1.004 | 0.006 | 0.94523832 | 0.96987016 | 0.0011 | 0.0063 | 0.0046 | 0.0046 | TRUE | 0.0413 |
| Cox17 | 3618 | 2604 | 2503 | 3105 | 2501 | 2942 | 2810 | 2770 | 2701 | 3091 | 2821 | 2904 | 2849.47 | 2760 | 2939 | 1.065 | 0.09 | 0.13784513 | 0.27835835 | 9.00E-04 | 0.0033 | 0.0023 | 0.0023 | TRUE | 0.0249 |
| Cox18 | 2305 | 1735 | 1968 | 2132 | 1906 | 2138 | 1790 | 1845 | 2123 | 2122 | 2150 | 2111 | 2023.67 | 1919 | 2128 | 1.109 | 0.15 | 0.04853921 | 0.13189562 | 0.0036 | 0.0036 | 0.0036 | 0.0036 | TRUE | 0.1033 |
| Cox19 | 729 | 491 | 423 | 473 | 403 | 466 | 566 | 522 | 456 | 471 | 455 | 460 | 488.39 | 515 | 462 | 0.895 | -0.16 | 0.13695028 | 0.27721715 | 0.0043 | 0.0073 | 0.0063 | 0.0063 | TRUE | 0.1136 |
| Cox20 | 257 | 158 | 148 | 188 | 143 | 156 | 200 | 168 | 160 | 187 | 161 | 154 | 171.63 | 176 | 167 | 0.949 | -0.075 | 0.64724743 | 0.78051363 | 0.0074 | 0.0162 | 0.0136 | 0.0136 | TRUE | 0.1629 |
| Cox4i1 | 27673 | 20038 | 20779 | 22663 | 20245 | 23141 | 21492 | 21314 | 22420 | 22559 | 22838 | 22844 | 22244.65 | 21742 | 22747 | 1.046 | 0.065 | 0.15554692 | 0.30322316 | 3.00E-04 | 0.0025 | 0.0015 | 0.0015 | TRUE | 0.0091 |
| Cox4i2 | 209 | 156 | 233 | 298 | 295 | 307 | 162 | 166 | 251 | 297 | 333 | 303 | 252.02 | 193 | 311 | 1.615 | 0.691 | 3.97E-05 | 0.00058341 | 0.0276 | 0.0118 | 0.0162 | 0.0162 | TRUE | 0.7331 |
| Cox5a | 13288 | 9599 | 10007 | 11203 | 9554 | 10902 | 10320 | 10210 | 10797 | 11151 | 10778 | 10762 | 10669.89 | 10442 | 10897 | 1.044 | 0.062 | 0.21460757 | 0.37733867 | 5.00E-04 | 0.0026 | 0.0017 | 0.0017 | TRUE | 0.0107 |
| Cox5b | 6509 | 4673 | 4990 | 5939 | 5130 | 5900 | 5055 | 4971 | 5384 | 5912 | 5787 | 5824 | 5488.85 | 5137 | 5841 | 1.137 | 0.186 | 0.00063261 | 0.00517269 | 7.00E-04 | 0.0029 | 0.0019 | 0.0019 | TRUE | 0.0214 |
| Cox6a1 | 7464 | 5336 | 5876 | 6361 | 5814 | 6703 | 5797 | 5676 | 6340 | 6332 | 6559 | 6617 | 6220.06 | 5938 | 6503 | 1.095 | 0.131 | 0.02765616 | 0.08722187 | 0.0018 | 0.0028 | 0.0024 | 0.0024 | TRUE | 0.0425 |
| Cox6a2 | 2 | 1 | 0 | 0 | 2 | 0 | 2 | 1 | 0 | 0 | 2 | 0 | 0.81 | 1 | 1 | 0.816 | -0.293 | 0.89959898 | NA | 0 | 2.9205 | 2.6456 | 2.6456 | TRUE | 0.9748 |
| Cox6b1 | 4463 | 3116 | 3499 | 3843 | 3353 | 4260 | 3466 | 3314 | 3775 | 3825 | 3783 | 4205 | 3728.2 | 3518 | 3938 | 1.119 | 0.163 | 0.0202921 | 0.06992978 | 0.0036 | 0.0031 | 0.0033 | 0.0033 | TRUE | 0.0489 |
| Cox6b2 | 1244 | 1200 | 1453 | 1719 | 1446 | 1703 | 966 | 1276 | 1568 | 1711 | 1631 | 1681 | 1472.3 | 1270 | 1674 | 1.321 | 0.401 | 0.00181885 | 0.01123942 | 0.0273 | 0.004 | 0.0112 | 0.0112 | TRUE | 0.2733 |
| Cox6c | 7223 | 5286 | 5926 | 6649 | 5579 | 6454 | 5610 | 5623 | 6394 | 6618 | 6294 | 6371 | 6151.62 | 5876 | 6428 | 1.094 | 0.13 | 0.0457894 | 0.12632075 | 0.003 | 0.0028 | 0.0029 | 0.0029 | TRUE | 0.0723 |
| Cox7a1 | 576 | 445 | 513 | 629 | 539 | 603 | 447 | 473 | 554 | 626 | 608 | 595 | 550.6 | 491 | 610 | 1.244 | 0.315 | 0.00256315 | 0.01465407 | 0.0045 | 0.0067 | 0.006 | 0.006 | TRUE | 0.1402 |
| Cox7a2 | 6287 | 4382 | 4769 | 5536 | 4912 | 5578 | 4883 | 4661 | 5146 | 5511 | 5541 | 5506 | 5207.96 | 4897 | 5519 | 1.127 | 0.173 | 0.00210738 | 0.01255935 | 9.00E-04 | 0.0029 | 0.0021 | 0.0021 | TRUE | 0.0237 |
| Cox7a2l | 5775 | 4198 | 4679 | 5360 | 4970 | 5607 | 4485 | 4465 | 5049 | 5335 | 5607 | 5535 | 5079.36 | 4666 | 5492 | 1.177 | 0.235 | 0.0002363 | 0.00242655 | 0.0025 | 0.0029 | 0.0028 | 0.0028 | TRUE | 0.0621 |
| Cox7b | 7440 | 5501 | 5920 | 6798 | 5948 | 7003 | 5778 | 5851 | 6388 | 6767 | 6710 | 6913 | 6401.17 | 6006 | 6797 | 1.132 | 0.179 | 0.00187105 | 0.01149738 | 0.0014 | 0.0028 | 0.0022 | 0.0022 | TRUE | 0.0375 |
| Cox7b2 | 264 | 167 | 154 | 67 | 58 | 52 | 205 | 178 | 166 | 67 | 65 | 51 | 122.05 | 183 | 61 | 0.333 | -1.588 | 1.89E-15 | 3.37E-13 | 0 | 0.0218 | 0.0176 | 0.0176 | TRUE | 0.1772 |
| Cox7c | 8346 | 5817 | 6339 | 7252 | 6259 | 6873 | 6482 | 6187 | 6840 | 7219 | 7061 | 6785 | 6762.22 | 6503 | 7022 | 1.08 | 0.111 | 0.05627048 | 0.14681891 | 0.0015 | 0.0028 | 0.0023 | 0.0023 | TRUE | 0.0247 |
| Cox8a | 10312 | 7524 | 8480 | 8908 | 8831 | 10143 | 8009 | 8003 | 9150 | 8867 | 9962 | 10013 | 9000.65 | 8387 | 9614 | 1.146 | 0.197 | 0.0063781 | 0.02952089 | 0.0052 | 0.0027 | 0.0037 | 0.0037 | TRUE | 0.077 |
| Cox8b | 0 | 0 | 0 | 1 | 0 | 1 | 0 | 0 | 0 | 1 | 0 | 1 | 0.33 | 0 | 1 | 3.763 | 1.912 | 0.57887588 | NA | 0 | 7.1752 | 6.7756 | 6.7756 | TRUE | 0.2094 |
| Cox8c | 97 | 85 | 70 | 87 | 63 | 67 | 75 | 90 | 76 | 87 | 71 | 66 | 77.51 | 80 | 75 | 0.93 | -0.105 | 0.64691368 | 0.78037811 | 0 | 0.033 | 0.0246 | 0.0246 | TRUE | 0.1854 |
| Cp | 1 | 1 | 0 | 0 | 0 | 1 | 1 | 1 | 0 | 0 | 0 | 1 | 0.47 | 1 | 0 | 0.599 | -0.74 | 0.79209856 | NA | 0 | 5.0317 | 3.8925 | 3.8925 | TRUE | 0.3835 |
| Cpa1 | 7 | 3 | 5 | 2 | 6 | 4 | 5 | 3 | 5 | 2 | 7 | 4 | 4.46 | 4 | 4 | 0.892 | -0.165 | 0.85698167 | NA | 0 | 0.5344 | 0.3755 | 0.3755 | TRUE | 0.3009 |
| Cpa2 | 78 | 58 | 56 | 52 | 53 | 62 | 61 | 62 | 60 | 52 | 60 | 61 | 59.24 | 61 | 58 | 0.945 | -0.081 | 0.74371062 | 0.84754073 | 0 | 0.0424 | 0.0276 | 0.0276 | TRUE | 0.0683 |
| Cpa3 | 0 | 0 | 0 | 0 | 0 | 0 | 0 | 0 | 0 | 0 | 0 | 0 | 0 | 0 | 0 | NA | NA | NA | NA | NA | NA | NA | NA | NA | NA |
| Cpa4 | 1 | 0 | 0 | 0 | 0 | 0 | 1 | 0 | 0 | 0 | 0 | 0 | 0.13 | 0 | 0 | 0.552 | -0.857 | 0.8336548 | NA | 0 | 18.3121 | 10 | 10 | TRUE | 0.2566 |
| Cpa5 | 1 | 0 | 0 | 0 | 0 | 2 | 1 | 0 | 0 | 0 | 0 | 2 | 0.46 | 0 | 1 | 1.93 | 0.948 | 0.7608301 | NA | 0 | 5.1717 | 5.216 | 5.216 | TRUE | 0.9737 |
| Cpa6 | 0 | 0 | 0 | 0 | 0 | 0 | 0 | 0 | 0 | 0 | 0 | 0 | 0 | 0 | 0 | NA | NA | NA | NA | NA | NA | NA | NA | NA | NA |

| | | | | | | | | | | | | | | | | | | | | | | | | | |
|---|---|---|---|---|---|---|---|---|---|---|---|---|---|---|---|---|---|---|---|---|---|---|---|---|---|
| Cpb1 | 25 | 16 | 12 | 16 | 22 | 18 | 19 | 17 | 13 | 16 | 25 | 18 | 17.98 | 16 | 20 | 1.171 | 0.228 | 0.61909537 | 0.75961286 | 0 | 0.1342 | 0.096 | 0.096 | TRUE | 0.2772 |
| Cpb2 | 41 | 34 | 10 | 10 | 12 | 8 | 32 | 36 | 11 | 10 | 14 | 8 | 18.36 | 26 | 11 | 0.394 | -1.343 | 0.01309011 | 0.05045765 | 0.1716 | 0.1315 | 0.143 | 0.143 | TRUE | 1.2597 |
| Cpd | 3818 | 2719 | 2880 | 3575 | 2788 | 3222 | 2965 | 2892 | 3107 | 3559 | 3145 | 3181 | 3141.54 | 2988 | 3295 | 1.103 | 0.141 | 0.0373775 | 0.10905013 | 0.0026 | 0.0032 | 0.003 | 0.003 | TRUE | 0.0598 |
| Cpe | 1075 | 793 | 873 | 1022 | 986 | 1150 | 835 | 844 | 942 | 1017 | 1112 | 1135 | 980.87 | 874 | 1088 | 1.247 | 0.318 | 0.00017825 | 0.00195377 | 0.0029 | 0.0048 | 0.0042 | 0.0042 | TRUE | 0.0549 |
| Cpeb1 | 492 | 347 | 391 | 556 | 400 | 519 | 382 | 369 | 422 | 553 | 451 | 512 | 448.35 | 391 | 505 | 1.295 | 0.373 | 0.00097453 | 0.00711026 | 0.0054 | 0.0077 | 0.007 | 0.007 | TRUE | 0.1006 |
| Cpeb2 | 1164 | 802 | 800 | 907 | 814 | 811 | 904 | 853 | 863 | 903 | 918 | 801 | 873.66 | 873 | 874 | 0.999 | -0.001 | 0.98960914 | 0.99454953 | 0.0021 | 0.0051 | 0.0041 | 0.0041 | TRUE | 0.0648 |
| Cpeb3 | 632 | 462 | 502 | 591 | 498 | 557 | 491 | 491 | 542 | 588 | 562 | 550 | 537.31 | 508 | 567 | 1.117 | 0.159 | 0.09645949 | 0.21562332 | 0 | 0.0068 | 0.0048 | 0.0048 | TRUE | 0.038 |
| Cpeb4 | 820 | 560 | 506 | 560 | 490 | 594 | 637 | 596 | 546 | 557 | 553 | 586 | 579.18 | 593 | 565 | 0.952 | -0.07 | 0.46679906 | 0.63566683 | 0.002 | 0.0065 | 0.005 | 0.005 | TRUE | 0.0555 |
| Cped1 | 8 | 6 | 1 | 2 | 1 | 0 | 6 | 6 | 1 | 2 | 1 | 0 | 2.8 | 4 | 1 | 0.225 | -2.151 | 0.11319757 | NA | 0 | 0.8492 | 0.724 | 0.724 | TRUE | 0.7609 |
| Cphx1 | 0 | 0 | 0 | 0 | 0 | 0 | 0 | 0 | 0 | 0 | 0 | 0 | 0 | 0 | 0 | NA | NA | NA | NA | NA | NA | NA | NA | NA | NA |
| Cphx2 | 0 | 0 | 0 | 0 | 0 | 0 | 0 | 0 | 0 | 0 | 0 | 0 | 0 | 0 | 0 | NA | NA | NA | NA | NA | NA | NA | NA | NA | NA |
| Cplx1 | 135 | 80 | 73 | 176 | 141 | 176 | 105 | 85 | 79 | 175 | 159 | 174 | 129.45 | 90 | 169 | 1.878 | 0.909 | 7.64E-07 | 2.33E-05 | 1.00E-04 | 0.0207 | 0.016 | 0.016 | TRUE | 0.2398 |
| Cplx2 | 10 | 12 | 11 | 17 | 5 | 11 | 8 | 13 | 12 | 17 | 6 | 11 | 10.97 | 11 | 11 | 1.049 | 0.069 | 0.9098911 | NA | 0.0707 | 0.2185 | 0.1781 | 0.1781 | TRUE | 0.2168 |
| Cplx3 | 1 | 0 | 0 | 0 | 0 | 0 | 1 | 0 | 0 | 0 | 0 | 0 | 0.13 | 0 | 0 | 0.552 | -0.857 | 0.8336548 | NA | 0 | 18.3121 | 10 | 10 | TRUE | 0.2566 |
| Cplx4 | 0 | 4 | 0 | 0 | 0 | 0 | 0 | 4 | 0 | 0 | 0 | 0 | 0.71 | 1 | 0 | 0.136 | -2.881 | 0.33173343 | NA | 10 | 3.3447 | 5.0011 | 5.0011 | TRUE | 1.8998 |
| Cpm | 377 | 275 | 250 | 267 | 240 | 268 | 293 | 293 | 270 | 266 | 271 | 265 | 276.02 | 285 | 267 | 0.936 | -0.096 | 0.43157409 | 0.60360153 | 0 | 0.011 | 0.0071 | 0.0071 | TRUE | 0.0237 |
| Cpn1 | 1100 | 831 | 880 | 902 | 808 | 914 | 854 | 884 | 950 | 898 | 912 | 902 | 899.9 | 896 | 904 | 1.01 | 0.014 | 0.85715033 | 0.91983255 | 0 | 0.0051 | 0.0035 | 0.0035 | TRUE | 0.0322 |
| Cpn2 | 16 | 6 | 4 | 6 | 9 | 13 | 12 | 6 | 4 | 6 | 10 | 13 | 8.68 | 7 | 10 | 1.224 | 0.291 | 0.67196352 | NA | 0.1037 | 0.2755 | 0.2263 | 0.2263 | TRUE | 0.8107 |
| Cpne1 | 4291 | 2949 | 3226 | 3220 | 3036 | 3487 | 3333 | 3137 | 3481 | 3205 | 3425 | 3442 | 3337.1 | 3317 | 3357 | 1.012 | 0.017 | 0.78394771 | 0.87279675 | 0.0018 | 0.0031 | 0.0026 | 0.0026 | TRUE | 0.0269 |
| Cpne2 | 353 | 212 | 190 | 200 | 180 | 173 | 274 | 226 | 205 | 199 | 203 | 171 | 212.93 | 235 | 191 | 0.809 | -0.306 | 0.05157905 | 0.13792691 | 0.0119 | 0.0136 | 0.013 | 0.013 | TRUE | 0.2519 |
| Cpne3 | 4799 | 3626 | 4239 | 4883 | 4243 | 4792 | 3727 | 3857 | 4574 | 4861 | 4787 | 4730 | 4422.58 | 4053 | 4793 | 1.183 | 0.243 | 0.00165119 | 0.0103967 | 0.0061 | 0.003 | 0.0041 | 0.0041 | TRUE | 0.1534 |
| Cpne4 | 4 | 4 | 8 | 4 | 6 | 6 | 3 | 4 | 9 | 4 | 7 | 6 | 5.44 | 5 | 6 | 1.057 | 0.08 | 0.92340996 | NA | 0 | 0.4377 | 0.3134 | 0.3134 | TRUE | 0.5874 |
| Cpne5 | 1052 | 812 | 868 | 1162 | 1053 | 1292 | 817 | 864 | 937 | 1157 | 1188 | 1275 | 1039.55 | 873 | 1207 | 1.385 | 0.47 | 1.59E-08 | 7.66E-07 | 0.0027 | 0.0047 | 0.004 | 0.004 | TRUE | 0.0488 |
| Cpne6 | 3 | 5 | 5 | 7 | 2 | 6 | 2 | 5 | 5 | 7 | 2 | 6 | 4.7 | 4 | 5 | 1.193 | 0.255 | 0.77733425 | NA | 0 | 0.5069 | 0.3704 | 0.3704 | TRUE | 0.418 |
| Cpne7 | 265 | 263 | 286 | 404 | 342 | 416 | 206 | 280 | 309 | 402 | 386 | 411 | 332.13 | 265 | 400 | 1.518 | 0.602 | 3.48E-05 | 0.00052523 | 0.0183 | 0.0096 | 0.0121 | 0.0121 | TRUE | 0.4579 |
| Cpne8 | 284 | 236 | 233 | 309 | 295 | 326 | 221 | 251 | 251 | 308 | 333 | 322 | 280.86 | 241 | 321 | 1.334 | 0.416 | 0.00088502 | 0.00663675 | 0 | 0.0109 | 0.0077 | 0.0077 | TRUE | 0.0673 |
| Cpne9 | 290 | 199 | 241 | 250 | 254 | 297 | 225 | 212 | 260 | 249 | 287 | 293 | 254.25 | 232 | 276 | 1.19 | 0.251 | 0.07004342 | 0.17161517 | 0.0052 | 0.0117 | 0.0098 | 0.0098 | TRUE | 0.1142 |
| Cpox | 4258 | 3191 | 3221 | 3622 | 3093 | 3458 | 3307 | 3394 | 3475 | 3605 | 3489 | 3414 | 3447.47 | 3392 | 3503 | 1.033 | 0.047 | 0.40462271 | 0.57734576 | 4.00E-04 | 0.0031 | 0.002 | 0.002 | TRUE | 0.0081 |
| Cpped1 | 1526 | 1080 | 953 | 978 | 845 | 925 | 1185 | 1149 | 1028 | 974 | 953 | 913 | 1033.68 | 1121 | 947 | 0.844 | -0.245 | 0.00289973 | 0.01610797 | 0.0024 | 0.0047 | 0.0039 | 0.0039 | TRUE | 0.0614 |
| Cpq | 52 | 39 | 44 | 27 | 25 | 25 | 40 | 41 | 47 | 27 | 28 | 25 | 34.85 | 43 | 27 | 0.617 | -0.696 | 0.03295512 | 0.0992891 | 0 | 0.0704 | 0.046 | 0.046 | TRUE | 0.0575 |
| Cps1 | 147 | 89 | 73 | 32 | 35 | 36 | 114 | 95 | 79 | 32 | 39 | 36 | 65.75 | 96 | 36 | 0.369 | -1.438 | 6.25E-08 | 2.55E-06 | 0 | 0.0385 | 0.0313 | 0.0313 | TRUE | 0.1266 |
| Cpsf1 | 9191 | 7251 | 7637 | 7809 | 7167 | 8279 | 7138 | 7713 | 8240 | 7773 | 8085 | 8173 | 7853.69 | 7697 | 8010 | 1.041 | 0.058 | 0.35960532 | 0.53425891 | 0.0026 | 0.0027 | 0.0028 | 0.0028 | TRUE | 0.0498 |
| Cpsf2 | 6735 | 4857 | 4697 | 4857 | 4166 | 4884 | 5231 | 5166 | 5068 | 4835 | 4700 | 4821 | 4970.12 | 5155 | 4785 | 0.928 | -0.108 | 0.03286243 | 0.0990725 | 0 | 0.0029 | 0.0016 | 0.0016 | TRUE | 0.0029 |
| Cpsf3 | 7122 | 5346 | 5443 | 5409 | 4787 | 5714 | 5531 | 5686 | 5873 | 5384 | 5400 | 5641 | 5585.95 | 5697 | 5475 | 0.961 | -0.057 | 0.28657714 | 0.45858118 | 5.00E-04 | 0.0029 | 0.0019 | 0.0019 | TRUE | 0.0089 |
| Cpsf3l | 5734 | 4167 | 4062 | 4246 | 3815 | 4555 | 4453 | 4432 | 4383 | 4226 | 4304 | 4496 | 4382.54 | 4423 | 4342 | 0.982 | -0.027 | 0.61776088 | 0.75859223 | 3.00E-04 | 0.003 | 0.0018 | 0.0018 | TRUE | 0.0119 |
| Cpsf4 | 2760 | 2001 | 1904 | 1697 | 1511 | 1757 | 2144 | 2128 | 2054 | 1689 | 1705 | 1734 | 1909.1 | 2109 | 1709 | 0.81 | -0.303 | 7.28E-07 | 2.23E-05 | 0 | 0.0037 | 0.0022 | 0.0022 | TRUE | 0.0061 |
| Cpsf4l | 738 | 601 | 677 | 567 | 588 | 663 | 573 | 639 | 730 | 564 | 663 | 654 | 637.52 | 647 | 627 | 0.97 | -0.043 | 0.696305 | 0.81509315 | 0.0099 | 0.0061 | 0.0073 | 0.0073 | TRUE | 0.147 |
| Cpsf6 | 11842 | 8806 | 8143 | 9115 | 7806 | 8942 | 9197 | 9367 | 8786 | 9073 | 8806 | 8827 | 9009.4 | 9117 | 8902 | 0.976 | -0.034 | 0.49476089 | 0.65985046 | 5.00E-04 | 0.0027 | 0.0017 | 0.0017 | TRUE | 0.0122 |
| Cpsf7 | 4814 | 3594 | 3389 | 3412 | 3066 | 3705 | 3739 | 3823 | 3657 | 3396 | 3459 | 3657 | 3621.82 | 3740 | 3504 | 0.937 | -0.094 | 0.102412 | 0.22502301 | 6.00E-04 | 0.0031 | 0.0021 | 0.0021 | TRUE | 0.0179 |
| Cpt1a | 6236 | 5127 | 6039 | 6933 | 6727 | 7662 | 4843 | 5454 | 6516 | 6901 | 7589 | 7564 | 6477.7 | 5604 | 7351 | 1.312 | 0.392 | 2.29E-05 | 0.00037234 | 0.0126 | 0.0028 | 0.006 | 0.006 | TRUE | 0.2469 |
| Cpt1b | 1 | 2 | 6 | 2 | 6 | 6 | 1 | 2 | 6 | 2 | 7 | 6 | 4.01 | 3 | 5 | 1.609 | 0.686 | 0.51256168 | NA | 0.3517 | 0.5935 | 0.5252 | 0.5252 | TRUE | 1.1243 |
| Cpt1c | 1402 | 1157 | 1395 | 1402 | 1293 | 1427 | 1089 | 1231 | 1505 | 1396 | 1459 | 1409 | 1347.93 | 1275 | 1421 | 1.116 | 0.158 | 0.12787177 | 0.26392365 | 0.0129 | 0.0042 | 0.0071 | 0.0071 | TRUE | 0.299 |
| Cpt2 | 1894 | 1414 | 1577 | 1836 | 1556 | 1868 | 1471 | 1504 | 1702 | 1828 | 1755 | 1844 | 1683.92 | 1559 | 1809 | 1.162 | 0.216 | 0.00395087 | 0.02041567 | 0.0026 | 0.0038 | 0.0035 | 0.0035 | TRUE | 0.0763 |
| Cpvl | 10 | 4 | 7 | 5 | 11 | 8 | 8 | 4 | 8 | 5 | 12 | 8 | 7.48 | 7 | 8 | 1.274 | 0.349 | 0.62663001 | NA | 0 | 0.3194 | 0.2353 | 0.2353 | TRUE | 0.2634 |
| Cpxcr1 | 0 | 0 | 0 | 0 | 0 | 0 | 0 | 0 | 0 | 0 | 0 | 0 | 0 | 0 | 0 | NA | NA | NA | NA | NA | NA | NA | NA | NA | NA |
| Cpxm1 | 1516 | 1114 | 1021 | 1106 | 1075 | 1102 | 1177 | 1185 | 1102 | 1101 | 1213 | 1088 | 1144.25 | 1155 | 1134 | 0.981 | -0.028 | 0.72096768 | 0.83257583 | 0.0016 | 0.0045 | 0.0035 | 0.0035 | TRUE | 0.0435 |
| Cpxm2 | 177 | 138 | 127 | 161 | 152 | 156 | 137 | 147 | 137 | 160 | 171 | 154 | 151.17 | 140 | 162 | 1.153 | 0.205 | 0.20007224 | 0.36070036 | 0 | 0.0181 | 0.0118 | 0.0118 | TRUE | 0.0271 |
| Cpz | 69 | 47 | 56 | 29 | 18 | 31 | 54 | 50 | 60 | 29 | 20 | 31 | 40.63 | 55 | 27 | 0.489 | -1.032 | 0.00109161 | 0.00770108 | 0 | 0.0608 | 0.0437 | 0.0437 | TRUE | 0.2422 |
| Cr1l | 1010 | 700 | 655 | 663 | 575 | 639 | 784 | 745 | 707 | 660 | 649 | 631 | 695.86 | 745 | 647 | 0.866 | -0.207 | 0.0171448 | 0.06177226 | 0 | 0.0058 | 0.004 | 0.004 | TRUE | 0.0264 |
| Cr2 | 100 | 61 | 56 | 74 | 61 | 54 | 78 | 65 | 60 | 74 | 69 | 53 | 66.46 | 68 | 65 | 0.959 | -0.061 | 0.80762666 | 0.88877485 | 0 | 0.0381 | 0.0298 | 0.0298 | TRUE | 0.2338 |
| Crabp1 | 2201 | 1643 | 1880 | 2333 | 2077 | 2306 | 1709 | 1748 | 2029 | 2322 | 2343 | 2276 | 2071.21 | 1829 | 2314 | 1.266 | 0.341 | 7.77E-06 | 0.00015243 | 0.0039 | 0.0036 | 0.0037 | 0.0037 | TRUE | 0.1094 |
| Crabp2 | 192 | 181 | 223 | 361 | 320 | 373 | 149 | 193 | 241 | 359 | 361 | 368 | 278.47 | 194 | 363 | 1.882 | 0.912 | 8.73E-09 | 4.42E-07 | 0.022 | 0.0109 | 0.0142 | 0.0142 | TRUE | 0.4294 |
| Cradd | 516 | 376 | 374 | 353 | 330 | 380 | 401 | 400 | 404 | 351 | 372 | 375 | 383.84 | 402 | 366 | 0.912 | -0.132 | 0.20877958 | 0.37050988 | 0 | 0.0086 | 0.0054 | 0.0054 | TRUE | 0.0147 |
| Cramp1l | 2435 | 1935 | 1931 | 1983 | 1820 | 2186 | 1891 | 2058 | 2084 | 1974 | 2053 | 2158 | 2036.31 | 2011 | 2062 | 1.026 | 0.037 | 0.59100875 | 0.73920352 | 0.002 | 0.0036 | 0.003 | 0.003 | TRUE | 0.034 |
| Crat | 798 | 583 | 528 | 538 | 477 | 559 | 620 | 620 | 570 | 536 | 538 | 552 | 572.51 | 603 | 542 | 0.898 | -0.156 | 0.08951708 | 0.20399806 | 0 | 0.0066 | 0.0043 | 0.0043 | TRUE | 0.0268 |
| Crb1 | 4 | 2 | 6 | 3 | 1 | 2 | 3 | 2 | 6 | 3 | 1 | 2 | 2.97 | 4 | 2 | 0.529 | -0.92 | 0.41817257 | NA | 0 | 0.8015 | 0.5526 | 0.5526 | TRUE | 0.4947 |
| Crb2 | 108 | 58 | 55 | 82 | 84 | 102 | 84 | 62 | 59 | 82 | 95 | 101 | 80.33 | 68 | 93 | 1.341 | 0.423 | 0.06949373 | 0.17074751 | 0.0127 | 0.0319 | 0.0265 | 0.0265 | TRUE | 0.401 |
| Crb3 | 224 | 196 | 338 | 329 | 352 | 389 | 174 | 208 | 365 | 327 | 397 | 384 | 309.29 | 249 | 369 | 1.486 | 0.571 | 0.08753659 | 0.20083668 | 0.0772 | 0.0101 | 0.0285 | 0.0772 | TRUE | 1.8223 |
| Crbn | 5133 | 3697 | 3915 | 3926 | 3607 | 4165 | 3987 | 3932 | 4224 | 3908 | 4069 | 4111 | 4038.63 | 4048 | 4029 | 0.996 | -0.006 | 0.91169893 | 0.9513752 | 8.00E-04 | 0.003 | 0.0021 | 0.0021 | TRUE | 0.0175 |
| Crcp | 2338 | 1585 | 1320 | 1548 | 1318 | 1617 | 1816 | 1686 | 1424 | 1541 | 1487 | 1596 | 1591.66 | 1642 | 1541 | 0.938 | -0.093 | 0.3029872 | 0.47552416 | 0.0071 | 0.0039 | 0.0052 | 0.0052 | TRUE | 0.161 |
| Crct1 | 7 | 5 | 3 | 1 | 3 | 2 | 5 | 5 | 3 | 1 | 3 | 2 | 3.39 | 4 | 2 | 0.446 | -1.166 | 0.2742636 | NA | 0 | 0.7014 | 0.4687 | 0.4687 | TRUE | 0.2266 |
| Creb1 | 2540 | 1722 | 1793 | 1853 | 1654 | 1934 | 1973 | 1832 | 1935 | 1844 | 1866 | 1909 | 1893.09 | 1913 | 1873 | 0.979 | -0.031 | 0.62602253 | 0.76437212 | 4.00E-04 | 0.0037 | 0.0024 | 0.0024 | TRUE | 0.0166 |
| Creb3 | 357 | 242 | 181 | 220 | 193 | 214 | 277 | 257 | 195 | 219 | 218 | 211 | 229.66 | 243 | 216 | 0.884 | -0.177 | 0.24958566 | 0.4184058 | 0.0126 | 0.0127 | 0.0127 | 0.0127 | TRUE | 0.321 |
| Creb3l1 | 613 | 459 | 418 | 561 | 515 | 620 | 476 | 488 | 451 | 558 | 581 | 612 | 527.79 | 472 | 584 | 1.237 | 0.307 | 0.00135748 | 0.00898265 | 0 | 0.0069 | 0.0047 | 0.0047 | TRUE | 0.0215 |
| Creb3l2 | 667 | 558 | 585 | 702 | 672 | 666 | 518 | 594 | 631 | 699 | 758 | 657 | 642.85 | 581 | 705 | 1.215 | 0.281 | 0.00642677 | 0.02970823 | 0.0061 | 0.0061 | 0.0061 | 0.0061 | TRUE | 0.1115 |
| Creb3l3 | 87 | 64 | 49 | 119 | 107 | 128 | 68 | 68 | 53 | 118 | 121 | 126 | 92.34 | 63 | 122 | 1.932 | 0.95 | 5.71E-06 | 0.00011774 | 0 | 0.0281 | 0.0197 | 0.0197 | TRUE | 0.1555 |
| Creb3l4 | 133 | 94 | 119 | 104 | 92 | 133 | 103 | 100 | 128 | 104 | 104 | 131 | 111.71 | 110 | 113 | 1.024 | 0.035 | 0.86151792 | 0.92334101 | 0.01 | 0.0236 | 0.0198 | 0.0198 | TRUE | 0.2077 |
| Creb5 | 17 | 10 | 9 | 11 | 2 | 8 | 13 | 11 | 10 | 11 | 2 | 8 | 9.11 | 11 | 7 | 0.633 | -0.66 | 0.32927742 | NA | 0.0319 | 0.2626 | 0.2142 | 0.2142 | TRUE | 0.5869 |
| Crebbp | 2472 | 1721 | 1304 | 1502 | 1408 | 1464 | 1920 | 1831 | 1407 | 1495 | 1588 | 1445 | 1614.36 | 1719 | 1509 | 0.877 | -0.189 | 0.06929509 | 0.17044136 | 0.0129 | 0.0039 | 0.0072 | 0.0072 | TRUE | 0.302 |

| | | | | | | | | | | | | | | | | | | | | | | | | | |
|---|---|---|---|---|---|---|---|---|---|---|---|---|---|---|---|---|---|---|---|---|---|---|---|---|---|
| Crebl2 | 1011 | 787 | 773 | 901 | 834 | 880 | 785 | 837 | 834 | 897 | 941 | 869 | 860.46 | 819 | 902 | 1.102 | 0.141 | 0.08086122 | 0.19052909 | 0 | 0.0052 | 0.0035 | 0.0035 | TRUE | 0.0164 |
| Crebrf | 2019 | 1734 | 2208 | 2589 | 2286 | 2901 | 1568 | 1844 | 2382 | 2577 | 2579 | 2864 | 2302.43 | 1931 | 2673 | 1.385 | 0.47 | 6.62E-05 | 0.00088016 | 0.0242 | 0.0035 | 0.0096 | 0.0096 | TRUE | 0.5033 |
| Crebzf | 2372 | 2143 | 2045 | 1762 | 1559 | 1895 | 1842 | 2279 | 2207 | 1754 | 1759 | 1871 | 1951.92 | 2109 | 1795 | 0.852 | -0.232 | 0.00678813 | 0.03104726 | 0.0061 | 0.0036 | 0.0048 | 0.0048 | TRUE | 0.1521 |
| Creg1 | 858 | 699 | 748 | 804 | 784 | 962 | 666 | 744 | 807 | 800 | 884 | 950 | 808.56 | 739 | 878 | 1.19 | 0.252 | 0.01181063 | 0.04678752 | 0.0071 | 0.0054 | 0.0059 | 0.0059 | TRUE | 0.0918 |
| Creg2 | 127 | 148 | 159 | 110 | 128 | 161 | 99 | 157 | 172 | 109 | 144 | 159 | 140.07 | 143 | 137 | 0.971 | -0.042 | 0.850003 | 0.91532381 | 0.0529 | 0.0193 | 0.0292 | 0.0292 | TRUE | 0.8226 |
| Creld1 | 1471 | 1223 | 1320 | 1533 | 1393 | 1574 | 1142 | 1301 | 1424 | 1526 | 1571 | 1554 | 1419.8 | 1289 | 1550 | 1.205 | 0.268 | 0.00174137 | 0.01086399 | 0.005 | 0.0041 | 0.0046 | 0.0046 | TRUE | 0.1235 |
| Creld2 | 4550 | 3130 | 2948 | 3155 | 2741 | 3350 | 3534 | 3329 | 3181 | 3140 | 3092 | 3307 | 3263.93 | 3348 | 3180 | 0.949 | -0.075 | 0.23688447 | 0.40350253 | 0.0016 | 0.0032 | 0.0026 | 0.0026 | TRUE | 0.0292 |
| Crem | 642 | 412 | 397 | 442 | 356 | 495 | 499 | 438 | 428 | 440 | 402 | 489 | 449.24 | 455 | 444 | 0.973 | -0.039 | 0.73153827 | 0.83929626 | 0.0061 | 0.0077 | 0.0072 | 0.0072 | TRUE | 0.0925 |
| Crh | 0 | 0 | 0 | 0 | 0 | 0 | 0 | 0 | 0 | 0 | 0 | 0 | 0 | 0 | 0 | NA | NA | NA | NA | NA | NA | NA | NA | NA | NA |
| Crhbp | 23 | 23 | 17 | 12 | 17 | 13 | 18 | 24 | 18 | 12 | 19 | 13 | 17.44 | 20 | 15 | 0.723 | -0.468 | 0.31770713 | 0.49160854 | 0 | 0.1383 | 0.0983 | 0.0983 | TRUE | 0.2969 |
| Crhr1 | 23 | 16 | 10 | 21 | 21 | 6 | 18 | 17 | 11 | 21 | 24 | 6 | 16.03 | 15 | 17 | 1.094 | 0.13 | 0.81472237 | NA | 0.1837 | 0.1503 | 0.1592 | 0.1592 | TRUE | 1.6263 |
| Crhr2 | 103 | 77 | 67 | 70 | 87 | 101 | 80 | 82 | 72 | 70 | 98 | 100 | 83.62 | 78 | 89 | 1.14 | 0.189 | 0.40425299 | 0.57716392 | 0.0097 | 0.0308 | 0.0249 | 0.0249 | TRUE | 0.356 |
| Crim1 | 1020 | 724 | 707 | 845 | 719 | 871 | 792 | 770 | 763 | 841 | 811 | 860 | 806.2 | 775 | 837 | 1.08 | 0.111 | 0.16337038 | 0.31398857 | 0 | 0.0054 | 0.0033 | 0.0033 | TRUE | 0.0087 |
| Crip1 | 1141 | 953 | 1217 | 1302 | 1176 | 1499 | 886 | 1014 | 1313 | 1296 | 1327 | 1480 | 1219.23 | 1071 | 1368 | 1.279 | 0.355 | 0.0033468 | 0.01801047 | 0.0223 | 0.0044 | 0.0097 | 0.0097 | TRUE | 0.4665 |
| Crip2 | 2240 | 1746 | 2031 | 2018 | 1779 | 2186 | 1740 | 1857 | 2191 | 2009 | 2007 | 2158 | 1993.65 | 1929 | 2058 | 1.068 | 0.094 | 0.27905036 | 0.45067422 | 0.007 | 0.0036 | 0.005 | 0.005 | TRUE | 0.1698 |
| Crip3 | 4 | 4 | 7 | 7 | 4 | 9 | 3 | 4 | 8 | 7 | 5 | 9 | 5.88 | 5 | 7 | 1.396 | 0.481 | 0.54820354 | NA | 0 | 0.4055 | 0.2883 | 0.2883 | TRUE | 0.38 |
| Cript | 2136 | 1476 | 1485 | 1639 | 1392 | 1604 | 1659 | 1570 | 1602 | 1631 | 1570 | 1583 | 1602.74 | 1610 | 1595 | 0.99 | -0.015 | 0.82335371 | 0.89863229 | 0 | 0.0039 | 0.0024 | 0.0024 | TRUE | 0.0088 |
| Crisp1 | 6 | 6 | 3 | 2 | 6 | 3 | 5 | 6 | 3 | 2 | 7 | 3 | 4.33 | 5 | 4 | 0.811 | -0.303 | 0.74582415 | NA | 0 | 0.5494 | 0.3939 | 0.3939 | TRUE | 0.5988 |
| Crisp2 | 1 | 3 | 1 | 0 | 1 | 0 | 1 | 3 | 1 | 0 | 1 | 0 | 1.03 | 2 | 0 | 0.223 | -2.162 | 0.29067121 | NA | 0 | 2.3054 | 1.7266 | 1.7266 | TRUE | 0.5075 |
| Crisp3 | 0 | 0 | 0 | 0 | 0 | 0 | 0 | 0 | 0 | 0 | 0 | 0 | 0 | 0 | 0 | NA | NA | NA | NA | NA | NA | NA | NA | NA | NA |
| Crisp4 | 0 | 0 | 0 | 0 | 0 | 0 | 0 | 0 | 0 | 0 | 0 | 0 | 0 | 0 | 0 | NA | NA | NA | NA | NA | NA | NA | NA | NA | NA |
| Crispld1 | 99 | 63 | 63 | 61 | 46 | 34 | 77 | 67 | 68 | 61 | 52 | 34 | 59.68 | 71 | 49 | 0.687 | -0.541 | 0.04980089 | 0.13445487 | 0.0229 | 0.0421 | 0.0373 | 0.0373 | TRUE | 0.6198 |
| Crispld2 | 549 | 383 | 488 | 593 | 492 | 566 | 426 | 407 | 527 | 590 | 555 | 559 | 510.73 | 453 | 568 | 1.255 | 0.328 | 0.00379437 | 0.01979508 | 0.0076 | 0.0071 | 0.0072 | 0.0072 | TRUE | 0.2235 |
| Crk | 13285 | 9505 | 9544 | 10633 | 9041 | 10735 | 10318 | 10110 | 10298 | 10584 | 10199 | 10597 | 10351.08 | 10242 | 10460 | 1.021 | 0.03 | 0.51707048 | 0.67923873 | 2.00E-04 | 0.0027 | 0.0015 | 0.0015 | TRUE | 0.0058 |
| Crkl | 4404 | 3098 | 3161 | 3216 | 2766 | 3129 | 3420 | 3295 | 3411 | 3201 | 3120 | 3089 | 3256.12 | 3375 | 3137 | 0.929 | -0.106 | 0.05304421 | 0.14071287 | 0 | 0.0032 | 0.0019 | 0.0019 | TRUE | 0.0052 |
| Crlf1 | 178 | 142 | 193 | 209 | 176 | 226 | 138 | 151 | 208 | 208 | 199 | 223 | 187.87 | 166 | 210 | 1.272 | 0.348 | 0.04539771 | 0.12550666 | 0.0189 | 0.015 | 0.0163 | 0.0163 | TRUE | 0.5203 |
| Crlf2 | 492 | 425 | 394 | 320 | 305 | 394 | 382 | 452 | 425 | 319 | 344 | 389 | 385.14 | 420 | 351 | 0.837 | -0.257 | 0.03338945 | 0.10028062 | 0.0063 | 0.0086 | 0.0079 | 0.0079 | TRUE | 0.1074 |
| Crlf3 | 6445 | 4670 | 4877 | 5056 | 4553 | 5072 | 5006 | 4967 | 5262 | 5033 | 5136 | 5007 | 5068.51 | 5078 | 5059 | 0.996 | -0.006 | 0.91646194 | 0.95388891 | 4.00E-04 | 0.0029 | 0.0018 | 0.0018 | TRUE | 0.0121 |
| Crls1 | 1850 | 1450 | 1573 | 1654 | 1461 | 1743 | 1437 | 1542 | 1697 | 1646 | 1648 | 1721 | 1615.26 | 1559 | 1672 | 1.074 | 0.103 | 0.18230833 | 0.33792413 | 0.0029 | 0.0039 | 0.0037 | 0.0037 | TRUE | 0.0725 |
| Crmp1 | 2078 | 1540 | 1601 | 2068 | 1708 | 2234 | 1614 | 1638 | 1727 | 2058 | 1927 | 2205 | 1861.67 | 1660 | 2063 | 1.244 | 0.315 | 1.27E-05 | 0.00022872 | 0.0024 | 0.0037 | 0.0032 | 0.0032 | TRUE | 0.0437 |
| Crnkl1 | 3497 | 2475 | 2288 | 2428 | 2205 | 2500 | 2716 | 2633 | 2469 | 2417 | 2487 | 2468 | 2531.59 | 2606 | 2457 | 0.942 | -0.086 | 0.16867694 | 0.32029553 | 9.00E-04 | 0.0034 | 0.0024 | 0.0024 | TRUE | 0.0255 |
| Crnn | 1 | 0 | 0 | 0 | 0 | 0 | 1 | 0 | 0 | 0 | 0 | 0 | 0.13 | 0 | 0 | 0.552 | -0.857 | 0.8336548 | NA | 0 | 18.3121 | 10 | 10 | TRUE | 0.2566 |
| Crocc | 1804 | 1617 | 1441 | 1353 | 1235 | 1439 | 1401 | 1720 | 1555 | 1347 | 1393 | 1421 | 1472.73 | 1559 | 1387 | 0.891 | -0.167 | 0.04757563 | 0.1299443 | 0.0051 | 0.004 | 0.0044 | 0.0044 | TRUE | 0.0991 |
| Crot | 2568 | 1970 | 1981 | 2013 | 1782 | 1926 | 1994 | 2095 | 2137 | 2004 | 2010 | 1901 | 2023.78 | 2075 | 1972 | 0.95 | -0.073 | 0.25061716 | 0.41949538 | 7.00E-04 | 0.0036 | 0.0025 | 0.0025 | TRUE | 0.0147 |
| Crp | 0 | 0 | 0 | 0 | 0 | 0 | 0 | 0 | 0 | 0 | 0 | 0 | 0 | 0 | 0 | NA | NA | NA | NA | NA | NA | NA | NA | NA | NA |
| Crtac1 | 743 | 562 | 748 | 693 | 718 | 804 | 577 | 598 | 807 | 690 | 810 | 794 | 712.57 | 661 | 765 | 1.159 | 0.213 | 0.09149081 | 0.20723901 | 0.0201 | 0.0058 | 0.01 | 0.01 | TRUE | 0.4386 |
| Crtam | 51 | 40 | 23 | 29 | 21 | 17 | 40 | 43 | 25 | 29 | 24 | 17 | 29.39 | 36 | 23 | 0.645 | -0.632 | 0.10121864 | 0.22315564 | 0.0382 | 0.0831 | 0.0711 | 0.0711 | TRUE | 0.472 |
| Crtap | 2022 | 1389 | 1270 | 1424 | 1116 | 1331 | 1570 | 1477 | 1370 | 1417 | 1259 | 1314 | 1401.42 | 1472 | 1330 | 0.903 | -0.148 | 0.06411937 | 0.16100811 | 0.0035 | 0.0041 | 0.0039 | 0.0039 | TRUE | 0.0443 |
| Crtc1 | 1045 | 755 | 673 | 745 | 653 | 753 | 812 | 803 | 726 | 742 | 737 | 743 | 760.4 | 780 | 741 | 0.948 | -0.077 | 0.36486006 | 0.53957008 | 0 | 0.0055 | 0.0039 | 0.0039 | TRUE | 0.0425 |
| Crtc2 | 1087 | 860 | 654 | 614 | 538 | 644 | 844 | 915 | 706 | 611 | 607 | 636 | 719.75 | 822 | 618 | 0.752 | -0.411 | 7.75E-05 | 0.00098796 | 0.0077 | 0.0057 | 0.0064 | 0.0064 | TRUE | 0.1757 |
| Crtc3 | 334 | 236 | 206 | 203 | 171 | 203 | 259 | 251 | 222 | 202 | 193 | 200 | 221.35 | 244 | 198 | 0.811 | -0.302 | 0.02866462 | 0.08962839 | 0 | 0.0131 | 0.0091 | 0.0091 | TRUE | 0.0665 |
| Crx | 4 | 9 | 6 | 1 | 1 | 1 | 3 | 10 | 6 | 1 | 1 | 1 | 3.71 | 6 | 1 | 0.164 | -2.607 | 0.03031362 | NA | 0 | 0.6411 | 0.4675 | 0.4675 | TRUE | 0.0836 |
| Crxos1 | 597 | 500 | 516 | 478 | 463 | 504 | 464 | 532 | 557 | 476 | 522 | 498 | 507.98 | 518 | 499 | 0.966 | -0.051 | 0.63158884 | 0.76886971 | 0.0039 | 0.0071 | 0.0061 | 0.0061 | TRUE | 0.1028 |
| Cry1 | 3094 | 2338 | 2276 | 2571 | 2312 | 2570 | 2403 | 2487 | 2456 | 2559 | 2608 | 2537 | 2508.33 | 2449 | 2568 | 1.049 | 0.069 | 0.22475459 | 0.3892273 | 0 | 0.0034 | 0.0019 | 0.0019 | TRUE | 0.0033 |
| Cry2 | 1720 | 1414 | 1410 | 1661 | 1520 | 1824 | 1336 | 1504 | 1521 | 1653 | 1715 | 1801 | 1588.32 | 1454 | 1723 | 1.187 | 0.247 | 0.00118657 | 0.00819866 | 0.0029 | 0.0039 | 0.0035 | 0.0035 | TRUE | 0.063 |
| Cryaa | 31 | 18 | 4 | 19 | 7 | 14 | 24 | 19 | 4 | 19 | 8 | 14 | 14.69 | 16 | 14 | 0.848 | -0.238 | 0.6982274 | NA | 0.3114 | 0.1637 | 0.2036 | 0.2036 | TRUE | 0.78 |
| Cryab | 47 | 40 | 46 | 36 | 33 | 32 | 37 | 43 | 50 | 36 | 37 | 32 | 38.89 | 43 | 35 | 0.817 | -0.292 | 0.34904495 | 0.52360406 | 0 | 0.0634 | 0.044 | 0.044 | TRUE | 0.1547 |
| Cryba1 | 4 | 1 | 2 | 4 | 3 | 4 | 3 | 1 | 2 | 4 | 3 | 4 | 2.94 | 2 | 4 | 1.751 | 0.808 | 0.46324143 | NA | 0 | 0.8084 | 0.514 | 0.514 | TRUE | 0.2207 |
| Cryba2 | 0 | 0 | 0 | 0 | 1 | 2 | 0 | 0 | 0 | 0 | 1 | 2 | 0.52 | 0 | 1 | 5.84 | 2.546 | 0.38239357 | NA | 0 | 4.5861 | 4.6218 | 4.6218 | TRUE | 0.0847 |
| Cryba4 | 3 | 1 | 6 | 3 | 4 | 5 | 2 | 1 | 6 | 3 | 5 | 5 | 3.72 | 3 | 4 | 1.276 | 0.351 | 0.73125039 | NA | 0 | 0.6401 | 0.4802 | 0.4802 | TRUE | 0.9135 |
| Crybb1 | 1 | 1 | 0 | 1 | 1 | 0 | 1 | 1 | 0 | 1 | 1 | 0 | 0.66 | 1 | 1 | 1.128 | 0.173 | 0.94106052 | NA | 0 | 3.5899 | 2.4471 | 2.4471 | TRUE | 0.2681 |
| Crybb2 | 0 | 0 | 0 | 0 | 0 | 0 | 0 | 0 | 0 | 0 | 0 | 0 | 0 | 0 | 0 | NA | NA | NA | NA | NA | NA | NA | NA | NA | NA |
| Crybb3 | 63 | 51 | 68 | 47 | 65 | 57 | 49 | 54 | 73 | 47 | 73 | 56 | 58.82 | 59 | 59 | 1.001 | 0.002 | 0.99496948 | 0.99706224 | 0.0313 | 0.0427 | 0.0395 | 0.0395 | TRUE | 0.3862 |
| Crybg3 | 515 | 319 | 285 | 420 | 358 | 431 | 400 | 339 | 308 | 418 | 404 | 425 | 382.37 | 349 | 416 | 1.187 | 0.248 | 0.04178112 | 0.11801244 | 0.0067 | 0.0086 | 0.008 | 0.008 | TRUE | 0.1985 |
| Cryga | 63 | 43 | 30 | 34 | 45 | 38 | 49 | 46 | 32 | 34 | 51 | 38 | 41.53 | 42 | 41 | 0.953 | -0.07 | 0.82591936 | 0.90019138 | 0.0202 | 0.0595 | 0.0489 | 0.0489 | TRUE | 0.3298 |
| Crygb | 10 | 4 | 3 | 1 | 2 | 2 | 8 | 4 | 3 | 1 | 2 | 2 | 3.41 | 5 | 2 | 0.335 | -1.578 | 0.15130305 | NA | 0 | 0.6966 | 0.4787 | 0.4787 | TRUE | 0.5613 |
| Crygc | 37 | 23 | 9 | 14 | 15 | 11 | 29 | 24 | 10 | 14 | 17 | 11 | 17.44 | 21 | 14 | 0.654 | -0.613 | 0.23454395 | 0.40061384 | 0.1132 | 0.1383 | 0.1313 | 0.1313 | TRUE | 0.8955 |
| Crygd | 91 | 71 | 61 | 60 | 53 | 64 | 71 | 76 | 66 | 60 | 60 | 63 | 65.78 | 71 | 61 | 0.862 | -0.214 | 0.36381293 | 0.53841025 | 0 | 0.0385 | 0.0249 | 0.0249 | TRUE | 0.0298 |
| Cryge | 96 | 51 | 41 | 55 | 53 | 48 | 75 | 54 | 44 | 55 | 60 | 47 | 55.83 | 58 | 54 | 0.926 | -0.111 | 0.69395645 | 0.81322833 | 0.0273 | 0.0449 | 0.0397 | 0.0397 | TRUE | 0.5481 |
| Crygf | 1 | 2 | 0 | 0 | 0 | 0 | 1 | 2 | 0 | 0 | 0 | 0 | 0.48 | 1 | 0 | 0.197 | -2.343 | 0.43249452 | NA | 0 | 4.8992 | 4.9172 | 4.9172 | TRUE | 0.1226 |
| Crygn | 15 | 25 | 20 | 18 | 23 | 16 | 12 | 27 | 22 | 18 | 26 | 16 | 19.91 | 20 | 20 | 1.006 | 0.009 | 0.98416991 | 0.9918029 | 0.0722 | 0.1214 | 0.1073 | 0.1073 | TRUE | 0.5996 |
| Crygs | 9 | 8 | 10 | 4 | 14 | 7 | 7 | 9 | 11 | 4 | 16 | 7 | 8.83 | 9 | 9 | 1.011 | 0.016 | 0.98205913 | NA | 0.134 | 0.2709 | 0.2323 | 0.2323 | TRUE | 0.9349 |
| Cryl1 | 827 | 562 | 544 | 612 | 537 | 545 | 642 | 598 | 587 | 609 | 606 | 538 | 596.67 | 609 | 584 | 0.958 | -0.062 | 0.51549616 | 0.67795239 | 0.002 | 0.0064 | 0.005 | 0.005 | TRUE | 0.0573 |
| Crym | 302 | 247 | 254 | 292 | 276 | 318 | 235 | 263 | 274 | 291 | 311 | 314 | 281.21 | 257 | 305 | 1.191 | 0.252 | 0.0452554 | 0.1252101 | 0 | 0.0109 | 0.0078 | 0.0078 | TRUE | 0.0721 |
| Cryz | 2461 | 1665 | 1859 | 1977 | 1704 | 1833 | 1911 | 1771 | 2006 | 1968 | 1922 | 1809 | 1897.98 | 1896 | 1900 | 1.002 | 0.003 | 0.97044054 | 0.98397988 | 0.0023 | 0.0037 | 0.0032 | 0.0032 | TRUE | 0.0392 |
| Cryzl1 | 889 | 624 | 652 | 720 | 590 | 652 | 690 | 664 | 704 | 717 | 666 | 644 | 680.6 | 686 | 676 | 0.985 | -0.022 | 0.79851129 | 0.88227596 | 0 | 0.0059 | 0.0041 | 0.0041 | TRUE | 0.0345 |
| Cs | 20965 | 15133 | 16117 | 17890 | 16051 | 18383 | 16283 | 16097 | 17390 | 17808 | 18107 | 18147 | 17305.21 | 16590 | 18021 | 1.086 | 0.119 | 0.01812968 | 0.06423588 | 9.00E-04 | 0.0026 | 0.0018 | 0.0018 | TRUE | 0.0217 |
| Csad | 1589 | 1233 | 1137 | 1708 | 1561 | 1683 | 1234 | 1312 | 1227 | 1700 | 1761 | 1661 | 1482.49 | 1258 | 1707 | 1.358 | 0.442 | 1.07E-10 | 8.63E-09 | 4.00E-04 | 0.004 | 0.0027 | 0.0027 | TRUE | 0.0166 |
| Csda | 38925 | 29221 | 27246 | 29187 | 25311 | 29718 | 30231 | 31082 | 29398 | 29053 | 28553 | 29336 | 29609 | 30237 | 28981 | 0.958 | -0.061 | 0.18595195 | 0.34228091 | 4.00E-04 | 0.0025 | 0.0015 | 0.0015 | TRUE | 0.0073 |

| | | | | | | | | | | | | | | | | | | | | | | | | | |
|---|---|---|---|---|---|---|---|---|---|---|---|---|---|---|---|---|---|---|---|---|---|---|---|---|---|
| Csdc2 | 86 | 51 | 47 | 48 | 42 | 62 | 67 | 54 | 51 | 48 | 47 | 61 | 54.69 | 57 | 52 | 0.907 | -0.141 | 0.60095574 | 0.74639037 | 0 | 0.0458 | 0.0344 | 0.0344 | TRUE | 0.2056 |
| Csde1 | 32625 | 23392 | 23542 | 26294 | 22657 | 25317 | 25338 | 24882 | 25402 | 26173 | 25559 | 24992 | 25391.02 | 25207 | 25575 | 1.015 | 0.021 | 0.64390043 | 0.77815751 | 3.00E-04 | 0.0025 | 0.0014 | 0.0014 | TRUE | 0.0051 |
| Cse1l | 22260 | 15386 | 15497 | 15646 | 13832 | 15457 | 17288 | 16366 | 16721 | 15574 | 15604 | 15258 | 16135.3 | 16792 | 15479 | 0.922 | -0.118 | 0.01257618 | 0.0490454 | 4.00E-04 | 0.0026 | 0.0015 | 0.0015 | TRUE | 0.0082 |
| Csf1 | 43 | 21 | 22 | 29 | 25 | 20 | 33 | 22 | 24 | 29 | 28 | 20 | 26.05 | 26 | 26 | 0.956 | -0.065 | 0.86741493 | 0.92681182 | 0 | 0.0934 | 0.0706 | 0.0706 | TRUE | 0.3883 |
| Csf1r | 1 | 1 | 0 | 3 | 0 | 1 | 1 | 1 | 0 | 3 | 0 | 1 | 0.97 | 1 | 1 | 2.153 | 1.107 | 0.59058006 | NA | 0 | 2.4484 | 1.889 | 1.889 | TRUE | 0.3164 |
| Csf2 | 0 | 0 | 0 | 0 | 0 | 0 | 0 | 0 | 0 | 0 | 0 | 0 | 0 | 0 | 0 | NA | NA | NA | NA | NA | NA | NA | NA | NA | NA |
| Csf2ra | 318 | 241 | 249 | 221 | 224 | 237 | 247 | 256 | 269 | 220 | 253 | 234 | 246.44 | 257 | 236 | 0.916 | -0.127 | 0.33140891 | 0.50569314 | 0 | 0.012 | 0.0083 | 0.0083 | TRUE | 0.0431 |
| Csf2rb | 0 | 3 | 0 | 0 | 0 | 0 | 0 | 3 | 0 | 0 | 0 | 0 | 0.53 | 1 | 0 | 0.181 | -2.465 | 0.45184603 | NA | 10 | 4.4587 | 6.272 | 6.272 | TRUE | 1.443 |
| Csf2rb2 | 0 | 1 | 0 | 0 | 0 | 0 | 0 | 1 | 0 | 0 | 0 | 0 | 0.18 | 0 | 0 | 0.552 | -0.857 | 0.8336548 | NA | 0 | 13.3714 | 10 | 10 | TRUE | 0.5216 |
| Csf3 | 0 | 0 | 0 | 0 | 1 | 0 | 0 | 0 | 0 | 0 | 1 | 0 | 0.19 | 0 | 0 | 2.094 | 1.067 | 0.79379329 | NA | 0 | 12.608 | 10 | 10 | TRUE | 0.5075 |
| Csf3r | 18 | 12 | 2 | 9 | 7 | 13 | 14 | 13 | 2 | 9 | 8 | 13 | 9.77 | 10 | 10 | 1.015 | 0.022 | 0.97452774 | NA | 0.2336 | 0.2451 | 0.2423 | 0.2423 | TRUE | 1.4183 |
| Csgalnact1 | 28 | 21 | 24 | 17 | 18 | 16 | 22 | 22 | 26 | 17 | 20 | 16 | 20.5 | 23 | 18 | 0.758 | -0.401 | 0.34047102 | 0.51463117 | 0 | 0.118 | 0.0772 | 0.0772 | TRUE | 0.0774 |
| Csgalnact2 | 1614 | 1116 | 1013 | 1198 | 981 | 1093 | 1254 | 1187 | 1093 | 1192 | 1107 | 1079 | 1151.96 | 1178 | 1126 | 0.955 | -0.066 | 0.41575363 | 0.58852794 | 0.0029 | 0.0045 | 0.0039 | 0.0039 | TRUE | 0.0472 |
| Csk | 3260 | 2438 | 2479 | 2131 | 1885 | 2310 | 2532 | 2593 | 2675 | 2121 | 2126 | 2280 | 2388 | 2600 | 2176 | 0.837 | -0.256 | 4.12E-05 | 0.00059971 | 8.00E-04 | 0.0034 | 0.0024 | 0.0024 | TRUE | 0.0215 |
| Csl | 3 | 2 | 4 | 3 | 2 | 4 | 2 | 2 | 4 | 3 | 2 | 4 | 2.99 | 3 | 3 | 1.062 | 0.087 | 0.93544572 | NA | 0 | 0.794 | 0.5038 | 0.5038 | TRUE | 0.1991 |
| Csmd1 | 14 | 7 | 3 | 4 | 3 | 1 | 11 | 7 | 3 | 4 | 3 | 1 | 4.98 | 7 | 3 | 0.38 | -1.396 | 0.13775231 | NA | 0 | 0.4779 | 0.3839 | 0.3839 | TRUE | 0.1452 |
| Csmd3 | 2 | 0 | 0 | 1 | 0 | 0 | 2 | 0 | 0 | 1 | 0 | 0 | 0.42 | 1 | 0 | 0.681 | -0.553 | 0.8627148 | NA | 0 | 5.5819 | 5.4906 | 5.4906 | TRUE | 0.9727 |
| Csn1s1 | 2 | 0 | 0 | 0 | 0 | 0 | 2 | 0 | 0 | 0 | 0 | 0 | 0.26 | 1 | 0 | 0.356 | -1.491 | 0.71362796 | NA | 0 | 9.1573 | 10 | 10 | TRUE | 0.968 |
| Csn1s2a | 0 | 0 | 0 | 0 | 0 | 0 | 0 | 0 | 0 | 0 | 0 | 0 | 0 | 0 | 0 | NA | NA | NA | NA | NA | NA | NA | NA | NA | NA |
| Csn1s2b | 0 | 0 | 0 | 0 | 0 | 0 | 0 | 0 | 0 | 0 | 0 | 0 | 0 | 0 | 0 | NA | NA | NA | NA | NA | NA | NA | NA | NA | NA |
| Csn2 | 1 | 0 | 0 | 0 | 0 | 0 | 1 | 0 | 0 | 0 | 0 | 0 | 0.13 | 0 | 0 | 0.552 | -0.857 | 0.8336548 | NA | 0 | 18.3121 | 10 | 10 | TRUE | 0.2566 |
| Csn3 | 4 | 0 | 1 | 0 | 0 | 0 | 3 | 0 | 1 | 0 | 0 | 0 | 0.7 | 1 | 0 | 0.134 | -2.902 | 0.26925366 | NA | 6.5513 | 3.3999 | 3.6307 | 3.6307 | TRUE | 0.1404 |
| Csnk1a1 | 18778 | 13974 | 15556 | 17572 | 15877 | 17951 | 14584 | 14864 | 16785 | 17491 | 17911 | 17720 | 16559.22 | 15411 | 17707 | 1.149 | 0.201 | 0.00125404 | 0.00849884 | 0.003 | 0.0026 | 0.0027 | 0.0027 | TRUE | 0.0742 |
| Csnk1d | 11345 | 8153 | 8170 | 8356 | 7284 | 8461 | 8811 | 8672 | 8815 | 8318 | 8217 | 8352 | 8530.96 | 8766 | 8296 | 0.946 | -0.08 | 0.08277103 | 0.19371971 | 0 | 0.0027 | 0.0014 | 0.0014 | TRUE | 0.0011 |
| Csnk1e | 7522 | 5880 | 7326 | 9293 | 8108 | 9304 | 5842 | 6254 | 7905 | 9250 | 9147 | 9184 | 7930.43 | 6667 | 9194 | 1.379 | 0.464 | 5.18E-07 | 1.68E-05 | 0.0132 | 0.0027 | 0.006 | 0.006 | TRUE | 0.3214 |
| Csnk1g1 | 2780 | 2074 | 1983 | 2129 | 1866 | 1993 | 2159 | 2206 | 2140 | 2119 | 2105 | 1967 | 2116.08 | 2168 | 2064 | 0.952 | -0.071 | 0.2537801 | 0.4228898 | 5.00E-04 | 0.0035 | 0.0023 | 0.0023 | TRUE | 0.0204 |
| Csnk1g2 | 7716 | 6101 | 5956 | 5952 | 5469 | 6528 | 5993 | 6490 | 6426 | 5925 | 6170 | 6444 | 6241.18 | 6303 | 6180 | 0.981 | -0.028 | 0.63301829 | 0.7701507 | 0.0014 | 0.0028 | 0.0023 | 0.0023 | TRUE | 0.0229 |
| Csnk1g3 | 2306 | 1538 | 1516 | 1643 | 1368 | 1612 | 1791 | 1636 | 1636 | 1635 | 1543 | 1591 | 1638.78 | 1688 | 1590 | 0.941 | -0.087 | 0.21208128 | 0.37437048 | 0.0013 | 0.0039 | 0.0029 | 0.0029 | TRUE | 0.0359 |
| Csnk2a1 | 14822 | 10476 | 10955 | 12007 | 10330 | 11862 | 11512 | 11143 | 11820 | 11952 | 11653 | 11710 | 11631.64 | 11492 | 11772 | 1.024 | 0.035 | 0.47339028 | 0.64189046 | 4.00E-04 | 0.0026 | 0.0016 | 0.0016 | TRUE | 0.0085 |
| Csnk2a2 | 8983 | 6759 | 6364 | 7705 | 6703 | 7431 | 6977 | 7189 | 6867 | 7670 | 7562 | 7336 | 7266.61 | 7011 | 7523 | 1.073 | 0.102 | 0.04320357 | 0.1210564 | 3.00E-04 | 0.0028 | 0.0017 | 0.0017 | TRUE | 0.006 |
| Csnk2b | 5211 | 3524 | 3415 | 3470 | 3111 | 3695 | 4047 | 3748 | 3685 | 3454 | 3510 | 3648 | 3681.91 | 3827 | 3537 | 0.924 | -0.114 | 0.06032278 | 0.1538675 | 0.0013 | 0.0031 | 0.0024 | 0.0024 | TRUE | 0.0315 |
| Csnka2ip | 0 | 0 | 0 | 0 | 0 | 0 | 0 | 0 | 0 | 0 | 0 | 0 | 0 | 0 | 0 | NA | NA | NA | NA | NA | NA | NA | NA | NA | NA |
| Cspg4 | 56 | 50 | 58 | 60 | 63 | 66 | 43 | 53 | 63 | 60 | 71 | 65 | 59.2 | 53 | 65 | 1.238 | 0.308 | 0.23747537 | 0.40417188 | 0 | 0.0425 | 0.0318 | 0.0318 | TRUE | 0.244 |
| Cspg5 | 175 | 135 | 165 | 225 | 192 | 237 | 136 | 144 | 178 | 224 | 217 | 234 | 188.68 | 153 | 225 | 1.481 | 0.567 | 0.00028415 | 0.00278744 | 0.0043 | 0.015 | 0.0121 | 0.0121 | TRUE | 0.2183 |
| Cspp1 | 2533 | 1845 | 1676 | 1599 | 1489 | 1690 | 1967 | 1963 | 1808 | 1592 | 1680 | 1668 | 1779.64 | 1913 | 1647 | 0.86 | -0.217 | 0.00129946 | 0.00870843 | 0.001 | 0.0038 | 0.0027 | 0.0027 | TRUE | 0.027 |
| Csprs | 0 | 0 | 0 | 0 | 0 | 1 | 0 | 0 | 0 | 0 | 0 | 1 | 0.16 | 0 | 0 | 2.094 | 1.067 | 0.79379329 | NA | 0 | 14.4078 | 10 | 10 | TRUE | 0.3835 |
| Csrnp1 | 1286 | 779 | 710 | 685 | 623 | 776 | 999 | 829 | 766 | 682 | 703 | 766 | 790.69 | 865 | 717 | 0.828 | -0.273 | 0.01119646 | 0.04486432 | 0.0097 | 0.0054 | 0.0071 | 0.0071 | TRUE | 0.2264 |
| Csrnp2 | 2685 | 1903 | 1472 | 1755 | 1743 | 1763 | 2085 | 2024 | 1588 | 1747 | 1966 | 1740 | 1858.56 | 1899 | 1818 | 0.956 | -0.065 | 0.51668561 | 0.67889202 | 0.0127 | 0.0037 | 0.0066 | 0.0066 | TRUE | 0.2459 |
| Csrnp3 | 902 | 694 | 741 | 884 | 735 | 923 | 701 | 738 | 800 | 880 | 829 | 911 | 809.75 | 746 | 873 | 1.173 | 0.23 | 0.00842971 | 0.03645564 | 0.0021 | 0.0054 | 0.0043 | 0.0043 | TRUE | 0.0461 |
| Csrp1 | 2225 | 1569 | 1519 | 1771 | 1519 | 1728 | 1728 | 1669 | 1639 | 1763 | 1714 | 1706 | 1703.04 | 1679 | 1728 | 1.028 | 0.041 | 0.52482187 | 0.68598801 | 0 | 0.0038 | 0.0023 | 0.0023 | TRUE | 0.0083 |
| Csrp2 | 7767 | 5934 | 6243 | 6429 | 5824 | 6894 | 6032 | 6312 | 6736 | 6399 | 6570 | 6805 | 6475.88 | 6360 | 6591 | 1.037 | 0.052 | 0.38267282 | 0.55658084 | 0.0018 | 0.0028 | 0.0024 | 0.0024 | TRUE | 0.0326 |
| Csrp2bp | 3617 | 2734 | 2606 | 2933 | 2540 | 2920 | 2809 | 2908 | 2812 | 2920 | 2865 | 2882 | 2866.09 | 2843 | 2889 | 1.016 | 0.024 | 0.67019283 | 0.79708707 | 0 | 0.0033 | 0.0019 | 0.0019 | TRUE | 0.0048 |
| Csrp3 | 1 | 0 | 0 | 0 | 0 | 1 | 1 | 0 | 0 | 0 | 0 | 1 | 0.29 | 0 | 0 | 1.075 | 0.105 | 0.97702352 | NA | 0 | 8.0647 | 7.5458 | 7.5458 | TRUE | 0.3835 |
| Cst10 | 0 | 0 | 0 | 0 | 1 | 0 | 0 | 0 | 0 | 0 | 1 | 0 | 0.19 | 0 | 0 | 2.094 | 1.067 | 0.79379329 | NA | 0 | 12.608 | 10 | 10 | TRUE | 0.5075 |
| Cst11 | 0 | 0 | 0 | 0 | 0 | 0 | 0 | 0 | 0 | 0 | 0 | 0 | 0 | 0 | 0 | NA | NA | NA | NA | NA | NA | NA | NA | NA | NA |
| Cst12 | 0 | 0 | 0 | 0 | 0 | 0 | 0 | 0 | 0 | 0 | 0 | 0 | 0 | 0 | 0 | NA | NA | NA | NA | NA | NA | NA | NA | NA | NA |
| Cst13 | 3 | 2 | 2 | 1 | 2 | 1 | 2 | 2 | 2 | 1 | 2 | 1 | 1.81 | 2 | 1 | 0.631 | -0.663 | 0.63172155 | NA | 0 | 1.3126 | 0.7908 | 0.7908 | TRUE | 0.1547 |
| Cst3 | 2428 | 1753 | 2075 | 2017 | 1954 | 2121 | 1886 | 1865 | 2239 | 2008 | 2204 | 2094 | 2049.18 | 1997 | 2102 | 1.053 | 0.075 | 0.36381408 | 0.53841025 | 0.0054 | 0.0036 | 0.0044 | 0.0044 | TRUE | 0.135 |
| Cst6 | 6 | 3 | 3 | 3 | 3 | 2 | 5 | 3 | 3 | 3 | 3 | 2 | 3.24 | 4 | 3 | 0.742 | -0.431 | 0.67764279 | NA | 0 | 0.7342 | 0.4574 | 0.4574 | TRUE | 0.1085 |
| Cst7 | 0 | 1 | 1 | 0 | 0 | 0 | 0 | 1 | 1 | 0 | 0 | 0 | 0.36 | 1 | 0 | 0.271 | -1.881 | 0.5755355 | NA | 0 | 6.6391 | 6.4428 | 6.4428 | TRUE | 0.3513 |
| Cst8 | 0 | 0 | 0 | 0 | 0 | 0 | 0 | 0 | 0 | 0 | 0 | 0 | 0 | 0 | 0 | NA | NA | NA | NA | NA | NA | NA | NA | NA | NA |
| Cst9 | 0 | 0 | 0 | 0 | 0 | 1 | 0 | 0 | 0 | 0 | 0 | 1 | 0.16 | 0 | 0 | 2.094 | 1.067 | 0.79379329 | NA | 0 | 14.4078 | 10 | 10 | TRUE | 0.3835 |
| Csta | 0 | 0 | 0 | 0 | 0 | 0 | 0 | 0 | 0 | 0 | 0 | 0 | 0 | 0 | 0 | NA | NA | NA | NA | NA | NA | NA | NA | NA | NA |
| Cstad | 1 | 0 | 0 | 0 | 0 | 0 | 1 | 0 | 0 | 0 | 0 | 0 | 0.13 | 0 | 0 | 0.552 | -0.857 | 0.8336548 | NA | 0 | 18.3121 | 10 | 10 | TRUE | 0.2566 |
| Cstb | 2008 | 1530 | 1841 | 2169 | 1948 | 2232 | 1560 | 1627 | 1986 | 2159 | 2198 | 2203 | 1955.55 | 1724 | 2187 | 1.269 | 0.344 | 9.97E-05 | 0.0012072 | 0.0081 | 0.0036 | 0.0051 | 0.0051 | TRUE | 0.2116 |
| Cstf1 | 4065 | 2967 | 2941 | 3009 | 2749 | 3153 | 3157 | 3156 | 3173 | 2995 | 3101 | 3112 | 3115.87 | 3162 | 3069 | 0.971 | -0.043 | 0.42937354 | 0.60143435 | 0 | 0.0032 | 0.0018 | 0.0018 | TRUE | 0.0055 |
| Cstf2 | 7940 | 5773 | 5843 | 6037 | 5028 | 5914 | 6167 | 6141 | 6305 | 6009 | 5672 | 5838 | 6021.87 | 6204 | 5840 | 0.941 | -0.087 | 0.08822872 | 0.20181091 | 3.00E-04 | 0.0028 | 0.0017 | 0.0017 | TRUE | 0.0079 |
| Cstf2t | 2990 | 2243 | 1835 | 1670 | 1499 | 1763 | 2322 | 2386 | 1980 | 1662 | 1691 | 1740 | 1963.62 | 2229 | 1698 | 0.761 | -0.393 | 7.99E-07 | 2.43E-05 | 0.0043 | 0.0036 | 0.0041 | 0.0041 | TRUE | 0.1143 |
| Cstf3 | 6064 | 3900 | 3026 | 3068 | 2933 | 3507 | 4710 | 4148 | 3265 | 3054 | 3309 | 3462 | 3657.94 | 4041 | 3275 | 0.81 | -0.304 | 0.00448335 | 0.0225954 | 0.0178 | 0.0031 | 0.008 | 0.008 | TRUE | 0.2873 |
| Cstl1 | 0 | 0 | 0 | 0 | 0 | 0 | 0 | 0 | 0 | 0 | 0 | 0 | 0 | 0 | 0 | NA | NA | NA | NA | NA | NA | NA | NA | NA | NA |
| Ctag2 | 17 | 3 | 5 | 3 | 5 | 2 | 13 | 3 | 5 | 3 | 6 | 2 | 5.4 | 7 | 4 | 0.471 | -1.087 | 0.23188312 | NA | 0.2524 | 0.4415 | 0.382 | 0.382 | TRUE | 0.9761 |
| Ctage5 | 2980 | 2138 | 1995 | 2081 | 1831 | 2028 | 2314 | 2274 | 2153 | 2071 | 2066 | 2002 | 2146.69 | 2247 | 2046 | 0.91 | -0.136 | 0.02875986 | 0.08983064 | 5.00E-04 | 0.0035 | 0.0023 | 0.0023 | TRUE | 0.0162 |
| Ctbp1 | 6636 | 4817 | 4847 | 4931 | 4472 | 5163 | 5154 | 5124 | 5230 | 4908 | 5045 | 5097 | 5092.9 | 5169 | 5017 | 0.971 | -0.043 | 0.3892958 | 0.5633074 | 0 | 0.0029 | 0.0016 | 0.0016 | TRUE | 0.0044 |
| Ctbp2 | 14329 | 10024 | 9579 | 11709 | 10064 | 11689 | 11129 | 10662 | 10336 | 11655 | 11353 | 11539 | 11112.33 | 10709 | 11516 | 1.075 | 0.105 | 0.03849862 | 0.11138883 | 6.00E-04 | 0.0026 | 0.0017 | 0.0017 | TRUE | 0.0145 |
| Ctbs | 741 | 513 | 636 | 630 | 545 | 654 | 576 | 546 | 686 | 627 | 615 | 646 | 615.82 | 603 | 629 | 1.046 | 0.064 | 0.53436798 | 0.69345575 | 0.0058 | 0.0063 | 0.0061 | 0.0061 | TRUE | 0.1686 |
| Ctc1 | 13877 | 10634 | 11845 | 12712 | 11286 | 13408 | 10778 | 11311 | 12781 | 12654 | 12732 | 13236 | 12248.44 | 11623 | 12874 | 1.108 | 0.148 | 0.02877812 | 0.08983762 | 0.0042 | 0.0026 | 0.0032 | 0.0032 | TRUE | 0.0926 |
| Ctcf | 12495 | 8913 | 8457 | 8287 | 7496 | 8768 | 9704 | 9481 | 9125 | 8249 | 8456 | 8655 | 8945.09 | 9437 | 8453 | 0.896 | -0.159 | 0.00189074 | 0.01157726 | 6.00E-04 | 0.0027 | 0.0018 | 0.0018 | TRUE | 0.0102 |
| Ctcfl | 150 | 134 | 92 | 98 | 60 | 78 | 116 | 143 | 99 | 98 | 68 | 77 | 100.09 | 119 | 81 | 0.678 | -0.561 | 0.01160739 | 0.04616152 | 0.0226 | 0.0261 | 0.0251 | 0.0251 | TRUE | 0.2588 |
| Ctdnep1 | 2989 | 2053 | 2070 | 2129 | 1976 | 2261 | 2321 | 2184 | 2234 | 2119 | 2229 | 2232 | 2219.83 | 2246 | 2193 | 0.976 | -0.035 | 0.56865337 | 0.72227518 | 5.00E-04 | 0.0035 | 0.0023 | 0.0023 | TRUE | 0.0107 |

| | | | | | | | | | | | | | | | | | | | | | | | | | |
|---|---|---|---|---|---|---|---|---|---|---|---|---|---|---|---|---|---|---|---|---|---|---|---|---|---|
| Ctdp1 | 3564 | 2457 | 1993 | 2043 | 1815 | 2077 | 2768 | 2613 | 2150 | 2034 | 2048 | 2050 | 2277.22 | 2510 | 2044 | 0.813 | -0.298 | 0.00070025 | 0.00557273 | 0.0075 | 0.0035 | 0.0051 | 0.0051 | TRUE | 0.1897 |
| Ctdsp1 | 966 | 714 | 504 | 486 | 425 | 504 | 750 | 759 | 544 | 484 | 479 | 498 | 585.71 | 684 | 487 | 0.71 | -0.493 | 7.83E-05 | 0.00099324 | 0.016 | 0.0065 | 0.0095 | 0.0095 | TRUE | 0.3743 |
| Ctdsp2 | 7687 | 5758 | 5985 | 6126 | 5705 | 6649 | 5970 | 6125 | 6458 | 6098 | 6436 | 6564 | 6274.98 | 6184 | 6366 | 1.03 | 0.042 | 0.46189107 | 0.6311481 | 0.0013 | 0.0028 | 0.0022 | 0.0022 | TRUE | 0.0182 |
| Ctdspl | 713 | 588 | 489 | 484 | 430 | 499 | 554 | 625 | 528 | 482 | 485 | 493 | 527.71 | 569 | 487 | 0.856 | -0.225 | 0.02540853 | 0.08229415 | 0.0022 | 0.0069 | 0.0054 | 0.0054 | TRUE | 0.0854 |
| Ctdspl2 | 3689 | 2809 | 2905 | 2947 | 2476 | 2786 | 2865 | 2988 | 3134 | 2933 | 2793 | 2750 | 2910.72 | 2996 | 2825 | 0.944 | -0.084 | 0.17961349 | 0.33446994 | 0.0013 | 0.0032 | 0.0024 | 0.0024 | TRUE | 0.0199 |
| Ctf1 | 114 | 74 | 98 | 106 | 83 | 125 | 89 | 79 | 106 | 106 | 94 | 123 | 99.26 | 91 | 108 | 1.185 | 0.245 | 0.24391998 | 0.41185449 | 0.0095 | 0.0263 | 0.0217 | 0.0217 | TRUE | 0.1792 |
| Ctf2 | 29 | 11 | 6 | 7 | 4 | 10 | 23 | 12 | 6 | 7 | 5 | 10 | 10.34 | 14 | 7 | 0.519 | -0.945 | 0.15874098 | NA | 0.1836 | 0.2316 | 0.217 | 0.217 | TRUE | 0.3453 |
| Ctgf | 1634 | 967 | 837 | 491 | 361 | 348 | 1269 | 1029 | 903 | 489 | 407 | 344 | 740.05 | 1067 | 413 | 0.387 | -1.371 | 1.12E-22 | 4.46E-20 | 0.0282 | 0.0056 | 0.0124 | 0.0124 | TRUE | 0.2346 |
| Cth | 964 | 638 | 636 | 919 | 793 | 840 | 749 | 679 | 686 | 915 | 895 | 829 | 792.02 | 705 | 880 | 1.246 | 0.317 | 0.00024836 | 0.00252114 | 0.0017 | 0.0054 | 0.0041 | 0.0041 | TRUE | 0.0376 |
| Cthrc1 | 126 | 73 | 110 | 113 | 118 | 140 | 98 | 78 | 119 | 112 | 133 | 138 | 113 | 98 | 128 | 1.305 | 0.384 | 0.06237858 | 0.15791304 | 0.0164 | 0.0234 | 0.0215 | 0.0215 | TRUE | 0.2303 |
| Ctif | 631 | 471 | 464 | 475 | 469 | 481 | 490 | 501 | 501 | 473 | 529 | 475 | 494.74 | 497 | 492 | 0.989 | -0.015 | 0.87585006 | 0.93088377 | 0 | 0.0072 | 0.0049 | 0.0049 | TRUE | 0.0485 |
| Ctla2a | 0 | 0 | 0 | 0 | 0 | 0 | 0 | 0 | 0 | 0 | 0 | 0 | 0 | 0 | 0 | NA | NA | NA | NA | NA | NA | NA | NA | NA | NA |
| Ctla2b | 0 | 0 | 1 | 0 | 0 | 0 | 0 | 0 | 1 | 0 | 0 | 0 | 0.18 | 0 | 0 | 0.552 | -0.857 | 0.8336548 | NA | 0 | 13.1817 | 10 | 10 | TRUE | 0.5365 |
| Ctla4 | 0 | 0 | 0 | 0 | 0 | 0 | 0 | 0 | 0 | 0 | 0 | 0 | 0 | 0 | 0 | NA | NA | NA | NA | NA | NA | NA | NA | NA | NA |
| Ctnna1 | 32693 | 22845 | 23381 | 24356 | 21545 | 24520 | 25391 | 24300 | 25228 | 24244 | 24305 | 24205 | 24612.18 | 24973 | 24251 | 0.971 | -0.042 | 0.34338184 | 0.51766234 | 2.00E-04 | 0.0025 | 0.0014 | 0.0014 | TRUE | 0.0068 |
| Ctnna2 | 17 | 14 | 10 | 10 | 14 | 7 | 13 | 15 | 11 | 10 | 16 | 7 | 11.92 | 13 | 11 | 0.833 | -0.264 | 0.64185402 | NA | 0 | 0.2012 | 0.1478 | 0.1478 | TRUE | 0.4874 |
| Ctnna3 | 1 | 0 | 0 | 0 | 1 | 0 | 1 | 0 | 0 | 0 | 1 | 0 | 0.32 | 0 | 0 | 1.075 | 0.105 | 0.97638847 | NA | 0 | 7.4681 | 7.0392 | 7.0392 | TRUE | 0.5075 |
| Ctnnal1 | 7639 | 5730 | 5888 | 4768 | 4508 | 5399 | 5933 | 6095 | 6353 | 4746 | 5085 | 5330 | 5590.35 | 6127 | 5054 | 0.825 | -0.278 | 6.35E-06 | 0.00012929 | 0.002 | 0.0029 | 0.0025 | 0.0025 | TRUE | 0.0348 |
| Ctnnb1 | 59889 | 43788 | 42299 | 40691 | 37722 | 42647 | 46513 | 46577 | 45640 | 40504 | 42554 | 42099 | 43981.25 | 46243 | 41719 | 0.902 | -0.149 | 0.00099889 | 0.00723765 | 4.00E-04 | 0.0025 | 0.0014 | 0.0014 | TRUE | 0.008 |
| Ctnnbip1 | 2266 | 1663 | 1649 | 1780 | 1646 | 1830 | 1760 | 1769 | 1779 | 1772 | 1857 | 1806 | 1790.54 | 1769 | 1812 | 1.024 | 0.034 | 0.58231737 | 0.73253195 | 0 | 0.0038 | 0.0022 | 0.0022 | TRUE | 0.0056 |
| Ctnnbl1 | 4245 | 3042 | 2857 | 3211 | 3073 | 3528 | 3297 | 3236 | 3083 | 3196 | 3467 | 3483 | 3293.48 | 3205 | 3382 | 1.055 | 0.077 | 0.2151466 | 0.37765193 | 0.0013 | 0.0031 | 0.0025 | 0.0025 | TRUE | 0.0282 |
| Ctnnd1 | 14116 | 9932 | 8904 | 10375 | 9081 | 10165 | 10963 | 10565 | 9607 | 10327 | 10244 | 10034 | 10290.2 | 10378 | 10202 | 0.983 | -0.025 | 0.67723321 | 0.80217727 | 0.0019 | 0.0027 | 0.0025 | 0.0025 | TRUE | 0.0515 |
| Ctnnd2 | 324 | 290 | 288 | 272 | 259 | 262 | 252 | 308 | 311 | 271 | 292 | 259 | 282.07 | 290 | 274 | 0.946 | -0.08 | 0.54985952 | 0.70618583 | 0.006 | 0.0108 | 0.0093 | 0.0093 | TRUE | 0.1655 |
| Ctns | 1410 | 1109 | 1180 | 1297 | 1186 | 1381 | 1095 | 1180 | 1273 | 1291 | 1338 | 1363 | 1256.69 | 1183 | 1331 | 1.127 | 0.172 | 0.02839871 | 0.08893575 | 0.0025 | 0.0043 | 0.0037 | 0.0037 | TRUE | 0.0537 |
| Ctps | 11652 | 8410 | 8737 | 9133 | 8068 | 9411 | 9050 | 8946 | 9427 | 9091 | 9102 | 9290 | 9150.83 | 9141 | 9161 | 1.002 | 0.003 | 0.94598153 | 0.97029019 | 3.00E-04 | 0.0027 | 0.0016 | 0.0016 | TRUE | 0.0091 |
| Ctps2 | 1814 | 1256 | 1317 | 1400 | 1206 | 1362 | 1409 | 1336 | 1421 | 1394 | 1360 | 1345 | 1377.41 | 1389 | 1366 | 0.984 | -0.024 | 0.72574163 | 0.83572689 | 0 | 0.0041 | 0.0026 | 0.0026 | TRUE | 0.0129 |
| Ctr9 | 8425 | 5875 | 5026 | 4990 | 4687 | 5225 | 6543 | 6249 | 5423 | 4967 | 5287 | 5158 | 5604.64 | 6072 | 5137 | 0.846 | -0.242 | 0.00096302 | 0.00705871 | 0.0048 | 0.0028 | 0.0037 | 0.0037 | TRUE | 0.1062 |
| Ctrb1 | 10 | 0 | 5 | 11 | 7 | 7 | 8 | 0 | 5 | 11 | 8 | 7 | 6.49 | 4 | 9 | 1.903 | 0.929 | 0.27978794 | NA | 0.3459 | 0.3678 | 0.3647 | 0.3647 | TRUE | 0.9843 |
| Ctrc | 1 | 1 | 1 | 1 | 0 | 2 | 1 | 1 | 1 | 1 | 0 | 2 | 0.98 | 1 | 1 | 1.044 | 0.062 | 0.97363744 | NA | 0 | 2.4171 | 1.5475 | 1.5475 | TRUE | 0.1802 |
| Ctrl | 9 | 16 | 6 | 6 | 2 | 9 | 7 | 17 | 6 | 6 | 2 | 9 | 7.93 | 10 | 6 | 0.572 | -0.806 | 0.28510182 | NA | 0.1975 | 0.3012 | 0.2723 | 0.2723 | TRUE | 0.6637 |
| Cts3 | 2 | 0 | 1 | 3 | 2 | 1 | 2 | 0 | 1 | 3 | 2 | 1 | 1.48 | 1 | 2 | 2.274 | 1.185 | 0.46249813 | NA | 0 | 1.6071 | 1.0982 | 1.0982 | TRUE | 0.2816 |
| Cts6 | 0 | 0 | 0 | 0 | 0 | 0 | 0 | 0 | 0 | 0 | 0 | 0 | 0 | 0 | 0 | NA | NA | NA | NA | NA | NA | NA | NA | NA | NA |
| Cts7 | 0 | 1 | 0 | 0 | 0 | 0 | 0 | 1 | 0 | 0 | 0 | 0 | 0.18 | 0 | 0 | 0.552 | -0.857 | 0.8336548 | NA | 0 | 13.3714 | 10 | 10 | TRUE | 0.5216 |
| Cts8 | 0 | 0 | 0 | 0 | 0 | 0 | 0 | 0 | 0 | 0 | 0 | 0 | 0 | 0 | 0 | NA | NA | NA | NA | NA | NA | NA | NA | NA | NA |
| Cts8-ps | 0 | 0 | 0 | 0 | 0 | 0 | 0 | 0 | 0 | 0 | 0 | 0 | 0 | 0 | 0 | NA | NA | NA | NA | NA | NA | NA | NA | NA | NA |
| Ctsa | 2205 | 1753 | 2039 | 1894 | 1772 | 2064 | 1713 | 1865 | 2200 | 1885 | 1999 | 2037 | 1949.83 | 1926 | 1974 | 1.026 | 0.037 | 0.68070421 | 0.80400837 | 0.0085 | 0.0036 | 0.0053 | 0.0053 | TRUE | 0.1871 |
| Ctsb | 7089 | 5168 | 5741 | 6414 | 5835 | 6246 | 5506 | 5497 | 6194 | 6384 | 6582 | 6166 | 6055.02 | 5732 | 6377 | 1.113 | 0.154 | 0.0159749 | 0.05850095 | 0.0027 | 0.0028 | 0.0028 | 0.0028 | TRUE | 0.0603 |
| Ctsc | 1086 | 777 | 846 | 1027 | 818 | 780 | 843 | 826 | 913 | 1022 | 923 | 770 | 882.97 | 861 | 905 | 1.051 | 0.072 | 0.49187664 | 0.65736783 | 0.0104 | 0.0051 | 0.0068 | 0.0068 | TRUE | 0.2056 |
| Ctsd | 15707 | 11273 | 12627 | 14073 | 12474 | 13951 | 12199 | 11991 | 13624 | 14008 | 14072 | 13772 | 13277.72 | 12605 | 13951 | 1.107 | 0.147 | 0.01467104 | 0.05486476 | 0.0024 | 0.0026 | 0.0025 | 0.0025 | TRUE | 0.061 |
| Ctse | 0 | 0 | 0 | 0 | 0 | 0 | 0 | 0 | 0 | 0 | 0 | 0 | 0 | 0 | 0 | NA | NA | NA | NA | NA | NA | NA | NA | NA | NA |
| Ctsf | 289 | 182 | 220 | 223 | 208 | 271 | 224 | 194 | 237 | 222 | 235 | 268 | 229.93 | 218 | 242 | 1.105 | 0.143 | 0.31973527 | 0.49360512 | 0.0054 | 0.0127 | 0.0106 | 0.0106 | TRUE | 0.1035 |
| Ctsg | 1 | 3 | 0 | 0 | 1 | 2 | 1 | 3 | 0 | 0 | 1 | 2 | 1.18 | 1 | 1 | 0.79 | -0.339 | 0.85764261 | NA | 0 | 2.0138 | 1.7083 | 1.7083 | TRUE | 0.7212 |
| Ctsh | 574 | 434 | 504 | 713 | 614 | 714 | 446 | 462 | 544 | 710 | 693 | 705 | 593.08 | 484 | 703 | 1.455 | 0.542 | 7.12E-08 | 2.89E-06 | 0.0034 | 0.0064 | 0.0055 | 0.0055 | TRUE | 0.1344 |
| Ctsj | 1 | 0 | 0 | 0 | 0 | 0 | 1 | 0 | 0 | 0 | 0 | 0 | 0.13 | 0 | 0 | 0.552 | -0.857 | 0.8336548 | NA | 0 | 18.3121 | 10 | 10 | TRUE | 0.2566 |
| Ctsk | 12 | 7 | 4 | 4 | 9 | 2 | 9 | 7 | 4 | 4 | 10 | 2 | 6.2 | 7 | 5 | 0.745 | -0.424 | 0.60868752 | NA | 0.1764 | 0.3848 | 0.3287 | 0.3287 | TRUE | 1.1161 |
| Ctsl | 8228 | 5642 | 5605 | 5862 | 5100 | 5822 | 6390 | 6001 | 6048 | 5835 | 5753 | 5747 | 5962.5 | 6146 | 5778 | 0.94 | -0.09 | 0.08527138 | 0.19721623 | 5.00E-04 | 0.0028 | 0.0018 | 0.0018 | TRUE | 0.0149 |
| Ctsll3 | 1 | 0 | 0 | 0 | 0 | 0 | 1 | 0 | 0 | 0 | 0 | 0 | 0.13 | 0 | 0 | 0.552 | -0.857 | 0.8336548 | NA | 0 | 18.3121 | 10 | 10 | TRUE | 0.2566 |
| Ctsm | 2 | 2 | 1 | 2 | 1 | 4 | 2 | 2 | 1 | 2 | 1 | 4 | 1.97 | 2 | 2 | 1.498 | 0.583 | 0.66331476 | NA | 0 | 1.2048 | 0.7709 | 0.7709 | TRUE | 0.3732 |
| Ctso | 96 | 73 | 71 | 71 | 72 | 70 | 75 | 78 | 77 | 71 | 81 | 69 | 74.97 | 77 | 74 | 0.965 | -0.051 | 0.81718226 | 0.89501809 | 0 | 0.034 | 0.0223 | 0.0223 | TRUE | 0.0692 |
| Ctsq | 0 | 0 | 0 | 0 | 0 | 0 | 0 | 0 | 0 | 0 | 0 | 0 | 0 | 0 | 0 | NA | NA | NA | NA | NA | NA | NA | NA | NA | NA |
| Ctsr | 0 | 0 | 0 | 0 | 0 | 0 | 0 | 0 | 0 | 0 | 0 | 0 | 0 | 0 | 0 | NA | NA | NA | NA | NA | NA | NA | NA | NA | NA |
| Ctss | 13 | 6 | 4 | 3 | 1 | 7 | 10 | 6 | 4 | 3 | 1 | 7 | 5.3 | 7 | 4 | 0.53 | -0.915 | 0.30688271 | NA | 0.1417 | 0.4493 | 0.37 | 0.37 | TRUE | 0.74 |
| Ctsw | 19 | 10 | 18 | 14 | 20 | 18 | 15 | 11 | 19 | 14 | 23 | 18 | 16.51 | 15 | 18 | 1.208 | 0.272 | 0.57436682 | NA | 0 | 0.1459 | 0.1084 | 0.1084 | TRUE | 0.2257 |
| Ctsz | 4254 | 3269 | 3506 | 3641 | 3559 | 4263 | 3304 | 3477 | 3783 | 3624 | 4015 | 4208 | 3735.24 | 3521 | 3949 | 1.122 | 0.166 | 0.02674687 | 0.08537137 | 0.0048 | 0.0031 | 0.0038 | 0.0038 | TRUE | 0.0633 |
| Cttn | 10460 | 7439 | 7073 | 7032 | 6224 | 7045 | 8124 | 7913 | 7632 | 7000 | 7021 | 6955 | 7440.63 | 7890 | 6992 | 0.886 | -0.175 | 0.00048245 | 0.00420721 | 4.00E-04 | 0.0027 | 0.0017 | 0.0017 | TRUE | 0.0099 |
| Cttnbp2 | 143 | 112 | 108 | 110 | 95 | 118 | 111 | 119 | 117 | 109 | 107 | 116 | 113.31 | 116 | 111 | 0.963 | -0.055 | 0.76373589 | 0.85969854 | 0 | 0.0233 | 0.015 | 0.015 | TRUE | 0.0186 |
| Cttnbp2nl | 2054 | 1470 | 1398 | 1413 | 1254 | 1556 | 1595 | 1564 | 1508 | 1406 | 1415 | 1536 | 1504.08 | 1556 | 1452 | 0.933 | -0.1 | 0.15572725 | 0.30349189 | 0.001 | 0.004 | 0.0029 | 0.0029 | TRUE | 0.0309 |
| Ctu1 | 803 | 611 | 602 | 716 | 618 | 704 | 624 | 650 | 650 | 713 | 697 | 695 | 671.33 | 641 | 702 | 1.095 | 0.132 | 0.11891744 | 0.25043274 | 0 | 0.006 | 0.0036 | 0.0036 | TRUE | 0.0071 |
| Ctu2 | 4143 | 3076 | 2604 | 2881 | 2622 | 2925 | 3218 | 3272 | 2810 | 2868 | 2958 | 2887 | 3002.06 | 3100 | 2904 | 0.937 | -0.095 | 0.17760817 | 0.33188045 | 0.003 | 0.0032 | 0.0032 | 0.0032 | TRUE | 0.0804 |
| Ctxn1 | 493 | 389 | 468 | 469 | 413 | 562 | 383 | 414 | 505 | 467 | 466 | 555 | 464.86 | 434 | 496 | 1.146 | 0.197 | 0.1190277 | 0.25057473 | 0.0128 | 0.0075 | 0.0093 | 0.0093 | TRUE | 0.2349 |
| Ctxn2 | 0 | 0 | 0 | 0 | 0 | 0 | 0 | 0 | 0 | 0 | 0 | 0 | 0 | 0 | 0 | NA | NA | NA | NA | NA | NA | NA | NA | NA | NA |
| Ctxn3 | 0 | 0 | 3 | 1 | 1 | 3 | 0 | 0 | 3 | 1 | 1 | 3 | 1.39 | 1 | 2 | 1.648 | 0.72 | 0.68888556 | NA | 1.0211 | 1.7112 | 1.5615 | 1.5615 | TRUE | 1.4446 |
| Cubn | 240 | 107 | 38 | 246 | 144 | 78 | 186 | 114 | 41 | 245 | 162 | 77 | 137.59 | 114 | 161 | 1.417 | 0.503 | 0.49073587 | 0.65650391 | 0.3766 | 0.0197 | 0.1102 | 0.3766 | TRUE | 0.2095 |
| Cuedc1 | 1409 | 1043 | 1082 | 937 | 817 | 973 | 1094 | 1109 | 1167 | 933 | 922 | 961 | 1031.01 | 1123 | 939 | 0.836 | -0.259 | 0.0005046 | 0.00434472 | 0 | 0.0047 | 0.003 | 0.003 | TRUE | 0.0136 |
| Cuedc2 | 2720 | 2029 | 1960 | 2253 | 2109 | 2455 | 2113 | 2158 | 2115 | 2243 | 2379 | 2423 | 2238.47 | 2129 | 2348 | 1.103 | 0.142 | 0.02120569 | 0.07219936 | 5.00E-04 | 0.0035 | 0.0023 | 0.0023 | TRUE | 0.019 |
| Cul1 | 7529 | 4999 | 4540 | 4280 | 3686 | 4338 | 5847 | 5317 | 4899 | 4260 | 4158 | 4282 | 4794.04 | 5354 | 4233 | 0.79 | -0.34 | 1.19E-06 | 3.38E-05 | 0.0037 | 0.0029 | 0.0033 | 0.0033 | TRUE | 0.0799 |
| Cul2 | 7880 | 5567 | 5690 | 5999 | 4942 | 5798 | 6120 | 5922 | 6139 | 5971 | 5575 | 5724 | 5908.51 | 6060 | 5757 | 0.95 | -0.074 | 0.16250971 | 0.31278634 | 5.00E-04 | 0.0028 | 0.0019 | 0.0019 | TRUE | 0.013 |
| Cul3 | 8969 | 6248 | 6060 | 6149 | 5234 | 5986 | 6966 | 6646 | 6539 | 6121 | 5904 | 5909 | 6347.46 | 6717 | 5978 | 0.89 | -0.168 | 0.00134388 | 0.00891479 | 5.00E-04 | 0.0028 | 0.0018 | 0.0018 | TRUE | 0.013 |
| Cul4a | 6531 | 4741 | 5256 | 5431 | 4979 | 5543 | 5072 | 5043 | 5671 | 5406 | 5617 | 5472 | 5380.18 | 5262 | 5498 | 1.045 | 0.064 | 0.30620738 | 0.47873792 | 0.0022 | 0.0029 | 0.0026 | 0.0026 | TRUE | 0.0559 |

| | | | | | | | | | | | | | | | | | | | | | | | | | |
|---|---|---|---|---|---|---|---|---|---|---|---|---|---|---|---|---|---|---|---|---|---|---|---|---|---|
| Cul4b | 23846 | 16653 | 17426 | 17109 | 14824 | 16966 | 18520 | 17714 | 18803 | 17030 | 16723 | 16748 | 17589.6 | 18346 | 16834 | 0.918 | -0.124 | 0.00900373 | 0.03835777 | 4.00E-04 | 0.0026 | 0.0016 | 0.0016 | TRUE | 0.0111 |
| Cul5 | 7037 | 4899 | 4487 | 5083 | 4165 | 4824 | 5465 | 5211 | 4841 | 5060 | 4699 | 4762 | 5006.33 | 5172 | 4840 | 0.935 | -0.096 | 0.13033617 | 0.26723495 | 0.0023 | 0.0029 | 0.0027 | 0.0027 | TRUE | 0.0381 |
| Cul7 | 4532 | 3373 | 3503 | 3686 | 3356 | 3885 | 3520 | 3588 | 3780 | 3669 | 3786 | 3835 | 3696.23 | 3629 | 3763 | 1.037 | 0.053 | 0.35279016 | 0.52767159 | 7.00E-04 | 0.0031 | 0.0021 | 0.0021 | TRUE | 0.0159 |
| Cul9 | 302 | 250 | 304 | 358 | 325 | 341 | 235 | 266 | 328 | 356 | 367 | 337 | 314.68 | 276 | 353 | 1.284 | 0.36 | 0.0091601 | 0.03878534 | 0.0119 | 0.01 | 0.0105 | 0.0105 | TRUE | 0.2995 |
| Cuta | 1508 | 1193 | 1183 | 1152 | 1042 | 1315 | 1171 | 1269 | 1276 | 1147 | 1175 | 1298 | 1222.82 | 1239 | 1207 | 0.975 | -0.036 | 0.64921938 | 0.7820068 | 0.0026 | 0.0044 | 0.0037 | 0.0037 | TRUE | 0.0532 |
| Cutc | 484 | 344 | 295 | 376 | 318 | 330 | 376 | 366 | 318 | 374 | 359 | 326 | 353.15 | 353 | 353 | 0.997 | -0.005 | 0.96725137 | 0.98237219 | 0.0036 | 0.0091 | 0.0074 | 0.0074 | TRUE | 0.0835 |
| Cux1 | 1885 | 1428 | 1422 | 1582 | 1338 | 1496 | 1464 | 1519 | 1534 | 1575 | 1509 | 1477 | 1513.03 | 1506 | 1520 | 1.01 | 0.015 | 0.82543726 | 0.8999218 | 0 | 0.004 | 0.0026 | 0.0026 | TRUE | 0.012 |
| Cux2 | 394 | 310 | 325 | 306 | 326 | 302 | 306 | 330 | 351 | 305 | 368 | 298 | 326.15 | 329 | 324 | 0.984 | -0.023 | 0.85553958 | 0.91884279 | 0.0059 | 0.0097 | 0.0086 | 0.0086 | TRUE | 0.1582 |
| Cuzd1 | 15 | 3 | 0 | 3 | 2 | 5 | 12 | 3 | 0 | 3 | 2 | 5 | 4.17 | 5 | 3 | 0.668 | -0.582 | 0.60396397 | NA | 0.9279 | 0.5708 | 0.661 | 0.661 | TRUE | 0.6109 |
| Cwc15 | 4111 | 2955 | 2712 | 2753 | 2260 | 2708 | 3193 | 3143 | 2926 | 2740 | 2550 | 2673 | 2870.89 | 3087 | 2654 | 0.86 | -0.218 | 0.00053151 | 0.00451121 | 0.0012 | 0.0033 | 0.0025 | 0.0025 | TRUE | 0.0251 |
| Cwc22 | 5344 | 3815 | 3423 | 3392 | 3043 | 3439 | 4150 | 4058 | 3693 | 3376 | 3433 | 3395 | 3684.31 | 3967 | 3401 | 0.857 | -0.223 | 0.00033953 | 0.00319064 | 0.0015 | 0.0031 | 0.0025 | 0.0025 | TRUE | 0.044 |
| Cwc25 | 2410 | 1752 | 1539 | 1347 | 1362 | 1437 | 1872 | 1864 | 1661 | 1341 | 1536 | 1419 | 1615.29 | 1799 | 1432 | 0.795 | -0.33 | 3.15E-05 | 0.00048409 | 0.0036 | 0.0039 | 0.0039 | 0.0039 | TRUE | 0.0537 |
| Cwc27 | 2849 | 2199 | 2303 | 2272 | 2026 | 2455 | 2213 | 2339 | 2485 | 2262 | 2286 | 2423 | 2334.54 | 2346 | 2324 | 0.991 | -0.012 | 0.85511246 | 0.91861122 | 0.002 | 0.0034 | 0.0029 | 0.0029 | TRUE | 0.0327 |
| Cwf19l1 | 1586 | 1172 | 1138 | 1227 | 1097 | 1269 | 1232 | 1247 | 1228 | 1221 | 1238 | 1253 | 1236.32 | 1236 | 1237 | 1.001 | 0.002 | 0.9747074 | 0.98637021 | 0 | 0.0043 | 0.0025 | 0.0025 | TRUE | 0.0015 |
| Cwf19l2 | 3164 | 2272 | 2460 | 2498 | 2057 | 2391 | 2457 | 2417 | 2654 | 2487 | 2321 | 2360 | 2449.28 | 2509 | 2389 | 0.952 | -0.07 | 0.28119624 | 0.45275857 | 0.0013 | 0.0034 | 0.0027 | 0.0027 | TRUE | 0.0304 |
| Cwh43 | 29 | 21 | 40 | 51 | 42 | 46 | 23 | 22 | 43 | 51 | 47 | 45 | 38.6 | 29 | 48 | 1.647 | 0.72 | 0.03601837 | 0.10613172 | 0.0383 | 0.0638 | 0.0575 | 0.0575 | TRUE | 1.0537 |
| Cx3cl1 | 514 | 338 | 225 | 297 | 247 | 251 | 399 | 360 | 243 | 296 | 279 | 248 | 303.93 | 334 | 274 | 0.818 | -0.29 | 0.08565752 | 0.19785729 | 0.0334 | 0.0102 | 0.0171 | 0.0171 | TRUE | 0.6353 |
| Cx3cr1 | 0 | 0 | 0 | 0 | 0 | 1 | 0 | 0 | 0 | 0 | 0 | 1 | 0.16 | 0 | 0 | 2.094 | 1.067 | 0.79379329 | NA | 0 | 14.4078 | 10 | 10 | TRUE | 0.3835 |
| Cxadr | 1443 | 1043 | 1111 | 1117 | 969 | 1052 | 1121 | 1109 | 1199 | 1112 | 1093 | 1038 | 1112.06 | 1143 | 1081 | 0.946 | -0.08 | 0.28813572 | 0.46025077 | 6.00E-04 | 0.0046 | 0.0032 | 0.0032 | TRUE | 0.0212 |
| Cxcl1 | 4 | 0 | 1 | 0 | 0 | 0 | 3 | 0 | 1 | 0 | 0 | 0 | 0.7 | 1 | 0 | 0.134 | -2.902 | 0.26925366 | NA | 6.5513 | 3.3999 | 3.6307 | 3.6307 | TRUE | 0.1404 |
| Cxcl10 | 57 | 35 | 24 | 19 | 16 | 21 | 44 | 37 | 26 | 19 | 18 | 21 | 27.51 | 36 | 19 | 0.534 | -0.905 | 0.01915275 | 0.06691525 | 0 | 0.0886 | 0.0673 | 0.0673 | TRUE | 0.2197 |
| Cxcl11 | 51 | 38 | 40 | 47 | 59 | 79 | 40 | 40 | 43 | 47 | 67 | 78 | 52.42 | 41 | 64 | 1.555 | 0.637 | 0.02671983 | 0.08530413 | 0.0204 | 0.0476 | 0.0396 | 0.0396 | TRUE | 0.2908 |
| Cxcl12 | 128 | 76 | 114 | 94 | 107 | 134 | 99 | 81 | 123 | 94 | 121 | 132 | 108.3 | 101 | 116 | 1.143 | 0.193 | 0.37638769 | 0.55097391 | 0.027 | 0.0243 | 0.0251 | 0.0251 | TRUE | 0.272 |
| Cxcl13 | 0 | 0 | 0 | 0 | 0 | 0 | 0 | 0 | 0 | 0 | 0 | 0 | 0 | 0 | 0 | NA | NA | NA | NA | NA | NA | NA | NA | NA | NA |
| Cxcl14 | 333 | 238 | 256 | 200 | 198 | 227 | 259 | 253 | 276 | 199 | 223 | 224 | 239.09 | 263 | 215 | 0.82 | -0.285 | 0.03149863 | 0.0959936 | 0 | 0.0123 | 0.0084 | 0.0084 | TRUE | 0.0496 |
| Cxcl16 | 166 | 139 | 161 | 234 | 180 | 228 | 129 | 148 | 174 | 233 | 203 | 225 | 185.26 | 150 | 220 | 1.476 | 0.562 | 0.00047405 | 0.00415173 | 0.0075 | 0.0152 | 0.013 | 0.013 | TRUE | 0.1951 |
| Cxcl17 | 0 | 0 | 0 | 0 | 0 | 0 | 0 | 0 | 0 | 0 | 0 | 0 | 0 | 0 | 0 | NA | NA | NA | NA | NA | NA | NA | NA | NA | NA |
| Cxcl2 | 0 | 0 | 0 | 0 | 0 | 0 | 0 | 0 | 0 | 0 | 0 | 0 | 0 | 0 | 0 | NA | NA | NA | NA | NA | NA | NA | NA | NA | NA |
| Cxcl3 | 1 | 2 | 3 | 1 | 0 | 1 | 1 | 2 | 3 | 1 | 0 | 1 | 1.35 | 2 | 1 | 0.337 | -1.569 | 0.36866479 | NA | 0 | 1.7529 | 1.1898 | 1.1898 | TRUE | 0.1561 |
| Cxcl5 | 6 | 6 | 5 | 5 | 2 | 5 | 5 | 6 | 5 | 5 | 2 | 5 | 4.77 | 5 | 4 | 0.751 | -0.414 | 0.6329431 | NA | 0 | 0.4995 | 0.3262 | 0.3262 | TRUE | 0.2213 |
| Cxcl9 | 2 | 3 | 1 | 1 | 0 | 0 | 2 | 3 | 1 | 1 | 0 | 0 | 1.14 | 2 | 0 | 0.192 | -2.382 | 0.22414484 | NA | 0 | 2.0879 | 1.5163 | 1.5163 | TRUE | 0.3906 |
| Cxcr1 | 3 | 2 | 4 | 3 | 3 | 2 | 2 | 2 | 4 | 3 | 3 | 2 | 2.85 | 3 | 3 | 0.956 | -0.065 | 0.95301137 | NA | 0 | 0.8331 | 0.5258 | 0.5258 | TRUE | 0.1994 |
| Cxcr2 | 1 | 1 | 1 | 0 | 0 | 0 | 1 | 1 | 1 | 0 | 0 | 0 | 0.49 | 1 | 0 | 0.196 | -2.347 | 0.38405676 | NA | 0 | 4.8735 | 3.7415 | 3.7415 | TRUE | 0.0718 |
| Cxcr3 | 2 | 0 | 0 | 1 | 0 | 0 | 2 | 0 | 0 | 1 | 0 | 0 | 0.42 | 1 | 0 | 0.681 | -0.553 | 0.8627148 | NA | 0 | 5.5819 | 5.4906 | 5.4906 | TRUE | 0.9727 |
| Cxcr4 | 81 | 65 | 59 | 95 | 82 | 85 | 63 | 69 | 64 | 95 | 93 | 84 | 77.78 | 65 | 91 | 1.386 | 0.471 | 0.03172506 | 0.09647793 | 0 | 0.0329 | 0.0215 | 0.0215 | TRUE | 0.038 |
| Cxcr5 | 3 | 2 | 2 | 1 | 1 | 2 | 2 | 2 | 2 | 1 | 1 | 2 | 1.79 | 2 | 1 | 0.621 | -0.688 | 0.6197856 | NA | 0 | 1.3298 | 0.7901 | 0.7901 | TRUE | 0.0998 |
| Cxcr6 | 0 | 0 | 0 | 0 | 0 | 0 | 0 | 0 | 0 | 0 | 0 | 0 | 0 | 0 | 0 | NA | NA | NA | NA | NA | NA | NA | NA | NA | NA |
| Cxcr7 | 198 | 158 | 179 | 173 | 162 | 176 | 154 | 168 | 193 | 172 | 183 | 174 | 173.95 | 172 | 176 | 1.03 | 0.042 | 0.78670679 | 0.87477829 | 0 | 0.0161 | 0.0119 | 0.0119 | TRUE | 0.1258 |
| Cxx1a | 141 | 90 | 100 | 102 | 75 | 99 | 110 | 96 | 108 | 102 | 85 | 98 | 99.5 | 105 | 95 | 0.907 | -0.142 | 0.47469918 | 0.64293333 | 0 | 0.0262 | 0.0182 | 0.0182 | TRUE | 0.078 |
| Cxx1b | 127 | 89 | 110 | 128 | 101 | 120 | 99 | 95 | 119 | 127 | 114 | 118 | 111.97 | 104 | 120 | 1.157 | 0.21 | 0.26849407 | 0.43908303 | 0 | 0.0236 | 0.0169 | 0.0169 | TRUE | 0.1418 |
| Cxx1c | 23 | 18 | 12 | 25 | 22 | 17 | 18 | 19 | 13 | 25 | 25 | 17 | 19.41 | 17 | 22 | 1.324 | 0.405 | 0.36053445 | 0.5354446 | 0 | 0.1246 | 0.0888 | 0.0888 | TRUE | 0.2665 |
| Cxxc1 | 2488 | 1884 | 1725 | 1623 | 1408 | 1780 | 1932 | 2004 | 1861 | 1616 | 1588 | 1757 | 1793.1 | 1932 | 1654 | 0.856 | -0.224 | 0.00126081 | 0.0085164 | 0.0013 | 0.0037 | 0.0029 | 0.0029 | TRUE | 0.0364 |
| Cxxc4 | 14 | 3 | 6 | 8 | 8 | 8 | 11 | 3 | 6 | 8 | 9 | 8 | 7.57 | 7 | 8 | 1.184 | 0.243 | 0.73371927 | NA | 0 | 0.3155 | 0.2366 | 0.2366 | TRUE | 0.3696 |
| Cxxc5 | 2005 | 1665 | 1792 | 2058 | 1698 | 1996 | 1557 | 1771 | 1934 | 2049 | 1916 | 1970 | 1866.03 | 1754 | 1978 | 1.129 | 0.176 | 0.03562688 | 0.10523145 | 0.0056 | 0.0037 | 0.0045 | 0.0045 | TRUE | 0.1195 |
| Cyb5 | 2906 | 2024 | 2054 | 2238 | 1919 | 2407 | 2257 | 2153 | 2216 | 2228 | 2165 | 2376 | 2232.46 | 2209 | 2256 | 1.021 | 0.031 | 0.63225819 | 0.76942242 | 0.001 | 0.0035 | 0.0025 | 0.0025 | TRUE | 0.0263 |
| Cyb561 | 2489 | 1834 | 2107 | 2371 | 2166 | 2333 | 1933 | 1951 | 2273 | 2360 | 2443 | 2303 | 2210.66 | 2052 | 2369 | 1.155 | 0.208 | 0.00647245 | 0.02989038 | 0.0041 | 0.0035 | 0.0037 | 0.0037 | TRUE | 0.1062 |
| Cyb561d1 | 282 | 207 | 215 | 236 | 194 | 225 | 219 | 220 | 232 | 235 | 219 | 222 | 224.51 | 224 | 225 | 1.008 | 0.012 | 0.92936861 | 0.9609898 | 0 | 0.013 | 0.0083 | 0.0083 | TRUE | 0.0155 |
| Cyb561d2 | 665 | 442 | 412 | 474 | 428 | 540 | 516 | 470 | 445 | 472 | 483 | 533 | 486.48 | 477 | 496 | 1.038 | 0.053 | 0.6143748 | 0.75628792 | 0.0032 | 0.0073 | 0.006 | 0.006 | TRUE | 0.0644 |
| Cyb5b | 14225 | 9632 | 8226 | 8563 | 7239 | 8622 | 11048 | 10245 | 8876 | 8524 | 8166 | 8511 | 9228.39 | 10056 | 8400 | 0.835 | -0.26 | 0.00060433 | 0.00499371 | 0.0061 | 0.0027 | 0.004 | 0.004 | TRUE | 0.1287 |
| Cyb5d1 | 437 | 267 | 201 | 176 | 168 | 214 | 339 | 284 | 217 | 175 | 190 | 211 | 236.04 | 280 | 192 | 0.683 | -0.55 | 0.00129538 | 0.00870059 | 0.0261 | 0.0125 | 0.0167 | 0.0167 | TRUE | 0.1686 |
| Cyb5d2 | 252 | 195 | 182 | 203 | 194 | 182 | 196 | 207 | 196 | 202 | 219 | 180 | 200.02 | 200 | 200 | 1.001 | 0.001 | 0.99194757 | 0.99565418 | 0 | 0.0143 | 0.0102 | 0.0102 | TRUE | 0.0884 |
| Cyb5r1 | 2846 | 2252 | 1941 | 1911 | 1612 | 1966 | 2210 | 2395 | 2094 | 1902 | 1819 | 1941 | 2060.26 | 2233 | 1887 | 0.845 | -0.243 | 0.00060861 | 0.00502449 | 0.0021 | 0.0036 | 0.0031 | 0.0031 | TRUE | 0.0478 |
| Cyb5r2 | 27 | 14 | 18 | 10 | 9 | 14 | 21 | 15 | 19 | 10 | 10 | 14 | 14.87 | 18 | 11 | 0.612 | -0.708 | 0.15861189 | NA | 0 | 0.1618 | 0.1098 | 0.1098 | TRUE | 0.1482 |
| Cyb5r3 | 9134 | 7073 | 7879 | 8828 | 8255 | 9501 | 7094 | 7523 | 8501 | 8787 | 9312 | 9379 | 8432.94 | 7706 | 9159 | 1.189 | 0.25 | 0.00043599 | 0.00388262 | 0.0048 | 0.0027 | 0.0035 | 0.0035 | TRUE | 0.0993 |
| Cyb5r4 | 3078 | 2257 | 2188 | 2290 | 1962 | 2360 | 2391 | 2401 | 2361 | 2279 | 2213 | 2330 | 2329.1 | 2384 | 2274 | 0.954 | -0.068 | 0.24530089 | 0.41364861 | 0 | 0.0034 | 0.002 | 0.002 | TRUE | 0.0065 |
| Cyb5rl | 169 | 121 | 82 | 71 | 80 | 80 | 131 | 129 | 88 | 71 | 90 | 79 | 98.06 | 116 | 80 | 0.685 | -0.546 | 0.01413394 | 0.05340113 | 0.0213 | 0.0266 | 0.0251 | 0.0251 | TRUE | 0.418 |
| Cyba | 758 | 592 | 660 | 739 | 738 | 783 | 589 | 630 | 712 | 736 | 833 | 773 | 711.94 | 644 | 781 | 1.214 | 0.28 | 0.00443347 | 0.0223993 | 0.0052 | 0.0058 | 0.0056 | 0.0056 | TRUE | 0.1015 |
| Cybasc3 | 1193 | 844 | 915 | 953 | 791 | 908 | 927 | 898 | 987 | 949 | 892 | 896 | 924.81 | 937 | 912 | 0.974 | -0.038 | 0.63467582 | 0.77160293 | 6.00E-04 | 0.005 | 0.0035 | 0.0035 | TRUE | 0.025 |
| Cybb | 614 | 579 | 566 | 357 | 344 | 440 | 477 | 616 | 611 | 355 | 388 | 434 | 480.2 | 568 | 392 | 0.693 | -0.528 | 3.02E-05 | 0.0004674 | 0.0137 | 0.0074 | 0.0094 | 0.0094 | TRUE | 0.2412 |
| Cybrd1 | 146 | 80 | 74 | 118 | 95 | 119 | 113 | 85 | 80 | 117 | 107 | 117 | 103.41 | 93 | 114 | 1.221 | 0.288 | 0.16407638 | 0.3149958 | 0.0105 | 0.0253 | 0.0211 | 0.0211 | TRUE | 0.4033 |
| Cyc1 | 16322 | 11668 | 11561 | 13059 | 11479 | 13317 | 12677 | 12411 | 12474 | 12999 | 12950 | 13146 | 12776.05 | 12521 | 13032 | 1.041 | 0.058 | 0.19455562 | 0.35374424 | 0 | 0.0026 | 0.0013 | 0.0013 | TRUE | 0.0015 |
| Cycs | 4283 | 2729 | 2413 | 2801 | 2187 | 2602 | 3326 | 2903 | 2604 | 2788 | 2467 | 2569 | 2776.11 | 2944 | 2608 | 0.885 | -0.176 | 0.04780758 | 0.13045318 | 0.009 | 0.0033 | 0.0053 | 0.0053 | TRUE | 0.1585 |
| Cyct | 826 | 646 | 612 | 587 | 509 | 500 | 642 | 687 | 660 | 584 | 574 | 494 | 606.85 | 663 | 551 | 0.831 | -0.268 | 0.00633214 | 0.02934103 | 0.0029 | 0.0063 | 0.0052 | 0.0052 | TRUE | 0.098 |
| Cyfip1 | 7140 | 4851 | 4911 | 5242 | 4517 | 5251 | 5545 | 5160 | 5299 | 5218 | 5096 | 5184 | 5250.21 | 5335 | 5166 | 0.968 | -0.047 | 0.38040141 | 0.55491707 | 6.00E-04 | 0.0029 | 0.0019 | 0.0019 | TRUE | 0.0148 |
| Cyfip2 | 6105 | 4767 | 5510 | 6640 | 6301 | 7118 | 4741 | 5071 | 5945 | 6609 | 7108 | 7027 | 6083.59 | 5252 | 6915 | 1.317 | 0.397 | 7.13E-07 | 2.20E-05 | 0.0075 | 0.0028 | 0.0045 | 0.0045 | TRUE | 0.162 |
| Cygb | 25 | 27 | 18 | 21 | 17 | 27 | 19 | 29 | 19 | 21 | 19 | 27 | 22.38 | 22 | 22 | 0.995 | -0.007 | 0.98575846 | 0.99284375 | 0 | 0.1083 | 0.0783 | 0.0783 | TRUE | 0.3168 |
| Cyhr1 | 2964 | 2420 | 2697 | 2679 | 2485 | 2856 | 2302 | 2574 | 2910 | 2667 | 2803 | 2819 | 2679.25 | 2595 | 2763 | 1.066 | 0.092 | 0.27151799 | 0.44248176 | 0.0064 | 0.0033 | 0.0046 | 0.0046 | TRUE | 0.1364 |
| Cylc1 | 0 | 0 | 0 | 0 | 0 | 0 | 0 | 0 | 0 | 0 | 0 | 0 | 0 | 0 | 0 | NA | NA | NA | NA | NA | NA | NA | NA | NA | NA |
| Cylc2 | 0 | 0 | 0 | 0 | 1 | 0 | 0 | 0 | 0 | 0 | 1 | 0 | 0.19 | 0 | 0 | 2.094 | 1.067 | 0.79379329 | NA | 0 | 12.608 | 10 | 10 | TRUE | 0.5075 |

| | | | | | | | | | | | | | | | | | | | | | | | | | |
|---|---|---|---|---|---|---|---|---|---|---|---|---|---|---|---|---|---|---|---|---|---|---|---|---|---|
| Cyld | 2196 | 1732 | 1629 | 1992 | 1809 | 1945 | 1706 | 1842 | 1758 | 1983 | 2041 | 1920 | 1874.85 | 1769 | 1981 | 1.121 | 0.164 | 0.01184092 | 0.04686853 | 7.00E-04 | 0.0037 | 0.0025 | 0.0025 | TRUE | 0.016 |
| Cym | 0 | 0 | 0 | 0 | 1 | 2 | 0 | 0 | 0 | 0 | 1 | 2 | 0.52 | 0 | 1 | 5.84 | 2.546 | 0.38239357 | NA | 0 | 4.5861 | 4.6218 | 4.6218 | TRUE | 0.0847 |
| Cyp11a1 | 393 | 254 | 200 | 415 | 340 | 341 | 305 | 270 | 216 | 413 | 384 | 337 | 320.74 | 264 | 378 | 1.426 | 0.512 | 0.00039446 | 0.00360716 | 0.0168 | 0.0098 | 0.0119 | 0.0119 | TRUE | 0.2762 |
| Cyp11b1 | 0 | 0 | 0 | 0 | 0 | 0 | 0 | 0 | 0 | 0 | 0 | 0 | 0 | 0 | 0 | NA | NA | NA | NA | NA | NA | NA | NA | NA | NA |
| Cyp11b2 | 0 | 0 | 0 | 0 | 0 | 0 | 0 | 0 | 0 | 0 | 0 | 0 | 0 | 0 | 0 | NA | NA | NA | NA | NA | NA | NA | NA | NA | NA |
| Cyp17a1 | 0 | 1 | 1 | 0 | 0 | 1 | 0 | 1 | 1 | 0 | 0 | 1 | 0.52 | 1 | 0 | 0.535 | -0.904 | 0.7403423 | NA | 0 | 4.5459 | 3.6616 | 3.6616 | TRUE | 0.3835 |
| Cyp19a1 | 3 | 1 | 3 | 1 | 3 | 5 | 2 | 1 | 3 | 1 | 3 | 5 | 2.66 | 2 | 3 | 1.402 | 0.487 | 0.68207671 | NA | 0 | 0.8942 | 0.6357 | 0.6357 | TRUE | 0.2778 |
| Cyp1a1 | 11 | 7 | 4 | 8 | 8 | 7 | 9 | 7 | 4 | 8 | 9 | 7 | 7.37 | 7 | 8 | 1.161 | 0.216 | 0.75770325 | NA | 0 | 0.3241 | 0.2161 | 0.2161 | TRUE | 0.2368 |
| Cyp1a2 | 6 | 6 | 2 | 1 | 2 | 5 | 5 | 6 | 2 | 1 | 2 | 5 | 3.56 | 4 | 3 | 0.621 | -0.687 | 0.51476198 | NA | 0 | 0.6673 | 0.5029 | 0.5029 | TRUE | 0.451 |
| Cyp1b1 | 85 | 37 | 76 | 18 | 15 | 17 | 66 | 39 | 82 | 18 | 17 | 17 | 39.83 | 62 | 17 | 0.275 | -1.861 | 5.39E-07 | 1.74E-05 | 0.0606 | 0.0619 | 0.0615 | 0.0615 | TRUE | 0.1544 |
| Cyp20a1 | 844 | 620 | 684 | 730 | 595 | 690 | 655 | 659 | 738 | 727 | 671 | 681 | 688.67 | 684 | 693 | 1.014 | 0.02 | 0.82380207 | 0.89868745 | 0.0017 | 0.0059 | 0.0045 | 0.0045 | TRUE | 0.0543 |
| Cyp21a1 | 2 | 0 | 1 | 1 | 0 | 0 | 2 | 0 | 1 | 1 | 0 | 0 | 0.6 | 1 | 0 | 0.414 | -1.271 | 0.61961437 | NA | 0 | 3.9224 | 3.1813 | 3.1813 | TRUE | 0.3906 |
| Cyp24a1 | 57 | 33 | 33 | 34 | 28 | 36 | 44 | 35 | 36 | 34 | 32 | 36 | 35.99 | 38 | 34 | 0.874 | -0.194 | 0.54375939 | 0.70124992 | 0 | 0.0683 | 0.0458 | 0.0458 | TRUE | 0.1593 |
| Cyp26a1 | 40 | 31 | 14 | 38 | 26 | 24 | 31 | 33 | 15 | 38 | 29 | 24 | 28.33 | 26 | 30 | 1.141 | 0.19 | 0.63535726 | 0.77174822 | 0.0651 | 0.0861 | 0.0806 | 0.0806 | TRUE | 0.7538 |
| Cyp26b1 | 19 | 10 | 15 | 15 | 19 | 13 | 15 | 11 | 16 | 15 | 21 | 13 | 15.13 | 14 | 16 | 1.175 | 0.233 | 0.64172865 | NA | 0 | 0.1591 | 0.1136 | 0.1136 | TRUE | 0.317 |
| Cyp26c1 | 25 | 18 | 14 | 7 | 17 | 8 | 19 | 19 | 15 | 7 | 19 | 8 | 14.62 | 18 | 11 | 0.625 | -0.678 | 0.21292546 | NA | 0.0648 | 0.1646 | 0.1402 | 0.1402 | TRUE | 1.2556 |
| Cyp27a1 | 114 | 96 | 86 | 107 | 99 | 101 | 89 | 102 | 93 | 107 | 112 | 100 | 100.22 | 95 | 106 | 1.123 | 0.168 | 0.38995502 | 0.56381035 | 0 | 0.0261 | 0.0174 | 0.0174 | TRUE | 0.0472 |
| Cyp27b1 | 34 | 26 | 14 | 28 | 18 | 19 | 26 | 28 | 15 | 28 | 20 | 19 | 22.68 | 23 | 22 | 0.964 | -0.054 | 0.89936082 | 0.94442467 | 0.0268 | 0.1069 | 0.0851 | 0.0851 | TRUE | 0.4883 |
| Cyp2a12 | 0 | 0 | 0 | 0 | 0 | 1 | 0 | 0 | 0 | 0 | 0 | 1 | 0.16 | 0 | 0 | 2.094 | 1.067 | 0.79379329 | NA | 0 | 14.4078 | 10 | 10 | TRUE | 0.3835 |
| Cyp2a22 | 0 | 0 | 0 | 0 | 0 | 0 | 0 | 0 | 0 | 0 | 0 | 0 | 0 | 0 | 0 | NA | NA | NA | NA | NA | NA | NA | NA | NA | NA |
| Cyp2a4 | 4 | 0 | 0 | 0 | 0 | 0 | 3 | 0 | 0 | 0 | 0 | 0 | 0.52 | 1 | 0 | 0.179 | -2.48 | 0.45030014 | NA | 10 | 4.5798 | 6.3235 | 6.3235 | TRUE | 1.8962 |
| Cyp2a5 | 0 | 0 | 0 | 0 | 0 | 0 | 0 | 0 | 0 | 0 | 0 | 0 | 0 | 0 | 0 | NA | NA | NA | NA | NA | NA | NA | NA | NA | NA |
| Cyp2ab1 | 17 | 9 | 9 | 16 | 7 | 19 | 13 | 10 | 10 | 16 | 8 | 19 | 12.51 | 11 | 14 | 1.308 | 0.387 | 0.49100113 | NA | 0 | 0.1919 | 0.1472 | 0.1472 | TRUE | 0.5951 |
| Cyp2b10 | 1 | 0 | 0 | 0 | 0 | 0 | 1 | 0 | 0 | 0 | 0 | 0 | 0.13 | 0 | 0 | 0.552 | -0.857 | 0.8336548 | NA | 0 | 18.3121 | 10 | 10 | TRUE | 0.2566 |
| Cyp2b13 | 0 | 0 | 0 | 0 | 0 | 0 | 0 | 0 | 0 | 0 | 0 | 0 | 0 | 0 | 0 | NA | NA | NA | NA | NA | NA | NA | NA | NA | NA |
| Cyp2b19 | 2 | 3 | 0 | 0 | 0 | 0 | 2 | 3 | 0 | 0 | 0 | 0 | 0.79 | 2 | 0 | 0.12 | -3.055 | 0.22090216 | NA | 5.3026 | 2.9997 | 3.1827 | 3.1827 | TRUE | 0.0516 |
| Cyp2b23 | 549 | 411 | 360 | 193 | 198 | 226 | 426 | 437 | 388 | 192 | 223 | 223 | 315.1 | 417 | 213 | 0.509 | -0.973 | 5.17E-15 | 8.70E-13 | 0.0014 | 0.0099 | 0.0076 | 0.0076 | TRUE | 0.0811 |
| Cyp2b9 | 0 | 0 | 0 | 0 | 0 | 0 | 0 | 0 | 0 | 0 | 0 | 0 | 0 | 0 | 0 | NA | NA | NA | NA | NA | NA | NA | NA | NA | NA |
| Cyp2c29 | 8 | 0 | 4 | 6 | 2 | 4 | 6 | 0 | 4 | 6 | 2 | 4 | 3.78 | 3 | 4 | 1.135 | 0.183 | 0.86583876 | NA | 0.4163 | 0.6287 | 0.5845 | 0.5845 | TRUE | 0.6478 |
| Cyp2c37 | 0 | 0 | 0 | 1 | 0 | 0 | 0 | 0 | 0 | 1 | 0 | 0 | 0.17 | 0 | 0 | 2.094 | 1.067 | 0.79379329 | NA | 0 | 14.2885 | 10 | 10 | TRUE | 0.3906 |
| Cyp2c38 | 0 | 0 | 0 | 0 | 0 | 0 | 0 | 0 | 0 | 0 | 0 | 0 | 0 | 0 | 0 | NA | NA | NA | NA | NA | NA | NA | NA | NA | NA |
| Cyp2c39 | 0 | 3 | 0 | 0 | 0 | 0 | 0 | 3 | 0 | 0 | 0 | 0 | 0.53 | 1 | 0 | 0.181 | -2.465 | 0.45184603 | NA | 10 | 4.4587 | 6.272 | 6.272 | TRUE | 1.443 |
| Cyp2c40 | 1 | 1 | 0 | 0 | 0 | 0 | 1 | 1 | 0 | 0 | 0 | 0 | 0.31 | 1 | 0 | 0.309 | -1.696 | 0.63183309 | NA | 0 | 7.7294 | 7.2472 | 7.2472 | TRUE | 0.2015 |
| Cyp2c44 | 61 | 40 | 60 | 57 | 55 | 53 | 47 | 43 | 65 | 57 | 62 | 52 | 54.29 | 52 | 57 | 1.108 | 0.149 | 0.58796189 | 0.73712285 | 0.0075 | 0.0461 | 0.0357 | 0.0357 | TRUE | 0.3842 |
| Cyp2c50 | 0 | 0 | 0 | 0 | 0 | 0 | 0 | 0 | 0 | 0 | 0 | 0 | 0 | 0 | 0 | NA | NA | NA | NA | NA | NA | NA | NA | NA | NA |
| Cyp2c53-ps | 1 | 2 | 3 | 0 | 0 | 1 | 1 | 2 | 3 | 0 | 0 | 1 | 1.19 | 2 | 0 | 0.184 | -2.442 | 0.2099393 | NA | 0 | 1.9974 | 1.4928 | 1.4928 | TRUE | 0.2878 |
| Cyp2c54 | 0 | 0 | 0 | 0 | 0 | 0 | 0 | 0 | 0 | 0 | 0 | 0 | 0 | 0 | 0 | NA | NA | NA | NA | NA | NA | NA | NA | NA | NA |
| Cyp2c55 | 249 | 196 | 181 | 31 | 36 | 50 | 193 | 208 | 195 | 31 | 41 | 49 | 119.67 | 199 | 40 | 0.202 | -2.304 | 4.52E-27 | 2.94E-24 | 0 | 0.0222 | 0.0177 | 0.0177 | TRUE | 0.3321 |
| Cyp2c65 | 1 | 1 | 0 | 0 | 1 | 1 | 1 | 1 | 0 | 0 | 1 | 1 | 0.66 | 1 | 1 | 1.125 | 0.17 | 0.94233031 | NA | 0 | 3.5974 | 2.4494 | 2.4494 | TRUE | 0.2637 |
| Cyp2c66 | 1 | 0 | 0 | 0 | 0 | 0 | 1 | 0 | 0 | 0 | 0 | 0 | 0.13 | 0 | 0 | 0.552 | -0.857 | 0.8336548 | NA | 0 | 18.3121 | 10 | 10 | TRUE | 0.2566 |
| Cyp2c67 | 0 | 0 | 0 | 0 | 1 | 0 | 0 | 0 | 0 | 0 | 1 | 0 | 0.19 | 0 | 0 | 2.094 | 1.067 | 0.79379329 | NA | 0 | 12.608 | 10 | 10 | TRUE | 0.5075 |
| Cyp2c68 | 0 | 0 | 0 | 0 | 0 | 0 | 0 | 0 | 0 | 0 | 0 | 0 | 0 | 0 | 0 | NA | NA | NA | NA | NA | NA | NA | NA | NA | NA |
| Cyp2c69 | 30 | 10 | 16 | 10 | 13 | 11 | 23 | 11 | 17 | 10 | 15 | 11 | 14.45 | 17 | 12 | 0.682 | -0.551 | 0.29749704 | NA | 0 | 0.1665 | 0.1299 | 0.1299 | TRUE | 0.1569 |
| Cyp2c70 | 1 | 2 | 2 | 6 | 2 | 4 | 1 | 2 | 2 | 6 | 2 | 4 | 2.87 | 2 | 4 | 2.493 | 1.318 | 0.25909016 | NA | 0 | 0.8273 | 0.5623 | 0.5623 | TRUE | 0.1543 |
| Cyp2d10 | 0 | 0 | 0 | 0 | 0 | 0 | 0 | 0 | 0 | 0 | 0 | 0 | 0 | 0 | 0 | NA | NA | NA | NA | NA | NA | NA | NA | NA | NA |
| Cyp2d11 | 0 | 0 | 0 | 0 | 0 | 0 | 0 | 0 | 0 | 0 | 0 | 0 | 0 | 0 | 0 | NA | NA | NA | NA | NA | NA | NA | NA | NA | NA |
| Cyp2d12 | 2 | 2 | 0 | 0 | 0 | 0 | 2 | 2 | 0 | 0 | 0 | 0 | 0.61 | 1 | 0 | 0.154 | -2.696 | 0.32099274 | NA | 0 | 3.8659 | 3.9247 | 3.9247 | TRUE | 0.3505 |
| Cyp2d13 | 0 | 0 | 0 | 0 | 0 | 0 | 0 | 0 | 0 | 0 | 0 | 0 | 0 | 0 | 0 | NA | NA | NA | NA | NA | NA | NA | NA | NA | NA |
| Cyp2d22 | 138 | 140 | 153 | 182 | 173 | 203 | 107 | 149 | 165 | 181 | 195 | 200 | 166.32 | 140 | 192 | 1.379 | 0.464 | 0.01033336 | 0.04229436 | 0.0191 | 0.0167 | 0.0174 | 0.0174 | TRUE | 0.4947 |
| Cyp2d26 | 0 | 2 | 2 | 0 | 3 | 5 | 0 | 2 | 2 | 0 | 3 | 5 | 2.1 | 1 | 3 | 2.019 | 1.014 | 0.50720333 | NA | 0 | 1.1305 | 1.145 | 1.145 | TRUE | 0.3839 |
| Cyp2d34 | 0 | 0 | 0 | 0 | 0 | 0 | 0 | 0 | 0 | 0 | 0 | 0 | 0 | 0 | 0 | NA | NA | NA | NA | NA | NA | NA | NA | NA | NA |
| Cyp2d37-ps | 0 | 0 | 0 | 0 | 0 | 0 | 0 | 0 | 0 | 0 | 0 | 0 | 0 | 0 | 0 | NA | NA | NA | NA | NA | NA | NA | NA | NA | NA |
| Cyp2d40 | 0 | 0 | 0 | 0 | 0 | 0 | 0 | 0 | 0 | 0 | 0 | 0 | 0 | 0 | 0 | NA | NA | NA | NA | NA | NA | NA | NA | NA | NA |
| Cyp2d9 | 0 | 1 | 0 | 0 | 0 | 0 | 0 | 1 | 0 | 0 | 0 | 0 | 0.18 | 0 | 0 | 0.552 | -0.857 | 0.8336548 | NA | 0 | 13.3714 | 10 | 10 | TRUE | 0.5216 |
| Cyp2e1 | 0 | 0 | 0 | 1 | 1 | 0 | 0 | 0 | 0 | 1 | 1 | 0 | 0.35 | 0 | 1 | 3.978 | 1.992 | 0.55384016 | NA | 0 | 6.6991 | 6.4234 | 6.4234 | TRUE | 0.2647 |
| Cyp2f2 | 3 | 1 | 1 | 4 | 2 | 3 | 2 | 1 | 1 | 4 | 2 | 3 | 2.28 | 1 | 3 | 2.006 | 1.004 | 0.42441461 | NA | 0 | 1.0425 | 0.6567 | 0.6567 | TRUE | 0.2267 |
| Cyp2g1 | 0 | 0 | 0 | 0 | 0 | 0 | 0 | 0 | 0 | 0 | 0 | 0 | 0 | 0 | 0 | NA | NA | NA | NA | NA | NA | NA | NA | NA | NA |
| Cyp2j11 | 0 | 0 | 0 | 0 | 0 | 0 | 0 | 0 | 0 | 0 | 0 | 0 | 0 | 0 | 0 | NA | NA | NA | NA | NA | NA | NA | NA | NA | NA |
| Cyp2j12 | 1 | 0 | 0 | 0 | 0 | 0 | 1 | 0 | 0 | 0 | 0 | 0 | 0.13 | 0 | 0 | 0.552 | -0.857 | 0.8336548 | NA | 0 | 18.3121 | 10 | 10 | TRUE | 0.2566 |
| Cyp2j13 | 60 | 45 | 52 | 44 | 36 | 53 | 47 | 48 | 56 | 44 | 41 | 52 | 47.88 | 50 | 46 | 0.912 | -0.132 | 0.63772027 | 0.77376365 | 0 | 0.0519 | 0.036 | 0.036 | TRUE | 0.1341 |
| Cyp2j5 | 0 | 0 | 0 | 0 | 0 | 0 | 0 | 0 | 0 | 0 | 0 | 0 | 0 | 0 | 0 | NA | NA | NA | NA | NA | NA | NA | NA | NA | NA |
| Cyp2j6 | 103 | 64 | 53 | 60 | 44 | 49 | 80 | 68 | 57 | 60 | 50 | 48 | 60.5 | 68 | 53 | 0.765 | -0.387 | 0.13652032 | 0.2764643 | 0 | 0.0416 | 0.0318 | 0.0318 | TRUE | 0.1854 |
| Cyp2j8 | 0 | 0 | 0 | 0 | 0 | 0 | 0 | 0 | 0 | 0 | 0 | 0 | 0 | 0 | 0 | NA | NA | NA | NA | NA | NA | NA | NA | NA | NA |
| Cyp2j9 | 135 | 94 | 91 | 115 | 111 | 94 | 105 | 100 | 98 | 114 | 125 | 93 | 105.92 | 101 | 111 | 1.094 | 0.129 | 0.5134665 | 0.67602954 | 0 | 0.0248 | 0.0186 | 0.0186 | TRUE | 0.2053 |
| Cyp2r1 | 33 | 14 | 9 | 11 | 15 | 5 | 26 | 15 | 10 | 11 | 17 | 5 | 13.84 | 17 | 11 | 0.639 | -0.646 | 0.27953253 | NA | 0.1997 | 0.1737 | 0.181 | 0.181 | TRUE | 0.3053 |
| Cyp2s1 | 2101 | 1897 | 2584 | 3187 | 3019 | 3263 | 1632 | 2018 | 2788 | 3172 | 3406 | 3221 | 2706.14 | 2146 | 3266 | 1.523 | 0.607 | 0.00838501 | 0.03631726 | 0.0378 | 0.0033 | 0.0127 | 0.0378 | TRUE | 0.8064 |
| Cyp2t4 | 6 | 4 | 7 | 10 | 4 | 4 | 4 | 5 | 4 | 8 | 10 | 5 | 4 | 5.81 | 6 | 6 | 1.13 | 0.176 | 0.82787279 | NA | 0 | 0.4101 | 0.2989 | 0.2989 | TRUE | 0.7254 |
| Cyp2u1 | 98 | 42 | 37 | 64 | 66 | 63 | 76 | 45 | 40 | 64 | 74 | 62 | 60.18 | 54 | 67 | 1.229 | 0.298 | 0.30607074 | 0.47857664 | 0.051 | 0.0418 | 0.0444 | 0.0444 | TRUE | 1.2631 |
| Cyp2w1 | 0 | 0 | 1 | 2 | 2 | 2 | 0 | 0 | 1 | 2 | 2 | 2 | 1.22 | 0 | 2 | 6.008 | 2.587 | 0.17406446 | NA | 0 | 1.9503 | 1.3605 | 1.3605 | TRUE | 0.5365 |

| | | | | | | | | | | | | | | | | | | | | | | | | | |
|---|---|---|---|---|---|---|---|---|---|---|---|---|---|---|---|---|---|---|---|---|---|---|---|---|---|
| Cyp39a1 | 187 | 135 | 150 | 174 | 161 | 214 | 145 | 144 | 162 | 173 | 182 | 211 | 169.46 | 150 | 189 | 1.258 | 0.332 | 0.03727401 | 0.10886143 | 0 | 0.0164 | 0.0123 | 0.0123 | TRUE | 0.1207 |
| Cyp3a11 | 0 | 0 | 0 | 0 | 0 | 0 | 0 | 0 | 0 | 0 | 0 | 0 | 0 | 0 | 0 | NA | NA | NA | NA | NA | NA | NA | NA | NA | NA |
| Cyp3a13 | 5 | 6 | 2 | 2 | 2 | 3 | 4 | 6 | 2 | 2 | 2 | 3 | 3.27 | 4 | 2 | 0.582 | -0.781 | 0.46242058 | NA | 0 | 0.7268 | 0.4826 | 0.4826 | TRUE | 0.2001 |
| Cyp3a16 | 0 | 0 | 0 | 0 | 0 | 0 | 0 | 0 | 0 | 0 | 0 | 0 | 0 | 0 | 0 | NA | NA | NA | NA | NA | NA | NA | NA | NA | NA |
| Cyp3a25 | 0 | 0 | 0 | 0 | 0 | 0 | 0 | 0 | 0 | 0 | 0 | 0 | 0 | 0 | 0 | NA | NA | NA | NA | NA | NA | NA | NA | NA | NA |
| Cyp3a41a | 0 | 0 | 0 | 0 | 0 | 0 | 0 | 0 | 0 | 0 | 0 | 0 | 0 | 0 | 0 | NA | NA | NA | NA | NA | NA | NA | NA | NA | NA |
| Cyp3a41b | 0 | 0 | 0 | 0 | 0 | 0 | 0 | 0 | 0 | 0 | 0 | 0 | 0 | 0 | 0 | NA | NA | NA | NA | NA | NA | NA | NA | NA | NA |
| Cyp3a44 | 0 | 0 | 0 | 0 | 0 | 0 | 0 | 0 | 0 | 0 | 0 | 0 | 0 | 0 | 0 | NA | NA | NA | NA | NA | NA | NA | NA | NA | NA |
| Cyp3a57 | 0 | 0 | 0 | 0 | 0 | 0 | 0 | 0 | 0 | 0 | 0 | 0 | 0 | 0 | 0 | NA | NA | NA | NA | NA | NA | NA | NA | NA | NA |
| Cyp3a59 | 0 | 0 | 0 | 1 | 0 | 0 | 0 | 0 | 0 | 1 | 0 | 0 | 0.17 | 0 | 0 | 2.094 | 1.067 | 0.79379329 | NA | 0 | 14.2885 | 10 | 10 | TRUE | 0.3906 |
| Cyp46a1 | 129 | 113 | 140 | 183 | 166 | 200 | 100 | 120 | 151 | 182 | 187 | 197 | 156.38 | 124 | 189 | 1.536 | 0.62 | 0.00056319 | 0.004724 | 0.014 | 0.0176 | 0.0166 | 0.0166 | TRUE | 0.3723 |
| Cyp4a10 | 0 | 1 | 0 | 0 | 0 | 0 | 0 | 1 | 0 | 0 | 0 | 0 | 0.18 | 0 | 0 | 0.552 | -0.857 | 0.8336548 | NA | 0 | 13.3714 | 10 | 10 | TRUE | 0.5216 |
| Cyp4a12a | 0 | 0 | 0 | 0 | 0 | 0 | 0 | 0 | 0 | 0 | 0 | 0 | 0 | 0 | 0 | NA | NA | NA | NA | NA | NA | NA | NA | NA | NA |
| Cyp4a12b | 12 | 7 | 3 | 4 | 3 | 2 | 9 | 7 | 3 | 4 | 3 | 2 | 4.89 | 6 | 3 | 0.46 | -1.12 | 0.21593338 | NA | 0 | 0.4871 | 0.352 | 0.352 | TRUE | 0.3734 |
| Cyp4a14 | 1 | 1 | 0 | 0 | 0 | 0 | 1 | 1 | 0 | 0 | 0 | 0 | 0.31 | 1 | 0 | 0.309 | -1.696 | 0.63183309 | NA | 0 | 7.7294 | 7.2472 | 7.2472 | TRUE | 0.2015 |
| Cyp4a29-ps | 0 | 0 | 0 | 0 | 0 | 0 | 0 | 0 | 0 | 0 | 0 | 0 | 0 | 0 | 0 | NA | NA | NA | NA | NA | NA | NA | NA | NA | NA |
| Cyp4a30b-ps | 0 | 0 | 1 | 0 | 1 | 0 | 0 | 0 | 1 | 0 | 1 | 0 | 0.37 | 0 | 0 | 1.075 | 0.105 | 0.97514588 | NA | 0 | 6.4454 | 6.1578 | 6.1578 | TRUE | 0.5365 |
| Cyp4a31 | 2 | 0 | 0 | 0 | 0 | 0 | 2 | 0 | 0 | 0 | 0 | 0 | 0.26 | 1 | 0 | 0.356 | -1.491 | 0.71362796 | NA | 0 | 9.1573 | 10 | 10 | TRUE | 0.968 |
| Cyp4a32 | 0 | 0 | 0 | 0 | 0 | 1 | 0 | 0 | 0 | 0 | 0 | 1 | 0.16 | 0 | 0 | 2.094 | 1.067 | 0.79379329 | NA | 0 | 14.4078 | 10 | 10 | TRUE | 0.3835 |
| Cyp4b1 | 0 | 2 | 0 | 0 | 1 | 0 | 0 | 2 | 0 | 0 | 1 | 0 | 0.54 | 1 | 0 | 0.535 | -0.903 | 0.76014553 | NA | 0 | 4.3706 | 4.6038 | 4.6038 | TRUE | 0.9746 |
| Cyp4f13 | 384 | 274 | 361 | 478 | 442 | 460 | 298 | 291 | 390 | 476 | 499 | 454 | 401.29 | 326 | 476 | 1.462 | 0.548 | 1.68E-05 | 0.00029134 | 0.0113 | 0.0083 | 0.0091 | 0.0091 | TRUE | 0.3155 |
| Cyp4f14 | 9 | 2 | 5 | 10 | 6 | 4 | 7 | 2 | 5 | 10 | 7 | 4 | 5.86 | 5 | 7 | 1.4 | 0.485 | 0.5547741 | NA | 0 | 0.4066 | 0.3131 | 0.3131 | TRUE | 0.2165 |
| Cyp4f15 | 4 | 2 | 0 | 2 | 1 | 2 | 3 | 2 | 0 | 2 | 1 | 2 | 1.72 | 2 | 2 | 0.95 | -0.074 | 0.96016809 | NA | 0 | 1.3794 | 0.9994 | 0.9994 | TRUE | 0.5096 |
| Cyp4f16 | 882 | 664 | 778 | 813 | 692 | 799 | 685 | 706 | 839 | 809 | 781 | 789 | 768.23 | 743 | 793 | 1.068 | 0.095 | 0.32111262 | 0.49500245 | 0.005 | 0.0055 | 0.0053 | 0.0053 | TRUE | 0.1482 |
| Cyp4f17 | 222 | 185 | 210 | 162 | 147 | 198 | 172 | 197 | 227 | 161 | 166 | 195 | 186.39 | 199 | 174 | 0.881 | -0.183 | 0.26114836 | 0.43111498 | 0.0102 | 0.0151 | 0.0137 | 0.0137 | TRUE | 0.164 |
| Cyp4f18 | 2 | 2 | 4 | 3 | 2 | 6 | 2 | 2 | 4 | 3 | 2 | 6 | 3.19 | 3 | 4 | 1.433 | 0.519 | 0.63087731 | NA | 0 | 0.7445 | 0.5171 | 0.5171 | TRUE | 0.4316 |
| Cyp4f39 | 163 | 108 | 112 | 115 | 107 | 130 | 127 | 115 | 121 | 114 | 121 | 128 | 120.97 | 121 | 121 | 1.002 | 0.002 | 0.98941411 | 0.99442345 | 0 | 0.022 | 0.0145 | 0.0145 | TRUE | 0.0281 |
| Cyp4f40 | 1 | 3 | 4 | 3 | 1 | 1 | 1 | 3 | 4 | 3 | 1 | 1 | 2.23 | 3 | 2 | 0.635 | -0.656 | 0.61870122 | NA | 0 | 1.0648 | 0.769 | 0.769 | TRUE | 0.6045 |
| Cyp4f41-ps | 11 | 8 | 11 | 7 | 9 | 11 | 9 | 9 | 12 | 7 | 10 | 11 | 9.48 | 10 | 9 | 0.971 | -0.043 | 0.94477085 | NA | 0 | 0.2523 | 0.1678 | 0.1678 | TRUE | 0.1672 |
| Cyp4v3 | 159 | 94 | 120 | 98 | 87 | 91 | 123 | 100 | 129 | 98 | 98 | 90 | 106.41 | 117 | 95 | 0.807 | -0.309 | 0.11545195 | 0.24530321 | 0 | 0.0247 | 0.0182 | 0.0182 | TRUE | 0.1649 |
| Cyp4x1 | 2 | 2 | 0 | 0 | 0 | 0 | 2 | 2 | 0 | 0 | 0 | 0 | 0.61 | 1 | 0 | 0.154 | -2.696 | 0.32099274 | NA | 0 | 3.8659 | 3.9247 | 3.9247 | TRUE | 0.3505 |
| Cyp51 | 7908 | 5913 | 7014 | 7979 | 6386 | 7659 | 6142 | 6290 | 7568 | 7942 | 7204 | 7561 | 7117.73 | 6667 | 7569 | 1.136 | 0.184 | 0.02248151 | 0.07512935 | 0.0079 | 0.0028 | 0.0045 | 0.0045 | TRUE | 0.1704 |
| Cyp7a1 | 1 | 0 | 0 | 0 | 0 | 0 | 1 | 0 | 0 | 0 | 0 | 0 | 0.13 | 0 | 0 | 0.552 | -0.857 | 0.8336548 | NA | 0 | 18.3121 | 10 | 10 | TRUE | 0.2566 |
| Cyp7b1 | 105 | 90 | 39 | 56 | 35 | 54 | 82 | 96 | 42 | 56 | 39 | 53 | 61.32 | 73 | 49 | 0.677 | -0.563 | 0.07185188 | 0.17495383 | 0.085 | 0.0411 | 0.0534 | 0.0534 | TRUE | 0.6056 |
| Cyp8b1 | 0 | 0 | 0 | 0 | 0 | 0 | 0 | 0 | 0 | 0 | 0 | 0 | 0 | 0 | 0 | NA | NA | NA | NA | NA | NA | NA | NA | NA | NA |
| Cypt1 | 0 | 0 | 0 | 0 | 0 | 0 | 0 | 0 | 0 | 0 | 0 | 0 | 0 | 0 | 0 | NA | NA | NA | NA | NA | NA | NA | NA | NA | NA |
| Cypt10 | 0 | 0 | 0 | 0 | 0 | 0 | 0 | 0 | 0 | 0 | 0 | 0 | 0 | 0 | 0 | NA | NA | NA | NA | NA | NA | NA | NA | NA | NA |
| Cypt12 | 0 | 0 | 0 | 0 | 0 | 0 | 0 | 0 | 0 | 0 | 0 | 0 | 0 | 0 | 0 | NA | NA | NA | NA | NA | NA | NA | NA | NA | NA |
| Cypt14 | 1 | 0 | 0 | 0 | 0 | 0 | 1 | 0 | 0 | 0 | 0 | 0 | 0.13 | 0 | 0 | 0.552 | -0.857 | 0.8336548 | NA | 0 | 18.3121 | 10 | 10 | TRUE | 0.2566 |
| Cypt15 | 0 | 0 | 0 | 0 | 0 | 0 | 0 | 0 | 0 | 0 | 0 | 0 | 0 | 0 | 0 | NA | NA | NA | NA | NA | NA | NA | NA | NA | NA |
| Cypt2 | 5 | 3 | 1 | 3 | 0 | 3 | 4 | 3 | 1 | 3 | 0 | 3 | 2.35 | 3 | 2 | 0.731 | -0.451 | 0.72945774 | NA | 0 | 1.0109 | 0.7951 | 0.7951 | TRUE | 0.5842 |
| Cypt3 | 48 | 36 | 36 | 25 | 28 | 24 | 37 | 38 | 39 | 25 | 32 | 24 | 32.43 | 38 | 27 | 0.699 | -0.517 | 0.12576117 | 0.26082056 | 0 | 0.0755 | 0.0501 | 0.0501 | TRUE | 0.1463 |
| Cypt4 | 0 | 0 | 0 | 0 | 0 | 0 | 0 | 0 | 0 | 0 | 0 | 0 | 0 | 0 | 0 | NA | NA | NA | NA | NA | NA | NA | NA | NA | NA |
| Cypt7 | 0 | 0 | 0 | 0 | 0 | 0 | 0 | 0 | 0 | 0 | 0 | 0 | 0 | 0 | 0 | NA | NA | NA | NA | NA | NA | NA | NA | NA | NA |
| Cypt8 | 0 | 0 | 0 | 0 | 0 | 0 | 0 | 0 | 0 | 0 | 0 | 0 | 0 | 0 | 0 | NA | NA | NA | NA | NA | NA | NA | NA | NA | NA |
| Cypt9 | 0 | 0 | 0 | 0 | 0 | 0 | 0 | 0 | 0 | 0 | 0 | 0 | 0 | 0 | 0 | NA | NA | NA | NA | NA | NA | NA | NA | NA | NA |
| Cyr61 | 1252 | 804 | 636 | 444 | 395 | 397 | 972 | 855 | 686 | 442 | 446 | 392 | 632.21 | 838 | 427 | 0.508 | -0.978 | 7.47E-15 | 1.23E-12 | 0.0168 | 0.0062 | 0.0096 | 0.0096 | TRUE | 0.1749 |
| Cys1 | 1 | 3 | 2 | 1 | 0 | 3 | 1 | 3 | 2 | 1 | 0 | 3 | 1.68 | 2 | 1 | 0.671 | -0.575 | 0.70851352 | NA | 0 | 1.4128 | 1.0772 | 1.0772 | TRUE | 0.6207 |
| Cysltr1 | 2 | 3 | 1 | 0 | 1 | 2 | 2 | 3 | 1 | 0 | 1 | 2 | 1.49 | 2 | 1 | 0.535 | -0.901 | 0.57596528 | NA | 0 | 1.5956 | 1.12 | 1.12 | TRUE | 0.3878 |
| Cysltr2 | 98 | 77 | 46 | 13 | 21 | 34 | 76 | 82 | 50 | 13 | 24 | 34 | 46.31 | 69 | 24 | 0.337 | -1.567 | 1.22E-05 | 0.00022062 | 0.0958 | 0.0536 | 0.0634 | 0.0634 | TRUE | 0.681 |
| Cystm1 | 725 | 478 | 485 | 662 | 568 | 662 | 563 | 508 | 523 | 659 | 641 | 653 | 591.34 | 531 | 651 | 1.223 | 0.29 | 0.00148404 | 0.00957626 | 0 | 0.0064 | 0.0043 | 0.0043 | TRUE | 0.0336 |
| Cyth1 | 533 | 385 | 421 | 479 | 436 | 426 | 414 | 410 | 454 | 477 | 492 | 421 | 444.49 | 426 | 463 | 1.087 | 0.121 | 0.26448208 | 0.43476048 | 0.0027 | 0.0078 | 0.0062 | 0.0062 | TRUE | 0.0758 |
| Cyth2 | 2424 | 1764 | 1717 | 1838 | 1632 | 1905 | 1883 | 1876 | 1853 | 1830 | 1841 | 1881 | 1860.45 | 1871 | 1851 | 0.989 | -0.016 | 0.79460772 | 0.87979922 | 0 | 0.0037 | 0.0021 | 0.0021 | TRUE | 0.0025 |
| Cyth3 | 1539 | 1102 | 1260 | 1248 | 1049 | 1220 | 1195 | 1172 | 1360 | 1242 | 1183 | 1204 | 1226.16 | 1242 | 1210 | 0.975 | -0.037 | 0.64590205 | 0.77978528 | 0.0028 | 0.0044 | 0.0038 | 0.0038 | TRUE | 0.08 |
| Cyth4 | 200 | 122 | 108 | 167 | 148 | 173 | 155 | 130 | 117 | 166 | 167 | 171 | 150.93 | 134 | 168 | 1.248 | 0.319 | 0.06028406 | 0.15380938 | 0.0042 | 0.0181 | 0.0142 | 0.0142 | TRUE | 0.2225 |
| Cytip | 0 | 1 | 0 | 1 | 0 | 0 | 0 | 1 | 0 | 1 | 0 | 0 | 0.34 | 0 | 0 | 1.075 | 0.105 | 0.97573409 | NA | 0 | 6.9086 | 6.5582 | 6.5582 | TRUE | 0.5216 |
| Cytl1 | 5 | 1 | 1 | 2 | 0 | 1 | 4 | 1 | 1 | 2 | 0 | 1 | 1.5 | 2 | 1 | 0.49 | -1.029 | 0.52752158 | NA | 0 | 1.5818 | 1.1652 | 1.1652 | TRUE | 0.5529 |
| Cyyr1 | 62 | 37 | 39 | 28 | 22 | 32 | 48 | 39 | 42 | 28 | 25 | 32 | 35.64 | 43 | 28 | 0.649 | -0.623 | 0.0552286 | 0.14470804 | 0 | 0.0689 | 0.0466 | 0.0466 | TRUE | 0.0904 |
| D030018L15I | 2 | 0 | 0 | 3 | 1 | 5 | 2 | 0 | 0 | 3 | 1 | 5 | 1.77 | 1 | 3 | 5.192 | 2.376 | 0.16250196 | NA | 0 | 1.3435 | 1.0946 | 1.0946 | TRUE | 0.4401 |
| D030024E09 | 0 | 0 | 0 | 0 | 0 | 0 | 0 | 0 | 0 | 0 | 0 | 0 | 0 | 0 | 0 | NA | NA | NA | NA | NA | NA | NA | NA | NA | NA |
| D030025E07 | 0 | 0 | 0 | 0 | 0 | 0 | 0 | 0 | 0 | 0 | 0 | 0 | 0 | 0 | 0 | NA | NA | NA | NA | NA | NA | NA | NA | NA | NA |
| D030025P21 | 32 | 36 | 18 | 28 | 24 | 16 | 25 | 38 | 19 | 28 | 27 | 16 | 25.55 | 27 | 24 | 0.857 | -0.223 | 0.59421219 | 0.74089299 | 0.064 | 0.0952 | 0.0866 | 0.0866 | TRUE | 0.6018 |
| D030028A08 | 130 | 78 | 72 | 96 | 73 | 77 | 101 | 83 | 78 | 96 | 82 | 76 | 85.92 | 87 | 85 | 0.966 | -0.05 | 0.81881027 | 0.89590869 | 0 | 0.03 | 0.0232 | 0.0232 | TRUE | 0.2016 |
| D030040B21 | 2 | 2 | 1 | 1 | 1 | 0 | 2 | 2 | 1 | 1 | 1 | 0 | 1.15 | 2 | 1 | 0.44 | -1.185 | 0.51270808 | NA | 0 | 2.0684 | 1.3164 | 1.3164 | TRUE | 0.2709 |
| D030045P18 | 2 | 1 | 2 | 0 | 0 | 0 | 2 | 1 | 2 | 0 | 0 | 0 | 0.8 | 2 | 0 | 0.12 | -3.062 | 0.17528122 | NA | 0 | 2.9805 | 2.3743 | 2.3743 | TRUE | 0.0718 |
| D030047H15 | 3 | 5 | 5 | 7 | 4 | 5 | 2 | 5 | 5 | 7 | 5 | 5 | 4.91 | 4 | 6 | 1.288 | 0.365 | 0.67214496 | NA | 0 | 0.4851 | 0.3279 | 0.3279 | TRUE | 0.4111 |
| D030056L22I | 3244 | 2369 | 2426 | 2594 | 2139 | 2458 | 2519 | 2520 | 2618 | 2582 | 2413 | 2426 | 2513.08 | 2552 | 2474 | 0.97 | -0.045 | 0.46161471 | 0.63095143 | 5.00E-04 | 0.0034 | 0.0022 | 0.0022 | TRUE | 0.0179 |
| D10Bwg1379 | 584 | 391 | 338 | 348 | 339 | 347 | 454 | 416 | 365 | 346 | 382 | 343 | 384.26 | 412 | 357 | 0.865 | -0.209 | 0.07970648 | 0.18867914 | 0.0055 | 0.0086 | 0.0076 | 0.0076 | TRUE | 0.1119 |
| D10Jhu81e | 2901 | 2164 | 2079 | 2683 | 2388 | 2730 | 2253 | 2302 | 2243 | 2671 | 2694 | 2695 | 2476.27 | 2266 | 2687 | 1.186 | 0.246 | 1.26E-05 | 0.00022872 | 0 | 0.0034 | 0.0019 | 0.0019 | TRUE | 0.0023 |

| | | | | | | | | | | | | | | | | | | | | | | | | | |
|---|---|---|---|---|---|---|---|---|---|---|---|---|---|---|---|---|---|---|---|---|---|---|---|---|---|
| D10Wsu102e | 4554 | 3314 | 3014 | 3246 | 2896 | 3292 | 3537 | 3525 | 3252 | 3231 | 3267 | 3250 | 3343.63 | 3438 | 3249 | 0.945 | -0.082 | 0.16273195 | 0.31312974 | 8.00E-04 | 0.0031 | 0.0022 | 0.0022 | TRUE | 0.0269 |
| D10Wsu52e | 12462 | 8800 | 8408 | 8472 | 7567 | 8684 | 9679 | 9360 | 9072 | 8433 | 8536 | 8572 | 8942.19 | 9370 | 8514 | 0.908 | -0.139 | 0.00527525 | 0.02553307 | 4.00E-04 | 0.0027 | 0.0017 | 0.0017 | TRUE | 0.0102 |
| D11Wsu47e | 3297 | 2517 | 2173 | 2365 | 2227 | 2426 | 2561 | 2677 | 2345 | 2354 | 2512 | 2395 | 2473.97 | 2528 | 2420 | 0.957 | -0.063 | 0.35933822 | 0.53411202 | 0.0022 | 0.0034 | 0.003 | 0.003 | TRUE | 0.0476 |
| D130009I18 | 0 | 0 | 0 | 0 | 0 | 0 | 0 | 0 | 0 | 0 | 0 | 0 | 0 | 0 | 0 | NA | NA | NA | NA | NA | NA | NA | NA | NA | NA |
| D130017N08 | 67 | 46 | 46 | 87 | 86 | 84 | 52 | 49 | 50 | 87 | 97 | 83 | 69.52 | 50 | 89 | 1.765 | 0.819 | 0.00046962 | 0.00412896 | 0 | 0.0365 | 0.0241 | 0.0241 | TRUE | 0.0586 |
| D130020L05 | 81 | 58 | 52 | 60 | 40 | 72 | 63 | 62 | 56 | 60 | 45 | 71 | 59.44 | 60 | 59 | 0.975 | -0.036 | 0.89151111 | 0.94018803 | 0.0075 | 0.0423 | 0.0331 | 0.0331 | TRUE | 0.2502 |
| D130040H23 | 169 | 125 | 91 | 120 | 95 | 95 | 131 | 133 | 98 | 119 | 107 | 94 | 113.8 | 121 | 107 | 0.882 | -0.182 | 0.36452826 | 0.53929671 | 0.0122 | 0.0233 | 0.0202 | 0.0202 | TRUE | 0.2582 |
| D130043K22 | 52 | 41 | 30 | 35 | 28 | 25 | 40 | 44 | 32 | 35 | 32 | 25 | 34.58 | 39 | 31 | 0.781 | -0.357 | 0.28590615 | 0.45786629 | 0 | 0.071 | 0.051 | 0.051 | TRUE | 0.187 |
| D130058E03 | 0 | 0 | 0 | 0 | 0 | 0 | 0 | 0 | 0 | 0 | 0 | 0 | 0 | 0 | 0 | NA | NA | NA | NA | NA | NA | NA | NA | NA | NA |
| D14Abb1e | 7200 | 5148 | 4442 | 4644 | 4114 | 4698 | 5592 | 5476 | 4793 | 4623 | 4641 | 4638 | 4960.33 | 5287 | 4634 | 0.876 | -0.191 | 0.00448903 | 0.02261602 | 0.003 | 0.0029 | 0.003 | 0.003 | TRUE | 0.081 |
| D15Ertd621e | 12406 | 8982 | 8511 | 8861 | 7714 | 8872 | 9635 | 9554 | 9183 | 8820 | 8702 | 8758 | 9108.83 | 9457 | 8760 | 0.926 | -0.111 | 0.02093969 | 0.07161785 | 2.00E-04 | 0.0027 | 0.0015 | 0.0015 | TRUE | 0.0078 |
| D16Ertd472e | 1488 | 1071 | 942 | 988 | 903 | 1051 | 1156 | 1139 | 1016 | 983 | 1019 | 1037 | 1058.49 | 1104 | 1013 | 0.917 | -0.125 | 0.11951027 | 0.25127395 | 0.0019 | 0.0047 | 0.0037 | 0.0037 | TRUE | 0.0562 |
| D16Ertd519e | 0 | 0 | 0 | 0 | 0 | 0 | 0 | 0 | 0 | 0 | 0 | 0 | 0 | 0 | 0 | NA | NA | NA | NA | NA | NA | NA | NA | NA | NA |
| D17Ertd648e | 3 | 0 | 3 | 2 | 0 | 0 | 2 | 0 | 3 | 2 | 0 | 0 | 1.26 | 2 | 1 | 0.362 | -1.465 | 0.46277884 | NA | 0 | 1.884 | 1.8435 | 1.8435 | TRUE | 0.8482 |
| D17H6S53E | 136 | 105 | 95 | 96 | 84 | 72 | 106 | 112 | 103 | 96 | 95 | 71 | 96.87 | 107 | 87 | 0.816 | -0.293 | 0.15623892 | 0.30432309 | 0 | 0.0269 | 0.0203 | 0.0203 | TRUE | 0.2533 |
| D17Wsu104e | 2263 | 1624 | 1706 | 2055 | 1877 | 2174 | 1758 | 1727 | 1841 | 2046 | 2117 | 2146 | 1939.14 | 1775 | 2103 | 1.185 | 0.245 | 0.00010605 | 0.00126949 | 3.00E-04 | 0.0036 | 0.0024 | 0.0024 | TRUE | 0.0122 |
| D17Wsu92e | 11257 | 8069 | 8442 | 8590 | 7739 | 8879 | 8743 | 8583 | 9109 | 8550 | 8730 | 8765 | 8746.73 | 8812 | 8682 | 0.985 | -0.021 | 0.66760943 | 0.79557625 | 4.00E-04 | 0.0027 | 0.0017 | 0.0017 | TRUE | 0.0105 |
| D19Bwg135 | 10675 | 7416 | 6854 | 7143 | 6312 | 7262 | 8291 | 7888 | 7395 | 7110 | 7121 | 7169 | 7495.66 | 7858 | 7133 | 0.907 | -0.14 | 0.01436423 | 0.05399997 | 0.0014 | 0.0027 | 0.0022 | 0.0022 | TRUE | 0.0323 |
| D19Ertd737e | 1414 | 980 | 943 | 934 | 759 | 871 | 1098 | 1042 | 1017 | 930 | 856 | 860 | 967.31 | 1052 | 882 | 0.837 | -0.256 | 0.00121314 | 0.00832028 | 9.00E-04 | 0.0049 | 0.0035 | 0.0035 | TRUE | 0.027 |
| D1Ertd622e | 888 | 659 | 655 | 773 | 695 | 758 | 690 | 701 | 707 | 769 | 784 | 748 | 733.19 | 699 | 767 | 1.098 | 0.134 | 0.09987995 | 0.22082633 | 0 | 0.0057 | 0.0034 | 0.0034 | TRUE | 0.0056 |
| D1Pas1 | 1765 | 1238 | 778 | 627 | 574 | 585 | 1371 | 1317 | 839 | 624 | 648 | 577 | 896.04 | 1176 | 616 | 0.524 | -0.933 | 1.58E-10 | 1.22E-08 | 0.0357 | 0.0051 | 0.0141 | 0.0141 | TRUE | 0.7443 |
| D230025D16 | 5108 | 3603 | 3551 | 3320 | 2901 | 3215 | 3967 | 3832 | 3831 | 3305 | 3273 | 3174 | 3563.7 | 3877 | 3251 | 0.838 | -0.255 | 2.65E-06 | 6.29E-05 | 0 | 0.0031 | 0.0018 | 0.0018 | TRUE | 0.0052 |
| D230030E09 | 0 | 0 | 1 | 0 | 0 | 0 | 0 | 0 | 1 | 0 | 0 | 0 | 0.18 | 0 | 0 | 0.552 | -0.857 | 0.8336548 | NA | 0 | 13.1817 | 10 | 10 | TRUE | 0.5365 |
| D2hgdh | 447 | 362 | 329 | 355 | 282 | 379 | 347 | 385 | 355 | 353 | 318 | 374 | 355.47 | 362 | 348 | 0.964 | -0.053 | 0.64880077 | 0.781832 | 0.0018 | 0.0091 | 0.0069 | 0.0069 | TRUE | 0.0642 |
| D2Wsu81e | 2602 | 2088 | 2090 | 2049 | 1941 | 2365 | 2021 | 2221 | 2255 | 2040 | 2190 | 2335 | 2176.8 | 2166 | 2188 | 1.011 | 0.016 | 0.83107823 | 0.9033735 | 0.0036 | 0.0035 | 0.0036 | 0.0036 | TRUE | 0.043 |
| D330022K07 | 1 | 1 | 0 | 0 | 0 | 0 | 1 | 1 | 0 | 0 | 0 | 0 | 0.31 | 1 | 0 | 0.309 | -1.696 | 0.63183309 | NA | 0 | 7.7294 | 7.2472 | 7.2472 | TRUE | 0.2015 |
| D330023K18 | 31 | 18 | 22 | 18 | 16 | 16 | 24 | 19 | 24 | 18 | 18 | 16 | 19.79 | 22 | 17 | 0.77 | -0.377 | 0.3770662 | 0.55151527 | 0 | 0.1222 | 0.0795 | 0.0795 | TRUE | 0.0835 |
| D330041H03 | 59 | 36 | 39 | 45 | 37 | 31 | 46 | 38 | 42 | 45 | 42 | 31 | 40.56 | 42 | 39 | 0.924 | -0.114 | 0.7122133 | 0.82701728 | 0 | 0.0609 | 0.044 | 0.044 | TRUE | 0.2756 |
| D330045A20 | 143 | 87 | 93 | 127 | 84 | 118 | 111 | 93 | 100 | 126 | 95 | 116 | 106.93 | 101 | 112 | 1.11 | 0.15 | 0.45087647 | 0.62090542 | 0.0049 | 0.0246 | 0.0192 | 0.0192 | TRUE | 0.1813 |
| D330050G23 | 17 | 5 | 13 | 9 | 10 | 8 | 13 | 5 | 14 | 9 | 11 | 8 | 10.11 | 11 | 9 | 0.856 | -0.225 | 0.71856751 | NA | 0 | 0.2368 | 0.1809 | 0.1809 | TRUE | 0.674 |
| D330050I16 | 74 | 45 | 51 | 38 | 37 | 36 | 57 | 48 | 55 | 38 | 42 | 36 | 45.91 | 53 | 39 | 0.715 | -0.484 | 0.08906896 | 0.20326724 | 0 | 0.054 | 0.036 | 0.036 | TRUE | 0.067 |
| D3Bwg0562e | 24 | 19 | 10 | 9 | 7 | 11 | 19 | 20 | 11 | 9 | 8 | 11 | 12.89 | 17 | 9 | 0.557 | -0.843 | 0.12642717 | NA | 0 | 0.1863 | 0.1343 | 0.1343 | TRUE | 0.4109 |
| D3Ertd254e | 1921 | 1479 | 1227 | 1287 | 1165 | 1247 | 1492 | 1573 | 1324 | 1281 | 1314 | 1231 | 1369.23 | 1463 | 1275 | 0.871 | -0.199 | 0.01395753 | 0.05292096 | 0.0036 | 0.0042 | 0.004 | 0.004 | TRUE | 0.0812 |
| D3Ertd751e | 481 | 339 | 294 | 352 | 336 | 345 | 374 | 361 | 317 | 350 | 379 | 341 | 353.56 | 351 | 357 | 1.015 | 0.021 | 0.85612078 | 0.91907385 | 0.0023 | 0.0091 | 0.0071 | 0.0071 | TRUE | 0.0764 |
| D430019H16 | 426 | 368 | 368 | 444 | 424 | 451 | 331 | 391 | 397 | 442 | 478 | 445 | 414.14 | 373 | 455 | 1.223 | 0.291 | 0.01008727 | 0.04160552 | 0.0037 | 0.0081 | 0.0068 | 0.0068 | TRUE | 0.1217 |
| D430020J02 | 431 | 335 | 272 | 208 | 215 | 190 | 335 | 356 | 293 | 207 | 243 | 188 | 270.28 | 328 | 213 | 0.645 | -0.632 | 7.75E-06 | 0.00015215 | 0.0086 | 0.0112 | 0.0105 | 0.0105 | TRUE | 0.163 |
| D430036J16 | 0 | 0 | 3 | 2 | 2 | 4 | 0 | 0 | 3 | 2 | 2 | 4 | 1.91 | 1 | 3 | 2.676 | 1.42 | 0.36641779 | NA | 0.455 | 1.2463 | 1.1216 | 1.1216 | TRUE | 1.4452 |
| D430041D05 | 93 | 79 | 51 | 72 | 54 | 64 | 72 | 84 | 55 | 72 | 61 | 63 | 67.84 | 70 | 65 | 0.926 | -0.11 | 0.65750382 | 0.78780301 | 0.0096 | 0.0374 | 0.0299 | 0.0299 | TRUE | 0.2704 |
| D430042O09 | 801 | 571 | 569 | 718 | 613 | 729 | 622 | 607 | 614 | 715 | 692 | 720 | 661.55 | 614 | 709 | 1.153 | 0.206 | 0.01511883 | 0.05614276 | 0 | 0.006 | 0.0036 | 0.0036 | TRUE | 0.0051 |
| D4Ertd617e | 0 | 0 | 0 | 0 | 0 | 0 | 0 | 0 | 0 | 0 | 0 | 0 | 0 | 0 | 0 | NA | NA | NA | NA | NA | NA | NA | NA | NA | NA |
| D4Wsu53e | 3445 | 3762 | 4090 | 3887 | 3222 | 4451 | 2676 | 4002 | 4413 | 3869 | 3635 | 4394 | 3831.32 | 3697 | 3966 | 1.073 | 0.102 | 0.65713927 | 0.78749824 | 0.0376 | 0.003 | 0.0127 | 0.0376 | TRUE | 0.7129 |
| D530049I02 | 0 | 0 | 0 | 0 | 0 | 0 | 0 | 0 | 0 | 0 | 0 | 0 | 0 | 0 | 0 | NA | NA | NA | NA | NA | NA | NA | NA | NA | NA |
| D5Ertd577e | 0 | 0 | 0 | 0 | 0 | 0 | 0 | 0 | 0 | 0 | 0 | 0 | 0 | 0 | 0 | NA | NA | NA | NA | NA | NA | NA | NA | NA | NA |
| D5Ertd579e | 2794 | 2076 | 2030 | 2227 | 1948 | 2203 | 2170 | 2208 | 2190 | 2217 | 2198 | 2175 | 2192.92 | 2189 | 2197 | 1.003 | 0.005 | 0.93499792 | 0.96412113 | 0 | 0.0035 | 0.0019 | 0.0019 | TRUE | 9.00E-04 |
| D5Ertd605e | 232 | 166 | 124 | 114 | 117 | 125 | 180 | 177 | 134 | 113 | 132 | 123 | 143.23 | 164 | 123 | 0.749 | -0.417 | 0.02079391 | 0.07122151 | 0.0098 | 0.019 | 0.0163 | 0.0163 | TRUE | 0.2574 |
| D630003M2 | 87 | 84 | 70 | 65 | 51 | 42 | 68 | 89 | 76 | 65 | 58 | 41 | 66.02 | 78 | 55 | 0.706 | -0.502 | 0.0530762 | 0.14074549 | 0.018 | 0.0383 | 0.0328 | 0.0328 | TRUE | 0.3827 |
| D630010B17 | 0 | 0 | 0 | 0 | 0 | 0 | 0 | 0 | 0 | 0 | 0 | 0 | 0 | 0 | 0 | NA | NA | NA | NA | NA | NA | NA | NA | NA | NA |
| D630013N20 | 0 | 0 | 0 | 0 | 0 | 0 | 0 | 0 | 0 | 0 | 0 | 0 | 0 | 0 | 0 | NA | NA | NA | NA | NA | NA | NA | NA | NA | NA |
| D630023F18 | 450 | 265 | 339 | 269 | 238 | 288 | 349 | 282 | 366 | 268 | 268 | 284 | 302.95 | 332 | 273 | 0.822 | -0.282 | 0.03239316 | 0.09796771 | 0.0068 | 0.0102 | 0.0092 | 0.0092 | TRUE | 0.1938 |
| D630029K05 | 1 | 4 | 4 | 5 | 1 | 4 | 1 | 4 | 4 | 5 | 1 | 4 | 3.23 | 3 | 3 | 1.117 | 0.16 | 0.88609177 | NA | 0 | 0.7354 | 0.585 | 0.585 | TRUE | 0.8105 |
| D630032N06 | 35 | 30 | 19 | 32 | 22 | 27 | 27 | 32 | 21 | 32 | 25 | 27 | 27.15 | 27 | 28 | 1.047 | 0.066 | 0.85886619 | 0.92132802 | 0 | 0.0897 | 0.0639 | 0.0639 | TRUE | 0.2234 |
| D630033O11 | 10 | 8 | 9 | 15 | 12 | 12 | 8 | 9 | 10 | 15 | 14 | 12 | 11.05 | 9 | 14 | 1.559 | 0.64 | 0.26046491 | NA | 0 | 0.2169 | 0.139 | 0.139 | TRUE | 0.0478 |
| D630037F22 | 1381 | 1058 | 1122 | 1266 | 1044 | 1167 | 1073 | 1125 | 1211 | 1260 | 1178 | 1152 | 1166.42 | 1136 | 1197 | 1.055 | 0.077 | 0.33258729 | 0.50692343 | 0.0022 | 0.0045 | 0.0036 | 0.0036 | TRUE | 0.0391 |
| D630039A03 | 126 | 78 | 81 | 65 | 82 | 92 | 98 | 83 | 87 | 65 | 93 | 91 | 86.04 | 89 | 83 | 0.92 | -0.12 | 0.59378079 | 0.74083813 | 0.01 | 0.03 | 0.0246 | 0.0246 | TRUE | 0.3456 |
| D630041G03 | 1 | 4 | 5 | 2 | 2 | 2 | 1 | 4 | 5 | 2 | 2 | 2 | 2.77 | 3 | 2 | 0.611 | -0.71 | 0.55051858 | NA | 0 | 0.8566 | 0.626 | 0.626 | TRUE | 0.9417 |
| D630045J12 | 2258 | 1761 | 1864 | 2072 | 1904 | 2090 | 1754 | 1873 | 2011 | 2062 | 2148 | 2063 | 1985.27 | 1879 | 2091 | 1.114 | 0.155 | 0.02727692 | 0.086388 | 0.0021 | 0.0036 | 0.0031 | 0.0031 | TRUE | 0.0455 |
| D630045M0 | 546 | 363 | 282 | 191 | 164 | 180 | 424 | 386 | 304 | 190 | 185 | 178 | 277.88 | 371 | 184 | 0.495 | -1.016 | 6.67E-12 | 7.17E-10 | 0.0132 | 0.011 | 0.0117 | 0.0117 | TRUE | 0.2817 |
| D6Ertd474e | 0 | 1 | 2 | 1 | 3 | 0 | 0 | 1 | 2 | 1 | 3 | 0 | 1.27 | 1 | 1 | 1.389 | 0.474 | 0.79905437 | NA | 0.0026 | 1.8732 | 1.6733 | 1.6733 | TRUE | 0.4582 |
| D6Ertd527e | 6 | 4 | 4 | 14 | 9 | 6 | 5 | 4 | 4 | 14 | 10 | 6 | 7.21 | 4 | 10 | 2.261 | 1.177 | 0.11194873 | NA | 0 | 0.3313 | 0.2343 | 0.2343 | TRUE | 0.1229 |
| D6Wsu163e | 1304 | 879 | 906 | 1004 | 842 | 1084 | 1013 | 935 | 978 | 999 | 950 | 1070 | 990.77 | 975 | 1006 | 1.032 | 0.045 | 0.57552453 | 0.72731849 | 0.0016 | 0.0048 | 0.0037 | 0.0037 | TRUE | 0.0367 |
| D730001G18 | 4 | 1 | 1 | 0 | 0 | 0 | 3 | 1 | 1 | 0 | 0 | 0 | 0.87 | 2 | 0 | 0.107 | -3.222 | 0.14630442 | NA | 0 | 2.7114 | 2.2736 | 2.2736 | TRUE | 0.4767 |
| D730005E14 | 54 | 52 | 65 | 60 | 59 | 59 | 42 | 55 | 70 | 60 | 67 | 58 | 58.65 | 56 | 62 | 1.111 | 0.152 | 0.57750948 | 0.72898905 | 0.0194 | 0.0428 | 0.0362 | 0.0362 | TRUE | 0.2436 |
| D730039F16 | 34 | 15 | 28 | 25 | 33 | 22 | 26 | 16 | 30 | 25 | 37 | 22 | 26.07 | 24 | 28 | 1.147 | 0.198 | 0.62785287 | 0.76543728 | 0.0474 | 0.0933 | 0.0811 | 0.0811 | TRUE | 0.5046 |
| D730045A05 | 7 | 3 | 2 | 0 | 6 | 3 | 5 | 3 | 2 | 0 | 7 | 3 | 3.42 | 3 | 3 | 0.875 | -0.193 | 0.86640743 | NA | 0.5118 | 0.6956 | 0.6545 | 0.6545 | TRUE | 0.2787 |
| D730048I06 | 0 | 1 | 0 | 0 | 0 | 0 | 0 | 1 | 0 | 0 | 0 | 0 | 0.18 | 0 | 0 | 0.552 | -0.857 | 0.8336548 | NA | 0 | 13.3714 | 10 | 10 | TRUE | 0.5216 |
| D730050B12 | 0 | 0 | 0 | 0 | 0 | 0 | 0 | 0 | 0 | 0 | 0 | 0 | 0 | 0 | 0 | NA | NA | NA | NA | NA | NA | NA | NA | NA | NA |
| D7Ertd143e | 1142 | 1053 | 529 | 448 | 386 | 414 | 887 | 1120 | 571 | 446 | 435 | 409 | 644.64 | 859 | 430 | 0.5 | -0.999 | 0.00039403 | 0.00360716 | 0.0555 | 0.0061 | 0.02 | 0.0555 | TRUE | 0.158 |
| D7Ertd443e | 0 | 2 | 1 | 1 | 3 | 2 | 0 | 2 | 1 | 1 | 3 | 2 | 1.59 | 1 | 2 | 2.066 | 1.047 | 0.51028291 | NA | 0 | 1.4899 | 1.101 | 1.101 | TRUE | 0.4472 |
| D7Ertd715e | 641 | 585 | 526 | 524 | 422 | 551 | 498 | 622 | 568 | 522 | 476 | 544 | 538.2 | 563 | 514 | 0.916 | -0.126 | 0.2518606 | 0.42083745 | 0.0069 | 0.0068 | 0.0069 | 0.0069 | TRUE | 0.1256 |

| | | | | | | | | | | | | | | | | | | | | | | | | | |
|---|---|---|---|---|---|---|---|---|---|---|---|---|---|---|---|---|---|---|---|---|---|---|---|---|---|
| D830005E20 | 0 | 0 | 0 | 0 | 0 | 0 | 0 | 0 | 0 | 0 | 0 | 0 | 0 | 0 | 0 | NA | NA | NA | NA | NA | NA | NA | NA | NA | NA |
| D830013O20 | 0 | 0 | 0 | 0 | 0 | 0 | 0 | 0 | 0 | 0 | 0 | 0 | 0 | 0 | 0 | NA | NA | NA | NA | NA | NA | NA | NA | NA | NA |
| D830015G02 | 1 | 0 | 0 | 0 | 0 | 0 | 1 | 0 | 0 | 0 | 0 | 0 | 0.13 | 0 | 0 | 0.552 | -0.857 | 0.8336548 | NA | 0 | 18.3121 | 10 | 10 | TRUE | 0.2566 |
| D830030K20 | 0 | 0 | 1 | 0 | 0 | 1 | 0 | 0 | 1 | 0 | 0 | 1 | 0.34 | 0 | 0 | 1.075 | 0.105 | 0.97570524 | NA | 0 | 6.885 | 6.5379 | 6.5379 | TRUE | 0.5365 |
| D830031N03 | 1401 | 999 | 983 | 1374 | 1148 | 1203 | 1088 | 1063 | 1061 | 1368 | 1295 | 1188 | 1176.94 | 1071 | 1284 | 1.198 | 0.261 | 0.00081328 | 0.00622615 | 0.0019 | 0.0044 | 0.0035 | 0.0035 | TRUE | 0.0521 |
| D830046C22 | 0 | 0 | 0 | 0 | 0 | 0 | 0 | 0 | 0 | 0 | 0 | 0 | 0 | 0 | 0 | NA | NA | NA | NA | NA | NA | NA | NA | NA | NA |
| D8Ertd738e | 4303 | 2894 | 2532 | 2382 | 2091 | 2585 | 3342 | 3078 | 2732 | 2371 | 2359 | 2552 | 2739 | 3051 | 2427 | 0.795 | -0.331 | 3.30E-05 | 0.00050347 | 0.0055 | 0.0033 | 0.0042 | 0.0042 | TRUE | 0.1011 |
| D8Ertd82e | 1085 | 949 | 956 | 778 | 766 | 874 | 843 | 1009 | 1032 | 774 | 864 | 863 | 897.49 | 961 | 834 | 0.869 | -0.203 | 0.03798235 | 0.11025219 | 0.0071 | 0.0051 | 0.0058 | 0.0058 | TRUE | 0.1446 |
| D930007P13 | 0 | 1 | 0 | 0 | 0 | 0 | 0 | 1 | 0 | 0 | 0 | 0 | 0.18 | 0 | 0 | 0.552 | -0.857 | 0.8336548 | NA | 0 | 13.3714 | 10 | 10 | TRUE | 0.5216 |
| D930015E06 | 2237 | 1835 | 1912 | 2126 | 2003 | 2155 | 1737 | 1952 | 2063 | 2116 | 2260 | 2127 | 2042.57 | 1917 | 2168 | 1.132 | 0.178 | 0.02001048 | 0.06920149 | 0.0037 | 0.0036 | 0.0037 | 0.0037 | TRUE | 0.084 |
| D930015M05 | 0 | 0 | 0 | 0 | 1 | 0 | 0 | 0 | 0 | 0 | 1 | 0 | 0.19 | 0 | 0 | 2.094 | 1.067 | 0.79379329 | NA | 0 | 12.608 | 10 | 10 | TRUE | 0.5075 |
| D930016D06 | 606 | 448 | 347 | 395 | 346 | 465 | 471 | 477 | 374 | 393 | 390 | 459 | 427.35 | 441 | 414 | 0.939 | -0.09 | 0.46804733 | 0.63700253 | 0.0107 | 0.008 | 0.0088 | 0.0088 | TRUE | 0.1937 |
| D930020B18 | 15 | 9 | 6 | 13 | 11 | 14 | 12 | 10 | 6 | 13 | 12 | 14 | 11.14 | 9 | 13 | 1.399 | 0.484 | 0.39680435 | NA | 0 | 0.2151 | 0.1439 | 0.1439 | TRUE | 0.2214 |
| D930028M14 | 95 | 53 | 82 | 83 | 81 | 90 | 74 | 56 | 88 | 83 | 91 | 89 | 80.25 | 73 | 88 | 1.201 | 0.265 | 0.25335969 | 0.42242648 | 0.0092 | 0.032 | 0.0262 | 0.0262 | TRUE | 0.2553 |
| D930032P07 | 0 | 0 | 0 | 0 | 0 | 0 | 0 | 0 | 0 | 0 | 0 | 0 | 0 | 0 | 0 | NA | NA | NA | NA | NA | NA | NA | NA | NA | NA |
| D930048N14 | 430 | 508 | 467 | 451 | 433 | 497 | 334 | 540 | 504 | 449 | 488 | 491 | 467.7 | 459 | 476 | 1.04 | 0.056 | 0.71320861 | 0.82762773 | 0.0312 | 0.0075 | 0.0146 | 0.0146 | TRUE | 0.6929 |
| Daam1 | 2410 | 1634 | 1596 | 1689 | 1472 | 1758 | 1872 | 1738 | 1722 | 1681 | 1661 | 1735 | 1734.85 | 1777 | 1692 | 0.952 | -0.072 | 0.28429123 | 0.45635383 | 9.00E-04 | 0.0038 | 0.0027 | 0.0027 | TRUE | 0.0271 |
| Daam2 | 43 | 29 | 18 | 21 | 18 | 22 | 33 | 31 | 19 | 21 | 20 | 22 | 24.43 | 28 | 21 | 0.747 | -0.42 | 0.29302659 | 0.46515206 | 0 | 0.0994 | 0.0732 | 0.0732 | TRUE | 0.4231 |
| Dab1 | 868 | 599 | 550 | 773 | 646 | 677 | 674 | 637 | 593 | 769 | 729 | 668 | 678.54 | 635 | 722 | 1.136 | 0.183 | 0.05329168 | 0.14101245 | 0.0032 | 0.0059 | 0.005 | 0.005 | TRUE | 0.0513 |
| Dab2 | 961 | 608 | 424 | 610 | 435 | 467 | 746 | 647 | 457 | 607 | 491 | 461 | 568.25 | 617 | 520 | 0.841 | -0.25 | 0.11238857 | 0.24076286 | 0.0391 | 0.0066 | 0.0161 | 0.0161 | TRUE | 0.3958 |
| Dab2ip | 4843 | 3991 | 3995 | 4703 | 4266 | 4693 | 3761 | 4245 | 4311 | 4681 | 4812 | 4633 | 4407.28 | 4106 | 4709 | 1.148 | 0.199 | 0.00246621 | 0.01421923 | 0.0025 | 0.003 | 0.0029 | 0.0029 | TRUE | 0.0667 |
| Dach1 | 31 | 22 | 30 | 24 | 14 | 14 | 24 | 23 | 32 | 24 | 16 | 14 | 22.23 | 26 | 18 | 0.673 | -0.57 | 0.17875086 | 0.33335655 | 0 | 0.1091 | 0.0825 | 0.0825 | TRUE | 0.4601 |
| Dach2 | 94 | 70 | 57 | 111 | 61 | 62 | 73 | 74 | 62 | 110 | 69 | 61 | 74.91 | 70 | 80 | 1.151 | 0.203 | 0.44386276 | 0.61487977 | 0.0445 | 0.0341 | 0.0371 | 0.0371 | TRUE | 1.0343 |
| Dact1 | 497 | 421 | 439 | 420 | 439 | 500 | 386 | 448 | 474 | 418 | 495 | 494 | 452.4 | 436 | 469 | 1.078 | 0.109 | 0.35577803 | 0.53025395 | 0.0081 | 0.0077 | 0.0078 | 0.0078 | TRUE | 0.1233 |
| Dact2 | 1253 | 982 | 1029 | 970 | 925 | 1068 | 973 | 1045 | 1110 | 966 | 1043 | 1054 | 1031.88 | 1043 | 1021 | 0.98 | -0.028 | 0.73011666 | 0.83880375 | 0.0025 | 0.0047 | 0.0039 | 0.0039 | TRUE | 0.0426 |
| Dact3 | 1218 | 1047 | 1236 | 1600 | 1386 | 1684 | 946 | 1114 | 1334 | 1593 | 1564 | 1662 | 1368.64 | 1131 | 1606 | 1.423 | 0.509 | 1.74E-06 | 4.51E-05 | 0.0141 | 0.0042 | 0.0074 | 0.0074 | TRUE | 0.2938 |
| Dad1 | 3843 | 2728 | 2452 | 2719 | 2457 | 2886 | 2985 | 2902 | 2646 | 2706 | 2772 | 2849 | 2809.88 | 2844 | 2776 | 0.975 | -0.036 | 0.58483205 | 0.734344 | 0.002 | 0.0033 | 0.0028 | 0.0028 | TRUE | 0.0448 |
| Daf2 | 9 | 8 | 8 | 7 | 8 | 8 | 7 | 9 | 9 | 7 | 9 | 8 | 8 | 8 | 8 | 0.993 | -0.01 | 0.98792848 | NA | 0 | 0.2986 | 0.1887 | 0.1887 | TRUE | 0.0514 |
| Dag1 | 19491 | 14081 | 13835 | 14674 | 13123 | 14925 | 15138 | 14978 | 14928 | 14606 | 14804 | 14733 | 14864.55 | 15015 | 14714 | 0.98 | -0.029 | 0.50256485 | 0.66680359 | 0 | 0.0026 | 0.0013 | 0.0013 | TRUE | 6.00E-04 |
| Dagla | 221 | 160 | 141 | 194 | 214 | 190 | 172 | 170 | 152 | 193 | 241 | 188 | 186.01 | 165 | 207 | 1.255 | 0.328 | 0.0391279 | 0.11268489 | 0.0069 | 0.0152 | 0.0127 | 0.0127 | TRUE | 0.218 |
| Daglb | 718 | 507 | 439 | 447 | 398 | 493 | 558 | 539 | 474 | 445 | 449 | 487 | 491.87 | 524 | 460 | 0.878 | -0.188 | 0.07315319 | 0.17706665 | 0.003 | 0.0072 | 0.0059 | 0.0059 | TRUE | 0.0791 |
| Dak | 1137 | 813 | 880 | 1085 | 963 | 1011 | 883 | 865 | 950 | 1080 | 1086 | 998 | 976.96 | 899 | 1055 | 1.173 | 0.231 | 0.00402933 | 0.02073115 | 0.0012 | 0.0049 | 0.0036 | 0.0036 | TRUE | 0.0277 |
| Dalrd3 | 2253 | 1989 | 1893 | 1965 | 1801 | 2122 | 1750 | 2116 | 2043 | 1956 | 2032 | 2095 | 1998.4 | 1970 | 2028 | 1.031 | 0.044 | 0.58797577 | 0.73712285 | 0.0052 | 0.0036 | 0.0042 | 0.0042 | TRUE | 0.1181 |
| Dancr | 262 | 185 | 191 | 173 | 129 | 170 | 203 | 197 | 206 | 172 | 146 | 168 | 181.98 | 202 | 162 | 0.802 | -0.318 | 0.03344008 | 0.1003694 | 0 | 0.0154 | 0.0106 | 0.0106 | TRUE | 0.0783 |
| Dand5 | 101 | 77 | 56 | 97 | 83 | 98 | 78 | 82 | 60 | 97 | 94 | 97 | 84.62 | 73 | 96 | 1.296 | 0.374 | 0.08434795 | 0.19580726 | 0 | 0.0304 | 0.022 | 0.022 | TRUE | 0.2073 |
| Dao | 1 | 0 | 0 | 0 | 0 | 2 | 1 | 0 | 0 | 0 | 0 | 2 | 0.46 | 0 | 1 | 1.93 | 0.948 | 0.7608301 | NA | 0 | 5.1717 | 5.216 | 5.216 | TRUE | 0.9737 |
| Dap | 697 | 522 | 582 | 641 | 619 | 694 | 541 | 555 | 628 | 638 | 698 | 685 | 624.33 | 575 | 674 | 1.174 | 0.231 | 0.01637551 | 0.0596471 | 0.0026 | 0.0062 | 0.0051 | 0.0051 | TRUE | 0.0751 |
| Dap3 | 8565 | 6198 | 6687 | 7318 | 6356 | 7058 | 6652 | 6593 | 7215 | 7284 | 7170 | 6967 | 6980.32 | 6820 | 7140 | 1.047 | 0.067 | 0.23935757 | 0.4063109 | 0.0013 | 0.0028 | 0.0022 | 0.0022 | TRUE | 0.0311 |
| Dapk1 | 3648 | 2713 | 3070 | 3776 | 3340 | 4003 | 2833 | 2886 | 3313 | 3759 | 3768 | 3952 | 3418.27 | 3011 | 3826 | 1.272 | 0.347 | 1.12E-06 | 3.22E-05 | 0.0038 | 0.0031 | 0.0034 | 0.0034 | TRUE | 0.0928 |
| Dapk2 | 272 | 172 | 175 | 225 | 161 | 169 | 211 | 183 | 189 | 224 | 182 | 167 | 192.57 | 194 | 191 | 0.98 | -0.029 | 0.85461878 | 0.91835698 | 0.0098 | 0.0147 | 0.0133 | 0.0133 | TRUE | 0.2546 |
| Dapk3 | 2168 | 1481 | 1211 | 1429 | 1164 | 1368 | 1684 | 1575 | 1307 | 1422 | 1313 | 1350 | 1441.96 | 1522 | 1362 | 0.894 | -0.162 | 0.08454215 | 0.19609882 | 0.0079 | 0.0041 | 0.0056 | 0.0056 | TRUE | 0.1827 |
| Dapl1 | 3 | 4 | 1 | 3 | 4 | 4 | 2 | 4 | 1 | 3 | 5 | 4 | 3.19 | 2 | 4 | 1.494 | 0.579 | 0.5877854 | NA | 0 | 0.7465 | 0.4969 | 0.4969 | TRUE | 0.3377 |
| Dapp1 | 427 | 339 | 315 | 384 | 359 | 444 | 332 | 361 | 340 | 382 | 405 | 438 | 376.27 | 344 | 408 | 1.189 | 0.25 | 0.0251818 | 0.08168941 | 0 | 0.0087 | 0.0063 | 0.0063 | TRUE | 0.0481 |
| Darc | 2 | 0 | 0 | 2 | 2 | 1 | 2 | 0 | 0 | 2 | 2 | 1 | 1.13 | 1 | 2 | 3.021 | 1.595 | 0.40823467 | NA | 0 | 2.0975 | 1.5463 | 1.5463 | TRUE | 0.9712 |
| Dars | 13063 | 9558 | 10004 | 10652 | 9513 | 10951 | 10145 | 10167 | 10794 | 10603 | 10732 | 10810 | 10541.9 | 10369 | 10715 | 1.034 | 0.048 | 0.34025738 | 0.51440079 | 5.00E-04 | 0.0027 | 0.0017 | 0.0017 | TRUE | 0.0157 |
| Dars2 | 4905 | 3352 | 3009 | 3726 | 3342 | 3712 | 3810 | 3565 | 3247 | 3709 | 3770 | 3664 | 3627.49 | 3541 | 3714 | 1.048 | 0.068 | 0.3170954 | 0.4911405 | 0.0029 | 0.0031 | 0.0031 | 0.0031 | TRUE | 0.0639 |
| Daxx | 3611 | 2492 | 2111 | 2159 | 1950 | 2318 | 2805 | 2651 | 2278 | 2149 | 2200 | 2288 | 2395.01 | 2578 | 2212 | 0.858 | -0.222 | 0.00616012 | 0.02877995 | 0.0054 | 0.0034 | 0.0043 | 0.0043 | TRUE | 0.1248 |
| Dazap1 | 9703 | 7448 | 7814 | 7442 | 6654 | 7729 | 7536 | 7922 | 8431 | 7408 | 7506 | 7630 | 7738.89 | 7963 | 7515 | 0.944 | -0.083 | 0.14680775 | 0.29115071 | 0.0015 | 0.0027 | 0.0022 | 0.0022 | TRUE | 0.0323 |
| Dazap2 | 3666 | 2814 | 3208 | 3170 | 3080 | 3420 | 2847 | 2993 | 3461 | 3155 | 3475 | 3376 | 3217.98 | 3100 | 3335 | 1.076 | 0.106 | 0.18200985 | 0.33758977 | 0.0061 | 0.0032 | 0.0042 | 0.0042 | TRUE | 0.1254 |
| Dazl | 8472 | 6561 | 5270 | 3620 | 3003 | 3109 | 6580 | 6979 | 5686 | 3603 | 3388 | 3069 | 4884.18 | 6415 | 3353 | 0.523 | -0.936 | 1.04E-28 | 7.42E-26 | 0.0082 | 0.0029 | 0.0049 | 0.0049 | TRUE | 0.1199 |
| Dbc1 | 250 | 194 | 145 | 179 | 166 | 174 | 194 | 206 | 156 | 178 | 187 | 172 | 182.36 | 185 | 179 | 0.962 | -0.055 | 0.72612395 | 0.83589639 | 0.005 | 0.0154 | 0.0125 | 0.0125 | TRUE | 0.1929 |
| Dbf4 | 8474 | 6140 | 6366 | 6748 | 6048 | 7102 | 6581 | 6531 | 6869 | 6717 | 6823 | 7011 | 6755.3 | 6660 | 6850 | 1.029 | 0.041 | 0.42585463 | 0.59841355 | 4.00E-04 | 0.0028 | 0.0017 | 0.0017 | TRUE | 0.0091 |
| Dbh | 0 | 0 | 0 | 0 | 0 | 0 | 0 | 0 | 0 | 0 | 0 | 0 | 0 | 0 | 0 | NA | NA | NA | NA | NA | NA | NA | NA | NA | NA |
| Dbi | 5010 | 3438 | 3276 | 3891 | 3377 | 3871 | 3891 | 3657 | 3535 | 3873 | 3810 | 3821 | 3764.46 | 3694 | 3835 | 1.037 | 0.053 | 0.36463785 | 0.53933831 | 8.00E-04 | 0.0031 | 0.0022 | 0.0022 | TRUE | 0.027 |
| Dbil5 | 27 | 20 | 24 | 23 | 13 | 23 | 21 | 21 | 26 | 23 | 15 | 23 | 21.4 | 23 | 20 | 0.891 | -0.166 | 0.69121135 | 0.81119007 | 0 | 0.1132 | 0.0793 | 0.0793 | TRUE | 0.2727 |
| Dbn1 | 1772 | 1356 | 1239 | 1751 | 1641 | 1768 | 1376 | 1442 | 1337 | 1743 | 1851 | 1745 | 1582.49 | 1385 | 1780 | 1.285 | 0.361 | 1.12E-07 | 4.34E-06 | 6.00E-04 | 0.0039 | 0.0027 | 0.0027 | TRUE | 0.0152 |
| Dbndd1 | 88 | 70 | 44 | 49 | 46 | 37 | 68 | 74 | 47 | 49 | 52 | 37 | 54.58 | 63 | 46 | 0.718 | -0.479 | 0.09394293 | 0.21115369 | 0.024 | 0.0459 | 0.0399 | 0.0399 | TRUE | 0.3972 |
| Dbndd2 | 234 | 158 | 195 | 245 | 257 | 283 | 182 | 168 | 210 | 244 | 290 | 279 | 228.89 | 187 | 271 | 1.452 | 0.538 | 0.00021087 | 0.00221448 | 0.0054 | 0.0128 | 0.0107 | 0.0107 | TRUE | 0.1246 |
| Dbnl | 4024 | 2819 | 2234 | 2387 | 2029 | 2196 | 3125 | 2999 | 2410 | 2376 | 2289 | 2168 | 2561.17 | 2845 | 2278 | 0.8 | -0.322 | 0.00045884 | 0.00405076 | 0.0094 | 0.0034 | 0.0057 | 0.0057 | TRUE | 0.2152 |
| Dbp | 131 | 88 | 105 | 131 | 123 | 134 | 102 | 94 | 113 | 130 | 139 | 132 | 118.35 | 103 | 134 | 1.301 | 0.379 | 0.03659902 | 0.10750303 | 0 | 0.0225 | 0.0152 | 0.0152 | TRUE | 0.0707 |
| Dbpht2 | 1 | 0 | 0 | 2 | 0 | 0 | 1 | 0 | 0 | 2 | 0 | 0 | 0.46 | 0 | 1 | 1.942 | 0.958 | 0.7580687 | NA | 0 | 5.1409 | 5.1921 | 5.1921 | TRUE | 0.9738 |
| Dbr1 | 2569 | 1760 | 1473 | 1408 | 1383 | 1466 | 1995 | 1872 | 1589 | 1402 | 1560 | 1447 | 1644.26 | 1819 | 1470 | 0.807 | -0.309 | 0.00057838 | 0.00482873 | 0.0072 | 0.0039 | 0.0052 | 0.0052 | TRUE | 0.1461 |
| Dbt | 3840 | 2753 | 3088 | 3448 | 3046 | 3609 | 2982 | 2928 | 3332 | 3432 | 3436 | 3563 | 3278.93 | 3081 | 3477 | 1.129 | 0.175 | 0.00799298 | 0.03499042 | 0.0023 | 0.0031 | 0.0028 | 0.0028 | TRUE | 0.0609 |
| Dbx1 | 2 | 1 | 0 | 1 | 0 | 0 | 2 | 1 | 0 | 1 | 0 | 0 | 0.6 | 1 | 0 | 0.416 | -1.265 | 0.62159733 | NA | 0 | 3.939 | 3.1896 | 3.1896 | TRUE | 0.3906 |
| Dbx2 | 216 | 172 | 222 | 269 | 252 | 282 | 168 | 183 | 240 | 268 | 284 | 278 | 236.78 | 197 | 277 | 1.413 | 0.499 | 0.00112112 | 0.00788539 | 0.0124 | 0.0124 | 0.0125 | 0.0125 | TRUE | 0.3847 |
| Dcaf10 | 1118 | 833 | 759 | 813 | 719 | 810 | 868 | 886 | 819 | 809 | 811 | 800 | 832.21 | 858 | 807 | 0.94 | -0.089 | 0.26191892 | 0.43223728 | 0 | 0.0053 | 0.0034 | 0.0034 | TRUE | 0.0179 |
| Dcaf11 | 1832 | 1429 | 1294 | 1399 | 1286 | 1502 | 1423 | 1520 | 1396 | 1393 | 1451 | 1483 | 1444.18 | 1446 | 1442 | 0.997 | -0.004 | 0.95462285 | 0.97522867 | 8.00E-04 | 0.0041 | 0.0029 | 0.0029 | TRUE | 0.0233 |
| Dcaf12 | 5881 | 4087 | 4390 | 4828 | 4250 | 4932 | 4568 | 4347 | 4737 | 4806 | 4794 | 4869 | 4686.74 | 4551 | 4823 | 1.06 | 0.084 | 0.12903074 | 0.26569804 | 6.00E-04 | 0.0029 | 0.002 | 0.002 | TRUE | 0.0183 |
| Dcaf12l1 | 1775 | 1067 | 974 | 744 | 592 | 710 | 1379 | 1135 | 1051 | 741 | 668 | 701 | 945.63 | 1188 | 703 | 0.591 | -0.759 | 6.69E-13 | 8.94E-11 | 0.0099 | 0.0049 | 0.0069 | 0.0069 | TRUE | 0.2411 |

| | | | | | | | | | | | | | | | | | | | | | | | | | |
|---|---|---|---|---|---|---|---|---|---|---|---|---|---|---|---|---|---|---|---|---|---|---|---|---|---|
| Dcaf12l2 | 13 | 15 | 7 | 11 | 10 | 3 | 10 | 16 | 8 | 11 | 11 | 3 | 9.8 | 11 | 8 | 0.747 | -0.421 | 0.53002095 | NA | 0.1411 | 0.2443 | 0.2182 | 0.2182 | TRUE | 1.0211 |
| Dcaf13 | 4741 | 3500 | 3498 | 3557 | 3103 | 3778 | 3682 | 3723 | 3774 | 3541 | 3501 | 3729 | 3658.33 | 3726 | 3590 | 0.964 | -0.053 | 0.33514767 | 0.50922352 | 4.00E-04 | 0.0031 | 0.0019 | 0.0019 | TRUE | 0.0141 |
| Dcaf15 | 5303 | 4093 | 4152 | 4444 | 3824 | 4503 | 4119 | 4354 | 4480 | 4424 | 4314 | 4445 | 4355.81 | 4318 | 4394 | 1.018 | 0.026 | 0.64185919 | 0.77660995 | 8.00E-04 | 0.003 | 0.0021 | 0.0021 | TRUE | 0.0202 |
| Dcaf17 | 3220 | 2368 | 2714 | 2793 | 2614 | 2916 | 2501 | 2519 | 2928 | 2780 | 2949 | 2879 | 2759.26 | 2649 | 2869 | 1.083 | 0.116 | 0.11793332 | 0.24916845 | 0.0041 | 0.0033 | 0.0036 | 0.0036 | TRUE | 0.1021 |
| Dcaf4 | 4203 | 3177 | 3823 | 3978 | 3802 | 4221 | 3264 | 3379 | 4125 | 3960 | 4289 | 4167 | 3864.02 | 3589 | 4139 | 1.153 | 0.206 | 0.01592955 | 0.05838933 | 0.0089 | 0.003 | 0.005 | 0.005 | TRUE | 0.2063 |
| Dcaf5 | 2334 | 1715 | 1608 | 1725 | 1526 | 1727 | 1813 | 1824 | 1735 | 1717 | 1721 | 1705 | 1752.55 | 1791 | 1714 | 0.957 | -0.063 | 0.31251291 | 0.48572717 | 0 | 0.0038 | 0.0022 | 0.0022 | TRUE | 0.0087 |
| Dcaf6 | 1379 | 1053 | 1009 | 968 | 919 | 954 | 1071 | 1120 | 1089 | 964 | 1037 | 942 | 1036.97 | 1093 | 981 | 0.897 | -0.157 | 0.03976542 | 0.11406125 | 4.00E-04 | 0.0047 | 0.0032 | 0.0032 | TRUE | 0.0292 |
| Dcaf7 | 11843 | 8591 | 8798 | 10799 | 9188 | 10799 | 9198 | 9138 | 9493 | 10749 | 10365 | 10660 | 9933.94 | 9276 | 10591 | 1.142 | 0.191 | 6.36E-05 | 0.00085333 | 2.00E-04 | 0.0027 | 0.0016 | 0.0016 | TRUE | 0.0051 |
| Dcaf8 | 4709 | 3447 | 3680 | 3678 | 3299 | 3669 | 3657 | 3667 | 3971 | 3661 | 3722 | 3622 | 3716.51 | 3765 | 3668 | 0.975 | -0.037 | 0.52372587 | 0.68511933 | 8.00E-04 | 0.0031 | 0.0022 | 0.0022 | TRUE | 0.0275 |
| Dcakd | 12821 | 9160 | 8411 | 7883 | 7257 | 8430 | 9958 | 9743 | 9075 | 7847 | 8187 | 8322 | 8855.23 | 9592 | 8119 | 0.846 | -0.241 | 2.02E-05 | 0.00033388 | 0.0014 | 0.0027 | 0.0022 | 0.0022 | TRUE | 0.027 |
| Dcbld1 | 2134 | 1491 | 1303 | 1328 | 1174 | 1250 | 1657 | 1586 | 1406 | 1322 | 1324 | 1234 | 1421.58 | 1550 | 1293 | 0.834 | -0.263 | 0.00113364 | 0.00794658 | 0.0038 | 0.0041 | 0.004 | 0.004 | TRUE | 0.0785 |
| Dcbld2 | 4859 | 3453 | 3319 | 3533 | 2977 | 3612 | 3774 | 3673 | 3581 | 3517 | 3358 | 3566 | 3578.1 | 3676 | 3480 | 0.947 | -0.079 | 0.16177703 | 0.311796 | 5.00E-04 | 0.0031 | 0.002 | 0.002 | TRUE | 0.0113 |
| Dcc | 116 | 92 | 29 | 35 | 28 | 21 | 90 | 98 | 31 | 35 | 32 | 21 | 51.07 | 73 | 29 | 0.396 | -1.338 | 0.00069259 | 0.00553646 | 0.1845 | 0.0488 | 0.0876 | 0.0876 | TRUE | 2.1883 |
| Dcdc2a | 60 | 46 | 50 | 49 | 61 | 45 | 47 | 49 | 54 | 49 | 69 | 44 | 51.91 | 50 | 54 | 1.082 | 0.114 | 0.68640965 | 0.80774354 | 0.0108 | 0.0481 | 0.0379 | 0.0379 | TRUE | 0.4528 |
| Dcdc2b | 4 | 5 | 3 | 7 | 2 | 7 | 3 | 5 | 3 | 7 | 2 | 7 | 4.63 | 4 | 5 | 1.407 | 0.492 | 0.58510812 | NA | 0 | 0.514 | 0.3658 | 0.3658 | TRUE | 0.486 |
| Dcdc2c | 1 | 2 | 1 | 2 | 3 | 0 | 1 | 2 | 1 | 2 | 3 | 0 | 1.56 | 1 | 2 | 1.353 | 0.437 | 0.78270562 | NA | 0 | 1.522 | 1.1441 | 1.1441 | TRUE | 0.2505 |
| Dchs1 | 135 | 84 | 68 | 83 | 65 | 72 | 105 | 89 | 73 | 83 | 73 | 71 | 82.43 | 89 | 76 | 0.844 | -0.245 | 0.28040107 | 0.45193645 | 0.0093 | 0.0312 | 0.0248 | 0.0248 | TRUE | 0.1767 |
| Dck | 3556 | 2558 | 2385 | 2414 | 2090 | 2205 | 2762 | 2721 | 2573 | 2403 | 2358 | 2177 | 2498.9 | 2685 | 2313 | 0.861 | -0.216 | 0.00097179 | 0.00709387 | 0.0014 | 0.0034 | 0.0027 | 0.0027 | TRUE | 0.0323 |
| Dclk1 | 41 | 34 | 58 | 68 | 69 | 79 | 32 | 36 | 63 | 68 | 78 | 78 | 59.02 | 44 | 75 | 1.728 | 0.789 | 0.00689129 | 0.03139855 | 0.0456 | 0.0426 | 0.0433 | 0.0433 | TRUE | 1.0924 |
| Dclk2 | 1797 | 1702 | 1831 | 1743 | 1698 | 1989 | 1396 | 1810 | 1976 | 1735 | 1916 | 1963 | 1799.27 | 1727 | 1871 | 1.085 | 0.117 | 0.28404604 | 0.45601147 | 0.0174 | 0.0037 | 0.0081 | 0.0081 | TRUE | 0.347 |
| Dclk3 | 130 | 102 | 93 | 119 | 121 | 135 | 101 | 108 | 100 | 118 | 137 | 133 | 116.34 | 103 | 129 | 1.253 | 0.326 | 0.07378961 | 0.17818463 | 0 | 0.0228 | 0.0152 | 0.0152 | TRUE | 0.0569 |
| Dclre1a | 923 | 653 | 583 | 597 | 587 | 614 | 717 | 695 | 629 | 594 | 662 | 606 | 650.51 | 680 | 621 | 0.911 | -0.135 | 0.15386037 | 0.30092994 | 0.0024 | 0.0061 | 0.0049 | 0.0049 | TRUE | 0.05 |
| Dclre1b | 763 | 485 | 426 | 470 | 414 | 437 | 593 | 516 | 460 | 468 | 467 | 431 | 489.06 | 523 | 455 | 0.869 | -0.203 | 0.07573894 | 0.1814206 | 0.0076 | 0.0073 | 0.0074 | 0.0074 | TRUE | 0.1672 |
| Dclre1c | 1125 | 897 | 868 | 945 | 790 | 980 | 874 | 954 | 937 | 941 | 891 | 967 | 927.28 | 922 | 933 | 1.014 | 0.02 | 0.79860754 | 0.8823141 | 0.001 | 0.005 | 0.0036 | 0.0036 | TRUE | 0.0259 |
| Dcn | 0 | 0 | 0 | 0 | 0 | 0 | 0 | 0 | 0 | 0 | 0 | 0 | 0 | 0 | 0 | NA | NA | NA | NA | NA | NA | NA | NA | NA | NA |
| Dcp1a | 4214 | 2962 | 2991 | 3045 | 2722 | 3039 | 3273 | 3151 | 3227 | 3031 | 3071 | 3000 | 3125.4 | 3217 | 3034 | 0.943 | -0.085 | 0.11842665 | 0.24987864 | 0 | 0.0032 | 0.0018 | 0.0018 | TRUE | 0.0039 |
| Dcp1b | 420 | 320 | 359 | 370 | 358 | 373 | 326 | 340 | 387 | 368 | 404 | 368 | 365.72 | 351 | 380 | 1.084 | 0.116 | 0.31978977 | 0.49360512 | 0.0028 | 0.0089 | 0.007 | 0.007 | TRUE | 0.0901 |
| Dcp2 | 13447 | 9462 | 8194 | 8350 | 7400 | 8324 | 10444 | 10065 | 8841 | 8312 | 8348 | 8217 | 9037.7 | 9783 | 8292 | 0.847 | -0.239 | 0.00034113 | 0.00319934 | 0.0035 | 0.0027 | 0.0031 | 0.0031 | TRUE | 0.0864 |
| Dcpp1 | 0 | 0 | 0 | 0 | 0 | 0 | 0 | 0 | 0 | 0 | 0 | 0 | 0 | 0 | 0 | NA | NA | NA | NA | NA | NA | NA | NA | NA | NA |
| Dcpp2 | 0 | 0 | 0 | 0 | 0 | 0 | 0 | 0 | 0 | 0 | 0 | 0 | 0 | 0 | 0 | NA | NA | NA | NA | NA | NA | NA | NA | NA | NA |
| Dcpp3 | 0 | 0 | 0 | 0 | 0 | 0 | 0 | 0 | 0 | 0 | 0 | 0 | 0 | 0 | 0 | NA | NA | NA | NA | NA | NA | NA | NA | NA | NA |
| Dcps | 3158 | 2279 | 2534 | 2416 | 2330 | 2718 | 2453 | 2424 | 2734 | 2405 | 2628 | 2683 | 2554.57 | 2537 | 2572 | 1.014 | 0.02 | 0.78012545 | 0.87046309 | 0.0035 | 0.0034 | 0.0034 | 0.0034 | TRUE | 0.0554 |
| Dcst1 | 8 | 1 | 1 | 3 | 0 | 3 | 6 | 1 | 1 | 3 | 0 | 3 | 2.38 | 3 | 2 | 0.699 | -0.517 | 0.70524109 | NA | 0.7074 | 0.9966 | 0.9205 | 0.9205 | TRUE | 1.8106 |
| Dcstamp | 93 | 63 | 53 | 58 | 53 | 44 | 72 | 67 | 57 | 58 | 60 | 43 | 59.56 | 65 | 54 | 0.815 | -0.295 | 0.25661478 | 0.42653739 | 0 | 0.0422 | 0.0318 | 0.0318 | TRUE | 0.2371 |
| Dct | 7 | 8 | 3 | 1 | 2 | 2 | 5 | 9 | 3 | 1 | 2 | 2 | 3.73 | 6 | 2 | 0.302 | -1.726 | 0.10713211 | NA | 0 | 0.637 | 0.4434 | 0.4434 | TRUE | 0.1347 |
| Dctd | 5965 | 4654 | 4818 | 4236 | 4001 | 4848 | 4633 | 4950 | 5199 | 4217 | 4514 | 4786 | 4716.25 | 4927 | 4506 | 0.915 | -0.129 | 0.05937416 | 0.15219419 | 0.0033 | 0.0029 | 0.0031 | 0.0031 | TRUE | 0.0385 |
| Dctn1 | 7365 | 5412 | 4728 | 5087 | 4467 | 5292 | 5720 | 5757 | 5101 | 5064 | 5039 | 5224 | 5317.51 | 5526 | 5109 | 0.924 | -0.114 | 0.06957868 | 0.17090359 | 0.0021 | 0.0029 | 0.0026 | 0.0026 | TRUE | 0.0546 |
| Dctn2 | 6514 | 4704 | 4533 | 4897 | 4316 | 5191 | 5059 | 5004 | 4891 | 4874 | 4869 | 5124 | 4970.25 | 4985 | 4956 | 0.994 | -0.008 | 0.87275339 | 0.92945862 | 4.00E-04 | 0.0029 | 0.0018 | 0.0018 | TRUE | 0.0108 |
| Dctn3 | 2258 | 1652 | 1709 | 1854 | 1624 | 1815 | 1754 | 1757 | 1844 | 1845 | 1832 | 1792 | 1804.02 | 1785 | 1823 | 1.022 | 0.031 | 0.6220306 | 0.76142749 | 0 | 0.0037 | 0.0023 | 0.0023 | TRUE | 0.0099 |
| Dctn4 | 2967 | 2277 | 2397 | 2450 | 2285 | 2631 | 2304 | 2422 | 2586 | 2439 | 2578 | 2597 | 2487.72 | 2437 | 2538 | 1.042 | 0.059 | 0.37527901 | 0.54963243 | 0.0017 | 0.0034 | 0.0028 | 0.0028 | TRUE | 0.0345 |
| Dctn5 | 1971 | 1342 | 1257 | 1396 | 1097 | 1394 | 1531 | 1427 | 1356 | 1390 | 1238 | 1376 | 1386.29 | 1438 | 1335 | 0.927 | -0.109 | 0.16938516 | 0.32083515 | 0.0032 | 0.0041 | 0.0038 | 0.0038 | TRUE | 0.0476 |
| Dctn6 | 2084 | 1535 | 1657 | 1858 | 1633 | 1875 | 1619 | 1633 | 1788 | 1849 | 1842 | 1851 | 1763.63 | 1680 | 1847 | 1.101 | 0.138 | 0.03988365 | 0.1142955 | 9.00E-04 | 0.0038 | 0.0027 | 0.0027 | TRUE | 0.0376 |
| Dctpp1 | 7570 | 5424 | 5730 | 6213 | 5676 | 6319 | 5879 | 5769 | 6183 | 6184 | 6403 | 6238 | 6109.45 | 5944 | 6275 | 1.056 | 0.078 | 0.13880024 | 0.27989163 | 6.00E-04 | 0.0028 | 0.0019 | 0.0019 | TRUE | 0.0149 |
| Dcun1d1 | 1622 | 1150 | 1197 | 1363 | 1139 | 1385 | 1260 | 1223 | 1292 | 1357 | 1285 | 1367 | 1297.23 | 1258 | 1336 | 1.062 | 0.087 | 0.21111562 | 0.37312669 | 0 | 0.0043 | 0.0028 | 0.0028 | TRUE | 0.0133 |
| Dcun1d2 | 1704 | 1178 | 1005 | 1035 | 963 | 1089 | 1323 | 1253 | 1084 | 1030 | 1086 | 1075 | 1142.07 | 1220 | 1064 | 0.871 | -0.2 | 0.02210414 | 0.07418045 | 0.0048 | 0.0045 | 0.0046 | 0.0046 | TRUE | 0.1125 |
| Dcun1d3 | 1704 | 1137 | 1189 | 1195 | 999 | 1169 | 1323 | 1209 | 1283 | 1190 | 1127 | 1154 | 1214.37 | 1272 | 1157 | 0.909 | -0.138 | 0.05970333 | 0.15275796 | 7.00E-04 | 0.0044 | 0.003 | 0.003 | TRUE | 0.022 |
| Dcun1d4 | 3814 | 3113 | 3047 | 3355 | 2808 | 3290 | 2962 | 3311 | 3288 | 3340 | 3168 | 3248 | 3219.36 | 3187 | 3252 | 1.021 | 0.03 | 0.64071896 | 0.77555861 | 0.002 | 0.0032 | 0.0027 | 0.0027 | TRUE | 0.0474 |
| Dcun1d5 | 4265 | 2952 | 2623 | 2738 | 2268 | 2741 | 3312 | 3140 | 2830 | 2725 | 2559 | 2706 | 2878.73 | 3094 | 2663 | 0.86 | -0.217 | 0.00238689 | 0.01386258 | 0.0032 | 0.0032 | 0.0033 | 0.0033 | TRUE | 0.0673 |
| Dcx | 0 | 0 | 1 | 0 | 0 | 0 | 0 | 0 | 1 | 0 | 0 | 0 | 0.18 | 0 | 0 | 0.552 | -0.857 | 0.8336548 | NA | 0 | 13.1817 | 10 | 10 | TRUE | 0.5365 |
| Dcxr | 598 | 462 | 643 | 699 | 673 | 762 | 464 | 491 | 694 | 696 | 759 | 752 | 642.81 | 550 | 736 | 1.341 | 0.423 | 0.00156722 | 0.00996895 | 0.0247 | 0.0061 | 0.0113 | 0.0113 | TRUE | 0.6072 |
| Dda1 | 4267 | 3016 | 2545 | 3020 | 2659 | 2985 | 3314 | 3208 | 2746 | 3006 | 3000 | 2947 | 3036.75 | 3089 | 2984 | 0.965 | -0.051 | 0.49900803 | 0.66359526 | 0.0047 | 0.0032 | 0.0038 | 0.0038 | TRUE | 0.1139 |
| Ddah1 | 7712 | 5938 | 5947 | 6331 | 5701 | 6537 | 5990 | 6316 | 6417 | 6302 | 6431 | 6453 | 6318.12 | 6241 | 6395 | 1.025 | 0.036 | 0.49489502 | 0.65988266 | 5.00E-04 | 0.0028 | 0.0018 | 0.0018 | TRUE | 0.0154 |
| Ddah2 | 3694 | 3133 | 4000 | 3152 | 3183 | 3666 | 2869 | 3333 | 4316 | 3137 | 3591 | 3619 | 3477.44 | 3506 | 3449 | 0.984 | -0.023 | 0.89994532 | 0.94496903 | 0.0244 | 0.0031 | 0.0095 | 0.0244 | TRUE | 0.4967 |
| Ddb1 | 79130 | 56262 | 59584 | 56802 | 51817 | 59230 | 61457 | 59845 | 64291 | 56541 | 58455 | 58469 | 59842.91 | 61864 | 57822 | 0.935 | -0.097 | 0.04517845 | 0.12502142 | 8.00E-04 | 0.0025 | 0.0017 | 0.0017 | TRUE | 0.0144 |
| Ddb2 | 684 | 597 | 713 | 858 | 695 | 807 | 531 | 635 | 769 | 854 | 784 | 797 | 728.38 | 645 | 812 | 1.261 | 0.335 | 0.00549354 | 0.02638414 | 0.0161 | 0.0057 | 0.0091 | 0.0091 | TRUE | 0.3331 |
| Ddc | 1116 | 815 | 734 | 585 | 545 | 617 | 867 | 867 | 792 | 582 | 615 | 609 | 721.97 | 842 | 602 | 0.714 | -0.485 | 2.04E-08 | 9.43E-07 | 0 | 0.0057 | 0.004 | 0.004 | TRUE | 0.0311 |
| Ddhd1 | 1498 | 1166 | 1111 | 1065 | 904 | 1101 | 1163 | 1240 | 1199 | 1060 | 1020 | 1087 | 1128.2 | 1201 | 1056 | 0.88 | -0.184 | 0.01168145 | 0.04635285 | 0 | 0.0045 | 0.003 | 0.003 | TRUE | 0.0103 |
| Ddhd2 | 4599 | 3264 | 3190 | 3567 | 2930 | 3585 | 3572 | 3472 | 3442 | 3551 | 3305 | 3539 | 3480.1 | 3495 | 3465 | 0.991 | -0.013 | 0.82734223 | 0.90110512 | 7.00E-04 | 0.0031 | 0.0021 | 0.0021 | TRUE | 0.0195 |
| Ddi1 | 0 | 0 | 0 | 0 | 0 | 0 | 0 | 0 | 0 | 0 | 0 | 0 | 0 | 0 | 0 | NA | NA | NA | NA | NA | NA | NA | NA | NA | NA |
| Ddi2 | 2738 | 1865 | 1526 | 1669 | 1497 | 1752 | 2126 | 1984 | 1647 | 1661 | 1689 | 1729 | 1806.07 | 1919 | 1693 | 0.881 | -0.182 | 0.04369176 | 0.12204243 | 0.0076 | 0.0037 | 0.0053 | 0.0053 | TRUE | 0.185 |
| Ddit3 | 209 | 165 | 198 | 215 | 163 | 260 | 162 | 176 | 214 | 214 | 184 | 257 | 201 | 184 | 218 | 1.193 | 0.254 | 0.13343932 | 0.27193109 | 0.0192 | 0.0142 | 0.0157 | 0.0157 | TRUE | 0.257 |
| Ddit4 | 4329 | 3529 | 4108 | 4050 | 3415 | 3990 | 3362 | 3754 | 4432 | 4031 | 3852 | 3939 | 3895.16 | 3849 | 3941 | 1.024 | 0.035 | 0.69351256 | 0.81291342 | 0.0098 | 0.003 | 0.0054 | 0.0054 | TRUE | 0.2134 |
| Ddit4l | 1136 | 731 | 612 | 785 | 684 | 726 | 882 | 778 | 660 | 781 | 772 | 717 | 764.98 | 773 | 757 | 0.976 | -0.035 | 0.74466217 | 0.84821935 | 0.0104 | 0.0055 | 0.0071 | 0.0071 | TRUE | 0.1922 |
| Ddn | 6 | 11 | 6 | 8 | 7 | 11 | 5 | 12 | 6 | 8 | 8 | 11 | 8.26 | 8 | 9 | 1.19 | 0.251 | 0.71231424 | NA | 0 | 0.2894 | 0.2114 | 0.2114 | TRUE | 0.601 |
| Ddo | 5 | 11 | 9 | 8 | 6 | 12 | 4 | 12 | 10 | 8 | 7 | 12 | 8.65 | 9 | 9 | 1.076 | 0.105 | 0.87787196 | NA | 0 | 0.2766 | 0.2204 | 0.2204 | TRUE | 0.9097 |
| Ddost | 10521 | 7728 | 7896 | 8742 | 7782 | 9013 | 8171 | 8220 | 8520 | 8702 | 8779 | 8897 | 8548.17 | 8304 | 8793 | 1.059 | 0.083 | 0.08420325 | 0.19569384 | 1.00E-04 | 0.0027 | 0.0015 | 0.0015 | TRUE | 0.0063 |
| Ddr1 | 1411 | 999 | 1065 | 1099 | 979 | 1251 | 1096 | 1063 | 1149 | 1094 | 1104 | 1235 | 1123.48 | 1103 | 1144 | 1.039 | 0.055 | 0.49514024 | 0.65996429 | 0.0022 | 0.0045 | 0.0037 | 0.0037 | TRUE | 0.0579 |
| Ddr2 | 1168 | 964 | 552 | 245 | 285 | 268 | 907 | 1025 | 596 | 244 | 322 | 265 | 559.68 | 843 | 277 | 0.328 | -1.609 | 1.93E-21 | 6.91E-19 | 0.047 | 0.0067 | 0.0182 | 0.0182 | TRUE | 0.3523 |
| Ddrgk1 | 1653 | 1272 | 1392 | 1363 | 1298 | 1373 | 1284 | 1353 | 1502 | 1357 | 1464 | 1355 | 1385.86 | 1380 | 1392 | 1.01 | 0.014 | 0.85997922 | 0.92203758 | 0.0035 | 0.0041 | 0.0039 | 0.0039 | TRUE | 0.0719 |

| | | | | | | | | | | | | | | | | | | | | | | | | | |
|---|---|---|---|---|---|---|---|---|---|---|---|---|---|---|---|---|---|---|---|---|---|---|---|---|---|
| Ddt | 2501 | 1865 | 2118 | 2418 | 2184 | 2665 | 1942 | 1984 | 2285 | 2407 | 2464 | 2631 | 2285.49 | 2070 | 2501 | 1.209 | 0.273 | 0.00040234 | 0.00366517 | 0.0045 | 0.0035 | 0.0039 | 0.0039 | TRUE | 0.0988 |
| Ddx1 | 18945 | 13517 | 13373 | 14694 | 12904 | 14836 | 14714 | 14378 | 14429 | 14626 | 14557 | 14645 | 14558.31 | 14507 | 14609 | 1.007 | 0.01 | 0.81845278 | 0.89565467 | 0 | 0.0026 | 0.0013 | 0.0013 | TRUE | 0.0019 |
| Ddx10 | 9516 | 6863 | 6306 | 6529 | 5789 | 6720 | 7391 | 7300 | 6804 | 6499 | 6531 | 6634 | 6859.68 | 7165 | 6555 | 0.915 | -0.129 | 0.01703615 | 0.06148931 | 8.00E-04 | 0.0028 | 0.002 | 0.002 | TRUE | 0.0235 |
| Ddx11 | 3166 | 2599 | 2241 | 2196 | 1954 | 2449 | 2459 | 2765 | 2418 | 2186 | 2204 | 2418 | 2408.2 | 2547 | 2269 | 0.891 | -0.166 | 0.02580861 | 0.08315654 | 0.0037 | 0.0034 | 0.0036 | 0.0036 | TRUE | 0.0669 |
| Ddx17 | 22877 | 17471 | 18535 | 18096 | 15533 | 18453 | 17768 | 18584 | 19999 | 18013 | 17523 | 18216 | 18350.32 | 18784 | 17917 | 0.954 | -0.068 | 0.23392507 | 0.40008269 | 0.0019 | 0.0026 | 0.0023 | 0.0023 | TRUE | 0.0392 |
| Ddx18 | 9192 | 6221 | 5987 | 5721 | 4985 | 5921 | 7139 | 6617 | 6460 | 5695 | 5624 | 5845 | 6229.9 | 6739 | 5721 | 0.849 | -0.237 | 3.95E-05 | 0.00058202 | 0.0013 | 0.0028 | 0.0022 | 0.0022 | TRUE | 0.0334 |
| Ddx19a | 4961 | 3553 | 3436 | 3499 | 3130 | 3756 | 3853 | 3779 | 3707 | 3483 | 3531 | 3708 | 3676.89 | 3780 | 3574 | 0.945 | -0.081 | 0.14594343 | 0.29012092 | 5.00E-04 | 0.0031 | 0.002 | 0.002 | TRUE | 0.0132 |
| Ddx19b | 6716 | 4798 | 5525 | 5579 | 4601 | 5365 | 5216 | 5104 | 5961 | 5553 | 5190 | 5296 | 5386.81 | 5427 | 5346 | 0.985 | -0.021 | 0.76157704 | 0.85859131 | 0.0034 | 0.0029 | 0.0033 | 0.0033 | TRUE | 0.0899 |
| Ddx20 | 4043 | 2826 | 2535 | 2800 | 2389 | 2718 | 3140 | 3006 | 2735 | 2787 | 2695 | 2683 | 2841.08 | 2960 | 2722 | 0.919 | -0.122 | 0.06966429 | 0.17108449 | 0.0021 | 0.0033 | 0.0029 | 0.0029 | TRUE | 0.0534 |
| Ddx21 | 32670 | 23610 | 22152 | 22352 | 20648 | 23938 | 25373 | 25114 | 23902 | 22249 | 23293 | 23630 | 23926.94 | 24796 | 23057 | 0.93 | -0.105 | 0.03801121 | 0.11026877 | 9.00E-04 | 0.0025 | 0.0018 | 0.0018 | TRUE | 0.0122 |
| Ddx23 | 8514 | 6452 | 5901 | 5661 | 5184 | 6098 | 6612 | 6863 | 6367 | 5635 | 5848 | 6020 | 6224.2 | 6614 | 5834 | 0.882 | -0.181 | 0.00112561 | 0.00790192 | 0.001 | 0.0028 | 0.0021 | 0.0021 | TRUE | 0.013 |
| Ddx24 | 9646 | 7069 | 6341 | 6769 | 6096 | 7036 | 7492 | 7519 | 6842 | 6738 | 6877 | 6946 | 7068.85 | 7284 | 6854 | 0.941 | -0.088 | 0.11969063 | 0.25135735 | 0.0013 | 0.0028 | 0.0022 | 0.0022 | TRUE | 0.0342 |
| Ddx25 | 931 | 669 | 608 | 701 | 559 | 596 | 723 | 712 | 656 | 698 | 631 | 588 | 667.91 | 697 | 639 | 0.916 | -0.126 | 0.18885096 | 0.34643323 | 0.0035 | 0.006 | 0.0052 | 0.0052 | TRUE | 0.0779 |
| Ddx26b | 619 | 595 | 666 | 599 | 512 | 620 | 481 | 633 | 719 | 596 | 578 | 612 | 603.02 | 611 | 595 | 0.978 | -0.032 | 0.80252029 | 0.88561013 | 0.0196 | 0.0064 | 0.0105 | 0.0105 | TRUE | 0.425 |
| Ddx27 | 8638 | 6586 | 6401 | 6113 | 5812 | 6887 | 6709 | 7005 | 6907 | 6085 | 6557 | 6799 | 6676.8 | 6874 | 6480 | 0.943 | -0.085 | 0.14669392 | 0.29108653 | 0.0016 | 0.0028 | 0.0023 | 0.0023 | TRUE | 0.0349 |
| Ddx28 | 1363 | 951 | 837 | 881 | 896 | 912 | 1059 | 1012 | 903 | 877 | 1011 | 900 | 960.21 | 991 | 929 | 0.936 | -0.096 | 0.29570371 | 0.46753243 | 0.0051 | 0.0049 | 0.005 | 0.005 | TRUE | 0.0711 |
| Ddx31 | 1764 | 1238 | 1179 | 1132 | 999 | 1143 | 1370 | 1317 | 1272 | 1127 | 1127 | 1128 | 1223.51 | 1320 | 1127 | 0.854 | -0.228 | 0.00119481 | 0.00822911 | 0 | 0.0044 | 0.0028 | 0.0028 | TRUE | 0.014 |
| Ddx39 | 15154 | 10984 | 9463 | 9919 | 9118 | 10355 | 11769 | 11684 | 10210 | 9873 | 10286 | 10222 | 10674.15 | 11221 | 10127 | 0.902 | -0.148 | 0.02168044 | 0.07325247 | 0.0031 | 0.0026 | 0.0029 | 0.0029 | TRUE | 0.0756 |
| Ddx39b | 13896 | 9893 | 7964 | 7790 | 7007 | 7585 | 10792 | 10523 | 8593 | 7754 | 7905 | 7488 | 8842.49 | 9969 | 7716 | 0.774 | -0.37 | 4.31E-06 | 9.37E-05 | 0.0074 | 0.0027 | 0.0046 | 0.0046 | TRUE | 0.1778 |
| Ddx3x | 49395 | 33487 | 30352 | 31075 | 27641 | 31872 | 38363 | 35620 | 32750 | 30932 | 31182 | 31463 | 33384.79 | 35578 | 31192 | 0.877 | -0.19 | 0.00228122 | 0.01336839 | 0.0031 | 0.0025 | 0.0028 | 0.0028 | TRUE | 0.0592 |
| Ddx3y | 6416 | 4598 | 3956 | 3938 | 3249 | 3559 | 4983 | 4891 | 4268 | 3920 | 3665 | 3513 | 4206.79 | 4714 | 3699 | 0.785 | -0.35 | 2.06E-06 | 5.18E-05 | 0.0046 | 0.003 | 0.0037 | 0.0037 | TRUE | 0.0829 |
| Ddx4 | 1152 | 819 | 873 | 1169 | 935 | 1059 | 895 | 871 | 942 | 1164 | 1055 | 1045 | 995.27 | 903 | 1088 | 1.206 | 0.27 | 0.00082655 | 0.00629899 | 0.0016 | 0.0048 | 0.0037 | 0.0037 | TRUE | 0.0448 |
| Ddx41 | 5893 | 4587 | 4600 | 4797 | 4490 | 5180 | 4577 | 4879 | 4963 | 4775 | 5065 | 5113 | 4895.49 | 4806 | 4984 | 1.037 | 0.053 | 0.36679911 | 0.54148519 | 0.0013 | 0.0029 | 0.0023 | 0.0023 | TRUE | 0.0217 |
| Ddx42 | 11777 | 8388 | 7511 | 7291 | 6493 | 7286 | 9147 | 8922 | 8104 | 7257 | 7325 | 7192 | 7991.31 | 8724 | 7258 | 0.832 | -0.266 | 7.16E-06 | 0.00014236 | 0.0018 | 0.0027 | 0.0024 | 0.0024 | TRUE | 0.047 |
| Ddx43 | 475 | 411 | 255 | 325 | 272 | 312 | 369 | 437 | 275 | 324 | 307 | 308 | 336.6 | 360 | 313 | 0.867 | -0.205 | 0.17877128 | 0.33335655 | 0.0239 | 0.0095 | 0.0138 | 0.0138 | TRUE | 0.2421 |
| Ddx46 | 15559 | 12113 | 11587 | 12258 | 11128 | 12990 | 12084 | 12884 | 12502 | 12202 | 12554 | 12823 | 12508.17 | 12490 | 12526 | 1.003 | 0.004 | 0.93174025 | 0.96216772 | 6.00E-04 | 0.0026 | 0.0018 | 0.0018 | TRUE | 0.01 |
| Ddx47 | 7826 | 5653 | 5635 | 5957 | 5535 | 6483 | 6078 | 6013 | 6080 | 5930 | 6244 | 6400 | 6124.1 | 6057 | 6191 | 1.022 | 0.032 | 0.5503674 | 0.7064132 | 5.00E-04 | 0.0028 | 0.0018 | 0.0018 | TRUE | 0.0168 |
| Ddx49 | 7131 | 5468 | 4874 | 5722 | 4946 | 5626 | 5538 | 5816 | 5259 | 5696 | 5580 | 5554 | 5573.77 | 5538 | 5610 | 1.013 | 0.019 | 0.74287632 | 0.84692759 | 9.00E-04 | 0.0029 | 0.0021 | 0.0021 | TRUE | 0.0233 |
| Ddx5 | 79025 | 56409 | 54393 | 60037 | 53616 | 60941 | 61375 | 60002 | 58690 | 59761 | 60484 | 60158 | 60078.3 | 60022 | 60134 | 1.002 | 0.003 | 0.95153767 | 0.97323401 | 2.00E-04 | 0.0025 | 0.0014 | 0.0014 | TRUE | 0.0048 |
| Ddx50 | 4750 | 3471 | 3994 | 4266 | 3777 | 4371 | 3689 | 3692 | 4309 | 4246 | 4261 | 4315 | 4085.46 | 3897 | 4274 | 1.097 | 0.134 | 0.05712132 | 0.14849672 | 0.0039 | 0.003 | 0.0033 | 0.0033 | TRUE | 0.1037 |
| Ddx51 | 2173 | 1637 | 1290 | 1316 | 1210 | 1266 | 1688 | 1741 | 1392 | 1310 | 1365 | 1250 | 1457.59 | 1607 | 1308 | 0.813 | -0.298 | 0.00101518 | 0.00732084 | 0.0068 | 0.0041 | 0.0052 | 0.0052 | TRUE | 0.1626 |
| Ddx52 | 5341 | 3675 | 3228 | 3435 | 2991 | 3165 | 4148 | 3909 | 3483 | 3419 | 3374 | 3124 | 3576.31 | 3847 | 3306 | 0.859 | -0.219 | 0.00331942 | 0.01787554 | 0.0046 | 0.0031 | 0.0037 | 0.0037 | TRUE | 0.0829 |
| Ddx54 | 14947 | 11357 | 11395 | 12041 | 11016 | 12803 | 11609 | 12080 | 12295 | 11986 | 12427 | 12639 | 12172.57 | 11995 | 12351 | 1.03 | 0.042 | 0.4007589 | 0.57383761 | 7.00E-04 | 0.0026 | 0.0017 | 0.0017 | TRUE | 0.0098 |
| Ddx55 | 1120 | 802 | 656 | 724 | 665 | 717 | 870 | 853 | 708 | 721 | 750 | 708 | 768.23 | 810 | 726 | 0.895 | -0.16 | 0.09915347 | 0.21977395 | 0.0055 | 0.0055 | 0.0055 | 0.0055 | TRUE | 0.1422 |
| Ddx56 | 6870 | 4846 | 4154 | 4497 | 3980 | 4475 | 5336 | 5155 | 4482 | 4476 | 4490 | 4418 | 4726.01 | 4991 | 4461 | 0.894 | -0.162 | 0.02095602 | 0.07163942 | 0.0038 | 0.0029 | 0.0034 | 0.0034 | TRUE | 0.0964 |
| Ddx58 | 963 | 787 | 673 | 975 | 948 | 1180 | 748 | 837 | 726 | 971 | 1069 | 1165 | 919.33 | 770 | 1068 | 1.387 | 0.472 | 5.70E-07 | 1.82E-05 | 0.0059 | 0.005 | 0.0053 | 0.0053 | TRUE | 0.0777 |
| Ddx59 | 1502 | 1023 | 1067 | 1134 | 995 | 1173 | 1167 | 1088 | 1151 | 1129 | 1122 | 1158 | 1135.86 | 1135 | 1136 | 1 | 0.001 | 0.99272148 | 0.99592003 | 0 | 0.0045 | 0.0029 | 0.0029 | TRUE | 0.0156 |
| Ddx6 | 13911 | 9624 | 9386 | 10686 | 8965 | 10144 | 10804 | 10237 | 10127 | 10637 | 10113 | 10014 | 10322.07 | 10389 | 10255 | 0.987 | -0.019 | 0.71955275 | 0.83168102 | 9.00E-04 | 0.0027 | 0.0019 | 0.0019 | TRUE | 0.015 |
| Ddx60 | 302 | 221 | 327 | 322 | 422 | 440 | 235 | 235 | 353 | 321 | 476 | 434 | 342.23 | 274 | 410 | 1.499 | 0.584 | 0.00101429 | 0.00731812 | 0.0456 | 0.0094 | 0.0197 | 0.0197 | TRUE | 0.6944 |
| Deaf1 | 1589 | 1421 | 1258 | 1024 | 901 | 1115 | 1234 | 1511 | 1357 | 1019 | 1016 | 1101 | 1206.56 | 1367 | 1045 | 0.766 | -0.385 | 1.38E-05 | 0.00024633 | 0.0056 | 0.0044 | 0.0048 | 0.0048 | TRUE | 0.1019 |
| Dear1 | 0 | 0 | 0 | 0 | 0 | 0 | 0 | 0 | 0 | 0 | 0 | 0 | 0 | 0 | 0 | NA | NA | NA | NA | NA | NA | NA | NA | NA | NA |
| Deb1 | 385 | 279 | 215 | 216 | 183 | 234 | 299 | 297 | 232 | 215 | 206 | 231 | 246.7 | 276 | 217 | 0.787 | -0.346 | 0.01638978 | 0.05968387 | 0.008 | 0.012 | 0.0108 | 0.0108 | TRUE | 0.21 |
| Decr1 | 1682 | 1153 | 1194 | 1468 | 1288 | 1390 | 1306 | 1226 | 1288 | 1461 | 1453 | 1372 | 1351.25 | 1273 | 1429 | 1.121 | 0.165 | 0.01875285 | 0.06595335 | 4.00E-04 | 0.0042 | 0.0028 | 0.0028 | TRUE | 0.0147 |
| Decr2 | 1041 | 776 | 728 | 672 | 652 | 672 | 809 | 825 | 786 | 669 | 736 | 663 | 747.87 | 807 | 689 | 0.854 | -0.228 | 0.00757788 | 0.03364767 | 0 | 0.0056 | 0.0039 | 0.0039 | TRUE | 0.0396 |
| Dedd | 1674 | 1199 | 1217 | 1320 | 1200 | 1266 | 1300 | 1275 | 1313 | 1314 | 1354 | 1250 | 1301 | 1296 | 1306 | 1.007 | 0.01 | 0.88590901 | 0.93662974 | 0 | 0.0042 | 0.0027 | 0.0027 | TRUE | 0.017 |
| Dedd2 | 1028 | 871 | 814 | 716 | 667 | 815 | 798 | 926 | 878 | 713 | 752 | 805 | 812.14 | 867 | 757 | 0.873 | -0.195 | 0.03223276 | 0.0975652 | 0.0036 | 0.0053 | 0.0048 | 0.0048 | TRUE | 0.0611 |
| Def6 | 1468 | 1091 | 1091 | 1200 | 1027 | 1165 | 1140 | 1160 | 1177 | 1194 | 1159 | 1150 | 1163.48 | 1159 | 1168 | 1.008 | 0.011 | 0.87331691 | 0.9297124 | 0 | 0.0045 | 0.0027 | 0.0027 | TRUE | 0.0049 |
| Def8 | 5396 | 3869 | 3560 | 3623 | 3257 | 3732 | 4191 | 4115 | 3841 | 3606 | 3674 | 3684 | 3852.01 | 4049 | 3655 | 0.902 | -0.148 | 0.01005551 | 0.04151454 | 9.00E-04 | 0.003 | 0.0021 | 0.0021 | TRUE | 0.0244 |
| Defa-ps1 | 0 | 0 | 0 | 0 | 0 | 0 | 0 | 0 | 0 | 0 | 0 | 0 | 0 | 0 | 0 | NA | NA | NA | NA | NA | NA | NA | NA | NA | NA |
| Defa-ps12 | 0 | 0 | 0 | 0 | 0 | 0 | 0 | 0 | 0 | 0 | 0 | 0 | 0 | 0 | 0 | NA | NA | NA | NA | NA | NA | NA | NA | NA | NA |
| Defa-ps13 | 0 | 0 | 0 | 0 | 0 | 0 | 0 | 0 | 0 | 0 | 0 | 0 | 0 | 0 | 0 | NA | NA | NA | NA | NA | NA | NA | NA | NA | NA |
| Defa-rs1 | 2 | 1 | 0 | 0 | 0 | 0 | 2 | 1 | 0 | 0 | 0 | 0 | 0.44 | 1 | 0 | 0.215 | -2.217 | 0.47268385 | NA | 0 | 5.4363 | 5.3181 | 5.3181 | TRUE | 0.2411 |
| Defa-rs7 | 0 | 0 | 0 | 0 | 0 | 0 | 0 | 0 | 0 | 0 | 0 | 0 | 0 | 0 | 0 | NA | NA | NA | NA | NA | NA | NA | NA | NA | NA |
| Defa17 | 0 | 0 | 0 | 0 | 0 | 0 | 0 | 0 | 0 | 0 | 0 | 0 | 0 | 0 | 0 | NA | NA | NA | NA | NA | NA | NA | NA | NA | NA |
| Defa20 | 0 | 0 | 0 | 0 | 0 | 0 | 0 | 0 | 0 | 0 | 0 | 0 | 0 | 0 | 0 | NA | NA | NA | NA | NA | NA | NA | NA | NA | NA |
| Defa21 | 0 | 0 | 0 | 0 | 0 | 0 | 0 | 0 | 0 | 0 | 0 | 0 | 0 | 0 | 0 | NA | NA | NA | NA | NA | NA | NA | NA | NA | NA |
| Defa22 | 0 | 0 | 0 | 0 | 0 | 0 | 0 | 0 | 0 | 0 | 0 | 0 | 0 | 0 | 0 | NA | NA | NA | NA | NA | NA | NA | NA | NA | NA |
| Defa23 | 0 | 0 | 0 | 0 | 0 | 0 | 0 | 0 | 0 | 0 | 0 | 0 | 0 | 0 | 0 | NA | NA | NA | NA | NA | NA | NA | NA | NA | NA |
| Defa24 | 0 | 0 | 0 | 0 | 0 | 0 | 0 | 0 | 0 | 0 | 0 | 0 | 0 | 0 | 0 | NA | NA | NA | NA | NA | NA | NA | NA | NA | NA |
| Defa25 | 0 | 0 | 0 | 0 | 0 | 0 | 0 | 0 | 0 | 0 | 0 | 0 | 0 | 0 | 0 | NA | NA | NA | NA | NA | NA | NA | NA | NA | NA |
| Defa26 | 0 | 0 | 0 | 0 | 0 | 0 | 0 | 0 | 0 | 0 | 0 | 0 | 0 | 0 | 0 | NA | NA | NA | NA | NA | NA | NA | NA | NA | NA |
| Defa3 | 0 | 0 | 0 | 0 | 0 | 0 | 0 | 0 | 0 | 0 | 0 | 0 | 0 | 0 | 0 | NA | NA | NA | NA | NA | NA | NA | NA | NA | NA |
| Defa4 | 0 | 0 | 0 | 0 | 0 | 0 | 0 | 0 | 0 | 0 | 0 | 0 | 0 | 0 | 0 | NA | NA | NA | NA | NA | NA | NA | NA | NA | NA |
| Defa5 | 0 | 0 | 0 | 0 | 0 | 0 | 0 | 0 | 0 | 0 | 0 | 0 | 0 | 0 | 0 | NA | NA | NA | NA | NA | NA | NA | NA | NA | NA |
| Defa6 | 0 | 0 | 0 | 0 | 0 | 0 | 0 | 0 | 0 | 0 | 0 | 0 | 0 | 0 | 0 | NA | NA | NA | NA | NA | NA | NA | NA | NA | NA |
| Defb1 | 0 | 0 | 1 | 0 | 0 | 0 | 0 | 0 | 1 | 0 | 0 | 0 | 0.18 | 0 | 0 | 0.552 | -0.857 | 0.8336548 | NA | 0 | 13.1817 | 10 | 10 | TRUE | 0.5365 |
| Defb10 | 0 | 0 | 0 | 0 | 0 | 0 | 0 | 0 | 0 | 0 | 0 | 0 | 0 | 0 | 0 | NA | NA | NA | NA | NA | NA | NA | NA | NA | NA |
| Defb11 | 0 | 0 | 0 | 0 | 0 | 0 | 0 | 0 | 0 | 0 | 0 | 0 | 0 | 0 | 0 | NA | NA | NA | NA | NA | NA | NA | NA | NA | NA |
| Defb12 | 0 | 0 | 0 | 0 | 0 | 0 | 0 | 0 | 0 | 0 | 0 | 0 | 0 | 0 | 0 | NA | NA | NA | NA | NA | NA | NA | NA | NA | NA |

| | | | | | | | | | | | | | | | | | | | | | | | | | |
|---|---|---|---|---|---|---|---|---|---|---|---|---|---|---|---|---|---|---|---|---|---|---|---|---|---|
| Defb13 | 0 | 1 | 0 | 0 | 0 | 0 | 0 | 1 | 0 | 0 | 0 | 0 | 0.18 | 0 | 0 | 0.552 | -0.857 | 0.8336548 | NA | 0 | 13.3714 | 10 | 10 | TRUE | 0.5216 |
| Defb14 | 0 | 0 | 0 | 0 | 0 | 0 | 0 | 0 | 0 | 0 | 0 | 0 | 0 | 0 | 0 | NA | NA | NA | NA | NA | NA | NA | NA | NA | NA |
| Defb15 | 0 | 0 | 0 | 0 | 0 | 0 | 0 | 0 | 0 | 0 | 0 | 0 | 0 | 0 | 0 | NA | NA | NA | NA | NA | NA | NA | NA | NA | NA |
| Defb18 | 0 | 0 | 0 | 0 | 0 | 0 | 0 | 0 | 0 | 0 | 0 | 0 | 0 | 0 | 0 | NA | NA | NA | NA | NA | NA | NA | NA | NA | NA |
| Defb19 | 0 | 0 | 0 | 0 | 0 | 0 | 0 | 0 | 0 | 0 | 0 | 0 | 0 | 0 | 0 | NA | NA | NA | NA | NA | NA | NA | NA | NA | NA |
| Defb2 | 0 | 0 | 0 | 0 | 0 | 0 | 0 | 0 | 0 | 0 | 0 | 0 | 0 | 0 | 0 | NA | NA | NA | NA | NA | NA | NA | NA | NA | NA |
| Defb20 | 6 | 4 | 1 | 6 | 2 | 5 | 5 | 4 | 1 | 6 | 2 | 5 | 3.86 | 3 | 4 | 1.304 | 0.383 | 0.70007967 | NA | 0 | 0.6165 | 0.4536 | 0.4536 | TRUE | 0.4452 |
| Defb21 | 0 | 0 | 0 | 0 | 0 | 0 | 0 | 0 | 0 | 0 | 0 | 0 | 0 | 0 | 0 | NA | NA | NA | NA | NA | NA | NA | NA | NA | NA |
| Defb22 | 0 | 0 | 0 | 0 | 0 | 0 | 0 | 0 | 0 | 0 | 0 | 0 | 0 | 0 | 0 | NA | NA | NA | NA | NA | NA | NA | NA | NA | NA |
| Defb23 | 4 | 1 | 0 | 0 | 0 | 0 | 3 | 1 | 0 | 0 | 0 | 0 | 0.7 | 1 | 0 | 0.134 | -2.897 | 0.27044302 | NA | 6.5513 | 3.4123 | 3.6399 | 3.6399 | TRUE | 0.1458 |
| Defb25 | 14 | 11 | 1 | 7 | 8 | 1 | 11 | 12 | 1 | 7 | 9 | 1 | 6.77 | 8 | 6 | 0.706 | -0.503 | 0.58284142 | NA | 0.7392 | 0.3524 | 0.4506 | 0.4506 | TRUE | 1.5167 |
| Defb26 | 0 | 0 | 0 | 0 | 0 | 0 | 0 | 0 | 0 | 0 | 0 | 0 | 0 | 0 | 0 | NA | NA | NA | NA | NA | NA | NA | NA | NA | NA |
| Defb28 | 0 | 0 | 1 | 0 | 0 | 0 | 0 | 0 | 1 | 0 | 0 | 0 | 0.18 | 0 | 0 | 0.552 | -0.857 | 0.8336548 | NA | 0 | 13.1817 | 10 | 10 | TRUE | 0.5365 |
| Defb29 | 3 | 4 | 3 | 0 | 2 | 2 | 2 | 4 | 3 | 0 | 2 | 2 | 2.34 | 3 | 1 | 0.429 | -1.22 | 0.35357125 | NA | 0 | 1.0144 | 0.7266 | 0.7266 | TRUE | 0.5169 |
| Defb3 | 0 | 0 | 0 | 0 | 0 | 0 | 0 | 0 | 0 | 0 | 0 | 0 | 0 | 0 | 0 | NA | NA | NA | NA | NA | NA | NA | NA | NA | NA |
| Defb30 | 40 | 22 | 22 | 24 | 22 | 15 | 31 | 23 | 24 | 24 | 25 | 15 | 23.62 | 26 | 21 | 0.804 | -0.314 | 0.44119667 | 0.61245438 | 0 | 0.1028 | 0.0769 | 0.0769 | TRUE | 0.3993 |
| Defb33 | 0 | 0 | 0 | 0 | 0 | 0 | 0 | 0 | 0 | 0 | 0 | 0 | 0 | 0 | 0 | NA | NA | NA | NA | NA | NA | NA | NA | NA | NA |
| Defb34 | 0 | 0 | 0 | 0 | 0 | 0 | 0 | 0 | 0 | 0 | 0 | 0 | 0 | 0 | 0 | NA | NA | NA | NA | NA | NA | NA | NA | NA | NA |
| Defb35 | 0 | 0 | 0 | 0 | 0 | 0 | 0 | 0 | 0 | 0 | 0 | 0 | 0 | 0 | 0 | NA | NA | NA | NA | NA | NA | NA | NA | NA | NA |
| Defb36 | 6 | 10 | 6 | 4 | 5 | 5 | 5 | 11 | 6 | 4 | 6 | 5 | 6.05 | 7 | 5 | 0.675 | -0.567 | 0.47067103 | NA | 0 | 0.3939 | 0.2716 | 0.2716 | TRUE | 0.4362 |
| Defb37 | 0 | 0 | 0 | 0 | 0 | 0 | 0 | 0 | 0 | 0 | 0 | 0 | 0 | 0 | 0 | NA | NA | NA | NA | NA | NA | NA | NA | NA | NA |
| Defb38 | 0 | 0 | 0 | 0 | 0 | 0 | 0 | 0 | 0 | 0 | 0 | 0 | 0 | 0 | 0 | NA | NA | NA | NA | NA | NA | NA | NA | NA | NA |
| Defb39 | 0 | 0 | 0 | 0 | 0 | 0 | 0 | 0 | 0 | 0 | 0 | 0 | 0 | 0 | 0 | NA | NA | NA | NA | NA | NA | NA | NA | NA | NA |
| Defb4 | 0 | 0 | 0 | 0 | 0 | 0 | 0 | 0 | 0 | 0 | 0 | 0 | 0 | 0 | 0 | NA | NA | NA | NA | NA | NA | NA | NA | NA | NA |
| Defb40 | 0 | 0 | 0 | 0 | 0 | 0 | 0 | 0 | 0 | 0 | 0 | 0 | 0 | 0 | 0 | NA | NA | NA | NA | NA | NA | NA | NA | NA | NA |
| Defb41 | 0 | 0 | 0 | 0 | 0 | 0 | 0 | 0 | 0 | 0 | 0 | 0 | 0 | 0 | 0 | NA | NA | NA | NA | NA | NA | NA | NA | NA | NA |
| Defb42 | 118 | 113 | 125 | 163 | 146 | 166 | 92 | 120 | 135 | 162 | 165 | 164 | 139.59 | 116 | 164 | 1.426 | 0.512 | 0.00542535 | 0.02612686 | 0.0109 | 0.0194 | 0.017 | 0.017 | TRUE | 0.3725 |
| Defb43 | 0 | 0 | 0 | 0 | 0 | 0 | 0 | 0 | 0 | 0 | 0 | 0 | 0 | 0 | 0 | NA | NA | NA | NA | NA | NA | NA | NA | NA | NA |
| Defb44-ps | 0 | 0 | 0 | 0 | 0 | 0 | 0 | 0 | 0 | 0 | 0 | 0 | 0 | 0 | 0 | NA | NA | NA | NA | NA | NA | NA | NA | NA | NA |
| Defb45 | 0 | 0 | 0 | 0 | 0 | 0 | 0 | 0 | 0 | 0 | 0 | 0 | 0 | 0 | 0 | NA | NA | NA | NA | NA | NA | NA | NA | NA | NA |
| Defb46 | 0 | 0 | 0 | 0 | 0 | 0 | 0 | 0 | 0 | 0 | 0 | 0 | 0 | 0 | 0 | NA | NA | NA | NA | NA | NA | NA | NA | NA | NA |
| Defb47 | 1 | 1 | 0 | 3 | 4 | 5 | 1 | 1 | 0 | 3 | 5 | 5 | 2.38 | 1 | 4 | 6.6 | 2.722 | 0.0647184 | NA | 0 | 0.9986 | 0.6768 | 0.6768 | TRUE | 0.1812 |
| Defb48 | 0 | 0 | 0 | 0 | 0 | 0 | 0 | 0 | 0 | 0 | 0 | 0 | 0 | 0 | 0 | NA | NA | NA | NA | NA | NA | NA | NA | NA | NA |
| Defb5 | 0 | 0 | 0 | 0 | 0 | 0 | 0 | 0 | 0 | 0 | 0 | 0 | 0 | 0 | 0 | NA | NA | NA | NA | NA | NA | NA | NA | NA | NA |
| Defb50 | 0 | 0 | 0 | 0 | 0 | 0 | 0 | 0 | 0 | 0 | 0 | 0 | 0 | 0 | 0 | NA | NA | NA | NA | NA | NA | NA | NA | NA | NA |
| Defb6 | 0 | 0 | 0 | 0 | 0 | 0 | 0 | 0 | 0 | 0 | 0 | 0 | 0 | 0 | 0 | NA | NA | NA | NA | NA | NA | NA | NA | NA | NA |
| Defb7 | 0 | 1 | 0 | 0 | 0 | 0 | 0 | 1 | 0 | 0 | 0 | 0 | 0.18 | 0 | 0 | 0.552 | -0.857 | 0.8336548 | NA | 0 | 13.3714 | 10 | 10 | TRUE | 0.5216 |
| Defb8 | 0 | 0 | 0 | 0 | 0 | 0 | 0 | 0 | 0 | 0 | 0 | 0 | 0 | 0 | 0 | NA | NA | NA | NA | NA | NA | NA | NA | NA | NA |
| Defb9 | 0 | 0 | 0 | 0 | 0 | 0 | 0 | 0 | 0 | 0 | 0 | 0 | 0 | 0 | 0 | NA | NA | NA | NA | NA | NA | NA | NA | NA | NA |
| Degs1 | 5933 | 4198 | 4285 | 4570 | 4232 | 4826 | 4608 | 4465 | 4623 | 4549 | 4774 | 4764 | 4630.64 | 4565 | 4696 | 1.028 | 0.04 | 0.44640897 | 0.61713158 | 3.00E-04 | 0.0029 | 0.0018 | 0.0018 | TRUE | 0.0091 |
| Degs2 | 32 | 18 | 23 | 37 | 20 | 33 | 25 | 19 | 25 | 37 | 23 | 33 | 26.8 | 23 | 31 | 1.337 | 0.419 | 0.2723339 | 0.44323256 | 0 | 0.0909 | 0.0673 | 0.0673 | TRUE | 0.3285 |
| Dek | 14250 | 10436 | 8415 | 7644 | 6757 | 7469 | 11067 | 11101 | 9080 | 7609 | 7623 | 7373 | 8975.37 | 10416 | 7535 | 0.723 | -0.467 | 1.06E-09 | 6.99E-08 | 0.0062 | 0.0027 | 0.0041 | 0.0041 | TRUE | 0.1535 |
| Dennd1a | 1238 | 854 | 813 | 798 | 751 | 849 | 962 | 908 | 877 | 794 | 847 | 838 | 871.12 | 916 | 826 | 0.901 | -0.15 | 0.06452489 | 0.16177062 | 0 | 0.0051 | 0.0036 | 0.0036 | TRUE | 0.024 |
| Dennd1b | 2032 | 1511 | 1459 | 1718 | 1413 | 1602 | 1578 | 1607 | 1574 | 1710 | 1594 | 1581 | 1607.53 | 1586 | 1628 | 1.027 | 0.038 | 0.56797609 | 0.72167146 | 4.00E-04 | 0.0039 | 0.0026 | 0.0026 | TRUE | 0.0234 |
| Dennd1c | 93 | 65 | 66 | 53 | 64 | 75 | 72 | 69 | 71 | 53 | 72 | 74 | 68.6 | 71 | 66 | 0.934 | -0.098 | 0.68461161 | 0.80669033 | 0 | 0.037 | 0.0274 | 0.0274 | TRUE | 0.2908 |
| Dennd2a | 2875 | 2103 | 1980 | 2118 | 1910 | 2122 | 2233 | 2237 | 2136 | 2108 | 2155 | 2095 | 2160.65 | 2202 | 2119 | 0.962 | -0.056 | 0.34976097 | 0.52434797 | 0 | 0.0035 | 0.0021 | 0.0021 | TRUE | 0.0081 |
| Dennd2c | 4489 | 3421 | 3874 | 4714 | 4187 | 4923 | 3486 | 3639 | 4180 | 4692 | 4723 | 4860 | 4263.46 | 3768 | 4758 | 1.263 | 0.337 | 3.14E-06 | 7.21E-05 | 0.0045 | 0.003 | 0.0035 | 0.0035 | TRUE | 0.1106 |
| Dennd2d | 101 | 85 | 97 | 107 | 86 | 76 | 78 | 90 | 105 | 107 | 97 | 75 | 92.01 | 91 | 93 | 1.022 | 0.032 | 0.88648015 | 0.93690919 | 0.016 | 0.0282 | 0.0247 | 0.0247 | TRUE | 0.2793 |
| Dennd3 | 909 | 753 | 802 | 910 | 820 | 934 | 706 | 801 | 865 | 906 | 925 | 922 | 854.19 | 791 | 918 | 1.163 | 0.218 | 0.01730223 | 0.06212025 | 0.0042 | 0.0052 | 0.0049 | 0.0049 | TRUE | 0.11 |
| Dennd4a | 2464 | 1858 | 2152 | 2373 | 1987 | 2515 | 1914 | 1976 | 2322 | 2362 | 2242 | 2483 | 2216.39 | 2071 | 2362 | 1.142 | 0.191 | 0.02071587 | 0.07102228 | 0.0063 | 0.0035 | 0.0045 | 0.0045 | TRUE | 0.1354 |
| Dennd4b | 1955 | 1604 | 1786 | 1942 | 1821 | 2097 | 1518 | 1706 | 1927 | 1933 | 2054 | 2070 | 1868.17 | 1717 | 2019 | 1.177 | 0.235 | 0.00692388 | 0.03148598 | 0.007 | 0.0037 | 0.0049 | 0.0049 | TRUE | 0.1379 |
| Dennd4c | 3075 | 2210 | 2021 | 2466 | 2203 | 2620 | 2388 | 2351 | 2181 | 2455 | 2485 | 2586 | 2407.64 | 2307 | 2509 | 1.087 | 0.121 | 0.05853642 | 0.1507769 | 0.0011 | 0.0034 | 0.0025 | 0.0025 | TRUE | 0.0273 |
| Dennd5a | 6481 | 4535 | 4678 | 4936 | 4281 | 4883 | 5034 | 4824 | 5048 | 4913 | 4829 | 4820 | 4911.31 | 4969 | 4854 | 0.977 | -0.034 | 0.51259156 | 0.67504942 | 2.00E-04 | 0.0029 | 0.0017 | 0.0017 | TRUE | 0.0078 |
| Dennd5b | 668 | 422 | 353 | 394 | 371 | 455 | 519 | 449 | 381 | 392 | 419 | 449 | 434.74 | 450 | 420 | 0.931 | -0.102 | 0.41824277 | 0.59082268 | 0.0122 | 0.0079 | 0.0092 | 0.0092 | TRUE | 0.1863 |
| Dennd6a | 4537 | 3219 | 3391 | 3930 | 3221 | 3767 | 3524 | 3424 | 3659 | 3912 | 3634 | 3719 | 3645.12 | 3536 | 3755 | 1.062 | 0.087 | 0.13809916 | 0.27871382 | 9.00E-04 | 0.0031 | 0.0022 | 0.0022 | TRUE | 0.0163 |
| Dennd6b | 470 | 486 | 385 | 358 | 311 | 364 | 365 | 517 | 415 | 356 | 351 | 359 | 393.98 | 432 | 355 | 0.824 | -0.278 | 0.03652949 | 0.10740917 | 0.0139 | 0.0084 | 0.0102 | 0.0102 | TRUE | 0.3368 |
| Denr | 7711 | 5578 | 5071 | 5061 | 4389 | 5299 | 5989 | 5933 | 5472 | 5038 | 4951 | 5231 | 5435.59 | 5798 | 5073 | 0.875 | -0.193 | 0.00094081 | 0.00694221 | 0.0013 | 0.0029 | 0.0023 | 0.0023 | TRUE | 0.0294 |
| Depdc1a | 1379 | 1078 | 1158 | 1120 | 902 | 1048 | 1071 | 1147 | 1249 | 1115 | 1018 | 1035 | 1105.68 | 1156 | 1056 | 0.915 | -0.128 | 0.1257366 | 0.26082056 | 0.0034 | 0.0046 | 0.0042 | 0.0042 | TRUE | 0.0601 |
| Depdc1b | 1852 | 1295 | 1258 | 1373 | 1231 | 1478 | 1438 | 1377 | 1357 | 1367 | 1389 | 1459 | 1397.93 | 1391 | 1405 | 1.009 | 0.013 | 0.84744446 | 0.91346333 | 0 | 0.0041 | 0.0027 | 0.0027 | TRUE | 0.0139 |
| Depdc5 | 5738 | 4064 | 4042 | 4038 | 3705 | 4217 | 4456 | 4323 | 4361 | 4019 | 4180 | 4163 | 4250.41 | 4380 | 4121 | 0.94 | -0.089 | 0.09038943 | 0.20529734 | 0 | 0.003 | 0.0017 | 0.0017 | TRUE | 0.0056 |
| Depdc7 | 1626 | 1034 | 920 | 969 | 850 | 960 | 1263 | 1100 | 993 | 965 | 959 | 948 | 1037.74 | 1119 | 957 | 0.854 | -0.228 | 0.01550216 | 0.05725384 | 0.0067 | 0.0047 | 0.0054 | 0.0054 | TRUE | 0.1573 |
| Deptor | 1635 | 1240 | 1250 | 1607 | 1403 | 1597 | 1270 | 1319 | 1349 | 1600 | 1583 | 1576 | 1449.39 | 1313 | 1586 | 1.21 | 0.274 | 3.49E-05 | 0.00052628 | 0 | 0.0041 | 0.0025 | 0.0025 | TRUE | 0.0102 |
| Dera | 5420 | 4094 | 4199 | 4519 | 4064 | 4544 | 4209 | 4355 | 4531 | 4498 | 4585 | 4486 | 4443.89 | 4365 | 4523 | 1.037 | 0.052 | 0.34257998 | 0.51674488 | 5.00E-04 | 0.003 | 0.0019 | 0.0019 | TRUE | 0.0134 |
| Derl1 | 3299 | 2377 | 2369 | 2620 | 2276 | 2538 | 2562 | 2528 | 2556 | 2608 | 2568 | 2505 | 2554.6 | 2549 | 2560 | 1.004 | 0.006 | 0.91168769 | 0.9513752 | 0 | 0.0034 | 0.0019 | 0.0019 | TRUE | 0.0043 |
| Derl2 | 3006 | 2241 | 2206 | 2456 | 2266 | 2523 | 2335 | 2384 | 2380 | 2445 | 2556 | 2491 | 2431.7 | 2366 | 2497 | 1.055 | 0.078 | 0.17686956 | 0.33067387 | 0 | 0.0034 | 0.002 | 0.002 | TRUE | 0.0051 |
| Derl3 | 374 | 259 | 260 | 382 | 375 | 437 | 290 | 275 | 281 | 380 | 423 | 431 | 346.86 | 282 | 411 | 1.457 | 0.543 | 2.11E-06 | 5.27E-05 | 0 | 0.0093 | 0.0065 | 0.0065 | TRUE | 0.0519 |
| Des | 1147 | 901 | 1142 | 1531 | 1405 | 1603 | 891 | 958 | 1232 | 1524 | 1585 | 1582 | 1295.46 | 1027 | 1564 | 1.525 | 0.609 | 9.85E-09 | 4.92E-07 | 0.0145 | 0.0043 | 0.0073 | 0.0073 | TRUE | 0.3621 |
| Desi1 | 3593 | 2560 | 2550 | 2670 | 2291 | 2772 | 2791 | 2723 | 2751 | 2658 | 2584 | 2736 | 2707.27 | 2755 | 2659 | 0.965 | -0.051 | 0.37416941 | 0.54868199 | 0 | 0.0033 | 0.002 | 0.002 | TRUE | 0.0078 |
| Desi2 | 4280 | 2909 | 3079 | 3584 | 2980 | 3435 | 3324 | 3094 | 3322 | 3568 | 3362 | 3391 | 3343.45 | 3247 | 3440 | 1.059 | 0.083 | 0.16479127 | 0.31552065 | 9.00E-04 | 0.0031 | 0.0023 | 0.0023 | TRUE | 0.0203 |

| | | | | | | | | | | | | | | | | | | | | | | | | | |
|---|---|---|---|---|---|---|---|---|---|---|---|---|---|---|---|---|---|---|---|---|---|---|---|---|---|
| Det1 | 907 | 700 | 706 | 671 | 708 | 831 | 704 | 745 | 762 | 668 | 799 | 820 | 749.62 | 737 | 762 | 1.035 | 0.05 | 0.61276381 | 0.75508227 | 0.0056 | 0.0056 | 0.0056 | 0.0056 | TRUE | 0.1407 |
| Dexi | 229 | 186 | 180 | 203 | 160 | 189 | 178 | 198 | 194 | 202 | 180 | 187 | 189.84 | 190 | 190 | 1.002 | 0.002 | 0.98779614 | 0.99380852 | 0 | 0.0149 | 0.0101 | 0.0101 | TRUE | 0.0372 |
| Dffa | 1407 | 949 | 894 | 1080 | 991 | 1121 | 1093 | 1009 | 965 | 1075 | 1118 | 1107 | 1061.07 | 1022 | 1100 | 1.074 | 0.103 | 0.18873591 | 0.34629041 | 0.0014 | 0.0047 | 0.0035 | 0.0035 | TRUE | 0.0455 |
| Dffb | 383 | 230 | 230 | 258 | 212 | 248 | 297 | 245 | 248 | 257 | 239 | 245 | 255.18 | 263 | 247 | 0.934 | -0.098 | 0.46799422 | 0.6369909 | 0.004 | 0.0117 | 0.0093 | 0.0093 | TRUE | 0.1533 |
| Dfna5 | 118 | 70 | 76 | 71 | 68 | 67 | 92 | 74 | 82 | 71 | 77 | 66 | 76.94 | 83 | 71 | 0.856 | -0.224 | 0.31717391 | 0.49116318 | 0 | 0.0332 | 0.023 | 0.023 | TRUE | 0.0945 |
| Dfnb59 | 3 | 2 | 4 | 0 | 1 | 1 | 2 | 2 | 4 | 0 | 1 | 1 | 1.81 | 3 | 1 | 0.239 | -2.065 | 0.19102188 | NA | 0 | 1.3084 | 0.886 | 0.886 | TRUE | 0.2672 |
| Dgat1 | 1070 | 749 | 623 | 746 | 688 | 738 | 831 | 797 | 672 | 743 | 776 | 729 | 757.86 | 767 | 749 | 0.975 | -0.036 | 0.70849887 | 0.82410273 | 0.0054 | 0.0056 | 0.0055 | 0.0055 | TRUE | 0.135 |
| Dgat2 | 774 | 545 | 491 | 564 | 451 | 528 | 601 | 580 | 530 | 561 | 509 | 521 | 550.34 | 570 | 530 | 0.929 | -0.106 | 0.28093045 | 0.45253453 | 0.0017 | 0.0067 | 0.0051 | 0.0051 | TRUE | 0.0442 |
| Dgat2l6 | 0 | 0 | 0 | 0 | 0 | 0 | 0 | 0 | 0 | 0 | 0 | 0 | 0 | 0 | 0 | NA | NA | NA | NA | NA | NA | NA | NA | NA | NA |
| Dgcr14 | 1866 | 1396 | 1330 | 1302 | 1270 | 1463 | 1449 | 1485 | 1435 | 1296 | 1433 | 1444 | 1423.69 | 1456 | 1391 | 0.955 | -0.067 | 0.3559418 | 0.53038743 | 0.0012 | 0.0041 | 0.003 | 0.003 | TRUE | 0.0433 |
| Dgcr2 | 6670 | 4885 | 5098 | 5360 | 4992 | 5916 | 5180 | 5196 | 5501 | 5335 | 5631 | 5840 | 5447.33 | 5292 | 5602 | 1.059 | 0.082 | 0.15711461 | 0.3056696 | 0.0013 | 0.0029 | 0.0023 | 0.0023 | TRUE | 0.0213 |
| Dgcr6 | 1343 | 976 | 1023 | 1110 | 978 | 1101 | 1043 | 1038 | 1104 | 1105 | 1103 | 1087 | 1080.01 | 1062 | 1098 | 1.035 | 0.05 | 0.49430777 | 0.65952267 | 0 | 0.0046 | 0.0029 | 0.0029 | TRUE | 0.014 |
| Dgcr8 | 3900 | 2794 | 1901 | 1854 | 1562 | 1703 | 3029 | 2972 | 2051 | 1845 | 1762 | 1681 | 2223.46 | 2684 | 1763 | 0.656 | -0.607 | 2.96E-07 | 1.03E-05 | 0.0213 | 0.0035 | 0.0096 | 0.0096 | TRUE | 0.5136 |
| Dgka | 1331 | 1110 | 1035 | 1181 | 1042 | 1202 | 1034 | 1181 | 1117 | 1176 | 1175 | 1187 | 1144.8 | 1111 | 1179 | 1.064 | 0.089 | 0.24919858 | 0.41810537 | 0.0015 | 0.0045 | 0.0034 | 0.0034 | TRUE | 0.0459 |
| Dgkb | 30 | 19 | 7 | 12 | 16 | 9 | 23 | 20 | 8 | 12 | 18 | 9 | 14.99 | 17 | 13 | 0.749 | -0.417 | 0.45458637 | NA | 0.1439 | 0.1605 | 0.156 | 0.156 | TRUE | 1.0824 |
| Dgkd | 7503 | 5389 | 6204 | 6597 | 5917 | 6630 | 5827 | 5732 | 6694 | 6567 | 6675 | 6545 | 6340 | 6084 | 6596 | 1.084 | 0.117 | 0.08273516 | 0.19369545 | 0.0036 | 0.0028 | 0.0031 | 0.0031 | TRUE | 0.0931 |
| Dgke | 2080 | 1580 | 1427 | 1832 | 1412 | 1574 | 1615 | 1681 | 1540 | 1824 | 1593 | 1554 | 1634.34 | 1612 | 1657 | 1.028 | 0.04 | 0.61769169 | 0.75859223 | 0.0041 | 0.0039 | 0.004 | 0.004 | TRUE | 0.0943 |
| Dgkg | 28 | 33 | 28 | 36 | 21 | 28 | 22 | 35 | 30 | 36 | 24 | 28 | 29.04 | 29 | 29 | 1.013 | 0.018 | 0.96104714 | 0.97845108 | 0 | 0.084 | 0.066 | 0.066 | TRUE | 0.3836 |
| Dgkh | 936 | 695 | 836 | 1187 | 999 | 1273 | 727 | 739 | 902 | 1182 | 1127 | 1257 | 988.9 | 789 | 1189 | 1.508 | 0.593 | 8.87E-10 | 6.07E-08 | 0.0076 | 0.0048 | 0.0057 | 0.0057 | TRUE | 0.1818 |
| Dgki | 30 | 37 | 20 | 31 | 27 | 31 | 23 | 39 | 22 | 31 | 30 | 31 | 29.36 | 28 | 31 | 1.097 | 0.134 | 0.71910124 | 0.83149535 | 0.0203 | 0.0832 | 0.0661 | 0.0661 | TRUE | 0.7529 |
| Dgkk | 168 | 116 | 74 | 80 | 75 | 69 | 130 | 123 | 80 | 80 | 85 | 68 | 94.34 | 111 | 78 | 0.692 | -0.531 | 0.02337724 | 0.0774527 | 0.0306 | 0.0275 | 0.0284 | 0.0284 | TRUE | 0.5837 |
| Dgkq | 1155 | 829 | 750 | 767 | 722 | 796 | 897 | 882 | 809 | 763 | 814 | 786 | 825.3 | 863 | 788 | 0.912 | -0.133 | 0.11092592 | 0.23852327 | 8.00E-04 | 0.0053 | 0.0038 | 0.0038 | TRUE | 0.0342 |
| Dgkz | 5175 | 3874 | 3605 | 4254 | 3795 | 4267 | 4019 | 4121 | 3890 | 4234 | 4281 | 4212 | 4126.24 | 4010 | 4242 | 1.058 | 0.081 | 0.12579481 | 0.26082056 | 2.00E-04 | 0.003 | 0.0018 | 0.0018 | TRUE | 0.0082 |
| Dguok | 662 | 497 | 539 | 507 | 507 | 515 | 514 | 529 | 582 | 505 | 572 | 508 | 534.9 | 542 | 528 | 0.976 | -0.035 | 0.73115511 | 0.83927567 | 0.0027 | 0.0069 | 0.0055 | 0.0055 | TRUE | 0.0594 |
| Dhcr24 | 18190 | 12885 | 11868 | 11524 | 10251 | 11950 | 14127 | 13706 | 12805 | 11471 | 11564 | 11796 | 12578.36 | 13546 | 11610 | 0.857 | -0.223 | 3.86E-05 | 0.0005731 | 0.0012 | 0.0026 | 0.002 | 0.002 | TRUE | 0.0279 |
| Dhcr7 | 9540 | 6842 | 7229 | 8927 | 7552 | 8577 | 7409 | 7278 | 7800 | 8886 | 8519 | 8467 | 8059.88 | 7496 | 8624 | 1.151 | 0.202 | 0.00013216 | 0.00153576 | 8.00E-04 | 0.0027 | 0.0019 | 0.0019 | TRUE | 0.0153 |
| Dhdds | 1277 | 969 | 921 | 927 | 843 | 971 | 992 | 1031 | 994 | 923 | 951 | 959 | 974.75 | 1006 | 944 | 0.939 | -0.09 | 0.22383052 | 0.38825359 | 0 | 0.0049 | 0.003 | 0.003 | TRUE | 0.0057 |
| Dhdh | 1040 | 829 | 891 | 976 | 848 | 1019 | 808 | 882 | 961 | 972 | 957 | 1006 | 930.83 | 884 | 978 | 1.109 | 0.149 | 0.08477615 | 0.19648216 | 0.0031 | 0.005 | 0.0043 | 0.0043 | TRUE | 0.0707 |
| Dhfr | 4533 | 3149 | 3042 | 2912 | 2597 | 2877 | 3521 | 3350 | 3282 | 2899 | 2930 | 2840 | 3136.79 | 3384 | 2890 | 0.853 | -0.229 | 7.07E-05 | 0.00092492 | 5.00E-04 | 0.0032 | 0.0021 | 0.0021 | TRUE | 0.0155 |
| Dhh | 62 | 43 | 58 | 76 | 78 | 65 | 48 | 46 | 63 | 76 | 88 | 64 | 64.05 | 52 | 76 | 1.458 | 0.544 | 0.03434519 | 0.10248347 | 0.0099 | 0.0394 | 0.0314 | 0.0314 | TRUE | 0.2024 |
| Dhodh | 4303 | 3431 | 3573 | 3911 | 3356 | 4390 | 3342 | 3650 | 3855 | 3893 | 3786 | 4334 | 3809.87 | 3616 | 4004 | 1.108 | 0.148 | 0.04678618 | 0.12832755 | 0.0048 | 0.003 | 0.0037 | 0.0037 | TRUE | 0.0632 |
| Dhps | 1871 | 1234 | 1236 | 1262 | 1053 | 1177 | 1453 | 1313 | 1334 | 1256 | 1188 | 1162 | 1284.22 | 1367 | 1202 | 0.879 | -0.187 | 0.01353166 | 0.05181034 | 0.0017 | 0.0043 | 0.0033 | 0.0033 | TRUE | 0.0385 |
| Dhrs1 | 1449 | 1040 | 1102 | 1149 | 1086 | 1230 | 1125 | 1106 | 1189 | 1144 | 1225 | 1214 | 1167.28 | 1140 | 1194 | 1.048 | 0.067 | 0.3625227 | 0.53722519 | 5.00E-04 | 0.0045 | 0.003 | 0.003 | TRUE | 0.0166 |
| Dhrs11 | 312 | 232 | 231 | 222 | 208 | 232 | 242 | 247 | 249 | 221 | 235 | 229 | 237.16 | 246 | 228 | 0.927 | -0.109 | 0.39968213 | 0.5730421 | 0 | 0.0124 | 0.0078 | 0.0078 | TRUE | 0.0085 |
| Dhrs13 | 1132 | 803 | 711 | 872 | 723 | 775 | 879 | 854 | 767 | 868 | 816 | 765 | 824.86 | 833 | 816 | 0.978 | -0.032 | 0.7228289 | 0.83413959 | 0.0033 | 0.0053 | 0.0046 | 0.0046 | TRUE | 0.0565 |
| Dhrs2 | 2 | 0 | 1 | 2 | 1 | 3 | 2 | 0 | 1 | 2 | 1 | 3 | 1.45 | 1 | 2 | 2.238 | 1.162 | 0.47239142 | NA | 0 | 1.6346 | 1.1021 | 1.1021 | TRUE | 0.2816 |
| Dhrs3 | 54 | 32 | 35 | 51 | 49 | 46 | 42 | 34 | 38 | 51 | 55 | 45 | 44.2 | 38 | 50 | 1.324 | 0.405 | 0.16353484 | 0.31417234 | 0 | 0.056 | 0.0378 | 0.0378 | TRUE | 0.0754 |
| Dhrs4 | 648 | 389 | 355 | 336 | 313 | 353 | 503 | 414 | 383 | 334 | 353 | 348 | 389.35 | 433 | 345 | 0.794 | -0.333 | 0.00772698 | 0.0341291 | 0.0089 | 0.0085 | 0.0086 | 0.0086 | TRUE | 0.2417 |
| Dhrs7 | 1776 | 1404 | 2197 | 2547 | 2309 | 2572 | 1379 | 1493 | 2371 | 2535 | 2605 | 2539 | 2153.72 | 1748 | 2560 | 1.465 | 0.551 | 0.03285593 | 0.0990725 | 0.0475 | 0.0035 | 0.0146 | 0.0475 | TRUE | 1.1716 |
| Dhrs7b | 3026 | 2170 | 2233 | 2630 | 2314 | 2653 | 2350 | 2308 | 2409 | 2618 | 2610 | 2619 | 2485.83 | 2356 | 2616 | 1.11 | 0.151 | 0.00788711 | 0.03470765 | 0 | 0.0034 | 0.0019 | 0.0019 | TRUE | 0.0046 |
| Dhrs7c | 0 | 0 | 0 | 0 | 0 | 0 | 0 | 0 | 0 | 0 | 0 | 0 | 0 | 0 | 0 | NA | NA | NA | NA | NA | NA | NA | NA | NA | NA |
| Dhrs9 | 0 | 3 | 0 | 0 | 0 | 0 | 0 | 3 | 0 | 0 | 0 | 0 | 0.53 | 1 | 0 | 0.181 | -2.465 | 0.45184603 | NA | 10 | 4.4587 | 6.272 | 6.272 | TRUE | 1.443 |
| Dhrsx | 419 | 321 | 308 | 214 | 192 | 203 | 325 | 341 | 332 | 213 | 217 | 200 | 271.53 | 333 | 210 | 0.631 | -0.665 | 7.96E-08 | 3.18E-06 | 0 | 0.0112 | 0.0071 | 0.0071 | TRUE | 0.0178 |
| Dhtkd1 | 2509 | 1977 | 2004 | 2451 | 2292 | 2668 | 1949 | 2103 | 2162 | 2440 | 2586 | 2634 | 2312.15 | 2071 | 2553 | 1.234 | 0.303 | 6.72E-06 | 0.00013595 | 0.0018 | 0.0035 | 0.0028 | 0.0028 | TRUE | 0.0336 |
| Dhx15 | 22466 | 15834 | 16128 | 16014 | 14176 | 16409 | 17448 | 16842 | 17402 | 15940 | 15992 | 16198 | 16637.22 | 17231 | 16043 | 0.931 | -0.103 | 0.02197293 | 0.0738615 | 1.00E-04 | 0.0026 | 0.0014 | 0.0014 | TRUE | 0.0047 |
| Dhx16 | 9533 | 7052 | 6775 | 6588 | 6050 | 6698 | 7404 | 7501 | 7310 | 6558 | 6825 | 6612 | 7034.97 | 7405 | 6665 | 0.9 | -0.152 | 0.00178936 | 0.01109301 | 1.00E-04 | 0.0028 | 0.0016 | 0.0016 | TRUE | 0.0053 |
| Dhx29 | 5511 | 3930 | 3710 | 3859 | 3399 | 4028 | 4280 | 4180 | 4003 | 3841 | 3834 | 3976 | 4019.24 | 4154 | 3884 | 0.935 | -0.098 | 0.07853307 | 0.1866744 | 6.00E-04 | 0.003 | 0.002 | 0.002 | TRUE | 0.0123 |
| Dhx30 | 10124 | 7486 | 7327 | 8366 | 7609 | 8596 | 7863 | 7963 | 7906 | 8328 | 8584 | 8486 | 8188.04 | 7911 | 8466 | 1.07 | 0.098 | 0.03570431 | 0.10543837 | 0 | 0.0027 | 0.0014 | 0.0014 | TRUE | 0.0025 |
| Dhx32 | 4486 | 3029 | 3120 | 3534 | 3002 | 3437 | 3484 | 3222 | 3366 | 3518 | 3387 | 3393 | 3394.93 | 3357 | 3433 | 1.022 | 0.031 | 0.58941547 | 0.7382741 | 7.00E-04 | 0.0031 | 0.0021 | 0.0021 | TRUE | 0.0151 |
| Dhx33 | 5011 | 3461 | 3058 | 3141 | 2694 | 2974 | 3892 | 3681 | 3300 | 3127 | 3039 | 2936 | 3329.04 | 3624 | 3034 | 0.837 | -0.257 | 0.00031264 | 0.00300055 | 0.0035 | 0.0031 | 0.0034 | 0.0034 | TRUE | 0.0744 |
| Dhx34 | 912 | 714 | 736 | 801 | 747 | 792 | 708 | 759 | 794 | 797 | 843 | 782 | 780.63 | 754 | 807 | 1.072 | 0.1 | 0.24331201 | 0.41108106 | 0.0012 | 0.0055 | 0.004 | 0.004 | TRUE | 0.0353 |
| Dhx35 | 3022 | 2132 | 2154 | 2224 | 1947 | 2325 | 2347 | 2268 | 2324 | 2214 | 2196 | 2295 | 2274.05 | 2313 | 2235 | 0.966 | -0.05 | 0.40152167 | 0.57469951 | 0 | 0.0035 | 0.0021 | 0.0021 | TRUE | 0.0067 |
| Dhx36 | 11027 | 8193 | 8168 | 8369 | 7295 | 8160 | 8564 | 8715 | 8813 | 8330 | 8230 | 8055 | 8451.23 | 8697 | 8205 | 0.943 | -0.084 | 0.07663875 | 0.18308501 | 1.00E-04 | 0.0027 | 0.0015 | 0.0015 | TRUE | 0.0031 |
| Dhx37 | 3321 | 2364 | 1939 | 2061 | 1884 | 2085 | 2579 | 2515 | 2092 | 2052 | 2125 | 2058 | 2236.85 | 2395 | 2078 | 0.867 | -0.206 | 0.01241457 | 0.0485343 | 0.0056 | 0.0035 | 0.0044 | 0.0044 | TRUE | 0.1472 |
| Dhx38 | 7675 | 5469 | 4606 | 4665 | 4243 | 4911 | 5961 | 5817 | 4970 | 4644 | 4787 | 4848 | 5171 | 5583 | 4760 | 0.852 | -0.231 | 0.00153855 | 0.00983036 | 0.0046 | 0.0029 | 0.0036 | 0.0036 | TRUE | 0.1119 |
| Dhx40 | 7145 | 5374 | 5549 | 5785 | 5132 | 5800 | 5549 | 5716 | 5987 | 5758 | 5789 | 5725 | 5754.35 | 5751 | 5757 | 1.001 | 0.002 | 0.96792497 | 0.98270717 | 6.00E-04 | 0.0028 | 0.0018 | 0.0018 | TRUE | 0.0157 |
| Dhx57 | 1838 | 1362 | 1268 | 1260 | 1251 | 1444 | 1427 | 1449 | 1368 | 1254 | 1411 | 1425 | 1389.22 | 1415 | 1363 | 0.963 | -0.054 | 0.47865092 | 0.64580449 | 0.0022 | 0.0041 | 0.0034 | 0.0034 | TRUE | 0.0598 |
| Dhx58 | 147 | 116 | 132 | 184 | 162 | 208 | 114 | 123 | 142 | 183 | 183 | 205 | 158.54 | 126 | 190 | 1.51 | 0.595 | 0.00030687 | 0.00295953 | 0 | 0.0174 | 0.013 | 0.013 | TRUE | 0.1181 |
| Dhx8 | 10975 | 7601 | 6503 | 6764 | 5894 | 6760 | 8524 | 8085 | 7017 | 6733 | 6649 | 6673 | 7280.11 | 7875 | 6685 | 0.849 | -0.237 | 0.00096401 | 0.00706233 | 0.0047 | 0.0028 | 0.0036 | 0.0036 | TRUE | 0.1108 |
| Dhx9 | 32232 | 22223 | 16543 | 16520 | 14657 | 16628 | 25033 | 23638 | 17850 | 16444 | 16535 | 16414 | 19319.05 | 22174 | 16464 | 0.742 | -0.43 | 1.05E-05 | 0.00019453 | 0.0147 | 0.0025 | 0.0068 | 0.0068 | TRUE | 0.3558 |
| Diablo | 2861 | 2153 | 2045 | 2163 | 2059 | 2302 | 2222 | 2290 | 2207 | 2153 | 2323 | 2272 | 2244.49 | 2240 | 2249 | 1.004 | 0.006 | 0.92038252 | 0.95638148 | 5.00E-04 | 0.0035 | 0.0023 | 0.0023 | TRUE | 0.0171 |
| Diap1 | 9681 | 7157 | 7948 | 8519 | 7749 | 8961 | 7519 | 7613 | 8576 | 8480 | 8742 | 8846 | 8295.8 | 7903 | 8689 | 1.1 | 0.137 | 0.02937609 | 0.09119732 | 0.0028 | 0.0027 | 0.0027 | 0.0027 | TRUE | 0.0676 |
| Diap2 | 1944 | 1472 | 1554 | 1644 | 1557 | 1882 | 1510 | 1566 | 1677 | 1636 | 1756 | 1858 | 1667.17 | 1584 | 1750 | 1.106 | 0.145 | 0.0544425 | 0.14306797 | 0.0029 | 0.0038 | 0.0035 | 0.0035 | TRUE | 0.0395 |
| Diap3 | 3042 | 2151 | 2250 | 2494 | 2249 | 2514 | 2363 | 2288 | 2428 | 2483 | 2537 | 2482 | 2429.94 | 2360 | 2501 | 1.06 | 0.084 | 0.15386525 | 0.30092994 | 0 | 0.0034 | 0.0021 | 0.0021 | TRUE | 0.0082 |
| Dicer1 | 13420 | 9349 | 9320 | 9775 | 8720 | 10506 | 10423 | 9944 | 10056 | 9730 | 9837 | 10371 | 10060.25 | 10141 | 9979 | 0.984 | -0.023 | 0.65087204 | 0.78320543 | 7.00E-04 | 0.0027 | 0.0018 | 0.0018 | TRUE | 0.0145 |
| Dido1 | 8578 | 6253 | 5208 | 5885 | 5087 | 5553 | 6662 | 6651 | 5619 | 5858 | 5739 | 5482 | 6001.84 | 6311 | 5693 | 0.902 | -0.149 | 0.0412409 | 0.11683968 | 0.0048 | 0.0028 | 0.0037 | 0.0037 | TRUE | 0.1116 |
| Diexf | 2797 | 1979 | 1721 | 1914 | 1708 | 2065 | 2172 | 2105 | 1857 | 1905 | 1927 | 2038 | 2000.79 | 2045 | 1957 | 0.956 | -0.064 | 0.39291904 | 0.5665031 | 0.0032 | 0.0036 | 0.0036 | 0.0036 | TRUE | 0.0773 |
| Dimt1 | 2098 | 1427 | 1402 | 1463 | 1365 | 1418 | 1629 | 1518 | 1513 | 1456 | 1540 | 1400 | 1509.33 | 1553 | 1465 | 0.942 | -0.086 | 0.22998813 | 0.39566928 | 0.0014 | 0.004 | 0.003 | 0.003 | TRUE | 0.0234 |
| Dio1 | 0 | 1 | 0 | 0 | 1 | 3 | 0 | 1 | 0 | 0 | 1 | 3 | 0.86 | 0 | 1 | 3.963 | 1.987 | 0.3906091 | NA | 0 | 2.7619 | 2.4803 | 2.4803 | TRUE | 0.2997 |

| | | | | | | | | | | | | | | | | | | | | | | | | | |
|---|---|---|---|---|---|---|---|---|---|---|---|---|---|---|---|---|---|---|---|---|---|---|---|---|---|
| Dio2 | 0 | 1 | 0 | 0 | 0 | 0 | 0 | 1 | 0 | 0 | 0 | 0 | 0.18 | 0 | 0 | 0.552 | -0.857 | 0.8336548 | NA | 0 | 13.3714 | 10 | 10 | TRUE | 0.5216 |
| Dio3 | 54 | 33 | 22 | 39 | 24 | 42 | 42 | 35 | 24 | 39 | 27 | 41 | 34.69 | 34 | 36 | 1.06 | 0.084 | 0.81188612 | 0.89135487 | 0.0334 | 0.0707 | 0.0607 | 0.0607 | TRUE | 0.3801 |
| Dio3os | 5 | 4 | 3 | 0 | 2 | 6 | 4 | 4 | 3 | 0 | 2 | 6 | 3.26 | 4 | 3 | 0.721 | -0.472 | 0.68079661 | NA | 0.2419 | 0.7297 | 0.6303 | 0.6303 | TRUE | 0.5103 |
| Dip2a | 2945 | 2219 | 2264 | 2536 | 2243 | 2562 | 2287 | 2360 | 2443 | 2524 | 2530 | 2529 | 2445.7 | 2363 | 2528 | 1.07 | 0.098 | 0.09599482 | 0.21471894 | 0 | 0.0034 | 0.0021 | 0.0021 | TRUE | 0.0104 |
| Dip2b | 5602 | 4022 | 3998 | 4647 | 4262 | 5220 | 4351 | 4278 | 4314 | 4626 | 4808 | 5153 | 4588.22 | 4314 | 4862 | 1.127 | 0.172 | 0.00336781 | 0.01810307 | 0.0012 | 0.0029 | 0.0023 | 0.0023 | TRUE | 0.0336 |
| Dip2c | 2631 | 1834 | 1812 | 2386 | 2100 | 2317 | 2043 | 1951 | 1955 | 2375 | 2369 | 2287 | 2163.43 | 1983 | 2344 | 1.181 | 0.24 | 7.17E-05 | 0.00093338 | 0 | 0.0035 | 0.0022 | 0.0022 | TRUE | 0.0089 |
| Diras1 | 89 | 87 | 58 | 76 | 82 | 70 | 69 | 93 | 63 | 76 | 93 | 69 | 76.92 | 75 | 79 | 1.058 | 0.081 | 0.7375553 | 0.84368771 | 0.0184 | 0.0332 | 0.0291 | 0.0291 | TRUE | 0.3769 |
| Diras2 | 64 | 49 | 32 | 16 | 24 | 23 | 50 | 52 | 35 | 16 | 27 | 23 | 33.68 | 46 | 22 | 0.478 | -1.064 | 0.00319894 | 0.01735791 | 0.0204 | 0.0728 | 0.0595 | 0.0595 | TRUE | 0.3296 |
| Dirc2 | 2104 | 1548 | 1558 | 1784 | 1574 | 1863 | 1634 | 1647 | 1681 | 1776 | 1776 | 1839 | 1725.37 | 1654 | 1797 | 1.087 | 0.12 | 0.05464854 | 0.14350389 | 0 | 0.0038 | 0.0022 | 0.0022 | TRUE | 0.0052 |
| Dis3 | 9226 | 6687 | 6184 | 6051 | 5719 | 6961 | 7165 | 7113 | 6672 | 6023 | 6452 | 6872 | 6716.19 | 6983 | 6449 | 0.923 | -0.115 | 0.07088763 | 0.17307769 | 0.0027 | 0.0028 | 0.0028 | 0.0028 | TRUE | 0.0409 |
| Dis3l | 3334 | 2727 | 2530 | 2142 | 1965 | 2424 | 2589 | 2901 | 2730 | 2132 | 2217 | 2393 | 2493.61 | 2740 | 2247 | 0.821 | -0.285 | 5.92E-05 | 0.00080215 | 0.0028 | 0.0034 | 0.0032 | 0.0032 | TRUE | 0.0393 |
| Dis3l2 | 1393 | 1056 | 1174 | 1492 | 1290 | 1358 | 1082 | 1123 | 1267 | 1485 | 1455 | 1341 | 1292.14 | 1157 | 1427 | 1.234 | 0.304 | 0.000259 | 0.00260109 | 0.0038 | 0.0043 | 0.0042 | 0.0042 | TRUE | 0.0811 |
| Disc1 | 61 | 47 | 28 | 37 | 32 | 24 | 47 | 50 | 30 | 37 | 36 | 24 | 37.37 | 42 | 32 | 0.753 | -0.409 | 0.23304848 | 0.39925229 | 0.0345 | 0.0659 | 0.0575 | 0.0575 | TRUE | 0.4603 |
| Disp1 | 1003 | 719 | 739 | 721 | 662 | 772 | 779 | 765 | 797 | 718 | 747 | 762 | 761.29 | 780 | 742 | 0.951 | -0.072 | 0.37440874 | 0.54897663 | 0 | 0.0055 | 0.0035 | 0.0035 | TRUE | 0.0101 |
| Disp2 | 51 | 21 | 14 | 29 | 28 | 23 | 40 | 22 | 15 | 29 | 32 | 23 | 26.7 | 26 | 28 | 1.06 | 0.084 | 0.84320825 | 0.91102695 | 0.0953 | 0.0912 | 0.0924 | 0.0924 | TRUE | 0.7961 |
| Dixdc1 | 762 | 563 | 690 | 710 | 691 | 755 | 592 | 599 | 745 | 707 | 780 | 745 | 694.45 | 645 | 744 | 1.155 | 0.208 | 0.04925076 | 0.13329694 | 0.008 | 0.0058 | 0.0066 | 0.0066 | TRUE | 0.2104 |
| Dkc1 | 14091 | 10092 | 11027 | 11388 | 10025 | 11525 | 10944 | 10735 | 11898 | 11336 | 11309 | 11377 | 11266.41 | 11192 | 11341 | 1.013 | 0.019 | 0.72840434 | 0.8372403 | 0.0012 | 0.0026 | 0.0021 | 0.0021 | TRUE | 0.037 |
| Dkk1 | 23 | 7 | 10 | 8 | 6 | 10 | 18 | 7 | 11 | 8 | 7 | 10 | 10.12 | 12 | 8 | 0.673 | -0.57 | 0.36240462 | NA | 0 | 0.2367 | 0.1801 | 0.1801 | TRUE | 0.4847 |
| Dkk2 | 0 | 0 | 0 | 0 | 0 | 0 | 0 | 0 | 0 | 0 | 0 | 0 | 0 | 0 | 0 | NA | NA | NA | NA | NA | NA | NA | NA | NA | NA |
| Dkk3 | 786 | 624 | 767 | 1029 | 986 | 975 | 610 | 664 | 828 | 1024 | 1112 | 962 | 866.81 | 701 | 1033 | 1.477 | 0.563 | 5.18E-07 | 1.68E-05 | 0.0139 | 0.0052 | 0.0079 | 0.0079 | TRUE | 0.2947 |
| Dkk4 | 0 | 1 | 1 | 1 | 1 | 1 | 0 | 1 | 1 | 1 | 1 | 1 | 0.88 | 1 | 1 | 1.524 | 0.607 | 0.7620868 | NA | 0 | 2.7093 | 1.693 | 1.693 | TRUE | 0.3841 |
| Dkkl1 | 123 | 72 | 107 | 155 | 137 | 154 | 96 | 77 | 115 | 154 | 155 | 152 | 124.74 | 96 | 154 | 1.603 | 0.681 | 0.00035939 | 0.00334858 | 0.006 | 0.0214 | 0.0178 | 0.0178 | TRUE | 0.2834 |
| Dlat | 7280 | 5086 | 5154 | 6178 | 5002 | 5676 | 5654 | 5410 | 5561 | 6150 | 5643 | 5603 | 5670.09 | 5542 | 5799 | 1.046 | 0.065 | 0.26070879 | 0.43077006 | 0.0014 | 0.0028 | 0.0022 | 0.0022 | TRUE | 0.0343 |
| Dlc1 | 13652 | 10952 | 12050 | 14239 | 12964 | 14280 | 10603 | 11650 | 13002 | 14173 | 14625 | 14097 | 13024.84 | 11752 | 14298 | 1.217 | 0.283 | 8.08E-05 | 0.00101689 | 0.0053 | 0.0026 | 0.0036 | 0.0036 | TRUE | 0.1059 |
| Dld | 8919 | 6280 | 6296 | 6898 | 5934 | 6812 | 6927 | 6680 | 6793 | 6866 | 6694 | 6724 | 6780.87 | 6800 | 6761 | 0.994 | -0.008 | 0.86335286 | 0.92466653 | 1.00E-04 | 0.0028 | 0.0016 | 0.0016 | TRUE | 0.0033 |
| Dlec1 | 65 | 47 | 46 | 56 | 51 | 50 | 50 | 50 | 50 | 56 | 58 | 49 | 52.12 | 50 | 54 | 1.082 | 0.114 | 0.66678392 | 0.79491786 | 0 | 0.0479 | 0.0309 | 0.0309 | TRUE | 0.0524 |
| Dleu2 | 16 | 10 | 12 | 14 | 9 | 8 | 12 | 11 | 13 | 14 | 10 | 8 | 11.33 | 12 | 11 | 0.888 | -0.172 | 0.76157602 | NA | 0 | 0.2116 | 0.1428 | 0.1428 | TRUE | 0.2725 |
| Dleu7 | 56 | 46 | 45 | 56 | 36 | 40 | 43 | 49 | 49 | 56 | 41 | 39 | 46.14 | 47 | 45 | 0.968 | -0.048 | 0.87000898 | 0.92811502 | 0 | 0.0538 | 0.0391 | 0.0391 | TRUE | 0.3275 |
| Dlg1 | 8013 | 5744 | 5799 | 6199 | 5446 | 6327 | 6223 | 6110 | 6257 | 6170 | 6144 | 6246 | 6191.68 | 6197 | 6187 | 0.998 | -0.002 | 0.96062128 | 0.97828661 | 0 | 0.0028 | 0.0015 | 0.0015 | TRUE | 0.0018 |
| Dlg2 | 1 | 2 | 1 | 1 | 0 | 1 | 1 | 2 | 1 | 1 | 0 | 1 | 0.99 | 1 | 1 | 0.516 | -0.956 | 0.61758228 | NA | 0 | 2.3862 | 1.5048 | 1.5048 | TRUE | 0.2008 |
| Dlg3 | 1915 | 1460 | 1453 | 1732 | 1499 | 1855 | 1487 | 1553 | 1568 | 1724 | 1691 | 1831 | 1642.38 | 1536 | 1749 | 1.14 | 0.188 | 0.0052053 | 0.02530588 | 7.00E-04 | 0.0039 | 0.0027 | 0.0027 | TRUE | 0.0206 |
| Dlg4 | 250 | 183 | 211 | 324 | 237 | 391 | 194 | 195 | 228 | 323 | 267 | 386 | 265.39 | 206 | 325 | 1.588 | 0.667 | 1.60E-05 | 0.00028025 | 0.0177 | 0.0114 | 0.0133 | 0.0133 | TRUE | 0.1493 |
| Dlg5 | 4328 | 3161 | 3393 | 3671 | 3420 | 3936 | 3361 | 3362 | 3661 | 3654 | 3858 | 3885 | 3630.4 | 3461 | 3799 | 1.098 | 0.135 | 0.02815247 | 0.08841646 | 0.0014 | 0.0031 | 0.0024 | 0.0024 | TRUE | 0.0306 |
| Dlgap1 | 86 | 70 | 55 | 59 | 61 | 79 | 67 | 74 | 59 | 59 | 69 | 78 | 67.69 | 67 | 69 | 1.025 | 0.035 | 0.88544355 | 0.93634542 | 0 | 0.0374 | 0.0276 | 0.0276 | TRUE | 0.1442 |
| Dlgap2 | 73 | 42 | 35 | 42 | 47 | 48 | 57 | 45 | 38 | 42 | 53 | 47 | 46.89 | 47 | 47 | 1.012 | 0.018 | 0.95229645 | 0.97361754 | 0 | 0.053 | 0.0406 | 0.0406 | TRUE | 0.3451 |
| Dlgap3 | 2219 | 1737 | 1607 | 2110 | 1935 | 2178 | 1723 | 1848 | 1734 | 2100 | 2183 | 2150 | 1956.36 | 1768 | 2144 | 1.213 | 0.279 | 1.11E-05 | 0.00020328 | 4.00E-04 | 0.0036 | 0.0024 | 0.0024 | TRUE | 0.0183 |
| Dlgap4 | 3593 | 2638 | 2694 | 3062 | 2701 | 3222 | 2791 | 2806 | 2907 | 3048 | 3047 | 3181 | 2963.15 | 2835 | 3092 | 1.091 | 0.126 | 0.02636543 | 0.08447458 | 2.00E-04 | 0.0032 | 0.002 | 0.002 | TRUE | 0.0077 |
| Dlgap5 | 9654 | 7116 | 7859 | 8331 | 7426 | 8297 | 7498 | 7569 | 8480 | 8293 | 8377 | 8190 | 8067.87 | 7849 | 8287 | 1.056 | 0.079 | 0.19681239 | 0.35644191 | 0.0023 | 0.0027 | 0.0025 | 0.0025 | TRUE | 0.0601 |
| Dlk1 | 443 | 301 | 242 | 450 | 328 | 372 | 344 | 320 | 261 | 448 | 370 | 367 | 351.75 | 308 | 395 | 1.278 | 0.354 | 0.00931871 | 0.03933618 | 0.0134 | 0.0092 | 0.0105 | 0.0105 | TRUE | 0.199 |
| Dlk2 | 9 | 6 | 7 | 5 | 6 | 9 | 7 | 6 | 8 | 5 | 7 | 9 | 6.93 | 7 | 7 | 0.987 | -0.02 | 0.97819345 | NA | 0 | 0.3446 | 0.2236 | 0.2236 | TRUE | 0.18 |
| Dll1 | 114 | 100 | 165 | 508 | 348 | 418 | 89 | 106 | 178 | 506 | 393 | 413 | 280.64 | 124 | 437 | 3.543 | 1.825 | 2.90E-19 | 7.98E-17 | 0.0674 | 0.0109 | 0.0247 | 0.0247 | TRUE | 1.4307 |
| Dll3 | 216 | 137 | 160 | 195 | 145 | 208 | 168 | 146 | 173 | 194 | 164 | 205 | 174.85 | 162 | 188 | 1.159 | 0.213 | 0.18414821 | 0.34028957 | 0.0045 | 0.016 | 0.0128 | 0.0128 | TRUE | 0.1304 |
| Dll4 | 10 | 10 | 9 | 18 | 23 | 15 | 8 | 11 | 10 | 18 | 26 | 15 | 14.46 | 10 | 20 | 2.095 | 1.067 | 0.04190841 | NA | 0 | 0.1663 | 0.1198 | 0.1198 | TRUE | 0.4015 |
| Dlst | 13097 | 9326 | 9490 | 10740 | 9398 | 10604 | 10172 | 9920 | 10240 | 10691 | 10602 | 10468 | 10348.62 | 10111 | 10587 | 1.047 | 0.066 | 0.15007833 | 0.29566776 | 1.00E-04 | 0.0027 | 0.0014 | 0.0014 | TRUE | 0.0033 |
| Dlx1 | 2 | 9 | 4 | 5 | 4 | 2 | 2 | 10 | 4 | 5 | 5 | 2 | 4.48 | 5 | 4 | 0.756 | -0.404 | 0.6782689 | NA | 0.276 | 0.5309 | 0.4518 | 0.4518 | TRUE | 0.3757 |
| Dlx1as | 0 | 0 | 0 | 0 | 0 | 0 | 0 | 0 | 0 | 0 | 0 | 0 | 0 | 0 | 0 | NA | NA | NA | NA | NA | NA | NA | NA | NA | NA |
| Dlx2 | 23 | 26 | 16 | 11 | 11 | 10 | 18 | 28 | 17 | 11 | 12 | 10 | 16 | 21 | 11 | 0.53 | -0.915 | 0.06478224 | NA | 0 | 0.1505 | 0.107 | 0.107 | TRUE | 0.4156 |
| Dlx3 | 185 | 125 | 143 | 170 | 134 | 161 | 144 | 133 | 154 | 169 | 151 | 159 | 151.71 | 144 | 160 | 1.113 | 0.155 | 0.3390924 | 0.51314427 | 0 | 0.018 | 0.0123 | 0.0123 | TRUE | 0.0407 |
| Dlx4 | 20 | 17 | 19 | 32 | 25 | 33 | 16 | 18 | 21 | 32 | 28 | 33 | 24.46 | 18 | 31 | 1.726 | 0.787 | 0.04316644 | 0.12099979 | 0 | 0.0993 | 0.0656 | 0.0656 | TRUE | 0.0962 |
| Dlx5 | 1 | 1 | 0 | 0 | 1 | 0 | 1 | 1 | 0 | 0 | 1 | 0 | 0.49 | 1 | 0 | 0.599 | -0.74 | 0.78905321 | NA | 0 | 4.7929 | 3.7323 | 3.7323 | TRUE | 0.5075 |
| Dlx6 | 9 | 4 | 1 | 1 | 0 | 0 | 7 | 4 | 1 | 1 | 0 | 0 | 2.22 | 4 | 0 | 0.089 | -3.491 | 0.03802992 | NA | 0 | 1.0701 | 0.9248 | 0.9248 | TRUE | 0.1996 |
| Dlx6as1 | 0 | 0 | 0 | 0 | 0 | 0 | 0 | 0 | 0 | 0 | 0 | 0 | 0 | 0 | 0 | NA | NA | NA | NA | NA | NA | NA | NA | NA | NA |
| Dlx6as2 | 0 | 0 | 0 | 0 | 0 | 0 | 0 | 0 | 0 | 0 | 0 | 0 | 0 | 0 | 0 | NA | NA | NA | NA | NA | NA | NA | NA | NA | NA |
| Dmap1 | 2388 | 1746 | 1727 | 1716 | 1577 | 1853 | 1855 | 1857 | 1863 | 1708 | 1779 | 1829 | 1815.27 | 1858 | 1772 | 0.954 | -0.069 | 0.27615469 | 0.44722484 | 0 | 0.0037 | 0.0023 | 0.0023 | TRUE | 0.0122 |
| Dmbt1 | 2 | 5 | 2 | 2 | 0 | 1 | 2 | 5 | 2 | 2 | 0 | 1 | 2 | 3 | 1 | 0.342 | -1.55 | 0.29145816 | NA | 0 | 1.1867 | 0.8828 | 0.8828 | TRUE | 0.5581 |
| Dmbx1 | 118 | 90 | 111 | 125 | 68 | 81 | 92 | 96 | 120 | 124 | 77 | 80 | 98.04 | 103 | 94 | 0.92 | -0.121 | 0.60715354 | 0.75124163 | 0.0374 | 0.0266 | 0.0297 | 0.0297 | TRUE | 0.8013 |
| Dmc1 | 526 | 409 | 462 | 529 | 451 | 492 | 409 | 435 | 498 | 527 | 509 | 486 | 477.18 | 447 | 507 | 1.136 | 0.184 | 0.08874146 | 0.20281128 | 0.004 | 0.0074 | 0.0063 | 0.0063 | TRUE | 0.1143 |
| Dmd | 2653 | 1919 | 2095 | 2459 | 2138 | 2489 | 2060 | 2041 | 2260 | 2448 | 2412 | 2457 | 2279.8 | 2120 | 2439 | 1.151 | 0.202 | 0.00173659 | 0.01084361 | 0.001 | 0.0035 | 0.0026 | 0.0026 | TRUE | 0.0395 |
| Dmgdh | 86 | 63 | 81 | 113 | 65 | 66 | 67 | 67 | 87 | 112 | 73 | 65 | 78.69 | 74 | 83 | 1.139 | 0.188 | 0.46858865 | 0.63749644 | 0.0433 | 0.0325 | 0.0357 | 0.0357 | TRUE | 0.8678 |
| Dmkn | 10 | 4 | 9 | 7 | 4 | 3 | 8 | 4 | 10 | 7 | 5 | 3 | 6.03 | 7 | 5 | 0.664 | -0.591 | 0.45883018 | NA | 0 | 0.3955 | 0.2851 | 0.2851 | TRUE | 0.3127 |
| Dmp1 | 3 | 0 | 0 | 0 | 1 | 0 | 2 | 0 | 0 | 0 | 1 | 0 | 0.58 | 1 | 0 | 0.457 | -1.13 | 0.69406848 | NA | 0 | 4.1146 | 4.3478 | 4.3478 | TRUE | 1.4398 |
| Dmpk | 124 | 101 | 67 | 68 | 70 | 95 | 96 | 107 | 72 | 68 | 79 | 94 | 86.08 | 92 | 80 | 0.87 | -0.201 | 0.3886879 | 0.56275488 | 0.021 | 0.03 | 0.0275 | 0.0275 | TRUE | 0.3167 |
| Dmrt1 | 3091 | 2396 | 2501 | 2046 | 1965 | 2262 | 2401 | 2549 | 2699 | 2037 | 2217 | 2233 | 2355.68 | 2550 | 2162 | 0.849 | -0.237 | 0.00073934 | 0.00580305 | 0.0023 | 0.0034 | 0.0031 | 0.0031 | TRUE | 0.0324 |
| Dmrt2 | 79 | 43 | 46 | 60 | 44 | 55 | 61 | 46 | 50 | 60 | 50 | 54 | 53.4 | 52 | 55 | 1.039 | 0.055 | 0.8378831 | 0.90753737 | 0 | 0.0468 | 0.0336 | 0.0336 | TRUE | 0.2211 |
| Dmrt3 | 204 | 167 | 170 | 143 | 142 | 121 | 158 | 178 | 183 | 142 | 160 | 119 | 156.91 | 173 | 140 | 0.813 | -0.299 | 0.07880497 | 0.1871029 | 0.0064 | 0.0175 | 0.0144 | 0.0144 | TRUE | 0.1828 |
| Dmrta1 | 13 | 6 | 1 | 14 | 5 | 8 | 10 | 6 | 1 | 14 | 6 | 8 | 7.51 | 6 | 9 | 1.531 | 0.614 | 0.43823593 | NA | 0.3083 | 0.3182 | 0.3158 | 0.3158 | TRUE | 0.4283 |
| Dmrta2 | 175 | 113 | 80 | 119 | 117 | 106 | 136 | 120 | 86 | 118 | 132 | 105 | 116.25 | 114 | 118 | 1.03 | 0.042 | 0.8400374 | 0.90890647 | 0.024 | 0.0228 | 0.0231 | 0.0231 | TRUE | 0.4394 |
| Dmrtb1 | 365 | 304 | 218 | 228 | 211 | 243 | 283 | 323 | 235 | 227 | 238 | 240 | 257.82 | 280 | 235 | 0.837 | -0.257 | 0.07275292 | 0.17648633 | 0.0092 | 0.0116 | 0.0109 | 0.0109 | TRUE | 0.1921 |
| Dmrtc1a | 2 | 0 | 0 | 0 | 0 | 0 | 2 | 0 | 0 | 0 | 0 | 0 | 0.26 | 1 | 0 | 0.356 | -1.491 | 0.71362796 | NA | 0 | 9.1573 | 10 | 10 | TRUE | 0.968 |

| | | | | | | | | | | | | | | | | | | | | | | | | | |
|---|---|---|---|---|---|---|---|---|---|---|---|---|---|---|---|---|---|---|---|---|---|---|---|---|---|
| Dmrtc1b | 169 | 171 | 149 | 91 | 79 | 95 | 131 | 182 | 161 | 91 | 89 | 94 | 124.57 | 158 | 91 | 0.58 | -0.786 | 4.29E-05 | 0.00061859 | 0.0101 | 0.0215 | 0.0179 | 0.0179 | TRUE | 0.2054 |
| Dmrtc1c1 | 0 | 0 | 0 | 0 | 0 | 0 | 0 | 0 | 0 | 0 | 0 | 0 | 0 | 0 | 0 | NA | NA | NA | NA | NA | NA | NA | NA | NA | NA |
| Dmrtc1c2 | 0 | 0 | 0 | 0 | 0 | 0 | 0 | 0 | 0 | 0 | 0 | 0 | 0 | 0 | 0 | NA | NA | NA | NA | NA | NA | NA | NA | NA | NA |
| Dmrtc2 | 95 | 88 | 135 | 193 | 136 | 160 | 74 | 94 | 146 | 192 | 153 | 158 | 136.09 | 105 | 168 | 1.621 | 0.697 | 0.00247125 | 0.01423107 | 0.0596 | 0.0198 | 0.0305 | 0.0305 | TRUE | 1.1655 |
| Dmtf1 | 5672 | 4222 | 4072 | 4241 | 3654 | 4390 | 4405 | 4491 | 4394 | 4221 | 4122 | 4334 | 4327.82 | 4430 | 4226 | 0.954 | -0.068 | 0.19485959 | 0.35420681 | 1.00E-04 | 0.003 | 0.0017 | 0.0017 | TRUE | 0.0061 |
| Dmwd | 3777 | 2960 | 2803 | 3002 | 2716 | 3060 | 2933 | 3149 | 3024 | 2988 | 3064 | 3021 | 3029.86 | 3035 | 3024 | 0.997 | -0.005 | 0.93352226 | 0.96302394 | 4.00E-04 | 0.0032 | 0.002 | 0.002 | TRUE | 0.0129 |
| Dmxl1 | 5887 | 4171 | 4080 | 4697 | 4042 | 4785 | 4572 | 4437 | 4402 | 4675 | 4560 | 4724 | 4561.64 | 4470 | 4653 | 1.041 | 0.058 | 0.26718291 | 0.43769936 | 2.00E-04 | 0.0029 | 0.0017 | 0.0017 | TRUE | 0.0049 |
| Dmxl2 | 4195 | 2991 | 3030 | 3479 | 3077 | 3626 | 3258 | 3181 | 3269 | 3463 | 3471 | 3579 | 3370.42 | 3236 | 3504 | 1.083 | 0.115 | 0.0328383 | 0.09904154 | 0 | 0.0031 | 0.0018 | 0.0018 | TRUE | 0.0043 |
| Dna2 | 5237 | 3692 | 3611 | 3568 | 3053 | 3575 | 4067 | 3927 | 3896 | 3552 | 3444 | 3529 | 3735.91 | 3963 | 3508 | 0.885 | -0.176 | 0.00101392 | 0.00731812 | 0 | 0.0031 | 0.0018 | 0.0018 | TRUE | 0.0065 |
| Dnaaf1 | 44 | 42 | 35 | 37 | 42 | 32 | 34 | 45 | 38 | 37 | 47 | 32 | 38.74 | 39 | 39 | 0.994 | -0.009 | 0.9769023 | 0.98766016 | 0 | 0.0636 | 0.0475 | 0.0475 | TRUE | 0.2733 |
| Dnaaf2 | 1566 | 1175 | 1190 | 1211 | 1098 | 1152 | 1216 | 1250 | 1284 | 1205 | 1239 | 1137 | 1221.89 | 1250 | 1194 | 0.955 | -0.066 | 0.36161501 | 0.53649194 | 5.00E-04 | 0.0044 | 0.003 | 0.003 | TRUE | 0.0207 |
| Dnaaf3 | 462 | 378 | 461 | 503 | 437 | 565 | 359 | 402 | 497 | 501 | 493 | 558 | 468.29 | 419 | 517 | 1.237 | 0.307 | 0.01565663 | 0.05765882 | 0.0134 | 0.0075 | 0.0095 | 0.0095 | TRUE | 0.3029 |
| Dnahc1 | 252 | 216 | 248 | 308 | 288 | 329 | 196 | 230 | 268 | 307 | 325 | 325 | 274.89 | 231 | 319 | 1.386 | 0.471 | 0.00074663 | 0.00585063 | 0.0086 | 0.011 | 0.0103 | 0.0103 | TRUE | 0.2158 |
| Dnahc10 | 269 | 191 | 194 | 268 | 225 | 276 | 209 | 203 | 209 | 267 | 254 | 272 | 235.74 | 207 | 264 | 1.277 | 0.352 | 0.00676052 | 0.03095072 | 0 | 0.0125 | 0.0079 | 0.0079 | TRUE | 0.013 |
| Dnahc11 | 149 | 91 | 63 | 29 | 28 | 36 | 116 | 97 | 68 | 29 | 32 | 36 | 62.75 | 94 | 32 | 0.341 | -1.554 | 6.63E-08 | 2.70E-06 | 0.0342 | 0.0402 | 0.0384 | 0.0384 | TRUE | 0.1514 |
| Dnahc17 | 238 | 197 | 208 | 185 | 194 | 225 | 185 | 210 | 224 | 184 | 219 | 222 | 207.32 | 206 | 208 | 1.013 | 0.018 | 0.90254526 | 0.94610044 | 0.0057 | 0.0139 | 0.0114 | 0.0114 | TRUE | 0.1149 |
| Dnahc2 | 557 | 470 | 458 | 474 | 458 | 489 | 433 | 500 | 494 | 472 | 517 | 483 | 482.99 | 476 | 491 | 1.033 | 0.047 | 0.652775 | 0.78431896 | 0.0025 | 0.0073 | 0.0058 | 0.0058 | TRUE | 0.0779 |
| Dnahc5 | 45 | 37 | 27 | 25 | 17 | 27 | 35 | 39 | 29 | 25 | 19 | 27 | 29.03 | 34 | 24 | 0.686 | -0.545 | 0.13292006 | 0.27105528 | 0 | 0.0841 | 0.0588 | 0.0588 | TRUE | 0.1422 |
| Dnahc6 | 158 | 99 | 96 | 116 | 93 | 107 | 123 | 105 | 104 | 115 | 105 | 106 | 109.6 | 111 | 109 | 0.98 | -0.03 | 0.87573471 | 0.93088377 | 0 | 0.0241 | 0.0167 | 0.0167 | TRUE | 0.103 |
| Dnahc7a | 6 | 9 | 1 | 8 | 1 | 3 | 5 | 10 | 1 | 8 | 1 | 3 | 4.56 | 5 | 4 | 0.796 | -0.33 | 0.74737946 | NA | 0.5596 | 0.5221 | 0.5326 | 0.5326 | TRUE | 0.278 |
| Dnahc7b | 25 | 24 | 16 | 6 | 6 | 8 | 19 | 26 | 17 | 6 | 7 | 8 | 13.81 | 21 | 7 | 0.332 | -1.589 | 0.00387834 | NA | 0 | 0.1741 | 0.1196 | 0.1196 | TRUE | 0.2111 |
| Dnahc8 | 2769 | 2336 | 3094 | 6439 | 5898 | 6700 | 2151 | 2485 | 3338 | 6409 | 6654 | 6614 | 4608.43 | 2658 | 6559 | 2.468 | 1.304 | 1.18E-11 | 1.20E-09 | 0.0264 | 0.0029 | 0.0095 | 0.0264 | TRUE | 0.6082 |
| Dnahc9 | 13 | 7 | 7 | 9 | 8 | 5 | 10 | 7 | 8 | 9 | 9 | 5 | 8 | 8 | 8 | 0.903 | -0.147 | 0.82701494 | NA | 0 | 0.2986 | 0.2016 | 0.2016 | TRUE | 0.284 |
| Dnaic1 | 182 | 143 | 137 | 160 | 141 | 199 | 141 | 152 | 148 | 159 | 159 | 196 | 159.34 | 147 | 171 | 1.169 | 0.226 | 0.16834345 | 0.31985417 | 0 | 0.0173 | 0.0131 | 0.0131 | TRUE | 0.1742 |
| Dnaic2 | 340 | 250 | 309 | 421 | 401 | 396 | 264 | 266 | 333 | 419 | 452 | 391 | 354.29 | 288 | 421 | 1.465 | 0.551 | 1.61E-05 | 0.00028108 | 0.0077 | 0.0091 | 0.0088 | 0.0088 | TRUE | 0.2099 |
| Dnaja1 | 10878 | 7543 | 6228 | 5709 | 4709 | 5687 | 8448 | 8023 | 6720 | 5683 | 5312 | 5614 | 6633.46 | 7730 | 5536 | 0.716 | -0.482 | 2.13E-09 | 1.31E-07 | 0.0072 | 0.0028 | 0.0045 | 0.0045 | TRUE | 0.1593 |
| Dnaja2 | 20189 | 14456 | 14144 | 14847 | 12907 | 14538 | 15680 | 15377 | 15261 | 14779 | 14560 | 14351 | 15001.37 | 15439 | 14563 | 0.943 | -0.084 | 0.06105844 | 0.15523094 | 1.00E-04 | 0.0026 | 0.0014 | 0.0014 | TRUE | 0.0023 |
| Dnaja3 | 6510 | 4812 | 4789 | 5136 | 4508 | 5413 | 5056 | 5118 | 5167 | 5112 | 5085 | 5343 | 5147.19 | 5114 | 5180 | 1.013 | 0.019 | 0.71259374 | 0.82728023 | 2.00E-04 | 0.0029 | 0.0017 | 0.0017 | TRUE | 0.0093 |
| Dnaja4 | 526 | 402 | 375 | 397 | 379 | 457 | 409 | 428 | 405 | 395 | 428 | 451 | 419.1 | 414 | 425 | 1.027 | 0.038 | 0.71626986 | 0.82949644 | 0 | 0.0081 | 0.0057 | 0.0057 | TRUE | 0.0436 |
| Dnajb1 | 4535 | 3457 | 3443 | 2903 | 2591 | 3042 | 3522 | 3677 | 3715 | 2890 | 2923 | 3003 | 3288.29 | 3638 | 2939 | 0.808 | -0.307 | 4.23E-08 | 1.83E-06 | 4.00E-04 | 0.0031 | 0.002 | 0.002 | TRUE | 0.0097 |
| Dnajb11 | 4339 | 3060 | 2652 | 3056 | 2524 | 3013 | 3370 | 3255 | 2861 | 3042 | 2847 | 2974 | 3058.31 | 3162 | 2954 | 0.934 | -0.099 | 0.17433006 | 0.32746741 | 0.0036 | 0.0032 | 0.0035 | 0.0035 | TRUE | 0.0834 |
| Dnajb12 | 1440 | 1134 | 1010 | 895 | 880 | 949 | 1118 | 1206 | 1090 | 891 | 993 | 937 | 1039.14 | 1138 | 940 | 0.826 | -0.276 | 0.00061697 | 0.00507387 | 0.0018 | 0.0047 | 0.0037 | 0.0037 | TRUE | 0.032 |
| Dnajb13 | 105 | 61 | 60 | 116 | 86 | 70 | 82 | 65 | 65 | 115 | 97 | 69 | 82.13 | 71 | 94 | 1.326 | 0.407 | 0.09859431 | 0.21878722 | 0.0318 | 0.0313 | 0.0314 | 0.0314 | TRUE | 0.2632 |
| Dnajb14 | 299 | 232 | 244 | 235 | 168 | 200 | 232 | 247 | 263 | 234 | 190 | 197 | 227.19 | 247 | 207 | 0.839 | -0.253 | 0.07541367 | 0.1809447 | 0.0036 | 0.0129 | 0.0102 | 0.0102 | TRUE | 0.1429 |
| Dnajb2 | 772 | 557 | 547 | 582 | 507 | 605 | 600 | 592 | 590 | 579 | 572 | 597 | 588.46 | 594 | 583 | 0.981 | -0.028 | 0.7544369 | 0.85445957 | 0 | 0.0065 | 0.0039 | 0.0039 | TRUE | 0.0055 |
| Dnajb3 | 73 | 42 | 31 | 40 | 20 | 31 | 57 | 45 | 33 | 40 | 23 | 31 | 37.97 | 45 | 31 | 0.687 | -0.541 | 0.11945253 | 0.25122646 | 0.0458 | 0.0649 | 0.0596 | 0.0596 | TRUE | 0.1637 |
| Dnajb4 | 459 | 346 | 322 | 380 | 321 | 357 | 356 | 368 | 347 | 378 | 362 | 352 | 360.79 | 357 | 364 | 1.02 | 0.028 | 0.79768651 | 0.88189654 | 0 | 0.009 | 0.0058 | 0.0058 | TRUE | 0.0132 |
| Dnajb5 | 904 | 650 | 721 | 897 | 771 | 867 | 702 | 691 | 778 | 893 | 870 | 856 | 798.33 | 724 | 873 | 1.207 | 0.271 | 0.00133361 | 0.00886573 | 8.00E-04 | 0.0054 | 0.0039 | 0.0039 | TRUE | 0.049 |
| Dnajb6 | 5107 | 3609 | 3312 | 3020 | 2758 | 2943 | 3966 | 3839 | 3574 | 3006 | 3111 | 2905 | 3400.24 | 3793 | 3007 | 0.792 | -0.336 | 1.01E-07 | 3.98E-06 | 0.0015 | 0.0031 | 0.0026 | 0.0026 | TRUE | 0.031 |
| Dnajb7 | 0 | 0 | 0 | 0 | 0 | 0 | 0 | 0 | 0 | 0 | 0 | 0 | 0 | 0 | 0 | NA | NA | NA | NA | NA | NA | NA | NA | NA | NA |
| Dnajb8 | 0 | 0 | 0 | 0 | 0 | 0 | 0 | 0 | 0 | 0 | 0 | 0 | 0 | 0 | 0 | NA | NA | NA | NA | NA | NA | NA | NA | NA | NA |
| Dnajb9 | 920 | 700 | 672 | 652 | 531 | 562 | 715 | 745 | 725 | 649 | 599 | 555 | 664.5 | 728 | 601 | 0.826 | -0.276 | 0.00267868 | 0.01510692 | 0.0016 | 0.006 | 0.0046 | 0.0046 | TRUE | 0.0589 |
| Dnajc1 | 1249 | 907 | 962 | 1074 | 941 | 1078 | 970 | 965 | 1038 | 1069 | 1062 | 1064 | 1027.93 | 991 | 1065 | 1.075 | 0.105 | 0.15888652 | 0.30807401 | 0 | 0.0047 | 0.003 | 0.003 | TRUE | 0.02 |
| Dnajc10 | 7911 | 5450 | 5174 | 5166 | 4468 | 4968 | 6144 | 5797 | 5583 | 5142 | 5040 | 4904 | 5435.12 | 5841 | 5029 | 0.861 | -0.217 | 0.0001666 | 0.00185303 | 0.0012 | 0.0029 | 0.0022 | 0.0022 | TRUE | 0.0254 |
| Dnajc11 | 6386 | 4401 | 3496 | 3532 | 3180 | 3418 | 4960 | 4681 | 3772 | 3516 | 3587 | 3374 | 3981.73 | 4471 | 3492 | 0.781 | -0.357 | 6.96E-05 | 0.00091284 | 0.0097 | 0.003 | 0.0056 | 0.0056 | TRUE | 0.2269 |
| Dnajc12 | 265 | 168 | 165 | 195 | 160 | 179 | 206 | 179 | 178 | 194 | 180 | 177 | 185.64 | 188 | 184 | 0.977 | -0.034 | 0.82247667 | 0.89813272 | 0 | 0.0152 | 0.0108 | 0.0108 | TRUE | 0.0858 |
| Dnajc13 | 14070 | 10195 | 9497 | 9971 | 9066 | 10377 | 10928 | 10844 | 10247 | 9925 | 10227 | 10244 | 10402.54 | 10673 | 10132 | 0.949 | -0.075 | 0.13725006 | 0.27759363 | 6.00E-04 | 0.0027 | 0.0018 | 0.0018 | TRUE | 0.0148 |
| Dnajc14 | 2749 | 1936 | 1834 | 1855 | 1567 | 1803 | 2135 | 2059 | 1979 | 1846 | 1768 | 1780 | 1927.87 | 2058 | 1798 | 0.873 | -0.195 | 0.00230386 | 0.01347897 | 6.00E-04 | 0.0037 | 0.0024 | 0.0024 | TRUE | 0.0135 |
| Dnajc15 | 1585 | 1132 | 1023 | 1218 | 985 | 1156 | 1231 | 1204 | 1104 | 1212 | 1111 | 1141 | 1167.27 | 1180 | 1155 | 0.978 | -0.031 | 0.68684262 | 0.80798786 | 0.0018 | 0.0045 | 0.0035 | 0.0035 | TRUE | 0.0373 |
| Dnajc16 | 1456 | 1086 | 1063 | 1073 | 1044 | 1138 | 1131 | 1155 | 1147 | 1068 | 1178 | 1123 | 1133.69 | 1144 | 1123 | 0.981 | -0.027 | 0.71028876 | 0.82554796 | 0 | 0.0045 | 0.003 | 0.003 | TRUE | 0.022 |
| Dnajc17 | 802 | 618 | 569 | 649 | 561 | 635 | 623 | 657 | 614 | 646 | 633 | 627 | 633.32 | 631 | 635 | 1.006 | 0.009 | 0.91481726 | 0.95295409 | 0 | 0.0062 | 0.0039 | 0.0039 | TRUE | 0.0147 |
| Dnajc18 | 1124 | 840 | 901 | 1054 | 860 | 981 | 873 | 893 | 972 | 1049 | 970 | 968 | 954.39 | 913 | 996 | 1.092 | 0.127 | 0.11958304 | 0.25134212 | 0.0016 | 0.0049 | 0.0038 | 0.0038 | TRUE | 0.0378 |
| Dnajc19 | 2034 | 1370 | 1439 | 1797 | 1442 | 1715 | 1580 | 1457 | 1553 | 1789 | 1627 | 1693 | 1616.34 | 1530 | 1703 | 1.113 | 0.154 | 0.02894696 | 0.0902375 | 0.0014 | 0.0039 | 0.003 | 0.003 | TRUE | 0.0235 |
| Dnajc2 | 9310 | 7059 | 6703 | 7167 | 6413 | 7481 | 7231 | 7509 | 7232 | 7134 | 7235 | 7385 | 7287.53 | 7324 | 7251 | 0.99 | -0.014 | 0.77189159 | 0.86503422 | 2.00E-04 | 0.0028 | 0.0016 | 0.0016 | TRUE | 0.0059 |
| Dnajc21 | 7047 | 5436 | 5660 | 5949 | 5227 | 6362 | 5473 | 5782 | 6107 | 5922 | 5897 | 6280 | 5910.15 | 5787 | 6033 | 1.043 | 0.061 | 0.31742207 | 0.49148669 | 0.0019 | 0.0028 | 0.0025 | 0.0025 | TRUE | 0.0285 |
| Dnajc22 | 366 | 278 | 299 | 380 | 379 | 440 | 284 | 296 | 323 | 378 | 428 | 434 | 357.12 | 301 | 413 | 1.376 | 0.461 | 8.10E-05 | 0.00101851 | 0.0023 | 0.0091 | 0.007 | 0.007 | TRUE | 0.0647 |
| Dnajc24 | 830 | 577 | 554 | 615 | 538 | 588 | 645 | 614 | 598 | 612 | 607 | 580 | 609.28 | 619 | 600 | 0.968 | -0.047 | 0.60172422 | 0.74676082 | 0 | 0.0063 | 0.0041 | 0.0041 | TRUE | 0.0168 |
| Dnajc25 | 335 | 231 | 209 | 225 | 173 | 237 | 260 | 246 | 226 | 224 | 195 | 234 | 230.75 | 244 | 218 | 0.892 | -0.164 | 0.23951107 | 0.40642666 | 0.0021 | 0.0127 | 0.0097 | 0.0097 | TRUE | 0.0862 |
| Dnajc27 | 1641 | 1202 | 1142 | 1192 | 1118 | 1287 | 1274 | 1279 | 1232 | 1187 | 1261 | 1270 | 1250.58 | 1262 | 1239 | 0.982 | -0.026 | 0.70871926 | 0.82415787 | 0 | 0.0043 | 0.0028 | 0.0028 | TRUE | 0.0169 |
| Dnajc28 | 232 | 172 | 179 | 183 | 176 | 215 | 180 | 183 | 193 | 182 | 199 | 212 | 191.54 | 185 | 198 | 1.067 | 0.094 | 0.52131394 | 0.68258888 | 0 | 0.0148 | 0.0101 | 0.0101 | TRUE | 0.0525 |
| Dnajc3 | 2947 | 2065 | 2229 | 2797 | 2324 | 2696 | 2289 | 2197 | 2405 | 2784 | 2622 | 2661 | 2492.93 | 2297 | 2689 | 1.171 | 0.228 | 0.00031005 | 0.00298619 | 9.00E-04 | 0.0034 | 0.0025 | 0.0025 | TRUE | 0.02 |
| Dnajc30 | 477 | 346 | 295 | 316 | 264 | 278 | 370 | 368 | 318 | 315 | 298 | 274 | 323.93 | 352 | 296 | 0.838 | -0.255 | 0.03687545 | 0.10809288 | 0.0029 | 0.0097 | 0.0077 | 0.0077 | TRUE | 0.0785 |
| Dnajc4 | 530 | 420 | 475 | 514 | 438 | 528 | 412 | 447 | 513 | 512 | 494 | 521 | 482.98 | 457 | 509 | 1.117 | 0.16 | 0.14169196 | 0.28399988 | 0.0045 | 0.0073 | 0.0064 | 0.0064 | TRUE | 0.1299 |
| Dnajc5 | 6217 | 4680 | 4486 | 4912 | 4330 | 4885 | 4828 | 4978 | 4840 | 4889 | 4885 | 4822 | 4873.87 | 4882 | 4865 | 0.997 | -0.005 | 0.92266857 | 0.95719768 | 0 | 0.0029 | 0.0016 | 0.0016 | TRUE | 0.0036 |
| Dnajc5b | 2 | 2 | 1 | 0 | 0 | 0 | 2 | 2 | 1 | 0 | 0 | 0 | 0.79 | 2 | 0 | 0.12 | -3.059 | 0.17609562 | NA | 0 | 2.9901 | 2.379 | 2.379 | TRUE | 0.0718 |
| Dnajc5g | 147 | 115 | 112 | 124 | 124 | 129 | 114 | 122 | 121 | 123 | 140 | 127 | 124.67 | 119 | 130 | 1.094 | 0.129 | 0.4607321 | 0.63028752 | 0 | 0.0214 | 0.0141 | 0.0141 | TRUE | 0.0412 |
| Dnajc6 | 3931 | 3029 | 3932 | 3703 | 3807 | 4167 | 3053 | 3222 | 4243 | 3686 | 4295 | 4113 | 3768.61 | 3506 | 4031 | 1.15 | 0.202 | 0.0585645 | 0.15077703 | 0.0195 | 0.0031 | 0.008 | 0.008 | TRUE | 0.41 |
| Dnajc7 | 18414 | 13055 | 12915 | 13603 | 12190 | 14376 | 14301 | 13886 | 13935 | 13540 | 13752 | 14191 | 13934.38 | 14041 | 13828 | 0.985 | -0.022 | 0.63786834 | 0.77387365 | 3.00E-04 | 0.0026 | 0.0015 | 0.0015 | TRUE | 0.0065 |
| Dnajc8 | 12702 | 9226 | 9246 | 9643 | 8655 | 10205 | 9865 | 9814 | 9976 | 9599 | 9764 | 10074 | 9848.55 | 9885 | 9812 | 0.993 | -0.011 | 0.8228382 | 0.89832647 | 2.00E-04 | 0.0027 | 0.0015 | 0.0015 | TRUE | 0.0067 |

| | | | | | | | | | | | | | | | | | | | | | | | | | |
|---|---|---|---|---|---|---|---|---|---|---|---|---|---|---|---|---|---|---|---|---|---|---|---|---|---|
| Dnajc9 | 2950 | 2015 | 2071 | 2148 | 1818 | 2142 | 2291 | 2143 | 2235 | 2138 | 2051 | 2114 | 2162.09 | 2223 | 2101 | 0.945 | -0.082 | 0.18415587 | 0.34028957 | 4.00E-04 | 0.0035 | 0.0023 | 0.0023 | TRUE | 0.0119 |
| Dnalc1 | 708 | 536 | 495 | 580 | 421 | 485 | 550 | 570 | 534 | 577 | 475 | 479 | 530.86 | 551 | 510 | 0.926 | -0.111 | 0.29721372 | 0.46928978 | 0.0047 | 0.0069 | 0.0062 | 0.0062 | TRUE | 0.1552 |
| Dnalc4 | 789 | 565 | 562 | 549 | 515 | 553 | 613 | 601 | 606 | 546 | 581 | 546 | 582.25 | 607 | 558 | 0.919 | -0.123 | 0.17141317 | 0.32330542 | 0 | 0.0065 | 0.0041 | 0.0041 | TRUE | 0.015 |
| Dnali1 | 7 | 3 | 6 | 8 | 4 | 7 | 5 | 3 | 6 | 8 | 5 | 7 | 5.75 | 5 | 7 | 1.286 | 0.363 | 0.647243 | NA | 0 | 0.4148 | 0.2782 | 0.2782 | TRUE | 0.1867 |
| Dnase1 | 0 | 0 | 0 | 1 | 0 | 0 | 0 | 0 | 0 | 1 | 0 | 0 | 0.17 | 0 | 0 | 2.094 | 1.067 | 0.79379329 | NA | 0 | 14.2885 | 10 | 10 | TRUE | 0.3906 |
| Dnase1l1 | 52 | 46 | 45 | 56 | 30 | 43 | 40 | 49 | 49 | 56 | 34 | 42 | 44.98 | 46 | 44 | 0.965 | -0.052 | 0.86460347 | 0.92532964 | 0.0129 | 0.0551 | 0.0437 | 0.0437 | TRUE | 0.3538 |
| Dnase1l2 | 7 | 10 | 11 | 6 | 7 | 8 | 5 | 11 | 12 | 6 | 8 | 8 | 8.28 | 9 | 7 | 0.79 | -0.341 | 0.61378171 | NA | 0 | 0.2885 | 0.2042 | 0.2042 | TRUE | 0.5755 |
| Dnase1l3 | 2 | 4 | 3 | 7 | 3 | 7 | 2 | 4 | 3 | 7 | 3 | 7 | 4.38 | 3 | 6 | 1.966 | 0.975 | 0.2976912 | NA | 0 | 0.543 | 0.3787 | 0.3787 | TRUE | 0.3349 |
| Dnase2a | 539 | 324 | 302 | 337 | 295 | 320 | 419 | 345 | 326 | 335 | 333 | 316 | 345.54 | 363 | 328 | 0.9 | -0.152 | 0.23043146 | 0.39614588 | 0.0073 | 0.0093 | 0.0087 | 0.0087 | TRUE | 0.2171 |
| Dnase2b | 0 | 0 | 0 | 0 | 0 | 0 | 0 | 0 | 0 | 0 | 0 | 0 | 0 | 0 | 0 | NA | NA | NA | NA | NA | NA | NA | NA | NA | NA |
| Dnd1 | 1032 | 835 | 854 | 848 | 766 | 890 | 802 | 888 | 921 | 844 | 864 | 879 | 866.32 | 870 | 862 | 0.992 | -0.011 | 0.89730991 | 0.94337995 | 0.0019 | 0.0052 | 0.004 | 0.004 | TRUE | 0.0602 |
| Dner | 118 | 122 | 136 | 170 | 180 | 179 | 92 | 130 | 147 | 169 | 203 | 177 | 152.86 | 123 | 183 | 1.501 | 0.586 | 0.00241044 | 0.01396986 | 0.0255 | 0.0179 | 0.0201 | 0.0201 | TRUE | 0.5571 |
| Dnlz | 544 | 332 | 304 | 339 | 326 | 340 | 423 | 353 | 328 | 337 | 368 | 336 | 357.42 | 368 | 347 | 0.939 | -0.091 | 0.4714939 | 0.6403518 | 0.0078 | 0.0091 | 0.0086 | 0.0086 | TRUE | 0.2042 |
| Dnm1 | 706 | 542 | 631 | 733 | 656 | 723 | 548 | 577 | 681 | 730 | 740 | 714 | 664.84 | 602 | 728 | 1.212 | 0.277 | 0.00558905 | 0.02671713 | 0.005 | 0.006 | 0.0057 | 0.0057 | TRUE | 0.1524 |
| Dnm1l | 10234 | 7239 | 8030 | 8697 | 7561 | 8749 | 7948 | 7700 | 8664 | 8657 | 8530 | 8637 | 8355.97 | 8104 | 8608 | 1.062 | 0.087 | 0.13362962 | 0.27218333 | 0.0018 | 0.0027 | 0.0023 | 0.0023 | TRUE | 0.0443 |
| Dnm2 | 7473 | 5574 | 5148 | 5351 | 4925 | 5631 | 5804 | 5929 | 5555 | 5326 | 5556 | 5559 | 5621.43 | 5763 | 5480 | 0.951 | -0.073 | 0.17573793 | 0.32921205 | 6.00E-04 | 0.0028 | 0.0019 | 0.0019 | TRUE | 0.012 |
| Dnm3 | 19 | 13 | 11 | 6 | 8 | 3 | 15 | 14 | 12 | 6 | 9 | 3 | 9.74 | 14 | 6 | 0.438 | -1.192 | 0.06575198 | NA | 0 | 0.2459 | 0.1792 | 0.1792 | TRUE | 0.3785 |
| Dnm3os | 0 | 0 | 0 | 0 | 0 | 0 | 0 | 0 | 0 | 0 | 0 | 0 | 0 | 0 | 0 | NA | NA | NA | NA | NA | NA | NA | NA | NA | NA |
| Dnmbp | 2077 | 1609 | 1648 | 1543 | 1505 | 1862 | 1613 | 1711 | 1778 | 1536 | 1698 | 1838 | 1695.76 | 1701 | 1691 | 0.995 | -0.008 | 0.92288397 | 0.95719768 | 0.0047 | 0.0038 | 0.0041 | 0.0041 | TRUE | 0.0782 |
| Dnmt1 | 22370 | 17014 | 16833 | 17621 | 16219 | 19198 | 17374 | 18098 | 18163 | 17540 | 18297 | 18951 | 18070.35 | 17878 | 18263 | 1.022 | 0.031 | 0.54848291 | 0.70511479 | 9.00E-04 | 0.0026 | 0.0018 | 0.0018 | TRUE | 0.0147 |
| Dnmt3a | 10505 | 7339 | 6156 | 8113 | 6946 | 8148 | 8159 | 7806 | 6642 | 8076 | 7836 | 8043 | 7760.38 | 7536 | 7985 | 1.059 | 0.083 | 0.2635205 | 0.43387842 | 0.0052 | 0.0027 | 0.0039 | 0.0039 | TRUE | 0.131 |
| Dnmt3b | 6706 | 4921 | 3624 | 3659 | 3243 | 3933 | 5208 | 5234 | 3910 | 3642 | 3658 | 3882 | 4256 | 4784 | 3727 | 0.779 | -0.36 | 0.00020493 | 0.00217125 | 0.0128 | 0.003 | 0.0065 | 0.0065 | TRUE | 0.3099 |
| Dnmt3l | 18463 | 12574 | 12687 | 16152 | 13017 | 15369 | 14339 | 13375 | 13689 | 16078 | 14685 | 15172 | 14556.2 | 13801 | 15312 | 1.109 | 0.15 | 0.00719473 | 0.03240714 | 0.0016 | 0.0026 | 0.0022 | 0.0022 | TRUE | 0.0235 |
| Dnpep | 4213 | 3035 | 3536 | 3818 | 3519 | 3911 | 3272 | 3228 | 3815 | 3800 | 3970 | 3861 | 3657.78 | 3438 | 3877 | 1.128 | 0.174 | 0.01696243 | 0.06131615 | 0.0044 | 0.0031 | 0.0035 | 0.0035 | TRUE | 0.1108 |
| Dnph1 | 3167 | 2352 | 2600 | 2747 | 2485 | 2906 | 2460 | 2502 | 2805 | 2734 | 2803 | 2869 | 2695.54 | 2589 | 2802 | 1.083 | 0.115 | 0.09232774 | 0.2087378 | 0.0024 | 0.0033 | 0.003 | 0.003 | TRUE | 0.0643 |
| Dntt | 2 | 1 | 0 | 1 | 0 | 0 | 2 | 1 | 0 | 1 | 0 | 0 | 0.6 | 1 | 0 | 0.416 | -1.265 | 0.62159733 | NA | 0 | 3.939 | 3.1896 | 3.1896 | TRUE | 0.3906 |
| Dnttip1 | 1343 | 923 | 858 | 963 | 897 | 949 | 1043 | 982 | 926 | 959 | 1012 | 937 | 976.32 | 984 | 969 | 0.984 | -0.024 | 0.76843001 | 0.86304967 | 0.0016 | 0.0049 | 0.0037 | 0.0037 | TRUE | 0.035 |
| Dnttip2 | 8572 | 6059 | 5741 | 6198 | 5658 | 6434 | 6658 | 6445 | 6194 | 6169 | 6383 | 6351 | 6366.75 | 6432 | 6301 | 0.979 | -0.03 | 0.56844172 | 0.72207857 | 6.00E-04 | 0.0028 | 0.0019 | 0.0019 | TRUE | 0.0127 |
| Doc2a | 359 | 287 | 282 | 329 | 285 | 334 | 279 | 305 | 304 | 327 | 322 | 330 | 311.18 | 296 | 326 | 1.104 | 0.143 | 0.2192496 | 0.3823839 | 0 | 0.01 | 0.0065 | 0.0065 | TRUE | 0.0323 |
| Doc2b | 60 | 41 | 17 | 40 | 31 | 25 | 47 | 44 | 18 | 40 | 35 | 25 | 34.67 | 36 | 33 | 0.908 | -0.138 | 0.72356077 | 0.83442425 | 0.1105 | 0.0708 | 0.0816 | 0.0816 | TRUE | 1.2599 |
| Doc2g | 55 | 59 | 73 | 63 | 60 | 51 | 43 | 63 | 79 | 63 | 68 | 50 | 60.83 | 62 | 60 | 0.988 | -0.017 | 0.95205669 | 0.97351168 | 0.043 | 0.0414 | 0.0418 | 0.0418 | TRUE | 0.2716 |
| Dock1 | 4006 | 3021 | 3417 | 2882 | 3067 | 3424 | 3111 | 3213 | 3687 | 2869 | 3460 | 3380 | 3286.71 | 3337 | 3236 | 0.97 | -0.044 | 0.61275038 | 0.75508227 | 0.0089 | 0.0031 | 0.0051 | 0.0051 | TRUE | 0.1202 |
| Dock10 | 35 | 27 | 13 | 29 | 21 | 27 | 27 | 29 | 14 | 29 | 24 | 27 | 24.86 | 23 | 27 | 1.127 | 0.172 | 0.67070832 | 0.79734126 | 0.0217 | 0.0978 | 0.0784 | 0.0784 | TRUE | 0.6525 |
| Dock11 | 1682 | 1206 | 1166 | 1108 | 996 | 1072 | 1306 | 1283 | 1258 | 1103 | 1124 | 1058 | 1188.66 | 1282 | 1095 | 0.853 | -0.229 | 0.0011909 | 0.00821106 | 0 | 0.0044 | 0.0028 | 0.0028 | TRUE | 0.0104 |
| Dock2 | 70 | 43 | 27 | 23 | 30 | 39 | 54 | 46 | 29 | 23 | 34 | 38 | 37.41 | 43 | 32 | 0.731 | -0.452 | 0.20224189 | 0.36364868 | 0.0562 | 0.0658 | 0.0631 | 0.0631 | TRUE | 0.3584 |
| Dock3 | 1628 | 1071 | 911 | 1056 | 926 | 1108 | 1264 | 1139 | 983 | 1051 | 1045 | 1094 | 1096.02 | 1129 | 1063 | 0.94 | -0.089 | 0.349263 | 0.52376624 | 0.0074 | 0.0046 | 0.0056 | 0.0056 | TRUE | 0.1519 |
| Dock4 | 1118 | 850 | 880 | 1006 | 888 | 988 | 868 | 904 | 950 | 1001 | 1002 | 975 | 950.06 | 907 | 993 | 1.095 | 0.131 | 0.08876984 | 0.20283283 | 0 | 0.0049 | 0.0032 | 0.0032 | TRUE | 0.0196 |
| Dock5 | 2409 | 1719 | 1870 | 1932 | 1701 | 1945 | 1871 | 1828 | 2018 | 1923 | 1919 | 1920 | 1913.2 | 1906 | 1921 | 1.008 | 0.012 | 0.85691521 | 0.91964928 | 8.00E-04 | 0.0037 | 0.0026 | 0.0026 | TRUE | 0.0312 |
| Dock6 | 9718 | 8594 | 8686 | 8386 | 8193 | 9698 | 7548 | 9141 | 9372 | 8347 | 9243 | 9573 | 8870.73 | 8687 | 9054 | 1.043 | 0.06 | 0.47467346 | 0.64293333 | 0.0075 | 0.0027 | 0.005 | 0.005 | TRUE | 0.1617 |
| Dock7 | 1945 | 1418 | 1151 | 1265 | 1129 | 1276 | 1511 | 1508 | 1242 | 1259 | 1274 | 1260 | 1342.21 | 1420 | 1264 | 0.889 | -0.169 | 0.05024335 | 0.13529166 | 0.0055 | 0.0042 | 0.0046 | 0.0046 | TRUE | 0.1428 |
| Dock8 | 143 | 108 | 111 | 110 | 115 | 114 | 111 | 115 | 120 | 109 | 130 | 113 | 116.24 | 115 | 117 | 1.017 | 0.025 | 0.89283399 | 0.94109706 | 0 | 0.0228 | 0.0155 | 0.0155 | TRUE | 0.0831 |
| Dock9 | 3269 | 2299 | 2397 | 2477 | 2337 | 2521 | 2539 | 2445 | 2586 | 2466 | 2636 | 2489 | 2526.88 | 2523 | 2530 | 1.002 | 0.003 | 0.9568741 | 0.97620282 | 6.00E-04 | 0.0034 | 0.0023 | 0.0023 | TRUE | 0.0161 |
| Dohh | 5305 | 3761 | 3660 | 4000 | 3604 | 4176 | 4120 | 4001 | 3949 | 3982 | 4066 | 4122 | 4039.91 | 4023 | 4057 | 1.008 | 0.011 | 0.82889773 | 0.90190585 | 2.00E-04 | 0.003 | 0.0018 | 0.0018 | TRUE | 0.0055 |
| Dok1 | 454 | 360 | 327 | 377 | 328 | 414 | 353 | 383 | 353 | 375 | 370 | 409 | 373.72 | 363 | 385 | 1.062 | 0.086 | 0.43272075 | 0.60464148 | 0 | 0.0088 | 0.0061 | 0.0061 | TRUE | 0.0347 |
| Dok2 | 94 | 52 | 40 | 83 | 68 | 74 | 73 | 55 | 43 | 83 | 77 | 73 | 67.31 | 57 | 78 | 1.342 | 0.424 | 0.09953745 | 0.2204352 | 0.0197 | 0.0376 | 0.0327 | 0.0327 | TRUE | 0.4643 |
| Dok3 | 35 | 33 | 21 | 29 | 27 | 32 | 27 | 35 | 23 | 29 | 30 | 32 | 29.31 | 28 | 30 | 1.071 | 0.1 | 0.78072534 | 0.87070126 | 0 | 0.0833 | 0.0582 | 0.0582 | TRUE | 0.2612 |
| Dok4 | 478 | 372 | 391 | 458 | 432 | 538 | 371 | 396 | 422 | 456 | 487 | 531 | 443.86 | 396 | 491 | 1.243 | 0.313 | 0.00393286 | 0.02033732 | 0.003 | 0.0078 | 0.0062 | 0.0062 | TRUE | 0.0593 |
| Dok5 | 13 | 13 | 3 | 12 | 12 | 17 | 10 | 14 | 3 | 12 | 14 | 17 | 11.57 | 9 | 14 | 1.548 | 0.631 | 0.29944591 | NA | 0.0645 | 0.2073 | 0.1765 | 0.1765 | TRUE | 0.5857 |
| Dok6 | 1 | 0 | 0 | 2 | 0 | 0 | 1 | 0 | 0 | 2 | 0 | 0 | 0.46 | 0 | 1 | 1.942 | 0.958 | 0.7580687 | NA | 0 | 5.1409 | 5.1921 | 5.1921 | TRUE | 0.9738 |
| Dok7 | 35 | 31 | 33 | 57 | 42 | 55 | 27 | 33 | 36 | 57 | 47 | 54 | 42.36 | 32 | 53 | 1.668 | 0.738 | 0.01406185 | 0.05325542 | 0 | 0.0584 | 0.0401 | 0.0401 | TRUE | 0.1416 |
| Dolk | 447 | 296 | 220 | 254 | 245 | 215 | 347 | 315 | 237 | 253 | 276 | 212 | 273.48 | 300 | 247 | 0.821 | -0.285 | 0.07687017 | 0.18348453 | 0.024 | 0.0111 | 0.0149 | 0.0149 | TRUE | 0.3668 |
| Dolpp1 | 1582 | 1171 | 1208 | 1387 | 1281 | 1485 | 1229 | 1246 | 1303 | 1381 | 1445 | 1466 | 1344.89 | 1259 | 1431 | 1.137 | 0.185 | 0.00785868 | 0.034594 | 0 | 0.0042 | 0.0027 | 0.0027 | TRUE | 0.0114 |
| Dom3z | 894 | 695 | 535 | 482 | 512 | 521 | 694 | 739 | 577 | 480 | 578 | 514 | 597.09 | 670 | 524 | 0.78 | -0.358 | 0.0017886 | 0.01109301 | 0.0107 | 0.0064 | 0.0077 | 0.0077 | TRUE | 0.1691 |
| Donson | 3689 | 2722 | 2584 | 2781 | 2432 | 2833 | 2865 | 2895 | 2788 | 2768 | 2744 | 2797 | 2809.49 | 2849 | 2770 | 0.972 | -0.041 | 0.45776548 | 0.62749036 | 0 | 0.0033 | 0.0019 | 0.0019 | TRUE | 0.0042 |
| Dopey1 | 1749 | 1210 | 1031 | 1232 | 972 | 1188 | 1358 | 1287 | 1112 | 1226 | 1097 | 1173 | 1208.91 | 1252 | 1165 | 0.929 | -0.106 | 0.2383007 | 0.40514237 | 0.006 | 0.0044 | 0.0049 | 0.0049 | TRUE | 0.1137 |
| Dopey2 | 3438 | 2576 | 2908 | 3439 | 3363 | 3842 | 2670 | 2740 | 3138 | 3423 | 3794 | 3793 | 3259.59 | 2849 | 3670 | 1.289 | 0.366 | 1.74E-06 | 4.51E-05 | 0.0051 | 0.0032 | 0.0039 | 0.0039 | TRUE | 0.0946 |
| Dos | 1056 | 880 | 541 | 706 | 746 | 663 | 820 | 936 | 584 | 703 | 842 | 654 | 756.45 | 780 | 733 | 0.939 | -0.091 | 0.53334582 | 0.69282122 | 0.0353 | 0.0056 | 0.0142 | 0.0142 | TRUE | 0.4879 |
| Dot1l | 7406 | 5584 | 5242 | 5539 | 5066 | 5733 | 5752 | 5940 | 5656 | 5514 | 5715 | 5659 | 5705.91 | 5783 | 5629 | 0.973 | -0.039 | 0.44794933 | 0.61800577 | 2.00E-04 | 0.0028 | 0.0017 | 0.0017 | TRUE | 0.0068 |
| Doxl2 | 0 | 0 | 0 | 0 | 0 | 0 | 0 | 0 | 0 | 0 | 0 | 0 | 0 | 0 | 0 | NA | NA | NA | NA | NA | NA | NA | NA | NA | NA |
| Dpagt1 | 843 | 594 | 579 | 665 | 597 | 694 | 655 | 632 | 625 | 662 | 673 | 685 | 655.3 | 637 | 673 | 1.056 | 0.079 | 0.35458786 | 0.52897654 | 0 | 0.006 | 0.0037 | 0.0037 | TRUE | 0.0074 |
| Dpcd | 994 | 801 | 725 | 772 | 705 | 803 | 772 | 852 | 782 | 768 | 795 | 793 | 793.79 | 802 | 785 | 0.98 | -0.029 | 0.72643608 | 0.83598639 | 0 | 0.0054 | 0.0037 | 0.0037 | TRUE | 0.0346 |
| Dpcr1 | 66 | 48 | 39 | 53 | 48 | 46 | 51 | 51 | 42 | 53 | 54 | 45 | 49.45 | 48 | 51 | 1.051 | 0.071 | 0.7947605 | 0.87990022 | 0 | 0.0504 | 0.0342 | 0.0342 | TRUE | 0.0907 |
| Dpep1 | 2 | 0 | 0 | 2 | 0 | 0 | 2 | 0 | 0 | 2 | 0 | 0 | 0.59 | 1 | 1 | 1.22 | 0.287 | 0.92000925 | NA | 0 | 4.0148 | 4.2481 | 4.2481 | TRUE | 0.9738 |
| Dpep2 | 4 | 4 | 6 | 7 | 7 | 5 | 3 | 4 | 6 | 7 | 8 | 5 | 5.61 | 4 | 7 | 1.449 | 0.535 | 0.50751289 | NA | 0 | 0.4252 | 0.2845 | 0.2845 | TRUE | 0.2219 |
| Dpep3 | 201 | 162 | 197 | 229 | 235 | 236 | 156 | 172 | 213 | 228 | 265 | 233 | 211.17 | 180 | 242 | 1.346 | 0.429 | 0.00609285 | 0.0285525 | 0.0107 | 0.0137 | 0.0128 | 0.0128 | TRUE | 0.2578 |
| Dpf1 | 892 | 713 | 637 | 715 | 587 | 733 | 693 | 758 | 687 | 712 | 662 | 724 | 706 | 713 | 699 | 0.982 | -0.026 | 0.7681462 | 0.86284342 | 0.0011 | 0.0058 | 0.0042 | 0.0042 | TRUE | 0.036 |
| Dpf2 | 5613 | 4130 | 4187 | 4401 | 3838 | 4596 | 4359 | 4393 | 4518 | 4381 | 4330 | 4537 | 4419.59 | 4423 | 4416 | 0.999 | -0.002 | 0.96854623 | 0.9829888 | 2.00E-04 | 0.003 | 0.0018 | 0.0018 | TRUE | 0.007 |
| Dpf3 | 107 | 96 | 67 | 74 | 51 | 61 | 83 | 102 | 72 | 74 | 58 | 60 | 74.82 | 86 | 64 | 0.745 | -0.425 | 0.07615916 | 0.18206102 | 0.0106 | 0.0341 | 0.0276 | 0.0276 | TRUE | 0.2454 |
| Dph1 | 1839 | 1383 | 1225 | 1334 | 1171 | 1391 | 1428 | 1471 | 1322 | 1328 | 1321 | 1373 | 1373.85 | 1407 | 1341 | 0.953 | -0.07 | 0.33013995 | 0.50451089 | 9.00E-04 | 0.0042 | 0.003 | 0.003 | TRUE | 0.0328 |

| | | | | | | | | | | | | | | | | | | | | | | | | | |
|---|---|---|---|---|---|---|---|---|---|---|---|---|---|---|---|---|---|---|---|---|---|---|---|---|---|
| Dph2 | 1103 | 838 | 757 | 737 | 649 | 797 | 857 | 891 | 817 | 734 | 732 | 787 | 802.89 | 855 | 751 | 0.878 | -0.187 | 0.02466789 | 0.08058931 | 0 | 0.0054 | 0.0037 | 0.0037 | TRUE | 0.021 |
| Dph3 | 2366 | 1517 | 1321 | 1379 | 1154 | 1276 | 1838 | 1614 | 1425 | 1373 | 1302 | 1260 | 1468.44 | 1626 | 1312 | 0.806 | -0.312 | 0.0008358 | 0.00634877 | 0.008 | 0.004 | 0.0056 | 0.0056 | TRUE | 0.1603 |
| Dph5 | 1201 | 899 | 957 | 999 | 884 | 973 | 933 | 956 | 1033 | 994 | 997 | 961 | 978.96 | 974 | 984 | 1.011 | 0.016 | 0.83493465 | 0.90619796 | 7.00E-04 | 0.0048 | 0.0034 | 0.0034 | TRUE | 0.0327 |
| Dpm1 | 1836 | 1349 | 1543 | 1783 | 1496 | 1713 | 1426 | 1435 | 1665 | 1775 | 1688 | 1691 | 1613.2 | 1509 | 1718 | 1.14 | 0.189 | 0.01596714 | 0.05850095 | 0.0035 | 0.0039 | 0.0038 | 0.0038 | TRUE | 0.0975 |
| Dpm2 | 1042 | 692 | 534 | 557 | 524 | 547 | 809 | 736 | 576 | 554 | 591 | 540 | 634.51 | 707 | 562 | 0.793 | -0.335 | 0.00543203 | 0.02614143 | 0.0147 | 0.0062 | 0.0089 | 0.0089 | TRUE | 0.3063 |
| Dpm3 | 613 | 439 | 536 | 655 | 538 | 636 | 476 | 467 | 578 | 652 | 607 | 628 | 568.02 | 507 | 629 | 1.243 | 0.313 | 0.00308096 | 0.01687857 | 0.0054 | 0.0066 | 0.0063 | 0.0063 | TRUE | 0.171 |
| Dpp10 | 4 | 2 | 2 | 6 | 0 | 2 | 3 | 2 | 2 | 6 | 0 | 2 | 2.56 | 2 | 3 | 1.08 | 0.111 | 0.93060479 | NA | 0.3289 | 0.9295 | 0.7821 | 0.7821 | TRUE | 0.4822 |
| Dpp3 | 8690 | 6332 | 6283 | 6542 | 5917 | 6858 | 6749 | 6735 | 6779 | 6512 | 6675 | 6770 | 6703.42 | 6754 | 6652 | 0.985 | -0.022 | 0.64730665 | 0.78051363 | 0 | 0.0028 | 0.0015 | 0.0015 | TRUE | 0.0042 |
| Dpp4 | 2524 | 1787 | 1503 | 1457 | 1222 | 1379 | 1960 | 1901 | 1622 | 1450 | 1379 | 1361 | 1612.16 | 1828 | 1397 | 0.764 | -0.389 | 2.82E-06 | 6.56E-05 | 0.0051 | 0.0039 | 0.0043 | 0.0043 | TRUE | 0.116 |
| Dpp6 | 66 | 70 | 67 | 49 | 66 | 53 | 51 | 74 | 72 | 49 | 74 | 52 | 62.26 | 66 | 58 | 0.889 | -0.17 | 0.53601993 | 0.69493874 | 0.0315 | 0.0405 | 0.0379 | 0.0379 | TRUE | 0.4591 |
| Dpp7 | 762 | 591 | 632 | 568 | 495 | 594 | 592 | 629 | 682 | 565 | 558 | 586 | 602.09 | 634 | 570 | 0.901 | -0.151 | 0.10991573 | 0.23714529 | 0.0015 | 0.0064 | 0.0047 | 0.0047 | TRUE | 0.0508 |
| Dpp8 | 2082 | 1538 | 1458 | 1698 | 1504 | 1628 | 1617 | 1636 | 1573 | 1690 | 1697 | 1607 | 1636.68 | 1609 | 1665 | 1.034 | 0.049 | 0.45100063 | 0.62101657 | 0 | 0.0039 | 0.0024 | 0.0024 | TRUE | 0.0111 |
| Dpp9 | 3029 | 2116 | 1982 | 2151 | 1922 | 2338 | 2352 | 2251 | 2139 | 2141 | 2168 | 2308 | 2226.52 | 2247 | 2206 | 0.981 | -0.028 | 0.67690881 | 0.80191112 | 0.0015 | 0.0035 | 0.0027 | 0.0027 | TRUE | 0.0216 |
| Dppa1 | 14 | 2 | 4 | 0 | 0 | 0 | 11 | 2 | 4 | 0 | 0 | 0 | 2.89 | 6 | 0 | 0.032 | -4.944 | 0.00226966 | NA | 0.7227 | 0.8236 | 0.808 | 0.808 | TRUE | 0.3067 |
| Dppa2 | 7012 | 4520 | 2992 | 3030 | 2659 | 3060 | 5446 | 4808 | 3228 | 3016 | 3000 | 3021 | 3753.09 | 4494 | 3012 | 0.67 | -0.577 | 0.0062467 | 0.02906383 | 0.0319 | 0.0031 | 0.0122 | 0.0319 | TRUE | 0.5044 |
| Dppa3 | 1243 | 989 | 1163 | 308 | 335 | 398 | 965 | 1052 | 1255 | 307 | 378 | 393 | 724.94 | 1091 | 359 | 0.329 | -1.602 | 1.83E-39 | 2.91E-36 | 0.016 | 0.0057 | 0.0088 | 0.0088 | TRUE | 0.2065 |
| Dppa4 | 7858 | 5654 | 4988 | 5478 | 4862 | 5490 | 6103 | 6014 | 5382 | 5453 | 5485 | 5419 | 5642.7 | 5833 | 5452 | 0.934 | -0.098 | 0.11323216 | 0.24206212 | 0.002 | 0.0028 | 0.0026 | 0.0026 | TRUE | 0.0555 |
| Dppa5a | 60945 | 41970 | 49764 | 55485 | 48540 | 54870 | 47333 | 44643 | 53695 | 55230 | 54758 | 54165 | 51637.38 | 48557 | 54718 | 1.127 | 0.172 | 0.01080946 | 0.04374283 | 0.0046 | 0.0025 | 0.0033 | 0.0033 | TRUE | 0.1048 |
| Dpt | 1 | 0 | 0 | 0 | 1 | 0 | 1 | 0 | 0 | 0 | 1 | 0 | 0.32 | 0 | 0 | 1.075 | 0.105 | 0.97638847 | NA | 0 | 7.4681 | 7.0392 | 7.0392 | TRUE | 0.5075 |
| Dpy19l1 | 6812 | 5013 | 5194 | 5118 | 4503 | 5079 | 5291 | 5332 | 5604 | 5094 | 5080 | 5014 | 5235.87 | 5409 | 5063 | 0.936 | -0.095 | 0.06721443 | 0.16635426 | 4.00E-04 | 0.0029 | 0.0018 | 0.0018 | TRUE | 0.0121 |
| Dpy19l2 | 2 | 3 | 3 | 2 | 2 | 1 | 2 | 3 | 3 | 2 | 2 | 1 | 2.2 | 3 | 2 | 0.662 | -0.595 | 0.63892692 | NA | 0 | 1.0785 | 0.6774 | 0.6774 | TRUE | 0.2138 |
| Dpy19l3 | 1524 | 1129 | 1055 | 1261 | 1154 | 1305 | 1184 | 1201 | 1138 | 1255 | 1302 | 1288 | 1228.02 | 1174 | 1282 | 1.091 | 0.126 | 0.07069033 | 0.17280261 | 0 | 0.0044 | 0.0027 | 0.0027 | TRUE | 0.0083 |
| Dpy19l4 | 1383 | 957 | 1100 | 1337 | 1125 | 1271 | 1074 | 1018 | 1187 | 1331 | 1269 | 1255 | 1188.93 | 1093 | 1285 | 1.176 | 0.234 | 0.00342841 | 0.01835841 | 0.0023 | 0.0044 | 0.0038 | 0.0038 | TRUE | 0.0654 |
| Dpy30 | 5520 | 3872 | 3968 | 3956 | 3502 | 4163 | 4287 | 4119 | 4281 | 3938 | 3951 | 4110 | 4114.19 | 4229 | 4000 | 0.946 | -0.081 | 0.13316119 | 0.27146954 | 3.00E-04 | 0.003 | 0.0018 | 0.0018 | TRUE | 0.0071 |
| Dpyd | 40 | 31 | 36 | 30 | 22 | 21 | 31 | 33 | 39 | 30 | 25 | 21 | 29.72 | 34 | 25 | 0.735 | -0.443 | 0.21473519 | 0.37747018 | 0 | 0.0822 | 0.0573 | 0.0573 | TRUE | 0.172 |
| Dpys | 112 | 100 | 115 | 122 | 96 | 120 | 87 | 106 | 124 | 121 | 108 | 118 | 110.94 | 106 | 116 | 1.105 | 0.144 | 0.47165287 | 0.64050684 | 0.0093 | 0.0238 | 0.0196 | 0.0196 | TRUE | 0.2541 |
| Dpysl2 | 686 | 533 | 563 | 727 | 609 | 758 | 533 | 567 | 607 | 724 | 687 | 748 | 644.36 | 569 | 720 | 1.268 | 0.342 | 0.00021938 | 0.00229043 | 0.0015 | 0.0061 | 0.0046 | 0.0046 | TRUE | 0.0406 |
| Dpysl3 | 776 | 682 | 649 | 728 | 829 | 867 | 603 | 725 | 700 | 725 | 935 | 856 | 757.35 | 676 | 839 | 1.242 | 0.312 | 0.00495459 | 0.02436048 | 0.0118 | 0.0056 | 0.0076 | 0.0076 | TRUE | 0.1684 |
| Dpysl4 | 249 | 188 | 168 | 170 | 141 | 170 | 193 | 200 | 181 | 169 | 159 | 168 | 178.45 | 191 | 165 | 0.864 | -0.212 | 0.15329597 | 0.30015187 | 0 | 0.0157 | 0.0102 | 0.0102 | TRUE | 0.0223 |
| Dpysl5 | 1675 | 1380 | 1393 | 1365 | 1360 | 1609 | 1301 | 1468 | 1503 | 1359 | 1534 | 1588 | 1458.85 | 1424 | 1494 | 1.05 | 0.071 | 0.41085314 | 0.58367199 | 0.0057 | 0.0041 | 0.0046 | 0.0046 | TRUE | 0.076 |
| DQ267100 | 1 | 1 | 1 | 3 | 4 | 1 | 1 | 1 | 1 | 3 | 5 | 1 | 1.9 | 1 | 3 | 2.917 | 1.545 | 0.28614718 | NA | 0 | 1.2493 | 0.8269 | 0.8269 | TRUE | 0.1422 |
| DQ267101 | 0 | 2 | 0 | 1 | 0 | 0 | 0 | 2 | 0 | 1 | 0 | 0 | 0.52 | 1 | 0 | 0.534 | -0.904 | 0.76303124 | NA | 0 | 4.5562 | 4.7774 | 4.7774 | TRUE | 0.9746 |
| DQ267102 | 5 | 3 | 9 | 8 | 6 | 5 | 4 | 3 | 10 | 8 | 7 | 5 | 6.08 | 6 | 7 | 1.186 | 0.247 | 0.7575244 | NA | 0 | 0.3925 | 0.2934 | 0.2934 | TRUE | 0.8424 |
| Dqx1 | 101 | 73 | 59 | 96 | 71 | 79 | 78 | 78 | 64 | 96 | 80 | 78 | 78.9 | 73 | 85 | 1.152 | 0.204 | 0.36384924 | 0.53841025 | 0 | 0.0325 | 0.0236 | 0.0236 | TRUE | 0.1247 |
| Dr1 | 5491 | 3907 | 3526 | 3766 | 3244 | 3719 | 4265 | 4156 | 3805 | 3749 | 3660 | 3671 | 3884.07 | 4075 | 3693 | 0.906 | -0.143 | 0.02032484 | 0.07001757 | 0.0014 | 0.003 | 0.0025 | 0.0025 | TRUE | 0.0407 |
| Dram1 | 124 | 72 | 68 | 77 | 49 | 75 | 96 | 77 | 73 | 77 | 55 | 74 | 75.37 | 82 | 69 | 0.834 | -0.261 | 0.27562292 | 0.44669297 | 0.013 | 0.0339 | 0.028 | 0.028 | TRUE | 0.2434 |
| Dram2 | 1341 | 932 | 1129 | 1280 | 1180 | 1214 | 1041 | 991 | 1218 | 1274 | 1331 | 1198 | 1175.79 | 1083 | 1268 | 1.17 | 0.227 | 0.01212951 | 0.04772009 | 0.0061 | 0.0044 | 0.005 | 0.005 | TRUE | 0.138 |
| Drap1 | 2711 | 2018 | 2061 | 2033 | 1964 | 2280 | 2106 | 2147 | 2224 | 2024 | 2216 | 2251 | 2160.97 | 2159 | 2164 | 1.002 | 0.003 | 0.95861295 | 0.9773909 | 0.0016 | 0.0035 | 0.0028 | 0.0028 | TRUE | 0.0389 |
| Draxin | 161 | 116 | 89 | 91 | 98 | 121 | 125 | 123 | 96 | 91 | 111 | 119 | 110.84 | 115 | 107 | 0.928 | -0.108 | 0.59113279 | 0.73926512 | 0.0111 | 0.0238 | 0.0203 | 0.0203 | TRUE | 0.1961 |
| Drd1a | 1 | 2 | 2 | 10 | 4 | 5 | 1 | 2 | 2 | 10 | 5 | 5 | 4.08 | 2 | 7 | 3.981 | 1.993 | 0.05774433 | NA | 0 | 0.5837 | 0.4205 | 0.4205 | TRUE | 0.565 |
| Drd2 | 31 | 30 | 23 | 30 | 25 | 27 | 24 | 32 | 25 | 30 | 28 | 27 | 27.59 | 27 | 28 | 1.053 | 0.074 | 0.83874673 | 0.9080766 | 0 | 0.0883 | 0.0589 | 0.0589 | TRUE | 0.1569 |
| Drd3 | 16 | 7 | 2 | 2 | 4 | 4 | 12 | 7 | 2 | 2 | 5 | 4 | 5.41 | 7 | 4 | 0.463 | -1.11 | 0.22195641 | NA | 0 | 0.4402 | 0.3813 | 0.3813 | TRUE | 0.1106 |
| Drd4 | 54 | 34 | 35 | 52 | 51 | 46 | 42 | 36 | 38 | 52 | 58 | 45 | 45.1 | 39 | 52 | 1.328 | 0.409 | 0.15495192 | 0.3024758 | 0 | 0.055 | 0.0371 | 0.0371 | TRUE | 0.0909 |
| Drd5 | 27 | 22 | 23 | 13 | 20 | 16 | 21 | 23 | 25 | 13 | 23 | 16 | 20.08 | 23 | 17 | 0.74 | -0.435 | 0.31749292 | 0.49148985 | 0 | 0.1205 | 0.0852 | 0.0852 | TRUE | 0.3507 |
| Drg1 | 9623 | 6941 | 6465 | 6849 | 5950 | 6756 | 7474 | 7383 | 6976 | 6817 | 6712 | 6669 | 7005.24 | 7278 | 6733 | 0.925 | -0.112 | 0.03020086 | 0.09309055 | 5.00E-04 | 0.0028 | 0.0018 | 0.0018 | TRUE | 0.016 |
| Drg2 | 4931 | 3369 | 3132 | 3132 | 2715 | 3333 | 3830 | 3584 | 3379 | 3118 | 3063 | 3290 | 3377.21 | 3598 | 3157 | 0.877 | -0.189 | 0.00417008 | 0.02132485 | 0.0022 | 0.0031 | 0.0028 | 0.0028 | TRUE | 0.0395 |
| Drosha | 16347 | 11853 | 11400 | 10501 | 9862 | 11206 | 12696 | 12608 | 12301 | 10453 | 11125 | 11062 | 11707.42 | 12535 | 10880 | 0.868 | -0.204 | 4.30E-05 | 0.00061859 | 6.00E-04 | 0.0026 | 0.0017 | 0.0017 | TRUE | 0.0145 |
| Drp2 | 98 | 87 | 56 | 50 | 38 | 34 | 76 | 93 | 60 | 50 | 43 | 34 | 59.21 | 76 | 42 | 0.551 | -0.861 | 0.00201156 | 0.01215697 | 0.0238 | 0.0425 | 0.0374 | 0.0374 | TRUE | 0.1141 |
| Dsc1 | 0 | 0 | 0 | 0 | 0 | 0 | 0 | 0 | 0 | 0 | 0 | 0 | 0 | 0 | 0 | NA | NA | NA | NA | NA | NA | NA | NA | NA | NA |
| Dsc2 | 570 | 407 | 382 | 546 | 440 | 430 | 443 | 433 | 412 | 543 | 496 | 424 | 458.69 | 429 | 488 | 1.136 | 0.184 | 0.10761402 | 0.23347408 | 0.0064 | 0.0076 | 0.0072 | 0.0072 | TRUE | 0.1545 |
| Dsc3 | 62 | 43 | 49 | 31 | 42 | 30 | 48 | 46 | 53 | 31 | 47 | 30 | 42.44 | 49 | 36 | 0.731 | -0.451 | 0.14609219 | 0.29037626 | 0.0046 | 0.0583 | 0.0451 | 0.0451 | TRUE | 0.5175 |
| Dscam | 28 | 24 | 19 | 18 | 13 | 13 | 22 | 26 | 21 | 18 | 15 | 13 | 18.87 | 23 | 15 | 0.671 | -0.575 | 0.19314137 | 0.3518442 | 0 | 0.1281 | 0.0853 | 0.0853 | TRUE | 0.1231 |
| Dscaml1 | 78 | 75 | 96 | 133 | 106 | 130 | 61 | 80 | 104 | 132 | 120 | 128 | 104.04 | 82 | 127 | 1.574 | 0.654 | 0.00302391 | 0.01664257 | 0.0245 | 0.0252 | 0.025 | 0.025 | TRUE | 0.3875 |
| Dscc1 | 721 | 471 | 407 | 411 | 343 | 429 | 560 | 501 | 439 | 409 | 387 | 423 | 453.28 | 500 | 406 | 0.811 | -0.302 | 0.00887676 | 0.0379867 | 0.0067 | 0.0077 | 0.0073 | 0.0073 | TRUE | 0.1344 |
| Dscr3 | 2339 | 1719 | 1771 | 1994 | 1832 | 2016 | 1817 | 1828 | 1911 | 1985 | 2067 | 1990 | 1932.93 | 1852 | 2014 | 1.088 | 0.121 | 0.05153403 | 0.13783233 | 0 | 0.0037 | 0.0023 | 0.0023 | TRUE | 0.0092 |
| Dse | 4334 | 2881 | 2799 | 3054 | 2551 | 3004 | 3366 | 3064 | 3020 | 3040 | 2878 | 2965 | 3055.63 | 3150 | 2961 | 0.939 | -0.09 | 0.16196213 | 0.31205405 | 0.0017 | 0.0032 | 0.0027 | 0.0027 | TRUE | 0.0446 |
| Dsel | 643 | 450 | 410 | 382 | 311 | 368 | 499 | 479 | 442 | 380 | 351 | 363 | 419.13 | 473 | 365 | 0.77 | -0.378 | 0.00038362 | 0.00352154 | 0 | 0.0081 | 0.0057 | 0.0057 | TRUE | 0.0377 |
| Dsg1a | 0 | 0 | 1 | 0 | 1 | 1 | 0 | 0 | 1 | 0 | 1 | 1 | 0.53 | 0 | 1 | 2.03 | 1.022 | 0.7058734 | NA | 0 | 4.4543 | 3.5426 | 3.5426 | TRUE | 0.5365 |
| Dsg1b | 4 | 3 | 2 | 0 | 1 | 0 | 3 | 3 | 2 | 0 | 1 | 0 | 1.6 | 3 | 0 | 0.131 | -2.93 | 0.09577042 | NA | 0 | 1.4862 | 1.0592 | 1.0592 | TRUE | 0.5075 |
| Dsg1c | 0 | 0 | 0 | 0 | 0 | 0 | 0 | 0 | 0 | 0 | 0 | 0 | 0 | 0 | 0 | NA | NA | NA | NA | NA | NA | NA | NA | NA | NA |
| Dsg2 | 7918 | 5883 | 6004 | 6338 | 5881 | 6716 | 6150 | 6258 | 6478 | 6309 | 6634 | 6630 | 6409.74 | 6295 | 6524 | 1.037 | 0.052 | 0.32376037 | 0.4977955 | 5.00E-04 | 0.0028 | 0.0018 | 0.0018 | TRUE | 0.0102 |
| Dsg3 | 3 | 3 | 0 | 2 | 2 | 1 | 2 | 3 | 0 | 2 | 2 | 1 | 1.79 | 2 | 2 | 0.933 | -0.1 | 0.94508481 | NA | 0 | 1.3246 | 0.9748 | 0.9748 | TRUE | 0.5593 |
| Dsg4 | 1 | 1 | 0 | 0 | 0 | 0 | 1 | 1 | 0 | 0 | 0 | 0 | 0.31 | 1 | 0 | 0.309 | -1.696 | 0.63183309 | NA | 0 | 7.7294 | 7.2472 | 7.2472 | TRUE | 0.2015 |
| Dsn1 | 1714 | 1223 | 1216 | 1352 | 1213 | 1411 | 1331 | 1301 | 1312 | 1346 | 1368 | 1393 | 1341.86 | 1315 | 1369 | 1.041 | 0.058 | 0.38389667 | 0.55785229 | 0 | 0.0042 | 0.0024 | 0.0024 | TRUE | 0.0029 |
| Dsp | 4352 | 3179 | 3283 | 3372 | 3140 | 3436 | 3380 | 3381 | 3542 | 3356 | 3542 | 3392 | 3432.4 | 3434 | 3430 | 0.999 | -0.002 | 0.97449935 | 0.98624254 | 5.00E-04 | 0.0031 | 0.002 | 0.002 | TRUE | 0.0098 |
| Dspp | 0 | 0 | 0 | 0 | 0 | 0 | 0 | 0 | 0 | 0 | 0 | 0 | 0 | 0 | 0 | NA | NA | NA | NA | NA | NA | NA | NA | NA | NA |
| Dst | 11200 | 7844 | 8028 | 9505 | 9507 | 10979 | 8699 | 8344 | 8662 | 9461 | 10725 | 10838 | 9454.73 | 8568 | 10341 | 1.207 | 0.271 | 1.95E-05 | 0.00032626 | 0.0028 | 0.0027 | 0.0028 | 0.0028 | TRUE | 0.0679 |
| Dstn | 10191 | 7489 | 7919 | 7710 | 6735 | 8071 | 7915 | 7966 | 8545 | 7675 | 7598 | 7967 | 7944.17 | 8142 | 7747 | 0.952 | -0.071 | 0.18954301 | 0.3472809 | 0.0011 | 0.0027 | 0.002 | 0.002 | TRUE | 0.0227 |
| Dstyk | 1439 | 1004 | 1054 | 1101 | 963 | 948 | 1118 | 1068 | 1137 | 1096 | 1086 | 936 | 1073.49 | 1108 | 1039 | 0.938 | -0.093 | 0.27307449 | 0.44411731 | 0.0034 | 0.0046 | 0.0042 | 0.0042 | TRUE | 0.0919 |

| | | | | | | | | | | | | | | | | | | | | | | | | |
|---|---|---|---|---|---|---|---|---|---|---|---|---|---|---|---|---|---|---|---|---|---|---|---|---|
| Dtd1 | 1135 | 860 | 921 | 984 | 842 | 1054 | 882 | 915 | 994 | 979 | 950 | 1040 | 959.97 | 930 | 990 | 1.066 | 0.092 | 0.26370037 | 0.43402457 | 0.002 | 0.0049 | 0.0039 | 0.0039 | TRUE | 0.0424 |
| Dtd2 | 1419 | 1010 | 976 | 924 | 850 | 1011 | 1102 | 1074 | 1053 | 920 | 959 | 998 | 1017.69 | 1076 | 959 | 0.89 | -0.168 | 0.0264654 | 0.08469995 | 0 | 0.0048 | 0.0031 | 0.0031 | TRUE | 0.0156 |
| Dtl | 5836 | 4353 | 4528 | 4822 | 4235 | 4563 | 4533 | 4630 | 4886 | 4800 | 4778 | 4504 | 4688.37 | 4683 | 4694 | 1.003 | 0.004 | 0.94859871 | 0.97192267 | 0.0012 | 0.0029 | 0.0022 | 0.0022 | TRUE | 0.0174 |
| Dtna | 382 | 261 | 340 | 233 | 278 | 328 | 297 | 278 | 367 | 232 | 314 | 324 | 301.75 | 314 | 290 | 0.924 | -0.114 | 0.46481527 | 0.63387125 | 0.0234 | 0.0103 | 0.0141 | 0.0141 | TRUE | 0.3478 |
| Dtnb | 2732 | 2058 | 2134 | 2122 | 1931 | 2214 | 2122 | 2189 | 2303 | 2112 | 2178 | 2186 | 2181.6 | 2205 | 2159 | 0.98 | -0.029 | 0.63800925 | 0.77391719 | 6.00E-04 | 0.0035 | 0.0024 | 0.0024 | TRUE | 0.0182 |
| Dtnbp1 | 1323 | 946 | 897 | 946 | 772 | 922 | 1028 | 1006 | 968 | 942 | 871 | 910 | 954.05 | 1001 | 908 | 0.907 | -0.141 | 0.06831069 | 0.16830972 | 0 | 0.0049 | 0.0033 | 0.0033 | TRUE | 0.0145 |
| Dtwd1 | 774 | 553 | 607 | 617 | 522 | 595 | 601 | 588 | 655 | 614 | 589 | 587 | 605.78 | 615 | 597 | 0.972 | -0.042 | 0.64906704 | 0.7820068 | 0 | 0.0063 | 0.0044 | 0.0044 | TRUE | 0.037 |
| Dtwd2 | 903 | 632 | 710 | 745 | 633 | 779 | 701 | 672 | 766 | 742 | 714 | 769 | 727.39 | 713 | 742 | 1.041 | 0.058 | 0.5161934 | 0.67849478 | 0.0014 | 0.0057 | 0.0043 | 0.0043 | TRUE | 0.0477 |
| Dtx1 | 872 | 832 | 1106 | 1165 | 1274 | 1338 | 677 | 885 | 1193 | 1160 | 1437 | 1321 | 1112.21 | 918 | 1306 | 1.423 | 0.508 | 0.04232871 | 0.1192735 | 0.0443 | 0.0046 | 0.0157 | 0.0443 | TRUE | 0.4701 |
| Dtx2 | 2153 | 1561 | 1494 | 1679 | 1481 | 1699 | 1672 | 1660 | 1612 | 1671 | 1671 | 1677 | 1660.62 | 1648 | 1673 | 1.015 | 0.021 | 0.73541486 | 0.84204795 | 0 | 0.0039 | 0.0022 | 0.0022 | TRUE | 0.0044 |
| Dtx3 | 935 | 899 | 852 | 1143 | 954 | 1116 | 726 | 956 | 919 | 1138 | 1076 | 1102 | 986.22 | 867 | 1105 | 1.278 | 0.354 | 0.00057161 | 0.00478341 | 0.0101 | 0.0048 | 0.0066 | 0.0066 | TRUE | 0.2512 |
| Dtx3l | 1090 | 795 | 627 | 672 | 661 | 762 | 847 | 846 | 677 | 669 | 746 | 752 | 755.92 | 790 | 722 | 0.913 | -0.131 | 0.21346285 | 0.37574193 | 0.0088 | 0.0056 | 0.0066 | 0.0066 | TRUE | 0.1822 |
| Dtx4 | 173 | 128 | 130 | 133 | 101 | 146 | 134 | 136 | 140 | 132 | 114 | 144 | 133.54 | 137 | 130 | 0.953 | -0.069 | 0.68993374 | 0.81028948 | 0 | 0.0202 | 0.0143 | 0.0143 | TRUE | 0.1155 |
| Dtymk | 5315 | 3649 | 3279 | 3423 | 3026 | 3511 | 4128 | 3881 | 3538 | 3407 | 3414 | 3466 | 3639.02 | 3849 | 3429 | 0.89 | -0.168 | 0.01258593 | 0.04905665 | 0.0026 | 0.0031 | 0.003 | 0.003 | TRUE | 0.0606 |
| Duox1 | 47 | 44 | 24 | 28 | 27 | 37 | 37 | 47 | 26 | 28 | 30 | 37 | 34.01 | 37 | 32 | 0.869 | -0.203 | 0.56146771 | 0.71613939 | 0.0212 | 0.0721 | 0.0584 | 0.0584 | TRUE | 0.1558 |
| Duox2 | 14 | 5 | 13 | 12 | 14 | 16 | 11 | 5 | 14 | 12 | 16 | 16 | 12.29 | 10 | 15 | 1.435 | 0.521 | 0.3566429 | NA | 0 | 0.1952 | 0.1471 | 0.1471 | TRUE | 0.491 |
| Duoxa1 | 13 | 13 | 15 | 17 | 12 | 19 | 10 | 14 | 16 | 17 | 14 | 19 | 14.89 | 13 | 17 | 1.243 | 0.314 | 0.5285591 | NA | 0 | 0.1616 | 0.1113 | 0.1113 | TRUE | 0.2516 |
| Duoxa2 | 15 | 19 | 9 | 9 | 7 | 5 | 12 | 20 | 10 | 9 | 8 | 5 | 10.56 | 14 | 7 | 0.525 | -0.929 | 0.13610516 | NA | 0 | 0.2269 | 0.1733 | 0.1733 | TRUE | 0.6539 |
| Dupd1 | 0 | 2 | 0 | 0 | 0 | 2 | 0 | 2 | 0 | 0 | 0 | 2 | 0.68 | 1 | 1 | 0.965 | -0.051 | 0.98509429 | NA | 8.094 | 3.4694 | 3.9051 | 3.9051 | TRUE | 0.9747 |
| Dus1l | 5814 | 4373 | 3746 | 4209 | 3906 | 4412 | 4515 | 4652 | 4042 | 4190 | 4406 | 4355 | 4360.04 | 4403 | 4317 | 0.98 | -0.029 | 0.66221115 | 0.79149275 | 0.0028 | 0.003 | 0.0029 | 0.0029 | TRUE | 0.062 |
| Dus2l | 1287 | 893 | 760 | 967 | 807 | 862 | 1000 | 950 | 820 | 963 | 910 | 851 | 915.55 | 923 | 908 | 0.982 | -0.026 | 0.7824324 | 0.87192471 | 0.006 | 0.005 | 0.0054 | 0.0054 | TRUE | 0.1122 |
| Dus3l | 2677 | 2027 | 1618 | 1594 | 1500 | 1751 | 2079 | 2156 | 1746 | 1587 | 1692 | 1729 | 1831.39 | 1994 | 1669 | 0.837 | -0.257 | 0.00273749 | 0.01535594 | 0.0061 | 0.0037 | 0.0048 | 0.0048 | TRUE | 0.141 |
| Dus4l | 700 | 505 | 436 | 567 | 468 | 544 | 544 | 537 | 470 | 564 | 528 | 537 | 530.1 | 517 | 543 | 1.049 | 0.069 | 0.48780866 | 0.65375052 | 0.0017 | 0.0069 | 0.0053 | 0.0053 | TRUE | 0.0704 |
| Dusp1 | 1557 | 1239 | 1373 | 1549 | 1457 | 1620 | 1209 | 1318 | 1481 | 1542 | 1644 | 1599 | 1465.55 | 1336 | 1595 | 1.195 | 0.257 | 0.00218345 | 0.0129119 | 0.0047 | 0.004 | 0.0044 | 0.0044 | TRUE | 0.1083 |
| Dusp10 | 158 | 109 | 86 | 123 | 88 | 90 | 123 | 116 | 93 | 122 | 99 | 89 | 107 | 111 | 103 | 0.934 | -0.098 | 0.63630918 | 0.77264185 | 0.0147 | 0.0246 | 0.0218 | 0.0218 | TRUE | 0.2573 |
| Dusp11 | 4074 | 2857 | 2752 | 3371 | 2720 | 3101 | 3164 | 3039 | 2969 | 3355 | 3068 | 3061 | 3109.59 | 3057 | 3161 | 1.034 | 0.048 | 0.44751455 | 0.61788255 | 0.0014 | 0.0032 | 0.0026 | 0.0026 | TRUE | 0.0351 |
| Dusp12 | 1661 | 1162 | 1154 | 1142 | 1053 | 1089 | 1290 | 1236 | 1245 | 1137 | 1188 | 1075 | 1195.14 | 1257 | 1133 | 0.901 | -0.151 | 0.03991005 | 0.11433842 | 6.00E-04 | 0.0044 | 0.0031 | 0.0031 | TRUE | 0.0243 |
| Dusp13 | 3 | 9 | 6 | 4 | 1 | 3 | 2 | 10 | 6 | 4 | 1 | 3 | 4.41 | 6 | 3 | 0.452 | -1.146 | 0.24620453 | NA | 0 | 0.5401 | 0.4329 | 0.4329 | TRUE | 0.1731 |
| Dusp14 | 536 | 373 | 291 | 365 | 276 | 301 | 416 | 397 | 314 | 363 | 311 | 297 | 349.81 | 376 | 324 | 0.861 | -0.217 | 0.11364411 | 0.24268868 | 0.0139 | 0.0092 | 0.0106 | 0.0106 | TRUE | 0.2312 |
| Dusp15 | 17 | 19 | 13 | 32 | 20 | 16 | 13 | 20 | 14 | 32 | 23 | 16 | 19.61 | 16 | 24 | 1.49 | 0.575 | 0.21308849 | 0.37540408 | 0.0449 | 0.1233 | 0.1008 | 0.1008 | TRUE | 0.2132 |
| Dusp16 | 4677 | 3132 | 3092 | 3496 | 3017 | 3523 | 3632 | 3331 | 3336 | 3480 | 3403 | 3478 | 3443.55 | 3433 | 3454 | 1.005 | 0.008 | 0.8975337 | 0.94349536 | 0.0011 | 0.0031 | 0.0023 | 0.0023 | TRUE | 0.032 |
| Dusp18 | 314 | 216 | 245 | 261 | 249 | 341 | 244 | 230 | 264 | 260 | 281 | 337 | 269.22 | 246 | 293 | 1.19 | 0.251 | 0.0719976 | 0.17521909 | 0.0081 | 0.0112 | 0.0103 | 0.0103 | TRUE | 0.2006 |
| Dusp19 | 633 | 489 | 507 | 633 | 553 | 571 | 492 | 520 | 547 | 630 | 624 | 564 | 562.73 | 520 | 606 | 1.167 | 0.223 | 0.02156378 | 0.0730012 | 0.0017 | 0.0066 | 0.005 | 0.005 | TRUE | 0.0443 |
| Dusp2 | 38 | 19 | 22 | 47 | 41 | 49 | 30 | 20 | 24 | 47 | 46 | 48 | 35.81 | 25 | 47 | 1.907 | 0.931 | 0.00456778 | 0.02290783 | 0 | 0.0686 | 0.0472 | 0.0472 | TRUE | 0.2382 |
| Dusp21 | 0 | 0 | 0 | 0 | 0 | 0 | 0 | 0 | 0 | 0 | 0 | 0 | 0 | 0 | 0 | NA | NA | NA | NA | NA | NA | NA | NA | NA | NA |
| Dusp22 | 861 | 666 | 660 | 754 | 619 | 748 | 669 | 708 | 712 | 751 | 698 | 738 | 712.75 | 696 | 729 | 1.049 | 0.068 | 0.42148251 | 0.59375611 | 0 | 0.0058 | 0.0038 | 0.0038 | TRUE | 0.0157 |
| Dusp23 | 13 | 14 | 13 | 10 | 10 | 6 | 10 | 15 | 14 | 10 | 11 | 6 | 11.03 | 13 | 9 | 0.698 | -0.519 | 0.37624494 | NA | 0 | 0.2173 | 0.1526 | 0.1526 | TRUE | 0.3064 |
| Dusp26 | 285 | 234 | 269 | 278 | 281 | 350 | 221 | 249 | 290 | 277 | 317 | 346 | 283.29 | 253 | 313 | 1.239 | 0.309 | 0.03054756 | 0.09378286 | 0.0121 | 0.0108 | 0.0112 | 0.0112 | TRUE | 0.1775 |
| Dusp27 | 652 | 508 | 324 | 285 | 309 | 333 | 506 | 540 | 350 | 284 | 349 | 329 | 392.89 | 465 | 321 | 0.687 | -0.542 | 0.00047551 | 0.00415688 | 0.0293 | 0.0085 | 0.0147 | 0.0147 | TRUE | 0.5382 |
| Dusp28 | 236 | 175 | 173 | 155 | 122 | 160 | 183 | 186 | 187 | 154 | 138 | 158 | 167.66 | 185 | 150 | 0.81 | -0.303 | 0.04799566 | 0.13077845 | 0 | 0.0166 | 0.0109 | 0.0109 | TRUE | 0.0514 |
| Dusp3 | 2836 | 2298 | 2668 | 3590 | 3150 | 3826 | 2203 | 2444 | 2879 | 3573 | 3554 | 3777 | 3071.59 | 2509 | 3635 | 1.45 | 0.536 | 1.29E-09 | 8.37E-08 | 0.0094 | 0.0032 | 0.0053 | 0.0053 | TRUE | 0.2017 |
| Dusp4 | 1072 | 839 | 811 | 1109 | 994 | 1039 | 833 | 892 | 875 | 1104 | 1121 | 1026 | 975.16 | 867 | 1084 | 1.251 | 0.323 | 4.12E-05 | 0.00059971 | 8.00E-04 | 0.0049 | 0.0034 | 0.0034 | TRUE | 0.0265 |
| Dusp5 | 292 | 239 | 188 | 205 | 190 | 199 | 227 | 254 | 203 | 204 | 214 | 196 | 216.45 | 228 | 205 | 0.898 | -0.154 | 0.28240301 | 0.45408778 | 0.0026 | 0.0134 | 0.0102 | 0.0102 | TRUE | 0.1053 |
| Dusp6 | 2603 | 1832 | 1601 | 1899 | 1736 | 1908 | 2022 | 1949 | 1727 | 1890 | 1958 | 1883 | 1904.99 | 1899 | 1910 | 1.005 | 0.007 | 0.92297617 | 0.95719768 | 0.003 | 0.0037 | 0.0034 | 0.0034 | TRUE | 0.0748 |
| Dusp7 | 827 | 699 | 722 | 732 | 707 | 793 | 642 | 744 | 779 | 729 | 798 | 783 | 745.64 | 722 | 770 | 1.069 | 0.096 | 0.32105379 | 0.49496514 | 0.005 | 0.0056 | 0.0054 | 0.0054 | TRUE | 0.1153 |
| Dusp8 | 159 | 84 | 62 | 89 | 42 | 44 | 123 | 89 | 67 | 89 | 47 | 43 | 76.52 | 93 | 60 | 0.639 | -0.646 | 0.03677074 | 0.10783016 | 0.1102 | 0.0334 | 0.0551 | 0.0551 | TRUE | 0.4885 |
| Dusp9 | 14043 | 8705 | 7441 | 10636 | 8197 | 9350 | 10907 | 9259 | 8029 | 10587 | 9247 | 9230 | 9543.13 | 9398 | 9688 | 1.031 | 0.043 | 0.65243166 | 0.78420564 | 0.012 | 0.0027 | 0.0066 | 0.0066 | TRUE | 0.2416 |
| Dut | 9478 | 6799 | 6569 | 6126 | 5456 | 6494 | 7361 | 7232 | 7088 | 6098 | 6155 | 6411 | 6724.07 | 7227 | 6221 | 0.861 | -0.216 | 1.95E-05 | 0.00032626 | 3.00E-04 | 0.0028 | 0.0017 | 0.0017 | TRUE | 0.0087 |
| Dux | 39 | 30 | 14 | 22 | 17 | 12 | 30 | 32 | 15 | 22 | 19 | 12 | 21.7 | 26 | 18 | 0.68 | -0.557 | 0.22178386 | 0.38587421 | 0.0744 | 0.1116 | 0.1016 | 0.1016 | TRUE | 0.7516 |
| Duxbl1 | 0 | 0 | 0 | 0 | 0 | 0 | 0 | 0 | 0 | 0 | 0 | 0 | 0 | 0 | 0 | NA | NA | NA | NA | NA | NA | NA | NA | NA | NA |
| Duxbl2 | 0 | 0 | 0 | 0 | 0 | 0 | 0 | 0 | 0 | 0 | 0 | 0 | 0 | 0 | 0 | NA | NA | NA | NA | NA | NA | NA | NA | NA | NA |
| Duxbl3 | 0 | 0 | 0 | 0 | 0 | 0 | 0 | 0 | 0 | 0 | 0 | 0 | 0 | 0 | 0 | NA | NA | NA | NA | NA | NA | NA | NA | NA | NA |
| Dvl1 | 2454 | 1944 | 1795 | 1993 | 1768 | 1879 | 1906 | 2068 | 1937 | 1984 | 1994 | 1855 | 1957.28 | 1970 | 1944 | 0.987 | -0.019 | 0.78146119 | 0.87115299 | 0.0012 | 0.0036 | 0.0027 | 0.0027 | TRUE | 0.0225 |
| Dvl2 | 7416 | 5947 | 5799 | 5686 | 5187 | 6160 | 5760 | 6326 | 6257 | 5660 | 5851 | 6081 | 5989.12 | 6114 | 5864 | 0.959 | -0.06 | 0.31681545 | 0.49084801 | 0.0017 | 0.0028 | 0.0024 | 0.0024 | TRUE | 0.0318 |
| Dvl3 | 1305 | 981 | 862 | 808 | 811 | 794 | 1014 | 1043 | 930 | 804 | 915 | 784 | 915.01 | 996 | 834 | 0.837 | -0.257 | 0.00409097 | 0.02099542 | 0.0037 | 0.005 | 0.0047 | 0.0047 | TRUE | 0.0832 |
| DXBay18 | 0 | 0 | 0 | 0 | 0 | 0 | 0 | 0 | 0 | 0 | 0 | 0 | 0 | 0 | 0 | NA | NA | NA | NA | NA | NA | NA | NA | NA | NA |
| Dydc1 | 1 | 0 | 0 | 1 | 0 | 0 | 1 | 0 | 0 | 1 | 0 | 0 | 0.3 | 0 | 0 | 1.075 | 0.105 | 0.97698525 | NA | 0 | 8.0272 | 7.5141 | 7.5141 | TRUE | 0.3906 |
| Dydc2 | 3 | 2 | 1 | 3 | 1 | 4 | 2 | 2 | 1 | 3 | 1 | 4 | 2.27 | 2 | 3 | 1.451 | 0.538 | 0.66793691 | NA | 0 | 1.0481 | 0.6827 | 0.6827 | TRUE | 0.2636 |
| Dym | 3026 | 2298 | 2510 | 2615 | 2314 | 2788 | 2350 | 2444 | 2708 | 2603 | 2610 | 2752 | 2578.06 | 2501 | 2655 | 1.063 | 0.088 | 0.21151662 | 0.37365061 | 0.0027 | 0.0033 | 0.0031 | 0.0031 | TRUE | 0.0634 |
| Dynap | 0 | 0 | 0 | 0 | 0 | 0 | 0 | 0 | 0 | 0 | 0 | 0 | 0 | 0 | 0 | NA | NA | NA | NA | NA | NA | NA | NA | NA | NA |
| Dync1h1 | 39462 | 28028 | 27642 | 34814 | 33099 | 37939 | 30648 | 29813 | 29825 | 34654 | 37339 | 37452 | 33288.59 | 30095 | 36482 | 1.212 | 0.278 | 4.84E-08 | 2.05E-06 | 0.001 | 0.0025 | 0.0018 | 0.0018 | TRUE | 0.0235 |
| Dync1i1 | 80 | 65 | 51 | 58 | 68 | 50 | 62 | 69 | 55 | 58 | 77 | 49 | 61.68 | 62 | 61 | 0.983 | -0.025 | 0.92374567 | 0.9577958 | 0.0137 | 0.0408 | 0.0335 | 0.0335 | TRUE | 0.3987 |
| Dync1i2 | 6930 | 5096 | 5345 | 5231 | 4713 | 5526 | 5382 | 5421 | 5767 | 5207 | 5317 | 5455 | 5424.78 | 5523 | 5326 | 0.965 | -0.052 | 0.34184425 | 0.5158885 | 7.00E-04 | 0.0029 | 0.002 | 0.002 | TRUE | 0.0181 |
| Dync1li1 | 4043 | 3084 | 3056 | 3342 | 2914 | 3259 | 3140 | 3280 | 3297 | 3327 | 3287 | 3217 | 3258.15 | 3239 | 3277 | 1.012 | 0.017 | 0.7556065 | 0.85511255 | 3.00E-04 | 0.0032 | 0.0019 | 0.0019 | TRUE | 0.009 |
| Dync1li2 | 4588 | 3485 | 3424 | 3654 | 3255 | 3607 | 3563 | 3707 | 3694 | 3637 | 3672 | 3561 | 3639.09 | 3655 | 3623 | 0.992 | -0.012 | 0.82034869 | 0.89651557 | 0 | 0.0031 | 0.0018 | 0.0018 | TRUE | 0.006 |
| Dync2h1 | 893 | 606 | 473 | 634 | 582 | 583 | 694 | 645 | 510 | 631 | 657 | 576 | 618.61 | 616 | 621 | 1.005 | 0.008 | 0.94797249 | 0.97160938 | 0.013 | 0.0063 | 0.0084 | 0.0084 | TRUE | 0.261 |
| Dync2li1 | 810 | 566 | 673 | 736 | 692 | 903 | 629 | 602 | 726 | 733 | 781 | 891 | 726.99 | 652 | 802 | 1.23 | 0.298 | 0.00459045 | 0.02299766 | 0.0081 | 0.0057 | 0.0066 | 0.0066 | TRUE | 0.1155 |
| Dynll1 | 10893 | 7683 | 7305 | 7028 | 6229 | 7108 | 8460 | 8172 | 7882 | 6996 | 7027 | 7017 | 7592.3 | 8171 | 7013 | 0.858 | -0.221 | 1.41E-05 | 0.00025103 | 4.00E-04 | 0.0027 | 0.0017 | 0.0017 | TRUE | 0.0118 |
| Dynll2 | 10993 | 7869 | 7086 | 7082 | 6274 | 6661 | 8538 | 8370 | 7646 | 7049 | 7078 | 6575 | 7542.71 | 8185 | 6901 | 0.843 | -0.247 | 6.57E-05 | 0.00087614 | 0.0023 | 0.0027 | 0.0026 | 0.0026 | TRUE | 0.0403 |

| | | | | | | | | | | | | | | | | | | | | | | | | | |
|---|---|---|---|---|---|---|---|---|---|---|---|---|---|---|---|---|---|---|---|---|---|---|---|---|---|
| Dynlrb1 | 2943 | 1908 | 1920 | 2211 | 1793 | 2153 | 2286 | 2030 | 2072 | 2201 | 2023 | 2125 | 2122.62 | 2129 | 2116 | 0.993 | -0.01 | 0.89229297 | 0.94080432 | 0.0025 | 0.0035 | 0.0032 | 0.0032 | TRUE | 0.0516 |
| Dynlrb2 | 3 | 2 | 0 | 0 | 0 | 0 | 2 | 2 | 0 | 0 | 0 | 0 | 0.74 | 1 | 0 | 0.127 | -2.979 | 0.24053699 | NA | 0 | 3.1927 | 3.3198 | 3.3198 | TRUE | 0.4766 |
| Dynlt1a | 617 | 440 | 513 | 604 | 565 | 687 | 479 | 468 | 554 | 601 | 637 | 678 | 569.59 | 500 | 639 | 1.279 | 0.355 | 0.00052896 | 0.00449759 | 0.004 | 0.0066 | 0.0058 | 0.0058 | TRUE | 0.0978 |
| Dynlt1b | 599 | 483 | 468 | 498 | 505 | 575 | 465 | 514 | 505 | 496 | 570 | 568 | 519.49 | 495 | 545 | 1.102 | 0.14 | 0.17184694 | 0.32390983 | 0.0027 | 0.007 | 0.0056 | 0.0056 | TRUE | 0.0723 |
| Dynlt1c | 0 | 0 | 0 | 0 | 0 | 0 | 0 | 0 | 0 | 0 | 0 | 0 | 0 | 0 | 0 | NA | NA | NA | NA | NA | NA | NA | NA | NA | NA |
| Dynlt1f | 151 | 131 | 135 | 165 | 143 | 154 | 117 | 139 | 146 | 164 | 161 | 152 | 146.64 | 134 | 159 | 1.192 | 0.254 | 0.13113314 | 0.26856081 | 0 | 0.0186 | 0.0135 | 0.0135 | TRUE | 0.1394 |
| Dynlt3 | 1637 | 1206 | 1391 | 1543 | 1217 | 1364 | 1271 | 1283 | 1501 | 1536 | 1373 | 1346 | 1385.06 | 1352 | 1418 | 1.05 | 0.071 | 0.42041829 | 0.59260736 | 0.0057 | 0.0041 | 0.0049 | 0.0049 | TRUE | 0.1107 |
| Dyrk1a | 4105 | 3056 | 2822 | 2945 | 2600 | 2879 | 3188 | 3251 | 3045 | 2931 | 2933 | 2842 | 3031.71 | 3161 | 2902 | 0.918 | -0.124 | 0.0312059 | 0.09536583 | 4.00E-04 | 0.0032 | 0.002 | 0.002 | TRUE | 0.0123 |
| Dyrk1b | 723 | 577 | 514 | 414 | 401 | 520 | 562 | 614 | 555 | 412 | 452 | 513 | 517.94 | 577 | 459 | 0.797 | -0.327 | 0.00256814 | 0.01467083 | 0.0055 | 0.007 | 0.0065 | 0.0065 | TRUE | 0.1257 |
| Dyrk2 | 809 | 629 | 574 | 583 | 560 | 565 | 628 | 669 | 619 | 580 | 632 | 558 | 614.42 | 639 | 590 | 0.923 | -0.116 | 0.21395993 | 0.37643148 | 0.001 | 0.0063 | 0.0046 | 0.0046 | TRUE | 0.0439 |
| Dyrk3 | 2458 | 1733 | 1583 | 1639 | 1438 | 1665 | 1909 | 1843 | 1708 | 1631 | 1622 | 1644 | 1726.29 | 1820 | 1632 | 0.896 | -0.158 | 0.02061469 | 0.07074329 | 0.0011 | 0.0038 | 0.0028 | 0.0028 | TRUE | 0.0347 |
| Dyrk4 | 42 | 31 | 14 | 21 | 21 | 9 | 33 | 33 | 15 | 21 | 24 | 9 | 22.36 | 27 | 18 | 0.656 | -0.609 | 0.20458595 | 0.36606747 | 0.1479 | 0.1084 | 0.1189 | 0.1189 | TRUE | 1.0075 |
| Dysf | 606 | 462 | 464 | 458 | 467 | 554 | 471 | 491 | 501 | 456 | 527 | 547 | 498.72 | 488 | 510 | 1.046 | 0.065 | 0.53201185 | 0.6918429 | 0.0031 | 0.0072 | 0.0059 | 0.0059 | TRUE | 0.1017 |
| Dytn | 0 | 0 | 0 | 0 | 0 | 0 | 0 | 0 | 0 | 0 | 0 | 0 | 0 | 0 | 0 | NA | NA | NA | NA | NA | NA | NA | NA | NA | NA |
| Dyx1c1 | 235 | 202 | 219 | 315 | 256 | 316 | 183 | 215 | 236 | 314 | 289 | 312 | 257.99 | 211 | 305 | 1.45 | 0.536 | 0.00010158 | 0.00122517 | 0.0051 | 0.0116 | 0.0097 | 0.0097 | TRUE | 0.1706 |
| Dzank1 | 171 | 177 | 136 | 152 | 120 | 150 | 133 | 188 | 147 | 151 | 135 | 148 | 150.43 | 156 | 145 | 0.934 | -0.099 | 0.58225362 | 0.73253195 | 0.0126 | 0.0182 | 0.0165 | 0.0165 | TRUE | 0.3419 |
| Dzip1 | 875 | 716 | 751 | 797 | 663 | 782 | 680 | 762 | 810 | 793 | 748 | 772 | 760.79 | 751 | 771 | 1.03 | 0.042 | 0.64379362 | 0.77809422 | 0.0033 | 0.0055 | 0.0048 | 0.0048 | TRUE | 0.0855 |
| Dzip1l | 903 | 650 | 521 | 582 | 490 | 575 | 701 | 691 | 562 | 579 | 553 | 568 | 609.1 | 651 | 567 | 0.868 | -0.204 | 0.05203908 | 0.13879494 | 0.0062 | 0.0063 | 0.0063 | 0.0063 | TRUE | 0.1659 |
| Dzip3 | 5605 | 3848 | 3200 | 4354 | 3553 | 3799 | 4353 | 4093 | 3453 | 4334 | 4008 | 3750 | 3998.55 | 3966 | 4031 | 1.016 | 0.023 | 0.79573867 | 0.88057393 | 0.0094 | 0.003 | 0.0052 | 0.0052 | TRUE | 0.1556 |
| E030002O03 | 0 | 0 | 0 | 0 | 0 | 0 | 0 | 0 | 0 | 0 | 0 | 0 | 0 | 0 | 0 | NA | NA | NA | NA | NA | NA | NA | NA | NA | NA |
| E030003E18I | 0 | 0 | 0 | 0 | 0 | 0 | 0 | 0 | 0 | 0 | 0 | 0 | 0 | 0 | 0 | NA | NA | NA | NA | NA | NA | NA | NA | NA | NA |
| E030011O05 | 23 | 24 | 20 | 11 | 14 | 25 | 18 | 26 | 22 | 11 | 16 | 25 | 19.4 | 22 | 17 | 0.797 | -0.327 | 0.47980896 | 0.64660659 | 0.04 | 0.1246 | 0.1018 | 0.1018 | TRUE | 0.5581 |
| E030013I19R | 0 | 0 | 0 | 0 | 0 | 1 | 0 | 0 | 0 | 0 | 0 | 1 | 0.16 | 0 | 0 | 2.094 | 1.067 | 0.79379329 | NA | 0 | 14.4078 | 10 | 10 | TRUE | 0.3835 |
| E030018B13 | 0 | 0 | 0 | 0 | 0 | 0 | 0 | 0 | 0 | 0 | 0 | 0 | 0 | 0 | 0 | NA | NA | NA | NA | NA | NA | NA | NA | NA | NA |
| E030019B06 | 0 | 1 | 0 | 1 | 2 | 0 | 0 | 1 | 0 | 1 | 2 | 0 | 0.72 | 0 | 1 | 3.105 | 1.634 | 0.50365406 | NA | 0 | 3.2978 | 2.8211 | 2.8211 | TRUE | 0.291 |
| E030019B13 | 1 | 0 | 0 | 1 | 0 | 0 | 1 | 0 | 0 | 1 | 0 | 0 | 0.3 | 0 | 0 | 1.075 | 0.105 | 0.97698525 | NA | 0 | 8.0272 | 7.5141 | 7.5141 | TRUE | 0.3906 |
| E030024N20 | 1 | 2 | 1 | 9 | 1 | 2 | 1 | 2 | 1 | 9 | 1 | 2 | 2.67 | 1 | 4 | 3.116 | 1.64 | 0.21292634 | NA | 0 | 0.8888 | 0.7508 | 0.7508 | TRUE | 1.9667 |
| E030025P04I | 0 | 0 | 0 | 1 | 0 | 0 | 0 | 0 | 0 | 1 | 0 | 0 | 0.17 | 0 | 0 | 2.094 | 1.067 | 0.79379329 | NA | 0 | 14.2885 | 10 | 10 | TRUE | 0.3906 |
| E030030I06R | 37 | 43 | 45 | 66 | 47 | 54 | 29 | 46 | 49 | 66 | 53 | 53 | 49.18 | 41 | 57 | 1.415 | 0.501 | 0.09371824 | 0.21078123 | 0.0245 | 0.0506 | 0.0435 | 0.0435 | TRUE | 0.6237 |
| E030044B06 | 0 | 1 | 1 | 2 | 3 | 4 | 0 | 1 | 1 | 2 | 3 | 4 | 1.91 | 1 | 3 | 4.664 | 2.222 | 0.15365924 | NA | 0 | 1.2426 | 0.86 | 0.86 | TRUE | 0.4062 |
| E130003G02 | 2 | 1 | 1 | 1 | 2 | 1 | 2 | 1 | 1 | 1 | 2 | 1 | 1.32 | 1 | 1 | 1.122 | 0.165 | 0.91688508 | NA | 0 | 1.7946 | 1.0548 | 1.0548 | TRUE | 0.1554 |
| E130006D01 | 8 | 0 | 3 | 7 | 9 | 2 | 6 | 0 | 3 | 7 | 10 | 2 | 4.76 | 3 | 6 | 1.94 | 0.956 | 0.35598002 | NA | 0.6966 | 0.5006 | 0.5427 | 0.5427 | TRUE | 0.3779 |
| E130008D07 | 37 | 44 | 45 | 47 | 37 | 47 | 29 | 47 | 49 | 47 | 42 | 46 | 43.17 | 42 | 45 | 1.1 | 0.138 | 0.65943612 | 0.78925812 | 0.0211 | 0.0573 | 0.0471 | 0.0471 | TRUE | 0.6478 |
| E130012A19 | 25387 | 15281 | 8948 | 10177 | 8298 | 9539 | 19717 | 16254 | 9655 | 10130 | 9361 | 9416 | 12422.28 | 15209 | 9636 | 0.634 | -0.658 | 0.01968122 | 0.06831076 | 0.0574 | 0.0026 | 0.0181 | 0.0574 | TRUE | 0.3154 |
| E130018N17 | 17 | 17 | 26 | 20 | 12 | 17 | 13 | 18 | 28 | 20 | 14 | 17 | 18.26 | 20 | 17 | 0.859 | -0.22 | 0.6436629 | 0.77809422 | 0.0461 | 0.1322 | 0.1071 | 0.1071 | TRUE | 0.4517 |
| E130102H24 | 41 | 53 | 50 | 51 | 34 | 42 | 32 | 56 | 54 | 51 | 38 | 41 | 45.46 | 47 | 43 | 0.929 | -0.106 | 0.73797721 | 0.84383266 | 0.0394 | 0.0546 | 0.0501 | 0.0501 | TRUE | 0.7725 |
| E130112N10 | 48 | 44 | 36 | 47 | 30 | 41 | 37 | 47 | 39 | 47 | 34 | 40 | 40.67 | 41 | 40 | 0.991 | -0.013 | 0.96532205 | 0.98104011 | 0 | 0.0607 | 0.0431 | 0.0431 | TRUE | 0.1455 |
| E130114P18I | 0 | 0 | 0 | 0 | 0 | 0 | 0 | 0 | 0 | 0 | 0 | 0 | 0 | 0 | 0 | NA | NA | NA | NA | NA | NA | NA | NA | NA | NA |
| E130201H02 | 8 | 2 | 2 | 2 | 4 | 2 | 6 | 2 | 2 | 2 | 5 | 2 | 3.16 | 3 | 3 | 0.778 | -0.362 | 0.74038317 | NA | 0 | 0.7518 | 0.5413 | 0.5413 | TRUE | 0.893 |
| E130215H24 | 0 | 0 | 0 | 0 | 0 | 0 | 0 | 0 | 0 | 0 | 0 | 0 | 0 | 0 | 0 | NA | NA | NA | NA | NA | NA | NA | NA | NA | NA |
| E130218I03R | 3 | 1 | 2 | 0 | 2 | 0 | 2 | 1 | 2 | 0 | 2 | 0 | 1.3 | 2 | 1 | 0.386 | -1.372 | 0.45337271 | NA | 0 | 1.8237 | 1.4355 | 1.4355 | TRUE | 0.9751 |
| E130304I02R | 0 | 0 | 1 | 1 | 1 | 1 | 0 | 0 | 1 | 1 | 1 | 1 | 0.7 | 0 | 1 | 3.004 | 1.587 | 0.48859459 | NA | 0 | 3.3966 | 2.2833 | 2.2833 | TRUE | 0.5365 |
| E130307A14 | 51 | 37 | 40 | 46 | 29 | 35 | 40 | 39 | 43 | 46 | 33 | 35 | 39.2 | 41 | 38 | 0.929 | -0.107 | 0.73066013 | 0.8390909 | 0 | 0.0629 | 0.0439 | 0.0439 | TRUE | 0.2631 |
| E130308A19 | 1467 | 1140 | 1117 | 1191 | 1035 | 1100 | 1139 | 1213 | 1205 | 1186 | 1168 | 1086 | 1166.03 | 1186 | 1147 | 0.967 | -0.048 | 0.52210133 | 0.68332002 | 9.00E-04 | 0.0045 | 0.0032 | 0.0032 | TRUE | 0.0259 |
| E130309D02 | 2049 | 1420 | 1336 | 1329 | 1224 | 1369 | 1591 | 1510 | 1442 | 1323 | 1381 | 1351 | 1433.07 | 1514 | 1352 | 0.891 | -0.166 | 0.0187276 | 0.06588076 | 9.00E-04 | 0.0041 | 0.0029 | 0.0029 | TRUE | 0.0247 |
| E130309D14 | 1067 | 842 | 847 | 1344 | 1307 | 1456 | 829 | 896 | 914 | 1338 | 1474 | 1437 | 1147.96 | 880 | 1416 | 1.613 | 0.689 | 1.59E-18 | 4.20E-16 | 0.0018 | 0.0045 | 0.0035 | 0.0035 | TRUE | 0.0321 |
| E130309F12I | 30 | 16 | 23 | 20 | 26 | 12 | 23 | 17 | 25 | 20 | 29 | 12 | 21.04 | 22 | 20 | 0.93 | -0.105 | 0.8165226 | 0.89470696 | 0.0596 | 0.1151 | 0.1004 | 0.1004 | TRUE | 0.2327 |
| E130310I04R | 0 | 1 | 0 | 0 | 0 | 0 | 0 | 1 | 0 | 0 | 0 | 0 | 0.18 | 0 | 0 | 0.552 | -0.857 | 0.8336548 | NA | 0 | 13.3714 | 10 | 10 | TRUE | 0.5216 |
| E130311K13I | 260 | 242 | 260 | 283 | 215 | 255 | 202 | 257 | 281 | 282 | 243 | 252 | 252.64 | 247 | 259 | 1.054 | 0.076 | 0.61294749 | 0.75508227 | 0.0138 | 0.0118 | 0.0124 | 0.0124 | TRUE | 0.3022 |
| E130317F20I | 111 | 94 | 62 | 73 | 73 | 62 | 86 | 100 | 67 | 73 | 82 | 61 | 78.22 | 84 | 72 | 0.852 | -0.232 | 0.33354666 | 0.50773953 | 0.0169 | 0.0327 | 0.0284 | 0.0284 | TRUE | 0.2398 |
| E230008N13 | 6 | 5 | 7 | 3 | 1 | 3 | 5 | 5 | 8 | 3 | 1 | 3 | 4.1 | 6 | 2 | 0.411 | -1.282 | 0.1888379 | NA | 0 | 0.5803 | 0.3862 | 0.3862 | TRUE | 0.1856 |
| E230016K23I | 28 | 25 | 19 | 19 | 12 | 16 | 22 | 27 | 21 | 19 | 14 | 16 | 19.51 | 23 | 16 | 0.704 | -0.506 | 0.24450835 | 0.41270168 | 0 | 0.1239 | 0.0833 | 0.0833 | TRUE | 0.1137 |
| E230016M11 | 12 | 4 | 10 | 9 | 6 | 6 | 9 | 4 | 11 | 9 | 7 | 6 | 7.67 | 8 | 7 | 0.885 | -0.176 | 0.80196907 | NA | 0 | 0.3115 | 0.2239 | 0.2239 | TRUE | 0.471 |
| E230019M04 | 0 | 0 | 4 | 2 | 0 | 3 | 0 | 0 | 4 | 2 | 0 | 3 | 1.54 | 1 | 2 | 1.197 | 0.259 | 0.8891316 | NA | 3.5742 | 1.5367 | 1.837 | 1.837 | TRUE | 1.6606 |
| E230025N22 | 30 | 19 | 27 | 23 | 22 | 26 | 23 | 20 | 29 | 23 | 25 | 26 | 24.34 | 24 | 25 | 1.012 | 0.017 | 0.96520311 | 0.98098898 | 0 | 0.0998 | 0.0671 | 0.0671 | TRUE | 0.1716 |
| E230029C05I | 0 | 0 | 0 | 0 | 0 | 0 | 0 | 0 | 0 | 0 | 0 | 0 | 0 | 0 | 0 | NA | NA | NA | NA | NA | NA | NA | NA | NA | NA |
| E2f1 | 3057 | 2265 | 2428 | 2845 | 2607 | 2863 | 2374 | 2409 | 2620 | 2832 | 2941 | 2826 | 2667.06 | 2468 | 2866 | 1.162 | 0.217 | 0.00061682 | 0.00507387 | 0.0011 | 0.0033 | 0.0025 | 0.0025 | TRUE | 0.0348 |
| E2f2 | 471 | 347 | 358 | 582 | 493 | 574 | 366 | 369 | 386 | 579 | 556 | 567 | 470.55 | 374 | 567 | 1.52 | 0.604 | 6.40E-10 | 4.57E-08 | 0 | 0.0075 | 0.0047 | 0.0047 | TRUE | 0.0096 |
| E2f3 | 2671 | 1816 | 1510 | 1369 | 1334 | 1460 | 2074 | 1932 | 1629 | 1363 | 1505 | 1441 | 1657.37 | 1878 | 1436 | 0.764 | -0.389 | 2.05E-05 | 0.00033804 | 0.0076 | 0.0039 | 0.0054 | 0.0054 | TRUE | 0.1616 |
| E2f4 | 10174 | 7510 | 7150 | 6624 | 6645 | 7614 | 7902 | 7988 | 7715 | 6594 | 7496 | 7516 | 7535.13 | 7868 | 7202 | 0.915 | -0.128 | 0.04336725 | 0.12137225 | 0.0026 | 0.0027 | 0.0028 | 0.0028 | TRUE | 0.0668 |
| E2f5 | 1768 | 1357 | 1592 | 1636 | 1380 | 1465 | 1373 | 1443 | 1718 | 1628 | 1557 | 1446 | 1527.62 | 1511 | 1544 | 1.022 | 0.032 | 0.72736965 | 0.83661541 | 0.0082 | 0.004 | 0.0054 | 0.0054 | TRUE | 0.1706 |
| E2f6 | 1937 | 1281 | 1171 | 1420 | 1110 | 1258 | 1504 | 1363 | 1263 | 1413 | 1252 | 1242 | 1339.66 | 1377 | 1302 | 0.945 | -0.082 | 0.35136831 | 0.52609545 | 0.0055 | 0.0042 | 0.0048 | 0.0048 | TRUE | 0.0812 |
| E2f7 | 1726 | 1242 | 1219 | 1164 | 1110 | 1311 | 1341 | 1321 | 1315 | 1159 | 1252 | 1294 | 1280.32 | 1326 | 1235 | 0.931 | -0.103 | 0.1586339 | 0.30789419 | 8.00E-04 | 0.0043 | 0.003 | 0.003 | TRUE | 0.0356 |
| E2f8 | 3954 | 2815 | 2858 | 2355 | 2277 | 2788 | 3071 | 2994 | 3084 | 2344 | 2569 | 2752 | 2802.33 | 3050 | 2555 | 0.838 | -0.255 | 0.00025076 | 0.00253476 | 0.0028 | 0.0033 | 0.0031 | 0.0031 | TRUE | 0.0638 |
| E330009J07F | 324 | 245 | 262 | 292 | 241 | 240 | 252 | 261 | 283 | 291 | 272 | 237 | 265.73 | 265 | 267 | 1.007 | 0.01 | 0.94047543 | 0.96671811 | 0.0035 | 0.0113 | 0.009 | 0.009 | TRUE | 0.1081 |
| E330011O21 | 1 | 1 | 0 | 0 | 0 | 0 | 1 | 1 | 0 | 0 | 0 | 0 | 0.31 | 1 | 0 | 0.309 | -1.696 | 0.63183309 | NA | 0 | 7.7294 | 7.2472 | 7.2472 | TRUE | 0.2015 |
| E330012B07 | 0 | 0 | 0 | 0 | 0 | 0 | 0 | 0 | 0 | 0 | 0 | 0 | 0 | 0 | 0 | NA | NA | NA | NA | NA | NA | NA | NA | NA | NA |
| E330013P04I | 0 | 1 | 1 | 0 | 1 | 0 | 0 | 1 | 1 | 0 | 1 | 0 | 0.55 | 1 | 0 | 0.535 | -0.902 | 0.73748201 | NA | 0 | 4.3501 | 3.5264 | 3.5264 | TRUE | 0.5075 |
| E330014E10I | 0 | 0 | 0 | 0 | 0 | 0 | 0 | 0 | 0 | 0 | 0 | 0 | 0 | 0 | 0 | NA | NA | NA | NA | NA | NA | NA | NA | NA | NA |

| | | | | | | | | | | | | | | | | | | | | | | | | | |
|---|---|---|---|---|---|---|---|---|---|---|---|---|---|---|---|---|---|---|---|---|---|---|---|---|---|
| E330017A01 | 1 | 0 | 0 | 0 | 0 | 0 | 1 | 0 | 0 | 0 | 0 | 0 | 0.13 | 0 | 0 | 0.552 | -0.857 | 0.8336548 | NA | 0 | 18.3121 | 10 | 10 | TRUE | 0.2566 |
| E330017L17 | 1 | 0 | 0 | 1 | 0 | 0 | 1 | 0 | 0 | 1 | 0 | 0 | 0.3 | 0 | 0 | 1.075 | 0.105 | 0.97698525 | NA | 0 | 8.0272 | 7.5141 | 7.5141 | TRUE | 0.3906 |
| E330020D12 | 112 | 83 | 73 | 102 | 92 | 115 | 87 | 88 | 79 | 102 | 104 | 114 | 95.48 | 85 | 107 | 1.254 | 0.327 | 0.10066271 | 0.22210129 | 0 | 0.0272 | 0.018 | 0.018 | TRUE | 0.0361 |
| E330021D16 | 7 | 1 | 3 | 3 | 2 | 0 | 5 | 1 | 3 | 3 | 2 | 0 | 2.5 | 3 | 2 | 0.525 | -0.929 | 0.47921948 | NA | 0 | 0.9517 | 0.7976 | 0.7976 | TRUE | 0.2278 |
| E330023G01 | 5 | 3 | 3 | 0 | 1 | 1 | 4 | 3 | 3 | 0 | 1 | 1 | 2.07 | 3 | 1 | 0.201 | -2.313 | 0.12598527 | NA | 0 | 1.1468 | 0.7619 | 0.7619 | TRUE | 0.2676 |
| E330033B04 | 20 | 21 | 14 | 11 | 18 | 19 | 16 | 22 | 15 | 11 | 20 | 19 | 17.16 | 18 | 17 | 0.945 | -0.082 | 0.86374757 | 0.92475985 | 0 | 0.1405 | 0.1062 | 0.1062 | TRUE | 0.4494 |
| E330034G19 | 1 | 0 | 0 | 0 | 0 | 0 | 1 | 0 | 0 | 0 | 0 | 0 | 0.13 | 0 | 0 | 0.552 | -0.857 | 0.8336548 | NA | 0 | 18.3121 | 10 | 10 | TRUE | 0.2566 |
| E430016F16 | 0 | 0 | 0 | 0 | 0 | 0 | 0 | 0 | 0 | 0 | 0 | 0 | 0 | 0 | 0 | NA | NA | NA | NA | NA | NA | NA | NA | NA | NA |
| E430018J23 | 104 | 77 | 50 | 79 | 72 | 64 | 81 | 82 | 54 | 79 | 81 | 63 | 73.28 | 72 | 74 | 1.024 | 0.034 | 0.88997726 | 0.93898109 | 0.0197 | 0.0348 | 0.0307 | 0.0307 | TRUE | 0.4179 |
| E430025E21 | 9789 | 6911 | 6463 | 6246 | 5422 | 6400 | 7603 | 7351 | 6974 | 6217 | 6117 | 6318 | 6763.17 | 7309 | 6217 | 0.85 | -0.234 | 1.67E-05 | 0.00029072 | 8.00E-04 | 0.0028 | 0.002 | 0.002 | TRUE | 0.0196 |
| E4f1 | 1052 | 850 | 897 | 969 | 873 | 972 | 817 | 904 | 968 | 965 | 985 | 960 | 932.99 | 896 | 970 | 1.084 | 0.116 | 0.17527888 | 0.32873626 | 0.0028 | 0.005 | 0.0042 | 0.0042 | TRUE | 0.0751 |
| E530001F21 | 1 | 1 | 0 | 0 | 0 | 0 | 1 | 1 | 0 | 0 | 0 | 0 | 0.31 | 1 | 0 | 0.309 | -1.696 | 0.63183309 | NA | 0 | 7.7294 | 7.2472 | 7.2472 | TRUE | 0.2015 |
| E530011L22 | 6 | 15 | 6 | 18 | 7 | 19 | 5 | 16 | 6 | 18 | 8 | 19 | 11.94 | 9 | 15 | 1.682 | 0.75 | 0.23808193 | NA | 0.2068 | 0.2009 | 0.2025 | 0.2025 | TRUE | 1.3926 |
| Eaf1 | 5296 | 3816 | 3388 | 3880 | 3463 | 4033 | 4113 | 4059 | 3656 | 3862 | 3907 | 3981 | 3929.63 | 3943 | 3917 | 0.993 | -0.01 | 0.87215058 | 0.92903341 | 0.0019 | 0.003 | 0.0026 | 0.0026 | TRUE | 0.0489 |
| Eaf2 | 86 | 60 | 69 | 87 | 68 | 87 | 67 | 64 | 74 | 87 | 77 | 86 | 75.71 | 68 | 83 | 1.218 | 0.284 | 0.20136681 | 0.36221191 | 0 | 0.0337 | 0.0224 | 0.0224 | TRUE | 0.0503 |
| Eapp | 1648 | 1222 | 1125 | 1195 | 1085 | 1296 | 1280 | 1300 | 1214 | 1190 | 1224 | 1279 | 1247.74 | 1265 | 1231 | 0.973 | -0.039 | 0.58701617 | 0.73617813 | 5.00E-04 | 0.0043 | 0.0029 | 0.0029 | TRUE | 0.0145 |
| Ear1 | 0 | 0 | 0 | 0 | 0 | 0 | 0 | 0 | 0 | 0 | 0 | 0 | 0 | 0 | 0 | NA | NA | NA | NA | NA | NA | NA | NA | NA | NA |
| Ear10 | 0 | 0 | 0 | 0 | 0 | 0 | 0 | 0 | 0 | 0 | 0 | 0 | 0 | 0 | 0 | NA | NA | NA | NA | NA | NA | NA | NA | NA | NA |
| Ear11 | 0 | 0 | 0 | 0 | 0 | 0 | 0 | 0 | 0 | 0 | 0 | 0 | 0 | 0 | 0 | NA | NA | NA | NA | NA | NA | NA | NA | NA | NA |
| Ear12 | 0 | 0 | 0 | 0 | 0 | 0 | 0 | 0 | 0 | 0 | 0 | 0 | 0 | 0 | 0 | NA | NA | NA | NA | NA | NA | NA | NA | NA | NA |
| Ear14 | 0 | 0 | 0 | 0 | 0 | 0 | 0 | 0 | 0 | 0 | 0 | 0 | 0 | 0 | 0 | NA | NA | NA | NA | NA | NA | NA | NA | NA | NA |
| Ear2 | 0 | 0 | 0 | 0 | 0 | 0 | 0 | 0 | 0 | 0 | 0 | 0 | 0 | 0 | 0 | NA | NA | NA | NA | NA | NA | NA | NA | NA | NA |
| Ear3 | 0 | 0 | 0 | 0 | 0 | 0 | 0 | 0 | 0 | 0 | 0 | 0 | 0 | 0 | 0 | NA | NA | NA | NA | NA | NA | NA | NA | NA | NA |
| Ear4 | 0 | 0 | 0 | 0 | 0 | 0 | 0 | 0 | 0 | 0 | 0 | 0 | 0 | 0 | 0 | NA | NA | NA | NA | NA | NA | NA | NA | NA | NA |
| Ear5 | 0 | 0 | 0 | 0 | 0 | 0 | 0 | 0 | 0 | 0 | 0 | 0 | 0 | 0 | 0 | NA | NA | NA | NA | NA | NA | NA | NA | NA | NA |
| Ear6 | 0 | 0 | 0 | 0 | 0 | 0 | 0 | 0 | 0 | 0 | 0 | 0 | 0 | 0 | 0 | NA | NA | NA | NA | NA | NA | NA | NA | NA | NA |
| Ear7 | 0 | 0 | 0 | 0 | 0 | 0 | 0 | 0 | 0 | 0 | 0 | 0 | 0 | 0 | 0 | NA | NA | NA | NA | NA | NA | NA | NA | NA | NA |
| Ears2 | 1564 | 1099 | 1102 | 1185 | 1057 | 1166 | 1215 | 1169 | 1189 | 1180 | 1192 | 1151 | 1182.62 | 1191 | 1174 | 0.985 | -0.021 | 0.76125862 | 0.85830003 | 0 | 0.0044 | 0.0027 | 0.0027 | TRUE | 0.0039 |
| Ebag9 | 962 | 741 | 713 | 654 | 614 | 702 | 747 | 788 | 769 | 651 | 693 | 693 | 723.55 | 768 | 679 | 0.884 | -0.178 | 0.03415292 | 0.10205888 | 0 | 0.0057 | 0.0037 | 0.0037 | TRUE | 0.0154 |
| Ebf1 | 59 | 44 | 47 | 69 | 72 | 50 | 46 | 47 | 51 | 69 | 81 | 49 | 57.1 | 48 | 66 | 1.388 | 0.473 | 0.08484944 | 0.19652158 | 0.0176 | 0.0439 | 0.0364 | 0.0364 | TRUE | 0.2612 |
| Ebf2 | 6 | 5 | 6 | 8 | 6 | 1 | 5 | 5 | 6 | 8 | 7 | 1 | 5.36 | 5 | 5 | 0.956 | -0.065 | 0.94052087 | NA | 0 | 0.4444 | 0.3669 | 0.3669 | TRUE | 1.1107 |
| Ebf3 | 84 | 51 | 47 | 52 | 34 | 34 | 65 | 54 | 51 | 52 | 38 | 34 | 48.98 | 57 | 41 | 0.724 | -0.466 | 0.11042461 | 0.23780306 | 0.0125 | 0.0508 | 0.0404 | 0.0404 | TRUE | 0.3899 |
| Ebf4 | 40 | 35 | 43 | 50 | 39 | 51 | 31 | 37 | 46 | 50 | 44 | 50 | 43.13 | 38 | 48 | 1.268 | 0.342 | 0.25491281 | 0.42444884 | 0 | 0.0574 | 0.0417 | 0.0417 | TRUE | 0.2501 |
| Ebi3 | 11 | 14 | 10 | 12 | 22 | 24 | 9 | 15 | 11 | 12 | 25 | 24 | 15.78 | 12 | 20 | 1.78 | 0.832 | 0.10876143 | NA | 0 | 0.1526 | 0.1255 | 0.1255 | TRUE | 0.7052 |
| Ebna1bp2 | 9218 | 6397 | 5980 | 5937 | 5537 | 6373 | 7159 | 6804 | 6452 | 5910 | 6246 | 6291 | 6477.19 | 6805 | 6149 | 0.903 | -0.147 | 0.01320026 | 0.0507727 | 0.0017 | 0.0028 | 0.0024 | 0.0024 | TRUE | 0.0255 |
| Ebp | 939 | 655 | 647 | 787 | 680 | 754 | 729 | 697 | 698 | 783 | 767 | 744 | 736.48 | 708 | 765 | 1.079 | 0.11 | 0.1813723 | 0.33693193 | 0 | 0.0056 | 0.0035 | 0.0035 | TRUE | 0.0087 |
| Ebpl | 1608 | 1056 | 1145 | 1285 | 1131 | 1169 | 1249 | 1123 | 1235 | 1279 | 1276 | 1154 | 1219.42 | 1202 | 1236 | 1.027 | 0.039 | 0.62677001 | 0.76482743 | 0.0025 | 0.0044 | 0.0037 | 0.0037 | TRUE | 0.0412 |
| Ecd | 4551 | 3237 | 3069 | 3197 | 2881 | 3231 | 3535 | 3443 | 3311 | 3182 | 3250 | 3189 | 3318.5 | 3430 | 3207 | 0.935 | -0.097 | 0.0816878 | 0.19203351 | 3.00E-04 | 0.0031 | 0.002 | 0.002 | TRUE | 0.011 |
| Ece1 | 4093 | 3059 | 3511 | 4182 | 3780 | 4599 | 3179 | 3254 | 3788 | 4163 | 4264 | 4540 | 3864.66 | 3407 | 4322 | 1.269 | 0.344 | 5.66E-06 | 0.00011731 | 0.0054 | 0.003 | 0.0039 | 0.0039 | TRUE | 0.1159 |
| Ece2 | 998 | 755 | 711 | 858 | 783 | 845 | 775 | 803 | 767 | 854 | 883 | 834 | 819.48 | 782 | 857 | 1.096 | 0.133 | 0.09533766 | 0.21361674 | 0 | 0.0053 | 0.0033 | 0.0033 | TRUE | 0.0083 |
| Ecel1 | 234 | 143 | 134 | 209 | 193 | 175 | 182 | 152 | 145 | 208 | 218 | 173 | 179.49 | 160 | 200 | 1.244 | 0.315 | 0.05533024 | 0.14489466 | 0.0096 | 0.0156 | 0.0139 | 0.0139 | TRUE | 0.1716 |
| Ech1 | 2469 | 1936 | 2384 | 2549 | 2426 | 2775 | 1918 | 2059 | 2572 | 2537 | 2737 | 2739 | 2427.1 | 2183 | 2671 | 1.225 | 0.292 | 0.002629 | 0.01489843 | 0.0128 | 0.0034 | 0.0064 | 0.0064 | TRUE | 0.2935 |
| Echdc1 | 1189 | 867 | 936 | 1252 | 1096 | 1182 | 923 | 922 | 1010 | 1246 | 1236 | 1167 | 1084.18 | 952 | 1216 | 1.279 | 0.355 | 4.23E-06 | 9.25E-05 | 0.001 | 0.0046 | 0.0034 | 0.0034 | TRUE | 0.0331 |
| Echdc2 | 1770 | 1286 | 1258 | 1339 | 1112 | 1269 | 1375 | 1368 | 1357 | 1333 | 1254 | 1253 | 1323.32 | 1367 | 1280 | 0.937 | -0.094 | 0.16825255 | 0.31979172 | 0 | 0.0042 | 0.0026 | 0.0026 | TRUE | 0.0158 |
| Echdc3 | 78 | 53 | 59 | 63 | 59 | 66 | 61 | 56 | 64 | 63 | 67 | 65 | 62.51 | 60 | 65 | 1.076 | 0.105 | 0.66176918 | 0.79124932 | 0 | 0.0403 | 0.0259 | 0.0259 | TRUE | 0.0245 |
| Echs1 | 3173 | 2501 | 2792 | 3179 | 2826 | 3393 | 2464 | 2660 | 3013 | 3164 | 3188 | 3349 | 2973.16 | 2712 | 3234 | 1.193 | 0.255 | 0.00099459 | 0.00721977 | 0.0052 | 0.0032 | 0.004 | 0.004 | TRUE | 0.1134 |
| Eci1 | 941 | 657 | 670 | 707 | 677 | 747 | 731 | 699 | 723 | 704 | 764 | 737 | 726.25 | 718 | 735 | 1.023 | 0.033 | 0.69201604 | 0.81166791 | 0 | 0.0057 | 0.0037 | 0.0037 | TRUE | 0.0164 |
| Eci2 | 1634 | 1128 | 1234 | 1147 | 989 | 1180 | 1269 | 1200 | 1331 | 1142 | 1116 | 1165 | 1203.77 | 1267 | 1141 | 0.901 | -0.151 | 0.04077913 | 0.11608368 | 7.00E-04 | 0.0044 | 0.0031 | 0.0031 | TRUE | 0.0247 |
| Eci3 | 27 | 13 | 23 | 26 | 24 | 26 | 21 | 14 | 25 | 26 | 27 | 26 | 23.04 | 20 | 26 | 1.316 | 0.396 | 0.32979064 | 0.50423161 | 0 | 0.1053 | 0.075 | 0.075 | TRUE | 0.3463 |
| Ecm1 | 49 | 32 | 28 | 46 | 30 | 41 | 38 | 34 | 30 | 46 | 34 | 40 | 37.07 | 34 | 40 | 1.171 | 0.228 | 0.47438026 | 0.64280192 | 0 | 0.0664 | 0.0462 | 0.0462 | TRUE | 0.1269 |
| Ecm2 | 17 | 16 | 6 | 1 | 3 | 12 | 13 | 17 | 6 | 1 | 3 | 12 | 8.82 | 12 | 5 | 0.444 | -1.17 | 0.14022714 | NA | 0.4722 | 0.2711 | 0.319 | 0.319 | TRUE | 1.1924 |
| Ecscr | 129 | 96 | 148 | 214 | 232 | 258 | 100 | 102 | 160 | 213 | 262 | 255 | 181.9 | 121 | 243 | 2.025 | 1.018 | 1.02E-07 | 4.01E-06 | 0.0333 | 0.0155 | 0.0202 | 0.0202 | TRUE | 0.7847 |
| Ecsit | 2505 | 1833 | 1994 | 2258 | 2017 | 2564 | 1946 | 1950 | 2152 | 2248 | 2275 | 2531 | 2183.47 | 2016 | 2351 | 1.167 | 0.223 | 0.00248161 | 0.01427351 | 0.0032 | 0.0035 | 0.0035 | 0.0035 | TRUE | 0.0545 |
| Ect2 | 15847 | 11331 | 12453 | 12842 | 11275 | 12884 | 12308 | 12053 | 13437 | 12783 | 12719 | 12718 | 12669.63 | 12599 | 12740 | 1.011 | 0.016 | 0.77167643 | 0.86486091 | 0.0013 | 0.0026 | 0.0022 | 0.0022 | TRUE | 0.0412 |
| Ect2l | 7 | 2 | 2 | 0 | 3 | 3 | 5 | 2 | 2 | 0 | 3 | 3 | 2.68 | 3 | 2 | 0.634 | -0.658 | 0.59770032 | NA | 0 | 0.8875 | 0.7287 | 0.7287 | TRUE | 0.7512 |
| Eda | 436 | 297 | 266 | 298 | 260 | 303 | 339 | 316 | 287 | 297 | 293 | 299 | 305.1 | 314 | 296 | 0.942 | -0.086 | 0.47618642 | 0.64403222 | 0 | 0.0102 | 0.0073 | 0.0073 | TRUE | 0.0623 |
| Eda2r | 3704 | 2655 | 3272 | 4069 | 3351 | 3831 | 2877 | 2824 | 3530 | 4050 | 3780 | 3782 | 3473.94 | 3077 | 3871 | 1.259 | 0.332 | 9.34E-05 | 0.00114591 | 0.0085 | 0.0031 | 0.0049 | 0.0049 | TRUE | 0.2004 |
| Edar | 9 | 9 | 12 | 16 | 11 | 10 | 7 | 10 | 13 | 16 | 12 | 10 | 11.29 | 10 | 13 | 1.311 | 0.391 | 0.49965367 | NA | 0 | 0.2124 | 0.1514 | 0.1514 | TRUE | 0.2903 |
| Edaradd | 73 | 70 | 78 | 57 | 49 | 56 | 57 | 74 | 84 | 57 | 55 | 55 | 63.77 | 72 | 56 | 0.782 | -0.354 | 0.16476948 | 0.31552065 | 0 | 0.0396 | 0.0307 | 0.0307 | TRUE | 0.3503 |
| Edc3 | 4217 | 3033 | 2960 | 3194 | 2812 | 3373 | 3275 | 3226 | 3194 | 3179 | 3172 | 3330 | 3229.39 | 3232 | 3227 | 0.998 | -0.002 | 0.9684824 | 0.9829888 | 0 | 0.0032 | 0.0019 | 0.0019 | TRUE | 0.0095 |
| Edc4 | 11661 | 9091 | 8529 | 8679 | 8181 | 9151 | 9057 | 9670 | 9203 | 8639 | 9229 | 9033 | 9138.47 | 9310 | 8967 | 0.963 | -0.054 | 0.3134678 | 0.48694656 | 0.001 | 0.0027 | 0.002 | 0.002 | TRUE | 0.014 |
| Eddm3b | 0 | 1 | 0 | 0 | 0 | 0 | 0 | 1 | 0 | 0 | 0 | 0 | 0.18 | 0 | 0 | 0.552 | -0.857 | 0.8336548 | NA | 0 | 13.3714 | 10 | 10 | TRUE | 0.5216 |
| Edem1 | 6951 | 5117 | 5274 | 6127 | 5214 | 5643 | 5399 | 5443 | 5691 | 6099 | 5882 | 5571 | 5680.55 | 5511 | 5851 | 1.062 | 0.087 | 0.12986806 | 0.2665811 | 0.0012 | 0.0028 | 0.0022 | 0.0022 | TRUE | 0.0215 |
| Edem2 | 3439 | 2709 | 2857 | 3322 | 3004 | 3245 | 2671 | 2882 | 3083 | 3307 | 3389 | 3203 | 3089 | 2879 | 3300 | 1.147 | 0.198 | 0.00347818 | 0.01853312 | 0.0025 | 0.0032 | 0.003 | 0.003 | TRUE | 0.0494 |
| Edem3 | 5770 | 4171 | 4057 | 4832 | 4267 | 4812 | 4481 | 4437 | 4377 | 4810 | 4814 | 4750 | 4611.5 | 4432 | 4791 | 1.081 | 0.112 | 0.02434204 | 0.07981666 | 0 | 0.0029 | 0.0016 | 0.0016 | TRUE | 0.0014 |
| Edf1 | 4047 | 2828 | 2637 | 2793 | 2382 | 2771 | 3143 | 3008 | 2845 | 2780 | 2687 | 2735 | 2866.54 | 2999 | 2734 | 0.911 | -0.134 | 0.02934122 | 0.0911671 | 0.0011 | 0.0033 | 0.0024 | 0.0024 | TRUE | 0.0243 |
| Edil3 | 12 | 23 | 11 | 7 | 4 | 5 | 9 | 24 | 12 | 7 | 5 | 5 | 10.34 | 15 | 6 | 0.364 | -1.457 | 0.02918766 | NA | 0 | 0.2315 | 0.1953 | 0.1953 | TRUE | 1.2548 |
| Edn1 | 45 | 28 | 21 | 13 | 18 | 10 | 35 | 30 | 23 | 13 | 20 | 10 | 21.75 | 29 | 14 | 0.487 | -1.039 | 0.01956232 | 0.06798384 | 0.0264 | 0.1114 | 0.09 | 0.09 | TRUE | 0.4215 |
| Edn2 | 20 | 8 | 14 | 18 | 22 | 13 | 16 | 9 | 15 | 18 | 25 | 13 | 15.79 | 13 | 19 | 1.403 | 0.488 | 0.33641784 | NA | 3.00E-04 | 0.1526 | 0.121 | 0.121 | TRUE | 0.2032 |

| | | | | | | | | | | | | | | | | | | | | | | | | | |
|---|---|---|---|---|---|---|---|---|---|---|---|---|---|---|---|---|---|---|---|---|---|---|---|---|---|
| Edn3 | 9 | 14 | 14 | 14 | 12 | 9 | 7 | 15 | 15 | 14 | 14 | 9 | 12.22 | 12 | 12 | 0.997 | -0.005 | 0.99302414 | NA | 0 | 0.1963 | 0.1523 | 0.1523 | TRUE | 0.7482 |
| Ednra | 73 | 53 | 55 | 57 | 48 | 40 | 57 | 56 | 59 | 57 | 54 | 39 | 53.8 | 57 | 50 | 0.871 | -0.199 | 0.46084424 | 0.63032026 | 0 | 0.0465 | 0.0336 | 0.0336 | TRUE | 0.2899 |
| Ednrb | 8 | 4 | 10 | 10 | 9 | 5 | 6 | 4 | 11 | 10 | 10 | 5 | 7.72 | 7 | 8 | 1.18 | 0.239 | 0.73694093 | NA | 0 | 0.3096 | 0.2353 | 0.2353 | TRUE | 0.5042 |
| Eea1 | 2107 | 1537 | 1589 | 1722 | 1477 | 1654 | 1636 | 1635 | 1715 | 1714 | 1666 | 1633 | 1666.48 | 1662 | 1671 | 1.006 | 0.008 | 0.89716199 | 0.94337809 | 0 | 0.0038 | 0.0024 | 0.0024 | TRUE | 0.009 |
| Eed | 6743 | 4904 | 4704 | 4809 | 4086 | 4930 | 5237 | 5216 | 5076 | 4787 | 4609 | 4867 | 4965.31 | 5176 | 4754 | 0.919 | -0.123 | 0.01901068 | 0.06663062 | 2.00E-04 | 0.0029 | 0.0018 | 0.0018 | TRUE | 0.0086 |
| Eef1a1 | 643842 | 450583 | 481952 | 542430 | 482214 | 548892 | 500045 | 479279 | 520022 | 539934 | 543987 | 541841 | 520851.49 | 499782 | 541921 | 1.084 | 0.117 | 0.01534705 | 0.05684256 | 8.00E-04 | 0.0024 | 0.0017 | 0.0017 | TRUE | 0.0158 |
| Eef1a2 | 214 | 124 | 147 | 220 | 179 | 205 | 166 | 132 | 159 | 219 | 202 | 202 | 180 | 152 | 208 | 1.361 | 0.444 | 0.00425114 | 0.02165413 | 0 | 0.0156 | 0.0118 | 0.0118 | TRUE | 0.1387 |
| Eef1b2 | 48161 | 32851 | 34221 | 38336 | 32854 | 38416 | 37405 | 34943 | 36924 | 38160 | 37063 | 37923 | 37069.47 | 36424 | 37715 | 1.035 | 0.05 | 0.29968075 | 0.47188905 | 7.00E-04 | 0.0025 | 0.0017 | 0.0017 | TRUE | 0.0155 |
| Eef1d | 17947 | 13151 | 12685 | 13282 | 11739 | 13596 | 13939 | 13989 | 13687 | 13221 | 13243 | 13421 | 13583.21 | 13872 | 13295 | 0.958 | -0.061 | 0.16688733 | 0.31800034 | 0 | 0.0026 | 0.0013 | 0.0013 | TRUE | 0.0016 |
| Eef1e1 | 4545 | 3196 | 3369 | 3590 | 3163 | 3549 | 3530 | 3400 | 3635 | 3573 | 3568 | 3503 | 3534.94 | 3522 | 3548 | 1.008 | 0.011 | 0.84416309 | 0.9118518 | 3.00E-04 | 0.0031 | 0.0019 | 0.0019 | TRUE | 0.0109 |
| Eef1g | 59818 | 44135 | 44846 | 50056 | 44799 | 51231 | 46458 | 46946 | 48388 | 49826 | 50538 | 50573 | 48788.15 | 47264 | 50312 | 1.065 | 0.09 | 0.03900529 | 0.11246774 | 2.00E-04 | 0.0025 | 0.0014 | 0.0014 | TRUE | 0.0053 |
| Eef2 | 293149 | 224883 | 245332 | 262987 | 240714 | 275198 | 227676 | 239205 | 264711 | 261777 | 271550 | 271663 | 256097.15 | 243864 | 268330 | 1.1 | 0.138 | 0.02564351 | 0.08279257 | 0.0032 | 0.0024 | 0.0027 | 0.0027 | TRUE | 0.0685 |
| Eef2k | 1064 | 779 | 801 | 927 | 885 | 956 | 826 | 829 | 864 | 923 | 998 | 944 | 897.35 | 840 | 955 | 1.137 | 0.185 | 0.01827035 | 0.06459019 | 0 | 0.0051 | 0.0033 | 0.0033 | TRUE | 0.0184 |
| Eefsec | 2914 | 2125 | 2078 | 1996 | 1884 | 2228 | 2263 | 2260 | 2242 | 1987 | 2125 | 2199 | 2179.53 | 2255 | 2104 | 0.933 | -0.1 | 0.11615017 | 0.24664008 | 9.00E-04 | 0.0035 | 0.0025 | 0.0025 | TRUE | 0.029 |
| Eepd1 | 329 | 198 | 249 | 219 | 172 | 220 | 256 | 211 | 269 | 218 | 194 | 217 | 227.33 | 245 | 210 | 0.856 | -0.224 | 0.12296959 | 0.25650968 | 0.0056 | 0.0129 | 0.0107 | 0.0107 | TRUE | 0.1597 |
| Efcab1 | 90 | 81 | 72 | 80 | 58 | 68 | 70 | 86 | 78 | 80 | 65 | 67 | 74.32 | 78 | 71 | 0.912 | -0.133 | 0.56280642 | 0.71733477 | 0 | 0.0343 | 0.0244 | 0.0244 | TRUE | 0.1109 |
| Efcab10 | 56 | 50 | 69 | 67 | 67 | 78 | 43 | 53 | 74 | 67 | 76 | 77 | 65.07 | 57 | 73 | 1.292 | 0.369 | 0.16215578 | 0.31223125 | 0.0238 | 0.0389 | 0.0347 | 0.0347 | TRUE | 0.5898 |
| Efcab11 | 220 | 160 | 193 | 207 | 174 | 184 | 171 | 170 | 208 | 206 | 196 | 182 | 188.88 | 183 | 195 | 1.065 | 0.091 | 0.55091195 | 0.70690228 | 0.0034 | 0.015 | 0.0116 | 0.0116 | TRUE | 0.1492 |
| Efcab12 | 302 | 249 | 185 | 172 | 168 | 198 | 235 | 265 | 200 | 171 | 190 | 195 | 209.2 | 233 | 185 | 0.795 | -0.331 | 0.03047181 | 0.09360275 | 0.0076 | 0.0138 | 0.012 | 0.012 | TRUE | 0.1636 |
| Efcab14 | 2852 | 2030 | 2133 | 2517 | 2155 | 2415 | 2215 | 2159 | 2301 | 2505 | 2431 | 2384 | 2332.71 | 2225 | 2440 | 1.097 | 0.133 | 0.02822646 | 0.08855154 | 4.00E-04 | 0.0034 | 0.0022 | 0.0022 | TRUE | 0.0105 |
| Efcab2 | 281 | 223 | 186 | 222 | 160 | 199 | 218 | 237 | 201 | 221 | 180 | 196 | 209.01 | 219 | 199 | 0.912 | -0.132 | 0.36858036 | 0.54321707 | 0.0032 | 0.0138 | 0.0108 | 0.0108 | TRUE | 0.0985 |
| Efcab3 | 17 | 20 | 8 | 9 | 9 | 7 | 13 | 21 | 9 | 9 | 10 | 7 | 11.52 | 14 | 9 | 0.603 | -0.73 | 0.21979591 | NA | 0 | 0.2081 | 0.1617 | 0.1617 | TRUE | 0.268 |
| Efcab4a | 262 | 241 | 228 | 279 | 285 | 328 | 203 | 256 | 246 | 278 | 322 | 324 | 271.48 | 235 | 308 | 1.312 | 0.392 | 0.00463496 | 0.02313916 | 0.0076 | 0.0112 | 0.0101 | 0.0101 | TRUE | 0.1692 |
| Efcab4b | 36 | 32 | 16 | 33 | 25 | 31 | 28 | 34 | 17 | 33 | 28 | 31 | 28.49 | 26 | 31 | 1.154 | 0.207 | 0.58214869 | 0.73253195 | 0.0093 | 0.0856 | 0.0671 | 0.0671 | TRUE | 0.4265 |
| Efcab5 | 24 | 15 | 25 | 23 | 19 | 24 | 19 | 16 | 27 | 23 | 21 | 24 | 21.6 | 21 | 23 | 1.11 | 0.151 | 0.71784659 | 0.83038049 | 0 | 0.1122 | 0.0793 | 0.0793 | TRUE | 0.3797 |
| Efcab6 | 35 | 40 | 32 | 50 | 35 | 43 | 27 | 43 | 35 | 50 | 39 | 42 | 39.33 | 35 | 44 | 1.276 | 0.351 | 0.2706824 | 0.44170151 | 0 | 0.0627 | 0.0475 | 0.0475 | TRUE | 0.2863 |
| Efcab7 | 312 | 162 | 121 | 156 | 122 | 135 | 242 | 172 | 131 | 155 | 138 | 133 | 161.89 | 182 | 142 | 0.777 | -0.363 | 0.08583145 | 0.19802889 | 0.0466 | 0.0171 | 0.026 | 0.026 | TRUE | 0.3184 |
| Efcab8 | 8 | 10 | 6 | 6 | 10 | 8 | 6 | 11 | 6 | 6 | 11 | 8 | 8.08 | 8 | 8 | 1.08 | 0.112 | 0.86928015 | NA | 0 | 0.2958 | 0.2072 | 0.2072 | TRUE | 0.2804 |
| Efcab9 | 3 | 1 | 1 | 3 | 7 | 5 | 2 | 1 | 1 | 3 | 8 | 5 | 3.38 | 1 | 5 | 3.4 | 1.766 | 0.11443615 | NA | 0 | 0.7032 | 0.4909 | 0.4909 | TRUE | 0.1533 |
| Efcc1 | 3 | 3 | 1 | 1 | 3 | 1 | 2 | 3 | 1 | 1 | 3 | 1 | 1.99 | 2 | 2 | 0.797 | -0.327 | 0.80864887 | NA | 0 | 1.1907 | 0.7976 | 0.7976 | TRUE | 0.436 |
| Efemp1 | 61 | 43 | 30 | 50 | 32 | 29 | 47 | 46 | 32 | 50 | 36 | 29 | 40 | 42 | 38 | 0.909 | -0.138 | 0.67753469 | 0.80217909 | 0.0336 | 0.0617 | 0.054 | 0.054 | TRUE | 0.4903 |
| Efemp2 | 243 | 178 | 190 | 205 | 203 | 187 | 189 | 189 | 205 | 204 | 229 | 185 | 200.12 | 194 | 206 | 1.059 | 0.082 | 0.57684689 | 0.72821698 | 0 | 0.0143 | 0.0107 | 0.0107 | TRUE | 0.1014 |
| Efha1 | 923 | 634 | 665 | 728 | 637 | 686 | 717 | 674 | 718 | 725 | 719 | 677 | 704.87 | 703 | 707 | 1.005 | 0.007 | 0.93543726 | 0.96438292 | 0 | 0.0058 | 0.0038 | 0.0038 | TRUE | 0.0162 |
| Efha2 | 940 | 599 | 744 | 705 | 699 | 759 | 730 | 637 | 803 | 702 | 789 | 749 | 734.92 | 723 | 747 | 1.032 | 0.045 | 0.65811271 | 0.78826824 | 0.0068 | 0.0057 | 0.006 | 0.006 | TRUE | 0.1241 |
| Efhb | 138 | 118 | 124 | 106 | 97 | 116 | 107 | 126 | 134 | 106 | 109 | 115 | 115.99 | 122 | 110 | 0.903 | -0.147 | 0.4287768 | 0.60086404 | 0 | 0.0229 | 0.0163 | 0.0163 | TRUE | 0.131 |
| Efhc1 | 235 | 157 | 181 | 215 | 174 | 188 | 183 | 167 | 195 | 214 | 196 | 186 | 190.12 | 182 | 199 | 1.094 | 0.129 | 0.3847523 | 0.55881157 | 0 | 0.0149 | 0.0107 | 0.0107 | TRUE | 0.0511 |
| Efhc2 | 2025 | 1520 | 1304 | 827 | 688 | 629 | 1573 | 1617 | 1407 | 823 | 776 | 621 | 1136.13 | 1532 | 740 | 0.483 | -1.051 | 6.16E-24 | 2.84E-21 | 0.0113 | 0.0045 | 0.0068 | 0.0068 | TRUE | 0.238 |
| Efhd1 | 118 | 91 | 99 | 149 | 114 | 149 | 92 | 97 | 107 | 148 | 129 | 147 | 119.88 | 99 | 141 | 1.442 | 0.528 | 0.0038189 | 0.0198722 | 0 | 0.0222 | 0.0154 | 0.0154 | TRUE | 0.0615 |
| Efhd2 | 1742 | 1310 | 1497 | 1584 | 1566 | 1829 | 1353 | 1393 | 1615 | 1577 | 1767 | 1806 | 1585.07 | 1454 | 1717 | 1.181 | 0.241 | 0.00573395 | 0.02725484 | 0.0063 | 0.0039 | 0.0048 | 0.0048 | TRUE | 0.1124 |
| Efna1 | 136 | 106 | 136 | 129 | 112 | 133 | 106 | 113 | 147 | 128 | 126 | 131 | 125.19 | 122 | 128 | 1.062 | 0.087 | 0.64279847 | 0.7773516 | 0.0081 | 0.0214 | 0.0175 | 0.0175 | TRUE | 0.3188 |
| Efna2 | 519 | 363 | 367 | 393 | 295 | 447 | 403 | 386 | 396 | 391 | 333 | 441 | 391.74 | 395 | 388 | 0.984 | -0.023 | 0.85060665 | 0.91562892 | 0.0072 | 0.0085 | 0.0081 | 0.0081 | TRUE | 0.1749 |
| Efna3 | 269 | 223 | 245 | 297 | 230 | 248 | 209 | 237 | 264 | 296 | 259 | 245 | 251.73 | 237 | 267 | 1.13 | 0.176 | 0.21630922 | 0.37897863 | 0.008 | 0.0118 | 0.0107 | 0.0107 | TRUE | 0.1279 |
| Efna4 | 424 | 312 | 302 | 276 | 272 | 305 | 329 | 332 | 326 | 275 | 307 | 301 | 311.61 | 329 | 294 | 0.894 | -0.162 | 0.16470389 | 0.31551292 | 0 | 0.01 | 0.0066 | 0.0066 | TRUE | 0.0383 |
| Efna5 | 74 | 55 | 60 | 56 | 64 | 63 | 57 | 59 | 65 | 56 | 72 | 62 | 61.81 | 60 | 63 | 1.052 | 0.073 | 0.76916197 | 0.86332904 | 0 | 0.0408 | 0.0282 | 0.0282 | TRUE | 0.122 |
| Efnb1 | 171 | 123 | 131 | 151 | 109 | 131 | 133 | 131 | 141 | 150 | 123 | 129 | 134.6 | 135 | 134 | 0.996 | -0.006 | 0.97263701 | 0.98514693 | 0 | 0.02 | 0.0141 | 0.0141 | TRUE | 0.1146 |
| Efnb2 | 629 | 408 | 312 | 384 | 367 | 370 | 489 | 434 | 337 | 382 | 414 | 365 | 403.44 | 420 | 387 | 0.919 | -0.122 | 0.37489141 | 0.5492899 | 0.0173 | 0.0083 | 0.0111 | 0.0111 | TRUE | 0.3403 |
| Efnb3 | 1008 | 735 | 846 | 814 | 807 | 902 | 783 | 782 | 913 | 810 | 910 | 890 | 848.09 | 826 | 870 | 1.055 | 0.077 | 0.40864269 | 0.58139857 | 0.0047 | 0.0052 | 0.005 | 0.005 | TRUE | 0.0988 |
| Efr3a | 9223 | 6878 | 6958 | 7097 | 6284 | 7127 | 7163 | 7316 | 7508 | 7064 | 7089 | 7035 | 7195.93 | 7329 | 7063 | 0.964 | -0.053 | 0.27333229 | 0.44425045 | 2.00E-04 | 0.0028 | 0.0016 | 0.0016 | TRUE | 0.0055 |
| Efr3b | 1237 | 815 | 808 | 912 | 778 | 864 | 961 | 867 | 872 | 908 | 878 | 853 | 889.64 | 900 | 880 | 0.976 | -0.035 | 0.66803878 | 0.79602153 | 0.0013 | 0.0051 | 0.0037 | 0.0037 | TRUE | 0.0439 |
| Efs | 807 | 581 | 653 | 712 | 625 | 720 | 627 | 618 | 705 | 709 | 705 | 711 | 678.98 | 650 | 708 | 1.091 | 0.126 | 0.16095312 | 0.31066887 | 0.001 | 0.0059 | 0.0043 | 0.0043 | TRUE | 0.0621 |
| Eftud1 | 2785 | 1966 | 1774 | 2064 | 1805 | 2013 | 2163 | 2091 | 1914 | 2055 | 2036 | 1987 | 2041.03 | 2056 | 2026 | 0.985 | -0.023 | 0.74088374 | 0.84593796 | 0.0017 | 0.0036 | 0.0028 | 0.0028 | TRUE | 0.0437 |
| Eftud2 | 34017 | 24952 | 23625 | 25124 | 22589 | 26075 | 26420 | 26541 | 25491 | 25008 | 25483 | 25740 | 25780.51 | 26151 | 25410 | 0.972 | -0.041 | 0.35977122 | 0.53442216 | 3.00E-04 | 0.0025 | 0.0014 | 0.0014 | TRUE | 0.0059 |
| Egf | 89 | 62 | 65 | 76 | 60 | 62 | 69 | 66 | 70 | 76 | 68 | 61 | 68.29 | 68 | 68 | 0.996 | -0.005 | 0.98235673 | 0.99089635 | 0 | 0.0371 | 0.0248 | 0.0248 | TRUE | 0.0849 |
| Egfbp2 | 0 | 0 | 0 | 0 | 0 | 0 | 0 | 0 | 0 | 0 | 0 | 0 | 0 | 0 | 0 | NA | NA | NA | NA | NA | NA | NA | NA | NA | NA |
| Egfem1 | 0 | 1 | 0 | 1 | 0 | 0 | 0 | 1 | 0 | 1 | 0 | 0 | 0.34 | 0 | 0 | 1.075 | 0.105 | 0.97573409 | NA | 0 | 6.9086 | 6.5582 | 6.5582 | TRUE | 0.5216 |
| Egfl6 | 128 | 72 | 85 | 115 | 106 | 103 | 99 | 77 | 92 | 114 | 120 | 102 | 100.57 | 89 | 112 | 1.248 | 0.319 | 0.11377793 | 0.2428431 | 0 | 0.026 | 0.0193 | 0.0193 | TRUE | 0.1423 |
| Egfl7 | 2204 | 1610 | 1643 | 2045 | 1897 | 2192 | 1712 | 1713 | 1773 | 2036 | 2140 | 2164 | 1922.75 | 1733 | 2113 | 1.22 | 0.287 | 4.99E-06 | 0.00010575 | 0 | 0.0037 | 0.0023 | 0.0023 | TRUE | 0.0126 |
| Egfl8 | 1 | 6 | 4 | 5 | 3 | 3 | 1 | 6 | 4 | 5 | 3 | 3 | 3.8 | 4 | 4 | 1.017 | 0.024 | 0.98128231 | NA | 0 | 0.6262 | 0.4963 | 0.4963 | TRUE | 0.4933 |
| Egflam | 50 | 19 | 21 | 24 | 23 | 15 | 39 | 20 | 23 | 24 | 26 | 15 | 24.39 | 27 | 22 | 0.779 | -0.36 | 0.40184294 | 0.57492905 | 0.0714 | 0.0996 | 0.0914 | 0.0914 | TRUE | 1.0304 |
| Egfr | 208 | 149 | 172 | 142 | 124 | 121 | 162 | 158 | 186 | 141 | 140 | 119 | 151.05 | 169 | 133 | 0.793 | -0.335 | 0.04515963 | 0.12499353 | 0 | 0.0181 | 0.0134 | 0.0134 | TRUE | 0.0902 |
| Egln1 | 5009 | 4089 | 4706 | 4556 | 4044 | 4230 | 3890 | 4349 | 5078 | 4535 | 4562 | 4176 | 4431.7 | 4439 | 4424 | 0.997 | -0.004 | 0.96198866 | 0.97911443 | 0.01 | 0.003 | 0.0054 | 0.0054 | TRUE | 0.1927 |
| Egln2 | 2419 | 1817 | 1586 | 1612 | 1499 | 1736 | 1879 | 1933 | 1711 | 1605 | 1691 | 1714 | 1755.34 | 1841 | 1670 | 0.907 | -0.141 | 0.04835033 | 0.13145735 | 0.002 | 0.0038 | 0.0031 | 0.0031 | TRUE | 0.0448 |
| Egln3 | 2435 | 1857 | 2487 | 2993 | 2832 | 3364 | 1891 | 1975 | 2683 | 2979 | 3195 | 3321 | 2674.11 | 2183 | 3165 | 1.451 | 0.537 | 1.11E-06 | 3.21E-05 | 0.0208 | 0.0033 | 0.0084 | 0.0084 | TRUE | 0.4844 |
| Egr1 | 766 | 613 | 325 | 404 | 612 | 576 | 595 | 652 | 351 | 402 | 690 | 569 | 543.13 | 533 | 554 | 1.039 | 0.056 | 0.87266553 | 0.92943431 | 0.0846 | 0.0068 | 0.0274 | 0.0846 | TRUE | 0.4241 |
| Egr2 | 12 | 9 | 8 | 12 | 16 | 10 | 9 | 10 | 9 | 12 | 18 | 10 | 11.23 | 9 | 13 | 1.439 | 0.525 | 0.36104811 | NA | 0 | 0.2134 | 0.1467 | 0.1467 | TRUE | 0.3767 |
| Egr3 | 1 | 2 | 1 | 1 | 1 | 1 | 1 | 2 | 1 | 1 | 1 | 1 | 1.18 | 1 | 1 | 0.792 | -0.336 | 0.84173106 | NA | 0 | 2.0071 | 1.1654 | 1.1654 | TRUE | 0.1607 |
| Egr4 | 33 | 22 | 21 | 26 | 19 | 26 | 26 | 23 | 23 | 26 | 21 | 26 | 24.11 | 24 | 24 | 1.017 | 0.024 | 0.94936019 | 0.97225491 | 0 | 0.1007 | 0.0651 | 0.0651 | TRUE | 0.0591 |
| Ehbp1 | 1863 | 1379 | 1304 | 1399 | 1285 | 1473 | 1447 | 1467 | 1407 | 1393 | 1450 | 1454 | 1436.17 | 1440 | 1432 | 0.994 | -0.009 | 0.89784244 | 0.94349536 | 0 | 0.0041 | 0.0025 | 0.0025 | TRUE | 0.0071 |
| Ehbp1l1 | 701 | 619 | 656 | 825 | 779 | 859 | 544 | 658 | 708 | 821 | 879 | 848 | 743.11 | 637 | 849 | 1.337 | 0.419 | 5.65E-05 | 0.000773 | 0.0081 | 0.0056 | 0.0064 | 0.0064 | TRUE | 0.2004 |

| | | | | | | | | | | | | | | | | | | | | | | | | | |
|---|---|---|---|---|---|---|---|---|---|---|---|---|---|---|---|---|---|---|---|---|---|---|---|---|---|
| Ehd1 | 9474 | 6545 | 5999 | 6196 | 5601 | 6460 | 7358 | 6962 | 6473 | 6167 | 6319 | 6377 | 6609.3 | 6931 | 6288 | 0.907 | -0.141 | 0.01992419 | 0.06895944 | 0.002 | 0.0028 | 0.0025 | 0.0025 | TRUE | 0.0407 |
| Ehd2 | 142 | 88 | 52 | 51 | 46 | 61 | 110 | 94 | 56 | 51 | 52 | 60 | 70.48 | 87 | 54 | 0.623 | -0.683 | 0.01409169 | 0.05332515 | 0.0516 | 0.0361 | 0.0407 | 0.0407 | TRUE | 0.4043 |
| Ehd3 | 584 | 403 | 349 | 380 | 309 | 347 | 454 | 429 | 377 | 378 | 349 | 343 | 388.03 | 420 | 357 | 0.848 | -0.238 | 0.03912553 | 0.11268489 | 0.0036 | 0.0085 | 0.007 | 0.007 | TRUE | 0.091 |
| Ehd4 | 4385 | 2951 | 2784 | 3074 | 2646 | 3021 | 3406 | 3139 | 3004 | 3060 | 2985 | 2982 | 3095.92 | 3183 | 3009 | 0.945 | -0.082 | 0.20327936 | 0.36472149 | 0.0017 | 0.0032 | 0.0027 | 0.0027 | TRUE | 0.0465 |
| Ehf | 13 | 7 | 6 | 0 | 3 | 6 | 10 | 7 | 6 | 0 | 3 | 6 | 5.55 | 8 | 3 | 0.385 | -1.378 | 0.13520891 | NA | 0.1135 | 0.4292 | 0.386 | 0.386 | TRUE | 0.6854 |
| Ehhadh | 548 | 386 | 358 | 374 | 336 | 378 | 426 | 411 | 386 | 372 | 379 | 373 | 391.16 | 408 | 375 | 0.918 | -0.123 | 0.24591253 | 0.4145333 | 0 | 0.0085 | 0.0055 | 0.0055 | TRUE | 0.0235 |
| Ehmt1 | 5966 | 4384 | 4444 | 4545 | 4132 | 4763 | 4634 | 4663 | 4795 | 4524 | 4661 | 4702 | 4663.17 | 4697 | 4629 | 0.986 | -0.021 | 0.68578465 | 0.80738108 | 2.00E-04 | 0.0029 | 0.0017 | 0.0017 | TRUE | 0.0048 |
| Ehmt2 | 7288 | 5505 | 5767 | 6937 | 5948 | 6974 | 5660 | 5856 | 6223 | 6905 | 6710 | 6884 | 6372.98 | 5913 | 6833 | 1.156 | 0.209 | 0.00016976 | 0.00188033 | 0.0011 | 0.0028 | 0.0021 | 0.0021 | TRUE | 0.0255 |
| Ei24 | 8830 | 6411 | 7035 | 7733 | 6780 | 7805 | 6858 | 6819 | 7591 | 7697 | 7649 | 7705 | 7386.43 | 7089 | 7684 | 1.084 | 0.116 | 0.04532569 | 0.12535606 | 0.0017 | 0.0027 | 0.0023 | 0.0023 | TRUE | 0.0464 |
| Eid1 | 1575 | 1188 | 911 | 1001 | 830 | 897 | 1223 | 1264 | 983 | 996 | 936 | 885 | 1048.01 | 1157 | 939 | 0.812 | -0.301 | 0.0028983 | 0.0161063 | 0.0099 | 0.0047 | 0.0064 | 0.0064 | TRUE | 0.203 |
| Eid2 | 1458 | 1071 | 890 | 906 | 793 | 918 | 1132 | 1139 | 960 | 902 | 895 | 906 | 989.08 | 1077 | 901 | 0.835 | -0.259 | 0.00288536 | 0.01604689 | 0.0037 | 0.0048 | 0.0044 | 0.0044 | TRUE | 0.1055 |
| Eid2b | 352 | 260 | 246 | 302 | 247 | 275 | 273 | 277 | 265 | 301 | 279 | 271 | 277.68 | 272 | 284 | 1.043 | 0.061 | 0.61676059 | 0.75811482 | 0 | 0.011 | 0.0072 | 0.0072 | TRUE | 0.0317 |
| Eid3 | 17 | 10 | 12 | 14 | 17 | 11 | 13 | 11 | 13 | 14 | 19 | 11 | 13.46 | 12 | 15 | 1.186 | 0.246 | 0.63911075 | NA | 0 | 0.1785 | 0.1234 | 0.1234 | TRUE | 0.2975 |
| Eif1 | 23869 | 16435 | 16892 | 19028 | 15812 | 18935 | 18538 | 17482 | 18226 | 18940 | 17838 | 18692 | 18285.98 | 18082 | 18490 | 1.023 | 0.032 | 0.52527465 | 0.68634104 | 8.00E-04 | 0.0026 | 0.0018 | 0.0018 | TRUE | 0.0116 |
| Eif1a | 13108 | 9521 | 9052 | 8793 | 7841 | 8981 | 10180 | 10127 | 9767 | 8753 | 8845 | 8866 | 9423.08 | 10025 | 8821 | 0.88 | -0.185 | 9.10E-05 | 0.00111986 | 2.00E-04 | 0.0027 | 0.0015 | 0.0015 | TRUE | 0.0062 |
| Eif1ad | 3239 | 2126 | 1856 | 1840 | 1678 | 1948 | 2516 | 2261 | 2003 | 1832 | 1893 | 1923 | 2071.18 | 2260 | 1883 | 0.832 | -0.265 | 0.00160295 | 0.01013759 | 0.006 | 0.0036 | 0.0046 | 0.0046 | TRUE | 0.1208 |
| Eif1ax | 8802 | 6239 | 6210 | 7121 | 5872 | 7056 | 6836 | 6636 | 6701 | 7088 | 6624 | 6965 | 6808.47 | 6724 | 6892 | 1.025 | 0.036 | 0.49201459 | 0.65741649 | 5.00E-04 | 0.0028 | 0.0018 | 0.0018 | TRUE | 0.0141 |
| Eif1b | 1695 | 1327 | 1346 | 1431 | 1203 | 1450 | 1316 | 1412 | 1452 | 1424 | 1357 | 1431 | 1398.86 | 1393 | 1404 | 1.009 | 0.013 | 0.85375777 | 0.91756973 | 0.0011 | 0.0041 | 0.003 | 0.003 | TRUE | 0.0294 |
| Eif2a | 9987 | 7080 | 7500 | 7680 | 7052 | 7981 | 7756 | 7531 | 8092 | 7645 | 7955 | 7878 | 7809.73 | 7793 | 7826 | 1.004 | 0.006 | 0.90747349 | 0.94923397 | 6.00E-04 | 0.0027 | 0.0018 | 0.0018 | TRUE | 0.0136 |
| Eif2ak1 | 2934 | 1972 | 1777 | 1833 | 1732 | 1886 | 2279 | 2098 | 1917 | 1825 | 1954 | 1862 | 1988.98 | 2098 | 1880 | 0.895 | -0.16 | 0.03704148 | 0.10840169 | 0.0036 | 0.0036 | 0.0037 | 0.0037 | TRUE | 0.0704 |
| Eif2ak2 | 2078 | 1437 | 1419 | 1482 | 1323 | 1441 | 1614 | 1529 | 1531 | 1475 | 1492 | 1422 | 1510.61 | 1558 | 1463 | 0.939 | -0.092 | 0.17093694 | 0.32259143 | 0 | 0.004 | 0.0026 | 0.0026 | TRUE | 0.0125 |
| Eif2ak3 | 2698 | 2118 | 2017 | 2492 | 2168 | 2549 | 2095 | 2253 | 2176 | 2481 | 2446 | 2516 | 2327.86 | 2175 | 2481 | 1.142 | 0.191 | 0.00159033 | 0.01007559 | 4.00E-04 | 0.0034 | 0.0022 | 0.0022 | TRUE | 0.0129 |
| Eif2ak4 | 2436 | 1870 | 1836 | 1939 | 1671 | 1915 | 1892 | 1989 | 1981 | 1930 | 1885 | 1890 | 1927.93 | 1954 | 1902 | 0.974 | -0.038 | 0.53650557 | 0.69528281 | 0 | 0.0037 | 0.0022 | 0.0022 | TRUE | 0.0097 |
| Eif2b1 | 5132 | 3725 | 3437 | 3739 | 3448 | 3978 | 3986 | 3962 | 3708 | 3722 | 3890 | 3927 | 3865.82 | 3885 | 3846 | 0.99 | -0.015 | 0.79348628 | 0.87903421 | 9.00E-04 | 0.003 | 0.0022 | 0.0022 | TRUE | 0.0191 |
| Eif2b2 | 3194 | 2241 | 1952 | 1889 | 1720 | 1997 | 2481 | 2384 | 2106 | 1880 | 1940 | 1971 | 2127.09 | 2324 | 1930 | 0.83 | -0.268 | 0.00031373 | 0.0030075 | 0.0031 | 0.0035 | 0.0035 | 0.0035 | TRUE | 0.0804 |
| Eif2b3 | 5027 | 3409 | 3220 | 3850 | 3342 | 3732 | 3904 | 3626 | 3474 | 3832 | 3770 | 3684 | 3715.2 | 3668 | 3762 | 1.025 | 0.036 | 0.56762718 | 0.72138115 | 0.0015 | 0.0031 | 0.0025 | 0.0025 | TRUE | 0.0393 |
| Eif2b4 | 3175 | 2258 | 1958 | 1992 | 1804 | 2031 | 2466 | 2402 | 2113 | 1983 | 2035 | 2005 | 2167.2 | 2327 | 2008 | 0.862 | -0.214 | 0.0032389 | 0.01751557 | 0.003 | 0.0035 | 0.0033 | 0.0033 | TRUE | 0.0777 |
| Eif2b5 | 10660 | 7928 | 7682 | 8092 | 7205 | 8103 | 8279 | 8433 | 8289 | 8055 | 8128 | 7999 | 8197.09 | 8334 | 8061 | 0.967 | -0.048 | 0.29772829 | 0.46979802 | 0 | 0.0027 | 0.0014 | 0.0014 | TRUE | 0.0013 |
| Eif2c1 | 6048 | 4538 | 4661 | 4523 | 4123 | 4710 | 4697 | 4827 | 5029 | 4502 | 4651 | 4649 | 4726.05 | 4851 | 4601 | 0.949 | -0.076 | 0.16097736 | 0.31067369 | 6.00E-04 | 0.0029 | 0.0019 | 0.0019 | TRUE | 0.0125 |
| Eif2c2 | 7880 | 5937 | 6442 | 7727 | 7031 | 8061 | 6120 | 6315 | 6951 | 7691 | 7932 | 7957 | 7161.11 | 6462 | 7860 | 1.217 | 0.283 | 3.53E-06 | 7.93E-05 | 0.0022 | 0.0028 | 0.0025 | 0.0025 | TRUE | 0.0533 |
| Eif2c3 | 792 | 495 | 497 | 640 | 538 | 625 | 615 | 527 | 536 | 637 | 607 | 617 | 589.81 | 559 | 620 | 1.106 | 0.146 | 0.13441203 | 0.27321014 | 0.0026 | 0.0064 | 0.0052 | 0.0052 | TRUE | 0.0945 |
| Eif2c4 | 197 | 167 | 170 | 187 | 191 | 201 | 153 | 178 | 183 | 186 | 215 | 198 | 185.68 | 171 | 200 | 1.17 | 0.227 | 0.13754599 | 0.27787207 | 0 | 0.0152 | 0.0114 | 0.0114 | TRUE | 0.104 |
| Eif2d | 8263 | 5692 | 5495 | 5470 | 4945 | 5744 | 6418 | 6055 | 5929 | 5445 | 5578 | 5670 | 5849.1 | 6134 | 5564 | 0.907 | -0.141 | 0.01015066 | 0.04174403 | 8.00E-04 | 0.0028 | 0.002 | 0.002 | TRUE | 0.0203 |
| Eif2s1 | 17699 | 12393 | 12467 | 13279 | 11443 | 13330 | 13746 | 13182 | 13452 | 13218 | 12909 | 13159 | 13277.61 | 13460 | 13095 | 0.973 | -0.04 | 0.3897859 | 0.56374383 | 2.00E-04 | 0.0026 | 0.0015 | 0.0015 | TRUE | 0.0043 |
| Eif2s2 | 47256 | 32648 | 31374 | 24160 | 21717 | 25044 | 36702 | 34727 | 33852 | 24049 | 24499 | 24722 | 29758.57 | 35094 | 24423 | 0.696 | -0.523 | 1.68E-25 | 9.24E-23 | 9.00E-04 | 0.0025 | 0.0018 | 0.0018 | TRUE | 0.0197 |
| Eif2s3x | 6105 | 4332 | 3948 | 4565 | 4015 | 4584 | 4741 | 4608 | 4260 | 4544 | 4529 | 4525 | 4534.62 | 4536 | 4533 | 0.999 | -0.002 | 0.97637558 | 0.98728943 | 0.0013 | 0.0029 | 0.0023 | 0.0023 | TRUE | 0.0344 |
| Eif2s3y | 4330 | 3380 | 3247 | 3105 | 2681 | 3377 | 3363 | 3595 | 3503 | 3091 | 3024 | 3334 | 3318.41 | 3487 | 3150 | 0.904 | -0.146 | 0.01943751 | 0.06767228 | 0.0014 | 0.0031 | 0.0025 | 0.0025 | TRUE | 0.0319 |
| Eif3a | 51977 | 38815 | 40436 | 41889 | 37576 | 44528 | 40368 | 41287 | 43630 | 41696 | 42390 | 43956 | 42221.22 | 41762 | 42681 | 1.022 | 0.031 | 0.53913648 | 0.69753507 | 0.0011 | 0.0025 | 0.0019 | 0.0019 | TRUE | 0.0187 |
| Eif3b | 28104 | 20507 | 18852 | 18141 | 16111 | 18784 | 21827 | 21813 | 20341 | 18058 | 18175 | 18543 | 19792.75 | 21327 | 18259 | 0.856 | -0.224 | 8.09E-06 | 0.00015774 | 8.00E-04 | 0.0025 | 0.0018 | 0.0018 | TRUE | 0.02 |
| Eif3c | 26841 | 19551 | 20070 | 21747 | 19137 | 22512 | 20846 | 20796 | 21655 | 21647 | 21589 | 22223 | 21459.34 | 21099 | 21820 | 1.034 | 0.049 | 0.29081528 | 0.46287559 | 3.00E-04 | 0.0025 | 0.0015 | 0.0015 | TRUE | 0.0065 |
| Eif3d | 25070 | 18226 | 17997 | 18629 | 17246 | 19103 | 19471 | 19387 | 19419 | 18543 | 19455 | 18858 | 19188.72 | 19426 | 18952 | 0.976 | -0.036 | 0.43125976 | 0.60334273 | 2.00E-04 | 0.0025 | 0.0014 | 0.0014 | TRUE | 0.0066 |
| Eif3e | 17596 | 12201 | 11882 | 12842 | 11297 | 12444 | 13666 | 12978 | 12821 | 12783 | 12744 | 12284 | 12879.32 | 13155 | 12604 | 0.958 | -0.062 | 0.21919222 | 0.3823839 | 7.00E-04 | 0.0026 | 0.0018 | 0.0018 | TRUE | 0.0142 |
| Eif3f | 18158 | 13666 | 14504 | 15093 | 13525 | 16195 | 14103 | 14536 | 15650 | 15024 | 15258 | 15987 | 15092.79 | 14763 | 15423 | 1.045 | 0.063 | 0.2684599 | 0.4390774 | 0.0019 | 0.0026 | 0.0023 | 0.0023 | TRUE | 0.0337 |
| Eif3g | 11215 | 8340 | 7846 | 8143 | 7326 | 8603 | 8710 | 8871 | 8466 | 8106 | 8264 | 8492 | 8484.94 | 8682 | 8287 | 0.955 | -0.067 | 0.17686992 | 0.33067387 | 4.00E-04 | 0.0027 | 0.0017 | 0.0017 | TRUE | 0.0058 |
| Eif3h | 13746 | 9892 | 10704 | 11147 | 10106 | 11761 | 10676 | 10522 | 11550 | 11096 | 11401 | 11610 | 11142.28 | 10916 | 11369 | 1.042 | 0.059 | 0.28739649 | 0.45953216 | 0.0014 | 0.0026 | 0.0021 | 0.0021 | TRUE | 0.0314 |
| Eif3i | 11911 | 8519 | 8948 | 9741 | 8744 | 9926 | 9251 | 9062 | 9655 | 9696 | 9864 | 9798 | 9554.32 | 9323 | 9786 | 1.05 | 0.07 | 0.15535047 | 0.3030416 | 4.00E-04 | 0.0027 | 0.0016 | 0.0016 | TRUE | 0.0118 |
| Eif3j1 | 0 | 0 | 0 | 0 | 0 | 0 | 0 | 0 | 0 | 0 | 0 | 0 | 0 | 0 | 0 | NA | NA | NA | NA | NA | NA | NA | NA | NA | NA |
| Eif3j2 | 4 | 7 | 2 | 3 | 2 | 3 | 3 | 7 | 2 | 3 | 2 | 3 | 3.49 | 4 | 3 | 0.653 | -0.615 | 0.55382404 | NA | 0 | 0.6824 | 0.4741 | 0.4741 | TRUE | 0.7149 |
| Eif3k | 10089 | 7234 | 7592 | 8132 | 7106 | 8368 | 7836 | 7695 | 8192 | 8095 | 8016 | 8261 | 8015.59 | 7908 | 8124 | 1.027 | 0.039 | 0.43909795 | 0.61048068 | 5.00E-04 | 0.0027 | 0.0017 | 0.0017 | TRUE | 0.012 |
| Eif3l | 18593 | 13299 | 13380 | 14182 | 12896 | 15397 | 14440 | 14146 | 14437 | 14117 | 14548 | 15199 | 14481.21 | 14341 | 14621 | 1.02 | 0.028 | 0.5746605 | 0.72661204 | 7.00E-04 | 0.0026 | 0.0017 | 0.0017 | TRUE | 0.0147 |
| Eif3m | 12115 | 8428 | 9226 | 9794 | 8291 | 9640 | 9409 | 8965 | 9955 | 9749 | 9353 | 9516 | 9491.16 | 9443 | 9539 | 1.01 | 0.015 | 0.79211234 | 0.87812469 | 0.0012 | 0.0027 | 0.0022 | 0.0022 | TRUE | 0.0272 |
| Eif4a1 | 138527 | 92700 | 86310 | 96568 | 80604 | 92557 | 107588 | 98604 | 93128 | 96124 | 90930 | 91368 | 96290.17 | 99773 | 92807 | 0.93 | -0.104 | 0.09019876 | 0.20499458 | 0.0031 | 0.0025 | 0.0027 | 0.0027 | TRUE | 0.0575 |
| Eif4a2 | 48510 | 38066 | 45529 | 60159 | 49417 | 55879 | 37676 | 40490 | 49125 | 59882 | 55748 | 55161 | 49680.38 | 42430 | 56930 | 1.342 | 0.424 | 7.08E-07 | 2.19E-05 | 0.0109 | 0.0025 | 0.0053 | 0.0053 | TRUE | 0.2332 |
| Eif4a3 | 16361 | 11568 | 10424 | 10948 | 9964 | 11096 | 12707 | 12305 | 11247 | 10898 | 11240 | 10953 | 11558.43 | 12086 | 11030 | 0.912 | -0.132 | 0.02389037 | 0.0787314 | 0.002 | 0.0026 | 0.0024 | 0.0024 | TRUE | 0.045 |
| Eif4b | 57130 | 43386 | 48818 | 52083 | 47715 | 55328 | 44370 | 46149 | 52674 | 51843 | 53827 | 54617 | 50580.31 | 47731 | 53429 | 1.119 | 0.163 | 0.01558924 | 0.05750117 | 0.0045 | 0.0025 | 0.0032 | 0.0032 | TRUE | 0.1004 |
| Eif4e | 11506 | 8226 | 8496 | 9127 | 7858 | 9330 | 8936 | 8750 | 9167 | 9085 | 8865 | 9210 | 9002.17 | 8951 | 9053 | 1.011 | 0.016 | 0.7353234 | 0.84201069 | 3.00E-04 | 0.0027 | 0.0016 | 0.0016 | TRUE | 0.0054 |
| Eif4e1b | 7 | 2 | 1 | 0 | 0 | 1 | 5 | 2 | 1 | 0 | 0 | 1 | 1.61 | 3 | 0 | 0.126 | -2.987 | 0.09904477 | NA | 0 | 1.4791 | 1.1977 | 1.1977 | TRUE | 0.5788 |
| Eif4e2 | 6500 | 4466 | 4068 | 4282 | 3595 | 4126 | 5048 | 4750 | 4389 | 4262 | 4056 | 4073 | 4429.81 | 4729 | 4130 | 0.873 | -0.196 | 0.00264034 | 0.01493404 | 0.0025 | 0.003 | 0.0028 | 0.0028 | TRUE | 0.0481 |
| Eif4e3 | 562 | 460 | 584 | 613 | 573 | 672 | 436 | 489 | 630 | 610 | 646 | 663 | 579.31 | 518 | 640 | 1.237 | 0.307 | 0.01493681 | 0.05569836 | 0.0169 | 0.0065 | 0.0097 | 0.0097 | TRUE | 0.4098 |
| Eif4ebp1 | 13999 | 10305 | 9983 | 9713 | 8469 | 10044 | 10872 | 10961 | 10772 | 9668 | 9554 | 9915 | 10290.42 | 10868 | 9712 | 0.894 | -0.162 | 0.00046318 | 0.00408148 | 1.00E-04 | 0.0027 | 0.0014 | 0.0014 | TRUE | 0.0041 |
| Eif4ebp2 | 13420 | 9767 | 8637 | 9570 | 8536 | 9454 | 10423 | 10389 | 9319 | 9526 | 9629 | 9333 | 9769.84 | 10044 | 9496 | 0.945 | -0.081 | 0.16820496 | 0.31974379 | 0.0019 | 0.0027 | 0.0024 | 0.0024 | TRUE | 0.0484 |
| Eif4ebp3 | 429 | 288 | 272 | 308 | 253 | 349 | 333 | 306 | 293 | 307 | 285 | 345 | 311.59 | 311 | 312 | 1.003 | 0.004 | 0.97442599 | 0.98624254 | 0.0035 | 0.01 | 0.0081 | 0.0081 | TRUE | 0.0936 |
| Eif4enif1 | 6605 | 4722 | 4975 | 5086 | 4690 | 5574 | 5130 | 5023 | 5368 | 5063 | 5291 | 5502 | 5229.39 | 5174 | 5285 | 1.022 | 0.031 | 0.59034662 | 0.73882577 | 0.0011 | 0.0029 | 0.0022 | 0.0022 | TRUE | 0.0167 |
| Eif4g1 | 58422 | 41444 | 35693 | 36824 | 32329 | 36981 | 45374 | 44083 | 38512 | 36655 | 36470 | 36506 | 39600.13 | 42656 | 36544 | 0.857 | -0.223 | 0.00056627 | 0.00474424 | 0.0036 | 0.0025 | 0.003 | 0.003 | TRUE | 0.0884 |
| Eif4g2 | 106706 | 71782 | 64684 | 69123 | 60850 | 69817 | 82874 | 76354 | 69793 | 68805 | 68645 | 68920 | 72565.22 | 76340 | 68790 | 0.901 | -0.15 | 0.01912339 | 0.06686168 | 0.0037 | 0.0025 | 0.003 | 0.003 | TRUE | 0.0689 |
| Eif4g3 | 1566 | 1290 | 1337 | 1330 | 1304 | 1509 | 1216 | 1372 | 1443 | 1324 | 1471 | 1490 | 1385.93 | 1344 | 1428 | 1.064 | 0.09 | 0.29299579 | 0.46515206 | 0.0052 | 0.0041 | 0.0045 | 0.0045 | TRUE | 0.0857 |
| Eif4h | 24357 | 17340 | 16065 | 17631 | 15144 | 17301 | 18917 | 18444 | 17334 | 17550 | 17084 | 17079 | 17734.67 | 18232 | 17238 | 0.945 | -0.081 | 0.11850708 | 0.24997369 | 0.001 | 0.0026 | 0.0019 | 0.0019 | TRUE | 0.0227 |
| Eif5 | 31297 | 21855 | 19928 | 19826 | 17706 | 20209 | 24307 | 23247 | 21502 | 19735 | 19974 | 19949 | 21452.41 | 23019 | 19886 | 0.864 | -0.211 | 0.00018945 | 0.00205289 | 0.0018 | 0.0025 | 0.0023 | 0.0023 | TRUE | 0.0406 |
| Eif5a | 81094 | 56410 | 53660 | 56426 | 48101 | 57140 | 62982 | 60003 | 57899 | 56166 | 54263 | 56406 | 57953.14 | 60295 | 55612 | 0.922 | -0.117 | 0.0214115 | 0.07264297 | 0.0011 | 0.0025 | 0.0018 | 0.0018 | TRUE | 0.0187 |

| | | | | | | | | | | | | | | | | | | | | | | | | | |
|---|---|---|---|---|---|---|---|---|---|---|---|---|---|---|---|---|---|---|---|---|---|---|---|---|---|
| Eif5a2 | 2045 | 1601 | 1587 | 1682 | 1439 | 1563 | 1588 | 1703 | 1712 | 1674 | 1623 | 1543 | 1640.69 | 1668 | 1613 | 0.968 | -0.047 | 0.5007417 | 0.66509628 | 0.0012 | 0.0039 | 0.0029 | 0.0029 | TRUE | 0.0219 |
| Eif5b | 16855 | 12143 | 11333 | 12287 | 10169 | 12054 | 13091 | 12916 | 12228 | 12230 | 11472 | 11899 | 12306.07 | 12745 | 11867 | 0.931 | -0.103 | 0.05114018 | 0.1371382 | 0.001 | 0.0026 | 0.0019 | 0.0019 | TRUE | 0.0153 |
| Eif6 | 7429 | 5213 | 4236 | 4289 | 4012 | 4429 | 5770 | 5545 | 4571 | 4269 | 4526 | 4372 | 4842.12 | 5295 | 4389 | 0.829 | -0.271 | 0.00097079 | 0.00709378 | 0.0074 | 0.0029 | 0.0047 | 0.0047 | TRUE | 0.174 |
| Elac1 | 689 | 469 | 533 | 579 | 469 | 572 | 535 | 499 | 575 | 576 | 529 | 565 | 546.53 | 536 | 557 | 1.039 | 0.055 | 0.57668285 | 0.7280742 | 0.0014 | 0.0068 | 0.0051 | 0.0051 | TRUE | 0.0441 |
| Elac2 | 7446 | 5258 | 4640 | 4778 | 4438 | 5018 | 5783 | 5593 | 5007 | 4756 | 5007 | 4954 | 5183.08 | 5461 | 4906 | 0.898 | -0.155 | 0.01829178 | 0.06463398 | 0.0028 | 0.0029 | 0.0029 | 0.0029 | TRUE | 0.0643 |
| Elane | 0 | 0 | 0 | 0 | 0 | 0 | 0 | 0 | 0 | 0 | 0 | 0 | 0 | 0 | 0 | NA | NA | NA | NA | NA | NA | NA | NA | NA | NA |
| Elavl1 | 8135 | 5950 | 5994 | 6120 | 5202 | 6197 | 6318 | 6329 | 6467 | 6092 | 5868 | 6117 | 6198.69 | 6371 | 6026 | 0.946 | -0.08 | 0.10638279 | 0.23148179 | 2.00E-04 | 0.0028 | 0.0016 | 0.0016 | TRUE | 0.0063 |
| Elavl2 | 2547 | 1923 | 2150 | 2087 | 1912 | 2135 | 1978 | 2045 | 2320 | 2077 | 2157 | 2108 | 2114.23 | 2114 | 2114 | 1 | 0.001 | 0.99279326 | 0.99592003 | 0.0033 | 0.0035 | 0.0035 | 0.0035 | TRUE | 0.0866 |
| Elavl3 | 1354 | 1156 | 1444 | 1871 | 1720 | 2075 | 1052 | 1230 | 1558 | 1862 | 1940 | 2048 | 1615.06 | 1280 | 1950 | 1.526 | 0.61 | 1.07E-07 | 4.16E-06 | 0.0203 | 0.0039 | 0.0088 | 0.0088 | TRUE | 0.4339 |
| Elavl4 | 232 | 189 | 182 | 148 | 132 | 160 | 180 | 201 | 196 | 147 | 149 | 158 | 171.96 | 192 | 151 | 0.788 | -0.343 | 0.02399125 | 0.07895624 | 0 | 0.0162 | 0.0107 | 0.0107 | TRUE | 0.0375 |
| Elf1 | 840 | 579 | 540 | 501 | 462 | 537 | 652 | 616 | 583 | 499 | 521 | 530 | 566.82 | 617 | 517 | 0.836 | -0.258 | 0.00618407 | 0.02885699 | 0 | 0.0066 | 0.0046 | 0.0046 | TRUE | 0.0314 |
| Elf2 | 2919 | 2068 | 2127 | 2135 | 1956 | 2253 | 2267 | 2200 | 2295 | 2125 | 2207 | 2224 | 2219.6 | 2254 | 2185 | 0.969 | -0.045 | 0.45376379 | 0.62349989 | 0 | 0.0035 | 0.0021 | 0.0021 | TRUE | 0.0071 |
| Elf3 | 1627 | 1391 | 1384 | 1706 | 1703 | 1837 | 1264 | 1480 | 1493 | 1698 | 1921 | 1813 | 1611.54 | 1412 | 1811 | 1.284 | 0.36 | 1.95E-05 | 0.00032626 | 0.0051 | 0.0039 | 0.0045 | 0.0045 | TRUE | 0.1054 |
| Elf4 | 1421 | 1159 | 1192 | 1218 | 1109 | 1350 | 1104 | 1233 | 1286 | 1212 | 1251 | 1333 | 1236.45 | 1208 | 1265 | 1.05 | 0.07 | 0.39622369 | 0.56958266 | 0.0036 | 0.0043 | 0.0041 | 0.0041 | TRUE | 0.0708 |
| Elf5 | 15 | 18 | 12 | 7 | 5 | 7 | 12 | 19 | 13 | 7 | 6 | 7 | 10.54 | 15 | 7 | 0.45 | -1.151 | 0.05922713 | NA | 0 | 0.2272 | 0.1558 | 0.1558 | TRUE | 0.3214 |
| Elfn1 | 17 | 8 | 8 | 12 | 11 | 11 | 13 | 9 | 9 | 12 | 12 | 11 | 10.93 | 10 | 12 | 1.145 | 0.195 | 0.73373864 | NA | 0 | 0.2193 | 0.1458 | 0.1458 | TRUE | 0.3188 |
| Elfn2 | 62 | 43 | 54 | 74 | 63 | 74 | 48 | 46 | 58 | 74 | 71 | 73 | 61.66 | 51 | 73 | 1.435 | 0.521 | 0.03520454 | 0.10436408 | 0 | 0.0409 | 0.0275 | 0.0275 | TRUE | 0.128 |
| Elk1 | 1682 | 1199 | 1232 | 1326 | 1156 | 1293 | 1306 | 1275 | 1329 | 1320 | 1304 | 1276 | 1301.9 | 1303 | 1300 | 0.997 | -0.004 | 0.95258172 | 0.97381985 | 0 | 0.0042 | 0.0025 | 0.0025 | TRUE | 0.0042 |
| Elk3 | 784 | 644 | 696 | 543 | 567 | 623 | 609 | 685 | 751 | 541 | 640 | 615 | 640 | 682 | 599 | 0.879 | -0.186 | 0.08279228 | 0.19373772 | 0.0079 | 0.0061 | 0.0067 | 0.0067 | TRUE | 0.1079 |
| Elk4 | 367 | 283 | 238 | 298 | 231 | 300 | 285 | 301 | 257 | 297 | 261 | 296 | 282.7 | 281 | 285 | 1.013 | 0.019 | 0.88375707 | 0.93547084 | 0.0017 | 0.0108 | 0.0082 | 0.0082 | TRUE | 0.0599 |
| Ell | 3172 | 2336 | 2204 | 2202 | 2002 | 2326 | 2464 | 2485 | 2378 | 2192 | 2258 | 2296 | 2345.48 | 2442 | 2249 | 0.921 | -0.119 | 0.04420002 | 0.12307634 | 0 | 0.0034 | 0.0021 | 0.0021 | TRUE | 0.0062 |
| Ell2 | 4572 | 3531 | 3672 | 4716 | 4057 | 4574 | 3551 | 3756 | 3962 | 4694 | 4577 | 4515 | 4175.85 | 3756 | 4595 | 1.224 | 0.292 | 1.20E-06 | 3.40E-05 | 0.0013 | 0.003 | 0.0024 | 0.0024 | TRUE | 0.0285 |
| Ell3 | 775 | 615 | 683 | 846 | 700 | 888 | 602 | 654 | 737 | 842 | 790 | 877 | 750.23 | 664 | 836 | 1.262 | 0.336 | 0.00055206 | 0.00464725 | 0.0052 | 0.0056 | 0.0054 | 0.0054 | TRUE | 0.1071 |
| Elmo1 | 629 | 475 | 628 | 590 | 649 | 778 | 489 | 505 | 678 | 587 | 732 | 768 | 626.47 | 557 | 696 | 1.251 | 0.323 | 0.01808016 | 0.06412401 | 0.0247 | 0.0062 | 0.0118 | 0.0118 | TRUE | 0.4144 |
| Elmo2 | 3073 | 2355 | 2764 | 2623 | 2534 | 2921 | 2387 | 2505 | 2982 | 2611 | 2859 | 2883 | 2704.5 | 2625 | 2784 | 1.061 | 0.086 | 0.31788597 | 0.49181173 | 0.0074 | 0.0033 | 0.005 | 0.005 | TRUE | 0.1716 |
| Elmo3 | 466 | 349 | 467 | 389 | 413 | 450 | 362 | 371 | 504 | 387 | 466 | 444 | 422.4 | 412 | 432 | 1.051 | 0.071 | 0.60874414 | 0.75233722 | 0.0196 | 0.008 | 0.0115 | 0.0115 | TRUE | 0.4282 |
| Elmod1 | 6 | 5 | 2 | 3 | 0 | 3 | 5 | 5 | 2 | 3 | 0 | 3 | 3.01 | 4 | 2 | 0.497 | -1.008 | 0.38905923 | NA | 0 | 0.7888 | 0.6121 | 0.6121 | TRUE | 0.5794 |
| Elmod2 | 2354 | 1916 | 2262 | 2886 | 2519 | 2850 | 1828 | 2038 | 2441 | 2873 | 2842 | 2813 | 2472.46 | 2102 | 2843 | 1.354 | 0.437 | 2.16E-06 | 5.35E-05 | 0.0104 | 0.0034 | 0.0057 | 0.0057 | TRUE | 0.2394 |
| Elmod3 | 551 | 377 | 365 | 400 | 344 | 416 | 428 | 401 | 394 | 398 | 388 | 411 | 403.28 | 408 | 399 | 0.978 | -0.033 | 0.75569949 | 0.85511255 | 0 | 0.0083 | 0.0054 | 0.0054 | TRUE | 0.0236 |
| Elmsan1 | 3049 | 2591 | 2855 | 3192 | 3266 | 3506 | 2368 | 2756 | 3081 | 3177 | 3684 | 3461 | 3087.87 | 2735 | 3441 | 1.259 | 0.332 | 0.00032911 | 0.00312143 | 0.0107 | 0.0032 | 0.0058 | 0.0058 | TRUE | 0.1697 |
| Eln | 27 | 39 | 46 | 118 | 89 | 95 | 21 | 41 | 50 | 117 | 100 | 94 | 70.62 | 37 | 104 | 2.831 | 1.501 | 2.75E-07 | 9.65E-06 | 0.0686 | 0.036 | 0.0434 | 0.0434 | TRUE | 1.3193 |
| Elof1 | 1326 | 940 | 964 | 935 | 844 | 1019 | 1030 | 1000 | 1040 | 931 | 952 | 1006 | 993.1 | 1023 | 963 | 0.941 | -0.088 | 0.24705855 | 0.41572968 | 0 | 0.0048 | 0.0031 | 0.0031 | TRUE | 0.0185 |
| Elovl1 | 1224 | 837 | 942 | 1068 | 938 | 1144 | 951 | 890 | 1016 | 1063 | 1058 | 1129 | 1017.98 | 952 | 1083 | 1.138 | 0.186 | 0.02092569 | 0.07159746 | 0.0017 | 0.0048 | 0.0037 | 0.0037 | TRUE | 0.0393 |
| Elovl2 | 1200 | 863 | 939 | 1043 | 929 | 1089 | 932 | 918 | 1013 | 1038 | 1048 | 1075 | 1004.06 | 954 | 1054 | 1.105 | 0.144 | 0.06249888 | 0.15815058 | 5.00E-04 | 0.0048 | 0.0033 | 0.0033 | TRUE | 0.0336 |
| Elovl3 | 2 | 1 | 3 | 2 | 4 | 3 | 2 | 1 | 3 | 2 | 5 | 3 | 2.55 | 2 | 3 | 1.624 | 0.7 | 0.5575732 | NA | 0 | 0.9307 | 0.6113 | 0.6113 | TRUE | 0.259 |
| Elovl4 | 1597 | 1177 | 1124 | 1485 | 1216 | 1396 | 1240 | 1252 | 1213 | 1478 | 1372 | 1378 | 1322.18 | 1235 | 1409 | 1.141 | 0.191 | 0.00648694 | 0.02993794 | 0 | 0.0042 | 0.0028 | 0.0028 | TRUE | 0.0221 |
| Elovl5 | 8964 | 6461 | 6899 | 7886 | 6781 | 7748 | 6962 | 6872 | 7444 | 7850 | 7650 | 7648 | 7404.38 | 7093 | 7716 | 1.088 | 0.122 | 0.02297184 | 0.07646402 | 8.00E-04 | 0.0027 | 0.0019 | 0.0019 | TRUE | 0.0227 |
| Elovl6 | 19517 | 13067 | 13463 | 15938 | 13869 | 15258 | 15158 | 13899 | 14526 | 15865 | 15646 | 15062 | 15026 | 14528 | 15524 | 1.068 | 0.095 | 0.07337312 | 0.17747877 | 0.0012 | 0.0026 | 0.002 | 0.002 | TRUE | 0.0177 |
| Elovl7 | 518 | 456 | 515 | 641 | 531 | 642 | 402 | 485 | 556 | 638 | 599 | 634 | 552.31 | 481 | 624 | 1.301 | 0.38 | 0.00132628 | 0.008833 | 0.0116 | 0.0067 | 0.0083 | 0.0083 | TRUE | 0.2525 |
| Elp2 | 10789 | 8178 | 8600 | 9268 | 8258 | 9498 | 8379 | 8699 | 9279 | 9225 | 9316 | 9376 | 9045.79 | 8786 | 9306 | 1.059 | 0.083 | 0.12958954 | 0.26635365 | 0.0012 | 0.0027 | 0.0021 | 0.0021 | TRUE | 0.0294 |
| Elp3 | 8960 | 6396 | 6704 | 6912 | 6231 | 7291 | 6959 | 6803 | 7234 | 6880 | 7029 | 7197 | 7017.08 | 6999 | 7035 | 1.005 | 0.008 | 0.88196054 | 0.93404431 | 5.00E-04 | 0.0028 | 0.0018 | 0.0018 | TRUE | 0.0104 |
| Elp4 | 1418 | 1015 | 918 | 993 | 909 | 1025 | 1101 | 1080 | 991 | 988 | 1025 | 1012 | 1032.86 | 1057 | 1008 | 0.953 | -0.07 | 0.36891797 | 0.5435465 | 8.00E-04 | 0.0047 | 0.0033 | 0.0033 | TRUE | 0.0356 |
| Elp5 | 4107 | 2851 | 2538 | 2765 | 2471 | 2871 | 3190 | 3033 | 2738 | 2752 | 2788 | 2834 | 2889.12 | 2987 | 2791 | 0.934 | -0.099 | 0.15185468 | 0.29826288 | 0.0025 | 0.0032 | 0.0031 | 0.0031 | TRUE | 0.0639 |
| Elp6 | 666 | 512 | 415 | 406 | 361 | 460 | 517 | 545 | 448 | 404 | 407 | 454 | 462.52 | 503 | 422 | 0.838 | -0.255 | 0.02207757 | 0.07414349 | 0.005 | 0.0076 | 0.0068 | 0.0068 | TRUE | 0.104 |
| Eltd1 | 1 | 0 | 1 | 1 | 2 | 1 | 1 | 0 | 1 | 1 | 2 | 1 | 1.02 | 1 | 1 | 2.238 | 1.162 | 0.54175048 | NA | 0 | 2.3358 | 1.4761 | 1.4761 | TRUE | 0.1945 |
| Emb | 3600 | 2535 | 3086 | 3130 | 2825 | 3427 | 2796 | 2696 | 3330 | 3116 | 3187 | 3383 | 3084.61 | 2941 | 3229 | 1.098 | 0.135 | 0.09996208 | 0.22086228 | 0.0071 | 0.0032 | 0.0045 | 0.0045 | TRUE | 0.1612 |
| Emc1 | 3778 | 2887 | 2587 | 2979 | 2682 | 2906 | 2934 | 3071 | 2791 | 2965 | 3026 | 2869 | 2942.66 | 2932 | 2953 | 1.007 | 0.01 | 0.87023456 | 0.92815928 | 0.0011 | 0.0032 | 0.0024 | 0.0024 | TRUE | 0.0209 |
| Emc10 | 3966 | 2927 | 3037 | 3308 | 2983 | 3435 | 3080 | 3113 | 3277 | 3293 | 3365 | 3391 | 3253.22 | 3157 | 3350 | 1.061 | 0.086 | 0.12755168 | 0.26346117 | 4.00E-04 | 0.0032 | 0.002 | 0.002 | TRUE | 0.0133 |
| Emc2 | 3717 | 2621 | 2812 | 3024 | 2538 | 2948 | 2887 | 2788 | 3034 | 3010 | 2863 | 2910 | 2915.37 | 2903 | 2928 | 1.009 | 0.013 | 0.83451147 | 0.90597926 | 8.00E-04 | 0.0032 | 0.0023 | 0.0023 | TRUE | 0.0185 |
| Emc3 | 2940 | 2190 | 2217 | 2274 | 2052 | 2252 | 2283 | 2329 | 2392 | 2264 | 2315 | 2223 | 2301.07 | 2335 | 2267 | 0.971 | -0.042 | 0.4755956 | 0.64353763 | 0 | 0.0035 | 0.0021 | 0.0021 | TRUE | 0.0055 |
| Emc4 | 1996 | 1353 | 1249 | 1357 | 1145 | 1437 | 1550 | 1439 | 1348 | 1351 | 1292 | 1419 | 1399.67 | 1446 | 1354 | 0.936 | -0.096 | 0.21902966 | 0.38215222 | 0.0029 | 0.0041 | 0.0037 | 0.0037 | TRUE | 0.0497 |
| Emc6 | 5269 | 3899 | 3784 | 4081 | 3562 | 4019 | 4092 | 4147 | 4083 | 4062 | 4018 | 3967 | 4061.72 | 4107 | 4016 | 0.978 | -0.032 | 0.52423151 | 0.68559254 | 0 | 0.003 | 0.0016 | 0.0016 | TRUE | 0.0014 |
| Emc7 | 1512 | 1027 | 1074 | 1228 | 1044 | 1154 | 1174 | 1092 | 1159 | 1222 | 1178 | 1139 | 1160.8 | 1142 | 1180 | 1.033 | 0.046 | 0.53037023 | 0.69033619 | 5.00E-04 | 0.0045 | 0.003 | 0.003 | TRUE | 0.0169 |
| Emc8 | 6121 | 4440 | 3655 | 3173 | 2911 | 3303 | 4754 | 4723 | 3944 | 3158 | 3284 | 3261 | 3853.88 | 4474 | 3234 | 0.723 | -0.469 | 9.36E-10 | 6.31E-08 | 0.0051 | 0.003 | 0.004 | 0.004 | TRUE | 0.1299 |
| Emc9 | 187 | 152 | 133 | 137 | 121 | 144 | 145 | 162 | 144 | 136 | 137 | 142 | 144.24 | 150 | 138 | 0.923 | -0.116 | 0.47806134 | 0.64546861 | 0 | 0.0189 | 0.0124 | 0.0124 | TRUE | 0.0453 |
| Emcn | 1 | 0 | 0 | 0 | 0 | 0 | 1 | 0 | 0 | 0 | 0 | 0 | 0.13 | 0 | 0 | 0.552 | -0.857 | 0.8336548 | NA | 0 | 18.3121 | 10 | 10 | TRUE | 0.2566 |
| Emd | 3635 | 2631 | 2534 | 2783 | 2355 | 2684 | 2823 | 2799 | 2734 | 2770 | 2657 | 2650 | 2738.71 | 2785 | 2692 | 0.966 | -0.049 | 0.38696584 | 0.56071448 | 0 | 0.0033 | 0.002 | 0.002 | TRUE | 0.0079 |
| Eme1 | 2830 | 1993 | 1770 | 1910 | 1707 | 2106 | 2198 | 2120 | 1910 | 1901 | 1926 | 2079 | 2022.25 | 2076 | 1969 | 0.948 | -0.077 | 0.29855821 | 0.47091152 | 0.003 | 0.0036 | 0.0035 | 0.0035 | TRUE | 0.0587 |
| Eme2 | 110 | 77 | 48 | 37 | 38 | 49 | 85 | 82 | 52 | 37 | 43 | 48 | 57.87 | 73 | 43 | 0.582 | -0.781 | 0.00611286 | 0.02861813 | 0.031 | 0.0434 | 0.0398 | 0.0398 | TRUE | 0.5613 |
| Emg1 | 9784 | 6817 | 6432 | 6752 | 6210 | 6950 | 7599 | 7251 | 6940 | 6721 | 7006 | 6861 | 7062.87 | 7263 | 6863 | 0.944 | -0.082 | 0.13464319 | 0.27347663 | 0.001 | 0.0028 | 0.002 | 0.002 | TRUE | 0.0202 |
| Emid1 | 297 | 199 | 217 | 357 | 300 | 333 | 231 | 212 | 234 | 355 | 338 | 329 | 283.17 | 226 | 341 | 1.51 | 0.595 | 1.27E-06 | 3.57E-05 | 0 | 0.0108 | 0.0072 | 0.0072 | TRUE | 0.03 |
| Emilin1 | 277 | 178 | 141 | 218 | 188 | 198 | 215 | 189 | 152 | 217 | 212 | 195 | 196.86 | 185 | 208 | 1.116 | 0.159 | 0.32281994 | 0.49667012 | 0.0113 | 0.0145 | 0.0135 | 0.0135 | TRUE | 0.2602 |
| Emilin2 | 310 | 231 | 205 | 259 | 217 | 236 | 241 | 246 | 221 | 258 | 245 | 233 | 240.54 | 236 | 245 | 1.039 | 0.055 | 0.67794517 | 0.80231161 | 0 | 0.0123 | 0.0083 | 0.0083 | TRUE | 0.031 |
| Emilin3 | 428 | 323 | 279 | 345 | 282 | 310 | 332 | 344 | 301 | 343 | 318 | 306 | 324.1 | 326 | 322 | 0.99 | -0.015 | 0.90014439 | 0.94510863 | 0 | 0.0097 | 0.0071 | 0.0071 | TRUE | 0.0472 |
| Eml1 | 681 | 517 | 585 | 628 | 614 | 611 | 529 | 550 | 631 | 625 | 693 | 603 | 605.16 | 570 | 640 | 1.124 | 0.169 | 0.10079502 | 0.22232456 | 0.0053 | 0.0063 | 0.006 | 0.006 | TRUE | 0.1017 |
| Eml2 | 2439 | 1916 | 2476 | 2593 | 2401 | 2761 | 1894 | 2038 | 2672 | 2581 | 2709 | 2726 | 2436.51 | 2201 | 2672 | 1.215 | 0.281 | 0.00793601 | 0.03485564 | 0.0175 | 0.0034 | 0.0076 | 0.0076 | TRUE | 0.4216 |
| Eml3 | 1212 | 930 | 788 | 890 | 811 | 839 | 941 | 989 | 850 | 886 | 915 | 828 | 901.63 | 927 | 876 | 0.945 | -0.082 | 0.34630036 | 0.52063438 | 0.003 | 0.0051 | 0.0044 | 0.0044 | TRUE | 0.0602 |
| Eml4 | 10339 | 7820 | 7407 | 8182 | 7394 | 8606 | 8030 | 8318 | 7992 | 8144 | 8341 | 8495 | 8220.16 | 8113 | 8327 | 1.026 | 0.038 | 0.44485863 | 0.61576258 | 3.00E-04 | 0.0027 | 0.0016 | 0.0016 | TRUE | 0.0059 |
| Eml5 | 3404 | 2842 | 2856 | 3102 | 2716 | 3158 | 2644 | 3023 | 3082 | 3088 | 3064 | 3117 | 3002.91 | 2916 | 3090 | 1.061 | 0.085 | 0.2264663 | 0.39149527 | 0.0029 | 0.0032 | 0.0032 | 0.0032 | TRUE | 0.0829 |

| | | | | | | | | | | | | | | | | | | | | | | | | | |
|---|---|---|---|---|---|---|---|---|---|---|---|---|---|---|---|---|---|---|---|---|---|---|---|---|---|
| Eml6 | 144 | 92 | 61 | 66 | 64 | 75 | 112 | 98 | 66 | 66 | 72 | 74 | 81.24 | 92 | 71 | 0.764 | -0.388 | 0.11277213 | 0.24140364 | 0.0278 | 0.0316 | 0.0305 | 0.0305 | TRUE | 0.4901 |
| Emp1 | 2043 | 1456 | 1646 | 1624 | 1391 | 1593 | 1587 | 1549 | 1776 | 1617 | 1569 | 1573 | 1611.62 | 1637 | 1586 | 0.969 | -0.045 | 0.54203534 | 0.6999739 | 0.002 | 0.0039 | 0.0033 | 0.0033 | TRUE | 0.065 |
| Emp2 | 354 | 225 | 313 | 390 | 336 | 331 | 275 | 239 | 338 | 388 | 379 | 327 | 324.33 | 284 | 365 | 1.285 | 0.362 | 0.01087666 | 0.04395254 | 0.0148 | 0.0097 | 0.0114 | 0.0114 | TRUE | 0.2946 |
| Emp3 | 149 | 105 | 152 | 167 | 149 | 202 | 116 | 112 | 164 | 166 | 168 | 199 | 154.19 | 131 | 178 | 1.369 | 0.454 | 0.01536434 | 0.05687241 | 0.0204 | 0.0178 | 0.0186 | 0.0186 | TRUE | 0.4984 |
| Emr1 | 2 | 0 | 1 | 0 | 2 | 4 | 2 | 0 | 1 | 0 | 2 | 4 | 1.47 | 1 | 2 | 2.279 | 1.188 | 0.49415736 | NA | 0 | 1.6116 | 1.3949 | 1.3949 | TRUE | 0.1534 |
| Emr4 | 3 | 3 | 1 | 1 | 1 | 3 | 2 | 3 | 1 | 1 | 1 | 3 | 1.95 | 2 | 2 | 0.773 | -0.371 | 0.78399338 | NA | 0 | 1.2194 | 0.7954 | 0.7954 | TRUE | 0.3414 |
| Emx1 | 21 | 15 | 16 | 36 | 34 | 38 | 16 | 16 | 17 | 36 | 38 | 38 | 26.87 | 16 | 37 | 2.256 | 1.174 | 0.00173145 | 0.01081628 | 0 | 0.0906 | 0.0581 | 0.0581 | TRUE | 0.008 |
| Emx2 | 0 | 1 | 1 | 0 | 0 | 0 | 0 | 1 | 1 | 0 | 0 | 0 | 0.36 | 1 | 0 | 0.271 | -1.881 | 0.5755355 | NA | 0 | 6.6391 | 6.4428 | 6.4428 | TRUE | 0.3513 |
| Emx2os | 1 | 0 | 0 | 0 | 0 | 0 | 1 | 0 | 0 | 0 | 0 | 0 | 0.13 | 0 | 0 | 0.552 | -0.857 | 0.8336548 | NA | 0 | 18.3121 | 10 | 10 | TRUE | 0.2566 |
| En1 | 30 | 18 | 13 | 13 | 11 | 7 | 23 | 19 | 14 | 13 | 12 | 7 | 14.79 | 19 | 11 | 0.565 | -0.823 | 0.11453839 | NA | 0 | 0.1627 | 0.1224 | 0.1224 | TRUE | 0.2662 |
| En2 | 777 | 611 | 568 | 622 | 569 | 647 | 603 | 650 | 613 | 619 | 642 | 639 | 627.66 | 622 | 633 | 1.019 | 0.027 | 0.76207723 | 0.85895197 | 0 | 0.0062 | 0.004 | 0.004 | TRUE | 0.0177 |
| Enah | 16915 | 12220 | 12083 | 12790 | 11474 | 13239 | 13137 | 12998 | 13037 | 12731 | 12944 | 13069 | 12986.14 | 13057 | 12915 | 0.989 | -0.016 | 0.72005986 | 0.83202375 | 0 | 0.0026 | 0.0013 | 0.0013 | TRUE | 0.0019 |
| Enam | 0 | 0 | 0 | 0 | 0 | 0 | 0 | 0 | 0 | 0 | 0 | 0 | 0 | 0 | 0 | NA | NA | NA | NA | NA | NA | NA | NA | NA | NA |
| Enc1 | 5037 | 3813 | 3494 | 4395 | 3673 | 4122 | 3912 | 4056 | 3770 | 4375 | 4144 | 4069 | 4054.2 | 3913 | 4196 | 1.072 | 0.101 | 0.08527225 | 0.19721623 | 0.001 | 0.003 | 0.0022 | 0.0022 | TRUE | 0.0171 |
| Endod1 | 466 | 327 | 265 | 333 | 311 | 316 | 362 | 348 | 286 | 331 | 351 | 312 | 331.65 | 332 | 331 | 0.995 | -0.007 | 0.95799765 | 0.97700039 | 0.0063 | 0.0096 | 0.0086 | 0.0086 | TRUE | 0.1621 |
| Endog | 937 | 681 | 533 | 459 | 437 | 513 | 728 | 724 | 575 | 457 | 493 | 506 | 580.58 | 676 | 485 | 0.717 | -0.479 | 1.80E-05 | 0.00030868 | 0.0087 | 0.0065 | 0.0072 | 0.0072 | TRUE | 0.1958 |
| Endou | 7 | 3 | 2 | 1 | 2 | 2 | 5 | 3 | 2 | 1 | 2 | 2 | 2.67 | 3 | 2 | 0.474 | -1.078 | 0.36419283 | NA | 0 | 0.8906 | 0.5865 | 0.5865 | TRUE | 0.401 |
| Endov | 435 | 350 | 266 | 367 | 345 | 359 | 338 | 372 | 287 | 365 | 389 | 354 | 351.01 | 332 | 369 | 1.111 | 0.152 | 0.22289573 | 0.38719599 | 0.0061 | 0.0092 | 0.0083 | 0.0083 | TRUE | 0.1536 |
| Eng | 2403 | 1831 | 1788 | 2117 | 1944 | 2359 | 1866 | 1948 | 1929 | 2107 | 2193 | 2329 | 2062.02 | 1914 | 2210 | 1.155 | 0.208 | 0.00153416 | 0.00981986 | 0.0011 | 0.0036 | 0.0026 | 0.0026 | TRUE | 0.0272 |
| Engase | 1381 | 1087 | 1110 | 1085 | 1045 | 1267 | 1073 | 1156 | 1198 | 1080 | 1179 | 1251 | 1156.01 | 1142 | 1170 | 1.025 | 0.036 | 0.66460593 | 0.79331991 | 0.0035 | 0.0045 | 0.0041 | 0.0041 | TRUE | 0.0549 |
| Enho | 175 | 144 | 125 | 172 | 174 | 168 | 136 | 153 | 135 | 171 | 196 | 166 | 159.55 | 141 | 178 | 1.258 | 0.331 | 0.04055028 | 0.11561195 | 0 | 0.0173 | 0.0125 | 0.0125 | TRUE | 0.0855 |
| Enkd1 | 1036 | 856 | 826 | 778 | 641 | 741 | 805 | 911 | 891 | 774 | 723 | 731 | 805.9 | 869 | 743 | 0.857 | -0.223 | 0.01007092 | 0.04156617 | 0.0019 | 0.0054 | 0.0042 | 0.0042 | TRUE | 0.0524 |
| Enkur | 18 | 17 | 14 | 19 | 21 | 20 | 14 | 18 | 15 | 19 | 24 | 20 | 18.25 | 16 | 21 | 1.325 | 0.406 | 0.36144509 | 0.53638908 | 0 | 0.1323 | 0.0869 | 0.0869 | TRUE | 0.0773 |
| Eno1 | 843 | 629 | 871 | 1006 | 948 | 989 | 655 | 669 | 940 | 1001 | 1069 | 976 | 885.12 | 755 | 1015 | 1.348 | 0.431 | 0.00049601 | 0.00428797 | 0.022 | 0.0051 | 0.0099 | 0.0099 | TRUE | 0.5411 |
| Eno2 | 298 | 257 | 340 | 571 | 521 | 637 | 231 | 273 | 367 | 568 | 588 | 629 | 442.77 | 290 | 595 | 2.057 | 1.041 | 7.03E-13 | 9.13E-11 | 0.026 | 0.0078 | 0.0126 | 0.0126 | TRUE | 0.5896 |
| Eno3 | 858 | 603 | 579 | 683 | 608 | 680 | 666 | 641 | 625 | 680 | 686 | 671 | 661.59 | 644 | 679 | 1.053 | 0.074 | 0.38433371 | 0.55837383 | 0 | 0.006 | 0.0037 | 0.0037 | TRUE | 0.0114 |
| Eno4 | 99 | 67 | 74 | 51 | 61 | 68 | 77 | 71 | 80 | 51 | 69 | 67 | 69.12 | 76 | 62 | 0.817 | -0.291 | 0.22381487 | 0.38825359 | 0 | 0.0367 | 0.0266 | 0.0266 | TRUE | 0.231 |
| Enoph1 | 3346 | 2568 | 2618 | 2514 | 2363 | 2753 | 2599 | 2732 | 2825 | 2502 | 2666 | 2718 | 2673.47 | 2719 | 2629 | 0.967 | -0.048 | 0.45585159 | 0.62546672 | 0.0015 | 0.0033 | 0.0026 | 0.0026 | TRUE | 0.0216 |
| Enox1 | 1264 | 914 | 741 | 852 | 695 | 882 | 982 | 972 | 800 | 848 | 784 | 871 | 876.04 | 918 | 834 | 0.908 | -0.139 | 0.15449061 | 0.30190516 | 0.0068 | 0.0051 | 0.0057 | 0.0057 | TRUE | 0.1485 |
| Enox2 | 814 | 594 | 550 | 738 | 699 | 730 | 632 | 632 | 593 | 735 | 789 | 721 | 683.54 | 619 | 748 | 1.206 | 0.271 | 0.00197947 | 0.01201891 | 0 | 0.0059 | 0.004 | 0.004 | TRUE | 0.0261 |
| Enpep | 22 | 13 | 10 | 12 | 8 | 3 | 17 | 14 | 11 | 12 | 9 | 3 | 10.94 | 14 | 8 | 0.568 | -0.815 | 0.19515798 | NA | 0.0725 | 0.2191 | 0.1857 | 0.1857 | TRUE | 0.8082 |
| Enpp1 | 236 | 158 | 175 | 181 | 156 | 171 | 183 | 168 | 189 | 180 | 176 | 169 | 177.52 | 180 | 175 | 0.971 | -0.042 | 0.77710167 | 0.8682181 | 0 | 0.0158 | 0.0104 | 0.0104 | TRUE | 0.0344 |
| Enpp2 | 5 | 8 | 11 | 10 | 5 | 3 | 4 | 9 | 12 | 10 | 6 | 3 | 7.14 | 8 | 6 | 0.778 | -0.362 | 0.64069703 | NA | 0.1769 | 0.3345 | 0.2885 | 0.2885 | TRUE | 0.3017 |
| Enpp3 | 1355 | 1040 | 1586 | 1089 | 1052 | 1406 | 1052 | 1106 | 1711 | 1084 | 1187 | 1388 | 1254.76 | 1290 | 1220 | 0.946 | -0.08 | 0.74696912 | 0.84993112 | 0.044 | 0.0043 | 0.0155 | 0.044 | TRUE | 0.9787 |
| Enpp4 | 1419 | 935 | 864 | 933 | 780 | 982 | 1102 | 995 | 932 | 929 | 880 | 969 | 967.81 | 1010 | 926 | 0.916 | -0.127 | 0.15018855 | 0.2957626 | 0.0039 | 0.0049 | 0.0045 | 0.0045 | TRUE | 0.0796 |
| Enpp5 | 650 | 463 | 681 | 806 | 713 | 796 | 505 | 492 | 735 | 802 | 804 | 786 | 687.42 | 577 | 797 | 1.384 | 0.468 | 0.00042777 | 0.00382814 | 0.0258 | 0.0059 | 0.0113 | 0.0113 | TRUE | 0.658 |
| Enpp6 | 3 | 3 | 0 | 1 | 0 | 0 | 2 | 3 | 0 | 1 | 0 | 0 | 1.09 | 2 | 0 | 0.2 | -2.321 | 0.27406904 | NA | 0 | 2.1847 | 1.9836 | 1.9836 | TRUE | 0.3792 |
| Enpp7 | 0 | 0 | 0 | 0 | 0 | 0 | 0 | 0 | 0 | 0 | 0 | 0 | 0 | 0 | 0 | NA | NA | NA | NA | NA | NA | NA | NA | NA | NA |
| Ensa | 5381 | 3850 | 4234 | 4839 | 4177 | 4831 | 4179 | 4095 | 4568 | 4817 | 4712 | 4769 | 4523.43 | 4281 | 4766 | 1.114 | 0.155 | 0.00947998 | 0.03982872 | 0.0015 | 0.003 | 0.0024 | 0.0024 | TRUE | 0.0415 |
| Enthd1 | 28 | 37 | 50 | 25 | 33 | 36 | 22 | 39 | 54 | 25 | 37 | 36 | 35.45 | 38 | 33 | 0.857 | -0.223 | 0.56228317 | 0.71682025 | 0.0975 | 0.0693 | 0.0776 | 0.0776 | TRUE | 0.2235 |
| Enthd2 | 1965 | 1599 | 1392 | 1388 | 1259 | 1540 | 1526 | 1701 | 1502 | 1382 | 1420 | 1520 | 1508.51 | 1576 | 1441 | 0.914 | -0.129 | 0.09344437 | 0.21029765 | 0.0026 | 0.004 | 0.0036 | 0.0036 | TRUE | 0.0566 |
| Entpd1 | 26 | 14 | 16 | 10 | 20 | 22 | 20 | 15 | 17 | 10 | 23 | 22 | 17.76 | 17 | 18 | 1.027 | 0.038 | 0.93634423 | 0.96471383 | 0.0409 | 0.1358 | 0.1104 | 0.1104 | TRUE | 0.8099 |
| Entpd2 | 202 | 171 | 218 | 283 | 298 | 335 | 157 | 182 | 235 | 282 | 336 | 331 | 253.76 | 191 | 316 | 1.66 | 0.731 | 5.13E-06 | 0.00010806 | 0.0201 | 0.0118 | 0.0144 | 0.0144 | TRUE | 0.4301 |
| Entpd3 | 42 | 40 | 58 | 53 | 54 | 49 | 33 | 43 | 63 | 53 | 61 | 48 | 49.97 | 46 | 54 | 1.186 | 0.246 | 0.42142719 | 0.59373669 | 0.0407 | 0.0499 | 0.0473 | 0.0473 | TRUE | 0.5879 |
| Entpd4 | 0 | 0 | 0 | 0 | 0 | 1 | 0 | 0 | 0 | 0 | 0 | 1 | 0.16 | 0 | 0 | 2.094 | 1.067 | 0.79379329 | NA | 0 | 14.4078 | 10 | 10 | TRUE | 0.3835 |
| Entpd5 | 1401 | 1002 | 1040 | 1205 | 991 | 1174 | 1088 | 1066 | 1122 | 1199 | 1118 | 1159 | 1125.4 | 1092 | 1159 | 1.062 | 0.086 | 0.2359592 | 0.40240602 | 0 | 0.0045 | 0.0029 | 0.0029 | TRUE | 0.0113 |
| Entpd6 | 2942 | 2022 | 1991 | 2019 | 1850 | 2141 | 2285 | 2151 | 2148 | 2010 | 2087 | 2113 | 2132.36 | 2195 | 2070 | 0.943 | -0.085 | 0.17314682 | 0.32562993 | 6.00E-04 | 0.0035 | 0.0024 | 0.0024 | TRUE | 0.0162 |
| Entpd7 | 463 | 363 | 318 | 353 | 307 | 356 | 360 | 386 | 343 | 351 | 346 | 351 | 356.33 | 363 | 349 | 0.964 | -0.053 | 0.63148038 | 0.76880316 | 0 | 0.0091 | 0.006 | 0.006 | TRUE | 0.0338 |
| Entpd8 | 0 | 0 | 0 | 0 | 0 | 0 | 0 | 0 | 0 | 0 | 0 | 0 | 0 | 0 | 0 | NA | NA | NA | NA | NA | NA | NA | NA | NA | NA |
| Eny2 | 2179 | 1522 | 1345 | 1490 | 1187 | 1344 | 1692 | 1619 | 1451 | 1483 | 1339 | 1327 | 1485.24 | 1587 | 1383 | 0.871 | -0.2 | 0.01459452 | 0.05464647 | 0.004 | 0.004 | 0.0042 | 0.0042 | TRUE | 0.0673 |
| Eogt | 1272 | 929 | 975 | 1026 | 879 | 990 | 988 | 988 | 1052 | 1021 | 992 | 977 | 1003.04 | 1009 | 997 | 0.988 | -0.017 | 0.81798786 | 0.89536894 | 0 | 0.0048 | 0.0031 | 0.0031 | TRUE | 0.0159 |
| Eomes | 1038 | 736 | 669 | 714 | 657 | 693 | 806 | 783 | 722 | 711 | 741 | 684 | 741.14 | 770 | 712 | 0.923 | -0.116 | 0.18311718 | 0.338896 | 0.0011 | 0.0056 | 0.0041 | 0.0041 | TRUE | 0.0352 |
| Ep300 | 10770 | 7446 | 7027 | 7795 | 7033 | 7830 | 8365 | 7920 | 7582 | 7759 | 7934 | 7729 | 7881.57 | 7956 | 7807 | 0.981 | -0.028 | 0.61639158 | 0.75779489 | 0.001 | 0.0027 | 0.0021 | 0.0021 | TRUE | 0.025 |
| Ep400 | 14061 | 10369 | 10098 | 10980 | 10505 | 11996 | 10921 | 11029 | 10896 | 10929 | 11851 | 11842 | 11244.62 | 10949 | 11541 | 1.054 | 0.076 | 0.14887641 | 0.29411535 | 9.00E-04 | 0.0026 | 0.0019 | 0.0019 | TRUE | 0.0263 |
| Epas1 | 4024 | 2924 | 3129 | 4123 | 3672 | 4370 | 3125 | 3110 | 3376 | 4104 | 4142 | 4314 | 3695.32 | 3204 | 4187 | 1.307 | 0.387 | 7.57E-11 | 6.29E-09 | 0.001 | 0.0031 | 0.0023 | 0.0023 | TRUE | 0.0265 |
| Epb4.1 | 4401 | 3295 | 3327 | 3543 | 3145 | 3675 | 3418 | 3505 | 3590 | 3527 | 3548 | 3628 | 3535.85 | 3504 | 3568 | 1.018 | 0.026 | 0.62707793 | 0.76496691 | 0 | 0.0031 | 0.0018 | 0.0018 | TRUE | 0.0058 |
| Epb4.1l1 | 3100 | 2157 | 2293 | 2760 | 2498 | 2744 | 2408 | 2294 | 2474 | 2747 | 2818 | 2709 | 2575.03 | 2392 | 2758 | 1.153 | 0.205 | 0.0006302 | 0.00515888 | 5.00E-04 | 0.0033 | 0.0022 | 0.0022 | TRUE | 0.0151 |
| Epb4.1l2 | 9763 | 7215 | 6909 | 7821 | 6888 | 8072 | 7583 | 7675 | 7455 | 7785 | 7770 | 7968 | 7705.91 | 7571 | 7841 | 1.036 | 0.051 | 0.2852544 | 0.45727025 | 1.00E-04 | 0.0027 | 0.0015 | 0.0015 | TRUE | 0.0025 |
| Epb4.1l3 | 1079 | 878 | 810 | 716 | 709 | 735 | 838 | 934 | 874 | 713 | 800 | 726 | 814 | 882 | 746 | 0.846 | -0.241 | 0.00586648 | 0.02775557 | 0.0022 | 0.0053 | 0.0043 | 0.0043 | TRUE | 0.046 |
| Epb4.1l4a | 3168 | 2276 | 2340 | 2421 | 2385 | 2734 | 2460 | 2421 | 2525 | 2410 | 2691 | 2699 | 2534.25 | 2469 | 2600 | 1.053 | 0.074 | 0.2626181 | 0.43314107 | 0.0016 | 0.0034 | 0.0028 | 0.0028 | TRUE | 0.0498 |
| Epb4.1l4b | 1035 | 698 | 730 | 867 | 681 | 850 | 804 | 742 | 788 | 863 | 768 | 839 | 800.71 | 778 | 823 | 1.058 | 0.082 | 0.33951031 | 0.5135592 | 0.0013 | 0.0054 | 0.004 | 0.004 | TRUE | 0.0398 |
| Epb4.1l5 | 2367 | 1828 | 1992 | 1998 | 1807 | 2164 | 1838 | 1944 | 2149 | 1989 | 2038 | 2136 | 2015.94 | 1977 | 2054 | 1.04 | 0.057 | 0.44787854 | 0.61796775 | 0.0034 | 0.0036 | 0.0035 | 0.0035 | TRUE | 0.0697 |
| Epb4.2 | 1 | 0 | 0 | 2 | 1 | 1 | 1 | 0 | 0 | 2 | 1 | 1 | 0.81 | 0 | 1 | 3.977 | 1.992 | 0.35812725 | NA | 0 | 2.9148 | 2.0074 | 2.0074 | TRUE | 0.2566 |
| Epb4.9 | 2058 | 1618 | 2035 | 1983 | 1756 | 2144 | 1598 | 1721 | 2196 | 1974 | 1981 | 2116 | 1931.07 | 1838 | 2024 | 1.102 | 0.14 | 0.17347718 | 0.32616539 | 0.0136 | 0.0037 | 0.0071 | 0.0071 | TRUE | 0.348 |
| Epc1 | 1179 | 767 | 684 | 648 | 595 | 720 | 916 | 816 | 738 | 645 | 671 | 711 | 749.42 | 823 | 676 | 0.819 | -0.288 | 0.00372655 | 0.01953912 | 0.006 | 0.0056 | 0.0057 | 0.0057 | TRUE | 0.1191 |
| Epc2 | 2199 | 1528 | 1544 | 1581 | 1356 | 1715 | 1708 | 1625 | 1666 | 1574 | 1530 | 1693 | 1632.59 | 1666 | 1599 | 0.959 | -0.06 | 0.38656338 | 0.56041691 | 0.0011 | 0.0039 | 0.0028 | 0.0028 | TRUE | 0.0322 |
| Epcam | 5400 | 4419 | 5368 | 5390 | 4838 | 5457 | 4194 | 4700 | 5792 | 5365 | 5458 | 5387 | 5149.38 | 4895 | 5403 | 1.104 | 0.143 | 0.13344467 | 0.27193109 | 0.0128 | 0.0029 | 0.0064 | 0.0064 | TRUE | 0.3125 |
| Epdr1 | 857 | 594 | 649 | 837 | 742 | 811 | 666 | 632 | 700 | 833 | 837 | 801 | 744.75 | 666 | 824 | 1.237 | 0.306 | 0.0002865 | 0.00280669 | 0 | 0.0056 | 0.0038 | 0.0038 | TRUE | 0.0226 |
| Epg5 | 1909 | 1284 | 1170 | 1420 | 1155 | 1397 | 1483 | 1366 | 1262 | 1413 | 1303 | 1379 | 1367.72 | 1370 | 1365 | 0.995 | -0.007 | 0.93194142 | 0.96225044 | 0.0034 | 0.0042 | 0.0039 | 0.0039 | TRUE | 0.0639 |

| | | | | | | | | | | | | | | | | | | | | | | | | | |
|---|---|---|---|---|---|---|---|---|---|---|---|---|---|---|---|---|---|---|---|---|---|---|---|---|---|
| Epgn | 4 | 3 | 3 | 2 | 1 | 2 | 3 | 3 | 3 | 2 | 1 | 2 | 2.44 | 3 | 2 | 0.539 | -0.893 | 0.45828655 | NA | 0 | 0.9746 | 0.5899 | 0.5899 | TRUE | 0.0569 |
| Epha1 | 522 | 393 | 434 | 483 | 463 | 516 | 405 | 418 | 468 | 481 | 522 | 509 | 467.36 | 430 | 504 | 1.173 | 0.23 | 0.0273152 | 0.08641623 | 0.0016 | 0.0075 | 0.0057 | 0.0057 | TRUE | 0.0664 |
| Epha10 | 73 | 58 | 72 | 113 | 103 | 104 | 57 | 62 | 78 | 112 | 116 | 103 | 87.9 | 66 | 110 | 1.699 | 0.764 | 0.00044511 | 0.00394477 | 0 | 0.0294 | 0.022 | 0.022 | TRUE | 0.2314 |
| Epha2 | 10229 | 7093 | 5759 | 10789 | 8433 | 8966 | 7944 | 7545 | 6214 | 10739 | 9513 | 8851 | 8467.76 | 7234 | 9701 | 1.341 | 0.423 | 5.07E-06 | 0.00010681 | 0.0125 | 0.0027 | 0.0061 | 0.0061 | TRUE | 0.1855 |
| Epha3 | 2 | 5 | 2 | 3 | 1 | 0 | 2 | 5 | 2 | 3 | 1 | 0 | 2.19 | 3 | 1 | 0.463 | -1.112 | 0.42673636 | NA | 0.2746 | 1.0843 | 0.8714 | 0.8714 | TRUE | 0.6612 |
| Epha4 | 4430 | 3312 | 3617 | 4817 | 4395 | 4889 | 3441 | 3523 | 3903 | 4795 | 4958 | 4826 | 4240.88 | 3622 | 4860 | 1.342 | 0.425 | 1.93E-11 | 1.82E-09 | 0.0021 | 0.003 | 0.0026 | 0.0026 | TRUE | 0.0555 |
| Epha5 | 4 | 2 | 2 | 0 | 0 | 0 | 3 | 2 | 2 | 0 | 0 | 0 | 1.23 | 2 | 0 | 0.077 | -3.702 | 0.05731828 | NA | 0 | 1.9262 | 1.5379 | 1.5379 | TRUE | 0.0744 |
| Epha6 | 3 | 1 | 0 | 0 | 2 | 3 | 2 | 1 | 0 | 0 | 2 | 3 | 1.44 | 1 | 2 | 1.472 | 0.557 | 0.74618635 | NA | 0 | 1.6538 | 1.4284 | 1.4284 | TRUE | 0.3854 |
| Epha7 | 266 | 188 | 211 | 213 | 181 | 217 | 207 | 200 | 228 | 212 | 204 | 214 | 210.78 | 212 | 210 | 0.995 | -0.007 | 0.96113155 | 0.97845108 | 0 | 0.0137 | 0.0091 | 0.0091 | TRUE | 0.0468 |
| Epha8 | 253 | 219 | 226 | 414 | 316 | 434 | 196 | 233 | 244 | 412 | 356 | 428 | 311.72 | 224 | 399 | 1.787 | 0.838 | 3.05E-10 | 2.27E-08 | 0.0076 | 0.01 | 0.0093 | 0.0093 | TRUE | 0.1437 |
| Ephb1 | 44 | 53 | 29 | 15 | 22 | 21 | 34 | 56 | 31 | 15 | 25 | 21 | 30.39 | 40 | 20 | 0.496 | -1.011 | 0.00996549 | 0.04126208 | 0.055 | 0.0804 | 0.0732 | 0.0732 | TRUE | 0.8341 |
| Ephb2 | 1494 | 1113 | 1132 | 1261 | 1188 | 1370 | 1160 | 1184 | 1221 | 1255 | 1340 | 1352 | 1252.24 | 1188 | 1316 | 1.108 | 0.147 | 0.03928698 | 0.11302915 | 0 | 0.0043 | 0.0029 | 0.0029 | TRUE | 0.0198 |
| Ephb3 | 1997 | 1668 | 1782 | 1878 | 1827 | 2040 | 1551 | 1774 | 1923 | 1869 | 2061 | 2014 | 1865.36 | 1749 | 1981 | 1.134 | 0.181 | 0.03392992 | 0.1015695 | 0.0062 | 0.0037 | 0.0047 | 0.0047 | TRUE | 0.122 |
| Ephb4 | 3185 | 2652 | 3285 | 3161 | 3181 | 3591 | 2474 | 2821 | 3544 | 3146 | 3588 | 3545 | 3186.48 | 2946 | 3426 | 1.164 | 0.219 | 0.04315214 | 0.12099979 | 0.0192 | 0.0032 | 0.0081 | 0.0081 | TRUE | 0.3826 |
| Ephb6 | 174 | 106 | 125 | 127 | 112 | 139 | 135 | 113 | 135 | 126 | 126 | 137 | 128.79 | 128 | 130 | 1.017 | 0.025 | 0.88817941 | 0.93792008 | 0 | 0.0208 | 0.0146 | 0.0146 | TRUE | 0.1011 |
| Ephx1 | 1219 | 952 | 1390 | 1427 | 1349 | 1554 | 947 | 1013 | 1500 | 1420 | 1522 | 1534 | 1322.58 | 1153 | 1492 | 1.295 | 0.373 | 0.09006702 | 0.20486665 | 0.034 | 0.0042 | 0.0122 | 0.034 | TRUE | 0.8273 |
| Ephx2 | 691 | 583 | 810 | 767 | 836 | 985 | 537 | 620 | 874 | 763 | 943 | 972 | 784.95 | 677 | 893 | 1.321 | 0.402 | 0.00759938 | 0.03371555 | 0.0398 | 0.0054 | 0.0151 | 0.0151 | TRUE | 0.7623 |
| Ephx3 | 7 | 2 | 3 | 4 | 6 | 3 | 5 | 2 | 3 | 4 | 7 | 3 | 4.09 | 3 | 5 | 1.234 | 0.303 | 0.74940097 | NA | 0 | 0.5826 | 0.4045 | 0.4045 | TRUE | 0.4039 |
| Ephx4 | 94 | 60 | 54 | 74 | 43 | 60 | 73 | 64 | 58 | 74 | 49 | 59 | 62.75 | 65 | 61 | 0.929 | -0.107 | 0.67975044 | 0.80328012 | 0.0112 | 0.0402 | 0.0323 | 0.0323 | TRUE | 0.296 |
| Epm2a | 134 | 120 | 89 | 61 | 51 | 60 | 104 | 128 | 96 | 61 | 58 | 59 | 84.2 | 109 | 59 | 0.542 | -0.882 | 7.53E-05 | 0.00097057 | 0 | 0.0306 | 0.0226 | 0.0226 | TRUE | 0.2057 |
| Epm2aip1 | 1752 | 1418 | 1323 | 1432 | 1163 | 1345 | 1361 | 1508 | 1428 | 1425 | 1312 | 1328 | 1393.61 | 1432 | 1355 | 0.947 | -0.078 | 0.29053185 | 0.46257901 | 0.0017 | 0.0041 | 0.0032 | 0.0032 | TRUE | 0.026 |
| Epn1 | 5145 | 3866 | 3772 | 3897 | 3545 | 4080 | 3996 | 4112 | 4070 | 3879 | 3999 | 4028 | 4013.98 | 4059 | 3969 | 0.978 | -0.032 | 0.53644161 | 0.69526297 | 0 | 0.003 | 0.0017 | 0.0017 | TRUE | 0.0048 |
| Epn2 | 6486 | 4768 | 4999 | 5633 | 4971 | 5688 | 5037 | 5072 | 5394 | 5607 | 5608 | 5615 | 5388.79 | 5168 | 5610 | 1.086 | 0.119 | 0.0249556 | 0.08130624 | 4.00E-04 | 0.0029 | 0.0018 | 0.0018 | TRUE | 0.0177 |
| Epn3 | 14 | 17 | 5 | 12 | 9 | 6 | 11 | 18 | 5 | 12 | 10 | 6 | 10.4 | 11 | 9 | 0.815 | -0.295 | 0.64640286 | NA | 0.1219 | 0.2304 | 0.2006 | 0.2006 | TRUE | 0.2117 |
| Epo | 24 | 9 | 10 | 15 | 13 | 23 | 19 | 10 | 11 | 15 | 15 | 23 | 15.22 | 13 | 18 | 1.322 | 0.403 | 0.43383307 | NA | 0.0409 | 0.1582 | 0.1246 | 0.1246 | TRUE | 0.745 |
| Epor | 167 | 156 | 121 | 201 | 176 | 227 | 130 | 166 | 131 | 200 | 199 | 224 | 174.82 | 142 | 208 | 1.466 | 0.552 | 0.00067031 | 0.00538544 | 0.0054 | 0.016 | 0.0131 | 0.0131 | TRUE | 0.2169 |
| Eppin | 0 | 2 | 3 | 1 | 5 | 3 | 0 | 2 | 3 | 1 | 6 | 3 | 2.49 | 2 | 3 | 1.852 | 0.889 | 0.50625405 | NA | 0.499 | 0.9529 | 0.8452 | 0.8452 | TRUE | 0.3357 |
| Eppk1 | 113 | 97 | 97 | 113 | 112 | 165 | 88 | 103 | 105 | 112 | 126 | 163 | 116.22 | 99 | 134 | 1.365 | 0.449 | 0.02690795 | 0.08567505 | 0.0163 | 0.0228 | 0.0209 | 0.0209 | TRUE | 0.377 |
| Eprs | 26271 | 18713 | 18778 | 19697 | 17121 | 20166 | 20404 | 19905 | 20261 | 19606 | 19314 | 19907 | 19899.54 | 20190 | 19609 | 0.971 | -0.042 | 0.34137241 | 0.51533965 | 1.00E-04 | 0.0025 | 0.0014 | 0.0014 | TRUE | 0.0022 |
| Eps15 | 5127 | 3640 | 3618 | 4046 | 3453 | 3810 | 3982 | 3872 | 3904 | 4027 | 3895 | 3761 | 3906.89 | 3919 | 3894 | 0.994 | -0.009 | 0.86498418 | 0.92551298 | 4.00E-04 | 0.003 | 0.0019 | 0.0019 | TRUE | 0.0111 |
| Eps15l1 | 7104 | 5458 | 5170 | 5528 | 4891 | 5693 | 5517 | 5806 | 5578 | 5503 | 5518 | 5620 | 5590.23 | 5634 | 5547 | 0.985 | -0.022 | 0.66298399 | 0.7920772 | 3.00E-04 | 0.0029 | 0.0017 | 0.0017 | TRUE | 0.0086 |
| Eps8 | 5988 | 4268 | 3586 | 4097 | 3818 | 4114 | 4651 | 4540 | 3869 | 4078 | 4307 | 4061 | 4251.02 | 4353 | 4149 | 0.953 | -0.07 | 0.34918443 | 0.52375613 | 0.0048 | 0.003 | 0.0038 | 0.0038 | TRUE | 0.1145 |
| Eps8l1 | 4 | 4 | 3 | 4 | 3 | 0 | 3 | 4 | 3 | 4 | 3 | 0 | 2.99 | 3 | 2 | 0.692 | -0.531 | 0.64851591 | NA | 0 | 0.794 | 0.6265 | 0.6265 | TRUE | 0.8753 |
| Eps8l2 | 726 | 569 | 707 | 842 | 736 | 858 | 564 | 605 | 763 | 838 | 830 | 847 | 741.22 | 644 | 838 | 1.305 | 0.384 | 0.00048972 | 0.00424988 | 0.0113 | 0.0056 | 0.0074 | 0.0074 | TRUE | 0.3039 |
| Eps8l3 | 1 | 0 | 0 | 1 | 0 | 1 | 1 | 0 | 0 | 1 | 0 | 1 | 0.46 | 0 | 1 | 1.94 | 0.956 | 0.73503911 | NA | 0 | 5.1562 | 3.9771 | 3.9771 | TRUE | 0.2566 |
| Epsti1 | 0 | 0 | 0 | 0 | 1 | 0 | 0 | 0 | 0 | 0 | 1 | 0 | 0.19 | 0 | 0 | 2.094 | 1.067 | 0.79379329 | NA | 0 | 12.608 | 10 | 10 | TRUE | 0.5075 |
| Ept1 | 4164 | 2802 | 2702 | 2504 | 2189 | 2414 | 3234 | 2980 | 2915 | 2492 | 2469 | 2383 | 2745.8 | 3043 | 2448 | 0.804 | -0.315 | 8.59E-07 | 2.57E-05 | 0.0015 | 0.0033 | 0.0026 | 0.0026 | TRUE | 0.0373 |
| Epx | 51 | 33 | 37 | 76 | 55 | 61 | 40 | 35 | 40 | 76 | 62 | 60 | 52.09 | 38 | 66 | 1.725 | 0.787 | 0.00389645 | 0.02022222 | 0 | 0.0479 | 0.0331 | 0.0331 | TRUE | 0.1486 |
| Epyc | 0 | 0 | 0 | 0 | 0 | 0 | 0 | 0 | 0 | 0 | 0 | 0 | 0 | 0 | 0 | NA | NA | NA | NA | NA | NA | NA | NA | NA | NA |
| Eqtn | 0 | 0 | 0 | 0 | 0 | 0 | 0 | 0 | 0 | 0 | 0 | 0 | 0 | 0 | 0 | NA | NA | NA | NA | NA | NA | NA | NA | NA | NA |
| Eral1 | 2662 | 2011 | 2115 | 2087 | 1877 | 2167 | 2067 | 2139 | 2282 | 2077 | 2117 | 2139 | 2137.1 | 2163 | 2111 | 0.977 | -0.034 | 0.60005825 | 0.74566446 | 0.001 | 0.0035 | 0.0025 | 0.0025 | TRUE | 0.0279 |
| Erap1 | 174 | 119 | 103 | 125 | 116 | 141 | 135 | 127 | 111 | 124 | 131 | 139 | 127.89 | 124 | 131 | 1.055 | 0.077 | 0.66429584 | 0.79308223 | 0 | 0.021 | 0.0148 | 0.0148 | TRUE | 0.0843 |
| Eras | 1747 | 1251 | 1269 | 1145 | 1077 | 1324 | 1357 | 1331 | 1369 | 1140 | 1215 | 1307 | 1286.4 | 1352 | 1221 | 0.903 | -0.148 | 0.05031058 | 0.13539839 | 0.0016 | 0.0043 | 0.0033 | 0.0033 | TRUE | 0.0469 |
| Erbb2 | 1454 | 995 | 893 | 834 | 799 | 843 | 1129 | 1058 | 964 | 830 | 901 | 832 | 952.48 | 1050 | 854 | 0.812 | -0.3 | 0.00054595 | 0.00461185 | 0.0034 | 0.0049 | 0.0044 | 0.0044 | TRUE | 0.062 |
| Erbb2ip | 2310 | 1678 | 1515 | 1749 | 1491 | 1619 | 1794 | 1785 | 1635 | 1741 | 1682 | 1598 | 1705.8 | 1738 | 1674 | 0.962 | -0.055 | 0.43588755 | 0.60716701 | 0.0017 | 0.0038 | 0.003 | 0.003 | TRUE | 0.032 |
| Erbb3 | 1769 | 1226 | 1242 | 1146 | 997 | 1145 | 1374 | 1304 | 1340 | 1141 | 1125 | 1130 | 1235.64 | 1339 | 1132 | 0.845 | -0.244 | 0.00041819 | 0.00376159 | 0 | 0.0043 | 0.0026 | 0.0026 | TRUE | 0.0064 |
| Erbb4 | 1 | 0 | 0 | 0 | 0 | 0 | 1 | 0 | 0 | 0 | 0 | 0 | 0.13 | 0 | 0 | 0.552 | -0.857 | 0.8336548 | NA | 0 | 18.3121 | 10 | 10 | TRUE | 0.2566 |
| Erc1 | 10575 | 7323 | 6340 | 6440 | 5681 | 6273 | 8213 | 7789 | 6841 | 6410 | 6409 | 6192 | 6975.81 | 7614 | 6337 | 0.832 | -0.265 | 0.00015983 | 0.00179315 | 0.0042 | 0.0028 | 0.0034 | 0.0034 | TRUE | 0.0962 |
| Erc2 | 633 | 436 | 419 | 497 | 411 | 477 | 492 | 464 | 452 | 495 | 464 | 471 | 472.79 | 469 | 477 | 1.014 | 0.02 | 0.83616441 | 0.90650015 | 0 | 0.0074 | 0.0049 | 0.0049 | TRUE | 0.0218 |
| Ercc1 | 876 | 645 | 514 | 546 | 453 | 580 | 680 | 686 | 555 | 543 | 511 | 573 | 591.35 | 640 | 542 | 0.846 | -0.241 | 0.02510214 | 0.08154202 | 0.007 | 0.0064 | 0.0066 | 0.0066 | TRUE | 0.1574 |
| Ercc2 | 2710 | 2045 | 1752 | 1835 | 1563 | 1832 | 2105 | 2175 | 1890 | 1827 | 1763 | 1808 | 1928.1 | 2057 | 1799 | 0.875 | -0.193 | 0.0068105 | 0.03111733 | 0.0023 | 0.0037 | 0.0031 | 0.0031 | TRUE | 0.0593 |
| Ercc3 | 5416 | 3862 | 3464 | 3639 | 3310 | 3577 | 4206 | 4108 | 3738 | 3622 | 3734 | 3531 | 3823.22 | 4017 | 3629 | 0.903 | -0.147 | 0.02055693 | 0.07057897 | 0.0019 | 0.003 | 0.0027 | 0.0027 | TRUE | 0.0448 |
| Ercc4 | 1829 | 1254 | 997 | 1044 | 902 | 974 | 1421 | 1334 | 1076 | 1039 | 1018 | 961 | 1141.39 | 1277 | 1006 | 0.787 | -0.346 | 0.00063175 | 0.00516861 | 0.0104 | 0.0045 | 0.0065 | 0.0065 | TRUE | 0.2253 |
| Ercc5 | 2801 | 1959 | 2143 | 2294 | 1917 | 2178 | 2175 | 2084 | 2312 | 2283 | 2163 | 2150 | 2194.58 | 2190 | 2199 | 1.004 | 0.006 | 0.93123601 | 0.9621336 | 0.0014 | 0.0035 | 0.0027 | 0.0027 | TRUE | 0.0279 |
| Ercc6 | 2627 | 1979 | 1901 | 2381 | 2034 | 2364 | 2040 | 2105 | 2051 | 2370 | 2295 | 2334 | 2199.12 | 2065 | 2333 | 1.13 | 0.176 | 0.0027034 | 0.01520053 | 0 | 0.0035 | 0.002 | 0.002 | TRUE | 0.0034 |
| Ercc6l | 4260 | 2719 | 2233 | 2582 | 2252 | 2571 | 3309 | 2892 | 2409 | 2570 | 2540 | 2538 | 2709.78 | 2870 | 2549 | 0.888 | -0.172 | 0.07351976 | 0.17774321 | 0.0118 | 0.0033 | 0.0063 | 0.0063 | TRUE | 0.2 |
| Ercc6l2 | 4860 | 3368 | 3360 | 3526 | 3096 | 3565 | 3775 | 3582 | 3625 | 3510 | 3493 | 3519 | 3584.01 | 3661 | 3507 | 0.958 | -0.062 | 0.24775345 | 0.41665236 | 0 | 0.0031 | 0.0018 | 0.0018 | TRUE | 0.0092 |
| Ercc8 | 1348 | 1030 | 948 | 1084 | 989 | 1172 | 1047 | 1096 | 1023 | 1079 | 1116 | 1157 | 1086.18 | 1055 | 1117 | 1.059 | 0.083 | 0.26663496 | 0.43705861 | 0 | 0.0046 | 0.0031 | 0.0031 | TRUE | 0.013 |
| Erdr1 | 1319 | 1214 | 1268 | 1138 | 956 | 1330 | 1024 | 1291 | 1368 | 1133 | 1078 | 1313 | 1201.34 | 1228 | 1175 | 0.958 | -0.061 | 0.58256121 | 0.73263262 | 0.0159 | 0.0044 | 0.0081 | 0.0081 | TRUE | 0.2589 |
| Ereg | 0 | 2 | 0 | 1 | 1 | 0 | 0 | 2 | 0 | 1 | 1 | 0 | 0.71 | 1 | 1 | 1.021 | 0.03 | 0.99038186 | NA | 0 | 3.3477 | 2.9808 | 2.9808 | TRUE | 0.9748 |
| Erf | 13433 | 8521 | 5801 | 6784 | 5851 | 6181 | 10433 | 9064 | 6259 | 6753 | 6601 | 6102 | 7535.11 | 8585 | 6485 | 0.755 | -0.405 | 0.05474924 | 0.14366273 | 0.0319 | 0.0027 | 0.0117 | 0.0319 | TRUE | 0.4086 |
| Erg | 27 | 22 | 10 | 9 | 8 | 16 | 21 | 23 | 11 | 9 | 9 | 16 | 14.82 | 18 | 11 | 0.612 | -0.709 | 0.18634995 | NA | 0.0621 | 0.1623 | 0.1353 | 0.1353 | TRUE | 0.6189 |
| Ergic1 | 3507 | 2658 | 3227 | 3571 | 3267 | 3889 | 2724 | 2827 | 3482 | 3555 | 3686 | 3839 | 3352.01 | 3011 | 3693 | 1.227 | 0.295 | 0.00077302 | 0.00600154 | 0.0096 | 0.0031 | 0.0053 | 0.0053 | TRUE | 0.2264 |
| Ergic2 | 2477 | 1770 | 1683 | 1740 | 1453 | 1757 | 1924 | 1883 | 1816 | 1732 | 1639 | 1734 | 1788 | 1874 | 1702 | 0.908 | -0.139 | 0.03127601 | 0.095478 | 4.00E-04 | 0.0038 | 0.0025 | 0.0025 | TRUE | 0.0123 |
| Ergic3 | 4208 | 3193 | 3253 | 3742 | 3378 | 3804 | 3268 | 3396 | 3510 | 3725 | 3811 | 3755 | 3577.52 | 3391 | 3764 | 1.11 | 0.151 | 0.00695285 | 0.03158841 | 4.00E-04 | 0.0031 | 0.002 | 0.002 | TRUE | 0.0127 |
| Erh | 6014 | 3896 | 3813 | 3580 | 3071 | 3652 | 4671 | 4144 | 4114 | 3564 | 3464 | 3605 | 3927.03 | 4310 | 3544 | 0.822 | -0.283 | 1.74E-05 | 0.00029854 | 0.0025 | 0.003 | 0.0029 | 0.0029 | TRUE | 0.0664 |
| Eri1 | 9792 | 7116 | 6877 | 7012 | 6238 | 7190 | 7605 | 7569 | 7420 | 6980 | 7037 | 7098 | 7284.82 | 7531 | 7038 | 0.934 | -0.098 | 0.03753764 | 0.10938338 | 0 | 0.0028 | 0.0015 | 0.0015 | TRUE | 0.002 |
| Eri2 | 1798 | 1322 | 1266 | 1290 | 1168 | 1255 | 1396 | 1406 | 1366 | 1284 | 1318 | 1239 | 1334.87 | 1389 | 1280 | 0.921 | -0.119 | 0.08092077 | 0.19063798 | 0 | 0.0042 | 0.0026 | 0.0026 | TRUE | 0.0096 |
| Eri3 | 2470 | 1833 | 1961 | 2158 | 1903 | 2264 | 1918 | 1950 | 2116 | 2148 | 2147 | 2235 | 2085.63 | 1995 | 2177 | 1.092 | 0.127 | 0.0536139 | 0.14156329 | 0.0012 | 0.0036 | 0.0026 | 0.0026 | TRUE | 0.0338 |

| | | | | | | | | | | | | | | | | | | | | | | | | | |
|---|---|---|---|---|---|---|---|---|---|---|---|---|---|---|---|---|---|---|---|---|---|---|---|---|---|
| Erich1 | 1783 | 1366 | 1354 | 1452 | 1239 | 1547 | 1385 | 1453 | 1461 | 1445 | 1398 | 1527 | 1444.82 | 1433 | 1457 | 1.018 | 0.025 | 0.71823853 | 0.83076664 | 8.00E-04 | 0.0041 | 0.0028 | 0.0028 | TRUE | 0.0216 |
| Erich2 | 7 | 6 | 10 | 3 | 7 | 5 | 5 | 6 | 11 | 3 | 8 | 5 | 6.4 | 7 | 5 | 0.7 | -0.514 | 0.50917896 | NA | 0 | 0.3725 | 0.2727 | 0.2727 | TRUE | 0.3758 |
| Erlec1 | 1981 | 1389 | 1560 | 1831 | 1505 | 1688 | 1539 | 1477 | 1683 | 1823 | 1698 | 1666 | 1647.66 | 1566 | 1729 | 1.104 | 0.143 | 0.05650676 | 0.14725635 | 0.0025 | 0.0039 | 0.0034 | 0.0034 | TRUE | 0.05 |
| Erlin1 | 3242 | 2308 | 2385 | 2656 | 2381 | 2629 | 2518 | 2455 | 2573 | 2644 | 2686 | 2595 | 2578.56 | 2515 | 2642 | 1.05 | 0.07 | 0.21994087 | 0.38308736 | 0 | 0.0033 | 0.002 | 0.002 | TRUE | 0.0052 |
| Erlin2 | 6085 | 4424 | 4233 | 4210 | 4072 | 4421 | 4726 | 4706 | 4567 | 4191 | 4594 | 4364 | 4524.59 | 4666 | 4383 | 0.939 | -0.091 | 0.11108676 | 0.2388332 | 9.00E-04 | 0.003 | 0.0021 | 0.0021 | TRUE | 0.0214 |
| Ermap | 1 | 0 | 1 | 2 | 2 | 3 | 1 | 0 | 1 | 2 | 2 | 3 | 1.51 | 1 | 2 | 3.834 | 1.939 | 0.2434068 | NA | 0 | 1.5713 | 1.0101 | 1.0101 | TRUE | 0.1911 |
| Ermn | 0 | 0 | 0 | 0 | 0 | 0 | 0 | 0 | 0 | 0 | 0 | 0 | 0 | 0 | 0 | NA | NA | NA | NA | NA | NA | NA | NA | NA | NA |
| Ermp1 | 6835 | 4764 | 5122 | 5693 | 5048 | 5594 | 5308 | 5067 | 5527 | 5667 | 5695 | 5522 | 5464.34 | 5301 | 5628 | 1.062 | 0.086 | 0.11727767 | 0.2483701 | 8.00E-04 | 0.0029 | 0.002 | 0.002 | TRUE | 0.0178 |
| Ern1 | 1182 | 847 | 830 | 1042 | 903 | 1054 | 918 | 901 | 896 | 1037 | 1019 | 1040 | 968.48 | 905 | 1032 | 1.14 | 0.189 | 0.01000201 | 0.04137733 | 0 | 0.0049 | 0.0029 | 0.0029 | TRUE | 0.0021 |
| Ern2 | 11 | 6 | 6 | 9 | 7 | 6 | 9 | 6 | 6 | 9 | 8 | 6 | 7.36 | 7 | 8 | 1.055 | 0.077 | 0.9112771 | NA | 0 | 0.3243 | 0.2098 | 0.2098 | TRUE | 0.11 |
| Ero1l | 6326 | 4798 | 6331 | 6546 | 5521 | 6248 | 4913 | 5104 | 6831 | 6516 | 6228 | 6168 | 5959.95 | 5616 | 6304 | 1.123 | 0.167 | 0.1007055 | 0.2221614 | 0.0168 | 0.0028 | 0.0073 | 0.0073 | TRUE | 0.4362 |
| Ero1lb | 586 | 452 | 472 | 567 | 485 | 575 | 455 | 481 | 509 | 564 | 547 | 568 | 520.72 | 482 | 560 | 1.164 | 0.219 | 0.02235308 | 0.07477009 | 0 | 0.007 | 0.0047 | 0.0047 | TRUE | 0.0293 |
| Erp27 | 1 | 0 | 0 | 0 | 0 | 1 | 1 | 0 | 0 | 0 | 0 | 1 | 0.29 | 0 | 0 | 1.075 | 0.105 | 0.97702352 | NA | 0 | 8.0647 | 7.5458 | 7.5458 | TRUE | 0.3835 |
| Erp29 | 5948 | 4347 | 4550 | 4778 | 4353 | 4883 | 4620 | 4624 | 4909 | 4756 | 4911 | 4820 | 4773.29 | 4718 | 4829 | 1.024 | 0.034 | 0.5297221 | 0.68986952 | 4.00E-04 | 0.0029 | 0.0019 | 0.0019 | TRUE | 0.0153 |
| Erp44 | 3180 | 2263 | 2481 | 2803 | 2475 | 2694 | 2470 | 2407 | 2677 | 2790 | 2792 | 2659 | 2632.57 | 2518 | 2747 | 1.091 | 0.126 | 0.05080663 | 0.13644854 | 0.0014 | 0.0033 | 0.0026 | 0.0026 | TRUE | 0.0363 |
| Errfi1 | 1617 | 1168 | 1106 | 1139 | 1007 | 1112 | 1256 | 1242 | 1193 | 1134 | 1136 | 1098 | 1176.51 | 1230 | 1123 | 0.912 | -0.134 | 0.0584301 | 0.15071711 | 0 | 0.0044 | 0.0027 | 0.0027 | TRUE | 0.0082 |
| Erv3 | 0 | 0 | 0 | 0 | 0 | 0 | 0 | 0 | 0 | 0 | 0 | 0 | 0 | 0 | 0 | NA | NA | NA | NA | NA | NA | NA | NA | NA | NA |
| Esam | 245 | 164 | 206 | 233 | 156 | 218 | 190 | 174 | 222 | 232 | 176 | 215 | 201.68 | 195 | 208 | 1.064 | 0.09 | 0.57395243 | 0.72603789 | 0.0118 | 0.0142 | 0.0135 | 0.0135 | TRUE | 0.1873 |
| Esco1 | 2967 | 2133 | 2061 | 2355 | 2032 | 2220 | 2304 | 2269 | 2224 | 2344 | 2292 | 2191 | 2270.82 | 2266 | 2276 | 1.004 | 0.006 | 0.92077132 | 0.95643782 | 3.00E-04 | 0.0035 | 0.0022 | 0.0022 | TRUE | 0.0129 |
| Esco2 | 8195 | 6284 | 7076 | 7338 | 6064 | 7295 | 6365 | 6684 | 7635 | 7304 | 6841 | 7201 | 7005.04 | 6895 | 7115 | 1.032 | 0.046 | 0.52255454 | 0.68379176 | 0.0042 | 0.0028 | 0.0036 | 0.0036 | TRUE | 0.1075 |
| Esd | 3835 | 2602 | 3096 | 3252 | 2769 | 3161 | 2978 | 2768 | 3341 | 3237 | 3124 | 3120 | 3094.65 | 3029 | 3160 | 1.044 | 0.062 | 0.40523572 | 0.57800733 | 0.0039 | 0.0032 | 0.0036 | 0.0036 | TRUE | 0.097 |
| Esf1 | 7519 | 5405 | 4507 | 5026 | 4127 | 4839 | 5840 | 5749 | 4863 | 5003 | 4656 | 4777 | 5147.89 | 5484 | 4812 | 0.877 | -0.189 | 0.01182748 | 0.04684127 | 0.0052 | 0.0029 | 0.0039 | 0.0039 | TRUE | 0.119 |
| Esm1 | 0 | 0 | 0 | 0 | 0 | 0 | 0 | 0 | 0 | 0 | 0 | 0 | 0 | 0 | 0 | NA | NA | NA | NA | NA | NA | NA | NA | NA | NA |
| Espl1 | 16605 | 12473 | 12522 | 14261 | 12973 | 14740 | 12896 | 13267 | 13511 | 14195 | 14635 | 14551 | 13842.64 | 13225 | 14460 | 1.094 | 0.129 | 0.00609932 | 0.02856408 | 3.00E-04 | 0.0026 | 0.0015 | 0.0015 | TRUE | 0.0058 |
| Espn | 511 | 450 | 583 | 658 | 600 | 603 | 397 | 479 | 629 | 655 | 677 | 595 | 571.95 | 502 | 642 | 1.284 | 0.361 | 0.01096569 | 0.04416248 | 0.0271 | 0.0066 | 0.0127 | 0.0127 | TRUE | 0.5749 |
| Espnl | 7 | 3 | 8 | 5 | 6 | 7 | 5 | 3 | 9 | 5 | 7 | 7 | 5.99 | 6 | 6 | 1.083 | 0.115 | 0.88356269 | NA | 0 | 0.3984 | 0.2754 | 0.2754 | TRUE | 0.2848 |
| Esr1 | 3 | 10 | 3 | 7 | 4 | 1 | 2 | 11 | 3 | 7 | 5 | 1 | 4.78 | 5 | 4 | 0.78 | -0.359 | 0.71601593 | NA | 0.455 | 0.4984 | 0.4854 | 0.4854 | TRUE | 0.6716 |
| Esr2 | 46 | 25 | 25 | 30 | 26 | 31 | 36 | 27 | 27 | 30 | 29 | 31 | 29.85 | 30 | 30 | 0.998 | -0.003 | 0.9941941 | 0.99663461 | 0 | 0.0818 | 0.0554 | 0.0554 | TRUE | 0.2423 |
| Esrp1 | 961 | 842 | 1061 | 889 | 912 | 1027 | 746 | 896 | 1145 | 885 | 1029 | 1014 | 952.39 | 929 | 976 | 1.052 | 0.073 | 0.57296114 | 0.72548197 | 0.0255 | 0.0049 | 0.0112 | 0.0112 | TRUE | 0.4033 |
| Esrp2 | 354 | 302 | 201 | 187 | 211 | 214 | 275 | 321 | 217 | 186 | 238 | 211 | 241.41 | 271 | 212 | 0.78 | -0.358 | 0.02919103 | 0.09080031 | 0.0223 | 0.0122 | 0.0152 | 0.0152 | TRUE | 0.1923 |
| Esrra | 1510 | 1136 | 1037 | 1261 | 1034 | 1170 | 1173 | 1208 | 1119 | 1255 | 1166 | 1155 | 1179.44 | 1167 | 1192 | 1.022 | 0.031 | 0.67575597 | 0.80101013 | 9.00E-04 | 0.0044 | 0.0032 | 0.0032 | TRUE | 0.0259 |
| Esrrb | 35661 | 27283 | 29821 | 33102 | 30890 | 34371 | 27696 | 29021 | 32177 | 32950 | 34847 | 33929 | 31769.98 | 29631 | 33909 | 1.144 | 0.195 | 0.0020389 | 0.01226001 | 0.0034 | 0.0025 | 0.0028 | 0.0028 | TRUE | 0.0691 |
| Esrrg | 24 | 33 | 16 | 19 | 17 | 32 | 19 | 35 | 17 | 19 | 19 | 32 | 23.45 | 24 | 23 | 0.99 | -0.015 | 0.97337091 | 0.98545797 | 0.0836 | 0.1035 | 0.0978 | 0.0978 | TRUE | 1.0057 |
| Esx1 | 641 | 407 | 370 | 727 | 493 | 505 | 498 | 433 | 399 | 724 | 556 | 499 | 518.05 | 443 | 593 | 1.335 | 0.417 | 0.00281368 | 0.01570932 | 0.0229 | 0.007 | 0.0121 | 0.0121 | TRUE | 0.4417 |
| Esyt1 | 3876 | 2756 | 2827 | 3044 | 2670 | 3075 | 3010 | 2932 | 3050 | 3030 | 3012 | 3035 | 3011.61 | 2997 | 3026 | 1.009 | 0.014 | 0.80297395 | 0.88603607 | 0 | 0.0032 | 0.0018 | 0.0018 | TRUE | 0.0044 |
| Esyt2 | 3777 | 2715 | 2887 | 3231 | 2811 | 3107 | 2933 | 2888 | 3115 | 3216 | 3171 | 3067 | 3065.12 | 2979 | 3151 | 1.058 | 0.082 | 0.16868962 | 0.32029553 | 6.00E-04 | 0.0032 | 0.0022 | 0.0022 | TRUE | 0.0191 |
| Esyt3 | 1513 | 1138 | 1205 | 1566 | 1388 | 1413 | 1175 | 1210 | 1300 | 1559 | 1566 | 1395 | 1367.53 | 1228 | 1507 | 1.227 | 0.295 | 0.00016032 | 0.00179722 | 0.0028 | 0.0042 | 0.0037 | 0.0037 | TRUE | 0.051 |
| Etaa1 | 3714 | 2726 | 2706 | 2605 | 2505 | 2860 | 2885 | 2900 | 2920 | 2593 | 2826 | 2823 | 2824.34 | 2902 | 2747 | 0.947 | -0.079 | 0.19391507 | 0.35280398 | 8.00E-04 | 0.0033 | 0.0023 | 0.0023 | TRUE | 0.0295 |
| Etd | 0 | 0 | 0 | 0 | 0 | 0 | 0 | 0 | 0 | 0 | 0 | 0 | 0 | 0 | 0 | NA | NA | NA | NA | NA | NA | NA | NA | NA | NA |
| Etf1 | 32341 | 22472 | 19362 | 21481 | 18562 | 20942 | 25118 | 23903 | 20891 | 21382 | 20940 | 20673 | 22151.25 | 23304 | 20998 | 0.901 | -0.15 | 0.02804184 | 0.0881271 | 0.0044 | 0.0025 | 0.0033 | 0.0033 | TRUE | 0.1003 |
| Etfa | 9986 | 6969 | 7011 | 8286 | 6949 | 7702 | 7756 | 7413 | 7565 | 8248 | 7839 | 7603 | 7737.25 | 7578 | 7897 | 1.042 | 0.059 | 0.27112369 | 0.44203202 | 9.00E-04 | 0.0027 | 0.002 | 0.002 | TRUE | 0.0186 |
| Etfb | 3556 | 2488 | 2533 | 2442 | 2134 | 2518 | 2762 | 2646 | 2733 | 2431 | 2407 | 2486 | 2577.52 | 2714 | 2441 | 0.899 | -0.153 | 0.00763193 | 0.03379281 | 0 | 0.0033 | 0.002 | 0.002 | TRUE | 0.0057 |
| Etfdh | 2069 | 1409 | 1402 | 1850 | 1503 | 1774 | 1607 | 1499 | 1513 | 1841 | 1696 | 1751 | 1651.1 | 1540 | 1763 | 1.144 | 0.195 | 0.0046112 | 0.02306122 | 0.001 | 0.0039 | 0.0028 | 0.0028 | TRUE | 0.0185 |
| Ethe1 | 483 | 289 | 332 | 453 | 398 | 452 | 375 | 307 | 358 | 451 | 449 | 446 | 397.81 | 347 | 449 | 1.29 | 0.368 | 0.00107772 | 0.00764457 | 0.0022 | 0.0084 | 0.0066 | 0.0066 | TRUE | 0.1107 |
| Etl4 | 9575 | 6711 | 6307 | 6716 | 6274 | 7373 | 7436 | 7138 | 6805 | 6685 | 7078 | 7278 | 7070.2 | 7126 | 7014 | 0.984 | -0.023 | 0.69019851 | 0.81040066 | 0.0015 | 0.0028 | 0.0024 | 0.0024 | TRUE | 0.0206 |
| Etnk1 | 10930 | 7917 | 7651 | 9663 | 7913 | 9262 | 8489 | 8421 | 8255 | 9619 | 8927 | 9143 | 8808.95 | 8388 | 9230 | 1.1 | 0.138 | 0.00756618 | 0.03362667 | 7.00E-04 | 0.0027 | 0.0018 | 0.0018 | TRUE | 0.0166 |
| Etnk2 | 661 | 467 | 545 | 570 | 542 | 672 | 513 | 497 | 588 | 567 | 611 | 663 | 573.39 | 533 | 614 | 1.154 | 0.207 | 0.04859389 | 0.131994 | 0.0053 | 0.0066 | 0.0061 | 0.0061 | TRUE | 0.0933 |
| Etohd2 | 24 | 20 | 22 | 25 | 26 | 35 | 19 | 21 | 24 | 25 | 29 | 35 | 25.4 | 21 | 30 | 1.403 | 0.488 | 0.20306135 | 0.36448021 | 0 | 0.0957 | 0.0656 | 0.0656 | TRUE | 0.148 |
| Etohi1 | 483 | 384 | 290 | 308 | 233 | 308 | 375 | 408 | 313 | 307 | 263 | 304 | 328.33 | 365 | 291 | 0.797 | -0.327 | 0.01362678 | 0.05200737 | 0.0095 | 0.0096 | 0.0096 | 0.0096 | TRUE | 0.1737 |
| Ets1 | 348 | 271 | 243 | 163 | 156 | 173 | 270 | 288 | 262 | 162 | 176 | 171 | 221.62 | 273 | 170 | 0.62 | -0.69 | 4.59E-07 | 1.52E-05 | 0 | 0.0131 | 0.0087 | 0.0087 | TRUE | 0.0235 |
| Ets2 | 1268 | 944 | 888 | 822 | 765 | 802 | 985 | 1004 | 958 | 818 | 863 | 792 | 903.33 | 982 | 824 | 0.839 | -0.254 | 0.00122638 | 0.00839094 | 0 | 0.0051 | 0.0033 | 0.0033 | TRUE | 0.0195 |
| Etv1 | 448 | 323 | 284 | 347 | 294 | 319 | 348 | 344 | 306 | 345 | 332 | 315 | 331.65 | 333 | 331 | 0.992 | -0.011 | 0.92518924 | 0.95858861 | 0 | 0.0096 | 0.0068 | 0.0068 | TRUE | 0.0522 |
| Etv2 | 47 | 33 | 16 | 19 | 14 | 28 | 37 | 35 | 17 | 19 | 16 | 28 | 25.2 | 30 | 21 | 0.7 | -0.515 | 0.22939765 | 0.39484352 | 0.0784 | 0.0965 | 0.0915 | 0.0915 | TRUE | 0.8232 |
| Etv3 | 1635 | 1088 | 1035 | 1049 | 961 | 1020 | 1270 | 1157 | 1117 | 1044 | 1084 | 1007 | 1113.18 | 1181 | 1045 | 0.883 | -0.179 | 0.02491256 | 0.08122152 | 0.0022 | 0.0046 | 0.0037 | 0.0037 | TRUE | 0.0536 |
| Etv4 | 10885 | 7885 | 7067 | 11296 | 9370 | 10694 | 8454 | 8387 | 7625 | 11244 | 10570 | 10557 | 9472.88 | 8155 | 10790 | 1.323 | 0.404 | 1.36E-11 | 1.31E-09 | 0.0021 | 0.0027 | 0.0025 | 0.0025 | TRUE | 0.0393 |
| Etv5 | 19360 | 14294 | 14042 | 14761 | 13361 | 15345 | 15036 | 15204 | 15151 | 14693 | 15073 | 15148 | 15050.87 | 15130 | 14971 | 0.989 | -0.015 | 0.73166526 | 0.83929626 | 0 | 0.0026 | 0.0014 | 0.0014 | TRUE | 0.0032 |
| Etv6 | 2115 | 1609 | 1502 | 1379 | 1347 | 1584 | 1643 | 1711 | 1621 | 1373 | 1520 | 1564 | 1571.77 | 1658 | 1486 | 0.896 | -0.159 | 0.03026537 | 0.09322909 | 0.002 | 0.0039 | 0.0032 | 0.0032 | TRUE | 0.0536 |
| EU599041 | 1081 | 899 | 749 | 814 | 722 | 825 | 840 | 956 | 808 | 810 | 814 | 814 | 840.52 | 868 | 813 | 0.937 | -0.094 | 0.28883078 | 0.4609489 | 0.0028 | 0.0052 | 0.0044 | 0.0044 | TRUE | 0.0916 |
| Eva1a | 129 | 109 | 76 | 104 | 65 | 76 | 100 | 116 | 82 | 104 | 73 | 75 | 91.67 | 99 | 84 | 0.846 | -0.242 | 0.29032018 | 0.4624223 | 0.0222 | 0.0283 | 0.0266 | 0.0266 | TRUE | 0.3355 |
| Eva1b | 730 | 556 | 593 | 762 | 607 | 751 | 567 | 591 | 640 | 758 | 685 | 741 | 663.8 | 599 | 728 | 1.217 | 0.284 | 0.00209233 | 0.01250758 | 0.0018 | 0.006 | 0.0046 | 0.0046 | TRUE | 0.0404 |
| Eva1c | 35 | 24 | 34 | 32 | 19 | 25 | 27 | 26 | 37 | 32 | 21 | 25 | 27.89 | 30 | 26 | 0.877 | -0.189 | 0.61408855 | 0.75613092 | 0 | 0.0874 | 0.0646 | 0.0646 | TRUE | 0.2522 |
| Evc | 711 | 601 | 533 | 551 | 490 | 566 | 552 | 639 | 575 | 548 | 553 | 559 | 571.09 | 589 | 553 | 0.941 | -0.088 | 0.36177857 | 0.53660525 | 0.0015 | 0.0066 | 0.0049 | 0.0049 | TRUE | 0.0651 |
| Evc2 | 1715 | 1226 | 1211 | 1221 | 1185 | 1297 | 1332 | 1304 | 1307 | 1215 | 1337 | 1280 | 1295.87 | 1314 | 1277 | 0.971 | -0.042 | 0.5544127 | 0.70999991 | 3.00E-04 | 0.0043 | 0.0029 | 0.0029 | TRUE | 0.0218 |
| Evi2a | 0 | 0 | 1 | 0 | 0 | 0 | 0 | 0 | 1 | 0 | 0 | 0 | 0.18 | 0 | 0 | 0.552 | -0.857 | 0.8336548 | NA | 0 | 13.1817 | 10 | 10 | TRUE | 0.5365 |
| Evi2b | 0 | 0 | 0 | 0 | 0 | 0 | 0 | 0 | 0 | 0 | 0 | 0 | 0 | 0 | 0 | NA | NA | NA | NA | NA | NA | NA | NA | NA | NA |
| Evi5 | 5474 | 3722 | 3844 | 4511 | 3815 | 4376 | 4251 | 3959 | 4148 | 4490 | 4304 | 4320 | 4245.31 | 4119 | 4371 | 1.061 | 0.085 | 0.12702043 | 0.26250766 | 6.00E-04 | 0.003 | 0.002 | 0.002 | TRUE | 0.0141 |
| Evi5l | 214 | 156 | 199 | 220 | 194 | 260 | 166 | 166 | 215 | 219 | 219 | 257 | 206.89 | 182 | 232 | 1.275 | 0.35 | 0.0256785 | 0.08285449 | 0.0103 | 0.0139 | 0.0128 | 0.0128 | TRUE | 0.2513 |
| Evl | 2454 | 1772 | 1761 | 2044 | 1823 | 2165 | 1906 | 1885 | 1900 | 2035 | 2057 | 2137 | 1986.53 | 1897 | 2076 | 1.094 | 0.13 | 0.03156786 | 0.09614307 | 0 | 0.0036 | 0.0021 | 0.0021 | TRUE | 0.0081 |
| Evpl | 760 | 526 | 538 | 799 | 661 | 759 | 590 | 559 | 580 | 795 | 746 | 749 | 670.08 | 576 | 763 | 1.323 | 0.404 | 2.92E-06 | 6.76E-05 | 0 | 0.006 | 0.0039 | 0.0039 | TRUE | 0.016 |

| | | | | | | | | | | | | | | | | | | | | | | | | | |
|---|---|---|---|---|---|---|---|---|---|---|---|---|---|---|---|---|---|---|---|---|---|---|---|---|---|
| Evx1 | 22 | 14 | 7 | 12 | 12 | 5 | 17 | 15 | 8 | 12 | 14 | 5 | 11.66 | 13 | 10 | 0.758 | -0.4 | 0.51102797 | NA | 0.1031 | 0.2057 | 0.1788 | 0.1788 | TRUE | 0.7334 |
| Evx2 | 1 | 0 | 0 | 0 | 0 | 0 | 1 | 0 | 0 | 0 | 0 | 0 | 0.13 | 0 | 0 | 0.552 | -0.857 | 0.8336548 | NA | 0 | 18.3121 | 10 | 10 | TRUE | 0.2566 |
| Ewsr1 | 23830 | 17995 | 17826 | 17717 | 16146 | 18343 | 18508 | 19141 | 19234 | 17635 | 18214 | 18107 | 18473.35 | 18961 | 17985 | 0.949 | -0.076 | 0.0983209 | 0.21845182 | 3.00E-04 | 0.0026 | 0.0015 | 0.0015 | TRUE | 0.0054 |
| Exd1 | 208 | 163 | 131 | 163 | 134 | 157 | 162 | 173 | 141 | 162 | 151 | 155 | 157.45 | 159 | 156 | 0.983 | -0.025 | 0.87913904 | 0.93262644 | 0 | 0.0175 | 0.0124 | 0.0124 | TRUE | 0.0906 |
| Exd2 | 2421 | 1754 | 1864 | 2075 | 1913 | 2082 | 1880 | 1866 | 2011 | 2065 | 2158 | 2055 | 2006 | 1919 | 2093 | 1.091 | 0.125 | 0.05132578 | 0.13742962 | 7.00E-04 | 0.0036 | 0.0025 | 0.0025 | TRUE | 0.0209 |
| Exo1 | 4506 | 3155 | 2963 | 2866 | 2717 | 3071 | 3500 | 3356 | 3197 | 2853 | 3065 | 3032 | 3167 | 3351 | 2983 | 0.89 | -0.169 | 0.00688397 | 0.0313852 | 0.0013 | 0.0032 | 0.0025 | 0.0025 | TRUE | 0.0195 |
| Exo5 | 1195 | 766 | 706 | 768 | 721 | 813 | 928 | 815 | 762 | 764 | 813 | 803 | 814.17 | 835 | 793 | 0.948 | -0.077 | 0.4126441 | 0.58551794 | 0.0047 | 0.0053 | 0.0051 | 0.0051 | TRUE | 0.118 |
| Exoc1 | 3232 | 2477 | 2245 | 2408 | 2174 | 2500 | 2510 | 2635 | 2422 | 2397 | 2452 | 2468 | 2480.76 | 2522 | 2439 | 0.967 | -0.048 | 0.42619907 | 0.59868927 | 6.00E-04 | 0.0034 | 0.0023 | 0.0023 | TRUE | 0.0179 |
| Exoc2 | 3333 | 2506 | 2646 | 2933 | 2535 | 2901 | 2589 | 2666 | 2855 | 2920 | 2860 | 2864 | 2792.03 | 2703 | 2881 | 1.067 | 0.093 | 0.13009855 | 0.26686673 | 0.001 | 0.0033 | 0.0024 | 0.0024 | TRUE | 0.0291 |
| Exoc3 | 4008 | 2797 | 2730 | 2980 | 2560 | 2911 | 3113 | 2975 | 2946 | 2966 | 2888 | 2874 | 2960.24 | 3011 | 2909 | 0.966 | -0.05 | 0.37704381 | 0.55151527 | 3.00E-04 | 0.0032 | 0.002 | 0.002 | TRUE | 0.0109 |
| Exoc3l | 235 | 155 | 163 | 240 | 201 | 228 | 183 | 165 | 176 | 239 | 227 | 225 | 202.33 | 175 | 230 | 1.318 | 0.398 | 0.00460917 | 0.02306122 | 0 | 0.0141 | 0.0092 | 0.0092 | TRUE | 0.0241 |
| Exoc3l4 | 36 | 19 | 26 | 50 | 31 | 30 | 28 | 20 | 28 | 50 | 35 | 30 | 31.76 | 25 | 38 | 1.496 | 0.581 | 0.10938172 | 0.23626915 | 0.0257 | 0.077 | 0.0625 | 0.0625 | TRUE | 0.5389 |
| Exoc4 | 7228 | 5295 | 6151 | 6937 | 6215 | 6521 | 5614 | 5632 | 6637 | 6905 | 7011 | 6437 | 6372.71 | 5961 | 6784 | 1.138 | 0.187 | 0.01175563 | 0.0466084 | 0.0056 | 0.0028 | 0.0038 | 0.0038 | TRUE | 0.1195 |
| Exoc5 | 3827 | 2677 | 2594 | 2864 | 2338 | 2675 | 2972 | 2847 | 2799 | 2851 | 2638 | 2641 | 2791.27 | 2873 | 2710 | 0.943 | -0.085 | 0.17191749 | 0.32394063 | 0.001 | 0.0033 | 0.0024 | 0.0024 | TRUE | 0.0253 |
| Exoc6 | 2034 | 1438 | 1502 | 1590 | 1442 | 1678 | 1580 | 1530 | 1621 | 1583 | 1627 | 1656 | 1599.3 | 1577 | 1622 | 1.029 | 0.041 | 0.53171173 | 0.69157269 | 0 | 0.0039 | 0.0024 | 0.0024 | TRUE | 0.0079 |
| Exoc6b | 2497 | 1775 | 1897 | 1981 | 1811 | 1960 | 1939 | 1888 | 2047 | 1972 | 2043 | 1935 | 1970.65 | 1958 | 1983 | 1.013 | 0.018 | 0.77599957 | 0.86759714 | 7.00E-04 | 0.0036 | 0.0025 | 0.0025 | TRUE | 0.0185 |
| Exoc7 | 1569 | 1226 | 1150 | 1257 | 1116 | 1250 | 1219 | 1304 | 1241 | 1251 | 1259 | 1234 | 1251.27 | 1255 | 1248 | 0.995 | -0.007 | 0.92382096 | 0.9577958 | 0 | 0.0043 | 0.0027 | 0.0027 | TRUE | 0.0142 |
| Exoc8 | 2352 | 1683 | 1297 | 1346 | 1263 | 1271 | 1827 | 1790 | 1399 | 1340 | 1425 | 1255 | 1505.94 | 1672 | 1340 | 0.8 | -0.321 | 0.0014347 | 0.00934219 | 0.011 | 0.004 | 0.0066 | 0.0066 | TRUE | 0.243 |
| Exog | 1139 | 862 | 791 | 815 | 745 | 948 | 885 | 917 | 853 | 811 | 840 | 936 | 873.75 | 885 | 862 | 0.975 | -0.037 | 0.66856803 | 0.79617284 | 0.0023 | 0.0051 | 0.0042 | 0.0042 | TRUE | 0.0668 |
| Exosc1 | 1883 | 1332 | 1232 | 1340 | 1219 | 1375 | 1462 | 1417 | 1329 | 1334 | 1375 | 1357 | 1379.15 | 1403 | 1355 | 0.965 | -0.051 | 0.46781972 | 0.63687468 | 6.00E-04 | 0.0041 | 0.0028 | 0.0028 | TRUE | 0.025 |
| Exosc10 | 5461 | 4003 | 3647 | 3809 | 3584 | 4133 | 4241 | 4258 | 3935 | 3791 | 4043 | 4080 | 4058.14 | 4145 | 3971 | 0.958 | -0.062 | 0.30588194 | 0.47853155 | 0.0014 | 0.003 | 0.0024 | 0.0024 | TRUE | 0.0236 |
| Exosc2 | 5193 | 3681 | 3499 | 3757 | 3390 | 3991 | 4033 | 3915 | 3775 | 3740 | 3824 | 3940 | 3871.29 | 3908 | 3835 | 0.981 | -0.028 | 0.62078124 | 0.76065413 | 7.00E-04 | 0.003 | 0.002 | 0.002 | TRUE | 0.0107 |
| Exosc3 | 2226 | 1601 | 1490 | 1412 | 1285 | 1493 | 1729 | 1703 | 1608 | 1406 | 1450 | 1474 | 1561.41 | 1680 | 1443 | 0.858 | -0.22 | 0.00106635 | 0.00758278 | 5.00E-04 | 0.0039 | 0.0026 | 0.0026 | TRUE | 0.0168 |
| Exosc4 | 1522 | 1100 | 950 | 909 | 792 | 884 | 1182 | 1170 | 1025 | 905 | 893 | 873 | 1008.02 | 1126 | 890 | 0.79 | -0.34 | 4.20E-05 | 0.00060804 | 0.0024 | 0.0048 | 0.004 | 0.004 | TRUE | 0.0718 |
| Exosc5 | 4444 | 3322 | 2984 | 3573 | 3293 | 3686 | 3451 | 3534 | 3220 | 3557 | 3715 | 3639 | 3519.13 | 3402 | 3637 | 1.069 | 0.096 | 0.10713622 | 0.23265586 | 0.001 | 0.0031 | 0.0023 | 0.0023 | TRUE | 0.0261 |
| Exosc6 | 2238 | 1625 | 1446 | 1508 | 1354 | 1619 | 1738 | 1728 | 1560 | 1501 | 1527 | 1598 | 1608.93 | 1675 | 1542 | 0.92 | -0.12 | 0.09374717 | 0.21081313 | 0.0017 | 0.0039 | 0.0031 | 0.0031 | TRUE | 0.043 |
| Exosc7 | 3151 | 2364 | 2182 | 2264 | 1970 | 2216 | 2447 | 2515 | 2354 | 2254 | 2222 | 2188 | 2329.94 | 2439 | 2221 | 0.911 | -0.135 | 0.02427954 | 0.07966656 | 2.00E-04 | 0.0034 | 0.0022 | 0.0022 | TRUE | 0.0108 |
| Exosc8 | 3582 | 2578 | 2872 | 2878 | 2667 | 2958 | 2782 | 2742 | 3099 | 2865 | 3009 | 2920 | 2902.74 | 2874 | 2931 | 1.02 | 0.029 | 0.66683448 | 0.79491786 | 0.0022 | 0.0032 | 0.0028 | 0.0028 | TRUE | 0.0559 |
| Exosc9 | 3404 | 2375 | 2218 | 2399 | 2136 | 2456 | 2644 | 2526 | 2393 | 2388 | 2410 | 2424 | 2464.21 | 2521 | 2407 | 0.954 | -0.068 | 0.27863388 | 0.4501033 | 9.00E-04 | 0.0034 | 0.0024 | 0.0024 | TRUE | 0.0238 |
| Exph5 | 16 | 6 | 12 | 13 | 11 | 8 | 12 | 6 | 13 | 13 | 12 | 8 | 10.83 | 10 | 11 | 1.038 | 0.054 | 0.92701843 | NA | 0 | 0.2212 | 0.1623 | 0.1623 | TRUE | 0.4011 |
| Ext1 | 1932 | 1337 | 1425 | 1667 | 1523 | 1737 | 1501 | 1422 | 1538 | 1659 | 1718 | 1715 | 1592.06 | 1487 | 1697 | 1.141 | 0.191 | 0.00413625 | 0.02119737 | 2.00E-04 | 0.0039 | 0.0026 | 0.0026 | TRUE | 0.0168 |
| Ext2 | 3526 | 2534 | 2615 | 3219 | 2796 | 3135 | 2738 | 2695 | 2822 | 3204 | 3154 | 3095 | 2951.42 | 2752 | 3151 | 1.145 | 0.196 | 0.00047462 | 0.00415413 | 0 | 0.0032 | 0.0019 | 0.0019 | TRUE | 0.0058 |
| Extl1 | 796 | 610 | 611 | 950 | 804 | 882 | 618 | 649 | 659 | 946 | 907 | 871 | 774.94 | 642 | 908 | 1.415 | 0.501 | 2.00E-09 | 1.25E-07 | 0 | 0.0055 | 0.0037 | 0.0037 | TRUE | 0.0162 |
| Extl2 | 451 | 327 | 335 | 382 | 338 | 390 | 350 | 348 | 361 | 380 | 381 | 385 | 367.68 | 353 | 382 | 1.082 | 0.114 | 0.28277129 | 0.45452655 | 0 | 0.0089 | 0.0054 | 0.0054 | TRUE | 0.0046 |
| Extl3 | 3343 | 2497 | 2576 | 2754 | 2482 | 2752 | 2596 | 2656 | 2779 | 2741 | 2800 | 2717 | 2714.97 | 2677 | 2753 | 1.029 | 0.041 | 0.48844429 | 0.65435695 | 4.00E-04 | 0.0033 | 0.0021 | 0.0021 | TRUE | 0.0134 |
| Eya1 | 158 | 118 | 111 | 66 | 83 | 106 | 123 | 126 | 120 | 66 | 94 | 105 | 105.33 | 123 | 88 | 0.717 | -0.48 | 0.02279415 | 0.07596103 | 0.0144 | 0.0249 | 0.0223 | 0.0223 | TRUE | 0.4798 |
| Eya2 | 77 | 77 | 101 | 119 | 88 | 95 | 60 | 82 | 109 | 118 | 99 | 94 | 93.7 | 84 | 104 | 1.253 | 0.326 | 0.17865632 | 0.33327261 | 0.0409 | 0.0277 | 0.0315 | 0.0315 | TRUE | 0.3023 |
| Eya3 | 3126 | 2395 | 2085 | 2248 | 2022 | 2323 | 2428 | 2548 | 2250 | 2238 | 2281 | 2293 | 2339.48 | 2409 | 2271 | 0.943 | -0.085 | 0.19749781 | 0.35727582 | 0.0014 | 0.0034 | 0.0027 | 0.0027 | TRUE | 0.0393 |
| Eya4 | 13 | 10 | 14 | 2 | 4 | 2 | 10 | 11 | 15 | 2 | 5 | 2 | 7.39 | 12 | 3 | 0.234 | -2.096 | 0.00843029 | NA | 0 | 0.3233 | 0.2297 | 0.2297 | TRUE | 0.2931 |
| Ezh1 | 4554 | 3585 | 4830 | 5664 | 5404 | 6466 | 3537 | 3813 | 5212 | 5638 | 6096 | 6383 | 5113.15 | 4187 | 6039 | 1.442 | 0.529 | 0.00439785 | 0.02228234 | 0.0246 | 0.0029 | 0.009 | 0.0246 | TRUE | 0.5571 |
| Ezh2 | 10960 | 8066 | 8253 | 7311 | 6531 | 7600 | 8512 | 8580 | 8905 | 7277 | 7368 | 7502 | 8024.03 | 8666 | 7382 | 0.852 | -0.231 | 2.35E-06 | 5.73E-05 | 2.00E-04 | 0.0027 | 0.0016 | 0.0016 | TRUE | 0.0071 |
| Ezr | 16531 | 12808 | 12744 | 13481 | 12096 | 14128 | 12839 | 13624 | 13751 | 13419 | 13646 | 13947 | 13537.39 | 13405 | 13671 | 1.02 | 0.029 | 0.57303032 | 0.72548197 | 7.00E-04 | 0.0026 | 0.0018 | 0.0018 | TRUE | 0.0168 |
| F10 | 0 | 0 | 0 | 0 | 0 | 0 | 0 | 0 | 0 | 0 | 0 | 0 | 0 | 0 | 0 | NA | NA | NA | NA | NA | NA | NA | NA | NA | NA |
| F11 | 1 | 0 | 0 | 0 | 0 | 0 | 1 | 0 | 0 | 0 | 0 | 0 | 0.13 | 0 | 0 | 0.552 | -0.857 | 0.8336548 | NA | 0 | 18.3121 | 10 | 10 | TRUE | 0.2566 |
| F11r | 3098 | 2170 | 2180 | 2117 | 1973 | 2221 | 2406 | 2308 | 2352 | 2107 | 2226 | 2192 | 2265.33 | 2355 | 2175 | 0.923 | -0.116 | 0.05411229 | 0.14242712 | 0 | 0.0035 | 0.0022 | 0.0022 | TRUE | 0.0091 |
| F12 | 1 | 2 | 3 | 1 | 1 | 0 | 1 | 2 | 3 | 1 | 1 | 0 | 1.38 | 2 | 1 | 0.347 | -1.526 | 0.37918523 | NA | 0 | 1.7231 | 1.1882 | 1.1882 | TRUE | 0.1962 |
| F13a1 | 0 | 0 | 0 | 0 | 0 | 0 | 0 | 0 | 0 | 0 | 0 | 0 | 0 | 0 | 0 | NA | NA | NA | NA | NA | NA | NA | NA | NA | NA |
| F13b | 0 | 0 | 0 | 0 | 0 | 0 | 0 | 0 | 0 | 0 | 0 | 0 | 0 | 0 | 0 | NA | NA | NA | NA | NA | NA | NA | NA | NA | NA |
| F2 | 75 | 64 | 81 | 74 | 52 | 63 | 58 | 68 | 87 | 74 | 59 | 62 | 68.04 | 71 | 65 | 0.917 | -0.125 | 0.6194914 | 0.75986884 | 0.0158 | 0.0373 | 0.0311 | 0.0311 | TRUE | 0.3477 |
| F2r | 1542 | 1045 | 909 | 1110 | 871 | 907 | 1198 | 1112 | 981 | 1105 | 983 | 895 | 1045.46 | 1097 | 994 | 0.906 | -0.143 | 0.15538501 | 0.3030416 | 0.0091 | 0.0047 | 0.0063 | 0.0063 | TRUE | 0.1144 |
| F2rl1 | 5312 | 3901 | 4021 | 4770 | 4026 | 4656 | 4126 | 4149 | 4339 | 4748 | 4542 | 4596 | 4416.61 | 4205 | 4629 | 1.101 | 0.139 | 0.00980376 | 0.04078145 | 4.00E-04 | 0.003 | 0.0019 | 0.0019 | TRUE | 0.0094 |
| F2rl2 | 5 | 1 | 1 | 2 | 0 | 3 | 4 | 1 | 1 | 2 | 0 | 3 | 1.83 | 2 | 2 | 0.811 | -0.303 | 0.83698691 | NA | 0 | 1.2978 | 1.0202 | 1.0202 | TRUE | 0.7894 |
| F2rl3 | 4 | 1 | 3 | 2 | 1 | 2 | 3 | 1 | 3 | 2 | 1 | 2 | 2.08 | 2 | 2 | 0.686 | -0.543 | 0.67617761 | NA | 0 | 1.14 | 0.7231 | 0.7231 | TRUE | 0.2461 |
| F3 | 326 | 301 | 352 | 392 | 352 | 357 | 253 | 320 | 380 | 390 | 397 | 352 | 348.81 | 318 | 380 | 1.2 | 0.263 | 0.07012604 | 0.17162869 | 0.0196 | 0.0092 | 0.0124 | 0.0124 | TRUE | 0.274 |
| F420014N23 | 0 | 5 | 0 | 3 | 1 | 2 | 0 | 5 | 0 | 3 | 1 | 2 | 1.9 | 2 | 2 | 1.185 | 0.245 | 0.87932499 | NA | 1.6537 | 1.2491 | 1.3354 | 1.3354 | TRUE | 2.3479 |
| F5 | 224 | 163 | 124 | 121 | 122 | 128 | 174 | 173 | 134 | 120 | 138 | 126 | 144.26 | 160 | 128 | 0.796 | -0.329 | 0.06184596 | 0.15678687 | 0.0066 | 0.0189 | 0.0153 | 0.0153 | TRUE | 0.2127 |
| F630028O10 | 0 | 0 | 0 | 0 | 0 | 0 | 0 | 0 | 0 | 0 | 0 | 0 | 0 | 0 | 0 | NA | NA | NA | NA | NA | NA | NA | NA | NA | NA |
| F630042J09F | 45 | 49 | 62 | 67 | 57 | 65 | 35 | 52 | 67 | 67 | 64 | 64 | 58.19 | 51 | 65 | 1.282 | 0.358 | 0.20668519 | 0.368394 | 0.0334 | 0.0432 | 0.0405 | 0.0405 | TRUE | 0.3206 |
| F630111L10F | 0 | 0 | 0 | 0 | 0 | 0 | 0 | 0 | 0 | 0 | 0 | 0 | 0 | 0 | 0 | NA | NA | NA | NA | NA | NA | NA | NA | NA | NA |
| F630206G17 | 0 | 0 | 0 | 0 | 0 | 0 | 0 | 0 | 0 | 0 | 0 | 0 | 0 | 0 | 0 | NA | NA | NA | NA | NA | NA | NA | NA | NA | NA |
| F7 | 5 | 2 | 4 | 2 | 3 | 0 | 4 | 2 | 4 | 2 | 3 | 0 | 2.62 | 3 | 2 | 0.509 | -0.974 | 0.43681749 | NA | 0 | 0.9081 | 0.6954 | 0.6954 | TRUE | 0.5169 |
| F730035M05 | 0 | 0 | 1 | 0 | 1 | 0 | 0 | 0 | 1 | 0 | 1 | 0 | 0.37 | 0 | 0 | 1.075 | 0.105 | 0.97514588 | NA | 0 | 6.4454 | 6.1578 | 6.1578 | TRUE | 0.5365 |
| F730043M19 | 69 | 76 | 66 | 91 | 67 | 81 | 54 | 81 | 71 | 91 | 76 | 80 | 75.29 | 69 | 82 | 1.207 | 0.272 | 0.25745571 | 0.42741156 | 0.0132 | 0.0339 | 0.0281 | 0.0281 | TRUE | 0.3779 |
| F8 | 75 | 71 | 31 | 48 | 25 | 31 | 58 | 76 | 33 | 48 | 28 | 31 | 45.63 | 56 | 36 | 0.638 | -0.648 | 0.06658793 | 0.16517551 | 0.0984 | 0.0544 | 0.0667 | 0.0667 | TRUE | 0.1985 |
| F830002L21F | 1 | 0 | 0 | 0 | 0 | 0 | 1 | 0 | 0 | 0 | 0 | 0 | 0.13 | 0 | 0 | 0.552 | -0.857 | 0.8336548 | NA | 0 | 18.3121 | 10 | 10 | TRUE | 0.2566 |
| F830016B08 | 0 | 0 | 0 | 0 | 0 | 0 | 0 | 0 | 0 | 0 | 0 | 0 | 0 | 0 | 0 | NA | NA | NA | NA | NA | NA | NA | NA | NA | NA |
| F830045P16F | 0 | 1 | 0 | 0 | 0 | 0 | 0 | 1 | 0 | 0 | 0 | 0 | 0.18 | 0 | 0 | 0.552 | -0.857 | 0.8336548 | NA | 0 | 13.3714 | 10 | 10 | TRUE | 0.5216 |
| F8a | 356 | 240 | 228 | 266 | 233 | 241 | 276 | 255 | 246 | 265 | 263 | 238 | 257.22 | 259 | 255 | 0.982 | -0.027 | 0.83592226 | 0.90645327 | 0 | 0.0116 | 0.0082 | 0.0082 | TRUE | 0.0408 |
| F9 | 0 | 0 | 0 | 0 | 0 | 0 | 0 | 0 | 0 | 0 | 0 | 0 | 0 | 0 | 0 | NA | NA | NA | NA | NA | NA | NA | NA | NA | NA |

| | | | | | | | | | | | | | | | | | | | | | | | | |
|---|---|---|---|---|---|---|---|---|---|---|---|---|---|---|---|---|---|---|---|---|---|---|---|---|
| F930015N05 | 1 | 2 | 0 | 4 | 1 | 1 | 1 | 2 | 0 | 4 | 1 | 1 | 1.5 | 1 | 2 | 2.126 | 1.088 | 0.50711268 | NA | 0 | 1.5823 | 1.1838 | 1.1838 | TRUE | 0.6559 |
| Fa2h | 98 | 67 | 89 | 120 | 88 | 101 | 76 | 71 | 96 | 119 | 99 | 100 | 93.64 | 81 | 106 | 1.313 | 0.392 | 0.06556755 | 0.16363837 | 0.0061 | 0.0277 | 0.0219 | 0.0219 | TRUE | 0.226 |
| Faah | 537 | 425 | 414 | 464 | 441 | 531 | 417 | 452 | 447 | 462 | 497 | 524 | 466.56 | 439 | 494 | 1.129 | 0.175 | 0.08788184 | 0.20135202 | 0 | 0.0075 | 0.0054 | 0.0054 | TRUE | 0.0397 |
| Fabp1 | 0 | 0 | 0 | 0 | 0 | 0 | 0 | 0 | 0 | 0 | 0 | 0 | 0 | 0 | 0 | NA | NA | NA | NA | NA | NA | NA | NA | NA | NA |
| Fabp12 | 2 | 4 | 4 | 0 | 0 | 2 | 2 | 4 | 4 | 0 | 0 | 2 | 2.02 | 3 | 1 | 0.203 | -2.298 | 0.15301186 | NA | 0 | 1.1778 | 0.9549 | 0.9549 | TRUE | 0.9733 |
| Fabp2 | 0 | 0 | 0 | 0 | 0 | 0 | 0 | 0 | 0 | 0 | 0 | 0 | 0 | 0 | 0 | NA | NA | NA | NA | NA | NA | NA | NA | NA | NA |
| Fabp3 | 5032 | 4463 | 7246 | 6755 | 6695 | 7786 | 3908 | 4747 | 7818 | 6724 | 7553 | 7686 | 6406.05 | 5491 | 7321 | 1.333 | 0.415 | 0.17805462 | 0.33250145 | 0.0683 | 0.0028 | 0.0191 | 0.0683 | TRUE | 1.6754 |
| Fabp4 | 5 | 3 | 3 | 1 | 1 | 1 | 4 | 3 | 3 | 1 | 1 | 1 | 2.24 | 3 | 1 | 0.3 | -1.738 | 0.19353651 | NA | 0 | 1.0619 | 0.6479 | 0.6479 | TRUE | 0.0251 |
| Fabp5 | 1897 | 1521 | 1524 | 1074 | 997 | 1080 | 1473 | 1618 | 1644 | 1069 | 1125 | 1066 | 1332.58 | 1578 | 1087 | 0.689 | -0.538 | 7.24E-13 | 9.32E-11 | 0.0017 | 0.0042 | 0.0033 | 0.0033 | TRUE | 0.0426 |
| Fabp6 | 15 | 3 | 6 | 3 | 2 | 7 | 12 | 3 | 6 | 3 | 2 | 7 | 5.58 | 7 | 4 | 0.565 | -0.824 | 0.34712036 | NA | 0.1987 | 0.4273 | 0.3597 | 0.3597 | TRUE | 0.2468 |
| Fabp7 | 4 | 0 | 1 | 0 | 0 | 1 | 3 | 0 | 1 | 0 | 0 | 1 | 0.86 | 1 | 0 | 0.259 | -1.95 | 0.39665209 | NA | 0 | 2.7515 | 2.4763 | 2.4763 | TRUE | 0.2206 |
| Fabp9 | 16 | 10 | 6 | 6 | 0 | 4 | 12 | 11 | 6 | 6 | 0 | 4 | 6.58 | 10 | 3 | 0.341 | -1.554 | 0.07031959 | NA | 0.1209 | 0.3628 | 0.3292 | 0.3292 | TRUE | 0.9187 |
| Fadd | 423 | 272 | 191 | 221 | 234 | 214 | 329 | 289 | 206 | 220 | 264 | 211 | 253.19 | 275 | 232 | 0.84 | -0.252 | 0.14482036 | 0.28828933 | 0.0311 | 0.0118 | 0.0175 | 0.0175 | TRUE | 0.4704 |
| Fads1 | 7914 | 5894 | 6849 | 7361 | 6502 | 7616 | 6146 | 6269 | 7390 | 7327 | 7335 | 7518 | 6997.68 | 6602 | 7393 | 1.12 | 0.164 | 0.02429131 | 0.07968687 | 0.0053 | 0.0028 | 0.0037 | 0.0037 | TRUE | 0.1326 |
| Fads2 | 4621 | 3196 | 3279 | 3878 | 3211 | 3775 | 3589 | 3400 | 3538 | 3860 | 3622 | 3727 | 3622.58 | 3509 | 3736 | 1.065 | 0.09 | 0.11139406 | 0.23927793 | 6.00E-04 | 0.0031 | 0.0021 | 0.0021 | TRUE | 0.0102 |
| Fads3 | 201 | 119 | 128 | 147 | 149 | 147 | 156 | 127 | 138 | 146 | 168 | 145 | 146.72 | 140 | 153 | 1.086 | 0.12 | 0.4820623 | 0.64862314 | 0 | 0.0186 | 0.014 | 0.014 | TRUE | 0.1116 |
| Fads6 | 288 | 237 | 267 | 365 | 319 | 324 | 224 | 252 | 288 | 363 | 360 | 320 | 301.15 | 255 | 348 | 1.37 | 0.454 | 0.00062507 | 0.00512869 | 0.007 | 0.0103 | 0.0093 | 0.0093 | TRUE | 0.1463 |
| Faf1 | 8696 | 6076 | 6667 | 7252 | 6436 | 7278 | 6754 | 6463 | 7194 | 7219 | 7260 | 7185 | 7012.34 | 6804 | 7221 | 1.061 | 0.086 | 0.12613353 | 0.26129144 | 0.0013 | 0.0028 | 0.0021 | 0.0021 | TRUE | 0.0304 |
| Faf2 | 4091 | 3016 | 2740 | 2853 | 2558 | 2892 | 3177 | 3208 | 2956 | 2840 | 2886 | 2855 | 2987.04 | 3114 | 2860 | 0.918 | -0.123 | 0.037315 | 0.10894998 | 7.00E-04 | 0.0032 | 0.0022 | 0.0022 | TRUE | 0.0234 |
| Fah | 971 | 788 | 1051 | 1422 | 1304 | 1571 | 754 | 838 | 1134 | 1415 | 1471 | 1551 | 1193.94 | 909 | 1479 | 1.631 | 0.705 | 8.14E-09 | 4.17E-07 | 0.0238 | 0.0044 | 0.0099 | 0.0099 | TRUE | 0.5572 |
| Fahd1 | 349 | 295 | 312 | 392 | 358 | 381 | 271 | 314 | 337 | 390 | 404 | 376 | 348.61 | 307 | 390 | 1.274 | 0.349 | 0.00375677 | 0.0196471 | 0.0038 | 0.0092 | 0.0076 | 0.0076 | TRUE | 0.1302 |
| Fahd2a | 388 | 327 | 307 | 351 | 260 | 304 | 301 | 348 | 331 | 349 | 293 | 300 | 320.53 | 327 | 314 | 0.964 | -0.052 | 0.67586574 | 0.80105861 | 0.0043 | 0.0098 | 0.0081 | 0.0081 | TRUE | 0.1093 |
| Faim | 1568 | 1059 | 1024 | 1066 | 907 | 1091 | 1218 | 1126 | 1105 | 1061 | 1023 | 1077 | 1101.73 | 1150 | 1054 | 0.916 | -0.127 | 0.09485804 | 0.21277558 | 0.001 | 0.0046 | 0.0033 | 0.0033 | TRUE | 0.0336 |
| Faim2 | 127 | 74 | 75 | 81 | 77 | 76 | 99 | 79 | 81 | 81 | 87 | 75 | 83.46 | 86 | 81 | 0.933 | -0.1 | 0.64631696 | 0.78008853 | 0 | 0.0308 | 0.0222 | 0.0222 | TRUE | 0.1723 |
| Faim3 | 3 | 2 | 2 | 1 | 0 | 0 | 2 | 2 | 2 | 1 | 0 | 0 | 1.27 | 2 | 0 | 0.168 | -2.574 | 0.16863336 | NA | 0 | 1.8709 | 1.3012 | 1.3012 | TRUE | 0.3906 |
| Fam101a | 35 | 23 | 19 | 23 | 23 | 21 | 27 | 24 | 21 | 23 | 26 | 21 | 23.62 | 24 | 23 | 0.957 | -0.063 | 0.8724393 | 0.9292626 | 0 | 0.1028 | 0.0686 | 0.0686 | TRUE | 0.0941 |
| Fam101b | 748 | 534 | 472 | 475 | 391 | 444 | 581 | 568 | 509 | 473 | 441 | 438 | 501.74 | 553 | 451 | 0.815 | -0.296 | 0.00341067 | 0.0182954 | 0.0015 | 0.0072 | 0.0053 | 0.0053 | TRUE | 0.054 |
| Fam102a | 919 | 752 | 865 | 813 | 887 | 859 | 714 | 800 | 933 | 809 | 1001 | 848 | 850.8 | 816 | 886 | 1.087 | 0.121 | 0.28726501 | 0.45937333 | 0.0133 | 0.0052 | 0.0081 | 0.0081 | TRUE | 0.1886 |
| Fam102b | 1022 | 791 | 731 | 882 | 668 | 753 | 794 | 841 | 789 | 878 | 754 | 743 | 799.78 | 808 | 792 | 0.98 | -0.028 | 0.75698869 | 0.85564566 | 0.0037 | 0.0054 | 0.0048 | 0.0048 | TRUE | 0.109 |
| Fam103a1 | 1756 | 1236 | 1368 | 1523 | 1254 | 1477 | 1364 | 1315 | 1476 | 1516 | 1415 | 1458 | 1423.87 | 1385 | 1463 | 1.057 | 0.08 | 0.27652613 | 0.44771047 | 0.0016 | 0.0041 | 0.0032 | 0.0032 | TRUE | 0.0387 |
| Fam104a | 3853 | 2816 | 2765 | 2914 | 2621 | 3211 | 2992 | 2995 | 2983 | 2901 | 2957 | 3170 | 2999.72 | 2990 | 3009 | 1.006 | 0.009 | 0.87828554 | 0.93196497 | 8.00E-04 | 0.0032 | 0.0022 | 0.0022 | TRUE | 0.0267 |
| Fam105a | 124 | 105 | 102 | 130 | 90 | 127 | 96 | 112 | 110 | 129 | 102 | 125 | 112.39 | 106 | 119 | 1.126 | 0.172 | 0.3722525 | 0.54682477 | 0 | 0.0235 | 0.0177 | 0.0177 | TRUE | 0.1544 |
| Fam105b | 1457 | 1002 | 837 | 817 | 698 | 807 | 1132 | 1066 | 903 | 813 | 787 | 797 | 916.3 | 1034 | 799 | 0.772 | -0.373 | 8.22E-05 | 0.00103137 | 0.006 | 0.005 | 0.0054 | 0.0054 | TRUE | 0.1437 |
| Fam107a | 3 | 10 | 3 | 7 | 6 | 5 | 2 | 11 | 3 | 7 | 7 | 5 | 5.81 | 5 | 6 | 1.174 | 0.231 | 0.78452238 | NA | 0.1736 | 0.4102 | 0.3408 | 0.3408 | TRUE | 1.4795 |
| Fam107b | 1104 | 725 | 726 | 721 | 582 | 691 | 857 | 771 | 783 | 718 | 657 | 682 | 744.72 | 804 | 686 | 0.852 | -0.232 | 0.00841728 | 0.03642392 | 0.0016 | 0.0056 | 0.0042 | 0.0042 | TRUE | 0.0422 |
| Fam108a | 3321 | 2452 | 2285 | 2573 | 2401 | 2594 | 2579 | 2608 | 2465 | 2561 | 2709 | 2561 | 2580.56 | 2551 | 2610 | 1.023 | 0.032 | 0.59005255 | 0.73868977 | 5.00E-04 | 0.0033 | 0.0022 | 0.0022 | TRUE | 0.013 |
| Fam108b | 2227 | 1551 | 1566 | 1558 | 1314 | 1464 | 1730 | 1650 | 1690 | 1551 | 1482 | 1445 | 1591.24 | 1690 | 1493 | 0.883 | -0.179 | 0.00698902 | 0.03171242 | 0 | 0.0039 | 0.0026 | 0.0026 | TRUE | 0.0141 |
| Fam108c | 4111 | 3143 | 3445 | 3434 | 3061 | 3727 | 3193 | 3343 | 3717 | 3418 | 3453 | 3679 | 3467.26 | 3418 | 3517 | 1.03 | 0.042 | 0.55004955 | 0.70633575 | 0.0036 | 0.0031 | 0.0033 | 0.0033 | TRUE | 0.0711 |
| Fam109a | 696 | 521 | 466 | 515 | 460 | 523 | 541 | 554 | 503 | 513 | 519 | 516 | 524.23 | 533 | 516 | 0.968 | -0.046 | 0.62320428 | 0.76236381 | 0 | 0.0069 | 0.0045 | 0.0045 | TRUE | 0.0265 |
| Fam109b | 75 | 53 | 44 | 73 | 55 | 65 | 58 | 56 | 47 | 73 | 62 | 64 | 60.16 | 54 | 66 | 1.224 | 0.292 | 0.24465037 | 0.41284388 | 0 | 0.0418 | 0.0285 | 0.0285 | TRUE | 0.0886 |
| Fam110a | 658 | 450 | 568 | 501 | 489 | 530 | 511 | 479 | 613 | 499 | 552 | 523 | 529.35 | 534 | 525 | 0.982 | -0.026 | 0.81912223 | 0.89618142 | 0.0076 | 0.0069 | 0.0071 | 0.0071 | TRUE | 0.188 |
| Fam110b | 710 | 526 | 669 | 583 | 579 | 673 | 551 | 559 | 722 | 580 | 653 | 664 | 621.77 | 611 | 632 | 1.037 | 0.053 | 0.6545111 | 0.78533835 | 0.0128 | 0.0062 | 0.0083 | 0.0083 | TRUE | 0.292 |
| Fam110c | 42 | 41 | 54 | 43 | 50 | 45 | 33 | 44 | 58 | 43 | 56 | 44 | 46.35 | 45 | 48 | 1.075 | 0.105 | 0.73611097 | 0.8425688 | 0.0331 | 0.0536 | 0.0477 | 0.0477 | TRUE | 0.2891 |
| Fam111a | 2577 | 1731 | 1560 | 1876 | 1465 | 1848 | 2001 | 1841 | 1683 | 1867 | 1653 | 1824 | 1811.7 | 1842 | 1781 | 0.967 | -0.049 | 0.54906964 | 0.70561521 | 0.0053 | 0.0037 | 0.0043 | 0.0043 | TRUE | 0.0709 |
| Fam114a1 | 379 | 324 | 404 | 639 | 568 | 658 | 294 | 345 | 436 | 636 | 641 | 650 | 500.21 | 358 | 642 | 1.8 | 0.848 | 5.79E-11 | 4.95E-09 | 0.0169 | 0.0072 | 0.0099 | 0.0099 | TRUE | 0.4081 |
| Fam114a2 | 7743 | 5974 | 5894 | 6593 | 6178 | 6686 | 6014 | 6354 | 6360 | 6563 | 6969 | 6600 | 6476.65 | 6243 | 6711 | 1.075 | 0.105 | 0.05435325 | 0.14288597 | 8.00E-04 | 0.0028 | 0.002 | 0.002 | TRUE | 0.0138 |
| Fam115a | 534 | 355 | 341 | 385 | 379 | 343 | 415 | 378 | 368 | 383 | 428 | 339 | 384.94 | 387 | 383 | 0.987 | -0.018 | 0.87976412 | 0.9329625 | 0.0061 | 0.0086 | 0.0078 | 0.0078 | TRUE | 0.1194 |
| Fam115c | 624 | 477 | 397 | 406 | 397 | 468 | 485 | 507 | 428 | 404 | 448 | 462 | 455.73 | 473 | 438 | 0.924 | -0.114 | 0.29994553 | 0.47214993 | 0.0038 | 0.0076 | 0.0065 | 0.0065 | TRUE | 0.0773 |
| Fam115e | 0 | 0 | 0 | 0 | 0 | 0 | 0 | 0 | 0 | 0 | 0 | 0 | 0 | 0 | 0 | NA | NA | NA | NA | NA | NA | NA | NA | NA | NA |
| Fam117a | 3236 | 2348 | 2344 | 2365 | 2284 | 2690 | 2513 | 2498 | 2529 | 2354 | 2577 | 2655 | 2521.02 | 2513 | 2529 | 1.006 | 0.009 | 0.893914 | 0.94181872 | 0.0016 | 0.0034 | 0.0027 | 0.0027 | TRUE | 0.0446 |
| Fam117b | 5286 | 4022 | 3957 | 4580 | 3992 | 4383 | 4105 | 4278 | 4270 | 4559 | 4503 | 4327 | 4340.36 | 4218 | 4463 | 1.058 | 0.082 | 0.12936194 | 0.26608271 | 4.00E-04 | 0.003 | 0.0019 | 0.0019 | TRUE | 0.0088 |
| Fam118a | 1993 | 1400 | 1350 | 1513 | 1266 | 1470 | 1548 | 1489 | 1457 | 1506 | 1428 | 1451 | 1479.84 | 1498 | 1462 | 0.975 | -0.036 | 0.59397932 | 0.74083813 | 0 | 0.004 | 0.0026 | 0.0026 | TRUE | 0.0107 |
| Fam118b | 1748 | 1222 | 1009 | 1026 | 949 | 1040 | 1358 | 1300 | 1089 | 1021 | 1071 | 1027 | 1144.1 | 1249 | 1040 | 0.831 | -0.267 | 0.00344993 | 0.01842671 | 0.0063 | 0.0045 | 0.0051 | 0.0051 | TRUE | 0.1488 |
| Fam120a | 11034 | 8695 | 8638 | 9416 | 8134 | 9415 | 8570 | 9249 | 9320 | 9373 | 9176 | 9294 | 9163.58 | 9046 | 9281 | 1.026 | 0.037 | 0.48435617 | 0.65044468 | 9.00E-04 | 0.0027 | 0.002 | 0.002 | TRUE | 0.0263 |
| Fam120aos | 174 | 160 | 158 | 176 | 159 | 148 | 135 | 170 | 170 | 175 | 179 | 146 | 162.74 | 158 | 167 | 1.056 | 0.079 | 0.64470193 | 0.77879688 | 0.0093 | 0.017 | 0.0147 | 0.0147 | TRUE | 0.1957 |
| Fam120b | 938 | 622 | 557 | 616 | 609 | 696 | 729 | 662 | 601 | 613 | 687 | 687 | 663.06 | 664 | 662 | 0.996 | -0.006 | 0.95048928 | 0.97274404 | 0.0054 | 0.006 | 0.0058 | 0.0058 | TRUE | 0.09 |
| Fam120c | 193 | 123 | 108 | 114 | 90 | 121 | 150 | 131 | 117 | 113 | 102 | 119 | 121.95 | 133 | 111 | 0.84 | -0.252 | 0.17581877 | 0.32931171 | 0 | 0.0219 | 0.0168 | 0.0168 | TRUE | 0.15 |
| Fam122a | 346 | 223 | 209 | 182 | 169 | 166 | 269 | 237 | 226 | 181 | 191 | 164 | 211.19 | 244 | 179 | 0.729 | -0.455 | 0.00181503 | 0.01122068 | 0 | 0.0136 | 0.0105 | 0.0105 | TRUE | 0.0952 |
| Fam122b | 3155 | 2320 | 2521 | 2813 | 2219 | 2603 | 2450 | 2468 | 2720 | 2800 | 2503 | 2570 | 2585.19 | 2546 | 2624 | 1.031 | 0.045 | 0.52952364 | 0.68979962 | 0.0027 | 0.0033 | 0.0032 | 0.0032 | TRUE | 0.0429 |
| Fam122c | 0 | 0 | 0 | 0 | 0 | 0 | 0 | 0 | 0 | 0 | 0 | 0 | 0 | 0 | 0 | NA | NA | NA | NA | NA | NA | NA | NA | NA | NA |
| Fam124a | 485 | 341 | 168 | 152 | 156 | 142 | 377 | 363 | 181 | 151 | 176 | 140 | 231.35 | 307 | 156 | 0.506 | -0.983 | 1.29E-05 | 0.00023158 | 0.0793 | 0.0127 | 0.0316 | 0.0316 | TRUE | 1.4248 |
| Fam124b | 10 | 9 | 7 | 10 | 6 | 6 | 8 | 10 | 8 | 10 | 7 | 6 | 7.92 | 9 | 8 | 0.913 | -0.131 | 0.84526923 | NA | 0 | 0.3016 | 0.1979 | 0.1979 | TRUE | 0.2262 |
| Fam126a | 1798 | 1310 | 1213 | 1621 | 1307 | 1626 | 1396 | 1393 | 1309 | 1614 | 1474 | 1605 | 1465.29 | 1366 | 1564 | 1.145 | 0.195 | 0.00635128 | 0.02941634 | 0.0012 | 0.004 | 0.003 | 0.003 | TRUE | 0.0299 |
| Fam126b | 1842 | 1418 | 1436 | 1664 | 1450 | 1539 | 1431 | 1508 | 1549 | 1656 | 1636 | 1519 | 1549.94 | 1496 | 1604 | 1.073 | 0.101 | 0.15308527 | 0.30002026 | 0.0013 | 0.004 | 0.003 | 0.003 | TRUE | 0.0259 |
| Fam129a | 1850 | 1425 | 1315 | 1536 | 1324 | 1555 | 1437 | 1516 | 1419 | 1529 | 1494 | 1535 | 1488.17 | 1457 | 1519 | 1.043 | 0.061 | 0.36204085 | 0.5367894 | 0 | 0.004 | 0.0025 | 0.0025 | TRUE | 0.0145 |
| Fam129b | 7576 | 6242 | 6998 | 8303 | 7981 | 9342 | 5884 | 6640 | 7551 | 8265 | 9003 | 9222 | 7760.74 | 6692 | 8830 | 1.32 | 0.401 | 2.12E-06 | 5.30E-05 | 0.0092 | 0.0027 | 0.005 | 0.005 | TRUE | 0.154 |
| Fam129c | 150 | 142 | 120 | 141 | 128 | 145 | 116 | 151 | 129 | 140 | 144 | 143 | 137.48 | 132 | 142 | 1.082 | 0.114 | 0.51434729 | 0.6770449 | 0 | 0.0197 | 0.0146 | 0.0146 | TRUE | 0.1557 |
| Fam131a | 222 | 130 | 171 | 226 | 206 | 205 | 172 | 138 | 185 | 225 | 232 | 202 | 192.49 | 165 | 220 | 1.33 | 0.411 | 0.00896234 | 0.03822702 | 0.0066 | 0.0147 | 0.0125 | 0.0125 | TRUE | 0.2025 |
| Fam131b | 110 | 74 | 63 | 72 | 74 | 70 | 85 | 79 | 68 | 72 | 83 | 69 | 76.06 | 77 | 75 | 0.961 | -0.058 | 0.79837471 | 0.88226146 | 0 | 0.0336 | 0.024 | 0.024 | TRUE | 0.1004 |

| | | | | | | | | | | | | | | | | | | | | | | | | | |
|---|---|---|---|---|---|---|---|---|---|---|---|---|---|---|---|---|---|---|---|---|---|---|---|---|---|
| Fam131c | 913 | 685 | 713 | 926 | 820 | 991 | 709 | 729 | 769 | 922 | 925 | 978 | 838.68 | 736 | 942 | 1.282 | 0.358 | 1.04E-05 | 0.00019326 | 0 | 0.0053 | 0.0035 | 0.0035 | TRUE | 0.0187 |
| Fam132a | 522 | 459 | 615 | 702 | 708 | 836 | 405 | 488 | 664 | 699 | 799 | 825 | 646.66 | 519 | 774 | 1.496 | 0.581 | 7.67E-05 | 0.00098376 | 0.0335 | 0.0061 | 0.0139 | 0.0139 | TRUE | 0.69 |
| Fam132b | 268 | 240 | 226 | 207 | 177 | 184 | 208 | 255 | 244 | 206 | 200 | 182 | 215.77 | 236 | 196 | 0.833 | -0.264 | 0.07033888 | 0.17206121 | 0.0043 | 0.0134 | 0.0106 | 0.0106 | TRUE | 0.1267 |
| Fam133b | 1569 | 1224 | 975 | 983 | 872 | 999 | 1219 | 1302 | 1052 | 978 | 984 | 986 | 1086.82 | 1191 | 983 | 0.825 | -0.278 | 0.00174871 | 0.01090028 | 0.005 | 0.0046 | 0.0047 | 0.0047 | TRUE | 0.1214 |
| Fam134a | 1967 | 1444 | 1412 | 1458 | 1332 | 1539 | 1528 | 1536 | 1524 | 1451 | 1503 | 1519 | 1510.06 | 1529 | 1491 | 0.975 | -0.036 | 0.57371581 | 0.7259313 | 0 | 0.004 | 0.0024 | 0.0024 | TRUE | 0.0066 |
| Fam134b | 1184 | 955 | 1315 | 1505 | 1466 | 1482 | 920 | 1016 | 1419 | 1498 | 1654 | 1463 | 1328.18 | 1118 | 1538 | 1.377 | 0.462 | 0.00032599 | 0.00309797 | 0.0291 | 0.0042 | 0.0111 | 0.0111 | TRUE | 0.6605 |
| Fam134c | 5033 | 3587 | 3524 | 3717 | 3418 | 3871 | 3909 | 3815 | 3802 | 3700 | 3856 | 3821 | 3817.29 | 3842 | 3792 | 0.987 | -0.019 | 0.71617728 | 0.82949644 | 0 | 0.003 | 0.0018 | 0.0018 | TRUE | 0.0056 |
| Fam135a | 2529 | 1844 | 1739 | 1809 | 1565 | 1799 | 1964 | 1961 | 1876 | 1801 | 1765 | 1776 | 1857.34 | 1934 | 1781 | 0.921 | -0.12 | 0.05283275 | 0.14030817 | 0 | 0.0037 | 0.0022 | 0.0022 | TRUE | 0.008 |
| Fam135b | 0 | 1 | 0 | 2 | 2 | 0 | 0 | 1 | 0 | 2 | 2 | 0 | 0.89 | 0 | 1 | 4.079 | 2.028 | 0.37173703 | NA | 0 | 2.6801 | 2.3469 | 2.3469 | TRUE | 0.5217 |
| Fam136a | 4148 | 2899 | 2287 | 2247 | 1851 | 2201 | 3222 | 3084 | 2468 | 2237 | 2088 | 2173 | 2545.06 | 2925 | 2166 | 0.74 | -0.434 | 2.41E-06 | 5.86E-05 | 0.0094 | 0.0034 | 0.0057 | 0.0057 | TRUE | 0.2252 |
| Fam13a | 85 | 68 | 62 | 62 | 52 | 64 | 66 | 72 | 67 | 62 | 59 | 63 | 64.8 | 68 | 61 | 0.896 | -0.158 | 0.50435532 | 0.66826706 | 0 | 0.039 | 0.025 | 0.025 | TRUE | 0.0216 |
| Fam13b | 1075 | 800 | 715 | 811 | 702 | 747 | 835 | 851 | 771 | 807 | 792 | 737 | 798.99 | 819 | 779 | 0.95 | -0.074 | 0.3866492 | 0.56048448 | 0.0011 | 0.0054 | 0.004 | 0.004 | TRUE | 0.0296 |
| Fam13c | 529 | 385 | 452 | 472 | 423 | 477 | 411 | 410 | 488 | 470 | 477 | 471 | 454.33 | 436 | 473 | 1.086 | 0.119 | 0.27000533 | 0.44094906 | 0.0027 | 0.0076 | 0.0061 | 0.0061 | TRUE | 0.1207 |
| Fam149a | 247 | 183 | 172 | 158 | 132 | 138 | 192 | 195 | 186 | 157 | 149 | 136 | 169.08 | 191 | 147 | 0.773 | -0.372 | 0.01544909 | 0.0571021 | 0 | 0.0164 | 0.0109 | 0.0109 | TRUE | 0.0482 |
| Fam149b | 1116 | 805 | 858 | 866 | 758 | 864 | 867 | 856 | 926 | 862 | 855 | 853 | 869.8 | 883 | 857 | 0.971 | -0.043 | 0.58622269 | 0.73550569 | 0 | 0.0052 | 0.0033 | 0.0033 | TRUE | 0.0207 |
| Fam150a | 23 | 17 | 14 | 22 | 14 | 18 | 18 | 18 | 15 | 22 | 16 | 18 | 17.75 | 17 | 19 | 1.086 | 0.12 | 0.79029479 | 0.87673234 | 0 | 0.1359 | 0.0897 | 0.0897 | TRUE | 0.1363 |
| Fam150b | 2 | 2 | 1 | 2 | 2 | 0 | 2 | 2 | 1 | 2 | 2 | 0 | 1.5 | 2 | 1 | 0.883 | -0.18 | 0.90853046 | NA | 0 | 1.5813 | 1.0992 | 1.0992 | TRUE | 0.5228 |
| Fam151a | 27 | 23 | 15 | 23 | 17 | 14 | 21 | 24 | 16 | 23 | 19 | 14 | 19.59 | 20 | 19 | 0.905 | -0.143 | 0.7460659 | 0.8492092 | 0 | 0.1234 | 0.09 | 0.09 | TRUE | 0.2775 |
| Fam151b | 44 | 17 | 22 | 21 | 13 | 21 | 34 | 18 | 24 | 21 | 15 | 21 | 22.05 | 25 | 19 | 0.736 | -0.443 | 0.30513056 | 0.47773372 | 0.0331 | 0.1099 | 0.0881 | 0.0881 | TRUE | 0.3992 |
| Fam154a | 37 | 26 | 41 | 31 | 31 | 28 | 29 | 28 | 44 | 31 | 35 | 28 | 32.35 | 34 | 31 | 0.932 | -0.102 | 0.77316975 | 0.86585562 | 0 | 0.0757 | 0.0584 | 0.0584 | TRUE | 0.5111 |
| Fam154b | 12 | 4 | 6 | 7 | 5 | 7 | 9 | 4 | 6 | 7 | 6 | 7 | 6.59 | 6 | 7 | 0.961 | -0.057 | 0.93871309 | NA | 0 | 0.3618 | 0.2442 | 0.2442 | TRUE | 0.3155 |
| Fam155a | 3 | 0 | 2 | 0 | 0 | 1 | 2 | 0 | 2 | 0 | 0 | 1 | 0.91 | 1 | 0 | 0.245 | -2.03 | 0.36330583 | NA | 0 | 2.5997 | 2.2719 | 2.2719 | TRUE | 0.484 |
| Fam159a | 105 | 66 | 78 | 129 | 94 | 154 | 82 | 70 | 84 | 128 | 106 | 152 | 103.73 | 79 | 129 | 1.64 | 0.713 | 0.00066685 | 0.00536071 | 0.0123 | 0.0253 | 0.0215 | 0.0215 | TRUE | 0.1307 |
| Fam159b | 61 | 45 | 57 | 48 | 38 | 49 | 47 | 48 | 62 | 48 | 43 | 48 | 49.29 | 52 | 46 | 0.891 | -0.166 | 0.54974391 | 0.70613719 | 0 | 0.0505 | 0.0351 | 0.0351 | TRUE | 0.1879 |
| Fam160a1 | 283 | 195 | 213 | 220 | 196 | 173 | 220 | 207 | 230 | 219 | 221 | 171 | 211.32 | 219 | 204 | 0.929 | -0.107 | 0.47707272 | 0.64471288 | 0.0062 | 0.0136 | 0.0115 | 0.0115 | TRUE | 0.2211 |
| Fam160a2 | 3956 | 2962 | 2971 | 3187 | 2808 | 3157 | 3072 | 3151 | 3206 | 3172 | 3168 | 3116 | 3147.55 | 3143 | 3152 | 1.003 | 0.005 | 0.93168446 | 0.96216772 | 0 | 0.0032 | 0.0018 | 0.0018 | TRUE | 0.0048 |
| Fam160b1 | 2230 | 1691 | 1835 | 1913 | 1669 | 1962 | 1732 | 1799 | 1980 | 1904 | 1883 | 1937 | 1872.4 | 1837 | 1908 | 1.04 | 0.056 | 0.4256134 | 0.59821932 | 0.002 | 0.0037 | 0.0031 | 0.0031 | TRUE | 0.0556 |
| Fam160b2 | 1888 | 1390 | 1225 | 1134 | 1099 | 1261 | 1466 | 1479 | 1322 | 1129 | 1240 | 1245 | 1313.33 | 1422 | 1205 | 0.846 | -0.241 | 0.00214621 | 0.01273909 | 0.0026 | 0.0042 | 0.0037 | 0.0037 | TRUE | 0.045 |
| Fam161a | 192 | 128 | 141 | 137 | 102 | 123 | 149 | 136 | 152 | 136 | 115 | 121 | 135.04 | 146 | 124 | 0.853 | -0.23 | 0.18135789 | 0.33693193 | 0 | 0.02 | 0.0138 | 0.0138 | TRUE | 0.0741 |
| Fam161b | 269 | 173 | 183 | 199 | 167 | 192 | 209 | 184 | 197 | 198 | 188 | 190 | 194.4 | 197 | 192 | 0.974 | -0.038 | 0.78994943 | 0.87647471 | 0 | 0.0146 | 0.0097 | 0.0097 | TRUE | 0.0346 |
| Fam162a | 4802 | 3418 | 4235 | 4969 | 4374 | 5026 | 3730 | 3636 | 4570 | 4946 | 4934 | 4961 | 4462.77 | 3979 | 4947 | 1.244 | 0.315 | 0.00012644 | 0.00147888 | 0.008 | 0.003 | 0.0046 | 0.0046 | TRUE | 0.2046 |
| Fam162b | 7 | 2 | 2 | 5 | 4 | 2 | 5 | 2 | 2 | 5 | 5 | 2 | 3.53 | 3 | 4 | 1.144 | 0.195 | 0.84946061 | NA | 0 | 0.6737 | 0.4765 | 0.4765 | TRUE | 0.6449 |
| Fam163a | 15 | 8 | 6 | 5 | 4 | 1 | 12 | 9 | 6 | 5 | 5 | 1 | 6.18 | 9 | 4 | 0.387 | -1.369 | 0.10001517 | NA | 0 | 0.3856 | 0.2961 | 0.2961 | TRUE | 0.5848 |
| Fam163b | 150 | 143 | 163 | 200 | 163 | 214 | 116 | 152 | 176 | 199 | 184 | 211 | 173.12 | 148 | 198 | 1.347 | 0.429 | 0.01489089 | 0.05557064 | 0.0175 | 0.0161 | 0.0165 | 0.0165 | TRUE | 0.4055 |
| Fam166a | 4 | 3 | 2 | 4 | 3 | 12 | 3 | 3 | 2 | 4 | 3 | 12 | 4.61 | 3 | 6 | 2.274 | 1.185 | 0.21212324 | NA | 0 | 0.5164 | 0.4001 | 0.4001 | TRUE | 1.4012 |
| Fam166b | 38 | 23 | 25 | 47 | 46 | 68 | 30 | 24 | 27 | 47 | 52 | 67 | 41.13 | 27 | 55 | 2.041 | 1.029 | 0.00108977 | 0.00769531 | 0 | 0.0601 | 0.0444 | 0.0444 | TRUE | 0.3038 |
| Fam167a | 176 | 170 | 200 | 166 | 177 | 205 | 137 | 181 | 216 | 165 | 200 | 202 | 183.43 | 178 | 189 | 1.069 | 0.096 | 0.60021489 | 0.74579426 | 0.0271 | 0.0153 | 0.0189 | 0.0189 | TRUE | 0.2897 |
| Fam167b | 1 | 0 | 0 | 0 | 1 | 2 | 1 | 0 | 0 | 0 | 1 | 2 | 0.65 | 0 | 1 | 2.998 | 1.584 | 0.52889475 | NA | 0 | 3.6683 | 3.06 | 3.06 | TRUE | 0.14 |
| Fam168a | 3566 | 2487 | 2478 | 2671 | 2429 | 2643 | 2770 | 2645 | 2674 | 2659 | 2740 | 2609 | 2682.77 | 2696 | 2669 | 0.989 | -0.015 | 0.79132371 | 0.87744725 | 3.00E-04 | 0.0033 | 0.0021 | 0.0021 | TRUE | 0.0071 |
| Fam168b | 5183 | 3944 | 3830 | 3525 | 3235 | 3590 | 4025 | 4195 | 4133 | 3509 | 3649 | 3544 | 3842.54 | 4118 | 3567 | 0.866 | -0.207 | 0.00011404 | 0.00135376 | 2.00E-04 | 0.003 | 0.0018 | 0.0018 | TRUE | 0.0049 |
| Fam169a | 14279 | 9697 | 9017 | 10666 | 8633 | 10221 | 11090 | 10315 | 9729 | 10617 | 9739 | 10090 | 10263.21 | 10378 | 10149 | 0.978 | -0.032 | 0.60837743 | 0.7520791 | 0.0026 | 0.0027 | 0.0028 | 0.0028 | TRUE | 0.0443 |
| Fam169b | 256 | 176 | 244 | 272 | 224 | 264 | 199 | 187 | 263 | 271 | 253 | 261 | 238.89 | 216 | 262 | 1.211 | 0.276 | 0.06883842 | 0.16943475 | 0.0122 | 0.0123 | 0.0124 | 0.0124 | TRUE | 0.3781 |
| Fam170a | 1 | 0 | 0 | 0 | 0 | 0 | 1 | 0 | 0 | 0 | 0 | 0 | 0.13 | 0 | 0 | 0.552 | -0.857 | 0.8336548 | NA | 0 | 18.3121 | 10 | 10 | TRUE | 0.2566 |
| Fam170b | 5 | 7 | 1 | 1 | 2 | 1 | 4 | 7 | 1 | 1 | 2 | 1 | 2.77 | 4 | 1 | 0.337 | -1.568 | 0.2136068 | NA | 0 | 0.8567 | 0.6578 | 0.6578 | TRUE | 0.1155 |
| Fam171a1 | 1147 | 891 | 845 | 793 | 685 | 737 | 891 | 948 | 912 | 789 | 773 | 728 | 839.99 | 917 | 763 | 0.833 | -0.264 | 0.00113218 | 0.00794416 | 0 | 0.0052 | 0.0035 | 0.0035 | TRUE | 0.0202 |
| Fam171a2 | 612 | 473 | 409 | 459 | 388 | 421 | 475 | 503 | 441 | 457 | 438 | 416 | 454.99 | 473 | 437 | 0.923 | -0.116 | 0.26298005 | 0.43333801 | 0 | 0.0076 | 0.0056 | 0.0056 | TRUE | 0.0383 |
| Fam171b | 506 | 373 | 320 | 388 | 293 | 353 | 393 | 397 | 345 | 386 | 331 | 348 | 366.71 | 378 | 355 | 0.938 | -0.092 | 0.43229149 | 0.6041598 | 0.0032 | 0.0089 | 0.0072 | 0.0072 | TRUE | 0.0675 |
| Fam172a | 1488 | 1028 | 1065 | 1154 | 913 | 1051 | 1156 | 1093 | 1149 | 1149 | 1030 | 1037 | 1102.4 | 1133 | 1072 | 0.946 | -0.08 | 0.30713167 | 0.4798156 | 0.0014 | 0.0046 | 0.0035 | 0.0035 | TRUE | 0.0472 |
| Fam173a | 356 | 288 | 299 | 295 | 276 | 324 | 276 | 306 | 323 | 294 | 311 | 320 | 305.05 | 302 | 308 | 1.024 | 0.034 | 0.77833937 | 0.86919326 | 0 | 0.0102 | 0.0074 | 0.0074 | TRUE | 0.0662 |
| Fam173b | 174 | 119 | 93 | 110 | 92 | 115 | 135 | 127 | 100 | 109 | 104 | 114 | 114.81 | 121 | 109 | 0.9 | -0.153 | 0.42701992 | 0.5991237 | 0 | 0.0231 | 0.0179 | 0.0179 | TRUE | 0.2113 |
| Fam174a | 438 | 360 | 382 | 456 | 367 | 422 | 340 | 383 | 412 | 454 | 414 | 417 | 403.3 | 378 | 428 | 1.135 | 0.183 | 0.1084601 | 0.23477513 | 0.0037 | 0.0083 | 0.0069 | 0.0069 | TRUE | 0.0967 |
| Fam174b | 389 | 285 | 213 | 240 | 199 | 224 | 302 | 303 | 230 | 239 | 224 | 221 | 253.27 | 278 | 228 | 0.818 | -0.29 | 0.04482031 | 0.12440291 | 0.0093 | 0.0118 | 0.011 | 0.011 | TRUE | 0.2517 |
| Fam175a | 1631 | 1185 | 1118 | 1146 | 1020 | 1158 | 1267 | 1260 | 1206 | 1141 | 1151 | 1143 | 1194.67 | 1244 | 1145 | 0.919 | -0.121 | 0.0816777 | 0.19203351 | 0 | 0.0044 | 0.0026 | 0.0026 | TRUE | 0.0084 |
| Fam175b | 1985 | 1520 | 1732 | 1963 | 1704 | 2122 | 1542 | 1617 | 1869 | 1954 | 1922 | 2095 | 1833.05 | 1676 | 1990 | 1.189 | 0.25 | 0.00248999 | 0.01431016 | 0.0055 | 0.0037 | 0.0044 | 0.0044 | TRUE | 0.1217 |
| Fam177a | 0 | 0 | 0 | 0 | 0 | 0 | 0 | 0 | 0 | 0 | 0 | 0 | 0 | 0 | 0 | NA | NA | NA | NA | NA | NA | NA | NA | NA | NA |
| Fam178a | 7526 | 5625 | 5447 | 5967 | 5088 | 6142 | 5845 | 5983 | 5877 | 5940 | 5740 | 6063 | 5908.01 | 5902 | 5914 | 1.002 | 0.003 | 0.94822314 | 0.97174686 | 2.00E-04 | 0.0028 | 0.0017 | 0.0017 | TRUE | 0.008 |
| Fam178b | 652 | 549 | 540 | 540 | 520 | 580 | 506 | 584 | 583 | 538 | 587 | 573 | 561.61 | 558 | 566 | 1.016 | 0.023 | 0.81685503 | 0.89487832 | 0.0028 | 0.0066 | 0.0054 | 0.0054 | TRUE | 0.0809 |
| Fam179a | 11 | 9 | 13 | 9 | 3 | 7 | 9 | 10 | 14 | 9 | 3 | 7 | 8.57 | 11 | 6 | 0.608 | -0.717 | 0.28913879 | NA | 0 | 0.2791 | 0.2043 | 0.2043 | TRUE | 0.3694 |
| Fam179b | 4604 | 3203 | 2843 | 3027 | 2757 | 3051 | 3576 | 3407 | 3068 | 3013 | 3110 | 3012 | 3197.56 | 3350 | 3045 | 0.908 | -0.139 | 0.04308027 | 0.12085305 | 0.0027 | 0.0032 | 0.0031 | 0.0031 | TRUE | 0.0658 |
| Fam180a | 3 | 1 | 0 | 0 | 0 | 0 | 2 | 1 | 0 | 0 | 0 | 0 | 0.57 | 1 | 0 | 0.165 | -2.597 | 0.35609012 | NA | 0 | 4.1927 | 4.2873 | 4.2873 | TRUE | 0.0685 |
| Fam181a | 3 | 7 | 4 | 4 | 4 | 5 | 2 | 7 | 4 | 4 | 5 | 5 | 4.59 | 4 | 5 | 0.971 | -0.042 | 0.96279266 | NA | 0 | 0.5191 | 0.3601 | 0.3601 | TRUE | 0.3346 |
| Fam181b | 79 | 87 | 73 | 67 | 56 | 90 | 61 | 93 | 79 | 67 | 63 | 89 | 75.23 | 78 | 73 | 0.948 | -0.078 | 0.7561484 | 0.85520897 | 0.0265 | 0.0339 | 0.0318 | 0.0318 | TRUE | 0.2694 |
| Fam183b | 71 | 68 | 71 | 72 | 66 | 87 | 55 | 72 | 77 | 72 | 74 | 86 | 72.68 | 68 | 77 | 1.145 | 0.196 | 0.41138321 | 0.58419276 | 0 | 0.035 | 0.0272 | 0.0272 | TRUE | 0.2839 |
| Fam184a | 631 | 452 | 420 | 428 | 406 | 449 | 490 | 481 | 453 | 426 | 458 | 443 | 458.55 | 475 | 442 | 0.931 | -0.104 | 0.29946806 | 0.47176203 | 0 | 0.0076 | 0.005 | 0.005 | TRUE | 0.0178 |
| Fam184b | 131 | 88 | 80 | 88 | 70 | 80 | 102 | 94 | 86 | 88 | 79 | 79 | 87.87 | 94 | 82 | 0.87 | -0.202 | 0.33270418 | 0.50702003 | 0 | 0.0294 | 0.0198 | 0.0198 | TRUE | 0.0578 |
| Fam185a | 1545 | 1125 | 1225 | 1463 | 1273 | 1429 | 1200 | 1197 | 1322 | 1456 | 1436 | 1411 | 1336.89 | 1240 | 1434 | 1.158 | 0.212 | 0.00341255 | 0.0182954 | 9.00E-04 | 0.0042 | 0.003 | 0.003 | TRUE | 0.0396 |
| Fam186b | 14 | 6 | 8 | 8 | 12 | 9 | 11 | 6 | 9 | 8 | 14 | 9 | 9.38 | 9 | 10 | 1.156 | 0.209 | 0.73967346 | NA | 0 | 0.2551 | 0.1774 | 0.1774 | TRUE | 0.2735 |
| Fam187a | 0 | 0 | 0 | 1 | 0 | 0 | 0 | 0 | 0 | 1 | 0 | 0 | 0.17 | 0 | 0 | 2.094 | 1.067 | 0.79379329 | NA | 0 | 14.2885 | 10 | 10 | TRUE | 0.3906 |
| Fam187b | 14 | 9 | 6 | 14 | 12 | 9 | 11 | 10 | 6 | 14 | 14 | 9 | 10.55 | 9 | 12 | 1.335 | 0.417 | 0.48139974 | NA | 0 | 0.2272 | 0.157 | 0.157 | TRUE | 0.2246 |

| | | | | | | | | | | | | | | | | | | | | | | | | | |
|---|---|---|---|---|---|---|---|---|---|---|---|---|---|---|---|---|---|---|---|---|---|---|---|---|---|
| Fam188a | 2957 | 2212 | 2299 | 2833 | 2359 | 2774 | 2297 | 2353 | 2481 | 2820 | 2661 | 2738 | 2558.26 | 2377 | 2740 | 1.154 | 0.206 | 0.00080609 | 0.00618439 | 8.00E-04 | 0.0034 | 0.0023 | 0.0023 | TRUE | 0.0176 |
| Fam188b | 159 | 134 | 155 | 147 | 132 | 141 | 123 | 143 | 167 | 146 | 149 | 139 | 144.61 | 144 | 145 | 1.007 | 0.01 | 0.9535522 | 0.97448316 | 0.0063 | 0.0188 | 0.0151 | 0.0151 | TRUE | 0.1971 |
| Fam189a1 | 27 | 17 | 13 | 16 | 10 | 10 | 21 | 18 | 14 | 16 | 11 | 10 | 15.03 | 18 | 12 | 0.695 | -0.525 | 0.29615285 | NA | 0 | 0.1602 | 0.113 | 0.113 | TRUE | 0.2649 |
| Fam189a2 | 413 | 320 | 381 | 430 | 456 | 489 | 321 | 340 | 411 | 428 | 514 | 483 | 416.23 | 357 | 475 | 1.332 | 0.413 | 0.00092353 | 0.00684293 | 0.0102 | 0.0081 | 0.0088 | 0.0088 | TRUE | 0.1941 |
| Fam189b | 1022 | 773 | 771 | 843 | 709 | 917 | 794 | 822 | 832 | 839 | 800 | 905 | 832.01 | 816 | 848 | 1.041 | 0.058 | 0.49162074 | 0.65729264 | 0.001 | 0.0053 | 0.0038 | 0.0038 | TRUE | 0.0415 |
| Fam192a | 4452 | 3111 | 2949 | 3068 | 2653 | 3057 | 3458 | 3309 | 3182 | 3054 | 2993 | 3018 | 3168.87 | 3316 | 3022 | 0.911 | -0.135 | 0.02039943 | 0.07017307 | 7.00E-04 | 0.0032 | 0.0021 | 0.0021 | TRUE | 0.0173 |
| Fam193a | 3417 | 2480 | 2405 | 2456 | 2410 | 2759 | 2654 | 2638 | 2595 | 2445 | 2719 | 2724 | 2628.96 | 2629 | 2629 | 1 | 0 | 0.99517655 | 0.99719984 | 0.0016 | 0.0033 | 0.0026 | 0.0026 | TRUE | 0.0459 |
| Fam193b | 2299 | 2172 | 1953 | 1890 | 1648 | 2024 | 1786 | 2310 | 2107 | 1881 | 1859 | 1998 | 1990.26 | 2068 | 1913 | 0.926 | -0.111 | 0.22200702 | 0.38612148 | 0.0082 | 0.0036 | 0.0054 | 0.0054 | TRUE | 0.1762 |
| Fam194a | 4 | 5 | 3 | 2 | 2 | 1 | 3 | 5 | 3 | 2 | 2 | 1 | 2.82 | 4 | 2 | 0.448 | -1.157 | 0.31702548 | NA | 0 | 0.8441 | 0.5398 | 0.5398 | TRUE | 0.1586 |
| Fam195a | 324 | 242 | 295 | 362 | 296 | 379 | 252 | 257 | 318 | 360 | 334 | 374 | 315.96 | 276 | 356 | 1.296 | 0.374 | 0.00398663 | 0.02054848 | 0.0065 | 0.0099 | 0.0089 | 0.0089 | TRUE | 0.1984 |
| Fam195b | 1956 | 1454 | 1678 | 2001 | 1768 | 2107 | 1519 | 1547 | 1811 | 1992 | 1994 | 2080 | 1823.75 | 1626 | 2022 | 1.245 | 0.316 | 6.44E-05 | 0.00086141 | 0.0044 | 0.0037 | 0.004 | 0.004 | TRUE | 0.1183 |
| Fam196a | 56 | 31 | 28 | 40 | 29 | 26 | 43 | 33 | 30 | 40 | 33 | 26 | 34.15 | 35 | 33 | 0.913 | -0.131 | 0.70423068 | 0.82094194 | 0 | 0.0718 | 0.0563 | 0.0563 | TRUE | 0.2473 |
| Fam196b | 20 | 19 | 18 | 18 | 26 | 26 | 16 | 20 | 19 | 18 | 29 | 26 | 21.35 | 18 | 24 | 1.326 | 0.407 | 0.33619639 | 0.51049134 | 0 | 0.1135 | 0.0816 | 0.0816 | TRUE | 0.3228 |
| Fam198a | 6 | 2 | 3 | 6 | 3 | 4 | 5 | 2 | 3 | 6 | 3 | 4 | 3.89 | 3 | 4 | 1.31 | 0.389 | 0.68400036 | NA | 0 | 0.612 | 0.4011 | 0.4011 | TRUE | 0.2291 |
| Fam198b | 471 | 332 | 366 | 289 | 252 | 277 | 366 | 353 | 395 | 288 | 284 | 273 | 326.54 | 371 | 282 | 0.759 | -0.397 | 0.00058676 | 0.00488177 | 0 | 0.0097 | 0.0065 | 0.0065 | TRUE | 0.0338 |
| Fam199x | 1380 | 969 | 1061 | 1390 | 1157 | 1273 | 1072 | 1031 | 1145 | 1384 | 1305 | 1257 | 1198.8 | 1083 | 1315 | 1.215 | 0.281 | 0.00026468 | 0.00264547 | 0.0017 | 0.0044 | 0.0034 | 0.0034 | TRUE | 0.0293 |
| Fam19a1 | 6 | 6 | 1 | 1 | 0 | 1 | 5 | 6 | 1 | 1 | 0 | 1 | 2.35 | 4 | 1 | 0.167 | -2.578 | 0.09031141 | NA | 0 | 1.0107 | 0.7897 | 0.7897 | TRUE | 0.2051 |
| Fam19a2 | 1 | 2 | 1 | 0 | 0 | 1 | 1 | 2 | 1 | 0 | 0 | 1 | 0.83 | 1 | 0 | 0.282 | -1.824 | 0.39899776 | NA | 0 | 2.8636 | 2.0027 | 2.0027 | TRUE | 0.3835 |
| Fam19a3 | 20 | 22 | 8 | 31 | 19 | 30 | 16 | 23 | 9 | 31 | 21 | 30 | 21.58 | 16 | 27 | 1.726 | 0.788 | 0.08183499 | 0.19228464 | 0.0514 | 0.1123 | 0.0983 | 0.0983 | TRUE | 0.3432 |
| Fam19a4 | 225 | 157 | 122 | 230 | 169 | 189 | 175 | 167 | 132 | 229 | 191 | 187 | 179.92 | 158 | 202 | 1.277 | 0.352 | 0.03464132 | 0.10308547 | 0.0116 | 0.0156 | 0.0144 | 0.0144 | TRUE | 0.2132 |
| Fam19a5 | 532 | 397 | 348 | 350 | 267 | 342 | 413 | 422 | 375 | 348 | 301 | 338 | 366.36 | 403 | 329 | 0.816 | -0.294 | 0.01045282 | 0.04264977 | 0.0015 | 0.0089 | 0.0067 | 0.0067 | TRUE | 0.06 |
| Fam203a | 1153 | 867 | 649 | 764 | 730 | 821 | 895 | 922 | 700 | 760 | 824 | 810 | 818.74 | 839 | 798 | 0.95 | -0.074 | 0.48654019 | 0.65247878 | 0.0106 | 0.0053 | 0.007 | 0.007 | TRUE | 0.2443 |
| Fam207a | 2375 | 1722 | 1505 | 1544 | 1398 | 1595 | 1845 | 1832 | 1624 | 1537 | 1577 | 1575 | 1664.77 | 1767 | 1563 | 0.884 | -0.178 | 0.01412747 | 0.05339076 | 0.0021 | 0.0039 | 0.0032 | 0.0032 | TRUE | 0.0594 |
| Fam209 | 0 | 0 | 0 | 0 | 0 | 0 | 0 | 0 | 0 | 0 | 0 | 0 | 0 | 0 | 0 | NA | NA | NA | NA | NA | NA | NA | NA | NA | NA |
| Fam20a | 22 | 25 | 23 | 26 | 22 | 33 | 17 | 27 | 25 | 26 | 25 | 33 | 25.3 | 23 | 28 | 1.23 | 0.299 | 0.44422622 | 0.6151255 | 0 | 0.0961 | 0.07 | 0.07 | TRUE | 0.3521 |
| Fam20b | 2995 | 2091 | 2236 | 2334 | 1992 | 2423 | 2326 | 2224 | 2413 | 2323 | 2247 | 2392 | 2320.87 | 2321 | 2321 | 1 | 0 | 0.99749236 | 0.99855014 | 8.00E-04 | 0.0034 | 0.0024 | 0.0024 | TRUE | 0.0157 |
| Fam20c | 127 | 113 | 117 | 130 | 130 | 118 | 99 | 120 | 126 | 129 | 147 | 116 | 122.94 | 115 | 131 | 1.141 | 0.191 | 0.3136531 | 0.48713345 | 0.0075 | 0.0217 | 0.0176 | 0.0176 | TRUE | 0.1754 |
| Fam21 | 10660 | 7691 | 7152 | 7460 | 6455 | 7275 | 8279 | 8181 | 7717 | 7426 | 7282 | 7182 | 7677.68 | 8059 | 7297 | 0.905 | -0.144 | 0.00594465 | 0.02804346 | 6.00E-04 | 0.0027 | 0.0018 | 0.0018 | TRUE | 0.0167 |
| Fam210a | 3626 | 2565 | 2811 | 2986 | 2577 | 3040 | 2816 | 2728 | 3033 | 2972 | 2907 | 3001 | 2909.65 | 2859 | 2960 | 1.036 | 0.051 | 0.41661389 | 0.58939545 | 0.0011 | 0.0032 | 0.0024 | 0.0024 | TRUE | 0.0337 |
| Fam210b | 903 | 686 | 646 | 733 | 681 | 750 | 701 | 730 | 697 | 730 | 768 | 740 | 727.71 | 709 | 746 | 1.052 | 0.073 | 0.37877997 | 0.55362533 | 0 | 0.0057 | 0.0035 | 0.0035 | TRUE | 0.0078 |
| Fam211a | 801 | 601 | 624 | 608 | 556 | 583 | 622 | 639 | 673 | 605 | 627 | 576 | 623.77 | 645 | 603 | 0.935 | -0.097 | 0.28137831 | 0.45294967 | 0 | 0.0062 | 0.0042 | 0.0042 | TRUE | 0.0183 |
| Fam211b | 50 | 45 | 41 | 53 | 53 | 60 | 39 | 48 | 44 | 53 | 60 | 59 | 50.45 | 44 | 57 | 1.317 | 0.397 | 0.14350942 | 0.28647768 | 0 | 0.0494 | 0.033 | 0.033 | TRUE | 0.0864 |
| Fam212a | 74 | 57 | 56 | 63 | 59 | 77 | 57 | 61 | 60 | 63 | 67 | 76 | 63.97 | 59 | 69 | 1.152 | 0.204 | 0.39687099 | 0.57027014 | 0 | 0.0395 | 0.0262 | 0.0262 | TRUE | 0.0867 |
| Fam212b | 83 | 34 | 20 | 33 | 25 | 20 | 64 | 36 | 22 | 33 | 28 | 20 | 33.83 | 41 | 27 | 0.653 | -0.615 | 0.14363202 | 0.28656232 | 0.153 | 0.0725 | 0.0966 | 0.0966 | TRUE | 0.3506 |
| Fam213a | 1626 | 1215 | 1450 | 1552 | 1388 | 1679 | 1263 | 1292 | 1565 | 1545 | 1566 | 1657 | 1481.31 | 1373 | 1589 | 1.159 | 0.213 | 0.01711656 | 0.06170509 | 0.007 | 0.004 | 0.005 | 0.005 | TRUE | 0.1767 |
| Fam213b | 580 | 454 | 456 | 542 | 441 | 496 | 450 | 483 | 492 | 540 | 497 | 490 | 492 | 475 | 509 | 1.073 | 0.102 | 0.30773226 | 0.48028142 | 0 | 0.0072 | 0.0051 | 0.0051 | TRUE | 0.0328 |
| Fam214a | 587 | 434 | 409 | 434 | 371 | 447 | 456 | 462 | 441 | 432 | 419 | 441 | 441.77 | 453 | 431 | 0.951 | -0.073 | 0.46455604 | 0.63371271 | 0 | 0.0078 | 0.0049 | 0.0049 | TRUE | 0.0067 |
| Fam214b | 505 | 377 | 373 | 434 | 357 | 440 | 392 | 401 | 402 | 432 | 403 | 434 | 410.8 | 398 | 423 | 1.063 | 0.088 | 0.3951452 | 0.56864784 | 0 | 0.0082 | 0.0052 | 0.0052 | TRUE | 0.0196 |
| Fam216a | 1224 | 848 | 946 | 974 | 808 | 898 | 951 | 902 | 1021 | 970 | 912 | 886 | 940.14 | 958 | 923 | 0.963 | -0.054 | 0.51734874 | 0.67945103 | 0.0019 | 0.0049 | 0.0039 | 0.0039 | TRUE | 0.0379 |
| Fam216b | 0 | 0 | 0 | 0 | 0 | 0 | 0 | 0 | 0 | 0 | 0 | 0 | 0 | 0 | 0 | NA | NA | NA | NA | NA | NA | NA | NA | NA | NA |
| Fam217a | 15 | 12 | 10 | 18 | 17 | 14 | 12 | 13 | 11 | 18 | 19 | 14 | 14.35 | 12 | 17 | 1.443 | 0.529 | 0.29107335 | NA | 0 | 0.1676 | 0.1096 | 0.1096 | TRUE | 0.1336 |
| Fam217b | 359 | 277 | 332 | 334 | 326 | 353 | 279 | 295 | 358 | 332 | 368 | 348 | 330.06 | 311 | 349 | 1.128 | 0.174 | 0.17473117 | 0.32804842 | 0.007 | 0.0096 | 0.0088 | 0.0088 | TRUE | 0.2001 |
| Fam219a | 230 | 161 | 132 | 124 | 143 | 141 | 179 | 171 | 142 | 123 | 161 | 139 | 152.71 | 164 | 141 | 0.857 | -0.222 | 0.2034347 | 0.36473811 | 0.009 | 0.0179 | 0.0154 | 0.0154 | TRUE | 0.1527 |
| Fam219b | 1017 | 790 | 861 | 951 | 883 | 930 | 790 | 840 | 929 | 947 | 996 | 918 | 903.33 | 853 | 954 | 1.119 | 0.163 | 0.0630062 | 0.15893886 | 0.0031 | 0.0051 | 0.0044 | 0.0044 | TRUE | 0.0716 |
| Fam220a | 1465 | 1083 | 1053 | 1212 | 1090 | 1271 | 1138 | 1152 | 1136 | 1206 | 1230 | 1255 | 1186.11 | 1142 | 1230 | 1.077 | 0.108 | 0.11983059 | 0.25153801 | 0 | 0.0044 | 0.0026 | 0.0026 | TRUE | 0.0037 |
| Fam221a | 364 | 242 | 283 | 350 | 283 | 355 | 283 | 257 | 305 | 348 | 319 | 350 | 310.59 | 282 | 339 | 1.205 | 0.269 | 0.0272189 | 0.08625771 | 9.00E-04 | 0.0101 | 0.0075 | 0.0075 | TRUE | 0.0604 |
| Fam221b | 6 | 4 | 6 | 8 | 6 | 7 | 5 | 4 | 6 | 8 | 7 | 7 | 6.17 | 5 | 7 | 1.414 | 0.5 | 0.50781918 | NA | 0 | 0.3864 | 0.2448 | 0.2448 | TRUE | 0.097 |
| Fam222a | 187 | 146 | 109 | 120 | 121 | 107 | 145 | 155 | 118 | 119 | 137 | 106 | 129.95 | 139 | 121 | 0.862 | -0.215 | 0.25342733 | 0.42246755 | 0.0098 | 0.0207 | 0.0177 | 0.0177 | TRUE | 0.1827 |
| Fam222b | 3299 | 2240 | 1816 | 1957 | 1747 | 2019 | 2562 | 2383 | 1959 | 1948 | 1971 | 1993 | 2136.03 | 2301 | 1971 | 0.856 | -0.225 | 0.012975 | 0.05009498 | 0.0084 | 0.0035 | 0.0054 | 0.0054 | TRUE | 0.2034 |
| Fam227a | 44 | 42 | 33 | 41 | 36 | 51 | 34 | 45 | 36 | 41 | 41 | 50 | 41.04 | 38 | 44 | 1.157 | 0.211 | 0.48916551 | 0.65489346 | 0 | 0.0602 | 0.0424 | 0.0424 | TRUE | 0.1579 |
| Fam227b | 1 | 0 | 0 | 0 | 0 | 0 | 1 | 0 | 0 | 0 | 0 | 0 | 0.13 | 0 | 0 | 0.552 | -0.857 | 0.8336548 | NA | 0 | 18.3121 | 10 | 10 | TRUE | 0.2566 |
| Fam228a | 11 | 10 | 10 | 20 | 15 | 10 | 9 | 11 | 11 | 20 | 17 | 10 | 12.78 | 10 | 16 | 1.566 | 0.647 | 0.24001068 | NA | 0 | 0.1879 | 0.1366 | 0.1366 | TRUE | 0.4976 |
| Fam228b | 231 | 200 | 164 | 226 | 165 | 204 | 179 | 213 | 177 | 225 | 186 | 201 | 196.93 | 190 | 204 | 1.079 | 0.11 | 0.47048043 | 0.63958271 | 0.0046 | 0.0145 | 0.0116 | 0.0116 | TRUE | 0.1176 |
| Fam229a | 8 | 8 | 4 | 9 | 4 | 7 | 6 | 9 | 4 | 9 | 5 | 7 | 6.57 | 6 | 7 | 1.076 | 0.106 | 0.88683349 | NA | 0 | 0.3632 | 0.2502 | 0.2502 | TRUE | 0.1956 |
| Fam229b | 160 | 103 | 85 | 178 | 116 | 153 | 124 | 110 | 92 | 177 | 131 | 151 | 130.77 | 109 | 153 | 1.405 | 0.491 | 0.0109524 | 0.04413998 | 0.0152 | 0.0206 | 0.019 | 0.019 | TRUE | 0.1966 |
| Fam24a | 0 | 0 | 0 | 0 | 0 | 0 | 0 | 0 | 0 | 0 | 0 | 0 | 0 | 0 | 0 | NA | NA | NA | NA | NA | NA | NA | NA | NA | NA |
| Fam25c | 1052 | 743 | 709 | 849 | 711 | 823 | 817 | 790 | 765 | 845 | 802 | 812 | 805.33 | 791 | 820 | 1.036 | 0.051 | 0.52632652 | 0.68732617 | 0 | 0.0054 | 0.0034 | 0.0034 | TRUE | 0.0106 |
| Fam26d | 2 | 2 | 2 | 2 | 2 | 0 | 2 | 2 | 2 | 2 | 2 | 0 | 1.68 | 2 | 1 | 0.724 | -0.466 | 0.75415884 | NA | 0 | 1.4124 | 0.9761 | 0.9761 | TRUE | 0.5238 |
| Fam26e | 184 | 116 | 94 | 105 | 81 | 98 | 143 | 123 | 101 | 105 | 91 | 97 | 110.06 | 122 | 98 | 0.793 | -0.335 | 0.09459417 | 0.21228363 | 0.0099 | 0.024 | 0.0198 | 0.0198 | TRUE | 0.176 |
| Fam26f | 149 | 101 | 65 | 82 | 67 | 75 | 116 | 107 | 70 | 82 | 76 | 74 | 87.42 | 98 | 77 | 0.785 | -0.35 | 0.13726433 | 0.27759363 | 0.0254 | 0.0295 | 0.0283 | 0.0283 | TRUE | 0.5691 |
| Fam32a | 4356 | 3199 | 2704 | 2590 | 2264 | 2759 | 3383 | 3403 | 2918 | 2578 | 2554 | 2724 | 2926.52 | 3235 | 2619 | 0.809 | -0.305 | 3.19E-05 | 0.00048948 | 0.0037 | 0.0032 | 0.0035 | 0.0035 | TRUE | 0.0884 |
| Fam35a | 1813 | 1200 | 1319 | 1514 | 1346 | 1462 | 1408 | 1276 | 1423 | 1507 | 1518 | 1443 | 1429.4 | 1369 | 1489 | 1.087 | 0.121 | 0.09700371 | 0.21653507 | 0.0014 | 0.0041 | 0.0031 | 0.0031 | TRUE | 0.0413 |
| Fam3a | 565 | 390 | 453 | 478 | 447 | 454 | 439 | 415 | 489 | 476 | 504 | 448 | 461.78 | 448 | 476 | 1.064 | 0.089 | 0.40441547 | 0.57722147 | 0.0028 | 0.0076 | 0.0061 | 0.0061 | TRUE | 0.0721 |
| Fam3b | 160 | 89 | 105 | 81 | 82 | 75 | 124 | 95 | 113 | 81 | 93 | 74 | 96.57 | 111 | 83 | 0.74 | -0.435 | 0.03819849 | 0.11069971 | 0 | 0.027 | 0.0211 | 0.0211 | TRUE | 0.157 |
| Fam3c | 2820 | 2146 | 2253 | 2004 | 1751 | 1992 | 2190 | 2283 | 2431 | 1995 | 1975 | 1966 | 2140.05 | 2301 | 1979 | 0.861 | -0.217 | 0.00081857 | 0.00624654 | 0.0011 | 0.0035 | 0.0025 | 0.0025 | TRUE | 0.0293 |
| Fam43a | 1244 | 862 | 773 | 850 | 748 | 731 | 966 | 917 | 834 | 846 | 844 | 722 | 854.77 | 906 | 804 | 0.886 | -0.175 | 0.06657057 | 0.16516111 | 0.0056 | 0.0052 | 0.0053 | 0.0053 | TRUE | 0.0962 |
| Fam43b | 108 | 83 | 99 | 120 | 110 | 112 | 84 | 88 | 107 | 119 | 124 | 111 | 105.51 | 93 | 118 | 1.273 | 0.349 | 0.07505286 | 0.18035146 | 0 | 0.0249 | 0.018 | 0.018 | TRUE | 0.1569 |
| Fam45a | 2520 | 1684 | 1810 | 2076 | 1679 | 2001 | 1957 | 1791 | 1953 | 2066 | 1894 | 1975 | 1939.54 | 1900 | 1978 | 1.041 | 0.058 | 0.40022305 | 0.573415 | 0.0017 | 0.0036 | 0.0029 | 0.0029 | TRUE | 0.0301 |
| Fam46a | 217 | 154 | 159 | 147 | 154 | 163 | 169 | 164 | 172 | 146 | 174 | 161 | 164.14 | 168 | 160 | 0.953 | -0.069 | 0.65761675 | 0.78781677 | 0 | 0.0169 | 0.0114 | 0.0114 | TRUE | 0.0619 |

| | | | | | | | | | | | | | | | | | | | | | | | | | |
|---|---|---|---|---|---|---|---|---|---|---|---|---|---|---|---|---|---|---|---|---|---|---|---|---|---|
| Fam46b | 293 | 220 | 206 | 260 | 229 | 262 | 228 | 234 | 222 | 259 | 258 | 259 | 243.27 | 228 | 259 | 1.134 | 0.182 | 0.15284916 | 0.2996397 | 0 | 0.0122 | 0.0076 | 0.0076 | TRUE | 0.0055 |
| Fam46c | 250 | 207 | 225 | 232 | 219 | 243 | 194 | 220 | 243 | 231 | 247 | 240 | 229.16 | 219 | 239 | 1.096 | 0.132 | 0.34527672 | 0.51936857 | 0.0032 | 0.0128 | 0.0098 | 0.0098 | TRUE | 0.119 |
| Fam46d | 9 | 5 | 4 | 5 | 8 | 2 | 7 | 5 | 4 | 5 | 9 | 2 | 5.43 | 5 | 5 | 0.941 | -0.088 | 0.91792509 | NA | 0 | 0.4386 | 0.3355 | 0.3355 | TRUE | 0.2746 |
| Fam47c | 0 | 0 | 0 | 0 | 0 | 0 | 0 | 0 | 0 | 0 | 0 | 0 | 0 | 0 | 0 | NA | NA | NA | NA | NA | NA | NA | NA | NA | NA |
| Fam47e | 0 | 1 | 3 | 0 | 0 | 0 | 0 | 1 | 3 | 0 | 0 | 0 | 0.72 | 1 | 0 | 0.135 | -2.888 | 0.27208013 | NA | 7.0304 | 3.309 | 3.6393 | 3.6393 | TRUE | 0.1576 |
| Fam49a | 737 | 538 | 465 | 622 | 529 | 551 | 572 | 572 | 502 | 619 | 597 | 544 | 567.7 | 549 | 587 | 1.067 | 0.094 | 0.34897562 | 0.52355501 | 0.0032 | 0.0066 | 0.0055 | 0.0055 | TRUE | 0.0638 |
| Fam49b | 8612 | 6285 | 6667 | 7412 | 6463 | 7379 | 6689 | 6685 | 7194 | 7378 | 7291 | 7284 | 7086.75 | 6856 | 7318 | 1.068 | 0.094 | 0.07500956 | 0.18030806 | 7.00E-04 | 0.0028 | 0.0019 | 0.0019 | TRUE | 0.0225 |
| Fam50a | 3164 | 2328 | 2215 | 2263 | 1941 | 2367 | 2457 | 2476 | 2390 | 2253 | 2190 | 2337 | 2350.4 | 2441 | 2260 | 0.926 | -0.111 | 0.06342999 | 0.15972602 | 2.00E-04 | 0.0034 | 0.0022 | 0.0022 | TRUE | 0.0108 |
| Fam50b | 2 | 0 | 2 | 1 | 2 | 0 | 2 | 0 | 2 | 1 | 2 | 0 | 1.16 | 1 | 1 | 0.854 | -0.227 | 0.90319719 | NA | 0 | 2.0447 | 1.641 | 1.641 | TRUE | 0.2593 |
| Fam53a | 2054 | 1361 | 1338 | 1342 | 1216 | 1383 | 1595 | 1448 | 1444 | 1336 | 1372 | 1365 | 1426.58 | 1496 | 1358 | 0.907 | -0.141 | 0.04876734 | 0.13233939 | 0.0013 | 0.0041 | 0.003 | 0.003 | TRUE | 0.0425 |
| Fam53b | 1059 | 862 | 917 | 1169 | 1041 | 1180 | 822 | 917 | 989 | 1164 | 1174 | 1165 | 1038.61 | 909 | 1168 | 1.286 | 0.363 | 1.90E-05 | 0.00032065 | 0.0033 | 0.0047 | 0.0042 | 0.0042 | TRUE | 0.088 |
| Fam53c | 3411 | 2407 | 1948 | 2003 | 1683 | 2030 | 2649 | 2560 | 2102 | 1994 | 1899 | 2004 | 2201.28 | 2437 | 1966 | 0.806 | -0.311 | 0.000334 | 0.00315315 | 0.007 | 0.0035 | 0.005 | 0.005 | TRUE | 0.1741 |
| Fam57a | 371 | 282 | 297 | 325 | 268 | 326 | 288 | 300 | 320 | 324 | 302 | 322 | 309.37 | 303 | 316 | 1.045 | 0.064 | 0.58641959 | 0.73568816 | 0 | 0.0101 | 0.0068 | 0.0068 | TRUE | 0.0288 |
| Fam57b | 122 | 88 | 110 | 185 | 151 | 160 | 95 | 94 | 119 | 184 | 170 | 158 | 136.58 | 103 | 171 | 1.674 | 0.744 | 3.19E-05 | 0.00048948 | 0.0032 | 0.0198 | 0.0153 | 0.0153 | TRUE | 0.1815 |
| Fam58b | 1402 | 1065 | 1025 | 1147 | 1033 | 1179 | 1089 | 1133 | 1106 | 1142 | 1165 | 1164 | 1133.1 | 1109 | 1157 | 1.043 | 0.061 | 0.38274912 | 0.55658084 | 0 | 0.0045 | 0.0027 | 0.0027 | TRUE | 0.0041 |
| Fam5b | 97 | 64 | 52 | 55 | 47 | 58 | 75 | 68 | 56 | 55 | 53 | 57 | 60.76 | 66 | 55 | 0.824 | -0.28 | 0.26726341 | 0.43772691 | 0 | 0.0414 | 0.0292 | 0.0292 | TRUE | 0.1594 |
| Fam5c | 1 | 0 | 0 | 0 | 0 | 0 | 1 | 0 | 0 | 0 | 0 | 0 | 0.13 | 0 | 0 | 0.552 | -0.857 | 0.8336548 | NA | 0 | 18.3121 | 10 | 10 | TRUE | 0.2566 |
| Fam60a | 1016 | 734 | 836 | 899 | 811 | 931 | 789 | 781 | 902 | 895 | 915 | 919 | 866.78 | 824 | 910 | 1.105 | 0.144 | 0.09260262 | 0.2091938 | 0.0021 | 0.0052 | 0.0041 | 0.0041 | TRUE | 0.0795 |
| Fam63a | 2121 | 1563 | 1645 | 1714 | 1570 | 1808 | 1647 | 1663 | 1775 | 1706 | 1771 | 1785 | 1724.46 | 1695 | 1754 | 1.035 | 0.05 | 0.44771498 | 0.61788255 | 5.00E-04 | 0.0038 | 0.0026 | 0.0026 | TRUE | 0.0202 |
| Fam63b | 961 | 832 | 897 | 1499 | 1292 | 1487 | 746 | 885 | 968 | 1492 | 1458 | 1468 | 1169.45 | 866 | 1473 | 1.704 | 0.769 | 2.57E-16 | 5.04E-14 | 0.007 | 0.0045 | 0.0054 | 0.0054 | TRUE | 0.183 |
| Fam64a | 3142 | 2352 | 2427 | 2571 | 2405 | 2662 | 2440 | 2502 | 2619 | 2559 | 2713 | 2628 | 2576.8 | 2520 | 2633 | 1.045 | 0.064 | 0.29455839 | 0.46650881 | 7.00E-04 | 0.0033 | 0.0023 | 0.0023 | TRUE | 0.0141 |
| Fam65a | 3012 | 2266 | 2189 | 2484 | 2376 | 2519 | 2339 | 2410 | 2362 | 2473 | 2680 | 2487 | 2458.52 | 2370 | 2547 | 1.074 | 0.103 | 0.09419779 | 0.21159343 | 7.00E-04 | 0.0034 | 0.0023 | 0.0023 | TRUE | 0.0253 |
| Fam65b | 74 | 66 | 85 | 78 | 62 | 93 | 57 | 70 | 92 | 78 | 70 | 92 | 76.46 | 73 | 80 | 1.101 | 0.138 | 0.57602199 | 0.72769249 | 0.0248 | 0.0334 | 0.031 | 0.031 | TRUE | 0.4153 |
| Fam65c | 19 | 11 | 10 | 16 | 10 | 12 | 15 | 12 | 11 | 16 | 11 | 12 | 12.72 | 13 | 13 | 1.043 | 0.06 | 0.90975545 | NA | 0 | 0.1888 | 0.126 | 0.126 | TRUE | 0.1622 |
| Fam69a | 658 | 452 | 405 | 457 | 364 | 406 | 511 | 481 | 437 | 455 | 411 | 401 | 449.19 | 476 | 422 | 0.885 | -0.177 | 0.10522674 | 0.22966724 | 0.0033 | 0.0077 | 0.0063 | 0.0063 | TRUE | 0.0596 |
| Fam69b | 645 | 559 | 537 | 645 | 710 | 697 | 501 | 595 | 579 | 642 | 801 | 688 | 634.33 | 558 | 710 | 1.274 | 0.349 | 0.00152905 | 0.00979155 | 0.0092 | 0.0062 | 0.0071 | 0.0071 | TRUE | 0.1448 |
| Fam69c | 7 | 0 | 4 | 7 | 3 | 3 | 5 | 0 | 4 | 7 | 3 | 3 | 3.84 | 3 | 4 | 1.338 | 0.42 | 0.69442281 | NA | 0.3456 | 0.6189 | 0.5627 | 0.5627 | TRUE | 0.9747 |
| Fam71a | 0 | 0 | 0 | 0 | 0 | 0 | 0 | 0 | 0 | 0 | 0 | 0 | 0 | 0 | 0 | NA | NA | NA | NA | NA | NA | NA | NA | NA | NA |
| Fam71b | 2 | 2 | 0 | 1 | 0 | 0 | 2 | 2 | 0 | 1 | 0 | 0 | 0.78 | 1 | 0 | 0.3 | -1.737 | 0.45896026 | NA | 0 | 3.0435 | 2.5735 | 2.5735 | TRUE | 0.3906 |
| Fam71d | 2 | 1 | 1 | 2 | 1 | 0 | 2 | 1 | 1 | 2 | 1 | 0 | 1.14 | 1 | 1 | 0.833 | -0.263 | 0.88205626 | NA | 0 | 2.0891 | 1.3766 | 1.3766 | TRUE | 0.3125 |
| Fam71e1 | 12 | 12 | 15 | 19 | 17 | 12 | 9 | 13 | 16 | 19 | 19 | 12 | 14.7 | 13 | 17 | 1.317 | 0.397 | 0.43957553 | NA | 0 | 0.1636 | 0.1205 | 0.1205 | TRUE | 0.3123 |
| Fam71e2 | 3 | 3 | 2 | 4 | 3 | 3 | 2 | 3 | 2 | 4 | 3 | 3 | 3 | 2 | 3 | 1.35 | 0.433 | 0.68508988 | NA | 0 | 0.7922 | 0.4826 | 0.4826 | TRUE | 0.0475 |
| Fam71f1 | 0 | 0 | 2 | 1 | 0 | 0 | 0 | 0 | 2 | 1 | 0 | 0 | 0.53 | 1 | 0 | 0.529 | -0.919 | 0.75834888 | NA | 0 | 4.512 | 4.7468 | 4.7468 | TRUE | 0.9746 |
| Fam71f2 | 15 | 4 | 8 | 12 | 12 | 19 | 12 | 4 | 9 | 12 | 14 | 19 | 11.46 | 8 | 15 | 1.773 | 0.826 | 0.16525295 | NA | 0 | 0.2092 | 0.1625 | 0.1625 | TRUE | 0.4821 |
| Fam72a | 582 | 368 | 387 | 446 | 338 | 413 | 452 | 391 | 418 | 444 | 381 | 408 | 415.66 | 420 | 411 | 0.977 | -0.034 | 0.75991709 | 0.85731129 | 0.0032 | 0.0081 | 0.0066 | 0.0066 | TRUE | 0.0568 |
| Fam73a | 4137 | 2961 | 3004 | 3556 | 3137 | 3522 | 3213 | 3150 | 3241 | 3540 | 3539 | 3477 | 3359.86 | 3201 | 3519 | 1.099 | 0.136 | 0.01032808 | 0.04228568 | 0 | 0.0031 | 0.0017 | 0.0017 | TRUE | 0.0024 |
| Fam73b | 885 | 686 | 585 | 646 | 567 | 610 | 687 | 730 | 631 | 643 | 640 | 602 | 655.51 | 683 | 628 | 0.92 | -0.121 | 0.19186458 | 0.35018776 | 0.0017 | 0.006 | 0.0046 | 0.0046 | TRUE | 0.0491 |
| Fam76a | 2419 | 1692 | 1777 | 1926 | 1641 | 1907 | 1879 | 1800 | 1917 | 1917 | 1851 | 1883 | 1874.45 | 1865 | 1884 | 1.01 | 0.014 | 0.82162297 | 0.89761195 | 0 | 0.0037 | 0.0023 | 0.0023 | TRUE | 0.0111 |
| Fam76b | 3090 | 2365 | 2430 | 2514 | 2186 | 2625 | 2400 | 2516 | 2622 | 2502 | 2466 | 2591 | 2516.2 | 2513 | 2520 | 1.004 | 0.005 | 0.93075943 | 0.96192556 | 0.001 | 0.0034 | 0.0024 | 0.0024 | TRUE | 0.0192 |
| Fam78a | 637 | 524 | 534 | 652 | 580 | 663 | 495 | 557 | 576 | 649 | 654 | 654 | 597.68 | 543 | 652 | 1.205 | 0.269 | 0.00459052 | 0.02299766 | 0.0016 | 0.0064 | 0.0048 | 0.0048 | TRUE | 0.0752 |
| Fam78b | 29 | 14 | 12 | 8 | 7 | 3 | 23 | 15 | 13 | 8 | 8 | 3 | 11.53 | 17 | 6 | 0.368 | -1.442 | 0.02015213 | NA | 0 | 0.208 | 0.166 | 0.166 | TRUE | 0.5464 |
| Fam81a | 2088 | 1616 | 1802 | 2255 | 2114 | 2412 | 1622 | 1719 | 1944 | 2245 | 2385 | 2381 | 2049.23 | 1762 | 2337 | 1.328 | 0.409 | 1.46E-07 | 5.46E-06 | 0.0043 | 0.0036 | 0.0039 | 0.0039 | TRUE | 0.0988 |
| Fam83a | 35 | 23 | 19 | 20 | 26 | 27 | 27 | 24 | 21 | 20 | 29 | 27 | 24.67 | 24 | 25 | 1.044 | 0.062 | 0.87363732 | 0.92995497 | 0 | 0.0985 | 0.069 | 0.069 | TRUE | 0.2166 |
| Fam83b | 119 | 104 | 96 | 84 | 62 | 71 | 92 | 111 | 104 | 84 | 70 | 70 | 88.38 | 102 | 75 | 0.732 | -0.449 | 0.03421743 | 0.10216624 | 0 | 0.0292 | 0.0208 | 0.0208 | TRUE | 0.1054 |
| Fam83c | 1 | 4 | 0 | 3 | 0 | 3 | 1 | 4 | 0 | 3 | 0 | 3 | 1.83 | 2 | 2 | 1.216 | 0.282 | 0.86003557 | NA | 1.2594 | 1.2977 | 1.2904 | 1.2904 | TRUE | 1.2824 |
| Fam83d | 3049 | 2229 | 2213 | 2397 | 2114 | 2260 | 2368 | 2371 | 2388 | 2386 | 2385 | 2231 | 2354.76 | 2376 | 2334 | 0.982 | -0.026 | 0.66939653 | 0.79684222 | 3.00E-04 | 0.0034 | 0.0022 | 0.0022 | TRUE | 0.0182 |
| Fam83e | 7 | 5 | 4 | 5 | 6 | 4 | 5 | 5 | 4 | 5 | 7 | 4 | 5.13 | 5 | 5 | 1.032 | 0.046 | 0.95581091 | NA | 0 | 0.4646 | 0.2951 | 0.2951 | TRUE | 0.1231 |
| Fam83f | 83 | 63 | 59 | 93 | 71 | 104 | 64 | 67 | 64 | 93 | 80 | 103 | 78.41 | 65 | 92 | 1.414 | 0.5 | 0.02470899 | 0.08070513 | 0 | 0.0327 | 0.0227 | 0.0227 | TRUE | 0.1125 |
| Fam83g | 897 | 682 | 650 | 826 | 707 | 772 | 697 | 725 | 701 | 822 | 798 | 762 | 750.88 | 708 | 794 | 1.122 | 0.166 | 0.04456024 | 0.1238294 | 0 | 0.0056 | 0.0036 | 0.0036 | TRUE | 0.015 |
| Fam83h | 325 | 242 | 189 | 249 | 224 | 211 | 252 | 257 | 204 | 248 | 253 | 208 | 237.1 | 238 | 236 | 0.99 | -0.014 | 0.92496258 | 0.95844943 | 0.009 | 0.0124 | 0.0114 | 0.0114 | TRUE | 0.1656 |
| Fam84a | 127 | 79 | 71 | 77 | 60 | 70 | 99 | 84 | 77 | 77 | 68 | 69 | 78.78 | 87 | 71 | 0.82 | -0.287 | 0.20070401 | 0.36140779 | 0 | 0.0325 | 0.0233 | 0.0233 | TRUE | 0.1616 |
| Fam84b | 91 | 58 | 56 | 62 | 53 | 55 | 71 | 62 | 60 | 62 | 60 | 54 | 61.43 | 64 | 59 | 0.908 | -0.139 | 0.57188324 | 0.72496988 | 0 | 0.041 | 0.0273 | 0.0273 | TRUE | 0.0765 |
| Fam86 | 1027 | 791 | 764 | 783 | 802 | 864 | 798 | 841 | 824 | 779 | 905 | 853 | 833.4 | 821 | 846 | 1.03 | 0.042 | 0.6216845 | 0.76141702 | 0.0019 | 0.0053 | 0.0041 | 0.0041 | TRUE | 0.0561 |
| Fam89a | 427 | 315 | 315 | 385 | 358 | 424 | 332 | 335 | 340 | 383 | 404 | 419 | 368.7 | 336 | 402 | 1.198 | 0.261 | 0.01578136 | 0.05798534 | 0 | 0.0089 | 0.0057 | 0.0057 | TRUE | 0.0195 |
| Fam89b | 515 | 357 | 380 | 363 | 338 | 382 | 400 | 380 | 410 | 361 | 381 | 377 | 384.91 | 397 | 373 | 0.941 | -0.088 | 0.40610971 | 0.5788896 | 0 | 0.0086 | 0.0055 | 0.0055 | TRUE | 0.0151 |
| Fam92a | 1124 | 817 | 911 | 844 | 680 | 809 | 873 | 869 | 983 | 840 | 767 | 799 | 855.13 | 908 | 802 | 0.884 | -0.178 | 0.04013709 | 0.11471294 | 0.0023 | 0.0052 | 0.0042 | 0.0042 | TRUE | 0.0601 |
| Fam92b | 7 | 4 | 7 | 9 | 5 | 8 | 5 | 4 | 8 | 9 | 6 | 8 | 6.62 | 6 | 8 | 1.313 | 0.393 | 0.59328213 | NA | 0 | 0.3603 | 0.2381 | 0.2381 | TRUE | 0.1511 |
| Fam96a | 2178 | 1523 | 1429 | 1631 | 1380 | 1518 | 1692 | 1620 | 1542 | 1623 | 1557 | 1499 | 1588.7 | 1618 | 1560 | 0.963 | -0.054 | 0.44240091 | 0.61332152 | 0.0013 | 0.0039 | 0.0029 | 0.0029 | TRUE | 0.0202 |
| Fam96b | 3246 | 2198 | 2137 | 2228 | 1884 | 2191 | 2521 | 2338 | 2306 | 2218 | 2125 | 2163 | 2278.46 | 2388 | 2169 | 0.907 | -0.14 | 0.02839012 | 0.08892833 | 0.0011 | 0.0035 | 0.0025 | 0.0025 | TRUE | 0.0295 |
| Fam98a | 5068 | 3711 | 3816 | 3959 | 3490 | 4015 | 3936 | 3947 | 4117 | 3941 | 3937 | 3963 | 3973.69 | 4000 | 3947 | 0.987 | -0.019 | 0.71831048 | 0.83078262 | 0 | 0.003 | 0.0017 | 0.0017 | TRUE | 0.0079 |
| Fam98b | 2494 | 1705 | 1541 | 1475 | 1279 | 1562 | 1937 | 1814 | 1663 | 1468 | 1443 | 1542 | 1644.38 | 1805 | 1484 | 0.822 | -0.283 | 0.00020151 | 0.00214459 | 0.003 | 0.0039 | 0.0036 | 0.0036 | TRUE | 0.0567 |
| Fam98c | 957 | 743 | 593 | 812 | 706 | 750 | 743 | 790 | 640 | 808 | 796 | 740 | 753.08 | 724 | 781 | 1.078 | 0.108 | 0.26479923 | 0.43523176 | 0.0053 | 0.0056 | 0.0055 | 0.0055 | TRUE | 0.1192 |
| Fan1 | 695 | 518 | 451 | 522 | 426 | 474 | 540 | 551 | 487 | 520 | 481 | 468 | 507.58 | 526 | 490 | 0.93 | -0.104 | 0.30092414 | 0.47332548 | 0.0016 | 0.0071 | 0.0054 | 0.0054 | TRUE | 0.0477 |
| Fanca | 7037 | 5485 | 5292 | 5799 | 5237 | 6042 | 5465 | 5834 | 5710 | 5772 | 5908 | 5964 | 5775.71 | 5670 | 5881 | 1.038 | 0.053 | 0.31071926 | 0.48365521 | 5.00E-04 | 0.0028 | 0.0018 | 0.0018 | TRUE | 0.0124 |
| Fancb | 1061 | 719 | 655 | 747 | 598 | 772 | 824 | 765 | 707 | 744 | 675 | 762 | 745.97 | 765 | 727 | 0.949 | -0.076 | 0.41845225 | 0.59106018 | 0.0037 | 0.0056 | 0.005 | 0.005 | TRUE | 0.0561 |
| Fancc | 995 | 793 | 791 | 864 | 816 | 884 | 773 | 844 | 853 | 860 | 921 | 873 | 853.83 | 823 | 885 | 1.075 | 0.105 | 0.20488513 | 0.36644055 | 0.0011 | 0.0052 | 0.0038 | 0.0038 | TRUE | 0.0362 |
| Fancd2 | 8427 | 6419 | 6773 | 7372 | 6429 | 7253 | 6545 | 6828 | 7308 | 7338 | 7253 | 7160 | 7071.87 | 6894 | 7250 | 1.052 | 0.073 | 0.20100046 | 0.36180686 | 0.0014 | 0.0028 | 0.0022 | 0.0022 | TRUE | 0.0337 |
| Fance | 2439 | 1863 | 1864 | 2073 | 1959 | 2162 | 1894 | 1982 | 2011 | 2063 | 2210 | 2134 | 2049.13 | 1962 | 2136 | 1.089 | 0.123 | 0.05332926 | 0.14101745 | 6.00E-04 | 0.0036 | 0.0024 | 0.0024 | TRUE | 0.0116 |
| Fancf | 504 | 407 | 324 | 300 | 293 | 303 | 391 | 433 | 350 | 299 | 331 | 299 | 350.37 | 391 | 310 | 0.79 | -0.34 | 0.00518991 | 0.02525685 | 0.0045 | 0.0092 | 0.0078 | 0.0078 | TRUE | 0.0947 |

| | | | | | | | | | | | | | | | | | | | | | | | | | |
|---|---|---|---|---|---|---|---|---|---|---|---|---|---|---|---|---|---|---|---|---|---|---|---|---|---|
| Fancg | 1103 | 800 | 804 | 787 | 728 | 839 | 857 | 851 | 868 | 783 | 821 | 828 | 834.66 | 859 | 811 | 0.945 | -0.082 | 0.29525241 | 0.46705464 | 0 | 0.0053 | 0.0032 | 0.0032 | TRUE | 0.0107 |
| Fanci | 5223 | 4110 | 4312 | 4735 | 4103 | 4509 | 4056 | 4372 | 4653 | 4713 | 4629 | 4451 | 4478.96 | 4360 | 4598 | 1.055 | 0.077 | 0.23359906 | 0.3997643 | 0.0024 | 0.003 | 0.0028 | 0.0028 | TRUE | 0.046 |
| Fancl | 3061 | 2179 | 2028 | 1950 | 1791 | 2076 | 2377 | 2318 | 2188 | 1941 | 2020 | 2049 | 2149.02 | 2294 | 2003 | 0.873 | -0.196 | 0.00215073 | 0.01275006 | 9.00E-04 | 0.0035 | 0.0025 | 0.0025 | TRUE | 0.0197 |
| Fancm | 6074 | 4243 | 3637 | 3598 | 3133 | 3568 | 4717 | 4513 | 3924 | 3581 | 3534 | 3522 | 3965.48 | 4385 | 3546 | 0.808 | -0.307 | 2.34E-05 | 0.00037797 | 0.0041 | 0.003 | 0.0035 | 0.0035 | TRUE | 0.1023 |
| Fank1 | 332 | 269 | 308 | 253 | 225 | 209 | 258 | 286 | 332 | 252 | 254 | 206 | 264.71 | 292 | 237 | 0.814 | -0.297 | 0.04107125 | 0.11659015 | 0.0113 | 0.0114 | 0.0114 | 0.0114 | TRUE | 0.1621 |
| Fap | 0 | 1 | 1 | 0 | 0 | 0 | 0 | 1 | 1 | 0 | 0 | 0 | 0.36 | 1 | 0 | 0.271 | -1.881 | 0.5755355 | NA | 0 | 6.6391 | 6.4428 | 6.4428 | TRUE | 0.3513 |
| Far1 | 6480 | 4797 | 4938 | 5514 | 4668 | 5316 | 5033 | 5103 | 5328 | 5489 | 5266 | 5248 | 5244.27 | 5155 | 5334 | 1.035 | 0.05 | 0.35144396 | 0.52609837 | 5.00E-04 | 0.0029 | 0.0019 | 0.0019 | TRUE | 0.0105 |
| Far2 | 18 | 11 | 8 | 7 | 5 | 3 | 14 | 12 | 9 | 7 | 6 | 3 | 8.31 | 12 | 5 | 0.45 | -1.153 | 0.09611715 | NA | 0 | 0.2875 | 0.2043 | 0.2043 | TRUE | 0.3121 |
| Farp1 | 2295 | 1796 | 1703 | 1934 | 1829 | 2036 | 1782 | 1910 | 1838 | 1925 | 2063 | 2010 | 1921.43 | 1843 | 1999 | 1.085 | 0.118 | 0.07011592 | 0.17162869 | 7.00E-04 | 0.0037 | 0.0025 | 0.0025 | TRUE | 0.0128 |
| Farp2 | 156 | 118 | 106 | 176 | 115 | 168 | 121 | 126 | 114 | 175 | 130 | 166 | 138.64 | 120 | 157 | 1.306 | 0.386 | 0.03123043 | 0.09539999 | 0.0063 | 0.0195 | 0.0158 | 0.0158 | TRUE | 0.2315 |
| Fars2 | 1257 | 902 | 896 | 1061 | 911 | 1010 | 976 | 959 | 967 | 1056 | 1028 | 997 | 997.22 | 967 | 1027 | 1.061 | 0.086 | 0.24621144 | 0.41487681 | 0 | 0.0048 | 0.0029 | 0.0029 | TRUE | 0.0079 |
| Farsa | 7858 | 5900 | 5340 | 5678 | 5239 | 6171 | 6103 | 6276 | 5762 | 5652 | 5910 | 6092 | 5965.71 | 6047 | 5885 | 0.973 | -0.039 | 0.49722919 | 0.66159903 | 0.0014 | 0.0028 | 0.0023 | 0.0023 | TRUE | 0.0205 |
| Farsb | 9283 | 6570 | 6654 | 7168 | 6247 | 7367 | 7210 | 6988 | 7180 | 7135 | 7047 | 7272 | 7138.73 | 7126 | 7151 | 1.004 | 0.005 | 0.91547703 | 0.95336344 | 1.00E-04 | 0.0028 | 0.0015 | 0.0015 | TRUE | 0.0035 |
| Fas | 48 | 31 | 45 | 39 | 27 | 37 | 37 | 33 | 49 | 39 | 30 | 37 | 37.44 | 40 | 35 | 0.895 | -0.161 | 0.61847417 | 0.75908957 | 0 | 0.0657 | 0.0481 | 0.0481 | TRUE | 0.2707 |
| Fasl | 0 | 0 | 0 | 0 | 0 | 0 | 0 | 0 | 0 | 0 | 0 | 0 | 0 | 0 | 0 | NA | NA | NA | NA | NA | NA | NA | NA | NA | NA |
| Fasn | 62124 | 43193 | 41106 | 46489 | 40103 | 45623 | 48249 | 45944 | 44353 | 46275 | 45240 | 45037 | 45849.72 | 46182 | 45517 | 0.986 | -0.021 | 0.67487804 | 0.80032886 | 9.00E-04 | 0.0025 | 0.0018 | 0.0018 | TRUE | 0.0188 |
| Fastk | 2818 | 2329 | 2321 | 2395 | 2142 | 2436 | 2189 | 2477 | 2504 | 2384 | 2416 | 2405 | 2395.9 | 2390 | 2402 | 1.006 | 0.008 | 0.90345564 | 0.9465688 | 0.0025 | 0.0034 | 0.003 | 0.003 | TRUE | 0.0678 |
| Fastkd1 | 3464 | 2540 | 2617 | 3046 | 2670 | 3090 | 2690 | 2702 | 2824 | 3032 | 3012 | 3050 | 2885.03 | 2739 | 3031 | 1.107 | 0.147 | 0.00859853 | 0.03698427 | 0 | 0.0032 | 0.0019 | 0.0019 | TRUE | 0.0088 |
| Fastkd2 | 2800 | 1989 | 1964 | 1992 | 1732 | 2050 | 2175 | 2116 | 2119 | 1983 | 1954 | 2024 | 2061.64 | 2137 | 1987 | 0.93 | -0.105 | 0.07815313 | 0.18592588 | 0 | 0.0036 | 0.0021 | 0.0021 | TRUE | 0.0032 |
| Fastkd3 | 1231 | 899 | 774 | 696 | 666 | 765 | 956 | 956 | 835 | 693 | 751 | 755 | 824.46 | 916 | 733 | 0.8 | -0.323 | 0.00030419 | 0.00294076 | 0.003 | 0.0053 | 0.0045 | 0.0045 | TRUE | 0.0691 |
| Fastkd5 | 1259 | 880 | 787 | 801 | 730 | 872 | 978 | 936 | 849 | 797 | 824 | 861 | 874.11 | 921 | 827 | 0.897 | -0.157 | 0.0677863 | 0.16750067 | 0.0023 | 0.0051 | 0.0042 | 0.0042 | TRUE | 0.0547 |
| Fat1 | 13496 | 9133 | 8226 | 10034 | 9166 | 10599 | 10482 | 9715 | 8876 | 9988 | 10340 | 10463 | 9977.18 | 9691 | 10264 | 1.059 | 0.082 | 0.21279147 | 0.37511451 | 0.0036 | 0.0027 | 0.0031 | 0.0031 | TRUE | 0.066 |
| Fat2 | 75 | 54 | 39 | 54 | 47 | 44 | 58 | 57 | 42 | 54 | 53 | 43 | 51.33 | 52 | 50 | 0.947 | -0.078 | 0.77862512 | 0.86937652 | 0 | 0.0486 | 0.0362 | 0.0362 | TRUE | 0.2371 |
| Fat3 | 76 | 56 | 32 | 52 | 39 | 37 | 59 | 60 | 35 | 52 | 44 | 37 | 47.57 | 51 | 44 | 0.86 | -0.218 | 0.48163712 | 0.64849796 | 0.0374 | 0.0523 | 0.0482 | 0.0482 | TRUE | 0.6162 |
| Fat4 | 2 | 1 | 1 | 1 | 3 | 0 | 2 | 1 | 1 | 1 | 3 | 0 | 1.35 | 1 | 1 | 1.148 | 0.199 | 0.90546773 | NA | 0 | 1.7633 | 1.2833 | 1.2833 | TRUE | 0.567 |
| Fate1 | 0 | 0 | 0 | 0 | 0 | 0 | 0 | 0 | 0 | 0 | 0 | 0 | 0 | 0 | 0 | NA | NA | NA | NA | NA | NA | NA | NA | NA | NA |
| Fau | 16681 | 12639 | 13675 | 13941 | 12732 | 14761 | 12955 | 13444 | 14755 | 13877 | 14363 | 14571 | 13994.31 | 13718 | 14270 | 1.04 | 0.057 | 0.34407905 | 0.51833072 | 0.0024 | 0.0026 | 0.0026 | 0.0026 | TRUE | 0.0534 |
| Faxc | 987 | 790 | 827 | 903 | 750 | 958 | 767 | 840 | 892 | 899 | 846 | 946 | 864.97 | 833 | 897 | 1.079 | 0.11 | 0.21808455 | 0.38110802 | 0.0034 | 0.0052 | 0.0046 | 0.0046 | TRUE | 0.061 |
| Fbf1 | 2549 | 2013 | 1814 | 2016 | 1848 | 2016 | 1980 | 2141 | 1957 | 2007 | 2085 | 1990 | 2026.63 | 2026 | 2027 | 1.001 | 0.001 | 0.98806725 | 0.99383851 | 0.001 | 0.0036 | 0.0026 | 0.0026 | TRUE | 0.0294 |
| Fbl | 10951 | 7863 | 7509 | 7374 | 6457 | 7937 | 8505 | 8364 | 8102 | 7340 | 7284 | 7835 | 7905.06 | 8324 | 7486 | 0.899 | -0.153 | 0.00445099 | 0.02246398 | 9.00E-04 | 0.0027 | 0.002 | 0.002 | TRUE | 0.0203 |
| Fblim1 | 9026 | 7066 | 7298 | 6995 | 7227 | 8567 | 7010 | 7516 | 7874 | 6963 | 8153 | 8457 | 7662.2 | 7467 | 7858 | 1.052 | 0.074 | 0.34259249 | 0.51674488 | 0.0062 | 0.0027 | 0.0042 | 0.0042 | TRUE | 0.1214 |
| Fbll1 | 16 | 13 | 3 | 11 | 11 | 5 | 12 | 14 | 3 | 11 | 12 | 5 | 9.63 | 10 | 9 | 0.947 | -0.079 | 0.90859571 | NA | 0.2082 | 0.2485 | 0.2381 | 0.2381 | TRUE | 1.0711 |
| Fbln1 | 2194 | 1785 | 2359 | 2484 | 2384 | 2754 | 1704 | 1899 | 2545 | 2473 | 2689 | 2719 | 2338.1 | 2049 | 2627 | 1.283 | 0.359 | 0.00201156 | 0.01215697 | 0.0238 | 0.0034 | 0.0093 | 0.0093 | TRUE | 0.5411 |
| Fbln2 | 396 | 262 | 317 | 361 | 302 | 372 | 308 | 279 | 342 | 359 | 341 | 367 | 332.59 | 310 | 356 | 1.151 | 0.203 | 0.09268717 | 0.20921947 | 0.0022 | 0.0096 | 0.0074 | 0.0074 | TRUE | 0.0903 |
| Fbln5 | 23 | 21 | 7 | 18 | 8 | 14 | 18 | 22 | 8 | 18 | 9 | 14 | 14.75 | 16 | 14 | 0.854 | -0.228 | 0.6786491 | NA | 0.1121 | 0.1631 | 0.1496 | 0.1496 | TRUE | 0.5222 |
| Fbln7 | 25 | 17 | 21 | 30 | 18 | 23 | 19 | 18 | 23 | 30 | 20 | 23 | 22.17 | 20 | 24 | 1.216 | 0.282 | 0.49025031 | 0.65603854 | 0 | 0.1093 | 0.0753 | 0.0753 | TRUE | 0.2444 |
| Fbn1 | 1088 | 637 | 533 | 761 | 600 | 646 | 845 | 678 | 575 | 757 | 677 | 638 | 694.96 | 699 | 691 | 0.985 | -0.021 | 0.86977675 | 0.92801725 | 0.0213 | 0.0058 | 0.0106 | 0.0106 | TRUE | 0.377 |
| Fbn2 | 551 | 375 | 304 | 396 | 378 | 384 | 428 | 399 | 328 | 394 | 426 | 379 | 392.42 | 385 | 400 | 1.035 | 0.05 | 0.68590972 | 0.80742115 | 0.0084 | 0.0085 | 0.0085 | 0.0085 | TRUE | 0.1879 |
| Fbp1 | 23 | 11 | 6 | 27 | 16 | 20 | 18 | 12 | 6 | 27 | 18 | 20 | 16.78 | 12 | 22 | 1.762 | 0.817 | 0.10711618 | NA | 0.0574 | 0.1436 | 0.1219 | 0.1219 | TRUE | 0.4579 |
| Fbp2 | 95 | 55 | 46 | 55 | 63 | 46 | 74 | 59 | 50 | 55 | 71 | 45 | 58.86 | 61 | 57 | 0.932 | -0.102 | 0.7140066 | 0.82815031 | 0.0289 | 0.0427 | 0.0388 | 0.0388 | TRUE | 0.3691 |
| Fbrs | 1338 | 1061 | 988 | 932 | 897 | 980 | 1039 | 1129 | 1066 | 928 | 1012 | 967 | 1023.47 | 1078 | 969 | 0.899 | -0.153 | 0.04899656 | 0.13273461 | 9.00E-04 | 0.0047 | 0.0034 | 0.0034 | TRUE | 0.0201 |
| Fbrsl1 | 2238 | 1854 | 1637 | 1636 | 1529 | 1661 | 1738 | 1972 | 1766 | 1628 | 1725 | 1640 | 1744.93 | 1825 | 1664 | 0.912 | -0.133 | 0.06848294 | 0.1686469 | 0.0021 | 0.0038 | 0.0033 | 0.0033 | TRUE | 0.059 |
| Fbxl12 | 622 | 517 | 586 | 669 | 571 | 612 | 483 | 550 | 632 | 666 | 644 | 604 | 596.58 | 555 | 638 | 1.152 | 0.205 | 0.06407969 | 0.16093673 | 0.0087 | 0.0064 | 0.0071 | 0.0071 | TRUE | 0.1729 |
| Fbxl13 | 9 | 5 | 7 | 7 | 9 | 17 | 7 | 5 | 8 | 7 | 10 | 17 | 8.96 | 7 | 11 | 1.706 | 0.771 | 0.24460902 | NA | 0 | 0.2669 | 0.1979 | 0.1979 | TRUE | 0.3853 |
| Fbxl14 | 3560 | 2786 | 2975 | 2631 | 2537 | 2971 | 2765 | 2963 | 3210 | 2619 | 2862 | 2933 | 2892.01 | 2979 | 2805 | 0.942 | -0.086 | 0.24363919 | 0.4114776 | 0.004 | 0.0032 | 0.0036 | 0.0036 | TRUE | 0.0556 |
| Fbxl15 | 237 | 170 | 136 | 121 | 108 | 112 | 184 | 181 | 147 | 120 | 122 | 111 | 144.08 | 171 | 118 | 0.687 | -0.541 | 0.00177729 | 0.01105434 | 0 | 0.0189 | 0.0143 | 0.0143 | TRUE | 0.1526 |
| Fbxl16 | 429 | 315 | 302 | 317 | 249 | 308 | 333 | 335 | 326 | 316 | 281 | 304 | 315.76 | 331 | 300 | 0.906 | -0.142 | 0.22271091 | 0.386969 | 0 | 0.0099 | 0.0065 | 0.0065 | TRUE | 0.0342 |
| Fbxl17 | 702 | 602 | 510 | 379 | 407 | 432 | 545 | 640 | 550 | 377 | 459 | 426 | 499.78 | 578 | 421 | 0.728 | -0.459 | 4.59E-05 | 0.00065104 | 0.0068 | 0.0072 | 0.0071 | 0.0071 | TRUE | 0.1005 |
| Fbxl18 | 1222 | 938 | 769 | 845 | 783 | 843 | 949 | 998 | 830 | 841 | 883 | 832 | 888.86 | 926 | 852 | 0.92 | -0.12 | 0.17900845 | 0.333601 | 0.0038 | 0.0051 | 0.0047 | 0.0047 | TRUE | 0.0946 |
| Fbxl19 | 5563 | 4074 | 3602 | 4042 | 3649 | 4169 | 4321 | 4333 | 3887 | 4023 | 4116 | 4115 | 4132.64 | 4180 | 4085 | 0.977 | -0.034 | 0.58381343 | 0.73344866 | 0.0015 | 0.003 | 0.0025 | 0.0025 | TRUE | 0.0455 |
| Fbxl2 | 503 | 360 | 443 | 526 | 439 | 550 | 391 | 383 | 478 | 524 | 495 | 543 | 468.89 | 417 | 521 | 1.251 | 0.323 | 0.004362 | 0.02211637 | 0.0058 | 0.0075 | 0.0071 | 0.0071 | TRUE | 0.1819 |
| Fbxl20 | 4130 | 3199 | 3581 | 3849 | 3449 | 3914 | 3208 | 3403 | 3864 | 3831 | 3891 | 3864 | 3676.67 | 3492 | 3862 | 1.107 | 0.146 | 0.04425867 | 0.12321565 | 0.0043 | 0.0031 | 0.0035 | 0.0035 | TRUE | 0.1056 |
| Fbxl21 | 89 | 80 | 78 | 59 | 59 | 81 | 69 | 85 | 84 | 59 | 67 | 80 | 73.94 | 79 | 69 | 0.865 | -0.209 | 0.37657084 | 0.55112975 | 0 | 0.0345 | 0.0265 | 0.0265 | TRUE | 0.2014 |
| Fbxl22 | 5 | 6 | 1 | 4 | 2 | 1 | 4 | 6 | 1 | 4 | 2 | 1 | 3.09 | 4 | 2 | 0.636 | -0.653 | 0.56702125 | NA | 0 | 0.7682 | 0.5954 | 0.5954 | TRUE | 0.1388 |
| Fbxl3 | 2083 | 1780 | 1984 | 2324 | 1911 | 2094 | 1618 | 1893 | 2141 | 2313 | 2156 | 2067 | 2031.35 | 1884 | 2179 | 1.158 | 0.211 | 0.02652869 | 0.08482652 | 0.0105 | 0.0036 | 0.0061 | 0.0061 | TRUE | 0.1886 |
| Fbxl4 | 691 | 466 | 463 | 542 | 427 | 508 | 537 | 496 | 500 | 540 | 482 | 501 | 509.1 | 511 | 508 | 0.993 | -0.01 | 0.91987531 | 0.95606295 | 0 | 0.0071 | 0.0051 | 0.0051 | TRUE | 0.0355 |
| Fbxl5 | 6007 | 4378 | 4257 | 5111 | 4228 | 4744 | 4665 | 4657 | 4593 | 5087 | 4770 | 4683 | 4742.61 | 4638 | 4847 | 1.045 | 0.063 | 0.25373488 | 0.4228898 | 7.00E-04 | 0.0029 | 0.002 | 0.002 | TRUE | 0.0231 |
| Fbxl6 | 2036 | 1610 | 1536 | 1670 | 1536 | 1781 | 1581 | 1713 | 1657 | 1662 | 1733 | 1758 | 1684.06 | 1650 | 1718 | 1.042 | 0.059 | 0.38008809 | 0.55468645 | 7.00E-04 | 0.0038 | 0.0026 | 0.0026 | TRUE | 0.0169 |
| Fbxl7 | 39 | 26 | 13 | 13 | 16 | 20 | 30 | 28 | 14 | 13 | 18 | 20 | 20.45 | 24 | 17 | 0.699 | -0.518 | 0.25870421 | 0.42861473 | 0.0546 | 0.1183 | 0.1007 | 0.1007 | TRUE | 0.7337 |
| Fbxl8 | 145 | 98 | 115 | 153 | 124 | 149 | 113 | 104 | 124 | 152 | 140 | 147 | 130.03 | 114 | 146 | 1.29 | 0.367 | 0.03454265 | 0.10285792 | 0 | 0.0207 | 0.014 | 0.014 | TRUE | 0.0594 |
| Fbxo10 | 837 | 678 | 642 | 515 | 475 | 548 | 650 | 721 | 693 | 513 | 536 | 541 | 608.9 | 688 | 530 | 0.771 | -0.375 | 4.19E-05 | 0.00060804 | 0 | 0.0063 | 0.0044 | 0.0044 | TRUE | 0.0292 |
| Fbxo11 | 3180 | 2356 | 2252 | 2545 | 2096 | 2427 | 2470 | 2506 | 2430 | 2533 | 2365 | 2396 | 2449.89 | 2469 | 2431 | 0.985 | -0.022 | 0.71659861 | 0.82974271 | 4.00E-04 | 0.0034 | 0.0022 | 0.0022 | TRUE | 0.0164 |
| Fbxo15 | 23227 | 16618 | 14233 | 9083 | 9366 | 9621 | 18039 | 17676 | 15357 | 9041 | 10566 | 9497 | 13362.92 | 17024 | 9701 | 0.57 | -0.812 | 4.38E-26 | 2.51E-23 | 0.0068 | 0.0026 | 0.0042 | 0.0042 | TRUE | 0.0896 |
| Fbxo16 | 226 | 181 | 152 | 212 | 198 | 234 | 176 | 193 | 164 | 211 | 223 | 231 | 199.57 | 178 | 222 | 1.251 | 0.323 | 0.02507695 | 0.08149724 | 0 | 0.0143 | 0.0099 | 0.0099 | TRUE | 0.056 |
| Fbxo17 | 414 | 296 | 268 | 306 | 266 | 325 | 322 | 315 | 289 | 305 | 300 | 321 | 308.51 | 309 | 309 | 0.999 | -0.002 | 0.98913911 | 0.99429134 | 0 | 0.0101 | 0.0068 | 0.0068 | TRUE | 0.033 |
| Fbxo18 | 5198 | 3841 | 3964 | 4726 | 4264 | 4800 | 4037 | 4086 | 4277 | 4704 | 4810 | 4738 | 4442.11 | 4133 | 4751 | 1.15 | 0.201 | 0.00014928 | 0.00170016 | 3.00E-04 | 0.003 | 0.0018 | 0.0018 | TRUE | 0.0112 |
| Fbxo2 | 1631 | 1344 | 1281 | 1631 | 1606 | 1838 | 1267 | 1430 | 1382 | 1623 | 1812 | 1814 | 1554.69 | 1360 | 1750 | 1.288 | 0.366 | 2.80E-06 | 6.54E-05 | 0.0034 | 0.004 | 0.0037 | 0.0037 | TRUE | 0.0485 |
| Fbxo21 | 5464 | 4126 | 4255 | 4265 | 3923 | 4520 | 4244 | 4389 | 4591 | 4245 | 4426 | 4462 | 4392.73 | 4408 | 4378 | 0.993 | -0.009 | 0.86796847 | 0.92704068 | 9.00E-04 | 0.003 | 0.0021 | 0.0021 | TRUE | 0.0161 |
| Fbxo22 | 6378 | 4696 | 5329 | 5086 | 4793 | 5764 | 4954 | 4995 | 5750 | 5063 | 5407 | 5690 | 5309.68 | 5233 | 5387 | 1.03 | 0.042 | 0.56619059 | 0.72043806 | 0.0043 | 0.0029 | 0.0037 | 0.0037 | TRUE | 0.0907 |

| | | | | | | | | | | | | | | | | | | | | | | | | | |
|---|---|---|---|---|---|---|---|---|---|---|---|---|---|---|---|---|---|---|---|---|---|---|---|---|---|
| Fbxo24 | 58 | 51 | 43 | 50 | 42 | 68 | 45 | 54 | 46 | 50 | 47 | 67 | 51.66 | 48 | 55 | 1.133 | 0.18 | 0.5177921 | 0.6797764 | 0 | 0.0483 | 0.0364 | 0.0364 | TRUE | 0.3329 |
| Fbxo25 | 1866 | 1384 | 1369 | 1612 | 1281 | 1594 | 1449 | 1472 | 1477 | 1605 | 1445 | 1574 | 1503.62 | 1466 | 1541 | 1.052 | 0.073 | 0.29620635 | 0.46812001 | 8.00E-04 | 0.004 | 0.0028 | 0.0028 | TRUE | 0.0351 |
| Fbxo27 | 321 | 278 | 251 | 255 | 254 | 274 | 249 | 296 | 271 | 254 | 287 | 270 | 271.11 | 272 | 270 | 0.996 | -0.006 | 0.9609021 | 0.97845108 | 0 | 0.0112 | 0.0084 | 0.0084 | TRUE | 0.0654 |
| Fbxo28 | 3958 | 2837 | 2990 | 3313 | 2927 | 3355 | 3074 | 3018 | 3226 | 3298 | 3302 | 3312 | 3204.92 | 3106 | 3304 | 1.064 | 0.089 | 0.1112306 | 0.23899865 | 2.00E-04 | 0.0032 | 0.002 | 0.002 | TRUE | 0.0136 |
| Fbxo3 | 1965 | 1443 | 1636 | 1699 | 1567 | 1752 | 1526 | 1535 | 1765 | 1691 | 1768 | 1729 | 1669.11 | 1609 | 1729 | 1.076 | 0.105 | 0.16545895 | 0.3164599 | 0.0029 | 0.0038 | 0.0036 | 0.0036 | TRUE | 0.086 |
| Fbxo30 | 3145 | 2308 | 2321 | 2553 | 2075 | 2509 | 2443 | 2455 | 2504 | 2541 | 2341 | 2477 | 2460.13 | 2467 | 2453 | 0.995 | -0.008 | 0.89853959 | 0.94384643 | 5.00E-04 | 0.0034 | 0.0022 | 0.0022 | TRUE | 0.0191 |
| Fbxo31 | 3574 | 2836 | 2797 | 2737 | 2657 | 3094 | 2776 | 3017 | 3018 | 2724 | 2997 | 3054 | 2931.06 | 2937 | 2925 | 0.997 | -0.005 | 0.94202336 | 0.9676829 | 0.0027 | 0.0032 | 0.003 | 0.003 | TRUE | 0.0441 |
| Fbxo32 | 131 | 73 | 83 | 66 | 78 | 85 | 102 | 78 | 90 | 66 | 88 | 84 | 84.42 | 90 | 79 | 0.878 | -0.187 | 0.40556985 | 0.5782354 | 0.0092 | 0.0305 | 0.0246 | 0.0246 | TRUE | 0.2098 |
| Fbxo33 | 1752 | 1305 | 1108 | 1138 | 1017 | 1162 | 1361 | 1388 | 1196 | 1133 | 1147 | 1147 | 1228.58 | 1315 | 1142 | 0.868 | -0.204 | 0.00996234 | 0.04126099 | 0.0025 | 0.0044 | 0.0037 | 0.0037 | TRUE | 0.0738 |
| Fbxo34 | 2211 | 1622 | 1564 | 1632 | 1361 | 1567 | 1717 | 1725 | 1688 | 1624 | 1535 | 1547 | 1639.46 | 1710 | 1569 | 0.918 | -0.124 | 0.0536096 | 0.14156329 | 0 | 0.0039 | 0.0024 | 0.0024 | TRUE | 0.0117 |
| Fbxo36 | 193 | 136 | 113 | 148 | 119 | 82 | 150 | 145 | 122 | 147 | 134 | 81 | 129.83 | 139 | 121 | 0.868 | -0.205 | 0.35328374 | 0.52780934 | 0.0445 | 0.0207 | 0.0273 | 0.0273 | TRUE | 0.863 |
| Fbxo38 | 6044 | 4322 | 4596 | 4904 | 4204 | 4919 | 4694 | 4597 | 4959 | 4881 | 4743 | 4856 | 4788.37 | 4750 | 4827 | 1.016 | 0.023 | 0.66994397 | 0.79708707 | 6.00E-04 | 0.0029 | 0.002 | 0.002 | TRUE | 0.0178 |
| Fbxo39 | 31 | 21 | 16 | 30 | 30 | 26 | 24 | 22 | 17 | 30 | 34 | 26 | 25.51 | 21 | 30 | 1.392 | 0.477 | 0.21255086 | 0.3748753 | 0 | 0.0953 | 0.0656 | 0.0656 | TRUE | 0.1385 |
| Fbxo4 | 71 | 44 | 53 | 53 | 44 | 40 | 55 | 47 | 57 | 53 | 50 | 39 | 50.17 | 53 | 47 | 0.89 | -0.169 | 0.54240322 | 0.70032243 | 0 | 0.0497 | 0.0353 | 0.0353 | TRUE | 0.1723 |
| Fbxo40 | 4 | 4 | 2 | 6 | 1 | 1 | 3 | 4 | 2 | 6 | 1 | 1 | 2.93 | 3 | 3 | 0.857 | -0.223 | 0.84790437 | NA | 0.1773 | 0.8101 | 0.628 | 0.628 | TRUE | 1.3618 |
| Fbxo41 | 243 | 185 | 205 | 245 | 241 | 257 | 189 | 197 | 221 | 244 | 272 | 254 | 229.36 | 202 | 257 | 1.27 | 0.345 | 0.01194631 | 0.04718632 | 0 | 0.0128 | 0.0092 | 0.0092 | TRUE | 0.0713 |
| Fbxo42 | 5488 | 4147 | 3859 | 4283 | 3658 | 4150 | 4262 | 4411 | 4164 | 4263 | 4127 | 4097 | 4220.64 | 4279 | 4162 | 0.973 | -0.04 | 0.46182114 | 0.63114231 | 4.00E-04 | 0.003 | 0.0019 | 0.0019 | TRUE | 0.0087 |
| Fbxo43 | 59 | 52 | 62 | 89 | 68 | 85 | 46 | 55 | 67 | 89 | 77 | 84 | 69.54 | 56 | 83 | 1.496 | 0.581 | 0.01707851 | 0.06162663 | 0 | 0.0365 | 0.0279 | 0.0279 | TRUE | 0.2517 |
| Fbxo44 | 257 | 212 | 237 | 252 | 258 | 349 | 200 | 226 | 256 | 251 | 291 | 345 | 261.21 | 227 | 296 | 1.306 | 0.386 | 0.01212627 | 0.04772009 | 0.0169 | 0.0115 | 0.0131 | 0.0131 | TRUE | 0.2196 |
| Fbxo45 | 2215 | 1584 | 1588 | 1748 | 1458 | 1696 | 1720 | 1685 | 1713 | 1740 | 1645 | 1674 | 1696.26 | 1706 | 1686 | 0.988 | -0.017 | 0.79079652 | 0.8770061 | 0 | 0.0038 | 0.0023 | 0.0023 | TRUE | 0.0094 |
| Fbxo46 | 748 | 531 | 489 | 461 | 401 | 443 | 581 | 565 | 528 | 459 | 452 | 437 | 503.66 | 558 | 449 | 0.805 | -0.313 | 0.00123941 | 0.00844449 | 0 | 0.0071 | 0.0048 | 0.0048 | TRUE | 0.0257 |
| Fbxo47 | 367 | 281 | 314 | 424 | 416 | 407 | 285 | 299 | 339 | 422 | 469 | 402 | 369.31 | 308 | 431 | 1.403 | 0.489 | 4.19E-05 | 0.00060778 | 0.0044 | 0.0088 | 0.0075 | 0.0075 | TRUE | 0.0874 |
| Fbxo48 | 130 | 108 | 99 | 68 | 74 | 90 | 101 | 115 | 107 | 68 | 83 | 89 | 93.78 | 108 | 80 | 0.745 | -0.425 | 0.04103955 | 0.11654644 | 0 | 0.0277 | 0.0202 | 0.0202 | TRUE | 0.1736 |
| Fbxo5 | 4360 | 3165 | 3447 | 3466 | 3147 | 3675 | 3386 | 3367 | 3719 | 3450 | 3550 | 3628 | 3516.68 | 3491 | 3543 | 1.015 | 0.022 | 0.7256839 | 0.83572689 | 0.0016 | 0.0031 | 0.0025 | 0.0025 | TRUE | 0.0394 |
| Fbxo6 | 2422 | 1754 | 1974 | 2044 | 1916 | 2108 | 1881 | 1866 | 2130 | 2035 | 2161 | 2081 | 2025.61 | 1959 | 2092 | 1.069 | 0.096 | 0.18587964 | 0.34223596 | 0.0026 | 0.0036 | 0.0033 | 0.0033 | TRUE | 0.0695 |
| Fbxo7 | 1535 | 1179 | 1289 | 1327 | 1148 | 1465 | 1192 | 1254 | 1391 | 1321 | 1295 | 1446 | 1316.54 | 1279 | 1354 | 1.06 | 0.084 | 0.30931223 | 0.48202942 | 0.0038 | 0.0042 | 0.0042 | 0.0042 | TRUE | 0.0697 |
| Fbxo8 | 1492 | 1113 | 980 | 1099 | 923 | 1139 | 1159 | 1184 | 1057 | 1094 | 1041 | 1124 | 1109.94 | 1133 | 1086 | 0.959 | -0.061 | 0.43533185 | 0.60698451 | 0.0015 | 0.0046 | 0.0035 | 0.0035 | TRUE | 0.0399 |
| Fbxo9 | 1392 | 939 | 798 | 763 | 638 | 718 | 1081 | 999 | 861 | 759 | 720 | 709 | 854.82 | 980 | 729 | 0.743 | -0.429 | 1.03E-05 | 0.00019238 | 0.0064 | 0.0052 | 0.0056 | 0.0056 | TRUE | 0.1338 |
| Fbxw10 | 111 | 78 | 62 | 84 | 76 | 89 | 86 | 83 | 67 | 84 | 86 | 88 | 82.21 | 79 | 86 | 1.085 | 0.118 | 0.58612856 | 0.73545215 | 0 | 0.0313 | 0.0219 | 0.0219 | TRUE | 0.151 |
| Fbxw11 | 8995 | 6429 | 6258 | 6455 | 5911 | 6497 | 6986 | 6838 | 6752 | 6425 | 6668 | 6414 | 6680.65 | 6859 | 6502 | 0.948 | -0.077 | 0.11885222 | 0.25040606 | 2.00E-04 | 0.0028 | 0.0016 | 0.0016 | TRUE | 0.006 |
| Fbxw13 | 1 | 2 | 0 | 0 | 0 | 0 | 1 | 2 | 0 | 0 | 0 | 0 | 0.48 | 1 | 0 | 0.197 | -2.343 | 0.43249452 | NA | 0 | 4.8992 | 4.9172 | 4.9172 | TRUE | 0.1226 |
| Fbxw14 | 0 | 0 | 0 | 0 | 0 | 0 | 0 | 0 | 0 | 0 | 0 | 0 | 0 | 0 | 0 | NA | NA | NA | NA | NA | NA | NA | NA | NA | NA |
| Fbxw15 | 1 | 0 | 0 | 0 | 0 | 0 | 1 | 0 | 0 | 0 | 0 | 0 | 0.13 | 0 | 0 | 0.552 | -0.857 | 0.8336548 | NA | 0 | 18.3121 | 10 | 10 | TRUE | 0.2566 |
| Fbxw16 | 0 | 0 | 0 | 0 | 0 | 0 | 0 | 0 | 0 | 0 | 0 | 0 | 0 | 0 | 0 | NA | NA | NA | NA | NA | NA | NA | NA | NA | NA |
| Fbxw17 | 366 | 277 | 261 | 222 | 221 | 225 | 284 | 295 | 282 | 221 | 249 | 222 | 258.82 | 287 | 231 | 0.804 | -0.315 | 0.01341327 | 0.05143974 | 0 | 0.0116 | 0.0078 | 0.0078 | TRUE | 0.0522 |
| Fbxw18 | 0 | 0 | 0 | 0 | 0 | 0 | 0 | 0 | 0 | 0 | 0 | 0 | 0 | 0 | 0 | NA | NA | NA | NA | NA | NA | NA | NA | NA | NA |
| Fbxw19 | 0 | 0 | 1 | 0 | 0 | 0 | 0 | 0 | 1 | 0 | 0 | 0 | 0.18 | 0 | 0 | 0.552 | -0.857 | 0.8336548 | NA | 0 | 13.1817 | 10 | 10 | TRUE | 0.5365 |
| Fbxw2 | 3905 | 2774 | 2776 | 3204 | 2686 | 2915 | 3033 | 2951 | 2995 | 3189 | 3030 | 2878 | 3012.62 | 2993 | 3032 | 1.013 | 0.019 | 0.76071474 | 0.85788984 | 0.0011 | 0.0032 | 0.0024 | 0.0024 | TRUE | 0.0252 |
| Fbxw20 | 1 | 0 | 1 | 0 | 0 | 1 | 1 | 0 | 1 | 0 | 0 | 1 | 0.47 | 1 | 0 | 0.596 | -0.748 | 0.78959815 | NA | 0 | 5.0047 | 3.8792 | 3.8792 | TRUE | 0.3835 |
| Fbxw21 | 1 | 0 | 0 | 0 | 0 | 0 | 1 | 0 | 0 | 0 | 0 | 0 | 0.13 | 0 | 0 | 0.552 | -0.857 | 0.8336548 | NA | 0 | 18.3121 | 10 | 10 | TRUE | 0.2566 |
| Fbxw22 | 0 | 0 | 0 | 0 | 0 | 0 | 0 | 0 | 0 | 0 | 0 | 0 | 0 | 0 | 0 | NA | NA | NA | NA | NA | NA | NA | NA | NA | NA |
| Fbxw24 | 0 | 0 | 0 | 0 | 0 | 0 | 0 | 0 | 0 | 0 | 0 | 0 | 0 | 0 | 0 | NA | NA | NA | NA | NA | NA | NA | NA | NA | NA |
| Fbxw26 | 0 | 0 | 0 | 0 | 0 | 0 | 0 | 0 | 0 | 0 | 0 | 0 | 0 | 0 | 0 | NA | NA | NA | NA | NA | NA | NA | NA | NA | NA |
| Fbxw28 | 0 | 0 | 0 | 0 | 0 | 0 | 0 | 0 | 0 | 0 | 0 | 0 | 0 | 0 | 0 | NA | NA | NA | NA | NA | NA | NA | NA | NA | NA |
| Fbxw4 | 236 | 169 | 196 | 242 | 236 | 223 | 183 | 180 | 211 | 241 | 266 | 220 | 216.97 | 191 | 242 | 1.266 | 0.341 | 0.01909893 | 0.06680651 | 0.0036 | 0.0134 | 0.0105 | 0.0105 | TRUE | 0.0859 |
| Fbxw5 | 1241 | 966 | 932 | 1046 | 835 | 1061 | 964 | 1028 | 1006 | 1041 | 942 | 1047 | 1004.58 | 999 | 1010 | 1.013 | 0.018 | 0.81955755 | 0.8964401 | 0.0012 | 0.0048 | 0.0036 | 0.0036 | TRUE | 0.0407 |
| Fbxw7 | 2211 | 1513 | 1275 | 1247 | 1136 | 1285 | 1717 | 1609 | 1376 | 1241 | 1282 | 1268 | 1415.59 | 1567 | 1264 | 0.805 | -0.312 | 0.00033864 | 0.00318645 | 0.006 | 0.0041 | 0.0048 | 0.0048 | TRUE | 0.1367 |
| Fbxw8 | 7416 | 5137 | 4931 | 5656 | 5191 | 5725 | 5760 | 5464 | 5321 | 5630 | 5856 | 5651 | 5613.63 | 5515 | 5712 | 1.035 | 0.05 | 0.36184156 | 0.53660525 | 8.00E-04 | 0.0028 | 0.002 | 0.002 | TRUE | 0.0187 |
| Fbxw9 | 2368 | 1870 | 1625 | 1655 | 1528 | 1783 | 1839 | 1989 | 1753 | 1647 | 1724 | 1760 | 1785.47 | 1860 | 1710 | 0.919 | -0.121 | 0.08950945 | 0.20399806 | 0.0018 | 0.0038 | 0.0031 | 0.0031 | TRUE | 0.0431 |
| Fcamr | 2 | 0 | 0 | 1 | 1 | 1 | 2 | 0 | 0 | 1 | 1 | 1 | 0.78 | 1 | 1 | 1.828 | 0.87 | 0.69255718 | NA | 0 | 3.0514 | 2.1415 | 2.1415 | TRUE | 0.9716 |
| Fcer1a | 0 | 0 | 0 | 0 | 0 | 0 | 0 | 0 | 0 | 0 | 0 | 0 | 0 | 0 | 0 | NA | NA | NA | NA | NA | NA | NA | NA | NA | NA |
| Fcer1g | 15 | 10 | 21 | 27 | 23 | 21 | 12 | 11 | 23 | 27 | 26 | 21 | 19.75 | 15 | 25 | 1.652 | 0.724 | 0.11544699 | 0.24530321 | 0.0275 | 0.1224 | 0.0988 | 0.0988 | TRUE | 0.8467 |
| Fcer2a | 0 | 0 | 0 | 0 | 0 | 0 | 0 | 0 | 0 | 0 | 0 | 0 | 0 | 0 | 0 | NA | NA | NA | NA | NA | NA | NA | NA | NA | NA |
| Fcf1 | 3556 | 2504 | 2419 | 2383 | 2104 | 2615 | 2762 | 2663 | 2610 | 2372 | 2374 | 2581 | 2560.39 | 2678 | 2442 | 0.912 | -0.133 | 0.03543606 | 0.10482488 | 0.001 | 0.0034 | 0.0025 | 0.0025 | TRUE | 0.0303 |
| Fcgbp | 339 | 236 | 234 | 323 | 268 | 317 | 263 | 251 | 252 | 322 | 302 | 313 | 283.93 | 255 | 312 | 1.221 | 0.288 | 0.01641601 | 0.05973653 | 0 | 0.0108 | 0.0068 | 0.0068 | TRUE | 0.0085 |
| Fcgr1 | 0 | 0 | 0 | 1 | 1 | 1 | 0 | 0 | 0 | 1 | 1 | 1 | 0.52 | 0 | 1 | 5.852 | 2.549 | 0.3348364 | NA | 0 | 4.5739 | 3.5345 | 3.5345 | TRUE | 0.0847 |
| Fcgr2b | 25 | 30 | 27 | 33 | 29 | 47 | 19 | 32 | 29 | 33 | 33 | 46 | 32.07 | 27 | 37 | 1.409 | 0.495 | 0.17065055 | 0.32232463 | 0.0247 | 0.0763 | 0.0619 | 0.0619 | TRUE | 0.4547 |
| Fcgr3 | 10 | 4 | 11 | 4 | 5 | 11 | 8 | 4 | 12 | 4 | 6 | 11 | 7.4 | 8 | 7 | 0.861 | -0.215 | 0.77250421 | NA | 0.1073 | 0.3229 | 0.263 | 0.263 | TRUE | 0.2837 |
| Fcgr4 | 0 | 0 | 0 | 0 | 0 | 0 | 0 | 0 | 0 | 0 | 0 | 0 | 0 | 0 | 0 | NA | NA | NA | NA | NA | NA | NA | NA | NA | NA |
| Fcgrt | 359 | 250 | 268 | 380 | 307 | 376 | 279 | 266 | 289 | 378 | 346 | 371 | 321.61 | 278 | 365 | 1.315 | 0.395 | 0.00064739 | 0.00525355 | 0 | 0.0098 | 0.0065 | 0.0065 | TRUE | 0.0227 |
| Fcho1 | 1955 | 1687 | 1532 | 1701 | 1520 | 1840 | 1518 | 1794 | 1653 | 1693 | 1715 | 1816 | 1698.35 | 1655 | 1741 | 1.053 | 0.075 | 0.33504625 | 0.50912355 | 0.0033 | 0.0038 | 0.0038 | 0.0038 | TRUE | 0.0654 |
| Fcho2 | 3403 | 2647 | 2724 | 3102 | 2905 | 3137 | 2643 | 2816 | 2939 | 3088 | 3277 | 3097 | 2976.55 | 2799 | 3154 | 1.127 | 0.173 | 0.0068395 | 0.03120235 | 0.0015 | 0.0032 | 0.0026 | 0.0026 | TRUE | 0.0297 |
| Fchsd1 | 941 | 760 | 720 | 731 | 690 | 792 | 731 | 808 | 777 | 728 | 778 | 782 | 767.33 | 772 | 763 | 0.989 | -0.016 | 0.85036772 | 0.91550962 | 9.00E-04 | 0.0055 | 0.0039 | 0.0039 | TRUE | 0.0274 |
| Fchsd2 | 1548 | 1168 | 1233 | 1245 | 1163 | 1372 | 1202 | 1242 | 1330 | 1239 | 1312 | 1354 | 1280.11 | 1258 | 1302 | 1.035 | 0.05 | 0.5046109 | 0.66826706 | 0.0016 | 0.0043 | 0.0033 | 0.0033 | TRUE | 0.0298 |
| Fcna | 1 | 1 | 2 | 0 | 1 | 1 | 1 | 1 | 2 | 0 | 1 | 1 | 1.02 | 1 | 1 | 0.531 | -0.914 | 0.6308001 | NA | 0 | 2.3285 | 1.4951 | 1.4951 | TRUE | 0.2654 |
| Fcnb | 1 | 0 | 1 | 0 | 0 | 1 | 1 | 0 | 1 | 0 | 0 | 1 | 0.47 | 1 | 0 | 0.596 | -0.748 | 0.78959815 | NA | 0 | 5.0047 | 3.8792 | 3.8792 | TRUE | 0.3835 |
| Fcrl1 | 1 | 0 | 0 | 1 | 2 | 1 | 1 | 0 | 0 | 1 | 2 | 1 | 0.84 | 0 | 1 | 4.069 | 2.025 | 0.34772519 | NA | 0 | 2.8378 | 1.9813 | 1.9813 | TRUE | 0.2566 |
| Fcrl5 | 0 | 1 | 1 | 0 | 1 | 4 | 0 | 1 | 1 | 0 | 1 | 4 | 1.2 | 1 | 2 | 2.501 | 1.322 | 0.49688351 | NA | 0 | 1.9722 | 1.7166 | 1.7166 | TRUE | 0.4359 |
| Fcrl6 | 0 | 0 | 0 | 0 | 0 | 0 | 0 | 0 | 0 | 0 | 0 | 0 | 0 | 0 | 0 | NA | NA | NA | NA | NA | NA | NA | NA | NA | NA |

| | | | | | | | | | | | | | | | | | | | | | | | | | |
|---|---|---|---|---|---|---|---|---|---|---|---|---|---|---|---|---|---|---|---|---|---|---|---|---|---|
| Fcrla | 0 | 0 | 1 | 4 | 1 | 1 | 0 | 0 | 1 | 4 | 1 | 1 | 1.2 | 0 | 2 | 5.92 | 2.565 | 0.19250712 | NA | 0 | 1.9841 | 1.5392 | 1.5392 | TRUE | 0.6551 |
| Fcrlb | 1 | 3 | 0 | 4 | 2 | 5 | 1 | 3 | 0 | 4 | 2 | 5 | 2.52 | 1 | 4 | 2.885 | 1.529 | 0.24435158 | NA | 0 | 0.9416 | 0.7333 | 0.7333 | TRUE | 0.8403 |
| Fcrls | 2 | 0 | 0 | 0 | 1 | 0 | 2 | 0 | 0 | 0 | 1 | 0 | 0.45 | 1 | 0 | 0.68 | -0.556 | 0.85993441 | NA | 0 | 5.3057 | 5.2576 | 5.2576 | TRUE | 0.9726 |
| Fdft1 | 9400 | 7193 | 8185 | 8768 | 7786 | 9289 | 7301 | 7651 | 8832 | 8728 | 8783 | 9170 | 8410.66 | 7928 | 8894 | 1.122 | 0.166 | 0.02168089 | 0.07325247 | 0.0053 | 0.0027 | 0.0037 | 0.0037 | TRUE | 0.1212 |
| Fdps | 10996 | 7966 | 8607 | 10369 | 8791 | 10861 | 8540 | 8473 | 9287 | 10321 | 9917 | 10721 | 9543.38 | 8767 | 10320 | 1.177 | 0.236 | 5.74E-05 | 0.00078022 | 0.0019 | 0.0027 | 0.0024 | 0.0024 | TRUE | 0.0327 |
| Fdx1 | 1483 | 1008 | 911 | 1080 | 877 | 1031 | 1152 | 1072 | 983 | 1075 | 989 | 1018 | 1048.18 | 1069 | 1027 | 0.96 | -0.059 | 0.48414366 | 0.65044468 | 0.0032 | 0.0047 | 0.0042 | 0.0042 | TRUE | 0.0588 |
| Fdx1l | 164 | 113 | 122 | 163 | 136 | 170 | 127 | 120 | 132 | 162 | 153 | 168 | 143.78 | 126 | 161 | 1.276 | 0.351 | 0.03189465 | 0.09680829 | 0 | 0.0189 | 0.0123 | 0.0123 | TRUE | 0.0178 |
| Fdxacb1 | 691 | 473 | 459 | 533 | 437 | 496 | 537 | 503 | 495 | 531 | 493 | 490 | 508.03 | 512 | 505 | 0.985 | -0.022 | 0.81770179 | 0.89531267 | 0 | 0.0071 | 0.0049 | 0.0049 | TRUE | 0.0243 |
| Fdxr | 2027 | 1527 | 1650 | 1946 | 1701 | 2031 | 1574 | 1624 | 1780 | 1937 | 1919 | 2005 | 1806.62 | 1659 | 1954 | 1.178 | 0.237 | 0.00074177 | 0.00581572 | 0.0015 | 0.0037 | 0.003 | 0.003 | TRUE | 0.0483 |
| Fech | 2746 | 2054 | 1971 | 2163 | 1890 | 2109 | 2133 | 2185 | 2127 | 2153 | 2132 | 2082 | 2135.21 | 2148 | 2122 | 0.988 | -0.017 | 0.76990446 | 0.86389107 | 0 | 0.0035 | 0.002 | 0.002 | TRUE | 0.0034 |
| Fem1a | 1974 | 1403 | 1083 | 1127 | 1070 | 1159 | 1533 | 1492 | 1169 | 1122 | 1207 | 1144 | 1277.84 | 1398 | 1158 | 0.827 | -0.274 | 0.00629483 | 0.0292592 | 0.0108 | 0.0043 | 0.0065 | 0.0065 | TRUE | 0.2451 |
| Fem1b | 18422 | 13577 | 13969 | 15469 | 14086 | 15959 | 14308 | 14442 | 15072 | 15398 | 15890 | 15754 | 15144 | 14607 | 15681 | 1.074 | 0.102 | 0.03191718 | 0.09685611 | 4.00E-04 | 0.0026 | 0.0016 | 0.0016 | TRUE | 0.0095 |
| Fem1c | 2851 | 2185 | 2203 | 2342 | 2043 | 2441 | 2214 | 2324 | 2377 | 2331 | 2305 | 2410 | 2326.83 | 2305 | 2349 | 1.02 | 0.028 | 0.64766359 | 0.78065911 | 6.00E-04 | 0.0034 | 0.0023 | 0.0023 | TRUE | 0.0149 |
| Fen1 | 6019 | 4068 | 3329 | 3499 | 3096 | 3438 | 4675 | 4327 | 3592 | 3483 | 3493 | 3394 | 3827.18 | 4198 | 3457 | 0.823 | -0.281 | 0.00111183 | 0.0078321 | 0.0084 | 0.003 | 0.0051 | 0.0051 | TRUE | 0.1934 |
| Fendrr | 0 | 0 | 1 | 2 | 1 | 1 | 0 | 0 | 1 | 2 | 1 | 1 | 0.86 | 0 | 1 | 3.977 | 1.992 | 0.35112459 | NA | 0 | 2.745 | 1.9107 | 1.9107 | TRUE | 0.5365 |
| Fer1l4 | 25 | 23 | 15 | 24 | 31 | 18 | 19 | 24 | 16 | 24 | 35 | 18 | 22.78 | 20 | 26 | 1.27 | 0.345 | 0.41907576 | 0.59168417 | 0.0352 | 0.1065 | 0.0868 | 0.0868 | TRUE | 0.3399 |
| Ferd3l | 0 | 0 | 0 | 0 | 0 | 0 | 0 | 0 | 0 | 0 | 0 | 0 | 0 | 0 | 0 | NA | NA | NA | NA | NA | NA | NA | NA | NA | NA |
| Fermt1 | 152 | 96 | 103 | 89 | 86 | 102 | 118 | 102 | 111 | 89 | 97 | 101 | 102.93 | 110 | 96 | 0.862 | -0.214 | 0.26774178 | 0.43830412 | 0 | 0.0255 | 0.0172 | 0.0172 | TRUE | 0.0425 |
| Fermt2 | 14800 | 10887 | 10679 | 11328 | 10193 | 11782 | 11495 | 11580 | 11523 | 11276 | 11499 | 11631 | 11500.46 | 11533 | 11469 | 0.994 | -0.008 | 0.85903727 | 0.92137324 | 0 | 0.0026 | 0.0014 | 0.0014 | TRUE | 0.0026 |
| Fermt3 | 44 | 41 | 59 | 48 | 50 | 47 | 34 | 44 | 64 | 48 | 56 | 46 | 48.67 | 47 | 50 | 1.073 | 0.101 | 0.74110634 | 0.84598937 | 0.0363 | 0.0511 | 0.0469 | 0.0469 | TRUE | 0.5808 |
| Fert2 | 1903 | 1321 | 1397 | 1412 | 1271 | 1475 | 1478 | 1405 | 1507 | 1406 | 1434 | 1456 | 1447.64 | 1463 | 1432 | 0.978 | -0.032 | 0.63712791 | 0.77324184 | 0 | 0.0041 | 0.0026 | 0.0026 | TRUE | 0.0142 |
| Fes | 136 | 114 | 120 | 152 | 142 | 184 | 106 | 121 | 129 | 151 | 160 | 182 | 141.58 | 119 | 164 | 1.39 | 0.475 | 0.00631746 | 0.02932781 | 0 | 0.0192 | 0.0146 | 0.0146 | TRUE | 0.1071 |
| Fetub | 505 | 374 | 350 | 416 | 344 | 406 | 392 | 398 | 378 | 414 | 388 | 401 | 395.1 | 389 | 401 | 1.03 | 0.043 | 0.67988207 | 0.80336925 | 0 | 0.0084 | 0.0053 | 0.0053 | TRUE | 0.0094 |
| Fev | 4 | 0 | 0 | 2 | 0 | 1 | 3 | 0 | 0 | 2 | 0 | 1 | 1.01 | 1 | 1 | 0.918 | -0.123 | 0.95416679 | NA | 2.3137 | 2.3395 | 2.3364 | 2.3364 | TRUE | 0.8566 |
| Fez1 | 414 | 265 | 237 | 378 | 294 | 344 | 322 | 282 | 256 | 376 | 332 | 340 | 317.77 | 287 | 349 | 1.215 | 0.282 | 0.02802471 | 0.0881271 | 0.0059 | 0.0099 | 0.0087 | 0.0087 | TRUE | 0.1389 |
| Fez2 | 2157 | 1631 | 1563 | 1650 | 1503 | 1614 | 1675 | 1735 | 1686 | 1642 | 1696 | 1593 | 1671.3 | 1699 | 1644 | 0.968 | -0.048 | 0.46009898 | 0.62988358 | 0 | 0.0038 | 0.0024 | 0.0024 | TRUE | 0.009 |
| Fezf1 | 2 | 0 | 0 | 1 | 1 | 2 | 2 | 0 | 0 | 1 | 1 | 2 | 0.94 | 1 | 1 | 2.399 | 1.262 | 0.53916972 | NA | 0 | 2.5188 | 1.8235 | 1.8235 | TRUE | 0.9714 |
| Fezf2 | 0 | 0 | 0 | 0 | 0 | 0 | 0 | 0 | 0 | 0 | 0 | 0 | 0 | 0 | 0 | NA | NA | NA | NA | NA | NA | NA | NA | NA | NA |
| Ffar1 | 0 | 1 | 1 | 0 | 0 | 3 | 0 | 1 | 1 | 0 | 0 | 3 | 0.85 | 1 | 1 | 1.453 | 0.539 | 0.81683464 | NA | 2.1834 | 2.7885 | 2.6825 | 2.6825 | TRUE | 1.4416 |
| Ffar2 | 6 | 4 | 1 | 1 | 1 | 1 | 5 | 4 | 1 | 1 | 1 | 1 | 2.18 | 3 | 1 | 0.307 | -1.705 | 0.21841454 | NA | 0 | 1.0876 | 0.739 | 0.739 | TRUE | 0.4524 |
| Ffar3 | 16 | 8 | 10 | 14 | 13 | 7 | 12 | 9 | 11 | 14 | 15 | 7 | 11.21 | 11 | 12 | 1.108 | 0.147 | 0.80087631 | NA | 0 | 0.2139 | 0.1568 | 0.1568 | TRUE | 0.5371 |
| Ffar4 | 14 | 16 | 15 | 24 | 10 | 17 | 11 | 17 | 16 | 24 | 11 | 17 | 16 | 15 | 17 | 1.196 | 0.258 | 0.60679305 | NA | 0 | 0.1505 | 0.1186 | 0.1186 | TRUE | 0.227 |
| Fga | 1 | 0 | 0 | 0 | 0 | 0 | 1 | 0 | 0 | 0 | 0 | 0 | 0.13 | 0 | 0 | 0.552 | -0.857 | 0.8336548 | NA | 0 | 18.3121 | 10 | 10 | TRUE | 0.2566 |
| Fgb | 0 | 0 | 0 | 0 | 0 | 0 | 0 | 0 | 0 | 0 | 1 | 0 | 0.19 | 0 | 0 | 2.094 | 1.067 | 0.79379329 | NA | 0 | 12.608 | 10 | 10 | TRUE | 0.5075 |
| Fgd1 | 7720 | 5620 | 5607 | 5362 | 5015 | 5621 | 5996 | 5978 | 6050 | 5337 | 5657 | 5549 | 5761.2 | 6008 | 5514 | 0.918 | -0.124 | 0.01500749 | 0.05588904 | 2.00E-04 | 0.0028 | 0.0017 | 0.0017 | TRUE | 0.0096 |
| Fgd2 | 58 | 45 | 45 | 59 | 55 | 61 | 45 | 48 | 49 | 59 | 62 | 60 | 53.74 | 47 | 60 | 1.281 | 0.358 | 0.16754074 | 0.31903274 | 0 | 0.0465 | 0.0296 | 0.0296 | TRUE | 0.0145 |
| Fgd3 | 8 | 2 | 5 | 1 | 5 | 2 | 6 | 2 | 5 | 1 | 6 | 2 | 3.72 | 4 | 3 | 0.61 | -0.714 | 0.49578674 | NA | 0.1456 | 0.6388 | 0.504 | 0.504 | TRUE | 0.7922 |
| Fgd4 | 208 | 153 | 108 | 81 | 81 | 89 | 162 | 163 | 117 | 81 | 91 | 88 | 116.78 | 147 | 87 | 0.587 | -0.768 | 0.00013357 | 0.00154965 | 0.013 | 0.0227 | 0.0198 | 0.0198 | TRUE | 0.3278 |
| Fgd5 | 2817 | 2274 | 2291 | 2923 | 2928 | 3147 | 2188 | 2419 | 2472 | 2910 | 3303 | 3107 | 2732.98 | 2360 | 3107 | 1.317 | 0.398 | 5.18E-08 | 2.15E-06 | 0.0035 | 0.0033 | 0.0035 | 0.0035 | TRUE | 0.0504 |
| Fgd6 | 1991 | 1344 | 1348 | 1506 | 1332 | 1603 | 1546 | 1430 | 1454 | 1499 | 1503 | 1582 | 1502.42 | 1477 | 1528 | 1.034 | 0.048 | 0.48652652 | 0.65247878 | 8.00E-04 | 0.004 | 0.0028 | 0.0028 | TRUE | 0.0213 |
| Fgf1 | 236 | 168 | 84 | 92 | 91 | 105 | 183 | 179 | 91 | 92 | 103 | 104 | 125.09 | 151 | 100 | 0.656 | -0.609 | 0.01264987 | 0.04923273 | 0.067 | 0.0214 | 0.0346 | 0.0346 | TRUE | 1.239 |
| Fgf10 | 353 | 316 | 434 | 322 | 312 | 359 | 274 | 336 | 468 | 321 | 352 | 354 | 350.91 | 359 | 342 | 0.955 | -0.066 | 0.6918862 | 0.81164884 | 0.0359 | 0.0092 | 0.0172 | 0.0172 | TRUE | 0.5378 |
| Fgf11 | 39 | 30 | 47 | 70 | 53 | 76 | 30 | 32 | 51 | 70 | 60 | 75 | 52.9 | 38 | 68 | 1.829 | 0.871 | 0.00279166 | 0.01561082 | 0.021 | 0.0472 | 0.0407 | 0.0407 | TRUE | 0.6418 |
| Fgf12 | 4 | 4 | 2 | 3 | 1 | 5 | 3 | 4 | 2 | 3 | 1 | 5 | 3.09 | 3 | 3 | 0.961 | -0.057 | 0.95819929 | NA | 0 | 0.7682 | 0.5272 | 0.5272 | TRUE | 0.2022 |
| Fgf13 | 603 | 434 | 427 | 460 | 375 | 424 | 468 | 462 | 461 | 458 | 423 | 419 | 448.36 | 464 | 433 | 0.934 | -0.098 | 0.32874245 | 0.50323614 | 0 | 0.0077 | 0.005 | 0.005 | TRUE | 0.0293 |
| Fgf14 | 15 | 8 | 4 | 8 | 10 | 6 | 12 | 9 | 4 | 8 | 11 | 6 | 8.27 | 8 | 8 | 1.006 | 0.009 | 0.98917209 | NA | 0 | 0.2889 | 0.22 | 0.22 | TRUE | 0.267 |
| Fgf15 | 243 | 184 | 257 | 405 | 346 | 375 | 189 | 196 | 277 | 403 | 390 | 370 | 304.23 | 221 | 388 | 1.766 | 0.82 | 4.44E-08 | 1.92E-06 | 0.019 | 0.0102 | 0.0126 | 0.0126 | TRUE | 0.5423 |
| Fgf16 | 0 | 0 | 1 | 0 | 0 | 0 | 0 | 0 | 1 | 0 | 0 | 0 | 0.18 | 0 | 0 | 0.552 | -0.857 | 0.8336548 | NA | 0 | 13.1817 | 10 | 10 | TRUE | 0.5365 |
| Fgf17 | 134 | 73 | 104 | 138 | 118 | 130 | 104 | 78 | 112 | 137 | 133 | 128 | 115.46 | 98 | 133 | 1.354 | 0.437 | 0.02492514 | 0.08122695 | 0.0053 | 0.023 | 0.0186 | 0.0186 | TRUE | 0.3012 |
| Fgf18 | 72 | 55 | 51 | 52 | 53 | 48 | 56 | 59 | 55 | 52 | 60 | 47 | 54.73 | 57 | 53 | 0.936 | -0.095 | 0.71450139 | 0.82843992 | 0 | 0.0457 | 0.0304 | 0.0304 | TRUE | 0.0981 |
| Fgf2 | 5 | 2 | 3 | 3 | 5 | 7 | 4 | 2 | 3 | 3 | 6 | 7 | 4.13 | 3 | 5 | 1.659 | 0.73 | 0.43846808 | NA | 0 | 0.5762 | 0.3853 | 0.3853 | TRUE | 0.2964 |
| Fgf20 | 3 | 0 | 0 | 0 | 2 | 2 | 2 | 0 | 0 | 0 | 2 | 2 | 1.09 | 1 | 1 | 1.679 | 0.748 | 0.71689736 | NA | 1.8421 | 2.17 | 2.1184 | 2.1184 | TRUE | 1.4382 |
| Fgf21 | 2 | 1 | 1 | 0 | 1 | 0 | 2 | 1 | 1 | 0 | 1 | 0 | 0.8 | 1 | 0 | 0.299 | -1.743 | 0.42212658 | NA | 0 | 2.9502 | 2.0092 | 2.0092 | TRUE | 0.5075 |
| Fgf22 | 18 | 16 | 14 | 15 | 13 | 12 | 14 | 17 | 15 | 15 | 15 | 12 | 14.59 | 15 | 14 | 0.901 | -0.151 | 0.76006512 | NA | 0 | 0.1649 | 0.1063 | 0.1063 | TRUE | 0.0699 |
| Fgf23 | 7 | 4 | 4 | 5 | 8 | 10 | 5 | 4 | 4 | 5 | 9 | 10 | 6.31 | 4 | 8 | 1.688 | 0.756 | 0.32302972 | NA | 0 | 0.3778 | 0.2537 | 0.2537 | TRUE | 0.325 |
| Fgf3 | 98 | 94 | 99 | 120 | 133 | 137 | 76 | 100 | 107 | 119 | 150 | 135 | 114.61 | 94 | 135 | 1.439 | 0.525 | 0.00924984 | 0.03910322 | 0.0131 | 0.0231 | 0.0203 | 0.0203 | TRUE | 0.3136 |
| Fgf4 | 12069 | 8773 | 11072 | 13053 | 11594 | 12574 | 9373 | 9332 | 11947 | 12993 | 13079 | 12412 | 11522.74 | 10217 | 12828 | 1.256 | 0.329 | 0.00015275 | 0.00172865 | 0.011 | 0.0026 | 0.0053 | 0.0053 | TRUE | 0.2674 |
| Fgf5 | 8 | 6 | 6 | 5 | 4 | 2 | 6 | 6 | 6 | 5 | 5 | 2 | 5.09 | 6 | 4 | 0.599 | -0.739 | 0.38449702 | NA | 0 | 0.4682 | 0.3084 | 0.3084 | TRUE | 0.3031 |
| Fgf6 | 0 | 0 | 0 | 1 | 1 | 1 | 0 | 0 | 0 | 1 | 1 | 1 | 0.52 | 0 | 1 | 5.852 | 2.549 | 0.3348364 | NA | 0 | 4.5739 | 3.5345 | 3.5345 | TRUE | 0.0847 |
| Fgf7 | 2 | 0 | 4 | 1 | 1 | 1 | 2 | 0 | 4 | 1 | 1 | 1 | 1.5 | 2 | 1 | 0.533 | -0.907 | 0.58130398 | NA | 0 | 1.586 | 1.2002 | 1.2002 | TRUE | 0.2521 |
| Fgf8 | 38 | 28 | 33 | 58 | 60 | 86 | 30 | 30 | 36 | 58 | 68 | 85 | 50.87 | 32 | 70 | 2.225 | 1.154 | 7.13E-05 | 0.00093103 | 0 | 0.049 | 0.0381 | 0.0381 | TRUE | 0.2348 |
| Fgf9 | 2 | 8 | 3 | 6 | 4 | 2 | 2 | 9 | 3 | 6 | 5 | 2 | 4.29 | 5 | 4 | 0.955 | -0.066 | 0.94667519 | NA | 0.2677 | 0.5545 | 0.4667 | 0.4667 | TRUE | 0.997 |
| Fgfbp1 | 908 | 479 | 478 | 280 | 220 | 248 | 705 | 510 | 516 | 279 | 248 | 245 | 417.03 | 577 | 257 | 0.445 | -1.169 | 2.05E-16 | 4.19E-14 | 0.0192 | 0.0081 | 0.0117 | 0.0117 | TRUE | 0.4583 |
| Fgfbp3 | 56 | 44 | 59 | 97 | 58 | 91 | 43 | 47 | 64 | 97 | 65 | 90 | 67.63 | 51 | 84 | 1.65 | 0.722 | 0.00600099 | 0.02821451 | 0.0254 | 0.0375 | 0.0342 | 0.0342 | TRUE | 0.3479 |
| Fgfr1 | 8988 | 6823 | 6588 | 7577 | 6671 | 7800 | 6981 | 7258 | 7108 | 7542 | 7526 | 7700 | 7352.34 | 7116 | 7589 | 1.067 | 0.093 | 0.05315415 | 0.14082154 | 1.00E-04 | 0.0027 | 0.0015 | 0.0015 | TRUE | 0.0037 |
| Fgfr1op | 3032 | 2202 | 2469 | 2159 | 1816 | 2363 | 2355 | 2342 | 2664 | 2149 | 2049 | 2333 | 2315.24 | 2454 | 2177 | 0.888 | -0.172 | 0.02505137 | 0.08146041 | 0.0043 | 0.0035 | 0.0038 | 0.0038 | TRUE | 0.0675 |
| Fgfr1op2 | 1240 | 996 | 937 | 997 | 889 | 987 | 963 | 1059 | 1011 | 992 | 1003 | 974 | 1000.52 | 1011 | 990 | 0.98 | -0.029 | 0.70166492 | 0.81928737 | 0 | 0.0048 | 0.0032 | 0.0032 | TRUE | 0.0219 |
| Fgfr2 | 338 | 181 | 156 | 169 | 114 | 153 | 263 | 193 | 168 | 168 | 129 | 151 | 178.54 | 208 | 149 | 0.716 | -0.482 | 0.01143882 | 0.04568177 | 0.0312 | 0.0157 | 0.0204 | 0.0204 | TRUE | 0.6112 |
| Fgfr3 | 672 | 559 | 586 | 651 | 558 | 615 | 522 | 595 | 632 | 648 | 629 | 607 | 605.57 | 583 | 628 | 1.08 | 0.112 | 0.26364138 | 0.43397745 | 0.0039 | 0.0063 | 0.0055 | 0.0055 | TRUE | 0.1047 |

| | | | | | | | | | | | | | | | | | | | | | | | | | |
|---|---|---|---|---|---|---|---|---|---|---|---|---|---|---|---|---|---|---|---|---|---|---|---|---|---|
| Fgfr4 | 387 | 257 | 355 | 410 | 365 | 437 | 301 | 273 | 383 | 408 | 412 | 431 | 368.04 | 319 | 417 | 1.31 | 0.389 | 0.00320003 | 0.01735791 | 0.0123 | 0.0089 | 0.0098 | 0.0098 | TRUE | 0.337 |
| Fgfrl1 | 968 | 772 | 915 | 1164 | 1018 | 1056 | 752 | 821 | 987 | 1159 | 1148 | 1042 | 984.96 | 853 | 1116 | 1.31 | 0.39 | 0.00015365 | 0.00173742 | 0.0102 | 0.0048 | 0.0066 | 0.0066 | TRUE | 0.2224 |
| Fgg | 17 | 16 | 9 | 20 | 16 | 19 | 13 | 17 | 10 | 20 | 18 | 19 | 16.11 | 13 | 19 | 1.421 | 0.507 | 0.29008201 | NA | 0 | 0.1496 | 0.1018 | 0.1018 | TRUE | 0.2231 |
| Fggy | 724 | 527 | 573 | 759 | 629 | 701 | 562 | 561 | 618 | 756 | 710 | 692 | 649.7 | 580 | 719 | 1.24 | 0.311 | 0.00064523 | 0.00523993 | 0.001 | 0.0061 | 0.0044 | 0.0044 | TRUE | 0.037 |
| Fgl1 | 26 | 24 | 26 | 16 | 10 | 9 | 20 | 26 | 28 | 16 | 11 | 9 | 18.31 | 25 | 12 | 0.492 | -1.022 | 0.02931604 | 0.09110984 | 0 | 0.1319 | 0.0958 | 0.0958 | TRUE | 0.3302 |
| Fgl2 | 170 | 109 | 140 | 158 | 124 | 165 | 132 | 116 | 151 | 157 | 140 | 163 | 143.18 | 133 | 153 | 1.155 | 0.208 | 0.23203251 | 0.39789331 | 0.0032 | 0.019 | 0.0148 | 0.0148 | TRUE | 0.1342 |
| Fgr | 120 | 94 | 77 | 125 | 109 | 128 | 93 | 100 | 83 | 124 | 123 | 126 | 108.34 | 92 | 124 | 1.352 | 0.435 | 0.02076069 | 0.0711589 | 0 | 0.0243 | 0.0161 | 0.0161 | TRUE | 0.0642 |
| Fh1 | 10969 | 7999 | 8526 | 9583 | 8460 | 9596 | 8519 | 8508 | 9199 | 9539 | 9544 | 9473 | 9130.41 | 8742 | 9519 | 1.089 | 0.123 | 0.01960895 | 0.06807645 | 9.00E-04 | 0.0027 | 0.0019 | 0.0019 | TRUE | 0.0254 |
| Fhad1 | 168 | 165 | 186 | 215 | 191 | 229 | 130 | 176 | 201 | 214 | 215 | 226 | 193.7 | 169 | 218 | 1.302 | 0.381 | 0.02560715 | 0.08271253 | 0.0185 | 0.0147 | 0.0158 | 0.0158 | TRUE | 0.4645 |
| Fhdc1 | 301 | 223 | 263 | 279 | 262 | 268 | 234 | 237 | 284 | 278 | 296 | 265 | 265.43 | 252 | 280 | 1.112 | 0.153 | 0.25158614 | 0.42057552 | 0.0035 | 0.0114 | 0.009 | 0.009 | TRUE | 0.135 |
| Fhit | 29 | 19 | 19 | 22 | 29 | 22 | 23 | 20 | 21 | 22 | 33 | 22 | 23.26 | 21 | 26 | 1.198 | 0.261 | 0.5159836 | 0.67828139 | 0 | 0.1043 | 0.0732 | 0.0732 | TRUE | 0.3542 |
| Fhl1 | 2225 | 1795 | 1823 | 2085 | 1874 | 2133 | 1728 | 1909 | 1967 | 2075 | 2114 | 2106 | 1983.24 | 1868 | 2098 | 1.125 | 0.169 | 0.01457487 | 0.05459327 | 0.0019 | 0.0036 | 0.003 | 0.003 | TRUE | 0.0539 |
| Fhl2 | 64 | 54 | 62 | 51 | 44 | 48 | 50 | 57 | 67 | 51 | 50 | 47 | 53.64 | 58 | 49 | 0.853 | -0.229 | 0.39195731 | 0.56559776 | 0 | 0.0466 | 0.0325 | 0.0325 | TRUE | 0.1555 |
| Fhl3 | 352 | 297 | 247 | 295 | 262 | 334 | 273 | 316 | 267 | 294 | 296 | 330 | 295.79 | 285 | 307 | 1.076 | 0.105 | 0.40747046 | 0.58030842 | 0.0029 | 0.0104 | 0.0082 | 0.0082 | TRUE | 0.0957 |
| Fhl4 | 290 | 178 | 134 | 194 | 153 | 150 | 225 | 189 | 145 | 193 | 173 | 148 | 178.82 | 186 | 171 | 0.914 | -0.129 | 0.48760292 | 0.65353606 | 0.0279 | 0.0157 | 0.0193 | 0.0193 | TRUE | 0.2256 |
| Fhl5 | 2 | 0 | 0 | 0 | 0 | 0 | 2 | 0 | 0 | 0 | 0 | 0 | 0.26 | 1 | 0 | 0.356 | -1.491 | 0.71362796 | NA | 0 | 9.1573 | 10 | 10 | TRUE | 0.968 |
| Fhod1 | 1497 | 1069 | 896 | 853 | 773 | 984 | 1163 | 1137 | 967 | 849 | 872 | 971 | 993.16 | 1089 | 897 | 0.824 | -0.28 | 0.00302634 | 0.01664953 | 0.0066 | 0.0048 | 0.0054 | 0.0054 | TRUE | 0.1133 |
| Fhod3 | 1574 | 1251 | 1271 | 1610 | 1492 | 1702 | 1222 | 1331 | 1371 | 1603 | 1683 | 1680 | 1481.73 | 1308 | 1655 | 1.267 | 0.341 | 2.42E-06 | 5.87E-05 | 0.0015 | 0.004 | 0.0031 | 0.0031 | TRUE | 0.0413 |
| Fibcd1 | 4 | 5 | 4 | 9 | 5 | 8 | 3 | 5 | 4 | 9 | 6 | 8 | 5.87 | 4 | 8 | 1.795 | 0.844 | 0.2852799 | NA | 0 | 0.406 | 0.2662 | 0.2662 | TRUE | 0.1356 |
| Fibin | 0 | 0 | 1 | 0 | 0 | 1 | 0 | 0 | 1 | 0 | 0 | 1 | 0.34 | 0 | 0 | 1.075 | 0.105 | 0.97570524 | NA | 0 | 6.885 | 6.5379 | 6.5379 | TRUE | 0.5365 |
| Fibp | 1052 | 771 | 760 | 750 | 759 | 860 | 817 | 820 | 820 | 747 | 856 | 849 | 818.15 | 819 | 817 | 0.997 | -0.004 | 0.96491875 | 0.9807735 | 0.0016 | 0.0053 | 0.0041 | 0.0041 | TRUE | 0.0687 |
| Ficd | 537 | 336 | 304 | 408 | 355 | 374 | 417 | 357 | 328 | 406 | 400 | 369 | 379.71 | 367 | 392 | 1.062 | 0.087 | 0.47145133 | 0.6403518 | 0.0065 | 0.0087 | 0.008 | 0.008 | TRUE | 0.1691 |
| Fig4 | 5673 | 4246 | 4242 | 4487 | 4157 | 4753 | 4406 | 4516 | 4577 | 4466 | 4690 | 4692 | 4557.88 | 4500 | 4616 | 1.026 | 0.037 | 0.48754871 | 0.65352468 | 4.00E-04 | 0.0029 | 0.0018 | 0.0018 | TRUE | 0.0098 |
| Figf | 208 | 150 | 159 | 234 | 207 | 220 | 162 | 160 | 172 | 233 | 234 | 217 | 196.05 | 165 | 228 | 1.388 | 0.473 | 0.00089458 | 0.00668736 | 0 | 0.0145 | 0.0094 | 0.0094 | TRUE | 0.0189 |
| Figla | 7 | 3 | 4 | 6 | 6 | 4 | 5 | 3 | 4 | 6 | 7 | 4 | 4.94 | 4 | 6 | 1.27 | 0.345 | 0.68343084 | NA | 0 | 0.4823 | 0.3131 | 0.3131 | TRUE | 0.1473 |
| Fign | 155 | 104 | 93 | 92 | 75 | 95 | 120 | 111 | 100 | 92 | 85 | 94 | 100.22 | 110 | 90 | 0.813 | -0.299 | 0.12919386 | 0.26592278 | 0 | 0.0261 | 0.0178 | 0.0178 | TRUE | 0.0684 |
| Fignl1 | 7157 | 4974 | 4882 | 5065 | 4243 | 4824 | 5559 | 5291 | 5268 | 5042 | 4787 | 4762 | 5117.87 | 5373 | 4864 | 0.905 | -0.144 | 0.00871522 | 0.03738495 | 7.00E-04 | 0.0029 | 0.002 | 0.002 | TRUE | 0.0126 |
| Fignl2 | 750 | 566 | 749 | 951 | 833 | 989 | 582 | 602 | 808 | 947 | 940 | 976 | 809.22 | 664 | 954 | 1.44 | 0.526 | 5.65E-06 | 0.00011729 | 0.016 | 0.0054 | 0.0084 | 0.0084 | TRUE | 0.4185 |
| Filip1 | 66 | 50 | 32 | 35 | 40 | 27 | 51 | 53 | 35 | 35 | 45 | 27 | 40.93 | 46 | 36 | 0.761 | -0.393 | 0.23156908 | 0.3975582 | 0.0324 | 0.0603 | 0.0529 | 0.0529 | TRUE | 0.2997 |
| Filip1l | 28 | 13 | 12 | 13 | 12 | 15 | 22 | 14 | 13 | 13 | 14 | 15 | 14.97 | 16 | 14 | 0.841 | -0.25 | 0.61672112 | NA | 0 | 0.1608 | 0.1132 | 0.1132 | TRUE | 0.5323 |
| Fip1l1 | 4684 | 3589 | 3578 | 3596 | 3354 | 3600 | 3638 | 3818 | 3861 | 3579 | 3784 | 3554 | 3705.49 | 3772 | 3639 | 0.965 | -0.052 | 0.37188076 | 0.54649756 | 7.00E-04 | 0.0031 | 0.0021 | 0.0021 | TRUE | 0.0146 |
| Fis1 | 2379 | 1626 | 1657 | 1704 | 1425 | 1634 | 1848 | 1730 | 1788 | 1696 | 1608 | 1613 | 1713.64 | 1789 | 1639 | 0.916 | -0.127 | 0.05415419 | 0.14249372 | 5.00E-04 | 0.0038 | 0.0025 | 0.0025 | TRUE | 0.0113 |
| Fitm1 | 43 | 19 | 48 | 58 | 44 | 55 | 33 | 20 | 52 | 58 | 50 | 54 | 44.51 | 35 | 54 | 1.539 | 0.622 | 0.06572815 | 0.1637818 | 0.0695 | 0.0557 | 0.0589 | 0.0589 | TRUE | 0.425 |
| Fitm2 | 668 | 470 | 459 | 546 | 484 | 501 | 519 | 500 | 495 | 543 | 546 | 495 | 516.34 | 505 | 528 | 1.045 | 0.063 | 0.5112392 | 0.67440166 | 0 | 0.007 | 0.0048 | 0.0048 | TRUE | 0.0365 |
| Fiz1 | 3686 | 2730 | 2349 | 2595 | 2519 | 2896 | 2863 | 2904 | 2535 | 2583 | 2842 | 2859 | 2764.12 | 2767 | 2761 | 0.997 | -0.004 | 0.95955733 | 0.97761443 | 0.004 | 0.0033 | 0.0035 | 0.0035 | TRUE | 0.0648 |
| Fjx1 | 364 | 287 | 183 | 285 | 249 | 275 | 283 | 305 | 197 | 284 | 281 | 271 | 270.25 | 262 | 279 | 1.062 | 0.087 | 0.57888051 | 0.72988172 | 0.0213 | 0.0112 | 0.0141 | 0.0141 | TRUE | 0.4962 |
| Fkbp10 | 920 | 607 | 767 | 915 | 854 | 979 | 715 | 646 | 828 | 911 | 963 | 966 | 838.06 | 730 | 947 | 1.299 | 0.377 | 0.00011928 | 0.00140662 | 0.0067 | 0.0053 | 0.0057 | 0.0057 | TRUE | 0.159 |
| Fkbp11 | 1150 | 912 | 952 | 1195 | 971 | 1141 | 893 | 970 | 1027 | 1190 | 1095 | 1126 | 1050.28 | 963 | 1137 | 1.182 | 0.242 | 0.0032295 | 0.01747795 | 0.0024 | 0.0047 | 0.0039 | 0.0039 | TRUE | 0.0512 |
| Fkbp14 | 249 | 189 | 198 | 257 | 195 | 253 | 193 | 201 | 214 | 256 | 220 | 250 | 222.27 | 203 | 242 | 1.197 | 0.259 | 0.06133902 | 0.15580569 | 0 | 0.0131 | 0.0093 | 0.0093 | TRUE | 0.0666 |
| Fkbp15 | 2178 | 1581 | 1566 | 1635 | 1511 | 1687 | 1692 | 1682 | 1690 | 1627 | 1705 | 1665 | 1676.72 | 1688 | 1666 | 0.987 | -0.019 | 0.75982315 | 0.8573099 | 0 | 0.0038 | 0.0022 | 0.0022 | TRUE | 0.0049 |
| Fkbp1a | 12196 | 8477 | 8515 | 9043 | 8005 | 9321 | 9472 | 9017 | 9188 | 9001 | 9030 | 9201 | 9151.62 | 9226 | 9077 | 0.984 | -0.024 | 0.62527569 | 0.76373375 | 2.00E-04 | 0.0027 | 0.0016 | 0.0016 | TRUE | 0.0068 |
| Fkbp1b | 463 | 341 | 420 | 494 | 434 | 481 | 360 | 363 | 453 | 492 | 490 | 475 | 438.61 | 392 | 486 | 1.241 | 0.312 | 0.00701207 | 0.03177665 | 0.0059 | 0.0078 | 0.0073 | 0.0073 | TRUE | 0.2102 |
| Fkbp2 | 916 | 723 | 716 | 803 | 735 | 841 | 711 | 769 | 773 | 799 | 829 | 830 | 785.28 | 751 | 819 | 1.093 | 0.128 | 0.1212139 | 0.25384766 | 0 | 0.0054 | 0.0036 | 0.0036 | TRUE | 0.0269 |
| Fkbp3 | 12287 | 8997 | 9869 | 9968 | 8655 | 10154 | 9543 | 9570 | 10649 | 9922 | 9764 | 10024 | 9911.8 | 9921 | 9903 | 0.998 | -0.002 | 0.96943866 | 0.9834812 | 0.0016 | 0.0027 | 0.0024 | 0.0024 | TRUE | 0.0502 |
| Fkbp4 | 54481 | 41130 | 40629 | 39465 | 35976 | 42347 | 42313 | 43749 | 43838 | 39283 | 40585 | 41803 | 41928.66 | 43300 | 40557 | 0.937 | -0.094 | 0.04750487 | 0.12980063 | 6.00E-04 | 0.0025 | 0.0016 | 0.0016 | TRUE | 0.0093 |
| Fkbp5 | 13569 | 9798 | 9705 | 10809 | 9737 | 11079 | 10538 | 10422 | 10472 | 10759 | 10984 | 10937 | 10685.4 | 10477 | 10893 | 1.04 | 0.056 | 0.21136073 | 0.37342137 | 0 | 0.0026 | 0.0014 | 0.0014 | TRUE | 0.0014 |
| Fkbp6 | 727 | 562 | 515 | 676 | 539 | 561 | 565 | 598 | 556 | 673 | 608 | 554 | 592.14 | 573 | 612 | 1.068 | 0.095 | 0.34411729 | 0.5183337 | 0.0039 | 0.0064 | 0.0056 | 0.0056 | TRUE | 0.0922 |
| Fkbp7 | 54 | 23 | 33 | 45 | 60 | 42 | 42 | 24 | 36 | 45 | 68 | 41 | 42.66 | 34 | 51 | 1.491 | 0.577 | 0.08227384 | 0.19306189 | 0.0478 | 0.058 | 0.0552 | 0.0552 | TRUE | 0.6054 |
| Fkbp8 | 11578 | 8485 | 8003 | 8464 | 7609 | 8725 | 8992 | 9025 | 8635 | 8425 | 8584 | 8613 | 8712.4 | 8884 | 8541 | 0.961 | -0.057 | 0.23680041 | 0.40340743 | 2.00E-04 | 0.0027 | 0.0016 | 0.0016 | TRUE | 0.0073 |
| Fkbp9 | 10088 | 6636 | 6533 | 7681 | 6285 | 7229 | 7835 | 7059 | 7049 | 7646 | 7090 | 7136 | 7302.42 | 7314 | 7291 | 0.996 | -0.005 | 0.93569552 | 0.96457959 | 0.0026 | 0.0028 | 0.0027 | 0.0027 | TRUE | 0.0478 |
| Fkbpl | 268 | 195 | 153 | 172 | 174 | 160 | 208 | 207 | 165 | 171 | 196 | 158 | 184.35 | 193 | 175 | 0.901 | -0.15 | 0.35476097 | 0.52912434 | 0.0088 | 0.0153 | 0.0134 | 0.0134 | TRUE | 0.172 |
| Fkrp | 1080 | 747 | 699 | 779 | 681 | 773 | 839 | 795 | 754 | 775 | 768 | 763 | 782.38 | 796 | 769 | 0.965 | -0.052 | 0.53364388 | 0.69295648 | 0 | 0.0055 | 0.0037 | 0.0037 | TRUE | 0.0279 |
| Fktn | 1267 | 900 | 946 | 1098 | 947 | 1087 | 984 | 957 | 1021 | 1093 | 1068 | 1073 | 1032.73 | 987 | 1078 | 1.092 | 0.127 | 0.08321149 | 0.19416945 | 0 | 0.0047 | 0.0029 | 0.0029 | TRUE | 0.0099 |
| Flad1 | 1732 | 1364 | 1335 | 1425 | 1263 | 1445 | 1345 | 1451 | 1440 | 1418 | 1425 | 1426 | 1417.7 | 1412 | 1423 | 1.009 | 0.013 | 0.85263942 | 0.9167369 | 0 | 0.0041 | 0.0027 | 0.0027 | TRUE | 0.0217 |
| Flcn | 2938 | 2244 | 2125 | 2232 | 1974 | 2317 | 2282 | 2387 | 2293 | 2222 | 2227 | 2287 | 2282.91 | 2321 | 2245 | 0.968 | -0.047 | 0.42557363 | 0.59821932 | 0 | 0.0035 | 0.0021 | 0.0021 | TRUE | 0.0075 |
| Flg2 | 0 | 0 | 0 | 0 | 0 | 0 | 0 | 0 | 0 | 0 | 0 | 0 | 0 | 0 | 0 | NA | NA | NA | NA | NA | NA | NA | NA | NA | NA |
| Fli1 | 5 | 5 | 4 | 2 | 1 | 3 | 4 | 5 | 4 | 2 | 1 | 3 | 3.27 | 4 | 2 | 0.456 | -1.131 | 0.29044076 | NA | 0 | 0.728 | 0.4652 | 0.4652 | TRUE | 0.1502 |
| Flii | 10325 | 7877 | 8073 | 8705 | 8053 | 9381 | 8019 | 8379 | 8711 | 8665 | 9085 | 9260 | 8686.4 | 8370 | 9003 | 1.076 | 0.106 | 0.05697042 | 0.14821227 | 0.0013 | 0.0027 | 0.0021 | 0.0021 | TRUE | 0.0166 |
| Flna | 23848 | 16203 | 15111 | 18052 | 15120 | 17634 | 18522 | 17235 | 16305 | 17969 | 17057 | 17407 | 17415.77 | 17354 | 17478 | 1.007 | 0.01 | 0.86475521 | 0.92542274 | 0.0019 | 0.0026 | 0.0025 | 0.0025 | TRUE | 0.0426 |
| Flnb | 26235 | 18847 | 21842 | 22561 | 21468 | 24079 | 20376 | 20047 | 23567 | 22457 | 24218 | 23770 | 22405.88 | 21330 | 23482 | 1.101 | 0.139 | 0.04457322 | 0.1238294 | 0.0048 | 0.0025 | 0.0034 | 0.0034 | TRUE | 0.1029 |
| Flnc | 2682 | 1988 | 1945 | 2956 | 2638 | 3058 | 2083 | 2115 | 2099 | 2942 | 2976 | 3019 | 2538.88 | 2099 | 2979 | 1.42 | 0.506 | 2.16E-19 | 6.06E-17 | 0 | 0.0034 | 0.0019 | 0.0019 | TRUE | 0.0017 |
| Flot1 | 1849 | 1572 | 1926 | 2085 | 2070 | 2279 | 1436 | 1672 | 2078 | 2075 | 2335 | 2250 | 1974.43 | 1729 | 2220 | 1.286 | 0.362 | 0.00104885 | 0.00749187 | 0.0187 | 0.0036 | 0.0083 | 0.0083 | TRUE | 0.3768 |
| Flot2 | 5711 | 4182 | 4912 | 5097 | 4729 | 5324 | 4435 | 4448 | 5300 | 5074 | 5335 | 5256 | 4974.63 | 4728 | 5222 | 1.105 | 0.144 | 0.05487683 | 0.1439711 | 0.0055 | 0.0029 | 0.0038 | 0.0038 | TRUE | 0.1358 |
| Flrt1 | 177 | 117 | 99 | 159 | 139 | 161 | 137 | 124 | 107 | 158 | 157 | 159 | 140.46 | 123 | 158 | 1.28 | 0.356 | 0.03800987 | 0.11026877 | 0 | 0.0193 | 0.014 | 0.014 | TRUE | 0.1291 |
| Flrt2 | 154 | 97 | 76 | 55 | 65 | 56 | 120 | 103 | 82 | 55 | 73 | 55 | 81.36 | 102 | 61 | 0.597 | -0.745 | 0.00193234 | 0.01177631 | 0.0203 | 0.0316 | 0.0283 | 0.0283 | TRUE | 0.1683 |
| Flrt3 | 317 | 182 | 165 | 226 | 174 | 171 | 246 | 194 | 178 | 225 | 196 | 169 | 201.31 | 206 | 197 | 0.951 | -0.073 | 0.66902708 | 0.79660147 | 0.0205 | 0.0142 | 0.0161 | 0.0161 | TRUE | 0.3395 |
| Flt1 | 98 | 91 | 75 | 113 | 107 | 114 | 76 | 97 | 81 | 112 | 121 | 113 | 99.93 | 85 | 115 | 1.366 | 0.45 | 0.02399759 | 0.07895892 | 0 | 0.0261 | 0.0185 | 0.0185 | TRUE | 0.1457 |
| Flt3 | 29 | 14 | 9 | 12 | 17 | 17 | 23 | 15 | 10 | 12 | 19 | 17 | 15.84 | 16 | 16 | 0.998 | -0.002 | 0.99659276 | NA | 0.0548 | 0.1521 | 0.1246 | 0.1246 | TRUE | 0.3586 |
| Flt3l | 17 | 12 | 11 | 11 | 9 | 22 | 13 | 13 | 12 | 11 | 10 | 22 | 13.44 | 13 | 14 | 1.135 | 0.183 | 0.73559608 | NA | 0 | 0.1787 | 0.1364 | 0.1364 | TRUE | 0.9423 |

| | | | | | | | | | | | | | | | | | | | | | | | | | |
|---|---|---|---|---|---|---|---|---|---|---|---|---|---|---|---|---|---|---|---|---|---|---|---|---|---|
| Flt4 | 1275 | 990 | 761 | 1002 | 890 | 866 | 990 | 1053 | 821 | 997 | 1004 | 855 | 953.45 | 955 | 952 | 0.996 | -0.005 | 0.95879957 | 0.97739513 | 0.0111 | 0.0049 | 0.0069 | 0.0069 | TRUE | 0.175 |
| Flywch1 | 1052 | 716 | 609 | 654 | 596 | 708 | 817 | 762 | 657 | 651 | 672 | 699 | 709.67 | 745 | 674 | 0.903 | -0.147 | 0.13700486 | 0.27724912 | 0.0055 | 0.0058 | 0.0057 | 0.0057 | TRUE | 0.1249 |
| Flywch2 | 77 | 50 | 64 | 85 | 72 | 87 | 60 | 53 | 69 | 85 | 81 | 86 | 72.29 | 61 | 84 | 1.385 | 0.469 | 0.0412853 | 0.11689596 | 0 | 0.0352 | 0.024 | 0.024 | TRUE | 0.1145 |
| Fmn1 | 131 | 121 | 88 | 88 | 83 | 80 | 102 | 129 | 95 | 88 | 94 | 79 | 97.6 | 109 | 87 | 0.8 | -0.322 | 0.1240323 | 0.25831176 | 0.0071 | 0.0267 | 0.0211 | 0.0211 | TRUE | 0.2527 |
| Fmn2 | 473 | 328 | 377 | 374 | 341 | 464 | 367 | 349 | 407 | 372 | 385 | 458 | 389.67 | 374 | 405 | 1.083 | 0.115 | 0.34121782 | 0.51526954 | 0.0069 | 0.0085 | 0.008 | 0.008 | TRUE | 0.1535 |
| Fmnl1 | 901 | 746 | 839 | 1181 | 1042 | 1410 | 700 | 794 | 905 | 1176 | 1175 | 1392 | 1023.58 | 800 | 1248 | 1.564 | 0.645 | 1.25E-09 | 8.12E-08 | 0.0118 | 0.0047 | 0.0071 | 0.0071 | TRUE | 0.1587 |
| Fmnl2 | 2386 | 1666 | 1583 | 1767 | 1535 | 1850 | 1853 | 1772 | 1708 | 1759 | 1732 | 1826 | 1775 | 1778 | 1772 | 0.996 | -0.005 | 0.93593428 | 0.96464483 | 7.00E-04 | 0.0038 | 0.0026 | 0.0026 | TRUE | 0.0172 |
| Fmnl3 | 1693 | 1194 | 986 | 984 | 858 | 1064 | 1315 | 1270 | 1064 | 979 | 968 | 1050 | 1107.75 | 1216 | 999 | 0.821 | -0.285 | 0.00202091 | 0.01219286 | 0.0065 | 0.0046 | 0.0052 | 0.0052 | TRUE | 0.1421 |
| Fmo1 | 19 | 14 | 7 | 5 | 4 | 2 | 15 | 15 | 8 | 5 | 5 | 2 | 8.11 | 13 | 4 | 0.305 | -1.713 | 0.02112972 | NA | 0 | 0.2946 | 0.223 | 0.223 | TRUE | 0.4347 |
| Fmo2 | 0 | 0 | 0 | 0 | 0 | 0 | 0 | 0 | 0 | 0 | 0 | 0 | 0 | 0 | 0 | NA | NA | NA | NA | NA | NA | NA | NA | NA | NA |
| Fmo3 | 0 | 0 | 0 | 0 | 0 | 0 | 0 | 0 | 0 | 0 | 0 | 0 | 0 | 0 | 0 | NA | NA | NA | NA | NA | NA | NA | NA | NA | NA |
| Fmo4 | 19 | 8 | 4 | 5 | 6 | 6 | 15 | 9 | 4 | 5 | 7 | 6 | 7.54 | 9 | 6 | 0.627 | -0.673 | 0.35980383 | NA | 0 | 0.3167 | 0.2497 | 0.2497 | TRUE | 0.2412 |
| Fmo5 | 175 | 143 | 137 | 138 | 127 | 137 | 136 | 152 | 148 | 137 | 143 | 135 | 141.95 | 145 | 138 | 0.956 | -0.065 | 0.69273584 | 0.81225511 | 0 | 0.0191 | 0.0125 | 0.0125 | TRUE | 0.0371 |
| Fmo6 | 0 | 0 | 0 | 0 | 0 | 0 | 0 | 0 | 0 | 0 | 0 | 0 | 0 | 0 | 0 | NA | NA | NA | NA | NA | NA | NA | NA | NA | NA |
| Fmo9 | 0 | 0 | 0 | 0 | 0 | 0 | 0 | 0 | 0 | 0 | 0 | 0 | 0 | 0 | 0 | NA | NA | NA | NA | NA | NA | NA | NA | NA | NA |
| Fmod | 1 | 0 | 0 | 0 | 0 | 0 | 1 | 0 | 0 | 0 | 0 | 0 | 0.13 | 0 | 0 | 0.552 | -0.857 | 0.8336548 | NA | 0 | 18.3121 | 10 | 10 | TRUE | 0.2566 |
| Fmr1 | 6635 | 4856 | 5184 | 5134 | 4279 | 5010 | 5153 | 5165 | 5593 | 5110 | 4827 | 4946 | 5132.51 | 5304 | 4961 | 0.936 | -0.096 | 0.09785995 | 0.21800379 | 0.0012 | 0.0029 | 0.0022 | 0.0022 | TRUE | 0.0276 |
| Fmr1nb | 2191 | 1496 | 1640 | 1607 | 1378 | 1507 | 1702 | 1591 | 1770 | 1600 | 1555 | 1488 | 1617.38 | 1688 | 1548 | 0.917 | -0.125 | 0.07652132 | 0.18286562 | 0.0014 | 0.0039 | 0.003 | 0.003 | TRUE | 0.0292 |
| Fn1 | 47322 | 36646 | 45048 | 47316 | 50313 | 57335 | 36753 | 38980 | 48606 | 47098 | 56758 | 56598 | 47465.72 | 41446 | 53485 | 1.291 | 0.368 | 0.00018905 | 0.00205172 | 0.0169 | 0.0025 | 0.007 | 0.007 | TRUE | 0.2796 |
| Fn3k | 33 | 20 | 46 | 61 | 61 | 69 | 26 | 21 | 50 | 61 | 69 | 68 | 49.03 | 32 | 66 | 2.064 | 1.045 | 0.00139275 | 0.00913848 | 0.0652 | 0.0508 | 0.0541 | 0.0541 | TRUE | 1.4458 |
| Fn3krp | 1010 | 759 | 823 | 994 | 811 | 976 | 784 | 807 | 888 | 989 | 915 | 963 | 891.26 | 826 | 956 | 1.158 | 0.212 | 0.01148442 | 0.04577433 | 0.0017 | 0.0051 | 0.0039 | 0.0039 | TRUE | 0.0494 |
| Fnbp1 | 2591 | 2034 | 2043 | 2163 | 1986 | 2343 | 2012 | 2164 | 2204 | 2153 | 2240 | 2313 | 2181.1 | 2127 | 2235 | 1.052 | 0.073 | 0.26858614 | 0.43913312 | 0.0014 | 0.0035 | 0.0027 | 0.0027 | TRUE | 0.0277 |
| Fnbp1l | 8412 | 6151 | 6131 | 6284 | 5671 | 6639 | 6533 | 6543 | 6615 | 6255 | 6397 | 6554 | 6482.93 | 6564 | 6402 | 0.975 | -0.036 | 0.46432612 | 0.63356183 | 1.00E-04 | 0.0028 | 0.0016 | 0.0016 | TRUE | 0.0053 |
| Fnbp4 | 5876 | 4477 | 4655 | 4827 | 4005 | 4967 | 4564 | 4762 | 5023 | 4805 | 4518 | 4903 | 4762.42 | 4783 | 4742 | 0.992 | -0.012 | 0.84958552 | 0.91494318 | 0.0018 | 0.0029 | 0.0025 | 0.0025 | TRUE | 0.0234 |
| Fnd3c2 | 0 | 0 | 0 | 1 | 0 | 0 | 0 | 0 | 0 | 1 | 0 | 0 | 0.17 | 0 | 0 | 2.094 | 1.067 | 0.79379329 | NA | 0 | 14.2885 | 10 | 10 | TRUE | 0.3906 |
| Fndc1 | 204 | 186 | 182 | 155 | 131 | 142 | 158 | 198 | 196 | 154 | 148 | 140 | 165.82 | 184 | 147 | 0.804 | -0.316 | 0.05322952 | 0.14094286 | 0 | 0.0167 | 0.0131 | 0.0131 | TRUE | 0.1775 |
| Fndc3a | 5349 | 3773 | 3882 | 4245 | 3730 | 4165 | 4154 | 4013 | 4189 | 4225 | 4208 | 4111 | 4150.18 | 4119 | 4181 | 1.015 | 0.022 | 0.67999978 | 0.80344191 | 0 | 0.003 | 0.0017 | 0.0017 | TRUE | 0.006 |
| Fndc3b | 2212 | 1598 | 1608 | 1830 | 1776 | 1880 | 1718 | 1700 | 1735 | 1822 | 2004 | 1856 | 1805.62 | 1718 | 1894 | 1.102 | 0.14 | 0.03437337 | 0.10254615 | 8.00E-04 | 0.0037 | 0.0026 | 0.0026 | TRUE | 0.0306 |
| Fndc3c1 | 136 | 81 | 77 | 74 | 64 | 93 | 106 | 86 | 83 | 74 | 72 | 92 | 85.42 | 92 | 79 | 0.862 | -0.215 | 0.33296571 | 0.50720228 | 0.0075 | 0.0302 | 0.0237 | 0.0237 | TRUE | 0.1905 |
| Fndc4 | 359 | 277 | 258 | 251 | 219 | 246 | 279 | 295 | 278 | 250 | 247 | 243 | 265.26 | 284 | 247 | 0.869 | -0.203 | 0.09984986 | 0.22082633 | 0 | 0.0114 | 0.0071 | 0.0071 | TRUE | 0.0118 |
| Fndc5 | 270 | 162 | 168 | 324 | 260 | 287 | 210 | 172 | 181 | 323 | 293 | 283 | 243.74 | 188 | 300 | 1.59 | 0.669 | 1.45E-06 | 3.93E-05 | 0.0035 | 0.0122 | 0.0096 | 0.0096 | TRUE | 0.1235 |
| Fndc7 | 7 | 11 | 8 | 6 | 6 | 5 | 5 | 12 | 9 | 6 | 7 | 5 | 7.24 | 9 | 6 | 0.693 | -0.53 | 0.46131042 | NA | 0 | 0.3297 | 0.2276 | 0.2276 | TRUE | 0.1737 |
| Fndc8 | 1 | 0 | 1 | 2 | 0 | 1 | 1 | 0 | 1 | 2 | 0 | 1 | 0.81 | 1 | 1 | 1.611 | 0.688 | 0.75133267 | NA | 0 | 2.9444 | 2.113 | 2.113 | TRUE | 0.131 |
| Fndc9 | 1 | 1 | 2 | 0 | 0 | 0 | 1 | 1 | 2 | 0 | 0 | 0 | 0.67 | 1 | 0 | 0.144 | -2.794 | 0.24876697 | NA | 0 | 3.559 | 2.8631 | 2.8631 | TRUE | 0.1682 |
| Fnip1 | 4283 | 3356 | 3939 | 4621 | 3988 | 4510 | 3326 | 3570 | 4250 | 4600 | 4499 | 4452 | 4116.16 | 3715 | 4517 | 1.216 | 0.283 | 0.00067923 | 0.00544481 | 0.0081 | 0.003 | 0.0047 | 0.0047 | TRUE | 0.1923 |
| Fnip2 | 4186 | 2989 | 3071 | 3264 | 3051 | 3524 | 3251 | 3179 | 3314 | 3249 | 3442 | 3479 | 3318.93 | 3248 | 3390 | 1.044 | 0.062 | 0.28399397 | 0.45597908 | 6.00E-04 | 0.0031 | 0.0021 | 0.0021 | TRUE | 0.0162 |
| Fnta | 9824 | 7198 | 7676 | 7601 | 7125 | 8171 | 7630 | 7656 | 8282 | 7566 | 8038 | 8066 | 7873.07 | 7856 | 7890 | 1.004 | 0.006 | 0.9117997 | 0.95140554 | 0.0016 | 0.0027 | 0.0022 | 0.0022 | TRUE | 0.0273 |
| Fntb | 3234 | 2378 | 2311 | 2308 | 2145 | 2472 | 2512 | 2529 | 2494 | 2297 | 2420 | 2440 | 2448.69 | 2512 | 2386 | 0.95 | -0.074 | 0.20639048 | 0.3682718 | 0 | 0.0034 | 0.0021 | 0.0021 | TRUE | 0.0129 |
| Focad | 4556 | 3569 | 3774 | 4182 | 3751 | 4045 | 3538 | 3796 | 4072 | 4163 | 4232 | 3993 | 3965.7 | 3802 | 4129 | 1.086 | 0.12 | 0.06973356 | 0.17116637 | 0.0025 | 0.003 | 0.0029 | 0.0029 | TRUE | 0.0469 |
| Folh1 | 20 | 14 | 16 | 9 | 8 | 8 | 16 | 15 | 17 | 9 | 9 | 8 | 12.26 | 16 | 9 | 0.543 | -0.882 | 0.10642384 | NA | 0 | 0.1957 | 0.1246 | 0.1246 | TRUE | 0.0244 |
| Folr1 | 1371 | 809 | 715 | 635 | 475 | 560 | 1065 | 861 | 771 | 632 | 536 | 553 | 736.26 | 899 | 574 | 0.637 | -0.65 | 8.13E-08 | 3.24E-06 | 0.0159 | 0.0056 | 0.0092 | 0.0092 | TRUE | 0.3178 |
| Folr2 | 15 | 7 | 1 | 4 | 2 | 6 | 12 | 7 | 1 | 4 | 2 | 6 | 5.39 | 7 | 4 | 0.596 | -0.748 | 0.42205394 | NA | 0.3886 | 0.4422 | 0.4274 | 0.4274 | TRUE | 0.2076 |
| Folr4 | 2 | 0 | 1 | 0 | 0 | 0 | 2 | 0 | 1 | 0 | 0 | 0 | 0.44 | 1 | 0 | 0.214 | -2.223 | 0.47051571 | NA | 0 | 5.4047 | 5.2946 | 5.2946 | TRUE | 0.2788 |
| Fopnl | 1696 | 1206 | 1190 | 1419 | 1200 | 1368 | 1317 | 1283 | 1284 | 1412 | 1354 | 1350 | 1333.44 | 1295 | 1372 | 1.06 | 0.083 | 0.21678788 | 0.37953812 | 0 | 0.0042 | 0.0025 | 0.0025 | TRUE | 0.008 |
| Fos | 131 | 92 | 106 | 99 | 111 | 125 | 102 | 98 | 114 | 99 | 125 | 123 | 110.19 | 105 | 116 | 1.106 | 0.145 | 0.45499728 | 0.62476716 | 0 | 0.0239 | 0.018 | 0.018 | TRUE | 0.1722 |
| Fosb | 8 | 11 | 3 | 4 | 6 | 11 | 6 | 12 | 3 | 4 | 7 | 11 | 7.13 | 7 | 7 | 1.026 | 0.037 | 0.96126047 | NA | 0.1532 | 0.335 | 0.2842 | 0.2842 | TRUE | 0.4238 |
| Fosl1 | 18 | 8 | 4 | 7 | 5 | 9 | 14 | 9 | 4 | 7 | 6 | 9 | 8.05 | 9 | 7 | 0.79 | -0.34 | 0.62934759 | NA | 0 | 0.2969 | 0.2333 | 0.2333 | TRUE | 0.2381 |
| Fosl2 | 113 | 121 | 111 | 68 | 72 | 73 | 88 | 129 | 120 | 68 | 81 | 72 | 92.87 | 112 | 74 | 0.66 | -0.599 | 0.00740368 | 0.0331311 | 0.0174 | 0.0279 | 0.0247 | 0.0247 | TRUE | 0.401 |
| Foxa1 | 266 | 187 | 160 | 189 | 178 | 192 | 207 | 199 | 173 | 188 | 201 | 190 | 192.77 | 193 | 193 | 0.997 | -0.004 | 0.97977411 | 0.98939603 | 0 | 0.0147 | 0.0105 | 0.0105 | TRUE | 0.0867 |
| Foxa2 | 18 | 12 | 1 | 7 | 3 | 2 | 14 | 13 | 1 | 7 | 3 | 2 | 6.69 | 9 | 4 | 0.439 | -1.187 | 0.19215761 | NA | 0.5899 | 0.3566 | 0.4196 | 0.4196 | TRUE | 1.8122 |
| Foxa3 | 125 | 100 | 81 | 122 | 107 | 124 | 97 | 106 | 87 | 121 | 121 | 122 | 109.23 | 97 | 121 | 1.253 | 0.326 | 0.08268041 | 0.19363001 | 0 | 0.0241 | 0.0161 | 0.0161 | TRUE | 0.0656 |
| Foxb1 | 0 | 0 | 0 | 0 | 0 | 0 | 0 | 0 | 0 | 0 | 0 | 0 | 0 | 0 | 0 | NA | NA | NA | NA | NA | NA | NA | NA | NA | NA |
| Foxb2 | 0 | 0 | 0 | 1 | 1 | 0 | 0 | 0 | 0 | 1 | 1 | 0 | 0.35 | 0 | 1 | 3.978 | 1.992 | 0.55384016 | NA | 0 | 6.6991 | 6.4234 | 6.4234 | TRUE | 0.2647 |
| Foxc1 | 46 | 27 | 30 | 35 | 37 | 30 | 36 | 29 | 32 | 35 | 42 | 30 | 33.83 | 32 | 36 | 1.089 | 0.123 | 0.71171829 | 0.82677093 | 0 | 0.0725 | 0.0507 | 0.0507 | TRUE | 0.163 |
| Foxc2 | 102 | 64 | 95 | 81 | 75 | 95 | 79 | 68 | 103 | 81 | 85 | 94 | 84.8 | 83 | 87 | 1.039 | 0.056 | 0.80646721 | 0.88818896 | 0.0115 | 0.0304 | 0.0252 | 0.0252 | TRUE | 0.3592 |
| Foxd1 | 40 | 40 | 39 | 60 | 45 | 68 | 31 | 43 | 42 | 60 | 51 | 67 | 48.88 | 39 | 59 | 1.551 | 0.633 | 0.02777512 | 0.08753969 | 0 | 0.0509 | 0.0383 | 0.0383 | TRUE | 0.2591 |
| Foxd2 | 45 | 27 | 21 | 26 | 17 | 19 | 35 | 29 | 23 | 26 | 19 | 19 | 25.02 | 29 | 21 | 0.735 | -0.444 | 0.26072845 | 0.43077006 | 0 | 0.0971 | 0.0715 | 0.0715 | TRUE | 0.1701 |
| Foxd3 | 928 | 690 | 501 | 693 | 676 | 678 | 721 | 734 | 541 | 690 | 763 | 669 | 686.16 | 665 | 707 | 1.061 | 0.086 | 0.46732158 | 0.63625712 | 0.0148 | 0.0059 | 0.0086 | 0.0086 | TRUE | 0.3094 |
| Foxd4 | 13 | 14 | 5 | 11 | 10 | 6 | 10 | 15 | 5 | 11 | 11 | 6 | 9.76 | 10 | 9 | 0.923 | -0.116 | 0.85710372 | NA | 0.0625 | 0.2454 | 0.1954 | 0.1954 | TRUE | 0.1627 |
| Foxe1 | 29 | 17 | 16 | 10 | 13 | 9 | 23 | 18 | 17 | 10 | 15 | 9 | 15.23 | 19 | 11 | 0.572 | -0.806 | 0.1080648 | NA | 0 | 0.1581 | 0.1095 | 0.1095 | TRUE | 0.2526 |
| Foxe3 | 3 | 5 | 2 | 2 | 3 | 1 | 2 | 5 | 2 | 2 | 3 | 1 | 2.69 | 3 | 2 | 0.647 | -0.628 | 0.59352499 | NA | 0 | 0.8819 | 0.6019 | 0.6019 | TRUE | 0.3933 |
| Foxf1 | 15 | 5 | 18 | 12 | 13 | 9 | 12 | 5 | 19 | 12 | 15 | 9 | 11.98 | 12 | 12 | 0.974 | -0.038 | 0.94976058 | NA | 0.1152 | 0.2002 | 0.1776 | 0.1776 | TRUE | 0.2222 |
| Foxf2 | 5 | 4 | 0 | 1 | 2 | 3 | 4 | 4 | 0 | 1 | 2 | 3 | 2.39 | 3 | 2 | 0.752 | -0.412 | 0.75419048 | NA | 0 | 0.9933 | 0.8207 | 0.8207 | TRUE | 0.8114 |
| Foxg1 | 148 | 96 | 86 | 83 | 73 | 90 | 115 | 102 | 93 | 83 | 82 | 89 | 93.94 | 103 | 85 | 0.816 | -0.293 | 0.15116906 | 0.2972839 | 0 | 0.0277 | 0.0192 | 0.0192 | TRUE | 0.1066 |
| Foxh1 | 1787 | 1308 | 1392 | 1639 | 1422 | 1645 | 1388 | 1391 | 1502 | 1631 | 1604 | 1624 | 1523.44 | 1427 | 1620 | 1.136 | 0.184 | 0.00651895 | 0.03004687 | 3.00E-04 | 0.004 | 0.0026 | 0.0026 | TRUE | 0.0248 |
| Foxi1 | 0 | 1 | 0 | 0 | 1 | 0 | 0 | 1 | 0 | 0 | 1 | 0 | 0.37 | 0 | 0 | 1.075 | 0.105 | 0.97520514 | NA | 0 | 6.4904 | 6.1969 | 6.1969 | TRUE | 0.5216 |
| Foxi2 | 4 | 2 | 2 | 1 | 1 | 3 | 3 | 2 | 2 | 1 | 1 | 3 | 2.08 | 2 | 2 | 0.686 | -0.543 | 0.67648408 | NA | 0 | 1.1422 | 0.723 | 0.723 | TRUE | 0.3417 |
| Foxi3 | 27 | 24 | 11 | 20 | 15 | 22 | 21 | 26 | 12 | 20 | 17 | 22 | 19.49 | 20 | 20 | 1.002 | 0.003 | 0.99539904 | 0.99730837 | 0 | 0.1241 | 0.0963 | 0.0963 | TRUE | 0.4616 |
| Foxj1 | 134 | 104 | 152 | 107 | 129 | 136 | 104 | 111 | 164 | 107 | 146 | 134 | 127.5 | 126 | 129 | 1.023 | 0.033 | 0.87737036 | 0.93155072 | 0.0364 | 0.021 | 0.0255 | 0.0255 | TRUE | 0.682 |

| | | | | | | | | | | | | | | | | | | | | | | | | | |
|---|---|---|---|---|---|---|---|---|---|---|---|---|---|---|---|---|---|---|---|---|---|---|---|---|---|
| Foxj2 | 1299 | 983 | 1102 | 1222 | 1163 | 1384 | 1009 | 1046 | 1189 | 1216 | 1312 | 1366 | 1189.69 | 1081 | 1298 | 1.202 | 0.266 | 0.0020424 | 0.01227055 | 0.0043 | 0.0044 | 0.0045 | 0.0045 | TRUE | 0.0901 |
| Foxj3 | 3834 | 2870 | 2887 | 3023 | 2774 | 3200 | 2978 | 3053 | 3115 | 3009 | 3129 | 3159 | 3073.81 | 3049 | 3099 | 1.017 | 0.024 | 0.66912819 | 0.79664374 | 3.00E-04 | 0.0032 | 0.002 | 0.002 | TRUE | 0.0079 |
| Foxk1 | 1930 | 1497 | 1570 | 1711 | 1543 | 1799 | 1499 | 1592 | 1694 | 1703 | 1741 | 1776 | 1667.5 | 1595 | 1740 | 1.092 | 0.127 | 0.07178019 | 0.17483883 | 0.0015 | 0.0038 | 0.003 | 0.003 | TRUE | 0.0354 |
| Foxk2 | 9063 | 6709 | 6817 | 8358 | 7261 | 8395 | 7039 | 7136 | 7355 | 8320 | 8191 | 8287 | 7721.41 | 7177 | 8266 | 1.152 | 0.204 | 2.21E-05 | 0.00036186 | 2.00E-04 | 0.0027 | 0.0015 | 0.0015 | TRUE | 0.0058 |
| Foxl1 | 1 | 0 | 0 | 1 | 0 | 1 | 1 | 0 | 0 | 1 | 0 | 1 | 0.46 | 0 | 1 | 1.94 | 0.956 | 0.73503911 | NA | 0 | 5.1562 | 3.9771 | 3.9771 | TRUE | 0.2566 |
| Foxl2 | 30 | 29 | 19 | 28 | 17 | 15 | 23 | 31 | 21 | 28 | 19 | 15 | 22.75 | 25 | 21 | 0.831 | -0.267 | 0.52790429 | 0.68874916 | 0.0253 | 0.1066 | 0.0848 | 0.0848 | TRUE | 0.538 |
| Foxl2os | 49 | 44 | 34 | 39 | 34 | 19 | 38 | 47 | 37 | 39 | 38 | 19 | 36.25 | 41 | 32 | 0.788 | -0.344 | 0.33249669 | 0.50686596 | 0.0468 | 0.0678 | 0.0625 | 0.0625 | TRUE | 0.9244 |
| Foxm1 | 5842 | 4297 | 4004 | 4366 | 4196 | 4664 | 4537 | 4571 | 4320 | 4346 | 4734 | 4604 | 4518.62 | 4476 | 4561 | 1.019 | 0.027 | 0.64487916 | 0.77893761 | 0.001 | 0.003 | 0.0022 | 0.0022 | TRUE | 0.0208 |
| Foxn1 | 0 | 1 | 2 | 2 | 1 | 1 | 0 | 1 | 2 | 2 | 1 | 1 | 1.22 | 1 | 1 | 1.337 | 0.419 | 0.81057392 | NA | 0 | 1.9431 | 1.3535 | 1.3535 | TRUE | 0.2998 |
| Foxn2 | 3827 | 2855 | 2807 | 3260 | 2908 | 3145 | 2972 | 3037 | 3029 | 3245 | 3281 | 3105 | 3111.33 | 3013 | 3210 | 1.066 | 0.092 | 0.10090894 | 0.22254151 | 2.00E-04 | 0.0032 | 0.0019 | 0.0019 | TRUE | 0.0101 |
| Foxn3 | 660 | 527 | 625 | 656 | 552 | 651 | 513 | 561 | 674 | 653 | 623 | 643 | 610.98 | 583 | 640 | 1.101 | 0.138 | 0.20735266 | 0.36908856 | 0.008 | 0.0063 | 0.007 | 0.007 | TRUE | 0.2213 |
| Foxn4 | 281 | 211 | 214 | 193 | 203 | 214 | 218 | 224 | 231 | 192 | 229 | 211 | 217.66 | 224 | 211 | 0.939 | -0.092 | 0.510218 | 0.67348965 | 0 | 0.0133 | 0.0093 | 0.0093 | TRUE | 0.0659 |
| Foxo1 | 2582 | 2062 | 2012 | 2147 | 1904 | 2253 | 2005 | 2193 | 2171 | 2137 | 2148 | 2224 | 2146.45 | 2123 | 2170 | 1.023 | 0.033 | 0.6141631 | 0.75615756 | 0.0011 | 0.0035 | 0.0025 | 0.0025 | TRUE | 0.0296 |
| Foxo3 | 2987 | 2371 | 2423 | 2590 | 2373 | 2628 | 2320 | 2522 | 2614 | 2578 | 2677 | 2594 | 2550.93 | 2485 | 2616 | 1.053 | 0.075 | 0.25226254 | 0.42118886 | 0.0017 | 0.0034 | 0.0027 | 0.0027 | TRUE | 0.0424 |
| Foxo4 | 2695 | 1975 | 2093 | 2266 | 2008 | 2387 | 2093 | 2101 | 2258 | 2256 | 2265 | 2356 | 2221.56 | 2151 | 2292 | 1.066 | 0.093 | 0.14142742 | 0.28362875 | 8.00E-04 | 0.0035 | 0.0024 | 0.0024 | TRUE | 0.0228 |
| Foxo6 | 387 | 319 | 332 | 379 | 357 | 434 | 301 | 339 | 358 | 377 | 403 | 428 | 367.75 | 333 | 403 | 1.214 | 0.28 | 0.01711752 | 0.06170509 | 0.0036 | 0.0089 | 0.0072 | 0.0072 | TRUE | 0.088 |
| Foxp1 | 2655 | 1943 | 2026 | 2522 | 2384 | 2852 | 2062 | 2067 | 2186 | 2510 | 2689 | 2815 | 2388.33 | 2105 | 2671 | 1.27 | 0.344 | 2.62E-07 | 9.26E-06 | 0.0018 | 0.0034 | 0.0028 | 0.0028 | TRUE | 0.0342 |
| Foxp2 | 11 | 5 | 3 | 1 | 2 | 0 | 9 | 5 | 3 | 1 | 2 | 0 | 3.39 | 6 | 1 | 0.182 | -2.457 | 0.04928886 | NA | 0 | 0.7012 | 0.5495 | 0.5495 | TRUE | 0.2856 |
| Foxp3 | 5 | 3 | 0 | 2 | 1 | 0 | 4 | 3 | 0 | 2 | 1 | 0 | 1.7 | 2 | 1 | 0.432 | -1.21 | 0.46294224 | NA | 0 | 1.3975 | 1.2578 | 1.2578 | TRUE | 0.7251 |
| Foxp4 | 2159 | 1719 | 1561 | 1753 | 1762 | 1917 | 1677 | 1828 | 1684 | 1745 | 1988 | 1892 | 1802.44 | 1730 | 1875 | 1.084 | 0.116 | 0.11573749 | 0.24580029 | 0.0025 | 0.0037 | 0.0034 | 0.0034 | TRUE | 0.0446 |
| Foxq1 | 208 | 192 | 172 | 183 | 160 | 175 | 162 | 204 | 186 | 182 | 180 | 173 | 181.13 | 184 | 178 | 0.975 | -0.037 | 0.81077607 | 0.8908004 | 0 | 0.0155 | 0.0117 | 0.0117 | TRUE | 0.1334 |
| Foxr1 | 146 | 129 | 122 | 106 | 91 | 143 | 113 | 137 | 132 | 106 | 103 | 141 | 121.93 | 127 | 117 | 0.918 | -0.123 | 0.53030768 | 0.69033619 | 0.013 | 0.0219 | 0.0193 | 0.0193 | TRUE | 0.3521 |
| Foxr2 | 186 | 126 | 128 | 166 | 138 | 192 | 144 | 134 | 138 | 165 | 156 | 190 | 154.51 | 139 | 170 | 1.225 | 0.293 | 0.07260171 | 0.1762691 | 0 | 0.0178 | 0.0127 | 0.0127 | TRUE | 0.1069 |
| Foxred1 | 2620 | 1864 | 1808 | 2038 | 1801 | 2072 | 2035 | 1983 | 1951 | 2029 | 2032 | 2045 | 2012.35 | 1990 | 2035 | 1.023 | 0.032 | 0.59036795 | 0.73882577 | 0 | 0.0036 | 0.0021 | 0.0021 | TRUE | 0.005 |
| Foxred2 | 1075 | 821 | 818 | 926 | 801 | 863 | 835 | 873 | 883 | 922 | 904 | 852 | 878.01 | 864 | 893 | 1.034 | 0.048 | 0.54332979 | 0.70095114 | 0 | 0.0051 | 0.0034 | 0.0034 | TRUE | 0.0191 |
| Foxs1 | 2 | 1 | 2 | 4 | 1 | 1 | 2 | 1 | 2 | 4 | 1 | 1 | 1.81 | 2 | 2 | 1.287 | 0.364 | 0.79546803 | NA | 0 | 1.3104 | 0.8662 | 0.8662 | TRUE | 0.6569 |
| Fpgs | 4815 | 3862 | 3966 | 3822 | 3669 | 4263 | 3740 | 4108 | 4279 | 3804 | 4139 | 4208 | 4046.42 | 4042 | 4050 | 1.003 | 0.004 | 0.95832707 | 0.97719689 | 0.0036 | 0.003 | 0.0032 | 0.0032 | TRUE | 0.0531 |
| Fpgt | 1352 | 913 | 872 | 927 | 786 | 868 | 1050 | 971 | 941 | 923 | 887 | 857 | 938.06 | 987 | 889 | 0.899 | -0.154 | 0.05900383 | 0.1515439 | 0.0015 | 0.005 | 0.0037 | 0.0037 | TRUE | 0.0385 |
| Fpr-rs3 | 0 | 0 | 0 | 0 | 0 | 0 | 0 | 0 | 0 | 0 | 0 | 0 | 0 | 0 | 0 | NA | NA | NA | NA | NA | NA | NA | NA | NA | NA |
| Fpr-rs4 | 0 | 0 | 0 | 0 | 0 | 0 | 0 | 0 | 0 | 0 | 0 | 0 | 0 | 0 | 0 | NA | NA | NA | NA | NA | NA | NA | NA | NA | NA |
| Fpr-rs6 | 0 | 0 | 0 | 0 | 0 | 0 | 0 | 0 | 0 | 0 | 0 | 0 | 0 | 0 | 0 | NA | NA | NA | NA | NA | NA | NA | NA | NA | NA |
| Fpr1 | 0 | 0 | 0 | 0 | 0 | 0 | 0 | 0 | 0 | 0 | 0 | 0 | 0 | 0 | 0 | NA | NA | NA | NA | NA | NA | NA | NA | NA | NA |
| Fpr2 | 0 | 0 | 0 | 0 | 0 | 0 | 0 | 0 | 0 | 0 | 0 | 0 | 0 | 0 | 0 | NA | NA | NA | NA | NA | NA | NA | NA | NA | NA |
| Fpr3 | 0 | 0 | 0 | 0 | 0 | 0 | 0 | 0 | 0 | 0 | 0 | 0 | 0 | 0 | 0 | NA | NA | NA | NA | NA | NA | NA | NA | NA | NA |
| Fra10ac1 | 1418 | 1064 | 1029 | 1124 | 936 | 1092 | 1101 | 1132 | 1110 | 1119 | 1056 | 1078 | 1099.34 | 1114 | 1084 | 0.973 | -0.039 | 0.58874079 | 0.73762904 | 0 | 0.0046 | 0.0028 | 0.0028 | TRUE | 0.0093 |
| Fras1 | 64 | 62 | 42 | 57 | 47 | 47 | 50 | 66 | 45 | 57 | 53 | 46 | 52.85 | 54 | 52 | 0.972 | -0.041 | 0.88155937 | 0.93382702 | 0 | 0.0473 | 0.0359 | 0.0359 | TRUE | 0.3154 |
| Frat1 | 214 | 129 | 124 | 175 | 134 | 159 | 166 | 137 | 134 | 174 | 151 | 157 | 153.26 | 146 | 161 | 1.099 | 0.136 | 0.41952083 | 0.59180924 | 0.0049 | 0.0179 | 0.0139 | 0.0139 | TRUE | 0.1726 |
| Frat2 | 493 | 433 | 450 | 453 | 424 | 505 | 383 | 461 | 486 | 451 | 478 | 499 | 459.46 | 443 | 476 | 1.078 | 0.108 | 0.3487954 | 0.52344945 | 0.007 | 0.0076 | 0.0074 | 0.0074 | TRUE | 0.1742 |
| Frem1 | 12 | 16 | 14 | 16 | 13 | 19 | 9 | 17 | 15 | 16 | 15 | 19 | 15.13 | 14 | 17 | 1.209 | 0.273 | 0.58452006 | NA | 0 | 0.1591 | 0.1133 | 0.1133 | TRUE | 0.4549 |
| Frem2 | 770 | 603 | 686 | 984 | 1069 | 1172 | 598 | 641 | 740 | 979 | 1206 | 1157 | 887 | 660 | 1114 | 1.691 | 0.758 | 9.35E-13 | 1.17E-10 | 0.01 | 0.0051 | 0.0069 | 0.0069 | TRUE | 0.1344 |
| Frem3 | 5 | 7 | 2 | 1 | 3 | 3 | 4 | 7 | 2 | 1 | 3 | 3 | 3.47 | 4 | 2 | 0.542 | -0.883 | 0.40865374 | NA | 0 | 0.6852 | 0.5018 | 0.5018 | TRUE | 0.3418 |
| Frg1 | 4391 | 3361 | 3268 | 3241 | 2868 | 3230 | 3410 | 3575 | 3526 | 3226 | 3235 | 3189 | 3360.25 | 3504 | 3217 | 0.918 | -0.123 | 0.02334602 | 0.07740444 | 0 | 0.0031 | 0.0018 | 0.0018 | TRUE | 0.0068 |
| Frk | 102 | 50 | 37 | 59 | 30 | 27 | 79 | 53 | 40 | 59 | 34 | 27 | 48.59 | 57 | 40 | 0.688 | -0.54 | 0.13179008 | 0.26948149 | 0.121 | 0.0512 | 0.0712 | 0.0712 | TRUE | 0.5541 |
| Frmd3 | 16 | 7 | 5 | 4 | 6 | 3 | 12 | 7 | 5 | 4 | 7 | 3 | 6.5 | 8 | 5 | 0.53 | -0.917 | 0.24290974 | NA | 0 | 0.3672 | 0.2734 | 0.2734 | TRUE | 0.6223 |
| Frmd4a | 1197 | 873 | 829 | 950 | 801 | 968 | 930 | 929 | 894 | 946 | 904 | 956 | 926.26 | 918 | 935 | 1.019 | 0.027 | 0.72283747 | 0.83413959 | 0 | 0.005 | 0.0031 | 0.0031 | TRUE | 0.01 |
| Frmd4b | 645 | 478 | 517 | 457 | 386 | 509 | 501 | 508 | 558 | 455 | 435 | 502 | 493.34 | 522 | 464 | 0.89 | -0.167 | 0.10584228 | 0.23064549 | 0.0023 | 0.0072 | 0.0057 | 0.0057 | TRUE | 0.0606 |
| Frmd5 | 614 | 397 | 413 | 460 | 382 | 431 | 477 | 422 | 446 | 458 | 431 | 425 | 443.18 | 448 | 438 | 0.976 | -0.035 | 0.73258515 | 0.84001922 | 0 | 0.0078 | 0.0055 | 0.0055 | TRUE | 0.0387 |
| Frmd6 | 1923 | 1392 | 1404 | 1324 | 1150 | 1382 | 1494 | 1481 | 1515 | 1318 | 1297 | 1364 | 1411.42 | 1497 | 1326 | 0.887 | -0.173 | 0.00895609 | 0.0382179 | 0 | 0.0041 | 0.0025 | 0.0025 | TRUE | 0.0075 |
| Frmd7 | 0 | 0 | 0 | 1 | 0 | 0 | 0 | 0 | 0 | 1 | 0 | 0 | 0.17 | 0 | 0 | 2.094 | 1.067 | 0.79379329 | NA | 0 | 14.2885 | 10 | 10 | TRUE | 0.3906 |
| Frmd8 | 3220 | 2151 | 1930 | 1956 | 1827 | 1847 | 2501 | 2288 | 2082 | 1947 | 2061 | 1823 | 2117.1 | 2290 | 1944 | 0.848 | -0.238 | 0.00340515 | 0.0182763 | 0.0053 | 0.0035 | 0.0043 | 0.0043 | TRUE | 0.0797 |
| Frmpd1 | 215 | 176 | 157 | 174 | 187 | 186 | 167 | 187 | 169 | 173 | 211 | 184 | 181.89 | 174 | 189 | 1.084 | 0.117 | 0.44762293 | 0.61788255 | 0 | 0.0155 | 0.0115 | 0.0115 | TRUE | 0.1045 |
| Frmpd3 | 2 | 1 | 0 | 3 | 3 | 5 | 2 | 1 | 0 | 3 | 3 | 5 | 2.32 | 1 | 4 | 4.112 | 2.04 | 0.13816039 | NA | 0 | 1.0238 | 0.7038 | 0.7038 | TRUE | 0.2685 |
| Frmpd4 | 2 | 1 | 0 | 1 | 1 | 0 | 2 | 1 | 0 | 1 | 1 | 0 | 0.79 | 1 | 1 | 0.78 | -0.358 | 0.86941444 | NA | 0 | 3.0022 | 2.1438 | 2.1438 | TRUE | 0.2798 |
| Frrs1 | 3194 | 2196 | 2225 | 2578 | 2381 | 2662 | 2481 | 2336 | 2401 | 2566 | 2686 | 2628 | 2516.2 | 2406 | 2627 | 1.091 | 0.126 | 0.03399688 | 0.10167764 | 3.00E-04 | 0.0034 | 0.0021 | 0.0021 | TRUE | 0.0093 |
| Frrs1l | 171 | 111 | 103 | 155 | 83 | 113 | 133 | 118 | 111 | 154 | 94 | 112 | 120.25 | 121 | 120 | 0.993 | -0.011 | 0.96008853 | 0.97787681 | 0.028 | 0.0221 | 0.0238 | 0.0238 | TRUE | 0.6421 |
| Frs2 | 3187 | 2351 | 2425 | 2474 | 2311 | 2751 | 2475 | 2501 | 2617 | 2463 | 2607 | 2716 | 2562.97 | 2531 | 2595 | 1.026 | 0.037 | 0.56477879 | 0.71920734 | 0.0013 | 0.0034 | 0.0025 | 0.0025 | TRUE | 0.0245 |
| Frs3 | 416 | 366 | 480 | 489 | 406 | 522 | 323 | 389 | 518 | 487 | 458 | 515 | 448.39 | 410 | 487 | 1.191 | 0.252 | 0.09018789 | 0.20499458 | 0.0275 | 0.0077 | 0.0137 | 0.0137 | TRUE | 0.5767 |
| Fry | 2057 | 1599 | 1650 | 2052 | 1928 | 2278 | 1598 | 1701 | 1780 | 2043 | 2175 | 2249 | 1924.17 | 1693 | 2156 | 1.274 | 0.35 | 8.22E-07 | 2.49E-05 | 0.0022 | 0.0037 | 0.0031 | 0.0031 | TRUE | 0.0304 |
| Fryl | 3169 | 2253 | 2368 | 2629 | 2425 | 2710 | 2461 | 2396 | 2555 | 2617 | 2736 | 2675 | 2573.42 | 2471 | 2676 | 1.083 | 0.115 | 0.05233019 | 0.13928405 | 3.00E-04 | 0.0033 | 0.0021 | 0.0021 | TRUE | 0.0104 |
| Frzb | 19 | 8 | 12 | 6 | 5 | 10 | 15 | 9 | 13 | 6 | 6 | 10 | 9.62 | 12 | 7 | 0.592 | -0.757 | 0.23178827 | NA | 0 | 0.2489 | 0.177 | 0.177 | TRUE | 0.306 |
| Fsbp | 51 | 33 | 39 | 36 | 33 | 34 | 40 | 35 | 42 | 36 | 37 | 34 | 37.24 | 39 | 36 | 0.912 | -0.133 | 0.6683077 | 0.79609449 | 0 | 0.0661 | 0.0429 | 0.0429 | TRUE | 0.0506 |
| Fscb | 2 | 0 | 0 | 1 | 0 | 0 | 2 | 0 | 0 | 1 | 0 | 0 | 0.42 | 1 | 0 | 0.681 | -0.553 | 0.8627148 | NA | 0 | 5.5819 | 5.4906 | 5.4906 | TRUE | 0.9727 |
| Fscn1 | 24741 | 19052 | 18451 | 18500 | 17322 | 20768 | 19215 | 20265 | 19908 | 18415 | 19541 | 20501 | 19641.04 | 19796 | 19486 | 0.984 | -0.023 | 0.68446069 | 0.80664285 | 0.0015 | 0.0025 | 0.0022 | 0.0022 | TRUE | 0.0283 |
| Fscn2 | 11 | 7 | 8 | 8 | 9 | 10 | 9 | 7 | 9 | 8 | 10 | 10 | 8.77 | 8 | 9 | 1.133 | 0.181 | 0.77458582 | NA | 0 | 0.2727 | 0.1723 | 0.1723 | TRUE | 0.0555 |
| Fscn3 | 1 | 0 | 0 | 0 | 0 | 0 | 1 | 0 | 0 | 0 | 0 | 0 | 0.13 | 0 | 0 | 0.552 | -0.857 | 0.8336548 | NA | 0 | 18.3121 | 10 | 10 | TRUE | 0.2566 |
| Fsd1 | 978 | 833 | 908 | 967 | 885 | 1070 | 760 | 886 | 980 | 963 | 998 | 1056 | 940.42 | 875 | 1006 | 1.152 | 0.204 | 0.04010811 | 0.11465304 | 0.0082 | 0.0049 | 0.006 | 0.006 | TRUE | 0.1656 |
| Fsd1l | 1702 | 1314 | 1158 | 1234 | 1211 | 1329 | 1322 | 1398 | 1249 | 1228 | 1366 | 1312 | 1312.57 | 1323 | 1302 | 0.984 | -0.024 | 0.75932347 | 0.8571376 | 0.0021 | 0.0042 | 0.0035 | 0.0035 | TRUE | 0.0296 |
| Fsd2 | 25 | 16 | 24 | 22 | 22 | 18 | 19 | 17 | 26 | 22 | 25 | 18 | 21.14 | 21 | 22 | 1.035 | 0.049 | 0.90663285 | 0.94912712 | 0 | 0.1146 | 0.0807 | 0.0807 | TRUE | 0.2343 |
| Fshb | 0 | 0 | 0 | 0 | 0 | 0 | 0 | 0 | 0 | 0 | 0 | 0 | 0 | 0 | 0 | NA | NA | NA | NA | NA | NA | NA | NA | NA | NA |

| | | | | | | | | | | | | | | | | | | | | | | | | | |
|---|---|---|---|---|---|---|---|---|---|---|---|---|---|---|---|---|---|---|---|---|---|---|---|---|---|
| Fshr | 1 | 1 | 1 | 0 | 0 | 0 | 1 | 1 | 1 | 0 | 0 | 0 | 0.49 | 1 | 0 | 0.196 | -2.347 | 0.38405676 | NA | 0 | 4.8735 | 3.7415 | 3.7415 | TRUE | 0.0718 |
| Fsip1 | 129 | 122 | 98 | 83 | 82 | 87 | 100 | 130 | 106 | 83 | 93 | 86 | 99.45 | 112 | 87 | 0.779 | -0.36 | 0.07689546 | 0.18351423 | 0 | 0.0263 | 0.0195 | 0.0195 | TRUE | 0.1904 |
| Fst | 38 | 32 | 32 | 31 | 40 | 23 | 30 | 34 | 35 | 31 | 45 | 23 | 32.79 | 33 | 33 | 1.004 | 0.006 | 0.98744712 | 0.99373352 | 0.0296 | 0.0747 | 0.0625 | 0.0625 | TRUE | 0.3913 |
| Fstl1 | 2262 | 1683 | 1893 | 2749 | 2365 | 2727 | 1757 | 1790 | 2043 | 2736 | 2668 | 2692 | 2280.97 | 1863 | 2699 | 1.45 | 0.536 | 8.19E-14 | 1.24E-11 | 0.0029 | 0.0035 | 0.0033 | 0.0033 | TRUE | 0.0847 |
| Fstl3 | 458 | 342 | 439 | 641 | 591 | 626 | 356 | 364 | 474 | 638 | 667 | 618 | 519.31 | 398 | 641 | 1.615 | 0.692 | 5.79E-09 | 3.17E-07 | 0.0114 | 0.007 | 0.0081 | 0.0081 | TRUE | 0.3144 |
| Fstl4 | 88 | 56 | 65 | 69 | 68 | 53 | 68 | 60 | 70 | 69 | 77 | 52 | 65.96 | 66 | 66 | 0.995 | -0.008 | 0.97519862 | 0.98653073 | 0 | 0.0384 | 0.0296 | 0.0296 | TRUE | 0.2934 |
| Fstl5 | 88 | 72 | 78 | 66 | 93 | 79 | 68 | 77 | 84 | 66 | 105 | 78 | 79.61 | 76 | 83 | 1.085 | 0.118 | 0.62426769 | 0.76326288 | 0.0212 | 0.0322 | 0.0291 | 0.0291 | TRUE | 0.4849 |
| Ftcd | 8 | 7 | 12 | 20 | 15 | 8 | 6 | 7 | 13 | 20 | 17 | 8 | 11.89 | 9 | 15 | 1.7 | 0.765 | 0.20542429 | NA | 0.1008 | 0.2018 | 0.1732 | 0.1732 | TRUE | 0.7647 |
| Fth1 | 30630 | 22000 | 20307 | 26133 | 22480 | 26724 | 23789 | 23401 | 21911 | 26013 | 25360 | 26381 | 24475.74 | 23034 | 25918 | 1.125 | 0.17 | 0.00095949 | 0.00704368 | 0.0011 | 0.0025 | 0.0019 | 0.0019 | TRUE | 0.0222 |
| Fthl17 | 39 | 32 | 19 | 11 | 14 | 17 | 30 | 34 | 21 | 11 | 16 | 17 | 21.39 | 28 | 15 | 0.511 | -0.968 | 0.02718054 | 0.08619736 | 0 | 0.1132 | 0.0857 | 0.0857 | TRUE | 0.3451 |
| Ftl1 | 111474 | 79120 | 79943 | 107376 | 89543 | 103182 | 86577 | 84159 | 86258 | 106882 | 101014 | 101857 | 94457.7 | 85665 | 103251 | 1.205 | 0.269 | 6.32E-09 | 3.41E-07 | 6.00E-04 | 0.0025 | 0.0015 | 0.0015 | TRUE | 0.0116 |
| Ftmt | 0 | 0 | 0 | 0 | 0 | 0 | 0 | 0 | 0 | 0 | 0 | 0 | 0 | 0 | 0 | NA | NA | NA | NA | NA | NA | NA | NA | NA | NA |
| Fto | 6974 | 4802 | 5173 | 5704 | 5048 | 5685 | 5416 | 5108 | 5582 | 5678 | 5695 | 5612 | 5515.04 | 5369 | 5662 | 1.054 | 0.077 | 0.16307629 | 0.31362345 | 8.00E-04 | 0.0029 | 0.002 | 0.002 | TRUE | 0.0217 |
| Ftsj1 | 1236 | 834 | 787 | 906 | 804 | 905 | 960 | 887 | 849 | 902 | 907 | 893 | 899.74 | 899 | 901 | 1.001 | 0.001 | 0.99282026 | 0.99592003 | 0.0011 | 0.0051 | 0.0037 | 0.0037 | TRUE | 0.0444 |
| Ftsj2 | 769 | 503 | 412 | 421 | 392 | 429 | 597 | 535 | 445 | 419 | 442 | 423 | 476.93 | 526 | 428 | 0.812 | -0.3 | 0.01206802 | 0.04755673 | 0.0097 | 0.0074 | 0.0082 | 0.0082 | TRUE | 0.2096 |
| Ftsj3 | 21409 | 15285 | 12864 | 13346 | 11775 | 13615 | 16627 | 16258 | 13880 | 13285 | 13283 | 13440 | 14462.36 | 15588 | 13336 | 0.855 | -0.225 | 0.00118878 | 0.00820835 | 0.0045 | 0.0026 | 0.0034 | 0.0034 | TRUE | 0.1122 |
| Ftsjd1 | 895 | 679 | 680 | 852 | 770 | 904 | 695 | 722 | 734 | 848 | 869 | 892 | 793.36 | 717 | 870 | 1.214 | 0.28 | 0.00051333 | 0.00439381 | 0 | 0.0054 | 0.0034 | 0.0034 | TRUE | 0.009 |
| Ftsjd2 | 5780 | 4343 | 4262 | 4356 | 4020 | 4573 | 4489 | 4620 | 4599 | 4336 | 4535 | 4514 | 4515.42 | 4569 | 4462 | 0.977 | -0.034 | 0.51358099 | 0.676118 | 2.00E-04 | 0.003 | 0.0017 | 0.0017 | TRUE | 0.0074 |
| Ftx | 129 | 126 | 95 | 93 | 88 | 79 | 100 | 134 | 103 | 93 | 99 | 78 | 101.09 | 112 | 90 | 0.802 | -0.318 | 0.13143144 | 0.26886355 | 0.0118 | 0.0259 | 0.0218 | 0.0218 | TRUE | 0.2801 |
| Fubp1 | 15785 | 12598 | 14279 | 12678 | 11555 | 13253 | 12260 | 13400 | 15407 | 12620 | 13035 | 13083 | 13300.74 | 13689 | 12913 | 0.943 | -0.084 | 0.27286022 | 0.44388699 | 0.0068 | 0.0026 | 0.0042 | 0.0042 | TRUE | 0.1473 |
| Fubp3 | 9978 | 6905 | 5865 | 5850 | 5163 | 6177 | 7749 | 7345 | 6328 | 5823 | 5824 | 6098 | 6527.94 | 7141 | 5915 | 0.828 | -0.272 | 0.00026676 | 0.00265888 | 0.0054 | 0.0028 | 0.0039 | 0.0039 | TRUE | 0.1206 |
| Fuca1 | 958 | 722 | 766 | 811 | 698 | 827 | 744 | 768 | 827 | 807 | 787 | 816 | 791.6 | 780 | 803 | 1.033 | 0.046 | 0.57883021 | 0.72988172 | 0 | 0.0054 | 0.0037 | 0.0037 | TRUE | 0.0325 |
| Fuca2 | 865 | 615 | 747 | 765 | 680 | 728 | 672 | 654 | 806 | 761 | 767 | 719 | 729.87 | 711 | 749 | 1.055 | 0.077 | 0.43451765 | 0.60626326 | 0.0057 | 0.0057 | 0.0057 | 0.0057 | TRUE | 0.1589 |
| Fuk | 3790 | 2994 | 3110 | 3574 | 3297 | 3666 | 2944 | 3185 | 3356 | 3558 | 3719 | 3619 | 3396.62 | 3162 | 3632 | 1.15 | 0.201 | 0.00191569 | 0.0116863 | 0.002 | 0.0031 | 0.0027 | 0.0027 | TRUE | 0.0452 |
| Fundc1 | 1169 | 845 | 775 | 809 | 646 | 752 | 908 | 899 | 836 | 805 | 729 | 742 | 819.89 | 881 | 759 | 0.861 | -0.216 | 0.01062719 | 0.04315307 | 0.0011 | 0.0053 | 0.0039 | 0.0039 | TRUE | 0.0343 |
| Fundc2 | 2143 | 1496 | 1761 | 1840 | 1633 | 1799 | 1664 | 1591 | 1900 | 1832 | 1842 | 1776 | 1767.56 | 1718 | 1817 | 1.057 | 0.081 | 0.3007335 | 0.47312977 | 0.0038 | 0.0038 | 0.0038 | 0.0038 | TRUE | 0.1011 |
| Fuom | 140 | 95 | 101 | 99 | 83 | 109 | 109 | 101 | 109 | 99 | 94 | 108 | 103.09 | 106 | 100 | 0.941 | -0.088 | 0.64527041 | 0.77909941 | 0 | 0.0254 | 0.0167 | 0.0167 | TRUE | 0.0446 |
| Furin | 1851 | 1390 | 1519 | 1683 | 1445 | 1712 | 1438 | 1479 | 1639 | 1675 | 1630 | 1690 | 1591.75 | 1519 | 1665 | 1.098 | 0.135 | 0.06421781 | 0.16114205 | 0.0019 | 0.0039 | 0.0032 | 0.0032 | TRUE | 0.0575 |
| Fus | 22140 | 15645 | 13017 | 11657 | 11430 | 12480 | 17195 | 16641 | 14045 | 11603 | 12894 | 12320 | 14116.52 | 15960 | 12272 | 0.769 | -0.379 | 9.50E-07 | 2.81E-05 | 0.0068 | 0.0026 | 0.0042 | 0.0042 | TRUE | 0.1346 |
| Fut1 | 1 | 1 | 0 | 2 | 1 | 6 | 1 | 1 | 0 | 2 | 1 | 6 | 1.81 | 1 | 3 | 4.853 | 2.279 | 0.16321302 | NA | 0 | 1.3092 | 0.9871 | 0.9871 | TRUE | 0.9512 |
| Fut10 | 503 | 400 | 413 | 520 | 448 | 531 | 391 | 425 | 446 | 518 | 505 | 524 | 468.16 | 421 | 516 | 1.229 | 0.298 | 0.00324359 | 0.01753427 | 0 | 0.0075 | 0.0052 | 0.0052 | TRUE | 0.0484 |
| Fut11 | 399 | 312 | 251 | 268 | 232 | 238 | 310 | 332 | 271 | 267 | 262 | 235 | 279.34 | 304 | 255 | 0.836 | -0.259 | 0.0487851 | 0.13235738 | 0.0038 | 0.0109 | 0.0088 | 0.0088 | TRUE | 0.0987 |
| Fut2 | 16 | 7 | 7 | 10 | 6 | 11 | 12 | 7 | 8 | 10 | 7 | 11 | 9.17 | 9 | 9 | 0.996 | -0.005 | 0.99341165 | NA | 0 | 0.2609 | 0.1813 | 0.1813 | TRUE | 0.4098 |
| Fut4 | 226 | 165 | 130 | 177 | 150 | 168 | 176 | 176 | 140 | 176 | 169 | 166 | 167.09 | 164 | 170 | 1.038 | 0.054 | 0.73707725 | 0.84327585 | 0 | 0.0166 | 0.0125 | 0.0125 | TRUE | 0.159 |
| Fut4-ps1 | 0 | 1 | 0 | 0 | 0 | 0 | 0 | 1 | 0 | 0 | 0 | 0 | 0.18 | 0 | 0 | 0.552 | -0.857 | 0.8336548 | NA | 0 | 13.3714 | 10 | 10 | TRUE | 0.5216 |
| Fut7 | 4 | 2 | 10 | 8 | 4 | 7 | 3 | 2 | 11 | 8 | 5 | 7 | 5.9 | 5 | 7 | 1.235 | 0.305 | 0.71963076 | NA | 0.2039 | 0.404 | 0.3477 | 0.3477 | TRUE | 1.5703 |
| Fut8 | 1667 | 1104 | 1099 | 1114 | 1009 | 1159 | 1295 | 1174 | 1186 | 1109 | 1138 | 1144 | 1174.34 | 1218 | 1130 | 0.927 | -0.11 | 0.14089233 | 0.28279373 | 0.001 | 0.0044 | 0.0032 | 0.0032 | TRUE | 0.0377 |
| Fut9 | 1612 | 1412 | 1956 | 2045 | 1816 | 2320 | 1252 | 1502 | 2111 | 2036 | 2049 | 2290 | 1873.14 | 1622 | 2125 | 1.311 | 0.391 | 0.0935067 | 0.21037444 | 0.0385 | 0.0037 | 0.0132 | 0.0385 | TRUE | 0.8387 |
| Fuz | 210 | 188 | 195 | 229 | 178 | 195 | 163 | 200 | 210 | 228 | 201 | 192 | 199.12 | 191 | 207 | 1.088 | 0.122 | 0.43375506 | 0.60567224 | 0.0084 | 0.0143 | 0.0126 | 0.0126 | TRUE | 0.196 |
| Fv1 | 20 | 16 | 11 | 6 | 8 | 16 | 16 | 17 | 12 | 6 | 9 | 16 | 12.54 | 15 | 10 | 0.694 | -0.528 | 0.35369047 | NA | 0.0268 | 0.1915 | 0.1503 | 0.1503 | TRUE | 0.8125 |
| Fxn | 853 | 585 | 532 | 516 | 478 | 569 | 662 | 622 | 574 | 514 | 539 | 562 | 578.89 | 619 | 538 | 0.867 | -0.206 | 0.03422916 | 0.10217989 | 0.0021 | 0.0065 | 0.0051 | 0.0051 | TRUE | 0.0481 |
| Fxr1 | 25985 | 20343 | 22335 | 24227 | 21059 | 24110 | 20181 | 21639 | 24099 | 24116 | 23757 | 23800 | 22931.98 | 21973 | 23891 | 1.087 | 0.121 | 0.06793377 | 0.16775697 | 0.004 | 0.0025 | 0.0031 | 0.0031 | TRUE | 0.0876 |
| Fxr2 | 18922 | 13777 | 14785 | 16615 | 14479 | 16247 | 14696 | 14654 | 15953 | 16539 | 16334 | 16038 | 15702.31 | 15101 | 16304 | 1.08 | 0.111 | 0.0373203 | 0.10894998 | 0.0012 | 0.0026 | 0.002 | 0.002 | TRUE | 0.0297 |
| Fxyd1 | 11 | 9 | 16 | 22 | 14 | 13 | 9 | 10 | 17 | 22 | 16 | 13 | 14.32 | 12 | 17 | 1.446 | 0.532 | 0.31834143 | NA | 0.04 | 0.168 | 0.1328 | 0.1328 | TRUE | 0.6005 |
| Fxyd2 | 23 | 14 | 18 | 21 | 20 | 22 | 18 | 15 | 19 | 21 | 23 | 22 | 19.56 | 17 | 22 | 1.247 | 0.319 | 0.45485422 | 0.6246379 | 0 | 0.1236 | 0.0797 | 0.0797 | TRUE | 0.0715 |
| Fxyd3 | 9 | 9 | 18 | 16 | 7 | 20 | 7 | 10 | 19 | 16 | 8 | 20 | 13.26 | 12 | 15 | 1.236 | 0.305 | 0.60513128 | NA | 0.1594 | 0.1812 | 0.175 | 0.175 | TRUE | 0.7806 |
| Fxyd4 | 14 | 8 | 18 | 14 | 9 | 18 | 11 | 9 | 19 | 14 | 10 | 18 | 13.44 | 13 | 14 | 1.09 | 0.125 | 0.82078452 | NA | 0.0487 | 0.1787 | 0.1434 | 0.1434 | TRUE | 0.5302 |
| Fxyd5 | 188 | 147 | 173 | 171 | 130 | 161 | 146 | 156 | 187 | 170 | 147 | 159 | 160.81 | 163 | 159 | 0.977 | -0.033 | 0.84194119 | 0.91013957 | 0.0051 | 0.0172 | 0.0137 | 0.0137 | TRUE | 0.1668 |
| Fxyd6 | 1435 | 1358 | 1991 | 1779 | 2009 | 2206 | 1115 | 1444 | 2148 | 1771 | 2266 | 2178 | 1820.35 | 1569 | 2072 | 1.321 | 0.401 | 0.17981507 | 0.33464802 | 0.0639 | 0.0037 | 0.0191 | 0.0639 | TRUE | 0.7607 |
| Fxyd7 | 5 | 4 | 1 | 7 | 4 | 10 | 4 | 4 | 1 | 7 | 5 | 10 | 5.09 | 3 | 7 | 2.295 | 1.198 | 0.17839396 | NA | 0 | 0.4676 | 0.3445 | 0.3445 | TRUE | 0.2306 |
| Fyb | 7 | 4 | 5 | 5 | 3 | 3 | 5 | 4 | 5 | 5 | 3 | 3 | 4.4 | 5 | 4 | 0.75 | -0.416 | 0.64120091 | NA | 0 | 0.5409 | 0.341 | 0.341 | TRUE | 0.1282 |
| Fyco1 | 1723 | 1282 | 1287 | 1448 | 1338 | 1422 | 1338 | 1364 | 1389 | 1441 | 1509 | 1404 | 1407.49 | 1364 | 1451 | 1.065 | 0.09 | 0.18541828 | 0.3417746 | 0 | 0.0041 | 0.0026 | 0.0026 | TRUE | 0.0144 |
| Fyn | 1853 | 1367 | 1378 | 1445 | 1338 | 1533 | 1439 | 1454 | 1487 | 1438 | 1509 | 1513 | 1473.52 | 1460 | 1487 | 1.019 | 0.027 | 0.68636915 | 0.80774354 | 0 | 0.004 | 0.0025 | 0.0025 | TRUE | 0.01 |
| Fyttd1 | 11975 | 8996 | 9271 | 9596 | 8335 | 9435 | 9300 | 9569 | 10003 | 9552 | 9403 | 9314 | 9523.52 | 9624 | 9423 | 0.979 | -0.03 | 0.55211777 | 0.70787759 | 6.00E-04 | 0.0027 | 0.0018 | 0.0018 | TRUE | 0.0145 |
| Fzd1 | 188 | 143 | 129 | 119 | 114 | 108 | 146 | 152 | 139 | 118 | 129 | 107 | 131.83 | 146 | 118 | 0.807 | -0.309 | 0.07526185 | 0.18072513 | 0 | 0.0204 | 0.014 | 0.014 | TRUE | 0.0714 |
| Fzd10 | 885 | 655 | 602 | 687 | 667 | 716 | 687 | 697 | 650 | 684 | 752 | 707 | 696.12 | 678 | 714 | 1.053 | 0.074 | 0.39442547 | 0.56801242 | 0 | 0.0058 | 0.004 | 0.004 | TRUE | 0.0251 |
| Fzd2 | 501 | 393 | 300 | 424 | 382 | 352 | 389 | 418 | 324 | 422 | 431 | 347 | 388.55 | 377 | 400 | 1.06 | 0.084 | 0.52051842 | 0.68185957 | 0.0125 | 0.0085 | 0.0097 | 0.0097 | TRUE | 0.1683 |
| Fzd3 | 1672 | 1219 | 1246 | 1645 | 1367 | 1540 | 1299 | 1297 | 1344 | 1637 | 1542 | 1520 | 1439.9 | 1313 | 1566 | 1.193 | 0.255 | 0.00019529 | 0.00209556 | 0 | 0.0041 | 0.0027 | 0.0027 | TRUE | 0.0191 |
| Fzd4 | 314 | 239 | 220 | 311 | 265 | 309 | 244 | 254 | 237 | 310 | 299 | 305 | 274.84 | 245 | 305 | 1.243 | 0.313 | 0.00987292 | 0.04098567 | 0 | 0.0111 | 0.007 | 0.007 | TRUE | 0.0109 |
| Fzd5 | 10620 | 8335 | 8276 | 9286 | 8633 | 9715 | 8248 | 8866 | 8930 | 9243 | 9739 | 9590 | 9102.68 | 8681 | 9524 | 1.097 | 0.134 | 0.01417591 | 0.05351723 | 0.0011 | 0.0027 | 0.002 | 0.002 | TRUE | 0.0235 |
| Fzd6 | 915 | 584 | 505 | 502 | 465 | 475 | 711 | 621 | 545 | 500 | 525 | 469 | 561.65 | 626 | 498 | 0.793 | -0.334 | 0.00316604 | 0.01721272 | 0.009 | 0.0066 | 0.0074 | 0.0074 | TRUE | 0.1731 |
| Fzd7 | 3102 | 2543 | 2865 | 2862 | 2719 | 3422 | 2409 | 2705 | 3091 | 2849 | 3067 | 3378 | 2916.61 | 2735 | 3098 | 1.134 | 0.181 | 0.05198753 | 0.13878554 | 0.011 | 0.0032 | 0.0059 | 0.0059 | TRUE | 0.1574 |
| Fzd8 | 216 | 145 | 136 | 125 | 125 | 143 | 168 | 154 | 147 | 124 | 141 | 141 | 145.89 | 156 | 135 | 0.865 | -0.21 | 0.20626746 | 0.36818458 | 0 | 0.0187 | 0.0129 | 0.0129 | TRUE | 0.0541 |
| Fzd9 | 547 | 433 | 418 | 541 | 547 | 543 | 425 | 461 | 451 | 539 | 617 | 536 | 504.67 | 446 | 564 | 1.266 | 0.34 | 0.00090023 | 0.00671049 | 0.0023 | 0.0071 | 0.0056 | 0.0056 | TRUE | 0.0779 |
| Fzr1 | 5991 | 4682 | 4528 | 4502 | 4137 | 4739 | 4653 | 4980 | 4886 | 4481 | 4667 | 4678 | 4724.2 | 4840 | 4609 | 0.953 | -0.07 | 0.20336777 | 0.36472149 | 7.00E-04 | 0.0029 | 0.002 | 0.002 | TRUE | 0.0142 |
| G0s2 | 57 | 27 | 39 | 69 | 56 | 62 | 44 | 29 | 42 | 69 | 63 | 61 | 51.35 | 38 | 64 | 1.668 | 0.738 | 0.00867286 | 0.03723674 | 0 | 0.0486 | 0.0364 | 0.0364 | TRUE | 0.3354 |
| G2e3 | 7798 | 6608 | 8767 | 10138 | 8987 | 10298 | 6056 | 7029 | 9460 | 10091 | 10138 | 10166 | 8823.35 | 7515 | 10132 | 1.348 | 0.431 | 0.02631672 | 0.08440938 | 0.027 | 0.0027 | 0.0096 | 0.027 | TRUE | 0.6254 |
| G3bp1 | 44403 | 33335 | 31941 | 31700 | 28825 | 32585 | 34486 | 35458 | 34464 | 31554 | 32518 | 32166 | 33441.03 | 34803 | 32079 | 0.922 | -0.118 | 0.00749874 | 0.0334309 | 2.00E-04 | 0.0025 | 0.0014 | 0.0014 | TRUE | 0.0033 |
| G3bp2 | 34321 | 23406 | 19308 | 18626 | 16426 | 18951 | 26656 | 24897 | 20833 | 18540 | 18530 | 18708 | 21360.6 | 24129 | 18593 | 0.77 | -0.376 | 2.07E-06 | 5.19E-05 | 0.0076 | 0.0025 | 0.0045 | 0.0045 | TRUE | 0.1745 |

| | | | | | | | | | | | | | | | | | | | | | | | | | |
|---|---|---|---|---|---|---|---|---|---|---|---|---|---|---|---|---|---|---|---|---|---|---|---|---|---|
| G530011O06 | 0 | 0 | 0 | 0 | 0 | 0 | 0 | 0 | 0 | 0 | 0 | 0 | 0 | 0 | 0 | NA | NA | NA | NA | NA | NA | NA | NA | NA | NA |
| G630025P09 | 24 | 7 | 4 | 16 | 11 | 7 | 19 | 7 | 4 | 16 | 12 | 7 | 10.94 | 10 | 12 | 1.129 | 0.176 | 0.79222773 | NA | 0.2541 | 0.219 | 0.2294 | 0.2294 | TRUE | 1.3711 |
| G630055G22 | 0 | 0 | 0 | 0 | 0 | 0 | 0 | 0 | 0 | 0 | 0 | 0 | 0 | 0 | 0 | NA | NA | NA | NA | NA | NA | NA | NA | NA | NA |
| G630071F17 | 0 | 1 | 0 | 0 | 0 | 0 | 0 | 1 | 0 | 0 | 0 | 0 | 0.18 | 0 | 0 | 0.552 | -0.857 | 0.8336548 | NA | 0 | 13.3714 | 10 | 10 | TRUE | 0.5216 |
| G630090E17 | 6 | 4 | 5 | 1 | 7 | 4 | 5 | 4 | 5 | 1 | 8 | 4 | 4.53 | 5 | 4 | 0.886 | -0.174 | 0.85305759 | NA | 0.0023 | 0.5262 | 0.4129 | 0.4129 | TRUE | 0.2906 |
| G630093K05 | 5 | 2 | 3 | 5 | 8 | 5 | 4 | 2 | 3 | 5 | 9 | 5 | 4.7 | 3 | 6 | 2.01 | 1.007 | 0.26020371 | NA | 0 | 0.507 | 0.3397 | 0.3397 | TRUE | 0.3098 |
| G6b | 2 | 1 | 1 | 2 | 1 | 3 | 2 | 1 | 1 | 2 | 1 | 3 | 1.63 | 1 | 2 | 1.636 | 0.71 | 0.6249077 | NA | 0 | 1.457 | 0.8805 | 0.8805 | TRUE | 0.1506 |
| G6pc | 1 | 0 | 0 | 1 | 3 | 1 | 1 | 0 | 0 | 1 | 3 | 1 | 1.02 | 0 | 2 | 5.137 | 2.361 | 0.24864342 | NA | 0 | 2.3171 | 1.7262 | 1.7262 | TRUE | 0.4395 |
| G6pc2 | 3 | 4 | 1 | 2 | 4 | 1 | 2 | 4 | 1 | 2 | 5 | 1 | 2.53 | 2 | 3 | 0.966 | -0.05 | 0.96769273 | NA | 0 | 0.9408 | 0.678 | 0.678 | TRUE | 0.4319 |
| G6pc3 | 1262 | 995 | 1269 | 1295 | 1225 | 1246 | 980 | 1058 | 1369 | 1289 | 1382 | 1230 | 1218.12 | 1136 | 1300 | 1.146 | 0.197 | 0.07903309 | 0.18745766 | 0.0169 | 0.0044 | 0.0082 | 0.0082 | TRUE | 0.3846 |
| G6pd2 | 1 | 0 | 0 | 0 | 0 | 1 | 1 | 0 | 0 | 0 | 0 | 1 | 0.29 | 0 | 0 | 1.075 | 0.105 | 0.97702352 | NA | 0 | 8.0647 | 7.5458 | 7.5458 | TRUE | 0.3835 |
| G6pdx | 2973 | 2081 | 2192 | 2156 | 1937 | 2250 | 2309 | 2214 | 2365 | 2146 | 2185 | 2221 | 2240 | 2296 | 2184 | 0.951 | -0.072 | 0.23421372 | 0.40038472 | 2.00E-04 | 0.0035 | 0.0022 | 0.0022 | TRUE | 0.0116 |
| G730013B05 | 267 | 204 | 229 | 257 | 227 | 265 | 207 | 217 | 247 | 256 | 256 | 262 | 240.82 | 224 | 258 | 1.155 | 0.208 | 0.11941927 | 0.25119347 | 0 | 0.0123 | 0.0087 | 0.0087 | TRUE | 0.0887 |
| Gaa | 6768 | 5162 | 5551 | 6163 | 6166 | 6676 | 5256 | 5491 | 5989 | 6135 | 6956 | 6590 | 6069.57 | 5579 | 6560 | 1.176 | 0.234 | 0.00072307 | 0.00570673 | 0.004 | 0.0028 | 0.0033 | 0.0033 | TRUE | 0.0504 |
| Gab1 | 28806 | 20368 | 19851 | 20990 | 18702 | 21604 | 22372 | 21665 | 21419 | 20893 | 21098 | 21326 | 21462.39 | 21819 | 21106 | 0.967 | -0.048 | 0.28803176 | 0.46018756 | 2.00E-04 | 0.0025 | 0.0014 | 0.0014 | TRUE | 0.0061 |
| Gab2 | 186 | 141 | 135 | 173 | 160 | 166 | 144 | 150 | 146 | 172 | 180 | 164 | 159.45 | 147 | 172 | 1.174 | 0.231 | 0.1373121 | 0.27763243 | 0 | 0.0173 | 0.0111 | 0.0111 | TRUE | 0.0192 |
| Gab3 | 54 | 31 | 34 | 34 | 17 | 29 | 42 | 33 | 37 | 34 | 19 | 29 | 32.21 | 37 | 27 | 0.732 | -0.45 | 0.20361368 | 0.36492167 | 0 | 0.076 | 0.0582 | 0.0582 | TRUE | 0.3848 |
| Gabarap | 3486 | 2475 | 2835 | 2824 | 2507 | 3044 | 2707 | 2633 | 3059 | 2811 | 2828 | 3005 | 2840.51 | 2800 | 2881 | 1.03 | 0.042 | 0.55740808 | 0.71234106 | 0.0035 | 0.0033 | 0.0034 | 0.0034 | TRUE | 0.0787 |
| Gabarapl1 | 3845 | 2919 | 3183 | 3164 | 2832 | 3321 | 2986 | 3105 | 3434 | 3149 | 3195 | 3278 | 3191.36 | 3175 | 3207 | 1.011 | 0.016 | 0.81692943 | 0.89487832 | 0.0025 | 0.0032 | 0.0029 | 0.0029 | TRUE | 0.0617 |
| Gabarapl2 | 6680 | 4135 | 3264 | 3570 | 2780 | 2923 | 5188 | 4398 | 3522 | 3554 | 3136 | 2885 | 3780.57 | 4369 | 3192 | 0.73 | -0.453 | 0.01233115 | 0.0483404 | 0.0234 | 0.0031 | 0.0093 | 0.0234 | TRUE | 0.1585 |
| Gabbr1 | 2926 | 2676 | 2762 | 2977 | 2431 | 2794 | 2273 | 2846 | 2980 | 2963 | 2742 | 2758 | 2760.49 | 2700 | 2821 | 1.046 | 0.065 | 0.48402975 | 0.65044468 | 0.0109 | 0.0033 | 0.0059 | 0.0059 | TRUE | 0.2362 |
| Gabbr2 | 8 | 8 | 9 | 7 | 3 | 4 | 6 | 9 | 10 | 7 | 3 | 4 | 6.46 | 8 | 5 | 0.593 | -0.753 | 0.32349519 | NA | 0 | 0.3696 | 0.2517 | 0.2517 | TRUE | 0.3233 |
| Gabpa | 12395 | 9059 | 9177 | 9499 | 8220 | 9551 | 9627 | 9636 | 9902 | 9455 | 9273 | 9428 | 9553.52 | 9722 | 9385 | 0.966 | -0.051 | 0.27470576 | 0.44572249 | 1.00E-04 | 0.0027 | 0.0014 | 0.0014 | TRUE | 0.0032 |
| Gabpb1 | 3788 | 2787 | 2936 | 2985 | 2440 | 2971 | 2942 | 2964 | 3168 | 2971 | 2753 | 2933 | 2955.18 | 3025 | 2886 | 0.955 | -0.067 | 0.28148524 | 0.45301905 | 0.0012 | 0.0032 | 0.0025 | 0.0025 | TRUE | 0.0206 |
| Gabpb2 | 5147 | 3999 | 3754 | 3499 | 3076 | 3430 | 3997 | 4254 | 4051 | 3483 | 3470 | 3386 | 3773.43 | 4101 | 3446 | 0.841 | -0.25 | 5.75E-06 | 0.00011824 | 4.00E-04 | 0.0031 | 0.0019 | 0.0019 | TRUE | 0.0129 |
| Gabra1 | 153 | 115 | 38 | 26 | 31 | 39 | 119 | 122 | 41 | 26 | 35 | 38 | 63.58 | 94 | 33 | 0.351 | -1.512 | 4.18E-05 | 0.00060674 | 0.1701 | 0.0397 | 0.0773 | 0.0773 | TRUE | 2.2723 |
| Gabra2 | 3 | 2 | 2 | 1 | 3 | 2 | 2 | 2 | 2 | 1 | 3 | 2 | 2.16 | 2 | 2 | 0.948 | -0.077 | 0.95156343 | NA | 0 | 1.0989 | 0.6862 | 0.6862 | TRUE | 0.228 |
| Gabra3 | 11 | 3 | 2 | 5 | 6 | 2 | 9 | 3 | 2 | 5 | 7 | 2 | 4.6 | 5 | 5 | 0.953 | -0.07 | 0.94128222 | NA | 0.2221 | 0.5174 | 0.4288 | 0.4288 | TRUE | 1.0587 |
| Gabra4 | 181 | 135 | 140 | 193 | 164 | 185 | 141 | 144 | 151 | 192 | 185 | 183 | 165.83 | 145 | 187 | 1.288 | 0.365 | 0.01666576 | 0.06044244 | 0 | 0.0167 | 0.0106 | 0.0106 | TRUE | 0.0131 |
| Gabra5 | 44 | 26 | 14 | 25 | 17 | 10 | 34 | 28 | 15 | 25 | 19 | 10 | 21.81 | 26 | 18 | 0.694 | -0.527 | 0.26882186 | 0.4393438 | 0.1293 | 0.1111 | 0.1161 | 0.1161 | TRUE | 0.4101 |
| Gabra6 | 0 | 0 | 0 | 0 | 0 | 0 | 0 | 0 | 0 | 0 | 0 | 0 | 0 | 0 | 0 | NA | NA | NA | NA | NA | NA | NA | NA | NA | NA |
| Gabrb1 | 3 | 1 | 1 | 1 | 0 | 0 | 2 | 1 | 1 | 1 | 0 | 0 | 0.91 | 1 | 0 | 0.245 | -2.029 | 0.33198448 | NA | 0 | 2.6031 | 1.8334 | 1.8334 | TRUE | 0.3906 |
| Gabrb2 | 6 | 13 | 15 | 10 | 7 | 4 | 5 | 14 | 16 | 10 | 8 | 4 | 9.41 | 12 | 7 | 0.638 | -0.648 | 0.35255681 | NA | 0.1935 | 0.2542 | 0.2362 | 0.2362 | TRUE | 1.3446 |
| Gabrb3 | 816 | 674 | 689 | 680 | 628 | 724 | 634 | 717 | 743 | 677 | 708 | 715 | 699.02 | 698 | 700 | 1.005 | 0.007 | 0.93930665 | 0.966328 | 0.0027 | 0.0058 | 0.0048 | 0.0048 | TRUE | 0.0813 |
| Gabrd | 2 | 5 | 4 | 6 | 5 | 2 | 2 | 5 | 4 | 6 | 6 | 2 | 4.13 | 4 | 5 | 1.238 | 0.308 | 0.75136737 | NA | 0 | 0.5764 | 0.4342 | 0.4342 | TRUE | 0.5711 |
| Gabre | 32 | 37 | 32 | 36 | 37 | 31 | 25 | 39 | 35 | 36 | 42 | 31 | 34.49 | 33 | 36 | 1.103 | 0.141 | 0.6791997 | 0.8028453 | 0 | 0.0712 | 0.0547 | 0.0547 | TRUE | 0.3857 |
| Gabrg1 | 5 | 5 | 7 | 4 | 1 | 4 | 4 | 5 | 8 | 4 | 1 | 4 | 4.3 | 6 | 3 | 0.553 | -0.854 | 0.36641733 | NA | 0 | 0.5533 | 0.3882 | 0.3882 | TRUE | 0.3368 |
| Gabrg2 | 18 | 12 | 8 | 8 | 13 | 7 | 14 | 13 | 9 | 8 | 15 | 7 | 10.82 | 12 | 10 | 0.823 | -0.282 | 0.63819297 | NA | 0 | 0.2215 | 0.1653 | 0.1653 | TRUE | 0.5676 |
| Gabrg3 | 23 | 11 | 5 | 11 | 10 | 10 | 18 | 12 | 5 | 11 | 11 | 10 | 11.18 | 12 | 11 | 0.901 | -0.15 | 0.80444675 | NA | 0.078 | 0.2145 | 0.1758 | 0.1758 | TRUE | 0.1835 |
| Gabrp | 31 | 34 | 44 | 64 | 59 | 63 | 24 | 36 | 47 | 64 | 67 | 62 | 50.03 | 36 | 64 | 1.81 | 0.856 | 0.00460206 | 0.02304741 | 0.0266 | 0.0498 | 0.0441 | 0.0441 | TRUE | 0.4248 |
| Gabrq | 10 | 5 | 5 | 8 | 4 | 4 | 8 | 5 | 5 | 8 | 5 | 4 | 5.82 | 6 | 6 | 0.882 | -0.18 | 0.81907044 | NA | 0 | 0.4098 | 0.2765 | 0.2765 | TRUE | 0.3552 |
| Gabrr1 | 79 | 57 | 26 | 55 | 47 | 64 | 61 | 61 | 28 | 55 | 53 | 63 | 53.5 | 50 | 57 | 1.133 | 0.18 | 0.56592715 | 0.72022053 | 0.0663 | 0.0467 | 0.0518 | 0.0518 | TRUE | 1.1269 |
| Gabrr2 | 1 | 0 | 2 | 0 | 0 | 2 | 1 | 0 | 2 | 0 | 0 | 2 | 0.82 | 1 | 1 | 0.69 | -0.536 | 0.81780122 | NA | 0 | 2.8993 | 2.6298 | 2.6298 | TRUE | 0.7053 |
| Gabrr3 | 3 | 3 | 1 | 0 | 0 | 0 | 2 | 3 | 1 | 0 | 0 | 0 | 1.1 | 2 | 0 | 0.086 | -3.533 | 0.0835999 | NA | 0 | 2.157 | 1.7853 | 1.7853 | TRUE | 0.1125 |
| Gad1 | 277 | 289 | 517 | 645 | 661 | 749 | 215 | 307 | 558 | 642 | 746 | 739 | 534.58 | 360 | 709 | 1.971 | 0.979 | 0.01809142 | 0.06414803 | 0.1215 | 0.0069 | 0.0341 | 0.1215 | TRUE | 1.8267 |
| Gad2 | 4 | 3 | 6 | 3 | 0 | 3 | 3 | 3 | 6 | 3 | 0 | 3 | 3.12 | 4 | 2 | 0.479 | -1.062 | 0.35799725 | NA | 0 | 0.7621 | 0.5911 | 0.5911 | TRUE | 0.5787 |
| Gadd45a | 1281 | 878 | 850 | 867 | 803 | 954 | 995 | 934 | 917 | 863 | 906 | 942 | 926.1 | 949 | 904 | 0.951 | -0.072 | 0.37024448 | 0.5449938 | 9.00E-04 | 0.005 | 0.0036 | 0.0036 | TRUE | 0.0228 |
| Gadd45b | 951 | 796 | 979 | 1096 | 899 | 1092 | 739 | 847 | 1056 | 1091 | 1014 | 1078 | 970.79 | 881 | 1061 | 1.208 | 0.272 | 0.01748059 | 0.0625094 | 0.016 | 0.0049 | 0.0084 | 0.0084 | TRUE | 0.3619 |
| Gadd45g | 70 | 64 | 115 | 72 | 68 | 83 | 54 | 68 | 124 | 72 | 77 | 82 | 79.47 | 82 | 77 | 0.941 | -0.087 | 0.76267194 | 0.8591534 | 0.0849 | 0.0322 | 0.0478 | 0.0478 | TRUE | 1.5878 |
| Gadd45gip1 | 4202 | 3152 | 3154 | 3304 | 3044 | 3486 | 3264 | 3353 | 3403 | 3289 | 3434 | 3441 | 3363.89 | 3340 | 3388 | 1.015 | 0.021 | 0.70489843 | 0.82131844 | 3.00E-04 | 0.0031 | 0.0019 | 0.0019 | TRUE | 0.008 |
| Gadl1 | 35 | 21 | 18 | 12 | 10 | 17 | 27 | 22 | 19 | 12 | 11 | 17 | 18.16 | 23 | 13 | 0.579 | -0.789 | 0.08636376 | 0.19896797 | 0 | 0.1329 | 0.093 | 0.093 | TRUE | 0.2225 |
| Gak | 6189 | 4679 | 4560 | 4613 | 4091 | 4814 | 4807 | 4977 | 4920 | 4592 | 4615 | 4752 | 4777.16 | 4901 | 4653 | 0.95 | -0.075 | 0.14536599 | 0.28921438 | 1.00E-04 | 0.0029 | 0.0017 | 0.0017 | TRUE | 0.0043 |
| Gal | 13 | 9 | 12 | 12 | 10 | 10 | 10 | 10 | 13 | 12 | 11 | 10 | 10.95 | 11 | 11 | 1.017 | 0.025 | 0.96487028 | NA | 0 | 0.2188 | 0.1407 | 0.1407 | TRUE | 0.0933 |
| Gal3st1 | 190 | 138 | 143 | 172 | 154 | 196 | 148 | 147 | 154 | 171 | 174 | 193 | 164.51 | 150 | 179 | 1.201 | 0.265 | 0.08737669 | 0.20058862 | 0 | 0.0168 | 0.0112 | 0.0112 | TRUE | 0.0511 |
| Gal3st2 | 1 | 0 | 1 | 0 | 0 | 0 | 1 | 0 | 1 | 0 | 0 | 0 | 0.31 | 1 | 0 | 0.307 | -1.706 | 0.62894649 | NA | 0 | 7.6657 | 7.2 | 7.2 | TRUE | 0.2067 |
| Gal3st3 | 138 | 97 | 99 | 83 | 81 | 80 | 107 | 103 | 107 | 83 | 91 | 79 | 95.02 | 106 | 84 | 0.796 | -0.329 | 0.09806862 | 0.21824375 | 0 | 0.0274 | 0.0178 | 0.0178 | TRUE | 0.0476 |
| Gal3st4 | 4 | 4 | 1 | 2 | 6 | 2 | 3 | 4 | 1 | 2 | 7 | 2 | 3.2 | 3 | 4 | 1.248 | 0.32 | 0.77241228 | NA | 0 | 0.7441 | 0.5658 | 0.5658 | TRUE | 0.8266 |
| Galc | 870 | 594 | 644 | 757 | 661 | 746 | 676 | 632 | 695 | 754 | 746 | 736 | 706.33 | 668 | 745 | 1.116 | 0.159 | 0.06214513 | 0.15737781 | 0 | 0.0058 | 0.0038 | 0.0038 | TRUE | 0.0246 |
| Gale | 742 | 591 | 429 | 564 | 501 | 580 | 576 | 629 | 463 | 561 | 565 | 573 | 561.16 | 556 | 566 | 1.018 | 0.026 | 0.82290552 | 0.89832647 | 0.01 | 0.0067 | 0.0077 | 0.0077 | TRUE | 0.2424 |
| Galk1 | 10541 | 8082 | 8936 | 9926 | 9050 | 10238 | 8187 | 8597 | 9642 | 9880 | 10209 | 10106 | 9436.91 | 8809 | 10065 | 1.143 | 0.193 | 0.00352344 | 0.01869359 | 0.0036 | 0.0027 | 0.003 | 0.003 | TRUE | 0.0835 |
| Galk2 | 1389 | 1025 | 963 | 1018 | 921 | 1174 | 1079 | 1090 | 1039 | 1013 | 1039 | 1159 | 1069.89 | 1069 | 1070 | 1.001 | 0.002 | 0.98473269 | 0.99216018 | 0.002 | 0.0046 | 0.0037 | 0.0037 | TRUE | 0.0634 |
| Galm | 660 | 517 | 521 | 595 | 535 | 699 | 513 | 550 | 562 | 592 | 604 | 690 | 585.08 | 542 | 629 | 1.163 | 0.218 | 0.02844254 | 0.08905351 | 0.0032 | 0.0065 | 0.0054 | 0.0054 | TRUE | 0.0871 |
| Galns | 664 | 492 | 484 | 567 | 554 | 587 | 516 | 523 | 522 | 564 | 625 | 579 | 555.01 | 520 | 589 | 1.132 | 0.179 | 0.05390625 | 0.14199817 | 0 | 0.0067 | 0.0044 | 0.0044 | TRUE | 0.0314 |
| Galnt1 | 5917 | 4308 | 4196 | 4308 | 3773 | 4280 | 4595 | 4582 | 4527 | 4288 | 4256 | 4225 | 4412.47 | 4568 | 4256 | 0.932 | -0.102 | 0.04081104 | 0.11612826 | 0 | 0.003 | 0.0016 | 0.0016 | TRUE | 7.00E-04 |
| Galnt10 | 1511 | 1077 | 957 | 951 | 880 | 998 | 1174 | 1146 | 1033 | 947 | 993 | 985 | 1046.04 | 1118 | 975 | 0.871 | -0.198 | 0.01312055 | 0.05051639 | 0.0017 | 0.0047 | 0.0036 | 0.0036 | TRUE | 0.0516 |
| Galnt11 | 3407 | 2533 | 2692 | 2870 | 2411 | 2758 | 2646 | 2694 | 2905 | 2857 | 2720 | 2723 | 2757.38 | 2748 | 2767 | 1.007 | 0.01 | 0.86986389 | 0.92804095 | 0.0013 | 0.0033 | 0.0025 | 0.0025 | TRUE | 0.0298 |
| Galnt12 | 653 | 516 | 632 | 642 | 635 | 689 | 507 | 549 | 682 | 639 | 716 | 680 | 628.91 | 579 | 678 | 1.174 | 0.231 | 0.04549939 | 0.12573913 | 0.0116 | 0.0062 | 0.008 | 0.008 | TRUE | 0.2788 |
| Galnt13 | 19 | 10 | 10 | 16 | 5 | 11 | 15 | 11 | 11 | 16 | 6 | 11 | 11.43 | 12 | 11 | 0.896 | -0.159 | 0.78808804 | NA | 4.00E-04 | 0.2097 | 0.1636 | 0.1636 | TRUE | 0.2042 |
| Galnt14 | 80 | 53 | 65 | 89 | 67 | 79 | 62 | 56 | 70 | 89 | 76 | 78 | 71.8 | 63 | 81 | 1.286 | 0.363 | 0.11716208 | 0.24816755 | 0 | 0.0354 | 0.0245 | 0.0245 | TRUE | 0.081 |

| | | | | | | | | | | | | | | | | | | | | | | | | |
|---|---|---|---|---|---|---|---|---|---|---|---|---|---|---|---|---|---|---|---|---|---|---|---|---|
| Galnt15 | 1 | 1 | 0 | 1 | 1 | 0 | 1 | 1 | 0 | 1 | 1 | 0 | 0.66 | 1 | 1 | 1.128 | 0.173 | 0.94106052 | NA | 0 | 3.5899 | 2.4471 | 2.4471 | TRUE | 0.2681 |
| Galnt16 | 272 | 169 | 176 | 254 | 172 | 173 | 211 | 180 | 190 | 253 | 194 | 171 | 199.76 | 194 | 206 | 1.061 | 0.086 | 0.61294664 | 0.75508227 | 0.0193 | 0.0143 | 0.0158 | 0.0158 | TRUE | 0.4411 |
| Galnt18 | 47 | 45 | 44 | 61 | 45 | 43 | 37 | 48 | 47 | 61 | 51 | 42 | 47.63 | 44 | 51 | 1.175 | 0.233 | 0.4246349 | 0.59737271 | 0 | 0.0522 | 0.04 | 0.04 | TRUE | 0.2136 |
| Galnt2 | 7787 | 5468 | 5526 | 6346 | 5575 | 5990 | 6048 | 5816 | 5963 | 6317 | 6289 | 5913 | 6057.6 | 5942 | 6173 | 1.039 | 0.055 | 0.30625336 | 0.47875744 | 6.00E-04 | 0.0028 | 0.0019 | 0.0019 | TRUE | 0.0166 |
| Galnt3 | 300 | 205 | 232 | 309 | 309 | 298 | 233 | 218 | 250 | 308 | 349 | 294 | 275.29 | 234 | 317 | 1.354 | 0.437 | 0.00079741 | 0.00613381 | 0.0025 | 0.011 | 0.0085 | 0.0085 | TRUE | 0.0849 |
| Galnt4 | 303 | 219 | 142 | 191 | 167 | 179 | 235 | 233 | 153 | 190 | 188 | 177 | 196.12 | 207 | 185 | 0.89 | -0.168 | 0.34071359 | 0.51478006 | 0.0237 | 0.0145 | 0.0172 | 0.0172 | TRUE | 0.5464 |
| Galnt5 | 0 | 0 | 0 | 0 | 0 | 0 | 0 | 0 | 0 | 0 | 0 | 0 | 0 | 0 | 0 | NA | NA | NA | NA | NA | NA | NA | NA | NA | NA |
| Galnt6 | 1164 | 830 | 818 | 865 | 832 | 954 | 904 | 883 | 883 | 861 | 939 | 942 | 901.81 | 890 | 914 | 1.026 | 0.037 | 0.63701451 | 0.77323554 | 0 | 0.0051 | 0.0034 | 0.0034 | TRUE | 0.0308 |
| Galnt7 | 2578 | 1716 | 1770 | 1805 | 1506 | 1634 | 2002 | 1825 | 1910 | 1797 | 1699 | 1613 | 1807.66 | 1912 | 1703 | 0.89 | -0.168 | 0.01788355 | 0.06358446 | 0.002 | 0.0037 | 0.0031 | 0.0031 | TRUE | 0.0284 |
| Galnt9 | 36 | 31 | 33 | 32 | 31 | 29 | 28 | 33 | 36 | 32 | 35 | 29 | 32 | 32 | 32 | 0.992 | -0.011 | 0.97327693 | 0.98545797 | 0 | 0.0765 | 0.0512 | 0.0512 | TRUE | 0.1092 |
| Galntl5 | 2 | 4 | 5 | 3 | 1 | 2 | 2 | 4 | 5 | 3 | 1 | 2 | 2.88 | 4 | 2 | 0.559 | -0.839 | 0.46692867 | NA | 0 | 0.8248 | 0.5733 | 0.5733 | TRUE | 0.5622 |
| Galntl6 | 11 | 7 | 2 | 3 | 1 | 0 | 9 | 7 | 2 | 3 | 1 | 0 | 3.71 | 6 | 1 | 0.225 | -2.151 | 0.07106971 | NA | 0.2912 | 0.6412 | 0.5682 | 0.5682 | TRUE | 0.7044 |
| Galp | 2 | 1 | 0 | 2 | 0 | 1 | 2 | 1 | 0 | 2 | 0 | 1 | 0.93 | 1 | 1 | 1.12 | 0.164 | 0.93546391 | NA | 0 | 2.5441 | 1.8977 | 1.8977 | TRUE | 0.1669 |
| Galr1 | 3 | 0 | 1 | 1 | 0 | 1 | 2 | 0 | 1 | 1 | 0 | 1 | 0.9 | 1 | 1 | 0.574 | -0.8 | 0.7018232 | NA | 0 | 2.64 | 1.9656 | 1.9656 | TRUE | 0.1659 |
| Galr2 | 67 | 31 | 44 | 60 | 45 | 84 | 52 | 33 | 47 | 60 | 51 | 83 | 54.32 | 44 | 65 | 1.455 | 0.541 | 0.06804536 | 0.16791649 | 0.0404 | 0.0461 | 0.0445 | 0.0445 | TRUE | 0.5568 |
| Galr3 | 2 | 1 | 2 | 1 | 2 | 1 | 2 | 1 | 2 | 1 | 2 | 1 | 1.5 | 2 | 1 | 0.879 | -0.186 | 0.90143055 | NA | 0 | 1.5801 | 0.9492 | 0.9492 | TRUE | 0.1551 |
| Galt | 1285 | 992 | 1056 | 1130 | 981 | 1116 | 998 | 1055 | 1139 | 1125 | 1107 | 1102 | 1087.62 | 1064 | 1111 | 1.045 | 0.064 | 0.41146655 | 0.58425307 | 0.0015 | 0.0046 | 0.0035 | 0.0035 | TRUE | 0.0455 |
| Gamt | 64 | 64 | 49 | 63 | 63 | 54 | 50 | 68 | 53 | 63 | 71 | 53 | 59.62 | 57 | 62 | 1.099 | 0.136 | 0.60446692 | 0.7493188 | 0 | 0.0422 | 0.0327 | 0.0327 | TRUE | 0.2432 |
| Gan | 633 | 423 | 303 | 361 | 313 | 290 | 492 | 450 | 327 | 359 | 353 | 286 | 377.87 | 423 | 333 | 0.785 | -0.349 | 0.02284784 | 0.07607432 | 0.0273 | 0.0087 | 0.0143 | 0.0143 | TRUE | 0.447 |
| Ganab | 19416 | 13886 | 13912 | 15820 | 14164 | 15525 | 15080 | 14770 | 15011 | 15747 | 15978 | 15326 | 15318.69 | 14954 | 15684 | 1.049 | 0.069 | 0.13271575 | 0.27079318 | 2.00E-04 | 0.0026 | 0.0014 | 0.0014 | TRUE | 0.0049 |
| Ganc | 361 | 247 | 201 | 234 | 206 | 234 | 280 | 263 | 217 | 233 | 232 | 231 | 242.71 | 253 | 232 | 0.913 | -0.131 | 0.35057507 | 0.52523789 | 0.005 | 0.0122 | 0.01 | 0.01 | TRUE | 0.1713 |
| Gap43 | 180 | 100 | 107 | 155 | 131 | 205 | 140 | 106 | 115 | 154 | 148 | 202 | 144.34 | 120 | 168 | 1.39 | 0.475 | 0.01231397 | 0.0483128 | 0.019 | 0.0188 | 0.0189 | 0.0189 | TRUE | 0.341 |
| Gapdh | 173020 | 128554 | 144585 | 150707 | 134521 | 158712 | 134377 | 136741 | 156006 | 150014 | 151754 | 156673 | 147594.16 | 142375 | 152814 | 1.073 | 0.102 | 0.10925447 | 0.23606563 | 0.0037 | 0.0024 | 0.0029 | 0.0029 | TRUE | 0.0859 |
| Gapdhs | 0 | 0 | 0 | 0 | 0 | 0 | 0 | 0 | 0 | 0 | 0 | 0 | 0 | 0 | 0 | NA | NA | NA | NA | NA | NA | NA | NA | NA | NA |
| Gapt | 0 | 0 | 0 | 0 | 0 | 0 | 0 | 0 | 0 | 0 | 0 | 0 | 0 | 0 | 0 | NA | NA | NA | NA | NA | NA | NA | NA | NA | NA |
| Gapvd1 | 8340 | 5820 | 5793 | 6683 | 5727 | 6427 | 6477 | 6191 | 6251 | 6652 | 6461 | 6344 | 6395.99 | 6306 | 6486 | 1.028 | 0.04 | 0.43237188 | 0.60421307 | 3.00E-04 | 0.0028 | 0.0017 | 0.0017 | TRUE | 0.007 |
| Gar1 | 5040 | 3657 | 3217 | 3238 | 2746 | 3291 | 3914 | 3890 | 3471 | 3223 | 3098 | 3249 | 3474.16 | 3758 | 3190 | 0.849 | -0.237 | 0.00028982 | 0.00283207 | 0.0021 | 0.0031 | 0.0028 | 0.0028 | TRUE | 0.0539 |
| Garem | 1831 | 1336 | 1342 | 1404 | 1365 | 1637 | 1422 | 1421 | 1448 | 1398 | 1540 | 1616 | 1474.09 | 1430 | 1518 | 1.061 | 0.086 | 0.25135119 | 0.42028106 | 0.0021 | 0.004 | 0.0033 | 0.0033 | TRUE | 0.0586 |
| Gareml | 515 | 404 | 369 | 391 | 335 | 385 | 400 | 430 | 398 | 389 | 378 | 380 | 395.84 | 409 | 382 | 0.935 | -0.097 | 0.35571563 | 0.53023831 | 0 | 0.0084 | 0.0054 | 0.0054 | TRUE | 0.0213 |
| Garnl3 | 1052 | 787 | 804 | 989 | 889 | 996 | 817 | 837 | 868 | 984 | 1003 | 983 | 915.37 | 841 | 990 | 1.179 | 0.237 | 0.00183336 | 0.01130464 | 0 | 0.005 | 0.0031 | 0.0031 | TRUE | 0.0093 |
| Gars | 18367 | 12852 | 12514 | 12987 | 11447 | 13124 | 14265 | 13671 | 13503 | 12927 | 12913 | 12955 | 13372.32 | 13813 | 12932 | 0.936 | -0.095 | 0.04389726 | 0.12249581 | 3.00E-04 | 0.0026 | 0.0015 | 0.0015 | TRUE | 0.0101 |
| Gart | 33796 | 25111 | 24951 | 28211 | 25774 | 29843 | 26248 | 26710 | 26922 | 28081 | 29076 | 29460 | 27749.45 | 26627 | 28872 | 1.084 | 0.117 | 0.01032391 | 0.04228069 | 3.00E-04 | 0.0025 | 0.0015 | 0.0015 | TRUE | 0.007 |
| Gas1 | 258 | 215 | 182 | 197 | 185 | 193 | 200 | 229 | 196 | 196 | 209 | 191 | 203.46 | 208 | 199 | 0.952 | -0.071 | 0.62239351 | 0.76150235 | 0 | 0.0141 | 0.0099 | 0.0099 | TRUE | 0.0754 |
| Gas2 | 933 | 672 | 658 | 689 | 581 | 633 | 725 | 715 | 710 | 686 | 655 | 625 | 685.92 | 717 | 655 | 0.914 | -0.129 | 0.13201671 | 0.26982268 | 0 | 0.0059 | 0.0038 | 0.0038 | TRUE | 0.0201 |
| Gas2l1 | 1631 | 1177 | 805 | 811 | 835 | 837 | 1267 | 1252 | 869 | 807 | 942 | 826 | 993.79 | 1129 | 858 | 0.759 | -0.397 | 0.00189796 | 0.0116057 | 0.0244 | 0.0048 | 0.0108 | 0.0108 | TRUE | 0.4827 |
| Gas2l2 | 8 | 2 | 1 | 1 | 4 | 4 | 6 | 2 | 1 | 1 | 5 | 4 | 3.15 | 3 | 3 | 0.965 | -0.051 | 0.96432694 | NA | 0.2943 | 0.7558 | 0.6184 | 0.6184 | TRUE | 1.2853 |
| Gas2l3 | 3158 | 2255 | 2683 | 3116 | 2699 | 2981 | 2453 | 2399 | 2895 | 3102 | 3045 | 2943 | 2805.89 | 2582 | 3030 | 1.174 | 0.231 | 0.00295319 | 0.01632249 | 0.0054 | 0.0033 | 0.004 | 0.004 | TRUE | 0.1349 |
| Gas5 | 11498 | 9257 | 10327 | 12331 | 9842 | 11846 | 8930 | 9847 | 11143 | 12274 | 11103 | 11694 | 10831.7 | 9973 | 11690 | 1.173 | 0.23 | 0.00343292 | 0.01836329 | 0.0073 | 0.0026 | 0.0043 | 0.0043 | TRUE | 0.1286 |
| Gas6 | 2162 | 1710 | 1807 | 2216 | 2082 | 2338 | 1679 | 1819 | 1950 | 2206 | 2349 | 2308 | 2051.71 | 1816 | 2288 | 1.261 | 0.334 | 3.97E-06 | 8.78E-05 | 0.0026 | 0.0036 | 0.0033 | 0.0033 | TRUE | 0.0542 |
| Gas7 | 1170 | 854 | 1016 | 1143 | 1029 | 1205 | 909 | 908 | 1096 | 1138 | 1161 | 1190 | 1066.9 | 971 | 1163 | 1.199 | 0.262 | 0.00342944 | 0.01835841 | 0.0051 | 0.0046 | 0.0048 | 0.0048 | TRUE | 0.149 |
| Gas8 | 513 | 389 | 222 | 261 | 187 | 237 | 398 | 414 | 240 | 260 | 211 | 234 | 292.74 | 351 | 235 | 0.669 | -0.579 | 0.00159698 | 0.0101088 | 0.0454 | 0.0105 | 0.0207 | 0.0207 | TRUE | 0.8576 |
| Gast | 2 | 2 | 0 | 5 | 4 | 2 | 2 | 2 | 0 | 5 | 5 | 2 | 2.52 | 1 | 4 | 3.054 | 1.611 | 0.21384392 | NA | 0 | 0.9414 | 0.6888 | 0.6888 | TRUE | 0.3671 |
| Gata1 | 182 | 156 | 177 | 195 | 171 | 205 | 141 | 166 | 191 | 194 | 193 | 202 | 181.27 | 166 | 196 | 1.189 | 0.25 | 0.1175224 | 0.24870412 | 0.0063 | 0.0155 | 0.0128 | 0.0128 | TRUE | 0.2 |
| Gata2 | 138 | 123 | 103 | 126 | 110 | 112 | 107 | 131 | 111 | 125 | 124 | 111 | 118.2 | 116 | 120 | 1.033 | 0.048 | 0.79722907 | 0.88160891 | 0 | 0.0225 | 0.0162 | 0.0162 | TRUE | 0.1154 |
| Gata3 | 112 | 67 | 40 | 33 | 19 | 21 | 87 | 71 | 43 | 33 | 21 | 21 | 46.07 | 67 | 25 | 0.371 | -1.43 | 4.28E-05 | 0.00061771 | 0.0762 | 0.0539 | 0.0603 | 0.0603 | TRUE | 0.2021 |
| Gata4 | 260 | 184 | 130 | 233 | 170 | 156 | 202 | 196 | 140 | 232 | 192 | 154 | 185.94 | 179 | 193 | 1.071 | 0.099 | 0.60420532 | 0.74918943 | 0.0344 | 0.0152 | 0.0208 | 0.0208 | TRUE | 0.2042 |
| Gata5 | 9 | 6 | 3 | 6 | 5 | 11 | 7 | 6 | 3 | 6 | 6 | 11 | 6.51 | 5 | 8 | 1.343 | 0.425 | 0.5760474 | NA | 0 | 0.3663 | 0.262 | 0.262 | TRUE | 0.4457 |
| Gata6 | 358 | 200 | 180 | 304 | 202 | 202 | 278 | 213 | 194 | 303 | 228 | 199 | 235.81 | 228 | 243 | 1.062 | 0.086 | 0.63614819 | 0.77255774 | 0.037 | 0.0125 | 0.0198 | 0.0198 | TRUE | 0.5105 |
| Gatad1 | 5580 | 3993 | 3934 | 4633 | 3947 | 4301 | 4334 | 4247 | 4245 | 4612 | 4453 | 4246 | 4355.98 | 4275 | 4437 | 1.038 | 0.053 | 0.33932035 | 0.51338051 | 6.00E-04 | 0.003 | 0.002 | 0.002 | TRUE | 0.0174 |
| Gatad2a | 22929 | 16759 | 15420 | 15636 | 14068 | 16263 | 17808 | 17826 | 16638 | 15564 | 15870 | 16054 | 16626.78 | 17424 | 15829 | 0.908 | -0.139 | 0.00599778 | 0.02820871 | 8.00E-04 | 0.0026 | 0.0018 | 0.0018 | TRUE | 0.019 |
| Gatad2b | 2150 | 1556 | 1582 | 1747 | 1571 | 1724 | 1670 | 1655 | 1707 | 1739 | 1772 | 1702 | 1707.49 | 1677 | 1738 | 1.036 | 0.051 | 0.4168122 | 0.58950096 | 0 | 0.0038 | 0.0022 | 0.0022 | TRUE | 0.0039 |
| Gatc | 511 | 409 | 384 | 362 | 321 | 419 | 397 | 435 | 414 | 360 | 362 | 414 | 397.05 | 415 | 379 | 0.913 | -0.131 | 0.2376187 | 0.4042714 | 0.0017 | 0.0084 | 0.0063 | 0.0063 | TRUE | 0.0759 |
| Gatm | 557 | 382 | 413 | 498 | 387 | 420 | 433 | 406 | 446 | 496 | 437 | 415 | 438.57 | 428 | 449 | 1.049 | 0.068 | 0.53220615 | 0.69203262 | 0.0031 | 0.0078 | 0.0064 | 0.0064 | TRUE | 0.0978 |
| Gatsl2 | 427 | 303 | 253 | 445 | 342 | 400 | 332 | 322 | 273 | 443 | 386 | 395 | 358.42 | 309 | 408 | 1.318 | 0.398 | 0.00106753 | 0.00758734 | 0.005 | 0.009 | 0.0078 | 0.0078 | TRUE | 0.1136 |
| Gatsl3 | 2039 | 1390 | 1657 | 1692 | 1489 | 1796 | 1584 | 1479 | 1788 | 1684 | 1680 | 1773 | 1664.48 | 1617 | 1712 | 1.06 | 0.084 | 0.2986198 | 0.47094274 | 0.0043 | 0.0039 | 0.0041 | 0.0041 | TRUE | 0.1012 |
| Gba | 1070 | 774 | 730 | 675 | 615 | 725 | 831 | 823 | 788 | 672 | 694 | 716 | 753.89 | 814 | 694 | 0.852 | -0.231 | 0.00505019 | 0.02473696 | 0 | 0.0056 | 0.0036 | 0.0036 | TRUE | 0.0093 |
| Gba2 | 1171 | 845 | 638 | 732 | 673 | 817 | 909 | 899 | 688 | 729 | 759 | 807 | 798.5 | 832 | 765 | 0.918 | -0.124 | 0.26162073 | 0.43179504 | 0.012 | 0.0054 | 0.0075 | 0.0075 | TRUE | 0.2668 |
| Gbas | 2594 | 1892 | 1950 | 2234 | 1962 | 2180 | 2015 | 2012 | 2104 | 2224 | 2213 | 2152 | 2120.04 | 2044 | 2196 | 1.075 | 0.104 | 0.08347485 | 0.19450702 | 0 | 0.0035 | 0.0021 | 0.0021 | TRUE | 0.0079 |
| Gbe1 | 683 | 424 | 441 | 469 | 377 | 446 | 530 | 451 | 476 | 467 | 425 | 440 | 464.95 | 486 | 444 | 0.913 | -0.132 | 0.2149599 | 0.37765193 | 0.0029 | 0.0075 | 0.006 | 0.006 | TRUE | 0.0799 |
| Gbf1 | 5186 | 3809 | 3852 | 4223 | 3813 | 4463 | 4028 | 4052 | 4156 | 4204 | 4301 | 4406 | 4191.05 | 4079 | 4304 | 1.055 | 0.078 | 0.14091851 | 0.28280654 | 2.00E-04 | 0.003 | 0.0018 | 0.0018 | TRUE | 0.0053 |
| Gbgt1 | 55 | 43 | 33 | 37 | 26 | 33 | 43 | 46 | 36 | 37 | 29 | 33 | 37.13 | 42 | 33 | 0.796 | -0.329 | 0.30009506 | 0.47228129 | 0 | 0.0663 | 0.0452 | 0.0452 | TRUE | 0.1033 |
| Gbp1 | 52 | 25 | 35 | 42 | 35 | 27 | 40 | 27 | 38 | 42 | 39 | 27 | 35.45 | 35 | 36 | 1.024 | 0.034 | 0.92168981 | 0.95683559 | 0.022 | 0.0693 | 0.0565 | 0.0565 | TRUE | 0.3821 |
| Gbp10 | 0 | 0 | 0 | 1 | 0 | 0 | 0 | 0 | 0 | 1 | 0 | 0 | 0.17 | 0 | 0 | 2.094 | 1.067 | 0.79379329 | NA | 0 | 14.2885 | 10 | 10 | TRUE | 0.3906 |
| Gbp11 | 2 | 0 | 0 | 0 | 0 | 1 | 2 | 0 | 0 | 0 | 0 | 1 | 0.42 | 1 | 0 | 0.682 | -0.553 | 0.86290097 | NA | 0 | 5.6 | 5.5066 | 5.5066 | TRUE | 0.9727 |
| Gbp2 | 79 | 41 | 29 | 49 | 35 | 34 | 61 | 44 | 31 | 49 | 39 | 34 | 43.01 | 45 | 41 | 0.885 | -0.176 | 0.59687858 | 0.74287579 | 0.0535 | 0.0575 | 0.0564 | 0.0564 | TRUE | 0.2966 |
| Gbp3 | 22 | 15 | 14 | 8 | 10 | 11 | 17 | 16 | 15 | 8 | 11 | 11 | 13.04 | 16 | 10 | 0.622 | -0.684 | 0.19541108 | NA | 0 | 0.1842 | 0.1195 | 0.1195 | TRUE | 0.1167 |
| Gbp4 | 87 | 63 | 51 | 64 | 62 | 66 | 68 | 67 | 55 | 64 | 70 | 65 | 64.74 | 63 | 66 | 1.045 | 0.063 | 0.79384946 | 0.87936839 | 0 | 0.039 | 0.0264 | 0.0264 | TRUE | 0.1051 |
| Gbp5 | 4 | 1 | 1 | 0 | 0 | 0 | 3 | 1 | 1 | 0 | 0 | 0 | 0.87 | 2 | 0 | 0.107 | -3.222 | 0.14630442 | NA | 0 | 2.7114 | 2.2736 | 2.2736 | TRUE | 0.4767 |
| Gbp6 | 3 | 0 | 1 | 2 | 3 | 0 | 2 | 0 | 1 | 2 | 3 | 0 | 1.46 | 1 | 2 | 1.499 | 0.584 | 0.73366154 | NA | 0 | 1.6213 | 1.4221 | 1.4221 | TRUE | 0.2266 |

| | | | | | | | | | | | | | | | | | | | | | | | | | |
|---|---|---|---|---|---|---|---|---|---|---|---|---|---|---|---|---|---|---|---|---|---|---|---|---|---|
| Gbp7 | 43 | 35 | 43 | 44 | 46 | 39 | 33 | 37 | 46 | 44 | 52 | 38 | 41.87 | 39 | 45 | 1.15 | 0.202 | 0.50889605 | 0.6721171 | 0 | 0.059 | 0.0435 | 0.0435 | TRUE | 0.1962 |
| Gbp8 | 9 | 2 | 3 | 1 | 3 | 4 | 7 | 2 | 3 | 1 | 3 | 4 | 3.45 | 4 | 3 | 0.657 | -0.605 | 0.57042739 | NA | 0 | 0.69 | 0.5186 | 0.5186 | TRUE | 0.8279 |
| Gbp9 | 64 | 60 | 65 | 34 | 36 | 32 | 50 | 64 | 70 | 34 | 41 | 32 | 48.28 | 61 | 36 | 0.579 | -0.788 | 0.00705155 | 0.03192517 | 0 | 0.0515 | 0.039 | 0.039 | TRUE | 0.2695 |
| Gbx1 | 26 | 16 | 14 | 36 | 25 | 38 | 20 | 17 | 15 | 36 | 28 | 38 | 25.64 | 17 | 34 | 1.93 | 0.949 | 0.01428324 | 0.05380873 | 0 | 0.0948 | 0.0654 | 0.0654 | TRUE | 0.1379 |
| Gbx2 | 1291 | 896 | 619 | 1514 | 1183 | 1324 | 1003 | 953 | 668 | 1507 | 1335 | 1307 | 1128.7 | 875 | 1383 | 1.579 | 0.659 | 1.68E-07 | 6.14E-06 | 0.023 | 0.0045 | 0.0105 | 0.0105 | TRUE | 0.5021 |
| Gc | 2 | 4 | 2 | 3 | 2 | 0 | 2 | 4 | 2 | 3 | 2 | 0 | 2.2 | 3 | 2 | 0.662 | -0.594 | 0.65983364 | NA | 0 | 1.079 | 0.8308 | 0.8308 | TRUE | 0.6411 |
| Gca | 2679 | 1853 | 1512 | 1699 | 1248 | 1479 | 2081 | 1971 | 1631 | 1691 | 1408 | 1460 | 1707.03 | 1894 | 1520 | 0.802 | -0.319 | 0.00139155 | 0.00913619 | 0.0117 | 0.0038 | 0.0066 | 0.0066 | TRUE | 0.1769 |
| Gcat | 3068 | 2383 | 3008 | 3667 | 3458 | 3989 | 2383 | 2535 | 3246 | 3650 | 3901 | 3938 | 3275.34 | 2721 | 3830 | 1.408 | 0.494 | 5.56E-07 | 1.79E-05 | 0.0147 | 0.0032 | 0.0067 | 0.0067 | TRUE | 0.344 |
| Gcc1 | 1092 | 829 | 837 | 856 | 681 | 850 | 848 | 882 | 903 | 852 | 768 | 839 | 848.73 | 878 | 820 | 0.935 | -0.096 | 0.24311939 | 0.41089104 | 7.00E-04 | 0.0052 | 0.0037 | 0.0037 | TRUE | 0.035 |
| Gcc2 | 2225 | 1539 | 1466 | 1351 | 1170 | 1360 | 1728 | 1637 | 1582 | 1345 | 1320 | 1343 | 1492.35 | 1649 | 1336 | 0.809 | -0.305 | 8.26E-06 | 0.00016011 | 6.00E-04 | 0.004 | 0.0027 | 0.0027 | TRUE | 0.0221 |
| Gcdh | 4305 | 3280 | 3436 | 3949 | 3436 | 3901 | 3344 | 3489 | 3707 | 3931 | 3876 | 3851 | 3699.62 | 3513 | 3886 | 1.107 | 0.146 | 0.01384003 | 0.05265252 | 0.001 | 0.0031 | 0.0023 | 0.0023 | TRUE | 0.0283 |
| Gcfc2 | 1278 | 1087 | 1094 | 1101 | 921 | 1145 | 993 | 1156 | 1180 | 1096 | 1039 | 1130 | 1099.07 | 1110 | 1088 | 0.983 | -0.025 | 0.77476276 | 0.86709608 | 0.0046 | 0.0046 | 0.0046 | 0.0046 | TRUE | 0.1065 |
| Gcg | 0 | 0 | 0 | 0 | 0 | 0 | 0 | 0 | 0 | 0 | 0 | 0 | 0 | 0 | 0 | NA | NA | NA | NA | NA | NA | NA | NA | NA | NA |
| Gcgr | 21 | 25 | 13 | 19 | 23 | 20 | 16 | 27 | 14 | 19 | 26 | 20 | 20.26 | 19 | 22 | 1.137 | 0.185 | 0.67732007 | 0.80217727 | 0.0176 | 0.1194 | 0.0925 | 0.0925 | TRUE | 0.6041 |
| Gch1 | 784 | 542 | 632 | 826 | 666 | 751 | 609 | 577 | 682 | 822 | 751 | 741 | 697.04 | 623 | 771 | 1.241 | 0.311 | 0.00111601 | 0.0078577 | 0.0036 | 0.0058 | 0.0051 | 0.0051 | TRUE | 0.079 |
| Gchfr | 311 | 207 | 304 | 358 | 316 | 367 | 242 | 220 | 328 | 356 | 356 | 362 | 310.81 | 263 | 358 | 1.364 | 0.448 | 0.00226491 | 0.01328915 | 0.0186 | 0.0101 | 0.0123 | 0.0123 | TRUE | 0.4998 |
| Gck | 22 | 30 | 43 | 41 | 44 | 49 | 17 | 32 | 46 | 41 | 50 | 48 | 39.04 | 32 | 46 | 1.476 | 0.562 | 0.1257983 | 0.26082056 | 0.0905 | 0.0631 | 0.0705 | 0.0705 | TRUE | 0.3137 |
| Gckr | 9 | 6 | 4 | 3 | 7 | 3 | 7 | 6 | 4 | 3 | 8 | 3 | 5.26 | 6 | 5 | 0.768 | -0.382 | 0.65318397 | NA | 0 | 0.4534 | 0.3248 | 0.3248 | TRUE | 0.6524 |
| Gclc | 4410 | 3336 | 2964 | 3272 | 2833 | 3177 | 3425 | 3548 | 3198 | 3257 | 3196 | 3136 | 3293.45 | 3390 | 3196 | 0.943 | -0.085 | 0.16199825 | 0.31205405 | 0.0011 | 0.0031 | 0.0024 | 0.0024 | TRUE | 0.0294 |
| Gclm | 3743 | 2835 | 2643 | 3092 | 2651 | 3076 | 2907 | 3016 | 2852 | 3078 | 2991 | 3036 | 2979.87 | 2925 | 3035 | 1.038 | 0.054 | 0.34137113 | 0.51533965 | 1.00E-04 | 0.0032 | 0.0019 | 0.0019 | TRUE | 0.0087 |
| Gcm1 | 42 | 28 | 19 | 12 | 19 | 16 | 33 | 30 | 21 | 12 | 21 | 16 | 22.01 | 28 | 16 | 0.587 | -0.77 | 0.07546 | 0.18096474 | 0 | 0.1101 | 0.086 | 0.086 | TRUE | 0.3163 |
| Gcm2 | 131 | 115 | 118 | 87 | 68 | 74 | 102 | 122 | 127 | 87 | 77 | 73 | 97.96 | 117 | 79 | 0.676 | -0.566 | 0.00593265 | 0.028013 | 0 | 0.0266 | 0.0197 | 0.0197 | TRUE | 0.1484 |
| Gcn1l1 | 14859 | 11450 | 10908 | 12560 | 11550 | 12733 | 11540 | 12179 | 11770 | 12502 | 13030 | 12569 | 12265.08 | 11830 | 12700 | 1.074 | 0.103 | 0.03675931 | 0.10783016 | 5.00E-04 | 0.0026 | 0.0017 | 0.0017 | TRUE | 0.0082 |
| Gcnt1 | 2117 | 1525 | 1602 | 1964 | 1719 | 1838 | 1644 | 1622 | 1729 | 1955 | 1939 | 1814 | 1783.9 | 1665 | 1903 | 1.143 | 0.193 | 0.00383449 | 0.01994408 | 8.00E-04 | 0.0038 | 0.0026 | 0.0026 | TRUE | 0.0202 |
| Gcnt2 | 2859 | 2432 | 2775 | 2593 | 2655 | 3037 | 2220 | 2587 | 2994 | 2581 | 2995 | 2998 | 2729.29 | 2600 | 2858 | 1.1 | 0.137 | 0.17217186 | 0.32430843 | 0.0134 | 0.0033 | 0.0069 | 0.0069 | TRUE | 0.2128 |
| Gcnt3 | 59 | 36 | 22 | 30 | 20 | 29 | 46 | 38 | 24 | 30 | 23 | 29 | 31.48 | 36 | 27 | 0.747 | -0.421 | 0.2552892 | 0.42502604 | 0.0367 | 0.0777 | 0.0661 | 0.0661 | TRUE | 0.3655 |
| Gcnt4 | 224 | 188 | 212 | 221 | 223 | 256 | 174 | 200 | 229 | 220 | 252 | 253 | 221.16 | 201 | 242 | 1.206 | 0.271 | 0.07099097 | 0.17330042 | 0.008 | 0.0131 | 0.0116 | 0.0116 | TRUE | 0.1641 |
| Gcnt7 | 0 | 0 | 0 | 0 | 0 | 0 | 0 | 0 | 0 | 0 | 0 | 0 | 0 | 0 | 0 | NA | NA | NA | NA | NA | NA | NA | NA | NA | NA |
| Gcsam | 0 | 0 | 0 | 0 | 0 | 0 | 0 | 0 | 0 | 0 | 0 | 0 | 0 | 0 | 0 | NA | NA | NA | NA | NA | NA | NA | NA | NA | NA |
| Gcsh | 10699 | 7456 | 7060 | 7743 | 6506 | 7753 | 8309 | 7931 | 7618 | 7707 | 7339 | 7653 | 7759.7 | 7953 | 7566 | 0.951 | -0.072 | 0.1906195 | 0.3487296 | 0.0011 | 0.0027 | 0.0021 | 0.0021 | TRUE | 0.019 |
| Gda | 75 | 39 | 34 | 43 | 21 | 44 | 58 | 41 | 37 | 43 | 24 | 43 | 41.06 | 45 | 37 | 0.804 | -0.316 | 0.3534731 | 0.52780934 | 0.0549 | 0.0602 | 0.0587 | 0.0587 | TRUE | 0.6384 |
| Gdap1 | 11 | 8 | 8 | 14 | 11 | 18 | 9 | 9 | 9 | 14 | 12 | 18 | 11.63 | 9 | 15 | 1.722 | 0.784 | 0.1612215 | NA | 0 | 0.2062 | 0.1343 | 0.1343 | TRUE | 0.1529 |
| Gdap10 | 185 | 196 | 217 | 182 | 161 | 203 | 144 | 208 | 234 | 181 | 182 | 200 | 191.58 | 195 | 188 | 0.966 | -0.049 | 0.78829785 | 0.87559381 | 0.0285 | 0.0148 | 0.0189 | 0.0189 | TRUE | 0.6304 |
| Gdap1l1 | 84 | 63 | 53 | 129 | 106 | 123 | 65 | 67 | 57 | 128 | 120 | 121 | 93.14 | 63 | 123 | 1.947 | 0.961 | 2.70E-06 | 6.38E-05 | 0 | 0.0279 | 0.0184 | 0.0184 | TRUE | 0.0546 |
| Gdap2 | 1147 | 830 | 874 | 955 | 870 | 882 | 891 | 883 | 943 | 951 | 981 | 871 | 919.91 | 906 | 934 | 1.032 | 0.045 | 0.58553777 | 0.7349044 | 0.0014 | 0.005 | 0.0038 | 0.0038 | TRUE | 0.0426 |
| Gde1 | 3302 | 2419 | 2673 | 2896 | 2444 | 2940 | 2565 | 2573 | 2884 | 2883 | 2757 | 2902 | 2760.62 | 2674 | 2847 | 1.066 | 0.092 | 0.17270372 | 0.32505324 | 0.0022 | 0.0033 | 0.0029 | 0.0029 | TRUE | 0.0568 |
| Gdf1 | 0 | 0 | 0 | 0 | 0 | 0 | 0 | 0 | 0 | 0 | 0 | 0 | 0 | 0 | 0 | NA | NA | NA | NA | NA | NA | NA | NA | NA | NA |
| Gdf10 | 13 | 6 | 2 | 10 | 8 | 5 | 10 | 6 | 2 | 10 | 9 | 5 | 7.09 | 6 | 8 | 1.251 | 0.323 | 0.67319488 | NA | 0.1298 | 0.3366 | 0.2818 | 0.2818 | TRUE | 0.2773 |
| Gdf11 | 66 | 76 | 42 | 56 | 41 | 37 | 51 | 81 | 45 | 56 | 46 | 37 | 52.66 | 59 | 46 | 0.783 | -0.353 | 0.251649 | 0.42058223 | 0.0524 | 0.0474 | 0.0489 | 0.0489 | TRUE | 0.7536 |
| Gdf15 | 84 | 58 | 96 | 119 | 111 | 121 | 65 | 62 | 104 | 118 | 125 | 119 | 98.94 | 77 | 121 | 1.583 | 0.663 | 0.00351353 | 0.0186618 | 0.0272 | 0.0264 | 0.0266 | 0.0266 | TRUE | 0.8146 |
| Gdf2 | 1 | 0 | 3 | 1 | 0 | 0 | 1 | 0 | 3 | 1 | 0 | 0 | 0.83 | 1 | 0 | 0.28 | -1.835 | 0.43260394 | NA | 0 | 2.8414 | 2.5763 | 2.5763 | TRUE | 0.5398 |
| Gdf3 | 4248 | 2977 | 2962 | 2701 | 2510 | 2942 | 3299 | 3167 | 3196 | 2689 | 2832 | 2904 | 3014.36 | 3221 | 2808 | 0.872 | -0.198 | 0.0007768 | 0.0060276 | 7.00E-04 | 0.0032 | 0.0022 | 0.0022 | TRUE | 0.017 |
| Gdf5 | 6 | 3 | 3 | 3 | 4 | 1 | 5 | 3 | 3 | 3 | 5 | 1 | 3.26 | 4 | 3 | 0.75 | -0.416 | 0.69519335 | NA | 0 | 0.7289 | 0.4973 | 0.4973 | TRUE | 0.3315 |
| Gdf6 | 4 | 1 | 1 | 1 | 5 | 2 | 3 | 1 | 1 | 1 | 6 | 2 | 2.31 | 2 | 3 | 1.566 | 0.647 | 0.61870373 | NA | 0 | 1.0285 | 0.7676 | 0.7676 | TRUE | 0.7956 |
| Gdf7 | 22 | 22 | 9 | 20 | 23 | 20 | 17 | 23 | 10 | 20 | 26 | 20 | 19.3 | 17 | 22 | 1.301 | 0.38 | 0.41050219 | 0.58328937 | 0.0259 | 0.1252 | 0.1011 | 0.1011 | TRUE | 0.2645 |
| Gdf9 | 29 | 10 | 17 | 9 | 15 | 12 | 23 | 11 | 18 | 9 | 17 | 12 | 14.87 | 17 | 13 | 0.722 | -0.471 | 0.37244374 | NA | 0.0522 | 0.1618 | 0.1315 | 0.1315 | TRUE | 0.3397 |
| Gdi1 | 3525 | 2714 | 2960 | 2684 | 2404 | 2928 | 2738 | 2887 | 3194 | 2672 | 2712 | 2890 | 2848.73 | 2940 | 2758 | 0.939 | -0.091 | 0.2083802 | 0.36993892 | 0.0035 | 0.0033 | 0.0034 | 0.0034 | TRUE | 0.0693 |
| Gdi2 | 30800 | 21972 | 22680 | 23929 | 20498 | 23520 | 23921 | 23371 | 24472 | 23819 | 23124 | 23218 | 23654.09 | 23921 | 23387 | 0.978 | -0.033 | 0.47706151 | 0.64471288 | 3.00E-04 | 0.0025 | 0.0015 | 0.0015 | TRUE | 0.0049 |
| Gdnf | 6 | 5 | 4 | 9 | 1 | 8 | 5 | 5 | 4 | 9 | 1 | 8 | 5.38 | 5 | 6 | 1.273 | 0.348 | 0.69093766 | NA | 0 | 0.443 | 0.3659 | 0.3659 | TRUE | 1.0393 |
| Gdpd1 | 993 | 662 | 731 | 809 | 656 | 744 | 771 | 704 | 789 | 805 | 740 | 734 | 757.31 | 755 | 760 | 1.007 | 0.01 | 0.91312812 | 0.9523745 | 0.0017 | 0.0056 | 0.0043 | 0.0043 | TRUE | 0.0394 |
| Gdpd2 | 47 | 35 | 29 | 35 | 15 | 37 | 37 | 37 | 31 | 35 | 17 | 37 | 32.22 | 35 | 30 | 0.845 | -0.243 | 0.50900458 | 0.67219832 | 0.0367 | 0.076 | 0.0666 | 0.0666 | TRUE | 0.8309 |
| Gdpd3 | 17 | 4 | 12 | 11 | 8 | 10 | 13 | 4 | 13 | 11 | 9 | 10 | 10.04 | 10 | 10 | 0.973 | -0.039 | 0.9506506 | NA | 0 | 0.2384 | 0.188 | 0.188 | TRUE | 0.8332 |
| Gdpd4 | 3 | 3 | 0 | 1 | 0 | 1 | 2 | 3 | 0 | 1 | 0 | 1 | 1.25 | 2 | 1 | 0.364 | -1.458 | 0.43728581 | NA | 0 | 1.8976 | 1.5133 | 1.5133 | TRUE | 0.4727 |
| Gdpd5 | 273 | 179 | 199 | 279 | 234 | 262 | 212 | 190 | 215 | 278 | 264 | 259 | 236.25 | 206 | 267 | 1.296 | 0.374 | 0.00484477 | 0.02392754 | 0 | 0.0125 | 0.0084 | 0.0084 | TRUE | 0.044 |
| Gdpgp1 | 812 | 544 | 450 | 526 | 409 | 430 | 631 | 579 | 486 | 524 | 461 | 424 | 517.38 | 565 | 470 | 0.83 | -0.269 | 0.0274246 | 0.08664451 | 0.0125 | 0.007 | 0.0087 | 0.0087 | TRUE | 0.1747 |
| Gem | 151 | 98 | 123 | 114 | 83 | 99 | 117 | 104 | 133 | 113 | 94 | 98 | 109.84 | 118 | 102 | 0.862 | -0.214 | 0.27099611 | 0.44196113 | 0 | 0.024 | 0.0182 | 0.0182 | TRUE | 0.1102 |
| Gemin2 | 1957 | 1428 | 1504 | 1675 | 1429 | 1601 | 1520 | 1519 | 1623 | 1667 | 1612 | 1580 | 1586.91 | 1554 | 1620 | 1.043 | 0.061 | 0.36559098 | 0.54031557 | 5.00E-04 | 0.0039 | 0.0026 | 0.0026 | TRUE | 0.0177 |
| Gemin4 | 7647 | 5522 | 4308 | 4488 | 4118 | 4606 | 5939 | 5874 | 4648 | 4467 | 4646 | 4547 | 5020.13 | 5487 | 4553 | 0.83 | -0.27 | 0.0017163 | 0.01074038 | 0.0087 | 0.0029 | 0.0051 | 0.0051 | TRUE | 0.2171 |
| Gemin5 | 23680 | 17197 | 15905 | 19091 | 16562 | 19430 | 18391 | 18292 | 17161 | 19003 | 18684 | 19180 | 18452.01 | 17948 | 18956 | 1.056 | 0.079 | 0.11321662 | 0.24206212 | 7.00E-04 | 0.0026 | 0.0017 | 0.0017 | TRUE | 0.0179 |
| Gemin6 | 2674 | 1813 | 1749 | 1976 | 1827 | 1937 | 2077 | 1928 | 1887 | 1967 | 2061 | 1912 | 1972.08 | 1964 | 1980 | 1.007 | 0.01 | 0.88383112 | 0.93547084 | 0.0015 | 0.0036 | 0.0028 | 0.0028 | TRUE | 0.0314 |
| Gemin7 | 1441 | 944 | 1029 | 1200 | 1084 | 1192 | 1119 | 1004 | 1110 | 1194 | 1223 | 1177 | 1137.93 | 1078 | 1198 | 1.11 | 0.151 | 0.0469808 | 0.12871316 | 0.001 | 0.0045 | 0.0033 | 0.0033 | TRUE | 0.042 |
| Gemin8 | 526 | 408 | 442 | 503 | 437 | 524 | 409 | 434 | 477 | 501 | 493 | 517 | 471.73 | 440 | 504 | 1.148 | 0.199 | 0.05301757 | 0.14069029 | 0.0014 | 0.0075 | 0.0055 | 0.0055 | TRUE | 0.0623 |
| Gen1 | 1203 | 863 | 586 | 562 | 454 | 519 | 934 | 918 | 632 | 559 | 512 | 512 | 678.08 | 828 | 528 | 0.637 | -0.651 | 1.07E-06 | 3.11E-05 | 0.0233 | 0.0059 | 0.0113 | 0.0113 | TRUE | 0.5018 |
| Get4 | 4791 | 3470 | 3366 | 3852 | 3317 | 3840 | 3721 | 3691 | 3632 | 3834 | 3742 | 3791 | 3735.12 | 3681 | 3789 | 1.029 | 0.041 | 0.42522049 | 0.59794435 | 0 | 0.0031 | 0.0017 | 0.0017 | TRUE | 0.0017 |
| Gfap | 54 | 40 | 30 | 79 | 70 | 66 | 42 | 43 | 32 | 79 | 79 | 65 | 56.6 | 39 | 74 | 1.897 | 0.924 | 0.00052346 | 0.00446411 | 0 | 0.0443 | 0.0318 | 0.0318 | TRUE | 0.1488 |
| Gfer | 3010 | 2379 | 2628 | 2533 | 2239 | 2637 | 2338 | 2531 | 2836 | 2521 | 2526 | 2603 | 2559.02 | 2568 | 2550 | 0.994 | -0.009 | 0.90852201 | 0.94963864 | 0.0046 | 0.0034 | 0.0038 | 0.0038 | TRUE | 0.1006 |
| Gfi1 | 3 | 1 | 2 | 5 | 3 | 5 | 2 | 1 | 2 | 5 | 3 | 5 | 3.14 | 2 | 4 | 2.371 | 1.245 | 0.25258603 | NA | 0 | 0.7569 | 0.4811 | 0.4811 | TRUE | 0.1043 |
| Gfi1b | 5 | 3 | 3 | 2 | 4 | 2 | 4 | 3 | 3 | 2 | 5 | 2 | 3.13 | 3 | 3 | 0.809 | -0.306 | 0.7729614 | NA | 0 | 0.7593 | 0.4852 | 0.4852 | TRUE | 0.2964 |
| Gfm1 | 6836 | 4615 | 4466 | 4743 | 4073 | 4877 | 5309 | 4909 | 4819 | 4721 | 4595 | 4814 | 4861.2 | 5012 | 4710 | 0.939 | -0.09 | 0.12564134 | 0.26077638 | 0.0013 | 0.0029 | 0.0023 | 0.0023 | TRUE | 0.0333 |

| | | | | | | | | | | | | | | | | | | | | | | | | |
|---|---|---|---|---|---|---|---|---|---|---|---|---|---|---|---|---|---|---|---|---|---|---|---|---|
| Gfm2 | 4929 | 3333 | 2856 | 3880 | 3322 | 3831 | 3828 | 3545 | 3082 | 3862 | 3748 | 3782 | 3641.09 | 3485 | 3797 | 1.089 | 0.123 | 0.1138788 | 0.24288051 | 0.0056 | 0.0031 | 0.0041 | 0.0041 | TRUE | 0.1242 |
| Gfod1 | 1035 | 898 | 1008 | 1198 | 1123 | 1336 | 804 | 955 | 1088 | 1192 | 1267 | 1319 | 1104.14 | 949 | 1259 | 1.33 | 0.411 | 7.70E-05 | 0.00098558 | 0.0111 | 0.0046 | 0.0069 | 0.0069 | TRUE | 0.2211 |
| Gfod2 | 864 | 589 | 542 | 554 | 479 | 636 | 671 | 627 | 585 | 551 | 540 | 628 | 600.33 | 628 | 573 | 0.913 | -0.132 | 0.19091614 | 0.34908077 | 0.0041 | 0.0064 | 0.0056 | 0.0056 | TRUE | 0.0823 |
| Gfpt1 | 3829 | 2667 | 2896 | 3748 | 3125 | 3591 | 2974 | 2837 | 3125 | 3731 | 3525 | 3545 | 3289.4 | 2979 | 3600 | 1.209 | 0.274 | 8.14E-06 | 0.00015845 | 0.0012 | 0.0031 | 0.0024 | 0.0024 | TRUE | 0.0219 |
| Gfpt2 | 5311 | 4012 | 5269 | 6500 | 5961 | 7498 | 4125 | 4268 | 5685 | 6470 | 6725 | 7402 | 5778.99 | 4693 | 6866 | 1.464 | 0.55 | 1.29E-07 | 4.90E-06 | 0.0191 | 0.0028 | 0.0076 | 0.0076 | TRUE | 0.4164 |
| Gfra1 | 137 | 90 | 106 | 123 | 107 | 108 | 106 | 96 | 114 | 122 | 121 | 107 | 111.04 | 105 | 117 | 1.104 | 0.143 | 0.44749571 | 0.61788255 | 0 | 0.0238 | 0.0165 | 0.0165 | TRUE | 0.0601 |
| Gfra2 | 7 | 7 | 18 | 14 | 11 | 17 | 5 | 7 | 19 | 14 | 12 | 17 | 12.57 | 10 | 14 | 1.363 | 0.446 | 0.45759111 | NA | 0.1464 | 0.1909 | 0.1786 | 0.1786 | TRUE | 1.7156 |
| Gfra3 | 49 | 31 | 25 | 28 | 27 | 25 | 38 | 33 | 27 | 28 | 30 | 25 | 30.17 | 33 | 28 | 0.84 | -0.251 | 0.47652094 | 0.64430176 | 0 | 0.081 | 0.0561 | 0.0561 | TRUE | 0.1695 |
| Gfra4 | 295 | 189 | 173 | 235 | 177 | 222 | 229 | 201 | 187 | 234 | 200 | 219 | 211.59 | 206 | 218 | 1.056 | 0.078 | 0.59588446 | 0.74235543 | 0.0043 | 0.0136 | 0.0108 | 0.0108 | TRUE | 0.1181 |
| Gfral | 0 | 0 | 0 | 0 | 0 | 1 | 0 | 0 | 0 | 0 | 0 | 1 | 0.16 | 0 | 0 | 2.094 | 1.067 | 0.79379329 | NA | 0 | 14.4078 | 10 | 10 | TRUE | 0.3835 |
| Gga1 | 3759 | 2819 | 2871 | 2995 | 2941 | 3259 | 2919 | 2999 | 3098 | 2981 | 3318 | 3217 | 3088.65 | 3005 | 3172 | 1.056 | 0.078 | 0.21758157 | 0.38036856 | 0.0014 | 0.0032 | 0.0026 | 0.0026 | TRUE | 0.0337 |
| Gga2 | 13987 | 10873 | 11517 | 12226 | 11205 | 13267 | 10863 | 11565 | 12427 | 12170 | 12640 | 13097 | 12127.01 | 11618 | 12636 | 1.088 | 0.121 | 0.05169117 | 0.138143 | 0.0028 | 0.0026 | 0.0027 | 0.0027 | TRUE | 0.0453 |
| Gga3 | 7647 | 5692 | 5334 | 5137 | 4956 | 5834 | 5939 | 6055 | 5755 | 5113 | 5591 | 5759 | 5702.04 | 5916 | 5488 | 0.927 | -0.109 | 0.07528423 | 0.18072513 | 0.0019 | 0.0028 | 0.0025 | 0.0025 | TRUE | 0.0436 |
| Ggact | 593 | 458 | 444 | 573 | 508 | 601 | 461 | 487 | 479 | 570 | 573 | 593 | 527.25 | 476 | 579 | 1.219 | 0.285 | 0.00218488 | 0.012915 | 0 | 0.0069 | 0.0043 | 0.0043 | TRUE | 0.0096 |
| Ggct | 847 | 605 | 549 | 640 | 537 | 594 | 658 | 644 | 592 | 637 | 606 | 586 | 620.49 | 631 | 610 | 0.965 | -0.052 | 0.57430152 | 0.72641519 | 0 | 0.0062 | 0.0045 | 0.0045 | TRUE | 0.0334 |
| Ggcx | 357 | 352 | 307 | 296 | 244 | 280 | 277 | 374 | 331 | 295 | 275 | 276 | 304.87 | 327 | 282 | 0.864 | -0.21 | 0.12218308 | 0.25539086 | 0.0097 | 0.0102 | 0.01 | 0.01 | TRUE | 0.2209 |
| Ggh | 489 | 299 | 303 | 264 | 224 | 313 | 380 | 318 | 327 | 263 | 253 | 309 | 308.2 | 342 | 275 | 0.803 | -0.316 | 0.01678359 | 0.06082348 | 0.0074 | 0.0101 | 0.0093 | 0.0093 | TRUE | 0.134 |
| Ggn | 10 | 7 | 0 | 6 | 4 | 6 | 8 | 7 | 0 | 6 | 5 | 6 | 5.27 | 5 | 6 | 1.062 | 0.087 | 0.92586577 | NA | 0.4242 | 0.4522 | 0.4474 | 0.4474 | TRUE | 1.4085 |
| Ggnbp1 | 308 | 280 | 247 | 254 | 237 | 305 | 239 | 298 | 267 | 253 | 267 | 301 | 270.8 | 268 | 274 | 1.025 | 0.036 | 0.79163186 | 0.87766011 | 0.0068 | 0.0112 | 0.0098 | 0.0098 | TRUE | 0.1074 |
| Ggnbp2 | 6901 | 5007 | 5583 | 6276 | 5389 | 6262 | 5360 | 5326 | 6024 | 6247 | 6079 | 6182 | 5869.61 | 5570 | 6169 | 1.108 | 0.148 | 0.01765569 | 0.06296227 | 0.0023 | 0.0028 | 0.0026 | 0.0026 | TRUE | 0.0616 |
| Ggps1 | 1437 | 1038 | 1041 | 1191 | 1000 | 1172 | 1116 | 1104 | 1123 | 1186 | 1128 | 1157 | 1135.66 | 1114 | 1157 | 1.038 | 0.054 | 0.44195535 | 0.61299056 | 0 | 0.0045 | 0.0027 | 0.0027 | TRUE | 0.0056 |
| Ggt1 | 593 | 467 | 549 | 780 | 698 | 847 | 461 | 497 | 592 | 776 | 787 | 836 | 658.27 | 517 | 800 | 1.553 | 0.635 | 1.62E-09 | 1.04E-07 | 0.0071 | 0.006 | 0.0064 | 0.0064 | TRUE | 0.1901 |
| Ggt5 | 33 | 29 | 37 | 30 | 30 | 38 | 26 | 31 | 40 | 30 | 34 | 38 | 32.94 | 32 | 34 | 1.058 | 0.082 | 0.81211308 | 0.89148699 | 0 | 0.0744 | 0.0548 | 0.0548 | TRUE | 0.298 |
| Ggt6 | 113 | 72 | 71 | 99 | 88 | 97 | 88 | 77 | 77 | 99 | 99 | 96 | 89.09 | 81 | 98 | 1.213 | 0.279 | 0.17357982 | 0.32631545 | 0 | 0.029 | 0.0191 | 0.0191 | TRUE | 0.0697 |
| Ggt7 | 226 | 182 | 211 | 240 | 200 | 257 | 176 | 194 | 228 | 239 | 226 | 254 | 219.17 | 199 | 240 | 1.209 | 0.274 | 0.0631412 | 0.15913899 | 0.0058 | 0.0132 | 0.0111 | 0.0111 | TRUE | 0.1709 |
| Ggta1 | 1736 | 1387 | 1604 | 1986 | 1675 | 1953 | 1348 | 1475 | 1731 | 1977 | 1890 | 1928 | 1724.78 | 1518 | 1932 | 1.274 | 0.349 | 8.69E-05 | 0.0010781 | 0.0076 | 0.0038 | 0.0051 | 0.0051 | TRUE | 0.1801 |
| Gh | 56 | 43 | 53 | 45 | 60 | 44 | 43 | 46 | 57 | 45 | 68 | 43 | 50.39 | 49 | 52 | 1.064 | 0.09 | 0.75877893 | 0.85685607 | 0.0229 | 0.0495 | 0.0421 | 0.0421 | TRUE | 0.5439 |
| Ghdc | 1173 | 871 | 966 | 1178 | 1073 | 1232 | 911 | 926 | 1042 | 1173 | 1210 | 1216 | 1079.83 | 960 | 1200 | 1.251 | 0.323 | 5.20E-05 | 0.00072392 | 0.0018 | 0.0046 | 0.0037 | 0.0037 | TRUE | 0.066 |
| Ghitm | 11931 | 8282 | 8433 | 9576 | 8134 | 8973 | 9266 | 8809 | 9099 | 9532 | 9176 | 8858 | 9123.43 | 9058 | 9189 | 1.014 | 0.02 | 0.69799009 | 0.81651154 | 8.00E-04 | 0.0027 | 0.0019 | 0.0019 | TRUE | 0.0131 |
| Ghr | 127 | 132 | 123 | 119 | 104 | 132 | 99 | 140 | 133 | 118 | 117 | 130 | 122.97 | 124 | 122 | 0.991 | -0.012 | 0.9489039 | 0.97209601 | 0.0126 | 0.0217 | 0.019 | 0.019 | TRUE | 0.3624 |
| Ghrh | 1 | 0 | 0 | 1 | 0 | 2 | 1 | 0 | 0 | 1 | 0 | 2 | 0.62 | 0 | 1 | 2.903 | 1.538 | 0.54465186 | NA | 0 | 3.7982 | 3.1252 | 3.1252 | TRUE | 0.1038 |
| Ghrhr | 0 | 1 | 0 | 0 | 0 | 0 | 0 | 1 | 0 | 0 | 0 | 0 | 0.18 | 0 | 0 | 0.552 | -0.857 | 0.8336548 | NA | 0 | 13.3714 | 10 | 10 | TRUE | 0.5216 |
| Ghrl | 1 | 0 | 0 | 0 | 0 | 0 | 1 | 0 | 0 | 0 | 0 | 0 | 0.13 | 0 | 0 | 0.552 | -0.857 | 0.8336548 | NA | 0 | 18.3121 | 10 | 10 | TRUE | 0.2566 |
| Ghsr | 8 | 7 | 3 | 8 | 9 | 7 | 6 | 7 | 3 | 8 | 10 | 7 | 6.99 | 5 | 8 | 1.468 | 0.554 | 0.44553068 | NA | 0 | 0.3416 | 0.2329 | 0.2329 | TRUE | 0.271 |
| Gid4 | 8936 | 6984 | 8062 | 8663 | 7845 | 9270 | 6940 | 7429 | 8699 | 8623 | 8850 | 9151 | 8281.98 | 7689 | 8875 | 1.155 | 0.207 | 0.00820452 | 0.03568766 | 0.0072 | 0.0027 | 0.0043 | 0.0043 | TRUE | 0.1609 |
| Gid8 | 4215 | 2945 | 2615 | 2644 | 2450 | 2880 | 3274 | 3133 | 2822 | 2632 | 2764 | 2843 | 2911.07 | 3076 | 2746 | 0.892 | -0.164 | 0.02034346 | 0.0700311 | 0.0031 | 0.0032 | 0.0033 | 0.0033 | TRUE | 0.0632 |
| Gif | 0 | 0 | 0 | 0 | 0 | 0 | 0 | 0 | 0 | 0 | 0 | 0 | 0 | 0 | 0 | NA | NA | NA | NA | NA | NA | NA | NA | NA | NA |
| Gigyf1 | 3804 | 3362 | 3492 | 3109 | 2895 | 3509 | 2954 | 3576 | 3768 | 3095 | 3266 | 3464 | 3353.81 | 3433 | 3275 | 0.955 | -0.067 | 0.44772414 | 0.61788255 | 0.0094 | 0.0031 | 0.0053 | 0.0053 | TRUE | 0.1832 |
| Gigyf2 | 9285 | 7440 | 7390 | 7477 | 7142 | 8151 | 7211 | 7914 | 7974 | 7443 | 8057 | 8046 | 7774.11 | 7700 | 7849 | 1.02 | 0.028 | 0.65014976 | 0.7827712 | 0.002 | 0.0027 | 0.0026 | 0.0026 | TRUE | 0.038 |
| Gimap1 | 7 | 3 | 2 | 0 | 3 | 3 | 5 | 3 | 2 | 0 | 3 | 3 | 2.86 | 3 | 2 | 0.574 | -0.802 | 0.50626361 | NA | 0 | 0.8325 | 0.6696 | 0.6696 | TRUE | 0.7521 |
| Gimap3 | 0 | 0 | 0 | 0 | 0 | 1 | 0 | 0 | 0 | 0 | 0 | 1 | 0.16 | 0 | 0 | 2.094 | 1.067 | 0.79379329 | NA | 0 | 14.4078 | 10 | 10 | TRUE | 0.3835 |
| Gimap4 | 0 | 0 | 1 | 0 | 0 | 1 | 0 | 0 | 1 | 0 | 0 | 1 | 0.34 | 0 | 0 | 1.075 | 0.105 | 0.97570524 | NA | 0 | 6.885 | 6.5379 | 6.5379 | TRUE | 0.5365 |
| Gimap5 | 1 | 1 | 1 | 2 | 0 | 0 | 1 | 1 | 1 | 2 | 0 | 0 | 0.82 | 1 | 1 | 0.698 | -0.519 | 0.8106088 | NA | 0 | 2.8986 | 2.1114 | 2.1114 | TRUE | 0.9736 |
| Gimap6 | 2 | 0 | 0 | 0 | 0 | 0 | 2 | 0 | 0 | 0 | 0 | 0 | 0.26 | 1 | 0 | 0.356 | -1.491 | 0.71362796 | NA | 0 | 9.1573 | 10 | 10 | TRUE | 0.968 |
| Gimap7 | 1 | 0 | 0 | 0 | 0 | 0 | 1 | 0 | 0 | 0 | 0 | 0 | 0.13 | 0 | 0 | 0.552 | -0.857 | 0.8336548 | NA | 0 | 18.3121 | 10 | 10 | TRUE | 0.2566 |
| Gimap8 | 21 | 17 | 12 | 6 | 7 | 14 | 16 | 18 | 13 | 6 | 8 | 14 | 12.5 | 16 | 9 | 0.586 | -0.771 | 0.17225728 | NA | 0 | 0.192 | 0.1434 | 0.1434 | TRUE | 0.6466 |
| Gimap9 | 172 | 111 | 111 | 133 | 108 | 130 | 134 | 118 | 120 | 132 | 122 | 128 | 125.66 | 124 | 127 | 1.028 | 0.039 | 0.82208519 | 0.89791102 | 0 | 0.0213 | 0.0141 | 0.0141 | TRUE | 0.054 |
| Gin1 | 597 | 424 | 329 | 382 | 300 | 364 | 464 | 451 | 355 | 380 | 338 | 359 | 391.28 | 423 | 359 | 0.848 | -0.238 | 0.05849623 | 0.15072007 | 0.0097 | 0.0085 | 0.0089 | 0.0089 | TRUE | 0.2238 |
| Ginm1 | 1186 | 842 | 819 | 935 | 847 | 899 | 921 | 896 | 884 | 931 | 956 | 887 | 912.35 | 900 | 925 | 1.026 | 0.037 | 0.63165224 | 0.76888141 | 0 | 0.005 | 0.0032 | 0.0032 | TRUE | 0.0148 |
| Gins1 | 2543 | 1809 | 1861 | 1895 | 1640 | 1941 | 1975 | 1924 | 2008 | 1886 | 1850 | 1916 | 1926.62 | 1969 | 1884 | 0.957 | -0.063 | 0.29913208 | 0.47138862 | 0 | 0.0037 | 0.0022 | 0.0022 | TRUE | 0.0047 |
| Gins2 | 4709 | 3353 | 3579 | 4474 | 3973 | 4486 | 3657 | 3567 | 3862 | 4453 | 4482 | 4428 | 4074.88 | 3695 | 4454 | 1.206 | 0.27 | 1.17E-06 | 3.33E-05 | 5.00E-04 | 0.003 | 0.002 | 0.002 | TRUE | 0.0185 |
| Gins3 | 1421 | 984 | 849 | 895 | 778 | 882 | 1104 | 1047 | 916 | 891 | 878 | 871 | 950.93 | 1022 | 880 | 0.859 | -0.219 | 0.01293004 | 0.04996189 | 0.0037 | 0.0049 | 0.0045 | 0.0045 | TRUE | 0.0971 |
| Gins4 | 6027 | 4245 | 4305 | 4272 | 3784 | 4282 | 4681 | 4515 | 4645 | 4252 | 4269 | 4227 | 4431.57 | 4614 | 4249 | 0.921 | -0.119 | 0.01933768 | 0.06744595 | 0 | 0.003 | 0.0016 | 0.0016 | TRUE | 0.0042 |
| Gip | 5 | 5 | 1 | 3 | 2 | 1 | 4 | 5 | 1 | 3 | 2 | 1 | 2.75 | 3 | 2 | 0.601 | -0.734 | 0.53533073 | NA | 0 | 0.8637 | 0.6193 | 0.6193 | TRUE | 0.3267 |
| Gipc1 | 6749 | 5080 | 4650 | 5602 | 4853 | 5415 | 5242 | 5404 | 5017 | 5576 | 5475 | 5345 | 5343.14 | 5221 | 5465 | 1.047 | 0.066 | 0.22438861 | 0.38882112 | 6.00E-04 | 0.0029 | 0.0019 | 0.0019 | TRUE | 0.0139 |
| Gipc2 | 160 | 118 | 125 | 146 | 155 | 171 | 124 | 126 | 135 | 145 | 175 | 169 | 145.61 | 128 | 163 | 1.272 | 0.347 | 0.03808588 | 0.11044056 | 0 | 0.0187 | 0.0132 | 0.0132 | TRUE | 0.0965 |
| Gipc3 | 18 | 8 | 11 | 11 | 10 | 6 | 14 | 9 | 12 | 11 | 11 | 6 | 10.42 | 12 | 9 | 0.81 | -0.304 | 0.61423833 | NA | 0 | 0.2299 | 0.1645 | 0.1645 | TRUE | 0.3593 |
| Gipr | 62 | 72 | 64 | 94 | 72 | 98 | 48 | 77 | 69 | 94 | 81 | 97 | 77.55 | 65 | 91 | 1.416 | 0.502 | 0.03830264 | 0.11091159 | 0.0184 | 0.033 | 0.029 | 0.029 | TRUE | 0.5093 |
| Git1 | 12138 | 9369 | 9432 | 8532 | 8419 | 9884 | 9427 | 9966 | 10177 | 8493 | 9498 | 9757 | 9552.85 | 9857 | 9249 | 0.938 | -0.092 | 0.15889922 | 0.30807401 | 0.0032 | 0.0027 | 0.0029 | 0.0029 | TRUE | 0.0627 |
| Git2 | 2121 | 1560 | 1584 | 1516 | 1445 | 1659 | 1647 | 1659 | 1709 | 1509 | 1630 | 1638 | 1632.1 | 1672 | 1592 | 0.952 | -0.07 | 0.29636303 | 0.46830626 | 6.00E-04 | 0.0039 | 0.0026 | 0.0026 | TRUE | 0.0255 |
| Gja1 | 3427 | 2408 | 2563 | 2563 | 2244 | 3117 | 2662 | 2561 | 2765 | 2551 | 2531 | 3077 | 2691.34 | 2663 | 2720 | 1.022 | 0.031 | 0.70556049 | 0.82174868 | 0.0066 | 0.0033 | 0.0045 | 0.0045 | TRUE | 0.1606 |
| Gja10 | 2 | 0 | 1 | 0 | 0 | 0 | 2 | 0 | 1 | 0 | 0 | 0 | 0.44 | 1 | 0 | 0.214 | -2.223 | 0.47051571 | NA | 0 | 5.4047 | 5.2946 | 5.2946 | TRUE | 0.2788 |
| Gja3 | 119 | 90 | 90 | 126 | 92 | 106 | 92 | 96 | 97 | 125 | 104 | 105 | 103.18 | 95 | 111 | 1.173 | 0.23 | 0.23736963 | 0.40415765 | 0 | 0.0254 | 0.0175 | 0.0175 | TRUE | 0.125 |
| Gja4 | 99 | 88 | 91 | 77 | 79 | 84 | 77 | 94 | 98 | 77 | 89 | 83 | 86.23 | 90 | 83 | 0.929 | -0.106 | 0.62236592 | 0.76150235 | 0 | 0.0299 | 0.0217 | 0.0217 | TRUE | 0.1664 |
| Gja5 | 14 | 11 | 13 | 14 | 10 | 13 | 11 | 12 | 14 | 14 | 11 | 13 | 12.44 | 12 | 13 | 1.046 | 0.065 | 0.90221639 | NA | 0 | 0.1929 | 0.1239 | 0.1239 | TRUE | 0.0641 |
| Gja6 | 6 | 2 | 7 | 8 | 5 | 6 | 5 | 2 | 8 | 8 | 6 | 6 | 5.64 | 5 | 7 | 1.367 | 0.451 | 0.58090653 | NA | 0 | 0.4223 | 0.3008 | 0.3008 | TRUE | 0.2535 |
| Gja8 | 6 | 3 | 2 | 0 | 0 | 2 | 5 | 3 | 2 | 0 | 0 | 2 | 2 | 3 | 1 | 0.201 | -2.315 | 0.1475297 | NA | 0 | 1.1891 | 0.9332 | 0.9332 | TRUE | 0.97 |
| Gjb1 | 62 | 60 | 47 | 58 | 59 | 73 | 48 | 64 | 51 | 58 | 67 | 72 | 59.84 | 54 | 66 | 1.212 | 0.277 | 0.28151649 | 0.45301905 | 0 | 0.042 | 0.0309 | 0.0309 | TRUE | 0.193 |
| Gjb2 | 61 | 51 | 39 | 41 | 37 | 29 | 47 | 54 | 42 | 41 | 42 | 29 | 42.48 | 48 | 37 | 0.772 | -0.373 | 0.22238619 | 0.38673389 | 0 | 0.0582 | 0.0434 | 0.0434 | TRUE | 0.2973 |

| | | | | | | | | | | | | | | | | | | | | | | | | | |
|---|---|---|---|---|---|---|---|---|---|---|---|---|---|---|---|---|---|---|---|---|---|---|---|---|---|
| Gjb3 | 1923 | 1455 | 1668 | 860 | 819 | 856 | 1494 | 1548 | 1800 | 856 | 924 | 845 | 1244.32 | 1614 | 875 | 0.542 | -0.882 | 2.65E-23 | 1.11E-20 | 0.0056 | 0.0043 | 0.0048 | 0.0048 | TRUE | 0.1218 |
| Gjb4 | 114 | 81 | 89 | 66 | 59 | 87 | 89 | 86 | 96 | 66 | 67 | 86 | 81.48 | 90 | 73 | 0.808 | -0.308 | 0.16435836 | 0.31522329 | 0 | 0.0315 | 0.023 | 0.023 | TRUE | 0.2334 |
| Gjb5 | 805 | 754 | 924 | 784 | 760 | 828 | 625 | 802 | 997 | 780 | 857 | 817 | 813.22 | 808 | 818 | 1.015 | 0.021 | 0.87542028 | 0.93077304 | 0.0269 | 0.0053 | 0.0119 | 0.0119 | TRUE | 0.2129 |
| Gjb6 | 0 | 0 | 0 | 1 | 0 | 1 | 0 | 0 | 0 | 1 | 0 | 1 | 0.33 | 0 | 1 | 3.763 | 1.912 | 0.57887588 | NA | 0 | 7.1752 | 6.7756 | 6.7756 | TRUE | 0.2094 |
| Gjc1 | 5552 | 4219 | 4286 | 4748 | 4097 | 4880 | 4312 | 4488 | 4625 | 4726 | 4622 | 4817 | 4598.26 | 4475 | 4722 | 1.056 | 0.078 | 0.1532612 | 0.30015187 | 6.00E-04 | 0.0029 | 0.0019 | 0.0019 | TRUE | 0.0126 |
| Gjc2 | 3 | 4 | 3 | 0 | 1 | 3 | 2 | 4 | 3 | 0 | 1 | 3 | 2.32 | 3 | 1 | 0.422 | -1.245 | 0.35227803 | NA | 0 | 1.0246 | 0.7637 | 0.7637 | TRUE | 0.6864 |
| Gjc3 | 1 | 1 | 1 | 1 | 0 | 1 | 1 | 1 | 1 | 1 | 0 | 1 | 0.82 | 1 | 1 | 0.698 | -0.518 | 0.79971927 | NA | 0 | 2.9034 | 1.738 | 1.738 | TRUE | 0.2018 |
| Gjd2 | 1 | 1 | 1 | 0 | 0 | 0 | 1 | 1 | 1 | 0 | 0 | 0 | 0.49 | 1 | 0 | 0.196 | -2.347 | 0.38405676 | NA | 0 | 4.8735 | 3.7415 | 3.7415 | TRUE | 0.0718 |
| Gjd3 | 16 | 13 | 16 | 21 | 16 | 21 | 12 | 14 | 17 | 21 | 18 | 21 | 17.2 | 14 | 20 | 1.383 | 0.467 | 0.30803836 | 0.48060171 | 0 | 0.1402 | 0.0921 | 0.0921 | TRUE | 0.1163 |
| Gjd4 | 1 | 2 | 2 | 1 | 1 | 0 | 1 | 2 | 2 | 1 | 1 | 0 | 1.2 | 2 | 1 | 0.421 | -1.249 | 0.48774863 | NA | 0 | 1.9815 | 1.3019 | 1.3019 | TRUE | 0.2709 |
| Gje1 | 0 | 0 | 0 | 0 | 0 | 0 | 0 | 0 | 0 | 0 | 0 | 0 | 0 | 0 | 0 | NA | NA | NA | NA | NA | NA | NA | NA | NA | NA |
| Gk2 | 0 | 0 | 0 | 0 | 0 | 0 | 0 | 0 | 0 | 0 | 0 | 0 | 0 | 0 | 0 | NA | NA | NA | NA | NA | NA | NA | NA | NA | NA |
| Gk5 | 989 | 717 | 709 | 832 | 705 | 820 | 768 | 763 | 765 | 828 | 795 | 809 | 788.12 | 765 | 811 | 1.06 | 0.084 | 0.2890831 | 0.46114562 | 0 | 0.0054 | 0.0032 | 0.0032 | TRUE | 0.0041 |
| Gkap1 | 1615 | 1148 | 1225 | 1256 | 1095 | 1232 | 1254 | 1221 | 1322 | 1250 | 1235 | 1216 | 1249.81 | 1266 | 1234 | 0.975 | -0.036 | 0.6057091 | 0.75027301 | 0 | 0.0043 | 0.0028 | 0.0028 | TRUE | 0.0173 |
| Gkn1 | 0 | 1 | 0 | 1 | 0 | 0 | 0 | 1 | 0 | 1 | 0 | 0 | 0.34 | 0 | 0 | 1.075 | 0.105 | 0.97573409 | NA | 0 | 6.9086 | 6.5582 | 6.5582 | TRUE | 0.5216 |
| Gkn2 | 0 | 1 | 0 | 0 | 0 | 0 | 0 | 1 | 0 | 0 | 0 | 0 | 0.18 | 0 | 0 | 0.552 | -0.857 | 0.8336548 | NA | 0 | 13.3714 | 10 | 10 | TRUE | 0.5216 |
| Gkn3 | 2 | 1 | 1 | 2 | 1 | 1 | 2 | 1 | 1 | 2 | 1 | 1 | 1.3 | 1 | 1 | 1.101 | 0.139 | 0.93032816 | NA | 0 | 1.8251 | 1.0578 | 1.0578 | TRUE | 0.1028 |
| Gla | 822 | 656 | 743 | 885 | 688 | 837 | 638 | 698 | 802 | 881 | 776 | 826 | 770.2 | 713 | 828 | 1.164 | 0.22 | 0.03024849 | 0.09321726 | 0.0068 | 0.0055 | 0.0061 | 0.0061 | TRUE | 0.1405 |
| Glb1 | 5076 | 3717 | 3988 | 4230 | 4100 | 4551 | 3942 | 3954 | 4303 | 4211 | 4625 | 4493 | 4254.56 | 4066 | 4443 | 1.093 | 0.128 | 0.04314606 | 0.12099979 | 0.0021 | 0.003 | 0.0026 | 0.0026 | TRUE | 0.0313 |
| Glb1l | 51 | 41 | 32 | 32 | 30 | 17 | 40 | 44 | 35 | 32 | 34 | 17 | 33.37 | 40 | 28 | 0.697 | -0.52 | 0.14901554 | 0.29422284 | 0.027 | 0.0734 | 0.0624 | 0.0624 | TRUE | 0.7657 |
| Glb1l2 | 390 | 323 | 378 | 412 | 386 | 362 | 303 | 344 | 408 | 410 | 435 | 357 | 376.2 | 352 | 401 | 1.144 | 0.194 | 0.14815256 | 0.29300464 | 0.0136 | 0.0087 | 0.0102 | 0.0102 | TRUE | 0.2238 |
| Glb1l3 | 1 | 0 | 1 | 0 | 2 | 1 | 1 | 0 | 1 | 0 | 2 | 1 | 0.85 | 1 | 1 | 1.705 | 0.77 | 0.7198077 | NA | 0 | 2.7913 | 2.0637 | 2.0637 | TRUE | 0.1683 |
| Glcci1 | 1147 | 785 | 522 | 597 | 529 | 580 | 891 | 835 | 563 | 594 | 597 | 573 | 675.44 | 763 | 588 | 0.769 | -0.379 | 0.00681215 | 0.03111733 | 0.0281 | 0.0059 | 0.0126 | 0.0126 | TRUE | 0.6141 |
| Glce | 3533 | 2379 | 2277 | 2486 | 2148 | 2257 | 2744 | 2531 | 2457 | 2475 | 2423 | 2228 | 2476.17 | 2577 | 2375 | 0.921 | -0.119 | 0.09137404 | 0.20704012 | 0.0026 | 0.0034 | 0.0032 | 0.0032 | TRUE | 0.0398 |
| Gldc | 14591 | 11466 | 12883 | 13899 | 12587 | 14663 | 11332 | 12196 | 13901 | 13835 | 14199 | 14475 | 13323.04 | 12476 | 14170 | 1.136 | 0.184 | 0.01136064 | 0.04543301 | 0.0056 | 0.0026 | 0.0037 | 0.0037 | TRUE | 0.1218 |
| Gldn | 5 | 1 | 2 | 1 | 0 | 1 | 4 | 1 | 2 | 1 | 0 | 1 | 1.51 | 2 | 1 | 0.281 | -1.831 | 0.2764636 | NA | 0 | 1.5673 | 1.0759 | 1.0759 | TRUE | 0.2155 |
| Gle1 | 4989 | 3701 | 3299 | 3342 | 3115 | 3598 | 3875 | 3937 | 3560 | 3327 | 3514 | 3552 | 3627.25 | 3791 | 3464 | 0.914 | -0.13 | 0.03781887 | 0.10992992 | 0.0016 | 0.0031 | 0.0026 | 0.0026 | TRUE | 0.0341 |
| Glg1 | 12335 | 9013 | 9712 | 10460 | 9076 | 10065 | 9580 | 9587 | 10479 | 10412 | 10239 | 9936 | 10038.75 | 9882 | 10196 | 1.032 | 0.045 | 0.4201115 | 0.59229576 | 0.0015 | 0.0027 | 0.0022 | 0.0022 | TRUE | 0.034 |
| Gli1 | 305 | 279 | 350 | 425 | 408 | 513 | 237 | 297 | 378 | 423 | 460 | 506 | 383.5 | 304 | 463 | 1.532 | 0.615 | 6.09E-05 | 0.00082292 | 0.027 | 0.0086 | 0.0142 | 0.0142 | TRUE | 0.4387 |
| Gli2 | 1528 | 1138 | 1501 | 1511 | 1511 | 1819 | 1187 | 1210 | 1620 | 1504 | 1705 | 1796 | 1503.5 | 1339 | 1668 | 1.247 | 0.318 | 0.0050807 | 0.02480985 | 0.0195 | 0.004 | 0.0086 | 0.0086 | TRUE | 0.4013 |
| Gli3 | 1038 | 647 | 595 | 515 | 471 | 544 | 806 | 688 | 642 | 513 | 531 | 537 | 619.56 | 712 | 527 | 0.738 | -0.438 | 2.95E-05 | 0.00045785 | 0.0063 | 0.0063 | 0.0063 | 0.0063 | TRUE | 0.1641 |
| Glipr1 | 12 | 9 | 2 | 3 | 4 | 7 | 9 | 10 | 2 | 3 | 5 | 7 | 5.91 | 7 | 5 | 0.678 | -0.56 | 0.50613218 | NA | 0.1456 | 0.4035 | 0.3357 | 0.3357 | TRUE | 0.8852 |
| Glipr1l1 | 0 | 0 | 0 | 0 | 0 | 0 | 0 | 0 | 0 | 0 | 0 | 0 | 0 | 0 | 0 | NA | NA | NA | NA | NA | NA | NA | NA | NA | NA |
| Glipr1l2 | 3 | 2 | 2 | 2 | 1 | 0 | 2 | 2 | 2 | 2 | 1 | 0 | 1.62 | 2 | 1 | 0.469 | -1.092 | 0.47727727 | NA | 0 | 1.4633 | 0.9833 | 0.9833 | TRUE | 0.3962 |
| Glipr2 | 1243 | 892 | 676 | 793 | 648 | 797 | 965 | 949 | 729 | 789 | 731 | 787 | 825.12 | 881 | 769 | 0.872 | -0.198 | 0.06950318 | 0.17074751 | 0.0116 | 0.0053 | 0.0073 | 0.0073 | TRUE | 0.2657 |
| Glis1 | 129 | 101 | 99 | 99 | 97 | 84 | 100 | 107 | 107 | 99 | 109 | 83 | 100.89 | 105 | 97 | 0.925 | -0.113 | 0.57075392 | 0.72416753 | 0 | 0.0259 | 0.0187 | 0.0187 | TRUE | 0.1581 |
| Glis2 | 1707 | 1172 | 1116 | 1162 | 1068 | 1268 | 1326 | 1247 | 1204 | 1157 | 1205 | 1252 | 1231.62 | 1259 | 1205 | 0.956 | -0.065 | 0.38307915 | 0.55694744 | 0.0013 | 0.0044 | 0.0032 | 0.0032 | TRUE | 0.027 |
| Glis3 | 216 | 176 | 181 | 122 | 146 | 166 | 168 | 187 | 195 | 121 | 165 | 164 | 166.71 | 183 | 150 | 0.819 | -0.289 | 0.09133145 | 0.20697644 | 0.0106 | 0.0166 | 0.0149 | 0.0149 | TRUE | 0.2958 |
| Glmn | 1946 | 1284 | 1202 | 1268 | 1126 | 1241 | 1511 | 1366 | 1297 | 1262 | 1270 | 1225 | 1321.93 | 1391 | 1252 | 0.899 | -0.154 | 0.04929362 | 0.13338766 | 0.0027 | 0.0042 | 0.0037 | 0.0037 | TRUE | 0.0708 |
| Glo1 | 12900 | 9580 | 10164 | 11178 | 10249 | 11559 | 10019 | 10190 | 10967 | 11127 | 11562 | 11411 | 10879.15 | 10392 | 11367 | 1.094 | 0.129 | 0.01710399 | 0.06169339 | 0.0012 | 0.0026 | 0.002 | 0.002 | TRUE | 0.0285 |
| Glod4 | 6271 | 4641 | 5098 | 6177 | 5522 | 6286 | 4870 | 4937 | 5501 | 6149 | 6229 | 6205 | 5648.49 | 5103 | 6194 | 1.214 | 0.28 | 4.79E-06 | 0.0001024 | 0.0021 | 0.0028 | 0.0025 | 0.0025 | TRUE | 0.0564 |
| Glod5 | 105 | 97 | 110 | 57 | 63 | 73 | 82 | 103 | 119 | 57 | 71 | 72 | 83.88 | 101 | 67 | 0.662 | -0.596 | 0.0107097 | 0.04343748 | 0.0172 | 0.0307 | 0.0267 | 0.0267 | TRUE | 0.2287 |
| Glp1r | 309 | 180 | 212 | 349 | 313 | 320 | 240 | 191 | 229 | 347 | 353 | 316 | 279.43 | 220 | 339 | 1.534 | 0.617 | 3.33E-06 | 7.59E-05 | 0.0041 | 0.0109 | 0.009 | 0.009 | TRUE | 0.1379 |
| Glp2r | 5 | 2 | 2 | 4 | 1 | 1 | 4 | 2 | 2 | 4 | 1 | 1 | 2.38 | 3 | 2 | 0.741 | -0.433 | 0.72758386 | NA | 0 | 0.9993 | 0.6867 | 0.6867 | TRUE | 0.6578 |
| Glra1 | 1 | 0 | 0 | 0 | 0 | 0 | 1 | 0 | 0 | 0 | 0 | 0 | 0.13 | 0 | 0 | 0.552 | -0.857 | 0.8336548 | NA | 0 | 18.3121 | 10 | 10 | TRUE | 0.2566 |
| Glra2 | 4 | 1 | 0 | 0 | 1 | 2 | 3 | 1 | 0 | 0 | 1 | 2 | 1.21 | 1 | 1 | 0.716 | -0.481 | 0.79574926 | NA | 0 | 1.9577 | 1.6521 | 1.6521 | TRUE | 0.4379 |
| Glra3 | 0 | 0 | 0 | 0 | 0 | 0 | 0 | 0 | 0 | 0 | 0 | 0 | 0 | 0 | 0 | NA | NA | NA | NA | NA | NA | NA | NA | NA | NA |
| Glra4 | 0 | 0 | 0 | 0 | 0 | 0 | 0 | 0 | 0 | 0 | 0 | 0 | 0 | 0 | 0 | NA | NA | NA | NA | NA | NA | NA | NA | NA | NA |
| Glrb | 259 | 155 | 176 | 218 | 177 | 191 | 201 | 165 | 190 | 217 | 200 | 189 | 193.52 | 185 | 202 | 1.086 | 0.119 | 0.43046423 | 0.60260776 | 0 | 0.0147 | 0.0112 | 0.0112 | TRUE | 0.0979 |
| Glrp1 | 2 | 0 | 0 | 0 | 0 | 0 | 2 | 0 | 0 | 0 | 0 | 0 | 0.26 | 1 | 0 | 0.356 | -1.491 | 0.71362796 | NA | 0 | 9.1573 | 10 | 10 | TRUE | 0.968 |
| Glrx | 1122 | 732 | 746 | 651 | 628 | 674 | 871 | 779 | 805 | 648 | 708 | 665 | 746.13 | 818 | 674 | 0.822 | -0.283 | 0.00131639 | 0.00878395 | 0.0016 | 0.0056 | 0.0042 | 0.0042 | TRUE | 0.0402 |
| Glrx2 | 851 | 581 | 564 | 599 | 503 | 581 | 661 | 618 | 609 | 596 | 567 | 574 | 604.12 | 629 | 579 | 0.919 | -0.121 | 0.17865511 | 0.33327261 | 0 | 0.0063 | 0.0042 | 0.0042 | TRUE | 0.0244 |
| Glrx3 | 1478 | 1009 | 1185 | 1181 | 1100 | 1261 | 1148 | 1073 | 1279 | 1176 | 1241 | 1245 | 1193.51 | 1167 | 1221 | 1.046 | 0.065 | 0.42880438 | 0.60086404 | 0.0032 | 0.0044 | 0.0041 | 0.0041 | TRUE | 0.082 |
| Glrx5 | 2364 | 1699 | 1566 | 1809 | 1592 | 1838 | 1836 | 1807 | 1690 | 1801 | 1796 | 1814 | 1790.66 | 1778 | 1804 | 1.014 | 0.02 | 0.75673251 | 0.85555908 | 4.00E-04 | 0.0038 | 0.0025 | 0.0025 | TRUE | 0.0223 |
| Gls | 3936 | 3140 | 3170 | 3268 | 2649 | 3136 | 3057 | 3340 | 3420 | 3253 | 2988 | 3096 | 3192.39 | 3272 | 3112 | 0.952 | -0.071 | 0.2830022 | 0.45474428 | 0.0021 | 0.0032 | 0.0029 | 0.0029 | TRUE | 0.0412 |
| Gls2 | 2247 | 1902 | 1977 | 2268 | 1995 | 2292 | 1745 | 2023 | 2133 | 2258 | 2251 | 2263 | 2112.02 | 1967 | 2257 | 1.149 | 0.2 | 0.01146251 | 0.04572163 | 0.0045 | 0.0035 | 0.004 | 0.004 | TRUE | 0.1211 |
| Glt1d1 | 1076 | 836 | 895 | 951 | 839 | 937 | 836 | 889 | 966 | 947 | 946 | 925 | 918.12 | 897 | 939 | 1.049 | 0.069 | 0.40605997 | 0.57887643 | 0.0018 | 0.005 | 0.0039 | 0.0039 | TRUE | 0.0534 |
| Glt25d1 | 11204 | 8478 | 9008 | 10082 | 8964 | 10320 | 8702 | 9018 | 9720 | 10036 | 10112 | 10187 | 9629.09 | 9147 | 10112 | 1.106 | 0.145 | 0.00990411 | 0.04109127 | 0.0015 | 0.0027 | 0.0022 | 0.0022 | TRUE | 0.0366 |
| Glt25d2 | 70 | 46 | 46 | 36 | 40 | 53 | 54 | 49 | 50 | 36 | 45 | 52 | 47.7 | 51 | 44 | 0.869 | -0.202 | 0.47873403 | 0.64580449 | 0 | 0.0521 | 0.0374 | 0.0374 | TRUE | 0.2315 |
| Glt28d2 | 49 | 33 | 26 | 38 | 27 | 21 | 38 | 35 | 28 | 38 | 30 | 21 | 31.7 | 34 | 30 | 0.875 | -0.193 | 0.59404924 | 0.74083813 | 0.0214 | 0.0772 | 0.0624 | 0.0624 | TRUE | 0.3139 |
| Glt6d1 | 0 | 0 | 0 | 0 | 0 | 0 | 0 | 0 | 0 | 0 | 0 | 0 | 0 | 0 | 0 | NA | NA | NA | NA | NA | NA | NA | NA | NA | NA |
| Glt8d1 | 644 | 503 | 564 | 547 | 511 | 550 | 500 | 535 | 609 | 544 | 576 | 543 | 551.27 | 548 | 554 | 1.014 | 0.02 | 0.8450311 | 0.91202932 | 0.0039 | 0.0067 | 0.0058 | 0.0058 | TRUE | 0.1085 |
| Glt8d2 | 33 | 15 | 22 | 34 | 36 | 15 | 26 | 16 | 24 | 34 | 41 | 15 | 25.76 | 22 | 30 | 1.353 | 0.436 | 0.32160812 | 0.49533888 | 0.1149 | 0.0944 | 0.1002 | 0.1002 | TRUE | 0.7107 |
| Gltp | 2194 | 1614 | 1707 | 1631 | 1436 | 1650 | 1704 | 1717 | 1842 | 1623 | 1620 | 1629 | 1689.15 | 1754 | 1624 | 0.926 | -0.11 | 0.09272219 | 0.20923244 | 4.00E-04 | 0.0038 | 0.0025 | 0.0025 | TRUE | 0.0227 |
| Gltpd1 | 403 | 265 | 222 | 240 | 197 | 205 | 313 | 282 | 240 | 239 | 222 | 202 | 249.65 | 278 | 221 | 0.793 | -0.335 | 0.02054122 | 0.07054196 | 0.009 | 0.0119 | 0.011 | 0.011 | TRUE | 0.1623 |
| Gltpd2 | 15 | 17 | 18 | 29 | 20 | 33 | 12 | 18 | 19 | 29 | 23 | 33 | 22.19 | 16 | 28 | 1.736 | 0.796 | 0.06053708 | 0.15417969 | 0 | 0.1092 | 0.0814 | 0.0814 | TRUE | 0.3977 |
| Gltscr1 | 614 | 458 | 468 | 413 | 399 | 388 | 477 | 487 | 505 | 411 | 450 | 383 | 452.21 | 490 | 415 | 0.847 | -0.24 | 0.02182013 | 0.07356156 | 9.00E-04 | 0.0077 | 0.0057 | 0.0057 | TRUE | 0.0623 |
| Gltscr1l | 3413 | 2530 | 2487 | 2517 | 2422 | 2736 | 2651 | 2691 | 2683 | 2505 | 2732 | 2701 | 2660.64 | 2675 | 2646 | 0.989 | -0.016 | 0.79503624 | 0.87993286 | 7.00E-04 | 0.0033 | 0.0023 | 0.0023 | TRUE | 0.0264 |
| Gltscr2 | 12618 | 9847 | 9859 | 10892 | 9803 | 11276 | 9800 | 10474 | 10638 | 10842 | 11059 | 11131 | 10657.27 | 10304 | 11011 | 1.069 | 0.096 | 0.06702382 | 0.16605504 | 9.00E-04 | 0.0026 | 0.0019 | 0.0019 | TRUE | 0.0226 |

| | | | | | | | | | | | | | | | | | | | | | | | | | |
|---|---|---|---|---|---|---|---|---|---|---|---|---|---|---|---|---|---|---|---|---|---|---|---|---|---|
| Glud1 | 33771 | 23740 | 25341 | 27932 | 24459 | 27410 | 26229 | 25252 | 27343 | 27803 | 27592 | 27058 | 26879.47 | 26275 | 27484 | 1.046 | 0.065 | 0.19062161 | 0.3487296 | 8.00E-04 | 0.0025 | 0.0017 | 0.0017 | TRUE | 0.0154 |
| Glul | 8903 | 6107 | 5678 | 6005 | 5493 | 6028 | 6915 | 6496 | 6127 | 5977 | 6197 | 5951 | 6276.94 | 6513 | 6042 | 0.927 | -0.109 | 0.07008821 | 0.17162425 | 0.0018 | 0.0028 | 0.0024 | 0.0024 | TRUE | 0.036 |
| Glyat | 0 | 0 | 0 | 0 | 0 | 0 | 0 | 0 | 0 | 0 | 0 | 0 | 0 | 0 | 0 | NA | NA | NA | NA | NA | NA | NA | NA | NA | NA |
| Glyatl3 | 0 | 0 | 1 | 0 | 0 | 0 | 0 | 0 | 1 | 0 | 0 | 0 | 0.18 | 0 | 0 | 0.552 | -0.857 | 0.8336548 | NA | 0 | 13.1817 | 10 | 10 | TRUE | 0.5365 |
| Glycam1 | 0 | 0 | 0 | 0 | 0 | 0 | 0 | 0 | 0 | 0 | 0 | 0 | 0 | 0 | 0 | NA | NA | NA | NA | NA | NA | NA | NA | NA | NA |
| Glyctk | 523 | 390 | 367 | 483 | 431 | 447 | 406 | 415 | 396 | 481 | 486 | 441 | 437.54 | 406 | 469 | 1.156 | 0.21 | 0.04002606 | 0.11446428 | 0 | 0.0078 | 0.0052 | 0.0052 | TRUE | 0.0325 |
| Glyr1 | 9635 | 6732 | 5694 | 5652 | 5145 | 5852 | 7483 | 7161 | 6144 | 5626 | 5804 | 5777 | 6332.42 | 6929 | 5736 | 0.827 | -0.273 | 0.0002014 | 0.00214459 | 0.005 | 0.0028 | 0.0037 | 0.0037 | TRUE | 0.1196 |
| Gm10007 | 0 | 0 | 0 | 0 | 0 | 0 | 0 | 0 | 0 | 0 | 0 | 0 | 0 | 0 | 0 | NA | NA | NA | NA | NA | NA | NA | NA | NA | NA |
| Gm10012 | 0 | 2 | 1 | 1 | 0 | 2 | 0 | 2 | 1 | 1 | 0 | 2 | 1.03 | 1 | 1 | 0.975 | -0.037 | 0.98512122 | NA | 0 | 2.3049 | 1.8419 | 1.8419 | TRUE | 0.2274 |
| Gm10024 | 0 | 0 | 0 | 0 | 0 | 0 | 0 | 0 | 0 | 0 | 0 | 0 | 0 | 0 | 0 | NA | NA | NA | NA | NA | NA | NA | NA | NA | NA |
| Gm10033 | 360 | 263 | 219 | 339 | 250 | 244 | 280 | 280 | 236 | 337 | 282 | 241 | 276 | 265 | 287 | 1.08 | 0.111 | 0.45601332 | 0.62562856 | 0.0153 | 0.011 | 0.0123 | 0.0123 | TRUE | 0.2633 |
| Gm10046 | 12 | 9 | 7 | 8 | 7 | 10 | 9 | 10 | 8 | 8 | 8 | 10 | 8.7 | 9 | 9 | 0.972 | -0.041 | 0.94845398 | NA | 0 | 0.275 | 0.1747 | 0.1747 | TRUE | 0.056 |
| Gm10052 | 0 | 0 | 0 | 0 | 0 | 0 | 0 | 0 | 0 | 0 | 0 | 0 | 0 | 0 | 0 | NA | NA | NA | NA | NA | NA | NA | NA | NA | NA |
| Gm10057 | 0 | 0 | 0 | 0 | 0 | 0 | 0 | 0 | 0 | 0 | 0 | 0 | 0 | 0 | 0 | NA | NA | NA | NA | NA | NA | NA | NA | NA | NA |
| Gm10058 | 0 | 0 | 0 | 0 | 0 | 0 | 0 | 0 | 0 | 0 | 0 | 0 | 0 | 0 | 0 | NA | NA | NA | NA | NA | NA | NA | NA | NA | NA |
| Gm10069 | 141 | 106 | 123 | 130 | 115 | 138 | 110 | 113 | 133 | 129 | 130 | 136 | 125.06 | 119 | 132 | 1.117 | 0.16 | 0.36858019 | 0.54321707 | 0 | 0.0214 | 0.0148 | 0.0148 | TRUE | 0.1099 |
| Gm10081 | 0 | 0 | 0 | 0 | 0 | 0 | 0 | 0 | 0 | 0 | 0 | 0 | 0 | 0 | 0 | NA | NA | NA | NA | NA | NA | NA | NA | NA | NA |
| Gm10094 | 0 | 0 | 0 | 0 | 0 | 0 | 0 | 0 | 0 | 0 | 0 | 0 | 0 | 0 | 0 | NA | NA | NA | NA | NA | NA | NA | NA | NA | NA |
| Gm10096 | 0 | 0 | 0 | 0 | 0 | 0 | 0 | 0 | 0 | 0 | 0 | 0 | 0 | 0 | 0 | NA | NA | NA | NA | NA | NA | NA | NA | NA | NA |
| Gm101 | 6 | 8 | 10 | 11 | 12 | 4 | 5 | 9 | 11 | 11 | 14 | 4 | 8.73 | 8 | 10 | 1.198 | 0.26 | 0.70982648 | NA | 0.1386 | 0.2738 | 0.236 | 0.236 | TRUE | 0.7694 |
| Gm10100 | 0 | 0 | 0 | 0 | 0 | 0 | 0 | 0 | 0 | 0 | 0 | 0 | 0 | 0 | 0 | NA | NA | NA | NA | NA | NA | NA | NA | NA | NA |
| Gm10104 | 0 | 0 | 0 | 0 | 0 | 0 | 0 | 0 | 0 | 0 | 0 | 0 | 0 | 0 | 0 | NA | NA | NA | NA | NA | NA | NA | NA | NA | NA |
| Gm10125 | 2 | 1 | 0 | 3 | 1 | 3 | 2 | 1 | 0 | 3 | 1 | 3 | 1.62 | 1 | 2 | 2.607 | 1.382 | 0.37575807 | NA | 0 | 1.4695 | 1.0091 | 1.0091 | TRUE | 0.2722 |
| Gm10142 | 0 | 0 | 0 | 0 | 0 | 0 | 0 | 0 | 0 | 0 | 0 | 0 | 0 | 0 | 0 | NA | NA | NA | NA | NA | NA | NA | NA | NA | NA |
| Gm10147 | 0 | 0 | 0 | 0 | 0 | 0 | 0 | 0 | 0 | 0 | 0 | 0 | 0 | 0 | 0 | NA | NA | NA | NA | NA | NA | NA | NA | NA | NA |
| Gm10190 | 6 | 5 | 0 | 2 | 2 | 2 | 5 | 5 | 0 | 2 | 2 | 2 | 2.7 | 3 | 2 | 0.615 | -0.702 | 0.57464914 | NA | 0 | 0.8802 | 0.7349 | 0.7349 | TRUE | 0.9772 |
| Gm10220 | 1 | 3 | 0 | 1 | 0 | 0 | 1 | 3 | 0 | 1 | 0 | 0 | 0.83 | 1 | 0 | 0.283 | -1.823 | 0.43666934 | NA | 0 | 2.8676 | 2.5891 | 2.5891 | TRUE | 0.5266 |
| Gm10228 | 0 | 0 | 0 | 0 | 0 | 0 | 0 | 0 | 0 | 0 | 0 | 0 | 0 | 0 | 0 | NA | NA | NA | NA | NA | NA | NA | NA | NA | NA |
| Gm10229 | 0 | 0 | 0 | 0 | 0 | 0 | 0 | 0 | 0 | 0 | 0 | 0 | 0 | 0 | 0 | NA | NA | NA | NA | NA | NA | NA | NA | NA | NA |
| Gm10230 | 0 | 0 | 0 | 0 | 0 | 0 | 0 | 0 | 0 | 0 | 0 | 0 | 0 | 0 | 0 | NA | NA | NA | NA | NA | NA | NA | NA | NA | NA |
| Gm10248 | 0 | 0 | 0 | 0 | 0 | 0 | 0 | 0 | 0 | 0 | 0 | 0 | 0 | 0 | 0 | NA | NA | NA | NA | NA | NA | NA | NA | NA | NA |
| Gm10256 | 0 | 0 | 0 | 0 | 0 | 0 | 0 | 0 | 0 | 0 | 0 | 0 | 0 | 0 | 0 | NA | NA | NA | NA | NA | NA | NA | NA | NA | NA |
| Gm10267 | 0 | 0 | 0 | 0 | 0 | 0 | 0 | 0 | 0 | 0 | 0 | 0 | 0 | 0 | 0 | NA | NA | NA | NA | NA | NA | NA | NA | NA | NA |
| Gm10272 | 0 | 0 | 0 | 0 | 0 | 0 | 0 | 0 | 0 | 0 | 0 | 0 | 0 | 0 | 0 | NA | NA | NA | NA | NA | NA | NA | NA | NA | NA |
| Gm10280 | 0 | 0 | 0 | 0 | 0 | 0 | 0 | 0 | 0 | 0 | 0 | 0 | 0 | 0 | 0 | NA | NA | NA | NA | NA | NA | NA | NA | NA | NA |
| Gm10318 | 0 | 0 | 0 | 0 | 1 | 0 | 0 | 0 | 0 | 0 | 1 | 0 | 0.19 | 0 | 0 | 2.094 | 1.067 | 0.79379329 | NA | 0 | 12.608 | 10 | 10 | TRUE | 0.5075 |
| Gm10319 | 0 | 0 | 0 | 0 | 0 | 0 | 0 | 0 | 0 | 0 | 0 | 0 | 0 | 0 | 0 | NA | NA | NA | NA | NA | NA | NA | NA | NA | NA |
| Gm10324 | 487 | 349 | 257 | 291 | 231 | 204 | 378 | 371 | 277 | 290 | 261 | 201 | 296.4 | 342 | 251 | 0.73 | -0.453 | 0.00520255 | 0.02530116 | 0.0277 | 0.0104 | 0.0155 | 0.0155 | TRUE | 0.334 |
| Gm10334 | 0 | 0 | 0 | 0 | 0 | 0 | 0 | 0 | 0 | 0 | 0 | 0 | 0 | 0 | 0 | NA | NA | NA | NA | NA | NA | NA | NA | NA | NA |
| Gm10336 | 1703 | 1036 | 1163 | 1390 | 1099 | 1305 | 1323 | 1102 | 1255 | 1384 | 1240 | 1288 | 1265.19 | 1227 | 1304 | 1.062 | 0.087 | 0.31336937 | 0.48684656 | 0.0046 | 0.0043 | 0.0046 | 0.0046 | TRUE | 0.0938 |
| Gm10352 | 0 | 0 | 0 | 0 | 0 | 0 | 0 | 0 | 0 | 0 | 0 | 0 | 0 | 0 | 0 | NA | NA | NA | NA | NA | NA | NA | NA | NA | NA |
| Gm10364 | 2 | 1 | 2 | 2 | 1 | 1 | 2 | 1 | 2 | 2 | 1 | 1 | 1.48 | 2 | 1 | 0.864 | -0.211 | 0.88836451 | NA | 0 | 1.6036 | 0.9503 | 0.9503 | TRUE | 0.1029 |
| Gm10373 | 4 | 1 | 1 | 0 | 0 | 0 | 3 | 1 | 1 | 0 | 0 | 0 | 0.87 | 2 | 0 | 0.107 | -3.222 | 0.14630442 | NA | 0 | 2.7114 | 2.2736 | 2.2736 | TRUE | 0.4767 |
| Gm10375 | 0 | 0 | 0 | 0 | 0 | 0 | 0 | 0 | 0 | 0 | 0 | 0 | 0 | 0 | 0 | NA | NA | NA | NA | NA | NA | NA | NA | NA | NA |
| Gm10377 | 0 | 0 | 0 | 0 | 0 | 0 | 0 | 0 | 0 | 0 | 0 | 0 | 0 | 0 | 0 | NA | NA | NA | NA | NA | NA | NA | NA | NA | NA |
| Gm10389 | 0 | 0 | 0 | 0 | 0 | 0 | 0 | 0 | 0 | 0 | 0 | 0 | 0 | 0 | 0 | NA | NA | NA | NA | NA | NA | NA | NA | NA | NA |
| Gm10390 | 1 | 2 | 0 | 1 | 0 | 0 | 1 | 2 | 0 | 1 | 0 | 0 | 0.65 | 1 | 0 | 0.384 | -1.379 | 0.58322298 | NA | 0 | 3.6492 | 3.0509 | 3.0509 | TRUE | 0.2622 |
| Gm10400 | 0 | 0 | 0 | 0 | 0 | 1 | 0 | 0 | 0 | 0 | 0 | 1 | 0.16 | 0 | 0 | 2.094 | 1.067 | 0.79379329 | NA | 0 | 14.4078 | 10 | 10 | TRUE | 0.3835 |
| Gm10406 | 87 | 54 | 54 | 64 | 59 | 61 | 68 | 57 | 58 | 64 | 67 | 60 | 62.29 | 61 | 64 | 1.034 | 0.049 | 0.84227949 | 0.91032497 | 0 | 0.0405 | 0.0268 | 0.0268 | TRUE | 0.0856 |
| Gm10408 | 0 | 0 | 0 | 0 | 0 | 0 | 0 | 0 | 0 | 0 | 0 | 0 | 0 | 0 | 0 | NA | NA | NA | NA | NA | NA | NA | NA | NA | NA |
| Gm10409 | 0 | 0 | 0 | 0 | 0 | 0 | 0 | 0 | 0 | 0 | 0 | 0 | 0 | 0 | 0 | NA | NA | NA | NA | NA | NA | NA | NA | NA | NA |
| Gm10413 | 0 | 0 | 0 | 0 | 0 | 0 | 0 | 0 | 0 | 0 | 0 | 0 | 0 | 0 | 0 | NA | NA | NA | NA | NA | NA | NA | NA | NA | NA |
| Gm10415 | 0 | 0 | 1 | 0 | 0 | 0 | 0 | 0 | 1 | 0 | 0 | 0 | 0.18 | 0 | 0 | 0.552 | -0.857 | 0.8336548 | NA | 0 | 13.1817 | 10 | 10 | TRUE | 0.5365 |
| Gm10416 | 130 | 86 | 56 | 67 | 73 | 53 | 101 | 91 | 60 | 67 | 82 | 52 | 75.7 | 84 | 67 | 0.79 | -0.34 | 0.1994492 | 0.3601222 | 0.0461 | 0.0337 | 0.0372 | 0.0372 | TRUE | 0.4004 |
| Gm10421 | 0 | 0 | 0 | 0 | 0 | 0 | 0 | 0 | 0 | 0 | 0 | 0 | 0 | 0 | 0 | NA | NA | NA | NA | NA | NA | NA | NA | NA | NA |
| Gm10432 | 24 | 15 | 16 | 18 | 15 | 17 | 19 | 16 | 17 | 18 | 17 | 17 | 17.25 | 17 | 17 | 0.993 | -0.011 | 0.98093008 | 0.9900737 | 0 | 0.1398 | 0.0886 | 0.0886 | TRUE | 0.0292 |
| Gm10433 | 26 | 12 | 9 | 10 | 5 | 8 | 20 | 13 | 10 | 10 | 6 | 8 | 11.03 | 14 | 8 | 0.546 | -0.872 | 0.15034395 | NA | 0 | 0.2174 | 0.1655 | 0.1655 | TRUE | 0.5704 |
| Gm10436 | 2 | 1 | 0 | 0 | 0 | 0 | 2 | 1 | 0 | 0 | 0 | 0 | 0.44 | 1 | 0 | 0.215 | -2.217 | 0.47268385 | NA | 0 | 5.4363 | 5.3181 | 5.3181 | TRUE | 0.2411 |
| Gm10439 | 77 | 54 | 51 | 19 | 18 | 15 | 60 | 57 | 55 | 19 | 20 | 15 | 37.72 | 57 | 18 | 0.312 | -1.679 | 5.79E-07 | 1.84E-05 | 0 | 0.0653 | 0.0441 | 0.0441 | TRUE | 0.1276 |
| Gm10440 | 0 | 0 | 0 | 0 | 0 | 0 | 0 | 0 | 0 | 0 | 0 | 0 | 0 | 0 | 0 | NA | NA | NA | NA | NA | NA | NA | NA | NA | NA |
| Gm10445 | 0 | 0 | 0 | 0 | 0 | 0 | 0 | 0 | 0 | 0 | 0 | 0 | 0 | 0 | 0 | NA | NA | NA | NA | NA | NA | NA | NA | NA | NA |
| Gm1045 | 32 | 10 | 13 | 11 | 7 | 6 | 25 | 11 | 14 | 11 | 8 | 6 | 12.38 | 17 | 8 | 0.494 | -1.018 | 0.0861806 | NA | 0.0901 | 0.1939 | 0.1623 | 0.1623 | TRUE | 0.8306 |
| Gm10451 | 368 | 266 | 239 | 260 | 188 | 245 | 286 | 283 | 258 | 259 | 212 | 242 | 256.56 | 276 | 238 | 0.863 | -0.213 | 0.10993666 | 0.23714529 | 0.0021 | 0.0117 | 0.0089 | 0.0089 | TRUE | 0.0934 |
| Gm10466 | 0 | 0 | 0 | 0 | 0 | 0 | 0 | 0 | 0 | 0 | 0 | 0 | 0 | 0 | 0 | NA | NA | NA | NA | NA | NA | NA | NA | NA | NA |
| Gm10471 | 2 | 1 | 2 | 2 | 1 | 1 | 2 | 1 | 2 | 2 | 1 | 1 | 1.48 | 2 | 1 | 0.864 | -0.211 | 0.88836451 | NA | 0 | 1.6036 | 0.9503 | 0.9503 | TRUE | 0.1029 |
| Gm10474 | 5 | 10 | 8 | 9 | 7 | 8 | 4 | 11 | 9 | 9 | 8 | 8 | 7.98 | 8 | 8 | 1.091 | 0.125 | 0.85612373 | NA | 0 | 0.2993 | 0.2172 | 0.2172 | TRUE | 0.728 |
| Gm10485 | 0 | 0 | 0 | 0 | 0 | 0 | 0 | 0 | 0 | 0 | 0 | 0 | 0 | 0 | 0 | NA | NA | NA | NA | NA | NA | NA | NA | NA | NA |
| Gm10486 | 0 | 0 | 0 | 0 | 0 | 0 | 0 | 0 | 0 | 0 | 0 | 0 | 0 | 0 | 0 | NA | NA | NA | NA | NA | NA | NA | NA | NA | NA |
| Gm10487 | 0 | 0 | 0 | 0 | 0 | 0 | 0 | 0 | 0 | 0 | 0 | 0 | 0 | 0 | 0 | NA | NA | NA | NA | NA | NA | NA | NA | NA | NA |

| | | | | | | | | | | | | | | | | | | | | | | | | | |
|---|---|---|---|---|---|---|---|---|---|---|---|---|---|---|---|---|---|---|---|---|---|---|---|---|---|
| Gm10488 | 0 | 0 | 0 | 0 | 0 | 0 | 0 | 0 | 0 | 0 | 0 | 0 | 0 | 0 | 0 | NA | NA | NA | NA | NA | NA | NA | NA | NA | NA |
| Gm10494 | 0 | 1 | 0 | 0 | 0 | 0 | 0 | 1 | 0 | 0 | 0 | 0 | 0.18 | 0 | 0 | 0.552 | -0.857 | 0.8336548 | NA | 0 | 13.3714 | 10 | 10 | TRUE | 0.5216 |
| Gm10509 | 197 | 138 | 118 | 167 | 121 | 121 | 153 | 147 | 127 | 166 | 137 | 119 | 141.55 | 142 | 141 | 0.986 | -0.02 | 0.91397264 | 0.95262969 | 0.0116 | 0.0192 | 0.017 | 0.017 | TRUE | 0.2671 |
| Gm10510 | 0 | 0 | 0 | 0 | 0 | 0 | 0 | 0 | 0 | 0 | 0 | 0 | 0 | 0 | 0 | NA | NA | NA | NA | NA | NA | NA | NA | NA | NA |
| Gm10512 | 0 | 1 | 0 | 0 | 0 | 0 | 0 | 1 | 0 | 0 | 0 | 0 | 0.18 | 0 | 0 | 0.552 | -0.857 | 0.8336548 | NA | 0 | 13.3714 | 10 | 10 | TRUE | 0.5216 |
| Gm10516 | 4 | 4 | 4 | 3 | 4 | 3 | 3 | 4 | 4 | 3 | 5 | 3 | 3.69 | 4 | 4 | 0.898 | -0.155 | 0.87294567 | NA | 0 | 0.6448 | 0.401 | 0.401 | TRUE | 0.0865 |
| Gm10532 | 77 | 48 | 37 | 32 | 31 | 37 | 60 | 51 | 40 | 32 | 35 | 37 | 42.36 | 50 | 35 | 0.681 | -0.554 | 0.0707339 | 0.17285 | 0 | 0.0584 | 0.0432 | 0.0432 | TRUE | 0.1833 |
| Gm10536 | 1 | 0 | 0 | 1 | 0 | 0 | 1 | 0 | 0 | 1 | 0 | 0 | 0.3 | 0 | 0 | 1.075 | 0.105 | 0.97698525 | NA | 0 | 8.0272 | 7.5141 | 7.5141 | TRUE | 0.3906 |
| Gm10538 | 0 | 0 | 0 | 0 | 0 | 0 | 0 | 0 | 0 | 0 | 0 | 0 | 0 | 0 | 0 | NA | NA | NA | NA | NA | NA | NA | NA | NA | NA |
| Gm10548 | 2 | 2 | 0 | 0 | 0 | 1 | 2 | 2 | 0 | 0 | 0 | 1 | 0.78 | 1 | 0 | 0.3 | -1.737 | 0.45918821 | NA | 0 | 3.0489 | 2.5775 | 2.5775 | TRUE | 0.3899 |
| Gm10549 | 1 | 0 | 0 | 0 | 0 | 1 | 1 | 0 | 0 | 0 | 0 | 1 | 0.29 | 0 | 0 | 1.075 | 0.105 | 0.97702352 | NA | 0 | 8.0647 | 7.5458 | 7.5458 | TRUE | 0.3835 |
| Gm10556 | 2 | 1 | 0 | 0 | 2 | 0 | 2 | 1 | 0 | 0 | 2 | 0 | 0.81 | 1 | 1 | 0.816 | -0.293 | 0.89959898 | NA | 0 | 2.9205 | 2.6456 | 2.6456 | TRUE | 0.9748 |
| Gm10560 | 47 | 28 | 34 | 51 | 32 | 36 | 37 | 30 | 37 | 51 | 36 | 36 | 37.56 | 35 | 41 | 1.188 | 0.248 | 0.44168663 | 0.61279625 | 0 | 0.0655 | 0.0484 | 0.0484 | TRUE | 0.3515 |
| Gm10578 | 0 | 0 | 0 | 0 | 0 | 0 | 0 | 0 | 0 | 0 | 0 | 0 | 0 | 0 | 0 | NA | NA | NA | NA | NA | NA | NA | NA | NA | NA |
| Gm10584 | 1 | 0 | 2 | 0 | 0 | 0 | 1 | 0 | 2 | 0 | 0 | 0 | 0.49 | 1 | 0 | 0.195 | -2.355 | 0.4284796 | NA | 0 | 4.8481 | 4.8763 | 4.8763 | TRUE | 0.127 |
| Gm10591 | 0 | 0 | 0 | 0 | 0 | 0 | 0 | 0 | 0 | 0 | 0 | 0 | 0 | 0 | 0 | NA | NA | NA | NA | NA | NA | NA | NA | NA | NA |
| Gm10619 | 0 | 0 | 0 | 0 | 0 | 0 | 0 | 0 | 0 | 0 | 0 | 0 | 0 | 0 | 0 | NA | NA | NA | NA | NA | NA | NA | NA | NA | NA |
| Gm10635 | 4 | 2 | 1 | 2 | 2 | 0 | 3 | 2 | 1 | 2 | 2 | 0 | 1.76 | 2 | 1 | 0.654 | -0.612 | 0.6787561 | NA | 0 | 1.3491 | 0.9788 | 0.9788 | TRUE | 0.5079 |
| Gm10636 | 0 | 0 | 0 | 0 | 0 | 0 | 0 | 0 | 0 | 0 | 0 | 0 | 0 | 0 | 0 | NA | NA | NA | NA | NA | NA | NA | NA | NA | NA |
| Gm10637 | 0 | 0 | 0 | 0 | 1 | 0 | 0 | 0 | 0 | 0 | 1 | 0 | 0.19 | 0 | 0 | 2.094 | 1.067 | 0.79379329 | NA | 0 | 12.608 | 10 | 10 | TRUE | 0.5075 |
| Gm10638 | 11 | 7 | 10 | 5 | 10 | 6 | 9 | 7 | 11 | 5 | 11 | 6 | 8.16 | 9 | 7 | 0.822 | -0.282 | 0.67783781 | NA | 0 | 0.2929 | 0.208 | 0.208 | TRUE | 0.5251 |
| Gm10639 | 2 | 1 | 0 | 0 | 1 | 0 | 2 | 1 | 0 | 0 | 1 | 0 | 0.62 | 1 | 0 | 0.416 | -1.266 | 0.61771935 | NA | 0 | 3.7995 | 3.0914 | 3.0914 | TRUE | 0.5075 |
| Gm10640 | 4 | 1 | 6 | 1 | 1 | 3 | 3 | 1 | 6 | 1 | 1 | 3 | 2.62 | 3 | 2 | 0.484 | -1.048 | 0.4015948 | NA | 0 | 0.9065 | 0.6851 | 0.6851 | TRUE | 0.21 |
| Gm10649 | 0 | 0 | 0 | 0 | 0 | 0 | 0 | 0 | 0 | 0 | 0 | 0 | 0 | 0 | 0 | NA | NA | NA | NA | NA | NA | NA | NA | NA | NA |
| Gm10653 | 4 | 3 | 2 | 8 | 1 | 4 | 3 | 3 | 2 | 8 | 1 | 4 | 3.58 | 3 | 4 | 1.55 | 0.632 | 0.5470063 | NA | 0 | 0.664 | 0.506 | 0.506 | TRUE | 0.2103 |
| Gm10658 | 74 | 43 | 91 | 65 | 60 | 76 | 57 | 46 | 98 | 65 | 68 | 75 | 68.14 | 67 | 69 | 1.034 | 0.049 | 0.86562063 | 0.92572508 | 0.0653 | 0.0372 | 0.0451 | 0.0451 | TRUE | 1.2587 |
| Gm10662 | 0 | 0 | 0 | 0 | 0 | 0 | 0 | 0 | 0 | 0 | 0 | 0 | 0 | 0 | 0 | NA | NA | NA | NA | NA | NA | NA | NA | NA | NA |
| Gm10664 | 921 | 731 | 946 | 1209 | 1018 | 1334 | 715 | 778 | 1021 | 1203 | 1148 | 1317 | 1030.38 | 838 | 1223 | 1.463 | 0.548 | 3.49E-06 | 7.87E-05 | 0.0195 | 0.0047 | 0.0091 | 0.0091 | TRUE | 0.4304 |
| Gm10665 | 0 | 0 | 0 | 0 | 0 | 0 | 0 | 0 | 0 | 0 | 0 | 0 | 0 | 0 | 0 | NA | NA | NA | NA | NA | NA | NA | NA | NA | NA |
| Gm10666 | 0 | 0 | 0 | 0 | 0 | 0 | 0 | 0 | 0 | 0 | 0 | 0 | 0 | 0 | 0 | NA | NA | NA | NA | NA | NA | NA | NA | NA | NA |
| Gm10670 | 0 | 0 | 0 | 0 | 0 | 0 | 0 | 0 | 0 | 0 | 0 | 0 | 0 | 0 | 0 | NA | NA | NA | NA | NA | NA | NA | NA | NA | NA |
| Gm10677 | 1 | 0 | 1 | 1 | 1 | 2 | 1 | 0 | 1 | 1 | 1 | 2 | 0.99 | 1 | 1 | 2.189 | 1.13 | 0.55451068 | NA | 0 | 2.391 | 1.4884 | 1.4884 | TRUE | 0.1946 |
| Gm10681 | 0 | 0 | 0 | 0 | 0 | 0 | 0 | 0 | 0 | 0 | 0 | 0 | 0 | 0 | 0 | NA | NA | NA | NA | NA | NA | NA | NA | NA | NA |
| Gm10684 | 0 | 0 | 0 | 0 | 0 | 0 | 0 | 0 | 0 | 0 | 0 | 0 | 0 | 0 | 0 | NA | NA | NA | NA | NA | NA | NA | NA | NA | NA |
| Gm10696 | 7 | 14 | 1 | 3 | 0 | 0 | 5 | 15 | 1 | 3 | 0 | 0 | 4.07 | 7 | 1 | 0.143 | -2.806 | 0.03929861 | NA | 1.594 | 0.5854 | 0.759 | 0.759 | TRUE | 0.4771 |
| Gm10697 | 0 | 0 | 0 | 0 | 0 | 0 | 0 | 0 | 0 | 0 | 0 | 0 | 0 | 0 | 0 | NA | NA | NA | NA | NA | NA | NA | NA | NA | NA |
| Gm10714 | 3 | 4 | 7 | 10 | 4 | 12 | 2 | 4 | 8 | 10 | 5 | 12 | 6.74 | 5 | 9 | 1.918 | 0.939 | 0.23432038 | NA | 0.0954 | 0.354 | 0.2861 | 0.2861 | TRUE | 0.523 |
| Gm10731 | 0 | 0 | 0 | 0 | 0 | 0 | 0 | 0 | 0 | 0 | 0 | 0 | 0 | 0 | 0 | NA | NA | NA | NA | NA | NA | NA | NA | NA | NA |
| Gm10745 | 0 | 1 | 0 | 0 | 0 | 0 | 0 | 1 | 0 | 0 | 0 | 0 | 0.18 | 0 | 0 | 0.552 | -0.857 | 0.8336548 | NA | 0 | 13.3714 | 10 | 10 | TRUE | 0.5216 |
| Gm10754 | 1 | 1 | 2 | 0 | 0 | 1 | 1 | 1 | 2 | 0 | 0 | 1 | 0.83 | 1 | 0 | 0.282 | -1.828 | 0.39771905 | NA | 0 | 2.8548 | 1.9998 | 1.9998 | TRUE | 0.3835 |
| Gm10767 | 6 | 12 | 9 | 9 | 4 | 8 | 5 | 13 | 10 | 9 | 5 | 8 | 8.08 | 9 | 7 | 0.805 | -0.313 | 0.65670356 | NA | 0 | 0.2956 | 0.2302 | 0.2302 | TRUE | 0.382 |
| Gm10768 | 95 | 90 | 103 | 98 | 88 | 95 | 74 | 96 | 111 | 98 | 99 | 94 | 95.21 | 94 | 97 | 1.042 | 0.06 | 0.78131 | 0.87115299 | 0.0126 | 0.0273 | 0.023 | 0.023 | TRUE | 0.3524 |
| Gm10778 | 14 | 14 | 10 | 13 | 13 | 11 | 11 | 15 | 11 | 13 | 15 | 11 | 12.5 | 12 | 13 | 1.054 | 0.076 | 0.8876536 | NA | 0 | 0.192 | 0.1267 | 0.1267 | TRUE | 0.1404 |
| Gm1078 | 1 | 1 | 0 | 0 | 1 | 0 | 1 | 1 | 0 | 0 | 1 | 0 | 0.49 | 1 | 0 | 0.599 | -0.74 | 0.78905321 | NA | 0 | 4.7929 | 3.7323 | 3.7323 | TRUE | 0.5075 |
| Gm10782 | 2 | 0 | 0 | 0 | 0 | 0 | 2 | 0 | 0 | 0 | 0 | 0 | 0.26 | 1 | 0 | 0.356 | -1.491 | 0.71362796 | NA | 0 | 9.1573 | 10 | 10 | TRUE | 0.968 |
| Gm10785 | 2 | 0 | 1 | 2 | 0 | 0 | 2 | 0 | 1 | 2 | 0 | 0 | 0.77 | 1 | 1 | 0.751 | -0.414 | 0.86074347 | NA | 0 | 3.0784 | 2.7066 | 2.7066 | TRUE | 0.9737 |
| Gm10787 | 1 | 0 | 3 | 0 | 1 | 0 | 1 | 0 | 3 | 0 | 1 | 0 | 0.86 | 1 | 0 | 0.28 | -1.835 | 0.4293679 | NA | 0 | 2.7681 | 2.5201 | 2.5201 | TRUE | 0.5637 |
| Gm10789 | 31 | 29 | 26 | 38 | 37 | 28 | 24 | 31 | 28 | 38 | 42 | 28 | 31.7 | 28 | 36 | 1.295 | 0.373 | 0.28611518 | 0.45811825 | 0 | 0.0772 | 0.0562 | 0.0562 | TRUE | 0.2883 |
| Gm10790 | 0 | 0 | 0 | 0 | 0 | 0 | 0 | 0 | 0 | 0 | 0 | 0 | 0 | 0 | 0 | NA | NA | NA | NA | NA | NA | NA | NA | NA | NA |
| Gm10791 | 1 | 0 | 0 | 0 | 0 | 0 | 1 | 0 | 0 | 0 | 0 | 0 | 0.13 | 0 | 0 | 0.552 | -0.857 | 0.8336548 | NA | 0 | 18.3121 | 10 | 10 | TRUE | 0.2566 |
| Gm10804 | 1 | 0 | 0 | 0 | 0 | 0 | 1 | 0 | 0 | 0 | 0 | 0 | 0.13 | 0 | 0 | 0.552 | -0.857 | 0.8336548 | NA | 0 | 18.3121 | 10 | 10 | TRUE | 0.2566 |
| Gm10814 | 1 | 1 | 1 | 3 | 2 | 1 | 1 | 1 | 1 | 3 | 2 | 1 | 1.52 | 1 | 2 | 2.154 | 1.107 | 0.47101675 | NA | 0 | 1.5568 | 0.9502 | 0.9502 | TRUE | 0.2059 |
| Gm10818 | 1 | 0 | 1 | 0 | 1 | 2 | 1 | 0 | 1 | 0 | 1 | 2 | 0.83 | 1 | 1 | 1.655 | 0.727 | 0.73646206 | NA | 0 | 2.8706 | 2.089 | 2.089 | TRUE | 0.2033 |
| Gm1082 | 33 | 26 | 26 | 38 | 35 | 44 | 26 | 28 | 28 | 38 | 39 | 43 | 33.68 | 27 | 40 | 1.489 | 0.574 | 0.08015609 | 0.18942973 | 0 | 0.0728 | 0.0469 | 0.0469 | TRUE | 0.0373 |
| Gm10823 | 0 | 0 | 0 | 0 | 0 | 0 | 0 | 0 | 0 | 0 | 0 | 0 | 0 | 0 | 0 | NA | NA | NA | NA | NA | NA | NA | NA | NA | NA |
| Gm10825 | 9 | 5 | 5 | 6 | 3 | 5 | 7 | 5 | 5 | 6 | 3 | 5 | 5.33 | 6 | 5 | 0.806 | -0.311 | 0.70156673 | NA | 0 | 0.4469 | 0.2863 | 0.2863 | TRUE | 0.1087 |
| Gm10845 | 0 | 0 | 0 | 0 | 0 | 1 | 0 | 0 | 0 | 0 | 0 | 1 | 0.16 | 0 | 0 | 2.094 | 1.067 | 0.79379329 | NA | 0 | 14.4078 | 10 | 10 | TRUE | 0.3835 |
| Gm10857 | 4 | 1 | 0 | 3 | 1 | 1 | 3 | 1 | 0 | 3 | 1 | 1 | 1.55 | 1 | 2 | 1.173 | 0.23 | 0.88452285 | NA | 0 | 1.5362 | 1.1721 | 1.1721 | TRUE | 0.6566 |
| Gm10863 | 30 | 26 | 19 | 22 | 30 | 27 | 23 | 28 | 21 | 22 | 34 | 27 | 25.64 | 24 | 28 | 1.15 | 0.201 | 0.60182224 | 0.74681762 | 0 | 0.0949 | 0.068 | 0.068 | TRUE | 0.2428 |
| Gm10865 | 1 | 0 | 0 | 0 | 0 | 0 | 1 | 0 | 0 | 0 | 0 | 0 | 0.13 | 0 | 0 | 0.552 | -0.857 | 0.8336548 | NA | 0 | 18.3121 | 10 | 10 | TRUE | 0.2566 |
| Gm10872 | 1 | 0 | 0 | 0 | 0 | 0 | 1 | 0 | 0 | 0 | 0 | 0 | 0.13 | 0 | 0 | 0.552 | -0.857 | 0.8336548 | NA | 0 | 18.3121 | 10 | 10 | TRUE | 0.2566 |
| Gm10921 | 0 | 0 | 0 | 0 | 0 | 0 | 0 | 0 | 0 | 0 | 0 | 0 | 0 | 0 | 0 | NA | NA | NA | NA | NA | NA | NA | NA | NA | NA |
| Gm10922 | 0 | 0 | 0 | 0 | 0 | 0 | 0 | 0 | 0 | 0 | 0 | 0 | 0 | 0 | 0 | NA | NA | NA | NA | NA | NA | NA | NA | NA | NA |
| Gm10941 | 17 | 3 | 14 | 24 | 10 | 12 | 13 | 3 | 15 | 24 | 11 | 12 | 13.09 | 10 | 16 | 1.485 | 0.571 | 0.35588319 | NA | 0.2362 | 0.1835 | 0.1969 | 0.1969 | TRUE | 1.2233 |
| Gm1110 | 0 | 0 | 1 | 0 | 0 | 0 | 0 | 0 | 1 | 0 | 0 | 0 | 0.18 | 0 | 0 | 0.552 | -0.857 | 0.8336548 | NA | 0 | 13.1817 | 10 | 10 | TRUE | 0.5365 |
| Gm11110 | 26 | 15 | 20 | 15 | 32 | 36 | 20 | 16 | 22 | 15 | 36 | 36 | 24.05 | 19 | 29 | 1.493 | 0.578 | 0.18792718 | 0.34507234 | 0.0833 | 0.101 | 0.096 | 0.096 | TRUE | 1.1892 |
| Gm11127 | 3 | 0 | 2 | 2 | 2 | 1 | 2 | 2 | 0 | 2 | 2 | 1 | 1.6 | 1 | 2 | 1.118 | 0.16 | 0.91577865 | NA | 0 | 1.4866 | 1.0373 | 1.0373 | TRUE | 0.4725 |
| Gm11128 | 0 | 0 | 0 | 0 | 0 | 0 | 0 | 0 | 0 | 0 | 0 | 0 | 0 | 0 | 0 | NA | NA | NA | NA | NA | NA | NA | NA | NA | NA |
| Gm11149 | 0 | 0 | 0 | 0 | 0 | 0 | 0 | 0 | 0 | 0 | 0 | 0 | 0 | 0 | 0 | NA | NA | NA | NA | NA | NA | NA | NA | NA | NA |

| | | | | | | | | | | | | | | | | | | | | | | | | | |
|---|---|---|---|---|---|---|---|---|---|---|---|---|---|---|---|---|---|---|---|---|---|---|---|---|---|
| Gm11166 | 2 | 0 | 0 | 0 | 0 | 0 | 2 | 0 | 0 | 0 | 0 | 0 | 0.26 | 1 | 0 | 0.356 | -1.491 | 0.71362796 | NA | 0 | 9.1573 | 10 | 10 | TRUE | 0.968 |
| Gm11186 | 0 | 0 | 0 | 0 | 0 | 0 | 0 | 0 | 0 | 0 | 0 | 0 | 0 | 0 | 0 | NA | NA | NA | NA | NA | NA | NA | NA | NA | NA |
| Gm11190 | 17 | 23 | 10 | 15 | 13 | 17 | 13 | 24 | 11 | 15 | 15 | 17 | 15.81 | 16 | 16 | 0.963 | -0.054 | 0.91508385 | NA | 0.0359 | 0.1524 | 0.1199 | 0.1199 | TRUE | 0.9054 |
| Gm11194 | 2 | 5 | 1 | 2 | 0 | 0 | 2 | 5 | 1 | 2 | 0 | 0 | 1.66 | 3 | 1 | 0.258 | -1.954 | 0.2622894 | NA | 0.5666 | 1.4328 | 1.2427 | 1.2427 | TRUE | 0.9735 |
| Gm11201 | 1 | 1 | 1 | 0 | 0 | 0 | 1 | 1 | 1 | 0 | 0 | 0 | 0.49 | 1 | 0 | 0.196 | -2.347 | 0.38405676 | NA | 0 | 4.8735 | 3.7415 | 3.7415 | TRUE | 0.0718 |
| Gm11202 | 0 | 2 | 1 | 0 | 0 | 0 | 0 | 2 | 1 | 0 | 0 | 0 | 0.53 | 1 | 0 | 0.181 | -2.465 | 0.39476468 | NA | 0 | 4.4375 | 4.5834 | 4.5834 | TRUE | 0.0598 |
| Gm11213 | 0 | 0 | 0 | 0 | 0 | 0 | 0 | 0 | 0 | 0 | 0 | 0 | 0 | 0 | 0 | NA | NA | NA | NA | NA | NA | NA | NA | NA | NA |
| Gm1123 | 3 | 1 | 2 | 0 | 4 | 2 | 2 | 1 | 2 | 0 | 5 | 2 | 2.01 | 2 | 2 | 1.14 | 0.189 | 0.89392924 | NA | 0 | 1.1837 | 0.9416 | 0.9416 | TRUE | 0.2209 |
| Gm11240 | 0 | 0 | 0 | 0 | 0 | 0 | 0 | 0 | 0 | 0 | 0 | 0 | 0 | 0 | 0 | NA | NA | NA | NA | NA | NA | NA | NA | NA | NA |
| Gm11346 | 7 | 5 | 6 | 5 | 2 | 8 | 5 | 5 | 6 | 5 | 2 | 8 | 5.39 | 5 | 5 | 0.889 | -0.17 | 0.83745761 | NA | 0 | 0.4419 | 0.3104 | 0.3104 | TRUE | 0.2237 |
| Gm11351 | 0 | 0 | 0 | 0 | 0 | 0 | 0 | 0 | 0 | 0 | 0 | 0 | 0 | 0 | 0 | NA | NA | NA | NA | NA | NA | NA | NA | NA | NA |
| Gm1140 | 0 | 0 | 0 | 0 | 0 | 0 | 0 | 0 | 0 | 0 | 0 | 0 | 0 | 0 | 0 | NA | NA | NA | NA | NA | NA | NA | NA | NA | NA |
| Gm1141 | 53 | 37 | 16 | 11 | 8 | 6 | 41 | 39 | 17 | 11 | 9 | 6 | 20.61 | 32 | 9 | 0.263 | -1.927 | 0.00017778 | 0.0019501 | 0.1113 | 0.1174 | 0.1157 | 0.1157 | TRUE | 1.1018 |
| Gm11413 | 72 | 47 | 65 | 77 | 62 | 63 | 56 | 50 | 70 | 77 | 70 | 62 | 64.14 | 59 | 70 | 1.188 | 0.249 | 0.32140216 | 0.49524911 | 0 | 0.0394 | 0.0297 | 0.0297 | TRUE | 0.2312 |
| Gm11426 | 5 | 2 | 3 | 1 | 0 | 2 | 4 | 2 | 3 | 1 | 0 | 2 | 2.04 | 3 | 1 | 0.327 | -1.614 | 0.25858225 | NA | 0 | 1.1664 | 0.8051 | 0.8051 | TRUE | 0.3493 |
| Gm11435 | 2 | 2 | 0 | 1 | 0 | 0 | 2 | 2 | 0 | 1 | 0 | 0 | 0.78 | 1 | 0 | 0.3 | -1.737 | 0.45896026 | NA | 0 | 3.0435 | 2.5735 | 2.5735 | TRUE | 0.3906 |
| Gm11437 | 0 | 0 | 0 | 0 | 0 | 0 | 0 | 0 | 0 | 0 | 0 | 0 | 0 | 0 | 0 | NA | NA | NA | NA | NA | NA | NA | NA | NA | NA |
| Gm11468 | 0 | 0 | 0 | 0 | 0 | 0 | 0 | 0 | 0 | 0 | 0 | 0 | 0 | 0 | 0 | NA | NA | NA | NA | NA | NA | NA | NA | NA | NA |
| Gm11487 | 11 | 9 | 3 | 3 | 15 | 0 | 9 | 10 | 3 | 3 | 17 | 0 | 6.88 | 7 | 7 | 0.914 | -0.129 | 0.89667015 | NA | 1.3028 | 0.3471 | 0.5661 | 0.5661 | TRUE | 2.2117 |
| Gm11517 | 41 | 17 | 5 | 3 | 3 | 1 | 32 | 18 | 5 | 3 | 3 | 1 | 10.45 | 18 | 2 | 0.13 | -2.943 | 0.00052041 | NA | 0.4062 | 0.2293 | 0.2793 | 0.2793 | TRUE | 0.0675 |
| Gm11529 | 3 | 1 | 4 | 3 | 0 | 3 | 2 | 1 | 4 | 3 | 0 | 3 | 2.28 | 2 | 2 | 0.787 | -0.345 | 0.79523816 | NA | 0 | 1.0437 | 0.8261 | 0.8261 | TRUE | 0.4667 |
| Gm11541 | 1 | 3 | 1 | 1 | 0 | 1 | 1 | 3 | 1 | 1 | 0 | 1 | 1.17 | 2 | 1 | 0.408 | -1.292 | 0.48108248 | NA | 0 | 2.0254 | 1.3674 | 1.3674 | TRUE | 0.4459 |
| Gm11544 | 31 | 20 | 5 | 12 | 5 | 5 | 24 | 21 | 5 | 12 | 6 | 5 | 12.21 | 17 | 8 | 0.441 | -1.18 | 0.0768981 | NA | 0.2927 | 0.1965 | 0.2237 | 0.2237 | TRUE | 1.6252 |
| Gm11545 | 276 | 181 | 98 | 126 | 101 | 91 | 214 | 193 | 106 | 125 | 114 | 90 | 140.3 | 171 | 110 | 0.639 | -0.645 | 0.00789537 | 0.03472264 | 0.0736 | 0.0193 | 0.035 | 0.035 | TRUE | 1.1017 |
| Gm11548 | 0 | 0 | 0 | 0 | 0 | 0 | 0 | 0 | 0 | 0 | 0 | 0 | 0 | 0 | 0 | NA | NA | NA | NA | NA | NA | NA | NA | NA | NA |
| Gm11549 | 3 | 3 | 0 | 2 | 2 | 5 | 2 | 3 | 0 | 2 | 2 | 5 | 2.45 | 2 | 3 | 1.648 | 0.721 | 0.57263325 | NA | 0 | 0.9695 | 0.749 | 0.749 | TRUE | 0.5523 |
| Gm11554 | 0 | 0 | 0 | 0 | 0 | 0 | 0 | 0 | 0 | 0 | 0 | 0 | 0 | 0 | 0 | NA | NA | NA | NA | NA | NA | NA | NA | NA | NA |
| Gm11559 | 0 | 0 | 0 | 0 | 0 | 0 | 0 | 0 | 0 | 0 | 0 | 0 | 0 | 0 | 0 | NA | NA | NA | NA | NA | NA | NA | NA | NA | NA |
| Gm11562 | 0 | 0 | 0 | 0 | 0 | 0 | 0 | 0 | 0 | 0 | 0 | 0 | 0 | 0 | 0 | NA | NA | NA | NA | NA | NA | NA | NA | NA | NA |
| Gm11563 | 0 | 0 | 0 | 0 | 0 | 0 | 0 | 0 | 0 | 0 | 0 | 0 | 0 | 0 | 0 | NA | NA | NA | NA | NA | NA | NA | NA | NA | NA |
| Gm11564 | 6 | 5 | 2 | 0 | 4 | 1 | 5 | 5 | 2 | 0 | 5 | 1 | 2.94 | 4 | 2 | 0.438 | -1.19 | 0.33810192 | NA | 0.2574 | 0.8088 | 0.7021 | 0.7021 | TRUE | 1.2101 |
| Gm11565 | 0 | 0 | 0 | 0 | 0 | 0 | 0 | 0 | 0 | 0 | 0 | 0 | 0 | 0 | 0 | NA | NA | NA | NA | NA | NA | NA | NA | NA | NA |
| Gm11567 | 0 | 0 | 0 | 0 | 0 | 0 | 0 | 0 | 0 | 0 | 0 | 0 | 0 | 0 | 0 | NA | NA | NA | NA | NA | NA | NA | NA | NA | NA |
| Gm11568 | 0 | 0 | 0 | 0 | 0 | 0 | 0 | 0 | 0 | 0 | 0 | 0 | 0 | 0 | 0 | NA | NA | NA | NA | NA | NA | NA | NA | NA | NA |
| Gm11569 | 0 | 0 | 0 | 0 | 0 | 0 | 0 | 0 | 0 | 0 | 0 | 0 | 0 | 0 | 0 | NA | NA | NA | NA | NA | NA | NA | NA | NA | NA |
| Gm11570 | 0 | 0 | 0 | 0 | 1 | 0 | 0 | 0 | 0 | 0 | 1 | 0 | 0.19 | 0 | 0 | 2.094 | 1.067 | 0.79379329 | NA | 0 | 12.608 | 10 | 10 | TRUE | 0.5075 |
| Gm11595 | 0 | 0 | 0 | 0 | 0 | 0 | 0 | 0 | 0 | 0 | 0 | 0 | 0 | 0 | 0 | NA | NA | NA | NA | NA | NA | NA | NA | NA | NA |
| Gm11596 | 0 | 0 | 0 | 0 | 0 | 0 | 0 | 0 | 0 | 0 | 0 | 0 | 0 | 0 | 0 | NA | NA | NA | NA | NA | NA | NA | NA | NA | NA |
| Gm11602 | 39 | 13 | 9 | 22 | 17 | 12 | 30 | 14 | 10 | 22 | 19 | 12 | 17.79 | 18 | 18 | 0.963 | -0.054 | 0.91881044 | 0.95538599 | 0.1695 | 0.1356 | 0.1457 | 0.1457 | TRUE | 1.8883 |
| Gm11627 | 87 | 68 | 41 | 54 | 53 | 76 | 68 | 72 | 44 | 54 | 60 | 75 | 62.12 | 61 | 63 | 1.021 | 0.03 | 0.91118327 | 0.95117897 | 0.0297 | 0.0406 | 0.0376 | 0.0376 | TRUE | 0.4843 |
| Gm11648 | 8 | 3 | 2 | 5 | 3 | 2 | 6 | 3 | 2 | 5 | 3 | 2 | 3.65 | 4 | 3 | 0.874 | -0.194 | 0.84711763 | NA | 0 | 0.6518 | 0.4593 | 0.4593 | TRUE | 0.6078 |
| Gm11696 | 43 | 33 | 39 | 42 | 35 | 44 | 33 | 35 | 42 | 42 | 39 | 43 | 39.22 | 37 | 41 | 1.133 | 0.181 | 0.55400553 | 0.70972493 | 0 | 0.0629 | 0.0416 | 0.0416 | TRUE | 0.1059 |
| Gm11710 | 0 | 0 | 0 | 0 | 0 | 0 | 0 | 0 | 0 | 0 | 0 | 0 | 0 | 0 | 0 | NA | NA | NA | NA | NA | NA | NA | NA | NA | NA |
| Gm11711 | 0 | 0 | 0 | 0 | 0 | 0 | 0 | 0 | 0 | 0 | 0 | 0 | 0 | 0 | 0 | NA | NA | NA | NA | NA | NA | NA | NA | NA | NA |
| Gm11744 | 49 | 27 | 33 | 41 | 23 | 36 | 38 | 29 | 36 | 41 | 26 | 36 | 34.11 | 34 | 34 | 0.999 | -0.002 | 0.9959794 | 0.99753911 | 0 | 0.0719 | 0.0539 | 0.0539 | TRUE | 0.2824 |
| Gm11747 | 0 | 0 | 0 | 0 | 0 | 0 | 0 | 0 | 0 | 0 | 0 | 0 | 0 | 0 | 0 | NA | NA | NA | NA | NA | NA | NA | NA | NA | NA |
| Gm11756 | 4 | 5 | 1 | 1 | 1 | 0 | 3 | 5 | 1 | 1 | 1 | 0 | 1.94 | 3 | 1 | 0.22 | -2.187 | 0.16703121 | NA | 0 | 1.2254 | 0.9097 | 0.9097 | TRUE | 0.1267 |
| Gm11757 | 0 | 0 | 0 | 0 | 0 | 0 | 0 | 0 | 0 | 0 | 0 | 0 | 0 | 0 | 0 | NA | NA | NA | NA | NA | NA | NA | NA | NA | NA |
| Gm11758 | 0 | 0 | 0 | 0 | 0 | 0 | 0 | 0 | 0 | 0 | 0 | 0 | 0 | 0 | 0 | NA | NA | NA | NA | NA | NA | NA | NA | NA | NA |
| Gm11762 | 1 | 0 | 0 | 0 | 0 | 0 | 1 | 0 | 0 | 0 | 0 | 0 | 0.13 | 0 | 0 | 0.552 | -0.857 | 0.8336548 | NA | 0 | 18.3121 | 10 | 10 | TRUE | 0.2566 |
| Gm11837 | 399 | 267 | 291 | 317 | 285 | 290 | 310 | 284 | 314 | 316 | 322 | 286 | 305.2 | 303 | 308 | 1.016 | 0.022 | 0.85249176 | 0.9167369 | 0 | 0.0102 | 0.0072 | 0.0072 | TRUE | 0.0429 |
| Gm11937 | 0 | 0 | 0 | 0 | 0 | 0 | 0 | 0 | 0 | 0 | 0 | 0 | 0 | 0 | 0 | NA | NA | NA | NA | NA | NA | NA | NA | NA | NA |
| Gm11938 | 0 | 0 | 0 | 0 | 0 | 0 | 0 | 0 | 0 | 0 | 0 | 0 | 0 | 0 | 0 | NA | NA | NA | NA | NA | NA | NA | NA | NA | NA |
| Gm11944 | 15 | 10 | 11 | 11 | 7 | 8 | 12 | 11 | 12 | 11 | 8 | 8 | 10.15 | 12 | 9 | 0.784 | -0.351 | 0.5524469 | NA | 0 | 0.2359 | 0.1516 | 0.1516 | TRUE | 0.1307 |
| Gm11961 | 22 | 14 | 18 | 20 | 10 | 18 | 17 | 15 | 19 | 20 | 11 | 18 | 16.73 | 17 | 16 | 0.958 | -0.062 | 0.89494802 | NA | 0 | 0.1441 | 0.1016 | 0.1016 | TRUE | 0.3143 |
| Gm11974 | 1683 | 1131 | 1000 | 1286 | 1022 | 1200 | 1307 | 1203 | 1079 | 1280 | 1153 | 1185 | 1201.12 | 1196 | 1206 | 1.007 | 0.01 | 0.91170406 | 0.9513752 | 0.0053 | 0.0044 | 0.0047 | 0.0047 | TRUE | 0.0877 |
| Gm11978 | 2 | 1 | 0 | 0 | 1 | 0 | 2 | 1 | 0 | 0 | 1 | 0 | 0.62 | 1 | 0 | 0.416 | -1.266 | 0.61771935 | NA | 0 | 3.7995 | 3.0914 | 3.0914 | TRUE | 0.5075 |
| Gm11981 | 0 | 0 | 0 | 1 | 1 | 0 | 0 | 0 | 0 | 1 | 1 | 0 | 0.35 | 0 | 1 | 3.978 | 1.992 | 0.55384016 | NA | 0 | 6.6991 | 6.4234 | 6.4234 | TRUE | 0.2647 |
| Gm11985 | 5 | 7 | 2 | 2 | 7 | 3 | 4 | 7 | 2 | 2 | 8 | 3 | 4.39 | 4 | 4 | 0.944 | -0.083 | 0.93113872 | NA | 0.1886 | 0.5424 | 0.4399 | 0.4399 | TRUE | 0.7494 |
| Gm11992 | 115 | 78 | 91 | 105 | 78 | 110 | 89 | 83 | 98 | 105 | 88 | 109 | 95.26 | 90 | 101 | 1.116 | 0.158 | 0.43783558 | 0.60928673 | 0 | 0.0273 | 0.0194 | 0.0194 | TRUE | 0.1076 |
| Gm12 | 0 | 0 | 0 | 0 | 0 | 1 | 0 | 0 | 0 | 0 | 0 | 1 | 0.16 | 0 | 0 | 2.094 | 1.067 | 0.79379329 | NA | 0 | 14.4078 | 10 | 10 | TRUE | 0.3835 |
| Gm12060 | 1 | 1 | 0 | 0 | 0 | 0 | 1 | 1 | 0 | 0 | 0 | 0 | 0.31 | 1 | 0 | 0.309 | -1.696 | 0.63183309 | NA | 0 | 7.7294 | 7.2472 | 7.2472 | TRUE | 0.2015 |
| Gm12070 | 8 | 7 | 8 | 14 | 17 | 12 | 6 | 7 | 9 | 14 | 19 | 12 | 11.21 | 7 | 15 | 2.026 | 1.019 | 0.08137949 | NA | 0 | 0.2139 | 0.1462 | 0.1462 | TRUE | 0.2488 |
| Gm12130 | 0 | 0 | 0 | 0 | 0 | 0 | 0 | 0 | 0 | 0 | 0 | 0 | 0 | 0 | 0 | NA | NA | NA | NA | NA | NA | NA | NA | NA | NA |
| Gm12159 | 0 | 0 | 0 | 0 | 0 | 0 | 0 | 0 | 0 | 0 | 0 | 0 | 0 | 0 | 0 | NA | NA | NA | NA | NA | NA | NA | NA | NA | NA |
| Gm12169 | 1 | 0 | 0 | 0 | 1 | 0 | 1 | 0 | 0 | 0 | 1 | 0 | 0.32 | 0 | 0 | 1.075 | 0.105 | 0.97638847 | NA | 0 | 7.4681 | 7.0392 | 7.0392 | TRUE | 0.5075 |
| Gm12171 | 0 | 0 | 0 | 0 | 0 | 0 | 0 | 0 | 0 | 0 | 0 | 0 | 0 | 0 | 0 | NA | NA | NA | NA | NA | NA | NA | NA | NA | NA |
| Gm12185 | 0 | 0 | 0 | 0 | 0 | 0 | 0 | 0 | 0 | 0 | 0 | 0 | 0 | 0 | 0 | NA | NA | NA | NA | NA | NA | NA | NA | NA | NA |
| Gm12191 | 0 | 0 | 0 | 0 | 0 | 0 | 0 | 0 | 0 | 0 | 0 | 0 | 0 | 0 | 0 | NA | NA | NA | NA | NA | NA | NA | NA | NA | NA |

| | | | | | | | | | | | | | | | | | | | | | | | | | |
|---|---|---|---|---|---|---|---|---|---|---|---|---|---|---|---|---|---|---|---|---|---|---|---|---|---|
| Gm12216 | 1 | 0 | 0 | 1 | 0 | 3 | 1 | 0 | 0 | 1 | 0 | 3 | 0.79 | 0 | 1 | 3.865 | 1.95 | 0.40970091 | NA | 0 | 3.0066 | 2.6451 | 2.6451 | TRUE | 0.2104 |
| Gm12228 | 0 | 1 | 0 | 0 | 1 | 2 | 0 | 1 | 0 | 0 | 1 | 2 | 0.69 | 0 | 1 | 2.998 | 1.584 | 0.52044796 | NA | 0 | 3.4158 | 2.8778 | 2.8778 | TRUE | 0.3487 |
| Gm12238 | 5 | 3 | 7 | 3 | 9 | 0 | 4 | 3 | 8 | 3 | 10 | 0 | 4.63 | 5 | 4 | 0.888 | -0.171 | 0.87154933 | NA | 0.8247 | 0.5146 | 0.5846 | 0.5846 | TRUE | 0.7343 |
| Gm12250 | 65 | 43 | 34 | 58 | 52 | 78 | 50 | 46 | 37 | 58 | 59 | 77 | 54.38 | 44 | 65 | 1.449 | 0.535 | 0.05205447 | 0.13880868 | 0 | 0.046 | 0.0358 | 0.0358 | TRUE | 0.2597 |
| Gm12253 | 0 | 0 | 0 | 0 | 0 | 0 | 0 | 0 | 0 | 0 | 0 | 0 | 0 | 0 | 0 | NA | NA | NA | NA | NA | NA | NA | NA | NA | NA |
| Gm12295 | 0 | 0 | 0 | 0 | 0 | 0 | 0 | 0 | 0 | 0 | 0 | 0 | 0 | 0 | 0 | NA | NA | NA | NA | NA | NA | NA | NA | NA | NA |
| Gm12298 | 1 | 0 | 1 | 0 | 0 | 0 | 1 | 0 | 1 | 0 | 0 | 0 | 0.31 | 1 | 0 | 0.307 | -1.706 | 0.62894649 | NA | 0 | 7.6657 | 7.2 | 7.2 | TRUE | 0.2067 |
| Gm12359 | 14 | 13 | 3 | 5 | 8 | 6 | 11 | 14 | 3 | 5 | 9 | 6 | 7.98 | 9 | 7 | 0.705 | -0.505 | 0.49026421 | NA | 0.1354 | 0.2995 | 0.2554 | 0.2554 | TRUE | 0.5582 |
| Gm12360 | 0 | 1 | 0 | 0 | 0 | 0 | 0 | 1 | 0 | 0 | 0 | 0 | 0.18 | 0 | 0 | 0.552 | -0.857 | 0.8336548 | NA | 0 | 13.3714 | 10 | 10 | TRUE | 0.5216 |
| Gm12409 | 2 | 1 | 0 | 0 | 0 | 0 | 2 | 1 | 0 | 0 | 0 | 0 | 0.44 | 1 | 0 | 0.215 | -2.217 | 0.47268385 | NA | 0 | 5.4363 | 5.3181 | 5.3181 | TRUE | 0.2411 |
| Gm12504 | 11 | 11 | 7 | 12 | 11 | 13 | 9 | 12 | 8 | 12 | 12 | 13 | 10.83 | 10 | 12 | 1.342 | 0.424 | 0.46007465 | NA | 0 | 0.2213 | 0.1438 | 0.1438 | TRUE | 0.1581 |
| Gm12505 | 1 | 2 | 2 | 0 | 0 | 1 | 1 | 2 | 2 | 0 | 0 | 1 | 1.01 | 2 | 0 | 0.223 | -2.165 | 0.286395 | NA | 0 | 2.3532 | 1.6816 | 1.6816 | TRUE | 0.3835 |
| Gm12522 | 29 | 25 | 34 | 55 | 48 | 37 | 23 | 27 | 37 | 55 | 54 | 37 | 38.54 | 29 | 49 | 1.707 | 0.772 | 0.02232694 | 0.07470012 | 0.0296 | 0.0639 | 0.0545 | 0.0545 | TRUE | 0.3838 |
| Gm12530 | 0 | 0 | 0 | 0 | 0 | 0 | 0 | 0 | 0 | 0 | 0 | 0 | 0 | 0 | 0 | NA | NA | NA | NA | NA | NA | NA | NA | NA | NA |
| Gm12597 | 0 | 0 | 0 | 0 | 0 | 0 | 0 | 0 | 0 | 0 | 0 | 0 | 0 | 0 | 0 | NA | NA | NA | NA | NA | NA | NA | NA | NA | NA |
| Gm12603 | 0 | 0 | 0 | 0 | 0 | 0 | 0 | 0 | 0 | 0 | 0 | 0 | 0 | 0 | 0 | NA | NA | NA | NA | NA | NA | NA | NA | NA | NA |
| Gm12633 | 0 | 0 | 0 | 0 | 0 | 0 | 0 | 0 | 0 | 0 | 0 | 0 | 0 | 0 | 0 | NA | NA | NA | NA | NA | NA | NA | NA | NA | NA |
| Gm12657 | 8 | 1 | 8 | 5 | 10 | 4 | 6 | 1 | 9 | 5 | 11 | 4 | 6.02 | 5 | 7 | 1.253 | 0.326 | 0.70680687 | NA | 0.3003 | 0.3962 | 0.3716 | 0.3716 | TRUE | 0.6208 |
| Gm12669 | 0 | 0 | 0 | 0 | 0 | 0 | 0 | 0 | 0 | 0 | 0 | 0 | 0 | 0 | 0 | NA | NA | NA | NA | NA | NA | NA | NA | NA | NA |
| Gm12695 | 0 | 0 | 1 | 0 | 0 | 0 | 0 | 0 | 1 | 0 | 0 | 0 | 0.18 | 0 | 0 | 0.552 | -0.857 | 0.8336548 | NA | 0 | 13.1817 | 10 | 10 | TRUE | 0.5365 |
| Gm12709 | 0 | 0 | 0 | 0 | 0 | 0 | 0 | 0 | 0 | 0 | 0 | 0 | 0 | 0 | 0 | NA | NA | NA | NA | NA | NA | NA | NA | NA | NA |
| Gm12718 | 3 | 0 | 1 | 1 | 1 | 0 | 2 | 0 | 1 | 1 | 1 | 0 | 0.92 | 1 | 1 | 0.596 | -0.746 | 0.7193098 | NA | 0 | 2.5728 | 1.9492 | 1.9492 | TRUE | 0.1743 |
| Gm12789 | 1 | 1 | 0 | 0 | 0 | 0 | 1 | 1 | 0 | 0 | 0 | 0 | 0.31 | 1 | 0 | 0.309 | -1.696 | 0.63183309 | NA | 0 | 7.7294 | 7.2472 | 7.2472 | TRUE | 0.2015 |
| Gm12794 | 384 | 272 | 114 | 124 | 139 | 88 | 298 | 289 | 123 | 123 | 157 | 87 | 179.61 | 237 | 122 | 0.516 | -0.955 | 0.03676625 | 0.10783016 | 0.1444 | 0.0156 | 0.0504 | 0.1444 | TRUE | 1.2425 |
| Gm128 | 107 | 101 | 95 | 137 | 98 | 98 | 83 | 107 | 103 | 136 | 111 | 97 | 106.12 | 98 | 115 | 1.179 | 0.237 | 0.25805525 | 0.42808539 | 0.0161 | 0.0248 | 0.0222 | 0.0222 | TRUE | 0.2847 |
| Gm12830 | 1 | 0 | 0 | 0 | 0 | 0 | 1 | 0 | 0 | 0 | 0 | 0 | 0.13 | 0 | 0 | 0.552 | -0.857 | 0.8336548 | NA | 0 | 18.3121 | 10 | 10 | TRUE | 0.2566 |
| Gm12839 | 0 | 0 | 0 | 0 | 0 | 0 | 0 | 0 | 0 | 0 | 0 | 0 | 0 | 0 | 0 | NA | NA | NA | NA | NA | NA | NA | NA | NA | NA |
| Gm12886 | 0 | 0 | 0 | 0 | 0 | 0 | 0 | 0 | 0 | 0 | 0 | 0 | 0 | 0 | 0 | NA | NA | NA | NA | NA | NA | NA | NA | NA | NA |
| Gm12887 | 0 | 0 | 0 | 0 | 0 | 0 | 0 | 0 | 0 | 0 | 0 | 0 | 0 | 0 | 0 | NA | NA | NA | NA | NA | NA | NA | NA | NA | NA |
| Gm12888 | 0 | 0 | 0 | 0 | 0 | 0 | 0 | 0 | 0 | 0 | 0 | 0 | 0 | 0 | 0 | NA | NA | NA | NA | NA | NA | NA | NA | NA | NA |
| Gm129 | 151 | 151 | 138 | 143 | 115 | 139 | 117 | 161 | 149 | 142 | 130 | 137 | 139.35 | 142 | 136 | 0.965 | -0.052 | 0.77431519 | 0.86673091 | 0.0086 | 0.0194 | 0.0162 | 0.0162 | TRUE | 0.2733 |
| Gm12942 | 119 | 101 | 188 | 230 | 195 | 270 | 92 | 107 | 203 | 229 | 220 | 267 | 186.36 | 134 | 239 | 1.788 | 0.838 | 0.00038937 | 0.0035675 | 0.0869 | 0.0151 | 0.0344 | 0.0344 | TRUE | 2.0298 |
| Gm13003 | 0 | 2 | 0 | 2 | 0 | 0 | 0 | 2 | 0 | 2 | 0 | 0 | 0.69 | 1 | 1 | 0.971 | -0.042 | 0.98762268 | NA | 8.094 | 3.4555 | 3.8966 | 3.8966 | TRUE | 0.9747 |
| Gm13011 | 0 | 0 | 0 | 0 | 0 | 0 | 0 | 0 | 0 | 0 | 0 | 0 | 0 | 0 | 0 | NA | NA | NA | NA | NA | NA | NA | NA | NA | NA |
| Gm13023 | 0 | 0 | 0 | 0 | 0 | 1 | 0 | 0 | 0 | 0 | 0 | 1 | 0.16 | 0 | 0 | 2.094 | 1.067 | 0.79379329 | NA | 0 | 14.4078 | 10 | 10 | TRUE | 0.3835 |
| Gm13031 | 4 | 2 | 4 | 1 | 3 | 4 | 3 | 2 | 4 | 1 | 3 | 4 | 2.98 | 3 | 3 | 0.871 | -0.2 | 0.85637666 | NA | 0 | 0.7978 | 0.5412 | 0.5412 | TRUE | 0.397 |
| Gm13032 | 11 | 13 | 19 | 11 | 17 | 13 | 9 | 14 | 21 | 11 | 19 | 13 | 14.31 | 15 | 14 | 1.011 | 0.016 | 0.97570646 | NA | 0.0712 | 0.1681 | 0.1403 | 0.1403 | TRUE | 0.2532 |
| Gm13034 | 57 | 43 | 25 | 20 | 35 | 26 | 44 | 46 | 27 | 20 | 39 | 26 | 33.67 | 39 | 28 | 0.72 | -0.474 | 0.20660131 | 0.36835958 | 0.0649 | 0.0728 | 0.0707 | 0.0707 | TRUE | 0.7087 |
| Gm13037 | 0 | 0 | 0 | 0 | 0 | 0 | 0 | 0 | 0 | 0 | 0 | 0 | 0 | 0 | 0 | NA | NA | NA | NA | NA | NA | NA | NA | NA | NA |
| Gm13040 | 0 | 0 | 0 | 0 | 0 | 0 | 0 | 0 | 0 | 0 | 0 | 0 | 0 | 0 | 0 | NA | NA | NA | NA | NA | NA | NA | NA | NA | NA |
| Gm13043 | 0 | 0 | 0 | 0 | 0 | 0 | 0 | 0 | 0 | 0 | 0 | 0 | 0 | 0 | 0 | NA | NA | NA | NA | NA | NA | NA | NA | NA | NA |
| Gm13051 | 534 | 397 | 329 | 333 | 294 | 370 | 415 | 422 | 355 | 331 | 332 | 365 | 370.06 | 397 | 343 | 0.862 | -0.214 | 0.06724417 | 0.16639905 | 0.0033 | 0.0088 | 0.0071 | 0.0071 | TRUE | 0.0962 |
| Gm13057 | 0 | 0 | 0 | 0 | 0 | 0 | 0 | 0 | 0 | 0 | 0 | 0 | 0 | 0 | 0 | NA | NA | NA | NA | NA | NA | NA | NA | NA | NA |
| Gm13078 | 48 | 43 | 19 | 46 | 32 | 18 | 37 | 46 | 21 | 46 | 36 | 18 | 33.86 | 35 | 33 | 0.959 | -0.06 | 0.88467217 | 0.93608375 | 0.1496 | 0.0724 | 0.0937 | 0.0937 | TRUE | 0.4953 |
| Gm13083 | 18 | 10 | 7 | 11 | 4 | 9 | 14 | 11 | 8 | 11 | 5 | 9 | 9.42 | 11 | 8 | 0.755 | -0.406 | 0.52797824 | NA | 0 | 0.254 | 0.1905 | 0.1905 | TRUE | 0.2805 |
| Gm13084 | 0 | 0 | 0 | 0 | 0 | 0 | 0 | 0 | 0 | 0 | 0 | 0 | 0 | 0 | 0 | NA | NA | NA | NA | NA | NA | NA | NA | NA | NA |
| Gm13088 | 0 | 0 | 0 | 0 | 0 | 0 | 0 | 0 | 0 | 0 | 0 | 0 | 0 | 0 | 0 | NA | NA | NA | NA | NA | NA | NA | NA | NA | NA |
| Gm13102 | 0 | 0 | 0 | 0 | 0 | 0 | 0 | 0 | 0 | 0 | 0 | 0 | 0 | 0 | 0 | NA | NA | NA | NA | NA | NA | NA | NA | NA | NA |
| Gm13103 | 0 | 0 | 0 | 0 | 0 | 0 | 0 | 0 | 0 | 0 | 0 | 0 | 0 | 0 | 0 | NA | NA | NA | NA | NA | NA | NA | NA | NA | NA |
| Gm13109 | 155 | 99 | 45 | 74 | 73 | 63 | 120 | 105 | 49 | 74 | 82 | 62 | 82.07 | 91 | 73 | 0.79 | -0.34 | 0.25249454 | 0.42140407 | 0.102 | 0.0313 | 0.0512 | 0.0512 | TRUE | 1.1758 |
| Gm13119 | 336 | 216 | 126 | 158 | 180 | 115 | 261 | 230 | 136 | 157 | 203 | 114 | 183.42 | 209 | 158 | 0.752 | -0.41 | 0.08754023 | 0.20083668 | 0.0895 | 0.0153 | 0.036 | 0.036 | TRUE | 0.3895 |
| Gm13124 | 4 | 5 | 0 | 1 | 1 | 0 | 3 | 5 | 0 | 1 | 1 | 0 | 1.76 | 3 | 1 | 0.247 | -2.015 | 0.24260422 | NA | 0 | 1.3505 | 1.2292 | 1.2292 | TRUE | 0.1174 |
| Gm13125 | 2 | 0 | 1 | 0 | 0 | 0 | 2 | 0 | 1 | 0 | 0 | 0 | 0.44 | 1 | 0 | 0.214 | -2.223 | 0.47051571 | NA | 0 | 5.4047 | 5.2946 | 5.2946 | TRUE | 0.2788 |
| Gm13128 | 28 | 18 | 13 | 11 | 8 | 7 | 22 | 19 | 14 | 11 | 9 | 7 | 13.63 | 18 | 9 | 0.486 | -1.04 | 0.0526292 | NA | 0 | 0.1763 | 0.1235 | 0.1235 | TRUE | 0.2022 |
| Gm13139 | 41 | 35 | 23 | 31 | 30 | 36 | 32 | 37 | 25 | 31 | 34 | 36 | 32.35 | 31 | 34 | 1.066 | 0.093 | 0.78557775 | 0.87390788 | 0 | 0.0757 | 0.0528 | 0.0528 | TRUE | 0.2002 |
| Gm13152 | 897 | 667 | 625 | 698 | 568 | 686 | 697 | 709 | 674 | 695 | 641 | 677 | 682.21 | 693 | 671 | 0.968 | -0.047 | 0.58229865 | 0.73253195 | 0 | 0.0059 | 0.0038 | 0.0038 | TRUE | 0.0177 |
| Gm13154 | 3155 | 2287 | 2206 | 2375 | 2007 | 2424 | 2450 | 2433 | 2380 | 2364 | 2264 | 2393 | 2380.72 | 2421 | 2340 | 0.967 | -0.049 | 0.40722244 | 0.58012135 | 0 | 0.0034 | 0.0021 | 0.0021 | TRUE | 0.0097 |
| Gm13157 | 2719 | 2060 | 2037 | 2408 | 2047 | 2501 | 2112 | 2191 | 2198 | 2397 | 2309 | 2469 | 2279.31 | 2167 | 2392 | 1.104 | 0.143 | 0.01866276 | 0.06569256 | 4.00E-04 | 0.0035 | 0.0022 | 0.0022 | TRUE | 0.0109 |
| Gm13177 | 0 | 0 | 0 | 0 | 0 | 0 | 0 | 0 | 0 | 0 | 0 | 0 | 0 | 0 | 0 | NA | NA | NA | NA | NA | NA | NA | NA | NA | NA |
| Gm13178 | 0 | 0 | 0 | 2 | 0 | 0 | 0 | 0 | 0 | 2 | 0 | 0 | 0.33 | 0 | 1 | 3.77 | 1.915 | 0.62344813 | NA | 0 | 7.1455 | 9.1834 | 9.1834 | TRUE | 0.9739 |
| Gm13212 | 491 | 354 | 297 | 358 | 369 | 384 | 381 | 377 | 320 | 356 | 416 | 379 | 371.67 | 359 | 384 | 1.065 | 0.091 | 0.44566849 | 0.61646566 | 0.0047 | 0.0088 | 0.0076 | 0.0076 | TRUE | 0.0996 |
| Gm1322 | 12 | 21 | 14 | 5 | 1 | 8 | 9 | 22 | 15 | 5 | 1 | 8 | 10.13 | 15 | 5 | 0.307 | -1.703 | 0.01671167 | NA | 0.1839 | 0.2364 | 0.2247 | 0.2247 | TRUE | 0.2485 |
| Gm13238 | 4 | 4 | 3 | 3 | 2 | 2 | 3 | 4 | 3 | 3 | 2 | 2 | 2.97 | 3 | 2 | 0.685 | -0.547 | 0.61290796 | NA | 0 | 0.8006 | 0.4896 | 0.4896 | TRUE | 0.0495 |
| Gm13242 | 2514 | 1884 | 1725 | 2182 | 1794 | 2381 | 1953 | 2004 | 1861 | 2172 | 2024 | 2350 | 2060.66 | 1939 | 2182 | 1.126 | 0.171 | 0.01956325 | 0.06798384 | 0.0028 | 0.0036 | 0.0034 | 0.0034 | TRUE | 0.0552 |
| Gm13247 | 4247 | 3149 | 2803 | 2975 | 2536 | 2865 | 3298 | 3350 | 3024 | 2961 | 2861 | 2828 | 3053.8 | 3224 | 2883 | 0.894 | -0.161 | 0.01003293 | 0.04145727 | 0.0013 | 0.0032 | 0.0025 | 0.0025 | TRUE | 0.0351 |
| Gm13251 | 125 | 75 | 42 | 33 | 45 | 49 | 97 | 80 | 45 | 33 | 51 | 48 | 59.03 | 74 | 44 | 0.589 | -0.763 | 0.01606986 | 0.05875839 | 0.0828 | 0.0426 | 0.0542 | 0.0542 | TRUE | 0.3193 |
| Gm13271 | 0 | 0 | 0 | 0 | 0 | 0 | 0 | 0 | 0 | 0 | 0 | 0 | 0 | 0 | 0 | NA | NA | NA | NA | NA | NA | NA | NA | NA | NA |
| Gm13272 | 0 | 0 | 0 | 0 | 0 | 0 | 0 | 0 | 0 | 0 | 0 | 0 | 0 | 0 | 0 | NA | NA | NA | NA | NA | NA | NA | NA | NA | NA |
| Gm13275 | 0 | 0 | 0 | 0 | 0 | 0 | 0 | 0 | 0 | 0 | 0 | 0 | 0 | 0 | 0 | NA | NA | NA | NA | NA | NA | NA | NA | NA | NA |

| | | | | | | | | | | | | | | | | | | | | | | | | | |
|---|---|---|---|---|---|---|---|---|---|---|---|---|---|---|---|---|---|---|---|---|---|---|---|---|---|
| Gm13276 | 0 | 0 | 0 | 0 | 0 | 0 | 0 | 0 | 0 | 0 | 0 | 0 | 0 | 0 | 0 | NA | NA | NA | NA | NA | NA | NA | NA | NA | NA |
| Gm13277 | 0 | 0 | 0 | 0 | 0 | 0 | 0 | 0 | 0 | 0 | 0 | 0 | 0 | 0 | 0 | NA | NA | NA | NA | NA | NA | NA | NA | NA | NA |
| Gm13278 | 0 | 0 | 0 | 0 | 0 | 0 | 0 | 0 | 0 | 0 | 0 | 0 | 0 | 0 | 0 | NA | NA | NA | NA | NA | NA | NA | NA | NA | NA |
| Gm13279 | 0 | 0 | 0 | 0 | 0 | 0 | 0 | 0 | 0 | 0 | 0 | 0 | 0 | 0 | 0 | NA | NA | NA | NA | NA | NA | NA | NA | NA | NA |
| Gm13280 | 0 | 0 | 0 | 0 | 0 | 0 | 0 | 0 | 0 | 0 | 0 | 0 | 0 | 0 | 0 | NA | NA | NA | NA | NA | NA | NA | NA | NA | NA |
| Gm13283 | 0 | 0 | 0 | 0 | 0 | 0 | 0 | 0 | 0 | 0 | 0 | 0 | 0 | 0 | 0 | NA | NA | NA | NA | NA | NA | NA | NA | NA | NA |
| Gm13285 | 0 | 0 | 0 | 0 | 0 | 0 | 0 | 0 | 0 | 0 | 0 | 0 | 0 | 0 | 0 | NA | NA | NA | NA | NA | NA | NA | NA | NA | NA |
| Gm13286 | 0 | 0 | 0 | 0 | 0 | 0 | 0 | 0 | 0 | 0 | 0 | 0 | 0 | 0 | 0 | NA | NA | NA | NA | NA | NA | NA | NA | NA | NA |
| Gm13287 | 0 | 0 | 0 | 0 | 0 | 0 | 0 | 0 | 0 | 0 | 0 | 0 | 0 | 0 | 0 | NA | NA | NA | NA | NA | NA | NA | NA | NA | NA |
| Gm13288 | 0 | 0 | 0 | 0 | 0 | 0 | 0 | 0 | 0 | 0 | 0 | 0 | 0 | 0 | 0 | NA | NA | NA | NA | NA | NA | NA | NA | NA | NA |
| Gm13289 | 0 | 0 | 0 | 0 | 0 | 0 | 0 | 0 | 0 | 0 | 0 | 0 | 0 | 0 | 0 | NA | NA | NA | NA | NA | NA | NA | NA | NA | NA |
| Gm13290 | 0 | 0 | 0 | 0 | 0 | 0 | 0 | 0 | 0 | 0 | 0 | 0 | 0 | 0 | 0 | NA | NA | NA | NA | NA | NA | NA | NA | NA | NA |
| Gm13293 | 0 | 0 | 0 | 0 | 0 | 0 | 0 | 0 | 0 | 0 | 0 | 0 | 0 | 0 | 0 | NA | NA | NA | NA | NA | NA | NA | NA | NA | NA |
| Gm13298 | 0 | 0 | 0 | 0 | 0 | 0 | 0 | 0 | 0 | 0 | 0 | 0 | 0 | 0 | 0 | NA | NA | NA | NA | NA | NA | NA | NA | NA | NA |
| Gm13304 | 0 | 0 | 0 | 0 | 0 | 0 | 0 | 0 | 0 | 0 | 0 | 0 | 0 | 0 | 0 | NA | NA | NA | NA | NA | NA | NA | NA | NA | NA |
| Gm13305 | 0 | 0 | 0 | 0 | 0 | 0 | 0 | 0 | 0 | 0 | 0 | 0 | 0 | 0 | 0 | NA | NA | NA | NA | NA | NA | NA | NA | NA | NA |
| Gm13306 | 0 | 0 | 0 | 0 | 0 | 0 | 0 | 0 | 0 | 0 | 0 | 0 | 0 | 0 | 0 | NA | NA | NA | NA | NA | NA | NA | NA | NA | NA |
| Gm13308 | 0 | 0 | 0 | 0 | 0 | 0 | 0 | 0 | 0 | 0 | 0 | 0 | 0 | 0 | 0 | NA | NA | NA | NA | NA | NA | NA | NA | NA | NA |
| Gm13315 | 1 | 0 | 0 | 1 | 0 | 1 | 1 | 0 | 0 | 1 | 0 | 1 | 0.46 | 0 | 1 | 1.94 | 0.956 | 0.73503911 | NA | 0 | 5.1562 | 3.9771 | 3.9771 | TRUE | 0.2566 |
| Gm1332 | 0 | 1 | 0 | 0 | 1 | 0 | 0 | 1 | 0 | 0 | 1 | 0 | 0.37 | 0 | 0 | 1.075 | 0.105 | 0.97520514 | NA | 0 | 6.4904 | 6.1969 | 6.1969 | TRUE | 0.5216 |
| Gm13363 | 0 | 0 | 0 | 0 | 0 | 0 | 0 | 0 | 0 | 0 | 0 | 0 | 0 | 0 | 0 | NA | NA | NA | NA | NA | NA | NA | NA | NA | NA |
| Gm13375 | 107 | 91 | 104 | 125 | 116 | 121 | 83 | 97 | 112 | 124 | 131 | 119 | 111.14 | 97 | 125 | 1.289 | 0.367 | 0.05882104 | 0.15121027 | 0 | 0.0238 | 0.018 | 0.018 | TRUE | 0.182 |
| Gm13399 | 0 | 0 | 0 | 0 | 0 | 0 | 0 | 0 | 0 | 0 | 0 | 0 | 0 | 0 | 0 | NA | NA | NA | NA | NA | NA | NA | NA | NA | NA |
| Gm13446 | 3 | 2 | 2 | 0 | 0 | 0 | 2 | 2 | 2 | 0 | 0 | 0 | 1.1 | 2 | 0 | 0.086 | -3.536 | 0.0788983 | NA | 0 | 2.152 | 1.6987 | 1.6987 | TRUE | 0.0718 |
| Gm13476 | 1 | 0 | 1 | 0 | 0 | 0 | 1 | 0 | 1 | 0 | 0 | 0 | 0.31 | 1 | 0 | 0.307 | -1.706 | 0.62894649 | NA | 0 | 7.6657 | 7.2 | 7.2 | TRUE | 0.2067 |
| Gm13483 | 0 | 0 | 0 | 0 | 0 | 0 | 0 | 0 | 0 | 0 | 0 | 0 | 0 | 0 | 0 | NA | NA | NA | NA | NA | NA | NA | NA | NA | NA |
| Gm13490 | 0 | 0 | 0 | 0 | 1 | 0 | 0 | 0 | 0 | 0 | 1 | 0 | 0.19 | 0 | 0 | 2.094 | 1.067 | 0.79379329 | NA | 0 | 12.608 | 10 | 10 | TRUE | 0.5075 |
| Gm13497 | 0 | 0 | 0 | 0 | 0 | 0 | 0 | 0 | 0 | 0 | 0 | 0 | 0 | 0 | 0 | NA | NA | NA | NA | NA | NA | NA | NA | NA | NA |
| Gm13498 | 74 | 71 | 20 | 19 | 19 | 17 | 57 | 76 | 22 | 19 | 21 | 17 | 35.28 | 52 | 19 | 0.368 | -1.442 | 0.00092884 | 0.00687517 | 0.1726 | 0.0696 | 0.1001 | 0.1001 | TRUE | 0.2331 |
| Gm13539 | 0 | 0 | 0 | 0 | 0 | 0 | 0 | 0 | 0 | 0 | 0 | 0 | 0 | 0 | 0 | NA | NA | NA | NA | NA | NA | NA | NA | NA | NA |
| Gm13544 | 14 | 2 | 7 | 3 | 1 | 3 | 11 | 2 | 8 | 3 | 1 | 3 | 4.6 | 7 | 2 | 0.343 | -1.543 | 0.12129019 | NA | 0 | 0.5171 | 0.4281 | 0.4281 | TRUE | 0.1728 |
| Gm13546 | 0 | 2 | 0 | 0 | 0 | 0 | 0 | 2 | 0 | 0 | 0 | 0 | 0.35 | 1 | 0 | 0.272 | -1.878 | 0.62190613 | NA | 10 | 6.6869 | 8.7633 | 8.7633 | TRUE | 0.9743 |
| Gm13547 | 0 | 0 | 0 | 0 | 0 | 0 | 0 | 0 | 0 | 0 | 0 | 0 | 0 | 0 | 0 | NA | NA | NA | NA | NA | NA | NA | NA | NA | NA |
| Gm13580 | 20 | 25 | 21 | 14 | 13 | 12 | 16 | 27 | 23 | 14 | 15 | 12 | 17.54 | 22 | 14 | 0.629 | -0.669 | 0.15541139 | 0.3030416 | 0 | 0.1376 | 0.0988 | 0.0988 | TRUE | 0.392 |
| Gm13582 | 0 | 0 | 0 | 0 | 0 | 0 | 0 | 0 | 0 | 0 | 0 | 0 | 0 | 0 | 0 | NA | NA | NA | NA | NA | NA | NA | NA | NA | NA |
| Gm136 | 0 | 2 | 0 | 0 | 0 | 0 | 0 | 2 | 0 | 0 | 0 | 0 | 0.35 | 1 | 0 | 0.272 | -1.878 | 0.62190613 | NA | 10 | 6.6869 | 8.7633 | 8.7633 | TRUE | 0.9743 |
| Gm13629 | 0 | 0 | 0 | 0 | 0 | 0 | 0 | 0 | 0 | 0 | 0 | 0 | 0 | 0 | 0 | NA | NA | NA | NA | NA | NA | NA | NA | NA | NA |
| Gm13704 | 5 | 5 | 2 | 1 | 4 | 1 | 4 | 5 | 2 | 1 | 5 | 1 | 2.98 | 4 | 2 | 0.559 | -0.839 | 0.46794111 | NA | 0 | 0.7989 | 0.5922 | 0.5922 | TRUE | 0.6592 |
| Gm13710 | 4 | 3 | 1 | 2 | 2 | 2 | 3 | 3 | 1 | 2 | 2 | 2 | 2.27 | 2 | 2 | 0.833 | -0.264 | 0.83080562 | NA | 0 | 1.0482 | 0.6565 | 0.6565 | TRUE | 0.233 |
| Gm13718 | 164 | 144 | 124 | 136 | 111 | 102 | 127 | 153 | 134 | 135 | 125 | 101 | 129.27 | 138 | 120 | 0.873 | -0.195 | 0.29383007 | 0.46585837 | 0.008 | 0.0208 | 0.0172 | 0.0172 | TRUE | 0.2141 |
| Gm13749 | 0 | 0 | 0 | 0 | 0 | 0 | 0 | 0 | 0 | 0 | 0 | 0 | 0 | 0 | 0 | NA | NA | NA | NA | NA | NA | NA | NA | NA | NA |
| Gm13752 | 12 | 8 | 9 | 1 | 5 | 4 | 9 | 9 | 10 | 1 | 6 | 4 | 6.35 | 9 | 4 | 0.379 | -1.399 | 0.08575378 | NA | 0 | 0.3754 | 0.2763 | 0.2763 | TRUE | 0.6113 |
| Gm13769 | 0 | 0 | 0 | 0 | 0 | 0 | 0 | 0 | 0 | 0 | 0 | 0 | 0 | 0 | 0 | NA | NA | NA | NA | NA | NA | NA | NA | NA | NA |
| Gm13807 | 0 | 0 | 0 | 0 | 0 | 0 | 0 | 0 | 0 | 0 | 0 | 0 | 0 | 0 | 0 | NA | NA | NA | NA | NA | NA | NA | NA | NA | NA |
| Gm13826 | 403 | 257 | 303 | 352 | 303 | 323 | 313 | 273 | 327 | 350 | 342 | 319 | 320.72 | 304 | 337 | 1.106 | 0.145 | 0.23115256 | 0.39705123 | 0.0018 | 0.0098 | 0.0075 | 0.0075 | TRUE | 0.0858 |
| Gm13845 | 5 | 1 | 3 | 4 | 4 | 4 | 4 | 1 | 3 | 4 | 5 | 4 | 3.44 | 3 | 4 | 1.491 | 0.576 | 0.57362854 | NA | 0 | 0.6918 | 0.4564 | 0.4564 | TRUE | 0.31 |
| Gm13871 | 0 | 0 | 0 | 0 | 0 | 0 | 0 | 0 | 0 | 0 | 0 | 0 | 0 | 0 | 0 | NA | NA | NA | NA | NA | NA | NA | NA | NA | NA |
| Gm13889 | 18 | 9 | 16 | 15 | 18 | 13 | 14 | 10 | 17 | 15 | 20 | 13 | 14.81 | 14 | 16 | 1.172 | 0.229 | 0.65182949 | NA | 0 | 0.1624 | 0.1179 | 0.1179 | TRUE | 0.2596 |
| Gm13939 | 35 | 26 | 27 | 30 | 26 | 32 | 27 | 28 | 29 | 30 | 29 | 32 | 29.13 | 28 | 30 | 1.083 | 0.115 | 0.74099125 | 0.84598937 | 0 | 0.0838 | 0.053 | 0.053 | TRUE | 0.0101 |
| Gm13944 | 0 | 0 | 0 | 0 | 0 | 0 | 0 | 0 | 0 | 0 | 0 | 0 | 0 | 0 | 0 | NA | NA | NA | NA | NA | NA | NA | NA | NA | NA |
| Gm14005 | 18 | 9 | 13 | 7 | 14 | 8 | 14 | 10 | 14 | 7 | 16 | 8 | 11.37 | 13 | 10 | 0.805 | -0.312 | 0.59631829 | NA | 2.00E-04 | 0.2108 | 0.1607 | 0.1607 | TRUE | 0.7251 |
| Gm14015 | 0 | 0 | 0 | 1 | 0 | 0 | 0 | 0 | 0 | 1 | 0 | 0 | 0.17 | 0 | 0 | 2.094 | 1.067 | 0.79379329 | NA | 0 | 14.2885 | 10 | 10 | TRUE | 0.3906 |
| Gm14023 | 36 | 24 | 25 | 38 | 27 | 46 | 28 | 26 | 27 | 38 | 30 | 45 | 32.36 | 27 | 38 | 1.415 | 0.501 | 0.14427018 | 0.28752025 | 0 | 0.0757 | 0.0531 | 0.0531 | TRUE | 0.1456 |
| Gm14047 | 0 | 2 | 2 | 1 | 2 | 3 | 0 | 2 | 2 | 1 | 2 | 3 | 1.75 | 1 | 2 | 1.514 | 0.599 | 0.69008791 | NA | 0 | 1.3569 | 1.0206 | 1.0206 | TRUE | 0.7318 |
| Gm14057 | 111 | 75 | 66 | 83 | 60 | 81 | 86 | 80 | 71 | 83 | 68 | 80 | 77.91 | 79 | 77 | 0.969 | -0.045 | 0.84038224 | 0.90907815 | 0 | 0.0328 | 0.0231 | 0.0231 | TRUE | 0.0923 |
| Gm14085 | 2 | 0 | 0 | 0 | 0 | 0 | 2 | 0 | 0 | 0 | 0 | 0 | 0.26 | 1 | 0 | 0.356 | -1.491 | 0.71362796 | NA | 0 | 9.1573 | 10 | 10 | TRUE | 0.968 |
| Gm14092 | 0 | 0 | 0 | 0 | 0 | 0 | 0 | 0 | 0 | 0 | 0 | 0 | 0 | 0 | 0 | NA | NA | NA | NA | NA | NA | NA | NA | NA | NA |
| Gm14124 | 927 | 644 | 429 | 495 | 388 | 459 | 720 | 685 | 463 | 493 | 438 | 453 | 541.9 | 623 | 461 | 0.74 | -0.435 | 0.00261865 | 0.0148584 | 0.0279 | 0.0068 | 0.0132 | 0.0132 | TRUE | 0.5838 |
| Gm14137 | 3 | 3 | 2 | 5 | 3 | 4 | 2 | 3 | 2 | 5 | 3 | 4 | 3.33 | 2 | 4 | 1.614 | 0.69 | 0.50029371 | NA | 0 | 0.7138 | 0.4412 | 0.4412 | TRUE | 0.0602 |
| Gm14139 | 637 | 433 | 273 | 312 | 241 | 292 | 495 | 461 | 295 | 311 | 272 | 288 | 353.43 | 417 | 290 | 0.695 | -0.524 | 0.00178361 | 0.01108305 | 0.0368 | 0.0091 | 0.0174 | 0.0174 | TRUE | 0.7449 |
| Gm14151 | 0 | 1 | 3 | 0 | 1 | 2 | 0 | 1 | 3 | 0 | 1 | 2 | 1.23 | 1 | 1 | 0.745 | -0.426 | 0.82060566 | NA | 0 | 1.9233 | 1.6887 | 1.6887 | TRUE | 0.42 |
| Gm14164 | 23 | 16 | 18 | 23 | 24 | 38 | 18 | 17 | 19 | 23 | 27 | 38 | 23.63 | 18 | 29 | 1.614 | 0.691 | 0.0890302 | 0.20321121 | 0 | 0.1027 | 0.0745 | 0.0745 | TRUE | 0.425 |
| Gm14169 | 8 | 5 | 10 | 15 | 11 | 7 | 6 | 5 | 11 | 15 | 12 | 7 | 9.43 | 7 | 11 | 1.544 | 0.627 | 0.33494522 | NA | 0 | 0.2538 | 0.1937 | 0.1937 | TRUE | 0.4736 |
| Gm14204 | 2 | 1 | 1 | 2 | 1 | 1 | 2 | 1 | 1 | 2 | 1 | 1 | 1.3 | 1 | 1 | 1.101 | 0.139 | 0.93032816 | NA | 0 | 1.8251 | 1.0578 | 1.0578 | TRUE | 0.1028 |
| Gm14207 | 1 | 1 | 2 | 2 | 5 | 4 | 1 | 1 | 2 | 2 | 6 | 4 | 2.6 | 1 | 4 | 2.954 | 1.563 | 0.21106935 | NA | 0 | 0.9153 | 0.6203 | 0.6203 | TRUE | 0.1201 |
| Gm14288 | 0 | 0 | 0 | 0 | 0 | 0 | 0 | 0 | 0 | 0 | 0 | 0 | 0 | 0 | 0 | NA | NA | NA | NA | NA | NA | NA | NA | NA | NA |
| Gm14295 | 238 | 129 | 195 | 241 | 194 | 204 | 185 | 137 | 210 | 240 | 219 | 201 | 198.76 | 177 | 220 | 1.239 | 0.309 | 0.07076566 | 0.17289804 | 0.0202 | 0.0144 | 0.016 | 0.016 | TRUE | 0.4019 |
| Gm14305 | 231 | 173 | 172 | 207 | 163 | 196 | 179 | 184 | 186 | 206 | 184 | 193 | 188.74 | 183 | 194 | 1.064 | 0.09 | 0.53372822 | 0.69300304 | 0 | 0.015 | 0.0097 | 0.0097 | TRUE | 0.0294 |
| Gm14306 | 0 | 0 | 0 | 0 | 0 | 0 | 0 | 0 | 0 | 0 | 0 | 0 | 0 | 0 | 0 | NA | NA | NA | NA | NA | NA | NA | NA | NA | NA |

| | | | | | | | | | | | | | | | | | | | | | | | | | |
|---|---|---|---|---|---|---|---|---|---|---|---|---|---|---|---|---|---|---|---|---|---|---|---|---|---|
| Gm14308 | 0 | 0 | 0 | 0 | 0 | 0 | 0 | 0 | 0 | 0 | 0 | 0 | 0 | 0 | 0 | NA | NA | NA | NA | NA | NA | NA | NA | NA | NA |
| Gm14318 | 0 | 0 | 0 | 0 | 0 | 0 | 0 | 0 | 0 | 0 | 0 | 0 | 0 | 0 | 0 | NA | NA | NA | NA | NA | NA | NA | NA | NA | NA |
| Gm14322 | 427 | 268 | 233 | 273 | 226 | 259 | 332 | 285 | 251 | 272 | 255 | 256 | 275.08 | 289 | 261 | 0.898 | -0.155 | 0.25822975 | 0.42827544 | 0.0075 | 0.011 | 0.0099 | 0.0099 | TRUE | 0.1955 |
| Gm14325 | 853 | 649 | 574 | 666 | 591 | 604 | 662 | 690 | 619 | 663 | 667 | 596 | 649.67 | 657 | 642 | 0.976 | -0.035 | 0.70415035 | 0.82094194 | 0.0018 | 0.0061 | 0.0047 | 0.0047 | TRUE | 0.0464 |
| Gm14326 | 1282 | 950 | 818 | 958 | 921 | 951 | 996 | 1011 | 883 | 954 | 1039 | 939 | 970.03 | 963 | 977 | 1.013 | 0.019 | 0.82232182 | 0.89810085 | 0.0031 | 0.0049 | 0.0043 | 0.0043 | TRUE | 0.0618 |
| Gm14327 | 1 | 2 | 2 | 2 | 1 | 1 | 1 | 2 | 2 | 2 | 1 | 1 | 1.53 | 2 | 1 | 0.832 | -0.266 | 0.85909569 | NA | 0 | 1.5535 | 0.9495 | 0.9495 | TRUE | 0.2415 |
| Gm14345 | 0 | 0 | 0 | 0 | 0 | 0 | 0 | 0 | 0 | 0 | 0 | 0 | 0 | 0 | 0 | NA | NA | NA | NA | NA | NA | NA | NA | NA | NA |
| Gm14346 | 0 | 0 | 0 | 0 | 0 | 0 | 0 | 0 | 0 | 0 | 0 | 0 | 0 | 0 | 0 | NA | NA | NA | NA | NA | NA | NA | NA | NA | NA |
| Gm14347 | 0 | 0 | 0 | 0 | 0 | 0 | 0 | 0 | 0 | 0 | 0 | 0 | 0 | 0 | 0 | NA | NA | NA | NA | NA | NA | NA | NA | NA | NA |
| Gm14351 | 0 | 0 | 0 | 0 | 0 | 0 | 0 | 0 | 0 | 0 | 0 | 0 | 0 | 0 | 0 | NA | NA | NA | NA | NA | NA | NA | NA | NA | NA |
| Gm14354 | 0 | 0 | 0 | 0 | 0 | 0 | 0 | 0 | 0 | 0 | 0 | 0 | 0 | 0 | 0 | NA | NA | NA | NA | NA | NA | NA | NA | NA | NA |
| Gm14374 | 0 | 0 | 0 | 0 | 0 | 0 | 0 | 0 | 0 | 0 | 0 | 0 | 0 | 0 | 0 | NA | NA | NA | NA | NA | NA | NA | NA | NA | NA |
| Gm14378 | 17 | 11 | 22 | 17 | 14 | 13 | 13 | 12 | 24 | 17 | 16 | 13 | 15.7 | 16 | 15 | 0.942 | -0.085 | 0.86483568 | NA | 0 | 0.1534 | 0.1176 | 0.1176 | TRUE | 0.7314 |
| Gm14379 | 5 | 0 | 2 | 2 | 0 | 0 | 4 | 0 | 2 | 2 | 0 | 0 | 1.34 | 2 | 1 | 0.329 | -1.604 | 0.41524811 | NA | 1.8833 | 1.7729 | 1.7902 | 1.7902 | TRUE | 0.3581 |
| Gm14391 | 0 | 0 | 0 | 0 | 0 | 0 | 0 | 0 | 0 | 0 | 0 | 0 | 0 | 0 | 0 | NA | NA | NA | NA | NA | NA | NA | NA | NA | NA |
| Gm14393 | 88 | 76 | 64 | 85 | 67 | 59 | 68 | 81 | 69 | 85 | 76 | 58 | 72.78 | 73 | 73 | 1.002 | 0.003 | 0.98998285 | 0.99485516 | 0.0075 | 0.035 | 0.0276 | 0.0276 | TRUE | 0.2878 |
| Gm14403 | 386 | 268 | 202 | 186 | 156 | 142 | 300 | 285 | 218 | 185 | 176 | 140 | 217.35 | 268 | 167 | 0.622 | -0.684 | 4.44E-05 | 0.00063252 | 0.0201 | 0.0133 | 0.0153 | 0.0153 | TRUE | 0.2872 |
| Gm14405 | 1 | 0 | 0 | 0 | 1 | 1 | 1 | 0 | 0 | 0 | 1 | 1 | 0.48 | 0 | 1 | 2.031 | 1.022 | 0.71376068 | NA | 0 | 4.9197 | 3.859 | 3.859 | TRUE | 0.2623 |
| Gm14420 | 393 | 290 | 251 | 238 | 211 | 219 | 305 | 308 | 271 | 237 | 238 | 216 | 262.61 | 295 | 230 | 0.78 | -0.358 | 0.00554999 | 0.02656599 | 0 | 0.0115 | 0.0082 | 0.0082 | TRUE | 0.0548 |
| Gm14430 | 0 | 0 | 0 | 0 | 0 | 0 | 0 | 0 | 0 | 0 | 0 | 0 | 0 | 0 | 0 | NA | NA | NA | NA | NA | NA | NA | NA | NA | NA |
| Gm14431 | 0 | 0 | 0 | 0 | 0 | 0 | 0 | 0 | 0 | 0 | 0 | 0 | 0 | 0 | 0 | NA | NA | NA | NA | NA | NA | NA | NA | NA | NA |
| Gm14434 | 0 | 0 | 0 | 0 | 0 | 0 | 0 | 0 | 0 | 0 | 0 | 0 | 0 | 0 | 0 | NA | NA | NA | NA | NA | NA | NA | NA | NA | NA |
| Gm14436 | 1 | 1 | 0 | 0 | 0 | 0 | 1 | 1 | 0 | 0 | 0 | 0 | 0.31 | 1 | 0 | 0.309 | -1.696 | 0.63183309 | NA | 0 | 7.7294 | 7.2472 | 7.2472 | TRUE | 0.2015 |
| Gm14440 | 4 | 2 | 0 | 3 | 1 | 1 | 3 | 2 | 0 | 3 | 1 | 1 | 1.72 | 2 | 2 | 0.952 | -0.071 | 0.96218513 | NA | 0 | 1.3783 | 1.0499 | 1.0499 | TRUE | 0.5109 |
| Gm14446 | 5 | 5 | 3 | 6 | 9 | 4 | 4 | 5 | 3 | 6 | 10 | 4 | 5.42 | 4 | 7 | 1.605 | 0.683 | 0.41343647 | NA | 0 | 0.4398 | 0.3078 | 0.3078 | TRUE | 0.3668 |
| Gm14458 | 203 | 162 | 114 | 74 | 60 | 58 | 158 | 172 | 123 | 74 | 68 | 57 | 108.6 | 151 | 66 | 0.438 | -1.192 | 2.00E-08 | 9.28E-07 | 0.0145 | 0.0243 | 0.0215 | 0.0215 | TRUE | 0.2625 |
| Gm14459 | 0 | 0 | 0 | 1 | 0 | 0 | 0 | 0 | 0 | 0 | 1 | 0 | 0.17 | 0 | 0 | 2.094 | 1.067 | 0.79379329 | NA | 0 | 14.2885 | 10 | 10 | TRUE | 0.3906 |
| Gm14461 | 0 | 0 | 0 | 0 | 0 | 0 | 0 | 0 | 0 | 0 | 0 | 0 | 0 | 0 | 0 | NA | NA | NA | NA | NA | NA | NA | NA | NA | NA |
| Gm14474 | 0 | 0 | 0 | 0 | 0 | 0 | 0 | 0 | 0 | 0 | 0 | 0 | 0 | 0 | 0 | NA | NA | NA | NA | NA | NA | NA | NA | NA | NA |
| Gm14475 | 0 | 0 | 0 | 0 | 0 | 0 | 0 | 0 | 0 | 0 | 0 | 0 | 0 | 0 | 0 | NA | NA | NA | NA | NA | NA | NA | NA | NA | NA |
| Gm14476 | 0 | 0 | 0 | 0 | 0 | 0 | 0 | 0 | 0 | 0 | 0 | 0 | 0 | 0 | 0 | NA | NA | NA | NA | NA | NA | NA | NA | NA | NA |
| Gm14477 | 0 | 0 | 0 | 0 | 0 | 0 | 0 | 0 | 0 | 0 | 0 | 0 | 0 | 0 | 0 | NA | NA | NA | NA | NA | NA | NA | NA | NA | NA |
| Gm14478 | 0 | 0 | 0 | 0 | 0 | 0 | 0 | 0 | 0 | 0 | 0 | 0 | 0 | 0 | 0 | NA | NA | NA | NA | NA | NA | NA | NA | NA | NA |
| Gm14479 | 0 | 0 | 0 | 0 | 0 | 0 | 0 | 0 | 0 | 0 | 0 | 0 | 0 | 0 | 0 | NA | NA | NA | NA | NA | NA | NA | NA | NA | NA |
| Gm14482 | 0 | 0 | 0 | 0 | 0 | 0 | 0 | 0 | 0 | 0 | 0 | 0 | 0 | 0 | 0 | NA | NA | NA | NA | NA | NA | NA | NA | NA | NA |
| Gm14483 | 0 | 0 | 0 | 0 | 0 | 0 | 0 | 0 | 0 | 0 | 0 | 0 | 0 | 0 | 0 | NA | NA | NA | NA | NA | NA | NA | NA | NA | NA |
| Gm14484 | 0 | 0 | 0 | 0 | 0 | 0 | 0 | 0 | 0 | 0 | 0 | 0 | 0 | 0 | 0 | NA | NA | NA | NA | NA | NA | NA | NA | NA | NA |
| Gm14492 | 0 | 2 | 0 | 1 | 1 | 0 | 0 | 2 | 0 | 1 | 1 | 0 | 0.71 | 1 | 1 | 1.021 | 0.03 | 0.99038186 | NA | 0 | 3.3477 | 2.9808 | 2.9808 | TRUE | 0.9748 |
| Gm14496 | 0 | 0 | 0 | 0 | 0 | 0 | 0 | 0 | 0 | 0 | 0 | 0 | 0 | 0 | 0 | NA | NA | NA | NA | NA | NA | NA | NA | NA | NA |
| Gm14501 | 0 | 0 | 2 | 1 | 2 | 2 | 0 | 0 | 2 | 1 | 2 | 2 | 1.23 | 1 | 2 | 2.553 | 1.352 | 0.47083384 | NA | 0 | 1.9283 | 1.5281 | 1.5281 | TRUE | 0.9754 |
| Gm14511 | 8 | 9 | 8 | 3 | 3 | 2 | 6 | 10 | 9 | 3 | 3 | 2 | 5.46 | 8 | 3 | 0.342 | -1.547 | 0.07251115 | NA | 0 | 0.4365 | 0.2877 | 0.2877 | TRUE | 0.1663 |
| Gm14525 | 0 | 1 | 0 | 0 | 0 | 0 | 0 | 1 | 0 | 0 | 0 | 0 | 0.18 | 0 | 0 | 0.552 | -0.857 | 0.8336548 | NA | 0 | 13.3714 | 10 | 10 | TRUE | 0.5216 |
| Gm14548 | 0 | 0 | 0 | 0 | 0 | 0 | 0 | 0 | 0 | 0 | 0 | 0 | 0 | 0 | 0 | NA | NA | NA | NA | NA | NA | NA | NA | NA | NA |
| Gm14625 | 0 | 1 | 0 | 0 | 0 | 0 | 0 | 1 | 0 | 0 | 0 | 0 | 0.18 | 0 | 0 | 0.552 | -0.857 | 0.8336548 | NA | 0 | 13.3714 | 10 | 10 | TRUE | 0.5216 |
| Gm14632 | 0 | 0 | 0 | 0 | 0 | 0 | 0 | 0 | 0 | 0 | 0 | 0 | 0 | 0 | 0 | NA | NA | NA | NA | NA | NA | NA | NA | NA | NA |
| Gm14634 | 5 | 7 | 3 | 4 | 3 | 3 | 4 | 7 | 3 | 4 | 3 | 3 | 4.15 | 5 | 3 | 0.714 | -0.486 | 0.6027993 | NA | 0 | 0.5737 | 0.3794 | 0.3794 | TRUE | 0.3975 |
| Gm14635 | 1 | 0 | 0 | 0 | 0 | 0 | 1 | 0 | 0 | 0 | 0 | 0 | 0.13 | 0 | 0 | 0.552 | -0.857 | 0.8336548 | NA | 0 | 18.3121 | 10 | 10 | TRUE | 0.2566 |
| Gm14685 | 0 | 0 | 0 | 0 | 0 | 0 | 0 | 0 | 0 | 0 | 0 | 0 | 0 | 0 | 0 | NA | NA | NA | NA | NA | NA | NA | NA | NA | NA |
| Gm14692 | 0 | 0 | 0 | 0 | 0 | 0 | 0 | 0 | 0 | 0 | 0 | 0 | 0 | 0 | 0 | NA | NA | NA | NA | NA | NA | NA | NA | NA | NA |
| Gm14718 | 0 | 0 | 0 | 0 | 0 | 0 | 0 | 0 | 0 | 0 | 0 | 0 | 0 | 0 | 0 | NA | NA | NA | NA | NA | NA | NA | NA | NA | NA |
| Gm14725 | 0 | 0 | 0 | 0 | 0 | 0 | 0 | 0 | 0 | 0 | 0 | 0 | 0 | 0 | 0 | NA | NA | NA | NA | NA | NA | NA | NA | NA | NA |
| Gm14743 | 0 | 0 | 0 | 0 | 0 | 0 | 0 | 0 | 0 | 0 | 0 | 0 | 0 | 0 | 0 | NA | NA | NA | NA | NA | NA | NA | NA | NA | NA |
| Gm14744 | 0 | 0 | 0 | 0 | 0 | 0 | 0 | 0 | 0 | 0 | 0 | 0 | 0 | 0 | 0 | NA | NA | NA | NA | NA | NA | NA | NA | NA | NA |
| Gm14781 | 0 | 0 | 0 | 0 | 0 | 0 | 0 | 0 | 0 | 0 | 0 | 0 | 0 | 0 | 0 | NA | NA | NA | NA | NA | NA | NA | NA | NA | NA |
| Gm14812 | 0 | 0 | 0 | 1 | 0 | 1 | 0 | 0 | 0 | 1 | 0 | 1 | 0.33 | 0 | 1 | 3.763 | 1.912 | 0.57887588 | NA | 0 | 7.1752 | 6.7756 | 6.7756 | TRUE | 0.2094 |
| Gm14819 | 0 | 0 | 0 | 0 | 0 | 0 | 0 | 0 | 0 | 0 | 0 | 0 | 0 | 0 | 0 | NA | NA | NA | NA | NA | NA | NA | NA | NA | NA |
| Gm14827 | 8 | 7 | 4 | 3 | 1 | 1 | 6 | 7 | 4 | 3 | 1 | 1 | 3.85 | 6 | 2 | 0.286 | -1.808 | 0.08762097 | NA | 0 | 0.6187 | 0.4251 | 0.4251 | TRUE | 0.3488 |
| Gm14850 | 0 | 0 | 0 | 0 | 0 | 0 | 0 | 0 | 0 | 0 | 0 | 0 | 0 | 0 | 0 | NA | NA | NA | NA | NA | NA | NA | NA | NA | NA |
| Gm14851 | 0 | 0 | 0 | 0 | 0 | 0 | 0 | 0 | 0 | 0 | 0 | 0 | 0 | 0 | 0 | NA | NA | NA | NA | NA | NA | NA | NA | NA | NA |
| Gm14858 | 1 | 0 | 0 | 0 | 0 | 0 | 1 | 0 | 0 | 0 | 0 | 0 | 0.13 | 0 | 0 | 0.552 | -0.857 | 0.8336548 | NA | 0 | 18.3121 | 10 | 10 | TRUE | 0.2566 |
| Gm14872 | 9 | 9 | 3 | 9 | 6 | 9 | 7 | 10 | 3 | 9 | 7 | 9 | 7.4 | 7 | 8 | 1.242 | 0.313 | 0.66075179 | NA | 0 | 0.3226 | 0.2305 | 0.2305 | TRUE | 0.3248 |
| Gm14873 | 65 | 62 | 45 | 73 | 54 | 63 | 50 | 66 | 49 | 73 | 61 | 62 | 60.13 | 55 | 65 | 1.191 | 0.252 | 0.32649762 | 0.50065771 | 0 | 0.0418 | 0.0309 | 0.0309 | TRUE | 0.2413 |
| Gm14920 | 0 | 0 | 0 | 0 | 0 | 0 | 0 | 0 | 0 | 0 | 0 | 0 | 0 | 0 | 0 | NA | NA | NA | NA | NA | NA | NA | NA | NA | NA |
| Gm15008 | 0 | 0 | 1 | 0 | 0 | 0 | 0 | 0 | 1 | 0 | 0 | 0 | 0.18 | 0 | 0 | 0.552 | -0.857 | 0.8336548 | NA | 0 | 13.1817 | 10 | 10 | TRUE | 0.5365 |
| Gm15023 | 0 | 0 | 0 | 0 | 0 | 0 | 0 | 0 | 0 | 0 | 0 | 0 | 0 | 0 | 0 | NA | NA | NA | NA | NA | NA | NA | NA | NA | NA |
| Gm15056 | 0 | 0 | 0 | 0 | 0 | 0 | 0 | 0 | 0 | 0 | 0 | 0 | 0 | 0 | 0 | NA | NA | NA | NA | NA | NA | NA | NA | NA | NA |
| Gm15070 | 13 | 13 | 6 | 5 | 5 | 13 | 10 | 14 | 6 | 5 | 6 | 13 | 8.97 | 10 | 8 | 0.775 | -0.368 | 0.58719728 | NA | 0.0853 | 0.2665 | 0.217 | 0.217 | TRUE | 0.4499 |
| Gm15080 | 0 | 0 | 0 | 0 | 0 | 0 | 0 | 0 | 0 | 0 | 0 | 0 | 0 | 0 | 0 | NA | NA | NA | NA | NA | NA | NA | NA | NA | NA |
| Gm15085 | 0 | 0 | 0 | 0 | 0 | 0 | 0 | 0 | 0 | 0 | 0 | 0 | 0 | 0 | 0 | NA | NA | NA | NA | NA | NA | NA | NA | NA | NA |

| | | | | | | | | | | | | | | | | | | | | | | | | | |
|---|---|---|---|---|---|---|---|---|---|---|---|---|---|---|---|---|---|---|---|---|---|---|---|---|---|
| Gm15091 | 24 | 21 | 19 | 7 | 10 | 8 | 19 | 22 | 21 | 7 | 11 | 8 | 14.6 | 21 | 9 | 0.424 | -1.239 | 0.01704692 | NA | 0 | 0.1647 | 0.1109 | 0.1109 | TRUE | 0.1846 |
| Gm15093 | 0 | 0 | 0 | 0 | 0 | 0 | 0 | 0 | 0 | 0 | 0 | 0 | 0 | 0 | 0 | NA | NA | NA | NA | NA | NA | NA | NA | NA | NA |
| Gm15097 | 83 | 67 | 114 | 48 | 32 | 39 | 64 | 71 | 123 | 48 | 36 | 38 | 63.52 | 86 | 41 | 0.476 | -1.07 | 0.00040643 | 0.00369071 | 0.0656 | 0.0397 | 0.0477 | 0.0477 | TRUE | 1.2863 |
| Gm15104 | 2 | 0 | 0 | 0 | 0 | 0 | 2 | 0 | 0 | 0 | 0 | 0 | 0.26 | 1 | 0 | 0.356 | -1.491 | 0.71362796 | NA | 0 | 9.1573 | 10 | 10 | TRUE | 0.968 |
| Gm15107 | 10 | 11 | 8 | 7 | 8 | 6 | 8 | 12 | 9 | 7 | 9 | 6 | 8.34 | 10 | 7 | 0.781 | -0.356 | 0.58842792 | NA | 0 | 0.2868 | 0.1893 | 0.1893 | TRUE | 0.1496 |
| Gm15114 | 2 | 0 | 1 | 0 | 0 | 1 | 2 | 0 | 1 | 0 | 0 | 1 | 0.6 | 1 | 0 | 0.414 | -1.271 | 0.61985863 | NA | 0 | 3.9313 | 3.1876 | 3.1876 | TRUE | 0.3835 |
| Gm15127 | 9 | 6 | 8 | 2 | 3 | 4 | 7 | 6 | 9 | 2 | 3 | 4 | 5.22 | 7 | 3 | 0.424 | -1.239 | 0.14858728 | NA | 0 | 0.4563 | 0.297 | 0.297 | TRUE | 0.1391 |
| Gm15133 | 0 | 0 | 0 | 0 | 0 | 0 | 0 | 0 | 0 | 0 | 0 | 0 | 0 | 0 | 0 | NA | NA | NA | NA | NA | NA | NA | NA | NA | NA |
| Gm15140 | 0 | 0 | 0 | 0 | 0 | 0 | 0 | 0 | 0 | 0 | 0 | 0 | 0 | 0 | 0 | NA | NA | NA | NA | NA | NA | NA | NA | NA | NA |
| Gm15179 | 0 | 0 | 1 | 1 | 1 | 0 | 0 | 0 | 1 | 1 | 1 | 0 | 0.53 | 0 | 1 | 2.036 | 1.026 | 0.70454668 | NA | 0 | 4.4428 | 3.5371 | 3.5371 | TRUE | 0.5365 |
| Gm15217 | 2 | 0 | 1 | 2 | 2 | 0 | 2 | 0 | 1 | 2 | 2 | 0 | 1.15 | 1 | 1 | 1.551 | 0.634 | 0.73541734 | NA | 0 | 2.0695 | 1.6309 | 1.6309 | TRUE | 0.5196 |
| Gm1527 | 1 | 1 | 0 | 0 | 1 | 1 | 1 | 1 | 0 | 0 | 1 | 1 | 0.66 | 1 | 1 | 1.125 | 0.17 | 0.94233031 | NA | 0 | 3.5974 | 2.4494 | 2.4494 | TRUE | 0.2637 |
| Gm15284 | 3 | 2 | 3 | 1 | 0 | 0 | 2 | 2 | 3 | 1 | 0 | 0 | 1.45 | 2 | 0 | 0.145 | -2.786 | 0.12270328 | NA | 0 | 1.6389 | 1.1594 | 1.1594 | TRUE | 0.3906 |
| Gm15292 | 0 | 0 | 0 | 0 | 0 | 0 | 0 | 0 | 0 | 0 | 0 | 0 | 0 | 0 | 0 | NA | NA | NA | NA | NA | NA | NA | NA | NA | NA |
| Gm15293 | 0 | 0 | 0 | 0 | 0 | 0 | 0 | 0 | 0 | 0 | 0 | 0 | 0 | 0 | 0 | NA | NA | NA | NA | NA | NA | NA | NA | NA | NA |
| Gm15299 | 0 | 0 | 0 | 0 | 0 | 0 | 0 | 0 | 0 | 0 | 0 | 0 | 0 | 0 | 0 | NA | NA | NA | NA | NA | NA | NA | NA | NA | NA |
| Gm15308 | 0 | 0 | 0 | 0 | 0 | 0 | 0 | 0 | 0 | 0 | 0 | 0 | 0 | 0 | 0 | NA | NA | NA | NA | NA | NA | NA | NA | NA | NA |
| Gm15315 | 0 | 0 | 0 | 0 | 0 | 0 | 0 | 0 | 0 | 0 | 0 | 0 | 0 | 0 | 0 | NA | NA | NA | NA | NA | NA | NA | NA | NA | NA |
| Gm15319 | 0 | 0 | 0 | 0 | 0 | 0 | 0 | 0 | 0 | 0 | 0 | 0 | 0 | 0 | 0 | NA | NA | NA | NA | NA | NA | NA | NA | NA | NA |
| Gm15328 | 1 | 0 | 0 | 0 | 0 | 0 | 1 | 0 | 0 | 0 | 0 | 0 | 0.13 | 0 | 0 | 0.552 | -0.857 | 0.8336548 | NA | 0 | 18.3121 | 10 | 10 | TRUE | 0.2566 |
| Gm15348 | 0 | 0 | 0 | 0 | 0 | 0 | 0 | 0 | 0 | 0 | 0 | 0 | 0 | 0 | 0 | NA | NA | NA | NA | NA | NA | NA | NA | NA | NA |
| Gm15350 | 0 | 0 | 0 | 0 | 0 | 0 | 0 | 0 | 0 | 0 | 0 | 0 | 0 | 0 | 0 | NA | NA | NA | NA | NA | NA | NA | NA | NA | NA |
| Gm15386 | 0 | 0 | 0 | 0 | 0 | 0 | 0 | 0 | 0 | 0 | 0 | 0 | 0 | 0 | 0 | NA | NA | NA | NA | NA | NA | NA | NA | NA | NA |
| Gm15401 | 0 | 0 | 2 | 1 | 2 | 4 | 0 | 0 | 2 | 1 | 2 | 4 | 1.56 | 1 | 2 | 3.553 | 1.829 | 0.29690172 | NA | 0 | 1.522 | 1.2891 | 1.2891 | TRUE | 0.6975 |
| Gm15408 | 16 | 6 | 14 | 14 | 8 | 13 | 12 | 6 | 15 | 14 | 9 | 13 | 11.62 | 11 | 12 | 1.056 | 0.079 | 0.89157115 | NA | 0 | 0.2064 | 0.155 | 0.155 | TRUE | 0.4995 |
| Gm15412 | 0 | 0 | 0 | 0 | 0 | 0 | 0 | 0 | 0 | 0 | 0 | 0 | 0 | 0 | 0 | NA | NA | NA | NA | NA | NA | NA | NA | NA | NA |
| Gm15413 | 0 | 1 | 0 | 0 | 0 | 0 | 0 | 1 | 0 | 0 | 0 | 0 | 0.18 | 0 | 0 | 0.552 | -0.857 | 0.8336548 | NA | 0 | 13.3714 | 10 | 10 | TRUE | 0.5216 |
| Gm15417 | 23 | 4 | 12 | 23 | 23 | 16 | 18 | 4 | 13 | 23 | 26 | 16 | 16.62 | 12 | 22 | 1.804 | 0.851 | 0.11358089 | NA | 0.1403 | 0.1451 | 0.1439 | 0.1439 | TRUE | 0.6956 |
| Gm15421 | 73 | 54 | 64 | 83 | 70 | 69 | 57 | 57 | 69 | 83 | 79 | 68 | 68.81 | 61 | 77 | 1.257 | 0.33 | 0.16454645 | 0.3153476 | 0 | 0.0369 | 0.0259 | 0.0259 | TRUE | 0.1075 |
| Gm15441 | 14 | 16 | 19 | 15 | 21 | 14 | 11 | 17 | 21 | 15 | 24 | 14 | 16.81 | 16 | 18 | 1.091 | 0.126 | 0.79806965 | NA | 0.0409 | 0.1435 | 0.114 | 0.114 | TRUE | 0.493 |
| Gm15446 | 44 | 33 | 19 | 40 | 34 | 42 | 34 | 35 | 21 | 40 | 38 | 41 | 34.9 | 30 | 40 | 1.325 | 0.406 | 0.22927939 | 0.39472864 | 0 | 0.0703 | 0.0533 | 0.0533 | TRUE | 0.4623 |
| Gm15455 | 153 | 108 | 57 | 47 | 48 | 50 | 119 | 115 | 62 | 47 | 54 | 49 | 74.25 | 99 | 50 | 0.506 | -0.982 | 0.00043326 | 0.00386795 | 0.0563 | 0.0343 | 0.0409 | 0.0409 | TRUE | 1.0066 |
| Gm15471 | 0 | 0 | 0 | 0 | 0 | 0 | 0 | 0 | 0 | 0 | 0 | 0 | 0 | 0 | 0 | NA | NA | NA | NA | NA | NA | NA | NA | NA | NA |
| Gm1553 | 1 | 0 | 0 | 0 | 0 | 0 | 1 | 0 | 0 | 0 | 0 | 0 | 0.13 | 0 | 0 | 0.552 | -0.857 | 0.8336548 | NA | 0 | 18.3121 | 10 | 10 | TRUE | 0.2566 |
| Gm15545 | 32 | 27 | 16 | 31 | 15 | 27 | 25 | 29 | 17 | 31 | 17 | 27 | 24.21 | 24 | 25 | 1.053 | 0.075 | 0.85612182 | 0.91907385 | 0.0269 | 0.1003 | 0.0815 | 0.0815 | TRUE | 0.4275 |
| Gm156 | 0 | 0 | 0 | 0 | 0 | 0 | 0 | 0 | 0 | 0 | 0 | 0 | 0 | 0 | 0 | NA | NA | NA | NA | NA | NA | NA | NA | NA | NA |
| Gm15612 | 4 | 3 | 1 | 3 | 1 | 1 | 3 | 3 | 1 | 3 | 1 | 1 | 2.08 | 2 | 2 | 0.689 | -0.538 | 0.6835725 | NA | 0 | 1.142 | 0.7599 | 0.7599 | TRUE | 0.3474 |
| Gm1564 | 3109 | 1990 | 2138 | 3363 | 2411 | 2717 | 2415 | 2117 | 2307 | 3348 | 2720 | 2682 | 2597.96 | 2280 | 2917 | 1.279 | 0.355 | 9.00E-05 | 0.00110778 | 0.0098 | 0.0033 | 0.0055 | 0.0055 | TRUE | 0.2035 |
| Gm15645 | 0 | 0 | 0 | 0 | 1 | 0 | 0 | 0 | 0 | 0 | 1 | 0 | 0.19 | 0 | 0 | 2.094 | 1.067 | 0.79379329 | NA | 0 | 12.608 | 10 | 10 | TRUE | 0.5075 |
| Gm15663 | 0 | 1 | 0 | 2 | 1 | 2 | 0 | 1 | 0 | 2 | 1 | 2 | 1.03 | 0 | 2 | 4.943 | 2.306 | 0.25230205 | NA | 0 | 2.3121 | 1.6193 | 1.6193 | TRUE | 0.5217 |
| Gm15698 | 34 | 17 | 14 | 17 | 14 | 10 | 26 | 18 | 15 | 17 | 16 | 10 | 17.03 | 20 | 14 | 0.706 | -0.503 | 0.3008021 | NA | 0 | 0.1416 | 0.1098 | 0.1098 | TRUE | 0.5379 |
| Gm15706 | 55 | 51 | 36 | 43 | 38 | 39 | 43 | 54 | 39 | 43 | 43 | 38 | 43.33 | 45 | 41 | 0.915 | -0.128 | 0.66671109 | 0.79491786 | 0 | 0.0571 | 0.0401 | 0.0401 | TRUE | 0.2215 |
| Gm15708 | 5 | 3 | 3 | 2 | 1 | 0 | 4 | 3 | 3 | 2 | 1 | 0 | 2.24 | 3 | 1 | 0.3 | -1.736 | 0.20858401 | NA | 0 | 1.0613 | 0.7336 | 0.7336 | TRUE | 0.3987 |
| Gm15713 | 0 | 0 | 0 | 0 | 0 | 0 | 0 | 0 | 0 | 0 | 0 | 0 | 0 | 0 | 0 | NA | NA | NA | NA | NA | NA | NA | NA | NA | NA |
| Gm15760 | 10 | 15 | 7 | 6 | 18 | 8 | 8 | 16 | 8 | 6 | 20 | 8 | 10.91 | 11 | 11 | 1.09 | 0.124 | 0.84999354 | NA | 0.2139 | 0.2197 | 0.218 | 0.218 | TRUE | 1.6297 |
| Gm15772 | 0 | 0 | 0 | 0 | 0 | 0 | 0 | 0 | 0 | 0 | 0 | 0 | 0 | 0 | 0 | NA | NA | NA | NA | NA | NA | NA | NA | NA | NA |
| Gm15787 | 6 | 5 | 3 | 9 | 3 | 4 | 5 | 5 | 3 | 9 | 3 | 4 | 4.92 | 4 | 5 | 1.236 | 0.306 | 0.72556714 | NA | 0 | 0.4844 | 0.342 | 0.342 | TRUE | 0.7124 |
| Gm15800 | 2663 | 1830 | 1624 | 1845 | 1730 | 2018 | 2068 | 1947 | 1752 | 1837 | 1952 | 1992 | 1924.55 | 1922 | 1927 | 1.001 | 0.002 | 0.98107297 | 0.990148 | 0.0039 | 0.0037 | 0.0037 | 0.0037 | TRUE | 0.0719 |
| Gm15850 | 0 | 0 | 0 | 0 | 0 | 2 | 0 | 0 | 0 | 0 | 0 | 2 | 0.33 | 0 | 1 | 3.743 | 1.904 | 0.62640105 | NA | 0 | 7.2051 | 9.2465 | 9.2465 | TRUE | 0.9739 |
| Gm1587 | 0 | 0 | 0 | 0 | 0 | 0 | 0 | 0 | 0 | 0 | 0 | 0 | 0 | 0 | 0 | NA | NA | NA | NA | NA | NA | NA | NA | NA | NA |
| Gm15880 | 0 | 0 | 0 | 0 | 0 | 0 | 0 | 0 | 0 | 0 | 0 | 0 | 0 | 0 | 0 | NA | NA | NA | NA | NA | NA | NA | NA | NA | NA |
| Gm15881 | 0 | 0 | 0 | 0 | 0 | 0 | 0 | 0 | 0 | 0 | 0 | 0 | 0 | 0 | 0 | NA | NA | NA | NA | NA | NA | NA | NA | NA | NA |
| Gm15910 | 66 | 39 | 34 | 33 | 25 | 35 | 51 | 41 | 37 | 33 | 28 | 35 | 37.51 | 43 | 32 | 0.735 | -0.443 | 0.16669487 | 0.31778393 | 0 | 0.0656 | 0.0467 | 0.0467 | TRUE | 0.2414 |
| Gm15915 | 73 | 55 | 73 | 94 | 76 | 94 | 57 | 59 | 79 | 94 | 86 | 93 | 77.68 | 65 | 91 | 1.411 | 0.497 | 0.03070528 | 0.09415803 | 0 | 0.0329 | 0.0249 | 0.0249 | TRUE | 0.3059 |
| Gm15941 | 0 | 0 | 0 | 0 | 0 | 0 | 0 | 0 | 0 | 0 | 0 | 0 | 0 | 0 | 0 | NA | NA | NA | NA | NA | NA | NA | NA | NA | NA |
| Gm15987 | 5 | 5 | 4 | 3 | 0 | 1 | 4 | 5 | 4 | 3 | 0 | 1 | 2.92 | 4 | 1 | 0.302 | -1.729 | 0.16012635 | NA | 0 | 0.8154 | 0.6017 | 0.6017 | TRUE | 0.733 |
| Gm15997 | 0 | 0 | 0 | 0 | 0 | 0 | 0 | 0 | 0 | 0 | 0 | 0 | 0 | 0 | 0 | NA | NA | NA | NA | NA | NA | NA | NA | NA | NA |
| Gm16023 | 22 | 11 | 9 | 11 | 5 | 12 | 17 | 12 | 10 | 11 | 6 | 12 | 11.16 | 13 | 10 | 0.736 | -0.443 | 0.45471908 | NA | 0 | 0.2149 | 0.1614 | 0.1614 | TRUE | 0.43 |
| Gm16039 | 126 | 63 | 76 | 85 | 88 | 72 | 98 | 67 | 82 | 85 | 99 | 71 | 83.64 | 82 | 85 | 1.024 | 0.035 | 0.88170987 | 0.93384803 | 0.0201 | 0.0308 | 0.0277 | 0.0277 | TRUE | 0.2241 |
| Gm1604b | 0 | 0 | 0 | 0 | 0 | 0 | 0 | 0 | 0 | 0 | 0 | 0 | 0 | 0 | 0 | NA | NA | NA | NA | NA | NA | NA | NA | NA | NA |
| Gm16062 | 49 | 31 | 35 | 53 | 40 | 31 | 38 | 33 | 38 | 53 | 45 | 31 | 39.55 | 36 | 43 | 1.178 | 0.236 | 0.46471542 | 0.63379556 | 0 | 0.0624 | 0.0497 | 0.0497 | TRUE | 0.4252 |
| Gm16063 | 51 | 35 | 35 | 33 | 27 | 31 | 40 | 37 | 38 | 33 | 30 | 31 | 34.75 | 38 | 31 | 0.819 | -0.289 | 0.36758197 | 0.54224885 | 0 | 0.0706 | 0.0448 | 0.0448 | TRUE | 0.0129 |
| Gm16065 | 23 | 19 | 23 | 19 | 16 | 27 | 18 | 20 | 25 | 19 | 18 | 27 | 21.08 | 21 | 21 | 1.02 | 0.029 | 0.94582474 | 0.97029019 | 0 | 0.1148 | 0.0814 | 0.0814 | TRUE | 0.2892 |
| Gm16119 | 2 | 2 | 3 | 3 | 2 | 0 | 2 | 2 | 3 | 3 | 2 | 0 | 2.03 | 2 | 2 | 0.762 | -0.392 | 0.77686614 | NA | 0 | 1.1719 | 0.8687 | 0.8687 | TRUE | 0.6406 |
| Gm16130 | 0 | 0 | 0 | 0 | 1 | 1 | 0 | 0 | 0 | 0 | 1 | 1 | 0.35 | 0 | 1 | 3.965 | 1.987 | 0.55529285 | NA | 0 | 6.7252 | 6.4426 | 6.4426 | TRUE | 0.261 |
| Gm16157 | 4 | 1 | 1 | 0 | 0 | 1 | 3 | 1 | 1 | 0 | 0 | 1 | 1.04 | 2 | 0 | 0.208 | -2.267 | 0.26260263 | NA | 0 | 2.2826 | 1.679 | 1.679 | TRUE | 0.4795 |
| Gm16197 | 49 | 39 | 21 | 23 | 34 | 25 | 38 | 41 | 23 | 23 | 38 | 25 | 31.35 | 34 | 29 | 0.833 | -0.263 | 0.4881889 | 0.65413743 | 0.0532 | 0.078 | 0.0713 | 0.0713 | TRUE | 0.5598 |
| Gm16287 | 0 | 0 | 0 | 0 | 0 | 0 | 0 | 0 | 0 | 0 | 0 | 0 | 0 | 0 | 0 | NA | NA | NA | NA | NA | NA | NA | NA | NA | NA |
| Gm16291 | 0 | 0 | 0 | 0 | 0 | 0 | 0 | 0 | 0 | 0 | 0 | 0 | 0 | 0 | 0 | NA | NA | NA | NA | NA | NA | NA | NA | NA | NA |

| | | | | | | | | | | | | | | | | | | | | | | | | | |
|---|---|---|---|---|---|---|---|---|---|---|---|---|---|---|---|---|---|---|---|---|---|---|---|---|---|
| Gm16294 | 0 | 0 | 0 | 0 | 0 | 0 | 0 | 0 | 0 | 0 | 0 | 0 | 0 | 0 | 0 | NA | NA | NA | NA | NA | NA | NA | NA | NA | NA |
| Gm1631 | 271 | 187 | 177 | 259 | 212 | 235 | 210 | 199 | 191 | 258 | 239 | 232 | 221.55 | 200 | 243 | 1.212 | 0.277 | 0.04179521 | 0.11801244 | 0 | 0.0131 | 0.0088 | 0.0088 | TRUE | 0.0323 |
| Gm16325 | 0 | 0 | 0 | 0 | 0 | 0 | 0 | 0 | 0 | 0 | 0 | 0 | 0 | 0 | 0 | NA | NA | NA | NA | NA | NA | NA | NA | NA | NA |
| Gm16336 | 0 | 0 | 0 | 0 | 0 | 0 | 0 | 0 | 0 | 0 | 0 | 0 | 0 | 0 | 0 | NA | NA | NA | NA | NA | NA | NA | NA | NA | NA |
| Gm16367 | 0 | 0 | 0 | 0 | 0 | 0 | 0 | 0 | 0 | 0 | 0 | 0 | 0 | 0 | 0 | NA | NA | NA | NA | NA | NA | NA | NA | NA | NA |
| Gm16381 | 0 | 0 | 0 | 0 | 0 | 0 | 0 | 0 | 0 | 0 | 0 | 0 | 0 | 0 | 0 | NA | NA | NA | NA | NA | NA | NA | NA | NA | NA |
| Gm16386 | 40 | 27 | 22 | 19 | 31 | 33 | 31 | 29 | 24 | 19 | 35 | 33 | 28.33 | 28 | 29 | 1.028 | 0.04 | 0.91724599 | 0.95437118 | 0.0217 | 0.0861 | 0.069 | 0.069 | TRUE | 0.603 |
| Gm16390 | 2 | 0 | 0 | 0 | 1 | 0 | 2 | 0 | 0 | 0 | 1 | 0 | 0.45 | 1 | 0 | 0.68 | -0.556 | 0.85993441 | NA | 0 | 5.3057 | 5.2576 | 5.2576 | TRUE | 0.9726 |
| Gm16404 | 0 | 0 | 0 | 0 | 0 | 0 | 0 | 0 | 0 | 0 | 0 | 0 | 0 | 0 | 0 | NA | NA | NA | NA | NA | NA | NA | NA | NA | NA |
| Gm16405 | 0 | 0 | 0 | 0 | 0 | 0 | 0 | 0 | 0 | 0 | 0 | 0 | 0 | 0 | 0 | NA | NA | NA | NA | NA | NA | NA | NA | NA | NA |
| Gm16430 | 0 | 0 | 0 | 0 | 0 | 0 | 0 | 0 | 0 | 0 | 0 | 0 | 0 | 0 | 0 | NA | NA | NA | NA | NA | NA | NA | NA | NA | NA |
| Gm16432 | 2 | 1 | 1 | 0 | 0 | 1 | 2 | 1 | 1 | 0 | 0 | 1 | 0.78 | 1 | 0 | 0.299 | -1.743 | 0.4255496 | NA | 0 | 3.0389 | 2.0589 | 2.0589 | TRUE | 0.3835 |
| Gm16442 | 0 | 0 | 0 | 0 | 0 | 0 | 0 | 0 | 0 | 0 | 0 | 0 | 0 | 0 | 0 | NA | NA | NA | NA | NA | NA | NA | NA | NA | NA |
| Gm16445 | 0 | 0 | 0 | 0 | 0 | 0 | 0 | 0 | 0 | 0 | 0 | 0 | 0 | 0 | 0 | NA | NA | NA | NA | NA | NA | NA | NA | NA | NA |
| Gm16451 | 0 | 0 | 0 | 0 | 0 | 0 | 0 | 0 | 0 | 0 | 0 | 0 | 0 | 0 | 0 | NA | NA | NA | NA | NA | NA | NA | NA | NA | NA |
| Gm1647 | 1 | 0 | 0 | 0 | 0 | 0 | 1 | 0 | 0 | 0 | 0 | 0 | 0.13 | 0 | 0 | 0.552 | -0.857 | 0.8336548 | NA | 0 | 18.3121 | 10 | 10 | TRUE | 0.2566 |
| Gm16497 | 0 | 0 | 0 | 0 | 0 | 0 | 0 | 0 | 0 | 0 | 0 | 0 | 0 | 0 | 0 | NA | NA | NA | NA | NA | NA | NA | NA | NA | NA |
| Gm16501 | 0 | 0 | 0 | 0 | 0 | 0 | 0 | 0 | 0 | 0 | 0 | 0 | 0 | 0 | 0 | NA | NA | NA | NA | NA | NA | NA | NA | NA | NA |
| Gm16515 | 3870 | 2729 | 2313 | 2561 | 2338 | 2479 | 3006 | 2903 | 2496 | 2549 | 2638 | 2447 | 2673.01 | 2802 | 2545 | 0.908 | -0.14 | 0.07216134 | 0.17552792 | 0.0048 | 0.0033 | 0.004 | 0.004 | TRUE | 0.1097 |
| Gm16516 | 271 | 196 | 236 | 314 | 256 | 332 | 210 | 208 | 255 | 313 | 289 | 328 | 267.11 | 224 | 310 | 1.384 | 0.468 | 0.00049719 | 0.00428926 | 0.0039 | 0.0113 | 0.0092 | 0.0092 | TRUE | 0.1454 |
| Gm16523 | 3 | 1 | 0 | 4 | 2 | 2 | 2 | 1 | 0 | 4 | 2 | 2 | 1.93 | 1 | 3 | 2.298 | 1.2 | 0.4008949 | NA | 0 | 1.2277 | 0.8793 | 0.8793 | TRUE | 0.5542 |
| Gm1653 | 14 | 9 | 10 | 7 | 9 | 13 | 11 | 10 | 11 | 7 | 10 | 13 | 10.2 | 11 | 10 | 0.957 | -0.063 | 0.91606353 | NA | 0 | 0.2348 | 0.1576 | 0.1576 | TRUE | 0.2478 |
| Gm16532 | 39 | 33 | 44 | 53 | 31 | 46 | 30 | 35 | 47 | 53 | 35 | 45 | 41 | 37 | 44 | 1.192 | 0.253 | 0.4317187 | 0.60365441 | 0.023 | 0.0602 | 0.05 | 0.05 | TRUE | 0.3646 |
| Gm16548 | 3 | 1 | 2 | 0 | 1 | 1 | 2 | 1 | 2 | 0 | 1 | 1 | 1.28 | 2 | 1 | 0.373 | -1.422 | 0.41672566 | NA | 0 | 1.8572 | 1.2076 | 1.2076 | TRUE | 0.2661 |
| Gm16551 | 2 | 4 | 4 | 2 | 1 | 6 | 2 | 4 | 4 | 2 | 1 | 6 | 3.19 | 3 | 3 | 0.92 | -0.12 | 0.91409796 | NA | 0 | 0.7444 | 0.5717 | 0.5717 | TRUE | 0.9514 |
| Gm16576 | 74 | 51 | 53 | 94 | 59 | 75 | 57 | 54 | 57 | 94 | 67 | 74 | 67.18 | 56 | 78 | 1.388 | 0.473 | 0.05357155 | 0.14156003 | 0 | 0.0377 | 0.0281 | 0.0281 | TRUE | 0.2814 |
| Gm16596 | 0 | 2 | 2 | 1 | 0 | 2 | 0 | 2 | 2 | 1 | 0 | 2 | 1.21 | 1 | 1 | 0.728 | -0.458 | 0.80519722 | NA | 0 | 1.9625 | 1.6239 | 1.6239 | TRUE | 0.3966 |
| Gm166 | 188 | 180 | 165 | 128 | 139 | 158 | 146 | 191 | 178 | 127 | 157 | 156 | 159.28 | 172 | 147 | 0.857 | -0.223 | 0.1993506 | 0.35998966 | 0.011 | 0.0173 | 0.0154 | 0.0154 | TRUE | 0.2026 |
| Gm1661 | 9 | 7 | 8 | 7 | 10 | 4 | 7 | 7 | 9 | 7 | 11 | 4 | 7.54 | 8 | 7 | 0.957 | -0.063 | 0.92877278 | NA | 0 | 0.3166 | 0.2274 | 0.2274 | TRUE | 0.2922 |
| Gm16617 | 0 | 0 | 0 | 0 | 0 | 0 | 0 | 0 | 0 | 0 | 0 | 0 | 0 | 0 | 0 | NA | NA | NA | NA | NA | NA | NA | NA | NA | NA |
| Gm16675 | 1 | 0 | 1 | 0 | 2 | 0 | 1 | 0 | 1 | 0 | 2 | 0 | 0.69 | 1 | 1 | 1.173 | 0.23 | 0.92656408 | NA | 0 | 3.4608 | 3.0217 | 3.0217 | TRUE | 0.9747 |
| Gm16677 | 0 | 0 | 0 | 0 | 0 | 0 | 0 | 0 | 0 | 0 | 0 | 0 | 0 | 0 | 0 | NA | NA | NA | NA | NA | NA | NA | NA | NA | NA |
| Gm16701 | 0 | 0 | 1 | 0 | 0 | 0 | 0 | 0 | 1 | 0 | 0 | 0 | 0.18 | 0 | 0 | 0.552 | -0.857 | 0.8336548 | NA | 0 | 13.1817 | 10 | 10 | TRUE | 0.5365 |
| Gm16702 | 5 | 4 | 10 | 3 | 4 | 5 | 4 | 4 | 11 | 3 | 5 | 5 | 5.23 | 6 | 4 | 0.665 | -0.589 | 0.49647979 | NA | 0 | 0.4558 | 0.337 | 0.337 | TRUE | 0.8809 |
| Gm1673 | 165 | 132 | 159 | 207 | 163 | 153 | 128 | 140 | 172 | 206 | 184 | 151 | 163.51 | 147 | 180 | 1.233 | 0.303 | 0.09229738 | 0.20873521 | 0.0176 | 0.0169 | 0.0171 | 0.0171 | TRUE | 0.2248 |
| Gm16793 | 2 | 1 | 0 | 2 | 1 | 1 | 2 | 1 | 0 | 2 | 1 | 1 | 1.12 | 1 | 1 | 1.516 | 0.6 | 0.73587335 | NA | 0 | 2.1176 | 1.3707 | 1.3707 | TRUE | 0.2754 |
| Gm16796 | 2 | 1 | 1 | 0 | 0 | 0 | 2 | 1 | 1 | 0 | 0 | 0 | 0.62 | 1 | 0 | 0.154 | -2.702 | 0.27412557 | NA | 0 | 3.8499 | 3.0088 | 3.0088 | TRUE | 0.0718 |
| Gm16833 | 0 | 0 | 0 | 0 | 0 | 0 | 0 | 0 | 0 | 0 | 0 | 0 | 0 | 0 | 0 | NA | NA | NA | NA | NA | NA | NA | NA | NA | NA |
| Gm16845 | 53 | 37 | 12 | 25 | 20 | 23 | 41 | 39 | 13 | 25 | 23 | 23 | 27.27 | 31 | 24 | 0.745 | -0.425 | 0.3339295 | 0.50810999 | 0.1355 | 0.0893 | 0.102 | 0.102 | TRUE | 1.6547 |
| Gm16853 | 2 | 1 | 0 | 1 | 0 | 1 | 2 | 1 | 0 | 1 | 0 | 1 | 0.77 | 1 | 1 | 0.75 | -0.415 | 0.8501676 | NA | 0 | 3.0941 | 2.17 | 2.17 | TRUE | 0.2799 |
| Gm16861 | 10 | 6 | 4 | 2 | 2 | 7 | 8 | 6 | 4 | 2 | 2 | 7 | 4.94 | 6 | 4 | 0.604 | -0.727 | 0.41683786 | NA | 0 | 0.4825 | 0.3623 | 0.3623 | TRUE | 0.9154 |
| Gm16863 | 0 | 0 | 1 | 3 | 4 | 0 | 0 | 0 | 1 | 3 | 5 | 0 | 1.43 | 0 | 3 | 7.195 | 2.847 | 0.15228139 | NA | 0 | 1.6603 | 1.6418 | 1.6418 | TRUE | 0.3576 |
| Gm16880 | 70 | 48 | 35 | 78 | 49 | 50 | 54 | 51 | 38 | 78 | 55 | 49 | 54.24 | 48 | 61 | 1.268 | 0.342 | 0.23521964 | 0.40157601 | 0.0294 | 0.0461 | 0.0414 | 0.0414 | TRUE | 0.5209 |
| Gm16894 | 0 | 0 | 0 | 0 | 0 | 1 | 0 | 0 | 0 | 0 | 0 | 1 | 0.16 | 0 | 0 | 2.094 | 1.067 | 0.79379329 | NA | 0 | 14.4078 | 10 | 10 | TRUE | 0.3835 |
| Gm16897 | 19 | 30 | 16 | 29 | 26 | 15 | 15 | 32 | 17 | 29 | 29 | 15 | 22.82 | 21 | 24 | 1.149 | 0.201 | 0.66153079 | 0.79103653 | 0.1096 | 0.1063 | 0.1072 | 0.1072 | TRUE | 1.0229 |
| Gm16907 | 5 | 1 | 0 | 2 | 3 | 1 | 4 | 1 | 0 | 2 | 3 | 1 | 1.88 | 2 | 2 | 1.213 | 0.279 | 0.84904649 | NA | 0.3783 | 1.2598 | 1.024 | 1.024 | TRUE | 1.1794 |
| Gm16938 | 38 | 33 | 27 | 34 | 30 | 19 | 30 | 35 | 29 | 34 | 34 | 19 | 30.03 | 31 | 29 | 0.921 | -0.119 | 0.74777452 | 0.85030594 | 0.0176 | 0.0813 | 0.0649 | 0.0649 | TRUE | 0.6286 |
| Gm16973 | 167 | 112 | 129 | 109 | 89 | 173 | 130 | 119 | 139 | 108 | 100 | 171 | 127.95 | 129 | 126 | 0.981 | -0.028 | 0.89687219 | 0.94326668 | 0.0377 | 0.0209 | 0.0258 | 0.0258 | TRUE | 0.9627 |
| Gm16982 | 3 | 0 | 0 | 1 | 0 | 0 | 2 | 0 | 0 | 1 | 0 | 0 | 0.55 | 1 | 0 | 0.458 | -1.128 | 0.69820437 | NA | 0 | 4.2787 | 4.502 | 4.502 | TRUE | 1.4399 |
| Gm16998 | 1 | 1 | 2 | 2 | 1 | 4 | 1 | 1 | 2 | 2 | 1 | 4 | 1.84 | 1 | 2 | 1.821 | 0.865 | 0.53903271 | NA | 0 | 1.2875 | 0.8419 | 0.8419 | TRUE | 0.3732 |
| Gm17019 | 0 | 0 | 0 | 0 | 0 | 0 | 0 | 0 | 0 | 0 | 0 | 0 | 0 | 0 | 0 | NA | NA | NA | NA | NA | NA | NA | NA | NA | NA |
| Gm17066 | 2163 | 1721 | 1682 | 1857 | 1595 | 1829 | 1680 | 1831 | 1815 | 1848 | 1799 | 1806 | 1796.44 | 1775 | 1818 | 1.025 | 0.036 | 0.58848687 | 0.73756952 | 8.00E-04 | 0.0037 | 0.0026 | 0.0026 | TRUE | 0.0277 |
| Gm1715 | 0 | 0 | 0 | 0 | 0 | 0 | 0 | 0 | 0 | 0 | 0 | 0 | 0 | 0 | 0 | NA | NA | NA | NA | NA | NA | NA | NA | NA | NA |
| Gm1720 | 0 | 0 | 0 | 0 | 0 | 0 | 0 | 0 | 0 | 0 | 0 | 0 | 0 | 0 | 0 | NA | NA | NA | NA | NA | NA | NA | NA | NA | NA |
| Gm17252 | 0 | 0 | 0 | 0 | 0 | 0 | 0 | 0 | 0 | 0 | 0 | 0 | 0 | 0 | 0 | NA | NA | NA | NA | NA | NA | NA | NA | NA | NA |
| Gm17296 | 2438 | 1884 | 1704 | 1794 | 1647 | 1716 | 1893 | 2004 | 1839 | 1786 | 1858 | 1694 | 1845.63 | 1912 | 1779 | 0.93 | -0.104 | 0.12957627 | 0.26635365 | 0.0014 | 0.0037 | 0.0029 | 0.0029 | TRUE | 0.0213 |
| Gm17359 | 0 | 1 | 0 | 1 | 0 | 0 | 0 | 1 | 0 | 1 | 0 | 0 | 0.34 | 0 | 0 | 1.075 | 0.105 | 0.97573409 | NA | 0 | 6.9086 | 6.5582 | 6.5582 | TRUE | 0.5216 |
| Gm17365 | 0 | 0 | 0 | 0 | 0 | 0 | 0 | 0 | 0 | 0 | 0 | 0 | 0 | 0 | 0 | NA | NA | NA | NA | NA | NA | NA | NA | NA | NA |
| Gm17384 | 0 | 0 | 0 | 0 | 0 | 0 | 0 | 0 | 0 | 0 | 0 | 0 | 0 | 0 | 0 | NA | NA | NA | NA | NA | NA | NA | NA | NA | NA |
| Gm17455 | 1 | 4 | 0 | 0 | 3 | 1 | 1 | 4 | 0 | 0 | 3 | 1 | 1.57 | 2 | 1 | 0.865 | -0.209 | 0.90409321 | NA | 1.5383 | 1.5148 | 1.5204 | 1.5204 | TRUE | 1.0786 |
| Gm17495 | 2 | 2 | 3 | 0 | 2 | 2 | 2 | 2 | 3 | 0 | 2 | 2 | 1.86 | 2 | 1 | 0.612 | -0.709 | 0.62204188 | NA | 0 | 1.278 | 0.9067 | 0.9067 | TRUE | 0.5154 |
| Gm17644 | 0 | 0 | 0 | 0 | 0 | 0 | 0 | 0 | 0 | 0 | 0 | 0 | 0 | 0 | 0 | NA | NA | NA | NA | NA | NA | NA | NA | NA | NA |
| Gm17660 | 0 | 0 | 0 | 0 | 0 | 0 | 0 | 0 | 0 | 0 | 0 | 0 | 0 | 0 | 0 | NA | NA | NA | NA | NA | NA | NA | NA | NA | NA |
| Gm17677 | 0 | 0 | 0 | 0 | 0 | 0 | 0 | 0 | 0 | 0 | 0 | 0 | 0 | 0 | 0 | NA | NA | NA | NA | NA | NA | NA | NA | NA | NA |
| Gm17689 | 0 | 0 | 0 | 0 | 0 | 0 | 0 | 0 | 0 | 0 | 0 | 0 | 0 | 0 | 0 | NA | NA | NA | NA | NA | NA | NA | NA | NA | NA |
| Gm17727 | 0 | 1 | 0 | 0 | 0 | 0 | 0 | 1 | 0 | 0 | 0 | 0 | 0.18 | 0 | 0 | 0.552 | -0.857 | 0.8336548 | NA | 0 | 13.3714 | 10 | 10 | TRUE | 0.5216 |
| Gm17745 | 20 | 12 | 8 | 3 | 2 | 0 | 16 | 13 | 9 | 3 | 2 | 0 | 7.03 | 13 | 2 | 0.14 | -2.84 | 0.00206575 | NA | 0 | 0.3396 | 0.2745 | 0.2745 | TRUE | 0.5367 |
| Gm17746 | 0 | 0 | 0 | 0 | 0 | 1 | 0 | 0 | 0 | 0 | 0 | 1 | 0.16 | 0 | 0 | 2.094 | 1.067 | 0.79379329 | NA | 0 | 14.4078 | 10 | 10 | TRUE | 0.3835 |
| Gm17751 | 0 | 0 | 0 | 0 | 0 | 0 | 0 | 0 | 0 | 0 | 0 | 0 | 0 | 0 | 0 | NA | NA | NA | NA | NA | NA | NA | NA | NA | NA |

| | | | | | | | | | | | | | | | | | | | | | | | | | |
|---|---|---|---|---|---|---|---|---|---|---|---|---|---|---|---|---|---|---|---|---|---|---|---|---|---|
| Gm17757 | 0 | 0 | 0 | 0 | 0 | 0 | 0 | 0 | 0 | 0 | 0 | 0 | 0 | 0 | 0 | NA | NA | NA | NA | NA | NA | NA | NA | NA | NA |
| Gm17762 | 124 | 99 | 125 | 135 | 111 | 147 | 96 | 105 | 135 | 134 | 125 | 145 | 123.53 | 112 | 135 | 1.21 | 0.275 | 0.15129181 | 0.29744345 | 0.0098 | 0.0216 | 0.0183 | 0.0183 | TRUE | 0.3053 |
| Gm17769 | 2 | 4 | 2 | 2 | 0 | 5 | 2 | 4 | 2 | 2 | 0 | 5 | 2.48 | 3 | 2 | 0.895 | -0.16 | 0.90166758 | NA | 0.3126 | 0.9573 | 0.799 | 0.799 | TRUE | 0.3554 |
| Gm17801 | 2 | 3 | 0 | 1 | 2 | 2 | 2 | 3 | 0 | 1 | 2 | 2 | 1.66 | 2 | 2 | 1.095 | 0.131 | 0.93053997 | NA | 0 | 1.4287 | 1.0365 | 1.0365 | TRUE | 0.1941 |
| Gm17821 | 0 | 0 | 0 | 0 | 0 | 0 | 0 | 0 | 0 | 0 | 0 | 0 | 0 | 0 | 0 | NA | NA | NA | NA | NA | NA | NA | NA | NA | NA |
| Gm17830 | 0 | 0 | 0 | 0 | 0 | 0 | 0 | 0 | 0 | 0 | 0 | 0 | 0 | 0 | 0 | NA | NA | NA | NA | NA | NA | NA | NA | NA | NA |
| Gm1821 | 436 | 335 | 340 | 381 | 339 | 339 | 339 | 356 | 367 | 379 | 382 | 335 | 359.69 | 354 | 365 | 1.033 | 0.047 | 0.67823754 | 0.80231161 | 0 | 0.009 | 0.0065 | 0.0065 | TRUE | 0.0633 |
| Gm18409 | 0 | 0 | 0 | 0 | 0 | 0 | 0 | 0 | 0 | 0 | 0 | 0 | 0 | 0 | 0 | NA | NA | NA | NA | NA | NA | NA | NA | NA | NA |
| Gm18853 | 0 | 0 | 0 | 0 | 0 | 0 | 0 | 0 | 0 | 0 | 0 | 0 | 0 | 0 | 0 | NA | NA | NA | NA | NA | NA | NA | NA | NA | NA |
| Gm19276 | 0 | 0 | 0 | 0 | 0 | 0 | 0 | 0 | 0 | 0 | 0 | 0 | 0 | 0 | 0 | NA | NA | NA | NA | NA | NA | NA | NA | NA | NA |
| Gm19277 | 0 | 0 | 0 | 0 | 0 | 0 | 0 | 0 | 0 | 0 | 0 | 0 | 0 | 0 | 0 | NA | NA | NA | NA | NA | NA | NA | NA | NA | NA |
| Gm19299 | 0 | 0 | 1 | 0 | 0 | 0 | 0 | 0 | 1 | 0 | 0 | 0 | 0.18 | 0 | 0 | 0.552 | -0.857 | 0.8336548 | NA | 0 | 13.1817 | 10 | 10 | TRUE | 0.5365 |
| Gm19303 | 0 | 0 | 0 | 0 | 0 | 0 | 0 | 0 | 0 | 0 | 0 | 0 | 0 | 0 | 0 | NA | NA | NA | NA | NA | NA | NA | NA | NA | NA |
| Gm19345 | 36 | 38 | 39 | 42 | 21 | 40 | 28 | 40 | 42 | 42 | 24 | 39 | 35.91 | 37 | 35 | 0.962 | -0.056 | 0.8736749 | 0.92995497 | 0.0378 | 0.0684 | 0.0601 | 0.0601 | TRUE | 0.5224 |
| Gm19395 | 0 | 0 | 0 | 0 | 0 | 0 | 0 | 0 | 0 | 0 | 0 | 0 | 0 | 0 | 0 | NA | NA | NA | NA | NA | NA | NA | NA | NA | NA |
| Gm19402 | 0 | 0 | 0 | 0 | 0 | 0 | 0 | 0 | 0 | 0 | 0 | 0 | 0 | 0 | 0 | NA | NA | NA | NA | NA | NA | NA | NA | NA | NA |
| Gm19424 | 0 | 0 | 0 | 0 | 0 | 0 | 0 | 0 | 0 | 0 | 0 | 0 | 0 | 0 | 0 | NA | NA | NA | NA | NA | NA | NA | NA | NA | NA |
| Gm1943 | 126 | 79 | 79 | 70 | 64 | 86 | 98 | 84 | 85 | 70 | 72 | 85 | 82.32 | 89 | 76 | 0.847 | -0.24 | 0.27087006 | 0.44195727 | 0 | 0.0312 | 0.022 | 0.022 | TRUE | 0.1094 |
| Gm19434 | 0 | 0 | 1 | 0 | 0 | 0 | 0 | 0 | 1 | 0 | 0 | 0 | 0.18 | 0 | 0 | 0.552 | -0.857 | 0.8336548 | NA | 0 | 13.1817 | 10 | 10 | TRUE | 0.5365 |
| Gm19461 | 3 | 1 | 2 | 4 | 0 | 1 | 2 | 1 | 2 | 4 | 0 | 1 | 1.75 | 2 | 2 | 0.901 | -0.151 | 0.91990282 | NA | 0 | 1.3541 | 1.05 | 1.05 | TRUE | 1.0812 |
| Gm19466 | 0 | 0 | 0 | 0 | 0 | 0 | 0 | 0 | 0 | 0 | 0 | 0 | 0 | 0 | 0 | NA | NA | NA | NA | NA | NA | NA | NA | NA | NA |
| Gm19510 | 4 | 1 | 0 | 0 | 1 | 0 | 3 | 1 | 0 | 0 | 1 | 0 | 0.88 | 1 | 0 | 0.259 | -1.947 | 0.39420209 | NA | 0 | 2.6863 | 2.4242 | 2.4242 | TRUE | 0.3212 |
| Gm19522 | 0 | 0 | 0 | 0 | 0 | 0 | 0 | 0 | 0 | 0 | 0 | 0 | 0 | 0 | 0 | NA | NA | NA | NA | NA | NA | NA | NA | NA | NA |
| Gm19557 | 0 | 0 | 0 | 0 | 0 | 0 | 0 | 0 | 0 | 0 | 0 | 0 | 0 | 0 | 0 | NA | NA | NA | NA | NA | NA | NA | NA | NA | NA |
| Gm19583 | 20 | 19 | 12 | 3 | 6 | 10 | 16 | 20 | 13 | 3 | 7 | 10 | 11.39 | 16 | 7 | 0.403 | -1.31 | 0.03404842 | NA | 0.0038 | 0.2106 | 0.1662 | 0.1662 | TRUE | 0.5331 |
| Gm19589 | 0 | 0 | 0 | 0 | 0 | 0 | 0 | 0 | 0 | 0 | 0 | 0 | 0 | 0 | 0 | NA | NA | NA | NA | NA | NA | NA | NA | NA | NA |
| Gm19619 | 0 | 0 | 0 | 0 | 0 | 0 | 0 | 0 | 0 | 0 | 0 | 0 | 0 | 0 | 0 | NA | NA | NA | NA | NA | NA | NA | NA | NA | NA |
| Gm1965 | 0 | 0 | 0 | 0 | 0 | 0 | 0 | 0 | 0 | 0 | 0 | 0 | 0 | 0 | 0 | NA | NA | NA | NA | NA | NA | NA | NA | NA | NA |
| Gm19668 | 0 | 0 | 0 | 0 | 0 | 0 | 0 | 0 | 0 | 0 | 0 | 0 | 0 | 0 | 0 | NA | NA | NA | NA | NA | NA | NA | NA | NA | NA |
| Gm1968 | 0 | 0 | 2 | 0 | 2 | 0 | 0 | 0 | 2 | 0 | 2 | 0 | 0.74 | 1 | 1 | 1.052 | 0.074 | 0.97802666 | NA | 9.2874 | 3.2239 | 3.7544 | 3.7544 | TRUE | 0.9748 |
| Gm19689 | 0 | 0 | 0 | 0 | 0 | 0 | 0 | 0 | 0 | 0 | 0 | 0 | 0 | 0 | 0 | NA | NA | NA | NA | NA | NA | NA | NA | NA | NA |
| Gm19705 | 107 | 87 | 63 | 63 | 43 | 55 | 83 | 93 | 68 | 63 | 49 | 54 | 68.19 | 81 | 55 | 0.68 | -0.556 | 0.0239882 | 0.07895624 | 0 | 0.0372 | 0.0284 | 0.0284 | TRUE | 0.1765 |
| Gm19710 | 4 | 3 | 4 | 6 | 2 | 3 | 3 | 3 | 4 | 6 | 2 | 3 | 3.63 | 3 | 4 | 1.067 | 0.093 | 0.92528323 | NA | 0 | 0.6546 | 0.4324 | 0.4324 | TRUE | 0.4428 |
| Gm19757 | 4 | 2 | 0 | 3 | 0 | 1 | 3 | 2 | 0 | 3 | 0 | 1 | 1.53 | 2 | 1 | 0.752 | -0.41 | 0.80700198 | NA | 0.6399 | 1.5469 | 1.3774 | 1.3774 | TRUE | 0.617 |
| Gm1976 | 60 | 50 | 56 | 100 | 68 | 79 | 47 | 53 | 60 | 100 | 77 | 78 | 69.07 | 53 | 85 | 1.597 | 0.676 | 0.00591926 | 0.02795903 | 0 | 0.0367 | 0.0282 | 0.0282 | TRUE | 0.2228 |
| Gm19782 | 0 | 0 | 0 | 0 | 0 | 0 | 0 | 0 | 0 | 0 | 0 | 0 | 0 | 0 | 0 | NA | NA | NA | NA | NA | NA | NA | NA | NA | NA |
| Gm19784 | 60 | 42 | 19 | 22 | 25 | 28 | 47 | 45 | 21 | 22 | 28 | 28 | 31.59 | 38 | 26 | 0.69 | -0.536 | 0.16857985 | 0.32020091 | 0.0729 | 0.0775 | 0.0762 | 0.0762 | TRUE | 1.0604 |
| Gm1987 | 0 | 0 | 0 | 0 | 0 | 0 | 0 | 0 | 0 | 0 | 0 | 0 | 0 | 0 | 0 | NA | NA | NA | NA | NA | NA | NA | NA | NA | NA |
| Gm19897 | 5 | 0 | 0 | 2 | 1 | 0 | 4 | 0 | 0 | 2 | 1 | 0 | 1.17 | 1 | 1 | 0.763 | -0.391 | 0.84922522 | NA | 2.9812 | 2.0333 | 2.1906 | 2.1906 | TRUE | 1.7221 |
| Gm1993 | 0 | 0 | 0 | 0 | 0 | 0 | 0 | 0 | 0 | 0 | 0 | 0 | 0 | 0 | 0 | NA | NA | NA | NA | NA | NA | NA | NA | NA | NA |
| Gm1995 | 302 | 269 | 90 | 94 | 113 | 80 | 235 | 286 | 97 | 94 | 127 | 79 | 152.97 | 206 | 100 | 0.485 | -1.044 | 0.03220847 | 0.09753299 | 0.1636 | 0.0179 | 0.0576 | 0.1636 | TRUE | 0.4516 |
| Gm19990 | 0 | 0 | 0 | 0 | 0 | 0 | 0 | 0 | 0 | 0 | 0 | 0 | 0 | 0 | 0 | NA | NA | NA | NA | NA | NA | NA | NA | NA | NA |
| Gm2002 | 0 | 0 | 0 | 0 | 0 | 0 | 0 | 0 | 0 | 0 | 0 | 0 | 0 | 0 | 0 | NA | NA | NA | NA | NA | NA | NA | NA | NA | NA |
| Gm20063 | 0 | 0 | 0 | 0 | 0 | 0 | 0 | 0 | 0 | 0 | 0 | 0 | 0 | 0 | 0 | NA | NA | NA | NA | NA | NA | NA | NA | NA | NA |
| Gm20098 | 0 | 1 | 0 | 0 | 0 | 0 | 0 | 1 | 0 | 0 | 0 | 0 | 0.18 | 0 | 0 | 0.552 | -0.857 | 0.8336548 | NA | 0 | 13.3714 | 10 | 10 | TRUE | 0.5216 |
| Gm2011 | 5 | 3 | 1 | 6 | 2 | 5 | 4 | 3 | 1 | 6 | 2 | 5 | 3.55 | 3 | 4 | 1.594 | 0.672 | 0.51381285 | NA | 0 | 0.6695 | 0.4733 | 0.4733 | TRUE | 0.2959 |
| Gm20110 | 0 | 0 | 0 | 0 | 0 | 0 | 0 | 0 | 0 | 0 | 0 | 0 | 0 | 0 | 0 | NA | NA | NA | NA | NA | NA | NA | NA | NA | NA |
| Gm2012 | 2 | 0 | 1 | 0 | 0 | 0 | 2 | 0 | 1 | 0 | 0 | 0 | 0.44 | 1 | 0 | 0.214 | -2.223 | 0.47051571 | NA | 0 | 5.4047 | 5.2946 | 5.2946 | TRUE | 0.2788 |
| Gm20125 | 1 | 0 | 0 | 0 | 0 | 0 | 1 | 0 | 0 | 0 | 0 | 0 | 0.13 | 0 | 0 | 0.552 | -0.857 | 0.8336548 | NA | 0 | 18.3121 | 10 | 10 | TRUE | 0.2566 |
| Gm20139 | 0 | 0 | 0 | 0 | 0 | 0 | 0 | 0 | 0 | 0 | 0 | 0 | 0 | 0 | 0 | NA | NA | NA | NA | NA | NA | NA | NA | NA | NA |
| Gm2016 | 1457 | 1038 | 447 | 424 | 421 | 354 | 1132 | 1104 | 482 | 422 | 475 | 349 | 660.74 | 906 | 415 | 0.458 | -1.125 | 0.00474979 | 0.02355614 | 0.1127 | 0.006 | 0.0332 | 0.1127 | TRUE | 1.9881 |
| Gm20172 | 0 | 0 | 0 | 0 | 0 | 0 | 0 | 0 | 0 | 0 | 0 | 0 | 0 | 0 | 0 | NA | NA | NA | NA | NA | NA | NA | NA | NA | NA |
| Gm20187 | 0 | 1 | 2 | 1 | 2 | 1 | 0 | 1 | 2 | 1 | 2 | 1 | 1.24 | 1 | 1 | 1.364 | 0.448 | 0.79696708 | NA | 0 | 1.9085 | 1.3477 | 1.3477 | TRUE | 0.3082 |
| Gm20199 | 2 | 5 | 3 | 9 | 3 | 5 | 2 | 5 | 3 | 9 | 3 | 5 | 4.56 | 3 | 6 | 1.76 | 0.816 | 0.38236396 | NA | 0 | 0.5217 | 0.3925 | 0.3925 | TRUE | 0.4996 |
| Gm2022 | 1328 | 1027 | 446 | 418 | 415 | 302 | 1031 | 1092 | 481 | 416 | 468 | 298 | 631.23 | 868 | 394 | 0.454 | -1.14 | 0.00479181 | 0.02371617 | 0.1159 | 0.0062 | 0.034 | 0.1159 | TRUE | 1.8042 |
| Gm20257 | 1 | 0 | 0 | 0 | 0 | 0 | 0 | 1 | 0 | 0 | 0 | 0 | 0.13 | 0 | 0 | 0.552 | -0.857 | 0.8336548 | NA | 0 | 18.3121 | 10 | 10 | TRUE | 0.2566 |
| Gm20268 | 0 | 0 | 0 | 0 | 0 | 0 | 0 | 0 | 0 | 0 | 0 | 0 | 0 | 0 | 0 | NA | NA | NA | NA | NA | NA | NA | NA | NA | NA |
| Gm2027 | 263 | 199 | 61 | 86 | 72 | 76 | 204 | 212 | 66 | 86 | 81 | 75 | 120.6 | 161 | 81 | 0.502 | -0.996 | 0.04911804 | 0.1329881 | 0.1751 | 0.0221 | 0.0644 | 0.1751 | TRUE | 2.7793 |
| Gm2030 | 1 | 2 | 0 | 0 | 0 | 0 | 0 | 1 | 2 | 0 | 0 | 0 | 0.48 | 1 | 0 | 0.197 | -2.343 | 0.43249452 | NA | 0 | 4.8992 | 4.9172 | 4.9172 | TRUE | 0.1226 |
| Gm20300 | 353 | 279 | 275 | 262 | 243 | 273 | 274 | 297 | 297 | 261 | 274 | 269 | 278.68 | 289 | 268 | 0.928 | -0.107 | 0.37933062 | 0.55390237 | 0 | 0.0109 | 0.0071 | 0.0071 | TRUE | 0.0255 |
| Gm20324 | 16 | 6 | 9 | 7 | 5 | 8 | 12 | 6 | 10 | 7 | 6 | 8 | 8.17 | 9 | 7 | 0.714 | -0.486 | 0.46945149 | NA | 0 | 0.2925 | 0.1998 | 0.1998 | TRUE | 0.2022 |
| Gm20337 | 37 | 28 | 21 | 21 | 20 | 27 | 29 | 30 | 23 | 21 | 23 | 27 | 25.22 | 27 | 24 | 0.862 | -0.214 | 0.57519813 | 0.72711578 | 0 | 0.0964 | 0.065 | 0.065 | TRUE | 0.1178 |
| Gm20356 | 0 | 0 | 0 | 0 | 0 | 0 | 0 | 0 | 0 | 0 | 0 | 0 | 0 | 0 | 0 | NA | NA | NA | NA | NA | NA | NA | NA | NA | NA |
| Gm20362 | 1 | 0 | 0 | 0 | 0 | 1 | 1 | 0 | 0 | 0 | 0 | 1 | 0.29 | 0 | 0 | 1.075 | 0.105 | 0.97702352 | NA | 0 | 8.0647 | 7.5458 | 7.5458 | TRUE | 0.3835 |
| Gm20408 | 0 | 0 | 0 | 0 | 0 | 0 | 0 | 0 | 0 | 0 | 0 | 0 | 0 | 0 | 0 | NA | NA | NA | NA | NA | NA | NA | NA | NA | NA |
| Gm20410 | 0 | 0 | 0 | 0 | 0 | 0 | 0 | 0 | 0 | 0 | 0 | 0 | 0 | 0 | 0 | NA | NA | NA | NA | NA | NA | NA | NA | NA | NA |
| Gm2042 | 0 | 0 | 0 | 0 | 0 | 0 | 0 | 0 | 0 | 0 | 0 | 0 | 0 | 0 | 0 | NA | NA | NA | NA | NA | NA | NA | NA | NA | NA |
| Gm20516 | 0 | 0 | 0 | 0 | 0 | 0 | 0 | 0 | 0 | 0 | 0 | 0 | 0 | 0 | 0 | NA | NA | NA | NA | NA | NA | NA | NA | NA | NA |
| Gm20554 | 16 | 13 | 9 | 11 | 7 | 19 | 12 | 14 | 10 | 11 | 8 | 19 | 12.26 | 12 | 13 | 1.05 | 0.07 | 0.90095383 | NA | 0 | 0.1957 | 0.1497 | 0.1497 | TRUE | 0.7482 |

| | | | | | | | | | | | | | | | | | | | | | | | | | |
|---|---|---|---|---|---|---|---|---|---|---|---|---|---|---|---|---|---|---|---|---|---|---|---|---|---|
| Gm20556 | 2 | 3 | 3 | 3 | 3 | 3 | 2 | 3 | 3 | 3 | 3 | 3 | 2.89 | 3 | 3 | 1.189 | 0.25 | 0.81908496 | NA | 0 | 0.8238 | 0.5119 | 0.5119 | TRUE | 0.2175 |
| Gm20579 | 0 | 0 | 0 | 0 | 0 | 0 | 0 | 0 | 0 | 0 | 0 | 0 | 0 | 0 | 0 | NA | NA | NA | NA | NA | NA | NA | NA | NA | NA |
| Gm20580 | 1 | 0 | 0 | 0 | 0 | 0 | 1 | 0 | 0 | 0 | 0 | 0 | 0.13 | 0 | 0 | 0.552 | -0.857 | 0.8336548 | NA | 0 | 18.3121 | 10 | 10 | TRUE | 0.2566 |
| Gm20581 | 0 | 0 | 0 | 0 | 0 | 0 | 0 | 0 | 0 | 0 | 0 | 0 | 0 | 0 | 0 | NA | NA | NA | NA | NA | NA | NA | NA | NA | NA |
| Gm20594 | 0 | 0 | 0 | 0 | 0 | 0 | 0 | 0 | 0 | 0 | 0 | 0 | 0 | 0 | 0 | NA | NA | NA | NA | NA | NA | NA | NA | NA | NA |
| Gm20597 | 2 | 6 | 6 | 2 | 0 | 3 | 2 | 6 | 6 | 2 | 0 | 3 | 3.23 | 5 | 2 | 0.357 | -1.485 | 0.21611287 | NA | 0 | 0.7369 | 0.629 | 0.629 | TRUE | 0.9135 |
| Gm20604 | 44 | 36 | 36 | 38 | 35 | 37 | 34 | 38 | 39 | 38 | 39 | 37 | 37.52 | 37 | 38 | 1.025 | 0.036 | 0.9073221 | 0.94923397 | 0 | 0.0656 | 0.0421 | 0.0421 | TRUE | 0.0417 |
| Gm20605 | 59 | 61 | 57 | 57 | 58 | 71 | 46 | 65 | 62 | 57 | 65 | 70 | 60.74 | 58 | 64 | 1.124 | 0.169 | 0.51463303 | 0.67719112 | 0 | 0.0414 | 0.032 | 0.032 | TRUE | 0.31 |
| Gm2061 | 25 | 17 | 29 | 30 | 16 | 20 | 19 | 18 | 31 | 30 | 18 | 20 | 22.74 | 23 | 23 | 0.992 | -0.012 | 0.977961 | 0.98819346 | 0.0407 | 0.1066 | 0.0883 | 0.0883 | TRUE | 0.5535 |
| Gm20611 | 0 | 0 | 0 | 0 | 0 | 0 | 0 | 0 | 0 | 0 | 0 | 0 | 0 | 0 | 0 | NA | NA | NA | NA | NA | NA | NA | NA | NA | NA |
| Gm20735 | 0 | 0 | 1 | 2 | 1 | 0 | 0 | 0 | 1 | 2 | 1 | 0 | 0.7 | 0 | 1 | 3.01 | 1.59 | 0.51808674 | NA | 0 | 3.3899 | 2.8624 | 2.8624 | TRUE | 0.3541 |
| Gm20736 | 0 | 0 | 0 | 0 | 0 | 0 | 0 | 0 | 0 | 0 | 0 | 0 | 0 | 0 | 0 | NA | NA | NA | NA | NA | NA | NA | NA | NA | NA |
| Gm20738 | 0 | 0 | 0 | 0 | 0 | 0 | 0 | 0 | 0 | 0 | 0 | 0 | 0 | 0 | 0 | NA | NA | NA | NA | NA | NA | NA | NA | NA | NA |
| Gm20740 | 0 | 0 | 0 | 0 | 0 | 0 | 0 | 0 | 0 | 0 | 0 | 0 | 0 | 0 | 0 | NA | NA | NA | NA | NA | NA | NA | NA | NA | NA |
| Gm20741 | 0 | 0 | 0 | 0 | 0 | 0 | 0 | 0 | 0 | 0 | 0 | 0 | 0 | 0 | 0 | NA | NA | NA | NA | NA | NA | NA | NA | NA | NA |
| Gm20743 | 0 | 0 | 0 | 0 | 0 | 0 | 0 | 0 | 0 | 0 | 0 | 0 | 0 | 0 | 0 | NA | NA | NA | NA | NA | NA | NA | NA | NA | NA |
| Gm20744 | 0 | 0 | 0 | 0 | 0 | 1 | 0 | 0 | 0 | 0 | 0 | 1 | 0.16 | 0 | 0 | 2.094 | 1.067 | 0.79379329 | NA | 0 | 14.4078 | 10 | 10 | TRUE | 0.3835 |
| Gm20745 | 0 | 0 | 0 | 0 | 0 | 0 | 0 | 0 | 0 | 0 | 0 | 0 | 0 | 0 | 0 | NA | NA | NA | NA | NA | NA | NA | NA | NA | NA |
| Gm20747 | 0 | 0 | 0 | 0 | 0 | 0 | 0 | 0 | 0 | 0 | 0 | 0 | 0 | 0 | 0 | NA | NA | NA | NA | NA | NA | NA | NA | NA | NA |
| Gm20748 | 194 | 146 | 176 | 197 | 172 | 227 | 151 | 155 | 190 | 196 | 194 | 224 | 185.01 | 165 | 205 | 1.243 | 0.314 | 0.04644527 | 0.12768653 | 0.0057 | 0.0152 | 0.0125 | 0.0125 | TRUE | 0.1743 |
| Gm20750 | 2 | 0 | 0 | 0 | 1 | 0 | 2 | 0 | 0 | 0 | 1 | 0 | 0.45 | 1 | 0 | 0.68 | -0.556 | 0.85993441 | NA | 0 | 5.3057 | 5.2576 | 5.2576 | TRUE | 0.9726 |
| Gm20751 | 10 | 10 | 17 | 11 | 8 | 13 | 8 | 11 | 18 | 11 | 9 | 13 | 11.59 | 12 | 11 | 0.908 | -0.14 | 0.81062546 | NA | 0 | 0.2069 | 0.1586 | 0.1586 | TRUE | 0.7212 |
| Gm20752 | 0 | 0 | 0 | 0 | 0 | 0 | 0 | 0 | 0 | 0 | 0 | 0 | 0 | 0 | 0 | NA | NA | NA | NA | NA | NA | NA | NA | NA | NA |
| Gm20753 | 0 | 0 | 0 | 0 | 0 | 1 | 0 | 0 | 0 | 0 | 0 | 1 | 0.16 | 0 | 0 | 2.094 | 1.067 | 0.79379329 | NA | 0 | 14.4078 | 10 | 10 | TRUE | 0.3835 |
| Gm20754 | 0 | 0 | 0 | 0 | 0 | 0 | 0 | 0 | 0 | 0 | 0 | 0 | 0 | 0 | 0 | NA | NA | NA | NA | NA | NA | NA | NA | NA | NA |
| Gm20755 | 1 | 0 | 0 | 0 | 0 | 0 | 1 | 0 | 0 | 0 | 0 | 0 | 0.13 | 0 | 0 | 0.552 | -0.857 | 0.8336548 | NA | 0 | 18.3121 | 10 | 10 | TRUE | 0.2566 |
| Gm20756 | 8 | 7 | 1 | 3 | 1 | 2 | 6 | 7 | 1 | 3 | 1 | 2 | 3.47 | 5 | 2 | 0.413 | -1.274 | 0.25321741 | NA | 0 | 0.6852 | 0.546 | 0.546 | TRUE | 0.8386 |
| Gm20757 | 1 | 0 | 0 | 0 | 0 | 0 | 1 | 0 | 0 | 0 | 0 | 0 | 0.13 | 0 | 0 | 0.552 | -0.857 | 0.8336548 | NA | 0 | 18.3121 | 10 | 10 | TRUE | 0.2566 |
| Gm20758 | 1 | 0 | 0 | 1 | 0 | 0 | 1 | 0 | 0 | 1 | 0 | 0 | 0.3 | 0 | 0 | 1.075 | 0.105 | 0.97698525 | NA | 0 | 8.0272 | 7.5141 | 7.5141 | TRUE | 0.3906 |
| Gm20759 | 0 | 0 | 0 | 0 | 1 | 0 | 0 | 0 | 0 | 0 | 1 | 0 | 0.19 | 0 | 0 | 2.094 | 1.067 | 0.79379329 | NA | 0 | 12.608 | 10 | 10 | TRUE | 0.5075 |
| Gm20765 | 0 | 0 | 0 | 0 | 0 | 0 | 0 | 0 | 0 | 0 | 0 | 0 | 0 | 0 | 0 | NA | NA | NA | NA | NA | NA | NA | NA | NA | NA |
| Gm20767 | 161 | 123 | 61 | 77 | 75 | 56 | 125 | 131 | 66 | 77 | 85 | 55 | 89.7 | 107 | 72 | 0.67 | -0.579 | 0.03538882 | 0.10474477 | 0.0783 | 0.0288 | 0.0428 | 0.0428 | TRUE | 1.0867 |
| Gm20806 | 0 | 0 | 0 | 0 | 0 | 0 | 0 | 0 | 0 | 0 | 0 | 0 | 0 | 0 | 0 | NA | NA | NA | NA | NA | NA | NA | NA | NA | NA |
| Gm20809 | 0 | 0 | 0 | 0 | 0 | 0 | 0 | 0 | 0 | 0 | 0 | 0 | 0 | 0 | 0 | NA | NA | NA | NA | NA | NA | NA | NA | NA | NA |
| Gm20815 | 0 | 0 | 0 | 0 | 0 | 0 | 0 | 0 | 0 | 0 | 0 | 0 | 0 | 0 | 0 | NA | NA | NA | NA | NA | NA | NA | NA | NA | NA |
| Gm20816 | 0 | 0 | 0 | 0 | 0 | 0 | 0 | 0 | 0 | 0 | 0 | 0 | 0 | 0 | 0 | NA | NA | NA | NA | NA | NA | NA | NA | NA | NA |
| Gm20822 | 0 | 0 | 0 | 0 | 0 | 0 | 0 | 0 | 0 | 0 | 0 | 0 | 0 | 0 | 0 | NA | NA | NA | NA | NA | NA | NA | NA | NA | NA |
| Gm20823 | 0 | 0 | 0 | 0 | 0 | 0 | 0 | 0 | 0 | 0 | 0 | 0 | 0 | 0 | 0 | NA | NA | NA | NA | NA | NA | NA | NA | NA | NA |
| Gm20826 | 0 | 0 | 0 | 0 | 0 | 0 | 0 | 0 | 0 | 0 | 0 | 0 | 0 | 0 | 0 | NA | NA | NA | NA | NA | NA | NA | NA | NA | NA |
| Gm2083 | 0 | 0 | 0 | 0 | 0 | 0 | 0 | 0 | 0 | 0 | 0 | 0 | 0 | 0 | 0 | NA | NA | NA | NA | NA | NA | NA | NA | NA | NA |
| Gm20831 | 0 | 0 | 0 | 0 | 0 | 0 | 0 | 0 | 0 | 0 | 0 | 0 | 0 | 0 | 0 | NA | NA | NA | NA | NA | NA | NA | NA | NA | NA |
| Gm20854 | 0 | 0 | 0 | 0 | 0 | 0 | 0 | 0 | 0 | 0 | 0 | 0 | 0 | 0 | 0 | NA | NA | NA | NA | NA | NA | NA | NA | NA | NA |
| Gm20857 | 0 | 0 | 0 | 0 | 0 | 0 | 0 | 0 | 0 | 0 | 0 | 0 | 0 | 0 | 0 | NA | NA | NA | NA | NA | NA | NA | NA | NA | NA |
| Gm20858 | 0 | 0 | 0 | 0 | 0 | 0 | 0 | 0 | 0 | 0 | 0 | 0 | 0 | 0 | 0 | NA | NA | NA | NA | NA | NA | NA | NA | NA | NA |
| Gm20865 | 0 | 0 | 0 | 0 | 0 | 0 | 0 | 0 | 0 | 0 | 0 | 0 | 0 | 0 | 0 | NA | NA | NA | NA | NA | NA | NA | NA | NA | NA |
| Gm20867 | 0 | 0 | 0 | 0 | 0 | 0 | 0 | 0 | 0 | 0 | 0 | 0 | 0 | 0 | 0 | NA | NA | NA | NA | NA | NA | NA | NA | NA | NA |
| Gm20871 | 0 | 0 | 0 | 0 | 0 | 0 | 0 | 0 | 0 | 0 | 0 | 0 | 0 | 0 | 0 | NA | NA | NA | NA | NA | NA | NA | NA | NA | NA |
| Gm20877 | 0 | 0 | 0 | 0 | 0 | 0 | 0 | 0 | 0 | 0 | 0 | 0 | 0 | 0 | 0 | NA | NA | NA | NA | NA | NA | NA | NA | NA | NA |
| Gm20878 | 0 | 0 | 0 | 0 | 0 | 0 | 0 | 0 | 0 | 0 | 0 | 0 | 0 | 0 | 0 | NA | NA | NA | NA | NA | NA | NA | NA | NA | NA |
| Gm20917 | 0 | 0 | 0 | 0 | 0 | 0 | 0 | 0 | 0 | 0 | 0 | 0 | 0 | 0 | 0 | NA | NA | NA | NA | NA | NA | NA | NA | NA | NA |
| Gm20939 | 761 | 577 | 450 | 459 | 393 | 514 | 591 | 614 | 486 | 457 | 443 | 507 | 516.33 | 564 | 469 | 0.832 | -0.265 | 0.02025752 | 0.06986986 | 0.0083 | 0.007 | 0.0074 | 0.0074 | TRUE | 0.1662 |
| Gm21002 | 0 | 0 | 0 | 0 | 0 | 0 | 0 | 0 | 0 | 0 | 0 | 0 | 0 | 0 | 0 | NA | NA | NA | NA | NA | NA | NA | NA | NA | NA |
| Gm21057 | 0 | 0 | 0 | 0 | 0 | 0 | 0 | 0 | 0 | 0 | 0 | 0 | 0 | 0 | 0 | NA | NA | NA | NA | NA | NA | NA | NA | NA | NA |
| Gm2109 | 4 | 3 | 1 | 1 | 1 | 3 | 3 | 3 | 1 | 1 | 1 | 3 | 2.08 | 2 | 2 | 0.687 | -0.541 | 0.68212909 | NA | 0 | 1.1436 | 0.7597 | 0.7597 | TRUE | 0.3415 |
| Gm21119 | 1 | 0 | 3 | 0 | 0 | 0 | 1 | 0 | 3 | 0 | 0 | 0 | 0.67 | 1 | 0 | 0.143 | -2.802 | 0.29586147 | NA | 6.534 | 3.5454 | 3.8112 | 3.8112 | TRUE | 0.3623 |
| Gm2115 | 0 | 0 | 0 | 1 | 0 | 0 | 0 | 0 | 0 | 1 | 0 | 0 | 0.17 | 0 | 0 | 2.094 | 1.067 | 0.79379329 | NA | 0 | 14.2885 | 10 | 10 | TRUE | 0.3906 |
| Gm21276 | 0 | 0 | 0 | 0 | 0 | 0 | 0 | 0 | 0 | 0 | 0 | 0 | 0 | 0 | 0 | NA | NA | NA | NA | NA | NA | NA | NA | NA | NA |
| Gm21293 | 0 | 0 | 0 | 0 | 0 | 0 | 0 | 0 | 0 | 0 | 0 | 0 | 0 | 0 | 0 | NA | NA | NA | NA | NA | NA | NA | NA | NA | NA |
| Gm21304 | 0 | 0 | 0 | 0 | 0 | 0 | 0 | 0 | 0 | 0 | 0 | 0 | 0 | 0 | 0 | NA | NA | NA | NA | NA | NA | NA | NA | NA | NA |
| Gm21312 | 0 | 0 | 0 | 0 | 0 | 0 | 0 | 0 | 0 | 0 | 0 | 0 | 0 | 0 | 0 | NA | NA | NA | NA | NA | NA | NA | NA | NA | NA |
| Gm21319 | 13 | 13 | 5 | 11 | 3 | 5 | 10 | 14 | 5 | 11 | 3 | 5 | 8.1 | 10 | 6 | 0.661 | -0.597 | 0.41129137 | NA | 0.1309 | 0.2951 | 0.2512 | 0.2512 | TRUE | 0.3995 |
| Gm21498 | 0 | 0 | 0 | 0 | 0 | 0 | 0 | 0 | 0 | 0 | 0 | 0 | 0 | 0 | 0 | NA | NA | NA | NA | NA | NA | NA | NA | NA | NA |
| Gm21541 | 0 | 0 | 0 | 0 | 0 | 0 | 0 | 0 | 0 | 0 | 0 | 0 | 0 | 0 | 0 | NA | NA | NA | NA | NA | NA | NA | NA | NA | NA |
| Gm21586 | 0 | 0 | 0 | 0 | 0 | 0 | 0 | 0 | 0 | 0 | 0 | 0 | 0 | 0 | 0 | NA | NA | NA | NA | NA | NA | NA | NA | NA | NA |
| Gm216 | 38 | 35 | 37 | 36 | 29 | 31 | 30 | 37 | 40 | 36 | 33 | 31 | 34.3 | 36 | 33 | 0.936 | -0.096 | 0.7711783 | 0.864506 | 0 | 0.0715 | 0.049 | 0.049 | TRUE | 0.1902 |
| Gm21637 | 0 | 0 | 0 | 0 | 0 | 0 | 0 | 0 | 0 | 0 | 0 | 0 | 0 | 0 | 0 | NA | NA | NA | NA | NA | NA | NA | NA | NA | NA |
| Gm21677 | 0 | 0 | 0 | 0 | 0 | 0 | 0 | 0 | 0 | 0 | 0 | 0 | 0 | 0 | 0 | NA | NA | NA | NA | NA | NA | NA | NA | NA | NA |
| Gm21693 | 0 | 0 | 0 | 0 | 0 | 0 | 0 | 0 | 0 | 0 | 0 | 0 | 0 | 0 | 0 | NA | NA | NA | NA | NA | NA | NA | NA | NA | NA |
| Gm21704 | 0 | 0 | 0 | 0 | 0 | 0 | 0 | 0 | 0 | 0 | 0 | 0 | 0 | 0 | 0 | NA | NA | NA | NA | NA | NA | NA | NA | NA | NA |

| | | | | | | | | | | | | | | | | | | | | | | | | | |
|---|---|---|---|---|---|---|---|---|---|---|---|---|---|---|---|---|---|---|---|---|---|---|---|---|---|
| Gm21708 | 0 | 0 | 0 | 0 | 0 | 0 | 0 | 0 | 0 | 0 | 0 | 0 | 0 | 0 | 0 | NA | NA | NA | NA | NA | NA | NA | NA | NA | NA |
| Gm2176 | 0 | 0 | 0 | 0 | 0 | 0 | 0 | 0 | 0 | 0 | 0 | 0 | 0 | 0 | 0 | NA | NA | NA | NA | NA | NA | NA | NA | NA | NA |
| Gm21943 | 0 | 0 | 0 | 0 | 0 | 0 | 0 | 0 | 0 | 0 | 0 | 0 | 0 | 0 | 0 | NA | NA | NA | NA | NA | NA | NA | NA | NA | NA |
| Gm21944 | 0 | 0 | 0 | 0 | 0 | 0 | 0 | 0 | 0 | 0 | 0 | 0 | 0 | 0 | 0 | NA | NA | NA | NA | NA | NA | NA | NA | NA | NA |
| Gm21945 | 0 | 0 | 0 | 0 | 0 | 0 | 0 | 0 | 0 | 0 | 0 | 0 | 0 | 0 | 0 | NA | NA | NA | NA | NA | NA | NA | NA | NA | NA |
| Gm21946 | 0 | 0 | 0 | 0 | 0 | 0 | 0 | 0 | 0 | 0 | 0 | 0 | 0 | 0 | 0 | NA | NA | NA | NA | NA | NA | NA | NA | NA | NA |
| Gm21947 | 8 | 8 | 1 | 3 | 4 | 1 | 6 | 9 | 1 | 3 | 5 | 1 | 4.05 | 5 | 3 | 0.528 | -0.921 | 0.3798418 | NA | 0.3186 | 0.5879 | 0.5171 | 0.5171 | TRUE | 0.3659 |
| Gm21949 | 0 | 0 | 0 | 0 | 0 | 0 | 0 | 0 | 0 | 0 | 0 | 0 | 0 | 0 | 0 | NA | NA | NA | NA | NA | NA | NA | NA | NA | NA |
| Gm21950 | 0 | 0 | 0 | 0 | 0 | 0 | 0 | 0 | 0 | 0 | 0 | 0 | 0 | 0 | 0 | NA | NA | NA | NA | NA | NA | NA | NA | NA | NA |
| Gm21951 | 0 | 0 | 0 | 0 | 0 | 0 | 0 | 0 | 0 | 0 | 0 | 0 | 0 | 0 | 0 | NA | NA | NA | NA | NA | NA | NA | NA | NA | NA |
| Gm2381 | 1151 | 930 | 721 | 731 | 613 | 759 | 894 | 989 | 778 | 728 | 692 | 749 | 804.92 | 887 | 723 | 0.815 | -0.295 | 0.00295209 | 0.01632249 | 0.0068 | 0.0054 | 0.0058 | 0.0058 | TRUE | 0.1333 |
| Gm2382 | 135 | 90 | 97 | 84 | 72 | 90 | 105 | 96 | 105 | 84 | 81 | 89 | 93.15 | 102 | 85 | 0.831 | -0.267 | 0.18186208 | 0.33745782 | 0 | 0.0279 | 0.018 | 0.018 | TRUE | 0.0251 |
| Gm2447 | 0 | 0 | 0 | 0 | 0 | 0 | 0 | 0 | 0 | 0 | 0 | 0 | 0 | 0 | 0 | NA | NA | NA | NA | NA | NA | NA | NA | NA | NA |
| Gm2506 | 0 | 0 | 0 | 0 | 0 | 0 | 0 | 0 | 0 | 0 | 0 | 0 | 0 | 0 | 0 | NA | NA | NA | NA | NA | NA | NA | NA | NA | NA |
| Gm2516 | 0 | 0 | 0 | 0 | 0 | 0 | 0 | 0 | 0 | 0 | 0 | 0 | 0 | 0 | 0 | NA | NA | NA | NA | NA | NA | NA | NA | NA | NA |
| Gm2518 | 9 | 5 | 2 | 5 | 3 | 3 | 7 | 5 | 2 | 5 | 3 | 3 | 4.3 | 5 | 4 | 0.771 | -0.375 | 0.68690504 | NA | 0 | 0.5538 | 0.3876 | 0.3876 | TRUE | 0.3678 |
| Gm266 | 35 | 35 | 42 | 62 | 36 | 53 | 27 | 37 | 45 | 62 | 41 | 52 | 44.06 | 36 | 52 | 1.427 | 0.513 | 0.10598618 | 0.23088866 | 0.0307 | 0.0562 | 0.0492 | 0.0492 | TRUE | 0.2956 |
| Gm2663 | 0 | 0 | 0 | 0 | 0 | 0 | 0 | 0 | 0 | 0 | 0 | 0 | 0 | 0 | 0 | NA | NA | NA | NA | NA | NA | NA | NA | NA | NA |
| Gm2694 | 847 | 635 | 677 | 699 | 560 | 661 | 658 | 675 | 730 | 696 | 632 | 653 | 673.96 | 688 | 660 | 0.961 | -0.057 | 0.52508573 | 0.68627014 | 0.0013 | 0.0059 | 0.0044 | 0.0044 | TRUE | 0.034 |
| Gm2696 | 0 | 0 | 0 | 0 | 0 | 0 | 0 | 0 | 0 | 0 | 0 | 0 | 0 | 0 | 0 | NA | NA | NA | NA | NA | NA | NA | NA | NA | NA |
| Gm2721 | 0 | 0 | 0 | 2 | 2 | 2 | 0 | 0 | 0 | 2 | 2 | 2 | 1.04 | 0 | 2 | 11.705 | 3.549 | 0.08512207 | NA | 0 | 2.2882 | 1.8123 | 1.8123 | TRUE | 0.0847 |
| Gm2762 | 50 | 29 | 26 | 28 | 32 | 21 | 39 | 31 | 28 | 28 | 36 | 21 | 30.41 | 33 | 28 | 0.857 | -0.222 | 0.54333154 | 0.70095114 | 0.0176 | 0.0804 | 0.0633 | 0.0633 | TRUE | 0.2376 |
| Gm2799 | 0 | 0 | 0 | 0 | 0 | 0 | 0 | 0 | 0 | 0 | 0 | 0 | 0 | 0 | 0 | NA | NA | NA | NA | NA | NA | NA | NA | NA | NA |
| Gm2825 | 0 | 0 | 0 | 0 | 0 | 0 | 0 | 0 | 0 | 0 | 0 | 0 | 0 | 0 | 0 | NA | NA | NA | NA | NA | NA | NA | NA | NA | NA |
| Gm2837 | 0 | 0 | 0 | 0 | 0 | 0 | 0 | 0 | 0 | 0 | 0 | 0 | 0 | 0 | 0 | NA | NA | NA | NA | NA | NA | NA | NA | NA | NA |
| Gm2848 | 9 | 7 | 9 | 6 | 1 | 6 | 7 | 7 | 10 | 6 | 1 | 6 | 6.2 | 8 | 4 | 0.549 | -0.864 | 0.2837082 | NA | 0 | 0.385 | 0.2904 | 0.2904 | TRUE | 0.6481 |
| Gm2863 | 0 | 0 | 0 | 0 | 0 | 0 | 0 | 0 | 0 | 0 | 0 | 0 | 0 | 0 | 0 | NA | NA | NA | NA | NA | NA | NA | NA | NA | NA |
| Gm2897 | 45 | 31 | 51 | 44 | 45 | 31 | 35 | 33 | 55 | 44 | 51 | 31 | 41.35 | 41 | 42 | 1.02 | 0.029 | 0.93208372 | 0.96225044 | 0.0477 | 0.0597 | 0.0564 | 0.0564 | TRUE | 0.6443 |
| Gm2913 | 0 | 0 | 0 | 0 | 0 | 0 | 0 | 0 | 0 | 0 | 0 | 0 | 0 | 0 | 0 | NA | NA | NA | NA | NA | NA | NA | NA | NA | NA |
| Gm2933 | 0 | 0 | 0 | 0 | 0 | 0 | 0 | 0 | 0 | 0 | 0 | 0 | 0 | 0 | 0 | NA | NA | NA | NA | NA | NA | NA | NA | NA | NA |
| Gm2a | 3509 | 2721 | 3398 | 4010 | 3455 | 3855 | 2725 | 2894 | 3666 | 3992 | 3898 | 3805 | 3496.77 | 3095 | 3898 | 1.26 | 0.334 | 0.00044452 | 0.00394382 | 0.013 | 0.0031 | 0.0062 | 0.0062 | TRUE | 0.3155 |
| Gm3002 | 11 | 8 | 2 | 3 | 3 | 3 | 9 | 9 | 2 | 3 | 3 | 3 | 4.76 | 7 | 3 | 0.48 | -1.06 | 0.25198484 | NA | 0 | 0.5006 | 0.3751 | 0.3751 | TRUE | 0.756 |
| Gm3020 | 0 | 0 | 0 | 0 | 0 | 0 | 0 | 0 | 0 | 0 | 0 | 0 | 0 | 0 | 0 | NA | NA | NA | NA | NA | NA | NA | NA | NA | NA |
| Gm3086 | 1 | 0 | 0 | 0 | 0 | 0 | 1 | 0 | 0 | 0 | 0 | 0 | 0.13 | 0 | 0 | 0.552 | -0.857 | 0.8336548 | NA | 0 | 18.3121 | 10 | 10 | TRUE | 0.2566 |
| Gm3139 | 0 | 0 | 0 | 0 | 0 | 0 | 0 | 0 | 0 | 0 | 0 | 0 | 0 | 0 | 0 | NA | NA | NA | NA | NA | NA | NA | NA | NA | NA |
| Gm3143 | 17 | 12 | 8 | 5 | 9 | 7 | 13 | 13 | 9 | 5 | 10 | 7 | 9.44 | 12 | 7 | 0.629 | -0.668 | 0.29246998 | NA | 0 | 0.2535 | 0.1779 | 0.1779 | TRUE | 0.274 |
| Gm3219 | 48 | 41 | 28 | 33 | 20 | 44 | 37 | 44 | 30 | 33 | 23 | 43 | 34.99 | 37 | 33 | 0.894 | -0.162 | 0.6443111 | 0.77858144 | 0.0332 | 0.0702 | 0.0605 | 0.0605 | TRUE | 0.2123 |
| Gm3230 | 2 | 2 | 2 | 8 | 3 | 5 | 2 | 2 | 2 | 8 | 3 | 5 | 3.69 | 2 | 5 | 2.835 | 1.503 | 0.1471976 | NA | 0 | 0.6452 | 0.4304 | 0.4304 | TRUE | 0.2985 |
| Gm3238 | 0 | 0 | 0 | 0 | 0 | 0 | 0 | 0 | 0 | 0 | 0 | 0 | 0 | 0 | 0 | NA | NA | NA | NA | NA | NA | NA | NA | NA | NA |
| Gm3258 | 10 | 3 | 8 | 7 | 12 | 7 | 8 | 3 | 9 | 7 | 14 | 7 | 7.83 | 7 | 9 | 1.38 | 0.465 | 0.51416101 | NA | 0 | 0.305 | 0.2359 | 0.2359 | TRUE | 0.521 |
| Gm3259 | 0 | 0 | 0 | 0 | 0 | 0 | 0 | 0 | 0 | 0 | 0 | 0 | 0 | 0 | 0 | NA | NA | NA | NA | NA | NA | NA | NA | NA | NA |
| Gm3264 | 1 | 1 | 0 | 1 | 0 | 2 | 1 | 1 | 0 | 1 | 0 | 2 | 0.8 | 1 | 1 | 1.615 | 0.692 | 0.75042087 | NA | 0 | 2.9588 | 2.1172 | 2.1172 | TRUE | 0.1299 |
| Gm3279 | 0 | 0 | 0 | 0 | 0 | 0 | 0 | 0 | 0 | 0 | 0 | 0 | 0 | 0 | 0 | NA | NA | NA | NA | NA | NA | NA | NA | NA | NA |
| Gm3285 | 0 | 0 | 0 | 0 | 0 | 1 | 0 | 0 | 0 | 0 | 0 | 1 | 0.16 | 0 | 0 | 2.094 | 1.067 | 0.79379329 | NA | 0 | 14.4078 | 10 | 10 | TRUE | 0.3835 |
| Gm3286 | 0 | 0 | 0 | 0 | 0 | 0 | 0 | 0 | 0 | 0 | 0 | 0 | 0 | 0 | 0 | NA | NA | NA | NA | NA | NA | NA | NA | NA | NA |
| Gm3317 | 0 | 0 | 0 | 0 | 0 | 0 | 0 | 0 | 0 | 0 | 0 | 0 | 0 | 0 | 0 | NA | NA | NA | NA | NA | NA | NA | NA | NA | NA |
| Gm3336 | 109 | 93 | 113 | 143 | 122 | 141 | 85 | 99 | 122 | 142 | 138 | 139 | 120.78 | 102 | 140 | 1.381 | 0.466 | 0.01520964 | 0.05640669 | 0.0074 | 0.022 | 0.0181 | 0.0181 | TRUE | 0.2874 |
| Gm3376 | 0 | 0 | 0 | 0 | 0 | 0 | 0 | 0 | 0 | 0 | 0 | 0 | 0 | 0 | 0 | NA | NA | NA | NA | NA | NA | NA | NA | NA | NA |
| Gm3383 | 0 | 0 | 0 | 0 | 0 | 0 | 0 | 0 | 0 | 0 | 0 | 0 | 0 | 0 | 0 | NA | NA | NA | NA | NA | NA | NA | NA | NA | NA |
| Gm3402 | 0 | 0 | 0 | 0 | 0 | 0 | 0 | 0 | 0 | 0 | 0 | 0 | 0 | 0 | 0 | NA | NA | NA | NA | NA | NA | NA | NA | NA | NA |
| Gm3404 | 0 | 0 | 0 | 0 | 0 | 0 | 0 | 0 | 0 | 0 | 0 | 0 | 0 | 0 | 0 | NA | NA | NA | NA | NA | NA | NA | NA | NA | NA |
| Gm3409 | 0 | 0 | 0 | 0 | 0 | 0 | 0 | 0 | 0 | 0 | 0 | 0 | 0 | 0 | 0 | NA | NA | NA | NA | NA | NA | NA | NA | NA | NA |
| Gm3414 | 184 | 117 | 109 | 131 | 117 | 149 | 143 | 124 | 118 | 130 | 132 | 147 | 132.41 | 128 | 136 | 1.06 | 0.084 | 0.63260163 | 0.76977482 | 0 | 0.0203 | 0.0147 | 0.0147 | TRUE | 0.1117 |
| Gm3415 | 0 | 0 | 0 | 0 | 0 | 0 | 0 | 0 | 0 | 0 | 0 | 0 | 0 | 0 | 0 | NA | NA | NA | NA | NA | NA | NA | NA | NA | NA |
| Gm3417 | 0 | 0 | 0 | 0 | 0 | 0 | 0 | 0 | 0 | 0 | 0 | 0 | 0 | 0 | 0 | NA | NA | NA | NA | NA | NA | NA | NA | NA | NA |
| Gm3428 | 0 | 0 | 0 | 0 | 0 | 0 | 0 | 0 | 0 | 0 | 0 | 0 | 0 | 0 | 0 | NA | NA | NA | NA | NA | NA | NA | NA | NA | NA |
| Gm3434 | 0 | 0 | 0 | 0 | 0 | 0 | 0 | 0 | 0 | 0 | 0 | 0 | 0 | 0 | 0 | NA | NA | NA | NA | NA | NA | NA | NA | NA | NA |
| Gm3435 | 18 | 13 | 8 | 18 | 13 | 20 | 14 | 14 | 9 | 18 | 15 | 20 | 14.79 | 12 | 18 | 1.429 | 0.515 | 0.30369451 | NA | 0 | 0.1626 | 0.1119 | 0.1119 | TRUE | 0.2306 |
| Gm3448 | 0 | 0 | 0 | 0 | 0 | 0 | 0 | 0 | 0 | 0 | 0 | 0 | 0 | 0 | 0 | NA | NA | NA | NA | NA | NA | NA | NA | NA | NA |
| Gm3494 | 0 | 0 | 0 | 0 | 0 | 0 | 0 | 0 | 0 | 0 | 0 | 0 | 0 | 0 | 0 | NA | NA | NA | NA | NA | NA | NA | NA | NA | NA |
| Gm3500 | 52 | 23 | 27 | 40 | 35 | 34 | 40 | 24 | 29 | 40 | 39 | 34 | 34.47 | 31 | 38 | 1.187 | 0.247 | 0.46703601 | 0.6359289 | 0 | 0.0712 | 0.0544 | 0.0544 | TRUE | 0.512 |
| Gm3558 | 282 | 189 | 221 | 232 | 176 | 187 | 219 | 201 | 238 | 231 | 199 | 185 | 212.1 | 219 | 205 | 0.933 | -0.1 | 0.50087272 | 0.66514669 | 0.0053 | 0.0136 | 0.0112 | 0.0112 | TRUE | 0.14 |
| Gm3604 | 654 | 548 | 461 | 443 | 375 | 458 | 508 | 583 | 497 | 441 | 423 | 452 | 484.06 | 529 | 439 | 0.83 | -0.269 | 0.00993381 | 0.04116669 | 0.0025 | 0.0073 | 0.0058 | 0.0058 | TRUE | 0.0887 |
| Gm362 | 0 | 0 | 0 | 0 | 0 | 0 | 0 | 0 | 0 | 0 | 0 | 0 | 0 | 0 | 0 | NA | NA | NA | NA | NA | NA | NA | NA | NA | NA |
| Gm364 | 854 | 544 | 471 | 218 | 169 | 139 | 663 | 579 | 508 | 217 | 191 | 137 | 382.5 | 583 | 182 | 0.311 | -1.687 | 9.93E-27 | 6.17E-24 | 0.0285 | 0.0086 | 0.0142 | 0.0142 | TRUE | 0.3183 |
| Gm3646 | 0 | 0 | 0 | 0 | 0 | 0 | 0 | 0 | 0 | 0 | 0 | 0 | 0 | 0 | 0 | NA | NA | NA | NA | NA | NA | NA | NA | NA | NA |
| Gm3696 | 22 | 25 | 23 | 22 | 16 | 12 | 17 | 27 | 25 | 22 | 18 | 12 | 20.05 | 23 | 17 | 0.762 | -0.393 | 0.38143193 | 0.55579288 | 0 | 0.1206 | 0.0939 | 0.0939 | TRUE | 0.3923 |
| Gm3701 | 0 | 0 | 0 | 0 | 0 | 0 | 0 | 0 | 0 | 0 | 0 | 0 | 0 | 0 | 0 | NA | NA | NA | NA | NA | NA | NA | NA | NA | NA |
| Gm3706 | 0 | 0 | 0 | 0 | 0 | 0 | 0 | 0 | 0 | 0 | 0 | 0 | 0 | 0 | 0 | NA | NA | NA | NA | NA | NA | NA | NA | NA | NA |

| | | | | | | | | | | | | | | | | | | | | | | | | | |
|---|---|---|---|---|---|---|---|---|---|---|---|---|---|---|---|---|---|---|---|---|---|---|---|---|---|
| Gm3716 | 247 | 187 | 239 | 265 | 205 | 259 | 192 | 199 | 258 | 264 | 231 | 256 | 233.22 | 216 | 250 | 1.162 | 0.217 | 0.15114298 | 0.29727351 | 0.0111 | 0.0126 | 0.0121 | 0.0121 | TRUE | 0.3036 |
| Gm3750 | 0 | 0 | 0 | 0 | 0 | 0 | 0 | 0 | 0 | 0 | 0 | 0 | 0 | 0 | 0 | NA | NA | NA | NA | NA | NA | NA | NA | NA | NA |
| Gm3763 | 0 | 0 | 0 | 0 | 0 | 0 | 0 | 0 | 0 | 0 | 0 | 0 | 0 | 0 | 0 | NA | NA | NA | NA | NA | NA | NA | NA | NA | NA |
| Gm3776 | 0 | 0 | 0 | 0 | 0 | 0 | 0 | 0 | 0 | 0 | 0 | 0 | 0 | 0 | 0 | NA | NA | NA | NA | NA | NA | NA | NA | NA | NA |
| Gm382 | 0 | 0 | 0 | 0 | 0 | 0 | 0 | 0 | 0 | 0 | 0 | 0 | 0 | 0 | 0 | NA | NA | NA | NA | NA | NA | NA | NA | NA | NA |
| Gm3833 | 2 | 1 | 5 | 8 | 7 | 5 | 2 | 1 | 5 | 8 | 8 | 5 | 4.8 | 3 | 7 | 2.654 | 1.408 | 0.13476353 | NA | 0 | 0.4961 | 0.3795 | 0.3795 | TRUE | 0.844 |
| Gm3893 | 0 | 0 | 0 | 0 | 0 | 0 | 0 | 0 | 0 | 0 | 0 | 0 | 0 | 0 | 0 | NA | NA | NA | NA | NA | NA | NA | NA | NA | NA |
| Gm3925 | 69 | 34 | 45 | 78 | 87 | 75 | 54 | 36 | 49 | 78 | 98 | 74 | 64.69 | 46 | 83 | 1.791 | 0.841 | 0.00124533 | 0.00846776 | 0.0133 | 0.0391 | 0.0322 | 0.0322 | TRUE | 0.2684 |
| Gm3985 | 2 | 4 | 3 | 1 | 0 | 1 | 2 | 4 | 3 | 1 | 0 | 1 | 1.84 | 3 | 1 | 0.228 | -2.132 | 0.17837465 | NA | 0 | 1.292 | 0.8837 | 0.8837 | TRUE | 0.2794 |
| Gm4013 | 19 | 12 | 17 | 14 | 10 | 16 | 15 | 13 | 18 | 14 | 11 | 16 | 14.48 | 15 | 14 | 0.898 | -0.154 | 0.75762741 | NA | 0 | 0.1661 | 0.1111 | 0.1111 | TRUE | 0.1271 |
| Gm4027 | 0 | 0 | 0 | 0 | 0 | 0 | 0 | 0 | 0 | 0 | 0 | 0 | 0 | 0 | 0 | NA | NA | NA | NA | NA | NA | NA | NA | NA | NA |
| Gm4064 | 0 | 0 | 0 | 0 | 0 | 0 | 0 | 0 | 0 | 0 | 0 | 0 | 0 | 0 | 0 | NA | NA | NA | NA | NA | NA | NA | NA | NA | NA |
| Gm4070 | 0 | 0 | 0 | 0 | 0 | 0 | 0 | 0 | 0 | 0 | 0 | 0 | 0 | 0 | 0 | NA | NA | NA | NA | NA | NA | NA | NA | NA | NA |
| Gm41 | 63 | 53 | 53 | 28 | 45 | 29 | 49 | 56 | 57 | 28 | 51 | 29 | 44.96 | 54 | 36 | 0.657 | -0.605 | 0.05663607 | 0.14753041 | 0.0309 | 0.0551 | 0.0491 | 0.0491 | TRUE | 0.9033 |
| Gm410 | 0 | 0 | 0 | 0 | 0 | 0 | 0 | 0 | 0 | 0 | 0 | 0 | 0 | 0 | 0 | NA | NA | NA | NA | NA | NA | NA | NA | NA | NA |
| Gm4133 | 0 | 0 | 0 | 0 | 0 | 0 | 0 | 0 | 0 | 0 | 0 | 0 | 0 | 0 | 0 | NA | NA | NA | NA | NA | NA | NA | NA | NA | NA |
| Gm4141 | 0 | 0 | 0 | 0 | 0 | 0 | 0 | 0 | 0 | 0 | 0 | 0 | 0 | 0 | 0 | NA | NA | NA | NA | NA | NA | NA | NA | NA | NA |
| Gm4172 | 0 | 0 | 0 | 0 | 0 | 0 | 0 | 0 | 0 | 0 | 0 | 0 | 0 | 0 | 0 | NA | NA | NA | NA | NA | NA | NA | NA | NA | NA |
| Gm4175 | 0 | 0 | 0 | 0 | 0 | 0 | 0 | 0 | 0 | 0 | 0 | 0 | 0 | 0 | 0 | NA | NA | NA | NA | NA | NA | NA | NA | NA | NA |
| Gm4177 | 0 | 0 | 0 | 0 | 0 | 0 | 0 | 0 | 0 | 0 | 0 | 0 | 0 | 0 | 0 | NA | NA | NA | NA | NA | NA | NA | NA | NA | NA |
| Gm4179 | 0 | 0 | 0 | 0 | 0 | 0 | 0 | 0 | 0 | 0 | 0 | 0 | 0 | 0 | 0 | NA | NA | NA | NA | NA | NA | NA | NA | NA | NA |
| Gm4187 | 0 | 0 | 0 | 0 | 0 | 0 | 0 | 0 | 0 | 0 | 0 | 0 | 0 | 0 | 0 | NA | NA | NA | NA | NA | NA | NA | NA | NA | NA |
| Gm4201 | 0 | 0 | 0 | 0 | 0 | 0 | 0 | 0 | 0 | 0 | 0 | 0 | 0 | 0 | 0 | NA | NA | NA | NA | NA | NA | NA | NA | NA | NA |
| Gm4214 | 0 | 0 | 0 | 0 | 0 | 0 | 0 | 0 | 0 | 0 | 0 | 0 | 0 | 0 | 0 | NA | NA | NA | NA | NA | NA | NA | NA | NA | NA |
| Gm4216 | 0 | 0 | 0 | 0 | 0 | 0 | 0 | 0 | 0 | 0 | 0 | 0 | 0 | 0 | 0 | NA | NA | NA | NA | NA | NA | NA | NA | NA | NA |
| Gm4220 | 0 | 0 | 0 | 0 | 0 | 0 | 0 | 0 | 0 | 0 | 0 | 0 | 0 | 0 | 0 | NA | NA | NA | NA | NA | NA | NA | NA | NA | NA |
| Gm4224 | 0 | 0 | 0 | 0 | 0 | 0 | 0 | 0 | 0 | 0 | 0 | 0 | 0 | 0 | 0 | NA | NA | NA | NA | NA | NA | NA | NA | NA | NA |
| Gm4251 | 0 | 0 | 0 | 0 | 0 | 0 | 0 | 0 | 0 | 0 | 0 | 0 | 0 | 0 | 0 | NA | NA | NA | NA | NA | NA | NA | NA | NA | NA |
| Gm4262 | 37 | 30 | 31 | 44 | 29 | 52 | 29 | 32 | 33 | 44 | 33 | 51 | 36.99 | 31 | 43 | 1.368 | 0.452 | 0.16573008 | 0.31676786 | 0 | 0.0665 | 0.0491 | 0.0491 | TRUE | 0.2326 |
| Gm4265 | 0 | 0 | 0 | 0 | 0 | 0 | 0 | 0 | 0 | 0 | 0 | 0 | 0 | 0 | 0 | NA | NA | NA | NA | NA | NA | NA | NA | NA | NA |
| Gm4278 | 0 | 0 | 0 | 0 | 0 | 0 | 0 | 0 | 0 | 0 | 0 | 0 | 0 | 0 | 0 | NA | NA | NA | NA | NA | NA | NA | NA | NA | NA |
| Gm428 | 1 | 2 | 2 | 0 | 0 | 3 | 1 | 2 | 2 | 0 | 0 | 3 | 1.34 | 2 | 1 | 0.606 | -0.723 | 0.68645422 | NA | 0.5179 | 1.7748 | 1.5067 | 1.5067 | TRUE | 1.4412 |
| Gm4285 | 17 | 11 | 14 | 10 | 10 | 11 | 13 | 12 | 15 | 10 | 11 | 11 | 12.02 | 13 | 11 | 0.802 | -0.319 | 0.55687234 | NA | 0 | 0.1996 | 0.1273 | 0.1273 | TRUE | 0.0508 |
| Gm4297 | 0 | 0 | 1 | 0 | 0 | 0 | 0 | 0 | 1 | 0 | 0 | 0 | 0.18 | 0 | 0 | 0.552 | -0.857 | 0.8336548 | NA | 0 | 13.1817 | 10 | 10 | TRUE | 0.5365 |
| Gm4301 | 0 | 0 | 0 | 0 | 0 | 0 | 0 | 0 | 0 | 0 | 0 | 0 | 0 | 0 | 0 | NA | NA | NA | NA | NA | NA | NA | NA | NA | NA |
| Gm4302 | 0 | 0 | 0 | 0 | 0 | 0 | 0 | 0 | 0 | 0 | 0 | 0 | 0 | 0 | 0 | NA | NA | NA | NA | NA | NA | NA | NA | NA | NA |
| Gm4303 | 0 | 0 | 0 | 0 | 0 | 0 | 0 | 0 | 0 | 0 | 0 | 0 | 0 | 0 | 0 | NA | NA | NA | NA | NA | NA | NA | NA | NA | NA |
| Gm4305 | 0 | 0 | 0 | 0 | 0 | 0 | 0 | 0 | 0 | 0 | 0 | 0 | 0 | 0 | 0 | NA | NA | NA | NA | NA | NA | NA | NA | NA | NA |
| Gm4307 | 0 | 0 | 0 | 0 | 0 | 0 | 0 | 0 | 0 | 0 | 0 | 0 | 0 | 0 | 0 | NA | NA | NA | NA | NA | NA | NA | NA | NA | NA |
| Gm4312 | 0 | 0 | 0 | 0 | 0 | 0 | 0 | 0 | 0 | 0 | 0 | 0 | 0 | 0 | 0 | NA | NA | NA | NA | NA | NA | NA | NA | NA | NA |
| Gm4340 | 0 | 0 | 0 | 0 | 0 | 0 | 0 | 0 | 0 | 0 | 0 | 0 | 0 | 0 | 0 | NA | NA | NA | NA | NA | NA | NA | NA | NA | NA |
| Gm4345 | 0 | 0 | 0 | 0 | 0 | 0 | 0 | 0 | 0 | 0 | 0 | 0 | 0 | 0 | 0 | NA | NA | NA | NA | NA | NA | NA | NA | NA | NA |
| Gm4349 | 134 | 107 | 143 | 125 | 111 | 143 | 104 | 114 | 154 | 124 | 125 | 141 | 127.16 | 124 | 130 | 1.056 | 0.078 | 0.68957613 | 0.81013576 | 0.017 | 0.0211 | 0.0199 | 0.0199 | TRUE | 0.4511 |
| Gm436 | 0 | 0 | 0 | 0 | 0 | 0 | 0 | 0 | 0 | 0 | 0 | 0 | 0 | 0 | 0 | NA | NA | NA | NA | NA | NA | NA | NA | NA | NA |
| Gm4371 | 121 | 86 | 52 | 90 | 67 | 97 | 94 | 91 | 56 | 90 | 76 | 96 | 83.75 | 80 | 87 | 1.076 | 0.106 | 0.66359662 | 0.79257847 | 0.0317 | 0.0307 | 0.031 | 0.031 | TRUE | 0.6216 |
| Gm438 | 0 | 0 | 2 | 1 | 0 | 1 | 0 | 0 | 2 | 1 | 0 | 1 | 0.69 | 1 | 1 | 0.966 | -0.049 | 0.98423749 | NA | 0 | 3.4369 | 3.0198 | 3.0198 | TRUE | 0.9749 |
| Gm44 | 94 | 56 | 44 | 30 | 32 | 39 | 73 | 60 | 47 | 30 | 36 | 38 | 47.42 | 60 | 35 | 0.576 | -0.795 | 0.00798661 | 0.03498395 | 0 | 0.0524 | 0.0419 | 0.0419 | TRUE | 0.1473 |
| Gm4461 | 3 | 1 | 0 | 1 | 1 | 0 | 2 | 1 | 0 | 1 | 1 | 0 | 0.92 | 1 | 1 | 0.598 | -0.741 | 0.72094137 | NA | 0 | 2.5799 | 1.9516 | 1.9516 | TRUE | 0.1813 |
| Gm4477 | 0 | 0 | 0 | 0 | 0 | 0 | 0 | 0 | 0 | 0 | 0 | 0 | 0 | 0 | 0 | NA | NA | NA | NA | NA | NA | NA | NA | NA | NA |
| Gm4489 | 5 | 3 | 1 | 2 | 1 | 1 | 4 | 3 | 1 | 2 | 1 | 1 | 2.04 | 3 | 1 | 0.498 | -1.006 | 0.45585313 | NA | 0 | 1.1624 | 0.7599 | 0.7599 | TRUE | 0.3002 |
| Gm4498 | 0 | 0 | 0 | 0 | 0 | 0 | 0 | 0 | 0 | 0 | 0 | 0 | 0 | 0 | 0 | NA | NA | NA | NA | NA | NA | NA | NA | NA | NA |
| Gm4532 | 29 | 30 | 34 | 28 | 15 | 30 | 23 | 32 | 37 | 28 | 17 | 30 | 27.59 | 31 | 25 | 0.828 | -0.273 | 0.48534707 | 0.65118424 | 0.034 | 0.0883 | 0.0735 | 0.0735 | TRUE | 0.426 |
| Gm4541 | 0 | 0 | 0 | 0 | 0 | 0 | 0 | 0 | 0 | 0 | 0 | 0 | 0 | 0 | 0 | NA | NA | NA | NA | NA | NA | NA | NA | NA | NA |
| Gm4559 | 0 | 0 | 0 | 0 | 0 | 0 | 0 | 0 | 0 | 0 | 0 | 0 | 0 | 0 | 0 | NA | NA | NA | NA | NA | NA | NA | NA | NA | NA |
| Gm4566 | 0 | 0 | 0 | 0 | 0 | 0 | 0 | 0 | 0 | 0 | 0 | 0 | 0 | 0 | 0 | NA | NA | NA | NA | NA | NA | NA | NA | NA | NA |
| Gm4567 | 0 | 0 | 0 | 0 | 0 | 0 | 0 | 0 | 0 | 0 | 0 | 0 | 0 | 0 | 0 | NA | NA | NA | NA | NA | NA | NA | NA | NA | NA |
| Gm4598 | 11 | 2 | 3 | 0 | 3 | 4 | 9 | 2 | 3 | 0 | 3 | 4 | 3.54 | 5 | 2 | 0.513 | -0.962 | 0.40087609 | NA | 0.4934 | 0.6719 | 0.6303 | 0.6303 | TRUE | 1.3159 |
| Gm4636 | 3206 | 2159 | 1890 | 1969 | 1684 | 2020 | 2490 | 2297 | 2039 | 1960 | 1900 | 1994 | 2113.25 | 2275 | 1951 | 0.857 | -0.223 | 0.00494596 | 0.02432645 | 0.0045 | 0.0035 | 0.0041 | 0.0041 | TRUE | 0.0992 |
| Gm4710 | 0 | 0 | 0 | 0 | 0 | 0 | 0 | 0 | 0 | 0 | 0 | 0 | 0 | 0 | 0 | NA | NA | NA | NA | NA | NA | NA | NA | NA | NA |
| Gm4719 | 0 | 0 | 0 | 0 | 0 | 0 | 0 | 0 | 0 | 0 | 0 | 0 | 0 | 0 | 0 | NA | NA | NA | NA | NA | NA | NA | NA | NA | NA |
| Gm4724 | 0 | 0 | 0 | 0 | 0 | 0 | 0 | 0 | 0 | 0 | 0 | 0 | 0 | 0 | 0 | NA | NA | NA | NA | NA | NA | NA | NA | NA | NA |
| Gm4736 | 0 | 0 | 0 | 0 | 0 | 0 | 0 | 0 | 0 | 0 | 0 | 0 | 0 | 0 | 0 | NA | NA | NA | NA | NA | NA | NA | NA | NA | NA |
| Gm4745 | 1 | 1 | 0 | 0 | 0 | 0 | 1 | 1 | 0 | 0 | 0 | 0 | 0.31 | 1 | 0 | 0.309 | -1.696 | 0.63183309 | NA | 0 | 7.7294 | 7.2472 | 7.2472 | TRUE | 0.2015 |
| Gm4759 | 0 | 0 | 0 | 0 | 0 | 0 | 0 | 0 | 0 | 0 | 0 | 0 | 0 | 0 | 0 | NA | NA | NA | NA | NA | NA | NA | NA | NA | NA |
| Gm4763 | 0 | 0 | 0 | 0 | 0 | 0 | 0 | 0 | 0 | 0 | 0 | 0 | 0 | 0 | 0 | NA | NA | NA | NA | NA | NA | NA | NA | NA | NA |
| Gm4776 | 0 | 0 | 0 | 0 | 0 | 0 | 0 | 0 | 0 | 0 | 0 | 0 | 0 | 0 | 0 | NA | NA | NA | NA | NA | NA | NA | NA | NA | NA |
| Gm4787 | 6 | 2 | 3 | 7 | 3 | 3 | 5 | 2 | 3 | 7 | 3 | 3 | 3.89 | 3 | 4 | 1.311 | 0.39 | 0.68748226 | NA | 0 | 0.6118 | 0.4206 | 0.4206 | TRUE | 0.4671 |
| Gm4788 | 0 | 0 | 0 | 0 | 0 | 0 | 0 | 0 | 0 | 0 | 0 | 0 | 0 | 0 | 0 | NA | NA | NA | NA | NA | NA | NA | NA | NA | NA |
| Gm4791 | 0 | 0 | 0 | 0 | 0 | 0 | 0 | 0 | 0 | 0 | 0 | 0 | 0 | 0 | 0 | NA | NA | NA | NA | NA | NA | NA | NA | NA | NA |

| | | | | | | | | | | | | | | | | | | | | | | | | | |
|---|---|---|---|---|---|---|---|---|---|---|---|---|---|---|---|---|---|---|---|---|---|---|---|---|---|
| Gm4792 | 0 | 0 | 0 | 0 | 0 | 0 | 0 | 0 | 0 | 0 | 0 | 0 | 0 | 0 | 0 | NA | NA | NA | NA | NA | NA | NA | NA | NA | NA |
| Gm4794 | 0 | 0 | 0 | 0 | 0 | 0 | 0 | 0 | 0 | 0 | 0 | 0 | 0 | 0 | 0 | NA | NA | NA | NA | NA | NA | NA | NA | NA | NA |
| Gm4814 | 0 | 0 | 0 | 0 | 0 | 0 | 0 | 0 | 0 | 0 | 0 | 0 | 0 | 0 | 0 | NA | NA | NA | NA | NA | NA | NA | NA | NA | NA |
| Gm4827 | 5 | 2 | 3 | 4 | 1 | 2 | 4 | 2 | 3 | 4 | 1 | 2 | 2.72 | 3 | 2 | 0.766 | -0.385 | 0.73746942 | NA | 0 | 0.8732 | 0.5763 | 0.5763 | TRUE | 0.3812 |
| Gm4832 | 1 | 0 | 0 | 1 | 1 | 0 | 1 | 0 | 0 | 1 | 1 | 0 | 0.48 | 0 | 1 | 2.037 | 1.027 | 0.71241723 | NA | 0 | 4.9057 | 3.8521 | 3.8521 | TRUE | 0.2664 |
| Gm4836 | 0 | 0 | 0 | 0 | 0 | 0 | 0 | 0 | 0 | 0 | 0 | 0 | 0 | 0 | 0 | NA | NA | NA | NA | NA | NA | NA | NA | NA | NA |
| Gm4841 | 0 | 0 | 0 | 0 | 0 | 0 | 0 | 0 | 0 | 0 | 0 | 0 | 0 | 0 | 0 | NA | NA | NA | NA | NA | NA | NA | NA | NA | NA |
| Gm4846 | 0 | 0 | 0 | 0 | 0 | 0 | 0 | 0 | 0 | 0 | 0 | 0 | 0 | 0 | 0 | NA | NA | NA | NA | NA | NA | NA | NA | NA | NA |
| Gm4847 | 0 | 0 | 0 | 0 | 0 | 0 | 0 | 0 | 0 | 0 | 0 | 0 | 0 | 0 | 0 | NA | NA | NA | NA | NA | NA | NA | NA | NA | NA |
| Gm4850 | 0 | 0 | 0 | 0 | 0 | 0 | 0 | 0 | 0 | 0 | 0 | 0 | 0 | 0 | 0 | NA | NA | NA | NA | NA | NA | NA | NA | NA | NA |
| Gm4858 | 644 | 390 | 118 | 84 | 71 | 60 | 500 | 415 | 127 | 84 | 80 | 59 | 210.88 | 347 | 74 | 0.214 | -2.226 | 0.00012344 | 0.00144771 | 0.2339 | 0.0137 | 0.0708 | 0.2339 | TRUE | 0.452 |
| Gm4861 | 0 | 0 | 0 | 0 | 0 | 0 | 0 | 0 | 0 | 0 | 0 | 0 | 0 | 0 | 0 | NA | NA | NA | NA | NA | NA | NA | NA | NA | NA |
| Gm4871 | 0 | 0 | 0 | 0 | 0 | 0 | 0 | 0 | 0 | 0 | 0 | 0 | 0 | 0 | 0 | NA | NA | NA | NA | NA | NA | NA | NA | NA | NA |
| Gm4872 | 1 | 0 | 0 | 0 | 2 | 3 | 1 | 0 | 0 | 0 | 2 | 3 | 1 | 0 | 2 | 5.033 | 2.331 | 0.28653323 | NA | 0 | 2.3747 | 2.1499 | 2.1499 | TRUE | 0.6217 |
| Gm4884 | 0 | 0 | 0 | 1 | 0 | 0 | 0 | 0 | 0 | 1 | 0 | 0 | 0.17 | 0 | 0 | 2.094 | 1.067 | 0.79379329 | NA | 0 | 14.2885 | 10 | 10 | TRUE | 0.3906 |
| Gm4890 | 35 | 23 | 6 | 10 | 9 | 8 | 27 | 24 | 6 | 10 | 10 | 8 | 14.35 | 19 | 9 | 0.476 | -1.07 | 0.07309259 | NA | 0.1979 | 0.1675 | 0.1764 | 0.1764 | TRUE | 1.6798 |
| Gm4894 | 0 | 0 | 0 | 0 | 0 | 0 | 0 | 0 | 0 | 0 | 0 | 0 | 0 | 0 | 0 | NA | NA | NA | NA | NA | NA | NA | NA | NA | NA |
| Gm4906 | 0 | 0 | 0 | 0 | 0 | 0 | 0 | 0 | 0 | 0 | 0 | 0 | 0 | 0 | 0 | NA | NA | NA | NA | NA | NA | NA | NA | NA | NA |
| Gm4907 | 0 | 0 | 0 | 0 | 0 | 0 | 0 | 0 | 0 | 0 | 0 | 0 | 0 | 0 | 0 | NA | NA | NA | NA | NA | NA | NA | NA | NA | NA |
| Gm4922 | 2 | 0 | 3 | 1 | 0 | 1 | 2 | 0 | 3 | 1 | 0 | 1 | 1.13 | 2 | 1 | 0.423 | -1.24 | 0.5235556 | NA | 0 | 2.102 | 1.6529 | 1.6529 | TRUE | 0.1028 |
| Gm4925 | 5 | 1 | 1 | 4 | 1 | 3 | 4 | 1 | 1 | 4 | 1 | 3 | 2.35 | 2 | 3 | 1.301 | 0.379 | 0.76349978 | NA | 0 | 1.0112 | 0.7237 | 0.7237 | TRUE | 0.7951 |
| Gm4926 | 1 | 1 | 1 | 0 | 1 | 0 | 1 | 1 | 1 | 0 | 1 | 0 | 0.67 | 1 | 0 | 0.383 | -1.383 | 0.54981825 | NA | 0 | 3.5158 | 2.3538 | 2.3538 | TRUE | 0.5075 |
| Gm4937 | 0 | 0 | 0 | 0 | 0 | 0 | 0 | 0 | 0 | 0 | 0 | 0 | 0 | 0 | 0 | NA | NA | NA | NA | NA | NA | NA | NA | NA | NA |
| Gm4944 | 289 | 209 | 149 | 181 | 142 | 135 | 224 | 222 | 161 | 180 | 160 | 133 | 180.19 | 202 | 158 | 0.777 | -0.363 | 0.04364214 | 0.12199825 | 0.0229 | 0.0156 | 0.0177 | 0.0177 | TRUE | 0.3419 |
| Gm4951 | 0 | 1 | 0 | 0 | 0 | 0 | 0 | 1 | 0 | 0 | 0 | 0 | 0.18 | 0 | 0 | 0.552 | -0.857 | 0.8336548 | NA | 0 | 13.3714 | 10 | 10 | TRUE | 0.5216 |
| Gm4952 | 1 | 1 | 0 | 1 | 1 | 2 | 1 | 1 | 0 | 1 | 1 | 2 | 0.99 | 1 | 1 | 2.197 | 1.135 | 0.55299242 | NA | 0 | 2.3972 | 1.4894 | 1.4894 | TRUE | 0.1884 |
| Gm4956 | 0 | 0 | 0 | 0 | 0 | 0 | 0 | 0 | 0 | 0 | 0 | 0 | 0 | 0 | 0 | NA | NA | NA | NA | NA | NA | NA | NA | NA | NA |
| Gm4961 | 216 | 163 | 152 | 180 | 162 | 190 | 168 | 173 | 164 | 179 | 183 | 188 | 175.77 | 168 | 183 | 1.088 | 0.122 | 0.41031371 | 0.58307953 | 0 | 0.0159 | 0.01 | 0.01 | TRUE | 0.0067 |
| Gm4971 | 443 | 404 | 139 | 126 | 149 | 136 | 344 | 430 | 150 | 125 | 168 | 134 | 225.25 | 308 | 142 | 0.463 | -1.112 | 0.01363215 | 0.05201396 | 0.1412 | 0.0129 | 0.0476 | 0.1412 | TRUE | 0.334 |
| Gm4975 | 207 | 225 | 282 | 129 | 164 | 234 | 161 | 239 | 304 | 128 | 185 | 231 | 208.13 | 235 | 181 | 0.776 | -0.367 | 0.11634936 | 0.24684301 | 0.0865 | 0.0138 | 0.0343 | 0.0343 | TRUE | 0.1945 |
| Gm4980 | 21 | 24 | 12 | 24 | 21 | 13 | 16 | 26 | 13 | 24 | 24 | 13 | 19.2 | 18 | 20 | 1.103 | 0.142 | 0.76395854 | 0.85985191 | 0.0601 | 0.1259 | 0.108 | 0.108 | TRUE | 0.5781 |
| Gm4981 | 9 | 2 | 2 | 2 | 1 | 4 | 7 | 2 | 2 | 2 | 1 | 4 | 3.06 | 4 | 2 | 0.617 | -0.698 | 0.53942765 | NA | 0 | 0.7777 | 0.5886 | 0.5886 | TRUE | 1.1579 |
| Gm4984 | 47 | 42 | 46 | 54 | 62 | 62 | 37 | 45 | 50 | 54 | 70 | 61 | 52.62 | 44 | 62 | 1.421 | 0.506 | 0.06581987 | 0.16392463 | 0 | 0.0475 | 0.0351 | 0.0351 | TRUE | 0.188 |
| Gm5 | 13 | 5 | 6 | 10 | 7 | 3 | 10 | 5 | 6 | 10 | 8 | 3 | 7.12 | 7 | 7 | 0.936 | -0.096 | 0.89820302 | NA | 0 | 0.3354 | 0.2626 | 0.2626 | TRUE | 0.6951 |
| Gm5039 | 1007 | 765 | 301 | 327 | 315 | 260 | 782 | 814 | 325 | 325 | 355 | 257 | 476.35 | 640 | 312 | 0.488 | -1.035 | 0.01468443 | 0.0549005 | 0.1273 | 0.0074 | 0.0382 | 0.1273 | TRUE | 2.1794 |
| Gm5065 | 1 | 0 | 0 | 0 | 0 | 0 | 1 | 0 | 0 | 0 | 0 | 0 | 0.13 | 0 | 0 | 0.552 | -0.857 | 0.8336548 | NA | 0 | 18.3121 | 10 | 10 | TRUE | 0.2566 |
| Gm5069 | 5 | 7 | 3 | 12 | 5 | 9 | 4 | 7 | 3 | 12 | 6 | 9 | 6.84 | 5 | 9 | 1.841 | 0.881 | 0.24517164 | NA | 0 | 0.349 | 0.2555 | 0.2555 | TRUE | 0.2124 |
| Gm5071 | 1 | 2 | 0 | 0 | 0 | 0 | 1 | 2 | 0 | 0 | 0 | 0 | 0.48 | 1 | 0 | 0.197 | -2.343 | 0.43249452 | NA | 0 | 4.8992 | 4.9172 | 4.9172 | TRUE | 0.1226 |
| Gm5072 | 0 | 0 | 0 | 0 | 0 | 0 | 0 | 0 | 0 | 0 | 0 | 0 | 0 | 0 | 0 | NA | NA | NA | NA | NA | NA | NA | NA | NA | NA |
| Gm5077 | 49 | 46 | 41 | 31 | 36 | 43 | 38 | 49 | 44 | 31 | 41 | 42 | 40.86 | 44 | 38 | 0.871 | -0.199 | 0.51791387 | 0.6797764 | 0 | 0.0604 | 0.0435 | 0.0435 | TRUE | 0.2031 |
| Gm5082 | 0 | 0 | 0 | 0 | 0 | 0 | 0 | 0 | 0 | 0 | 0 | 0 | 0 | 0 | 0 | NA | NA | NA | NA | NA | NA | NA | NA | NA | NA |
| Gm5083 | 10 | 12 | 10 | 9 | 8 | 22 | 8 | 13 | 11 | 9 | 9 | 22 | 11.84 | 11 | 13 | 1.285 | 0.362 | 0.54491774 | NA | 0.1016 | 0.2027 | 0.1725 | 0.1725 | TRUE | 1.3563 |
| Gm5084 | 0 | 0 | 0 | 0 | 0 | 0 | 0 | 0 | 0 | 0 | 0 | 0 | 0 | 0 | 0 | NA | NA | NA | NA | NA | NA | NA | NA | NA | NA |
| Gm5086 | 4 | 6 | 2 | 3 | 4 | 6 | 3 | 6 | 2 | 3 | 5 | 6 | 4.18 | 4 | 5 | 1.161 | 0.215 | 0.81905341 | NA | 0 | 0.5697 | 0.3971 | 0.3971 | TRUE | 0.4728 |
| Gm5087 | 0 | 0 | 0 | 0 | 0 | 0 | 0 | 0 | 0 | 0 | 0 | 0 | 0 | 0 | 0 | NA | NA | NA | NA | NA | NA | NA | NA | NA | NA |
| Gm5088 | 1 | 1 | 0 | 2 | 0 | 1 | 1 | 1 | 0 | 2 | 0 | 1 | 0.8 | 1 | 1 | 1.618 | 0.694 | 0.74938576 | NA | 0 | 2.9538 | 2.1156 | 2.1156 | TRUE | 0.1303 |
| Gm5089 | 3 | 4 | 4 | 4 | 5 | 3 | 2 | 4 | 4 | 4 | 6 | 3 | 3.91 | 3 | 4 | 1.166 | 0.221 | 0.81656682 | NA | 0 | 0.608 | 0.3953 | 0.3953 | TRUE | 0.2103 |
| Gm5091 | 521 | 428 | 534 | 568 | 528 | 613 | 405 | 455 | 576 | 565 | 596 | 605 | 533.7 | 479 | 589 | 1.233 | 0.303 | 0.01554763 | 0.0573773 | 0.0147 | 0.0069 | 0.0094 | 0.0094 | TRUE | 0.3663 |
| Gm5095 | 1 | 1 | 0 | 0 | 0 | 0 | 1 | 1 | 0 | 0 | 0 | 0 | 0.31 | 1 | 0 | 0.309 | -1.696 | 0.63183309 | NA | 0 | 7.7294 | 7.2472 | 7.2472 | TRUE | 0.2015 |
| Gm5105 | 0 | 1 | 1 | 0 | 0 | 0 | 0 | 1 | 1 | 0 | 0 | 0 | 0.36 | 1 | 0 | 0.271 | -1.881 | 0.5755355 | NA | 0 | 6.6391 | 6.4428 | 6.4428 | TRUE | 0.3513 |
| Gm5108 | 0 | 0 | 0 | 0 | 0 | 0 | 0 | 0 | 0 | 0 | 0 | 0 | 0 | 0 | 0 | NA | NA | NA | NA | NA | NA | NA | NA | NA | NA |
| Gm5111 | 0 | 0 | 0 | 0 | 0 | 0 | 0 | 0 | 0 | 0 | 0 | 0 | 0 | 0 | 0 | NA | NA | NA | NA | NA | NA | NA | NA | NA | NA |
| Gm5113 | 269 | 175 | 167 | 187 | 171 | 174 | 209 | 186 | 180 | 186 | 193 | 172 | 187.68 | 192 | 184 | 0.954 | -0.068 | 0.64742825 | 0.78051363 | 0 | 0.0151 | 0.0107 | 0.0107 | TRUE | 0.0724 |
| Gm5114 | 0 | 0 | 0 | 0 | 0 | 0 | 0 | 0 | 0 | 0 | 0 | 0 | 0 | 0 | 0 | NA | NA | NA | NA | NA | NA | NA | NA | NA | NA |
| Gm5122 | 4 | 5 | 1 | 0 | 2 | 2 | 3 | 5 | 1 | 0 | 2 | 2 | 2.29 | 3 | 1 | 0.44 | -1.185 | 0.38593705 | NA | 0 | 1.0378 | 0.8218 | 0.8218 | TRUE | 0.2072 |
| Gm5124 | 44 | 36 | 23 | 22 | 25 | 23 | 34 | 38 | 25 | 22 | 28 | 23 | 28.35 | 32 | 24 | 0.745 | -0.425 | 0.25030506 | 0.41918343 | 0 | 0.086 | 0.0624 | 0.0624 | TRUE | 0.2661 |
| Gm5126 | 14 | 5 | 8 | 12 | 7 | 10 | 11 | 5 | 9 | 12 | 8 | 10 | 9.09 | 8 | 10 | 1.186 | 0.246 | 0.69965406 | NA | 0 | 0.2632 | 0.1834 | 0.1834 | TRUE | 0.2948 |
| Gm5127 | 0 | 1 | 0 | 0 | 0 | 0 | 0 | 1 | 0 | 0 | 0 | 0 | 0.18 | 0 | 0 | 0.552 | -0.857 | 0.8336548 | NA | 0 | 13.3714 | 10 | 10 | TRUE | 0.5216 |
| Gm5128 | 0 | 0 | 0 | 0 | 0 | 0 | 0 | 0 | 0 | 0 | 0 | 0 | 0 | 0 | 0 | NA | NA | NA | NA | NA | NA | NA | NA | NA | NA |
| Gm5129 | 1 | 0 | 1 | 1 | 2 | 3 | 1 | 0 | 1 | 1 | 2 | 3 | 1.34 | 1 | 2 | 3.302 | 1.723 | 0.32219666 | NA | 0 | 1.7648 | 1.1667 | 1.1667 | TRUE | 0.1985 |
| Gm5132 | 0 | 0 | 0 | 0 | 0 | 0 | 0 | 0 | 0 | 0 | 0 | 0 | 0 | 0 | 0 | NA | NA | NA | NA | NA | NA | NA | NA | NA | NA |
| Gm5134 | 341 | 255 | 232 | 181 | 176 | 177 | 265 | 271 | 250 | 180 | 199 | 175 | 223.31 | 262 | 185 | 0.703 | -0.509 | 0.00019445 | 0.00208809 | 0 | 0.013 | 0.0088 | 0.0088 | TRUE | 0.0458 |
| Gm5136 | 19 | 14 | 12 | 4 | 3 | 3 | 15 | 15 | 13 | 4 | 3 | 3 | 8.82 | 14 | 3 | 0.242 | -2.046 | 0.00389166 | NA | 0 | 0.2711 | 0.1777 | 0.1777 | TRUE | 0.0293 |
| Gm514 | 28 | 29 | 22 | 23 | 17 | 19 | 22 | 31 | 24 | 23 | 19 | 19 | 22.86 | 26 | 20 | 0.802 | -0.319 | 0.42901428 | 0.60104893 | 0 | 0.1061 | 0.0727 | 0.0727 | TRUE | 0.204 |
| Gm5141 | 298 | 268 | 212 | 232 | 221 | 224 | 231 | 285 | 229 | 231 | 249 | 221 | 241.1 | 248 | 234 | 0.942 | -0.086 | 0.54256479 | 0.70040449 | 0.0056 | 0.0123 | 0.0103 | 0.0103 | TRUE | 0.1796 |
| Gm5142 | 2 | 0 | 0 | 0 | 1 | 0 | 2 | 0 | 0 | 0 | 1 | 0 | 0.45 | 1 | 0 | 0.68 | -0.556 | 0.85993441 | NA | 0 | 5.3057 | 5.2576 | 5.2576 | TRUE | 0.9726 |
| Gm5148 | 1 | 0 | 0 | 0 | 0 | 0 | 1 | 0 | 0 | 0 | 0 | 0 | 0.13 | 0 | 0 | 0.552 | -0.857 | 0.8336548 | NA | 0 | 18.3121 | 10 | 10 | TRUE | 0.2566 |
| Gm5150 | 1 | 0 | 0 | 0 | 0 | 0 | 1 | 0 | 0 | 0 | 0 | 0 | 0.13 | 0 | 0 | 0.552 | -0.857 | 0.8336548 | NA | 0 | 18.3121 | 10 | 10 | TRUE | 0.2566 |
| Gm5166 | 5 | 4 | 0 | 0 | 1 | 1 | 4 | 4 | 0 | 0 | 1 | 1 | 1.71 | 3 | 1 | 0.254 | -1.978 | 0.25274805 | NA | 0 | 1.3893 | 1.2362 | 1.2362 | TRUE | 0.8232 |

| | | | | | | | | | | | | | | | | | | | | | | | | | |
|---|---|---|---|---|---|---|---|---|---|---|---|---|---|---|---|---|---|---|---|---|---|---|---|---|---|
| Gm5168 | 0 | 0 | 0 | 0 | 0 | 0 | 0 | 0 | 0 | 0 | 0 | 0 | 0 | 0 | 0 | NA | NA | NA | NA | NA | NA | NA | NA | NA | NA |
| Gm5169 | 0 | 0 | 0 | 0 | 0 | 0 | 0 | 0 | 0 | 0 | 0 | 0 | 0 | 0 | 0 | NA | NA | NA | NA | NA | NA | NA | NA | NA | NA |
| Gm5176 | 11 | 10 | 1 | 2 | 3 | 6 | 9 | 11 | 1 | 2 | 3 | 6 | 5.26 | 7 | 4 | 0.555 | -0.85 | 0.3681351 | NA | 0.3852 | 0.453 | 0.435 | 0.435 | TRUE | 0.8614 |
| Gm5177 | 1 | 5 | 0 | 1 | 2 | 2 | 1 | 5 | 0 | 1 | 2 | 2 | 1.89 | 2 | 2 | 0.869 | -0.202 | 0.8931849 | NA | 0.6645 | 1.2585 | 1.0915 | 1.0915 | TRUE | 1.7092 |
| Gm525 | 18 | 6 | 5 | 7 | 11 | 7 | 14 | 6 | 5 | 7 | 12 | 7 | 8.67 | 8 | 9 | 0.992 | -0.011 | 0.98733286 | NA | 0.1007 | 0.2757 | 0.2235 | 0.2235 | TRUE | 1.179 |
| Gm527 | 221 | 165 | 191 | 221 | 170 | 219 | 172 | 176 | 206 | 220 | 192 | 216 | 196.86 | 185 | 209 | 1.139 | 0.188 | 0.2089125 | 0.37065372 | 0.0026 | 0.0145 | 0.011 | 0.011 | TRUE | 0.11 |
| Gm5294 | 0 | 0 | 0 | 1 | 2 | 1 | 0 | 0 | 0 | 1 | 2 | 1 | 0.71 | 0 | 1 | 7.937 | 2.989 | 0.20904898 | NA | 0 | 3.3573 | 2.7105 | 2.7105 | TRUE | 0.1573 |
| Gm53 | 192 | 176 | 143 | 116 | 111 | 124 | 149 | 187 | 154 | 115 | 125 | 122 | 142.29 | 163 | 121 | 0.742 | -0.431 | 0.01273427 | 0.04944604 | 0 | 0.0191 | 0.0144 | 0.0144 | TRUE | 0.166 |
| Gm5334 | 0 | 0 | 0 | 0 | 0 | 0 | 0 | 0 | 0 | 0 | 0 | 0 | 0 | 0 | 0 | NA | NA | NA | NA | NA | NA | NA | NA | NA | NA |
| Gm5346 | 0 | 0 | 0 | 0 | 0 | 0 | 0 | 0 | 0 | 0 | 0 | 0 | 0 | 0 | 0 | NA | NA | NA | NA | NA | NA | NA | NA | NA | NA |
| Gm5347 | 0 | 0 | 0 | 0 | 0 | 0 | 0 | 0 | 0 | 0 | 0 | 0 | 0 | 0 | 0 | NA | NA | NA | NA | NA | NA | NA | NA | NA | NA |
| Gm5382 | 0 | 0 | 0 | 0 | 0 | 0 | 0 | 0 | 0 | 0 | 0 | 0 | 0 | 0 | 0 | NA | NA | NA | NA | NA | NA | NA | NA | NA | NA |
| Gm5409 | 0 | 0 | 0 | 0 | 0 | 0 | 0 | 0 | 0 | 0 | 0 | 0 | 0 | 0 | 0 | NA | NA | NA | NA | NA | NA | NA | NA | NA | NA |
| Gm5414 | 0 | 0 | 0 | 0 | 0 | 0 | 0 | 0 | 0 | 0 | 0 | 0 | 0 | 0 | 0 | NA | NA | NA | NA | NA | NA | NA | NA | NA | NA |
| Gm5415 | 0 | 0 | 0 | 0 | 0 | 1 | 0 | 0 | 0 | 0 | 0 | 1 | 0.16 | 0 | 0 | 2.094 | 1.067 | 0.79379329 | NA | 0 | 14.4078 | 10 | 10 | TRUE | 0.3835 |
| Gm5416 | 0 | 0 | 0 | 0 | 0 | 0 | 0 | 0 | 0 | 0 | 0 | 0 | 0 | 0 | 0 | NA | NA | NA | NA | NA | NA | NA | NA | NA | NA |
| Gm5420 | 0 | 0 | 0 | 0 | 0 | 0 | 0 | 0 | 0 | 0 | 0 | 0 | 0 | 0 | 0 | NA | NA | NA | NA | NA | NA | NA | NA | NA | NA |
| Gm5424 | 0 | 0 | 0 | 0 | 0 | 0 | 0 | 0 | 0 | 0 | 0 | 0 | 0 | 0 | 0 | NA | NA | NA | NA | NA | NA | NA | NA | NA | NA |
| Gm5431 | 0 | 0 | 0 | 0 | 0 | 0 | 0 | 0 | 0 | 0 | 0 | 0 | 0 | 0 | 0 | NA | NA | NA | NA | NA | NA | NA | NA | NA | NA |
| Gm5434 | 0 | 0 | 0 | 2 | 0 | 1 | 0 | 0 | 0 | 2 | 0 | 1 | 0.5 | 0 | 1 | 5.65 | 2.498 | 0.39774729 | NA | 0 | 4.7776 | 4.7714 | 4.7714 | TRUE | 0.0529 |
| Gm5441 | 0 | 0 | 0 | 0 | 0 | 0 | 0 | 0 | 0 | 0 | 0 | 0 | 0 | 0 | 0 | NA | NA | NA | NA | NA | NA | NA | NA | NA | NA |
| Gm5458 | 5 | 7 | 10 | 7 | 6 | 8 | 4 | 7 | 11 | 7 | 7 | 8 | 7.29 | 7 | 7 | 0.998 | -0.004 | 0.9960003 | NA | 0 | 0.3274 | 0.2352 | 0.2352 | TRUE | 0.2347 |
| Gm5460 | 0 | 0 | 0 | 0 | 0 | 0 | 0 | 0 | 0 | 0 | 0 | 0 | 0 | 0 | 0 | NA | NA | NA | NA | NA | NA | NA | NA | NA | NA |
| Gm5464 | 2 | 1 | 2 | 2 | 1 | 1 | 2 | 1 | 2 | 2 | 1 | 1 | 1.48 | 2 | 1 | 0.864 | -0.211 | 0.88836451 | NA | 0 | 1.6036 | 0.9503 | 0.9503 | TRUE | 0.1029 |
| Gm5468 | 4 | 2 | 1 | 3 | 2 | 2 | 3 | 2 | 1 | 3 | 2 | 2 | 2.25 | 2 | 2 | 1.121 | 0.164 | 0.89407593 | NA | 0 | 1.0535 | 0.6584 | 0.6584 | TRUE | 0.2203 |
| Gm5475 | 1 | 1 | 1 | 2 | 1 | 1 | 1 | 1 | 1 | 2 | 1 | 1 | 1.17 | 1 | 1 | 1.424 | 0.51 | 0.76157012 | NA | 0 | 2.0266 | 1.1613 | 1.1613 | TRUE | 0.1027 |
| Gm5476 | 0 | 1 | 0 | 0 | 0 | 0 | 0 | 1 | 0 | 0 | 0 | 0 | 0.18 | 0 | 0 | 0.552 | -0.857 | 0.8336548 | NA | 0 | 13.3714 | 10 | 10 | TRUE | 0.5216 |
| Gm5477 | 0 | 0 | 0 | 0 | 0 | 0 | 0 | 0 | 0 | 0 | 0 | 0 | 0 | 0 | 0 | NA | NA | NA | NA | NA | NA | NA | NA | NA | NA |
| Gm5478 | 0 | 0 | 0 | 0 | 0 | 0 | 0 | 0 | 0 | 0 | 0 | 0 | 0 | 0 | 0 | NA | NA | NA | NA | NA | NA | NA | NA | NA | NA |
| Gm5480 | 43 | 40 | 49 | 45 | 40 | 50 | 33 | 43 | 53 | 45 | 45 | 49 | 44.68 | 43 | 46 | 1.091 | 0.125 | 0.6749462 | 0.80032886 | 0 | 0.0555 | 0.042 | 0.042 | TRUE | 0.2479 |
| Gm5483 | 0 | 0 | 0 | 0 | 0 | 0 | 0 | 0 | 0 | 0 | 0 | 0 | 0 | 0 | 0 | NA | NA | NA | NA | NA | NA | NA | NA | NA | NA |
| Gm5485 | 0 | 0 | 0 | 0 | 0 | 0 | 0 | 0 | 0 | 0 | 0 | 0 | 0 | 0 | 0 | NA | NA | NA | NA | NA | NA | NA | NA | NA | NA |
| Gm5494 | 5 | 2 | 1 | 0 | 0 | 0 | 4 | 2 | 1 | 0 | 0 | 0 | 1.18 | 2 | 0 | 0.08 | -3.652 | 0.06909248 | NA | 0 | 2.0082 | 1.7071 | 1.7071 | TRUE | 0.1449 |
| Gm5506 | 6788 | 4958 | 5956 | 6286 | 6000 | 6883 | 5272 | 5274 | 6426 | 6257 | 6769 | 6795 | 6132.08 | 5657 | 6607 | 1.168 | 0.224 | 0.00534996 | 0.02580728 | 0.0077 | 0.0028 | 0.0045 | 0.0045 | TRUE | 0.1718 |
| Gm5512 | 12 | 9 | 11 | 10 | 13 | 10 | 9 | 10 | 12 | 10 | 15 | 10 | 10.88 | 10 | 12 | 1.121 | 0.164 | 0.775231 | NA | 0 | 0.2204 | 0.1459 | 0.1459 | TRUE | 0.2005 |
| Gm5523 | 20 | 7 | 22 | 17 | 13 | 18 | 16 | 7 | 24 | 17 | 15 | 18 | 16.01 | 16 | 17 | 1.059 | 0.083 | 0.87289732 | NA | 0.0733 | 0.1504 | 0.1308 | 0.1308 | TRUE | 0.1828 |
| Gm5531 | 0 | 2 | 0 | 0 | 0 | 0 | 0 | 2 | 0 | 0 | 0 | 0 | 0.35 | 1 | 0 | 0.272 | -1.878 | 0.62190613 | NA | 10 | 6.6869 | 8.7633 | 8.7633 | TRUE | 0.9743 |
| Gm5535 | 1 | 0 | 1 | 0 | 2 | 2 | 1 | 0 | 1 | 0 | 2 | 2 | 1.01 | 1 | 1 | 2.24 | 1.163 | 0.56316354 | NA | 0 | 2.3389 | 1.7772 | 1.7772 | TRUE | 0.5111 |
| Gm5538 | 0 | 0 | 0 | 0 | 0 | 0 | 0 | 0 | 0 | 0 | 0 | 0 | 0 | 0 | 0 | NA | NA | NA | NA | NA | NA | NA | NA | NA | NA |
| Gm5544 | 0 | 1 | 0 | 0 | 0 | 0 | 0 | 1 | 0 | 0 | 0 | 0 | 0.18 | 0 | 0 | 0.552 | -0.857 | 0.8336548 | NA | 0 | 13.3714 | 10 | 10 | TRUE | 0.5216 |
| Gm5547 | 0 | 0 | 0 | 0 | 0 | 0 | 0 | 0 | 0 | 0 | 0 | 0 | 0 | 0 | 0 | NA | NA | NA | NA | NA | NA | NA | NA | NA | NA |
| Gm5549 | 0 | 0 | 0 | 0 | 0 | 0 | 0 | 0 | 0 | 0 | 0 | 0 | 0 | 0 | 0 | NA | NA | NA | NA | NA | NA | NA | NA | NA | NA |
| Gm5577 | 42 | 33 | 35 | 32 | 32 | 33 | 33 | 35 | 38 | 32 | 36 | 33 | 34.34 | 35 | 34 | 0.955 | -0.067 | 0.8354933 | 0.90637245 | 0 | 0.0715 | 0.0461 | 0.0461 | TRUE | 0.0342 |
| Gm5591 | 0 | 0 | 0 | 0 | 0 | 0 | 0 | 0 | 0 | 0 | 0 | 0 | 0 | 0 | 0 | NA | NA | NA | NA | NA | NA | NA | NA | NA | NA |
| Gm5592 | 0 | 0 | 0 | 0 | 0 | 0 | 0 | 0 | 0 | 0 | 0 | 0 | 0 | 0 | 0 | NA | NA | NA | NA | NA | NA | NA | NA | NA | NA |
| Gm5595 | 1279 | 973 | 931 | 859 | 866 | 892 | 993 | 1035 | 1005 | 855 | 977 | 881 | 957.56 | 1011 | 904 | 0.894 | -0.162 | 0.04734929 | 0.12952409 | 0.0014 | 0.0049 | 0.0037 | 0.0037 | TRUE | 0.0578 |
| Gm5607 | 8 | 1 | 3 | 4 | 5 | 6 | 6 | 1 | 3 | 4 | 6 | 6 | 4.34 | 3 | 5 | 1.432 | 0.518 | 0.58350551 | NA | 0 | 0.5481 | 0.4088 | 0.4088 | TRUE | 0.4611 |
| Gm561 | 385 | 258 | 265 | 374 | 296 | 374 | 299 | 274 | 286 | 372 | 334 | 369 | 322.46 | 286 | 358 | 1.251 | 0.323 | 0.00584036 | 0.02765032 | 0 | 0.0098 | 0.0067 | 0.0067 | TRUE | 0.0396 |
| Gm5615 | 0 | 0 | 0 | 0 | 0 | 0 | 0 | 0 | 0 | 0 | 0 | 0 | 0 | 0 | 0 | NA | NA | NA | NA | NA | NA | NA | NA | NA | NA |
| Gm5617 | 302 | 275 | 203 | 286 | 219 | 252 | 235 | 293 | 219 | 285 | 247 | 249 | 254.43 | 249 | 260 | 1.048 | 0.068 | 0.64299841 | 0.77752761 | 0.0107 | 0.0117 | 0.0114 | 0.0114 | TRUE | 0.2546 |
| Gm5622 | 3 | 1 | 0 | 2 | 0 | 0 | 2 | 1 | 0 | 2 | 0 | 0 | 0.9 | 1 | 1 | 0.579 | -0.789 | 0.7251066 | NA | 0 | 2.6434 | 2.4396 | 2.4396 | TRUE | 0.4576 |
| Gm5627 | 0 | 0 | 0 | 0 | 0 | 0 | 0 | 0 | 0 | 0 | 0 | 0 | 0 | 0 | 0 | NA | NA | NA | NA | NA | NA | NA | NA | NA | NA |
| Gm5634 | 1 | 0 | 1 | 1 | 0 | 0 | 1 | 0 | 1 | 1 | 0 | 0 | 0.48 | 1 | 0 | 0.596 | -0.748 | 0.78941713 | NA | 0 | 4.9902 | 3.8695 | 3.8695 | TRUE | 0.3906 |
| Gm5635 | 212 | 166 | 198 | 124 | 124 | 119 | 165 | 177 | 214 | 123 | 140 | 117 | 155.94 | 185 | 127 | 0.687 | -0.541 | 0.00172523 | 0.01078679 | 0.008 | 0.0176 | 0.0148 | 0.0148 | TRUE | 0.1943 |
| Gm5640 | 0 | 0 | 1 | 0 | 1 | 0 | 0 | 0 | 1 | 0 | 1 | 0 | 0.37 | 0 | 0 | 1.075 | 0.105 | 0.97514588 | NA | 0 | 6.4454 | 6.1578 | 6.1578 | TRUE | 0.5365 |
| Gm5643 | 27 | 11 | 17 | 16 | 17 | 16 | 21 | 12 | 18 | 16 | 19 | 16 | 16.99 | 17 | 17 | 0.987 | -0.018 | 0.9687293 | NA | 0 | 0.142 | 0.101 | 0.101 | TRUE | 0.3457 |
| Gm5662 | 2342 | 1528 | 665 | 668 | 638 | 572 | 1819 | 1625 | 718 | 665 | 720 | 565 | 1018.51 | 1387 | 650 | 0.468 | -1.095 | 0.00719882 | 0.03240714 | 0.1184 | 0.0048 | 0.0328 | 0.1184 | TRUE | 1.1878 |
| Gm5712 | 0 | 0 | 0 | 0 | 2 | 0 | 0 | 0 | 0 | 0 | 2 | 0 | 0.38 | 0 | 1 | 4.195 | 2.069 | 0.57837236 | NA | 10 | 6.3052 | 8.2866 | 8.2866 | TRUE | 0.9743 |
| Gm572 | 22 | 17 | 9 | 9 | 11 | 21 | 17 | 18 | 10 | 9 | 12 | 21 | 14.5 | 15 | 14 | 0.935 | -0.097 | 0.85741465 | NA | 0.0693 | 0.1659 | 0.1395 | 0.1395 | TRUE | 0.7458 |
| Gm5725 | 0 | 0 | 0 | 0 | 0 | 0 | 0 | 0 | 0 | 0 | 0 | 0 | 0 | 0 | 0 | NA | NA | NA | NA | NA | NA | NA | NA | NA | NA |
| Gm5726 | 0 | 0 | 0 | 0 | 0 | 0 | 0 | 0 | 0 | 0 | 0 | 0 | 0 | 0 | 0 | NA | NA | NA | NA | NA | NA | NA | NA | NA | NA |
| Gm5728 | 0 | 0 | 0 | 0 | 0 | 0 | 0 | 0 | 0 | 0 | 0 | 0 | 0 | 0 | 0 | NA | NA | NA | NA | NA | NA | NA | NA | NA | NA |
| Gm5741 | 7 | 6 | 1 | 4 | 3 | 5 | 5 | 6 | 1 | 4 | 3 | 5 | 4.2 | 4 | 4 | 0.947 | -0.079 | 0.93386336 | NA | 0 | 0.5667 | 0.421 | 0.421 | TRUE | 0.6796 |
| Gm5766 | 0 | 0 | 0 | 1 | 0 | 0 | 0 | 0 | 0 | 1 | 0 | 0 | 0.17 | 0 | 0 | 2.094 | 1.067 | 0.79379329 | NA | 0 | 14.2885 | 10 | 10 | TRUE | 0.3906 |
| Gm5771 | 0 | 0 | 0 | 0 | 0 | 0 | 0 | 0 | 0 | 0 | 0 | 0 | 0 | 0 | 0 | NA | NA | NA | NA | NA | NA | NA | NA | NA | NA |
| Gm5779 | 1 | 0 | 1 | 0 | 0 | 5 | 1 | 0 | 1 | 0 | 0 | 5 | 1.13 | 1 | 2 | 2.663 | 1.413 | 0.51229993 | NA | 3.0613 | 2.0963 | 2.2661 | 2.2661 | TRUE | 2.3422 |
| Gm5795 | 1 | 0 | 2 | 0 | 0 | 1 | 1 | 0 | 2 | 0 | 0 | 1 | 0.65 | 1 | 0 | 0.382 | -1.39 | 0.57974155 | NA | 0 | 3.6284 | 3.0424 | 3.0424 | TRUE | 0.2583 |
| Gm5796 | 59 | 52 | 29 | 48 | 29 | 46 | 46 | 55 | 31 | 48 | 33 | 45 | 43.05 | 44 | 42 | 0.952 | -0.071 | 0.82539104 | 0.8999218 | 0.0322 | 0.0575 | 0.0508 | 0.0508 | TRUE | 0.2986 |
| Gm5797 | 1 | 0 | 3 | 0 | 1 | 0 | 1 | 0 | 3 | 0 | 1 | 0 | 0.86 | 1 | 0 | 0.28 | -1.835 | 0.4293679 | NA | 0 | 2.7681 | 2.5201 | 2.5201 | TRUE | 0.5637 |

| | | | | | | | | | | | | | | | | | | | | | | | | | |
|---|---|---|---|---|---|---|---|---|---|---|---|---|---|---|---|---|---|---|---|---|---|---|---|---|---|
| Gm5800 | 0 | 0 | 0 | 0 | 0 | 0 | 0 | 0 | 0 | 0 | 0 | 0 | 0 | 0 | 0 | NA | NA | NA | NA | NA | NA | NA | NA | NA | NA |
| Gm5801 | 12 | 9 | 4 | 20 | 7 | 5 | 9 | 10 | 4 | 20 | 8 | 5 | 9.32 | 8 | 11 | 1.404 | 0.489 | 0.48897283 | NA | 0.2387 | 0.2566 | 0.2512 | 0.2512 | TRUE | 1.3573 |
| Gm5803 | 7 | 7 | 6 | 18 | 12 | 6 | 5 | 7 | 6 | 18 | 14 | 6 | 9.46 | 6 | 13 | 1.942 | 0.957 | 0.15029899 | NA | 0 | 0.2531 | 0.2023 | 0.2023 | TRUE | 0.2573 |
| Gm581 | 4 | 5 | 5 | 10 | 9 | 11 | 3 | 5 | 5 | 10 | 10 | 11 | 7.46 | 4 | 10 | 2.281 | 1.19 | 0.09487789 | NA | 0 | 0.3199 | 0.2093 | 0.2093 | TRUE | 0.2156 |
| Gm5820 | 2 | 3 | 2 | 5 | 10 | 4 | 2 | 3 | 2 | 5 | 11 | 4 | 4.52 | 2 | 7 | 2.955 | 1.563 | 0.10778377 | NA | 0 | 0.527 | 0.3904 | 0.3904 | TRUE | 0.811 |
| Gm5833 | 0 | 0 | 0 | 0 | 0 | 0 | 0 | 0 | 0 | 0 | 0 | 0 | 0 | 0 | 0 | NA | NA | NA | NA | NA | NA | NA | NA | NA | NA |
| Gm5860 | 0 | 0 | 0 | 0 | 0 | 0 | 0 | 0 | 0 | 0 | 0 | 0 | 0 | 0 | 0 | NA | NA | NA | NA | NA | NA | NA | NA | NA | NA |
| Gm5868 | 5 | 3 | 1 | 1 | 1 | 0 | 4 | 3 | 1 | 1 | 1 | 0 | 1.71 | 3 | 1 | 0.253 | -1.983 | 0.22167582 | NA | 0 | 1.3862 | 0.978 | 0.978 | TRUE | 0.3021 |
| Gm5878 | 39 | 35 | 22 | 36 | 27 | 25 | 30 | 37 | 24 | 36 | 30 | 25 | 30.37 | 30 | 30 | 0.996 | -0.005 | 0.98853211 | 0.99395634 | 0 | 0.0805 | 0.0608 | 0.0608 | TRUE | 0.1808 |
| Gm5885 | 0 | 0 | 0 | 0 | 0 | 0 | 0 | 0 | 0 | 0 | 0 | 0 | 0 | 0 | 0 | NA | NA | NA | NA | NA | NA | NA | NA | NA | NA |
| Gm5886 | 0 | 0 | 0 | 0 | 0 | 0 | 0 | 0 | 0 | 0 | 0 | 0 | 0 | 0 | 0 | NA | NA | NA | NA | NA | NA | NA | NA | NA | NA |
| Gm5891 | 0 | 0 | 0 | 0 | 0 | 0 | 0 | 0 | 0 | 0 | 0 | 0 | 0 | 0 | 0 | NA | NA | NA | NA | NA | NA | NA | NA | NA | NA |
| Gm590 | 11 | 9 | 3 | 14 | 7 | 6 | 9 | 10 | 3 | 14 | 8 | 6 | 8.18 | 7 | 9 | 1.291 | 0.369 | 0.60281213 | NA | 0.093 | 0.292 | 0.2385 | 0.2385 | TRUE | 0.662 |
| Gm5901 | 1 | 0 | 0 | 0 | 0 | 0 | 1 | 0 | 0 | 0 | 0 | 0 | 0.13 | 0 | 0 | 0.552 | -0.857 | 0.8336548 | NA | 0 | 18.3121 | 10 | 10 | TRUE | 0.2566 |
| Gm5916 | 0 | 0 | 0 | 0 | 0 | 0 | 0 | 0 | 0 | 0 | 0 | 0 | 0 | 0 | 0 | NA | NA | NA | NA | NA | NA | NA | NA | NA | NA |
| Gm5918 | 214 | 138 | 165 | 152 | 152 | 183 | 166 | 147 | 178 | 151 | 171 | 181 | 165.74 | 164 | 168 | 1.025 | 0.035 | 0.8275769 | 0.90122336 | 0 | 0.0167 | 0.0127 | 0.0127 | TRUE | 0.0812 |
| Gm5925 | 0 | 0 | 0 | 0 | 0 | 0 | 0 | 0 | 0 | 0 | 0 | 0 | 0 | 0 | 0 | NA | NA | NA | NA | NA | NA | NA | NA | NA | NA |
| Gm5934 | 0 | 0 | 0 | 0 | 0 | 0 | 0 | 0 | 0 | 0 | 0 | 0 | 0 | 0 | 0 | NA | NA | NA | NA | NA | NA | NA | NA | NA | NA |
| Gm5935 | 0 | 0 | 0 | 0 | 0 | 0 | 0 | 0 | 0 | 0 | 0 | 0 | 0 | 0 | 0 | NA | NA | NA | NA | NA | NA | NA | NA | NA | NA |
| Gm5936 | 0 | 0 | 0 | 2 | 0 | 0 | 0 | 0 | 0 | 2 | 0 | 0 | 0.33 | 0 | 1 | 3.77 | 1.915 | 0.62344813 | NA | 0 | 7.1455 | 9.1834 | 9.1834 | TRUE | 0.9739 |
| Gm5938 | 0 | 0 | 0 | 0 | 0 | 0 | 0 | 0 | 0 | 0 | 0 | 0 | 0 | 0 | 0 | NA | NA | NA | NA | NA | NA | NA | NA | NA | NA |
| Gm5941 | 0 | 0 | 0 | 1 | 0 | 1 | 0 | 0 | 0 | 1 | 0 | 1 | 0.33 | 0 | 1 | 3.763 | 1.912 | 0.57887588 | NA | 0 | 7.1752 | 6.7756 | 6.7756 | TRUE | 0.2094 |
| Gm595 | 233 | 159 | 149 | 140 | 125 | 84 | 181 | 169 | 161 | 139 | 141 | 83 | 145.69 | 170 | 121 | 0.709 | -0.497 | 0.01460386 | 0.05465643 | 0.0339 | 0.0187 | 0.0227 | 0.0227 | TRUE | 0.7867 |
| Gm597 | 0 | 0 | 0 | 0 | 0 | 0 | 0 | 0 | 0 | 0 | 0 | 0 | 0 | 0 | 0 | NA | NA | NA | NA | NA | NA | NA | NA | NA | NA |
| Gm6026 | 0 | 0 | 0 | 0 | 0 | 0 | 0 | 0 | 0 | 0 | 0 | 0 | 0 | 0 | 0 | NA | NA | NA | NA | NA | NA | NA | NA | NA | NA |
| Gm6034 | 1 | 0 | 0 | 0 | 0 | 1 | 1 | 0 | 0 | 0 | 0 | 1 | 0.29 | 0 | 0 | 1.075 | 0.105 | 0.97702352 | NA | 0 | 8.0647 | 7.5458 | 7.5458 | TRUE | 0.3835 |
| Gm6040 | 0 | 0 | 0 | 0 | 0 | 0 | 0 | 0 | 0 | 0 | 0 | 0 | 0 | 0 | 0 | NA | NA | NA | NA | NA | NA | NA | NA | NA | NA |
| Gm6042 | 0 | 0 | 0 | 0 | 0 | 0 | 0 | 0 | 0 | 0 | 0 | 0 | 0 | 0 | 0 | NA | NA | NA | NA | NA | NA | NA | NA | NA | NA |
| Gm608 | 2119 | 1492 | 1320 | 1462 | 1354 | 1463 | 1646 | 1587 | 1424 | 1455 | 1527 | 1444 | 1513.99 | 1552 | 1475 | 0.949 | -0.075 | 0.32407281 | 0.49809594 | 0.0026 | 0.004 | 0.0035 | 0.0035 | TRUE | 0.062 |
| Gm6084 | 0 | 0 | 0 | 0 | 0 | 0 | 0 | 0 | 0 | 0 | 0 | 0 | 0 | 0 | 0 | NA | NA | NA | NA | NA | NA | NA | NA | NA | NA |
| Gm6086 | 5 | 3 | 1 | 2 | 1 | 2 | 4 | 3 | 1 | 2 | 1 | 2 | 2.21 | 3 | 2 | 0.619 | -0.692 | 0.5883643 | NA | 0 | 1.0759 | 0.699 | 0.699 | TRUE | 0.2995 |
| Gm609 | 0 | 0 | 0 | 0 | 0 | 0 | 0 | 0 | 0 | 0 | 0 | 0 | 0 | 0 | 0 | NA | NA | NA | NA | NA | NA | NA | NA | NA | NA |
| Gm6116 | 0 | 0 | 0 | 0 | 0 | 0 | 0 | 0 | 0 | 0 | 0 | 0 | 0 | 0 | 0 | NA | NA | NA | NA | NA | NA | NA | NA | NA | NA |
| Gm6121 | 0 | 0 | 0 | 0 | 0 | 0 | 0 | 0 | 0 | 0 | 0 | 0 | 0 | 0 | 0 | NA | NA | NA | NA | NA | NA | NA | NA | NA | NA |
| Gm614 | 0 | 1 | 0 | 0 | 0 | 0 | 0 | 1 | 0 | 0 | 0 | 0 | 0.18 | 0 | 0 | 0.552 | -0.857 | 0.8336548 | NA | 0 | 13.3714 | 10 | 10 | TRUE | 0.5216 |
| Gm6150 | 0 | 0 | 0 | 0 | 0 | 0 | 0 | 0 | 0 | 0 | 0 | 0 | 0 | 0 | 0 | NA | NA | NA | NA | NA | NA | NA | NA | NA | NA |
| Gm6164 | 0 | 0 | 0 | 0 | 0 | 0 | 0 | 0 | 0 | 0 | 0 | 0 | 0 | 0 | 0 | NA | NA | NA | NA | NA | NA | NA | NA | NA | NA |
| Gm6194 | 0 | 0 | 2 | 1 | 1 | 1 | 0 | 0 | 2 | 1 | 1 | 1 | 0.88 | 1 | 1 | 1.509 | 0.594 | 0.77995955 | NA | 0 | 2.7015 | 2.0514 | 2.0514 | TRUE | 0.9752 |
| Gm6213 | 0 | 0 | 0 | 1 | 0 | 1 | 0 | 0 | 0 | 1 | 0 | 1 | 0.33 | 0 | 1 | 3.763 | 1.912 | 0.57887588 | NA | 0 | 7.1752 | 6.7756 | 6.7756 | TRUE | 0.2094 |
| Gm6225 | 2 | 2 | 0 | 0 | 0 | 0 | 2 | 2 | 0 | 0 | 0 | 0 | 0.61 | 1 | 0 | 0.154 | -2.696 | 0.32099274 | NA | 0 | 3.8659 | 3.9247 | 3.9247 | TRUE | 0.3505 |
| Gm6249 | 0 | 0 | 0 | 0 | 0 | 0 | 0 | 0 | 0 | 0 | 0 | 0 | 0 | 0 | 0 | NA | NA | NA | NA | NA | NA | NA | NA | NA | NA |
| Gm6251 | 25 | 14 | 28 | 33 | 19 | 28 | 19 | 15 | 30 | 33 | 21 | 28 | 24.41 | 21 | 27 | 1.279 | 0.355 | 0.39338463 | 0.56697051 | 0.0363 | 0.0995 | 0.083 | 0.083 | TRUE | 0.5711 |
| Gm6260 | 0 | 0 | 0 | 0 | 0 | 0 | 0 | 0 | 0 | 0 | 0 | 0 | 0 | 0 | 0 | NA | NA | NA | NA | NA | NA | NA | NA | NA | NA |
| Gm6268 | 2 | 0 | 1 | 0 | 0 | 0 | 2 | 0 | 1 | 0 | 0 | 0 | 0.44 | 1 | 0 | 0.214 | -2.223 | 0.47051571 | NA | 0 | 5.4047 | 5.2946 | 5.2946 | TRUE | 0.2788 |
| Gm6277 | 3 | 0 | 0 | 1 | 5 | 2 | 2 | 0 | 0 | 1 | 6 | 2 | 1.82 | 1 | 3 | 3.316 | 1.73 | 0.2916909 | NA | 0.7947 | 1.3023 | 1.1952 | 1.1952 | TRUE | 1.4366 |
| Gm6289 | 0 | 0 | 1 | 0 | 0 | 0 | 0 | 0 | 1 | 0 | 0 | 0 | 0.18 | 0 | 0 | 0.552 | -0.857 | 0.8336548 | NA | 0 | 13.1817 | 10 | 10 | TRUE | 0.5365 |
| Gm6297 | 49 | 37 | 54 | 59 | 33 | 45 | 38 | 39 | 58 | 59 | 37 | 44 | 46.01 | 45 | 47 | 1.043 | 0.061 | 0.84551556 | 0.9121221 | 0.034 | 0.0539 | 0.0484 | 0.0484 | TRUE | 0.477 |
| Gm6300 | 0 | 0 | 0 | 0 | 0 | 0 | 0 | 0 | 0 | 0 | 0 | 0 | 0 | 0 | 0 | NA | NA | NA | NA | NA | NA | NA | NA | NA | NA |
| Gm6307 | 0 | 0 | 0 | 0 | 0 | 0 | 0 | 0 | 0 | 0 | 0 | 0 | 0 | 0 | 0 | NA | NA | NA | NA | NA | NA | NA | NA | NA | NA |
| Gm6313 | 0 | 0 | 0 | 0 | 0 | 0 | 0 | 0 | 0 | 0 | 0 | 0 | 0 | 0 | 0 | NA | NA | NA | NA | NA | NA | NA | NA | NA | NA |
| Gm6329 | 162 | 140 | 146 | 101 | 90 | 115 | 126 | 149 | 158 | 101 | 102 | 114 | 124.64 | 144 | 106 | 0.733 | -0.447 | 0.01484577 | 0.0554167 | 0 | 0.0214 | 0.016 | 0.016 | TRUE | 0.142 |
| Gm6367 | 0 | 0 | 0 | 0 | 0 | 0 | 0 | 0 | 0 | 0 | 0 | 0 | 0 | 0 | 0 | NA | NA | NA | NA | NA | NA | NA | NA | NA | NA |
| Gm6370 | 0 | 0 | 0 | 0 | 0 | 0 | 0 | 0 | 0 | 0 | 0 | 0 | 0 | 0 | 0 | NA | NA | NA | NA | NA | NA | NA | NA | NA | NA |
| Gm6377 | 1 | 1 | 3 | 1 | 1 | 1 | 1 | 1 | 3 | 1 | 1 | 1 | 1.36 | 2 | 1 | 0.625 | -0.679 | 0.673999 | NA | 0 | 1.7392 | 1.0856 | 1.0856 | TRUE | 0.4559 |
| Gm6402 | 38 | 23 | 25 | 33 | 23 | 34 | 30 | 24 | 27 | 33 | 26 | 34 | 28.89 | 27 | 31 | 1.14 | 0.188 | 0.59614886 | 0.74235543 | 0 | 0.0845 | 0.0565 | 0.0565 | TRUE | 0.1166 |
| Gm6406 | 42 | 33 | 17 | 17 | 20 | 8 | 33 | 35 | 18 | 17 | 23 | 8 | 22.24 | 29 | 16 | 0.545 | -0.877 | 0.06317213 | 0.15918886 | 0.1158 | 0.109 | 0.1107 | 0.1107 | TRUE | 0.6415 |
| Gm6408 | 0 | 0 | 0 | 1 | 0 | 0 | 0 | 0 | 0 | 1 | 0 | 0 | 0.17 | 0 | 0 | 2.094 | 1.067 | 0.79379329 | NA | 0 | 14.2885 | 10 | 10 | TRUE | 0.3906 |
| Gm6416 | 0 | 0 | 0 | 0 | 0 | 0 | 0 | 0 | 0 | 0 | 0 | 0 | 0 | 0 | 0 | NA | NA | NA | NA | NA | NA | NA | NA | NA | NA |
| Gm6432 | 31 | 32 | 12 | 27 | 29 | 22 | 24 | 34 | 13 | 27 | 33 | 22 | 25.4 | 24 | 27 | 1.14 | 0.189 | 0.65741997 | 0.78776858 | 0.0766 | 0.0958 | 0.0908 | 0.0908 | TRUE | 0.2253 |
| Gm6455 | 0 | 0 | 0 | 0 | 0 | 0 | 0 | 0 | 0 | 0 | 0 | 0 | 0 | 0 | 0 | NA | NA | NA | NA | NA | NA | NA | NA | NA | NA |
| Gm6460 | 0 | 0 | 0 | 0 | 0 | 0 | 0 | 0 | 0 | 0 | 0 | 0 | 0 | 0 | 0 | NA | NA | NA | NA | NA | NA | NA | NA | NA | NA |
| Gm648 | 4 | 3 | 2 | 2 | 0 | 1 | 3 | 3 | 2 | 2 | 0 | 1 | 1.91 | 3 | 1 | 0.359 | -1.478 | 0.30981231 | NA | 0 | 1.2462 | 0.8456 | 0.8456 | TRUE | 0.354 |
| Gm6484 | 1 | 0 | 0 | 0 | 1 | 1 | 1 | 0 | 0 | 0 | 1 | 1 | 0.48 | 0 | 1 | 2.031 | 1.022 | 0.71376068 | NA | 0 | 4.9197 | 3.859 | 3.859 | TRUE | 0.2623 |
| Gm6498 | 5 | 1 | 0 | 1 | 3 | 0 | 4 | 1 | 0 | 1 | 3 | 0 | 1.55 | 2 | 1 | 0.836 | -0.259 | 0.88041448 | NA | 1.3629 | 1.5271 | 1.4907 | 1.4907 | TRUE | 0.8803 |
| Gm6524 | 4 | 2 | 6 | 2 | 2 | 5 | 3 | 2 | 6 | 2 | 2 | 5 | 3.48 | 4 | 3 | 0.795 | -0.331 | 0.75021889 | NA | 0 | 0.6831 | 0.4868 | 0.4868 | TRUE | 0.4929 |
| Gm6525 | 1 | 0 | 0 | 1 | 0 | 1 | 1 | 0 | 0 | 1 | 0 | 1 | 0.46 | 0 | 1 | 1.94 | 0.956 | 0.73503911 | NA | 0 | 5.1562 | 3.9771 | 3.9771 | TRUE | 0.2566 |
| Gm6531 | 0 | 0 | 0 | 0 | 0 | 0 | 0 | 0 | 0 | 0 | 0 | 0 | 0 | 0 | 0 | NA | NA | NA | NA | NA | NA | NA | NA | NA | NA |
| Gm6537 | 1 | 1 | 1 | 1 | 0 | 2 | 1 | 1 | 1 | 1 | 0 | 2 | 0.98 | 1 | 1 | 1.044 | 0.062 | 0.97363744 | NA | 0 | 2.4171 | 1.5475 | 1.5475 | TRUE | 0.1802 |
| Gm6548 | 671 | 448 | 349 | 444 | 364 | 477 | 521 | 477 | 377 | 442 | 411 | 471 | 449.62 | 458 | 441 | 0.961 | -0.057 | 0.65470578 | 0.78550405 | 0.0138 | 0.0077 | 0.0096 | 0.0096 | TRUE | 0.276 |

| | | | | | | | | | | | | | | | | | | | | | | | | | |
|---|---|---|---|---|---|---|---|---|---|---|---|---|---|---|---|---|---|---|---|---|---|---|---|---|---|
| Gm6567 | 0 | 0 | 0 | 0 | 0 | 0 | 0 | 0 | 0 | 0 | 0 | 0 | 0 | 0 | 0 | NA | NA | NA | NA | NA | NA | NA | NA | NA | NA |
| Gm6568 | 0 | 0 | 1 | 0 | 0 | 0 | 0 | 0 | 1 | 0 | 0 | 0 | 0.18 | 0 | 0 | 0.552 | -0.857 | 0.8336548 | NA | 0 | 13.1817 | 10 | 10 | TRUE | 0.5365 |
| Gm6578 | 0 | 0 | 0 | 0 | 0 | 0 | 0 | 0 | 0 | 0 | 0 | 0 | 0 | 0 | 0 | NA | NA | NA | NA | NA | NA | NA | NA | NA | NA |
| Gm6583 | 8 | 4 | 1 | 4 | 3 | 3 | 6 | 4 | 1 | 4 | 3 | 3 | 3.65 | 4 | 3 | 0.875 | -0.193 | 0.84920384 | NA | 0 | 0.6525 | 0.4721 | 0.4721 | TRUE | 0.3108 |
| Gm6588 | 12 | 12 | 7 | 13 | 5 | 9 | 9 | 13 | 8 | 13 | 6 | 9 | 9.52 | 10 | 9 | 0.934 | -0.099 | 0.87610371 | NA | 0 | 0.2515 | 0.1838 | 0.1838 | TRUE | 0.2639 |
| Gm6592 | 1 | 0 | 1 | 0 | 0 | 0 | 1 | 0 | 1 | 0 | 0 | 0 | 0.31 | 1 | 0 | 0.307 | -1.706 | 0.62894649 | NA | 0 | 7.6657 | 7.2 | 7.2 | TRUE | 0.2067 |
| Gm6602 | 0 | 0 | 0 | 0 | 0 | 0 | 0 | 0 | 0 | 0 | 0 | 0 | 0 | 0 | 0 | NA | NA | NA | NA | NA | NA | NA | NA | NA | NA |
| Gm6607 | 4 | 2 | 3 | 1 | 3 | 1 | 3 | 2 | 3 | 1 | 3 | 1 | 2.31 | 3 | 2 | 0.62 | -0.689 | 0.58373734 | NA | 0 | 1.0301 | 0.675 | 0.675 | TRUE | 0.4351 |
| Gm6614 | 0 | 0 | 0 | 0 | 0 | 0 | 0 | 0 | 0 | 0 | 0 | 0 | 0 | 0 | 0 | NA | NA | NA | NA | NA | NA | NA | NA | NA | NA |
| Gm6623 | 2 | 2 | 0 | 0 | 1 | 0 | 2 | 2 | 0 | 0 | 1 | 0 | 0.8 | 1 | 0 | 0.3 | -1.737 | 0.45535154 | NA | 0 | 2.9596 | 2.5115 | 2.5115 | TRUE | 0.5075 |
| Gm6634 | 0 | 0 | 0 | 0 | 0 | 0 | 0 | 0 | 0 | 0 | 0 | 0 | 0 | 0 | 0 | NA | NA | NA | NA | NA | NA | NA | NA | NA | NA |
| Gm6639 | 13 | 12 | 6 | 10 | 13 | 10 | 10 | 13 | 6 | 10 | 15 | 10 | 10.64 | 10 | 12 | 1.168 | 0.225 | 0.70494265 | NA | 0 | 0.2252 | 0.159 | 0.159 | TRUE | 0.2635 |
| Gm6642 | 0 | 1 | 0 | 0 | 0 | 0 | 0 | 0 | 1 | 0 | 0 | 0 | 0.18 | 0 | 0 | 0.552 | -0.857 | 0.8336548 | NA | 0 | 13.3714 | 10 | 10 | TRUE | 0.5216 |
| Gm6644 | 0 | 0 | 0 | 0 | 0 | 0 | 0 | 0 | 0 | 0 | 0 | 0 | 0 | 0 | 0 | NA | NA | NA | NA | NA | NA | NA | NA | NA | NA |
| Gm6654 | 1 | 0 | 0 | 1 | 1 | 1 | 1 | 0 | 0 | 1 | 1 | 1 | 0.65 | 0 | 1 | 3.004 | 1.587 | 0.49664754 | NA | 0 | 3.6606 | 2.4271 | 2.4271 | TRUE | 0.2566 |
| Gm6682 | 384 | 245 | 258 | 254 | 225 | 244 | 298 | 261 | 278 | 253 | 254 | 241 | 264.12 | 279 | 249 | 0.891 | -0.167 | 0.18802223 | 0.3451582 | 0 | 0.0114 | 0.0078 | 0.0078 | TRUE | 0.0439 |
| Gm6696 | 0 | 0 | 0 | 0 | 0 | 0 | 0 | 0 | 0 | 0 | 0 | 0 | 0 | 0 | 0 | NA | NA | NA | NA | NA | NA | NA | NA | NA | NA |
| Gm6710 | 0 | 0 | 0 | 0 | 0 | 0 | 0 | 0 | 0 | 0 | 0 | 0 | 0 | 0 | 0 | NA | NA | NA | NA | NA | NA | NA | NA | NA | NA |
| Gm6756 | 15 | 8 | 13 | 12 | 4 | 9 | 12 | 9 | 14 | 12 | 5 | 9 | 9.92 | 12 | 9 | 0.747 | -0.421 | 0.50223606 | NA | 0 | 0.2413 | 0.1806 | 0.1806 | TRUE | 0.3421 |
| Gm6760 | 0 | 0 | 0 | 0 | 0 | 1 | 0 | 0 | 0 | 0 | 0 | 1 | 0.16 | 0 | 0 | 2.094 | 1.067 | 0.79379329 | NA | 0 | 14.4078 | 10 | 10 | TRUE | 0.3835 |
| Gm6763 | 0 | 0 | 0 | 0 | 0 | 0 | 0 | 0 | 0 | 0 | 0 | 0 | 0 | 0 | 0 | NA | NA | NA | NA | NA | NA | NA | NA | NA | NA |
| Gm6772 | 21 | 23 | 15 | 29 | 15 | 24 | 16 | 24 | 16 | 29 | 17 | 24 | 21.07 | 19 | 23 | 1.231 | 0.3 | 0.48989261 | 0.65568265 | 0 | 0.1149 | 0.0878 | 0.0878 | TRUE | 0.2303 |
| Gm6787 | 11 | 4 | 5 | 5 | 6 | 7 | 9 | 4 | 5 | 5 | 7 | 7 | 6.14 | 6 | 6 | 1.008 | 0.011 | 0.98813714 | NA | 0 | 0.3883 | 0.2602 | 0.2602 | TRUE | 0.3949 |
| Gm6792 | 81 | 55 | 71 | 86 | 73 | 87 | 63 | 59 | 77 | 86 | 82 | 86 | 75.31 | 66 | 85 | 1.285 | 0.362 | 0.11005516 | 0.23724903 | 0 | 0.0339 | 0.0235 | 0.0235 | TRUE | 0.1622 |
| Gm6793 | 0 | 0 | 0 | 0 | 0 | 0 | 0 | 0 | 0 | 0 | 0 | 0 | 0 | 0 | 0 | NA | NA | NA | NA | NA | NA | NA | NA | NA | NA |
| Gm6812 | 7 | 1 | 5 | 6 | 4 | 2 | 5 | 1 | 5 | 6 | 5 | 2 | 4.06 | 4 | 4 | 1.033 | 0.046 | 0.96262047 | NA | 0 | 0.5863 | 0.4599 | 0.4599 | TRUE | 0.6211 |
| Gm684 | 63 | 30 | 34 | 77 | 63 | 44 | 49 | 32 | 37 | 77 | 71 | 43 | 51.45 | 39 | 64 | 1.61 | 0.687 | 0.02705455 | 0.08591216 | 0.0513 | 0.0485 | 0.0493 | 0.0493 | TRUE | 0.6965 |
| Gm6878 | 0 | 1 | 0 | 0 | 0 | 0 | 0 | 1 | 0 | 0 | 0 | 0 | 0.18 | 0 | 0 | 0.552 | -0.857 | 0.8336548 | NA | 0 | 13.3714 | 10 | 10 | TRUE | 0.5216 |
| Gm6880 | 349 | 225 | 219 | 207 | 188 | 212 | 271 | 239 | 236 | 206 | 212 | 209 | 229.02 | 249 | 209 | 0.838 | -0.256 | 0.05931208 | 0.15214423 | 0 | 0.0128 | 0.0088 | 0.0088 | TRUE | 0.0729 |
| Gm6890 | 22 | 19 | 14 | 18 | 12 | 7 | 17 | 20 | 15 | 18 | 14 | 7 | 15.13 | 17 | 13 | 0.731 | -0.451 | 0.38419498 | NA | 0.0217 | 0.1591 | 0.1256 | 0.1256 | TRUE | 0.4497 |
| Gm6902 | 1 | 0 | 0 | 0 | 0 | 0 | 1 | 0 | 0 | 0 | 0 | 0 | 0.13 | 0 | 0 | 0.552 | -0.857 | 0.8336548 | NA | 0 | 18.3121 | 10 | 10 | TRUE | 0.2566 |
| Gm6904 | 0 | 0 | 0 | 1 | 0 | 0 | 0 | 0 | 0 | 1 | 0 | 0 | 0.17 | 0 | 0 | 2.094 | 1.067 | 0.79379329 | NA | 0 | 14.2885 | 10 | 10 | TRUE | 0.3906 |
| Gm6927 | 0 | 0 | 0 | 0 | 0 | 0 | 0 | 0 | 0 | 0 | 0 | 0 | 0 | 0 | 0 | NA | NA | NA | NA | NA | NA | NA | NA | NA | NA |
| Gm6936 | 0 | 0 | 0 | 0 | 0 | 0 | 0 | 0 | 0 | 0 | 0 | 0 | 0 | 0 | 0 | NA | NA | NA | NA | NA | NA | NA | NA | NA | NA |
| Gm6938 | 3 | 3 | 3 | 2 | 2 | 3 | 2 | 3 | 3 | 2 | 2 | 3 | 2.66 | 3 | 2 | 0.831 | -0.267 | 0.81337489 | NA | 0 | 0.8931 | 0.5406 | 0.5406 | TRUE | 0.0545 |
| Gm694 | 2 | 2 | 3 | 0 | 0 | 1 | 2 | 2 | 3 | 0 | 0 | 1 | 1.32 | 2 | 0 | 0.162 | -2.624 | 0.15867857 | NA | 0 | 1.8014 | 1.286 | 1.286 | TRUE | 0.3835 |
| Gm6981 | 56 | 41 | 20 | 27 | 23 | 23 | 43 | 44 | 22 | 27 | 26 | 23 | 30.7 | 36 | 25 | 0.69 | -0.535 | 0.16017723 | 0.30958933 | 0.0474 | 0.0796 | 0.0706 | 0.0706 | TRUE | 0.8434 |
| Gm6994 | 0 | 0 | 0 | 0 | 0 | 0 | 0 | 0 | 0 | 0 | 0 | 0 | 0 | 0 | 0 | NA | NA | NA | NA | NA | NA | NA | NA | NA | NA |
| Gm7008 | 4 | 1 | 4 | 2 | 3 | 0 | 3 | 1 | 4 | 2 | 3 | 0 | 2.31 | 3 | 2 | 0.622 | -0.686 | 0.60863342 | NA | 0 | 1.0283 | 0.8295 | 0.8295 | TRUE | 0.4485 |
| Gm7030 | 3 | 1 | 0 | 1 | 0 | 0 | 2 | 1 | 0 | 1 | 0 | 0 | 0.73 | 1 | 0 | 0.319 | -1.646 | 0.49448041 | NA | 0 | 3.2424 | 2.7675 | 2.7675 | TRUE | 0.189 |
| Gm7056 | 0 | 0 | 0 | 0 | 0 | 0 | 0 | 0 | 0 | 0 | 0 | 0 | 0 | 0 | 0 | NA | NA | NA | NA | NA | NA | NA | NA | NA | NA |
| Gm7073 | 0 | 0 | 0 | 0 | 0 | 0 | 0 | 0 | 0 | 0 | 0 | 0 | 0 | 0 | 0 | NA | NA | NA | NA | NA | NA | NA | NA | NA | NA |
| Gm7092 | 0 | 0 | 1 | 1 | 0 | 0 | 0 | 0 | 1 | 1 | 0 | 0 | 0.35 | 0 | 0 | 1.075 | 0.105 | 0.97567165 | NA | 0 | 6.8576 | 6.5143 | 6.5143 | TRUE | 0.5365 |
| Gm7102 | 17 | 7 | 6 | 6 | 3 | 5 | 13 | 7 | 6 | 6 | 3 | 5 | 6.9 | 9 | 5 | 0.522 | -0.937 | 0.21277471 | NA | 0 | 0.3458 | 0.2467 | 0.2467 | TRUE | 0.6579 |
| Gm7104 | 48 | 37 | 22 | 14 | 14 | 11 | 37 | 39 | 24 | 14 | 16 | 11 | 23.49 | 33 | 14 | 0.401 | -1.317 | 0.00212285 | 0.0126162 | 0 | 0.1033 | 0.0795 | 0.0795 | TRUE | 0.4204 |
| Gm711 | 8 | 6 | 3 | 1 | 4 | 5 | 6 | 6 | 3 | 1 | 5 | 5 | 4.38 | 5 | 4 | 0.652 | -0.616 | 0.51480613 | NA | 0 | 0.5436 | 0.4043 | 0.4043 | TRUE | 0.6016 |
| Gm7120 | 95 | 57 | 68 | 49 | 51 | 84 | 74 | 61 | 73 | 49 | 58 | 83 | 66.17 | 69 | 63 | 0.91 | -0.135 | 0.60914609 | 0.75263876 | 0.0281 | 0.0382 | 0.0354 | 0.0354 | TRUE | 0.6793 |
| Gm7134 | 0 | 0 | 0 | 0 | 0 | 0 | 0 | 0 | 0 | 0 | 0 | 0 | 0 | 0 | 0 | NA | NA | NA | NA | NA | NA | NA | NA | NA | NA |
| Gm715 | 35 | 23 | 13 | 24 | 13 | 20 | 27 | 24 | 14 | 24 | 15 | 20 | 20.66 | 22 | 20 | 0.884 | -0.179 | 0.68875122 | 0.80956516 | 0.0332 | 0.1171 | 0.0948 | 0.0948 | TRUE | 0.5223 |
| Gm7157 | 1 | 0 | 0 | 0 | 0 | 1 | 1 | 0 | 0 | 0 | 0 | 1 | 0.29 | 0 | 0 | 1.075 | 0.105 | 0.97702352 | NA | 0 | 8.0647 | 7.5458 | 7.5458 | TRUE | 0.3835 |
| Gm7168 | 0 | 0 | 0 | 0 | 0 | 0 | 0 | 0 | 0 | 0 | 0 | 0 | 0 | 0 | 0 | NA | NA | NA | NA | NA | NA | NA | NA | NA | NA |
| Gm7173 | 33 | 17 | 22 | 17 | 23 | 20 | 26 | 18 | 24 | 17 | 26 | 20 | 21.68 | 23 | 21 | 0.92 | -0.12 | 0.77370579 | 0.86638803 | 0 | 0.1118 | 0.0797 | 0.0797 | TRUE | 0.2292 |
| Gm7244 | 3 | 0 | 0 | 1 | 0 | 0 | 2 | 0 | 0 | 1 | 0 | 0 | 0.55 | 1 | 0 | 0.458 | -1.128 | 0.69820437 | NA | 0 | 4.2787 | 4.502 | 4.502 | TRUE | 1.4399 |
| Gm7257 | 0 | 0 | 0 | 0 | 0 | 0 | 0 | 0 | 0 | 0 | 0 | 0 | 0 | 0 | 0 | NA | NA | NA | NA | NA | NA | NA | NA | NA | NA |
| Gm7271 | 0 | 0 | 0 | 0 | 0 | 0 | 0 | 0 | 0 | 0 | 0 | 0 | 0 | 0 | 0 | NA | NA | NA | NA | NA | NA | NA | NA | NA | NA |
| Gm732 | 0 | 0 | 0 | 0 | 0 | 0 | 0 | 0 | 0 | 0 | 0 | 0 | 0 | 0 | 0 | NA | NA | NA | NA | NA | NA | NA | NA | NA | NA |
| Gm7325 | 2943 | 2394 | 2520 | 3166 | 2913 | 3432 | 2286 | 2546 | 2719 | 3151 | 3286 | 3388 | 2896.12 | 2517 | 3275 | 1.302 | 0.381 | 2.37E-07 | 8.48E-06 | 0.0039 | 0.0032 | 0.0036 | 0.0036 | TRUE | 0.0803 |
| Gm7334 | 0 | 0 | 0 | 0 | 0 | 0 | 0 | 0 | 0 | 0 | 0 | 0 | 0 | 0 | 0 | NA | NA | NA | NA | NA | NA | NA | NA | NA | NA |
| Gm7337 | 0 | 0 | 0 | 0 | 0 | 0 | 0 | 0 | 0 | 0 | 0 | 0 | 0 | 0 | 0 | NA | NA | NA | NA | NA | NA | NA | NA | NA | NA |
| Gm7367 | 0 | 0 | 0 | 0 | 0 | 0 | 0 | 0 | 0 | 0 | 0 | 0 | 0 | 0 | 0 | NA | NA | NA | NA | NA | NA | NA | NA | NA | NA |
| Gm7444 | 2 | 0 | 0 | 2 | 1 | 0 | 2 | 0 | 0 | 2 | 1 | 0 | 0.78 | 1 | 1 | 1.851 | 0.888 | 0.70714822 | NA | 0 | 3.046 | 2.6704 | 2.6704 | TRUE | 0.5318 |
| Gm7457 | 0 | 0 | 0 | 0 | 0 | 0 | 0 | 0 | 0 | 0 | 0 | 0 | 0 | 0 | 0 | NA | NA | NA | NA | NA | NA | NA | NA | NA | NA |
| Gm7534 | 3 | 5 | 4 | 4 | 3 | 1 | 2 | 5 | 4 | 4 | 3 | 1 | 3.39 | 4 | 3 | 0.705 | -0.504 | 0.63280964 | NA | 0 | 0.7023 | 0.4933 | 0.4933 | TRUE | 0.4086 |
| Gm7538 | 2 | 1 | 0 | 0 | 0 | 0 | 2 | 1 | 0 | 0 | 0 | 0 | 0.44 | 1 | 0 | 0.215 | -2.217 | 0.47268385 | NA | 0 | 5.4363 | 5.3181 | 5.3181 | TRUE | 0.2411 |
| Gm7550 | 0 | 0 | 0 | 0 | 0 | 0 | 0 | 0 | 0 | 0 | 0 | 0 | 0 | 0 | 0 | NA | NA | NA | NA | NA | NA | NA | NA | NA | NA |
| Gm757 | 7 | 9 | 3 | 19 | 9 | 14 | 5 | 10 | 3 | 19 | 10 | 14 | 10.19 | 6 | 14 | 2.368 | 1.244 | 0.05432081 | NA | 0.0331 | 0.235 | 0.1854 | 0.1854 | TRUE | 0.3371 |
| Gm7609 | 0 | 0 | 0 | 0 | 0 | 0 | 0 | 0 | 0 | 0 | 0 | 0 | 0 | 0 | 0 | NA | NA | NA | NA | NA | NA | NA | NA | NA | NA |
| Gm7616 | 0 | 0 | 0 | 1 | 0 | 0 | 0 | 0 | 0 | 1 | 0 | 0 | 0.17 | 0 | 0 | 2.094 | 1.067 | 0.79379329 | NA | 0 | 14.2885 | 10 | 10 | TRUE | 0.3906 |
| Gm765 | 0 | 0 | 0 | 0 | 0 | 0 | 0 | 0 | 0 | 0 | 0 | 0 | 0 | 0 | 0 | NA | NA | NA | NA | NA | NA | NA | NA | NA | NA |

| | | | | | | | | | | | | | | | | | | | | | | | | | |
|---|---|---|---|---|---|---|---|---|---|---|---|---|---|---|---|---|---|---|---|---|---|---|---|---|---|
| Gm766 | 18 | 10 | 14 | 14 | 6 | 18 | 14 | 11 | 15 | 14 | 7 | 18 | 13.03 | 13 | 13 | 0.974 | -0.038 | 0.94546971 | NA | 0 | 0.1843 | 0.1438 | 0.1438 | TRUE | 0.427 |
| Gm7694 | 142 | 110 | 116 | 102 | 132 | 130 | 110 | 117 | 125 | 102 | 149 | 128 | 121.87 | 117 | 126 | 1.074 | 0.103 | 0.5945875 | 0.74105677 | 0.0117 | 0.0219 | 0.019 | 0.019 | TRUE | 0.2498 |
| Gm7714 | 0 | 0 | 0 | 0 | 1 | 0 | 0 | 0 | 0 | 0 | 1 | 0 | 0.19 | 0 | 0 | 2.094 | 1.067 | 0.79379329 | NA | 0 | 12.608 | 10 | 10 | TRUE | 0.5075 |
| Gm773 | 44 | 38 | 15 | 6 | 2 | 2 | 34 | 40 | 16 | 6 | 2 | 2 | 16.83 | 30 | 3 | 0.113 | -3.145 | 1.82E-06 | NA | 0.1564 | 0.1432 | 0.1463 | 0.1463 | TRUE | 0.4238 |
| Gm7849 | 0 | 0 | 0 | 0 | 0 | 0 | 0 | 0 | 0 | 0 | 0 | 0 | 0 | 0 | 0 | NA | NA | NA | NA | NA | NA | NA | NA | NA | NA |
| Gm7854 | 2 | 0 | 0 | 3 | 0 | 0 | 2 | 0 | 0 | 3 | 0 | 0 | 0.76 | 1 | 1 | 1.815 | 0.86 | 0.74367993 | NA | 8.431 | 3.135 | 3.5951 | 3.5951 | TRUE | 1.4418 |
| Gm7861 | 0 | 0 | 0 | 0 | 0 | 0 | 0 | 0 | 0 | 0 | 0 | 0 | 0 | 0 | 0 | NA | NA | NA | NA | NA | NA | NA | NA | NA | NA |
| Gm7903 | 0 | 0 | 0 | 0 | 0 | 0 | 0 | 0 | 0 | 0 | 0 | 0 | 0 | 0 | 0 | NA | NA | NA | NA | NA | NA | NA | NA | NA | NA |
| Gm7904 | 0 | 2 | 2 | 1 | 4 | 3 | 0 | 2 | 2 | 1 | 5 | 3 | 2.13 | 1 | 3 | 2.058 | 1.041 | 0.46084173 | NA | 0 | 1.1173 | 0.8985 | 0.8985 | TRUE | 0.3806 |
| Gm7977 | 0 | 1 | 0 | 0 | 1 | 2 | 0 | 1 | 0 | 0 | 1 | 2 | 0.69 | 0 | 1 | 2.998 | 1.584 | 0.52044796 | NA | 0 | 3.4158 | 2.8778 | 2.8778 | TRUE | 0.3487 |
| Gm7978 | 0 | 0 | 0 | 0 | 0 | 0 | 0 | 0 | 0 | 0 | 0 | 0 | 0 | 0 | 0 | NA | NA | NA | NA | NA | NA | NA | NA | NA | NA |
| Gm805 | 20 | 20 | 13 | 12 | 9 | 11 | 16 | 21 | 14 | 12 | 10 | 11 | 13.97 | 17 | 11 | 0.651 | -0.62 | 0.22888448 | NA | 0 | 0.1721 | 0.1154 | 0.1154 | TRUE | 0.2285 |
| Gm806 | 1 | 3 | 3 | 2 | 4 | 0 | 1 | 3 | 3 | 2 | 5 | 0 | 2.28 | 2 | 2 | 0.908 | -0.14 | 0.91848965 | NA | 0.4605 | 1.0398 | 0.8972 | 0.8972 | TRUE | 0.2685 |
| Gm8096 | 2 | 1 | 0 | 0 | 2 | 0 | 2 | 1 | 0 | 0 | 2 | 0 | 0.81 | 1 | 1 | 0.816 | -0.293 | 0.89959898 | NA | 0 | 2.9205 | 2.6456 | 2.6456 | TRUE | 0.9748 |
| Gm813 | 0 | 0 | 0 | 0 | 0 | 0 | 0 | 0 | 0 | 0 | 0 | 0 | 0 | 0 | 0 | NA | NA | NA | NA | NA | NA | NA | NA | NA | NA |
| Gm815 | 0 | 0 | 0 | 0 | 0 | 0 | 0 | 0 | 0 | 0 | 0 | 0 | 0 | 0 | 0 | NA | NA | NA | NA | NA | NA | NA | NA | NA | NA |
| Gm8179 | 0 | 0 | 0 | 0 | 0 | 0 | 0 | 0 | 0 | 0 | 0 | 0 | 0 | 0 | 0 | NA | NA | NA | NA | NA | NA | NA | NA | NA | NA |
| Gm8221 | 7 | 5 | 4 | 3 | 1 | 1 | 5 | 5 | 4 | 3 | 1 | 1 | 3.36 | 5 | 2 | 0.34 | -1.556 | 0.1558732 | NA | 0 | 0.7074 | 0.4686 | 0.4686 | TRUE | 0.3486 |
| Gm8234 | 2 | 2 | 4 | 0 | 0 | 1 | 2 | 2 | 4 | 0 | 0 | 1 | 1.5 | 3 | 0 | 0.141 | -2.83 | 0.11801785 | NA | 0 | 1.5853 | 1.1789 | 1.1789 | TRUE | 0.3835 |
| Gm826 | 0 | 1 | 0 | 1 | 0 | 0 | 0 | 1 | 0 | 1 | 0 | 0 | 0.34 | 0 | 0 | 1.075 | 0.105 | 0.97573409 | NA | 0 | 6.9086 | 6.5582 | 6.5582 | TRUE | 0.5216 |
| Gm8267 | 1 | 1 | 0 | 0 | 0 | 0 | 1 | 1 | 0 | 0 | 0 | 0 | 0.31 | 1 | 0 | 0.309 | -1.696 | 0.63183309 | NA | 0 | 7.7294 | 7.2472 | 7.2472 | TRUE | 0.2015 |
| Gm829 | 0 | 0 | 0 | 1 | 0 | 0 | 0 | 0 | 0 | 1 | 0 | 0 | 0.17 | 0 | 0 | 2.094 | 1.067 | 0.79379329 | NA | 0 | 14.2885 | 10 | 10 | TRUE | 0.3906 |
| Gm8298 | 0 | 0 | 0 | 0 | 0 | 0 | 0 | 0 | 0 | 0 | 0 | 0 | 0 | 0 | 0 | NA | NA | NA | NA | NA | NA | NA | NA | NA | NA |
| Gm8300 | 2212 | 1551 | 620 | 717 | 703 | 559 | 1718 | 1650 | 669 | 714 | 793 | 552 | 1015.88 | 1346 | 686 | 0.51 | -0.972 | 0.02632146 | 0.08440938 | 0.1368 | 0.0048 | 0.0366 | 0.1368 | TRUE | 2.3219 |
| Gm833 | 1 | 0 | 0 | 0 | 0 | 1 | 1 | 0 | 0 | 0 | 0 | 1 | 0.29 | 0 | 0 | 1.075 | 0.105 | 0.97702352 | NA | 0 | 8.0647 | 7.5458 | 7.5458 | TRUE | 0.3835 |
| Gm8363 | 0 | 0 | 0 | 0 | 0 | 0 | 0 | 0 | 0 | 0 | 0 | 0 | 0 | 0 | 0 | NA | NA | NA | NA | NA | NA | NA | NA | NA | NA |
| Gm8369 | 0 | 0 | 0 | 0 | 0 | 0 | 0 | 0 | 0 | 0 | 0 | 0 | 0 | 0 | 0 | NA | NA | NA | NA | NA | NA | NA | NA | NA | NA |
| Gm839 | 0 | 0 | 0 | 0 | 0 | 0 | 0 | 0 | 0 | 0 | 0 | 0 | 0 | 0 | 0 | NA | NA | NA | NA | NA | NA | NA | NA | NA | NA |
| Gm8439 | 0 | 0 | 0 | 0 | 0 | 0 | 0 | 0 | 0 | 0 | 0 | 0 | 0 | 0 | 0 | NA | NA | NA | NA | NA | NA | NA | NA | NA | NA |
| Gm8453 | 0 | 0 | 0 | 0 | 0 | 0 | 0 | 0 | 0 | 0 | 0 | 0 | 0 | 0 | 0 | NA | NA | NA | NA | NA | NA | NA | NA | NA | NA |
| Gm853 | 34 | 40 | 48 | 85 | 74 | 68 | 26 | 43 | 52 | 85 | 83 | 67 | 59.33 | 40 | 78 | 1.973 | 0.981 | 0.0007153 | 0.00565472 | 0.0399 | 0.0424 | 0.0418 | 0.0418 | TRUE | 0.8236 |
| Gm8579 | 1 | 1 | 0 | 0 | 0 | 0 | 1 | 1 | 0 | 0 | 0 | 0 | 0.31 | 1 | 0 | 0.309 | -1.696 | 0.63183309 | NA | 0 | 7.7294 | 7.2472 | 7.2472 | TRUE | 0.2015 |
| Gm8580 | 1 | 0 | 2 | 0 | 2 | 0 | 1 | 0 | 2 | 0 | 2 | 0 | 0.87 | 1 | 1 | 0.751 | -0.414 | 0.85669157 | NA | 0 | 2.7419 | 2.5715 | 2.5715 | TRUE | 0.7602 |
| Gm8615 | 535 | 408 | 449 | 472 | 435 | 517 | 416 | 434 | 484 | 470 | 491 | 510 | 467.48 | 445 | 490 | 1.105 | 0.144 | 0.16904232 | 0.32056811 | 0.0019 | 0.0075 | 0.0057 | 0.0057 | TRUE | 0.0698 |
| Gm8633 | 0 | 0 | 0 | 0 | 0 | 0 | 0 | 0 | 0 | 0 | 0 | 0 | 0 | 0 | 0 | NA | NA | NA | NA | NA | NA | NA | NA | NA | NA |
| Gm8660 | 0 | 0 | 0 | 0 | 0 | 0 | 0 | 0 | 0 | 0 | 0 | 0 | 0 | 0 | 0 | NA | NA | NA | NA | NA | NA | NA | NA | NA | NA |
| Gm8677 | 0 | 0 | 0 | 0 | 0 | 0 | 0 | 0 | 0 | 0 | 0 | 0 | 0 | 0 | 0 | NA | NA | NA | NA | NA | NA | NA | NA | NA | NA |
| Gm8693 | 0 | 0 | 0 | 0 | 0 | 0 | 0 | 0 | 0 | 0 | 0 | 0 | 0 | 0 | 0 | NA | NA | NA | NA | NA | NA | NA | NA | NA | NA |
| Gm8709 | 0 | 0 | 0 | 0 | 0 | 0 | 0 | 0 | 0 | 0 | 0 | 0 | 0 | 0 | 0 | NA | NA | NA | NA | NA | NA | NA | NA | NA | NA |
| Gm872 | 1 | 0 | 2 | 2 | 1 | 0 | 1 | 0 | 2 | 2 | 1 | 0 | 1.01 | 1 | 1 | 1.071 | 0.098 | 0.96051014 | NA | 0 | 2.3515 | 1.8491 | 1.8491 | TRUE | 0.3232 |
| Gm8720 | 0 | 0 | 0 | 0 | 0 | 0 | 0 | 0 | 0 | 0 | 0 | 0 | 0 | 0 | 0 | NA | NA | NA | NA | NA | NA | NA | NA | NA | NA |
| Gm8764 | 0 | 0 | 0 | 0 | 0 | 0 | 0 | 0 | 0 | 0 | 0 | 0 | 0 | 0 | 0 | NA | NA | NA | NA | NA | NA | NA | NA | NA | NA |
| Gm8765 | 0 | 0 | 0 | 1 | 0 | 1 | 0 | 0 | 0 | 1 | 0 | 1 | 0.33 | 0 | 1 | 3.763 | 1.912 | 0.57887588 | NA | 0 | 7.1752 | 6.7756 | 6.7756 | TRUE | 0.2094 |
| Gm8773 | 11 | 6 | 10 | 10 | 6 | 11 | 9 | 6 | 11 | 10 | 7 | 11 | 8.88 | 9 | 9 | 1.078 | 0.108 | 0.86537087 | NA | 0 | 0.2692 | 0.1817 | 0.1817 | TRUE | 0.1543 |
| Gm8787 | 13 | 12 | 17 | 11 | 6 | 12 | 10 | 13 | 18 | 11 | 7 | 12 | 11.79 | 14 | 10 | 0.728 | -0.457 | 0.42452274 | NA | 0 | 0.2034 | 0.1486 | 0.1486 | TRUE | 0.354 |
| Gm8801 | 220 | 137 | 95 | 124 | 105 | 104 | 171 | 146 | 103 | 123 | 118 | 103 | 127.27 | 140 | 115 | 0.817 | -0.291 | 0.16582877 | 0.31682805 | 0.0304 | 0.021 | 0.0238 | 0.0238 | TRUE | 0.3278 |
| Gm8817 | 13 | 5 | 3 | 7 | 3 | 3 | 10 | 5 | 3 | 7 | 3 | 3 | 5.33 | 6 | 4 | 0.702 | -0.51 | 0.55577767 | NA | 0 | 0.4473 | 0.3477 | 0.3477 | TRUE | 0.6058 |
| Gm884 | 31 | 24 | 16 | 21 | 12 | 13 | 24 | 26 | 17 | 21 | 14 | 13 | 19.02 | 22 | 16 | 0.706 | -0.501 | 0.2686823 | 0.43924008 | 0 | 0.127 | 0.0937 | 0.0937 | TRUE | 0.3936 |
| Gm8882 | 0 | 0 | 0 | 0 | 0 | 0 | 0 | 0 | 0 | 0 | 0 | 0 | 0 | 0 | 0 | NA | NA | NA | NA | NA | NA | NA | NA | NA | NA |
| Gm8883 | 0 | 0 | 0 | 0 | 0 | 0 | 0 | 0 | 0 | 0 | 0 | 0 | 0 | 0 | 0 | NA | NA | NA | NA | NA | NA | NA | NA | NA | NA |
| Gm8884 | 0 | 0 | 0 | 0 | 0 | 0 | 0 | 0 | 0 | 0 | 0 | 0 | 0 | 0 | 0 | NA | NA | NA | NA | NA | NA | NA | NA | NA | NA |
| Gm889 | 0 | 0 | 0 | 0 | 0 | 0 | 0 | 0 | 0 | 0 | 0 | 0 | 0 | 0 | 0 | NA | NA | NA | NA | NA | NA | NA | NA | NA | NA |
| Gm8898 | 0 | 0 | 0 | 0 | 0 | 0 | 0 | 0 | 0 | 0 | 0 | 0 | 0 | 0 | 0 | NA | NA | NA | NA | NA | NA | NA | NA | NA | NA |
| Gm8909 | 10 | 6 | 4 | 5 | 12 | 14 | 8 | 6 | 4 | 5 | 14 | 14 | 8.47 | 6 | 11 | 1.726 | 0.787 | 0.25832554 | NA | 0 | 0.2824 | 0.2239 | 0.2239 | TRUE | 0.8343 |
| Gm8979 | 0 | 0 | 0 | 0 | 0 | 0 | 0 | 0 | 0 | 0 | 0 | 0 | 0 | 0 | 0 | NA | NA | NA | NA | NA | NA | NA | NA | NA | NA |
| Gm8989 | 0 | 0 | 0 | 0 | 0 | 0 | 0 | 0 | 0 | 0 | 0 | 0 | 0 | 0 | 0 | NA | NA | NA | NA | NA | NA | NA | NA | NA | NA |
| Gm8994 | 127 | 120 | 45 | 67 | 76 | 65 | 99 | 128 | 49 | 67 | 86 | 64 | 81.9 | 92 | 72 | 0.785 | -0.348 | 0.24927377 | 0.41812816 | 0.1103 | 0.0314 | 0.0535 | 0.0535 | TRUE | 0.3138 |
| Gm9 | 548 | 388 | 284 | 291 | 218 | 248 | 426 | 413 | 306 | 290 | 246 | 245 | 320.86 | 382 | 260 | 0.681 | -0.555 | 0.00015203 | 0.00172595 | 0.0177 | 0.0098 | 0.0122 | 0.0122 | TRUE | 0.3327 |
| Gm904 | 3 | 2 | 3 | 1 | 2 | 0 | 2 | 2 | 3 | 1 | 2 | 0 | 1.82 | 2 | 1 | 0.413 | -1.274 | 0.38959591 | NA | 0 | 1.3016 | 0.9027 | 0.9027 | TRUE | 0.3965 |
| Gm9041 | 0 | 0 | 0 | 0 | 0 | 0 | 0 | 0 | 0 | 0 | 0 | 0 | 0 | 0 | 0 | NA | NA | NA | NA | NA | NA | NA | NA | NA | NA |
| Gm9047 | 0 | 0 | 1 | 0 | 0 | 0 | 0 | 0 | 1 | 0 | 0 | 0 | 0.18 | 0 | 0 | 0.552 | -0.857 | 0.8336548 | NA | 0 | 13.1817 | 10 | 10 | TRUE | 0.5365 |
| Gm9054 | 3 | 1 | 1 | 1 | 0 | 1 | 2 | 1 | 1 | 1 | 0 | 1 | 1.08 | 1 | 1 | 0.445 | -1.168 | 0.52935111 | NA | 0 | 2.2054 | 1.4011 | 1.4011 | TRUE | 0.2213 |
| Gm906 | 5 | 5 | 4 | 7 | 9 | 8 | 4 | 5 | 4 | 7 | 10 | 8 | 6.42 | 4 | 8 | 1.857 | 0.893 | 0.23543384 | NA | 0 | 0.3714 | 0.2386 | 0.2386 | TRUE | 0.0985 |
| Gm9079 | 31 | 27 | 21 | 40 | 24 | 23 | 24 | 29 | 23 | 40 | 27 | 23 | 27.51 | 25 | 30 | 1.19 | 0.251 | 0.51075399 | 0.6741349 | 0 | 0.0886 | 0.0688 | 0.0688 | TRUE | 0.5781 |
| Gm9112 | 195 | 175 | 200 | 126 | 144 | 140 | 151 | 186 | 216 | 125 | 162 | 138 | 163.24 | 184 | 142 | 0.772 | -0.373 | 0.03995299 | 0.11436973 | 0.0189 | 0.0169 | 0.0175 | 0.0175 | TRUE | 0.1939 |
| Gm9125 | 6 | 1 | 1 | 2 | 0 | 1 | 5 | 1 | 1 | 2 | 0 | 1 | 1.63 | 2 | 1 | 0.434 | -1.205 | 0.45086575 | NA | 0 | 1.4564 | 1.1239 | 1.1239 | TRUE | 0.8899 |
| Gm9159 | 0 | 0 | 0 | 1 | 0 | 0 | 0 | 0 | 0 | 1 | 0 | 0 | 0.17 | 0 | 0 | 2.094 | 1.067 | 0.79379329 | NA | 0 | 14.2885 | 10 | 10 | TRUE | 0.3906 |
| Gm9199 | 0 | 0 | 0 | 0 | 1 | 0 | 0 | 0 | 0 | 0 | 1 | 0 | 0.19 | 0 | 0 | 2.094 | 1.067 | 0.79379329 | NA | 0 | 12.608 | 10 | 10 | TRUE | 0.5075 |
| Gm9268 | 0 | 0 | 0 | 0 | 0 | 0 | 0 | 0 | 0 | 0 | 0 | 0 | 0 | 0 | 0 | NA | NA | NA | NA | NA | NA | NA | NA | NA | NA |

| | | | | | | | | | | | | | | | | | | | | | | | | | |
|---|---|---|---|---|---|---|---|---|---|---|---|---|---|---|---|---|---|---|---|---|---|---|---|---|---|
| Gm933 | 2 | 4 | 10 | 3 | 5 | 5 | 2 | 4 | 11 | 3 | 6 | 5 | 5.03 | 6 | 5 | 0.834 | -0.262 | 0.77748007 | NA | 0.2891 | 0.4739 | 0.4165 | 0.4165 | TRUE | 0.5907 |
| Gm9376 | 0 | 1 | 0 | 0 | 0 | 2 | 0 | 1 | 0 | 0 | 0 | 2 | 0.51 | 0 | 1 | 1.931 | 0.949 | 0.75306031 | NA | 0 | 4.6833 | 4.7842 | 4.7842 | TRUE | 0.9737 |
| Gm94 | 0 | 0 | 0 | 0 | 0 | 0 | 0 | 0 | 0 | 0 | 0 | 0 | 0 | 0 | 0 | NA | NA | NA | NA | NA | NA | NA | NA | NA | NA |
| Gm9513 | 0 | 0 | 0 | 0 | 0 | 0 | 0 | 0 | 0 | 0 | 0 | 0 | 0 | 0 | 0 | NA | NA | NA | NA | NA | NA | NA | NA | NA | NA |
| Gm9573 | 0 | 0 | 0 | 0 | 0 | 0 | 0 | 0 | 0 | 0 | 0 | 0 | 0 | 0 | 0 | NA | NA | NA | NA | NA | NA | NA | NA | NA | NA |
| Gm960 | 9 | 7 | 3 | 11 | 6 | 10 | 7 | 7 | 3 | 11 | 7 | 10 | 7.54 | 6 | 9 | 1.553 | 0.635 | 0.36837998 | NA | 0 | 0.3166 | 0.222 | 0.222 | TRUE | 0.3189 |
| Gm9696 | 0 | 0 | 0 | 0 | 0 | 0 | 0 | 0 | 0 | 0 | 0 | 0 | 0 | 0 | 0 | NA | NA | NA | NA | NA | NA | NA | NA | NA | NA |
| Gm97 | 0 | 0 | 0 | 0 | 0 | 0 | 0 | 0 | 0 | 0 | 0 | 0 | 0 | 0 | 0 | NA | NA | NA | NA | NA | NA | NA | NA | NA | NA |
| Gm9705 | 3 | 5 | 6 | 3 | 2 | 2 | 2 | 5 | 6 | 3 | 2 | 2 | 3.56 | 4 | 2 | 0.52 | -0.942 | 0.36214227 | NA | 0 | 0.6688 | 0.4515 | 0.4515 | TRUE | 0.5074 |
| Gm973 | 175 | 111 | 85 | 62 | 56 | 58 | 136 | 118 | 92 | 62 | 63 | 57 | 87.97 | 115 | 61 | 0.524 | -0.932 | 4.44E-05 | 0.00063252 | 0.0147 | 0.0294 | 0.0249 | 0.0249 | TRUE | 0.1871 |
| Gm9731 | 0 | 0 | 0 | 0 | 0 | 0 | 0 | 0 | 0 | 0 | 0 | 0 | 0 | 0 | 0 | NA | NA | NA | NA | NA | NA | NA | NA | NA | NA |
| Gm9733 | 0 | 0 | 0 | 0 | 0 | 0 | 0 | 0 | 0 | 0 | 0 | 0 | 0 | 0 | 0 | NA | NA | NA | NA | NA | NA | NA | NA | NA | NA |
| Gm9758 | 0 | 0 | 0 | 0 | 0 | 0 | 0 | 0 | 0 | 0 | 0 | 0 | 0 | 0 | 0 | NA | NA | NA | NA | NA | NA | NA | NA | NA | NA |
| Gm9767 | 0 | 1 | 0 | 1 | 0 | 0 | 0 | 1 | 0 | 1 | 0 | 0 | 0.34 | 0 | 0 | 1.075 | 0.105 | 0.97573409 | NA | 0 | 6.9086 | 6.5582 | 6.5582 | TRUE | 0.5216 |
| Gm9776 | 69 | 50 | 31 | 45 | 38 | 29 | 54 | 53 | 33 | 45 | 43 | 29 | 42.75 | 47 | 39 | 0.824 | -0.279 | 0.38921965 | 0.56329652 | 0.0365 | 0.0579 | 0.0521 | 0.0521 | TRUE | 0.4624 |
| Gm9833 | 878 | 643 | 588 | 559 | 484 | 555 | 682 | 684 | 634 | 556 | 546 | 548 | 608.43 | 667 | 550 | 0.825 | -0.278 | 0.00179127 | 0.0110978 | 0 | 0.0063 | 0.0041 | 0.0041 | TRUE | 0.0205 |
| Gm9839 | 0 | 0 | 0 | 0 | 0 | 0 | 0 | 0 | 0 | 0 | 0 | 0 | 0 | 0 | 0 | NA | NA | NA | NA | NA | NA | NA | NA | NA | NA |
| Gm9846 | 0 | 0 | 0 | 0 | 0 | 0 | 0 | 0 | 0 | 0 | 0 | 0 | 0 | 0 | 0 | NA | NA | NA | NA | NA | NA | NA | NA | NA | NA |
| Gm9855 | 0 | 0 | 0 | 0 | 0 | 0 | 0 | 0 | 0 | 0 | 0 | 0 | 0 | 0 | 0 | NA | NA | NA | NA | NA | NA | NA | NA | NA | NA |
| Gm9866 | 3 | 7 | 10 | 3 | 5 | 8 | 2 | 7 | 11 | 3 | 6 | 8 | 6.18 | 7 | 6 | 0.821 | -0.285 | 0.73273107 | NA | 0.2157 | 0.3858 | 0.3352 | 0.3352 | TRUE | 0.3286 |
| Gm9871 | 0 | 0 | 0 | 0 | 0 | 0 | 0 | 0 | 0 | 0 | 0 | 0 | 0 | 0 | 0 | NA | NA | NA | NA | NA | NA | NA | NA | NA | NA |
| Gm9895 | 55 | 33 | 75 | 53 | 64 | 81 | 43 | 35 | 81 | 53 | 72 | 80 | 60.61 | 53 | 68 | 1.297 | 0.376 | 0.2429913 | 0.41077169 | 0.1001 | 0.0415 | 0.0578 | 0.0578 | TRUE | 1.6972 |
| Gm9897 | 88 | 50 | 55 | 64 | 59 | 59 | 68 | 53 | 59 | 64 | 67 | 58 | 61.56 | 60 | 63 | 1.036 | 0.051 | 0.83738991 | 0.9072782 | 0 | 0.0409 | 0.0284 | 0.0284 | TRUE | 0.1351 |
| Gm9899 | 6 | 6 | 8 | 6 | 11 | 9 | 5 | 6 | 9 | 6 | 12 | 9 | 7.82 | 7 | 9 | 1.396 | 0.482 | 0.48883512 | NA | 0 | 0.3054 | 0.2173 | 0.2173 | TRUE | 0.1754 |
| Gm9920 | 19 | 10 | 6 | 13 | 14 | 21 | 15 | 11 | 6 | 13 | 16 | 21 | 13.56 | 11 | 17 | 1.528 | 0.612 | 0.26025333 | NA | 0 | 0.1773 | 0.1367 | 0.1367 | TRUE | 0.3185 |
| Gm9926 | 9 | 10 | 12 | 11 | 10 | 6 | 7 | 11 | 13 | 11 | 11 | 6 | 9.79 | 10 | 9 | 0.928 | -0.107 | 0.86380527 | NA | 0 | 0.2446 | 0.1784 | 0.1784 | TRUE | 0.3589 |
| Gm9958 | 32 | 43 | 31 | 29 | 25 | 29 | 25 | 46 | 33 | 29 | 28 | 29 | 31.62 | 35 | 29 | 0.831 | -0.266 | 0.46041287 | 0.63003171 | 0 | 0.0774 | 0.0617 | 0.0617 | TRUE | 0.2573 |
| Gm996 | 51 | 49 | 43 | 60 | 55 | 60 | 40 | 52 | 46 | 60 | 62 | 59 | 53.19 | 46 | 60 | 1.317 | 0.398 | 0.13497301 | 0.27399079 | 0 | 0.047 | 0.0319 | 0.0319 | TRUE | 0.1412 |
| Gm9961 | 0 | 0 | 0 | 0 | 0 | 0 | 0 | 0 | 0 | 0 | 0 | 0 | 0 | 0 | 0 | NA | NA | NA | NA | NA | NA | NA | NA | NA | NA |
| Gm9962 | 3 | 0 | 0 | 0 | 0 | 0 | 2 | 0 | 0 | 0 | 0 | 0 | 0.39 | 1 | 0 | 0.239 | -2.067 | 0.57190649 | NA | 10 | 6.1057 | 8.0283 | 8.0283 | TRUE | 1.4408 |
| Gm9992 | 24 | 21 | 18 | 15 | 10 | 7 | 19 | 22 | 19 | 15 | 11 | 7 | 15.59 | 20 | 11 | 0.549 | -0.865 | 0.08611067 | NA | 0 | 0.1545 | 0.112 | 0.112 | TRUE | 0.3982 |
| Gm9994 | 1 | 0 | 0 | 2 | 0 | 1 | 1 | 0 | 0 | 2 | 0 | 1 | 0.63 | 0 | 1 | 2.909 | 1.541 | 0.54363287 | NA | 0 | 3.7899 | 3.121 | 3.121 | TRUE | 0.1026 |
| Gm9999 | 0 | 0 | 0 | 0 | 0 | 0 | 0 | 0 | 0 | 0 | 0 | 0 | 0 | 0 | 0 | NA | NA | NA | NA | NA | NA | NA | NA | NA | NA |
| Gmcl1 | 3329 | 2500 | 2403 | 2816 | 2491 | 2835 | 2585 | 2659 | 2593 | 2803 | 2810 | 2799 | 2708.21 | 2612 | 2804 | 1.073 | 0.102 | 0.06414461 | 0.16104321 | 0 | 0.0033 | 0.0018 | 0.0018 | TRUE | 0.0029 |
| Gmcl1l | 0 | 0 | 0 | 0 | 0 | 0 | 0 | 0 | 0 | 0 | 0 | 0 | 0 | 0 | 0 | NA | NA | NA | NA | NA | NA | NA | NA | NA | NA |
| Gmds | 809 | 548 | 553 | 684 | 609 | 656 | 628 | 583 | 597 | 681 | 687 | 648 | 637.22 | 603 | 672 | 1.113 | 0.155 | 0.07997763 | 0.18919567 | 0 | 0.0061 | 0.004 | 0.004 | TRUE | 0.0174 |
| Gmeb1 | 1702 | 1175 | 1099 | 1128 | 986 | 1195 | 1322 | 1250 | 1186 | 1123 | 1112 | 1180 | 1195.38 | 1253 | 1138 | 0.908 | -0.139 | 0.06502605 | 0.16271312 | 0.0013 | 0.0044 | 0.0033 | 0.0033 | TRUE | 0.0293 |
| Gmeb2 | 1924 | 1490 | 1496 | 1640 | 1428 | 1591 | 1494 | 1585 | 1614 | 1632 | 1611 | 1571 | 1584.55 | 1564 | 1605 | 1.027 | 0.038 | 0.57206222 | 0.72498188 | 5.00E-04 | 0.0039 | 0.0026 | 0.0026 | TRUE | 0.0194 |
| Gmfb | 7084 | 5118 | 5186 | 5588 | 4852 | 5452 | 5502 | 5444 | 5596 | 5562 | 5474 | 5382 | 5493.21 | 5514 | 5473 | 0.993 | -0.011 | 0.82772051 | 0.90131108 | 0 | 0.0029 | 0.0016 | 0.0016 | TRUE | 0.0026 |
| Gmfg | 320 | 205 | 304 | 496 | 380 | 428 | 249 | 218 | 328 | 494 | 429 | 423 | 356.58 | 265 | 449 | 1.696 | 0.762 | 2.07E-07 | 7.50E-06 | 0.0223 | 0.0091 | 0.0125 | 0.0125 | TRUE | 0.4678 |
| Gmip | 216 | 168 | 180 | 215 | 167 | 192 | 168 | 179 | 194 | 214 | 188 | 190 | 188.77 | 180 | 197 | 1.098 | 0.135 | 0.36508143 | 0.53984159 | 0 | 0.015 | 0.0107 | 0.0107 | TRUE | 0.0603 |
| Gml | 23 | 12 | 15 | 2 | 5 | 4 | 18 | 13 | 16 | 2 | 6 | 4 | 9.73 | 16 | 4 | 0.244 | -2.037 | 0.00314342 | NA | 0 | 0.246 | 0.1765 | 0.1765 | TRUE | 0.2987 |
| Gmnc | 0 | 0 | 0 | 0 | 0 | 0 | 0 | 0 | 0 | 0 | 0 | 0 | 0 | 0 | 0 | NA | NA | NA | NA | NA | NA | NA | NA | NA | NA |
| Gmnn | 5880 | 4311 | 4454 | 4488 | 3928 | 4373 | 4567 | 4586 | 4806 | 4467 | 4431 | 4317 | 4528.92 | 4653 | 4405 | 0.947 | -0.079 | 0.13922876 | 0.28048626 | 3.00E-04 | 0.0029 | 0.0018 | 0.0018 | TRUE | 0.01 |
| Gmppa | 2260 | 1682 | 1698 | 2019 | 1857 | 2230 | 1755 | 1789 | 1832 | 2010 | 2095 | 2201 | 1947.07 | 1792 | 2102 | 1.173 | 0.231 | 0.00040085 | 0.00365623 | 8.00E-04 | 0.0036 | 0.0025 | 0.0025 | TRUE | 0.021 |
| Gmppb | 1060 | 708 | 456 | 493 | 435 | 507 | 823 | 753 | 492 | 491 | 491 | 500 | 591.72 | 689 | 494 | 0.715 | -0.484 | 0.00137636 | 0.00906143 | 0.0343 | 0.0064 | 0.0147 | 0.0147 | TRUE | 0.7324 |
| Gmpr | 1098 | 855 | 881 | 1016 | 883 | 991 | 853 | 909 | 951 | 1011 | 996 | 978 | 949.75 | 904 | 995 | 1.102 | 0.14 | 0.07523125 | 0.18071937 | 7.00E-04 | 0.0049 | 0.0034 | 0.0034 | TRUE | 0.0313 |
| Gmpr2 | 1339 | 971 | 942 | 1065 | 901 | 1085 | 1040 | 1033 | 1016 | 1060 | 1016 | 1071 | 1039.46 | 1030 | 1049 | 1.019 | 0.027 | 0.70940237 | 0.82475095 | 0 | 0.0047 | 0.0029 | 0.0029 | TRUE | 0.0087 |
| Gmps | 9745 | 6890 | 6938 | 7325 | 6201 | 7000 | 7569 | 7329 | 7486 | 7291 | 6995 | 6910 | 7263.35 | 7461 | 7065 | 0.947 | -0.079 | 0.11747912 | 0.24870412 | 3.00E-04 | 0.0028 | 0.0017 | 0.0017 | TRUE | 0.0096 |
| Gna11 | 5279 | 3824 | 3717 | 4396 | 3999 | 4320 | 4100 | 4068 | 4011 | 4376 | 4511 | 4265 | 4221.62 | 4060 | 4384 | 1.08 | 0.111 | 0.03698381 | 0.10832635 | 2.00E-04 | 0.003 | 0.0018 | 0.0018 | TRUE | 0.0078 |
| Gna12 | 2413 | 1702 | 1863 | 1962 | 1698 | 1966 | 1874 | 1810 | 2010 | 1953 | 1916 | 1941 | 1917.31 | 1898 | 1937 | 1.02 | 0.029 | 0.65762554 | 0.78781677 | 9.00E-04 | 0.0037 | 0.0026 | 0.0026 | TRUE | 0.0313 |
| Gna13 | 10646 | 7922 | 8959 | 8296 | 7920 | 9277 | 8268 | 8427 | 9667 | 8258 | 8935 | 9158 | 8785.29 | 8787 | 8784 | 1 | 0 | 0.99586885 | 0.99753911 | 0.0041 | 0.0027 | 0.0036 | 0.0036 | TRUE | 0.0934 |
| Gna14 | 40 | 35 | 87 | 78 | 87 | 91 | 31 | 37 | 94 | 78 | 98 | 90 | 71.3 | 54 | 89 | 1.652 | 0.725 | 0.03634141 | 0.10694417 | 0.1677 | 0.0357 | 0.0715 | 0.0715 | TRUE | 3.3803 |
| Gna15 | 600 | 486 | 675 | 884 | 799 | 894 | 466 | 517 | 728 | 880 | 901 | 883 | 729.18 | 570 | 888 | 1.56 | 0.642 | 1.96E-06 | 5.01E-05 | 0.0277 | 0.0057 | 0.0117 | 0.0117 | TRUE | 0.6824 |
| Gnai1 | 1755 | 1240 | 1138 | 1217 | 973 | 1142 | 1363 | 1319 | 1228 | 1211 | 1098 | 1127 | 1224.38 | 1303 | 1145 | 0.878 | -0.187 | 0.01565485 | 0.05765882 | 0.0019 | 0.0044 | 0.0035 | 0.0035 | TRUE | 0.0305 |
| Gnai2 | 9038 | 7083 | 7124 | 7267 | 6763 | 7941 | 7019 | 7534 | 7687 | 7234 | 7629 | 7839 | 7490.36 | 7413 | 7567 | 1.021 | 0.03 | 0.60993442 | 0.75322701 | 0.0015 | 0.0027 | 0.0024 | 0.0024 | TRUE | 0.0267 |
| Gnai3 | 10243 | 7292 | 7164 | 7722 | 6561 | 7562 | 7955 | 7756 | 7730 | 7686 | 7401 | 7465 | 7665.74 | 7814 | 7517 | 0.962 | -0.056 | 0.2484834 | 0.41748833 | 2.00E-04 | 0.0027 | 0.0016 | 0.0016 | TRUE | 0.0047 |
| Gnal | 521 | 342 | 383 | 227 | 225 | 218 | 405 | 364 | 413 | 226 | 254 | 215 | 312.78 | 394 | 232 | 0.587 | -0.769 | 6.66E-10 | 4.73E-08 | 0.002 | 0.01 | 0.0077 | 0.0077 | TRUE | 0.0743 |
| Gnao1 | 459 | 284 | 319 | 319 | 326 | 344 | 356 | 302 | 344 | 318 | 368 | 340 | 337.94 | 334 | 342 | 1.019 | 0.027 | 0.82043123 | 0.89651557 | 0.0034 | 0.0094 | 0.0076 | 0.0076 | TRUE | 0.0789 |
| Gnaq | 2440 | 1756 | 1751 | 1980 | 1827 | 2009 | 1895 | 1868 | 1889 | 1971 | 2061 | 1983 | 1944.55 | 1884 | 2005 | 1.064 | 0.089 | 0.14112347 | 0.28313837 | 0 | 0.0036 | 0.0021 | 0.0021 | TRUE | 0.0071 |
| Gnas | 9956 | 7132 | 7352 | 8755 | 7646 | 8732 | 7732 | 7586 | 7933 | 8715 | 8625 | 8620 | 8201.9 | 7750 | 8653 | 1.117 | 0.159 | 0.00084857 | 0.00642065 | 1.00E-04 | 0.0027 | 0.0015 | 0.0015 | TRUE | 0.0051 |
| Gnat1 | 23 | 13 | 13 | 11 | 15 | 11 | 18 | 14 | 14 | 11 | 17 | 11 | 14.07 | 15 | 13 | 0.837 | -0.256 | 0.61697192 | NA | 0 | 0.1708 | 0.1175 | 0.1175 | TRUE | 0.2762 |
| Gnat2 | 187 | 113 | 111 | 171 | 115 | 135 | 145 | 120 | 120 | 170 | 130 | 133 | 136.4 | 128 | 144 | 1.121 | 0.165 | 0.3717248 | 0.54649756 | 0.0114 | 0.0198 | 0.0173 | 0.0173 | TRUE | 0.2574 |
| Gnat3 | 0 | 0 | 0 | 0 | 0 | 0 | 0 | 0 | 0 | 0 | 0 | 0 | 0 | 0 | 0 | NA | NA | NA | NA | NA | NA | NA | NA | NA | NA |
| Gnaz | 478 | 413 | 415 | 433 | 375 | 408 | 371 | 439 | 448 | 431 | 423 | 403 | 419.19 | 419 | 419 | 1.002 | 0.002 | 0.9823701 | 0.99089635 | 0.004 | 0.0081 | 0.0067 | 0.0067 | TRUE | 0.1253 |
| Gnb1 | 23462 | 16771 | 17423 | 17643 | 15250 | 17794 | 18222 | 17839 | 18799 | 17562 | 17204 | 17565 | 17865.19 | 18287 | 17444 | 0.954 | -0.068 | 0.14369332 | 0.28660462 | 3.00E-04 | 0.0026 | 0.0015 | 0.0015 | TRUE | 0.0073 |
| Gnb1l | 744 | 563 | 550 | 713 | 550 | 604 | 578 | 599 | 593 | 710 | 620 | 596 | 616.09 | 590 | 642 | 1.089 | 0.123 | 0.20458125 | 0.36606747 | 0.0029 | 0.0063 | 0.0052 | 0.0052 | TRUE | 0.1012 |
| Gnb2 | 8292 | 6024 | 6159 | 6838 | 5862 | 6797 | 6440 | 6408 | 6646 | 6807 | 6613 | 6710 | 6603.73 | 6498 | 6710 | 1.033 | 0.047 | 0.34219714 | 0.51631201 | 1.00E-04 | 0.0028 | 0.0016 | 0.0016 | TRUE | 0.0048 |
| Gnb2l1 | 131223 | 95141 | 103031 | 111452 | 102589 | 116123 | 101915 | 101200 | 111170 | 110939 | 115731 | 114631 | 109264.45 | 104762 | 113767 | 1.086 | 0.119 | 0.0269673 | 0.08578747 | 0.0016 | 0.0024 | 0.0021 | 0.0021 | TRUE | 0.035 |

| | | | | | | | | | | | | | | | | | | | | | | | | | |
|---|---|---|---|---|---|---|---|---|---|---|---|---|---|---|---|---|---|---|---|---|---|---|---|---|---|
| Gnb3 | 1 | 2 | 1 | 2 | 0 | 4 | 1 | 2 | 1 | 2 | 0 | 4 | 1.65 | 1 | 2 | 1.539 | 0.622 | 0.68779711 | NA | 0 | 1.4356 | 1.0968 | 1.0968 | TRUE | 0.1314 |
| Gnb4 | 1256 | 878 | 901 | 1019 | 844 | 953 | 975 | 934 | 972 | 1014 | 952 | 941 | 964.79 | 960 | 969 | 1.009 | 0.012 | 0.87084013 | 0.92839601 | 0 | 0.0049 | 0.0032 | 0.0032 | TRUE | 0.0202 |
| Gnb5 | 305 | 235 | 234 | 248 | 214 | 258 | 237 | 250 | 252 | 247 | 241 | 255 | 247.05 | 246 | 248 | 1.007 | 0.01 | 0.93965297 | 0.96635918 | 0 | 0.012 | 0.0077 | 0.0077 | TRUE | 0.014 |
| Gne | 982 | 734 | 859 | 969 | 868 | 982 | 763 | 781 | 927 | 965 | 979 | 969 | 897.23 | 824 | 971 | 1.181 | 0.24 | 0.008596 | 0.03698427 | 0.0043 | 0.0051 | 0.0049 | 0.0049 | TRUE | 0.141 |
| Gng10 | 568 | 396 | 384 | 420 | 365 | 377 | 441 | 421 | 414 | 418 | 412 | 372 | 413.11 | 425 | 401 | 0.94 | -0.089 | 0.40008896 | 0.57339532 | 0 | 0.0082 | 0.0057 | 0.0057 | TRUE | 0.0455 |
| Gng11 | 33 | 21 | 26 | 24 | 17 | 8 | 26 | 22 | 28 | 24 | 19 | 8 | 21.16 | 25 | 17 | 0.669 | -0.581 | 0.20854252 | 0.3701811 | 0.07 | 0.1144 | 0.1041 | 0.1041 | TRUE | 0.9876 |
| Gng12 | 1731 | 1329 | 1685 | 1786 | 1425 | 1796 | 1344 | 1414 | 1818 | 1778 | 1608 | 1773 | 1622.4 | 1525 | 1720 | 1.129 | 0.175 | 0.09499933 | 0.21299223 | 0.0147 | 0.0039 | 0.0073 | 0.0073 | TRUE | 0.3376 |
| Gng13 | 17 | 11 | 6 | 13 | 9 | 13 | 13 | 12 | 6 | 13 | 10 | 13 | 11.22 | 10 | 12 | 1.135 | 0.183 | 0.75053848 | NA | 0 | 0.2137 | 0.1502 | 0.1502 | TRUE | 0.3642 |
| Gng2 | 1655 | 1242 | 1495 | 2026 | 1727 | 2107 | 1285 | 1321 | 1613 | 2017 | 1948 | 2080 | 1710.73 | 1406 | 2015 | 1.434 | 0.52 | 4.75E-09 | 2.66E-07 | 0.0078 | 0.0038 | 0.0051 | 0.0051 | TRUE | 0.1966 |
| Gng3 | 662 | 519 | 531 | 449 | 384 | 569 | 514 | 552 | 573 | 447 | 433 | 562 | 513.49 | 546 | 481 | 0.881 | -0.182 | 0.11925588 | 0.25096058 | 0.0093 | 0.007 | 0.0079 | 0.0079 | TRUE | 0.2555 |
| Gng4 | 3 | 6 | 2 | 3 | 1 | 6 | 2 | 6 | 2 | 3 | 1 | 6 | 3.48 | 3 | 3 | 0.944 | -0.082 | 0.93842383 | NA | 0.1695 | 0.6826 | 0.5336 | 0.5336 | TRUE | 0.3745 |
| Gng5 | 2808 | 1927 | 1983 | 2051 | 1714 | 2124 | 2181 | 2050 | 2140 | 2042 | 1934 | 2097 | 2073.68 | 2124 | 2024 | 0.953 | -0.069 | 0.28261325 | 0.45437472 | 8.00E-04 | 0.0036 | 0.0025 | 0.0025 | TRUE | 0.0182 |
| Gng7 | 242 | 165 | 145 | 257 | 175 | 192 | 188 | 176 | 156 | 256 | 197 | 190 | 193.78 | 173 | 214 | 1.234 | 0.303 | 0.06495191 | 0.16258452 | 0.0134 | 0.0147 | 0.0143 | 0.0143 | TRUE | 0.3194 |
| Gng8 | 26 | 35 | 26 | 37 | 35 | 40 | 20 | 37 | 28 | 37 | 39 | 39 | 33.55 | 28 | 38 | 1.37 | 0.454 | 0.1935143 | 0.3522542 | 0 | 0.0731 | 0.0573 | 0.0573 | TRUE | 0.3205 |
| Gngt1 | 0 | 0 | 0 | 0 | 0 | 0 | 0 | 0 | 0 | 0 | 0 | 0 | 0 | 0 | 0 | NA | NA | NA | NA | NA | NA | NA | NA | NA | NA |
| Gngt2 | 173 | 121 | 184 | 226 | 172 | 297 | 134 | 129 | 199 | 225 | 194 | 293 | 195.63 | 154 | 237 | 1.55 | 0.632 | 0.00155865 | 0.00992774 | 0.0466 | 0.0145 | 0.0234 | 0.0234 | TRUE | 0.5852 |
| Gnl1 | 6274 | 4527 | 4396 | 4262 | 3764 | 4336 | 4873 | 4815 | 4743 | 4242 | 4246 | 4280 | 4533.36 | 4810 | 4256 | 0.885 | -0.177 | 0.00041464 | 0.00373674 | 0 | 0.0029 | 0.0016 | 0.0016 | TRUE | 0.0018 |
| Gnl2 | 6710 | 4945 | 4857 | 4998 | 4565 | 5158 | 5211 | 5260 | 5241 | 4975 | 5150 | 5092 | 5154.75 | 5237 | 5072 | 0.968 | -0.046 | 0.35016034 | 0.52478155 | 0 | 0.0029 | 0.0016 | 0.0016 | TRUE | 0.0034 |
| Gnl3 | 11690 | 8566 | 8812 | 9400 | 8124 | 9890 | 9079 | 9112 | 9508 | 9357 | 9165 | 9763 | 9330.53 | 9233 | 9428 | 1.021 | 0.03 | 0.55476873 | 0.71019341 | 7.00E-04 | 0.0027 | 0.0018 | 0.0018 | TRUE | 0.0118 |
| Gnl3l | 12117 | 8817 | 9170 | 11052 | 9665 | 11110 | 9411 | 9379 | 9894 | 11001 | 10903 | 10967 | 10259.2 | 9561 | 10957 | 1.146 | 0.197 | 4.67E-05 | 0.00066008 | 3.00E-04 | 0.0027 | 0.0016 | 0.0016 | TRUE | 0.0113 |
| Gnmt | 127 | 101 | 144 | 181 | 146 | 198 | 99 | 107 | 155 | 180 | 165 | 195 | 150.29 | 120 | 180 | 1.505 | 0.59 | 0.00233994 | 0.01365653 | 0.0254 | 0.0182 | 0.0201 | 0.0201 | TRUE | 0.633 |
| Gnpat | 12949 | 9601 | 10098 | 10286 | 9252 | 10934 | 10057 | 10212 | 10896 | 10239 | 10437 | 10794 | 10439.08 | 10388 | 10490 | 1.01 | 0.014 | 0.79137847 | 0.87744725 | 0.001 | 0.0027 | 0.002 | 0.002 | TRUE | 0.0222 |
| Gnpda1 | 5723 | 4448 | 4966 | 5022 | 4740 | 5192 | 4445 | 4731 | 5358 | 4999 | 5347 | 5125 | 5000.96 | 4845 | 5157 | 1.065 | 0.091 | 0.21619925 | 0.37886288 | 0.0049 | 0.0029 | 0.0037 | 0.0037 | TRUE | 0.1045 |
| Gnpda2 | 1667 | 1114 | 1273 | 1186 | 981 | 1136 | 1295 | 1185 | 1374 | 1181 | 1107 | 1121 | 1210.3 | 1285 | 1136 | 0.885 | -0.177 | 0.02608513 | 0.08387734 | 0.0025 | 0.0044 | 0.0037 | 0.0037 | TRUE | 0.0534 |
| Gnpnat1 | 3014 | 2069 | 2225 | 2250 | 1900 | 2285 | 2341 | 2201 | 2401 | 2240 | 2143 | 2256 | 2263.51 | 2314 | 2213 | 0.956 | -0.064 | 0.31088083 | 0.48382183 | 9.00E-04 | 0.0035 | 0.0025 | 0.0025 | TRUE | 0.0218 |
| Gnptab | 3210 | 2286 | 2373 | 2493 | 2181 | 2639 | 2493 | 2432 | 2560 | 2482 | 2460 | 2605 | 2505.36 | 2495 | 2516 | 1.008 | 0.012 | 0.83876214 | 0.9080766 | 4.00E-04 | 0.0034 | 0.0022 | 0.0022 | TRUE | 0.0118 |
| Gnptg | 413 | 328 | 356 | 359 | 333 | 392 | 321 | 349 | 384 | 357 | 376 | 387 | 362.29 | 351 | 373 | 1.066 | 0.092 | 0.42909467 | 0.60110262 | 0.0024 | 0.009 | 0.0069 | 0.0069 | TRUE | 0.076 |
| Gnrh1 | 3 | 0 | 1 | 0 | 1 | 1 | 2 | 0 | 1 | 0 | 1 | 1 | 0.92 | 1 | 1 | 0.595 | -0.749 | 0.7182143 | NA | 0 | 2.5766 | 1.95 | 1.95 | TRUE | 0.1738 |
| Gnrhr | 0 | 0 | 0 | 0 | 0 | 0 | 0 | 0 | 0 | 0 | 0 | 0 | 0 | 0 | 0 | NA | NA | NA | NA | NA | NA | NA | NA | NA | NA |
| Gns | 4299 | 3313 | 3181 | 3278 | 3190 | 3529 | 3339 | 3524 | 3432 | 3263 | 3599 | 3484 | 3440.06 | 3432 | 3449 | 1.005 | 0.007 | 0.90681966 | 0.94912712 | 0.0013 | 0.0031 | 0.0024 | 0.0024 | TRUE | 0.027 |
| Golga1 | 661 | 508 | 516 | 447 | 359 | 480 | 513 | 540 | 557 | 445 | 405 | 474 | 489.04 | 537 | 441 | 0.824 | -0.28 | 0.00643919 | 0.02975599 | 0.0016 | 0.0073 | 0.0055 | 0.0055 | TRUE | 0.0579 |
| Golga2 | 2467 | 1919 | 1885 | 2124 | 1798 | 2103 | 1916 | 2041 | 2034 | 2114 | 2028 | 2076 | 2034.95 | 1997 | 2073 | 1.039 | 0.055 | 0.37959683 | 0.55419586 | 4.00E-04 | 0.0036 | 0.0023 | 0.0023 | TRUE | 0.0159 |
| Golga3 | 4065 | 2899 | 2742 | 3247 | 2846 | 3398 | 3157 | 3084 | 2959 | 3232 | 3211 | 3354 | 3166.06 | 3067 | 3266 | 1.065 | 0.09 | 0.11638121 | 0.24687394 | 5.00E-04 | 0.0032 | 0.0021 | 0.0021 | TRUE | 0.0114 |
| Golga4 | 8896 | 6623 | 6464 | 7808 | 6847 | 7906 | 6909 | 7045 | 6975 | 7772 | 7724 | 7804 | 7371.53 | 6976 | 7767 | 1.113 | 0.155 | 0.00084802 | 0.00641992 | 0 | 0.0027 | 0.0014 | 0.0014 | TRUE | 9.00E-04 |
| Golga5 | 3046 | 2268 | 2245 | 2418 | 2177 | 2457 | 2366 | 2412 | 2422 | 2407 | 2456 | 2425 | 2414.78 | 2400 | 2429 | 1.012 | 0.018 | 0.75469707 | 0.8545742 | 0 | 0.0034 | 0.0019 | 0.0019 | TRUE | 0.002 |
| Golga7 | 7044 | 5180 | 5170 | 5707 | 4848 | 5392 | 5471 | 5510 | 5578 | 5681 | 5469 | 5323 | 5505.26 | 5520 | 5491 | 0.995 | -0.007 | 0.88660236 | 0.93690919 | 4.00E-04 | 0.0029 | 0.0018 | 0.0018 | TRUE | 0.0112 |
| Golga7b | 19 | 13 | 14 | 21 | 17 | 19 | 15 | 14 | 15 | 21 | 19 | 19 | 17.09 | 15 | 20 | 1.346 | 0.429 | 0.34493008 | 0.51916211 | 0 | 0.1411 | 0.0893 | 0.0893 | TRUE | 0.0183 |
| Golgb1 | 5793 | 4184 | 4024 | 4359 | 4040 | 4759 | 4499 | 4450 | 4342 | 4339 | 4558 | 4698 | 4480.98 | 4430 | 4532 | 1.023 | 0.032 | 0.56279355 | 0.71733477 | 6.00E-04 | 0.003 | 0.002 | 0.002 | TRUE | 0.017 |
| Golim4 | 1790 | 1273 | 1347 | 1423 | 1355 | 1500 | 1390 | 1354 | 1453 | 1416 | 1529 | 1481 | 1437.24 | 1399 | 1475 | 1.054 | 0.076 | 0.27506095 | 0.44599502 | 6.00E-04 | 0.0041 | 0.0028 | 0.0028 | TRUE | 0.0147 |
| Golm1 | 6396 | 4459 | 4996 | 5409 | 4870 | 5475 | 4968 | 4743 | 5391 | 5384 | 5494 | 5405 | 5230.63 | 5034 | 5428 | 1.078 | 0.109 | 0.07482381 | 0.18004295 | 0.0019 | 0.0029 | 0.0025 | 0.0025 | TRUE | 0.0463 |
| Golph3 | 7562 | 5436 | 5513 | 5747 | 5057 | 5919 | 5873 | 5782 | 5948 | 5721 | 5705 | 5843 | 5812.02 | 5868 | 5756 | 0.981 | -0.028 | 0.57091993 | 0.724253 | 0 | 0.0028 | 0.0015 | 0.0015 | TRUE | 0.0021 |
| Golph3l | 1944 | 1355 | 1333 | 1608 | 1361 | 1594 | 1510 | 1441 | 1438 | 1601 | 1535 | 1574 | 1516.48 | 1463 | 1570 | 1.072 | 0.101 | 0.12629177 | 0.26153119 | 0 | 0.004 | 0.0025 | 0.0025 | TRUE | 0.0098 |
| Golt1a | 2 | 1 | 3 | 1 | 2 | 1 | 2 | 1 | 3 | 1 | 2 | 1 | 1.68 | 2 | 1 | 0.722 | -0.47 | 0.7453713 | NA | 0 | 1.4114 | 0.8904 | 0.8904 | TRUE | 0.2622 |
| Golt1b | 2378 | 1866 | 1758 | 1846 | 1542 | 1841 | 1847 | 1985 | 1897 | 1838 | 1740 | 1817 | 1853.83 | 1910 | 1798 | 0.942 | -0.086 | 0.18740226 | 0.34432968 | 6.00E-04 | 0.0037 | 0.0025 | 0.0025 | TRUE | 0.0143 |
| Gon4l | 4558 | 3140 | 2774 | 2867 | 2625 | 2966 | 3540 | 3340 | 2993 | 2854 | 2961 | 2928 | 3102.68 | 3291 | 2914 | 0.885 | -0.176 | 0.01300897 | 0.05021258 | 0.0032 | 0.0032 | 0.0033 | 0.0033 | TRUE | 0.0758 |
| Gopc | 2368 | 1661 | 1483 | 1595 | 1293 | 1508 | 1839 | 1767 | 1600 | 1588 | 1459 | 1489 | 1623.5 | 1735 | 1512 | 0.871 | -0.2 | 0.00852496 | 0.03674523 | 0.003 | 0.0039 | 0.0035 | 0.0035 | TRUE | 0.0555 |
| Gorab | 1087 | 731 | 717 | 794 | 667 | 733 | 844 | 778 | 774 | 790 | 752 | 724 | 776.97 | 799 | 755 | 0.945 | -0.082 | 0.33802742 | 0.51223769 | 0.0011 | 0.0055 | 0.004 | 0.004 | TRUE | 0.0314 |
| Gorasp1 | 1335 | 1067 | 1050 | 1164 | 1076 | 1292 | 1037 | 1135 | 1133 | 1159 | 1214 | 1275 | 1158.77 | 1102 | 1216 | 1.105 | 0.144 | 0.06313997 | 0.15913899 | 0.0017 | 0.0045 | 0.0035 | 0.0035 | TRUE | 0.0332 |
| Gorasp2 | 6020 | 4317 | 4390 | 4703 | 4080 | 4717 | 4675 | 4592 | 4737 | 4681 | 4603 | 4656 | 4657.44 | 4668 | 4647 | 0.995 | -0.007 | 0.89658038 | 0.94309857 | 0 | 0.0029 | 0.0016 | 0.0016 | TRUE | 0.0024 |
| Gosr1 | 2916 | 2080 | 2077 | 2209 | 1898 | 2213 | 2265 | 2212 | 2241 | 2199 | 2141 | 2185 | 2207.14 | 2239 | 2175 | 0.971 | -0.042 | 0.46631646 | 0.63525173 | 0 | 0.0035 | 0.002 | 0.002 | TRUE | 0.0022 |
| Gosr2 | 6526 | 4508 | 4341 | 4456 | 4000 | 4330 | 5068 | 4795 | 4684 | 4435 | 4512 | 4274 | 4628.29 | 4849 | 4407 | 0.908 | -0.139 | 0.01505567 | 0.0559955 | 9.00E-04 | 0.0029 | 0.0021 | 0.0021 | TRUE | 0.0194 |
| Got1 | 10127 | 7278 | 7925 | 8374 | 7902 | 9278 | 7865 | 7742 | 8551 | 8335 | 8914 | 9159 | 8427.72 | 8053 | 8803 | 1.093 | 0.129 | 0.03670102 | 0.10771411 | 0.0024 | 0.0027 | 0.0026 | 0.0026 | TRUE | 0.0356 |
| Got1l1 | 1744 | 1294 | 1548 | 2082 | 1661 | 1870 | 1354 | 1376 | 1670 | 2072 | 1874 | 1846 | 1698.89 | 1467 | 1931 | 1.317 | 0.398 | 1.15E-05 | 0.00020982 | 0.0084 | 0.0038 | 0.0053 | 0.0053 | TRUE | 0.175 |
| Got2 | 25879 | 18297 | 19030 | 20884 | 17913 | 20763 | 20099 | 19462 | 20533 | 20788 | 20208 | 20496 | 20264.42 | 20031 | 20497 | 1.023 | 0.033 | 0.47709565 | 0.64471288 | 4.00E-04 | 0.0025 | 0.0015 | 0.0015 | TRUE | 0.0075 |
| Gp1ba | 0 | 1 | 0 | 1 | 0 | 0 | 0 | 1 | 0 | 1 | 0 | 0 | 0.34 | 0 | 0 | 1.075 | 0.105 | 0.97573409 | NA | 0 | 6.9086 | 6.5582 | 6.5582 | TRUE | 0.5216 |
| Gp1bb | 6 | 3 | 3 | 0 | 2 | 2 | 5 | 3 | 3 | 0 | 2 | 2 | 2.55 | 4 | 1 | 0.374 | -1.419 | 0.26468136 | NA | 0 | 0.9307 | 0.6655 | 0.6655 | TRUE | 0.5175 |
| Gp2 | 0 | 0 | 0 | 0 | 0 | 0 | 0 | 0 | 0 | 0 | 0 | 0 | 0 | 0 | 0 | NA | NA | NA | NA | NA | NA | NA | NA | NA | NA |
| Gp49a | 0 | 0 | 0 | 0 | 0 | 0 | 0 | 0 | 0 | 0 | 0 | 0 | 0 | 0 | 0 | NA | NA | NA | NA | NA | NA | NA | NA | NA | NA |
| Gp5 | 8 | 2 | 3 | 4 | 6 | 1 | 6 | 2 | 3 | 4 | 7 | 1 | 3.89 | 4 | 4 | 0.982 | -0.027 | 0.9792124 | NA | 0.203 | 0.6124 | 0.4977 | 0.4977 | TRUE | 0.2053 |
| Gp6 | 1 | 1 | 2 | 2 | 3 | 2 | 1 | 1 | 2 | 2 | 3 | 2 | 1.89 | 1 | 2 | 1.87 | 0.903 | 0.51161833 | NA | 0 | 1.2556 | 0.7814 | 0.7814 | TRUE | 0.1665 |
| Gp9 | 0 | 0 | 0 | 0 | 0 | 0 | 0 | 0 | 0 | 0 | 0 | 0 | 0 | 0 | 0 | NA | NA | NA | NA | NA | NA | NA | NA | NA | NA |
| Gpa33 | 1056 | 974 | 1083 | 1061 | 1156 | 1327 | 820 | 1036 | 1169 | 1056 | 1304 | 1310 | 1115.81 | 1008 | 1223 | 1.215 | 0.281 | 0.02222306 | 0.07447842 | 0.0209 | 0.0046 | 0.01 | 0.01 | TRUE | 0.3269 |
| Gpaa1 | 1786 | 1275 | 1179 | 1189 | 1081 | 1323 | 1387 | 1356 | 1272 | 1184 | 1219 | 1306 | 1287.41 | 1338 | 1236 | 0.923 | -0.115 | 0.12429829 | 0.25875262 | 0.0015 | 0.0043 | 0.0033 | 0.0033 | TRUE | 0.0296 |
| Gpam | 1511 | 1138 | 1087 | 1277 | 1165 | 1202 | 1174 | 1210 | 1173 | 1271 | 1314 | 1187 | 1221.47 | 1186 | 1257 | 1.06 | 0.084 | 0.2491384 | 0.41809736 | 7.00E-04 | 0.0044 | 0.003 | 0.003 | TRUE | 0.0293 |
| Gpank1 | 1156 | 822 | 679 | 762 | 740 | 808 | 898 | 874 | 733 | 758 | 835 | 798 | 815.95 | 835 | 797 | 0.953 | -0.07 | 0.47069548 | 0.63963211 | 0.0059 | 0.0053 | 0.0055 | 0.0055 | TRUE | 0.1339 |
| Gpat2 | 1220 | 1073 | 979 | 1082 | 932 | 1081 | 948 | 1141 | 1056 | 1077 | 1051 | 1067 | 1056.79 | 1048 | 1065 | 1.018 | 0.026 | 0.7649314 | 0.86058711 | 0.0036 | 0.0047 | 0.0043 | 0.0043 | TRUE | 0.0887 |
| Gpatch1 | 2148 | 1673 | 1514 | 1534 | 1378 | 1614 | 1668 | 1780 | 1634 | 1527 | 1555 | 1593 | 1626.02 | 1694 | 1558 | 0.92 | -0.12 | 0.07499602 | 0.18030582 | 6.00E-04 | 0.0039 | 0.0027 | 0.0027 | TRUE | 0.0231 |
| Gpatch11 | 1276 | 890 | 888 | 882 | 777 | 939 | 991 | 947 | 958 | 878 | 877 | 927 | 929.54 | 965 | 894 | 0.926 | -0.112 | 0.14566674 | 0.28966556 | 0 | 0.005 | 0.0032 | 0.0032 | TRUE | 0.0127 |
| Gpatch2 | 2025 | 1451 | 1272 | 1257 | 1181 | 1303 | 1573 | 1543 | 1372 | 1251 | 1332 | 1286 | 1393.06 | 1496 | 1290 | 0.861 | -0.216 | 0.00506497 | 0.02476688 | 0.0025 | 0.0041 | 0.0035 | 0.0035 | TRUE | 0.062 |

| | | | | | | | | | | | | | | | | | | | | | | | | | |
|---|---|---|---|---|---|---|---|---|---|---|---|---|---|---|---|---|---|---|---|---|---|---|---|---|---|
| Gpatch2l | 1506 | 1093 | 1082 | 1174 | 985 | 1132 | 1170 | 1163 | 1167 | 1169 | 1111 | 1117 | 1149.49 | 1167 | 1132 | 0.971 | -0.043 | 0.54423483 | 0.70173644 | 0 | 0.0045 | 0.0027 | 0.0027 | TRUE | 0.0094 |
| Gpatch3 | 694 | 435 | 289 | 271 | 248 | 250 | 539 | 463 | 312 | 270 | 280 | 247 | 351.64 | 438 | 266 | 0.604 | -0.727 | 1.98E-05 | 0.00032822 | 0.038 | 0.0092 | 0.0178 | 0.0178 | TRUE | 0.3212 |
| Gpatch4 | 1425 | 1037 | 754 | 760 | 707 | 827 | 1107 | 1103 | 814 | 757 | 798 | 816 | 898.96 | 1008 | 790 | 0.783 | -0.353 | 0.00189842 | 0.0116057 | 0.0147 | 0.0051 | 0.0082 | 0.0082 | TRUE | 0.3345 |
| Gpatch8 | 6827 | 5040 | 4771 | 4962 | 4401 | 5250 | 5302 | 5361 | 5148 | 4939 | 4965 | 5183 | 5149.6 | 5270 | 5029 | 0.954 | -0.068 | 0.19599765 | 0.35550691 | 3.00E-04 | 0.0029 | 0.0018 | 0.0018 | TRUE | 0.0088 |
| Gpbar1 | 16 | 8 | 10 | 14 | 13 | 13 | 12 | 9 | 11 | 14 | 15 | 13 | 12.19 | 11 | 14 | 1.295 | 0.373 | 0.49014585 | NA | 0 | 0.1968 | 0.1277 | 0.1277 | TRUE | 0.1094 |
| Gpbp1 | 11779 | 8351 | 8549 | 8802 | 7769 | 8893 | 9148 | 8883 | 9224 | 8762 | 8764 | 8779 | 8926.65 | 9085 | 8768 | 0.965 | -0.051 | 0.27214214 | 0.44316998 | 1.00E-04 | 0.0027 | 0.0015 | 0.0015 | TRUE | 0.0046 |
| Gpbp1l1 | 7508 | 5605 | 5615 | 5443 | 5138 | 5684 | 5831 | 5962 | 6059 | 5418 | 5796 | 5611 | 5779.46 | 5951 | 5608 | 0.943 | -0.085 | 0.10638802 | 0.23148179 | 5.00E-04 | 0.0028 | 0.0018 | 0.0018 | TRUE | 0.0108 |
| Gpc1 | 1675 | 1155 | 1073 | 1102 | 923 | 1094 | 1301 | 1229 | 1158 | 1097 | 1041 | 1080 | 1150.89 | 1229 | 1073 | 0.872 | -0.198 | 0.00979927 | 0.04077467 | 0.0014 | 0.0045 | 0.0033 | 0.0033 | TRUE | 0.0327 |
| Gpc2 | 721 | 560 | 695 | 903 | 700 | 823 | 560 | 596 | 750 | 899 | 790 | 812 | 734.41 | 635 | 834 | 1.316 | 0.396 | 0.00049726 | 0.00428926 | 0.0129 | 0.0057 | 0.0079 | 0.0079 | TRUE | 0.2907 |
| Gpc3 | 3376 | 2656 | 3601 | 3163 | 2729 | 3319 | 2622 | 2825 | 3885 | 3148 | 3079 | 3276 | 3139.33 | 3111 | 3168 | 1.019 | 0.027 | 0.81299811 | 0.89199678 | 0.0226 | 0.0032 | 0.0091 | 0.0091 | TRUE | 0.5753 |
| Gpc4 | 585 | 351 | 357 | 285 | 223 | 304 | 454 | 373 | 385 | 284 | 252 | 300 | 341.37 | 404 | 279 | 0.688 | -0.54 | 2.30E-05 | 0.0003733 | 0.0072 | 0.0094 | 0.0087 | 0.0087 | TRUE | 0.1425 |
| Gpc5 | 7 | 3 | 2 | 4 | 7 | 5 | 5 | 3 | 2 | 4 | 8 | 5 | 4.6 | 3 | 6 | 1.514 | 0.599 | 0.50540605 | NA | 0 | 0.5177 | 0.3585 | 0.3585 | TRUE | 0.4072 |
| Gpc6 | 87 | 80 | 73 | 62 | 52 | 63 | 68 | 85 | 79 | 62 | 59 | 62 | 69 | 77 | 61 | 0.793 | -0.335 | 0.15481199 | 0.30245048 | 0 | 0.0368 | 0.0252 | 0.0252 | TRUE | 0.1245 |
| Gpcpd1 | 1270 | 908 | 773 | 736 | 671 | 679 | 986 | 966 | 834 | 733 | 757 | 670 | 824.35 | 929 | 720 | 0.774 | -0.37 | 8.75E-05 | 0.00108463 | 0.0049 | 0.0053 | 0.0052 | 0.0052 | TRUE | 0.0931 |
| Gpd1 | 68 | 53 | 50 | 42 | 49 | 44 | 53 | 56 | 54 | 42 | 55 | 43 | 50.61 | 54 | 47 | 0.86 | -0.218 | 0.42392402 | 0.59649005 | 0 | 0.0493 | 0.0337 | 0.0337 | TRUE | 0.1842 |
| Gpd1l | 2667 | 1794 | 1999 | 2060 | 1801 | 2144 | 2071 | 1908 | 2157 | 2051 | 2032 | 2116 | 2055.87 | 2045 | 2066 | 1.01 | 0.015 | 0.83082365 | 0.9033735 | 0.0016 | 0.0036 | 0.0028 | 0.0028 | TRUE | 0.0407 |
| Gpd2 | 8117 | 5701 | 5785 | 7430 | 6052 | 6903 | 6304 | 6064 | 6242 | 7396 | 6827 | 6814 | 6607.93 | 6203 | 7012 | 1.13 | 0.177 | 0.00150367 | 0.0096854 | 0.0011 | 0.0028 | 0.0021 | 0.0021 | TRUE | 0.028 |
| Gpha2 | 45 | 26 | 35 | 32 | 26 | 34 | 35 | 28 | 38 | 32 | 29 | 34 | 32.52 | 34 | 32 | 0.944 | -0.083 | 0.80416861 | 0.88656654 | 0 | 0.0753 | 0.0509 | 0.0509 | TRUE | 0.1483 |
| Gphb5 | 0 | 0 | 0 | 0 | 0 | 0 | 0 | 0 | 0 | 0 | 0 | 0 | 0 | 0 | 0 | NA | NA | NA | NA | NA | NA | NA | NA | NA | NA |
| Gphn | 795 | 514 | 511 | 576 | 536 | 572 | 617 | 547 | 551 | 573 | 605 | 565 | 576.37 | 572 | 581 | 1.013 | 0.019 | 0.84390769 | 0.91164482 | 0.0016 | 0.0065 | 0.0049 | 0.0049 | TRUE | 0.0605 |
| Gpi1 | 43970 | 33701 | 38104 | 37169 | 33772 | 39262 | 34150 | 35847 | 41114 | 36998 | 38098 | 38758 | 37494.13 | 37037 | 37951 | 1.025 | 0.035 | 0.61103074 | 0.75392458 | 0.0041 | 0.0025 | 0.0034 | 0.0034 | TRUE | 0.1135 |
| Gpihbp1 | 0 | 0 | 0 | 0 | 0 | 0 | 0 | 0 | 0 | 0 | 0 | 0 | 0 | 0 | 0 | NA | NA | NA | NA | NA | NA | NA | NA | NA | NA |
| Gpkow | 1297 | 1009 | 907 | 916 | 775 | 951 | 1007 | 1073 | 979 | 912 | 874 | 939 | 964.01 | 1020 | 908 | 0.891 | -0.166 | 0.03504045 | 0.10392886 | 6.00E-04 | 0.0049 | 0.0034 | 0.0034 | TRUE | 0.0244 |
| Gpld1 | 17 | 9 | 9 | 6 | 10 | 17 | 13 | 10 | 10 | 6 | 11 | 17 | 11.09 | 11 | 11 | 1.04 | 0.057 | 0.924931 | NA | 0 | 0.2162 | 0.1704 | 0.1704 | TRUE | 0.191 |
| Gpm6a | 36 | 13 | 12 | 17 | 20 | 18 | 28 | 14 | 13 | 17 | 23 | 18 | 18.66 | 18 | 19 | 1.025 | 0.036 | 0.94055776 | 0.96673321 | 0.0662 | 0.1294 | 0.1108 | 0.1108 | TRUE | 1.3379 |
| Gpm6b | 30 | 14 | 24 | 35 | 29 | 30 | 23 | 15 | 26 | 35 | 33 | 30 | 26.88 | 21 | 33 | 1.511 | 0.595 | 0.11758497 | 0.24876287 | 0 | 0.0906 | 0.0658 | 0.0658 | TRUE | 0.3597 |
| Gpn1 | 3398 | 2540 | 2513 | 2621 | 2375 | 2874 | 2639 | 2702 | 2712 | 2609 | 2679 | 2837 | 2696.27 | 2684 | 2708 | 1.009 | 0.013 | 0.82289263 | 0.89832647 | 7.00E-04 | 0.0033 | 0.0022 | 0.0022 | TRUE | 0.0212 |
| Gpn2 | 2193 | 1555 | 1400 | 1501 | 1348 | 1578 | 1703 | 1654 | 1511 | 1494 | 1521 | 1558 | 1573.39 | 1623 | 1524 | 0.939 | -0.092 | 0.20094347 | 0.36176911 | 0.0016 | 0.0039 | 0.0031 | 0.0031 | TRUE | 0.0434 |
| Gpn3 | 3477 | 2447 | 2533 | 2674 | 2337 | 2726 | 2700 | 2603 | 2733 | 2662 | 2636 | 2691 | 2670.9 | 2679 | 2663 | 0.994 | -0.009 | 0.88004702 | 0.93298562 | 0 | 0.0033 | 0.0019 | 0.0019 | TRUE | 0.0073 |
| Gpnmb | 67 | 51 | 53 | 48 | 54 | 58 | 52 | 54 | 57 | 48 | 61 | 57 | 54.9 | 54 | 55 | 1.015 | 0.022 | 0.93266142 | 0.96256262 | 0 | 0.0456 | 0.0308 | 0.0308 | TRUE | 0.1212 |
| Gpr1 | 10 | 10 | 5 | 6 | 4 | 14 | 8 | 11 | 5 | 6 | 5 | 14 | 8.02 | 8 | 8 | 1.029 | 0.041 | 0.95442843 | NA | 0.1033 | 0.298 | 0.2429 | 0.2429 | TRUE | 1.0673 |
| Gpr101 | 1 | 0 | 0 | 0 | 1 | 2 | 1 | 0 | 0 | 0 | 1 | 2 | 0.65 | 0 | 1 | 2.998 | 1.584 | 0.52889475 | NA | 0 | 3.6683 | 3.06 | 3.06 | TRUE | 0.14 |
| Gpr107 | 3407 | 2447 | 2525 | 2763 | 2566 | 2821 | 2646 | 2603 | 2724 | 2750 | 2895 | 2785 | 2733.86 | 2658 | 2810 | 1.057 | 0.08 | 0.16602069 | 0.31702522 | 2.00E-04 | 0.0033 | 0.002 | 0.002 | TRUE | 0.0083 |
| Gpr108 | 1196 | 959 | 1028 | 1032 | 1023 | 1162 | 929 | 1020 | 1109 | 1027 | 1154 | 1147 | 1064.42 | 1019 | 1109 | 1.09 | 0.124 | 0.16589854 | 0.31683429 | 0.0052 | 0.0047 | 0.0048 | 0.0048 | TRUE | 0.0751 |
| Gpr110 | 5 | 4 | 5 | 0 | 3 | 1 | 4 | 4 | 5 | 0 | 3 | 1 | 2.98 | 4 | 1 | 0.314 | -1.669 | 0.17361803 | NA | 0 | 0.7967 | 0.6107 | 0.6107 | TRUE | 0.7926 |
| Gpr111 | 3 | 2 | 1 | 1 | 1 | 2 | 2 | 2 | 1 | 1 | 1 | 2 | 1.61 | 2 | 1 | 0.735 | -0.444 | 0.76045804 | NA | 0 | 1.4785 | 0.8907 | 0.8907 | TRUE | 0.0997 |
| Gpr113 | 56 | 57 | 50 | 64 | 52 | 45 | 43 | 61 | 54 | 64 | 59 | 44 | 54.14 | 53 | 56 | 1.06 | 0.085 | 0.76066164 | 0.85788984 | 0.014 | 0.0462 | 0.0372 | 0.0372 | TRUE | 0.2686 |
| Gpr114 | 72 | 52 | 61 | 69 | 70 | 87 | 56 | 55 | 66 | 69 | 79 | 86 | 68.43 | 59 | 78 | 1.322 | 0.403 | 0.09111794 | 0.20660909 | 0 | 0.0371 | 0.0261 | 0.0261 | TRUE | 0.1009 |
| Gpr115 | 0 | 1 | 3 | 3 | 6 | 3 | 0 | 1 | 3 | 3 | 7 | 3 | 2.84 | 1 | 4 | 3.11 | 1.637 | 0.20095119 | NA | 0.1895 | 0.8381 | 0.7021 | 0.7021 | TRUE | 0.7722 |
| Gpr116 | 212 | 187 | 134 | 150 | 146 | 152 | 165 | 199 | 145 | 149 | 165 | 150 | 162.03 | 170 | 155 | 0.913 | -0.131 | 0.43826401 | 0.60976418 | 0.008 | 0.0171 | 0.0145 | 0.0145 | TRUE | 0.2349 |
| Gpr119 | 0 | 0 | 0 | 0 | 0 | 0 | 0 | 0 | 0 | 0 | 0 | 0 | 0 | 0 | 0 | NA | NA | NA | NA | NA | NA | NA | NA | NA | NA |
| Gpr12 | 13 | 19 | 17 | 13 | 19 | 17 | 10 | 20 | 18 | 13 | 21 | 17 | 16.63 | 16 | 17 | 1.062 | 0.087 | 0.86065705 | NA | 0 | 0.1449 | 0.1152 | 0.1152 | TRUE | 0.6056 |
| Gpr123 | 4 | 2 | 3 | 2 | 1 | 1 | 3 | 2 | 3 | 2 | 1 | 1 | 2.1 | 3 | 1 | 0.485 | -1.044 | 0.4261152 | NA | 0 | 1.1331 | 0.6938 | 0.6938 | TRUE | 0.1032 |
| Gpr124 | 398 | 245 | 181 | 294 | 224 | 187 | 309 | 261 | 195 | 293 | 253 | 185 | 249.16 | 255 | 244 | 0.951 | -0.073 | 0.70421178 | 0.82094194 | 0.0486 | 0.0119 | 0.0225 | 0.0225 | TRUE | 0.2859 |
| Gpr125 | 4528 | 3320 | 2979 | 3166 | 2686 | 3100 | 3517 | 3531 | 3214 | 3151 | 3030 | 3060 | 3250.69 | 3421 | 3080 | 0.9 | -0.151 | 0.01356843 | 0.05188163 | 0.0011 | 0.0032 | 0.0024 | 0.0024 | TRUE | 0.0335 |
| Gpr126 | 183 | 111 | 107 | 74 | 92 | 73 | 142 | 118 | 115 | 74 | 104 | 72 | 104.19 | 125 | 83 | 0.659 | -0.601 | 0.00492686 | 0.02424916 | 0.0165 | 0.0252 | 0.0228 | 0.0228 | TRUE | 0.4198 |
| Gpr128 | 0 | 0 | 0 | 0 | 0 | 0 | 0 | 0 | 0 | 0 | 0 | 0 | 0 | 0 | 0 | NA | NA | NA | NA | NA | NA | NA | NA | NA | NA |
| Gpr132 | 3 | 3 | 5 | 6 | 1 | 2 | 2 | 3 | 5 | 6 | 1 | 2 | 3.33 | 3 | 3 | 0.851 | -0.233 | 0.82904335 | NA | 0.1179 | 0.7138 | 0.5385 | 0.5385 | TRUE | 0.964 |
| Gpr133 | 79 | 48 | 51 | 61 | 56 | 57 | 61 | 51 | 55 | 61 | 63 | 56 | 57.93 | 56 | 60 | 1.071 | 0.099 | 0.69575672 | 0.81479785 | 0 | 0.0433 | 0.0288 | 0.0288 | TRUE | 0.0736 |
| Gpr135 | 65 | 70 | 52 | 69 | 62 | 63 | 50 | 74 | 56 | 69 | 70 | 62 | 63.64 | 60 | 67 | 1.115 | 0.157 | 0.53742826 | 0.69609988 | 0 | 0.0397 | 0.0307 | 0.0307 | TRUE | 0.3521 |
| Gpr137 | 671 | 494 | 480 | 519 | 414 | 487 | 521 | 525 | 518 | 517 | 467 | 481 | 504.82 | 521 | 488 | 0.937 | -0.095 | 0.3248569 | 0.49873037 | 0 | 0.0071 | 0.0047 | 0.0047 | TRUE | 0.0305 |
| Gpr137b | 687 | 551 | 646 | 586 | 514 | 630 | 534 | 586 | 697 | 583 | 580 | 622 | 600.29 | 606 | 595 | 0.985 | -0.022 | 0.84470188 | 0.91195135 | 0.0086 | 0.0064 | 0.0071 | 0.0071 | TRUE | 0.2036 |
| Gpr137b-ps | 68 | 52 | 54 | 61 | 49 | 65 | 53 | 55 | 58 | 61 | 55 | 64 | 57.76 | 55 | 60 | 1.086 | 0.119 | 0.63478014 | 0.77160293 | 0 | 0.0435 | 0.0283 | 0.0283 | TRUE | 0.039 |
| Gpr137c | 28 | 25 | 37 | 33 | 39 | 30 | 22 | 27 | 40 | 33 | 44 | 30 | 32.45 | 30 | 36 | 1.214 | 0.28 | 0.44528458 | 0.61613756 | 0.0397 | 0.0755 | 0.0655 | 0.0655 | TRUE | 0.6181 |
| Gpr139 | 2 | 0 | 0 | 1 | 3 | 0 | 2 | 0 | 0 | 1 | 3 | 0 | 0.99 | 1 | 1 | 2.55 | 1.35 | 0.54059712 | NA | 1.8421 | 2.3992 | 2.2957 | 2.2957 | TRUE | 0.5657 |
| Gpr141 | 1 | 0 | 0 | 1 | 0 | 0 | 1 | 0 | 0 | 1 | 0 | 0 | 0.3 | 0 | 0 | 1.075 | 0.105 | 0.97698525 | NA | 0 | 8.0272 | 7.5141 | 7.5141 | TRUE | 0.3906 |
| Gpr142 | 9 | 9 | 3 | 10 | 2 | 9 | 7 | 10 | 3 | 10 | 2 | 9 | 6.82 | 7 | 7 | 1.072 | 0.101 | 0.89841006 | NA | 0.1699 | 0.3502 | 0.3049 | 0.3049 | TRUE | 0.5403 |
| Gpr143 | 21 | 9 | 7 | 15 | 12 | 11 | 16 | 10 | 8 | 15 | 14 | 11 | 12.13 | 11 | 13 | 1.155 | 0.207 | 0.71318335 | NA | 0 | 0.1979 | 0.1474 | 0.1474 | TRUE | 0.7711 |
| Gpr146 | 64 | 66 | 97 | 93 | 116 | 128 | 50 | 70 | 105 | 93 | 131 | 126 | 95.73 | 75 | 117 | 1.57 | 0.651 | 0.01431534 | 0.05391544 | 0.0729 | 0.0272 | 0.0399 | 0.0399 | TRUE | 0.6131 |
| Gpr149 | 3 | 1 | 1 | 1 | 0 | 0 | 2 | 1 | 1 | 1 | 0 | 0 | 0.91 | 1 | 0 | 0.245 | -2.029 | 0.33198448 | NA | 0 | 2.6031 | 1.8334 | 1.8334 | TRUE | 0.3906 |
| Gpr15 | 0 | 0 | 0 | 0 | 0 | 0 | 0 | 0 | 0 | 0 | 0 | 0 | 0 | 0 | 0 | NA | NA | NA | NA | NA | NA | NA | NA | NA | NA |
| Gpr150 | 8 | 12 | 4 | 12 | 10 | 5 | 6 | 13 | 4 | 12 | 11 | 5 | 8.58 | 8 | 9 | 1.213 | 0.278 | 0.69124274 | NA | 0.125 | 0.2788 | 0.2364 | 0.2364 | TRUE | 0.8436 |
| Gpr151 | 0 | 0 | 0 | 0 | 0 | 0 | 0 | 0 | 0 | 0 | 0 | 0 | 0 | 0 | 0 | NA | NA | NA | NA | NA | NA | NA | NA | NA | NA |
| Gpr152 | 0 | 0 | 0 | 0 | 2 | 0 | 0 | 0 | 0 | 0 | 2 | 0 | 0.38 | 0 | 1 | 4.195 | 2.069 | 0.57837236 | NA | 10 | 6.3052 | 8.2866 | 8.2866 | TRUE | 0.9743 |
| Gpr153 | 261 | 197 | 216 | 358 | 343 | 350 | 203 | 210 | 233 | 356 | 387 | 346 | 289.02 | 215 | 363 | 1.69 | 0.757 | 1.75E-09 | 1.10E-07 | 0 | 0.0106 | 0.0077 | 0.0077 | TRUE | 0.0567 |
| Gpr155 | 279 | 199 | 156 | 212 | 166 | 207 | 217 | 212 | 168 | 211 | 187 | 204 | 199.89 | 199 | 201 | 1.008 | 0.011 | 0.94039393 | 0.96671811 | 0.0059 | 0.0143 | 0.0119 | 0.0119 | TRUE | 0.1886 |
| Gpr156 | 62 | 41 | 41 | 42 | 42 | 58 | 48 | 44 | 44 | 42 | 47 | 57 | 47.07 | 45 | 49 | 1.075 | 0.104 | 0.71354398 | 0.82781527 | 0 | 0.0528 | 0.0368 | 0.0368 | TRUE | 0.1912 |
| Gpr157 | 218 | 156 | 150 | 185 | 162 | 178 | 169 | 166 | 162 | 184 | 183 | 176 | 173.29 | 166 | 181 | 1.09 | 0.125 | 0.40054986 | 0.57368196 | 0 | 0.0161 | 0.0101 | 0.0101 | TRUE | 0.0069 |
| Gpr158 | 3 | 1 | 1 | 0 | 1 | 3 | 2 | 1 | 1 | 0 | 1 | 3 | 1.43 | 1 | 1 | 0.9 | -0.152 | 0.92546202 | NA | 0 | 1.6633 | 1.2161 | 1.2161 | TRUE | 0.4066 |

| | | | | | | | | | | | | | | | | | | | | | | | | | |
|---|---|---|---|---|---|---|---|---|---|---|---|---|---|---|---|---|---|---|---|---|---|---|---|---|---|
| Gpr160 | 875 | 632 | 673 | 726 | 690 | 747 | 680 | 672 | 726 | 723 | 778 | 737 | 719.41 | 693 | 746 | 1.077 | 0.107 | 0.2090536 | 0.37076598 | 0 | 0.0057 | 0.0039 | 0.0039 | TRUE | 0.0204 |
| Gpr161 | 141 | 118 | 94 | 73 | 80 | 91 | 110 | 126 | 101 | 73 | 90 | 90 | 98.2 | 112 | 84 | 0.751 | -0.413 | 0.04409309 | 0.1228504 | 0 | 0.0266 | 0.0199 | 0.0199 | TRUE | 0.139 |
| Gpr162 | 150 | 136 | 129 | 120 | 123 | 151 | 116 | 145 | 139 | 119 | 139 | 149 | 134.6 | 133 | 136 | 1.021 | 0.03 | 0.86641233 | 0.92629452 | 0.0063 | 0.02 | 0.016 | 0.016 | TRUE | 0.1419 |
| Gpr165 | 3 | 3 | 1 | 2 | 2 | 4 | 2 | 3 | 1 | 2 | 2 | 4 | 2.47 | 2 | 3 | 1.241 | 0.312 | 0.79384935 | NA | 0 | 0.9635 | 0.6195 | 0.6195 | TRUE | 0.1887 |
| Gpr17 | 0 | 1 | 0 | 0 | 0 | 0 | 0 | 1 | 0 | 0 | 0 | 0 | 0.18 | 0 | 0 | 0.552 | -0.857 | 0.8336548 | NA | 0 | 13.3714 | 10 | 10 | TRUE | 0.5216 |
| Gpr171 | 0 | 0 | 0 | 0 | 0 | 0 | 0 | 0 | 0 | 0 | 0 | 0 | 0 | 0 | 0 | NA | NA | NA | NA | NA | NA | NA | NA | NA | NA |
| Gpr173 | 29 | 18 | 16 | 26 | 18 | 18 | 23 | 19 | 17 | 26 | 20 | 18 | 20.48 | 20 | 21 | 1.079 | 0.11 | 0.79495079 | 0.87990642 | 0 | 0.1181 | 0.0812 | 0.0812 | TRUE | 0.2027 |
| Gpr174 | 0 | 0 | 0 | 0 | 0 | 0 | 0 | 0 | 0 | 0 | 0 | 0 | 0 | 0 | 0 | NA | NA | NA | NA | NA | NA | NA | NA | NA | NA |
| Gpr176 | 18 | 14 | 5 | 2 | 6 | 3 | 14 | 15 | 5 | 2 | 7 | 3 | 7.66 | 11 | 4 | 0.335 | -1.579 | 0.04464439 | NA | 0.1451 | 0.3117 | 0.269 | 0.269 | TRUE | 0.7473 |
| Gpr179 | 443 | 339 | 324 | 387 | 361 | 448 | 344 | 361 | 350 | 385 | 407 | 442 | 381.49 | 352 | 411 | 1.172 | 0.229 | 0.03694852 | 0.10826266 | 0 | 0.0086 | 0.0061 | 0.0061 | TRUE | 0.0502 |
| Gpr18 | 0 | 0 | 0 | 1 | 0 | 0 | 0 | 0 | 0 | 1 | 0 | 0 | 0.17 | 0 | 0 | 2.094 | 1.067 | 0.79379329 | NA | 0 | 14.2885 | 10 | 10 | TRUE | 0.3906 |
| Gpr180 | 1328 | 884 | 910 | 1006 | 865 | 929 | 1031 | 940 | 982 | 1001 | 976 | 917 | 974.64 | 984 | 965 | 0.979 | -0.031 | 0.69759457 | 0.81631335 | 0.0012 | 0.0049 | 0.0036 | 0.0036 | TRUE | 0.0227 |
| Gpr182 | 16 | 8 | 13 | 12 | 13 | 17 | 12 | 9 | 14 | 12 | 15 | 17 | 13.06 | 12 | 15 | 1.238 | 0.308 | 0.56107044 | NA | 0 | 0.1839 | 0.1255 | 0.1255 | TRUE | 0.195 |
| Gpr183 | 0 | 0 | 0 | 0 | 0 | 0 | 0 | 0 | 0 | 0 | 0 | 0 | 0 | 0 | 0 | NA | NA | NA | NA | NA | NA | NA | NA | NA | NA |
| Gpr19 | 1483 | 1085 | 1246 | 1431 | 1192 | 1441 | 1152 | 1154 | 1344 | 1424 | 1345 | 1422 | 1306.99 | 1217 | 1397 | 1.15 | 0.201 | 0.01377148 | 0.05248954 | 0.0035 | 0.0042 | 0.004 | 0.004 | TRUE | 0.0993 |
| Gpr20 | 119 | 72 | 77 | 116 | 106 | 97 | 92 | 77 | 83 | 115 | 120 | 96 | 97.15 | 84 | 110 | 1.306 | 0.385 | 0.06037677 | 0.15393599 | 0 | 0.0268 | 0.0198 | 0.0198 | TRUE | 0.1345 |
| Gpr21 | 12 | 9 | 8 | 6 | 11 | 6 | 9 | 10 | 9 | 6 | 12 | 6 | 8.64 | 9 | 8 | 0.874 | -0.194 | 0.76810572 | NA | 0 | 0.2768 | 0.1943 | 0.1943 | TRUE | 0.5775 |
| Gpr22 | 0 | 0 | 0 | 0 | 0 | 0 | 0 | 0 | 0 | 0 | 0 | 0 | 0 | 0 | 0 | NA | NA | NA | NA | NA | NA | NA | NA | NA | NA |
| Gpr25 | 21 | 18 | 21 | 36 | 22 | 25 | 16 | 19 | 23 | 36 | 25 | 25 | 23.91 | 19 | 29 | 1.481 | 0.567 | 0.15947861 | 0.30880403 | 0 | 0.1016 | 0.0737 | 0.0737 | TRUE | 0.3412 |
| Gpr26 | 0 | 1 | 1 | 1 | 0 | 3 | 0 | 1 | 1 | 1 | 0 | 3 | 1.02 | 1 | 1 | 1.961 | 0.971 | 0.63207857 | NA | 0 | 2.3338 | 1.8722 | 1.8722 | TRUE | 0.3332 |
| Gpr27 | 140 | 102 | 118 | 138 | 138 | 134 | 109 | 108 | 127 | 137 | 156 | 132 | 128.31 | 115 | 142 | 1.235 | 0.305 | 0.08726996 | 0.20044183 | 0 | 0.0209 | 0.015 | 0.015 | TRUE | 0.0862 |
| Gpr3 | 66 | 50 | 38 | 80 | 63 | 68 | 51 | 53 | 41 | 80 | 71 | 67 | 60.55 | 48 | 73 | 1.494 | 0.579 | 0.02223379 | 0.07448392 | 0 | 0.0416 | 0.0292 | 0.0292 | TRUE | 0.1348 |
| Gpr30 | 0 | 1 | 0 | 0 | 0 | 0 | 0 | 1 | 0 | 0 | 0 | 0 | 0.18 | 0 | 0 | 0.552 | -0.857 | 0.8336548 | NA | 0 | 13.3714 | 10 | 10 | TRUE | 0.5216 |
| Gpr31b | 0 | 0 | 0 | 0 | 0 | 0 | 0 | 0 | 0 | 0 | 0 | 0 | 0 | 0 | 0 | NA | NA | NA | NA | NA | NA | NA | NA | NA | NA |
| Gpr33 | 0 | 0 | 0 | 0 | 0 | 0 | 0 | 0 | 0 | 0 | 0 | 0 | 0 | 0 | 0 | NA | NA | NA | NA | NA | NA | NA | NA | NA | NA |
| Gpr34 | 0 | 0 | 0 | 0 | 0 | 0 | 0 | 0 | 0 | 0 | 0 | 0 | 0 | 0 | 0 | NA | NA | NA | NA | NA | NA | NA | NA | NA | NA |
| Gpr35 | 5 | 4 | 2 | 3 | 2 | 3 | 4 | 4 | 2 | 3 | 2 | 3 | 3.08 | 3 | 3 | 0.795 | -0.331 | 0.75473705 | NA | 0 | 0.7711 | 0.4808 | 0.4808 | TRUE | 0.1374 |
| Gpr37 | 2 | 2 | 0 | 1 | 1 | 0 | 2 | 2 | 0 | 1 | 1 | 0 | 0.97 | 1 | 1 | 0.564 | -0.825 | 0.68413901 | NA | 0 | 2.4524 | 1.8344 | 1.8344 | TRUE | 0.3849 |
| Gpr37l1 | 33 | 16 | 29 | 37 | 39 | 30 | 26 | 17 | 31 | 37 | 44 | 30 | 30.73 | 25 | 37 | 1.488 | 0.573 | 0.12053183 | 0.25260432 | 0.0205 | 0.0796 | 0.0647 | 0.0647 | TRUE | 0.2757 |
| Gpr39 | 18 | 19 | 9 | 9 | 8 | 7 | 14 | 20 | 10 | 9 | 9 | 7 | 11.47 | 15 | 8 | 0.566 | -0.82 | 0.1616324 | NA | 0 | 0.2091 | 0.1518 | 0.1518 | TRUE | 0.1925 |
| Gpr4 | 343 | 293 | 273 | 372 | 314 | 370 | 266 | 312 | 295 | 370 | 354 | 365 | 327.06 | 291 | 363 | 1.253 | 0.325 | 0.00568189 | 0.02707943 | 0 | 0.0097 | 0.0069 | 0.0069 | TRUE | 0.067 |
| Gpr44 | 0 | 0 | 1 | 2 | 0 | 1 | 0 | 0 | 1 | 2 | 0 | 1 | 0.68 | 0 | 1 | 2.911 | 1.541 | 0.53439701 | NA | 0 | 3.5076 | 2.9193 | 2.9193 | TRUE | 0.2835 |
| Gpr45 | 3 | 4 | 3 | 4 | 1 | 3 | 2 | 4 | 3 | 4 | 1 | 3 | 2.98 | 3 | 3 | 0.839 | -0.253 | 0.81758211 | NA | 0 | 0.7972 | 0.5267 | 0.5267 | TRUE | 0.2622 |
| Gpr50 | 26 | 18 | 14 | 6 | 11 | 7 | 20 | 19 | 15 | 6 | 12 | 7 | 13.29 | 18 | 8 | 0.458 | -1.125 | 0.04150317 | NA | 0 | 0.1808 | 0.1311 | 0.1311 | TRUE | 0.4658 |
| Gpr52 | 0 | 0 | 0 | 0 | 0 | 1 | 0 | 0 | 0 | 0 | 0 | 1 | 0.16 | 0 | 0 | 2.094 | 1.067 | 0.79379329 | NA | 0 | 14.4078 | 10 | 10 | TRUE | 0.3835 |
| Gpr55 | 3 | 3 | 2 | 1 | 2 | 3 | 2 | 3 | 2 | 1 | 2 | 3 | 2.32 | 2 | 2 | 0.811 | -0.303 | 0.80485573 | NA | 0 | 1.0261 | 0.6423 | 0.6423 | TRUE | 0.2004 |
| Gpr56 | 410 | 375 | 623 | 741 | 783 | 974 | 318 | 399 | 672 | 738 | 883 | 961 | 661.98 | 463 | 861 | 1.86 | 0.895 | 0.00820469 | 0.03568766 | 0.081 | 0.006 | 0.0256 | 0.081 | TRUE | 1.8041 |
| Gpr6 | 0 | 0 | 0 | 0 | 0 | 0 | 0 | 0 | 0 | 0 | 0 | 0 | 0 | 0 | 0 | NA | NA | NA | NA | NA | NA | NA | NA | NA | NA |
| Gpr61 | 8 | 11 | 8 | 13 | 12 | 12 | 6 | 12 | 9 | 13 | 14 | 12 | 10.81 | 9 | 13 | 1.461 | 0.547 | 0.34881134 | NA | 0 | 0.2216 | 0.1501 | 0.1501 | TRUE | 0.2894 |
| Gpr62 | 19 | 12 | 16 | 21 | 15 | 21 | 15 | 13 | 17 | 21 | 17 | 21 | 17.22 | 15 | 20 | 1.311 | 0.391 | 0.39260848 | 0.56630871 | 0 | 0.14 | 0.0919 | 0.0919 | TRUE | 0.0753 |
| Gpr63 | 83 | 41 | 43 | 42 | 41 | 39 | 64 | 44 | 46 | 42 | 46 | 38 | 46.84 | 51 | 42 | 0.811 | -0.302 | 0.3063345 | 0.47877557 | 0 | 0.053 | 0.0411 | 0.0411 | TRUE | 0.4541 |
| Gpr64 | 53 | 25 | 25 | 18 | 32 | 23 | 41 | 27 | 27 | 18 | 36 | 23 | 28.58 | 32 | 26 | 0.798 | -0.326 | 0.41318319 | 0.585992 | 0.0609 | 0.0854 | 0.0785 | 0.0785 | TRUE | 0.7391 |
| Gpr65 | 0 | 0 | 0 | 0 | 0 | 0 | 0 | 0 | 0 | 0 | 0 | 0 | 0 | 0 | 0 | NA | NA | NA | NA | NA | NA | NA | NA | NA | NA |
| Gpr68 | 34 | 38 | 47 | 57 | 58 | 83 | 26 | 40 | 51 | 57 | 65 | 82 | 53.61 | 39 | 68 | 1.759 | 0.815 | 0.00752719 | 0.0335368 | 0.0474 | 0.0466 | 0.0469 | 0.0469 | TRUE | 0.5712 |
| Gpr75 | 96 | 93 | 103 | 82 | 63 | 82 | 75 | 99 | 111 | 82 | 71 | 81 | 86.38 | 95 | 78 | 0.827 | -0.273 | 0.22873014 | 0.39402675 | 0.015 | 0.0299 | 0.0254 | 0.0254 | TRUE | 0.3824 |
| Gpr81 | 10 | 12 | 6 | 10 | 6 | 11 | 8 | 13 | 6 | 10 | 7 | 11 | 9.1 | 9 | 9 | 1.031 | 0.044 | 0.94581802 | NA | 0 | 0.2629 | 0.1856 | 0.1856 | TRUE | 0.3963 |
| Gpr82 | 0 | 0 | 0 | 0 | 0 | 0 | 0 | 0 | 0 | 0 | 0 | 0 | 0 | 0 | 0 | NA | NA | NA | NA | NA | NA | NA | NA | NA | NA |
| Gpr83 | 18 | 12 | 4 | 10 | 15 | 17 | 14 | 13 | 4 | 10 | 17 | 17 | 12.45 | 10 | 15 | 1.384 | 0.469 | 0.42498671 | NA | 0.0876 | 0.1927 | 0.167 | 0.167 | TRUE | 0.8401 |
| Gpr84 | 4 | 2 | 0 | 1 | 1 | 2 | 3 | 2 | 0 | 1 | 1 | 2 | 1.56 | 2 | 1 | 0.764 | -0.388 | 0.80391439 | NA | 0 | 1.5263 | 1.1091 | 1.1091 | TRUE | 0.4538 |
| Gpr85 | 39 | 23 | 25 | 9 | 10 | 15 | 30 | 24 | 27 | 9 | 11 | 15 | 19.46 | 27 | 12 | 0.428 | -1.225 | 0.0069008 | 0.03140804 | 0 | 0.1242 | 0.0862 | 0.0862 | TRUE | 0.2239 |
| Gpr87 | 0 | 0 | 1 | 1 | 0 | 0 | 0 | 0 | 1 | 1 | 0 | 0 | 0.35 | 0 | 0 | 1.075 | 0.105 | 0.97567165 | NA | 0 | 6.8576 | 6.5143 | 6.5143 | TRUE | 0.5365 |
| Gpr88 | 0 | 0 | 0 | 0 | 0 | 1 | 0 | 0 | 0 | 0 | 0 | 1 | 0.16 | 0 | 0 | 2.094 | 1.067 | 0.79379329 | NA | 0 | 14.4078 | 10 | 10 | TRUE | 0.3835 |
| Gpr89 | 1347 | 959 | 884 | 1034 | 961 | 1011 | 1046 | 1020 | 954 | 1029 | 1084 | 998 | 1021.9 | 1007 | 1037 | 1.029 | 0.041 | 0.60085775 | 0.74639037 | 0.001 | 0.0047 | 0.0034 | 0.0034 | TRUE | 0.0248 |
| Gpr97 | 9 | 0 | 0 | 12 | 8 | 3 | 7 | 0 | 0 | 12 | 9 | 3 | 5.15 | 2 | 8 | 3.175 | 1.667 | 0.14159786 | NA | 1.8476 | 0.4623 | 0.6695 | 0.6695 | TRUE | 1.808 |
| Gpr98 | 598 | 393 | 380 | 427 | 325 | 414 | 464 | 418 | 410 | 425 | 367 | 409 | 415.47 | 431 | 400 | 0.928 | -0.108 | 0.32981608 | 0.50423161 | 0.0029 | 0.0081 | 0.0065 | 0.0065 | TRUE | 0.0597 |
| Gprasp1 | 8821 | 6227 | 5794 | 6924 | 5992 | 6748 | 6851 | 6624 | 6252 | 6892 | 6760 | 6661 | 6673.2 | 6576 | 6771 | 1.029 | 0.042 | 0.4473662 | 0.61788255 | 0.001 | 0.0028 | 0.002 | 0.002 | TRUE | 0.0226 |
| Gprasp2 | 2514 | 2035 | 2059 | 2480 | 2120 | 2315 | 1953 | 2165 | 2222 | 2469 | 2392 | 2285 | 2247.37 | 2113 | 2382 | 1.128 | 0.174 | 0.01393111 | 0.05291453 | 0.0027 | 0.0035 | 0.0032 | 0.0032 | TRUE | 0.055 |
| Gprc5a | 2971 | 2578 | 3173 | 4292 | 4213 | 5210 | 2307 | 2742 | 3424 | 4272 | 4753 | 5143 | 3773.55 | 2824 | 4723 | 1.673 | 0.742 | 4.71E-05 | 0.00066505 | 0.0237 | 0.0031 | 0.0092 | 0.0237 | TRUE | 0.418 |
| Gprc5b | 960 | 693 | 785 | 933 | 780 | 907 | 746 | 737 | 847 | 929 | 880 | 895 | 838.95 | 777 | 901 | 1.162 | 0.217 | 0.01211884 | 0.04771753 | 0.002 | 0.0053 | 0.0042 | 0.0042 | TRUE | 0.0726 |
| Gprc5c | 573 | 431 | 453 | 468 | 399 | 512 | 445 | 458 | 489 | 466 | 450 | 505 | 468.94 | 464 | 474 | 1.023 | 0.033 | 0.74956687 | 0.8514989 | 0 | 0.0075 | 0.0054 | 0.0054 | TRUE | 0.04 |
| Gprc5d | 4 | 0 | 4 | 1 | 1 | 1 | 3 | 0 | 4 | 1 | 1 | 1 | 1.76 | 2 | 1 | 0.416 | -1.266 | 0.415765 | NA | 0 | 1.3525 | 1.0499 | 1.0499 | TRUE | 0.4309 |
| Gprc6a | 0 | 0 | 0 | 0 | 0 | 0 | 0 | 0 | 0 | 0 | 0 | 0 | 0 | 0 | 0 | NA | NA | NA | NA | NA | NA | NA | NA | NA | NA |
| Gprin1 | 66 | 57 | 38 | 56 | 56 | 57 | 51 | 61 | 41 | 56 | 63 | 56 | 54.68 | 51 | 58 | 1.144 | 0.194 | 0.47052489 | 0.63958271 | 0 | 0.0458 | 0.0336 | 0.0336 | TRUE | 0.2113 |
| Gprin2 | 105 | 99 | 98 | 109 | 95 | 107 | 82 | 105 | 106 | 108 | 107 | 106 | 102.31 | 98 | 107 | 1.105 | 0.144 | 0.47223146 | 0.64086802 | 0 | 0.0256 | 0.0189 | 0.0189 | TRUE | 0.2276 |
| Gprin3 | 19 | 10 | 4 | 7 | 1 | 5 | 15 | 11 | 4 | 7 | 1 | 5 | 7.12 | 10 | 4 | 0.439 | -1.187 | 0.14505432 | NA | 0.2443 | 0.3351 | 0.3126 | 0.3126 | TRUE | 0.2412 |
| Gps1 | 9202 | 6902 | 6597 | 6860 | 6037 | 6994 | 7147 | 7342 | 7118 | 6828 | 6810 | 6904 | 7024.91 | 7202 | 6847 | 0.951 | -0.073 | 0.12690154 | 0.26237578 | 0 | 0.0028 | 0.0015 | 0.0015 | TRUE | 0.0035 |
| Gps2 | 1514 | 1167 | 1010 | 892 | 855 | 1017 | 1176 | 1241 | 1090 | 888 | 965 | 1004 | 1060.55 | 1169 | 952 | 0.814 | -0.296 | 0.00041145 | 0.00371974 | 0.0031 | 0.0047 | 0.0041 | 0.0041 | TRUE | 0.0422 |
| Gpsm1 | 514 | 417 | 384 | 493 | 401 | 488 | 399 | 444 | 414 | 491 | 452 | 482 | 446.99 | 419 | 475 | 1.136 | 0.184 | 0.07364558 | 0.17789695 | 0 | 0.0077 | 0.0053 | 0.0053 | TRUE | 0.03 |
| Gpsm2 | 2987 | 2038 | 2311 | 2140 | 1826 | 2052 | 2320 | 2168 | 2494 | 2130 | 2060 | 2026 | 2199.49 | 2327 | 2072 | 0.89 | -0.167 | 0.01647747 | 0.05987196 | 0.0021 | 0.0035 | 0.0031 | 0.0031 | TRUE | 0.0463 |

| | | | | | | | | | | | | | | | | | | | | | | | | | |
|---|---|---|---|---|---|---|---|---|---|---|---|---|---|---|---|---|---|---|---|---|---|---|---|---|---|
| Gpsm3 | 192 | 147 | 132 | 141 | 119 | 139 | 149 | 156 | 142 | 140 | 134 | 137 | 143.29 | 149 | 137 | 0.92 | -0.121 | 0.45867345 | 0.62843363 | 0 | 0.019 | 0.0121 | 0.0121 | TRUE | 0.0166 |
| Gpt | 190 | 157 | 132 | 175 | 137 | 158 | 148 | 167 | 142 | 174 | 155 | 156 | 156.95 | 152 | 162 | 1.062 | 0.087 | 0.58861514 | 0.73762904 | 0 | 0.0175 | 0.0124 | 0.0124 | TRUE | 0.0708 |
| Gpt2 | 2277 | 1705 | 1746 | 2099 | 1813 | 1961 | 1768 | 1814 | 1884 | 2089 | 2045 | 1936 | 1922.73 | 1822 | 2023 | 1.111 | 0.152 | 0.01992601 | 0.06895944 | 8.00E-04 | 0.0037 | 0.0025 | 0.0025 | TRUE | 0.0176 |
| Gpx1 | 12428 | 8393 | 7564 | 7473 | 6124 | 7140 | 9652 | 8928 | 8161 | 7439 | 6909 | 7048 | 8022.79 | 8914 | 7132 | 0.8 | -0.322 | 2.81E-06 | 6.56E-05 | 0.0041 | 0.0027 | 0.0033 | 0.0033 | TRUE | 0.0665 |
| Gpx2 | 156 | 131 | 160 | 250 | 265 | 320 | 121 | 139 | 173 | 249 | 299 | 316 | 216.14 | 144 | 288 | 2.004 | 1.003 | 1.48E-09 | 9.50E-08 | 0.0179 | 0.0134 | 0.0147 | 0.0147 | TRUE | 0.3009 |
| Gpx2-ps1 | 5 | 4 | 3 | 3 | 1 | 3 | 4 | 4 | 3 | 3 | 1 | 3 | 3.08 | 4 | 2 | 0.628 | -0.671 | 0.5329569 | NA | 0 | 0.7731 | 0.4911 | 0.4911 | TRUE | 0.1868 |
| Gpx3 | 2111 | 1443 | 1617 | 1918 | 1680 | 2014 | 1640 | 1535 | 1745 | 1909 | 1895 | 1988 | 1785.28 | 1640 | 1931 | 1.178 | 0.236 | 0.00078251 | 0.00606152 | 0.0015 | 0.0038 | 0.003 | 0.003 | TRUE | 0.0365 |
| Gpx4 | 13454 | 9388 | 11040 | 10941 | 10218 | 11264 | 10449 | 9986 | 11912 | 10891 | 11527 | 11119 | 10980.67 | 10782 | 11179 | 1.037 | 0.052 | 0.45210319 | 0.62194332 | 0.004 | 0.0026 | 0.0034 | 0.0034 | TRUE | 0.1023 |
| Gpx5 | 0 | 1 | 0 | 0 | 0 | 0 | 0 | 1 | 0 | 0 | 0 | 0 | 0.18 | 0 | 0 | 0.552 | -0.857 | 0.8336548 | NA | 0 | 13.3714 | 10 | 10 | TRUE | 0.5216 |
| Gpx6 | 30 | 24 | 13 | 35 | 35 | 42 | 23 | 26 | 14 | 35 | 39 | 41 | 29.77 | 21 | 38 | 1.831 | 0.873 | 0.01776462 | 0.06324029 | 0 | 0.082 | 0.0614 | 0.0614 | TRUE | 0.4168 |
| Gpx7 | 338 | 233 | 243 | 345 | 264 | 296 | 263 | 248 | 262 | 343 | 298 | 292 | 284.33 | 258 | 311 | 1.208 | 0.273 | 0.03052216 | 0.09373727 | 0 | 0.0108 | 0.0079 | 0.0079 | TRUE | 0.0946 |
| Gpx8 | 161 | 111 | 108 | 136 | 144 | 144 | 125 | 118 | 117 | 135 | 162 | 142 | 133.27 | 120 | 146 | 1.219 | 0.286 | 0.09779628 | 0.2178959 | 0 | 0.0202 | 0.014 | 0.014 | TRUE | 0.0887 |
| Gramd1a | 1727 | 1451 | 1770 | 1472 | 1380 | 1599 | 1341 | 1543 | 1910 | 1465 | 1557 | 1578 | 1565.83 | 1598 | 1533 | 0.961 | -0.058 | 0.59404877 | 0.74083813 | 0.0162 | 0.0039 | 0.0079 | 0.0079 | TRUE | 0.3503 |
| Gramd1b | 907 | 676 | 740 | 743 | 733 | 826 | 704 | 719 | 798 | 740 | 827 | 815 | 767.3 | 740 | 794 | 1.073 | 0.101 | 0.26271637 | 0.43318169 | 0.0028 | 0.0055 | 0.0046 | 0.0046 | TRUE | 0.0542 |
| Gramd1c | 102 | 61 | 56 | 51 | 56 | 73 | 79 | 65 | 60 | 51 | 63 | 72 | 65.09 | 68 | 62 | 0.904 | -0.145 | 0.56844806 | 0.72207857 | 0.0113 | 0.0388 | 0.0311 | 0.0311 | TRUE | 0.2255 |
| Gramd2 | 105 | 74 | 82 | 76 | 63 | 60 | 82 | 79 | 88 | 76 | 71 | 59 | 75.78 | 83 | 69 | 0.828 | -0.272 | 0.2280983 | 0.39336499 | 0 | 0.0337 | 0.0234 | 0.0234 | TRUE | 0.1335 |
| Gramd3 | 466 | 304 | 343 | 374 | 309 | 336 | 362 | 323 | 370 | 372 | 349 | 332 | 351.32 | 352 | 351 | 0.997 | -0.005 | 0.96599481 | 0.98144468 | 0 | 0.0092 | 0.0068 | 0.0068 | TRUE | 0.0548 |
| Gramd4 | 1821 | 1456 | 1690 | 1848 | 1760 | 2085 | 1414 | 1549 | 1823 | 1839 | 1985 | 2058 | 1778.28 | 1595 | 1961 | 1.231 | 0.299 | 0.00129617 | 0.00870121 | 0.0093 | 0.0038 | 0.0057 | 0.0057 | TRUE | 0.1877 |
| Grap | 3 | 2 | 4 | 2 | 2 | 3 | 2 | 2 | 4 | 2 | 2 | 3 | 2.66 | 3 | 2 | 0.83 | -0.269 | 0.81375573 | NA | 0 | 0.8922 | 0.5567 | 0.5567 | TRUE | 0.1997 |
| Grap2 | 8 | 5 | 3 | 4 | 2 | 5 | 6 | 5 | 3 | 4 | 2 | 5 | 4.32 | 5 | 4 | 0.755 | -0.406 | 0.65652123 | NA | 0 | 0.5506 | 0.3654 | 0.3654 | TRUE | 0.1642 |
| Grasp | 1369 | 1047 | 1134 | 1525 | 1376 | 1600 | 1063 | 1114 | 1224 | 1518 | 1552 | 1579 | 1341.7 | 1134 | 1550 | 1.369 | 0.453 | 2.26E-09 | 1.37E-07 | 0.0019 | 0.0042 | 0.0034 | 0.0034 | TRUE | 0.0573 |
| Grb10 | 1664 | 1143 | 1030 | 1184 | 952 | 1018 | 1292 | 1216 | 1111 | 1179 | 1074 | 1005 | 1146.16 | 1206 | 1086 | 0.899 | -0.154 | 0.0824015 | 0.19326627 | 0.0053 | 0.0045 | 0.0048 | 0.0048 | TRUE | 0.0677 |
| Grb14 | 530 | 401 | 448 | 457 | 392 | 422 | 412 | 427 | 483 | 455 | 442 | 417 | 439.21 | 441 | 438 | 0.996 | -0.006 | 0.95497625 | 0.9753363 | 0.0025 | 0.0078 | 0.0061 | 0.0061 | TRUE | 0.0822 |
| Grb2 | 9970 | 7349 | 6698 | 6628 | 5955 | 6925 | 7743 | 7817 | 7227 | 6598 | 6718 | 6836 | 7156.47 | 7596 | 6717 | 0.884 | -0.178 | 0.00099358 | 0.00721607 | 8.00E-04 | 0.0028 | 0.002 | 0.002 | TRUE | 0.0218 |
| Grb7 | 3127 | 2509 | 3199 | 3066 | 2948 | 3451 | 2429 | 2669 | 3452 | 3052 | 3326 | 3407 | 3055.55 | 2850 | 3262 | 1.145 | 0.196 | 0.06736875 | 0.16667847 | 0.0188 | 0.0032 | 0.0079 | 0.0079 | TRUE | 0.4138 |
| Grcc10 | 1922 | 1458 | 1712 | 1970 | 1781 | 2055 | 1493 | 1551 | 1847 | 1961 | 2009 | 2029 | 1814.92 | 1630 | 2000 | 1.228 | 0.296 | 0.00043791 | 0.003896 | 0.0061 | 0.0037 | 0.0046 | 0.0046 | TRUE | 0.1621 |
| Greb1 | 455 | 339 | 355 | 584 | 524 | 606 | 353 | 361 | 383 | 581 | 591 | 598 | 477.94 | 366 | 590 | 1.616 | 0.693 | 1.54E-12 | 1.84E-10 | 0 | 0.0074 | 0.0047 | 0.0047 | TRUE | 0.0192 |
| Greb1l | 1672 | 1137 | 1008 | 1074 | 975 | 987 | 1299 | 1209 | 1088 | 1069 | 1100 | 974 | 1123.15 | 1199 | 1048 | 0.873 | -0.196 | 0.02636217 | 0.08447458 | 0.0051 | 0.0045 | 0.0047 | 0.0047 | TRUE | 0.078 |
| Grem1 | 2 | 0 | 2 | 1 | 4 | 3 | 2 | 0 | 2 | 1 | 5 | 3 | 2.03 | 1 | 3 | 2.241 | 1.164 | 0.41238461 | NA | 0 | 1.1699 | 0.8837 | 0.8837 | TRUE | 0.2206 |
| Grem2 | 30 | 9 | 11 | 19 | 8 | 11 | 23 | 10 | 12 | 19 | 9 | 11 | 13.92 | 15 | 13 | 0.856 | -0.224 | 0.69297988 | NA | 0.1285 | 0.1727 | 0.1595 | 0.1595 | TRUE | 1.333 |
| Grhl1 | 257 | 199 | 187 | 150 | 155 | 186 | 200 | 212 | 202 | 149 | 175 | 184 | 186.8 | 205 | 169 | 0.829 | -0.271 | 0.07162918 | 0.17450075 | 0 | 0.0151 | 0.0109 | 0.0109 | TRUE | 0.116 |
| Grhl2 | 625 | 473 | 487 | 283 | 257 | 332 | 485 | 503 | 525 | 282 | 290 | 328 | 402.23 | 504 | 300 | 0.595 | -0.749 | 1.34E-11 | 1.31E-09 | 9.00E-04 | 0.0083 | 0.0062 | 0.0062 | TRUE | 0.0768 |
| Grhl3 | 2629 | 2085 | 2378 | 2429 | 2389 | 2692 | 2042 | 2218 | 2566 | 2418 | 2695 | 2657 | 2432.62 | 2275 | 2590 | 1.139 | 0.188 | 0.03029704 | 0.09329138 | 0.0079 | 0.0034 | 0.005 | 0.005 | TRUE | 0.1509 |
| Grhpr | 3742 | 2800 | 2956 | 3013 | 2796 | 3140 | 2906 | 2978 | 3189 | 2999 | 3154 | 3100 | 3054.51 | 3024 | 3084 | 1.02 | 0.029 | 0.63966154 | 0.77477341 | 0.0012 | 0.0032 | 0.0024 | 0.0024 | TRUE | 0.0274 |
| Gria1 | 14 | 11 | 4 | 1 | 2 | 5 | 11 | 12 | 4 | 1 | 2 | 5 | 5.85 | 9 | 3 | 0.304 | -1.717 | 0.05595274 | NA | 0.1179 | 0.4078 | 0.339 | 0.339 | TRUE | 0.63 |
| Gria2 | 6 | 1 | 2 | 1 | 3 | 0 | 5 | 1 | 2 | 1 | 3 | 0 | 2.04 | 3 | 1 | 0.53 | -0.915 | 0.52709511 | NA | 0.3801 | 1.1622 | 0.9644 | 0.9644 | TRUE | 0.7434 |
| Gria3 | 91 | 57 | 56 | 41 | 37 | 26 | 71 | 61 | 60 | 41 | 42 | 26 | 49.99 | 64 | 36 | 0.561 | -0.834 | 0.00430589 | 0.02186838 | 0.0061 | 0.0498 | 0.0396 | 0.0396 | TRUE | 0.4741 |
| Gria4 | 67 | 36 | 34 | 36 | 22 | 25 | 52 | 38 | 37 | 36 | 25 | 25 | 35.39 | 42 | 29 | 0.668 | -0.581 | 0.08932977 | 0.20376483 | 0.0176 | 0.0694 | 0.0548 | 0.0548 | TRUE | 0.3536 |
| Grid1 | 125 | 105 | 101 | 71 | 76 | 72 | 97 | 112 | 109 | 71 | 86 | 71 | 90.87 | 106 | 76 | 0.716 | -0.481 | 0.02118064 | 0.07213125 | 0 | 0.0285 | 0.02 | 0.02 | TRUE | 0.113 |
| Grid2 | 13 | 11 | 13 | 5 | 1 | 3 | 10 | 12 | 14 | 5 | 1 | 3 | 7.48 | 12 | 3 | 0.258 | -1.954 | 0.0125207 | NA | 0 | 0.3192 | 0.232 | 0.232 | TRUE | 0.4166 |
| Grid2ip | 70 | 63 | 74 | 124 | 101 | 116 | 54 | 67 | 80 | 123 | 114 | 115 | 92.18 | 67 | 117 | 1.764 | 0.818 | 0.00015647 | 0.00176233 | 0.004 | 0.0281 | 0.0221 | 0.0221 | TRUE | 0.2864 |
| Grifin | 7 | 7 | 2 | 6 | 6 | 8 | 5 | 7 | 2 | 6 | 7 | 8 | 5.95 | 5 | 7 | 1.366 | 0.45 | 0.57048916 | NA | 0 | 0.401 | 0.2831 | 0.2831 | TRUE | 0.4006 |
| Grik1 | 0 | 1 | 1 | 0 | 0 | 0 | 0 | 1 | 1 | 0 | 0 | 0 | 0.36 | 1 | 0 | 0.271 | -1.881 | 0.5755355 | NA | 0 | 6.6391 | 6.4428 | 6.4428 | TRUE | 0.3513 |
| Grik2 | 38 | 26 | 14 | 21 | 22 | 15 | 30 | 28 | 15 | 21 | 25 | 15 | 22.13 | 24 | 20 | 0.828 | -0.272 | 0.53394655 | 0.69315568 | 0.0451 | 0.1095 | 0.0922 | 0.0922 | TRUE | 0.5917 |
| Grik3 | 2261 | 1720 | 1731 | 1705 | 1652 | 1780 | 1756 | 1830 | 1868 | 1697 | 1864 | 1757 | 1795.2 | 1818 | 1773 | 0.975 | -0.036 | 0.5938377 | 0.74083813 | 0.001 | 0.0037 | 0.0027 | 0.0027 | TRUE | 0.0239 |
| Grik4 | 80 | 60 | 64 | 88 | 74 | 96 | 62 | 64 | 69 | 88 | 83 | 95 | 76.81 | 65 | 89 | 1.367 | 0.451 | 0.04082966 | 0.11615809 | 0 | 0.0333 | 0.0218 | 0.0218 | TRUE | 0.0355 |
| Grik5 | 705 | 526 | 585 | 695 | 605 | 662 | 548 | 559 | 631 | 692 | 683 | 653 | 627.68 | 579 | 676 | 1.168 | 0.224 | 0.01687815 | 0.06107328 | 0.0017 | 0.0062 | 0.0048 | 0.0048 | TRUE | 0.0701 |
| Grin1 | 1590 | 1995 | 2988 | 1861 | 2547 | 2605 | 1235 | 2122 | 3224 | 1852 | 2873 | 2572 | 2313.04 | 2194 | 2432 | 1.109 | 0.149 | 0.72750948 | 0.83661541 | 0.1318 | 0.0035 | 0.0332 | 0.1318 | TRUE | 0.2757 |
| Grin2a | 7 | 13 | 8 | 9 | 13 | 12 | 5 | 14 | 9 | 9 | 15 | 12 | 10.56 | 9 | 12 | 1.288 | 0.365 | 0.55226281 | NA | 0 | 0.2268 | 0.1749 | 0.1749 | TRUE | 0.4368 |
| Grin2b | 12 | 5 | 2 | 12 | 10 | 9 | 9 | 5 | 2 | 12 | 11 | 9 | 8.15 | 5 | 11 | 1.854 | 0.891 | 0.20914896 | NA | 0.0268 | 0.2932 | 0.2306 | 0.2306 | TRUE | 0.5325 |
| Grin2c | 58 | 67 | 99 | 210 | 207 | 220 | 45 | 71 | 107 | 209 | 234 | 217 | 147.14 | 74 | 220 | 2.993 | 1.581 | 2.90E-11 | 2.68E-09 | 0.075 | 0.0185 | 0.0318 | 0.0318 | TRUE | 0.9968 |
| Grin2d | 98 | 70 | 69 | 95 | 89 | 90 | 76 | 74 | 74 | 95 | 100 | 89 | 84.8 | 75 | 95 | 1.259 | 0.333 | 0.11145273 | 0.2393036 | 0 | 0.0304 | 0.0196 | 0.0196 | TRUE | 0.0276 |
| Grin3a | 3 | 0 | 3 | 1 | 1 | 0 | 2 | 0 | 3 | 1 | 1 | 0 | 1.28 | 2 | 1 | 0.374 | -1.418 | 0.44694565 | NA | 0 | 1.8515 | 1.5066 | 1.5066 | TRUE | 0.4398 |
| Grin3b | 0 | 3 | 2 | 4 | 1 | 5 | 0 | 3 | 2 | 4 | 1 | 5 | 2.57 | 2 | 3 | 1.979 | 0.985 | 0.45148 | NA | 0 | 0.9262 | 0.7968 | 0.7968 | TRUE | 0.9011 |
| Grina | 2521 | 2078 | 2317 | 2843 | 2851 | 3286 | 1958 | 2210 | 2500 | 2830 | 3216 | 3244 | 2659.71 | 2223 | 3097 | 1.394 | 0.48 | 1.30E-07 | 4.93E-06 | 0.0097 | 0.0033 | 0.0056 | 0.0056 | TRUE | 0.1442 |
| Grip1 | 334 | 227 | 213 | 128 | 133 | 134 | 259 | 241 | 230 | 127 | 150 | 132 | 190.07 | 243 | 136 | 0.559 | -0.84 | 2.68E-08 | 1.20E-06 | 0 | 0.0149 | 0.0107 | 0.0107 | TRUE | 0.073 |
| Grip2 | 94 | 64 | 76 | 92 | 82 | 86 | 73 | 68 | 82 | 92 | 93 | 85 | 82.01 | 74 | 90 | 1.206 | 0.27 | 0.20753889 | 0.36913307 | 0 | 0.0313 | 0.0209 | 0.0209 | TRUE | 0.0679 |
| Gripap1 | 488 | 333 | 284 | 312 | 224 | 295 | 379 | 354 | 306 | 311 | 253 | 291 | 315.69 | 346 | 285 | 0.821 | -0.284 | 0.03116574 | 0.0953246 | 0.008 | 0.0099 | 0.0093 | 0.0093 | TRUE | 0.1147 |
| Grk1 | 12 | 11 | 9 | 10 | 7 | 4 | 9 | 12 | 10 | 10 | 8 | 4 | 8.76 | 10 | 7 | 0.71 | -0.494 | 0.45241098 | NA | 0 | 0.2731 | 0.193 | 0.193 | TRUE | 0.4598 |
| Grk4 | 18 | 13 | 13 | 5 | 7 | 9 | 14 | 14 | 14 | 5 | 8 | 9 | 10.6 | 14 | 7 | 0.519 | -0.945 | 0.11252827 | NA | 0 | 0.226 | 0.1497 | 0.1497 | TRUE | 0.2147 |
| Grk5 | 707 | 480 | 461 | 484 | 399 | 472 | 549 | 511 | 497 | 482 | 450 | 466 | 492.48 | 519 | 466 | 0.897 | -0.157 | 0.10992329 | 0.23714529 | 0 | 0.0072 | 0.0049 | 0.0049 | TRUE | 0.0319 |
| Grk6 | 5057 | 3904 | 4229 | 4421 | 4169 | 4747 | 3928 | 4153 | 4563 | 4401 | 4703 | 4686 | 4405.5 | 4215 | 4597 | 1.091 | 0.126 | 0.06482267 | 0.16231787 | 0.0033 | 0.003 | 0.0031 | 0.0031 | TRUE | 0.0636 |
| Grlf1 | 8275 | 5523 | 5053 | 5332 | 4720 | 5420 | 6427 | 5875 | 5452 | 5307 | 5325 | 5350 | 5622.7 | 5918 | 5327 | 0.9 | -0.152 | 0.0222242 | 0.07447842 | 0.0032 | 0.0028 | 0.003 | 0.003 | TRUE | 0.0698 |
| Grm1 | 0 | 0 | 0 | 1 | 0 | 0 | 0 | 0 | 0 | 1 | 0 | 0 | 0.17 | 0 | 0 | 2.094 | 1.067 | 0.79379329 | NA | 0 | 14.2885 | 10 | 10 | TRUE | 0.3906 |
| Grm2 | 21 | 9 | 7 | 15 | 9 | 8 | 16 | 10 | 8 | 15 | 10 | 8 | 11.07 | 11 | 11 | 0.971 | -0.042 | 0.94427534 | NA | 0.0625 | 0.2165 | 0.1701 | 0.1701 | TRUE | 0.7678 |
| Grm3 | 9 | 0 | 6 | 2 | 1 | 0 | 7 | 0 | 6 | 2 | 1 | 0 | 2.76 | 4 | 1 | 0.229 | -2.129 | 0.14611976 | NA | 0 | 0.86 | 0.9402 | 0.9402 | TRUE | 1.3177 |
| Grm4 | 56 | 43 | 49 | 42 | 34 | 42 | 43 | 46 | 53 | 42 | 38 | 41 | 43.95 | 47 | 40 | 0.859 | -0.219 | 0.44899131 | 0.61914451 | 0 | 0.0563 | 0.0371 | 0.0371 | TRUE | 0.079 |
| Grm5 | 3 | 3 | 0 | 1 | 0 | 0 | 2 | 3 | 0 | 1 | 0 | 0 | 1.09 | 2 | 0 | 0.2 | -2.321 | 0.27406904 | NA | 0 | 2.1847 | 1.9836 | 1.9836 | TRUE | 0.3792 |
| Grm6 | 3 | 0 | 0 | 1 | 1 | 0 | 2 | 0 | 0 | 1 | 1 | 0 | 0.74 | 1 | 1 | 0.851 | -0.233 | 0.92310399 | NA | 0 | 3.1955 | 2.8611 | 2.8611 | TRUE | 1.4389 |

| | | | | | | | | | | | | | | | | | | | | | | | | | |
|---|---|---|---|---|---|---|---|---|---|---|---|---|---|---|---|---|---|---|---|---|---|---|---|---|---|
| Grm7 | 191 | 127 | 133 | 136 | 113 | 143 | 148 | 135 | 144 | 135 | 127 | 141 | 138.49 | 142 | 134 | 0.946 | -0.08 | 0.6291173 | 0.76644848 | 0 | 0.0195 | 0.0128 | 0.0128 | TRUE | 0.0214 |
| Grm8 | 1 | 1 | 0 | 1 | 0 | 1 | 1 | 1 | 0 | 1 | 0 | 1 | 0.64 | 1 | 1 | 1.082 | 0.113 | 0.96183421 | NA | 0 | 3.7222 | 2.4899 | 2.4899 | TRUE | 0.2041 |
| Grn | 6717 | 4761 | 5008 | 5056 | 4620 | 5346 | 5217 | 5064 | 5404 | 5033 | 5212 | 5277 | 5201.09 | 5228 | 5174 | 0.99 | -0.015 | 0.78048446 | 0.87053897 | 5.00E-04 | 0.0029 | 0.0019 | 0.0019 | TRUE | 0.0103 |
| Grp | 6 | 4 | 7 | 6 | 2 | 2 | 5 | 4 | 8 | 6 | 2 | 2 | 4.45 | 6 | 3 | 0.627 | -0.675 | 0.46824816 | NA | 0 | 0.5356 | 0.3834 | 0.3834 | TRUE | 0.6527 |
| Grpel1 | 6771 | 4691 | 4218 | 5118 | 4488 | 5015 | 5259 | 4990 | 4551 | 5094 | 5063 | 4951 | 4984.61 | 4933 | 5036 | 1.02 | 0.029 | 0.65104593 | 0.78333581 | 0.0026 | 0.0029 | 0.0028 | 0.0028 | TRUE | 0.0557 |
| Grpel2 | 2621 | 2028 | 2324 | 2692 | 2468 | 2815 | 2036 | 2157 | 2508 | 2680 | 2784 | 2779 | 2490.49 | 2234 | 2748 | 1.231 | 0.3 | 0.00017202 | 0.0018997 | 0.0056 | 0.0034 | 0.0042 | 0.0042 | TRUE | 0.1389 |
| Grpr | 0 | 1 | 0 | 0 | 0 | 0 | 0 | 1 | 0 | 0 | 0 | 0 | 0.18 | 0 | 0 | 0.552 | -0.857 | 0.8336548 | NA | 0 | 13.3714 | 10 | 10 | TRUE | 0.5216 |
| Grrp1 | 43 | 54 | 46 | 47 | 42 | 55 | 33 | 57 | 50 | 47 | 47 | 54 | 48.15 | 47 | 49 | 1.068 | 0.095 | 0.75120356 | 0.85220887 | 0.0229 | 0.0516 | 0.0434 | 0.0434 | TRUE | 0.5922 |
| Grsf1 | 7512 | 5440 | 5978 | 6504 | 5560 | 6277 | 5834 | 5786 | 6450 | 6474 | 6272 | 6196 | 6168.94 | 6023 | 6314 | 1.048 | 0.068 | 0.2562077 | 0.42612273 | 0.0019 | 0.0028 | 0.0024 | 0.0024 | TRUE | 0.0465 |
| Grtp1 | 1829 | 1520 | 1553 | 1873 | 1791 | 1986 | 1421 | 1617 | 1676 | 1864 | 2020 | 1960 | 1759.71 | 1571 | 1948 | 1.242 | 0.312 | 6.76E-05 | 0.00089485 | 0.0036 | 0.0038 | 0.0039 | 0.0039 | TRUE | 0.0876 |
| Grwd1 | 6352 | 4429 | 3490 | 3763 | 3246 | 3767 | 4933 | 4711 | 3766 | 3746 | 3662 | 3719 | 4089.37 | 4470 | 3709 | 0.829 | -0.27 | 0.00231743 | 0.01354171 | 0.0093 | 0.003 | 0.0054 | 0.0054 | TRUE | 0.2305 |
| Grxcr1 | 0 | 0 | 0 | 0 | 0 | 0 | 0 | 0 | 0 | 0 | 0 | 0 | 0 | 0 | 0 | NA | NA | NA | NA | NA | NA | NA | NA | NA | NA |
| Grxcr2 | 1 | 0 | 0 | 0 | 1 | 0 | 1 | 0 | 0 | 0 | 1 | 0 | 0.32 | 0 | 0 | 1.075 | 0.105 | 0.97638847 | NA | 0 | 7.4681 | 7.0392 | 7.0392 | TRUE | 0.5075 |
| Gsc | 80 | 69 | 72 | 65 | 50 | 77 | 62 | 73 | 78 | 65 | 56 | 76 | 68.39 | 71 | 66 | 0.931 | -0.104 | 0.66963133 | 0.79697335 | 0 | 0.0371 | 0.0278 | 0.0278 | TRUE | 0.1678 |
| Gsc2 | 30 | 15 | 8 | 16 | 14 | 13 | 23 | 16 | 9 | 16 | 16 | 13 | 15.41 | 16 | 15 | 0.915 | -0.128 | 0.80539112 | NA | 0.0618 | 0.1563 | 0.1295 | 0.1295 | TRUE | 0.1842 |
| Gsdma | 32 | 21 | 20 | 20 | 17 | 14 | 25 | 22 | 22 | 20 | 19 | 14 | 20.28 | 23 | 18 | 0.766 | -0.385 | 0.3646656 | 0.53933831 | 0 | 0.1193 | 0.0799 | 0.0799 | TRUE | 0.1877 |
| Gsdma2 | 0 | 0 | 0 | 0 | 0 | 1 | 0 | 0 | 0 | 0 | 0 | 1 | 0.16 | 0 | 0 | 2.094 | 1.067 | 0.79379329 | NA | 0 | 14.4078 | 10 | 10 | TRUE | 0.3835 |
| Gsdma3 | 0 | 1 | 0 | 0 | 0 | 0 | 0 | 1 | 0 | 0 | 0 | 0 | 0.18 | 0 | 0 | 0.552 | -0.857 | 0.8336548 | NA | 0 | 13.3714 | 10 | 10 | TRUE | 0.5216 |
| Gsdmc | 2 | 3 | 1 | 2 | 2 | 0 | 2 | 3 | 1 | 2 | 2 | 0 | 1.68 | 2 | 1 | 0.725 | -0.464 | 0.75911796 | NA | 0 | 1.4145 | 1.0263 | 1.0263 | TRUE | 0.5234 |
| Gsdmc2 | 1 | 1 | 1 | 0 | 0 | 0 | 1 | 1 | 1 | 0 | 0 | 0 | 0.49 | 1 | 0 | 0.196 | -2.347 | 0.38405676 | NA | 0 | 4.8735 | 3.7415 | 3.7415 | TRUE | 0.0718 |
| Gsdmc3 | 0 | 0 | 0 | 1 | 0 | 0 | 0 | 0 | 0 | 1 | 0 | 0 | 0.17 | 0 | 0 | 2.094 | 1.067 | 0.79379329 | NA | 0 | 14.2885 | 10 | 10 | TRUE | 0.3906 |
| Gsdmc4 | 4 | 5 | 5 | 0 | 2 | 6 | 3 | 5 | 5 | 0 | 2 | 6 | 3.67 | 4 | 3 | 0.6 | -0.738 | 0.50299766 | NA | 0.3014 | 0.6488 | 0.5789 | 0.5789 | TRUE | 0.6126 |
| Gsdmcl-ps | 0 | 0 | 0 | 0 | 0 | 1 | 0 | 0 | 0 | 0 | 0 | 1 | 0.16 | 0 | 0 | 2.094 | 1.067 | 0.79379329 | NA | 0 | 14.4078 | 10 | 10 | TRUE | 0.3835 |
| Gsdmcl1 | 0 | 0 | 0 | 0 | 0 | 0 | 0 | 0 | 0 | 0 | 0 | 0 | 0 | 0 | 0 | NA | NA | NA | NA | NA | NA | NA | NA | NA | NA |
| Gsdmd | 156 | 104 | 127 | 119 | 96 | 98 | 121 | 111 | 137 | 118 | 108 | 97 | 115.38 | 123 | 108 | 0.878 | -0.188 | 0.31978584 | 0.49360512 | 0 | 0.023 | 0.0171 | 0.0171 | TRUE | 0.0953 |
| Gse1 | 6029 | 4977 | 4527 | 5532 | 5213 | 5666 | 4682 | 5294 | 4885 | 5507 | 5881 | 5593 | 5306.93 | 4954 | 5660 | 1.143 | 0.193 | 0.00218962 | 0.01293765 | 0.0023 | 0.0029 | 0.0027 | 0.0027 | TRUE | 0.0439 |
| Gsg1 | 57 | 49 | 38 | 61 | 41 | 39 | 44 | 52 | 41 | 61 | 46 | 38 | 47.14 | 46 | 48 | 1.06 | 0.085 | 0.77519558 | 0.86721285 | 0.0134 | 0.0527 | 0.0419 | 0.0419 | TRUE | 0.4028 |
| Gsg1l | 8 | 5 | 8 | 10 | 13 | 11 | 6 | 5 | 9 | 10 | 15 | 11 | 9.27 | 7 | 12 | 1.761 | 0.816 | 0.19741365 | NA | 0 | 0.258 | 0.1731 | 0.1731 | TRUE | 0.1556 |
| Gsg2 | 3043 | 2125 | 1660 | 1680 | 1524 | 1714 | 2363 | 2260 | 1791 | 1672 | 1719 | 1692 | 1916.39 | 2138 | 1694 | 0.792 | -0.337 | 0.00038148 | 0.00350865 | 0.0095 | 0.0037 | 0.0059 | 0.0059 | TRUE | 0.2422 |
| Gsk3a | 6057 | 4327 | 4305 | 4446 | 4045 | 4552 | 4704 | 4603 | 4645 | 4426 | 4563 | 4494 | 4572.35 | 4651 | 4494 | 0.966 | -0.05 | 0.32492938 | 0.49878806 | 0 | 0.0029 | 0.0016 | 0.0016 | TRUE | 0.0022 |
| Gsk3b | 4029 | 2784 | 2589 | 2704 | 2500 | 2737 | 3129 | 2961 | 2794 | 2692 | 2820 | 2702 | 2849.6 | 2961 | 2738 | 0.924 | -0.114 | 0.07460747 | 0.17970416 | 0.0014 | 0.0033 | 0.0026 | 0.0026 | TRUE | 0.0305 |
| Gskip | 451 | 291 | 284 | 349 | 305 | 336 | 350 | 310 | 306 | 347 | 344 | 332 | 331.56 | 322 | 341 | 1.056 | 0.078 | 0.50181539 | 0.6660891 | 0 | 0.0096 | 0.0068 | 0.0068 | TRUE | 0.0717 |
| Gsn | 690 | 453 | 539 | 652 | 621 | 644 | 536 | 482 | 582 | 649 | 701 | 636 | 597.43 | 533 | 662 | 1.241 | 0.311 | 0.00188376 | 0.01155564 | 0.0036 | 0.0064 | 0.0055 | 0.0055 | TRUE | 0.0787 |
| Gspt1 | 23059 | 16130 | 14398 | 16137 | 14170 | 16019 | 17909 | 17157 | 15535 | 16063 | 15985 | 15813 | 16410.46 | 16867 | 15954 | 0.946 | -0.081 | 0.18400479 | 0.34014232 | 0.0025 | 0.0026 | 0.0026 | 0.0026 | TRUE | 0.0583 |
| Gspt2 | 742 | 589 | 506 | 448 | 373 | 464 | 576 | 627 | 546 | 446 | 421 | 458 | 512.25 | 583 | 442 | 0.758 | -0.399 | 6.93E-05 | 0.00091049 | 0.0014 | 0.0071 | 0.0053 | 0.0053 | TRUE | 0.0481 |
| Gsr | 7275 | 5428 | 5545 | 5346 | 4727 | 5536 | 5650 | 5774 | 5983 | 5321 | 5333 | 5465 | 5587.62 | 5802 | 5373 | 0.926 | -0.111 | 0.03237488 | 0.09794411 | 4.00E-04 | 0.0029 | 0.0017 | 0.0017 | TRUE | 0.009 |
| Gss | 1682 | 1224 | 1189 | 1322 | 1100 | 1365 | 1306 | 1302 | 1283 | 1316 | 1241 | 1347 | 1299.25 | 1297 | 1301 | 1.004 | 0.005 | 0.93988781 | 0.9663921 | 0 | 0.0043 | 0.0027 | 0.0027 | TRUE | 0.0194 |
| Gsta1 | 0 | 0 | 0 | 0 | 1 | 0 | 0 | 0 | 0 | 0 | 1 | 0 | 0.19 | 0 | 0 | 2.094 | 1.067 | 0.79379329 | NA | 0 | 12.608 | 10 | 10 | TRUE | 0.5075 |
| Gsta2 | 0 | 1 | 3 | 1 | 1 | 3 | 0 | 1 | 3 | 1 | 1 | 3 | 1.56 | 1 | 2 | 1.238 | 0.309 | 0.84765071 | NA | 0 | 1.5175 | 1.2044 | 1.2044 | TRUE | 0.5799 |
| Gsta3 | 125 | 85 | 58 | 50 | 74 | 58 | 97 | 90 | 63 | 50 | 83 | 57 | 73.43 | 83 | 63 | 0.756 | -0.404 | 0.13278664 | 0.27086049 | 0.0473 | 0.0347 | 0.0382 | 0.0382 | TRUE | 0.6299 |
| Gsta4 | 13142 | 9675 | 9872 | 10672 | 9680 | 11759 | 10207 | 10291 | 10652 | 10623 | 10920 | 11608 | 10716.78 | 10383 | 11050 | 1.064 | 0.09 | 0.09571148 | 0.21438711 | 0.0012 | 0.0026 | 0.002 | 0.002 | TRUE | 0.0239 |
| Gstcd | 2275 | 1543 | 1477 | 1544 | 1418 | 1650 | 1767 | 1641 | 1594 | 1537 | 1600 | 1629 | 1627.86 | 1667 | 1589 | 0.952 | -0.071 | 0.30834041 | 0.48096797 | 0.0014 | 0.0039 | 0.0029 | 0.0029 | TRUE | 0.0341 |
| Gstk1 | 280 | 164 | 218 | 220 | 170 | 247 | 217 | 174 | 235 | 219 | 192 | 244 | 213.62 | 209 | 218 | 1.044 | 0.063 | 0.69350759 | 0.81291342 | 0.0134 | 0.0135 | 0.0135 | 0.0135 | TRUE | 0.2188 |
| Gstm1 | 1379 | 1189 | 1431 | 1446 | 1457 | 1636 | 1071 | 1265 | 1544 | 1439 | 1644 | 1615 | 1429.63 | 1293 | 1566 | 1.212 | 0.278 | 0.01457463 | 0.05459327 | 0.0183 | 0.0041 | 0.0086 | 0.0086 | TRUE | 0.3454 |
| Gstm2 | 252 | 212 | 266 | 183 | 171 | 207 | 196 | 226 | 287 | 182 | 193 | 204 | 214.61 | 236 | 193 | 0.821 | -0.284 | 0.08344508 | 0.19449859 | 0.0172 | 0.0135 | 0.0146 | 0.0146 | TRUE | 0.39 |
| Gstm3 | 109 | 84 | 152 | 159 | 155 | 182 | 85 | 89 | 164 | 158 | 175 | 180 | 141.8 | 113 | 171 | 1.526 | 0.61 | 0.00835102 | 0.03619195 | 0.0629 | 0.0191 | 0.0312 | 0.0312 | TRUE | 1.5521 |
| Gstm4 | 957 | 702 | 747 | 801 | 704 | 808 | 743 | 747 | 806 | 797 | 794 | 798 | 780.85 | 765 | 796 | 1.041 | 0.059 | 0.47343536 | 0.64189069 | 0 | 0.0055 | 0.0035 | 0.0035 | TRUE | 0.0249 |
| Gstm5 | 1574 | 1140 | 1216 | 1303 | 1138 | 1322 | 1222 | 1213 | 1312 | 1297 | 1284 | 1305 | 1272.15 | 1249 | 1295 | 1.038 | 0.053 | 0.44935791 | 0.61947074 | 0 | 0.0043 | 0.0028 | 0.0028 | TRUE | 0.0227 |
| Gstm6 | 143 | 103 | 121 | 101 | 90 | 101 | 111 | 110 | 131 | 101 | 102 | 100 | 108.82 | 117 | 101 | 0.861 | -0.216 | 0.25230301 | 0.42118886 | 0 | 0.0242 | 0.0164 | 0.0164 | TRUE | 0.0975 |
| Gstm7 | 172 | 112 | 147 | 184 | 113 | 189 | 134 | 119 | 159 | 183 | 127 | 187 | 151.42 | 137 | 166 | 1.213 | 0.278 | 0.14744532 | 0.29209092 | 0.0247 | 0.0181 | 0.0199 | 0.0199 | TRUE | 0.4162 |
| Gsto1 | 1165 | 815 | 868 | 903 | 866 | 922 | 905 | 867 | 937 | 899 | 977 | 910 | 915.7 | 903 | 929 | 1.028 | 0.04 | 0.61639427 | 0.75779489 | 5.00E-04 | 0.005 | 0.0035 | 0.0035 | TRUE | 0.0241 |
| Gsto2 | 31 | 20 | 13 | 24 | 21 | 19 | 24 | 21 | 14 | 24 | 24 | 19 | 20.95 | 20 | 22 | 1.106 | 0.146 | 0.73286895 | 0.84027723 | 0 | 0.1155 | 0.0836 | 0.0836 | TRUE | 0.3246 |
| Gstp1 | 2294 | 1626 | 1651 | 1641 | 1476 | 1727 | 1782 | 1730 | 1781 | 1633 | 1665 | 1705 | 1715.99 | 1764 | 1668 | 0.945 | -0.081 | 0.19593017 | 0.35547397 | 0 | 0.0038 | 0.0023 | 0.0023 | TRUE | 0.0046 |
| Gstp2 | 397 | 309 | 309 | 268 | 276 | 340 | 308 | 329 | 333 | 267 | 311 | 336 | 314.03 | 323 | 305 | 0.943 | -0.085 | 0.49962393 | 0.66410535 | 0.0043 | 0.01 | 0.0083 | 0.0083 | TRUE | 0.1363 |
| Gstt1 | 123 | 78 | 107 | 155 | 137 | 146 | 96 | 83 | 115 | 154 | 155 | 144 | 124.49 | 98 | 151 | 1.542 | 0.625 | 0.00078186 | 0.00606152 | 0.0016 | 0.0215 | 0.0166 | 0.0166 | TRUE | 0.2183 |
| Gstt2 | 245 | 188 | 216 | 331 | 241 | 308 | 190 | 200 | 233 | 329 | 272 | 304 | 254.79 | 208 | 302 | 1.458 | 0.544 | 0.00010153 | 0.00122517 | 0.0061 | 0.0117 | 0.0101 | 0.0101 | TRUE | 0.1207 |
| Gstt3 | 373 | 309 | 296 | 308 | 300 | 346 | 290 | 329 | 319 | 307 | 338 | 342 | 320.72 | 313 | 329 | 1.054 | 0.076 | 0.52434686 | 0.68568066 | 0 | 0.0098 | 0.0072 | 0.0072 | TRUE | 0.0506 |
| Gstt4 | 12 | 17 | 5 | 27 | 20 | 22 | 9 | 18 | 5 | 27 | 23 | 22 | 17.33 | 11 | 24 | 2.182 | 1.126 | 0.02705138 | 0.08591216 | 0.0488 | 0.1392 | 0.1208 | 0.1208 | TRUE | 1.0467 |
| Gstz1 | 1568 | 1077 | 1093 | 1357 | 1130 | 1270 | 1218 | 1146 | 1179 | 1351 | 1275 | 1254 | 1236.99 | 1181 | 1293 | 1.094 | 0.13 | 0.07132096 | 0.17389806 | 5.00E-04 | 0.0043 | 0.0029 | 0.0029 | TRUE | 0.0185 |
| Gsx1 | 24 | 16 | 12 | 13 | 9 | 15 | 19 | 17 | 13 | 13 | 10 | 15 | 14.42 | 16 | 13 | 0.778 | -0.362 | 0.47309886 | NA | 0 | 0.1668 | 0.1128 | 0.1128 | TRUE | 0.1377 |
| Gsx2 | 22 | 22 | 15 | 21 | 12 | 19 | 17 | 23 | 16 | 21 | 14 | 19 | 18.31 | 19 | 18 | 0.945 | -0.081 | 0.85798831 | 0.92052447 | 0 | 0.1319 | 0.0932 | 0.0932 | TRUE | 0.2136 |
| Gt(ROSA)26 | 307 | 253 | 271 | 319 | 252 | 302 | 238 | 269 | 292 | 318 | 284 | 298 | 283.31 | 266 | 300 | 1.129 | 0.176 | 0.17562727 | 0.3291865 | 0.0035 | 0.0108 | 0.0086 | 0.0086 | TRUE | 0.1045 |
| Gtdc1 | 1079 | 706 | 803 | 816 | 641 | 813 | 838 | 751 | 866 | 812 | 723 | 803 | 798.89 | 818 | 779 | 0.952 | -0.07 | 0.43984933 | 0.61125586 | 0.0034 | 0.0054 | 0.0047 | 0.0047 | TRUE | 0.06 |
| Gtdc2 | 467 | 338 | 310 | 377 | 351 | 343 | 363 | 360 | 334 | 375 | 396 | 339 | 361.09 | 352 | 370 | 1.048 | 0.068 | 0.55189925 | 0.70785139 | 0 | 0.009 | 0.0066 | 0.0066 | TRUE | 0.0636 |
| Gtf2a1 | 3768 | 2675 | 2579 | 2914 | 2295 | 2570 | 2926 | 2845 | 2783 | 2901 | 2589 | 2537 | 2763.52 | 2851 | 2676 | 0.938 | -0.092 | 0.17862951 | 0.33327261 | 0.0025 | 0.0033 | 0.003 | 0.003 | TRUE | 0.0661 |
| Gtf2a1l | 276 | 215 | 232 | 312 | 208 | 282 | 214 | 229 | 250 | 311 | 235 | 278 | 252.83 | 231 | 275 | 1.192 | 0.254 | 0.07797143 | 0.18558628 | 0.0089 | 0.0118 | 0.0109 | 0.0109 | TRUE | 0.1748 |
| Gtf2a2 | 2564 | 1750 | 1671 | 1878 | 1566 | 1725 | 1991 | 1861 | 1803 | 1869 | 1767 | 1703 | 1832.43 | 1885 | 1780 | 0.943 | -0.084 | 0.23169722 | 0.3976626 | 0.0019 | 0.0037 | 0.003 | 0.003 | TRUE | 0.0302 |
| Gtf2b | 2971 | 2027 | 1793 | 1691 | 1510 | 1655 | 2307 | 2156 | 1935 | 1683 | 1703 | 1634 | 1903.1 | 2133 | 1673 | 0.784 | -0.351 | 4.66E-06 | 0.00010026 | 0.0038 | 0.0037 | 0.0037 | 0.0037 | TRUE | 0.0794 |
| Gtf2e1 | 6899 | 4765 | 4580 | 4990 | 4278 | 4708 | 5358 | 5068 | 4942 | 4967 | 4826 | 4648 | 4968.17 | 5123 | 4814 | 0.939 | -0.09 | 0.11732274 | 0.24842873 | 0.0011 | 0.0029 | 0.0022 | 0.0022 | TRUE | 0.02 |

| | | | | | | | | | | | | | | | | | | | | | | | | | |
|---|---|---|---|---|---|---|---|---|---|---|---|---|---|---|---|---|---|---|---|---|---|---|---|---|---|
| Gtf2e2 | 8788 | 6647 | 6692 | 7121 | 6160 | 6806 | 6825 | 7070 | 7221 | 7088 | 6949 | 6719 | 6978.69 | 7039 | 6919 | 0.983 | -0.025 | 0.63791922 | 0.77387365 | 6.00E-04 | 0.0028 | 0.0018 | 0.0018 | TRUE | 0.0087 |
| Gtf2f1 | 5485 | 4069 | 3440 | 3410 | 3126 | 3578 | 4260 | 4328 | 3712 | 3394 | 3526 | 3532 | 3792.11 | 4100 | 3484 | 0.85 | -0.235 | 0.00072895 | 0.00574046 | 0.0032 | 0.0031 | 0.0032 | 0.0032 | TRUE | 0.0827 |
| Gtf2f2 | 3606 | 2647 | 2406 | 2740 | 2288 | 2740 | 2801 | 2816 | 2596 | 2727 | 2581 | 2705 | 2704.26 | 2738 | 2671 | 0.976 | -0.036 | 0.56710709 | 0.72102217 | 0.0011 | 0.0033 | 0.0024 | 0.0024 | TRUE | 0.0244 |
| Gtf2h1 | 4318 | 2860 | 2800 | 2598 | 2078 | 2584 | 3354 | 3042 | 3021 | 2586 | 2344 | 2551 | 2816.33 | 3139 | 2494 | 0.794 | -0.332 | 1.52E-06 | 4.10E-05 | 0.0026 | 0.0033 | 0.0031 | 0.0031 | TRUE | 0.0443 |
| Gtf2h2 | 2199 | 1598 | 1598 | 1707 | 1451 | 1817 | 1708 | 1700 | 1724 | 1699 | 1637 | 1794 | 1710.26 | 1711 | 1710 | 1 | 0 | 0.99970119 | 0.9998411 | 4.00E-04 | 0.0038 | 0.0025 | 0.0025 | TRUE | 0.0222 |
| Gtf2h3 | 3086 | 2219 | 2028 | 2207 | 1990 | 2243 | 2397 | 2360 | 2188 | 2197 | 2245 | 2214 | 2266.87 | 2315 | 2219 | 0.958 | -0.062 | 0.32299098 | 0.49687978 | 8.00E-04 | 0.0035 | 0.0024 | 0.0024 | TRUE | 0.0275 |
| Gtf2h4 | 2976 | 2143 | 2117 | 2306 | 2060 | 2302 | 2311 | 2279 | 2284 | 2295 | 2324 | 2272 | 2294.46 | 2291 | 2297 | 1.002 | 0.003 | 0.95644171 | 0.97583135 | 0 | 0.0035 | 0.0019 | 0.0019 | TRUE | 0.0012 |
| Gtf2h5 | 2718 | 1842 | 1738 | 2077 | 1669 | 1913 | 2111 | 1959 | 1875 | 2067 | 1883 | 1888 | 1964.04 | 1982 | 1946 | 0.981 | -0.027 | 0.70904968 | 0.82447503 | 0.0028 | 0.0036 | 0.0033 | 0.0033 | TRUE | 0.0404 |
| Gtf2i | 11370 | 7739 | 7412 | 7869 | 7029 | 8589 | 8831 | 8232 | 7997 | 7833 | 7929 | 8479 | 8216.81 | 8353 | 8080 | 0.967 | -0.048 | 0.42057974 | 0.59277648 | 0.002 | 0.0027 | 0.0025 | 0.0025 | TRUE | 0.0308 |
| Gtf2ird1 | 3895 | 2870 | 3067 | 2706 | 2481 | 2776 | 3025 | 3053 | 3309 | 2694 | 2799 | 2740 | 2936.64 | 3129 | 2744 | 0.877 | -0.189 | 0.00208668 | 0.01248428 | 0.0011 | 0.0032 | 0.0024 | 0.0024 | TRUE | 0.0305 |
| Gtf2ird2 | 161 | 120 | 68 | 64 | 55 | 69 | 125 | 128 | 73 | 64 | 62 | 68 | 86.65 | 109 | 65 | 0.593 | -0.754 | 0.00243799 | 0.01408179 | 0.0381 | 0.0298 | 0.0322 | 0.0322 | TRUE | 0.7676 |
| Gtf3a | 4664 | 3443 | 3641 | 3651 | 3151 | 3651 | 3622 | 3662 | 3929 | 3634 | 3555 | 3604 | 3667.7 | 3738 | 3598 | 0.963 | -0.054 | 0.34364938 | 0.51790179 | 6.00E-04 | 0.0031 | 0.0021 | 0.0021 | TRUE | 0.0241 |
| Gtf3c1 | 11042 | 7789 | 7628 | 8034 | 7003 | 7986 | 8576 | 8285 | 8231 | 7997 | 7900 | 7883 | 8145.34 | 8364 | 7927 | 0.948 | -0.078 | 0.10428327 | 0.2282574 | 2.00E-04 | 0.0027 | 0.0015 | 0.0015 | TRUE | 0.0061 |
| Gtf3c2 | 8685 | 6305 | 5376 | 5847 | 5120 | 5933 | 6745 | 6707 | 5801 | 5820 | 5776 | 5857 | 6117.54 | 6418 | 5818 | 0.906 | -0.142 | 0.03398811 | 0.10167268 | 0.0032 | 0.0028 | 0.0031 | 0.0031 | TRUE | 0.0858 |
| Gtf3c3 | 3769 | 2814 | 2586 | 2671 | 2301 | 2579 | 2927 | 2993 | 2790 | 2659 | 2596 | 2546 | 2751.84 | 2903 | 2600 | 0.895 | -0.159 | 0.00706079 | 0.0319468 | 5.00E-04 | 0.0033 | 0.0022 | 0.0022 | TRUE | 0.0138 |
| Gtf3c4 | 4982 | 3679 | 3854 | 4243 | 3687 | 4099 | 3869 | 3913 | 4158 | 4223 | 4159 | 4046 | 4061.7 | 3980 | 4143 | 1.041 | 0.058 | 0.302753 | 0.4753129 | 6.00E-04 | 0.003 | 0.0021 | 0.0021 | TRUE | 0.0185 |
| Gtf3c5 | 3142 | 2345 | 2029 | 2296 | 2043 | 2342 | 2440 | 2494 | 2189 | 2285 | 2305 | 2312 | 2337.66 | 2374 | 2301 | 0.968 | -0.046 | 0.49492354 | 0.65988266 | 0.0017 | 0.0034 | 0.0029 | 0.0029 | TRUE | 0.0555 |
| Gtf3c6 | 1470 | 931 | 785 | 787 | 720 | 711 | 1142 | 990 | 847 | 783 | 812 | 702 | 879.41 | 993 | 766 | 0.77 | -0.378 | 0.00065359 | 0.00528679 | 0.0121 | 0.0051 | 0.0077 | 0.0077 | TRUE | 0.1592 |
| Gtl3 | 4968 | 3788 | 4458 | 4424 | 3853 | 4549 | 3858 | 4029 | 4810 | 4404 | 4347 | 4491 | 4323.1 | 4232 | 4414 | 1.043 | 0.061 | 0.4443208 | 0.61519687 | 0.0059 | 0.003 | 0.0044 | 0.0044 | TRUE | 0.1729 |
| Gtpbp1 | 5301 | 4064 | 4213 | 4581 | 4165 | 4743 | 4117 | 4323 | 4546 | 4560 | 4699 | 4682 | 4487.7 | 4329 | 4647 | 1.074 | 0.103 | 0.07554235 | 0.18113183 | 0.001 | 0.003 | 0.0022 | 0.0022 | TRUE | 0.0235 |
| Gtpbp10 | 3350 | 2416 | 2476 | 2712 | 2372 | 2629 | 2602 | 2570 | 2672 | 2700 | 2676 | 2595 | 2635.64 | 2615 | 2657 | 1.016 | 0.023 | 0.68361293 | 0.80598552 | 0 | 0.0033 | 0.002 | 0.002 | TRUE | 0.0051 |
| Gtpbp2 | 988 | 805 | 856 | 857 | 814 | 901 | 767 | 856 | 924 | 853 | 918 | 889 | 868 | 849 | 887 | 1.046 | 0.065 | 0.47068731 | 0.63963211 | 0.004 | 0.0052 | 0.0048 | 0.0048 | TRUE | 0.0886 |
| Gtpbp3 | 3010 | 2297 | 2310 | 2526 | 2370 | 2724 | 2338 | 2443 | 2492 | 2514 | 2674 | 2689 | 2525.08 | 2424 | 2626 | 1.083 | 0.116 | 0.06159378 | 0.15634167 | 9.00E-04 | 0.0034 | 0.0024 | 0.0024 | TRUE | 0.0168 |
| Gtpbp4 | 15881 | 11068 | 10393 | 12536 | 10303 | 12011 | 12334 | 11773 | 11214 | 12478 | 11623 | 11857 | 11879.81 | 11774 | 11986 | 1.018 | 0.026 | 0.65207397 | 0.78392575 | 0.0015 | 0.0026 | 0.0022 | 0.0022 | TRUE | 0.0213 |
| Gtpbp5 | 744 | 535 | 431 | 441 | 400 | 421 | 578 | 569 | 465 | 439 | 451 | 416 | 486.29 | 537 | 435 | 0.808 | -0.307 | 0.00608074 | 0.02852379 | 0.0061 | 0.0073 | 0.0069 | 0.0069 | TRUE | 0.1577 |
| Gtpbp6 | 1008 | 857 | 706 | 770 | 721 | 741 | 783 | 912 | 762 | 766 | 813 | 731 | 794.59 | 819 | 770 | 0.941 | -0.087 | 0.35745058 | 0.53191475 | 0.0049 | 0.0054 | 0.0052 | 0.0052 | TRUE | 0.1144 |
| Gtpbp8 | 327 | 209 | 201 | 234 | 195 | 238 | 254 | 222 | 217 | 233 | 220 | 235 | 230.17 | 231 | 229 | 0.99 | -0.015 | 0.91270862 | 0.95207592 | 0 | 0.0127 | 0.0091 | 0.0091 | TRUE | 0.0901 |
| Gtse1 | 9937 | 7108 | 7225 | 7942 | 6754 | 7886 | 7718 | 7561 | 7796 | 7905 | 7619 | 7785 | 7730.57 | 7692 | 7770 | 1.01 | 0.015 | 0.7599618 | 0.85731129 | 1.00E-04 | 0.0027 | 0.0015 | 0.0015 | TRUE | 0.0035 |
| Gtsf1 | 713 | 551 | 494 | 356 | 307 | 319 | 554 | 586 | 533 | 354 | 346 | 315 | 448.08 | 558 | 338 | 0.607 | -0.72 | 6.59E-12 | 7.14E-10 | 0 | 0.0077 | 0.0055 | 0.0055 | TRUE | 0.0435 |
| Gtsf1l | 799 | 568 | 636 | 487 | 418 | 524 | 621 | 604 | 686 | 485 | 472 | 517 | 564.09 | 637 | 491 | 0.772 | -0.373 | 0.00012864 | 0.00150214 | 0.0017 | 0.0066 | 0.005 | 0.005 | TRUE | 0.052 |
| Guca1a | 459 | 395 | 368 | 467 | 428 | 485 | 356 | 420 | 397 | 465 | 483 | 479 | 433.36 | 391 | 476 | 1.218 | 0.285 | 0.00753745 | 0.03356162 | 0.0014 | 0.0079 | 0.0059 | 0.0059 | TRUE | 0.0753 |
| Guca1b | 16 | 2 | 3 | 7 | 5 | 8 | 12 | 2 | 3 | 7 | 6 | 8 | 6.38 | 6 | 7 | 1.115 | 0.158 | 0.8499131 | NA | 0.2802 | 0.3738 | 0.3459 | 0.3459 | TRUE | 2.686 |
| Guca2a | 1 | 2 | 1 | 2 | 0 | 0 | 1 | 2 | 1 | 2 | 0 | 0 | 1 | 1 | 1 | 0.515 | -0.958 | 0.63579447 | NA | 0 | 2.3829 | 1.8115 | 1.8115 | TRUE | 0.9736 |
| Guca2b | 1 | 0 | 0 | 0 | 0 | 0 | 1 | 0 | 0 | 0 | 0 | 0 | 0.13 | 0 | 0 | 0.552 | -0.857 | 0.8336548 | NA | 0 | 18.3121 | 10 | 10 | TRUE | 0.2566 |
| Gucd1 | 993 | 698 | 590 | 642 | 535 | 586 | 771 | 742 | 637 | 639 | 604 | 578 | 661.89 | 717 | 607 | 0.846 | -0.242 | 0.01536455 | 0.05687241 | 0.0049 | 0.006 | 0.0057 | 0.0057 | TRUE | 0.111 |
| Gucy1a2 | 130 | 93 | 66 | 77 | 81 | 98 | 101 | 99 | 71 | 77 | 91 | 97 | 89.31 | 90 | 88 | 0.972 | -0.04 | 0.85652423 | 0.91936774 | 0.0135 | 0.029 | 0.0247 | 0.0247 | TRUE | 0.3122 |
| Gucy1a3 | 8 | 7 | 10 | 6 | 4 | 2 | 6 | 7 | 11 | 6 | 5 | 2 | 6.15 | 8 | 4 | 0.512 | -0.965 | 0.22865874 | NA | 0 | 0.3877 | 0.2782 | 0.2782 | TRUE | 0.3861 |
| Gucy1b2 | 2 | 1 | 0 | 0 | 0 | 1 | 2 | 1 | 0 | 0 | 0 | 1 | 0.6 | 1 | 0 | 0.416 | -1.265 | 0.62184281 | NA | 0 | 3.948 | 3.1959 | 3.1959 | TRUE | 0.3835 |
| Gucy1b3 | 156 | 109 | 102 | 97 | 104 | 90 | 121 | 116 | 110 | 97 | 117 | 89 | 108.31 | 116 | 101 | 0.868 | -0.204 | 0.29524546 | 0.46705464 | 0 | 0.0243 | 0.018 | 0.018 | TRUE | 0.1888 |
| Gucy2c | 17 | 9 | 11 | 15 | 11 | 4 | 13 | 10 | 12 | 15 | 12 | 4 | 10.99 | 12 | 10 | 0.896 | -0.158 | 0.7984668 | NA | 0.0894 | 0.2181 | 0.1857 | 0.1857 | TRUE | 1.1044 |
| Gucy2d | 0 | 2 | 0 | 0 | 1 | 0 | 0 | 2 | 0 | 0 | 1 | 0 | 0.54 | 1 | 0 | 0.535 | -0.903 | 0.76014553 | NA | 0 | 4.3706 | 4.6038 | 4.6038 | TRUE | 0.9746 |
| Gucy2e | 37 | 35 | 37 | 32 | 31 | 40 | 29 | 37 | 40 | 32 | 35 | 39 | 35.37 | 35 | 35 | 1.012 | 0.017 | 0.95981392 | 0.97773647 | 0 | 0.0694 | 0.049 | 0.049 | TRUE | 0.2272 |
| Gucy2f | 0 | 0 | 0 | 0 | 1 | 0 | 0 | 0 | 0 | 0 | 1 | 0 | 0.19 | 0 | 0 | 2.094 | 1.067 | 0.79379329 | NA | 0 | 12.608 | 10 | 10 | TRUE | 0.5075 |
| Gucy2g | 34 | 34 | 29 | 35 | 32 | 30 | 26 | 36 | 31 | 35 | 36 | 30 | 32.4 | 31 | 34 | 1.076 | 0.106 | 0.75444482 | 0.85445957 | 0 | 0.0756 | 0.052 | 0.052 | TRUE | 0.1537 |
| Guf1 | 2571 | 1897 | 1888 | 2148 | 1781 | 2030 | 1997 | 2018 | 2037 | 2138 | 2009 | 2004 | 2033.82 | 2017 | 2050 | 1.017 | 0.024 | 0.69874352 | 0.8170087 | 0 | 0.0036 | 0.0023 | 0.0023 | TRUE | 0.0171 |
| Guk1 | 5253 | 3572 | 3170 | 3691 | 3031 | 3520 | 4080 | 3799 | 3420 | 3674 | 3419 | 3475 | 3644.63 | 3766 | 3523 | 0.935 | -0.097 | 0.18285426 | 0.33854092 | 0.0042 | 0.0031 | 0.0036 | 0.0036 | TRUE | 0.0784 |
| Gulo | 295 | 253 | 251 | 243 | 241 | 249 | 229 | 269 | 271 | 242 | 272 | 246 | 254.77 | 256 | 253 | 0.99 | -0.015 | 0.91334087 | 0.95246365 | 0.0031 | 0.0117 | 0.0091 | 0.0091 | TRUE | 0.1051 |
| Gulp1 | 282 | 200 | 144 | 197 | 170 | 171 | 219 | 213 | 155 | 196 | 192 | 169 | 190.63 | 196 | 186 | 0.945 | -0.082 | 0.6231914 | 0.76236381 | 0.015 | 0.0149 | 0.0149 | 0.0149 | TRUE | 0.3395 |
| Gusb | 1851 | 1354 | 1310 | 1613 | 1450 | 1506 | 1438 | 1440 | 1413 | 1606 | 1636 | 1487 | 1503.21 | 1430 | 1576 | 1.101 | 0.139 | 0.0436921 | 0.12204243 | 7.00E-04 | 0.004 | 0.0028 | 0.0028 | TRUE | 0.0299 |
| Gvin1 | 0 | 0 | 0 | 0 | 0 | 0 | 0 | 0 | 0 | 0 | 0 | 0 | 0 | 0 | 0 | NA | NA | NA | NA | NA | NA | NA | NA | NA | NA |
| Gxylt1 | 3430 | 2497 | 2457 | 2440 | 2203 | 2397 | 2664 | 2656 | 2651 | 2429 | 2485 | 2366 | 2541.87 | 2657 | 2427 | 0.913 | -0.131 | 0.02135887 | 0.07251363 | 0 | 0.0034 | 0.0019 | 0.0019 | TRUE | 0.0058 |
| Gxylt2 | 41 | 34 | 23 | 47 | 26 | 38 | 32 | 36 | 25 | 47 | 29 | 38 | 34.41 | 31 | 38 | 1.226 | 0.294 | 0.39144793 | 0.56513379 | 0 | 0.0713 | 0.0557 | 0.0557 | TRUE | 0.1832 |
| Gyg | 1516 | 1143 | 1255 | 1530 | 1280 | 1366 | 1177 | 1216 | 1354 | 1523 | 1444 | 1348 | 1343.79 | 1249 | 1438 | 1.153 | 0.205 | 0.01212493 | 0.04772009 | 0.0035 | 0.0042 | 0.0041 | 0.0041 | TRUE | 0.0642 |
| Gyk | 1865 | 1410 | 1684 | 1743 | 1557 | 1775 | 1448 | 1500 | 1817 | 1735 | 1756 | 1752 | 1668.15 | 1588 | 1748 | 1.102 | 0.139 | 0.11146952 | 0.2393036 | 0.007 | 0.0038 | 0.0049 | 0.0049 | TRUE | 0.1895 |
| Gykl1 | 0 | 0 | 0 | 0 | 0 | 0 | 0 | 0 | 0 | 0 | 0 | 0 | 0 | 0 | 0 | NA | NA | NA | NA | NA | NA | NA | NA | NA | NA |
| Gyltl1b | 1023 | 842 | 937 | 885 | 844 | 949 | 795 | 896 | 1011 | 881 | 952 | 937 | 911.84 | 901 | 923 | 1.027 | 0.039 | 0.69038447 | 0.81055242 | 0.0071 | 0.005 | 0.0057 | 0.0057 | TRUE | 0.1375 |
| Gypa | 2 | 1 | 0 | 0 | 3 | 0 | 2 | 1 | 0 | 0 | 3 | 0 | 1 | 1 | 1 | 1.229 | 0.297 | 0.89095244 | NA | 2.3937 | 2.372 | 2.3763 | 2.3763 | TRUE | 1.4437 |
| Gypc | 125 | 76 | 52 | 81 | 43 | 57 | 97 | 81 | 56 | 81 | 49 | 56 | 69.91 | 78 | 62 | 0.79 | -0.341 | 0.22477312 | 0.3892273 | 0.0574 | 0.0363 | 0.0423 | 0.0423 | TRUE | 0.3113 |
| Gys1 | 2107 | 1489 | 1500 | 1653 | 1461 | 1691 | 1636 | 1584 | 1618 | 1645 | 1648 | 1669 | 1633.59 | 1613 | 1654 | 1.025 | 0.036 | 0.56474315 | 0.71920734 | 0 | 0.0039 | 0.0022 | 0.0022 | TRUE | 0.003 |
| Gys2 | 1 | 1 | 1 | 0 | 0 | 0 | 1 | 1 | 1 | 0 | 0 | 0 | 0.49 | 1 | 0 | 0.196 | -2.347 | 0.38405676 | NA | 0 | 4.8735 | 3.7415 | 3.7415 | TRUE | 0.0718 |
| Gzf1 | 3350 | 2466 | 2847 | 3015 | 2724 | 2972 | 2602 | 2623 | 3072 | 3001 | 3073 | 2934 | 2884.11 | 2766 | 3003 | 1.086 | 0.119 | 0.10944329 | 0.23635187 | 0.0044 | 0.0032 | 0.0037 | 0.0037 | TRUE | 0.113 |
| Gzma | 0 | 0 | 0 | 0 | 0 | 0 | 0 | 0 | 0 | 0 | 0 | 0 | 0 | 0 | 0 | NA | NA | NA | NA | NA | NA | NA | NA | NA | NA |
| Gzmb | 2 | 0 | 0 | 0 | 0 | 0 | 2 | 0 | 0 | 0 | 0 | 0 | 0.26 | 1 | 0 | 0.356 | -1.491 | 0.71362796 | NA | 0 | 9.1573 | 10 | 10 | TRUE | 0.968 |
| Gzmc | 0 | 1 | 0 | 0 | 0 | 0 | 0 | 1 | 0 | 0 | 0 | 0 | 0.18 | 0 | 0 | 0.552 | -0.857 | 0.8336548 | NA | 0 | 13.3714 | 10 | 10 | TRUE | 0.5216 |
| Gzmd | 0 | 0 | 0 | 0 | 0 | 0 | 0 | 0 | 0 | 0 | 0 | 0 | 0 | 0 | 0 | NA | NA | NA | NA | NA | NA | NA | NA | NA | NA |
| Gzme | 0 | 0 | 0 | 0 | 0 | 0 | 0 | 0 | 0 | 0 | 0 | 0 | 0 | 0 | 0 | NA | NA | NA | NA | NA | NA | NA | NA | NA | NA |
| Gzmf | 0 | 0 | 0 | 0 | 0 | 0 | 0 | 0 | 0 | 0 | 0 | 0 | 0 | 0 | 0 | NA | NA | NA | NA | NA | NA | NA | NA | NA | NA |
| Gzmg | 0 | 0 | 0 | 0 | 0 | 0 | 0 | 0 | 0 | 0 | 0 | 0 | 0 | 0 | 0 | NA | NA | NA | NA | NA | NA | NA | NA | NA | NA |
| Gzmk | 0 | 0 | 0 | 0 | 0 | 0 | 0 | 0 | 0 | 0 | 0 | 0 | 0 | 0 | 0 | NA | NA | NA | NA | NA | NA | NA | NA | NA | NA |

| | | | | | | | | | | | | | | | | | | | | | | | | | |
|---|---|---|---|---|---|---|---|---|---|---|---|---|---|---|---|---|---|---|---|---|---|---|---|---|---|
| Gzmm | 29 | 35 | 49 | 46 | 34 | 56 | 23 | 37 | 53 | 46 | 38 | 55 | 42.01 | 38 | 46 | 1.256 | 0.329 | 0.34836974 | 0.52303033 | 0.0787 | 0.0588 | 0.0644 | 0.0644 | TRUE | 0.2635 |
| Gzmn | 1 | 1 | 0 | 1 | 1 | 0 | 1 | 1 | 0 | 1 | 1 | 0 | 0.66 | 1 | 1 | 1.128 | 0.173 | 0.94106052 | NA | 0 | 3.5899 | 2.4471 | 2.4471 | TRUE | 0.2681 |
| H13 | 1780 | 1325 | 1168 | 1094 | 980 | 1180 | 1382 | 1409 | 1260 | 1089 | 1106 | 1165 | 1235.24 | 1350 | 1120 | 0.829 | -0.271 | 0.00037477 | 0.00346252 | 0.0016 | 0.0043 | 0.0034 | 0.0034 | TRUE | 0.0402 |
| H19 | 4559 | 3491 | 2728 | 1314 | 1263 | 1363 | 3541 | 3713 | 2943 | 1308 | 1425 | 1345 | 2379.31 | 3399 | 1359 | 0.4 | -1.323 | 5.34E-51 | 2.54E-47 | 0.0073 | 0.0034 | 0.0051 | 0.0051 | TRUE | 0.1658 |
| H1f0 | 2419 | 1556 | 1036 | 1248 | 1119 | 1288 | 1879 | 1655 | 1118 | 1242 | 1262 | 1271 | 1404.62 | 1551 | 1258 | 0.811 | -0.301 | 0.17053841 | 0.32218543 | 0.0341 | 0.0041 | 0.0128 | 0.0341 | TRUE | 0.5639 |
| H1fnt | 1 | 0 | 0 | 0 | 0 | 0 | 1 | 0 | 0 | 0 | 0 | 0 | 0.13 | 0 | 0 | 0.552 | -0.857 | 0.8336548 | NA | 0 | 18.3121 | 10 | 10 | TRUE | 0.2566 |
| H1foo | 2 | 1 | 3 | 0 | 0 | 0 | 2 | 1 | 3 | 0 | 0 | 0 | 0.98 | 2 | 0 | 0.098 | -3.351 | 0.11532532 | NA | 0 | 2.4316 | 2.0136 | 2.0136 | TRUE | 0.2678 |
| H1fx | 4589 | 3668 | 2880 | 2972 | 2684 | 3171 | 3564 | 3902 | 3107 | 2958 | 3028 | 3130 | 3281.6 | 3524 | 3039 | 0.862 | -0.214 | 0.00801276 | 0.03506623 | 0.0063 | 0.0031 | 0.0044 | 0.0044 | TRUE | 0.1287 |
| H2-Aa | 8 | 8 | 8 | 14 | 9 | 11 | 6 | 9 | 9 | 14 | 10 | 11 | 9.72 | 8 | 12 | 1.513 | 0.598 | 0.32722254 | NA | 0 | 0.2463 | 0.1614 | 0.1614 | TRUE | 0.1209 |
| H2-Ab1 | 245 | 209 | 206 | 325 | 278 | 295 | 190 | 222 | 222 | 324 | 314 | 291 | 260.53 | 211 | 310 | 1.467 | 0.553 | 2.79E-05 | 0.0004368 | 0 | 0.0115 | 0.0086 | 0.0086 | TRUE | 0.0944 |
| H2-Bl | 231 | 163 | 153 | 125 | 119 | 140 | 179 | 173 | 165 | 124 | 134 | 138 | 152.46 | 172 | 132 | 0.765 | -0.386 | 0.01591881 | 0.05837035 | 0 | 0.018 | 0.0118 | 0.0118 | TRUE | 0.0286 |
| H2-D1 | 576 | 435 | 380 | 601 | 588 | 591 | 447 | 463 | 410 | 598 | 663 | 583 | 527.51 | 440 | 615 | 1.396 | 0.481 | 2.00E-06 | 5.07E-05 | 0.0022 | 0.0069 | 0.0054 | 0.0054 | TRUE | 0.0543 |
| H2-DMa | 262 | 174 | 202 | 243 | 238 | 246 | 203 | 185 | 218 | 242 | 268 | 243 | 226.62 | 202 | 251 | 1.241 | 0.311 | 0.02408425 | 0.07917117 | 0 | 0.0129 | 0.0092 | 0.0092 | TRUE | 0.0555 |
| H2-DMb1 | 82 | 70 | 89 | 113 | 105 | 155 | 64 | 74 | 96 | 112 | 118 | 153 | 103.02 | 78 | 128 | 1.652 | 0.724 | 0.00115016 | 0.00802303 | 0.0256 | 0.0254 | 0.0255 | 0.0255 | TRUE | 0.367 |
| H2-DMb2 | 0 | 1 | 3 | 3 | 2 | 1 | 0 | 1 | 3 | 3 | 2 | 1 | 1.76 | 1 | 2 | 1.512 | 0.596 | 0.69560704 | NA | 0 | 1.3529 | 1.0717 | 1.0717 | TRUE | 0.6562 |
| H2-Ea-ps | 0 | 0 | 0 | 0 | 0 | 0 | 0 | 0 | 0 | 0 | 0 | 0 | 0 | 0 | 0 | NA | NA | NA | NA | NA | NA | NA | NA | NA | NA |
| H2-Eb1 | 127 | 85 | 73 | 83 | 72 | 66 | 99 | 90 | 79 | 83 | 81 | 65 | 82.8 | 89 | 76 | 0.851 | -0.233 | 0.29395687 | 0.46590435 | 0 | 0.031 | 0.0233 | 0.0233 | TRUE | 0.1552 |
| H2-Eb2 | 6 | 4 | 3 | 7 | 3 | 8 | 5 | 4 | 3 | 7 | 3 | 8 | 5.07 | 4 | 6 | 1.501 | 0.586 | 0.48944956 | NA | 0 | 0.4702 | 0.3157 | 0.3157 | TRUE | 0.3151 |
| H2-K1 | 3067 | 2549 | 2375 | 3503 | 2998 | 3480 | 2382 | 2711 | 2563 | 3487 | 3382 | 3435 | 2993.36 | 2552 | 3435 | 1.347 | 0.43 | 3.22E-11 | 2.94E-09 | 0.0017 | 0.0032 | 0.0027 | 0.0027 | TRUE | 0.0424 |
| H2-K2 | 56 | 55 | 59 | 64 | 59 | 70 | 43 | 59 | 64 | 64 | 67 | 69 | 60.84 | 55 | 67 | 1.213 | 0.279 | 0.27827723 | 0.44973049 | 0 | 0.0414 | 0.0311 | 0.0311 | TRUE | 0.3411 |
| H2-Ke2 | 4933 | 3557 | 3165 | 2974 | 2683 | 3237 | 3831 | 3784 | 3415 | 2960 | 3027 | 3195 | 3368.71 | 3677 | 3061 | 0.832 | -0.265 | 6.45E-05 | 0.00086232 | 0.0023 | 0.0031 | 0.0029 | 0.0029 | TRUE | 0.0467 |
| H2-Ke6 | 612 | 606 | 643 | 748 | 767 | 914 | 475 | 645 | 694 | 745 | 865 | 902 | 720.96 | 605 | 837 | 1.389 | 0.474 | 0.00029255 | 0.00285425 | 0.0228 | 0.0057 | 0.0109 | 0.0109 | TRUE | 0.4284 |
| H2-L | 0 | 0 | 0 | 0 | 0 | 0 | 0 | 0 | 0 | 0 | 0 | 0 | 0 | 0 | 0 | NA | NA | NA | NA | NA | NA | NA | NA | NA | NA |
| H2-M1 | 0 | 0 | 0 | 0 | 0 | 0 | 0 | 0 | 0 | 0 | 0 | 0 | 0 | 0 | 0 | NA | NA | NA | NA | NA | NA | NA | NA | NA | NA |
| H2-M10.1 | 0 | 1 | 0 | 0 | 0 | 0 | 0 | 1 | 0 | 0 | 0 | 0 | 0.18 | 0 | 0 | 0.552 | -0.857 | 0.8336548 | NA | 0 | 13.3714 | 10 | 10 | TRUE | 0.5216 |
| H2-M10.2 | 0 | 0 | 0 | 0 | 0 | 0 | 0 | 0 | 0 | 0 | 0 | 0 | 0 | 0 | 0 | NA | NA | NA | NA | NA | NA | NA | NA | NA | NA |
| H2-M10.3 | 0 | 0 | 0 | 0 | 0 | 0 | 0 | 0 | 0 | 0 | 0 | 0 | 0 | 0 | 0 | NA | NA | NA | NA | NA | NA | NA | NA | NA | NA |
| H2-M10.4 | 0 | 1 | 0 | 0 | 0 | 1 | 0 | 1 | 0 | 0 | 0 | 1 | 0.34 | 0 | 0 | 1.075 | 0.105 | 0.97576789 | NA | 0 | 6.9363 | 6.5821 | 6.5821 | TRUE | 0.5216 |
| H2-M10.5 | 0 | 0 | 0 | 0 | 0 | 0 | 0 | 0 | 0 | 0 | 0 | 0 | 0 | 0 | 0 | NA | NA | NA | NA | NA | NA | NA | NA | NA | NA |
| H2-M10.6 | 0 | 1 | 0 | 0 | 0 | 0 | 0 | 1 | 0 | 0 | 0 | 0 | 0.18 | 0 | 0 | 0.552 | -0.857 | 0.8336548 | NA | 0 | 13.3714 | 10 | 10 | TRUE | 0.5216 |
| H2-M11 | 0 | 0 | 0 | 0 | 0 | 0 | 0 | 0 | 0 | 0 | 0 | 0 | 0 | 0 | 0 | NA | NA | NA | NA | NA | NA | NA | NA | NA | NA |
| H2-M2 | 1 | 1 | 0 | 0 | 0 | 0 | 1 | 1 | 0 | 0 | 0 | 0 | 0.31 | 1 | 0 | 0.309 | -1.696 | 0.63183309 | NA | 0 | 7.7294 | 7.2472 | 7.2472 | TRUE | 0.2015 |
| H2-M3 | 202 | 141 | 134 | 193 | 165 | 138 | 157 | 150 | 145 | 192 | 186 | 136 | 160.99 | 151 | 171 | 1.137 | 0.185 | 0.28913471 | 0.46117647 | 0.0122 | 0.0171 | 0.0157 | 0.0157 | TRUE | 0.3526 |
| H2-M5 | 394 | 261 | 190 | 247 | 179 | 200 | 306 | 278 | 205 | 246 | 202 | 197 | 238.98 | 263 | 215 | 0.816 | -0.294 | 0.07958733 | 0.18845952 | 0.0245 | 0.0123 | 0.016 | 0.016 | TRUE | 0.4046 |
| H2-M9 | 0 | 0 | 0 | 0 | 0 | 0 | 0 | 0 | 0 | 0 | 0 | 0 | 0 | 0 | 0 | NA | NA | NA | NA | NA | NA | NA | NA | NA | NA |
| H2-Oa | 41 | 42 | 42 | 77 | 59 | 75 | 32 | 45 | 45 | 77 | 67 | 74 | 56.51 | 41 | 73 | 1.802 | 0.85 | 0.00156537 | 0.0099616 | 0 | 0.0444 | 0.0329 | 0.0329 | TRUE | 0.3273 |
| H2-Ob | 1 | 0 | 1 | 0 | 0 | 0 | 1 | 0 | 1 | 0 | 0 | 0 | 0.31 | 1 | 0 | 0.307 | -1.706 | 0.62894649 | NA | 0 | 7.6657 | 7.2 | 7.2 | TRUE | 0.2067 |
| H2-Q1 | 5 | 8 | 3 | 1 | 5 | 6 | 4 | 9 | 3 | 1 | 6 | 6 | 4.7 | 5 | 4 | 0.805 | -0.312 | 0.74016426 | NA | 0.1785 | 0.5069 | 0.4203 | 0.4203 | TRUE | 0.8015 |
| H2-Q10 | 336 | 293 | 270 | 325 | 311 | 374 | 261 | 312 | 291 | 324 | 351 | 369 | 317.91 | 288 | 348 | 1.211 | 0.277 | 0.02545079 | 0.08237502 | 0.0033 | 0.0099 | 0.0079 | 0.0079 | TRUE | 0.0827 |
| H2-Q2 | 63 | 52 | 59 | 118 | 116 | 131 | 49 | 55 | 64 | 117 | 131 | 129 | 90.92 | 56 | 126 | 2.262 | 1.178 | 3.75E-08 | 1.64E-06 | 0 | 0.0285 | 0.0202 | 0.0202 | TRUE | 0.1219 |
| H2-Q4 | 332 | 288 | 249 | 324 | 316 | 326 | 258 | 306 | 269 | 323 | 356 | 322 | 305.61 | 278 | 334 | 1.203 | 0.267 | 0.0320001 | 0.09702533 | 0.0024 | 0.0102 | 0.0079 | 0.0079 | TRUE | 0.0903 |
| H2-Q5 | 4 | 3 | 0 | 3 | 6 | 7 | 3 | 3 | 0 | 3 | 7 | 7 | 3.83 | 2 | 6 | 2.574 | 1.364 | 0.20123931 | NA | 0 | 0.6217 | 0.5036 | 0.5036 | TRUE | 0.6222 |
| H2-Q6 | 0 | 0 | 0 | 0 | 0 | 0 | 0 | 0 | 0 | 0 | 0 | 0 | 0 | 0 | 0 | NA | NA | NA | NA | NA | NA | NA | NA | NA | NA |
| H2-Q7 | 0 | 0 | 0 | 0 | 0 | 0 | 0 | 0 | 0 | 0 | 0 | 0 | 0 | 0 | 0 | NA | NA | NA | NA | NA | NA | NA | NA | NA | NA |
| H2-Q8 | 0 | 0 | 0 | 0 | 0 | 0 | 0 | 0 | 0 | 0 | 0 | 0 | 0 | 0 | 0 | NA | NA | NA | NA | NA | NA | NA | NA | NA | NA |
| H2-Q9 | 0 | 0 | 0 | 0 | 0 | 0 | 0 | 0 | 0 | 0 | 0 | 0 | 0 | 0 | 0 | NA | NA | NA | NA | NA | NA | NA | NA | NA | NA |
| H2-T10 | 7 | 3 | 11 | 5 | 4 | 1 | 5 | 3 | 12 | 5 | 5 | 1 | 5.16 | 7 | 4 | 0.511 | -0.968 | 0.29692562 | NA | 0.2356 | 0.4615 | 0.3999 | 0.3999 | TRUE | 0.5638 |
| H2-T22 | 4 | 1 | 2 | 7 | 2 | 2 | 3 | 1 | 2 | 7 | 2 | 2 | 2.92 | 2 | 4 | 1.742 | 0.8 | 0.48749095 | NA | 0 | 0.8138 | 0.596 | 0.596 | TRUE | 0.926 |
| H2-T23 | 8 | 4 | 6 | 7 | 5 | 7 | 6 | 4 | 6 | 7 | 6 | 7 | 6.08 | 5 | 7 | 1.15 | 0.202 | 0.78986225 | NA | 0 | 0.3924 | 0.2489 | 0.2489 | TRUE | 0.0947 |
| H2-T24 | 0 | 1 | 1 | 0 | 0 | 0 | 0 | 1 | 1 | 0 | 0 | 0 | 0.36 | 1 | 0 | 0.271 | -1.881 | 0.5755355 | NA | 0 | 6.6391 | 6.4428 | 6.4428 | TRUE | 0.3513 |
| H2-T3 | 5 | 0 | 0 | 1 | 2 | 1 | 4 | 0 | 0 | 1 | 2 | 1 | 1.35 | 1 | 1 | 1.018 | 0.026 | 0.98864103 | NA | 1.145 | 1.7533 | 1.6084 | 1.6084 | TRUE | 2.3344 |
| H2-T9 | 0 | 0 | 0 | 0 | 0 | 0 | 0 | 0 | 0 | 0 | 0 | 0 | 0 | 0 | 0 | NA | NA | NA | NA | NA | NA | NA | NA | NA | NA |
| H2afb1 | 0 | 0 | 0 | 0 | 0 | 0 | 0 | 0 | 0 | 0 | 0 | 0 | 0 | 0 | 0 | NA | NA | NA | NA | NA | NA | NA | NA | NA | NA |
| H2afj | 2480 | 1951 | 1925 | 2215 | 1778 | 2178 | 1926 | 2075 | 2077 | 2205 | 2006 | 2150 | 2073.17 | 2026 | 2120 | 1.048 | 0.067 | 0.32339403 | 0.49739276 | 0.0017 | 0.0036 | 0.0028 | 0.0028 | TRUE | 0.0266 |
| H2afv | 3871 | 2840 | 3128 | 3370 | 2944 | 3386 | 3006 | 3021 | 3375 | 3354 | 3321 | 3343 | 3236.76 | 3134 | 3339 | 1.066 | 0.092 | 0.14948107 | 0.29501973 | 0.0018 | 0.0032 | 0.0026 | 0.0026 | TRUE | 0.0544 |
| H2afx | 4737 | 3545 | 4022 | 4272 | 3905 | 4282 | 3679 | 3771 | 4340 | 4252 | 4405 | 4227 | 4112.35 | 3930 | 4295 | 1.093 | 0.129 | 0.06993359 | 0.17145226 | 0.004 | 0.003 | 0.0034 | 0.0034 | TRUE | 0.1008 |
| H2afy | 9454 | 7662 | 9301 | 7054 | 7157 | 8022 | 7343 | 8150 | 10036 | 7022 | 8074 | 7919 | 8090.42 | 8510 | 7672 | 0.902 | -0.149 | 0.12912782 | 0.26582514 | 0.0157 | 0.0027 | 0.0068 | 0.0068 | TRUE | 0.3005 |
| H2afy2 | 2404 | 1814 | 1680 | 1606 | 1431 | 1696 | 1867 | 1930 | 1813 | 1599 | 1614 | 1674 | 1749.41 | 1870 | 1629 | 0.871 | -0.199 | 0.00203674 | 0.0122573 | 0 | 0.0038 | 0.0024 | 0.0024 | TRUE | 0.0091 |
| H2afy3 | 0 | 0 | 1 | 0 | 1 | 0 | 0 | 0 | 1 | 0 | 1 | 0 | 0.37 | 0 | 0 | 1.075 | 0.105 | 0.97514588 | NA | 0 | 6.4454 | 6.1578 | 6.1578 | TRUE | 0.5365 |
| H2afz | 23762 | 16157 | 14329 | 13426 | 11979 | 14079 | 18455 | 17186 | 15461 | 13364 | 13514 | 13898 | 15312.95 | 17034 | 13592 | 0.798 | -0.326 | 1.21E-06 | 3.41E-05 | 0.004 | 0.0026 | 0.0032 | 0.0032 | TRUE | 0.0798 |
| H2bfm | 49 | 23 | 15 | 6 | 4 | 7 | 38 | 24 | 16 | 6 | 5 | 7 | 16.02 | 26 | 6 | 0.22 | -2.185 | 0.00015019 | NA | 0 | 0.1504 | 0.1323 | 0.1323 | TRUE | 0.1521 |
| H3f3a | 8876 | 6407 | 6592 | 6718 | 5734 | 6822 | 6894 | 6815 | 7113 | 6687 | 6469 | 6734 | 6785.23 | 6941 | 6630 | 0.955 | -0.066 | 0.18944854 | 0.3471523 | 3.00E-04 | 0.0028 | 0.0017 | 0.0017 | TRUE | 0.0057 |
| H3f3b | 41503 | 28790 | 24976 | 26050 | 22686 | 24465 | 32234 | 30624 | 26949 | 25930 | 25592 | 24151 | 27579.85 | 29936 | 25224 | 0.843 | -0.247 | 0.00036605 | 0.00339407 | 0.0047 | 0.0025 | 0.0034 | 0.0034 | TRUE | 0.0932 |
| H60b | 80 | 56 | 67 | 92 | 68 | 84 | 62 | 60 | 72 | 92 | 77 | 83 | 74.2 | 65 | 84 | 1.299 | 0.377 | 0.09790169 | 0.2180288 | 0 | 0.0344 | 0.0238 | 0.0238 | TRUE | 0.0872 |
| H60c | 3 | 2 | 2 | 7 | 6 | 6 | 2 | 2 | 2 | 7 | 7 | 6 | 4.38 | 2 | 7 | 2.96 | 1.566 | 0.0958689 | NA | 0 | 0.5436 | 0.3408 | 0.3408 | TRUE | 0.0186 |
| H6pd | 2533 | 1860 | 1728 | 2062 | 1950 | 2073 | 1967 | 1978 | 1864 | 2053 | 2200 | 2046 | 2018.15 | 1936 | 2100 | 1.083 | 0.116 | 0.07509742 | 0.18042821 | 8.00E-04 | 0.0036 | 0.0025 | 0.0025 | TRUE | 0.0208 |
| Haao | 14 | 19 | 21 | 27 | 21 | 25 | 11 | 20 | 23 | 27 | 24 | 25 | 21.5 | 18 | 25 | 1.424 | 0.51 | 0.24036873 | 0.4077368 | 0 | 0.1127 | 0.0879 | 0.0879 | TRUE | 0.7689 |
| Habp2 | 0 | 3 | 1 | 5 | 8 | 2 | 0 | 3 | 1 | 5 | 9 | 2 | 3.37 | 1 | 5 | 3.927 | 1.973 | 0.11177783 | NA | 0.4763 | 0.7048 | 0.6546 | 0.6546 | TRUE | 0.3005 |
| Habp4 | 589 | 471 | 374 | 337 | 316 | 345 | 457 | 501 | 404 | 335 | 356 | 341 | 399.08 | 454 | 344 | 0.757 | -0.401 | 0.00051279 | 0.00439381 | 0.004 | 0.0084 | 0.007 | 0.007 | TRUE | 0.1043 |

| | | | | | | | | | | | | | | | | | | | | | | | | | |
|---|---|---|---|---|---|---|---|---|---|---|---|---|---|---|---|---|---|---|---|---|---|---|---|---|---|
| Hace1 | 1728 | 1189 | 1160 | 1294 | 1082 | 1243 | 1342 | 1265 | 1252 | 1288 | 1221 | 1227 | 1265.68 | 1286 | 1245 | 0.967 | -0.048 | 0.5040585 | 0.66813578 | 0 | 0.0043 | 0.0029 | 0.0029 | TRUE | 0.0182 |
| Hacl1 | 1561 | 1163 | 1305 | 1469 | 1385 | 1537 | 1212 | 1237 | 1408 | 1462 | 1562 | 1517 | 1399.91 | 1286 | 1514 | 1.178 | 0.237 | 0.00266571 | 0.0150522 | 0.0029 | 0.0041 | 0.0038 | 0.0038 | TRUE | 0.0819 |
| Hadh | 8916 | 6583 | 7034 | 6786 | 5946 | 6880 | 6925 | 7002 | 7590 | 6755 | 6708 | 6792 | 6961.78 | 7172 | 6752 | 0.942 | -0.087 | 0.11655768 | 0.24717492 | 0.0011 | 0.0028 | 0.0021 | 0.0021 | TRUE | 0.0315 |
| Hadha | 10627 | 7563 | 7284 | 9012 | 8150 | 9183 | 8254 | 8045 | 7859 | 8971 | 9194 | 9065 | 8564.53 | 8053 | 9077 | 1.127 | 0.172 | 0.00036577 | 0.00339407 | 2.00E-04 | 0.0027 | 0.0016 | 0.0016 | TRUE | 0.0059 |
| Hadhb | 2879 | 1991 | 1810 | 1978 | 1750 | 1925 | 2236 | 2118 | 1953 | 1969 | 1974 | 1900 | 2025.02 | 2102 | 1948 | 0.926 | -0.112 | 0.11081564 | 0.23850141 | 0.0021 | 0.0036 | 0.003 | 0.003 | TRUE | 0.0464 |
| Hagh | 1811 | 1340 | 1502 | 1864 | 1576 | 1860 | 1407 | 1425 | 1621 | 1855 | 1778 | 1836 | 1653.66 | 1484 | 1823 | 1.23 | 0.298 | 6.65E-05 | 0.0008827 | 0.0025 | 0.0039 | 0.0034 | 0.0034 | TRUE | 0.0768 |
| Haghl | 347 | 300 | 299 | 298 | 269 | 307 | 270 | 319 | 323 | 297 | 303 | 303 | 302.39 | 304 | 301 | 0.994 | -0.008 | 0.94841746 | 0.97180664 | 0 | 0.0103 | 0.0078 | 0.0078 | TRUE | 0.1195 |
| Hal | 1855 | 1416 | 1285 | 1626 | 1619 | 1754 | 1441 | 1506 | 1387 | 1619 | 1826 | 1731 | 1584.96 | 1445 | 1725 | 1.194 | 0.256 | 0.00047297 | 0.00414927 | 0.002 | 0.0039 | 0.0032 | 0.0032 | TRUE | 0.0355 |
| Hamp | 1 | 0 | 0 | 0 | 0 | 0 | 1 | 0 | 0 | 0 | 0 | 0 | 0.13 | 0 | 0 | 0.552 | -0.857 | 0.8336548 | NA | 0 | 18.3121 | 10 | 10 | TRUE | 0.2566 |
| Hamp2 | 0 | 0 | 0 | 0 | 0 | 0 | 0 | 0 | 0 | 0 | 0 | 0 | 0 | 0 | 0 | NA | NA | NA | NA | NA | NA | NA | NA | NA | NA |
| Hand1 | 574 | 343 | 403 | 267 | 215 | 261 | 446 | 365 | 435 | 266 | 243 | 258 | 335.24 | 415 | 256 | 0.614 | -0.703 | 1.64E-08 | 7.86E-07 | 0.0047 | 0.0095 | 0.008 | 0.008 | TRUE | 0.1269 |
| Hand2 | 10 | 8 | 5 | 2 | 2 | 5 | 8 | 9 | 5 | 2 | 2 | 5 | 5.14 | 7 | 3 | 0.425 | -1.233 | 0.15929139 | NA | 0 | 0.4633 | 0.3186 | 0.3186 | TRUE | 0.3959 |
| Hao1 | 0 | 0 | 0 | 0 | 0 | 0 | 0 | 0 | 0 | 0 | 0 | 0 | 0 | 0 | 0 | NA | NA | NA | NA | NA | NA | NA | NA | NA | NA |
| Hao2 | 0 | 1 | 1 | 1 | 0 | 0 | 0 | 1 | 1 | 1 | 0 | 0 | 0.52 | 1 | 0 | 0.535 | -0.904 | 0.74017109 | NA | 0 | 4.534 | 3.6534 | 3.6534 | TRUE | 0.3906 |
| Hap1 | 8102 | 6394 | 5039 | 4781 | 4653 | 5386 | 6292 | 6801 | 5437 | 4759 | 5249 | 5317 | 5642.6 | 6177 | 5108 | 0.827 | -0.274 | 0.00083867 | 0.00636255 | 0.0077 | 0.0028 | 0.0047 | 0.0047 | TRUE | 0.1331 |
| Hapln1 | 1 | 0 | 3 | 0 | 0 | 2 | 1 | 0 | 3 | 0 | 0 | 2 | 1 | 1 | 1 | 0.507 | -0.979 | 0.65508135 | NA | 2.0505 | 2.3773 | 2.3252 | 2.3252 | TRUE | 0.8631 |
| Hapln2 | 0 | 0 | 0 | 0 | 1 | 0 | 0 | 0 | 0 | 0 | 1 | 0 | 0.19 | 0 | 0 | 2.094 | 1.067 | 0.79379329 | NA | 0 | 12.608 | 10 | 10 | TRUE | 0.5075 |
| Hapln3 | 56 | 33 | 27 | 58 | 46 | 44 | 43 | 35 | 29 | 58 | 52 | 43 | 43.46 | 36 | 51 | 1.408 | 0.494 | 0.10517063 | 0.22960193 | 0 | 0.057 | 0.0435 | 0.0435 | TRUE | 0.2886 |
| Hapln4 | 621 | 486 | 429 | 648 | 607 | 611 | 482 | 517 | 463 | 645 | 685 | 603 | 565.85 | 487 | 644 | 1.321 | 0.402 | 3.70E-05 | 0.0005513 | 0.0017 | 0.0066 | 0.005 | 0.005 | TRUE | 0.0369 |
| Harbi1 | 1060 | 782 | 823 | 900 | 870 | 910 | 823 | 832 | 888 | 896 | 981 | 898 | 886.45 | 848 | 925 | 1.092 | 0.126 | 0.12310844 | 0.25676184 | 0.001 | 0.0051 | 0.0037 | 0.0037 | TRUE | 0.033 |
| Hars | 6670 | 4471 | 4252 | 4327 | 3708 | 4386 | 5180 | 4756 | 4588 | 4307 | 4183 | 4330 | 4557.28 | 4841 | 4273 | 0.882 | -0.181 | 0.00353835 | 0.01874489 | 0.0018 | 0.0029 | 0.0025 | 0.0025 | TRUE | 0.0464 |
| Hars2 | 1375 | 1054 | 1023 | 1065 | 940 | 1080 | 1068 | 1121 | 1104 | 1060 | 1060 | 1066 | 1079.91 | 1098 | 1062 | 0.968 | -0.046 | 0.5179294 | 0.6797764 | 0 | 0.0046 | 0.0028 | 0.0028 | TRUE | 0.0071 |
| Has1 | 8 | 10 | 2 | 7 | 6 | 8 | 6 | 11 | 2 | 7 | 7 | 8 | 6.77 | 6 | 7 | 1.139 | 0.187 | 0.80686798 | NA | 0.0409 | 0.3523 | 0.2757 | 0.2757 | TRUE | 0.2139 |
| Has2 | 17 | 7 | 10 | 25 | 12 | 19 | 13 | 7 | 11 | 25 | 14 | 19 | 14.77 | 10 | 19 | 1.805 | 0.852 | 0.10239188 | NA | 0 | 0.1629 | 0.1235 | 0.1235 | TRUE | 0.1454 |
| Has2as | 1 | 1 | 0 | 0 | 0 | 0 | 1 | 1 | 0 | 0 | 0 | 0 | 0.31 | 1 | 0 | 0.309 | -1.696 | 0.63183309 | NA | 0 | 7.7294 | 7.2472 | 7.2472 | TRUE | 0.2015 |
| Has3 | 88 | 58 | 46 | 50 | 52 | 52 | 68 | 62 | 50 | 50 | 59 | 51 | 56.57 | 60 | 53 | 0.883 | -0.179 | 0.49625054 | 0.66061075 | 0 | 0.0443 | 0.0322 | 0.0322 | TRUE | 0.1849 |
| Hat1 | 22729 | 16314 | 17975 | 16000 | 13768 | 16002 | 17653 | 17353 | 19395 | 15926 | 15532 | 15796 | 16942.51 | 18134 | 15751 | 0.869 | -0.203 | 0.00033593 | 0.00316721 | 0.0018 | 0.0026 | 0.0023 | 0.0023 | TRUE | 0.0452 |
| Haus1 | 2491 | 1716 | 1714 | 1893 | 1582 | 1814 | 1935 | 1825 | 1849 | 1884 | 1785 | 1791 | 1844.83 | 1870 | 1820 | 0.973 | -0.04 | 0.53729572 | 0.69599128 | 5.00E-04 | 0.0037 | 0.0024 | 0.0024 | TRUE | 0.0117 |
| Haus2 | 3676 | 2574 | 2534 | 2667 | 2304 | 2412 | 2855 | 2738 | 2734 | 2655 | 2599 | 2381 | 2660.33 | 2776 | 2545 | 0.916 | -0.126 | 0.05129214 | 0.13736529 | 0.0014 | 0.0033 | 0.0026 | 0.0026 | TRUE | 0.0389 |
| Haus3 | 2701 | 1954 | 1734 | 1873 | 1582 | 1806 | 2098 | 2078 | 1871 | 1864 | 1785 | 1783 | 1913.17 | 2016 | 1811 | 0.898 | -0.156 | 0.02543309 | 0.0823364 | 0.0018 | 0.0037 | 0.003 | 0.003 | TRUE | 0.047 |
| Haus4 | 1944 | 1326 | 1221 | 1245 | 1208 | 1417 | 1510 | 1410 | 1317 | 1239 | 1363 | 1399 | 1373.09 | 1412 | 1334 | 0.943 | -0.085 | 0.29448487 | 0.46649416 | 0.0037 | 0.0042 | 0.004 | 0.004 | TRUE | 0.0465 |
| Haus5 | 2485 | 1833 | 1613 | 1780 | 1599 | 1919 | 1930 | 1950 | 1740 | 1772 | 1804 | 1894 | 1848.36 | 1873 | 1823 | 0.973 | -0.04 | 0.57521151 | 0.72711578 | 0.002 | 0.0037 | 0.0031 | 0.0031 | TRUE | 0.0457 |
| Haus6 | 4667 | 3301 | 3405 | 3710 | 3243 | 3647 | 3625 | 3511 | 3674 | 3693 | 3658 | 3600 | 3626.9 | 3603 | 3650 | 1.013 | 0.019 | 0.72744234 | 0.83661541 | 0 | 0.0031 | 0.0018 | 0.0018 | TRUE | 0.006 |
| Haus7 | 2026 | 1512 | 1585 | 2037 | 1769 | 2085 | 1574 | 1608 | 1710 | 2028 | 1996 | 2058 | 1828.91 | 1631 | 2027 | 1.244 | 0.315 | 1.18E-06 | 3.37E-05 | 5.00E-04 | 0.0037 | 0.0025 | 0.0025 | TRUE | 0.0217 |
| Haus8 | 2783 | 2057 | 1953 | 1965 | 1707 | 1998 | 2161 | 2188 | 2107 | 1956 | 1926 | 1972 | 2051.78 | 2152 | 1951 | 0.907 | -0.141 | 0.01770294 | 0.06309929 | 0 | 0.0036 | 0.0021 | 0.0021 | TRUE | 0.0039 |
| Havcr1 | 8 | 3 | 5 | 12 | 11 | 7 | 6 | 3 | 5 | 12 | 12 | 7 | 7.68 | 5 | 10 | 2.081 | 1.057 | 0.1375144 | NA | 0 | 0.3111 | 0.2189 | 0.2189 | TRUE | 0.3213 |
| Havcr2 | 4 | 8 | 10 | 7 | 12 | 15 | 3 | 9 | 11 | 7 | 14 | 15 | 9.62 | 8 | 12 | 1.612 | 0.688 | 0.3068105 | NA | 0.1304 | 0.2488 | 0.2168 | 0.2168 | TRUE | 0.9882 |
| Hax1 | 3163 | 2233 | 2046 | 2236 | 2020 | 2255 | 2457 | 2375 | 2208 | 2226 | 2279 | 2226 | 2294.99 | 2347 | 2244 | 0.955 | -0.066 | 0.3069856 | 0.47963983 | 0.0012 | 0.0035 | 0.0026 | 0.0026 | TRUE | 0.0322 |
| Hba-a1 | 0 | 0 | 0 | 0 | 0 | 0 | 0 | 0 | 0 | 0 | 0 | 0 | 0 | 0 | 0 | NA | NA | NA | NA | NA | NA | NA | NA | NA | NA |
| Hba-a2 | 0 | 0 | 0 | 0 | 0 | 0 | 0 | 0 | 0 | 0 | 0 | 0 | 0 | 0 | 0 | NA | NA | NA | NA | NA | NA | NA | NA | NA | NA |
| Hba-x | 0 | 0 | 1 | 0 | 0 | 0 | 0 | 0 | 1 | 0 | 0 | 0 | 0.18 | 0 | 0 | 0.552 | -0.857 | 0.8336548 | NA | 0 | 13.1817 | 10 | 10 | TRUE | 0.5365 |
| Hbb-b1 | 0 | 0 | 0 | 0 | 0 | 0 | 0 | 0 | 0 | 0 | 0 | 0 | 0 | 0 | 0 | NA | NA | NA | NA | NA | NA | NA | NA | NA | NA |
| Hbb-b2 | 0 | 0 | 0 | 0 | 0 | 0 | 0 | 0 | 0 | 0 | 0 | 0 | 0 | 0 | 0 | NA | NA | NA | NA | NA | NA | NA | NA | NA | NA |
| Hbb-bh1 | 1 | 0 | 1 | 1 | 0 | 0 | 1 | 0 | 1 | 1 | 0 | 0 | 0.48 | 1 | 0 | 0.596 | -0.748 | 0.78941713 | NA | 0 | 4.9902 | 3.8695 | 3.8695 | TRUE | 0.3906 |
| Hbb-bh2 | 0 | 0 | 0 | 0 | 0 | 0 | 0 | 0 | 0 | 0 | 0 | 0 | 0 | 0 | 0 | NA | NA | NA | NA | NA | NA | NA | NA | NA | NA |
| Hbb-y | 1 | 1 | 0 | 1 | 1 | 0 | 1 | 1 | 0 | 1 | 1 | 0 | 0.66 | 1 | 1 | 1.128 | 0.173 | 0.94106052 | NA | 0 | 3.5899 | 2.4471 | 2.4471 | TRUE | 0.2681 |
| Hbegf | 1035 | 806 | 777 | 1020 | 798 | 944 | 804 | 857 | 838 | 1015 | 900 | 932 | 891.16 | 833 | 949 | 1.141 | 0.19 | 0.02190227 | 0.07375763 | 0.0014 | 0.0051 | 0.0038 | 0.0038 | TRUE | 0.0445 |
| Hbp1 | 4624 | 3233 | 3991 | 4193 | 3799 | 4303 | 3591 | 3439 | 4306 | 4174 | 4286 | 4248 | 4007.25 | 3779 | 4236 | 1.121 | 0.165 | 0.04101674 | 0.11654644 | 0.0072 | 0.003 | 0.0045 | 0.0045 | TRUE | 0.1801 |
| Hbq1a | 27 | 19 | 18 | 30 | 21 | 18 | 21 | 20 | 19 | 30 | 24 | 18 | 21.99 | 20 | 24 | 1.175 | 0.232 | 0.57462999 | 0.72661204 | 0 | 0.1102 | 0.078 | 0.078 | TRUE | 0.1756 |
| Hbq1b | 55 | 41 | 29 | 33 | 35 | 47 | 43 | 44 | 31 | 33 | 39 | 46 | 39.39 | 39 | 39 | 1.006 | 0.009 | 0.97780681 | 0.98810752 | 0 | 0.0626 | 0.047 | 0.047 | TRUE | 0.2156 |
| Hbs1l | 8114 | 5660 | 5943 | 6495 | 5728 | 6623 | 6302 | 6020 | 6412 | 6465 | 6462 | 6538 | 6366.59 | 6245 | 6488 | 1.039 | 0.055 | 0.27992274 | 0.45144608 | 3.00E-04 | 0.0028 | 0.0017 | 0.0017 | TRUE | 0.0119 |
| Hc | 18 | 6 | 6 | 9 | 7 | 14 | 14 | 6 | 6 | 9 | 8 | 14 | 9.59 | 9 | 10 | 1.124 | 0.168 | 0.79373701 | NA | 0.0639 | 0.2497 | 0.1958 | 0.1958 | TRUE | 0.9789 |
| Hccs | 2523 | 1797 | 1841 | 1965 | 1675 | 1860 | 1960 | 1911 | 1986 | 1956 | 1890 | 1836 | 1923.17 | 1952 | 1894 | 0.97 | -0.044 | 0.48180842 | 0.64850624 | 0 | 0.0037 | 0.0023 | 0.0023 | TRUE | 0.0101 |
| Hcfc1 | 30441 | 21885 | 20977 | 22363 | 20568 | 23546 | 23642 | 23279 | 22634 | 22260 | 23203 | 23244 | 23043.58 | 23185 | 22902 | 0.988 | -0.018 | 0.70561264 | 0.82174868 | 4.00E-04 | 0.0025 | 0.0016 | 0.0016 | TRUE | 0.0074 |
| Hcfc1r1 | 998 | 754 | 774 | 771 | 771 | 845 | 775 | 802 | 835 | 767 | 870 | 834 | 813.94 | 804 | 824 | 1.025 | 0.035 | 0.6786669 | 0.8026633 | 0.0015 | 0.0053 | 0.004 | 0.004 | TRUE | 0.0427 |
| Hcfc2 | 645 | 464 | 482 | 581 | 496 | 527 | 501 | 494 | 520 | 578 | 560 | 520 | 528.78 | 505 | 553 | 1.095 | 0.131 | 0.17149762 | 0.32342203 | 0 | 0.0069 | 0.0047 | 0.0047 | TRUE | 0.0316 |
| Hck | 209 | 148 | 158 | 251 | 211 | 223 | 162 | 157 | 170 | 250 | 238 | 220 | 199.71 | 163 | 236 | 1.444 | 0.531 | 0.00021237 | 0.00222866 | 0 | 0.0143 | 0.0096 | 0.0096 | TRUE | 0.0393 |
| Hcls1 | 7 | 5 | 0 | 1 | 3 | 1 | 5 | 5 | 0 | 1 | 3 | 1 | 2.69 | 3 | 2 | 0.484 | -1.046 | 0.42835487 | NA | 0.6482 | 0.8845 | 0.83 | 0.83 | TRUE | 1.0544 |
| Hcn1 | 36 | 23 | 9 | 14 | 10 | 11 | 28 | 24 | 10 | 14 | 11 | 11 | 16.37 | 21 | 12 | 0.575 | -0.797 | 0.12975914 | NA | 0.099 | 0.1472 | 0.1334 | 0.1334 | TRUE | 1.1092 |
| Hcn2 | 1937 | 1731 | 1709 | 2343 | 2279 | 2433 | 1504 | 1841 | 1844 | 2332 | 2571 | 2402 | 2082.42 | 1730 | 2435 | 1.409 | 0.495 | 1.08E-08 | 5.37E-07 | 0.0069 | 0.0036 | 0.0049 | 0.0049 | TRUE | 0.1613 |
| Hcn3 | 324 | 308 | 311 | 379 | 364 | 351 | 252 | 328 | 336 | 377 | 411 | 346 | 341.53 | 305 | 378 | 1.244 | 0.315 | 0.02106961 | 0.07189019 | 0.0131 | 0.0094 | 0.0105 | 0.0105 | TRUE | 0.2846 |
| Hcn4 | 28 | 23 | 33 | 32 | 43 | 40 | 22 | 24 | 36 | 32 | 49 | 39 | 33.61 | 27 | 40 | 1.473 | 0.559 | 0.11514156 | 0.24500795 | 0.0247 | 0.0729 | 0.0598 | 0.0598 | TRUE | 0.2993 |
| Hcrt | 0 | 1 | 3 | 0 | 0 | 2 | 0 | 1 | 3 | 0 | 0 | 2 | 1.05 | 1 | 1 | 0.48 | -1.059 | 0.62568106 | NA | 2.327 | 2.2686 | 2.2798 | 2.2798 | TRUE | 0.5605 |
| Hcrtr1 | 10 | 11 | 9 | 16 | 11 | 8 | 8 | 12 | 10 | 16 | 12 | 8 | 10.9 | 10 | 12 | 1.251 | 0.323 | 0.58381366 | NA | 0 | 0.2198 | 0.1577 | 0.1577 | TRUE | 0.2288 |
| Hcrtr2 | 10 | 0 | 3 | 3 | 2 | 1 | 8 | 0 | 3 | 3 | 2 | 1 | 2.87 | 4 | 2 | 0.55 | -0.863 | 0.49996505 | NA | 0.724 | 0.8276 | 0.8012 | 0.8012 | TRUE | 0.1904 |
| Hcst | 1 | 1 | 1 | 2 | 1 | 2 | 1 | 1 | 1 | 2 | 1 | 2 | 1.34 | 1 | 2 | 1.773 | 0.826 | 0.60444684 | NA | 0 | 1.7772 | 1.0296 | 1.0296 | TRUE | 0.0577 |
| Hdac1 | 8726 | 6047 | 5449 | 5502 | 4676 | 5598 | 6777 | 6432 | 5879 | 5477 | 5275 | 5526 | 5894.41 | 6363 | 5426 | 0.853 | -0.23 | 0.00032647 | 0.00310052 | 0.0026 | 0.0028 | 0.0028 | 0.0028 | TRUE | 0.0537 |
| Hdac10 | 258 | 200 | 169 | 184 | 168 | 181 | 200 | 213 | 182 | 183 | 190 | 179 | 191.14 | 198 | 184 | 0.925 | -0.112 | 0.4405555 | 0.61193459 | 0 | 0.0148 | 0.0101 | 0.0101 | TRUE | 0.0512 |
| Hdac11 | 502 | 485 | 580 | 669 | 612 | 755 | 390 | 516 | 626 | 666 | 690 | 745 | 605.53 | 511 | 700 | 1.377 | 0.461 | 0.00105429 | 0.00751941 | 0.0259 | 0.0063 | 0.0126 | 0.0126 | TRUE | 0.3224 |

| | | | | | | | | | | | | | | | | | | | | | | | | |
|---|---|---|---|---|---|---|---|---|---|---|---|---|---|---|---|---|---|---|---|---|---|---|---|---|
| Hdac2 | 8790 | 6444 | 6346 | 6550 | 5486 | 6548 | 6827 | 6854 | 6847 | 6520 | 6189 | 6464 | 6616.84 | 6843 | 6391 | 0.934 | -0.098 | 0.04741383 | 0.12960143 | 2.00E-04 | 0.0028 | 0.0016 | 0.0016 | TRUE | 0.0093 |
| Hdac3 | 3481 | 2511 | 2549 | 2276 | 2093 | 2483 | 2704 | 2671 | 2750 | 2266 | 2361 | 2451 | 2533.76 | 2708 | 2359 | 0.871 | -0.199 | 0.00092178 | 0.00683354 | 4.00E-04 | 0.0034 | 0.0022 | 0.0022 | TRUE | 0.0148 |
| Hdac4 | 1628 | 1249 | 1392 | 1670 | 1487 | 1785 | 1264 | 1329 | 1502 | 1662 | 1677 | 1762 | 1532.8 | 1365 | 1700 | 1.248 | 0.319 | 5.96E-05 | 0.00080738 | 0.0036 | 0.004 | 0.0039 | 0.0039 | TRUE | 0.0919 |
| Hdac5 | 4956 | 3773 | 4166 | 4085 | 3883 | 4530 | 3849 | 4013 | 4495 | 4066 | 4380 | 4472 | 4212.65 | 4119 | 4306 | 1.046 | 0.065 | 0.3678644 | 0.54249751 | 0.0039 | 0.003 | 0.0035 | 0.0035 | TRUE | 0.0773 |
| Hdac6 | 7440 | 5882 | 6307 | 5565 | 5137 | 6131 | 5778 | 6257 | 6805 | 5539 | 5795 | 6052 | 6037.81 | 6280 | 5795 | 0.923 | -0.115 | 0.09878862 | 0.21911634 | 0.0041 | 0.0028 | 0.0033 | 0.0033 | TRUE | 0.0653 |
| Hdac7 | 561 | 496 | 592 | 645 | 661 | 779 | 436 | 528 | 639 | 642 | 746 | 769 | 626.46 | 534 | 719 | 1.35 | 0.433 | 0.00095091 | 0.00699502 | 0.0201 | 0.0062 | 0.0107 | 0.0107 | TRUE | 0.3262 |
| Hdac8 | 307 | 215 | 228 | 213 | 200 | 211 | 238 | 229 | 246 | 212 | 226 | 208 | 226.51 | 238 | 215 | 0.905 | -0.144 | 0.27953875 | 0.45115712 | 0 | 0.0129 | 0.0083 | 0.0083 | TRUE | 0.0184 |
| Hdac9 | 251 | 215 | 207 | 202 | 174 | 185 | 195 | 229 | 223 | 201 | 196 | 183 | 204.49 | 216 | 193 | 0.899 | -0.154 | 0.28707571 | 0.45917336 | 0 | 0.014 | 0.0101 | 0.0101 | TRUE | 0.0851 |
| Hdc | 77 | 53 | 32 | 48 | 38 | 35 | 60 | 56 | 35 | 48 | 43 | 35 | 45.98 | 50 | 42 | 0.825 | -0.277 | 0.3752331 | 0.54962151 | 0.0334 | 0.054 | 0.0483 | 0.0483 | TRUE | 0.5754 |
| Hddc2 | 1854 | 1427 | 1371 | 1327 | 1149 | 1335 | 1440 | 1518 | 1479 | 1321 | 1296 | 1318 | 1395.34 | 1479 | 1312 | 0.887 | -0.172 | 0.00979508 | 0.04076911 | 0 | 0.0041 | 0.0025 | 0.0025 | TRUE | 0.0068 |
| Hddc3 | 384 | 273 | 299 | 318 | 277 | 331 | 298 | 290 | 323 | 317 | 312 | 327 | 311.17 | 304 | 319 | 1.05 | 0.07 | 0.54724584 | 0.7040944 | 0 | 0.01 | 0.0066 | 0.0066 | TRUE | 0.0316 |
| Hdgf | 31759 | 23168 | 21936 | 23985 | 21589 | 24929 | 24666 | 24644 | 23669 | 23875 | 24355 | 24609 | 24302.7 | 24326 | 24280 | 0.998 | -0.003 | 0.95064369 | 0.97276276 | 3.00E-04 | 0.0025 | 0.0015 | 0.0015 | TRUE | 0.0068 |
| Hdgfl1 | 95 | 98 | 63 | 53 | 60 | 52 | 74 | 104 | 68 | 53 | 68 | 51 | 69.63 | 82 | 57 | 0.698 | -0.519 | 0.04502541 | 0.12471863 | 0.0253 | 0.0365 | 0.0333 | 0.0333 | TRUE | 0.5061 |
| Hdgfrp2 | 6912 | 5603 | 4859 | 4787 | 4400 | 4954 | 5368 | 5960 | 5243 | 4765 | 4964 | 4890 | 5198.32 | 5524 | 4873 | 0.882 | -0.181 | 0.00421274 | 0.02147946 | 0.0023 | 0.0029 | 0.0027 | 0.0027 | TRUE | 0.0578 |
| Hdgfrp3 | 1566 | 1177 | 1027 | 1100 | 933 | 1035 | 1216 | 1252 | 1108 | 1095 | 1053 | 1022 | 1124.25 | 1192 | 1057 | 0.886 | -0.175 | 0.02570602 | 0.08285449 | 0.0017 | 0.0045 | 0.0035 | 0.0035 | TRUE | 0.0442 |
| Hdhd1a | 3 | 0 | 3 | 0 | 3 | 0 | 2 | 0 | 3 | 0 | 3 | 0 | 1.49 | 2 | 1 | 0.587 | -0.769 | 0.67979657 | NA | 2.9285 | 1.5911 | 1.7719 | 1.7719 | TRUE | 1.3873 |
| Hdhd2 | 1503 | 1058 | 995 | 1032 | 918 | 996 | 1167 | 1125 | 1074 | 1027 | 1036 | 983 | 1068.73 | 1122 | 1015 | 0.904 | -0.146 | 0.05231095 | 0.13925878 | 0 | 0.0046 | 0.0031 | 0.0031 | TRUE | 0.017 |
| Hdhd3 | 86 | 56 | 36 | 52 | 36 | 30 | 67 | 60 | 39 | 52 | 41 | 30 | 47.86 | 55 | 41 | 0.734 | -0.446 | 0.16416177 | 0.31507508 | 0.0544 | 0.0519 | 0.0526 | 0.0526 | TRUE | 0.2977 |
| Hdlbp | 41990 | 28437 | 25172 | 28074 | 24197 | 27220 | 32612 | 30248 | 27160 | 27945 | 27297 | 26870 | 28688.7 | 30007 | 27371 | 0.912 | -0.133 | 0.04888911 | 0.13246864 | 0.0043 | 0.0025 | 0.0032 | 0.0032 | TRUE | 0.0842 |
| Hdx | 234 | 143 | 114 | 96 | 90 | 106 | 182 | 152 | 123 | 96 | 102 | 105 | 126.43 | 152 | 101 | 0.657 | -0.606 | 0.00197754 | 0.0120123 | 0.0152 | 0.0212 | 0.0193 | 0.0193 | TRUE | 0.1385 |
| Heatr1 | 18966 | 13654 | 11998 | 13006 | 12076 | 13845 | 14730 | 14524 | 12946 | 12946 | 13623 | 13667 | 13739.28 | 14067 | 13412 | 0.953 | -0.069 | 0.2670126 | 0.4375113 | 0.0028 | 0.0026 | 0.0027 | 0.0027 | TRUE | 0.0592 |
| Heatr2 | 4182 | 3319 | 3319 | 3320 | 3010 | 3446 | 3248 | 3530 | 3581 | 3305 | 3396 | 3402 | 3410.26 | 3453 | 3368 | 0.976 | -0.035 | 0.55950025 | 0.71433113 | 0.0013 | 0.0031 | 0.0024 | 0.0024 | TRUE | 0.0337 |
| Heatr3 | 11052 | 7821 | 6017 | 6224 | 5496 | 6231 | 8584 | 8319 | 6492 | 6195 | 6200 | 6151 | 6990.23 | 7798 | 6182 | 0.793 | -0.335 | 0.00018108 | 0.00198181 | 0.0105 | 0.0028 | 0.0056 | 0.0056 | TRUE | 0.2614 |
| Heatr5a | 5232 | 3947 | 3881 | 4222 | 3864 | 4546 | 4063 | 4198 | 4188 | 4203 | 4359 | 4488 | 4249.76 | 4150 | 4350 | 1.048 | 0.068 | 0.21106763 | 0.37308801 | 5.00E-04 | 0.003 | 0.0019 | 0.0019 | TRUE | 0.0108 |
| Heatr5b | 3870 | 2791 | 3023 | 3770 | 3212 | 3694 | 3006 | 2969 | 3262 | 3753 | 3623 | 3647 | 3376.48 | 3079 | 3674 | 1.194 | 0.256 | 2.28E-05 | 0.0003722 | 0.0011 | 0.0031 | 0.0023 | 0.0023 | TRUE | 0.0324 |
| Heatr6 | 3664 | 2626 | 2787 | 2650 | 2472 | 2791 | 2846 | 2793 | 3007 | 2638 | 2789 | 2755 | 2804.62 | 2882 | 2727 | 0.946 | -0.08 | 0.18778335 | 0.3449412 | 8.00E-04 | 0.0033 | 0.0023 | 0.0023 | TRUE | 0.0172 |
| Heatr8 | 53 | 44 | 48 | 49 | 42 | 53 | 41 | 47 | 52 | 49 | 47 | 52 | 48.04 | 47 | 49 | 1.068 | 0.095 | 0.73141204 | 0.83929626 | 0 | 0.0518 | 0.0345 | 0.0345 | TRUE | 0.0974 |
| Hebp1 | 778 | 515 | 580 | 782 | 666 | 772 | 604 | 548 | 626 | 778 | 751 | 762 | 678.28 | 593 | 764 | 1.289 | 0.366 | 4.08E-05 | 0.00059527 | 5.00E-04 | 0.0059 | 0.0042 | 0.0042 | TRUE | 0.0492 |
| Hebp2 | 476 | 352 | 349 | 314 | 259 | 287 | 370 | 374 | 377 | 313 | 292 | 283 | 334.79 | 374 | 296 | 0.793 | -0.335 | 0.00300203 | 0.01655575 | 0 | 0.0095 | 0.0061 | 0.0061 | TRUE | 0.0276 |
| Heca | 2416 | 1786 | 1933 | 2041 | 1749 | 2133 | 1876 | 1900 | 2086 | 2032 | 1973 | 2106 | 1995.35 | 1954 | 2037 | 1.043 | 0.061 | 0.3731665 | 0.54766083 | 0.0015 | 0.0036 | 0.0029 | 0.0029 | TRUE | 0.0415 |
| Hectd1 | 8053 | 5412 | 5539 | 6655 | 5923 | 6678 | 6254 | 5757 | 5977 | 6624 | 6682 | 6592 | 6314.33 | 5996 | 6633 | 1.106 | 0.145 | 0.00653254 | 0.03009939 | 7.00E-04 | 0.0028 | 0.0019 | 0.0019 | TRUE | 0.0176 |
| Hectd2 | 748 | 531 | 552 | 659 | 558 | 584 | 581 | 565 | 596 | 656 | 629 | 576 | 600.55 | 581 | 620 | 1.069 | 0.096 | 0.30121427 | 0.47354972 | 0 | 0.0064 | 0.0046 | 0.0046 | TRUE | 0.0465 |
| Hectd3 | 2567 | 1847 | 1869 | 2102 | 1832 | 2238 | 1994 | 1965 | 2017 | 2092 | 2067 | 2209 | 2057.2 | 1992 | 2123 | 1.066 | 0.092 | 0.1353363 | 0.27461125 | 0 | 0.0036 | 0.0023 | 0.0023 | TRUE | 0.0155 |
| Hecw1 | 20 | 6 | 8 | 8 | 8 | 7 | 16 | 6 | 9 | 8 | 9 | 7 | 9.07 | 10 | 8 | 0.767 | -0.383 | 0.55820976 | NA | 0 | 0.2636 | 0.1961 | 0.1961 | TRUE | 0.9294 |
| Hecw2 | 317 | 187 | 191 | 94 | 81 | 80 | 246 | 199 | 206 | 94 | 91 | 79 | 152.52 | 217 | 88 | 0.404 | -1.309 | 1.41E-13 | 2.04E-11 | 0 | 0.018 | 0.0145 | 0.0145 | TRUE | 0.1616 |
| Heg1 | 537 | 379 | 413 | 422 | 429 | 465 | 417 | 403 | 446 | 420 | 484 | 459 | 438.14 | 422 | 454 | 1.076 | 0.106 | 0.31746237 | 0.49148985 | 0.0013 | 0.0078 | 0.0059 | 0.0059 | TRUE | 0.0508 |
| Helb | 5676 | 4623 | 4653 | 4760 | 4761 | 5605 | 4408 | 4917 | 5021 | 4738 | 5371 | 5533 | 4998.05 | 4782 | 5214 | 1.091 | 0.125 | 0.09653244 | 0.21575265 | 0.0053 | 0.0029 | 0.0039 | 0.0039 | TRUE | 0.0779 |
| Hells | 21184 | 15321 | 14463 | 14815 | 12805 | 14608 | 16453 | 16297 | 15605 | 14747 | 14445 | 14420 | 15327.91 | 16118 | 14537 | 0.902 | -0.149 | 0.00164828 | 0.0103875 | 4.00E-04 | 0.0026 | 0.0016 | 0.0016 | TRUE | 0.0094 |
| Helq | 1607 | 1120 | 1010 | 1075 | 897 | 1069 | 1248 | 1191 | 1090 | 1070 | 1012 | 1055 | 1111.07 | 1176 | 1046 | 0.888 | -0.171 | 0.03117974 | 0.09532751 | 0.002 | 0.0046 | 0.0036 | 0.0036 | TRUE | 0.0491 |
| Helt | 1 | 2 | 1 | 2 | 2 | 2 | 1 | 2 | 1 | 2 | 2 | 2 | 1.7 | 1 | 2 | 1.591 | 0.67 | 0.63785066 | NA | 0 | 1.396 | 0.8408 | 0.8408 | TRUE | 0.1605 |
| Helz | 8735 | 6253 | 6147 | 6661 | 6297 | 7497 | 6784 | 6651 | 6633 | 6630 | 7104 | 7401 | 6867.1 | 6689 | 7045 | 1.053 | 0.075 | 0.19283775 | 0.35133587 | 0.0014 | 0.0028 | 0.0022 | 0.0022 | TRUE | 0.0325 |
| Helz2 | 567 | 385 | 350 | 504 | 468 | 533 | 440 | 410 | 378 | 502 | 528 | 526 | 463.89 | 409 | 519 | 1.264 | 0.338 | 0.00108154 | 0.00766789 | 0.0011 | 0.0075 | 0.0055 | 0.0055 | TRUE | 0.0549 |
| Hemgn | 0 | 2 | 0 | 0 | 0 | 0 | 0 | 2 | 0 | 0 | 0 | 0 | 0.35 | 1 | 0 | 0.272 | -1.878 | 0.62190613 | NA | 10 | 6.6869 | 8.7633 | 8.7633 | TRUE | 0.9743 |
| Hemk1 | 1568 | 1084 | 1057 | 1195 | 1139 | 1205 | 1218 | 1153 | 1140 | 1190 | 1285 | 1190 | 1195.88 | 1170 | 1222 | 1.042 | 0.059 | 0.42241809 | 0.59472469 | 8.00E-04 | 0.0044 | 0.0031 | 0.0031 | TRUE | 0.0244 |
| Hemt1 | 41 | 21 | 20 | 13 | 6 | 9 | 32 | 22 | 22 | 13 | 7 | 9 | 17.39 | 25 | 10 | 0.377 | -1.408 | 0.00460803 | 0.02306122 | 0 | 0.1387 | 0.1049 | 0.1049 | TRUE | 0.3765 |
| Henmt1 | 88 | 48 | 49 | 48 | 31 | 38 | 68 | 51 | 53 | 48 | 35 | 38 | 48.76 | 57 | 40 | 0.695 | -0.524 | 0.07033447 | 0.17206121 | 0 | 0.051 | 0.0392 | 0.0392 | TRUE | 0.2666 |
| Hepacam | 34 | 18 | 16 | 28 | 25 | 20 | 26 | 19 | 17 | 28 | 28 | 20 | 23.11 | 21 | 25 | 1.192 | 0.254 | 0.5351776 | 0.69419125 | 0 | 0.105 | 0.0774 | 0.0774 | TRUE | 0.3518 |
| Hepacam2 | 22 | 9 | 12 | 17 | 12 | 15 | 17 | 10 | 13 | 17 | 14 | 15 | 14.15 | 13 | 15 | 1.132 | 0.178 | 0.72727971 | NA | 0 | 0.17 | 0.1181 | 0.1181 | TRUE | 0.3313 |
| Heph | 43 | 18 | 20 | 20 | 14 | 11 | 33 | 19 | 22 | 20 | 16 | 11 | 20.11 | 25 | 16 | 0.621 | -0.686 | 0.13488291 | 0.27384678 | 0.047 | 0.1203 | 0.0992 | 0.0992 | TRUE | 0.6767 |
| Hephl1 | 16 | 10 | 4 | 0 | 2 | 3 | 12 | 11 | 4 | 0 | 2 | 3 | 5.43 | 9 | 2 | 0.188 | -2.413 | 0.01692311 | NA | 0 | 0.4387 | 0.3836 | 0.3836 | TRUE | 0.6411 |
| Herc1 | 6835 | 4580 | 4089 | 5311 | 4770 | 5623 | 5308 | 4872 | 4412 | 5287 | 5381 | 5551 | 5135.09 | 4864 | 5406 | 1.111 | 0.152 | 0.03314644 | 0.09971839 | 0.0043 | 0.0029 | 0.0035 | 0.0035 | TRUE | 0.0804 |
| Herc2 | 9968 | 6897 | 6598 | 8242 | 7749 | 8823 | 7742 | 7336 | 7119 | 8204 | 8742 | 8710 | 7975.43 | 7399 | 8552 | 1.155 | 0.208 | 0.00021549 | 0.0022597 | 0.0013 | 0.0027 | 0.0022 | 0.0022 | TRUE | 0.0203 |
| Herc3 | 1481 | 1067 | 1118 | 1041 | 923 | 1083 | 1150 | 1135 | 1206 | 1036 | 1041 | 1069 | 1106.34 | 1164 | 1049 | 0.902 | -0.15 | 0.03849862 | 0.11138883 | 0 | 0.0046 | 0.0029 | 0.0029 | TRUE | 0.0118 |
| Herc4 | 12201 | 8795 | 8916 | 9214 | 8333 | 9307 | 9476 | 9355 | 9620 | 9172 | 9400 | 9187 | 9368.49 | 9484 | 9253 | 0.976 | -0.036 | 0.44387274 | 0.61487977 | 0 | 0.0027 | 0.0014 | 0.0014 | TRUE | 0.0024 |
| Herc6 | 159 | 125 | 104 | 65 | 44 | 61 | 123 | 133 | 112 | 65 | 50 | 60 | 90.54 | 123 | 58 | 0.475 | -1.075 | 6.30E-07 | 1.99E-05 | 0 | 0.0286 | 0.0207 | 0.0207 | TRUE | 0.1302 |
| Herpud1 | 501 | 338 | 348 | 410 | 321 | 362 | 389 | 360 | 375 | 408 | 362 | 357 | 375.28 | 375 | 376 | 1.002 | 0.003 | 0.9755677 | 0.98670326 | 0 | 0.0087 | 0.0063 | 0.0063 | TRUE | 0.0656 |
| Herpud2 | 2792 | 2080 | 1867 | 2046 | 1738 | 1896 | 2168 | 2212 | 2014 | 2037 | 1961 | 1872 | 2044.04 | 2131 | 1957 | 0.917 | -0.124 | 0.06680805 | 0.16561513 | 0.0016 | 0.0036 | 0.0028 | 0.0028 | TRUE | 0.0274 |
| Hes1 | 1075 | 898 | 927 | 1174 | 1047 | 1168 | 835 | 955 | 1000 | 1169 | 1181 | 1153 | 1048.84 | 930 | 1168 | 1.258 | 0.331 | 0.00010254 | 0.00123058 | 0.0035 | 0.0047 | 0.0043 | 0.0043 | TRUE | 0.1005 |
| Hes2 | 37 | 37 | 35 | 47 | 50 | 57 | 29 | 39 | 38 | 47 | 56 | 56 | 44.22 | 35 | 53 | 1.517 | 0.601 | 0.04298161 | 0.12069473 | 0 | 0.056 | 0.0401 | 0.0401 | TRUE | 0.2294 |
| Hes3 | 1 | 2 | 1 | 0 | 2 | 8 | 1 | 2 | 1 | 0 | 2 | 8 | 2.36 | 1 | 3 | 2.607 | 1.382 | 0.33811405 | NA | 0.9027 | 1.0084 | 0.9789 | 0.9789 | TRUE | 0.3869 |
| Hes5 | 26 | 19 | 22 | 12 | 22 | 41 | 20 | 20 | 24 | 12 | 25 | 40 | 23.56 | 21 | 26 | 1.207 | 0.272 | 0.55709046 | 0.71220875 | 0.1333 | 0.103 | 0.1117 | 0.1117 | TRUE | 0.2071 |
| Hes6 | 562 | 413 | 452 | 508 | 407 | 504 | 436 | 439 | 488 | 506 | 459 | 498 | 470.97 | 454 | 488 | 1.075 | 0.104 | 0.31180661 | 0.48510415 | 0.0011 | 0.0075 | 0.0055 | 0.0055 | TRUE | 0.046 |
| Hes7 | 15 | 19 | 15 | 32 | 24 | 34 | 12 | 20 | 16 | 32 | 27 | 34 | 23.42 | 16 | 31 | 1.953 | 0.966 | 0.01894597 | 0.06645269 | 0 | 0.1036 | 0.0748 | 0.0748 | TRUE | 0.3521 |
| Hesx1 | 90 | 59 | 69 | 21 | 20 | 28 | 70 | 63 | 74 | 21 | 23 | 28 | 46.37 | 69 | 24 | 0.344 | -1.541 | 3.53E-07 | 1.21E-05 | 0 | 0.0535 | 0.0372 | 0.0372 | TRUE | 0.131 |
| Hexa | 2549 | 2005 | 2600 | 3144 | 2766 | 3315 | 1980 | 2133 | 2805 | 3130 | 3120 | 3272 | 2740.01 | 2306 | 3174 | 1.378 | 0.462 | 1.19E-05 | 0.00021625 | 0.0179 | 0.0033 | 0.0077 | 0.0077 | TRUE | 0.4336 |
| Hexb | 4150 | 3270 | 4132 | 4699 | 4314 | 5184 | 3223 | 3478 | 4458 | 4677 | 4867 | 5117 | 4303.53 | 3720 | 4887 | 1.314 | 0.394 | 8.05E-05 | 0.001015 | 0.016 | 0.003 | 0.007 | 0.007 | TRUE | 0.3662 |
| Hexdc | 692 | 508 | 443 | 528 | 446 | 508 | 537 | 540 | 478 | 526 | 503 | 501 | 514.33 | 518 | 510 | 0.983 | -0.025 | 0.79687135 | 0.88159845 | 0 | 0.007 | 0.005 | 0.005 | TRUE | 0.0528 |
| Hexim1 | 4170 | 3005 | 2319 | 2171 | 2011 | 2261 | 3239 | 3196 | 2502 | 2161 | 2269 | 2232 | 2599.8 | 2979 | 2221 | 0.745 | -0.425 | 3.74E-06 | 8.37E-05 | 0.0093 | 0.0033 | 0.0057 | 0.0057 | TRUE | 0.2365 |
| Hexim2 | 472 | 363 | 344 | 381 | 359 | 382 | 367 | 386 | 371 | 379 | 405 | 377 | 380.87 | 375 | 387 | 1.033 | 0.047 | 0.65709801 | 0.78749824 | 0 | 0.0086 | 0.0056 | 0.0056 | TRUE | 0.0181 |

| | | | | | | | | | | | | | | | | | | | | | | | | | |
|---|---|---|---|---|---|---|---|---|---|---|---|---|---|---|---|---|---|---|---|---|---|---|---|---|---|
| Hey1 | 271 | 253 | 248 | 226 | 232 | 274 | 210 | 269 | 268 | 225 | 262 | 270 | 250.72 | 249 | 252 | 1.017 | 0.025 | 0.86757215 | 0.92684288 | 0.0108 | 0.0119 | 0.0116 | 0.0116 | TRUE | 0.2214 |
| Hey2 | 69 | 55 | 65 | 69 | 75 | 80 | 54 | 59 | 70 | 69 | 85 | 79 | 69.08 | 61 | 78 | 1.279 | 0.355 | 0.13898793 | 0.28023058 | 0 | 0.0367 | 0.0268 | 0.0268 | TRUE | 0.1529 |
| Heyl | 4 | 3 | 3 | 2 | 0 | 2 | 3 | 3 | 3 | 2 | 0 | 2 | 2.25 | 3 | 1 | 0.425 | -1.233 | 0.35005735 | NA | 0 | 1.0558 | 0.7197 | 0.7197 | TRUE | 0.392 |
| Hfe | 43 | 24 | 21 | 22 | 20 | 26 | 33 | 26 | 23 | 22 | 23 | 26 | 25.29 | 27 | 24 | 0.853 | -0.229 | 0.55124407 | 0.70713799 | 0 | 0.0962 | 0.067 | 0.067 | TRUE | 0.3008 |
| Hfe2 | 149 | 92 | 79 | 107 | 87 | 94 | 116 | 98 | 85 | 107 | 98 | 93 | 99.38 | 100 | 99 | 0.99 | -0.015 | 0.94351874 | 0.96859249 | 0 | 0.0263 | 0.0203 | 0.0203 | TRUE | 0.2167 |
| Hfm1 | 163 | 148 | 115 | 172 | 100 | 116 | 127 | 157 | 124 | 171 | 113 | 115 | 134.44 | 136 | 133 | 0.98 | -0.03 | 0.88475163 | 0.93609243 | 0.0305 | 0.0201 | 0.0231 | 0.0231 | TRUE | 0.663 |
| Hgd | 3 | 0 | 1 | 2 | 0 | 1 | 2 | 0 | 1 | 2 | 0 | 1 | 1.06 | 1 | 1 | 0.858 | -0.222 | 0.90857918 | NA | 0 | 2.2289 | 1.7585 | 1.7585 | TRUE | 0.2274 |
| Hgf | 4 | 0 | 2 | 3 | 0 | 3 | 3 | 0 | 2 | 3 | 0 | 3 | 1.87 | 2 | 2 | 1.117 | 0.16 | 0.91656777 | NA | 0 | 1.2707 | 1.1556 | 1.1556 | TRUE | 0.5908 |
| Hgfac | 2 | 2 | 6 | 0 | 1 | 0 | 2 | 2 | 6 | 0 | 1 | 0 | 1.88 | 3 | 0 | 0.111 | -3.172 | 0.06811502 | NA | 0 | 1.2628 | 1.0334 | 1.0334 | TRUE | 0.8653 |
| Hgs | 5826 | 4356 | 4303 | 4189 | 3844 | 4454 | 4525 | 4633 | 4643 | 4170 | 4336 | 4397 | 4450.68 | 4600 | 4301 | 0.935 | -0.097 | 0.06630409 | 0.16472872 | 3.00E-04 | 0.003 | 0.0018 | 0.0018 | TRUE | 0.0087 |
| Hgsnat | 2208 | 1668 | 1844 | 1881 | 1657 | 1893 | 1715 | 1774 | 1990 | 1872 | 1869 | 1869 | 1848.17 | 1826 | 1870 | 1.025 | 0.036 | 0.62528597 | 0.76373375 | 0.0025 | 0.0037 | 0.0033 | 0.0033 | TRUE | 0.0733 |
| Hhat | 93 | 53 | 64 | 78 | 76 | 85 | 72 | 56 | 69 | 78 | 86 | 84 | 74.16 | 66 | 83 | 1.246 | 0.317 | 0.16439337 | 0.31522329 | 0 | 0.0344 | 0.0239 | 0.0239 | TRUE | 0.135 |
| Hhatl | 0 | 0 | 0 | 1 | 0 | 0 | 0 | 0 | 0 | 1 | 0 | 0 | 0.17 | 0 | 0 | 2.094 | 1.067 | 0.79379329 | NA | 0 | 14.2885 | 10 | 10 | TRUE | 0.3906 |
| Hhex | 467 | 396 | 412 | 406 | 346 | 403 | 363 | 421 | 445 | 404 | 390 | 398 | 403.46 | 410 | 397 | 0.974 | -0.038 | 0.73579733 | 0.84235092 | 0.0037 | 0.0083 | 0.0068 | 0.0068 | TRUE | 0.1238 |
| Hhip | 27 | 20 | 9 | 12 | 7 | 6 | 21 | 21 | 10 | 12 | 8 | 6 | 12.95 | 17 | 9 | 0.493 | -1.019 | 0.07872823 | NA | 0.07 | 0.1854 | 0.1543 | 0.1543 | TRUE | 0.6797 |
| Hhipl1 | 22 | 16 | 8 | 36 | 19 | 18 | 17 | 17 | 9 | 36 | 21 | 18 | 19.63 | 14 | 25 | 1.744 | 0.802 | 0.09448257 | 0.21206649 | 0.081 | 0.1232 | 0.1115 | 0.1115 | TRUE | 0.8909 |
| Hhipl2 | 12 | 7 | 8 | 12 | 6 | 10 | 9 | 7 | 9 | 12 | 7 | 10 | 9 | 8 | 10 | 1.126 | 0.171 | 0.78587592 | NA | 0 | 0.2659 | 0.1762 | 0.1762 | TRUE | 0.1908 |
| Hhla1 | 1 | 0 | 1 | 0 | 0 | 1 | 1 | 0 | 1 | 0 | 0 | 1 | 0.47 | 1 | 0 | 0.596 | -0.748 | 0.78959815 | NA | 0 | 5.0047 | 3.8792 | 3.8792 | TRUE | 0.3835 |
| Hiat1 | 6476 | 4718 | 4734 | 4588 | 3988 | 4702 | 5030 | 5018 | 5108 | 4567 | 4499 | 4642 | 4810.57 | 5052 | 4569 | 0.905 | -0.145 | 0.00381928 | 0.0198722 | 0 | 0.0029 | 0.0016 | 0.0016 | TRUE | 0.0023 |
| Hiatl1 | 3965 | 3020 | 3123 | 3260 | 2853 | 3233 | 3079 | 3212 | 3370 | 3245 | 3218 | 3191 | 3219.4 | 3220 | 3218 | 1 | 0 | 0.99771382 | 0.99869204 | 8.00E-04 | 0.0032 | 0.0022 | 0.0022 | TRUE | 0.0199 |
| Hibadh | 4504 | 3511 | 4293 | 4973 | 4403 | 4994 | 3498 | 3735 | 4632 | 4950 | 4967 | 4930 | 4451.96 | 3955 | 4949 | 1.252 | 0.324 | 0.00030968 | 0.00298466 | 0.0111 | 0.003 | 0.0056 | 0.0056 | TRUE | 0.2724 |
| Hibch | 1876 | 1284 | 1351 | 1763 | 1563 | 1742 | 1457 | 1366 | 1458 | 1755 | 1763 | 1720 | 1586.37 | 1427 | 1746 | 1.223 | 0.29 | 1.02E-05 | 0.00019222 | 0 | 0.0039 | 0.0025 | 0.0025 | TRUE | 0.0165 |
| Hic1 | 250 | 186 | 167 | 217 | 200 | 192 | 194 | 198 | 180 | 216 | 226 | 190 | 200.56 | 191 | 211 | 1.101 | 0.139 | 0.33885439 | 0.51299541 | 0 | 0.0142 | 0.0102 | 0.0102 | TRUE | 0.0833 |
| Hic2 | 5019 | 3642 | 3305 | 3287 | 3068 | 3490 | 3898 | 3874 | 3566 | 3272 | 3461 | 3445 | 3586.02 | 3779 | 3393 | 0.897 | -0.156 | 0.01055945 | 0.04296222 | 0.0013 | 0.0031 | 0.0024 | 0.0024 | TRUE | 0.0293 |
| Hid1 | 798 | 670 | 655 | 704 | 619 | 739 | 620 | 713 | 707 | 701 | 698 | 730 | 694.62 | 680 | 710 | 1.046 | 0.065 | 0.47523563 | 0.64341602 | 0.0022 | 0.0058 | 0.0046 | 0.0046 | TRUE | 0.0747 |
| Hif1a | 8326 | 5567 | 4800 | 5406 | 4613 | 5087 | 6466 | 5922 | 5179 | 5381 | 5204 | 5022 | 5528.98 | 5856 | 5202 | 0.888 | -0.171 | 0.02867632 | 0.08962839 | 0.0064 | 0.0029 | 0.0042 | 0.0042 | TRUE | 0.1243 |
| Hif1an | 5241 | 3815 | 3768 | 3995 | 3541 | 3994 | 4070 | 4058 | 4066 | 3977 | 3995 | 3943 | 4018 | 4065 | 3972 | 0.977 | -0.034 | 0.5055148 | 0.66894945 | 0 | 0.003 | 0.0016 | 0.0016 | TRUE | 5.00E-04 |
| Hif3a | 937 | 836 | 1128 | 1460 | 1392 | 1634 | 728 | 889 | 1217 | 1453 | 1570 | 1613 | 1245.11 | 945 | 1545 | 1.637 | 0.711 | 0.00142698 | 0.00931319 | 0.035 | 0.0043 | 0.0129 | 0.035 | TRUE | 0.7578 |
| Higd1a | 2547 | 1922 | 2060 | 1985 | 1778 | 2050 | 1978 | 2044 | 2223 | 1976 | 2006 | 2024 | 2041.76 | 2082 | 2002 | 0.962 | -0.055 | 0.40947916 | 0.58224091 | 0.0015 | 0.0036 | 0.0028 | 0.0028 | TRUE | 0.0421 |
| Higd1b | 2 | 2 | 1 | 3 | 3 | 3 | 2 | 2 | 1 | 3 | 3 | 3 | 2.35 | 2 | 3 | 1.959 | 0.97 | 0.42894269 | NA | 0 | 1.0116 | 0.6133 | 0.6133 | TRUE | 0.0556 |
| Higd1c | 0 | 0 | 0 | 0 | 0 | 0 | 0 | 0 | 0 | 0 | 0 | 0 | 0 | 0 | 0 | NA | NA | NA | NA | NA | NA | NA | NA | NA | NA |
| Higd2a | 2426 | 1747 | 2114 | 2384 | 2239 | 2628 | 1884 | 1858 | 2281 | 2373 | 2526 | 2594 | 2252.75 | 2008 | 2498 | 1.245 | 0.316 | 0.00020602 | 0.00217956 | 0.0073 | 0.0035 | 0.0048 | 0.0048 | TRUE | 0.1696 |
| Hilpda | 460 | 341 | 317 | 392 | 282 | 334 | 357 | 363 | 342 | 390 | 318 | 330 | 350.01 | 354 | 346 | 0.978 | -0.032 | 0.78910347 | 0.87619699 | 0.0035 | 0.0092 | 0.0075 | 0.0075 | TRUE | 0.1435 |
| Hils1 | 1 | 2 | 0 | 1 | 0 | 0 | 1 | 2 | 0 | 1 | 0 | 0 | 0.65 | 1 | 0 | 0.384 | -1.379 | 0.58322298 | NA | 0 | 3.6492 | 3.0509 | 3.0509 | TRUE | 0.2622 |
| Hinfp | 936 | 735 | 713 | 796 | 610 | 746 | 727 | 782 | 769 | 792 | 688 | 736 | 749.16 | 759 | 739 | 0.975 | -0.037 | 0.67825727 | 0.80231161 | 0.0018 | 0.0056 | 0.0043 | 0.0043 | TRUE | 0.0472 |
| Hint1 | 11300 | 7859 | 7970 | 8815 | 7313 | 8650 | 8776 | 8360 | 8600 | 8774 | 8250 | 8539 | 8549.74 | 8579 | 8521 | 0.993 | -0.01 | 0.84920808 | 0.91474347 | 5.00E-04 | 0.0027 | 0.0018 | 0.0018 | TRUE | 0.0094 |
| Hint2 | 807 | 557 | 628 | 656 | 602 | 632 | 627 | 592 | 678 | 653 | 679 | 624 | 642.14 | 632 | 652 | 1.031 | 0.044 | 0.63409356 | 0.7710651 | 0.0014 | 0.0061 | 0.0046 | 0.0046 | TRUE | 0.0441 |
| Hint3 | 345 | 272 | 284 | 306 | 258 | 287 | 268 | 289 | 306 | 305 | 291 | 283 | 290.44 | 288 | 293 | 1.02 | 0.029 | 0.81445455 | 0.89319411 | 0 | 0.0106 | 0.0073 | 0.0073 | TRUE | 0.0451 |
| Hip1 | 2355 | 1675 | 1549 | 1458 | 1398 | 1449 | 1829 | 1782 | 1671 | 1451 | 1577 | 1430 | 1623.47 | 1761 | 1486 | 0.843 | -0.246 | 0.00064307 | 0.0052283 | 0.0018 | 0.0039 | 0.0031 | 0.0031 | TRUE | 0.0339 |
| Hip1r | 2533 | 2045 | 2104 | 2205 | 2211 | 2588 | 1967 | 2175 | 2270 | 2195 | 2494 | 2555 | 2276.09 | 2137 | 2415 | 1.13 | 0.177 | 0.02794979 | 0.08795383 | 0.0052 | 0.0035 | 0.0042 | 0.0042 | TRUE | 0.0773 |
| Hipk1 | 3931 | 2793 | 2810 | 3193 | 2815 | 3212 | 3053 | 2971 | 3032 | 3178 | 3176 | 3171 | 3096.76 | 3019 | 3175 | 1.052 | 0.073 | 0.17563676 | 0.3291865 | 0 | 0.0032 | 0.0017 | 0.0017 | TRUE | 0.0023 |
| Hipk2 | 1163 | 793 | 722 | 683 | 585 | 680 | 903 | 844 | 779 | 680 | 660 | 671 | 756.14 | 842 | 670 | 0.795 | -0.331 | 0.00018259 | 0.0019938 | 0.0019 | 0.0056 | 0.0043 | 0.0043 | TRUE | 0.0505 |
| Hipk3 | 1578 | 1178 | 1292 | 1206 | 1039 | 1218 | 1226 | 1253 | 1394 | 1200 | 1172 | 1202 | 1241.26 | 1291 | 1191 | 0.924 | -0.114 | 0.13708286 | 0.27736768 | 0.0018 | 0.0043 | 0.0034 | 0.0034 | TRUE | 0.0579 |
| Hipk4 | 37 | 38 | 19 | 41 | 22 | 30 | 29 | 40 | 21 | 41 | 25 | 30 | 30.82 | 30 | 32 | 1.066 | 0.092 | 0.80827796 | 0.8891493 | 0.0517 | 0.0793 | 0.0719 | 0.0719 | TRUE | 0.2963 |
| Hira | 7493 | 5128 | 4239 | 4628 | 4041 | 4664 | 5819 | 5455 | 4574 | 4607 | 4559 | 4604 | 4936.23 | 5283 | 4590 | 0.868 | -0.203 | 0.01218496 | 0.04787235 | 0.0071 | 0.0029 | 0.0045 | 0.0045 | TRUE | 0.1674 |
| Hirip3 | 4436 | 3442 | 3104 | 2553 | 2440 | 2931 | 3445 | 3661 | 3349 | 2541 | 2753 | 2893 | 3107.14 | 3485 | 2729 | 0.783 | -0.353 | 2.51E-07 | 8.92E-06 | 0.0027 | 0.0032 | 0.003 | 0.003 | TRUE | 0.0444 |
| Hist1h1a | 51 | 39 | 29 | 39 | 34 | 28 | 40 | 41 | 31 | 39 | 38 | 28 | 36.2 | 37 | 35 | 0.929 | -0.106 | 0.74516202 | 0.84852395 | 0 | 0.0679 | 0.0495 | 0.0495 | TRUE | 0.2449 |
| Hist1h1b | 14 | 9 | 5 | 8 | 8 | 14 | 11 | 10 | 5 | 8 | 9 | 14 | 9.44 | 9 | 10 | 1.182 | 0.241 | 0.70346046 | NA | 0 | 0.2534 | 0.1834 | 0.1834 | TRUE | 0.3355 |
| Hist1h1c | 211 | 145 | 125 | 128 | 101 | 114 | 164 | 154 | 135 | 127 | 114 | 113 | 134.48 | 151 | 118 | 0.779 | -0.36 | 0.03988708 | 0.1142955 | 0 | 0.0201 | 0.0145 | 0.0145 | TRUE | 0.0887 |
| Hist1h1d | 19 | 11 | 14 | 14 | 20 | 9 | 15 | 12 | 15 | 14 | 23 | 9 | 14.49 | 14 | 15 | 1.081 | 0.113 | 0.83092873 | NA | 0.0417 | 0.166 | 0.1323 | 0.1323 | TRUE | 0.3515 |
| Hist1h1e | 15 | 9 | 7 | 13 | 7 | 11 | 12 | 10 | 8 | 13 | 8 | 11 | 10.08 | 10 | 11 | 1.096 | 0.132 | 0.82526726 | NA | 0 | 0.2376 | 0.1609 | 0.1609 | TRUE | 0.1567 |
| Hist1h1t | 9 | 4 | 1 | 1 | 1 | 1 | 7 | 4 | 1 | 1 | 1 | 1 | 2.57 | 4 | 1 | 0.247 | -2.016 | 0.13380856 | NA | 0 | 0.9238 | 0.6879 | 0.6879 | TRUE | 0.0898 |
| Hist1h2aa | 2 | 3 | 0 | 0 | 0 | 0 | 2 | 3 | 0 | 0 | 0 | 0 | 0.79 | 2 | 0 | 0.12 | -3.055 | 0.22090216 | NA | 5.3026 | 2.9997 | 3.1827 | 3.1827 | TRUE | 0.0516 |
| Hist1h2ab | 3 | 9 | 3 | 9 | 5 | 14 | 2 | 10 | 3 | 9 | 6 | 14 | 7.26 | 5 | 10 | 1.928 | 0.947 | 0.22881153 | NA | 0.2013 | 0.3289 | 0.2941 | 0.2941 | TRUE | 0.6059 |
| Hist1h2ac | 9 | 8 | 7 | 9 | 5 | 5 | 7 | 9 | 8 | 9 | 6 | 5 | 7.1 | 8 | 7 | 0.852 | -0.231 | 0.74492103 | NA | 0 | 0.3363 | 0.2211 | 0.2211 | TRUE | 0.2799 |
| Hist1h2ad | 7 | 1 | 2 | 2 | 1 | 1 | 5 | 1 | 2 | 2 | 1 | 1 | 2.13 | 3 | 1 | 0.463 | -1.11 | 0.41265486 | NA | 0 | 1.1165 | 0.783 | 0.783 | TRUE | 0.8892 |
| Hist1h2ae | 56 | 38 | 22 | 25 | 39 | 29 | 43 | 40 | 24 | 25 | 44 | 29 | 34.19 | 36 | 33 | 0.896 | -0.159 | 0.66750483 | 0.79551793 | 0.063 | 0.0717 | 0.0694 | 0.0694 | TRUE | 0.6148 |
| Hist1h2af | 30 | 26 | 24 | 17 | 17 | 21 | 23 | 28 | 26 | 17 | 19 | 21 | 22.28 | 26 | 19 | 0.742 | -0.431 | 0.28462228 | 0.45655853 | 0 | 0.1088 | 0.0706 | 0.0706 | TRUE | 0.0476 |
| Hist1h2ag | 9 | 4 | 4 | 8 | 6 | 7 | 7 | 4 | 4 | 8 | 7 | 7 | 6.2 | 5 | 7 | 1.371 | 0.455 | 0.54742963 | NA | 0 | 0.3847 | 0.2488 | 0.2488 | TRUE | 0.2555 |
| Hist1h2ah | 5 | 12 | 3 | 7 | 5 | 7 | 4 | 13 | 3 | 7 | 6 | 7 | 6.57 | 7 | 7 | 0.997 | -0.005 | 0.99508016 | NA | 0.1554 | 0.3633 | 0.3017 | 0.3017 | TRUE | 1.5707 |
| Hist1h2ai | 7 | 9 | 7 | 4 | 9 | 9 | 5 | 10 | 8 | 4 | 10 | 9 | 7.6 | 8 | 8 | 1.026 | 0.037 | 0.95871302 | NA | 0 | 0.3144 | 0.2304 | 0.2304 | TRUE | 0.5189 |
| Hist1h2ak | 15 | 9 | 9 | 7 | 7 | 13 | 12 | 10 | 10 | 7 | 8 | 13 | 9.77 | 11 | 9 | 0.894 | -0.162 | 0.79073178 | NA | 0 | 0.245 | 0.1672 | 0.1672 | TRUE | 0.3958 |
| Hist1h2an | 3 | 2 | 0 | 0 | 1 | 2 | 2 | 2 | 0 | 0 | 1 | 2 | 1.26 | 1 | 1 | 0.683 | -0.55 | 0.7607184 | NA | 0 | 1.8835 | 1.5306 | 1.5306 | TRUE | 0.4181 |
| Hist1h2ao | 0 | 0 | 0 | 0 | 0 | 0 | 0 | 0 | 0 | 0 | 0 | 0 | 0 | 0 | 0 | NA | NA | NA | NA | NA | NA | NA | NA | NA | NA |
| Hist1h2ap | 0 | 0 | 0 | 0 | 0 | 0 | 0 | 0 | 0 | 0 | 0 | 0 | 0 | 0 | 0 | NA | NA | NA | NA | NA | NA | NA | NA | NA | NA |
| Hist1h2ba | 1 | 1 | 0 | 0 | 0 | 1 | 1 | 1 | 0 | 0 | 0 | 1 | 0.47 | 1 | 0 | 0.599 | -0.74 | 0.79209856 | NA | 0 | 5.0317 | 3.8925 | 3.8925 | TRUE | 0.3835 |
| Hist1h2bb | 4 | 7 | 1 | 3 | 4 | 8 | 3 | 7 | 1 | 3 | 5 | 8 | 4.5 | 4 | 5 | 1.335 | 0.416 | 0.664908 | NA | 0.2033 | 0.5286 | 0.4399 | 0.4399 | TRUE | 0.6493 |
| Hist1h2bc | 93 | 46 | 39 | 61 | 43 | 54 | 72 | 49 | 42 | 61 | 49 | 53 | 54.3 | 54 | 54 | 0.987 | -0.019 | 0.94770835 | 0.97142825 | 0.0316 | 0.0461 | 0.0418 | 0.0418 | TRUE | 0.7723 |
| Hist1h2be | 6 | 9 | 9 | 6 | 7 | 4 | 5 | 10 | 10 | 6 | 8 | 4 | 6.96 | 8 | 6 | 0.75 | -0.414 | 0.57634346 | NA | 0 | 0.3429 | 0.2467 | 0.2467 | TRUE | 0.5156 |

| | | | | | | | | | | | | | | | | | | | | | | | | | |
|---|---|---|---|---|---|---|---|---|---|---|---|---|---|---|---|---|---|---|---|---|---|---|---|---|---|
| Hist1h2bf | 3 | 3 | 5 | 4 | 2 | 6 | 2 | 3 | 5 | 4 | 2 | 6 | 3.85 | 3 | 4 | 1.14 | 0.189 | 0.84661831 | NA | 0 | 0.6186 | 0.4246 | 0.4246 | TRUE | 0.2009 |
| Hist1h2bg | 24 | 15 | 13 | 17 | 12 | 20 | 19 | 16 | 14 | 17 | 14 | 20 | 16.47 | 16 | 17 | 1.03 | 0.042 | 0.92846416 | NA | 0 | 0.1463 | 0.0984 | 0.0984 | TRUE | 0.1227 |
| Hist1h2bh | 9 | 3 | 3 | 4 | 5 | 1 | 7 | 3 | 3 | 4 | 6 | 1 | 4 | 4 | 4 | 0.768 | -0.381 | 0.70187737 | NA | 0 | 0.5943 | 0.4591 | 0.4591 | TRUE | 0.5244 |
| Hist1h2bj | 12 | 15 | 11 | 20 | 12 | 20 | 9 | 16 | 12 | 20 | 14 | 20 | 15.06 | 12 | 18 | 1.452 | 0.538 | 0.28555359 | NA | 0 | 0.1598 | 0.1144 | 0.1144 | TRUE | 0.249 |
| Hist1h2bk | 16 | 11 | 6 | 7 | 8 | 9 | 12 | 12 | 6 | 7 | 9 | 9 | 9.25 | 10 | 8 | 0.805 | -0.313 | 0.62028378 | NA | 0 | 0.2588 | 0.1779 | 0.1779 | TRUE | 0.3287 |
| Hist1h2bl | 5 | 6 | 4 | 9 | 6 | 6 | 4 | 6 | 4 | 9 | 7 | 6 | 6.04 | 5 | 7 | 1.499 | 0.584 | 0.44926061 | NA | 0 | 0.3949 | 0.2579 | 0.2579 | TRUE | 0.1391 |
| Hist1h2bm | 0 | 1 | 3 | 4 | 4 | 3 | 0 | 1 | 3 | 4 | 5 | 3 | 2.63 | 1 | 4 | 2.813 | 1.492 | 0.24816847 | NA | 0 | 0.905 | 0.7118 | 0.7118 | TRUE | 0.772 |
| Hist1h2bn | 2 | 1 | 1 | 4 | 3 | 1 | 2 | 1 | 1 | 4 | 3 | 1 | 2.01 | 1 | 3 | 2.221 | 1.151 | 0.39950538 | NA | 0 | 1.1826 | 0.774 | 0.774 | TRUE | 0.4055 |
| Hist1h2bp | 13 | 11 | 13 | 12 | 8 | 7 | 10 | 12 | 14 | 12 | 9 | 7 | 10.62 | 12 | 9 | 0.783 | -0.353 | 0.54823324 | NA | 0 | 0.2257 | 0.1527 | 0.1527 | TRUE | 0.2141 |
| Hist1h2bq | 0 | 0 | 0 | 0 | 0 | 0 | 0 | 0 | 0 | 0 | 0 | 0 | 0 | 0 | 0 | NA | NA | NA | NA | NA | NA | NA | NA | NA | NA |
| Hist1h2br | 0 | 0 | 0 | 0 | 0 | 0 | 0 | 0 | 0 | 0 | 0 | 0 | 0 | 0 | 0 | NA | NA | NA | NA | NA | NA | NA | NA | NA | NA |
| Hist1h3a | 14 | 19 | 10 | 12 | 6 | 5 | 11 | 20 | 11 | 12 | 7 | 5 | 10.92 | 14 | 8 | 0.569 | -0.813 | 0.19193311 | NA | 0.0747 | 0.2195 | 0.1791 | 0.1791 | TRUE | 0.6348 |
| Hist1h3b | 4 | 4 | 3 | 2 | 2 | 5 | 3 | 4 | 3 | 2 | 2 | 5 | 3.3 | 3 | 3 | 0.875 | -0.193 | 0.852462 | NA | 0 | 0.7213 | 0.4691 | 0.4691 | TRUE | 0.3944 |
| Hist1h3c | 44 | 42 | 40 | 44 | 32 | 32 | 34 | 45 | 43 | 44 | 36 | 32 | 38.92 | 41 | 37 | 0.919 | -0.121 | 0.70054043 | 0.81850931 | 0 | 0.0633 | 0.046 | 0.046 | TRUE | 0.1837 |
| Hist1h3d | 14 | 5 | 6 | 3 | 10 | 9 | 11 | 5 | 6 | 3 | 11 | 9 | 7.64 | 7 | 8 | 1 | 0 | 0.99988115 | NA | 0.1211 | 0.3128 | 0.2604 | 0.2604 | TRUE | 0.7185 |
| Hist1h3e | 12 | 21 | 14 | 14 | 10 | 24 | 9 | 22 | 15 | 14 | 11 | 24 | 15.95 | 15 | 16 | 1.064 | 0.09 | 0.86549234 | NA | 0.1087 | 0.1511 | 0.1388 | 0.1388 | TRUE | 0.2897 |
| Hist1h3f | 3 | 10 | 7 | 6 | 3 | 9 | 2 | 11 | 8 | 6 | 3 | 9 | 6.46 | 7 | 6 | 0.914 | -0.129 | 0.87392985 | NA | 0.2075 | 0.3693 | 0.3221 | 0.3221 | TRUE | 0.4256 |
| Hist1h3g | 28 | 17 | 9 | 14 | 16 | 11 | 22 | 18 | 10 | 14 | 18 | 11 | 15.4 | 17 | 14 | 0.852 | -0.231 | 0.65344914 | NA | 0.0409 | 0.1564 | 0.1259 | 0.1259 | TRUE | 0.5304 |
| Hist1h3h | 6 | 8 | 4 | 9 | 5 | 13 | 5 | 9 | 4 | 9 | 6 | 13 | 7.49 | 6 | 9 | 1.59 | 0.669 | 0.3535672 | NA | 0 | 0.319 | 0.2355 | 0.2355 | TRUE | 0.1705 |
| Hist1h3i | 6 | 5 | 5 | 5 | 6 | 9 | 5 | 5 | 5 | 5 | 7 | 9 | 6 | 5 | 7 | 1.348 | 0.431 | 0.57527473 | NA | 0 | 0.3974 | 0.2567 | 0.2567 | TRUE | 0.1796 |
| Hist1h4a | 10 | 7 | 9 | 3 | 4 | 6 | 8 | 7 | 10 | 3 | 5 | 6 | 6.39 | 8 | 5 | 0.54 | -0.889 | 0.24381073 | NA | 0 | 0.3733 | 0.2454 | 0.2454 | TRUE | 0.1642 |
| Hist1h4b | 21 | 19 | 16 | 13 | 18 | 12 | 16 | 20 | 17 | 13 | 20 | 12 | 16.48 | 18 | 15 | 0.836 | -0.259 | 0.5884245 | NA | 0 | 0.1462 | 0.103 | 0.103 | TRUE | 0.3908 |
| Hist1h4c | 4 | 2 | 2 | 2 | 2 | 1 | 3 | 2 | 2 | 2 | 2 | 1 | 2.1 | 2 | 2 | 0.697 | -0.521 | 0.68400953 | NA | 0 | 1.1287 | 0.6894 | 0.6894 | TRUE | 0.1192 |
| Hist1h4d | 26 | 20 | 22 | 11 | 8 | 23 | 20 | 21 | 24 | 11 | 9 | 23 | 17.98 | 22 | 14 | 0.66 | -0.599 | 0.21857666 | 0.381688 | 0.0454 | 0.1342 | 0.1122 | 0.1122 | TRUE | 1.2244 |
| Hist1h4f | 2 | 3 | 1 | 3 | 5 | 6 | 2 | 3 | 1 | 3 | 6 | 6 | 3.4 | 2 | 5 | 2.518 | 1.332 | 0.21486089 | NA | 0 | 0.7004 | 0.4742 | 0.4742 | TRUE | 0.2482 |
| Hist1h4h | 13 | 6 | 6 | 9 | 7 | 8 | 10 | 6 | 6 | 9 | 8 | 8 | 7.95 | 7 | 8 | 1.065 | 0.091 | 0.89148377 | NA | 0 | 0.3005 | 0.1965 | 0.1965 | TRUE | 0.2881 |
| Hist1h4i | 214 | 144 | 61 | 90 | 66 | 64 | 166 | 153 | 66 | 90 | 74 | 63 | 102.07 | 128 | 76 | 0.588 | -0.767 | 0.00947198 | 0.03982292 | 0.1198 | 0.0256 | 0.0524 | 0.0524 | TRUE | 1.8064 |
| Hist1h4j | 3 | 2 | 3 | 1 | 7 | 3 | 2 | 2 | 3 | 1 | 8 | 3 | 3.26 | 2 | 4 | 1.526 | 0.61 | 0.58307599 | NA | 0 | 0.7299 | 0.568 | 0.568 | TRUE | 0.5149 |
| Hist1h4k | 11 | 16 | 13 | 18 | 9 | 9 | 9 | 17 | 14 | 18 | 10 | 9 | 12.76 | 13 | 12 | 0.947 | -0.079 | 0.88952132 | NA | 0.0619 | 0.1882 | 0.1519 | 0.1519 | TRUE | 0.6506 |
| Hist1h4m | 0 | 0 | 0 | 0 | 0 | 0 | 0 | 0 | 0 | 0 | 0 | 0 | 0 | 0 | 0 | NA | NA | NA | NA | NA | NA | NA | NA | NA | NA |
| Hist1h4n | 10 | 7 | 6 | 7 | 8 | 8 | 8 | 7 | 6 | 7 | 9 | 8 | 7.6 | 7 | 8 | 1.096 | 0.132 | 0.8456151 | NA | 0 | 0.3144 | 0.1978 | 0.1978 | TRUE | 0.0355 |
| Hist2h2aa1 | 0 | 0 | 0 | 0 | 0 | 0 | 0 | 0 | 0 | 0 | 0 | 0 | 0 | 0 | 0 | NA | NA | NA | NA | NA | NA | NA | NA | NA | NA |
| Hist2h2aa2 | 0 | 1 | 0 | 0 | 0 | 0 | 0 | 1 | 0 | 0 | 0 | 0 | 0.18 | 0 | 0 | 0.552 | -0.857 | 0.8336548 | NA | 0 | 13.3714 | 10 | 10 | TRUE | 0.5216 |
| Hist2h2ab | 0 | 0 | 0 | 0 | 0 | 1 | 0 | 0 | 0 | 0 | 0 | 1 | 0.16 | 0 | 0 | 2.094 | 1.067 | 0.79379329 | NA | 0 | 14.4078 | 10 | 10 | TRUE | 0.3835 |
| Hist2h2ac | 3 | 4 | 0 | 1 | 1 | 1 | 2 | 4 | 0 | 1 | 1 | 1 | 1.62 | 2 | 1 | 0.47 | -1.089 | 0.49461224 | NA | 0 | 1.4691 | 1.1128 | 1.1128 | TRUE | 0.1272 |
| Hist2h2bb | 12 | 8 | 11 | 8 | 4 | 11 | 9 | 9 | 12 | 8 | 5 | 11 | 8.84 | 10 | 8 | 0.794 | -0.334 | 0.60864326 | NA | 0 | 0.2706 | 0.1909 | 0.1909 | TRUE | 0.2517 |
| Hist2h2be | 44 | 19 | 32 | 26 | 33 | 27 | 34 | 20 | 35 | 26 | 37 | 27 | 29.78 | 30 | 30 | 1.001 | 0.002 | 0.99668306 | 0.99800974 | 0.0257 | 0.082 | 0.0671 | 0.0671 | TRUE | 0.4799 |
| Hist2h3b | 5 | 4 | 2 | 3 | 4 | 3 | 4 | 4 | 2 | 3 | 5 | 3 | 3.46 | 3 | 4 | 1.005 | 0.007 | 0.99468822 | NA | 0 | 0.6875 | 0.4357 | 0.4357 | TRUE | 0.1369 |
| Hist2h3c1 | 19 | 11 | 20 | 16 | 9 | 16 | 15 | 12 | 22 | 16 | 10 | 16 | 14.99 | 16 | 14 | 0.879 | -0.187 | 0.71337579 | NA | 0 | 0.1606 | 0.1193 | 0.1193 | TRUE | 0.4005 |
| Hist2h3c2 | 6 | 4 | 4 | 5 | 1 | 3 | 5 | 4 | 4 | 5 | 1 | 3 | 3.72 | 4 | 3 | 0.692 | -0.532 | 0.59286423 | NA | 0 | 0.6402 | 0.4339 | 0.4339 | TRUE | 0.2768 |
| Hist2h4 | 3 | 2 | 2 | 1 | 0 | 1 | 2 | 2 | 2 | 1 | 0 | 1 | 1.43 | 2 | 1 | 0.307 | -1.705 | 0.30957199 | NA | 0 | 1.6564 | 1.0462 | 1.0462 | TRUE | 0.1985 |
| Hist3h2a | 44 | 22 | 24 | 29 | 24 | 33 | 34 | 23 | 26 | 29 | 27 | 33 | 28.66 | 28 | 30 | 1.053 | 0.074 | 0.83768612 | 0.90746155 | 0 | 0.0851 | 0.0598 | 0.0598 | TRUE | 0.3057 |
| Hist3h2ba | 46 | 37 | 26 | 28 | 28 | 31 | 36 | 39 | 28 | 28 | 32 | 31 | 32.2 | 34 | 30 | 0.871 | -0.199 | 0.55727625 | 0.7122577 | 0 | 0.076 | 0.0518 | 0.0518 | TRUE | 0.1669 |
| Hist3h2bb-ps | 0 | 1 | 0 | 0 | 0 | 0 | 0 | 1 | 0 | 0 | 0 | 0 | 0.18 | 0 | 0 | 0.552 | -0.857 | 0.8336548 | NA | 0 | 13.3714 | 10 | 10 | TRUE | 0.5216 |
| Hist4h4 | 15 | 9 | 12 | 8 | 5 | 8 | 12 | 10 | 13 | 8 | 6 | 8 | 9.28 | 12 | 7 | 0.632 | -0.663 | 0.28896609 | NA | 0 | 0.2579 | 0.1675 | 0.1675 | TRUE | 0.0823 |
| Hivep1 | 1413 | 1061 | 1063 | 1017 | 966 | 1088 | 1097 | 1129 | 1147 | 1012 | 1090 | 1074 | 1091.51 | 1124 | 1059 | 0.942 | -0.086 | 0.24115622 | 0.40878154 | 0 | 0.0046 | 0.003 | 0.003 | TRUE | 0.0177 |
| Hivep2 | 67 | 36 | 38 | 33 | 37 | 29 | 52 | 38 | 41 | 33 | 42 | 29 | 39.09 | 44 | 35 | 0.778 | -0.362 | 0.25828192 | 0.42831227 | 0 | 0.0631 | 0.0477 | 0.0477 | TRUE | 0.244 |
| Hivep3 | 60 | 43 | 27 | 23 | 23 | 39 | 47 | 46 | 29 | 23 | 26 | 38 | 34.8 | 41 | 29 | 0.717 | -0.48 | 0.1792926 | 0.33398007 | 0.0403 | 0.0705 | 0.0623 | 0.0623 | TRUE | 0.5375 |
| Hjurp | 891 | 736 | 737 | 636 | 589 | 638 | 692 | 783 | 795 | 633 | 664 | 630 | 699.57 | 757 | 642 | 0.85 | -0.234 | 0.01060898 | 0.04311465 | 0.0023 | 0.0058 | 0.0046 | 0.0046 | TRUE | 0.0701 |
| Hk1 | 6472 | 5571 | 7680 | 8330 | 8140 | 9101 | 5027 | 5926 | 8287 | 8292 | 9183 | 8984 | 7616.25 | 6413 | 8820 | 1.375 | 0.46 | 0.03912035 | 0.11268489 | 0.0357 | 0.0027 | 0.0116 | 0.0357 | TRUE | 0.7969 |
| Hk2 | 21413 | 18706 | 21561 | 20473 | 19616 | 23004 | 16631 | 19897 | 23264 | 20379 | 22129 | 22708 | 20834.7 | 19931 | 21739 | 1.091 | 0.125 | 0.19567176 | 0.35532161 | 0.0137 | 0.0025 | 0.0067 | 0.0067 | TRUE | 0.2092 |
| Hk3 | 8 | 6 | 5 | 5 | 10 | 4 | 6 | 6 | 5 | 5 | 11 | 4 | 6.37 | 6 | 7 | 1.109 | 0.15 | 0.84623056 | NA | 0 | 0.3747 | 0.2721 | 0.2721 | TRUE | 0.8077 |
| Hkdc1 | 37 | 30 | 16 | 35 | 25 | 41 | 29 | 32 | 17 | 35 | 28 | 40 | 30.24 | 26 | 34 | 1.326 | 0.407 | 0.27223834 | 0.44317796 | 0.0195 | 0.0808 | 0.0656 | 0.0656 | TRUE | 0.486 |
| Hlcs | 964 | 732 | 676 | 707 | 670 | 735 | 749 | 779 | 729 | 704 | 756 | 726 | 740.31 | 752 | 729 | 0.968 | -0.047 | 0.5735967 | 0.72584485 | 0 | 0.0056 | 0.0037 | 0.0037 | TRUE | 0.0125 |
| Hlf | 55 | 35 | 36 | 50 | 25 | 45 | 43 | 37 | 39 | 50 | 28 | 44 | 40.2 | 40 | 41 | 1.032 | 0.046 | 0.88655692 | 0.93690919 | 0.0142 | 0.0614 | 0.0491 | 0.0491 | TRUE | 0.5089 |
| Hltf | 4946 | 3743 | 4380 | 4258 | 4025 | 4678 | 3841 | 3981 | 4726 | 4238 | 4541 | 4618 | 4324.27 | 4183 | 4466 | 1.068 | 0.095 | 0.23795731 | 0.40470298 | 0.0067 | 0.003 | 0.0044 | 0.0044 | TRUE | 0.1565 |
| Hlx | 12 | 14 | 11 | 14 | 10 | 9 | 9 | 15 | 12 | 14 | 11 | 9 | 11.7 | 12 | 11 | 0.953 | -0.069 | 0.90166936 | NA | 0 | 0.205 | 0.1411 | 0.1411 | TRUE | 0.1983 |
| Hmbox1 | 871 | 673 | 654 | 705 | 600 | 729 | 676 | 716 | 706 | 702 | 677 | 720 | 699.37 | 699 | 700 | 1.001 | 0.002 | 0.98026568 | 0.98968266 | 0 | 0.0058 | 0.0037 | 0.0037 | TRUE | 0.0103 |
| Hmbs | 4190 | 3055 | 3422 | 3778 | 3308 | 3845 | 3254 | 3250 | 3692 | 3761 | 3732 | 3796 | 3580.68 | 3399 | 3763 | 1.108 | 0.147 | 0.02497856 | 0.08134396 | 0.0024 | 0.0031 | 0.0028 | 0.0028 | TRUE | 0.0688 |
| Hmcn1 | 139 | 60 | 61 | 66 | 89 | 68 | 108 | 64 | 66 | 66 | 100 | 67 | 78.47 | 79 | 78 | 0.97 | -0.043 | 0.87465721 | 0.93043734 | 0.0637 | 0.0326 | 0.0418 | 0.0418 | TRUE | 1.0234 |
| Hmg20a | 3138 | 2434 | 2519 | 2747 | 2478 | 2842 | 2437 | 2589 | 2718 | 2734 | 2795 | 2805 | 2679.91 | 2581 | 2778 | 1.077 | 0.107 | 0.08885393 | 0.20287367 | 0.0013 | 0.0033 | 0.0025 | 0.0025 | TRUE | 0.0299 |
| Hmg20b | 1327 | 935 | 886 | 866 | 715 | 851 | 1031 | 995 | 956 | 862 | 807 | 840 | 914.97 | 994 | 836 | 0.841 | -0.25 | 0.00150107 | 0.00967301 | 0 | 0.005 | 0.0034 | 0.0034 | TRUE | 0.0132 |
| Hmga1 | 234 | 185 | 178 | 191 | 139 | 204 | 182 | 197 | 192 | 190 | 157 | 201 | 186.48 | 190 | 183 | 0.964 | -0.053 | 0.73319429 | 0.84058281 | 0.0026 | 0.0151 | 0.0117 | 0.0117 | TRUE | 0.1587 |
| Hmga1-rs1 | 0 | 0 | 0 | 0 | 0 | 0 | 0 | 0 | 0 | 0 | 0 | 0 | 0 | 0 | 0 | NA | NA | NA | NA | NA | NA | NA | NA | NA | NA |
| Hmga2 | 2787 | 1952 | 1599 | 1573 | 1301 | 1491 | 2165 | 2076 | 1725 | 1566 | 1468 | 1472 | 1745.24 | 1989 | 1502 | 0.755 | -0.406 | 4.16E-06 | 9.13E-05 | 0.0066 | 0.0038 | 0.005 | 0.005 | TRUE | 0.1609 |
| Hmga2-ps1 | 45 | 55 | 46 | 67 | 59 | 60 | 35 | 59 | 50 | 67 | 67 | 59 | 55.93 | 48 | 64 | 1.358 | 0.441 | 0.11136378 | 0.23924884 | 0.0163 | 0.0448 | 0.0371 | 0.0371 | TRUE | 0.5208 |
| Hmgb1 | 7194 | 5005 | 4702 | 4943 | 4180 | 4732 | 5587 | 5324 | 5073 | 4920 | 4715 | 4671 | 5048.57 | 5328 | 4769 | 0.895 | -0.16 | 0.00598106 | 0.02817645 | 0.0013 | 0.0029 | 0.0023 | 0.0023 | TRUE | 0.0224 |
| Hmgb1-rs17 | 2 | 0 | 0 | 1 | 0 | 0 | 2 | 0 | 0 | 1 | 0 | 0 | 0.42 | 1 | 0 | 0.681 | -0.553 | 0.8627148 | NA | 0 | 5.5819 | 5.4906 | 5.4906 | TRUE | 0.9727 |
| Hmgb2 | 25104 | 17825 | 16796 | 16987 | 13999 | 15886 | 19497 | 18960 | 18123 | 16909 | 15792 | 15682 | 17493.88 | 18860 | 16128 | 0.855 | -0.226 | 3.51E-05 | 0.0005286 | 0.0015 | 0.0026 | 0.0021 | 0.0021 | TRUE | 0.022 |

| | | | | | | | | | | | | | | | | | | | | | | | | | |
|---|---|---|---|---|---|---|---|---|---|---|---|---|---|---|---|---|---|---|---|---|---|---|---|---|---|
| Hmgb3 | 1884 | 1320 | 1201 | 1067 | 958 | 1012 | 1463 | 1404 | 1296 | 1062 | 1081 | 999 | 1217.5 | 1388 | 1047 | 0.754 | -0.408 | 1.61E-07 | 5.94E-06 | 0.002 | 0.0044 | 0.0035 | 0.0035 | TRUE | 0.0399 |
| Hmgb4 | 1 | 0 | 0 | 0 | 0 | 1 | 1 | 0 | 0 | 0 | 0 | 1 | 0.29 | 0 | 0 | 1.075 | 0.105 | 0.97702352 | NA | 0 | 8.0647 | 7.5458 | 7.5458 | TRUE | 0.3835 |
| Hmgcl | 1872 | 1333 | 1384 | 1661 | 1429 | 1679 | 1454 | 1418 | 1493 | 1653 | 1612 | 1657 | 1548 | 1455 | 1641 | 1.128 | 0.174 | 0.00737833 | 0.03305908 | 0 | 0.004 | 0.0024 | 0.0024 | TRUE | 0.0061 |
| Hmgcll1 | 303 | 211 | 211 | 279 | 199 | 225 | 235 | 224 | 228 | 278 | 224 | 222 | 235.29 | 229 | 241 | 1.054 | 0.075 | 0.59318889 | 0.74070371 | 0.0045 | 0.0125 | 0.0101 | 0.0101 | TRUE | 0.194 |
| Hmgcr | 16315 | 12222 | 12902 | 14185 | 11977 | 14170 | 12671 | 13000 | 13921 | 14120 | 13511 | 13988 | 13535.28 | 13197 | 13873 | 1.051 | 0.072 | 0.18453422 | 0.34072312 | 0.0014 | 0.0026 | 0.0021 | 0.0021 | TRUE | 0.0281 |
| Hmgcs1 | 22117 | 16201 | 17109 | 18741 | 15860 | 18388 | 17177 | 17233 | 18460 | 18655 | 17892 | 18152 | 17928.15 | 17623 | 18233 | 1.035 | 0.049 | 0.33878667 | 0.51295317 | 0.001 | 0.0026 | 0.0018 | 0.0018 | TRUE | 0.0211 |
| Hmgcs2 | 130 | 90 | 92 | 84 | 78 | 60 | 101 | 96 | 99 | 84 | 88 | 59 | 87.8 | 99 | 77 | 0.778 | -0.363 | 0.10063805 | 0.22208115 | 0.006 | 0.0294 | 0.0236 | 0.0236 | TRUE | 0.3829 |
| Hmgn1 | 30708 | 21843 | 20235 | 22512 | 19092 | 22219 | 23850 | 23234 | 21833 | 22408 | 21538 | 21934 | 22466.15 | 22972 | 21960 | 0.956 | -0.065 | 0.21201485 | 0.37430505 | 0.0011 | 0.0025 | 0.0019 | 0.0019 | TRUE | 0.023 |
| Hmgn2 | 3000 | 2080 | 1686 | 1819 | 1419 | 1590 | 2330 | 2212 | 1819 | 1811 | 1601 | 1570 | 1890.43 | 2120 | 1661 | 0.782 | -0.354 | 0.00022525 | 0.00234147 | 0.0103 | 0.0037 | 0.0061 | 0.0061 | TRUE | 0.1858 |
| Hmgn3 | 614 | 491 | 419 | 411 | 390 | 420 | 477 | 522 | 452 | 409 | 440 | 415 | 452.48 | 484 | 421 | 0.871 | -0.2 | 0.05552032 | 0.14525936 | 0.0011 | 0.0077 | 0.0056 | 0.0056 | TRUE | 0.054 |
| Hmgn5 | 1818 | 1219 | 1054 | 1208 | 998 | 1136 | 1412 | 1297 | 1137 | 1202 | 1126 | 1121 | 1215.93 | 1282 | 1150 | 0.896 | -0.159 | 0.07567027 | 0.1812865 | 0.006 | 0.0044 | 0.0049 | 0.0049 | TRUE | 0.1161 |
| Hmgxb3 | 5674 | 4071 | 4069 | 4272 | 3700 | 4521 | 4407 | 4330 | 4390 | 4252 | 4174 | 4463 | 4336.12 | 4376 | 4296 | 0.982 | -0.026 | 0.62579326 | 0.76415746 | 4.00E-04 | 0.003 | 0.0019 | 0.0019 | TRUE | 0.014 |
| Hmgxb4 | 4397 | 3226 | 3255 | 3452 | 3241 | 3764 | 3415 | 3431 | 3512 | 3436 | 3656 | 3716 | 3527.75 | 3453 | 3603 | 1.043 | 0.061 | 0.28484732 | 0.45683603 | 7.00E-04 | 0.0031 | 0.0021 | 0.0021 | TRUE | 0.02 |
| Hmha1 | 873 | 832 | 1162 | 1274 | 1293 | 1566 | 678 | 885 | 1254 | 1268 | 1459 | 1546 | 1181.58 | 939 | 1424 | 1.518 | 0.602 | 0.02613081 | 0.08392982 | 0.0519 | 0.0044 | 0.0172 | 0.0519 | TRUE | 0.6707 |
| Hmmr | 16353 | 11889 | 13620 | 15275 | 13445 | 14975 | 12701 | 12646 | 14696 | 15205 | 15167 | 14783 | 14199.57 | 13348 | 15052 | 1.128 | 0.174 | 0.00842409 | 0.03644234 | 0.0038 | 0.0026 | 0.0031 | 0.0031 | TRUE | 0.0952 |
| Hmox1 | 4716 | 3352 | 2875 | 2924 | 2537 | 2853 | 3663 | 3565 | 3102 | 2911 | 2862 | 2816 | 3153.2 | 3443 | 2863 | 0.831 | -0.267 | 0.00020332 | 0.00215841 | 0.0034 | 0.0032 | 0.0034 | 0.0034 | TRUE | 0.0907 |
| Hmox2 | 2313 | 1603 | 1515 | 1569 | 1408 | 1640 | 1796 | 1705 | 1635 | 1562 | 1588 | 1619 | 1650.88 | 1712 | 1590 | 0.928 | -0.108 | 0.10907921 | 0.23575823 | 8.00E-04 | 0.0039 | 0.0027 | 0.0027 | TRUE | 0.0233 |
| Hmx1 | 120 | 98 | 74 | 97 | 82 | 80 | 93 | 104 | 80 | 97 | 93 | 79 | 90.89 | 92 | 90 | 0.966 | -0.051 | 0.81146113 | 0.89134752 | 0 | 0.0285 | 0.0214 | 0.0214 | TRUE | 0.1257 |
| Hmx2 | 6 | 10 | 12 | 13 | 5 | 5 | 5 | 11 | 13 | 13 | 6 | 5 | 8.63 | 10 | 8 | 0.849 | -0.236 | 0.73946625 | NA | 0.1634 | 0.2772 | 0.2436 | 0.2436 | TRUE | 0.9618 |
| Hmx3 | 4 | 5 | 3 | 7 | 3 | 3 | 3 | 5 | 3 | 7 | 3 | 3 | 4.16 | 4 | 4 | 1.154 | 0.207 | 0.82458111 | NA | 0 | 0.5718 | 0.3887 | 0.3887 | TRUE | 0.4674 |
| Hn1 | 9769 | 6917 | 6728 | 7494 | 6568 | 7485 | 7587 | 7358 | 7259 | 7460 | 7409 | 7389 | 7410.32 | 7401 | 7419 | 1.002 | 0.003 | 0.94722501 | 0.97107433 | 2.00E-04 | 0.0027 | 0.0015 | 0.0015 | TRUE | 0.006 |
| Hn1l | 14067 | 9874 | 10256 | 11051 | 9583 | 11223 | 10925 | 10503 | 11066 | 11000 | 10811 | 11079 | 10897.3 | 10831 | 10963 | 1.012 | 0.017 | 0.71655823 | 0.82974271 | 3.00E-04 | 0.0026 | 0.0016 | 0.0016 | TRUE | 0.0085 |
| Hnf1a | 299 | 252 | 253 | 293 | 296 | 354 | 232 | 268 | 273 | 292 | 334 | 349 | 291.38 | 258 | 325 | 1.265 | 0.339 | 0.00979176 | 0.04076717 | 0.0052 | 0.0106 | 0.0089 | 0.0089 | TRUE | 0.0932 |
| Hnf1b | 105 | 82 | 50 | 76 | 75 | 63 | 82 | 87 | 54 | 76 | 85 | 62 | 74.19 | 74 | 74 | 0.994 | -0.009 | 0.97097279 | 0.98433322 | 0.0278 | 0.0344 | 0.0326 | 0.0326 | TRUE | 0.4906 |
| Hnf4a | 252 | 238 | 315 | 252 | 260 | 279 | 196 | 253 | 340 | 251 | 293 | 275 | 268.05 | 263 | 273 | 1.043 | 0.061 | 0.73210236 | 0.83960034 | 0.0375 | 0.0113 | 0.0191 | 0.0191 | TRUE | 0.3425 |
| Hnf4g | 0 | 1 | 0 | 0 | 0 | 0 | 0 | 1 | 0 | 0 | 0 | 0 | 0.18 | 0 | 0 | 0.552 | -0.857 | 0.8336548 | NA | 0 | 13.3714 | 10 | 10 | TRUE | 0.5216 |
| Hnmt | 2 | 8 | 3 | 0 | 0 | 1 | 2 | 9 | 3 | 0 | 0 | 1 | 2.38 | 5 | 0 | 0.085 | -3.561 | 0.03189355 | NA | 0 | 0.9978 | 0.874 | 0.874 | TRUE | 0.3819 |
| Hnrnpa0 | 21582 | 16561 | 14137 | 14154 | 12758 | 14137 | 16762 | 17616 | 15254 | 14089 | 14392 | 13955 | 15344.65 | 16544 | 14145 | 0.855 | -0.226 | 0.00022272 | 0.00231857 | 0.0026 | 0.0026 | 0.0026 | 0.0026 | TRUE | 0.0567 |
| Hnrnpa1 | 2252 | 1741 | 1940 | 1901 | 1681 | 2069 | 1749 | 1852 | 2093 | 1892 | 1896 | 2042 | 1920.86 | 1898 | 1943 | 1.025 | 0.036 | 0.65263539 | 0.78431896 | 0.0047 | 0.0037 | 0.004 | 0.004 | TRUE | 0.0973 |
| Hnrnpa2b1 | 80246 | 58021 | 55252 | 52639 | 45613 | 54201 | 62324 | 61716 | 59616 | 52397 | 51456 | 53505 | 56835.67 | 61219 | 52453 | 0.857 | -0.223 | 9.74E-07 | 2.88E-05 | 4.00E-04 | 0.0025 | 0.0015 | 0.0015 | TRUE | 0.0064 |
| Hnrnpa3 | 2642 | 1816 | 2047 | 2195 | 2020 | 2144 | 2052 | 1932 | 2209 | 2185 | 2279 | 2116 | 2128.73 | 2064 | 2193 | 1.063 | 0.087 | 0.214824 | 0.37748702 | 0.0022 | 0.0035 | 0.0031 | 0.0031 | TRUE | 0.0443 |
| Hnrnpab | 60083 | 42991 | 38621 | 37168 | 33268 | 38442 | 46664 | 45729 | 41672 | 36997 | 37530 | 37948 | 41089.92 | 44688 | 37492 | 0.839 | -0.253 | 5.38E-06 | 0.00011279 | 0.0018 | 0.0025 | 0.0022 | 0.0022 | TRUE | 0.0427 |
| Hnrnpc | 33098 | 22633 | 21619 | 24332 | 20516 | 22905 | 25706 | 24074 | 23327 | 24220 | 23144 | 22611 | 23846.99 | 24369 | 23325 | 0.957 | -0.063 | 0.25782663 | 0.42780543 | 0.0018 | 0.0025 | 0.0022 | 0.0022 | TRUE | 0.0283 |
| Hnrnpd | 21747 | 16441 | 13121 | 11899 | 11371 | 13197 | 16890 | 17488 | 14157 | 11844 | 12828 | 13027 | 14372.48 | 16178 | 12566 | 0.777 | -0.365 | 3.42E-06 | 7.76E-05 | 0.0072 | 0.0026 | 0.0044 | 0.0044 | TRUE | 0.1458 |
| Hnrnpf | 25410 | 18768 | 17536 | 17795 | 15989 | 18496 | 19735 | 19963 | 18921 | 17713 | 18037 | 18258 | 18771.35 | 19540 | 18003 | 0.921 | -0.118 | 0.0123182 | 0.04831616 | 4.00E-04 | 0.0026 | 0.0016 | 0.0016 | TRUE | 0.0093 |
| Hnrnph1 | 15090 | 11293 | 9155 | 8880 | 7366 | 9248 | 11720 | 12012 | 9878 | 8839 | 8310 | 9129 | 9981.35 | 11203 | 8759 | 0.782 | -0.355 | 3.40E-06 | 7.71E-05 | 0.0063 | 0.0027 | 0.0041 | 0.0041 | TRUE | 0.1305 |
| Hnrnph2 | 4224 | 2992 | 3001 | 3224 | 2779 | 3209 | 3281 | 3183 | 3238 | 3209 | 3135 | 3168 | 3202.19 | 3234 | 3171 | 0.98 | -0.029 | 0.59440136 | 0.7409266 | 0 | 0.0032 | 0.0018 | 0.0018 | TRUE | 0.0023 |
| Hnrnph3 | 3706 | 2977 | 2905 | 3326 | 2833 | 3325 | 2878 | 3167 | 3134 | 3311 | 3196 | 3282 | 3161.38 | 3060 | 3263 | 1.067 | 0.094 | 0.12567887 | 0.26079096 | 0.0013 | 0.0032 | 0.0024 | 0.0024 | TRUE | 0.0336 |
| Hnrnpk | 45246 | 32727 | 31826 | 33024 | 29387 | 33831 | 35141 | 34811 | 34340 | 32872 | 33152 | 33396 | 33952 | 34764 | 33140 | 0.953 | -0.069 | 0.10325194 | 0.22645082 | 1.00E-04 | 0.0025 | 0.0013 | 0.0013 | TRUE | 0.0014 |
| Hnrnpl | 33792 | 25772 | 26097 | 25246 | 23523 | 28129 | 26245 | 27413 | 28158 | 25130 | 26536 | 27768 | 26875.08 | 27272 | 26478 | 0.971 | -0.043 | 0.44715859 | 0.61780935 | 0.0017 | 0.0025 | 0.0022 | 0.0022 | TRUE | 0.0243 |
| Hnrnpm | 25887 | 19357 | 16867 | 15810 | 14631 | 16778 | 20105 | 20590 | 18199 | 15737 | 16505 | 16562 | 17949.92 | 19631 | 16268 | 0.829 | -0.271 | 5.74E-06 | 0.00011823 | 0.0024 | 0.0026 | 0.0025 | 0.0025 | TRUE | 0.0497 |
| Hnrnpr | 5150 | 3800 | 3387 | 3435 | 2892 | 3409 | 4000 | 4042 | 3655 | 3419 | 3262 | 3365 | 3623.87 | 3899 | 3349 | 0.859 | -0.22 | 0.0003602 | 0.00335182 | 0.0014 | 0.0031 | 0.0024 | 0.0024 | TRUE | 0.0361 |
| Hnrnpu | 68929 | 52065 | 47556 | 49890 | 44344 | 51342 | 53534 | 55381 | 51313 | 49660 | 50025 | 50682 | 51765.86 | 53409 | 50122 | 0.938 | -0.092 | 0.05772932 | 0.14953649 | 8.00E-04 | 0.0025 | 0.0017 | 0.0017 | TRUE | 0.0144 |
| Hnrnpul1 | 11866 | 9263 | 9522 | 9332 | 8478 | 10122 | 9216 | 9853 | 10274 | 9289 | 9564 | 9992 | 9698 | 9781 | 9615 | 0.983 | -0.024 | 0.68116762 | 0.80435631 | 0.0017 | 0.0027 | 0.0024 | 0.0024 | TRUE | 0.0315 |
| Hnrnpul2 | 18976 | 13820 | 13815 | 13737 | 12664 | 14164 | 14738 | 14700 | 14906 | 13674 | 14286 | 13982 | 14381.07 | 14781 | 13981 | 0.946 | -0.08 | 0.07847755 | 0.18657345 | 2.00E-04 | 0.0026 | 0.0014 | 0.0014 | TRUE | 0.0045 |
| Hnrpdl | 9319 | 7124 | 6515 | 6019 | 5373 | 6214 | 7238 | 7578 | 7030 | 5991 | 6061 | 6134 | 6671.97 | 7282 | 6062 | 0.833 | -0.264 | 4.68E-07 | 1.53E-05 | 6.00E-04 | 0.0028 | 0.0018 | 0.0018 | TRUE | 0.0153 |
| Hnrpll | 5769 | 4357 | 4364 | 4285 | 3935 | 4365 | 4481 | 4634 | 4709 | 4265 | 4439 | 4309 | 4472.84 | 4608 | 4338 | 0.942 | -0.087 | 0.10185923 | 0.22408403 | 3.00E-04 | 0.003 | 0.0018 | 0.0018 | TRUE | 0.0073 |
| Hoga1 | 38 | 21 | 18 | 20 | 14 | 19 | 30 | 22 | 19 | 20 | 16 | 19 | 20.95 | 24 | 18 | 0.759 | -0.397 | 0.34860226 | 0.52326949 | 0 | 0.1155 | 0.0808 | 0.0808 | TRUE | 0.3194 |
| Homer1 | 2445 | 1834 | 1755 | 1867 | 1497 | 1764 | 1899 | 1951 | 1894 | 1858 | 1689 | 1741 | 1838.65 | 1915 | 1763 | 0.921 | -0.118 | 0.07255341 | 0.17618176 | 7.00E-04 | 0.0037 | 0.0026 | 0.0026 | TRUE | 0.0272 |
| Homer2 | 1328 | 1019 | 974 | 1264 | 1037 | 1171 | 1031 | 1084 | 1051 | 1258 | 1170 | 1156 | 1125.04 | 1055 | 1195 | 1.133 | 0.18 | 0.01575002 | 0.05789994 | 6.00E-04 | 0.0045 | 0.0031 | 0.0031 | TRUE | 0.0262 |
| Homer3 | 894 | 607 | 635 | 499 | 477 | 515 | 694 | 646 | 685 | 497 | 538 | 508 | 594.72 | 675 | 514 | 0.761 | -0.394 | 1.64E-05 | 0.00028516 | 0 | 0.0064 | 0.0043 | 0.0043 | TRUE | 0.0184 |
| Homez | 407 | 283 | 200 | 245 | 250 | 244 | 316 | 301 | 216 | 244 | 282 | 241 | 266.61 | 278 | 256 | 0.917 | -0.125 | 0.42605602 | 0.59860599 | 0.0206 | 0.0113 | 0.0141 | 0.0141 | TRUE | 0.4142 |
| Hook1 | 10083 | 7249 | 7193 | 8024 | 6902 | 7708 | 7831 | 7711 | 7761 | 7987 | 7786 | 7609 | 7780.86 | 7768 | 7794 | 1.003 | 0.005 | 0.92073838 | 0.95643782 | 2.00E-04 | 0.0027 | 0.0016 | 0.0016 | TRUE | 0.0058 |
| Hook2 | 3644 | 3109 | 3584 | 3329 | 3294 | 3888 | 2830 | 3307 | 3867 | 3314 | 3716 | 3838 | 3478.66 | 3335 | 3623 | 1.087 | 0.12 | 0.22850102 | 0.39382191 | 0.0132 | 0.0031 | 0.0069 | 0.0069 | TRUE | 0.2371 |
| Hook3 | 1998 | 1561 | 1419 | 1410 | 1196 | 1449 | 1552 | 1660 | 1531 | 1404 | 1349 | 1430 | 1487.73 | 1581 | 1394 | 0.882 | -0.18 | 0.00932655 | 0.03934605 | 7.00E-04 | 0.004 | 0.0028 | 0.0028 | TRUE | 0.0227 |
| Hopx | 159 | 137 | 152 | 201 | 184 | 220 | 123 | 146 | 164 | 200 | 208 | 217 | 176.34 | 144 | 208 | 1.45 | 0.536 | 0.00082676 | 0.00629899 | 0.0045 | 0.0159 | 0.0127 | 0.0127 | TRUE | 0.1876 |
| Hormad1 | 1056 | 805 | 896 | 653 | 518 | 632 | 820 | 856 | 967 | 650 | 584 | 624 | 750.24 | 881 | 619 | 0.704 | -0.506 | 8.69E-08 | 3.45E-06 | 0.004 | 0.0056 | 0.005 | 0.005 | TRUE | 0.0852 |
| Hormad2 | 329 | 225 | 246 | 214 | 186 | 170 | 256 | 239 | 265 | 213 | 210 | 168 | 225.16 | 253 | 197 | 0.776 | -0.366 | 0.01150356 | 0.04583785 | 0.0043 | 0.013 | 0.0106 | 0.0106 | TRUE | 0.1851 |
| Hotair | 0 | 0 | 0 | 0 | 0 | 0 | 0 | 0 | 0 | 0 | 0 | 0 | 0 | 0 | 0 | NA | NA | NA | NA | NA | NA | NA | NA | NA | NA |
| Hoxa1 | 293 | 190 | 150 | 162 | 147 | 121 | 228 | 202 | 162 | 161 | 166 | 119 | 173.01 | 197 | 149 | 0.751 | -0.413 | 0.02501778 | 0.08143454 | 0.0244 | 0.0161 | 0.0185 | 0.0185 | TRUE | 0.3181 |
| Hoxa10 | 16 | 12 | 6 | 18 | 3 | 7 | 12 | 13 | 6 | 18 | 3 | 7 | 9.98 | 10 | 9 | 0.893 | -0.163 | 0.81416168 | NA | 0.2596 | 0.2399 | 0.2453 | 0.2453 | TRUE | 1.1731 |
| Hoxa11 | 3 | 3 | 1 | 2 | 1 | 0 | 2 | 3 | 1 | 2 | 1 | 0 | 1.62 | 2 | 1 | 0.47 | -1.09 | 0.48478046 | NA | 0 | 1.4656 | 1.0344 | 1.0344 | TRUE | 0.3957 |
| Hoxa11as | 4 | 2 | 1 | 2 | 2 | 0 | 3 | 2 | 1 | 2 | 2 | 0 | 1.76 | 2 | 1 | 0.654 | -0.612 | 0.6787561 | NA | 0 | 1.3491 | 0.9788 | 0.9788 | TRUE | 0.5079 |
| Hoxa13 | 16 | 13 | 13 | 19 | 8 | 15 | 12 | 14 | 14 | 19 | 9 | 15 | 13.84 | 13 | 14 | 1.07 | 0.098 | 0.85117519 | NA | 0 | 0.1737 | 0.1244 | 0.1244 | TRUE | 0.2592 |
| Hoxa2 | 1 | 2 | 1 | 2 | 0 | 0 | 1 | 2 | 1 | 2 | 0 | 0 | 1 | 1 | 1 | 0.515 | -0.958 | 0.63579447 | NA | 0 | 2.3829 | 1.8115 | 1.8115 | TRUE | 0.9736 |
| Hoxa3 | 27 | 10 | 5 | 9 | 5 | 5 | 21 | 11 | 5 | 9 | 6 | 5 | 9.42 | 12 | 7 | 0.52 | -0.945 | 0.17736146 | NA | 0.1948 | 0.254 | 0.2356 | 0.2356 | TRUE | 0.3173 |
| Hoxa4 | 25 | 15 | 15 | 31 | 19 | 16 | 19 | 16 | 16 | 31 | 21 | 16 | 19.94 | 17 | 23 | 1.315 | 0.395 | 0.37701224 | 0.55151527 | 0 | 0.1213 | 0.093 | 0.093 | TRUE | 0.3161 |
| Hoxa5 | 21 | 7 | 7 | 12 | 9 | 8 | 16 | 7 | 8 | 12 | 10 | 8 | 10.22 | 10 | 10 | 0.939 | -0.091 | 0.88407431 | NA | 0 | 0.2344 | 0.1813 | 0.1813 | TRUE | 1.082 |
| Hoxa6 | 1 | 1 | 0 | 0 | 0 | 0 | 1 | 1 | 0 | 0 | 0 | 0 | 0.31 | 1 | 0 | 0.309 | -1.696 | 0.63183309 | NA | 0 | 7.7294 | 7.2472 | 7.2472 | TRUE | 0.2015 |

| | | | | | | | | | | | | | | | | | | | | | | | | | |
|---|---|---|---|---|---|---|---|---|---|---|---|---|---|---|---|---|---|---|---|---|---|---|---|---|---|
| Hoxa7 | 24 | 18 | 8 | 16 | 10 | 8 | 19 | 19 | 9 | 16 | 11 | 8 | 13.59 | 16 | 12 | 0.751 | -0.413 | 0.45818281 | NA | 0.0679 | 0.1769 | 0.1476 | 0.1476 | TRUE | 0.5709 |
| Hoxa9 | 13 | 9 | 9 | 12 | 5 | 7 | 10 | 10 | 10 | 12 | 6 | 7 | 8.98 | 10 | 8 | 0.837 | -0.257 | 0.6886658 | NA | 0 | 0.2664 | 0.1833 | 0.1833 | TRUE | 0.4979 |
| Hoxb1 | 6 | 6 | 2 | 4 | 2 | 2 | 5 | 6 | 2 | 4 | 2 | 2 | 3.57 | 4 | 3 | 0.622 | -0.684 | 0.50358736 | NA | 0 | 0.6666 | 0.4566 | 0.4566 | TRUE | 0.2071 |
| Hoxb13 | 79 | 49 | 41 | 61 | 51 | 51 | 61 | 52 | 44 | 61 | 58 | 50 | 54.39 | 52 | 56 | 1.062 | 0.087 | 0.74711545 | 0.84993112 | 0 | 0.046 | 0.0335 | 0.0335 | TRUE | 0.2028 |
| Hoxb2 | 12 | 2 | 5 | 2 | 7 | 3 | 9 | 2 | 5 | 2 | 8 | 3 | 4.95 | 5 | 4 | 0.738 | -0.437 | 0.63952428 | NA | 0.277 | 0.4814 | 0.4215 | 0.4215 | TRUE | 0.2781 |
| Hoxb3 | 2 | 2 | 1 | 0 | 1 | 2 | 2 | 2 | 1 | 0 | 1 | 2 | 1.31 | 2 | 1 | 0.652 | -0.618 | 0.7119827 | NA | 0 | 1.8112 | 1.2143 | 1.2143 | TRUE | 0.3697 |
| Hoxb4 | 53 | 17 | 18 | 24 | 19 | 20 | 41 | 18 | 19 | 24 | 21 | 20 | 23.96 | 26 | 22 | 0.814 | -0.297 | 0.49981659 | 0.66429966 | 0.0884 | 0.1014 | 0.0974 | 0.0974 | TRUE | 1.8019 |
| Hoxb5 | 4 | 0 | 0 | 0 | 0 | 0 | 3 | 0 | 0 | 0 | 0 | 0 | 0.52 | 1 | 0 | 0.179 | -2.48 | 0.45030014 | NA | 10 | 4.5798 | 6.3235 | 6.3235 | TRUE | 1.8962 |
| Hoxb6 | 1 | 1 | 0 | 0 | 0 | 0 | 1 | 1 | 0 | 0 | 0 | 0 | 0.31 | 1 | 0 | 0.309 | -1.696 | 0.63183309 | NA | 0 | 7.7294 | 7.2472 | 7.2472 | TRUE | 0.2015 |
| Hoxb7 | 64 | 21 | 30 | 29 | 24 | 38 | 50 | 22 | 32 | 29 | 27 | 38 | 32.98 | 35 | 31 | 0.886 | -0.175 | 0.64455576 | 0.77868612 | 0.0692 | 0.0743 | 0.0728 | 0.0728 | TRUE | 0.3934 |
| Hoxb8 | 1 | 0 | 2 | 4 | 0 | 0 | 1 | 0 | 2 | 4 | 0 | 0 | 1.15 | 1 | 1 | 1.382 | 0.467 | 0.8220587 | NA | 3.0477 | 2.0585 | 2.2215 | 2.2215 | TRUE | 1.8538 |
| Hoxb9 | 102 | 71 | 77 | 67 | 77 | 69 | 79 | 76 | 83 | 67 | 87 | 68 | 76.58 | 79 | 74 | 0.93 | -0.105 | 0.64158123 | 0.77633937 | 0 | 0.0334 | 0.0238 | 0.0238 | TRUE | 0.2034 |
| Hoxc10 | 7 | 4 | 6 | 10 | 8 | 3 | 5 | 4 | 6 | 10 | 9 | 3 | 6.35 | 5 | 7 | 1.352 | 0.435 | 0.57890737 | NA | 0 | 0.3756 | 0.2825 | 0.2825 | TRUE | 0.7769 |
| Hoxc11 | 0 | 0 | 0 | 0 | 0 | 0 | 0 | 0 | 0 | 0 | 0 | 0 | 0 | 0 | 0 | NA | NA | NA | NA | NA | NA | NA | NA | NA | NA |
| Hoxc12 | 9 | 9 | 12 | 8 | 2 | 7 | 7 | 10 | 13 | 8 | 2 | 7 | 7.77 | 10 | 6 | 0.592 | -0.755 | 0.29592111 | NA | 0 | 0.3073 | 0.2373 | 0.2373 | TRUE | 0.4422 |
| Hoxc13 | 193 | 183 | 157 | 180 | 141 | 168 | 150 | 195 | 169 | 179 | 159 | 166 | 169.67 | 171 | 168 | 0.985 | -0.021 | 0.8950818 | 0.94242388 | 0.0054 | 0.0164 | 0.0131 | 0.0131 | TRUE | 0.1512 |
| Hoxc4 | 55 | 39 | 38 | 47 | 45 | 42 | 43 | 41 | 41 | 47 | 51 | 41 | 44.04 | 42 | 46 | 1.107 | 0.147 | 0.60826758 | 0.7520791 | 0 | 0.0562 | 0.0367 | 0.0367 | TRUE | 0.0682 |
| Hoxc5 | 4 | 2 | 3 | 1 | 2 | 2 | 3 | 2 | 3 | 1 | 2 | 2 | 2.28 | 3 | 2 | 0.611 | -0.71 | 0.56717131 | NA | 0 | 1.0406 | 0.6402 | 0.6402 | TRUE | 0.1145 |
| Hoxc6 | 6 | 3 | 4 | 1 | 1 | 1 | 5 | 3 | 4 | 1 | 1 | 1 | 2.55 | 4 | 1 | 0.254 | -1.977 | 0.12576373 | NA | 0 | 0.9332 | 0.5834 | 0.5834 | TRUE | 0.0552 |
| Hoxc8 | 1 | 0 | 0 | 1 | 0 | 0 | 1 | 0 | 0 | 1 | 0 | 0 | 0.3 | 0 | 0 | 1.075 | 0.105 | 0.97698525 | NA | 0 | 8.0272 | 7.5141 | 7.5141 | TRUE | 0.3906 |
| Hoxc9 | 5 | 5 | 3 | 3 | 1 | 2 | 4 | 5 | 3 | 3 | 1 | 2 | 3.09 | 4 | 2 | 0.495 | -1.015 | 0.35415639 | NA | 0 | 0.77 | 0.4951 | 0.4951 | TRUE | 0.1554 |
| Hoxd1 | 96 | 63 | 47 | 95 | 68 | 76 | 75 | 67 | 51 | 95 | 77 | 75 | 73.1 | 64 | 82 | 1.274 | 0.35 | 0.15048311 | 0.29622022 | 0.0127 | 0.0349 | 0.0288 | 0.0288 | TRUE | 0.2791 |
| Hoxd10 | 6 | 4 | 4 | 4 | 1 | 4 | 5 | 4 | 4 | 4 | 1 | 4 | 3.71 | 4 | 3 | 0.691 | -0.532 | 0.59007676 | NA | 0 | 0.6404 | 0.4248 | 0.4248 | TRUE | 0.3376 |
| Hoxd11 | 13 | 3 | 6 | 9 | 6 | 9 | 10 | 3 | 6 | 9 | 7 | 9 | 7.4 | 6 | 8 | 1.223 | 0.291 | 0.68590607 | NA | 0 | 0.3229 | 0.2381 | 0.2381 | TRUE | 0.289 |
| Hoxd12 | 1 | 0 | 1 | 0 | 0 | 0 | 1 | 0 | 1 | 0 | 0 | 0 | 0.31 | 1 | 0 | 0.307 | -1.706 | 0.62894649 | NA | 0 | 7.6657 | 7.2 | 7.2 | TRUE | 0.2067 |
| Hoxd13 | 203 | 139 | 136 | 148 | 108 | 129 | 158 | 148 | 147 | 147 | 122 | 127 | 141.46 | 151 | 132 | 0.876 | -0.19 | 0.25912558 | 0.42901446 | 0 | 0.0192 | 0.0134 | 0.0134 | TRUE | 0.1044 |
| Hoxd3 | 1 | 0 | 1 | 0 | 0 | 0 | 1 | 0 | 1 | 0 | 0 | 0 | 0.31 | 1 | 0 | 0.307 | -1.706 | 0.62894649 | NA | 0 | 7.6657 | 7.2 | 7.2 | TRUE | 0.2067 |
| Hoxd4 | 9 | 10 | 19 | 13 | 17 | 21 | 7 | 11 | 21 | 13 | 19 | 21 | 15.16 | 13 | 18 | 1.409 | 0.495 | 0.36104089 | NA | 0.103 | 0.1587 | 0.1434 | 0.1434 | TRUE | 1.0293 |
| Hoxd8 | 113 | 74 | 66 | 117 | 74 | 81 | 88 | 79 | 71 | 116 | 83 | 80 | 86.26 | 79 | 93 | 1.175 | 0.232 | 0.31119856 | 0.48421087 | 0.0175 | 0.0299 | 0.0263 | 0.0263 | TRUE | 0.4614 |
| Hoxd9 | 40 | 44 | 32 | 33 | 36 | 17 | 31 | 47 | 35 | 33 | 41 | 17 | 33.77 | 38 | 30 | 0.802 | -0.318 | 0.40449522 | 0.57722147 | 0.0794 | 0.0726 | 0.0744 | 0.0744 | TRUE | 0.6708 |
| Hp | 3 | 1 | 1 | 0 | 0 | 2 | 2 | 1 | 1 | 0 | 0 | 2 | 1.07 | 1 | 1 | 0.444 | -1.171 | 0.55043018 | NA | 0 | 2.2082 | 1.6818 | 1.6818 | TRUE | 0.9734 |
| Hp1bp3 | 2311 | 1849 | 1802 | 1780 | 1608 | 1791 | 1795 | 1967 | 1944 | 1772 | 1814 | 1768 | 1843.29 | 1902 | 1785 | 0.939 | -0.091 | 0.17094434 | 0.32259143 | 9.00E-04 | 0.0037 | 0.0026 | 0.0026 | TRUE | 0.0305 |
| Hpca | 264 | 206 | 160 | 181 | 165 | 180 | 205 | 219 | 173 | 180 | 186 | 178 | 190.13 | 199 | 181 | 0.91 | -0.136 | 0.36991564 | 0.54467954 | 0 | 0.0149 | 0.0113 | 0.0113 | TRUE | 0.1368 |
| Hpcal1 | 2164 | 1761 | 2490 | 3032 | 2811 | 3179 | 1681 | 1873 | 2687 | 3018 | 3171 | 3138 | 2594.64 | 2080 | 3109 | 1.495 | 0.58 | 0.00677041 | 0.03097615 | 0.0327 | 0.0033 | 0.0113 | 0.0327 | TRUE | 0.7864 |
| Hpcal4 | 230 | 161 | 138 | 182 | 171 | 219 | 179 | 171 | 149 | 181 | 193 | 216 | 181.51 | 166 | 197 | 1.181 | 0.24 | 0.12354249 | 0.25751682 | 0.0032 | 0.0155 | 0.0119 | 0.0119 | TRUE | 0.0855 |
| Hpd | 5 | 6 | 2 | 5 | 8 | 7 | 4 | 6 | 2 | 5 | 9 | 7 | 5.56 | 4 | 7 | 1.679 | 0.748 | 0.36603839 | NA | 0 | 0.429 | 0.3022 | 0.3022 | TRUE | 0.3137 |
| Hpdl | 499 | 345 | 329 | 341 | 329 | 423 | 388 | 367 | 355 | 339 | 371 | 418 | 372.94 | 370 | 376 | 1.016 | 0.022 | 0.84860759 | 0.91423444 | 0.0039 | 0.0088 | 0.0073 | 0.0073 | TRUE | 0.1096 |
| Hpgd | 274 | 197 | 143 | 148 | 110 | 137 | 213 | 210 | 154 | 147 | 124 | 135 | 163.88 | 192 | 135 | 0.704 | -0.506 | 0.00403427 | 0.02074911 | 0.0139 | 0.0169 | 0.016 | 0.016 | TRUE | 0.3104 |
| Hpgds | 0 | 0 | 0 | 0 | 0 | 0 | 0 | 0 | 0 | 0 | 0 | 0 | 0 | 0 | 0 | NA | NA | NA | NA | NA | NA | NA | NA | NA | NA |
| Hpn | 865 | 676 | 1061 | 1283 | 1175 | 1449 | 672 | 719 | 1145 | 1277 | 1326 | 1430 | 1094.78 | 845 | 1344 | 1.591 | 0.67 | 0.00719822 | 0.03240714 | 0.0439 | 0.0046 | 0.0155 | 0.0439 | TRUE | 1.1399 |
| Hprt | 9447 | 6903 | 7691 | 8139 | 7004 | 8258 | 7337 | 7343 | 8299 | 8102 | 7901 | 8152 | 7855.49 | 7660 | 8052 | 1.051 | 0.072 | 0.24378672 | 0.41167811 | 0.0026 | 0.0027 | 0.0027 | 0.0027 | TRUE | 0.0647 |
| Hps1 | 2547 | 2054 | 1970 | 1777 | 1691 | 1772 | 1978 | 2185 | 2126 | 1769 | 1908 | 1749 | 1952.38 | 2096 | 1809 | 0.863 | -0.212 | 0.0023615 | 0.01375423 | 0.002 | 0.0036 | 0.003 | 0.003 | TRUE | 0.0303 |
| Hps3 | 1543 | 1062 | 1182 | 1355 | 1193 | 1344 | 1198 | 1130 | 1275 | 1349 | 1346 | 1327 | 1270.79 | 1201 | 1341 | 1.116 | 0.158 | 0.03112054 | 0.09520709 | 9.00E-04 | 0.0043 | 0.0031 | 0.0031 | TRUE | 0.0336 |
| Hps4 | 1103 | 885 | 870 | 980 | 849 | 1028 | 857 | 941 | 939 | 975 | 958 | 1015 | 947.46 | 912 | 983 | 1.079 | 0.11 | 0.16849116 | 0.32007499 | 0.001 | 0.0049 | 0.0035 | 0.0035 | TRUE | 0.0358 |
| Hps5 | 1132 | 713 | 629 | 687 | 562 | 646 | 879 | 758 | 679 | 684 | 634 | 638 | 711.97 | 772 | 652 | 0.842 | -0.247 | 0.01920385 | 0.0670446 | 0.0083 | 0.0058 | 0.0066 | 0.0066 | TRUE | 0.1807 |
| Hps6 | 280 | 197 | 147 | 186 | 180 | 166 | 217 | 210 | 159 | 185 | 203 | 164 | 189.62 | 195 | 184 | 0.939 | -0.091 | 0.58367907 | 0.73334431 | 0.0145 | 0.0149 | 0.0148 | 0.0148 | TRUE | 0.2813 |
| Hpse | 161 | 130 | 124 | 150 | 138 | 139 | 125 | 138 | 134 | 149 | 156 | 137 | 139.89 | 132 | 147 | 1.115 | 0.157 | 0.3477414 | 0.52236276 | 0 | 0.0194 | 0.0129 | 0.0129 | TRUE | 0.0387 |
| Hpse2 | 13 | 14 | 5 | 8 | 13 | 6 | 10 | 15 | 5 | 8 | 15 | 6 | 9.82 | 10 | 10 | 0.931 | -0.103 | 0.87580312 | NA | 0.1111 | 0.2437 | 0.2075 | 0.2075 | TRUE | 0.2231 |
| Hpx | 6 | 3 | 2 | 2 | 3 | 0 | 5 | 3 | 2 | 2 | 3 | 0 | 2.56 | 3 | 2 | 0.521 | -0.941 | 0.45673589 | NA | 0 | 0.9268 | 0.7136 | 0.7136 | TRUE | 0.5058 |
| Hr | 781 | 651 | 652 | 717 | 715 | 773 | 607 | 692 | 704 | 714 | 807 | 763 | 714.32 | 668 | 761 | 1.142 | 0.192 | 0.04392215 | 0.12254135 | 0.0039 | 0.0057 | 0.0051 | 0.0051 | TRUE | 0.0797 |
| Hras1 | 2425 | 1619 | 1356 | 1393 | 1248 | 1381 | 1883 | 1722 | 1463 | 1387 | 1408 | 1363 | 1537.72 | 1689 | 1386 | 0.819 | -0.288 | 0.00151438 | 0.00973686 | 0.007 | 0.004 | 0.0053 | 0.0053 | TRUE | 0.1647 |
| Hrasls | 19 | 18 | 12 | 19 | 11 | 9 | 15 | 19 | 13 | 19 | 12 | 9 | 14.51 | 16 | 13 | 0.86 | -0.217 | 0.67656168 | NA | 0 | 0.1658 | 0.1257 | 0.1257 | TRUE | 0.5167 |
| Hrasls5 | 1 | 0 | 1 | 0 | 0 | 0 | 1 | 0 | 1 | 0 | 0 | 0 | 0.31 | 1 | 0 | 0.307 | -1.706 | 0.62894649 | NA | 0 | 7.6657 | 7.2 | 7.2 | TRUE | 0.2067 |
| Hrc | 52 | 45 | 69 | 121 | 115 | 97 | 40 | 48 | 74 | 120 | 130 | 96 | 84.77 | 54 | 115 | 2.144 | 1.101 | 1.91E-05 | 0.00032172 | 0.0457 | 0.0304 | 0.0343 | 0.0343 | TRUE | 0.8608 |
| Hrct1 | 12 | 5 | 7 | 2 | 2 | 4 | 9 | 5 | 8 | 2 | 2 | 4 | 5.06 | 7 | 3 | 0.368 | -1.441 | 0.10549802 | NA | 0 | 0.4704 | 0.318 | 0.318 | TRUE | 0.185 |
| Hrg | 11 | 9 | 8 | 7 | 6 | 7 | 9 | 10 | 9 | 7 | 7 | 7 | 7.9 | 9 | 7 | 0.773 | -0.371 | 0.57641896 | NA | 0 | 0.3025 | 0.1883 | 0.1883 | TRUE | 0.0124 |
| Hrh1 | 186 | 159 | 169 | 154 | 153 | 189 | 144 | 169 | 182 | 153 | 173 | 187 | 168.07 | 165 | 171 | 1.037 | 0.053 | 0.74738532 | 0.85003409 | 0.0066 | 0.0165 | 0.0136 | 0.0136 | TRUE | 0.1429 |
| Hrh2 | 38 | 18 | 18 | 33 | 30 | 37 | 30 | 19 | 19 | 33 | 34 | 37 | 28.55 | 23 | 35 | 1.497 | 0.582 | 0.11393631 | 0.24294968 | 0 | 0.0854 | 0.0618 | 0.0618 | TRUE | 0.4856 |
| Hrh3 | 19 | 16 | 22 | 50 | 32 | 45 | 15 | 17 | 24 | 50 | 36 | 44 | 30.97 | 19 | 43 | 2.377 | 1.249 | 0.00069957 | 0.00557069 | 0 | 0.079 | 0.0596 | 0.0596 | TRUE | 0.2992 |
| Hrh4 | 0 | 0 | 0 | 0 | 0 | 0 | 0 | 0 | 0 | 0 | 0 | 0 | 0 | 0 | 0 | NA | NA | NA | NA | NA | NA | NA | NA | NA | NA |
| Hrk | 140 | 107 | 110 | 171 | 123 | 152 | 109 | 114 | 119 | 170 | 139 | 150 | 133.38 | 114 | 153 | 1.349 | 0.432 | 0.01329568 | 0.051071 | 0 | 0.0202 | 0.0143 | 0.0143 | TRUE | 0.1021 |
| Hrnr | 0 | 0 | 0 | 0 | 0 | 0 | 0 | 0 | 0 | 0 | 0 | 0 | 0 | 0 | 0 | NA | NA | NA | NA | NA | NA | NA | NA | NA | NA |
| Hrsp12 | 1534 | 1061 | 1134 | 1405 | 1119 | 1341 | 1191 | 1129 | 1224 | 1399 | 1262 | 1324 | 1254.7 | 1181 | 1328 | 1.125 | 0.17 | 0.02345748 | 0.07762856 | 0.0013 | 0.0043 | 0.0032 | 0.0032 | TRUE | 0.0258 |
| Hs1bp3 | 2851 | 2148 | 2219 | 2221 | 2139 | 2411 | 2214 | 2285 | 2394 | 2211 | 2413 | 2380 | 2316.19 | 2298 | 2335 | 1.016 | 0.023 | 0.72120344 | 0.8327808 | 0.0014 | 0.0034 | 0.0027 | 0.0027 | TRUE | 0.0262 |
| Hs2st1 | 3308 | 2234 | 2043 | 1892 | 1713 | 1952 | 2569 | 2376 | 2204 | 1883 | 1932 | 1927 | 2148.75 | 2383 | 1914 | 0.802 | -0.318 | 9.06E-06 | 0.00017317 | 0.0027 | 0.0035 | 0.0032 | 0.0032 | TRUE | 0.0577 |
| Hs3st1 | 62 | 37 | 33 | 34 | 26 | 15 | 48 | 39 | 36 | 34 | 29 | 15 | 33.52 | 41 | 26 | 0.629 | -0.669 | 0.07213291 | 0.17548862 | 0.0524 | 0.0731 | 0.0681 | 0.0681 | TRUE | 0.9154 |
| Hs3st2 | 4 | 2 | 0 | 2 | 2 | 0 | 3 | 2 | 0 | 2 | 2 | 0 | 1.58 | 2 | 1 | 0.784 | -0.351 | 0.83082926 | NA | 0 | 1.5023 | 1.3079 | 1.3079 | TRUE | 0.4656 |
| Hs3st3a1 | 48 | 21 | 45 | 35 | 35 | 31 | 37 | 22 | 49 | 35 | 39 | 31 | 35.52 | 36 | 35 | 0.968 | -0.047 | 0.89420427 | 0.94184685 | 0.0468 | 0.0692 | 0.0632 | 0.0632 | TRUE | 0.248 |
| Hs3st3b1 | 1166 | 1038 | 1156 | 1152 | 1121 | 1316 | 906 | 1104 | 1247 | 1147 | 1265 | 1299 | 1161.23 | 1086 | 1237 | 1.141 | 0.191 | 0.07864051 | 0.18686764 | 0.0133 | 0.0045 | 0.0076 | 0.0076 | TRUE | 0.2597 |

| | | | | | | | | | | | | | | | | | | | | | | | | | |
|---|---|---|---|---|---|---|---|---|---|---|---|---|---|---|---|---|---|---|---|---|---|---|---|---|---|
| Hs3st4 | 11 | 4 | 3 | 4 | 1 | 1 | 9 | 4 | 3 | 4 | 1 | 1 | 3.69 | 5 | 2 | 0.377 | -1.406 | 0.19223918 | NA | 0 | 0.645 | 0.4968 | 0.4968 | TRUE | 0.7489 |
| Hs3st5 | 7 | 6 | 3 | 0 | 0 | 2 | 5 | 6 | 3 | 0 | 0 | 2 | 2.84 | 5 | 1 | 0.135 | -2.888 | 0.04673294 | NA | 0 | 0.8374 | 0.6625 | 0.6625 | TRUE | 0.9732 |
| Hs3st6 | 117 | 84 | 96 | 108 | 103 | 142 | 91 | 89 | 104 | 108 | 116 | 140 | 107.95 | 95 | 121 | 1.285 | 0.362 | 0.06682823 | 0.16562786 | 0 | 0.0244 | 0.0187 | 0.0187 | TRUE | 0.1926 |
| Hs6st1 | 4230 | 3413 | 3106 | 2894 | 2955 | 3131 | 3285 | 3630 | 3351 | 2881 | 3334 | 3091 | 3262 | 3422 | 3102 | 0.906 | -0.142 | 0.04717257 | 0.12915939 | 0.0036 | 0.0032 | 0.0034 | 0.0034 | TRUE | 0.0515 |
| Hs6st2 | 200 | 130 | 106 | 118 | 82 | 117 | 155 | 138 | 114 | 117 | 93 | 115 | 122.24 | 136 | 108 | 0.796 | -0.329 | 0.09148094 | 0.20723901 | 0.0122 | 0.0218 | 0.0191 | 0.0191 | TRUE | 0.1944 |
| Hs6st3 | 20 | 12 | 5 | 6 | 4 | 8 | 16 | 13 | 5 | 6 | 5 | 8 | 8.68 | 11 | 6 | 0.542 | -0.883 | 0.20128159 | NA | 0 | 0.2755 | 0.2179 | 0.2179 | TRUE | 0.5182 |
| Hsbp1 | 4175 | 2978 | 3125 | 3111 | 2683 | 3375 | 3243 | 3168 | 3372 | 3097 | 3027 | 3332 | 3206.18 | 3261 | 3152 | 0.967 | -0.049 | 0.433894 | 0.60572701 | 0.0013 | 0.0032 | 0.0025 | 0.0025 | TRUE | 0.0304 |
| Hsbp1l1 | 25 | 12 | 11 | 19 | 17 | 14 | 19 | 13 | 12 | 19 | 19 | 14 | 15.99 | 15 | 17 | 1.161 | 0.216 | 0.6571096 | NA | 0 | 0.1506 | 0.108 | 0.108 | TRUE | 0.4444 |
| Hscb | 669 | 501 | 567 | 691 | 601 | 607 | 520 | 533 | 612 | 688 | 678 | 599 | 604.88 | 555 | 655 | 1.182 | 0.241 | 0.01871621 | 0.06585693 | 0.0051 | 0.0063 | 0.0059 | 0.0059 | TRUE | 0.0929 |
| Hsd11b1 | 1 | 0 | 0 | 0 | 0 | 0 | 1 | 0 | 0 | 0 | 0 | 0 | 0.13 | 0 | 0 | 0.552 | -0.857 | 0.8336548 | NA | 0 | 18.3121 | 10 | 10 | TRUE | 0.2566 |
| Hsd11b2 | 150 | 107 | 93 | 133 | 143 | 145 | 116 | 114 | 100 | 132 | 161 | 143 | 127.92 | 110 | 145 | 1.316 | 0.396 | 0.0268955 | 0.0856545 | 0 | 0.021 | 0.0152 | 0.0152 | TRUE | 0.089 |
| Hsd17b1 | 23 | 16 | 17 | 38 | 13 | 20 | 18 | 17 | 18 | 38 | 15 | 20 | 20.91 | 18 | 24 | 1.363 | 0.447 | 0.33465959 | 0.50875235 | 0.0827 | 0.1158 | 0.106 | 0.106 | TRUE | 1.0729 |
| Hsd17b10 | 2548 | 1891 | 2180 | 2455 | 2294 | 2540 | 1979 | 2011 | 2352 | 2444 | 2588 | 2507 | 2313.58 | 2114 | 2513 | 1.189 | 0.25 | 0.00117074 | 0.00812808 | 0.0045 | 0.0035 | 0.0038 | 0.0038 | TRUE | 0.1163 |
| Hsd17b11 | 1481 | 1070 | 1169 | 1373 | 1239 | 1496 | 1150 | 1138 | 1261 | 1367 | 1398 | 1477 | 1298.48 | 1183 | 1414 | 1.196 | 0.258 | 0.00060443 | 0.00499371 | 0.0016 | 0.0043 | 0.0033 | 0.0033 | TRUE | 0.039 |
| Hsd17b12 | 6681 | 4716 | 5335 | 6286 | 5475 | 6307 | 5189 | 5016 | 5756 | 6257 | 6176 | 6226 | 5770.17 | 5320 | 6220 | 1.169 | 0.226 | 0.00032201 | 0.00307037 | 0.0024 | 0.0028 | 0.0027 | 0.0027 | TRUE | 0.062 |
| Hsd17b13 | 2 | 3 | 2 | 1 | 0 | 2 | 2 | 3 | 2 | 1 | 0 | 2 | 1.65 | 2 | 1 | 0.444 | -1.17 | 0.44674399 | NA | 0 | 1.4429 | 0.9809 | 0.9809 | TRUE | 0.3261 |
| Hsd17b14 | 3520 | 2505 | 2743 | 3083 | 2869 | 3397 | 2734 | 2665 | 2960 | 3069 | 3237 | 3353 | 3002.79 | 2786 | 3220 | 1.156 | 0.209 | 0.00152192 | 0.00975698 | 0.002 | 0.0032 | 0.0028 | 0.0028 | TRUE | 0.0355 |
| Hsd17b2 | 2 | 0 | 1 | 3 | 1 | 1 | 2 | 0 | 1 | 3 | 1 | 1 | 1.29 | 1 | 2 | 1.878 | 0.909 | 0.59350075 | NA | 0 | 1.8412 | 1.2591 | 1.2591 | TRUE | 0.3462 |
| Hsd17b3 | 2 | 4 | 2 | 2 | 1 | 2 | 2 | 4 | 2 | 2 | 1 | 2 | 2.18 | 3 | 2 | 0.653 | -0.615 | 0.63125515 | NA | 0 | 1.0913 | 0.6943 | 0.6943 | TRUE | 0.3073 |
| Hsd17b4 | 5684 | 3926 | 4282 | 4617 | 4170 | 4556 | 4415 | 4176 | 4620 | 4596 | 4704 | 4497 | 4501.37 | 4404 | 4599 | 1.044 | 0.063 | 0.28637134 | 0.45837651 | 0.0012 | 0.003 | 0.0023 | 0.0023 | TRUE | 0.0245 |
| Hsd17b6 | 2 | 2 | 2 | 2 | 1 | 1 | 2 | 2 | 2 | 2 | 1 | 1 | 1.66 | 2 | 1 | 0.711 | -0.492 | 0.73096965 | NA | 0 | 1.4324 | 0.8486 | 0.8486 | TRUE | 0.103 |
| Hsd17b7 | 1888 | 1388 | 1333 | 1614 | 1363 | 1448 | 1466 | 1476 | 1438 | 1607 | 1538 | 1429 | 1492.43 | 1460 | 1525 | 1.044 | 0.062 | 0.38438861 | 0.55839683 | 0.0012 | 0.004 | 0.003 | 0.003 | TRUE | 0.0365 |
| Hsd3b1 | 0 | 0 | 0 | 0 | 0 | 0 | 0 | 0 | 0 | 0 | 0 | 0 | 0 | 0 | 0 | NA | NA | NA | NA | NA | NA | NA | NA | NA | NA |
| Hsd3b2 | 1 | 0 | 0 | 0 | 1 | 0 | 1 | 0 | 0 | 0 | 1 | 0 | 0.32 | 0 | 0 | 1.075 | 0.105 | 0.97638847 | NA | 0 | 7.4681 | 7.0392 | 7.0392 | TRUE | 0.5075 |
| Hsd3b3 | 0 | 0 | 0 | 0 | 0 | 0 | 0 | 0 | 0 | 0 | 0 | 0 | 0 | 0 | 0 | NA | NA | NA | NA | NA | NA | NA | NA | NA | NA |
| Hsd3b4 | 0 | 0 | 0 | 0 | 0 | 0 | 0 | 0 | 0 | 0 | 0 | 0 | 0 | 0 | 0 | NA | NA | NA | NA | NA | NA | NA | NA | NA | NA |
| Hsd3b5 | 0 | 0 | 0 | 0 | 0 | 0 | 0 | 0 | 0 | 0 | 0 | 0 | 0 | 0 | 0 | NA | NA | NA | NA | NA | NA | NA | NA | NA | NA |
| Hsd3b6 | 0 | 0 | 0 | 0 | 0 | 0 | 0 | 0 | 0 | 0 | 0 | 0 | 0 | 0 | 0 | NA | NA | NA | NA | NA | NA | NA | NA | NA | NA |
| Hsd3b7 | 425 | 322 | 366 | 444 | 409 | 479 | 330 | 343 | 395 | 442 | 461 | 473 | 407.28 | 356 | 459 | 1.292 | 0.37 | 0.00090843 | 0.00675553 | 0.0023 | 0.0082 | 0.0065 | 0.0065 | TRUE | 0.103 |
| Hsdl1 | 2583 | 1896 | 2030 | 1992 | 1793 | 1924 | 2006 | 2017 | 2190 | 1983 | 2023 | 1899 | 2019.67 | 2071 | 1968 | 0.951 | -0.073 | 0.27268454 | 0.44367209 | 0.0012 | 0.0036 | 0.0027 | 0.0027 | TRUE | 0.0303 |
| Hsdl2 | 1264 | 901 | 880 | 1055 | 898 | 994 | 982 | 958 | 950 | 1050 | 1013 | 981 | 989 | 963 | 1015 | 1.053 | 0.075 | 0.32003223 | 0.49392297 | 0 | 0.0048 | 0.003 | 0.003 | TRUE | 0.0113 |
| Hsf1 | 2321 | 1843 | 1686 | 1581 | 1459 | 1796 | 1803 | 1960 | 1819 | 1574 | 1646 | 1773 | 1762.46 | 1861 | 1664 | 0.895 | -0.16 | 0.02738248 | 0.08656884 | 0.0021 | 0.0038 | 0.0032 | 0.0032 | TRUE | 0.0399 |
| Hsf2 | 3243 | 2332 | 2348 | 2289 | 1951 | 2353 | 2519 | 2481 | 2533 | 2278 | 2201 | 2323 | 2389.14 | 2511 | 2267 | 0.903 | -0.147 | 0.01188698 | 0.0470118 | 0 | 0.0034 | 0.002 | 0.002 | TRUE | 0.0078 |
| Hsf2bp | 3747 | 2797 | 2971 | 2724 | 2379 | 2656 | 2910 | 2975 | 3206 | 2711 | 2684 | 2622 | 2851.34 | 3030 | 2672 | 0.882 | -0.181 | 0.00346767 | 0.01850069 | 0.0011 | 0.0033 | 0.0024 | 0.0024 | TRUE | 0.0309 |
| Hsf3 | 35 | 22 | 25 | 5 | 6 | 3 | 27 | 23 | 27 | 5 | 7 | 3 | 15.38 | 26 | 5 | 0.187 | -2.417 | 2.42E-05 | NA | 0 | 0.1566 | 0.1107 | 0.1107 | TRUE | 0.2479 |
| Hsf4 | 97 | 91 | 110 | 178 | 157 | 184 | 75 | 97 | 119 | 177 | 177 | 182 | 137.79 | 97 | 179 | 1.859 | 0.894 | 2.78E-06 | 6.50E-05 | 0.0145 | 0.0196 | 0.0183 | 0.0183 | TRUE | 0.404 |
| Hsf5 | 129 | 81 | 114 | 178 | 117 | 133 | 100 | 86 | 123 | 177 | 132 | 131 | 124.97 | 103 | 147 | 1.427 | 0.513 | 0.0118362 | 0.04686283 | 0.0225 | 0.0214 | 0.0217 | 0.0217 | TRUE | 0.3456 |
| Hsfy2 | 0 | 0 | 0 | 0 | 0 | 0 | 0 | 0 | 0 | 0 | 0 | 0 | 0 | 0 | 0 | NA | NA | NA | NA | NA | NA | NA | NA | NA | NA |
| Hsh2d | 86 | 64 | 93 | 93 | 81 | 86 | 67 | 68 | 100 | 93 | 91 | 85 | 84.01 | 78 | 90 | 1.149 | 0.2 | 0.38687145 | 0.56071448 | 0.0158 | 0.0306 | 0.0265 | 0.0265 | TRUE | 0.5326 |
| Hsp90aa1 | 220315 | 153880 | 148792 | 144769 | 120961 | 138903 | 171109 | 163680 | 160545 | 144103 | 136457 | 137119 | 152168.83 | 165111 | 139226 | 0.843 | -0.246 | 6.68E-07 | 2.09E-05 | 0.001 | 0.0024 | 0.0018 | 0.0018 | TRUE | 0.0124 |
| Hsp90ab1 | 401502 | 287955 | 275022 | 292132 | 259974 | 300011 | 311830 | 306294 | 296746 | 290788 | 293278 | 296157 | 299182.11 | 304957 | 293408 | 0.962 | -0.056 | 0.20547137 | 0.36719209 | 3.00E-04 | 0.0024 | 0.0014 | 0.0014 | TRUE | 0.0068 |
| Hsp90b1 | 50164 | 37581 | 39391 | 43944 | 36755 | 43025 | 38960 | 39974 | 42503 | 43742 | 41463 | 42472 | 41519.13 | 40479 | 42559 | 1.051 | 0.072 | 0.16827854 | 0.31979859 | 0.0013 | 0.0025 | 0.002 | 0.002 | TRUE | 0.0234 |
| Hspa12a | 780 | 605 | 711 | 918 | 857 | 980 | 606 | 644 | 767 | 914 | 967 | 967 | 810.74 | 672 | 949 | 1.415 | 0.501 | 3.92E-07 | 1.32E-05 | 0.0065 | 0.0053 | 0.0058 | 0.0058 | TRUE | 0.1782 |
| Hspa12b | 64 | 41 | 36 | 63 | 49 | 46 | 50 | 44 | 39 | 63 | 55 | 45 | 49.26 | 44 | 54 | 1.23 | 0.298 | 0.29138394 | 0.46351736 | 0 | 0.0505 | 0.0372 | 0.0372 | TRUE | 0.1835 |
| Hspa13 | 1596 | 1067 | 947 | 1126 | 872 | 1043 | 1240 | 1135 | 1022 | 1121 | 984 | 1030 | 1088.41 | 1132 | 1045 | 0.922 | -0.117 | 0.19869382 | 0.35898506 | 0.006 | 0.0046 | 0.0051 | 0.0051 | TRUE | 0.0865 |
| Hspa14 | 9031 | 6414 | 5951 | 5909 | 5261 | 5926 | 7014 | 6822 | 6421 | 5882 | 5935 | 5850 | 6320.7 | 6752 | 5889 | 0.872 | -0.198 | 0.00027408 | 0.00271107 | 8.00E-04 | 0.0028 | 0.002 | 0.002 | TRUE | 0.0224 |
| Hspa1a | 72 | 76 | 51 | 41 | 56 | 60 | 56 | 81 | 55 | 41 | 63 | 59 | 59.17 | 64 | 54 | 0.852 | -0.231 | 0.41129409 | 0.58412425 | 0.0329 | 0.0425 | 0.0398 | 0.0398 | TRUE | 0.4528 |
| Hspa1b | 341 | 310 | 239 | 185 | 165 | 156 | 265 | 330 | 258 | 184 | 186 | 154 | 229.46 | 284 | 175 | 0.615 | -0.701 | 4.56E-06 | 9.83E-05 | 0.0108 | 0.0128 | 0.0122 | 0.0122 | TRUE | 0.2158 |
| Hspa1l | 7 | 3 | 2 | 4 | 4 | 1 | 5 | 3 | 2 | 4 | 5 | 1 | 3.38 | 3 | 3 | 0.856 | -0.224 | 0.83279233 | NA | 0 | 0.7041 | 0.5122 | 0.5122 | TRUE | 0.515 |
| Hspa2 | 576 | 436 | 451 | 414 | 376 | 486 | 447 | 464 | 487 | 412 | 424 | 480 | 452.29 | 466 | 439 | 0.943 | -0.085 | 0.42503774 | 0.59781097 | 0.002 | 0.0077 | 0.0059 | 0.0059 | TRUE | 0.0791 |
| Hspa4 | 50412 | 35317 | 30785 | 30842 | 27100 | 31566 | 39153 | 37566 | 33217 | 30700 | 30572 | 31161 | 33728.01 | 36645 | 30811 | 0.841 | -0.25 | 0.00010124 | 0.00122375 | 0.0035 | 0.0025 | 0.003 | 0.003 | TRUE | 0.0819 |
| Hspa4l | 2135 | 1451 | 1626 | 1468 | 1095 | 1368 | 1658 | 1543 | 1754 | 1461 | 1235 | 1350 | 1500.49 | 1652 | 1349 | 0.817 | -0.292 | 0.00043466 | 0.00387561 | 0.0045 | 0.004 | 0.0043 | 0.0043 | TRUE | 0.0643 |
| Hspa5 | 28473 | 19735 | 17446 | 20846 | 16900 | 18596 | 22114 | 20992 | 18824 | 20750 | 19065 | 18357 | 20016.98 | 20643 | 19391 | 0.939 | -0.09 | 0.20326403 | 0.36472149 | 0.0052 | 0.0025 | 0.0036 | 0.0036 | TRUE | 0.0726 |
| Hspa8 | 125417 | 86703 | 78463 | 76193 | 69678 | 79278 | 97406 | 92225 | 84661 | 75842 | 78604 | 78260 | 84499.65 | 91431 | 77569 | 0.848 | -0.237 | 6.78E-05 | 0.00089719 | 0.0026 | 0.0025 | 0.0025 | 0.0025 | TRUE | 0.0514 |
| Hspa9 | 57160 | 40908 | 41525 | 45855 | 40276 | 45544 | 44394 | 43513 | 44805 | 45644 | 45436 | 44959 | 44791.78 | 44237 | 45346 | 1.025 | 0.036 | 0.40129297 | 0.57443865 | 1.00E-04 | 0.0025 | 0.0013 | 0.0013 | TRUE | 0.0025 |
| Hspb1 | 18080 | 13272 | 12120 | 13809 | 13111 | 14419 | 14042 | 14117 | 13077 | 13745 | 14791 | 14234 | 14001.07 | 13745 | 14257 | 1.037 | 0.052 | 0.34064751 | 0.51473462 | 0.0015 | 0.0026 | 0.0021 | 0.0021 | TRUE | 0.022 |
| Hspb11 | 317 | 212 | 249 | 389 | 263 | 333 | 246 | 226 | 269 | 387 | 297 | 329 | 292.17 | 247 | 338 | 1.37 | 0.454 | 0.00101148 | 0.00730522 | 0.0095 | 0.0105 | 0.0102 | 0.0102 | TRUE | 0.1891 |
| Hspb2 | 79 | 43 | 56 | 87 | 73 | 90 | 61 | 46 | 60 | 87 | 82 | 89 | 70.89 | 56 | 86 | 1.534 | 0.617 | 0.00889353 | 0.03803567 | 0 | 0.0359 | 0.0255 | 0.0255 | TRUE | 0.199 |
| Hspb3 | 6 | 2 | 5 | 7 | 2 | 2 | 5 | 2 | 5 | 7 | 2 | 2 | 3.9 | 4 | 4 | 0.92 | -0.12 | 0.90438521 | NA | 0 | 0.6106 | 0.4613 | 0.4613 | TRUE | 0.9273 |
| Hspb6 | 88 | 66 | 73 | 88 | 78 | 79 | 68 | 70 | 79 | 88 | 88 | 78 | 78.48 | 72 | 85 | 1.169 | 0.225 | 0.30403952 | 0.47654753 | 0 | 0.0326 | 0.0217 | 0.0217 | TRUE | 0.0507 |
| Hspb7 | 13 | 11 | 5 | 18 | 9 | 9 | 10 | 12 | 5 | 18 | 10 | 9 | 10.69 | 9 | 12 | 1.355 | 0.438 | 0.47453752 | NA | 6.00E-04 | 0.2241 | 0.1757 | 0.1757 | TRUE | 0.6497 |
| Hspb8 | 1394 | 1076 | 1464 | 1439 | 1393 | 1464 | 1083 | 1145 | 1580 | 1432 | 1571 | 1445 | 1375.98 | 1269 | 1483 | 1.17 | 0.226 | 0.05849292 | 0.15072007 | 0.023 | 0.0041 | 0.0096 | 0.0096 | TRUE | 0.5479 |
| Hspb9 | 9 | 8 | 10 | 8 | 5 | 9 | 7 | 9 | 11 | 8 | 6 | 9 | 8.13 | 9 | 8 | 0.866 | -0.207 | 0.75524723 | NA | 0 | 0.294 | 0.1943 | 0.1943 | TRUE | 0.1296 |
| Hspbap1 | 5196 | 3918 | 3839 | 4028 | 3625 | 4351 | 4036 | 4168 | 4142 | 4009 | 4089 | 4295 | 4123.21 | 4115 | 4131 | 1.004 | 0.006 | 0.91241368 | 0.95183773 | 6.00E-04 | 0.003 | 0.002 | 0.002 | TRUE | 0.0148 |
| Hspbp1 | 2871 | 2199 | 1961 | 2083 | 1845 | 2067 | 2230 | 2339 | 2116 | 2073 | 2081 | 2040 | 2146.66 | 2228 | 2065 | 0.927 | -0.11 | 0.08572939 | 0.19785729 | 8.00E-04 | 0.0035 | 0.0025 | 0.0025 | TRUE | 0.0228 |
| Hspd1 | 54063 | 36548 | 32767 | 34152 | 28356 | 32808 | 41988 | 38876 | 35355 | 33995 | 31989 | 32387 | 35764.89 | 38740 | 32790 | 0.846 | -0.241 | 0.00030156 | 0.00291821 | 0.0042 | 0.0025 | 0.0032 | 0.0032 | TRUE | 0.0715 |
| Hspe1 | 11974 | 8536 | 8212 | 8564 | 7381 | 8171 | 9300 | 9080 | 8861 | 8525 | 8327 | 8066 | 8692.86 | 9080 | 8306 | 0.915 | -0.129 | 0.01089229 | 0.0439894 | 5.00E-04 | 0.0027 | 0.0017 | 0.0017 | TRUE | 0.0078 |
| Hspg2 | 19013 | 12944 | 13534 | 18155 | 16605 | 17655 | 14767 | 13768 | 14603 | 18071 | 18732 | 17428 | 16228.31 | 14379 | 18077 | 1.257 | 0.33 | 6.85E-10 | 4.85E-08 | 0.0012 | 0.0026 | 0.002 | 0.002 | TRUE | 0.0169 |
| Hsph1 | 7562 | 5405 | 3850 | 3589 | 2954 | 3625 | 5873 | 5749 | 4154 | 3572 | 3332 | 3578 | 4376.63 | 5259 | 3494 | 0.664 | -0.59 | 2.81E-08 | 1.26E-06 | 0.0171 | 0.003 | 0.0079 | 0.0079 | TRUE | 0.4104 |

| | | | | | | | | | | | | | | | | | | | | | | | | | |
|---|---|---|---|---|---|---|---|---|---|---|---|---|---|---|---|---|---|---|---|---|---|---|---|---|---|
| Htatip2 | 836 | 526 | 553 | 605 | 549 | 584 | 649 | 559 | 597 | 602 | 619 | 576 | 600.59 | 602 | 599 | 0.994 | -0.009 | 0.92206404 | 0.95711845 | 0.002 | 0.0064 | 0.0049 | 0.0049 | TRUE | 0.0592 |
| Htatsf1 | 4930 | 3553 | 3596 | 4034 | 3593 | 3976 | 3829 | 3779 | 3880 | 4015 | 4053 | 3925 | 3913.65 | 3829 | 3998 | 1.044 | 0.062 | 0.23367239 | 0.39984191 | 0 | 0.003 | 0.0017 | 0.0017 | TRUE | 0.0031 |
| Htr1a | 0 | 0 | 0 | 0 | 0 | 0 | 0 | 0 | 0 | 0 | 0 | 0 | 0 | 0 | 0 | NA | NA | NA | NA | NA | NA | NA | NA | NA | NA |
| Htr1b | 10 | 9 | 6 | 9 | 11 | 4 | 8 | 10 | 6 | 9 | 12 | 4 | 8.19 | 8 | 8 | 1.056 | 0.078 | 0.910147 | NA | 0 | 0.2919 | 0.221 | 0.221 | TRUE | 0.3027 |
| Htr1d | 10 | 16 | 8 | 11 | 6 | 11 | 8 | 17 | 9 | 11 | 7 | 11 | 10.33 | 11 | 10 | 0.868 | -0.205 | 0.74086965 | NA | 0 | 0.2318 | 0.1781 | 0.1781 | TRUE | 0.7599 |
| Htr1f | 1 | 0 | 0 | 0 | 0 | 0 | 1 | 0 | 0 | 0 | 0 | 0 | 0.13 | 0 | 0 | 0.552 | -0.857 | 0.8336548 | NA | 0 | 18.3121 | 10 | 10 | TRUE | 0.2566 |
| Htr2a | 0 | 1 | 1 | 0 | 0 | 0 | 0 | 1 | 1 | 0 | 0 | 0 | 0.36 | 1 | 0 | 0.271 | -1.881 | 0.5755355 | NA | 0 | 6.6391 | 6.4428 | 6.4428 | TRUE | 0.3513 |
| Htr2b | 0 | 0 | 0 | 0 | 0 | 0 | 0 | 0 | 0 | 0 | 0 | 0 | 0 | 0 | 0 | NA | NA | NA | NA | NA | NA | NA | NA | NA | NA |
| Htr2c | 0 | 0 | 0 | 0 | 0 | 0 | 0 | 0 | 0 | 0 | 0 | 0 | 0 | 0 | 0 | NA | NA | NA | NA | NA | NA | NA | NA | NA | NA |
| Htr3a | 74 | 43 | 63 | 74 | 58 | 70 | 57 | 46 | 68 | 74 | 65 | 69 | 63.23 | 57 | 69 | 1.217 | 0.284 | 0.2603869 | 0.43057079 | 0 | 0.0399 | 0.0298 | 0.0298 | TRUE | 0.2325 |
| Htr3b | 3 | 0 | 1 | 1 | 0 | 1 | 2 | 0 | 1 | 1 | 0 | 1 | 0.9 | 1 | 1 | 0.574 | -0.8 | 0.7018232 | NA | 0 | 2.64 | 1.9656 | 1.9656 | TRUE | 0.1659 |
| Htr4 | 26 | 15 | 17 | 28 | 16 | 19 | 20 | 16 | 18 | 28 | 18 | 19 | 19.86 | 18 | 22 | 1.184 | 0.244 | 0.5744703 | 0.72653815 | 0 | 0.1218 | 0.086 | 0.086 | TRUE | 0.3763 |
| Htr5a | 1 | 2 | 1 | 1 | 0 | 0 | 1 | 2 | 1 | 1 | 0 | 0 | 0.83 | 1 | 0 | 0.282 | -1.824 | 0.39881264 | NA | 0 | 2.8588 | 2 | 2 | TRUE | 0.3906 |
| Htr5b | 92 | 64 | 46 | 48 | 37 | 53 | 71 | 68 | 50 | 48 | 42 | 52 | 55.17 | 63 | 47 | 0.748 | -0.419 | 0.12449623 | 0.25908921 | 0 | 0.0454 | 0.035 | 0.035 | TRUE | 0.2874 |
| Htr6 | 48 | 37 | 37 | 55 | 43 | 49 | 37 | 39 | 40 | 55 | 49 | 48 | 44.7 | 39 | 51 | 1.304 | 0.383 | 0.17870525 | 0.33332039 | 0 | 0.0555 | 0.0357 | 0.0357 | TRUE | 0.0441 |
| Htr7 | 7 | 5 | 0 | 1 | 0 | 6 | 5 | 5 | 0 | 1 | 0 | 6 | 2.95 | 3 | 2 | 0.645 | -0.633 | 0.64704855 | NA | 1.993 | 0.807 | 1.0243 | 1.0243 | TRUE | 1.9621 |
| Htra1 | 2511 | 1753 | 1727 | 1602 | 1452 | 1594 | 1950 | 1865 | 1863 | 1595 | 1638 | 1574 | 1747.4 | 1893 | 1602 | 0.846 | -0.242 | 0.0001419 | 0.00162905 | 0 | 0.0038 | 0.0023 | 0.0023 | TRUE | 0.0089 |
| Htra2 | 2325 | 1725 | 1344 | 1346 | 1176 | 1396 | 1806 | 1835 | 1450 | 1340 | 1327 | 1378 | 1522.55 | 1697 | 1348 | 0.794 | -0.333 | 0.0002753 | 0.0027174 | 0.008 | 0.004 | 0.0054 | 0.0054 | TRUE | 0.1926 |
| Htra3 | 48 | 29 | 32 | 63 | 69 | 79 | 37 | 31 | 35 | 63 | 78 | 78 | 53.53 | 34 | 73 | 2.118 | 1.083 | 7.58E-05 | 0.00097514 | 0 | 0.0467 | 0.0328 | 0.0328 | TRUE | 0.1371 |
| Htra4 | 18 | 7 | 9 | 17 | 10 | 14 | 14 | 7 | 10 | 17 | 11 | 14 | 12.19 | 10 | 14 | 1.334 | 0.416 | 0.45443013 | NA | 0 | 0.1968 | 0.1404 | 0.1404 | TRUE | 0.4184 |
| Htt | 7124 | 5262 | 5426 | 6665 | 6042 | 6524 | 5533 | 5597 | 5855 | 6634 | 6816 | 6440 | 6145.86 | 5662 | 6630 | 1.171 | 0.228 | 1.82E-05 | 0.00031011 | 6.00E-04 | 0.0028 | 0.0019 | 0.0019 | TRUE | 0.0108 |
| Hunk | 4437 | 3083 | 3167 | 2783 | 2667 | 2911 | 3446 | 3279 | 3417 | 2770 | 3009 | 2874 | 3132.5 | 3381 | 2884 | 0.853 | -0.23 | 0.00011425 | 0.00135515 | 7.00E-04 | 0.0032 | 0.0022 | 0.0022 | TRUE | 0.0171 |
| Hus1 | 5048 | 3691 | 3704 | 4147 | 3702 | 4242 | 3921 | 3926 | 3997 | 4128 | 4176 | 4188 | 4055.81 | 3948 | 4164 | 1.055 | 0.077 | 0.12971612 | 0.26638397 | 0 | 0.003 | 0.0016 | 0.0016 | TRUE | 0.0014 |
| Hus1b | 173 | 151 | 147 | 166 | 127 | 151 | 134 | 161 | 159 | 165 | 143 | 149 | 151.86 | 151 | 152 | 1.013 | 0.019 | 0.90769522 | 0.94923397 | 0 | 0.018 | 0.0133 | 0.0133 | TRUE | 0.1108 |
| Huwe1 | 46341 | 31794 | 29072 | 34554 | 32633 | 36512 | 35991 | 33819 | 31368 | 34395 | 36813 | 36043 | 34738.3 | 33726 | 35750 | 1.06 | 0.084 | 0.17066765 | 0.32232463 | 0.0029 | 0.0025 | 0.0027 | 0.0027 | TRUE | 0.0458 |
| Hvcn1 | 1020 | 662 | 615 | 756 | 700 | 790 | 792 | 704 | 664 | 753 | 790 | 780 | 747 | 720 | 774 | 1.072 | 0.101 | 0.27493013 | 0.44593467 | 0.0033 | 0.0056 | 0.0048 | 0.0048 | TRUE | 0.0955 |
| Hyal1 | 210 | 188 | 199 | 224 | 198 | 231 | 163 | 200 | 215 | 223 | 223 | 228 | 208.69 | 193 | 225 | 1.173 | 0.23 | 0.1242512 | 0.25869226 | 0.0056 | 0.0138 | 0.0114 | 0.0114 | TRUE | 0.2145 |
| Hyal2 | 1195 | 939 | 853 | 995 | 848 | 1061 | 928 | 999 | 920 | 990 | 957 | 1047 | 973.62 | 949 | 998 | 1.053 | 0.074 | 0.35198075 | 0.52657116 | 0.001 | 0.0049 | 0.0035 | 0.0035 | TRUE | 0.0244 |
| Hyal3 | 8 | 12 | 11 | 14 | 11 | 13 | 6 | 13 | 12 | 14 | 12 | 13 | 11.67 | 10 | 13 | 1.292 | 0.37 | 0.51521792 | NA | 0 | 0.2055 | 0.1456 | 0.1456 | TRUE | 0.5635 |
| Hyal4 | 0 | 0 | 0 | 0 | 0 | 0 | 0 | 0 | 0 | 0 | 0 | 0 | 0 | 0 | 0 | NA | NA | NA | NA | NA | NA | NA | NA | NA | NA |
| Hyal5 | 0 | 0 | 0 | 0 | 0 | 0 | 0 | 0 | 0 | 0 | 0 | 0 | 0 | 0 | 0 | NA | NA | NA | NA | NA | NA | NA | NA | NA | NA |
| Hyal6 | 2 | 1 | 1 | 2 | 1 | 1 | 2 | 1 | 1 | 2 | 1 | 1 | 1.3 | 1 | 1 | 1.101 | 0.139 | 0.93032816 | NA | 0 | 1.8251 | 1.0578 | 1.0578 | TRUE | 0.1028 |
| Hydin | 2396 | 1923 | 1846 | 2141 | 1930 | 2208 | 1861 | 2045 | 1992 | 2131 | 2177 | 2180 | 2064.36 | 1966 | 2163 | 1.101 | 0.139 | 0.03112065 | 0.09520709 | 8.00E-04 | 0.0036 | 0.0025 | 0.0025 | TRUE | 0.0275 |
| Hyi | 196 | 131 | 148 | 187 | 162 | 173 | 152 | 139 | 160 | 186 | 183 | 171 | 165.15 | 150 | 180 | 1.195 | 0.257 | 0.09720186 | 0.21687577 | 0 | 0.0168 | 0.0112 | 0.0112 | TRUE | 0.0406 |
| Hyls1 | 1278 | 883 | 825 | 916 | 786 | 852 | 993 | 939 | 890 | 912 | 887 | 841 | 910.25 | 941 | 880 | 0.934 | -0.098 | 0.23026701 | 0.39605372 | 0.0014 | 0.005 | 0.0037 | 0.0037 | TRUE | 0.0291 |
| Hyou1 | 15326 | 11463 | 11692 | 13783 | 11878 | 13785 | 11903 | 12193 | 12616 | 13720 | 13400 | 13608 | 12906.46 | 12237 | 13576 | 1.11 | 0.15 | 0.00177111 | 0.01102549 | 4.00E-04 | 0.0026 | 0.0016 | 0.0016 | TRUE | 0.0089 |
| Hypk | 5011 | 3531 | 3379 | 3526 | 2908 | 3707 | 3892 | 3756 | 3646 | 3510 | 3281 | 3659 | 3623.88 | 3765 | 3483 | 0.925 | -0.112 | 0.07322733 | 0.17715611 | 0.0016 | 0.0031 | 0.0025 | 0.0025 | TRUE | 0.0312 |
| I730028E13R | 0 | 0 | 0 | 0 | 0 | 0 | 0 | 0 | 0 | 0 | 0 | 0 | 0 | 0 | 0 | NA | NA | NA | NA | NA | NA | NA | NA | NA | NA |
| I730030J21R | 0 | 0 | 0 | 1 | 0 | 0 | 0 | 0 | 0 | 1 | 0 | 0 | 0.17 | 0 | 0 | 2.094 | 1.067 | 0.79379329 | NA | 0 | 14.2885 | 10 | 10 | TRUE | 0.3906 |
| I830012O16F | 0 | 0 | 0 | 0 | 0 | 0 | 0 | 0 | 0 | 0 | 0 | 0 | 0 | 0 | 0 | NA | NA | NA | NA | NA | NA | NA | NA | NA | NA |
| I830077J02R | 1 | 5 | 2 | 1 | 3 | 1 | 1 | 5 | 2 | 1 | 3 | 1 | 2.27 | 3 | 2 | 0.654 | -0.612 | 0.64586985 | NA | 0 | 1.0465 | 0.8044 | 0.8044 | TRUE | 0.4748 |
| Iah1 | 772 | 567 | 507 | 583 | 522 | 568 | 600 | 603 | 547 | 580 | 589 | 561 | 579.94 | 583 | 577 | 0.988 | -0.018 | 0.8446447 | 0.91195135 | 0 | 0.0065 | 0.0044 | 0.0044 | TRUE | 0.0333 |
| Iapp | 1 | 0 | 0 | 0 | 0 | 0 | 1 | 0 | 0 | 0 | 0 | 0 | 0.13 | 0 | 0 | 0.552 | -0.857 | 0.8336548 | NA | 0 | 18.3121 | 10 | 10 | TRUE | 0.2566 |
| Iars | 17806 | 12774 | 12099 | 12030 | 10348 | 12423 | 13829 | 13588 | 13055 | 11975 | 11674 | 12263 | 12730.52 | 13491 | 11971 | 0.887 | -0.173 | 0.00053858 | 0.00455764 | 6.00E-04 | 0.0026 | 0.0017 | 0.0017 | TRUE | 0.0097 |
| Iars2 | 10510 | 7615 | 8366 | 8574 | 7786 | 8549 | 8163 | 8100 | 9027 | 8535 | 8783 | 8439 | 8507.77 | 8430 | 8586 | 1.019 | 0.027 | 0.65151549 | 0.78364982 | 0.0016 | 0.0027 | 0.0024 | 0.0024 | TRUE | 0.0466 |
| Iba57 | 919 | 638 | 562 | 683 | 545 | 645 | 714 | 679 | 606 | 680 | 615 | 637 | 655.03 | 666 | 644 | 0.965 | -0.051 | 0.59398553 | 0.74083813 | 0.0033 | 0.006 | 0.0051 | 0.0051 | TRUE | 0.0716 |
| Ibsp | 0 | 0 | 0 | 0 | 0 | 0 | 0 | 0 | 0 | 0 | 0 | 0 | 0 | 0 | 0 | NA | NA | NA | NA | NA | NA | NA | NA | NA | NA |
| Ibtk | 9787 | 6867 | 6698 | 7791 | 6753 | 7663 | 7601 | 7304 | 7227 | 7755 | 7618 | 7565 | 7511.73 | 7377 | 7646 | 1.036 | 0.051 | 0.30024648 | 0.47244031 | 3.00E-04 | 0.0027 | 0.0016 | 0.0016 | TRUE | 0.0087 |
| Ica1 | 515 | 420 | 411 | 467 | 373 | 458 | 400 | 447 | 443 | 465 | 421 | 452 | 437.99 | 430 | 446 | 1.04 | 0.056 | 0.59354991 | 0.74083813 | 0 | 0.0078 | 0.0057 | 0.0057 | TRUE | 0.0467 |
| Ica1l | 35 | 21 | 14 | 18 | 9 | 16 | 27 | 22 | 15 | 18 | 10 | 16 | 18.08 | 21 | 15 | 0.677 | -0.564 | 0.23403471 | 0.40022231 | 0 | 0.1335 | 0.1042 | 0.1042 | TRUE | 0.231 |
| Icam1 | 4769 | 3788 | 4529 | 6214 | 5383 | 6540 | 3704 | 4029 | 4887 | 6185 | 6073 | 6456 | 5222.31 | 4207 | 6238 | 1.484 | 0.569 | 1.33E-10 | 1.05E-08 | 0.0108 | 0.0029 | 0.0055 | 0.0055 | TRUE | 0.2431 |
| Icam2 | 8 | 1 | 2 | 7 | 3 | 2 | 6 | 1 | 2 | 7 | 3 | 2 | 3.63 | 3 | 4 | 1.264 | 0.338 | 0.75158089 | NA | 0.292 | 0.6559 | 0.5457 | 0.5457 | TRUE | 1.2842 |
| Icam4 | 51 | 40 | 36 | 27 | 27 | 43 | 40 | 43 | 39 | 27 | 30 | 42 | 36.8 | 41 | 33 | 0.826 | -0.275 | 0.39897136 | 0.57224403 | 0 | 0.0668 | 0.0491 | 0.0491 | TRUE | 0.4197 |
| Icam5 | 99 | 82 | 103 | 104 | 101 | 110 | 77 | 87 | 111 | 104 | 114 | 109 | 100.22 | 92 | 109 | 1.191 | 0.252 | 0.22666544 | 0.39165004 | 0.0088 | 0.0261 | 0.0212 | 0.0212 | TRUE | 0.3203 |
| Ick | 1393 | 1093 | 1030 | 991 | 975 | 1061 | 1082 | 1163 | 1111 | 986 | 1100 | 1047 | 1081.59 | 1119 | 1044 | 0.934 | -0.099 | 0.2040077 | 0.36539875 | 0.0012 | 0.0046 | 0.0034 | 0.0034 | TRUE | 0.0284 |
| Icmt | 3816 | 2830 | 2956 | 3299 | 2794 | 3307 | 2964 | 3010 | 3189 | 3284 | 3152 | 3265 | 3143.95 | 3054 | 3234 | 1.059 | 0.083 | 0.15648955 | 0.30468671 | 7.00E-04 | 0.0032 | 0.0021 | 0.0021 | TRUE | 0.018 |
| Icos | 0 | 0 | 1 | 0 | 0 | 1 | 0 | 0 | 1 | 0 | 0 | 1 | 0.34 | 0 | 0 | 1.075 | 0.105 | 0.97570524 | NA | 0 | 6.885 | 6.5379 | 6.5379 | TRUE | 0.5365 |
| Icosl | 824 | 585 | 540 | 725 | 552 | 649 | 640 | 622 | 583 | 722 | 623 | 641 | 638.32 | 615 | 662 | 1.075 | 0.105 | 0.27233194 | 0.44323256 | 0.0027 | 0.0061 | 0.005 | 0.005 | TRUE | 0.0746 |
| Ict1 | 2667 | 1883 | 1869 | 2005 | 1751 | 2047 | 2071 | 2003 | 2017 | 1996 | 1975 | 2021 | 2013.78 | 2030 | 1997 | 0.983 | -0.024 | 0.68677087 | 0.80798786 | 0 | 0.0036 | 0.0021 | 0.0021 | TRUE | 0.0039 |
| Id1 | 1488 | 874 | 880 | 947 | 925 | 894 | 1156 | 930 | 950 | 943 | 1043 | 883 | 983.92 | 1012 | 956 | 0.943 | -0.085 | 0.40912011 | 0.58184614 | 0.0101 | 0.0048 | 0.0066 | 0.0066 | TRUE | 0.1907 |
| Id2 | 417 | 338 | 373 | 286 | 305 | 353 | 324 | 360 | 402 | 285 | 344 | 348 | 343.85 | 362 | 326 | 0.902 | -0.149 | 0.25256559 | 0.42147344 | 0.0094 | 0.0093 | 0.0093 | 0.0093 | TRUE | 0.1401 |
| Id3 | 688 | 485 | 303 | 243 | 226 | 209 | 534 | 516 | 327 | 242 | 255 | 206 | 346.72 | 459 | 234 | 0.509 | -0.974 | 1.53E-08 | 7.38E-07 | 0.0389 | 0.0093 | 0.0181 | 0.0181 | TRUE | 0.7235 |
| Id4 | 142 | 143 | 104 | 88 | 85 | 96 | 110 | 152 | 112 | 88 | 96 | 95 | 108.81 | 125 | 93 | 0.745 | -0.425 | 0.03667159 | 0.10767194 | 0.0114 | 0.0242 | 0.0204 | 0.0204 | TRUE | 0.3607 |
| Ide | 11627 | 8321 | 8985 | 9485 | 8328 | 9886 | 9030 | 8851 | 9695 | 9441 | 9395 | 9759 | 9361.85 | 9192 | 9532 | 1.037 | 0.053 | 0.33724521 | 0.51143059 | 0.0012 | 0.0027 | 0.0021 | 0.0021 | TRUE | 0.0278 |
| Idh1 | 11663 | 7894 | 8040 | 8861 | 7593 | 8626 | 9058 | 8397 | 8675 | 8820 | 8566 | 8515 | 8671.85 | 8710 | 8634 | 0.991 | -0.013 | 0.80338387 | 0.88615262 | 6.00E-04 | 0.0027 | 0.0018 | 0.0018 | TRUE | 0.0151 |
| Idh2 | 8691 | 6707 | 7470 | 8240 | 7809 | 8651 | 6750 | 7134 | 8060 | 8202 | 8809 | 8540 | 7915.91 | 7315 | 8517 | 1.165 | 0.22 | 0.00184968 | 0.01139544 | 0.0047 | 0.0027 | 0.0035 | 0.0035 | TRUE | 0.0968 |
| Idh3a | 10452 | 7363 | 7746 | 8572 | 7494 | 8869 | 8118 | 7832 | 8358 | 8533 | 8454 | 8755 | 8341.51 | 8103 | 8581 | 1.059 | 0.083 | 0.1032186 | 0.22641245 | 5.00E-04 | 0.0027 | 0.0017 | 0.0017 | TRUE | 0.0103 |
| Idh3b | 5894 | 4139 | 4473 | 5016 | 4369 | 4971 | 4578 | 4403 | 4826 | 4993 | 4929 | 4907 | 4772.55 | 4602 | 4943 | 1.074 | 0.103 | 0.06547181 | 0.16342798 | 8.00E-04 | 0.0029 | 0.0021 | 0.0021 | TRUE | 0.0217 |
| Idh3g | 7618 | 5662 | 5766 | 6493 | 5666 | 6549 | 5917 | 6023 | 6221 | 6463 | 6392 | 6465 | 6246.75 | 6054 | 6440 | 1.064 | 0.09 | 0.07113777 | 0.17352385 | 2.00E-04 | 0.0028 | 0.0016 | 0.0016 | TRUE | 0.0072 |

| | | | | | | | | | | | | | | | | | | | | | | | | | |
|---|---|---|---|---|---|---|---|---|---|---|---|---|---|---|---|---|---|---|---|---|---|---|---|---|---|
| Idi1 | 2204 | 1525 | 1579 | 1744 | 1476 | 1730 | 1712 | 1622 | 1704 | 1736 | 1665 | 1708 | 1691.07 | 1679 | 1703 | 1.014 | 0.02 | 0.75580259 | 0.85511255 | 0 | 0.0038 | 0.0024 | 0.0024 | TRUE | 0.0105 |
| Idi2 | 0 | 0 | 0 | 0 | 0 | 0 | 0 | 0 | 0 | 0 | 0 | 0 | 0 | 0 | 0 | NA | NA | NA | NA | NA | NA | NA | NA | NA | NA |
| Idnk | 557 | 398 | 368 | 386 | 348 | 462 | 433 | 423 | 397 | 384 | 393 | 456 | 414.31 | 418 | 411 | 0.984 | -0.024 | 0.82995226 | 0.90264098 | 0.003 | 0.0081 | 0.0066 | 0.0066 | TRUE | 0.1081 |
| Ido1 | 11 | 0 | 2 | 2 | 4 | 3 | 9 | 0 | 2 | 2 | 5 | 3 | 3.36 | 4 | 3 | 0.847 | -0.239 | 0.84068954 | NA | 0.7784 | 0.7076 | 0.7258 | 0.7258 | TRUE | 0.9493 |
| Ido2 | 57 | 41 | 35 | 55 | 55 | 45 | 44 | 44 | 38 | 55 | 62 | 44 | 47.81 | 42 | 54 | 1.277 | 0.353 | 0.21515565 | 0.37765193 | 0 | 0.052 | 0.0372 | 0.0372 | TRUE | 0.1948 |
| Ids | 1345 | 1013 | 1005 | 1044 | 937 | 1070 | 1045 | 1078 | 1084 | 1039 | 1057 | 1056 | 1059.83 | 1069 | 1051 | 0.984 | -0.024 | 0.7406065 | 0.84588493 | 0 | 0.0047 | 0.0028 | 0.0028 | TRUE | 0.005 |
| Idua | 274 | 242 | 336 | 444 | 368 | 408 | 213 | 257 | 363 | 442 | 415 | 403 | 348.77 | 278 | 420 | 1.52 | 0.604 | 0.00023152 | 0.00239098 | 0.035 | 0.0092 | 0.0164 | 0.0164 | TRUE | 0.7987 |
| Ier2 | 6941 | 4830 | 3916 | 4444 | 4070 | 4203 | 5391 | 5138 | 4225 | 4424 | 4591 | 4149 | 4652.94 | 4918 | 4388 | 0.892 | -0.165 | 0.05497821 | 0.14413124 | 0.0087 | 0.0029 | 0.0051 | 0.0051 | TRUE | 0.1843 |
| Ier3 | 907 | 934 | 1196 | 1111 | 1221 | 1386 | 704 | 993 | 1290 | 1106 | 1377 | 1368 | 1139.98 | 996 | 1284 | 1.29 | 0.367 | 0.16384789 | 0.31462634 | 0.0491 | 0.0045 | 0.0173 | 0.0491 | TRUE | 0.2495 |
| Ier3ip1 | 1927 | 1340 | 1442 | 1519 | 1295 | 1537 | 1497 | 1425 | 1556 | 1512 | 1461 | 1517 | 1494.67 | 1493 | 1497 | 1.003 | 0.004 | 0.95042489 | 0.97274404 | 4.00E-04 | 0.004 | 0.0027 | 0.0027 | TRUE | 0.018 |
| Ier5 | 213 | 175 | 187 | 181 | 167 | 207 | 165 | 186 | 202 | 180 | 188 | 204 | 187.71 | 184 | 191 | 1.039 | 0.056 | 0.71475693 | 0.82854995 | 0 | 0.0151 | 0.0113 | 0.0113 | TRUE | 0.0967 |
| Ier5l | 733 | 570 | 483 | 524 | 483 | 477 | 569 | 606 | 521 | 522 | 545 | 471 | 539.01 | 565 | 513 | 0.905 | -0.144 | 0.16358151 | 0.31417234 | 0.0036 | 0.0068 | 0.0058 | 0.0058 | TRUE | 0.0593 |
| Iffo1 | 616 | 510 | 478 | 470 | 413 | 468 | 478 | 542 | 516 | 468 | 466 | 462 | 488.73 | 512 | 465 | 0.91 | -0.136 | 0.16977855 | 0.32136735 | 0 | 0.0073 | 0.005 | 0.005 | TRUE | 0.0418 |
| Iffo2 | 1614 | 1336 | 1512 | 1442 | 1397 | 1591 | 1254 | 1421 | 1631 | 1435 | 1576 | 1571 | 1481.32 | 1435 | 1527 | 1.065 | 0.091 | 0.33728012 | 0.51143059 | 0.0094 | 0.004 | 0.0059 | 0.0059 | TRUE | 0.1717 |
| Ifi202b | 0 | 0 | 0 | 0 | 0 | 0 | 0 | 0 | 0 | 0 | 0 | 0 | 0 | 0 | 0 | NA | NA | NA | NA | NA | NA | NA | NA | NA | NA |
| Ifi203 | 0 | 0 | 1 | 0 | 0 | 0 | 0 | 0 | 1 | 0 | 0 | 0 | 0.18 | 0 | 0 | 0.552 | -0.857 | 0.8336548 | NA | 0 | 13.1817 | 10 | 10 | TRUE | 0.5365 |
| Ifi204 | 0 | 0 | 0 | 0 | 0 | 0 | 0 | 0 | 0 | 0 | 0 | 0 | 0 | 0 | 0 | NA | NA | NA | NA | NA | NA | NA | NA | NA | NA |
| Ifi205 | 0 | 0 | 0 | 0 | 0 | 0 | 0 | 0 | 0 | 0 | 0 | 0 | 0 | 0 | 0 | NA | NA | NA | NA | NA | NA | NA | NA | NA | NA |
| Ifi27l1 | 45 | 40 | 35 | 25 | 29 | 22 | 35 | 43 | 38 | 25 | 33 | 22 | 32.43 | 39 | 27 | 0.687 | -0.541 | 0.11668601 | 0.24741036 | 0 | 0.0755 | 0.0535 | 0.0535 | TRUE | 0.248 |
| Ifi27l2a | 10 | 6 | 12 | 10 | 3 | 10 | 8 | 6 | 13 | 10 | 3 | 10 | 8.38 | 9 | 8 | 0.869 | -0.203 | 0.77014108 | NA | 0.0625 | 0.2851 | 0.2271 | 0.2271 | TRUE | 0.6116 |
| Ifi27l2b | 0 | 0 | 0 | 0 | 0 | 0 | 0 | 0 | 0 | 0 | 0 | 0 | 0 | 0 | 0 | NA | NA | NA | NA | NA | NA | NA | NA | NA | NA |
| Ifi30 | 2990 | 2327 | 3144 | 3648 | 3336 | 4140 | 2322 | 2475 | 3392 | 3631 | 3763 | 4087 | 3278.52 | 2730 | 3827 | 1.402 | 0.488 | 0.00776315 | 0.03425 | 0.0239 | 0.0031 | 0.009 | 0.0239 | TRUE | 0.5462 |
| Ifi35 | 286 | 179 | 135 | 155 | 145 | 171 | 222 | 190 | 146 | 154 | 164 | 169 | 174.14 | 186 | 162 | 0.868 | -0.205 | 0.24628083 | 0.41490946 | 0.0184 | 0.016 | 0.0167 | 0.0167 | TRUE | 0.2564 |
| Ifi44 | 0 | 0 | 0 | 0 | 0 | 0 | 0 | 0 | 0 | 0 | 0 | 0 | 0 | 0 | 0 | NA | NA | NA | NA | NA | NA | NA | NA | NA | NA |
| Ifi44l | 0 | 0 | 0 | 0 | 0 | 0 | 0 | 0 | 0 | 0 | 0 | 0 | 0 | 0 | 0 | NA | NA | NA | NA | NA | NA | NA | NA | NA | NA |
| Ifi47 | 3 | 1 | 0 | 1 | 2 | 0 | 2 | 1 | 0 | 1 | 2 | 0 | 1.11 | 1 | 1 | 0.908 | -0.139 | 0.94226433 | NA | 0 | 2.1424 | 1.738 | 1.738 | TRUE | 0.2546 |
| Ifih1 | 7 | 8 | 6 | 3 | 5 | 3 | 5 | 9 | 6 | 3 | 6 | 3 | 5.33 | 7 | 4 | 0.566 | -0.821 | 0.32590242 | NA | 0 | 0.4467 | 0.2972 | 0.2972 | TRUE | 0.2319 |
| Ifit1 | 2 | 3 | 4 | 1 | 2 | 4 | 2 | 3 | 4 | 1 | 2 | 4 | 2.71 | 3 | 2 | 0.811 | -0.303 | 0.795789 | NA | 0 | 0.877 | 0.6066 | 0.6066 | TRUE | 0.333 |
| Ifit2 | 116 | 80 | 106 | 117 | 119 | 134 | 90 | 85 | 114 | 116 | 134 | 132 | 112.09 | 96 | 127 | 1.325 | 0.406 | 0.03778742 | 0.10988719 | 0.0045 | 0.0236 | 0.0185 | 0.0185 | TRUE | 0.2396 |
| Ifit3 | 3 | 1 | 1 | 0 | 0 | 0 | 2 | 1 | 1 | 0 | 0 | 0 | 0.75 | 1 | 0 | 0.126 | -2.986 | 0.19880652 | NA | 0 | 3.1818 | 2.5705 | 2.5705 | TRUE | 0.2178 |
| Ifitm1 | 938 | 963 | 1541 | 622 | 866 | 1186 | 729 | 1024 | 1663 | 619 | 977 | 1171 | 1030.4 | 1139 | 922 | 0.81 | -0.304 | 0.47243734 | 0.64096316 | 0.1276 | 0.0047 | 0.0357 | 0.1276 | TRUE | 0.4729 |
| Ifitm10 | 1 | 3 | 3 | 3 | 3 | 4 | 1 | 3 | 3 | 3 | 3 | 4 | 2.92 | 2 | 3 | 1.479 | 0.564 | 0.6143525 | NA | 0 | 0.8139 | 0.5462 | 0.5462 | TRUE | 0.5339 |
| Ifitm2 | 8259 | 6786 | 8228 | 6296 | 6424 | 8126 | 6414 | 7218 | 8878 | 6267 | 7247 | 8022 | 7341.02 | 7503 | 7179 | 0.957 | -0.064 | 0.55891687 | 0.7137776 | 0.0179 | 0.0027 | 0.0084 | 0.0084 | TRUE | 0.3133 |
| Ifitm3 | 1324 | 1104 | 1414 | 816 | 901 | 1174 | 1028 | 1174 | 1526 | 812 | 1016 | 1159 | 1119.31 | 1243 | 996 | 0.802 | -0.319 | 0.15366058 | 0.30072858 | 0.0351 | 0.0045 | 0.0135 | 0.0351 | TRUE | 0.4753 |
| Ifitm5 | 12 | 23 | 16 | 15 | 14 | 12 | 9 | 24 | 17 | 15 | 16 | 12 | 15.6 | 17 | 14 | 0.844 | -0.244 | 0.63939936 | NA | 0.0733 | 0.1543 | 0.1302 | 0.1302 | TRUE | 0.2019 |
| Ifitm6 | 2 | 1 | 6 | 1 | 1 | 0 | 2 | 1 | 6 | 1 | 1 | 0 | 1.87 | 3 | 1 | 0.233 | -2.102 | 0.19787936 | NA | 0 | 1.2705 | 1.0164 | 1.0164 | TRUE | 1.2033 |
| Ifitm7 | 9 | 5 | 10 | 16 | 3 | 6 | 7 | 5 | 11 | 16 | 3 | 6 | 8.06 | 8 | 8 | 1.105 | 0.143 | 0.84757897 | NA | 0.2295 | 0.2966 | 0.2771 | 0.2771 | TRUE | 1.5164 |
| Ifltd1 | 0 | 0 | 0 | 0 | 0 | 0 | 0 | 0 | 0 | 0 | 0 | 0 | 0 | 0 | 0 | NA | NA | NA | NA | NA | NA | NA | NA | NA | NA |
| Ifna1 | 0 | 0 | 0 | 0 | 0 | 0 | 0 | 0 | 0 | 0 | 0 | 0 | 0 | 0 | 0 | NA | NA | NA | NA | NA | NA | NA | NA | NA | NA |
| Ifna11 | 0 | 0 | 0 | 0 | 0 | 0 | 0 | 0 | 0 | 0 | 0 | 0 | 0 | 0 | 0 | NA | NA | NA | NA | NA | NA | NA | NA | NA | NA |
| Ifna12 | 0 | 0 | 0 | 0 | 0 | 0 | 0 | 0 | 0 | 0 | 0 | 0 | 0 | 0 | 0 | NA | NA | NA | NA | NA | NA | NA | NA | NA | NA |
| Ifna13 | 0 | 0 | 0 | 0 | 0 | 0 | 0 | 0 | 0 | 0 | 0 | 0 | 0 | 0 | 0 | NA | NA | NA | NA | NA | NA | NA | NA | NA | NA |
| Ifna14 | 0 | 0 | 0 | 0 | 0 | 0 | 0 | 0 | 0 | 0 | 0 | 0 | 0 | 0 | 0 | NA | NA | NA | NA | NA | NA | NA | NA | NA | NA |
| Ifna2 | 0 | 0 | 0 | 0 | 0 | 0 | 0 | 0 | 0 | 0 | 0 | 0 | 0 | 0 | 0 | NA | NA | NA | NA | NA | NA | NA | NA | NA | NA |
| Ifna4 | 0 | 0 | 0 | 0 | 0 | 0 | 0 | 0 | 0 | 0 | 0 | 0 | 0 | 0 | 0 | NA | NA | NA | NA | NA | NA | NA | NA | NA | NA |
| Ifna5 | 0 | 0 | 0 | 0 | 0 | 0 | 0 | 0 | 0 | 0 | 0 | 0 | 0 | 0 | 0 | NA | NA | NA | NA | NA | NA | NA | NA | NA | NA |
| Ifna6 | 0 | 0 | 0 | 0 | 0 | 0 | 0 | 0 | 0 | 0 | 0 | 0 | 0 | 0 | 0 | NA | NA | NA | NA | NA | NA | NA | NA | NA | NA |
| Ifna7 | 0 | 0 | 0 | 0 | 0 | 0 | 0 | 0 | 0 | 0 | 0 | 0 | 0 | 0 | 0 | NA | NA | NA | NA | NA | NA | NA | NA | NA | NA |
| Ifna9 | 0 | 0 | 0 | 0 | 0 | 0 | 0 | 0 | 0 | 0 | 0 | 0 | 0 | 0 | 0 | NA | NA | NA | NA | NA | NA | NA | NA | NA | NA |
| Ifnab | 0 | 0 | 0 | 0 | 0 | 0 | 0 | 0 | 0 | 0 | 0 | 0 | 0 | 0 | 0 | NA | NA | NA | NA | NA | NA | NA | NA | NA | NA |
| Ifnar1 | 2737 | 1924 | 1917 | 2213 | 1949 | 2130 | 2126 | 2047 | 2068 | 2203 | 2199 | 2103 | 2124.13 | 2080 | 2168 | 1.042 | 0.059 | 0.33043739 | 0.50485747 | 0 | 0.0035 | 0.0022 | 0.0022 | TRUE | 0.0085 |
| Ifnar2 | 523 | 387 | 463 | 542 | 472 | 527 | 406 | 412 | 500 | 540 | 532 | 520 | 484.94 | 439 | 531 | 1.211 | 0.276 | 0.01125769 | 0.04506203 | 0.0046 | 0.0073 | 0.0065 | 0.0065 | TRUE | 0.1636 |
| Ifnb1 | 0 | 0 | 0 | 0 | 0 | 0 | 0 | 0 | 0 | 0 | 0 | 0 | 0 | 0 | 0 | NA | NA | NA | NA | NA | NA | NA | NA | NA | NA |
| Ifne | 0 | 0 | 0 | 0 | 0 | 0 | 0 | 0 | 0 | 0 | 0 | 0 | 0 | 0 | 0 | NA | NA | NA | NA | NA | NA | NA | NA | NA | NA |
| Ifng | 0 | 1 | 0 | 0 | 1 | 0 | 0 | 1 | 0 | 0 | 1 | 0 | 0.37 | 0 | 0 | 1.075 | 0.105 | 0.97520514 | NA | 0 | 6.4904 | 6.1969 | 6.1969 | TRUE | 0.5216 |
| Ifngr1 | 526 | 399 | 426 | 451 | 364 | 452 | 409 | 424 | 460 | 449 | 411 | 446 | 433.06 | 431 | 435 | 1.013 | 0.018 | 0.86447312 | 0.92525942 | 0 | 0.0079 | 0.0057 | 0.0057 | TRUE | 0.0389 |
| Ifngr2 | 308 | 277 | 279 | 337 | 323 | 399 | 239 | 295 | 301 | 335 | 364 | 394 | 321.43 | 278 | 364 | 1.315 | 0.395 | 0.00262987 | 0.01489843 | 0.0081 | 0.0098 | 0.0093 | 0.0093 | TRUE | 0.1841 |
| Ifnk | 0 | 0 | 0 | 1 | 0 | 0 | 0 | 0 | 0 | 1 | 0 | 0 | 0.17 | 0 | 0 | 2.094 | 1.067 | 0.79379329 | NA | 0 | 14.2885 | 10 | 10 | TRUE | 0.3906 |
| Ifnl2 | 1 | 0 | 0 | 0 | 2 | 0 | 1 | 0 | 0 | 0 | 2 | 0 | 0.51 | 0 | 1 | 2.139 | 1.097 | 0.71688146 | NA | 0 | 4.6912 | 4.8901 | 4.8901 | TRUE | 0.9745 |
| Ifnl3 | 0 | 0 | 0 | 0 | 0 | 0 | 0 | 0 | 0 | 0 | 0 | 0 | 0 | 0 | 0 | NA | NA | NA | NA | NA | NA | NA | NA | NA | NA |
| Ifnlr1 | 448 | 297 | 355 | 376 | 350 | 417 | 348 | 316 | 383 | 374 | 395 | 412 | 371.27 | 349 | 394 | 1.128 | 0.174 | 0.13318637 | 0.27148214 | 0.0026 | 0.0088 | 0.007 | 0.007 | TRUE | 0.0782 |
| Ifnz | 0 | 0 | 0 | 0 | 0 | 0 | 0 | 0 | 0 | 0 | 0 | 0 | 0 | 0 | 0 | NA | NA | NA | NA | NA | NA | NA | NA | NA | NA |
| Ifrd1 | 7980 | 5541 | 5012 | 5121 | 4902 | 5754 | 6198 | 5894 | 5408 | 5097 | 5530 | 5680 | 5634.5 | 5833 | 5436 | 0.932 | -0.102 | 0.13548139 | 0.27482764 | 0.0036 | 0.0028 | 0.0032 | 0.0032 | TRUE | 0.0495 |
| Ifrd2 | 4643 | 3363 | 3242 | 3406 | 3308 | 3902 | 3606 | 3577 | 3498 | 3390 | 3732 | 3852 | 3609.21 | 3560 | 3658 | 1.027 | 0.039 | 0.54453753 | 0.70202845 | 0.0018 | 0.0031 | 0.0027 | 0.0027 | TRUE | 0.0502 |
| Ift122 | 1256 | 933 | 1047 | 1009 | 914 | 1042 | 975 | 992 | 1130 | 1004 | 1031 | 1029 | 1026.94 | 1032 | 1021 | 0.99 | -0.014 | 0.86395301 | 0.92491052 | 0.0024 | 0.0047 | 0.0039 | 0.0039 | TRUE | 0.0796 |
| Ift140 | 2089 | 1544 | 1781 | 1589 | 1537 | 1722 | 1622 | 1642 | 1922 | 1582 | 1734 | 1700 | 1700.32 | 1729 | 1672 | 0.968 | -0.048 | 0.56314483 | 0.71753952 | 0.0051 | 0.0038 | 0.0043 | 0.0043 | TRUE | 0.1137 |
| Ift172 | 1534 | 1132 | 1171 | 1317 | 1149 | 1332 | 1191 | 1204 | 1263 | 1311 | 1296 | 1315 | 1263.5 | 1219 | 1307 | 1.073 | 0.101 | 0.14241725 | 0.28505388 | 0 | 0.0043 | 0.0026 | 0.0026 | TRUE | 0.0116 |
| Ift20 | 2168 | 1599 | 1436 | 1488 | 1395 | 1614 | 1684 | 1701 | 1549 | 1481 | 1574 | 1593 | 1597.03 | 1645 | 1549 | 0.942 | -0.087 | 0.21948904 | 0.3825944 | 0.0014 | 0.0039 | 0.003 | 0.003 | TRUE | 0.0303 |
| Ift27 | 575 | 398 | 366 | 401 | 367 | 373 | 447 | 423 | 395 | 399 | 414 | 368 | 407.7 | 422 | 394 | 0.932 | -0.102 | 0.34890327 | 0.52351256 | 0 | 0.0082 | 0.0061 | 0.0061 | TRUE | 0.0375 |

| | | | | | | | | | | | | | | | | | | | | | | | | | |
|---|---|---|---|---|---|---|---|---|---|---|---|---|---|---|---|---|---|---|---|---|---|---|---|---|---|
| Ift43 | 484 | 361 | 413 | 348 | 334 | 383 | 376 | 384 | 446 | 346 | 377 | 378 | 384.46 | 402 | 367 | 0.915 | -0.128 | 0.26433533 | 0.43456923 | 0.0032 | 0.0086 | 0.0069 | 0.0069 | TRUE | 0.1021 |
| Ift46 | 4562 | 3189 | 3065 | 3060 | 2882 | 3182 | 3543 | 3392 | 3307 | 3046 | 3251 | 3141 | 3280.09 | 3414 | 3146 | 0.921 | -0.119 | 0.04393704 | 0.12255312 | 9.00E-04 | 0.0031 | 0.0022 | 0.0022 | TRUE | 0.0136 |
| Ift52 | 2714 | 2006 | 2225 | 2634 | 2330 | 2538 | 2108 | 2134 | 2401 | 2622 | 2628 | 2505 | 2399.69 | 2214 | 2585 | 1.168 | 0.224 | 0.0012686 | 0.0085529 | 0.0024 | 0.0034 | 0.0031 | 0.0031 | TRUE | 0.0651 |
| Ift57 | 1054 | 807 | 874 | 865 | 766 | 857 | 819 | 858 | 943 | 861 | 864 | 846 | 865.19 | 873 | 857 | 0.983 | -0.025 | 0.76497301 | 0.86058711 | 0.0017 | 0.0052 | 0.0039 | 0.0039 | TRUE | 0.0574 |
| Ift74 | 1129 | 733 | 741 | 626 | 553 | 626 | 877 | 780 | 800 | 623 | 624 | 618 | 720.16 | 819 | 622 | 0.758 | -0.4 | 4.91E-06 | 0.00010455 | 0 | 0.0057 | 0.0041 | 0.0041 | TRUE | 0.0482 |
| Ift80 | 1641 | 1129 | 1214 | 1096 | 1042 | 1093 | 1274 | 1201 | 1310 | 1091 | 1175 | 1079 | 1188.45 | 1262 | 1115 | 0.883 | -0.179 | 0.01764676 | 0.06296079 | 0.0011 | 0.0044 | 0.0033 | 0.0033 | TRUE | 0.0263 |
| Ift81 | 1339 | 917 | 930 | 1007 | 907 | 1061 | 1040 | 975 | 1003 | 1002 | 1023 | 1047 | 1015.29 | 1006 | 1024 | 1.017 | 0.024 | 0.74248033 | 0.84667875 | 0 | 0.0048 | 0.003 | 0.003 | TRUE | 0.0108 |
| Ift88 | 876 | 720 | 616 | 747 | 616 | 714 | 680 | 766 | 665 | 744 | 695 | 705 | 709.03 | 704 | 715 | 1.016 | 0.023 | 0.79691122 | 0.88159845 | 0.0021 | 0.0058 | 0.0046 | 0.0046 | TRUE | 0.0688 |
| Igbp1 | 3117 | 2237 | 2501 | 2580 | 2326 | 2830 | 2421 | 2379 | 2699 | 2568 | 2624 | 2794 | 2580.77 | 2500 | 2662 | 1.065 | 0.091 | 0.19368834 | 0.35249463 | 0.0028 | 0.0033 | 0.0032 | 0.0032 | TRUE | 0.058 |
| Igbp1b | 80 | 60 | 64 | 27 | 28 | 34 | 62 | 64 | 69 | 27 | 32 | 34 | 47.84 | 65 | 31 | 0.472 | -1.082 | 0.00015514 | 0.00174873 | 0 | 0.052 | 0.0346 | 0.0346 | TRUE | 0.0819 |
| Igdcc3 | 855 | 633 | 521 | 796 | 644 | 675 | 664 | 673 | 562 | 792 | 726 | 666 | 680.78 | 633 | 728 | 1.149 | 0.201 | 0.05333662 | 0.14101745 | 0.0071 | 0.0059 | 0.0063 | 0.0063 | TRUE | 0.1101 |
| Igdcc4 | 968 | 699 | 744 | 1079 | 903 | 1061 | 752 | 744 | 803 | 1074 | 1019 | 1047 | 906.36 | 766 | 1047 | 1.368 | 0.452 | 9.00E-09 | 4.53E-07 | 0 | 0.005 | 0.0033 | 0.0033 | TRUE | 0.0201 |
| Igf1 | 3 | 3 | 6 | 1 | 1 | 2 | 2 | 3 | 6 | 1 | 1 | 2 | 2.68 | 4 | 1 | 0.348 | -1.523 | 0.21728577 | NA | 0 | 0.8861 | 0.5993 | 0.5993 | TRUE | 0.4603 |
| Igf1r | 3666 | 2659 | 2738 | 3053 | 2743 | 2974 | 2847 | 2828 | 2954 | 3039 | 3094 | 2936 | 2949.83 | 2876 | 3023 | 1.051 | 0.072 | 0.20989368 | 0.3717485 | 3.00E-04 | 0.0032 | 0.002 | 0.002 | TRUE | 0.0078 |
| Igf2 | 59 | 49 | 57 | 71 | 44 | 54 | 46 | 52 | 62 | 71 | 50 | 53 | 55.51 | 53 | 58 | 1.096 | 0.133 | 0.62717327 | 0.76496691 | 0.0121 | 0.0451 | 0.0359 | 0.0359 | TRUE | 0.3225 |
| Igf2as | 4 | 0 | 1 | 7 | 2 | 1 | 3 | 0 | 1 | 7 | 2 | 1 | 2.4 | 1 | 3 | 2.322 | 1.215 | 0.38046069 | NA | 0.6683 | 0.9902 | 0.9043 | 0.9043 | TRUE | 0.9303 |
| Igf2bp1 | 34752 | 23738 | 20038 | 20867 | 17691 | 19966 | 26990 | 25250 | 21621 | 20771 | 19957 | 19710 | 22383.14 | 24620 | 20146 | 0.818 | -0.289 | 0.00012996 | 0.00151481 | 0.0065 | 0.0025 | 0.0041 | 0.0041 | TRUE | 0.1389 |
| Igf2bp2 | 7393 | 5980 | 6880 | 5534 | 5197 | 6197 | 5742 | 6361 | 7423 | 5509 | 5863 | 6117 | 6169.14 | 6509 | 5830 | 0.896 | -0.158 | 0.06594094 | 0.16411964 | 0.0097 | 0.0028 | 0.0052 | 0.0052 | TRUE | 0.1843 |
| Igf2bp3 | 5287 | 4110 | 3787 | 4113 | 3789 | 4403 | 4106 | 4372 | 4086 | 4094 | 4274 | 4346 | 4213.16 | 4188 | 4238 | 1.012 | 0.017 | 0.76115347 | 0.85829385 | 9.00E-04 | 0.003 | 0.0021 | 0.0021 | TRUE | 0.0177 |
| Igf2r | 17191 | 12724 | 12928 | 15350 | 14503 | 16131 | 13352 | 13534 | 13949 | 15279 | 16361 | 15924 | 14733.19 | 13612 | 15855 | 1.165 | 0.22 | 1.20E-05 | 0.0002178 | 7.00E-04 | 0.0026 | 0.0018 | 0.0018 | TRUE | 0.0123 |
| Igfals | 82 | 63 | 72 | 85 | 92 | 75 | 64 | 67 | 78 | 85 | 104 | 74 | 78.47 | 70 | 88 | 1.26 | 0.334 | 0.14953303 | 0.2950473 | 0.0075 | 0.0326 | 0.0257 | 0.0257 | TRUE | 0.2409 |
| Igfbp1 | 0 | 1 | 0 | 0 | 0 | 1 | 0 | 1 | 0 | 0 | 0 | 1 | 0.34 | 0 | 0 | 1.075 | 0.105 | 0.97576789 | NA | 0 | 6.9363 | 6.5821 | 6.5821 | TRUE | 0.5216 |
| Igfbp2 | 20904 | 16035 | 19064 | 22885 | 19582 | 22083 | 16235 | 17056 | 20570 | 22780 | 22091 | 21799 | 20088.49 | 17954 | 22223 | 1.238 | 0.308 | 0.00011494 | 0.00136222 | 0.0084 | 0.0025 | 0.0045 | 0.0045 | TRUE | 0.1986 |
| Igfbp3 | 1235 | 855 | 642 | 756 | 661 | 764 | 959 | 909 | 693 | 753 | 746 | 754 | 802.29 | 854 | 751 | 0.878 | -0.188 | 0.09738951 | 0.21719273 | 0.0135 | 0.0054 | 0.008 | 0.008 | TRUE | 0.3192 |
| Igfbp4 | 1462 | 1233 | 1913 | 2317 | 2606 | 2799 | 1135 | 1312 | 2064 | 2306 | 2940 | 2763 | 2086.72 | 1504 | 2670 | 1.776 | 0.829 | 0.00441205 | 0.02233053 | 0.0605 | 0.0036 | 0.0176 | 0.0605 | TRUE | 1.2788 |
| Igfbp5 | 76 | 60 | 77 | 98 | 83 | 82 | 59 | 64 | 83 | 98 | 94 | 81 | 79.68 | 69 | 91 | 1.328 | 0.409 | 0.07564898 | 0.1812865 | 0.0076 | 0.0322 | 0.0255 | 0.0255 | TRUE | 0.292 |
| Igfbp6 | 28 | 8 | 12 | 10 | 11 | 16 | 22 | 9 | 13 | 10 | 12 | 16 | 13.56 | 15 | 13 | 0.87 | -0.202 | 0.71588281 | NA | 0.0762 | 0.1772 | 0.1475 | 0.1475 | TRUE | 0.5087 |
| Igfbp7 | 96 | 46 | 40 | 70 | 47 | 56 | 75 | 49 | 43 | 70 | 53 | 55 | 57.44 | 56 | 59 | 1.058 | 0.082 | 0.7762497 | 0.86766476 | 0.0389 | 0.0437 | 0.0423 | 0.0423 | TRUE | 0.8469 |
| Igfbpl1 | 313 | 189 | 201 | 215 | 186 | 232 | 243 | 201 | 217 | 214 | 210 | 229 | 218.98 | 220 | 218 | 0.985 | -0.022 | 0.87710369 | 0.93138507 | 0 | 0.0132 | 0.0098 | 0.0098 | TRUE | 0.0973 |
| Igfl3 | 0 | 0 | 0 | 0 | 0 | 0 | 0 | 0 | 0 | 0 | 0 | 0 | 0 | 0 | 0 | NA | NA | NA | NA | NA | NA | NA | NA | NA | NA |
| Igflr1 | 83 | 65 | 78 | 61 | 50 | 70 | 64 | 69 | 84 | 61 | 56 | 69 | 67.33 | 72 | 62 | 0.86 | -0.218 | 0.37028591 | 0.5449938 | 0 | 0.0376 | 0.0275 | 0.0275 | TRUE | 0.1708 |
| Igfn1 | 4 | 5 | 3 | 9 | 4 | 13 | 3 | 5 | 3 | 9 | 5 | 13 | 6.33 | 4 | 9 | 2.292 | 1.196 | 0.1361173 | NA | 0 | 0.377 | 0.2806 | 0.2806 | TRUE | 0.1158 |
| Ighmbp2 | 1073 | 813 | 742 | 798 | 666 | 773 | 833 | 865 | 801 | 794 | 751 | 763 | 801.24 | 833 | 769 | 0.924 | -0.114 | 0.16212246 | 0.31220914 | 0 | 0.0054 | 0.0035 | 0.0035 | TRUE | 0.013 |
| Igj | 1 | 1 | 2 | 0 | 0 | 0 | 1 | 1 | 2 | 0 | 0 | 0 | 0.67 | 1 | 0 | 0.144 | -2.794 | 0.24876697 | NA | 0 | 3.559 | 2.8631 | 2.8631 | TRUE | 0.1682 |
| Igll1 | 8 | 5 | 8 | 7 | 5 | 5 | 6 | 5 | 9 | 7 | 6 | 5 | 6.28 | 7 | 6 | 0.873 | -0.195 | 0.79472047 | NA | 0 | 0.3795 | 0.2452 | 0.2452 | TRUE | 0.1429 |
| Iglon5 | 264 | 199 | 209 | 234 | 239 | 228 | 205 | 212 | 226 | 233 | 270 | 225 | 228.31 | 214 | 243 | 1.133 | 0.18 | 0.19483568 | 0.35420681 | 0 | 0.0128 | 0.0095 | 0.0095 | TRUE | 0.1022 |
| Igsf1 | 340 | 260 | 279 | 304 | 277 | 303 | 264 | 277 | 301 | 303 | 312 | 299 | 292.64 | 281 | 305 | 1.088 | 0.122 | 0.31450268 | 0.48807674 | 0 | 0.0105 | 0.0071 | 0.0071 | TRUE | 0.045 |
| Igsf10 | 581 | 362 | 326 | 353 | 319 | 307 | 451 | 385 | 352 | 351 | 360 | 303 | 367.06 | 396 | 338 | 0.851 | -0.234 | 0.069934 | 0.17145226 | 0.01 | 0.0089 | 0.0092 | 0.0092 | TRUE | 0.1801 |
| Igsf11 | 150 | 105 | 133 | 107 | 76 | 115 | 116 | 112 | 144 | 107 | 86 | 114 | 112.91 | 124 | 102 | 0.826 | -0.275 | 0.16695587 | 0.31808854 | 0.0097 | 0.0234 | 0.0196 | 0.0196 | TRUE | 0.1861 |
| Igsf21 | 672 | 574 | 632 | 962 | 890 | 1015 | 522 | 611 | 682 | 958 | 1004 | 1002 | 796.32 | 605 | 988 | 1.638 | 0.712 | 2.39E-12 | 2.78E-10 | 0.0071 | 0.0054 | 0.0061 | 0.0061 | TRUE | 0.1788 |
| Igsf23 | 101 | 65 | 60 | 56 | 44 | 53 | 78 | 69 | 65 | 56 | 50 | 52 | 61.67 | 71 | 53 | 0.741 | -0.433 | 0.08001504 | 0.18922445 | 0 | 0.0409 | 0.0275 | 0.0275 | TRUE | 0.0917 |
| Igsf3 | 3684 | 2832 | 2932 | 2804 | 2606 | 3045 | 2861 | 3012 | 3164 | 2791 | 2940 | 3006 | 2962.33 | 3012 | 2912 | 0.967 | -0.048 | 0.45312062 | 0.62297547 | 0.0015 | 0.0032 | 0.0026 | 0.0026 | TRUE | 0.024 |
| Igsf5 | 34 | 26 | 14 | 26 | 19 | 13 | 26 | 28 | 15 | 26 | 21 | 13 | 21.55 | 23 | 20 | 0.864 | -0.211 | 0.63841009 | 0.77423614 | 0.0586 | 0.1124 | 0.0981 | 0.0981 | TRUE | 0.5567 |
| Igsf6 | 0 | 0 | 0 | 0 | 0 | 0 | 0 | 0 | 0 | 0 | 0 | 0 | 0 | 0 | 0 | NA | NA | NA | NA | NA | NA | NA | NA | NA | NA |
| Igsf8 | 3133 | 2486 | 2836 | 3751 | 3172 | 3771 | 2433 | 2644 | 3060 | 3734 | 3578 | 3723 | 3195.38 | 2712 | 3678 | 1.357 | 0.441 | 5.55E-08 | 2.29E-06 | 0.0067 | 0.0032 | 0.0044 | 0.0044 | TRUE | 0.152 |
| Igsf9 | 1460 | 1234 | 1242 | 1243 | 1219 | 1403 | 1134 | 1313 | 1340 | 1237 | 1375 | 1385 | 1297.34 | 1262 | 1332 | 1.057 | 0.08 | 0.3559088 | 0.53038743 | 0.0054 | 0.0043 | 0.0046 | 0.0046 | TRUE | 0.0984 |
| Igsf9b | 592 | 459 | 474 | 529 | 450 | 547 | 460 | 488 | 511 | 527 | 508 | 540 | 505.61 | 486 | 525 | 1.081 | 0.112 | 0.24906341 | 0.41806702 | 0 | 0.0071 | 0.0049 | 0.0049 | TRUE | 0.0289 |
| Igtp | 187 | 133 | 125 | 245 | 183 | 254 | 145 | 141 | 135 | 244 | 206 | 251 | 187.11 | 140 | 234 | 1.663 | 0.734 | 1.52E-06 | 4.10E-05 | 0 | 0.0151 | 0.0111 | 0.0111 | TRUE | 0.1106 |
| Ihh | 9 | 2 | 4 | 11 | 4 | 5 | 7 | 2 | 4 | 11 | 5 | 5 | 5.64 | 4 | 7 | 1.491 | 0.577 | 0.49497657 | NA | 0.1179 | 0.4228 | 0.3331 | 0.3331 | TRUE | 0.5078 |
| Iigp1 | 0 | 0 | 0 | 1 | 0 | 0 | 0 | 0 | 0 | 1 | 0 | 0 | 0.17 | 0 | 0 | 2.094 | 1.067 | 0.79379329 | NA | 0 | 14.2885 | 10 | 10 | TRUE | 0.3906 |
| Ik | 7252 | 5310 | 5385 | 5579 | 5042 | 5750 | 5632 | 5648 | 5810 | 5553 | 5688 | 5676 | 5668.04 | 5697 | 5639 | 0.99 | -0.015 | 0.76762299 | 0.86257577 | 0 | 0.0028 | 0.0016 | 0.0016 | TRUE | 0.0037 |
| Ikbip | 1155 | 802 | 916 | 1076 | 907 | 1058 | 897 | 853 | 988 | 1071 | 1023 | 1044 | 979.52 | 913 | 1046 | 1.147 | 0.198 | 0.01601421 | 0.05862991 | 0.0018 | 0.0048 | 0.0038 | 0.0038 | TRUE | 0.0601 |
| Ikbkap | 10670 | 7324 | 6749 | 6872 | 6084 | 6847 | 8287 | 7790 | 7282 | 6840 | 6863 | 6759 | 7303.72 | 7786 | 6821 | 0.876 | -0.191 | 0.00137236 | 0.00904524 | 0.0019 | 0.0028 | 0.0024 | 0.0024 | TRUE | 0.0392 |
| Ikbkb | 2036 | 1587 | 1465 | 1501 | 1403 | 1667 | 1581 | 1688 | 1581 | 1494 | 1583 | 1646 | 1595.41 | 1617 | 1574 | 0.974 | -0.038 | 0.58864754 | 0.73762904 | 0.0013 | 0.0039 | 0.0029 | 0.0029 | TRUE | 0.0242 |
| Ikbke | 151 | 98 | 97 | 77 | 63 | 60 | 117 | 104 | 105 | 77 | 71 | 59 | 88.85 | 109 | 69 | 0.632 | -0.661 | 0.00194049 | 0.01181238 | 0 | 0.0291 | 0.0208 | 0.0208 | TRUE | 0.1424 |
| Ikbkg | 1847 | 1363 | 1364 | 1562 | 1409 | 1475 | 1434 | 1450 | 1472 | 1555 | 1589 | 1456 | 1492.73 | 1452 | 1533 | 1.056 | 0.079 | 0.24931638 | 0.41813278 | 4.00E-04 | 0.004 | 0.0027 | 0.0027 | TRUE | 0.0237 |
| Ikzf1 | 184 | 132 | 115 | 134 | 102 | 107 | 143 | 140 | 124 | 133 | 115 | 106 | 126.91 | 136 | 118 | 0.868 | -0.205 | 0.25889907 | 0.42878846 | 0 | 0.0211 | 0.0157 | 0.0157 | TRUE | 0.1342 |
| Ikzf2 | 79 | 42 | 62 | 63 | 63 | 55 | 61 | 45 | 67 | 63 | 71 | 54 | 60.17 | 58 | 63 | 1.083 | 0.115 | 0.66193018 | 0.79124932 | 0.0107 | 0.0418 | 0.0336 | 0.0336 | TRUE | 0.3085 |
| Ikzf3 | 50 | 33 | 20 | 39 | 43 | 36 | 39 | 35 | 22 | 39 | 49 | 36 | 36.4 | 32 | 41 | 1.272 | 0.348 | 0.30862926 | 0.48126096 | 0.0243 | 0.0675 | 0.0563 | 0.0563 | TRUE | 0.5061 |
| Ikzf4 | 132 | 97 | 93 | 125 | 119 | 134 | 103 | 103 | 100 | 124 | 134 | 132 | 116.17 | 102 | 130 | 1.277 | 0.352 | 0.04962982 | 0.13406899 | 0 | 0.0228 | 0.0145 | 0.0145 | TRUE | 0.0162 |
| Ikzf5 | 1435 | 1116 | 1055 | 1031 | 935 | 1118 | 1115 | 1187 | 1138 | 1026 | 1055 | 1104 | 1104.1 | 1147 | 1062 | 0.926 | -0.11 | 0.13727221 | 0.27759363 | 0 | 0.0046 | 0.0031 | 0.0031 | TRUE | 0.0147 |
| Il10 | 0 | 0 | 0 | 0 | 0 | 0 | 0 | 0 | 0 | 0 | 0 | 0 | 0 | 0 | 0 | NA | NA | NA | NA | NA | NA | NA | NA | NA | NA |
| Il10ra | 7 | 11 | 6 | 6 | 5 | 3 | 5 | 12 | 6 | 6 | 6 | 3 | 6.36 | 8 | 5 | 0.621 | -0.687 | 0.37885646 | NA | 0 | 0.3748 | 0.2703 | 0.2703 | TRUE | 0.5019 |
| Il10rb | 144 | 115 | 129 | 131 | 120 | 133 | 112 | 122 | 139 | 130 | 135 | 131 | 128.4 | 124 | 132 | 1.067 | 0.094 | 0.59533161 | 0.74186165 | 0 | 0.0209 | 0.0147 | 0.0147 | TRUE | 0.1072 |
| Il11 | 19 | 12 | 14 | 26 | 15 | 24 | 15 | 13 | 15 | 26 | 17 | 24 | 18.19 | 14 | 22 | 1.563 | 0.644 | 0.15486284 | 0.3024725 | 0 | 0.1327 | 0.0906 | 0.0906 | TRUE | 0.2229 |
| Il11ra1 | 152 | 117 | 103 | 138 | 121 | 155 | 118 | 124 | 111 | 137 | 137 | 153 | 130.09 | 118 | 142 | 1.208 | 0.272 | 0.11523519 | 0.24514202 | 0 | 0.0206 | 0.0138 | 0.0138 | TRUE | 0.046 |
| Il11ra2 | 0 | 0 | 0 | 0 | 0 | 0 | 0 | 0 | 0 | 0 | 0 | 0 | 0 | 0 | 0 | NA | NA | NA | NA | NA | NA | NA | NA | NA | NA |
| Il12a | 0 | 0 | 0 | 0 | 0 | 0 | 0 | 0 | 0 | 0 | 0 | 0 | 0 | 0 | 0 | NA | NA | NA | NA | NA | NA | NA | NA | NA | NA |
| Il12b | 1 | 1 | 1 | 0 | 0 | 0 | 1 | 1 | 1 | 0 | 0 | 0 | 0.49 | 1 | 0 | 0.196 | -2.347 | 0.38405676 | NA | 0 | 4.8735 | 3.7415 | 3.7415 | TRUE | 0.0718 |

| | | | | | | | | | | | | | | | | | | | | | | | | | |
|---|---|---|---|---|---|---|---|---|---|---|---|---|---|---|---|---|---|---|---|---|---|---|---|---|---|
| Il12rb1 | 197 | 168 | 196 | 228 | 215 | 299 | 153 | 179 | 211 | 227 | 243 | 295 | 217.97 | 181 | 255 | 1.415 | 0.501 | 0.00231668 | 0.01354171 | 0.0182 | 0.0133 | 0.0147 | 0.0147 | TRUE | 0.2305 |
| Il12rb2 | 6 | 3 | 5 | 7 | 9 | 7 | 5 | 3 | 5 | 7 | 10 | 7 | 6.21 | 4 | 8 | 1.803 | 0.85 | 0.26810839 | NA | 0 | 0.3839 | 0.2513 | 0.2513 | TRUE | 0.1447 |
| Il13 | 0 | 0 | 0 | 1 | 3 | 1 | 0 | 0 | 0 | 1 | 3 | 1 | 0.89 | 0 | 2 | 10.025 | 3.326 | 0.13427406 | NA | 0 | 2.6521 | 2.2624 | 2.2624 | TRUE | 0.4405 |
| Il13ra1 | 354 | 249 | 318 | 302 | 244 | 265 | 275 | 265 | 343 | 301 | 275 | 262 | 286.73 | 294 | 279 | 0.95 | -0.073 | 0.59424874 | 0.74089299 | 0.009 | 0.0107 | 0.0102 | 0.0102 | TRUE | 0.2295 |
| Il13ra2 | 0 | 0 | 0 | 0 | 0 | 0 | 0 | 0 | 0 | 0 | 0 | 0 | 0 | 0 | 0 | NA | NA | NA | NA | NA | NA | NA | NA | NA | NA |
| Il15 | 28 | 14 | 24 | 17 | 20 | 20 | 22 | 15 | 26 | 17 | 23 | 20 | 20.29 | 21 | 20 | 0.944 | -0.083 | 0.84767172 | 0.91363938 | 0 | 0.1192 | 0.0861 | 0.0861 | TRUE | 0.3157 |
| Il15ra | 1 | 3 | 3 | 0 | 6 | 7 | 1 | 3 | 3 | 0 | 7 | 7 | 3.48 | 2 | 5 | 1.94 | 0.956 | 0.42578908 | NA | 0.8359 | 0.6834 | 0.7186 | 0.7186 | TRUE | 1.4739 |
| Il16 | 6 | 6 | 3 | 4 | 4 | 7 | 5 | 6 | 3 | 4 | 5 | 7 | 4.95 | 5 | 5 | 1.082 | 0.113 | 0.89408639 | NA | 0 | 0.4815 | 0.3183 | 0.3183 | TRUE | 0.197 |
| Il17a | 0 | 0 | 0 | 0 | 0 | 0 | 0 | 0 | 0 | 0 | 0 | 0 | 0 | 0 | 0 | NA | NA | NA | NA | NA | NA | NA | NA | NA | NA |
| Il17b | 87 | 71 | 81 | 90 | 89 | 96 | 68 | 76 | 87 | 90 | 100 | 95 | 85.87 | 77 | 95 | 1.24 | 0.311 | 0.14680606 | 0.29115071 | 0 | 0.03 | 0.0213 | 0.0213 | TRUE | 0.1305 |
| Il17c | 2 | 1 | 1 | 0 | 0 | 0 | 2 | 1 | 1 | 0 | 0 | 0 | 0.62 | 1 | 0 | 0.154 | -2.702 | 0.27412557 | NA | 0 | 3.8499 | 3.0088 | 3.0088 | TRUE | 0.0718 |
| Il17d | 178 | 123 | 101 | 136 | 107 | 132 | 138 | 131 | 109 | 135 | 121 | 130 | 127.41 | 126 | 129 | 1.019 | 0.028 | 0.8776312 | 0.93172788 | 0 | 0.021 | 0.0155 | 0.0155 | TRUE | 0.1371 |
| Il17f | 40 | 28 | 31 | 37 | 42 | 39 | 31 | 30 | 33 | 37 | 47 | 38 | 36.17 | 31 | 41 | 1.299 | 0.377 | 0.23757811 | 0.40425047 | 0 | 0.068 | 0.0455 | 0.0455 | TRUE | 0.1353 |
| Il17ra | 1632 | 1426 | 1368 | 1679 | 1533 | 1809 | 1268 | 1517 | 1476 | 1671 | 1729 | 1786 | 1574.47 | 1420 | 1729 | 1.219 | 0.286 | 0.00045449 | 0.00401487 | 0.0045 | 0.0039 | 0.0042 | 0.0042 | TRUE | 0.1103 |
| Il17rb | 8 | 3 | 1 | 2 | 8 | 7 | 6 | 3 | 1 | 2 | 9 | 7 | 4.73 | 3 | 6 | 1.648 | 0.721 | 0.4536787 | NA | 0.294 | 0.503 | 0.4459 | 0.4459 | TRUE | 0.5592 |
| Il17rc | 656 | 444 | 501 | 606 | 508 | 583 | 509 | 472 | 541 | 603 | 573 | 576 | 545.69 | 507 | 584 | 1.151 | 0.203 | 0.03494987 | 0.10376785 | 0 | 0.0068 | 0.0048 | 0.0048 | TRUE | 0.0405 |
| Il17rd | 2991 | 2203 | 2505 | 2611 | 2688 | 3335 | 2323 | 2343 | 2703 | 2599 | 3032 | 3292 | 2715.44 | 2456 | 2974 | 1.211 | 0.277 | 0.00260867 | 0.01482319 | 0.0102 | 0.0033 | 0.0057 | 0.0057 | TRUE | 0.1488 |
| Il17re | 25 | 25 | 17 | 24 | 20 | 10 | 19 | 27 | 18 | 24 | 23 | 10 | 20.11 | 21 | 19 | 0.875 | -0.193 | 0.67729581 | 0.80217727 | 0.0616 | 0.1203 | 0.1051 | 0.1051 | TRUE | 0.9323 |
| Il18 | 2 | 0 | 2 | 0 | 0 | 0 | 2 | 0 | 2 | 0 | 0 | 0 | 0.62 | 1 | 0 | 0.153 | -2.705 | 0.31809589 | NA | 0 | 3.8341 | 3.9014 | 3.9014 | TRUE | 0.3054 |
| Il18bp | 50 | 58 | 45 | 36 | 49 | 52 | 39 | 62 | 49 | 36 | 55 | 51 | 48.59 | 50 | 47 | 0.96 | -0.059 | 0.84556201 | 0.9121221 | 0.0323 | 0.0512 | 0.0459 | 0.0459 | TRUE | 0.2781 |
| Il18r1 | 3 | 1 | 6 | 8 | 3 | 1 | 2 | 1 | 6 | 8 | 3 | 1 | 3.7 | 3 | 4 | 1.267 | 0.341 | 0.75682474 | NA | 0.4958 | 0.6429 | 0.5992 | 0.5992 | TRUE | 0.4043 |
| Il18rap | 2 | 0 | 0 | 0 | 0 | 0 | 2 | 0 | 0 | 0 | 0 | 0 | 0.26 | 1 | 0 | 0.356 | -1.491 | 0.71362796 | NA | 0 | 9.1573 | 10 | 10 | TRUE | 0.968 |
| Il19 | 1 | 0 | 2 | 3 | 0 | 0 | 1 | 0 | 2 | 3 | 0 | 0 | 0.99 | 1 | 1 | 1.038 | 0.054 | 0.98034809 | NA | 2.347 | 2.4042 | 2.3958 | 2.3958 | TRUE | 1.1969 |
| Il1a | 0 | 0 | 0 | 0 | 0 | 0 | 0 | 0 | 0 | 0 | 0 | 0 | 0 | 0 | 0 | NA | NA | NA | NA | NA | NA | NA | NA | NA | NA |
| Il1b | 0 | 0 | 0 | 0 | 0 | 0 | 0 | 0 | 0 | 0 | 0 | 0 | 0 | 0 | 0 | NA | NA | NA | NA | NA | NA | NA | NA | NA | NA |
| Il1f10 | 0 | 0 | 0 | 0 | 0 | 0 | 0 | 0 | 0 | 0 | 0 | 0 | 0 | 0 | 0 | NA | NA | NA | NA | NA | NA | NA | NA | NA | NA |
| Il1f5 | 0 | 0 | 0 | 0 | 0 | 0 | 0 | 0 | 0 | 0 | 0 | 0 | 0 | 0 | 0 | NA | NA | NA | NA | NA | NA | NA | NA | NA | NA |
| Il1f6 | 0 | 0 | 0 | 0 | 0 | 0 | 0 | 0 | 0 | 0 | 0 | 0 | 0 | 0 | 0 | NA | NA | NA | NA | NA | NA | NA | NA | NA | NA |
| Il1f8 | 0 | 0 | 0 | 0 | 1 | 0 | 0 | 0 | 0 | 0 | 1 | 0 | 0.19 | 0 | 0 | 2.094 | 1.067 | 0.79379329 | NA | 0 | 12.608 | 10 | 10 | TRUE | 0.5075 |
| Il1f9 | 0 | 0 | 0 | 0 | 0 | 0 | 0 | 0 | 0 | 0 | 0 | 0 | 0 | 0 | 0 | NA | NA | NA | NA | NA | NA | NA | NA | NA | NA |
| Il1r1 | 225 | 127 | 126 | 130 | 98 | 107 | 175 | 135 | 136 | 129 | 111 | 106 | 131.9 | 149 | 115 | 0.772 | -0.374 | 0.04639949 | 0.1275922 | 0.0113 | 0.0204 | 0.0176 | 0.0176 | TRUE | 0.2687 |
| Il1r2 | 117 | 92 | 80 | 82 | 73 | 94 | 91 | 98 | 86 | 82 | 82 | 93 | 88.64 | 92 | 86 | 0.934 | -0.098 | 0.63527883 | 0.77174822 | 0 | 0.0292 | 0.0194 | 0.0194 | TRUE | 0.0525 |
| Il1rap | 573 | 347 | 344 | 394 | 330 | 335 | 445 | 369 | 371 | 392 | 372 | 331 | 380.08 | 395 | 365 | 0.921 | -0.119 | 0.3339882 | 0.50810999 | 0.0074 | 0.0087 | 0.0083 | 0.0083 | TRUE | 0.1486 |
| Il1rapl1 | 0 | 0 | 0 | 0 | 0 | 0 | 0 | 0 | 0 | 0 | 0 | 0 | 0 | 0 | 0 | NA | NA | NA | NA | NA | NA | NA | NA | NA | NA |
| Il1rapl2 | 0 | 0 | 0 | 1 | 0 | 0 | 0 | 0 | 0 | 1 | 0 | 0 | 0.17 | 0 | 0 | 2.094 | 1.067 | 0.79379329 | NA | 0 | 14.2885 | 10 | 10 | TRUE | 0.3906 |
| Il1rl1 | 3 | 5 | 3 | 4 | 3 | 1 | 2 | 5 | 3 | 4 | 3 | 1 | 3.21 | 3 | 3 | 0.773 | -0.371 | 0.73131012 | NA | 0 | 0.7416 | 0.5178 | 0.5178 | TRUE | 0.4083 |
| Il1rl2 | 56 | 29 | 31 | 23 | 18 | 20 | 43 | 31 | 33 | 23 | 20 | 20 | 28.46 | 36 | 21 | 0.58 | -0.785 | 0.03336483 | 0.10026986 | 0 | 0.0857 | 0.0601 | 0.0601 | TRUE | 0.2841 |
| Il1rn | 0 | 0 | 1 | 0 | 0 | 0 | 0 | 0 | 1 | 0 | 0 | 0 | 0.18 | 0 | 0 | 0.552 | -0.857 | 0.8336548 | NA | 0 | 13.1817 | 10 | 10 | TRUE | 0.5365 |
| Il2 | 0 | 0 | 0 | 0 | 0 | 0 | 0 | 0 | 0 | 0 | 0 | 0 | 0 | 0 | 0 | NA | NA | NA | NA | NA | NA | NA | NA | NA | NA |
| Il20 | 0 | 0 | 0 | 0 | 0 | 1 | 0 | 0 | 0 | 0 | 0 | 1 | 0.16 | 0 | 0 | 2.094 | 1.067 | 0.79379329 | NA | 0 | 14.4078 | 10 | 10 | TRUE | 0.3835 |
| Il20ra | 6 | 7 | 1 | 11 | 6 | 6 | 5 | 7 | 1 | 11 | 7 | 6 | 6.14 | 4 | 8 | 1.789 | 0.839 | 0.31017964 | NA | 0.0735 | 0.3886 | 0.3175 | 0.3175 | TRUE | 0.3886 |
| Il20rb | 50 | 14 | 10 | 14 | 8 | 10 | 39 | 15 | 11 | 14 | 9 | 10 | 16.22 | 22 | 11 | 0.502 | -0.995 | 0.0854884 | NA | 0.2223 | 0.1485 | 0.1721 | 0.1721 | TRUE | 2.3523 |
| Il21 | 1 | 1 | 0 | 0 | 0 | 1 | 1 | 1 | 0 | 0 | 0 | 1 | 0.47 | 1 | 0 | 0.599 | -0.74 | 0.79209856 | NA | 0 | 5.0317 | 3.8925 | 3.8925 | TRUE | 0.3835 |
| Il21r | 0 | 0 | 4 | 3 | 2 | 2 | 0 | 0 | 4 | 3 | 2 | 2 | 1.92 | 1 | 2 | 1.746 | 0.804 | 0.60701262 | NA | 1.0157 | 1.2355 | 1.195 | 1.195 | TRUE | 1.9027 |
| Il22 | 0 | 1 | 0 | 0 | 0 | 0 | 0 | 1 | 0 | 0 | 0 | 0 | 0.18 | 0 | 0 | 0.552 | -0.857 | 0.8336548 | NA | 0 | 13.3714 | 10 | 10 | TRUE | 0.5216 |
| Il22ra1 | 73 | 52 | 37 | 46 | 37 | 39 | 57 | 55 | 40 | 46 | 42 | 38 | 46.33 | 51 | 42 | 0.826 | -0.276 | 0.34348745 | 0.51776694 | 0 | 0.0536 | 0.0393 | 0.0393 | TRUE | 0.2633 |
| Il22ra2 | 1 | 0 | 0 | 1 | 0 | 0 | 1 | 0 | 0 | 1 | 0 | 0 | 0.3 | 0 | 0 | 1.075 | 0.105 | 0.97698525 | NA | 0 | 8.0272 | 7.5141 | 7.5141 | TRUE | 0.3906 |
| Il23a | 38 | 30 | 48 | 58 | 53 | 63 | 30 | 32 | 52 | 58 | 60 | 62 | 48.82 | 38 | 60 | 1.602 | 0.68 | 0.02336448 | 0.07742837 | 0.0208 | 0.051 | 0.0432 | 0.0432 | TRUE | 0.7415 |
| Il23r | 0 | 0 | 0 | 1 | 0 | 0 | 0 | 0 | 0 | 1 | 0 | 0 | 0.17 | 0 | 0 | 2.094 | 1.067 | 0.79379329 | NA | 0 | 14.2885 | 10 | 10 | TRUE | 0.3906 |
| Il24 | 0 | 2 | 0 | 0 | 0 | 0 | 0 | 2 | 0 | 0 | 0 | 0 | 0.35 | 1 | 0 | 0.272 | -1.878 | 0.62190613 | NA | 10 | 6.6869 | 8.7633 | 8.7633 | TRUE | 0.9743 |
| Il25 | 1 | 2 | 5 | 3 | 6 | 4 | 1 | 2 | 5 | 3 | 7 | 4 | 3.67 | 3 | 5 | 1.696 | 0.762 | 0.47018006 | NA | 0.1456 | 0.6487 | 0.5088 | 0.5088 | TRUE | 0.7951 |
| Il27 | 0 | 0 | 0 | 0 | 0 | 0 | 0 | 0 | 0 | 0 | 0 | 0 | 0 | 0 | 0 | NA | NA | NA | NA | NA | NA | NA | NA | NA | NA |
| Il27ra | 205 | 156 | 190 | 246 | 221 | 295 | 159 | 166 | 205 | 245 | 249 | 291 | 219.26 | 177 | 262 | 1.487 | 0.573 | 0.00017082 | 0.00188831 | 0.009 | 0.0132 | 0.012 | 0.012 | TRUE | 0.2039 |
| Il2ra | 1 | 2 | 1 | 2 | 1 | 0 | 1 | 2 | 1 | 2 | 1 | 0 | 1.18 | 1 | 1 | 0.793 | -0.335 | 0.84888331 | NA | 0 | 2.0048 | 1.3645 | 1.3645 | TRUE | 0.3277 |
| Il2rb | 1 | 2 | 3 | 5 | 8 | 7 | 1 | 2 | 3 | 5 | 9 | 7 | 4.51 | 2 | 7 | 3.529 | 1.819 | 0.0640173 | NA | 0 | 0.5281 | 0.3766 | 0.3766 | TRUE | 0.319 |
| Il2rg | 93 | 57 | 52 | 70 | 50 | 76 | 72 | 61 | 56 | 70 | 56 | 75 | 65.01 | 63 | 67 | 1.061 | 0.086 | 0.73083716 | 0.83911492 | 0 | 0.0389 | 0.0293 | 0.0293 | TRUE | 0.1636 |
| Il3 | 0 | 0 | 0 | 0 | 0 | 1 | 0 | 0 | 0 | 0 | 0 | 1 | 0.16 | 0 | 0 | 2.094 | 1.067 | 0.79379329 | NA | 0 | 14.4078 | 10 | 10 | TRUE | 0.3835 |
| Il31 | 0 | 0 | 0 | 0 | 0 | 0 | 0 | 0 | 0 | 0 | 0 | 0 | 0 | 0 | 0 | NA | NA | NA | NA | NA | NA | NA | NA | NA | NA |
| Il31ra | 2 | 3 | 1 | 1 | 3 | 3 | 2 | 3 | 1 | 1 | 3 | 3 | 2.19 | 2 | 2 | 1.263 | 0.337 | 0.79263965 | NA | 0 | 1.0826 | 0.72 | 0.72 | TRUE | 0.3001 |
| Il33 | 17 | 18 | 6 | 12 | 11 | 5 | 13 | 19 | 6 | 12 | 12 | 5 | 11.35 | 13 | 10 | 0.75 | -0.415 | 0.50797114 | NA | 0.1412 | 0.2112 | 0.1925 | 0.1925 | TRUE | 0.1641 |
| Il34 | 73 | 69 | 43 | 45 | 43 | 48 | 57 | 73 | 46 | 45 | 49 | 47 | 52.86 | 59 | 47 | 0.797 | -0.327 | 0.24172644 | 0.40937375 | 0 | 0.0473 | 0.0368 | 0.0368 | TRUE | 0.2848 |
| Il3ra | 401 | 259 | 283 | 285 | 278 | 283 | 311 | 275 | 305 | 284 | 314 | 279 | 294.83 | 297 | 292 | 0.98 | -0.029 | 0.81403328 | 0.89286914 | 0 | 0.0105 | 0.0075 | 0.0075 | TRUE | 0.0457 |
| Il4 | 0 | 0 | 0 | 0 | 0 | 1 | 0 | 0 | 0 | 0 | 0 | 1 | 0.16 | 0 | 0 | 2.094 | 1.067 | 0.79379329 | NA | 0 | 14.4078 | 10 | 10 | TRUE | 0.3835 |
| Il4i1 | 1 | 0 | 0 | 0 | 0 | 1 | 1 | 0 | 0 | 0 | 0 | 1 | 0.29 | 0 | 0 | 1.075 | 0.105 | 0.97702352 | NA | 0 | 8.0647 | 7.5458 | 7.5458 | TRUE | 0.3835 |
| Il4ra | 327 | 223 | 241 | 268 | 293 | 273 | 254 | 237 | 260 | 267 | 331 | 269 | 269.67 | 250 | 289 | 1.151 | 0.203 | 0.13065553 | 0.26773612 | 0.0049 | 0.0112 | 0.0093 | 0.0093 | TRUE | 0.1734 |
| Il5 | 0 | 1 | 0 | 0 | 0 | 0 | 0 | 1 | 0 | 0 | 0 | 0 | 0.18 | 0 | 0 | 0.552 | -0.857 | 0.8336548 | NA | 0 | 13.3714 | 10 | 10 | TRUE | 0.5216 |
| Il5ra | 0 | 0 | 0 | 0 | 0 | 0 | 0 | 0 | 0 | 0 | 0 | 0 | 0 | 0 | 0 | NA | NA | NA | NA | NA | NA | NA | NA | NA | NA |
| Il6 | 6 | 3 | 1 | 2 | 6 | 6 | 5 | 3 | 1 | 2 | 7 | 6 | 3.94 | 3 | 5 | 1.596 | 0.675 | 0.50221434 | NA | 0 | 0.6047 | 0.4643 | 0.4643 | TRUE | 0.5498 |
| Il6ra | 139 | 112 | 64 | 59 | 45 | 62 | 108 | 119 | 69 | 59 | 51 | 61 | 77.81 | 99 | 57 | 0.576 | -0.796 | 0.00185516 | 0.01142352 | 0.0339 | 0.0329 | 0.0332 | 0.0332 | TRUE | 0.6394 |

| | | | | | | | | | | | | | | | | | | | | | | | | | |
|---|---|---|---|---|---|---|---|---|---|---|---|---|---|---|---|---|---|---|---|---|---|---|---|---|---|
| Il6st | 5378 | 4209 | 4575 | 4567 | 4727 | 5535 | 4177 | 4477 | 4936 | 4546 | 5333 | 5464 | 4822.12 | 4530 | 5114 | 1.129 | 0.175 | 0.03530924 | 0.10461754 | 0.008 | 0.0029 | 0.0048 | 0.0048 | TRUE | 0.1154 |
| Il7 | 22 | 18 | 9 | 21 | 16 | 14 | 17 | 19 | 10 | 21 | 18 | 14 | 16.45 | 15 | 18 | 1.142 | 0.192 | 0.69368549 | NA | 0 | 0.1465 | 0.1104 | 0.1104 | TRUE | 0.4273 |
| Il7r | 0 | 0 | 0 | 0 | 0 | 0 | 0 | 0 | 0 | 0 | 0 | 0 | 0 | 0 | 0 | NA | NA | NA | NA | NA | NA | NA | NA | NA | NA |
| Il8 | 0 | 0 | 0 | 0 | 0 | 0 | 0 | 0 | 0 | 0 | 0 | 0 | 0 | 0 | 0 | NA | NA | NA | NA | NA | NA | NA | NA | NA | NA |
| Il9 | 0 | 0 | 0 | 0 | 0 | 0 | 0 | 0 | 0 | 0 | 0 | 0 | 0 | 0 | 0 | NA | NA | NA | NA | NA | NA | NA | NA | NA | NA |
| Il9r | 26 | 14 | 17 | 19 | 12 | 11 | 20 | 15 | 18 | 19 | 14 | 11 | 16.12 | 18 | 15 | 0.808 | -0.308 | 0.52389764 | NA | 0 | 0.1494 | 0.1059 | 0.1059 | TRUE | 0.3395 |
| Ildr1 | 1197 | 968 | 1078 | 1459 | 1299 | 1334 | 930 | 1030 | 1163 | 1452 | 1465 | 1317 | 1226.17 | 1041 | 1411 | 1.358 | 0.442 | 1.70E-06 | 4.47E-05 | 0.0068 | 0.0044 | 0.0053 | 0.0053 | TRUE | 0.1261 |
| Ildr2 | 260 | 189 | 142 | 184 | 188 | 215 | 202 | 201 | 153 | 183 | 212 | 212 | 193.94 | 185 | 202 | 1.089 | 0.122 | 0.44334655 | 0.61433424 | 0.0097 | 0.0146 | 0.0132 | 0.0132 | TRUE | 0.2377 |
| Ilf2 | 11171 | 7839 | 7644 | 8370 | 7421 | 8599 | 8676 | 8338 | 8248 | 8331 | 8372 | 8489 | 8408.96 | 8421 | 8397 | 0.997 | -0.004 | 0.92942463 | 0.9609898 | 3.00E-04 | 0.0027 | 0.0016 | 0.0016 | TRUE | 0.0087 |
| Ilf3 | 17882 | 14054 | 14475 | 14952 | 13676 | 15742 | 13888 | 14949 | 15618 | 14883 | 15428 | 15540 | 15051.1 | 14818 | 15284 | 1.032 | 0.045 | 0.43564236 | 0.60705138 | 0.0018 | 0.0026 | 0.0023 | 0.0023 | TRUE | 0.0371 |
| Ilk | 2248 | 1501 | 1359 | 1351 | 1144 | 1338 | 1746 | 1597 | 1466 | 1345 | 1291 | 1321 | 1460.84 | 1603 | 1319 | 0.822 | -0.283 | 0.00036719 | 0.00339914 | 0.0035 | 0.004 | 0.0039 | 0.0039 | TRUE | 0.0755 |
| Ilkap | 2216 | 1676 | 1743 | 1577 | 1462 | 1688 | 1721 | 1783 | 1881 | 1570 | 1649 | 1666 | 1711.64 | 1795 | 1628 | 0.908 | -0.139 | 0.039948 | 0.11436973 | 0.001 | 0.0038 | 0.0027 | 0.0027 | TRUE | 0.021 |
| Iltifb | 0 | 0 | 0 | 0 | 0 | 0 | 0 | 0 | 0 | 0 | 0 | 0 | 0 | 0 | 0 | NA | NA | NA | NA | NA | NA | NA | NA | NA | NA |
| Ilvbl | 460 | 353 | 395 | 391 | 355 | 390 | 357 | 375 | 426 | 389 | 400 | 385 | 388.94 | 386 | 391 | 1.016 | 0.022 | 0.84230754 | 0.91032497 | 0.002 | 0.0085 | 0.0065 | 0.0065 | TRUE | 0.0921 |
| Immp1l | 703 | 464 | 502 | 571 | 502 | 506 | 546 | 494 | 542 | 568 | 566 | 500 | 535.9 | 527 | 545 | 1.032 | 0.046 | 0.65018258 | 0.7827712 | 0.0023 | 0.0068 | 0.0054 | 0.0054 | TRUE | 0.0627 |
| Immp2l | 103 | 65 | 85 | 96 | 86 | 74 | 80 | 69 | 92 | 96 | 97 | 73 | 84.41 | 80 | 89 | 1.102 | 0.14 | 0.53524826 | 0.69421991 | 0.0091 | 0.0305 | 0.0247 | 0.0247 | TRUE | 0.2282 |
| Immt | 17585 | 12391 | 12278 | 13814 | 11791 | 14039 | 13658 | 13180 | 13248 | 13750 | 13301 | 13859 | 13499.35 | 13362 | 13637 | 1.021 | 0.029 | 0.53468972 | 0.69374729 | 3.00E-04 | 0.0026 | 0.0015 | 0.0015 | TRUE | 0.0056 |
| Imp3 | 2796 | 2143 | 2224 | 2505 | 2266 | 2597 | 2172 | 2279 | 2400 | 2493 | 2556 | 2564 | 2410.68 | 2284 | 2538 | 1.112 | 0.153 | 0.01491447 | 0.0556441 | 0.001 | 0.0034 | 0.0024 | 0.0024 | TRUE | 0.0239 |
| Imp4 | 4153 | 2846 | 2319 | 2364 | 2096 | 2361 | 3225 | 3027 | 2502 | 2353 | 2365 | 2331 | 2633.87 | 2918 | 2350 | 0.804 | -0.314 | 0.000305 | 0.00294556 | 0.0077 | 0.0033 | 0.0051 | 0.0051 | TRUE | 0.1879 |
| Impa1 | 5127 | 3684 | 3457 | 3229 | 2787 | 3176 | 3982 | 3919 | 3730 | 3214 | 3144 | 3135 | 3520.67 | 3877 | 3164 | 0.816 | -0.293 | 1.54E-07 | 5.71E-06 | 4.00E-04 | 0.0031 | 0.002 | 0.002 | TRUE | 0.0132 |
| Impa2 | 4639 | 3789 | 3965 | 3528 | 3302 | 3847 | 3603 | 4030 | 4278 | 3512 | 3725 | 3798 | 3824.3 | 3970 | 3678 | 0.927 | -0.11 | 0.1324866 | 0.27048008 | 0.0041 | 0.003 | 0.0036 | 0.0036 | TRUE | 0.0811 |
| Impact | 9768 | 6652 | 6704 | 6811 | 6095 | 7039 | 7586 | 7076 | 7234 | 6780 | 6876 | 6949 | 7083.27 | 7299 | 6868 | 0.941 | -0.088 | 0.08858418 | 0.20258138 | 5.00E-04 | 0.0028 | 0.0018 | 0.0018 | TRUE | 0.0147 |
| Impad1 | 5068 | 3742 | 3545 | 3969 | 3316 | 3486 | 3936 | 3980 | 3825 | 3951 | 3741 | 3441 | 3812.37 | 3914 | 3711 | 0.948 | -0.077 | 0.23537299 | 0.40178982 | 0.0022 | 0.003 | 0.0028 | 0.0028 | TRUE | 0.0496 |
| Impdh1 | 6294 | 4485 | 4284 | 4759 | 4143 | 4918 | 4888 | 4771 | 4622 | 4737 | 4674 | 4855 | 4757.83 | 4760 | 4755 | 0.999 | -0.002 | 0.97177666 | 0.98486874 | 4.00E-04 | 0.0029 | 0.0018 | 0.0018 | TRUE | 0.0078 |
| Impdh2 | 28064 | 20870 | 22582 | 23032 | 20708 | 24753 | 21796 | 22199 | 24366 | 22926 | 23361 | 24435 | 23180.48 | 22787 | 23574 | 1.035 | 0.049 | 0.40202221 | 0.5750704 | 0.0022 | 0.0025 | 0.0024 | 0.0024 | TRUE | 0.0449 |
| Impg1 | 0 | 1 | 0 | 0 | 1 | 0 | 0 | 1 | 0 | 0 | 1 | 0 | 0.37 | 0 | 0 | 1.075 | 0.105 | 0.97520514 | NA | 0 | 6.4904 | 6.1969 | 6.1969 | TRUE | 0.5216 |
| Impg2 | 1 | 1 | 1 | 0 | 1 | 0 | 1 | 1 | 1 | 0 | 1 | 0 | 0.67 | 1 | 0 | 0.383 | -1.383 | 0.54981825 | NA | 0 | 3.5158 | 2.3538 | 2.3538 | TRUE | 0.5075 |
| Ina | 1806 | 1238 | 908 | 1188 | 1021 | 1321 | 1403 | 1317 | 980 | 1183 | 1152 | 1304 | 1222.93 | 1233 | 1213 | 0.983 | -0.025 | 0.82705607 | 0.9008621 | 0.0189 | 0.0044 | 0.0089 | 0.0089 | TRUE | 0.384 |
| Inadl | 774 | 552 | 667 | 581 | 558 | 604 | 601 | 587 | 720 | 578 | 629 | 596 | 618.67 | 636 | 601 | 0.946 | -0.08 | 0.44020683 | 0.61163373 | 0.0056 | 0.0063 | 0.006 | 0.006 | TRUE | 0.1521 |
| Inca1 | 171 | 121 | 161 | 230 | 183 | 206 | 133 | 129 | 174 | 229 | 206 | 203 | 179 | 145 | 213 | 1.473 | 0.558 | 0.00069319 | 0.00553814 | 0.0082 | 0.0157 | 0.0138 | 0.0138 | TRUE | 0.2971 |
| Incenp | 12047 | 8986 | 8390 | 8804 | 7810 | 9198 | 9356 | 9558 | 9053 | 8763 | 8810 | 9080 | 9103.54 | 9322 | 8884 | 0.953 | -0.069 | 0.16091598 | 0.31065866 | 4.00E-04 | 0.0027 | 0.0017 | 0.0017 | TRUE | 0.0077 |
| Inf2 | 1541 | 1320 | 1247 | 1504 | 1536 | 1665 | 1197 | 1404 | 1346 | 1497 | 1733 | 1644 | 1469.98 | 1316 | 1625 | 1.236 | 0.306 | 0.00032976 | 0.00312554 | 0.0049 | 0.004 | 0.0045 | 0.0045 | TRUE | 0.0775 |
| Ing1 | 4011 | 3000 | 2961 | 2900 | 2543 | 3038 | 3115 | 3191 | 3195 | 2887 | 2869 | 2999 | 3042.59 | 3167 | 2918 | 0.922 | -0.118 | 0.03419238 | 0.10211276 | 0 | 0.0032 | 0.0019 | 0.0019 | TRUE | 0.0072 |
| Ing2 | 1940 | 1499 | 1494 | 1426 | 1260 | 1436 | 1507 | 1594 | 1612 | 1419 | 1421 | 1418 | 1495.27 | 1571 | 1419 | 0.904 | -0.145 | 0.02922203 | 0.09085718 | 0 | 0.004 | 0.0025 | 0.0025 | TRUE | 0.0162 |
| Ing3 | 1563 | 1168 | 1074 | 1134 | 1021 | 1198 | 1214 | 1242 | 1159 | 1129 | 1152 | 1183 | 1179.72 | 1205 | 1155 | 0.958 | -0.062 | 0.38556258 | 0.55936317 | 0 | 0.0044 | 0.0029 | 0.0029 | TRUE | 0.013 |
| Ing4 | 2671 | 1985 | 2157 | 1977 | 1850 | 2203 | 2074 | 2111 | 2327 | 1968 | 2087 | 2175 | 2123.81 | 2171 | 2077 | 0.957 | -0.063 | 0.37705103 | 0.55151527 | 0.0025 | 0.0035 | 0.0032 | 0.0032 | TRUE | 0.0475 |
| Ing5 | 5268 | 3875 | 3691 | 3954 | 3422 | 3977 | 4091 | 4122 | 3983 | 3936 | 3860 | 3926 | 3986.31 | 4065 | 3907 | 0.961 | -0.057 | 0.27033142 | 0.44122953 | 0 | 0.003 | 0.0017 | 0.0017 | TRUE | 0.0038 |
| Inha | 18 | 17 | 11 | 28 | 26 | 33 | 14 | 18 | 12 | 28 | 29 | 33 | 22.28 | 15 | 30 | 2.048 | 1.034 | 0.01265178 | 0.04923273 | 0 | 0.1088 | 0.0737 | 0.0737 | TRUE | 0.17 |
| Inhba | 0 | 0 | 0 | 0 | 0 | 1 | 0 | 0 | 0 | 0 | 0 | 1 | 0.16 | 0 | 0 | 2.094 | 1.067 | 0.79379329 | NA | 0 | 14.4078 | 10 | 10 | TRUE | 0.3835 |
| Inhbb | 1499 | 1168 | 1125 | 1499 | 1197 | 1357 | 1164 | 1242 | 1214 | 1492 | 1350 | 1340 | 1300.41 | 1207 | 1394 | 1.156 | 0.209 | 0.00543646 | 0.02614512 | 0.0017 | 0.0042 | 0.0033 | 0.0033 | TRUE | 0.046 |
| Inhbc | 39 | 25 | 7 | 19 | 24 | 28 | 30 | 27 | 8 | 19 | 27 | 28 | 23.01 | 22 | 25 | 1.128 | 0.174 | 0.71860386 | 0.8310547 | 0.1809 | 0.1054 | 0.1246 | 0.1246 | TRUE | 1.6787 |
| Inhbe | 2 | 2 | 4 | 1 | 1 | 4 | 2 | 2 | 4 | 1 | 1 | 4 | 2.34 | 3 | 2 | 0.778 | -0.362 | 0.77476777 | NA | 0 | 1.0132 | 0.7155 | 0.7155 | TRUE | 0.6498 |
| Inip | 2097 | 1453 | 1364 | 1465 | 1346 | 1522 | 1629 | 1546 | 1472 | 1458 | 1518 | 1502 | 1520.84 | 1549 | 1493 | 0.963 | -0.054 | 0.43500601 | 0.60670774 | 0.001 | 0.004 | 0.0028 | 0.0028 | TRUE | 0.0256 |
| Inmt | 0 | 0 | 0 | 0 | 0 | 1 | 0 | 0 | 0 | 0 | 0 | 1 | 0.16 | 0 | 0 | 2.094 | 1.067 | 0.79379329 | NA | 0 | 14.4078 | 10 | 10 | TRUE | 0.3835 |
| Ino80 | 5491 | 3757 | 3493 | 3448 | 3095 | 3614 | 4265 | 3996 | 3769 | 3432 | 3491 | 3568 | 3753.5 | 4010 | 3497 | 0.872 | -0.198 | 0.00163125 | 0.01028927 | 0.0017 | 0.0031 | 0.0026 | 0.0026 | TRUE | 0.0382 |
| Ino80b | 585 | 424 | 315 | 318 | 300 | 346 | 454 | 451 | 340 | 317 | 338 | 342 | 373.63 | 415 | 332 | 0.799 | -0.325 | 0.01320925 | 0.05077996 | 0.0116 | 0.0088 | 0.0096 | 0.0096 | TRUE | 0.2818 |
| Ino80c | 1307 | 935 | 796 | 796 | 720 | 771 | 1015 | 995 | 859 | 792 | 812 | 761 | 872.36 | 956 | 788 | 0.823 | -0.28 | 0.00174116 | 0.01086399 | 0.0036 | 0.0051 | 0.0046 | 0.0046 | TRUE | 0.0927 |
| Ino80d | 2250 | 1545 | 1289 | 1483 | 1237 | 1521 | 1747 | 1643 | 1391 | 1476 | 1395 | 1501 | 1525.8 | 1594 | 1457 | 0.914 | -0.13 | 0.14242182 | 0.28508327 | 0.0064 | 0.004 | 0.005 | 0.005 | TRUE | 0.1484 |
| Ino80e | 1807 | 1239 | 1010 | 999 | 980 | 1172 | 1403 | 1318 | 1090 | 994 | 1106 | 1157 | 1178 | 1270 | 1086 | 0.854 | -0.229 | 0.02370434 | 0.07826428 | 0.0098 | 0.0044 | 0.0065 | 0.0065 | TRUE | 0.1838 |
| Inpp1 | 182 | 158 | 145 | 173 | 154 | 159 | 141 | 168 | 156 | 172 | 174 | 157 | 161.46 | 155 | 168 | 1.082 | 0.114 | 0.47398867 | 0.64249276 | 0 | 0.0171 | 0.0121 | 0.0121 | TRUE | 0.0725 |
| Inpp4a | 974 | 743 | 745 | 890 | 794 | 915 | 756 | 790 | 804 | 886 | 896 | 903 | 839.25 | 783 | 895 | 1.143 | 0.193 | 0.01380817 | 0.05261536 | 0 | 0.0053 | 0.0032 | 0.0032 | TRUE | 0.0116 |
| Inpp4b | 87 | 53 | 17 | 23 | 20 | 14 | 68 | 56 | 18 | 23 | 23 | 14 | 33.59 | 47 | 20 | 0.413 | -1.276 | 0.00517314 | 0.02518381 | 0.2139 | 0.073 | 0.1137 | 0.1137 | TRUE | 0.6272 |
| Inpp5a | 3548 | 2477 | 2350 | 2703 | 2381 | 2621 | 2756 | 2635 | 2536 | 2691 | 2686 | 2587 | 2648.31 | 2642 | 2655 | 1.004 | 0.006 | 0.92311709 | 0.95727437 | 8.00E-04 | 0.0033 | 0.0023 | 0.0023 | TRUE | 0.0177 |
| Inpp5b | 2431 | 1730 | 1632 | 1720 | 1611 | 1856 | 1888 | 1840 | 1761 | 1712 | 1817 | 1832 | 1808.46 | 1830 | 1787 | 0.976 | -0.035 | 0.59767668 | 0.74367484 | 8.00E-04 | 0.0037 | 0.0026 | 0.0026 | TRUE | 0.0165 |
| Inpp5d | 14370 | 10830 | 12210 | 11729 | 10344 | 11966 | 11161 | 11520 | 13174 | 11675 | 11669 | 11812 | 11835.2 | 11952 | 11719 | 0.981 | -0.028 | 0.67343905 | 0.79945717 | 0.0033 | 0.0026 | 0.0031 | 0.0031 | TRUE | 0.0978 |
| Inpp5e | 712 | 483 | 473 | 458 | 416 | 463 | 553 | 514 | 510 | 456 | 469 | 457 | 493.22 | 526 | 461 | 0.875 | -0.193 | 0.04633997 | 0.12749512 | 0 | 0.0072 | 0.0047 | 0.0047 | TRUE | 0.0257 |
| Inpp5f | 2628 | 1919 | 1894 | 2223 | 2042 | 2356 | 2041 | 2041 | 2044 | 2213 | 2304 | 2326 | 2161.33 | 2042 | 2281 | 1.117 | 0.159 | 0.00722945 | 0.03250409 | 0 | 0.0035 | 0.0021 | 0.0021 | TRUE | 0.0083 |
| Inpp5j | 6 | 8 | 2 | 7 | 6 | 6 | 5 | 9 | 2 | 7 | 7 | 6 | 5.83 | 5 | 7 | 1.285 | 0.362 | 0.6542378 | NA | 0 | 0.4089 | 0.2966 | 0.2966 | TRUE | 0.3772 |
| Inpp5k | 1457 | 1159 | 1133 | 1120 | 1049 | 1158 | 1132 | 1233 | 1222 | 1115 | 1183 | 1143 | 1171.38 | 1196 | 1147 | 0.96 | -0.059 | 0.43106637 | 0.60321438 | 9.00E-04 | 0.0044 | 0.0031 | 0.0031 | TRUE | 0.0277 |
| Inppl1 | 5187 | 3923 | 4118 | 4057 | 3884 | 4409 | 4029 | 4173 | 4443 | 4038 | 4382 | 4352 | 4236.15 | 4215 | 4257 | 1.01 | 0.015 | 0.81229066 | 0.89151249 | 0.002 | 0.003 | 0.0026 | 0.0026 | TRUE | 0.0273 |
| Ins1 | 0 | 0 | 0 | 0 | 0 | 0 | 0 | 0 | 0 | 0 | 0 | 0 | 0 | 0 | 0 | NA | NA | NA | NA | NA | NA | NA | NA | NA | NA |
| Ins2 | 0 | 0 | 0 | 0 | 0 | 0 | 0 | 0 | 0 | 0 | 0 | 0 | 0 | 0 | 0 | NA | NA | NA | NA | NA | NA | NA | NA | NA | NA |
| Insc | 90 | 79 | 96 | 125 | 85 | 112 | 70 | 84 | 104 | 124 | 96 | 111 | 98.06 | 86 | 110 | 1.295 | 0.373 | 0.08741437 | 0.20061232 | 0.0174 | 0.0266 | 0.024 | 0.024 | TRUE | 0.3049 |
| Insig1 | 8965 | 7143 | 8067 | 9168 | 7609 | 8784 | 6963 | 7598 | 8704 | 9126 | 8584 | 8671 | 8274.27 | 7755 | 8794 | 1.134 | 0.182 | 0.01890772 | 0.06638369 | 0.0068 | 0.0027 | 0.0042 | 0.0042 | TRUE | 0.1399 |
| Insig2 | 1719 | 1232 | 1296 | 1456 | 1269 | 1405 | 1335 | 1310 | 1398 | 1449 | 1432 | 1387 | 1385.29 | 1348 | 1423 | 1.056 | 0.078 | 0.25337314 | 0.42242648 | 0 | 0.0041 | 0.0026 | 0.0026 | TRUE | 0.0124 |
| Insl3 | 12 | 16 | 20 | 10 | 20 | 12 | 9 | 17 | 22 | 10 | 23 | 12 | 15.38 | 16 | 15 | 0.931 | -0.103 | 0.84936824 | NA | 0.1191 | 0.1565 | 0.1457 | 0.1457 | TRUE | 0.5345 |
| Insl5 | 1 | 2 | 0 | 0 | 1 | 0 | 1 | 2 | 0 | 0 | 1 | 0 | 0.67 | 1 | 0 | 0.385 | -1.379 | 0.57963618 | NA | 0 | 3.5292 | 2.9645 | 2.9645 | TRUE | 0.3656 |
| Insl6 | 142 | 65 | 113 | 142 | 117 | 148 | 110 | 69 | 122 | 141 | 132 | 146 | 120.13 | 100 | 140 | 1.389 | 0.474 | 0.02620783 | 0.08413938 | 0.0305 | 0.0222 | 0.0243 | 0.0243 | TRUE | 0.6895 |
| Insm1 | 42 | 23 | 22 | 29 | 28 | 29 | 33 | 24 | 24 | 29 | 32 | 29 | 28.32 | 27 | 30 | 1.092 | 0.127 | 0.72540409 | 0.83572689 | 0 | 0.0861 | 0.0587 | 0.0587 | TRUE | 0.2578 |

| | | | | | | | | | | | | | | | | | | | | | | | | | |
|---|---|---|---|---|---|---|---|---|---|---|---|---|---|---|---|---|---|---|---|---|---|---|---|---|---|
| Insm2 | 2 | 2 | 2 | 1 | 1 | 2 | 2 | 2 | 2 | 1 | 1 | 2 | 1.66 | 2 | 1 | 0.71 | -0.494 | 0.73016878 | NA | 0 | 1.4336 | 0.8486 | 0.8486 | TRUE | 0.0997 |
| Insr | 1824 | 1248 | 1361 | 1585 | 1296 | 1450 | 1417 | 1327 | 1469 | 1578 | 1462 | 1431 | 1447.29 | 1404 | 1490 | 1.061 | 0.086 | 0.24697399 | 0.41568896 | 0.0019 | 0.0041 | 0.0033 | 0.0033 | TRUE | 0.032 |
| Insrr | 13 | 4 | 7 | 6 | 6 | 5 | 10 | 4 | 8 | 6 | 7 | 5 | 6.6 | 7 | 6 | 0.794 | -0.333 | 0.65656131 | NA | 0 | 0.3617 | 0.2498 | 0.2498 | TRUE | 0.224 |
| Ints1 | 10278 | 7614 | 7185 | 7780 | 6979 | 8049 | 7982 | 8099 | 7753 | 7744 | 7873 | 7946 | 7899.47 | 7945 | 7854 | 0.989 | -0.017 | 0.73171247 | 0.83929626 | 1.00E-04 | 0.0027 | 0.0016 | 0.0016 | TRUE | 0.0054 |
| Ints10 | 6003 | 4313 | 4348 | 4411 | 3985 | 4669 | 4662 | 4588 | 4691 | 4391 | 4495 | 4609 | 4572.77 | 4647 | 4498 | 0.968 | -0.047 | 0.36429652 | 0.53901555 | 1.00E-04 | 0.0029 | 0.0017 | 0.0017 | TRUE | 0.0057 |
| Ints12 | 1817 | 1289 | 1197 | 1238 | 1081 | 1256 | 1411 | 1371 | 1292 | 1232 | 1219 | 1240 | 1294.25 | 1358 | 1230 | 0.906 | -0.143 | 0.04266803 | 0.11997918 | 0 | 0.0043 | 0.0028 | 0.0028 | TRUE | 0.0217 |
| Ints2 | 8920 | 6244 | 6182 | 6880 | 5917 | 6689 | 6928 | 6642 | 6670 | 6848 | 6675 | 6603 | 6727.7 | 6747 | 6709 | 0.994 | -0.008 | 0.86677211 | 0.92654056 | 3.00E-04 | 0.0028 | 0.0017 | 0.0017 | TRUE | 0.0069 |
| Ints3 | 11850 | 8453 | 8074 | 7992 | 7437 | 8568 | 9203 | 8991 | 8712 | 7955 | 8390 | 8458 | 8618.23 | 8969 | 8268 | 0.922 | -0.118 | 0.02451788 | 0.08020922 | 7.00E-04 | 0.0027 | 0.0019 | 0.0019 | TRUE | 0.0134 |
| Ints4 | 6482 | 4745 | 4808 | 5241 | 4826 | 5642 | 5034 | 5047 | 5188 | 5217 | 5444 | 5570 | 5249.99 | 5090 | 5410 | 1.063 | 0.088 | 0.09623971 | 0.21523302 | 4.00E-04 | 0.0029 | 0.0018 | 0.0018 | TRUE | 0.012 |
| Ints5 | 4158 | 2833 | 2015 | 2002 | 1797 | 2014 | 3229 | 3013 | 2174 | 1993 | 2027 | 1988 | 2404.18 | 2805 | 2003 | 0.713 | -0.487 | 1.82E-05 | 0.00031011 | 0.0192 | 0.0034 | 0.0089 | 0.0089 | TRUE | 0.4685 |
| Ints6 | 2620 | 1694 | 1476 | 1447 | 1326 | 1443 | 2035 | 1802 | 1593 | 1440 | 1496 | 1424 | 1631.67 | 1810 | 1453 | 0.802 | -0.318 | 0.00036074 | 0.00335464 | 0.0069 | 0.0039 | 0.0051 | 0.0051 | TRUE | 0.1458 |
| Ints7 | 4679 | 3442 | 3255 | 3276 | 3185 | 3564 | 3634 | 3661 | 3512 | 3261 | 3593 | 3518 | 3529.91 | 3602 | 3457 | 0.959 | -0.06 | 0.32205535 | 0.49587577 | 0.0011 | 0.0031 | 0.0023 | 0.0023 | TRUE | 0.0301 |
| Ints8 | 6145 | 4476 | 4464 | 4794 | 4197 | 4828 | 4773 | 4761 | 4817 | 4772 | 4735 | 4766 | 4770.47 | 4784 | 4758 | 0.995 | -0.008 | 0.87451953 | 0.93036901 | 0 | 0.0029 | 0.0015 | 0.0015 | TRUE | 4.00E-04 |
| Ints9 | 3627 | 2543 | 2298 | 2215 | 2122 | 2521 | 2817 | 2705 | 2480 | 2205 | 2394 | 2489 | 2514.78 | 2667 | 2363 | 0.885 | -0.176 | 0.01629402 | 0.05939566 | 0.0033 | 0.0034 | 0.0035 | 0.0035 | TRUE | 0.0456 |
| Intu | 1468 | 1097 | 1086 | 966 | 927 | 1099 | 1140 | 1167 | 1172 | 962 | 1046 | 1085 | 1095.16 | 1160 | 1031 | 0.889 | -0.17 | 0.02740669 | 0.08660707 | 0.001 | 0.0046 | 0.0033 | 0.0033 | TRUE | 0.0419 |
| Invs | 265 | 148 | 78 | 113 | 110 | 110 | 206 | 157 | 84 | 112 | 124 | 109 | 132.09 | 149 | 115 | 0.767 | -0.383 | 0.13929137 | 0.28048626 | 0.0913 | 0.0204 | 0.0407 | 0.0407 | TRUE | 0.2562 |
| Ip6k1 | 4843 | 3797 | 4028 | 4280 | 3938 | 4245 | 3761 | 4039 | 4346 | 4260 | 4442 | 4190 | 4173.27 | 4049 | 4297 | 1.062 | 0.087 | 0.19120754 | 0.34937075 | 0.0027 | 0.003 | 0.0029 | 0.0029 | TRUE | 0.0502 |
| Ip6k2 | 591 | 365 | 410 | 396 | 324 | 398 | 459 | 388 | 442 | 394 | 366 | 393 | 407.03 | 430 | 384 | 0.893 | -0.164 | 0.14049527 | 0.28231392 | 0.0024 | 0.0082 | 0.0064 | 0.0064 | TRUE | 0.0813 |
| Ip6k3 | 2 | 2 | 0 | 0 | 0 | 0 | 2 | 2 | 0 | 0 | 0 | 0 | 0.61 | 1 | 0 | 0.154 | -2.696 | 0.32099274 | NA | 0 | 3.8659 | 3.9247 | 3.9247 | TRUE | 0.3505 |
| Ipcef1 | 18 | 5 | 5 | 4 | 2 | 5 | 14 | 5 | 5 | 4 | 2 | 5 | 5.98 | 8 | 4 | 0.447 | -1.16 | 0.16838167 | NA | 0 | 0.3989 | 0.3158 | 0.3158 | TRUE | 1.4018 |
| Ipmk | 5398 | 4073 | 3979 | 4261 | 3899 | 4409 | 4192 | 4332 | 4293 | 4241 | 4398 | 4352 | 4301.72 | 4272 | 4330 | 1.014 | 0.02 | 0.70488706 | 0.82131844 | 0 | 0.003 | 0.0017 | 0.0017 | TRUE | 0.004 |
| Ipo11 | 7362 | 5318 | 5387 | 6246 | 5476 | 5938 | 5718 | 5657 | 5813 | 6217 | 6177 | 5862 | 5907.24 | 5729 | 6085 | 1.062 | 0.087 | 0.09326835 | 0.2100007 | 4.00E-04 | 0.0028 | 0.0018 | 0.0018 | TRUE | 0.0127 |
| Ipo13 | 2423 | 1518 | 1104 | 1166 | 1029 | 1142 | 1882 | 1615 | 1191 | 1161 | 1161 | 1127 | 1356.08 | 1563 | 1150 | 0.735 | -0.445 | 0.00043263 | 0.00386476 | 0.0238 | 0.0042 | 0.0107 | 0.0107 | TRUE | 0.2665 |
| Ipo4 | 9276 | 6694 | 5774 | 5707 | 5066 | 6145 | 7204 | 7120 | 6230 | 5681 | 5715 | 6066 | 6336.08 | 6851 | 5821 | 0.849 | -0.236 | 0.00051961 | 0.00444188 | 0.0035 | 0.0028 | 0.0032 | 0.0032 | TRUE | 0.0765 |
| Ipo5 | 47801 | 35496 | 35959 | 37712 | 33559 | 39352 | 37125 | 37757 | 38799 | 37538 | 37858 | 38846 | 37987.35 | 37894 | 38081 | 1.005 | 0.007 | 0.87433248 | 0.93023925 | 3.00E-04 | 0.0025 | 0.0015 | 0.0015 | TRUE | 0.0053 |
| Ipo7 | 22247 | 16348 | 17415 | 19162 | 16375 | 19143 | 17278 | 17389 | 18791 | 19074 | 18473 | 18897 | 18316.95 | 17819 | 18815 | 1.056 | 0.079 | 0.13483928 | 0.27379711 | 0.0012 | 0.0026 | 0.0019 | 0.0019 | TRUE | 0.0277 |
| Ipo8 | 2907 | 2017 | 2067 | 2245 | 1965 | 2256 | 2258 | 2145 | 2230 | 2235 | 2217 | 2227 | 2218.65 | 2211 | 2226 | 1.006 | 0.009 | 0.87534008 | 0.93075701 | 0 | 0.0035 | 0.0021 | 0.0021 | TRUE | 0.0081 |
| Ipo9 | 12546 | 8759 | 8432 | 8190 | 7696 | 8838 | 9744 | 9317 | 9098 | 8152 | 8682 | 8724 | 8952.92 | 9386 | 8519 | 0.907 | -0.14 | 0.01041675 | 0.04253902 | 0.0011 | 0.0027 | 0.002 | 0.002 | TRUE | 0.0174 |
| Ipp | 948 | 700 | 705 | 845 | 708 | 791 | 736 | 745 | 761 | 841 | 799 | 781 | 777.03 | 747 | 807 | 1.08 | 0.112 | 0.17069288 | 0.32232967 | 0 | 0.0055 | 0.0035 | 0.0035 | TRUE | 0.0166 |
| Ippk | 1757 | 1239 | 1198 | 1326 | 1168 | 1386 | 1365 | 1318 | 1293 | 1320 | 1318 | 1368 | 1330.14 | 1325 | 1335 | 1.007 | 0.01 | 0.88047185 | 0.93336677 | 0 | 0.0042 | 0.0026 | 0.0026 | TRUE | 0.0086 |
| Ipw | 1 | 1 | 0 | 1 | 0 | 0 | 1 | 1 | 0 | 1 | 0 | 0 | 0.47 | 1 | 0 | 0.599 | -0.74 | 0.79191656 | NA | 0 | 5.0171 | 3.8827 | 3.8827 | TRUE | 0.3906 |
| Iqca | 59 | 66 | 75 | 44 | 40 | 69 | 46 | 70 | 81 | 44 | 45 | 68 | 59 | 66 | 52 | 0.805 | -0.313 | 0.29533782 | 0.46709939 | 0.0581 | 0.0426 | 0.0471 | 0.0471 | TRUE | 0.5845 |
| Iqcb1 | 2424 | 1864 | 2170 | 2142 | 1991 | 2114 | 1883 | 1983 | 2341 | 2132 | 2246 | 2087 | 2111.96 | 2069 | 2155 | 1.042 | 0.06 | 0.48050556 | 0.64736224 | 0.0068 | 0.0035 | 0.0047 | 0.0047 | TRUE | 0.1597 |
| Iqcc | 965 | 711 | 677 | 702 | 683 | 764 | 749 | 756 | 730 | 699 | 770 | 754 | 743.28 | 745 | 741 | 0.994 | -0.009 | 0.91356446 | 0.95246365 | 0 | 0.0056 | 0.0038 | 0.0038 | TRUE | 0.0299 |
| Iqcd | 4 | 5 | 4 | 3 | 1 | 4 | 3 | 5 | 4 | 3 | 1 | 4 | 3.47 | 4 | 3 | 0.646 | -0.63 | 0.54028165 | NA | 0 | 0.686 | 0.4578 | 0.4578 | TRUE | 0.2608 |
| Iqce | 525 | 376 | 362 | 352 | 313 | 360 | 408 | 400 | 391 | 350 | 353 | 355 | 376.19 | 400 | 353 | 0.883 | -0.179 | 0.0893623 | 0.20380651 | 0 | 0.0087 | 0.0054 | 0.0054 | TRUE | 0.0043 |
| Iqcf1 | 0 | 0 | 0 | 0 | 1 | 0 | 0 | 0 | 0 | 0 | 1 | 0 | 0.19 | 0 | 0 | 2.094 | 1.067 | 0.79379329 | NA | 0 | 12.608 | 10 | 10 | TRUE | 0.5075 |
| Iqcf3 | 0 | 0 | 0 | 0 | 0 | 0 | 0 | 0 | 0 | 0 | 0 | 0 | 0 | 0 | 0 | NA | NA | NA | NA | NA | NA | NA | NA | NA | NA |
| Iqcf4 | 0 | 0 | 0 | 0 | 0 | 0 | 0 | 0 | 0 | 0 | 0 | 0 | 0 | 0 | 0 | NA | NA | NA | NA | NA | NA | NA | NA | NA | NA |
| Iqcf5 | 9 | 1 | 2 | 3 | 1 | 2 | 7 | 1 | 2 | 3 | 1 | 2 | 2.72 | 3 | 2 | 0.583 | -0.779 | 0.52278842 | NA | 0 | 0.8748 | 0.6811 | 0.6811 | TRUE | 1.5885 |
| Iqcf6 | 0 | 0 | 0 | 0 | 0 | 1 | 0 | 0 | 0 | 0 | 0 | 1 | 0.16 | 0 | 0 | 2.094 | 1.067 | 0.79379329 | NA | 0 | 14.4078 | 10 | 10 | TRUE | 0.3835 |
| Iqcg | 784 | 648 | 719 | 712 | 600 | 692 | 609 | 689 | 776 | 709 | 677 | 683 | 690.44 | 691 | 690 | 1 | 0 | 0.99993023 | 0.99993023 | 0.0064 | 0.0059 | 0.006 | 0.006 | TRUE | 0.1352 |
| Iqch | 14 | 8 | 5 | 3 | 7 | 5 | 11 | 9 | 5 | 3 | 8 | 5 | 6.77 | 8 | 5 | 0.626 | -0.676 | 0.37340759 | NA | 0 | 0.3527 | 0.2584 | 0.2584 | TRUE | 0.2868 |
| Iqcj | 0 | 1 | 0 | 0 | 0 | 0 | 0 | 1 | 0 | 0 | 0 | 0 | 0.18 | 0 | 0 | 0.552 | -0.857 | 0.8336548 | NA | 0 | 13.3714 | 10 | 10 | TRUE | 0.5216 |
| Iqck | 11 | 10 | 2 | 7 | 5 | 4 | 9 | 11 | 2 | 7 | 6 | 4 | 6.32 | 7 | 6 | 0.77 | -0.377 | 0.64013607 | NA | 0 | 0.3777 | 0.3084 | 0.3084 | TRUE | 0.79 |
| Iqgap1 | 17033 | 12215 | 12286 | 13320 | 12172 | 13924 | 13229 | 12993 | 13256 | 13259 | 13731 | 13745 | 13368.89 | 13159 | 13578 | 1.032 | 0.045 | 0.32558531 | 0.49968762 | 2.00E-04 | 0.0026 | 0.0014 | 0.0014 | TRUE | 0.0052 |
| Iqgap2 | 2541 | 1773 | 1759 | 1705 | 1523 | 1749 | 1973 | 1886 | 1898 | 1697 | 1718 | 1727 | 1816.52 | 1919 | 1714 | 0.893 | -0.164 | 0.00814869 | 0.03554475 | 0 | 0.0037 | 0.0022 | 0.0022 | TRUE | 0.0077 |
| Iqgap3 | 3899 | 2874 | 2865 | 2877 | 2715 | 2976 | 3028 | 3057 | 3091 | 2864 | 3063 | 2938 | 3006.81 | 3059 | 2955 | 0.966 | -0.05 | 0.37900443 | 0.55372691 | 3.00E-04 | 0.0032 | 0.002 | 0.002 | TRUE | 0.0123 |
| Iqsec1 | 2468 | 1825 | 1608 | 1840 | 1665 | 1847 | 1917 | 1941 | 1735 | 1832 | 1878 | 1823 | 1854.36 | 1864 | 1844 | 0.989 | -0.016 | 0.81033817 | 0.89066161 | 0.0014 | 0.0037 | 0.0028 | 0.0028 | TRUE | 0.0435 |
| Iqsec2 | 427 | 332 | 333 | 427 | 368 | 391 | 332 | 353 | 359 | 425 | 415 | 386 | 378.37 | 348 | 409 | 1.176 | 0.234 | 0.0319843 | 0.09699802 | 0 | 0.0087 | 0.0059 | 0.0059 | TRUE | 0.0279 |
| Iqsec3 | 84 | 59 | 73 | 129 | 105 | 86 | 65 | 63 | 79 | 128 | 118 | 85 | 89.75 | 69 | 110 | 1.606 | 0.684 | 0.00300234 | 0.01655575 | 0.021 | 0.0288 | 0.0266 | 0.0266 | TRUE | 0.4213 |
| Iqub | 50 | 41 | 37 | 26 | 22 | 21 | 39 | 44 | 40 | 26 | 25 | 21 | 32.3 | 41 | 24 | 0.584 | -0.776 | 0.02258464 | 0.07540347 | 0 | 0.0758 | 0.0498 | 0.0498 | TRUE | 0.0794 |
| Irak1 | 1710 | 1372 | 1160 | 1081 | 994 | 1169 | 1328 | 1459 | 1252 | 1076 | 1121 | 1154 | 1231.74 | 1346 | 1117 | 0.83 | -0.269 | 0.00079799 | 0.00613381 | 0.0028 | 0.0044 | 0.0038 | 0.0038 | TRUE | 0.0631 |
| Irak1bp1 | 659 | 533 | 563 | 462 | 421 | 499 | 512 | 567 | 607 | 460 | 475 | 493 | 518.94 | 562 | 476 | 0.848 | -0.237 | 0.02069652 | 0.07097298 | 0.0029 | 0.007 | 0.0056 | 0.0056 | TRUE | 0.0763 |
| Irak2 | 818 | 641 | 643 | 708 | 613 | 755 | 635 | 682 | 694 | 705 | 692 | 745 | 692.08 | 670 | 714 | 1.067 | 0.093 | 0.28537156 | 0.45732168 | 0 | 0.0059 | 0.0041 | 0.0041 | TRUE | 0.0263 |
| Irak3 | 910 | 655 | 559 | 791 | 742 | 798 | 707 | 697 | 603 | 787 | 837 | 788 | 736.47 | 669 | 804 | 1.2 | 0.263 | 0.00421384 | 0.02147946 | 0.0028 | 0.0056 | 0.0047 | 0.0047 | TRUE | 0.0848 |
| Irak4 | 231 | 170 | 160 | 165 | 148 | 164 | 179 | 181 | 173 | 164 | 167 | 162 | 170.99 | 178 | 164 | 0.925 | -0.113 | 0.44967689 | 0.61973116 | 0 | 0.0163 | 0.0102 | 0.0102 | TRUE | 0.0061 |
| Ireb2 | 20964 | 15225 | 15271 | 16609 | 14450 | 16459 | 16282 | 16195 | 16477 | 16533 | 16301 | 16248 | 16339.17 | 16318 | 16361 | 1.003 | 0.004 | 0.93092114 | 0.96192556 | 0 | 0.0026 | 0.0013 | 0.0013 | TRUE | 0.001 |
| Irf1 | 1752 | 1456 | 1314 | 1337 | 1326 | 1545 | 1361 | 1549 | 1418 | 1331 | 1496 | 1525 | 1446.52 | 1443 | 1451 | 1.006 | 0.009 | 0.9115872 | 0.9513752 | 0.0042 | 0.0041 | 0.0041 | 0.0041 | TRUE | 0.0633 |
| Irf2 | 56 | 44 | 59 | 65 | 62 | 44 | 43 | 47 | 64 | 65 | 70 | 43 | 55.34 | 51 | 59 | 1.16 | 0.214 | 0.45883652 | 0.62859679 | 0.0334 | 0.0453 | 0.042 | 0.042 | TRUE | 0.4833 |
| Irf2bp1 | 1267 | 993 | 802 | 801 | 714 | 787 | 984 | 1056 | 865 | 797 | 805 | 777 | 880.88 | 968 | 793 | 0.819 | -0.289 | 0.00146106 | 0.0094621 | 0.0041 | 0.0051 | 0.0048 | 0.0048 | TRUE | 0.1003 |
| Irf2bp2 | 2587 | 2272 | 2214 | 1729 | 1609 | 1848 | 2009 | 2417 | 2389 | 1721 | 1815 | 1824 | 2029.2 | 2272 | 1787 | 0.787 | -0.345 | 2.27E-05 | 0.00037071 | 0.0054 | 0.0036 | 0.0043 | 0.0043 | TRUE | 0.1266 |
| Irf2bpl | 5468 | 3913 | 3804 | 4740 | 4006 | 4384 | 4247 | 4162 | 4104 | 4718 | 4519 | 4328 | 4346.42 | 4171 | 4522 | 1.084 | 0.116 | 0.04022493 | 0.11489506 | 7.00E-04 | 0.003 | 0.0021 | 0.0021 | TRUE | 0.0178 |
| Irf3 | 1342 | 1049 | 845 | 835 | 743 | 858 | 1042 | 1116 | 912 | 831 | 838 | 847 | 931.02 | 1023 | 839 | 0.819 | -0.287 | 0.00137264 | 0.00904524 | 0.0042 | 0.005 | 0.0047 | 0.0047 | TRUE | 0.1053 |
| Irf4 | 7 | 5 | 2 | 1 | 5 | 4 | 5 | 5 | 2 | 1 | 6 | 4 | 3.92 | 4 | 4 | 0.804 | -0.315 | 0.75358397 | NA | 0 | 0.6076 | 0.4641 | 0.4641 | TRUE | 0.4819 |
| Irf5 | 253 | 181 | 179 | 280 | 217 | 273 | 196 | 193 | 193 | 279 | 245 | 269 | 229.19 | 194 | 264 | 1.363 | 0.447 | 0.00086969 | 0.00654239 | 0 | 0.0128 | 0.0085 | 0.0085 | TRUE | 0.0449 |
| Irf6 | 827 | 658 | 855 | 1082 | 914 | 968 | 642 | 700 | 923 | 1077 | 1031 | 956 | 888.07 | 755 | 1021 | 1.355 | 0.439 | 0.00026936 | 0.00267544 | 0.0193 | 0.0051 | 0.0093 | 0.0093 | TRUE | 0.4438 |
| Irf7 | 10 | 10 | 5 | 4 | 8 | 11 | 8 | 11 | 5 | 4 | 9 | 11 | 7.94 | 8 | 8 | 1.001 | 0.002 | 0.99784301 | NA | 0 | 0.3008 | 0.2286 | 0.2286 | TRUE | 0.5292 |
| Irf8 | 474 | 356 | 419 | 512 | 511 | 544 | 368 | 379 | 452 | 510 | 576 | 537 | 470.34 | 400 | 541 | 1.356 | 0.439 | 9.29E-05 | 0.00114202 | 0.0057 | 0.0075 | 0.0069 | 0.0069 | TRUE | 0.1484 |

| | | | | | | | | | | | | | | | | | | | | | | | | | |
|---|---|---|---|---|---|---|---|---|---|---|---|---|---|---|---|---|---|---|---|---|---|---|---|---|---|
| Irf9 | 451 | 364 | 422 | 385 | 323 | 411 | 350 | 387 | 455 | 383 | 364 | 406 | 391.02 | 397 | 384 | 0.97 | -0.044 | 0.72351628 | 0.83442425 | 0.008 | 0.0085 | 0.0083 | 0.0083 | TRUE | 0.1841 |
| Irg1 | 0 | 1 | 0 | 0 | 0 | 0 | 0 | 1 | 0 | 0 | 0 | 0 | 0.18 | 0 | 0 | 0.552 | -0.857 | 0.8336548 | NA | 0 | 13.3714 | 10 | 10 | TRUE | 0.5216 |
| Irgc1 | 1 | 2 | 1 | 1 | 2 | 2 | 1 | 2 | 1 | 1 | 2 | 2 | 1.53 | 1 | 2 | 1.331 | 0.412 | 0.78295172 | NA | 0 | 1.5466 | 0.9442 | 0.9442 | TRUE | 0.1606 |
| Irgm1 | 144 | 75 | 39 | 76 | 76 | 105 | 112 | 80 | 42 | 76 | 86 | 104 | 83.12 | 78 | 89 | 1.125 | 0.169 | 0.57222895 | 0.72501271 | 0.1099 | 0.0309 | 0.0528 | 0.0528 | TRUE | 0.2542 |
| Irgm2 | 169 | 116 | 92 | 211 | 169 | 182 | 131 | 123 | 99 | 210 | 191 | 180 | 155.71 | 118 | 194 | 1.633 | 0.707 | 3.55E-05 | 0.00053401 | 0.0053 | 0.0176 | 0.0143 | 0.0143 | TRUE | 0.1853 |
| Irgq | 546 | 376 | 431 | 621 | 560 | 573 | 424 | 400 | 465 | 618 | 632 | 566 | 517.43 | 430 | 605 | 1.408 | 0.494 | 1.35E-06 | 3.73E-05 | 0.0023 | 0.007 | 0.0056 | 0.0056 | TRUE | 0.0567 |
| Irs1 | 104 | 61 | 83 | 83 | 78 | 70 | 81 | 65 | 90 | 83 | 88 | 69 | 79.15 | 79 | 80 | 1.017 | 0.024 | 0.91703939 | 0.95426832 | 0.0061 | 0.0324 | 0.0254 | 0.0254 | TRUE | 0.1968 |
| Irs2 | 642 | 502 | 387 | 331 | 350 | 346 | 499 | 534 | 418 | 329 | 395 | 342 | 419.34 | 484 | 355 | 0.734 | -0.447 | 0.00032045 | 0.00305953 | 0.01 | 0.0081 | 0.0087 | 0.0087 | TRUE | 0.1593 |
| Irs3 | 10 | 6 | 10 | 8 | 8 | 8 | 8 | 6 | 11 | 8 | 9 | 8 | 8.3 | 8 | 8 | 0.999 | -0.001 | 0.99909841 | NA | 0 | 0.2878 | 0.1875 | 0.1875 | TRUE | 0.1843 |
| Irs4 | 129 | 126 | 104 | 118 | 114 | 117 | 100 | 134 | 112 | 117 | 129 | 115 | 118 | 115 | 120 | 1.047 | 0.067 | 0.72559549 | 0.83572689 | 0 | 0.0225 | 0.0174 | 0.0174 | TRUE | 0.1962 |
| Irx1 | 41 | 25 | 31 | 25 | 23 | 28 | 32 | 27 | 33 | 25 | 26 | 28 | 28.39 | 31 | 26 | 0.853 | -0.229 | 0.52029569 | 0.68175525 | 0 | 0.0859 | 0.0561 | 0.0561 | TRUE | 0.0826 |
| Irx2 | 321 | 279 | 258 | 281 | 244 | 249 | 249 | 297 | 278 | 280 | 275 | 246 | 270.87 | 275 | 267 | 0.973 | -0.039 | 0.76764597 | 0.86257577 | 0.0031 | 0.0112 | 0.0087 | 0.0087 | TRUE | 0.0805 |
| Irx3 | 209 | 193 | 209 | 209 | 194 | 201 | 162 | 205 | 226 | 208 | 219 | 198 | 203.07 | 198 | 208 | 1.059 | 0.083 | 0.6012016 | 0.74656598 | 0.0109 | 0.0141 | 0.0131 | 0.0131 | TRUE | 0.2916 |
| Irx4 | 31 | 20 | 20 | 11 | 14 | 13 | 24 | 21 | 22 | 11 | 16 | 13 | 17.75 | 22 | 13 | 0.587 | -0.768 | 0.09313256 | 0.20989335 | 0 | 0.1359 | 0.0896 | 0.0896 | TRUE | 0.112 |
| Irx5 | 76 | 55 | 72 | 66 | 40 | 61 | 59 | 59 | 78 | 66 | 45 | 60 | 61.04 | 65 | 57 | 0.882 | -0.182 | 0.49152233 | 0.65729264 | 0.0133 | 0.0413 | 0.0338 | 0.0338 | TRUE | 0.263 |
| Irx6 | 14 | 6 | 5 | 10 | 3 | 7 | 11 | 6 | 5 | 10 | 3 | 7 | 7.15 | 7 | 7 | 0.887 | -0.172 | 0.81586573 | NA | 0 | 0.3339 | 0.2561 | 0.2561 | TRUE | 0.2274 |
| Isca1 | 2257 | 1712 | 1624 | 1698 | 1445 | 1685 | 1753 | 1821 | 1752 | 1690 | 1630 | 1663 | 1718.31 | 1775 | 1661 | 0.936 | -0.095 | 0.13025099 | 0.26709862 | 0 | 0.0038 | 0.0023 | 0.0023 | TRUE | 0.006 |
| Isca2 | 867 | 626 | 746 | 812 | 691 | 821 | 673 | 666 | 805 | 808 | 780 | 810 | 757.07 | 715 | 799 | 1.12 | 0.164 | 0.08558857 | 0.19785175 | 0.0042 | 0.0056 | 0.0052 | 0.0052 | TRUE | 0.1407 |
| Iscu | 1622 | 1159 | 984 | 1113 | 1001 | 1110 | 1260 | 1233 | 1062 | 1108 | 1129 | 1096 | 1147.86 | 1185 | 1111 | 0.937 | -0.095 | 0.25719177 | 0.42719836 | 0.0035 | 0.0045 | 0.0042 | 0.0042 | TRUE | 0.0972 |
| Isg15 | 1 | 2 | 0 | 0 | 0 | 1 | 1 | 2 | 0 | 0 | 0 | 1 | 0.65 | 1 | 0 | 0.384 | -1.379 | 0.58345004 | NA | 0 | 3.6569 | 3.0564 | 3.0564 | TRUE | 0.2561 |
| Isg20 | 9 | 6 | 6 | 1 | 4 | 6 | 7 | 6 | 6 | 1 | 5 | 6 | 5.21 | 6 | 4 | 0.573 | -0.802 | 0.35700249 | NA | 0 | 0.4571 | 0.3378 | 0.3378 | TRUE | 0.7004 |
| Isg20l2 | 4714 | 3340 | 3135 | 3005 | 2803 | 3210 | 3661 | 3553 | 3383 | 2991 | 3162 | 3169 | 3319.76 | 3532 | 3107 | 0.879 | -0.186 | 0.00193641 | 0.01179256 | 0.001 | 0.0031 | 0.0023 | 0.0023 | TRUE | 0.0166 |
| Isl1 | 38 | 26 | 11 | 20 | 23 | 11 | 30 | 28 | 12 | 20 | 26 | 11 | 20.96 | 23 | 19 | 0.811 | -0.303 | 0.53498883 | 0.69400937 | 0.143 | 0.1155 | 0.123 | 0.123 | TRUE | 0.5024 |
| Isl2 | 32 | 36 | 29 | 36 | 29 | 47 | 25 | 38 | 31 | 36 | 33 | 46 | 34.9 | 31 | 38 | 1.23 | 0.298 | 0.3810034 | 0.55559628 | 0 | 0.0703 | 0.0545 | 0.0545 | TRUE | 0.2773 |
| Islr | 26 | 20 | 24 | 48 | 29 | 35 | 20 | 21 | 26 | 48 | 33 | 35 | 30.4 | 22 | 39 | 1.719 | 0.782 | 0.03038028 | 0.09343351 | 0 | 0.0804 | 0.0589 | 0.0589 | TRUE | 0.3461 |
| Islr2 | 2054 | 1553 | 1462 | 1903 | 1698 | 1887 | 1595 | 1652 | 1577 | 1894 | 1916 | 1863 | 1749.53 | 1608 | 1891 | 1.176 | 0.234 | 0.00018921 | 0.00205187 | 0 | 0.0038 | 0.0022 | 0.0022 | TRUE | 0.0066 |
| Ism1 | 9 | 5 | 1 | 3 | 6 | 7 | 7 | 5 | 1 | 3 | 7 | 7 | 5.01 | 4 | 6 | 1.215 | 0.281 | 0.75607072 | NA | 0.1456 | 0.4756 | 0.3897 | 0.3897 | TRUE | 0.7294 |
| Isoc1 | 4792 | 3679 | 4132 | 3838 | 3485 | 3930 | 3722 | 3913 | 4458 | 3820 | 3931 | 3880 | 3954.12 | 4031 | 3877 | 0.962 | -0.056 | 0.44179471 | 0.61286955 | 0.004 | 0.003 | 0.0035 | 0.0035 | TRUE | 0.1043 |
| Isoc2a | 2950 | 2053 | 2254 | 2517 | 2238 | 2485 | 2291 | 2184 | 2432 | 2505 | 2525 | 2453 | 2398.36 | 2302 | 2494 | 1.083 | 0.116 | 0.0694647 | 0.17071391 | 9.00E-04 | 0.0034 | 0.0025 | 0.0025 | TRUE | 0.0286 |
| Isoc2b | 40 | 37 | 28 | 49 | 39 | 35 | 31 | 39 | 30 | 49 | 44 | 35 | 37.99 | 33 | 43 | 1.268 | 0.342 | 0.28519681 | 0.45727025 | 0 | 0.0648 | 0.0473 | 0.0473 | TRUE | 0.2103 |
| Ispd | 71 | 50 | 48 | 35 | 37 | 46 | 55 | 53 | 52 | 35 | 42 | 45 | 47.02 | 53 | 41 | 0.761 | -0.395 | 0.1611013 | 0.31078701 | 0 | 0.0528 | 0.0354 | 0.0354 | TRUE | 0.1228 |
| Ist1 | 5302 | 3880 | 3864 | 3794 | 3526 | 4061 | 4118 | 4127 | 4169 | 3777 | 3978 | 4009 | 4029.54 | 4138 | 3921 | 0.948 | -0.078 | 0.14913372 | 0.29437485 | 3.00E-04 | 0.003 | 0.0018 | 0.0018 | TRUE | 0.0127 |
| Isx | 0 | 0 | 0 | 0 | 1 | 0 | 0 | 0 | 0 | 0 | 1 | 0 | 0.19 | 0 | 0 | 2.094 | 1.067 | 0.79379329 | NA | 0 | 12.608 | 10 | 10 | TRUE | 0.5075 |
| Isy1 | 2620 | 1904 | 1686 | 1780 | 1619 | 1847 | 2035 | 2025 | 1819 | 1772 | 1826 | 1823 | 1883.46 | 1960 | 1807 | 0.922 | -0.118 | 0.08739 | 0.20058862 | 0.0016 | 0.0037 | 0.0029 | 0.0029 | TRUE | 0.0468 |
| Isyna1 | 8673 | 6608 | 6272 | 7382 | 6555 | 8308 | 6736 | 7029 | 6767 | 7348 | 7395 | 8201 | 7246.04 | 6844 | 7648 | 1.118 | 0.16 | 0.00762347 | 0.03376582 | 0.002 | 0.0028 | 0.0025 | 0.0025 | TRUE | 0.049 |
| Itch | 7557 | 5148 | 5849 | 6753 | 5578 | 6438 | 5869 | 5476 | 6311 | 6722 | 6293 | 6355 | 6170.98 | 5885 | 6457 | 1.097 | 0.134 | 0.03922092 | 0.11288455 | 0.0029 | 0.0028 | 0.0029 | 0.0029 | TRUE | 0.0483 |
| Itfg1 | 3204 | 2272 | 2598 | 2689 | 2386 | 2551 | 2488 | 2417 | 2803 | 2677 | 2692 | 2518 | 2599.14 | 2569 | 2629 | 1.023 | 0.033 | 0.64373557 | 0.77809422 | 0.0034 | 0.0033 | 0.0034 | 0.0034 | TRUE | 0.0758 |
| Itfg2 | 2157 | 1623 | 1593 | 1514 | 1369 | 1656 | 1675 | 1726 | 1719 | 1507 | 1544 | 1635 | 1634.43 | 1707 | 1562 | 0.916 | -0.127 | 0.05532683 | 0.14489466 | 4.00E-04 | 0.0039 | 0.0026 | 0.0026 | TRUE | 0.0203 |
| Itfg3 | 2941 | 2016 | 2175 | 2275 | 2103 | 2359 | 2284 | 2144 | 2347 | 2265 | 2372 | 2329 | 2290.16 | 2258 | 2322 | 1.028 | 0.04 | 0.53167248 | 0.69157269 | 8.00E-04 | 0.0035 | 0.0024 | 0.0024 | TRUE | 0.0231 |
| Itga1 | 143 | 119 | 114 | 40 | 44 | 52 | 111 | 127 | 123 | 40 | 50 | 51 | 83.57 | 120 | 47 | 0.391 | -1.356 | 2.15E-09 | 1.32E-07 | 0 | 0.0308 | 0.0219 | 0.0219 | TRUE | 0.1445 |
| Itga10 | 78 | 93 | 55 | 57 | 57 | 59 | 61 | 99 | 59 | 57 | 64 | 58 | 66.35 | 73 | 60 | 0.822 | -0.283 | 0.29319015 | 0.46521877 | 0.0339 | 0.0381 | 0.0369 | 0.0369 | TRUE | 0.8539 |
| Itga11 | 49 | 29 | 31 | 22 | 26 | 32 | 38 | 31 | 33 | 22 | 29 | 32 | 30.86 | 34 | 28 | 0.805 | -0.313 | 0.37187252 | 0.54649756 | 0 | 0.0792 | 0.0554 | 0.0554 | TRUE | 0.2159 |
| Itga2 | 6 | 4 | 2 | 5 | 3 | 4 | 5 | 4 | 2 | 5 | 3 | 4 | 3.9 | 4 | 4 | 1.102 | 0.14 | 0.88233733 | NA | 0 | 0.6106 | 0.3926 | 0.3926 | TRUE | 0.1855 |
| Itga2b | 488 | 323 | 382 | 498 | 395 | 416 | 379 | 344 | 412 | 496 | 446 | 411 | 414.45 | 378 | 451 | 1.192 | 0.253 | 0.03123763 | 0.09540158 | 0.0061 | 0.0081 | 0.0075 | 0.0075 | TRUE | 0.0903 |
| Itga3 | 8749 | 6313 | 6255 | 7286 | 6275 | 6996 | 6795 | 6715 | 6749 | 7252 | 7079 | 6906 | 6916.1 | 6753 | 7079 | 1.048 | 0.068 | 0.16465379 | 0.31546872 | 2.00E-04 | 0.0028 | 0.0016 | 0.0016 | TRUE | 0.0056 |
| Itga4 | 36 | 28 | 18 | 30 | 22 | 16 | 28 | 30 | 19 | 30 | 25 | 16 | 24.61 | 26 | 24 | 0.909 | -0.137 | 0.73796189 | 0.84383266 | 0.0287 | 0.0987 | 0.0803 | 0.0803 | TRUE | 0.4755 |
| Itga5 | 3786 | 2554 | 2396 | 3519 | 2760 | 3260 | 2940 | 2717 | 2585 | 3503 | 3114 | 3218 | 3012.81 | 2747 | 3278 | 1.193 | 0.254 | 0.0004059 | 0.00368822 | 0.0035 | 0.0032 | 0.0034 | 0.0034 | TRUE | 0.0467 |
| Itga6 | 8961 | 6399 | 6820 | 7819 | 6759 | 7684 | 6960 | 6807 | 7359 | 7783 | 7625 | 7585 | 7353.01 | 7042 | 7664 | 1.089 | 0.122 | 0.02016832 | 0.06961261 | 7.00E-04 | 0.0027 | 0.0019 | 0.0019 | TRUE | 0.0187 |
| Itga7 | 288 | 224 | 170 | 164 | 136 | 170 | 224 | 238 | 183 | 163 | 153 | 168 | 188.31 | 215 | 161 | 0.75 | -0.414 | 0.00798319 | 0.03498395 | 0.0052 | 0.015 | 0.0121 | 0.0121 | TRUE | 0.1728 |
| Itga8 | 126 | 76 | 81 | 68 | 58 | 73 | 98 | 81 | 87 | 68 | 65 | 72 | 78.55 | 89 | 68 | 0.769 | -0.379 | 0.08588627 | 0.19812337 | 0 | 0.0326 | 0.0221 | 0.0221 | TRUE | 0.0872 |
| Itga9 | 1270 | 926 | 959 | 1201 | 1078 | 1308 | 986 | 985 | 1035 | 1195 | 1216 | 1291 | 1118.14 | 1002 | 1234 | 1.232 | 0.302 | 4.67E-05 | 0.00065966 | 0 | 0.0045 | 0.0031 | 0.0031 | TRUE | 0.0198 |
| Itgad | 1 | 0 | 0 | 0 | 0 | 0 | 1 | 0 | 0 | 0 | 0 | 0 | 0.13 | 0 | 0 | 0.552 | -0.857 | 0.8336548 | NA | 0 | 18.3121 | 10 | 10 | TRUE | 0.2566 |
| Itgae | 277 | 202 | 209 | 254 | 194 | 212 | 215 | 215 | 226 | 253 | 219 | 209 | 222.74 | 219 | 227 | 1.04 | 0.056 | 0.68630149 | 0.80774354 | 0 | 0.0131 | 0.0095 | 0.0095 | TRUE | 0.1114 |
| Itgal | 769 | 582 | 513 | 441 | 445 | 446 | 597 | 619 | 554 | 439 | 502 | 440 | 525.18 | 590 | 460 | 0.779 | -0.36 | 0.00040493 | 0.00368176 | 0.0023 | 0.0069 | 0.0055 | 0.0055 | TRUE | 0.0704 |
| Itgam | 229 | 168 | 90 | 164 | 120 | 111 | 178 | 179 | 97 | 163 | 135 | 110 | 143.64 | 151 | 136 | 0.897 | -0.157 | 0.5032296 | 0.6672846 | 0.0676 | 0.0189 | 0.0327 | 0.0327 | TRUE | 0.7101 |
| Itgav | 3568 | 2565 | 2712 | 3403 | 2876 | 3316 | 2771 | 2728 | 2926 | 3387 | 3244 | 3273 | 3055.14 | 2808 | 3301 | 1.176 | 0.234 | 6.25E-05 | 0.00083977 | 5.00E-04 | 0.0032 | 0.0021 | 0.0021 | TRUE | 0.016 |
| Itgax | 0 | 0 | 0 | 1 | 0 | 0 | 0 | 0 | 0 | 1 | 0 | 0 | 0.17 | 0 | 0 | 2.094 | 1.067 | 0.79379329 | NA | 0 | 14.2885 | 10 | 10 | TRUE | 0.3906 |
| Itgb1 | 22102 | 15606 | 16944 | 19430 | 16445 | 18694 | 17166 | 16600 | 18282 | 19341 | 18552 | 18454 | 18065.69 | 17349 | 18782 | 1.083 | 0.115 | 0.03532781 | 0.10462432 | 0.0015 | 0.0026 | 0.0021 | 0.0021 | TRUE | 0.027 |
| Itgb1bp1 | 937 | 620 | 630 | 654 | 584 | 670 | 728 | 659 | 680 | 651 | 659 | 661 | 673.03 | 689 | 657 | 0.952 | -0.071 | 0.41669393 | 0.58945035 | 0 | 0.0059 | 0.004 | 0.004 | TRUE | 0.0303 |
| Itgb1bp2 | 26 | 19 | 21 | 22 | 17 | 17 | 20 | 20 | 23 | 22 | 19 | 17 | 20.15 | 21 | 19 | 0.919 | -0.122 | 0.77212845 | 0.86523183 | 0 | 0.12 | 0.0779 | 0.0779 | TRUE | 0.0776 |
| Itgb2 | 16 | 6 | 6 | 5 | 6 | 5 | 12 | 6 | 6 | 5 | 7 | 5 | 6.99 | 8 | 6 | 0.647 | -0.629 | 0.39203144 | NA | 0 | 0.3413 | 0.2389 | 0.2389 | TRUE | 0.668 |
| Itgb2l | 0 | 0 | 0 | 0 | 0 | 0 | 0 | 0 | 0 | 0 | 0 | 0 | 0 | 0 | 0 | NA | NA | NA | NA | NA | NA | NA | NA | NA | NA |
| Itgb3 | 1323 | 974 | 956 | 1165 | 1006 | 1015 | 1028 | 1036 | 1032 | 1160 | 1135 | 1002 | 1065.26 | 1032 | 1099 | 1.065 | 0.091 | 0.2629378 | 0.43331833 | 0.0022 | 0.0047 | 0.0038 | 0.0038 | TRUE | 0.0721 |
| Itgb3bp | 470 | 336 | 311 | 337 | 299 | 364 | 365 | 357 | 336 | 335 | 337 | 359 | 348.35 | 353 | 344 | 0.975 | -0.037 | 0.73986605 | 0.84538261 | 0 | 0.0092 | 0.0061 | 0.0061 | TRUE | 0.02 |
| Itgb4 | 602 | 459 | 508 | 624 | 523 | 625 | 468 | 488 | 548 | 621 | 590 | 617 | 555.33 | 501 | 609 | 1.218 | 0.285 | 0.00377217 | 0.01971318 | 0.0018 | 0.0067 | 0.0052 | 0.0052 | TRUE | 0.0765 |
| Itgb5 | 1949 | 1460 | 1677 | 1533 | 1454 | 1707 | 1514 | 1553 | 1809 | 1526 | 1640 | 1685 | 1621.24 | 1625 | 1617 | 0.996 | -0.006 | 0.94103848 | 0.96681409 | 0.0054 | 0.0039 | 0.0044 | 0.0044 | TRUE | 0.1172 |
| Itgb6 | 4 | 2 | 0 | 1 | 0 | 1 | 3 | 2 | 0 | 1 | 0 | 1 | 1.2 | 2 | 1 | 0.379 | -1.401 | 0.46100567 | NA | 0 | 1.9729 | 1.5651 | 1.5651 | TRUE | 0.3143 |
| Itgb7 | 1017 | 823 | 848 | 1288 | 1251 | 1302 | 790 | 875 | 915 | 1282 | 1411 | 1285 | 1093.14 | 860 | 1326 | 1.544 | 0.627 | 1.09E-13 | 1.61E-11 | 0.0034 | 0.0046 | 0.0042 | 0.0042 | TRUE | 0.064 |
| Itgb8 | 2 | 1 | 0 | 0 | 0 | 0 | 2 | 1 | 0 | 0 | 0 | 0 | 0.44 | 1 | 0 | 0.215 | -2.217 | 0.47268385 | NA | 0 | 5.4363 | 5.3181 | 5.3181 | TRUE | 0.2411 |

| | | | | | | | | | | | | | | | | | | | | | | | | | |
|---|---|---|---|---|---|---|---|---|---|---|---|---|---|---|---|---|---|---|---|---|---|---|---|---|---|
| Itgbl1 | 4 | 2 | 5 | 1 | 3 | 1 | 3 | 2 | 5 | 1 | 3 | 1 | 2.67 | 3 | 2 | 0.498 | -1.006 | 0.40496846 | NA | 0 | 0.8914 | 0.6207 | 0.6207 | TRUE | 0.4347 |
| Itih1 | 0 | 0 | 1 | 0 | 0 | 0 | 0 | 0 | 1 | 0 | 0 | 0 | 0.18 | 0 | 0 | 0.552 | -0.857 | 0.8336548 | NA | 0 | 13.1817 | 10 | 10 | TRUE | 0.5365 |
| Itih2 | 4 | 1 | 4 | 1 | 0 | 2 | 3 | 1 | 4 | 1 | 0 | 2 | 1.91 | 3 | 1 | 0.357 | -1.484 | 0.31853402 | NA | 0 | 1.2437 | 0.9144 | 0.9144 | TRUE | 0.2795 |
| Itih3 | 2 | 1 | 3 | 2 | 1 | 2 | 2 | 1 | 3 | 2 | 1 | 2 | 1.82 | 2 | 2 | 0.884 | -0.179 | 0.89668509 | NA | 0 | 1.3014 | 0.8096 | 0.8096 | TRUE | 0.2615 |
| Itih4 | 7 | 1 | 4 | 10 | 5 | 13 | 5 | 1 | 4 | 10 | 6 | 13 | 6.54 | 3 | 10 | 2.578 | 1.366 | 0.0933969 | NA | 0 | 0.3648 | 0.2931 | 0.2931 | TRUE | 0.3263 |
| Itih5 | 356 | 226 | 206 | 222 | 197 | 193 | 276 | 240 | 222 | 221 | 222 | 191 | 228.82 | 246 | 211 | 0.854 | -0.228 | 0.11631056 | 0.24684301 | 0.0062 | 0.0128 | 0.0108 | 0.0108 | TRUE | 0.1359 |
| Itk | 2 | 1 | 0 | 1 | 0 | 0 | 2 | 1 | 0 | 1 | 0 | 0 | 0.6 | 1 | 0 | 0.416 | -1.265 | 0.62159733 | NA | 0 | 3.939 | 3.1896 | 3.1896 | TRUE | 0.3906 |
| Itln1 | 0 | 1 | 0 | 0 | 0 | 0 | 0 | 1 | 0 | 0 | 0 | 0 | 0.18 | 0 | 0 | 0.552 | -0.857 | 0.8336548 | NA | 0 | 13.3714 | 10 | 10 | TRUE | 0.5216 |
| Itm2a | 894 | 612 | 651 | 790 | 692 | 701 | 694 | 651 | 702 | 786 | 781 | 692 | 717.79 | 682 | 753 | 1.102 | 0.141 | 0.11984705 | 0.25153801 | 0.002 | 0.0057 | 0.0045 | 0.0045 | TRUE | 0.0604 |
| Itm2b | 4949 | 3632 | 3916 | 4317 | 3911 | 4329 | 3844 | 3863 | 4225 | 4297 | 4412 | 4273 | 4152.48 | 3977 | 4327 | 1.088 | 0.122 | 0.03997973 | 0.11440046 | 0.0012 | 0.003 | 0.0023 | 0.0023 | TRUE | 0.0358 |
| Itm2c | 2319 | 1683 | 1853 | 2094 | 1912 | 2236 | 1801 | 1790 | 1999 | 2084 | 2157 | 2207 | 2006.53 | 1863 | 2149 | 1.154 | 0.207 | 0.00273696 | 0.01535594 | 0.0017 | 0.0036 | 0.0029 | 0.0029 | TRUE | 0.0483 |
| Itpa | 4149 | 2678 | 2478 | 2431 | 2039 | 2457 | 3222 | 2849 | 2674 | 2420 | 2300 | 2425 | 2648.35 | 2915 | 2382 | 0.817 | -0.292 | 0.00013557 | 0.00156774 | 0.0045 | 0.0033 | 0.0038 | 0.0038 | TRUE | 0.1051 |
| Itpk1 | 7535 | 5365 | 5203 | 6158 | 5701 | 6611 | 5852 | 5707 | 5614 | 6130 | 6431 | 6526 | 6043.31 | 5724 | 6362 | 1.111 | 0.152 | 0.00390412 | 0.02023994 | 5.00E-04 | 0.0028 | 0.0018 | 0.0018 | TRUE | 0.0126 |
| Itpka | 1123 | 917 | 915 | 1057 | 954 | 1063 | 872 | 975 | 987 | 1052 | 1076 | 1049 | 1002.09 | 945 | 1059 | 1.123 | 0.167 | 0.0373576 | 0.10903659 | 0.0015 | 0.0048 | 0.0036 | 0.0036 | TRUE | 0.0572 |
| Itpkb | 106 | 103 | 84 | 146 | 129 | 123 | 82 | 110 | 91 | 145 | 146 | 121 | 115.8 | 94 | 137 | 1.465 | 0.551 | 0.00472143 | 0.0234725 | 0.0069 | 0.0229 | 0.0185 | 0.0185 | TRUE | 0.1937 |
| Itpkc | 812 | 642 | 618 | 616 | 529 | 551 | 631 | 683 | 667 | 613 | 597 | 544 | 622.37 | 660 | 585 | 0.886 | -0.174 | 0.06079793 | 0.15473376 | 0.0012 | 0.0062 | 0.0046 | 0.0046 | TRUE | 0.0444 |
| Itpr1 | 10475 | 7846 | 8139 | 9380 | 9012 | 10494 | 8135 | 8346 | 8782 | 9337 | 10166 | 10359 | 9187.6 | 8421 | 9954 | 1.182 | 0.241 | 5.46E-05 | 0.00075395 | 0.0021 | 0.0027 | 0.0025 | 0.0025 | TRUE | 0.036 |
| Itpr2 | 175 | 117 | 122 | 195 | 164 | 185 | 136 | 124 | 132 | 194 | 185 | 183 | 158.96 | 131 | 187 | 1.431 | 0.517 | 0.00095025 | 0.00699502 | 0 | 0.0173 | 0.0112 | 0.0112 | TRUE | 0.0174 |
| Itpr3 | 7646 | 5903 | 6308 | 7171 | 6713 | 7763 | 5938 | 6279 | 6806 | 7138 | 7573 | 7663 | 6899.63 | 6341 | 7458 | 1.177 | 0.235 | 0.0002664 | 0.00265709 | 0.0029 | 0.0028 | 0.0028 | 0.0028 | TRUE | 0.0502 |
| Itprip | 209 | 166 | 145 | 163 | 116 | 151 | 162 | 177 | 156 | 162 | 131 | 149 | 156.25 | 165 | 147 | 0.895 | -0.161 | 0.32573976 | 0.49986916 | 0 | 0.0176 | 0.0129 | 0.0129 | TRUE | 0.0956 |
| Itpripl1 | 798 | 604 | 621 | 646 | 567 | 654 | 620 | 642 | 670 | 643 | 640 | 646 | 643.43 | 644 | 643 | 0.999 | -0.001 | 0.9881683 | 0.99387021 | 0 | 0.0061 | 0.0039 | 0.0039 | TRUE | 0.0145 |
| Itpripl2 | 180 | 129 | 104 | 167 | 140 | 138 | 140 | 137 | 112 | 166 | 158 | 136 | 141.6 | 130 | 153 | 1.179 | 0.237 | 0.17672496 | 0.33053256 | 0.0049 | 0.0192 | 0.0152 | 0.0152 | TRUE | 0.1364 |
| Itsn1 | 4257 | 3161 | 3226 | 3183 | 2967 | 3238 | 3306 | 3362 | 3481 | 3168 | 3347 | 3196 | 3310.2 | 3383 | 3237 | 0.957 | -0.063 | 0.26500432 | 0.43545523 | 5.00E-04 | 0.0031 | 0.002 | 0.002 | TRUE | 0.0106 |
| Itsn2 | 2634 | 1888 | 1635 | 1660 | 1539 | 1835 | 2046 | 2008 | 1764 | 1652 | 1736 | 1811 | 1836.34 | 1939 | 1733 | 0.893 | -0.163 | 0.03417083 | 0.10206975 | 0.0033 | 0.0037 | 0.0037 | 0.0037 | TRUE | 0.0746 |
| Ivd | 1570 | 1112 | 1107 | 1389 | 1243 | 1258 | 1219 | 1183 | 1194 | 1383 | 1402 | 1242 | 1270.55 | 1199 | 1342 | 1.119 | 0.162 | 0.03168906 | 0.09638896 | 0.0016 | 0.0043 | 0.0033 | 0.0033 | TRUE | 0.0519 |
| Ivl | 0 | 0 | 0 | 0 | 0 | 0 | 0 | 0 | 0 | 0 | 0 | 0 | 0 | 0 | 0 | NA | NA | NA | NA | NA | NA | NA | NA | NA | NA |
| Ivns1abp | 9561 | 6884 | 7025 | 8280 | 6736 | 7992 | 7426 | 7322 | 7580 | 8242 | 7599 | 7889 | 7676.35 | 7443 | 7910 | 1.063 | 0.088 | 0.09689223 | 0.2163876 | 8.00E-04 | 0.0027 | 0.0019 | 0.0019 | TRUE | 0.0164 |
| Iws1 | 5778 | 4253 | 3783 | 3854 | 3436 | 3782 | 4488 | 4524 | 4082 | 3836 | 3876 | 3733 | 4089.84 | 4365 | 3815 | 0.874 | -0.194 | 0.00137216 | 0.00904524 | 0.0014 | 0.003 | 0.0024 | 0.0024 | TRUE | 0.0386 |
| Iyd | 4 | 1 | 3 | 3 | 5 | 2 | 3 | 1 | 3 | 3 | 6 | 2 | 3 | 2 | 4 | 1.403 | 0.489 | 0.65953597 | NA | 0 | 0.7921 | 0.5461 | 0.5461 | TRUE | 0.3636 |
| Izumo1 | 25 | 45 | 36 | 50 | 43 | 60 | 19 | 48 | 39 | 50 | 49 | 59 | 43.94 | 35 | 53 | 1.507 | 0.592 | 0.08601595 | 0.19829451 | 0.0771 | 0.0564 | 0.0619 | 0.0619 | TRUE | 0.979 |
| Izumo2 | 13 | 12 | 10 | 13 | 17 | 14 | 10 | 13 | 11 | 13 | 19 | 14 | 13.26 | 11 | 15 | 1.366 | 0.45 | 0.3912413 | NA | 0 | 0.1811 | 0.121 | 0.121 | TRUE | 0.2047 |
| Izumo3 | 0 | 0 | 0 | 0 | 0 | 0 | 0 | 0 | 0 | 0 | 0 | 0 | 0 | 0 | 0 | NA | NA | NA | NA | NA | NA | NA | NA | NA | NA |
| Izumo4 | 78 | 90 | 57 | 92 | 71 | 78 | 61 | 96 | 62 | 92 | 80 | 77 | 77.75 | 73 | 83 | 1.148 | 0.199 | 0.42084832 | 0.5929796 | 0.026 | 0.0329 | 0.031 | 0.031 | TRUE | 0.6824 |
| Jag1 | 1114 | 873 | 1087 | 1084 | 1014 | 1258 | 865 | 929 | 1173 | 1079 | 1144 | 1242 | 1071.9 | 989 | 1155 | 1.17 | 0.226 | 0.03968285 | 0.11391518 | 0.0146 | 0.0046 | 0.0078 | 0.0078 | TRUE | 0.3141 |
| Jag2 | 2519 | 1984 | 1935 | 2326 | 1923 | 2195 | 1956 | 2110 | 2088 | 2315 | 2169 | 2167 | 2134.34 | 2051 | 2217 | 1.082 | 0.113 | 0.08274711 | 0.19369545 | 0.0012 | 0.0035 | 0.0026 | 0.0026 | TRUE | 0.0206 |
| Jagn1 | 2395 | 1570 | 1469 | 1563 | 1419 | 1605 | 1860 | 1670 | 1585 | 1556 | 1601 | 1584 | 1642.68 | 1705 | 1580 | 0.926 | -0.112 | 0.14202479 | 0.28446754 | 0.003 | 0.0039 | 0.0036 | 0.0036 | TRUE | 0.0786 |
| Jak1 | 5998 | 4329 | 4358 | 4843 | 4331 | 4874 | 4658 | 4605 | 4702 | 4821 | 4886 | 4811 | 4747.21 | 4655 | 4839 | 1.04 | 0.056 | 0.25922295 | 0.42912598 | 0 | 0.0029 | 0.0016 | 0.0016 | TRUE | 0.0011 |
| Jak2 | 985 | 671 | 756 | 854 | 733 | 792 | 765 | 714 | 816 | 850 | 827 | 782 | 792.21 | 765 | 820 | 1.072 | 0.1 | 0.25179279 | 0.42077333 | 0.0017 | 0.0054 | 0.0042 | 0.0042 | TRUE | 0.0386 |
| Jak3 | 2408 | 2311 | 2272 | 1946 | 1806 | 2104 | 1870 | 2458 | 2451 | 1937 | 2037 | 2077 | 2138.54 | 2260 | 2017 | 0.894 | -0.162 | 0.09726066 | 0.21697309 | 0.0111 | 0.0035 | 0.0064 | 0.0064 | TRUE | 0.2806 |
| Jakmip1 | 489 | 412 | 505 | 542 | 535 | 610 | 380 | 438 | 545 | 540 | 604 | 602 | 518.02 | 454 | 582 | 1.284 | 0.361 | 0.00497208 | 0.02440449 | 0.016 | 0.007 | 0.0099 | 0.0099 | TRUE | 0.351 |
| Jakmip2 | 24 | 14 | 16 | 13 | 3 | 10 | 19 | 15 | 17 | 13 | 3 | 10 | 12.83 | 17 | 9 | 0.52 | -0.943 | 0.10145359 | NA | 0 | 0.1871 | 0.1519 | 0.1519 | TRUE | 0.7884 |
| Jakmip3 | 6 | 4 | 3 | 5 | 2 | 2 | 5 | 4 | 3 | 5 | 2 | 2 | 3.56 | 4 | 3 | 0.757 | -0.402 | 0.6885566 | NA | 0 | 0.6682 | 0.4394 | 0.4394 | TRUE | 0.4037 |
| Jam2 | 5659 | 3808 | 3042 | 3323 | 3169 | 3310 | 4395 | 4051 | 3282 | 3308 | 3575 | 3267 | 3646.35 | 3909 | 3383 | 0.865 | -0.209 | 0.02569898 | 0.08285449 | 0.0114 | 0.0031 | 0.0061 | 0.0061 | TRUE | 0.2388 |
| Jam3 | 957 | 662 | 564 | 591 | 531 | 623 | 743 | 704 | 609 | 588 | 599 | 615 | 643.05 | 685 | 601 | 0.875 | -0.193 | 0.05124715 | 0.13732119 | 0.0042 | 0.0061 | 0.0055 | 0.0055 | TRUE | 0.1114 |
| Jarid2 | 26688 | 17518 | 14495 | 18108 | 15325 | 16893 | 20727 | 18634 | 15640 | 18025 | 17288 | 16676 | 17831.66 | 18334 | 17330 | 0.945 | -0.081 | 0.34678896 | 0.52120448 | 0.0097 | 0.0026 | 0.0053 | 0.0053 | TRUE | 0.202 |
| Jazf1 | 111 | 80 | 89 | 107 | 84 | 103 | 86 | 85 | 96 | 107 | 95 | 102 | 95.05 | 89 | 101 | 1.136 | 0.184 | 0.35640175 | 0.53085142 | 0 | 0.0274 | 0.0181 | 0.0181 | TRUE | 0.0414 |
| Jdp2 | 321 | 282 | 386 | 390 | 369 | 383 | 249 | 300 | 416 | 388 | 416 | 378 | 358.05 | 322 | 394 | 1.229 | 0.297 | 0.06533848 | 0.16323816 | 0.0325 | 0.009 | 0.0159 | 0.0159 | TRUE | 0.7451 |
| Jhdm1d | 2235 | 1587 | 1726 | 2407 | 1953 | 2200 | 1736 | 1688 | 1862 | 2396 | 2203 | 2172 | 2009.52 | 1762 | 2257 | 1.281 | 0.358 | 3.87E-07 | 1.31E-05 | 0.002 | 0.0036 | 0.0031 | 0.0031 | TRUE | 0.0354 |
| Jkamp | 2348 | 1745 | 1694 | 1855 | 1750 | 1831 | 1824 | 1856 | 1828 | 1846 | 1974 | 1807 | 1855.94 | 1836 | 1876 | 1.022 | 0.031 | 0.63489103 | 0.77160293 | 5.00E-04 | 0.0037 | 0.0025 | 0.0025 | TRUE | 0.0249 |
| Jmjd1c | 10399 | 6951 | 6028 | 7861 | 6771 | 8148 | 8076 | 7394 | 6504 | 7825 | 7638 | 8043 | 7580.14 | 7325 | 7835 | 1.069 | 0.097 | 0.19995844 | 0.36058617 | 0.0058 | 0.0027 | 0.004 | 0.004 | TRUE | 0.1171 |
| Jmjd4 | 2108 | 1400 | 1117 | 1203 | 1130 | 1149 | 1637 | 1489 | 1205 | 1197 | 1275 | 1134 | 1323.01 | 1444 | 1202 | 0.831 | -0.266 | 0.0107974 | 0.04371878 | 0.0121 | 0.0042 | 0.0071 | 0.0071 | TRUE | 0.2497 |
| Jmjd6 | 4878 | 3706 | 3849 | 3704 | 3403 | 4038 | 3789 | 3942 | 4153 | 3687 | 3839 | 3986 | 3899.27 | 3961 | 3837 | 0.969 | -0.045 | 0.4600744 | 0.62988358 | 0.0014 | 0.003 | 0.0024 | 0.0024 | TRUE | 0.0218 |
| Jmjd7 | 790 | 652 | 675 | 773 | 699 | 831 | 614 | 694 | 728 | 769 | 789 | 820 | 735.62 | 679 | 793 | 1.171 | 0.228 | 0.01372024 | 0.0523361 | 0.0031 | 0.0056 | 0.0048 | 0.0048 | TRUE | 0.0877 |
| Jmjd8 | 764 | 532 | 488 | 389 | 405 | 464 | 593 | 566 | 527 | 387 | 457 | 458 | 497.99 | 562 | 434 | 0.771 | -0.375 | 0.00046181 | 0.00407194 | 0.0041 | 0.0072 | 0.0062 | 0.0062 | TRUE | 0.1045 |
| Jmy | 2759 | 2116 | 2034 | 2014 | 1769 | 2101 | 2143 | 2251 | 2195 | 2005 | 1996 | 2074 | 2110.43 | 2196 | 2025 | 0.923 | -0.116 | 0.05471966 | 0.14362257 | 0 | 0.0035 | 0.0022 | 0.0022 | TRUE | 0.0058 |
| Josd1 | 2195 | 1695 | 1695 | 1864 | 1663 | 1819 | 1705 | 1803 | 1829 | 1855 | 1876 | 1796 | 1810.62 | 1779 | 1842 | 1.036 | 0.052 | 0.42587549 | 0.59841355 | 5.00E-04 | 0.0037 | 0.0025 | 0.0025 | TRUE | 0.0167 |
| Josd2 | 265 | 184 | 118 | 170 | 154 | 152 | 206 | 196 | 127 | 169 | 174 | 150 | 170.31 | 176 | 164 | 0.928 | -0.108 | 0.57002423 | 0.72356628 | 0.0294 | 0.0163 | 0.0201 | 0.0201 | TRUE | 0.6106 |
| Jph1 | 546 | 381 | 355 | 337 | 302 | 350 | 424 | 405 | 383 | 335 | 341 | 346 | 372.33 | 404 | 341 | 0.841 | -0.249 | 0.02168925 | 0.07325247 | 0 | 0.0088 | 0.0058 | 0.0058 | TRUE | 0.0237 |
| Jph2 | 98 | 52 | 59 | 74 | 53 | 66 | 76 | 55 | 64 | 74 | 60 | 65 | 65.61 | 65 | 66 | 1.013 | 0.019 | 0.93798495 | 0.96554692 | 0 | 0.0385 | 0.029 | 0.029 | TRUE | 0.2226 |
| Jph3 | 240 | 155 | 186 | 243 | 195 | 243 | 186 | 165 | 201 | 242 | 220 | 240 | 208.95 | 184 | 234 | 1.272 | 0.347 | 0.01572937 | 0.05785381 | 0 | 0.0138 | 0.01 | 0.01 | TRUE | 0.0831 |
| Jph4 | 131 | 93 | 81 | 132 | 108 | 134 | 102 | 99 | 87 | 131 | 122 | 132 | 112.26 | 96 | 128 | 1.336 | 0.418 | 0.02421243 | 0.07951937 | 0 | 0.0235 | 0.0157 | 0.0157 | TRUE | 0.0569 |
| Jpx | 10 | 5 | 3 | 3 | 6 | 8 | 8 | 5 | 3 | 3 | 7 | 8 | 5.66 | 5 | 6 | 1.062 | 0.087 | 0.91605312 | NA | 0 | 0.421 | 0.3103 | 0.3103 | TRUE | 0.347 |
| Jrk | 946 | 685 | 502 | 599 | 564 | 490 | 735 | 729 | 542 | 596 | 636 | 484 | 620.2 | 669 | 572 | 0.854 | -0.227 | 0.08884536 | 0.20287367 | 0.0227 | 0.0062 | 0.0112 | 0.0112 | TRUE | 0.3191 |
| Jrkl | 257 | 192 | 153 | 181 | 159 | 145 | 200 | 204 | 165 | 180 | 179 | 143 | 178.6 | 190 | 167 | 0.881 | -0.182 | 0.26620224 | 0.436634 | 0.0088 | 0.0157 | 0.0138 | 0.0138 | TRUE | 0.1765 |
| Jsrp1 | 66 | 69 | 86 | 107 | 72 | 93 | 51 | 73 | 93 | 107 | 81 | 92 | 82.83 | 72 | 93 | 1.299 | 0.377 | 0.13441147 | 0.27321014 | 0.0397 | 0.031 | 0.0335 | 0.0335 | TRUE | 0.3132 |
| Jtb | 1224 | 817 | 815 | 800 | 763 | 873 | 951 | 869 | 879 | 796 | 861 | 862 | 869.65 | 900 | 840 | 0.932 | -0.102 | 0.21820281 | 0.38122145 | 0.0012 | 0.0052 | 0.0038 | 0.0038 | TRUE | 0.0307 |
| Jun | 766 | 505 | 374 | 439 | 323 | 289 | 595 | 537 | 404 | 437 | 364 | 285 | 437.04 | 512 | 362 | 0.706 | -0.502 | 0.00220948 | 0.01303884 | 0.0394 | 0.0078 | 0.017 | 0.017 | TRUE | 0.302 |
| Junb | 1088 | 1040 | 1029 | 930 | 902 | 1185 | 845 | 1106 | 1110 | 926 | 1018 | 1170 | 1029.09 | 1020 | 1038 | 1.019 | 0.027 | 0.81961762 | 0.8964401 | 0.0179 | 0.0047 | 0.0089 | 0.0089 | TRUE | 0.2784 |
| Jund | 6758 | 4504 | 3824 | 4044 | 3185 | 3502 | 5249 | 4791 | 4126 | 4025 | 3593 | 3457 | 4206.83 | 4722 | 3692 | 0.782 | -0.356 | 6.58E-05 | 0.0008768 | 0.01 | 0.003 | 0.0055 | 0.0055 | TRUE | 0.1481 |

| | | | | | | | | | | | | | | | | | | | | | | | | | |
|---|---|---|---|---|---|---|---|---|---|---|---|---|---|---|---|---|---|---|---|---|---|---|---|---|---|
| Jup | 7855 | 6127 | 6611 | 6619 | 6565 | 8023 | 6101 | 6517 | 7133 | 6589 | 7406 | 7920 | 6944.26 | 6584 | 7305 | 1.11 | 0.15 | 0.05810986 | 0.1502762 | 0.0071 | 0.0028 | 0.0044 | 0.0044 | TRUE | 0.09 |
| Kalrn | 1083 | 812 | 847 | 831 | 742 | 856 | 841 | 864 | 914 | 827 | 837 | 845 | 854.66 | 873 | 836 | 0.959 | -0.06 | 0.44775904 | 0.61788255 | 0 | 0.0052 | 0.0034 | 0.0034 | TRUE | 0.0198 |
| Kank1 | 2512 | 1796 | 1802 | 1806 | 1625 | 1883 | 1951 | 1910 | 1944 | 1798 | 1833 | 1859 | 1882.56 | 1935 | 1830 | 0.945 | -0.081 | 0.18166078 | 0.33724867 | 0 | 0.0037 | 0.0021 | 0.0021 | TRUE | 0.0029 |
| Kank2 | 5909 | 4836 | 6206 | 6956 | 6463 | 7326 | 4589 | 5144 | 6696 | 6924 | 7291 | 7232 | 6312.72 | 5476 | 7149 | 1.306 | 0.385 | 0.00027736 | 0.00272835 | 0.02 | 0.0028 | 0.0079 | 0.0079 | TRUE | 0.4625 |
| Kank3 | 2816 | 2284 | 2236 | 2555 | 2175 | 2474 | 2187 | 2429 | 2413 | 2543 | 2454 | 2442 | 2411.37 | 2343 | 2480 | 1.059 | 0.083 | 0.20597135 | 0.36784941 | 0.0016 | 0.0034 | 0.0027 | 0.0027 | TRUE | 0.0424 |
| Kank4 | 1082 | 886 | 724 | 752 | 704 | 735 | 840 | 942 | 781 | 749 | 794 | 726 | 805.37 | 854 | 756 | 0.884 | -0.177 | 0.05725773 | 0.14866207 | 0.0043 | 0.0054 | 0.005 | 0.005 | TRUE | 0.093 |
| Kansl1 | 8041 | 5939 | 5991 | 6360 | 5608 | 6463 | 6245 | 6317 | 6464 | 6331 | 6326 | 6380 | 6343.95 | 6342 | 6346 | 1.001 | 0.001 | 0.98288064 | 0.99091459 | 0 | 0.0028 | 0.0015 | 0.0015 | TRUE | 0.0034 |
| Kansl1l | 301 | 245 | 292 | 270 | 230 | 292 | 234 | 261 | 315 | 269 | 259 | 288 | 270.99 | 270 | 272 | 1.013 | 0.019 | 0.89585571 | 0.94272381 | 0.0097 | 0.0112 | 0.0107 | 0.0107 | TRUE | 0.2381 |
| Kansl2 | 2717 | 1928 | 1876 | 1710 | 1658 | 1954 | 2110 | 2051 | 2024 | 1702 | 1870 | 1929 | 1947.76 | 2062 | 1834 | 0.889 | -0.17 | 0.01432155 | 0.05392465 | 0.0018 | 0.0036 | 0.0029 | 0.0029 | TRUE | 0.0482 |
| Kansl3 | 4336 | 3220 | 3403 | 3460 | 3009 | 3551 | 3368 | 3425 | 3672 | 3444 | 3394 | 3505 | 3468.07 | 3488 | 3448 | 0.989 | -0.016 | 0.78554627 | 0.87390788 | 9.00E-04 | 0.0031 | 0.0022 | 0.0022 | TRUE | 0.0256 |
| Kap | 0 | 0 | 0 | 0 | 0 | 0 | 0 | 0 | 0 | 0 | 0 | 0 | 0 | 0 | 0 | NA | NA | NA | NA | NA | NA | NA | NA | NA | NA |
| Kars | 36242 | 26374 | 26810 | 29087 | 26476 | 31071 | 28148 | 28054 | 28928 | 28953 | 29868 | 30672 | 29103.63 | 28377 | 29831 | 1.051 | 0.072 | 0.12452897 | 0.25911961 | 5.00E-04 | 0.0025 | 0.0016 | 0.0016 | TRUE | 0.0081 |
| Kat2a | 14568 | 10729 | 11018 | 11317 | 10070 | 11994 | 11314 | 11412 | 11888 | 11265 | 11360 | 11840 | 11513.31 | 11538 | 11488 | 0.996 | -0.006 | 0.90286192 | 0.94622418 | 5.00E-04 | 0.0026 | 0.0017 | 0.0017 | TRUE | 0.0088 |
| Kat2b | 4131 | 2926 | 3141 | 3209 | 2701 | 3126 | 3208 | 3112 | 3389 | 3194 | 3047 | 3086 | 3172.82 | 3236 | 3109 | 0.961 | -0.058 | 0.33349781 | 0.50773953 | 8.00E-04 | 0.0032 | 0.0022 | 0.0022 | TRUE | 0.0202 |
| Kat5 | 2394 | 1798 | 1847 | 1927 | 1786 | 2053 | 1859 | 1913 | 1993 | 1918 | 2015 | 2027 | 1954.05 | 1922 | 1987 | 1.034 | 0.049 | 0.44776628 | 0.61788255 | 6.00E-04 | 0.0036 | 0.0024 | 0.0024 | TRUE | 0.0127 |
| Kat6a | 8204 | 6231 | 6263 | 6070 | 5782 | 6602 | 6372 | 6628 | 6758 | 6042 | 6523 | 6517 | 6473.2 | 6586 | 6361 | 0.966 | -0.05 | 0.37510547 | 0.54949088 | 0.0012 | 0.0028 | 0.0021 | 0.0021 | TRUE | 0.0235 |
| Kat6b | 11974 | 8900 | 7910 | 8475 | 7743 | 8415 | 9300 | 9467 | 8535 | 8436 | 8735 | 8307 | 8796.53 | 9101 | 8493 | 0.933 | -0.1 | 0.0830942 | 0.19410862 | 0.0017 | 0.0027 | 0.0023 | 0.0023 | TRUE | 0.0359 |
| Kat7 | 19676 | 14390 | 14777 | 15222 | 13944 | 16957 | 15282 | 15306 | 15944 | 15152 | 15730 | 16739 | 15692.27 | 15511 | 15874 | 1.023 | 0.033 | 0.54157105 | 0.69973309 | 0.0014 | 0.0026 | 0.0021 | 0.0021 | TRUE | 0.0279 |
| Kat8 | 1608 | 1156 | 1232 | 1255 | 1241 | 1432 | 1249 | 1230 | 1329 | 1249 | 1400 | 1414 | 1311.77 | 1269 | 1354 | 1.067 | 0.094 | 0.22776874 | 0.39299951 | 0.0024 | 0.0042 | 0.0036 | 0.0036 | TRUE | 0.0559 |
| Katna1 | 3163 | 2310 | 2628 | 2808 | 2591 | 3107 | 2457 | 2457 | 2836 | 2795 | 2923 | 3067 | 2755.73 | 2583 | 2928 | 1.134 | 0.182 | 0.01440215 | 0.05407144 | 0.0041 | 0.0033 | 0.0036 | 0.0036 | TRUE | 0.0878 |
| Katnal1 | 501 | 356 | 453 | 595 | 510 | 603 | 389 | 379 | 489 | 592 | 575 | 595 | 503.24 | 419 | 587 | 1.406 | 0.492 | 1.46E-05 | 0.00025853 | 0.0076 | 0.0071 | 0.0072 | 0.0072 | TRUE | 0.2389 |
| Katnal2 | 178 | 117 | 154 | 194 | 182 | 186 | 138 | 124 | 166 | 193 | 205 | 184 | 168.48 | 143 | 194 | 1.358 | 0.442 | 0.00696131 | 0.03160672 | 0.0042 | 0.0165 | 0.0132 | 0.0132 | TRUE | 0.1965 |
| Katnb1 | 3220 | 2396 | 2317 | 2356 | 2175 | 2487 | 2501 | 2549 | 2500 | 2345 | 2454 | 2455 | 2467.22 | 2517 | 2418 | 0.961 | -0.058 | 0.31944055 | 0.49335473 | 0 | 0.0034 | 0.002 | 0.002 | TRUE | 0.0085 |
| Katnbl1 | 2700 | 1914 | 1912 | 1969 | 1635 | 1926 | 2097 | 2036 | 2063 | 1960 | 1844 | 1901 | 1983.59 | 2065 | 1902 | 0.921 | -0.119 | 0.05330461 | 0.14101245 | 0 | 0.0036 | 0.0022 | 0.0022 | TRUE | 0.0086 |
| Kazald1 | 19 | 19 | 14 | 12 | 17 | 15 | 15 | 20 | 15 | 12 | 19 | 15 | 16 | 17 | 15 | 0.917 | -0.125 | 0.79593418 | NA | 0 | 0.1506 | 0.1047 | 0.1047 | TRUE | 0.2044 |
| Kazn | 3160 | 2646 | 3147 | 3326 | 3404 | 3803 | 2454 | 2815 | 3396 | 3311 | 3840 | 3754 | 3261.54 | 2888 | 3635 | 1.259 | 0.333 | 0.00121093 | 0.00831274 | 0.0161 | 0.0032 | 0.0073 | 0.0073 | TRUE | 0.2867 |
| Kbtbd11 | 569 | 452 | 509 | 600 | 559 | 722 | 442 | 481 | 549 | 597 | 631 | 713 | 568.75 | 491 | 647 | 1.322 | 0.402 | 0.00031201 | 0.00299704 | 0.0085 | 0.0066 | 0.0072 | 0.0072 | TRUE | 0.1259 |
| Kbtbd12 | 4 | 3 | 2 | 1 | 1 | 0 | 3 | 3 | 2 | 1 | 1 | 0 | 1.76 | 3 | 1 | 0.246 | -2.023 | 0.20085498 | NA | 0 | 1.3466 | 0.8883 | 0.8883 | TRUE | 0.2726 |
| Kbtbd13 | 109 | 67 | 50 | 84 | 46 | 49 | 85 | 71 | 54 | 84 | 52 | 48 | 65.62 | 70 | 61 | 0.873 | -0.196 | 0.49244103 | 0.65761558 | 0.0561 | 0.0385 | 0.0435 | 0.0435 | TRUE | 0.5847 |
| Kbtbd2 | 1678 | 1265 | 1193 | 1453 | 1198 | 1285 | 1303 | 1346 | 1287 | 1446 | 1351 | 1268 | 1333.72 | 1312 | 1355 | 1.033 | 0.047 | 0.52940019 | 0.68976453 | 0.0017 | 0.0042 | 0.0033 | 0.0033 | TRUE | 0.042 |
| Kbtbd3 | 119 | 104 | 76 | 92 | 92 | 102 | 92 | 111 | 82 | 92 | 104 | 101 | 96.85 | 95 | 99 | 1.039 | 0.055 | 0.79059179 | 0.87698319 | 0 | 0.0269 | 0.0202 | 0.0202 | TRUE | 0.1869 |
| Kbtbd4 | 1000 | 626 | 599 | 638 | 592 | 637 | 777 | 666 | 646 | 635 | 668 | 629 | 670.09 | 696 | 644 | 0.922 | -0.117 | 0.23314014 | 0.39931361 | 0.0044 | 0.006 | 0.0054 | 0.0054 | TRUE | 0.126 |
| Kbtbd7 | 1186 | 914 | 733 | 764 | 660 | 738 | 921 | 972 | 791 | 760 | 745 | 729 | 819.63 | 895 | 745 | 0.832 | -0.266 | 0.00455951 | 0.02288582 | 0.0047 | 0.0053 | 0.0051 | 0.0051 | TRUE | 0.119 |
| Kbtbd8 | 3904 | 2758 | 2516 | 2660 | 2278 | 2570 | 3032 | 2934 | 2715 | 2648 | 2570 | 2537 | 2739.17 | 2894 | 2585 | 0.893 | -0.164 | 0.0106275 | 0.04315307 | 0.0015 | 0.0033 | 0.0026 | 0.0026 | TRUE | 0.0352 |
| Kcmf1 | 6838 | 4599 | 4671 | 5152 | 4332 | 4961 | 5311 | 4892 | 5040 | 5128 | 4887 | 4897 | 5025.86 | 5081 | 4971 | 0.978 | -0.032 | 0.57189082 | 0.72496988 | 0.001 | 0.0029 | 0.0021 | 0.0021 | TRUE | 0.0194 |
| Kcna1 | 34 | 16 | 8 | 18 | 16 | 9 | 26 | 17 | 9 | 18 | 18 | 9 | 16.15 | 17 | 15 | 0.846 | -0.241 | 0.65747599 | NA | 0.153 | 0.1492 | 0.1502 | 0.1502 | TRUE | 0.1964 |
| Kcna10 | 0 | 0 | 0 | 0 | 0 | 0 | 0 | 0 | 0 | 0 | 0 | 0 | 0 | 0 | 0 | NA | NA | NA | NA | NA | NA | NA | NA | NA | NA |
| Kcna2 | 14 | 5 | 9 | 17 | 14 | 11 | 11 | 5 | 10 | 17 | 16 | 11 | 11.58 | 9 | 15 | 1.661 | 0.732 | 0.20539283 | NA | 0 | 0.2071 | 0.1501 | 0.1501 | TRUE | 0.3253 |
| Kcna3 | 28 | 24 | 11 | 17 | 13 | 13 | 22 | 26 | 12 | 17 | 15 | 13 | 17.26 | 20 | 15 | 0.748 | -0.419 | 0.3858065 | 0.55960344 | 0 | 0.1397 | 0.1085 | 0.1085 | TRUE | 0.5927 |
| Kcna4 | 1 | 2 | 2 | 0 | 3 | 3 | 1 | 2 | 2 | 0 | 3 | 3 | 1.9 | 2 | 2 | 1.271 | 0.345 | 0.81151642 | NA | 0 | 1.249 | 0.9737 | 0.9737 | TRUE | 0.7485 |
| Kcna5 | 15 | 8 | 5 | 5 | 3 | 11 | 12 | 9 | 5 | 5 | 3 | 11 | 7.46 | 9 | 6 | 0.749 | -0.416 | 0.57419729 | NA | 0.0988 | 0.32 | 0.259 | 0.259 | TRUE | 0.4819 |
| Kcna6 | 56 | 32 | 34 | 34 | 27 | 28 | 43 | 34 | 37 | 34 | 30 | 28 | 34.36 | 38 | 31 | 0.801 | -0.32 | 0.33008836 | 0.50448598 | 0 | 0.0714 | 0.0484 | 0.0484 | TRUE | 0.1339 |
| Kcna7 | 49 | 40 | 25 | 39 | 43 | 29 | 38 | 43 | 27 | 39 | 49 | 29 | 37.26 | 36 | 39 | 1.071 | 0.099 | 0.77012962 | 0.86394025 | 0.0302 | 0.066 | 0.0565 | 0.0565 | TRUE | 0.1862 |
| Kcnab1 | 6 | 2 | 0 | 0 | 0 | 1 | 5 | 2 | 0 | 0 | 0 | 1 | 1.3 | 2 | 0 | 0.16 | -2.64 | 0.19533709 | NA | 0 | 1.8315 | 1.7847 | 1.7847 | TRUE | 0.1402 |
| Kcnab2 | 236 | 166 | 201 | 184 | 175 | 182 | 183 | 177 | 217 | 183 | 197 | 180 | 189.5 | 192 | 187 | 0.972 | -0.041 | 0.78802318 | 0.87535675 | 0 | 0.0149 | 0.0112 | 0.0112 | TRUE | 0.1296 |
| Kcnab3 | 465 | 412 | 353 | 463 | 426 | 446 | 361 | 438 | 381 | 461 | 481 | 440 | 427 | 393 | 461 | 1.173 | 0.23 | 0.03948949 | 0.11349754 | 0.0036 | 0.008 | 0.0066 | 0.0066 | TRUE | 0.1128 |
| Kcnb1 | 250 | 213 | 198 | 266 | 209 | 254 | 194 | 227 | 214 | 265 | 236 | 251 | 230.94 | 212 | 251 | 1.189 | 0.249 | 0.06842381 | 0.16853034 | 0 | 0.0127 | 0.0091 | 0.0091 | TRUE | 0.0619 |
| Kcnb2 | 2 | 2 | 2 | 1 | 1 | 2 | 2 | 2 | 2 | 1 | 1 | 2 | 1.66 | 2 | 1 | 0.71 | -0.494 | 0.73016878 | NA | 0 | 1.4336 | 0.8486 | 0.8486 | TRUE | 0.0997 |
| Kcnc1 | 21 | 10 | 16 | 22 | 21 | 20 | 16 | 11 | 17 | 22 | 24 | 20 | 18.26 | 15 | 22 | 1.469 | 0.555 | 0.21882229 | 0.38193028 | 0 | 0.1322 | 0.0906 | 0.0906 | TRUE | 0.2441 |
| Kcnc2 | 13 | 7 | 15 | 8 | 9 | 8 | 10 | 7 | 16 | 8 | 10 | 8 | 9.96 | 11 | 9 | 0.772 | -0.373 | 0.54603333 | NA | 0 | 0.2405 | 0.172 | 0.172 | TRUE | 0.4916 |
| Kcnc3 | 374 | 326 | 235 | 336 | 221 | 274 | 290 | 347 | 254 | 334 | 249 | 270 | 290.84 | 297 | 284 | 0.96 | -0.058 | 0.7025432 | 0.81985628 | 0.0203 | 0.0106 | 0.0134 | 0.0134 | TRUE | 0.2633 |
| Kcnc4 | 43 | 38 | 41 | 36 | 52 | 40 | 33 | 40 | 44 | 36 | 59 | 39 | 42.01 | 39 | 45 | 1.136 | 0.184 | 0.56230243 | 0.71682025 | 0.023 | 0.0588 | 0.0488 | 0.0488 | TRUE | 0.5526 |
| Kcnd1 | 395 | 346 | 404 | 370 | 364 | 450 | 307 | 368 | 436 | 368 | 411 | 444 | 388.98 | 370 | 408 | 1.105 | 0.144 | 0.29827121 | 0.47055082 | 0.0173 | 0.0085 | 0.0112 | 0.0112 | TRUE | 0.2323 |
| Kcnd2 | 20 | 22 | 16 | 9 | 16 | 7 | 16 | 23 | 17 | 9 | 18 | 7 | 15.02 | 19 | 11 | 0.6 | -0.737 | 0.16792221 | NA | 0.0555 | 0.1602 | 0.1337 | 0.1337 | TRUE | 0.9337 |
| Kcnd3 | 15 | 9 | 10 | 7 | 8 | 3 | 12 | 10 | 11 | 7 | 9 | 3 | 8.49 | 11 | 6 | 0.585 | -0.773 | 0.25406184 | NA | 0 | 0.2814 | 0.2031 | 0.2031 | TRUE | 0.5502 |
| Kcne1 | 7 | 9 | 9 | 7 | 4 | 6 | 5 | 10 | 10 | 7 | 5 | 6 | 7.02 | 8 | 6 | 0.716 | -0.483 | 0.50497145 | NA | 0 | 0.34 | 0.2298 | 0.2298 | TRUE | 0.352 |
| Kcne1l | 15 | 10 | 4 | 13 | 8 | 8 | 12 | 11 | 4 | 13 | 9 | 8 | 9.41 | 9 | 10 | 1.11 | 0.151 | 0.81552922 | NA | 0 | 0.2543 | 0.195 | 0.195 | TRUE | 0.5823 |
| Kcne2 | 2 | 1 | 0 | 2 | 1 | 1 | 2 | 1 | 0 | 2 | 1 | 1 | 1.12 | 1 | 1 | 1.516 | 0.6 | 0.73587335 | NA | 0 | 2.1176 | 1.3707 | 1.3707 | TRUE | 0.2754 |
| Kcne3 | 4 | 0 | 0 | 5 | 0 | 2 | 3 | 0 | 0 | 5 | 0 | 2 | 1.68 | 1 | 2 | 2.105 | 1.074 | 0.54907079 | NA | 3.1419 | 1.4163 | 1.6586 | 1.6586 | TRUE | 0.477 |
| Kcne4 | 0 | 0 | 0 | 0 | 0 | 0 | 0 | 0 | 0 | 0 | 0 | 0 | 0 | 0 | 0 | NA | NA | NA | NA | NA | NA | NA | NA | NA | NA |
| Kcnf1 | 103 | 71 | 66 | 68 | 35 | 51 | 80 | 76 | 71 | 68 | 39 | 50 | 64.04 | 76 | 52 | 0.696 | -0.522 | 0.04616136 | 0.12710159 | 0.0142 | 0.0394 | 0.0333 | 0.0333 | TRUE | 0.5035 |
| Kcng1 | 229 | 183 | 196 | 222 | 198 | 202 | 178 | 195 | 211 | 221 | 223 | 199 | 204.62 | 195 | 214 | 1.105 | 0.144 | 0.3201854 | 0.49405267 | 0 | 0.014 | 0.0102 | 0.0102 | TRUE | 0.0682 |
| Kcng2 | 34 | 34 | 20 | 44 | 33 | 46 | 26 | 36 | 22 | 44 | 37 | 45 | 35.1 | 28 | 42 | 1.508 | 0.592 | 0.07905057 | 0.18746802 | 0 | 0.07 | 0.0525 | 0.0525 | TRUE | 0.3786 |
| Kcng3 | 22 | 23 | 12 | 16 | 21 | 22 | 17 | 24 | 13 | 16 | 24 | 22 | 19.31 | 18 | 21 | 1.125 | 0.17 | 0.70753581 | 0.82345161 | 0 | 0.1252 | 0.0953 | 0.0953 | TRUE | 0.4231 |
| Kcng4 | 18 | 12 | 6 | 10 | 11 | 10 | 14 | 13 | 6 | 10 | 12 | 10 | 10.91 | 11 | 11 | 0.958 | -0.062 | 0.91624669 | NA | 0 | 0.2197 | 0.1574 | 0.1574 | TRUE | 0.4501 |
| Kcnh1 | 162 | 135 | 112 | 113 | 124 | 115 | 126 | 144 | 121 | 112 | 140 | 114 | 126.03 | 130 | 122 | 0.936 | -0.095 | 0.6049271 | 0.74963243 | 0.0032 | 0.0212 | 0.0162 | 0.0162 | TRUE | 0.1601 |
| Kcnh2 | 1159 | 939 | 1174 | 1347 | 1388 | 1570 | 900 | 999 | 1267 | 1341 | 1566 | 1550 | 1270.35 | 1055 | 1486 | 1.41 | 0.495 | 1.54E-05 | 0.00027247 | 0.0185 | 0.0043 | 0.0087 | 0.0087 | TRUE | 0.3661 |
| Kcnh3 | 291 | 316 | 346 | 401 | 455 | 526 | 226 | 336 | 373 | 399 | 513 | 519 | 394.52 | 312 | 477 | 1.537 | 0.62 | 0.00023142 | 0.00239098 | 0.0408 | 0.0084 | 0.0178 | 0.0178 | TRUE | 0.6945 |
| Kcnh4 | 118 | 104 | 108 | 138 | 117 | 157 | 92 | 111 | 117 | 137 | 132 | 155 | 123.86 | 107 | 141 | 1.339 | 0.421 | 0.02280782 | 0.07598884 | 0 | 0.0216 | 0.0165 | 0.0165 | TRUE | 0.1628 |

| | | | | | | | | | | | | | | | | | | | | | | | | | |
|---|---|---|---|---|---|---|---|---|---|---|---|---|---|---|---|---|---|---|---|---|---|---|---|---|---|
| Kcnh5 | 13 | 9 | 10 | 9 | 14 | 10 | 10 | 10 | 11 | 9 | 16 | 10 | 10.85 | 10 | 12 | 1.13 | 0.177 | 0.76067461 | NA | 0 | 0.2209 | 0.15 | 0.15 | TRUE | 0.3602 |
| Kcnh6 | 262 | 204 | 198 | 233 | 240 | 218 | 203 | 217 | 214 | 232 | 271 | 215 | 225.33 | 211 | 239 | 1.131 | 0.178 | 0.20939616 | 0.37114327 | 0.0032 | 0.0129 | 0.0101 | 0.0101 | TRUE | 0.1416 |
| Kcnh7 | 0 | 3 | 0 | 5 | 1 | 2 | 0 | 3 | 0 | 5 | 1 | 2 | 1.88 | 1 | 3 | 2.661 | 1.412 | 0.38347802 | NA | 1.0671 | 1.2642 | 1.2268 | 1.2268 | TRUE | 1.4447 |
| Kcnh8 | 3 | 3 | 4 | 7 | 3 | 10 | 2 | 3 | 4 | 7 | 3 | 10 | 5.01 | 3 | 7 | 2.101 | 1.071 | 0.22684196 | NA | 0 | 0.4755 | 0.3407 | 0.3407 | TRUE | 0.137 |
| Kcnip1 | 2 | 6 | 5 | 1 | 2 | 1 | 2 | 6 | 5 | 1 | 2 | 1 | 2.93 | 4 | 1 | 0.319 | -1.647 | 0.17817349 | NA | 0 | 0.8118 | 0.5963 | 0.5963 | TRUE | 0.7979 |
| Kcnip2 | 299 | 270 | 352 | 529 | 489 | 562 | 232 | 287 | 380 | 527 | 552 | 555 | 422.04 | 300 | 545 | 1.824 | 0.867 | 6.16E-09 | 3.35E-07 | 0.0275 | 0.008 | 0.0135 | 0.0135 | TRUE | 0.614 |
| Kcnip3 | 335 | 218 | 230 | 296 | 259 | 281 | 260 | 232 | 248 | 295 | 292 | 277 | 267.41 | 247 | 288 | 1.165 | 0.22 | 0.07828389 | 0.18617497 | 0 | 0.0113 | 0.0075 | 0.0075 | TRUE | 0.0304 |
| Kcnip4 | 0 | 0 | 0 | 1 | 0 | 0 | 0 | 0 | 0 | 1 | 0 | 0 | 0.17 | 0 | 0 | 2.094 | 1.067 | 0.79379329 | NA | 0 | 14.2885 | 10 | 10 | TRUE | 0.3906 |
| Kcnj1 | 0 | 0 | 0 | 0 | 0 | 1 | 0 | 0 | 0 | 0 | 0 | 1 | 0.16 | 0 | 0 | 2.094 | 1.067 | 0.79379329 | NA | 0 | 14.4078 | 10 | 10 | TRUE | 0.3835 |
| Kcnj10 | 233 | 176 | 147 | 221 | 186 | 211 | 181 | 187 | 159 | 220 | 210 | 208 | 194.15 | 176 | 213 | 1.21 | 0.275 | 0.05877241 | 0.1511267 | 0 | 0.0146 | 0.0101 | 0.0101 | TRUE | 0.0719 |
| Kcnj11 | 34 | 32 | 37 | 47 | 38 | 35 | 26 | 34 | 40 | 47 | 43 | 35 | 37.43 | 33 | 42 | 1.247 | 0.319 | 0.32930243 | 0.50366182 | 0 | 0.0657 | 0.0498 | 0.0498 | TRUE | 0.2873 |
| Kcnj12 | 679 | 480 | 498 | 513 | 527 | 576 | 527 | 511 | 537 | 511 | 595 | 569 | 541.5 | 525 | 558 | 1.062 | 0.086 | 0.37920406 | 0.55384873 | 0.0014 | 0.0068 | 0.0051 | 0.0051 | TRUE | 0.0648 |
| Kcnj13 | 0 | 0 | 0 | 0 | 0 | 0 | 0 | 0 | 0 | 0 | 0 | 0 | 0 | 0 | 0 | NA | NA | NA | NA | NA | NA | NA | NA | NA | NA |
| Kcnj14 | 16 | 16 | 12 | 9 | 17 | 8 | 12 | 17 | 13 | 9 | 19 | 8 | 13.07 | 14 | 12 | 0.845 | -0.243 | 0.66376915 | NA | 0.0482 | 0.1837 | 0.1473 | 0.1473 | TRUE | 0.9771 |
| Kcnj15 | 13 | 6 | 4 | 2 | 2 | 0 | 10 | 6 | 4 | 2 | 2 | 0 | 4.17 | 7 | 1 | 0.199 | -2.333 | 0.03583086 | NA | 0 | 0.5703 | 0.4495 | 0.4495 | TRUE | 0.41 |
| Kcnj16 | 3 | 5 | 2 | 2 | 6 | 1 | 2 | 5 | 2 | 2 | 7 | 1 | 3.26 | 3 | 3 | 0.986 | -0.02 | 0.98586486 | NA | 0.2541 | 0.7297 | 0.5872 | 0.5872 | TRUE | 1.1227 |
| Kcnj2 | 36 | 27 | 19 | 25 | 30 | 18 | 28 | 29 | 21 | 25 | 34 | 18 | 25.61 | 26 | 26 | 0.983 | -0.025 | 0.95032515 | 0.97271537 | 0.0227 | 0.095 | 0.076 | 0.076 | TRUE | 0.2417 |
| Kcnj3 | 172 | 120 | 98 | 107 | 106 | 117 | 134 | 128 | 106 | 107 | 120 | 115 | 118.09 | 123 | 114 | 0.927 | -0.109 | 0.55727824 | 0.7122577 | 0 | 0.0225 | 0.0165 | 0.0165 | TRUE | 0.1368 |
| Kcnj4 | 321 | 236 | 240 | 332 | 310 | 322 | 249 | 251 | 259 | 330 | 350 | 318 | 292.89 | 253 | 333 | 1.314 | 0.394 | 0.00096967 | 0.00708925 | 0 | 0.0105 | 0.0068 | 0.0068 | TRUE | 0.0221 |
| Kcnj5 | 1 | 3 | 2 | 1 | 2 | 1 | 1 | 3 | 2 | 1 | 2 | 1 | 1.73 | 2 | 1 | 0.7 | -0.515 | 0.72125608 | NA | 0 | 1.3745 | 0.8913 | 0.8913 | TRUE | 0.3002 |
| Kcnj6 | 0 | 0 | 0 | 1 | 1 | 2 | 0 | 0 | 0 | 1 | 1 | 2 | 0.68 | 0 | 1 | 7.73 | 2.951 | 0.21871019 | NA | 0 | 3.4727 | 2.7687 | 2.7687 | TRUE | 0.0985 |
| Kcnj8 | 1 | 1 | 0 | 0 | 2 | 0 | 1 | 1 | 0 | 0 | 2 | 0 | 0.68 | 1 | 1 | 1.179 | 0.238 | 0.92432435 | NA | 0 | 3.4737 | 3.0272 | 3.0272 | TRUE | 0.9747 |
| Kcnj9 | 14 | 7 | 13 | 8 | 6 | 9 | 11 | 7 | 14 | 8 | 7 | 9 | 9.33 | 11 | 8 | 0.732 | -0.45 | 0.47422941 | NA | 0 | 0.2565 | 0.1742 | 0.1742 | TRUE | 0.1493 |
| Kcnk1 | 1455 | 1136 | 1281 | 1263 | 1223 | 1317 | 1130 | 1208 | 1382 | 1257 | 1380 | 1300 | 1276.25 | 1240 | 1312 | 1.059 | 0.083 | 0.34399374 | 0.51831147 | 0.0052 | 0.0043 | 0.0048 | 0.0048 | TRUE | 0.1196 |
| Kcnk10 | 0 | 0 | 0 | 0 | 1 | 1 | 0 | 0 | 0 | 0 | 1 | 1 | 0.35 | 0 | 1 | 3.965 | 1.987 | 0.55529285 | NA | 0 | 6.7252 | 6.4426 | 6.4426 | TRUE | 0.261 |
| Kcnk12 | 34 | 30 | 21 | 21 | 24 | 15 | 26 | 32 | 23 | 21 | 27 | 15 | 23.96 | 27 | 21 | 0.772 | -0.374 | 0.35944738 | 0.53419275 | 0 | 0.1013 | 0.0772 | 0.0772 | TRUE | 0.2357 |
| Kcnk13 | 2 | 1 | 1 | 2 | 0 | 0 | 2 | 1 | 1 | 2 | 0 | 0 | 0.95 | 1 | 1 | 0.544 | -0.879 | 0.6666706 | NA | 0 | 2.503 | 1.8451 | 1.8451 | TRUE | 0.9736 |
| Kcnk15 | 0 | 0 | 0 | 0 | 0 | 0 | 0 | 0 | 0 | 0 | 0 | 0 | 0 | 0 | 0 | NA | NA | NA | NA | NA | NA | NA | NA | NA | NA |
| Kcnk16 | 1 | 0 | 1 | 1 | 3 | 3 | 1 | 0 | 1 | 1 | 3 | 3 | 1.53 | 1 | 2 | 3.884 | 1.958 | 0.24418512 | NA | 0 | 1.5487 | 1.0582 | 1.0582 | TRUE | 0.299 |
| Kcnk18 | 7 | 3 | 7 | 9 | 11 | 8 | 5 | 3 | 8 | 9 | 12 | 8 | 7.57 | 5 | 10 | 1.802 | 0.85 | 0.23066705 | NA | 0 | 0.3153 | 0.2197 | 0.2197 | TRUE | 0.2406 |
| Kcnk2 | 49 | 19 | 14 | 11 | 9 | 11 | 38 | 20 | 15 | 11 | 10 | 11 | 17.56 | 24 | 11 | 0.43 | -1.218 | 0.01866438 | 0.06569256 | 0.099 | 0.1374 | 0.125 | 0.125 | TRUE | 0.8059 |
| Kcnk3 | 6 | 0 | 8 | 7 | 9 | 3 | 5 | 0 | 9 | 7 | 10 | 3 | 5.56 | 5 | 7 | 1.502 | 0.587 | 0.5407204 | NA | 0.6722 | 0.4285 | 0.4762 | 0.4762 | TRUE | 0.3205 |
| Kcnk4 | 11 | 12 | 12 | 15 | 8 | 11 | 9 | 13 | 13 | 15 | 9 | 11 | 11.51 | 12 | 12 | 1.03 | 0.043 | 0.94001815 | NA | 0 | 0.2083 | 0.1447 | 0.1447 | TRUE | 0.2451 |
| Kcnk5 | 4038 | 3124 | 2903 | 3003 | 3064 | 3259 | 3136 | 3323 | 3132 | 2989 | 3457 | 3217 | 3209.04 | 3197 | 3221 | 1.007 | 0.011 | 0.87775246 | 0.93172788 | 0.0029 | 0.0032 | 0.0031 | 0.0031 | TRUE | 0.0493 |
| Kcnk6 | 659 | 494 | 555 | 828 | 766 | 880 | 512 | 525 | 599 | 824 | 864 | 869 | 698.86 | 545 | 852 | 1.566 | 0.647 | 2.46E-12 | 2.81E-10 | 0.0018 | 0.0058 | 0.0046 | 0.0046 | TRUE | 0.0841 |
| Kcnk7 | 5 | 10 | 6 | 5 | 6 | 6 | 4 | 11 | 6 | 5 | 7 | 6 | 6.44 | 7 | 6 | 0.853 | -0.23 | 0.76440194 | NA | 0 | 0.3702 | 0.2646 | 0.2646 | TRUE | 0.3154 |
| Kcnk9 | 0 | 0 | 0 | 0 | 0 | 0 | 0 | 0 | 0 | 0 | 0 | 0 | 0 | 0 | 0 | NA | NA | NA | NA | NA | NA | NA | NA | NA | NA |
| Kcnma1 | 135 | 95 | 82 | 102 | 119 | 117 | 105 | 101 | 88 | 102 | 134 | 115 | 107.61 | 98 | 117 | 1.188 | 0.248 | 0.20922474 | 0.37093389 | 0 | 0.0245 | 0.0189 | 0.0189 | TRUE | 0.1581 |
| Kcnmb1 | 28 | 13 | 11 | 26 | 22 | 31 | 22 | 14 | 12 | 26 | 25 | 31 | 21.46 | 16 | 27 | 1.688 | 0.755 | 0.07943784 | 0.1881679 | 0 | 0.1129 | 0.0845 | 0.0845 | TRUE | 0.6285 |
| Kcnmb2 | 2 | 2 | 3 | 1 | 2 | 1 | 2 | 2 | 3 | 1 | 2 | 1 | 1.86 | 2 | 1 | 0.613 | -0.707 | 0.60891049 | NA | 0 | 1.2771 | 0.7922 | 0.7922 | TRUE | 0.1547 |
| Kcnmb3 | 0 | 1 | 1 | 0 | 0 | 0 | 0 | 1 | 1 | 0 | 0 | 0 | 0.36 | 1 | 0 | 0.271 | -1.881 | 0.5755355 | NA | 0 | 6.6391 | 6.4428 | 6.4428 | TRUE | 0.3513 |
| Kcnmb4 | 31 | 23 | 15 | 30 | 22 | 22 | 24 | 24 | 16 | 30 | 25 | 22 | 23.52 | 21 | 26 | 1.175 | 0.232 | 0.56315137 | 0.71753952 | 0 | 0.1032 | 0.0739 | 0.0739 | TRUE | 0.2466 |
| Kcnn1 | 386 | 291 | 308 | 408 | 372 | 409 | 300 | 310 | 332 | 406 | 420 | 404 | 361.86 | 314 | 410 | 1.307 | 0.387 | 0.00043122 | 0.00385451 | 0 | 0.009 | 0.0059 | 0.0059 | TRUE | 0.0293 |
| Kcnn2 | 404 | 309 | 301 | 303 | 262 | 306 | 314 | 329 | 325 | 302 | 296 | 302 | 311.08 | 323 | 300 | 0.931 | -0.104 | 0.3662573 | 0.54090882 | 0 | 0.01 | 0.0062 | 0.0062 | TRUE | 0.0068 |
| Kcnn3 | 53 | 41 | 42 | 55 | 52 | 55 | 41 | 44 | 45 | 55 | 59 | 54 | 49.63 | 43 | 56 | 1.291 | 0.368 | 0.17219834 | 0.32431557 | 0 | 0.0502 | 0.0321 | 0.0321 | TRUE | 0.0182 |
| Kcnn4 | 41 | 35 | 39 | 68 | 70 | 52 | 32 | 37 | 42 | 68 | 79 | 51 | 51.52 | 37 | 66 | 1.788 | 0.838 | 0.00379476 | 0.01979508 | 0.0176 | 0.0484 | 0.0394 | 0.0394 | TRUE | 0.2001 |
| Kcnq1 | 1 | 2 | 2 | 16 | 17 | 9 | 1 | 2 | 2 | 16 | 19 | 9 | 8.18 | 2 | 15 | 9.046 | 3.177 | 0.00032413 | NA | 0 | 0.2923 | 0.231 | 0.231 | TRUE | 0.22 |
| Kcnq1ot1 | 579 | 473 | 459 | 694 | 550 | 721 | 450 | 503 | 495 | 691 | 620 | 712 | 578.51 | 483 | 674 | 1.401 | 0.486 | 9.21E-07 | 2.74E-05 | 0.0028 | 0.0065 | 0.0053 | 0.0053 | TRUE | 0.0562 |
| Kcnq2 | 329 | 215 | 224 | 286 | 247 | 249 | 256 | 229 | 242 | 285 | 279 | 246 | 255.84 | 242 | 270 | 1.112 | 0.153 | 0.24309403 | 0.41089104 | 0 | 0.0117 | 0.0085 | 0.0085 | TRUE | 0.0688 |
| Kcnq3 | 6 | 4 | 4 | 2 | 1 | 3 | 5 | 4 | 4 | 2 | 1 | 3 | 3.22 | 4 | 2 | 0.463 | -1.11 | 0.3011144 | NA | 0 | 0.7388 | 0.4678 | 0.4678 | TRUE | 0.1502 |
| Kcnq4 | 51 | 36 | 39 | 59 | 56 | 68 | 40 | 38 | 42 | 59 | 63 | 67 | 51.5 | 40 | 63 | 1.576 | 0.657 | 0.01423379 | 0.05367784 | 0 | 0.0484 | 0.0314 | 0.0314 | TRUE | 0.0319 |
| Kcnq5 | 76 | 55 | 55 | 60 | 54 | 46 | 59 | 59 | 59 | 60 | 61 | 45 | 57.15 | 59 | 55 | 0.937 | -0.094 | 0.7170145 | 0.82988811 | 0 | 0.0439 | 0.0306 | 0.0306 | TRUE | 0.2132 |
| Kcnrg | 19 | 7 | 8 | 22 | 19 | 16 | 15 | 7 | 9 | 22 | 21 | 16 | 14.99 | 10 | 20 | 1.878 | 0.909 | 0.07867473 | NA | 0 | 0.1605 | 0.1203 | 0.1203 | TRUE | 0.6555 |
| Kcns1 | 122 | 89 | 83 | 81 | 73 | 84 | 95 | 95 | 90 | 81 | 82 | 83 | 87.48 | 93 | 82 | 0.881 | -0.183 | 0.37032859 | 0.5449938 | 0 | 0.0295 | 0.0187 | 0.0187 | TRUE | 0.0095 |
| Kcns2 | 42 | 31 | 20 | 26 | 28 | 25 | 33 | 33 | 22 | 26 | 32 | 25 | 28.22 | 29 | 28 | 0.935 | -0.096 | 0.79459814 | 0.87979922 | 0 | 0.0864 | 0.0629 | 0.0629 | TRUE | 0.3068 |
| Kcns3 | 449 | 361 | 336 | 276 | 262 | 306 | 349 | 384 | 363 | 275 | 296 | 302 | 327.94 | 365 | 291 | 0.797 | -0.327 | 0.00483824 | 0.02390355 | 0 | 0.0097 | 0.0066 | 0.0066 | TRUE | 0.0269 |
| Kcnt1 | 102 | 86 | 80 | 69 | 70 | 80 | 79 | 91 | 86 | 69 | 79 | 79 | 80.61 | 85 | 76 | 0.884 | -0.179 | 0.41024014 | 0.58307953 | 0 | 0.0318 | 0.0214 | 0.0214 | TRUE | 0.0591 |
| Kcnt2 | 19 | 16 | 11 | 17 | 23 | 27 | 15 | 17 | 12 | 17 | 26 | 27 | 18.86 | 15 | 23 | 1.589 | 0.668 | 0.1398359 | 0.28110754 | 0 | 0.1281 | 0.0921 | 0.0921 | TRUE | 0.3346 |
| Kcnu1 | 125 | 121 | 149 | 118 | 122 | 114 | 97 | 129 | 161 | 117 | 138 | 113 | 125.7 | 129 | 123 | 0.956 | -0.065 | 0.75618051 | 0.85520897 | 0.0303 | 0.0213 | 0.024 | 0.024 | TRUE | 0.1903 |
| Kcnv1 | 9 | 10 | 6 | 3 | 3 | 9 | 7 | 11 | 6 | 3 | 3 | 9 | 6.56 | 8 | 5 | 0.64 | -0.645 | 0.40836719 | NA | 0.0331 | 0.3638 | 0.2766 | 0.2766 | TRUE | 0.9066 |
| Kcnv2 | 29 | 24 | 15 | 23 | 15 | 15 | 23 | 26 | 16 | 23 | 17 | 15 | 19.81 | 22 | 18 | 0.849 | -0.236 | 0.5929396 | 0.74048805 | 0 | 0.1221 | 0.0891 | 0.0891 | TRUE | 0.2682 |
| Kcp | 253 | 241 | 181 | 255 | 261 | 306 | 196 | 256 | 195 | 254 | 294 | 302 | 249.75 | 216 | 283 | 1.315 | 0.395 | 0.00819267 | 0.03566794 | 0.0122 | 0.0119 | 0.012 | 0.012 | TRUE | 0.2839 |
| Kctd1 | 305 | 213 | 179 | 186 | 180 | 187 | 237 | 227 | 193 | 185 | 203 | 185 | 204.9 | 219 | 191 | 0.869 | -0.202 | 0.16888153 | 0.32052596 | 0 | 0.014 | 0.0106 | 0.0106 | TRUE | 0.1123 |
| Kctd10 | 2624 | 1854 | 1778 | 1796 | 1595 | 1867 | 2038 | 1972 | 1918 | 1788 | 1799 | 1843 | 1893.09 | 1976 | 1810 | 0.915 | -0.127 | 0.04127866 | 0.11689596 | 0 | 0.0037 | 0.0023 | 0.0023 | TRUE | 0.0094 |
| Kctd11 | 72 | 56 | 50 | 63 | 51 | 75 | 56 | 60 | 54 | 63 | 58 | 74 | 60.62 | 57 | 65 | 1.149 | 0.2 | 0.42246133 | 0.59472469 | 0 | 0.0415 | 0.0285 | 0.0285 | TRUE | 0.14 |
| Kctd12 | 687 | 562 | 404 | 381 | 339 | 366 | 534 | 598 | 436 | 379 | 382 | 361 | 448.37 | 523 | 374 | 0.716 | -0.482 | 0.00010222 | 0.00122979 | 0.0111 | 0.0077 | 0.0088 | 0.0088 | TRUE | 0.2364 |
| Kctd12b | 8 | 8 | 5 | 2 | 2 | 3 | 6 | 9 | 5 | 2 | 2 | 3 | 4.55 | 7 | 2 | 0.36 | -1.475 | 0.11323946 | NA | 0 | 0.5228 | 0.3382 | 0.3382 | TRUE | 0.131 |
| Kctd13 | 1158 | 908 | 1044 | 1264 | 1037 | 1190 | 899 | 966 | 1126 | 1258 | 1170 | 1175 | 1099.07 | 997 | 1201 | 1.206 | 0.271 | 0.00343786 | 0.01838284 | 0.0063 | 0.0046 | 0.0053 | 0.0053 | TRUE | 0.1524 |
| Kctd14 | 31 | 16 | 14 | 9 | 18 | 20 | 24 | 17 | 15 | 9 | 20 | 20 | 17.53 | 19 | 16 | 0.861 | -0.216 | 0.66030272 | 0.78996458 | 0.0577 | 0.1376 | 0.1163 | 0.1163 | TRUE | 0.771 |

| | | | | | | | | | | | | | | | | | | | | | | | | | |
|---|---|---|---|---|---|---|---|---|---|---|---|---|---|---|---|---|---|---|---|---|---|---|---|---|---|
| Kctd15 | 831 | 646 | 653 | 415 | 394 | 462 | 645 | 687 | 705 | 413 | 444 | 456 | 558.46 | 679 | 438 | 0.646 | -0.631 | 4.34E-11 | 3.80E-09 | 0 | 0.0067 | 0.0047 | 0.0047 | TRUE | 0.029 |
| Kctd16 | 0 | 1 | 0 | 0 | 0 | 0 | 0 | 1 | 0 | 0 | 0 | 0 | 0.18 | 0 | 0 | 0.552 | -0.857 | 0.8336548 | NA | 0 | 13.3714 | 10 | 10 | TRUE | 0.5216 |
| Kctd17 | 956 | 721 | 691 | 757 | 659 | 829 | 742 | 767 | 746 | 754 | 743 | 818 | 761.71 | 752 | 772 | 1.028 | 0.039 | 0.6403055 | 0.77532078 | 0 | 0.0055 | 0.0037 | 0.0037 | TRUE | 0.0334 |
| Kctd18 | 831 | 612 | 629 | 615 | 578 | 637 | 645 | 651 | 679 | 612 | 652 | 629 | 644.68 | 658 | 631 | 0.959 | -0.061 | 0.48266054 | 0.64916412 | 0 | 0.0061 | 0.0039 | 0.0039 | TRUE | 0.0097 |
| Kctd19 | 57 | 41 | 51 | 97 | 89 | 81 | 44 | 44 | 55 | 97 | 100 | 80 | 69.97 | 48 | 92 | 1.942 | 0.958 | 8.07E-05 | 0.00101576 | 0 | 0.0363 | 0.0268 | 0.0268 | TRUE | 0.1373 |
| Kctd2 | 2311 | 1780 | 2030 | 1992 | 1920 | 1957 | 1795 | 1893 | 2190 | 1983 | 2166 | 1932 | 1993.2 | 1959 | 2027 | 1.035 | 0.05 | 0.55958755 | 0.71437878 | 0.0067 | 0.0036 | 0.0047 | 0.0047 | TRUE | 0.1277 |
| Kctd20 | 4573 | 3147 | 3089 | 3187 | 2871 | 3298 | 3552 | 3347 | 3333 | 3172 | 3239 | 3256 | 3316.47 | 3411 | 3222 | 0.944 | -0.083 | 0.14475549 | 0.2882527 | 5.00E-04 | 0.0031 | 0.002 | 0.002 | TRUE | 0.0163 |
| Kctd21 | 234 | 177 | 158 | 179 | 155 | 168 | 182 | 188 | 170 | 178 | 175 | 166 | 176.56 | 180 | 173 | 0.959 | -0.06 | 0.68826698 | 0.80906325 | 0 | 0.0158 | 0.0103 | 0.0103 | TRUE | 0.0221 |
| Kctd3 | 5257 | 4037 | 3851 | 4192 | 3793 | 4330 | 4083 | 4294 | 4155 | 4173 | 4279 | 4274 | 4209.7 | 4177 | 4242 | 1.016 | 0.022 | 0.67100281 | 0.79747076 | 2.00E-04 | 0.003 | 0.0018 | 0.0018 | TRUE | 0.0072 |
| Kctd4 | 0 | 0 | 0 | 0 | 0 | 0 | 0 | 0 | 0 | 0 | 0 | 0 | 0 | 0 | 0 | NA | NA | NA | NA | NA | NA | NA | NA | NA | NA |
| Kctd5 | 4792 | 3486 | 3308 | 3645 | 3151 | 3535 | 3722 | 3708 | 3569 | 3628 | 3555 | 3490 | 3611.92 | 3666 | 3558 | 0.97 | -0.044 | 0.41973944 | 0.59200077 | 2.00E-04 | 0.0031 | 0.0018 | 0.0018 | TRUE | 0.0064 |
| Kctd6 | 839 | 618 | 664 | 681 | 641 | 761 | 652 | 657 | 716 | 678 | 723 | 751 | 696.27 | 675 | 717 | 1.064 | 0.089 | 0.32048718 | 0.49441158 | 0.0013 | 0.0058 | 0.0043 | 0.0043 | TRUE | 0.033 |
| Kctd7 | 349 | 218 | 220 | 292 | 218 | 241 | 271 | 232 | 237 | 291 | 246 | 238 | 252.47 | 247 | 258 | 1.044 | 0.062 | 0.65803131 | 0.7882368 | 0.006 | 0.0118 | 0.0101 | 0.0101 | TRUE | 0.1373 |
| Kctd8 | 8 | 9 | 5 | 7 | 5 | 5 | 6 | 10 | 5 | 7 | 6 | 5 | 6.45 | 7 | 6 | 0.832 | -0.265 | 0.72191042 | NA | 0 | 0.3696 | 0.2434 | 0.2434 | TRUE | 0.2385 |
| Kctd9 | 2078 | 1426 | 1528 | 1690 | 1426 | 1519 | 1614 | 1517 | 1649 | 1682 | 1609 | 1499 | 1594.97 | 1593 | 1597 | 1.002 | 0.003 | 0.96789075 | 0.98270717 | 0.002 | 0.0039 | 0.0032 | 0.0032 | TRUE | 0.0348 |
| Kdelc1 | 1621 | 1121 | 1340 | 1565 | 1279 | 1652 | 1259 | 1192 | 1446 | 1558 | 1443 | 1631 | 1421.44 | 1299 | 1544 | 1.189 | 0.25 | 0.0040695 | 0.02091528 | 0.0059 | 0.0041 | 0.0048 | 0.0048 | TRUE | 0.1149 |
| Kdelc2 | 477 | 380 | 379 | 489 | 409 | 474 | 370 | 404 | 409 | 487 | 461 | 468 | 433.28 | 394 | 472 | 1.199 | 0.262 | 0.01094375 | 0.04413628 | 0 | 0.0079 | 0.0053 | 0.0053 | TRUE | 0.0356 |
| Kdelr1 | 3379 | 2440 | 2830 | 2881 | 2652 | 3058 | 2624 | 2595 | 3054 | 2868 | 2992 | 3019 | 2858.58 | 2758 | 2960 | 1.074 | 0.102 | 0.16637577 | 0.31740641 | 0.0042 | 0.0033 | 0.0036 | 0.0036 | TRUE | 0.1058 |
| Kdelr2 | 6252 | 4362 | 4466 | 4683 | 4237 | 4513 | 4856 | 4640 | 4819 | 4661 | 4780 | 4455 | 4701.75 | 4772 | 4632 | 0.971 | -0.043 | 0.43340521 | 0.60536115 | 6.00E-04 | 0.0029 | 0.002 | 0.002 | TRUE | 0.0137 |
| Kdelr3 | 2022 | 1519 | 2108 | 2828 | 2569 | 2834 | 1570 | 1616 | 2275 | 2815 | 2898 | 2798 | 2328.56 | 1820 | 2837 | 1.56 | 0.641 | 1.81E-08 | 8.47E-07 | 0.0229 | 0.0034 | 0.0089 | 0.0089 | TRUE | 0.5732 |
| Kdm1a | 23239 | 17310 | 17725 | 19043 | 17134 | 19730 | 18049 | 18412 | 19125 | 18955 | 19329 | 19477 | 18891.19 | 18529 | 19254 | 1.039 | 0.055 | 0.24173764 | 0.40937375 | 5.00E-04 | 0.0026 | 0.0016 | 0.0016 | TRUE | 0.0097 |
| Kdm1b | 2226 | 1713 | 1581 | 1629 | 1447 | 1717 | 1729 | 1822 | 1706 | 1622 | 1632 | 1695 | 1700.94 | 1752 | 1650 | 0.942 | -0.087 | 0.18446688 | 0.34064408 | 3.00E-04 | 0.0038 | 0.0025 | 0.0025 | TRUE | 0.0143 |
| Kdm2a | 5850 | 4380 | 4034 | 4089 | 3608 | 4314 | 4543 | 4659 | 4353 | 4070 | 4070 | 4259 | 4325.67 | 4518 | 4133 | 0.915 | -0.129 | 0.02098145 | 0.0716608 | 6.00E-04 | 0.003 | 0.002 | 0.002 | TRUE | 0.0123 |
| Kdm2b | 5965 | 4351 | 4151 | 4522 | 3976 | 4625 | 4633 | 4628 | 4479 | 4501 | 4485 | 4566 | 4548.65 | 4580 | 4517 | 0.986 | -0.02 | 0.69445079 | 0.81365564 | 0 | 0.0029 | 0.0016 | 0.0016 | TRUE | 0.0045 |
| Kdm3a | 7444 | 5380 | 5007 | 4441 | 3959 | 4958 | 5781 | 5723 | 5403 | 4421 | 4466 | 4894 | 5114.6 | 5636 | 4594 | 0.815 | -0.295 | 1.72E-06 | 4.50E-05 | 0.002 | 0.0029 | 0.0025 | 0.0025 | TRUE | 0.0401 |
| Kdm3b | 10251 | 7532 | 7486 | 8501 | 7690 | 8284 | 7962 | 8012 | 8077 | 8462 | 8675 | 8178 | 8227.52 | 8017 | 8438 | 1.053 | 0.074 | 0.13349343 | 0.27199168 | 3.00E-04 | 0.0027 | 0.0016 | 0.0016 | TRUE | 0.0089 |
| Kdm4a | 4828 | 3467 | 3210 | 3220 | 2908 | 3447 | 3750 | 3688 | 3464 | 3205 | 3281 | 3403 | 3464.92 | 3634 | 3296 | 0.907 | -0.141 | 0.01772473 | 0.06316121 | 9.00E-04 | 0.0031 | 0.0023 | 0.0023 | TRUE | 0.0202 |
| Kdm4b | 2625 | 2201 | 2460 | 2379 | 2201 | 2664 | 2039 | 2341 | 2654 | 2368 | 2483 | 2630 | 2419.17 | 2345 | 2494 | 1.065 | 0.09 | 0.32153529 | 0.49528009 | 0.0097 | 0.0034 | 0.0056 | 0.0056 | TRUE | 0.1617 |
| Kdm4c | 6434 | 4780 | 6064 | 6892 | 6323 | 7756 | 4997 | 5084 | 6543 | 6860 | 7133 | 7656 | 6379.02 | 5541 | 7216 | 1.303 | 0.381 | 4.73E-05 | 0.00066704 | 0.0136 | 0.0028 | 0.0062 | 0.0062 | TRUE | 0.3041 |
| Kdm4d | 6 | 11 | 5 | 3 | 1 | 5 | 5 | 12 | 5 | 3 | 1 | 5 | 5.13 | 7 | 3 | 0.426 | -1.232 | 0.17776103 | NA | 0 | 0.464 | 0.3661 | 0.3661 | TRUE | 0.6383 |
| Kdm5a | 10227 | 7248 | 7100 | 7700 | 6859 | 7617 | 7943 | 7710 | 7661 | 7665 | 7738 | 7519 | 7705.79 | 7771 | 7641 | 0.983 | -0.025 | 0.60723307 | 0.75124899 | 2.00E-04 | 0.0027 | 0.0015 | 0.0015 | TRUE | 0.0046 |
| Kdm5b | 30468 | 22305 | 21403 | 22090 | 20129 | 23065 | 23663 | 23726 | 23094 | 21988 | 22708 | 22769 | 22991.18 | 23494 | 22488 | 0.957 | -0.063 | 0.1601182 | 0.3095171 | 2.00E-04 | 0.0025 | 0.0014 | 0.0014 | TRUE | 0.0046 |
| Kdm5c | 13314 | 9696 | 9836 | 10258 | 9216 | 10832 | 10340 | 10314 | 10613 | 10211 | 10397 | 10693 | 10427.86 | 10422 | 10434 | 1.001 | 0.002 | 0.97279652 | 0.98520376 | 2.00E-04 | 0.0027 | 0.0015 | 0.0015 | TRUE | 0.0058 |
| Kdm5d | 729 | 515 | 430 | 196 | 186 | 177 | 566 | 548 | 464 | 195 | 210 | 175 | 359.6 | 526 | 193 | 0.366 | -1.449 | 3.13E-29 | 2.36E-26 | 0.007 | 0.009 | 0.0084 | 0.0084 | TRUE | 0.1218 |
| Kdm6a | 3937 | 2806 | 2710 | 2891 | 2644 | 3110 | 3058 | 2985 | 2924 | 2878 | 2983 | 3070 | 2982.82 | 2989 | 2977 | 0.996 | -0.006 | 0.91364499 | 0.95246365 | 5.00E-04 | 0.0032 | 0.0021 | 0.0021 | TRUE | 0.0104 |
| Kdm6b | 4312 | 3592 | 3474 | 3564 | 3442 | 4027 | 3349 | 3821 | 3748 | 3548 | 3883 | 3975 | 3720.66 | 3639 | 3802 | 1.045 | 0.064 | 0.37386687 | 0.54846343 | 0.0036 | 0.0031 | 0.0034 | 0.0034 | TRUE | 0.0603 |
| Kdm8 | 250 | 177 | 143 | 161 | 158 | 164 | 194 | 188 | 154 | 160 | 178 | 162 | 172.85 | 179 | 167 | 0.929 | -0.107 | 0.50356443 | 0.66766664 | 0.0032 | 0.0161 | 0.0125 | 0.0125 | TRUE | 0.1493 |
| Kdr | 106 | 75 | 58 | 54 | 54 | 60 | 82 | 80 | 63 | 54 | 61 | 59 | 66.43 | 75 | 58 | 0.771 | -0.376 | 0.122574 | 0.25590858 | 0 | 0.0381 | 0.0273 | 0.0273 | TRUE | 0.1797 |
| Kdsr | 2344 | 1841 | 1988 | 2283 | 1837 | 2260 | 1820 | 1958 | 2145 | 2272 | 2072 | 2231 | 2083.26 | 1974 | 2192 | 1.111 | 0.152 | 0.04690039 | 0.1285668 | 0.0038 | 0.0036 | 0.0038 | 0.0038 | TRUE | 0.0688 |
| Keap1 | 1041 | 669 | 680 | 672 | 643 | 683 | 809 | 712 | 734 | 669 | 725 | 674 | 720.39 | 752 | 689 | 0.916 | -0.127 | 0.15921233 | 0.30843343 | 0.0021 | 0.0057 | 0.0045 | 0.0045 | TRUE | 0.0553 |
| Keg1 | 0 | 0 | 0 | 0 | 0 | 0 | 0 | 0 | 0 | 0 | 0 | 0 | 0 | 0 | 0 | NA | NA | NA | NA | NA | NA | NA | NA | NA | NA |
| Kel | 1 | 0 | 0 | 1 | 0 | 1 | 1 | 0 | 0 | 1 | 0 | 1 | 0.46 | 0 | 1 | 1.94 | 0.956 | 0.73503911 | NA | 0 | 5.1562 | 3.9771 | 3.9771 | TRUE | 0.2566 |
| Kera | 0 | 0 | 0 | 0 | 0 | 0 | 0 | 0 | 0 | 0 | 0 | 0 | 0 | 0 | 0 | NA | NA | NA | NA | NA | NA | NA | NA | NA | NA |
| Khdc1a | 0 | 0 | 1 | 0 | 2 | 0 | 0 | 0 | 1 | 0 | 2 | 0 | 0.56 | 0 | 1 | 2.136 | 1.095 | 0.70886258 | NA | 0 | 4.2661 | 4.493 | 4.493 | TRUE | 0.9746 |
| Khdc1b | 0 | 0 | 0 | 0 | 0 | 0 | 0 | 0 | 0 | 0 | 0 | 0 | 0 | 0 | 0 | NA | NA | NA | NA | NA | NA | NA | NA | NA | NA |
| Khdc1c | 0 | 0 | 0 | 0 | 0 | 0 | 0 | 0 | 0 | 0 | 0 | 0 | 0 | 0 | 0 | NA | NA | NA | NA | NA | NA | NA | NA | NA | NA |
| Khdc3 | 6388 | 5386 | 5522 | 4308 | 4125 | 4630 | 4961 | 5729 | 5958 | 4288 | 4653 | 4571 | 5026.77 | 5549 | 4504 | 0.812 | -0.301 | 5.71E-05 | 0.00077769 | 0.005 | 0.0029 | 0.0038 | 0.0038 | TRUE | 0.106 |
| Khdrbs1 | 14651 | 10556 | 9887 | 10214 | 8651 | 10032 | 11379 | 11228 | 10668 | 10167 | 9759 | 9903 | 10517.41 | 11092 | 9943 | 0.896 | -0.158 | 0.00187871 | 0.01153949 | 6.00E-04 | 0.0027 | 0.0018 | 0.0018 | TRUE | 0.0136 |
| Khdrbs2 | 20 | 13 | 11 | 10 | 11 | 12 | 16 | 14 | 12 | 10 | 12 | 12 | 12.57 | 14 | 11 | 0.824 | -0.279 | 0.59994024 | NA | 0 | 0.1909 | 0.1233 | 0.1233 | TRUE | 0.0703 |
| Khdrbs3 | 2764 | 1984 | 1949 | 1974 | 1762 | 2009 | 2147 | 2110 | 2103 | 1965 | 1988 | 1983 | 2049.3 | 2120 | 1979 | 0.933 | -0.1 | 0.08879413 | 0.20283283 | 0 | 0.0036 | 0.002 | 0.002 | TRUE | 0.0015 |
| Khk | 175 | 124 | 136 | 162 | 149 | 163 | 136 | 132 | 147 | 161 | 168 | 161 | 150.8 | 138 | 163 | 1.183 | 0.242 | 0.12904749 | 0.26569804 | 0 | 0.0181 | 0.0117 | 0.0117 | TRUE | 0.0283 |
| Khnyn | 2073 | 1685 | 2014 | 2082 | 1830 | 2037 | 1610 | 1792 | 2173 | 2072 | 2064 | 2011 | 1953.85 | 1858 | 2049 | 1.104 | 0.143 | 0.13960656 | 0.28083355 | 0.0111 | 0.0036 | 0.0062 | 0.0062 | TRUE | 0.2643 |
| Khsrp | 17029 | 13006 | 12144 | 12050 | 11070 | 12581 | 13226 | 13834 | 13103 | 11995 | 12488 | 12419 | 12844.22 | 13388 | 12301 | 0.919 | -0.122 | 0.01312302 | 0.05051639 | 5.00E-04 | 0.0026 | 0.0017 | 0.0017 | TRUE | 0.0104 |
| Kidins220 | 4226 | 3186 | 3107 | 3422 | 3220 | 3649 | 3282 | 3389 | 3352 | 3406 | 3632 | 3602 | 3444.06 | 3341 | 3547 | 1.062 | 0.086 | 0.1253879 | 0.26060334 | 5.00E-04 | 0.0031 | 0.002 | 0.002 | TRUE | 0.0147 |
| Kif11 | 11985 | 8519 | 9114 | 9446 | 8373 | 9897 | 9308 | 9062 | 9834 | 9403 | 9446 | 9770 | 9470.29 | 9401 | 9540 | 1.015 | 0.021 | 0.68991765 | 0.81028948 | 8.00E-04 | 0.0027 | 0.0019 | 0.0019 | TRUE | 0.0196 |
| Kif12 | 0 | 0 | 0 | 1 | 0 | 2 | 0 | 0 | 0 | 1 | 0 | 2 | 0.49 | 0 | 1 | 5.637 | 2.495 | 0.3989397 | NA | 0 | 4.7909 | 4.7853 | 4.7853 | TRUE | 0.0526 |
| Kif13a | 4382 | 3056 | 3113 | 3093 | 2794 | 3336 | 3403 | 3251 | 3359 | 3079 | 3152 | 3293 | 3256.11 | 3338 | 3175 | 0.951 | -0.072 | 0.20772494 | 0.3693261 | 6.00E-04 | 0.0032 | 0.0021 | 0.0021 | TRUE | 0.0131 |
| Kif13b | 4897 | 3579 | 3707 | 4476 | 4107 | 4470 | 3803 | 3807 | 4000 | 4455 | 4633 | 4413 | 4185.19 | 3870 | 4500 | 1.163 | 0.218 | 7.17E-05 | 0.00093338 | 4.00E-04 | 0.003 | 0.0019 | 0.0019 | TRUE | 0.0103 |
| Kif14 | 3885 | 2827 | 2962 | 3266 | 2947 | 3199 | 3017 | 3007 | 3196 | 3251 | 3325 | 3158 | 3158.96 | 3073 | 3245 | 1.056 | 0.078 | 0.17743361 | 0.33159762 | 6.00E-04 | 0.0032 | 0.0021 | 0.0021 | TRUE | 0.0146 |
| Kif15 | 5012 | 3762 | 3908 | 3811 | 3337 | 3938 | 3893 | 4002 | 4217 | 3793 | 3764 | 3887 | 3926.04 | 4037 | 3815 | 0.945 | -0.081 | 0.15376679 | 0.3008197 | 7.00E-04 | 0.003 | 0.0021 | 0.0021 | TRUE | 0.0184 |
| Kif16b | 2475 | 1680 | 1857 | 1840 | 1545 | 1779 | 1922 | 1787 | 2004 | 1832 | 1743 | 1756 | 1840.59 | 1904 | 1777 | 0.933 | -0.1 | 0.14641409 | 0.29077347 | 0.0014 | 0.0037 | 0.0029 | 0.0029 | TRUE | 0.0342 |
| Kif17 | 212 | 196 | 170 | 235 | 269 | 244 | 165 | 208 | 183 | 234 | 303 | 241 | 222.47 | 185 | 259 | 1.4 | 0.486 | 0.00194375 | 0.01182716 | 0.0132 | 0.0131 | 0.0131 | 0.0131 | TRUE | 0.2395 |
| Kif18a | 3177 | 2359 | 2730 | 2918 | 2482 | 2908 | 2467 | 2509 | 2946 | 2905 | 2800 | 2871 | 2749.58 | 2641 | 2859 | 1.083 | 0.115 | 0.12902175 | 0.26569804 | 0.0047 | 0.0033 | 0.0038 | 0.0038 | TRUE | 0.1228 |
| Kif18b | 6989 | 5142 | 5094 | 5072 | 4748 | 5351 | 5428 | 5469 | 5496 | 5049 | 5356 | 5282 | 5346.85 | 5464 | 5229 | 0.957 | -0.064 | 0.21751887 | 0.38030547 | 3.00E-04 | 0.0029 | 0.0017 | 0.0017 | TRUE | 0.0111 |
| Kif19a | 127 | 108 | 106 | 154 | 146 | 155 | 99 | 115 | 114 | 153 | 165 | 153 | 133.15 | 109 | 157 | 1.441 | 0.527 | 0.0022741 | 0.01333759 | 0 | 0.0202 | 0.0138 | 0.0138 | TRUE | 0.0826 |
| Kif1a | 768 | 569 | 682 | 1152 | 1111 | 1170 | 596 | 605 | 736 | 1147 | 1253 | 1155 | 915.43 | 646 | 1185 | 1.837 | 0.878 | 8.11E-20 | 2.57E-17 | 0.0068 | 0.005 | 0.0055 | 0.0055 | TRUE | 0.1716 |
| Kif1b | 7583 | 5074 | 4832 | 6046 | 5194 | 5891 | 5889 | 5397 | 5214 | 6018 | 5859 | 5815 | 5698.85 | 5500 | 5897 | 1.072 | 0.1 | 0.10030519 | 0.2214834 | 0.0019 | 0.0028 | 0.0025 | 0.0025 | TRUE | 0.0474 |
| Kif1c | 5960 | 4339 | 4635 | 5733 | 4962 | 5519 | 4629 | 4615 | 5001 | 5707 | 5598 | 5448 | 5166.29 | 4748 | 5584 | 1.176 | 0.234 | 3.73E-05 | 0.00055524 | 0.001 | 0.0029 | 0.0021 | 0.0021 | TRUE | 0.0261 |
| Kif20a | 13295 | 9338 | 9844 | 9440 | 8829 | 10229 | 10326 | 9933 | 10622 | 9397 | 9960 | 10098 | 10055.69 | 10294 | 9818 | 0.954 | -0.068 | 0.20735501 | 0.36908856 | 0.0011 | 0.0027 | 0.002 | 0.002 | TRUE | 0.0173 |

| | | | | | | | | | | | | | | | | | | | | | | | | | |
|---|---|---|---|---|---|---|---|---|---|---|---|---|---|---|---|---|---|---|---|---|---|---|---|---|---|
| Kif20b | 11224 | 8768 | 9270 | 10735 | 9379 | 11077 | 8717 | 9326 | 10002 | 10686 | 10580 | 10935 | 10041.11 | 9348 | 10734 | 1.148 | 0.2 | 0.00100162 | 0.00724482 | 0.0024 | 0.0027 | 0.0026 | 0.0026 | TRUE | 0.0457 |
| Kif21a | 1361 | 982 | 928 | 994 | 922 | 1023 | 1057 | 1045 | 1001 | 989 | 1040 | 1010 | 1023.71 | 1034 | 1013 | 0.979 | -0.031 | 0.67512136 | 0.80045704 | 0 | 0.0047 | 0.003 | 0.003 | TRUE | 0.0091 |
| Kif21b | 2110 | 1714 | 2163 | 2452 | 2333 | 2586 | 1639 | 1823 | 2334 | 2441 | 2632 | 2553 | 2236.86 | 1932 | 2542 | 1.317 | 0.397 | 0.00019329 | 0.00207725 | 0.0174 | 0.0035 | 0.0077 | 0.0077 | TRUE | 0.3994 |
| Kif22 | 7167 | 5228 | 5386 | 5675 | 4994 | 5912 | 5566 | 5561 | 5811 | 5649 | 5634 | 5836 | 5676.23 | 5646 | 5706 | 1.011 | 0.016 | 0.76234333 | 0.85900698 | 3.00E-04 | 0.0028 | 0.0017 | 0.0017 | TRUE | 0.0079 |
| Kif23 | 9364 | 7414 | 8396 | 8339 | 8189 | 9269 | 7273 | 7886 | 9059 | 8301 | 9238 | 9150 | 8484.44 | 8073 | 8896 | 1.102 | 0.14 | 0.08070217 | 0.19031119 | 0.0078 | 0.0027 | 0.0045 | 0.0045 | TRUE | 0.1394 |
| Kif24 | 2059 | 1464 | 1400 | 1491 | 1352 | 1678 | 1599 | 1557 | 1511 | 1484 | 1525 | 1656 | 1555.46 | 1556 | 1555 | 0.999 | -0.001 | 0.99063998 | 0.99533061 | 0.0014 | 0.004 | 0.003 | 0.003 | TRUE | 0.0395 |
| Kif26a | 2468 | 2175 | 2346 | 2691 | 2703 | 3003 | 1917 | 2314 | 2531 | 2679 | 3049 | 2964 | 2575.66 | 2254 | 2897 | 1.287 | 0.364 | 0.00013626 | 0.00157192 | 0.0112 | 0.0033 | 0.0062 | 0.0062 | TRUE | 0.2111 |
| Kif26b | 62 | 42 | 40 | 55 | 47 | 52 | 48 | 45 | 43 | 55 | 53 | 51 | 49.18 | 45 | 53 | 1.167 | 0.223 | 0.41058326 | 0.58334658 | 0 | 0.0506 | 0.0324 | 0.0324 | TRUE | 0.0274 |
| Kif27 | 93 | 72 | 47 | 63 | 60 | 52 | 72 | 77 | 51 | 63 | 68 | 51 | 63.54 | 67 | 61 | 0.906 | -0.142 | 0.58717429 | 0.73631182 | 0.0159 | 0.0397 | 0.0333 | 0.0333 | TRUE | 0.3586 |
| Kif2a | 7204 | 5105 | 4922 | 5290 | 4445 | 5294 | 5595 | 5430 | 5311 | 5266 | 5014 | 5226 | 5307.01 | 5445 | 5169 | 0.949 | -0.075 | 0.15454068 | 0.30196171 | 4.00E-04 | 0.0029 | 0.0018 | 0.0018 | TRUE | 0.0082 |
| Kif2b | 2 | 0 | 0 | 0 | 0 | 2 | 2 | 0 | 0 | 0 | 0 | 2 | 0.59 | 1 | 1 | 1.213 | 0.279 | 0.92253239 | NA | 0 | 4.0336 | 4.2602 | 4.2602 | TRUE | 0.9737 |
| Kif2c | 7579 | 5511 | 5599 | 5912 | 5243 | 6037 | 5886 | 5862 | 6041 | 5885 | 5915 | 5959 | 5924.74 | 5930 | 5920 | 0.998 | -0.002 | 0.9611296 | 0.97845108 | 0 | 0.0028 | 0.0015 | 0.0015 | TRUE | 0.0033 |
| Kif3a | 4328 | 3082 | 2864 | 3114 | 2793 | 3208 | 3361 | 3278 | 3090 | 3100 | 3151 | 3167 | 3191.19 | 3243 | 3139 | 0.967 | -0.048 | 0.413127 | 0.58597045 | 7.00E-04 | 0.0032 | 0.0021 | 0.0021 | TRUE | 0.0205 |
| Kif3b | 2910 | 2071 | 1973 | 2095 | 1929 | 2261 | 2260 | 2203 | 2129 | 2085 | 2176 | 2232 | 2180.87 | 2197 | 2164 | 0.985 | -0.022 | 0.72025246 | 0.83208603 | 6.00E-04 | 0.0035 | 0.0024 | 0.0024 | TRUE | 0.0125 |
| Kif3c | 887 | 723 | 806 | 813 | 738 | 826 | 689 | 769 | 870 | 809 | 833 | 815 | 797.47 | 776 | 819 | 1.058 | 0.081 | 0.40300495 | 0.57589979 | 0.0058 | 0.0054 | 0.0055 | 0.0055 | TRUE | 0.1323 |
| Kif4 | 4346 | 3096 | 3386 | 3646 | 3281 | 3778 | 3375 | 3293 | 3653 | 3629 | 3701 | 3729 | 3563.67 | 3440 | 3686 | 1.072 | 0.1 | 0.09807141 | 0.21824375 | 0.0012 | 0.0031 | 0.0023 | 0.0023 | TRUE | 0.035 |
| Kif4-ps | 7 | 2 | 4 | 4 | 5 | 0 | 5 | 2 | 4 | 4 | 6 | 0 | 3.58 | 4 | 3 | 0.792 | -0.337 | 0.76097999 | NA | 0.3523 | 0.6638 | 0.6014 | 0.6014 | TRUE | 0.8255 |
| Kif5a | 507 | 459 | 573 | 506 | 500 | 613 | 394 | 488 | 618 | 504 | 564 | 605 | 528.85 | 500 | 558 | 1.118 | 0.161 | 0.26521368 | 0.43561108 | 0.0277 | 0.0069 | 0.0132 | 0.0132 | TRUE | 0.3209 |
| Kif5b | 16762 | 11898 | 11500 | 12228 | 10725 | 12479 | 13018 | 12656 | 12408 | 12172 | 12099 | 12319 | 12445.31 | 12694 | 12197 | 0.961 | -0.058 | 0.21554152 | 0.37814348 | 2.00E-04 | 0.0026 | 0.0015 | 0.0015 | TRUE | 0.0062 |
| Kif5c | 1842 | 1373 | 1396 | 1360 | 1205 | 1359 | 1431 | 1460 | 1506 | 1354 | 1359 | 1342 | 1408.66 | 1466 | 1352 | 0.923 | -0.116 | 0.07934649 | 0.18801381 | 0 | 0.0041 | 0.0025 | 0.0025 | TRUE | 0.007 |
| Kif6 | 70 | 61 | 47 | 46 | 49 | 56 | 54 | 65 | 51 | 46 | 55 | 55 | 54.38 | 57 | 52 | 0.92 | -0.12 | 0.65125872 | 0.78347285 | 0 | 0.046 | 0.0323 | 0.0323 | TRUE | 0.1283 |
| Kif7 | 1273 | 932 | 815 | 1100 | 934 | 1107 | 989 | 991 | 879 | 1095 | 1054 | 1093 | 1016.8 | 953 | 1081 | 1.133 | 0.18 | 0.02423297 | 0.07955026 | 0.0014 | 0.0048 | 0.0036 | 0.0036 | TRUE | 0.0531 |
| Kif9 | 136 | 125 | 81 | 108 | 105 | 83 | 106 | 133 | 87 | 108 | 118 | 82 | 105.65 | 109 | 103 | 0.943 | -0.085 | 0.70339562 | 0.82070478 | 0.0291 | 0.0249 | 0.026 | 0.026 | TRUE | 0.2939 |
| Kifap3 | 3839 | 2811 | 2697 | 2721 | 2333 | 2743 | 2982 | 2990 | 2910 | 2708 | 2632 | 2708 | 2821.63 | 2961 | 2683 | 0.906 | -0.142 | 0.01037866 | 0.04242206 | 0 | 0.0033 | 0.0019 | 0.0019 | TRUE | 0.0033 |
| Kifc1 | 6119 | 4415 | 4453 | 4606 | 4224 | 4690 | 4752 | 4696 | 4805 | 4585 | 4765 | 4630 | 4705.49 | 4751 | 4660 | 0.981 | -0.028 | 0.58033991 | 0.73107688 | 0 | 0.0029 | 0.0017 | 0.0017 | TRUE | 0.0047 |
| Kifc2 | 370 | 315 | 303 | 314 | 246 | 334 | 287 | 335 | 327 | 313 | 278 | 330 | 311.52 | 316 | 307 | 0.973 | -0.04 | 0.74991092 | 0.85155134 | 0.0041 | 0.01 | 0.0082 | 0.0082 | TRUE | 0.0793 |
| Kifc3 | 3608 | 2933 | 2879 | 2694 | 2797 | 3319 | 2802 | 3120 | 3106 | 2682 | 3155 | 3276 | 3023.61 | 3009 | 3038 | 1.01 | 0.014 | 0.86514725 | 0.92551298 | 0.0071 | 0.0032 | 0.0046 | 0.0046 | TRUE | 0.1283 |
| Kifc5b | 1764 | 1417 | 1418 | 1567 | 1396 | 1485 | 1370 | 1507 | 1530 | 1560 | 1575 | 1466 | 1501.3 | 1469 | 1534 | 1.045 | 0.063 | 0.39007573 | 0.56385036 | 0.002 | 0.004 | 0.0032 | 0.0032 | TRUE | 0.0436 |
| Kin | 858 | 564 | 463 | 464 | 458 | 512 | 666 | 600 | 500 | 462 | 517 | 505 | 541.64 | 589 | 495 | 0.838 | -0.255 | 0.02963944 | 0.09179557 | 0.0106 | 0.0068 | 0.008 | 0.008 | TRUE | 0.2029 |
| Kir3dl1 | 0 | 0 | 0 | 0 | 0 | 0 | 0 | 0 | 0 | 0 | 0 | 0 | 0 | 0 | 0 | NA | NA | NA | NA | NA | NA | NA | NA | NA | NA |
| Kir3dl2 | 0 | 0 | 0 | 0 | 0 | 0 | 0 | 0 | 0 | 0 | 0 | 0 | 0 | 0 | 0 | NA | NA | NA | NA | NA | NA | NA | NA | NA | NA |
| Kirrel | 2476 | 1594 | 1322 | 1128 | 991 | 1120 | 1923 | 1696 | 1426 | 1123 | 1118 | 1106 | 1398.55 | 1682 | 1116 | 0.662 | -0.594 | 2.31E-09 | 1.40E-07 | 0.0098 | 0.0041 | 0.0064 | 0.0064 | TRUE | 0.1596 |
| Kirrel2 | 300 | 266 | 308 | 507 | 487 | 590 | 233 | 283 | 332 | 505 | 549 | 582 | 414.12 | 283 | 545 | 1.938 | 0.955 | 7.00E-13 | 9.13E-11 | 0.0148 | 0.0081 | 0.0101 | 0.0101 | TRUE | 0.2883 |
| Kirrel3 | 13 | 5 | 8 | 12 | 8 | 13 | 10 | 5 | 9 | 12 | 9 | 13 | 9.64 | 8 | 11 | 1.395 | 0.48 | 0.43735491 | NA | 0 | 0.2482 | 0.17 | 0.17 | TRUE | 0.2383 |
| Kis2 | 2 | 2 | 0 | 0 | 1 | 1 | 2 | 2 | 0 | 0 | 1 | 1 | 0.97 | 1 | 1 | 0.563 | -0.828 | 0.68307268 | NA | 0 | 2.4559 | 1.835 | 1.835 | TRUE | 0.3849 |
| Kiss1 | 3 | 8 | 1 | 7 | 1 | 2 | 2 | 9 | 1 | 7 | 1 | 2 | 3.66 | 4 | 3 | 0.861 | -0.215 | 0.8485852 | NA | 0.6163 | 0.6491 | 0.6389 | 0.6389 | TRUE | 1.5525 |
| Kiss1r | 29 | 27 | 36 | 43 | 30 | 41 | 23 | 29 | 39 | 43 | 34 | 40 | 34.53 | 30 | 39 | 1.316 | 0.396 | 0.24957931 | 0.4184058 | 0.0142 | 0.0711 | 0.0557 | 0.0557 | TRUE | 0.4251 |
| Kit | 8755 | 6781 | 7333 | 7463 | 7046 | 7580 | 6800 | 7213 | 7912 | 7429 | 7949 | 7483 | 7464.11 | 7308 | 7620 | 1.043 | 0.061 | 0.36135848 | 0.53636467 | 0.0032 | 0.0027 | 0.003 | 0.003 | TRUE | 0.0637 |
| Kitl | 549 | 420 | 453 | 388 | 390 | 433 | 426 | 447 | 489 | 386 | 440 | 427 | 435.92 | 454 | 418 | 0.921 | -0.118 | 0.27690346 | 0.4480676 | 0.0025 | 0.0079 | 0.0062 | 0.0062 | TRUE | 0.0515 |
| Kl | 5 | 8 | 10 | 5 | 0 | 3 | 4 | 9 | 11 | 5 | 0 | 3 | 5.19 | 8 | 3 | 0.354 | -1.5 | 0.12114144 | NA | 0.2558 | 0.4594 | 0.4196 | 0.4196 | TRUE | 0.5573 |
| Klb | 43 | 27 | 20 | 35 | 22 | 24 | 33 | 29 | 22 | 35 | 25 | 24 | 27.84 | 28 | 28 | 0.99 | -0.014 | 0.9697974 | 0.98349044 | 0 | 0.0876 | 0.0669 | 0.0669 | TRUE | 0.3236 |
| Klc1 | 2254 | 1854 | 1759 | 1675 | 1535 | 1879 | 1751 | 1972 | 1898 | 1667 | 1732 | 1855 | 1812.4 | 1874 | 1751 | 0.936 | -0.096 | 0.1949267 | 0.35428374 | 0.0028 | 0.0037 | 0.0034 | 0.0034 | TRUE | 0.0411 |
| Klc2 | 2592 | 2061 | 2137 | 2275 | 2125 | 2476 | 2013 | 2192 | 2306 | 2265 | 2397 | 2444 | 2269.52 | 2170 | 2369 | 1.092 | 0.127 | 0.07267774 | 0.17639378 | 0.0024 | 0.0035 | 0.0032 | 0.0032 | TRUE | 0.0501 |
| Klc3 | 562 | 516 | 591 | 684 | 630 | 663 | 436 | 549 | 638 | 681 | 711 | 654 | 611.51 | 541 | 682 | 1.264 | 0.338 | 0.00683152 | 0.03118585 | 0.017 | 0.0063 | 0.0096 | 0.0096 | TRUE | 0.3393 |
| Klc4 | 1482 | 1104 | 1155 | 1212 | 1097 | 1295 | 1151 | 1174 | 1246 | 1206 | 1238 | 1278 | 1215.65 | 1190 | 1241 | 1.043 | 0.061 | 0.40173462 | 0.5748316 | 5.00E-04 | 0.0044 | 0.003 | 0.003 | TRUE | 0.0198 |
| Klf1 | 32 | 20 | 19 | 21 | 15 | 12 | 25 | 21 | 21 | 21 | 17 | 12 | 19.38 | 22 | 17 | 0.742 | -0.431 | 0.33096713 | 0.50534272 | 0 | 0.1247 | 0.0885 | 0.0885 | TRUE | 0.3157 |
| Klf10 | 5483 | 4431 | 5244 | 5855 | 5033 | 5954 | 4258 | 4713 | 5658 | 5828 | 5678 | 5878 | 5335.53 | 4876 | 5795 | 1.189 | 0.25 | 0.00481637 | 0.02382845 | 0.0106 | 0.0029 | 0.0055 | 0.0055 | TRUE | 0.2393 |
| Klf11 | 2857 | 2039 | 2015 | 2259 | 1950 | 2189 | 2219 | 2169 | 2174 | 2249 | 2200 | 2161 | 2195.2 | 2187 | 2203 | 1.007 | 0.01 | 0.86497537 | 0.92551298 | 0 | 0.0035 | 0.002 | 0.002 | TRUE | 0.004 |
| Klf12 | 950 | 680 | 614 | 630 | 651 | 646 | 738 | 723 | 663 | 627 | 734 | 638 | 687.14 | 708 | 666 | 0.94 | -0.09 | 0.3499423 | 0.52455531 | 0.0038 | 0.0059 | 0.0052 | 0.0052 | TRUE | 0.0919 |
| Klf13 | 3376 | 2532 | 1905 | 2248 | 2243 | 2031 | 2622 | 2693 | 2055 | 2238 | 2530 | 2005 | 2357.27 | 2457 | 2258 | 0.918 | -0.123 | 0.24752377 | 0.41631645 | 0.0155 | 0.0034 | 0.0077 | 0.0077 | TRUE | 0.2458 |
| Klf14 | 25 | 15 | 9 | 16 | 15 | 19 | 19 | 16 | 10 | 16 | 17 | 19 | 16.11 | 15 | 17 | 1.132 | 0.179 | 0.71328284 | NA | 0 | 0.1495 | 0.1083 | 0.1083 | TRUE | 0.4057 |
| Klf15 | 882 | 808 | 953 | 1042 | 1038 | 1070 | 685 | 859 | 1028 | 1037 | 1171 | 1056 | 972.86 | 857 | 1088 | 1.271 | 0.346 | 0.00507026 | 0.02477916 | 0.0207 | 0.0049 | 0.01 | 0.01 | TRUE | 0.2538 |
| Klf16 | 3103 | 2444 | 2349 | 2179 | 2102 | 2324 | 2410 | 2600 | 2535 | 2169 | 2371 | 2294 | 2396.43 | 2515 | 2278 | 0.906 | -0.142 | 0.02821633 | 0.08853919 | 0.0013 | 0.0034 | 0.0026 | 0.0026 | TRUE | 0.0213 |
| Klf17 | 290 | 245 | 204 | 192 | 151 | 182 | 225 | 261 | 220 | 191 | 170 | 180 | 207.85 | 235 | 180 | 0.768 | -0.381 | 0.00856844 | 0.03689928 | 0 | 0.0138 | 0.0102 | 0.0102 | TRUE | 0.0942 |
| Klf2 | 12469 | 9791 | 8795 | 9984 | 9863 | 10475 | 9684 | 10415 | 9490 | 9938 | 11126 | 10340 | 10165.57 | 9863 | 10468 | 1.061 | 0.086 | 0.16760188 | 0.31906415 | 0.0027 | 0.0027 | 0.0027 | 0.0027 | TRUE | 0.0368 |
| Klf3 | 5268 | 3972 | 4473 | 4719 | 4494 | 4902 | 4091 | 4225 | 4826 | 4697 | 5070 | 4839 | 4624.79 | 4381 | 4869 | 1.112 | 0.153 | 0.03326215 | 0.10000334 | 0.0044 | 0.0029 | 0.0035 | 0.0035 | TRUE | 0.0959 |
| Klf4 | 26157 | 20226 | 18311 | 18869 | 17013 | 19777 | 20315 | 21514 | 19757 | 18782 | 19192 | 19523 | 19847.36 | 20529 | 19166 | 0.934 | -0.099 | 0.05563446 | 0.14545151 | 0.0011 | 0.0025 | 0.0019 | 0.0019 | TRUE | 0.0215 |
| Klf5 | 5279 | 4250 | 3706 | 4015 | 3628 | 4187 | 4100 | 4521 | 3999 | 3997 | 4093 | 4133 | 4140.32 | 4207 | 4074 | 0.969 | -0.046 | 0.46564895 | 0.63463117 | 0.002 | 0.003 | 0.0026 | 0.0026 | TRUE | 0.0515 |
| Klf6 | 3326 | 2199 | 2216 | 2498 | 2103 | 2266 | 2583 | 2339 | 2391 | 2487 | 2372 | 2237 | 2401.51 | 2438 | 2365 | 0.97 | -0.045 | 0.51878221 | 0.68026262 | 0.0024 | 0.0034 | 0.003 | 0.003 | TRUE | 0.0338 |
| Klf7 | 344 | 254 | 249 | 253 | 220 | 252 | 267 | 270 | 269 | 252 | 248 | 249 | 259.13 | 269 | 250 | 0.929 | -0.106 | 0.39121942 | 0.56492733 | 0 | 0.0116 | 0.0071 | 0.0071 | TRUE | 7.00E-04 |
| Klf8 | 1950 | 1378 | 1410 | 1525 | 1327 | 1502 | 1514 | 1466 | 1521 | 1518 | 1497 | 1483 | 1499.88 | 1500 | 1499 | 0.999 | -0.002 | 0.98123638 | 0.99018074 | 0 | 0.004 | 0.0023 | 0.0023 | TRUE | 0.0048 |
| Klf9 | 10485 | 7604 | 6773 | 7033 | 6343 | 6658 | 8143 | 8088 | 7308 | 7001 | 7156 | 6572 | 7378.04 | 7846 | 6910 | 0.88 | -0.184 | 0.00341178 | 0.0182954 | 0.0025 | 0.0027 | 0.0027 | 0.0027 | TRUE | 0.0438 |
| Klhdc1 | 108 | 80 | 95 | 155 | 111 | 129 | 84 | 85 | 103 | 154 | 125 | 127 | 113.05 | 91 | 135 | 1.505 | 0.589 | 0.00245131 | 0.01414475 | 0 | 0.0234 | 0.0181 | 0.0181 | TRUE | 0.1514 |
| Klhdc10 | 8969 | 6102 | 5423 | 6426 | 5450 | 6257 | 6966 | 6491 | 5851 | 6396 | 6148 | 6177 | 6338.18 | 6436 | 6240 | 0.969 | -0.045 | 0.51174283 | 0.67469683 | 0.0034 | 0.0028 | 0.0032 | 0.0032 | TRUE | 0.0769 |
| Klhdc2 | 2939 | 2163 | 2401 | 3055 | 2644 | 2926 | 2283 | 2301 | 2591 | 3041 | 2983 | 2888 | 2681.01 | 2392 | 2971 | 1.243 | 0.314 | 4.29E-06 | 9.35E-05 | 0.0024 | 0.0033 | 0.003 | 0.003 | TRUE | 0.0637 |
| Klhdc3 | 6959 | 5090 | 5140 | 5603 | 5438 | 6075 | 5405 | 5414 | 5546 | 5577 | 6135 | 5997 | 5678.96 | 5455 | 5903 | 1.082 | 0.114 | 0.04395209 | 0.12255312 | 0.0011 | 0.0028 | 0.0021 | 0.0021 | TRUE | 0.0285 |
| Klhdc4 | 2686 | 2005 | 1631 | 1613 | 1315 | 1554 | 2086 | 2133 | 1760 | 1606 | 1483 | 1534 | 1766.95 | 1993 | 1541 | 0.773 | -0.371 | 8.63E-06 | 0.00016598 | 0.0051 | 0.0038 | 0.0044 | 0.0044 | TRUE | 0.1247 |
| Klhdc5 | 326 | 299 | 218 | 258 | 209 | 248 | 253 | 318 | 235 | 257 | 236 | 245 | 257.31 | 269 | 246 | 0.916 | -0.127 | 0.37978315 | 0.55435463 | 0.0097 | 0.0116 | 0.0111 | 0.0111 | TRUE | 0.2773 |
| Klhdc7a | 109 | 105 | 131 | 174 | 174 | 165 | 85 | 112 | 141 | 173 | 196 | 163 | 145.01 | 113 | 177 | 1.587 | 0.666 | 0.00083634 | 0.00634877 | 0.0283 | 0.0188 | 0.0214 | 0.0214 | TRUE | 0.3492 |

| | | | | | | | | | | | | | | | | | | | | | | | | | |
|---|---|---|---|---|---|---|---|---|---|---|---|---|---|---|---|---|---|---|---|---|---|---|---|---|---|
| Klhdc7b | 1 | 2 | 2 | 4 | 1 | 1 | 1 | 2 | 2 | 4 | 1 | 1 | 1.86 | 2 | 2 | 1.241 | 0.311 | 0.82436628 | NA | 0 | 1.2768 | 0.8697 | 0.8697 | TRUE | 0.6569 |
| Klhdc8a | 22 | 7 | 18 | 15 | 14 | 11 | 17 | 7 | 19 | 15 | 16 | 11 | 14.26 | 14 | 14 | 0.939 | -0.09 | 0.866146 | NA | 0.0506 | 0.1687 | 0.1373 | 0.1373 | TRUE | 0.7663 |
| Klhdc8b | 277 | 188 | 188 | 211 | 174 | 204 | 215 | 200 | 203 | 210 | 196 | 201 | 204.28 | 206 | 202 | 0.982 | -0.026 | 0.85299941 | 0.91687758 | 0 | 0.014 | 0.009 | 0.009 | TRUE | 0.0181 |
| Klhdc9 | 60 | 49 | 75 | 83 | 88 | 109 | 47 | 52 | 81 | 83 | 99 | 108 | 78.19 | 60 | 97 | 1.624 | 0.7 | 0.0063326 | 0.02934103 | 0.0358 | 0.0327 | 0.0338 | 0.0338 | TRUE | 0.7862 |
| Klhl1 | 1 | 1 | 0 | 0 | 0 | 0 | 1 | 1 | 0 | 0 | 0 | 0 | 0.31 | 1 | 0 | 0.309 | -1.696 | 0.63183309 | NA | 0 | 7.7294 | 7.2472 | 7.2472 | TRUE | 0.2015 |
| Klhl10 | 81 | 66 | 78 | 82 | 63 | 74 | 63 | 70 | 84 | 82 | 71 | 73 | 73.84 | 72 | 75 | 1.045 | 0.063 | 0.78431772 | 0.87314067 | 0 | 0.0345 | 0.0252 | 0.0252 | TRUE | 0.1777 |
| Klhl11 | 2317 | 1679 | 1647 | 1763 | 1604 | 1855 | 1800 | 1786 | 1777 | 1755 | 1809 | 1831 | 1793.01 | 1788 | 1798 | 1.006 | 0.009 | 0.88926309 | 0.93844044 | 0 | 0.0037 | 0.0022 | 0.0022 | TRUE | 0.0055 |
| Klhl12 | 3902 | 2846 | 2717 | 3099 | 2667 | 3146 | 3031 | 3027 | 2932 | 3085 | 3009 | 3106 | 3031.4 | 2997 | 3067 | 1.023 | 0.033 | 0.54697845 | 0.70406727 | 0 | 0.0032 | 0.0019 | 0.0019 | TRUE | 0.0043 |
| Klhl13 | 6847 | 5160 | 5150 | 2764 | 2481 | 2599 | 5318 | 5489 | 5557 | 2751 | 2799 | 2566 | 4079.82 | 5455 | 2705 | 0.496 | -1.012 | 1.88E-67 | 1.34E-63 | 9.00E-04 | 0.003 | 0.0022 | 0.0022 | TRUE | 0.0249 |
| Klhl14 | 11 | 1 | 4 | 3 | 8 | 8 | 9 | 1 | 4 | 3 | 9 | 8 | 5.64 | 5 | 7 | 1.383 | 0.468 | 0.59921523 | NA | 0.3068 | 0.4228 | 0.392 | 0.392 | TRUE | 0.4043 |
| Klhl15 | 386 | 287 | 337 | 365 | 280 | 331 | 300 | 305 | 364 | 363 | 316 | 327 | 329.1 | 323 | 335 | 1.041 | 0.058 | 0.64249427 | 0.77724677 | 0.0054 | 0.0096 | 0.0084 | 0.0084 | TRUE | 0.1346 |
| Klhl17 | 491 | 449 | 376 | 300 | 284 | 345 | 381 | 478 | 406 | 299 | 320 | 341 | 370.7 | 422 | 320 | 0.76 | -0.395 | 0.00136073 | 0.00899579 | 0.0069 | 0.0088 | 0.0082 | 0.0082 | TRUE | 0.1548 |
| Klhl18 | 616 | 416 | 326 | 375 | 310 | 361 | 478 | 442 | 352 | 373 | 350 | 356 | 392 | 424 | 360 | 0.846 | -0.241 | 0.05913594 | 0.15180136 | 0.0108 | 0.0085 | 0.0092 | 0.0092 | TRUE | 0.2527 |
| Klhl2 | 3407 | 2482 | 2636 | 2696 | 2409 | 2791 | 2646 | 2640 | 2844 | 2684 | 2718 | 2755 | 2714.45 | 2710 | 2719 | 1.004 | 0.005 | 0.93159347 | 0.96216772 | 6.00E-04 | 0.0033 | 0.0022 | 0.0022 | TRUE | 0.0224 |
| Klhl20 | 1955 | 1565 | 1368 | 1500 | 1370 | 1507 | 1518 | 1665 | 1476 | 1493 | 1546 | 1488 | 1530.89 | 1553 | 1509 | 0.972 | -0.042 | 0.56387391 | 0.71824702 | 0.0015 | 0.004 | 0.0031 | 0.0031 | TRUE | 0.0466 |
| Klhl21 | 2108 | 1541 | 1662 | 1506 | 1531 | 1650 | 1637 | 1639 | 1793 | 1499 | 1727 | 1629 | 1654.1 | 1690 | 1618 | 0.958 | -0.062 | 0.4164786 | 0.58932072 | 0.003 | 0.0039 | 0.0036 | 0.0036 | TRUE | 0.0502 |
| Klhl22 | 3439 | 2334 | 2004 | 2004 | 1807 | 1963 | 2671 | 2483 | 2162 | 1995 | 2038 | 1938 | 2214.49 | 2439 | 1990 | 0.815 | -0.294 | 0.00028147 | 0.00276308 | 0.0052 | 0.0035 | 0.0043 | 0.0043 | TRUE | 0.1184 |
| Klhl23 | 454 | 305 | 248 | 344 | 268 | 320 | 353 | 324 | 268 | 342 | 302 | 316 | 317.54 | 315 | 320 | 1.014 | 0.021 | 0.87661044 | 0.93106882 | 0.0087 | 0.0099 | 0.0095 | 0.0095 | TRUE | 0.191 |
| Klhl24 | 4392 | 3588 | 4463 | 4777 | 4566 | 5300 | 3411 | 3817 | 4816 | 4755 | 5151 | 5232 | 4530.16 | 4015 | 5046 | 1.258 | 0.331 | 0.00123953 | 0.00844449 | 0.0172 | 0.0029 | 0.0073 | 0.0073 | TRUE | 0.3706 |
| Klhl25 | 870 | 668 | 648 | 648 | 593 | 736 | 676 | 711 | 699 | 645 | 669 | 727 | 687.66 | 695 | 680 | 0.979 | -0.03 | 0.73442765 | 0.8412894 | 7.00E-04 | 0.0059 | 0.0042 | 0.0042 | TRUE | 0.0428 |
| Klhl26 | 2201 | 1628 | 1886 | 1897 | 1591 | 1754 | 1709 | 1732 | 2035 | 1888 | 1795 | 1731 | 1815.11 | 1825 | 1805 | 0.99 | -0.015 | 0.8527263 | 0.9167369 | 0.0052 | 0.0037 | 0.0043 | 0.0043 | TRUE | 0.1207 |
| Klhl28 | 656 | 475 | 493 | 537 | 462 | 567 | 509 | 505 | 532 | 535 | 521 | 560 | 527.02 | 515 | 539 | 1.045 | 0.064 | 0.49545008 | 0.6600921 | 0 | 0.0069 | 0.0044 | 0.0044 | TRUE | 0.014 |
| Klhl29 | 170 | 109 | 106 | 64 | 55 | 64 | 132 | 116 | 114 | 64 | 62 | 63 | 91.88 | 121 | 63 | 0.52 | -0.943 | 5.65E-06 | 0.00011729 | 0 | 0.0282 | 0.0189 | 0.0189 | TRUE | 0.074 |
| Klhl3 | 199 | 153 | 130 | 114 | 112 | 125 | 155 | 163 | 140 | 113 | 126 | 123 | 136.8 | 153 | 121 | 0.793 | -0.335 | 0.04902268 | 0.1327802 | 0 | 0.0198 | 0.0134 | 0.0134 | TRUE | 0.0485 |
| Klhl30 | 18 | 3 | 11 | 12 | 9 | 10 | 14 | 3 | 12 | 12 | 10 | 10 | 10.17 | 10 | 11 | 1.086 | 0.119 | 0.85409545 | NA | 0.1018 | 0.2355 | 0.2025 | 0.2025 | TRUE | 1.0807 |
| Klhl31 | 3 | 2 | 4 | 9 | 4 | 2 | 2 | 2 | 4 | 9 | 5 | 2 | 4.04 | 3 | 5 | 1.78 | 0.832 | 0.40454169 | NA | 0 | 0.5896 | 0.4509 | 0.4509 | TRUE | 0.2784 |
| Klhl32 | 43 | 38 | 35 | 25 | 29 | 29 | 33 | 40 | 38 | 25 | 33 | 29 | 32.97 | 37 | 29 | 0.774 | -0.37 | 0.2699583 | 0.44092264 | 0 | 0.0743 | 0.0501 | 0.0501 | TRUE | 0.0905 |
| Klhl33 | 8 | 4 | 4 | 9 | 4 | 7 | 6 | 4 | 4 | 9 | 5 | 7 | 5.86 | 5 | 7 | 1.37 | 0.454 | 0.56289497 | NA | 0 | 0.4068 | 0.2712 | 0.2712 | TRUE | 0.1673 |
| Klhl34 | 39 | 39 | 18 | 30 | 13 | 23 | 30 | 41 | 19 | 30 | 15 | 23 | 26.4 | 30 | 23 | 0.741 | -0.433 | 0.30593101 | 0.47853155 | 0.0847 | 0.0922 | 0.0902 | 0.0902 | TRUE | 0.138 |
| Klhl35 | 3 | 2 | 0 | 3 | 2 | 2 | 2 | 2 | 0 | 3 | 2 | 2 | 1.95 | 1 | 2 | 1.577 | 0.658 | 0.63508714 | NA | 0 | 1.2205 | 0.8526 | 0.8526 | TRUE | 0.4555 |
| Klhl36 | 852 | 608 | 525 | 576 | 645 | 651 | 662 | 647 | 566 | 573 | 728 | 643 | 636.42 | 625 | 648 | 1.034 | 0.049 | 0.65451273 | 0.78533835 | 0.0089 | 0.0062 | 0.007 | 0.007 | TRUE | 0.1341 |
| Klhl38 | 2 | 1 | 0 | 0 | 0 | 0 | 2 | 1 | 0 | 0 | 0 | 0 | 0.44 | 1 | 0 | 0.215 | -2.217 | 0.47268385 | NA | 0 | 5.4363 | 5.3181 | 5.3181 | TRUE | 0.2411 |
| Klhl4 | 1 | 1 | 1 | 1 | 2 | 0 | 1 | 1 | 1 | 1 | 2 | 0 | 1.03 | 1 | 1 | 1.102 | 0.141 | 0.94001409 | NA | 0 | 2.3068 | 1.5256 | 1.5256 | TRUE | 0.2204 |
| Klhl40 | 7 | 5 | 4 | 2 | 1 | 0 | 5 | 5 | 4 | 2 | 1 | 0 | 3.03 | 5 | 1 | 0.206 | -2.282 | 0.07082482 | NA | 0 | 0.7842 | 0.5542 | 0.5542 | TRUE | 0.4009 |
| Klhl41 | 5 | 10 | 4 | 7 | 6 | 8 | 4 | 11 | 4 | 7 | 7 | 8 | 6.74 | 6 | 7 | 1.166 | 0.222 | 0.76846873 | NA | 0 | 0.3538 | 0.2596 | 0.2596 | TRUE | 0.8486 |
| Klhl5 | 1124 | 714 | 689 | 748 | 695 | 723 | 873 | 759 | 743 | 745 | 784 | 714 | 769.69 | 792 | 748 | 0.941 | -0.087 | 0.35021184 | 0.52480371 | 0.004 | 0.0055 | 0.005 | 0.005 | TRUE | 0.0992 |
| Klhl6 | 1 | 0 | 0 | 0 | 0 | 2 | 1 | 0 | 0 | 0 | 0 | 2 | 0.46 | 0 | 1 | 1.93 | 0.948 | 0.7608301 | NA | 0 | 5.1717 | 5.216 | 5.216 | TRUE | 0.9737 |
| Klhl7 | 2137 | 1559 | 1691 | 1850 | 1641 | 1850 | 1660 | 1658 | 1825 | 1841 | 1851 | 1826 | 1776.92 | 1714 | 1839 | 1.074 | 0.103 | 0.12688162 | 0.26237254 | 9.00E-04 | 0.0038 | 0.0027 | 0.0027 | TRUE | 0.0376 |
| Klhl8 | 1792 | 1290 | 1377 | 1341 | 1150 | 1470 | 1392 | 1372 | 1486 | 1335 | 1297 | 1451 | 1388.83 | 1417 | 1361 | 0.962 | -0.057 | 0.44959066 | 0.61967207 | 0.0018 | 0.0041 | 0.0033 | 0.0033 | TRUE | 0.0405 |
| Klhl9 | 2884 | 2130 | 1828 | 1716 | 1510 | 1721 | 2240 | 2266 | 1972 | 1708 | 1703 | 1699 | 1931.39 | 2159 | 1703 | 0.788 | -0.343 | 1.92E-06 | 4.95E-05 | 0.0025 | 0.0037 | 0.0032 | 0.0032 | TRUE | 0.0683 |
| Klk1 | 13 | 12 | 28 | 29 | 37 | 42 | 10 | 13 | 30 | 29 | 42 | 41 | 27.52 | 18 | 37 | 2.146 | 1.102 | 0.01292108 | 0.04996189 | 0.1336 | 0.0885 | 0.0999 | 0.0999 | TRUE | 1.8425 |
| Klk10 | 297 | 244 | 246 | 320 | 305 | 286 | 231 | 260 | 265 | 319 | 344 | 282 | 283.43 | 252 | 315 | 1.252 | 0.325 | 0.01347124 | 0.05162047 | 0.0045 | 0.0108 | 0.0089 | 0.0089 | TRUE | 0.094 |
| Klk11 | 61 | 59 | 71 | 81 | 66 | 90 | 47 | 63 | 77 | 81 | 74 | 89 | 71.78 | 62 | 81 | 1.319 | 0.4 | 0.10889162 | 0.2354596 | 0.0178 | 0.0354 | 0.0306 | 0.0306 | TRUE | 0.3121 |
| Klk12 | 2 | 3 | 0 | 1 | 1 | 2 | 2 | 3 | 0 | 1 | 1 | 2 | 1.47 | 2 | 1 | 0.866 | -0.208 | 0.8963711 | NA | 0 | 1.6107 | 1.156 | 1.156 | TRUE | 0.16 |
| Klk13 | 105 | 115 | 122 | 177 | 179 | 240 | 82 | 122 | 132 | 176 | 202 | 237 | 158.42 | 112 | 205 | 1.848 | 0.886 | 1.04E-05 | 0.00019326 | 0.0347 | 0.0174 | 0.0222 | 0.0222 | TRUE | 0.5421 |
| Klk14 | 12 | 22 | 11 | 22 | 21 | 20 | 9 | 23 | 12 | 22 | 24 | 20 | 18.32 | 15 | 22 | 1.484 | 0.57 | 0.23858324 | 0.40547804 | 0.0549 | 0.1318 | 0.1116 | 0.1116 | TRUE | 1.102 |
| Klk15 | 12 | 11 | 8 | 10 | 3 | 5 | 9 | 12 | 9 | 10 | 3 | 5 | 7.99 | 10 | 6 | 0.622 | -0.685 | 0.32800561 | NA | 0 | 0.2991 | 0.2199 | 0.2199 | TRUE | 0.741 |
| Klk1b1 | 0 | 0 | 0 | 0 | 0 | 0 | 0 | 0 | 0 | 0 | 0 | 0 | 0 | 0 | 0 | NA | NA | NA | NA | NA | NA | NA | NA | NA | NA |
| Klk1b11 | 0 | 0 | 0 | 0 | 0 | 0 | 0 | 0 | 0 | 0 | 0 | 0 | 0 | 0 | 0 | NA | NA | NA | NA | NA | NA | NA | NA | NA | NA |
| Klk1b16 | 0 | 0 | 0 | 0 | 0 | 0 | 0 | 0 | 0 | 0 | 0 | 0 | 0 | 0 | 0 | NA | NA | NA | NA | NA | NA | NA | NA | NA | NA |
| Klk1b21 | 0 | 0 | 0 | 0 | 0 | 1 | 0 | 0 | 0 | 0 | 0 | 1 | 0.16 | 0 | 0 | 2.094 | 1.067 | 0.79379329 | NA | 0 | 14.4078 | 10 | 10 | TRUE | 0.3835 |
| Klk1b22 | 0 | 0 | 0 | 0 | 0 | 1 | 0 | 0 | 0 | 0 | 0 | 1 | 0.16 | 0 | 0 | 2.094 | 1.067 | 0.79379329 | NA | 0 | 14.4078 | 10 | 10 | TRUE | 0.3835 |
| Klk1b24 | 0 | 0 | 0 | 0 | 0 | 0 | 0 | 0 | 0 | 0 | 0 | 0 | 0 | 0 | 0 | NA | NA | NA | NA | NA | NA | NA | NA | NA | NA |
| Klk1b26 | 0 | 0 | 0 | 0 | 0 | 0 | 0 | 0 | 0 | 0 | 0 | 0 | 0 | 0 | 0 | NA | NA | NA | NA | NA | NA | NA | NA | NA | NA |
| Klk1b27 | 0 | 0 | 0 | 0 | 0 | 0 | 0 | 0 | 0 | 0 | 0 | 0 | 0 | 0 | 0 | NA | NA | NA | NA | NA | NA | NA | NA | NA | NA |
| Klk1b3 | 0 | 0 | 0 | 0 | 0 | 0 | 0 | 0 | 0 | 0 | 0 | 0 | 0 | 0 | 0 | NA | NA | NA | NA | NA | NA | NA | NA | NA | NA |
| Klk1b4 | 0 | 0 | 0 | 0 | 0 | 0 | 0 | 0 | 0 | 0 | 0 | 0 | 0 | 0 | 0 | NA | NA | NA | NA | NA | NA | NA | NA | NA | NA |
| Klk1b5 | 0 | 1 | 2 | 0 | 1 | 7 | 0 | 1 | 2 | 0 | 1 | 7 | 1.88 | 1 | 3 | 2.624 | 1.392 | 0.41129161 | NA | 1.8889 | 1.2654 | 1.4064 | 1.4064 | TRUE | 1.6394 |
| Klk1b7-ps | 0 | 0 | 0 | 0 | 0 | 0 | 0 | 0 | 0 | 0 | 0 | 0 | 0 | 0 | 0 | NA | NA | NA | NA | NA | NA | NA | NA | NA | NA |
| Klk1b8 | 0 | 0 | 0 | 0 | 0 | 0 | 0 | 0 | 0 | 0 | 0 | 0 | 0 | 0 | 0 | NA | NA | NA | NA | NA | NA | NA | NA | NA | NA |
| Klk1b9 | 2 | 1 | 1 | 0 | 0 | 1 | 2 | 1 | 1 | 0 | 0 | 1 | 0.78 | 1 | 0 | 0.299 | -1.743 | 0.4255496 | NA | 0 | 3.0389 | 2.0589 | 2.0589 | TRUE | 0.3835 |
| Klk4 | 9 | 9 | 4 | 7 | 3 | 13 | 7 | 10 | 4 | 7 | 3 | 13 | 7.34 | 7 | 8 | 1.119 | 0.162 | 0.82874218 | NA | 0.12 | 0.3251 | 0.2689 | 0.2689 | TRUE | 0.3398 |
| Klk5 | 13 | 10 | 4 | 14 | 15 | 8 | 10 | 11 | 4 | 14 | 17 | 8 | 10.63 | 8 | 13 | 1.526 | 0.61 | 0.32683704 | NA | 0.0506 | 0.2253 | 0.1814 | 0.1814 | TRUE | 0.4195 |
| Klk6 | 3 | 1 | 0 | 3 | 3 | 3 | 2 | 1 | 0 | 3 | 3 | 3 | 2.12 | 1 | 3 | 2.595 | 1.376 | 0.31499094 | NA | 0 | 1.1199 | 0.7821 | 0.7821 | TRUE | 0.6022 |
| Klk7 | 17 | 10 | 17 | 17 | 14 | 21 | 13 | 11 | 18 | 17 | 16 | 21 | 15.94 | 14 | 18 | 1.273 | 0.349 | 0.47164211 | NA | 0 | 0.1511 | 0.1055 | 0.1055 | TRUE | 0.279 |
| Klk8 | 1028 | 766 | 884 | 1028 | 947 | 1127 | 798 | 815 | 954 | 1023 | 1068 | 1113 | 961.85 | 856 | 1068 | 1.25 | 0.322 | 0.00034008 | 0.00319364 | 0.0043 | 0.0049 | 0.0048 | 0.0048 | TRUE | 0.1177 |
| Klk9 | 1 | 0 | 2 | 2 | 0 | 5 | 1 | 0 | 2 | 2 | 0 | 5 | 1.64 | 1 | 2 | 2.415 | 1.272 | 0.45128121 | NA | 0.912 | 1.4445 | 1.3346 | 1.3346 | TRUE | 0.1697 |
| Klkb1 | 0 | 0 | 0 | 0 | 0 | 0 | 0 | 0 | 0 | 0 | 0 | 0 | 0 | 0 | 0 | NA | NA | NA | NA | NA | NA | NA | NA | NA | NA |
| Klra1 | 0 | 0 | 0 | 0 | 0 | 0 | 0 | 0 | 0 | 0 | 0 | 0 | 0 | 0 | 0 | NA | NA | NA | NA | NA | NA | NA | NA | NA | NA |

| | | | | | | | | | | | | | | | | | | | | | | | | | |
|---|---|---|---|---|---|---|---|---|---|---|---|---|---|---|---|---|---|---|---|---|---|---|---|---|---|
| Klra10 | 0 | 0 | 0 | 0 | 0 | 0 | 0 | 0 | 0 | 0 | 0 | 0 | 0 | 0 | 0 | NA | NA | NA | NA | NA | NA | NA | NA | NA | NA |
| Klra12 | 0 | 0 | 0 | 0 | 0 | 0 | 0 | 0 | 0 | 0 | 0 | 0 | 0 | 0 | 0 | NA | NA | NA | NA | NA | NA | NA | NA | NA | NA |
| Klra13-ps | 0 | 0 | 0 | 0 | 0 | 0 | 0 | 0 | 0 | 0 | 0 | 0 | 0 | 0 | 0 | NA | NA | NA | NA | NA | NA | NA | NA | NA | NA |
| Klra14 | 0 | 0 | 0 | 0 | 0 | 0 | 0 | 0 | 0 | 0 | 0 | 0 | 0 | 0 | 0 | NA | NA | NA | NA | NA | NA | NA | NA | NA | NA |
| Klra15 | 0 | 0 | 0 | 0 | 0 | 0 | 0 | 0 | 0 | 0 | 0 | 0 | 0 | 0 | 0 | NA | NA | NA | NA | NA | NA | NA | NA | NA | NA |
| Klra17 | 0 | 0 | 1 | 0 | 0 | 0 | 0 | 0 | 1 | 0 | 0 | 0 | 0.18 | 0 | 0 | 0.552 | -0.857 | 0.8336548 | NA | 0 | 13.1817 | 10 | 10 | TRUE | 0.5365 |
| Klra18 | 0 | 0 | 0 | 0 | 0 | 0 | 0 | 0 | 0 | 0 | 0 | 0 | 0 | 0 | 0 | NA | NA | NA | NA | NA | NA | NA | NA | NA | NA |
| Klra19 | 0 | 0 | 0 | 0 | 0 | 0 | 0 | 0 | 0 | 0 | 0 | 0 | 0 | 0 | 0 | NA | NA | NA | NA | NA | NA | NA | NA | NA | NA |
| Klra2 | 0 | 0 | 0 | 0 | 0 | 0 | 0 | 0 | 0 | 0 | 0 | 0 | 0 | 0 | 0 | NA | NA | NA | NA | NA | NA | NA | NA | NA | NA |
| Klra21 | 0 | 0 | 0 | 0 | 0 | 0 | 0 | 0 | 0 | 0 | 0 | 0 | 0 | 0 | 0 | NA | NA | NA | NA | NA | NA | NA | NA | NA | NA |
| Klra22 | 0 | 0 | 0 | 0 | 0 | 0 | 0 | 0 | 0 | 0 | 0 | 0 | 0 | 0 | 0 | NA | NA | NA | NA | NA | NA | NA | NA | NA | NA |
| Klra23 | 0 | 0 | 0 | 0 | 0 | 0 | 0 | 0 | 0 | 0 | 0 | 0 | 0 | 0 | 0 | NA | NA | NA | NA | NA | NA | NA | NA | NA | NA |
| Klra3 | 0 | 0 | 0 | 0 | 0 | 0 | 0 | 0 | 0 | 0 | 0 | 0 | 0 | 0 | 0 | NA | NA | NA | NA | NA | NA | NA | NA | NA | NA |
| Klra33 | 0 | 0 | 0 | 0 | 0 | 0 | 0 | 0 | 0 | 0 | 0 | 0 | 0 | 0 | 0 | NA | NA | NA | NA | NA | NA | NA | NA | NA | NA |
| Klra4 | 0 | 0 | 0 | 0 | 0 | 0 | 0 | 0 | 0 | 0 | 0 | 0 | 0 | 0 | 0 | NA | NA | NA | NA | NA | NA | NA | NA | NA | NA |
| Klra5 | 0 | 0 | 0 | 0 | 0 | 0 | 0 | 0 | 0 | 0 | 0 | 0 | 0 | 0 | 0 | NA | NA | NA | NA | NA | NA | NA | NA | NA | NA |
| Klra6 | 0 | 0 | 0 | 0 | 0 | 0 | 0 | 0 | 0 | 0 | 0 | 0 | 0 | 0 | 0 | NA | NA | NA | NA | NA | NA | NA | NA | NA | NA |
| Klra7 | 0 | 0 | 0 | 0 | 0 | 0 | 0 | 0 | 0 | 0 | 0 | 0 | 0 | 0 | 0 | NA | NA | NA | NA | NA | NA | NA | NA | NA | NA |
| Klra8 | 0 | 0 | 0 | 0 | 0 | 0 | 0 | 0 | 0 | 0 | 0 | 0 | 0 | 0 | 0 | NA | NA | NA | NA | NA | NA | NA | NA | NA | NA |
| Klra9 | 0 | 0 | 0 | 0 | 0 | 0 | 0 | 0 | 0 | 0 | 0 | 0 | 0 | 0 | 0 | NA | NA | NA | NA | NA | NA | NA | NA | NA | NA |
| Klrb1 | 0 | 0 | 0 | 0 | 0 | 0 | 0 | 0 | 0 | 0 | 0 | 0 | 0 | 0 | 0 | NA | NA | NA | NA | NA | NA | NA | NA | NA | NA |
| Klrb1-ps1 | 0 | 0 | 0 | 0 | 0 | 0 | 0 | 0 | 0 | 0 | 0 | 0 | 0 | 0 | 0 | NA | NA | NA | NA | NA | NA | NA | NA | NA | NA |
| Klrb1a | 1 | 2 | 2 | 0 | 2 | 0 | 1 | 2 | 2 | 0 | 2 | 0 | 1.22 | 2 | 1 | 0.434 | -1.205 | 0.52273097 | NA | 0 | 1.9456 | 1.5507 | 1.5507 | TRUE | 0.9751 |
| Klrb1b | 0 | 0 | 0 | 0 | 0 | 0 | 0 | 0 | 0 | 0 | 0 | 0 | 0 | 0 | 0 | NA | NA | NA | NA | NA | NA | NA | NA | NA | NA |
| Klrb1c | 0 | 0 | 0 | 0 | 1 | 0 | 0 | 0 | 0 | 0 | 1 | 0 | 0.19 | 0 | 0 | 2.094 | 1.067 | 0.79379329 | NA | 0 | 12.608 | 10 | 10 | TRUE | 0.5075 |
| Klrb1f | 7 | 5 | 3 | 5 | 5 | 4 | 5 | 5 | 3 | 5 | 6 | 4 | 4.76 | 4 | 5 | 1.03 | 0.043 | 0.95964823 | NA | 0 | 0.5004 | 0.3185 | 0.3185 | TRUE | 0.123 |
| Klrc1 | 0 | 0 | 0 | 0 | 0 | 0 | 0 | 0 | 0 | 0 | 0 | 0 | 0 | 0 | 0 | NA | NA | NA | NA | NA | NA | NA | NA | NA | NA |
| Klrc2 | 0 | 0 | 0 | 0 | 0 | 0 | 0 | 0 | 0 | 0 | 0 | 0 | 0 | 0 | 0 | NA | NA | NA | NA | NA | NA | NA | NA | NA | NA |
| Klrc3 | 0 | 0 | 0 | 0 | 0 | 0 | 0 | 0 | 0 | 0 | 0 | 0 | 0 | 0 | 0 | NA | NA | NA | NA | NA | NA | NA | NA | NA | NA |
| Klrd1 | 0 | 0 | 0 | 0 | 0 | 0 | 0 | 0 | 0 | 0 | 0 | 0 | 0 | 0 | 0 | NA | NA | NA | NA | NA | NA | NA | NA | NA | NA |
| Klre1 | 0 | 0 | 0 | 0 | 0 | 1 | 0 | 0 | 0 | 0 | 0 | 1 | 0.16 | 0 | 0 | 2.094 | 1.067 | 0.79379329 | NA | 0 | 14.4078 | 10 | 10 | TRUE | 0.3835 |
| Klrg1 | 3 | 5 | 0 | 0 | 1 | 0 | 2 | 5 | 0 | 0 | 1 | 0 | 1.46 | 2 | 0 | 0.146 | -2.781 | 0.15926534 | NA | 1.491 | 1.6227 | 1.6222 | 1.6222 | TRUE | 0.2076 |
| Klrg2 | 178 | 112 | 104 | 150 | 121 | 145 | 138 | 119 | 112 | 149 | 137 | 143 | 133.09 | 123 | 143 | 1.156 | 0.209 | 0.23086994 | 0.39662875 | 0 | 0.0202 | 0.0145 | 0.0145 | TRUE | 0.1285 |
| Klri1 | 0 | 0 | 0 | 0 | 0 | 0 | 0 | 0 | 0 | 0 | 0 | 0 | 0 | 0 | 0 | NA | NA | NA | NA | NA | NA | NA | NA | NA | NA |
| Klri2 | 0 | 0 | 0 | 0 | 0 | 0 | 0 | 0 | 0 | 0 | 0 | 0 | 0 | 0 | 0 | NA | NA | NA | NA | NA | NA | NA | NA | NA | NA |
| Klrk1 | 0 | 0 | 0 | 0 | 0 | 0 | 0 | 0 | 0 | 0 | 0 | 0 | 0 | 0 | 0 | NA | NA | NA | NA | NA | NA | NA | NA | NA | NA |
| Kmo | 0 | 0 | 0 | 0 | 0 | 0 | 0 | 0 | 0 | 0 | 0 | 0 | 0 | 0 | 0 | NA | NA | NA | NA | NA | NA | NA | NA | NA | NA |
| Kncn | 0 | 0 | 0 | 0 | 0 | 0 | 0 | 0 | 0 | 0 | 0 | 0 | 0 | 0 | 0 | NA | NA | NA | NA | NA | NA | NA | NA | NA | NA |
| Kndc1 | 474 | 422 | 440 | 427 | 366 | 484 | 368 | 449 | 475 | 425 | 413 | 478 | 434.58 | 431 | 439 | 1.022 | 0.032 | 0.79402702 | 0.87942876 | 0.0098 | 0.0079 | 0.0085 | 0.0085 | TRUE | 0.1981 |
| Kng1 | 17 | 21 | 12 | 16 | 10 | 11 | 13 | 22 | 13 | 16 | 11 | 11 | 14.43 | 16 | 13 | 0.791 | -0.338 | 0.51460846 | NA | 0 | 0.1667 | 0.1234 | 0.1234 | TRUE | 0.504 |
| Kng2 | 4 | 5 | 3 | 0 | 2 | 1 | 3 | 5 | 3 | 0 | 2 | 1 | 2.48 | 4 | 1 | 0.272 | -1.878 | 0.16305492 | NA | 0 | 0.9565 | 0.6893 | 0.6893 | TRUE | 0.3988 |
| Knstrn | 3119 | 2276 | 2286 | 2513 | 2344 | 2602 | 2422 | 2421 | 2467 | 2501 | 2644 | 2569 | 2504.03 | 2437 | 2571 | 1.055 | 0.078 | 0.18052863 | 0.33560668 | 0 | 0.0034 | 0.002 | 0.002 | TRUE | 0.0073 |
| Kntc1 | 9440 | 6722 | 6940 | 7932 | 7079 | 7621 | 7332 | 7150 | 7488 | 7896 | 7986 | 7523 | 7562.4 | 7323 | 7802 | 1.065 | 0.091 | 0.0779633 | 0.18558628 | 6.00E-04 | 0.0027 | 0.0018 | 0.0018 | TRUE | 0.0119 |
| Kpna1 | 8483 | 5741 | 5975 | 6493 | 5568 | 6220 | 6588 | 6107 | 6447 | 6463 | 6281 | 6140 | 6337.75 | 6381 | 6295 | 0.986 | -0.02 | 0.71605085 | 0.82948564 | 8.00E-04 | 0.0028 | 0.002 | 0.002 | TRUE | 0.0172 |
| Kpna2 | 21228 | 14442 | 15057 | 17112 | 14841 | 16907 | 16487 | 15362 | 16246 | 17033 | 16742 | 16690 | 16426.72 | 16032 | 16822 | 1.049 | 0.069 | 0.1606068 | 0.31025179 | 7.00E-04 | 0.0026 | 0.0017 | 0.0017 | TRUE | 0.0163 |
| Kpna3 | 14789 | 10939 | 10900 | 11699 | 10089 | 11764 | 11486 | 11636 | 11761 | 11645 | 11381 | 11613 | 11587.03 | 11628 | 11546 | 0.993 | -0.01 | 0.82630691 | 0.90029631 | 1.00E-04 | 0.0026 | 0.0014 | 0.0014 | TRUE | 0.0019 |
| Kpna4 | 8317 | 5663 | 5196 | 5637 | 4730 | 5540 | 6459 | 6024 | 5606 | 5611 | 5336 | 5469 | 5750.9 | 6030 | 5472 | 0.907 | -0.141 | 0.02796343 | 0.08797739 | 0.0026 | 0.0028 | 0.0028 | 0.0028 | TRUE | 0.0479 |
| Kpna6 | 12453 | 8869 | 8659 | 9819 | 8284 | 9415 | 9672 | 9434 | 9343 | 9774 | 9345 | 9294 | 9476.94 | 9483 | 9471 | 0.999 | -0.002 | 0.96882396 | 0.98315003 | 3.00E-04 | 0.0027 | 0.0016 | 0.0016 | TRUE | 0.0096 |
| Kpna7 | 45 | 28 | 34 | 39 | 33 | 23 | 35 | 30 | 37 | 39 | 37 | 23 | 33.36 | 34 | 33 | 0.971 | -0.043 | 0.90146915 | 0.94568228 | 0 | 0.0735 | 0.0575 | 0.0575 | TRUE | 0.5275 |
| Kpnb1 | 107874 | 77220 | 76385 | 83481 | 73156 | 86375 | 83781 | 82138 | 82419 | 83097 | 82528 | 85265 | 83204.63 | 82779 | 83630 | 1.01 | 0.015 | 0.73039006 | 0.83891556 | 2.00E-04 | 0.0025 | 0.0013 | 0.0013 | TRUE | 0.0036 |
| Kprp | 0 | 0 | 0 | 0 | 0 | 0 | 0 | 0 | 0 | 0 | 0 | 0 | 0 | 0 | 0 | NA | NA | NA | NA | NA | NA | NA | NA | NA | NA |
| Kptn | 729 | 608 | 513 | 606 | 511 | 574 | 566 | 647 | 554 | 603 | 576 | 567 | 585.45 | 589 | 582 | 0.99 | -0.015 | 0.87660329 | 0.93106882 | 0.0024 | 0.0065 | 0.0052 | 0.0052 | TRUE | 0.0844 |
| Kras | 3258 | 2530 | 2628 | 2883 | 2390 | 2827 | 2530 | 2691 | 2836 | 2870 | 2696 | 2791 | 2735.61 | 2686 | 2786 | 1.038 | 0.054 | 0.40782673 | 0.5805844 | 0.0016 | 0.0033 | 0.0027 | 0.0027 | TRUE | 0.0319 |
| Krba1 | 1798 | 1357 | 1193 | 1072 | 967 | 1071 | 1396 | 1443 | 1287 | 1067 | 1091 | 1057 | 1223.71 | 1375 | 1072 | 0.779 | -0.361 | 1.33E-06 | 3.69E-05 | 0.0011 | 0.0044 | 0.0032 | 0.0032 | TRUE | 0.0369 |
| Krcc1 | 1947 | 1364 | 1127 | 1221 | 1127 | 1228 | 1512 | 1451 | 1216 | 1215 | 1271 | 1212 | 1313 | 1393 | 1233 | 0.884 | -0.178 | 0.04502411 | 0.12471863 | 0.0062 | 0.0042 | 0.0049 | 0.0049 | TRUE | 0.1467 |
| Kremen1 | 2257 | 1593 | 1651 | 1750 | 1596 | 1775 | 1753 | 1694 | 1781 | 1742 | 1800 | 1752 | 1753.9 | 1743 | 1765 | 1.012 | 0.018 | 0.77905217 | 0.86964954 | 0 | 0.0038 | 0.0023 | 0.0023 | TRUE | 0.0069 |
| Kremen2 | 84 | 60 | 61 | 93 | 70 | 79 | 65 | 64 | 66 | 93 | 79 | 78 | 74.07 | 65 | 83 | 1.281 | 0.357 | 0.11251351 | 0.24092219 | 0 | 0.0344 | 0.0229 | 0.0229 | TRUE | 0.0934 |
| Kri1 | 3739 | 2823 | 2544 | 2644 | 2474 | 2763 | 2904 | 3003 | 2745 | 2632 | 2791 | 2728 | 2800.32 | 2884 | 2717 | 0.942 | -0.087 | 0.1616657 | 0.31166548 | 9.00E-04 | 0.0033 | 0.0024 | 0.0024 | TRUE | 0.0211 |
| Krit1 | 2711 | 1847 | 1874 | 1956 | 1740 | 2011 | 2106 | 1965 | 2022 | 1947 | 1963 | 1985 | 1997.87 | 2031 | 1965 | 0.967 | -0.048 | 0.435444 | 0.60702244 | 0 | 0.0036 | 0.0023 | 0.0023 | TRUE | 0.013 |
| Krr1 | 6304 | 4478 | 4313 | 4630 | 3995 | 4754 | 4896 | 4763 | 4654 | 4609 | 4507 | 4693 | 4686.89 | 4771 | 4603 | 0.965 | -0.052 | 0.32386117 | 0.49789692 | 3.00E-04 | 0.0029 | 0.0018 | 0.0018 | TRUE | 0.0065 |
| Krt1 | 11 | 16 | 12 | 13 | 11 | 11 | 9 | 17 | 13 | 13 | 12 | 11 | 12.45 | 13 | 12 | 0.951 | -0.072 | 0.89598354 | NA | 0 | 0.1927 | 0.1367 | 0.1367 | TRUE | 0.2328 |
| Krt10 | 88 | 63 | 55 | 69 | 53 | 54 | 68 | 67 | 59 | 69 | 60 | 53 | 62.75 | 65 | 61 | 0.932 | -0.102 | 0.67974097 | 0.80328012 | 0 | 0.0402 | 0.0279 | 0.0279 | TRUE | 0.1219 |
| Krt12 | 96 | 69 | 54 | 50 | 45 | 43 | 75 | 73 | 58 | 50 | 51 | 42 | 58.2 | 69 | 48 | 0.69 | -0.534 | 0.039568 | 0.11367747 | 0 | 0.0431 | 0.0307 | 0.0307 | TRUE | 0.1506 |
| Krt13 | 3 | 1 | 2 | 2 | 1 | 0 | 2 | 1 | 2 | 2 | 1 | 0 | 1.45 | 2 | 1 | 0.556 | -0.848 | 0.59994311 | NA | 0 | 1.6425 | 1.1248 | 1.1248 | TRUE | 0.395 |
| Krt14 | 29 | 20 | 11 | 25 | 19 | 23 | 23 | 21 | 12 | 25 | 21 | 23 | 20.78 | 19 | 23 | 1.23 | 0.299 | 0.48882694 | 0.65456738 | 0 | 0.1165 | 0.0861 | 0.0861 | TRUE | 0.4726 |
| Krt15 | 5 | 1 | 4 | 2 | 4 | 4 | 4 | 1 | 4 | 2 | 5 | 4 | 3.29 | 3 | 4 | 1.113 | 0.154 | 0.88425494 | NA | 0 | 0.7237 | 0.5059 | 0.5059 | TRUE | 0.3959 |
| Krt16 | 35 | 25 | 25 | 36 | 25 | 24 | 27 | 27 | 27 | 36 | 28 | 24 | 28.08 | 27 | 29 | 1.086 | 0.12 | 0.74286379 | 0.84692759 | 0 | 0.0868 | 0.0601 | 0.0601 | TRUE | 0.2639 |
| Krt17 | 242 | 209 | 217 | 294 | 262 | 284 | 188 | 222 | 234 | 293 | 296 | 280 | 252.16 | 215 | 290 | 1.353 | 0.437 | 0.00126347 | 0.00852839 | 0.0027 | 0.0118 | 0.0092 | 0.0092 | TRUE | 0.1448 |
| Krt18 | 749 | 526 | 608 | 520 | 469 | 555 | 582 | 559 | 656 | 518 | 529 | 548 | 565.3 | 599 | 532 | 0.888 | -0.171 | 0.08131893 | 0.19138671 | 0.0022 | 0.0066 | 0.0052 | 0.0052 | TRUE | 0.0783 |

| | | | | | | | | | | | | | | | | | | | | | | | | | |
|---|---|---|---|---|---|---|---|---|---|---|---|---|---|---|---|---|---|---|---|---|---|---|---|---|---|
| Krt19 | 1455 | 1254 | 1615 | 1677 | 1593 | 1883 | 1130 | 1334 | 1743 | 1669 | 1797 | 1859 | 1588.61 | 1402 | 1775 | 1.268 | 0.342 | 0.005328 | 0.02571872 | 0.0253 | 0.0039 | 0.0102 | 0.0102 | TRUE | 0.5418 |
| Krt2 | 1 | 0 | 0 | 1 | 0 | 0 | 1 | 0 | 0 | 1 | 0 | 0 | 0.3 | 0 | 0 | 1.075 | 0.105 | 0.97698525 | NA | 0 | 8.0272 | 7.5141 | 7.5141 | TRUE | 0.3906 |
| Krt20 | 41 | 25 | 19 | 19 | 19 | 16 | 32 | 27 | 21 | 19 | 21 | 16 | 22.51 | 27 | 19 | 0.704 | -0.506 | 0.22057115 | 0.38409156 | 0 | 0.1077 | 0.077 | 0.077 | TRUE | 0.1815 |
| Krt222 | 66 | 42 | 47 | 51 | 44 | 47 | 51 | 45 | 51 | 51 | 50 | 46 | 48.91 | 49 | 49 | 0.999 | -0.002 | 0.9954942 | 0.99730837 | 0 | 0.0509 | 0.0329 | 0.0329 | TRUE | 0.0434 |
| Krt23 | 175 | 121 | 144 | 115 | 78 | 105 | 136 | 129 | 155 | 114 | 88 | 104 | 121.02 | 140 | 102 | 0.731 | -0.452 | 0.01641127 | 0.05973653 | 0.0032 | 0.022 | 0.017 | 0.017 | TRUE | 0.1341 |
| Krt24 | 3 | 0 | 1 | 1 | 2 | 0 | 2 | 0 | 1 | 1 | 2 | 0 | 1.11 | 1 | 1 | 0.906 | -0.142 | 0.94060664 | NA | 0 | 2.1374 | 1.7366 | 1.7366 | TRUE | 0.2449 |
| Krt25 | 4 | 3 | 0 | 1 | 0 | 0 | 3 | 3 | 0 | 1 | 0 | 0 | 1.22 | 2 | 0 | 0.175 | -2.517 | 0.22079851 | NA | 0 | 1.9523 | 1.8163 | 1.8163 | TRUE | 0.6537 |
| Krt26 | 2 | 3 | 0 | 1 | 1 | 2 | 2 | 3 | 0 | 1 | 1 | 2 | 1.47 | 2 | 1 | 0.866 | -0.208 | 0.8963711 | NA | 0 | 1.6107 | 1.156 | 1.156 | TRUE | 0.16 |
| Krt27 | 23 | 17 | 12 | 18 | 15 | 18 | 18 | 18 | 13 | 18 | 17 | 18 | 16.92 | 16 | 18 | 1.072 | 0.1 | 0.82712691 | NA | 0 | 0.1425 | 0.0933 | 0.0933 | TRUE | 0.1415 |
| Krt28 | 8 | 9 | 5 | 5 | 5 | 6 | 6 | 10 | 5 | 5 | 6 | 6 | 6.29 | 7 | 6 | 0.784 | -0.351 | 0.64101911 | NA | 0 | 0.3794 | 0.247 | 0.247 | TRUE | 0.2386 |
| Krt31 | 0 | 0 | 0 | 0 | 0 | 0 | 0 | 0 | 0 | 0 | 0 | 0 | 0 | 0 | 0 | NA | NA | NA | NA | NA | NA | NA | NA | NA | NA |
| Krt32 | 0 | 0 | 0 | 0 | 0 | 0 | 0 | 0 | 0 | 0 | 0 | 0 | 0 | 0 | 0 | NA | NA | NA | NA | NA | NA | NA | NA | NA | NA |
| Krt33a | 8 | 1 | 3 | 0 | 7 | 1 | 6 | 1 | 3 | 0 | 8 | 1 | 3.23 | 3 | 3 | 0.812 | -0.3 | 0.81461451 | NA | 1.2592 | 0.7355 | 0.8686 | 0.8686 | TRUE | 1.0573 |
| Krt33b | 0 | 0 | 0 | 0 | 0 | 0 | 0 | 0 | 0 | 0 | 0 | 0 | 0 | 0 | 0 | NA | NA | NA | NA | NA | NA | NA | NA | NA | NA |
| Krt34 | 0 | 0 | 1 | 2 | 1 | 0 | 0 | 0 | 1 | 2 | 1 | 0 | 0.7 | 0 | 1 | 3.01 | 1.59 | 0.51808674 | NA | 0 | 3.3899 | 2.8624 | 2.8624 | TRUE | 0.3541 |
| Krt35 | 0 | 1 | 0 | 0 | 0 | 0 | 0 | 1 | 0 | 0 | 0 | 0 | 0.18 | 0 | 0 | 0.552 | -0.857 | 0.8336548 | NA | 0 | 13.3714 | 10 | 10 | TRUE | 0.5216 |
| Krt36 | 2 | 0 | 0 | 1 | 0 | 0 | 2 | 0 | 0 | 1 | 0 | 0 | 0.42 | 1 | 0 | 0.681 | -0.553 | 0.8627148 | NA | 0 | 5.5819 | 5.4906 | 5.4906 | TRUE | 0.9727 |
| Krt39 | 0 | 0 | 0 | 0 | 0 | 0 | 0 | 0 | 0 | 0 | 0 | 0 | 0 | 0 | 0 | NA | NA | NA | NA | NA | NA | NA | NA | NA | NA |
| Krt4 | 5 | 2 | 2 | 0 | 0 | 3 | 4 | 2 | 2 | 0 | 0 | 3 | 1.86 | 3 | 1 | 0.367 | -1.447 | 0.35674647 | NA | 0.0772 | 1.2801 | 1.0874 | 1.0874 | TRUE | 1.441 |
| Krt40 | 4 | 4 | 2 | 1 | 2 | 1 | 3 | 4 | 2 | 1 | 2 | 1 | 2.29 | 3 | 1 | 0.439 | -1.187 | 0.35409322 | NA | 0 | 1.036 | 0.6584 | 0.6584 | TRUE | 0.1543 |
| Krt42 | 431 | 362 | 374 | 375 | 382 | 475 | 335 | 385 | 404 | 373 | 431 | 469 | 399.41 | 375 | 424 | 1.136 | 0.184 | 0.13736703 | 0.27766029 | 0.009 | 0.0084 | 0.0086 | 0.0086 | TRUE | 0.1305 |
| Krt5 | 9 | 14 | 8 | 9 | 11 | 5 | 7 | 15 | 9 | 9 | 12 | 5 | 9.47 | 10 | 9 | 0.865 | -0.21 | 0.74896877 | NA | 0.0712 | 0.2527 | 0.2016 | 0.2016 | TRUE | 0.2982 |
| Krt6a | 0 | 0 | 0 | 0 | 0 | 0 | 0 | 0 | 0 | 0 | 0 | 0 | 0 | 0 | 0 | NA | NA | NA | NA | NA | NA | NA | NA | NA | NA |
| Krt6b | 2 | 2 | 0 | 1 | 0 | 0 | 2 | 2 | 0 | 1 | 0 | 0 | 0.78 | 1 | 0 | 0.3 | -1.737 | 0.45896026 | NA | 0 | 3.0435 | 2.5735 | 2.5735 | TRUE | 0.3906 |
| Krt7 | 125 | 141 | 219 | 286 | 265 | 288 | 97 | 150 | 236 | 285 | 299 | 284 | 225.22 | 161 | 289 | 1.807 | 0.854 | 0.00020624 | 0.00218036 | 0.0883 | 0.013 | 0.0333 | 0.0333 | TRUE | 0.7157 |
| Krt71 | 24 | 12 | 13 | 5 | 13 | 10 | 19 | 13 | 14 | 5 | 15 | 10 | 12.49 | 15 | 10 | 0.64 | -0.645 | 0.26215118 | NA | 0.0331 | 0.1922 | 0.1533 | 0.1533 | TRUE | 0.2881 |
| Krt72 | 3 | 0 | 0 | 0 | 0 | 0 | 2 | 0 | 0 | 0 | 0 | 0 | 0.39 | 1 | 0 | 0.239 | -2.067 | 0.57190649 | NA | 10 | 6.1057 | 8.0283 | 8.0283 | TRUE | 1.4408 |
| Krt73 | 3 | 2 | 0 | 0 | 0 | 1 | 2 | 2 | 0 | 0 | 0 | 1 | 0.91 | 1 | 0 | 0.246 | -2.022 | 0.36558656 | NA | 0 | 2.6143 | 2.2789 | 2.2789 | TRUE | 0.472 |
| Krt74 | 2 | 0 | 1 | 1 | 1 | 1 | 2 | 0 | 1 | 1 | 1 | 1 | 0.96 | 1 | 1 | 1.143 | 0.193 | 0.91871967 | NA | 0 | 2.4786 | 1.5559 | 1.5559 | TRUE | 0.2848 |
| Krt75 | 8 | 7 | 6 | 1 | 8 | 3 | 6 | 7 | 6 | 1 | 9 | 3 | 5.52 | 6 | 4 | 0.634 | -0.657 | 0.45745703 | NA | 0.1616 | 0.4319 | 0.3692 | 0.3692 | TRUE | 1.2896 |
| Krt76 | 0 | 0 | 0 | 0 | 0 | 0 | 0 | 0 | 0 | 0 | 0 | 0 | 0 | 0 | 0 | NA | NA | NA | NA | NA | NA | NA | NA | NA | NA |
| Krt77 | 0 | 0 | 0 | 2 | 0 | 0 | 0 | 0 | 0 | 2 | 0 | 0 | 0.33 | 0 | 1 | 3.77 | 1.915 | 0.62344813 | NA | 0 | 7.1455 | 9.1834 | 9.1834 | TRUE | 0.9739 |
| Krt78 | 7 | 5 | 7 | 13 | 7 | 8 | 5 | 5 | 8 | 13 | 8 | 8 | 7.84 | 6 | 10 | 1.582 | 0.662 | 0.33480586 | NA | 0 | 0.3047 | 0.2057 | 0.2057 | TRUE | 0.3244 |
| Krt79 | 107 | 78 | 81 | 100 | 89 | 101 | 83 | 83 | 87 | 100 | 100 | 100 | 92.19 | 84 | 100 | 1.183 | 0.242 | 0.22391079 | 0.38829859 | 0 | 0.0281 | 0.0177 | 0.0177 | TRUE | 0.008 |
| Krt8 | 682 | 570 | 532 | 655 | 563 | 602 | 530 | 606 | 574 | 652 | 635 | 594 | 598.56 | 570 | 627 | 1.102 | 0.14 | 0.14270579 | 0.28539162 | 0.002 | 0.0064 | 0.0049 | 0.0049 | TRUE | 0.048 |
| Krt80 | 7 | 11 | 8 | 12 | 34 | 19 | 5 | 12 | 9 | 12 | 38 | 19 | 15.8 | 9 | 23 | 2.705 | 1.436 | 0.01361887 | NA | 0.1925 | 0.1524 | 0.1644 | 0.1644 | TRUE | 0.453 |
| Krt81 | 0 | 1 | 2 | 2 | 3 | 3 | 0 | 1 | 2 | 2 | 3 | 3 | 1.93 | 1 | 3 | 2.736 | 1.452 | 0.32179261 | NA | 0 | 1.2327 | 0.8859 | 0.8859 | TRUE | 0.5369 |
| Krt82 | 0 | 1 | 1 | 1 | 0 | 0 | 0 | 1 | 1 | 1 | 0 | 0 | 0.52 | 1 | 0 | 0.535 | -0.904 | 0.74017109 | NA | 0 | 4.534 | 3.6534 | 3.6534 | TRUE | 0.3906 |
| Krt83 | 5 | 1 | 3 | 1 | 5 | 1 | 4 | 1 | 3 | 1 | 6 | 1 | 2.63 | 3 | 3 | 0.901 | -0.15 | 0.90329478 | NA | 0.2874 | 0.902 | 0.7218 | 0.7218 | TRUE | 1.1402 |
| Krt84 | 2 | 0 | 3 | 1 | 0 | 1 | 2 | 0 | 3 | 1 | 0 | 1 | 1.13 | 2 | 1 | 0.423 | -1.24 | 0.5235556 | NA | 0 | 2.102 | 1.6529 | 1.6529 | TRUE | 0.1028 |
| Krt85 | 2 | 0 | 1 | 0 | 0 | 1 | 2 | 0 | 1 | 0 | 0 | 1 | 0.6 | 1 | 0 | 0.414 | -1.271 | 0.61985863 | NA | 0 | 3.9313 | 3.1876 | 3.1876 | TRUE | 0.3835 |
| Krt86 | 3 | 0 | 0 | 2 | 0 | 2 | 2 | 0 | 0 | 2 | 0 | 2 | 1.05 | 1 | 1 | 1.603 | 0.681 | 0.74347332 | NA | 0 | 2.2614 | 2.1558 | 2.1558 | TRUE | 1.4382 |
| Krt9 | 4 | 3 | 0 | 2 | 0 | 1 | 3 | 3 | 0 | 2 | 0 | 1 | 1.55 | 2 | 1 | 0.475 | -1.074 | 0.52519413 | NA | 0 | 1.5355 | 1.3203 | 1.3203 | TRUE | 0.5117 |
| Krtap1-3 | 0 | 0 | 0 | 0 | 0 | 0 | 0 | 0 | 0 | 0 | 0 | 0 | 0 | 0 | 0 | NA | NA | NA | NA | NA | NA | NA | NA | NA | NA |
| Krtap1-4 | 0 | 0 | 0 | 0 | 0 | 1 | 0 | 0 | 0 | 0 | 0 | 1 | 0.16 | 0 | 0 | 2.094 | 1.067 | 0.79379329 | NA | 0 | 14.4078 | 10 | 10 | TRUE | 0.3835 |
| Krtap1-5 | 0 | 0 | 0 | 0 | 0 | 0 | 0 | 0 | 0 | 0 | 0 | 0 | 0 | 0 | 0 | NA | NA | NA | NA | NA | NA | NA | NA | NA | NA |
| Krtap10-10 | 4 | 2 | 6 | 3 | 3 | 3 | 3 | 2 | 6 | 3 | 3 | 3 | 3.51 | 4 | 3 | 0.803 | -0.317 | 0.75496514 | NA | 0 | 0.6783 | 0.4525 | 0.4525 | TRUE | 0.4918 |
| Krtap10-4 | 0 | 0 | 0 | 0 | 0 | 0 | 0 | 0 | 0 | 0 | 0 | 0 | 0 | 0 | 0 | NA | NA | NA | NA | NA | NA | NA | NA | NA | NA |
| Krtap11-1 | 0 | 0 | 0 | 0 | 0 | 0 | 0 | 0 | 0 | 0 | 0 | 0 | 0 | 0 | 0 | NA | NA | NA | NA | NA | NA | NA | NA | NA | NA |
| Krtap12-1 | 0 | 0 | 0 | 0 | 0 | 0 | 0 | 0 | 0 | 0 | 0 | 0 | 0 | 0 | 0 | NA | NA | NA | NA | NA | NA | NA | NA | NA | NA |
| Krtap13 | 0 | 0 | 0 | 0 | 0 | 0 | 0 | 0 | 0 | 0 | 0 | 0 | 0 | 0 | 0 | NA | NA | NA | NA | NA | NA | NA | NA | NA | NA |
| Krtap13-1 | 0 | 0 | 0 | 0 | 0 | 0 | 0 | 0 | 0 | 0 | 0 | 0 | 0 | 0 | 0 | NA | NA | NA | NA | NA | NA | NA | NA | NA | NA |
| Krtap14 | 0 | 0 | 0 | 0 | 0 | 0 | 0 | 0 | 0 | 0 | 0 | 0 | 0 | 0 | 0 | NA | NA | NA | NA | NA | NA | NA | NA | NA | NA |
| Krtap15 | 0 | 0 | 0 | 0 | 0 | 0 | 0 | 0 | 0 | 0 | 0 | 0 | 0 | 0 | 0 | NA | NA | NA | NA | NA | NA | NA | NA | NA | NA |
| Krtap16-1 | 0 | 0 | 0 | 0 | 0 | 0 | 0 | 0 | 0 | 0 | 0 | 0 | 0 | 0 | 0 | NA | NA | NA | NA | NA | NA | NA | NA | NA | NA |
| Krtap16-3 | 0 | 0 | 0 | 0 | 0 | 0 | 0 | 0 | 0 | 0 | 0 | 0 | 0 | 0 | 0 | NA | NA | NA | NA | NA | NA | NA | NA | NA | NA |
| Krtap17-1 | 0 | 0 | 0 | 0 | 0 | 0 | 0 | 0 | 0 | 0 | 0 | 0 | 0 | 0 | 0 | NA | NA | NA | NA | NA | NA | NA | NA | NA | NA |
| Krtap19-1 | 0 | 0 | 0 | 0 | 0 | 0 | 0 | 0 | 0 | 0 | 0 | 0 | 0 | 0 | 0 | NA | NA | NA | NA | NA | NA | NA | NA | NA | NA |
| Krtap19-3 | 0 | 0 | 0 | 0 | 0 | 0 | 0 | 0 | 0 | 0 | 0 | 0 | 0 | 0 | 0 | NA | NA | NA | NA | NA | NA | NA | NA | NA | NA |
| Krtap19-4 | 0 | 0 | 0 | 0 | 0 | 0 | 0 | 0 | 0 | 0 | 0 | 0 | 0 | 0 | 0 | NA | NA | NA | NA | NA | NA | NA | NA | NA | NA |
| Krtap19-5 | 0 | 0 | 0 | 0 | 0 | 0 | 0 | 0 | 0 | 0 | 0 | 0 | 0 | 0 | 0 | NA | NA | NA | NA | NA | NA | NA | NA | NA | NA |
| Krtap19-9b | 0 | 0 | 0 | 0 | 0 | 0 | 0 | 0 | 0 | 0 | 0 | 0 | 0 | 0 | 0 | NA | NA | NA | NA | NA | NA | NA | NA | NA | NA |
| Krtap2-4 | 0 | 0 | 0 | 0 | 0 | 0 | 0 | 0 | 0 | 0 | 0 | 0 | 0 | 0 | 0 | NA | NA | NA | NA | NA | NA | NA | NA | NA | NA |
| Krtap20-2 | 0 | 0 | 0 | 0 | 0 | 0 | 0 | 0 | 0 | 0 | 0 | 0 | 0 | 0 | 0 | NA | NA | NA | NA | NA | NA | NA | NA | NA | NA |
| Krtap21-1 | 0 | 0 | 0 | 0 | 0 | 0 | 0 | 0 | 0 | 0 | 0 | 0 | 0 | 0 | 0 | NA | NA | NA | NA | NA | NA | NA | NA | NA | NA |
| Krtap22-2 | 0 | 0 | 0 | 0 | 0 | 0 | 0 | 0 | 0 | 0 | 0 | 0 | 0 | 0 | 0 | NA | NA | NA | NA | NA | NA | NA | NA | NA | NA |
| Krtap24-1 | 0 | 0 | 0 | 0 | 0 | 0 | 0 | 0 | 0 | 0 | 0 | 0 | 0 | 0 | 0 | NA | NA | NA | NA | NA | NA | NA | NA | NA | NA |
| Krtap26-1 | 0 | 0 | 0 | 0 | 0 | 0 | 0 | 0 | 0 | 0 | 0 | 0 | 0 | 0 | 0 | NA | NA | NA | NA | NA | NA | NA | NA | NA | NA |
| Krtap27-1 | 0 | 0 | 0 | 0 | 0 | 0 | 0 | 0 | 0 | 0 | 0 | 0 | 0 | 0 | 0 | NA | NA | NA | NA | NA | NA | NA | NA | NA | NA |

| | | | | | | | | | | | | | | | | | | | | | | | | | |
|---|---|---|---|---|---|---|---|---|---|---|---|---|---|---|---|---|---|---|---|---|---|---|---|---|---|
| Krtap3-1 | 0 | 0 | 0 | 0 | 0 | 0 | 0 | 0 | 0 | 0 | 0 | 0 | 0 | 0 | 0 | NA | NA | NA | NA | NA | NA | NA | NA | NA | NA |
| Krtap3-2 | 0 | 0 | 0 | 0 | 0 | 0 | 0 | 0 | 0 | 0 | 0 | 0 | 0 | 0 | 0 | NA | NA | NA | NA | NA | NA | NA | NA | NA | NA |
| Krtap3-3 | 0 | 0 | 0 | 0 | 0 | 0 | 0 | 0 | 0 | 0 | 0 | 0 | 0 | 0 | 0 | NA | NA | NA | NA | NA | NA | NA | NA | NA | NA |
| Krtap31-1 | 0 | 0 | 0 | 0 | 0 | 0 | 0 | 0 | 0 | 0 | 0 | 0 | 0 | 0 | 0 | NA | NA | NA | NA | NA | NA | NA | NA | NA | NA |
| Krtap31-2 | 0 | 0 | 0 | 0 | 0 | 0 | 0 | 0 | 0 | 0 | 0 | 0 | 0 | 0 | 0 | NA | NA | NA | NA | NA | NA | NA | NA | NA | NA |
| Krtap4-1 | 0 | 0 | 0 | 0 | 0 | 0 | 0 | 0 | 0 | 0 | 0 | 0 | 0 | 0 | 0 | NA | NA | NA | NA | NA | NA | NA | NA | NA | NA |
| Krtap4-13 | 5 | 0 | 1 | 0 | 2 | 1 | 4 | 0 | 1 | 0 | 2 | 1 | 1.37 | 2 | 1 | 0.619 | -0.691 | 0.70135126 | NA | 1.0128 | 1.7354 | 1.5703 | 1.5703 | TRUE | 0.6748 |
| Krtap4-16 | 1 | 1 | 0 | 0 | 0 | 0 | 1 | 1 | 0 | 0 | 0 | 0 | 0.31 | 1 | 0 | 0.309 | -1.696 | 0.63183309 | NA | 0 | 7.7294 | 7.2472 | 7.2472 | TRUE | 0.2015 |
| Krtap4-2 | 0 | 0 | 0 | 0 | 0 | 0 | 0 | 0 | 0 | 0 | 0 | 0 | 0 | 0 | 0 | NA | NA | NA | NA | NA | NA | NA | NA | NA | NA |
| Krtap4-6 | 0 | 0 | 0 | 0 | 0 | 0 | 0 | 0 | 0 | 0 | 0 | 0 | 0 | 0 | 0 | NA | NA | NA | NA | NA | NA | NA | NA | NA | NA |
| Krtap4-7 | 0 | 0 | 0 | 0 | 0 | 0 | 0 | 0 | 0 | 0 | 0 | 0 | 0 | 0 | 0 | NA | NA | NA | NA | NA | NA | NA | NA | NA | NA |
| Krtap4-8 | 0 | 0 | 0 | 0 | 0 | 0 | 0 | 0 | 0 | 0 | 0 | 0 | 0 | 0 | 0 | NA | NA | NA | NA | NA | NA | NA | NA | NA | NA |
| Krtap4-9 | 0 | 0 | 0 | 0 | 0 | 0 | 0 | 0 | 0 | 0 | 0 | 0 | 0 | 0 | 0 | NA | NA | NA | NA | NA | NA | NA | NA | NA | NA |
| Krtap5-1 | 0 | 0 | 0 | 0 | 0 | 0 | 0 | 0 | 0 | 0 | 0 | 0 | 0 | 0 | 0 | NA | NA | NA | NA | NA | NA | NA | NA | NA | NA |
| Krtap5-2 | 0 | 0 | 0 | 0 | 0 | 0 | 0 | 0 | 0 | 0 | 0 | 0 | 0 | 0 | 0 | NA | NA | NA | NA | NA | NA | NA | NA | NA | NA |
| Krtap5-3 | 0 | 0 | 0 | 0 | 0 | 0 | 0 | 0 | 0 | 0 | 0 | 0 | 0 | 0 | 0 | NA | NA | NA | NA | NA | NA | NA | NA | NA | NA |
| Krtap5-4 | 0 | 0 | 0 | 0 | 0 | 0 | 0 | 0 | 0 | 0 | 0 | 0 | 0 | 0 | 0 | NA | NA | NA | NA | NA | NA | NA | NA | NA | NA |
| Krtap5-5 | 0 | 0 | 0 | 0 | 0 | 0 | 0 | 0 | 0 | 0 | 0 | 0 | 0 | 0 | 0 | NA | NA | NA | NA | NA | NA | NA | NA | NA | NA |
| Krtap6-1 | 0 | 0 | 0 | 0 | 0 | 0 | 0 | 0 | 0 | 0 | 0 | 0 | 0 | 0 | 0 | NA | NA | NA | NA | NA | NA | NA | NA | NA | NA |
| Krtap6-2 | 0 | 0 | 0 | 0 | 0 | 0 | 0 | 0 | 0 | 0 | 0 | 0 | 0 | 0 | 0 | NA | NA | NA | NA | NA | NA | NA | NA | NA | NA |
| Krtap6-5 | 1 | 0 | 0 | 0 | 1 | 0 | 1 | 0 | 0 | 0 | 1 | 0 | 0.32 | 0 | 0 | 1.075 | 0.105 | 0.97638847 | NA | 0 | 7.4681 | 7.0392 | 7.0392 | TRUE | 0.5075 |
| Krtap7-1 | 0 | 0 | 0 | 0 | 0 | 0 | 0 | 0 | 0 | 0 | 0 | 0 | 0 | 0 | 0 | NA | NA | NA | NA | NA | NA | NA | NA | NA | NA |
| Krtap8-1 | 0 | 0 | 0 | 0 | 0 | 0 | 0 | 0 | 0 | 0 | 0 | 0 | 0 | 0 | 0 | NA | NA | NA | NA | NA | NA | NA | NA | NA | NA |
| Krtap9-1 | 0 | 0 | 0 | 0 | 0 | 0 | 0 | 0 | 0 | 0 | 0 | 0 | 0 | 0 | 0 | NA | NA | NA | NA | NA | NA | NA | NA | NA | NA |
| Krtap9-3 | 0 | 0 | 0 | 0 | 0 | 0 | 0 | 0 | 0 | 0 | 0 | 0 | 0 | 0 | 0 | NA | NA | NA | NA | NA | NA | NA | NA | NA | NA |
| Krtap9-5 | 0 | 0 | 0 | 0 | 0 | 0 | 0 | 0 | 0 | 0 | 0 | 0 | 0 | 0 | 0 | NA | NA | NA | NA | NA | NA | NA | NA | NA | NA |
| Krtcap2 | 2421 | 1719 | 1681 | 1916 | 1614 | 1870 | 1880 | 1828 | 1814 | 1907 | 1821 | 1846 | 1849.41 | 1841 | 1858 | 1.009 | 0.013 | 0.83445624 | 0.90597926 | 0 | 0.0037 | 0.0022 | 0.0022 | TRUE | 0.0065 |
| Krtcap3 | 226 | 196 | 225 | 260 | 254 | 317 | 176 | 208 | 243 | 259 | 287 | 313 | 247.51 | 209 | 286 | 1.376 | 0.46 | 0.00239611 | 0.01390485 | 0.0134 | 0.012 | 0.0124 | 0.0124 | TRUE | 0.2334 |
| Krtdap | 7 | 2 | 0 | 3 | 0 | 2 | 5 | 2 | 0 | 3 | 0 | 2 | 2.09 | 2 | 2 | 0.646 | -0.63 | 0.67560238 | NA | 1.1341 | 1.1378 | 1.1391 | 1.1391 | TRUE | 0.2559 |
| Ksr1 | 1522 | 1065 | 980 | 980 | 959 | 983 | 1182 | 1133 | 1057 | 975 | 1082 | 970 | 1066.67 | 1124 | 1009 | 0.896 | -0.158 | 0.05423511 | 0.14265097 | 0.0025 | 0.0046 | 0.0039 | 0.0039 | TRUE | 0.0464 |
| Ksr2 | 75 | 59 | 52 | 55 | 49 | 51 | 58 | 63 | 56 | 55 | 55 | 50 | 56.25 | 59 | 53 | 0.905 | -0.144 | 0.57210317 | 0.72498188 | 0 | 0.0446 | 0.0287 | 0.0287 | TRUE | 0.0242 |
| Kti12 | 1387 | 1021 | 951 | 1020 | 860 | 1027 | 1077 | 1086 | 1026 | 1015 | 970 | 1014 | 1031.44 | 1063 | 1000 | 0.94 | -0.089 | 0.23182367 | 0.39772736 | 0 | 0.0047 | 0.003 | 0.003 | TRUE | 0.0107 |
| Ktn1 | 7599 | 5249 | 5155 | 5610 | 4935 | 5635 | 5902 | 5583 | 5562 | 5584 | 5567 | 5563 | 5626.89 | 5682 | 5571 | 0.98 | -0.029 | 0.57606772 | 0.72769249 | 4.00E-04 | 0.0028 | 0.0018 | 0.0018 | TRUE | 0.0142 |
| Kxd1 | 2960 | 2346 | 2485 | 2866 | 2572 | 3076 | 2299 | 2495 | 2681 | 2853 | 2901 | 3036 | 2711.07 | 2492 | 2930 | 1.177 | 0.235 | 0.00088476 | 0.00663675 | 0.0029 | 0.0033 | 0.0032 | 0.0032 | TRUE | 0.0571 |
| Ky | 114 | 90 | 70 | 120 | 101 | 106 | 89 | 96 | 76 | 119 | 114 | 105 | 99.64 | 87 | 113 | 1.299 | 0.378 | 0.05856812 | 0.15077703 | 0 | 0.0262 | 0.0186 | 0.0186 | TRUE | 0.1089 |
| Kynu | 2 | 2 | 1 | 2 | 0 | 0 | 2 | 2 | 1 | 2 | 0 | 0 | 1.13 | 2 | 1 | 0.426 | -1.23 | 0.52142546 | NA | 0 | 2.109 | 1.582 | 1.582 | TRUE | 0.9735 |
| L1cam | 766 | 520 | 545 | 868 | 774 | 859 | 595 | 553 | 588 | 864 | 873 | 848 | 720.2 | 579 | 862 | 1.488 | 0.573 | 8.19E-12 | 8.67E-10 | 0 | 0.0057 | 0.0036 | 0.0036 | TRUE | 0.0171 |
| L1td1 | 31694 | 21781 | 16206 | 17298 | 15184 | 17328 | 24615 | 23168 | 17486 | 17218 | 17129 | 17105 | 19453.77 | 21756 | 17151 | 0.788 | -0.343 | 0.00045981 | 0.00405686 | 0.0149 | 0.0025 | 0.0069 | 0.0069 | TRUE | 0.3605 |
| L2hgdh | 3981 | 3154 | 3276 | 2949 | 2632 | 2831 | 3092 | 3355 | 3535 | 2935 | 2969 | 2795 | 3113.46 | 3327 | 2900 | 0.872 | -0.198 | 0.00326428 | 0.01761949 | 0.0023 | 0.0032 | 0.0029 | 0.0029 | TRUE | 0.0475 |
| L3hypdh | 137 | 108 | 109 | 155 | 114 | 127 | 106 | 115 | 118 | 154 | 129 | 125 | 124.52 | 113 | 136 | 1.208 | 0.273 | 0.13203245 | 0.26982268 | 0 | 0.0215 | 0.0156 | 0.0156 | TRUE | 0.1438 |
| L3mbtl1 | 51 | 68 | 56 | 62 | 52 | 57 | 40 | 72 | 60 | 62 | 59 | 56 | 58.17 | 57 | 59 | 1.035 | 0.049 | 0.86120151 | 0.92314033 | 0.0318 | 0.0432 | 0.0399 | 0.0399 | TRUE | 0.4787 |
| L3mbtl2 | 10443 | 7605 | 7131 | 7463 | 6775 | 7849 | 8111 | 8089 | 7694 | 7429 | 7643 | 7748 | 7785.67 | 7965 | 7607 | 0.955 | -0.067 | 0.19113818 | 0.34930802 | 5.00E-04 | 0.0027 | 0.0017 | 0.0017 | TRUE | 0.0107 |
| L3mbtl3 | 407 | 316 | 298 | 259 | 225 | 244 | 316 | 336 | 322 | 258 | 254 | 241 | 287.71 | 325 | 251 | 0.773 | -0.371 | 0.00200997 | 0.01215697 | 0 | 0.0107 | 0.0068 | 0.0068 | TRUE | 0.0138 |
| L3mbtl4 | 5 | 6 | 3 | 1 | 5 | 2 | 4 | 6 | 3 | 1 | 6 | 2 | 3.69 | 4 | 3 | 0.629 | -0.669 | 0.51891889 | NA | 0.0042 | 0.6455 | 0.4846 | 0.4846 | TRUE | 0.7918 |
| l7Rn6 | 3087 | 2335 | 2445 | 2694 | 2349 | 2772 | 2398 | 2484 | 2638 | 2682 | 2650 | 2736 | 2597.88 | 2507 | 2689 | 1.074 | 0.103 | 0.09736289 | 0.21716725 | 9.00E-04 | 0.0033 | 0.0024 | 0.0024 | TRUE | 0.0255 |
| Lacc1 | 774 | 475 | 461 | 482 | 386 | 496 | 601 | 505 | 497 | 480 | 435 | 490 | 501.44 | 534 | 468 | 0.874 | -0.194 | 0.07913695 | 0.18764174 | 0.006 | 0.0072 | 0.0068 | 0.0068 | TRUE | 0.1454 |
| Lace1 | 722 | 526 | 632 | 699 | 690 | 704 | 561 | 559 | 682 | 696 | 778 | 695 | 661.88 | 601 | 723 | 1.205 | 0.269 | 0.01005003 | 0.04150393 | 0.0068 | 0.006 | 0.0063 | 0.0063 | TRUE | 0.1607 |
| Lactb | 1127 | 786 | 686 | 881 | 642 | 772 | 875 | 836 | 740 | 877 | 724 | 762 | 802.47 | 817 | 788 | 0.963 | -0.054 | 0.59243393 | 0.74024462 | 0.0074 | 0.0054 | 0.006 | 0.006 | TRUE | 0.1168 |
| Lactb2 | 4926 | 3836 | 3976 | 4561 | 3916 | 4458 | 3826 | 4080 | 4290 | 4540 | 4418 | 4401 | 4259.1 | 4065 | 4453 | 1.096 | 0.132 | 0.02890192 | 0.09013641 | 0.0014 | 0.003 | 0.0024 | 0.0024 | TRUE | 0.033 |
| Lactbl1 | 0 | 1 | 0 | 1 | 0 | 1 | 0 | 1 | 0 | 1 | 0 | 1 | 0.51 | 0 | 1 | 1.941 | 0.957 | 0.72720315 | NA | 0 | 4.6706 | 3.6514 | 3.6514 | TRUE | 0.5217 |
| Lad1 | 298 | 250 | 359 | 345 | 384 | 450 | 231 | 266 | 387 | 343 | 433 | 444 | 350.92 | 295 | 407 | 1.384 | 0.469 | 0.00692811 | 0.03148598 | 0.0434 | 0.0092 | 0.0188 | 0.0188 | TRUE | 0.8404 |
| Lag3 | 240 | 168 | 244 | 322 | 322 | 361 | 186 | 179 | 263 | 321 | 363 | 356 | 278.08 | 209 | 347 | 1.66 | 0.731 | 2.34E-06 | 5.72E-05 | 0.0201 | 0.0109 | 0.0135 | 0.0135 | TRUE | 0.5339 |
| Lage3 | 1598 | 1091 | 1143 | 1081 | 1034 | 1149 | 1241 | 1160 | 1233 | 1076 | 1166 | 1134 | 1168.6 | 1211 | 1125 | 0.928 | -0.107 | 0.14659083 | 0.29093505 | 6.00E-04 | 0.0045 | 0.0031 | 0.0031 | TRUE | 0.0178 |
| Lair1 | 3 | 7 | 3 | 3 | 3 | 2 | 2 | 7 | 3 | 3 | 3 | 2 | 3.56 | 4 | 3 | 0.648 | -0.625 | 0.54456458 | NA | 0 | 0.6682 | 0.4679 | 0.4679 | TRUE | 0.6717 |
| Lalba | 0 | 1 | 2 | 1 | 2 | 0 | 0 | 1 | 2 | 1 | 2 | 0 | 1.08 | 1 | 1 | 1.03 | 0.043 | 0.98266035 | NA | 0 | 2.1992 | 1.8152 | 1.8152 | TRUE | 0.2391 |
| Lama1 | 8208 | 5787 | 6067 | 6437 | 6499 | 7081 | 6375 | 6156 | 6546 | 6407 | 7332 | 6990 | 6634.26 | 6359 | 6910 | 1.086 | 0.12 | 0.0581207 | 0.1502762 | 0.0025 | 0.0028 | 0.0027 | 0.0027 | TRUE | 0.0494 |
| Lama2 | 29 | 20 | 14 | 11 | 11 | 16 | 23 | 21 | 15 | 11 | 12 | 16 | 16.34 | 20 | 13 | 0.662 | -0.595 | 0.21581019 | NA | 0 | 0.1474 | 0.1025 | 0.1025 | TRUE | 0.2028 |
| Lama3 | 1063 | 773 | 765 | 787 | 631 | 863 | 826 | 822 | 825 | 783 | 712 | 852 | 803.4 | 824 | 782 | 0.95 | -0.074 | 0.40412902 | 0.57710222 | 0.0026 | 0.0054 | 0.0045 | 0.0045 | TRUE | 0.072 |
| Lama4 | 173 | 104 | 58 | 69 | 44 | 36 | 134 | 111 | 63 | 69 | 50 | 36 | 76.9 | 103 | 52 | 0.498 | -1.006 | 0.00123711 | 0.00843611 | 0.1109 | 0.0332 | 0.0551 | 0.0551 | TRUE | 0.2916 |
| Lama5 | 9269 | 7062 | 6910 | 11638 | 10005 | 11040 | 7199 | 7512 | 7456 | 11584 | 11287 | 10898 | 9322.62 | 7389 | 11256 | 1.524 | 0.608 | 5.67E-33 | 5.78E-30 | 6.00E-04 | 0.0027 | 0.0017 | 0.0017 | TRUE | 0.0095 |
| Lamb1 | 6223 | 4265 | 4025 | 6033 | 4679 | 5163 | 4833 | 4537 | 4343 | 6005 | 5278 | 5097 | 5015.5 | 4571 | 5460 | 1.194 | 0.256 | 0.00051052 | 0.0043825 | 0.005 | 0.0029 | 0.0037 | 0.0037 | TRUE | 0.0933 |
| Lamb2 | 1886 | 1353 | 1367 | 1583 | 1252 | 1535 | 1465 | 1439 | 1475 | 1576 | 1412 | 1515 | 1480.39 | 1460 | 1501 | 1.029 | 0.041 | 0.55834857 | 0.71311815 | 8.00E-04 | 0.004 | 0.0029 | 0.0029 | TRUE | 0.0316 |
| Lamb3 | 220 | 144 | 208 | 313 | 278 | 300 | 171 | 153 | 224 | 312 | 314 | 296 | 244.96 | 183 | 307 | 1.683 | 0.751 | 8.14E-07 | 2.47E-05 | 0.0132 | 0.0121 | 0.0124 | 0.0124 | TRUE | 0.4055 |
| Lamc1 | 23175 | 16316 | 16795 | 18164 | 17185 | 19258 | 17999 | 17355 | 18122 | 18080 | 19386 | 19011 | 18325.55 | 17825 | 18826 | 1.056 | 0.079 | 0.11804837 | 0.24930044 | 8.00E-04 | 0.0026 | 0.0018 | 0.0018 | TRUE | 0.0147 |
| Lamc2 | 2179 | 1663 | 1577 | 1656 | 1509 | 1780 | 1692 | 1769 | 1702 | 1648 | 1702 | 1757 | 1711.77 | 1721 | 1702 | 0.99 | -0.015 | 0.81712266 | 0.89501809 | 0 | 0.0038 | 0.0024 | 0.0024 | TRUE | 0.0096 |
| Lamc3 | 12 | 11 | 10 | 20 | 10 | 15 | 9 | 12 | 11 | 20 | 11 | 15 | 12.97 | 11 | 15 | 1.459 | 0.545 | 0.30964116 | NA | 0 | 0.1852 | 0.1281 | 0.1281 | TRUE | 0.2399 |
| Lamp1 | 34142 | 24049 | 24146 | 25644 | 22749 | 25080 | 26517 | 25581 | 26053 | 25526 | 25663 | 24758 | 25682.94 | 26050 | 25316 | 0.972 | -0.041 | 0.36112708 | 0.53610193 | 3.00E-04 | 0.0025 | 0.0014 | 0.0014 | TRUE | 0.0046 |
| Lamp2 | 2790 | 2180 | 3166 | 3443 | 3110 | 3578 | 2167 | 2319 | 3416 | 3427 | 3508 | 3532 | 3061.57 | 2634 | 3489 | 1.325 | 0.406 | 0.05947863 | 0.1523527 | 0.0331 | 0.0032 | 0.0112 | 0.0331 | TRUE | 0.8178 |
| Lamp3 | 6 | 2 | 3 | 7 | 3 | 3 | 5 | 2 | 3 | 7 | 3 | 3 | 3.89 | 3 | 4 | 1.311 | 0.39 | 0.68748226 | NA | 0 | 0.6118 | 0.4206 | 0.4206 | TRUE | 0.4671 |

| | | | | | | | | | | | | | | | | | | | | | | | | | |
|---|---|---|---|---|---|---|---|---|---|---|---|---|---|---|---|---|---|---|---|---|---|---|---|---|---|
| Lamp5 | 1 | 0 | 0 | 1 | 0 | 0 | 1 | 0 | 0 | 1 | 0 | 0 | 0.3 | 0 | 0 | 1.075 | 0.105 | 0.97698525 | NA | 0 | 8.0272 | 7.5141 | 7.5141 | TRUE | 0.3906 |
| Lamtor1 | 4061 | 2891 | 2538 | 2669 | 2357 | 2681 | 3154 | 3075 | 2738 | 2657 | 2659 | 2647 | 2821.64 | 2989 | 2654 | 0.887 | -0.172 | 0.0110604 | 0.04445772 | 0.0022 | 0.0033 | 0.003 | 0.003 | TRUE | 0.0648 |
| Lamtor2 | 3765 | 2639 | 2848 | 3353 | 2881 | 3334 | 2924 | 2807 | 3073 | 3338 | 3250 | 3291 | 3113.83 | 2935 | 3293 | 1.122 | 0.166 | 0.00487055 | 0.02402993 | 6.00E-04 | 0.0032 | 0.0022 | 0.0022 | TRUE | 0.0201 |
| Lamtor3 | 1431 | 985 | 1024 | 1162 | 1105 | 1240 | 1111 | 1048 | 1105 | 1157 | 1247 | 1224 | 1148.55 | 1088 | 1209 | 1.11 | 0.151 | 0.03996572 | 0.11438327 | 0 | 0.0045 | 0.003 | 0.003 | TRUE | 0.0174 |
| Lamtor4 | 991 | 643 | 567 | 625 | 566 | 610 | 770 | 684 | 612 | 622 | 639 | 602 | 654.7 | 689 | 621 | 0.9 | -0.153 | 0.13439645 | 0.27321014 | 0.0058 | 0.006 | 0.006 | 0.006 | TRUE | 0.1311 |
| Lamtor5 | 2210 | 1573 | 1629 | 1705 | 1433 | 1656 | 1716 | 1673 | 1758 | 1697 | 1617 | 1635 | 1682.62 | 1716 | 1650 | 0.962 | -0.057 | 0.37875386 | 0.55362533 | 0 | 0.0038 | 0.0024 | 0.0024 | TRUE | 0.0077 |
| Lancl1 | 2768 | 2015 | 2148 | 2206 | 1964 | 2257 | 2150 | 2143 | 2318 | 2196 | 2216 | 2228 | 2208.37 | 2204 | 2213 | 1.005 | 0.007 | 0.91235037 | 0.95183773 | 6.00E-04 | 0.0035 | 0.0023 | 0.0023 | TRUE | 0.0244 |
| Lancl2 | 1921 | 1402 | 1298 | 1307 | 1139 | 1188 | 1492 | 1491 | 1401 | 1301 | 1285 | 1173 | 1357.07 | 1461 | 1253 | 0.857 | -0.223 | 0.00256096 | 0.01464736 | 0.0014 | 0.0042 | 0.0032 | 0.0032 | TRUE | 0.0381 |
| Lancl3 | 38 | 26 | 9 | 18 | 15 | 20 | 30 | 28 | 10 | 18 | 17 | 20 | 20.24 | 23 | 18 | 0.809 | -0.306 | 0.52411604 | 0.68550426 | 0.106 | 0.1195 | 0.1159 | 0.1159 | TRUE | 1.2974 |
| Lao1 | 0 | 0 | 0 | 0 | 0 | 0 | 0 | 0 | 0 | 0 | 0 | 0 | 0 | 0 | 0 | NA | NA | NA | NA | NA | NA | NA | NA | NA | NA |
| Lap3 | 16623 | 12693 | 13788 | 13990 | 13165 | 15271 | 12910 | 13501 | 14877 | 13926 | 14851 | 15075 | 14190.14 | 13763 | 14617 | 1.062 | 0.087 | 0.17801938 | 0.33250145 | 0.0035 | 0.0026 | 0.0029 | 0.0029 | TRUE | 0.0613 |
| Laptm4a | 9960 | 7085 | 7663 | 8230 | 7401 | 8064 | 7736 | 7536 | 8268 | 8192 | 8349 | 7960 | 8006.95 | 7847 | 8167 | 1.041 | 0.058 | 0.29890991 | 0.47129832 | 0.0013 | 0.0027 | 0.0021 | 0.0021 | TRUE | 0.0267 |
| Laptm4b | 6758 | 5192 | 5816 | 6285 | 5793 | 6650 | 5249 | 5523 | 6275 | 6256 | 6535 | 6565 | 6067.08 | 5682 | 6452 | 1.136 | 0.184 | 0.0094378 | 0.03972157 | 0.0045 | 0.0028 | 0.0034 | 0.0034 | TRUE | 0.1014 |
| Laptm5 | 1517 | 1203 | 1262 | 1680 | 1518 | 1723 | 1178 | 1280 | 1362 | 1672 | 1712 | 1701 | 1484.18 | 1273 | 1695 | 1.333 | 0.415 | 2.21E-08 | 1.01E-06 | 0.002 | 0.004 | 0.0033 | 0.0033 | TRUE | 0.0535 |
| Large | 1432 | 995 | 1007 | 1042 | 958 | 1180 | 1112 | 1058 | 1087 | 1037 | 1081 | 1165 | 1089.98 | 1086 | 1094 | 1.008 | 0.011 | 0.88872747 | 0.93815227 | 0.0012 | 0.0046 | 0.0034 | 0.0034 | TRUE | 0.0388 |
| Larp1 | 36478 | 26578 | 25314 | 28399 | 25753 | 30704 | 28331 | 28271 | 27314 | 28268 | 29052 | 30310 | 28590.86 | 27972 | 29210 | 1.044 | 0.062 | 0.20368489 | 0.36500353 | 8.00E-04 | 0.0025 | 0.0017 | 0.0017 | TRUE | 0.0133 |
| Larp1b | 1413 | 1073 | 1149 | 1400 | 1179 | 1378 | 1097 | 1141 | 1240 | 1394 | 1330 | 1360 | 1260.4 | 1159 | 1361 | 1.176 | 0.233 | 0.00188058 | 0.01154106 | 0.0014 | 0.0043 | 0.0033 | 0.0033 | TRUE | 0.0435 |
| Larp4 | 12678 | 9101 | 9179 | 10586 | 8860 | 10226 | 9846 | 9681 | 9904 | 10537 | 9995 | 10095 | 10009.68 | 9810 | 10209 | 1.041 | 0.058 | 0.2355711 | 0.40203197 | 3.00E-04 | 0.0027 | 0.0016 | 0.0016 | TRUE | 0.0097 |
| Larp4b | 7209 | 5220 | 4900 | 5227 | 4514 | 5215 | 5599 | 5552 | 5287 | 5203 | 5092 | 5148 | 5313.61 | 5479 | 5148 | 0.939 | -0.09 | 0.08205176 | 0.19260401 | 4.00E-04 | 0.0029 | 0.0018 | 0.0018 | TRUE | 0.0114 |
| Larp6 | 37 | 11 | 16 | 27 | 19 | 14 | 29 | 12 | 17 | 27 | 21 | 14 | 19.97 | 19 | 21 | 1.06 | 0.084 | 0.86143041 | 0.92331645 | 0.1071 | 0.1211 | 0.1171 | 0.1171 | TRUE | 0.6381 |
| Larp7 | 4953 | 3691 | 3134 | 3162 | 2745 | 3097 | 3847 | 3926 | 3382 | 3147 | 3097 | 3057 | 3409.29 | 3718 | 3100 | 0.834 | -0.263 | 0.00013004 | 0.00151481 | 0.0028 | 0.0031 | 0.0031 | 0.0031 | TRUE | 0.0754 |
| Lars | 14367 | 10079 | 9643 | 10160 | 8908 | 10378 | 11158 | 10721 | 10405 | 10113 | 10049 | 10245 | 10448.49 | 10761 | 10136 | 0.942 | -0.087 | 0.08253897 | 0.19342383 | 5.00E-04 | 0.0027 | 0.0017 | 0.0017 | TRUE | 0.0129 |
| Lars2 | 18302 | 13496 | 16662 | 19232 | 15260 | 18906 | 14214 | 14356 | 17978 | 19144 | 17215 | 18663 | 16928.26 | 15516 | 18341 | 1.182 | 0.241 | 0.00505491 | 0.02474042 | 0.0109 | 0.0026 | 0.0053 | 0.0053 | TRUE | 0.2354 |
| Las1l | 8218 | 5986 | 5793 | 5775 | 5336 | 6254 | 6383 | 6367 | 6251 | 5748 | 6020 | 6174 | 6157.01 | 6334 | 5981 | 0.944 | -0.083 | 0.11345059 | 0.24238405 | 5.00E-04 | 0.0028 | 0.0018 | 0.0018 | TRUE | 0.0142 |
| Lasp1 | 18844 | 13473 | 13008 | 15978 | 14382 | 16570 | 14635 | 14331 | 14036 | 15904 | 16224 | 16357 | 15247.99 | 14334 | 16162 | 1.127 | 0.173 | 0.00017244 | 0.00190175 | 2.00E-04 | 0.0026 | 0.0015 | 0.0015 | TRUE | 0.0042 |
| Lat | 363 | 254 | 223 | 326 | 268 | 341 | 282 | 270 | 241 | 325 | 302 | 337 | 292.69 | 264 | 321 | 1.213 | 0.279 | 0.02513045 | 0.08157836 | 0 | 0.0105 | 0.0078 | 0.0078 | TRUE | 0.0663 |
| Lat2 | 41 | 34 | 28 | 46 | 32 | 38 | 32 | 36 | 30 | 46 | 36 | 38 | 36.27 | 33 | 40 | 1.218 | 0.285 | 0.37200975 | 0.54658053 | 0 | 0.0678 | 0.0457 | 0.0457 | TRUE | 0.1317 |
| Lats1 | 1397 | 1032 | 1123 | 1268 | 1081 | 1186 | 1085 | 1098 | 1212 | 1262 | 1219 | 1171 | 1174.47 | 1132 | 1217 | 1.077 | 0.107 | 0.16764151 | 0.3190971 | 0.0017 | 0.0044 | 0.0035 | 0.0035 | TRUE | 0.0453 |
| Lats2 | 419 | 282 | 202 | 214 | 176 | 160 | 325 | 300 | 218 | 213 | 199 | 158 | 235.47 | 281 | 190 | 0.673 | -0.571 | 0.00104402 | 0.00746594 | 0.0292 | 0.0125 | 0.0174 | 0.0174 | TRUE | 0.4244 |
| Lax1 | 25 | 17 | 13 | 10 | 11 | 8 | 19 | 18 | 14 | 10 | 12 | 8 | 13.63 | 17 | 10 | 0.582 | -0.781 | 0.13728954 | NA | 0 | 0.1763 | 0.1196 | 0.1196 | TRUE | 0.1289 |
| Layn | 93 | 90 | 83 | 110 | 86 | 96 | 72 | 96 | 90 | 109 | 97 | 95 | 93.13 | 86 | 100 | 1.177 | 0.235 | 0.26312895 | 0.43335254 | 0 | 0.0279 | 0.0211 | 0.0211 | TRUE | 0.2087 |
| Lbh | 345 | 291 | 292 | 258 | 252 | 253 | 268 | 310 | 315 | 257 | 284 | 250 | 280.57 | 298 | 264 | 0.888 | -0.172 | 0.18455736 | 0.34072312 | 0.0032 | 0.0109 | 0.0085 | 0.0085 | TRUE | 0.0924 |
| Lbp | 35 | 42 | 31 | 51 | 53 | 46 | 27 | 45 | 33 | 51 | 60 | 45 | 43.55 | 35 | 52 | 1.491 | 0.576 | 0.06379195 | 0.16046785 | 0.0156 | 0.0569 | 0.0458 | 0.0458 | TRUE | 0.3975 |
| Lbr | 16117 | 11660 | 12371 | 13900 | 12086 | 13577 | 12517 | 12403 | 13348 | 13836 | 13634 | 13403 | 13190.18 | 12756 | 13624 | 1.068 | 0.095 | 0.06281338 | 0.15859241 | 8.00E-04 | 0.0026 | 0.0018 | 0.0018 | TRUE | 0.0201 |
| Lbx1 | 29 | 14 | 13 | 26 | 12 | 10 | 23 | 15 | 14 | 26 | 14 | 10 | 16.79 | 17 | 17 | 0.951 | -0.073 | 0.88754841 | NA | 0.1019 | 0.1436 | 0.1316 | 0.1316 | TRUE | 1.2527 |
| Lbx2 | 8 | 1 | 1 | 2 | 2 | 2 | 6 | 1 | 1 | 2 | 2 | 2 | 2.43 | 3 | 2 | 0.715 | -0.484 | 0.70238072 | NA | 0 | 0.9779 | 0.7412 | 0.7412 | TRUE | 1.8152 |
| Lca5 | 1054 | 723 | 819 | 851 | 792 | 901 | 819 | 769 | 884 | 847 | 893 | 889 | 850.22 | 824 | 876 | 1.064 | 0.09 | 0.28823887 | 0.46036409 | 0.0015 | 0.0052 | 0.004 | 0.004 | TRUE | 0.0459 |
| Lca5l | 9 | 7 | 10 | 8 | 3 | 2 | 7 | 7 | 11 | 8 | 3 | 2 | 6.42 | 8 | 4 | 0.533 | -0.908 | 0.25587408 | NA | 0.0219 | 0.3713 | 0.2863 | 0.2863 | TRUE | 0.9123 |
| Lcat | 13 | 20 | 16 | 19 | 15 | 25 | 10 | 21 | 17 | 19 | 17 | 25 | 18.19 | 16 | 20 | 1.265 | 0.339 | 0.47268912 | 0.64119729 | 0 | 0.1327 | 0.1053 | 0.1053 | TRUE | 0.6383 |
| Lce1a1 | 0 | 0 | 0 | 0 | 0 | 0 | 0 | 0 | 0 | 0 | 0 | 0 | 0 | 0 | 0 | NA | NA | NA | NA | NA | NA | NA | NA | NA | NA |
| Lce1a2 | 0 | 0 | 0 | 0 | 0 | 0 | 0 | 0 | 0 | 0 | 0 | 0 | 0 | 0 | 0 | NA | NA | NA | NA | NA | NA | NA | NA | NA | NA |
| Lce1b | 0 | 0 | 0 | 0 | 0 | 0 | 0 | 0 | 0 | 0 | 0 | 0 | 0 | 0 | 0 | NA | NA | NA | NA | NA | NA | NA | NA | NA | NA |
| Lce1c | 0 | 0 | 0 | 0 | 0 | 0 | 0 | 0 | 0 | 0 | 0 | 0 | 0 | 0 | 0 | NA | NA | NA | NA | NA | NA | NA | NA | NA | NA |
| Lce1d | 0 | 0 | 0 | 0 | 0 | 0 | 0 | 0 | 0 | 0 | 0 | 0 | 0 | 0 | 0 | NA | NA | NA | NA | NA | NA | NA | NA | NA | NA |
| Lce1e | 0 | 0 | 0 | 0 | 0 | 0 | 0 | 0 | 0 | 0 | 0 | 0 | 0 | 0 | 0 | NA | NA | NA | NA | NA | NA | NA | NA | NA | NA |
| Lce1f | 0 | 1 | 0 | 0 | 0 | 0 | 0 | 1 | 0 | 0 | 0 | 0 | 0.18 | 0 | 0 | 0.552 | -0.857 | 0.8336548 | NA | 0 | 13.3714 | 10 | 10 | TRUE | 0.5216 |
| Lce1g | 0 | 0 | 0 | 0 | 0 | 0 | 0 | 0 | 0 | 0 | 0 | 0 | 0 | 0 | 0 | NA | NA | NA | NA | NA | NA | NA | NA | NA | NA |
| Lce1h | 0 | 0 | 0 | 0 | 0 | 0 | 0 | 0 | 0 | 0 | 0 | 0 | 0 | 0 | 0 | NA | NA | NA | NA | NA | NA | NA | NA | NA | NA |
| Lce1i | 0 | 0 | 0 | 0 | 0 | 0 | 0 | 0 | 0 | 0 | 0 | 0 | 0 | 0 | 0 | NA | NA | NA | NA | NA | NA | NA | NA | NA | NA |
| Lce1k | 0 | 0 | 0 | 0 | 0 | 0 | 0 | 0 | 0 | 0 | 0 | 0 | 0 | 0 | 0 | NA | NA | NA | NA | NA | NA | NA | NA | NA | NA |
| Lce1l | 0 | 0 | 0 | 0 | 0 | 0 | 0 | 0 | 0 | 0 | 0 | 0 | 0 | 0 | 0 | NA | NA | NA | NA | NA | NA | NA | NA | NA | NA |
| Lce1m | 0 | 0 | 0 | 0 | 0 | 0 | 0 | 0 | 0 | 0 | 0 | 0 | 0 | 0 | 0 | NA | NA | NA | NA | NA | NA | NA | NA | NA | NA |
| Lce3a | 0 | 0 | 0 | 0 | 0 | 0 | 0 | 0 | 0 | 0 | 0 | 0 | 0 | 0 | 0 | NA | NA | NA | NA | NA | NA | NA | NA | NA | NA |
| Lce3b | 0 | 0 | 0 | 0 | 0 | 0 | 0 | 0 | 0 | 0 | 0 | 0 | 0 | 0 | 0 | NA | NA | NA | NA | NA | NA | NA | NA | NA | NA |
| Lce3c | 0 | 0 | 0 | 0 | 0 | 0 | 0 | 0 | 0 | 0 | 0 | 0 | 0 | 0 | 0 | NA | NA | NA | NA | NA | NA | NA | NA | NA | NA |
| Lce3d | 0 | 0 | 0 | 0 | 0 | 0 | 0 | 0 | 0 | 0 | 0 | 0 | 0 | 0 | 0 | NA | NA | NA | NA | NA | NA | NA | NA | NA | NA |
| Lce3e | 0 | 0 | 0 | 0 | 0 | 0 | 0 | 0 | 0 | 0 | 0 | 0 | 0 | 0 | 0 | NA | NA | NA | NA | NA | NA | NA | NA | NA | NA |
| Lce3f | 0 | 0 | 0 | 0 | 0 | 0 | 0 | 0 | 0 | 0 | 0 | 0 | 0 | 0 | 0 | NA | NA | NA | NA | NA | NA | NA | NA | NA | NA |
| Lce6a | 0 | 0 | 0 | 0 | 0 | 0 | 0 | 0 | 0 | 0 | 0 | 0 | 0 | 0 | 0 | NA | NA | NA | NA | NA | NA | NA | NA | NA | NA |
| Lck | 1062 | 836 | 665 | 675 | 543 | 697 | 825 | 889 | 718 | 672 | 613 | 688 | 734.01 | 811 | 658 | 0.811 | -0.302 | 0.00263549 | 0.01492239 | 0.0063 | 0.0057 | 0.0059 | 0.0059 | TRUE | 0.1157 |
| Lclat1 | 1313 | 988 | 1062 | 1025 | 895 | 1063 | 1020 | 1051 | 1146 | 1020 | 1010 | 1049 | 1049.31 | 1072 | 1026 | 0.959 | -0.061 | 0.43412046 | 0.60582736 | 0.0012 | 0.0047 | 0.0034 | 0.0034 | TRUE | 0.0427 |
| Lcmt1 | 2835 | 2230 | 2257 | 2440 | 2123 | 2573 | 2202 | 2372 | 2435 | 2429 | 2395 | 2540 | 2395.47 | 2336 | 2455 | 1.052 | 0.073 | 0.26573906 | 0.43612452 | 0.0015 | 0.0034 | 0.0026 | 0.0026 | TRUE | 0.0317 |
| Lcmt2 | 638 | 447 | 285 | 305 | 265 | 248 | 496 | 475 | 308 | 304 | 299 | 245 | 354.31 | 426 | 283 | 0.661 | -0.597 | 0.00038525 | 0.00353203 | 0.0372 | 0.0091 | 0.0174 | 0.0174 | TRUE | 0.6734 |
| Lcn10 | 0 | 0 | 0 | 0 | 0 | 0 | 0 | 0 | 0 | 0 | 0 | 0 | 0 | 0 | 0 | NA | NA | NA | NA | NA | NA | NA | NA | NA | NA |
| Lcn11 | 0 | 0 | 0 | 0 | 0 | 0 | 0 | 0 | 0 | 0 | 0 | 0 | 0 | 0 | 0 | NA | NA | NA | NA | NA | NA | NA | NA | NA | NA |
| Lcn12 | 0 | 0 | 0 | 0 | 0 | 0 | 0 | 0 | 0 | 0 | 0 | 0 | 0 | 0 | 0 | NA | NA | NA | NA | NA | NA | NA | NA | NA | NA |
| Lcn2 | 0 | 0 | 0 | 2 | 0 | 0 | 0 | 0 | 0 | 2 | 0 | 0 | 0.33 | 0 | 1 | 3.77 | 1.915 | 0.62344813 | NA | 0 | 7.1455 | 9.1834 | 9.1834 | TRUE | 0.9739 |
| Lcn3 | 0 | 0 | 0 | 0 | 0 | 0 | 0 | 0 | 0 | 0 | 0 | 0 | 0 | 0 | 0 | NA | NA | NA | NA | NA | NA | NA | NA | NA | NA |
| Lcn4 | 0 | 0 | 0 | 0 | 0 | 0 | 0 | 0 | 0 | 0 | 0 | 0 | 0 | 0 | 0 | NA | NA | NA | NA | NA | NA | NA | NA | NA | NA |

| | | | | | | | | | | | | | | | | | | | | | | | | | |
|---|---|---|---|---|---|---|---|---|---|---|---|---|---|---|---|---|---|---|---|---|---|---|---|---|---|
| Lcn5 | 0 | 0 | 0 | 0 | 0 | 0 | 0 | 0 | 0 | 0 | 0 | 0 | 0 | 0 | 0 | NA | NA | NA | NA | NA | NA | NA | NA | NA | NA |
| Lcn6 | 0 | 0 | 0 | 0 | 0 | 0 | 0 | 0 | 0 | 0 | 0 | 0 | 0 | 0 | 0 | NA | NA | NA | NA | NA | NA | NA | NA | NA | NA |
| Lcn8 | 0 | 0 | 0 | 0 | 0 | 0 | 0 | 0 | 0 | 0 | 0 | 0 | 0 | 0 | 0 | NA | NA | NA | NA | NA | NA | NA | NA | NA | NA |
| Lcn9 | 2 | 0 | 0 | 0 | 1 | 0 | 2 | 0 | 0 | 0 | 1 | 0 | 0.45 | 1 | 0 | 0.68 | -0.556 | 0.85993441 | NA | 0 | 5.3057 | 5.2576 | 5.2576 | TRUE | 0.9726 |
| Lcor | 248 | 179 | 182 | 170 | 167 | 182 | 193 | 190 | 196 | 169 | 188 | 180 | 186.11 | 193 | 179 | 0.927 | -0.11 | 0.4488155 | 0.61896178 | 0 | 0.0152 | 0.0098 | 0.0098 | TRUE | 0.025 |
| Lcorl | 821 | 723 | 869 | 1130 | 963 | 1229 | 638 | 769 | 938 | 1125 | 1086 | 1213 | 961.45 | 782 | 1141 | 1.464 | 0.55 | 3.97E-06 | 8.78E-05 | 0.0185 | 0.0049 | 0.0092 | 0.0092 | TRUE | 0.3625 |
| Lcp1 | 178 | 158 | 125 | 48 | 58 | 71 | 138 | 168 | 135 | 48 | 65 | 70 | 104.08 | 147 | 61 | 0.416 | -1.267 | 4.74E-09 | 2.66E-07 | 0.0122 | 0.0252 | 0.022 | 0.022 | TRUE | 0.3248 |
| Lcp2 | 5 | 2 | 1 | 0 | 0 | 2 | 4 | 2 | 1 | 0 | 0 | 2 | 1.51 | 2 | 1 | 0.281 | -1.831 | 0.29858965 | NA | 0 | 1.5713 | 1.2683 | 1.2683 | TRUE | 0.8202 |
| Lct | 42 | 41 | 30 | 32 | 31 | 30 | 33 | 44 | 32 | 32 | 35 | 30 | 34.17 | 36 | 32 | 0.89 | -0.169 | 0.61098974 | 0.75392458 | 0 | 0.0718 | 0.0497 | 0.0497 | TRUE | 0.2197 |
| Lctl | 91 | 71 | 65 | 90 | 47 | 70 | 71 | 76 | 70 | 90 | 53 | 69 | 71.34 | 72 | 71 | 0.982 | -0.026 | 0.91860543 | 0.95536837 | 0.0185 | 0.0356 | 0.031 | 0.031 | TRUE | 0.261 |
| Ldb1 | 5905 | 4748 | 4977 | 4618 | 4489 | 5453 | 4586 | 5050 | 5370 | 4597 | 5064 | 5383 | 5008.41 | 5002 | 5015 | 1.003 | 0.004 | 0.95748559 | 0.97670491 | 0.0061 | 0.0029 | 0.0041 | 0.0041 | TRUE | 0.0652 |
| Ldb2 | 0 | 0 | 1 | 0 | 0 | 0 | 0 | 0 | 1 | 0 | 0 | 0 | 0.18 | 0 | 0 | 0.552 | -0.857 | 0.8336548 | NA | 0 | 13.1817 | 10 | 10 | TRUE | 0.5365 |
| Ldb3 | 583 | 414 | 565 | 719 | 707 | 810 | 453 | 440 | 610 | 716 | 798 | 800 | 635.94 | 501 | 771 | 1.542 | 0.624 | 3.69E-07 | 1.25E-05 | 0.0172 | 0.0062 | 0.0092 | 0.0092 | TRUE | 0.4115 |
| Ldha | 83026 | 64933 | 76682 | 75474 | 69839 | 81044 | 64483 | 69068 | 82739 | 75127 | 78786 | 80003 | 75034.28 | 72097 | 77972 | 1.082 | 0.113 | 0.16438648 | 0.31522329 | 0.0087 | 0.0025 | 0.0048 | 0.0048 | TRUE | 0.2042 |
| Ldhal6b | 21 | 19 | 25 | 29 | 30 | 25 | 16 | 20 | 27 | 29 | 34 | 25 | 25.15 | 21 | 29 | 1.387 | 0.472 | 0.23247265 | 0.39840907 | 0 | 0.0967 | 0.0716 | 0.0716 | TRUE | 0.3091 |
| Ldhb | 3194 | 2763 | 3516 | 3564 | 3413 | 4075 | 2481 | 2939 | 3794 | 3548 | 3850 | 4023 | 3438.97 | 3071 | 3807 | 1.24 | 0.31 | 0.09844693 | 0.21859593 | 0.0251 | 0.0031 | 0.0096 | 0.0251 | TRUE | 0.5145 |
| Ldhc | 118 | 84 | 94 | 57 | 50 | 69 | 92 | 89 | 101 | 57 | 56 | 68 | 77.28 | 94 | 60 | 0.643 | -0.636 | 0.00462237 | 0.0230937 | 0 | 0.0331 | 0.0227 | 0.0227 | TRUE | 0.1098 |
| Ldhd | 108 | 82 | 126 | 100 | 105 | 119 | 84 | 87 | 136 | 100 | 118 | 117 | 107.09 | 102 | 112 | 1.098 | 0.134 | 0.5491833 | 0.70563439 | 0.0326 | 0.0246 | 0.0269 | 0.0269 | TRUE | 0.7857 |
| Ldlr | 8226 | 6196 | 6304 | 7003 | 6399 | 7171 | 6389 | 6591 | 6802 | 6971 | 7219 | 7079 | 6841.63 | 6594 | 7090 | 1.075 | 0.105 | 0.04117833 | 0.11674104 | 4.00E-04 | 0.0028 | 0.0018 | 0.0018 | TRUE | 0.0093 |
| Ldlrad1 | 0 | 0 | 0 | 0 | 0 | 0 | 0 | 0 | 0 | 0 | 0 | 0 | 0 | 0 | 0 | NA | NA | NA | NA | NA | NA | NA | NA | NA | NA |
| Ldlrad2 | 0 | 0 | 0 | 0 | 0 | 0 | 0 | 0 | 0 | 0 | 0 | 0 | 0 | 0 | 0 | NA | NA | NA | NA | NA | NA | NA | NA | NA | NA |
| Ldlrad3 | 4280 | 3125 | 3382 | 4009 | 3556 | 4207 | 3324 | 3324 | 3649 | 3991 | 4012 | 4153 | 3742.05 | 3432 | 4052 | 1.181 | 0.24 | 7.73E-05 | 0.00098723 | 0.0013 | 0.0031 | 0.0024 | 0.0024 | TRUE | 0.0367 |
| Ldlrad4 | 50 | 28 | 30 | 43 | 49 | 50 | 39 | 30 | 32 | 43 | 55 | 49 | 41.4 | 34 | 49 | 1.448 | 0.534 | 0.07936481 | 0.18802607 | 0 | 0.0597 | 0.042 | 0.042 | TRUE | 0.1508 |
| Ldlrap1 | 1574 | 1211 | 1234 | 1496 | 1327 | 1435 | 1222 | 1288 | 1331 | 1489 | 1497 | 1417 | 1374.12 | 1280 | 1468 | 1.147 | 0.198 | 0.0052246 | 0.02534801 | 7.00E-04 | 0.0042 | 0.0029 | 0.0029 | TRUE | 0.0199 |
| Ldoc1 | 186 | 152 | 126 | 115 | 114 | 132 | 144 | 162 | 136 | 114 | 129 | 130 | 135.91 | 147 | 124 | 0.845 | -0.244 | 0.15795796 | 0.30687686 | 0 | 0.0199 | 0.014 | 0.014 | TRUE | 0.0713 |
| Ldoc1l | 515 | 389 | 339 | 325 | 307 | 349 | 400 | 414 | 366 | 324 | 346 | 345 | 365.65 | 393 | 338 | 0.859 | -0.219 | 0.04938669 | 0.13358893 | 0 | 0.0089 | 0.0062 | 0.0062 | TRUE | 0.0406 |
| Leap2 | 0 | 0 | 0 | 0 | 0 | 0 | 0 | 0 | 0 | 0 | 0 | 0 | 0 | 0 | 0 | NA | NA | NA | NA | NA | NA | NA | NA | NA | NA |
| Lect1 | 0 | 2 | 0 | 0 | 2 | 3 | 0 | 2 | 0 | 0 | 2 | 3 | 1.22 | 1 | 2 | 2.552 | 1.352 | 0.50518172 | NA | 2.2758 | 1.9385 | 1.9582 | 1.9582 | TRUE | 0.9751 |
| Lect2 | 0 | 0 | 0 | 0 | 0 | 0 | 0 | 0 | 0 | 0 | 0 | 0 | 0 | 0 | 0 | NA | NA | NA | NA | NA | NA | NA | NA | NA | NA |
| Lef1 | 891 | 709 | 896 | 888 | 822 | 1211 | 692 | 754 | 967 | 884 | 927 | 1195 | 903.27 | 804 | 1002 | 1.248 | 0.32 | 0.01710564 | 0.06169339 | 0.0285 | 0.0051 | 0.0118 | 0.0118 | TRUE | 0.3686 |
| Lefty1 | 1382 | 829 | 641 | 637 | 669 | 746 | 1073 | 882 | 692 | 634 | 755 | 736 | 795.33 | 882 | 708 | 0.801 | -0.32 | 0.01844935 | 0.06507812 | 0.0257 | 0.0054 | 0.012 | 0.012 | TRUE | 0.1533 |
| Lefty2 | 5136 | 3940 | 2876 | 2832 | 2662 | 2606 | 3989 | 4191 | 3103 | 2819 | 3003 | 2573 | 3279.59 | 3761 | 2798 | 0.744 | -0.427 | 2.40E-05 | 0.0003854 | 0.0143 | 0.0031 | 0.0071 | 0.0071 | TRUE | 0.2832 |
| Lekr1 | 65 | 72 | 63 | 62 | 59 | 61 | 50 | 77 | 68 | 62 | 67 | 60 | 63.92 | 65 | 63 | 0.973 | -0.04 | 0.87622303 | 0.93096567 | 0 | 0.0395 | 0.0313 | 0.0313 | TRUE | 0.3907 |
| Lelp1 | 0 | 0 | 0 | 0 | 0 | 0 | 0 | 0 | 0 | 0 | 0 | 0 | 0 | 0 | 0 | NA | NA | NA | NA | NA | NA | NA | NA | NA | NA |
| Lemd1 | 9 | 2 | 3 | 3 | 2 | 3 | 7 | 2 | 3 | 3 | 2 | 3 | 3.43 | 4 | 3 | 0.651 | -0.619 | 0.55203938 | NA | 0 | 0.6941 | 0.4783 | 0.4783 | TRUE | 0.8296 |
| Lemd2 | 1459 | 1255 | 1274 | 1434 | 1307 | 1422 | 1133 | 1335 | 1375 | 1427 | 1474 | 1404 | 1358.05 | 1281 | 1435 | 1.122 | 0.166 | 0.05010561 | 0.13499707 | 0.005 | 0.0042 | 0.0045 | 0.0045 | TRUE | 0.127 |
| Lemd3 | 2449 | 1744 | 1674 | 1718 | 1494 | 1579 | 1902 | 1855 | 1806 | 1710 | 1685 | 1559 | 1752.92 | 1854 | 1651 | 0.89 | -0.168 | 0.01286992 | 0.04983737 | 0.001 | 0.0038 | 0.0027 | 0.0027 | TRUE | 0.0294 |
| Lenep | 69 | 52 | 46 | 44 | 42 | 52 | 54 | 55 | 50 | 44 | 47 | 51 | 50.17 | 53 | 47 | 0.898 | -0.154 | 0.56664532 | 0.72074944 | 0 | 0.0497 | 0.0323 | 0.0323 | TRUE | 0.0415 |
| Leng1 | 414 | 295 | 289 | 295 | 296 | 343 | 322 | 314 | 312 | 294 | 334 | 339 | 318.88 | 316 | 322 | 1.019 | 0.027 | 0.81812904 | 0.89536894 | 0 | 0.0099 | 0.0069 | 0.0069 | TRUE | 0.0688 |
| Leng8 | 6172 | 6066 | 5610 | 5833 | 4984 | 6089 | 4794 | 6452 | 6053 | 5806 | 5622 | 6011 | 5789.74 | 5766 | 5813 | 1.009 | 0.012 | 0.89357459 | 0.9417388 | 0.0123 | 0.0028 | 0.006 | 0.006 | TRUE | 0.2677 |
| Leng9 | 15 | 18 | 18 | 12 | 12 | 16 | 12 | 19 | 19 | 12 | 14 | 16 | 15.25 | 17 | 14 | 0.831 | -0.266 | 0.59288654 | NA | 0 | 0.1578 | 0.1121 | 0.1121 | TRUE | 0.437 |
| Leo1 | 4828 | 3492 | 3039 | 3316 | 2937 | 3183 | 3750 | 3714 | 3279 | 3301 | 3313 | 3142 | 3416.54 | 3581 | 3252 | 0.908 | -0.14 | 0.03979512 | 0.11412351 | 0.0027 | 0.0031 | 0.003 | 0.003 | TRUE | 0.0656 |
| Lep | 0 | 0 | 0 | 0 | 0 | 0 | 0 | 0 | 0 | 0 | 0 | 0 | 0 | 0 | 0 | NA | NA | NA | NA | NA | NA | NA | NA | NA | NA |
| Lepr | 285 | 244 | 263 | 321 | 248 | 244 | 221 | 260 | 284 | 320 | 280 | 241 | 267.47 | 255 | 280 | 1.102 | 0.141 | 0.34542243 | 0.51953308 | 0.0144 | 0.0113 | 0.0122 | 0.0122 | TRUE | 0.1744 |
| Lepre1 | 2814 | 2020 | 2096 | 2391 | 2213 | 2436 | 2186 | 2149 | 2262 | 2380 | 2496 | 2405 | 2312.82 | 2199 | 2427 | 1.104 | 0.143 | 0.01740235 | 0.06230756 | 2.00E-04 | 0.0035 | 0.0022 | 0.0022 | TRUE | 0.0075 |
| Leprel1 | 70 | 37 | 21 | 22 | 18 | 29 | 54 | 39 | 23 | 22 | 20 | 29 | 31.2 | 39 | 24 | 0.604 | -0.728 | 0.06779482 | 0.16750067 | 0.0843 | 0.0784 | 0.0801 | 0.0801 | TRUE | 0.1353 |
| Leprel2 | 277 | 232 | 155 | 207 | 176 | 219 | 215 | 247 | 167 | 206 | 199 | 216 | 208.32 | 210 | 207 | 0.986 | -0.02 | 0.90162957 | 0.94568228 | 0.0143 | 0.0138 | 0.0139 | 0.0139 | TRUE | 0.3091 |
| Leprel4 | 477 | 374 | 398 | 561 | 485 | 596 | 370 | 398 | 429 | 558 | 547 | 588 | 481.94 | 399 | 564 | 1.418 | 0.504 | 8.71E-07 | 2.60E-05 | 0.0011 | 0.0073 | 0.0054 | 0.0054 | TRUE | 0.0501 |
| Leprot | 4447 | 3399 | 3870 | 3519 | 3302 | 3743 | 3454 | 3615 | 4176 | 3503 | 3725 | 3695 | 3694.62 | 3748 | 3641 | 0.972 | -0.041 | 0.58682296 | 0.736065 | 0.0046 | 0.0031 | 0.0039 | 0.0039 | TRUE | 0.1206 |
| Leprotl1 | 2484 | 1788 | 1831 | 1808 | 1629 | 1931 | 1929 | 1902 | 1976 | 1800 | 1838 | 1906 | 1891.71 | 1936 | 1848 | 0.955 | -0.067 | 0.28527802 | 0.45727025 | 0 | 0.0037 | 0.0023 | 0.0023 | TRUE | 0.0094 |
| Letm1 | 4575 | 3264 | 3001 | 3249 | 2876 | 3385 | 3553 | 3472 | 3238 | 3234 | 3244 | 3342 | 3347.19 | 3421 | 3273 | 0.956 | -0.064 | 0.28229076 | 0.45400943 | 0.001 | 0.0031 | 0.0023 | 0.0023 | TRUE | 0.0264 |
| Letm2 | 128 | 92 | 92 | 142 | 120 | 120 | 99 | 98 | 99 | 141 | 135 | 118 | 115.29 | 99 | 131 | 1.332 | 0.413 | 0.0245618 | 0.08028517 | 0 | 0.023 | 0.0155 | 0.0155 | TRUE | 0.082 |
| Letmd1 | 482 | 352 | 349 | 408 | 326 | 318 | 374 | 374 | 377 | 406 | 368 | 314 | 368.86 | 375 | 363 | 0.966 | -0.049 | 0.68417427 | 0.80650746 | 0.0055 | 0.0089 | 0.0078 | 0.0078 | TRUE | 0.1618 |
| Lfng | 106 | 88 | 109 | 123 | 116 | 133 | 82 | 94 | 118 | 122 | 131 | 131 | 113.02 | 98 | 128 | 1.318 | 0.398 | 0.04356126 | 0.12184367 | 0.0075 | 0.0234 | 0.019 | 0.019 | TRUE | 0.2971 |
| Lgals1 | 1367 | 975 | 1015 | 1207 | 1179 | 1396 | 1062 | 1037 | 1095 | 1201 | 1330 | 1378 | 1183.92 | 1065 | 1303 | 1.224 | 0.292 | 0.00020848 | 0.00219585 | 0.0021 | 0.0044 | 0.0036 | 0.0036 | TRUE | 0.0567 |
| Lgals12 | 53 | 40 | 42 | 78 | 65 | 62 | 41 | 43 | 45 | 78 | 73 | 61 | 56.87 | 43 | 71 | 1.647 | 0.72 | 0.00569204 | 0.02710973 | 0 | 0.0441 | 0.0302 | 0.0302 | TRUE | 0.1288 |
| Lgals2 | 2 | 1 | 2 | 2 | 1 | 3 | 2 | 1 | 2 | 2 | 1 | 3 | 1.81 | 2 | 2 | 1.285 | 0.362 | 0.79210957 | NA | 0 | 1.3124 | 0.8048 | 0.8048 | TRUE | 0.1505 |
| Lgals3 | 444 | 312 | 339 | 460 | 422 | 484 | 345 | 332 | 366 | 458 | 476 | 478 | 409.04 | 348 | 471 | 1.354 | 0.438 | 2.75E-05 | 0.00043257 | 0 | 0.0082 | 0.0054 | 0.0054 | TRUE | 0.0226 |
| Lgals3bp | 39 | 38 | 26 | 25 | 31 | 25 | 30 | 40 | 28 | 25 | 35 | 25 | 30.55 | 33 | 28 | 0.855 | -0.227 | 0.52823538 | 0.68874916 | 0 | 0.08 | 0.0599 | 0.0599 | TRUE | 0.2649 |
| Lgals4 | 5 | 10 | 14 | 8 | 4 | 10 | 4 | 11 | 15 | 8 | 5 | 10 | 8.66 | 10 | 8 | 0.773 | -0.371 | 0.6028302 | NA | 0.1775 | 0.276 | 0.2469 | 0.2469 | TRUE | 0.299 |
| Lgals6 | 0 | 0 | 0 | 0 | 0 | 0 | 0 | 0 | 0 | 0 | 0 | 0 | 0 | 0 | 0 | NA | NA | NA | NA | NA | NA | NA | NA | NA | NA |
| Lgals7 | 28 | 27 | 23 | 17 | 26 | 24 | 22 | 29 | 25 | 17 | 29 | 24 | 24.2 | 25 | 23 | 0.93 | -0.105 | 0.79293907 | 0.87869058 | 0 | 0.1003 | 0.0741 | 0.0741 | TRUE | 0.2531 |
| Lgals8 | 548 | 393 | 449 | 451 | 401 | 508 | 426 | 418 | 484 | 449 | 452 | 501 | 455.15 | 443 | 467 | 1.058 | 0.081 | 0.4513889 | 0.62125196 | 0.0029 | 0.0076 | 0.0062 | 0.0062 | TRUE | 0.0762 |
| Lgals9 | 163 | 136 | 103 | 143 | 121 | 127 | 127 | 145 | 111 | 142 | 137 | 125 | 131.1 | 128 | 135 | 1.057 | 0.08 | 0.65607128 | 0.78648216 | 0 | 0.0205 | 0.0155 | 0.0155 | TRUE | 0.1324 |
| Lgalsl | 1451 | 1012 | 1166 | 1249 | 1023 | 1108 | 1127 | 1076 | 1258 | 1243 | 1154 | 1094 | 1158.76 | 1154 | 1164 | 1.009 | 0.013 | 0.8799873 | 0.93298562 | 0.0044 | 0.0045 | 0.0044 | 0.0044 | TRUE | 0.073 |
| Lgi1 | 4 | 2 | 0 | 0 | 0 | 0 | 3 | 2 | 0 | 0 | 0 | 0 | 0.87 | 2 | 0 | 0.108 | -3.215 | 0.18197869 | NA | 0 | 2.7193 | 2.9075 | 2.9075 | TRUE | 0.1779 |
| Lgi2 | 574 | 406 | 448 | 477 | 457 | 521 | 446 | 432 | 483 | 475 | 516 | 514 | 477.62 | 454 | 502 | 1.106 | 0.145 | 0.15083287 | 0.29674524 | 0 | 0.0074 | 0.0053 | 0.0053 | TRUE | 0.0363 |
| Lgi3 | 68 | 51 | 52 | 92 | 66 | 74 | 53 | 54 | 56 | 92 | 74 | 73 | 67.04 | 54 | 80 | 1.468 | 0.554 | 0.02096385 | 0.07164621 | 0 | 0.0378 | 0.0262 | 0.0262 | TRUE | 0.161 |
| Lgi4 | 4 | 3 | 2 | 8 | 6 | 4 | 3 | 3 | 2 | 8 | 7 | 4 | 4.52 | 3 | 6 | 2.193 | 1.133 | 0.21450995 | NA | 0 | 0.5265 | 0.3473 | 0.3473 | TRUE | 0.2591 |

| | | | | | | | | | | | | | | | | | | | | | | | | | |
|---|---|---|---|---|---|---|---|---|---|---|---|---|---|---|---|---|---|---|---|---|---|---|---|---|---|
| Lgmn | 3691 | 2586 | 2729 | 3202 | 2838 | 3144 | 2867 | 2751 | 2945 | 3187 | 3202 | 3104 | 3009.06 | 2854 | 3164 | 1.109 | 0.149 | 0.00960088 | 0.04018312 | 3.00E-04 | 0.0032 | 0.002 | 0.002 | TRUE | 0.0119 |
| Lgr4 | 5793 | 4323 | 4282 | 4788 | 4123 | 4688 | 4499 | 4598 | 4620 | 4766 | 4651 | 4628 | 4627.11 | 4572 | 4682 | 1.024 | 0.034 | 0.4988419 | 0.66349779 | 0 | 0.0029 | 0.0016 | 0.0016 | TRUE | 0.003 |
| Lgr5 | 10 | 8 | 3 | 5 | 2 | 1 | 8 | 9 | 3 | 5 | 2 | 1 | 4.62 | 7 | 3 | 0.42 | -1.252 | 0.19346859 | NA | 0 | 0.5152 | 0.4056 | 0.4056 | TRUE | 0.6372 |
| Lgr6 | 175 | 120 | 141 | 189 | 158 | 186 | 136 | 128 | 152 | 188 | 178 | 184 | 160.95 | 139 | 183 | 1.325 | 0.406 | 0.01018864 | 0.04187069 | 0 | 0.0172 | 0.0117 | 0.0117 | TRUE | 0.0706 |
| Lgsn | 2 | 0 | 0 | 0 | 0 | 1 | 2 | 0 | 0 | 0 | 0 | 1 | 0.42 | 1 | 0 | 0.682 | -0.553 | 0.86290097 | NA | 0 | 5.6 | 5.5066 | 5.5066 | TRUE | 0.9727 |
| Lhb | 0 | 1 | 0 | 0 | 3 | 4 | 0 | 1 | 0 | 0 | 3 | 4 | 1.4 | 0 | 2 | 7.068 | 2.821 | 0.15741364 | NA | 0 | 1.696 | 1.6573 | 1.6573 | TRUE | 0.8498 |
| Lhcgr | 0 | 0 | 0 | 0 | 0 | 0 | 0 | 0 | 0 | 0 | 0 | 0 | 0 | 0 | 0 | NA | NA | NA | NA | NA | NA | NA | NA | NA | NA |
| Lhfp | 346 | 233 | 213 | 278 | 246 | 255 | 269 | 248 | 230 | 277 | 278 | 252 | 258.72 | 249 | 269 | 1.076 | 0.106 | 0.41607662 | 0.58881021 | 0 | 0.0116 | 0.0084 | 0.0084 | TRUE | 0.0591 |
| Lhfpl1 | 0 | 2 | 2 | 1 | 0 | 0 | 0 | 2 | 2 | 1 | 0 | 0 | 0.88 | 1 | 0 | 0.266 | -1.912 | 0.40263886 | NA | 0 | 2.6953 | 2.4144 | 2.4144 | TRUE | 0.6892 |
| Lhfpl2 | 257 | 183 | 231 | 223 | 204 | 209 | 200 | 195 | 249 | 222 | 230 | 206 | 216.99 | 215 | 219 | 1.025 | 0.035 | 0.81166379 | 0.89135487 | 0.0062 | 0.0133 | 0.0113 | 0.0113 | TRUE | 0.2116 |
| Lhfpl3 | 1 | 0 | 1 | 2 | 2 | 0 | 1 | 0 | 1 | 2 | 2 | 0 | 1.02 | 1 | 1 | 2.246 | 1.167 | 0.56159335 | NA | 0 | 2.3326 | 1.7754 | 1.7754 | TRUE | 0.519 |
| Lhfpl4 | 82 | 49 | 98 | 132 | 97 | 132 | 64 | 52 | 106 | 131 | 109 | 130 | 98.78 | 74 | 123 | 1.684 | 0.752 | 0.00261385 | 0.01484299 | 0.0567 | 0.0264 | 0.0342 | 0.0342 | TRUE | 1.238 |
| Lhfpl5 | 84 | 60 | 60 | 95 | 101 | 86 | 65 | 64 | 65 | 95 | 114 | 85 | 81.2 | 65 | 98 | 1.51 | 0.594 | 0.00764043 | 0.03381996 | 0 | 0.0316 | 0.023 | 0.023 | TRUE | 0.1936 |
| Lhpp | 889 | 626 | 571 | 800 | 701 | 830 | 690 | 666 | 616 | 796 | 791 | 819 | 729.81 | 657 | 802 | 1.218 | 0.285 | 0.00089791 | 0.00670525 | 0 | 0.0057 | 0.0039 | 0.0039 | TRUE | 0.035 |
| Lhx1 | 4 | 8 | 1 | 3 | 4 | 6 | 3 | 9 | 1 | 3 | 5 | 6 | 4.35 | 4 | 5 | 1.066 | 0.092 | 0.92470055 | NA | 0.208 | 0.5469 | 0.4528 | 0.4528 | TRUE | 0.794 |
| Lhx2 | 126 | 92 | 73 | 80 | 64 | 61 | 98 | 98 | 79 | 80 | 72 | 60 | 81.09 | 92 | 71 | 0.77 | -0.376 | 0.09586964 | 0.21460686 | 0 | 0.0317 | 0.0242 | 0.0242 | TRUE | 0.157 |
| Lhx3 | 3 | 10 | 2 | 7 | 2 | 13 | 2 | 11 | 2 | 7 | 2 | 13 | 6.2 | 5 | 7 | 1.494 | 0.579 | 0.52351936 | NA | 0.5269 | 0.3849 | 0.4267 | 0.4267 | TRUE | 0.3752 |
| Lhx4 | 6 | 3 | 9 | 3 | 4 | 6 | 5 | 3 | 10 | 3 | 5 | 6 | 5.16 | 6 | 5 | 0.771 | -0.376 | 0.66334345 | NA | 0 | 0.4614 | 0.3391 | 0.3391 | TRUE | 0.6966 |
| Lhx5 | 14 | 13 | 13 | 15 | 12 | 24 | 11 | 14 | 14 | 15 | 14 | 24 | 15.15 | 13 | 18 | 1.361 | 0.444 | 0.37661272 | NA | 0 | 0.1589 | 0.1146 | 0.1146 | TRUE | 0.5136 |
| Lhx6 | 179 | 157 | 131 | 154 | 114 | 143 | 139 | 167 | 141 | 153 | 129 | 141 | 145.07 | 149 | 141 | 0.949 | -0.076 | 0.65710083 | 0.78749824 | 0 | 0.0188 | 0.0141 | 0.0141 | TRUE | 0.1117 |
| Lhx8 | 8 | 4 | 6 | 10 | 8 | 3 | 6 | 4 | 6 | 10 | 9 | 3 | 6.48 | 5 | 7 | 1.285 | 0.362 | 0.64092009 | NA | 0 | 0.3682 | 0.2782 | 0.2782 | TRUE | 0.7771 |
| Lhx9 | 2 | 1 | 3 | 0 | 0 | 0 | 2 | 1 | 3 | 0 | 0 | 0 | 0.98 | 2 | 0 | 0.098 | -3.351 | 0.11532532 | NA | 0 | 2.4316 | 2.0136 | 2.0136 | TRUE | 0.2678 |
| Lias | 2826 | 2058 | 2098 | 2399 | 2118 | 2310 | 2195 | 2189 | 2264 | 2388 | 2389 | 2280 | 2284.21 | 2216 | 2352 | 1.062 | 0.086 | 0.14644514 | 0.29079472 | 0 | 0.0035 | 0.0021 | 0.0021 | TRUE | 0.0088 |
| Lif | 148 | 81 | 80 | 120 | 67 | 98 | 115 | 86 | 86 | 119 | 76 | 97 | 96.53 | 96 | 97 | 1.012 | 0.017 | 0.93950572 | 0.966328 | 0.0298 | 0.027 | 0.0278 | 0.0278 | TRUE | 0.1983 |
| Lifr | 7023 | 5383 | 5706 | 6685 | 6421 | 7296 | 5454 | 5726 | 6157 | 6654 | 7244 | 7202 | 6406.18 | 5779 | 7033 | 1.217 | 0.284 | 9.67E-06 | 0.00018328 | 0.0027 | 0.0028 | 0.0028 | 0.0028 | TRUE | 0.0398 |
| Lig1 | 20084 | 15438 | 15197 | 15387 | 13860 | 15960 | 15598 | 16421 | 16397 | 15316 | 15636 | 15755 | 15853.96 | 16139 | 15569 | 0.965 | -0.052 | 0.27911753 | 0.45068084 | 4.00E-04 | 0.0026 | 0.0016 | 0.0016 | TRUE | 0.0106 |
| Lig3 | 9727 | 6961 | 6462 | 6044 | 5706 | 6249 | 7555 | 7404 | 6972 | 6016 | 6437 | 6169 | 6758.87 | 7310 | 6207 | 0.849 | -0.236 | 2.77E-05 | 0.00043417 | 0.0012 | 0.0028 | 0.0021 | 0.0021 | TRUE | 0.0199 |
| Lig4 | 1840 | 1349 | 1223 | 1286 | 1109 | 1235 | 1429 | 1435 | 1320 | 1280 | 1251 | 1219 | 1322.31 | 1395 | 1250 | 0.896 | -0.159 | 0.0265034 | 0.08476462 | 7.00E-04 | 0.0042 | 0.0029 | 0.0029 | TRUE | 0.0259 |
| Lilra5 | 1 | 3 | 0 | 0 | 0 | 1 | 1 | 3 | 0 | 0 | 0 | 1 | 0.83 | 1 | 0 | 0.283 | -1.823 | 0.43687644 | NA | 0 | 2.8724 | 2.5927 | 2.5927 | TRUE | 0.5251 |
| Lilra6 | 0 | 0 | 0 | 0 | 0 | 0 | 0 | 0 | 0 | 0 | 0 | 0 | 0 | 0 | 0 | NA | NA | NA | NA | NA | NA | NA | NA | NA | NA |
| Lilrb3 | 1 | 0 | 1 | 0 | 0 | 0 | 1 | 0 | 1 | 0 | 0 | 0 | 0.31 | 1 | 0 | 0.307 | -1.706 | 0.62894649 | NA | 0 | 7.6657 | 7.2 | 7.2 | TRUE | 0.2067 |
| Lilrb4 | 0 | 0 | 0 | 0 | 0 | 0 | 0 | 0 | 0 | 0 | 0 | 0 | 0 | 0 | 0 | NA | NA | NA | NA | NA | NA | NA | NA | NA | NA |
| Lim2 | 4 | 6 | 4 | 3 | 0 | 6 | 3 | 6 | 4 | 3 | 0 | 6 | 3.79 | 4 | 3 | 0.663 | -0.594 | 0.57873892 | NA | 0.2206 | 0.6285 | 0.5472 | 0.5472 | TRUE | 0.2782 |
| Lima1 | 8920 | 7083 | 9713 | 10171 | 10295 | 11631 | 6928 | 7534 | 10480 | 10124 | 11614 | 11482 | 9693.62 | 8314 | 11073 | 1.332 | 0.414 | 0.03871282 | 0.11182746 | 0.0287 | 0.0027 | 0.0099 | 0.0287 | TRUE | 0.6342 |
| Limch1 | 319 | 221 | 212 | 193 | 197 | 211 | 248 | 235 | 229 | 192 | 222 | 208 | 222.37 | 237 | 207 | 0.873 | -0.196 | 0.15171428 | 0.29811001 | 0 | 0.0131 | 0.009 | 0.009 | TRUE | 0.0464 |
| Limd1 | 1857 | 1359 | 1321 | 1195 | 1086 | 1266 | 1442 | 1446 | 1425 | 1190 | 1225 | 1250 | 1329.59 | 1438 | 1222 | 0.849 | -0.235 | 0.00047022 | 0.00413085 | 0 | 0.0042 | 0.0025 | 0.0025 | TRUE | 0.0064 |
| Limd2 | 2368 | 1559 | 1681 | 2016 | 1811 | 2024 | 1839 | 1658 | 1814 | 2007 | 2043 | 1998 | 1893.15 | 1770 | 2016 | 1.138 | 0.186 | 0.00526591 | 0.02550512 | 0.001 | 0.0037 | 0.0027 | 0.0027 | TRUE | 0.0365 |
| Lime1 | 545 | 475 | 362 | 326 | 296 | 358 | 423 | 505 | 391 | 325 | 334 | 353 | 388.49 | 440 | 337 | 0.768 | -0.381 | 0.00186763 | 0.01148133 | 0.0075 | 0.0085 | 0.0082 | 0.0082 | TRUE | 0.1906 |
| Limk1 | 3113 | 2250 | 2090 | 2185 | 1870 | 2298 | 2418 | 2393 | 2255 | 2175 | 2110 | 2268 | 2269.85 | 2355 | 2184 | 0.927 | -0.109 | 0.08644686 | 0.19909524 | 9.00E-04 | 0.0035 | 0.0025 | 0.0025 | TRUE | 0.0166 |
| Limk2 | 4697 | 3489 | 3865 | 3836 | 3776 | 4210 | 3648 | 3711 | 4170 | 3818 | 4260 | 4156 | 3960.58 | 3843 | 4078 | 1.061 | 0.086 | 0.22846617 | 0.39382191 | 0.004 | 0.003 | 0.0034 | 0.0034 | TRUE | 0.0671 |
| Lims1 | 4717 | 3503 | 3995 | 4519 | 3883 | 4587 | 3663 | 3726 | 4311 | 4498 | 4380 | 4528 | 4184.48 | 3900 | 4469 | 1.146 | 0.197 | 0.0052546 | 0.02545899 | 0.004 | 0.003 | 0.0034 | 0.0034 | TRUE | 0.1026 |
| Lims2 | 72 | 56 | 61 | 106 | 86 | 84 | 56 | 60 | 66 | 106 | 97 | 83 | 77.79 | 61 | 95 | 1.579 | 0.659 | 0.00364703 | 0.01920668 | 0 | 0.0329 | 0.0236 | 0.0236 | TRUE | 0.1261 |
| Lin28a | 10619 | 7992 | 8824 | 11369 | 10489 | 13252 | 8247 | 8501 | 9521 | 11317 | 11833 | 13082 | 10416.75 | 8756 | 12077 | 1.379 | 0.464 | 2.20E-10 | 1.66E-08 | 0.0056 | 0.0027 | 0.0038 | 0.0038 | TRUE | 0.0711 |
| Lin28b | 2150 | 1487 | 1312 | 1332 | 1177 | 1372 | 1670 | 1582 | 1416 | 1326 | 1328 | 1354 | 1445.86 | 1556 | 1336 | 0.858 | -0.221 | 0.00456019 | 0.02288582 | 0.003 | 0.0041 | 0.0037 | 0.0037 | TRUE | 0.074 |
| Lin37 | 621 | 426 | 401 | 406 | 398 | 445 | 482 | 453 | 433 | 404 | 449 | 439 | 443.42 | 456 | 431 | 0.943 | -0.085 | 0.41419827 | 0.58679114 | 0 | 0.0078 | 0.0056 | 0.0056 | TRUE | 0.0342 |
| Lin52 | 716 | 586 | 667 | 856 | 665 | 799 | 556 | 623 | 720 | 852 | 750 | 789 | 715.01 | 633 | 797 | 1.263 | 0.336 | 0.0016159 | 0.01020301 | 0.0084 | 0.0057 | 0.0068 | 0.0068 | TRUE | 0.1682 |
| Lin54 | 2618 | 2000 | 2212 | 2377 | 2132 | 2525 | 2033 | 2127 | 2387 | 2366 | 2405 | 2493 | 2301.86 | 2182 | 2421 | 1.11 | 0.151 | 0.03899254 | 0.11246774 | 0.0032 | 0.0035 | 0.0034 | 0.0034 | TRUE | 0.0806 |
| Lin7a | 4 | 7 | 9 | 4 | 3 | 7 | 3 | 7 | 10 | 4 | 3 | 7 | 5.76 | 7 | 5 | 0.721 | -0.471 | 0.57034265 | NA | 0 | 0.4141 | 0.3154 | 0.3154 | TRUE | 0.4661 |
| Lin7b | 46 | 38 | 35 | 34 | 30 | 34 | 36 | 40 | 38 | 34 | 34 | 34 | 35.86 | 38 | 34 | 0.891 | -0.167 | 0.59683941 | 0.74287579 | 0 | 0.0685 | 0.0436 | 0.0436 | TRUE | 0.0235 |
| Lin7c | 8659 | 6103 | 6156 | 6791 | 5644 | 6454 | 6725 | 6492 | 6642 | 6760 | 6367 | 6371 | 6559.48 | 6620 | 6499 | 0.982 | -0.027 | 0.6115481 | 0.75404219 | 5.00E-04 | 0.0028 | 0.0018 | 0.0018 | TRUE | 0.015 |
| Lin9 | 1281 | 934 | 875 | 855 | 745 | 866 | 995 | 993 | 944 | 851 | 840 | 855 | 913.15 | 977 | 849 | 0.868 | -0.204 | 0.00735769 | 0.03301836 | 0 | 0.005 | 0.0031 | 0.0031 | TRUE | 0.0104 |
| Lingo1 | 240 | 210 | 198 | 279 | 220 | 247 | 186 | 223 | 214 | 278 | 248 | 244 | 232.19 | 208 | 257 | 1.239 | 0.31 | 0.02688405 | 0.0856371 | 0.0032 | 0.0126 | 0.0098 | 0.0098 | TRUE | 0.0979 |
| Lingo2 | 0 | 1 | 0 | 0 | 0 | 0 | 0 | 1 | 0 | 0 | 0 | 0 | 0.18 | 0 | 0 | 0.552 | -0.857 | 0.8336548 | NA | 0 | 13.3714 | 10 | 10 | TRUE | 0.5216 |
| Lingo3 | 78 | 56 | 79 | 137 | 108 | 130 | 61 | 60 | 85 | 136 | 122 | 128 | 98.65 | 69 | 129 | 1.893 | 0.921 | 1.49E-05 | 0.00026441 | 0.0056 | 0.0265 | 0.0215 | 0.0215 | TRUE | 0.3883 |
| Lingo4 | 0 | 0 | 0 | 0 | 0 | 0 | 0 | 0 | 0 | 0 | 0 | 0 | 0 | 0 | 0 | NA | NA | NA | NA | NA | NA | NA | NA | NA | NA |
| Lins | 610 | 456 | 432 | 506 | 407 | 463 | 474 | 485 | 466 | 504 | 459 | 457 | 474.13 | 475 | 473 | 0.997 | -0.005 | 0.96336862 | 0.97975151 | 0 | 0.0074 | 0.005 | 0.005 | TRUE | 0.0372 |
| Lipa | 1531 | 1064 | 1231 | 1530 | 1336 | 1479 | 1189 | 1132 | 1328 | 1523 | 1507 | 1460 | 1356.53 | 1216 | 1497 | 1.231 | 0.3 | 0.00012347 | 0.00144771 | 0.0026 | 0.0042 | 0.0037 | 0.0037 | TRUE | 0.0754 |
| Lipc | 23 | 10 | 11 | 9 | 17 | 8 | 18 | 11 | 12 | 9 | 19 | 8 | 12.73 | 14 | 12 | 0.875 | -0.192 | 0.737475 | NA | 0.0813 | 0.1886 | 0.1584 | 0.1584 | TRUE | 0.978 |
| Lipe | 686 | 544 | 484 | 496 | 467 | 495 | 533 | 579 | 522 | 494 | 527 | 489 | 523.81 | 545 | 503 | 0.924 | -0.114 | 0.23786334 | 0.40459129 | 0 | 0.007 | 0.0049 | 0.0049 | TRUE | 0.0339 |
| Lipf | 0 | 1 | 0 | 0 | 0 | 0 | 0 | 1 | 0 | 0 | 0 | 0 | 0.18 | 0 | 0 | 0.552 | -0.857 | 0.8336548 | NA | 0 | 13.3714 | 10 | 10 | TRUE | 0.5216 |
| Lipg | 93 | 94 | 80 | 113 | 115 | 97 | 72 | 100 | 86 | 112 | 130 | 96 | 99.42 | 86 | 113 | 1.312 | 0.392 | 0.0684001 | 0.16850098 | 0.0149 | 0.0263 | 0.023 | 0.023 | TRUE | 0.1874 |
| Liph | 2239 | 1870 | 2054 | 1887 | 1932 | 2183 | 1739 | 1989 | 2216 | 1878 | 2179 | 2155 | 2026.18 | 1981 | 2071 | 1.046 | 0.065 | 0.48883035 | 0.65456738 | 0.0102 | 0.0036 | 0.0058 | 0.0058 | TRUE | 0.1415 |
| Lipi | 0 | 0 | 0 | 0 | 0 | 0 | 0 | 0 | 0 | 0 | 0 | 0 | 0 | 0 | 0 | NA | NA | NA | NA | NA | NA | NA | NA | NA | NA |
| Lipk | 1 | 0 | 0 | 0 | 0 | 0 | 1 | 0 | 0 | 0 | 0 | 0 | 0.13 | 0 | 0 | 0.552 | -0.857 | 0.8336548 | NA | 0 | 18.3121 | 10 | 10 | TRUE | 0.2566 |
| Lipm | 1 | 1 | 0 | 2 | 3 | 1 | 1 | 1 | 0 | 2 | 3 | 1 | 1.37 | 1 | 2 | 3.367 | 1.751 | 0.31352097 | NA | 0 | 1.7361 | 1.1614 | 1.1614 | TRUE | 0.1569 |
| Lipn | 0 | 0 | 0 | 0 | 0 | 0 | 0 | 0 | 0 | 0 | 0 | 0 | 0 | 0 | 0 | NA | NA | NA | NA | NA | NA | NA | NA | NA | NA |
| Lipo1 | 533 | 349 | 297 | 220 | 185 | 174 | 414 | 371 | 320 | 219 | 209 | 172 | 284.18 | 368 | 200 | 0.54 | -0.888 | 1.03E-09 | 6.82E-08 | 0.0127 | 0.0108 | 0.0113 | 0.0113 | TRUE | 0.1588 |
| Lipt1 | 534 | 465 | 466 | 535 | 480 | 582 | 415 | 495 | 503 | 533 | 541 | 575 | 510.12 | 471 | 550 | 1.171 | 0.228 | 0.03313457 | 0.09970366 | 0.0046 | 0.0071 | 0.0063 | 0.0063 | TRUE | 0.1347 |
| Lipt2 | 611 | 424 | 382 | 481 | 416 | 491 | 475 | 451 | 412 | 479 | 469 | 485 | 461.75 | 446 | 478 | 1.069 | 0.096 | 0.34483707 | 0.51910603 | 0 | 0.0076 | 0.0054 | 0.0054 | TRUE | 0.0498 |

| | | | | | | | | | | | | | | | | | | | | | | | | | |
|---|---|---|---|---|---|---|---|---|---|---|---|---|---|---|---|---|---|---|---|---|---|---|---|---|---|
| Litaf | 2371 | 1746 | 1946 | 2083 | 1956 | 2466 | 1841 | 1857 | 2100 | 2073 | 2207 | 2434 | 2085.45 | 1933 | 2238 | 1.159 | 0.212 | 0.00891549 | 0.03811064 | 0.0054 | 0.0036 | 0.0043 | 0.0043 | TRUE | 0.0718 |
| Lix1 | 30 | 21 | 20 | 12 | 19 | 16 | 23 | 22 | 22 | 12 | 21 | 16 | 19.4 | 22 | 16 | 0.727 | -0.461 | 0.29662474 | 0.4685222 | 0 | 0.1246 | 0.0873 | 0.0873 | TRUE | 0.3164 |
| Lix1l | 979 | 737 | 863 | 1080 | 909 | 1038 | 760 | 784 | 931 | 1075 | 1025 | 1025 | 933.43 | 825 | 1042 | 1.265 | 0.339 | 0.00023946 | 0.00245 | 0.005 | 0.005 | 0.005 | 0.005 | TRUE | 0.1478 |
| Llgl1 | 10560 | 7798 | 7824 | 7675 | 6921 | 7988 | 8202 | 8295 | 8442 | 7640 | 7808 | 7885 | 8045.14 | 8313 | 7778 | 0.936 | -0.096 | 0.04367732 | 0.12204243 | 1.00E-04 | 0.0027 | 0.0015 | 0.0015 | TRUE | 0.003 |
| Llgl2 | 16334 | 12007 | 11706 | 14207 | 12699 | 14078 | 12686 | 12772 | 12631 | 14142 | 14326 | 13897 | 13408.81 | 12696 | 14122 | 1.112 | 0.153 | 0.00058678 | 0.00488177 | 1.00E-04 | 0.0026 | 0.0014 | 0.0014 | TRUE | 0.0024 |
| Llph | 1355 | 989 | 870 | 871 | 777 | 903 | 1052 | 1052 | 939 | 867 | 877 | 891 | 946.33 | 1014 | 878 | 0.865 | -0.209 | 0.00977736 | 0.04071906 | 0.0013 | 0.0049 | 0.0036 | 0.0036 | TRUE | 0.0495 |
| Lman1 | 9698 | 7210 | 7789 | 8334 | 7379 | 8521 | 7532 | 7669 | 8404 | 8296 | 8324 | 8412 | 8106.16 | 7868 | 8344 | 1.061 | 0.085 | 0.13951116 | 0.2807311 | 0.0016 | 0.0027 | 0.0023 | 0.0023 | TRUE | 0.0431 |
| Lman1l | 0 | 1 | 0 | 2 | 3 | 2 | 0 | 1 | 0 | 2 | 3 | 2 | 1.4 | 0 | 2 | 7.074 | 2.823 | 0.12385354 | NA | 0 | 1.6927 | 1.2125 | 1.2125 | TRUE | 0.5217 |
| Lman2 | 7012 | 5078 | 5123 | 5444 | 4759 | 5582 | 5446 | 5401 | 5528 | 5419 | 5369 | 5510 | 5445.48 | 5458 | 5433 | 0.995 | -0.007 | 0.89167028 | 0.94021713 | 0 | 0.0029 | 0.0015 | 0.0015 | TRUE | 0.0019 |
| Lman2l | 751 | 585 | 559 | 630 | 513 | 661 | 583 | 622 | 603 | 627 | 579 | 653 | 611.17 | 603 | 620 | 1.029 | 0.041 | 0.65278516 | 0.78431896 | 0 | 0.0063 | 0.0045 | 0.0045 | TRUE | 0.0378 |
| Lmbr1 | 998 | 681 | 678 | 685 | 585 | 680 | 775 | 724 | 732 | 682 | 660 | 671 | 707.35 | 744 | 671 | 0.901 | -0.15 | 0.07526641 | 0.18072513 | 0 | 0.0058 | 0.0037 | 0.0037 | TRUE | 0.0172 |
| Lmbr1l | 1945 | 1497 | 1396 | 1500 | 1385 | 1522 | 1511 | 1592 | 1506 | 1493 | 1562 | 1502 | 1527.86 | 1536 | 1519 | 0.989 | -0.016 | 0.80942067 | 0.88999535 | 0 | 0.004 | 0.0025 | 0.0025 | TRUE | 0.012 |
| Lmbrd1 | 1054 | 752 | 740 | 757 | 674 | 771 | 819 | 800 | 798 | 754 | 760 | 761 | 781.98 | 806 | 758 | 0.941 | -0.088 | 0.2658213 | 0.4362094 | 0 | 0.0055 | 0.0032 | 0.0032 | TRUE | 0.0025 |
| Lmbrd2 | 1277 | 935 | 899 | 1167 | 1003 | 1122 | 992 | 995 | 970 | 1162 | 1131 | 1108 | 1059.51 | 986 | 1134 | 1.15 | 0.202 | 0.00522269 | 0.02534732 | 0 | 0.0047 | 0.0028 | 0.0028 | TRUE | 0.0057 |
| Lmcd1 | 181 | 122 | 195 | 277 | 241 | 320 | 141 | 130 | 210 | 276 | 272 | 316 | 224.04 | 160 | 288 | 1.803 | 0.851 | 1.57E-06 | 4.19E-05 | 0.0308 | 0.013 | 0.0178 | 0.0178 | TRUE | 0.7646 |
| Lmf1 | 520 | 407 | 482 | 531 | 442 | 539 | 404 | 433 | 520 | 529 | 499 | 532 | 486.02 | 452 | 520 | 1.153 | 0.205 | 0.07162411 | 0.17450075 | 0.0072 | 0.0073 | 0.0073 | 0.0073 | TRUE | 0.1966 |
| Lmf2 | 2244 | 1799 | 1662 | 1635 | 1466 | 1735 | 1743 | 1914 | 1793 | 1627 | 1654 | 1713 | 1740.61 | 1817 | 1665 | 0.917 | -0.125 | 0.06465595 | 0.16204234 | 0.001 | 0.0038 | 0.0027 | 0.0027 | TRUE | 0.0262 |
| Lmln | 452 | 337 | 371 | 381 | 339 | 393 | 351 | 358 | 400 | 379 | 382 | 388 | 376.57 | 370 | 383 | 1.038 | 0.053 | 0.62726262 | 0.76497138 | 0 | 0.0087 | 0.006 | 0.006 | TRUE | 0.0575 |
| Lmna | 1113 | 798 | 936 | 994 | 912 | 1132 | 864 | 849 | 1010 | 989 | 1029 | 1117 | 976.48 | 908 | 1045 | 1.153 | 0.205 | 0.02570394 | 0.08285449 | 0.0052 | 0.0049 | 0.0051 | 0.0051 | TRUE | 0.1131 |
| Lmnb1 | 10454 | 7760 | 7149 | 6984 | 6224 | 7145 | 8119 | 8254 | 7714 | 6952 | 7021 | 7053 | 7518.92 | 8029 | 7009 | 0.873 | -0.196 | 0.00011805 | 0.00139624 | 4.00E-04 | 0.0027 | 0.0017 | 0.0017 | TRUE | 0.0143 |
| Lmnb2 | 5456 | 4277 | 3911 | 3933 | 3551 | 4317 | 4237 | 4549 | 4220 | 3915 | 4006 | 4262 | 4198.19 | 4335 | 4061 | 0.937 | -0.094 | 0.12224521 | 0.25544602 | 0.0015 | 0.003 | 0.0024 | 0.0024 | TRUE | 0.0229 |
| Lmo1 | 90 | 89 | 80 | 117 | 79 | 99 | 70 | 95 | 86 | 116 | 89 | 98 | 92.37 | 84 | 101 | 1.218 | 0.284 | 0.19205004 | 0.35037387 | 0.0112 | 0.0281 | 0.0233 | 0.0233 | TRUE | 0.2219 |
| Lmo2 | 121 | 92 | 52 | 95 | 92 | 98 | 94 | 98 | 56 | 95 | 104 | 97 | 90.51 | 83 | 99 | 1.184 | 0.244 | 0.30084778 | 0.47325746 | 0.0298 | 0.0286 | 0.0289 | 0.0289 | TRUE | 0.697 |
| Lmo3 | 0 | 1 | 0 | 0 | 0 | 0 | 0 | 1 | 0 | 0 | 0 | 0 | 0.18 | 0 | 0 | 0.552 | -0.857 | 0.8336548 | NA | 0 | 13.3714 | 10 | 10 | TRUE | 0.5216 |
| Lmo4 | 508 | 328 | 242 | 263 | 233 | 276 | 395 | 349 | 261 | 262 | 263 | 272 | 300.27 | 335 | 266 | 0.791 | -0.339 | 0.02512761 | 0.08157836 | 0.0193 | 0.0103 | 0.0131 | 0.0131 | TRUE | 0.3804 |
| Lmo7 | 2275 | 1656 | 1464 | 1951 | 1694 | 1825 | 1767 | 1761 | 1580 | 1942 | 1911 | 1802 | 1793.77 | 1703 | 1885 | 1.106 | 0.146 | 0.04289748 | 0.12050585 | 0.0022 | 0.0037 | 0.0032 | 0.0032 | TRUE | 0.0473 |
| Lmod1 | 99 | 75 | 79 | 113 | 68 | 99 | 77 | 80 | 85 | 112 | 77 | 98 | 88.14 | 81 | 96 | 1.191 | 0.252 | 0.25095041 | 0.41975825 | 0.0082 | 0.0293 | 0.0235 | 0.0235 | TRUE | 0.2467 |
| Lmod2 | 1 | 2 | 2 | 0 | 0 | 3 | 1 | 2 | 2 | 0 | 0 | 3 | 1.34 | 2 | 1 | 0.606 | -0.723 | 0.68645422 | NA | 0.5179 | 1.7748 | 1.5067 | 1.5067 | TRUE | 1.4412 |
| Lmod3 | 11 | 3 | 5 | 4 | 1 | 4 | 9 | 3 | 5 | 4 | 1 | 4 | 4.36 | 6 | 3 | 0.527 | -0.923 | 0.33367873 | NA | 0 | 0.5454 | 0.4039 | 0.4039 | TRUE | 0.2241 |
| Lmtk2 | 6946 | 4791 | 4527 | 5042 | 4438 | 5215 | 5395 | 5096 | 4885 | 5019 | 5007 | 5148 | 5091.45 | 5125 | 5058 | 0.986 | -0.02 | 0.73164929 | 0.83929626 | 0.0012 | 0.0029 | 0.0022 | 0.0022 | TRUE | 0.0262 |
| Lmtk3 | 417 | 326 | 326 | 307 | 293 | 349 | 324 | 347 | 352 | 306 | 331 | 345 | 333.84 | 341 | 327 | 0.961 | -0.058 | 0.61564355 | 0.75732793 | 0 | 0.0095 | 0.0066 | 0.0066 | TRUE | 0.0378 |
| Lmx1a | 18 | 24 | 18 | 24 | 15 | 22 | 14 | 26 | 19 | 24 | 17 | 22 | 20.24 | 20 | 21 | 1.075 | 0.105 | 0.81183177 | 0.89135487 | 0 | 0.1195 | 0.0902 | 0.0902 | TRUE | 0.2485 |
| Lmx1b | 94 | 77 | 61 | 72 | 65 | 65 | 73 | 82 | 66 | 72 | 73 | 64 | 71.65 | 74 | 70 | 0.947 | -0.079 | 0.73244538 | 0.83992634 | 0 | 0.0355 | 0.0243 | 0.0243 | TRUE | 0.0825 |
| Lnp | 3245 | 2363 | 2535 | 2658 | 2347 | 2692 | 2520 | 2513 | 2735 | 2646 | 2648 | 2657 | 2619.97 | 2589 | 2650 | 1.024 | 0.034 | 0.57793401 | 0.7293317 | 8.00E-04 | 0.0033 | 0.0023 | 0.0023 | TRUE | 0.0289 |
| Lnpep | 1740 | 1213 | 1169 | 1173 | 907 | 1157 | 1351 | 1290 | 1261 | 1168 | 1023 | 1142 | 1205.98 | 1301 | 1111 | 0.854 | -0.228 | 0.00368094 | 0.0193283 | 0.0021 | 0.0044 | 0.0036 | 0.0036 | TRUE | 0.056 |
| Lnx1 | 63 | 75 | 97 | 89 | 100 | 100 | 49 | 80 | 105 | 89 | 113 | 99 | 88.91 | 78 | 100 | 1.297 | 0.375 | 0.15712269 | 0.3056696 | 0.0648 | 0.0291 | 0.0392 | 0.0392 | TRUE | 0.348 |
| Lnx2 | 1294 | 1003 | 1017 | 1095 | 1073 | 1189 | 1005 | 1067 | 1097 | 1090 | 1210 | 1174 | 1107.23 | 1056 | 1158 | 1.097 | 0.134 | 0.08807534 | 0.20167587 | 0.0016 | 0.0046 | 0.0035 | 0.0035 | TRUE | 0.0318 |
| LOC1000389 | 2 | 2 | 0 | 2 | 2 | 0 | 2 | 2 | 0 | 2 | 2 | 0 | 1.32 | 1 | 1 | 1.131 | 0.177 | 0.91971667 | NA | 0 | 1.7962 | 1.4631 | 1.4631 | TRUE | 0.5204 |
| LOC1000407 | 0 | 0 | 0 | 0 | 0 | 0 | 0 | 0 | 0 | 0 | 0 | 0 | 0 | 0 | 0 | NA | NA | NA | NA | NA | NA | NA | NA | NA | NA |
| LOC1000433 | 0 | 0 | 0 | 0 | 0 | 0 | 0 | 0 | 0 | 0 | 0 | 0 | 0 | 0 | 0 | NA | NA | NA | NA | NA | NA | NA | NA | NA | NA |
| LOC1000488 | 0 | 0 | 0 | 0 | 0 | 0 | 0 | 0 | 0 | 0 | 0 | 0 | 0 | 0 | 0 | NA | NA | NA | NA | NA | NA | NA | NA | NA | NA |
| LOC1005029 | 0 | 0 | 0 | 0 | 0 | 0 | 0 | 0 | 0 | 0 | 0 | 0 | 0 | 0 | 0 | NA | NA | NA | NA | NA | NA | NA | NA | NA | NA |
| LOC1005032 | 62 | 31 | 27 | 28 | 27 | 16 | 48 | 33 | 29 | 28 | 30 | 16 | 30.73 | 37 | 25 | 0.664 | -0.591 | 0.12610711 | 0.26129144 | 0.0573 | 0.0795 | 0.0734 | 0.0734 | TRUE | 0.6218 |
| LOC1005034 | 7 | 4 | 7 | 7 | 2 | 3 | 5 | 4 | 8 | 7 | 2 | 3 | 4.9 | 6 | 4 | 0.714 | -0.487 | 0.580401 | NA | 0 | 0.4856 | 0.3482 | 0.3482 | TRUE | 0.6755 |
| LOC1005046 | 7 | 4 | 3 | 5 | 3 | 5 | 5 | 4 | 3 | 5 | 3 | 5 | 4.37 | 4 | 4 | 1.021 | 0.031 | 0.97253601 | NA | 0 | 0.5447 | 0.3455 | 0.3455 | TRUE | 0.1243 |
| LOC1005047 | 21 | 2 | 10 | 10 | 11 | 7 | 16 | 2 | 11 | 10 | 12 | 7 | 9.75 | 10 | 10 | 0.98 | -0.03 | 0.96646902 | NA | 0.2893 | 0.2455 | 0.257 | 0.257 | TRUE | 0.3332 |
| LOC1008616 | 36 | 22 | 18 | 21 | 25 | 24 | 28 | 23 | 19 | 21 | 28 | 24 | 23.93 | 23 | 24 | 1.018 | 0.026 | 0.94687208 | 0.97098886 | 0 | 0.1015 | 0.0707 | 0.0707 | TRUE | 0.1872 |
| LOC1008619 | 0 | 0 | 0 | 0 | 0 | 0 | 0 | 0 | 0 | 0 | 0 | 0 | 0 | 0 | 0 | NA | NA | NA | NA | NA | NA | NA | NA | NA | NA |
| LOC106740 | 257 | 213 | 204 | 266 | 228 | 249 | 200 | 227 | 220 | 265 | 257 | 246 | 235.68 | 216 | 256 | 1.191 | 0.252 | 0.05807958 | 0.1502762 | 0 | 0.0125 | 0.0085 | 0.0085 | TRUE | 0.0501 |
| LOC171588 | 6 | 2 | 1 | 1 | 3 | 0 | 5 | 2 | 1 | 1 | 3 | 0 | 2.04 | 3 | 1 | 0.531 | -0.913 | 0.5279756 | NA | 0.3758 | 1.1636 | 0.9645 | 0.9645 | TRUE | 0.7853 |
| LOC666331 | 69 | 62 | 93 | 77 | 62 | 124 | 54 | 66 | 100 | 77 | 70 | 122 | 81.48 | 73 | 90 | 1.234 | 0.303 | 0.29287037 | 0.46503064 | 0.0843 | 0.0315 | 0.0474 | 0.0474 | TRUE | 0.9842 |
| Loh12cr1 | 608 | 426 | 372 | 427 | 342 | 409 | 472 | 453 | 401 | 425 | 386 | 404 | 423.55 | 442 | 405 | 0.914 | -0.129 | 0.2376903 | 0.40434509 | 0.0024 | 0.008 | 0.0063 | 0.0063 | TRUE | 0.0739 |
| Lonp1 | 16708 | 13404 | 13921 | 15248 | 14336 | 16114 | 12976 | 14258 | 15021 | 15178 | 16172 | 15907 | 14918.67 | 14085 | 15752 | 1.119 | 0.162 | 0.01088683 | 0.04398119 | 0.0031 | 0.0026 | 0.0028 | 0.0028 | TRUE | 0.0583 |
| Lonp2 | 4548 | 3249 | 3214 | 3367 | 2939 | 3375 | 3532 | 3456 | 3468 | 3352 | 3315 | 3332 | 3409.11 | 3485 | 3333 | 0.956 | -0.065 | 0.21723895 | 0.380002 | 0 | 0.0031 | 0.0017 | 0.0017 | TRUE | 0.0017 |
| Lonrf1 | 2859 | 2139 | 2186 | 2119 | 1861 | 2116 | 2220 | 2275 | 2359 | 2109 | 2099 | 2089 | 2191.97 | 2285 | 2099 | 0.919 | -0.122 | 0.04234197 | 0.1192735 | 0 | 0.0035 | 0.0021 | 0.0021 | TRUE | 0.0096 |
| Lonrf2 | 121 | 81 | 46 | 72 | 74 | 73 | 94 | 86 | 50 | 72 | 83 | 72 | 76.16 | 77 | 76 | 0.981 | -0.027 | 0.91766703 | 0.95467028 | 0.0436 | 0.0335 | 0.0363 | 0.0363 | TRUE | 0.8312 |
| Lonrf3 | 1365 | 1096 | 1028 | 1133 | 966 | 1034 | 1060 | 1166 | 1109 | 1128 | 1090 | 1021 | 1095.57 | 1112 | 1080 | 0.972 | -0.041 | 0.59779264 | 0.74375096 | 0.0016 | 0.0046 | 0.0035 | 0.0035 | TRUE | 0.0276 |
| Lor | 16 | 20 | 9 | 9 | 6 | 7 | 12 | 21 | 10 | 9 | 7 | 7 | 11.01 | 14 | 8 | 0.524 | -0.931 | 0.12490169 | NA | 0 | 0.2177 | 0.1635 | 0.1635 | TRUE | 0.7045 |
| Lox | 17 | 13 | 14 | 16 | 17 | 16 | 13 | 14 | 15 | 16 | 19 | 16 | 15.51 | 14 | 17 | 1.209 | 0.273 | 0.56709806 | NA | 0 | 0.1553 | 0.0993 | 0.0993 | TRUE | 0.058 |
| Loxhd1 | 7 | 5 | 6 | 2 | 4 | 5 | 5 | 5 | 6 | 2 | 5 | 5 | 4.78 | 5 | 4 | 0.663 | -0.592 | 0.49702453 | NA | 0 | 0.4985 | 0.3278 | 0.3278 | TRUE | 0.2919 |
| Loxl1 | 303 | 227 | 220 | 330 | 272 | 235 | 235 | 241 | 237 | 328 | 307 | 232 | 263.58 | 238 | 289 | 1.214 | 0.28 | 0.05864905 | 0.15095819 | 0.0131 | 0.0114 | 0.0119 | 0.0119 | TRUE | 0.3427 |
| Loxl2 | 2240 | 1672 | 1884 | 2209 | 2001 | 2353 | 1740 | 1778 | 2033 | 2199 | 2257 | 2323 | 2054.99 | 1850 | 2260 | 1.222 | 0.29 | 0.00010459 | 0.00125413 | 0.0033 | 0.0036 | 0.0035 | 0.0035 | TRUE | 0.089 |
| Loxl3 | 55 | 39 | 40 | 61 | 54 | 56 | 43 | 41 | 43 | 61 | 61 | 55 | 50.71 | 42 | 59 | 1.388 | 0.473 | 0.0769648 | 0.18362731 | 0 | 0.0492 | 0.0314 | 0.0314 | TRUE | 0.0263 |
| Loxl4 | 85 | 62 | 61 | 77 | 61 | 67 | 66 | 66 | 66 | 77 | 69 | 66 | 68.23 | 66 | 71 | 1.07 | 0.098 | 0.67309551 | 0.79918211 | 0 | 0.0372 | 0.0241 | 0.0241 | TRUE | 0.0532 |
| Lpar1 | 1855 | 1234 | 1007 | 992 | 827 | 973 | 1441 | 1313 | 1087 | 987 | 933 | 961 | 1120.12 | 1280 | 960 | 0.749 | -0.416 | 3.36E-05 | 0.00051093 | 0.0099 | 0.0045 | 0.0063 | 0.0063 | TRUE | 0.208 |
| Lpar2 | 1597 | 1259 | 1570 | 1781 | 1638 | 1951 | 1240 | 1339 | 1694 | 1773 | 1848 | 1926 | 1636.68 | 1424 | 1849 | 1.3 | 0.378 | 0.00023916 | 0.00244866 | 0.014 | 0.0039 | 0.007 | 0.007 | TRUE | 0.328 |
| Lpar3 | 94 | 80 | 63 | 56 | 59 | 58 | 73 | 85 | 68 | 56 | 67 | 57 | 67.61 | 75 | 60 | 0.793 | -0.334 | 0.16356807 | 0.31417234 | 0 | 0.0375 | 0.0263 | 0.0263 | TRUE | 0.1101 |
| Lpar4 | 237 | 144 | 105 | 111 | 94 | 118 | 184 | 153 | 113 | 110 | 106 | 116 | 130.59 | 150 | 111 | 0.736 | -0.443 | 0.02909669 | 0.09058571 | 0.0248 | 0.0206 | 0.0218 | 0.0218 | TRUE | 0.1955 |
| Lpar5 | 313 | 229 | 211 | 229 | 216 | 229 | 243 | 244 | 228 | 228 | 244 | 226 | 235.34 | 238 | 233 | 0.975 | -0.036 | 0.78180141 | 0.87142537 | 0 | 0.0125 | 0.0081 | 0.0081 | TRUE | 0.0185 |

| | | | | | | | | | | | | | | | | | | | | | | | |
|---|---|---|---|---|---|---|---|---|---|---|---|---|---|---|---|---|---|---|---|---|---|---|---|
| Lpar6 | 193 | 131 | 109 | 70 | 72 | 85 | 150 | 139 | 118 | 70 | 81 | 84 | 106.94 | 136 | 78 | 0.575 | -0.798 | 6.84E-05 | 0.00090277 | 0 | 0.0246 | 0.0188 | 0.0188 | TRUE | 0.135 |
| Lpcat1 | 3201 | 2323 | 2333 | 2361 | 2156 | 2367 | 2486 | 2471 | 2517 | 2350 | 2432 | 2337 | 2432.21 | 2491 | 2373 | 0.952 | -0.071 | 0.2192178 | 0.3823839 | 0 | 0.0034 | 0.002 | 0.002 | TRUE | 0.0057 |
| Lpcat2 | 333 | 222 | 318 | 347 | 330 | 316 | 259 | 236 | 343 | 345 | 372 | 312 | 311.25 | 279 | 343 | 1.23 | 0.299 | 0.04491774 | 0.12451691 | 0.0194 | 0.01 | 0.0128 | 0.0128 | TRUE | 0.4341 |
| Lpcat2b | 0 | 0 | 0 | 0 | 0 | 0 | 0 | 0 | 0 | 0 | 0 | 0 | 0 | 0 | 0 | NA | NA | NA | NA | NA | NA | NA | NA | NA | NA |
| Lpcat3 | 6556 | 4758 | 4993 | 5664 | 5214 | 5930 | 5092 | 5061 | 5387 | 5638 | 5882 | 5854 | 5485.65 | 5180 | 5791 | 1.118 | 0.161 | 0.0027825 | 0.01557176 | 6.00E-04 | 0.0029 | 0.0019 | 0.0019 | TRUE | 0.0148 |
| Lpcat4 | 1151 | 884 | 948 | 1001 | 872 | 1074 | 894 | 940 | 1023 | 996 | 984 | 1060 | 982.9 | 952 | 1013 | 1.066 | 0.092 | 0.26587516 | 0.4362477 | 0.0022 | 0.0048 | 0.0039 | 0.0039 | TRUE | 0.0496 |
| Lpgat1 | 5151 | 3882 | 3959 | 4382 | 3810 | 4203 | 4001 | 4129 | 4272 | 4362 | 4298 | 4149 | 4201.74 | 4134 | 4270 | 1.033 | 0.047 | 0.39699391 | 0.57038943 | 7.00E-04 | 0.003 | 0.002 | 0.002 | TRUE | 0.0104 |
| Lphn1 | 12483 | 9818 | 9364 | 12149 | 11431 | 13010 | 9695 | 10443 | 10104 | 12093 | 12895 | 12843 | 11345.55 | 10081 | 12610 | 1.251 | 0.323 | 2.37E-09 | 1.42E-07 | 0.0012 | 0.0026 | 0.002 | 0.002 | TRUE | 0.0158 |
| Lphn2 | 1501 | 1059 | 1100 | 1023 | 963 | 1136 | 1166 | 1126 | 1187 | 1018 | 1086 | 1121 | 1117.53 | 1160 | 1075 | 0.927 | -0.109 | 0.14563607 | 0.28966556 | 6.00E-04 | 0.0045 | 0.0032 | 0.0032 | TRUE | 0.0262 |
| Lphn3 | 216 | 129 | 78 | 59 | 63 | 51 | 168 | 137 | 84 | 59 | 71 | 50 | 94.88 | 130 | 60 | 0.459 | -1.122 | 2.55E-05 | 0.00040446 | 0.066 | 0.0274 | 0.0388 | 0.0388 | TRUE | 0.2173 |
| Lpin1 | 2239 | 1868 | 2462 | 3324 | 2831 | 3502 | 1739 | 1987 | 2656 | 3309 | 3194 | 3457 | 2723.63 | 2127 | 3320 | 1.561 | 0.642 | 0.0006338 | 0.00517655 | 0.0251 | 0.0033 | 0.0095 | 0.0251 | TRUE | 0.5722 |
| Lpin2 | 1900 | 1382 | 1359 | 1326 | 1154 | 1366 | 1476 | 1470 | 1466 | 1320 | 1302 | 1348 | 1397.03 | 1471 | 1323 | 0.9 | -0.152 | 0.0204304 | 0.0702458 | 0 | 0.0041 | 0.0024 | 0.0024 | TRUE | 0.0033 |
| Lpin3 | 1161 | 925 | 887 | 978 | 871 | 905 | 902 | 984 | 957 | 974 | 983 | 893 | 948.69 | 948 | 950 | 1.003 | 0.004 | 0.95621152 | 0.9757358 | 0.0014 | 0.0049 | 0.0037 | 0.0037 | TRUE | 0.0326 |
| Lpl | 2226 | 1674 | 1646 | 1796 | 1422 | 1635 | 1729 | 1781 | 1776 | 1788 | 1604 | 1614 | 1715.23 | 1762 | 1669 | 0.948 | -0.078 | 0.26527383 | 0.43561108 | 0.0014 | 0.0038 | 0.0029 | 0.0029 | TRUE | 0.0473 |
| Lpo | 19 | 5 | 6 | 7 | 4 | 5 | 15 | 5 | 6 | 7 | 5 | 5 | 7.16 | 9 | 6 | 0.608 | -0.718 | 0.34036725 | NA | 0 | 0.3334 | 0.261 | 0.261 | TRUE | 1.3516 |
| Lpp | 1221 | 833 | 845 | 1171 | 968 | 1099 | 948 | 886 | 912 | 1166 | 1092 | 1085 | 1014.77 | 915 | 1114 | 1.216 | 0.282 | 0.00022928 | 0.00237818 | 0 | 0.0048 | 0.0032 | 0.0032 | TRUE | 0.0197 |
| Lpxn | 20 | 14 | 9 | 5 | 15 | 8 | 16 | 15 | 10 | 5 | 17 | 8 | 11.66 | 14 | 10 | 0.73 | -0.453 | 0.45741085 | NA | 0.1042 | 0.2058 | 0.1794 | 0.1794 | TRUE | 1.1695 |
| Lrat | 29 | 24 | 31 | 13 | 21 | 25 | 23 | 26 | 33 | 13 | 24 | 25 | 23.8 | 27 | 21 | 0.755 | -0.406 | 0.33180172 | 0.50596842 | 0.0287 | 0.102 | 0.0828 | 0.0828 | TRUE | 0.5805 |
| Lrba | 4397 | 3035 | 2945 | 3642 | 3156 | 3403 | 3415 | 3228 | 3178 | 3625 | 3560 | 3359 | 3394.28 | 3274 | 3515 | 1.073 | 0.102 | 0.09326605 | 0.2100007 | 0.0011 | 0.0031 | 0.0024 | 0.0024 | TRUE | 0.0184 |
| Lrch1 | 1498 | 1140 | 1175 | 1206 | 1136 | 1332 | 1163 | 1213 | 1268 | 1200 | 1282 | 1315 | 1240.12 | 1215 | 1266 | 1.043 | 0.061 | 0.4186999 | 0.59135154 | 0.0013 | 0.0043 | 0.0032 | 0.0032 | TRUE | 0.0247 |
| Lrch2 | 1107 | 778 | 916 | 939 | 863 | 1016 | 860 | 828 | 988 | 935 | 974 | 1003 | 931.14 | 892 | 971 | 1.089 | 0.123 | 0.16608361 | 0.31707019 | 0.0036 | 0.005 | 0.0046 | 0.0046 | TRUE | 0.1037 |
| Lrch3 | 3074 | 2264 | 2426 | 2732 | 2389 | 2944 | 2387 | 2408 | 2618 | 2719 | 2695 | 2906 | 2622.32 | 2471 | 2773 | 1.123 | 0.167 | 0.01069334 | 0.04338347 | 0.0016 | 0.0033 | 0.0027 | 0.0027 | TRUE | 0.0322 |
| Lrch4 | 0 | 0 | 0 | 0 | 0 | 0 | 0 | 0 | 0 | 0 | 0 | 0 | 0 | 0 | 0 | NA | NA | NA | NA | NA | NA | NA | NA | NA | NA |
| Lrcol1 | 52 | 57 | 63 | 72 | 80 | 81 | 40 | 61 | 68 | 72 | 90 | 80 | 68.48 | 56 | 81 | 1.444 | 0.53 | 0.04221651 | 0.11903741 | 0.0258 | 0.037 | 0.034 | 0.034 | TRUE | 0.6085 |
| Lrdd | 1383 | 1047 | 932 | 1007 | 897 | 1097 | 1074 | 1114 | 1006 | 1002 | 1012 | 1083 | 1048.43 | 1065 | 1032 | 0.97 | -0.044 | 0.57321096 | 0.72554943 | 0.0012 | 0.0047 | 0.0035 | 0.0035 | TRUE | 0.0269 |
| Lrfn1 | 1024 | 737 | 746 | 849 | 786 | 855 | 795 | 784 | 805 | 845 | 887 | 844 | 826.66 | 795 | 859 | 1.08 | 0.111 | 0.15807695 | 0.30702457 | 0 | 0.0053 | 0.0032 | 0.0032 | TRUE | 0.0095 |
| Lrfn2 | 3 | 1 | 5 | 3 | 2 | 3 | 2 | 1 | 5 | 3 | 2 | 3 | 2.83 | 3 | 3 | 0.945 | -0.082 | 0.94274495 | NA | 0 | 0.8393 | 0.5774 | 0.5774 | TRUE | 0.5968 |
| Lrfn3 | 293 | 191 | 275 | 270 | 247 | 293 | 228 | 203 | 297 | 269 | 279 | 289 | 260.68 | 243 | 279 | 1.152 | 0.204 | 0.17602481 | 0.32965443 | 0.0149 | 0.0115 | 0.0126 | 0.0126 | TRUE | 0.4077 |
| Lrfn4 | 1057 | 922 | 924 | 1236 | 1107 | 1182 | 821 | 981 | 997 | 1230 | 1249 | 1167 | 1074.1 | 933 | 1215 | 1.305 | 0.384 | 1.93E-05 | 0.00032336 | 0.0054 | 0.0046 | 0.0049 | 0.0049 | TRUE | 0.1376 |
| Lrfn5 | 18 | 5 | 4 | 2 | 2 | 3 | 14 | 5 | 4 | 2 | 2 | 3 | 5.14 | 8 | 2 | 0.3 | -1.738 | 0.065949 | NA | 0 | 0.4638 | 0.3686 | 0.3686 | TRUE | 1.6686 |
| Lrg1 | 0 | 0 | 0 | 0 | 0 | 0 | 0 | 0 | 0 | 0 | 0 | 0 | 0 | 0 | 0 | NA | NA | NA | NA | NA | NA | NA | NA | NA | NA |
| Lrguk | 87 | 80 | 51 | 79 | 73 | 90 | 68 | 85 | 55 | 79 | 82 | 89 | 76.25 | 69 | 83 | 1.204 | 0.268 | 0.25749954 | 0.42741156 | 0.0084 | 0.0335 | 0.027 | 0.027 | TRUE | 0.3224 |
| Lrif1 | 725 | 529 | 298 | 364 | 297 | 295 | 563 | 563 | 322 | 362 | 335 | 291 | 405.98 | 483 | 329 | 0.682 | -0.552 | 0.0021248 | 0.01262252 | 0.0518 | 0.0083 | 0.0207 | 0.0207 | TRUE | 0.9749 |
| Lrig1 | 2546 | 1969 | 1958 | 1977 | 1969 | 2044 | 1977 | 2094 | 2113 | 1968 | 2221 | 2018 | 2065.22 | 2061 | 2069 | 1.004 | 0.005 | 0.93873197 | 0.96617672 | 0.0022 | 0.0036 | 0.0031 | 0.0031 | TRUE | 0.0495 |
| Lrig2 | 1888 | 1375 | 1382 | 1909 | 1580 | 1808 | 1466 | 1463 | 1491 | 1900 | 1782 | 1785 | 1647.91 | 1473 | 1822 | 1.237 | 0.307 | 2.48E-06 | 5.96E-05 | 0 | 0.0039 | 0.0025 | 0.0025 | TRUE | 0.0169 |
| Lrig3 | 1036 | 778 | 780 | 903 | 803 | 974 | 805 | 828 | 842 | 899 | 906 | 961 | 873.33 | 825 | 922 | 1.119 | 0.162 | 0.03900425 | 0.11246774 | 0 | 0.0051 | 0.0033 | 0.0033 | TRUE | 0.0169 |
| Lrit1 | 0 | 0 | 0 | 1 | 0 | 0 | 0 | 0 | 0 | 1 | 0 | 0 | 0.17 | 0 | 0 | 2.094 | 1.067 | 0.79379329 | NA | 0 | 14.2885 | 10 | 10 | TRUE | 0.3906 |
| Lrit2 | 0 | 0 | 0 | 0 | 0 | 0 | 0 | 0 | 0 | 0 | 0 | 0 | 0 | 0 | 0 | NA | NA | NA | NA | NA | NA | NA | NA | NA | NA |
| Lrit3 | 0 | 2 | 0 | 0 | 1 | 0 | 0 | 2 | 0 | 0 | 1 | 0 | 0.54 | 1 | 0 | 0.535 | -0.903 | 0.76014553 | NA | 0 | 4.3706 | 4.6038 | 4.6038 | TRUE | 0.9746 |
| Lrmp | 181 | 207 | 178 | 184 | 171 | 224 | 141 | 220 | 192 | 183 | 193 | 221 | 191.67 | 184 | 199 | 1.087 | 0.12 | 0.50514603 | 0.66864718 | 0.0257 | 0.0148 | 0.0181 | 0.0181 | TRUE | 0.505 |
| Lrp1 | 4672 | 3531 | 4496 | 5801 | 5466 | 6558 | 3629 | 3756 | 4851 | 5774 | 6166 | 6474 | 5108.31 | 4079 | 6138 | 1.506 | 0.59 | 1.17E-09 | 7.66E-08 | 0.0149 | 0.0029 | 0.0066 | 0.0066 | TRUE | 0.3334 |
| Lrp10 | 3462 | 2532 | 2716 | 2952 | 2730 | 3287 | 2689 | 2693 | 2931 | 2938 | 3080 | 3245 | 2929.25 | 2771 | 3088 | 1.115 | 0.157 | 0.01755321 | 0.06272201 | 0.002 | 0.0032 | 0.0028 | 0.0028 | TRUE | 0.0305 |
| Lrp11 | 4846 | 3525 | 4029 | 4399 | 3898 | 4451 | 3764 | 3749 | 4347 | 4379 | 4397 | 4394 | 4171.73 | 3953 | 4390 | 1.111 | 0.152 | 0.02660105 | 0.08500085 | 0.0034 | 0.003 | 0.0031 | 0.0031 | TRUE | 0.0917 |
| Lrp12 | 881 | 637 | 587 | 658 | 549 | 577 | 684 | 678 | 633 | 655 | 619 | 570 | 639.84 | 665 | 615 | 0.923 | -0.115 | 0.21738939 | 0.38017209 | 0.0017 | 0.0061 | 0.0047 | 0.0047 | TRUE | 0.0495 |
| Lrp1b | 25 | 12 | 7 | 6 | 3 | 2 | 19 | 13 | 8 | 6 | 3 | 2 | 8.51 | 13 | 4 | 0.282 | -1.824 | 0.01541911 | NA | 0 | 0.2809 | 0.2359 | 0.2359 | TRUE | 0.1438 |
| Lrp2 | 2862 | 1685 | 1254 | 2138 | 1954 | 2232 | 2223 | 1792 | 1353 | 2128 | 2204 | 2203 | 1983.99 | 1789 | 2178 | 1.217 | 0.283 | 0.1613522 | 0.31118702 | 0.029 | 0.0036 | 0.0113 | 0.029 | TRUE | 0.2317 |
| Lrp2bp | 38 | 38 | 33 | 38 | 30 | 26 | 30 | 40 | 36 | 38 | 34 | 26 | 33.81 | 35 | 33 | 0.927 | -0.11 | 0.74657574 | 0.8496542 | 0 | 0.0725 | 0.0536 | 0.0536 | TRUE | 0.2391 |
| Lrp3 | 619 | 491 | 496 | 575 | 553 | 581 | 481 | 522 | 535 | 572 | 624 | 574 | 551.32 | 513 | 590 | 1.152 | 0.204 | 0.03464791 | 0.10308547 | 0.0011 | 0.0067 | 0.0049 | 0.0049 | TRUE | 0.0373 |
| Lrp4 | 1308 | 1046 | 1179 | 1340 | 1169 | 1251 | 1016 | 1113 | 1272 | 1334 | 1319 | 1235 | 1214.69 | 1134 | 1296 | 1.145 | 0.195 | 0.03134181 | 0.09565845 | 0.0061 | 0.0044 | 0.0051 | 0.0051 | TRUE | 0.1365 |
| Lrp5 | 4212 | 3325 | 3575 | 4230 | 4065 | 4610 | 3271 | 3537 | 3857 | 4211 | 4586 | 4551 | 4002.08 | 3555 | 4449 | 1.252 | 0.324 | 6.69E-06 | 0.00013534 | 0.0041 | 0.003 | 0.0035 | 0.0035 | TRUE | 0.0672 |
| Lrp6 | 9421 | 6775 | 7009 | 7941 | 7237 | 8386 | 7317 | 7206 | 7563 | 7904 | 8164 | 8278 | 7738.81 | 7362 | 8115 | 1.102 | 0.141 | 0.00521141 | 0.02530978 | 4.00E-04 | 0.0027 | 0.0017 | 0.0017 | TRUE | 0.0069 |
| Lrp8 | 1033 | 800 | 953 | 1156 | 992 | 1129 | 802 | 851 | 1028 | 1151 | 1119 | 1114 | 1010.96 | 894 | 1128 | 1.264 | 0.338 | 0.00044386 | 0.00394041 | 0.0075 | 0.0048 | 0.0057 | 0.0057 | TRUE | 0.2041 |
| Lrpap1 | 3521 | 2357 | 2139 | 1945 | 1880 | 2018 | 2735 | 2507 | 2308 | 1936 | 2121 | 1992 | 2266.44 | 2517 | 2016 | 0.8 | -0.321 | 2.80E-05 | 0.00043761 | 0.004 | 0.0035 | 0.0038 | 0.0038 | TRUE | 0.071 |
| Lrpprc | 16132 | 12059 | 10607 | 11942 | 10443 | 11858 | 12529 | 12827 | 11445 | 11887 | 11781 | 11706 | 12029.07 | 12267 | 11791 | 0.961 | -0.057 | 0.31385966 | 0.48723755 | 0.0017 | 0.0026 | 0.0022 | 0.0022 | TRUE | 0.0418 |
| Lrr1 | 858 | 537 | 446 | 545 | 523 | 567 | 666 | 571 | 481 | 542 | 590 | 560 | 568.5 | 573 | 564 | 0.981 | -0.027 | 0.8200262 | 0.89651557 | 0.0125 | 0.0066 | 0.0085 | 0.0085 | TRUE | 0.2022 |
| Lrrc1 | 1514 | 1029 | 1043 | 1073 | 1034 | 1123 | 1176 | 1095 | 1125 | 1068 | 1166 | 1109 | 1123.15 | 1132 | 1114 | 0.983 | -0.024 | 0.74572803 | 0.84889223 | 8.00E-04 | 0.0045 | 0.0032 | 0.0032 | TRUE | 0.0196 |
| Lrrc10 | 6 | 10 | 5 | 5 | 5 | 5 | 5 | 11 | 5 | 5 | 6 | 5 | 6.04 | 7 | 5 | 0.759 | -0.398 | 0.61167501 | NA | 0 | 0.3948 | 0.2725 | 0.2725 | TRUE | 0.56 |
| Lrrc10b | 20 | 7 | 13 | 17 | 7 | 12 | 16 | 7 | 14 | 17 | 8 | 12 | 12.28 | 12 | 12 | 0.987 | -0.018 | 0.97457745 | NA | 0.0506 | 0.1955 | 0.1561 | 0.1561 | TRUE | 0.2634 |
| Lrrc14 | 605 | 446 | 293 | 272 | 241 | 236 | 470 | 474 | 316 | 271 | 272 | 233 | 339.34 | 420 | 259 | 0.614 | -0.704 | 7.57E-06 | 0.00014967 | 0.0267 | 0.0094 | 0.0147 | 0.0147 | TRUE | 0.5295 |
| Lrrc14b | 0 | 0 | 0 | 1 | 1 | 2 | 0 | 0 | 0 | 1 | 1 | 2 | 0.68 | 0 | 1 | 7.73 | 2.951 | 0.21871019 | NA | 0 | 3.4727 | 2.7687 | 2.7687 | TRUE | 0.0985 |
| Lrrc15 | 249 | 159 | 144 | 105 | 110 | 122 | 193 | 169 | 155 | 105 | 124 | 120 | 144.49 | 172 | 116 | 0.671 | -0.576 | 0.00094665 | 0.00697444 | 0 | 0.0188 | 0.0146 | 0.0146 | TRUE | 0.1279 |
| Lrrc16a | 1923 | 1378 | 1329 | 1432 | 1309 | 1370 | 1494 | 1466 | 1434 | 1425 | 1477 | 1352 | 1441.29 | 1465 | 1418 | 0.968 | -0.047 | 0.49179004 | 0.65736783 | 4.00E-04 | 0.0041 | 0.0027 | 0.0027 | TRUE | 0.0199 |
| Lrrc16b | 118 | 133 | 151 | 151 | 146 | 163 | 92 | 141 | 163 | 150 | 165 | 161 | 145.33 | 132 | 159 | 1.21 | 0.276 | 0.18291064 | 0.33860145 | 0.0365 | 0.0187 | 0.0239 | 0.0239 | TRUE | 0.8127 |
| Lrrc17 | 8 | 3 | 1 | 1 | 1 | 0 | 6 | 3 | 1 | 1 | 1 | 0 | 2.1 | 3 | 1 | 0.195 | -2.356 | 0.1314555 | NA | 0 | 1.1304 | 0.8794 | 0.8794 | TRUE | 0.1324 |
| Lrrc18 | 2 | 4 | 8 | 3 | 0 | 5 | 2 | 4 | 9 | 3 | 0 | 5 | 3.73 | 5 | 3 | 0.568 | -0.816 | 0.47074222 | NA | 0.5904 | 0.6383 | 0.6271 | 0.6271 | TRUE | 0.2868 |
| Lrrc19 | 3 | 4 | 2 | 0 | 3 | 6 | 2 | 4 | 2 | 0 | 3 | 6 | 3.01 | 3 | 3 | 1.071 | 0.099 | 0.93419318 | NA | 0.3724 | 0.7903 | 0.6955 | 0.6955 | TRUE | 0.2669 |
| Lrrc2 | 1700 | 1230 | 1398 | 1447 | 1387 | 1665 | 1320 | 1308 | 1508 | 1440 | 1565 | 1644 | 1464.29 | 1379 | 1550 | 1.124 | 0.169 | 0.04164116 | 0.11771699 | 0.0045 | 0.004 | 0.0043 | 0.0043 | TRUE | 0.0797 |
| Lrrc20 | 736 | 479 | 383 | 433 | 365 | 406 | 572 | 510 | 413 | 431 | 412 | 401 | 456.32 | 498 | 415 | 0.83 | -0.269 | 0.03159603 | 0.09616738 | 0.0122 | 0.0076 | 0.0091 | 0.0091 | TRUE | 0.255 |
| Lrrc23 | 11 | 7 | 9 | 7 | 8 | 10 | 9 | 7 | 10 | 7 | 9 | 10 | 8.59 | 9 | 9 | 1.006 | 0.009 | 0.98902123 | NA | 0 | 0.2782 | 0.1785 | 0.1785 | TRUE | 0.091 |

| | | | | | | | | | | | | | | | | | | | | | | | | | |
|---|---|---|---|---|---|---|---|---|---|---|---|---|---|---|---|---|---|---|---|---|---|---|---|---|---|
| Lrrc24 | 17 | 19 | 22 | 24 | 19 | 31 | 13 | 20 | 24 | 24 | 21 | 31 | 22.18 | 19 | 25 | 1.348 | 0.431 | 0.30947876 | 0.48221737 | 0 | 0.1093 | 0.0837 | 0.0837 | TRUE | 0.4809 |
| Lrrc25 | 4 | 2 | 14 | 8 | 4 | 9 | 3 | 2 | 15 | 8 | 5 | 9 | 6.95 | 7 | 7 | 1.074 | 0.102 | 0.90340979 | NA | 0.426 | 0.3434 | 0.368 | 0.368 | TRUE | 2.7988 |
| Lrrc26 | 1 | 0 | 0 | 0 | 0 | 1 | 1 | 0 | 0 | 0 | 0 | 1 | 0.29 | 0 | 0 | 1.075 | 0.105 | 0.97702352 | NA | 0 | 8.0647 | 7.5458 | 7.5458 | TRUE | 0.3835 |
| Lrrc27 | 299 | 268 | 290 | 351 | 334 | 345 | 232 | 285 | 313 | 349 | 377 | 341 | 316.16 | 277 | 356 | 1.29 | 0.367 | 0.0062878 | 0.02923601 | 0.0095 | 0.0099 | 0.0098 | 0.0098 | TRUE | 0.241 |
| Lrrc28 | 1471 | 911 | 1029 | 1133 | 905 | 1134 | 1142 | 969 | 1110 | 1128 | 1021 | 1119 | 1081.65 | 1074 | 1089 | 1.014 | 0.02 | 0.8197977 | 0.8964401 | 0.0043 | 0.0046 | 0.0045 | 0.0045 | TRUE | 0.0862 |
| Lrrc29 | 184 | 147 | 133 | 194 | 147 | 180 | 143 | 156 | 144 | 193 | 166 | 178 | 163.23 | 148 | 179 | 1.215 | 0.28 | 0.07431282 | 0.17919794 | 0 | 0.0169 | 0.0116 | 0.0116 | TRUE | 0.052 |
| Lrrc3 | 327 | 232 | 208 | 277 | 241 | 267 | 254 | 247 | 224 | 276 | 272 | 264 | 256.06 | 242 | 271 | 1.116 | 0.159 | 0.21275112 | 0.37509708 | 0 | 0.0117 | 0.0078 | 0.0078 | TRUE | 0.042 |
| Lrrc30 | 1 | 0 | 0 | 0 | 0 | 0 | 1 | 0 | 0 | 0 | 0 | 0 | 0.13 | 0 | 0 | 0.552 | -0.857 | 0.8336548 | NA | 0 | 18.3121 | 10 | 10 | TRUE | 0.2566 |
| Lrrc32 | 63 | 37 | 45 | 46 | 37 | 50 | 49 | 39 | 49 | 46 | 42 | 49 | 45.62 | 46 | 46 | 0.999 | -0.001 | 0.99664955 | 0.99800974 | 0 | 0.0544 | 0.0368 | 0.0368 | TRUE | 0.1073 |
| Lrrc33 | 239 | 173 | 107 | 172 | 133 | 164 | 186 | 184 | 115 | 171 | 150 | 162 | 161.37 | 162 | 161 | 0.993 | -0.01 | 0.95808314 | 0.97701786 | 0.0294 | 0.0171 | 0.0206 | 0.0206 | TRUE | 0.6369 |
| Lrrc34 | 1623 | 1370 | 1991 | 1967 | 2052 | 2226 | 1261 | 1457 | 2148 | 1958 | 2315 | 2197 | 1889.38 | 1622 | 2157 | 1.33 | 0.411 | 0.09811879 | 0.21824375 | 0.0441 | 0.0037 | 0.0143 | 0.0441 | TRUE | 0.9704 |
| Lrrc36 | 13 | 13 | 8 | 24 | 20 | 20 | 10 | 14 | 9 | 24 | 23 | 20 | 16.46 | 11 | 22 | 2.038 | 1.027 | 0.03279933 | NA | 0 | 0.1464 | 0.099 | 0.099 | TRUE | 0.1906 |
| Lrrc38 | 9 | 9 | 25 | 24 | 13 | 20 | 7 | 10 | 27 | 24 | 15 | 20 | 16.97 | 15 | 20 | 1.365 | 0.449 | 0.41935948 | NA | 0.2147 | 0.1421 | 0.1626 | 0.1626 | TRUE | 2.0102 |
| Lrrc39 | 5 | 5 | 1 | 2 | 1 | 1 | 4 | 5 | 1 | 2 | 1 | 1 | 2.4 | 3 | 1 | 0.399 | -1.326 | 0.30560479 | NA | 0 | 0.9909 | 0.6906 | 0.6906 | TRUE | 0.2642 |
| Lrrc3b | 121 | 100 | 83 | 75 | 62 | 81 | 94 | 106 | 90 | 75 | 70 | 80 | 85.74 | 97 | 75 | 0.776 | -0.366 | 0.0842591 | 0.19572816 | 0 | 0.0301 | 0.0204 | 0.0204 | TRUE | 0.0712 |
| Lrrc4 | 74 | 64 | 42 | 60 | 48 | 78 | 57 | 68 | 45 | 60 | 54 | 77 | 60.29 | 57 | 64 | 1.119 | 0.162 | 0.54716212 | 0.7040944 | 0.0188 | 0.0417 | 0.0355 | 0.0355 | TRUE | 0.2686 |
| Lrrc40 | 5147 | 3623 | 3587 | 3772 | 3324 | 3715 | 3997 | 3854 | 3870 | 3755 | 3750 | 3667 | 3815.55 | 3907 | 3724 | 0.953 | -0.07 | 0.18602615 | 0.34228524 | 0 | 0.003 | 0.0017 | 0.0017 | TRUE | 0.0051 |
| Lrrc41 | 2126 | 1400 | 1572 | 1771 | 1520 | 1695 | 1651 | 1489 | 1696 | 1763 | 1715 | 1673 | 1664.55 | 1612 | 1717 | 1.065 | 0.09 | 0.21180999 | 0.37403017 | 0.002 | 0.0039 | 0.0032 | 0.0032 | TRUE | 0.0525 |
| Lrrc42 | 3768 | 2672 | 2716 | 2931 | 2438 | 2888 | 2926 | 2842 | 2931 | 2918 | 2750 | 2851 | 2869.65 | 2900 | 2840 | 0.979 | -0.03 | 0.5983021 | 0.74416177 | 2.00E-04 | 0.0033 | 0.002 | 0.002 | TRUE | 0.009 |
| Lrrc43 | 3 | 1 | 2 | 1 | 0 | 1 | 2 | 1 | 2 | 1 | 0 | 1 | 1.26 | 2 | 1 | 0.363 | -1.462 | 0.4062211 | NA | 0 | 1.8899 | 1.2105 | 1.2105 | TRUE | 0.1995 |
| Lrrc45 | 777 | 723 | 654 | 649 | 638 | 732 | 603 | 769 | 706 | 646 | 720 | 723 | 694.42 | 693 | 696 | 1.007 | 0.01 | 0.92484341 | 0.95843872 | 0.0082 | 0.0058 | 0.0066 | 0.0066 | TRUE | 0.1577 |
| Lrrc46 | 55 | 49 | 50 | 50 | 47 | 45 | 43 | 52 | 54 | 50 | 53 | 44 | 49.33 | 50 | 49 | 0.993 | -0.01 | 0.97211505 | 0.98493233 | 0 | 0.0505 | 0.0347 | 0.0347 | TRUE | 0.1407 |
| Lrrc47 | 6567 | 4934 | 4479 | 4418 | 4288 | 4784 | 5100 | 5248 | 4833 | 4398 | 4837 | 4723 | 4856.48 | 5060 | 4653 | 0.919 | -0.122 | 0.04668237 | 0.12814117 | 0.0018 | 0.0029 | 0.0025 | 0.0025 | TRUE | 0.028 |
| Lrrc48 | 282 | 220 | 292 | 333 | 290 | 319 | 219 | 234 | 315 | 331 | 327 | 315 | 290.27 | 256 | 324 | 1.272 | 0.347 | 0.01812736 | 0.06423588 | 0.0155 | 0.0106 | 0.0121 | 0.0121 | TRUE | 0.4448 |
| Lrrc49 | 422 | 265 | 257 | 273 | 259 | 297 | 328 | 282 | 277 | 272 | 292 | 293 | 290.67 | 296 | 286 | 0.963 | -0.054 | 0.66675953 | 0.79491786 | 0.0026 | 0.0106 | 0.0081 | 0.0081 | TRUE | 0.1093 |
| Lrrc4b | 9 | 5 | 5 | 16 | 16 | 16 | 7 | 5 | 5 | 16 | 18 | 16 | 11.25 | 6 | 17 | 2.778 | 1.474 | 0.0124308 | NA | 0 | 0.2132 | 0.1388 | 0.1388 | TRUE | 0.0819 |
| Lrrc4c | 0 | 0 | 0 | 0 | 0 | 0 | 0 | 0 | 0 | 0 | 0 | 0 | 0 | 0 | 0 | NA | NA | NA | NA | NA | NA | NA | NA | NA | NA |
| Lrrc51 | 116 | 84 | 117 | 216 | 181 | 204 | 90 | 89 | 126 | 215 | 204 | 201 | 154.38 | 102 | 207 | 2.04 | 1.029 | 7.47E-09 | 3.88E-07 | 0.0084 | 0.0178 | 0.0156 | 0.0156 | TRUE | 0.4083 |
| Lrrc52 | 1 | 5 | 2 | 1 | 1 | 4 | 1 | 5 | 2 | 1 | 1 | 4 | 2.39 | 3 | 2 | 0.759 | -0.398 | 0.75985723 | NA | 0.3416 | 0.9951 | 0.7866 | 0.7866 | TRUE | 0.5116 |
| Lrrc55 | 32 | 14 | 16 | 7 | 9 | 5 | 25 | 15 | 17 | 7 | 10 | 5 | 13.18 | 19 | 7 | 0.38 | -1.395 | 0.01442506 | NA | 0 | 0.1823 | 0.1379 | 0.1379 | TRUE | 0.4584 |
| Lrrc56 | 285 | 234 | 218 | 214 | 215 | 236 | 221 | 249 | 235 | 213 | 243 | 233 | 232.33 | 235 | 230 | 0.977 | -0.033 | 0.8044796 | 0.88681374 | 0 | 0.0126 | 0.0089 | 0.0089 | TRUE | 0.0439 |
| Lrrc57 | 1430 | 1006 | 975 | 1072 | 834 | 933 | 1111 | 1070 | 1052 | 1067 | 941 | 921 | 1026.94 | 1078 | 976 | 0.906 | -0.143 | 0.0855471 | 0.19778789 | 0.0026 | 0.0047 | 0.004 | 0.004 | TRUE | 0.0798 |
| Lrrc58 | 4740 | 3584 | 3444 | 3688 | 3349 | 3441 | 3681 | 3812 | 3716 | 3671 | 3778 | 3397 | 3675.92 | 3736 | 3615 | 0.967 | -0.048 | 0.43159642 | 0.60360153 | 0.0012 | 0.0031 | 0.0024 | 0.0024 | TRUE | 0.0341 |
| Lrrc59 | 28828 | 20574 | 18869 | 20917 | 18533 | 20847 | 22389 | 21884 | 20359 | 20821 | 20907 | 20579 | 21156.73 | 21544 | 20769 | 0.964 | -0.053 | 0.31234339 | 0.48562212 | 0.0012 | 0.0025 | 0.0019 | 0.0019 | TRUE | 0.0283 |
| Lrrc6 | 16 | 16 | 22 | 27 | 14 | 15 | 12 | 17 | 24 | 27 | 16 | 15 | 18.44 | 18 | 19 | 1.095 | 0.131 | 0.7846015 | 0.87338856 | 0.0598 | 0.1309 | 0.1105 | 0.1105 | TRUE | 0.4953 |
| Lrrc61 | 563 | 453 | 450 | 597 | 604 | 669 | 437 | 482 | 486 | 594 | 681 | 660 | 556.78 | 468 | 645 | 1.38 | 0.465 | 3.28E-06 | 7.50E-05 | 0.0026 | 0.0067 | 0.0054 | 0.0054 | TRUE | 0.0569 |
| Lrrc63 | 1 | 1 | 2 | 1 | 2 | 2 | 1 | 1 | 2 | 1 | 2 | 2 | 1.54 | 1 | 2 | 1.328 | 0.409 | 0.78439372 | NA | 0 | 1.5441 | 0.9441 | 0.9441 | TRUE | 0.1668 |
| Lrrc66 | 2 | 2 | 2 | 0 | 1 | 0 | 2 | 2 | 2 | 0 | 1 | 0 | 1.16 | 2 | 0 | 0.192 | -2.384 | 0.21556583 | NA | 0 | 2.0436 | 1.421 | 1.421 | TRUE | 0.5075 |
| Lrrc69 | 1 | 0 | 1 | 1 | 1 | 1 | 1 | 0 | 1 | 1 | 1 | 1 | 0.83 | 1 | 1 | 1.656 | 0.728 | 0.72085425 | NA | 0 | 2.8658 | 1.7182 | 1.7182 | TRUE | 0.1959 |
| Lrrc7 | 0 | 0 | 0 | 0 | 0 | 0 | 0 | 0 | 0 | 0 | 0 | 0 | 0 | 0 | 0 | NA | NA | NA | NA | NA | NA | NA | NA | NA | NA |
| Lrrc71 | 11 | 11 | 9 | 12 | 12 | 19 | 9 | 12 | 10 | 12 | 14 | 19 | 12.37 | 10 | 15 | 1.488 | 0.574 | 0.29444993 | NA | 0 | 0.1941 | 0.1321 | 0.1321 | TRUE | 0.2646 |
| Lrrc72 | 1 | 1 | 1 | 0 | 0 | 0 | 1 | 1 | 1 | 0 | 0 | 0 | 0.49 | 1 | 0 | 0.196 | -2.347 | 0.38405676 | NA | 0 | 4.8735 | 3.7415 | 3.7415 | TRUE | 0.0718 |
| Lrrc73 | 41 | 29 | 19 | 23 | 21 | 23 | 32 | 31 | 21 | 23 | 24 | 23 | 25.41 | 28 | 23 | 0.828 | -0.273 | 0.47959184 | 0.64649686 | 0 | 0.0957 | 0.0673 | 0.0673 | TRUE | 0.3095 |
| Lrrc8a | 1223 | 951 | 874 | 991 | 876 | 923 | 950 | 1012 | 943 | 986 | 988 | 911 | 965.04 | 968 | 962 | 0.994 | -0.009 | 0.90757962 | 0.94923397 | 7.00E-04 | 0.0049 | 0.0035 | 0.0035 | TRUE | 0.0258 |
| Lrrc8b | 616 | 468 | 489 | 607 | 550 | 624 | 478 | 498 | 528 | 604 | 620 | 616 | 557.42 | 501 | 613 | 1.226 | 0.294 | 0.00153577 | 0.00982212 | 0 | 0.0067 | 0.0044 | 0.0044 | TRUE | 0.0243 |
| Lrrc8c | 444 | 305 | 334 | 324 | 319 | 346 | 345 | 324 | 360 | 323 | 360 | 342 | 342.26 | 343 | 342 | 0.994 | -0.009 | 0.9359479 | 0.96464483 | 0 | 0.0094 | 0.0065 | 0.0065 | TRUE | 0.0265 |
| Lrrc8d | 2606 | 1870 | 1786 | 1806 | 1656 | 1952 | 2024 | 1989 | 1927 | 1798 | 1868 | 1927 | 1922.15 | 1980 | 1864 | 0.941 | -0.088 | 0.16901753 | 0.32056364 | 4.00E-04 | 0.0037 | 0.0024 | 0.0024 | TRUE | 0.012 |
| Lrrc8e | 166 | 121 | 144 | 169 | 135 | 183 | 129 | 129 | 155 | 168 | 152 | 181 | 152.36 | 138 | 167 | 1.218 | 0.284 | 0.0899662 | 0.20469387 | 0 | 0.018 | 0.0137 | 0.0137 | TRUE | 0.1252 |
| Lrrc9 | 266 | 218 | 197 | 167 | 169 | 211 | 207 | 232 | 213 | 166 | 191 | 208 | 202.7 | 217 | 188 | 0.869 | -0.202 | 0.17371866 | 0.32644765 | 0.0029 | 0.0141 | 0.0109 | 0.0109 | TRUE | 0.1175 |
| Lrrcc1 | 1125 | 887 | 783 | 768 | 694 | 771 | 874 | 943 | 845 | 764 | 783 | 761 | 828.42 | 887 | 769 | 0.867 | -0.205 | 0.01262164 | 0.049169 | 0 | 0.0053 | 0.0037 | 0.0037 | TRUE | 0.0352 |
| Lrrd1 | 2 | 2 | 2 | 1 | 0 | 3 | 2 | 2 | 2 | 1 | 0 | 3 | 1.63 | 2 | 1 | 0.696 | -0.524 | 0.73027627 | NA | 0 | 1.4541 | 1.0268 | 1.0268 | TRUE | 0.6417 |
| Lrrfip1 | 2523 | 1993 | 1888 | 2198 | 1961 | 2368 | 1960 | 2120 | 2037 | 2188 | 2212 | 2338 | 2142.38 | 2039 | 2246 | 1.102 | 0.141 | 0.02934722 | 0.0911671 | 0.001 | 0.0035 | 0.0025 | 0.0025 | TRUE | 0.0156 |
| Lrrfip2 | 3159 | 2248 | 2076 | 2312 | 2057 | 2396 | 2453 | 2391 | 2240 | 2301 | 2321 | 2365 | 2345.28 | 2361 | 2329 | 0.986 | -0.021 | 0.73879331 | 0.84449558 | 8.00E-04 | 0.0034 | 0.0024 | 0.0024 | TRUE | 0.0243 |
| Lrriq1 | 45 | 25 | 10 | 20 | 13 | 16 | 35 | 27 | 11 | 20 | 15 | 16 | 20.45 | 24 | 17 | 0.689 | -0.537 | 0.27175282 | 0.44274058 | 0.1285 | 0.1183 | 0.1213 | 0.1213 | TRUE | 0.3285 |
| Lrriq3 | 23 | 24 | 18 | 27 | 20 | 16 | 18 | 26 | 19 | 27 | 23 | 16 | 21.34 | 21 | 22 | 1.043 | 0.061 | 0.88669773 | 0.93690919 | 0 | 0.1135 | 0.0845 | 0.0845 | TRUE | 0.3385 |
| Lrriq4 | 247 | 232 | 267 | 267 | 257 | 295 | 192 | 247 | 288 | 266 | 290 | 291 | 262.27 | 242 | 282 | 1.171 | 0.228 | 0.14199165 | 0.28444102 | 0.0182 | 0.0115 | 0.0135 | 0.0135 | TRUE | 0.343 |
| Lrrk1 | 263 | 224 | 195 | 195 | 163 | 224 | 204 | 238 | 210 | 194 | 184 | 221 | 208.67 | 217 | 200 | 0.92 | -0.12 | 0.41390856 | 0.58649698 | 0.0031 | 0.0138 | 0.0107 | 0.0107 | TRUE | 0.0966 |
| Lrrk2 | 125 | 123 | 75 | 89 | 75 | 98 | 97 | 131 | 81 | 89 | 85 | 97 | 96.46 | 103 | 90 | 0.876 | -0.191 | 0.39229356 | 0.56596869 | 0.0219 | 0.027 | 0.0255 | 0.0255 | TRUE | 0.4619 |
| Lrrn1 | 413 | 312 | 337 | 349 | 325 | 342 | 321 | 332 | 364 | 347 | 367 | 338 | 344.65 | 339 | 351 | 1.036 | 0.051 | 0.65563697 | 0.78615933 | 0 | 0.0093 | 0.0065 | 0.0065 | TRUE | 0.046 |
| Lrrn2 | 70 | 50 | 51 | 77 | 58 | 59 | 54 | 53 | 55 | 77 | 65 | 58 | 60.48 | 54 | 67 | 1.232 | 0.301 | 0.23001701 | 0.39567133 | 0 | 0.0416 | 0.0287 | 0.0287 | TRUE | 0.1531 |
| Lrrn3 | 4 | 0 | 0 | 1 | 0 | 1 | 3 | 0 | 0 | 1 | 0 | 1 | 0.85 | 1 | 1 | 0.616 | -0.699 | 0.76265936 | NA | 1.9552 | 2.7967 | 2.6428 | 2.6428 | TRUE | 1.8937 |
| Lrrn4 | 707 | 494 | 368 | 329 | 261 | 323 | 549 | 525 | 397 | 327 | 294 | 319 | 402.07 | 490 | 313 | 0.638 | -0.648 | 1.31E-06 | 3.65E-05 | 0.0145 | 0.0083 | 0.0103 | 0.0103 | TRUE | 0.3169 |
| Lrrn4cl | 105 | 79 | 86 | 77 | 79 | 94 | 82 | 84 | 93 | 77 | 89 | 93 | 86.16 | 86 | 86 | 1.002 | 0.003 | 0.98731439 | 0.99373352 | 0 | 0.0299 | 0.0206 | 0.0206 | TRUE | 0.0911 |
| Lrrtm1 | 0 | 1 | 0 | 1 | 0 | 0 | 0 | 1 | 0 | 1 | 0 | 0 | 0.34 | 0 | 0 | 1.075 | 0.105 | 0.97573409 | NA | 0 | 6.9086 | 6.5582 | 6.5582 | TRUE | 0.5216 |
| Lrrtm2 | 0 | 0 | 0 | 0 | 0 | 0 | 0 | 0 | 0 | 0 | 0 | 0 | 0 | 0 | 0 | NA | NA | NA | NA | NA | NA | NA | NA | NA | NA |
| Lrrtm3 | 2 | 1 | 0 | 1 | 4 | 0 | 2 | 1 | 0 | 1 | 5 | 0 | 1.35 | 1 | 2 | 2 | 1 | 0.5837664 | NA | 0.0033 | 1.7527 | 1.5776 | 1.5776 | TRUE | 0.815 |
| Lrrtm4 | 0 | 0 | 1 | 0 | 0 | 1 | 0 | 0 | 1 | 0 | 0 | 1 | 0.34 | 0 | 0 | 1.075 | 0.105 | 0.97570524 | NA | 0 | 6.885 | 6.5379 | 6.5379 | TRUE | 0.5365 |
| Lrsam1 | 958 | 775 | 683 | 699 | 653 | 721 | 744 | 824 | 737 | 696 | 737 | 712 | 741.59 | 768 | 715 | 0.93 | -0.104 | 0.23118032 | 0.39705123 | 0.001 | 0.0056 | 0.0041 | 0.0041 | TRUE | 0.0468 |
| Lrtm1 | 1 | 0 | 0 | 0 | 0 | 0 | 1 | 0 | 0 | 0 | 0 | 0 | 0.13 | 0 | 0 | 0.552 | -0.857 | 0.8336548 | NA | 0 | 18.3121 | 10 | 10 | TRUE | 0.2566 |

| | | | | | | | | | | | | | | | | | | | | | | | | | |
|---|---|---|---|---|---|---|---|---|---|---|---|---|---|---|---|---|---|---|---|---|---|---|---|---|---|
| Lrtm2 | 37 | 32 | 34 | 63 | 46 | 55 | 29 | 34 | 37 | 63 | 52 | 54 | 44.73 | 33 | 56 | 1.711 | 0.774 | 0.00823709 | 0.03580676 | 0 | 0.0554 | 0.0381 | 0.0381 | TRUE | 0.1156 |
| Lrwd1 | 3828 | 2833 | 2712 | 2830 | 2636 | 3042 | 2973 | 3013 | 2926 | 2817 | 2974 | 3003 | 2951.05 | 2971 | 2931 | 0.987 | -0.02 | 0.73378835 | 0.84079179 | 3.00E-04 | 0.0032 | 0.002 | 0.002 | TRUE | 0.0142 |
| Lsamp | 0 | 0 | 0 | 0 | 0 | 0 | 0 | 0 | 0 | 0 | 0 | 0 | 0 | 0 | 0 | NA | NA | NA | NA | NA | NA | NA | NA | NA | NA |
| Lsg1 | 5614 | 3840 | 3525 | 3895 | 3442 | 4002 | 4360 | 4085 | 3803 | 3877 | 3883 | 3951 | 3993.13 | 4083 | 3904 | 0.956 | -0.066 | 0.30392658 | 0.476475 | 0.002 | 0.003 | 0.0027 | 0.0027 | TRUE | 0.0437 |
| Lsm1 | 2552 | 1708 | 1750 | 1698 | 1460 | 1788 | 1982 | 1817 | 1888 | 1690 | 1647 | 1765 | 1798.22 | 1896 | 1701 | 0.897 | -0.157 | 0.01976379 | 0.06854739 | 0.0011 | 0.0037 | 0.0027 | 0.0027 | TRUE | 0.0199 |
| Lsm10 | 675 | 482 | 476 | 478 | 468 | 482 | 524 | 513 | 514 | 476 | 528 | 476 | 505.02 | 517 | 493 | 0.953 | -0.07 | 0.47401429 | 0.64249276 | 0 | 0.0071 | 0.0048 | 0.0048 | TRUE | 0.043 |
| Lsm11 | 2301 | 1770 | 1919 | 2204 | 2075 | 2419 | 1787 | 1883 | 2071 | 2194 | 2341 | 2388 | 2110.5 | 1914 | 2308 | 1.207 | 0.271 | 0.00023982 | 0.00245022 | 0.0031 | 0.0035 | 0.0035 | 0.0035 | TRUE | 0.062 |
| Lsm12 | 4317 | 3267 | 3322 | 3427 | 2849 | 3424 | 3353 | 3475 | 3584 | 3411 | 3214 | 3380 | 3402.92 | 3471 | 3335 | 0.961 | -0.057 | 0.3309092 | 0.50532178 | 8.00E-04 | 0.0031 | 0.0022 | 0.0022 | TRUE | 0.0121 |
| Lsm14a | 11595 | 8344 | 8773 | 8530 | 7551 | 9025 | 9005 | 8875 | 9466 | 8491 | 8518 | 8909 | 8877.48 | 9115 | 8639 | 0.948 | -0.077 | 0.13945272 | 0.28069219 | 8.00E-04 | 0.0027 | 0.0019 | 0.0019 | TRUE | 0.0137 |
| Lsm14b | 5428 | 4300 | 4706 | 4365 | 4082 | 4633 | 4216 | 4574 | 5078 | 4345 | 4605 | 4573 | 4565.1 | 4623 | 4508 | 0.976 | -0.036 | 0.62509789 | 0.76369981 | 0.004 | 0.0029 | 0.0036 | 0.0036 | TRUE | 0.0904 |
| Lsm2 | 3064 | 2243 | 2190 | 2145 | 1911 | 2220 | 2380 | 2386 | 2363 | 2135 | 2156 | 2191 | 2268.49 | 2376 | 2161 | 0.909 | -0.137 | 0.01691755 | 0.06116936 | 0 | 0.0035 | 0.0019 | 0.0019 | TRUE | 0.0019 |
| Lsm3 | 3214 | 2124 | 2066 | 2168 | 1800 | 2105 | 2496 | 2259 | 2229 | 2158 | 2031 | 2078 | 2208.54 | 2328 | 2089 | 0.897 | -0.158 | 0.02170892 | 0.07328425 | 0.0021 | 0.0035 | 0.0029 | 0.0029 | TRUE | 0.0495 |
| Lsm4 | 9010 | 6945 | 7210 | 8351 | 7509 | 8496 | 6998 | 7387 | 7780 | 8313 | 8471 | 8387 | 7889.15 | 7388 | 8390 | 1.136 | 0.184 | 0.00098809 | 0.00718351 | 0.0013 | 0.0027 | 0.0021 | 0.0021 | TRUE | 0.0264 |
| Lsm5 | 2164 | 1402 | 1460 | 1707 | 1398 | 1500 | 1681 | 1491 | 1575 | 1699 | 1577 | 1481 | 1584.05 | 1582 | 1586 | 1.001 | 0.002 | 0.98274864 | 0.99089635 | 0.0036 | 0.0039 | 0.0038 | 0.0038 | TRUE | 0.0478 |
| Lsm6 | 4561 | 3019 | 2954 | 3016 | 2723 | 2890 | 3542 | 3211 | 3187 | 3002 | 3072 | 2853 | 3144.63 | 3313 | 2976 | 0.897 | -0.156 | 0.01734948 | 0.06222392 | 0.002 | 0.0032 | 0.0028 | 0.0028 | TRUE | 0.0452 |
| Lsm7 | 1261 | 892 | 912 | 953 | 882 | 996 | 979 | 949 | 984 | 949 | 995 | 983 | 973.17 | 971 | 976 | 1.005 | 0.007 | 0.92945659 | 0.9609898 | 0 | 0.0049 | 0.003 | 0.003 | TRUE | 0.007 |
| Lsmd1 | 3142 | 2217 | 2152 | 2232 | 1962 | 2309 | 2440 | 2358 | 2322 | 2222 | 2213 | 2279 | 2305.81 | 2373 | 2238 | 0.943 | -0.085 | 0.14897918 | 0.29422284 | 0 | 0.0035 | 0.0021 | 0.0021 | TRUE | 0.0076 |
| Lsp1 | 36 | 11 | 14 | 36 | 13 | 21 | 28 | 12 | 15 | 36 | 15 | 21 | 21 | 18 | 24 | 1.285 | 0.362 | 0.46313879 | 0.63236939 | 0.1581 | 0.1153 | 0.1278 | 0.1278 | TRUE | 0.6626 |
| Lsr | 2582 | 2012 | 1992 | 1992 | 1930 | 2259 | 2005 | 2140 | 2149 | 1983 | 2177 | 2230 | 2114.15 | 2098 | 2130 | 1.016 | 0.022 | 0.74869774 | 0.85098503 | 0.0022 | 0.0035 | 0.003 | 0.003 | TRUE | 0.0446 |
| Lss | 7702 | 5649 | 6009 | 7190 | 6293 | 7569 | 5982 | 6009 | 6484 | 7157 | 7099 | 7472 | 6700.35 | 6158 | 7243 | 1.176 | 0.234 | 3.01E-05 | 0.00046621 | 0.0012 | 0.0028 | 0.0021 | 0.0021 | TRUE | 0.0259 |
| Lst1 | 8 | 3 | 8 | 6 | 3 | 14 | 6 | 3 | 9 | 6 | 3 | 14 | 6.87 | 6 | 8 | 1.294 | 0.372 | 0.63834251 | NA | 0.2016 | 0.3475 | 0.3056 | 0.3056 | TRUE | 0.856 |
| Lta | 5 | 3 | 1 | 6 | 8 | 4 | 4 | 3 | 1 | 6 | 9 | 4 | 4.52 | 3 | 6 | 2.259 | 1.176 | 0.21072036 | NA | 0 | 0.5272 | 0.3802 | 0.3802 | TRUE | 0.2183 |
| Lta4h | 6821 | 4669 | 4698 | 5336 | 4539 | 5375 | 5298 | 4966 | 5069 | 5311 | 5120 | 5306 | 5178.48 | 5111 | 5246 | 1.026 | 0.037 | 0.48727765 | 0.65322261 | 5.00E-04 | 0.0029 | 0.0019 | 0.0019 | TRUE | 0.0127 |
| Ltb | 58 | 41 | 57 | 76 | 76 | 92 | 45 | 44 | 62 | 76 | 86 | 91 | 67.06 | 50 | 84 | 1.687 | 0.754 | 0.00246929 | 0.01422555 | 0 | 0.0378 | 0.0289 | 0.0289 | TRUE | 0.3051 |
| Ltb4r1 | 13 | 7 | 4 | 9 | 4 | 10 | 10 | 7 | 4 | 9 | 5 | 10 | 7.53 | 7 | 8 | 1.059 | 0.083 | 0.90744141 | NA | 0 | 0.317 | 0.236 | 0.236 | TRUE | 0.2585 |
| Ltb4r2 | 25 | 12 | 11 | 15 | 9 | 13 | 19 | 13 | 12 | 15 | 10 | 13 | 13.66 | 15 | 13 | 0.854 | -0.228 | 0.66317876 | NA | 0 | 0.1759 | 0.1247 | 0.1247 | TRUE | 0.4422 |
| Ltbp1 | 1199 | 835 | 1032 | 1161 | 1007 | 1141 | 931 | 888 | 1114 | 1156 | 1136 | 1126 | 1058.49 | 978 | 1139 | 1.166 | 0.222 | 0.01586091 | 0.05821777 | 0.0061 | 0.0047 | 0.0052 | 0.0052 | TRUE | 0.1725 |
| Ltbp2 | 96 | 93 | 94 | 115 | 113 | 95 | 75 | 99 | 101 | 114 | 127 | 94 | 101.77 | 92 | 112 | 1.227 | 0.295 | 0.1689678 | 0.32056364 | 0.0167 | 0.0257 | 0.0231 | 0.0231 | TRUE | 0.2895 |
| Ltbp3 | 206 | 191 | 251 | 301 | 290 | 380 | 160 | 203 | 271 | 300 | 327 | 375 | 272.64 | 211 | 334 | 1.589 | 0.668 | 0.00014309 | 0.00163871 | 0.0357 | 0.0111 | 0.0184 | 0.0184 | TRUE | 0.5663 |
| Ltbp4 | 5337 | 3964 | 3802 | 5058 | 4312 | 4684 | 4145 | 4216 | 4102 | 5035 | 4864 | 4624 | 4497.79 | 4154 | 4841 | 1.165 | 0.221 | 8.21E-05 | 0.00103137 | 7.00E-04 | 0.003 | 0.002 | 0.002 | TRUE | 0.0189 |
| Ltbr | 306 | 240 | 219 | 267 | 221 | 267 | 238 | 255 | 236 | 266 | 249 | 264 | 251.32 | 243 | 260 | 1.069 | 0.096 | 0.44828686 | 0.6182924 | 0 | 0.0119 | 0.0077 | 0.0077 | TRUE | 0.0205 |
| Ltc4s | 9 | 10 | 5 | 13 | 3 | 3 | 7 | 11 | 5 | 13 | 3 | 3 | 7.05 | 8 | 6 | 0.845 | -0.243 | 0.75941847 | NA | 0.2446 | 0.3385 | 0.3116 | 0.3116 | TRUE | 1.9879 |
| Ltf | 13 | 12 | 10 | 9 | 11 | 5 | 10 | 13 | 11 | 9 | 12 | 5 | 9.99 | 11 | 9 | 0.778 | -0.362 | 0.55842865 | NA | 0 | 0.2396 | 0.1724 | 0.1724 | TRUE | 0.2844 |
| Ltk | 177 | 168 | 170 | 192 | 129 | 196 | 137 | 179 | 183 | 191 | 146 | 193 | 171.62 | 166 | 177 | 1.068 | 0.095 | 0.59328851 | 0.74072962 | 0.0189 | 0.0162 | 0.017 | 0.017 | TRUE | 0.2717 |
| Ltn1 | 7136 | 5043 | 4827 | 5468 | 4627 | 5382 | 5542 | 5364 | 5208 | 5443 | 5220 | 5313 | 5348.36 | 5371 | 5325 | 0.991 | -0.013 | 0.80885737 | 0.88964974 | 5.00E-04 | 0.0029 | 0.0018 | 0.0018 | TRUE | 0.0096 |
| Ltv1 | 6438 | 4539 | 4342 | 4286 | 3856 | 4587 | 5000 | 4828 | 4685 | 4266 | 4350 | 4528 | 4609.58 | 4838 | 4381 | 0.905 | -0.143 | 0.01015198 | 0.04174403 | 7.00E-04 | 0.0029 | 0.002 | 0.002 | TRUE | 0.0107 |
| Luc7l | 6002 | 4713 | 4693 | 4430 | 4000 | 4647 | 4661 | 5013 | 5064 | 4410 | 4512 | 4587 | 4707.95 | 4913 | 4503 | 0.917 | -0.125 | 0.02819023 | 0.08849615 | 9.00E-04 | 0.0029 | 0.0021 | 0.0021 | TRUE | 0.0249 |
| Luc7l2 | 7573 | 5681 | 5401 | 5544 | 4660 | 5891 | 5882 | 6043 | 5828 | 5518 | 5257 | 5815 | 5723.81 | 5918 | 5530 | 0.935 | -0.097 | 0.08802506 | 0.20159303 | 0.0012 | 0.0028 | 0.0022 | 0.0022 | TRUE | 0.0248 |
| Luc7l3 | 7085 | 6015 | 5443 | 5152 | 4392 | 5411 | 5503 | 6398 | 5873 | 5128 | 4955 | 5341 | 5533.01 | 5925 | 5141 | 0.868 | -0.204 | 0.00244192 | 0.01409629 | 0.0033 | 0.0029 | 0.0031 | 0.0031 | TRUE | 0.0596 |
| Lum | 0 | 1 | 0 | 0 | 0 | 0 | 0 | 1 | 0 | 0 | 0 | 0 | 0.18 | 0 | 0 | 0.552 | -0.857 | 0.8336548 | NA | 0 | 13.3714 | 10 | 10 | TRUE | 0.5216 |
| Lurap1 | 113 | 61 | 77 | 100 | 76 | 86 | 88 | 65 | 83 | 100 | 86 | 85 | 84.32 | 79 | 90 | 1.142 | 0.192 | 0.38507011 | 0.55910271 | 0 | 0.0305 | 0.0233 | 0.0233 | TRUE | 0.2066 |
| Lurap1l | 164 | 129 | 125 | 156 | 122 | 150 | 127 | 137 | 135 | 155 | 138 | 148 | 140.07 | 133 | 147 | 1.107 | 0.146 | 0.37772881 | 0.55231667 | 0 | 0.0193 | 0.0127 | 0.0127 | TRUE | 0.0309 |
| Luzp1 | 13230 | 9980 | 9732 | 11956 | 10645 | 12107 | 10275 | 10616 | 10501 | 11901 | 12009 | 11951 | 11208.78 | 10464 | 11954 | 1.142 | 0.192 | 2.30E-05 | 0.0003733 | 1.00E-04 | 0.0026 | 0.0014 | 0.0014 | TRUE | 0.0031 |
| Luzp2 | 2 | 2 | 0 | 2 | 0 | 1 | 2 | 2 | 0 | 2 | 0 | 1 | 1.11 | 1 | 1 | 0.814 | -0.297 | 0.87469488 | NA | 0 | 2.1381 | 1.6614 | 1.6614 | TRUE | 0.2204 |
| Luzp4 | 69 | 42 | 54 | 37 | 28 | 34 | 54 | 45 | 58 | 37 | 32 | 34 | 43.08 | 52 | 34 | 0.652 | -0.617 | 0.03791581 | 0.11014851 | 0 | 0.0574 | 0.0393 | 0.0393 | TRUE | 0.1216 |
| Lxn | 106 | 91 | 69 | 72 | 50 | 61 | 82 | 97 | 74 | 72 | 56 | 60 | 73.64 | 84 | 63 | 0.744 | -0.426 | 0.0698302 | 0.17134473 | 0 | 0.0346 | 0.0259 | 0.0259 | TRUE | 0.143 |
| Ly6a | 624 | 552 | 514 | 175 | 173 | 178 | 485 | 587 | 555 | 174 | 195 | 176 | 361.91 | 542 | 182 | 0.335 | -1.576 | 1.14E-35 | 1.36E-32 | 0.0053 | 0.009 | 0.0078 | 0.0078 | TRUE | 0.1062 |
| Ly6c1 | 4 | 6 | 6 | 8 | 4 | 1 | 3 | 6 | 6 | 8 | 5 | 1 | 4.9 | 5 | 5 | 0.854 | -0.227 | 0.80603317 | NA | 0.2025 | 0.4857 | 0.4102 | 0.4102 | TRUE | 0.2299 |
| Ly6c2 | 2 | 1 | 0 | 0 | 0 | 1 | 2 | 1 | 0 | 0 | 0 | 1 | 0.6 | 1 | 0 | 0.416 | -1.265 | 0.62184281 | NA | 0 | 3.948 | 3.1959 | 3.1959 | TRUE | 0.3835 |
| Ly6d | 0 | 0 | 1 | 0 | 0 | 0 | 0 | 0 | 1 | 0 | 0 | 0 | 0.18 | 0 | 0 | 0.552 | -0.857 | 0.8336548 | NA | 0 | 13.1817 | 10 | 10 | TRUE | 0.5365 |
| Ly6e | 4947 | 3796 | 4350 | 4138 | 3788 | 4776 | 3842 | 4038 | 4694 | 4119 | 4273 | 4715 | 4280.06 | 4191 | 4369 | 1.043 | 0.061 | 0.46248194 | 0.63177429 | 0.0067 | 0.003 | 0.0047 | 0.0047 | TRUE | 0.1335 |
| Ly6f | 3 | 2 | 2 | 3 | 0 | 0 | 2 | 2 | 2 | 3 | 0 | 0 | 1.6 | 2 | 1 | 0.459 | -1.124 | 0.49412827 | NA | 0.0089 | 1.4835 | 1.2185 | 1.2185 | TRUE | 1.4413 |
| Ly6g5b | 6 | 8 | 2 | 2 | 1 | 2 | 5 | 9 | 2 | 2 | 1 | 2 | 3.4 | 5 | 2 | 0.335 | -1.576 | 0.15900042 | NA | 0 | 0.6988 | 0.5049 | 0.5049 | TRUE | 0.1562 |
| Ly6g5c | 2 | 0 | 0 | 0 | 1 | 0 | 2 | 0 | 0 | 0 | 1 | 0 | 0.45 | 1 | 0 | 0.68 | -0.556 | 0.85993441 | NA | 0 | 5.3057 | 5.2576 | 5.2576 | TRUE | 0.9726 |
| Ly6g6c | 21 | 30 | 41 | 29 | 38 | 30 | 16 | 32 | 44 | 29 | 43 | 30 | 32.3 | 31 | 34 | 1.108 | 0.148 | 0.7145461 | 0.82843992 | 0.1117 | 0.0758 | 0.0861 | 0.0861 | TRUE | 0.3089 |
| Ly6g6d | 82 | 95 | 80 | 54 | 63 | 70 | 64 | 101 | 86 | 54 | 71 | 69 | 74.16 | 84 | 65 | 0.776 | -0.365 | 0.14900568 | 0.29422284 | 0.0273 | 0.0344 | 0.0323 | 0.0323 | TRUE | 0.3002 |
| Ly6g6e | 1650 | 1482 | 2164 | 1464 | 1756 | 1858 | 1281 | 1576 | 2335 | 1457 | 1981 | 1834 | 1744.19 | 1731 | 1757 | 1.016 | 0.022 | 0.93809391 | 0.96558954 | 0.0585 | 0.0038 | 0.0184 | 0.0585 | TRUE | 0.7823 |
| Ly6g6f | 0 | 0 | 4 | 1 | 0 | 3 | 0 | 0 | 4 | 1 | 0 | 3 | 1.38 | 1 | 1 | 0.955 | -0.066 | 0.97310535 | NA | 3.7875 | 1.7214 | 2.0529 | 2.0529 | TRUE | 1.3909 |
| Ly6h | 67 | 30 | 31 | 72 | 50 | 61 | 52 | 32 | 33 | 72 | 56 | 60 | 50.95 | 39 | 63 | 1.586 | 0.665 | 0.0242403 | 0.07955607 | 0.0248 | 0.0489 | 0.0424 | 0.0424 | TRUE | 0.7187 |
| Ly6i | 0 | 0 | 0 | 0 | 0 | 1 | 0 | 0 | 0 | 0 | 0 | 1 | 0.16 | 0 | 0 | 2.094 | 1.067 | 0.79379329 | NA | 0 | 14.4078 | 10 | 10 | TRUE | 0.3835 |
| Ly6k | 386 | 281 | 290 | 287 | 269 | 297 | 300 | 299 | 313 | 286 | 303 | 293 | 298.99 | 304 | 294 | 0.968 | -0.047 | 0.68795598 | 0.80889723 | 0 | 0.0104 | 0.0065 | 0.0065 | TRUE | 0.0084 |
| Ly75 | 2426 | 1870 | 2324 | 2403 | 2327 | 2705 | 1884 | 1989 | 2508 | 2392 | 2625 | 2670 | 2344.69 | 2127 | 2562 | 1.206 | 0.27 | 0.00623165 | 0.02903161 | 0.0134 | 0.0034 | 0.0066 | 0.0066 | TRUE | 0.2954 |
| Ly86 | 3 | 1 | 1 | 0 | 0 | 0 | 2 | 1 | 1 | 0 | 0 | 0 | 0.75 | 1 | 0 | 0.126 | -2.986 | 0.19880652 | NA | 0 | 3.1818 | 2.5705 | 2.5705 | TRUE | 0.2178 |
| Ly9 | 1 | 4 | 1 | 0 | 0 | 2 | 1 | 4 | 1 | 0 | 0 | 2 | 1.35 | 2 | 1 | 0.336 | -1.575 | 0.39970923 | NA | 0.6948 | 1.7613 | 1.5108 | 1.5108 | TRUE | 0.9734 |
| Ly96 | 7 | 4 | 0 | 4 | 2 | 1 | 5 | 4 | 0 | 4 | 2 | 1 | 2.82 | 3 | 2 | 0.731 | -0.453 | 0.71769575 | NA | 0.5122 | 0.8431 | 0.7678 | 0.7678 | TRUE | 0.9133 |
| Lyar | 11115 | 8512 | 7896 | 8441 | 7325 | 8822 | 8633 | 9054 | 8520 | 8402 | 8263 | 8709 | 8596.76 | 8736 | 8458 | 0.968 | -0.046 | 0.3720992 | 0.54665576 | 7.00E-04 | 0.0027 | 0.0018 | 0.0018 | TRUE | 0.0123 |
| Lyg1 | 0 | 0 | 0 | 0 | 0 | 0 | 0 | 0 | 0 | 0 | 0 | 0 | 0 | 0 | 0 | NA | NA | NA | NA | NA | NA | NA | NA | NA | NA |
| Lyg2 | 3 | 4 | 1 | 2 | 1 | 2 | 2 | 4 | 1 | 2 | 1 | 2 | 2.13 | 2 | 2 | 0.67 | -0.577 | 0.65722385 | NA | 0 | 1.1171 | 0.7259 | 0.7259 | TRUE | 0.2403 |
| Lyl1 | 3 | 6 | 5 | 9 | 14 | 14 | 2 | 6 | 5 | 9 | 16 | 14 | 8.78 | 4 | 13 | 2.803 | 1.487 | 0.03316506 | NA | 0 | 0.2724 | 0.2056 | 0.2056 | TRUE | 0.5333 |

| | | | | | | | | | | | | | | | | | | | | | | | | | |
|---|---|---|---|---|---|---|---|---|---|---|---|---|---|---|---|---|---|---|---|---|---|---|---|---|---|
| Lyn | 445 | 290 | 264 | 319 | 264 | 280 | 346 | 308 | 285 | 318 | 298 | 276 | 305.12 | 313 | 297 | 0.947 | -0.079 | 0.53592699 | 0.69491105 | 0.0044 | 0.0102 | 0.0084 | 0.0084 | TRUE | 0.1007 |
| Lynx1 | 99 | 58 | 90 | 106 | 88 | 104 | 77 | 62 | 97 | 106 | 99 | 103 | 90.52 | 79 | 103 | 1.306 | 0.386 | 0.08049473 | 0.18996968 | 0.0098 | 0.0286 | 0.0239 | 0.0239 | TRUE | 0.3254 |
| Lypd1 | 53 | 41 | 25 | 11 | 22 | 17 | 41 | 44 | 27 | 11 | 25 | 17 | 27.38 | 37 | 18 | 0.465 | -1.104 | 0.00765461 | 0.03386176 | 0.0547 | 0.089 | 0.0807 | 0.0807 | TRUE | 0.5265 |
| Lypd2 | 1 | 3 | 1 | 4 | 2 | 5 | 1 | 3 | 1 | 4 | 2 | 5 | 2.7 | 2 | 4 | 2.287 | 1.194 | 0.320512 | NA | 0 | 0.8791 | 0.6083 | 0.6083 | TRUE | 0.4438 |
| Lypd3 | 8 | 7 | 7 | 11 | 17 | 9 | 6 | 7 | 8 | 11 | 19 | 9 | 10.04 | 7 | 13 | 1.838 | 0.878 | 0.16124079 | NA | 0 | 0.2386 | 0.1743 | 0.1743 | TRUE | 0.6545 |
| Lypd4 | 30 | 23 | 23 | 13 | 18 | 23 | 23 | 24 | 25 | 13 | 20 | 23 | 21.42 | 24 | 19 | 0.771 | -0.375 | 0.37266715 | 0.54720892 | 0 | 0.1131 | 0.08 | 0.08 | TRUE | 0.3862 |
| Lypd5 | 4 | 1 | 1 | 3 | 2 | 0 | 3 | 1 | 1 | 3 | 2 | 0 | 1.75 | 2 | 2 | 0.967 | -0.049 | 0.97390668 | NA | 0 | 1.3578 | 1.0492 | 1.0492 | TRUE | 0.6383 |
| Lypd6 | 217 | 185 | 130 | 146 | 144 | 147 | 169 | 197 | 140 | 145 | 162 | 145 | 159.75 | 169 | 151 | 0.895 | -0.161 | 0.35134601 | 0.52609545 | 0.0096 | 0.0173 | 0.0151 | 0.0151 | TRUE | 0.156 |
| Lypd6b | 9 | 11 | 9 | 10 | 11 | 7 | 7 | 12 | 10 | 10 | 12 | 7 | 9.61 | 10 | 10 | 1.036 | 0.052 | 0.93386745 | NA | 0 | 0.249 | 0.1732 | 0.1732 | TRUE | 0.2304 |
| Lypd8 | 0 | 0 | 0 | 0 | 0 | 0 | 0 | 0 | 0 | 0 | 0 | 0 | 0 | 0 | 0 | NA | NA | NA | NA | NA | NA | NA | NA | NA | NA |
| Lypla1 | 7896 | 5547 | 5745 | 5722 | 4902 | 5523 | 6132 | 5900 | 6199 | 5696 | 5530 | 5452 | 5818.21 | 6077 | 5559 | 0.915 | -0.129 | 0.01295939 | 0.05006176 | 3.00E-04 | 0.0028 | 0.0018 | 0.0018 | TRUE | 0.0078 |
| Lypla2 | 2356 | 1721 | 1715 | 1714 | 1647 | 1828 | 1830 | 1831 | 1850 | 1706 | 1858 | 1805 | 1813.25 | 1837 | 1790 | 0.974 | -0.038 | 0.55518808 | 0.71041211 | 3.00E-04 | 0.0037 | 0.0024 | 0.0024 | TRUE | 0.0201 |
| Lyplal1 | 1338 | 847 | 817 | 786 | 688 | 720 | 1039 | 901 | 882 | 782 | 776 | 711 | 848.49 | 941 | 756 | 0.803 | -0.317 | 0.00062642 | 0.00513384 | 0.0046 | 0.0052 | 0.005 | 0.005 | TRUE | 0.1041 |
| Lyrm1 | 96 | 74 | 63 | 77 | 68 | 62 | 75 | 79 | 68 | 77 | 77 | 61 | 72.63 | 74 | 72 | 0.968 | -0.047 | 0.83911752 | 0.90832375 | 0 | 0.0351 | 0.0246 | 0.0246 | TRUE | 0.1479 |
| Lyrm2 | 485 | 331 | 314 | 316 | 295 | 294 | 377 | 352 | 339 | 315 | 333 | 290 | 334.19 | 356 | 313 | 0.876 | -0.191 | 0.10283863 | 0.2257522 | 0 | 0.0095 | 0.0069 | 0.0069 | TRUE | 0.0445 |
| Lyrm4 | 656 | 479 | 466 | 544 | 445 | 501 | 509 | 510 | 503 | 541 | 502 | 495 | 509.98 | 507 | 513 | 1.011 | 0.015 | 0.87122446 | 0.9285222 | 0 | 0.0071 | 0.0046 | 0.0046 | TRUE | 0.0287 |
| Lyrm5 | 719 | 490 | 508 | 620 | 472 | 600 | 558 | 521 | 548 | 617 | 532 | 592 | 561.61 | 542 | 580 | 1.07 | 0.098 | 0.31371006 | 0.48713345 | 0.0016 | 0.0066 | 0.005 | 0.005 | TRUE | 0.06 |
| Lyrm7 | 199 | 118 | 125 | 192 | 169 | 245 | 155 | 126 | 135 | 191 | 191 | 242 | 173.09 | 139 | 208 | 1.498 | 0.584 | 0.0005244 | 0.00446945 | 0.0105 | 0.0161 | 0.0144 | 0.0144 | TRUE | 0.2262 |
| Lyrm9 | 1207 | 921 | 975 | 1205 | 979 | 1259 | 937 | 980 | 1052 | 1199 | 1104 | 1243 | 1085.97 | 990 | 1182 | 1.197 | 0.259 | 0.00156972 | 0.00997598 | 0.0026 | 0.0046 | 0.0039 | 0.0039 | TRUE | 0.039 |
| Lysmd1 | 42 | 27 | 27 | 23 | 32 | 26 | 33 | 29 | 29 | 23 | 36 | 26 | 29.19 | 30 | 28 | 0.929 | -0.107 | 0.768186 | 0.86284342 | 0 | 0.0836 | 0.06 | 0.06 | TRUE | 0.3558 |
| Lysmd2 | 245 | 164 | 137 | 177 | 126 | 185 | 190 | 174 | 148 | 176 | 142 | 183 | 168.92 | 171 | 167 | 0.976 | -0.035 | 0.83593091 | 0.90645327 | 0.0107 | 0.0165 | 0.0148 | 0.0148 | TRUE | 0.1724 |
| Lysmd3 | 1014 | 738 | 707 | 675 | 640 | 724 | 788 | 785 | 763 | 672 | 722 | 715 | 740.66 | 779 | 703 | 0.902 | -0.148 | 0.07339258 | 0.1774958 | 0 | 0.0056 | 0.0036 | 0.0036 | TRUE | 0.0177 |
| Lysmd4 | 344 | 274 | 284 | 359 | 328 | 393 | 267 | 291 | 306 | 357 | 370 | 388 | 330.06 | 288 | 372 | 1.293 | 0.37 | 0.00154578 | 0.00986774 | 0 | 0.0096 | 0.0068 | 0.0068 | TRUE | 0.0509 |
| Lyst | 1161 | 802 | 793 | 927 | 808 | 1036 | 902 | 853 | 856 | 923 | 912 | 1023 | 911.23 | 870 | 953 | 1.094 | 0.13 | 0.11618582 | 0.24667693 | 0.0015 | 0.005 | 0.0038 | 0.0038 | TRUE | 0.0503 |
| Lyve1 | 0 | 0 | 0 | 0 | 0 | 0 | 0 | 0 | 0 | 0 | 0 | 0 | 0 | 0 | 0 | NA | NA | NA | NA | NA | NA | NA | NA | NA | NA |
| Lyz1 | 0 | 0 | 0 | 0 | 1 | 0 | 0 | 0 | 0 | 0 | 1 | 0 | 0.19 | 0 | 0 | 2.094 | 1.067 | 0.79379329 | NA | 0 | 12.608 | 10 | 10 | TRUE | 0.5075 |
| Lyz2 | 0 | 0 | 0 | 0 | 0 | 0 | 0 | 0 | 0 | 0 | 0 | 0 | 0 | 0 | 0 | NA | NA | NA | NA | NA | NA | NA | NA | NA | NA |
| Lyzl1 | 0 | 0 | 0 | 0 | 0 | 0 | 0 | 0 | 0 | 0 | 0 | 0 | 0 | 0 | 0 | NA | NA | NA | NA | NA | NA | NA | NA | NA | NA |
| Lyzl4 | 6 | 0 | 0 | 3 | 2 | 1 | 5 | 0 | 0 | 3 | 2 | 1 | 1.81 | 2 | 2 | 1.254 | 0.327 | 0.83872587 | NA | 1.34 | 1.3083 | 1.3154 | 1.3154 | TRUE | 2.7646 |
| Lyzl6 | 0 | 0 | 0 | 0 | 0 | 0 | 0 | 0 | 0 | 0 | 0 | 0 | 0 | 0 | 0 | NA | NA | NA | NA | NA | NA | NA | NA | NA | NA |
| Lzic | 1531 | 1031 | 1156 | 1131 | 967 | 1153 | 1189 | 1097 | 1247 | 1126 | 1091 | 1138 | 1147.98 | 1178 | 1118 | 0.95 | -0.075 | 0.33356708 | 0.50773953 | 0.0014 | 0.0045 | 0.0034 | 0.0034 | TRUE | 0.0419 |
| Lztfl1 | 1046 | 762 | 880 | 863 | 732 | 893 | 812 | 811 | 950 | 859 | 826 | 882 | 856.46 | 858 | 856 | 0.999 | -0.001 | 0.98840112 | 0.99389455 | 0.0035 | 0.0052 | 0.0046 | 0.0046 | TRUE | 0.1026 |
| Lztr1 | 2052 | 1518 | 1527 | 1576 | 1413 | 1482 | 1594 | 1615 | 1648 | 1569 | 1594 | 1463 | 1580.29 | 1619 | 1542 | 0.953 | -0.07 | 0.29901576 | 0.47133998 | 5.00E-04 | 0.0039 | 0.0026 | 0.0026 | TRUE | 0.0244 |
| Lzts1 | 18 | 17 | 12 | 23 | 10 | 8 | 14 | 18 | 13 | 23 | 11 | 8 | 14.51 | 15 | 14 | 0.939 | -0.091 | 0.86724355 | NA | 0.0926 | 0.1657 | 0.1453 | 0.1453 | TRUE | 1.3489 |
| Lzts2 | 1769 | 1571 | 1688 | 1905 | 1754 | 2094 | 1374 | 1671 | 1821 | 1896 | 1979 | 2067 | 1801.39 | 1622 | 1981 | 1.223 | 0.29 | 0.00243338 | 0.01407738 | 0.0099 | 0.0037 | 0.006 | 0.006 | TRUE | 0.2214 |
| M1ap | 1004 | 749 | 798 | 806 | 616 | 777 | 780 | 797 | 861 | 802 | 695 | 767 | 783.62 | 813 | 755 | 0.93 | -0.104 | 0.24673017 | 0.41551954 | 0.0026 | 0.0055 | 0.0045 | 0.0045 | TRUE | 0.0555 |
| M6pr | 13662 | 9826 | 10014 | 11347 | 10048 | 11389 | 10611 | 10452 | 10805 | 11295 | 11335 | 11243 | 10956.7 | 10623 | 11291 | 1.063 | 0.088 | 0.05271707 | 0.14007596 | 0 | 0.0026 | 0.0014 | 0.0014 | TRUE | 0.0027 |
| Maats1 | 190 | 173 | 149 | 144 | 136 | 172 | 148 | 184 | 161 | 143 | 153 | 170 | 159.82 | 164 | 155 | 0.951 | -0.073 | 0.66027378 | 0.78996458 | 0.0044 | 0.0173 | 0.0135 | 0.0135 | TRUE | 0.1183 |
| Mab21l1 | 0 | 0 | 0 | 0 | 0 | 1 | 0 | 0 | 0 | 0 | 0 | 1 | 0.16 | 0 | 0 | 2.094 | 1.067 | 0.79379329 | NA | 0 | 14.4078 | 10 | 10 | TRUE | 0.3835 |
| Mab21l2 | 4 | 1 | 4 | 4 | 1 | 0 | 3 | 1 | 4 | 4 | 1 | 0 | 2.27 | 3 | 2 | 0.604 | -0.728 | 0.59672713 | NA | 0.3699 | 1.0484 | 0.8919 | 0.8919 | TRUE | 0.9446 |
| Mab21l3 | 370 | 316 | 378 | 375 | 351 | 414 | 287 | 336 | 408 | 373 | 396 | 409 | 368.21 | 344 | 393 | 1.146 | 0.197 | 0.1441882 | 0.28743123 | 0.0138 | 0.0089 | 0.0104 | 0.0104 | TRUE | 0.3018 |
| Macc1 | 0 | 0 | 0 | 0 | 1 | 0 | 0 | 0 | 0 | 0 | 1 | 0 | 0.19 | 0 | 0 | 2.094 | 1.067 | 0.79379329 | NA | 0 | 12.608 | 10 | 10 | TRUE | 0.5075 |
| Macf1 | 17729 | 11604 | 11859 | 16861 | 15079 | 17416 | 13769 | 12343 | 12796 | 16783 | 17011 | 17192 | 14982.42 | 12969 | 16995 | 1.31 | 0.39 | 1.86E-12 | 2.20E-10 | 0.0015 | 0.0026 | 0.0021 | 0.0021 | TRUE | 0.0358 |
| Macrod1 | 201 | 136 | 183 | 273 | 248 | 261 | 156 | 145 | 197 | 272 | 280 | 258 | 217.9 | 166 | 270 | 1.627 | 0.702 | 3.39E-06 | 7.71E-05 | 0.0069 | 0.0133 | 0.0116 | 0.0116 | TRUE | 0.275 |
| Macrod2 | 549 | 359 | 360 | 422 | 321 | 362 | 426 | 382 | 388 | 420 | 362 | 357 | 389.37 | 399 | 380 | 0.951 | -0.073 | 0.52801154 | 0.68874916 | 0.0035 | 0.0085 | 0.0069 | 0.0069 | TRUE | 0.0998 |
| Mad1l1 | 3116 | 2148 | 1789 | 1590 | 1438 | 1772 | 2420 | 2285 | 1930 | 1583 | 1622 | 1749 | 1931.55 | 2212 | 1651 | 0.746 | -0.423 | 1.66E-06 | 4.40E-05 | 0.0071 | 0.0037 | 0.0051 | 0.0051 | TRUE | 0.149 |
| Mad2l1 | 6817 | 4863 | 4871 | 5004 | 4348 | 5005 | 5294 | 5173 | 5256 | 4981 | 4905 | 4941 | 5091.61 | 5241 | 4942 | 0.943 | -0.085 | 0.08442541 | 0.19592342 | 0 | 0.0029 | 0.0015 | 0.0015 | TRUE | 0.0016 |
| Mad2l1bp | 824 | 515 | 421 | 461 | 408 | 442 | 640 | 548 | 454 | 459 | 460 | 436 | 499.58 | 547 | 452 | 0.823 | -0.281 | 0.02442156 | 0.07998564 | 0.0135 | 0.0072 | 0.0092 | 0.0092 | TRUE | 0.1601 |
| Mad2l2 | 2353 | 1622 | 1841 | 2005 | 1828 | 2214 | 1827 | 1725 | 1986 | 1996 | 2062 | 2186 | 1963.78 | 1846 | 2081 | 1.127 | 0.173 | 0.01885732 | 0.06625559 | 0.0028 | 0.0036 | 0.0034 | 0.0034 | TRUE | 0.0519 |
| Madcam1 | 42 | 38 | 33 | 58 | 59 | 77 | 33 | 40 | 36 | 58 | 67 | 76 | 51.49 | 36 | 67 | 1.852 | 0.889 | 0.00139633 | 0.00915491 | 0 | 0.0485 | 0.0347 | 0.0347 | TRUE | 0.1335 |
| Madd | 995 | 717 | 647 | 717 | 612 | 749 | 773 | 763 | 698 | 714 | 690 | 739 | 729.5 | 745 | 714 | 0.959 | -0.06 | 0.48615464 | 0.6521453 | 7.00E-04 | 0.0057 | 0.004 | 0.004 | TRUE | 0.0343 |
| Maea | 9592 | 7149 | 7327 | 8098 | 7414 | 8630 | 7450 | 7604 | 7906 | 8061 | 8364 | 8519 | 7983.9 | 7653 | 8315 | 1.087 | 0.12 | 0.02149724 | 0.07284496 | 7.00E-04 | 0.0027 | 0.0018 | 0.0018 | TRUE | 0.0101 |
| Mael | 699 | 509 | 519 | 707 | 539 | 570 | 543 | 541 | 560 | 704 | 608 | 563 | 586.46 | 548 | 625 | 1.14 | 0.19 | 0.06783849 | 0.16755428 | 0.0052 | 0.0065 | 0.0061 | 0.0061 | TRUE | 0.1458 |
| Maf | 2 | 4 | 3 | 5 | 4 | 8 | 2 | 4 | 3 | 5 | 5 | 8 | 4.41 | 3 | 6 | 1.976 | 0.982 | 0.29059171 | NA | 0 | 0.5404 | 0.3701 | 0.3701 | TRUE | 0.3354 |
| Maf1 | 3981 | 2914 | 3170 | 3072 | 2804 | 3253 | 3092 | 3100 | 3420 | 3058 | 3163 | 3211 | 3174.02 | 3204 | 3144 | 0.982 | -0.027 | 0.67383708 | 0.79979329 | 0.0014 | 0.0032 | 0.0026 | 0.0026 | TRUE | 0.042 |
| Mafa | 235 | 167 | 178 | 210 | 143 | 174 | 183 | 178 | 192 | 209 | 161 | 172 | 182.39 | 184 | 181 | 0.983 | -0.024 | 0.87772581 | 0.93172788 | 0.0043 | 0.0154 | 0.0122 | 0.0122 | TRUE | 0.2034 |
| Mafb | 37 | 21 | 29 | 22 | 30 | 28 | 29 | 22 | 31 | 22 | 34 | 28 | 27.62 | 27 | 28 | 1.007 | 0.011 | 0.97700569 | 0.98766016 | 0 | 0.0882 | 0.0645 | 0.0645 | TRUE | 0.1927 |
| Maff | 611 | 499 | 638 | 868 | 717 | 861 | 475 | 531 | 688 | 864 | 809 | 850 | 702.75 | 565 | 841 | 1.494 | 0.579 | 2.13E-06 | 5.30E-05 | 0.0176 | 0.0058 | 0.0093 | 0.0093 | TRUE | 0.4282 |
| Mafg | 1897 | 1405 | 1103 | 1229 | 1051 | 1209 | 1473 | 1494 | 1190 | 1223 | 1186 | 1193 | 1293.4 | 1386 | 1201 | 0.866 | -0.208 | 0.02348952 | 0.07771661 | 0.0073 | 0.0043 | 0.0053 | 0.0053 | TRUE | 0.1807 |
| Mafk | 687 | 492 | 491 | 518 | 481 | 567 | 534 | 523 | 530 | 516 | 543 | 560 | 534.11 | 529 | 540 | 1.019 | 0.028 | 0.76604248 | 0.8611802 | 0 | 0.0069 | 0.0044 | 0.0044 | TRUE | 0.0177 |
| Mag | 13 | 19 | 24 | 24 | 25 | 41 | 10 | 20 | 26 | 24 | 28 | 40 | 24.79 | 19 | 31 | 1.678 | 0.747 | 0.09094136 | 0.20625593 | 0.0963 | 0.098 | 0.0976 | 0.0976 | TRUE | 0.811 |
| Magea1 | 0 | 0 | 0 | 0 | 0 | 0 | 0 | 0 | 0 | 0 | 0 | 0 | 0 | 0 | 0 | NA | NA | NA | NA | NA | NA | NA | NA | NA | NA |
| Magea10 | 26 | 26 | 17 | 3 | 2 | 4 | 20 | 28 | 18 | 3 | 2 | 4 | 12.56 | 22 | 3 | 0.14 | -2.834 | 2.66E-05 | NA | 0 | 0.1911 | 0.1386 | 0.1386 | TRUE | 0.2652 |
| Magea2 | 21 | 20 | 12 | 5 | 1 | 6 | 16 | 21 | 13 | 5 | 1 | 6 | 10.43 | 17 | 4 | 0.242 | -2.044 | 0.00284572 | NA | 0 | 0.2297 | 0.1827 | 0.1827 | TRUE | 0.4524 |
| Magea3 | 27 | 26 | 11 | 7 | 4 | 0 | 21 | 28 | 12 | 7 | 5 | 0 | 12 | 20 | 4 | 0.189 | -2.407 | 0.00095064 | NA | 0.3942 | 0.2 | 0.2225 | 0.2225 | TRUE | 0.4877 |
| Magea4 | 168 | 109 | 111 | 21 | 17 | 21 | 130 | 116 | 120 | 21 | 19 | 21 | 71.17 | 122 | 20 | 0.166 | -2.59 | 3.12E-21 | 1.09E-18 | 0 | 0.0357 | 0.0246 | 0.0246 | TRUE | 0.0403 |
| Magea5 | 165 | 119 | 89 | 48 | 35 | 20 | 128 | 127 | 96 | 48 | 39 | 20 | 76.29 | 117 | 36 | 0.304 | -1.719 | 2.51E-09 | 1.50E-07 | 0.0716 | 0.0335 | 0.0412 | 0.0412 | TRUE | 1.1408 |
| Magea6 | 38 | 22 | 14 | 8 | 7 | 5 | 30 | 23 | 15 | 8 | 8 | 5 | 14.8 | 23 | 7 | 0.302 | -1.726 | 0.0019321 | NA | 0 | 0.1625 | 0.1274 | 0.1274 | TRUE | 0.1245 |
| Magea8 | 143 | 108 | 56 | 17 | 16 | 19 | 111 | 115 | 60 | 17 | 18 | 19 | 56.68 | 95 | 18 | 0.187 | -2.419 | 1.13E-12 | 1.39E-10 | 0.0612 | 0.0442 | 0.0494 | 0.0494 | TRUE | 0.9588 |

| | | | | | | | | | | | | | | | | | | | | | | | | | |
|---|---|---|---|---|---|---|---|---|---|---|---|---|---|---|---|---|---|---|---|---|---|---|---|---|---|
| Mageb1 | 0 | 0 | 0 | 0 | 0 | 0 | 0 | 0 | 0 | 0 | 0 | 0 | 0 | 0 | 0 | NA | NA | NA | NA | NA | NA | NA | NA | NA | NA |
| Mageb16 | 977 | 668 | 738 | 606 | 487 | 647 | 759 | 711 | 796 | 603 | 549 | 639 | 676.15 | 755 | 597 | 0.791 | -0.338 | 0.00034442 | 0.00322383 | 0.0027 | 0.0059 | 0.0049 | 0.0049 | TRUE | 0.0558 |
| Mageb16-ps | 0 | 1 | 0 | 0 | 1 | 0 | 0 | 1 | 0 | 0 | 1 | 0 | 0.37 | 0 | 0 | 1.075 | 0.105 | 0.97520514 | NA | 0 | 6.4904 | 6.1969 | 6.1969 | TRUE | 0.5216 |
| Mageb18 | 22 | 11 | 5 | 3 | 2 | 1 | 17 | 12 | 5 | 3 | 2 | 1 | 6.74 | 11 | 2 | 0.18 | -2.477 | 0.00612591 | NA | 0 | 0.3543 | 0.2969 | 0.2969 | TRUE | 0.0809 |
| Mageb2 | 0 | 0 | 0 | 0 | 0 | 0 | 0 | 0 | 0 | 0 | 0 | 0 | 0 | 0 | 0 | NA | NA | NA | NA | NA | NA | NA | NA | NA | NA |
| Mageb3 | 4 | 3 | 7 | 1 | 7 | 7 | 3 | 3 | 8 | 1 | 8 | 7 | 4.94 | 5 | 5 | 1.148 | 0.199 | 0.83234003 | NA | 0.2852 | 0.482 | 0.4291 | 0.4291 | TRUE | 1.0952 |
| Mageb4 | 153 | 109 | 91 | 121 | 86 | 81 | 119 | 116 | 98 | 120 | 97 | 80 | 105.06 | 111 | 99 | 0.891 | -0.166 | 0.4319858 | 0.60396881 | 0.0161 | 0.025 | 0.0225 | 0.0225 | TRUE | 0.3265 |
| Mageb5 | 0 | 0 | 0 | 0 | 0 | 0 | 0 | 0 | 0 | 0 | 0 | 0 | 0 | 0 | 0 | NA | NA | NA | NA | NA | NA | NA | NA | NA | NA |
| Maged1 | 9886 | 7127 | 6528 | 6306 | 5927 | 6866 | 7678 | 7581 | 7044 | 6277 | 6686 | 6778 | 7007.27 | 7434 | 6580 | 0.885 | -0.176 | 0.00268386 | 0.01512635 | 0.0017 | 0.0028 | 0.0023 | 0.0023 | TRUE | 0.0257 |
| Maged2 | 1000 | 732 | 730 | 836 | 728 | 811 | 777 | 779 | 788 | 832 | 821 | 801 | 799.49 | 781 | 818 | 1.048 | 0.067 | 0.39531284 | 0.56877455 | 0 | 0.0054 | 0.0032 | 0.0032 | TRUE | 0.0042 |
| Magee1 | 465 | 298 | 281 | 298 | 268 | 280 | 361 | 317 | 303 | 297 | 302 | 276 | 309.45 | 327 | 292 | 0.889 | -0.17 | 0.16993359 | 0.32144796 | 0.0029 | 0.0101 | 0.0078 | 0.0078 | TRUE | 0.1007 |
| Magee2 | 168 | 108 | 78 | 84 | 71 | 86 | 130 | 115 | 84 | 84 | 80 | 85 | 96.35 | 110 | 83 | 0.751 | -0.414 | 0.05945467 | 0.15231862 | 0.0171 | 0.027 | 0.0241 | 0.0241 | TRUE | 0.3781 |
| Mageh1 | 314 | 238 | 207 | 239 | 231 | 290 | 244 | 253 | 223 | 238 | 261 | 286 | 250.86 | 240 | 262 | 1.089 | 0.123 | 0.36147062 | 0.53638908 | 0.0021 | 0.0119 | 0.0091 | 0.0091 | TRUE | 0.0783 |
| Magel2 | 272 | 245 | 91 | 64 | 86 | 63 | 211 | 261 | 98 | 64 | 97 | 62 | 132.16 | 190 | 74 | 0.39 | -1.36 | 4.39E-06 | 9.53E-05 | 0.1405 | 0.0204 | 0.0537 | 0.0537 | TRUE | 0.3048 |
| Magi1 | 2363 | 1770 | 1522 | 1659 | 1558 | 1768 | 1835 | 1883 | 1642 | 1651 | 1758 | 1745 | 1752.41 | 1787 | 1718 | 0.961 | -0.057 | 0.43597995 | 0.60723655 | 0.0026 | 0.0038 | 0.0033 | 0.0033 | TRUE | 0.0592 |
| Magi2 | 85 | 81 | 68 | 63 | 56 | 65 | 66 | 86 | 73 | 63 | 63 | 64 | 69.27 | 75 | 63 | 0.846 | -0.241 | 0.30746071 | 0.48011973 | 0 | 0.0366 | 0.0256 | 0.0256 | TRUE | 0.1473 |
| Magi3 | 1740 | 1225 | 959 | 932 | 777 | 874 | 1351 | 1303 | 1035 | 928 | 877 | 863 | 1059.36 | 1230 | 889 | 0.722 | -0.47 | 3.98E-06 | 8.79E-05 | 0.0101 | 0.0047 | 0.0065 | 0.0065 | TRUE | 0.2279 |
| Magix | 3 | 4 | 5 | 5 | 5 | 3 | 2 | 4 | 5 | 5 | 6 | 3 | 4.26 | 4 | 5 | 1.149 | 0.2 | 0.82786231 | NA | 0 | 0.5588 | 0.3743 | 0.3743 | TRUE | 0.3066 |
| Magoh | 4084 | 2770 | 2317 | 2352 | 2000 | 2442 | 3172 | 2946 | 2500 | 2341 | 2256 | 2411 | 2604.39 | 2873 | 2336 | 0.813 | -0.299 | 0.0004341 | 0.00387307 | 0.007 | 0.0033 | 0.0048 | 0.0048 | TRUE | 0.1556 |
| Magohb | 1409 | 971 | 949 | 887 | 743 | 805 | 1094 | 1033 | 1024 | 883 | 838 | 795 | 944.48 | 1050 | 839 | 0.798 | -0.326 | 4.91E-05 | 0.00069137 | 0.001 | 0.0049 | 0.0036 | 0.0036 | TRUE | 0.026 |
| Magt1 | 2978 | 2110 | 2365 | 2920 | 2491 | 2723 | 2313 | 2244 | 2552 | 2907 | 2810 | 2688 | 2585.63 | 2370 | 2802 | 1.183 | 0.242 | 0.00046972 | 0.00412896 | 0.0025 | 0.0033 | 0.0031 | 0.0031 | TRUE | 0.0538 |
| Mak | 203 | 186 | 176 | 141 | 162 | 190 | 158 | 198 | 190 | 140 | 183 | 188 | 176.01 | 182 | 170 | 0.939 | -0.091 | 0.59440369 | 0.7409266 | 0.014 | 0.0159 | 0.0153 | 0.0153 | TRUE | 0.2556 |
| Mak16 | 10581 | 6953 | 6315 | 6667 | 5705 | 6493 | 8218 | 7396 | 6814 | 6636 | 6436 | 6410 | 6984.87 | 7476 | 6494 | 0.868 | -0.204 | 0.00385999 | 0.02006211 | 0.0044 | 0.0028 | 0.0034 | 0.0034 | TRUE | 0.0927 |
| Mal | 30 | 24 | 41 | 35 | 34 | 22 | 23 | 26 | 44 | 35 | 38 | 22 | 31.33 | 31 | 32 | 1.025 | 0.036 | 0.92695345 | 0.95969289 | 0.0737 | 0.0781 | 0.0768 | 0.0768 | TRUE | 0.8959 |
| Mal2 | 134 | 113 | 135 | 131 | 120 | 123 | 104 | 120 | 146 | 130 | 135 | 121 | 126.19 | 123 | 129 | 1.051 | 0.072 | 0.700776 | 0.8185838 | 0.0087 | 0.0212 | 0.0176 | 0.0176 | TRUE | 0.2519 |
| Malat1 | 13726 | 11817 | 9745 | 18527 | 13398 | 16079 | 10660 | 12570 | 10515 | 18442 | 15114 | 15872 | 13862.22 | 11248 | 16476 | 1.465 | 0.551 | 1.63E-10 | 1.25E-08 | 0.0107 | 0.0026 | 0.0053 | 0.0053 | TRUE | 0.1332 |
| Mall | 127 | 85 | 103 | 132 | 118 | 123 | 99 | 90 | 111 | 131 | 133 | 121 | 114.35 | 100 | 128 | 1.286 | 0.363 | 0.05072659 | 0.13631043 | 0 | 0.0232 | 0.016 | 0.016 | TRUE | 0.0843 |
| Malsu1 | 1440 | 1117 | 1102 | 1307 | 1093 | 1235 | 1118 | 1188 | 1189 | 1301 | 1233 | 1219 | 1208.12 | 1165 | 1251 | 1.075 | 0.104 | 0.15136149 | 0.29753952 | 0 | 0.0044 | 0.003 | 0.003 | TRUE | 0.0156 |
| Malt1 | 1042 | 776 | 966 | 1106 | 965 | 1103 | 809 | 825 | 1042 | 1101 | 1089 | 1089 | 992.56 | 892 | 1093 | 1.227 | 0.295 | 0.00316998 | 0.0172276 | 0.0091 | 0.0048 | 0.0062 | 0.0062 | TRUE | 0.2539 |
| Mamdc2 | 14 | 11 | 6 | 6 | 11 | 10 | 11 | 12 | 6 | 6 | 12 | 10 | 9.55 | 10 | 9 | 0.963 | -0.054 | 0.93122033 | NA | 0 | 0.2506 | 0.1821 | 0.1821 | TRUE | 0.3474 |
| Mamdc4 | 45 | 40 | 32 | 33 | 35 | 31 | 35 | 43 | 35 | 33 | 39 | 31 | 35.83 | 38 | 34 | 0.918 | -0.123 | 0.70372017 | 0.82088242 | 0 | 0.0686 | 0.0469 | 0.0469 | TRUE | 0.1135 |
| Maml1 | 2999 | 2057 | 1962 | 2059 | 1889 | 2088 | 2329 | 2188 | 2117 | 2050 | 2131 | 2061 | 2145.98 | 2211 | 2081 | 0.94 | -0.089 | 0.16642131 | 0.31740856 | 0.001 | 0.0035 | 0.0025 | 0.0025 | TRUE | 0.0271 |
| Maml2 | 25 | 35 | 30 | 32 | 23 | 21 | 19 | 37 | 32 | 32 | 26 | 21 | 27.92 | 29 | 26 | 0.892 | -0.165 | 0.67522221 | 0.8005102 | 0.0476 | 0.0873 | 0.0758 | 0.0758 | TRUE | 0.7331 |
| Maml3 | 92 | 35 | 43 | 45 | 57 | 52 | 71 | 37 | 46 | 45 | 64 | 51 | 52.58 | 51 | 53 | 1.021 | 0.03 | 0.92236406 | 0.95711845 | 0.056 | 0.0475 | 0.05 | 0.05 | TRUE | 0.9801 |
| Mamld1 | 8 | 2 | 3 | 7 | 5 | 9 | 6 | 2 | 3 | 7 | 6 | 9 | 5.51 | 4 | 7 | 1.811 | 0.857 | 0.30321368 | NA | 0 | 0.4324 | 0.3053 | 0.3053 | TRUE | 0.6131 |
| Mamstr | 49 | 60 | 75 | 69 | 79 | 89 | 38 | 64 | 81 | 69 | 89 | 88 | 71.41 | 61 | 82 | 1.357 | 0.441 | 0.12006381 | 0.25177113 | 0.0641 | 0.0356 | 0.0436 | 0.0436 | TRUE | 0.4917 |
| Man1a | 575 | 423 | 510 | 522 | 437 | 522 | 447 | 450 | 550 | 520 | 493 | 515 | 495.78 | 482 | 509 | 1.058 | 0.082 | 0.45567244 | 0.62528093 | 0.0055 | 0.0072 | 0.0067 | 0.0067 | TRUE | 0.1729 |
| Man1a2 | 4178 | 2834 | 2697 | 2694 | 2303 | 2639 | 3245 | 3014 | 2910 | 2682 | 2598 | 2605 | 2842.36 | 3056 | 2628 | 0.859 | -0.219 | 0.00054675 | 0.0046152 | 0.0015 | 0.0033 | 0.0025 | 0.0025 | TRUE | 0.0361 |
| Man1b1 | 5127 | 3879 | 3922 | 3869 | 3667 | 4055 | 3982 | 4126 | 4232 | 3851 | 4137 | 4003 | 4055.11 | 4113 | 3997 | 0.972 | -0.041 | 0.4718302 | 0.64057116 | 7.00E-04 | 0.003 | 0.0021 | 0.0021 | TRUE | 0.0124 |
| Man1c1 | 582 | 482 | 597 | 638 | 572 | 740 | 452 | 513 | 644 | 635 | 645 | 730 | 603.28 | 536 | 670 | 1.253 | 0.326 | 0.00982484 | 0.04085726 | 0.0173 | 0.0064 | 0.0098 | 0.0098 | TRUE | 0.3598 |
| Man2a1 | 5486 | 3495 | 3463 | 3727 | 3101 | 3590 | 4261 | 3718 | 3737 | 3710 | 3498 | 3544 | 3744.48 | 3905 | 3584 | 0.917 | -0.125 | 0.07082565 | 0.17298548 | 0.0032 | 0.0031 | 0.0032 | 0.0032 | TRUE | 0.0785 |
| Man2a2 | 2020 | 1707 | 1615 | 1846 | 1667 | 1927 | 1569 | 1816 | 1743 | 1838 | 1881 | 1902 | 1791.24 | 1709 | 1874 | 1.098 | 0.134 | 0.0657608 | 0.1638346 | 0.0025 | 0.0037 | 0.0033 | 0.0033 | TRUE | 0.0645 |
| Man2b1 | 4279 | 3227 | 3529 | 3747 | 3426 | 3931 | 3323 | 3433 | 3808 | 3730 | 3865 | 3881 | 3673.12 | 3521 | 3825 | 1.087 | 0.12 | 0.06809057 | 0.16796046 | 0.0024 | 0.0031 | 0.0029 | 0.0029 | TRUE | 0.0613 |
| Man2b2 | 1101 | 866 | 978 | 909 | 916 | 974 | 855 | 921 | 1055 | 905 | 1033 | 961 | 955.19 | 944 | 966 | 1.025 | 0.036 | 0.70856888 | 0.8241171 | 0.0069 | 0.0049 | 0.0056 | 0.0056 | TRUE | 0.1264 |
| Man2c1 | 2716 | 2154 | 1947 | 2274 | 2027 | 2355 | 2109 | 2291 | 2101 | 2264 | 2287 | 2325 | 2229.39 | 2167 | 2292 | 1.058 | 0.081 | 0.20051663 | 0.36124939 | 8.00E-04 | 0.0035 | 0.0024 | 0.0024 | TRUE | 0.0299 |
| Manba | 9755 | 7127 | 6741 | 6035 | 5852 | 6699 | 7576 | 7581 | 7273 | 6007 | 6602 | 6613 | 6942.09 | 7477 | 6407 | 0.857 | -0.223 | 0.00012109 | 0.00142444 | 0.0015 | 0.0028 | 0.0023 | 0.0023 | TRUE | 0.0365 |
| Manbal | 1182 | 770 | 787 | 829 | 654 | 750 | 918 | 819 | 849 | 825 | 738 | 740 | 814.92 | 862 | 768 | 0.89 | -0.169 | 0.05708174 | 0.14844785 | 0.0027 | 0.0053 | 0.0044 | 0.0044 | TRUE | 0.0512 |
| Manea | 1116 | 751 | 788 | 999 | 748 | 896 | 867 | 799 | 850 | 994 | 844 | 884 | 873.09 | 839 | 907 | 1.082 | 0.114 | 0.20000108 | 0.36061756 | 0.0034 | 0.0051 | 0.0045 | 0.0045 | TRUE | 0.0837 |
| Maneal | 421 | 319 | 347 | 378 | 337 | 369 | 327 | 339 | 374 | 376 | 380 | 364 | 360.23 | 347 | 373 | 1.079 | 0.109 | 0.32924181 | 0.50362565 | 0 | 0.009 | 0.0063 | 0.0063 | TRUE | 0.0538 |
| Manf | 6097 | 4440 | 4239 | 5072 | 4113 | 4834 | 4735 | 4723 | 4574 | 5049 | 4640 | 4772 | 4748.73 | 4677 | 4820 | 1.031 | 0.043 | 0.43929743 | 0.61060761 | 8.00E-04 | 0.0029 | 0.0021 | 0.0021 | TRUE | 0.021 |
| Mansc1 | 172 | 140 | 160 | 161 | 140 | 157 | 134 | 149 | 173 | 160 | 158 | 155 | 154.72 | 152 | 158 | 1.044 | 0.062 | 0.70837297 | 0.82410273 | 0 | 0.0177 | 0.0134 | 0.0134 | TRUE | 0.1506 |
| Mansc4 | 9 | 10 | 9 | 8 | 4 | 12 | 7 | 11 | 10 | 8 | 5 | 12 | 8.61 | 9 | 8 | 0.904 | -0.146 | 0.82699363 | NA | 0 | 0.2777 | 0.2029 | 0.2029 | TRUE | 0.237 |
| Maoa | 2892 | 2046 | 1962 | 2254 | 1882 | 2137 | 2246 | 2176 | 2117 | 2244 | 2123 | 2110 | 2169.27 | 2180 | 2159 | 0.99 | -0.015 | 0.81592992 | 0.89433177 | 6.00E-04 | 0.0035 | 0.0024 | 0.0024 | TRUE | 0.0144 |
| Maob | 179 | 98 | 119 | 154 | 145 | 118 | 139 | 104 | 128 | 153 | 164 | 116 | 134.17 | 124 | 144 | 1.16 | 0.214 | 0.27032314 | 0.44122953 | 0.0188 | 0.0201 | 0.0197 | 0.0197 | TRUE | 0.3036 |
| Map1a | 5104 | 3741 | 3291 | 4539 | 4237 | 4600 | 3964 | 3979 | 3551 | 4518 | 4780 | 4541 | 4222.18 | 3831 | 4613 | 1.203 | 0.267 | 2.86E-05 | 0.00044588 | 0.0021 | 0.003 | 0.0027 | 0.0027 | TRUE | 0.0494 |
| Map1b | 7246 | 5601 | 4933 | 6439 | 6138 | 7306 | 5628 | 5958 | 5323 | 6409 | 6924 | 7212 | 6242.31 | 5636 | 6848 | 1.215 | 0.281 | 1.97E-05 | 0.0003271 | 0.0031 | 0.0028 | 0.003 | 0.003 | TRUE | 0.0386 |
| Map1lc3a | 2043 | 1334 | 1383 | 1542 | 1302 | 1591 | 1587 | 1419 | 1492 | 1535 | 1469 | 1571 | 1512.03 | 1499 | 1525 | 1.016 | 0.023 | 0.74723532 | 0.84993112 | 0.0016 | 0.004 | 0.0031 | 0.0031 | TRUE | 0.0325 |
| Map1lc3b | 5677 | 4208 | 4545 | 4329 | 4036 | 4743 | 4409 | 4476 | 4904 | 4309 | 4553 | 4682 | 4555.55 | 4596 | 4515 | 0.983 | -0.025 | 0.6898295 | 0.81028948 | 0.0023 | 0.0029 | 0.0027 | 0.0027 | TRUE | 0.0415 |
| Map1s | 6452 | 4846 | 4306 | 3932 | 3683 | 4083 | 5011 | 5155 | 4646 | 3914 | 4155 | 4031 | 4485.17 | 4937 | 4033 | 0.817 | -0.292 | 1.41E-06 | 3.87E-05 | 0.0015 | 0.003 | 0.0024 | 0.0024 | TRUE | 0.032 |
| Map2 | 737 | 576 | 666 | 698 | 598 | 728 | 572 | 613 | 719 | 695 | 675 | 719 | 665.29 | 635 | 696 | 1.1 | 0.137 | 0.17900847 | 0.333601 | 0.0059 | 0.006 | 0.006 | 0.006 | TRUE | 0.1563 |
| Map2k1 | 9431 | 7015 | 7387 | 7493 | 6556 | 7491 | 7325 | 7462 | 7971 | 7459 | 7396 | 7395 | 7501.01 | 7586 | 7417 | 0.978 | -0.032 | 0.54603703 | 0.70317212 | 8.00E-04 | 0.0027 | 0.0019 | 0.0019 | TRUE | 0.0239 |
| Map2k2 | 4884 | 3596 | 3135 | 3411 | 3061 | 3311 | 3793 | 3825 | 3383 | 3395 | 3453 | 3268 | 3519.63 | 3667 | 3372 | 0.919 | -0.121 | 0.06488747 | 0.16245167 | 0.0022 | 0.0031 | 0.0028 | 0.0028 | TRUE | 0.0553 |
| Map2k3 | 4099 | 3127 | 2886 | 2945 | 2645 | 3132 | 3184 | 3326 | 3114 | 2931 | 2984 | 3092 | 3105.11 | 3208 | 3002 | 0.936 | -0.095 | 0.10196176 | 0.22420607 | 6.00E-04 | 0.0032 | 0.0021 | 0.0021 | TRUE | 0.0124 |
| Map2k4 | 7448 | 5249 | 5492 | 5637 | 5004 | 5967 | 5785 | 5583 | 5926 | 5611 | 5645 | 5890 | 5740.02 | 5765 | 5715 | 0.992 | -0.012 | 0.81747719 | 0.89520392 | 5.00E-04 | 0.0028 | 0.0019 | 0.0019 | TRUE | 0.0091 |
| Map2k5 | 759 | 478 | 551 | 557 | 535 | 571 | 589 | 508 | 595 | 554 | 604 | 564 | 569.01 | 564 | 574 | 1.016 | 0.022 | 0.82253984 | 0.89813308 | 0.0029 | 0.0066 | 0.0054 | 0.0054 | TRUE | 0.0849 |
| Map2k6 | 1285 | 952 | 1062 | 1153 | 1008 | 1144 | 998 | 1013 | 1146 | 1148 | 1137 | 1129 | 1095.11 | 1052 | 1138 | 1.083 | 0.115 | 0.15073611 | 0.29659571 | 0.002 | 0.0046 | 0.0037 | 0.0037 | TRUE | 0.0713 |
| Map2k7 | 4483 | 3456 | 3748 | 3996 | 3477 | 4280 | 3482 | 3676 | 4044 | 3978 | 3922 | 4225 | 3887.83 | 3734 | 4042 | 1.083 | 0.115 | 0.09577718 | 0.21444386 | 0.0033 | 0.003 | 0.0032 | 0.0032 | TRUE | 0.064 |
| Map3k1 | 3530 | 2988 | 3234 | 3327 | 3235 | 3939 | 2742 | 3178 | 3489 | 3312 | 3649 | 3888 | 3376.48 | 3136 | 3616 | 1.154 | 0.206 | 0.02216084 | 0.07433579 | 0.0099 | 0.0031 | 0.0056 | 0.0056 | TRUE | 0.1494 |
| Map3k10 | 1478 | 1081 | 1064 | 1106 | 876 | 976 | 1148 | 1150 | 1148 | 1101 | 988 | 963 | 1083.06 | 1149 | 1017 | 0.886 | -0.174 | 0.02678878 | 0.08542884 | 0.0015 | 0.0046 | 0.0035 | 0.0035 | TRUE | 0.0621 |

| | | | | | | | | | | | | | | | | | | | | | | | | |
|---|---|---|---|---|---|---|---|---|---|---|---|---|---|---|---|---|---|---|---|---|---|---|---|---|
| Map3k11 | 1284 | 924 | 826 | 864 | 775 | 815 | 997 | 983 | 891 | 860 | 874 | 805 | 901.69 | 957 | 846 | 0.883 | -0.179 | 0.03163621 | 0.09624868 | 0.0017 | 0.0051 | 0.0039 | 0.0039 | TRUE | 0.0423 |
| Map3k12 | 1732 | 1591 | 1748 | 1857 | 1672 | 2084 | 1345 | 1692 | 1886 | 1848 | 1886 | 2057 | 1785.91 | 1641 | 1930 | 1.178 | 0.236 | 0.02601144 | 0.08370189 | 0.0148 | 0.0038 | 0.0076 | 0.0076 | TRUE | 0.3066 |
| Map3k13 | 238 | 153 | 190 | 218 | 206 | 209 | 185 | 163 | 205 | 217 | 232 | 206 | 201.38 | 184 | 218 | 1.186 | 0.246 | 0.0981509 | 0.21824375 | 0.0021 | 0.0142 | 0.0109 | 0.0109 | TRUE | 0.1042 |
| Map3k14 | 255 | 201 | 203 | 184 | 226 | 235 | 198 | 214 | 219 | 183 | 255 | 232 | 216.83 | 210 | 223 | 1.062 | 0.087 | 0.5731069 | 0.72548197 | 0.0107 | 0.0134 | 0.0126 | 0.0126 | TRUE | 0.2752 |
| Map3k15 | 1205 | 917 | 780 | 849 | 656 | 727 | 936 | 975 | 842 | 845 | 740 | 718 | 842.61 | 918 | 768 | 0.836 | -0.258 | 0.00664403 | 0.03051514 | 0.0054 | 0.0052 | 0.0053 | 0.0053 | TRUE | 0.0935 |
| Map3k19 | 2 | 1 | 2 | 0 | 0 | 0 | 2 | 1 | 2 | 0 | 0 | 0 | 0.8 | 2 | 0 | 0.12 | -3.062 | 0.17528122 | NA | 0 | 2.9805 | 2.3743 | 2.3743 | TRUE | 0.0718 |
| Map3k2 | 1226 | 945 | 1015 | 1124 | 962 | 1080 | 952 | 1005 | 1095 | 1119 | 1085 | 1066 | 1053.79 | 1017 | 1090 | 1.073 | 0.101 | 0.20679783 | 0.36850285 | 0.0019 | 0.0047 | 0.0037 | 0.0037 | TRUE | 0.0529 |
| Map3k3 | 4626 | 3253 | 3278 | 3917 | 3318 | 3628 | 3593 | 3460 | 3537 | 3899 | 3743 | 3581 | 3635.56 | 3530 | 3741 | 1.06 | 0.084 | 0.14889937 | 0.29411535 | 8.00E-04 | 0.0031 | 0.0021 | 0.0021 | TRUE | 0.0172 |
| Map3k4 | 6868 | 5141 | 5766 | 5940 | 5372 | 6259 | 5334 | 5468 | 6221 | 5913 | 6060 | 6179 | 5862.57 | 5674 | 6051 | 1.067 | 0.093 | 0.16769493 | 0.31915627 | 0.0035 | 0.0028 | 0.0031 | 0.0031 | TRUE | 0.0863 |
| Map3k5 | 1201 | 914 | 846 | 954 | 836 | 891 | 933 | 972 | 913 | 950 | 943 | 880 | 931.68 | 939 | 924 | 0.984 | -0.024 | 0.76273683 | 0.8591534 | 0 | 0.005 | 0.0034 | 0.0034 | TRUE | 0.0215 |
| Map3k6 | 517 | 377 | 349 | 341 | 267 | 340 | 402 | 401 | 377 | 339 | 301 | 336 | 359.23 | 393 | 325 | 0.828 | -0.272 | 0.01531214 | 0.05672794 | 0 | 0.009 | 0.0062 | 0.0062 | TRUE | 0.0463 |
| Map3k7 | 6108 | 4174 | 3624 | 3498 | 3011 | 3441 | 4744 | 4440 | 3910 | 3482 | 3397 | 3397 | 3894.89 | 4365 | 3425 | 0.784 | -0.35 | 2.03E-06 | 5.12E-05 | 0.0044 | 0.003 | 0.0037 | 0.0037 | TRUE | 0.1006 |
| Map3k8 | 64 | 57 | 61 | 71 | 76 | 60 | 50 | 61 | 66 | 71 | 86 | 59 | 65.3 | 59 | 72 | 1.228 | 0.296 | 0.24623242 | 0.41487681 | 0.013 | 0.0387 | 0.0315 | 0.0315 | TRUE | 0.2162 |
| Map3k9 | 1771 | 1352 | 1472 | 1570 | 1478 | 1656 | 1375 | 1438 | 1588 | 1563 | 1667 | 1635 | 1544.45 | 1467 | 1622 | 1.106 | 0.146 | 0.05446669 | 0.14308376 | 0.0023 | 0.004 | 0.0035 | 0.0035 | TRUE | 0.0621 |
| Map4 | 21715 | 15674 | 15742 | 15820 | 13790 | 16136 | 16865 | 16672 | 16985 | 15747 | 15557 | 15929 | 16292.55 | 16841 | 15744 | 0.935 | -0.097 | 0.02699576 | 0.0858208 | 0 | 0.0026 | 0.0013 | 0.0013 | TRUE | 0.0013 |
| Map4k1 | 1490 | 1120 | 1111 | 1223 | 1059 | 1324 | 1157 | 1191 | 1199 | 1217 | 1195 | 1307 | 1211.06 | 1182 | 1240 | 1.049 | 0.069 | 0.33961084 | 0.51360255 | 5.00E-04 | 0.0044 | 0.003 | 0.003 | TRUE | 0.0273 |
| Map4k2 | 776 | 626 | 581 | 677 | 616 | 683 | 603 | 666 | 627 | 674 | 695 | 674 | 656.41 | 632 | 681 | 1.079 | 0.11 | 0.21163652 | 0.37381622 | 0 | 0.006 | 0.004 | 0.004 | TRUE | 0.026 |
| Map4k3 | 3052 | 2403 | 2437 | 2610 | 2374 | 2630 | 2370 | 2556 | 2630 | 2598 | 2678 | 2596 | 2571.37 | 2519 | 2624 | 1.043 | 0.06 | 0.34278795 | 0.51693059 | 0.0013 | 0.0033 | 0.0025 | 0.0025 | TRUE | 0.0332 |
| Map4k4 | 16224 | 12066 | 13212 | 12321 | 11523 | 13513 | 12600 | 12834 | 14256 | 12264 | 12999 | 13339 | 13048.91 | 13230 | 12867 | 0.973 | -0.04 | 0.52775905 | 0.68871694 | 0.0027 | 0.0026 | 0.0028 | 0.0028 | TRUE | 0.0561 |
| Map4k5 | 2470 | 1759 | 1793 | 1702 | 1484 | 1662 | 1918 | 1871 | 1935 | 1694 | 1674 | 1641 | 1788.82 | 1908 | 1670 | 0.875 | -0.193 | 0.0018259 | 0.01126836 | 0 | 0.0038 | 0.0022 | 0.0022 | TRUE | 0.0034 |
| Map6 | 682 | 466 | 453 | 510 | 417 | 576 | 530 | 496 | 489 | 508 | 470 | 569 | 510.14 | 505 | 516 | 1.021 | 0.03 | 0.77243898 | 0.86543807 | 0.0035 | 0.0071 | 0.0059 | 0.0059 | TRUE | 0.0949 |
| Map6d1 | 146 | 115 | 141 | 159 | 145 | 148 | 113 | 122 | 152 | 158 | 164 | 146 | 142.63 | 129 | 156 | 1.211 | 0.276 | 0.1178666 | 0.24913743 | 0.0062 | 0.019 | 0.0154 | 0.0154 | TRUE | 0.2384 |
| Map7 | 4311 | 3247 | 3209 | 2852 | 2767 | 3204 | 3348 | 3454 | 3462 | 2839 | 3121 | 3163 | 3231.27 | 3421 | 3041 | 0.889 | -0.17 | 0.00670643 | 0.03074338 | 0.0014 | 0.0032 | 0.0025 | 0.0025 | TRUE | 0.0414 |
| Map7d1 | 1875 | 1454 | 1465 | 1419 | 1376 | 1781 | 1456 | 1547 | 1581 | 1412 | 1552 | 1758 | 1551.07 | 1528 | 1574 | 1.031 | 0.044 | 0.61101741 | 0.75392458 | 0.0063 | 0.004 | 0.0048 | 0.0048 | TRUE | 0.1271 |
| Map7d2 | 3721 | 2777 | 2902 | 3695 | 3218 | 3645 | 2890 | 2954 | 3131 | 3678 | 3630 | 3598 | 3313.57 | 2992 | 3635 | 1.216 | 0.282 | 1.02E-06 | 2.98E-05 | 6.00E-04 | 0.0031 | 0.0021 | 0.0021 | TRUE | 0.0201 |
| Map9 | 802 | 687 | 699 | 874 | 746 | 924 | 623 | 731 | 754 | 870 | 842 | 912 | 788.59 | 703 | 875 | 1.248 | 0.32 | 0.00075033 | 0.00586996 | 0.0048 | 0.0054 | 0.0052 | 0.0052 | TRUE | 0.1232 |
| Mapk1 | 21747 | 15626 | 16123 | 16400 | 14643 | 16645 | 16890 | 16621 | 17397 | 16325 | 16519 | 16431 | 16697.05 | 16969 | 16425 | 0.968 | -0.047 | 0.30123236 | 0.47354972 | 2.00E-04 | 0.0026 | 0.0014 | 0.0014 | TRUE | 0.0059 |
| Mapk10 | 72 | 59 | 47 | 16 | 14 | 24 | 56 | 63 | 51 | 16 | 16 | 24 | 37.47 | 57 | 19 | 0.328 | -1.607 | 2.74E-06 | 6.44E-05 | 0 | 0.0657 | 0.0481 | 0.0481 | TRUE | 0.3269 |
| Mapk11 | 614 | 451 | 501 | 453 | 516 | 563 | 477 | 480 | 541 | 451 | 582 | 556 | 514.33 | 499 | 530 | 1.061 | 0.086 | 0.46022432 | 0.62988358 | 0.0095 | 0.007 | 0.0078 | 0.0078 | TRUE | 0.1989 |
| Mapk12 | 406 | 272 | 314 | 456 | 366 | 424 | 315 | 289 | 339 | 454 | 413 | 419 | 371.47 | 314 | 429 | 1.363 | 0.447 | 8.41E-05 | 0.00104864 | 0.0011 | 0.0088 | 0.0066 | 0.0066 | TRUE | 0.0519 |
| Mapk13 | 749 | 639 | 833 | 691 | 658 | 759 | 582 | 680 | 899 | 688 | 742 | 749 | 723.26 | 720 | 726 | 1.011 | 0.016 | 0.90618604 | 0.94880353 | 0.0244 | 0.0057 | 0.0114 | 0.0114 | TRUE | 0.5558 |
| Mapk14 | 9475 | 6355 | 5570 | 5189 | 4800 | 5307 | 7359 | 6760 | 6010 | 5165 | 5415 | 5239 | 5991.23 | 6710 | 5273 | 0.786 | -0.348 | 2.42E-06 | 5.86E-05 | 0.0051 | 0.0028 | 0.0038 | 0.0038 | TRUE | 0.1013 |
| Mapk15 | 57 | 72 | 43 | 48 | 53 | 59 | 44 | 77 | 46 | 48 | 60 | 58 | 55.51 | 56 | 55 | 0.996 | -0.006 | 0.98484926 | 0.99220769 | 0.0388 | 0.0451 | 0.0433 | 0.0433 | TRUE | 0.8729 |
| Mapk1ip1 | 493 | 393 | 362 | 442 | 339 | 466 | 383 | 418 | 391 | 440 | 382 | 460 | 412.32 | 397 | 427 | 1.079 | 0.109 | 0.32901315 | 0.50354267 | 0.003 | 0.0082 | 0.0066 | 0.0066 | TRUE | 0.0952 |
| Mapk1ip1l | 11571 | 8374 | 7829 | 8060 | 7158 | 8336 | 8987 | 8907 | 8447 | 8023 | 8075 | 8229 | 8444.71 | 8780 | 8109 | 0.923 | -0.115 | 0.02267633 | 0.07563893 | 5.00E-04 | 0.0027 | 0.0017 | 0.0017 | TRUE | 0.0134 |
| Mapk3 | 4233 | 3172 | 3537 | 3807 | 3734 | 4437 | 3288 | 3374 | 3816 | 3789 | 4212 | 4380 | 3809.97 | 3493 | 4127 | 1.182 | 0.241 | 0.00176683 | 0.01100364 | 0.0056 | 0.003 | 0.004 | 0.004 | TRUE | 0.0796 |
| Mapk4 | 700 | 552 | 466 | 510 | 467 | 522 | 544 | 587 | 503 | 508 | 527 | 515 | 530.57 | 545 | 517 | 0.948 | -0.076 | 0.4370583 | 0.60850129 | 0.0011 | 0.0069 | 0.0051 | 0.0051 | TRUE | 0.052 |
| Mapk6 | 12326 | 9141 | 9717 | 11350 | 9680 | 11406 | 9573 | 9723 | 10485 | 11298 | 10920 | 11259 | 10543.02 | 9927 | 11159 | 1.124 | 0.169 | 0.00190192 | 0.0116221 | 0.0012 | 0.0027 | 0.002 | 0.002 | TRUE | 0.0294 |
| Mapk7 | 4838 | 3562 | 3304 | 3222 | 2965 | 3388 | 3757 | 3789 | 3565 | 3207 | 3345 | 3344 | 3501.3 | 3704 | 3299 | 0.89 | -0.167 | 0.00317283 | 0.01723653 | 5.00E-04 | 0.0031 | 0.002 | 0.002 | TRUE | 0.0129 |
| Mapk8 | 3985 | 2753 | 3105 | 3404 | 2943 | 3442 | 3095 | 2928 | 3350 | 3388 | 3320 | 3398 | 3246.62 | 3124 | 3369 | 1.078 | 0.109 | 0.09239608 | 0.20885928 | 0.0019 | 0.0032 | 0.0027 | 0.0027 | TRUE | 0.0476 |
| Mapk8ip1 | 2454 | 1861 | 1941 | 2215 | 1870 | 2300 | 1906 | 1980 | 2094 | 2205 | 2110 | 2270 | 2094.1 | 1993 | 2195 | 1.102 | 0.14 | 0.03453077 | 0.10284397 | 0.0013 | 0.0036 | 0.0027 | 0.0027 | TRUE | 0.0237 |
| Mapk8ip2 | 521 | 487 | 494 | 528 | 549 | 581 | 405 | 518 | 533 | 526 | 619 | 574 | 529.02 | 485 | 573 | 1.184 | 0.243 | 0.0453026 | 0.12531644 | 0.0127 | 0.0069 | 0.0087 | 0.0087 | TRUE | 0.2598 |
| Mapk8ip3 | 2275 | 1791 | 1989 | 2217 | 1936 | 2324 | 1767 | 1905 | 2146 | 2207 | 2184 | 2294 | 2083.84 | 1939 | 2228 | 1.15 | 0.202 | 0.0101615 | 0.04177116 | 0.0045 | 0.0036 | 0.004 | 0.004 | TRUE | 0.1048 |
| Mapk9 | 6560 | 4761 | 5128 | 5790 | 4850 | 5839 | 5095 | 5064 | 5533 | 5763 | 5471 | 5764 | 5448.47 | 5231 | 5666 | 1.084 | 0.116 | 0.04771786 | 0.13023322 | 0.0014 | 0.0029 | 0.0023 | 0.0023 | TRUE | 0.0309 |
| Mapkap1 | 3342 | 2385 | 2193 | 2334 | 2091 | 2418 | 2596 | 2537 | 2366 | 2323 | 2359 | 2387 | 2427.96 | 2500 | 2356 | 0.942 | -0.086 | 0.16818377 | 0.31974379 | 9.00E-04 | 0.0034 | 0.0024 | 0.0024 | TRUE | 0.0261 |
| Mapkapk2 | 2614 | 1890 | 1832 | 1721 | 1550 | 1762 | 2030 | 2010 | 1977 | 1713 | 1749 | 1739 | 1869.71 | 2006 | 1734 | 0.864 | -0.211 | 0.00049621 | 0.00428797 | 0 | 0.0037 | 0.0021 | 0.0021 | TRUE | 0.0019 |
| Mapkapk3 | 2703 | 2213 | 2648 | 2943 | 2610 | 3289 | 2099 | 2354 | 2857 | 2929 | 2944 | 3247 | 2738.5 | 2437 | 3040 | 1.249 | 0.32 | 0.00116076 | 0.00807731 | 0.0137 | 0.0033 | 0.0066 | 0.0066 | TRUE | 0.2756 |
| Mapkapk5 | 2727 | 1906 | 1874 | 1870 | 1720 | 1867 | 2118 | 2027 | 2022 | 1861 | 1940 | 1843 | 1968.69 | 2056 | 1881 | 0.915 | -0.129 | 0.0389355 | 0.1123345 | 0 | 0.0036 | 0.0023 | 0.0023 | TRUE | 0.0089 |
| Mapkbp1 | 1447 | 1116 | 1142 | 1205 | 1145 | 1250 | 1124 | 1187 | 1232 | 1199 | 1292 | 1234 | 1211.36 | 1181 | 1242 | 1.052 | 0.073 | 0.32447928 | 0.49836475 | 0.001 | 0.0044 | 0.0032 | 0.0032 | TRUE | 0.0226 |
| Mapre1 | 11099 | 7536 | 6520 | 7205 | 5899 | 6800 | 8620 | 8016 | 7035 | 7172 | 6655 | 6713 | 7368.38 | 7890 | 6847 | 0.867 | -0.205 | 0.00644488 | 0.02977267 | 0.0058 | 0.0027 | 0.0039 | 0.0039 | TRUE | 0.1096 |
| Mapre2 | 4085 | 3089 | 2958 | 3630 | 3221 | 3801 | 3173 | 3286 | 3192 | 3613 | 3634 | 3752 | 3441.52 | 3217 | 3666 | 1.14 | 0.189 | 0.00049234 | 0.00426298 | 0 | 0.0031 | 0.0018 | 0.0018 | TRUE | 0.0051 |
| Mapre3 | 382 | 247 | 279 | 307 | 226 | 269 | 297 | 263 | 301 | 306 | 255 | 266 | 281.09 | 287 | 276 | 0.96 | -0.059 | 0.65217019 | 0.78397549 | 0.0035 | 0.0109 | 0.0087 | 0.0087 | TRUE | 0.1041 |
| Mapt | 1383 | 1025 | 1046 | 1447 | 1234 | 1405 | 1074 | 1090 | 1129 | 1440 | 1392 | 1387 | 1252.07 | 1098 | 1406 | 1.282 | 0.359 | 2.31E-07 | 8.26E-06 | 0 | 0.0043 | 0.0027 | 0.0027 | TRUE | 0.0072 |
| Mar-01 | 3 | 7 | 5 | 0 | 3 | 0 | 2 | 7 | 5 | 0 | 3 | 0 | 3.09 | 5 | 1 | 0.215 | -2.22 | 0.10102576 | NA | 0.5745 | 0.7688 | 0.7415 | 0.7415 | TRUE | 0.9703 |
| Mar-02 | 1969 | 1406 | 1602 | 1476 | 1531 | 1665 | 1529 | 1496 | 1729 | 1469 | 1727 | 1644 | 1598.88 | 1585 | 1613 | 1.018 | 0.026 | 0.75756594 | 0.85595969 | 0.0058 | 0.0039 | 0.0046 | 0.0046 | TRUE | 0.0751 |
| Mar-01 | 9 | 5 | 6 | 4 | 3 | 3 | 7 | 5 | 6 | 4 | 3 | 3 | 4.85 | 6 | 3 | 0.548 | -0.868 | 0.31347909 | NA | 0 | 0.4909 | 0.3077 | 0.3077 | TRUE | 0.0397 |
| Mar-10 | 4 | 2 | 8 | 3 | 5 | 8 | 3 | 2 | 9 | 3 | 6 | 8 | 5.06 | 5 | 6 | 1.208 | 0.273 | 0.76145742 | NA | 0.1621 | 0.4704 | 0.3819 | 0.3819 | TRUE | 0.6213 |
| Mar-11 | 83 | 89 | 88 | 114 | 107 | 117 | 64 | 95 | 95 | 113 | 121 | 115 | 100.63 | 85 | 116 | 1.387 | 0.472 | 0.02640555 | 0.08458415 | 0.0131 | 0.026 | 0.0224 | 0.0224 | TRUE | 0.4734 |
| Mar-02 | 1488 | 1071 | 1152 | 1313 | 1116 | 1281 | 1156 | 1139 | 1243 | 1307 | 1259 | 1265 | 1228.06 | 1179 | 1277 | 1.083 | 0.116 | 0.10999811 | 0.23719438 | 4.00E-04 | 0.0044 | 0.0029 | 0.0029 | TRUE | 0.026 |
| Mar-03 | 42 | 43 | 39 | 31 | 37 | 18 | 33 | 46 | 42 | 31 | 42 | 18 | 35.13 | 40 | 30 | 0.75 | -0.416 | 0.25857766 | 0.42850468 | 0.063 | 0.0699 | 0.0681 | 0.0681 | TRUE | 0.3198 |
| Mar-04 | 1 | 0 | 1 | 0 | 0 | 0 | 1 | 0 | 1 | 0 | 0 | 0 | 0.31 | 1 | 0 | 0.307 | -1.706 | 0.62894649 | NA | 0 | 7.6657 | 7.2 | 7.2 | TRUE | 0.2067 |
| Mar-05 | 5883 | 4333 | 4338 | 5072 | 4125 | 4734 | 4569 | 4609 | 4681 | 5049 | 4653 | 4673 | 4705.66 | 4620 | 4792 | 1.037 | 0.053 | 0.34627287 | 0.52063438 | 8.00E-04 | 0.0029 | 0.0021 | 0.0021 | TRUE | 0.0269 |
| Mar-06 | 7417 | 5321 | 5495 | 6581 | 5662 | 6433 | 5760 | 5660 | 5929 | 6551 | 6387 | 6350 | 6106.3 | 5783 | 6429 | 1.112 | 0.153 | 0.00232977 | 0.01360826 | 2.00E-04 | 0.0028 | 0.0017 | 0.0017 | TRUE | 0.0058 |
| Mar-07 | 6624 | 5002 | 4932 | 5172 | 4434 | 5231 | 5145 | 5321 | 5322 | 5148 | 5002 | 5164 | 5183.46 | 5263 | 5105 | 0.97 | -0.043 | 0.39286902 | 0.5665031 | 2.00E-04 | 0.0029 | 0.0017 | 0.0017 | TRUE | 0.0048 |
| Mar-08 | 2143 | 1620 | 1739 | 1766 | 1491 | 1686 | 1664 | 1723 | 1876 | 1758 | 1682 | 1664 | 1728.02 | 1754 | 1701 | 0.971 | -0.043 | 0.54353728 | 0.70106712 | 0.0018 | 0.0038 | 0.0031 | 0.0031 | TRUE | 0.0441 |
| Mar-09 | 1105 | 807 | 799 | 878 | 788 | 958 | 858 | 858 | 862 | 874 | 889 | 946 | 881.22 | 859 | 903 | 1.051 | 0.071 | 0.36230909 | 0.53696431 | 0 | 0.0051 | 0.0033 | 0.0033 | TRUE | 0.0209 |
| Marcks | 3864 | 3043 | 2531 | 2888 | 2427 | 2843 | 3001 | 3237 | 2731 | 2875 | 2738 | 2806 | 2897.97 | 2990 | 2806 | 0.939 | -0.091 | 0.20305427 | 0.36448021 | 0.0034 | 0.0032 | 0.0034 | 0.0034 | TRUE | 0.0683 |
| Marcksl1 | 14232 | 10389 | 9475 | 6221 | 5666 | 6857 | 11053 | 11051 | 10223 | 6192 | 6392 | 6769 | 8613.44 | 10776 | 6451 | 0.599 | -0.74 | 2.20E-36 | 2.86E-33 | 0.0018 | 0.0027 | 0.0024 | 0.0024 | TRUE | 0.0245 |
| Marcksl1-ps | 0 | 0 | 0 | 0 | 0 | 0 | 0 | 0 | 0 | 0 | 0 | 0 | 0 | 0 | 0 | NA | NA | NA | NA | NA | NA | NA | NA | NA | NA |

| | | | | | | | | | | | | | | | | | | | | | | | | | |
|---|---|---|---|---|---|---|---|---|---|---|---|---|---|---|---|---|---|---|---|---|---|---|---|---|---|
| Marco | 0 | 1 | 0 | 0 | 0 | 0 | 0 | 1 | 0 | 0 | 0 | 0 | 0.18 | 0 | 0 | 0.552 | -0.857 | 0.8336548 | NA | 0 | 13.3714 | 10 | 10 | TRUE | 0.5216 |
| Marf1 | 4844 | 4139 | 4332 | 4306 | 4034 | 4599 | 3762 | 4403 | 4674 | 4286 | 4551 | 4540 | 4369.3 | 4280 | 4459 | 1.043 | 0.06 | 0.44753066 | 0.61788255 | 0.0066 | 0.003 | 0.0043 | 0.0043 | TRUE | 0.138 |
| Mark1 | 934 | 701 | 694 | 764 | 688 | 798 | 725 | 746 | 749 | 760 | 776 | 788 | 757.37 | 740 | 775 | 1.048 | 0.067 | 0.40433961 | 0.5771907 | 0 | 0.0056 | 0.0034 | 0.0034 | TRUE | 0.0037 |
| Mark2 | 3687 | 2914 | 2657 | 2538 | 2318 | 2712 | 2864 | 3100 | 2867 | 2526 | 2615 | 2677 | 2774.74 | 2944 | 2606 | 0.886 | -0.175 | 0.00479202 | 0.02371617 | 0.0011 | 0.0033 | 0.0024 | 0.0024 | TRUE | 0.0259 |
| Mark3 | 6914 | 4938 | 4632 | 4843 | 4369 | 4976 | 5370 | 5252 | 4998 | 4821 | 4929 | 4912 | 5046.94 | 5207 | 4887 | 0.938 | -0.092 | 0.08691781 | 0.19988502 | 6.00E-04 | 0.0029 | 0.0019 | 0.0019 | TRUE | 0.0149 |
| Mark4 | 2411 | 2001 | 2360 | 2469 | 2292 | 2665 | 1873 | 2128 | 2546 | 2458 | 2586 | 2631 | 2370.23 | 2182 | 2558 | 1.173 | 0.23 | 0.01724868 | 0.06195913 | 0.0121 | 0.0034 | 0.0063 | 0.0063 | TRUE | 0.2574 |
| Mars | 11085 | 8147 | 7257 | 7377 | 6542 | 7640 | 8609 | 8666 | 7830 | 7343 | 7380 | 7542 | 7895.05 | 8368 | 7422 | 0.887 | -0.173 | 0.0024365 | 0.01408179 | 0.0015 | 0.0027 | 0.0022 | 0.0022 | TRUE | 0.0384 |
| Mars2 | 961 | 698 | 491 | 568 | 507 | 514 | 746 | 742 | 530 | 565 | 572 | 507 | 610.56 | 673 | 548 | 0.813 | -0.298 | 0.02027345 | 0.06990793 | 0.019 | 0.0063 | 0.0102 | 0.0102 | TRUE | 0.401 |
| Marveld1 | 804 | 611 | 488 | 459 | 400 | 448 | 624 | 650 | 527 | 457 | 451 | 442 | 525.21 | 600 | 450 | 0.749 | -0.417 | 8.76E-05 | 0.0010853 | 0.0046 | 0.0069 | 0.0062 | 0.0062 | TRUE | 0.1311 |
| Marveld2 | 589 | 412 | 437 | 428 | 360 | 422 | 457 | 438 | 472 | 426 | 406 | 417 | 435.99 | 456 | 416 | 0.914 | -0.13 | 0.19563403 | 0.35529825 | 0 | 0.0079 | 0.005 | 0.005 | TRUE | 0.0124 |
| Marveld3 | 183 | 165 | 146 | 166 | 164 | 139 | 142 | 176 | 158 | 165 | 185 | 137 | 160.44 | 159 | 162 | 1.027 | 0.038 | 0.82375367 | 0.89868745 | 0.0105 | 0.0172 | 0.0153 | 0.0153 | TRUE | 0.1984 |
| Mas1 | 3 | 3 | 1 | 6 | 1 | 2 | 2 | 3 | 1 | 6 | 1 | 2 | 2.61 | 2 | 3 | 1.382 | 0.467 | 0.70007773 | NA | 0 | 0.9096 | 0.6699 | 0.6699 | TRUE | 0.9641 |
| Masp1 | 2 | 5 | 1 | 3 | 2 | 1 | 2 | 5 | 1 | 3 | 2 | 1 | 2.36 | 3 | 2 | 0.792 | -0.336 | 0.79166591 | NA | 0 | 1.0053 | 0.729 | 0.729 | TRUE | 0.8398 |
| Masp2 | 24 | 27 | 41 | 44 | 36 | 45 | 19 | 29 | 44 | 44 | 41 | 44 | 36.74 | 31 | 43 | 1.427 | 0.513 | 0.15301595 | 0.29992553 | 0.0593 | 0.0669 | 0.0649 | 0.0649 | TRUE | 0.4727 |
| Mast1 | 684 | 517 | 527 | 549 | 486 | 541 | 531 | 550 | 569 | 546 | 548 | 534 | 546.43 | 550 | 543 | 0.988 | -0.017 | 0.85188392 | 0.91645167 | 0 | 0.0068 | 0.0042 | 0.0042 | TRUE | 0.0111 |
| Mast2 | 4646 | 3456 | 3253 | 3173 | 2985 | 3325 | 3608 | 3676 | 3510 | 3158 | 3367 | 3282 | 3433.75 | 3598 | 3269 | 0.908 | -0.139 | 0.01434814 | 0.05399226 | 5.00E-04 | 0.0031 | 0.002 | 0.002 | TRUE | 0.0107 |
| Mast3 | 2306 | 1932 | 1969 | 1824 | 1784 | 2188 | 1791 | 2055 | 2125 | 1816 | 2013 | 2160 | 1993.1 | 1990 | 1996 | 1.004 | 0.006 | 0.94912986 | 0.97218812 | 0.0074 | 0.0036 | 0.0049 | 0.0049 | TRUE | 0.0949 |
| Mast4 | 623 | 433 | 419 | 485 | 467 | 508 | 484 | 461 | 452 | 483 | 527 | 501 | 484.6 | 466 | 504 | 1.08 | 0.111 | 0.25747882 | 0.42741156 | 0 | 0.0073 | 0.0049 | 0.0049 | TRUE | 0.0183 |
| Mastl | 3438 | 2460 | 2475 | 2744 | 2256 | 2685 | 2670 | 2617 | 2671 | 2731 | 2545 | 2651 | 2647.37 | 2653 | 2642 | 0.996 | -0.005 | 0.92779231 | 0.960172 | 3.00E-04 | 0.0033 | 0.0021 | 0.0021 | TRUE | 0.0124 |
| Mat1a | 24 | 13 | 8 | 9 | 7 | 6 | 19 | 14 | 9 | 9 | 8 | 6 | 10.65 | 14 | 8 | 0.547 | -0.869 | 0.15541918 | NA | 0 | 0.225 | 0.1667 | 0.1667 | TRUE | 0.1334 |
| Mat2a | 20684 | 13793 | 9357 | 10494 | 9017 | 9034 | 16064 | 14671 | 10096 | 10446 | 10172 | 8918 | 11727.95 | 13610 | 9845 | 0.723 | -0.467 | 0.02132349 | 0.07242791 | 0.0296 | 0.0026 | 0.011 | 0.0296 | TRUE | 0.6237 |
| Mat2b | 6911 | 5329 | 5626 | 5731 | 5294 | 6086 | 5367 | 5668 | 6070 | 5705 | 5972 | 6008 | 5798.48 | 5702 | 5895 | 1.034 | 0.048 | 0.43045861 | 0.60260776 | 0.002 | 0.0028 | 0.0026 | 0.0026 | TRUE | 0.0389 |
| Matk | 53 | 60 | 63 | 84 | 63 | 91 | 41 | 64 | 68 | 84 | 71 | 90 | 69.58 | 58 | 82 | 1.43 | 0.516 | 0.04652379 | 0.12783363 | 0.0262 | 0.0365 | 0.0337 | 0.0337 | TRUE | 0.6248 |
| Matn1 | 15 | 18 | 15 | 18 | 14 | 19 | 12 | 19 | 16 | 18 | 16 | 19 | 16.57 | 16 | 18 | 1.13 | 0.176 | 0.70850766 | NA | 0 | 0.1454 | 0.0998 | 0.0998 | TRUE | 0.3016 |
| Matn2 | 167 | 102 | 107 | 158 | 133 | 150 | 130 | 108 | 115 | 157 | 150 | 148 | 134.84 | 118 | 152 | 1.283 | 0.359 | 0.0360224 | 0.10613172 | 0 | 0.02 | 0.0137 | 0.0137 | TRUE | 0.0867 |
| Matn3 | 743 | 544 | 821 | 930 | 928 | 1035 | 577 | 579 | 886 | 926 | 1047 | 1022 | 839.31 | 681 | 998 | 1.469 | 0.555 | 7.79E-05 | 0.00098921 | 0.0343 | 0.0053 | 0.013 | 0.013 | TRUE | 0.8147 |
| Matn4 | 13 | 9 | 12 | 13 | 8 | 10 | 10 | 10 | 13 | 13 | 9 | 10 | 10.74 | 11 | 11 | 0.981 | -0.027 | 0.96273104 | NA | 0 | 0.2231 | 0.1465 | 0.1465 | TRUE | 0.1357 |
| Matr3 | 33333 | 24853 | 25064 | 26477 | 22726 | 25919 | 25888 | 26436 | 27044 | 26355 | 25637 | 25586 | 26157.75 | 26456 | 25859 | 0.978 | -0.033 | 0.47103931 | 0.63985601 | 3.00E-04 | 0.0025 | 0.0015 | 0.0015 | TRUE | 0.0046 |
| Mau2 | 5467 | 4139 | 4083 | 3864 | 3704 | 4208 | 4246 | 4403 | 4406 | 3846 | 4178 | 4154 | 4205.46 | 4352 | 4059 | 0.933 | -0.1 | 0.0838555 | 0.19523484 | 9.00E-04 | 0.003 | 0.0022 | 0.0022 | TRUE | 0.0258 |
| Mavs | 1754 | 1278 | 1433 | 1531 | 1335 | 1698 | 1362 | 1359 | 1546 | 1524 | 1506 | 1676 | 1495.67 | 1422 | 1569 | 1.104 | 0.142 | 0.074822 | 0.18004295 | 0.0036 | 0.004 | 0.0039 | 0.0039 | TRUE | 0.0684 |
| Max | 4857 | 3513 | 3339 | 3303 | 3082 | 3793 | 3772 | 3737 | 3603 | 3288 | 3477 | 3744 | 3603.43 | 3704 | 3503 | 0.946 | -0.081 | 0.21041106 | 0.37232983 | 0.002 | 0.0031 | 0.0027 | 0.0027 | TRUE | 0.0445 |
| Maz | 10207 | 7852 | 7932 | 8408 | 7979 | 9466 | 7927 | 8352 | 8559 | 8369 | 9001 | 9344 | 8592.14 | 8279 | 8905 | 1.076 | 0.105 | 0.08038987 | 0.18982528 | 0.0021 | 0.0027 | 0.0025 | 0.0025 | TRUE | 0.0339 |
| Mb | 18 | 8 | 14 | 11 | 15 | 10 | 14 | 9 | 15 | 11 | 17 | 10 | 12.56 | 13 | 13 | 0.994 | -0.008 | 0.98844196 | NA | 0 | 0.1912 | 0.14 | 0.14 | TRUE | 0.3357 |
| Mb21d1 | 235 | 154 | 126 | 148 | 141 | 161 | 183 | 164 | 136 | 147 | 159 | 159 | 157.93 | 161 | 155 | 0.96 | -0.058 | 0.72909834 | 0.83790324 | 0.0062 | 0.0174 | 0.0141 | 0.0141 | TRUE | 0.1874 |
| Mb21d2 | 1209 | 961 | 939 | 697 | 585 | 706 | 939 | 1022 | 1013 | 694 | 660 | 697 | 837.5 | 991 | 684 | 0.691 | -0.534 | 8.18E-11 | 6.75E-09 | 0 | 0.0053 | 0.0036 | 0.0036 | TRUE | 0.027 |
| Mbd1 | 952 | 734 | 597 | 541 | 494 | 567 | 739 | 781 | 644 | 539 | 557 | 560 | 636.63 | 721 | 552 | 0.764 | -0.388 | 7.76E-05 | 0.00098796 | 0.0036 | 0.0061 | 0.0053 | 0.0053 | TRUE | 0.1001 |
| Mbd2 | 992 | 678 | 594 | 597 | 552 | 551 | 770 | 721 | 641 | 594 | 623 | 544 | 648.91 | 711 | 587 | 0.824 | -0.28 | 0.00564398 | 0.02693891 | 0.0052 | 0.0061 | 0.0058 | 0.0058 | TRUE | 0.0864 |
| Mbd3 | 7258 | 5239 | 4711 | 5081 | 4494 | 5036 | 5637 | 5573 | 5083 | 5058 | 5070 | 4971 | 5231.9 | 5431 | 5033 | 0.926 | -0.11 | 0.05995501 | 0.15313383 | 0.0013 | 0.0029 | 0.0023 | 0.0023 | TRUE | 0.038 |
| Mbd3l1 | 0 | 0 | 0 | 0 | 0 | 0 | 0 | 0 | 0 | 0 | 0 | 0 | 0 | 0 | 0 | NA | NA | NA | NA | NA | NA | NA | NA | NA | NA |
| Mbd3l2 | 1 | 0 | 0 | 0 | 0 | 1 | 1 | 0 | 0 | 0 | 0 | 1 | 0.29 | 0 | 0 | 1.075 | 0.105 | 0.97702352 | NA | 0 | 8.0647 | 7.5458 | 7.5458 | TRUE | 0.3835 |
| Mbd4 | 2027 | 1511 | 1420 | 1498 | 1351 | 1537 | 1574 | 1607 | 1532 | 1491 | 1524 | 1517 | 1541.02 | 1571 | 1511 | 0.961 | -0.057 | 0.37906416 | 0.55375757 | 0 | 0.004 | 0.0024 | 0.0024 | TRUE | 0.0055 |
| Mbd5 | 812 | 568 | 482 | 490 | 448 | 501 | 631 | 604 | 520 | 488 | 505 | 495 | 540.43 | 585 | 496 | 0.846 | -0.241 | 0.01906059 | 0.06674007 | 0.0036 | 0.0068 | 0.0058 | 0.0058 | TRUE | 0.1081 |
| Mbd6 | 2552 | 1956 | 1865 | 2046 | 1767 | 2098 | 1982 | 2081 | 2012 | 2037 | 1993 | 2071 | 2029.32 | 2025 | 2034 | 1.005 | 0.007 | 0.90891042 | 0.94963864 | 0 | 0.0036 | 0.0022 | 0.0022 | TRUE | 0.0069 |
| Mbip | 1236 | 858 | 868 | 799 | 760 | 812 | 960 | 913 | 937 | 795 | 857 | 802 | 877.23 | 937 | 818 | 0.873 | -0.197 | 0.01308825 | 0.05045765 | 0 | 0.0051 | 0.0034 | 0.0034 | TRUE | 0.0204 |
| Mbl1 | 1 | 0 | 0 | 0 | 0 | 0 | 1 | 0 | 0 | 0 | 0 | 0 | 0.13 | 0 | 0 | 0.552 | -0.857 | 0.8336548 | NA | 0 | 18.3121 | 10 | 10 | TRUE | 0.2566 |
| Mbl2 | 29 | 21 | 11 | 13 | 15 | 18 | 23 | 22 | 12 | 13 | 17 | 18 | 17.39 | 19 | 16 | 0.833 | -0.264 | 0.57973665 | 0.73056319 | 0 | 0.1387 | 0.1052 | 0.1052 | TRUE | 0.5115 |
| Mblac1 | 68 | 44 | 25 | 34 | 25 | 35 | 53 | 47 | 27 | 34 | 28 | 35 | 37.2 | 42 | 32 | 0.759 | -0.398 | 0.24953097 | 0.4184058 | 0.0401 | 0.0661 | 0.0588 | 0.0588 | TRUE | 0.7201 |
| Mblac2 | 99 | 67 | 51 | 47 | 58 | 48 | 77 | 71 | 55 | 47 | 65 | 47 | 60.46 | 68 | 53 | 0.779 | -0.36 | 0.17821367 | 0.33270743 | 0.0158 | 0.0416 | 0.0346 | 0.0346 | TRUE | 0.3156 |
| Mbnl1 | 712 | 495 | 384 | 470 | 381 | 360 | 553 | 527 | 414 | 468 | 430 | 355 | 457.81 | 498 | 418 | 0.837 | -0.257 | 0.05877487 | 0.1511267 | 0.019 | 0.0076 | 0.0111 | 0.0111 | TRUE | 0.2469 |
| Mbnl2 | 1775 | 1285 | 1330 | 1446 | 1195 | 1423 | 1379 | 1367 | 1435 | 1439 | 1348 | 1405 | 1395.44 | 1394 | 1397 | 1.003 | 0.005 | 0.94324332 | 0.96837931 | 0 | 0.0041 | 0.0027 | 0.0027 | TRUE | 0.0112 |
| Mbnl3 | 788 | 564 | 517 | 502 | 414 | 469 | 612 | 600 | 558 | 500 | 467 | 463 | 533.24 | 590 | 477 | 0.807 | -0.309 | 0.00130294 | 0.00871753 | 0 | 0.0069 | 0.0048 | 0.0048 | TRUE | 0.026 |
| Mboat1 | 323 | 240 | 245 | 235 | 171 | 201 | 251 | 255 | 264 | 234 | 193 | 198 | 232.62 | 257 | 208 | 0.813 | -0.299 | 0.0299326 | 0.09252306 | 0 | 0.0126 | 0.0093 | 0.0093 | TRUE | 0.1265 |
| Mboat2 | 2392 | 1810 | 2091 | 2287 | 1967 | 2377 | 1858 | 1925 | 2256 | 2276 | 2219 | 2346 | 2146.86 | 2013 | 2280 | 1.134 | 0.181 | 0.02376659 | 0.07842742 | 0.0053 | 0.0035 | 0.0042 | 0.0042 | TRUE | 0.134 |
| Mboat4 | 23 | 18 | 22 | 17 | 14 | 24 | 18 | 19 | 24 | 17 | 16 | 24 | 19.53 | 20 | 19 | 0.936 | -0.096 | 0.82601698 | 0.9002105 | 0 | 0.1238 | 0.0864 | 0.0864 | TRUE | 0.277 |
| Mboat7 | 1739 | 1316 | 1396 | 1536 | 1415 | 1562 | 1351 | 1400 | 1506 | 1529 | 1596 | 1542 | 1487.3 | 1419 | 1556 | 1.097 | 0.134 | 0.05932455 | 0.15214888 | 0.0012 | 0.004 | 0.003 | 0.003 | TRUE | 0.0346 |
| Mbp | 257 | 232 | 299 | 294 | 256 | 287 | 200 | 247 | 323 | 293 | 289 | 283 | 272.29 | 257 | 288 | 1.13 | 0.176 | 0.28003214 | 0.45149457 | 0.0255 | 0.0111 | 0.0154 | 0.0154 | TRUE | 0.4063 |
| Mbtd1 | 9953 | 7656 | 7765 | 8169 | 7418 | 8913 | 7730 | 8144 | 8378 | 8131 | 8368 | 8799 | 8258.37 | 8084 | 8433 | 1.043 | 0.061 | 0.28008351 | 0.45152646 | 0.0014 | 0.0027 | 0.0022 | 0.0022 | TRUE | 0.0181 |
| Mbtps1 | 20081 | 14224 | 13377 | 14340 | 12834 | 14146 | 15596 | 15130 | 14434 | 14274 | 14478 | 13964 | 14646 | 15053 | 14239 | 0.946 | -0.08 | 0.11378439 | 0.2428431 | 8.00E-04 | 0.0026 | 0.0018 | 0.0018 | TRUE | 0.0158 |
| Mbtps2 | 2817 | 2052 | 2336 | 2329 | 1910 | 2290 | 2188 | 2183 | 2521 | 2318 | 2155 | 2261 | 2270.77 | 2297 | 2245 | 0.978 | -0.032 | 0.66457587 | 0.79331991 | 0.0037 | 0.0035 | 0.0035 | 0.0035 | TRUE | 0.0868 |
| Mc1r | 10 | 3 | 3 | 2 | 4 | 2 | 8 | 3 | 3 | 2 | 5 | 2 | 3.78 | 5 | 3 | 0.578 | -0.79 | 0.43519221 | NA | 0 | 0.6296 | 0.4529 | 0.4529 | TRUE | 0.785 |
| Mc2r | 0 | 0 | 0 | 0 | 0 | 0 | 0 | 0 | 0 | 0 | 0 | 0 | 0 | 0 | 0 | NA | NA | NA | NA | NA | NA | NA | NA | NA | NA |
| Mc3r | 1 | 1 | 0 | 1 | 0 | 0 | 1 | 1 | 0 | 1 | 0 | 0 | 0.47 | 1 | 0 | 0.599 | -0.74 | 0.79191656 | NA | 0 | 5.0171 | 3.8827 | 3.8827 | TRUE | 0.3906 |
| Mc4r | 1 | 2 | 2 | 1 | 2 | 2 | 1 | 2 | 2 | 1 | 2 | 2 | 1.71 | 2 | 2 | 1.052 | 0.073 | 0.95914088 | NA | 0 | 1.3847 | 0.8587 | 0.8587 | TRUE | 0.2431 |
| Mc5r | 35 | 17 | 10 | 12 | 12 | 14 | 27 | 18 | 11 | 12 | 14 | 14 | 15.89 | 19 | 13 | 0.691 | -0.533 | 0.2989605 | NA | 0.0621 | 0.1516 | 0.1248 | 0.1248 | TRUE | 0.196 |
| Mcam | 3590 | 2975 | 4113 | 4199 | 3935 | 4607 | 2788 | 3164 | 4438 | 4180 | 4439 | 4548 | 3926.19 | 3463 | 4389 | 1.267 | 0.342 | 0.10447165 | 0.22844046 | 0.0317 | 0.003 | 0.0109 | 0.0317 | TRUE | 0.736 |
| Mcat | 2229 | 1581 | 1684 | 1891 | 1648 | 1873 | 1731 | 1682 | 1817 | 1882 | 1859 | 1849 | 1803.37 | 1743 | 1863 | 1.069 | 0.096 | 0.13137523 | 0.26878702 | 0 | 0.0037 | 0.0024 | 0.0024 | TRUE | 0.016 |
| Mcc | 448 | 307 | 256 | 326 | 307 | 342 | 348 | 327 | 276 | 325 | 346 | 338 | 326.53 | 317 | 336 | 1.057 | 0.08 | 0.51818355 | 0.68004751 | 0.0044 | 0.0097 | 0.0081 | 0.0081 | TRUE | 0.1393 |
| Mccc1 | 763 | 610 | 504 | 605 | 530 | 555 | 593 | 649 | 544 | 602 | 598 | 548 | 588.87 | 595 | 583 | 0.979 | -0.031 | 0.75706607 | 0.85566545 | 0.0034 | 0.0065 | 0.0055 | 0.0055 | TRUE | 0.0701 |
| Mccc2 | 1638 | 1142 | 1117 | 1351 | 1245 | 1329 | 1272 | 1215 | 1205 | 1345 | 1404 | 1312 | 1292.22 | 1231 | 1354 | 1.099 | 0.136 | 0.05394665 | 0.14207841 | 0 | 0.0043 | 0.0028 | 0.0028 | TRUE | 0.0127 |

| | | | | | | | | | | | | | | | | | | | | | | | | | |
|---|---|---|---|---|---|---|---|---|---|---|---|---|---|---|---|---|---|---|---|---|---|---|---|---|---|
| Mcee | 587 | 446 | 436 | 531 | 499 | 533 | 456 | 474 | 470 | 529 | 563 | 526 | 503.06 | 467 | 539 | 1.155 | 0.208 | 0.02919761 | 0.090801 | 0 | 0.0071 | 0.0046 | 0.0046 | TRUE | 0.0168 |
| Mcf2 | 185 | 122 | 121 | 213 | 157 | 190 | 144 | 130 | 131 | 212 | 177 | 188 | 163.45 | 135 | 192 | 1.425 | 0.511 | 0.00140502 | 0.00919927 | 0 | 0.0169 | 0.0122 | 0.0122 | TRUE | 0.0882 |
| Mcf2l | 1607 | 1291 | 1656 | 2014 | 1862 | 2226 | 1248 | 1373 | 1787 | 2005 | 2101 | 2197 | 1785.13 | 1469 | 2101 | 1.432 | 0.518 | 2.65E-06 | 6.29E-05 | 0.0186 | 0.0038 | 0.0082 | 0.0082 | TRUE | 0.4284 |
| Mcfd2 | 1798 | 1296 | 1294 | 1343 | 1165 | 1286 | 1396 | 1379 | 1396 | 1337 | 1314 | 1269 | 1348.62 | 1390 | 1307 | 0.94 | -0.09 | 0.1816396 | 0.33724867 | 0 | 0.0042 | 0.0025 | 0.0025 | TRUE | 0.0077 |
| Mchr1 | 3 | 0 | 3 | 3 | 1 | 3 | 2 | 0 | 3 | 3 | 1 | 3 | 2.11 | 2 | 2 | 1.27 | 0.345 | 0.79980855 | NA | 0 | 1.1272 | 0.8557 | 0.8557 | TRUE | 0.5723 |
| Mcin | 6 | 8 | 7 | 8 | 8 | 2 | 5 | 9 | 8 | 8 | 9 | 2 | 6.61 | 7 | 6 | 0.919 | -0.122 | 0.87633863 | NA | 0 | 0.3608 | 0.293 | 0.293 | TRUE | 0.9239 |
| Mcl1 | 36527 | 28672 | 30107 | 34427 | 30897 | 36326 | 28369 | 30498 | 32485 | 34269 | 34855 | 35859 | 32722.53 | 30451 | 34994 | 1.149 | 0.201 | 0.00072147 | 0.0057004 | 0.0025 | 0.0025 | 0.0025 | 0.0025 | TRUE | 0.0439 |
| Mcm10 | 8187 | 5879 | 5693 | 5789 | 5224 | 6038 | 6358 | 6253 | 6143 | 5762 | 5893 | 5960 | 6061.77 | 6251 | 5872 | 0.939 | -0.091 | 0.06707839 | 0.16613264 | 1.00E-04 | 0.0028 | 0.0016 | 0.0016 | TRUE | 0.0033 |
| Mcm2 | 24209 | 17741 | 17309 | 16992 | 15393 | 17850 | 18802 | 18871 | 18676 | 16914 | 17365 | 17621 | 18041.44 | 18783 | 17300 | 0.921 | -0.119 | 0.00805489 | 0.03522906 | 1.00E-04 | 0.0026 | 0.0014 | 0.0014 | TRUE | 0.0047 |
| Mcm3 | 21579 | 16302 | 16771 | 16748 | 15273 | 18014 | 16759 | 17340 | 18096 | 16671 | 17230 | 17783 | 17313.09 | 17398 | 17228 | 0.99 | -0.014 | 0.78986681 | 0.87645107 | 0.001 | 0.0026 | 0.002 | 0.002 | TRUE | 0.015 |
| Mcm3ap | 12310 | 8580 | 8496 | 8373 | 7375 | 8434 | 9561 | 9126 | 9167 | 8334 | 8320 | 8326 | 8805.68 | 9285 | 8327 | 0.897 | -0.157 | 0.00103212 | 0.00739827 | 3.00E-04 | 0.0027 | 0.0015 | 0.0015 | TRUE | 0.0084 |
| Mcm4 | 18312 | 13055 | 13165 | 12777 | 11215 | 12857 | 14222 | 13886 | 14205 | 12718 | 12652 | 12692 | 13395.88 | 14104 | 12687 | 0.899 | -0.153 | 0.0005589 | 0.00469354 | 0 | 0.0026 | 0.0013 | 0.0013 | TRUE | 0.0022 |
| Mcm5 | 31477 | 23301 | 22581 | 22487 | 19765 | 23175 | 24447 | 24785 | 24365 | 22384 | 22297 | 22877 | 23525.72 | 24532 | 22519 | 0.918 | -0.123 | 0.0044022 | 0.02229649 | 1.00E-04 | 0.0025 | 0.0013 | 0.0013 | TRUE | 0.0024 |
| Mcm6 | 27580 | 20570 | 22744 | 22507 | 20115 | 23686 | 21420 | 21880 | 24541 | 22403 | 22692 | 23382 | 22719.64 | 22614 | 22826 | 1.009 | 0.014 | 0.82511539 | 0.89977678 | 0.0024 | 0.0025 | 0.0027 | 0.0027 | TRUE | 0.0679 |
| Mcm7 | 23098 | 17441 | 18007 | 17327 | 16122 | 18727 | 17939 | 18552 | 19429 | 17247 | 18187 | 18486 | 18306.9 | 18640 | 17973 | 0.964 | -0.052 | 0.33157149 | 0.50583321 | 0.0014 | 0.0026 | 0.002 | 0.002 | TRUE | 0.0168 |
| Mcm8 | 1802 | 1289 | 1075 | 1269 | 1129 | 1193 | 1400 | 1371 | 1160 | 1263 | 1274 | 1178 | 1274.17 | 1310 | 1238 | 0.944 | -0.083 | 0.33707002 | 0.51127473 | 0.0053 | 0.0043 | 0.0046 | 0.0046 | TRUE | 0.1191 |
| Mcm9 | 1257 | 893 | 713 | 746 | 668 | 834 | 976 | 950 | 769 | 743 | 754 | 823 | 835.81 | 898 | 773 | 0.86 | -0.218 | 0.03343812 | 0.1003694 | 0.0087 | 0.0053 | 0.0064 | 0.0064 | TRUE | 0.1851 |
| Mcmbp | 9584 | 7009 | 7500 | 7460 | 6870 | 7715 | 7443 | 7455 | 8092 | 7426 | 7750 | 7616 | 7630.49 | 7663 | 7597 | 0.991 | -0.012 | 0.82337409 | 0.89863229 | 0.001 | 0.0027 | 0.0021 | 0.0021 | TRUE | 0.0291 |
| Mcmdc2 | 144 | 122 | 157 | 154 | 126 | 152 | 112 | 130 | 169 | 153 | 142 | 150 | 142.75 | 137 | 148 | 1.09 | 0.125 | 0.50524275 | 0.66871328 | 0.0166 | 0.019 | 0.0183 | 0.0183 | TRUE | 0.4354 |
| Mcoln1 | 2502 | 1909 | 2049 | 2023 | 1909 | 2124 | 1943 | 2031 | 2211 | 2014 | 2154 | 2097 | 2074.76 | 2062 | 2088 | 1.013 | 0.019 | 0.78380741 | 0.87273272 | 0.0023 | 0.0036 | 0.0031 | 0.0031 | TRUE | 0.0483 |
| Mcoln2 | 5 | 3 | 2 | 4 | 2 | 4 | 4 | 3 | 2 | 4 | 2 | 4 | 3.24 | 3 | 3 | 1.097 | 0.134 | 0.89705218 | NA | 0 | 0.7347 | 0.4635 | 0.4635 | TRUE | 0.1088 |
| Mcoln3 | 72 | 45 | 80 | 70 | 72 | 77 | 56 | 48 | 86 | 70 | 81 | 76 | 69.5 | 63 | 76 | 1.198 | 0.26 | 0.32431587 | 0.49820569 | 0.0334 | 0.0365 | 0.0357 | 0.0357 | TRUE | 0.8306 |
| Mcph1 | 7774 | 5513 | 5215 | 5237 | 4880 | 5528 | 6038 | 5864 | 5627 | 5213 | 5505 | 5457 | 5617.3 | 5843 | 5392 | 0.922 | -0.116 | 0.03405961 | 0.10184393 | 8.00E-04 | 0.0028 | 0.002 | 0.002 | TRUE | 0.0127 |
| Mcpt-ps1 | 0 | 0 | 0 | 0 | 0 | 0 | 0 | 0 | 0 | 0 | 0 | 0 | 0 | 0 | 0 | NA | NA | NA | NA | NA | NA | NA | NA | NA | NA |
| Mcpt1 | 0 | 0 | 0 | 0 | 0 | 0 | 0 | 0 | 0 | 0 | 0 | 0 | 0 | 0 | 0 | NA | NA | NA | NA | NA | NA | NA | NA | NA | NA |
| Mcpt2 | 0 | 0 | 0 | 0 | 0 | 0 | 0 | 0 | 0 | 0 | 0 | 0 | 0 | 0 | 0 | NA | NA | NA | NA | NA | NA | NA | NA | NA | NA |
| Mcpt4 | 0 | 0 | 0 | 0 | 0 | 0 | 0 | 0 | 0 | 0 | 0 | 0 | 0 | 0 | 0 | NA | NA | NA | NA | NA | NA | NA | NA | NA | NA |
| Mcpt8 | 0 | 0 | 0 | 0 | 0 | 0 | 0 | 0 | 0 | 0 | 0 | 0 | 0 | 0 | 0 | NA | NA | NA | NA | NA | NA | NA | NA | NA | NA |
| Mcpt9 | 0 | 0 | 0 | 0 | 0 | 0 | 0 | 0 | 0 | 0 | 0 | 0 | 0 | 0 | 0 | NA | NA | NA | NA | NA | NA | NA | NA | NA | NA |
| Mcrs1 | 5833 | 4535 | 4582 | 4595 | 4295 | 4970 | 4530 | 4824 | 4944 | 4574 | 4845 | 4906 | 4770.54 | 4766 | 4775 | 1.002 | 0.003 | 0.95617297 | 0.9757358 | 0.0015 | 0.0029 | 0.0023 | 0.0023 | TRUE | 0.0232 |
| Mctp1 | 4 | 2 | 1 | 3 | 1 | 0 | 3 | 2 | 1 | 3 | 1 | 0 | 1.74 | 2 | 1 | 0.643 | -0.637 | 0.67192813 | NA | 0 | 1.3662 | 1.0284 | 1.0284 | TRUE | 0.5289 |
| Mctp2 | 465 | 355 | 471 | 529 | 525 | 621 | 361 | 378 | 508 | 527 | 592 | 613 | 496.47 | 416 | 577 | 1.392 | 0.477 | 0.00031072 | 0.0029907 | 0.0187 | 0.0072 | 0.0106 | 0.0106 | TRUE | 0.4313 |
| Mcts1 | 920 | 613 | 654 | 700 | 551 | 735 | 715 | 652 | 706 | 697 | 622 | 726 | 686.02 | 691 | 682 | 0.987 | -0.02 | 0.83436819 | 0.90597926 | 0.0028 | 0.0059 | 0.0049 | 0.0049 | TRUE | 0.0676 |
| Mcts2 | 747 | 593 | 471 | 514 | 451 | 481 | 580 | 631 | 508 | 512 | 509 | 475 | 535.73 | 573 | 499 | 0.869 | -0.202 | 0.05622293 | 0.14672162 | 0.0048 | 0.0069 | 0.0062 | 0.0062 | TRUE | 0.11 |
| Mcu | 1591 | 1160 | 1318 | 1382 | 1309 | 1471 | 1236 | 1234 | 1422 | 1376 | 1477 | 1452 | 1366.01 | 1297 | 1435 | 1.107 | 0.146 | 0.06623788 | 0.16465009 | 0.003 | 0.0042 | 0.0038 | 0.0038 | TRUE | 0.0837 |
| Mdc1 | 15888 | 13034 | 15459 | 15776 | 14701 | 17137 | 12340 | 13864 | 16680 | 15703 | 16584 | 16917 | 15348.05 | 14295 | 16401 | 1.148 | 0.199 | 0.02826426 | 0.08861178 | 0.0125 | 0.0026 | 0.0058 | 0.0058 | TRUE | 0.2606 |
| Mdfi | 607 | 478 | 498 | 657 | 603 | 713 | 471 | 508 | 537 | 654 | 680 | 704 | 592.55 | 505 | 679 | 1.346 | 0.429 | 5.70E-06 | 0.00011773 | 0.0011 | 0.0064 | 0.0047 | 0.0047 | TRUE | 0.0442 |
| Mdfic | 434 | 331 | 300 | 332 | 281 | 332 | 337 | 352 | 324 | 330 | 317 | 328 | 331.34 | 338 | 325 | 0.963 | -0.054 | 0.63111442 | 0.76842307 | 0 | 0.0096 | 0.0061 | 0.0061 | TRUE | 0.015 |
| Mdga1 | 167 | 141 | 136 | 192 | 140 | 150 | 130 | 150 | 147 | 191 | 158 | 148 | 153.92 | 142 | 166 | 1.17 | 0.226 | 0.18501956 | 0.34127842 | 0.0065 | 0.0178 | 0.0145 | 0.0145 | TRUE | 0.1948 |
| Mdga2 | 63 | 43 | 14 | 15 | 16 | 12 | 49 | 46 | 15 | 15 | 18 | 12 | 25.77 | 37 | 15 | 0.404 | -1.307 | 0.00624198 | 0.02906078 | 0.1706 | 0.0944 | 0.1167 | 0.1167 | TRUE | 1.8082 |
| Mdh1 | 7212 | 5011 | 5046 | 5511 | 4685 | 5263 | 5601 | 5330 | 5445 | 5486 | 5285 | 5195 | 5390.36 | 5459 | 5322 | 0.975 | -0.037 | 0.48650575 | 0.65247878 | 4.00E-04 | 0.0029 | 0.0018 | 0.0018 | TRUE | 0.0089 |
| Mdh1b | 6 | 6 | 0 | 6 | 5 | 8 | 5 | 6 | 0 | 6 | 6 | 8 | 5.09 | 4 | 7 | 1.748 | 0.806 | 0.38671636 | NA | 0.0859 | 0.4679 | 0.4157 | 0.4157 | TRUE | 1.0429 |
| Mdh2 | 20381 | 13914 | 13037 | 13482 | 11347 | 13740 | 15829 | 14800 | 14067 | 13420 | 12801 | 13563 | 14080.01 | 14899 | 13261 | 0.89 | -0.168 | 0.00419892 | 0.02143089 | 0.0021 | 0.0026 | 0.0024 | 0.0024 | TRUE | 0.0367 |
| Mdk | 8078 | 5425 | 5125 | 6648 | 5400 | 6471 | 6274 | 5771 | 5530 | 6617 | 6092 | 6388 | 6111.87 | 5858 | 6366 | 1.086 | 0.119 | 0.06259897 | 0.158288 | 0.0027 | 0.0028 | 0.0028 | 0.0028 | TRUE | 0.0476 |
| Mdm1 | 2299 | 1745 | 1812 | 1715 | 1581 | 1889 | 1786 | 1856 | 1955 | 1707 | 1784 | 1865 | 1825.36 | 1866 | 1785 | 0.958 | -0.063 | 0.36456201 | 0.53929671 | 0.0015 | 0.0037 | 0.0029 | 0.0029 | TRUE | 0.0212 |
| Mdm2 | 8388 | 5844 | 6223 | 6186 | 5115 | 5772 | 6515 | 6216 | 6715 | 6158 | 5770 | 5698 | 6178.5 | 6482 | 5875 | 0.906 | -0.142 | 0.01406028 | 0.05325542 | 0.0014 | 0.0028 | 0.0022 | 0.0022 | TRUE | 0.0216 |
| Mdm4 | 5872 | 4955 | 5677 | 6452 | 5613 | 6418 | 4561 | 5271 | 6125 | 6422 | 6332 | 6336 | 5841.08 | 5319 | 6363 | 1.197 | 0.259 | 0.00351526 | 0.01866403 | 0.0106 | 0.0028 | 0.0055 | 0.0055 | TRUE | 0.2146 |
| Mdn1 | 19479 | 13224 | 10815 | 14340 | 13334 | 15357 | 15129 | 14066 | 11669 | 14274 | 15042 | 15160 | 14223.31 | 13621 | 14825 | 1.088 | 0.122 | 0.14298466 | 0.28582933 | 0.0087 | 0.0026 | 0.0049 | 0.0049 | TRUE | 0.1919 |
| Mdp1 | 1681 | 1254 | 1357 | 1402 | 1193 | 1410 | 1306 | 1334 | 1464 | 1396 | 1346 | 1392 | 1372.81 | 1368 | 1378 | 1.008 | 0.012 | 0.87006319 | 0.92811502 | 0.0014 | 0.0042 | 0.0031 | 0.0031 | TRUE | 0.045 |
| Me1 | 3715 | 2728 | 3091 | 3522 | 2929 | 3132 | 2885 | 2902 | 3335 | 3506 | 3304 | 3092 | 3170.66 | 3041 | 3301 | 1.086 | 0.119 | 0.11882889 | 0.25039384 | 0.005 | 0.0032 | 0.0039 | 0.0039 | TRUE | 0.0865 |
| Me2 | 8572 | 6205 | 6224 | 6671 | 5889 | 6852 | 6658 | 6600 | 6716 | 6640 | 6643 | 6764 | 6670.17 | 6658 | 6682 | 1.004 | 0.005 | 0.90899382 | 0.94965636 | 0 | 0.0028 | 0.0015 | 0.0015 | TRUE | 0.0014 |
| Me3 | 16 | 5 | 6 | 7 | 6 | 6 | 12 | 5 | 6 | 7 | 7 | 6 | 7.31 | 8 | 7 | 0.796 | -0.329 | 0.64658053 | NA | 0 | 0.3265 | 0.2329 | 0.2329 | TRUE | 0.8369 |
| Mea1 | 1771 | 1424 | 1452 | 1487 | 1385 | 1606 | 1375 | 1515 | 1567 | 1480 | 1562 | 1585 | 1514.13 | 1486 | 1542 | 1.04 | 0.056 | 0.4545087 | 0.62428338 | 0.0024 | 0.004 | 0.0034 | 0.0034 | TRUE | 0.0527 |
| Meaf6 | 859 | 606 | 586 | 660 | 559 | 665 | 667 | 645 | 632 | 657 | 631 | 656 | 648.01 | 648 | 648 | 0.999 | -0.001 | 0.9928242 | 0.99592003 | 0 | 0.0061 | 0.0038 | 0.0038 | TRUE | 0.0084 |
| Mecom | 7 | 7 | 7 | 5 | 7 | 4 | 5 | 7 | 8 | 5 | 8 | 4 | 6.21 | 7 | 6 | 0.823 | -0.281 | 0.71313799 | NA | 0 | 0.3841 | 0.2552 | 0.2552 | TRUE | 0.2521 |
| Mecp2 | 1427 | 1106 | 1070 | 1225 | 1033 | 1270 | 1108 | 1176 | 1155 | 1219 | 1165 | 1254 | 1179.61 | 1146 | 1213 | 1.059 | 0.083 | 0.25379897 | 0.4228898 | 0 | 0.0044 | 0.0029 | 0.0029 | TRUE | 0.0138 |
| Mecr | 1305 | 909 | 938 | 1027 | 919 | 960 | 1014 | 967 | 1012 | 1022 | 1037 | 948 | 999.87 | 998 | 1002 | 1.004 | 0.006 | 0.94070906 | 0.96674968 | 0 | 0.0048 | 0.0033 | 0.0033 | TRUE | 0.0274 |
| Med1 | 9952 | 6578 | 5942 | 7106 | 6234 | 6990 | 7729 | 6997 | 6411 | 7073 | 7033 | 6900 | 7023.95 | 7046 | 7002 | 0.993 | -0.01 | 0.89087208 | 0.93970406 | 0.0043 | 0.0028 | 0.0034 | 0.0034 | TRUE | 0.0886 |
| Med10 | 2181 | 1535 | 1377 | 1313 | 1231 | 1376 | 1694 | 1633 | 1486 | 1307 | 1389 | 1358 | 1477.73 | 1604 | 1351 | 0.841 | -0.249 | 0.00085219 | 0.00644459 | 0.0021 | 0.004 | 0.0033 | 0.0033 | TRUE | 0.0497 |
| Med11 | 1365 | 918 | 908 | 1018 | 886 | 934 | 1060 | 976 | 980 | 1013 | 999 | 922 | 991.86 | 1005 | 978 | 0.972 | -0.042 | 0.60492972 | 0.74963243 | 0.0015 | 0.0048 | 0.0036 | 0.0036 | TRUE | 0.0307 |
| Med12 | 3302 | 2390 | 2434 | 2862 | 2708 | 3211 | 2565 | 2542 | 2626 | 2849 | 3055 | 3170 | 2801.08 | 2578 | 3025 | 1.173 | 0.231 | 0.00023411 | 0.00241245 | 0.0013 | 0.0033 | 0.0025 | 0.0025 | TRUE | 0.0316 |
| Med12l | 1207 | 872 | 861 | 1028 | 886 | 874 | 937 | 928 | 929 | 1023 | 999 | 863 | 946.59 | 931 | 962 | 1.032 | 0.046 | 0.59627621 | 0.74244933 | 0.0032 | 0.0049 | 0.0043 | 0.0043 | TRUE | 0.0982 |
| Med13 | 8425 | 5719 | 5533 | 7214 | 6011 | 6572 | 6543 | 6083 | 5970 | 7181 | 6781 | 6488 | 6507.67 | 6199 | 6817 | 1.099 | 0.137 | 0.02736803 | 0.08654467 | 0.0023 | 0.0028 | 0.0026 | 0.0026 | TRUE | 0.0292 |
| Med13l | 9465 | 6494 | 6403 | 7392 | 6395 | 7083 | 7351 | 6908 | 6909 | 7358 | 7214 | 6992 | 7121.94 | 7056 | 7188 | 1.018 | 0.026 | 0.62186051 | 0.76142749 | 7.00E-04 | 0.0028 | 0.0019 | 0.0019 | TRUE | 0.0166 |
| Med14 | 2868 | 2019 | 1843 | 2007 | 1729 | 1932 | 2227 | 2148 | 1989 | 1998 | 1950 | 1907 | 2036.51 | 2121 | 1952 | 0.919 | -0.121 | 0.07232709 | 0.17575928 | 0.0015 | 0.0036 | 0.0028 | 0.0028 | TRUE | 0.0359 |
| Med15 | 4074 | 2990 | 2966 | 3139 | 2849 | 3437 | 3164 | 3180 | 3200 | 3125 | 3214 | 3393 | 3212.7 | 3181 | 3244 | 1.02 | 0.028 | 0.62674845 | 0.76482743 | 6.00E-04 | 0.0032 | 0.0021 | 0.0021 | TRUE | 0.0198 |
| Med16 | 3063 | 2255 | 2137 | 2282 | 2152 | 2280 | 2379 | 2399 | 2306 | 2272 | 2428 | 2251 | 2338.87 | 2361 | 2317 | 0.981 | -0.028 | 0.64912797 | 0.7820068 | 6.00E-04 | 0.0034 | 0.0023 | 0.0023 | TRUE | 0.021 |
| Med17 | 4526 | 3244 | 2989 | 3011 | 2790 | 3170 | 3515 | 3451 | 3225 | 2997 | 3147 | 3129 | 3244.12 | 3397 | 3091 | 0.91 | -0.137 | 0.02327507 | 0.07725745 | 0.0011 | 0.0032 | 0.0023 | 0.0023 | TRUE | 0.0236 |
| Med18 | 789 | 575 | 522 | 584 | 498 | 522 | 613 | 612 | 563 | 581 | 562 | 515 | 574.34 | 596 | 553 | 0.927 | -0.11 | 0.25163469 | 0.42058223 | 0.0012 | 0.0066 | 0.0049 | 0.0049 | TRUE | 0.0421 |

| | | | | | | | | | | | | | | | | | | | | | | | | | |
|---|---|---|---|---|---|---|---|---|---|---|---|---|---|---|---|---|---|---|---|---|---|---|---|---|---|
| Med19 | 3873 | 2724 | 2711 | 2764 | 2454 | 2807 | 3008 | 2897 | 2925 | 2751 | 2768 | 2771 | 2853.54 | 2943 | 2763 | 0.939 | -0.092 | 0.09677595 | 0.2161617 | 0 | 0.0033 | 0.0018 | 0.0018 | TRUE | 0.0046 |
| Med20 | 1897 | 1331 | 1209 | 1364 | 1166 | 1337 | 1473 | 1416 | 1305 | 1358 | 1315 | 1320 | 1364.42 | 1398 | 1331 | 0.951 | -0.072 | 0.32597684 | 0.50001404 | 0.0014 | 0.0042 | 0.0031 | 0.0031 | TRUE | 0.0406 |
| Med21 | 1285 | 791 | 807 | 875 | 718 | 808 | 998 | 841 | 871 | 871 | 810 | 798 | 864.78 | 903 | 826 | 0.913 | -0.131 | 0.15176458 | 0.29812687 | 0.0044 | 0.0052 | 0.0049 | 0.0049 | TRUE | 0.1041 |
| Med22 | 2468 | 1639 | 1415 | 1430 | 1294 | 1489 | 1917 | 1743 | 1527 | 1423 | 1460 | 1470 | 1590 | 1729 | 1451 | 0.838 | -0.255 | 0.00310463 | 0.01696923 | 0.0056 | 0.0039 | 0.0047 | 0.0047 | TRUE | 0.1256 |
| Med23 | 3736 | 2762 | 2576 | 2560 | 2431 | 2771 | 2902 | 2938 | 2779 | 2548 | 2742 | 2735 | 2774.17 | 2873 | 2675 | 0.931 | -0.103 | 0.09091762 | 0.2062348 | 9.00E-04 | 0.0033 | 0.0023 | 0.0023 | TRUE | 0.0211 |
| Med24 | 8696 | 6544 | 6330 | 6695 | 6294 | 7296 | 6754 | 6961 | 6830 | 6664 | 7100 | 7202 | 6918.56 | 6848 | 6989 | 1.021 | 0.029 | 0.58176518 | 0.73222699 | 7.00E-04 | 0.0028 | 0.0019 | 0.0019 | TRUE | 0.0202 |
| Med25 | 6280 | 4785 | 5070 | 4943 | 4599 | 5405 | 4877 | 5090 | 5470 | 4920 | 5188 | 5336 | 5146.94 | 5146 | 5148 | 1.001 | 0.001 | 0.98617135 | 0.99318962 | 0.0024 | 0.0029 | 0.0027 | 0.0027 | TRUE | 0.037 |
| Med26 | 2882 | 2242 | 2105 | 2046 | 1812 | 2097 | 2238 | 2385 | 2271 | 2037 | 2044 | 2070 | 2174.19 | 2298 | 2050 | 0.893 | -0.164 | 0.00676351 | 0.03095447 | 0 | 0.0035 | 0.0022 | 0.0022 | TRUE | 0.0131 |
| Med27 | 939 | 665 | 633 | 659 | 567 | 710 | 729 | 707 | 683 | 656 | 640 | 701 | 686.02 | 706 | 666 | 0.942 | -0.087 | 0.31839122 | 0.49229399 | 0 | 0.0059 | 0.004 | 0.004 | TRUE | 0.0257 |
| Med28 | 2990 | 1987 | 1977 | 2106 | 1829 | 2013 | 2322 | 2114 | 2133 | 2096 | 2063 | 1987 | 2119.28 | 2190 | 2049 | 0.935 | -0.097 | 0.14253003 | 0.28511991 | 0.0014 | 0.0035 | 0.0027 | 0.0027 | TRUE | 0.035 |
| Med29 | 959 | 672 | 530 | 559 | 505 | 581 | 745 | 715 | 572 | 556 | 570 | 574 | 621.86 | 677 | 567 | 0.835 | -0.26 | 0.01737706 | 0.0622638 | 0.0085 | 0.0062 | 0.007 | 0.007 | TRUE | 0.2141 |
| Med30 | 1601 | 1160 | 1288 | 1353 | 1166 | 1397 | 1243 | 1234 | 1390 | 1347 | 1315 | 1379 | 1318.04 | 1289 | 1347 | 1.046 | 0.065 | 0.38873702 | 0.56276899 | 0.0017 | 0.0042 | 0.0033 | 0.0033 | TRUE | 0.0549 |
| Med31 | 1284 | 851 | 906 | 932 | 746 | 872 | 997 | 905 | 978 | 928 | 842 | 861 | 918.34 | 960 | 877 | 0.913 | -0.132 | 0.11066548 | 0.23828588 | 0.0015 | 0.005 | 0.0038 | 0.0038 | TRUE | 0.031 |
| Med4 | 3392 | 2334 | 2202 | 2199 | 1986 | 2294 | 2634 | 2483 | 2376 | 2189 | 2240 | 2265 | 2364.47 | 2498 | 2231 | 0.893 | -0.164 | 0.01047521 | 0.04271676 | 0.0012 | 0.0034 | 0.0025 | 0.0025 | TRUE | 0.0286 |
| Med6 | 1756 | 1207 | 1121 | 1141 | 975 | 1086 | 1364 | 1284 | 1210 | 1136 | 1100 | 1072 | 1194.16 | 1286 | 1103 | 0.857 | -0.223 | 0.00348023 | 0.01853312 | 0.0016 | 0.0044 | 0.0034 | 0.0034 | TRUE | 0.0352 |
| Med7 | 2864 | 1978 | 1816 | 2003 | 1669 | 1971 | 2224 | 2104 | 1959 | 1994 | 1883 | 1946 | 2018.34 | 2096 | 1941 | 0.925 | -0.112 | 0.10738854 | 0.23309102 | 0.002 | 0.0036 | 0.003 | 0.003 | TRUE | 0.0391 |
| Med8 | 2320 | 1583 | 1489 | 1506 | 1405 | 1677 | 1802 | 1684 | 1607 | 1499 | 1585 | 1655 | 1638.63 | 1698 | 1580 | 0.93 | -0.105 | 0.15511902 | 0.30271931 | 0.0024 | 0.0039 | 0.0033 | 0.0033 | TRUE | 0.036 |
| Med9 | 3425 | 2388 | 2199 | 2436 | 2212 | 2477 | 2660 | 2540 | 2373 | 2425 | 2495 | 2445 | 2489.7 | 2524 | 2455 | 0.972 | -0.041 | 0.52211138 | 0.68332002 | 0.0014 | 0.0034 | 0.0026 | 0.0026 | TRUE | 0.0333 |
| Medag | 12 | 8 | 6 | 11 | 8 | 6 | 9 | 9 | 6 | 11 | 9 | 6 | 8.37 | 8 | 9 | 1.058 | 0.081 | 0.90193164 | NA | 0 | 0.2857 | 0.1929 | 0.1929 | TRUE | 0.2468 |
| Mef2a | 662 | 458 | 422 | 464 | 416 | 453 | 514 | 487 | 455 | 462 | 469 | 447 | 472.5 | 485 | 459 | 0.944 | -0.083 | 0.41029446 | 0.58307953 | 0 | 0.0074 | 0.0051 | 0.0051 | TRUE | 0.034 |
| Mef2b | 139 | 116 | 119 | 166 | 136 | 192 | 108 | 123 | 128 | 165 | 153 | 190 | 144.66 | 120 | 169 | 1.42 | 0.506 | 0.00343191 | 0.01836329 | 0 | 0.0188 | 0.0144 | 0.0144 | TRUE | 0.1161 |
| Mef2c | 8 | 2 | 4 | 4 | 5 | 4 | 6 | 2 | 4 | 4 | 6 | 4 | 4.37 | 4 | 5 | 1.049 | 0.068 | 0.94007783 | NA | 0 | 0.5446 | 0.369 | 0.369 | TRUE | 0.26 |
| Mef2d | 2036 | 1711 | 1804 | 1878 | 1843 | 1929 | 1581 | 1820 | 1947 | 1869 | 2079 | 1904 | 1866.74 | 1783 | 1951 | 1.095 | 0.131 | 0.12466055 | 0.25931788 | 0.0063 | 0.0037 | 0.0047 | 0.0047 | TRUE | 0.121 |
| Mefv | 0 | 0 | 0 | 1 | 0 | 0 | 0 | 0 | 0 | 1 | 0 | 0 | 0.17 | 0 | 0 | 2.094 | 1.067 | 0.79379329 | NA | 0 | 14.2885 | 10 | 10 | TRUE | 0.3906 |
| Meg3 | 7978 | 8266 | 6929 | 8331 | 7954 | 9554 | 6196 | 8792 | 7476 | 8293 | 8973 | 9431 | 8193.64 | 7488 | 8899 | 1.189 | 0.249 | 0.01395667 | 0.05292096 | 0.0169 | 0.0027 | 0.0073 | 0.0073 | TRUE | 0.2285 |
| Megf10 | 6 | 9 | 4 | 6 | 2 | 6 | 5 | 10 | 4 | 6 | 2 | 6 | 5.45 | 6 | 5 | 0.776 | -0.366 | 0.66350833 | NA | 0 | 0.4373 | 0.3227 | 0.3227 | TRUE | 0.5177 |
| Megf11 | 221 | 208 | 220 | 374 | 352 | 356 | 172 | 221 | 237 | 372 | 397 | 351 | 291.84 | 210 | 373 | 1.788 | 0.838 | 3.48E-09 | 2.00E-07 | 0.0114 | 0.0105 | 0.0108 | 0.0108 | TRUE | 0.3087 |
| Megf6 | 964 | 674 | 717 | 1027 | 976 | 919 | 749 | 717 | 774 | 1022 | 1101 | 907 | 878.29 | 747 | 1010 | 1.352 | 0.435 | 2.02E-06 | 5.11E-05 | 0.0044 | 0.0051 | 0.0049 | 0.0049 | TRUE | 0.0955 |
| Megf8 | 3474 | 2503 | 2342 | 2731 | 2178 | 2219 | 2698 | 2662 | 2527 | 2718 | 2457 | 2190 | 2542.24 | 2629 | 2455 | 0.934 | -0.099 | 0.21992475 | 0.38308736 | 0.0058 | 0.0034 | 0.0043 | 0.0043 | TRUE | 0.1088 |
| Megf9 | 199 | 125 | 136 | 142 | 110 | 130 | 155 | 133 | 147 | 141 | 124 | 128 | 138 | 145 | 131 | 0.905 | -0.143 | 0.39953113 | 0.57288306 | 0 | 0.0196 | 0.0136 | 0.0136 | TRUE | 0.0518 |
| Mei1 | 86 | 73 | 64 | 102 | 86 | 98 | 67 | 78 | 69 | 102 | 97 | 97 | 84.8 | 71 | 99 | 1.387 | 0.472 | 0.02500589 | 0.08141439 | 0 | 0.0304 | 0.0199 | 0.0199 | TRUE | 0.0554 |
| Mei4 | 168 | 103 | 100 | 99 | 98 | 90 | 130 | 110 | 108 | 99 | 111 | 89 | 107.65 | 116 | 100 | 0.851 | -0.232 | 0.23668757 | 0.40326325 | 0 | 0.0244 | 0.0184 | 0.0184 | TRUE | 0.1327 |
| Meig1 | 0 | 0 | 1 | 1 | 1 | 0 | 0 | 0 | 1 | 1 | 1 | 0 | 0.53 | 0 | 1 | 2.036 | 1.026 | 0.70454668 | NA | 0 | 4.4428 | 3.5371 | 3.5371 | TRUE | 0.5365 |
| Meiob | 70 | 50 | 100 | 162 | 136 | 144 | 54 | 53 | 108 | 161 | 153 | 142 | 112.05 | 72 | 152 | 2.136 | 1.095 | 1.26E-05 | 0.00022872 | 0.0698 | 0.0236 | 0.0352 | 0.0352 | TRUE | 1.6873 |
| Meis1 | 84 | 48 | 49 | 33 | 28 | 26 | 65 | 51 | 53 | 33 | 32 | 26 | 43.21 | 56 | 30 | 0.529 | -0.917 | 0.00254897 | 0.01460218 | 0 | 0.0573 | 0.0408 | 0.0408 | TRUE | 0.1809 |
| Meis2 | 130 | 55 | 51 | 53 | 31 | 30 | 101 | 59 | 55 | 53 | 35 | 30 | 55.31 | 72 | 39 | 0.543 | -0.88 | 0.00739134 | 0.03308625 | 0.088 | 0.0453 | 0.0578 | 0.0578 | TRUE | 1.2824 |
| Meis3 | 87 | 63 | 42 | 62 | 32 | 46 | 68 | 67 | 45 | 62 | 36 | 45 | 53.85 | 60 | 48 | 0.796 | -0.33 | 0.26625349 | 0.43666794 | 0.0392 | 0.0464 | 0.0445 | 0.0445 | TRUE | 0.5067 |
| Melk | 6660 | 4782 | 5107 | 5084 | 4473 | 5343 | 5173 | 5087 | 5510 | 5061 | 5046 | 5274 | 5191.75 | 5257 | 5127 | 0.976 | -0.036 | 0.52540884 | 0.68644136 | 9.00E-04 | 0.0029 | 0.0021 | 0.0021 | TRUE | 0.0215 |
| Memo1 | 2224 | 1594 | 1643 | 1775 | 1456 | 1664 | 1727 | 1696 | 1773 | 1767 | 1643 | 1643 | 1707.93 | 1732 | 1684 | 0.973 | -0.04 | 0.54521258 | 0.70230023 | 5.00E-04 | 0.0038 | 0.0026 | 0.0026 | TRUE | 0.0225 |
| Men1 | 4513 | 3278 | 2915 | 2842 | 2716 | 3046 | 3505 | 3487 | 3145 | 2829 | 3064 | 3007 | 3172.8 | 3379 | 2967 | 0.877 | -0.189 | 0.00455246 | 0.02287112 | 0.0022 | 0.0032 | 0.0029 | 0.0029 | TRUE | 0.0441 |
| Meox1 | 49 | 39 | 29 | 70 | 56 | 67 | 38 | 41 | 31 | 70 | 63 | 66 | 51.64 | 37 | 66 | 1.794 | 0.843 | 0.00198785 | 0.01205958 | 0 | 0.0483 | 0.0327 | 0.0327 | TRUE | 0.1169 |
| Meox2 | 5 | 2 | 3 | 3 | 2 | 5 | 4 | 2 | 3 | 3 | 2 | 5 | 3.24 | 3 | 3 | 1.096 | 0.132 | 0.89914175 | NA | 0 | 0.7345 | 0.473 | 0.473 | TRUE | 0.2353 |
| Mep1a | 0 | 0 | 0 | 0 | 0 | 0 | 0 | 0 | 0 | 0 | 0 | 0 | 0 | 0 | 0 | NA | NA | NA | NA | NA | NA | NA | NA | NA | NA |
| Mep1b | 1024 | 849 | 886 | 358 | 374 | 498 | 795 | 903 | 956 | 356 | 422 | 492 | 654.04 | 885 | 423 | 0.479 | -1.062 | 1.33E-18 | 3.58E-16 | 0.0144 | 0.0061 | 0.0087 | 0.0087 | TRUE | 0.2352 |
| Mepce | 2467 | 1834 | 1616 | 1631 | 1519 | 1693 | 1916 | 1951 | 1744 | 1623 | 1714 | 1671 | 1769.8 | 1870 | 1669 | 0.892 | -0.165 | 0.01841128 | 0.06499196 | 0.0016 | 0.0038 | 0.0029 | 0.0029 | TRUE | 0.0414 |
| Mepe | 0 | 0 | 0 | 0 | 0 | 0 | 0 | 0 | 0 | 0 | 0 | 0 | 0 | 0 | 0 | NA | NA | NA | NA | NA | NA | NA | NA | NA | NA |
| Mertk | 486 | 312 | 310 | 377 | 353 | 414 | 377 | 332 | 334 | 375 | 398 | 409 | 371 | 348 | 394 | 1.129 | 0.176 | 0.11912523 | 0.25072257 | 0 | 0.0088 | 0.0064 | 0.0064 | TRUE | 0.0676 |
| Mesdc1 | 1075 | 793 | 709 | 756 | 679 | 708 | 835 | 844 | 765 | 753 | 766 | 699 | 776.8 | 815 | 739 | 0.907 | -0.142 | 0.10062494 | 0.22208115 | 0.0012 | 0.0055 | 0.0041 | 0.0041 | TRUE | 0.0325 |
| Mesdc2 | 7188 | 5298 | 4899 | 5502 | 4885 | 5458 | 5583 | 5635 | 5286 | 5477 | 5511 | 5388 | 5479.89 | 5501 | 5459 | 0.992 | -0.012 | 0.82584893 | 0.90019138 | 4.00E-04 | 0.0029 | 0.0018 | 0.0018 | TRUE | 0.0141 |
| Mesp1 | 22 | 14 | 7 | 19 | 12 | 16 | 17 | 15 | 8 | 19 | 14 | 16 | 14.63 | 13 | 16 | 1.209 | 0.273 | 0.59768016 | NA | 0 | 0.1644 | 0.1252 | 0.1252 | TRUE | 0.5366 |
| Mesp2 | 197 | 125 | 98 | 120 | 100 | 124 | 153 | 133 | 106 | 119 | 113 | 122 | 124.39 | 131 | 118 | 0.901 | -0.15 | 0.43403696 | 0.60576998 | 0.0111 | 0.0215 | 0.0184 | 0.0184 | TRUE | 0.2501 |
| Mest | 11946 | 8513 | 7848 | 9122 | 8385 | 9708 | 9278 | 9055 | 8468 | 9080 | 9459 | 9583 | 9153.92 | 8934 | 9374 | 1.049 | 0.069 | 0.21454754 | 0.37727953 | 0.0013 | 0.0027 | 0.0021 | 0.0021 | TRUE | 0.0253 |
| Met | 1078 | 817 | 893 | 751 | 729 | 818 | 837 | 869 | 964 | 748 | 822 | 807 | 841.21 | 890 | 792 | 0.891 | -0.166 | 0.06015436 | 0.15353327 | 0.0028 | 0.0052 | 0.0044 | 0.0044 | TRUE | 0.0615 |
| Metap1 | 5256 | 3827 | 4022 | 3928 | 3712 | 4097 | 4082 | 4071 | 4340 | 3910 | 4188 | 4044 | 4105.73 | 4164 | 4047 | 0.972 | -0.041 | 0.47799896 | 0.64546861 | 9.00E-04 | 0.003 | 0.0022 | 0.0022 | TRUE | 0.0164 |
| Metap1d | 383 | 304 | 256 | 263 | 208 | 248 | 297 | 323 | 276 | 262 | 235 | 245 | 273.05 | 299 | 247 | 0.827 | -0.274 | 0.03217654 | 0.09751892 | 0 | 0.0111 | 0.0081 | 0.0081 | TRUE | 0.0541 |
| Metap2 | 17184 | 12544 | 11843 | 12101 | 10595 | 12047 | 13346 | 13343 | 12778 | 12045 | 11952 | 11892 | 12559.55 | 13156 | 11963 | 0.909 | -0.137 | 0.00328385 | 0.01770504 | 2.00E-04 | 0.0026 | 0.0015 | 0.0015 | TRUE | 0.0077 |
| Metrn | 668 | 505 | 387 | 556 | 454 | 543 | 519 | 537 | 418 | 553 | 512 | 536 | 512.53 | 491 | 534 | 1.086 | 0.119 | 0.29097406 | 0.46307674 | 0.0074 | 0.0071 | 0.0072 | 0.0072 | TRUE | 0.1932 |
| Metrnl | 109 | 89 | 73 | 112 | 81 | 115 | 85 | 95 | 79 | 111 | 91 | 114 | 95.75 | 86 | 105 | 1.229 | 0.297 | 0.14730616 | 0.29189615 | 0 | 0.0272 | 0.0198 | 0.0198 | TRUE | 0.1277 |
| Mettl1 | 1057 | 795 | 664 | 782 | 634 | 747 | 821 | 846 | 716 | 778 | 715 | 737 | 769.01 | 794 | 743 | 0.936 | -0.096 | 0.29856574 | 0.47091152 | 0.0034 | 0.0055 | 0.0048 | 0.0048 | TRUE | 0.0845 |
| Mettl10 | 646 | 474 | 507 | 502 | 388 | 501 | 502 | 504 | 547 | 500 | 438 | 495 | 497.49 | 518 | 478 | 0.924 | -0.114 | 0.2615426 | 0.43171595 | 0.0015 | 0.0072 | 0.0054 | 0.0054 | TRUE | 0.0593 |
| Mettl11b | 0 | 0 | 0 | 0 | 0 | 0 | 0 | 0 | 0 | 0 | 0 | 0 | 0 | 0 | 0 | NA | NA | NA | NA | NA | NA | NA | NA | NA | NA |
| Mettl13 | 2129 | 1461 | 1349 | 1471 | 1235 | 1430 | 1654 | 1554 | 1456 | 1464 | 1393 | 1412 | 1488.7 | 1555 | 1423 | 0.915 | -0.129 | 0.07896576 | 0.18732906 | 0.0018 | 0.004 | 0.0032 | 0.0032 | TRUE | 0.0388 |
| Mettl14 | 2850 | 2143 | 2177 | 2223 | 1966 | 2421 | 2213 | 2279 | 2349 | 2213 | 2218 | 2390 | 2277.07 | 2280 | 2274 | 0.998 | -0.004 | 0.95495292 | 0.9753363 | 0.001 | 0.0035 | 0.0025 | 0.0025 | TRUE | 0.0245 |
| Mettl15 | 391 | 286 | 284 | 347 | 299 | 290 | 304 | 304 | 306 | 345 | 337 | 286 | 313.88 | 305 | 323 | 1.059 | 0.083 | 0.49697373 | 0.66144385 | 0.0021 | 0.01 | 0.0076 | 0.0076 | TRUE | 0.1146 |
| Mettl16 | 7980 | 5803 | 5570 | 6110 | 5347 | 6367 | 6198 | 6173 | 6010 | 6082 | 6032 | 6285 | 6129.89 | 6127 | 6133 | 1.001 | 0.001 | 0.97737061 | 0.98786577 | 2.00E-04 | 0.0028 | 0.0016 | 0.0016 | TRUE | 0.0058 |
| Mettl17 | 625 | 505 | 415 | 369 | 329 | 386 | 485 | 537 | 448 | 367 | 371 | 381 | 431.64 | 490 | 373 | 0.762 | -0.393 | 0.00029894 | 0.00290271 | 0.0023 | 0.0079 | 0.0061 | 0.0061 | TRUE | 0.0784 |
| Mettl18 | 650 | 416 | 358 | 398 | 344 | 360 | 505 | 442 | 386 | 396 | 388 | 355 | 412.2 | 444 | 380 | 0.852 | -0.231 | 0.05846072 | 0.15071711 | 0.0087 | 0.0082 | 0.0083 | 0.0083 | TRUE | 0.1712 |
| Mettl2 | 3498 | 2571 | 2440 | 2839 | 2518 | 2838 | 2717 | 2735 | 2633 | 2826 | 2841 | 2802 | 2758.71 | 2695 | 2823 | 1.047 | 0.067 | 0.2307553 | 0.39651184 | 0 | 0.0033 | 0.0019 | 0.0019 | TRUE | 0.0048 |
| Mettl20 | 260 | 200 | 240 | 314 | 237 | 322 | 202 | 213 | 259 | 313 | 267 | 318 | 261.9 | 225 | 299 | 1.339 | 0.421 | 0.0031594 | 0.01718317 | 0.009 | 0.0115 | 0.0107 | 0.0107 | TRUE | 0.1932 |

| | | | | | | | | | | | | | | | | | | | | | | | | | |
|---|---|---|---|---|---|---|---|---|---|---|---|---|---|---|---|---|---|---|---|---|---|---|---|---|---|
| Mettl21a | 719 | 536 | 547 | 603 | 519 | 558 | 558 | 570 | 590 | 600 | 585 | 551 | 575.88 | 573 | 579 | 1.011 | 0.016 | 0.86406703 | 0.9249633 | 0 | 0.0065 | 0.0043 | 0.0043 | TRUE | 0.0215 |
| Mettl21c | 0 | 0 | 0 | 0 | 0 | 0 | 0 | 0 | 0 | 0 | 0 | 0 | 0 | 0 | 0 | NA | NA | NA | NA | NA | NA | NA | NA | NA | NA |
| Mettl21d | 1252 | 867 | 953 | 981 | 908 | 954 | 972 | 922 | 1028 | 976 | 1024 | 942 | 977.57 | 974 | 981 | 1.006 | 0.009 | 0.90886218 | 0.94963864 | 0.0012 | 0.0049 | 0.0036 | 0.0036 | TRUE | 0.0268 |
| Mettl21e | 2 | 0 | 1 | 3 | 1 | 3 | 2 | 0 | 1 | 3 | 1 | 3 | 1.62 | 1 | 2 | 2.601 | 1.379 | 0.37676984 | NA | 0 | 1.4672 | 1.0091 | 1.0091 | TRUE | 0.2808 |
| Mettl22 | 853 | 618 | 753 | 790 | 707 | 751 | 662 | 657 | 812 | 786 | 798 | 741 | 742.94 | 710 | 775 | 1.092 | 0.127 | 0.20832757 | 0.36989141 | 0.0064 | 0.0056 | 0.006 | 0.006 | TRUE | 0.1814 |
| Mettl23 | 780 | 588 | 617 | 617 | 515 | 677 | 606 | 625 | 666 | 614 | 581 | 668 | 626.74 | 632 | 621 | 0.984 | -0.023 | 0.80714022 | 0.88851318 | 0.002 | 0.0062 | 0.0048 | 0.0048 | TRUE | 0.0521 |
| Mettl24 | 8 | 10 | 6 | 8 | 6 | 5 | 6 | 11 | 6 | 8 | 7 | 5 | 7.17 | 8 | 7 | 0.849 | -0.237 | 0.73971343 | NA | 0 | 0.3332 | 0.225 | 0.225 | TRUE | 0.2808 |
| Mettl25 | 618 | 412 | 409 | 357 | 299 | 302 | 480 | 438 | 441 | 355 | 337 | 298 | 391.72 | 453 | 330 | 0.727 | -0.459 | 5.35E-05 | 0.00074001 | 0.0025 | 0.0085 | 0.0067 | 0.0067 | TRUE | 0.0846 |
| Mettl3 | 1859 | 1273 | 1180 | 1128 | 1018 | 1186 | 1444 | 1354 | 1273 | 1123 | 1148 | 1171 | 1252.18 | 1357 | 1147 | 0.844 | -0.244 | 0.00126006 | 0.0085164 | 0.0016 | 0.0043 | 0.0033 | 0.0033 | TRUE | 0.039 |
| Mettl4 | 1713 | 1256 | 1337 | 1444 | 1220 | 1419 | 1330 | 1336 | 1443 | 1437 | 1376 | 1401 | 1387.24 | 1370 | 1405 | 1.026 | 0.038 | 0.59061892 | 0.73901229 | 5.00E-04 | 0.0041 | 0.0028 | 0.0028 | TRUE | 0.0256 |
| Mettl5 | 2034 | 1516 | 1581 | 1850 | 1535 | 1757 | 1580 | 1613 | 1706 | 1841 | 1732 | 1734 | 1700.95 | 1633 | 1769 | 1.084 | 0.117 | 0.08305878 | 0.19410862 | 8.00E-04 | 0.0038 | 0.0027 | 0.0027 | TRUE | 0.0183 |
| Mettl6 | 1624 | 1077 | 983 | 961 | 865 | 883 | 1261 | 1146 | 1061 | 957 | 976 | 872 | 1045.26 | 1156 | 935 | 0.807 | -0.309 | 0.00050629 | 0.00435139 | 0.0048 | 0.0047 | 0.0047 | 0.0047 | TRUE | 0.0788 |
| Mettl7a1 | 466 | 354 | 417 | 611 | 580 | 582 | 362 | 377 | 450 | 608 | 654 | 575 | 504.24 | 396 | 612 | 1.549 | 0.631 | 1.66E-08 | 7.89E-07 | 0.0061 | 0.0071 | 0.0069 | 0.0069 | TRUE | 0.1591 |
| Mettl7a2 | 22 | 13 | 12 | 20 | 18 | 16 | 17 | 14 | 13 | 20 | 20 | 16 | 16.65 | 15 | 19 | 1.266 | 0.34 | 0.46552368 | NA | 0 | 0.1448 | 0.0959 | 0.0959 | TRUE | 0.123 |
| Mettl7a3 | 9 | 4 | 8 | 7 | 6 | 6 | 7 | 4 | 9 | 7 | 7 | 6 | 6.59 | 7 | 7 | 0.987 | -0.019 | 0.97929031 | NA | 0 | 0.3621 | 0.2381 | 0.2381 | TRUE | 0.2237 |
| Mettl7b | 3 | 3 | 2 | 0 | 0 | 0 | 2 | 3 | 2 | 0 | 0 | 0 | 1.28 | 2 | 0 | 0.074 | -3.747 | 0.0514076 | NA | 0 | 1.8543 | 1.4762 | 1.4762 | TRUE | 0.0718 |
| Mettl8 | 1411 | 901 | 732 | 852 | 740 | 836 | 1096 | 958 | 790 | 848 | 835 | 825 | 892.03 | 948 | 836 | 0.88 | -0.184 | 0.09271388 | 0.20923244 | 0.0125 | 0.0051 | 0.0075 | 0.0075 | TRUE | 0.214 |
| Mettl9 | 3294 | 2456 | 2738 | 2883 | 2448 | 2824 | 2558 | 2612 | 2954 | 2870 | 2762 | 2788 | 2757.34 | 2708 | 2807 | 1.037 | 0.052 | 0.45325366 | 0.62309845 | 0.0026 | 0.0033 | 0.0031 | 0.0031 | TRUE | 0.0759 |
| Mex3a | 3457 | 2720 | 2850 | 3022 | 2576 | 3001 | 2685 | 2893 | 3075 | 3008 | 2906 | 2962 | 2921.63 | 2884 | 2959 | 1.027 | 0.038 | 0.56579981 | 0.72012259 | 0.0022 | 0.0032 | 0.0028 | 0.0028 | TRUE | 0.0456 |
| Mex3b | 165 | 141 | 113 | 129 | 108 | 117 | 128 | 150 | 122 | 128 | 122 | 115 | 127.63 | 133 | 122 | 0.915 | -0.128 | 0.4732528 | 0.64176492 | 0 | 0.021 | 0.015 | 0.015 | TRUE | 0.1164 |
| Mex3c | 4143 | 3005 | 2994 | 3067 | 2592 | 2968 | 3218 | 3196 | 3231 | 3053 | 2924 | 2930 | 3091.9 | 3215 | 2969 | 0.924 | -0.115 | 0.0365522 | 0.10745384 | 0 | 0.0032 | 0.0018 | 0.0018 | TRUE | 0.0075 |
| Mex3d | 2139 | 1702 | 1886 | 1772 | 1625 | 1963 | 1661 | 1810 | 2035 | 1764 | 1833 | 1938 | 1840.24 | 1835 | 1845 | 1.006 | 0.009 | 0.91454778 | 0.9529197 | 0.0058 | 0.0037 | 0.0045 | 0.0045 | TRUE | 0.1089 |
| Mfap1a | 735 | 541 | 473 | 485 | 424 | 529 | 571 | 575 | 510 | 483 | 478 | 522 | 523.33 | 552 | 494 | 0.895 | -0.16 | 0.10714547 | 0.23265586 | 0.0014 | 0.007 | 0.0052 | 0.0052 | TRUE | 0.0497 |
| Mfap1b | 313 | 189 | 182 | 171 | 170 | 181 | 243 | 201 | 196 | 170 | 192 | 179 | 196.86 | 213 | 180 | 0.84 | -0.252 | 0.09859259 | 0.21878722 | 0.0052 | 0.0145 | 0.0116 | 0.0116 | TRUE | 0.1733 |
| Mfap2 | 65 | 47 | 56 | 65 | 47 | 92 | 50 | 50 | 60 | 65 | 53 | 91 | 61.57 | 53 | 70 | 1.303 | 0.382 | 0.16288941 | 0.3133243 | 0.0289 | 0.0409 | 0.0375 | 0.0375 | TRUE | 0.4942 |
| Mfap3 | 4294 | 3279 | 2886 | 3240 | 2809 | 3167 | 3335 | 3488 | 3114 | 3225 | 3169 | 3126 | 3242.84 | 3312 | 3173 | 0.958 | -0.062 | 0.31752727 | 0.49148985 | 0.0013 | 0.0032 | 0.0025 | 0.0025 | TRUE | 0.0327 |
| Mfap3l | 383 | 281 | 277 | 266 | 221 | 280 | 297 | 299 | 299 | 265 | 249 | 276 | 280.95 | 298 | 263 | 0.884 | -0.178 | 0.14138522 | 0.28358392 | 0 | 0.0109 | 0.007 | 0.007 | TRUE | 0.0238 |
| Mfap4 | 101 | 60 | 53 | 99 | 87 | 89 | 78 | 64 | 57 | 99 | 98 | 88 | 80.67 | 66 | 95 | 1.416 | 0.501 | 0.02570906 | 0.08285449 | 0 | 0.0318 | 0.0238 | 0.0238 | TRUE | 0.25 |
| Mfap5 | 282 | 188 | 172 | 194 | 195 | 213 | 219 | 200 | 186 | 193 | 220 | 210 | 204.65 | 202 | 208 | 1.027 | 0.039 | 0.78914741 | 0.87619699 | 0 | 0.014 | 0.0102 | 0.0102 | TRUE | 0.0682 |
| Mff | 3787 | 2684 | 3140 | 3287 | 2869 | 3333 | 2941 | 2855 | 3388 | 3272 | 3237 | 3290 | 3163.79 | 3061 | 3266 | 1.067 | 0.094 | 0.193696 | 0.35249463 | 0.0039 | 0.0032 | 0.0035 | 0.0035 | TRUE | 0.1047 |
| Mfge8 | 9504 | 7358 | 8897 | 8752 | 7694 | 8356 | 7381 | 7827 | 9600 | 8712 | 8680 | 8249 | 8407.96 | 8269 | 8547 | 1.034 | 0.048 | 0.57825386 | 0.72941333 | 0.0084 | 0.0027 | 0.0052 | 0.0052 | TRUE | 0.2416 |
| Mfhas1 | 775 | 629 | 561 | 534 | 459 | 468 | 602 | 669 | 605 | 532 | 518 | 462 | 564.6 | 625 | 504 | 0.806 | -0.311 | 0.00178972 | 0.01109301 | 0.0027 | 0.0066 | 0.0054 | 0.0054 | TRUE | 0.0626 |
| Mfi2 | 70 | 55 | 87 | 141 | 108 | 134 | 54 | 59 | 94 | 140 | 122 | 132 | 100.2 | 69 | 131 | 1.925 | 0.945 | 4.41E-05 | 0.0006306 | 0.0319 | 0.0261 | 0.0275 | 0.0275 | TRUE | 0.8667 |
| Mfn1 | 4480 | 3105 | 2936 | 3038 | 2619 | 3100 | 3479 | 3303 | 3168 | 3024 | 2955 | 3060 | 3164.8 | 3317 | 3013 | 0.908 | -0.139 | 0.02039206 | 0.07016459 | 0.001 | 0.0032 | 0.0023 | 0.0023 | TRUE | 0.0229 |
| Mfn2 | 2337 | 1672 | 1378 | 1489 | 1348 | 1594 | 1815 | 1778 | 1487 | 1482 | 1521 | 1574 | 1609.46 | 1693 | 1526 | 0.9 | -0.152 | 0.07279092 | 0.17652139 | 0.0052 | 0.0039 | 0.0045 | 0.0045 | TRUE | 0.1357 |
| Mfng | 853 | 623 | 717 | 779 | 673 | 848 | 662 | 663 | 774 | 775 | 759 | 837 | 745.09 | 700 | 790 | 1.132 | 0.179 | 0.05807294 | 0.1502762 | 0.004 | 0.0056 | 0.0051 | 0.0051 | TRUE | 0.0989 |
| Mfrp | 0 | 0 | 0 | 0 | 0 | 0 | 0 | 0 | 0 | 0 | 0 | 0 | 0 | 0 | 0 | NA | NA | NA | NA | NA | NA | NA | NA | NA | NA |
| Mfsd1 | 4942 | 3417 | 3253 | 3316 | 2915 | 3165 | 3838 | 3635 | 3510 | 3301 | 3288 | 3124 | 3449.39 | 3661 | 3238 | 0.884 | -0.178 | 0.00323483 | 0.01750017 | 0.0011 | 0.0031 | 0.0023 | 0.0023 | TRUE | 0.0223 |
| Mfsd10 | 2043 | 1448 | 1307 | 1289 | 1219 | 1377 | 1587 | 1540 | 1410 | 1283 | 1375 | 1359 | 1425.79 | 1512 | 1339 | 0.885 | -0.177 | 0.01735287 | 0.06222392 | 0.0019 | 0.0041 | 0.0033 | 0.0033 | TRUE | 0.0415 |
| Mfsd11 | 1001 | 751 | 745 | 768 | 707 | 855 | 777 | 799 | 804 | 764 | 798 | 844 | 797.69 | 793 | 802 | 1.012 | 0.017 | 0.84068881 | 0.90926714 | 0 | 0.0054 | 0.0036 | 0.0036 | TRUE | 0.0255 |
| Mfsd12 | 349 | 234 | 171 | 233 | 201 | 221 | 271 | 249 | 185 | 232 | 227 | 218 | 230.22 | 235 | 226 | 0.957 | -0.063 | 0.68794467 | 0.80889723 | 0.0157 | 0.0127 | 0.0136 | 0.0136 | TRUE | 0.3772 |
| Mfsd2a | 118 | 65 | 50 | 50 | 35 | 60 | 92 | 69 | 54 | 50 | 39 | 59 | 60.54 | 72 | 49 | 0.689 | -0.538 | 0.05876676 | 0.1511267 | 0.0407 | 0.0416 | 0.0413 | 0.0413 | TRUE | 0.2486 |
| Mfsd2b | 97 | 59 | 73 | 78 | 80 | 72 | 75 | 63 | 79 | 78 | 90 | 71 | 75.97 | 72 | 80 | 1.098 | 0.135 | 0.55506783 | 0.71032183 | 0 | 0.0336 | 0.0246 | 0.0246 | TRUE | 0.1193 |
| Mfsd3 | 379 | 275 | 264 | 338 | 297 | 297 | 294 | 293 | 285 | 336 | 335 | 293 | 306.07 | 291 | 321 | 1.105 | 0.145 | 0.22847353 | 0.39382191 | 0 | 0.0102 | 0.0071 | 0.0071 | TRUE | 0.0689 |
| Mfsd4 | 98 | 92 | 82 | 145 | 126 | 130 | 76 | 98 | 88 | 144 | 142 | 128 | 112.88 | 87 | 138 | 1.588 | 0.667 | 0.00049041 | 0.00425074 | 0 | 0.0234 | 0.0171 | 0.0171 | TRUE | 0.142 |
| Mfsd5 | 2448 | 1817 | 1593 | 1729 | 1541 | 1832 | 1901 | 1933 | 1719 | 1721 | 1738 | 1808 | 1803.45 | 1851 | 1756 | 0.948 | -0.077 | 0.27594975 | 0.4470764 | 0.0018 | 0.0037 | 0.003 | 0.003 | TRUE | 0.0461 |
| Mfsd6 | 228 | 155 | 216 | 235 | 213 | 271 | 177 | 165 | 233 | 234 | 240 | 268 | 219.46 | 192 | 247 | 1.293 | 0.371 | 0.01900608 | 0.06663062 | 0.0137 | 0.0132 | 0.0134 | 0.0134 | TRUE | 0.3702 |
| Mfsd6l | 20 | 28 | 28 | 25 | 41 | 36 | 16 | 30 | 30 | 25 | 46 | 36 | 30.37 | 25 | 36 | 1.426 | 0.512 | 0.19718236 | 0.35693103 | 0.0777 | 0.0805 | 0.0797 | 0.0797 | TRUE | 0.2421 |
| Mfsd7a | 45 | 44 | 31 | 42 | 28 | 51 | 35 | 47 | 33 | 42 | 32 | 50 | 39.82 | 38 | 41 | 1.081 | 0.112 | 0.72747364 | 0.83661541 | 0.0167 | 0.0619 | 0.0498 | 0.0498 | TRUE | 0.2069 |
| Mfsd7b | 490 | 349 | 257 | 262 | 243 | 257 | 381 | 371 | 277 | 261 | 274 | 254 | 302.95 | 343 | 263 | 0.764 | -0.388 | 0.00604345 | 0.02835818 | 0.0128 | 0.0102 | 0.011 | 0.011 | TRUE | 0.3113 |
| Mfsd7c | 333 | 204 | 277 | 413 | 322 | 374 | 259 | 217 | 299 | 411 | 363 | 369 | 319.67 | 258 | 381 | 1.477 | 0.563 | 3.23E-05 | 0.00049431 | 0.01 | 0.0098 | 0.01 | 0.01 | TRUE | 0.2047 |
| Mfsd8 | 224 | 141 | 171 | 200 | 148 | 157 | 174 | 150 | 185 | 199 | 167 | 155 | 171.58 | 170 | 174 | 1.024 | 0.035 | 0.83309738 | 0.90489102 | 0.008 | 0.0162 | 0.0139 | 0.0139 | TRUE | 0.1785 |
| Mfsd9 | 466 | 364 | 377 | 401 | 381 | 433 | 362 | 387 | 407 | 399 | 430 | 427 | 402.05 | 385 | 419 | 1.089 | 0.123 | 0.25290496 | 0.42184276 | 0 | 0.0083 | 0.0058 | 0.0058 | TRUE | 0.035 |
| Mga | 15219 | 11142 | 10559 | 14024 | 12300 | 14445 | 11820 | 11852 | 11393 | 13959 | 13876 | 14259 | 12859.87 | 11688 | 14031 | 1.2 | 0.264 | 1.64E-08 | 7.86E-07 | 2.00E-04 | 0.0026 | 0.0015 | 0.0015 | TRUE | 0.0059 |
| Mgam | 3 | 3 | 1 | 1 | 1 | 1 | 2 | 3 | 1 | 1 | 1 | 1 | 1.62 | 2 | 1 | 0.469 | -1.092 | 0.46568167 | NA | 0 | 1.4668 | 0.8963 | 0.8963 | TRUE | 0.1718 |
| Mgarp | 1096 | 884 | 961 | 991 | 863 | 1038 | 851 | 940 | 1037 | 986 | 974 | 1025 | 968.85 | 943 | 995 | 1.057 | 0.081 | 0.36572421 | 0.54034485 | 0.0043 | 0.0049 | 0.0047 | 0.0047 | TRUE | 0.0908 |
| Mgat1 | 1251 | 934 | 900 | 987 | 916 | 1001 | 972 | 993 | 971 | 982 | 1033 | 988 | 990.02 | 979 | 1001 | 1.023 | 0.033 | 0.65910784 | 0.78906344 | 0 | 0.0048 | 0.0029 | 0.0029 | TRUE | 0.0091 |
| Mgat2 | 2344 | 1653 | 1284 | 1283 | 1285 | 1282 | 1820 | 1758 | 1385 | 1277 | 1450 | 1266 | 1492.74 | 1654 | 1331 | 0.803 | -0.316 | 0.00210803 | 0.01255935 | 0.0121 | 0.004 | 0.0069 | 0.0069 | TRUE | 0.2423 |
| Mgat3 | 1187 | 832 | 1001 | 1159 | 1035 | 1210 | 922 | 885 | 1080 | 1154 | 1168 | 1194 | 1067.11 | 962 | 1172 | 1.219 | 0.286 | 0.00116309 | 0.00808954 | 0.0045 | 0.0046 | 0.0046 | 0.0046 | TRUE | 0.1333 |
| Mgat4a | 1257 | 919 | 893 | 1026 | 850 | 954 | 976 | 978 | 964 | 1021 | 959 | 942 | 973.21 | 973 | 974 | 1.002 | 0.002 | 0.97567923 | 0.98672491 | 0 | 0.0049 | 0.0031 | 0.0031 | TRUE | 0.0219 |
| Mgat4b | 3429 | 2495 | 2742 | 2883 | 2563 | 3022 | 2663 | 2654 | 2959 | 2870 | 2891 | 2983 | 2836.65 | 2759 | 2915 | 1.057 | 0.08 | 0.21614968 | 0.37885523 | 0.0017 | 0.0033 | 0.0027 | 0.0027 | TRUE | 0.0482 |
| Mgat4c | 10 | 14 | 7 | 3 | 8 | 5 | 8 | 15 | 8 | 3 | 9 | 5 | 7.86 | 10 | 6 | 0.558 | -0.841 | 0.2483414 | NA | 0.0752 | 0.304 | 0.2414 | 0.2414 | TRUE | 0.5777 |
| Mgat5 | 508 | 361 | 345 | 294 | 269 | 299 | 395 | 384 | 372 | 293 | 303 | 295 | 340.34 | 384 | 297 | 0.774 | -0.371 | 0.00085704 | 0.00647038 | 0 | 0.0094 | 0.0059 | 0.0059 | TRUE | 0.0077 |
| Mgat5b | 1418 | 1107 | 1135 | 1385 | 1180 | 1361 | 1101 | 1178 | 1225 | 1379 | 1331 | 1344 | 1259.46 | 1168 | 1351 | 1.159 | 0.212 | 0.00366749 | 0.01930231 | 9.00E-04 | 0.0043 | 0.0031 | 0.0031 | TRUE | 0.0313 |
| Mgea5 | 14549 | 11501 | 11367 | 11523 | 10296 | 12004 | 11300 | 12233 | 12265 | 11470 | 11615 | 11850 | 11788.78 | 11933 | 11645 | 0.976 | -0.035 | 0.51285383 | 0.6752851 | 0.0011 | 0.0026 | 0.002 | 0.002 | TRUE | 0.0266 |
| Mgl2 | 178 | 135 | 124 | 150 | 132 | 144 | 138 | 144 | 134 | 149 | 149 | 142 | 142.67 | 139 | 147 | 1.059 | 0.083 | 0.60999109 | 0.75322701 | 0 | 0.019 | 0.0121 | 0.0121 | TRUE | 0.0097 |
| Mgll | 305 | 231 | 194 | 170 | 178 | 173 | 237 | 246 | 209 | 169 | 201 | 171 | 205.45 | 231 | 180 | 0.78 | -0.359 | 0.01503576 | 0.05595057 | 0.0026 | 0.014 | 0.0108 | 0.0108 | TRUE | 0.1024 |
| Mgme1 | 2397 | 1806 | 2040 | 2086 | 1949 | 2167 | 1862 | 1921 | 2201 | 2076 | 2199 | 2139 | 2066.34 | 1995 | 2138 | 1.073 | 0.101 | 0.18440923 | 0.34062562 | 0.0038 | 0.0036 | 0.0037 | 0.0037 | TRUE | 0.0984 |
| Mgmt | 843 | 640 | 786 | 832 | 773 | 815 | 655 | 681 | 848 | 828 | 872 | 805 | 781.38 | 728 | 835 | 1.149 | 0.2 | 0.05775493 | 0.1495726 | 0.0093 | 0.0055 | 0.0067 | 0.0067 | TRUE | 0.2439 |

| | | | | | | | | | | | | | | | | | | | | | | | | | |
|---|---|---|---|---|---|---|---|---|---|---|---|---|---|---|---|---|---|---|---|---|---|---|---|---|---|
| Mgp | 1 | 2 | 3 | 0 | 0 | 1 | 1 | 2 | 3 | 0 | 0 | 1 | 1.19 | 2 | 0 | 0.184 | -2.442 | 0.2099393 | NA | 0 | 1.9974 | 1.4928 | 1.4928 | TRUE | 0.2878 |
| Mgrn1 | 2672 | 2005 | 2150 | 2102 | 2014 | 2339 | 2075 | 2133 | 2320 | 2092 | 2272 | 2309 | 2200.17 | 2176 | 2224 | 1.023 | 0.033 | 0.64667903 | 0.78033832 | 0.0026 | 0.0035 | 0.0032 | 0.0032 | TRUE | 0.0402 |
| Mgst1 | 301 | 197 | 222 | 227 | 199 | 238 | 234 | 210 | 240 | 226 | 224 | 235 | 228.04 | 228 | 228 | 1.003 | 0.005 | 0.9729811 | 0.98525108 | 0 | 0.0128 | 0.0086 | 0.0086 | TRUE | 0.0506 |
| Mgst2 | 12 | 4 | 6 | 8 | 4 | 6 | 9 | 4 | 6 | 8 | 5 | 6 | 6.41 | 6 | 6 | 0.909 | -0.138 | 0.85511066 | NA | 0 | 0.3723 | 0.2587 | 0.2587 | TRUE | 0.3006 |
| Mgst3 | 926 | 659 | 800 | 893 | 837 | 972 | 719 | 701 | 863 | 889 | 944 | 960 | 846 | 761 | 931 | 1.224 | 0.292 | 0.00237033 | 0.01378883 | 0.0058 | 0.0052 | 0.0055 | 0.0055 | TRUE | 0.1593 |
| Mia | 28 | 24 | 19 | 24 | 18 | 18 | 22 | 26 | 21 | 24 | 20 | 18 | 21.62 | 23 | 21 | 0.916 | -0.127 | 0.75591877 | 0.85511255 | 0 | 0.112 | 0.0745 | 0.0745 | TRUE | 0.1074 |
| Mia2 | 4 | 1 | 2 | 0 | 2 | 0 | 3 | 1 | 2 | 0 | 2 | 0 | 1.43 | 2 | 1 | 0.337 | -1.571 | 0.3778833 | NA | 0 | 1.6589 | 1.3408 | 1.3408 | TRUE | 0.888 |
| Mia3 | 5967 | 4317 | 4013 | 4129 | 3637 | 4254 | 4634 | 4592 | 4330 | 4110 | 4103 | 4199 | 4328.09 | 4519 | 4137 | 0.915 | -0.128 | 0.01981352 | 0.0686699 | 5.00E-04 | 0.003 | 0.0019 | 0.0019 | TRUE | 0.0161 |
| Miat | 62 | 74 | 40 | 67 | 40 | 77 | 48 | 79 | 43 | 67 | 45 | 76 | 59.64 | 57 | 63 | 1.112 | 0.153 | 0.61355411 | 0.75566815 | 0.0688 | 0.0422 | 0.0496 | 0.0496 | TRUE | 0.9408 |
| Mib1 | 3156 | 2292 | 2504 | 2901 | 2482 | 2746 | 2451 | 2438 | 2702 | 2888 | 2800 | 2711 | 2664.87 | 2530 | 2800 | 1.107 | 0.146 | 0.02547677 | 0.08242179 | 0.0017 | 0.0033 | 0.0027 | 0.0027 | TRUE | 0.042 |
| Mib2 | 2428 | 1961 | 1728 | 1763 | 1634 | 1848 | 1886 | 2086 | 1864 | 1755 | 1843 | 1824 | 1876.43 | 1945 | 1807 | 0.929 | -0.106 | 0.12887042 | 0.26544819 | 0.0017 | 0.0037 | 0.003 | 0.003 | TRUE | 0.0475 |
| Mical1 | 1497 | 1213 | 1180 | 1456 | 1366 | 1654 | 1163 | 1290 | 1273 | 1449 | 1541 | 1633 | 1391.53 | 1242 | 1541 | 1.242 | 0.313 | 5.66E-05 | 0.00077314 | 0.0027 | 0.0041 | 0.0036 | 0.0036 | TRUE | 0.0392 |
| Mical2 | 201 | 179 | 153 | 251 | 187 | 260 | 156 | 190 | 165 | 250 | 211 | 257 | 204.84 | 170 | 239 | 1.408 | 0.494 | 0.00114465 | 0.00799647 | 0.0058 | 0.014 | 0.0116 | 0.0116 | TRUE | 0.1136 |
| Mical3 | 5754 | 4191 | 4081 | 4900 | 4336 | 4770 | 4469 | 4458 | 4403 | 4877 | 4891 | 4709 | 4634.63 | 4443 | 4826 | 1.086 | 0.119 | 0.01957091 | 0.06799392 | 0 | 0.0029 | 0.0017 | 0.0017 | TRUE | 0.0055 |
| Micalcl | 5 | 6 | 2 | 3 | 3 | 3 | 4 | 6 | 2 | 3 | 3 | 3 | 3.63 | 4 | 3 | 0.751 | -0.413 | 0.67859634 | NA | 0 | 0.6561 | 0.433 | 0.433 | TRUE | 0.2331 |
| Micall1 | 1605 | 1229 | 1229 | 1755 | 1438 | 1683 | 1247 | 1307 | 1326 | 1747 | 1622 | 1661 | 1485.07 | 1293 | 1677 | 1.298 | 0.376 | 5.00E-08 | 2.10E-06 | 6.00E-04 | 0.004 | 0.0027 | 0.0027 | TRUE | 0.0162 |
| Micall2 | 406 | 317 | 454 | 656 | 576 | 705 | 315 | 337 | 490 | 653 | 650 | 696 | 523.51 | 381 | 666 | 1.755 | 0.812 | 1.28E-08 | 6.29E-07 | 0.0268 | 0.007 | 0.0126 | 0.0126 | TRUE | 0.7116 |
| Micu1 | 969 | 637 | 617 | 624 | 533 | 668 | 753 | 678 | 666 | 621 | 601 | 659 | 662.95 | 699 | 627 | 0.897 | -0.157 | 0.08971566 | 0.20422136 | 0.0021 | 0.006 | 0.0047 | 0.0047 | TRUE | 0.0568 |
| Mid1 | 14 | 12 | 6 | 5 | 8 | 9 | 11 | 13 | 6 | 5 | 9 | 9 | 8.83 | 10 | 8 | 0.755 | -0.405 | 0.53687986 | NA | 0 | 0.2707 | 0.194 | 0.194 | TRUE | 0.3018 |
| Mid1ip1 | 908 | 729 | 758 | 664 | 576 | 693 | 705 | 775 | 818 | 661 | 650 | 684 | 715.56 | 766 | 665 | 0.87 | -0.201 | 0.02600685 | 0.08370189 | 0.0021 | 0.0057 | 0.0045 | 0.0045 | TRUE | 0.0607 |
| Mid2 | 86 | 58 | 56 | 89 | 81 | 85 | 67 | 62 | 60 | 89 | 91 | 84 | 75.46 | 63 | 88 | 1.392 | 0.477 | 0.03118761 | 0.09533031 | 0 | 0.0338 | 0.0219 | 0.0219 | TRUE | 0.0286 |
| Midn | 4948 | 3629 | 3006 | 3434 | 2982 | 3296 | 3843 | 3860 | 3243 | 3418 | 3364 | 3254 | 3497.06 | 3649 | 3345 | 0.916 | -0.126 | 0.09265839 | 0.20921947 | 0.0045 | 0.0031 | 0.0038 | 0.0038 | TRUE | 0.1139 |
| Mien1 | 2218 | 1696 | 1703 | 1819 | 1595 | 1882 | 1723 | 1804 | 1838 | 1811 | 1799 | 1858 | 1805.32 | 1788 | 1823 | 1.02 | 0.029 | 0.65073638 | 0.78320543 | 0 | 0.0037 | 0.0024 | 0.0024 | TRUE | 0.0129 |
| Mier1 | 4215 | 3080 | 3185 | 3057 | 2588 | 3053 | 3274 | 3276 | 3437 | 3043 | 2920 | 3014 | 3160.43 | 3329 | 2992 | 0.899 | -0.153 | 0.00662307 | 0.03042866 | 3.00E-04 | 0.0032 | 0.002 | 0.002 | TRUE | 0.0096 |
| Mier2 | 3246 | 2542 | 2455 | 2496 | 2337 | 2677 | 2521 | 2704 | 2649 | 2485 | 2636 | 2643 | 2606.23 | 2625 | 2588 | 0.986 | -0.02 | 0.74903809 | 0.85119878 | 9.00E-04 | 0.0033 | 0.0024 | 0.0024 | TRUE | 0.0149 |
| Mier3 | 2113 | 1631 | 1529 | 1616 | 1455 | 1550 | 1641 | 1735 | 1650 | 1609 | 1641 | 1530 | 1634.3 | 1675 | 1593 | 0.951 | -0.072 | 0.27998926 | 0.45147636 | 5.00E-04 | 0.0039 | 0.0026 | 0.0026 | TRUE | 0.0146 |
| Mif | 22599 | 17136 | 20157 | 20797 | 19065 | 23259 | 17552 | 18227 | 21749 | 20701 | 21507 | 22960 | 20449.51 | 19176 | 21723 | 1.133 | 0.18 | 0.02343418 | 0.07758738 | 0.0082 | 0.0025 | 0.0045 | 0.0045 | TRUE | 0.1684 |
| Mif4gd | 3429 | 2310 | 2072 | 2053 | 1832 | 2090 | 2663 | 2457 | 2236 | 2044 | 2067 | 2063 | 2254.89 | 2452 | 2058 | 0.838 | -0.254 | 0.00058805 | 0.00488522 | 0.0031 | 0.0035 | 0.0035 | 0.0035 | TRUE | 0.0719 |
| Miip | 506 | 387 | 429 | 319 | 338 | 419 | 393 | 412 | 463 | 318 | 381 | 414 | 396.66 | 423 | 371 | 0.879 | -0.186 | 0.13866664 | 0.27970471 | 0.0099 | 0.0084 | 0.0089 | 0.0089 | TRUE | 0.1844 |
| Mill1 | 0 | 0 | 0 | 0 | 0 | 0 | 0 | 0 | 0 | 0 | 0 | 0 | 0 | 0 | 0 | NA | NA | NA | NA | NA | NA | NA | NA | NA | NA |
| Mill2 | 2 | 1 | 1 | 0 | 2 | 2 | 2 | 1 | 1 | 0 | 2 | 2 | 1.32 | 1 | 1 | 1.122 | 0.166 | 0.92013686 | NA | 0 | 1.7965 | 1.2293 | 1.2293 | TRUE | 0.5133 |
| Milr1 | 5 | 4 | 0 | 0 | 1 | 1 | 4 | 4 | 0 | 0 | 1 | 1 | 1.71 | 3 | 1 | 0.254 | -1.978 | 0.25274805 | NA | 0 | 1.3893 | 1.2362 | 1.2362 | TRUE | 0.8232 |
| Mina | 4252 | 3082 | 2860 | 2809 | 2549 | 2869 | 3302 | 3278 | 3086 | 2796 | 2876 | 2832 | 3028.38 | 3222 | 2835 | 0.879 | -0.186 | 0.00133569 | 0.00887497 | 5.00E-04 | 0.0032 | 0.0021 | 0.0021 | TRUE | 0.0164 |
| Mink1 | 6864 | 5164 | 4593 | 4816 | 4463 | 4980 | 5331 | 5493 | 4956 | 4794 | 5035 | 4916 | 5087.38 | 5260 | 4915 | 0.934 | -0.098 | 0.09560641 | 0.21418534 | 0.0014 | 0.0029 | 0.0023 | 0.0023 | TRUE | 0.0308 |
| Minos1 | 3684 | 2516 | 2526 | 2878 | 2427 | 2838 | 2861 | 2676 | 2726 | 2865 | 2738 | 2802 | 2777.86 | 2754 | 2802 | 1.017 | 0.024 | 0.68630228 | 0.80774354 | 5.00E-04 | 0.0033 | 0.0022 | 0.0022 | TRUE | 0.0144 |
| Minpp1 | 4601 | 3226 | 3197 | 3474 | 3019 | 3471 | 3573 | 3431 | 3450 | 3458 | 3406 | 3426 | 3457.43 | 3485 | 3430 | 0.984 | -0.023 | 0.66388742 | 0.79285953 | 0 | 0.0031 | 0.0018 | 0.0018 | TRUE | 0.0062 |
| Mios | 1789 | 1239 | 1354 | 1371 | 1222 | 1321 | 1389 | 1318 | 1461 | 1365 | 1379 | 1304 | 1369.26 | 1389 | 1349 | 0.971 | -0.043 | 0.55275471 | 0.70844002 | 0.001 | 0.0042 | 0.003 | 0.003 | TRUE | 0.0235 |
| Miox | 20 | 11 | 11 | 11 | 14 | 9 | 16 | 12 | 12 | 11 | 16 | 9 | 12.45 | 13 | 12 | 0.899 | -0.153 | 0.77936067 | NA | 0 | 0.1927 | 0.1339 | 0.1339 | TRUE | 0.2943 |
| Mip | 87 | 59 | 53 | 73 | 60 | 81 | 68 | 63 | 57 | 73 | 68 | 80 | 67.97 | 63 | 74 | 1.172 | 0.23 | 0.32924297 | 0.50362565 | 0 | 0.0373 | 0.0252 | 0.0252 | TRUE | 0.0555 |
| Mipep | 2612 | 1979 | 2224 | 2290 | 2050 | 2395 | 2029 | 2105 | 2400 | 2279 | 2313 | 2364 | 2248.28 | 2178 | 2319 | 1.066 | 0.092 | 0.21819342 | 0.38122145 | 0.0035 | 0.0035 | 0.0036 | 0.0036 | TRUE | 0.0955 |
| Mipol1 | 758 | 524 | 603 | 731 | 589 | 722 | 589 | 557 | 651 | 728 | 664 | 713 | 650.25 | 599 | 702 | 1.173 | 0.23 | 0.01504226 | 0.05596019 | 0.0024 | 0.0061 | 0.0049 | 0.0049 | TRUE | 0.0642 |
| Mir100 | 0 | 0 | 0 | 0 | 0 | 0 | 0 | 0 | 0 | 0 | 0 | 0 | 0 | 0 | 0 | NA | NA | NA | NA | NA | NA | NA | NA | NA | NA |
| Mir101a | 0 | 0 | 0 | 0 | 0 | 0 | 0 | 0 | 0 | 0 | 0 | 0 | 0 | 0 | 0 | NA | NA | NA | NA | NA | NA | NA | NA | NA | NA |
| Mir101b | 0 | 0 | 0 | 0 | 0 | 0 | 0 | 0 | 0 | 0 | 0 | 0 | 0 | 0 | 0 | NA | NA | NA | NA | NA | NA | NA | NA | NA | NA |
| Mir101c | 0 | 0 | 0 | 0 | 0 | 0 | 0 | 0 | 0 | 0 | 0 | 0 | 0 | 0 | 0 | NA | NA | NA | NA | NA | NA | NA | NA | NA | NA |
| Mir103-1 | 0 | 0 | 0 | 0 | 0 | 0 | 0 | 0 | 0 | 0 | 0 | 0 | 0 | 0 | 0 | NA | NA | NA | NA | NA | NA | NA | NA | NA | NA |
| Mir103-2 | 0 | 0 | 0 | 1 | 0 | 0 | 0 | 0 | 0 | 1 | 0 | 0 | 0.17 | 0 | 0 | 2.094 | 1.067 | 0.79379329 | NA | 0 | 14.2885 | 10 | 10 | TRUE | 0.3906 |
| Mir105 | 0 | 0 | 0 | 0 | 0 | 0 | 0 | 0 | 0 | 0 | 0 | 0 | 0 | 0 | 0 | NA | NA | NA | NA | NA | NA | NA | NA | NA | NA |
| Mir106a | 0 | 0 | 0 | 0 | 0 | 0 | 0 | 0 | 0 | 0 | 0 | 0 | 0 | 0 | 0 | NA | NA | NA | NA | NA | NA | NA | NA | NA | NA |
| Mir106b | 0 | 0 | 0 | 0 | 0 | 0 | 0 | 0 | 0 | 0 | 0 | 0 | 0 | 0 | 0 | NA | NA | NA | NA | NA | NA | NA | NA | NA | NA |
| Mir107 | 0 | 0 | 0 | 0 | 0 | 0 | 0 | 0 | 0 | 0 | 0 | 0 | 0 | 0 | 0 | NA | NA | NA | NA | NA | NA | NA | NA | NA | NA |
| Mir10a | 0 | 0 | 0 | 0 | 0 | 0 | 0 | 0 | 0 | 0 | 0 | 0 | 0 | 0 | 0 | NA | NA | NA | NA | NA | NA | NA | NA | NA | NA |
| Mir10b | 0 | 0 | 0 | 0 | 0 | 0 | 0 | 0 | 0 | 0 | 0 | 0 | 0 | 0 | 0 | NA | NA | NA | NA | NA | NA | NA | NA | NA | NA |
| Mir1186 | 0 | 0 | 0 | 0 | 0 | 0 | 0 | 0 | 0 | 0 | 0 | 0 | 0 | 0 | 0 | NA | NA | NA | NA | NA | NA | NA | NA | NA | NA |
| Mir1186b | 0 | 0 | 0 | 0 | 0 | 0 | 0 | 0 | 0 | 0 | 0 | 0 | 0 | 0 | 0 | NA | NA | NA | NA | NA | NA | NA | NA | NA | NA |
| Mir1187 | 0 | 0 | 0 | 0 | 0 | 0 | 0 | 0 | 0 | 0 | 0 | 0 | 0 | 0 | 0 | NA | NA | NA | NA | NA | NA | NA | NA | NA | NA |
| Mir1188 | 3 | 0 | 1 | 1 | 2 | 0 | 2 | 0 | 1 | 1 | 2 | 0 | 1.11 | 1 | 1 | 0.906 | -0.142 | 0.94060664 | NA | 0 | 2.1374 | 1.7366 | 1.7366 | TRUE | 0.2449 |
| Mir1190 | 0 | 0 | 0 | 0 | 0 | 0 | 0 | 0 | 0 | 0 | 0 | 0 | 0 | 0 | 0 | NA | NA | NA | NA | NA | NA | NA | NA | NA | NA |
| Mir1191 | 0 | 0 | 0 | 0 | 0 | 0 | 0 | 0 | 0 | 0 | 0 | 0 | 0 | 0 | 0 | NA | NA | NA | NA | NA | NA | NA | NA | NA | NA |
| Mir1192 | 1 | 1 | 1 | 3 | 0 | 2 | 1 | 1 | 1 | 3 | 0 | 2 | 1.31 | 1 | 2 | 1.739 | 0.799 | 0.63748462 | NA | 0 | 1.8071 | 1.2614 | 1.2614 | TRUE | 0.3155 |
| Mir1193 | 0 | 0 | 0 | 0 | 0 | 0 | 0 | 0 | 0 | 0 | 0 | 0 | 0 | 0 | 0 | NA | NA | NA | NA | NA | NA | NA | NA | NA | NA |
| Mir1195 | 0 | 0 | 0 | 0 | 0 | 0 | 0 | 0 | 0 | 0 | 0 | 0 | 0 | 0 | 0 | NA | NA | NA | NA | NA | NA | NA | NA | NA | NA |
| Mir1196 | 0 | 1 | 0 | 1 | 0 | 0 | 0 | 1 | 0 | 1 | 0 | 0 | 0.34 | 0 | 0 | 1.075 | 0.105 | 0.97573409 | NA | 0 | 6.9086 | 6.5582 | 6.5582 | TRUE | 0.5216 |
| Mir1197 | 0 | 0 | 0 | 0 | 0 | 0 | 0 | 0 | 0 | 0 | 0 | 0 | 0 | 0 | 0 | NA | NA | NA | NA | NA | NA | NA | NA | NA | NA |
| Mir1198 | 0 | 0 | 0 | 0 | 0 | 0 | 0 | 0 | 0 | 0 | 0 | 0 | 0 | 0 | 0 | NA | NA | NA | NA | NA | NA | NA | NA | NA | NA |
| Mir1199 | 21 | 36 | 33 | 25 | 33 | 37 | 16 | 38 | 36 | 25 | 37 | 37 | 31.47 | 30 | 33 | 1.107 | 0.146 | 0.71221966 | 0.82701728 | 0.091 | 0.0777 | 0.0815 | 0.0815 | TRUE | 1.2925 |
| Mir1224 | 0 | 0 | 0 | 0 | 0 | 0 | 0 | 0 | 0 | 0 | 0 | 0 | 0 | 0 | 0 | NA | NA | NA | NA | NA | NA | NA | NA | NA | NA |
| Mir122a | 0 | 0 | 0 | 0 | 0 | 0 | 0 | 0 | 0 | 0 | 0 | 0 | 0 | 0 | 0 | NA | NA | NA | NA | NA | NA | NA | NA | NA | NA |
| Mir1247 | 0 | 0 | 0 | 0 | 0 | 0 | 0 | 0 | 0 | 0 | 0 | 0 | 0 | 0 | 0 | NA | NA | NA | NA | NA | NA | NA | NA | NA | NA |

| | | | | | | | | | | | | | | | | | | | | | | | | | |
|---|---|---|---|---|---|---|---|---|---|---|---|---|---|---|---|---|---|---|---|---|---|---|---|---|---|
| Mir1249 | 0 | 2 | 2 | 2 | 0 | 3 | 0 | 2 | 2 | 2 | 0 | 3 | 1.54 | 1 | 2 | 1.215 | 0.281 | 0.86677387 | NA | | 0 | 1.5418 | 1.3615 | 1.3615 | TRUE | 0.5958 |
| Mir124a-1 | 0 | 0 | 0 | 0 | 0 | 0 | 0 | 0 | 0 | 0 | 0 | 0 | 0 | 0 | 0 | NA | NA | NA | NA | NA | NA | NA | NA | NA | NA |
| Mir124a-2 | 0 | 0 | 0 | 0 | 0 | 0 | 0 | 0 | 0 | 0 | 0 | 0 | 0 | 0 | 0 | NA | NA | NA | NA | NA | NA | NA | NA | NA | NA |
| Mir124a-3 | 0 | 0 | 0 | 0 | 0 | 0 | 0 | 0 | 0 | 0 | 0 | 0 | 0 | 0 | 0 | NA | NA | NA | NA | NA | NA | NA | NA | NA | NA |
| Mir1251 | 0 | 0 | 0 | 0 | 0 | 0 | 0 | 0 | 0 | 0 | 0 | 0 | 0 | 0 | 0 | NA | NA | NA | NA | NA | NA | NA | NA | NA | NA |
| Mir125a | 0 | 0 | 0 | 0 | 0 | 0 | 0 | 0 | 0 | 0 | 0 | 0 | 0 | 0 | 0 | NA | NA | NA | NA | NA | NA | NA | NA | NA | NA |
| Mir125b-1 | 0 | 0 | 0 | 0 | 0 | 0 | 0 | 0 | 0 | 0 | 0 | 0 | 0 | 0 | 0 | NA | NA | NA | NA | NA | NA | NA | NA | NA | NA |
| Mir125b-2 | 0 | 0 | 0 | 0 | 0 | 0 | 0 | 0 | 0 | 0 | 0 | 0 | 0 | 0 | 0 | NA | NA | NA | NA | NA | NA | NA | NA | NA | NA |
| Mir126 | 0 | 0 | 0 | 0 | 0 | 0 | 0 | 0 | 0 | 0 | 0 | 0 | 0 | 0 | 0 | NA | NA | NA | NA | NA | NA | NA | NA | NA | NA |
| Mir1264 | 0 | 0 | 0 | 0 | 0 | 0 | 0 | 0 | 0 | 0 | 0 | 0 | 0 | 0 | 0 | NA | NA | NA | NA | NA | NA | NA | NA | NA | NA |
| Mir127 | 0 | 0 | 0 | 1 | 0 | 0 | 0 | 0 | 0 | 1 | 0 | 0 | 0.17 | 0 | 0 | 2.094 | 1.067 | 0.79379329 | NA | | 0 | 14.2885 | 10 | 10 | TRUE | 0.3906 |
| Mir128-1 | 0 | 0 | 0 | 0 | 0 | 0 | 0 | 0 | 0 | 0 | 0 | 0 | 0 | 0 | 0 | NA | NA | NA | NA | NA | NA | NA | NA | NA | NA |
| Mir128-2 | 0 | 0 | 0 | 0 | 0 | 0 | 0 | 0 | 0 | 0 | 0 | 0 | 0 | 0 | 0 | NA | NA | NA | NA | NA | NA | NA | NA | NA | NA |
| Mir129-1 | 0 | 0 | 0 | 0 | 0 | 0 | 0 | 0 | 0 | 0 | 0 | 0 | 0 | 0 | 0 | NA | NA | NA | NA | NA | NA | NA | NA | NA | NA |
| Mir129-2 | 0 | 0 | 0 | 0 | 0 | 0 | 0 | 0 | 0 | 0 | 0 | 0 | 0 | 0 | 0 | NA | NA | NA | NA | NA | NA | NA | NA | NA | NA |
| Mir1298 | 0 | 0 | 0 | 0 | 0 | 0 | 0 | 0 | 0 | 0 | 0 | 0 | 0 | 0 | 0 | NA | NA | NA | NA | NA | NA | NA | NA | NA | NA |
| Mir1306 | 0 | 0 | 0 | 0 | 0 | 0 | 0 | 0 | 0 | 0 | 0 | 0 | 0 | 0 | 0 | NA | NA | NA | NA | NA | NA | NA | NA | NA | NA |
| Mir130a | 0 | 0 | 0 | 0 | 0 | 0 | 0 | 0 | 0 | 0 | 0 | 0 | 0 | 0 | 0 | NA | NA | NA | NA | NA | NA | NA | NA | NA | NA |
| Mir130b | 0 | 0 | 0 | 0 | 0 | 0 | 0 | 0 | 0 | 0 | 0 | 0 | 0 | 0 | 0 | NA | NA | NA | NA | NA | NA | NA | NA | NA | NA |
| Mir132 | 0 | 0 | 0 | 0 | 0 | 0 | 0 | 0 | 0 | 0 | 0 | 0 | 0 | 0 | 0 | NA | NA | NA | NA | NA | NA | NA | NA | NA | NA |
| Mir133a-1 | 0 | 0 | 0 | 0 | 0 | 0 | 0 | 0 | 0 | 0 | 0 | 0 | 0 | 0 | 0 | NA | NA | NA | NA | NA | NA | NA | NA | NA | NA |
| Mir133a-2 | 0 | 0 | 0 | 0 | 0 | 0 | 0 | 0 | 0 | 0 | 0 | 0 | 0 | 0 | 0 | NA | NA | NA | NA | NA | NA | NA | NA | NA | NA |
| Mir133b | 0 | 0 | 0 | 0 | 0 | 0 | 0 | 0 | 0 | 0 | 0 | 0 | 0 | 0 | 0 | NA | NA | NA | NA | NA | NA | NA | NA | NA | NA |
| Mir134 | 0 | 0 | 0 | 0 | 0 | 0 | 0 | 0 | 0 | 0 | 0 | 0 | 0 | 0 | 0 | NA | NA | NA | NA | NA | NA | NA | NA | NA | NA |
| Mir135a-1 | 0 | 0 | 0 | 0 | 0 | 0 | 0 | 0 | 0 | 0 | 0 | 0 | 0 | 0 | 0 | NA | NA | NA | NA | NA | NA | NA | NA | NA | NA |
| Mir135a-2 | 0 | 0 | 0 | 0 | 0 | 0 | 0 | 0 | 0 | 0 | 0 | 0 | 0 | 0 | 0 | NA | NA | NA | NA | NA | NA | NA | NA | NA | NA |
| Mir135b | 0 | 0 | 0 | 0 | 0 | 0 | 0 | 0 | 0 | 0 | 0 | 0 | 0 | 0 | 0 | NA | NA | NA | NA | NA | NA | NA | NA | NA | NA |
| Mir136 | 0 | 0 | 0 | 0 | 0 | 0 | 0 | 0 | 0 | 0 | 0 | 0 | 0 | 0 | 0 | NA | NA | NA | NA | NA | NA | NA | NA | NA | NA |
| Mir137 | 0 | 0 | 0 | 0 | 0 | 0 | 0 | 0 | 0 | 0 | 0 | 0 | 0 | 0 | 0 | NA | NA | NA | NA | NA | NA | NA | NA | NA | NA |
| Mir138-1 | 1 | 0 | 0 | 0 | 0 | 0 | 1 | 0 | 0 | 0 | 0 | 0 | 0.13 | 0 | 0 | 0.552 | -0.857 | 0.8336548 | NA | | 0 | 18.3121 | 10 | 10 | TRUE | 0.2566 |
| Mir138-2 | 0 | 0 | 0 | 0 | 0 | 0 | 0 | 0 | 0 | 0 | 0 | 0 | 0 | 0 | 0 | NA | NA | NA | NA | NA | NA | NA | NA | NA | NA |
| Mir139 | 0 | 0 | 0 | 0 | 0 | 0 | 0 | 0 | 0 | 0 | 0 | 0 | 0 | 0 | 0 | NA | NA | NA | NA | NA | NA | NA | NA | NA | NA |
| Mir140 | 0 | 0 | 0 | 0 | 0 | 0 | 0 | 0 | 0 | 0 | 0 | 0 | 0 | 0 | 0 | NA | NA | NA | NA | NA | NA | NA | NA | NA | NA |
| Mir141 | 0 | 0 | 0 | 0 | 0 | 0 | 0 | 0 | 0 | 0 | 0 | 0 | 0 | 0 | 0 | NA | NA | NA | NA | NA | NA | NA | NA | NA | NA |
| Mir142 | 0 | 0 | 0 | 0 | 0 | 0 | 0 | 0 | 0 | 0 | 0 | 0 | 0 | 0 | 0 | NA | NA | NA | NA | NA | NA | NA | NA | NA | NA |
| Mir143 | 1 | 0 | 0 | 0 | 0 | 0 | 1 | 0 | 0 | 0 | 0 | 0 | 0.13 | 0 | 0 | 0.552 | -0.857 | 0.8336548 | NA | | 0 | 18.3121 | 10 | 10 | TRUE | 0.2566 |
| Mir143hg | 6 | 3 | 3 | 0 | 2 | 6 | 5 | 3 | 3 | 0 | 2 | 6 | 3.21 | 4 | 3 | 0.735 | -0.445 | 0.7007535 | NA | | 0.2949 | 0.7405 | 0.6457 | 0.6457 | TRUE | 0.4982 |
| Mir144 | 0 | 0 | 0 | 0 | 0 | 0 | 0 | 0 | 0 | 0 | 0 | 0 | 0 | 0 | 0 | NA | NA | NA | NA | NA | NA | NA | NA | NA | NA |
| Mir145 | 0 | 0 | 0 | 0 | 0 | 0 | 0 | 0 | 0 | 0 | 0 | 0 | 0 | 0 | 0 | NA | NA | NA | NA | NA | NA | NA | NA | NA | NA |
| Mir146 | 0 | 0 | 0 | 0 | 0 | 0 | 0 | 0 | 0 | 0 | 0 | 0 | 0 | 0 | 0 | NA | NA | NA | NA | NA | NA | NA | NA | NA | NA |
| Mir146b | 0 | 0 | 0 | 0 | 0 | 0 | 0 | 0 | 0 | 0 | 0 | 0 | 0 | 0 | 0 | NA | NA | NA | NA | NA | NA | NA | NA | NA | NA |
| Mir147 | 0 | 0 | 0 | 0 | 0 | 0 | 0 | 0 | 0 | 0 | 0 | 0 | 0 | 0 | 0 | NA | NA | NA | NA | NA | NA | NA | NA | NA | NA |
| Mir148a | 0 | 0 | 0 | 0 | 0 | 0 | 0 | 0 | 0 | 0 | 0 | 0 | 0 | 0 | 0 | NA | NA | NA | NA | NA | NA | NA | NA | NA | NA |
| Mir148b | 0 | 0 | 0 | 0 | 1 | 0 | 0 | 0 | 0 | 0 | 1 | 0 | 0.19 | 0 | 0 | 2.094 | 1.067 | 0.79379329 | NA | | 0 | 12.608 | 10 | 10 | TRUE | 0.5075 |
| Mir149 | 0 | 0 | 0 | 0 | 0 | 0 | 0 | 0 | 0 | 0 | 0 | 0 | 0 | 0 | 0 | NA | NA | NA | NA | NA | NA | NA | NA | NA | NA |
| Mir150 | 0 | 0 | 0 | 0 | 0 | 0 | 0 | 0 | 0 | 0 | 0 | 0 | 0 | 0 | 0 | NA | NA | NA | NA | NA | NA | NA | NA | NA | NA |
| Mir152 | 0 | 0 | 0 | 0 | 0 | 0 | 0 | 0 | 0 | 0 | 0 | 0 | 0 | 0 | 0 | NA | NA | NA | NA | NA | NA | NA | NA | NA | NA |
| Mir153 | 0 | 0 | 0 | 0 | 0 | 0 | 0 | 0 | 0 | 0 | 0 | 0 | 0 | 0 | 0 | NA | NA | NA | NA | NA | NA | NA | NA | NA | NA |
| Mir154 | 0 | 0 | 0 | 0 | 0 | 0 | 0 | 0 | 0 | 0 | 0 | 0 | 0 | 0 | 0 | NA | NA | NA | NA | NA | NA | NA | NA | NA | NA |
| Mir155 | 0 | 0 | 0 | 0 | 0 | 0 | 0 | 0 | 0 | 0 | 0 | 0 | 0 | 0 | 0 | NA | NA | NA | NA | NA | NA | NA | NA | NA | NA |
| Mir15a | 0 | 0 | 0 | 0 | 0 | 0 | 0 | 0 | 0 | 0 | 0 | 0 | 0 | 0 | 0 | NA | NA | NA | NA | NA | NA | NA | NA | NA | NA |
| Mir15b | 0 | 0 | 0 | 0 | 0 | 0 | 0 | 0 | 0 | 0 | 0 | 0 | 0 | 0 | 0 | NA | NA | NA | NA | NA | NA | NA | NA | NA | NA |
| Mir16-1 | 0 | 0 | 0 | 0 | 0 | 0 | 0 | 0 | 0 | 0 | 0 | 0 | 0 | 0 | 0 | NA | NA | NA | NA | NA | NA | NA | NA | NA | NA |
| Mir16-2 | 0 | 0 | 0 | 0 | 0 | 0 | 0 | 0 | 0 | 0 | 0 | 0 | 0 | 0 | 0 | NA | NA | NA | NA | NA | NA | NA | NA | NA | NA |
| Mir17 | 1 | 0 | 0 | 0 | 0 | 0 | 1 | 0 | 0 | 0 | 0 | 0 | 0.13 | 0 | 0 | 0.552 | -0.857 | 0.8336548 | NA | | 0 | 18.3121 | 10 | 10 | TRUE | 0.2566 |
| Mir17hg | 676 | 786 | 506 | 326 | 326 | 349 | 525 | 836 | 546 | 325 | 368 | 345 | 490.64 | 636 | 346 | 0.544 | -0.878 | 3.00E-08 | 1.32E-06 | | 0.036 | 0.0073 | 0.0158 | 0.0158 | TRUE | 0.8901 |
| Mir18 | 1 | 1 | 0 | 1 | 0 | 1 | 1 | 1 | 0 | 1 | 0 | 1 | 0.64 | 1 | 1 | 1.082 | 0.113 | 0.96183421 | NA | | 0 | 3.7222 | 2.4899 | 2.4899 | TRUE | 0.2041 |
| Mir181a-1 | 0 | 0 | 0 | 0 | 0 | 0 | 0 | 0 | 0 | 0 | 0 | 0 | 0 | 0 | 0 | NA | NA | NA | NA | NA | NA | NA | NA | NA | NA |
| Mir181a-2 | 0 | 0 | 0 | 0 | 0 | 0 | 0 | 0 | 0 | 0 | 0 | 0 | 0 | 0 | 0 | NA | NA | NA | NA | NA | NA | NA | NA | NA | NA |
| Mir181b-1 | 0 | 0 | 0 | 0 | 0 | 0 | 0 | 0 | 0 | 0 | 0 | 0 | 0 | 0 | 0 | NA | NA | NA | NA | NA | NA | NA | NA | NA | NA |
| Mir181b-2 | 0 | 0 | 0 | 0 | 0 | 0 | 0 | 0 | 0 | 0 | 0 | 0 | 0 | 0 | 0 | NA | NA | NA | NA | NA | NA | NA | NA | NA | NA |
| Mir181c | 1 | 0 | 0 | 0 | 0 | 0 | 1 | 0 | 0 | 0 | 0 | 0 | 0.13 | 0 | 0 | 0.552 | -0.857 | 0.8336548 | NA | | 0 | 18.3121 | 10 | 10 | TRUE | 0.2566 |
| Mir181d | 0 | 0 | 0 | 0 | 1 | 1 | 0 | 0 | 0 | 0 | 1 | 1 | 0.35 | 0 | 1 | 3.965 | 1.987 | 0.55529285 | NA | | 0 | 6.7252 | 6.4426 | 6.4426 | TRUE | 0.261 |
| Mir182 | 0 | 0 | 0 | 0 | 0 | 0 | 0 | 0 | 0 | 0 | 0 | 0 | 0 | 0 | 0 | NA | NA | NA | NA | NA | NA | NA | NA | NA | NA |
| Mir183 | 0 | 0 | 0 | 0 | 0 | 0 | 0 | 0 | 0 | 0 | 0 | 0 | 0 | 0 | 0 | NA | NA | NA | NA | NA | NA | NA | NA | NA | NA |
| Mir1839 | 1 | 3 | 3 | 3 | 3 | 0 | 1 | 3 | 3 | 3 | 3 | 0 | 2.26 | 2 | 2 | 0.896 | -0.159 | 0.90676634 | NA | | 0 | 1.05 | 0.8726 | 0.8726 | TRUE | 0.7615 |
| Mir184 | 0 | 0 | 0 | 0 | 0 | 0 | 0 | 0 | 0 | 0 | 0 | 0 | 0 | 0 | 0 | NA | NA | NA | NA | NA | NA | NA | NA | NA | NA |
| Mir1843 | 0 | 0 | 0 | 0 | 0 | 0 | 0 | 0 | 0 | 0 | 0 | 0 | 0 | 0 | 0 | NA | NA | NA | NA | NA | NA | NA | NA | NA | NA |
| Mir1843b | 0 | 0 | 0 | 0 | 0 | 0 | 0 | 0 | 0 | 0 | 0 | 0 | 0 | 0 | 0 | NA | NA | NA | NA | NA | NA | NA | NA | NA | NA |

| | | | | | | | | | | | | | | | | | | | | | | | | | |
|---|---|---|---|---|---|---|---|---|---|---|---|---|---|---|---|---|---|---|---|---|---|---|---|---|---|
| Mir185 | 0 | 0 | 0 | 0 | 0 | 0 | 0 | 0 | 0 | 0 | 0 | 0 | 0 | 0 | 0 | NA | NA | NA | NA | NA | NA | NA | NA | NA | NA | |
| Mir186 | 2 | 2 | 1 | 2 | 3 | 3 | 2 | 2 | 1 | 2 | 3 | 3 | 2.18 | 2 | 3 | 1.746 | 0.804 | 0.52511666 | NA | | 0 | 1.0883 | 0.6642 | 0.6642 | TRUE | 0.0764 |
| Mir187 | 0 | 0 | 0 | 0 | 0 | 0 | 0 | 0 | 0 | 0 | 0 | 0 | 0 | 0 | 0 | NA | NA | NA | NA | NA | NA | NA | NA | NA | NA | |
| Mir188 | 0 | 0 | 0 | 0 | 0 | 0 | 0 | 0 | 0 | 0 | 0 | 0 | 0 | 0 | 0 | NA | NA | NA | NA | NA | NA | NA | NA | NA | NA | |
| Mir1892 | 0 | 0 | 0 | 0 | 0 | 0 | 0 | 0 | 0 | 0 | 0 | 0 | 0 | 0 | 0 | NA | NA | NA | NA | NA | NA | NA | NA | NA | NA | |
| Mir1893 | 0 | 0 | 0 | 0 | 0 | 0 | 0 | 0 | 0 | 0 | 0 | 0 | 0 | 0 | 0 | NA | NA | NA | NA | NA | NA | NA | NA | NA | NA | |
| Mir1894 | 2 | 4 | 1 | 0 | 2 | 1 | 2 | 4 | 1 | 0 | 2 | 1 | 1.69 | 2 | 1 | 0.465 | -1.106 | 0.47914146 | NA | | 0 | 1.4062 | 1.0576 | 1.0576 | TRUE | 0.4685 |
| Mir1895 | 0 | 0 | 0 | 0 | 0 | 0 | 0 | 0 | 0 | 0 | 0 | 0 | 0 | 0 | 0 | NA | NA | NA | NA | NA | NA | NA | NA | NA | NA | |
| Mir1896 | 0 | 0 | 0 | 0 | 0 | 0 | 0 | 0 | 0 | 0 | 0 | 0 | 0 | 0 | 0 | NA | NA | NA | NA | NA | NA | NA | NA | NA | NA | |
| Mir1897 | 0 | 0 | 0 | 0 | 0 | 0 | 0 | 0 | 0 | 0 | 0 | 0 | 0 | 0 | 0 | NA | NA | NA | NA | NA | NA | NA | NA | NA | NA | |
| Mir1898 | 3 | 1 | 0 | 3 | 0 | 1 | 2 | 1 | 0 | 3 | 0 | 1 | 1.23 | 1 | 1 | 1.143 | 0.193 | 0.91642119 | NA | | 0 | 1.9327 | 1.6311 | 1.6311 | TRUE | 0.3958 |
| Mir1899 | 0 | 0 | 0 | 0 | 0 | 0 | 0 | 0 | 0 | 0 | 0 | 0 | 0 | 0 | 0 | NA | NA | NA | NA | NA | NA | NA | NA | NA | NA | |
| Mir18b | 0 | 0 | 0 | 0 | 0 | 0 | 0 | 0 | 0 | 0 | 0 | 0 | 0 | 0 | 0 | NA | NA | NA | NA | NA | NA | NA | NA | NA | NA | |
| Mir190 | 0 | 0 | 0 | 0 | 0 | 0 | 0 | 0 | 0 | 0 | 0 | 0 | 0 | 0 | 0 | NA | NA | NA | NA | NA | NA | NA | NA | NA | NA | |
| Mir1900 | 0 | 0 | 0 | 1 | 0 | 0 | 0 | 0 | 0 | 1 | 0 | 0 | 0.17 | 0 | 0 | 2.094 | 1.067 | 0.79379329 | NA | | 0 | 14.2885 | 10 | 10 | TRUE | 0.3906 |
| Mir1901 | 0 | 0 | 0 | 0 | 0 | 0 | 0 | 0 | 0 | 0 | 0 | 0 | 0 | 0 | 0 | NA | NA | NA | NA | NA | NA | NA | NA | NA | NA | |
| Mir1902 | 0 | 0 | 0 | 0 | 0 | 0 | 0 | 0 | 0 | 0 | 0 | 0 | 0 | 0 | 0 | NA | NA | NA | NA | NA | NA | NA | NA | NA | NA | |
| Mir1903 | 0 | 0 | 0 | 0 | 0 | 0 | 0 | 0 | 0 | 0 | 0 | 0 | 0 | 0 | 0 | NA | NA | NA | NA | NA | NA | NA | NA | NA | NA | |
| Mir1904 | 0 | 0 | 0 | 0 | 0 | 0 | 0 | 0 | 0 | 0 | 0 | 0 | 0 | 0 | 0 | NA | NA | NA | NA | NA | NA | NA | NA | NA | NA | |
| Mir1905 | 0 | 0 | 0 | 0 | 0 | 0 | 0 | 0 | 0 | 0 | 0 | 0 | 0 | 0 | 0 | NA | NA | NA | NA | NA | NA | NA | NA | NA | NA | |
| Mir1906-1 | 0 | 0 | 0 | 0 | 0 | 0 | 0 | 0 | 0 | 0 | 0 | 0 | 0 | 0 | 0 | NA | NA | NA | NA | NA | NA | NA | NA | NA | NA | |
| Mir1906-2 | 0 | 0 | 0 | 0 | 0 | 0 | 0 | 0 | 0 | 0 | 0 | 0 | 0 | 0 | 0 | NA | NA | NA | NA | NA | NA | NA | NA | NA | NA | |
| Mir1907 | 2 | 2 | 0 | 1 | 2 | 0 | 2 | 2 | 0 | 1 | 2 | 0 | 1.16 | 1 | 1 | 0.859 | -0.22 | 0.90645413 | NA | | 0 | 2.0537 | 1.6433 | 1.6433 | TRUE | 0.2579 |
| Mir190b | 0 | 0 | 0 | 0 | 0 | 0 | 0 | 0 | 0 | 0 | 0 | 0 | 0 | 0 | 0 | NA | NA | NA | NA | NA | NA | NA | NA | NA | NA | |
| Mir191 | 1 | 0 | 0 | 0 | 0 | 0 | 1 | 0 | 0 | 0 | 0 | 0 | 0.13 | 0 | 0 | 0.552 | -0.857 | 0.8336548 | NA | | 0 | 18.3121 | 10 | 10 | TRUE | 0.2566 |
| Mir1912 | 0 | 0 | 0 | 0 | 0 | 0 | 0 | 0 | 0 | 0 | 0 | 0 | 0 | 0 | 0 | NA | NA | NA | NA | NA | NA | NA | NA | NA | NA | |
| Mir192 | 0 | 0 | 0 | 0 | 0 | 0 | 0 | 0 | 0 | 0 | 0 | 0 | 0 | 0 | 0 | NA | NA | NA | NA | NA | NA | NA | NA | NA | NA | |
| Mir1928 | 0 | 0 | 0 | 0 | 0 | 0 | 0 | 0 | 0 | 0 | 0 | 0 | 0 | 0 | 0 | NA | NA | NA | NA | NA | NA | NA | NA | NA | NA | |
| Mir1929 | 1 | 1 | 0 | 0 | 0 | 0 | 1 | 1 | 0 | 0 | 0 | 0 | 0.31 | 1 | 0 | 0.309 | -1.696 | 0.63183309 | NA | | 0 | 7.7294 | 7.2472 | 7.2472 | TRUE | 0.2015 |
| Mir193 | 0 | 0 | 0 | 0 | 0 | 0 | 0 | 0 | 0 | 0 | 0 | 0 | 0 | 0 | 0 | NA | NA | NA | NA | NA | NA | NA | NA | NA | NA | |
| Mir1930 | 0 | 0 | 0 | 0 | 0 | 0 | 0 | 0 | 0 | 0 | 0 | 0 | 0 | 0 | 0 | NA | NA | NA | NA | NA | NA | NA | NA | NA | NA | |
| Mir1931 | 0 | 0 | 0 | 0 | 1 | 0 | 0 | 0 | 0 | 0 | 1 | 0 | 0.19 | 0 | 0 | 2.094 | 1.067 | 0.79379329 | NA | | 0 | 12.608 | 10 | 10 | TRUE | 0.5075 |
| Mir1932 | 0 | 0 | 0 | 0 | 0 | 0 | 0 | 0 | 0 | 0 | 0 | 0 | 0 | 0 | 0 | NA | NA | NA | NA | NA | NA | NA | NA | NA | NA | |
| Mir1933 | 0 | 0 | 0 | 0 | 0 | 0 | 0 | 0 | 0 | 0 | 0 | 0 | 0 | 0 | 0 | NA | NA | NA | NA | NA | NA | NA | NA | NA | NA | |
| Mir1934 | 1 | 0 | 3 | 0 | 0 | 0 | 1 | 0 | 3 | 0 | 0 | 0 | 0.67 | 1 | 0 | 0.143 | -2.802 | 0.29586147 | NA | | 6.534 | 3.5454 | 3.8112 | 3.8112 | TRUE | 0.3623 |
| Mir1935 | 0 | 0 | 0 | 0 | 0 | 0 | 0 | 0 | 0 | 0 | 0 | 0 | 0 | 0 | 0 | NA | NA | NA | NA | NA | NA | NA | NA | NA | NA | |
| Mir1936 | 0 | 0 | 0 | 0 | 1 | 0 | 0 | 0 | 0 | 0 | 1 | 0 | 0.19 | 0 | 0 | 2.094 | 1.067 | 0.79379329 | NA | | 0 | 12.608 | 10 | 10 | TRUE | 0.5075 |
| Mir1938 | 2 | 0 | 2 | 0 | 0 | 1 | 2 | 0 | 2 | 0 | 0 | 1 | 0.78 | 1 | 0 | 0.298 | -1.746 | 0.45627758 | NA | | 0 | 3.029 | 2.568 | 2.568 | TRUE | 0.4004 |
| Mir193b | 0 | 0 | 0 | 0 | 0 | 0 | 0 | 0 | 0 | 0 | 0 | 0 | 0 | 0 | 0 | NA | NA | NA | NA | NA | NA | NA | NA | NA | NA | |
| Mir194-1 | 0 | 0 | 0 | 0 | 0 | 0 | 0 | 0 | 0 | 0 | 0 | 0 | 0 | 0 | 0 | NA | NA | NA | NA | NA | NA | NA | NA | NA | NA | |
| Mir194-2 | 0 | 0 | 0 | 0 | 0 | 0 | 0 | 0 | 0 | 0 | 0 | 0 | 0 | 0 | 0 | NA | NA | NA | NA | NA | NA | NA | NA | NA | NA | |
| Mir1940 | 0 | 0 | 0 | 0 | 0 | 0 | 0 | 0 | 0 | 0 | 0 | 0 | 0 | 0 | 0 | NA | NA | NA | NA | NA | NA | NA | NA | NA | NA | |
| Mir1941 | 0 | 0 | 0 | 0 | 0 | 0 | 0 | 0 | 0 | 0 | 0 | 0 | 0 | 0 | 0 | NA | NA | NA | NA | NA | NA | NA | NA | NA | NA | |
| Mir1942 | 0 | 0 | 0 | 0 | 0 | 0 | 0 | 0 | 0 | 0 | 0 | 0 | 0 | 0 | 0 | NA | NA | NA | NA | NA | NA | NA | NA | NA | NA | |
| Mir1943 | 2 | 2 | 1 | 1 | 2 | 4 | 2 | 2 | 1 | 1 | 2 | 4 | 1.99 | 2 | 2 | 1.515 | 0.6 | 0.65398804 | NA | | 0 | 1.1914 | 0.7712 | 0.7712 | TRUE | 0.221 |
| Mir1945 | 4 | 0 | 1 | 0 | 0 | 0 | 3 | 0 | 1 | 0 | 0 | 0 | 0.7 | 1 | 0 | 0.134 | -2.902 | 0.26925366 | NA | | 6.5513 | 3.3999 | 3.6307 | 3.6307 | TRUE | 0.1404 |
| Mir1946a | 0 | 0 | 0 | 0 | 0 | 0 | 0 | 0 | 0 | 0 | 0 | 0 | 0 | 0 | 0 | NA | NA | NA | NA | NA | NA | NA | NA | NA | NA | |
| Mir1946b | 1 | 0 | 0 | 0 | 0 | 0 | 1 | 0 | 0 | 0 | 0 | 0 | 0.13 | 0 | 0 | 0.552 | -0.857 | 0.8336548 | NA | | 0 | 18.3121 | 10 | 10 | TRUE | 0.2566 |
| Mir1947 | 0 | 0 | 0 | 0 | 0 | 0 | 0 | 0 | 0 | 0 | 0 | 0 | 0 | 0 | 0 | NA | NA | NA | NA | NA | NA | NA | NA | NA | NA | |
| Mir1948 | 0 | 0 | 0 | 0 | 0 | 0 | 0 | 0 | 0 | 0 | 0 | 0 | 0 | 0 | 0 | NA | NA | NA | NA | NA | NA | NA | NA | NA | NA | |
| Mir1949 | 10 | 16 | 6 | 10 | 7 | 8 | 8 | 17 | 6 | 10 | 8 | 8 | 9.5 | 10 | 9 | 0.831 | -0.267 | 0.68071178 | NA | | 0.0625 | 0.2519 | 0.196 | 0.196 | TRUE | 1.0288 |
| Mir195 | 0 | 0 | 0 | 0 | 0 | 0 | 0 | 0 | 0 | 0 | 0 | 0 | 0 | 0 | 0 | NA | NA | NA | NA | NA | NA | NA | NA | NA | NA | |
| Mir1950 | 0 | 0 | 0 | 0 | 0 | 0 | 0 | 0 | 0 | 0 | 0 | 0 | 0 | 0 | 0 | NA | NA | NA | NA | NA | NA | NA | NA | NA | NA | |
| Mir1951 | 0 | 0 | 0 | 0 | 0 | 0 | 0 | 0 | 0 | 0 | 0 | 0 | 0 | 0 | 0 | NA | NA | NA | NA | NA | NA | NA | NA | NA | NA | |
| Mir1952 | 0 | 0 | 0 | 0 | 0 | 0 | 0 | 0 | 0 | 0 | 0 | 0 | 0 | 0 | 0 | NA | NA | NA | NA | NA | NA | NA | NA | NA | NA | |
| Mir1953 | 0 | 0 | 0 | 0 | 0 | 0 | 0 | 0 | 0 | 0 | 0 | 0 | 0 | 0 | 0 | NA | NA | NA | NA | NA | NA | NA | NA | NA | NA | |
| Mir1954 | 0 | 0 | 0 | 0 | 0 | 0 | 0 | 0 | 0 | 0 | 0 | 0 | 0 | 0 | 0 | NA | NA | NA | NA | NA | NA | NA | NA | NA | NA | |
| Mir1955 | 0 | 0 | 0 | 0 | 0 | 0 | 0 | 0 | 0 | 0 | 0 | 0 | 0 | 0 | 0 | NA | NA | NA | NA | NA | NA | NA | NA | NA | NA | |
| Mir1956 | 0 | 0 | 0 | 0 | 0 | 0 | 0 | 0 | 0 | 0 | 0 | 0 | 0 | 0 | 0 | NA | NA | NA | NA | NA | NA | NA | NA | NA | NA | |
| Mir1957 | 0 | 0 | 0 | 0 | 0 | 0 | 0 | 0 | 0 | 0 | 0 | 0 | 0 | 0 | 0 | NA | NA | NA | NA | NA | NA | NA | NA | NA | NA | |
| Mir1958 | 0 | 0 | 0 | 0 | 0 | 0 | 0 | 0 | 0 | 0 | 0 | 0 | 0 | 0 | 0 | NA | NA | NA | NA | NA | NA | NA | NA | NA | NA | |
| Mir1960 | 0 | 0 | 0 | 0 | 0 | 0 | 0 | 0 | 0 | 0 | 0 | 0 | 0 | 0 | 0 | NA | NA | NA | NA | NA | NA | NA | NA | NA | NA | |
| Mir1961 | 0 | 0 | 0 | 0 | 0 | 0 | 0 | 0 | 0 | 0 | 0 | 0 | 0 | 0 | 0 | NA | NA | NA | NA | NA | NA | NA | NA | NA | NA | |
| Mir1962 | 0 | 0 | 0 | 0 | 0 | 0 | 0 | 0 | 0 | 0 | 0 | 0 | 0 | 0 | 0 | NA | NA | NA | NA | NA | NA | NA | NA | NA | NA | |
| Mir1963 | 0 | 1 | 0 | 0 | 0 | 1 | 0 | 1 | 0 | 0 | 0 | 1 | 0.34 | 0 | 0 | 1.075 | 0.105 | 0.97576789 | NA | | 0 | 6.9363 | 6.5821 | 6.5821 | TRUE | 0.5216 |
| Mir1964 | 0 | 1 | 0 | 0 | 0 | 0 | 0 | 1 | 0 | 0 | 0 | 0 | 0.18 | 0 | 0 | 0.552 | -0.857 | 0.8336548 | NA | | 0 | 13.3714 | 10 | 10 | TRUE | 0.5216 |
| Mir1965 | 0 | 0 | 0 | 0 | 0 | 0 | 0 | 0 | 0 | 0 | 0 | 0 | 0 | 0 | 0 | NA | NA | NA | NA | NA | NA | NA | NA | NA | NA | |
| Mir1966 | 0 | 0 | 1 | 0 | 0 | 0 | 0 | 0 | 1 | 0 | 0 | 0 | 0.18 | 0 | 0 | 0.552 | -0.857 | 0.8336548 | NA | | 0 | 13.1817 | 10 | 10 | TRUE | 0.5365 |
| Mir1967 | 0 | 0 | 0 | 0 | 0 | 0 | 0 | 0 | 0 | 0 | 0 | 0 | 0 | 0 | 0 | NA | NA | NA | NA | NA | NA | NA | NA | NA | NA | |

| | | | | | | | | | | | | | | | | | | | | | | | | | |
|---|---|---|---|---|---|---|---|---|---|---|---|---|---|---|---|---|---|---|---|---|---|---|---|---|---|
| Mir1968 | 5 | 3 | 3 | 2 | 4 | 3 | 4 | 3 | 3 | 2 | 5 | 3 | 3.3 | 3 | 3 | 0.906 | -0.142 | 0.88985342 | NA | 0 | 0.7215 | 0.4545 | 0.4545 | TRUE | 0.1696 |
| Mir1969 | 0 | 0 | 0 | 0 | 0 | 0 | 0 | 0 | 0 | 0 | 0 | 0 | 0 | 0 | 0 | NA | NA | NA | NA | NA | NA | NA | NA | NA | NA |
| Mir196a-1 | 7 | 13 | 6 | 9 | 4 | 7 | 5 | 14 | 6 | 9 | 5 | 7 | 7.69 | 8 | 7 | 0.806 | -0.312 | 0.6650617 | NA | 0 | 0.3108 | 0.2414 | 0.2414 | TRUE | 0.8469 |
| Mir196a-2 | 0 | 0 | 0 | 0 | 0 | 0 | 0 | 0 | 0 | 0 | 0 | 0 | 0 | 0 | 0 | NA | NA | NA | NA | NA | NA | NA | NA | NA | NA |
| Mir196b | 0 | 0 | 0 | 0 | 0 | 0 | 0 | 0 | 0 | 0 | 0 | 0 | 0 | 0 | 0 | NA | NA | NA | NA | NA | NA | NA | NA | NA | NA |
| Mir1970 | 0 | 0 | 0 | 0 | 0 | 0 | 0 | 0 | 0 | 0 | 0 | 0 | 0 | 0 | 0 | NA | NA | NA | NA | NA | NA | NA | NA | NA | NA |
| Mir1971 | 0 | 0 | 0 | 0 | 0 | 0 | 0 | 0 | 0 | 0 | 0 | 0 | 0 | 0 | 0 | NA | NA | NA | NA | NA | NA | NA | NA | NA | NA |
| Mir1981 | 0 | 0 | 0 | 0 | 0 | 0 | 0 | 0 | 0 | 0 | 0 | 0 | 0 | 0 | 0 | NA | NA | NA | NA | NA | NA | NA | NA | NA | NA |
| Mir1982 | 0 | 0 | 0 | 0 | 0 | 0 | 0 | 0 | 0 | 0 | 0 | 0 | 0 | 0 | 0 | NA | NA | NA | NA | NA | NA | NA | NA | NA | NA |
| Mir1983 | 0 | 1 | 0 | 0 | 0 | 0 | 0 | 1 | 0 | 0 | 0 | 0 | 0.18 | 0 | 0 | 0.552 | -0.857 | 0.8336548 | NA | 0 | 13.3714 | 10 | 10 | TRUE | 0.5216 |
| Mir199a-1 | 0 | 0 | 0 | 0 | 0 | 0 | 0 | 0 | 0 | 0 | 0 | 0 | 0 | 0 | 0 | NA | NA | NA | NA | NA | NA | NA | NA | NA | NA |
| Mir199a-2 | 0 | 0 | 0 | 0 | 0 | 0 | 0 | 0 | 0 | 0 | 0 | 0 | 0 | 0 | 0 | NA | NA | NA | NA | NA | NA | NA | NA | NA | NA |
| Mir199b | 0 | 0 | 0 | 0 | 0 | 0 | 0 | 0 | 0 | 0 | 0 | 0 | 0 | 0 | 0 | NA | NA | NA | NA | NA | NA | NA | NA | NA | NA |
| Mir19a | 0 | 0 | 0 | 0 | 0 | 1 | 0 | 0 | 0 | 0 | 0 | 1 | 0.16 | 0 | 0 | 2.094 | 1.067 | 0.79379329 | NA | 0 | 14.4078 | 10 | 10 | TRUE | 0.3835 |
| Mir19b-1 | 1 | 1 | 1 | 0 | 0 | 0 | 1 | 1 | 1 | 0 | 0 | 0 | 0.49 | 1 | 0 | 0.196 | -2.347 | 0.38405676 | NA | 0 | 4.8735 | 3.7415 | 3.7415 | TRUE | 0.0718 |
| Mir19b-2 | 0 | 0 | 0 | 0 | 0 | 0 | 0 | 0 | 0 | 0 | 0 | 0 | 0 | 0 | 0 | NA | NA | NA | NA | NA | NA | NA | NA | NA | NA |
| Mir1a-1 | 0 | 0 | 0 | 0 | 0 | 0 | 0 | 0 | 0 | 0 | 0 | 0 | 0 | 0 | 0 | NA | NA | NA | NA | NA | NA | NA | NA | NA | NA |
| Mir1a-2 | 0 | 0 | 0 | 0 | 0 | 0 | 0 | 0 | 0 | 0 | 0 | 0 | 0 | 0 | 0 | NA | NA | NA | NA | NA | NA | NA | NA | NA | NA |
| Mir1b | 0 | 0 | 0 | 0 | 0 | 0 | 0 | 0 | 0 | 0 | 0 | 0 | 0 | 0 | 0 | NA | NA | NA | NA | NA | NA | NA | NA | NA | NA |
| Mir200a | 0 | 0 | 0 | 0 | 0 | 0 | 0 | 0 | 0 | 0 | 0 | 0 | 0 | 0 | 0 | NA | NA | NA | NA | NA | NA | NA | NA | NA | NA |
| Mir200b | 1 | 0 | 0 | 0 | 0 | 0 | 1 | 0 | 0 | 0 | 0 | 0 | 0.13 | 0 | 0 | 0.552 | -0.857 | 0.8336548 | NA | 0 | 18.3121 | 10 | 10 | TRUE | 0.2566 |
| Mir200c | 0 | 0 | 0 | 0 | 0 | 0 | 0 | 0 | 0 | 0 | 0 | 0 | 0 | 0 | 0 | NA | NA | NA | NA | NA | NA | NA | NA | NA | NA |
| Mir201 | 0 | 0 | 0 | 0 | 0 | 0 | 0 | 0 | 0 | 0 | 0 | 0 | 0 | 0 | 0 | NA | NA | NA | NA | NA | NA | NA | NA | NA | NA |
| Mir202 | 0 | 0 | 0 | 0 | 0 | 0 | 0 | 0 | 0 | 0 | 0 | 0 | 0 | 0 | 0 | NA | NA | NA | NA | NA | NA | NA | NA | NA | NA |
| Mir203 | 0 | 0 | 0 | 0 | 0 | 0 | 0 | 0 | 0 | 0 | 0 | 0 | 0 | 0 | 0 | NA | NA | NA | NA | NA | NA | NA | NA | NA | NA |
| Mir204 | 0 | 0 | 0 | 0 | 0 | 0 | 0 | 0 | 0 | 0 | 0 | 0 | 0 | 0 | 0 | NA | NA | NA | NA | NA | NA | NA | NA | NA | NA |
| Mir205 | 0 | 0 | 0 | 0 | 0 | 0 | 0 | 0 | 0 | 0 | 0 | 0 | 0 | 0 | 0 | NA | NA | NA | NA | NA | NA | NA | NA | NA | NA |
| Mir206 | 0 | 0 | 0 | 0 | 0 | 0 | 0 | 0 | 0 | 0 | 0 | 0 | 0 | 0 | 0 | NA | NA | NA | NA | NA | NA | NA | NA | NA | NA |
| Mir207 | 0 | 0 | 0 | 0 | 0 | 0 | 0 | 0 | 0 | 0 | 0 | 0 | 0 | 0 | 0 | NA | NA | NA | NA | NA | NA | NA | NA | NA | NA |
| Mir208a | 0 | 0 | 0 | 0 | 0 | 0 | 0 | 0 | 0 | 0 | 0 | 0 | 0 | 0 | 0 | NA | NA | NA | NA | NA | NA | NA | NA | NA | NA |
| Mir208b | 0 | 0 | 0 | 0 | 0 | 0 | 0 | 0 | 0 | 0 | 0 | 0 | 0 | 0 | 0 | NA | NA | NA | NA | NA | NA | NA | NA | NA | NA |
| Mir20a | 1 | 4 | 0 | 0 | 0 | 0 | 1 | 4 | 0 | 0 | 0 | 0 | 0.84 | 2 | 0 | 0.114 | -3.128 | 0.21102852 | NA | 7.4731 | 2.8287 | 3.2172 | 3.2172 | TRUE | 0.737 |
| Mir20b | 0 | 0 | 0 | 0 | 0 | 0 | 0 | 0 | 0 | 0 | 0 | 0 | 0 | 0 | 0 | NA | NA | NA | NA | NA | NA | NA | NA | NA | NA |
| Mir21 | 0 | 0 | 0 | 0 | 0 | 0 | 0 | 0 | 0 | 0 | 0 | 0 | 0 | 0 | 0 | NA | NA | NA | NA | NA | NA | NA | NA | NA | NA |
| Mir210 | 0 | 0 | 0 | 1 | 0 | 0 | 0 | 0 | 0 | 1 | 0 | 0 | 0.17 | 0 | 0 | 2.094 | 1.067 | 0.79379329 | NA | 0 | 14.2885 | 10 | 10 | TRUE | 0.3906 |
| Mir211 | 0 | 0 | 0 | 0 | 0 | 0 | 0 | 0 | 0 | 0 | 0 | 0 | 0 | 0 | 0 | NA | NA | NA | NA | NA | NA | NA | NA | NA | NA |
| Mir212 | 0 | 0 | 0 | 0 | 0 | 0 | 0 | 0 | 0 | 0 | 0 | 0 | 0 | 0 | 0 | NA | NA | NA | NA | NA | NA | NA | NA | NA | NA |
| Mir2136 | 0 | 0 | 0 | 0 | 0 | 0 | 0 | 0 | 0 | 0 | 0 | 0 | 0 | 0 | 0 | NA | NA | NA | NA | NA | NA | NA | NA | NA | NA |
| Mir2137 | 0 | 0 | 0 | 0 | 0 | 0 | 0 | 0 | 0 | 0 | 0 | 0 | 0 | 0 | 0 | NA | NA | NA | NA | NA | NA | NA | NA | NA | NA |
| Mir214 | 0 | 0 | 0 | 0 | 0 | 0 | 0 | 0 | 0 | 0 | 0 | 0 | 0 | 0 | 0 | NA | NA | NA | NA | NA | NA | NA | NA | NA | NA |
| Mir215 | 0 | 0 | 0 | 0 | 0 | 0 | 0 | 0 | 0 | 0 | 0 | 0 | 0 | 0 | 0 | NA | NA | NA | NA | NA | NA | NA | NA | NA | NA |
| Mir216a | 0 | 0 | 0 | 0 | 0 | 0 | 0 | 0 | 0 | 0 | 0 | 0 | 0 | 0 | 0 | NA | NA | NA | NA | NA | NA | NA | NA | NA | NA |
| Mir216b | 0 | 0 | 0 | 0 | 0 | 0 | 0 | 0 | 0 | 0 | 0 | 0 | 0 | 0 | 0 | NA | NA | NA | NA | NA | NA | NA | NA | NA | NA |
| Mir217 | 0 | 0 | 0 | 0 | 0 | 0 | 0 | 0 | 0 | 0 | 0 | 0 | 0 | 0 | 0 | NA | NA | NA | NA | NA | NA | NA | NA | NA | NA |
| Mir218-1 | 0 | 0 | 0 | 0 | 0 | 0 | 0 | 0 | 0 | 0 | 0 | 0 | 0 | 0 | 0 | NA | NA | NA | NA | NA | NA | NA | NA | NA | NA |
| Mir218-2 | 0 | 0 | 0 | 0 | 0 | 0 | 0 | 0 | 0 | 0 | 0 | 0 | 0 | 0 | 0 | NA | NA | NA | NA | NA | NA | NA | NA | NA | NA |
| Mir219-1 | 0 | 0 | 0 | 0 | 0 | 0 | 0 | 0 | 0 | 0 | 0 | 0 | 0 | 0 | 0 | NA | NA | NA | NA | NA | NA | NA | NA | NA | NA |
| Mir219-2 | 0 | 0 | 0 | 0 | 0 | 0 | 0 | 0 | 0 | 0 | 0 | 0 | 0 | 0 | 0 | NA | NA | NA | NA | NA | NA | NA | NA | NA | NA |
| Mir22 | 0 | 0 | 0 | 0 | 0 | 0 | 0 | 0 | 0 | 0 | 0 | 0 | 0 | 0 | 0 | NA | NA | NA | NA | NA | NA | NA | NA | NA | NA |
| Mir221 | 0 | 0 | 0 | 0 | 0 | 0 | 0 | 0 | 0 | 0 | 0 | 0 | 0 | 0 | 0 | NA | NA | NA | NA | NA | NA | NA | NA | NA | NA |
| Mir222 | 0 | 0 | 0 | 0 | 0 | 0 | 0 | 0 | 0 | 0 | 0 | 0 | 0 | 0 | 0 | NA | NA | NA | NA | NA | NA | NA | NA | NA | NA |
| Mir223 | 0 | 0 | 0 | 0 | 0 | 0 | 0 | 0 | 0 | 0 | 0 | 0 | 0 | 0 | 0 | NA | NA | NA | NA | NA | NA | NA | NA | NA | NA |
| Mir22hg | 156 | 103 | 88 | 104 | 76 | 69 | 121 | 110 | 95 | 104 | 86 | 68 | 97.17 | 109 | 86 | 0.788 | -0.345 | 0.11693618 | 0.24783049 | 0.0179 | 0.0268 | 0.0243 | 0.0243 | TRUE | 0.2409 |
| Mir23a | 0 | 0 | 0 | 0 | 0 | 0 | 0 | 0 | 0 | 0 | 0 | 0 | 0 | 0 | 0 | NA | NA | NA | NA | NA | NA | NA | NA | NA | NA |
| Mir23b | 0 | 0 | 0 | 0 | 0 | 0 | 0 | 0 | 0 | 0 | 0 | 0 | 0 | 0 | 0 | NA | NA | NA | NA | NA | NA | NA | NA | NA | NA |
| Mir24-1 | 1 | 0 | 0 | 0 | 0 | 0 | 1 | 0 | 0 | 0 | 0 | 0 | 0.13 | 0 | 0 | 0.552 | -0.857 | 0.8336548 | NA | 0 | 18.3121 | 10 | 10 | TRUE | 0.2566 |
| Mir24-2 | 0 | 0 | 0 | 1 | 0 | 0 | 0 | 0 | 0 | 1 | 0 | 0 | 0.17 | 0 | 0 | 2.094 | 1.067 | 0.79379329 | NA | 0 | 14.2885 | 10 | 10 | TRUE | 0.3906 |
| Mir25 | 5 | 6 | 6 | 7 | 3 | 6 | 4 | 6 | 6 | 7 | 3 | 6 | 5.5 | 5 | 5 | 0.99 | -0.014 | 0.98624738 | NA | 0 | 0.4332 | 0.2906 | 0.2906 | TRUE | 0.2119 |
| Mir26a-1 | 0 | 0 | 0 | 0 | 0 | 0 | 0 | 0 | 0 | 0 | 0 | 0 | 0 | 0 | 0 | NA | NA | NA | NA | NA | NA | NA | NA | NA | NA |
| Mir26a-2 | 0 | 0 | 0 | 0 | 0 | 0 | 0 | 0 | 0 | 0 | 0 | 0 | 0 | 0 | 0 | NA | NA | NA | NA | NA | NA | NA | NA | NA | NA |
| Mir26b | 0 | 0 | 0 | 0 | 0 | 0 | 0 | 0 | 0 | 0 | 0 | 0 | 0 | 0 | 0 | NA | NA | NA | NA | NA | NA | NA | NA | NA | NA |
| Mir27a | 0 | 0 | 0 | 0 | 0 | 0 | 0 | 0 | 0 | 0 | 0 | 0 | 0 | 0 | 0 | NA | NA | NA | NA | NA | NA | NA | NA | NA | NA |
| Mir27b | 0 | 0 | 0 | 0 | 0 | 0 | 0 | 0 | 0 | 0 | 0 | 0 | 0 | 0 | 0 | NA | NA | NA | NA | NA | NA | NA | NA | NA | NA |
| Mir28 | 0 | 0 | 0 | 0 | 0 | 0 | 0 | 0 | 0 | 0 | 0 | 0 | 0 | 0 | 0 | NA | NA | NA | NA | NA | NA | NA | NA | NA | NA |
| Mir2861 | 4 | 5 | 2 | 5 | 3 | 4 | 3 | 5 | 2 | 5 | 3 | 4 | 3.82 | 3 | 4 | 1.171 | 0.228 | 0.81277008 | NA | 0 | 0.6236 | 0.4056 | 0.4056 | TRUE | 0.2668 |
| Mir28b | 0 | 0 | 0 | 0 | 0 | 0 | 0 | 0 | 0 | 0 | 0 | 0 | 0 | 0 | 0 | NA | NA | NA | NA | NA | NA | NA | NA | NA | NA |
| Mir28c | 0 | 0 | 0 | 0 | 0 | 0 | 0 | 0 | 0 | 0 | 0 | 0 | 0 | 0 | 0 | NA | NA | NA | NA | NA | NA | NA | NA | NA | NA |
| Mir290 | 2 | 1 | 1 | 1 | 0 | 0 | 2 | 1 | 1 | 1 | 0 | 0 | 0.78 | 1 | 0 | 0.299 | -1.743 | 0.425346 | NA | 0 | 3.0336 | 2.0559 | 2.0559 | TRUE | 0.3906 |
| Mir291a | 0 | 0 | 0 | 0 | 0 | 0 | 0 | 0 | 0 | 0 | 0 | 0 | 0 | 0 | 0 | NA | NA | NA | NA | NA | NA | NA | NA | NA | NA |

| | | | | | | | | | | | | | | | | | | | | | | | | | |
|---|---|---|---|---|---|---|---|---|---|---|---|---|---|---|---|---|---|---|---|---|---|---|---|---|---|
| Mir291b | 0 | 0 | 0 | 0 | 0 | 0 | 0 | 0 | 0 | 0 | 0 | 0 | 0 | 0 | 0 | NA | NA | NA | NA | NA | NA | NA | NA | NA | NA |
| Mir292 | 0 | 0 | 0 | 0 | 0 | 0 | 0 | 0 | 0 | 0 | 0 | 0 | 0 | 0 | 0 | NA | NA | NA | NA | NA | NA | NA | NA | NA | NA |
| Mir293 | 0 | 0 | 0 | 0 | 0 | 0 | 0 | 0 | 0 | 0 | 0 | 0 | 0 | 0 | 0 | NA | NA | NA | NA | NA | NA | NA | NA | NA | NA |
| Mir294 | 0 | 0 | 0 | 0 | 0 | 0 | 0 | 0 | 0 | 0 | 0 | 0 | 0 | 0 | 0 | NA | NA | NA | NA | NA | NA | NA | NA | NA | NA |
| Mir295 | 0 | 0 | 0 | 0 | 0 | 0 | 0 | 0 | 0 | 0 | 0 | 0 | 0 | 0 | 0 | NA | NA | NA | NA | NA | NA | NA | NA | NA | NA |
| Mir296 | 0 | 0 | 0 | 0 | 0 | 0 | 0 | 0 | 0 | 0 | 0 | 0 | 0 | 0 | 0 | NA | NA | NA | NA | NA | NA | NA | NA | NA | NA |
| Mir297-1 | 0 | 0 | 0 | 0 | 0 | 0 | 0 | 0 | 0 | 0 | 0 | 0 | 0 | 0 | 0 | NA | NA | NA | NA | NA | NA | NA | NA | NA | NA |
| Mir297-2 | 0 | 0 | 0 | 0 | 0 | 0 | 0 | 0 | 0 | 0 | 0 | 0 | 0 | 0 | 0 | NA | NA | NA | NA | NA | NA | NA | NA | NA | NA |
| Mir297a-3 | 0 | 0 | 0 | 0 | 0 | 0 | 0 | 0 | 0 | 0 | 0 | 0 | 0 | 0 | 0 | NA | NA | NA | NA | NA | NA | NA | NA | NA | NA |
| Mir297a-4 | 0 | 0 | 0 | 0 | 0 | 0 | 0 | 0 | 0 | 0 | 0 | 0 | 0 | 0 | 0 | NA | NA | NA | NA | NA | NA | NA | NA | NA | NA |
| Mir297b | 0 | 0 | 0 | 0 | 0 | 0 | 0 | 0 | 0 | 0 | 0 | 0 | 0 | 0 | 0 | NA | NA | NA | NA | NA | NA | NA | NA | NA | NA |
| Mir297c | 0 | 0 | 0 | 0 | 0 | 0 | 0 | 0 | 0 | 0 | 0 | 0 | 0 | 0 | 0 | NA | NA | NA | NA | NA | NA | NA | NA | NA | NA |
| Mir298 | 0 | 0 | 0 | 0 | 0 | 0 | 0 | 0 | 0 | 0 | 0 | 0 | 0 | 0 | 0 | NA | NA | NA | NA | NA | NA | NA | NA | NA | NA |
| Mir299 | 0 | 0 | 0 | 0 | 0 | 0 | 0 | 0 | 0 | 0 | 0 | 0 | 0 | 0 | 0 | NA | NA | NA | NA | NA | NA | NA | NA | NA | NA |
| Mir29a | 0 | 0 | 0 | 0 | 0 | 0 | 0 | 0 | 0 | 0 | 0 | 0 | 0 | 0 | 0 | NA | NA | NA | NA | NA | NA | NA | NA | NA | NA |
| Mir29b-1 | 0 | 0 | 0 | 0 | 0 | 0 | 0 | 0 | 0 | 0 | 0 | 0 | 0 | 0 | 0 | NA | NA | NA | NA | NA | NA | NA | NA | NA | NA |
| Mir29b-2 | 0 | 0 | 0 | 0 | 0 | 0 | 0 | 0 | 0 | 0 | 0 | 0 | 0 | 0 | 0 | NA | NA | NA | NA | NA | NA | NA | NA | NA | NA |
| Mir29c | 0 | 0 | 0 | 0 | 0 | 0 | 0 | 0 | 0 | 0 | 0 | 0 | 0 | 0 | 0 | NA | NA | NA | NA | NA | NA | NA | NA | NA | NA |
| Mir300 | 0 | 0 | 0 | 0 | 0 | 0 | 0 | 0 | 0 | 0 | 0 | 0 | 0 | 0 | 0 | NA | NA | NA | NA | NA | NA | NA | NA | NA | NA |
| Mir301 | 0 | 0 | 0 | 0 | 1 | 0 | 0 | 0 | 0 | 0 | 1 | 0 | 0.19 | 0 | 0 | 2.094 | 1.067 | 0.79379329 | NA | 0 | 12.608 | 10 | 10 | TRUE | 0.5075 |
| Mir301b | 0 | 0 | 0 | 0 | 0 | 0 | 0 | 0 | 0 | 0 | 0 | 0 | 0 | 0 | 0 | NA | NA | NA | NA | NA | NA | NA | NA | NA | NA |
| Mir302a | 0 | 0 | 0 | 0 | 0 | 0 | 0 | 0 | 0 | 0 | 0 | 0 | 0 | 0 | 0 | NA | NA | NA | NA | NA | NA | NA | NA | NA | NA |
| Mir302b | 0 | 0 | 0 | 0 | 0 | 0 | 0 | 0 | 0 | 0 | 0 | 0 | 0 | 0 | 0 | NA | NA | NA | NA | NA | NA | NA | NA | NA | NA |
| Mir302c | 0 | 0 | 0 | 0 | 0 | 0 | 0 | 0 | 0 | 0 | 0 | 0 | 0 | 0 | 0 | NA | NA | NA | NA | NA | NA | NA | NA | NA | NA |
| Mir302d | 0 | 0 | 0 | 0 | 0 | 0 | 0 | 0 | 0 | 0 | 0 | 0 | 0 | 0 | 0 | NA | NA | NA | NA | NA | NA | NA | NA | NA | NA |
| Mir3057 | 1 | 2 | 3 | 3 | 2 | 0 | 1 | 2 | 3 | 3 | 2 | 0 | 1.9 | 2 | 2 | 0.868 | -0.204 | 0.8879786 | NA | 0 | 1.2516 | 0.9783 | 0.9783 | TRUE | 0.5261 |
| Mir3058 | 3 | 1 | 4 | 4 | 4 | 2 | 2 | 1 | 4 | 4 | 5 | 2 | 3.03 | 2 | 4 | 1.359 | 0.442 | 0.6887743 | NA | 0 | 0.7847 | 0.5402 | 0.5402 | TRUE | 0.3502 |
| Mir3059 | 0 | 0 | 0 | 0 | 0 | 0 | 0 | 0 | 0 | 0 | 0 | 0 | 0 | 0 | 0 | NA | NA | NA | NA | NA | NA | NA | NA | NA | NA |
| Mir3060 | 2 | 0 | 0 | 1 | 0 | 1 | 2 | 0 | 0 | 1 | 0 | 1 | 0.59 | 1 | 1 | 1.206 | 0.27 | 0.91756405 | NA | 0 | 4.0242 | 3.277 | 3.277 | TRUE | 0.9721 |
| Mir3061 | 7 | 8 | 7 | 2 | 9 | 9 | 5 | 9 | 8 | 2 | 10 | 9 | 7.09 | 7 | 7 | 0.98 | -0.029 | 0.96956758 | NA | 0.1179 | 0.3368 | 0.2801 | 0.2801 | TRUE | 1.0746 |
| Mir3062 | 2 | 1 | 1 | 0 | 0 | 2 | 2 | 1 | 1 | 0 | 0 | 2 | 0.95 | 1 | 1 | 0.541 | -0.886 | 0.66458129 | NA | 0 | 2.5103 | 1.8467 | 1.8467 | TRUE | 0.9735 |
| Mir3063 | 0 | 0 | 0 | 0 | 0 | 0 | 0 | 0 | 0 | 0 | 0 | 0 | 0 | 0 | 0 | NA | NA | NA | NA | NA | NA | NA | NA | NA | NA |
| Mir3064 | 105 | 124 | 85 | 104 | 92 | 103 | 82 | 132 | 92 | 104 | 104 | 102 | 102.36 | 102 | 103 | 1.018 | 0.026 | 0.90699225 | 0.94914932 | 0.0237 | 0.0256 | 0.025 | 0.025 | TRUE | 0.6492 |
| Mir3065 | 0 | 0 | 0 | 0 | 0 | 0 | 0 | 0 | 0 | 0 | 0 | 0 | 0 | 0 | 0 | NA | NA | NA | NA | NA | NA | NA | NA | NA | NA |
| Mir3066 | 0 | 0 | 0 | 0 | 0 | 0 | 0 | 0 | 0 | 0 | 0 | 0 | 0 | 0 | 0 | NA | NA | NA | NA | NA | NA | NA | NA | NA | NA |
| Mir3067 | 0 | 0 | 0 | 0 | 0 | 0 | 0 | 0 | 0 | 0 | 0 | 0 | 0 | 0 | 0 | NA | NA | NA | NA | NA | NA | NA | NA | NA | NA |
| Mir3068 | 8 | 1 | 1 | 2 | 2 | 4 | 6 | 1 | 1 | 2 | 2 | 4 | 2.76 | 3 | 3 | 0.944 | -0.083 | 0.94461314 | NA | 0.3032 | 0.8616 | 0.6824 | 0.6824 | TRUE | 1.8171 |
| Mir3069 | 0 | 0 | 0 | 0 | 0 | 0 | 0 | 0 | 0 | 0 | 0 | 0 | 0 | 0 | 0 | NA | NA | NA | NA | NA | NA | NA | NA | NA | NA |
| Mir3070a | 0 | 0 | 0 | 0 | 0 | 0 | 0 | 0 | 0 | 0 | 0 | 0 | 0 | 0 | 0 | NA | NA | NA | NA | NA | NA | NA | NA | NA | NA |
| Mir3070b | 1 | 0 | 0 | 2 | 0 | 0 | 1 | 0 | 0 | 2 | 0 | 0 | 0.46 | 0 | 1 | 1.942 | 0.958 | 0.7580687 | NA | 0 | 5.1409 | 5.1921 | 5.1921 | TRUE | 0.9738 |
| Mir3071 | 0 | 0 | 0 | 0 | 0 | 0 | 0 | 0 | 0 | 0 | 0 | 0 | 0 | 0 | 0 | NA | NA | NA | NA | NA | NA | NA | NA | NA | NA |
| Mir3072 | 4 | 6 | 5 | 6 | 1 | 3 | 3 | 6 | 5 | 6 | 1 | 3 | 4.16 | 5 | 3 | 0.692 | -0.531 | 0.58434504 | NA | 0 | 0.5725 | 0.427 | 0.427 | TRUE | 0.4984 |
| Mir3073 | 0 | 0 | 0 | 0 | 0 | 0 | 0 | 0 | 0 | 0 | 0 | 0 | 0 | 0 | 0 | NA | NA | NA | NA | NA | NA | NA | NA | NA | NA |
| Mir3074-1 | 0 | 0 | 0 | 0 | 0 | 0 | 0 | 0 | 0 | 0 | 0 | 0 | 0 | 0 | 0 | NA | NA | NA | NA | NA | NA | NA | NA | NA | NA |
| Mir3074-2 | 0 | 0 | 0 | 0 | 0 | 0 | 0 | 0 | 0 | 0 | 0 | 0 | 0 | 0 | 0 | NA | NA | NA | NA | NA | NA | NA | NA | NA | NA |
| Mir3075 | 0 | 0 | 0 | 0 | 0 | 0 | 0 | 0 | 0 | 0 | 0 | 0 | 0 | 0 | 0 | NA | NA | NA | NA | NA | NA | NA | NA | NA | NA |
| Mir3076 | 0 | 0 | 1 | 0 | 0 | 0 | 0 | 0 | 1 | 0 | 0 | 0 | 0.18 | 0 | 0 | 0.552 | -0.857 | 0.8336548 | NA | 0 | 13.1817 | 10 | 10 | TRUE | 0.5365 |
| Mir3077 | 0 | 0 | 0 | 0 | 0 | 0 | 0 | 0 | 0 | 0 | 0 | 0 | 0 | 0 | 0 | NA | NA | NA | NA | NA | NA | NA | NA | NA | NA |
| Mir3078 | 0 | 0 | 0 | 0 | 0 | 0 | 0 | 0 | 0 | 0 | 0 | 0 | 0 | 0 | 0 | NA | NA | NA | NA | NA | NA | NA | NA | NA | NA |
| Mir3079 | 0 | 0 | 0 | 0 | 0 | 0 | 0 | 0 | 0 | 0 | 0 | 0 | 0 | 0 | 0 | NA | NA | NA | NA | NA | NA | NA | NA | NA | NA |
| Mir3081 | 0 | 0 | 0 | 0 | 0 | 0 | 0 | 0 | 0 | 0 | 0 | 0 | 0 | 0 | 0 | NA | NA | NA | NA | NA | NA | NA | NA | NA | NA |
| Mir3082 | 20 | 19 | 16 | 18 | 11 | 13 | 16 | 20 | 17 | 18 | 12 | 13 | 16.03 | 18 | 14 | 0.82 | -0.287 | 0.54836521 | NA | 0 | 0.1503 | 0.1014 | 0.1014 | TRUE | 0.2091 |
| Mir3083 | 0 | 0 | 0 | 0 | 0 | 0 | 0 | 0 | 0 | 0 | 0 | 0 | 0 | 0 | 0 | NA | NA | NA | NA | NA | NA | NA | NA | NA | NA |
| Mir3085 | 0 | 0 | 0 | 0 | 0 | 0 | 0 | 0 | 0 | 0 | 0 | 0 | 0 | 0 | 0 | NA | NA | NA | NA | NA | NA | NA | NA | NA | NA |
| Mir3086 | 0 | 0 | 0 | 0 | 0 | 0 | 0 | 0 | 0 | 0 | 0 | 0 | 0 | 0 | 0 | NA | NA | NA | NA | NA | NA | NA | NA | NA | NA |
| Mir3087 | 0 | 0 | 0 | 0 | 0 | 0 | 0 | 0 | 0 | 0 | 0 | 0 | 0 | 0 | 0 | NA | NA | NA | NA | NA | NA | NA | NA | NA | NA |
| Mir3088 | 0 | 0 | 0 | 0 | 0 | 0 | 0 | 0 | 0 | 0 | 0 | 0 | 0 | 0 | 0 | NA | NA | NA | NA | NA | NA | NA | NA | NA | NA |
| Mir3089 | 0 | 0 | 0 | 0 | 0 | 0 | 0 | 0 | 0 | 0 | 0 | 0 | 0 | 0 | 0 | NA | NA | NA | NA | NA | NA | NA | NA | NA | NA |
| Mir3091 | 72 | 50 | 28 | 57 | 31 | 44 | 56 | 53 | 30 | 57 | 35 | 43 | 45.74 | 46 | 45 | 0.967 | -0.049 | 0.88141212 | 0.93380946 | 0.0576 | 0.0542 | 0.0552 | 0.0552 | TRUE | 0.6199 |
| Mir3092 | 0 | 0 | 0 | 0 | 0 | 0 | 0 | 0 | 0 | 0 | 0 | 0 | 0 | 0 | 0 | NA | NA | NA | NA | NA | NA | NA | NA | NA | NA |
| Mir3093 | 0 | 0 | 0 | 0 | 0 | 0 | 0 | 0 | 0 | 0 | 0 | 0 | 0 | 0 | 0 | NA | NA | NA | NA | NA | NA | NA | NA | NA | NA |
| Mir3094 | 0 | 0 | 0 | 0 | 0 | 0 | 0 | 0 | 0 | 0 | 0 | 0 | 0 | 0 | 0 | NA | NA | NA | NA | NA | NA | NA | NA | NA | NA |
| Mir3095 | 0 | 0 | 0 | 0 | 0 | 0 | 0 | 0 | 0 | 0 | 0 | 0 | 0 | 0 | 0 | NA | NA | NA | NA | NA | NA | NA | NA | NA | NA |
| Mir3096 | 1 | 0 | 0 | 0 | 1 | 1 | 1 | 0 | 0 | 0 | 1 | 1 | 0.48 | 0 | 1 | 2.031 | 1.022 | 0.71376068 | NA | 0 | 4.9197 | 3.859 | 3.859 | TRUE | 0.2623 |
| Mir3096b | 1 | 2 | 0 | 0 | 0 | 3 | 1 | 2 | 0 | 0 | 0 | 3 | 0.98 | 1 | 1 | 1.039 | 0.055 | 0.9797912 | NA | 2.2872 | 2.4268 | 2.4046 | 2.4046 | TRUE | 1.1825 |
| Mir3097 | 0 | 0 | 0 | 0 | 0 | 1 | 0 | 0 | 0 | 0 | 0 | 1 | 0.16 | 0 | 0 | 2.094 | 1.067 | 0.79379329 | NA | 0 | 14.4078 | 10 | 10 | TRUE | 0.3835 |
| Mir3098 | 0 | 0 | 0 | 0 | 0 | 0 | 0 | 0 | 0 | 0 | 0 | 0 | 0 | 0 | 0 | NA | NA | NA | NA | NA | NA | NA | NA | NA | NA |
| Mir3099 | 0 | 0 | 0 | 0 | 0 | 0 | 0 | 0 | 0 | 0 | 0 | 0 | 0 | 0 | 0 | NA | NA | NA | NA | NA | NA | NA | NA | NA | NA |
| Mir30a | 0 | 0 | 0 | 0 | 0 | 0 | 0 | 0 | 0 | 0 | 0 | 0 | 0 | 0 | 0 | NA | NA | NA | NA | NA | NA | NA | NA | NA | NA |

| | | | | | | | | | | | | | | | | | | | | | | | | | |
|---|---|---|---|---|---|---|---|---|---|---|---|---|---|---|---|---|---|---|---|---|---|---|---|---|---|
| Mir30b | 0 | 0 | 0 | 0 | 0 | 0 | 0 | 0 | 0 | 0 | 0 | 0 | 0 | 0 | 0 | NA | NA | NA | NA | NA | NA | NA | NA | NA | NA |
| Mir30c-1 | 0 | 0 | 0 | 0 | 0 | 0 | 0 | 0 | 0 | 0 | 0 | 0 | 0 | 0 | 0 | NA | NA | NA | NA | NA | NA | NA | NA | NA | NA |
| Mir30c-2 | 0 | 0 | 0 | 0 | 0 | 0 | 0 | 0 | 0 | 0 | 0 | 0 | 0 | 0 | 0 | NA | NA | NA | NA | NA | NA | NA | NA | NA | NA |
| Mir30d | 0 | 0 | 0 | 1 | 0 | 0 | 0 | 0 | 0 | 1 | 0 | 0 | 0.17 | 0 | 0 | 2.094 | 1.067 | 0.79379329 | NA | 0 | 14.2885 | 10 | 10 | TRUE | 0.3906 |
| Mir30e | 0 | 0 | 0 | 0 | 0 | 0 | 0 | 0 | 0 | 0 | 0 | 0 | 0 | 0 | 0 | NA | NA | NA | NA | NA | NA | NA | NA | NA | NA |
| Mir31 | 0 | 0 | 0 | 0 | 0 | 0 | 0 | 0 | 0 | 0 | 0 | 0 | 0 | 0 | 0 | NA | NA | NA | NA | NA | NA | NA | NA | NA | NA |
| Mir3100 | 0 | 0 | 0 | 0 | 1 | 0 | 0 | 0 | 0 | 0 | 1 | 0 | 0.19 | 0 | 0 | 2.094 | 1.067 | 0.79379329 | NA | 0 | 12.608 | 10 | 10 | TRUE | 0.5075 |
| Mir3101 | 4 | 9 | 10 | 5 | 2 | 3 | 3 | 10 | 11 | 5 | 2 | 3 | 5.61 | 8 | 3 | 0.445 | -1.169 | 0.1824873 | NA | 0 | 0.4248 | 0.3398 | 0.3398 | TRUE | 1.0566 |
| Mir3102 | 0 | 1 | 0 | 0 | 3 | 0 | 0 | 1 | 0 | 0 | 3 | 0 | 0.74 | 0 | 1 | 3.214 | 1.684 | 0.52836502 | NA | 8.5607 | 3.1995 | 3.6756 | 3.6756 | TRUE | 1.4433 |
| Mir3103 | 0 | 0 | 0 | 0 | 0 | 0 | 0 | 0 | 0 | 0 | 0 | 0 | 0 | 0 | 0 | NA | NA | NA | NA | NA | NA | NA | NA | NA | NA |
| Mir3104 | 0 | 0 | 0 | 0 | 0 | 0 | 0 | 0 | 0 | 0 | 0 | 0 | 0 | 0 | 0 | NA | NA | NA | NA | NA | NA | NA | NA | NA | NA |
| Mir3106 | 0 | 0 | 0 | 0 | 0 | 0 | 0 | 0 | 0 | 0 | 0 | 0 | 0 | 0 | 0 | NA | NA | NA | NA | NA | NA | NA | NA | NA | NA |
| Mir3107 | 0 | 0 | 0 | 0 | 0 | 0 | 0 | 0 | 0 | 0 | 0 | 0 | 0 | 0 | 0 | NA | NA | NA | NA | NA | NA | NA | NA | NA | NA |
| Mir3108 | 0 | 0 | 0 | 0 | 1 | 0 | 0 | 0 | 0 | 0 | 1 | 0 | 0.19 | 0 | 0 | 2.094 | 1.067 | 0.79379329 | NA | 0 | 12.608 | 10 | 10 | TRUE | 0.5075 |
| Mir3109 | 0 | 0 | 0 | 0 | 0 | 0 | 0 | 0 | 0 | 0 | 0 | 0 | 0 | 0 | 0 | NA | NA | NA | NA | NA | NA | NA | NA | NA | NA |
| Mir3110 | 0 | 0 | 0 | 0 | 0 | 0 | 0 | 0 | 0 | 0 | 0 | 0 | 0 | 0 | 0 | NA | NA | NA | NA | NA | NA | NA | NA | NA | NA |
| Mir3112 | 18 | 16 | 22 | 12 | 12 | 19 | 14 | 17 | 24 | 12 | 14 | 19 | 16.5 | 18 | 15 | 0.816 | -0.293 | 0.54667389 | NA | 0 | 0.1461 | 0.1082 | 0.1082 | TRUE | 0.3401 |
| Mir32 | 0 | 0 | 0 | 0 | 0 | 0 | 0 | 0 | 0 | 0 | 0 | 0 | 0 | 0 | 0 | NA | NA | NA | NA | NA | NA | NA | NA | NA | NA |
| Mir320 | 0 | 0 | 1 | 0 | 0 | 0 | 0 | 0 | 1 | 0 | 0 | 0 | 0.18 | 0 | 0 | 0.552 | -0.857 | 0.8336548 | NA | 0 | 13.1817 | 10 | 10 | TRUE | 0.5365 |
| Mir322 | 0 | 0 | 0 | 0 | 0 | 0 | 0 | 0 | 0 | 0 | 0 | 0 | 0 | 0 | 0 | NA | NA | NA | NA | NA | NA | NA | NA | NA | NA |
| Mir323 | 0 | 0 | 0 | 0 | 0 | 0 | 0 | 0 | 0 | 0 | 0 | 0 | 0 | 0 | 0 | NA | NA | NA | NA | NA | NA | NA | NA | NA | NA |
| Mir324 | 0 | 1 | 0 | 0 | 0 | 0 | 0 | 1 | 0 | 0 | 0 | 0 | 0.18 | 0 | 0 | 0.552 | -0.857 | 0.8336548 | NA | 0 | 13.3714 | 10 | 10 | TRUE | 0.5216 |
| Mir325 | 0 | 0 | 0 | 0 | 1 | 0 | 0 | 0 | 0 | 0 | 1 | 0 | 0.19 | 0 | 0 | 2.094 | 1.067 | 0.79379329 | NA | 0 | 12.608 | 10 | 10 | TRUE | 0.5075 |
| Mir326 | 0 | 0 | 0 | 0 | 0 | 0 | 0 | 0 | 0 | 0 | 0 | 0 | 0 | 0 | 0 | NA | NA | NA | NA | NA | NA | NA | NA | NA | NA |
| Mir328 | 0 | 0 | 0 | 0 | 0 | 0 | 0 | 0 | 0 | 0 | 0 | 0 | 0 | 0 | 0 | NA | NA | NA | NA | NA | NA | NA | NA | NA | NA |
| Mir329 | 0 | 0 | 0 | 0 | 0 | 0 | 0 | 0 | 0 | 0 | 0 | 0 | 0 | 0 | 0 | NA | NA | NA | NA | NA | NA | NA | NA | NA | NA |
| Mir33 | 0 | 0 | 0 | 0 | 0 | 0 | 0 | 0 | 0 | 0 | 0 | 0 | 0 | 0 | 0 | NA | NA | NA | NA | NA | NA | NA | NA | NA | NA |
| Mir330 | 0 | 0 | 1 | 0 | 0 | 0 | 0 | 0 | 1 | 0 | 0 | 0 | 0.18 | 0 | 0 | 0.552 | -0.857 | 0.8336548 | NA | 0 | 13.1817 | 10 | 10 | TRUE | 0.5365 |
| Mir331 | 1 | 0 | 0 | 0 | 0 | 0 | 1 | 0 | 0 | 0 | 0 | 0 | 0.13 | 0 | 0 | 0.552 | -0.857 | 0.8336548 | NA | 0 | 18.3121 | 10 | 10 | TRUE | 0.2566 |
| Mir335 | 0 | 0 | 0 | 0 | 0 | 0 | 0 | 0 | 0 | 0 | 0 | 0 | 0 | 0 | 0 | NA | NA | NA | NA | NA | NA | NA | NA | NA | NA |
| Mir337 | 0 | 0 | 0 | 0 | 0 | 0 | 0 | 0 | 0 | 0 | 0 | 0 | 0 | 0 | 0 | NA | NA | NA | NA | NA | NA | NA | NA | NA | NA |
| Mir338 | 0 | 0 | 0 | 0 | 0 | 0 | 0 | 0 | 0 | 0 | 0 | 0 | 0 | 0 | 0 | NA | NA | NA | NA | NA | NA | NA | NA | NA | NA |
| Mir339 | 1 | 0 | 0 | 0 | 1 | 1 | 1 | 0 | 0 | 0 | 1 | 1 | 0.48 | 0 | 1 | 2.031 | 1.022 | 0.71376068 | NA | 0 | 4.9197 | 3.859 | 3.859 | TRUE | 0.2623 |
| Mir340 | 0 | 0 | 0 | 0 | 0 | 0 | 0 | 0 | 0 | 0 | 0 | 0 | 0 | 0 | 0 | NA | NA | NA | NA | NA | NA | NA | NA | NA | NA |
| Mir341 | 0 | 0 | 0 | 0 | 0 | 0 | 0 | 0 | 0 | 0 | 0 | 0 | 0 | 0 | 0 | NA | NA | NA | NA | NA | NA | NA | NA | NA | NA |
| Mir343 | 1 | 2 | 0 | 4 | 2 | 2 | 1 | 2 | 0 | 4 | 2 | 2 | 1.85 | 1 | 3 | 2.857 | 1.515 | 0.30818879 | NA | 0 | 1.2817 | 0.9024 | 0.9024 | TRUE | 0.4346 |
| Mir344 | 0 | 0 | 0 | 0 | 0 | 0 | 0 | 0 | 0 | 0 | 0 | 0 | 0 | 0 | 0 | NA | NA | NA | NA | NA | NA | NA | NA | NA | NA |
| Mir344-2 | 1 | 0 | 0 | 0 | 0 | 0 | 1 | 0 | 0 | 0 | 0 | 0 | 0.13 | 0 | 0 | 0.552 | -0.857 | 0.8336548 | NA | 0 | 18.3121 | 10 | 10 | TRUE | 0.2566 |
| Mir344b | 0 | 0 | 0 | 0 | 0 | 0 | 0 | 0 | 0 | 0 | 0 | 0 | 0 | 0 | 0 | NA | NA | NA | NA | NA | NA | NA | NA | NA | NA |
| Mir344c | 0 | 0 | 0 | 0 | 0 | 0 | 0 | 0 | 0 | 0 | 0 | 0 | 0 | 0 | 0 | NA | NA | NA | NA | NA | NA | NA | NA | NA | NA |
| Mir344d-1 | 0 | 0 | 0 | 0 | 0 | 0 | 0 | 0 | 0 | 0 | 0 | 0 | 0 | 0 | 0 | NA | NA | NA | NA | NA | NA | NA | NA | NA | NA |
| Mir344d-2 | 0 | 0 | 0 | 0 | 0 | 0 | 0 | 0 | 0 | 0 | 0 | 0 | 0 | 0 | 0 | NA | NA | NA | NA | NA | NA | NA | NA | NA | NA |
| Mir344d-3 | 0 | 0 | 0 | 0 | 0 | 0 | 0 | 0 | 0 | 0 | 0 | 0 | 0 | 0 | 0 | NA | NA | NA | NA | NA | NA | NA | NA | NA | NA |
| Mir344e | 0 | 0 | 0 | 0 | 0 | 0 | 0 | 0 | 0 | 0 | 0 | 0 | 0 | 0 | 0 | NA | NA | NA | NA | NA | NA | NA | NA | NA | NA |
| Mir344f | 0 | 0 | 0 | 0 | 0 | 0 | 0 | 0 | 0 | 0 | 0 | 0 | 0 | 0 | 0 | NA | NA | NA | NA | NA | NA | NA | NA | NA | NA |
| Mir344g | 0 | 0 | 0 | 0 | 0 | 0 | 0 | 0 | 0 | 0 | 0 | 0 | 0 | 0 | 0 | NA | NA | NA | NA | NA | NA | NA | NA | NA | NA |
| Mir345 | 0 | 0 | 0 | 0 | 0 | 0 | 0 | 0 | 0 | 0 | 0 | 0 | 0 | 0 | 0 | NA | NA | NA | NA | NA | NA | NA | NA | NA | NA |
| Mir346 | 0 | 0 | 0 | 0 | 0 | 0 | 0 | 0 | 0 | 0 | 0 | 0 | 0 | 0 | 0 | NA | NA | NA | NA | NA | NA | NA | NA | NA | NA |
| Mir3470a | 0 | 0 | 0 | 0 | 0 | 0 | 0 | 0 | 0 | 0 | 0 | 0 | 0 | 0 | 0 | NA | NA | NA | NA | NA | NA | NA | NA | NA | NA |
| Mir3470b | 0 | 0 | 0 | 0 | 0 | 0 | 0 | 0 | 0 | 0 | 0 | 0 | 0 | 0 | 0 | NA | NA | NA | NA | NA | NA | NA | NA | NA | NA |
| Mir3471-1 | 0 | 0 | 0 | 0 | 0 | 0 | 0 | 0 | 0 | 0 | 0 | 0 | 0 | 0 | 0 | NA | NA | NA | NA | NA | NA | NA | NA | NA | NA |
| Mir3473 | 0 | 0 | 0 | 0 | 0 | 0 | 0 | 0 | 0 | 0 | 0 | 0 | 0 | 0 | 0 | NA | NA | NA | NA | NA | NA | NA | NA | NA | NA |
| Mir3473c | 0 | 0 | 0 | 0 | 0 | 0 | 0 | 0 | 0 | 0 | 0 | 0 | 0 | 0 | 0 | NA | NA | NA | NA | NA | NA | NA | NA | NA | NA |
| Mir3473d | 0 | 1 | 1 | 1 | 0 | 1 | 0 | 1 | 1 | 1 | 0 | 1 | 0.69 | 1 | 1 | 0.979 | -0.031 | 0.98943171 | NA | 0 | 3.4496 | 2.4101 | 2.4101 | TRUE | 0.3734 |
| Mir3474 | 1 | 0 | 0 | 0 | 0 | 0 | 1 | 0 | 0 | 0 | 0 | 0 | 0.13 | 0 | 0 | 0.552 | -0.857 | 0.8336548 | NA | 0 | 18.3121 | 10 | 10 | TRUE | 0.2566 |
| Mir3475 | 0 | 0 | 0 | 0 | 0 | 0 | 0 | 0 | 0 | 0 | 0 | 0 | 0 | 0 | 0 | NA | NA | NA | NA | NA | NA | NA | NA | NA | NA |
| Mir34a | 0 | 0 | 0 | 1 | 0 | 0 | 0 | 0 | 0 | 1 | 0 | 0 | 0.17 | 0 | 0 | 2.094 | 1.067 | 0.79379329 | NA | 0 | 14.2885 | 10 | 10 | TRUE | 0.3906 |
| Mir34b | 0 | 0 | 0 | 0 | 0 | 0 | 0 | 0 | 0 | 0 | 0 | 0 | 0 | 0 | 0 | NA | NA | NA | NA | NA | NA | NA | NA | NA | NA |
| Mir34c | 0 | 0 | 0 | 0 | 1 | 0 | 0 | 0 | 0 | 0 | 1 | 0 | 0.19 | 0 | 0 | 2.094 | 1.067 | 0.79379329 | NA | 0 | 12.608 | 10 | 10 | TRUE | 0.5075 |
| Mir350 | 0 | 0 | 0 | 0 | 0 | 0 | 0 | 0 | 0 | 0 | 0 | 0 | 0 | 0 | 0 | NA | NA | NA | NA | NA | NA | NA | NA | NA | NA |
| Mir351 | 0 | 0 | 0 | 0 | 0 | 0 | 0 | 0 | 0 | 0 | 0 | 0 | 0 | 0 | 0 | NA | NA | NA | NA | NA | NA | NA | NA | NA | NA |
| Mir3572 | 0 | 0 | 0 | 0 | 0 | 0 | 0 | 0 | 0 | 0 | 0 | 0 | 0 | 0 | 0 | NA | NA | NA | NA | NA | NA | NA | NA | NA | NA |
| Mir362 | 0 | 0 | 0 | 0 | 0 | 0 | 0 | 0 | 0 | 0 | 0 | 0 | 0 | 0 | 0 | NA | NA | NA | NA | NA | NA | NA | NA | NA | NA |
| Mir363 | 4 | 2 | 1 | 1 | 2 | 2 | 3 | 2 | 1 | 1 | 2 | 2 | 1.92 | 2 | 2 | 0.808 | -0.307 | 0.8191094 | NA | 0 | 1.2348 | 0.7762 | 0.7762 | TRUE | 0.2191 |
| Mir365-1 | 0 | 0 | 0 | 0 | 0 | 0 | 0 | 0 | 0 | 0 | 0 | 0 | 0 | 0 | 0 | NA | NA | NA | NA | NA | NA | NA | NA | NA | NA |
| Mir365-2 | 0 | 0 | 0 | 0 | 0 | 0 | 0 | 0 | 0 | 0 | 0 | 0 | 0 | 0 | 0 | NA | NA | NA | NA | NA | NA | NA | NA | NA | NA |
| Mir367 | 0 | 0 | 0 | 0 | 0 | 0 | 0 | 0 | 0 | 0 | 0 | 0 | 0 | 0 | 0 | NA | NA | NA | NA | NA | NA | NA | NA | NA | NA |
| Mir369 | 0 | 0 | 0 | 0 | 0 | 0 | 0 | 0 | 0 | 0 | 0 | 0 | 0 | 0 | 0 | NA | NA | NA | NA | NA | NA | NA | NA | NA | NA |
| Mir370 | 0 | 0 | 0 | 0 | 0 | 0 | 0 | 0 | 0 | 0 | 0 | 0 | 0 | 0 | 0 | NA | NA | NA | NA | NA | NA | NA | NA | NA | NA |

| | | | | | | | | | | | | | | | | | | | | | | | | | | |
|---|---|---|---|---|---|---|---|---|---|---|---|---|---|---|---|---|---|---|---|---|---|---|---|---|---|---|
| Mir374 | 0 | 0 | 0 | 0 | 0 | 0 | 0 | 0 | 0 | 0 | 0 | 0 | 0 | 0 | 0 | NA | NA | NA | NA | NA | NA | NA | NA | NA | NA | |
| Mir374c | 0 | 0 | 0 | 0 | 0 | 0 | 0 | 0 | 0 | 0 | 0 | 0 | 0 | 0 | 0 | NA | NA | NA | NA | NA | NA | NA | NA | NA | NA | |
| Mir375 | 0 | 0 | 0 | 0 | 0 | 0 | 0 | 0 | 0 | 0 | 0 | 0 | 0 | 0 | 0 | NA | NA | NA | NA | NA | NA | NA | NA | NA | NA | |
| Mir376a | 0 | 0 | 0 | 0 | 0 | 0 | 0 | 0 | 0 | 0 | 0 | 0 | 0 | 0 | 0 | NA | NA | NA | NA | NA | NA | NA | NA | NA | NA | |
| Mir376b | 0 | 0 | 0 | 0 | 0 | 0 | 0 | 0 | 0 | 0 | 0 | 0 | 0 | 0 | 0 | NA | NA | NA | NA | NA | NA | NA | NA | NA | NA | |
| Mir376c | 0 | 0 | 0 | 0 | 0 | 1 | 0 | 0 | 0 | 0 | 0 | 1 | 0.16 | 0 | 0 | 2.094 | 1.067 | 0.79379329 | NA | | 0 | 14.4078 | 10 | 10 | TRUE | 0.3835 |
| Mir377 | 1 | 1 | 1 | 1 | 0 | 0 | 1 | 1 | 1 | 1 | 0 | 0 | 0.65 | 1 | 0 | 0.383 | -1.383 | 0.55308337 | NA | | 0 | 3.6349 | 2.4186 | 2.4186 | TRUE | 0.3906 |
| Mir378 | 0 | 0 | 0 | 0 | 0 | 0 | 0 | 0 | 0 | 0 | 0 | 0 | 0 | 0 | 0 | NA | NA | NA | NA | NA | NA | NA | NA | NA | NA | |
| Mir378b | 0 | 0 | 0 | 0 | 0 | 0 | 0 | 0 | 0 | 0 | 0 | 0 | 0 | 0 | 0 | NA | NA | NA | NA | NA | NA | NA | NA | NA | NA | |
| Mir379 | 0 | 0 | 0 | 0 | 0 | 0 | 0 | 0 | 0 | 0 | 0 | 0 | 0 | 0 | 0 | NA | NA | NA | NA | NA | NA | NA | NA | NA | NA | |
| Mir380 | 0 | 0 | 0 | 0 | 0 | 0 | 0 | 0 | 0 | 0 | 0 | 0 | 0 | 0 | 0 | NA | NA | NA | NA | NA | NA | NA | NA | NA | NA | |
| Mir381 | 0 | 0 | 0 | 0 | 0 | 0 | 0 | 0 | 0 | 0 | 0 | 0 | 0 | 0 | 0 | NA | NA | NA | NA | NA | NA | NA | NA | NA | NA | |
| Mir382 | 0 | 0 | 0 | 0 | 0 | 0 | 0 | 0 | 0 | 0 | 0 | 0 | 0 | 0 | 0 | NA | NA | NA | NA | NA | NA | NA | NA | NA | NA | |
| Mir383 | 0 | 0 | 0 | 0 | 0 | 0 | 0 | 0 | 0 | 0 | 0 | 0 | 0 | 0 | 0 | NA | NA | NA | NA | NA | NA | NA | NA | NA | NA | |
| Mir384 | 0 | 0 | 0 | 0 | 0 | 0 | 0 | 0 | 0 | 0 | 0 | 0 | 0 | 0 | 0 | NA | NA | NA | NA | NA | NA | NA | NA | NA | NA | |
| Mir3960 | 7 | 5 | 3 | 3 | 2 | 7 | 5 | 5 | 3 | 3 | 2 | 7 | 4.36 | 4 | 4 | 0.87 | -0.201 | 0.82774225 | NA | | 0 | 0.5463 | 0.3848 | 0.3848 | TRUE | 0.6619 |
| Mir3962 | 0 | 0 | 0 | 0 | 0 | 0 | 0 | 0 | 0 | 0 | 0 | 0 | 0 | 0 | 0 | NA | NA | NA | NA | NA | NA | NA | NA | NA | NA | |
| Mir3963 | 0 | 0 | 0 | 0 | 0 | 0 | 0 | 0 | 0 | 0 | 0 | 0 | 0 | 0 | 0 | NA | NA | NA | NA | NA | NA | NA | NA | NA | NA | |
| Mir3964 | 0 | 0 | 0 | 0 | 0 | 0 | 0 | 0 | 0 | 0 | 0 | 0 | 0 | 0 | 0 | NA | NA | NA | NA | NA | NA | NA | NA | NA | NA | |
| Mir3965 | 0 | 0 | 0 | 0 | 0 | 0 | 0 | 0 | 0 | 0 | 0 | 0 | 0 | 0 | 0 | NA | NA | NA | NA | NA | NA | NA | NA | NA | NA | |
| Mir3966 | 0 | 0 | 0 | 0 | 0 | 0 | 0 | 0 | 0 | 0 | 0 | 0 | 0 | 0 | 0 | NA | NA | NA | NA | NA | NA | NA | NA | NA | NA | |
| Mir3967 | 0 | 0 | 0 | 0 | 0 | 0 | 0 | 0 | 0 | 0 | 0 | 0 | 0 | 0 | 0 | NA | NA | NA | NA | NA | NA | NA | NA | NA | NA | |
| Mir3968 | 0 | 0 | 0 | 0 | 0 | 0 | 0 | 0 | 0 | 0 | 0 | 0 | 0 | 0 | 0 | NA | NA | NA | NA | NA | NA | NA | NA | NA | NA | |
| Mir3969 | 0 | 0 | 0 | 0 | 0 | 0 | 0 | 0 | 0 | 0 | 0 | 0 | 0 | 0 | 0 | NA | NA | NA | NA | NA | NA | NA | NA | NA | NA | |
| Mir3970 | 0 | 0 | 0 | 0 | 0 | 0 | 0 | 0 | 0 | 0 | 0 | 0 | 0 | 0 | 0 | NA | NA | NA | NA | NA | NA | NA | NA | NA | NA | |
| Mir3971 | 2 | 0 | 0 | 0 | 0 | 0 | 2 | 0 | 0 | 0 | 0 | 0 | 0.26 | 1 | 0 | 0.356 | -1.491 | 0.71362796 | NA | | 0 | 9.1573 | 10 | 10 | TRUE | 0.968 |
| Mir409 | 2 | 0 | 0 | 0 | 0 | 0 | 2 | 0 | 0 | 0 | 0 | 0 | 0.26 | 1 | 0 | 0.356 | -1.491 | 0.71362796 | NA | | 0 | 9.1573 | 10 | 10 | TRUE | 0.968 |
| Mir410 | 0 | 0 | 0 | 0 | 0 | 0 | 0 | 0 | 0 | 0 | 0 | 0 | 0 | 0 | 0 | NA | NA | NA | NA | NA | NA | NA | NA | NA | NA | |
| Mir411 | 0 | 0 | 0 | 0 | 0 | 0 | 0 | 0 | 0 | 0 | 0 | 0 | 0 | 0 | 0 | NA | NA | NA | NA | NA | NA | NA | NA | NA | NA | |
| Mir412 | 0 | 0 | 0 | 0 | 1 | 0 | 0 | 0 | 0 | 0 | 1 | 0 | 0.19 | 0 | 0 | 2.094 | 1.067 | 0.79379329 | NA | | 0 | 12.608 | 10 | 10 | TRUE | 0.5075 |
| Mir421 | 0 | 0 | 0 | 0 | 0 | 0 | 0 | 0 | 0 | 0 | 0 | 0 | 0 | 0 | 0 | NA | NA | NA | NA | NA | NA | NA | NA | NA | NA | |
| Mir423 | 1 | 0 | 0 | 0 | 0 | 0 | 1 | 0 | 0 | 0 | 0 | 0 | 0.13 | 0 | 0 | 0.552 | -0.857 | 0.8336548 | NA | | 0 | 18.3121 | 10 | 10 | TRUE | 0.2566 |
| Mir425 | 2 | 0 | 1 | 1 | 0 | 0 | 2 | 0 | 1 | 1 | 0 | 0 | 0.6 | 1 | 0 | 0.414 | -1.271 | 0.61961437 | NA | | 0 | 3.9224 | 3.1813 | 3.1813 | TRUE | 0.3906 |
| Mir429 | 0 | 0 | 0 | 0 | 0 | 0 | 0 | 0 | 0 | 0 | 0 | 0 | 0 | 0 | 0 | NA | NA | NA | NA | NA | NA | NA | NA | NA | NA | |
| Mir431 | 0 | 0 | 0 | 0 | 0 | 0 | 0 | 0 | 0 | 0 | 0 | 0 | 0 | 0 | 0 | NA | NA | NA | NA | NA | NA | NA | NA | NA | NA | |
| Mir432 | 0 | 0 | 0 | 0 | 0 | 0 | 0 | 0 | 0 | 0 | 0 | 0 | 0 | 0 | 0 | NA | NA | NA | NA | NA | NA | NA | NA | NA | NA | |
| Mir433 | 0 | 0 | 0 | 0 | 0 | 0 | 0 | 0 | 0 | 0 | 0 | 0 | 0 | 0 | 0 | NA | NA | NA | NA | NA | NA | NA | NA | NA | NA | |
| Mir434 | 0 | 0 | 0 | 0 | 0 | 0 | 0 | 0 | 0 | 0 | 0 | 0 | 0 | 0 | 0 | NA | NA | NA | NA | NA | NA | NA | NA | NA | NA | |
| Mir448 | 0 | 0 | 0 | 0 | 0 | 0 | 0 | 0 | 0 | 0 | 0 | 0 | 0 | 0 | 0 | NA | NA | NA | NA | NA | NA | NA | NA | NA | NA | |
| Mir449a | 0 | 0 | 0 | 0 | 0 | 0 | 0 | 0 | 0 | 0 | 0 | 0 | 0 | 0 | 0 | NA | NA | NA | NA | NA | NA | NA | NA | NA | NA | |
| Mir449b | 0 | 0 | 0 | 0 | 0 | 0 | 0 | 0 | 0 | 0 | 0 | 0 | 0 | 0 | 0 | NA | NA | NA | NA | NA | NA | NA | NA | NA | NA | |
| Mir449c | 0 | 0 | 0 | 0 | 0 | 0 | 0 | 0 | 0 | 0 | 0 | 0 | 0 | 0 | 0 | NA | NA | NA | NA | NA | NA | NA | NA | NA | NA | |
| Mir450-1 | 0 | 0 | 0 | 0 | 0 | 0 | 0 | 0 | 0 | 0 | 0 | 0 | 0 | 0 | 0 | NA | NA | NA | NA | NA | NA | NA | NA | NA | NA | |
| Mir450-2 | 0 | 0 | 0 | 0 | 0 | 0 | 0 | 0 | 0 | 0 | 0 | 0 | 0 | 0 | 0 | NA | NA | NA | NA | NA | NA | NA | NA | NA | NA | |
| Mir450b | 0 | 0 | 0 | 0 | 0 | 1 | 0 | 0 | 0 | 0 | 0 | 1 | 0.16 | 0 | 0 | 2.094 | 1.067 | 0.79379329 | NA | | 0 | 14.4078 | 10 | 10 | TRUE | 0.3835 |
| Mir451 | 0 | 0 | 0 | 0 | 0 | 0 | 0 | 0 | 0 | 0 | 0 | 0 | 0 | 0 | 0 | NA | NA | NA | NA | NA | NA | NA | NA | NA | NA | |
| Mir452 | 0 | 0 | 0 | 0 | 0 | 0 | 0 | 0 | 0 | 0 | 0 | 0 | 0 | 0 | 0 | NA | NA | NA | NA | NA | NA | NA | NA | NA | NA | |
| Mir453 | 2 | 0 | 0 | 0 | 0 | 1 | 2 | 0 | 0 | 0 | 0 | 1 | 0.42 | 1 | 0 | 0.682 | -0.553 | 0.86290097 | NA | | 0 | 5.6 | 5.5066 | 5.5066 | TRUE | 0.9727 |
| Mir455 | 0 | 0 | 0 | 0 | 0 | 0 | 0 | 0 | 0 | 0 | 0 | 0 | 0 | 0 | 0 | NA | NA | NA | NA | NA | NA | NA | NA | NA | NA | |
| Mir463 | 0 | 0 | 0 | 0 | 0 | 0 | 0 | 0 | 0 | 0 | 0 | 0 | 0 | 0 | 0 | NA | NA | NA | NA | NA | NA | NA | NA | NA | NA | |
| Mir465 | 0 | 0 | 0 | 0 | 0 | 0 | 0 | 0 | 0 | 0 | 0 | 0 | 0 | 0 | 0 | NA | NA | NA | NA | NA | NA | NA | NA | NA | NA | |
| Mir465b-1 | 0 | 0 | 0 | 0 | 0 | 0 | 0 | 0 | 0 | 0 | 0 | 0 | 0 | 0 | 0 | NA | NA | NA | NA | NA | NA | NA | NA | NA | NA | |
| Mir465b-2 | 0 | 0 | 0 | 0 | 0 | 0 | 0 | 0 | 0 | 0 | 0 | 0 | 0 | 0 | 0 | NA | NA | NA | NA | NA | NA | NA | NA | NA | NA | |
| Mir465c-1 | 0 | 0 | 0 | 0 | 0 | 0 | 0 | 0 | 0 | 0 | 0 | 0 | 0 | 0 | 0 | NA | NA | NA | NA | NA | NA | NA | NA | NA | NA | |
| Mir465c-2 | 0 | 0 | 0 | 0 | 0 | 0 | 0 | 0 | 0 | 0 | 0 | 0 | 0 | 0 | 0 | NA | NA | NA | NA | NA | NA | NA | NA | NA | NA | |
| Mir466 | 0 | 0 | 0 | 0 | 0 | 0 | 0 | 0 | 0 | 0 | 0 | 0 | 0 | 0 | 0 | NA | NA | NA | NA | NA | NA | NA | NA | NA | NA | |
| Mir4660 | 0 | 0 | 0 | 0 | 0 | 0 | 0 | 0 | 0 | 0 | 0 | 0 | 0 | 0 | 0 | NA | NA | NA | NA | NA | NA | NA | NA | NA | NA | |
| Mir466b-2 | 0 | 0 | 0 | 0 | 0 | 0 | 0 | 0 | 0 | 0 | 0 | 0 | 0 | 0 | 0 | NA | NA | NA | NA | NA | NA | NA | NA | NA | NA | |
| Mir466b-3 | 0 | 0 | 0 | 0 | 0 | 0 | 0 | 0 | 0 | 0 | 0 | 0 | 0 | 0 | 0 | NA | NA | NA | NA | NA | NA | NA | NA | NA | NA | |
| Mir466d | 0 | 0 | 0 | 0 | 0 | 0 | 0 | 0 | 0 | 0 | 0 | 0 | 0 | 0 | 0 | NA | NA | NA | NA | NA | NA | NA | NA | NA | NA | |
| Mir466f-1 | 0 | 0 | 0 | 0 | 0 | 0 | 0 | 0 | 0 | 0 | 0 | 0 | 0 | 0 | 0 | NA | NA | NA | NA | NA | NA | NA | NA | NA | NA | |
| Mir466f-2 | 0 | 0 | 0 | 1 | 0 | 0 | 0 | 0 | 0 | 1 | 0 | 0 | 0.17 | 0 | 0 | 2.094 | 1.067 | 0.79379329 | NA | | 0 | 14.2885 | 10 | 10 | TRUE | 0.3906 |
| Mir466f-3 | 0 | 0 | 0 | 0 | 0 | 0 | 0 | 0 | 0 | 0 | 0 | 0 | 0 | 0 | 0 | NA | NA | NA | NA | NA | NA | NA | NA | NA | NA | |
| Mir466g | 0 | 0 | 0 | 0 | 0 | 0 | 0 | 0 | 0 | 0 | 0 | 0 | 0 | 0 | 0 | NA | NA | NA | NA | NA | NA | NA | NA | NA | NA | |
| Mir466h | 0 | 0 | 0 | 0 | 0 | 0 | 0 | 0 | 0 | 0 | 0 | 0 | 0 | 0 | 0 | NA | NA | NA | NA | NA | NA | NA | NA | NA | NA | |
| Mir466i | 0 | 0 | 0 | 0 | 0 | 0 | 0 | 0 | 0 | 0 | 0 | 0 | 0 | 0 | 0 | NA | NA | NA | NA | NA | NA | NA | NA | NA | NA | |
| Mir466n | 0 | 0 | 0 | 0 | 0 | 0 | 0 | 0 | 0 | 0 | 0 | 0 | 0 | 0 | 0 | NA | NA | NA | NA | NA | NA | NA | NA | NA | NA | |
| Mir467a-1 | 0 | 0 | 0 | 0 | 0 | 0 | 0 | 0 | 0 | 0 | 0 | 0 | 0 | 0 | 0 | NA | NA | NA | NA | NA | NA | NA | NA | NA | NA | |
| Mir467a-10 | 0 | 0 | 0 | 0 | 0 | 0 | 0 | 0 | 0 | 0 | 0 | 0 | 0 | 0 | 0 | NA | NA | NA | NA | NA | NA | NA | NA | NA | NA | |

| | | | | | | | | | | | | | | | | | | | | | | | | | |
|---|---|---|---|---|---|---|---|---|---|---|---|---|---|---|---|---|---|---|---|---|---|---|---|---|---|
| Mir467a-2 | 0 | 0 | 0 | 0 | 0 | 0 | 0 | 0 | 0 | 0 | 0 | 0 | 0 | 0 | 0 | NA | NA | NA | NA | NA | NA | NA | NA | NA | NA |
| Mir467a-3 | 0 | 0 | 0 | 0 | 0 | 0 | 0 | 0 | 0 | 0 | 0 | 0 | 0 | 0 | 0 | NA | NA | NA | NA | NA | NA | NA | NA | NA | NA |
| Mir467a-4 | 0 | 0 | 0 | 0 | 0 | 0 | 0 | 0 | 0 | 0 | 0 | 0 | 0 | 0 | 0 | NA | NA | NA | NA | NA | NA | NA | NA | NA | NA |
| Mir467a-5 | 0 | 0 | 0 | 0 | 0 | 0 | 0 | 0 | 0 | 0 | 0 | 0 | 0 | 0 | 0 | NA | NA | NA | NA | NA | NA | NA | NA | NA | NA |
| Mir467a-6 | 0 | 0 | 0 | 0 | 0 | 0 | 0 | 0 | 0 | 0 | 0 | 0 | 0 | 0 | 0 | NA | NA | NA | NA | NA | NA | NA | NA | NA | NA |
| Mir467a-7 | 0 | 0 | 0 | 0 | 0 | 0 | 0 | 0 | 0 | 0 | 0 | 0 | 0 | 0 | 0 | NA | NA | NA | NA | NA | NA | NA | NA | NA | NA |
| Mir467a-8 | 0 | 0 | 0 | 0 | 0 | 0 | 0 | 0 | 0 | 0 | 0 | 0 | 0 | 0 | 0 | NA | NA | NA | NA | NA | NA | NA | NA | NA | NA |
| Mir467a-9 | 0 | 0 | 0 | 0 | 0 | 0 | 0 | 0 | 0 | 0 | 0 | 0 | 0 | 0 | 0 | NA | NA | NA | NA | NA | NA | NA | NA | NA | NA |
| Mir467b | 0 | 0 | 0 | 0 | 0 | 0 | 0 | 0 | 0 | 0 | 0 | 0 | 0 | 0 | 0 | NA | NA | NA | NA | NA | NA | NA | NA | NA | NA |
| Mir467c | 0 | 0 | 0 | 0 | 0 | 0 | 0 | 0 | 0 | 0 | 0 | 0 | 0 | 0 | 0 | NA | NA | NA | NA | NA | NA | NA | NA | NA | NA |
| Mir467d | 0 | 0 | 0 | 0 | 0 | 0 | 0 | 0 | 0 | 0 | 0 | 0 | 0 | 0 | 0 | NA | NA | NA | NA | NA | NA | NA | NA | NA | NA |
| Mir467e | 0 | 0 | 0 | 0 | 0 | 0 | 0 | 0 | 0 | 0 | 0 | 0 | 0 | 0 | 0 | NA | NA | NA | NA | NA | NA | NA | NA | NA | NA |
| Mir467f | 0 | 0 | 0 | 0 | 0 | 0 | 0 | 0 | 0 | 0 | 0 | 0 | 0 | 0 | 0 | NA | NA | NA | NA | NA | NA | NA | NA | NA | NA |
| Mir468 | 0 | 0 | 0 | 0 | 0 | 0 | 0 | 0 | 0 | 0 | 0 | 0 | 0 | 0 | 0 | NA | NA | NA | NA | NA | NA | NA | NA | NA | NA |
| Mir470 | 0 | 0 | 0 | 0 | 0 | 0 | 0 | 0 | 0 | 0 | 0 | 0 | 0 | 0 | 0 | NA | NA | NA | NA | NA | NA | NA | NA | NA | NA |
| Mir471 | 0 | 0 | 0 | 0 | 0 | 0 | 0 | 0 | 0 | 0 | 0 | 0 | 0 | 0 | 0 | NA | NA | NA | NA | NA | NA | NA | NA | NA | NA |
| Mir483 | 0 | 0 | 0 | 0 | 0 | 0 | 0 | 0 | 0 | 0 | 0 | 0 | 0 | 0 | 0 | NA | NA | NA | NA | NA | NA | NA | NA | NA | NA |
| Mir484 | 1 | 0 | 0 | 0 | 0 | 0 | 1 | 0 | 0 | 0 | 0 | 0 | 0.13 | 0 | 0 | 0.552 | -0.857 | 0.8336548 | NA | 0 | 18.3121 | 10 | 10 | TRUE | 0.2566 |
| Mir485 | 1 | 0 | 0 | 0 | 1 | 0 | 1 | 0 | 0 | 0 | 1 | 0 | 0.32 | 0 | 0 | 1.075 | 0.105 | 0.97638847 | NA | 0 | 7.4681 | 7.0392 | 7.0392 | TRUE | 0.5075 |
| Mir486 | 0 | 0 | 0 | 0 | 0 | 0 | 0 | 0 | 0 | 0 | 0 | 0 | 0 | 0 | 0 | NA | NA | NA | NA | NA | NA | NA | NA | NA | NA |
| Mir487b | 0 | 0 | 0 | 0 | 0 | 0 | 0 | 0 | 0 | 0 | 0 | 0 | 0 | 0 | 0 | NA | NA | NA | NA | NA | NA | NA | NA | NA | NA |
| Mir488 | 0 | 0 | 0 | 0 | 0 | 0 | 0 | 0 | 0 | 0 | 0 | 0 | 0 | 0 | 0 | NA | NA | NA | NA | NA | NA | NA | NA | NA | NA |
| Mir489 | 0 | 0 | 0 | 0 | 0 | 0 | 0 | 0 | 0 | 0 | 0 | 0 | 0 | 0 | 0 | NA | NA | NA | NA | NA | NA | NA | NA | NA | NA |
| Mir490 | 0 | 0 | 0 | 0 | 0 | 0 | 0 | 0 | 0 | 0 | 0 | 0 | 0 | 0 | 0 | NA | NA | NA | NA | NA | NA | NA | NA | NA | NA |
| Mir491 | 0 | 0 | 0 | 0 | 0 | 0 | 0 | 0 | 0 | 0 | 0 | 0 | 0 | 0 | 0 | NA | NA | NA | NA | NA | NA | NA | NA | NA | NA |
| Mir493 | 0 | 0 | 0 | 0 | 0 | 0 | 0 | 0 | 0 | 0 | 0 | 0 | 0 | 0 | 0 | NA | NA | NA | NA | NA | NA | NA | NA | NA | NA |
| Mir494 | 0 | 0 | 0 | 0 | 0 | 0 | 0 | 0 | 0 | 0 | 0 | 0 | 0 | 0 | 0 | NA | NA | NA | NA | NA | NA | NA | NA | NA | NA |
| Mir495 | 0 | 0 | 0 | 0 | 0 | 0 | 0 | 0 | 0 | 0 | 0 | 0 | 0 | 0 | 0 | NA | NA | NA | NA | NA | NA | NA | NA | NA | NA |
| Mir496 | 0 | 1 | 0 | 0 | 1 | 0 | 0 | 1 | 0 | 0 | 1 | 0 | 0.37 | 0 | 0 | 1.075 | 0.105 | 0.97520514 | NA | 0 | 6.4904 | 6.1969 | 6.1969 | TRUE | 0.5216 |
| Mir497 | 0 | 0 | 0 | 0 | 0 | 0 | 0 | 0 | 0 | 0 | 0 | 0 | 0 | 0 | 0 | NA | NA | NA | NA | NA | NA | NA | NA | NA | NA |
| Mir499 | 0 | 0 | 0 | 0 | 0 | 0 | 0 | 0 | 0 | 0 | 0 | 0 | 0 | 0 | 0 | NA | NA | NA | NA | NA | NA | NA | NA | NA | NA |
| Mir500 | 0 | 0 | 0 | 0 | 0 | 0 | 0 | 0 | 0 | 0 | 0 | 0 | 0 | 0 | 0 | NA | NA | NA | NA | NA | NA | NA | NA | NA | NA |
| Mir501 | 0 | 0 | 0 | 0 | 0 | 0 | 0 | 0 | 0 | 0 | 0 | 0 | 0 | 0 | 0 | NA | NA | NA | NA | NA | NA | NA | NA | NA | NA |
| Mir503 | 0 | 0 | 0 | 0 | 0 | 0 | 0 | 0 | 0 | 0 | 0 | 0 | 0 | 0 | 0 | NA | NA | NA | NA | NA | NA | NA | NA | NA | NA |
| Mir504 | 0 | 0 | 0 | 0 | 0 | 0 | 0 | 0 | 0 | 0 | 0 | 0 | 0 | 0 | 0 | NA | NA | NA | NA | NA | NA | NA | NA | NA | NA |
| Mir5046 | 0 | 0 | 0 | 0 | 0 | 0 | 0 | 0 | 0 | 0 | 0 | 0 | 0 | 0 | 0 | NA | NA | NA | NA | NA | NA | NA | NA | NA | NA |
| Mir505 | 0 | 0 | 0 | 0 | 0 | 0 | 0 | 0 | 0 | 0 | 0 | 0 | 0 | 0 | 0 | NA | NA | NA | NA | NA | NA | NA | NA | NA | NA |
| Mir509 | 0 | 0 | 0 | 0 | 0 | 0 | 0 | 0 | 0 | 0 | 0 | 0 | 0 | 0 | 0 | NA | NA | NA | NA | NA | NA | NA | NA | NA | NA |
| Mir5097 | 0 | 0 | 0 | 0 | 0 | 0 | 0 | 0 | 0 | 0 | 0 | 0 | 0 | 0 | 0 | NA | NA | NA | NA | NA | NA | NA | NA | NA | NA |
| Mir5098 | 0 | 0 | 0 | 0 | 0 | 0 | 0 | 0 | 0 | 0 | 0 | 0 | 0 | 0 | 0 | NA | NA | NA | NA | NA | NA | NA | NA | NA | NA |
| Mir5100 | 0 | 0 | 0 | 0 | 0 | 0 | 0 | 0 | 0 | 0 | 0 | 0 | 0 | 0 | 0 | NA | NA | NA | NA | NA | NA | NA | NA | NA | NA |
| Mir5101 | 0 | 0 | 0 | 0 | 0 | 0 | 0 | 0 | 0 | 0 | 0 | 0 | 0 | 0 | 0 | NA | NA | NA | NA | NA | NA | NA | NA | NA | NA |
| Mir5103 | 0 | 0 | 0 | 0 | 0 | 0 | 0 | 0 | 0 | 0 | 0 | 0 | 0 | 0 | 0 | NA | NA | NA | NA | NA | NA | NA | NA | NA | NA |
| Mir5104 | 0 | 0 | 0 | 0 | 0 | 0 | 0 | 0 | 0 | 0 | 0 | 0 | 0 | 0 | 0 | NA | NA | NA | NA | NA | NA | NA | NA | NA | NA |
| Mir5106 | 1 | 0 | 0 | 0 | 0 | 1 | 1 | 0 | 0 | 0 | 0 | 1 | 0.29 | 0 | 0 | 1.075 | 0.105 | 0.97702352 | NA | 0 | 8.0647 | 7.5458 | 7.5458 | TRUE | 0.3835 |
| Mir5107 | 0 | 0 | 0 | 0 | 0 | 1 | 0 | 0 | 0 | 0 | 0 | 1 | 0.16 | 0 | 0 | 2.094 | 1.067 | 0.79379329 | NA | 0 | 14.4078 | 10 | 10 | TRUE | 0.3835 |
| Mir5108 | 0 | 0 | 0 | 0 | 0 | 0 | 0 | 0 | 0 | 0 | 0 | 0 | 0 | 0 | 0 | NA | NA | NA | NA | NA | NA | NA | NA | NA | NA |
| Mir5109 | 2 | 0 | 0 | 0 | 0 | 0 | 2 | 0 | 0 | 0 | 0 | 0 | 0.26 | 1 | 0 | 0.356 | -1.491 | 0.71362796 | NA | 0 | 9.1573 | 10 | 10 | TRUE | 0.968 |
| Mir511 | 0 | 0 | 0 | 0 | 0 | 0 | 0 | 0 | 0 | 0 | 0 | 0 | 0 | 0 | 0 | NA | NA | NA | NA | NA | NA | NA | NA | NA | NA |
| Mir5112 | 0 | 0 | 0 | 0 | 0 | 0 | 0 | 0 | 0 | 0 | 0 | 0 | 0 | 0 | 0 | NA | NA | NA | NA | NA | NA | NA | NA | NA | NA |
| Mir5113 | 0 | 0 | 0 | 0 | 0 | 0 | 0 | 0 | 0 | 0 | 0 | 0 | 0 | 0 | 0 | NA | NA | NA | NA | NA | NA | NA | NA | NA | NA |
| Mir5114 | 0 | 0 | 0 | 0 | 0 | 0 | 0 | 0 | 0 | 0 | 0 | 0 | 0 | 0 | 0 | NA | NA | NA | NA | NA | NA | NA | NA | NA | NA |
| Mir5115 | 0 | 0 | 0 | 0 | 0 | 0 | 0 | 0 | 0 | 0 | 0 | 0 | 0 | 0 | 0 | NA | NA | NA | NA | NA | NA | NA | NA | NA | NA |
| Mir5116 | 0 | 0 | 0 | 0 | 0 | 0 | 0 | 0 | 0 | 0 | 0 | 0 | 0 | 0 | 0 | NA | NA | NA | NA | NA | NA | NA | NA | NA | NA |
| Mir5117 | 0 | 0 | 0 | 0 | 0 | 0 | 0 | 0 | 0 | 0 | 0 | 0 | 0 | 0 | 0 | NA | NA | NA | NA | NA | NA | NA | NA | NA | NA |
| Mir5119 | 1 | 0 | 0 | 3 | 0 | 1 | 1 | 0 | 0 | 3 | 0 | 1 | 0.79 | 0 | 1 | 3.877 | 1.955 | 0.4082088 | NA | 0 | 2.9962 | 2.64 | 2.64 | TRUE | 0.219 |
| Mir5120 | 0 | 0 | 0 | 0 | 0 | 0 | 0 | 0 | 0 | 0 | 0 | 0 | 0 | 0 | 0 | NA | NA | NA | NA | NA | NA | NA | NA | NA | NA |
| Mir5121 | 5 | 5 | 0 | 0 | 0 | 0 | 4 | 5 | 0 | 0 | 0 | 0 | 1.53 | 3 | 0 | 0.062 | -4.017 | 0.04794395 | NA | 6.698 | 1.5478 | 1.8151 | 1.8151 | TRUE | 0.2445 |
| Mir5122 | 0 | 0 | 0 | 0 | 0 | 0 | 0 | 0 | 0 | 0 | 0 | 0 | 0 | 0 | 0 | NA | NA | NA | NA | NA | NA | NA | NA | NA | NA |
| Mir5123 | 0 | 0 | 0 | 0 | 0 | 0 | 0 | 0 | 0 | 0 | 0 | 0 | 0 | 0 | 0 | NA | NA | NA | NA | NA | NA | NA | NA | NA | NA |
| Mir5124 | 0 | 0 | 0 | 0 | 0 | 0 | 0 | 0 | 0 | 0 | 0 | 0 | 0 | 0 | 0 | NA | NA | NA | NA | NA | NA | NA | NA | NA | NA |
| Mir5125 | 0 | 0 | 0 | 0 | 0 | 0 | 0 | 0 | 0 | 0 | 0 | 0 | 0 | 0 | 0 | NA | NA | NA | NA | NA | NA | NA | NA | NA | NA |
| Mir5126 | 0 | 0 | 0 | 0 | 0 | 0 | 0 | 0 | 0 | 0 | 0 | 0 | 0 | 0 | 0 | NA | NA | NA | NA | NA | NA | NA | NA | NA | NA |
| Mir5127 | 0 | 0 | 0 | 0 | 0 | 0 | 0 | 0 | 0 | 0 | 0 | 0 | 0 | 0 | 0 | NA | NA | NA | NA | NA | NA | NA | NA | NA | NA |
| Mir5128 | 0 | 0 | 0 | 0 | 0 | 0 | 0 | 0 | 0 | 0 | 0 | 0 | 0 | 0 | 0 | NA | NA | NA | NA | NA | NA | NA | NA | NA | NA |
| Mir5129 | 0 | 0 | 0 | 0 | 0 | 0 | 0 | 0 | 0 | 0 | 0 | 0 | 0 | 0 | 0 | NA | NA | NA | NA | NA | NA | NA | NA | NA | NA |
| Mir5130 | 0 | 0 | 0 | 0 | 0 | 0 | 0 | 0 | 0 | 0 | 0 | 0 | 0 | 0 | 0 | NA | NA | NA | NA | NA | NA | NA | NA | NA | NA |
| Mir5131 | 0 | 0 | 0 | 0 | 0 | 0 | 0 | 0 | 0 | 0 | 0 | 0 | 0 | 0 | 0 | NA | NA | NA | NA | NA | NA | NA | NA | NA | NA |
| Mir5132 | 0 | 0 | 0 | 0 | 0 | 0 | 0 | 0 | 0 | 0 | 0 | 0 | 0 | 0 | 0 | NA | NA | NA | NA | NA | NA | NA | NA | NA | NA |

| | | | | | | | | | | | | | | | | | | | | | | | | | |
|---|---|---|---|---|---|---|---|---|---|---|---|---|---|---|---|---|---|---|---|---|---|---|---|---|---|
| Mir5133 | 32 | 13 | 16 | 18 | 19 | 13 | 25 | 14 | 17 | 18 | 21 | 13 | 18.02 | 19 | 17 | 0.919 | -0.122 | 0.79488101 | 0.87990506 | 0 | 0.1339 | 0.1039 | 0.1039 | TRUE | 0.5351 |
| Mir5134 | 0 | 0 | 1 | 1 | 1 | 0 | 0 | 0 | 1 | 1 | 1 | 0 | 0.53 | 0 | 1 | 2.036 | 1.026 | 0.70454668 | NA | 0 | 4.4428 | 3.5371 | 3.5371 | TRUE | 0.5365 |
| Mir5135 | 0 | 0 | 0 | 0 | 0 | 0 | 0 | 0 | 0 | 0 | 0 | 0 | 0 | 0 | 0 | NA | NA | NA | NA | NA | NA | NA | NA | NA | NA |
| Mir5136 | 0 | 0 | 0 | 0 | 0 | 0 | 0 | 0 | 0 | 0 | 0 | 0 | 0 | 0 | 0 | NA | NA | NA | NA | NA | NA | NA | NA | NA | NA |
| Mir532 | 0 | 0 | 0 | 0 | 0 | 0 | 0 | 0 | 0 | 0 | 0 | 0 | 0 | 0 | 0 | NA | NA | NA | NA | NA | NA | NA | NA | NA | NA |
| Mir539 | 0 | 0 | 0 | 0 | 0 | 0 | 0 | 0 | 0 | 0 | 0 | 0 | 0 | 0 | 0 | NA | NA | NA | NA | NA | NA | NA | NA | NA | NA |
| Mir540 | 0 | 0 | 0 | 0 | 0 | 0 | 0 | 0 | 0 | 0 | 0 | 0 | 0 | 0 | 0 | NA | NA | NA | NA | NA | NA | NA | NA | NA | NA |
| Mir541 | 0 | 0 | 0 | 0 | 0 | 1 | 0 | 0 | 0 | 0 | 0 | 1 | 0.16 | 0 | 0 | 2.094 | 1.067 | 0.79379329 | NA | 0 | 14.4078 | 10 | 10 | TRUE | 0.3835 |
| Mir542 | 0 | 0 | 0 | 0 | 0 | 0 | 0 | 0 | 0 | 0 | 0 | 0 | 0 | 0 | 0 | NA | NA | NA | NA | NA | NA | NA | NA | NA | NA |
| Mir543 | 0 | 0 | 0 | 0 | 0 | 0 | 0 | 0 | 0 | 0 | 0 | 0 | 0 | 0 | 0 | NA | NA | NA | NA | NA | NA | NA | NA | NA | NA |
| Mir544 | 0 | 0 | 0 | 0 | 0 | 0 | 0 | 0 | 0 | 0 | 0 | 0 | 0 | 0 | 0 | NA | NA | NA | NA | NA | NA | NA | NA | NA | NA |
| Mir546 | 0 | 0 | 0 | 0 | 0 | 0 | 0 | 0 | 0 | 0 | 0 | 0 | 0 | 0 | 0 | NA | NA | NA | NA | NA | NA | NA | NA | NA | NA |
| Mir547 | 0 | 0 | 0 | 0 | 0 | 0 | 0 | 0 | 0 | 0 | 0 | 0 | 0 | 0 | 0 | NA | NA | NA | NA | NA | NA | NA | NA | NA | NA |
| Mir551b | 0 | 0 | 0 | 0 | 0 | 0 | 0 | 0 | 0 | 0 | 0 | 0 | 0 | 0 | 0 | NA | NA | NA | NA | NA | NA | NA | NA | NA | NA |
| Mir568 | 0 | 0 | 2 | 0 | 0 | 1 | 0 | 0 | 2 | 0 | 0 | 1 | 0.52 | 1 | 0 | 0.529 | -0.92 | 0.75853134 | NA | 0 | 4.5238 | 4.7579 | 4.7579 | TRUE | 0.9746 |
| Mir574 | 0 | 0 | 0 | 0 | 1 | 0 | 0 | 0 | 0 | 0 | 1 | 0 | 0.19 | 0 | 0 | 2.094 | 1.067 | 0.79379329 | NA | 0 | 12.608 | 10 | 10 | TRUE | 0.5075 |
| Mir582 | 0 | 0 | 0 | 0 | 0 | 0 | 0 | 0 | 0 | 0 | 0 | 0 | 0 | 0 | 0 | NA | NA | NA | NA | NA | NA | NA | NA | NA | NA |
| Mir592 | 0 | 0 | 0 | 0 | 0 | 0 | 0 | 0 | 0 | 0 | 0 | 0 | 0 | 0 | 0 | NA | NA | NA | NA | NA | NA | NA | NA | NA | NA |
| Mir598 | 0 | 0 | 0 | 0 | 0 | 0 | 0 | 0 | 0 | 0 | 0 | 0 | 0 | 0 | 0 | NA | NA | NA | NA | NA | NA | NA | NA | NA | NA |
| Mir599 | 0 | 0 | 0 | 0 | 0 | 0 | 0 | 0 | 0 | 0 | 0 | 0 | 0 | 0 | 0 | NA | NA | NA | NA | NA | NA | NA | NA | NA | NA |
| Mir615 | 0 | 0 | 0 | 0 | 0 | 0 | 0 | 0 | 0 | 0 | 0 | 0 | 0 | 0 | 0 | NA | NA | NA | NA | NA | NA | NA | NA | NA | NA |
| Mir653 | 0 | 0 | 0 | 0 | 0 | 0 | 0 | 0 | 0 | 0 | 0 | 0 | 0 | 0 | 0 | NA | NA | NA | NA | NA | NA | NA | NA | NA | NA |
| Mir654 | 0 | 0 | 0 | 0 | 0 | 0 | 0 | 0 | 0 | 0 | 0 | 0 | 0 | 0 | 0 | NA | NA | NA | NA | NA | NA | NA | NA | NA | NA |
| Mir664 | 0 | 0 | 0 | 0 | 0 | 0 | 0 | 0 | 0 | 0 | 0 | 0 | 0 | 0 | 0 | NA | NA | NA | NA | NA | NA | NA | NA | NA | NA |
| Mir665 | 0 | 0 | 0 | 0 | 0 | 0 | 0 | 0 | 0 | 0 | 0 | 0 | 0 | 0 | 0 | NA | NA | NA | NA | NA | NA | NA | NA | NA | NA |
| Mir666 | 0 | 0 | 0 | 0 | 0 | 0 | 0 | 0 | 0 | 0 | 0 | 0 | 0 | 0 | 0 | NA | NA | NA | NA | NA | NA | NA | NA | NA | NA |
| Mir667 | 0 | 0 | 0 | 0 | 0 | 0 | 0 | 0 | 0 | 0 | 0 | 0 | 0 | 0 | 0 | NA | NA | NA | NA | NA | NA | NA | NA | NA | NA |
| Mir668 | 0 | 0 | 0 | 0 | 0 | 0 | 0 | 0 | 0 | 0 | 0 | 0 | 0 | 0 | 0 | NA | NA | NA | NA | NA | NA | NA | NA | NA | NA |
| Mir669a-1 | 0 | 0 | 0 | 0 | 0 | 0 | 0 | 0 | 0 | 0 | 0 | 0 | 0 | 0 | 0 | NA | NA | NA | NA | NA | NA | NA | NA | NA | NA |
| Mir669a-10 | 0 | 0 | 0 | 0 | 0 | 0 | 0 | 0 | 0 | 0 | 0 | 0 | 0 | 0 | 0 | NA | NA | NA | NA | NA | NA | NA | NA | NA | NA |
| Mir669a-11 | 0 | 0 | 0 | 0 | 0 | 0 | 0 | 0 | 0 | 0 | 0 | 0 | 0 | 0 | 0 | NA | NA | NA | NA | NA | NA | NA | NA | NA | NA |
| Mir669a-12 | 0 | 0 | 0 | 0 | 0 | 0 | 0 | 0 | 0 | 0 | 0 | 0 | 0 | 0 | 0 | NA | NA | NA | NA | NA | NA | NA | NA | NA | NA |
| Mir669a-2 | 0 | 0 | 0 | 0 | 0 | 0 | 0 | 0 | 0 | 0 | 0 | 0 | 0 | 0 | 0 | NA | NA | NA | NA | NA | NA | NA | NA | NA | NA |
| Mir669a-3 | 0 | 0 | 0 | 0 | 0 | 0 | 0 | 0 | 0 | 0 | 0 | 0 | 0 | 0 | 0 | NA | NA | NA | NA | NA | NA | NA | NA | NA | NA |
| Mir669a-4 | 0 | 0 | 0 | 0 | 0 | 0 | 0 | 0 | 0 | 0 | 0 | 0 | 0 | 0 | 0 | NA | NA | NA | NA | NA | NA | NA | NA | NA | NA |
| Mir669a-5 | 0 | 0 | 0 | 0 | 0 | 0 | 0 | 0 | 0 | 0 | 0 | 0 | 0 | 0 | 0 | NA | NA | NA | NA | NA | NA | NA | NA | NA | NA |
| Mir669a-6 | 0 | 0 | 0 | 0 | 0 | 0 | 0 | 0 | 0 | 0 | 0 | 0 | 0 | 0 | 0 | NA | NA | NA | NA | NA | NA | NA | NA | NA | NA |
| Mir669a-7 | 0 | 0 | 0 | 0 | 0 | 0 | 0 | 0 | 0 | 0 | 0 | 0 | 0 | 0 | 0 | NA | NA | NA | NA | NA | NA | NA | NA | NA | NA |
| Mir669a-8 | 0 | 0 | 0 | 0 | 0 | 0 | 0 | 0 | 0 | 0 | 0 | 0 | 0 | 0 | 0 | NA | NA | NA | NA | NA | NA | NA | NA | NA | NA |
| Mir669a-9 | 0 | 0 | 0 | 0 | 0 | 0 | 0 | 0 | 0 | 0 | 0 | 0 | 0 | 0 | 0 | NA | NA | NA | NA | NA | NA | NA | NA | NA | NA |
| Mir669b | 0 | 0 | 0 | 0 | 0 | 0 | 0 | 0 | 0 | 0 | 0 | 0 | 0 | 0 | 0 | NA | NA | NA | NA | NA | NA | NA | NA | NA | NA |
| Mir669c | 0 | 0 | 0 | 0 | 0 | 0 | 0 | 0 | 0 | 0 | 0 | 0 | 0 | 0 | 0 | NA | NA | NA | NA | NA | NA | NA | NA | NA | NA |
| Mir669e | 0 | 0 | 0 | 0 | 0 | 0 | 0 | 0 | 0 | 0 | 0 | 0 | 0 | 0 | 0 | NA | NA | NA | NA | NA | NA | NA | NA | NA | NA |
| Mir669g | 0 | 0 | 0 | 0 | 0 | 0 | 0 | 0 | 0 | 0 | 0 | 0 | 0 | 0 | 0 | NA | NA | NA | NA | NA | NA | NA | NA | NA | NA |
| Mir669h | 0 | 0 | 0 | 0 | 0 | 0 | 0 | 0 | 0 | 0 | 0 | 0 | 0 | 0 | 0 | NA | NA | NA | NA | NA | NA | NA | NA | NA | NA |
| Mir669i | 1 | 2 | 0 | 1 | 0 | 0 | 1 | 2 | 0 | 1 | 0 | 0 | 0.65 | 1 | 0 | 0.384 | -1.379 | 0.58322298 | NA | 0 | 3.6492 | 3.0509 | 3.0509 | TRUE | 0.2622 |
| Mir669j | 0 | 0 | 0 | 0 | 0 | 0 | 0 | 0 | 0 | 0 | 0 | 0 | 0 | 0 | 0 | NA | NA | NA | NA | NA | NA | NA | NA | NA | NA |
| Mir669k | 0 | 0 | 0 | 0 | 0 | 0 | 0 | 0 | 0 | 0 | 0 | 0 | 0 | 0 | 0 | NA | NA | NA | NA | NA | NA | NA | NA | NA | NA |
| Mir669m-1 | 0 | 0 | 0 | 0 | 0 | 0 | 0 | 0 | 0 | 0 | 0 | 0 | 0 | 0 | 0 | NA | NA | NA | NA | NA | NA | NA | NA | NA | NA |
| Mir669m-2 | 0 | 0 | 0 | 0 | 0 | 0 | 0 | 0 | 0 | 0 | 0 | 0 | 0 | 0 | 0 | NA | NA | NA | NA | NA | NA | NA | NA | NA | NA |
| Mir669p-1 | 0 | 0 | 0 | 0 | 0 | 0 | 0 | 0 | 0 | 0 | 0 | 0 | 0 | 0 | 0 | NA | NA | NA | NA | NA | NA | NA | NA | NA | NA |
| Mir669p-2 | 0 | 0 | 0 | 0 | 0 | 0 | 0 | 0 | 0 | 0 | 0 | 0 | 0 | 0 | 0 | NA | NA | NA | NA | NA | NA | NA | NA | NA | NA |
| Mir670 | 0 | 0 | 0 | 0 | 0 | 0 | 0 | 0 | 0 | 0 | 0 | 0 | 0 | 0 | 0 | NA | NA | NA | NA | NA | NA | NA | NA | NA | NA |
| Mir671 | 0 | 0 | 0 | 0 | 0 | 0 | 0 | 0 | 0 | 0 | 0 | 0 | 0 | 0 | 0 | NA | NA | NA | NA | NA | NA | NA | NA | NA | NA |
| Mir672 | 0 | 0 | 0 | 0 | 0 | 0 | 0 | 0 | 0 | 0 | 0 | 0 | 0 | 0 | 0 | NA | NA | NA | NA | NA | NA | NA | NA | NA | NA |
| Mir673 | 0 | 0 | 0 | 0 | 0 | 0 | 0 | 0 | 0 | 0 | 0 | 0 | 0 | 0 | 0 | NA | NA | NA | NA | NA | NA | NA | NA | NA | NA |
| Mir674 | 0 | 0 | 0 | 0 | 0 | 0 | 0 | 0 | 0 | 0 | 0 | 0 | 0 | 0 | 0 | NA | NA | NA | NA | NA | NA | NA | NA | NA | NA |
| Mir675 | 0 | 0 | 0 | 0 | 0 | 0 | 0 | 0 | 0 | 0 | 0 | 0 | 0 | 0 | 0 | NA | NA | NA | NA | NA | NA | NA | NA | NA | NA |
| Mir676 | 0 | 0 | 0 | 0 | 0 | 0 | 0 | 0 | 0 | 0 | 0 | 0 | 0 | 0 | 0 | NA | NA | NA | NA | NA | NA | NA | NA | NA | NA |
| Mir677 | 8 | 3 | 3 | 3 | 8 | 5 | 6 | 3 | 3 | 3 | 9 | 5 | 4.93 | 4 | 6 | 1.304 | 0.383 | 0.66430594 | NA | 0 | 0.483 | 0.3551 | 0.3551 | TRUE | 0.4163 |
| Mir678 | 0 | 0 | 0 | 0 | 0 | 0 | 0 | 0 | 0 | 0 | 0 | 0 | 0 | 0 | 0 | NA | NA | NA | NA | NA | NA | NA | NA | NA | NA |
| Mir679 | 0 | 0 | 1 | 0 | 0 | 0 | 0 | 0 | 1 | 0 | 0 | 0 | 0.18 | 0 | 0 | 0.552 | -0.857 | 0.8336548 | NA | 0 | 13.1817 | 10 | 10 | TRUE | 0.5365 |
| Mir680-2 | 0 | 3 | 1 | 0 | 0 | 1 | 0 | 3 | 1 | 0 | 0 | 1 | 0.88 | 1 | 0 | 0.266 | -1.909 | 0.40962126 | NA | 0 | 2.7074 | 2.5155 | 2.5155 | TRUE | 0.2246 |
| Mir680-3 | 0 | 0 | 0 | 0 | 0 | 0 | 0 | 0 | 0 | 0 | 0 | 0 | 0 | 0 | 0 | NA | NA | NA | NA | NA | NA | NA | NA | NA | NA |
| Mir681 | 0 | 0 | 0 | 0 | 0 | 0 | 0 | 0 | 0 | 0 | 0 | 0 | 0 | 0 | 0 | NA | NA | NA | NA | NA | NA | NA | NA | NA | NA |
| Mir682 | 58 | 47 | 30 | 59 | 35 | 42 | 45 | 50 | 32 | 59 | 39 | 41 | 44.51 | 42 | 46 | 1.096 | 0.132 | 0.67126682 | 0.79760717 | 0.0244 | 0.0557 | 0.0472 | 0.0472 | TRUE | 0.4204 |
| Mir683-1 | 0 | 0 | 0 | 0 | 0 | 0 | 0 | 0 | 0 | 0 | 0 | 0 | 0 | 0 | 0 | NA | NA | NA | NA | NA | NA | NA | NA | NA | NA |
| Mir683-2 | 0 | 0 | 0 | 0 | 0 | 0 | 0 | 0 | 0 | 0 | 0 | 0 | 0 | 0 | 0 | NA | NA | NA | NA | NA | NA | NA | NA | NA | NA |
| Mir684-1 | 10 | 5 | 1 | 3 | 1 | 4 | 8 | 5 | 1 | 3 | 1 | 4 | 3.7 | 5 | 3 | 0.565 | -0.825 | 0.43948866 | NA | 0 | 0.6422 | 0.5293 | 0.5293 | TRUE | 0.251 |

| | | | | | | | | | | | | | | | | | | | | | | | | | |
|---|---|---|---|---|---|---|---|---|---|---|---|---|---|---|---|---|---|---|---|---|---|---|---|---|---|
| Mir684-2 | 0 | 0 | 0 | 0 | 0 | 0 | 0 | 0 | 0 | 0 | 0 | 0 | 0 | 0 | 0 | NA | NA | NA | NA | NA | NA | NA | NA | NA | NA |
| Mir686 | 0 | 0 | 0 | 0 | 0 | 0 | 0 | 0 | 0 | 0 | 0 | 0 | 0 | 0 | 0 | NA | NA | NA | NA | NA | NA | NA | NA | NA | NA |
| Mir687 | 0 | 0 | 0 | 0 | 0 | 0 | 0 | 0 | 0 | 0 | 0 | 0 | 0 | 0 | 0 | NA | NA | NA | NA | NA | NA | NA | NA | NA | NA |
| Mir688 | 1 | 1 | 0 | 0 | 0 | 0 | 1 | 1 | 0 | 0 | 0 | 0 | 0.31 | 1 | 0 | 0.309 | -1.696 | 0.63183309 | NA | 0 | 7.7294 | 7.2472 | 7.2472 | TRUE | 0.2015 |
| Mir690 | 0 | 0 | 0 | 0 | 0 | 0 | 0 | 0 | 0 | 0 | 0 | 0 | 0 | 0 | 0 | NA | NA | NA | NA | NA | NA | NA | NA | NA | NA |
| Mir691 | 0 | 0 | 0 | 0 | 0 | 0 | 0 | 0 | 0 | 0 | 0 | 0 | 0 | 0 | 0 | NA | NA | NA | NA | NA | NA | NA | NA | NA | NA |
| Mir692-1 | 0 | 0 | 0 | 0 | 0 | 2 | 0 | 0 | 0 | 0 | 0 | 2 | 0.33 | 0 | 1 | 3.743 | 1.904 | 0.62640105 | NA | 0 | 7.2051 | 9.2465 | 9.2465 | TRUE | 0.9739 |
| Mir693 | 0 | 0 | 0 | 0 | 0 | 0 | 0 | 0 | 0 | 0 | 0 | 0 | 0 | 0 | 0 | NA | NA | NA | NA | NA | NA | NA | NA | NA | NA |
| Mir694 | 0 | 0 | 0 | 0 | 0 | 0 | 0 | 0 | 0 | 0 | 0 | 0 | 0 | 0 | 0 | NA | NA | NA | NA | NA | NA | NA | NA | NA | NA |
| Mir695 | 0 | 0 | 0 | 0 | 0 | 0 | 0 | 0 | 0 | 0 | 0 | 0 | 0 | 0 | 0 | NA | NA | NA | NA | NA | NA | NA | NA | NA | NA |
| Mir697 | 3 | 0 | 1 | 0 | 0 | 2 | 2 | 0 | 1 | 0 | 0 | 2 | 0.9 | 1 | 1 | 0.574 | -0.8 | 0.72123729 | NA | 0 | 2.644 | 2.4388 | 2.4388 | TRUE | 0.4452 |
| Mir698 | 0 | 0 | 0 | 0 | 0 | 0 | 0 | 0 | 0 | 0 | 0 | 0 | 0 | 0 | 0 | NA | NA | NA | NA | NA | NA | NA | NA | NA | NA |
| Mir7-1 | 3 | 0 | 1 | 4 | 3 | 1 | 2 | 0 | 1 | 4 | 3 | 1 | 1.96 | 1 | 3 | 2.324 | 1.217 | 0.40132493 | NA | 0 | 1.2114 | 0.9246 | 0.9246 | TRUE | 0.5442 |
| Mir7-2 | 0 | 0 | 0 | 0 | 0 | 0 | 0 | 0 | 0 | 0 | 0 | 0 | 0 | 0 | 0 | NA | NA | NA | NA | NA | NA | NA | NA | NA | NA |
| Mir700 | 0 | 0 | 0 | 0 | 0 | 0 | 0 | 0 | 0 | 0 | 0 | 0 | 0 | 0 | 0 | NA | NA | NA | NA | NA | NA | NA | NA | NA | NA |
| Mir701 | 0 | 0 | 0 | 0 | 0 | 0 | 0 | 0 | 0 | 0 | 0 | 0 | 0 | 0 | 0 | NA | NA | NA | NA | NA | NA | NA | NA | NA | NA |
| Mir702 | 0 | 0 | 0 | 0 | 0 | 0 | 0 | 0 | 0 | 0 | 0 | 0 | 0 | 0 | 0 | NA | NA | NA | NA | NA | NA | NA | NA | NA | NA |
| Mir703 | 373 | 266 | 289 | 401 | 276 | 336 | 290 | 283 | 312 | 399 | 311 | 332 | 321.11 | 295 | 347 | 1.18 | 0.239 | 0.06426204 | 0.16119645 | 0.0069 | 0.0098 | 0.0089 | 0.0089 | TRUE | 0.1948 |
| Mir704 | 1 | 0 | 0 | 0 | 0 | 1 | 1 | 0 | 0 | 0 | 0 | 1 | 0.29 | 0 | 0 | 1.075 | 0.105 | 0.97702352 | NA | 0 | 8.0647 | 7.5458 | 7.5458 | TRUE | 0.3835 |
| Mir705 | 0 | 0 | 0 | 0 | 0 | 0 | 0 | 0 | 0 | 0 | 0 | 0 | 0 | 0 | 0 | NA | NA | NA | NA | NA | NA | NA | NA | NA | NA |
| Mir706 | 0 | 0 | 0 | 0 | 0 | 0 | 0 | 0 | 0 | 0 | 0 | 0 | 0 | 0 | 0 | NA | NA | NA | NA | NA | NA | NA | NA | NA | NA |
| Mir707 | 3 | 1 | 2 | 3 | 3 | 0 | 2 | 1 | 2 | 3 | 3 | 0 | 1.99 | 2 | 2 | 1.126 | 0.171 | 0.90304644 | NA | 0 | 1.1952 | 0.9174 | 0.9174 | TRUE | 0.7609 |
| Mir708 | 0 | 0 | 0 | 0 | 0 | 0 | 0 | 0 | 0 | 0 | 0 | 0 | 0 | 0 | 0 | NA | NA | NA | NA | NA | NA | NA | NA | NA | NA |
| Mir709 | 0 | 1 | 0 | 0 | 0 | 0 | 0 | 1 | 0 | 0 | 0 | 0 | 0.18 | 0 | 0 | 0.552 | -0.857 | 0.8336548 | NA | 0 | 13.3714 | 10 | 10 | TRUE | 0.5216 |
| Mir710 | 0 | 0 | 0 | 0 | 0 | 0 | 0 | 0 | 0 | 0 | 0 | 0 | 0 | 0 | 0 | NA | NA | NA | NA | NA | NA | NA | NA | NA | NA |
| Mir711 | 0 | 0 | 0 | 0 | 0 | 0 | 0 | 0 | 0 | 0 | 0 | 0 | 0 | 0 | 0 | NA | NA | NA | NA | NA | NA | NA | NA | NA | NA |
| Mir713 | 0 | 0 | 0 | 0 | 0 | 0 | 0 | 0 | 0 | 0 | 0 | 0 | 0 | 0 | 0 | NA | NA | NA | NA | NA | NA | NA | NA | NA | NA |
| Mir717 | 0 | 1 | 0 | 0 | 0 | 0 | 0 | 1 | 0 | 0 | 0 | 0 | 0.18 | 0 | 0 | 0.552 | -0.857 | 0.8336548 | NA | 0 | 13.3714 | 10 | 10 | TRUE | 0.5216 |
| Mir718 | 0 | 0 | 0 | 0 | 0 | 0 | 0 | 0 | 0 | 0 | 0 | 0 | 0 | 0 | 0 | NA | NA | NA | NA | NA | NA | NA | NA | NA | NA |
| Mir719 | 0 | 0 | 0 | 0 | 0 | 0 | 0 | 0 | 0 | 0 | 0 | 0 | 0 | 0 | 0 | NA | NA | NA | NA | NA | NA | NA | NA | NA | NA |
| Mir721 | 0 | 0 | 0 | 0 | 0 | 0 | 0 | 0 | 0 | 0 | 0 | 0 | 0 | 0 | 0 | NA | NA | NA | NA | NA | NA | NA | NA | NA | NA |
| Mir741 | 0 | 0 | 0 | 0 | 0 | 0 | 0 | 0 | 0 | 0 | 0 | 0 | 0 | 0 | 0 | NA | NA | NA | NA | NA | NA | NA | NA | NA | NA |
| Mir742 | 0 | 0 | 0 | 0 | 0 | 0 | 0 | 0 | 0 | 0 | 0 | 0 | 0 | 0 | 0 | NA | NA | NA | NA | NA | NA | NA | NA | NA | NA |
| Mir743 | 0 | 0 | 0 | 0 | 0 | 0 | 0 | 0 | 0 | 0 | 0 | 0 | 0 | 0 | 0 | NA | NA | NA | NA | NA | NA | NA | NA | NA | NA |
| Mir743b | 0 | 0 | 0 | 0 | 0 | 0 | 0 | 0 | 0 | 0 | 0 | 0 | 0 | 0 | 0 | NA | NA | NA | NA | NA | NA | NA | NA | NA | NA |
| Mir744 | 0 | 0 | 0 | 0 | 0 | 0 | 0 | 0 | 0 | 0 | 0 | 0 | 0 | 0 | 0 | NA | NA | NA | NA | NA | NA | NA | NA | NA | NA |
| Mir758 | 0 | 0 | 0 | 0 | 0 | 0 | 0 | 0 | 0 | 0 | 0 | 0 | 0 | 0 | 0 | NA | NA | NA | NA | NA | NA | NA | NA | NA | NA |
| Mir759 | 0 | 0 | 0 | 0 | 0 | 0 | 0 | 0 | 0 | 0 | 0 | 0 | 0 | 0 | 0 | NA | NA | NA | NA | NA | NA | NA | NA | NA | NA |
| Mir760 | 0 | 0 | 0 | 0 | 0 | 1 | 0 | 0 | 0 | 0 | 0 | 1 | 0.16 | 0 | 0 | 2.094 | 1.067 | 0.79379329 | NA | 0 | 14.4078 | 10 | 10 | TRUE | 0.3835 |
| Mir761 | 2 | 1 | 0 | 2 | 1 | 1 | 2 | 1 | 0 | 2 | 1 | 1 | 1.12 | 1 | 1 | 1.516 | 0.6 | 0.73587335 | NA | 0 | 2.1176 | 1.3707 | 1.3707 | TRUE | 0.2754 |
| Mir762 | 1 | 1 | 1 | 3 | 0 | 1 | 1 | 1 | 1 | 3 | 0 | 1 | 1.15 | 1 | 1 | 1.393 | 0.478 | 0.78963432 | NA | 0 | 2.0655 | 1.4301 | 1.4301 | TRUE | 0.4184 |
| Mir764 | 0 | 0 | 0 | 0 | 0 | 0 | 0 | 0 | 0 | 0 | 0 | 0 | 0 | 0 | 0 | NA | NA | NA | NA | NA | NA | NA | NA | NA | NA |
| Mir767 | 0 | 0 | 0 | 0 | 0 | 0 | 0 | 0 | 0 | 0 | 0 | 0 | 0 | 0 | 0 | NA | NA | NA | NA | NA | NA | NA | NA | NA | NA |
| Mir770 | 0 | 0 | 0 | 0 | 0 | 0 | 0 | 0 | 0 | 0 | 0 | 0 | 0 | 0 | 0 | NA | NA | NA | NA | NA | NA | NA | NA | NA | NA |
| Mir7b | 0 | 0 | 0 | 0 | 0 | 0 | 0 | 0 | 0 | 0 | 0 | 0 | 0 | 0 | 0 | NA | NA | NA | NA | NA | NA | NA | NA | NA | NA |
| Mir802 | 0 | 0 | 0 | 0 | 0 | 0 | 0 | 0 | 0 | 0 | 0 | 0 | 0 | 0 | 0 | NA | NA | NA | NA | NA | NA | NA | NA | NA | NA |
| Mir804 | 0 | 0 | 0 | 0 | 0 | 0 | 0 | 0 | 0 | 0 | 0 | 0 | 0 | 0 | 0 | NA | NA | NA | NA | NA | NA | NA | NA | NA | NA |
| Mir871 | 0 | 0 | 0 | 0 | 0 | 0 | 0 | 0 | 0 | 0 | 0 | 0 | 0 | 0 | 0 | NA | NA | NA | NA | NA | NA | NA | NA | NA | NA |
| Mir872 | 0 | 0 | 0 | 0 | 0 | 0 | 0 | 0 | 0 | 0 | 0 | 0 | 0 | 0 | 0 | NA | NA | NA | NA | NA | NA | NA | NA | NA | NA |
| Mir873 | 0 | 0 | 0 | 0 | 0 | 0 | 0 | 0 | 0 | 0 | 0 | 0 | 0 | 0 | 0 | NA | NA | NA | NA | NA | NA | NA | NA | NA | NA |
| Mir874 | 0 | 0 | 0 | 0 | 0 | 0 | 0 | 0 | 0 | 0 | 0 | 0 | 0 | 0 | 0 | NA | NA | NA | NA | NA | NA | NA | NA | NA | NA |
| Mir875 | 0 | 0 | 0 | 0 | 0 | 0 | 0 | 0 | 0 | 0 | 0 | 0 | 0 | 0 | 0 | NA | NA | NA | NA | NA | NA | NA | NA | NA | NA |
| Mir876 | 0 | 0 | 0 | 0 | 0 | 0 | 0 | 0 | 0 | 0 | 0 | 0 | 0 | 0 | 0 | NA | NA | NA | NA | NA | NA | NA | NA | NA | NA |
| Mir877 | 1 | 0 | 0 | 1 | 0 | 0 | 1 | 0 | 0 | 1 | 0 | 0 | 0.3 | 0 | 0 | 1.075 | 0.105 | 0.97698525 | NA | 0 | 8.0272 | 7.5141 | 7.5141 | TRUE | 0.3906 |
| Mir878 | 0 | 0 | 0 | 0 | 0 | 0 | 0 | 0 | 0 | 0 | 0 | 0 | 0 | 0 | 0 | NA | NA | NA | NA | NA | NA | NA | NA | NA | NA |
| Mir879 | 0 | 0 | 0 | 0 | 0 | 0 | 0 | 0 | 0 | 0 | 0 | 0 | 0 | 0 | 0 | NA | NA | NA | NA | NA | NA | NA | NA | NA | NA |
| Mir880 | 0 | 0 | 0 | 0 | 0 | 0 | 0 | 0 | 0 | 0 | 0 | 0 | 0 | 0 | 0 | NA | NA | NA | NA | NA | NA | NA | NA | NA | NA |
| Mir881 | 0 | 0 | 0 | 0 | 0 | 0 | 0 | 0 | 0 | 0 | 0 | 0 | 0 | 0 | 0 | NA | NA | NA | NA | NA | NA | NA | NA | NA | NA |
| Mir882 | 0 | 0 | 0 | 0 | 0 | 0 | 0 | 0 | 0 | 0 | 0 | 0 | 0 | 0 | 0 | NA | NA | NA | NA | NA | NA | NA | NA | NA | NA |
| Mir883a | 0 | 0 | 0 | 0 | 0 | 0 | 0 | 0 | 0 | 0 | 0 | 0 | 0 | 0 | 0 | NA | NA | NA | NA | NA | NA | NA | NA | NA | NA |
| Mir883b | 0 | 0 | 0 | 0 | 0 | 0 | 0 | 0 | 0 | 0 | 0 | 0 | 0 | 0 | 0 | NA | NA | NA | NA | NA | NA | NA | NA | NA | NA |
| Mir9-1 | 0 | 0 | 0 | 0 | 0 | 0 | 0 | 0 | 0 | 0 | 0 | 0 | 0 | 0 | 0 | NA | NA | NA | NA | NA | NA | NA | NA | NA | NA |
| Mir9-2 | 0 | 0 | 0 | 0 | 0 | 0 | 0 | 0 | 0 | 0 | 0 | 0 | 0 | 0 | 0 | NA | NA | NA | NA | NA | NA | NA | NA | NA | NA |
| Mir9-3 | 0 | 0 | 0 | 0 | 0 | 0 | 0 | 0 | 0 | 0 | 0 | 0 | 0 | 0 | 0 | NA | NA | NA | NA | NA | NA | NA | NA | NA | NA |
| Mir92-1 | 0 | 0 | 0 | 0 | 0 | 0 | 0 | 0 | 0 | 0 | 0 | 0 | 0 | 0 | 0 | NA | NA | NA | NA | NA | NA | NA | NA | NA | NA |
| Mir92-2 | 0 | 1 | 1 | 0 | 0 | 0 | 0 | 1 | 1 | 0 | 0 | 0 | 0.36 | 1 | 0 | 0.271 | -1.881 | 0.5755355 | NA | 0 | 6.6391 | 6.4428 | 6.4428 | TRUE | 0.3513 |
| Mir92b | 0 | 0 | 0 | 0 | 0 | 0 | 0 | 0 | 0 | 0 | 0 | 0 | 0 | 0 | 0 | NA | NA | NA | NA | NA | NA | NA | NA | NA | NA |
| Mir93 | 0 | 0 | 0 | 1 | 0 | 1 | 0 | 0 | 0 | 1 | 0 | 1 | 0.33 | 0 | 1 | 3.763 | 1.912 | 0.57887588 | NA | 0 | 7.1752 | 6.7756 | 6.7756 | TRUE | 0.2094 |
| Mir96 | 0 | 0 | 0 | 0 | 0 | 0 | 0 | 0 | 0 | 0 | 0 | 0 | 0 | 0 | 0 | NA | NA | NA | NA | NA | NA | NA | NA | NA | NA |

| | | | | | | | | | | | | | | | | | | | | | | | | | |
|---|---|---|---|---|---|---|---|---|---|---|---|---|---|---|---|---|---|---|---|---|---|---|---|---|---|
| Mir98 | 0 | 0 | 0 | 0 | 0 | 0 | 0 | 0 | 0 | 0 | 0 | 0 | 0 | 0 | 0 | NA | NA | NA | NA | NA | NA | NA | NA | NA | NA |
| Mir99a | 0 | 0 | 0 | 0 | 0 | 0 | 0 | 0 | 0 | 0 | 0 | 0 | 0 | 0 | 0 | NA | NA | NA | NA | NA | NA | NA | NA | NA | NA |
| Mir99b | 0 | 0 | 0 | 0 | 0 | 0 | 0 | 0 | 0 | 0 | 0 | 0 | 0 | 0 | 0 | NA | NA | NA | NA | NA | NA | NA | NA | NA | NA |
| Mira | 19 | 11 | 4 | 6 | 4 | 7 | 15 | 12 | 4 | 6 | 5 | 7 | 8.03 | 10 | 6 | 0.56 | -0.836 | 0.24601645 | NA | 0 | 0.2977 | 0.2389 | 0.2389 | TRUE | 0.4381 |
| Mirg | 524 | 443 | 270 | 332 | 304 | 375 | 407 | 471 | 291 | 330 | 343 | 370 | 368.85 | 390 | 348 | 0.892 | -0.165 | 0.28442726 | 0.45640291 | 0.0279 | 0.0089 | 0.0145 | 0.0145 | TRUE | 0.3761 |
| Mirlet7a-1 | 0 | 0 | 0 | 1 | 0 | 0 | 0 | 0 | 0 | 1 | 0 | 0 | 0.17 | 0 | 0 | 2.094 | 1.067 | 0.79379329 | NA | 0 | 14.2885 | 10 | 10 | TRUE | 0.3906 |
| Mirlet7a-2 | 0 | 0 | 0 | 0 | 0 | 0 | 0 | 0 | 0 | 0 | 0 | 0 | 0 | 0 | 0 | NA | NA | NA | NA | NA | NA | NA | NA | NA | NA |
| Mirlet7b | 0 | 0 | 0 | 0 | 0 | 0 | 0 | 0 | 0 | 0 | 0 | 0 | 0 | 0 | 0 | NA | NA | NA | NA | NA | NA | NA | NA | NA | NA |
| Mirlet7c-1 | 0 | 0 | 0 | 0 | 0 | 0 | 0 | 0 | 0 | 0 | 0 | 0 | 0 | 0 | 0 | NA | NA | NA | NA | NA | NA | NA | NA | NA | NA |
| Mirlet7c-2 | 0 | 0 | 0 | 0 | 0 | 0 | 0 | 0 | 0 | 0 | 0 | 0 | 0 | 0 | 0 | NA | NA | NA | NA | NA | NA | NA | NA | NA | NA |
| Mirlet7d | 4 | 3 | 0 | 1 | 0 | 0 | 3 | 3 | 0 | 1 | 0 | 0 | 1.22 | 2 | 0 | 0.175 | -2.517 | 0.22079851 | NA | 0 | 1.9523 | 1.8163 | 1.8163 | TRUE | 0.6537 |
| Mirlet7e | 0 | 0 | 0 | 0 | 0 | 0 | 0 | 0 | 0 | 0 | 0 | 0 | 0 | 0 | 0 | NA | NA | NA | NA | NA | NA | NA | NA | NA | NA |
| Mirlet7f-1 | 0 | 0 | 0 | 0 | 0 | 0 | 0 | 0 | 0 | 0 | 0 | 0 | 0 | 0 | 0 | NA | NA | NA | NA | NA | NA | NA | NA | NA | NA |
| Mirlet7f-2 | 0 | 0 | 0 | 0 | 0 | 0 | 0 | 0 | 0 | 0 | 0 | 0 | 0 | 0 | 0 | NA | NA | NA | NA | NA | NA | NA | NA | NA | NA |
| Mirlet7g | 0 | 0 | 0 | 1 | 0 | 0 | 0 | 0 | 0 | 1 | 0 | 0 | 0.17 | 0 | 0 | 2.094 | 1.067 | 0.79379329 | NA | 0 | 14.2885 | 10 | 10 | TRUE | 0.3906 |
| Mirlet7i | 0 | 0 | 0 | 0 | 0 | 0 | 0 | 0 | 0 | 0 | 0 | 0 | 0 | 0 | 0 | NA | NA | NA | NA | NA | NA | NA | NA | NA | NA |
| Mis12 | 3778 | 2665 | 2457 | 2623 | 2298 | 2604 | 2934 | 2835 | 2651 | 2611 | 2592 | 2571 | 2698.98 | 2807 | 2591 | 0.923 | -0.116 | 0.06068767 | 0.15448066 | 0.001 | 0.0033 | 0.0024 | 0.0024 | TRUE | 0.0283 |
| Mis18a | 2174 | 1607 | 1596 | 1770 | 1584 | 1886 | 1688 | 1709 | 1722 | 1762 | 1787 | 1862 | 1755.07 | 1706 | 1804 | 1.057 | 0.08 | 0.2029231 | 0.36436932 | 0 | 0.0038 | 0.0023 | 0.0023 | TRUE | 0.0098 |
| Mis18bp1 | 6084 | 4219 | 4104 | 4410 | 3677 | 4123 | 4725 | 4488 | 4428 | 4390 | 4148 | 4070 | 4374.81 | 4547 | 4203 | 0.924 | -0.114 | 0.04987878 | 0.13453809 | 0.001 | 0.003 | 0.0022 | 0.0022 | TRUE | 0.0186 |
| Mitd1 | 1223 | 913 | 859 | 985 | 882 | 949 | 950 | 971 | 927 | 980 | 995 | 937 | 960.02 | 949 | 971 | 1.022 | 0.032 | 0.67464986 | 0.80024529 | 0 | 0.0049 | 0.0031 | 0.0031 | TRUE | 0.0113 |
| Mitf | 1559 | 1358 | 1537 | 1471 | 1517 | 1719 | 1211 | 1444 | 1658 | 1464 | 1711 | 1697 | 1531.03 | 1438 | 1624 | 1.131 | 0.177 | 0.09854035 | 0.2187354 | 0.0148 | 0.004 | 0.0077 | 0.0077 | TRUE | 0.2351 |
| Mixl1 | 79 | 47 | 47 | 43 | 34 | 35 | 61 | 50 | 51 | 43 | 38 | 35 | 46.29 | 54 | 39 | 0.711 | -0.493 | 0.08610379 | 0.198465 | 0 | 0.0536 | 0.0371 | 0.0371 | TRUE | 0.1348 |
| Mki67 | 18208 | 12896 | 13431 | 14872 | 13478 | 15752 | 14141 | 13717 | 14492 | 14804 | 15205 | 15550 | 14651.41 | 14117 | 15186 | 1.076 | 0.105 | 0.03184647 | 0.09670089 | 6.00E-04 | 0.0026 | 0.0017 | 0.0017 | TRUE | 0.0074 |
| Mki67ip | 4694 | 3173 | 2662 | 2868 | 2501 | 2873 | 3646 | 3375 | 2872 | 2855 | 2821 | 2836 | 3067.55 | 3298 | 2837 | 0.86 | -0.218 | 0.00827496 | 0.0359168 | 0.0066 | 0.0032 | 0.0046 | 0.0046 | TRUE | 0.1541 |
| Mkks | 2213 | 1548 | 1373 | 1473 | 1299 | 1449 | 1719 | 1647 | 1481 | 1466 | 1465 | 1430 | 1534.8 | 1616 | 1454 | 0.899 | -0.154 | 0.04063284 | 0.11580563 | 0.0024 | 0.004 | 0.0034 | 0.0034 | TRUE | 0.0629 |
| Mkl1 | 2734 | 2000 | 1757 | 1783 | 1651 | 1768 | 2123 | 2127 | 1896 | 1775 | 1862 | 1745 | 1921.52 | 2049 | 1794 | 0.875 | -0.193 | 0.00669575 | 0.03073292 | 0.0022 | 0.0037 | 0.0031 | 0.0031 | TRUE | 0.0508 |
| Mkl2 | 2462 | 1707 | 1625 | 1514 | 1420 | 1625 | 1912 | 1816 | 1753 | 1507 | 1602 | 1604 | 1699.05 | 1827 | 1571 | 0.859 | -0.219 | 0.00134411 | 0.00891479 | 0.0011 | 0.0038 | 0.0028 | 0.0028 | TRUE | 0.0207 |
| Mkln1 | 1917 | 1374 | 1581 | 1694 | 1513 | 1655 | 1489 | 1462 | 1706 | 1686 | 1707 | 1634 | 1613.84 | 1552 | 1676 | 1.08 | 0.111 | 0.14784952 | 0.29256724 | 0.003 | 0.0039 | 0.0036 | 0.0036 | TRUE | 0.0889 |
| Mknk1 | 1408 | 1043 | 1071 | 1228 | 1143 | 1218 | 1094 | 1109 | 1156 | 1222 | 1289 | 1202 | 1178.78 | 1120 | 1238 | 1.106 | 0.145 | 0.04449671 | 0.12375784 | 0 | 0.0044 | 0.0029 | 0.0029 | TRUE | 0.0155 |
| Mknk2 | 5271 | 3963 | 3305 | 3256 | 2956 | 3405 | 4094 | 4215 | 3566 | 3241 | 3335 | 3361 | 3635.36 | 3958 | 3312 | 0.837 | -0.257 | 0.000297 | 0.00288579 | 0.0036 | 0.0031 | 0.0034 | 0.0034 | TRUE | 0.0906 |
| Mkrn1 | 66866 | 48520 | 50888 | 66187 | 57423 | 66535 | 51932 | 51610 | 54908 | 65882 | 64779 | 65680 | 59131.95 | 52817 | 65447 | 1.239 | 0.309 | 4.30E-11 | 3.79E-09 | 6.00E-04 | 0.0025 | 0.0016 | 0.0016 | TRUE | 0.0147 |
| Mkrn2 | 4182 | 3124 | 3420 | 3890 | 3390 | 3866 | 3248 | 3323 | 3690 | 3872 | 3824 | 3816 | 3628.97 | 3420 | 3837 | 1.123 | 0.167 | 0.00921837 | 0.03900479 | 0.002 | 0.0031 | 0.0027 | 0.0027 | TRUE | 0.0575 |
| Mkrn3 | 370 | 274 | 281 | 292 | 221 | 257 | 287 | 291 | 303 | 291 | 249 | 254 | 279.28 | 294 | 265 | 0.901 | -0.15 | 0.23198001 | 0.39785101 | 0 | 0.0109 | 0.0078 | 0.0078 | TRUE | 0.0842 |
| Mks1 | 1639 | 1314 | 1403 | 1351 | 1313 | 1534 | 1273 | 1398 | 1514 | 1345 | 1481 | 1514 | 1420.79 | 1395 | 1447 | 1.038 | 0.054 | 0.52017018 | 0.68171579 | 0.0051 | 0.0041 | 0.0044 | 0.0044 | TRUE | 0.0727 |
| Mkx | 394 | 290 | 282 | 341 | 289 | 353 | 306 | 308 | 304 | 339 | 326 | 348 | 322.11 | 306 | 338 | 1.104 | 0.143 | 0.20648218 | 0.36830772 | 0 | 0.0098 | 0.0061 | 0.0061 | TRUE | 0.0105 |
| Mlana | 40 | 36 | 23 | 37 | 24 | 35 | 31 | 38 | 25 | 37 | 27 | 35 | 32.11 | 31 | 33 | 1.048 | 0.068 | 0.84523936 | 0.91204939 | 0 | 0.0762 | 0.0563 | 0.0563 | TRUE | 0.2113 |
| Mlc1 | 7 | 3 | 3 | 13 | 4 | 4 | 5 | 3 | 3 | 13 | 5 | 4 | 5.54 | 4 | 7 | 1.782 | 0.834 | 0.3340859 | NA | 0.1583 | 0.4299 | 0.3447 | 0.3447 | TRUE | 1.3945 |
| Mlec | 14502 | 10652 | 10302 | 10163 | 9196 | 10053 | 11263 | 11330 | 11116 | 10116 | 10374 | 9924 | 10687.23 | 11236 | 10138 | 0.902 | -0.149 | 0.00151869 | 0.00975142 | 2.00E-04 | 0.0026 | 0.0015 | 0.0015 | TRUE | 0.005 |
| Mlf1 | 476 | 348 | 341 | 376 | 272 | 343 | 370 | 370 | 368 | 374 | 307 | 339 | 354.58 | 369 | 340 | 0.922 | -0.118 | 0.31062138 | 0.48357602 | 0.0015 | 0.0091 | 0.0069 | 0.0069 | TRUE | 0.089 |
| Mlf1ip | 2330 | 1734 | 1625 | 1749 | 1647 | 1820 | 1810 | 1844 | 1753 | 1741 | 1858 | 1797 | 1800.49 | 1802 | 1799 | 0.997 | -0.004 | 0.95408544 | 0.97481901 | 2.00E-04 | 0.0037 | 0.0024 | 0.0024 | TRUE | 0.0099 |
| Mlf2 | 18762 | 13293 | 13467 | 13773 | 12229 | 14258 | 14572 | 14140 | 14531 | 13710 | 13796 | 14075 | 14137.01 | 14414 | 13860 | 0.962 | -0.057 | 0.21291207 | 0.3752326 | 1.00E-04 | 0.0026 | 0.0014 | 0.0014 | TRUE | 0.0034 |
| Mlh1 | 2719 | 1923 | 1778 | 1986 | 1729 | 1974 | 2112 | 2045 | 1918 | 1977 | 1950 | 1949 | 1991.94 | 2025 | 1959 | 0.966 | -0.049 | 0.44611322 | 0.61678238 | 8.00E-04 | 0.0036 | 0.0025 | 0.0025 | TRUE | 0.0255 |
| Mlh3 | 4471 | 3181 | 2772 | 3117 | 2890 | 3197 | 3472 | 3384 | 2991 | 3103 | 3260 | 3156 | 3227.63 | 3282 | 3173 | 0.966 | -0.05 | 0.4722327 | 0.64086802 | 0.0028 | 0.0032 | 0.0032 | 0.0032 | TRUE | 0.0727 |
| Mlip | 2 | 1 | 0 | 1 | 1 | 2 | 2 | 1 | 0 | 1 | 1 | 2 | 1.12 | 1 | 1 | 1.514 | 0.598 | 0.73673685 | NA | 0 | 2.1202 | 1.3712 | 1.3712 | TRUE | 0.2754 |
| Mlkl | 80 | 35 | 43 | 48 | 57 | 67 | 62 | 37 | 46 | 48 | 64 | 66 | 54 | 48 | 59 | 1.21 | 0.276 | 0.34173958 | 0.51578499 | 0.0308 | 0.0463 | 0.042 | 0.042 | TRUE | 0.5448 |
| Mll1 | 5120 | 3793 | 3472 | 4080 | 3899 | 4567 | 3976 | 4035 | 3746 | 4061 | 4398 | 4508 | 4120.89 | 3919 | 4322 | 1.103 | 0.141 | 0.02456357 | 0.08028517 | 0.0018 | 0.003 | 0.0026 | 0.0026 | TRUE | 0.0343 |
| Mll2 | 5573 | 3979 | 3301 | 4321 | 4182 | 4865 | 4328 | 4232 | 3562 | 4301 | 4718 | 4803 | 4323.97 | 4041 | 4607 | 1.14 | 0.189 | 0.01904303 | 0.06671127 | 0.0067 | 0.003 | 0.0044 | 0.0044 | TRUE | 0.1302 |
| Mll3 | 6040 | 3984 | 3802 | 5088 | 4632 | 5093 | 4691 | 4238 | 4102 | 5065 | 5225 | 5028 | 4724.77 | 4344 | 5106 | 1.175 | 0.232 | 0.00029248 | 0.00285425 | 0.0024 | 0.0029 | 0.0027 | 0.0027 | TRUE | 0.0605 |
| Mll5 | 1626 | 1274 | 1154 | 1258 | 1110 | 1210 | 1263 | 1355 | 1245 | 1252 | 1252 | 1194 | 1260.33 | 1288 | 1233 | 0.958 | -0.062 | 0.38770725 | 0.56161951 | 6.00E-04 | 0.0043 | 0.003 | 0.003 | TRUE | 0.0246 |
| Mllt1 | 2054 | 1506 | 1297 | 1435 | 1287 | 1327 | 1595 | 1602 | 1399 | 1428 | 1452 | 1310 | 1464.47 | 1532 | 1397 | 0.911 | -0.135 | 0.09170408 | 0.20759051 | 0.0037 | 0.004 | 0.0039 | 0.0039 | TRUE | 0.068 |
| Mllt10 | 6132 | 4734 | 4721 | 4540 | 4258 | 4803 | 4762 | 5035 | 5094 | 4519 | 4803 | 4741 | 4825.96 | 4964 | 4688 | 0.945 | -0.082 | 0.14530503 | 0.28917905 | 8.00E-04 | 0.0029 | 0.0021 | 0.0021 | TRUE | 0.0157 |
| Mllt11 | 437 | 328 | 301 | 447 | 387 | 457 | 339 | 349 | 325 | 445 | 437 | 451 | 390.95 | 338 | 444 | 1.315 | 0.396 | 0.00016434 | 0.00183222 | 0 | 0.0085 | 0.0053 | 0.0053 | TRUE | 0.0119 |
| Mllt3 | 1749 | 1271 | 1244 | 1230 | 1099 | 1216 | 1358 | 1352 | 1342 | 1224 | 1240 | 1200 | 1286.18 | 1351 | 1221 | 0.904 | -0.146 | 0.03016653 | 0.09304493 | 0 | 0.0043 | 0.0025 | 0.0025 | TRUE | 0.0028 |
| Mllt4 | 13668 | 10479 | 11200 | 11319 | 10148 | 12074 | 10615 | 11146 | 12085 | 11267 | 11448 | 11919 | 11413.38 | 11282 | 11545 | 1.023 | 0.033 | 0.58173878 | 0.73222699 | 0.0021 | 0.0026 | 0.0026 | 0.0026 | TRUE | 0.0473 |
| Mllt6 | 9904 | 8484 | 6018 | 7149 | 6127 | 6850 | 7692 | 9024 | 6493 | 7116 | 6912 | 6762 | 7333.29 | 7736 | 6930 | 0.896 | -0.159 | 0.09369094 | 0.21075301 | 0.0135 | 0.0027 | 0.0063 | 0.0063 | TRUE | 0.1899 |
| Mlph | 84 | 65 | 71 | 45 | 61 | 80 | 65 | 69 | 77 | 45 | 69 | 79 | 67.26 | 70 | 64 | 0.914 | -0.13 | 0.62198705 | 0.76142749 | 0.0272 | 0.0377 | 0.0348 | 0.0348 | TRUE | 0.6302 |
| Mlst8 | 1370 | 1005 | 1047 | 1063 | 960 | 1100 | 1064 | 1069 | 1130 | 1058 | 1083 | 1086 | 1081.61 | 1088 | 1076 | 0.99 | -0.015 | 0.83454299 | 0.90597926 | 0 | 0.0046 | 0.0029 | 0.0029 | TRUE | 0.0134 |
| Mlx | 3136 | 2047 | 1945 | 2207 | 1974 | 2307 | 2436 | 2177 | 2099 | 2197 | 2227 | 2277 | 2235.45 | 2237 | 2234 | 0.997 | -0.004 | 0.95782958 | 0.97691359 | 0.0029 | 0.0035 | 0.0032 | 0.0032 | TRUE | 0.0747 |
| Mlxip | 4260 | 3031 | 3312 | 3585 | 3336 | 3999 | 3309 | 3224 | 3574 | 3569 | 3763 | 3948 | 3564.28 | 3369 | 3760 | 1.116 | 0.159 | 0.01565618 | 0.05765882 | 0.0023 | 0.0031 | 0.0028 | 0.0028 | TRUE | 0.0339 |
| Mlxipl | 298 | 216 | 233 | 288 | 285 | 346 | 231 | 230 | 251 | 287 | 322 | 342 | 277.06 | 237 | 317 | 1.334 | 0.416 | 0.00123122 | 0.00840893 | 0 | 0.011 | 0.0083 | 0.0083 | TRUE | 0.0791 |
| Mlycd | 631 | 437 | 491 | 657 | 625 | 631 | 490 | 465 | 530 | 654 | 705 | 623 | 577.77 | 495 | 661 | 1.334 | 0.416 | 2.13E-05 | 0.00035113 | 0.0021 | 0.0065 | 0.0051 | 0.0051 | TRUE | 0.042 |
| Mmaa | 264 | 162 | 155 | 160 | 138 | 139 | 205 | 172 | 167 | 159 | 156 | 137 | 166.13 | 181 | 151 | 0.827 | -0.275 | 0.09194905 | 0.20804619 | 0.0047 | 0.0167 | 0.0131 | 0.0131 | TRUE | 0.1494 |
| Mmab | 916 | 817 | 842 | 1035 | 1005 | 1012 | 711 | 869 | 909 | 1030 | 1134 | 999 | 941.99 | 830 | 1054 | 1.273 | 0.348 | 0.0006331 | 0.00517378 | 0.0094 | 0.0049 | 0.0064 | 0.0064 | TRUE | 0.1928 |
| Mmachc | 1537 | 1128 | 1108 | 1363 | 1164 | 1327 | 1194 | 1200 | 1196 | 1357 | 1313 | 1310 | 1261.48 | 1197 | 1327 | 1.109 | 0.149 | 0.02759784 | 0.08708615 | 0 | 0.0043 | 0.0025 | 0.0025 | TRUE | 0.0048 |
| Mmadhc | 3519 | 2410 | 2461 | 2642 | 2241 | 2577 | 2733 | 2563 | 2655 | 2630 | 2528 | 2544 | 2608.96 | 2650 | 2567 | 0.968 | -0.047 | 0.43019706 | 0.60241075 | 4.00E-04 | 0.0033 | 0.0021 | 0.0021 | TRUE | 0.01 |
| Mmd | 1439 | 1021 | 1153 | 1069 | 939 | 1050 | 1118 | 1086 | 1244 | 1064 | 1059 | 1037 | 1101.27 | 1149 | 1053 | 0.917 | -0.125 | 0.11415157 | 0.24326351 | 0.0018 | 0.0046 | 0.0036 | 0.0036 | TRUE | 0.0608 |
| Mmd2 | 19 | 16 | 11 | 40 | 27 | 22 | 15 | 17 | 12 | 40 | 30 | 22 | 22.61 | 15 | 31 | 2.105 | 1.074 | 0.01289823 | 0.04991653 | 0 | 0.1073 | 0.0845 | 0.0845 | TRUE | 0.1166 |
| Mme | 3557 | 2845 | 2511 | 1648 | 1339 | 1432 | 2763 | 3026 | 2709 | 1640 | 1511 | 1414 | 2177.11 | 2833 | 1522 | 0.537 | -0.896 | 7.07E-32 | 6.74E-29 | 0.0038 | 0.0035 | 0.0037 | 0.0037 | TRUE | 0.057 |
| Mmel1 | 79 | 81 | 85 | 100 | 84 | 102 | 61 | 86 | 92 | 100 | 95 | 101 | 89.04 | 80 | 99 | 1.244 | 0.315 | 0.15641892 | 0.30459068 | 0.0118 | 0.029 | 0.0242 | 0.0242 | TRUE | 0.4353 |

| | | | | | | | | | | | | | | | | | | | | | | | | | |
|---|---|---|---|---|---|---|---|---|---|---|---|---|---|---|---|---|---|---|---|---|---|---|---|---|---|
| Mmgt1 | 1701 | 1285 | 1425 | 1658 | 1333 | 1636 | 1321 | 1367 | 1538 | 1650 | 1504 | 1615 | 1499.1 | 1409 | 1590 | 1.13 | 0.177 | 0.02647424 | 0.08470927 | 0.0034 | 0.004 | 0.0039 | 0.0039 | TRUE | 0.0765 |
| Mmgt2 | 401 | 302 | 333 | 360 | 329 | 355 | 311 | 321 | 359 | 358 | 371 | 350 | 345.32 | 330 | 360 | 1.09 | 0.125 | 0.27689756 | 0.4480676 | 0 | 0.0093 | 0.0066 | 0.0066 | TRUE | 0.0637 |
| Mmp10 | 0 | 0 | 0 | 0 | 0 | 0 | 0 | 0 | 0 | 0 | 0 | 0 | 0 | 0 | 0 | NA | NA | NA | NA | NA | NA | NA | NA | NA | NA |
| Mmp11 | 2144 | 1756 | 1809 | 2215 | 1910 | 2287 | 1665 | 1868 | 1952 | 2205 | 2155 | 2258 | 2017 | 1828 | 2206 | 1.208 | 0.273 | 0.00021876 | 0.00228562 | 0.0028 | 0.0036 | 0.0034 | 0.0034 | TRUE | 0.0761 |
| Mmp12 | 0 | 0 | 0 | 0 | 0 | 0 | 0 | 0 | 0 | 0 | 0 | 0 | 0 | 0 | 0 | NA | NA | NA | NA | NA | NA | NA | NA | NA | NA |
| Mmp13 | 0 | 0 | 0 | 0 | 0 | 0 | 0 | 0 | 0 | 0 | 0 | 0 | 0 | 0 | 0 | NA | NA | NA | NA | NA | NA | NA | NA | NA | NA |
| Mmp14 | 671 | 503 | 417 | 608 | 521 | 621 | 521 | 535 | 450 | 605 | 588 | 613 | 552.01 | 502 | 602 | 1.198 | 0.261 | 0.00886362 | 0.03794182 | 0.0021 | 0.0067 | 0.0053 | 0.0053 | TRUE | 0.0921 |
| Mmp15 | 1430 | 1056 | 1090 | 1423 | 1317 | 1434 | 1111 | 1123 | 1176 | 1416 | 1486 | 1416 | 1287.95 | 1137 | 1439 | 1.267 | 0.341 | 1.06E-06 | 3.08E-05 | 0 | 0.0043 | 0.0027 | 0.0027 | TRUE | 0.0108 |
| Mmp16 | 9 | 5 | 4 | 4 | 7 | 4 | 7 | 5 | 4 | 4 | 8 | 4 | 5.41 | 5 | 5 | 0.934 | -0.098 | 0.90470208 | NA | 0 | 0.4406 | 0.2982 | 0.2982 | TRUE | 0.3556 |
| Mmp17 | 1642 | 1393 | 1526 | 1949 | 1715 | 1841 | 1275 | 1482 | 1647 | 1940 | 1935 | 1817 | 1682.6 | 1468 | 1897 | 1.294 | 0.372 | 4.06E-05 | 0.0005945 | 0.0078 | 0.0038 | 0.0053 | 0.0053 | TRUE | 0.1633 |
| Mmp19 | 45 | 36 | 29 | 27 | 22 | 21 | 35 | 38 | 31 | 27 | 25 | 21 | 29.49 | 35 | 24 | 0.692 | -0.531 | 0.13446766 | 0.27321014 | 0 | 0.0828 | 0.0553 | 0.0553 | TRUE | 0.0956 |
| Mmp1a | 0 | 0 | 0 | 0 | 0 | 0 | 0 | 0 | 0 | 0 | 0 | 0 | 0 | 0 | 0 | NA | NA | NA | NA | NA | NA | NA | NA | NA | NA |
| Mmp1b | 1 | 1 | 0 | 0 | 1 | 1 | 1 | 1 | 0 | 0 | 1 | 1 | 0.66 | 1 | 1 | 1.125 | 0.17 | 0.94233031 | NA | 0 | 3.5974 | 2.4494 | 2.4494 | TRUE | 0.2637 |
| Mmp2 | 2654 | 1955 | 2465 | 3004 | 2651 | 2810 | 2061 | 2080 | 2660 | 2990 | 2991 | 2774 | 2592.53 | 2267 | 2918 | 1.288 | 0.365 | 9.86E-05 | 0.00119782 | 0.0116 | 0.0033 | 0.0059 | 0.0059 | TRUE | 0.2767 |
| Mmp20 | 11 | 2 | 3 | 3 | 0 | 1 | 9 | 2 | 3 | 3 | 0 | 1 | 2.98 | 5 | 1 | 0.286 | -1.805 | 0.15801655 | NA | 0.3876 | 0.7977 | 0.6933 | 0.6933 | TRUE | 1.3137 |
| Mmp21 | 4 | 5 | 4 | 3 | 3 | 2 | 3 | 5 | 4 | 3 | 3 | 2 | 3.51 | 4 | 3 | 0.658 | -0.603 | 0.54880118 | NA | 0 | 0.6768 | 0.4276 | 0.4276 | TRUE | 0.1319 |
| Mmp23 | 38 | 41 | 35 | 34 | 33 | 31 | 30 | 44 | 38 | 34 | 37 | 31 | 35.43 | 37 | 34 | 0.922 | -0.117 | 0.72227561 | 0.83375105 | 0 | 0.0693 | 0.0499 | 0.0499 | TRUE | 0.2714 |
| Mmp24 | 91 | 82 | 42 | 45 | 44 | 50 | 71 | 87 | 45 | 45 | 50 | 49 | 57.83 | 68 | 48 | 0.706 | -0.502 | 0.08193296 | 0.19242231 | 0.0387 | 0.0434 | 0.0421 | 0.0421 | TRUE | 0.2359 |
| Mmp25 | 387 | 271 | 275 | 426 | 422 | 497 | 301 | 288 | 297 | 424 | 476 | 491 | 379.38 | 295 | 464 | 1.569 | 0.65 | 6.58E-09 | 3.51E-07 | 0 | 0.0087 | 0.0063 | 0.0063 | TRUE | 0.0657 |
| Mmp27 | 0 | 0 | 0 | 0 | 0 | 0 | 0 | 0 | 0 | 0 | 0 | 0 | 0 | 0 | 0 | NA | NA | NA | NA | NA | NA | NA | NA | NA | NA |
| Mmp28 | 221 | 137 | 129 | 148 | 121 | 151 | 172 | 146 | 139 | 147 | 137 | 149 | 148.24 | 152 | 144 | 0.945 | -0.082 | 0.62574477 | 0.76415746 | 0 | 0.0184 | 0.0135 | 0.0135 | TRUE | 0.1437 |
| Mmp3 | 1 | 0 | 0 | 0 | 0 | 0 | 1 | 0 | 0 | 0 | 0 | 0 | 0.13 | 0 | 0 | 0.552 | -0.857 | 0.8336548 | NA | 0 | 18.3121 | 10 | 10 | TRUE | 0.2566 |
| Mmp7 | 0 | 0 | 1 | 0 | 0 | 0 | 0 | 0 | 1 | 0 | 0 | 0 | 0.18 | 0 | 0 | 0.552 | -0.857 | 0.8336548 | NA | 0 | 13.1817 | 10 | 10 | TRUE | 0.5365 |
| Mmp8 | 0 | 0 | 0 | 0 | 0 | 0 | 0 | 0 | 0 | 0 | 0 | 0 | 0 | 0 | 0 | NA | NA | NA | NA | NA | NA | NA | NA | NA | NA |
| Mmp9 | 170 | 115 | 118 | 135 | 121 | 141 | 132 | 122 | 127 | 134 | 137 | 139 | 131.96 | 127 | 137 | 1.073 | 0.102 | 0.54741544 | 0.70424923 | 0 | 0.0204 | 0.0129 | 0.0129 | TRUE | 0.0125 |
| Mmrn1 | 1 | 0 | 0 | 0 | 0 | 0 | 1 | 0 | 0 | 0 | 0 | 0 | 0.13 | 0 | 0 | 0.552 | -0.857 | 0.8336548 | NA | 0 | 18.3121 | 10 | 10 | TRUE | 0.2566 |
| Mmrn2 | 1962 | 1445 | 1461 | 1601 | 1453 | 1604 | 1524 | 1537 | 1576 | 1594 | 1639 | 1583 | 1575.57 | 1546 | 1605 | 1.039 | 0.055 | 0.39227174 | 0.56596869 | 0 | 0.0039 | 0.0023 | 0.0023 | TRUE | 0.004 |
| Mms19 | 8307 | 6616 | 7251 | 8171 | 7511 | 8871 | 6452 | 7037 | 7824 | 8133 | 8473 | 8757 | 7779.41 | 7104 | 8454 | 1.19 | 0.252 | 0.00053594 | 0.0045386 | 0.0052 | 0.0027 | 0.0037 | 0.0037 | TRUE | 0.0958 |
| Mms22l | 6052 | 4488 | 4345 | 4375 | 3869 | 4574 | 4700 | 4774 | 4688 | 4355 | 4365 | 4515 | 4566.19 | 4721 | 4412 | 0.935 | -0.098 | 0.05588246 | 0.14596637 | 0 | 0.0029 | 0.0017 | 0.0017 | TRUE | 0.0052 |
| Mn1 | 151 | 129 | 131 | 116 | 118 | 127 | 117 | 137 | 141 | 115 | 133 | 125 | 128.3 | 132 | 124 | 0.947 | -0.078 | 0.66188166 | 0.79124932 | 0 | 0.0209 | 0.0151 | 0.0151 | TRUE | 0.1087 |
| Mnat1 | 2953 | 2306 | 2290 | 2674 | 2435 | 2498 | 2293 | 2453 | 2471 | 2662 | 2747 | 2466 | 2515.29 | 2406 | 2625 | 1.091 | 0.126 | 0.05943815 | 0.15230361 | 0.0017 | 0.0034 | 0.0028 | 0.0028 | TRUE | 0.0342 |
| Mnd1 | 1185 | 854 | 984 | 1064 | 1007 | 1192 | 920 | 908 | 1062 | 1059 | 1136 | 1177 | 1043.71 | 963 | 1124 | 1.168 | 0.223 | 0.01042251 | 0.04253902 | 0.0039 | 0.0047 | 0.0045 | 0.0045 | TRUE | 0.0929 |
| Mnda | 0 | 0 | 0 | 0 | 0 | 0 | 0 | 0 | 0 | 0 | 0 | 0 | 0 | 0 | 0 | NA | NA | NA | NA | NA | NA | NA | NA | NA | NA |
| Mndal | 0 | 0 | 0 | 0 | 0 | 0 | 0 | 0 | 0 | 0 | 0 | 0 | 0 | 0 | 0 | NA | NA | NA | NA | NA | NA | NA | NA | NA | NA |
| Mnf1 | 2274 | 1733 | 1893 | 2185 | 1851 | 2169 | 1766 | 1843 | 2043 | 2175 | 2088 | 2141 | 2009.37 | 1884 | 2135 | 1.134 | 0.182 | 0.01116093 | 0.04474703 | 0.0023 | 0.0036 | 0.0032 | 0.0032 | TRUE | 0.065 |
| Mns1 | 1054 | 865 | 695 | 668 | 594 | 702 | 819 | 920 | 750 | 665 | 670 | 693 | 752.76 | 830 | 676 | 0.815 | -0.295 | 0.0018942 | 0.01158982 | 0.0043 | 0.0056 | 0.0051 | 0.0051 | TRUE | 0.1048 |
| Mnt | 2897 | 2431 | 2374 | 2468 | 2323 | 2769 | 2250 | 2586 | 2562 | 2457 | 2621 | 2733 | 2534.66 | 2466 | 2604 | 1.057 | 0.08 | 0.28630981 | 0.45835867 | 0.0037 | 0.0034 | 0.0036 | 0.0036 | TRUE | 0.0728 |
| Mnx1 | 72 | 66 | 54 | 57 | 57 | 86 | 56 | 70 | 58 | 57 | 64 | 85 | 65.05 | 61 | 69 | 1.121 | 0.165 | 0.52026825 | 0.68175525 | 0.0154 | 0.0389 | 0.0322 | 0.0322 | TRUE | 0.3941 |
| Moap1 | 10 | 8 | 10 | 10 | 3 | 8 | 8 | 9 | 11 | 10 | 3 | 8 | 8.05 | 9 | 7 | 0.796 | -0.33 | 0.6318472 | NA | 0 | 0.2968 | 0.2152 | 0.2152 | TRUE | 0.4889 |
| Mob1a | 3188 | 2157 | 2046 | 2126 | 1863 | 2062 | 2476 | 2294 | 2208 | 2116 | 2102 | 2036 | 2205.23 | 2326 | 2085 | 0.895 | -0.16 | 0.01676264 | 0.06076299 | 0.0016 | 0.0035 | 0.0027 | 0.0027 | TRUE | 0.0396 |
| Mob1b | 903 | 600 | 448 | 527 | 415 | 421 | 701 | 638 | 483 | 525 | 468 | 416 | 538.54 | 607 | 470 | 0.771 | -0.375 | 0.00651669 | 0.03004687 | 0.0231 | 0.0068 | 0.0118 | 0.0118 | TRUE | 0.3712 |
| Mob2 | 1073 | 747 | 807 | 855 | 723 | 847 | 833 | 795 | 871 | 851 | 816 | 836 | 833.58 | 833 | 834 | 1.002 | 0.003 | 0.97226787 | 0.98494753 | 0 | 0.0053 | 0.0035 | 0.0035 | TRUE | 0.0183 |
| Mob3a | 1115 | 780 | 803 | 942 | 850 | 1054 | 866 | 830 | 866 | 938 | 959 | 1040 | 916.52 | 854 | 979 | 1.146 | 0.197 | 0.01419569 | 0.05356361 | 8.00E-04 | 0.005 | 0.0036 | 0.0036 | TRUE | 0.0365 |
| Mob3b | 982 | 688 | 669 | 700 | 567 | 708 | 763 | 732 | 722 | 697 | 640 | 699 | 708.61 | 739 | 679 | 0.918 | -0.123 | 0.15259523 | 0.29927984 | 0 | 0.0058 | 0.0039 | 0.0039 | TRUE | 0.0287 |
| Mob3c | 307 | 267 | 289 | 293 | 283 | 301 | 238 | 284 | 312 | 292 | 319 | 297 | 290.38 | 278 | 303 | 1.092 | 0.127 | 0.34374689 | 0.51799413 | 0.0072 | 0.0106 | 0.0095 | 0.0095 | TRUE | 0.1894 |
| Mob4 | 2798 | 1957 | 2019 | 2280 | 1932 | 2197 | 2173 | 2082 | 2178 | 2270 | 2179 | 2169 | 2175.17 | 2144 | 2206 | 1.029 | 0.041 | 0.50306919 | 0.66713379 | 0 | 0.0035 | 0.0022 | 0.0022 | TRUE | 0.0078 |
| Mobp | 334 | 233 | 275 | 125 | 111 | 162 | 259 | 248 | 297 | 124 | 125 | 160 | 202.25 | 268 | 136 | 0.511 | -0.97 | 9.95E-10 | 6.65E-08 | 0.0083 | 0.0141 | 0.0126 | 0.0126 | TRUE | 0.2369 |
| Mocos | 167 | 108 | 159 | 205 | 171 | 188 | 130 | 115 | 172 | 204 | 193 | 186 | 166.45 | 139 | 194 | 1.403 | 0.489 | 0.00528223 | 0.02554953 | 0.0143 | 0.0167 | 0.016 | 0.016 | TRUE | 0.4211 |
| Mocs1 | 541 | 444 | 466 | 531 | 518 | 574 | 420 | 472 | 503 | 529 | 584 | 567 | 512.47 | 465 | 560 | 1.206 | 0.271 | 0.00972205 | 0.04054778 | 0.0036 | 0.0071 | 0.0059 | 0.0059 | TRUE | 0.0889 |
| Mocs2 | 1334 | 912 | 822 | 907 | 785 | 909 | 1036 | 970 | 887 | 903 | 886 | 897 | 929.8 | 964 | 895 | 0.927 | -0.109 | 0.19224328 | 0.35052088 | 0.0022 | 0.005 | 0.004 | 0.004 | TRUE | 0.0583 |
| Mocs3 | 567 | 433 | 271 | 366 | 320 | 344 | 440 | 461 | 292 | 364 | 361 | 340 | 376.37 | 398 | 355 | 0.891 | -0.167 | 0.27550956 | 0.44657044 | 0.0274 | 0.0087 | 0.0142 | 0.0142 | TRUE | 0.6024 |
| Mog | 6 | 7 | 3 | 3 | 7 | 7 | 5 | 7 | 3 | 3 | 8 | 7 | 5.52 | 5 | 6 | 1.157 | 0.211 | 0.79942314 | NA | 0 | 0.4316 | 0.3136 | 0.3136 | TRUE | 0.4512 |
| Mogat1 | 0 | 0 | 0 | 1 | 2 | 0 | 0 | 0 | 0 | 1 | 2 | 0 | 0.54 | 0 | 1 | 6.07 | 2.602 | 0.36554818 | NA | 0 | 4.3758 | 4.4761 | 4.4761 | TRUE | 0.0722 |
| Mogat2 | 134 | 62 | 88 | 129 | 112 | 115 | 104 | 66 | 95 | 128 | 126 | 114 | 105.54 | 88 | 123 | 1.381 | 0.466 | 0.02803961 | 0.0881271 | 0.0169 | 0.0249 | 0.0228 | 0.0228 | TRUE | 0.4495 |
| Mogs | 1996 | 1547 | 1511 | 1647 | 1647 | 1766 | 1550 | 1646 | 1630 | 1639 | 1858 | 1743 | 1677.8 | 1609 | 1747 | 1.086 | 0.119 | 0.09672648 | 0.21611827 | 0.0019 | 0.0038 | 0.0031 | 0.0031 | TRUE | 0.0368 |
| Mon1a | 382 | 229 | 230 | 223 | 210 | 240 | 297 | 244 | 248 | 222 | 237 | 237 | 247.37 | 263 | 232 | 0.879 | -0.186 | 0.17470953 | 0.32804842 | 0.004 | 0.012 | 0.0095 | 0.0095 | TRUE | 0.1524 |
| Mon1b | 2003 | 1443 | 1405 | 1525 | 1439 | 1657 | 1556 | 1535 | 1516 | 1518 | 1623 | 1636 | 1563.93 | 1536 | 1592 | 1.037 | 0.052 | 0.43732087 | 0.60879862 | 0 | 0.0039 | 0.0026 | 0.0026 | TRUE | 0.0203 |
| Mon2 | 6601 | 4713 | 4751 | 5487 | 4784 | 5605 | 5127 | 5013 | 5126 | 5462 | 5397 | 5533 | 5276.29 | 5089 | 5464 | 1.074 | 0.103 | 0.03747976 | 0.10925927 | 0 | 0.0029 | 0.0016 | 0.0016 | TRUE | 0.002 |
| Morc1 | 7765 | 6172 | 6599 | 6384 | 6111 | 7031 | 6031 | 6565 | 7120 | 6355 | 6894 | 6941 | 6650.87 | 6572 | 6730 | 1.024 | 0.035 | 0.62359176 | 0.76270726 | 0.0045 | 0.0028 | 0.0034 | 0.0034 | TRUE | 0.065 |
| Morc2a | 7268 | 5033 | 4146 | 4565 | 4028 | 4656 | 5645 | 5354 | 4473 | 4544 | 4544 | 4596 | 4859.33 | 5157 | 4561 | 0.884 | -0.178 | 0.02671229 | 0.08529911 | 0.0067 | 0.0029 | 0.0044 | 0.0044 | TRUE | 0.1634 |
| Morc2b | 34 | 33 | 22 | 22 | 23 | 15 | 26 | 35 | 24 | 22 | 26 | 15 | 24.65 | 28 | 21 | 0.733 | -0.448 | 0.26841536 | 0.4390774 | 0 | 0.0986 | 0.076 | 0.076 | TRUE | 0.3689 |
| Morc3 | 11185 | 8430 | 8372 | 8340 | 7440 | 8487 | 8687 | 8967 | 9033 | 8302 | 8393 | 8378 | 8626.64 | 8896 | 8358 | 0.94 | -0.09 | 0.05709845 | 0.14846428 | 1.00E-04 | 0.0027 | 0.0015 | 0.0015 | TRUE | 0.0052 |
| Morc4 | 1006 | 682 | 733 | 827 | 667 | 788 | 781 | 725 | 791 | 823 | 752 | 778 | 775.2 | 766 | 784 | 1.024 | 0.035 | 0.68350765 | 0.80598552 | 7.00E-04 | 0.0055 | 0.0039 | 0.0039 | TRUE | 0.0245 |
| Morf4l1 | 17374 | 12139 | 12582 | 12885 | 11185 | 12745 | 13494 | 12912 | 13576 | 12826 | 12618 | 12581 | 13001.08 | 13327 | 12675 | 0.951 | -0.072 | 0.12599191 | 0.2611751 | 3.00E-04 | 0.0026 | 0.0015 | 0.0015 | TRUE | 0.009 |
| Morf4l2 | 23146 | 16002 | 15452 | 17562 | 14678 | 16663 | 17977 | 17021 | 16673 | 17481 | 16558 | 16449 | 17026.44 | 17224 | 16829 | 0.977 | -0.034 | 0.52889495 | 0.68931326 | 0.0012 | 0.0026 | 0.002 | 0.002 | TRUE | 0.018 |
| Morn1 | 94 | 98 | 132 | 145 | 125 | 150 | 73 | 104 | 142 | 144 | 141 | 148 | 125.52 | 106 | 144 | 1.367 | 0.451 | 0.04429203 | 0.1232605 | 0.0456 | 0.0213 | 0.0281 | 0.0281 | TRUE | 0.3338 |
| Morn2 | 261 | 176 | 166 | 197 | 183 | 209 | 203 | 187 | 179 | 196 | 206 | 206 | 196.31 | 190 | 203 | 1.067 | 0.094 | 0.51258337 | 0.67504942 | 0 | 0.0145 | 0.0096 | 0.0096 | TRUE | 0.0428 |
| Morn3 | 6 | 5 | 7 | 5 | 6 | 3 | 5 | 5 | 8 | 5 | 7 | 3 | 5.37 | 6 | 5 | 0.84 | -0.252 | 0.75931183 | NA | 0 | 0.4435 | 0.2984 | 0.2984 | TRUE | 0.2456 |
| Morn4 | 1011 | 671 | 820 | 818 | 749 | 847 | 785 | 714 | 885 | 814 | 845 | 836 | 813.17 | 795 | 832 | 1.047 | 0.066 | 0.47856592 | 0.64580449 | 0.0045 | 0.0053 | 0.0051 | 0.0051 | TRUE | 0.1127 |

| | | | | | | | | | | | | | | | | | | | | | | | | | |
|---|---|---|---|---|---|---|---|---|---|---|---|---|---|---|---|---|---|---|---|---|---|---|---|---|---|
| Morn5 | 3 | 6 | 16 | 21 | 18 | 15 | 2 | 6 | 17 | 21 | 20 | 15 | 13.67 | 8 | 19 | 2.218 | 1.149 | 0.07087759 | NA | 0.3037 | 0.1759 | 0.2066 | 0.2066 | TRUE | 1.634 |
| Mos | 8 | 8 | 8 | 9 | 9 | 8 | 6 | 9 | 9 | 9 | 10 | 8 | 8.39 | 8 | 9 | 1.164 | 0.219 | 0.73532668 | NA | 0 | 0.2848 | 0.1827 | 0.1827 | TRUE | 0.1201 |
| Mospd1 | 573 | 369 | 364 | 447 | 414 | 435 | 445 | 393 | 393 | 445 | 467 | 429 | 428.61 | 410 | 447 | 1.087 | 0.12 | 0.26069846 | 0.43077006 | 0.0017 | 0.008 | 0.0059 | 0.0059 | TRUE | 0.0685 |
| Mospd2 | 1035 | 766 | 902 | 1023 | 870 | 1029 | 804 | 815 | 973 | 1018 | 981 | 1016 | 934.57 | 864 | 1005 | 1.165 | 0.221 | 0.01544787 | 0.0571021 | 0.0046 | 0.005 | 0.0049 | 0.0049 | TRUE | 0.1431 |
| Mospd3 | 1581 | 1086 | 1021 | 929 | 761 | 937 | 1228 | 1155 | 1102 | 925 | 858 | 925 | 1032.15 | 1162 | 903 | 0.777 | -0.365 | 4.82E-06 | 0.00010277 | 0.0015 | 0.0047 | 0.0036 | 0.0036 | TRUE | 0.0311 |
| Mospd4 | 1 | 1 | 4 | 4 | 3 | 5 | 1 | 1 | 4 | 4 | 3 | 5 | 3.08 | 2 | 4 | 2.07 | 1.05 | 0.35501848 | NA | 0 | 0.7728 | 0.5601 | 0.5601 | TRUE | 0.7963 |
| Mov10 | 7840 | 5956 | 7365 | 6979 | 6613 | 7680 | 6089 | 6335 | 7947 | 6947 | 7460 | 7581 | 7059.91 | 6790 | 7329 | 1.08 | 0.111 | 0.21956907 | 0.38267293 | 0.0107 | 0.0028 | 0.0057 | 0.0057 | TRUE | 0.2704 |
| Mov10l1 | 1668 | 1333 | 1400 | 1914 | 1653 | 1713 | 1295 | 1418 | 1511 | 1905 | 1865 | 1691 | 1614.15 | 1408 | 1820 | 1.294 | 0.372 | 4.04E-06 | 8.87E-05 | 0.0039 | 0.0039 | 0.0041 | 0.0041 | TRUE | 0.0609 |
| Moxd1 | 389 | 295 | 320 | 344 | 324 | 341 | 302 | 314 | 345 | 342 | 366 | 337 | 334.29 | 320 | 348 | 1.088 | 0.122 | 0.29497477 | 0.4669447 | 0 | 0.0095 | 0.0067 | 0.0067 | TRUE | 0.0513 |
| Moxd2 | 0 | 0 | 0 | 0 | 0 | 0 | 0 | 0 | 0 | 0 | 0 | 0 | 0 | 0 | 0 | NA | NA | NA | NA | NA | NA | NA | NA | NA | NA |
| Mpc1 | 576 | 409 | 474 | 619 | 467 | 543 | 447 | 435 | 511 | 616 | 527 | 536 | 512.14 | 464 | 560 | 1.206 | 0.271 | 0.01280514 | 0.04968075 | 0.0055 | 0.0071 | 0.0066 | 0.0066 | TRUE | 0.092 |
| Mpc2 | 582 | 328 | 394 | 389 | 328 | 344 | 452 | 349 | 425 | 387 | 370 | 340 | 387.14 | 409 | 366 | 0.892 | -0.164 | 0.18807812 | 0.34521647 | 0.0088 | 0.0085 | 0.0086 | 0.0086 | TRUE | 0.1856 |
| Mpdu1 | 4347 | 3121 | 2715 | 2688 | 2428 | 2710 | 3376 | 3320 | 2929 | 2676 | 2739 | 2675 | 2952.54 | 3208 | 2697 | 0.84 | -0.252 | 0.00024274 | 0.00247642 | 0.0025 | 0.0032 | 0.003 | 0.003 | TRUE | 0.0697 |
| Mpdz | 1332 | 937 | 1029 | 1011 | 917 | 1022 | 1035 | 997 | 1110 | 1006 | 1034 | 1009 | 1031.86 | 1047 | 1016 | 0.971 | -0.043 | 0.57977879 | 0.73056319 | 6.00E-04 | 0.0047 | 0.0033 | 0.0033 | TRUE | 0.032 |
| Mpeg1 | 5 | 6 | 2 | 3 | 0 | 3 | 4 | 6 | 2 | 3 | 0 | 3 | 3.06 | 4 | 2 | 0.489 | -1.033 | 0.37788221 | NA | 0 | 0.7765 | 0.6162 | 0.6162 | TRUE | 0.3942 |
| Mpg | 2767 | 2206 | 2349 | 2289 | 2141 | 2470 | 2149 | 2346 | 2535 | 2278 | 2415 | 2438 | 2360.34 | 2343 | 2377 | 1.015 | 0.022 | 0.76656827 | 0.86170352 | 0.0037 | 0.0034 | 0.0035 | 0.0035 | TRUE | 0.0654 |
| Mphosph10 | 5919 | 4137 | 3817 | 4127 | 3788 | 4337 | 4597 | 4400 | 4119 | 4108 | 4273 | 4281 | 4296.43 | 4372 | 4221 | 0.965 | -0.051 | 0.39463192 | 0.56811659 | 0.0014 | 0.003 | 0.0024 | 0.0024 | TRUE | 0.0312 |
| Mphosph6 | 3786 | 2750 | 2690 | 2704 | 2314 | 2716 | 2940 | 2925 | 2902 | 2692 | 2610 | 2681 | 2791.86 | 2922 | 2661 | 0.911 | -0.135 | 0.01407761 | 0.05328687 | 0 | 0.0033 | 0.0018 | 0.0018 | TRUE | 0.0033 |
| Mphosph8 | 9933 | 7566 | 7729 | 9343 | 7890 | 9233 | 7715 | 8048 | 8340 | 9300 | 8901 | 9114 | 8569.51 | 8034 | 9105 | 1.134 | 0.181 | 0.00061318 | 0.00505141 | 8.00E-04 | 0.0027 | 0.0019 | 0.0019 | TRUE | 0.015 |
| Mphosph9 | 2107 | 1544 | 1280 | 1461 | 1305 | 1492 | 1636 | 1642 | 1381 | 1454 | 1472 | 1473 | 1509.86 | 1553 | 1466 | 0.943 | -0.084 | 0.29772393 | 0.46979802 | 0.0041 | 0.004 | 0.004 | 0.004 | TRUE | 0.1113 |
| Mpi | 2085 | 1505 | 1580 | 1707 | 1526 | 1760 | 1619 | 1601 | 1705 | 1699 | 1721 | 1737 | 1680.5 | 1642 | 1719 | 1.048 | 0.067 | 0.29602606 | 0.46793856 | 0 | 0.0038 | 0.0024 | 0.0024 | TRUE | 0.0133 |
| Mpl | 1 | 4 | 1 | 0 | 1 | 1 | 1 | 4 | 1 | 0 | 1 | 1 | 1.37 | 2 | 1 | 0.348 | -1.525 | 0.38894729 | NA | 0 | 1.7312 | 1.2727 | 1.2727 | TRUE | 0.7869 |
| Mplkip | 1101 | 807 | 915 | 1073 | 869 | 1038 | 855 | 858 | 987 | 1068 | 980 | 1025 | 962.3 | 900 | 1024 | 1.139 | 0.188 | 0.02881464 | 0.08990343 | 0.0032 | 0.0049 | 0.0043 | 0.0043 | TRUE | 0.0834 |
| Mpnd | 3480 | 2618 | 2832 | 3374 | 3003 | 3416 | 2703 | 2785 | 3056 | 3358 | 3388 | 3372 | 3110.25 | 2848 | 3373 | 1.185 | 0.245 | 0.00012246 | 0.00143824 | 0.0016 | 0.0032 | 0.0026 | 0.0026 | TRUE | 0.0492 |
| Mpo | 1 | 3 | 3 | 3 | 1 | 1 | 1 | 3 | 3 | 3 | 1 | 1 | 2.05 | 2 | 2 | 0.73 | -0.455 | 0.73590261 | NA | 0 | 1.158 | 0.8012 | 0.8012 | TRUE | 0.52 |
| Mpp1 | 1615 | 1082 | 1115 | 947 | 828 | 929 | 1254 | 1151 | 1203 | 943 | 934 | 917 | 1067.01 | 1203 | 931 | 0.774 | -0.37 | 7.17E-07 | 2.20E-05 | 0 | 0.0046 | 0.0031 | 0.0031 | TRUE | 0.0177 |
| Mpp2 | 1269 | 887 | 979 | 1158 | 1074 | 1199 | 986 | 943 | 1056 | 1153 | 1212 | 1184 | 1088.88 | 995 | 1183 | 1.188 | 0.249 | 0.00114054 | 0.00797935 | 8.00E-04 | 0.0046 | 0.0033 | 0.0033 | TRUE | 0.0331 |
| Mpp3 | 412 | 410 | 420 | 461 | 404 | 521 | 320 | 436 | 453 | 459 | 456 | 514 | 439.7 | 403 | 476 | 1.187 | 0.247 | 0.06748073 | 0.16689774 | 0.0176 | 0.0078 | 0.0108 | 0.0108 | TRUE | 0.3974 |
| Mpp4 | 6 | 5 | 4 | 2 | 2 | 0 | 5 | 5 | 4 | 2 | 2 | 0 | 3.09 | 5 | 1 | 0.293 | -1.773 | 0.13664931 | NA | 0 | 0.7694 | 0.55 | 0.55 | TRUE | 0.5288 |
| Mpp5 | 3710 | 2630 | 2729 | 2894 | 2350 | 2724 | 2881 | 2797 | 2945 | 2881 | 2651 | 2689 | 2807.37 | 2874 | 2740 | 0.953 | -0.069 | 0.26094147 | 0.43087297 | 8.00E-04 | 0.0033 | 0.0023 | 0.0023 | TRUE | 0.0245 |
| Mpp6 | 5700 | 4178 | 4335 | 4515 | 3969 | 4423 | 4427 | 4444 | 4677 | 4494 | 4477 | 4366 | 4481.05 | 4516 | 4446 | 0.985 | -0.022 | 0.67612848 | 0.8011859 | 4.00E-04 | 0.003 | 0.0018 | 0.0018 | TRUE | 0.0118 |
| Mpp7 | 2649 | 1857 | 1681 | 1754 | 1524 | 1728 | 2057 | 1975 | 1814 | 1746 | 1719 | 1706 | 1836.23 | 1949 | 1724 | 0.884 | -0.178 | 0.01030137 | 0.04221497 | 0.0017 | 0.0037 | 0.0029 | 0.0029 | TRUE | 0.044 |
| Mppe1 | 378 | 235 | 221 | 180 | 161 | 161 | 294 | 250 | 238 | 179 | 182 | 159 | 216.95 | 261 | 173 | 0.662 | -0.595 | 5.60E-05 | 0.00076783 | 0.0055 | 0.0134 | 0.0109 | 0.0109 | TRUE | 0.1454 |
| Mpped1 | 116 | 68 | 66 | 93 | 70 | 89 | 90 | 72 | 71 | 93 | 79 | 88 | 82.17 | 78 | 87 | 1.105 | 0.144 | 0.51221018 | 0.67499724 | 0 | 0.0313 | 0.0228 | 0.0228 | TRUE | 0.1961 |
| Mpped2 | 432 | 272 | 255 | 228 | 188 | 219 | 336 | 289 | 275 | 227 | 212 | 216 | 259.2 | 300 | 218 | 0.726 | -0.462 | 0.00058822 | 0.00488522 | 0.0037 | 0.0116 | 0.009 | 0.009 | TRUE | 0.1293 |
| Mprip | 14918 | 10666 | 10053 | 11534 | 10242 | 11660 | 11586 | 11345 | 10847 | 11481 | 11554 | 11510 | 11387.29 | 11259 | 11515 | 1.023 | 0.032 | 0.51087077 | 0.67418335 | 4.00E-04 | 0.0026 | 0.0016 | 0.0016 | TRUE | 0.0125 |
| Mpst | 1363 | 1086 | 1366 | 1332 | 1320 | 1491 | 1059 | 1155 | 1474 | 1326 | 1489 | 1472 | 1329.08 | 1229 | 1429 | 1.164 | 0.219 | 0.04723534 | 0.12926535 | 0.0165 | 0.0042 | 0.008 | 0.008 | TRUE | 0.3621 |
| Mptx1 | 0 | 0 | 0 | 0 | 0 | 0 | 0 | 0 | 0 | 0 | 0 | 0 | 0 | 0 | 0 | NA | NA | NA | NA | NA | NA | NA | NA | NA | NA |
| Mptx2 | 0 | 0 | 0 | 0 | 0 | 0 | 0 | 0 | 0 | 0 | 0 | 0 | 0 | 0 | 0 | NA | NA | NA | NA | NA | NA | NA | NA | NA | NA |
| Mpv17 | 2133 | 1460 | 1569 | 1678 | 1488 | 1720 | 1657 | 1553 | 1693 | 1670 | 1679 | 1698 | 1658.22 | 1634 | 1682 | 1.029 | 0.042 | 0.52990456 | 0.68998141 | 3.00E-04 | 0.0039 | 0.0025 | 0.0025 | TRUE | 0.0222 |
| Mpv17l | 945 | 625 | 666 | 788 | 674 | 741 | 734 | 665 | 719 | 784 | 760 | 731 | 732.26 | 706 | 758 | 1.074 | 0.103 | 0.23223024 | 0.39813684 | 0 | 0.0057 | 0.004 | 0.004 | TRUE | 0.0299 |
| Mpv17l2 | 1600 | 1263 | 1190 | 1502 | 1284 | 1457 | 1243 | 1343 | 1284 | 1495 | 1448 | 1438 | 1375.32 | 1290 | 1460 | 1.133 | 0.18 | 0.00910557 | 0.03865336 | 0 | 0.0041 | 0.0027 | 0.0027 | TRUE | 0.0157 |
| Mpz | 18 | 12 | 12 | 14 | 18 | 16 | 14 | 13 | 13 | 14 | 20 | 16 | 14.95 | 13 | 17 | 1.253 | 0.326 | 0.50667883 | NA | 0 | 0.1609 | 0.1058 | 0.1058 | TRUE | 0.1602 |
| Mpzl1 | 3328 | 2464 | 2560 | 2444 | 2163 | 2477 | 2585 | 2621 | 2762 | 2433 | 2440 | 2445 | 2547.65 | 2656 | 2439 | 0.919 | -0.122 | 0.03766997 | 0.10964193 | 3.00E-04 | 0.0034 | 0.0021 | 0.0021 | TRUE | 0.0147 |
| Mpzl2 | 1647 | 1276 | 1445 | 1514 | 1315 | 1569 | 1279 | 1357 | 1559 | 1507 | 1483 | 1549 | 1455.82 | 1398 | 1513 | 1.083 | 0.116 | 0.16509218 | 0.31592751 | 0.0045 | 0.0041 | 0.0043 | 0.0043 | TRUE | 0.1206 |
| Mpzl3 | 70 | 48 | 57 | 54 | 47 | 63 | 54 | 51 | 62 | 54 | 53 | 62 | 55.98 | 56 | 56 | 1.014 | 0.021 | 0.93665632 | 0.96487414 | 0 | 0.0448 | 0.0302 | 0.0302 | TRUE | 0.0718 |
| Mr1 | 6 | 12 | 5 | 4 | 5 | 5 | 5 | 13 | 5 | 4 | 6 | 5 | 6.23 | 8 | 5 | 0.644 | -0.635 | 0.42513868 | NA | 0 | 0.3829 | 0.2862 | 0.2862 | TRUE | 0.9479 |
| Mrap | 36 | 20 | 14 | 24 | 17 | 23 | 28 | 21 | 15 | 24 | 19 | 23 | 21.69 | 21 | 22 | 1.013 | 0.018 | 0.96559642 | 0.98122801 | 0 | 0.1117 | 0.0835 | 0.0835 | TRUE | 0.2341 |
| Mrap2 | 2 | 0 | 3 | 3 | 4 | 2 | 2 | 0 | 3 | 3 | 5 | 2 | 2.38 | 2 | 3 | 1.973 | 0.98 | 0.45112441 | NA | 0 | 0.9994 | 0.7518 | 0.7518 | TRUE | 0.321 |
| Mras | 1484 | 1092 | 1012 | 1310 | 1227 | 1484 | 1153 | 1162 | 1092 | 1304 | 1384 | 1465 | 1259.86 | 1136 | 1384 | 1.219 | 0.286 | 0.0001526 | 0.00172865 | 0.0015 | 0.0043 | 0.0033 | 0.0033 | TRUE | 0.0317 |
| Mrc1 | 131 | 101 | 64 | 84 | 61 | 71 | 102 | 107 | 69 | 84 | 69 | 70 | 83.46 | 93 | 74 | 0.798 | -0.325 | 0.16901018 | 0.32056364 | 0.0212 | 0.0308 | 0.0281 | 0.0281 | TRUE | 0.4541 |
| Mrc2 | 610 | 354 | 367 | 418 | 323 | 359 | 474 | 377 | 396 | 416 | 364 | 354 | 396.86 | 416 | 378 | 0.908 | -0.139 | 0.26381695 | 0.43407614 | 0.0091 | 0.0084 | 0.0086 | 0.0086 | TRUE | 0.1833 |
| Mre11a | 5409 | 4000 | 4025 | 4186 | 3557 | 4261 | 4201 | 4255 | 4343 | 4167 | 4013 | 4206 | 4197.38 | 4266 | 4129 | 0.968 | -0.047 | 0.37657127 | 0.55112975 | 2.00E-04 | 0.003 | 0.0018 | 0.0018 | TRUE | 0.0073 |
| Mreg | 25099 | 16585 | 12245 | 11088 | 9632 | 10570 | 19493 | 17641 | 13212 | 11037 | 10866 | 10434 | 13780.66 | 16782 | 10779 | 0.642 | -0.639 | 1.56E-09 | 1.00E-07 | 0.0188 | 0.0026 | 0.008 | 0.008 | TRUE | 0.4233 |
| Mrfap1 | 15248 | 11469 | 10221 | 10239 | 9206 | 10428 | 11842 | 12199 | 11028 | 10192 | 10385 | 10294 | 10990.26 | 11690 | 10290 | 0.88 | -0.184 | 0.00072452 | 0.00571221 | 0.0012 | 0.0026 | 0.002 | 0.002 | TRUE | 0.0298 |
| Mrgbp | 725 | 570 | 524 | 614 | 502 | 550 | 563 | 606 | 565 | 611 | 566 | 543 | 575.86 | 578 | 573 | 0.992 | -0.011 | 0.90873719 | 0.94963864 | 9.00E-04 | 0.0065 | 0.0047 | 0.0047 | TRUE | 0.0396 |
| Mrgpra1 | 0 | 0 | 0 | 0 | 0 | 0 | 0 | 0 | 0 | 0 | 0 | 0 | 0 | 0 | 0 | NA | NA | NA | NA | NA | NA | NA | NA | NA | NA |
| Mrgpra2a | 0 | 0 | 0 | 0 | 0 | 0 | 0 | 0 | 0 | 0 | 0 | 0 | 0 | 0 | 0 | NA | NA | NA | NA | NA | NA | NA | NA | NA | NA |
| Mrgpra2b | 0 | 0 | 0 | 0 | 0 | 0 | 0 | 0 | 0 | 0 | 0 | 0 | 0 | 0 | 0 | NA | NA | NA | NA | NA | NA | NA | NA | NA | NA |
| Mrgpra3 | 0 | 0 | 0 | 0 | 0 | 0 | 0 | 0 | 0 | 0 | 0 | 0 | 0 | 0 | 0 | NA | NA | NA | NA | NA | NA | NA | NA | NA | NA |
| Mrgpra4 | 0 | 0 | 0 | 0 | 0 | 0 | 0 | 0 | 0 | 0 | 0 | 0 | 0 | 0 | 0 | NA | NA | NA | NA | NA | NA | NA | NA | NA | NA |
| Mrgpra6 | 0 | 0 | 0 | 0 | 0 | 0 | 0 | 0 | 0 | 0 | 0 | 0 | 0 | 0 | 0 | NA | NA | NA | NA | NA | NA | NA | NA | NA | NA |
| Mrgpra9 | 0 | 0 | 0 | 0 | 0 | 0 | 0 | 0 | 0 | 0 | 0 | 0 | 0 | 0 | 0 | NA | NA | NA | NA | NA | NA | NA | NA | NA | NA |
| Mrgprb1 | 1 | 0 | 0 | 0 | 0 | 0 | 1 | 0 | 0 | 0 | 0 | 0 | 0.13 | 0 | 0 | 0.552 | -0.857 | 0.8336548 | NA | 0 | 18.3121 | 10 | 10 | TRUE | 0.2566 |
| Mrgprb2 | 0 | 0 | 0 | 0 | 0 | 0 | 0 | 0 | 0 | 0 | 0 | 0 | 0 | 0 | 0 | NA | NA | NA | NA | NA | NA | NA | NA | NA | NA |
| Mrgprb3 | 0 | 0 | 0 | 0 | 0 | 0 | 0 | 0 | 0 | 0 | 0 | 0 | 0 | 0 | 0 | NA | NA | NA | NA | NA | NA | NA | NA | NA | NA |
| Mrgprb4 | 0 | 0 | 0 | 0 | 0 | 0 | 0 | 0 | 0 | 0 | 0 | 0 | 0 | 0 | 0 | NA | NA | NA | NA | NA | NA | NA | NA | NA | NA |
| Mrgprb5 | 0 | 0 | 0 | 0 | 0 | 0 | 0 | 0 | 0 | 0 | 0 | 0 | 0 | 0 | 0 | NA | NA | NA | NA | NA | NA | NA | NA | NA | NA |
| Mrgprb8 | 0 | 0 | 0 | 0 | 0 | 0 | 0 | 0 | 0 | 0 | 0 | 0 | 0 | 0 | 0 | NA | NA | NA | NA | NA | NA | NA | NA | NA | NA |

| | | | | | | | | | | | | | | | | | | | | | | | | | |
|---|---|---|---|---|---|---|---|---|---|---|---|---|---|---|---|---|---|---|---|---|---|---|---|---|---|
| Mrgprd | 0 | 0 | 0 | 0 | 0 | 0 | 0 | 0 | 0 | 0 | 0 | 0 | 0 | 0 | 0 | NA | NA | NA | NA | NA | NA | NA | NA | NA | NA |
| Mrgpre | 12 | 11 | 12 | 8 | 14 | 12 | 9 | 12 | 13 | 8 | 16 | 12 | 11.6 | 11 | 12 | 1.05 | 0.07 | 0.90142789 | NA | 0 | 0.2068 | 0.146 | 0.146 | TRUE | 0.1804 |
| Mrgprf | 44 | 16 | 18 | 34 | 25 | 32 | 34 | 17 | 19 | 34 | 28 | 32 | 27.37 | 23 | 31 | 1.306 | 0.385 | 0.32872444 | 0.50323614 | 0.0403 | 0.089 | 0.0754 | 0.0754 | TRUE | 1.0979 |
| Mrgprg | 0 | 0 | 0 | 0 | 0 | 0 | 0 | 0 | 0 | 0 | 0 | 0 | 0 | 0 | 0 | NA | NA | NA | NA | NA | NA | NA | NA | NA | NA |
| Mrgprh | 0 | 0 | 0 | 0 | 0 | 0 | 0 | 0 | 0 | 0 | 0 | 0 | 0 | 0 | 0 | NA | NA | NA | NA | NA | NA | NA | NA | NA | NA |
| Mrgprx1 | 0 | 0 | 0 | 0 | 0 | 0 | 0 | 0 | 0 | 0 | 0 | 0 | 0 | 0 | 0 | NA | NA | NA | NA | NA | NA | NA | NA | NA | NA |
| Mrgprx2 | 0 | 0 | 0 | 0 | 0 | 0 | 0 | 0 | 0 | 0 | 0 | 0 | 0 | 0 | 0 | NA | NA | NA | NA | NA | NA | NA | NA | NA | NA |
| Mri1 | 2116 | 1665 | 1811 | 1742 | 1568 | 1927 | 1643 | 1771 | 1954 | 1734 | 1769 | 1902 | 1795.6 | 1789 | 1802 | 1.008 | 0.011 | 0.88567856 | 0.93652464 | 0.0045 | 0.0037 | 0.004 | 0.004 | TRUE | 0.0779 |
| Mrm1 | 2264 | 1721 | 1623 | 1804 | 1589 | 1863 | 1758 | 1831 | 1751 | 1796 | 1793 | 1839 | 1794.58 | 1780 | 1809 | 1.017 | 0.024 | 0.70221291 | 0.81967939 | 0 | 0.0037 | 0.0022 | 0.0022 | TRUE | 0.0073 |
| Mro | 8 | 5 | 10 | 2 | 11 | 8 | 6 | 5 | 11 | 2 | 12 | 8 | 7.44 | 7 | 7 | 0.996 | -0.006 | 0.99361108 | NA | 0.2234 | 0.3211 | 0.2952 | 0.2952 | TRUE | 0.2725 |
| Mroh1 | 2458 | 1983 | 1991 | 2308 | 2158 | 2328 | 1909 | 2109 | 2148 | 2297 | 2434 | 2298 | 2199.42 | 2055 | 2343 | 1.141 | 0.19 | 0.00590412 | 0.02790178 | 0.0021 | 0.0035 | 0.003 | 0.003 | TRUE | 0.0486 |
| Mroh2a | 0 | 1 | 1 | 1 | 0 | 0 | 0 | 1 | 1 | 1 | 0 | 0 | 0.52 | 1 | 0 | 0.535 | -0.904 | 0.74017109 | NA | 0 | 4.534 | 3.6534 | 3.6534 | TRUE | 0.3906 |
| Mroh2b | 12 | 9 | 7 | 1 | 0 | 1 | 9 | 10 | 8 | 1 | 0 | 1 | 4.74 | 9 | 1 | 0.077 | -3.691 | 0.00367987 | NA | 0 | 0.5026 | 0.353 | 0.353 | TRUE | 0.193 |
| Mroh4 | 13 | 14 | 10 | 4 | 11 | 7 | 10 | 15 | 11 | 4 | 12 | 7 | 9.85 | 12 | 8 | 0.647 | -0.627 | 0.32876136 | NA | 0.0268 | 0.2431 | 0.1893 | 0.1893 | TRUE | 0.5841 |
| Mroh5 | 0 | 0 | 0 | 0 | 1 | 1 | 0 | 0 | 0 | 0 | 1 | 1 | 0.35 | 0 | 1 | 3.965 | 1.987 | 0.55529285 | NA | 0 | 6.7252 | 6.4426 | 6.4426 | TRUE | 0.261 |
| Mroh8 | 25 | 23 | 21 | 39 | 45 | 46 | 19 | 24 | 23 | 39 | 51 | 45 | 33.59 | 22 | 45 | 2.038 | 1.027 | 0.00257852 | 0.01470663 | 0 | 0.073 | 0.0503 | 0.0503 | TRUE | 0.1141 |
| Mroh9 | 0 | 0 | 0 | 0 | 0 | 0 | 0 | 0 | 0 | 0 | 0 | 0 | 0 | 0 | 0 | NA | NA | NA | NA | NA | NA | NA | NA | NA | NA |
| Mrp63 | 1636 | 1219 | 1232 | 1361 | 1196 | 1286 | 1271 | 1297 | 1329 | 1355 | 1349 | 1269 | 1311.67 | 1299 | 1324 | 1.02 | 0.029 | 0.68167563 | 0.80469794 | 0 | 0.0042 | 0.0027 | 0.0027 | TRUE | 0.0161 |
| Mrpl1 | 3825 | 2782 | 2646 | 2932 | 2616 | 2907 | 2971 | 2959 | 2855 | 2919 | 2951 | 2870 | 2920.7 | 2928 | 2913 | 0.994 | -0.008 | 0.88577902 | 0.93656158 | 0 | 0.0032 | 0.0019 | 0.0019 | TRUE | 0.0057 |
| Mrpl10 | 6924 | 4889 | 4749 | 5130 | 4574 | 5312 | 5378 | 5200 | 5124 | 5106 | 5160 | 5244 | 5202.03 | 5234 | 5170 | 0.988 | -0.018 | 0.72307502 | 0.83420061 | 2.00E-04 | 0.0029 | 0.0017 | 0.0017 | TRUE | 0.0072 |
| Mrpl11 | 3143 | 2264 | 2136 | 2322 | 2021 | 2436 | 2441 | 2408 | 2305 | 2311 | 2280 | 2405 | 2358.31 | 2385 | 2332 | 0.978 | -0.033 | 0.59151388 | 0.73948289 | 4.00E-04 | 0.0034 | 0.0022 | 0.0022 | TRUE | 0.0103 |
| Mrpl12 | 6613 | 4499 | 4074 | 4149 | 3817 | 4266 | 5136 | 4786 | 4396 | 4130 | 4306 | 4211 | 4494.08 | 4773 | 4216 | 0.883 | -0.18 | 0.00717132 | 0.03232409 | 0.0029 | 0.003 | 0.003 | 0.003 | TRUE | 0.0579 |
| Mrpl13 | 2955 | 1886 | 1769 | 1978 | 1550 | 1811 | 2295 | 2006 | 1909 | 1969 | 1749 | 1788 | 1952.51 | 2070 | 1835 | 0.886 | -0.175 | 0.03693817 | 0.10825451 | 0.006 | 0.0036 | 0.0046 | 0.0046 | TRUE | 0.1116 |
| Mrpl14 | 1108 | 821 | 918 | 895 | 775 | 961 | 861 | 873 | 991 | 891 | 874 | 949 | 906.36 | 908 | 905 | 0.998 | -0.003 | 0.96811001 | 0.98282523 | 0.0029 | 0.005 | 0.0043 | 0.0043 | TRUE | 0.0737 |
| Mrpl15 | 10382 | 7384 | 7398 | 8154 | 7340 | 8448 | 8063 | 7854 | 7982 | 8116 | 8280 | 8339 | 8106.02 | 7966 | 8245 | 1.035 | 0.049 | 0.29318426 | 0.46521877 | 0 | 0.0027 | 0.0015 | 0.0015 | TRUE | 0.0023 |
| Mrpl16 | 2431 | 1631 | 1500 | 1546 | 1246 | 1468 | 1888 | 1735 | 1618 | 1539 | 1406 | 1449 | 1605.84 | 1747 | 1465 | 0.838 | -0.256 | 0.00106371 | 0.00757153 | 0.0035 | 0.0039 | 0.0038 | 0.0038 | TRUE | 0.0617 |
| Mrpl17 | 5445 | 3773 | 3434 | 3525 | 3166 | 3530 | 4229 | 4013 | 3705 | 3509 | 3572 | 3485 | 3752.08 | 3982 | 3522 | 0.884 | -0.178 | 0.00514797 | 0.02510964 | 0.0019 | 0.0031 | 0.0027 | 0.0027 | TRUE | 0.0449 |
| Mrpl18 | 4526 | 3118 | 3155 | 3219 | 2691 | 3245 | 3515 | 3317 | 3404 | 3204 | 3036 | 3203 | 3279.86 | 3412 | 3148 | 0.922 | -0.117 | 0.04344933 | 0.12157816 | 6.00E-04 | 0.0031 | 0.0021 | 0.0021 | TRUE | 0.0116 |
| Mrpl19 | 3045 | 2107 | 2057 | 2219 | 1903 | 2269 | 2365 | 2241 | 2219 | 2209 | 2147 | 2240 | 2236.84 | 2275 | 2199 | 0.966 | -0.05 | 0.41340837 | 0.5860164 | 5.00E-04 | 0.0035 | 0.0023 | 0.0023 | TRUE | 0.0149 |
| Mrpl2 | 5442 | 3966 | 3991 | 4199 | 3896 | 4480 | 4227 | 4219 | 4306 | 4180 | 4395 | 4422 | 4291.44 | 4251 | 4332 | 1.019 | 0.028 | 0.60588991 | 0.75040841 | 3.00E-04 | 0.003 | 0.0018 | 0.0018 | TRUE | 0.0116 |
| Mrpl20 | 3538 | 2277 | 1899 | 2103 | 1685 | 2059 | 2748 | 2422 | 2049 | 2093 | 1901 | 2033 | 2207.59 | 2406 | 2009 | 0.834 | -0.261 | 0.00655814 | 0.03018856 | 0.0111 | 0.0035 | 0.0062 | 0.0062 | TRUE | 0.1855 |
| Mrpl21 | 2293 | 1623 | 1692 | 2021 | 1689 | 2099 | 1781 | 1726 | 1826 | 2012 | 1905 | 2072 | 1887 | 1778 | 1996 | 1.123 | 0.168 | 0.01028594 | 0.04217353 | 7.00E-04 | 0.0037 | 0.0026 | 0.0026 | TRUE | 0.0189 |
| Mrpl22 | 2935 | 2147 | 2137 | 2322 | 1950 | 2379 | 2279 | 2284 | 2306 | 2311 | 2200 | 2348 | 2288.1 | 2290 | 2286 | 0.999 | -0.001 | 0.98061516 | 0.9899403 | 0 | 0.0035 | 0.0021 | 0.0021 | TRUE | 0.0131 |
| Mrpl23 | 896 | 678 | 765 | 949 | 764 | 919 | 696 | 721 | 825 | 945 | 862 | 907 | 826.03 | 747 | 905 | 1.213 | 0.278 | 0.00238114 | 0.01384045 | 0.0038 | 0.0053 | 0.0048 | 0.0048 | TRUE | 0.097 |
| Mrpl24 | 5914 | 4467 | 4781 | 5394 | 4847 | 5476 | 4593 | 4751 | 5159 | 5369 | 5468 | 5406 | 5124.34 | 4834 | 5414 | 1.12 | 0.164 | 0.00598847 | 0.02819578 | 0.0016 | 0.0029 | 0.0024 | 0.0024 | TRUE | 0.0418 |
| Mrpl27 | 1851 | 1297 | 1389 | 1465 | 1356 | 1426 | 1438 | 1380 | 1499 | 1458 | 1530 | 1408 | 1451.93 | 1439 | 1465 | 1.018 | 0.026 | 0.71448869 | 0.82843992 | 0.001 | 0.0041 | 0.0029 | 0.0029 | TRUE | 0.0175 |
| Mrpl28 | 5676 | 4167 | 4317 | 4692 | 4165 | 4811 | 4408 | 4432 | 4658 | 4670 | 4699 | 4749 | 4602.81 | 4499 | 4706 | 1.046 | 0.065 | 0.21609939 | 0.3788432 | 3.00E-04 | 0.0029 | 0.0018 | 0.0018 | TRUE | 0.0114 |
| Mrpl3 | 3858 | 2542 | 2454 | 2627 | 2319 | 2565 | 2996 | 2704 | 2648 | 2615 | 2616 | 2532 | 2685.19 | 2783 | 2588 | 0.929 | -0.106 | 0.11181373 | 0.23992699 | 0.0019 | 0.0033 | 0.0028 | 0.0028 | TRUE | 0.056 |
| Mrpl30 | 2999 | 2114 | 2312 | 2731 | 2282 | 2667 | 2329 | 2249 | 2495 | 2718 | 2574 | 2633 | 2499.66 | 2358 | 2642 | 1.121 | 0.165 | 0.01030195 | 0.04221497 | 0.0012 | 0.0034 | 0.0026 | 0.0026 | TRUE | 0.0307 |
| Mrpl32 | 1859 | 1349 | 1451 | 1733 | 1446 | 1701 | 1444 | 1435 | 1566 | 1725 | 1631 | 1679 | 1579.96 | 1482 | 1678 | 1.134 | 0.181 | 0.00869226 | 0.03729763 | 9.00E-04 | 0.0039 | 0.0028 | 0.0028 | TRUE | 0.0291 |
| Mrpl33 | 2006 | 1369 | 1233 | 1408 | 1136 | 1253 | 1558 | 1456 | 1330 | 1402 | 1282 | 1237 | 1377.42 | 1448 | 1307 | 0.901 | -0.15 | 0.07279207 | 0.17652139 | 0.0041 | 0.0041 | 0.0043 | 0.0043 | TRUE | 0.0605 |
| Mrpl34 | 2453 | 1689 | 1527 | 1486 | 1214 | 1578 | 1905 | 1797 | 1648 | 1479 | 1370 | 1558 | 1625.96 | 1783 | 1469 | 0.823 | -0.28 | 0.00044844 | 0.00397121 | 0.0038 | 0.0039 | 0.004 | 0.004 | TRUE | 0.053 |
| Mrpl35 | 3369 | 2330 | 2360 | 2810 | 2353 | 2720 | 2617 | 2478 | 2546 | 2797 | 2654 | 2685 | 2629.65 | 2547 | 2712 | 1.065 | 0.09 | 0.12655765 | 0.26189207 | 4.00E-04 | 0.0033 | 0.0021 | 0.0021 | TRUE | 0.0091 |
| Mrpl36 | 1887 | 1267 | 1200 | 1219 | 1013 | 1265 | 1466 | 1348 | 1295 | 1213 | 1143 | 1249 | 1285.49 | 1370 | 1202 | 0.877 | -0.19 | 0.01478608 | 0.05523716 | 0.0024 | 0.0043 | 0.0036 | 0.0036 | TRUE | 0.047 |
| Mrpl37 | 4343 | 2958 | 2669 | 2780 | 2512 | 2827 | 3373 | 3146 | 2880 | 2767 | 2834 | 2791 | 2965.15 | 3133 | 2797 | 0.892 | -0.165 | 0.01728458 | 0.0620725 | 0.0027 | 0.0032 | 0.0031 | 0.0031 | TRUE | 0.0605 |
| Mrpl38 | 5616 | 4204 | 4173 | 4526 | 4183 | 4844 | 4362 | 4472 | 4503 | 4505 | 4719 | 4782 | 4556.98 | 4446 | 4669 | 1.05 | 0.071 | 0.18540922 | 0.3417746 | 4.00E-04 | 0.0029 | 0.0018 | 0.0018 | TRUE | 0.0115 |
| Mrpl39 | 5177 | 3613 | 3568 | 4058 | 3380 | 4080 | 4021 | 3843 | 3850 | 4039 | 3813 | 4028 | 3932.27 | 3905 | 3960 | 1.014 | 0.02 | 0.71922956 | 0.83150918 | 6.00E-04 | 0.003 | 0.002 | 0.002 | TRUE | 0.0127 |
| Mrpl4 | 3855 | 2775 | 2733 | 2949 | 2546 | 2948 | 2994 | 2952 | 2949 | 2935 | 2872 | 2910 | 2935.39 | 2965 | 2906 | 0.98 | -0.029 | 0.59103183 | 0.73920352 | 0 | 0.0032 | 0.0018 | 0.0018 | TRUE | 0.0012 |
| Mrpl40 | 3537 | 2501 | 2565 | 2489 | 2299 | 2551 | 2747 | 2660 | 2768 | 2478 | 2594 | 2518 | 2627.37 | 2725 | 2530 | 0.928 | -0.108 | 0.06155768 | 0.15627777 | 0 | 0.0033 | 0.002 | 0.002 | TRUE | 0.0058 |
| Mrpl41 | 876 | 574 | 586 | 661 | 519 | 654 | 680 | 611 | 632 | 658 | 585 | 646 | 635.37 | 641 | 630 | 0.982 | -0.027 | 0.77430909 | 0.86673091 | 0.0019 | 0.0062 | 0.0048 | 0.0048 | TRUE | 0.0431 |
| Mrpl42 | 6134 | 4274 | 4667 | 4805 | 4257 | 5245 | 4764 | 4546 | 5036 | 4783 | 4802 | 5178 | 4851.45 | 4782 | 4921 | 1.029 | 0.042 | 0.50431608 | 0.66826706 | 0.0019 | 0.0029 | 0.0026 | 0.0026 | TRUE | 0.0259 |
| Mrpl43 | 3101 | 2076 | 1917 | 2061 | 1907 | 2038 | 2408 | 2208 | 2068 | 2052 | 2151 | 2012 | 2149.95 | 2228 | 2072 | 0.929 | -0.107 | 0.14308742 | 0.28595477 | 0.0028 | 0.0035 | 0.0034 | 0.0034 | TRUE | 0.062 |
| Mrpl44 | 2996 | 2168 | 2015 | 2106 | 1816 | 2216 | 2327 | 2306 | 2174 | 2096 | 2049 | 2188 | 2189.93 | 2269 | 2111 | 0.93 | -0.105 | 0.09901188 | 0.2195434 | 8.00E-04 | 0.0035 | 0.0024 | 0.0024 | TRUE | 0.016 |
| Mrpl45 | 10558 | 7846 | 7877 | 8380 | 7251 | 8678 | 8200 | 8346 | 8499 | 8341 | 8180 | 8567 | 8355.45 | 8348 | 8363 | 1.002 | 0.003 | 0.95500116 | 0.9753363 | 3.00E-04 | 0.0027 | 0.0016 | 0.0016 | TRUE | 0.0056 |
| Mrpl46 | 2288 | 1336 | 1276 | 1349 | 1138 | 1351 | 1777 | 1421 | 1377 | 1343 | 1284 | 1334 | 1422.52 | 1525 | 1320 | 0.864 | -0.21 | 0.02960352 | 0.09174394 | 0.0096 | 0.0041 | 0.006 | 0.006 | TRUE | 0.2569 |
| Mrpl47 | 2864 | 2065 | 2195 | 2556 | 2153 | 2664 | 2224 | 2197 | 2368 | 2544 | 2429 | 2630 | 2398.68 | 2263 | 2534 | 1.12 | 0.164 | 0.01038217 | 0.04242206 | 0.001 | 0.0034 | 0.0025 | 0.0025 | TRUE | 0.0197 |
| Mrpl48 | 1725 | 1275 | 1320 | 1524 | 1369 | 1438 | 1340 | 1356 | 1424 | 1517 | 1544 | 1420 | 1433.52 | 1373 | 1494 | 1.088 | 0.121 | 0.08476127 | 0.19647955 | 8.00E-04 | 0.0041 | 0.0029 | 0.0029 | TRUE | 0.0229 |
| Mrpl49 | 3394 | 2509 | 2439 | 2489 | 2234 | 2580 | 2636 | 2669 | 2632 | 2478 | 2520 | 2547 | 2580.17 | 2646 | 2515 | 0.951 | -0.073 | 0.19125083 | 0.34937075 | 0 | 0.0033 | 0.0019 | 0.0019 | TRUE | 0.0021 |
| Mrpl50 | 3052 | 2244 | 2170 | 2286 | 1948 | 2227 | 2370 | 2387 | 2341 | 2275 | 2198 | 2198 | 2295.02 | 2366 | 2224 | 0.94 | -0.089 | 0.12283606 | 0.25626855 | 0 | 0.0035 | 0.002 | 0.002 | TRUE | 0.005 |
| Mrpl51 | 4286 | 3021 | 2797 | 2889 | 2510 | 2981 | 3329 | 3213 | 3018 | 2876 | 2832 | 2943 | 3035.01 | 3187 | 2884 | 0.904 | -0.145 | 0.0176841 | 0.06304787 | 0.0011 | 0.0032 | 0.0024 | 0.0024 | TRUE | 0.0259 |
| Mrpl52 | 1902 | 1367 | 1408 | 1528 | 1314 | 1658 | 1477 | 1454 | 1519 | 1521 | 1482 | 1637 | 1515.08 | 1483 | 1547 | 1.043 | 0.061 | 0.38322704 | 0.55704913 | 9.00E-04 | 0.004 | 0.0028 | 0.0028 | TRUE | 0.0314 |
| Mrpl53 | 1023 | 729 | 789 | 852 | 744 | 867 | 795 | 775 | 851 | 848 | 839 | 856 | 827.42 | 807 | 848 | 1.051 | 0.072 | 0.37386637 | 0.54846343 | 0 | 0.0053 | 0.0035 | 0.0035 | TRUE | 0.0262 |
| Mrpl54 | 1015 | 779 | 806 | 824 | 721 | 841 | 788 | 829 | 870 | 820 | 813 | 830 | 825.06 | 829 | 821 | 0.992 | -0.011 | 0.88961982 | 0.93874758 | 0 | 0.0053 | 0.0035 | 0.0035 | TRUE | 0.0231 |
| Mrpl55 | 1701 | 1077 | 1046 | 1122 | 978 | 1145 | 1321 | 1146 | 1129 | 1117 | 1103 | 1130 | 1157.62 | 1199 | 1117 | 0.93 | -0.104 | 0.20762515 | 0.36924054 | 0.0033 | 0.0045 | 0.0041 | 0.0041 | TRUE | 0.0996 |
| Mrpl9 | 4053 | 2828 | 3008 | 3263 | 3013 | 3340 | 3148 | 3008 | 3246 | 3248 | 3399 | 3297 | 3224.26 | 3134 | 3315 | 1.058 | 0.081 | 0.16574939 | 0.31676786 | 5.00E-04 | 0.0032 | 0.0021 | 0.0021 | TRUE | 0.0146 |
| Mrps10 | 3026 | 2230 | 2279 | 2484 | 2074 | 2448 | 2350 | 2372 | 2459 | 2473 | 2340 | 2417 | 2401.67 | 2394 | 2410 | 1.007 | 0.01 | 0.86301082 | 0.92445579 | 3.00E-04 | 0.0034 | 0.0021 | 0.0021 | TRUE | 0.0077 |
| Mrps11 | 1649 | 1069 | 900 | 1047 | 883 | 954 | 1281 | 1137 | 971 | 1042 | 996 | 942 | 1061.49 | 1130 | 993 | 0.878 | -0.188 | 0.0638327 | 0.16054209 | 0.0093 | 0.0047 | 0.0065 | 0.0065 | TRUE | 0.1793 |
| Mrps12 | 2022 | 1593 | 1453 | 1483 | 1248 | 1621 | 1570 | 1694 | 1568 | 1476 | 1408 | 1600 | 1552.81 | 1611 | 1495 | 0.929 | -0.107 | 0.15373008 | 0.30078905 | 0.0022 | 0.004 | 0.0034 | 0.0034 | TRUE | 0.046 |
| Mrps14 | 3434 | 2253 | 2328 | 2638 | 2240 | 2465 | 2667 | 2396 | 2512 | 2626 | 2527 | 2433 | 2526.93 | 2525 | 2529 | 1.001 | 0.001 | 0.98752986 | 0.99373352 | 0.0018 | 0.0034 | 0.0028 | 0.0028 | TRUE | 0.0301 |

| | | | | | | | | | | | | | | | | | | | | | | | | | |
|---|---|---|---|---|---|---|---|---|---|---|---|---|---|---|---|---|---|---|---|---|---|---|---|---|---|
| Mrps15 | 2357 | 1761 | 1620 | 1781 | 1557 | 1696 | 1831 | 1873 | 1748 | 1773 | 1756 | 1674 | 1775.86 | 1817 | 1734 | 0.954 | -0.068 | 0.30116428 | 0.47354686 | 5.00E-04 | 0.0038 | 0.0025 | 0.0025 | TRUE | 0.013 |
| Mrps16 | 2293 | 1682 | 1627 | 1849 | 1711 | 1905 | 1781 | 1789 | 1756 | 1840 | 1930 | 1881 | 1829.45 | 1775 | 1884 | 1.061 | 0.085 | 0.16674018 | 0.31778393 | 0 | 0.0037 | 0.0022 | 0.0022 | TRUE | 0.0055 |
| Mrps17 | 3011 | 1966 | 1886 | 2189 | 1877 | 2273 | 2339 | 2091 | 2035 | 2179 | 2117 | 2244 | 2167.48 | 2155 | 2180 | 1.011 | 0.015 | 0.8295227 | 0.90237978 | 0.0028 | 0.0035 | 0.0032 | 0.0032 | TRUE | 0.069 |
| Mrps18a | 2394 | 1740 | 1821 | 1900 | 1657 | 1943 | 1859 | 1851 | 1965 | 1891 | 1869 | 1918 | 1892.26 | 1892 | 1893 | 1.001 | 0.002 | 0.97991331 | 0.98946668 | 0 | 0.0037 | 0.0023 | 0.0023 | TRUE | 0.0135 |
| Mrps18b | 3494 | 2471 | 2272 | 2506 | 2358 | 2728 | 2714 | 2628 | 2451 | 2494 | 2660 | 2693 | 2606.83 | 2598 | 2616 | 1.006 | 0.009 | 0.89092071 | 0.93970406 | 0.0018 | 0.0033 | 0.0027 | 0.0027 | TRUE | 0.0292 |
| Mrps18c | 1620 | 1178 | 1127 | 1139 | 977 | 1130 | 1258 | 1253 | 1216 | 1134 | 1102 | 1115 | 1179.77 | 1242 | 1117 | 0.899 | -0.154 | 0.02706662 | 0.08591231 | 0 | 0.0044 | 0.0026 | 0.0026 | TRUE | 0.004 |
| Mrps2 | 5273 | 3783 | 3902 | 3844 | 3529 | 4076 | 4095 | 4024 | 4210 | 3826 | 3981 | 4024 | 4026.75 | 4110 | 3944 | 0.96 | -0.06 | 0.27298869 | 0.44404545 | 3.00E-04 | 0.003 | 0.0019 | 0.0019 | TRUE | 0.0083 |
| Mrps21 | 1138 | 811 | 789 | 795 | 653 | 792 | 884 | 863 | 851 | 791 | 737 | 782 | 817.94 | 866 | 770 | 0.889 | -0.17 | 0.03441353 | 0.1026231 | 0 | 0.0053 | 0.0034 | 0.0034 | TRUE | 0.0165 |
| Mrps22 | 2620 | 1915 | 1884 | 1860 | 1650 | 2024 | 2035 | 2037 | 2033 | 1851 | 1861 | 1998 | 1969.24 | 2035 | 1903 | 0.936 | -0.096 | 0.12932299 | 0.26608271 | 4.00E-04 | 0.0036 | 0.0024 | 0.0024 | TRUE | 0.023 |
| Mrps23 | 4278 | 3098 | 2908 | 3292 | 2877 | 3302 | 3323 | 3295 | 3138 | 3277 | 3246 | 3260 | 3256.26 | 3252 | 3261 | 1.002 | 0.003 | 0.94983072 | 0.97250044 | 2.00E-04 | 0.0032 | 0.0019 | 0.0019 | TRUE | 0.0113 |
| Mrps24 | 3588 | 2756 | 3043 | 3136 | 2804 | 3204 | 2787 | 2932 | 3283 | 3122 | 3163 | 3163 | 3074.86 | 3001 | 3149 | 1.05 | 0.071 | 0.31372444 | 0.48713345 | 0.0032 | 0.0032 | 0.0032 | 0.0032 | TRUE | 0.0822 |
| Mrps25 | 4962 | 3489 | 3174 | 3703 | 2967 | 3586 | 3854 | 3711 | 3425 | 3686 | 3347 | 3540 | 3593.78 | 3663 | 3524 | 0.962 | -0.056 | 0.40022083 | 0.573415 | 0.0025 | 0.0031 | 0.0029 | 0.0029 | TRUE | 0.0393 |
| Mrps26 | 2236 | 1534 | 1430 | 1480 | 1300 | 1525 | 1737 | 1632 | 1543 | 1473 | 1467 | 1505 | 1559.4 | 1637 | 1482 | 0.904 | -0.145 | 0.04001301 | 0.11444986 | 0.0014 | 0.0039 | 0.003 | 0.003 | TRUE | 0.0353 |
| Mrps27 | 5678 | 4106 | 4005 | 4320 | 3744 | 4580 | 4410 | 4367 | 4321 | 4300 | 4224 | 4521 | 4357.27 | 4366 | 4348 | 0.996 | -0.006 | 0.91344289 | 0.95246365 | 4.00E-04 | 0.003 | 0.0019 | 0.0019 | TRUE | 0.0148 |
| Mrps28 | 1708 | 1280 | 1294 | 1328 | 1301 | 1504 | 1327 | 1362 | 1396 | 1322 | 1468 | 1485 | 1393.08 | 1362 | 1425 | 1.047 | 0.066 | 0.37233998 | 0.54688296 | 0.0016 | 0.0041 | 0.0032 | 0.0032 | TRUE | 0.0485 |
| Mrps30 | 5342 | 3848 | 4074 | 4185 | 3681 | 4214 | 4149 | 4093 | 4396 | 4166 | 4153 | 4160 | 4185.99 | 4213 | 4160 | 0.988 | -0.018 | 0.74124677 | 0.8460229 | 4.00E-04 | 0.003 | 0.0019 | 0.0019 | TRUE | 0.0174 |
| Mrps31 | 4816 | 3570 | 3737 | 4525 | 3937 | 4467 | 3740 | 3797 | 4032 | 4504 | 4441 | 4410 | 4154.18 | 3856 | 4452 | 1.155 | 0.208 | 0.00018456 | 0.00200923 | 5.00E-04 | 0.003 | 0.002 | 0.002 | TRUE | 0.0191 |
| Mrps33 | 785 | 534 | 511 | 653 | 463 | 602 | 610 | 568 | 551 | 650 | 522 | 594 | 582.6 | 576 | 589 | 1.021 | 0.031 | 0.76975157 | 0.86378733 | 0.0055 | 0.0065 | 0.0062 | 0.0062 | TRUE | 0.1118 |
| Mrps34 | 1315 | 1003 | 882 | 1062 | 853 | 1000 | 1021 | 1067 | 952 | 1057 | 962 | 987 | 1007.73 | 1013 | 1002 | 0.989 | -0.016 | 0.84630598 | 0.91278686 | 0.0017 | 0.0048 | 0.0037 | 0.0037 | TRUE | 0.0325 |
| Mrps35 | 1298 | 947 | 1089 | 1085 | 923 | 1052 | 1008 | 1007 | 1175 | 1080 | 1041 | 1038 | 1058.36 | 1063 | 1053 | 0.991 | -0.012 | 0.88382497 | 0.93547084 | 0.0033 | 0.0047 | 0.0042 | 0.0042 | TRUE | 0.0988 |
| Mrps36 | 744 | 574 | 633 | 676 | 609 | 670 | 578 | 611 | 683 | 673 | 687 | 661 | 648.78 | 624 | 674 | 1.082 | 0.114 | 0.22775172 | 0.39299951 | 0.0024 | 0.0061 | 0.0049 | 0.0049 | TRUE | 0.0801 |
| Mrps5 | 6366 | 4658 | 4621 | 4773 | 4369 | 5059 | 4944 | 4955 | 4986 | 4751 | 4929 | 4994 | 4926.44 | 4962 | 4891 | 0.986 | -0.021 | 0.68689991 | 0.80798786 | 1.00E-04 | 0.0029 | 0.0017 | 0.0017 | TRUE | 0.0077 |
| Mrps6 | 986 | 707 | 616 | 593 | 613 | 696 | 766 | 752 | 665 | 590 | 692 | 687 | 691.89 | 728 | 656 | 0.901 | -0.151 | 0.1294586 | 0.26619936 | 0.0053 | 0.0059 | 0.0057 | 0.0057 | TRUE | 0.0922 |
| Mrps7 | 4484 | 3130 | 2886 | 2892 | 2671 | 3009 | 3483 | 3329 | 3114 | 2879 | 3013 | 2970 | 3131.34 | 3309 | 2954 | 0.892 | -0.164 | 0.00897469 | 0.03825684 | 0.0014 | 0.0032 | 0.0025 | 0.0025 | TRUE | 0.032 |
| Mrps9 | 4976 | 3591 | 3582 | 3506 | 3204 | 3742 | 3865 | 3820 | 3865 | 3490 | 3614 | 3694 | 3724.59 | 3850 | 3599 | 0.935 | -0.097 | 0.07135636 | 0.17395306 | 1.00E-04 | 0.0031 | 0.0018 | 0.0018 | TRUE | 0.0087 |
| Mrrf | 1603 | 1185 | 1190 | 1441 | 1155 | 1441 | 1245 | 1260 | 1284 | 1434 | 1303 | 1422 | 1324.88 | 1263 | 1386 | 1.099 | 0.136 | 0.05841324 | 0.15071711 | 7.00E-04 | 0.0042 | 0.003 | 0.003 | TRUE | 0.0328 |
| Mrs2 | 1519 | 1203 | 1193 | 1344 | 1157 | 1313 | 1180 | 1280 | 1287 | 1338 | 1305 | 1296 | 1280.96 | 1249 | 1313 | 1.053 | 0.074 | 0.30229088 | 0.474796 | 7.00E-04 | 0.0043 | 0.0029 | 0.0029 | TRUE | 0.0296 |
| Mrto4 | 6498 | 4362 | 3917 | 4287 | 3822 | 4313 | 5047 | 4640 | 4226 | 4267 | 4312 | 4258 | 4458.24 | 4638 | 4279 | 0.922 | -0.117 | 0.0929763 | 0.20964036 | 0.0036 | 0.003 | 0.0033 | 0.0033 | TRUE | 0.0734 |
| Mrvi1 | 31 | 15 | 18 | 16 | 15 | 16 | 24 | 16 | 19 | 16 | 17 | 16 | 18.02 | 20 | 16 | 0.811 | -0.302 | 0.50289635 | 0.66702835 | 0 | 0.134 | 0.0901 | 0.0901 | TRUE | 0.2323 |
| Ms4a1 | 1 | 0 | 1 | 0 | 0 | 1 | 1 | 0 | 1 | 0 | 0 | 1 | 0.47 | 1 | 0 | 0.596 | -0.748 | 0.78959815 | NA | 0 | 5.0047 | 3.8792 | 3.8792 | TRUE | 0.3835 |
| Ms4a10 | 4 | 3 | 5 | 2 | 3 | 2 | 3 | 3 | 5 | 2 | 3 | 2 | 3.17 | 4 | 2 | 0.629 | -0.67 | 0.52863039 | NA | 0 | 0.7492 | 0.4767 | 0.4767 | TRUE | 0.1698 |
| Ms4a13 | 1 | 0 | 0 | 1 | 1 | 0 | 1 | 0 | 0 | 1 | 1 | 0 | 0.48 | 0 | 1 | 2.037 | 1.027 | 0.71241723 | NA | 0 | 4.9057 | 3.8521 | 3.8521 | TRUE | 0.2664 |
| Ms4a15 | 68 | 67 | 77 | 62 | 65 | 52 | 53 | 71 | 83 | 62 | 73 | 51 | 65.59 | 69 | 62 | 0.904 | -0.146 | 0.5826513 | 0.73263262 | 0.0278 | 0.0386 | 0.0355 | 0.0355 | TRUE | 0.3576 |
| Ms4a18 | 0 | 0 | 0 | 0 | 0 | 0 | 0 | 0 | 0 | 0 | 0 | 0 | 0 | 0 | 0 | NA | NA | NA | NA | NA | NA | NA | NA | NA | NA |
| Ms4a2 | 0 | 0 | 0 | 0 | 0 | 0 | 0 | 0 | 0 | 0 | 0 | 0 | 0 | 0 | 0 | NA | NA | NA | NA | NA | NA | NA | NA | NA | NA |
| Ms4a3 | 0 | 0 | 0 | 0 | 0 | 0 | 0 | 0 | 0 | 0 | 0 | 0 | 0 | 0 | 0 | NA | NA | NA | NA | NA | NA | NA | NA | NA | NA |
| Ms4a4b | 0 | 0 | 0 | 0 | 0 | 0 | 0 | 0 | 0 | 0 | 0 | 0 | 0 | 0 | 0 | NA | NA | NA | NA | NA | NA | NA | NA | NA | NA |
| Ms4a4c | 0 | 0 | 0 | 0 | 0 | 0 | 0 | 0 | 0 | 0 | 0 | 0 | 0 | 0 | 0 | NA | NA | NA | NA | NA | NA | NA | NA | NA | NA |
| Ms4a4d | 44 | 27 | 19 | 28 | 5 | 21 | 34 | 29 | 21 | 28 | 6 | 21 | 22.94 | 28 | 18 | 0.653 | -0.615 | 0.21442761 | 0.37711504 | 0.2151 | 0.1057 | 0.1312 | 0.1312 | TRUE | 0.7861 |
| Ms4a5 | 3 | 3 | 0 | 3 | 4 | 2 | 2 | 3 | 0 | 3 | 5 | 2 | 2.5 | 2 | 3 | 1.683 | 0.751 | 0.55072791 | NA | 0 | 0.9508 | 0.7175 | 0.7175 | TRUE | 0.5512 |
| Ms4a6b | 0 | 0 | 0 | 0 | 0 | 0 | 0 | 0 | 0 | 0 | 0 | 0 | 0 | 0 | 0 | NA | NA | NA | NA | NA | NA | NA | NA | NA | NA |
| Ms4a6c | 0 | 0 | 0 | 0 | 0 | 0 | 0 | 0 | 0 | 0 | 0 | 0 | 0 | 0 | 0 | NA | NA | NA | NA | NA | NA | NA | NA | NA | NA |
| Ms4a6d | 8 | 8 | 15 | 18 | 7 | 14 | 6 | 9 | 16 | 18 | 8 | 14 | 11.76 | 10 | 13 | 1.309 | 0.388 | 0.52183174 | NA | 0.1119 | 0.204 | 0.1782 | 0.1782 | TRUE | 0.4564 |
| Ms4a7 | 0 | 0 | 1 | 0 | 0 | 0 | 0 | 0 | 1 | 0 | 0 | 0 | 0.18 | 0 | 0 | 0.552 | -0.857 | 0.8336548 | NA | 0 | 13.1817 | 10 | 10 | TRUE | 0.5365 |
| Ms4a8a | 1 | 0 | 0 | 0 | 0 | 0 | 1 | 0 | 0 | 0 | 0 | 0 | 0.13 | 0 | 0 | 0.552 | -0.857 | 0.8336548 | NA | 0 | 18.3121 | 10 | 10 | TRUE | 0.2566 |
| Msantd1 | 5 | 0 | 2 | 2 | 0 | 1 | 4 | 0 | 2 | 2 | 0 | 1 | 1.5 | 2 | 1 | 0.49 | -1.029 | 0.54883526 | NA | 0 | 1.5791 | 1.3764 | 1.3764 | TRUE | 0.1407 |
| Msantd2 | 1584 | 1307 | 1158 | 1243 | 1089 | 1391 | 1230 | 1390 | 1249 | 1237 | 1229 | 1373 | 1284.81 | 1290 | 1280 | 0.993 | -0.01 | 0.90089179 | 0.94547664 | 0.0034 | 0.0043 | 0.004 | 0.004 | TRUE | 0.0549 |
| Msantd3 | 1381 | 1066 | 1057 | 1450 | 1251 | 1459 | 1073 | 1134 | 1140 | 1443 | 1411 | 1440 | 1273.63 | 1116 | 1431 | 1.285 | 0.361 | 1.92E-07 | 6.96E-06 | 0 | 0.0043 | 0.0027 | 0.0027 | TRUE | 0.0145 |
| Msantd4 | 3201 | 2114 | 2141 | 2200 | 1887 | 2055 | 2486 | 2249 | 2310 | 2190 | 2129 | 2029 | 2232.01 | 2348 | 2116 | 0.9 | -0.152 | 0.02409096 | 0.07917502 | 0.0017 | 0.0035 | 0.0028 | 0.0028 | TRUE | 0.0328 |
| Msc | 433 | 335 | 224 | 206 | 191 | 240 | 336 | 356 | 242 | 205 | 215 | 237 | 265.29 | 311 | 219 | 0.703 | -0.509 | 0.0012604 | 0.0085164 | 0.0203 | 0.0114 | 0.014 | 0.014 | TRUE | 0.4205 |
| Msgn1 | 3 | 4 | 0 | 2 | 3 | 3 | 2 | 4 | 0 | 2 | 3 | 3 | 2.49 | 2 | 3 | 1.257 | 0.33 | 0.79333692 | NA | 0 | 0.9555 | 0.7373 | 0.7373 | TRUE | 0.2632 |
| Msh2 | 19899 | 15215 | 16533 | 16712 | 15206 | 17822 | 15455 | 16184 | 17839 | 16635 | 17154 | 17593 | 16809.97 | 16493 | 17127 | 1.039 | 0.055 | 0.38205252 | 0.55626553 | 0.0028 | 0.0026 | 0.0028 | 0.0028 | TRUE | 0.0623 |
| Msh3 | 3942 | 2763 | 2475 | 2301 | 2117 | 2590 | 3062 | 2939 | 2671 | 2290 | 2388 | 2557 | 2651.07 | 2891 | 2412 | 0.834 | -0.262 | 0.00032314 | 0.00307704 | 0.0034 | 0.0033 | 0.0034 | 0.0034 | TRUE | 0.0534 |
| Msh4 | 39 | 34 | 33 | 28 | 29 | 23 | 30 | 36 | 36 | 28 | 33 | 23 | 30.89 | 34 | 28 | 0.817 | -0.292 | 0.4021203 | 0.57507729 | 0 | 0.0791 | 0.0547 | 0.0547 | TRUE | 0.166 |
| Msh5 | 103 | 123 | 106 | 142 | 118 | 138 | 80 | 131 | 114 | 141 | 133 | 136 | 122.65 | 108 | 137 | 1.273 | 0.348 | 0.08964358 | 0.20415483 | 0.023 | 0.0218 | 0.0221 | 0.0221 | TRUE | 0.5879 |
| Msh6 | 45245 | 31754 | 31150 | 36807 | 31267 | 35989 | 35140 | 33776 | 33611 | 36638 | 35272 | 35527 | 34993.92 | 34176 | 35812 | 1.048 | 0.067 | 0.1453146 | 0.28917905 | 5.00E-04 | 0.0025 | 0.0015 | 0.0015 | TRUE | 0.0075 |
| Msi1 | 925 | 762 | 933 | 963 | 867 | 1033 | 718 | 811 | 1007 | 959 | 978 | 1020 | 915.33 | 845 | 986 | 1.168 | 0.224 | 0.04489621 | 0.12450552 | 0.014 | 0.005 | 0.0079 | 0.0079 | TRUE | 0.3307 |
| Msi2 | 6070 | 4793 | 4755 | 5901 | 5093 | 6107 | 4714 | 5098 | 5131 | 5874 | 5745 | 6029 | 5431.83 | 4981 | 5883 | 1.182 | 0.241 | 2.47E-05 | 0.00039545 | 0.0011 | 0.0029 | 0.0022 | 0.0022 | TRUE | 0.0273 |
| Msl1 | 4046 | 2934 | 2972 | 2803 | 2518 | 3050 | 3142 | 3121 | 3207 | 2790 | 2841 | 3011 | 3018.58 | 3157 | 2881 | 0.913 | -0.132 | 0.02389539 | 0.0787314 | 5.00E-04 | 0.0032 | 0.0021 | 0.0021 | TRUE | 0.0192 |
| Msl2 | 2806 | 2038 | 2040 | 2189 | 1871 | 2156 | 2179 | 2168 | 2201 | 2179 | 2111 | 2128 | 2161.03 | 2183 | 2139 | 0.98 | -0.029 | 0.62188815 | 0.76142749 | 0 | 0.0035 | 0.002 | 0.002 | TRUE | 0.0032 |
| Msl3 | 409 | 319 | 309 | 370 | 323 | 316 | 318 | 339 | 333 | 368 | 364 | 312 | 339.17 | 330 | 348 | 1.055 | 0.078 | 0.51197434 | 0.67487312 | 0.0021 | 0.0094 | 0.0071 | 0.0071 | TRUE | 0.0965 |
| Msl3l2 | 447 | 365 | 242 | 288 | 234 | 255 | 347 | 388 | 261 | 287 | 264 | 252 | 299.82 | 332 | 268 | 0.804 | -0.314 | 0.03703764 | 0.10840169 | 0.0191 | 0.0103 | 0.0129 | 0.0129 | TRUE | 0.3839 |
| Msln | 33 | 23 | 26 | 26 | 18 | 22 | 26 | 24 | 28 | 26 | 20 | 22 | 24.34 | 26 | 23 | 0.871 | -0.199 | 0.60434809 | 0.74930146 | 0 | 0.0998 | 0.0651 | 0.0651 | TRUE | 0.0923 |
| Mslnl | 20 | 16 | 19 | 15 | 16 | 8 | 16 | 17 | 21 | 15 | 18 | 8 | 15.66 | 18 | 14 | 0.769 | -0.378 | 0.45213793 | NA | 0 | 0.1538 | 0.1169 | 0.1169 | TRUE | 0.5998 |
| Msmb | 0 | 0 | 0 | 0 | 0 | 0 | 0 | 0 | 0 | 0 | 0 | 0 | 0 | 0 | 0 | NA | NA | NA | NA | NA | NA | NA | NA | NA | NA |
| Msmp | 0 | 0 | 0 | 0 | 0 | 0 | 0 | 0 | 0 | 0 | 0 | 0 | 0 | 0 | 0 | NA | NA | NA | NA | NA | NA | NA | NA | NA | NA |
| Msn | 8347 | 5990 | 6044 | 6805 | 5804 | 6530 | 6483 | 6371 | 6521 | 6774 | 6548 | 6446 | 6523.83 | 6458 | 6589 | 1.02 | 0.029 | 0.5621183 | 0.71677731 | 2.00E-04 | 0.0028 | 0.0016 | 0.0016 | TRUE | 0.0074 |
| Msr1 | 0 | 1 | 0 | 0 | 0 | 0 | 0 | 1 | 0 | 0 | 0 | 0 | 0.18 | 0 | 0 | 0.552 | -0.857 | 0.8336548 | NA | 0 | 13.3714 | 10 | 10 | TRUE | 0.5216 |
| Msra | 371 | 319 | 264 | 278 | 295 | 317 | 288 | 339 | 285 | 277 | 333 | 313 | 305.79 | 304 | 308 | 1.011 | 0.016 | 0.89894348 | 0.94405578 | 0.0058 | 0.0102 | 0.0089 | 0.0089 | TRUE | 0.1126 |

| | | | | | | | | | | | | | | | | | | | | | | | | | |
|---|---|---|---|---|---|---|---|---|---|---|---|---|---|---|---|---|---|---|---|---|---|---|---|---|---|
| Msrb1 | 1172 | 806 | 805 | 903 | 813 | 973 | 910 | 857 | 869 | 899 | 917 | 961 | 902.11 | 879 | 926 | 1.052 | 0.074 | 0.34712276 | 0.52154164 | 0 | 0.0051 | 0.0033 | 0.0033 | TRUE | 0.0133 |
| Msrb2 | 476 | 349 | 340 | 283 | 222 | 318 | 370 | 371 | 367 | 282 | 250 | 314 | 325.64 | 369 | 282 | 0.765 | -0.387 | 0.00151394 | 0.00973686 | 0.0017 | 0.0097 | 0.0076 | 0.0076 | TRUE | 0.1101 |
| Msrb3 | 527 | 312 | 277 | 302 | 245 | 273 | 409 | 332 | 299 | 301 | 276 | 269 | 314.42 | 347 | 282 | 0.811 | -0.302 | 0.02999633 | 0.09261991 | 0.0125 | 0.01 | 0.0108 | 0.0108 | TRUE | 0.2995 |
| Mss51 | 65 | 44 | 55 | 69 | 54 | 66 | 50 | 47 | 59 | 69 | 61 | 65 | 58.56 | 52 | 65 | 1.247 | 0.318 | 0.2096071 | 0.37145244 | 0 | 0.0429 | 0.029 | 0.029 | TRUE | 0.1084 |
| Mst1 | 49 | 27 | 45 | 37 | 41 | 50 | 38 | 29 | 49 | 37 | 46 | 49 | 41.3 | 39 | 44 | 1.149 | 0.2 | 0.52906109 | 0.68944841 | 0.016 | 0.0598 | 0.0484 | 0.0484 | TRUE | 0.2237 |
| Mst1r | 34 | 16 | 26 | 27 | 33 | 23 | 26 | 17 | 28 | 27 | 37 | 23 | 26.38 | 24 | 29 | 1.205 | 0.269 | 0.49522658 | 0.65996429 | 0.0234 | 0.0923 | 0.0741 | 0.0741 | TRUE | 0.3813 |
| Mstn | 0 | 0 | 0 | 0 | 0 | 0 | 0 | 0 | 0 | 0 | 0 | 0 | 0 | 0 | 0 | NA | NA | NA | NA | NA | NA | NA | NA | NA | NA |
| Msto1 | 5313 | 3946 | 3995 | 3780 | 3644 | 4221 | 4126 | 4197 | 4311 | 3763 | 4111 | 4167 | 4112.41 | 4211 | 4014 | 0.953 | -0.069 | 0.25147346 | 0.42043632 | 0.0014 | 0.003 | 0.0024 | 0.0024 | TRUE | 0.0366 |
| Msx1 | 36 | 34 | 28 | 42 | 28 | 29 | 28 | 36 | 30 | 42 | 32 | 29 | 32.73 | 31 | 34 | 1.087 | 0.12 | 0.72641126 | 0.83598639 | 0 | 0.0748 | 0.0544 | 0.0544 | TRUE | 0.2901 |
| Msx1as | 0 | 0 | 0 | 0 | 0 | 1 | 0 | 0 | 0 | 0 | 0 | 1 | 0.16 | 0 | 0 | 2.094 | 1.067 | 0.79379329 | NA | 0 | 14.4078 | 10 | 10 | TRUE | 0.3835 |
| Msx2 | 180 | 101 | 131 | 158 | 118 | 132 | 140 | 107 | 141 | 157 | 133 | 130 | 134.88 | 129 | 140 | 1.081 | 0.112 | 0.54191644 | 0.69994683 | 0.0089 | 0.02 | 0.0169 | 0.0169 | TRUE | 0.2181 |
| Msx3 | 32 | 22 | 21 | 29 | 15 | 19 | 25 | 23 | 23 | 29 | 17 | 19 | 22.58 | 24 | 22 | 0.911 | -0.134 | 0.74516704 | 0.84852395 | 0 | 0.1074 | 0.0777 | 0.0777 | TRUE | 0.5105 |
| Mt1 | 4608 | 4190 | 4682 | 4838 | 4527 | 5029 | 3579 | 4457 | 5052 | 4816 | 5107 | 4964 | 4662.43 | 4363 | 4962 | 1.138 | 0.187 | 0.06009653 | 0.15346787 | 0.0145 | 0.0029 | 0.0069 | 0.0069 | TRUE | 0.3034 |
| Mt2 | 6260 | 5052 | 5525 | 5994 | 5169 | 5575 | 4862 | 5374 | 5961 | 5966 | 5831 | 5503 | 5583 | 5399 | 5767 | 1.069 | 0.096 | 0.20750056 | 0.36911083 | 0.0057 | 0.0029 | 0.004 | 0.004 | TRUE | 0.1012 |
| Mt3 | 306 | 223 | 235 | 307 | 267 | 364 | 238 | 237 | 254 | 306 | 301 | 359 | 282.42 | 243 | 322 | 1.328 | 0.41 | 0.00143762 | 0.00934884 | 0.0027 | 0.0108 | 0.0083 | 0.0083 | TRUE | 0.1179 |
| Mt4 | 19 | 20 | 22 | 17 | 20 | 22 | 15 | 21 | 24 | 17 | 23 | 22 | 20.16 | 20 | 21 | 1.033 | 0.046 | 0.91475591 | 0.95295409 | 0 | 0.12 | 0.0858 | 0.0858 | TRUE | 0.3486 |
| Mta1 | 6478 | 5123 | 5605 | 5559 | 5309 | 6254 | 5031 | 5449 | 6048 | 5533 | 5989 | 6174 | 5704.07 | 5509 | 5899 | 1.071 | 0.099 | 0.18796093 | 0.34508998 | 0.0056 | 0.0028 | 0.0039 | 0.0039 | TRUE | 0.089 |
| Mta2 | 7416 | 5317 | 5025 | 4993 | 4532 | 5115 | 5760 | 5656 | 5422 | 4970 | 5113 | 5049 | 5328.19 | 5613 | 5044 | 0.898 | -0.155 | 0.00311716 | 0.01701864 | 4.00E-04 | 0.0029 | 0.0018 | 0.0018 | TRUE | 0.0107 |
| Mta3 | 5006 | 3582 | 3786 | 3571 | 3068 | 3592 | 3888 | 3810 | 4085 | 3555 | 3461 | 3546 | 3724.1 | 3928 | 3521 | 0.897 | -0.158 | 0.00473254 | 0.02350322 | 5.00E-04 | 0.0031 | 0.002 | 0.002 | TRUE | 0.0147 |
| Mtag2 | 4 | 5 | 2 | 1 | 1 | 3 | 3 | 5 | 2 | 1 | 1 | 3 | 2.61 | 3 | 2 | 0.486 | -1.041 | 0.3895365 | NA | 0 | 0.91 | 0.6154 | 0.6154 | TRUE | 0.3421 |
| Mtap | 7270 | 5417 | 5122 | 5238 | 4429 | 5230 | 5646 | 5762 | 5527 | 5214 | 4996 | 5163 | 5384.66 | 5645 | 5124 | 0.908 | -0.139 | 0.00660686 | 0.03038347 | 2.00E-04 | 0.0029 | 0.0017 | 0.0017 | TRUE | 0.0058 |
| Mtap7d3 | 343 | 262 | 241 | 266 | 285 | 334 | 266 | 279 | 260 | 265 | 322 | 330 | 286.85 | 268 | 306 | 1.137 | 0.186 | 0.15486558 | 0.3024725 | 0.0043 | 0.0107 | 0.0088 | 0.0088 | TRUE | 0.1552 |
| Mtbp | 1839 | 1278 | 1238 | 1261 | 1029 | 1187 | 1428 | 1359 | 1336 | 1255 | 1161 | 1172 | 1285.2 | 1374 | 1196 | 0.87 | -0.202 | 0.00542254 | 0.02612213 | 8.00E-04 | 0.0043 | 0.003 | 0.003 | TRUE | 0.0227 |
| Mtch1 | 3189 | 2226 | 2081 | 2138 | 1851 | 2220 | 2477 | 2368 | 2245 | 2128 | 2088 | 2191 | 2249.61 | 2363 | 2136 | 0.903 | -0.147 | 0.02283438 | 0.07605962 | 0.0011 | 0.0035 | 0.0025 | 0.0025 | TRUE | 0.023 |
| Mtch2 | 10670 | 7507 | 7714 | 9036 | 7654 | 8973 | 8287 | 7985 | 8323 | 8994 | 8635 | 8858 | 8513.67 | 8198 | 8829 | 1.077 | 0.107 | 0.0295164 | 0.09154428 | 3.00E-04 | 0.0027 | 0.0016 | 0.0016 | TRUE | 0.0063 |
| Mtcp1 | 236 | 214 | 219 | 237 | 180 | 249 | 183 | 228 | 236 | 236 | 203 | 246 | 222 | 216 | 228 | 1.064 | 0.089 | 0.55653047 | 0.71162015 | 0.0099 | 0.0131 | 0.0122 | 0.0122 | TRUE | 0.2071 |
| Mtdh | 9462 | 7006 | 7392 | 7824 | 6900 | 8135 | 7349 | 7452 | 7976 | 7788 | 7784 | 8030 | 7729.87 | 7592 | 7867 | 1.036 | 0.052 | 0.33800606 | 0.51223769 | 9.00E-04 | 0.0027 | 0.002 | 0.002 | TRUE | 0.0237 |
| Mterf | 200 | 140 | 136 | 131 | 123 | 129 | 155 | 149 | 147 | 130 | 139 | 127 | 141.25 | 150 | 132 | 0.877 | -0.189 | 0.25047345 | 0.41935606 | 0 | 0.0192 | 0.0123 | 0.0123 | TRUE | 0.0186 |
| Mterfd1 | 1162 | 803 | 845 | 906 | 790 | 889 | 902 | 854 | 912 | 902 | 891 | 878 | 889.83 | 889 | 890 | 1 | 0.001 | 0.99295806 | 0.99595436 | 0 | 0.0051 | 0.0032 | 0.0032 | TRUE | 0.0139 |
| Mterfd2 | 1220 | 819 | 698 | 736 | 688 | 768 | 948 | 871 | 753 | 733 | 776 | 758 | 806.45 | 857 | 756 | 0.88 | -0.185 | 0.05750927 | 0.14912554 | 0.006 | 0.0054 | 0.0056 | 0.0056 | TRUE | 0.1328 |
| Mterfd3 | 382 | 254 | 259 | 266 | 226 | 278 | 297 | 270 | 279 | 265 | 255 | 274 | 273.41 | 282 | 265 | 0.937 | -0.093 | 0.4492945 | 0.61944307 | 0 | 0.0111 | 0.0073 | 0.0073 | TRUE | 0.0249 |
| Mtf1 | 2343 | 1681 | 1682 | 1725 | 1636 | 1768 | 1820 | 1788 | 1815 | 1717 | 1846 | 1745 | 1788.43 | 1808 | 1769 | 0.978 | -0.032 | 0.62138642 | 0.76111715 | 0 | 0.0038 | 0.0024 | 0.0024 | TRUE | 0.0169 |
| Mtf2 | 20561 | 14747 | 13672 | 12061 | 10860 | 12618 | 15969 | 15686 | 14752 | 12006 | 12251 | 12456 | 13853.27 | 15469 | 12238 | 0.791 | -0.338 | 6.37E-11 | 5.39E-09 | 9.00E-04 | 0.0026 | 0.0019 | 0.0019 | TRUE | 0.0201 |
| Mtfmt | 969 | 696 | 637 | 723 | 655 | 734 | 753 | 740 | 687 | 720 | 739 | 725 | 727.23 | 727 | 728 | 1 | 0 | 0.99803848 | 0.99894706 | 0 | 0.0057 | 0.0037 | 0.0037 | TRUE | 0.026 |
| Mtfp1 | 469 | 330 | 301 | 312 | 282 | 316 | 364 | 351 | 325 | 311 | 318 | 312 | 330.11 | 347 | 314 | 0.903 | -0.148 | 0.19664091 | 0.35635711 | 0 | 0.0096 | 0.0064 | 0.0064 | TRUE | 0.0341 |
| Mtfr1 | 4991 | 3448 | 3584 | 4003 | 3526 | 3945 | 3876 | 3668 | 3867 | 3985 | 3978 | 3894 | 3877.93 | 3804 | 3952 | 1.039 | 0.055 | 0.31228241 | 0.48561942 | 3.00E-04 | 0.003 | 0.0019 | 0.0019 | TRUE | 0.0118 |
| Mtfr1l | 4001 | 3065 | 3156 | 3591 | 3095 | 3705 | 3107 | 3260 | 3405 | 3574 | 3491 | 3657 | 3416.05 | 3257 | 3574 | 1.098 | 0.135 | 0.02356592 | 0.07787923 | 9.00E-04 | 0.0031 | 0.0023 | 0.0023 | TRUE | 0.0203 |
| Mtfr2 | 1768 | 1299 | 1334 | 1432 | 1216 | 1414 | 1373 | 1382 | 1439 | 1425 | 1372 | 1396 | 1397.88 | 1398 | 1398 | 1 | 0 | 0.99467969 | 0.99691164 | 0 | 0.0041 | 0.0025 | 0.0025 | TRUE | 0.0079 |
| Mtg1 | 1546 | 1119 | 1069 | 1287 | 1178 | 1310 | 1201 | 1190 | 1153 | 1281 | 1329 | 1293 | 1241.26 | 1181 | 1301 | 1.101 | 0.138 | 0.04478629 | 0.12434546 | 0 | 0.0043 | 0.0026 | 0.0026 | TRUE | 0.0051 |
| Mthfd1 | 20474 | 14719 | 14330 | 15418 | 13788 | 15186 | 15901 | 15656 | 15462 | 15347 | 15554 | 14991 | 15485.32 | 15673 | 15297 | 0.976 | -0.035 | 0.44036211 | 0.61173055 | 2.00E-04 | 0.0026 | 0.0014 | 0.0014 | TRUE | 0.0038 |
| Mthfd1l | 7602 | 5769 | 6162 | 6271 | 5587 | 6433 | 5904 | 6136 | 6649 | 6242 | 6303 | 6350 | 6264.09 | 6230 | 6298 | 1.011 | 0.016 | 0.78259473 | 0.87203762 | 0.0017 | 0.0028 | 0.0024 | 0.0024 | TRUE | 0.0421 |
| Mthfd2 | 5756 | 3920 | 3457 | 4115 | 3287 | 4116 | 4470 | 4170 | 3730 | 4096 | 3708 | 4063 | 4039.57 | 4123 | 3956 | 0.959 | -0.06 | 0.42626528 | 0.59872344 | 0.0048 | 0.003 | 0.0039 | 0.0039 | TRUE | 0.0845 |
| Mthfd2l | 94 | 71 | 73 | 67 | 49 | 71 | 73 | 76 | 79 | 67 | 55 | 70 | 69.89 | 76 | 64 | 0.848 | -0.238 | 0.30763908 | 0.48024087 | 0 | 0.0363 | 0.0247 | 0.0247 | TRUE | 0.1168 |
| Mthfr | 1068 | 727 | 547 | 483 | 490 | 524 | 829 | 773 | 590 | 481 | 553 | 517 | 623.97 | 731 | 517 | 0.706 | -0.503 | 5.39E-05 | 0.00074567 | 0.0167 | 0.0062 | 0.0095 | 0.0095 | TRUE | 0.3313 |
| Mthfs | 191 | 130 | 150 | 174 | 154 | 190 | 148 | 138 | 162 | 173 | 174 | 188 | 163.83 | 149 | 178 | 1.193 | 0.254 | 0.10400404 | 0.22775084 | 0 | 0.0169 | 0.0115 | 0.0115 | TRUE | 0.0506 |
| Mthfsd | 586 | 413 | 385 | 375 | 305 | 349 | 455 | 439 | 415 | 373 | 344 | 345 | 395.28 | 436 | 354 | 0.81 | -0.304 | 0.00462101 | 0.0230937 | 0 | 0.0084 | 0.0057 | 0.0057 | TRUE | 0.0264 |
| Mtif2 | 5232 | 3795 | 3650 | 4123 | 3627 | 4010 | 4063 | 4037 | 3938 | 4104 | 4092 | 3958 | 4032.11 | 4013 | 4051 | 1.009 | 0.013 | 0.79779768 | 0.88189654 | 0 | 0.003 | 0.0017 | 0.0017 | TRUE | 0.0049 |
| Mtif3 | 1133 | 706 | 685 | 731 | 655 | 794 | 880 | 751 | 739 | 728 | 739 | 784 | 770.06 | 790 | 750 | 0.948 | -0.078 | 0.41210163 | 0.58508961 | 0.0046 | 0.0055 | 0.0052 | 0.0052 | TRUE | 0.1227 |
| Mtl5 | 524 | 402 | 444 | 494 | 453 | 374 | 407 | 428 | 479 | 492 | 511 | 369 | 447.6 | 438 | 457 | 1.045 | 0.063 | 0.63295814 | 0.77014308 | 0.0164 | 0.0077 | 0.0104 | 0.0104 | TRUE | 0.3332 |
| Mtm1 | 731 | 582 | 670 | 716 | 615 | 678 | 568 | 619 | 723 | 713 | 694 | 669 | 664.25 | 637 | 692 | 1.089 | 0.124 | 0.2323762 | 0.39829153 | 0.0066 | 0.006 | 0.0062 | 0.0062 | TRUE | 0.1645 |
| Mtmr1 | 1076 | 748 | 728 | 815 | 706 | 813 | 836 | 796 | 786 | 811 | 796 | 803 | 804.51 | 806 | 803 | 0.996 | -0.005 | 0.94722719 | 0.97107433 | 0 | 0.0054 | 0.0033 | 0.0033 | TRUE | 0.0134 |
| Mtmr10 | 1694 | 1125 | 1045 | 965 | 857 | 1002 | 1316 | 1197 | 1128 | 961 | 967 | 989 | 1092.72 | 1214 | 972 | 0.8 | -0.322 | 7.92E-05 | 0.00100314 | 0.0026 | 0.0046 | 0.0039 | 0.0039 | TRUE | 0.0678 |
| Mtmr11 | 85 | 69 | 81 | 95 | 72 | 97 | 66 | 73 | 87 | 95 | 81 | 96 | 83.06 | 75 | 91 | 1.205 | 0.269 | 0.22312981 | 0.38750843 | 0 | 0.031 | 0.023 | 0.023 | TRUE | 0.1665 |
| Mtmr12 | 4044 | 2966 | 2855 | 2554 | 2311 | 2689 | 3141 | 3155 | 3081 | 2542 | 2607 | 2654 | 2863.33 | 3126 | 2601 | 0.832 | -0.265 | 2.00E-06 | 5.07E-05 | 0 | 0.0033 | 0.0019 | 0.0019 | TRUE | 0.0048 |
| Mtmr14 | 5251 | 3876 | 4401 | 4233 | 4139 | 4595 | 4078 | 4123 | 4749 | 4214 | 4669 | 4536 | 4394.74 | 4317 | 4473 | 1.036 | 0.052 | 0.48174219 | 0.64849796 | 0.0042 | 0.003 | 0.0037 | 0.0037 | TRUE | 0.0929 |
| Mtmr2 | 4844 | 3476 | 3500 | 3877 | 3449 | 3967 | 3762 | 3697 | 3776 | 3859 | 3891 | 3916 | 3817 | 3745 | 3889 | 1.038 | 0.054 | 0.29254273 | 0.46479694 | 0 | 0.003 | 0.0016 | 0.0016 | TRUE | 0.0015 |
| Mtmr3 | 4803 | 3559 | 4029 | 4445 | 3907 | 4449 | 3730 | 3786 | 4347 | 4425 | 4408 | 4392 | 4181.18 | 3954 | 4408 | 1.115 | 0.157 | 0.0215789 | 0.07303509 | 0.0034 | 0.003 | 0.0031 | 0.0031 | TRUE | 0.0913 |
| Mtmr4 | 4915 | 3667 | 3434 | 3758 | 3352 | 3837 | 3817 | 3901 | 3705 | 3741 | 3781 | 3788 | 3788.82 | 3808 | 3770 | 0.99 | -0.014 | 0.78560022 | 0.87390788 | 0 | 0.0031 | 0.0018 | 0.0018 | TRUE | 0.0066 |
| Mtmr6 | 1646 | 1141 | 1163 | 1369 | 1153 | 1359 | 1278 | 1214 | 1255 | 1363 | 1301 | 1342 | 1291.98 | 1249 | 1335 | 1.069 | 0.096 | 0.16504224 | 0.31587424 | 0 | 0.0043 | 0.0026 | 0.0026 | TRUE | 0.0073 |
| Mtmr7 | 7531 | 5167 | 5114 | 5441 | 4835 | 5253 | 5849 | 5496 | 5518 | 5416 | 5454 | 5186 | 5486.49 | 5621 | 5352 | 0.952 | -0.071 | 0.19209737 | 0.35037387 | 7.00E-04 | 0.0029 | 0.002 | 0.002 | TRUE | 0.0156 |
| Mtmr9 | 4023 | 2812 | 2907 | 3061 | 2736 | 3148 | 3124 | 2991 | 3137 | 3047 | 3086 | 3108 | 3082.2 | 3084 | 3080 | 0.999 | -0.002 | 0.9709029 | 0.98433219 | 0 | 0.0032 | 0.0019 | 0.0019 | TRUE | 0.0083 |
| Mtnr1a | 8 | 3 | 6 | 4 | 5 | 9 | 6 | 3 | 6 | 4 | 6 | 9 | 5.73 | 5 | 6 | 1.16 | 0.214 | 0.79007207 | NA | 0 | 0.416 | 0.2923 | 0.2923 | TRUE | 0.3723 |
| Mtnr1b | 0 | 0 | 0 | 0 | 0 | 0 | 0 | 0 | 0 | 0 | 0 | 0 | 0 | 0 | 0 | NA | NA | NA | NA | NA | NA | NA | NA | NA | NA |
| Mto1 | 1941 | 1374 | 1309 | 1428 | 1334 | 1512 | 1507 | 1462 | 1412 | 1421 | 1505 | 1493 | 1466.72 | 1460 | 1473 | 1.008 | 0.011 | 0.87119071 | 0.9285222 | 0 | 0.004 | 0.0027 | 0.0027 | TRUE | 0.0114 |
| Mtor | 9422 | 6858 | 6817 | 7425 | 6877 | 8013 | 7318 | 7295 | 7355 | 7391 | 7758 | 7910 | 7504.47 | 7323 | 7686 | 1.05 | 0.07 | 0.16835254 | 0.31985417 | 4.00E-04 | 0.0027 | 0.0017 | 0.0017 | TRUE | 0.0139 |
| Mtpap | 3041 | 2115 | 2193 | 2309 | 1996 | 2410 | 2362 | 2250 | 2366 | 2298 | 2252 | 2379 | 2317.81 | 2326 | 2310 | 0.993 | -0.01 | 0.8670744 | 0.92677197 | 3.00E-04 | 0.0034 | 0.0022 | 0.0022 | TRUE | 0.0098 |
| Mtpn | 9562 | 6798 | 7558 | 8244 | 6997 | 8107 | 7426 | 7231 | 8155 | 8206 | 7893 | 8003 | 7819.11 | 7604 | 8034 | 1.057 | 0.08 | 0.18334707 | 0.33923362 | 0.0021 | 0.0027 | 0.0025 | 0.0025 | TRUE | 0.0487 |
| Mtr | 4222 | 2940 | 2631 | 2855 | 2585 | 2695 | 3279 | 3127 | 2839 | 2842 | 2916 | 2660 | 2943.92 | 3082 | 2806 | 0.91 | -0.136 | 0.05600937 | 0.14627115 | 0.0032 | 0.0032 | 0.0033 | 0.0033 | TRUE | 0.0574 |
| Mtrf1 | 773 | 530 | 602 | 595 | 569 | 647 | 600 | 564 | 650 | 592 | 642 | 639 | 614.42 | 605 | 624 | 1.033 | 0.046 | 0.62472514 | 0.76346484 | 0.0017 | 0.0063 | 0.0048 | 0.0048 | TRUE | 0.0474 |

| | | | | | | | | | | | | | | | | | | | | | | | | | |
|---|---|---|---|---|---|---|---|---|---|---|---|---|---|---|---|---|---|---|---|---|---|---|---|---|---|
| Mtrf1l | 190 | 139 | 119 | 123 | 133 | 123 | 148 | 148 | 128 | 122 | 150 | 121 | 136.28 | 141 | 131 | 0.926 | -0.111 | 0.52879918 | 0.68929563 | 0 | 0.0198 | 0.015 | 0.015 | TRUE | 0.1531 |
| Mtrr | 3114 | 2322 | 2149 | 2117 | 1975 | 2232 | 2419 | 2470 | 2319 | 2107 | 2228 | 2203 | 2290.96 | 2403 | 2179 | 0.907 | -0.141 | 0.02198155 | 0.07387311 | 5.00E-04 | 0.0035 | 0.0023 | 0.0023 | TRUE | 0.0109 |
| Mtss1 | 4805 | 3787 | 3493 | 4294 | 4003 | 4825 | 3732 | 4028 | 3769 | 4274 | 4516 | 4763 | 4180.33 | 3843 | 4518 | 1.176 | 0.234 | 0.00021592 | 0.00226095 | 0.002 | 0.003 | 0.0026 | 0.0026 | TRUE | 0.0277 |
| Mtss1l | 1759 | 1394 | 1238 | 1366 | 1285 | 1466 | 1366 | 1483 | 1336 | 1360 | 1450 | 1447 | 1406.87 | 1395 | 1419 | 1.017 | 0.025 | 0.73450066 | 0.8412894 | 0.0014 | 0.0041 | 0.0031 | 0.0031 | TRUE | 0.0357 |
| Mttp | 423 | 296 | 279 | 304 | 270 | 265 | 329 | 315 | 301 | 303 | 305 | 262 | 302.2 | 315 | 290 | 0.918 | -0.123 | 0.31485116 | 0.48851147 | 0 | 0.0103 | 0.0076 | 0.0076 | TRUE | 0.082 |
| Mtus1 | 3520 | 2668 | 2685 | 2866 | 2548 | 3185 | 2734 | 2838 | 2897 | 2853 | 2874 | 3144 | 2890.03 | 2823 | 2957 | 1.048 | 0.068 | 0.2879703 | 0.4601408 | 0.0014 | 0.0032 | 0.0026 | 0.0026 | TRUE | 0.0374 |
| Mtus2 | 947 | 665 | 523 | 592 | 608 | 659 | 735 | 707 | 564 | 589 | 686 | 651 | 655.48 | 669 | 642 | 0.957 | -0.063 | 0.57807171 | 0.72933171 | 0.0113 | 0.006 | 0.0077 | 0.0077 | TRUE | 0.2172 |
| Mtx1 | 3198 | 2212 | 1935 | 2146 | 1718 | 2047 | 2484 | 2353 | 2088 | 2136 | 1938 | 2021 | 2169.9 | 2308 | 2032 | 0.88 | -0.185 | 0.01798555 | 0.06387663 | 0.0043 | 0.0035 | 0.0039 | 0.0039 | TRUE | 0.0839 |
| Mtx2 | 3379 | 2398 | 2302 | 2480 | 2120 | 2385 | 2624 | 2551 | 2484 | 2469 | 2392 | 2354 | 2478.9 | 2553 | 2405 | 0.942 | -0.087 | 0.14325025 | 0.28616015 | 3.00E-04 | 0.0034 | 0.0021 | 0.0021 | TRUE | 0.0075 |
| Mtx3 | 605 | 550 | 459 | 357 | 330 | 370 | 470 | 585 | 495 | 355 | 372 | 365 | 440.51 | 517 | 364 | 0.706 | -0.502 | 1.10E-05 | 0.00020172 | 0.0055 | 0.0078 | 0.007 | 0.007 | TRUE | 0.1545 |
| Muc1 | 34 | 17 | 25 | 29 | 23 | 20 | 26 | 18 | 27 | 29 | 26 | 20 | 24.34 | 24 | 25 | 1.038 | 0.054 | 0.89105473 | 0.93977607 | 0 | 0.0998 | 0.0728 | 0.0728 | TRUE | 0.2458 |
| Muc13 | 14 | 6 | 5 | 7 | 1 | 3 | 11 | 6 | 5 | 7 | 1 | 3 | 5.62 | 7 | 4 | 0.488 | -1.034 | 0.23649006 | NA | 0.1202 | 0.4243 | 0.3479 | 0.3479 | TRUE | 0.9512 |
| Muc15 | 0 | 0 | 0 | 0 | 0 | 0 | 0 | 0 | 0 | 0 | 0 | 0 | 0 | 0 | 0 | NA | NA | NA | NA | NA | NA | NA | NA | NA | NA |
| Muc19 | 0 | 0 | 1 | 0 | 0 | 0 | 0 | 0 | 1 | 0 | 0 | 0 | 0.18 | 0 | 0 | 0.552 | -0.857 | 0.8336548 | NA | 0 | 13.1817 | 10 | 10 | TRUE | 0.5365 |
| Muc2 | 116 | 122 | 109 | 148 | 127 | 171 | 90 | 130 | 118 | 147 | 143 | 169 | 132.81 | 113 | 153 | 1.371 | 0.456 | 0.01652805 | 0.06001916 | 0.0135 | 0.0203 | 0.0183 | 0.0183 | TRUE | 0.3427 |
| Muc20 | 5 | 11 | 5 | 3 | 3 | 5 | 4 | 12 | 5 | 3 | 3 | 5 | 5.38 | 7 | 4 | 0.547 | -0.869 | 0.31477366 | NA | 0 | 0.4429 | 0.332 | 0.332 | TRUE | 0.8818 |
| Muc4 | 46 | 33 | 34 | 38 | 46 | 55 | 36 | 35 | 37 | 38 | 52 | 54 | 41.92 | 36 | 48 | 1.338 | 0.42 | 0.16588334 | 0.31683429 | 0 | 0.059 | 0.0421 | 0.0421 | TRUE | 0.2828 |
| Muc5ac | 4 | 1 | 1 | 2 | 0 | 0 | 3 | 1 | 1 | 2 | 0 | 0 | 1.21 | 2 | 1 | 0.379 | -1.401 | 0.46075088 | NA | 0 | 1.9665 | 1.5636 | 1.5636 | TRUE | 0.9735 |
| Muc5b | 2 | 0 | 0 | 0 | 0 | 0 | 2 | 0 | 0 | 0 | 0 | 0 | 0.26 | 1 | 0 | 0.356 | -1.491 | 0.71362796 | NA | 0 | 9.1573 | 10 | 10 | TRUE | 0.968 |
| Muc6 | 1 | 0 | 0 | 0 | 0 | 0 | 1 | 0 | 0 | 0 | 0 | 0 | 0.13 | 0 | 0 | 0.552 | -0.857 | 0.8336548 | NA | 0 | 18.3121 | 10 | 10 | TRUE | 0.2566 |
| Mucl1 | 0 | 0 | 0 | 0 | 0 | 0 | 0 | 0 | 0 | 0 | 0 | 0 | 0 | 0 | 0 | NA | NA | NA | NA | NA | NA | NA | NA | NA | NA |
| Mug-ps1 | 0 | 0 | 0 | 0 | 0 | 0 | 0 | 0 | 0 | 0 | 0 | 0 | 0 | 0 | 0 | NA | NA | NA | NA | NA | NA | NA | NA | NA | NA |
| Mug1 | 0 | 0 | 0 | 0 | 0 | 0 | 0 | 0 | 0 | 0 | 0 | 0 | 0 | 0 | 0 | NA | NA | NA | NA | NA | NA | NA | NA | NA | NA |
| Mug2 | 0 | 0 | 0 | 0 | 0 | 0 | 0 | 0 | 0 | 0 | 0 | 0 | 0 | 0 | 0 | NA | NA | NA | NA | NA | NA | NA | NA | NA | NA |
| Mul1 | 1855 | 1285 | 1097 | 1210 | 1073 | 1182 | 1441 | 1367 | 1184 | 1204 | 1210 | 1167 | 1262.15 | 1331 | 1194 | 0.896 | -0.158 | 0.06269719 | 0.15838299 | 0.0046 | 0.0043 | 0.0044 | 0.0044 | TRUE | 0.1105 |
| Mum1 | 4171 | 3185 | 3028 | 2810 | 2606 | 2995 | 3239 | 3388 | 3267 | 2797 | 2940 | 2957 | 3097.98 | 3298 | 2898 | 0.879 | -0.187 | 0.00114558 | 0.00799891 | 4.00E-04 | 0.0032 | 0.002 | 0.002 | TRUE | 0.0113 |
| Mum1l1 | 1423 | 990 | 838 | 1041 | 840 | 982 | 1105 | 1053 | 904 | 1036 | 948 | 969 | 1002.61 | 1021 | 984 | 0.963 | -0.054 | 0.55468073 | 0.71014436 | 0.0055 | 0.0048 | 0.005 | 0.005 | TRUE | 0.1176 |
| Mup1 | 0 | 0 | 0 | 0 | 0 | 0 | 0 | 0 | 0 | 0 | 0 | 0 | 0 | 0 | 0 | NA | NA | NA | NA | NA | NA | NA | NA | NA | NA |
| Mup10 | 0 | 0 | 0 | 0 | 0 | 0 | 0 | 0 | 0 | 0 | 0 | 0 | 0 | 0 | 0 | NA | NA | NA | NA | NA | NA | NA | NA | NA | NA |
| Mup11 | 0 | 0 | 0 | 0 | 0 | 0 | 0 | 0 | 0 | 0 | 0 | 0 | 0 | 0 | 0 | NA | NA | NA | NA | NA | NA | NA | NA | NA | NA |
| Mup12 | 0 | 0 | 0 | 0 | 0 | 0 | 0 | 0 | 0 | 0 | 0 | 0 | 0 | 0 | 0 | NA | NA | NA | NA | NA | NA | NA | NA | NA | NA |
| Mup13 | 0 | 0 | 0 | 0 | 0 | 0 | 0 | 0 | 0 | 0 | 0 | 0 | 0 | 0 | 0 | NA | NA | NA | NA | NA | NA | NA | NA | NA | NA |
| Mup14 | 0 | 0 | 0 | 0 | 0 | 0 | 0 | 0 | 0 | 0 | 0 | 0 | 0 | 0 | 0 | NA | NA | NA | NA | NA | NA | NA | NA | NA | NA |
| Mup15 | 0 | 0 | 0 | 0 | 0 | 0 | 0 | 0 | 0 | 0 | 0 | 0 | 0 | 0 | 0 | NA | NA | NA | NA | NA | NA | NA | NA | NA | NA |
| Mup16 | 0 | 0 | 0 | 0 | 0 | 0 | 0 | 0 | 0 | 0 | 0 | 0 | 0 | 0 | 0 | NA | NA | NA | NA | NA | NA | NA | NA | NA | NA |
| Mup17 | 0 | 0 | 0 | 0 | 0 | 0 | 0 | 0 | 0 | 0 | 0 | 0 | 0 | 0 | 0 | NA | NA | NA | NA | NA | NA | NA | NA | NA | NA |
| Mup19 | 0 | 0 | 0 | 0 | 0 | 0 | 0 | 0 | 0 | 0 | 0 | 0 | 0 | 0 | 0 | NA | NA | NA | NA | NA | NA | NA | NA | NA | NA |
| Mup2 | 0 | 0 | 0 | 0 | 0 | 0 | 0 | 0 | 0 | 0 | 0 | 0 | 0 | 0 | 0 | NA | NA | NA | NA | NA | NA | NA | NA | NA | NA |
| Mup20 | 0 | 0 | 0 | 0 | 0 | 0 | 0 | 0 | 0 | 0 | 0 | 0 | 0 | 0 | 0 | NA | NA | NA | NA | NA | NA | NA | NA | NA | NA |
| Mup21 | 0 | 0 | 0 | 0 | 0 | 0 | 0 | 0 | 0 | 0 | 0 | 0 | 0 | 0 | 0 | NA | NA | NA | NA | NA | NA | NA | NA | NA | NA |
| Mup3 | 17 | 13 | 6 | 1 | 3 | 4 | 13 | 14 | 6 | 1 | 3 | 4 | 6.97 | 11 | 3 | 0.246 | -2.022 | 0.01499238 | NA | 0 | 0.3424 | 0.2677 | 0.2677 | TRUE | 0.4673 |
| Mup4 | 0 | 0 | 0 | 0 | 0 | 0 | 0 | 0 | 0 | 0 | 0 | 0 | 0 | 0 | 0 | NA | NA | NA | NA | NA | NA | NA | NA | NA | NA |
| Mup5 | 0 | 0 | 0 | 0 | 0 | 0 | 0 | 0 | 0 | 0 | 0 | 0 | 0 | 0 | 0 | NA | NA | NA | NA | NA | NA | NA | NA | NA | NA |
| Mup6 | 60 | 22 | 37 | 17 | 13 | 27 | 47 | 23 | 40 | 17 | 15 | 27 | 28.03 | 37 | 20 | 0.529 | -0.919 | 0.02609346 | 0.08388525 | 0.074 | 0.087 | 0.0834 | 0.0834 | TRUE | 0.439 |
| Mup7 | 0 | 0 | 0 | 0 | 0 | 0 | 0 | 0 | 0 | 0 | 0 | 0 | 0 | 0 | 0 | NA | NA | NA | NA | NA | NA | NA | NA | NA | NA |
| Mup8 | 0 | 0 | 0 | 0 | 0 | 0 | 0 | 0 | 0 | 0 | 0 | 0 | 0 | 0 | 0 | NA | NA | NA | NA | NA | NA | NA | NA | NA | NA |
| Mup9 | 0 | 1 | 0 | 0 | 0 | 0 | 0 | 1 | 0 | 0 | 0 | 0 | 0.18 | 0 | 0 | 0.552 | -0.857 | 0.8336548 | NA | 0 | 13.3714 | 10 | 10 | TRUE | 0.5216 |
| Murc | 67 | 43 | 36 | 37 | 47 | 45 | 52 | 46 | 39 | 37 | 53 | 44 | 45.15 | 46 | 45 | 0.975 | -0.036 | 0.90208702 | 0.94574851 | 0 | 0.0549 | 0.0413 | 0.0413 | TRUE | 0.1906 |
| Mus81 | 471 | 333 | 249 | 299 | 268 | 296 | 366 | 354 | 269 | 298 | 302 | 292 | 313.47 | 330 | 297 | 0.9 | -0.152 | 0.26498876 | 0.43545523 | 0.0109 | 0.01 | 0.0103 | 0.0103 | TRUE | 0.2883 |
| Musk | 0 | 1 | 0 | 0 | 0 | 0 | 0 | 1 | 0 | 0 | 0 | 0 | 0.18 | 0 | 0 | 0.552 | -0.857 | 0.8336548 | NA | 0 | 13.3714 | 10 | 10 | TRUE | 0.5216 |
| Mustn1 | 3 | 3 | 5 | 8 | 2 | 4 | 2 | 3 | 5 | 8 | 2 | 4 | 4.18 | 3 | 5 | 1.328 | 0.409 | 0.66962972 | NA | 0 | 0.5693 | 0.4194 | 0.4194 | TRUE | 0.5747 |
| Mut | 5073 | 3556 | 3709 | 3942 | 3462 | 4093 | 3940 | 3782 | 4002 | 3924 | 3905 | 4040 | 3932.37 | 3908 | 3956 | 1.012 | 0.018 | 0.7437725 | 0.84754368 | 3.00E-04 | 0.003 | 0.0019 | 0.0019 | TRUE | 0.0095 |
| Mutyh | 744 | 606 | 610 | 699 | 666 | 703 | 578 | 645 | 658 | 696 | 751 | 694 | 670.28 | 627 | 714 | 1.14 | 0.189 | 0.040699 | 0.11594317 | 0.0021 | 0.006 | 0.0047 | 0.0047 | TRUE | 0.0587 |
| Mvb12a | 2315 | 1697 | 2075 | 2180 | 1961 | 2177 | 1798 | 1805 | 2239 | 2170 | 2212 | 2149 | 2062.19 | 1947 | 2177 | 1.119 | 0.162 | 0.06287082 | 0.15868471 | 0.0078 | 0.0036 | 0.005 | 0.005 | TRUE | 0.2056 |
| Mvb12b | 581 | 501 | 617 | 581 | 643 | 709 | 451 | 533 | 666 | 578 | 725 | 700 | 608.91 | 550 | 668 | 1.217 | 0.283 | 0.03863611 | 0.11169617 | 0.0238 | 0.0063 | 0.0119 | 0.0119 | TRUE | 0.3959 |
| Mvd | 5100 | 3777 | 3498 | 4016 | 3466 | 4107 | 3961 | 4018 | 3774 | 3998 | 3910 | 4054 | 3952.43 | 3918 | 3987 | 1.018 | 0.025 | 0.64436017 | 0.77858144 | 4.00E-04 | 0.003 | 0.0019 | 0.0019 | TRUE | 0.0122 |
| Mvk | 1629 | 1247 | 1158 | 1304 | 1061 | 1385 | 1265 | 1326 | 1249 | 1298 | 1197 | 1367 | 1283.87 | 1280 | 1287 | 1.006 | 0.009 | 0.90581292 | 0.94862134 | 0.0019 | 0.0043 | 0.0034 | 0.0034 | TRUE | 0.0445 |
| Mvp | 1763 | 1221 | 1347 | 1467 | 1305 | 1533 | 1369 | 1299 | 1453 | 1460 | 1472 | 1513 | 1427.86 | 1374 | 1482 | 1.079 | 0.11 | 0.12390415 | 0.25812009 | 9.00E-04 | 0.0041 | 0.003 | 0.003 | TRUE | 0.0297 |
| Mx1 | 16 | 10 | 9 | 14 | 8 | 9 | 12 | 11 | 10 | 14 | 9 | 9 | 10.77 | 11 | 11 | 0.969 | -0.046 | 0.93665019 | NA | 0 | 0.2225 | 0.1491 | 0.1491 | TRUE | 0.2779 |
| Mx2 | 13 | 13 | 4 | 4 | 0 | 4 | 10 | 14 | 4 | 4 | 0 | 4 | 6.03 | 9 | 3 | 0.286 | -1.804 | 0.05042433 | NA | 0.2535 | 0.3956 | 0.3701 | 0.3701 | TRUE | 0.3323 |
| Mxd1 | 399 | 308 | 277 | 295 | 276 | 331 | 310 | 328 | 299 | 294 | 311 | 327 | 311.36 | 312 | 311 | 0.995 | -0.007 | 0.95333552 | 0.97433139 | 0 | 0.01 | 0.0068 | 0.0068 | TRUE | 0.0265 |
| Mxd3 | 419 | 310 | 340 | 374 | 303 | 344 | 325 | 330 | 367 | 372 | 342 | 340 | 345.95 | 341 | 351 | 1.033 | 0.047 | 0.68561493 | 0.8072736 | 0 | 0.0093 | 0.0067 | 0.0067 | TRUE | 0.05 |
| Mxd4 | 344 | 229 | 213 | 265 | 272 | 333 | 267 | 244 | 230 | 264 | 307 | 329 | 273.32 | 247 | 300 | 1.211 | 0.276 | 0.04122129 | 0.11680727 | 0.0059 | 0.0111 | 0.0095 | 0.0095 | TRUE | 0.1271 |
| Mxi1 | 2878 | 2246 | 2550 | 2667 | 2443 | 2824 | 2235 | 2389 | 2751 | 2655 | 2756 | 2788 | 2595.68 | 2458 | 2733 | 1.112 | 0.154 | 0.05312077 | 0.14081141 | 0.0056 | 0.0033 | 0.0042 | 0.0042 | TRUE | 0.1311 |
| Mxra7 | 1180 | 919 | 1177 | 1438 | 1251 | 1507 | 916 | 978 | 1270 | 1431 | 1411 | 1488 | 1249.04 | 1055 | 1443 | 1.371 | 0.455 | 2.50E-05 | 0.00039933 | 0.0152 | 0.0043 | 0.0076 | 0.0076 | TRUE | 0.3788 |
| Mxra8 | 352 | 261 | 304 | 361 | 338 | 411 | 273 | 278 | 328 | 359 | 381 | 406 | 337.56 | 293 | 382 | 1.307 | 0.386 | 0.00156888 | 0.00997512 | 0.0038 | 0.0094 | 0.0078 | 0.0078 | TRUE | 0.1194 |
| Myadm | 6225 | 4309 | 4539 | 5332 | 4582 | 5291 | 4835 | 4583 | 4898 | 5307 | 5169 | 5223 | 5002.52 | 4772 | 5233 | 1.097 | 0.133 | 0.01261781 | 0.04916748 | 4.00E-04 | 0.0029 | 0.0018 | 0.0018 | TRUE | 0.0144 |
| Myadml2 | 8 | 8 | 1 | 3 | 2 | 4 | 6 | 9 | 1 | 3 | 2 | 4 | 4.17 | 5 | 3 | 0.581 | -0.784 | 0.43330746 | NA | 0 | 0.5714 | 0.4616 | 0.4616 | TRUE | 0.3826 |
| Myb | 178 | 170 | 229 | 281 | 234 | 328 | 138 | 181 | 247 | 280 | 264 | 324 | 238.94 | 189 | 289 | 1.542 | 0.625 | 0.00084078 | 0.00636848 | 0.0414 | 0.0123 | 0.0208 | 0.0208 | TRUE | 0.5425 |

| | | | | | | | | | | | | | | | | | | | | | | | | | |
|---|---|---|---|---|---|---|---|---|---|---|---|---|---|---|---|---|---|---|---|---|---|---|---|---|---|
| Mybbp1a | 67624 | 48486 | 44563 | 45928 | 41411 | 47841 | 52521 | 51574 | 48083 | 45717 | 46716 | 47226 | 48639.46 | 50726 | 46553 | 0.918 | -0.124 | 0.01605493 | 0.05871879 | 0.0012 | 0.0025 | 0.0019 | 0.0019 | TRUE | 0.0254 |
| Mybl1 | 1207 | 969 | 1079 | 1322 | 1030 | 1198 | 937 | 1031 | 1164 | 1316 | 1162 | 1183 | 1132.14 | 1044 | 1220 | 1.171 | 0.227 | 0.01552303 | 0.05731609 | 0.007 | 0.0045 | 0.0055 | 0.0055 | TRUE | 0.1207 |
| Mybl2 | 22322 | 17930 | 19155 | 17653 | 18166 | 20563 | 17337 | 19072 | 20668 | 17572 | 20493 | 20299 | 19240.05 | 19026 | 19455 | 1.023 | 0.032 | 0.67886826 | 0.80270219 | 0.006 | 0.0025 | 0.0043 | 0.0043 | TRUE | 0.0877 |
| Mybpc1 | 80 | 58 | 53 | 32 | 35 | 36 | 62 | 62 | 57 | 32 | 39 | 36 | 47.98 | 60 | 36 | 0.589 | -0.765 | 0.00640483 | 0.02961942 | 0 | 0.0518 | 0.0341 | 0.0341 | TRUE | 0.0618 |
| Mybpc2 | 45 | 22 | 35 | 46 | 52 | 56 | 35 | 23 | 38 | 46 | 59 | 55 | 42.64 | 32 | 53 | 1.654 | 0.726 | 0.01947746 | 0.06775159 | 0 | 0.058 | 0.0449 | 0.0449 | TRUE | 0.3508 |
| Mybpc3 | 77 | 74 | 55 | 75 | 69 | 72 | 60 | 79 | 59 | 75 | 78 | 71 | 70.24 | 66 | 75 | 1.133 | 0.18 | 0.44769472 | 0.61788255 | 0 | 0.0362 | 0.0262 | 0.0262 | TRUE | 0.2423 |
| Mybph | 38 | 31 | 52 | 49 | 30 | 63 | 30 | 33 | 56 | 49 | 34 | 62 | 43.9 | 40 | 48 | 1.235 | 0.304 | 0.38037292 | 0.55491707 | 0.0821 | 0.0564 | 0.0637 | 0.0637 | TRUE | 0.3342 |
| Mybphl | 5 | 4 | 2 | 1 | 3 | 2 | 4 | 4 | 2 | 1 | 3 | 2 | 2.78 | 3 | 2 | 0.607 | -0.721 | 0.53054353 | NA | 0 | 0.8565 | 0.5643 | 0.5643 | TRUE | 0.2273 |
| Myc | 644 | 431 | 235 | 444 | 338 | 353 | 500 | 458 | 254 | 442 | 381 | 348 | 397.32 | 404 | 390 | 0.966 | -0.05 | 0.87097615 | 0.92839601 | 0.0664 | 0.0084 | 0.0244 | 0.0664 | TRUE | 1.2136 |
| Mycbp | 1501 | 1100 | 975 | 1015 | 891 | 979 | 1166 | 1170 | 1052 | 1010 | 1005 | 966 | 1061.62 | 1129 | 994 | 0.879 | -0.185 | 0.01742939 | 0.06238876 | 0.0012 | 0.0047 | 0.0034 | 0.0034 | TRUE | 0.0418 |
| Mycbp2 | 8505 | 5838 | 5478 | 7171 | 6270 | 6896 | 6605 | 6210 | 5911 | 7138 | 7073 | 6807 | 6624.1 | 6242 | 7006 | 1.122 | 0.166 | 0.00473063 | 0.02350188 | 0.0016 | 0.0028 | 0.0023 | 0.0023 | TRUE | 0.0321 |
| Mycbpap | 16 | 17 | 7 | 8 | 5 | 8 | 12 | 18 | 8 | 8 | 6 | 8 | 9.93 | 13 | 7 | 0.567 | -0.818 | 0.19685855 | NA | 0 | 0.2412 | 0.1795 | 0.1795 | TRUE | 0.1348 |
| Mycl1 | 1012 | 803 | 742 | 713 | 624 | 698 | 786 | 854 | 801 | 710 | 704 | 689 | 757.24 | 814 | 701 | 0.862 | -0.214 | 0.00993286 | 0.04116669 | 0 | 0.0056 | 0.0036 | 0.0036 | TRUE | 0.0223 |
| Mycn | 5414 | 4575 | 5089 | 4665 | 4842 | 5631 | 4205 | 4866 | 5491 | 4644 | 5462 | 5559 | 5037.78 | 4854 | 5222 | 1.076 | 0.106 | 0.26845517 | 0.4390774 | 0.0119 | 0.0029 | 0.0064 | 0.0064 | TRUE | 0.1682 |
| Mycs | 1 | 0 | 1 | 0 | 0 | 0 | 1 | 0 | 1 | 0 | 0 | 0 | 0.31 | 1 | 0 | 0.307 | -1.706 | 0.62894649 | NA | 0 | 7.6657 | 7.2 | 7.2 | TRUE | 0.2067 |
| Myct1 | 0 | 0 | 0 | 0 | 0 | 0 | 0 | 0 | 0 | 0 | 0 | 0 | 0 | 0 | 0 | NA | NA | NA | NA | NA | NA | NA | NA | NA | NA |
| Myd88 | 506 | 338 | 329 | 307 | 259 | 317 | 393 | 360 | 355 | 306 | 292 | 313 | 336.37 | 369 | 304 | 0.821 | -0.284 | 0.01303249 | 0.05027622 | 0 | 0.0095 | 0.0064 | 0.0064 | TRUE | 0.0392 |
| Myef2 | 11200 | 7854 | 7447 | 7215 | 6311 | 7150 | 8699 | 8354 | 8035 | 7182 | 7119 | 7058 | 7741.24 | 8363 | 7120 | 0.851 | -0.232 | 8.27E-06 | 0.00016021 | 6.00E-04 | 0.0027 | 0.0018 | 0.0018 | TRUE | 0.0153 |
| Myeov2 | 1305 | 857 | 843 | 934 | 787 | 861 | 1014 | 912 | 910 | 930 | 888 | 850 | 917.03 | 945 | 889 | 0.94 | -0.09 | 0.28338918 | 0.45526374 | 0.0021 | 0.005 | 0.004 | 0.004 | TRUE | 0.0504 |
| Myf5 | 0 | 0 | 1 | 1 | 0 | 0 | 0 | 0 | 1 | 1 | 0 | 0 | 0.35 | 0 | 0 | 1.075 | 0.105 | 0.97567165 | NA | 0 | 6.8576 | 6.5143 | 6.5143 | TRUE | 0.5365 |
| Myf6 | 0 | 0 | 0 | 3 | 2 | 0 | 0 | 0 | 0 | 3 | 2 | 0 | 0.87 | 0 | 2 | 9.846 | 3.3 | 0.17284784 | NA | 0 | 2.715 | 2.9256 | 2.9256 | TRUE | 0.4506 |
| Myg1 | 2313 | 1631 | 1506 | 1545 | 1385 | 1643 | 1796 | 1735 | 1625 | 1538 | 1562 | 1622 | 1646.41 | 1719 | 1574 | 0.915 | -0.128 | 0.06404762 | 0.16088447 | 0.0011 | 0.0039 | 0.0028 | 0.0028 | TRUE | 0.0272 |
| Myh1 | 3 | 1 | 2 | 0 | 2 | 3 | 2 | 1 | 2 | 0 | 2 | 3 | 1.79 | 2 | 2 | 0.928 | -0.108 | 0.94087722 | NA | 0 | 1.3229 | 0.9746 | 0.9746 | TRUE | 0.6285 |
| Myh10 | 15665 | 10654 | 11167 | 11904 | 10299 | 11856 | 12166 | 11333 | 12049 | 11849 | 11618 | 11704 | 11786.54 | 11849 | 11724 | 0.989 | -0.016 | 0.75823583 | 0.85642898 | 6.00E-04 | 0.0026 | 0.0017 | 0.0017 | TRUE | 0.0177 |
| Myh11 | 88 | 61 | 41 | 51 | 38 | 39 | 68 | 65 | 44 | 51 | 43 | 38 | 51.6 | 59 | 44 | 0.741 | -0.432 | 0.13367641 | 0.27220928 | 0.0184 | 0.0484 | 0.04 | 0.04 | TRUE | 0.3956 |
| Myh13 | 560 | 489 | 357 | 498 | 538 | 574 | 435 | 520 | 385 | 496 | 607 | 567 | 501.59 | 447 | 557 | 1.245 | 0.316 | 0.01161235 | 0.04616838 | 0.0138 | 0.0072 | 0.0093 | 0.0093 | TRUE | 0.2315 |
| Myh14 | 1823 | 1427 | 1546 | 1936 | 1846 | 2095 | 1416 | 1518 | 1668 | 1927 | 2082 | 2068 | 1779.92 | 1534 | 2026 | 1.322 | 0.403 | 1.92E-07 | 6.96E-06 | 0.0035 | 0.0038 | 0.0037 | 0.0037 | TRUE | 0.0703 |
| Myh15 | 1 | 2 | 0 | 0 | 0 | 0 | 1 | 2 | 0 | 0 | 0 | 0 | 0.48 | 1 | 0 | 0.197 | -2.343 | 0.43249452 | NA | 0 | 4.8992 | 4.9172 | 4.9172 | TRUE | 0.1226 |
| Myh2 | 0 | 1 | 5 | 0 | 1 | 1 | 0 | 1 | 5 | 0 | 1 | 1 | 1.43 | 2 | 1 | 0.332 | -1.592 | 0.40020276 | NA | 1.4414 | 1.661 | 1.6069 | 1.6069 | TRUE | 0.9293 |
| Myh3 | 1779 | 1379 | 1273 | 1347 | 1274 | 1412 | 1382 | 1467 | 1374 | 1341 | 1437 | 1394 | 1398.99 | 1408 | 1391 | 0.988 | -0.017 | 0.80563878 | 0.88774921 | 5.00E-04 | 0.0041 | 0.0028 | 0.0028 | TRUE | 0.0161 |
| Myh4 | 17 | 25 | 12 | 14 | 5 | 9 | 13 | 27 | 13 | 14 | 6 | 9 | 13.53 | 18 | 10 | 0.547 | -0.871 | 0.13095392 | NA | 0.1077 | 0.1775 | 0.1579 | 0.1579 | TRUE | 0.9327 |
| Myh6 | 465 | 391 | 352 | 391 | 346 | 395 | 361 | 416 | 380 | 389 | 390 | 390 | 387.72 | 386 | 390 | 1.013 | 0.018 | 0.86613097 | 0.9261042 | 0 | 0.0085 | 0.006 | 0.006 | TRUE | 0.0537 |
| Myh7 | 159 | 126 | 103 | 83 | 60 | 86 | 123 | 134 | 111 | 83 | 68 | 85 | 100.64 | 123 | 79 | 0.639 | -0.645 | 0.00143738 | 0.00934844 | 0 | 0.026 | 0.019 | 0.019 | TRUE | 0.1237 |
| Myh7b | 868 | 799 | 876 | 887 | 911 | 995 | 674 | 850 | 945 | 883 | 1028 | 982 | 893.68 | 823 | 964 | 1.174 | 0.231 | 0.04749443 | 0.12979692 | 0.0165 | 0.0051 | 0.0087 | 0.0087 | TRUE | 0.3085 |
| Myh8 | 12 | 4 | 5 | 4 | 1 | 4 | 9 | 4 | 5 | 4 | 1 | 4 | 4.67 | 6 | 3 | 0.477 | -1.068 | 0.24845455 | NA | 0 | 0.5098 | 0.3727 | 0.3727 | TRUE | 0.5392 |
| Myh9 | 29104 | 22084 | 22874 | 23798 | 22926 | 25305 | 22604 | 23490 | 24681 | 23689 | 25863 | 24980 | 24217.75 | 23592 | 24844 | 1.053 | 0.075 | 0.18600194 | 0.34228524 | 0.0019 | 0.0025 | 0.0023 | 0.0023 | TRUE | 0.0203 |
| Myl1 | 0 | 1 | 0 | 0 | 0 | 0 | 0 | 1 | 0 | 0 | 0 | 0 | 0.18 | 0 | 0 | 0.552 | -0.857 | 0.8336548 | NA | 0 | 13.3714 | 10 | 10 | TRUE | 0.5216 |
| Myl10 | 4 | 3 | 3 | 1 | 8 | 8 | 3 | 3 | 3 | 1 | 9 | 8 | 4.58 | 3 | 6 | 1.87 | 0.903 | 0.35274328 | NA | 0 | 0.5204 | 0.4415 | 0.4415 | TRUE | 1.2794 |
| Myl12a | 7839 | 5102 | 4760 | 4705 | 3918 | 4674 | 6088 | 5427 | 5136 | 4683 | 4420 | 4614 | 5061.4 | 5550 | 4572 | 0.823 | -0.28 | 6.79E-05 | 0.00089831 | 0.004 | 0.0029 | 0.0034 | 0.0034 | TRUE | 0.0885 |
| Myl12b | 4801 | 3395 | 3251 | 3354 | 3106 | 3477 | 3729 | 3611 | 3508 | 3339 | 3504 | 3432 | 3520.42 | 3616 | 3425 | 0.947 | -0.079 | 0.16064626 | 0.31028609 | 5.00E-04 | 0.0031 | 0.002 | 0.002 | TRUE | 0.0093 |
| Myl2 | 10 | 5 | 6 | 16 | 12 | 20 | 8 | 5 | 6 | 16 | 14 | 20 | 11.46 | 6 | 17 | 2.497 | 1.32 | 0.02384508 | NA | 0 | 0.2092 | 0.1413 | 0.1413 | TRUE | 0.1556 |
| Myl3 | 12 | 7 | 24 | 43 | 55 | 46 | 9 | 7 | 26 | 43 | 62 | 45 | 32.15 | 14 | 50 | 3.576 | 1.838 | 2.62E-05 | 0.00041452 | 0.1591 | 0.0761 | 0.0929 | 0.0929 | TRUE | 2.1096 |
| Myl4 | 290 | 218 | 208 | 347 | 284 | 351 | 225 | 232 | 224 | 345 | 320 | 346 | 282.3 | 227 | 337 | 1.487 | 0.572 | 2.46E-06 | 5.92E-05 | 0 | 0.0108 | 0.007 | 0.007 | TRUE | 0.0215 |
| Myl6 | 7910 | 5423 | 5205 | 6048 | 5512 | 6204 | 6143 | 5768 | 5616 | 6020 | 6218 | 6124 | 5981.75 | 5842 | 6121 | 1.047 | 0.066 | 0.23149284 | 0.39754021 | 0.001 | 0.0028 | 0.0021 | 0.0021 | TRUE | 0.0251 |
| Myl6b | 369 | 258 | 264 | 312 | 247 | 322 | 287 | 274 | 285 | 311 | 279 | 318 | 292.16 | 282 | 303 | 1.073 | 0.102 | 0.40214773 | 0.57507729 | 0 | 0.0105 | 0.0072 | 0.0072 | TRUE | 0.0512 |
| Myl7 | 32 | 48 | 28 | 49 | 32 | 40 | 25 | 51 | 30 | 49 | 36 | 39 | 38.41 | 35 | 41 | 1.184 | 0.244 | 0.48609643 | 0.65212843 | 0.0577 | 0.0641 | 0.0623 | 0.0623 | TRUE | 1.0468 |
| Myl9 | 90 | 49 | 31 | 46 | 35 | 43 | 70 | 52 | 33 | 46 | 39 | 42 | 47.2 | 52 | 42 | 0.814 | -0.296 | 0.35376895 | 0.52808559 | 0.0509 | 0.0526 | 0.0521 | 0.0521 | TRUE | 0.1665 |
| Mylip | 491 | 389 | 393 | 418 | 394 | 458 | 381 | 414 | 424 | 416 | 444 | 452 | 421.97 | 406 | 437 | 1.079 | 0.109 | 0.29994419 | 0.47214993 | 0 | 0.008 | 0.0056 | 0.0056 | TRUE | 0.0362 |
| Mylk | 121 | 83 | 65 | 60 | 56 | 76 | 94 | 88 | 70 | 60 | 63 | 75 | 75.05 | 84 | 66 | 0.782 | -0.355 | 0.13068004 | 0.26774796 | 0 | 0.034 | 0.0262 | 0.0262 | TRUE | 0.1909 |
| Mylk2 | 28 | 16 | 13 | 28 | 20 | 25 | 22 | 17 | 14 | 28 | 23 | 25 | 21.32 | 18 | 25 | 1.41 | 0.496 | 0.23621008 | 0.4025937 | 0 | 0.1136 | 0.0785 | 0.0785 | TRUE | 0.2647 |
| Mylk3 | 21 | 9 | 5 | 11 | 6 | 8 | 16 | 10 | 5 | 11 | 7 | 8 | 9.48 | 10 | 9 | 0.807 | -0.31 | 0.636475 | NA | 0.0891 | 0.2524 | 0.2042 | 0.2042 | TRUE | 0.3365 |
| Mylk4 | 2 | 0 | 0 | 1 | 2 | 1 | 2 | 0 | 0 | 1 | 2 | 1 | 0.97 | 1 | 1 | 2.455 | 1.296 | 0.5265091 | NA | 0 | 2.4576 | 1.8045 | 1.8045 | TRUE | 0.9714 |
| Mylpf | 4491 | 2616 | 2079 | 2577 | 1961 | 2370 | 3488 | 2783 | 2243 | 2565 | 2212 | 2340 | 2605.12 | 2838 | 2372 | 0.836 | -0.259 | 0.17967968 | 0.33448315 | 0.0264 | 0.0033 | 0.0103 | 0.0264 | TRUE | 0.2257 |
| Mynn | 2233 | 1620 | 1684 | 1907 | 1563 | 1781 | 1734 | 1723 | 1817 | 1898 | 1763 | 1758 | 1782.34 | 1758 | 1806 | 1.028 | 0.04 | 0.55034696 | 0.7064132 | 8.00E-04 | 0.0038 | 0.0026 | 0.0026 | TRUE | 0.024 |
| Myo10 | 12073 | 9566 | 9694 | 9244 | 8797 | 10708 | 9377 | 10175 | 10460 | 9201 | 9924 | 10570 | 9951.23 | 10004 | 9898 | 0.99 | -0.015 | 0.82370555 | 0.89868745 | 0.0031 | 0.0027 | 0.0032 | 0.0032 | TRUE | 0.0465 |
| Myo15 | 43 | 34 | 29 | 39 | 42 | 40 | 33 | 36 | 31 | 39 | 47 | 39 | 37.76 | 33 | 42 | 1.244 | 0.315 | 0.31203166 | 0.48529581 | 0 | 0.0652 | 0.0431 | 0.0431 | TRUE | 0.0933 |
| Myo16 | 22 | 9 | 26 | 20 | 23 | 14 | 17 | 10 | 28 | 20 | 26 | 14 | 19.06 | 18 | 20 | 1.089 | 0.123 | 0.80360754 | 0.88633094 | 0.1184 | 0.1267 | 0.1245 | 0.1245 | TRUE | 0.3187 |
| Myo18a | 11850 | 9451 | 10268 | 13402 | 12954 | 14774 | 9203 | 10053 | 11079 | 13340 | 14613 | 14584 | 12145.57 | 10112 | 14179 | 1.403 | 0.488 | 1.96E-11 | 1.83E-09 | 0.0055 | 0.0026 | 0.0037 | 0.0037 | TRUE | 0.0856 |
| Myo18b | 199 | 194 | 198 | 276 | 269 | 292 | 155 | 206 | 214 | 275 | 303 | 288 | 240.16 | 192 | 289 | 1.516 | 0.601 | 6.92E-05 | 0.00091049 | 0.0116 | 0.0123 | 0.0121 | 0.0121 | TRUE | 0.3404 |
| Myo19 | 2029 | 1581 | 1384 | 1199 | 1079 | 1290 | 1576 | 1682 | 1493 | 1193 | 1217 | 1273 | 1405.83 | 1584 | 1228 | 0.776 | -0.367 | 7.05E-07 | 2.18E-05 | 0.0016 | 0.0041 | 0.0032 | 0.0032 | TRUE | 0.0343 |
| Myo1a | 164 | 156 | 140 | 153 | 133 | 182 | 127 | 166 | 151 | 152 | 150 | 180 | 154.39 | 148 | 161 | 1.09 | 0.125 | 0.46899951 | 0.63789455 | 0.0082 | 0.0178 | 0.015 | 0.015 | TRUE | 0.1747 |
| Myo1b | 7029 | 5125 | 5171 | 6148 | 5620 | 6205 | 5459 | 5451 | 5579 | 6120 | 6340 | 6125 | 5845.82 | 5496 | 6195 | 1.127 | 0.173 | 0.00050529 | 0.00434542 | 1.00E-04 | 0.0028 | 0.0016 | 0.0016 | TRUE | 0.0051 |
| Myo1c | 7275 | 5495 | 5368 | 6118 | 5541 | 6170 | 5650 | 5845 | 5792 | 6090 | 6251 | 6091 | 5953.1 | 5762 | 6144 | 1.066 | 0.093 | 0.06051113 | 0.15414109 | 1.00E-04 | 0.0028 | 0.0016 | 0.0016 | TRUE | 0.0036 |
| Myo1d | 1217 | 826 | 717 | 748 | 731 | 796 | 945 | 879 | 774 | 745 | 825 | 786 | 825.4 | 866 | 785 | 0.905 | -0.144 | 0.12864213 | 0.26509922 | 0.0053 | 0.0053 | 0.0053 | 0.0053 | TRUE | 0.102 |
| Myo1e | 4366 | 3085 | 3217 | 3493 | 3269 | 3671 | 3391 | 3281 | 3471 | 3477 | 3688 | 3624 | 3488.67 | 3381 | 3596 | 1.063 | 0.089 | 0.11833907 | 0.24976694 | 5.00E-04 | 0.0031 | 0.002 | 0.002 | TRUE | 0.0103 |
| Myo1f | 307 | 266 | 221 | 252 | 231 | 258 | 238 | 283 | 238 | 251 | 261 | 255 | 254.33 | 253 | 256 | 1.01 | 0.014 | 0.91557354 | 0.95339447 | 0 | 0.0117 | 0.0086 | 0.0086 | TRUE | 0.1133 |
| Myo1g | 360 | 242 | 226 | 386 | 358 | 380 | 280 | 257 | 244 | 384 | 404 | 375 | 324.01 | 260 | 388 | 1.485 | 0.57 | 1.29E-06 | 3.60E-05 | 0 | 0.0097 | 0.0068 | 0.0068 | TRUE | 0.0512 |
| Myo1h | 66 | 75 | 64 | 63 | 65 | 76 | 51 | 80 | 69 | 63 | 73 | 75 | 68.53 | 67 | 70 | 1.062 | 0.087 | 0.72921266 | 0.83796724 | 0.0166 | 0.037 | 0.0312 | 0.0312 | TRUE | 0.4212 |
| Myo3a | 7 | 3 | 10 | 8 | 4 | 4 | 5 | 3 | 11 | 8 | 5 | 4 | 5.97 | 6 | 6 | 0.853 | -0.229 | 0.7791946 | NA | 0.0954 | 0.3992 | 0.3098 | 0.3098 | TRUE | 0.5545 |

| | | | | | | | | | | | | | | | | | | | | | | | | |
|---|---|---|---|---|---|---|---|---|---|---|---|---|---|---|---|---|---|---|---|---|---|---|---|---|
| Myo3b | 3 | 5 | 4 | 6 | 5 | 7 | 2 | 5 | 4 | 6 | 6 | 7 | 5.08 | 4 | 6 | 1.581 | 0.661 | 0.43431237 | NA | 0 | 0.4689 | 0.3079 | 0.3079 | TRUE | 0.3097 |
| Myo5a | 7268 | 5114 | 4751 | 5511 | 4941 | 5554 | 5645 | 5440 | 5126 | 5486 | 5574 | 5483 | 5458.83 | 5404 | 5514 | 1.02 | 0.029 | 0.60993234 | 0.75322701 | 9.00E-04 | 0.0029 | 0.0021 | 0.0021 | TRUE | 0.0245 |
| Myo5b | 717 | 539 | 583 | 696 | 661 | 689 | 557 | 573 | 629 | 693 | 746 | 680 | 646.31 | 586 | 706 | 1.205 | 0.269 | 0.00367399 | 0.01930601 | 0.0016 | 0.0061 | 0.0046 | 0.0046 | TRUE | 0.0466 |
| Myo5c | 1195 | 888 | 783 | 924 | 720 | 905 | 928 | 945 | 845 | 920 | 812 | 893 | 890.48 | 906 | 875 | 0.966 | -0.05 | 0.56569626 | 0.72005492 | 0.0026 | 0.0051 | 0.0043 | 0.0043 | TRUE | 0.0459 |
| Myo6 | 1843 | 1360 | 1428 | 1585 | 1403 | 1615 | 1431 | 1447 | 1541 | 1578 | 1583 | 1594 | 1528.91 | 1473 | 1585 | 1.077 | 0.107 | 0.10936235 | 0.23626302 | 0 | 0.004 | 0.0025 | 0.0025 | TRUE | 0.0192 |
| Myo7a | 244 | 171 | 174 | 200 | 222 | 203 | 190 | 182 | 188 | 199 | 250 | 200 | 201.51 | 187 | 216 | 1.159 | 0.213 | 0.15537489 | 0.3030416 | 0.0042 | 0.0142 | 0.0113 | 0.0113 | TRUE | 0.1975 |
| Myo7b | 11 | 12 | 13 | 19 | 10 | 14 | 9 | 13 | 14 | 19 | 11 | 14 | 13.22 | 12 | 15 | 1.264 | 0.338 | 0.52800616 | NA | 0 | 0.1816 | 0.1304 | 0.1304 | TRUE | 0.2933 |
| Myo9a | 3537 | 2587 | 2081 | 2440 | 2070 | 2274 | 2747 | 2752 | 2245 | 2429 | 2335 | 2245 | 2458.82 | 2581 | 2336 | 0.905 | -0.145 | 0.08484599 | 0.19652158 | 0.0065 | 0.0034 | 0.0047 | 0.0047 | TRUE | 0.1557 |
| Myo9b | 8901 | 6797 | 5819 | 5986 | 5430 | 5972 | 6913 | 7230 | 6279 | 5958 | 6126 | 5895 | 6400.15 | 6807 | 5993 | 0.88 | -0.184 | 0.00359754 | 0.01900207 | 0.0025 | 0.0028 | 0.0027 | 0.0027 | TRUE | 0.0558 |
| Myoc | 2 | 0 | 1 | 1 | 0 | 0 | 2 | 0 | 1 | 1 | 0 | 0 | 0.6 | 1 | 0 | 0.414 | -1.271 | 0.61961437 | NA | 0 | 3.9224 | 3.1813 | 3.1813 | TRUE | 0.3906 |
| Myocd | 10 | 3 | 3 | 8 | 5 | 3 | 8 | 3 | 3 | 8 | 6 | 3 | 5.13 | 5 | 6 | 1.138 | 0.187 | 0.82983704 | NA | 0 | 0.4647 | 0.3517 | 0.3517 | TRUE | 0.4122 |
| Myod1 | 7 | 8 | 6 | 12 | 15 | 14 | 5 | 9 | 6 | 12 | 17 | 14 | 10.52 | 7 | 14 | 2.106 | 1.075 | 0.07397532 | NA | 0 | 0.2278 | 0.1527 | 0.1527 | TRUE | 0.1189 |
| Myof | 3593 | 2726 | 2652 | 2881 | 2790 | 3163 | 2791 | 2900 | 2861 | 2868 | 3147 | 3122 | 2948.19 | 2851 | 3046 | 1.069 | 0.096 | 0.12065103 | 0.25281707 | 0.0012 | 0.0032 | 0.0024 | 0.0024 | TRUE | 0.0319 |
| Myog | 9 | 6 | 8 | 8 | 7 | 4 | 7 | 6 | 9 | 8 | 8 | 4 | 6.97 | 7 | 7 | 0.899 | -0.154 | 0.83086589 | NA | 0 | 0.3425 | 0.2305 | 0.2305 | TRUE | 0.3275 |
| Myom1 | 148 | 127 | 76 | 95 | 83 | 70 | 115 | 135 | 82 | 95 | 94 | 69 | 98.22 | 111 | 86 | 0.773 | -0.372 | 0.11227259 | 0.24062261 | 0.0354 | 0.0266 | 0.029 | 0.029 | TRUE | 0.282 |
| Myom2 | 540 | 537 | 654 | 468 | 648 | 655 | 419 | 571 | 706 | 466 | 731 | 647 | 589.95 | 565 | 615 | 1.087 | 0.121 | 0.67249636 | 0.79866975 | 0.0572 | 0.0064 | 0.0206 | 0.0572 | TRUE | 0.2385 |
| Myom3 | 137 | 97 | 80 | 99 | 80 | 83 | 106 | 103 | 86 | 99 | 90 | 82 | 94.44 | 98 | 90 | 0.912 | -0.133 | 0.51970229 | 0.68129 | 0 | 0.0275 | 0.0199 | 0.0199 | TRUE | 0.1109 |
| Myot | 51 | 28 | 29 | 34 | 18 | 18 | 40 | 30 | 31 | 34 | 20 | 18 | 28.77 | 34 | 24 | 0.712 | -0.49 | 0.20302984 | 0.36448021 | 0.0328 | 0.0848 | 0.0709 | 0.0709 | TRUE | 0.7924 |
| Myoz1 | 1 | 1 | 1 | 1 | 3 | 2 | 1 | 1 | 1 | 1 | 3 | 2 | 1.55 | 1 | 2 | 2.182 | 1.126 | 0.46296369 | NA | 0 | 1.5359 | 0.9483 | 0.9483 | TRUE | 0.1866 |
| Myoz2 | 2 | 1 | 1 | 2 | 0 | 0 | 2 | 1 | 1 | 2 | 0 | 0 | 0.95 | 1 | 1 | 0.544 | -0.879 | 0.6666706 | NA | 0 | 2.503 | 1.8451 | 1.8451 | TRUE | 0.9736 |
| Myoz3 | 57 | 67 | 52 | 67 | 47 | 48 | 44 | 71 | 56 | 67 | 53 | 47 | 56.46 | 57 | 56 | 0.981 | -0.028 | 0.92101898 | 0.95655604 | 0.0282 | 0.0444 | 0.0397 | 0.0397 | TRUE | 0.2521 |
| Mypn | 914 | 604 | 522 | 452 | 407 | 460 | 710 | 642 | 563 | 450 | 459 | 454 | 546.45 | 638 | 454 | 0.71 | -0.494 | 3.98E-06 | 8.78E-05 | 0.0056 | 0.0068 | 0.0064 | 0.0064 | TRUE | 0.1239 |
| Mypop | 193 | 138 | 110 | 123 | 85 | 96 | 150 | 147 | 119 | 122 | 96 | 95 | 121.41 | 139 | 104 | 0.753 | -0.41 | 0.03452896 | 0.10284397 | 0.0101 | 0.0219 | 0.0186 | 0.0186 | TRUE | 0.2298 |
| Myrf | 3308 | 2554 | 2554 | 3941 | 3527 | 4282 | 2569 | 2717 | 2756 | 3923 | 3979 | 4227 | 3361.71 | 2681 | 4043 | 1.509 | 0.594 | 1.31E-22 | 5.06E-20 | 0.0012 | 0.0031 | 0.0023 | 0.0023 | TRUE | 0.0194 |
| Myrfl | 20 | 15 | 30 | 18 | 23 | 30 | 16 | 16 | 32 | 18 | 26 | 30 | 22.89 | 21 | 25 | 1.161 | 0.216 | 0.62785886 | 0.76543728 | 0.0791 | 0.106 | 0.0984 | 0.0984 | TRUE | 1.0976 |
| Myrip | 233 | 167 | 146 | 185 | 165 | 173 | 181 | 178 | 158 | 184 | 186 | 171 | 176.2 | 172 | 180 | 1.046 | 0.065 | 0.67007857 | 0.79708707 | 0 | 0.0159 | 0.0108 | 0.0108 | TRUE | 0.0557 |
| Mysm1 | 3541 | 2630 | 2310 | 2561 | 2393 | 2585 | 2750 | 2797 | 2492 | 2549 | 2700 | 2552 | 2640.11 | 2680 | 2600 | 0.97 | -0.044 | 0.50571171 | 0.66914807 | 0.0019 | 0.0033 | 0.0028 | 0.0028 | TRUE | 0.0447 |
| Myt1 | 5 | 4 | 2 | 4 | 5 | 1 | 4 | 4 | 2 | 4 | 6 | 1 | 3.48 | 3 | 4 | 1.014 | 0.021 | 0.984146 | NA | 0 | 0.6827 | 0.4978 | 0.4978 | TRUE | 0.4378 |
| Myt1l | 48 | 46 | 25 | 18 | 13 | 20 | 37 | 49 | 27 | 18 | 15 | 20 | 27.58 | 38 | 18 | 0.464 | -1.109 | 0.00517114 | 0.02518381 | 0 | 0.0883 | 0.0709 | 0.0709 | TRUE | 0.1134 |
| Myzap | 212 | 163 | 168 | 153 | 104 | 142 | 165 | 173 | 181 | 152 | 117 | 140 | 154.85 | 173 | 136 | 0.792 | -0.336 | 0.0432911 | 0.12124421 | 0 | 0.0177 | 0.0134 | 0.0134 | TRUE | 0.1492 |
| Mzb1 | 39 | 32 | 37 | 40 | 39 | 36 | 30 | 34 | 40 | 40 | 44 | 36 | 37.27 | 35 | 40 | 1.149 | 0.2 | 0.52700552 | 0.68806493 | 0 | 0.066 | 0.0453 | 0.0453 | TRUE | 0.116 |
| Mzf1 | 104 | 92 | 64 | 76 | 50 | 63 | 81 | 98 | 69 | 76 | 56 | 62 | 73.66 | 83 | 65 | 0.786 | -0.347 | 0.15174675 | 0.29812687 | 0.0121 | 0.0346 | 0.0285 | 0.0285 | TRUE | 0.2312 |
| Mzt1 | 3957 | 2998 | 3414 | 3797 | 3291 | 3774 | 3073 | 3189 | 3684 | 3780 | 3713 | 3726 | 3527.25 | 3315 | 3740 | 1.129 | 0.174 | 0.01723033 | 0.06190876 | 0.0044 | 0.0031 | 0.0036 | 0.0036 | TRUE | 0.1143 |
| Mzt2 | 723 | 491 | 522 | 529 | 443 | 545 | 562 | 522 | 563 | 527 | 500 | 538 | 535.22 | 549 | 522 | 0.95 | -0.074 | 0.43156281 | 0.60360153 | 0 | 0.0069 | 0.0046 | 0.0046 | TRUE | 0.0206 |
| N28178 | 2 | 2 | 2 | 3 | 1 | 4 | 2 | 2 | 2 | 3 | 1 | 4 | 2.32 | 2 | 3 | 1.407 | 0.493 | 0.68947576 | NA | 0 | 1.0254 | 0.6549 | 0.6549 | TRUE | 0.2633 |
| N4bp1 | 18929 | 14025 | 15349 | 15765 | 14503 | 16742 | 14701 | 14918 | 16561 | 15692 | 16361 | 16527 | 15793.55 | 15393 | 16193 | 1.052 | 0.073 | 0.22136846 | 0.3852634 | 0.0025 | 0.0026 | 0.0025 | 0.0025 | TRUE | 0.0538 |
| N4bp2 | 5028 | 3545 | 3250 | 3651 | 3239 | 3746 | 3905 | 3771 | 3507 | 3634 | 3654 | 3698 | 3694.76 | 3728 | 3662 | 0.982 | -0.026 | 0.66192348 | 0.79124932 | 0.0013 | 0.0031 | 0.0023 | 0.0023 | TRUE | 0.0325 |
| N4bp2l1 | 43 | 32 | 32 | 43 | 42 | 47 | 33 | 34 | 35 | 43 | 47 | 46 | 39.76 | 34 | 45 | 1.34 | 0.422 | 0.16021824 | 0.30962672 | 0 | 0.062 | 0.0395 | 0.0395 | TRUE | 0.0219 |
| N4bp2l2 | 4318 | 3225 | 3350 | 3667 | 3036 | 3630 | 3354 | 3430 | 3615 | 3650 | 3425 | 3583 | 3509.51 | 3466 | 3553 | 1.026 | 0.036 | 0.53771916 | 0.69625259 | 9.00E-04 | 0.0031 | 0.0022 | 0.0022 | TRUE | 0.017 |
| N4bp3 | 3199 | 2802 | 2983 | 3114 | 3005 | 3571 | 2485 | 2980 | 3219 | 3100 | 3390 | 3525 | 3116.39 | 2895 | 3338 | 1.154 | 0.207 | 0.02340693 | 0.07751511 | 0.01 | 0.0032 | 0.0057 | 0.0057 | TRUE | 0.1892 |
| N6amt1 | 723 | 598 | 510 | 486 | 469 | 550 | 562 | 636 | 550 | 484 | 529 | 543 | 550.61 | 583 | 519 | 0.891 | -0.167 | 0.09916206 | 0.21977395 | 0.0031 | 0.0067 | 0.0056 | 0.0056 | TRUE | 0.0735 |
| N6amt2 | 1060 | 671 | 567 | 592 | 509 | 620 | 823 | 714 | 612 | 589 | 574 | 612 | 654.05 | 716 | 592 | 0.825 | -0.278 | 0.01282402 | 0.04972699 | 0.0104 | 0.0061 | 0.0075 | 0.0075 | TRUE | 0.1748 |
| Naa10 | 3468 | 2286 | 2237 | 2436 | 2085 | 2515 | 2693 | 2432 | 2414 | 2425 | 2352 | 2483 | 2466.39 | 2513 | 2420 | 0.962 | -0.055 | 0.40725878 | 0.58012135 | 0.0017 | 0.0034 | 0.0028 | 0.0028 | TRUE | 0.0491 |
| Naa11 | 1157 | 888 | 939 | 795 | 708 | 794 | 899 | 945 | 1013 | 791 | 799 | 784 | 871.69 | 952 | 791 | 0.832 | -0.265 | 0.00121644 | 0.00833887 | 0.001 | 0.0051 | 0.0037 | 0.0037 | TRUE | 0.0373 |
| Naa15 | 14804 | 10656 | 10418 | 10906 | 9243 | 10813 | 11498 | 11335 | 11241 | 10856 | 10427 | 10674 | 11005.03 | 11358 | 10652 | 0.938 | -0.093 | 0.04674683 | 0.12826885 | 1.00E-04 | 0.0026 | 0.0015 | 0.0015 | TRUE | 0.0042 |
| Naa16 | 2186 | 1760 | 1985 | 1995 | 1713 | 2125 | 1698 | 1872 | 2142 | 1986 | 1932 | 2098 | 1954.6 | 1904 | 2005 | 1.055 | 0.077 | 0.37625166 | 0.55083119 | 0.0072 | 0.0036 | 0.0049 | 0.0049 | TRUE | 0.1441 |
| Naa20 | 2880 | 2093 | 2013 | 2080 | 1783 | 2040 | 2237 | 2226 | 2172 | 2070 | 2011 | 2014 | 2121.79 | 2212 | 2032 | 0.919 | -0.123 | 0.03826808 | 0.11085643 | 0 | 0.0035 | 0.002 | 0.002 | TRUE | 0.0034 |
| Naa25 | 8549 | 6041 | 6202 | 6750 | 5905 | 6861 | 6640 | 6426 | 6692 | 6719 | 6661 | 6773 | 6651.76 | 6586 | 6718 | 1.02 | 0.029 | 0.5570099 | 0.71216947 | 1.00E-04 | 0.0028 | 0.0016 | 0.0016 | TRUE | 0.0055 |
| Naa30 | 4538 | 3358 | 3478 | 4021 | 3368 | 3923 | 3524 | 3572 | 3753 | 4002 | 3799 | 3873 | 3753.94 | 3616 | 3891 | 1.077 | 0.106 | 0.05975165 | 0.1528329 | 6.00E-04 | 0.0031 | 0.002 | 0.002 | TRUE | 0.0131 |
| Naa35 | 5408 | 3973 | 3796 | 3529 | 3199 | 3705 | 4200 | 4226 | 4096 | 3513 | 3609 | 3657 | 3883.5 | 4174 | 3593 | 0.861 | -0.216 | 4.41E-05 | 0.0006306 | 0 | 0.003 | 0.0018 | 0.0018 | TRUE | 0.0047 |
| Naa38 | 2988 | 2217 | 2262 | 2252 | 1856 | 2194 | 2321 | 2358 | 2441 | 2242 | 2094 | 2166 | 2270.12 | 2373 | 2167 | 0.914 | -0.13 | 0.03414242 | 0.10204885 | 5.00E-04 | 0.0035 | 0.0023 | 0.0023 | TRUE | 0.011 |
| Naa40 | 5132 | 3848 | 3483 | 3303 | 3116 | 3463 | 3986 | 4093 | 3758 | 3288 | 3515 | 3419 | 3676.42 | 3946 | 3407 | 0.863 | -0.212 | 0.0004022 | 0.00366517 | 0.0011 | 0.0031 | 0.0023 | 0.0023 | TRUE | 0.0207 |
| Naa50 | 20334 | 14610 | 14734 | 15937 | 13305 | 15234 | 15793 | 15540 | 15898 | 15864 | 15009 | 15038 | 15523.71 | 15744 | 15304 | 0.972 | -0.041 | 0.3951963 | 0.56866412 | 5.00E-04 | 0.0026 | 0.0016 | 0.0016 | TRUE | 0.0125 |
| Naa60 | 2098 | 1396 | 1415 | 1469 | 1281 | 1564 | 1629 | 1485 | 1527 | 1462 | 1445 | 1544 | 1515.39 | 1547 | 1484 | 0.958 | -0.061 | 0.38621929 | 0.56003169 | 0.0012 | 0.004 | 0.0029 | 0.0029 | TRUE | 0.0272 |
| Naaa | 527 | 359 | 345 | 384 | 331 | 405 | 409 | 382 | 372 | 382 | 373 | 400 | 386.47 | 388 | 385 | 0.992 | -0.012 | 0.91220624 | 0.95176026 | 0 | 0.0086 | 0.0057 | 0.0057 | TRUE | 0.029 |
| Naalad2 | 66 | 55 | 36 | 19 | 9 | 10 | 51 | 59 | 39 | 19 | 10 | 10 | 31.26 | 50 | 13 | 0.263 | -1.927 | 1.65E-06 | 4.38E-05 | 0.0281 | 0.0783 | 0.0671 | 0.0671 | TRUE | 0.5159 |
| Naaladl1 | 42 | 44 | 37 | 27 | 47 | 33 | 33 | 47 | 40 | 27 | 53 | 33 | 38.64 | 40 | 38 | 0.942 | -0.086 | 0.8039625 | 0.88644882 | 0.0517 | 0.0638 | 0.0604 | 0.0604 | TRUE | 0.9006 |
| Nab1 | 187 | 134 | 158 | 194 | 141 | 156 | 145 | 143 | 170 | 193 | 159 | 154 | 160.74 | 153 | 169 | 1.107 | 0.147 | 0.38164544 | 0.55593746 | 0.0065 | 0.0172 | 0.0141 | 0.0141 | TRUE | 0.173 |
| Nab2 | 393 | 293 | 261 | 283 | 234 | 317 | 305 | 312 | 282 | 282 | 264 | 313 | 292.85 | 300 | 286 | 0.956 | -0.065 | 0.60379459 | 0.74881007 | 0 | 0.0105 | 0.0078 | 0.0078 | TRUE | 0.0756 |
| Nabp1 | 5746 | 4210 | 4203 | 4186 | 3659 | 4068 | 4463 | 4478 | 4535 | 4167 | 4128 | 4016 | 4297.67 | 4492 | 4104 | 0.914 | -0.13 | 0.01095393 | 0.04413998 | 0 | 0.003 | 0.0017 | 0.0017 | TRUE | 0.0043 |
| Nabp2 | 829 | 633 | 637 | 670 | 552 | 722 | 644 | 673 | 687 | 667 | 623 | 713 | 667.81 | 668 | 668 | 1.001 | 0.001 | 0.99158814 | 0.99565418 | 0.0013 | 0.006 | 0.0044 | 0.0044 | TRUE | 0.0416 |
| Naca | 29176 | 20361 | 21563 | 22329 | 19673 | 23287 | 22660 | 21658 | 23266 | 22226 | 22193 | 22988 | 22498.52 | 22528 | 22469 | 0.997 | -0.004 | 0.93949848 | 0.966328 | 6.00E-04 | 0.0025 | 0.0017 | 0.0017 | TRUE | 0.0139 |
| Nacad | 148 | 66 | 85 | 107 | 83 | 96 | 115 | 70 | 92 | 107 | 94 | 95 | 95.3 | 92 | 99 | 1.058 | 0.082 | 0.71671955 | 0.82975813 | 0.0226 | 0.0273 | 0.026 | 0.026 | TRUE | 0.2323 |
| Nacc1 | 19610 | 14670 | 14530 | 16118 | 14529 | 16547 | 15230 | 15604 | 15678 | 16044 | 16390 | 16334 | 15880.13 | 15504 | 16256 | 1.049 | 0.068 | 0.12580263 | 0.26082056 | 1.00E-04 | 0.0026 | 0.0014 | 0.0014 | TRUE | 0.0029 |
| Nacc2 | 842 | 632 | 667 | 761 | 671 | 815 | 654 | 672 | 720 | 757 | 757 | 805 | 727.48 | 682 | 773 | 1.135 | 0.183 | 0.03345179 | 0.10038345 | 0 | 0.0057 | 0.0039 | 0.0039 | TRUE | 0.0272 |
| Nadk | 11510 | 8049 | 8351 | 8984 | 8242 | 9322 | 8939 | 8562 | 9011 | 8943 | 9298 | 9202 | 8992.39 | 8837 | 9148 | 1.035 | 0.05 | 0.31790727 | 0.49181173 | 4.00E-04 | 0.0027 | 0.0017 | 0.0017 | TRUE | 0.009 |
| Nadkd1 | 7986 | 6343 | 7258 | 7836 | 6826 | 7967 | 6202 | 6747 | 7831 | 7800 | 7700 | 7865 | 7357.62 | 6927 | 7788 | 1.125 | 0.17 | 0.03019001 | 0.09309055 | 0.007 | 0.0027 | 0.0043 | 0.0043 | TRUE | 0.1591 |
| Nadsyn1 | 823 | 620 | 590 | 768 | 674 | 770 | 639 | 659 | 637 | 764 | 760 | 760 | 703.37 | 645 | 761 | 1.181 | 0.24 | 0.00359387 | 0.01898969 | 0 | 0.0058 | 0.0035 | 0.0035 | TRUE | 0.0043 |
| Nae1 | 6583 | 4759 | 4986 | 5323 | 4494 | 5125 | 5113 | 5062 | 5380 | 5299 | 5070 | 5059 | 5163.67 | 5185 | 5143 | 0.992 | -0.012 | 0.83112448 | 0.9033735 | 7.00E-04 | 0.0029 | 0.0019 | 0.0019 | TRUE | 0.013 |

| | | | | | | | | | | | | | | | | | | | | | | | | | |
|---|---|---|---|---|---|---|---|---|---|---|---|---|---|---|---|---|---|---|---|---|---|---|---|---|---|
| Naf1 | 5371 | 4144 | 4243 | 4057 | 3626 | 4299 | 4171 | 4408 | 4578 | 4038 | 4091 | 4244 | 4255.02 | 4386 | 4124 | 0.941 | -0.088 | 0.1337225 | 0.27225664 | 0.0011 | 0.003 | 0.0022 | 0.0022 | TRUE | 0.0227 |
| Naga | 716 | 602 | 836 | 986 | 930 | 988 | 556 | 640 | 902 | 981 | 1049 | 975 | 850.73 | 699 | 1002 | 1.435 | 0.521 | 0.00017526 | 0.00192687 | 0.0323 | 0.0052 | 0.0127 | 0.0127 | TRUE | 0.7549 |
| Nagk | 866 | 652 | 602 | 638 | 587 | 652 | 673 | 694 | 650 | 635 | 662 | 644 | 659.42 | 672 | 647 | 0.963 | -0.055 | 0.52110106 | 0.68243518 | 0 | 0.006 | 0.0038 | 0.0038 | TRUE | 0.0094 |
| Naglu | 273 | 178 | 184 | 338 | 321 | 303 | 212 | 189 | 199 | 336 | 362 | 299 | 266.26 | 200 | 332 | 1.658 | 0.729 | 4.51E-08 | 1.93E-06 | 0.0032 | 0.0113 | 0.0088 | 0.0088 | TRUE | 0.0888 |
| Nagpa | 795 | 534 | 464 | 416 | 389 | 488 | 617 | 568 | 501 | 414 | 439 | 482 | 503.46 | 562 | 445 | 0.79 | -0.34 | 0.00238631 | 0.01386258 | 0.0067 | 0.0071 | 0.007 | 0.007 | TRUE | 0.1056 |
| Nags | 141 | 146 | 177 | 240 | 216 | 267 | 110 | 155 | 191 | 239 | 244 | 264 | 200.32 | 152 | 249 | 1.649 | 0.722 | 9.58E-05 | 0.00116869 | 0.0328 | 0.0143 | 0.0194 | 0.0194 | TRUE | 0.4804 |
| Naif1 | 246 | 164 | 153 | 166 | 127 | 134 | 191 | 174 | 165 | 165 | 143 | 132 | 161.9 | 177 | 147 | 0.829 | -0.271 | 0.09813591 | 0.21824375 | 0 | 0.0171 | 0.0131 | 0.0131 | TRUE | 0.1273 |
| Naip1 | 1 | 4 | 1 | 3 | 3 | 0 | 1 | 4 | 1 | 3 | 3 | 0 | 2.08 | 2 | 2 | 1.054 | 0.076 | 0.95777985 | NA | 0.573 | 1.1418 | 0.9961 | 0.9961 | TRUE | 0.7861 |
| Naip2 | 2 | 2 | 0 | 1 | 4 | 0 | 2 | 2 | 0 | 1 | 5 | 0 | 1.53 | 1 | 2 | 1.453 | 0.539 | 0.75353212 | NA | 1.064 | 1.5501 | 1.4453 | 1.4453 | TRUE | 0.9663 |
| Naip5 | 46 | 25 | 8 | 27 | 9 | 12 | 36 | 27 | 9 | 27 | 10 | 12 | 19.97 | 24 | 16 | 0.685 | -0.547 | 0.32885253 | 0.50335074 | 0.3058 | 0.1211 | 0.174 | 0.174 | TRUE | 0.3619 |
| Naip6 | 13 | 5 | 2 | 6 | 6 | 7 | 10 | 5 | 2 | 6 | 7 | 7 | 6.2 | 6 | 7 | 1.088 | 0.122 | 0.87834061 | NA | 0 | 0.3844 | 0.3 | 0.3 | TRUE | 0.4013 |
| Naip7 | 0 | 0 | 2 | 0 | 0 | 0 | 0 | 0 | 2 | 0 | 0 | 0 | 0.36 | 1 | 0 | 0.269 | -1.896 | 0.61680529 | NA | 10 | 6.5921 | 8.6623 | 8.6623 | TRUE | 0.9744 |
| Nalcn | 54 | 36 | 33 | 40 | 38 | 36 | 42 | 38 | 36 | 40 | 43 | 36 | 39.01 | 39 | 40 | 1.015 | 0.022 | 0.94213757 | 0.96773067 | 0 | 0.0632 | 0.0417 | 0.0417 | TRUE | 0.056 |
| Nampt | 13183 | 10207 | 12010 | 12399 | 10790 | 12279 | 10239 | 10857 | 12959 | 12342 | 12172 | 12121 | 11781.65 | 11352 | 12212 | 1.076 | 0.106 | 0.18144814 | 0.33698521 | 0.0074 | 0.0026 | 0.0044 | 0.0044 | TRUE | 0.1873 |
| Nanog | 31506 | 21763 | 20265 | 18336 | 15487 | 16713 | 24469 | 23149 | 21866 | 18252 | 17471 | 16498 | 20284.17 | 23161 | 17407 | 0.751 | -0.412 | 1.73E-11 | 1.64E-09 | 0.0028 | 0.0025 | 0.0027 | 0.0027 | TRUE | 0.03 |
| Nanos1 | 688 | 470 | 461 | 611 | 488 | 551 | 534 | 500 | 497 | 608 | 551 | 544 | 539.05 | 510 | 568 | 1.11 | 0.151 | 0.11982024 | 0.25153801 | 0 | 0.0068 | 0.0049 | 0.0049 | TRUE | 0.0466 |
| Nanos2 | 0 | 1 | 1 | 0 | 0 | 0 | 0 | 1 | 1 | 0 | 0 | 0 | 0.36 | 1 | 0 | 0.271 | -1.881 | 0.5755355 | NA | 0 | 6.6391 | 6.4428 | 6.4428 | TRUE | 0.3513 |
| Nanos3 | 62 | 43 | 16 | 20 | 15 | 15 | 48 | 46 | 17 | 20 | 17 | 15 | 27.13 | 37 | 17 | 0.461 | -1.116 | 0.01229704 | 0.04825963 | 0.1251 | 0.0898 | 0.1001 | 0.1001 | TRUE | 1.4997 |
| Nanp | 19 | 9 | 9 | 13 | 12 | 11 | 15 | 10 | 10 | 13 | 14 | 11 | 11.9 | 12 | 13 | 1.082 | 0.114 | 0.83673443 | NA | 0 | 0.2017 | 0.1359 | 0.1359 | TRUE | 0.3319 |
| Nans | 6295 | 4371 | 4465 | 4861 | 4121 | 4966 | 4889 | 4649 | 4818 | 4839 | 4649 | 4902 | 4790.98 | 4785 | 4797 | 1.002 | 0.003 | 0.95132203 | 0.97323401 | 5.00E-04 | 0.0029 | 0.0019 | 0.0019 | TRUE | 0.0088 |
| Nap1l1 | 47298 | 32752 | 28799 | 31202 | 25732 | 30488 | 36734 | 34838 | 31074 | 31058 | 29028 | 30096 | 32138.21 | 34215 | 30061 | 0.879 | -0.187 | 0.00498992 | 0.02447524 | 0.0041 | 0.0025 | 0.0032 | 0.0032 | TRUE | 0.0789 |
| Nap1l2 | 66 | 49 | 27 | 37 | 20 | 21 | 51 | 52 | 29 | 37 | 23 | 21 | 35.44 | 44 | 27 | 0.603 | -0.729 | 0.04983241 | 0.13451457 | 0.0693 | 0.0693 | 0.0693 | 0.0693 | TRUE | 0.7067 |
| Nap1l3 | 173 | 137 | 112 | 136 | 129 | 126 | 134 | 146 | 121 | 135 | 146 | 124 | 134.37 | 134 | 135 | 1.009 | 0.013 | 0.9389335 | 0.96631456 | 0 | 0.0201 | 0.0144 | 0.0144 | TRUE | 0.0668 |
| Nap1l4 | 10774 | 7760 | 7260 | 7757 | 6627 | 7455 | 8368 | 8254 | 7833 | 7721 | 7476 | 7359 | 7835.31 | 8152 | 7519 | 0.922 | -0.117 | 0.02612997 | 0.08392982 | 7.00E-04 | 0.0027 | 0.0019 | 0.0019 | TRUE | 0.0142 |
| Nap1l5 | 119 | 75 | 54 | 80 | 54 | 81 | 92 | 80 | 58 | 80 | 61 | 80 | 75.16 | 77 | 74 | 0.953 | -0.069 | 0.78051892 | 0.87053897 | 0.0247 | 0.034 | 0.0314 | 0.0314 | TRUE | 0.3652 |
| Napa | 4344 | 2908 | 2672 | 2976 | 2586 | 2799 | 3374 | 3093 | 2883 | 2962 | 2917 | 2763 | 2998.78 | 3117 | 2881 | 0.924 | -0.115 | 0.10765121 | 0.23350461 | 0.0033 | 0.0032 | 0.0033 | 0.0033 | TRUE | 0.0644 |
| Napb | 223 | 177 | 171 | 182 | 152 | 188 | 173 | 188 | 185 | 181 | 171 | 186 | 180.7 | 182 | 179 | 0.988 | -0.017 | 0.90752144 | 0.94923397 | 0 | 0.0155 | 0.0101 | 0.0101 | TRUE | 0.0213 |
| Napepld | 1329 | 1051 | 1115 | 1196 | 1169 | 1330 | 1032 | 1118 | 1203 | 1190 | 1319 | 1313 | 1195.89 | 1118 | 1274 | 1.141 | 0.191 | 0.02261639 | 0.07547423 | 0.0038 | 0.0044 | 0.0042 | 0.0042 | TRUE | 0.056 |
| Napg | 2860 | 1979 | 1975 | 2283 | 1969 | 2207 | 2221 | 2105 | 2131 | 2272 | 2221 | 2179 | 2188.28 | 2152 | 2224 | 1.033 | 0.046 | 0.44317668 | 0.61426729 | 0 | 0.0035 | 0.0022 | 0.0022 | TRUE | 0.0098 |
| Naprt1 | 313 | 244 | 235 | 357 | 261 | 322 | 243 | 260 | 254 | 355 | 294 | 318 | 287.31 | 252 | 322 | 1.282 | 0.359 | 0.00457027 | 0.02291225 | 0 | 0.0107 | 0.008 | 0.008 | TRUE | 0.09 |
| Napsa | 11 | 3 | 12 | 21 | 19 | 20 | 9 | 3 | 13 | 21 | 21 | 20 | 14.46 | 8 | 21 | 2.513 | 1.329 | 0.01599767 | NA | 0 | 0.1663 | 0.1359 | 0.1359 | TRUE | 0.5142 |
| Narf | 3239 | 2373 | 2460 | 2980 | 2636 | 2977 | 2516 | 2524 | 2654 | 2966 | 2974 | 2939 | 2762.13 | 2565 | 2960 | 1.154 | 0.207 | 0.00027673 | 0.00272499 | 0 | 0.0033 | 0.002 | 0.002 | TRUE | 0.0112 |
| Narfl | 870 | 642 | 514 | 501 | 473 | 531 | 676 | 683 | 555 | 499 | 534 | 524 | 578.27 | 638 | 519 | 0.812 | -0.3 | 0.00443262 | 0.0223993 | 0.0057 | 0.0065 | 0.0063 | 0.0063 | TRUE | 0.1489 |
| Narg2 | 1948 | 1448 | 1472 | 1554 | 1305 | 1573 | 1513 | 1540 | 1588 | 1547 | 1472 | 1553 | 1535.54 | 1547 | 1524 | 0.986 | -0.021 | 0.75509536 | 0.85485766 | 0 | 0.004 | 0.0025 | 0.0025 | TRUE | 0.0104 |
| Nars | 14059 | 9543 | 8245 | 8308 | 7056 | 8204 | 10919 | 10151 | 8896 | 8270 | 7960 | 8099 | 9049.06 | 9989 | 8110 | 0.812 | -0.301 | 3.72E-05 | 0.00055438 | 0.0052 | 0.0027 | 0.0037 | 0.0037 | TRUE | 0.1117 |
| Nars2 | 1638 | 1196 | 1193 | 1411 | 1191 | 1326 | 1272 | 1272 | 1287 | 1405 | 1344 | 1309 | 1314.77 | 1277 | 1353 | 1.059 | 0.083 | 0.22901352 | 0.39446736 | 0 | 0.0042 | 0.0026 | 0.0026 | TRUE | 0.0139 |
| Nasp | 43924 | 31839 | 28936 | 29517 | 25404 | 29968 | 34114 | 33867 | 31222 | 29381 | 28658 | 29583 | 31137.49 | 33068 | 29207 | 0.883 | -0.179 | 0.00065981 | 0.00532508 | 0.0013 | 0.0025 | 0.002 | 0.002 | TRUE | 0.0291 |
| Nat1 | 59 | 44 | 52 | 44 | 27 | 24 | 46 | 47 | 56 | 44 | 30 | 24 | 41.11 | 50 | 33 | 0.661 | -0.598 | 0.06540517 | 0.1633472 | 0.0216 | 0.0601 | 0.0503 | 0.0503 | TRUE | 0.6355 |
| Nat10 | 7781 | 6001 | 5554 | 5753 | 5208 | 5890 | 6043 | 6383 | 5993 | 5727 | 5875 | 5814 | 5972.52 | 6140 | 5805 | 0.946 | -0.081 | 0.122237 | 0.25544602 | 4.00E-04 | 0.0028 | 0.0018 | 0.0018 | TRUE | 0.0145 |
| Nat14 | 27 | 21 | 20 | 25 | 23 | 33 | 21 | 22 | 22 | 25 | 26 | 33 | 24.72 | 22 | 28 | 1.289 | 0.366 | 0.33959949 | 0.51360255 | 0 | 0.0983 | 0.0648 | 0.0648 | TRUE | 0.1492 |
| Nat2 | 406 | 273 | 340 | 307 | 250 | 304 | 315 | 290 | 367 | 306 | 282 | 300 | 310.05 | 324 | 296 | 0.914 | -0.13 | 0.30885679 | 0.48156323 | 0.0046 | 0.0101 | 0.0084 | 0.0084 | TRUE | 0.1436 |
| Nat3 | 1 | 0 | 5 | 2 | 2 | 2 | 1 | 0 | 5 | 2 | 2 | 2 | 2.07 | 2 | 2 | 1.031 | 0.044 | 0.97545497 | NA | 0.5428 | 1.1499 | 0.9864 | 0.9864 | TRUE | 1.7146 |
| Nat6 | 288 | 202 | 193 | 231 | 196 | 256 | 224 | 215 | 208 | 230 | 221 | 253 | 225.09 | 216 | 235 | 1.088 | 0.121 | 0.37191505 | 0.54649756 | 0 | 0.013 | 0.0088 | 0.0088 | TRUE | 0.051 |
| Nat8 | 0 | 1 | 2 | 1 | 1 | 3 | 0 | 1 | 2 | 1 | 1 | 3 | 1.38 | 1 | 2 | 1.664 | 0.735 | 0.66159436 | NA | 0 | 1.7144 | 1.2556 | 1.2556 | TRUE | 0.362 |
| Nat8l | 527 | 372 | 342 | 489 | 387 | 423 | 409 | 396 | 369 | 487 | 437 | 418 | 419.15 | 391 | 447 | 1.141 | 0.19 | 0.08309326 | 0.19410862 | 0.0024 | 0.0081 | 0.0063 | 0.0063 | TRUE | 0.0715 |
| Nat9 | 812 | 678 | 630 | 702 | 597 | 732 | 631 | 721 | 680 | 699 | 673 | 723 | 687.74 | 677 | 698 | 1.033 | 0.047 | 0.60091497 | 0.74639037 | 0.0016 | 0.0059 | 0.0044 | 0.0044 | TRUE | 0.0455 |
| Nav1 | 3336 | 2614 | 2578 | 3145 | 3049 | 3417 | 2591 | 2780 | 2782 | 3131 | 3440 | 3373 | 3016.05 | 2718 | 3315 | 1.22 | 0.287 | 7.67E-06 | 0.00015076 | 0.0015 | 0.0032 | 0.0026 | 0.0026 | TRUE | 0.0287 |
| Nav2 | 1794 | 1293 | 1502 | 1524 | 1461 | 1797 | 1393 | 1375 | 1621 | 1517 | 1648 | 1774 | 1554.73 | 1463 | 1646 | 1.126 | 0.171 | 0.05042772 | 0.13568588 | 0.0064 | 0.004 | 0.0049 | 0.0049 | TRUE | 0.1053 |
| Nav3 | 7 | 5 | 3 | 2 | 1 | 0 | 5 | 5 | 3 | 2 | 1 | 0 | 2.85 | 4 | 1 | 0.221 | -2.178 | 0.09170083 | NA | 0 | 0.8335 | 0.5992 | 0.5992 | TRUE | 0.4003 |
| Nbas | 2844 | 2096 | 2030 | 2014 | 1822 | 2072 | 2209 | 2229 | 2190 | 2005 | 2055 | 2045 | 2122.36 | 2209 | 2035 | 0.921 | -0.119 | 0.04224676 | 0.11909921 | 0 | 0.0035 | 0.002 | 0.002 | TRUE | 0.0021 |
| Nbea | 1101 | 691 | 625 | 726 | 618 | 673 | 855 | 735 | 674 | 723 | 697 | 664 | 724.78 | 755 | 695 | 0.918 | -0.123 | 0.2291145 | 0.39449868 | 0.0071 | 0.0057 | 0.0062 | 0.0062 | TRUE | 0.1661 |
| Nbeal1 | 2178 | 1579 | 1512 | 2069 | 1644 | 1962 | 1692 | 1680 | 1631 | 2059 | 1855 | 1937 | 1808.91 | 1668 | 1950 | 1.17 | 0.226 | 0.00078329 | 0.00606152 | 0.001 | 0.0037 | 0.0027 | 0.0027 | TRUE | 0.029 |
| Nbeal2 | 2198 | 1681 | 1491 | 1785 | 1655 | 1819 | 1707 | 1788 | 1609 | 1777 | 1867 | 1796 | 1757.23 | 1701 | 1813 | 1.065 | 0.091 | 0.17962578 | 0.33446994 | 0.0011 | 0.0038 | 0.0028 | 0.0028 | TRUE | 0.0264 |
| Nbl1 | 92 | 58 | 48 | 69 | 67 | 84 | 71 | 62 | 52 | 69 | 76 | 83 | 68.69 | 62 | 76 | 1.221 | 0.288 | 0.23381369 | 0.39994006 | 0 | 0.0369 | 0.0276 | 0.0276 | TRUE | 0.1917 |
| Nbn | 1983 | 1521 | 1470 | 1510 | 1269 | 1547 | 1540 | 1618 | 1586 | 1503 | 1432 | 1527 | 1534.31 | 1581 | 1487 | 0.941 | -0.087 | 0.19131804 | 0.34937075 | 0 | 0.004 | 0.0026 | 0.0026 | TRUE | 0.0127 |
| Nbr1 | 15962 | 11524 | 11205 | 11508 | 10069 | 11477 | 12397 | 12258 | 12090 | 11455 | 11359 | 11330 | 11814.76 | 12248 | 11381 | 0.929 | -0.106 | 0.01773545 | 0.06318364 | 0 | 0.0026 | 0.0014 | 0.0014 | TRUE | 0.0016 |
| Ncald | 51 | 24 | 36 | 26 | 28 | 28 | 40 | 26 | 39 | 26 | 32 | 28 | 31.51 | 35 | 29 | 0.813 | -0.298 | 0.39745614 | 0.57076667 | 0 | 0.0776 | 0.0574 | 0.0574 | TRUE | 0.355 |
| Ncam1 | 1090 | 804 | 911 | 1091 | 1014 | 1075 | 847 | 855 | 983 | 1086 | 1144 | 1061 | 995.97 | 895 | 1097 | 1.227 | 0.295 | 0.00054699 | 0.0046152 | 0.0029 | 0.0048 | 0.0042 | 0.0042 | TRUE | 0.0865 |
| Ncam2 | 2 | 1 | 1 | 3 | 0 | 2 | 2 | 1 | 1 | 3 | 0 | 2 | 1.44 | 1 | 2 | 1.347 | 0.43 | 0.78837588 | NA | 0 | 1.6452 | 1.1526 | 1.1526 | TRUE | 0.3591 |
| Ncan | 355 | 243 | 257 | 406 | 368 | 371 | 276 | 258 | 277 | 404 | 415 | 366 | 332.83 | 270 | 395 | 1.459 | 0.545 | 2.89E-06 | 6.70E-05 | 0 | 0.0095 | 0.0067 | 0.0067 | TRUE | 0.0478 |
| Ncapd2 | 29092 | 21764 | 25014 | 25247 | 22608 | 25906 | 22595 | 23150 | 26990 | 25131 | 25504 | 25573 | 24823.79 | 24245 | 25403 | 1.048 | 0.067 | 0.32759612 | 0.5019575 | 0.0044 | 0.0025 | 0.0034 | 0.0034 | TRUE | 0.1199 |
| Ncapd3 | 7814 | 5654 | 5961 | 6477 | 5781 | 6824 | 6069 | 6014 | 6432 | 6447 | 6522 | 6736 | 6369.98 | 6172 | 6568 | 1.064 | 0.09 | 0.09203093 | 0.20816554 | 7.00E-04 | 0.0028 | 0.0019 | 0.0019 | TRUE | 0.0164 |
| Ncapg | 13326 | 9557 | 10084 | 9949 | 8692 | 9960 | 10350 | 10166 | 10881 | 9903 | 9805 | 9832 | 10156.12 | 10466 | 9847 | 0.941 | -0.088 | 0.07760276 | 0.18492435 | 5.00E-04 | 0.0027 | 0.0017 | 0.0017 | TRUE | 0.0146 |
| Ncapg2 | 6512 | 4530 | 4227 | 4375 | 3700 | 4443 | 5058 | 4819 | 4561 | 4355 | 4174 | 4386 | 4558.63 | 4813 | 4305 | 0.894 | -0.161 | 0.00690217 | 0.03140804 | 0.0014 | 0.0029 | 0.0023 | 0.0023 | TRUE | 0.0254 |
| Ncaph | 6704 | 4845 | 5076 | 5250 | 4583 | 5325 | 5207 | 5154 | 5477 | 5226 | 5170 | 5257 | 5248.3 | 5279 | 5218 | 0.989 | -0.017 | 0.74977946 | 0.85153057 | 3.00E-04 | 0.0029 | 0.0018 | 0.0018 | TRUE | 0.0129 |
| Ncaph2 | 12829 | 9712 | 10351 | 10336 | 9266 | 10292 | 9964 | 10331 | 11169 | 10288 | 10453 | 10160 | 10394.03 | 10488 | 10300 | 0.982 | -0.026 | 0.65247138 | 0.78420564 | 0.0015 | 0.0027 | 0.0023 | 0.0023 | TRUE | 0.0394 |
| Ncbp1 | 8465 | 6222 | 6040 | 5470 | 4927 | 5782 | 6574 | 6618 | 6517 | 5445 | 5558 | 5708 | 6070.08 | 6570 | 5570 | 0.848 | -0.238 | 1.54E-06 | 4.13E-05 | 1.00E-04 | 0.0028 | 0.0016 | 0.0016 | TRUE | 0.0057 |
| Ncbp2 | 5515 | 3997 | 4016 | 4359 | 3668 | 4279 | 4283 | 4252 | 4333 | 4339 | 4138 | 4224 | 4261.49 | 4289 | 4234 | 0.987 | -0.019 | 0.71991525 | 0.83196545 | 0 | 0.003 | 0.0017 | 0.0017 | TRUE | 0.0058 |
| Nccrp1 | 147 | 108 | 149 | 229 | 264 | 282 | 114 | 115 | 161 | 228 | 298 | 278 | 198.99 | 130 | 268 | 2.07 | 1.05 | 2.21E-09 | 1.35E-07 | 0.0216 | 0.0143 | 0.0165 | 0.0165 | TRUE | 0.4254 |

| | | | | | | | | | | | | | | | | | | | | | | | | | |
|---|---|---|---|---|---|---|---|---|---|---|---|---|---|---|---|---|---|---|---|---|---|---|---|---|---|
| Ncdn | 2604 | 1915 | 1494 | 1583 | 1466 | 1643 | 2022 | 2037 | 1612 | 1576 | 1654 | 1622 | 1753.8 | 1890 | 1617 | 0.855 | -0.226 | 0.01247151 | 0.04870363 | 0.0076 | 0.0038 | 0.0053 | 0.0053 | TRUE | 0.1982 |
| Nceh1 | 2385 | 1665 | 1647 | 1881 | 1597 | 1706 | 1852 | 1771 | 1777 | 1872 | 1802 | 1684 | 1793.08 | 1800 | 1786 | 0.992 | -0.012 | 0.85775151 | 0.92033949 | 0.0012 | 0.0037 | 0.0028 | 0.0028 | TRUE | 0.0305 |
| Ncf1 | 20 | 16 | 17 | 10 | 7 | 18 | 16 | 17 | 18 | 10 | 8 | 18 | 14.42 | 17 | 12 | 0.706 | -0.502 | 0.33693173 | NA | 0 | 0.1668 | 0.1248 | 0.1248 | TRUE | 0.7545 |
| Ncf2 | 54 | 51 | 41 | 43 | 58 | 62 | 42 | 54 | 44 | 43 | 65 | 61 | 51.64 | 47 | 56 | 1.208 | 0.273 | 0.33837467 | 0.51249222 | 0.015 | 0.0483 | 0.0391 | 0.0391 | TRUE | 0.386 |
| Ncf4 | 0 | 1 | 2 | 1 | 0 | 1 | 0 | 1 | 2 | 1 | 0 | 1 | 0.87 | 1 | 1 | 0.648 | -0.625 | 0.77012475 | NA | 0 | 2.7349 | 2.0642 | 2.0642 | TRUE | 0.1602 |
| Nck1 | 1800 | 1268 | 1239 | 1534 | 1232 | 1428 | 1398 | 1349 | 1337 | 1527 | 1390 | 1410 | 1401.67 | 1361 | 1442 | 1.059 | 0.083 | 0.24346297 | 0.41122859 | 9.00E-04 | 0.0041 | 0.0029 | 0.0029 | TRUE | 0.032 |
| Nck2 | 845 | 686 | 674 | 682 | 613 | 638 | 656 | 730 | 727 | 679 | 692 | 630 | 685.57 | 704 | 667 | 0.948 | -0.078 | 0.39363287 | 0.5672139 | 0.0018 | 0.0059 | 0.0045 | 0.0045 | TRUE | 0.0448 |
| Nckap1 | 11179 | 8247 | 8700 | 9506 | 8141 | 9553 | 8682 | 8772 | 9387 | 9462 | 9184 | 9430 | 9153.02 | 8947 | 9359 | 1.046 | 0.065 | 0.2171974 | 0.380002 | 9.00E-04 | 0.0027 | 0.0019 | 0.0019 | TRUE | 0.0225 |
| Nckap1l | 1531 | 1023 | 835 | 943 | 783 | 848 | 1189 | 1088 | 901 | 939 | 883 | 837 | 972.87 | 1059 | 886 | 0.835 | -0.26 | 0.01263159 | 0.04919436 | 0.0106 | 0.0049 | 0.0068 | 0.0068 | TRUE | 0.2027 |
| Nckap5 | 75 | 45 | 31 | 20 | 21 | 19 | 58 | 48 | 33 | 20 | 24 | 19 | 33.65 | 46 | 21 | 0.442 | -1.176 | 0.00118994 | 0.00821106 | 0 | 0.0729 | 0.0596 | 0.0596 | TRUE | 0.1544 |
| Nckap5l | 902 | 686 | 556 | 551 | 572 | 610 | 701 | 730 | 600 | 548 | 645 | 602 | 637.68 | 677 | 598 | 0.883 | -0.179 | 0.088119 | 0.20174353 | 0.0068 | 0.0061 | 0.0064 | 0.0064 | TRUE | 0.1128 |
| Nckipsd | 1020 | 803 | 801 | 752 | 739 | 817 | 792 | 854 | 864 | 749 | 834 | 807 | 816.55 | 837 | 797 | 0.952 | -0.071 | 0.40697905 | 0.57994402 | 0.0015 | 0.0053 | 0.004 | 0.004 | TRUE | 0.0328 |
| Ncl | 154973 | 112136 | 101933 | 104555 | 93209 | 109600 | 120361 | 119278 | 109985 | 104074 | 105149 | 108192 | 111173.15 | 116541 | 105805 | 0.908 | -0.139 | 0.00792622 | 0.03484743 | 0.0014 | 0.0024 | 0.002 | 0.002 | TRUE | 0.0297 |
| Ncln | 5159 | 3848 | 3284 | 3497 | 3240 | 3593 | 4007 | 4093 | 3543 | 3481 | 3655 | 3547 | 3721.01 | 3881 | 3561 | 0.917 | -0.125 | 0.06570415 | 0.1637818 | 0.0028 | 0.0031 | 0.003 | 0.003 | TRUE | 0.0697 |
| Ncmap | 118 | 70 | 47 | 75 | 47 | 47 | 92 | 74 | 51 | 75 | 53 | 46 | 65.15 | 72 | 58 | 0.798 | -0.325 | 0.26088168 | 0.4308495 | 0.0591 | 0.0388 | 0.0446 | 0.0446 | TRUE | 0.231 |
| Ncoa1 | 2376 | 1731 | 1687 | 1882 | 1737 | 1952 | 1845 | 1841 | 1820 | 1873 | 1960 | 1927 | 1877.77 | 1835 | 1920 | 1.046 | 0.064 | 0.29141307 | 0.46351736 | 0 | 0.0037 | 0.0021 | 0.0021 | TRUE | 0.0055 |
| Ncoa2 | 3619 | 2670 | 2496 | 2536 | 2379 | 2734 | 2811 | 2840 | 2693 | 2524 | 2684 | 2699 | 2708.48 | 2781 | 2636 | 0.947 | -0.078 | 0.19595462 | 0.35547397 | 7.00E-04 | 0.0033 | 0.0023 | 0.0023 | TRUE | 0.0167 |
| Ncoa3 | 15501 | 10530 | 9903 | 11070 | 9522 | 10952 | 12039 | 11201 | 10685 | 11019 | 10742 | 10811 | 11082.84 | 11308 | 10857 | 0.96 | -0.059 | 0.304551 | 0.47713992 | 0.0018 | 0.0026 | 0.0023 | 0.0023 | TRUE | 0.0393 |
| Ncoa4 | 2706 | 1810 | 1574 | 1685 | 1463 | 1651 | 2102 | 1925 | 1698 | 1677 | 1650 | 1630 | 1780.45 | 1908 | 1652 | 0.865 | -0.209 | 0.01110326 | 0.04455329 | 0.0049 | 0.0038 | 0.0043 | 0.0043 | TRUE | 0.1115 |
| Ncoa5 | 3259 | 2179 | 1604 | 1827 | 1625 | 1840 | 2531 | 2318 | 1731 | 1819 | 1833 | 1816 | 2007.95 | 2193 | 1823 | 0.83 | -0.268 | 0.01547585 | 0.05718623 | 0.0171 | 0.0036 | 0.0083 | 0.0083 | TRUE | 0.4098 |
| Ncoa6 | 2939 | 2264 | 2396 | 2416 | 2117 | 2496 | 2283 | 2408 | 2585 | 2405 | 2388 | 2464 | 2422.18 | 2425 | 2419 | 0.998 | -0.003 | 0.96963042 | 0.9834812 | 0.0017 | 0.0034 | 0.0027 | 0.0027 | TRUE | 0.0402 |
| Ncoa7 | 529 | 377 | 277 | 307 | 287 | 295 | 411 | 401 | 299 | 306 | 324 | 291 | 338.55 | 370 | 307 | 0.826 | -0.275 | 0.04608188 | 0.12698058 | 0.0137 | 0.0094 | 0.0107 | 0.0107 | TRUE | 0.317 |
| Ncor1 | 18123 | 12708 | 11759 | 11935 | 10768 | 12086 | 14075 | 13517 | 12688 | 11880 | 12147 | 11931 | 12706.47 | 13427 | 11986 | 0.893 | -0.164 | 0.00261568 | 0.01484747 | 0.0013 | 0.0026 | 0.0021 | 0.0021 | TRUE | 0.0283 |
| Ncor2 | 8076 | 6098 | 6587 | 6977 | 6965 | 8363 | 6272 | 6486 | 7107 | 6945 | 7857 | 8256 | 7153.95 | 6622 | 7686 | 1.161 | 0.215 | 0.00416175 | 0.02128986 | 0.0058 | 0.0028 | 0.0039 | 0.0039 | TRUE | 0.0871 |
| Ncr1 | 0 | 0 | 0 | 0 | 0 | 0 | 0 | 0 | 0 | 0 | 0 | 0 | 0 | 0 | 0 | NA | NA | NA | NA | NA | NA | NA | NA | NA | NA |
| Ncrna00085 | 46 | 46 | 34 | 45 | 41 | 39 | 36 | 49 | 37 | 45 | 46 | 38 | 41.81 | 41 | 43 | 1.071 | 0.099 | 0.7446423 | 0.84821935 | 0 | 0.0591 | 0.0422 | 0.0422 | TRUE | 0.2426 |
| Ncrna00086 | 740 | 522 | 532 | 567 | 492 | 548 | 575 | 555 | 574 | 564 | 555 | 541 | 560.73 | 568 | 553 | 0.974 | -0.038 | 0.67400197 | 0.79979329 | 0 | 0.0067 | 0.0041 | 0.0041 | TRUE | 0.0047 |
| Ncs1 | 898 | 716 | 655 | 763 | 702 | 778 | 697 | 762 | 707 | 759 | 792 | 768 | 747.53 | 722 | 773 | 1.072 | 0.1 | 0.23434656 | 0.4005639 | 0 | 0.0056 | 0.0037 | 0.0037 | TRUE | 0.0268 |
| Ncstn | 2376 | 1751 | 1598 | 1600 | 1445 | 1674 | 1845 | 1863 | 1724 | 1593 | 1630 | 1652 | 1717.89 | 1811 | 1625 | 0.897 | -0.157 | 0.01739095 | 0.06229794 | 5.00E-04 | 0.0038 | 0.0025 | 0.0025 | TRUE | 0.0206 |
| Nctc1 | 0 | 0 | 0 | 0 | 0 | 0 | 0 | 0 | 0 | 0 | 0 | 0 | 0 | 0 | 0 | NA | NA | NA | NA | NA | NA | NA | NA | NA | NA |
| Ndc80 | 3159 | 2183 | 2329 | 2438 | 2168 | 2546 | 2453 | 2322 | 2513 | 2427 | 2446 | 2513 | 2445.71 | 2429 | 2462 | 1.013 | 0.019 | 0.75454033 | 0.85450004 | 5.00E-04 | 0.0034 | 0.0023 | 0.0023 | TRUE | 0.0177 |
| Nde1 | 4195 | 3122 | 3314 | 3537 | 3055 | 3592 | 3258 | 3321 | 3576 | 3521 | 3446 | 3546 | 3444.6 | 3385 | 3504 | 1.036 | 0.051 | 0.39333481 | 0.56695587 | 9.00E-04 | 0.0031 | 0.0023 | 0.0023 | TRUE | 0.0293 |
| Ndel1 | 3367 | 2470 | 2907 | 3214 | 2862 | 3268 | 2615 | 2627 | 3137 | 3199 | 3229 | 3226 | 3005.47 | 2793 | 3218 | 1.153 | 0.205 | 0.00748183 | 0.03338676 | 0.0052 | 0.0032 | 0.0039 | 0.0039 | TRUE | 0.1395 |
| Ndfip1 | 3258 | 2532 | 2700 | 2970 | 2596 | 2781 | 2530 | 2693 | 2913 | 2956 | 2929 | 2745 | 2794.51 | 2712 | 2877 | 1.061 | 0.086 | 0.21926946 | 0.3823839 | 0.0028 | 0.0033 | 0.0032 | 0.0032 | TRUE | 0.0509 |
| Ndfip2 | 2498 | 1693 | 1753 | 2019 | 1636 | 1895 | 1940 | 1801 | 1891 | 2010 | 1846 | 1871 | 1893.06 | 1877 | 1909 | 1.016 | 0.023 | 0.72955427 | 0.8382924 | 0.0013 | 0.0037 | 0.0028 | 0.0028 | TRUE | 0.0261 |
| Ndn | 725 | 490 | 407 | 308 | 281 | 290 | 563 | 521 | 439 | 307 | 317 | 286 | 405.55 | 508 | 303 | 0.596 | -0.748 | 8.85E-10 | 6.07E-08 | 0.0076 | 0.0083 | 0.0081 | 0.0081 | TRUE | 0.1605 |
| Ndnf | 44 | 32 | 19 | 30 | 22 | 34 | 34 | 34 | 21 | 30 | 25 | 34 | 29.49 | 30 | 30 | 0.991 | -0.013 | 0.97288932 | 0.98522793 | 0 | 0.0828 | 0.0642 | 0.0642 | TRUE | 0.4401 |
| Ndnl2 | 1295 | 968 | 916 | 835 | 772 | 900 | 1006 | 1030 | 988 | 831 | 871 | 888 | 935.71 | 1008 | 863 | 0.857 | -0.223 | 0.00352583 | 0.01869932 | 0 | 0.005 | 0.0031 | 0.0031 | TRUE | 0.013 |
| Ndor1 | 2267 | 1798 | 1444 | 1429 | 1335 | 1493 | 1761 | 1913 | 1558 | 1422 | 1506 | 1474 | 1605.59 | 1744 | 1467 | 0.841 | -0.249 | 0.00260933 | 0.01482319 | 0.0047 | 0.0039 | 0.0043 | 0.0043 | TRUE | 0.1024 |
| Ndp | 0 | 1 | 0 | 0 | 0 | 0 | 0 | 1 | 0 | 0 | 0 | 0 | 0.18 | 0 | 0 | 0.552 | -0.857 | 0.8336548 | NA | 0 | 13.3714 | 10 | 10 | TRUE | 0.5216 |
| Ndrg1 | 4105 | 3475 | 4610 | 5341 | 4693 | 5535 | 3188 | 3696 | 4974 | 5316 | 5294 | 5464 | 4655.53 | 3953 | 5358 | 1.356 | 0.439 | 0.02363826 | 0.07806414 | 0.0269 | 0.0029 | 0.0097 | 0.0269 | TRUE | 0.6214 |
| Ndrg2 | 2416 | 1944 | 2943 | 3649 | 3577 | 4118 | 1876 | 2068 | 3175 | 3632 | 4035 | 4065 | 3142.04 | 2373 | 3911 | 1.648 | 0.721 | 0.00407747 | 0.02094872 | 0.0451 | 0.0032 | 0.0138 | 0.0451 | TRUE | 1.0589 |
| Ndrg3 | 3442 | 2423 | 2792 | 3476 | 3028 | 3509 | 2673 | 2577 | 3013 | 3460 | 3416 | 3464 | 3100.49 | 2754 | 3447 | 1.252 | 0.324 | 2.67E-06 | 6.33E-05 | 0.003 | 0.0032 | 0.0031 | 0.0031 | TRUE | 0.0804 |
| Ndrg4 | 3209 | 2291 | 2092 | 2713 | 2310 | 2610 | 2492 | 2437 | 2257 | 2701 | 2606 | 2576 | 2511.56 | 2395 | 2628 | 1.096 | 0.133 | 0.03738998 | 0.10905974 | 0.0012 | 0.0034 | 0.0025 | 0.0025 | TRUE | 0.0305 |
| Ndst1 | 1453 | 1108 | 1090 | 1109 | 1023 | 1149 | 1128 | 1179 | 1176 | 1104 | 1154 | 1134 | 1145.89 | 1161 | 1131 | 0.974 | -0.038 | 0.59784193 | 0.74375096 | 0 | 0.0045 | 0.0028 | 0.0028 | TRUE | 0.0076 |
| Ndst2 | 637 | 472 | 536 | 557 | 578 | 573 | 495 | 502 | 578 | 554 | 652 | 566 | 557.88 | 525 | 591 | 1.126 | 0.171 | 0.10995367 | 0.23714618 | 0.0059 | 0.0067 | 0.0064 | 0.0064 | TRUE | 0.0948 |
| Ndst3 | 5 | 3 | 5 | 4 | 1 | 2 | 4 | 3 | 5 | 4 | 1 | 2 | 3.26 | 4 | 2 | 0.575 | -0.797 | 0.45520304 | NA | 0 | 0.7297 | 0.4884 | 0.4884 | TRUE | 0.3811 |
| Ndst4 | 0 | 3 | 0 | 1 | 0 | 1 | 0 | 3 | 0 | 1 | 0 | 1 | 0.86 | 1 | 1 | 0.647 | -0.627 | 0.7874777 | NA | 2.2711 | 2.7511 | 2.6648 | 2.6648 | TRUE | 1.4437 |
| Ndufa1 | 1994 | 1352 | 1430 | 1767 | 1423 | 1643 | 1549 | 1438 | 1543 | 1759 | 1605 | 1622 | 1585.96 | 1510 | 1662 | 1.101 | 0.138 | 0.05227082 | 0.1392038 | 0.0015 | 0.0039 | 0.003 | 0.003 | TRUE | 0.0315 |
| Ndufa10 | 5501 | 4011 | 3649 | 4175 | 3638 | 4128 | 4272 | 4266 | 3937 | 4156 | 4104 | 4075 | 4135.15 | 4158 | 4112 | 0.988 | -0.017 | 0.76805378 | 0.86284342 | 9.00E-04 | 0.003 | 0.0021 | 0.0021 | TRUE | 0.0261 |
| Ndufa11 | 4020 | 2721 | 2739 | 3068 | 2722 | 3130 | 3122 | 2894 | 2955 | 3054 | 3071 | 3090 | 3031.03 | 2990 | 3072 | 1.026 | 0.038 | 0.51573983 | 0.67802331 | 5.00E-04 | 0.0032 | 0.0021 | 0.0021 | TRUE | 0.0185 |
| Ndufa12 | 3096 | 2133 | 2228 | 2489 | 2159 | 2590 | 2405 | 2269 | 2404 | 2478 | 2436 | 2557 | 2424.54 | 2359 | 2490 | 1.055 | 0.078 | 0.19848898 | 0.35875101 | 4.00E-04 | 0.0034 | 0.0022 | 0.0022 | TRUE | 0.0134 |
| Ndufa13 | 3009 | 2125 | 2212 | 2292 | 2003 | 2389 | 2337 | 2260 | 2387 | 2281 | 2260 | 2358 | 2313.9 | 2328 | 2300 | 0.988 | -0.017 | 0.77036431 | 0.86413569 | 0 | 0.0035 | 0.0021 | 0.0021 | TRUE | 0.0076 |
| Ndufa2 | 2394 | 1642 | 1579 | 1664 | 1486 | 1717 | 1859 | 1747 | 1704 | 1656 | 1676 | 1695 | 1722.88 | 1770 | 1676 | 0.946 | -0.08 | 0.22622579 | 0.39122144 | 7.00E-04 | 0.0038 | 0.0026 | 0.0026 | TRUE | 0.0245 |
| Ndufa3 | 1903 | 1356 | 1444 | 1684 | 1353 | 1638 | 1478 | 1442 | 1558 | 1676 | 1526 | 1617 | 1549.66 | 1493 | 1606 | 1.077 | 0.107 | 0.12998726 | 0.26674916 | 0.0012 | 0.004 | 0.0029 | 0.0029 | TRUE | 0.0226 |
| Ndufa4 | 7263 | 4882 | 5037 | 5849 | 4529 | 5476 | 5641 | 5193 | 5435 | 5822 | 5109 | 5406 | 5434.27 | 5423 | 5446 | 1.004 | 0.006 | 0.92703566 | 0.95969289 | 0.0028 | 0.0029 | 0.0028 | 0.0028 | TRUE | 0.0446 |
| Ndufa4l2 | 1168 | 935 | 1480 | 2054 | 1921 | 2276 | 907 | 995 | 1597 | 2045 | 2167 | 2247 | 1659.5 | 1166 | 2153 | 1.847 | 0.885 | 0.00105891 | 0.00754488 | 0.052 | 0.0039 | 0.0157 | 0.052 | TRUE | 1.2491 |
| Ndufa5 | 1781 | 1244 | 1170 | 1329 | 1023 | 1289 | 1383 | 1323 | 1262 | 1323 | 1154 | 1272 | 1286.38 | 1323 | 1250 | 0.945 | -0.082 | 0.29596127 | 0.46788789 | 0.0026 | 0.0043 | 0.0037 | 0.0037 | TRUE | 0.0529 |
| Ndufa6 | 3495 | 2668 | 3140 | 3320 | 2986 | 3430 | 2714 | 2838 | 3388 | 3305 | 3369 | 3386 | 3166.59 | 2980 | 3353 | 1.126 | 0.171 | 0.03545078 | 0.10484131 | 0.0068 | 0.0032 | 0.0044 | 0.0044 | TRUE | 0.1733 |
| Ndufa7 | 6572 | 4906 | 5517 | 6011 | 5219 | 6147 | 5104 | 5218 | 5953 | 5983 | 5888 | 6068 | 5702.4 | 5425 | 5980 | 1.103 | 0.141 | 0.03579386 | 0.10561553 | 0.0035 | 0.0028 | 0.0031 | 0.0031 | TRUE | 0.0879 |
| Ndufa8 | 3164 | 2346 | 2316 | 2503 | 2174 | 2604 | 2457 | 2495 | 2499 | 2491 | 2452 | 2571 | 2494.37 | 2484 | 2505 | 1.009 | 0.013 | 0.82593645 | 0.90019138 | 0 | 0.0034 | 0.002 | 0.002 | TRUE | 0.0064 |
| Ndufa9 | 9526 | 6759 | 6848 | 7660 | 6675 | 7672 | 7398 | 7189 | 7389 | 7625 | 7530 | 7573 | 7450.86 | 7325 | 7576 | 1.034 | 0.048 | 0.30454792 | 0.47713992 | 0 | 0.0027 | 0.0015 | 0.0015 | TRUE | 0.0032 |
| Ndufab1 | 2298 | 1543 | 1654 | 1895 | 1525 | 1867 | 1785 | 1641 | 1785 | 1886 | 1720 | 1843 | 1776.72 | 1737 | 1816 | 1.046 | 0.065 | 0.35721239 | 0.53167101 | 0.0017 | 0.0038 | 0.003 | 0.003 | TRUE | 0.0275 |
| Ndufaf1 | 1636 | 1169 | 1223 | 1412 | 1193 | 1348 | 1271 | 1243 | 1320 | 1406 | 1346 | 1331 | 1319.28 | 1278 | 1361 | 1.065 | 0.091 | 0.19027754 | 0.34831415 | 0 | 0.0042 | 0.0027 | 0.0027 | TRUE | 0.0101 |
| Ndufaf2 | 1260 | 839 | 803 | 739 | 763 | 822 | 979 | 892 | 866 | 736 | 861 | 811 | 857.54 | 912 | 803 | 0.878 | -0.188 | 0.03877525 | 0.11196256 | 0.004 | 0.0052 | 0.0048 | 0.0048 | TRUE | 0.0635 |
| Ndufaf3 | 717 | 565 | 555 | 663 | 536 | 705 | 557 | 601 | 599 | 660 | 605 | 696 | 619.54 | 586 | 654 | 1.118 | 0.161 | 0.08737262 | 0.20058862 | 0.0018 | 0.0063 | 0.0048 | 0.0048 | TRUE | 0.0491 |
| Ndufaf4 | 1928 | 1345 | 1216 | 1253 | 1036 | 1186 | 1497 | 1431 | 1312 | 1247 | 1169 | 1171 | 1304.47 | 1413 | 1196 | 0.845 | -0.243 | 0.00167823 | 0.01053442 | 0.0023 | 0.0042 | 0.0035 | 0.0035 | TRUE | 0.0468 |
| Ndufaf5 | 899 | 601 | 627 | 723 | 603 | 745 | 698 | 639 | 677 | 720 | 680 | 735 | 691.56 | 671 | 712 | 1.06 | 0.084 | 0.33762359 | 0.5118428 | 0 | 0.0059 | 0.004 | 0.004 | TRUE | 0.0204 |
| Ndufaf6 | 1097 | 840 | 886 | 1033 | 901 | 1033 | 852 | 893 | 956 | 1028 | 1016 | 1020 | 960.98 | 900 | 1021 | 1.136 | 0.184 | 0.01951614 | 0.06786962 | 7.00E-04 | 0.0049 | 0.0034 | 0.0034 | TRUE | 0.0343 |

| | | | | | | | | | | | | | | | | | | | | | | | | | |
|---|---|---|---|---|---|---|---|---|---|---|---|---|---|---|---|---|---|---|---|---|---|---|---|---|---|
| Ndufaf7 | 1847 | 1344 | 1535 | 1937 | 1719 | 1955 | 1434 | 1430 | 1656 | 1928 | 1939 | 1930 | 1719.59 | 1507 | 1932 | 1.284 | 0.36 | 1.56E-06 | 4.17E-05 | 0.0028 | 0.0038 | 0.0035 | 0.0035 | TRUE | 0.0892 |
| Ndufb10 | 3402 | 2317 | 2324 | 2582 | 2340 | 2688 | 2642 | 2465 | 2508 | 2570 | 2640 | 2653 | 2579.61 | 2538 | 2621 | 1.032 | 0.045 | 0.44564153 | 0.61646566 | 5.00E-04 | 0.0033 | 0.0022 | 0.0022 | TRUE | 0.0161 |
| Ndufb11 | 3463 | 2411 | 2634 | 3068 | 2758 | 3127 | 2690 | 2565 | 2842 | 3054 | 3111 | 3087 | 2891.37 | 2699 | 3084 | 1.143 | 0.193 | 0.001578 | 0.01001524 | 8.00E-04 | 0.0032 | 0.0023 | 0.0023 | TRUE | 0.0255 |
| Ndufb2 | 1515 | 1082 | 1078 | 1347 | 1183 | 1268 | 1177 | 1151 | 1163 | 1341 | 1335 | 1252 | 1236.29 | 1164 | 1309 | 1.124 | 0.169 | 0.01633085 | 0.05949957 | 0 | 0.0043 | 0.0028 | 0.0028 | TRUE | 0.0178 |
| Ndufb3 | 2356 | 1679 | 1627 | 2101 | 1725 | 1927 | 1830 | 1786 | 1756 | 2091 | 1946 | 1902 | 1885.14 | 1791 | 1980 | 1.105 | 0.145 | 0.02913648 | 0.09067009 | 0.001 | 0.0037 | 0.0026 | 0.0026 | TRUE | 0.0296 |
| Ndufb4 | 684 | 554 | 568 | 734 | 611 | 737 | 531 | 589 | 613 | 731 | 689 | 728 | 646.8 | 578 | 716 | 1.242 | 0.313 | 0.00075664 | 0.00589998 | 0.0017 | 0.0061 | 0.0047 | 0.0047 | TRUE | 0.0624 |
| Ndufb5 | 4495 | 3359 | 3586 | 3881 | 3488 | 4128 | 3491 | 3573 | 3869 | 3863 | 3935 | 4075 | 3801.03 | 3644 | 3958 | 1.086 | 0.12 | 0.05117135 | 0.13719604 | 0.0015 | 0.0031 | 0.0024 | 0.0024 | TRUE | 0.0352 |
| Ndufb6 | 3783 | 2562 | 2766 | 3253 | 2735 | 3000 | 2938 | 2725 | 2984 | 3238 | 3085 | 2961 | 2988.77 | 2882 | 3095 | 1.073 | 0.102 | 0.11310484 | 0.24189848 | 0.0017 | 0.0032 | 0.0027 | 0.0027 | TRUE | 0.0273 |
| Ndufb7 | 4325 | 3054 | 3062 | 3317 | 3000 | 3326 | 3359 | 3249 | 3304 | 3302 | 3384 | 3283 | 3313.46 | 3304 | 3323 | 1.005 | 0.008 | 0.88365576 | 0.93547084 | 0 | 0.0031 | 0.0018 | 0.0018 | TRUE | 0.0031 |
| Ndufb8 | 6380 | 4729 | 5276 | 5683 | 4923 | 5553 | 4955 | 5030 | 5693 | 5657 | 5554 | 5482 | 5395.03 | 5226 | 5564 | 1.065 | 0.091 | 0.16245134 | 0.31272783 | 0.0029 | 0.0029 | 0.0029 | 0.0029 | TRUE | 0.074 |
| Ndufb9 | 5494 | 3950 | 4274 | 4654 | 4294 | 4754 | 4267 | 4202 | 4612 | 4633 | 4844 | 4693 | 4541.62 | 4360 | 4723 | 1.083 | 0.116 | 0.04846148 | 0.13170943 | 0.0012 | 0.0029 | 0.0023 | 0.0023 | TRUE | 0.0307 |
| Ndufc1 | 2511 | 1698 | 1754 | 2071 | 1628 | 1899 | 1950 | 1806 | 1893 | 2061 | 1837 | 1875 | 1903.58 | 1883 | 1924 | 1.022 | 0.031 | 0.66278452 | 0.7919394 | 0.0021 | 0.0037 | 0.0031 | 0.0031 | TRUE | 0.0473 |
| Ndufc2 | 3568 | 2594 | 2597 | 2880 | 2476 | 2792 | 2771 | 2759 | 2802 | 2867 | 2793 | 2756 | 2791.42 | 2777 | 2805 | 1.01 | 0.014 | 0.79393524 | 0.87939525 | 0 | 0.0033 | 0.0019 | 0.0019 | TRUE | 0.0045 |
| Ndufs1 | 9780 | 6879 | 6619 | 7110 | 6228 | 6891 | 7596 | 7317 | 7142 | 7077 | 7026 | 6802 | 7160.04 | 7352 | 6968 | 0.948 | -0.078 | 0.13267719 | 0.27075316 | 5.00E-04 | 0.0028 | 0.0018 | 0.0018 | TRUE | 0.0105 |
| Ndufs2 | 9164 | 6740 | 7450 | 7683 | 6854 | 7932 | 7117 | 7169 | 8038 | 7648 | 7732 | 7830 | 7589.13 | 7441 | 7737 | 1.04 | 0.056 | 0.35546371 | 0.53000655 | 0.0022 | 0.0027 | 0.0026 | 0.0026 | TRUE | 0.0598 |
| Ndufs3 | 3028 | 2103 | 2188 | 2393 | 2210 | 2515 | 2352 | 2237 | 2361 | 2382 | 2493 | 2483 | 2384.55 | 2317 | 2453 | 1.058 | 0.082 | 0.17329926 | 0.32587375 | 3.00E-04 | 0.0034 | 0.0022 | 0.0022 | TRUE | 0.0107 |
| Ndufs4 | 3253 | 2352 | 2537 | 2799 | 2431 | 2757 | 2526 | 2502 | 2737 | 2786 | 2742 | 2722 | 2669.3 | 2588 | 2750 | 1.063 | 0.088 | 0.15315041 | 0.3000524 | 8.00E-04 | 0.0033 | 0.0024 | 0.0024 | TRUE | 0.0302 |
| Ndufs5 | 0 | 0 | 0 | 0 | 0 | 0 | 0 | 0 | 0 | 0 | 0 | 0 | 0 | 0 | 0 | NA | NA | NA | NA | NA | NA | NA | NA | NA | NA |
| Ndufs6 | 3720 | 2568 | 2679 | 2943 | 2492 | 2891 | 2889 | 2732 | 2891 | 2929 | 2811 | 2854 | 2850.98 | 2837 | 2865 | 1.01 | 0.014 | 0.81153721 | 0.89135487 | 4.00E-04 | 0.0033 | 0.0021 | 0.0021 | TRUE | 0.0127 |
| Ndufs7 | 2637 | 1878 | 1954 | 2074 | 1840 | 2011 | 2048 | 1998 | 2108 | 2064 | 2076 | 1985 | 2046.56 | 2051 | 2042 | 0.995 | -0.007 | 0.9111258 | 0.95117897 | 0 | 0.0036 | 0.0022 | 0.0022 | TRUE | 0.0072 |
| Ndufs8 | 2780 | 1993 | 2128 | 2308 | 2061 | 2350 | 2159 | 2120 | 2296 | 2297 | 2325 | 2320 | 2252.89 | 2192 | 2314 | 1.056 | 0.079 | 0.19972403 | 0.36043632 | 4.00E-04 | 0.0035 | 0.0023 | 0.0023 | TRUE | 0.0205 |
| Ndufv1 | 8908 | 6644 | 6686 | 7544 | 6621 | 7582 | 6918 | 7067 | 7214 | 7509 | 7469 | 7485 | 7277.13 | 7066 | 7488 | 1.06 | 0.084 | 0.08026802 | 0.18960021 | 1.00E-04 | 0.0028 | 0.0015 | 0.0015 | TRUE | 0.0042 |
| Ndufv2 | 6560 | 4771 | 5255 | 5808 | 4920 | 5870 | 5095 | 5075 | 5670 | 5781 | 5550 | 5795 | 5494.33 | 5280 | 5709 | 1.082 | 0.113 | 0.06592225 | 0.16411964 | 0.0021 | 0.0029 | 0.0025 | 0.0025 | TRUE | 0.0506 |
| Ndufv3 | 3807 | 3004 | 3263 | 3451 | 3203 | 3790 | 2957 | 3195 | 3521 | 3435 | 3613 | 3741 | 3410.43 | 3224 | 3596 | 1.116 | 0.159 | 0.03158669 | 0.09616738 | 0.0043 | 0.0031 | 0.0036 | 0.0036 | TRUE | 0.0785 |
| Neat1 | 5620 | 5931 | 4195 | 4853 | 3913 | 4916 | 4365 | 6309 | 4526 | 4831 | 4414 | 4853 | 4882.95 | 5067 | 4699 | 0.928 | -0.108 | 0.53725257 | 0.69599128 | 0.022 | 0.0029 | 0.0089 | 0.022 | TRUE | 0.5602 |
| Neb | 338 | 271 | 219 | 271 | 251 | 235 | 263 | 288 | 236 | 270 | 283 | 232 | 261.99 | 262 | 262 | 0.996 | -0.006 | 0.9665622 | 0.98188155 | 0.006 | 0.0115 | 0.0099 | 0.0099 | TRUE | 0.1111 |
| Nebl | 32 | 26 | 27 | 28 | 27 | 34 | 25 | 28 | 29 | 28 | 30 | 34 | 28.92 | 27 | 31 | 1.13 | 0.176 | 0.61733968 | 0.75838678 | 0 | 0.0844 | 0.055 | 0.055 | TRUE | 0.0491 |
| Necab1 | 152 | 122 | 87 | 64 | 57 | 67 | 118 | 130 | 94 | 64 | 64 | 66 | 89.31 | 114 | 65 | 0.568 | -0.817 | 0.00018469 | 0.00200923 | 0 | 0.029 | 0.0222 | 0.0222 | TRUE | 0.2229 |
| Necab2 | 1205 | 887 | 810 | 1082 | 886 | 1067 | 936 | 943 | 874 | 1077 | 999 | 1053 | 980.53 | 918 | 1043 | 1.137 | 0.185 | 0.01764952 | 0.06296079 | 5.00E-04 | 0.0048 | 0.0033 | 0.0033 | TRUE | 0.0201 |
| Necab3 | 566 | 467 | 502 | 703 | 660 | 713 | 440 | 497 | 542 | 700 | 745 | 704 | 604.36 | 493 | 716 | 1.457 | 0.543 | 7.68E-08 | 3.08E-06 | 0.0042 | 0.0063 | 0.0057 | 0.0057 | TRUE | 0.1108 |
| Necap1 | 9471 | 6727 | 7180 | 7800 | 6437 | 7716 | 7356 | 7155 | 7747 | 7764 | 7262 | 7617 | 7483.48 | 7419 | 7548 | 1.017 | 0.025 | 0.6544856 | 0.78533835 | 0.0011 | 0.0027 | 0.0021 | 0.0021 | TRUE | 0.018 |
| Necap2 | 2706 | 2037 | 2001 | 2016 | 1917 | 2230 | 2102 | 2167 | 2159 | 2007 | 2163 | 2201 | 2133.01 | 2143 | 2124 | 0.991 | -0.013 | 0.84520577 | 0.91204939 | 9.00E-04 | 0.0035 | 0.0025 | 0.0025 | TRUE | 0.0283 |
| Nedd1 | 3060 | 2258 | 2407 | 2438 | 2055 | 2361 | 2377 | 2402 | 2597 | 2427 | 2318 | 2331 | 2408.54 | 2459 | 2359 | 0.96 | -0.059 | 0.35344849 | 0.52780934 | 0.0011 | 0.0034 | 0.0025 | 0.0025 | TRUE | 0.0291 |
| Nedd4 | 95375 | 69566 | 77682 | 90904 | 81380 | 93259 | 74074 | 73996 | 83818 | 90486 | 91805 | 92061 | 84373.37 | 77296 | 91451 | 1.183 | 0.243 | 4.35E-05 | 0.00062491 | 0.0027 | 0.0025 | 0.0025 | 0.0025 | TRUE | 0.0667 |
| Nedd4l | 3375 | 2560 | 2694 | 2736 | 2531 | 2848 | 2621 | 2723 | 2907 | 2723 | 2855 | 2811 | 2773.52 | 2750 | 2796 | 1.017 | 0.025 | 0.69353293 | 0.81291342 | 0.0013 | 0.0033 | 0.0025 | 0.0025 | TRUE | 0.0299 |
| Nedd8 | 4947 | 3273 | 3035 | 3222 | 2841 | 3379 | 3842 | 3481 | 3275 | 3207 | 3205 | 3336 | 3391 | 3533 | 3249 | 0.919 | -0.122 | 0.08055752 | 0.19006416 | 0.0031 | 0.0031 | 0.0032 | 0.0032 | TRUE | 0.0725 |
| Nedd9 | 180 | 102 | 80 | 82 | 85 | 80 | 140 | 108 | 86 | 82 | 96 | 79 | 98.52 | 111 | 86 | 0.76 | -0.395 | 0.08081941 | 0.19046197 | 0.0269 | 0.0265 | 0.0266 | 0.0266 | TRUE | 0.2491 |
| Nefh | 3419 | 2356 | 1448 | 1669 | 1483 | 1564 | 2655 | 2506 | 1562 | 1661 | 1673 | 1544 | 1933.67 | 2241 | 1626 | 0.725 | -0.463 | 0.05063053 | 0.13610347 | 0.0399 | 0.0037 | 0.0139 | 0.0399 | TRUE | 0.8488 |
| Nefl | 944 | 712 | 671 | 878 | 699 | 784 | 733 | 757 | 724 | 874 | 789 | 774 | 775.16 | 738 | 812 | 1.101 | 0.138 | 0.1077859 | 0.23352795 | 0.0012 | 0.0055 | 0.004 | 0.004 | TRUE | 0.0533 |
| Nefm | 153 | 136 | 92 | 138 | 99 | 138 | 119 | 145 | 99 | 137 | 112 | 136 | 124.67 | 121 | 128 | 1.064 | 0.09 | 0.64727143 | 0.78051363 | 0.0148 | 0.0214 | 0.0196 | 0.0196 | TRUE | 0.2688 |
| Negr1 | 7 | 3 | 3 | 5 | 2 | 5 | 5 | 3 | 3 | 5 | 2 | 5 | 4.01 | 4 | 4 | 1.018 | 0.026 | 0.97808966 | NA | 0 | 0.5941 | 0.3942 | 0.3942 | TRUE | 0.2364 |
| Neil1 | 280 | 218 | 212 | 254 | 244 | 243 | 217 | 232 | 229 | 253 | 275 | 240 | 241.01 | 226 | 256 | 1.133 | 0.18 | 0.1723903 | 0.32459157 | 0 | 0.0123 | 0.0084 | 0.0084 | TRUE | 0.0466 |
| Neil2 | 399 | 376 | 485 | 443 | 491 | 538 | 310 | 400 | 523 | 441 | 554 | 531 | 459.85 | 411 | 509 | 1.241 | 0.311 | 0.05440001 | 0.1429826 | 0.0376 | 0.0076 | 0.0167 | 0.0167 | TRUE | 0.3563 |
| Neil3 | 964 | 707 | 723 | 834 | 700 | 800 | 749 | 752 | 780 | 830 | 790 | 790 | 781.73 | 760 | 803 | 1.057 | 0.08 | 0.32087356 | 0.49474065 | 0 | 0.0055 | 0.0034 | 0.0034 | TRUE | 0.0104 |
| Nek1 | 4710 | 3417 | 3535 | 4053 | 3443 | 4015 | 3658 | 3635 | 3814 | 4034 | 3884 | 3963 | 3831.46 | 3702 | 3960 | 1.07 | 0.098 | 0.07117806 | 0.17357925 | 2.00E-04 | 0.003 | 0.0019 | 0.0019 | TRUE | 0.0084 |
| Nek10 | 85 | 65 | 57 | 33 | 30 | 32 | 66 | 69 | 62 | 33 | 34 | 32 | 49.16 | 66 | 33 | 0.499 | -1.002 | 0.0003226 | 0.00307394 | 0 | 0.0506 | 0.0328 | 0.0328 | TRUE | 0.0239 |
| Nek11 | 1 | 4 | 9 | 2 | 3 | 8 | 1 | 4 | 10 | 2 | 3 | 8 | 4.67 | 5 | 4 | 0.927 | -0.11 | 0.9141909 | NA | 0.5763 | 0.51 | 0.5301 | 0.5301 | TRUE | 0.3251 |
| Nek2 | 8855 | 6306 | 6581 | 6894 | 6168 | 7051 | 6877 | 6708 | 7101 | 6862 | 6958 | 6960 | 6911.1 | 6895 | 6927 | 1.005 | 0.007 | 0.89387312 | 0.94181872 | 2.00E-04 | 0.0028 | 0.0016 | 0.0016 | TRUE | 0.0082 |
| Nek3 | 2302 | 1701 | 1617 | 1396 | 1283 | 1548 | 1788 | 1809 | 1745 | 1390 | 1447 | 1528 | 1617.83 | 1781 | 1455 | 0.817 | -0.291 | 1.69E-05 | 0.00029185 | 6.00E-04 | 0.0039 | 0.0027 | 0.0027 | TRUE | 0.0237 |
| Nek4 | 1712 | 1284 | 1208 | 1200 | 996 | 1159 | 1330 | 1366 | 1303 | 1194 | 1124 | 1144 | 1243.5 | 1333 | 1154 | 0.866 | -0.207 | 0.0031188 | 0.01702061 | 0 | 0.0043 | 0.0027 | 0.0027 | TRUE | 0.0113 |
| Nek5 | 15 | 8 | 15 | 13 | 12 | 17 | 12 | 9 | 16 | 13 | 14 | 17 | 13.27 | 12 | 15 | 1.194 | 0.256 | 0.62928042 | NA | 0 | 0.1811 | 0.1268 | 0.1268 | TRUE | 0.3052 |
| Nek6 | 2101 | 1481 | 1658 | 2170 | 1926 | 2219 | 1632 | 1575 | 1789 | 2160 | 2173 | 2190 | 1919.88 | 1665 | 2174 | 1.306 | 0.385 | 1.92E-08 | 8.93E-07 | 0.0014 | 0.0037 | 0.0029 | 0.0029 | TRUE | 0.0495 |
| Nek7 | 1779 | 1282 | 1419 | 1485 | 1323 | 1443 | 1382 | 1364 | 1531 | 1478 | 1492 | 1424 | 1445.25 | 1426 | 1465 | 1.028 | 0.04 | 0.5833858 | 0.7331692 | 0.0016 | 0.0041 | 0.0032 | 0.0032 | TRUE | 0.0494 |
| Nek8 | 955 | 848 | 806 | 885 | 886 | 901 | 742 | 902 | 870 | 881 | 999 | 889 | 880.54 | 838 | 923 | 1.104 | 0.142 | 0.14367314 | 0.28660435 | 0.0068 | 0.0051 | 0.0057 | 0.0057 | TRUE | 0.1253 |
| Nek9 | 6651 | 4804 | 4995 | 5994 | 5394 | 5800 | 5166 | 5110 | 5390 | 5966 | 6085 | 5725 | 5573.66 | 5222 | 5925 | 1.135 | 0.182 | 0.00070551 | 0.005599 | 6.00E-04 | 0.0029 | 0.0019 | 0.0019 | TRUE | 0.0107 |
| Nelfa | 5567 | 4310 | 4111 | 3801 | 3474 | 4273 | 4324 | 4584 | 4436 | 3784 | 3919 | 4218 | 4210.76 | 4448 | 3974 | 0.894 | -0.162 | 0.00823959 | 0.03580676 | 0.0016 | 0.003 | 0.0025 | 0.0025 | TRUE | 0.0355 |
| Nelfb | 5575 | 4069 | 4222 | 4258 | 3950 | 4627 | 4330 | 4328 | 4556 | 4238 | 4456 | 4568 | 4412.58 | 4405 | 4421 | 1.004 | 0.006 | 0.92296347 | 0.95719768 | 9.00E-04 | 0.003 | 0.0021 | 0.0021 | TRUE | 0.016 |
| Nelfcd | 4673 | 3584 | 3344 | 3508 | 3198 | 3761 | 3629 | 3812 | 3608 | 3492 | 3608 | 3713 | 3643.66 | 3683 | 3604 | 0.979 | -0.031 | 0.58477321 | 0.734344 | 7.00E-04 | 0.0031 | 0.0021 | 0.0021 | TRUE | 0.0113 |
| Nelfe | 3757 | 2772 | 2631 | 2741 | 2608 | 2959 | 2918 | 2949 | 2839 | 2728 | 2942 | 2921 | 2882.79 | 2902 | 2864 | 0.987 | -0.019 | 0.74547415 | 0.84872461 | 7.00E-04 | 0.0032 | 0.0022 | 0.0022 | TRUE | 0.0209 |
| Nell1 | 15 | 11 | 15 | 8 | 13 | 12 | 12 | 12 | 16 | 8 | 15 | 12 | 12.33 | 13 | 12 | 0.872 | -0.198 | 0.71957271 | NA | 0 | 0.1946 | 0.1363 | 0.1363 | TRUE | 0.2812 |
| Nell2 | 1006 | 727 | 667 | 915 | 901 | 868 | 781 | 773 | 720 | 911 | 1016 | 857 | 843.06 | 758 | 928 | 1.222 | 0.29 | 0.00135166 | 0.00895498 | 0.0036 | 0.0052 | 0.0047 | 0.0047 | TRUE | 0.0813 |
| Nemf | 5426 | 3991 | 3859 | 4237 | 3593 | 4202 | 4214 | 4245 | 4164 | 4218 | 4053 | 4148 | 4173.66 | 4208 | 4140 | 0.984 | -0.023 | 0.65040098 | 0.78296818 | 0 | 0.003 | 0.0017 | 0.0017 | TRUE | 0.004 |
| Nenf | 401 | 341 | 356 | 379 | 320 | 377 | 311 | 363 | 384 | 377 | 361 | 372 | 361.45 | 353 | 370 | 1.053 | 0.075 | 0.52813003 | 0.68874916 | 0.0037 | 0.009 | 0.0073 | 0.0073 | TRUE | 0.1296 |
| Neo1 | 5854 | 4081 | 4218 | 5216 | 4490 | 5011 | 4547 | 4341 | 4551 | 5192 | 5065 | 4947 | 4773.74 | 4480 | 5068 | 1.131 | 0.178 | 0.0008681 | 0.00653727 | 4.00E-04 | 0.0029 | 0.0018 | 0.0018 | TRUE | 0.0088 |
| Nepn | 1 | 0 | 1 | 1 | 1 | 2 | 1 | 0 | 1 | 1 | 1 | 2 | 0.99 | 1 | 1 | 2.189 | 1.13 | 0.55451068 | NA | 0 | 2.391 | 1.4884 | 1.4884 | TRUE | 0.1946 |
| Nes | 1892 | 1276 | 1073 | 1010 | 894 | 1098 | 1469 | 1357 | 1158 | 1005 | 1009 | 1084 | 1180.37 | 1328 | 1033 | 0.777 | -0.365 | 0.00010251 | 0.00123058 | 0.0075 | 0.0044 | 0.0055 | 0.0055 | TRUE | 0.15 |
| Nespas | 147 | 124 | 93 | 138 | 101 | 115 | 114 | 132 | 100 | 137 | 114 | 114 | 118.54 | 115 | 122 | 1.054 | 0.076 | 0.6908478 | 0.81102978 | 0.0062 | 0.0224 | 0.0179 | 0.0179 | TRUE | 0.1453 |
| Net1 | 4882 | 3544 | 3612 | 3216 | 2829 | 3293 | 3792 | 3770 | 3897 | 3201 | 3191 | 3251 | 3517 | 3820 | 3214 | 0.842 | -0.249 | 2.73E-06 | 6.42E-05 | 0 | 0.0031 | 0.0017 | 0.0017 | TRUE | 0.0038 |
| Neto1 | 0 | 3 | 0 | 0 | 1 | 1 | 0 | 3 | 0 | 0 | 1 | 1 | 0.88 | 1 | 1 | 0.674 | -0.569 | 0.80551363 | NA | 2.4361 | 2.6823 | 2.6397 | 2.6397 | TRUE | 1.4437 |

| | | | | | | | | | | | | | | | | | | | | | | | | |
|---|---|---|---|---|---|---|---|---|---|---|---|---|---|---|---|---|---|---|---|---|---|---|---|---|
| Neto2 | 652 | 403 | 224 | 253 | 177 | 196 | 506 | 429 | 242 | 252 | 200 | 193 | 303.62 | 392 | 215 | 0.547 | -0.87 | 4.32E-05 | 0.00062047 | 0.0768 | 0.0102 | 0.029 | 0.029 | TRUE | 0.4111 |
| Neu1 | 855 | 712 | 838 | 1028 | 896 | 1090 | 664 | 757 | 904 | 1023 | 1011 | 1076 | 905.94 | 775 | 1037 | 1.341 | 0.423 | 7.24E-05 | 0.00093961 | 0.0111 | 0.005 | 0.0071 | 0.0071 | TRUE | 0.2505 |
| Neu2 | 2 | 3 | 2 | 4 | 3 | 3 | 2 | 3 | 2 | 4 | 3 | 3 | 2.87 | 2 | 3 | 1.519 | 0.604 | 0.58419498 | NA | 0 | 0.8278 | 0.5135 | 0.5135 | TRUE | 0.1173 |
| Neu3 | 598 | 378 | 455 | 521 | 397 | 492 | 464 | 402 | 491 | 519 | 448 | 486 | 468.27 | 452 | 484 | 1.07 | 0.098 | 0.38241965 | 0.55640702 | 0.0054 | 0.0075 | 0.0069 | 0.0069 | TRUE | 0.1059 |
| Neu4 | 1 | 0 | 0 | 0 | 0 | 0 | 1 | 0 | 0 | 0 | 0 | 0 | 0.13 | 0 | 0 | 0.552 | -0.857 | 0.8336548 | NA | 0 | 18.3121 | 10 | 10 | TRUE | 0.2566 |
| Neurl1a | 1613 | 1300 | 1623 | 1824 | 1634 | 1863 | 1253 | 1383 | 1751 | 1816 | 1843 | 1839 | 1647.46 | 1462 | 1833 | 1.255 | 0.328 | 0.00171177 | 0.01071673 | 0.0147 | 0.0039 | 0.0072 | 0.0072 | TRUE | 0.3584 |
| Neurl1b | 314 | 240 | 294 | 385 | 307 | 343 | 244 | 255 | 317 | 383 | 346 | 339 | 314.09 | 272 | 356 | 1.313 | 0.393 | 0.00325906 | 0.01759878 | 0.0087 | 0.01 | 0.0096 | 0.0096 | TRUE | 0.2301 |
| Neurl2 | 82 | 69 | 71 | 71 | 92 | 79 | 64 | 73 | 77 | 71 | 104 | 78 | 77.69 | 71 | 84 | 1.182 | 0.241 | 0.3072407 | 0.47991901 | 0.013 | 0.0329 | 0.0273 | 0.0273 | TRUE | 0.374 |
| Neurl3 | 32 | 32 | 24 | 32 | 19 | 16 | 25 | 34 | 26 | 32 | 21 | 16 | 25.64 | 28 | 23 | 0.818 | -0.289 | 0.47484549 | 0.64299637 | 0.0339 | 0.0948 | 0.0784 | 0.0784 | TRUE | 0.5481 |
| Neurl4 | 5545 | 4199 | 3816 | 4126 | 3801 | 4287 | 4307 | 4466 | 4117 | 4107 | 4288 | 4232 | 4252.88 | 4297 | 4209 | 0.979 | -0.03 | 0.59611156 | 0.74235543 | 7.00E-04 | 0.003 | 0.0021 | 0.0021 | TRUE | 0.0159 |
| Neurod1 | 104 | 84 | 78 | 107 | 95 | 98 | 81 | 89 | 84 | 107 | 107 | 97 | 94.12 | 85 | 104 | 1.222 | 0.29 | 0.14700311 | 0.29136488 | 0 | 0.0276 | 0.018 | 0.018 | TRUE | 0.0326 |
| Neurod2 | 2 | 1 | 0 | 1 | 2 | 0 | 2 | 1 | 0 | 1 | 2 | 0 | 0.98 | 1 | 1 | 1.185 | 0.245 | 0.9025703 | NA | 0 | 2.4255 | 1.8643 | 1.8643 | TRUE | 0.205 |
| Neurod4 | 3 | 0 | 0 | 1 | 0 | 1 | 2 | 0 | 0 | 1 | 0 | 1 | 0.72 | 1 | 1 | 0.815 | -0.296 | 0.90325409 | NA | 0 | 3.2999 | 2.9046 | 2.9046 | TRUE | 1.439 |
| Neurod6 | 0 | 0 | 0 | 0 | 0 | 0 | 0 | 0 | 0 | 0 | 0 | 0 | 0 | 0 | 0 | NA | NA | NA | NA | NA | NA | NA | NA | NA | NA |
| Neurog1 | 50 | 36 | 23 | 10 | 4 | 6 | 39 | 38 | 25 | 10 | 5 | 6 | 20.39 | 34 | 7 | 0.201 | -2.315 | 5.16E-06 | 0.0001085 | 0 | 0.1187 | 0.0962 | 0.0962 | TRUE | 0.4346 |
| Neurog2 | 2 | 4 | 2 | 3 | 1 | 2 | 2 | 4 | 2 | 3 | 1 | 2 | 2.34 | 3 | 2 | 0.782 | -0.355 | 0.77421602 | NA | 0 | 1.0142 | 0.6632 | 0.6632 | TRUE | 0.3073 |
| Neurog3 | 34 | 53 | 66 | 45 | 54 | 89 | 26 | 56 | 71 | 45 | 61 | 88 | 57.93 | 51 | 65 | 1.271 | 0.346 | 0.33547555 | 0.50961335 | 0.155 | 0.0433 | 0.0753 | 0.0753 | TRUE | 0.6021 |
| Nexn | 164 | 136 | 136 | 199 | 170 | 212 | 127 | 145 | 147 | 198 | 192 | 209 | 169.65 | 140 | 200 | 1.437 | 0.523 | 0.00073981 | 0.00580359 | 0 | 0.0164 | 0.0112 | 0.0112 | TRUE | 0.0686 |
| Nf1 | 3394 | 2343 | 2318 | 3140 | 2820 | 3337 | 2636 | 2492 | 2501 | 3126 | 3181 | 3294 | 2871.71 | 2543 | 3200 | 1.258 | 0.331 | 1.80E-08 | 8.47E-07 | 6.00E-04 | 0.0033 | 0.0021 | 0.0021 | TRUE | 0.0128 |
| Nf2 | 7254 | 5024 | 4787 | 4996 | 4537 | 5234 | 5634 | 5344 | 5165 | 4973 | 5118 | 5167 | 5233.49 | 5381 | 5086 | 0.945 | -0.082 | 0.1435919 | 0.28656173 | 9.00E-04 | 0.0029 | 0.0021 | 0.0021 | TRUE | 0.0209 |
| Nfam1 | 354 | 287 | 283 | 227 | 209 | 216 | 275 | 305 | 305 | 226 | 236 | 213 | 260.09 | 295 | 225 | 0.763 | -0.39 | 0.00243843 | 0.01408179 | 0 | 0.0115 | 0.008 | 0.008 | TRUE | 0.0441 |
| Nfasc | 567 | 420 | 480 | 523 | 461 | 544 | 440 | 447 | 518 | 521 | 520 | 537 | 497.12 | 468 | 526 | 1.125 | 0.17 | 0.09833945 | 0.21845908 | 0.0021 | 0.0072 | 0.0056 | 0.0056 | TRUE | 0.097 |
| Nfat5 | 5629 | 3942 | 3312 | 3707 | 3228 | 4256 | 4372 | 4193 | 3574 | 3690 | 3642 | 4201 | 3945.21 | 4046 | 3844 | 0.95 | -0.074 | 0.38149983 | 0.55579288 | 0.0075 | 0.003 | 0.0049 | 0.0049 | TRUE | 0.1265 |
| Nfatc1 | 215 | 191 | 168 | 226 | 211 | 198 | 167 | 203 | 181 | 225 | 238 | 195 | 201.64 | 184 | 219 | 1.196 | 0.259 | 0.08789111 | 0.20135202 | 0.0052 | 0.0142 | 0.0115 | 0.0115 | TRUE | 0.1019 |
| Nfatc2 | 530 | 335 | 344 | 394 | 367 | 438 | 412 | 356 | 371 | 392 | 414 | 432 | 396.29 | 380 | 413 | 1.085 | 0.117 | 0.29033669 | 0.4624223 | 0.0018 | 0.0084 | 0.0063 | 0.0063 | TRUE | 0.0665 |
| Nfatc2ip | 5401 | 3749 | 3356 | 3326 | 2968 | 3323 | 4195 | 3988 | 3621 | 3311 | 3348 | 3280 | 3623.8 | 3935 | 3313 | 0.842 | -0.249 | 0.00017494 | 0.00192487 | 0.0024 | 0.0031 | 0.0029 | 0.0029 | TRUE | 0.0588 |
| Nfatc3 | 10977 | 7925 | 8657 | 9959 | 8733 | 10288 | 8525 | 8430 | 9341 | 9913 | 9852 | 10156 | 9369.45 | 8765 | 9974 | 1.138 | 0.187 | 0.00102395 | 0.00735812 | 0.0016 | 0.0027 | 0.0022 | 0.0022 | TRUE | 0.0401 |
| Nfatc4 | 635 | 565 | 770 | 841 | 819 | 993 | 493 | 601 | 831 | 837 | 924 | 980 | 777.71 | 642 | 914 | 1.427 | 0.513 | 0.0005117 | 0.00438999 | 0.0372 | 0.0055 | 0.0144 | 0.0144 | TRUE | 0.7814 |
| Nfe2 | 74 | 44 | 38 | 69 | 46 | 39 | 57 | 47 | 41 | 69 | 52 | 38 | 50.73 | 48 | 53 | 1.089 | 0.122 | 0.68516401 | 0.8070222 | 0.038 | 0.0492 | 0.046 | 0.046 | TRUE | 0.2368 |
| Nfe2l1 | 17467 | 12830 | 12770 | 14167 | 12693 | 14771 | 13566 | 13647 | 13779 | 14102 | 14319 | 14581 | 13998.97 | 13664 | 14334 | 1.049 | 0.069 | 0.12356069 | 0.2575172 | 1.00E-04 | 0.0026 | 0.0014 | 0.0014 | TRUE | 0.0028 |
| Nfe2l2 | 3078 | 2223 | 2319 | 2714 | 2424 | 2765 | 2391 | 2365 | 2502 | 2702 | 2735 | 2729 | 2570.47 | 2419 | 2722 | 1.125 | 0.17 | 0.00315781 | 0.01718106 | 0 | 0.0033 | 0.002 | 0.002 | TRUE | 0.0107 |
| Nfe2l3 | 215 | 169 | 214 | 212 | 236 | 245 | 167 | 180 | 231 | 211 | 266 | 242 | 216.13 | 193 | 240 | 1.248 | 0.32 | 0.04887036 | 0.13246864 | 0.0166 | 0.0134 | 0.0143 | 0.0143 | TRUE | 0.3218 |
| Nfia | 53 | 46 | 48 | 53 | 41 | 57 | 41 | 49 | 52 | 53 | 46 | 56 | 49.53 | 47 | 52 | 1.102 | 0.14 | 0.61169073 | 0.75415299 | 0 | 0.0503 | 0.0346 | 0.0346 | TRUE | 0.1215 |
| Nfib | 535 | 353 | 310 | 317 | 268 | 349 | 416 | 375 | 334 | 316 | 302 | 345 | 347.98 | 375 | 321 | 0.853 | -0.229 | 0.06424984 | 0.16119414 | 0.0057 | 0.0092 | 0.0081 | 0.0081 | TRUE | 0.1076 |
| Nfic | 1392 | 1173 | 1351 | 1531 | 1412 | 1588 | 1081 | 1248 | 1458 | 1524 | 1593 | 1568 | 1411.83 | 1262 | 1562 | 1.239 | 0.309 | 0.00168824 | 0.01058333 | 0.0104 | 0.0041 | 0.0063 | 0.0063 | TRUE | 0.2204 |
| Nfil3 | 429 | 361 | 397 | 394 | 402 | 440 | 333 | 384 | 428 | 392 | 453 | 434 | 404.26 | 382 | 426 | 1.12 | 0.164 | 0.18003739 | 0.33493094 | 0.0083 | 0.0083 | 0.0083 | 0.0083 | TRUE | 0.1527 |
| Nfix | 180 | 166 | 133 | 157 | 159 | 149 | 140 | 177 | 144 | 156 | 179 | 147 | 157.1 | 154 | 161 | 1.051 | 0.072 | 0.67466282 | 0.80024529 | 0.0072 | 0.0175 | 0.0145 | 0.0145 | TRUE | 0.1806 |
| Nfkb1 | 2800 | 1861 | 1802 | 2166 | 1885 | 2093 | 2175 | 1980 | 1944 | 2156 | 2126 | 2066 | 2074.52 | 2033 | 2116 | 1.04 | 0.056 | 0.40542484 | 0.57815607 | 0.0017 | 0.0036 | 0.0028 | 0.0028 | TRUE | 0.0464 |
| Nfkb2 | 946 | 719 | 831 | 939 | 801 | 911 | 735 | 765 | 897 | 935 | 904 | 899 | 855.62 | 799 | 913 | 1.145 | 0.195 | 0.0354692 | 0.10487409 | 0.0043 | 0.0052 | 0.005 | 0.005 | TRUE | 0.1346 |
| Nfkbia | 742 | 526 | 517 | 606 | 591 | 714 | 576 | 559 | 558 | 603 | 667 | 705 | 611.39 | 564 | 658 | 1.165 | 0.221 | 0.01982559 | 0.06869508 | 0.0018 | 0.0063 | 0.0048 | 0.0048 | TRUE | 0.0643 |
| Nfkbib | 2301 | 1726 | 1638 | 1933 | 1677 | 1979 | 1787 | 1836 | 1767 | 1924 | 1892 | 1954 | 1859.99 | 1797 | 1923 | 1.071 | 0.098 | 0.1085191 | 0.23486729 | 0 | 0.0037 | 0.0022 | 0.0022 | TRUE | 0.0043 |
| Nfkbid | 243 | 173 | 194 | 152 | 130 | 188 | 189 | 184 | 209 | 151 | 147 | 186 | 177.6 | 194 | 161 | 0.833 | -0.264 | 0.09712924 | 0.21675325 | 0.0043 | 0.0158 | 0.0126 | 0.0126 | TRUE | 0.1884 |
| Nfkbie | 195 | 152 | 124 | 137 | 128 | 132 | 151 | 162 | 134 | 136 | 144 | 130 | 143 | 149 | 137 | 0.918 | -0.123 | 0.46423608 | 0.63356183 | 0 | 0.019 | 0.0133 | 0.0133 | TRUE | 0.0775 |
| Nfkbil1 | 453 | 334 | 362 | 371 | 366 | 336 | 352 | 355 | 391 | 369 | 413 | 332 | 368.59 | 366 | 371 | 1.014 | 0.021 | 0.86253901 | 0.92408891 | 0.0048 | 0.0089 | 0.0076 | 0.0076 | TRUE | 0.1072 |
| Nfkbiz | 174 | 222 | 157 | 77 | 109 | 113 | 135 | 236 | 169 | 77 | 123 | 112 | 141.97 | 180 | 104 | 0.577 | -0.795 | 0.00065267 | 0.00528341 | 0.0618 | 0.0191 | 0.0315 | 0.0315 | TRUE | 0.3075 |
| Nfrkb | 2761 | 2126 | 1963 | 2113 | 2010 | 2225 | 2144 | 2261 | 2118 | 2103 | 2267 | 2196 | 2181.83 | 2174 | 2189 | 1.007 | 0.009 | 0.8816698 | 0.93384803 | 8.00E-04 | 0.0035 | 0.0025 | 0.0025 | TRUE | 0.0145 |
| Nfs1 | 3483 | 2435 | 2623 | 2993 | 2603 | 2791 | 2705 | 2590 | 2830 | 2979 | 2936 | 2755 | 2799.37 | 2708 | 2890 | 1.067 | 0.094 | 0.14081113 | 0.28268406 | 0.0013 | 0.0033 | 0.0026 | 0.0026 | TRUE | 0.0205 |
| Nfu1 | 3552 | 2615 | 2461 | 2972 | 2576 | 3099 | 2759 | 2782 | 2655 | 2958 | 2906 | 3059 | 2853.19 | 2732 | 2974 | 1.089 | 0.123 | 0.03294299 | 0.0992735 | 3.00E-04 | 0.0033 | 0.002 | 0.002 | TRUE | 0.0076 |
| Nfx1 | 9380 | 6885 | 6777 | 7243 | 6532 | 7601 | 7285 | 7323 | 7312 | 7210 | 7369 | 7503 | 7333.78 | 7307 | 7361 | 1.007 | 0.011 | 0.82428833 | 0.89901206 | 0 | 0.0027 | 0.0015 | 0.0015 | TRUE | 0.004 |
| Nfxl1 | 3032 | 2255 | 2276 | 2559 | 2266 | 2502 | 2355 | 2399 | 2456 | 2547 | 2556 | 2470 | 2463.77 | 2403 | 2524 | 1.051 | 0.071 | 0.21728859 | 0.38004232 | 0 | 0.0034 | 0.002 | 0.002 | TRUE | 0.0044 |
| Nfya | 3533 | 2834 | 3050 | 2653 | 2481 | 2704 | 2744 | 3014 | 3291 | 2641 | 2799 | 2669 | 2859.7 | 3016 | 2703 | 0.897 | -0.157 | 0.03480382 | 0.10340545 | 0.0041 | 0.0033 | 0.0037 | 0.0037 | TRUE | 0.0774 |
| Nfyb | 6278 | 4776 | 4650 | 4968 | 4376 | 4926 | 4876 | 5080 | 5017 | 4945 | 4937 | 4863 | 4952.96 | 4991 | 4915 | 0.985 | -0.022 | 0.66481242 | 0.79350012 | 0 | 0.0029 | 0.0016 | 0.0016 | TRUE | 0.0051 |
| Nfyc | 2432 | 1746 | 1270 | 1205 | 1072 | 1330 | 1889 | 1857 | 1370 | 1199 | 1209 | 1313 | 1473.01 | 1705 | 1240 | 0.727 | -0.46 | 2.73E-05 | 0.0004303 | 0.0163 | 0.004 | 0.008 | 0.008 | TRUE | 0.3534 |
| Ngb | 42 | 46 | 33 | 60 | 38 | 49 | 33 | 49 | 36 | 60 | 43 | 48 | 44.69 | 39 | 50 | 1.299 | 0.377 | 0.21645716 | 0.37914488 | 0.0151 | 0.0555 | 0.0444 | 0.0444 | TRUE | 0.3515 |
| Ngdn | 3198 | 2145 | 2068 | 2143 | 1913 | 2171 | 2484 | 2282 | 2231 | 2133 | 2158 | 2143 | 2238.5 | 2332 | 2145 | 0.919 | -0.122 | 0.06102733 | 0.15519395 | 0.0013 | 0.0035 | 0.0026 | 0.0026 | TRUE | 0.0403 |
| Ngef | 183 | 146 | 146 | 162 | 136 | 159 | 142 | 155 | 158 | 161 | 153 | 157 | 154.43 | 152 | 157 | 1.039 | 0.056 | 0.72390507 | 0.83455196 | 0 | 0.0178 | 0.0115 | 0.0115 | TRUE | 0.0354 |
| Ngf | 23 | 20 | 21 | 15 | 18 | 23 | 18 | 21 | 23 | 15 | 20 | 23 | 19.96 | 21 | 19 | 0.942 | -0.086 | 0.84088861 | 0.90934561 | 0 | 0.1212 | 0.0833 | 0.0833 | TRUE | 0.2165 |
| Ngfr | 1600 | 1258 | 1057 | 1663 | 1621 | 1678 | 1243 | 1338 | 1140 | 1655 | 1829 | 1656 | 1476.95 | 1240 | 1713 | 1.381 | 0.465 | 8.44E-09 | 4.31E-07 | 0.0038 | 0.004 | 0.004 | 0.004 | TRUE | 0.0576 |
| Ngfrap1 | 12215 | 8290 | 7415 | 7675 | 6829 | 7705 | 9487 | 8818 | 8001 | 7640 | 7704 | 7606 | 8209.18 | 8769 | 7650 | 0.872 | -0.197 | 0.00294634 | 0.01630355 | 0.0034 | 0.0027 | 0.0031 | 0.0031 | TRUE | 0.0716 |
| Ngly1 | 810 | 522 | 542 | 529 | 492 | 572 | 629 | 555 | 585 | 527 | 555 | 565 | 569.23 | 590 | 549 | 0.929 | -0.107 | 0.26403552 | 0.43412627 | 0 | 0.0066 | 0.0048 | 0.0048 | TRUE | 0.0425 |
| Ngp | 0 | 1 | 1 | 0 | 1 | 0 | 0 | 1 | 1 | 0 | 1 | 0 | 0.55 | 1 | 0 | 0.535 | -0.902 | 0.73748201 | NA | 0 | 4.3501 | 3.5264 | 3.5264 | TRUE | 0.5075 |
| Ngrn | 2583 | 1945 | 1796 | 1987 | 1778 | 2017 | 2006 | 2069 | 1938 | 1978 | 2006 | 1991 | 1997.93 | 2004 | 1992 | 0.994 | -0.009 | 0.87918622 | 0.93262644 | 0 | 0.0036 | 0.0022 | 0.0022 | TRUE | 0.0098 |
| Nhej1 | 259 | 192 | 180 | 168 | 169 | 175 | 201 | 204 | 194 | 167 | 191 | 173 | 188.37 | 200 | 177 | 0.884 | -0.178 | 0.2205189 | 0.38404735 | 0 | 0.015 | 0.0099 | 0.0099 | TRUE | 0.0475 |
| Nhlh1 | 0 | 0 | 0 | 2 | 0 | 0 | 0 | 0 | 0 | 2 | 0 | 0 | 0.33 | 0 | 1 | 3.77 | 1.915 | 0.62344813 | NA | 0 | 7.1455 | 9.1834 | 9.1834 | TRUE | 0.9739 |
| Nhlh2 | 0 | 1 | 1 | 3 | 0 | 0 | 0 | 1 | 1 | 3 | 0 | 0 | 0.85 | 1 | 1 | 1.462 | 0.548 | 0.81368289 | NA | 2.2127 | 2.775 | 2.6765 | 2.6765 | TRUE | 1.4417 |
| Nhlrc1 | 422 | 326 | 298 | 283 | 274 | 285 | 328 | 347 | 322 | 282 | 309 | 281 | 311.36 | 332 | 291 | 0.875 | -0.193 | 0.10155686 | 0.22365982 | 0 | 0.01 | 0.0067 | 0.0067 | TRUE | 0.0332 |
| Nhlrc2 | 1863 | 1343 | 1303 | 1328 | 1170 | 1343 | 1447 | 1429 | 1406 | 1322 | 1320 | 1326 | 1374.82 | 1427 | 1323 | 0.926 | -0.11 | 0.09330812 | 0.21005717 | 0 | 0.0042 | 0.0024 | 0.0024 | TRUE | 0.002 |
| Nhlrc3 | 370 | 264 | 316 | 351 | 355 | 361 | 287 | 281 | 341 | 349 | 400 | 356 | 335.89 | 303 | 368 | 1.218 | 0.284 | 0.02256063 | 0.0753409 | 0.0051 | 0.0095 | 0.0082 | 0.0082 | TRUE | 0.1303 |
| Nhlrc4 | 7 | 2 | 1 | 2 | 5 | 4 | 5 | 2 | 1 | 2 | 6 | 4 | 3.37 | 3 | 4 | 1.285 | 0.362 | 0.73607998 | NA | 0 | 0.7056 | 0.5356 | 0.5356 | TRUE | 0.6213 |

| | | | | | | | | | | | | | | | | | | | | | | | | | |
|---|---|---|---|---|---|---|---|---|---|---|---|---|---|---|---|---|---|---|---|---|---|---|---|---|---|
| Nhp2 | 12660 | 8822 | 8293 | 8387 | 7488 | 8821 | 9832 | 9384 | 8948 | 8348 | 8447 | 8708 | 8944.62 | 9388 | 8501 | 0.905 | -0.144 | 0.00896685 | 0.03823482 | 0.0012 | 0.0027 | 0.0021 | 0.0021 | TRUE | 0.0211 |
| Nhp2l1 | 7279 | 5178 | 4795 | 5213 | 4674 | 5451 | 5653 | 5508 | 5174 | 5189 | 5273 | 5381 | 5362.93 | 5445 | 5281 | 0.97 | -0.045 | 0.42638089 | 0.5987801 | 9.00E-04 | 0.0029 | 0.0021 | 0.0021 | TRUE | 0.023 |
| Nhs | 229 | 137 | 167 | 178 | 171 | 220 | 178 | 146 | 180 | 177 | 193 | 217 | 181.84 | 168 | 196 | 1.164 | 0.219 | 0.16990452 | 0.32144133 | 0.0063 | 0.0155 | 0.0128 | 0.0128 | TRUE | 0.1363 |
| Nhsl1 | 1334 | 950 | 1011 | 943 | 921 | 1128 | 1036 | 1011 | 1091 | 939 | 1039 | 1114 | 1038.1 | 1046 | 1031 | 0.985 | -0.021 | 0.80389436 | 0.88644746 | 0.0034 | 0.0047 | 0.0043 | 0.0043 | TRUE | 0.0736 |
| Nhsl2 | 194 | 123 | 143 | 151 | 146 | 185 | 151 | 131 | 154 | 150 | 165 | 183 | 155.57 | 145 | 166 | 1.141 | 0.19 | 0.25054145 | 0.41941777 | 0 | 0.0177 | 0.0133 | 0.0133 | TRUE | 0.0854 |
| Niacr1 | 0 | 0 | 0 | 0 | 0 | 0 | 0 | 0 | 0 | 0 | 0 | 0 | 0 | 0 | 0 | NA | NA | NA | NA | NA | NA | NA | NA | NA | NA |
| Nicn1 | 975 | 660 | 784 | 780 | 724 | 848 | 757 | 702 | 846 | 776 | 817 | 837 | 789.24 | 768 | 810 | 1.055 | 0.077 | 0.40547376 | 0.57815607 | 0.0037 | 0.0054 | 0.0049 | 0.0049 | TRUE | 0.0888 |
| Nid1 | 6731 | 5033 | 6553 | 7220 | 6798 | 7742 | 5228 | 5354 | 7071 | 7187 | 7669 | 7643 | 6691.67 | 5884 | 7500 | 1.275 | 0.35 | 0.00033173 | 0.00314095 | 0.0157 | 0.0028 | 0.0067 | 0.0067 | TRUE | 0.3789 |
| Nid2 | 37220 | 29319 | 32217 | 42360 | 39466 | 44666 | 28907 | 31186 | 34762 | 42165 | 44522 | 44092 | 37605.73 | 31618 | 43593 | 1.379 | 0.463 | 1.31E-11 | 1.30E-09 | 0.0047 | 0.0025 | 0.0034 | 0.0034 | TRUE | 0.0926 |
| Nif3l1 | 2441 | 1666 | 1480 | 1454 | 1347 | 1558 | 1896 | 1772 | 1597 | 1447 | 1520 | 1538 | 1628.28 | 1755 | 1502 | 0.855 | -0.227 | 0.00391899 | 0.0202876 | 0.0037 | 0.0039 | 0.0038 | 0.0038 | TRUE | 0.0744 |
| Nim1 | 147 | 88 | 102 | 169 | 139 | 151 | 114 | 94 | 110 | 168 | 157 | 149 | 131.99 | 106 | 158 | 1.487 | 0.573 | 0.00113891 | 0.00797572 | 0 | 0.0204 | 0.0145 | 0.0145 | TRUE | 0.0986 |
| Nin | 2287 | 1668 | 1332 | 1002 | 856 | 908 | 1776 | 1774 | 1437 | 997 | 966 | 896 | 1307.84 | 1662 | 953 | 0.573 | -0.804 | 1.59E-17 | 3.49E-15 | 0.0082 | 0.0042 | 0.0056 | 0.0056 | TRUE | 0.1675 |
| Ninj1 | 747 | 625 | 765 | 872 | 772 | 992 | 580 | 665 | 825 | 868 | 871 | 979 | 798.09 | 690 | 906 | 1.316 | 0.396 | 0.00089604 | 0.0066948 | 0.0168 | 0.0054 | 0.009 | 0.009 | TRUE | 0.346 |
| Ninj2 | 66 | 19 | 14 | 12 | 17 | 16 | 51 | 20 | 15 | 12 | 19 | 16 | 22.25 | 29 | 16 | 0.532 | -0.91 | 0.07675015 | 0.18328987 | 0.2102 | 0.109 | 0.1408 | 0.1408 | TRUE | 2.561 |
| Ninl | 3004 | 2415 | 2252 | 2111 | 1958 | 2168 | 2333 | 2569 | 2430 | 2101 | 2209 | 2140 | 2297.01 | 2444 | 2150 | 0.88 | -0.184 | 0.00411664 | 0.02111198 | 0.0011 | 0.0035 | 0.0025 | 0.0025 | TRUE | 0.0242 |
| Nip7 | 6552 | 4494 | 4078 | 4274 | 3899 | 4470 | 5089 | 4780 | 4400 | 4254 | 4398 | 4413 | 4555.73 | 4756 | 4355 | 0.915 | -0.128 | 0.04952296 | 0.13388155 | 0.0025 | 0.0029 | 0.0028 | 0.0028 | TRUE | 0.0521 |
| Nipa1 | 324 | 206 | 184 | 289 | 195 | 236 | 252 | 219 | 199 | 288 | 220 | 233 | 234.98 | 223 | 247 | 1.104 | 0.143 | 0.35312074 | 0.52780934 | 0.0134 | 0.0125 | 0.0128 | 0.0128 | TRUE | 0.2335 |
| Nipa2 | 2868 | 1980 | 1861 | 2063 | 1730 | 1890 | 2227 | 2106 | 2008 | 2054 | 1952 | 1866 | 2035.4 | 2114 | 1957 | 0.925 | -0.112 | 0.10701015 | 0.2324462 | 0.0021 | 0.0036 | 0.003 | 0.003 | TRUE | 0.0275 |
| Nipal1 | 89 | 60 | 49 | 110 | 83 | 101 | 69 | 64 | 53 | 109 | 94 | 100 | 81.44 | 62 | 101 | 1.623 | 0.699 | 0.00166406 | 0.01045928 | 0 | 0.0315 | 0.0227 | 0.0227 | TRUE | 0.1356 |
| Nipal2 | 25 | 14 | 26 | 14 | 15 | 16 | 19 | 15 | 28 | 14 | 17 | 16 | 18.17 | 21 | 16 | 0.749 | -0.417 | 0.36765494 | 0.54230052 | 0 | 0.1329 | 0.0977 | 0.0977 | TRUE | 0.3522 |
| Nipal3 | 1413 | 1020 | 1084 | 1379 | 1225 | 1413 | 1097 | 1085 | 1170 | 1373 | 1382 | 1395 | 1250.24 | 1117 | 1383 | 1.239 | 0.309 | 1.13E-05 | 0.00020732 | 0 | 0.0043 | 0.0028 | 0.0028 | TRUE | 0.0194 |
| Nipal4 | 150 | 93 | 113 | 137 | 120 | 143 | 116 | 99 | 122 | 136 | 135 | 141 | 125.04 | 112 | 137 | 1.223 | 0.29 | 0.10278832 | 0.22567642 | 0 | 0.0214 | 0.0148 | 0.0148 | TRUE | 0.1039 |
| Nipbl | 9090 | 6546 | 6370 | 7623 | 6603 | 7775 | 7060 | 6963 | 6873 | 7588 | 7449 | 7675 | 7267.97 | 6965 | 7571 | 1.087 | 0.12 | 0.01185397 | 0.04689422 | 0 | 0.0028 | 0.0015 | 0.0015 | TRUE | 0.0024 |
| Nipsnap1 | 3955 | 2905 | 3268 | 3776 | 3267 | 3962 | 3072 | 3090 | 3526 | 3759 | 3686 | 3911 | 3507.18 | 3229 | 3785 | 1.173 | 0.23 | 0.00086649 | 0.00652862 | 0.0032 | 0.0031 | 0.0031 | 0.0031 | TRUE | 0.0779 |
| Nipsnap3a | 0 | 0 | 2 | 0 | 0 | 2 | 0 | 0 | 2 | 0 | 0 | 2 | 0.69 | 1 | 1 | 0.955 | -0.066 | 0.98070217 | NA | 8.094 | 3.4437 | 3.8891 | 3.8891 | TRUE | 0.9748 |
| Nipsnap3b | 2058 | 1504 | 1609 | 1787 | 1583 | 1829 | 1598 | 1600 | 1736 | 1779 | 1786 | 1806 | 1717.39 | 1645 | 1790 | 1.089 | 0.123 | 0.0632158 | 0.15927084 | 5.00E-04 | 0.0038 | 0.0026 | 0.0026 | TRUE | 0.0279 |
| Nisch | 20519 | 15332 | 15058 | 14113 | 13572 | 15820 | 15936 | 16308 | 16247 | 14048 | 15311 | 15617 | 15577.94 | 16164 | 14992 | 0.927 | -0.109 | 0.04954175 | 0.13390701 | 0.0015 | 0.0026 | 0.0021 | 0.0021 | TRUE | 0.0372 |
| Nit1 | 1380 | 931 | 893 | 975 | 809 | 993 | 1072 | 990 | 964 | 971 | 913 | 980 | 981.5 | 1009 | 955 | 0.946 | -0.081 | 0.31398026 | 0.48737184 | 0.0014 | 0.0048 | 0.0036 | 0.0036 | TRUE | 0.0376 |
| Nit2 | 4461 | 3152 | 3549 | 4515 | 3851 | 4552 | 3465 | 3353 | 3829 | 4494 | 4344 | 4494 | 3996.47 | 3549 | 4444 | 1.253 | 0.325 | 4.37E-07 | 1.45E-05 | 0.0023 | 0.003 | 0.0027 | 0.0027 | TRUE | 0.0573 |
| Nkain1 | 4749 | 3426 | 3592 | 3606 | 3304 | 3842 | 3688 | 3644 | 3876 | 3589 | 3727 | 3793 | 3719.6 | 3736 | 3703 | 0.991 | -0.013 | 0.82445397 | 0.89912411 | 6.00E-04 | 0.0031 | 0.0021 | 0.0021 | TRUE | 0.0128 |
| Nkain2 | 0 | 0 | 0 | 0 | 0 | 1 | 0 | 0 | 0 | 0 | 0 | 1 | 0.16 | 0 | 0 | 2.094 | 1.067 | 0.79379329 | NA | 0 | 14.4078 | 10 | 10 | TRUE | 0.3835 |
| Nkain3 | 1 | 1 | 0 | 3 | 0 | 0 | 1 | 1 | 0 | 3 | 0 | 0 | 0.8 | 1 | 1 | 1.621 | 0.697 | 0.76811698 | NA | 1.7582 | 2.9487 | 2.7352 | 2.7352 | TRUE | 1.4417 |
| Nkain4 | 62 | 37 | 61 | 25 | 28 | 21 | 48 | 39 | 66 | 25 | 32 | 21 | 38.42 | 51 | 26 | 0.502 | -0.993 | 0.00374209 | 0.01959917 | 0.031 | 0.0641 | 0.0549 | 0.0549 | TRUE | 0.2922 |
| Nkap | 2143 | 1501 | 1644 | 1673 | 1443 | 1709 | 1664 | 1597 | 1774 | 1665 | 1628 | 1687 | 1669.17 | 1678 | 1660 | 0.989 | -0.015 | 0.82338368 | 0.89863229 | 9.00E-04 | 0.0038 | 0.0028 | 0.0028 | TRUE | 0.029 |
| Nkapl | 51 | 22 | 15 | 24 | 15 | 27 | 40 | 23 | 16 | 24 | 17 | 27 | 24.44 | 26 | 23 | 0.843 | -0.247 | 0.57347713 | 0.7257767 | 0.0927 | 0.0994 | 0.0974 | 0.0974 | TRUE | 0.5809 |
| Nkd1 | 492 | 358 | 380 | 362 | 339 | 374 | 382 | 381 | 410 | 360 | 382 | 369 | 380.81 | 391 | 370 | 0.948 | -0.077 | 0.47439379 | 0.64280192 | 0 | 0.0087 | 0.0056 | 0.0056 | TRUE | 0.0201 |
| Nkd2 | 632 | 520 | 422 | 452 | 413 | 478 | 491 | 553 | 455 | 450 | 466 | 472 | 481.16 | 500 | 463 | 0.926 | -0.111 | 0.29331731 | 0.46521877 | 0.003 | 0.0074 | 0.006 | 0.006 | TRUE | 0.0974 |
| Nkg7 | 2 | 3 | 6 | 0 | 2 | 3 | 2 | 3 | 6 | 0 | 2 | 3 | 2.74 | 4 | 2 | 0.471 | -1.087 | 0.39073285 | NA | 0 | 0.8676 | 0.7249 | 0.7249 | TRUE | 0.4136 |
| Nkiras1 | 574 | 455 | 426 | 454 | 390 | 382 | 446 | 484 | 460 | 452 | 440 | 377 | 443.07 | 463 | 423 | 0.914 | -0.13 | 0.23328911 | 0.39938195 | 0.0031 | 0.0078 | 0.0063 | 0.0063 | TRUE | 0.1061 |
| Nkiras2 | 3542 | 2431 | 2440 | 2801 | 2438 | 2617 | 2751 | 2586 | 2633 | 2788 | 2750 | 2583 | 2681.88 | 2657 | 2707 | 1.019 | 0.026 | 0.6683228 | 0.79609449 | 0.001 | 0.0033 | 0.0024 | 0.0024 | TRUE | 0.0196 |
| Nkpd1 | 8 | 4 | 5 | 13 | 8 | 9 | 6 | 4 | 5 | 13 | 9 | 9 | 7.79 | 5 | 10 | 1.931 | 0.95 | 0.16861688 | NA | 0 | 0.3069 | 0.2023 | 0.2023 | TRUE | 0.1852 |
| Nkrf | 1717 | 1093 | 921 | 1021 | 830 | 948 | 1334 | 1163 | 994 | 1016 | 936 | 936 | 1063.06 | 1164 | 963 | 0.826 | -0.275 | 0.00809068 | 0.03536396 | 0.0104 | 0.0047 | 0.0068 | 0.0068 | TRUE | 0.1597 |
| Nktr | 4050 | 2993 | 2367 | 2218 | 1941 | 2309 | 3145 | 3184 | 2554 | 2208 | 2190 | 2279 | 2593.31 | 2961 | 2226 | 0.751 | -0.412 | 1.10E-06 | 3.18E-05 | 0.0067 | 0.0033 | 0.0048 | 0.0048 | TRUE | 0.1741 |
| Nkx1-2 | 2 | 3 | 2 | 3 | 6 | 5 | 2 | 3 | 2 | 3 | 7 | 5 | 3.6 | 2 | 5 | 2.153 | 1.106 | 0.2810354 | NA | 0 | 0.661 | 0.4399 | 0.4399 | TRUE | 0.1265 |
| Nkx2-1 | 4 | 0 | 0 | 2 | 0 | 1 | 3 | 0 | 0 | 2 | 0 | 1 | 1.01 | 1 | 1 | 0.918 | -0.123 | 0.95416679 | NA | 2.3137 | 2.3395 | 2.3364 | 2.3364 | TRUE | 0.8566 |
| Nkx2-2 | 8 | 9 | 5 | 5 | 3 | 4 | 6 | 10 | 5 | 5 | 3 | 4 | 5.58 | 7 | 4 | 0.585 | -0.773 | 0.34118098 | NA | 0 | 0.427 | 0.2807 | 0.2807 | TRUE | 0.2395 |
| Nkx2-2as | 1 | 0 | 0 | 0 | 0 | 0 | 1 | 0 | 0 | 0 | 0 | 0 | 0.13 | 0 | 0 | 0.552 | -0.857 | 0.8336548 | NA | 0 | 18.3121 | 10 | 10 | TRUE | 0.2566 |
| Nkx2-3 | 17 | 13 | 7 | 17 | 7 | 12 | 13 | 14 | 8 | 17 | 8 | 12 | 11.87 | 12 | 12 | 1.06 | 0.083 | 0.88477297 | NA | 0 | 0.202 | 0.1549 | 0.1549 | TRUE | 0.2469 |
| Nkx2-4 | 106 | 77 | 81 | 118 | 82 | 120 | 82 | 82 | 87 | 117 | 93 | 118 | 96.67 | 84 | 109 | 1.309 | 0.389 | 0.05678721 | 0.14778944 | 0 | 0.0269 | 0.0195 | 0.0195 | TRUE | 0.1734 |
| Nkx2-5 | 29 | 16 | 18 | 42 | 39 | 32 | 23 | 17 | 19 | 42 | 44 | 32 | 29.39 | 20 | 39 | 1.974 | 0.981 | 0.0074085 | 0.03313193 | 0 | 0.0831 | 0.0591 | 0.0591 | TRUE | 0.2165 |
| Nkx2-6 | 1 | 0 | 0 | 0 | 0 | 1 | 1 | 0 | 0 | 0 | 0 | 1 | 0.29 | 0 | 0 | 1.075 | 0.105 | 0.97702352 | NA | 0 | 8.0647 | 7.5458 | 7.5458 | TRUE | 0.3835 |
| Nkx2-9 | 112 | 98 | 102 | 88 | 62 | 69 | 87 | 104 | 110 | 88 | 70 | 68 | 87.82 | 100 | 75 | 0.753 | -0.41 | 0.06198774 | 0.15706272 | 0 | 0.0294 | 0.023 | 0.023 | TRUE | 0.1937 |
| Nkx3-1 | 222 | 160 | 181 | 219 | 183 | 180 | 172 | 170 | 195 | 218 | 206 | 178 | 190 | 179 | 201 | 1.12 | 0.164 | 0.28267545 | 0.45442359 | 0.0032 | 0.0149 | 0.0115 | 0.0115 | TRUE | 0.1123 |
| Nkx3-2 | 0 | 0 | 1 | 5 | 3 | 1 | 0 | 0 | 1 | 5 | 3 | 1 | 1.74 | 0 | 3 | 9.026 | 3.174 | 0.07343958 | NA | 0 | 1.3662 | 1.0995 | 1.0995 | TRUE | 0.3985 |
| Nkx6-1 | 9 | 8 | 6 | 10 | 5 | 2 | 7 | 9 | 6 | 10 | 6 | 2 | 6.59 | 7 | 6 | 0.801 | -0.321 | 0.68170441 | NA | 0.0625 | 0.3621 | 0.287 | 0.287 | TRUE | 0.2914 |
| Nkx6-2 | 289 | 215 | 214 | 284 | 258 | 331 | 224 | 229 | 231 | 283 | 291 | 327 | 264.09 | 228 | 300 | 1.318 | 0.398 | 0.0018403 | 0.01134255 | 0 | 0.0114 | 0.0079 | 0.0079 | TRUE | 0.0691 |
| Nkx6-3 | 11 | 13 | 10 | 20 | 16 | 17 | 9 | 14 | 11 | 20 | 18 | 17 | 14.65 | 11 | 18 | 1.668 | 0.738 | 0.14268381 | NA | 0 | 0.1642 | 0.1104 | 0.1104 | TRUE | 0.1953 |
| Nle1 | 3426 | 2627 | 2514 | 2327 | 2151 | 2614 | 2661 | 2794 | 2713 | 2316 | 2427 | 2580 | 2581.83 | 2723 | 2441 | 0.897 | -0.157 | 0.01432964 | 0.05394087 | 0.0012 | 0.0033 | 0.0026 | 0.0026 | TRUE | 0.0306 |
| Nlgn1 | 2 | 5 | 2 | 2 | 4 | 0 | 2 | 5 | 2 | 2 | 5 | 0 | 2.59 | 3 | 2 | 0.717 | -0.48 | 0.7102909 | NA | 0.4236 | 0.9179 | 0.7965 | 0.7965 | TRUE | 0.3263 |
| Nlgn2 | 1427 | 1185 | 1270 | 1500 | 1297 | 1461 | 1108 | 1260 | 1370 | 1493 | 1463 | 1442 | 1356.26 | 1246 | 1466 | 1.178 | 0.237 | 0.00549768 | 0.02639513 | 0.0046 | 0.0042 | 0.0045 | 0.0045 | TRUE | 0.117 |
| Nlgn3 | 29 | 21 | 31 | 36 | 29 | 41 | 23 | 22 | 33 | 36 | 33 | 40 | 31.22 | 26 | 36 | 1.403 | 0.488 | 0.16640945 | 0.31740856 | 0 | 0.0783 | 0.057 | 0.057 | TRUE | 0.3517 |
| Nlk | 5837 | 4424 | 4184 | 4613 | 4046 | 4811 | 4533 | 4706 | 4515 | 4592 | 4564 | 4749 | 4609.81 | 4585 | 4635 | 1.011 | 0.016 | 0.75939837 | 0.8571376 | 3.00E-04 | 0.0029 | 0.0018 | 0.0018 | TRUE | 0.0064 |
| Nln | 5211 | 3875 | 4186 | 4625 | 4301 | 4720 | 4047 | 4122 | 4517 | 4604 | 4852 | 4659 | 4466.78 | 4229 | 4705 | 1.113 | 0.154 | 0.01236109 | 0.04843121 | 0.0018 | 0.003 | 0.0025 | 0.0025 | TRUE | 0.043 |
| Nlrc3 | 32 | 8 | 17 | 16 | 12 | 22 | 25 | 9 | 18 | 16 | 14 | 22 | 17.15 | 17 | 17 | 0.98 | -0.03 | 0.9536511 | 0.97451456 | 0.0994 | 0.1406 | 0.1293 | 0.1293 | TRUE | 0.3136 |
| Nlrc4 | 71 | 58 | 38 | 44 | 33 | 36 | 55 | 62 | 41 | 44 | 37 | 36 | 45.73 | 53 | 39 | 0.738 | -0.439 | 0.13946695 | 0.28069219 | 0 | 0.0542 | 0.0412 | 0.0412 | TRUE | 0.2855 |
| Nlrc5 | 95 | 87 | 76 | 86 | 77 | 106 | 74 | 93 | 82 | 86 | 87 | 105 | 87.57 | 83 | 93 | 1.121 | 0.165 | 0.4422824 | 0.61332152 | 0 | 0.0295 | 0.0219 | 0.0219 | TRUE | 0.1326 |
| Nlrp10 | 3 | 4 | 4 | 5 | 1 | 0 | 2 | 4 | 4 | 5 | 1 | 0 | 2.83 | 3 | 2 | 0.569 | -0.812 | 0.51372228 | NA | 0.327 | 0.8386 | 0.7278 | 0.7278 | TRUE | 1.505 |
| Nlrp12 | 20 | 16 | 22 | 27 | 13 | 28 | 16 | 17 | 24 | 27 | 15 | 28 | 20.91 | 19 | 23 | 1.244 | 0.315 | 0.47715894 | 0.64473744 | 0.0331 | 0.1158 | 0.093 | 0.093 | TRUE | 0.5392 |
| Nlrp14 | 1548 | 1238 | 1230 | 1163 | 1004 | 1040 | 1202 | 1317 | 1327 | 1158 | 1133 | 1027 | 1193.86 | 1282 | 1106 | 0.863 | -0.212 | 0.00826707 | 0.03589665 | 0.0027 | 0.0044 | 0.0038 | 0.0038 | TRUE | 0.0474 |

| | | | | | | | | | | | | | | | | | | | | | | | | | |
|---|---|---|---|---|---|---|---|---|---|---|---|---|---|---|---|---|---|---|---|---|---|---|---|---|---|
| Nlrp1a | 1 | 1 | 1 | 1 | 0 | 0 | 1 | 1 | 1 | 1 | 0 | 0 | 0.65 | 1 | 0 | 0.383 | -1.383 | 0.55308337 | NA | 0 | 3.6349 | 2.4186 | 2.4186 | TRUE | 0.3906 |
| Nlrp1b | 0 | 1 | 0 | 0 | 0 | 0 | 0 | 1 | 0 | 0 | 0 | 0 | 0.18 | 0 | 0 | 0.552 | -0.857 | 0.8336548 | NA | 0 | 13.3714 | 10 | 10 | TRUE | 0.5216 |
| Nlrp1c-ps | 0 | 0 | 0 | 0 | 0 | 0 | 0 | 0 | 0 | 0 | 0 | 0 | 0 | 0 | 0 | NA | NA | NA | NA | NA | NA | NA | NA | NA | NA |
| Nlrp2 | 3 | 3 | 1 | 1 | 2 | 2 | 2 | 3 | 1 | 1 | 2 | 2 | 1.97 | 2 | 2 | 0.785 | -0.349 | 0.79311622 | NA | 0 | 1.2049 | 0.7583 | 0.7583 | TRUE | 0.1705 |
| Nlrp3 | 139 | 107 | 173 | 122 | 138 | 150 | 108 | 114 | 187 | 121 | 156 | 148 | 138.94 | 136 | 142 | 1.045 | 0.063 | 0.77390586 | 0.86654419 | 0.0487 | 0.0195 | 0.028 | 0.028 | TRUE | 1.0646 |
| Nlrp4a | 105 | 69 | 79 | 75 | 43 | 38 | 82 | 73 | 85 | 75 | 49 | 38 | 66.81 | 80 | 54 | 0.669 | -0.579 | 0.03594593 | 0.10599859 | 0.0429 | 0.0379 | 0.0392 | 0.0392 | TRUE | 1.0069 |
| Nlrp4b | 0 | 0 | 0 | 0 | 0 | 0 | 0 | 0 | 0 | 0 | 0 | 0 | 0 | 0 | 0 | NA | NA | NA | NA | NA | NA | NA | NA | NA | NA |
| Nlrp4c | 2484 | 1640 | 1209 | 909 | 767 | 664 | 1929 | 1744 | 1305 | 905 | 865 | 655 | 1233.95 | 1659 | 808 | 0.487 | -1.039 | 5.34E-14 | 8.21E-12 | 0.0311 | 0.0043 | 0.0128 | 0.0128 | TRUE | 0.3748 |
| Nlrp4e | 14 | 9 | 7 | 2 | 3 | 4 | 11 | 10 | 8 | 2 | 3 | 4 | 6.22 | 10 | 3 | 0.331 | -1.596 | 0.0490759 | NA | 0 | 0.3834 | 0.2558 | 0.2558 | TRUE | 0.1395 |
| Nlrp4f | 1902 | 1249 | 1022 | 1206 | 897 | 916 | 1477 | 1329 | 1103 | 1200 | 1012 | 904 | 1170.85 | 1303 | 1039 | 0.797 | -0.328 | 0.00580202 | 0.02750521 | 0.0196 | 0.0045 | 0.0093 | 0.0093 | TRUE | 0.223 |
| Nlrp4g | 7 | 4 | 7 | 7 | 8 | 4 | 5 | 4 | 8 | 7 | 9 | 4 | 6.2 | 6 | 7 | 1.153 | 0.205 | 0.79047715 | NA | 0 | 0.3848 | 0.2664 | 0.2664 | TRUE | 0.2955 |
| Nlrp5 | 6 | 6 | 5 | 1 | 4 | 5 | 5 | 6 | 5 | 1 | 5 | 5 | 4.48 | 5 | 4 | 0.635 | -0.654 | 0.47989608 | NA | 0 | 0.5314 | 0.3805 | 0.3805 | TRUE | 0.6021 |
| Nlrp5-ps | 70 | 54 | 34 | 35 | 30 | 25 | 54 | 57 | 37 | 35 | 34 | 25 | 40.31 | 49 | 31 | 0.626 | -0.676 | 0.03781699 | 0.10992992 | 0.0198 | 0.0612 | 0.0501 | 0.0501 | TRUE | 0.3891 |
| Nlrp6 | 22 | 11 | 22 | 13 | 11 | 22 | 17 | 12 | 24 | 13 | 12 | 22 | 16.6 | 18 | 16 | 0.9 | -0.151 | 0.76156283 | NA | 0.0477 | 0.1452 | 0.1187 | 0.1187 | TRUE | 0.257 |
| Nlrp9a | 0 | 0 | 0 | 0 | 0 | 0 | 0 | 0 | 0 | 0 | 0 | 0 | 0 | 0 | 0 | NA | NA | NA | NA | NA | NA | NA | NA | NA | NA |
| Nlrp9b | 236 | 179 | 192 | 126 | 125 | 164 | 183 | 190 | 207 | 125 | 141 | 162 | 168.2 | 193 | 143 | 0.739 | -0.437 | 0.00713958 | 0.03222167 | 0.0032 | 0.0165 | 0.0129 | 0.0129 | TRUE | 0.1475 |
| Nlrp9c | 1 | 1 | 1 | 1 | 0 | 1 | 1 | 1 | 1 | 1 | 0 | 1 | 0.82 | 1 | 1 | 0.698 | -0.518 | 0.79971927 | NA | 0 | 2.9034 | 1.738 | 1.738 | TRUE | 0.2018 |
| Nlrx1 | 828 | 597 | 601 | 729 | 702 | 813 | 643 | 635 | 648 | 726 | 792 | 803 | 707.78 | 642 | 774 | 1.204 | 0.268 | 0.00186721 | 0.01148133 | 0 | 0.0058 | 0.0039 | 0.0039 | TRUE | 0.0351 |
| Nmb | 187 | 134 | 138 | 132 | 111 | 159 | 145 | 143 | 149 | 131 | 125 | 157 | 141.71 | 146 | 138 | 0.949 | -0.076 | 0.65484297 | 0.7856027 | 0 | 0.0192 | 0.0138 | 0.0138 | TRUE | 0.154 |
| Nmbr | 4 | 6 | 1 | 1 | 4 | 2 | 3 | 6 | 1 | 1 | 5 | 2 | 3.01 | 3 | 3 | 0.7 | -0.515 | 0.65775688 | NA | 0.2223 | 0.7903 | 0.624 | 0.624 | TRUE | 0.2626 |
| Nmd3 | 6794 | 4609 | 4466 | 4563 | 3903 | 4517 | 5277 | 4903 | 4819 | 4542 | 4403 | 4459 | 4733.65 | 5000 | 4468 | 0.893 | -0.163 | 0.00461128 | 0.02306122 | 0.001 | 0.0029 | 0.0022 | 0.0022 | TRUE | 0.0292 |
| Nme1 | 11067 | 7587 | 7280 | 7562 | 6448 | 7810 | 8595 | 8070 | 7855 | 7527 | 7274 | 7710 | 7838.57 | 8173 | 7504 | 0.918 | -0.124 | 0.0278584 | 0.08772418 | 0.0013 | 0.0027 | 0.0021 | 0.0021 | TRUE | 0.0252 |
| Nme2 | 29057 | 20569 | 21324 | 22640 | 19874 | 23288 | 22567 | 21879 | 23008 | 22536 | 22420 | 22989 | 22566.56 | 22485 | 22648 | 1.007 | 0.01 | 0.8203787 | 0.89651557 | 3.00E-04 | 0.0025 | 0.0015 | 0.0015 | TRUE | 0.0068 |
| Nme3 | 685 | 503 | 552 | 572 | 506 | 556 | 532 | 535 | 596 | 569 | 571 | 549 | 558.62 | 554 | 563 | 1.017 | 0.024 | 0.79848727 | 0.88227596 | 0 | 0.0067 | 0.0047 | 0.0047 | TRUE | 0.0486 |
| Nme4 | 1765 | 1349 | 1529 | 1773 | 1498 | 1842 | 1371 | 1435 | 1650 | 1765 | 1690 | 1818 | 1621.43 | 1485 | 1758 | 1.185 | 0.245 | 0.00270255 | 0.01520053 | 0.0046 | 0.0039 | 0.0042 | 0.0042 | TRUE | 0.1123 |
| Nme5 | 19 | 17 | 14 | 8 | 11 | 20 | 15 | 18 | 15 | 8 | 12 | 20 | 14.68 | 16 | 13 | 0.84 | -0.251 | 0.63039107 | NA | 0.0268 | 0.1639 | 0.1277 | 0.1277 | TRUE | 0.4531 |
| Nme6 | 1462 | 1019 | 1044 | 1026 | 891 | 1136 | 1135 | 1084 | 1126 | 1021 | 1005 | 1121 | 1082.28 | 1115 | 1049 | 0.941 | -0.088 | 0.2558829 | 0.42581608 | 0.0011 | 0.0046 | 0.0034 | 0.0034 | TRUE | 0.0436 |
| Nme7 | 1943 | 1327 | 1496 | 1387 | 1250 | 1535 | 1509 | 1412 | 1614 | 1381 | 1410 | 1515 | 1473.46 | 1512 | 1435 | 0.95 | -0.074 | 0.33289926 | 0.5071551 | 0.0024 | 0.004 | 0.0036 | 0.0036 | TRUE | 0.0411 |
| Nme8 | 2 | 3 | 2 | 3 | 4 | 1 | 2 | 3 | 2 | 3 | 5 | 1 | 2.56 | 2 | 3 | 1.234 | 0.303 | 0.79946558 | NA | 0 | 0.9265 | 0.6285 | 0.6285 | TRUE | 0.2377 |
| Nme9 | 0 | 0 | 1 | 0 | 0 | 1 | 0 | 0 | 1 | 0 | 0 | 1 | 0.34 | 0 | 0 | 1.075 | 0.105 | 0.97570524 | NA | 0 | 6.885 | 6.5379 | 6.5379 | TRUE | 0.5365 |
| Nmi | 102 | 76 | 50 | 49 | 52 | 47 | 79 | 81 | 54 | 49 | 59 | 46 | 61.31 | 71 | 51 | 0.715 | -0.485 | 0.06870933 | 0.16914612 | 0.0164 | 0.0411 | 0.0342 | 0.0342 | TRUE | 0.3879 |
| Nmnat1 | 347 | 241 | 232 | 264 | 230 | 275 | 270 | 256 | 250 | 263 | 259 | 271 | 261.65 | 259 | 264 | 1.021 | 0.031 | 0.80608864 | 0.88797109 | 0 | 0.0115 | 0.0073 | 0.0073 | TRUE | 0.0161 |
| Nmnat2 | 2180 | 1658 | 1949 | 2751 | 2359 | 2587 | 1693 | 1764 | 2103 | 2738 | 2661 | 2554 | 2252.16 | 1853 | 2651 | 1.432 | 0.518 | 7.12E-10 | 4.96E-08 | 0.0069 | 0.0035 | 0.0046 | 0.0046 | TRUE | 0.1667 |
| Nmnat3 | 152 | 112 | 129 | 131 | 150 | 171 | 118 | 119 | 139 | 130 | 169 | 169 | 140.8 | 125 | 156 | 1.246 | 0.317 | 0.07762064 | 0.18493612 | 0.0083 | 0.0193 | 0.0161 | 0.0161 | TRUE | 0.2227 |
| Nmral1 | 992 | 681 | 663 | 636 | 536 | 612 | 770 | 724 | 715 | 633 | 605 | 604 | 675.34 | 736 | 614 | 0.833 | -0.264 | 0.00224925 | 0.0132244 | 0 | 0.0059 | 0.0039 | 0.0039 | TRUE | 0.0201 |
| Nmrk1 | 453 | 367 | 336 | 439 | 341 | 391 | 352 | 390 | 363 | 437 | 385 | 386 | 385.4 | 368 | 403 | 1.095 | 0.131 | 0.24161638 | 0.40930928 | 0.0017 | 0.0086 | 0.0064 | 0.0064 | TRUE | 0.0652 |
| Nmrk2 | 1 | 1 | 0 | 0 | 1 | 5 | 1 | 1 | 0 | 0 | 1 | 5 | 1.32 | 1 | 2 | 3.261 | 1.705 | 0.37271727 | NA | 0 | 1.8015 | 1.6101 | 1.6101 | TRUE | 0.6406 |
| Nms | 0 | 1 | 0 | 0 | 0 | 0 | 0 | 1 | 0 | 0 | 0 | 0 | 0.18 | 0 | 0 | 0.552 | -0.857 | 0.8336548 | NA | 0 | 13.3714 | 10 | 10 | TRUE | 0.5216 |
| Nmt1 | 15335 | 11053 | 10933 | 11877 | 10693 | 12256 | 11910 | 11757 | 11797 | 11822 | 12063 | 12099 | 11907.89 | 11821 | 11995 | 1.015 | 0.021 | 0.64001035 | 0.77502905 | 0 | 0.0026 | 0.0014 | 0.0014 | TRUE | 0.0019 |
| Nmt2 | 5918 | 4375 | 4250 | 4154 | 3517 | 3957 | 4596 | 4654 | 4586 | 4135 | 3968 | 3906 | 4307.37 | 4612 | 4003 | 0.868 | -0.204 | 0.00011256 | 0.00133841 | 2.00E-04 | 0.003 | 0.0018 | 0.0018 | TRUE | 0.0102 |
| Nmu | 5 | 5 | 3 | 5 | 7 | 10 | 4 | 5 | 3 | 5 | 8 | 10 | 5.86 | 4 | 8 | 1.833 | 0.874 | 0.27348062 | NA | 0 | 0.4066 | 0.2742 | 0.2742 | TRUE | 0.2361 |
| Nmur1 | 3 | 0 | 0 | 4 | 2 | 0 | 2 | 0 | 0 | 4 | 2 | 0 | 1.43 | 1 | 2 | 2.468 | 1.304 | 0.49193104 | NA | 2.4168 | 1.6622 | 1.7699 | 1.7699 | TRUE | 0.4326 |
| Nmur2 | 5 | 2 | 4 | 4 | 3 | 4 | 4 | 2 | 4 | 4 | 3 | 4 | 3.61 | 3 | 4 | 1.092 | 0.127 | 0.89643617 | NA | 0 | 0.6595 | 0.4138 | 0.4138 | TRUE | 0.1476 |
| Nnat | 708 | 491 | 555 | 627 | 527 | 713 | 550 | 522 | 599 | 624 | 595 | 704 | 598.91 | 557 | 641 | 1.152 | 0.204 | 0.04486282 | 0.12444233 | 0.0045 | 0.0064 | 0.0058 | 0.0058 | TRUE | 0.0877 |
| Nnmt | 10 | 6 | 5 | 1 | 1 | 3 | 8 | 6 | 5 | 1 | 1 | 3 | 4.1 | 6 | 2 | 0.261 | -1.937 | 0.06119013 | NA | 0 | 0.5798 | 0.395 | 0.395 | TRUE | 0.3433 |
| Nnt | 2097 | 1397 | 1483 | 1758 | 1581 | 1820 | 1629 | 1486 | 1600 | 1750 | 1784 | 1797 | 1674.14 | 1572 | 1777 | 1.13 | 0.176 | 0.00863961 | 0.03712747 | 6.00E-04 | 0.0038 | 0.0026 | 0.0026 | TRUE | 0.027 |
| Noa1 | 2466 | 1884 | 1803 | 2016 | 1829 | 2108 | 1915 | 2004 | 1945 | 2007 | 2063 | 2081 | 2002.6 | 1955 | 2050 | 1.049 | 0.069 | 0.25533864 | 0.42505884 | 0 | 0.0036 | 0.0022 | 0.0022 | TRUE | 0.0058 |
| Nob1 | 5564 | 3949 | 4107 | 3971 | 3699 | 4511 | 4321 | 4201 | 4431 | 3953 | 4173 | 4453 | 4255.31 | 4318 | 4193 | 0.971 | -0.042 | 0.49597058 | 0.66060083 | 0.0017 | 0.003 | 0.0025 | 0.0025 | TRUE | 0.0362 |
| Nobox | 22 | 14 | 22 | 19 | 14 | 11 | 17 | 15 | 24 | 19 | 16 | 11 | 16.88 | 19 | 15 | 0.82 | -0.287 | 0.5498453 | NA | 0 | 0.1428 | 0.1058 | 0.1058 | TRUE | 0.2996 |
| Noc2l | 14912 | 11370 | 12230 | 12128 | 11412 | 13393 | 11582 | 12094 | 13196 | 12072 | 12874 | 13221 | 12506.46 | 12291 | 12722 | 1.035 | 0.05 | 0.43392291 | 0.60572701 | 0.0029 | 0.0026 | 0.0029 | 0.0029 | TRUE | 0.0507 |
| Noc3l | 5216 | 3738 | 3191 | 3308 | 2906 | 3381 | 4051 | 3976 | 3443 | 3293 | 3278 | 3338 | 3563.13 | 3823 | 3303 | 0.863 | -0.212 | 0.00273465 | 0.01535206 | 0.0034 | 0.0031 | 0.0033 | 0.0033 | TRUE | 0.0914 |
| Noc4l | 5020 | 3619 | 3340 | 3336 | 3054 | 3582 | 3899 | 3849 | 3604 | 3321 | 3445 | 3536 | 3609 | 3784 | 3434 | 0.907 | -0.141 | 0.01814484 | 0.06426088 | 0.001 | 0.0031 | 0.0023 | 0.0023 | TRUE | 0.0209 |
| Nod1 | 87 | 67 | 49 | 63 | 63 | 63 | 68 | 71 | 53 | 63 | 71 | 62 | 64.61 | 64 | 65 | 1.018 | 0.026 | 0.9146509 | 0.9529197 | 0 | 0.0391 | 0.0281 | 0.0281 | TRUE | 0.1858 |
| Nod2 | 141 | 116 | 138 | 147 | 151 | 160 | 110 | 123 | 149 | 146 | 170 | 158 | 142.73 | 127 | 158 | 1.248 | 0.32 | 0.07303066 | 0.17694985 | 0.0077 | 0.019 | 0.0158 | 0.0158 | TRUE | 0.2219 |
| Nodal | 3059 | 1975 | 1630 | 2400 | 1912 | 2092 | 2376 | 2101 | 1759 | 2389 | 2157 | 2065 | 2141.06 | 2079 | 2204 | 1.059 | 0.083 | 0.40908899 | 0.58184614 | 0.0136 | 0.0035 | 0.0068 | 0.0068 | TRUE | 0.2182 |
| Nog | 53 | 31 | 30 | 69 | 52 | 48 | 41 | 33 | 32 | 69 | 59 | 47 | 46.87 | 35 | 58 | 1.629 | 0.704 | 0.017218 | 0.061905 | 0 | 0.053 | 0.0406 | 0.0406 | TRUE | 0.1507 |
| Nol10 | 6740 | 4679 | 4123 | 4636 | 4107 | 4585 | 5235 | 4977 | 4449 | 4615 | 4633 | 4526 | 4739.04 | 4887 | 4591 | 0.939 | -0.091 | 0.17966876 | 0.33448315 | 0.0031 | 0.0029 | 0.0031 | 0.0031 | TRUE | 0.0746 |
| Nol11 | 12814 | 9185 | 9298 | 10083 | 8843 | 10556 | 9952 | 9770 | 10032 | 10037 | 9976 | 10420 | 10031.22 | 9918 | 10144 | 1.023 | 0.033 | 0.49490405 | 0.65988266 | 2.00E-04 | 0.0027 | 0.0015 | 0.0015 | TRUE | 0.0069 |
| Nol12 | 1598 | 1165 | 1005 | 980 | 985 | 1138 | 1241 | 1239 | 1084 | 975 | 1111 | 1123 | 1129.12 | 1188 | 1070 | 0.9 | -0.153 | 0.08331581 | 0.19429481 | 0.005 | 0.0045 | 0.0047 | 0.0047 | TRUE | 0.0721 |
| Nol3 | 282 | 223 | 220 | 270 | 254 | 276 | 219 | 237 | 237 | 269 | 287 | 272 | 253.56 | 231 | 276 | 1.195 | 0.257 | 0.04319839 | 0.1210564 | 0 | 0.0118 | 0.0077 | 0.0077 | TRUE | 0.0259 |
| Nol4 | 34 | 25 | 27 | 16 | 15 | 17 | 26 | 27 | 29 | 16 | 17 | 17 | 21.96 | 27 | 17 | 0.605 | -0.725 | 0.07447907 | 0.1794554 | 0 | 0.1104 | 0.0702 | 0.0702 | TRUE | 0.0188 |
| Nol6 | 15588 | 11037 | 9514 | 10295 | 9160 | 10947 | 12107 | 11740 | 10266 | 10248 | 10333 | 10806 | 10916.57 | 11371 | 10462 | 0.92 | -0.12 | 0.07587781 | 0.18166189 | 0.0039 | 0.0026 | 0.0032 | 0.0032 | TRUE | 0.0881 |
| Nol7 | 3582 | 2572 | 2536 | 2565 | 2304 | 2564 | 2782 | 2736 | 2736 | 2553 | 2599 | 2531 | 2656.25 | 2751 | 2561 | 0.931 | -0.104 | 0.06174188 | 0.15657854 | 0 | 0.0033 | 0.0018 | 0.0018 | TRUE | 0.002 |
| Nol8 | 5399 | 3934 | 3593 | 4116 | 3676 | 4206 | 4193 | 4185 | 3877 | 4097 | 4147 | 4152 | 4108.41 | 4085 | 4132 | 1.011 | 0.016 | 0.77584453 | 0.86757979 | 8.00E-04 | 0.003 | 0.0021 | 0.0021 | TRUE | 0.0239 |
| Nol9 | 13712 | 10094 | 9846 | 10619 | 9834 | 11749 | 10650 | 10737 | 10624 | 10570 | 11094 | 11598 | 10878.69 | 10670 | 11087 | 1.039 | 0.055 | 0.29350152 | 0.46538908 | 0.001 | 0.0026 | 0.0019 | 0.0019 | TRUE | 0.0204 |
| Nolc1 | 15907 | 12048 | 10676 | 10504 | 9909 | 11603 | 12354 | 12815 | 11519 | 10456 | 11178 | 11454 | 11629.48 | 12229 | 11029 | 0.902 | -0.149 | 0.0139435 | 0.0529194 | 0.0024 | 0.0026 | 0.0026 | 0.0026 | TRUE | 0.0314 |
| Nom1 | 3371 | 2366 | 2045 | 2106 | 1909 | 2158 | 2618 | 2517 | 2207 | 2096 | 2154 | 2130 | 2286.91 | 2447 | 2127 | 0.868 | -0.204 | 0.00612822 | 0.02867126 | 0.0033 | 0.0035 | 0.0035 | 0.0035 | TRUE | 0.0889 |
| Nomo1 | 14386 | 9937 | 10176 | 10869 | 9486 | 10775 | 11173 | 10570 | 10980 | 10819 | 10701 | 10637 | 10813.24 | 10908 | 10719 | 0.983 | -0.025 | 0.59764735 | 0.74367484 | 3.00E-04 | 0.0026 | 0.0016 | 0.0016 | TRUE | 0.0089 |
| Nono | 21391 | 15360 | 14948 | 15455 | 14143 | 16389 | 16613 | 16338 | 16129 | 15384 | 15955 | 16178 | 16099.6 | 16360 | 15839 | 0.968 | -0.047 | 0.31946129 | 0.49335473 | 4.00E-04 | 0.0026 | 0.0015 | 0.0015 | TRUE | 0.0077 |

| | | | | | | | | | | | | | | | | | | | | | | | | | |
|---|---|---|---|---|---|---|---|---|---|---|---|---|---|---|---|---|---|---|---|---|---|---|---|---|---|
| Nop10 | 5737 | 3986 | 4112 | 4441 | 3776 | 4654 | 4456 | 4240 | 4437 | 4421 | 4260 | 4594 | 4401.14 | 4378 | 4425 | 1.011 | 0.016 | 0.78322131 | 0.87246374 | 8.00E-04 | 0.003 | 0.0021 | 0.0021 | TRUE | 0.0137 |
| Nop14 | 7086 | 4870 | 4761 | 4862 | 3993 | 4763 | 5503 | 5180 | 5137 | 4840 | 4505 | 4702 | 4977.77 | 5273 | 4682 | 0.888 | -0.172 | 0.00272982 | 0.01533096 | 0.0011 | 0.0029 | 0.0022 | 0.0022 | TRUE | 0.018 |
| Nop16 | 3225 | 2298 | 2028 | 1773 | 1568 | 1917 | 2505 | 2444 | 2188 | 1765 | 1769 | 1892 | 2093.89 | 2379 | 1809 | 0.76 | -0.396 | 5.15E-08 | 2.15E-06 | 0.0029 | 0.0036 | 0.0033 | 0.0033 | TRUE | 0.0591 |
| Nop2 | 16777 | 11559 | 9612 | 9948 | 9032 | 10461 | 13030 | 12295 | 10371 | 9902 | 10189 | 10327 | 11019.05 | 11899 | 10139 | 0.852 | -0.231 | 0.00291262 | 0.01617331 | 0.0067 | 0.0026 | 0.0043 | 0.0043 | TRUE | 0.1539 |
| Nop56 | 15814 | 11417 | 9247 | 8675 | 8034 | 9368 | 12282 | 12144 | 9977 | 8635 | 9063 | 9248 | 10224.92 | 11468 | 8982 | 0.783 | -0.353 | 6.49E-06 | 0.0001318 | 0.0068 | 0.0027 | 0.0043 | 0.0043 | TRUE | 0.1576 |
| Nop58 | 17650 | 12499 | 11168 | 11892 | 10625 | 12160 | 13708 | 13295 | 12050 | 11837 | 11986 | 12004 | 12480.07 | 13018 | 11942 | 0.917 | -0.125 | 0.03511817 | 0.10413775 | 0.0021 | 0.0026 | 0.0024 | 0.0024 | TRUE | 0.0516 |
| Nop9 | 5837 | 4175 | 3998 | 4295 | 3787 | 4351 | 4533 | 4441 | 4314 | 4275 | 4272 | 4295 | 4355.09 | 4429 | 4281 | 0.966 | -0.05 | 0.33888635 | 0.51299541 | 0 | 0.003 | 0.0017 | 0.0017 | TRUE | 0.0063 |
| Nos1 | 451 | 354 | 293 | 431 | 332 | 397 | 350 | 377 | 316 | 429 | 375 | 392 | 373.07 | 348 | 399 | 1.147 | 0.197 | 0.09043852 | 0.20537001 | 0.0031 | 0.0088 | 0.0071 | 0.0071 | TRUE | 0.0676 |
| Nos1ap | 109 | 91 | 99 | 121 | 105 | 128 | 85 | 97 | 107 | 120 | 118 | 126 | 108.92 | 96 | 121 | 1.274 | 0.349 | 0.06761878 | 0.16718133 | 0 | 0.0242 | 0.017 | 0.017 | TRUE | 0.12 |
| Nos2 | 39 | 40 | 42 | 51 | 40 | 45 | 30 | 43 | 45 | 51 | 45 | 44 | 43.08 | 39 | 47 | 1.199 | 0.262 | 0.38700401 | 0.56071448 | 0 | 0.0574 | 0.0425 | 0.0425 | TRUE | 0.3661 |
| Nos3 | 523 | 440 | 474 | 685 | 653 | 816 | 406 | 468 | 511 | 682 | 737 | 806 | 601.61 | 462 | 742 | 1.61 | 0.687 | 4.75E-10 | 3.46E-08 | 0.0084 | 0.0064 | 0.007 | 0.007 | TRUE | 0.1377 |
| Nosip | 2525 | 1838 | 1740 | 1770 | 1601 | 1903 | 1961 | 1955 | 1877 | 1762 | 1806 | 1879 | 1873.34 | 1931 | 1816 | 0.94 | -0.089 | 0.15915091 | 0.30839584 | 3.00E-04 | 0.0037 | 0.0024 | 0.0024 | TRUE | 0.0113 |
| Nostrin | 31 | 21 | 13 | 16 | 18 | 14 | 24 | 22 | 14 | 16 | 20 | 14 | 18.42 | 20 | 17 | 0.819 | -0.288 | 0.53054756 | 0.69038787 | 0 | 0.1311 | 0.0967 | 0.0967 | TRUE | 0.3559 |
| Notch1 | 4047 | 3047 | 3485 | 4088 | 3723 | 4198 | 3143 | 3241 | 3760 | 4069 | 4200 | 4144 | 3759.61 | 3381 | 4138 | 1.224 | 0.292 | 6.62E-05 | 0.00088016 | 0.0045 | 0.0031 | 0.0036 | 0.0036 | TRUE | 0.1161 |
| Notch2 | 3823 | 2684 | 2384 | 2436 | 2183 | 2568 | 2969 | 2855 | 2572 | 2425 | 2463 | 2535 | 2636.48 | 2799 | 2474 | 0.883 | -0.179 | 0.00970522 | 0.04048939 | 0.0023 | 0.0033 | 0.0031 | 0.0031 | TRUE | 0.0604 |
| Notch3 | 4969 | 3619 | 3706 | 4770 | 4053 | 4172 | 3859 | 3849 | 3999 | 4748 | 4572 | 4118 | 4191.02 | 3902 | 4479 | 1.148 | 0.199 | 0.00246891 | 0.01422555 | 0.0025 | 0.003 | 0.0029 | 0.0029 | TRUE | 0.0609 |
| Notch4 | 2767 | 2238 | 2263 | 3613 | 3393 | 3524 | 2149 | 2381 | 2442 | 3596 | 3828 | 3479 | 2979.01 | 2324 | 3634 | 1.565 | 0.646 | 2.39E-20 | 7.96E-18 | 0.0029 | 0.0032 | 0.0032 | 0.0032 | TRUE | 0.0539 |
| Noto | 3 | 4 | 0 | 0 | 0 | 0 | 2 | 4 | 0 | 0 | 0 | 0 | 1.1 | 2 | 0 | 0.087 | -3.53 | 0.1154815 | NA | 6.0628 | 2.162 | 2.4038 | 2.4038 | TRUE | 0.0619 |
| Notum | 453 | 357 | 268 | 377 | 416 | 493 | 352 | 380 | 289 | 375 | 469 | 487 | 391.99 | 340 | 444 | 1.303 | 0.381 | 0.00520736 | 0.02530731 | 0.0164 | 0.0085 | 0.0108 | 0.0108 | TRUE | 0.2135 |
| Nov | 43 | 19 | 21 | 19 | 20 | 16 | 33 | 20 | 23 | 19 | 23 | 16 | 22.26 | 25 | 19 | 0.741 | -0.433 | 0.30584953 | 0.47853155 | 0 | 0.1089 | 0.0826 | 0.0826 | TRUE | 0.5488 |
| Nova1 | 227 | 151 | 96 | 110 | 92 | 111 | 176 | 161 | 104 | 109 | 104 | 110 | 127.23 | 147 | 108 | 0.73 | -0.454 | 0.03196535 | 0.09696111 | 0.0318 | 0.0211 | 0.0242 | 0.0242 | TRUE | 0.6703 |
| Nova2 | 909 | 789 | 801 | 928 | 844 | 904 | 706 | 839 | 864 | 924 | 952 | 892 | 862.96 | 803 | 923 | 1.151 | 0.203 | 0.03086648 | 0.09457119 | 0.0053 | 0.0052 | 0.0052 | 0.0052 | TRUE | 0.1398 |
| Nox1 | 3 | 2 | 2 | 0 | 1 | 0 | 2 | 2 | 2 | 0 | 1 | 0 | 1.29 | 2 | 0 | 0.168 | -2.574 | 0.16701241 | NA | 0 | 1.8389 | 1.2821 | 1.2821 | TRUE | 0.5075 |
| Nox3 | 0 | 0 | 0 | 0 | 0 | 0 | 0 | 0 | 0 | 0 | 0 | 0 | 0 | 0 | 0 | NA | NA | NA | NA | NA | NA | NA | NA | NA | NA |
| Nox4 | 357 | 237 | 245 | 264 | 199 | 245 | 277 | 252 | 264 | 263 | 224 | 242 | 253.81 | 264 | 243 | 0.918 | -0.123 | 0.34683518 | 0.52121914 | 0 | 0.0118 | 0.0084 | 0.0084 | TRUE | 0.056 |
| Noxa1 | 15 | 10 | 8 | 13 | 11 | 4 | 12 | 11 | 9 | 13 | 12 | 4 | 10.04 | 11 | 10 | 0.939 | -0.091 | 0.88632264 | NA | 0 | 0.2386 | 0.1923 | 0.1923 | TRUE | 0.9662 |
| Noxo1 | 124 | 106 | 105 | 89 | 78 | 86 | 96 | 113 | 113 | 89 | 88 | 85 | 97.3 | 107 | 87 | 0.814 | -0.297 | 0.13544478 | 0.27479237 | 0 | 0.0268 | 0.0181 | 0.0181 | TRUE | 0.092 |
| Noxred1 | 377 | 271 | 255 | 272 | 224 | 297 | 293 | 288 | 275 | 271 | 253 | 293 | 278.81 | 285 | 272 | 0.954 | -0.068 | 0.5850447 | 0.73447904 | 0 | 0.0109 | 0.0076 | 0.0076 | TRUE | 0.0511 |
| Npas1 | 11 | 7 | 6 | 13 | 15 | 12 | 9 | 7 | 6 | 13 | 17 | 12 | 10.7 | 7 | 14 | 1.838 | 0.878 | 0.13518998 | NA | 0 | 0.224 | 0.1476 | 0.1476 | TRUE | 0.1424 |
| Npas2 | 40 | 32 | 33 | 21 | 26 | 23 | 31 | 34 | 36 | 21 | 29 | 23 | 28.94 | 34 | 24 | 0.723 | -0.467 | 0.19320087 | 0.35190773 | 0 | 0.0843 | 0.0571 | 0.0571 | TRUE | 0.1789 |
| Npas3 | 26 | 20 | 13 | 23 | 25 | 19 | 20 | 21 | 14 | 23 | 28 | 19 | 20.89 | 18 | 23 | 1.25 | 0.321 | 0.45187908 | 0.62180684 | 0 | 0.1159 | 0.0832 | 0.0832 | TRUE | 0.2099 |
| Npas4 | 3 | 0 | 0 | 1 | 0 | 0 | 2 | 0 | 0 | 1 | 0 | 0 | 0.55 | 1 | 0 | 0.458 | -1.128 | 0.69820437 | NA | 0 | 4.2787 | 4.502 | 4.502 | TRUE | 1.4399 |
| Npat | 4400 | 3261 | 3184 | 3462 | 3005 | 3265 | 3417 | 3469 | 3436 | 3446 | 3390 | 3223 | 3396.76 | 3441 | 3353 | 0.975 | -0.037 | 0.50602237 | 0.66949715 | 3.00E-04 | 0.0031 | 0.002 | 0.002 | TRUE | 0.0141 |
| Npb | 14 | 2 | 12 | 7 | 9 | 12 | 11 | 2 | 13 | 7 | 10 | 12 | 9.15 | 9 | 10 | 1.108 | 0.147 | 0.83327684 | NA | 0.1886 | 0.2614 | 0.244 | 0.244 | TRUE | 1.2437 |
| Npbwr1 | 3 | 3 | 0 | 3 | 0 | 1 | 2 | 3 | 0 | 3 | 0 | 1 | 1.58 | 2 | 1 | 0.724 | -0.466 | 0.77897413 | NA | 0.6399 | 1.5002 | 1.3405 | 1.3405 | TRUE | 0.6377 |
| Npc1 | 3423 | 2474 | 2275 | 2436 | 2529 | 2793 | 2658 | 2632 | 2455 | 2425 | 2853 | 2757 | 2629.94 | 2582 | 2678 | 1.037 | 0.052 | 0.48397853 | 0.65044468 | 0.0037 | 0.0033 | 0.0036 | 0.0036 | TRUE | 0.0834 |
| Npc1l1 | 69 | 23 | 41 | 43 | 35 | 36 | 54 | 24 | 44 | 43 | 39 | 36 | 40.02 | 41 | 39 | 0.955 | -0.067 | 0.84534866 | 0.91209847 | 0.0541 | 0.0616 | 0.0596 | 0.0596 | TRUE | 0.5191 |
| Npc2 | 5374 | 3979 | 4816 | 5053 | 4397 | 5243 | 4174 | 4232 | 5196 | 5030 | 4960 | 5176 | 4794.71 | 4534 | 5055 | 1.115 | 0.158 | 0.05420458 | 0.14260005 | 0.0079 | 0.0029 | 0.0046 | 0.0046 | TRUE | 0.1981 |
| Npcd | 1 | 3 | 5 | 7 | 4 | 8 | 1 | 3 | 5 | 7 | 5 | 8 | 4.79 | 3 | 7 | 2.157 | 1.109 | 0.23481289 | NA | 0 | 0.4972 | 0.388 | 0.388 | TRUE | 0.5182 |
| Npdc1 | 1102 | 852 | 1030 | 1172 | 1122 | 1336 | 856 | 906 | 1111 | 1167 | 1266 | 1319 | 1104.11 | 958 | 1251 | 1.307 | 0.387 | 0.00013933 | 0.00160088 | 0.0105 | 0.0046 | 0.0065 | 0.0065 | TRUE | 0.2325 |
| Npepl1 | 1981 | 1516 | 1518 | 1546 | 1398 | 1635 | 1539 | 1613 | 1638 | 1539 | 1577 | 1614 | 1586.5 | 1597 | 1577 | 0.988 | -0.017 | 0.79921449 | 0.88284819 | 0 | 0.0039 | 0.0025 | 0.0025 | TRUE | 0.0126 |
| Npepps | 22597 | 17256 | 18107 | 18996 | 16882 | 19909 | 17550 | 18355 | 19537 | 18909 | 19045 | 19653 | 18841.49 | 18481 | 19202 | 1.039 | 0.055 | 0.31507363 | 0.48870278 | 0.0016 | 0.0026 | 0.0021 | 0.0021 | TRUE | 0.0306 |
| Npff | 18 | 11 | 1 | 3 | 1 | 1 | 14 | 12 | 1 | 3 | 1 | 1 | 5.31 | 9 | 2 | 0.19 | -2.397 | 0.02710758 | NA | 0 | 0.4488 | 0.4918 | 0.4918 | TRUE | 0.9092 |
| Npffr1 | 2 | 2 | 0 | 1 | 3 | 1 | 2 | 2 | 0 | 1 | 3 | 1 | 1.51 | 1 | 2 | 1.42 | 0.506 | 0.75038374 | NA | 0 | 1.5742 | 1.1441 | 1.1441 | TRUE | 0.4377 |
| Npffr2 | 3 | 2 | 0 | 1 | 2 | 0 | 2 | 2 | 0 | 1 | 2 | 0 | 1.28 | 1 | 1 | 0.703 | -0.509 | 0.77722559 | NA | 0 | 1.8471 | 1.525 | 1.525 | TRUE | 0.3027 |
| Nphp1 | 1181 | 853 | 845 | 933 | 727 | 938 | 917 | 907 | 912 | 929 | 820 | 926 | 901.85 | 912 | 892 | 0.978 | -0.032 | 0.69879881 | 0.8170087 | 0.0011 | 0.0051 | 0.0038 | 0.0038 | TRUE | 0.0571 |
| Nphp3 | 2423 | 1896 | 2151 | 2521 | 2181 | 2476 | 1882 | 2017 | 2321 | 2509 | 2460 | 2444 | 2272.25 | 2073 | 2471 | 1.193 | 0.255 | 0.00142391 | 0.0092974 | 0.0053 | 0.0035 | 0.0042 | 0.0042 | TRUE | 0.1317 |
| Nphp4 | 927 | 759 | 733 | 807 | 727 | 830 | 720 | 807 | 791 | 803 | 820 | 819 | 793.49 | 773 | 814 | 1.055 | 0.078 | 0.35445987 | 0.52886383 | 0 | 0.0054 | 0.0038 | 0.0038 | TRUE | 0.045 |
| Nphs1 | 1441 | 1345 | 1596 | 2945 | 2775 | 3309 | 1119 | 1431 | 1722 | 2931 | 3130 | 3266 | 2266.72 | 1424 | 3109 | 2.187 | 1.129 | 8.02E-22 | 2.94E-19 | 0.0231 | 0.0035 | 0.0095 | 0.0095 | TRUE | 0.4307 |
| Nphs1as | 50 | 55 | 46 | 55 | 59 | 64 | 39 | 59 | 50 | 55 | 67 | 63 | 55.24 | 49 | 62 | 1.264 | 0.338 | 0.21499918 | 0.37765193 | 0 | 0.0453 | 0.0351 | 0.0351 | TRUE | 0.3152 |
| Nphs2 | 8 | 3 | 10 | 5 | 3 | 3 | 6 | 3 | 11 | 5 | 3 | 3 | 5.25 | 7 | 4 | 0.563 | -0.828 | 0.34100227 | NA | 0 | 0.4536 | 0.3355 | 0.3355 | TRUE | 0.2007 |
| Npl | 1832 | 1187 | 1297 | 1329 | 1130 | 1295 | 1423 | 1263 | 1399 | 1323 | 1275 | 1278 | 1326.82 | 1362 | 1292 | 0.948 | -0.077 | 0.30350803 | 0.47603761 | 0.0016 | 0.0042 | 0.0032 | 0.0032 | TRUE | 0.0479 |
| Nploc4 | 11778 | 8111 | 7674 | 8448 | 7213 | 8412 | 9147 | 8628 | 8280 | 8409 | 8137 | 8304 | 8484.22 | 8685 | 8283 | 0.954 | -0.069 | 0.21481225 | 0.37748702 | 0.0012 | 0.0027 | 0.0021 | 0.0021 | TRUE | 0.0268 |
| Npm1 | 255733 | 183020 | 179515 | 185646 | 159292 | 188641 | 198617 | 194676 | 193695 | 184792 | 179698 | 186218 | 189615.94 | 195663 | 183569 | 0.938 | -0.092 | 0.03282646 | 0.09902672 | 3.00E-04 | 0.0024 | 0.0013 | 0.0013 | TRUE | 0.0042 |
| Npm2 | 37 | 27 | 19 | 28 | 23 | 17 | 29 | 29 | 21 | 28 | 26 | 17 | 24.76 | 26 | 24 | 0.9 | -0.151 | 0.70447761 | 0.82107628 | 0 | 0.0982 | 0.0743 | 0.0743 | TRUE | 0.3861 |
| Npm3 | 5542 | 4148 | 4351 | 5111 | 4424 | 5417 | 4304 | 4412 | 4695 | 5087 | 4991 | 5347 | 4806.12 | 4470 | 5142 | 1.151 | 0.203 | 0.00059858 | 0.00495683 | 0.0013 | 0.0029 | 0.0023 | 0.0023 | TRUE | 0.0234 |
| Npm3-ps1 | 541 | 369 | 369 | 462 | 407 | 465 | 420 | 393 | 398 | 460 | 459 | 459 | 431.48 | 404 | 459 | 1.137 | 0.185 | 0.06647425 | 0.1650274 | 0 | 0.0079 | 0.005 | 0.005 | TRUE | 0.0161 |
| Npnt | 740 | 571 | 634 | 697 | 620 | 679 | 575 | 607 | 684 | 694 | 699 | 670 | 654.94 | 622 | 688 | 1.108 | 0.148 | 0.12001987 | 0.25171593 | 0.0028 | 0.006 | 0.005 | 0.005 | TRUE | 0.0884 |
| Nppa | 0 | 0 | 0 | 0 | 0 | 0 | 0 | 0 | 0 | 0 | 0 | 0 | 0 | 0 | 0 | NA | NA | NA | NA | NA | NA | NA | NA | NA | NA |
| Nppb | 28 | 23 | 19 | 25 | 13 | 20 | 22 | 24 | 21 | 25 | 15 | 20 | 21 | 22 | 20 | 0.893 | -0.163 | 0.6997064 | 0.81780224 | 0 | 0.1153 | 0.0809 | 0.0809 | TRUE | 0.2044 |
| Nppc | 66 | 32 | 32 | 54 | 43 | 44 | 51 | 34 | 35 | 54 | 49 | 43 | 44.25 | 40 | 49 | 1.203 | 0.267 | 0.38064793 | 0.55516627 | 0.0142 | 0.056 | 0.0442 | 0.0442 | TRUE | 0.5346 |
| Npr1 | 1155 | 692 | 593 | 546 | 432 | 556 | 897 | 736 | 640 | 543 | 487 | 549 | 642.11 | 758 | 526 | 0.694 | -0.527 | 1.64E-05 | 0.00028514 | 0.0156 | 0.0061 | 0.0092 | 0.0092 | TRUE | 0.2858 |
| Npr2 | 182 | 152 | 132 | 153 | 189 | 154 | 141 | 162 | 142 | 152 | 213 | 152 | 160.5 | 148 | 172 | 1.16 | 0.214 | 0.23273537 | 0.39881149 | 0.0165 | 0.0172 | 0.017 | 0.017 | TRUE | 0.4405 |
| Npr3 | 252 | 171 | 172 | 315 | 272 | 348 | 196 | 182 | 186 | 314 | 307 | 344 | 254.52 | 188 | 322 | 1.71 | 0.774 | 2.65E-09 | 1.57E-07 | 0 | 0.0117 | 0.008 | 0.008 | TRUE | 0.042 |
| Nprl2 | 1150 | 815 | 970 | 963 | 856 | 1094 | 893 | 867 | 1047 | 959 | 966 | 1080 | 968.48 | 936 | 1002 | 1.072 | 0.1 | 0.29036872 | 0.4624223 | 0.0061 | 0.0049 | 0.0054 | 0.0054 | TRUE | 0.1258 |
| Nprl3 | 2837 | 2054 | 1761 | 1951 | 1822 | 1965 | 2203 | 2185 | 1900 | 1942 | 2055 | 1940 | 2037.58 | 2096 | 1979 | 0.943 | -0.084 | 0.26041749 | 0.43057079 | 0.0031 | 0.0036 | 0.0035 | 0.0035 | TRUE | 0.0799 |
| Nps | 5 | 2 | 4 | 2 | 1 | 2 | 4 | 2 | 4 | 2 | 1 | 2 | 2.57 | 3 | 2 | 0.495 | -1.015 | 0.39481061 | NA | 0 | 0.9246 | 0.5824 | 0.5824 | TRUE | 0.1495 |
| Npsr1 | 2 | 1 | 2 | 2 | 2 | 5 | 2 | 1 | 2 | 2 | 2 | 5 | 2.33 | 2 | 3 | 1.935 | 0.953 | 0.44703515 | NA | 0 | 1.0212 | 0.6594 | 0.6594 | TRUE | 0.3933 |
| Nptn | 5213 | 3686 | 3628 | 3812 | 3229 | 3793 | 4049 | 3921 | 3915 | 3794 | 3643 | 3744 | 3844.24 | 3962 | 3727 | 0.941 | -0.088 | 0.09875979 | 0.2190864 | 2.00E-04 | 0.003 | 0.0018 | 0.0018 | TRUE | 0.0047 |

| | | | | | | | | | | | | | | | | | | | | | | | | | |
|---|---|---|---|---|---|---|---|---|---|---|---|---|---|---|---|---|---|---|---|---|---|---|---|---|---|
| Nptx1 | 185 | 142 | 124 | 161 | 122 | 177 | 144 | 151 | 134 | 160 | 138 | 175 | 150.19 | 143 | 158 | 1.105 | 0.143 | 0.3926889 | 0.56631041 | 0 | 0.0182 | 0.0136 | 0.0136 | TRUE | 0.1231 |
| Nptx2 | 2383 | 1866 | 1681 | 2231 | 2120 | 2228 | 1851 | 1985 | 1814 | 2221 | 2392 | 2199 | 2076.85 | 1883 | 2271 | 1.206 | 0.27 | 7.17E-05 | 0.00093338 | 0.0015 | 0.0036 | 0.0028 | 0.0028 | TRUE | 0.0264 |
| Nptxr | 100 | 70 | 63 | 112 | 98 | 78 | 78 | 74 | 68 | 111 | 111 | 77 | 86.52 | 73 | 100 | 1.354 | 0.437 | 0.05369201 | 0.1415904 | 0.0137 | 0.0298 | 0.0252 | 0.0252 | TRUE | 0.3958 |
| Npvf | 1 | 0 | 0 | 0 | 0 | 0 | 1 | 0 | 0 | 0 | 0 | 0 | 0.13 | 0 | 0 | 0.552 | -0.857 | 0.8336548 | NA | 0 | 18.3121 | 10 | 10 | TRUE | 0.2566 |
| Npw | 66 | 49 | 49 | 75 | 54 | 59 | 51 | 52 | 53 | 75 | 61 | 58 | 58.34 | 52 | 65 | 1.242 | 0.313 | 0.21940498 | 0.38257355 | 0 | 0.043 | 0.0295 | 0.0295 | TRUE | 0.1665 |
| Npy | 16 | 5 | 9 | 7 | 12 | 14 | 12 | 5 | 10 | 7 | 14 | 14 | 10.3 | 9 | 12 | 1.231 | 0.3 | 0.63045654 | NA | 0 | 0.2326 | 0.1823 | 0.1823 | TRUE | 0.4426 |
| Npy1r | 12 | 1 | 4 | 9 | 13 | 16 | 9 | 1 | 4 | 9 | 15 | 16 | 9.02 | 5 | 13 | 2.575 | 1.364 | 0.06209863 | NA | 0.1994 | 0.2652 | 0.2515 | 0.2515 | TRUE | 1.0045 |
| Npy2r | 0 | 0 | 1 | 0 | 0 | 0 | 0 | 0 | 1 | 0 | 0 | 0 | 0.18 | 0 | 0 | 0.552 | -0.857 | 0.8336548 | NA | 0 | 13.1817 | 10 | 10 | TRUE | 0.5365 |
| Npy5r | 3 | 2 | 2 | 2 | 3 | 1 | 2 | 2 | 2 | 2 | 3 | 1 | 2.16 | 2 | 2 | 0.949 | -0.076 | 0.95228531 | NA | 0 | 1.0982 | 0.6863 | 0.6863 | TRUE | 0.2272 |
| Npy6r | 0 | 0 | 0 | 0 | 0 | 0 | 0 | 0 | 0 | 0 | 0 | 0 | 0 | 0 | 0 | NA | NA | NA | NA | NA | NA | NA | NA | NA | NA |
| Nqo1 | 1250 | 887 | 693 | 845 | 713 | 798 | 971 | 943 | 748 | 841 | 804 | 788 | 849.21 | 887 | 811 | 0.913 | -0.132 | 0.20398118 | 0.36539704 | 0.0093 | 0.0052 | 0.0066 | 0.0066 | TRUE | 0.2217 |
| Nqo2 | 1782 | 1301 | 1576 | 1753 | 1414 | 1620 | 1384 | 1384 | 1700 | 1745 | 1595 | 1599 | 1567.94 | 1489 | 1646 | 1.106 | 0.146 | 0.10698382 | 0.23242434 | 0.0079 | 0.0039 | 0.0053 | 0.0053 | TRUE | 0.1829 |
| Nr0b1 | 1275 | 882 | 772 | 823 | 733 | 945 | 990 | 938 | 833 | 819 | 827 | 933 | 890.06 | 920 | 860 | 0.933 | -0.1 | 0.28612759 | 0.45811825 | 0.0055 | 0.0051 | 0.0052 | 0.0052 | TRUE | 0.0813 |
| Nr0b2 | 6 | 3 | 5 | 9 | 12 | 11 | 5 | 3 | 5 | 9 | 14 | 11 | 7.77 | 4 | 11 | 2.505 | 1.325 | 0.06112259 | NA | 0 | 0.3076 | 0.2052 | 0.2052 | TRUE | 0.1225 |
| Nr1d1 | 537 | 399 | 523 | 594 | 558 | 672 | 417 | 424 | 564 | 591 | 629 | 663 | 548.32 | 468 | 628 | 1.343 | 0.426 | 0.0005311 | 0.00451094 | 0.0145 | 0.0067 | 0.009 | 0.009 | TRUE | 0.3643 |
| Nr1d2 | 1651 | 1331 | 1374 | 1476 | 1201 | 1502 | 1282 | 1416 | 1483 | 1469 | 1355 | 1483 | 1414.56 | 1394 | 1436 | 1.032 | 0.045 | 0.56711196 | 0.72102217 | 0.0033 | 0.0041 | 0.0038 | 0.0038 | TRUE | 0.0609 |
| Nr1h2 | 3188 | 2567 | 2366 | 2069 | 1967 | 2223 | 2476 | 2730 | 2553 | 2059 | 2219 | 2194 | 2372.04 | 2586 | 2157 | 0.834 | -0.261 | 8.28E-05 | 0.00103701 | 0.0017 | 0.0034 | 0.0027 | 0.0027 | TRUE | 0.0287 |
| Nr1h3 | 28 | 24 | 12 | 17 | 13 | 25 | 22 | 26 | 13 | 17 | 15 | 25 | 19.41 | 20 | 19 | 0.935 | -0.098 | 0.8318235 | 0.90385079 | 0.0361 | 0.1245 | 0.1011 | 0.1011 | TRUE | 0.4753 |
| Nr1h4 | 2 | 0 | 0 | 0 | 1 | 2 | 2 | 0 | 0 | 0 | 1 | 2 | 0.78 | 1 | 1 | 1.844 | 0.883 | 0.7091433 | NA | 0 | 3.0568 | 2.6753 | 2.6753 | TRUE | 0.5491 |
| Nr1h5 | 3 | 0 | 2 | 2 | 4 | 0 | 2 | 0 | 2 | 2 | 5 | 0 | 1.83 | 1 | 2 | 1.403 | 0.488 | 0.75595802 | NA | 0.888 | 1.2962 | 1.2195 | 1.2195 | TRUE | 0.1841 |
| Nr1i2 | 34 | 22 | 15 | 15 | 20 | 14 | 26 | 23 | 16 | 15 | 23 | 14 | 19.55 | 22 | 17 | 0.768 | -0.381 | 0.39777829 | 0.57100066 | 0 | 0.1237 | 0.0939 | 0.0939 | TRUE | 0.3508 |
| Nr1i3 | 2 | 3 | 0 | 0 | 0 | 2 | 2 | 3 | 0 | 0 | 0 | 2 | 1.12 | 2 | 1 | 0.424 | -1.237 | 0.551706 | NA | 1.491 | 2.119 | 2.0351 | 2.0351 | TRUE | 0.3265 |
| Nr2c1 | 573 | 427 | 399 | 473 | 431 | 425 | 445 | 454 | 431 | 471 | 486 | 420 | 451.05 | 443 | 459 | 1.034 | 0.049 | 0.64118524 | 0.77592588 | 0 | 0.0077 | 0.0056 | 0.0056 | TRUE | 0.066 |
| Nr2c2 | 3868 | 2914 | 2648 | 2880 | 2683 | 2981 | 3004 | 3100 | 2857 | 2867 | 3027 | 2943 | 2966.17 | 2987 | 2946 | 0.986 | -0.021 | 0.73173418 | 0.83929626 | 7.00E-04 | 0.0032 | 0.0023 | 0.0023 | TRUE | 0.0171 |
| Nr2c2ap | 1536 | 1056 | 964 | 1043 | 964 | 1092 | 1193 | 1123 | 1040 | 1038 | 1087 | 1078 | 1093.34 | 1119 | 1068 | 0.953 | -0.069 | 0.38198641 | 0.55626553 | 0.0019 | 0.0046 | 0.0036 | 0.0036 | TRUE | 0.0449 |
| Nr2e1 | 1 | 0 | 0 | 0 | 2 | 2 | 1 | 0 | 0 | 0 | 2 | 2 | 0.83 | 0 | 1 | 4.067 | 2.024 | 0.3803708 | NA | 0 | 2.8424 | 2.4672 | 2.4672 | TRUE | 0.5093 |
| Nr2e3 | 16 | 16 | 12 | 14 | 18 | 12 | 12 | 17 | 13 | 14 | 20 | 12 | 14.75 | 14 | 15 | 1.086 | 0.119 | 0.81286776 | NA | 0 | 0.1631 | 0.1153 | 0.1153 | TRUE | 0.3338 |
| Nr2f1 | 38 | 34 | 20 | 26 | 24 | 27 | 30 | 36 | 22 | 26 | 27 | 27 | 27.81 | 29 | 27 | 0.911 | -0.134 | 0.71736302 | 0.8300325 | 0 | 0.0876 | 0.063 | 0.063 | TRUE | 0.1971 |
| Nr2f2 | 11 | 6 | 6 | 7 | 1 | 2 | 9 | 6 | 6 | 7 | 1 | 2 | 5.24 | 7 | 3 | 0.474 | -1.078 | 0.23027479 | NA | 0.0217 | 0.4543 | 0.3622 | 0.3622 | TRUE | 1.2887 |
| Nr2f6 | 1633 | 1349 | 1279 | 1527 | 1390 | 1642 | 1268 | 1435 | 1380 | 1520 | 1568 | 1621 | 1465.36 | 1361 | 1570 | 1.155 | 0.208 | 0.00495794 | 0.02436856 | 0.0019 | 0.004 | 0.0032 | 0.0032 | TRUE | 0.0447 |
| Nr3c1 | 1877 | 1343 | 1360 | 1580 | 1324 | 1530 | 1458 | 1429 | 1467 | 1573 | 1494 | 1510 | 1488.41 | 1451 | 1526 | 1.051 | 0.072 | 0.27153204 | 0.44248176 | 0 | 0.004 | 0.0024 | 0.0024 | TRUE | 0.0089 |
| Nr3c2 | 271 | 202 | 154 | 172 | 153 | 168 | 210 | 215 | 166 | 171 | 173 | 166 | 183.53 | 197 | 170 | 0.86 | -0.218 | 0.16433346 | 0.31522329 | 0.0047 | 0.0153 | 0.0123 | 0.0123 | TRUE | 0.1958 |
| Nr4a1 | 1497 | 1234 | 1444 | 1680 | 1577 | 1813 | 1163 | 1313 | 1558 | 1672 | 1779 | 1790 | 1545.72 | 1345 | 1747 | 1.301 | 0.38 | 0.00010242 | 0.00123058 | 0.0107 | 0.004 | 0.0062 | 0.0062 | TRUE | 0.2317 |
| Nr4a2 | 57 | 42 | 30 | 25 | 23 | 17 | 44 | 45 | 32 | 25 | 26 | 17 | 31.49 | 40 | 23 | 0.554 | -0.852 | 0.01815594 | 0.0642652 | 0 | 0.0777 | 0.0588 | 0.0588 | TRUE | 0.2983 |
| Nr4a3 | 2 | 1 | 1 | 1 | 2 | 3 | 2 | 1 | 1 | 1 | 2 | 3 | 1.65 | 1 | 2 | 1.658 | 0.729 | 0.61498707 | NA | 0 | 1.4375 | 0.8795 | 0.8795 | TRUE | 0.1994 |
| Nr5a1 | 2 | 1 | 3 | 3 | 3 | 5 | 2 | 1 | 3 | 3 | 3 | 5 | 2.86 | 2 | 4 | 1.958 | 0.969 | 0.39326716 | NA | 0 | 0.8311 | 0.5419 | 0.5419 | TRUE | 0.258 |
| Nr5a2 | 3893 | 2754 | 2883 | 2759 | 2620 | 3007 | 3024 | 2929 | 3111 | 2746 | 2956 | 2968 | 2955.66 | 3021 | 2890 | 0.956 | -0.064 | 0.29327864 | 0.46521877 | 9.00E-04 | 0.0032 | 0.0023 | 0.0023 | TRUE | 0.0231 |
| Nr6a1 | 6288 | 4382 | 4250 | 4732 | 3922 | 4596 | 4884 | 4661 | 4586 | 4710 | 4424 | 4537 | 4633.67 | 4710 | 4557 | 0.967 | -0.048 | 0.39125556 | 0.56492733 | 7.00E-04 | 0.0029 | 0.002 | 0.002 | TRUE | 0.0129 |
| Nradd | 945 | 746 | 978 | 1196 | 1048 | 1291 | 734 | 794 | 1055 | 1190 | 1182 | 1274 | 1038.31 | 861 | 1215 | 1.415 | 0.501 | 1.93E-05 | 0.00032336 | 0.0191 | 0.0047 | 0.0089 | 0.0089 | TRUE | 0.4607 |
| Nrap | 31 | 37 | 23 | 25 | 23 | 19 | 24 | 39 | 25 | 25 | 26 | 19 | 26.31 | 29 | 23 | 0.791 | -0.338 | 0.3902546 | 0.56405188 | 0 | 0.0925 | 0.0725 | 0.0725 | TRUE | 0.5511 |
| Nrarp | 400 | 313 | 312 | 376 | 353 | 395 | 311 | 333 | 337 | 374 | 398 | 390 | 357.11 | 327 | 387 | 1.187 | 0.248 | 0.02440797 | 0.07998445 | 0 | 0.0091 | 0.0059 | 0.0059 | TRUE | 0.023 |
| Nras | 9806 | 6958 | 7036 | 8091 | 7034 | 7830 | 7616 | 7401 | 7592 | 8054 | 7935 | 7729 | 7721.18 | 7536 | 7906 | 1.049 | 0.069 | 0.1551865 | 0.30280965 | 2.00E-04 | 0.0027 | 0.0016 | 0.0016 | TRUE | 0.0047 |
| Nrbf2 | 989 | 743 | 679 | 741 | 641 | 749 | 768 | 790 | 733 | 738 | 723 | 739 | 748.53 | 764 | 733 | 0.96 | -0.059 | 0.47679114 | 0.64457056 | 0 | 0.0056 | 0.0035 | 0.0035 | TRUE | 0.0143 |
| Nrbp1 | 2911 | 2199 | 2383 | 2368 | 2178 | 2539 | 2261 | 2339 | 2571 | 2357 | 2457 | 2506 | 2415.27 | 2390 | 2440 | 1.022 | 0.031 | 0.65543954 | 0.78605448 | 0.0023 | 0.0034 | 0.003 | 0.003 | TRUE | 0.0527 |
| Nrbp2 | 351 | 307 | 438 | 338 | 346 | 370 | 273 | 327 | 473 | 336 | 390 | 365 | 360.63 | 358 | 364 | 1.022 | 0.031 | 0.85488715 | 0.91850192 | 0.0401 | 0.009 | 0.0182 | 0.0182 | TRUE | 0.8501 |
| Nrcam | 113 | 90 | 81 | 179 | 173 | 182 | 88 | 96 | 87 | 178 | 195 | 180 | 137.32 | 90 | 184 | 2.042 | 1.03 | 1.85E-09 | 1.16E-07 | 0 | 0.0197 | 0.013 | 0.013 | TRUE | 0.0274 |
| Nrd1 | 11038 | 7872 | 7820 | 8363 | 6840 | 8094 | 8573 | 8373 | 8438 | 8325 | 7716 | 7990 | 8235.76 | 8461 | 8010 | 0.947 | -0.079 | 0.12541354 | 0.26061874 | 6.00E-04 | 0.0027 | 0.0018 | 0.0018 | TRUE | 0.0144 |
| Nrde2 | 2929 | 2039 | 1855 | 1934 | 1641 | 2009 | 2275 | 2169 | 2002 | 1925 | 1851 | 1983 | 2034.12 | 2149 | 1920 | 0.893 | -0.163 | 0.02050547 | 0.07046997 | 0.0022 | 0.0036 | 0.0031 | 0.0031 | TRUE | 0.043 |
| Nrep | 402 | 324 | 366 | 344 | 326 | 333 | 312 | 345 | 395 | 342 | 368 | 329 | 348.44 | 351 | 346 | 0.99 | -0.015 | 0.90680984 | 0.94912712 | 0.006 | 0.0092 | 0.0082 | 0.0082 | TRUE | 0.1385 |
| Nrf1 | 3845 | 2683 | 2345 | 2280 | 2129 | 2419 | 2986 | 2854 | 2530 | 2270 | 2402 | 2388 | 2571.59 | 2790 | 2353 | 0.843 | -0.247 | 0.00079095 | 0.00609439 | 0.0034 | 0.0033 | 0.0035 | 0.0035 | TRUE | 0.08 |
| Nrg1 | 73 | 49 | 45 | 45 | 35 | 44 | 57 | 52 | 49 | 45 | 39 | 43 | 47.51 | 53 | 42 | 0.81 | -0.304 | 0.27382486 | 0.44484868 | 0 | 0.0523 | 0.0342 | 0.0342 | TRUE | 0.0476 |
| Nrg2 | 324 | 234 | 232 | 292 | 199 | 276 | 252 | 249 | 250 | 291 | 224 | 272 | 256.41 | 250 | 262 | 1.051 | 0.071 | 0.60200515 | 0.74697974 | 0.0043 | 0.0117 | 0.0096 | 0.0096 | TRUE | 0.1726 |
| Nrg3 | 1 | 0 | 0 | 0 | 0 | 2 | 1 | 0 | 0 | 0 | 0 | 2 | 0.46 | 0 | 1 | 1.93 | 0.948 | 0.7608301 | NA | 0 | 5.1717 | 5.216 | 5.216 | TRUE | 0.9737 |
| Nrg4 | 177 | 107 | 110 | 93 | 78 | 117 | 137 | 114 | 119 | 93 | 88 | 115 | 111.01 | 123 | 99 | 0.799 | -0.324 | 0.10110379 | 0.22293681 | 0.0069 | 0.0238 | 0.019 | 0.019 | TRUE | 0.2197 |
| Nrgn | 280 | 230 | 186 | 222 | 171 | 210 | 217 | 245 | 201 | 221 | 193 | 207 | 214 | 221 | 207 | 0.938 | -0.092 | 0.5244921 | 0.6856824 | 0.0021 | 0.0135 | 0.0103 | 0.0103 | TRUE | 0.0916 |
| Nrip1 | 733 | 544 | 576 | 553 | 541 | 558 | 569 | 579 | 621 | 550 | 610 | 551 | 580.17 | 590 | 570 | 0.967 | -0.048 | 0.6149587 | 0.75681115 | 0.001 | 0.0065 | 0.0048 | 0.0048 | TRUE | 0.0424 |
| Nrip2 | 22 | 11 | 16 | 26 | 20 | 19 | 17 | 12 | 17 | 26 | 23 | 19 | 18.87 | 15 | 23 | 1.452 | 0.538 | 0.22702864 | 0.39199328 | 0 | 0.128 | 0.0887 | 0.0887 | TRUE | 0.1845 |
| Nrip3 | 227 | 155 | 128 | 205 | 141 | 229 | 176 | 165 | 138 | 204 | 159 | 226 | 178.08 | 160 | 196 | 1.228 | 0.296 | 0.0895027 | 0.20399806 | 0.0177 | 0.0157 | 0.0163 | 0.0163 | TRUE | 0.2889 |
| Nrk | 686 | 514 | 618 | 424 | 450 | 556 | 533 | 547 | 667 | 422 | 508 | 549 | 537.48 | 582 | 493 | 0.848 | -0.238 | 0.05549601 | 0.14522235 | 0.0138 | 0.0068 | 0.0093 | 0.0093 | TRUE | 0.1875 |
| Nrl | 1 | 0 | 1 | 0 | 0 | 1 | 1 | 0 | 1 | 0 | 0 | 1 | 0.47 | 1 | 0 | 0.596 | -0.748 | 0.78959815 | NA | 0 | 5.0047 | 3.8792 | 3.8792 | TRUE | 0.3835 |
| Nrm | 1332 | 1043 | 1181 | 1346 | 1266 | 1430 | 1035 | 1109 | 1274 | 1340 | 1428 | 1412 | 1266.31 | 1139 | 1393 | 1.224 | 0.292 | 0.00080707 | 0.00618853 | 0.0052 | 0.0043 | 0.0047 | 0.0047 | TRUE | 0.1276 |
| Nrn1 | 11 | 12 | 9 | 13 | 12 | 8 | 9 | 13 | 10 | 13 | 14 | 8 | 10.9 | 11 | 12 | 1.112 | 0.153 | 0.79232725 | NA | 0 | 0.2199 | 0.1523 | 0.1523 | TRUE | 0.2926 |
| Nrn1l | 51 | 49 | 54 | 98 | 90 | 93 | 40 | 52 | 58 | 98 | 102 | 92 | 73.48 | 50 | 97 | 1.957 | 0.968 | 5.34E-05 | 0.00073993 | 0 | 0.0347 | 0.0263 | 0.0263 | TRUE | 0.3227 |
| Nron | 1 | 0 | 0 | 0 | 0 | 0 | 1 | 0 | 0 | 0 | 0 | 0 | 0.13 | 0 | 0 | 0.552 | -0.857 | 0.8336548 | NA | 0 | 18.3121 | 10 | 10 | TRUE | 0.2566 |
| Nrp | 44 | 42 | 30 | 24 | 27 | 31 | 34 | 45 | 32 | 24 | 30 | 31 | 32.69 | 37 | 28 | 0.765 | -0.387 | 0.25907672 | 0.42899676 | 0 | 0.0749 | 0.0533 | 0.0533 | TRUE | 0.2212 |
| Nrp1 | 83 | 56 | 30 | 42 | 24 | 42 | 64 | 60 | 32 | 42 | 27 | 41 | 44.46 | 52 | 37 | 0.704 | -0.507 | 0.13444514 | 0.27321014 | 0.0688 | 0.0557 | 0.0594 | 0.0594 | TRUE | 0.8607 |
| Nrp2 | 1006 | 761 | 1065 | 842 | 957 | 1085 | 781 | 809 | 1149 | 838 | 1080 | 1071 | 954.78 | 913 | 996 | 1.092 | 0.127 | 0.36309762 | 0.5378541 | 0.0323 | 0.0049 | 0.013 | 0.013 | TRUE | 0.604 |
| Nrsn1 | 71 | 45 | 57 | 60 | 42 | 55 | 55 | 48 | 62 | 60 | 47 | 54 | 54.32 | 55 | 54 | 0.983 | -0.025 | 0.92540152 | 0.95873904 | 0 | 0.0461 | 0.0324 | 0.0324 | TRUE | 0.0952 |
| Nrsn2 | 212 | 153 | 162 | 183 | 150 | 204 | 165 | 163 | 175 | 182 | 169 | 201 | 175.82 | 168 | 184 | 1.103 | 0.141 | 0.35463769 | 0.52899567 | 0 | 0.0159 | 0.0111 | 0.0111 | TRUE | 0.0716 |

| | | | | | | | | | | | | | | | | | | | | | | | | |
|---|---|---|---|---|---|---|---|---|---|---|---|---|---|---|---|---|---|---|---|---|---|---|---|---|
| Nrtn | 650 | 550 | 523 | 691 | 603 | 655 | 505 | 585 | 564 | 688 | 680 | 647 | 611.47 | 551 | 672 | 1.221 | 0.288 | 0.00249968 | 0.01435514 | 0.002 | 0.0063 | 0.0049 | 0.0049 | TRUE | 0.0685 |
| Nrxn1 | 10 | 10 | 15 | 11 | 8 | 4 | 8 | 11 | 16 | 11 | 9 | 4 | 9.75 | 12 | 8 | 0.697 | -0.521 | 0.42481646 | NA | 0.0762 | 0.2455 | 0.1991 | 0.1991 | TRUE | 0.4976 |
| Nrxn2 | 491 | 363 | 304 | 423 | 397 | 457 | 381 | 386 | 328 | 421 | 448 | 451 | 402.59 | 365 | 440 | 1.203 | 0.267 | 0.01615573 | 0.05902551 | 0.0017 | 0.0083 | 0.0064 | 0.0064 | TRUE | 0.0868 |
| Nrxn3 | 9 | 7 | 8 | 5 | 2 | 5 | 7 | 7 | 9 | 5 | 2 | 5 | 5.87 | 8 | 4 | 0.534 | -0.905 | 0.25595627 | NA | 0 | 0.406 | 0.2677 | 0.2677 | TRUE | 0.2201 |
| Nsa2 | 5848 | 4378 | 4280 | 4882 | 4161 | 4880 | 4542 | 4657 | 4618 | 4860 | 4694 | 4817 | 4697.95 | 4606 | 4790 | 1.04 | 0.057 | 0.26046348 | 0.43057079 | 0 | 0.0029 | 0.0016 | 0.0016 | TRUE | 0.0037 |
| Nsd1 | 14756 | 10320 | 9597 | 9860 | 9013 | 10422 | 11460 | 10977 | 10355 | 9815 | 10168 | 10288 | 10510.51 | 10931 | 10090 | 0.923 | -0.116 | 0.03822821 | 0.11076338 | 0.0014 | 0.0027 | 0.0022 | 0.0022 | TRUE | 0.0259 |
| Nsdhl | 3224 | 2162 | 2104 | 2595 | 2172 | 2584 | 2504 | 2300 | 2270 | 2583 | 2450 | 2551 | 2442.99 | 2358 | 2528 | 1.071 | 0.1 | 0.12552562 | 0.26066216 | 0.0015 | 0.0034 | 0.0026 | 0.0026 | TRUE | 0.0366 |
| Nsf | 14573 | 9873 | 8960 | 9144 | 7934 | 9149 | 11318 | 10502 | 9668 | 9102 | 8950 | 9031 | 9761.93 | 10496 | 9028 | 0.86 | -0.218 | 0.00064029 | 0.00521479 | 0.003 | 0.0027 | 0.0028 | 0.0028 | TRUE | 0.0582 |
| Nsfl1c | 6589 | 4756 | 4113 | 4355 | 4043 | 4562 | 5117 | 5059 | 4438 | 4335 | 4561 | 4503 | 4668.91 | 4871 | 4466 | 0.917 | -0.126 | 0.06150327 | 0.1561674 | 0.003 | 0.0029 | 0.003 | 0.003 | TRUE | 0.0733 |
| Nsg1 | 67 | 45 | 39 | 48 | 42 | 41 | 52 | 48 | 42 | 48 | 47 | 40 | 46.27 | 47 | 45 | 0.951 | -0.073 | 0.79700277 | 0.88160891 | 0 | 0.0536 | 0.0362 | 0.0362 | TRUE | 0.072 |
| Nsg2 | 101 | 100 | 96 | 66 | 88 | 101 | 78 | 106 | 104 | 66 | 99 | 100 | 92.18 | 96 | 88 | 0.921 | -0.119 | 0.61202064 | 0.75449465 | 0.0295 | 0.0281 | 0.0285 | 0.0285 | TRUE | 0.4836 |
| Nsl1 | 1859 | 1413 | 1551 | 1635 | 1455 | 1797 | 1444 | 1503 | 1674 | 1627 | 1641 | 1774 | 1610.52 | 1540 | 1681 | 1.093 | 0.128 | 0.10047531 | 0.22179053 | 0.0032 | 0.0039 | 0.0037 | 0.0037 | TRUE | 0.0686 |
| Nsmaf | 1341 | 936 | 981 | 962 | 884 | 979 | 1041 | 996 | 1058 | 958 | 997 | 966 | 1002.81 | 1032 | 974 | 0.943 | -0.084 | 0.25833584 | 0.42834726 | 0 | 0.0048 | 0.003 | 0.003 | TRUE | 0.0109 |
| Nsmce1 | 1830 | 1269 | 1538 | 1447 | 1304 | 1454 | 1421 | 1350 | 1659 | 1440 | 1471 | 1435 | 1462.88 | 1477 | 1449 | 0.982 | -0.027 | 0.75021847 | 0.85168585 | 0.0052 | 0.004 | 0.0045 | 0.0045 | TRUE | 0.1383 |
| Nsmce2 | 2097 | 1486 | 1515 | 1461 | 1329 | 1471 | 1629 | 1581 | 1635 | 1454 | 1499 | 1452 | 1541.6 | 1615 | 1468 | 0.909 | -0.137 | 0.03312319 | 0.09970366 | 0 | 0.004 | 0.0023 | 0.0023 | TRUE | 0.004 |
| Nsmce4a | 5485 | 4000 | 4341 | 4219 | 3809 | 4339 | 4260 | 4255 | 4684 | 4200 | 4297 | 4283 | 4329.74 | 4400 | 4260 | 0.969 | -0.046 | 0.4376694 | 0.60911477 | 0.0013 | 0.003 | 0.0023 | 0.0023 | TRUE | 0.0386 |
| Nsmf | 539 | 409 | 390 | 533 | 498 | 505 | 419 | 435 | 421 | 531 | 562 | 499 | 477.55 | 425 | 531 | 1.248 | 0.319 | 0.00136232 | 0.00899799 | 0 | 0.0074 | 0.0051 | 0.0051 | TRUE | 0.0323 |
| Nsun2 | 31463 | 23296 | 24548 | 24865 | 22558 | 26191 | 24436 | 24780 | 26487 | 24751 | 25448 | 25855 | 25292.61 | 25234 | 25351 | 1.005 | 0.007 | 0.89618365 | 0.94275064 | 9.00E-04 | 0.0025 | 0.0019 | 0.0019 | TRUE | 0.023 |
| Nsun3 | 298 | 182 | 135 | 159 | 158 | 162 | 231 | 194 | 146 | 158 | 178 | 160 | 177.85 | 190 | 165 | 0.865 | -0.21 | 0.24923093 | 0.41810537 | 0.0239 | 0.0158 | 0.0182 | 0.0182 | TRUE | 0.2191 |
| Nsun4 | 1276 | 990 | 947 | 931 | 828 | 1024 | 991 | 1053 | 1022 | 927 | 934 | 1011 | 989.58 | 1022 | 957 | 0.938 | -0.093 | 0.23230404 | 0.3982156 | 6.00E-04 | 0.0048 | 0.0034 | 0.0034 | TRUE | 0.0291 |
| Nsun5 | 1598 | 1108 | 997 | 1293 | 1073 | 1273 | 1241 | 1179 | 1076 | 1287 | 1210 | 1257 | 1208.26 | 1165 | 1251 | 1.073 | 0.102 | 0.1968096 | 0.35644191 | 0.0023 | 0.0044 | 0.0036 | 0.0036 | TRUE | 0.0534 |
| Nsun6 | 619 | 524 | 542 | 590 | 529 | 592 | 481 | 557 | 585 | 587 | 597 | 584 | 565.23 | 541 | 589 | 1.093 | 0.128 | 0.20815913 | 0.36975192 | 0.0037 | 0.0066 | 0.0056 | 0.0056 | TRUE | 0.1184 |
| Nsun7 | 9 | 5 | 6 | 8 | 13 | 3 | 7 | 5 | 6 | 8 | 15 | 3 | 7.4 | 6 | 9 | 1.343 | 0.426 | 0.57340641 | NA | 0.1515 | 0.3229 | 0.2739 | 0.2739 | TRUE | 0.2069 |
| Nt5c | 1428 | 954 | 1039 | 1037 | 909 | 1076 | 1109 | 1015 | 1121 | 1032 | 1025 | 1062 | 1060.79 | 1082 | 1040 | 0.961 | -0.057 | 0.45403731 | 0.62381576 | 7.00E-04 | 0.0047 | 0.0033 | 0.0033 | TRUE | 0.0341 |
| Nt5c1a | 5 | 1 | 1 | 3 | 0 | 2 | 4 | 1 | 1 | 3 | 0 | 2 | 1.83 | 2 | 2 | 0.811 | -0.301 | 0.83777068 | NA | 0 | 1.2968 | 1.0203 | 1.0203 | TRUE | 0.725 |
| Nt5c1b | 0 | 0 | 0 | 0 | 0 | 0 | 0 | 0 | 0 | 0 | 0 | 0 | 0 | 0 | 0 | NA | NA | NA | NA | NA | NA | NA | NA | NA | NA |
| Nt5c2 | 2909 | 1995 | 1704 | 1743 | 1631 | 1815 | 2259 | 2122 | 1839 | 1735 | 1840 | 1792 | 1931.09 | 2073 | 1789 | 0.862 | -0.214 | 0.00884699 | 0.03789331 | 0.005 | 0.0037 | 0.0043 | 0.0043 | TRUE | 0.1178 |
| Nt5c3 | 918 | 718 | 716 | 810 | 694 | 836 | 713 | 764 | 773 | 806 | 783 | 825 | 777.28 | 750 | 805 | 1.075 | 0.105 | 0.20600725 | 0.36784941 | 0 | 0.0055 | 0.0036 | 0.0036 | TRUE | 0.0232 |
| Nt5c3l | 2362 | 1595 | 1579 | 1556 | 1455 | 1725 | 1834 | 1697 | 1704 | 1549 | 1641 | 1703 | 1687.97 | 1745 | 1631 | 0.934 | -0.099 | 0.16177037 | 0.311796 | 0.0016 | 0.0038 | 0.003 | 0.003 | TRUE | 0.0252 |
| Nt5dc1 | 907 | 591 | 570 | 621 | 567 | 623 | 704 | 629 | 615 | 618 | 640 | 615 | 636.81 | 649 | 624 | 0.959 | -0.06 | 0.51567612 | 0.67800192 | 0.0018 | 0.0061 | 0.0046 | 0.0046 | TRUE | 0.0684 |
| Nt5dc2 | 7713 | 5816 | 6308 | 6353 | 5642 | 6709 | 5990 | 6186 | 6806 | 6324 | 6365 | 6623 | 6382.4 | 6327 | 6437 | 1.018 | 0.025 | 0.68499509 | 0.80694783 | 0.002 | 0.0028 | 0.0026 | 0.0026 | TRUE | 0.0532 |
| Nt5dc3 | 1314 | 937 | 816 | 1123 | 942 | 1000 | 1021 | 997 | 880 | 1118 | 1063 | 987 | 1010.89 | 966 | 1056 | 1.092 | 0.127 | 0.14693554 | 0.2913233 | 0.004 | 0.0048 | 0.0045 | 0.0045 | TRUE | 0.0701 |
| Nt5e | 93 | 42 | 28 | 53 | 38 | 52 | 72 | 45 | 30 | 53 | 43 | 51 | 49.01 | 49 | 49 | 0.988 | -0.017 | 0.95888124 | 0.97739513 | 0.0812 | 0.0508 | 0.0596 | 0.0596 | TRUE | 0.4972 |
| Nt5m | 293 | 194 | 195 | 176 | 190 | 187 | 228 | 206 | 210 | 175 | 214 | 185 | 203.07 | 215 | 191 | 0.888 | -0.172 | 0.24176337 | 0.40937375 | 0 | 0.0141 | 0.0105 | 0.0105 | TRUE | 0.1145 |
| Ntan1 | 2114 | 1500 | 1844 | 1945 | 1728 | 2041 | 1642 | 1596 | 1990 | 1936 | 1949 | 2015 | 1854.54 | 1743 | 1967 | 1.13 | 0.176 | 0.04163466 | 0.11771699 | 0.0071 | 0.0037 | 0.0048 | 0.0048 | TRUE | 0.1838 |
| Ntf3 | 10 | 3 | 12 | 6 | 2 | 5 | 8 | 3 | 13 | 6 | 2 | 5 | 6.18 | 8 | 4 | 0.556 | -0.846 | 0.30892525 | NA | 0.1484 | 0.386 | 0.3204 | 0.3204 | TRUE | 0.127 |
| Ntf5 | 88 | 68 | 67 | 77 | 85 | 69 | 68 | 72 | 72 | 77 | 96 | 68 | 75.6 | 71 | 80 | 1.128 | 0.174 | 0.45158026 | 0.62145549 | 0 | 0.0338 | 0.0252 | 0.0252 | TRUE | 0.2584 |
| Nthl1 | 1152 | 806 | 845 | 1047 | 854 | 953 | 895 | 857 | 912 | 1042 | 963 | 941 | 935.02 | 888 | 982 | 1.106 | 0.145 | 0.06943737 | 0.17071391 | 9.00E-04 | 0.005 | 0.0036 | 0.0036 | TRUE | 0.0346 |
| Ntm | 22 | 11 | 10 | 9 | 12 | 14 | 17 | 12 | 11 | 9 | 14 | 14 | 12.65 | 13 | 12 | 0.906 | -0.142 | 0.79335803 | NA | 0 | 0.1898 | 0.1338 | 0.1338 | TRUE | 0.3493 |
| Ntmt1 | 1459 | 1055 | 1066 | 1145 | 993 | 1078 | 1133 | 1122 | 1150 | 1140 | 1120 | 1064 | 1121.6 | 1135 | 1108 | 0.976 | -0.035 | 0.62682431 | 0.7648284 | 0 | 0.0045 | 0.0028 | 0.0028 | TRUE | 0.0146 |
| Ntn1 | 539 | 416 | 461 | 1080 | 990 | 1355 | 419 | 442 | 497 | 1075 | 1117 | 1338 | 814.66 | 453 | 1177 | 2.604 | 1.381 | 7.73E-38 | 1.11E-34 | 0.0098 | 0.0053 | 0.0068 | 0.0068 | TRUE | 0.1733 |
| Ntn3 | 45 | 23 | 14 | 21 | 27 | 18 | 35 | 24 | 15 | 21 | 30 | 18 | 23.94 | 25 | 23 | 0.912 | -0.133 | 0.76167288 | 0.85863163 | 0.0826 | 0.1014 | 0.0961 | 0.0961 | TRUE | 0.2132 |
| Ntn4 | 23 | 18 | 19 | 29 | 12 | 20 | 18 | 19 | 21 | 29 | 14 | 20 | 19.94 | 19 | 21 | 1.089 | 0.123 | 0.78350141 | 0.87257173 | 0 | 0.1213 | 0.0938 | 0.0938 | TRUE | 0.2949 |
| Ntn5 | 17 | 20 | 14 | 26 | 30 | 29 | 13 | 21 | 15 | 26 | 34 | 29 | 22.99 | 16 | 30 | 1.794 | 0.843 | 0.04088084 | 0.11628056 | 0 | 0.1055 | 0.0756 | 0.0756 | TRUE | 0.2924 |
| Ntng1 | 0 | 1 | 0 | 0 | 0 | 0 | 0 | 1 | 0 | 0 | 0 | 0 | 0.18 | 0 | 0 | 0.552 | -0.857 | 0.8336548 | NA | 0 | 13.3714 | 10 | 10 | TRUE | 0.5216 |
| Ntng2 | 30 | 25 | 30 | 51 | 39 | 28 | 23 | 27 | 32 | 51 | 44 | 28 | 34.11 | 27 | 41 | 1.495 | 0.58 | 0.10466039 | 0.22862769 | 0.0362 | 0.0719 | 0.0617 | 0.0617 | TRUE | 0.5303 |
| Ntpcr | 1839 | 1361 | 1322 | 1599 | 1496 | 1744 | 1428 | 1448 | 1426 | 1592 | 1688 | 1722 | 1550.54 | 1434 | 1667 | 1.162 | 0.217 | 0.00114044 | 0.00797935 | 0 | 0.004 | 0.0026 | 0.0026 | TRUE | 0.0191 |
| Ntrk1 | 256 | 256 | 199 | 262 | 260 | 267 | 199 | 272 | 215 | 261 | 293 | 264 | 250.59 | 229 | 273 | 1.195 | 0.257 | 0.08254988 | 0.19342383 | 0.0117 | 0.0119 | 0.0118 | 0.0118 | TRUE | 0.3039 |
| Ntrk2 | 276 | 232 | 207 | 207 | 205 | 229 | 214 | 247 | 223 | 206 | 231 | 226 | 224.64 | 228 | 221 | 0.97 | -0.043 | 0.75323412 | 0.85352858 | 0 | 0.013 | 0.0092 | 0.0092 | TRUE | 0.0552 |
| Ntrk3 | 30 | 27 | 35 | 59 | 49 | 62 | 23 | 29 | 38 | 59 | 55 | 61 | 44.17 | 30 | 58 | 1.974 | 0.981 | 0.00129229 | 0.00868394 | 0 | 0.0561 | 0.0418 | 0.0418 | TRUE | 0.3367 |
| Nts | 0 | 1 | 1 | 0 | 0 | 0 | 0 | 1 | 1 | 0 | 0 | 0 | 0.36 | 1 | 0 | 0.271 | -1.881 | 0.5755355 | NA | 0 | 6.6391 | 6.4428 | 6.4428 | TRUE | 0.3513 |
| Ntsr1 | 3 | 1 | 4 | 4 | 4 | 5 | 2 | 1 | 4 | 4 | 5 | 5 | 3.52 | 2 | 5 | 1.752 | 0.809 | 0.42929632 | NA | 0 | 0.6751 | 0.4512 | 0.4512 | TRUE | 0.3474 |
| Ntsr2 | 0 | 0 | 0 | 0 | 1 | 0 | 0 | 0 | 0 | 0 | 1 | 0 | 0.19 | 0 | 0 | 2.094 | 1.067 | 0.79379329 | NA | 0 | 12.608 | 10 | 10 | TRUE | 0.5075 |
| Nuak1 | 1107 | 874 | 906 | 848 | 788 | 927 | 860 | 930 | 978 | 844 | 889 | 915 | 902.52 | 923 | 883 | 0.958 | -0.061 | 0.46619 | 0.63513999 | 0.002 | 0.0051 | 0.004 | 0.004 | TRUE | 0.0442 |
| Nuak2 | 436 | 339 | 436 | 437 | 375 | 503 | 339 | 361 | 470 | 435 | 423 | 497 | 420.7 | 390 | 452 | 1.162 | 0.216 | 0.10892732 | 0.23550117 | 0.0166 | 0.0081 | 0.0107 | 0.0107 | TRUE | 0.3703 |
| Nub1 | 6499 | 4900 | 5154 | 5418 | 4884 | 5786 | 5047 | 5212 | 5561 | 5393 | 5510 | 5712 | 5405.85 | 5273 | 5538 | 1.051 | 0.071 | 0.22368853 | 0.38816531 | 0.0014 | 0.0029 | 0.0023 | 0.0023 | TRUE | 0.0276 |
| Nubp1 | 3514 | 2643 | 2522 | 2650 | 2487 | 2719 | 2729 | 2811 | 2721 | 2638 | 2806 | 2684 | 2731.53 | 2754 | 2709 | 0.984 | -0.024 | 0.68206384 | 0.80494909 | 3.00E-04 | 0.0033 | 0.0021 | 0.0021 | TRUE | 0.0116 |
| Nubp2 | 3945 | 2990 | 2684 | 2837 | 2515 | 2982 | 3064 | 3180 | 2896 | 2824 | 2837 | 2944 | 2957.53 | 3047 | 2868 | 0.941 | -0.087 | 0.15256528 | 0.29927984 | 9.00E-04 | 0.0032 | 0.0023 | 0.0023 | TRUE | 0.0222 |
| Nubpl | 1978 | 1476 | 1754 | 2038 | 1718 | 1883 | 1536 | 1570 | 1893 | 2029 | 1938 | 1859 | 1804.05 | 1666 | 1942 | 1.166 | 0.222 | 0.01060311 | 0.04310301 | 0.0071 | 0.0037 | 0.0049 | 0.0049 | TRUE | 0.1687 |
| Nucb1 | 3402 | 2550 | 2651 | 3120 | 2714 | 3039 | 2642 | 2712 | 2860 | 3106 | 3062 | 3000 | 2897.05 | 2738 | 3056 | 1.116 | 0.159 | 0.00722617 | 0.03250409 | 6.00E-04 | 0.0032 | 0.0022 | 0.0022 | TRUE | 0.0183 |
| Nucb2 | 724 | 495 | 497 | 587 | 521 | 610 | 562 | 527 | 536 | 584 | 588 | 602 | 566.55 | 542 | 591 | 1.091 | 0.125 | 0.16784417 | 0.31927026 | 0 | 0.0066 | 0.0042 | 0.0042 | TRUE | 0.0139 |
| Nucks1 | 23753 | 16583 | 14816 | 17722 | 14430 | 16588 | 18448 | 17639 | 15986 | 17640 | 16279 | 16375 | 17061.22 | 17358 | 16765 | 0.966 | -0.05 | 0.4390082 | 0.61048068 | 0.0033 | 0.0026 | 0.003 | 0.003 | TRUE | 0.0584 |
| Nudc | 20096 | 15601 | 14141 | 13999 | 12751 | 15172 | 15608 | 16595 | 15258 | 13935 | 14384 | 14977 | 15126.08 | 15820 | 14432 | 0.912 | -0.132 | 0.0162826 | 0.05938433 | 0.0015 | 0.0026 | 0.0021 | 0.0021 | TRUE | 0.0223 |
| Nudcd1 | 4923 | 3626 | 3584 | 3793 | 3476 | 4134 | 3823 | 3857 | 3867 | 3776 | 3921 | 4081 | 3887.54 | 3849 | 3926 | 1.02 | 0.029 | 0.60668504 | 0.75109132 | 5.00E-04 | 0.003 | 0.002 | 0.002 | TRUE | 0.0147 |
| Nudcd2 | 9741 | 6807 | 6763 | 7379 | 6449 | 7079 | 7565 | 7241 | 7297 | 7345 | 7275 | 6988 | 7285.24 | 7368 | 7203 | 0.977 | -0.033 | 0.51705266 | 0.67923873 | 4.00E-04 | 0.0028 | 0.0017 | 0.0017 | TRUE | 0.0083 |
| Nudcd3 | 6458 | 4753 | 4364 | 4357 | 4125 | 4619 | 5016 | 5056 | 4709 | 4337 | 4653 | 4560 | 4721.69 | 4927 | 4517 | 0.917 | -0.126 | 0.02958934 | 0.0917199 | 0.0011 | 0.0029 | 0.0022 | 0.0022 | TRUE | 0.0181 |
| Nudt1 | 1504 | 992 | 811 | 921 | 840 | 920 | 1168 | 1055 | 875 | 917 | 948 | 908 | 978.48 | 1033 | 924 | 0.893 | -0.163 | 0.11147299 | 0.2393036 | 0.0099 | 0.0048 | 0.0065 | 0.0065 | TRUE | 0.2113 |
| Nudt10 | 145 | 77 | 71 | 77 | 58 | 54 | 113 | 82 | 77 | 77 | 65 | 53 | 77.75 | 91 | 65 | 0.716 | -0.482 | 0.05267754 | 0.1400521 | 0.0278 | 0.0329 | 0.0314 | 0.0314 | TRUE | 0.486 |
| Nudt11 | 552 | 368 | 353 | 343 | 270 | 290 | 429 | 391 | 381 | 341 | 305 | 286 | 355.55 | 400 | 311 | 0.775 | -0.368 | 0.00195129 | 0.01186801 | 0.0032 | 0.0091 | 0.0073 | 0.0073 | TRUE | 0.0862 |

| | | | | | | | | | | | | | | | | | | | | | | | | | |
|---|---|---|---|---|---|---|---|---|---|---|---|---|---|---|---|---|---|---|---|---|---|---|---|---|---|
| Nudt12 | 652 | 437 | 432 | 554 | 439 | 538 | 506 | 465 | 466 | 551 | 495 | 531 | 502.52 | 479 | 526 | 1.097 | 0.133 | 0.18016432 | 0.33507985 | 0 | 0.0071 | 0.0051 | 0.0051 | TRUE | 0.0308 |
| Nudt13 | 247 | 202 | 154 | 209 | 167 | 216 | 192 | 215 | 166 | 208 | 188 | 213 | 197.09 | 191 | 203 | 1.065 | 0.091 | 0.55111046 | 0.70703005 | 0.0042 | 0.0145 | 0.0116 | 0.0116 | TRUE | 0.1299 |
| Nudt14 | 846 | 667 | 808 | 899 | 824 | 930 | 657 | 709 | 872 | 895 | 930 | 918 | 830.14 | 746 | 914 | 1.228 | 0.296 | 0.00479069 | 0.02371617 | 0.0096 | 0.0053 | 0.0067 | 0.0067 | TRUE | 0.255 |
| Nudt15 | 79 | 66 | 60 | 49 | 57 | 46 | 61 | 70 | 65 | 49 | 64 | 45 | 59.13 | 65 | 53 | 0.806 | -0.311 | 0.23040539 | 0.39614588 | 0 | 0.0425 | 0.0312 | 0.0312 | TRUE | 0.2804 |
| Nudt16 | 455 | 338 | 390 | 349 | 319 | 380 | 353 | 360 | 421 | 347 | 360 | 375 | 369.35 | 378 | 361 | 0.957 | -0.064 | 0.58171558 | 0.73222699 | 0.0029 | 0.0088 | 0.007 | 0.007 | TRUE | 0.1102 |
| Nudt16l1 | 3696 | 2672 | 2547 | 2870 | 2521 | 2878 | 2871 | 2842 | 2748 | 2857 | 2844 | 2841 | 2833.78 | 2820 | 2847 | 1.009 | 0.013 | 0.81060099 | 0.89074498 | 0 | 0.0033 | 0.0019 | 0.0019 | TRUE | 0.006 |
| Nudt17 | 474 | 381 | 446 | 517 | 510 | 567 | 368 | 405 | 481 | 515 | 575 | 560 | 484.05 | 418 | 550 | 1.318 | 0.399 | 0.00064801 | 0.00525355 | 0.0087 | 0.0073 | 0.0077 | 0.0077 | TRUE | 0.1986 |
| Nudt18 | 517 | 340 | 399 | 402 | 371 | 362 | 402 | 362 | 431 | 400 | 419 | 357 | 394.95 | 398 | 392 | 0.984 | -0.023 | 0.84348305 | 0.91125496 | 0.0043 | 0.0084 | 0.0072 | 0.0072 | TRUE | 0.0696 |
| Nudt19 | 1534 | 1028 | 959 | 1007 | 791 | 999 | 1191 | 1093 | 1035 | 1002 | 892 | 986 | 1033.41 | 1106 | 960 | 0.867 | -0.206 | 0.01651057 | 0.05997093 | 0.0036 | 0.0047 | 0.0043 | 0.0043 | TRUE | 0.0559 |
| Nudt2 | 833 | 573 | 644 | 766 | 598 | 721 | 647 | 609 | 695 | 762 | 675 | 712 | 683.36 | 650 | 716 | 1.102 | 0.14 | 0.13156298 | 0.26908385 | 0.0024 | 0.0059 | 0.0048 | 0.0048 | TRUE | 0.0401 |
| Nudt21 | 5577 | 3789 | 4269 | 4731 | 3987 | 4709 | 4331 | 4030 | 4606 | 4709 | 4498 | 4649 | 4470.57 | 4322 | 4619 | 1.069 | 0.096 | 0.13123006 | 0.2686689 | 0.0022 | 0.003 | 0.0027 | 0.0027 | TRUE | 0.042 |
| Nudt22 | 366 | 295 | 263 | 285 | 247 | 264 | 284 | 314 | 284 | 284 | 279 | 261 | 284.13 | 294 | 275 | 0.934 | -0.099 | 0.42175152 | 0.59401798 | 0 | 0.0108 | 0.0073 | 0.0073 | TRUE | 0.0381 |
| Nudt3 | 4517 | 3379 | 3700 | 3869 | 3490 | 4226 | 3508 | 3594 | 3992 | 3851 | 3937 | 4172 | 3842.44 | 3698 | 3987 | 1.079 | 0.109 | 0.10602111 | 0.23092955 | 0.0029 | 0.003 | 0.003 | 0.003 | TRUE | 0.0585 |
| Nudt4 | 23901 | 15632 | 13785 | 12405 | 11068 | 12263 | 18563 | 16628 | 14874 | 12348 | 12486 | 12105 | 14500.6 | 16688 | 12313 | 0.738 | -0.439 | 4.04E-09 | 2.31E-07 | 0.0061 | 0.0026 | 0.0039 | 0.0039 | TRUE | 0.1185 |
| Nudt5 | 2864 | 2005 | 1639 | 1892 | 1603 | 1652 | 2224 | 2133 | 1768 | 1883 | 1808 | 1631 | 1907.99 | 2042 | 1774 | 0.868 | -0.204 | 0.02760094 | 0.08708615 | 0.0089 | 0.0037 | 0.0056 | 0.0056 | TRUE | 0.1646 |
| Nudt6 | 143 | 101 | 76 | 88 | 99 | 84 | 111 | 107 | 82 | 88 | 112 | 83 | 97.12 | 100 | 94 | 0.933 | -0.099 | 0.64662049 | 0.78033832 | 0.015 | 0.0268 | 0.0236 | 0.0236 | TRUE | 0.2471 |
| Nudt7 | 136 | 95 | 103 | 115 | 107 | 133 | 106 | 101 | 111 | 114 | 121 | 131 | 114.05 | 106 | 122 | 1.154 | 0.206 | 0.260456 | 0.43057079 | 0 | 0.0232 | 0.0154 | 0.0154 | TRUE | 0.0448 |
| Nudt8 | 646 | 430 | 462 | 548 | 486 | 530 | 502 | 457 | 498 | 545 | 548 | 523 | 512.42 | 486 | 539 | 1.108 | 0.148 | 0.12385983 | 0.25806538 | 0 | 0.0071 | 0.0047 | 0.0047 | TRUE | 0.0297 |
| Nudt9 | 2164 | 1613 | 1579 | 1428 | 1262 | 1485 | 1681 | 1716 | 1704 | 1421 | 1424 | 1466 | 1568.53 | 1700 | 1437 | 0.846 | -0.242 | 0.00014564 | 0.00166527 | 0 | 0.0039 | 0.0023 | 0.0023 | TRUE | 0.0038 |
| Nuf2 | 3237 | 2299 | 2270 | 2254 | 1875 | 2204 | 2514 | 2445 | 2449 | 2244 | 2115 | 2176 | 2323.88 | 2469 | 2178 | 0.882 | -0.181 | 0.00228515 | 0.01338122 | 0 | 0.0034 | 0.0021 | 0.0021 | TRUE | 0.0084 |
| Nufip1 | 4497 | 3175 | 3393 | 3724 | 3136 | 3637 | 3493 | 3377 | 3661 | 3707 | 3538 | 3590 | 3560.96 | 3510 | 3612 | 1.029 | 0.041 | 0.47632533 | 0.64415914 | 7.00E-04 | 0.0031 | 0.0021 | 0.0021 | TRUE | 0.0168 |
| Nufip2 | 13192 | 10190 | 8918 | 9861 | 8346 | 10262 | 10246 | 10839 | 9622 | 9816 | 9415 | 10130 | 10011.34 | 10236 | 9787 | 0.956 | -0.065 | 0.28444953 | 0.45640291 | 0.0023 | 0.0027 | 0.0025 | 0.0025 | TRUE | 0.0333 |
| Nuggc | 57 | 52 | 48 | 40 | 27 | 35 | 44 | 55 | 52 | 40 | 30 | 35 | 42.7 | 50 | 35 | 0.697 | -0.521 | 0.08199578 | 0.19253583 | 0 | 0.0579 | 0.0403 | 0.0403 | TRUE | 0.1096 |
| Numa1 | 12469 | 9419 | 9418 | 10151 | 9691 | 11100 | 9684 | 10019 | 10162 | 10104 | 10932 | 10957 | 10309.85 | 9955 | 10664 | 1.071 | 0.099 | 0.06810601 | 0.16796046 | 0.0012 | 0.0027 | 0.002 | 0.002 | TRUE | 0.0259 |
| Numb | 1922 | 1446 | 1282 | 1319 | 1265 | 1413 | 1493 | 1538 | 1383 | 1313 | 1427 | 1395 | 1424.82 | 1471 | 1378 | 0.936 | -0.095 | 0.19655884 | 0.3563115 | 0.0016 | 0.0041 | 0.0032 | 0.0032 | TRUE | 0.0321 |
| Numbl | 1429 | 1087 | 1045 | 1092 | 1041 | 1091 | 1110 | 1156 | 1128 | 1087 | 1174 | 1077 | 1121.99 | 1131 | 1113 | 0.984 | -0.024 | 0.74719336 | 0.84993112 | 4.00E-04 | 0.0045 | 0.0031 | 0.0031 | TRUE | 0.0275 |
| Nup107 | 10766 | 7805 | 7574 | 7999 | 6979 | 8050 | 8361 | 8302 | 8172 | 7962 | 7873 | 7947 | 8102.95 | 8278 | 7927 | 0.958 | -0.063 | 0.17521167 | 0.32873463 | 0 | 0.0027 | 0.0014 | 0.0014 | TRUE | 0.0015 |
| Nup133 | 15894 | 11601 | 12415 | 13006 | 11960 | 13839 | 12344 | 12340 | 13396 | 12946 | 13492 | 13661 | 13029.87 | 12693 | 13366 | 1.053 | 0.075 | 0.17374867 | 0.32646111 | 0.0014 | 0.0026 | 0.0021 | 0.0021 | TRUE | 0.0285 |
| Nup153 | 14819 | 10902 | 9924 | 10597 | 9403 | 10578 | 11509 | 11596 | 10708 | 10548 | 10608 | 10442 | 10901.91 | 11271 | 10533 | 0.934 | -0.098 | 0.05971166 | 0.15275796 | 8.00E-04 | 0.0026 | 0.0019 | 0.0019 | TRUE | 0.0232 |
| Nup155 | 9123 | 6609 | 6525 | 7166 | 6186 | 7227 | 7085 | 7030 | 7040 | 7133 | 6978 | 7134 | 7066.9 | 7052 | 7082 | 1.004 | 0.006 | 0.89591604 | 0.94272381 | 0 | 0.0028 | 0.0014 | 0.0014 | TRUE | 0.002 |
| Nup160 | 15465 | 11089 | 11551 | 13146 | 11590 | 13590 | 12011 | 11795 | 12463 | 13086 | 13075 | 13415 | 12640.89 | 12090 | 13192 | 1.091 | 0.126 | 0.0087418 | 0.03748772 | 4.00E-04 | 0.0026 | 0.0016 | 0.0016 | TRUE | 0.0089 |
| Nup188 | 19714 | 14988 | 16413 | 18068 | 16237 | 18672 | 15311 | 15943 | 17709 | 17985 | 18317 | 18432 | 17282.85 | 16321 | 18245 | 1.118 | 0.161 | 0.00912589 | 0.03867728 | 0.0029 | 0.0026 | 0.0027 | 0.0027 | TRUE | 0.0677 |
| Nup205 | 17456 | 12263 | 12239 | 13388 | 11546 | 13788 | 13557 | 13044 | 13206 | 13326 | 13025 | 13611 | 13294.91 | 13269 | 13321 | 1.004 | 0.006 | 0.90688469 | 0.94912712 | 3.00E-04 | 0.0026 | 0.0015 | 0.0015 | TRUE | 0.0046 |
| Nup210 | 7820 | 6017 | 6372 | 6667 | 6092 | 6822 | 6073 | 6400 | 6875 | 6636 | 6872 | 6734 | 6598.68 | 6449 | 6747 | 1.047 | 0.066 | 0.27378824 | 0.44483975 | 0.0019 | 0.0028 | 0.0024 | 0.0024 | TRUE | 0.0406 |
| Nup210l | 77 | 90 | 119 | 59 | 48 | 66 | 60 | 96 | 128 | 59 | 54 | 65 | 76.99 | 95 | 59 | 0.632 | -0.661 | 0.01944045 | 0.06767228 | 0.0676 | 0.0332 | 0.0438 | 0.0438 | TRUE | 0.1478 |
| Nup214 | 7637 | 5303 | 5002 | 4965 | 4690 | 5333 | 5931 | 5641 | 5397 | 4942 | 5291 | 5264 | 5411.11 | 5656 | 5166 | 0.913 | -0.131 | 0.02702769 | 0.08588411 | 0.0015 | 0.0029 | 0.0024 | 0.0024 | TRUE | 0.0224 |
| Nup35 | 1404 | 919 | 727 | 832 | 745 | 886 | 1090 | 978 | 784 | 828 | 840 | 875 | 899.27 | 951 | 848 | 0.89 | -0.168 | 0.1285638 | 0.26504578 | 0.0132 | 0.0051 | 0.0077 | 0.0077 | TRUE | 0.2767 |
| Nup37 | 4183 | 2884 | 2922 | 3008 | 2644 | 3008 | 3249 | 3068 | 3153 | 2994 | 2983 | 2969 | 3069.25 | 3157 | 2982 | 0.944 | -0.083 | 0.13805498 | 0.27866401 | 0 | 0.0032 | 0.0019 | 0.0019 | TRUE | 0.0082 |
| Nup43 | 5393 | 3873 | 4086 | 4335 | 3805 | 4510 | 4189 | 4120 | 4409 | 4315 | 4292 | 4452 | 4296.08 | 4239 | 4353 | 1.027 | 0.039 | 0.48271478 | 0.64917598 | 5.00E-04 | 0.003 | 0.002 | 0.002 | TRUE | 0.0147 |
| Nup50 | 9606 | 6815 | 6321 | 6480 | 5929 | 6684 | 7461 | 7249 | 6820 | 6450 | 6689 | 6598 | 6877.79 | 7177 | 6579 | 0.916 | -0.126 | 0.02208324 | 0.07414511 | 0.001 | 0.0028 | 0.002 | 0.002 | TRUE | 0.0229 |
| Nup54 | 5865 | 4355 | 4292 | 4118 | 3657 | 4343 | 4555 | 4632 | 4631 | 4099 | 4125 | 4287 | 4388.37 | 4606 | 4170 | 0.906 | -0.143 | 0.00589636 | 0.02787849 | 0 | 0.003 | 0.0017 | 0.0017 | TRUE | 0.0073 |
| Nup62 | 10988 | 7737 | 6984 | 6864 | 6155 | 6981 | 8534 | 8230 | 7536 | 6832 | 6943 | 6891 | 7494.43 | 8100 | 6889 | 0.85 | -0.234 | 8.24E-05 | 0.00103268 | 0.0018 | 0.0027 | 0.0024 | 0.0024 | TRUE | 0.0452 |
| Nup62-il4i1 | 90 | 84 | 85 | 102 | 79 | 86 | 70 | 89 | 92 | 102 | 89 | 85 | 87.75 | 84 | 92 | 1.105 | 0.144 | 0.50759861 | 0.67089947 | 0 | 0.0294 | 0.0225 | 0.0225 | TRUE | 0.2227 |
| Nup62cl | 1054 | 686 | 632 | 579 | 461 | 477 | 819 | 730 | 682 | 576 | 520 | 471 | 632.91 | 744 | 522 | 0.702 | -0.511 | 2.08E-06 | 5.20E-05 | 0.0079 | 0.0062 | 0.0067 | 0.0067 | TRUE | 0.0971 |
| Nup85 | 15187 | 10905 | 10622 | 10600 | 9568 | 11560 | 11795 | 11600 | 11461 | 10551 | 10794 | 11412 | 11268.68 | 11619 | 10919 | 0.94 | -0.09 | 0.08140295 | 0.19155291 | 8.00E-04 | 0.0026 | 0.0018 | 0.0018 | TRUE | 0.0191 |
| Nup88 | 10244 | 7189 | 7147 | 6814 | 6003 | 6876 | 7956 | 7647 | 7712 | 6783 | 6772 | 6788 | 7276.13 | 7772 | 6781 | 0.872 | -0.197 | 3.97E-05 | 0.00058341 | 1.00E-04 | 0.0028 | 0.0015 | 0.0015 | TRUE | 0.0054 |
| Nup93 | 19736 | 14168 | 13530 | 14388 | 12425 | 14754 | 15328 | 15070 | 14599 | 14322 | 14017 | 14564 | 14650.03 | 14999 | 14301 | 0.953 | -0.069 | 0.1480364 | 0.29281542 | 4.00E-04 | 0.0026 | 0.0016 | 0.0016 | TRUE | 0.0067 |
| Nup98 | 8381 | 5860 | 5789 | 6428 | 5483 | 6539 | 6509 | 6233 | 6246 | 6398 | 6185 | 6455 | 6337.91 | 6329 | 6346 | 1.003 | 0.004 | 0.94374143 | 0.96875153 | 4.00E-04 | 0.0028 | 0.0017 | 0.0017 | TRUE | 0.0076 |
| Nupl1 | 5310 | 3928 | 3899 | 3780 | 3435 | 3882 | 4124 | 4178 | 4207 | 3763 | 3875 | 3832 | 3996.5 | 4170 | 3823 | 0.917 | -0.125 | 0.01515041 | 0.0562308 | 0 | 0.003 | 0.0017 | 0.0017 | TRUE | 0.0023 |
| Nupl2 | 651 | 445 | 362 | 428 | 357 | 381 | 506 | 473 | 391 | 426 | 403 | 376 | 429.07 | 457 | 402 | 0.878 | -0.188 | 0.1186847 | 0.25012686 | 0.0086 | 0.008 | 0.0082 | 0.0082 | TRUE | 0.1813 |
| Nupr1 | 145 | 105 | 90 | 106 | 80 | 119 | 113 | 112 | 97 | 106 | 90 | 117 | 105.77 | 107 | 104 | 0.975 | -0.037 | 0.8516248 | 0.91640026 | 0 | 0.0248 | 0.0185 | 0.0185 | TRUE | 0.1313 |
| Nupr1l | 25 | 9 | 17 | 18 | 26 | 26 | 19 | 10 | 18 | 18 | 29 | 26 | 20.04 | 16 | 24 | 1.523 | 0.607 | 0.18338207 | 0.33925446 | 0.0322 | 0.1207 | 0.0984 | 0.0984 | TRUE | 0.517 |
| Nus1 | 6230 | 4660 | 4800 | 5138 | 4436 | 5251 | 4839 | 4957 | 5179 | 5114 | 5004 | 5184 | 5046.12 | 4992 | 5101 | 1.022 | 0.032 | 0.55386801 | 0.70963062 | 5.00E-04 | 0.0029 | 0.0019 | 0.0019 | TRUE | 0.0131 |
| Nusap1 | 5816 | 4313 | 5105 | 5300 | 4635 | 5350 | 4517 | 4588 | 5508 | 5276 | 5229 | 5281 | 5066.44 | 4871 | 5262 | 1.081 | 0.112 | 0.14438829 | 0.28770972 | 0.0062 | 0.0029 | 0.004 | 0.004 | TRUE | 0.159 |
| Nutf2 | 1384 | 894 | 878 | 906 | 814 | 963 | 1075 | 951 | 947 | 902 | 918 | 951 | 957.32 | 991 | 924 | 0.93 | -0.104 | 0.21091379 | 0.37295445 | 0.0022 | 0.0049 | 0.0039 | 0.0039 | TRUE | 0.0683 |
| Nutf2-ps1 | 0 | 0 | 0 | 0 | 0 | 0 | 0 | 0 | 0 | 0 | 0 | 0 | 0 | 0 | 0 | NA | NA | NA | NA | NA | NA | NA | NA | NA | NA |
| Nvl | 6722 | 4833 | 4988 | 5148 | 4688 | 5411 | 5221 | 5141 | 5382 | 5124 | 5289 | 5341 | 5249.64 | 5248 | 5251 | 1.001 | 0.001 | 0.98388172 | 0.99165231 | 3.00E-04 | 0.0029 | 0.0017 | 0.0017 | TRUE | 0.006 |
| Nwd1 | 2 | 1 | 4 | 0 | 2 | 2 | 2 | 1 | 4 | 0 | 2 | 2 | 1.86 | 2 | 1 | 0.611 | -0.711 | 0.62893176 | NA | 0 | 1.2763 | 0.9781 | 0.9781 | TRUE | 0.5407 |
| Nxf1 | 6308 | 4408 | 3589 | 3114 | 2810 | 3421 | 4899 | 4689 | 3873 | 3100 | 3170 | 3377 | 3851.18 | 4487 | 3216 | 0.716 | -0.481 | 1.51E-08 | 7.29E-07 | 0.0079 | 0.003 | 0.0049 | 0.0049 | TRUE | 0.174 |
| Nxf2 | 48 | 36 | 33 | 29 | 22 | 27 | 37 | 38 | 36 | 29 | 25 | 27 | 31.92 | 37 | 27 | 0.723 | -0.467 | 0.16454507 | 0.3153476 | 0 | 0.0767 | 0.0491 | 0.0491 | TRUE | 0.0294 |
| Nxf3 | 598 | 428 | 420 | 252 | 177 | 191 | 464 | 455 | 453 | 251 | 200 | 189 | 335.32 | 457 | 213 | 0.466 | -1.102 | 1.48E-17 | 3.31E-15 | 0.0057 | 0.0095 | 0.0086 | 0.0086 | TRUE | 0.2692 |
| Nxf7 | 134 | 104 | 63 | 85 | 72 | 67 | 104 | 111 | 68 | 85 | 81 | 66 | 85.77 | 94 | 77 | 0.817 | -0.291 | 0.22654099 | 0.39151432 | 0.0292 | 0.0301 | 0.0298 | 0.0298 | TRUE | 0.543 |
| Nxn | 4731 | 3520 | 3754 | 4721 | 4199 | 5007 | 3674 | 3744 | 4051 | 4699 | 4737 | 4943 | 4307.99 | 3823 | 4793 | 1.254 | 0.327 | 4.97E-08 | 2.10E-06 | 0.0014 | 0.003 | 0.0024 | 0.0024 | TRUE | 0.0327 |
| Nxnl1 | 1 | 1 | 2 | 1 | 2 | 5 | 1 | 1 | 2 | 1 | 2 | 5 | 2.03 | 1 | 3 | 2.1 | 1.071 | 0.43764313 | NA | 0 | 1.1694 | 0.8125 | 0.8125 | TRUE | 0.3809 |
| Nxnl2 | 41 | 42 | 53 | 51 | 53 | 67 | 32 | 45 | 57 | 51 | 60 | 66 | 51.73 | 45 | 59 | 1.335 | 0.417 | 0.15889045 | 0.30807401 | 0.0307 | 0.0482 | 0.0434 | 0.0434 | TRUE | 0.2836 |
| Nxpe2 | 1 | 0 | 1 | 1 | 0 | 0 | 1 | 0 | 1 | 1 | 0 | 0 | 0.48 | 1 | 0 | 0.596 | -0.748 | 0.78941713 | NA | 0 | 4.9902 | 3.8695 | 3.8695 | TRUE | 0.3906 |
| Nxpe3 | 655 | 458 | 520 | 616 | 553 | 647 | 509 | 487 | 561 | 613 | 624 | 639 | 572.11 | 519 | 625 | 1.206 | 0.27 | 0.00452435 | 0.02275389 | 6.00E-04 | 0.0066 | 0.0047 | 0.0047 | TRUE | 0.056 |
| Nxpe4 | 46 | 37 | 25 | 18 | 22 | 18 | 36 | 39 | 27 | 18 | 25 | 18 | 27.09 | 34 | 20 | 0.589 | -0.763 | 0.04600433 | 0.12684026 | 0 | 0.0899 | 0.0655 | 0.0655 | TRUE | 0.2126 |
| Nxpe5 | 0 | 1 | 0 | 0 | 0 | 0 | 0 | 1 | 0 | 0 | 0 | 0 | 0.18 | 0 | 0 | 0.552 | -0.857 | 0.8336548 | NA | 0 | 13.3714 | 10 | 10 | TRUE | 0.5216 |

| | | | | | | | | | | | | | | | | | | | | | | | | | |
|---|---|---|---|---|---|---|---|---|---|---|---|---|---|---|---|---|---|---|---|---|---|---|---|---|---|
| Nxph1 | 0 | 0 | 0 | 1 | 0 | 0 | 0 | 0 | 0 | 1 | 0 | 0 | 0.17 | 0 | 0 | 2.094 | 1.067 | 0.79379329 | NA | 0 | 14.2885 | 10 | 10 | TRUE | 0.3906 |
| Nxph2 | 0 | 0 | 0 | 0 | 1 | 0 | 0 | 0 | 0 | 0 | 1 | 0 | 0.19 | 0 | 0 | 2.094 | 1.067 | 0.79379329 | NA | 0 | 12.608 | 10 | 10 | TRUE | 0.5075 |
| Nxph3 | 125 | 105 | 88 | 123 | 117 | 125 | 97 | 112 | 95 | 122 | 132 | 123 | 113.59 | 101 | 126 | 1.245 | 0.316 | 0.08695734 | 0.19988502 | 0 | 0.0233 | 0.0157 | 0.0157 | TRUE | 0.0757 |
| Nxph4 | 599 | 535 | 811 | 1082 | 995 | 1162 | 465 | 569 | 875 | 1077 | 1122 | 1147 | 875.99 | 636 | 1115 | 1.754 | 0.811 | 0.00361067 | 0.01905031 | 0.0547 | 0.0051 | 0.0178 | 0.0547 | TRUE | 1.2638 |
| Nxt1 | 2078 | 1571 | 1531 | 1545 | 1390 | 1469 | 1614 | 1671 | 1652 | 1538 | 1568 | 1450 | 1582.16 | 1646 | 1519 | 0.923 | -0.116 | 0.0828308 | 0.19379614 | 3.00E-04 | 0.0039 | 0.0026 | 0.0026 | TRUE | 0.0189 |
| Nxt2 | 2771 | 2075 | 2632 | 3114 | 2566 | 2876 | 2152 | 2207 | 2840 | 3100 | 2895 | 2839 | 2672.1 | 2400 | 2945 | 1.228 | 0.296 | 0.0022502 | 0.01322457 | 0.0132 | 0.0033 | 0.0064 | 0.0064 | TRUE | 0.3105 |
| Nyap1 | 725 | 479 | 631 | 786 | 664 | 793 | 563 | 510 | 681 | 782 | 749 | 783 | 677.95 | 585 | 771 | 1.321 | 0.402 | 0.00020134 | 0.00214459 | 0.0093 | 0.0059 | 0.0069 | 0.0069 | TRUE | 0.2363 |
| Nyap2 | 1 | 0 | 0 | 0 | 0 | 0 | 1 | 0 | 0 | 0 | 0 | 0 | 0.13 | 0 | 0 | 0.552 | -0.857 | 0.8336548 | NA | 0 | 18.3121 | 10 | 10 | TRUE | 0.2566 |
| Nynrin | 1218 | 934 | 836 | 1106 | 1062 | 1171 | 946 | 993 | 902 | 1101 | 1198 | 1156 | 1049.4 | 947 | 1152 | 1.215 | 0.281 | 0.00029975 | 0.00290664 | 0.001 | 0.0047 | 0.0034 | 0.0034 | TRUE | 0.0208 |
| Nyx | 12 | 11 | 6 | 7 | 6 | 15 | 9 | 12 | 6 | 7 | 7 | 15 | 9.34 | 9 | 10 | 1.042 | 0.059 | 0.92697754 | NA | 0 | 0.2562 | 0.1961 | 0.1961 | TRUE | 0.8153 |
| Oacyl | 59 | 55 | 51 | 69 | 70 | 60 | 46 | 59 | 55 | 69 | 79 | 59 | 61.04 | 53 | 69 | 1.302 | 0.381 | 0.13668999 | 0.27672946 | 0 | 0.0413 | 0.0306 | 0.0306 | TRUE | 0.1411 |
| Oaf | 140 | 97 | 120 | 242 | 176 | 199 | 109 | 103 | 129 | 241 | 199 | 196 | 162.88 | 114 | 212 | 1.867 | 0.901 | 1.26E-07 | 4.83E-06 | 0.0075 | 0.017 | 0.0143 | 0.0143 | TRUE | 0.1578 |
| Oard1 | 3198 | 2190 | 2266 | 2092 | 1833 | 2124 | 2484 | 2329 | 2445 | 2082 | 2068 | 2097 | 2250.85 | 2419 | 2082 | 0.86 | -0.217 | 0.00030536 | 0.00294703 | 0 | 0.0035 | 0.0022 | 0.0022 | TRUE | 0.0127 |
| Oas1a | 290 | 205 | 232 | 193 | 171 | 187 | 225 | 218 | 250 | 192 | 193 | 185 | 210.54 | 231 | 190 | 0.822 | -0.283 | 0.04274349 | 0.12012048 | 0 | 0.0137 | 0.0093 | 0.0093 | TRUE | 0.0549 |
| Oas1b | 33 | 29 | 28 | 26 | 20 | 35 | 26 | 31 | 30 | 26 | 23 | 35 | 28.28 | 29 | 28 | 0.964 | -0.052 | 0.88668124 | 0.93690919 | 0 | 0.0862 | 0.0613 | 0.0613 | TRUE | 0.3086 |
| Oas1c | 380 | 306 | 301 | 330 | 368 | 421 | 295 | 325 | 325 | 328 | 415 | 416 | 350.77 | 315 | 386 | 1.227 | 0.296 | 0.02013205 | 0.06952099 | 0.0079 | 0.0092 | 0.0088 | 0.0088 | TRUE | 0.2 |
| Oas1d | 30 | 24 | 26 | 35 | 27 | 29 | 23 | 26 | 28 | 35 | 30 | 29 | 28.47 | 26 | 31 | 1.227 | 0.295 | 0.40806117 | 0.58080247 | 0 | 0.0857 | 0.0562 | 0.0562 | TRUE | 0.0679 |
| Oas1e | 3 | 3 | 1 | 2 | 1 | 1 | 2 | 3 | 1 | 2 | 1 | 1 | 1.78 | 2 | 1 | 0.622 | -0.684 | 0.62649727 | NA | 0 | 1.3307 | 0.8298 | 0.8298 | TRUE | 0.1712 |
| Oas1f | 11 | 9 | 11 | 5 | 2 | 4 | 9 | 10 | 12 | 5 | 2 | 4 | 6.86 | 10 | 4 | 0.378 | -1.404 | 0.06594197 | NA | 0 | 0.3479 | 0.2337 | 0.2337 | TRUE | 0.1608 |
| Oas1g | 147 | 121 | 127 | 91 | 84 | 103 | 114 | 129 | 137 | 91 | 95 | 102 | 111.15 | 127 | 96 | 0.758 | -0.399 | 0.0346692 | 0.10312735 | 0 | 0.0237 | 0.0165 | 0.0165 | TRUE | 0.0844 |
| Oas1h | 6 | 5 | 2 | 4 | 3 | 1 | 5 | 5 | 2 | 4 | 3 | 1 | 3.41 | 4 | 3 | 0.681 | -0.554 | 0.59864073 | NA | 0 | 0.6965 | 0.493 | 0.493 | TRUE | 0.4086 |
| Oas2 | 98 | 98 | 85 | 84 | 94 | 121 | 76 | 104 | 92 | 84 | 106 | 119 | 96.86 | 91 | 103 | 1.142 | 0.191 | 0.38608022 | 0.55991846 | 0.019 | 0.0269 | 0.0246 | 0.0246 | TRUE | 0.2739 |
| Oas3 | 339 | 271 | 343 | 371 | 372 | 413 | 263 | 288 | 370 | 369 | 420 | 408 | 353.05 | 307 | 399 | 1.302 | 0.381 | 0.00576661 | 0.02738278 | 0.0144 | 0.0091 | 0.0109 | 0.0109 | TRUE | 0.3575 |
| Oasl1 | 72 | 66 | 58 | 68 | 54 | 73 | 56 | 70 | 63 | 68 | 61 | 72 | 64.9 | 63 | 67 | 1.069 | 0.097 | 0.69009404 | 0.81040066 | 0 | 0.0389 | 0.027 | 0.027 | TRUE | 0.0959 |
| Oasl2 | 21 | 15 | 11 | 2 | 7 | 6 | 16 | 16 | 12 | 2 | 8 | 6 | 9.99 | 15 | 5 | 0.353 | -1.501 | 0.02350047 | NA | 0 | 0.2397 | 0.1844 | 0.1844 | TRUE | 0.6462 |
| Oat | 4792 | 3263 | 3408 | 4033 | 3478 | 4151 | 3722 | 3471 | 3677 | 4014 | 3924 | 4098 | 3817.57 | 3623 | 4012 | 1.107 | 0.147 | 0.00947952 | 0.03982872 | 5.00E-04 | 0.003 | 0.002 | 0.002 | TRUE | 0.0163 |
| Oaz1 | 6 | 8 | 2 | 6 | 8 | 4 | 5 | 9 | 2 | 6 | 9 | 4 | 5.71 | 5 | 6 | 1.231 | 0.3 | 0.71930502 | NA | 0.0625 | 0.4173 | 0.3242 | 0.3242 | TRUE | 0.3677 |
| Oaz1-ps | 0 | 0 | 0 | 0 | 0 | 0 | 0 | 0 | 0 | 0 | 0 | 0 | 0 | 0 | 0 | NA | NA | NA | NA | NA | NA | NA | NA | NA | NA |
| Oaz2 | 3989 | 2665 | 2434 | 2597 | 2156 | 2379 | 3098 | 2835 | 2626 | 2585 | 2432 | 2348 | 2654.13 | 2853 | 2455 | 0.86 | -0.218 | 0.00375409 | 0.01964029 | 0.0041 | 0.0033 | 0.0037 | 0.0037 | TRUE | 0.0698 |
| Oaz3 | 3 | 0 | 1 | 2 | 1 | 2 | 2 | 0 | 1 | 2 | 1 | 2 | 1.42 | 1 | 2 | 1.437 | 0.523 | 0.74491962 | NA | 0 | 1.675 | 1.1519 | 1.1519 | TRUE | 0.365 |
| Obfc1 | 897 | 613 | 641 | 621 | 584 | 633 | 697 | 652 | 692 | 618 | 659 | 625 | 657.03 | 680 | 634 | 0.931 | -0.103 | 0.23738218 | 0.40415765 | 0 | 0.006 | 0.004 | 0.004 | TRUE | 0.015 |
| Obox1 | 0 | 1 | 0 | 0 | 0 | 0 | 0 | 0 | 1 | 0 | 0 | 0 | 0.18 | 0 | 0 | 0.552 | -0.857 | 0.8336548 | NA | 0 | 13.3714 | 10 | 10 | TRUE | 0.5216 |
| Obox2 | 3 | 1 | 1 | 1 | 0 | 0 | 2 | 1 | 1 | 1 | 0 | 0 | 0.91 | 1 | 0 | 0.245 | -2.029 | 0.33198448 | NA | 0 | 2.6031 | 1.8334 | 1.8334 | TRUE | 0.3906 |
| Obox3 | 2 | 5 | 1 | 0 | 4 | 1 | 2 | 5 | 1 | 0 | 5 | 1 | 2.24 | 3 | 2 | 0.683 | -0.55 | 0.6988922 | NA | 0.7706 | 1.0597 | 0.9838 | 0.9838 | TRUE | 1.2123 |
| Obox5 | 1 | 0 | 1 | 0 | 0 | 0 | 1 | 0 | 1 | 0 | 0 | 0 | 0.31 | 1 | 0 | 0.307 | -1.706 | 0.62894649 | NA | 0 | 7.6657 | 7.2 | 7.2 | TRUE | 0.2067 |
| Obox6 | 105 | 100 | 102 | 76 | 73 | 108 | 82 | 106 | 110 | 76 | 82 | 107 | 93.76 | 99 | 88 | 0.894 | -0.162 | 0.46901517 | 0.63789455 | 0.0198 | 0.0277 | 0.0254 | 0.0254 | TRUE | 0.3248 |
| Obp1a | 0 | 0 | 0 | 0 | 0 | 0 | 0 | 0 | 0 | 0 | 0 | 0 | 0 | 0 | 0 | NA | NA | NA | NA | NA | NA | NA | NA | NA | NA |
| Obp2a | 0 | 0 | 0 | 0 | 0 | 0 | 0 | 0 | 0 | 0 | 0 | 0 | 0 | 0 | 0 | NA | NA | NA | NA | NA | NA | NA | NA | NA | NA |
| Obp2b | 0 | 0 | 0 | 0 | 0 | 0 | 0 | 0 | 0 | 0 | 0 | 0 | 0 | 0 | 0 | NA | NA | NA | NA | NA | NA | NA | NA | NA | NA |
| Obscn | 105 | 82 | 62 | 77 | 75 | 66 | 82 | 87 | 67 | 77 | 85 | 65 | 77.01 | 79 | 76 | 0.957 | -0.063 | 0.7849565 | 0.87364767 | 0 | 0.0332 | 0.0252 | 0.0252 | TRUE | 0.1453 |
| Obsl1 | 1050 | 747 | 743 | 865 | 792 | 884 | 815 | 795 | 802 | 861 | 893 | 873 | 839.81 | 804 | 876 | 1.089 | 0.122 | 0.11435483 | 0.24366033 | 0 | 0.0052 | 0.0031 | 0.0031 | TRUE | 0.0036 |
| Oc90 | 8 | 4 | 5 | 0 | 2 | 1 | 6 | 4 | 5 | 0 | 2 | 1 | 3.18 | 5 | 1 | 0.198 | -2.339 | 0.06098556 | NA | 0 | 0.7467 | 0.5396 | 0.5396 | TRUE | 0.3987 |
| Oca2 | 0 | 3 | 1 | 2 | 2 | 0 | 0 | 3 | 1 | 2 | 2 | 0 | 1.42 | 1 | 1 | 1.018 | 0.026 | 0.98826537 | NA | 0 | 1.6721 | 1.5179 | 1.5179 | TRUE | 0.4899 |
| Ocel1 | 502 | 382 | 334 | 336 | 328 | 373 | 390 | 406 | 360 | 334 | 370 | 368 | 371.55 | 385 | 357 | 0.927 | -0.11 | 0.32600789 | 0.50001404 | 0 | 0.0088 | 0.0063 | 0.0063 | TRUE | 0.037 |
| Ociad1 | 6590 | 4939 | 5195 | 5518 | 4997 | 5726 | 5118 | 5254 | 5605 | 5493 | 5637 | 5652 | 5459.88 | 5326 | 5594 | 1.051 | 0.071 | 0.20326564 | 0.36472149 | 0.001 | 0.0029 | 0.0021 | 0.0021 | TRUE | 0.0256 |
| Ociad2 | 202 | 164 | 130 | 175 | 120 | 183 | 157 | 174 | 140 | 174 | 135 | 181 | 160.3 | 157 | 163 | 1.042 | 0.059 | 0.73361467 | 0.84079179 | 0.0104 | 0.0172 | 0.0153 | 0.0153 | TRUE | 0.2284 |
| Ocln | 777 | 581 | 602 | 761 | 686 | 721 | 603 | 618 | 650 | 757 | 774 | 712 | 685.69 | 624 | 748 | 1.2 | 0.262 | 0.00259851 | 0.01480007 | 0 | 0.0059 | 0.004 | 0.004 | TRUE | 0.0212 |
| Ocm | 4 | 5 | 0 | 2 | 3 | 0 | 3 | 5 | 0 | 2 | 3 | 0 | 2.3 | 3 | 2 | 0.626 | -0.675 | 0.6422665 | NA | 1.1623 | 1.0329 | 1.0557 | 1.0557 | TRUE | 0.1935 |
| Ocrl | 3164 | 2266 | 2391 | 2624 | 2208 | 2367 | 2457 | 2410 | 2580 | 2612 | 2491 | 2337 | 2481.15 | 2482 | 2480 | 0.999 | -0.001 | 0.98278505 | 0.99089635 | 0.0018 | 0.0034 | 0.0028 | 0.0028 | TRUE | 0.0313 |
| Ocstamp | 16 | 14 | 9 | 13 | 15 | 16 | 12 | 15 | 10 | 13 | 17 | 16 | 13.78 | 12 | 15 | 1.23 | 0.299 | 0.56005649 | NA | 0 | 0.1744 | 0.1162 | 0.1162 | TRUE | 0.1228 |
| Odam | 0 | 0 | 0 | 0 | 0 | 0 | 0 | 0 | 0 | 0 | 0 | 0 | 0 | 0 | 0 | NA | NA | NA | NA | NA | NA | NA | NA | NA | NA |
| Odc1 | 48438 | 33294 | 28670 | 30535 | 26087 | 30088 | 37620 | 35414 | 30935 | 30395 | 29429 | 29701 | 32248.95 | 34656 | 29842 | 0.861 | -0.216 | 0.00201491 | 0.01216793 | 0.0049 | 0.0025 | 0.0035 | 0.0035 | TRUE | 0.108 |
| Odf1 | 5 | 2 | 3 | 0 | 1 | 1 | 4 | 2 | 3 | 0 | 1 | 1 | 1.89 | 3 | 1 | 0.224 | -2.16 | 0.16477083 | NA | 0 | 1.2539 | 0.8424 | 0.8424 | TRUE | 0.2673 |
| Odf2 | 14211 | 11096 | 11319 | 11597 | 10622 | 12132 | 11037 | 11803 | 12213 | 11544 | 11983 | 11976 | 11759.23 | 11684 | 11834 | 1.013 | 0.019 | 0.7358866 | 0.84238565 | 0.0012 | 0.0026 | 0.0021 | 0.0021 | TRUE | 0.0289 |
| Odf2l | 1180 | 872 | 862 | 939 | 810 | 941 | 916 | 928 | 930 | 935 | 914 | 929 | 925.24 | 925 | 926 | 1.002 | 0.002 | 0.97558879 | 0.98670326 | 0 | 0.005 | 0.0029 | 0.0029 | TRUE | 0.0015 |
| Odf3 | 1 | 2 | 6 | 2 | 5 | 4 | 1 | 2 | 6 | 2 | 6 | 4 | 3.49 | 3 | 4 | 1.265 | 0.339 | 0.75671048 | NA | 0.2948 | 0.6809 | 0.5672 | 0.5672 | TRUE | 0.8116 |
| Odf3b | 20 | 8 | 14 | 16 | 16 | 22 | 16 | 9 | 15 | 16 | 18 | 22 | 15.81 | 13 | 19 | 1.413 | 0.498 | 0.31059309 | NA | 0 | 0.1524 | 0.1096 | 0.1096 | TRUE | 0.3566 |
| Odf3l1 | 9 | 6 | 5 | 4 | 3 | 6 | 7 | 6 | 5 | 4 | 3 | 6 | 5.34 | 6 | 4 | 0.708 | -0.497 | 0.54170864 | NA | 0 | 0.446 | 0.2857 | 0.2857 | TRUE | 0.1634 |
| Odf3l2 | 1 | 3 | 1 | 2 | 0 | 0 | 1 | 3 | 1 | 2 | 0 | 0 | 1.17 | 2 | 1 | 0.408 | -1.295 | 0.50272732 | NA | 0 | 2.0231 | 1.6328 | 1.6328 | TRUE | 0.9735 |
| Odf4 | 1 | 0 | 0 | 1 | 1 | 1 | 1 | 0 | 0 | 1 | 1 | 1 | 0.65 | 0 | 1 | 3.004 | 1.587 | 0.49664754 | NA | 0 | 3.6606 | 2.4271 | 2.4271 | TRUE | 0.2566 |
| Ofcc1 | 0 | 0 | 1 | 0 | 0 | 0 | 0 | 0 | 1 | 0 | 0 | 0 | 0.18 | 0 | 0 | 0.552 | -0.857 | 0.8336548 | NA | 0 | 13.1817 | 10 | 10 | TRUE | 0.5365 |
| Ofd1 | 2826 | 2053 | 1831 | 1897 | 1761 | 1845 | 2195 | 2184 | 1976 | 1888 | 1987 | 1821 | 2008.4 | 2118 | 1899 | 0.896 | -0.159 | 0.02396034 | 0.07889085 | 0.0019 | 0.0036 | 0.0031 | 0.0031 | TRUE | 0.0413 |
| Ogdh | 16822 | 11156 | 9789 | 11566 | 9682 | 10912 | 13065 | 11866 | 10562 | 11513 | 10922 | 10772 | 11450.1 | 11831 | 11069 | 0.935 | -0.096 | 0.19823432 | 0.35833605 | 0.0061 | 0.0026 | 0.004 | 0.004 | TRUE | 0.1075 |
| Ogdhl | 7967 | 6383 | 6831 | 6812 | 6496 | 7087 | 6188 | 6790 | 7371 | 6781 | 7328 | 6996 | 6908.75 | 6783 | 7035 | 1.038 | 0.053 | 0.44996119 | 0.62006318 | 0.0038 | 0.0028 | 0.0034 | 0.0034 | TRUE | 0.0726 |
| Ogfod1 | 6139 | 4260 | 3783 | 4357 | 3742 | 4305 | 4768 | 4531 | 4082 | 4337 | 4221 | 4250 | 4364.84 | 4460 | 4269 | 0.957 | -0.064 | 0.33903686 | 0.51311456 | 0.0028 | 0.003 | 0.003 | 0.003 | TRUE | 0.0668 |
| Ogfod2 | 342 | 325 | 287 | 278 | 266 | 295 | 266 | 346 | 310 | 277 | 300 | 291 | 298.17 | 307 | 289 | 0.945 | -0.081 | 0.54072099 | 0.69903447 | 0.0069 | 0.0104 | 0.0093 | 0.0093 | TRUE | 0.17 |
| Ogfod3 | 832 | 594 | 676 | 752 | 665 | 816 | 646 | 632 | 729 | 749 | 750 | 806 | 718.61 | 669 | 768 | 1.149 | 0.201 | 0.02810587 | 0.08830892 | 0.0023 | 0.0057 | 0.0046 | 0.0046 | TRUE | 0.0709 |
| Ogfr | 1766 | 1294 | 1232 | 1209 | 1087 | 1313 | 1372 | 1376 | 1329 | 1203 | 1226 | 1296 | 1300.52 | 1359 | 1242 | 0.914 | -0.13 | 0.06276667 | 0.15850248 | 0 | 0.0042 | 0.0028 | 0.0028 | TRUE | 0.0178 |
| Ogfrl1 | 1147 | 892 | 927 | 953 | 832 | 947 | 891 | 949 | 1000 | 949 | 939 | 935 | 943.65 | 947 | 941 | 0.995 | -0.007 | 0.92967139 | 0.96107285 | 9.00E-04 | 0.0049 | 0.0035 | 0.0035 | TRUE | 0.0335 |

| | | | | | | | | | | | | | | | | | | | | | | | | | |
|---|---|---|---|---|---|---|---|---|---|---|---|---|---|---|---|---|---|---|---|---|---|---|---|---|---|
| Ogg1 | 1344 | 869 | 807 | 924 | 857 | 882 | 1044 | 924 | 871 | 920 | 967 | 871 | 932.69 | 946 | 919 | 0.969 | -0.045 | 0.61769676 | 0.75859223 | 0.0048 | 0.005 | 0.0049 | 0.0049 | TRUE | 0.1006 |
| Ogn | 1 | 1 | 2 | 0 | 0 | 0 | 1 | 1 | 2 | 0 | 0 | 0 | 0.67 | 1 | 0 | 0.144 | -2.794 | 0.24876697 | NA | 0 | 3.559 | 2.8631 | 2.8631 | TRUE | 0.1682 |
| Ogt | 11608 | 9318 | 8282 | 8684 | 7267 | 8948 | 9015 | 9911 | 8936 | 8644 | 8198 | 8833 | 8923.02 | 9287 | 8558 | 0.922 | -0.118 | 0.05150592 | 0.13779485 | 0.0023 | 0.0027 | 0.0025 | 0.0025 | TRUE | 0.042 |
| Oip5 | 2299 | 1804 | 2183 | 2275 | 1999 | 2397 | 1786 | 1919 | 2355 | 2265 | 2255 | 2366 | 2157.61 | 2020 | 2295 | 1.137 | 0.186 | 0.04688225 | 0.12854171 | 0.0106 | 0.0035 | 0.0058 | 0.0058 | TRUE | 0.2543 |
| Oit1 | 12 | 7 | 7 | 4 | 1 | 0 | 9 | 7 | 8 | 4 | 1 | 0 | 4.9 | 8 | 2 | 0.211 | -2.246 | 0.02847014 | NA | 0 | 0.4856 | 0.3964 | 0.3964 | TRUE | 1.1065 |
| Oit3 | 4 | 1 | 0 | 0 | 1 | 0 | 3 | 1 | 0 | 0 | 1 | 0 | 0.88 | 1 | 0 | 0.259 | -1.947 | 0.39420209 | NA | 0 | 2.6863 | 2.4242 | 2.4242 | TRUE | 0.3212 |
| Ola1 | 8229 | 5870 | 6180 | 6346 | 5726 | 6630 | 6391 | 6244 | 6668 | 6317 | 6460 | 6545 | 6437.38 | 6434 | 6441 | 1.001 | 0.001 | 0.97805773 | 0.98822134 | 5.00E-04 | 0.0028 | 0.0018 | 0.0018 | TRUE | 0.0122 |
| Olah | 8 | 6 | 6 | 0 | 3 | 1 | 6 | 6 | 6 | 0 | 3 | 1 | 3.91 | 6 | 1 | 0.222 | -2.174 | 0.05326195 | NA | 0 | 0.6091 | 0.4683 | 0.4683 | TRUE | 0.791 |
| Olfm1 | 897 | 644 | 628 | 675 | 606 | 732 | 697 | 685 | 678 | 672 | 684 | 723 | 689.57 | 687 | 693 | 1.009 | 0.013 | 0.88002958 | 0.93298562 | 0 | 0.0059 | 0.0037 | 0.0037 | TRUE | 0.0172 |
| Olfm2 | 313 | 224 | 218 | 367 | 292 | 375 | 243 | 238 | 235 | 365 | 329 | 370 | 296.91 | 239 | 355 | 1.486 | 0.572 | 2.15E-06 | 5.33E-05 | 0 | 0.0104 | 0.007 | 0.007 | TRUE | 0.0438 |
| Olfm3 | 0 | 0 | 0 | 0 | 0 | 0 | 0 | 0 | 0 | 0 | 0 | 0 | 0 | 0 | 0 | NA | NA | NA | NA | NA | NA | NA | NA | NA | NA |
| Olfm4 | 4 | 0 | 2 | 1 | 1 | 2 | 3 | 0 | 2 | 1 | 1 | 2 | 1.56 | 2 | 1 | 0.762 | -0.393 | 0.80131751 | NA | 0 | 1.5213 | 1.1085 | 1.1085 | TRUE | 0.4995 |
| Olfml1 | 54 | 52 | 64 | 57 | 60 | 62 | 42 | 55 | 69 | 57 | 68 | 61 | 58.66 | 55 | 62 | 1.124 | 0.169 | 0.53345553 | 0.6928378 | 0.0187 | 0.0428 | 0.036 | 0.036 | TRUE | 0.234 |
| Olfml2a | 41 | 23 | 23 | 43 | 37 | 27 | 32 | 24 | 25 | 43 | 42 | 27 | 32.05 | 27 | 37 | 1.359 | 0.442 | 0.21480224 | 0.37748702 | 0 | 0.0764 | 0.0598 | 0.0598 | TRUE | 0.4528 |
| Olfml2b | 200 | 143 | 128 | 227 | 182 | 222 | 155 | 152 | 138 | 226 | 205 | 219 | 182.66 | 148 | 217 | 1.458 | 0.544 | 0.00025821 | 0.00259538 | 0 | 0.0154 | 0.0103 | 0.0103 | TRUE | 0.0376 |
| Olfml3 | 1 | 1 | 2 | 0 | 1 | 1 | 1 | 1 | 2 | 0 | 1 | 1 | 1.02 | 1 | 1 | 0.531 | -0.914 | 0.6308001 | NA | 0 | 2.3285 | 1.4951 | 1.4951 | TRUE | 0.2654 |
| Olfr1 | 0 | 0 | 0 | 0 | 0 | 0 | 0 | 0 | 0 | 0 | 0 | 0 | 0 | 0 | 0 | NA | NA | NA | NA | NA | NA | NA | NA | NA | NA |
| Olfr10 | 0 | 0 | 0 | 0 | 0 | 0 | 0 | 0 | 0 | 0 | 0 | 0 | 0 | 0 | 0 | NA | NA | NA | NA | NA | NA | NA | NA | NA | NA |
| Olfr100 | 0 | 0 | 0 | 0 | 0 | 0 | 0 | 0 | 0 | 0 | 0 | 0 | 0 | 0 | 0 | NA | NA | NA | NA | NA | NA | NA | NA | NA | NA |
| Olfr1000 | 0 | 0 | 0 | 0 | 0 | 0 | 0 | 0 | 0 | 0 | 0 | 0 | 0 | 0 | 0 | NA | NA | NA | NA | NA | NA | NA | NA | NA | NA |
| Olfr1002 | 0 | 0 | 0 | 0 | 0 | 0 | 0 | 0 | 0 | 0 | 0 | 0 | 0 | 0 | 0 | NA | NA | NA | NA | NA | NA | NA | NA | NA | NA |
| Olfr1006 | 0 | 0 | 0 | 0 | 0 | 0 | 0 | 0 | 0 | 0 | 0 | 0 | 0 | 0 | 0 | NA | NA | NA | NA | NA | NA | NA | NA | NA | NA |
| Olfr1008 | 0 | 0 | 0 | 0 | 0 | 0 | 0 | 0 | 0 | 0 | 0 | 0 | 0 | 0 | 0 | NA | NA | NA | NA | NA | NA | NA | NA | NA | NA |
| Olfr1009 | 0 | 0 | 0 | 0 | 0 | 0 | 0 | 0 | 0 | 0 | 0 | 0 | 0 | 0 | 0 | NA | NA | NA | NA | NA | NA | NA | NA | NA | NA |
| Olfr101 | 0 | 0 | 0 | 0 | 0 | 0 | 0 | 0 | 0 | 0 | 0 | 0 | 0 | 0 | 0 | NA | NA | NA | NA | NA | NA | NA | NA | NA | NA |
| Olfr1010 | 0 | 0 | 0 | 0 | 0 | 0 | 0 | 0 | 0 | 0 | 0 | 0 | 0 | 0 | 0 | NA | NA | NA | NA | NA | NA | NA | NA | NA | NA |
| Olfr1012 | 0 | 0 | 0 | 0 | 0 | 0 | 0 | 0 | 0 | 0 | 0 | 0 | 0 | 0 | 0 | NA | NA | NA | NA | NA | NA | NA | NA | NA | NA |
| Olfr1013 | 0 | 0 | 0 | 0 | 0 | 0 | 0 | 0 | 0 | 0 | 0 | 0 | 0 | 0 | 0 | NA | NA | NA | NA | NA | NA | NA | NA | NA | NA |
| Olfr1014 | 0 | 0 | 0 | 0 | 0 | 0 | 0 | 0 | 0 | 0 | 0 | 0 | 0 | 0 | 0 | NA | NA | NA | NA | NA | NA | NA | NA | NA | NA |
| Olfr1015 | 0 | 0 | 0 | 0 | 0 | 0 | 0 | 0 | 0 | 0 | 0 | 0 | 0 | 0 | 0 | NA | NA | NA | NA | NA | NA | NA | NA | NA | NA |
| Olfr1016 | 0 | 0 | 0 | 0 | 0 | 0 | 0 | 0 | 0 | 0 | 0 | 0 | 0 | 0 | 0 | NA | NA | NA | NA | NA | NA | NA | NA | NA | NA |
| Olfr1018 | 0 | 0 | 0 | 0 | 0 | 0 | 0 | 0 | 0 | 0 | 0 | 0 | 0 | 0 | 0 | NA | NA | NA | NA | NA | NA | NA | NA | NA | NA |
| Olfr1019 | 0 | 0 | 0 | 0 | 0 | 0 | 0 | 0 | 0 | 0 | 0 | 0 | 0 | 0 | 0 | NA | NA | NA | NA | NA | NA | NA | NA | NA | NA |
| Olfr102 | 0 | 0 | 0 | 0 | 0 | 0 | 0 | 0 | 0 | 0 | 0 | 0 | 0 | 0 | 0 | NA | NA | NA | NA | NA | NA | NA | NA | NA | NA |
| Olfr1020 | 0 | 0 | 0 | 0 | 0 | 0 | 0 | 0 | 0 | 0 | 0 | 0 | 0 | 0 | 0 | NA | NA | NA | NA | NA | NA | NA | NA | NA | NA |
| Olfr1022 | 0 | 0 | 0 | 0 | 0 | 0 | 0 | 0 | 0 | 0 | 0 | 0 | 0 | 0 | 0 | NA | NA | NA | NA | NA | NA | NA | NA | NA | NA |
| Olfr1023 | 0 | 0 | 0 | 0 | 0 | 0 | 0 | 0 | 0 | 0 | 0 | 0 | 0 | 0 | 0 | NA | NA | NA | NA | NA | NA | NA | NA | NA | NA |
| Olfr1024 | 0 | 0 | 0 | 0 | 0 | 0 | 0 | 0 | 0 | 0 | 0 | 0 | 0 | 0 | 0 | NA | NA | NA | NA | NA | NA | NA | NA | NA | NA |
| Olfr1026 | 0 | 0 | 0 | 0 | 0 | 0 | 0 | 0 | 0 | 0 | 0 | 0 | 0 | 0 | 0 | NA | NA | NA | NA | NA | NA | NA | NA | NA | NA |
| Olfr1028 | 0 | 0 | 0 | 0 | 0 | 0 | 0 | 0 | 0 | 0 | 0 | 0 | 0 | 0 | 0 | NA | NA | NA | NA | NA | NA | NA | NA | NA | NA |
| Olfr1029 | 0 | 0 | 0 | 0 | 0 | 0 | 0 | 0 | 0 | 0 | 0 | 0 | 0 | 0 | 0 | NA | NA | NA | NA | NA | NA | NA | NA | NA | NA |
| Olfr103 | 0 | 0 | 0 | 0 | 0 | 0 | 0 | 0 | 0 | 0 | 0 | 0 | 0 | 0 | 0 | NA | NA | NA | NA | NA | NA | NA | NA | NA | NA |
| Olfr1030 | 0 | 0 | 0 | 0 | 0 | 0 | 0 | 0 | 0 | 0 | 0 | 0 | 0 | 0 | 0 | NA | NA | NA | NA | NA | NA | NA | NA | NA | NA |
| Olfr1031 | 0 | 0 | 0 | 0 | 0 | 0 | 0 | 0 | 0 | 0 | 0 | 0 | 0 | 0 | 0 | NA | NA | NA | NA | NA | NA | NA | NA | NA | NA |
| Olfr1032 | 0 | 0 | 0 | 0 | 0 | 0 | 0 | 0 | 0 | 0 | 0 | 0 | 0 | 0 | 0 | NA | NA | NA | NA | NA | NA | NA | NA | NA | NA |
| Olfr1033 | 0 | 0 | 0 | 1 | 0 | 0 | 0 | 0 | 0 | 1 | 0 | 0 | 0.17 | 0 | 0 | 2.094 | 1.067 | 0.79379329 | NA | 0 | 14.2885 | 10 | 10 | TRUE | 0.3906 |
| Olfr1034 | 0 | 0 | 0 | 0 | 0 | 0 | 0 | 0 | 0 | 0 | 0 | 0 | 0 | 0 | 0 | NA | NA | NA | NA | NA | NA | NA | NA | NA | NA |
| Olfr1036 | 0 | 0 | 0 | 0 | 0 | 0 | 0 | 0 | 0 | 0 | 0 | 0 | 0 | 0 | 0 | NA | NA | NA | NA | NA | NA | NA | NA | NA | NA |
| Olfr1037 | 0 | 0 | 0 | 0 | 0 | 0 | 0 | 0 | 0 | 0 | 0 | 0 | 0 | 0 | 0 | NA | NA | NA | NA | NA | NA | NA | NA | NA | NA |
| Olfr1038-ps | 0 | 0 | 0 | 0 | 0 | 0 | 0 | 0 | 0 | 0 | 0 | 0 | 0 | 0 | 0 | NA | NA | NA | NA | NA | NA | NA | NA | NA | NA |
| Olfr1039 | 0 | 0 | 0 | 0 | 0 | 0 | 0 | 0 | 0 | 0 | 0 | 0 | 0 | 0 | 0 | NA | NA | NA | NA | NA | NA | NA | NA | NA | NA |
| Olfr1040 | 0 | 0 | 0 | 0 | 0 | 0 | 0 | 0 | 0 | 0 | 0 | 0 | 0 | 0 | 0 | NA | NA | NA | NA | NA | NA | NA | NA | NA | NA |
| Olfr1042 | 0 | 0 | 0 | 0 | 0 | 0 | 0 | 0 | 0 | 0 | 0 | 0 | 0 | 0 | 0 | NA | NA | NA | NA | NA | NA | NA | NA | NA | NA |
| Olfr1043 | 0 | 0 | 0 | 0 | 0 | 0 | 0 | 0 | 0 | 0 | 0 | 0 | 0 | 0 | 0 | NA | NA | NA | NA | NA | NA | NA | NA | NA | NA |
| Olfr1044 | 0 | 0 | 0 | 0 | 0 | 0 | 0 | 0 | 0 | 0 | 0 | 0 | 0 | 0 | 0 | NA | NA | NA | NA | NA | NA | NA | NA | NA | NA |
| Olfr1045 | 0 | 0 | 0 | 0 | 0 | 0 | 0 | 0 | 0 | 0 | 0 | 0 | 0 | 0 | 0 | NA | NA | NA | NA | NA | NA | NA | NA | NA | NA |
| Olfr1046 | 0 | 0 | 0 | 0 | 0 | 0 | 0 | 0 | 0 | 0 | 0 | 0 | 0 | 0 | 0 | NA | NA | NA | NA | NA | NA | NA | NA | NA | NA |
| Olfr1047 | 0 | 0 | 0 | 0 | 0 | 0 | 0 | 0 | 0 | 0 | 0 | 0 | 0 | 0 | 0 | NA | NA | NA | NA | NA | NA | NA | NA | NA | NA |
| Olfr1048 | 0 | 0 | 0 | 0 | 0 | 0 | 0 | 0 | 0 | 0 | 0 | 0 | 0 | 0 | 0 | NA | NA | NA | NA | NA | NA | NA | NA | NA | NA |
| Olfr1049 | 0 | 0 | 0 | 0 | 0 | 0 | 0 | 0 | 0 | 0 | 0 | 0 | 0 | 0 | 0 | NA | NA | NA | NA | NA | NA | NA | NA | NA | NA |
| Olfr1051 | 0 | 0 | 0 | 0 | 0 | 0 | 0 | 0 | 0 | 0 | 0 | 0 | 0 | 0 | 0 | NA | NA | NA | NA | NA | NA | NA | NA | NA | NA |
| Olfr1052 | 0 | 0 | 0 | 0 | 0 | 0 | 0 | 0 | 0 | 0 | 0 | 0 | 0 | 0 | 0 | NA | NA | NA | NA | NA | NA | NA | NA | NA | NA |
| Olfr1053 | 0 | 0 | 0 | 0 | 0 | 0 | 0 | 0 | 0 | 0 | 0 | 0 | 0 | 0 | 0 | NA | NA | NA | NA | NA | NA | NA | NA | NA | NA |
| Olfr1054 | 0 | 0 | 0 | 0 | 0 | 0 | 0 | 0 | 0 | 0 | 0 | 0 | 0 | 0 | 0 | NA | NA | NA | NA | NA | NA | NA | NA | NA | NA |
| Olfr1055 | 0 | 0 | 0 | 0 | 0 | 0 | 0 | 0 | 0 | 0 | 0 | 0 | 0 | 0 | 0 | NA | NA | NA | NA | NA | NA | NA | NA | NA | NA |
| Olfr1056 | 0 | 0 | 0 | 0 | 0 | 0 | 0 | 0 | 0 | 0 | 0 | 0 | 0 | 0 | 0 | NA | NA | NA | NA | NA | NA | NA | NA | NA | NA |
| Olfr1057 | 0 | 0 | 0 | 0 | 0 | 0 | 0 | 0 | 0 | 0 | 0 | 0 | 0 | 0 | 0 | NA | NA | NA | NA | NA | NA | NA | NA | NA | NA |
| Olfr1058 | 0 | 0 | 0 | 0 | 0 | 0 | 0 | 0 | 0 | 0 | 0 | 0 | 0 | 0 | 0 | NA | NA | NA | NA | NA | NA | NA | NA | NA | NA |
| Olfr1061 | 0 | 0 | 0 | 0 | 0 | 0 | 0 | 0 | 0 | 0 | 0 | 0 | 0 | 0 | 0 | NA | NA | NA | NA | NA | NA | NA | NA | NA | NA |

| | | | | | | | | | | | | | | | | | | | | | | | | | |
|---|---|---|---|---|---|---|---|---|---|---|---|---|---|---|---|---|---|---|---|---|---|---|---|---|---|
| Olfr1062 | 0 | 0 | 0 | 0 | 0 | 0 | 0 | 0 | 0 | 0 | 0 | 0 | 0 | 0 | 0 | NA | NA | NA | NA | NA | NA | NA | NA | NA | NA |
| Olfr1065 | 0 | 0 | 0 | 0 | 0 | 0 | 0 | 0 | 0 | 0 | 0 | 0 | 0 | 0 | 0 | NA | NA | NA | NA | NA | NA | NA | NA | NA | NA |
| Olfr1066 | 0 | 0 | 0 | 0 | 0 | 0 | 0 | 0 | 0 | 0 | 0 | 0 | 0 | 0 | 0 | NA | NA | NA | NA | NA | NA | NA | NA | NA | NA |
| Olfr107 | 0 | 0 | 0 | 0 | 0 | 0 | 0 | 0 | 0 | 0 | 0 | 0 | 0 | 0 | 0 | NA | NA | NA | NA | NA | NA | NA | NA | NA | NA |
| Olfr1076 | 0 | 0 | 0 | 0 | 0 | 0 | 0 | 0 | 0 | 0 | 0 | 0 | 0 | 0 | 0 | NA | NA | NA | NA | NA | NA | NA | NA | NA | NA |
| Olfr1077-ps1 | 0 | 0 | 0 | 0 | 0 | 0 | 0 | 0 | 0 | 0 | 0 | 0 | 0 | 0 | 0 | NA | NA | NA | NA | NA | NA | NA | NA | NA | NA |
| Olfr1079 | 0 | 0 | 0 | 0 | 0 | 0 | 0 | 0 | 0 | 0 | 0 | 0 | 0 | 0 | 0 | NA | NA | NA | NA | NA | NA | NA | NA | NA | NA |
| Olfr108 | 0 | 0 | 0 | 0 | 0 | 0 | 0 | 0 | 0 | 0 | 0 | 0 | 0 | 0 | 0 | NA | NA | NA | NA | NA | NA | NA | NA | NA | NA |
| Olfr1080 | 0 | 0 | 0 | 0 | 0 | 0 | 0 | 0 | 0 | 0 | 0 | 0 | 0 | 0 | 0 | NA | NA | NA | NA | NA | NA | NA | NA | NA | NA |
| Olfr1082 | 0 | 0 | 0 | 0 | 0 | 0 | 0 | 0 | 0 | 0 | 0 | 0 | 0 | 0 | 0 | NA | NA | NA | NA | NA | NA | NA | NA | NA | NA |
| Olfr1084 | 0 | 0 | 0 | 0 | 0 | 0 | 0 | 0 | 0 | 0 | 0 | 0 | 0 | 0 | 0 | NA | NA | NA | NA | NA | NA | NA | NA | NA | NA |
| Olfr1085 | 0 | 0 | 0 | 0 | 0 | 0 | 0 | 0 | 0 | 0 | 0 | 0 | 0 | 0 | 0 | NA | NA | NA | NA | NA | NA | NA | NA | NA | NA |
| Olfr1086 | 0 | 0 | 0 | 0 | 0 | 0 | 0 | 0 | 0 | 0 | 0 | 0 | 0 | 0 | 0 | NA | NA | NA | NA | NA | NA | NA | NA | NA | NA |
| Olfr1087 | 0 | 0 | 0 | 0 | 0 | 0 | 0 | 0 | 0 | 0 | 0 | 0 | 0 | 0 | 0 | NA | NA | NA | NA | NA | NA | NA | NA | NA | NA |
| Olfr1089 | 0 | 0 | 0 | 0 | 0 | 0 | 0 | 0 | 0 | 0 | 0 | 0 | 0 | 0 | 0 | NA | NA | NA | NA | NA | NA | NA | NA | NA | NA |
| Olfr109 | 0 | 0 | 0 | 0 | 0 | 0 | 0 | 0 | 0 | 0 | 0 | 0 | 0 | 0 | 0 | NA | NA | NA | NA | NA | NA | NA | NA | NA | NA |
| Olfr1090 | 0 | 0 | 0 | 0 | 0 | 0 | 0 | 0 | 0 | 0 | 0 | 0 | 0 | 0 | 0 | NA | NA | NA | NA | NA | NA | NA | NA | NA | NA |
| Olfr1093 | 0 | 0 | 0 | 0 | 0 | 0 | 0 | 0 | 0 | 0 | 0 | 0 | 0 | 0 | 0 | NA | NA | NA | NA | NA | NA | NA | NA | NA | NA |
| Olfr1094 | 0 | 0 | 0 | 0 | 0 | 0 | 0 | 0 | 0 | 0 | 0 | 0 | 0 | 0 | 0 | NA | NA | NA | NA | NA | NA | NA | NA | NA | NA |
| Olfr1095 | 0 | 0 | 0 | 0 | 0 | 0 | 0 | 0 | 0 | 0 | 0 | 0 | 0 | 0 | 0 | NA | NA | NA | NA | NA | NA | NA | NA | NA | NA |
| Olfr1097 | 0 | 0 | 0 | 0 | 0 | 0 | 0 | 0 | 0 | 0 | 0 | 0 | 0 | 0 | 0 | NA | NA | NA | NA | NA | NA | NA | NA | NA | NA |
| Olfr1098 | 0 | 0 | 0 | 0 | 0 | 0 | 0 | 0 | 0 | 0 | 0 | 0 | 0 | 0 | 0 | NA | NA | NA | NA | NA | NA | NA | NA | NA | NA |
| Olfr1099 | 0 | 0 | 0 | 0 | 0 | 0 | 0 | 0 | 0 | 0 | 0 | 0 | 0 | 0 | 0 | NA | NA | NA | NA | NA | NA | NA | NA | NA | NA |
| Olfr11 | 0 | 0 | 0 | 0 | 0 | 0 | 0 | 0 | 0 | 0 | 0 | 0 | 0 | 0 | 0 | NA | NA | NA | NA | NA | NA | NA | NA | NA | NA |
| Olfr110 | 0 | 0 | 0 | 0 | 0 | 0 | 0 | 0 | 0 | 0 | 0 | 0 | 0 | 0 | 0 | NA | NA | NA | NA | NA | NA | NA | NA | NA | NA |
| Olfr1100 | 0 | 0 | 0 | 0 | 0 | 0 | 0 | 0 | 0 | 0 | 0 | 0 | 0 | 0 | 0 | NA | NA | NA | NA | NA | NA | NA | NA | NA | NA |
| Olfr1101 | 0 | 0 | 0 | 0 | 0 | 0 | 0 | 0 | 0 | 0 | 0 | 0 | 0 | 0 | 0 | NA | NA | NA | NA | NA | NA | NA | NA | NA | NA |
| Olfr1102 | 0 | 0 | 0 | 0 | 0 | 0 | 0 | 0 | 0 | 0 | 0 | 0 | 0 | 0 | 0 | NA | NA | NA | NA | NA | NA | NA | NA | NA | NA |
| Olfr1104 | 0 | 0 | 0 | 0 | 0 | 0 | 0 | 0 | 0 | 0 | 0 | 0 | 0 | 0 | 0 | NA | NA | NA | NA | NA | NA | NA | NA | NA | NA |
| Olfr1105 | 0 | 0 | 0 | 0 | 0 | 0 | 0 | 0 | 0 | 0 | 0 | 0 | 0 | 0 | 0 | NA | NA | NA | NA | NA | NA | NA | NA | NA | NA |
| Olfr1106 | 0 | 0 | 0 | 0 | 0 | 0 | 0 | 0 | 0 | 0 | 0 | 0 | 0 | 0 | 0 | NA | NA | NA | NA | NA | NA | NA | NA | NA | NA |
| Olfr1107 | 0 | 0 | 0 | 0 | 0 | 0 | 0 | 0 | 0 | 0 | 0 | 0 | 0 | 0 | 0 | NA | NA | NA | NA | NA | NA | NA | NA | NA | NA |
| Olfr1109 | 0 | 0 | 0 | 0 | 0 | 0 | 0 | 0 | 0 | 0 | 0 | 0 | 0 | 0 | 0 | NA | NA | NA | NA | NA | NA | NA | NA | NA | NA |
| Olfr111 | 0 | 0 | 0 | 0 | 0 | 0 | 0 | 0 | 0 | 0 | 0 | 0 | 0 | 0 | 0 | NA | NA | NA | NA | NA | NA | NA | NA | NA | NA |
| Olfr1110 | 0 | 0 | 0 | 0 | 0 | 0 | 0 | 0 | 0 | 0 | 0 | 0 | 0 | 0 | 0 | NA | NA | NA | NA | NA | NA | NA | NA | NA | NA |
| Olfr1111 | 0 | 0 | 0 | 0 | 0 | 0 | 0 | 0 | 0 | 0 | 0 | 0 | 0 | 0 | 0 | NA | NA | NA | NA | NA | NA | NA | NA | NA | NA |
| Olfr1112 | 0 | 0 | 0 | 0 | 0 | 0 | 0 | 0 | 0 | 0 | 0 | 0 | 0 | 0 | 0 | NA | NA | NA | NA | NA | NA | NA | NA | NA | NA |
| Olfr1113 | 0 | 0 | 0 | 0 | 0 | 0 | 0 | 0 | 0 | 0 | 0 | 0 | 0 | 0 | 0 | NA | NA | NA | NA | NA | NA | NA | NA | NA | NA |
| Olfr1115 | 0 | 0 | 0 | 0 | 0 | 0 | 0 | 0 | 0 | 0 | 0 | 0 | 0 | 0 | 0 | NA | NA | NA | NA | NA | NA | NA | NA | NA | NA |
| Olfr1116-ps | 0 | 0 | 0 | 0 | 0 | 0 | 0 | 0 | 0 | 0 | 0 | 0 | 0 | 0 | 0 | NA | NA | NA | NA | NA | NA | NA | NA | NA | NA |
| Olfr1118 | 0 | 0 | 0 | 0 | 0 | 0 | 0 | 0 | 0 | 0 | 0 | 0 | 0 | 0 | 0 | NA | NA | NA | NA | NA | NA | NA | NA | NA | NA |
| Olfr112 | 0 | 0 | 0 | 0 | 0 | 0 | 0 | 0 | 0 | 0 | 0 | 0 | 0 | 0 | 0 | NA | NA | NA | NA | NA | NA | NA | NA | NA | NA |
| Olfr1120 | 0 | 0 | 0 | 0 | 0 | 0 | 0 | 0 | 0 | 0 | 0 | 0 | 0 | 0 | 0 | NA | NA | NA | NA | NA | NA | NA | NA | NA | NA |
| Olfr1121 | 0 | 0 | 0 | 0 | 0 | 0 | 0 | 0 | 0 | 0 | 0 | 0 | 0 | 0 | 0 | NA | NA | NA | NA | NA | NA | NA | NA | NA | NA |
| Olfr1122 | 0 | 0 | 0 | 0 | 0 | 0 | 0 | 0 | 0 | 0 | 0 | 0 | 0 | 0 | 0 | NA | NA | NA | NA | NA | NA | NA | NA | NA | NA |
| Olfr1123 | 0 | 0 | 0 | 0 | 0 | 0 | 0 | 0 | 0 | 0 | 0 | 0 | 0 | 0 | 0 | NA | NA | NA | NA | NA | NA | NA | NA | NA | NA |
| Olfr1124 | 0 | 0 | 0 | 0 | 0 | 0 | 0 | 0 | 0 | 0 | 0 | 0 | 0 | 0 | 0 | NA | NA | NA | NA | NA | NA | NA | NA | NA | NA |
| Olfr1126 | 0 | 0 | 0 | 0 | 0 | 0 | 0 | 0 | 0 | 0 | 0 | 0 | 0 | 0 | 0 | NA | NA | NA | NA | NA | NA | NA | NA | NA | NA |
| Olfr1128 | 0 | 0 | 0 | 0 | 0 | 0 | 0 | 0 | 0 | 0 | 0 | 0 | 0 | 0 | 0 | NA | NA | NA | NA | NA | NA | NA | NA | NA | NA |
| Olfr1129 | 0 | 0 | 0 | 0 | 0 | 0 | 0 | 0 | 0 | 0 | 0 | 0 | 0 | 0 | 0 | NA | NA | NA | NA | NA | NA | NA | NA | NA | NA |
| Olfr113 | 0 | 0 | 0 | 0 | 0 | 0 | 0 | 0 | 0 | 0 | 0 | 0 | 0 | 0 | 0 | NA | NA | NA | NA | NA | NA | NA | NA | NA | NA |
| Olfr1130 | 0 | 0 | 0 | 0 | 0 | 0 | 0 | 0 | 0 | 0 | 0 | 0 | 0 | 0 | 0 | NA | NA | NA | NA | NA | NA | NA | NA | NA | NA |
| Olfr1131 | 0 | 0 | 0 | 0 | 0 | 0 | 0 | 0 | 0 | 0 | 0 | 0 | 0 | 0 | 0 | NA | NA | NA | NA | NA | NA | NA | NA | NA | NA |
| Olfr1132 | 0 | 0 | 0 | 0 | 0 | 0 | 0 | 0 | 0 | 0 | 0 | 0 | 0 | 0 | 0 | NA | NA | NA | NA | NA | NA | NA | NA | NA | NA |
| Olfr1133 | 0 | 0 | 0 | 0 | 0 | 0 | 0 | 0 | 0 | 0 | 0 | 0 | 0 | 0 | 0 | NA | NA | NA | NA | NA | NA | NA | NA | NA | NA |
| Olfr1134 | 0 | 0 | 0 | 0 | 0 | 0 | 0 | 0 | 0 | 0 | 0 | 0 | 0 | 0 | 0 | NA | NA | NA | NA | NA | NA | NA | NA | NA | NA |
| Olfr1135 | 0 | 0 | 0 | 0 | 0 | 0 | 0 | 0 | 0 | 0 | 0 | 0 | 0 | 0 | 0 | NA | NA | NA | NA | NA | NA | NA | NA | NA | NA |
| Olfr1136 | 0 | 0 | 0 | 0 | 0 | 0 | 0 | 0 | 0 | 0 | 0 | 0 | 0 | 0 | 0 | NA | NA | NA | NA | NA | NA | NA | NA | NA | NA |
| Olfr1137 | 0 | 0 | 0 | 0 | 0 | 0 | 0 | 0 | 0 | 0 | 0 | 0 | 0 | 0 | 0 | NA | NA | NA | NA | NA | NA | NA | NA | NA | NA |
| Olfr1138 | 0 | 0 | 0 | 0 | 0 | 0 | 0 | 0 | 0 | 0 | 0 | 0 | 0 | 0 | 0 | NA | NA | NA | NA | NA | NA | NA | NA | NA | NA |
| Olfr114 | 0 | 0 | 0 | 0 | 0 | 0 | 0 | 0 | 0 | 0 | 0 | 0 | 0 | 0 | 0 | NA | NA | NA | NA | NA | NA | NA | NA | NA | NA |
| Olfr1140 | 0 | 0 | 0 | 0 | 0 | 0 | 0 | 0 | 0 | 0 | 0 | 0 | 0 | 0 | 0 | NA | NA | NA | NA | NA | NA | NA | NA | NA | NA |
| Olfr1141 | 0 | 0 | 0 | 0 | 0 | 0 | 0 | 0 | 0 | 0 | 0 | 0 | 0 | 0 | 0 | NA | NA | NA | NA | NA | NA | NA | NA | NA | NA |
| Olfr1143 | 0 | 0 | 0 | 0 | 0 | 0 | 0 | 0 | 0 | 0 | 0 | 0 | 0 | 0 | 0 | NA | NA | NA | NA | NA | NA | NA | NA | NA | NA |
| Olfr1145 | 0 | 0 | 0 | 0 | 0 | 0 | 0 | 0 | 0 | 0 | 0 | 0 | 0 | 0 | 0 | NA | NA | NA | NA | NA | NA | NA | NA | NA | NA |
| Olfr1148 | 0 | 0 | 0 | 0 | 0 | 0 | 0 | 0 | 0 | 0 | 0 | 0 | 0 | 0 | 0 | NA | NA | NA | NA | NA | NA | NA | NA | NA | NA |
| Olfr115 | 0 | 0 | 0 | 0 | 0 | 0 | 0 | 0 | 0 | 0 | 0 | 0 | 0 | 0 | 0 | NA | NA | NA | NA | NA | NA | NA | NA | NA | NA |
| Olfr1151 | 0 | 0 | 0 | 0 | 0 | 0 | 0 | 0 | 0 | 0 | 0 | 0 | 0 | 0 | 0 | NA | NA | NA | NA | NA | NA | NA | NA | NA | NA |
| Olfr1152 | 0 | 0 | 0 | 0 | 0 | 0 | 0 | 0 | 0 | 0 | 0 | 0 | 0 | 0 | 0 | NA | NA | NA | NA | NA | NA | NA | NA | NA | NA |

| | | | | | | | | | | | | | | | | | | | | | | | | | |
|---|---|---|---|---|---|---|---|---|---|---|---|---|---|---|---|---|---|---|---|---|---|---|---|---|---|
| Olfr1153 | 0 | 0 | 0 | 0 | 0 | 0 | 0 | 0 | 0 | 0 | 0 | 0 | 0 | 0 | 0 | NA | NA | NA | NA | NA | NA | NA | NA | NA | NA |
| Olfr1154 | 0 | 0 | 0 | 0 | 0 | 0 | 0 | 0 | 0 | 0 | 0 | 0 | 0 | 0 | 0 | NA | NA | NA | NA | NA | NA | NA | NA | NA | NA |
| Olfr1155 | 0 | 0 | 0 | 0 | 0 | 0 | 0 | 0 | 0 | 0 | 0 | 0 | 0 | 0 | 0 | NA | NA | NA | NA | NA | NA | NA | NA | NA | NA |
| Olfr1156 | 0 | 0 | 0 | 0 | 0 | 0 | 0 | 0 | 0 | 0 | 0 | 0 | 0 | 0 | 0 | NA | NA | NA | NA | NA | NA | NA | NA | NA | NA |
| Olfr1157 | 0 | 0 | 0 | 0 | 0 | 0 | 0 | 0 | 0 | 0 | 0 | 0 | 0 | 0 | 0 | NA | NA | NA | NA | NA | NA | NA | NA | NA | NA |
| Olfr1158 | 0 | 0 | 0 | 0 | 0 | 0 | 0 | 0 | 0 | 0 | 0 | 0 | 0 | 0 | 0 | NA | NA | NA | NA | NA | NA | NA | NA | NA | NA |
| Olfr116 | 0 | 0 | 0 | 0 | 0 | 0 | 0 | 0 | 0 | 0 | 0 | 0 | 0 | 0 | 0 | NA | NA | NA | NA | NA | NA | NA | NA | NA | NA |
| Olfr1160 | 0 | 0 | 0 | 0 | 0 | 0 | 0 | 0 | 0 | 0 | 0 | 0 | 0 | 0 | 0 | NA | NA | NA | NA | NA | NA | NA | NA | NA | NA |
| Olfr1161 | 0 | 0 | 0 | 0 | 0 | 0 | 0 | 0 | 0 | 0 | 0 | 0 | 0 | 0 | 0 | NA | NA | NA | NA | NA | NA | NA | NA | NA | NA |
| Olfr1162 | 0 | 0 | 0 | 0 | 0 | 0 | 0 | 0 | 0 | 0 | 0 | 0 | 0 | 0 | 0 | NA | NA | NA | NA | NA | NA | NA | NA | NA | NA |
| Olfr1163 | 0 | 0 | 0 | 0 | 0 | 0 | 0 | 0 | 0 | 0 | 0 | 0 | 0 | 0 | 0 | NA | NA | NA | NA | NA | NA | NA | NA | NA | NA |
| Olfr1164 | 0 | 0 | 0 | 0 | 0 | 0 | 0 | 0 | 0 | 0 | 0 | 0 | 0 | 0 | 0 | NA | NA | NA | NA | NA | NA | NA | NA | NA | NA |
| Olfr1166 | 0 | 0 | 0 | 0 | 0 | 0 | 0 | 0 | 0 | 0 | 0 | 0 | 0 | 0 | 0 | NA | NA | NA | NA | NA | NA | NA | NA | NA | NA |
| Olfr1167 | 0 | 0 | 0 | 0 | 0 | 0 | 0 | 0 | 0 | 0 | 0 | 0 | 0 | 0 | 0 | NA | NA | NA | NA | NA | NA | NA | NA | NA | NA |
| Olfr1168 | 0 | 0 | 0 | 0 | 0 | 0 | 0 | 0 | 0 | 0 | 0 | 0 | 0 | 0 | 0 | NA | NA | NA | NA | NA | NA | NA | NA | NA | NA |
| Olfr117 | 0 | 0 | 0 | 0 | 0 | 0 | 0 | 0 | 0 | 0 | 0 | 0 | 0 | 0 | 0 | NA | NA | NA | NA | NA | NA | NA | NA | NA | NA |
| Olfr1170 | 0 | 0 | 0 | 0 | 0 | 0 | 0 | 0 | 0 | 0 | 0 | 0 | 0 | 0 | 0 | NA | NA | NA | NA | NA | NA | NA | NA | NA | NA |
| Olfr1173 | 0 | 0 | 0 | 0 | 0 | 0 | 0 | 0 | 0 | 0 | 0 | 0 | 0 | 0 | 0 | NA | NA | NA | NA | NA | NA | NA | NA | NA | NA |
| Olfr1176 | 0 | 0 | 0 | 0 | 0 | 0 | 0 | 0 | 0 | 0 | 0 | 0 | 0 | 0 | 0 | NA | NA | NA | NA | NA | NA | NA | NA | NA | NA |
| Olfr1178 | 0 | 0 | 0 | 0 | 0 | 0 | 0 | 0 | 0 | 0 | 0 | 0 | 0 | 0 | 0 | NA | NA | NA | NA | NA | NA | NA | NA | NA | NA |
| Olfr1179 | 0 | 0 | 0 | 0 | 0 | 0 | 0 | 0 | 0 | 0 | 0 | 0 | 0 | 0 | 0 | NA | NA | NA | NA | NA | NA | NA | NA | NA | NA |
| Olfr118 | 0 | 1 | 1 | 1 | 1 | 0 | 0 | 1 | 1 | 1 | 1 | 0 | 0.71 | 1 | 1 | 1.021 | 0.029 | 0.98980508 | NA | 0 | 3.3357 | 2.3741 | 2.3741 | TRUE | 0.3739 |
| Olfr1180 | 0 | 0 | 0 | 0 | 0 | 0 | 0 | 0 | 0 | 0 | 0 | 0 | 0 | 0 | 0 | NA | NA | NA | NA | NA | NA | NA | NA | NA | NA |
| Olfr1181 | 0 | 0 | 0 | 0 | 0 | 0 | 0 | 0 | 0 | 0 | 0 | 0 | 0 | 0 | 0 | NA | NA | NA | NA | NA | NA | NA | NA | NA | NA |
| Olfr1182 | 0 | 0 | 0 | 0 | 0 | 0 | 0 | 0 | 0 | 0 | 0 | 0 | 0 | 0 | 0 | NA | NA | NA | NA | NA | NA | NA | NA | NA | NA |
| Olfr1183 | 0 | 0 | 0 | 0 | 0 | 0 | 0 | 0 | 0 | 0 | 0 | 0 | 0 | 0 | 0 | NA | NA | NA | NA | NA | NA | NA | NA | NA | NA |
| Olfr1184 | 0 | 0 | 0 | 0 | 0 | 0 | 0 | 0 | 0 | 0 | 0 | 0 | 0 | 0 | 0 | NA | NA | NA | NA | NA | NA | NA | NA | NA | NA |
| Olfr1186 | 0 | 0 | 0 | 0 | 0 | 0 | 0 | 0 | 0 | 0 | 0 | 0 | 0 | 0 | 0 | NA | NA | NA | NA | NA | NA | NA | NA | NA | NA |
| Olfr1188 | 0 | 0 | 0 | 0 | 0 | 0 | 0 | 0 | 0 | 0 | 0 | 0 | 0 | 0 | 0 | NA | NA | NA | NA | NA | NA | NA | NA | NA | NA |
| Olfr1189 | 0 | 0 | 0 | 0 | 0 | 0 | 0 | 0 | 0 | 0 | 0 | 0 | 0 | 0 | 0 | NA | NA | NA | NA | NA | NA | NA | NA | NA | NA |
| Olfr119 | 0 | 0 | 0 | 0 | 0 | 0 | 0 | 0 | 0 | 0 | 0 | 0 | 0 | 0 | 0 | NA | NA | NA | NA | NA | NA | NA | NA | NA | NA |
| Olfr1193 | 0 | 0 | 0 | 0 | 0 | 0 | 0 | 0 | 0 | 0 | 0 | 0 | 0 | 0 | 0 | NA | NA | NA | NA | NA | NA | NA | NA | NA | NA |
| Olfr1195 | 0 | 0 | 0 | 0 | 0 | 0 | 0 | 0 | 0 | 0 | 0 | 0 | 0 | 0 | 0 | NA | NA | NA | NA | NA | NA | NA | NA | NA | NA |
| Olfr1196 | 1 | 1 | 0 | 0 | 0 | 0 | 1 | 1 | 0 | 0 | 0 | 0 | 0.31 | 1 | 0 | 0.309 | -1.696 | 0.63183309 | NA | 0 | 7.7294 | 7.2472 | 7.2472 | TRUE | 0.2015 |
| Olfr1197 | 0 | 0 | 0 | 0 | 0 | 0 | 0 | 0 | 0 | 0 | 0 | 0 | 0 | 0 | 0 | NA | NA | NA | NA | NA | NA | NA | NA | NA | NA |
| Olfr1198 | 0 | 0 | 0 | 0 | 0 | 0 | 0 | 0 | 0 | 0 | 0 | 0 | 0 | 0 | 0 | NA | NA | NA | NA | NA | NA | NA | NA | NA | NA |
| Olfr1199 | 0 | 0 | 0 | 0 | 0 | 0 | 0 | 0 | 0 | 0 | 0 | 0 | 0 | 0 | 0 | NA | NA | NA | NA | NA | NA | NA | NA | NA | NA |
| Olfr12 | 0 | 0 | 0 | 0 | 0 | 0 | 0 | 0 | 0 | 0 | 0 | 0 | 0 | 0 | 0 | NA | NA | NA | NA | NA | NA | NA | NA | NA | NA |
| Olfr120 | 0 | 0 | 0 | 0 | 0 | 0 | 0 | 0 | 0 | 0 | 0 | 0 | 0 | 0 | 0 | NA | NA | NA | NA | NA | NA | NA | NA | NA | NA |
| Olfr1200 | 0 | 0 | 0 | 0 | 0 | 0 | 0 | 0 | 0 | 0 | 0 | 0 | 0 | 0 | 0 | NA | NA | NA | NA | NA | NA | NA | NA | NA | NA |
| Olfr1201 | 0 | 0 | 0 | 0 | 0 | 0 | 0 | 0 | 0 | 0 | 0 | 0 | 0 | 0 | 0 | NA | NA | NA | NA | NA | NA | NA | NA | NA | NA |
| Olfr1202 | 0 | 0 | 0 | 0 | 0 | 0 | 0 | 0 | 0 | 0 | 0 | 0 | 0 | 0 | 0 | NA | NA | NA | NA | NA | NA | NA | NA | NA | NA |
| Olfr1204 | 0 | 0 | 0 | 0 | 0 | 0 | 0 | 0 | 0 | 0 | 0 | 0 | 0 | 0 | 0 | NA | NA | NA | NA | NA | NA | NA | NA | NA | NA |
| Olfr1205 | 0 | 0 | 0 | 0 | 0 | 0 | 0 | 0 | 0 | 0 | 0 | 0 | 0 | 0 | 0 | NA | NA | NA | NA | NA | NA | NA | NA | NA | NA |
| Olfr1206 | 0 | 0 | 0 | 0 | 0 | 0 | 0 | 0 | 0 | 0 | 0 | 0 | 0 | 0 | 0 | NA | NA | NA | NA | NA | NA | NA | NA | NA | NA |
| Olfr1208 | 0 | 0 | 0 | 0 | 0 | 0 | 0 | 0 | 0 | 0 | 0 | 0 | 0 | 0 | 0 | NA | NA | NA | NA | NA | NA | NA | NA | NA | NA |
| Olfr1209 | 0 | 0 | 0 | 0 | 0 | 0 | 0 | 0 | 0 | 0 | 0 | 0 | 0 | 0 | 0 | NA | NA | NA | NA | NA | NA | NA | NA | NA | NA |
| Olfr121 | 1 | 0 | 0 | 0 | 0 | 0 | 1 | 0 | 0 | 0 | 0 | 0 | 0.13 | 0 | 0 | 0.552 | -0.857 | 0.8336548 | NA | 0 | 18.3121 | 10 | 10 | TRUE | 0.2566 |
| Olfr1211 | 0 | 0 | 0 | 0 | 0 | 0 | 0 | 0 | 0 | 0 | 0 | 0 | 0 | 0 | 0 | NA | NA | NA | NA | NA | NA | NA | NA | NA | NA |
| Olfr1212 | 0 | 0 | 0 | 0 | 0 | 0 | 0 | 0 | 0 | 0 | 0 | 0 | 0 | 0 | 0 | NA | NA | NA | NA | NA | NA | NA | NA | NA | NA |
| Olfr1213 | 0 | 0 | 0 | 0 | 0 | 0 | 0 | 0 | 0 | 0 | 0 | 0 | 0 | 0 | 0 | NA | NA | NA | NA | NA | NA | NA | NA | NA | NA |
| Olfr1214 | 0 | 0 | 0 | 0 | 0 | 0 | 0 | 0 | 0 | 0 | 0 | 0 | 0 | 0 | 0 | NA | NA | NA | NA | NA | NA | NA | NA | NA | NA |
| Olfr1215 | 0 | 0 | 0 | 0 | 0 | 0 | 0 | 0 | 0 | 0 | 0 | 0 | 0 | 0 | 0 | NA | NA | NA | NA | NA | NA | NA | NA | NA | NA |
| Olfr1216 | 0 | 0 | 0 | 0 | 0 | 0 | 0 | 0 | 0 | 0 | 0 | 0 | 0 | 0 | 0 | NA | NA | NA | NA | NA | NA | NA | NA | NA | NA |
| Olfr1217 | 0 | 0 | 0 | 0 | 0 | 0 | 0 | 0 | 0 | 0 | 0 | 0 | 0 | 0 | 0 | NA | NA | NA | NA | NA | NA | NA | NA | NA | NA |
| Olfr1218 | 0 | 0 | 0 | 0 | 0 | 0 | 0 | 0 | 0 | 0 | 0 | 0 | 0 | 0 | 0 | NA | NA | NA | NA | NA | NA | NA | NA | NA | NA |
| Olfr1219 | 0 | 0 | 0 | 0 | 0 | 0 | 0 | 0 | 0 | 0 | 0 | 0 | 0 | 0 | 0 | NA | NA | NA | NA | NA | NA | NA | NA | NA | NA |
| Olfr122 | 0 | 0 | 0 | 0 | 0 | 0 | 0 | 0 | 0 | 0 | 0 | 0 | 0 | 0 | 0 | NA | NA | NA | NA | NA | NA | NA | NA | NA | NA |
| Olfr1220 | 0 | 0 | 0 | 0 | 0 | 0 | 0 | 0 | 0 | 0 | 0 | 0 | 0 | 0 | 0 | NA | NA | NA | NA | NA | NA | NA | NA | NA | NA |
| Olfr1221 | 0 | 0 | 0 | 0 | 0 | 0 | 0 | 0 | 0 | 0 | 0 | 0 | 0 | 0 | 0 | NA | NA | NA | NA | NA | NA | NA | NA | NA | NA |
| Olfr1222 | 0 | 0 | 0 | 0 | 0 | 0 | 0 | 0 | 0 | 0 | 0 | 0 | 0 | 0 | 0 | NA | NA | NA | NA | NA | NA | NA | NA | NA | NA |
| Olfr1223 | 0 | 0 | 0 | 0 | 0 | 0 | 0 | 0 | 0 | 0 | 0 | 0 | 0 | 0 | 0 | NA | NA | NA | NA | NA | NA | NA | NA | NA | NA |
| Olfr1225 | 0 | 0 | 0 | 0 | 0 | 0 | 0 | 0 | 0 | 0 | 0 | 0 | 0 | 0 | 0 | NA | NA | NA | NA | NA | NA | NA | NA | NA | NA |
| Olfr1226 | 0 | 0 | 0 | 0 | 0 | 0 | 0 | 0 | 0 | 0 | 0 | 0 | 0 | 0 | 0 | NA | NA | NA | NA | NA | NA | NA | NA | NA | NA |
| Olfr1228 | 0 | 0 | 0 | 0 | 0 | 0 | 0 | 0 | 0 | 0 | 0 | 0 | 0 | 0 | 0 | NA | NA | NA | NA | NA | NA | NA | NA | NA | NA |
| Olfr1229 | 0 | 0 | 0 | 0 | 0 | 0 | 0 | 0 | 0 | 0 | 0 | 0 | 0 | 0 | 0 | NA | NA | NA | NA | NA | NA | NA | NA | NA | NA |
| Olfr123 | 0 | 0 | 0 | 0 | 0 | 0 | 0 | 0 | 0 | 0 | 0 | 0 | 0 | 0 | 0 | NA | NA | NA | NA | NA | NA | NA | NA | NA | NA |
| Olfr1230 | 0 | 0 | 0 | 0 | 0 | 0 | 0 | 0 | 0 | 0 | 0 | 0 | 0 | 0 | 0 | NA | NA | NA | NA | NA | NA | NA | NA | NA | NA |
| Olfr1231 | 0 | 0 | 0 | 0 | 0 | 0 | 0 | 0 | 0 | 0 | 0 | 0 | 0 | 0 | 0 | NA | NA | NA | NA | NA | NA | NA | NA | NA | NA |

| | | | | | | | | | | | | | | | | | | | | | | | | | | |
|---|---|---|---|---|---|---|---|---|---|---|---|---|---|---|---|---|---|---|---|---|---|---|---|---|---|---|
| Olfr1232 | 0 | 0 | 0 | 0 | 0 | 0 | 0 | 0 | 0 | 0 | 0 | 0 | 0 | 0 | 0 | NA | NA | NA | NA | NA | NA | NA | NA | NA | NA | |
| Olfr1233 | 0 | 0 | 0 | 0 | 0 | 0 | 0 | 0 | 0 | 0 | 0 | 0 | 0 | 0 | 0 | NA | NA | NA | NA | NA | NA | NA | NA | NA | NA | |
| Olfr1234 | 0 | 0 | 0 | 0 | 0 | 0 | 0 | 0 | 0 | 0 | 0 | 0 | 0 | 0 | 0 | NA | NA | NA | NA | NA | NA | NA | NA | NA | NA | |
| Olfr1238 | 0 | 0 | 0 | 0 | 0 | 0 | 0 | 0 | 0 | 0 | 0 | 0 | 0 | 0 | 0 | NA | NA | NA | NA | NA | NA | NA | NA | NA | NA | |
| Olfr1239 | 0 | 0 | 0 | 0 | 0 | 0 | 0 | 0 | 0 | 0 | 0 | 0 | 0 | 0 | 0 | NA | NA | NA | NA | NA | NA | NA | NA | NA | NA | |
| Olfr124 | 0 | 0 | 0 | 0 | 0 | 0 | 0 | 0 | 0 | 0 | 0 | 0 | 0 | 0 | 0 | NA | NA | NA | NA | NA | NA | NA | NA | NA | NA | |
| Olfr1240 | 0 | 0 | 0 | 0 | 0 | 0 | 0 | 0 | 0 | 0 | 0 | 0 | 0 | 0 | 0 | NA | NA | NA | NA | NA | NA | NA | NA | NA | NA | |
| Olfr1241 | 0 | 0 | 0 | 0 | 0 | 0 | 0 | 0 | 0 | 0 | 0 | 0 | 0 | 0 | 0 | NA | NA | NA | NA | NA | NA | NA | NA | NA | NA | |
| Olfr1242 | 0 | 0 | 0 | 0 | 0 | 0 | 0 | 0 | 0 | 0 | 0 | 0 | 0 | 0 | 0 | NA | NA | NA | NA | NA | NA | NA | NA | NA | NA | |
| Olfr1243 | 0 | 0 | 0 | 0 | 0 | 0 | 0 | 0 | 0 | 0 | 0 | 0 | 0 | 0 | 0 | NA | NA | NA | NA | NA | NA | NA | NA | NA | NA | |
| Olfr1245 | 0 | 0 | 0 | 0 | 0 | 0 | 0 | 0 | 0 | 0 | 0 | 0 | 0 | 0 | 0 | NA | NA | NA | NA | NA | NA | NA | NA | NA | NA | |
| Olfr1246 | 0 | 0 | 0 | 0 | 0 | 0 | 0 | 0 | 0 | 0 | 0 | 0 | 0 | 0 | 0 | NA | NA | NA | NA | NA | NA | NA | NA | NA | NA | |
| Olfr1247 | 0 | 0 | 0 | 0 | 0 | 0 | 0 | 0 | 0 | 0 | 0 | 0 | 0 | 0 | 0 | NA | NA | NA | NA | NA | NA | NA | NA | NA | NA | |
| Olfr1248 | 0 | 0 | 0 | 0 | 0 | 0 | 0 | 0 | 0 | 0 | 0 | 0 | 0 | 0 | 0 | NA | NA | NA | NA | NA | NA | NA | NA | NA | NA | |
| Olfr1249 | 0 | 0 | 0 | 0 | 0 | 0 | 0 | 0 | 0 | 0 | 0 | 0 | 0 | 0 | 0 | NA | NA | NA | NA | NA | NA | NA | NA | NA | NA | |
| Olfr125 | 0 | 0 | 0 | 0 | 0 | 0 | 0 | 0 | 0 | 0 | 0 | 0 | 0 | 0 | 0 | NA | NA | NA | NA | NA | NA | NA | NA | NA | NA | |
| Olfr1250 | 0 | 0 | 0 | 0 | 0 | 0 | 0 | 0 | 0 | 0 | 0 | 0 | 0 | 0 | 0 | NA | NA | NA | NA | NA | NA | NA | NA | NA | NA | |
| Olfr1251 | 0 | 0 | 0 | 0 | 0 | 0 | 0 | 0 | 0 | 0 | 0 | 0 | 0 | 0 | 0 | NA | NA | NA | NA | NA | NA | NA | NA | NA | NA | |
| Olfr1252 | 0 | 0 | 0 | 0 | 0 | 0 | 0 | 0 | 0 | 0 | 0 | 0 | 0 | 0 | 0 | NA | NA | NA | NA | NA | NA | NA | NA | NA | NA | |
| Olfr1253 | 0 | 0 | 0 | 0 | 0 | 0 | 0 | 0 | 0 | 0 | 0 | 0 | 0 | 0 | 0 | NA | NA | NA | NA | NA | NA | NA | NA | NA | NA | |
| Olfr1254 | 0 | 0 | 0 | 0 | 0 | 0 | 0 | 0 | 0 | 0 | 0 | 0 | 0 | 0 | 0 | NA | NA | NA | NA | NA | NA | NA | NA | NA | NA | |
| Olfr1255 | 0 | 0 | 0 | 0 | 0 | 0 | 0 | 0 | 0 | 0 | 0 | 0 | 0 | 0 | 0 | NA | NA | NA | NA | NA | NA | NA | NA | NA | NA | |
| Olfr1256 | 0 | 0 | 0 | 0 | 0 | 0 | 0 | 0 | 0 | 0 | 0 | 0 | 0 | 0 | 0 | NA | NA | NA | NA | NA | NA | NA | NA | NA | NA | |
| Olfr1257 | 0 | 0 | 0 | 0 | 0 | 0 | 0 | 0 | 0 | 0 | 0 | 0 | 0 | 0 | 0 | NA | NA | NA | NA | NA | NA | NA | NA | NA | NA | |
| Olfr1258 | 0 | 0 | 0 | 0 | 0 | 0 | 0 | 0 | 0 | 0 | 0 | 0 | 0 | 0 | 0 | NA | NA | NA | NA | NA | NA | NA | NA | NA | NA | |
| Olfr1259 | 2 | 4 | 4 | 0 | 0 | 0 | 2 | 4 | 4 | 0 | 0 | 0 | 1.69 | 3 | 0 | 0.057 | -4.132 | 0.02167482 | NA | | 0 | 1.407 | 1.1889 | 1.1889 | TRUE | 0.4261 |
| Olfr126 | 0 | 0 | 0 | 0 | 0 | 0 | 0 | 0 | 0 | 0 | 0 | 0 | 0 | 0 | 0 | NA | NA | NA | NA | NA | NA | NA | NA | NA | NA | |
| Olfr1260 | 0 | 0 | 0 | 0 | 0 | 0 | 0 | 0 | 0 | 0 | 0 | 0 | 0 | 0 | 0 | NA | NA | NA | NA | NA | NA | NA | NA | NA | NA | |
| Olfr1261 | 0 | 0 | 0 | 0 | 0 | 0 | 0 | 0 | 0 | 0 | 0 | 0 | 0 | 0 | 0 | NA | NA | NA | NA | NA | NA | NA | NA | NA | NA | |
| Olfr1262 | 0 | 0 | 0 | 0 | 0 | 0 | 0 | 0 | 0 | 0 | 0 | 0 | 0 | 0 | 0 | NA | NA | NA | NA | NA | NA | NA | NA | NA | NA | |
| Olfr1263 | 0 | 0 | 0 | 0 | 0 | 0 | 0 | 0 | 0 | 0 | 0 | 0 | 0 | 0 | 0 | NA | NA | NA | NA | NA | NA | NA | NA | NA | NA | |
| Olfr1264 | 0 | 0 | 0 | 0 | 0 | 0 | 0 | 0 | 0 | 0 | 0 | 0 | 0 | 0 | 0 | NA | NA | NA | NA | NA | NA | NA | NA | NA | NA | |
| Olfr1265 | 0 | 0 | 0 | 0 | 0 | 0 | 0 | 0 | 0 | 0 | 0 | 0 | 0 | 0 | 0 | NA | NA | NA | NA | NA | NA | NA | NA | NA | NA | |
| Olfr1269 | 0 | 0 | 0 | 0 | 0 | 0 | 0 | 0 | 0 | 0 | 0 | 0 | 0 | 0 | 0 | NA | NA | NA | NA | NA | NA | NA | NA | NA | NA | |
| Olfr127 | 0 | 0 | 0 | 0 | 0 | 0 | 0 | 0 | 0 | 0 | 0 | 0 | 0 | 0 | 0 | NA | NA | NA | NA | NA | NA | NA | NA | NA | NA | |
| Olfr1270 | 0 | 0 | 0 | 0 | 0 | 0 | 0 | 0 | 0 | 0 | 0 | 0 | 0 | 0 | 0 | NA | NA | NA | NA | NA | NA | NA | NA | NA | NA | |
| Olfr1271 | 0 | 0 | 0 | 0 | 0 | 0 | 0 | 0 | 0 | 0 | 0 | 0 | 0 | 0 | 0 | NA | NA | NA | NA | NA | NA | NA | NA | NA | NA | |
| Olfr1272 | 0 | 0 | 0 | 0 | 0 | 0 | 0 | 0 | 0 | 0 | 0 | 0 | 0 | 0 | 0 | NA | NA | NA | NA | NA | NA | NA | NA | NA | NA | |
| Olfr1273-ps | 0 | 0 | 0 | 0 | 0 | 0 | 0 | 0 | 0 | 0 | 0 | 0 | 0 | 0 | 0 | NA | NA | NA | NA | NA | NA | NA | NA | NA | NA | |
| Olfr1274-ps | 0 | 0 | 0 | 0 | 0 | 0 | 0 | 0 | 0 | 0 | 0 | 0 | 0 | 0 | 0 | NA | NA | NA | NA | NA | NA | NA | NA | NA | NA | |
| Olfr1275 | 0 | 0 | 0 | 0 | 0 | 0 | 0 | 0 | 0 | 0 | 0 | 0 | 0 | 0 | 0 | NA | NA | NA | NA | NA | NA | NA | NA | NA | NA | |
| Olfr1276 | 0 | 0 | 0 | 0 | 0 | 0 | 0 | 0 | 0 | 0 | 0 | 0 | 0 | 0 | 0 | NA | NA | NA | NA | NA | NA | NA | NA | NA | NA | |
| Olfr1277 | 0 | 0 | 0 | 0 | 4 | 0 | 0 | 0 | 0 | 0 | 5 | 0 | 0.75 | 0 | 2 | 8.401 | 3.071 | 0.29044532 | NA | | 10 | 3.1538 | 4.7319 | 4.7319 | TRUE | 1.8998 |
| Olfr1278 | 0 | 0 | 0 | 0 | 0 | 0 | 0 | 0 | 0 | 0 | 0 | 0 | 0 | 0 | 0 | NA | NA | NA | NA | NA | NA | NA | NA | NA | NA | |
| Olfr1279 | 0 | 0 | 0 | 0 | 0 | 0 | 0 | 0 | 0 | 0 | 0 | 0 | 0 | 0 | 0 | NA | NA | NA | NA | NA | NA | NA | NA | NA | NA | |
| Olfr128 | 0 | 0 | 0 | 0 | 0 | 0 | 0 | 0 | 0 | 0 | 0 | 0 | 0 | 0 | 0 | NA | NA | NA | NA | NA | NA | NA | NA | NA | NA | |
| Olfr1280 | 0 | 0 | 0 | 0 | 0 | 0 | 0 | 0 | 0 | 0 | 0 | 0 | 0 | 0 | 0 | NA | NA | NA | NA | NA | NA | NA | NA | NA | NA | |
| Olfr1281 | 0 | 0 | 0 | 0 | 0 | 0 | 0 | 0 | 0 | 0 | 0 | 0 | 0 | 0 | 0 | NA | NA | NA | NA | NA | NA | NA | NA | NA | NA | |
| Olfr1282 | 0 | 0 | 0 | 0 | 0 | 0 | 0 | 0 | 0 | 0 | 0 | 0 | 0 | 0 | 0 | NA | NA | NA | NA | NA | NA | NA | NA | NA | NA | |
| Olfr1283 | 0 | 0 | 0 | 0 | 0 | 0 | 0 | 0 | 0 | 0 | 0 | 0 | 0 | 0 | 0 | NA | NA | NA | NA | NA | NA | NA | NA | NA | NA | |
| Olfr1284 | 0 | 0 | 0 | 0 | 0 | 0 | 0 | 0 | 0 | 0 | 0 | 0 | 0 | 0 | 0 | NA | NA | NA | NA | NA | NA | NA | NA | NA | NA | |
| Olfr1286 | 0 | 0 | 0 | 0 | 0 | 0 | 0 | 0 | 0 | 0 | 0 | 0 | 0 | 0 | 0 | NA | NA | NA | NA | NA | NA | NA | NA | NA | NA | |
| Olfr1287 | 0 | 0 | 0 | 0 | 0 | 0 | 0 | 0 | 0 | 0 | 0 | 0 | 0 | 0 | 0 | NA | NA | NA | NA | NA | NA | NA | NA | NA | NA | |
| Olfr1288 | 0 | 0 | 0 | 0 | 0 | 0 | 0 | 0 | 0 | 0 | 0 | 0 | 0 | 0 | 0 | NA | NA | NA | NA | NA | NA | NA | NA | NA | NA | |
| Olfr1289 | 0 | 0 | 0 | 0 | 0 | 0 | 0 | 0 | 0 | 0 | 0 | 0 | 0 | 0 | 0 | NA | NA | NA | NA | NA | NA | NA | NA | NA | NA | |
| Olfr129 | 0 | 0 | 0 | 0 | 0 | 0 | 0 | 0 | 0 | 0 | 0 | 0 | 0 | 0 | 0 | NA | NA | NA | NA | NA | NA | NA | NA | NA | NA | |
| Olfr1290 | 0 | 0 | 0 | 0 | 0 | 0 | 0 | 0 | 0 | 0 | 0 | 0 | 0 | 0 | 0 | NA | NA | NA | NA | NA | NA | NA | NA | NA | NA | |
| Olfr1294 | 0 | 0 | 0 | 0 | 0 | 0 | 0 | 0 | 0 | 0 | 0 | 0 | 0 | 0 | 0 | NA | NA | NA | NA | NA | NA | NA | NA | NA | NA | |
| Olfr1295 | 0 | 0 | 0 | 0 | 0 | 0 | 0 | 0 | 0 | 0 | 0 | 0 | 0 | 0 | 0 | NA | NA | NA | NA | NA | NA | NA | NA | NA | NA | |
| Olfr1297 | 0 | 0 | 0 | 0 | 0 | 0 | 0 | 0 | 0 | 0 | 0 | 0 | 0 | 0 | 0 | NA | NA | NA | NA | NA | NA | NA | NA | NA | NA | |
| Olfr1298 | 0 | 0 | 0 | 0 | 0 | 0 | 0 | 0 | 0 | 0 | 0 | 0 | 0 | 0 | 0 | NA | NA | NA | NA | NA | NA | NA | NA | NA | NA | |
| Olfr1299 | 0 | 0 | 0 | 0 | 0 | 0 | 0 | 0 | 0 | 0 | 0 | 0 | 0 | 0 | 0 | NA | NA | NA | NA | NA | NA | NA | NA | NA | NA | |
| Olfr13 | 0 | 0 | 0 | 0 | 0 | 0 | 0 | 0 | 0 | 0 | 0 | 0 | 0 | 0 | 0 | NA | NA | NA | NA | NA | NA | NA | NA | NA | NA | |
| Olfr130 | 0 | 0 | 0 | 0 | 0 | 0 | 0 | 0 | 0 | 0 | 0 | 0 | 0 | 0 | 0 | NA | NA | NA | NA | NA | NA | NA | NA | NA | NA | |
| Olfr1300-ps1 | 0 | 0 | 0 | 0 | 0 | 0 | 0 | 0 | 0 | 0 | 0 | 0 | 0 | 0 | 0 | NA | NA | NA | NA | NA | NA | NA | NA | NA | NA | |
| Olfr1301 | 0 | 0 | 0 | 0 | 0 | 0 | 0 | 0 | 0 | 0 | 0 | 0 | 0 | 0 | 0 | NA | NA | NA | NA | NA | NA | NA | NA | NA | NA | |
| Olfr1302 | 0 | 0 | 0 | 0 | 0 | 0 | 0 | 0 | 0 | 0 | 0 | 0 | 0 | 0 | 0 | NA | NA | NA | NA | NA | NA | NA | NA | NA | NA | |
| Olfr1303 | 0 | 0 | 0 | 0 | 0 | 0 | 0 | 0 | 0 | 0 | 0 | 0 | 0 | 0 | 0 | NA | NA | NA | NA | NA | NA | NA | NA | NA | NA | |
| Olfr1305 | 0 | 0 | 0 | 0 | 0 | 0 | 0 | 0 | 0 | 0 | 0 | 0 | 0 | 0 | 0 | NA | NA | NA | NA | NA | NA | NA | NA | NA | NA | |

| | | | | | | | | | | | | | | | | | | | | | | | | | |
|---|---|---|---|---|---|---|---|---|---|---|---|---|---|---|---|---|---|---|---|---|---|---|---|---|---|
| Olfr1306 | 0 | 0 | 0 | 0 | 0 | 0 | 0 | 0 | 0 | 0 | 0 | 0 | 0 | 0 | 0 | NA | NA | NA | NA | NA | NA | NA | NA | NA | NA |
| Olfr1307 | 0 | 0 | 0 | 0 | 0 | 0 | 0 | 0 | 0 | 0 | 0 | 0 | 0 | 0 | 0 | NA | NA | NA | NA | NA | NA | NA | NA | NA | NA |
| Olfr1308 | 0 | 0 | 0 | 0 | 0 | 0 | 0 | 0 | 0 | 0 | 0 | 0 | 0 | 0 | 0 | NA | NA | NA | NA | NA | NA | NA | NA | NA | NA |
| Olfr1309 | 0 | 0 | 0 | 0 | 0 | 0 | 0 | 0 | 0 | 0 | 0 | 0 | 0 | 0 | 0 | NA | NA | NA | NA | NA | NA | NA | NA | NA | NA |
| Olfr131 | 0 | 0 | 0 | 0 | 0 | 0 | 0 | 0 | 0 | 0 | 0 | 0 | 0 | 0 | 0 | NA | NA | NA | NA | NA | NA | NA | NA | NA | NA |
| Olfr1310 | 0 | 0 | 0 | 0 | 0 | 0 | 0 | 0 | 0 | 0 | 0 | 0 | 0 | 0 | 0 | NA | NA | NA | NA | NA | NA | NA | NA | NA | NA |
| Olfr1311 | 0 | 0 | 0 | 0 | 0 | 0 | 0 | 0 | 0 | 0 | 0 | 0 | 0 | 0 | 0 | NA | NA | NA | NA | NA | NA | NA | NA | NA | NA |
| Olfr1312 | 0 | 0 | 0 | 0 | 0 | 0 | 0 | 0 | 0 | 0 | 0 | 0 | 0 | 0 | 0 | NA | NA | NA | NA | NA | NA | NA | NA | NA | NA |
| Olfr1313 | 0 | 0 | 0 | 0 | 0 | 0 | 0 | 0 | 0 | 0 | 0 | 0 | 0 | 0 | 0 | NA | NA | NA | NA | NA | NA | NA | NA | NA | NA |
| Olfr1314 | 0 | 0 | 0 | 0 | 0 | 0 | 0 | 0 | 0 | 0 | 0 | 0 | 0 | 0 | 0 | NA | NA | NA | NA | NA | NA | NA | NA | NA | NA |
| Olfr1316 | 0 | 0 | 0 | 0 | 0 | 0 | 0 | 0 | 0 | 0 | 0 | 0 | 0 | 0 | 0 | NA | NA | NA | NA | NA | NA | NA | NA | NA | NA |
| Olfr1317 | 0 | 0 | 0 | 0 | 0 | 0 | 0 | 0 | 0 | 0 | 0 | 0 | 0 | 0 | 0 | NA | NA | NA | NA | NA | NA | NA | NA | NA | NA |
| Olfr1318 | 0 | 0 | 0 | 0 | 0 | 0 | 0 | 0 | 0 | 0 | 0 | 0 | 0 | 0 | 0 | NA | NA | NA | NA | NA | NA | NA | NA | NA | NA |
| Olfr132 | 0 | 0 | 0 | 0 | 0 | 0 | 0 | 0 | 0 | 0 | 0 | 0 | 0 | 0 | 0 | NA | NA | NA | NA | NA | NA | NA | NA | NA | NA |
| Olfr1320 | 0 | 0 | 0 | 0 | 0 | 0 | 0 | 0 | 0 | 0 | 0 | 0 | 0 | 0 | 0 | NA | NA | NA | NA | NA | NA | NA | NA | NA | NA |
| Olfr1321 | 0 | 0 | 0 | 0 | 0 | 0 | 0 | 0 | 0 | 0 | 0 | 0 | 0 | 0 | 0 | NA | NA | NA | NA | NA | NA | NA | NA | NA | NA |
| Olfr1322 | 0 | 0 | 0 | 0 | 0 | 0 | 0 | 0 | 0 | 0 | 0 | 0 | 0 | 0 | 0 | NA | NA | NA | NA | NA | NA | NA | NA | NA | NA |
| Olfr1323 | 0 | 0 | 0 | 0 | 0 | 0 | 0 | 0 | 0 | 0 | 0 | 0 | 0 | 0 | 0 | NA | NA | NA | NA | NA | NA | NA | NA | NA | NA |
| Olfr1324 | 0 | 0 | 0 | 0 | 0 | 0 | 0 | 0 | 0 | 0 | 0 | 0 | 0 | 0 | 0 | NA | NA | NA | NA | NA | NA | NA | NA | NA | NA |
| Olfr1325 | 0 | 0 | 0 | 0 | 0 | 0 | 0 | 0 | 0 | 0 | 0 | 0 | 0 | 0 | 0 | NA | NA | NA | NA | NA | NA | NA | NA | NA | NA |
| Olfr1328 | 0 | 0 | 0 | 1 | 0 | 1 | 0 | 0 | 0 | 1 | 0 | 1 | 0.33 | 0 | 1 | 3.763 | 1.912 | 0.57887588 | NA | | 0 | 7.1752 | 6.7756 | 6.7756 | TRUE | 0.2094 |
| Olfr1329 | 0 | 0 | 0 | 0 | 0 | 0 | 0 | 0 | 0 | 0 | 0 | 0 | 0 | 0 | 0 | NA | NA | NA | NA | NA | NA | NA | NA | NA | NA |
| Olfr133 | 0 | 0 | 0 | 0 | 0 | 0 | 0 | 0 | 0 | 0 | 0 | 0 | 0 | 0 | 0 | NA | NA | NA | NA | NA | NA | NA | NA | NA | NA |
| Olfr1330 | 0 | 0 | 0 | 0 | 0 | 0 | 0 | 0 | 0 | 0 | 0 | 0 | 0 | 0 | 0 | NA | NA | NA | NA | NA | NA | NA | NA | NA | NA |
| Olfr1331 | 0 | 0 | 0 | 0 | 0 | 0 | 0 | 0 | 0 | 0 | 0 | 0 | 0 | 0 | 0 | NA | NA | NA | NA | NA | NA | NA | NA | NA | NA |
| Olfr1333 | 0 | 0 | 0 | 0 | 0 | 0 | 0 | 0 | 0 | 0 | 0 | 0 | 0 | 0 | 0 | NA | NA | NA | NA | NA | NA | NA | NA | NA | NA |
| Olfr1335 | 0 | 0 | 0 | 0 | 0 | 0 | 0 | 0 | 0 | 0 | 0 | 0 | 0 | 0 | 0 | NA | NA | NA | NA | NA | NA | NA | NA | NA | NA |
| Olfr1336 | 1 | 0 | 0 | 1 | 1 | 0 | 1 | 0 | 0 | 1 | 1 | 0 | 0.48 | 0 | 1 | 2.037 | 1.027 | 0.71241723 | NA | | 0 | 4.9057 | 3.8521 | 3.8521 | TRUE | 0.2664 |
| Olfr1337 | 0 | 0 | 0 | 0 | 0 | 0 | 0 | 0 | 0 | 0 | 0 | 0 | 0 | 0 | 0 | NA | NA | NA | NA | NA | NA | NA | NA | NA | NA |
| Olfr1338 | 0 | 0 | 0 | 0 | 0 | 0 | 0 | 0 | 0 | 0 | 0 | 0 | 0 | 0 | 0 | NA | NA | NA | NA | NA | NA | NA | NA | NA | NA |
| Olfr1339 | 0 | 0 | 0 | 0 | 0 | 0 | 0 | 0 | 0 | 0 | 0 | 0 | 0 | 0 | 0 | NA | NA | NA | NA | NA | NA | NA | NA | NA | NA |
| Olfr134 | 0 | 0 | 0 | 0 | 0 | 0 | 0 | 0 | 0 | 0 | 0 | 0 | 0 | 0 | 0 | NA | NA | NA | NA | NA | NA | NA | NA | NA | NA |
| Olfr1340 | 0 | 0 | 0 | 0 | 0 | 0 | 0 | 0 | 0 | 0 | 0 | 0 | 0 | 0 | 0 | NA | NA | NA | NA | NA | NA | NA | NA | NA | NA |
| Olfr1341 | 0 | 0 | 0 | 0 | 0 | 0 | 0 | 0 | 0 | 0 | 0 | 0 | 0 | 0 | 0 | NA | NA | NA | NA | NA | NA | NA | NA | NA | NA |
| Olfr1342 | 2 | 2 | 0 | 2 | 1 | 2 | 2 | 2 | 0 | 2 | 1 | 2 | 1.46 | 1 | 2 | 1.373 | 0.457 | 0.7717007 | NA | | 0 | 1.6232 | 1.0952 | 1.0952 | TRUE | 0.3759 |
| Olfr1344 | 25 | 13 | 25 | 32 | 28 | 39 | 19 | 14 | 27 | 32 | 32 | 38 | 27.03 | 20 | 34 | 1.698 | 0.764 | 0.04886713 | 0.13246864 | 0.0012 | 0.0901 | 0.0691 | 0.0691 | TRUE | | 0.3942 |
| Olfr1346 | 0 | 0 | 1 | 0 | 0 | 0 | 0 | 0 | 1 | 0 | 0 | 0 | 0.18 | 0 | 0 | 0.552 | -0.857 | 0.8336548 | NA | | 0 | 13.1817 | 10 | 10 | TRUE | 0.5365 |
| Olfr1347 | 1 | 1 | 0 | 0 | 0 | 0 | 1 | 1 | 0 | 0 | 0 | 0 | 0.31 | 1 | 0 | 0.309 | -1.696 | 0.63183309 | NA | | 0 | 7.7294 | 7.2472 | 7.2472 | TRUE | 0.2015 |
| Olfr1348 | 1 | 0 | 0 | 0 | 0 | 0 | 1 | 0 | 0 | 0 | 0 | 0 | 0.13 | 0 | 0 | 0.552 | -0.857 | 0.8336548 | NA | | 0 | 18.3121 | 10 | 10 | TRUE | 0.2566 |
| Olfr1349 | 6 | 6 | 1 | 0 | 0 | 1 | 5 | 6 | 1 | 0 | 0 | 1 | 2.18 | 4 | 0 | 0.091 | -3.452 | 0.04019666 | NA | | 0 | 1.0873 | 0.9201 | 0.9201 | TRUE | 0.2984 |
| Olfr135 | 0 | 0 | 0 | 0 | 0 | 0 | 0 | 0 | 0 | 0 | 0 | 0 | 0 | 0 | 0 | NA | NA | NA | NA | NA | NA | NA | NA | NA | NA |
| Olfr1350 | 0 | 1 | 0 | 1 | 0 | 1 | 0 | 1 | 0 | 1 | 0 | 1 | 0.51 | 0 | 1 | 1.941 | 0.957 | 0.72720315 | NA | | 0 | 4.6706 | 3.6514 | 3.6514 | TRUE | 0.5217 |
| Olfr1351 | 0 | 0 | 0 | 0 | 0 | 0 | 0 | 0 | 0 | 0 | 0 | 0 | 0 | 0 | 0 | NA | NA | NA | NA | NA | NA | NA | NA | NA | NA |
| Olfr1352 | 0 | 0 | 0 | 0 | 0 | 0 | 0 | 0 | 0 | 0 | 0 | 0 | 0 | 0 | 0 | NA | NA | NA | NA | NA | NA | NA | NA | NA | NA |
| Olfr1353 | 0 | 0 | 0 | 0 | 0 | 0 | 0 | 0 | 0 | 0 | 0 | 0 | 0 | 0 | 0 | NA | NA | NA | NA | NA | NA | NA | NA | NA | NA |
| Olfr1354 | 0 | 0 | 0 | 0 | 0 | 0 | 0 | 0 | 0 | 0 | 0 | 0 | 0 | 0 | 0 | NA | NA | NA | NA | NA | NA | NA | NA | NA | NA |
| Olfr1355 | 0 | 0 | 0 | 0 | 0 | 0 | 0 | 0 | 0 | 0 | 0 | 0 | 0 | 0 | 0 | NA | NA | NA | NA | NA | NA | NA | NA | NA | NA |
| Olfr1356 | 0 | 0 | 0 | 0 | 0 | 0 | 0 | 0 | 0 | 0 | 0 | 0 | 0 | 0 | 0 | NA | NA | NA | NA | NA | NA | NA | NA | NA | NA |
| Olfr1357 | 0 | 0 | 0 | 0 | 0 | 0 | 0 | 0 | 0 | 0 | 0 | 0 | 0 | 0 | 0 | NA | NA | NA | NA | NA | NA | NA | NA | NA | NA |
| Olfr1359 | 0 | 0 | 0 | 0 | 0 | 0 | 0 | 0 | 0 | 0 | 0 | 0 | 0 | 0 | 0 | NA | NA | NA | NA | NA | NA | NA | NA | NA | NA |
| Olfr136 | 0 | 0 | 0 | 0 | 0 | 0 | 0 | 0 | 0 | 0 | 0 | 0 | 0 | 0 | 0 | NA | NA | NA | NA | NA | NA | NA | NA | NA | NA |
| Olfr1360 | 0 | 0 | 0 | 0 | 0 | 0 | 0 | 0 | 0 | 0 | 0 | 0 | 0 | 0 | 0 | NA | NA | NA | NA | NA | NA | NA | NA | NA | NA |
| Olfr1361 | 0 | 0 | 0 | 0 | 0 | 0 | 0 | 0 | 0 | 0 | 0 | 0 | 0 | 0 | 0 | NA | NA | NA | NA | NA | NA | NA | NA | NA | NA |
| Olfr1362 | 0 | 0 | 0 | 0 | 0 | 0 | 0 | 0 | 0 | 0 | 0 | 0 | 0 | 0 | 0 | NA | NA | NA | NA | NA | NA | NA | NA | NA | NA |
| Olfr1364 | 0 | 0 | 0 | 0 | 0 | 0 | 0 | 0 | 0 | 0 | 0 | 0 | 0 | 0 | 0 | NA | NA | NA | NA | NA | NA | NA | NA | NA | NA |
| Olfr1366 | 0 | 0 | 0 | 0 | 0 | 0 | 0 | 0 | 0 | 0 | 0 | 0 | 0 | 0 | 0 | NA | NA | NA | NA | NA | NA | NA | NA | NA | NA |
| Olfr1367 | 0 | 0 | 0 | 0 | 0 | 0 | 0 | 0 | 0 | 0 | 0 | 0 | 0 | 0 | 0 | NA | NA | NA | NA | NA | NA | NA | NA | NA | NA |
| Olfr1368 | 0 | 0 | 0 | 0 | 0 | 0 | 0 | 0 | 0 | 0 | 0 | 0 | 0 | 0 | 0 | NA | NA | NA | NA | NA | NA | NA | NA | NA | NA |
| Olfr137 | 0 | 0 | 0 | 0 | 0 | 0 | 0 | 0 | 0 | 0 | 0 | 0 | 0 | 0 | 0 | NA | NA | NA | NA | NA | NA | NA | NA | NA | NA |
| Olfr1370 | 0 | 0 | 0 | 0 | 0 | 0 | 0 | 0 | 0 | 0 | 0 | 0 | 0 | 0 | 0 | NA | NA | NA | NA | NA | NA | NA | NA | NA | NA |
| Olfr1371 | 0 | 0 | 2 | 1 | 0 | 1 | 0 | 0 | 2 | 1 | 0 | 1 | 0.69 | 1 | 1 | 0.966 | -0.049 | 0.98423749 | NA | | 0 | 3.4369 | 3.0198 | 3.0198 | TRUE | 0.9749 |
| Olfr1372-ps1 | 89 | 62 | 64 | 79 | 75 | 90 | 69 | 66 | 69 | 79 | 85 | 89 | 76.04 | 68 | 84 | 1.234 | 0.304 | 0.16792678 | 0.3193849 | 0 | 0.0336 | 0.0217 | 0.0217 | TRUE | | 0.0306 |
| Olfr1373 | 0 | 0 | 0 | 0 | 0 | 0 | 0 | 0 | 0 | 0 | 0 | 0 | 0 | 0 | 0 | NA | NA | NA | NA | NA | NA | NA | NA | NA | NA |
| Olfr1377 | 0 | 0 | 0 | 0 | 0 | 0 | 0 | 0 | 0 | 0 | 0 | 0 | 0 | 0 | 0 | NA | NA | NA | NA | NA | NA | NA | NA | NA | NA |
| Olfr1378 | 0 | 0 | 0 | 0 | 0 | 0 | 0 | 0 | 0 | 0 | 0 | 0 | 0 | 0 | 0 | NA | NA | NA | NA | NA | NA | NA | NA | NA | NA |
| Olfr138 | 0 | 0 | 0 | 0 | 0 | 0 | 0 | 0 | 0 | 0 | 0 | 0 | 0 | 0 | 0 | NA | NA | NA | NA | NA | NA | NA | NA | NA | NA |
| Olfr1380 | 0 | 0 | 0 | 0 | 0 | 0 | 0 | 0 | 0 | 0 | 0 | 0 | 0 | 0 | 0 | NA | NA | NA | NA | NA | NA | NA | NA | NA | NA |
| Olfr1381 | 0 | 0 | 0 | 0 | 0 | 0 | 0 | 0 | 0 | 0 | 0 | 0 | 0 | 0 | 0 | NA | NA | NA | NA | NA | NA | NA | NA | NA | NA |
| Olfr1382 | 0 | 0 | 0 | 0 | 0 | 0 | 0 | 0 | 0 | 0 | 0 | 0 | 0 | 0 | 0 | NA | NA | NA | NA | NA | NA | NA | NA | NA | NA |

| | | | | | | | | | | | | | | | | | | | | | | | | | |
|---|---|---|---|---|---|---|---|---|---|---|---|---|---|---|---|---|---|---|---|---|---|---|---|---|---|
| Olfr1383 | 0 | 0 | 0 | 0 | 0 | 0 | 0 | 0 | 0 | 0 | 0 | 0 | 0 | 0 | 0 | NA | NA | NA | NA | NA | NA | NA | NA | NA | NA |
| Olfr1384 | 0 | 0 | 0 | 0 | 0 | 0 | 0 | 0 | 0 | 0 | 0 | 0 | 0 | 0 | 0 | NA | NA | NA | NA | NA | NA | NA | NA | NA | NA |
| Olfr1385 | 0 | 0 | 0 | 0 | 0 | 0 | 0 | 0 | 0 | 0 | 0 | 0 | 0 | 0 | 0 | NA | NA | NA | NA | NA | NA | NA | NA | NA | NA |
| Olfr1386 | 0 | 0 | 0 | 0 | 0 | 0 | 0 | 0 | 0 | 0 | 0 | 0 | 0 | 0 | 0 | NA | NA | NA | NA | NA | NA | NA | NA | NA | NA |
| Olfr1387 | 0 | 0 | 0 | 0 | 0 | 0 | 0 | 0 | 0 | 0 | 0 | 0 | 0 | 0 | 0 | NA | NA | NA | NA | NA | NA | NA | NA | NA | NA |
| Olfr1388 | 0 | 0 | 0 | 0 | 0 | 0 | 0 | 0 | 0 | 0 | 0 | 0 | 0 | 0 | 0 | NA | NA | NA | NA | NA | NA | NA | NA | NA | NA |
| Olfr1389 | 1 | 0 | 0 | 0 | 1 | 0 | 1 | 0 | 0 | 0 | 1 | 0 | 0.32 | 0 | 0 | 1.075 | 0.105 | 0.97638847 | NA | 0 | 7.4681 | 7.0392 | 7.0392 | TRUE | 0.5075 |
| Olfr139 | 0 | 0 | 0 | 0 | 0 | 0 | 0 | 0 | 0 | 0 | 0 | 0 | 0 | 0 | 0 | NA | NA | NA | NA | NA | NA | NA | NA | NA | NA |
| Olfr1390 | 0 | 0 | 0 | 0 | 0 | 0 | 0 | 0 | 0 | 0 | 0 | 0 | 0 | 0 | 0 | NA | NA | NA | NA | NA | NA | NA | NA | NA | NA |
| Olfr1391 | 0 | 0 | 0 | 0 | 0 | 0 | 0 | 0 | 0 | 0 | 0 | 0 | 0 | 0 | 0 | NA | NA | NA | NA | NA | NA | NA | NA | NA | NA |
| Olfr1392 | 0 | 1 | 0 | 0 | 0 | 0 | 0 | 1 | 0 | 0 | 0 | 0 | 0.18 | 0 | 0 | 0.552 | -0.857 | 0.8336548 | NA | 0 | 13.3714 | 10 | 10 | TRUE | 0.5216 |
| Olfr1393 | 3 | 1 | 1 | 1 | 1 | 0 | 2 | 1 | 1 | 1 | 1 | 0 | 1.1 | 1 | 1 | 0.46 | -1.122 | 0.54331629 | NA | 0 | 2.1583 | 1.3946 | 1.3946 | TRUE | 0.2706 |
| Olfr1394 | 0 | 0 | 0 | 0 | 0 | 0 | 0 | 0 | 0 | 0 | 0 | 0 | 0 | 0 | 0 | NA | NA | NA | NA | NA | NA | NA | NA | NA | NA |
| Olfr1395 | 0 | 0 | 0 | 0 | 0 | 0 | 0 | 0 | 0 | 0 | 0 | 0 | 0 | 0 | 0 | NA | NA | NA | NA | NA | NA | NA | NA | NA | NA |
| Olfr1396 | 0 | 0 | 0 | 0 | 0 | 0 | 0 | 0 | 0 | 0 | 0 | 0 | 0 | 0 | 0 | NA | NA | NA | NA | NA | NA | NA | NA | NA | NA |
| Olfr140 | 0 | 0 | 0 | 0 | 0 | 0 | 0 | 0 | 0 | 0 | 0 | 0 | 0 | 0 | 0 | NA | NA | NA | NA | NA | NA | NA | NA | NA | NA |
| Olfr1402 | 0 | 0 | 0 | 0 | 0 | 0 | 0 | 0 | 0 | 0 | 0 | 0 | 0 | 0 | 0 | NA | NA | NA | NA | NA | NA | NA | NA | NA | NA |
| Olfr1404 | 0 | 0 | 0 | 0 | 0 | 0 | 0 | 0 | 0 | 0 | 0 | 0 | 0 | 0 | 0 | NA | NA | NA | NA | NA | NA | NA | NA | NA | NA |
| Olfr1406 | 0 | 1 | 0 | 0 | 0 | 0 | 0 | 1 | 0 | 0 | 0 | 0 | 0.18 | 0 | 0 | 0.552 | -0.857 | 0.8336548 | NA | 0 | 13.3714 | 10 | 10 | TRUE | 0.5216 |
| Olfr1408 | 0 | 0 | 0 | 0 | 0 | 0 | 0 | 0 | 0 | 0 | 0 | 0 | 0 | 0 | 0 | NA | NA | NA | NA | NA | NA | NA | NA | NA | NA |
| Olfr141 | 0 | 0 | 0 | 0 | 0 | 0 | 0 | 0 | 0 | 0 | 0 | 0 | 0 | 0 | 0 | NA | NA | NA | NA | NA | NA | NA | NA | NA | NA |
| Olfr1410 | 0 | 0 | 0 | 0 | 0 | 0 | 0 | 0 | 0 | 0 | 0 | 0 | 0 | 0 | 0 | NA | NA | NA | NA | NA | NA | NA | NA | NA | NA |
| Olfr1411 | 0 | 0 | 0 | 0 | 0 | 0 | 0 | 0 | 0 | 0 | 0 | 0 | 0 | 0 | 0 | NA | NA | NA | NA | NA | NA | NA | NA | NA | NA |
| Olfr1412 | 0 | 0 | 0 | 0 | 0 | 0 | 0 | 0 | 0 | 0 | 0 | 0 | 0 | 0 | 0 | NA | NA | NA | NA | NA | NA | NA | NA | NA | NA |
| Olfr1413 | 0 | 0 | 0 | 0 | 0 | 0 | 0 | 0 | 0 | 0 | 0 | 0 | 0 | 0 | 0 | NA | NA | NA | NA | NA | NA | NA | NA | NA | NA |
| Olfr1414 | 0 | 0 | 0 | 0 | 0 | 0 | 0 | 0 | 0 | 0 | 0 | 0 | 0 | 0 | 0 | NA | NA | NA | NA | NA | NA | NA | NA | NA | NA |
| Olfr1415 | 0 | 0 | 0 | 0 | 0 | 0 | 0 | 0 | 0 | 0 | 0 | 0 | 0 | 0 | 0 | NA | NA | NA | NA | NA | NA | NA | NA | NA | NA |
| Olfr1416 | 0 | 0 | 0 | 0 | 0 | 0 | 0 | 0 | 0 | 0 | 0 | 0 | 0 | 0 | 0 | NA | NA | NA | NA | NA | NA | NA | NA | NA | NA |
| Olfr1417 | 0 | 0 | 0 | 0 | 0 | 0 | 0 | 0 | 0 | 0 | 0 | 0 | 0 | 0 | 0 | NA | NA | NA | NA | NA | NA | NA | NA | NA | NA |
| Olfr1418 | 0 | 0 | 0 | 0 | 0 | 0 | 0 | 0 | 0 | 0 | 0 | 0 | 0 | 0 | 0 | NA | NA | NA | NA | NA | NA | NA | NA | NA | NA |
| Olfr1419 | 0 | 1 | 0 | 0 | 0 | 1 | 0 | 1 | 0 | 0 | 0 | 1 | 0.34 | 0 | 0 | 1.075 | 0.105 | 0.97576789 | NA | 0 | 6.9363 | 6.5821 | 6.5821 | TRUE | 0.5216 |
| Olfr142 | 0 | 0 | 0 | 0 | 0 | 0 | 0 | 0 | 0 | 0 | 0 | 0 | 0 | 0 | 0 | NA | NA | NA | NA | NA | NA | NA | NA | NA | NA |
| Olfr1420 | 0 | 0 | 0 | 0 | 0 | 0 | 0 | 0 | 0 | 0 | 0 | 0 | 0 | 0 | 0 | NA | NA | NA | NA | NA | NA | NA | NA | NA | NA |
| Olfr1423 | 0 | 0 | 0 | 0 | 0 | 0 | 0 | 0 | 0 | 0 | 0 | 0 | 0 | 0 | 0 | NA | NA | NA | NA | NA | NA | NA | NA | NA | NA |
| Olfr1424 | 0 | 0 | 0 | 0 | 0 | 0 | 0 | 0 | 0 | 0 | 0 | 0 | 0 | 0 | 0 | NA | NA | NA | NA | NA | NA | NA | NA | NA | NA |
| Olfr1425 | 0 | 0 | 0 | 0 | 0 | 0 | 0 | 0 | 0 | 0 | 0 | 0 | 0 | 0 | 0 | NA | NA | NA | NA | NA | NA | NA | NA | NA | NA |
| Olfr1426 | 0 | 0 | 0 | 0 | 0 | 0 | 0 | 0 | 0 | 0 | 0 | 0 | 0 | 0 | 0 | NA | NA | NA | NA | NA | NA | NA | NA | NA | NA |
| Olfr1427 | 0 | 0 | 0 | 0 | 0 | 0 | 0 | 0 | 0 | 0 | 0 | 0 | 0 | 0 | 0 | NA | NA | NA | NA | NA | NA | NA | NA | NA | NA |
| Olfr1428 | 0 | 0 | 0 | 0 | 0 | 0 | 0 | 0 | 0 | 0 | 0 | 0 | 0 | 0 | 0 | NA | NA | NA | NA | NA | NA | NA | NA | NA | NA |
| Olfr143 | 0 | 0 | 0 | 0 | 0 | 0 | 0 | 0 | 0 | 0 | 0 | 0 | 0 | 0 | 0 | NA | NA | NA | NA | NA | NA | NA | NA | NA | NA |
| Olfr1431 | 0 | 0 | 0 | 0 | 0 | 0 | 0 | 0 | 0 | 0 | 0 | 0 | 0 | 0 | 0 | NA | NA | NA | NA | NA | NA | NA | NA | NA | NA |
| Olfr1433 | 0 | 0 | 0 | 0 | 0 | 0 | 0 | 0 | 0 | 0 | 0 | 0 | 0 | 0 | 0 | NA | NA | NA | NA | NA | NA | NA | NA | NA | NA |
| Olfr1434 | 0 | 0 | 0 | 0 | 0 | 0 | 0 | 0 | 0 | 0 | 0 | 0 | 0 | 0 | 0 | NA | NA | NA | NA | NA | NA | NA | NA | NA | NA |
| Olfr1436 | 0 | 0 | 0 | 0 | 0 | 0 | 0 | 0 | 0 | 0 | 0 | 0 | 0 | 0 | 0 | NA | NA | NA | NA | NA | NA | NA | NA | NA | NA |
| Olfr1437 | 0 | 0 | 0 | 0 | 0 | 0 | 0 | 0 | 0 | 0 | 0 | 0 | 0 | 0 | 0 | NA | NA | NA | NA | NA | NA | NA | NA | NA | NA |
| Olfr1440 | 0 | 0 | 0 | 0 | 0 | 0 | 0 | 0 | 0 | 0 | 0 | 0 | 0 | 0 | 0 | NA | NA | NA | NA | NA | NA | NA | NA | NA | NA |
| Olfr1441 | 0 | 0 | 0 | 0 | 0 | 0 | 0 | 0 | 0 | 0 | 0 | 0 | 0 | 0 | 0 | NA | NA | NA | NA | NA | NA | NA | NA | NA | NA |
| Olfr1442 | 0 | 0 | 0 | 0 | 0 | 0 | 0 | 0 | 0 | 0 | 0 | 0 | 0 | 0 | 0 | NA | NA | NA | NA | NA | NA | NA | NA | NA | NA |
| Olfr1443 | 0 | 0 | 0 | 0 | 0 | 0 | 0 | 0 | 0 | 0 | 0 | 0 | 0 | 0 | 0 | NA | NA | NA | NA | NA | NA | NA | NA | NA | NA |
| Olfr1444 | 0 | 0 | 0 | 0 | 0 | 0 | 0 | 0 | 0 | 0 | 0 | 0 | 0 | 0 | 0 | NA | NA | NA | NA | NA | NA | NA | NA | NA | NA |
| Olfr1445 | 0 | 0 | 0 | 0 | 0 | 0 | 0 | 0 | 0 | 0 | 0 | 0 | 0 | 0 | 0 | NA | NA | NA | NA | NA | NA | NA | NA | NA | NA |
| Olfr1446 | 0 | 0 | 0 | 0 | 0 | 0 | 0 | 0 | 0 | 0 | 0 | 0 | 0 | 0 | 0 | NA | NA | NA | NA | NA | NA | NA | NA | NA | NA |
| Olfr1447 | 0 | 0 | 0 | 0 | 0 | 1 | 0 | 0 | 0 | 0 | 0 | 1 | 0.16 | 0 | 0 | 2.094 | 1.067 | 0.79379329 | NA | 0 | 14.4078 | 10 | 10 | TRUE | 0.3835 |
| Olfr1448 | 0 | 0 | 0 | 0 | 0 | 0 | 0 | 0 | 0 | 0 | 0 | 0 | 0 | 0 | 0 | NA | NA | NA | NA | NA | NA | NA | NA | NA | NA |
| Olfr1449 | 0 | 0 | 0 | 0 | 0 | 0 | 0 | 0 | 0 | 0 | 0 | 0 | 0 | 0 | 0 | NA | NA | NA | NA | NA | NA | NA | NA | NA | NA |
| Olfr145 | 0 | 0 | 0 | 0 | 0 | 0 | 0 | 0 | 0 | 0 | 0 | 0 | 0 | 0 | 0 | NA | NA | NA | NA | NA | NA | NA | NA | NA | NA |
| Olfr1450 | 0 | 0 | 0 | 0 | 0 | 0 | 0 | 0 | 0 | 0 | 0 | 0 | 0 | 0 | 0 | NA | NA | NA | NA | NA | NA | NA | NA | NA | NA |
| Olfr1451 | 0 | 0 | 0 | 0 | 0 | 0 | 0 | 0 | 0 | 0 | 0 | 0 | 0 | 0 | 0 | NA | NA | NA | NA | NA | NA | NA | NA | NA | NA |
| Olfr1453 | 0 | 0 | 0 | 0 | 0 | 0 | 0 | 0 | 0 | 0 | 0 | 0 | 0 | 0 | 0 | NA | NA | NA | NA | NA | NA | NA | NA | NA | NA |
| Olfr1454 | 0 | 0 | 0 | 0 | 0 | 0 | 0 | 0 | 0 | 0 | 0 | 0 | 0 | 0 | 0 | NA | NA | NA | NA | NA | NA | NA | NA | NA | NA |
| Olfr1457 | 0 | 0 | 0 | 0 | 0 | 0 | 0 | 0 | 0 | 0 | 0 | 0 | 0 | 0 | 0 | NA | NA | NA | NA | NA | NA | NA | NA | NA | NA |
| Olfr1459 | 0 | 0 | 0 | 0 | 0 | 0 | 0 | 0 | 0 | 0 | 0 | 0 | 0 | 0 | 0 | NA | NA | NA | NA | NA | NA | NA | NA | NA | NA |
| Olfr146 | 0 | 0 | 0 | 0 | 0 | 0 | 0 | 0 | 0 | 0 | 0 | 0 | 0 | 0 | 0 | NA | NA | NA | NA | NA | NA | NA | NA | NA | NA |
| Olfr1461 | 0 | 0 | 0 | 0 | 0 | 0 | 0 | 0 | 0 | 0 | 0 | 0 | 0 | 0 | 0 | NA | NA | NA | NA | NA | NA | NA | NA | NA | NA |
| Olfr1462 | 0 | 0 | 0 | 0 | 0 | 0 | 0 | 0 | 0 | 0 | 0 | 0 | 0 | 0 | 0 | NA | NA | NA | NA | NA | NA | NA | NA | NA | NA |
| Olfr1463 | 0 | 0 | 0 | 0 | 0 | 0 | 0 | 0 | 0 | 0 | 0 | 0 | 0 | 0 | 0 | NA | NA | NA | NA | NA | NA | NA | NA | NA | NA |
| Olfr1465 | 0 | 0 | 0 | 0 | 0 | 0 | 0 | 0 | 0 | 0 | 0 | 0 | 0 | 0 | 0 | NA | NA | NA | NA | NA | NA | NA | NA | NA | NA |
| Olfr1466 | 0 | 0 | 0 | 0 | 0 | 0 | 0 | 0 | 0 | 0 | 0 | 0 | 0 | 0 | 0 | NA | NA | NA | NA | NA | NA | NA | NA | NA | NA |
| Olfr1467 | 0 | 0 | 0 | 0 | 0 | 0 | 0 | 0 | 0 | 0 | 0 | 0 | 0 | 0 | 0 | NA | NA | NA | NA | NA | NA | NA | NA | NA | NA |

| | | | | | | | | | | | | | | | | | | | | | | | | | | |
|---|---|---|---|---|---|---|---|---|---|---|---|---|---|---|---|---|---|---|---|---|---|---|---|---|---|---|
| Olfr1469 | 0 | 0 | 0 | 0 | 0 | 0 | 0 | 0 | 0 | 0 | 0 | 0 | 0 | 0 | 0 | NA | NA | NA | NA | NA | NA | NA | NA | NA | NA | |
| Olfr147 | 0 | 3 | 0 | 2 | 0 | 0 | 0 | 3 | 0 | 2 | 0 | 0 | 0.86 | 1 | 1 | 0.645 | -0.633 | 0.80250836 | NA | 10 | 2.7467 | 3.3778 | 3.3778 | TRUE | | 1.4434 |
| Olfr1471 | 0 | 0 | 0 | 0 | 0 | 0 | 0 | 0 | 0 | 0 | 0 | 0 | 0 | 0 | 0 | NA | NA | NA | NA | NA | NA | NA | NA | NA | NA | |
| Olfr1472 | 0 | 0 | 0 | 0 | 0 | 0 | 0 | 0 | 0 | 0 | 0 | 0 | 0 | 0 | 0 | NA | NA | NA | NA | NA | NA | NA | NA | NA | NA | |
| Olfr1474 | 0 | 0 | 0 | 0 | 0 | 0 | 0 | 0 | 0 | 0 | 0 | 0 | 0 | 0 | 0 | NA | NA | NA | NA | NA | NA | NA | NA | NA | NA | |
| Olfr1475 | 0 | 0 | 0 | 0 | 0 | 0 | 0 | 0 | 0 | 0 | 0 | 0 | 0 | 0 | 0 | NA | NA | NA | NA | NA | NA | NA | NA | NA | NA | |
| Olfr1477 | 0 | 0 | 0 | 0 | 0 | 0 | 0 | 0 | 0 | 0 | 0 | 0 | 0 | 0 | 0 | NA | NA | NA | NA | NA | NA | NA | NA | NA | NA | |
| Olfr148 | 0 | 0 | 0 | 0 | 0 | 0 | 0 | 0 | 0 | 0 | 0 | 0 | 0 | 0 | 0 | NA | NA | NA | NA | NA | NA | NA | NA | NA | NA | |
| Olfr1480 | 0 | 0 | 0 | 0 | 0 | 0 | 0 | 0 | 0 | 0 | 0 | 0 | 0 | 0 | 0 | NA | NA | NA | NA | NA | NA | NA | NA | NA | NA | |
| Olfr1484 | 0 | 0 | 0 | 0 | 0 | 0 | 0 | 0 | 0 | 0 | 0 | 0 | 0 | 0 | 0 | NA | NA | NA | NA | NA | NA | NA | NA | NA | NA | |
| Olfr1487 | 0 | 0 | 0 | 0 | 0 | 0 | 0 | 0 | 0 | 0 | 0 | 0 | 0 | 0 | 0 | NA | NA | NA | NA | NA | NA | NA | NA | NA | NA | |
| Olfr1489 | 0 | 0 | 0 | 0 | 0 | 0 | 0 | 0 | 0 | 0 | 0 | 0 | 0 | 0 | 0 | NA | NA | NA | NA | NA | NA | NA | NA | NA | NA | |
| Olfr149 | 0 | 0 | 0 | 0 | 0 | 0 | 0 | 0 | 0 | 0 | 0 | 0 | 0 | 0 | 0 | NA | NA | NA | NA | NA | NA | NA | NA | NA | NA | |
| Olfr1490 | 0 | 0 | 0 | 0 | 0 | 0 | 0 | 0 | 0 | 0 | 0 | 0 | 0 | 0 | 0 | NA | NA | NA | NA | NA | NA | NA | NA | NA | NA | |
| Olfr1491 | 0 | 0 | 0 | 0 | 0 | 0 | 0 | 0 | 0 | 0 | 0 | 0 | 0 | 0 | 0 | NA | NA | NA | NA | NA | NA | NA | NA | NA | NA | |
| Olfr1494 | 0 | 0 | 0 | 0 | 0 | 0 | 0 | 0 | 0 | 0 | 0 | 0 | 0 | 0 | 0 | NA | NA | NA | NA | NA | NA | NA | NA | NA | NA | |
| Olfr1495 | 0 | 0 | 0 | 0 | 0 | 0 | 0 | 0 | 0 | 0 | 0 | 0 | 0 | 0 | 0 | NA | NA | NA | NA | NA | NA | NA | NA | NA | NA | |
| Olfr1496 | 0 | 0 | 0 | 0 | 0 | 0 | 0 | 0 | 0 | 0 | 0 | 0 | 0 | 0 | 0 | NA | NA | NA | NA | NA | NA | NA | NA | NA | NA | |
| Olfr1497 | 0 | 0 | 0 | 0 | 0 | 0 | 0 | 0 | 0 | 0 | 0 | 0 | 0 | 0 | 0 | NA | NA | NA | NA | NA | NA | NA | NA | NA | NA | |
| Olfr1499 | 0 | 0 | 0 | 0 | 0 | 0 | 0 | 0 | 0 | 0 | 0 | 0 | 0 | 0 | 0 | NA | NA | NA | NA | NA | NA | NA | NA | NA | NA | |
| Olfr15 | 0 | 0 | 0 | 0 | 0 | 0 | 0 | 0 | 0 | 0 | 0 | 0 | 0 | 0 | 0 | NA | NA | NA | NA | NA | NA | NA | NA | NA | NA | |
| Olfr150 | 0 | 0 | 0 | 0 | 0 | 0 | 0 | 0 | 0 | 0 | 0 | 0 | 0 | 0 | 0 | NA | NA | NA | NA | NA | NA | NA | NA | NA | NA | |
| Olfr1500 | 0 | 0 | 0 | 0 | 0 | 0 | 0 | 0 | 0 | 0 | 0 | 0 | 0 | 0 | 0 | NA | NA | NA | NA | NA | NA | NA | NA | NA | NA | |
| Olfr1501 | 0 | 0 | 0 | 0 | 0 | 0 | 0 | 0 | 0 | 0 | 0 | 0 | 0 | 0 | 0 | NA | NA | NA | NA | NA | NA | NA | NA | NA | NA | |
| Olfr1502 | 0 | 0 | 0 | 0 | 0 | 0 | 0 | 0 | 0 | 0 | 0 | 0 | 0 | 0 | 0 | NA | NA | NA | NA | NA | NA | NA | NA | NA | NA | |
| Olfr1504 | 0 | 0 | 0 | 0 | 0 | 0 | 0 | 0 | 0 | 0 | 0 | 0 | 0 | 0 | 0 | NA | NA | NA | NA | NA | NA | NA | NA | NA | NA | |
| Olfr1505 | 0 | 0 | 0 | 0 | 0 | 0 | 0 | 0 | 0 | 0 | 0 | 0 | 0 | 0 | 0 | NA | NA | NA | NA | NA | NA | NA | NA | NA | NA | |
| Olfr1506 | 0 | 0 | 0 | 0 | 0 | 0 | 0 | 0 | 0 | 0 | 0 | 0 | 0 | 0 | 0 | NA | NA | NA | NA | NA | NA | NA | NA | NA | NA | |
| Olfr1507 | 0 | 0 | 0 | 0 | 0 | 0 | 0 | 0 | 0 | 0 | 0 | 0 | 0 | 0 | 0 | NA | NA | NA | NA | NA | NA | NA | NA | NA | NA | |
| Olfr1508 | 0 | 0 | 0 | 0 | 0 | 0 | 0 | 0 | 0 | 0 | 0 | 0 | 0 | 0 | 0 | NA | NA | NA | NA | NA | NA | NA | NA | NA | NA | |
| Olfr1509 | 0 | 0 | 0 | 0 | 0 | 0 | 0 | 0 | 0 | 0 | 0 | 0 | 0 | 0 | 0 | NA | NA | NA | NA | NA | NA | NA | NA | NA | NA | |
| Olfr151 | 0 | 0 | 0 | 0 | 0 | 0 | 0 | 0 | 0 | 0 | 0 | 0 | 0 | 0 | 0 | NA | NA | NA | NA | NA | NA | NA | NA | NA | NA | |
| Olfr1510 | 0 | 0 | 0 | 0 | 0 | 0 | 0 | 0 | 0 | 0 | 0 | 0 | 0 | 0 | 0 | NA | NA | NA | NA | NA | NA | NA | NA | NA | NA | |
| Olfr1511 | 0 | 0 | 0 | 0 | 0 | 0 | 0 | 0 | 0 | 0 | 0 | 0 | 0 | 0 | 0 | NA | NA | NA | NA | NA | NA | NA | NA | NA | NA | |
| Olfr1512 | 0 | 0 | 0 | 0 | 0 | 0 | 0 | 0 | 0 | 0 | 0 | 0 | 0 | 0 | 0 | NA | NA | NA | NA | NA | NA | NA | NA | NA | NA | |
| Olfr1513 | 0 | 0 | 0 | 0 | 0 | 0 | 0 | 0 | 0 | 0 | 0 | 0 | 0 | 0 | 0 | NA | NA | NA | NA | NA | NA | NA | NA | NA | NA | |
| Olfr152 | 0 | 0 | 0 | 0 | 0 | 0 | 0 | 0 | 0 | 0 | 0 | 0 | 0 | 0 | 0 | NA | NA | NA | NA | NA | NA | NA | NA | NA | NA | |
| Olfr153 | 9 | 5 | 4 | 6 | 6 | 2 | 7 | 5 | 4 | 6 | 7 | 2 | 5.22 | 5 | 5 | 0.871 | -0.2 | 0.81405367 | NA | | 0 | 0.4562 | 0.3275 | 0.3275 | TRUE | 0.5673 |
| Olfr1532-ps1 | 0 | 0 | 0 | 0 | 0 | 0 | 0 | 0 | 0 | 0 | 0 | 0 | 0 | 0 | 0 | NA | NA | NA | NA | NA | NA | NA | NA | NA | NA | |
| Olfr1535 | 0 | 0 | 0 | 0 | 0 | 0 | 0 | 0 | 0 | 0 | 0 | 0 | 0 | 0 | 0 | NA | NA | NA | NA | NA | NA | NA | NA | NA | NA | |
| Olfr1537 | 0 | 0 | 0 | 0 | 0 | 0 | 0 | 0 | 0 | 0 | 0 | 0 | 0 | 0 | 0 | NA | NA | NA | NA | NA | NA | NA | NA | NA | NA | |
| Olfr154 | 0 | 0 | 0 | 0 | 0 | 0 | 0 | 0 | 0 | 0 | 0 | 0 | 0 | 0 | 0 | NA | NA | NA | NA | NA | NA | NA | NA | NA | NA | |
| Olfr155 | 0 | 0 | 0 | 0 | 0 | 0 | 0 | 0 | 0 | 0 | 0 | 0 | 0 | 0 | 0 | NA | NA | NA | NA | NA | NA | NA | NA | NA | NA | |
| Olfr156 | 0 | 0 | 0 | 0 | 0 | 0 | 0 | 0 | 0 | 0 | 0 | 0 | 0 | 0 | 0 | NA | NA | NA | NA | NA | NA | NA | NA | NA | NA | |
| Olfr157 | 1 | 1 | 0 | 0 | 1 | 0 | 1 | 1 | 0 | 0 | 1 | 0 | 0.49 | 1 | 0 | 0.599 | -0.74 | 0.78905321 | NA | | 0 | 4.7929 | 3.7323 | 3.7323 | TRUE | 0.5075 |
| Olfr159 | 0 | 0 | 0 | 0 | 0 | 0 | 0 | 0 | 0 | 0 | 0 | 0 | 0 | 0 | 0 | NA | NA | NA | NA | NA | NA | NA | NA | NA | NA | |
| Olfr16 | 0 | 0 | 0 | 0 | 0 | 0 | 0 | 0 | 0 | 0 | 0 | 0 | 0 | 0 | 0 | NA | NA | NA | NA | NA | NA | NA | NA | NA | NA | |
| Olfr160 | 0 | 0 | 0 | 0 | 0 | 1 | 0 | 0 | 0 | 0 | 0 | 1 | 0.16 | 0 | 0 | 2.094 | 1.067 | 0.79379329 | NA | | 0 | 14.4078 | 10 | 10 | TRUE | 0.3835 |
| Olfr161 | 0 | 0 | 2 | 2 | 2 | 1 | 0 | 0 | 2 | 2 | 2 | 1 | 1.23 | 1 | 2 | 2.556 | 1.354 | 0.47019769 | NA | | 0 | 1.9261 | 1.5276 | 1.5276 | TRUE | 0.9754 |
| Olfr164 | 0 | 0 | 0 | 0 | 0 | 0 | 0 | 0 | 0 | 0 | 0 | 0 | 0 | 0 | 0 | NA | NA | NA | NA | NA | NA | NA | NA | NA | NA | |
| Olfr165 | 0 | 0 | 0 | 0 | 0 | 0 | 0 | 0 | 0 | 0 | 0 | 0 | 0 | 0 | 0 | NA | NA | NA | NA | NA | NA | NA | NA | NA | NA | |
| Olfr166 | 0 | 0 | 0 | 0 | 0 | 0 | 0 | 0 | 0 | 0 | 0 | 0 | 0 | 0 | 0 | NA | NA | NA | NA | NA | NA | NA | NA | NA | NA | |
| Olfr167 | 0 | 0 | 0 | 0 | 0 | 0 | 0 | 0 | 0 | 0 | 0 | 0 | 0 | 0 | 0 | NA | NA | NA | NA | NA | NA | NA | NA | NA | NA | |
| Olfr168 | 0 | 0 | 0 | 0 | 0 | 0 | 0 | 0 | 0 | 0 | 0 | 0 | 0 | 0 | 0 | NA | NA | NA | NA | NA | NA | NA | NA | NA | NA | |
| Olfr169 | 0 | 0 | 0 | 0 | 0 | 0 | 0 | 0 | 0 | 0 | 0 | 0 | 0 | 0 | 0 | NA | NA | NA | NA | NA | NA | NA | NA | NA | NA | |
| Olfr17 | 0 | 0 | 0 | 0 | 0 | 0 | 0 | 0 | 0 | 0 | 0 | 0 | 0 | 0 | 0 | NA | NA | NA | NA | NA | NA | NA | NA | NA | NA | |
| Olfr170 | 0 | 0 | 0 | 0 | 0 | 0 | 0 | 0 | 0 | 0 | 0 | 0 | 0 | 0 | 0 | NA | NA | NA | NA | NA | NA | NA | NA | NA | NA | |
| Olfr171 | 0 | 0 | 0 | 0 | 0 | 0 | 0 | 0 | 0 | 0 | 0 | 0 | 0 | 0 | 0 | NA | NA | NA | NA | NA | NA | NA | NA | NA | NA | |
| Olfr172 | 0 | 0 | 0 | 0 | 0 | 0 | 0 | 0 | 0 | 0 | 0 | 0 | 0 | 0 | 0 | NA | NA | NA | NA | NA | NA | NA | NA | NA | NA | |
| Olfr173 | 0 | 0 | 0 | 0 | 0 | 0 | 0 | 0 | 0 | 0 | 0 | 0 | 0 | 0 | 0 | NA | NA | NA | NA | NA | NA | NA | NA | NA | NA | |
| Olfr175-ps1 | 0 | 0 | 0 | 0 | 0 | 0 | 0 | 0 | 0 | 0 | 0 | 0 | 0 | 0 | 0 | NA | NA | NA | NA | NA | NA | NA | NA | NA | NA | |
| Olfr176 | 0 | 0 | 0 | 0 | 0 | 0 | 0 | 0 | 0 | 0 | 0 | 0 | 0 | 0 | 0 | NA | NA | NA | NA | NA | NA | NA | NA | NA | NA | |
| Olfr177 | 0 | 0 | 0 | 0 | 0 | 0 | 0 | 0 | 0 | 0 | 0 | 0 | 0 | 0 | 0 | NA | NA | NA | NA | NA | NA | NA | NA | NA | NA | |
| Olfr178 | 0 | 0 | 0 | 0 | 0 | 0 | 0 | 0 | 0 | 0 | 0 | 0 | 0 | 0 | 0 | NA | NA | NA | NA | NA | NA | NA | NA | NA | NA | |
| Olfr179 | 1 | 0 | 0 | 0 | 0 | 0 | 1 | 0 | 0 | 0 | 0 | 0 | 0.13 | 0 | 0 | 0.552 | -0.857 | 0.8336548 | NA | | 0 | 18.3121 | 10 | 10 | TRUE | 0.2566 |
| Olfr18 | 3 | 0 | 0 | 1 | 0 | 0 | 2 | 0 | 0 | 1 | 0 | 0 | 0.55 | 1 | 0 | 0.458 | -1.128 | 0.69820437 | NA | | 0 | 4.2787 | 4.502 | 4.502 | TRUE | 1.4399 |
| Olfr180 | 0 | 0 | 0 | 0 | 0 | 0 | 0 | 0 | 0 | 0 | 0 | 0 | 0 | 0 | 0 | NA | NA | NA | NA | NA | NA | NA | NA | NA | NA | |
| Olfr181 | 0 | 0 | 0 | 0 | 0 | 0 | 0 | 0 | 0 | 0 | 0 | 0 | 0 | 0 | 0 | NA | NA | NA | NA | NA | NA | NA | NA | NA | NA | |
| Olfr183 | 0 | 0 | 0 | 0 | 0 | 0 | 0 | 0 | 0 | 0 | 0 | 0 | 0 | 0 | 0 | NA | NA | NA | NA | NA | NA | NA | NA | NA | NA | |

| | | | | | | | | | | | | | | | | | | | | | | | | | |
|---|---|---|---|---|---|---|---|---|---|---|---|---|---|---|---|---|---|---|---|---|---|---|---|---|---|
| Olfr186 | 0 | 0 | 0 | 0 | 0 | 0 | 0 | 0 | 0 | 0 | 0 | 0 | 0 | 0 | 0 | NA | NA | NA | NA | NA | NA | NA | NA | NA | NA |
| Olfr187 | 0 | 0 | 0 | 0 | 0 | 0 | 0 | 0 | 0 | 0 | 0 | 0 | 0 | 0 | 0 | NA | NA | NA | NA | NA | NA | NA | NA | NA | NA |
| Olfr19 | 0 | 0 | 0 | 0 | 0 | 0 | 0 | 0 | 0 | 0 | 0 | 0 | 0 | 0 | 0 | NA | NA | NA | NA | NA | NA | NA | NA | NA | NA |
| Olfr190 | 0 | 0 | 0 | 0 | 0 | 0 | 0 | 0 | 0 | 0 | 0 | 0 | 0 | 0 | 0 | NA | NA | NA | NA | NA | NA | NA | NA | NA | NA |
| Olfr191 | 0 | 0 | 0 | 0 | 0 | 0 | 0 | 0 | 0 | 0 | 0 | 0 | 0 | 0 | 0 | NA | NA | NA | NA | NA | NA | NA | NA | NA | NA |
| Olfr192 | 0 | 0 | 0 | 0 | 0 | 0 | 0 | 0 | 0 | 0 | 0 | 0 | 0 | 0 | 0 | NA | NA | NA | NA | NA | NA | NA | NA | NA | NA |
| Olfr193 | 0 | 0 | 0 | 0 | 0 | 0 | 0 | 0 | 0 | 0 | 0 | 0 | 0 | 0 | 0 | NA | NA | NA | NA | NA | NA | NA | NA | NA | NA |
| Olfr194 | 0 | 0 | 0 | 0 | 0 | 0 | 0 | 0 | 0 | 0 | 0 | 0 | 0 | 0 | 0 | NA | NA | NA | NA | NA | NA | NA | NA | NA | NA |
| Olfr195 | 0 | 0 | 0 | 0 | 0 | 0 | 0 | 0 | 0 | 0 | 0 | 0 | 0 | 0 | 0 | NA | NA | NA | NA | NA | NA | NA | NA | NA | NA |
| Olfr196 | 0 | 0 | 0 | 0 | 0 | 0 | 0 | 0 | 0 | 0 | 0 | 0 | 0 | 0 | 0 | NA | NA | NA | NA | NA | NA | NA | NA | NA | NA |
| Olfr197 | 0 | 0 | 0 | 0 | 0 | 0 | 0 | 0 | 0 | 0 | 0 | 0 | 0 | 0 | 0 | NA | NA | NA | NA | NA | NA | NA | NA | NA | NA |
| Olfr198 | 0 | 0 | 0 | 0 | 0 | 0 | 0 | 0 | 0 | 0 | 0 | 0 | 0 | 0 | 0 | NA | NA | NA | NA | NA | NA | NA | NA | NA | NA |
| Olfr199 | 0 | 0 | 0 | 0 | 0 | 0 | 0 | 0 | 0 | 0 | 0 | 0 | 0 | 0 | 0 | NA | NA | NA | NA | NA | NA | NA | NA | NA | NA |
| Olfr2 | 0 | 0 | 0 | 0 | 0 | 0 | 0 | 0 | 0 | 0 | 0 | 0 | 0 | 0 | 0 | NA | NA | NA | NA | NA | NA | NA | NA | NA | NA |
| Olfr20 | 0 | 1 | 0 | 0 | 0 | 0 | 0 | 1 | 0 | 0 | 0 | 0 | 0.18 | 0 | 0 | 0.552 | -0.857 | 0.8336548 | NA | | 0 | 13.3714 | 10 | 10 | TRUE | 0.5216 |
| Olfr201 | 0 | 0 | 0 | 0 | 0 | 0 | 0 | 0 | 0 | 0 | 0 | 0 | 0 | 0 | 0 | NA | NA | NA | NA | NA | NA | NA | NA | NA | NA |
| Olfr202 | 0 | 0 | 0 | 0 | 0 | 0 | 0 | 0 | 0 | 0 | 0 | 0 | 0 | 0 | 0 | NA | NA | NA | NA | NA | NA | NA | NA | NA | NA |
| Olfr203 | 0 | 0 | 0 | 0 | 0 | 0 | 0 | 0 | 0 | 0 | 0 | 0 | 0 | 0 | 0 | NA | NA | NA | NA | NA | NA | NA | NA | NA | NA |
| Olfr204 | 0 | 0 | 0 | 0 | 0 | 0 | 0 | 0 | 0 | 0 | 0 | 0 | 0 | 0 | 0 | NA | NA | NA | NA | NA | NA | NA | NA | NA | NA |
| Olfr205 | 0 | 0 | 0 | 0 | 0 | 0 | 0 | 0 | 0 | 0 | 0 | 0 | 0 | 0 | 0 | NA | NA | NA | NA | NA | NA | NA | NA | NA | NA |
| Olfr206 | 0 | 0 | 0 | 0 | 0 | 0 | 0 | 0 | 0 | 0 | 0 | 0 | 0 | 0 | 0 | NA | NA | NA | NA | NA | NA | NA | NA | NA | NA |
| Olfr207 | 0 | 0 | 0 | 0 | 0 | 0 | 0 | 0 | 0 | 0 | 0 | 0 | 0 | 0 | 0 | NA | NA | NA | NA | NA | NA | NA | NA | NA | NA |
| Olfr208 | 0 | 0 | 0 | 0 | 0 | 0 | 0 | 0 | 0 | 0 | 0 | 0 | 0 | 0 | 0 | NA | NA | NA | NA | NA | NA | NA | NA | NA | NA |
| Olfr209 | 0 | 0 | 0 | 0 | 0 | 0 | 0 | 0 | 0 | 0 | 0 | 0 | 0 | 0 | 0 | NA | NA | NA | NA | NA | NA | NA | NA | NA | NA |
| Olfr211 | 0 | 0 | 0 | 0 | 0 | 0 | 0 | 0 | 0 | 0 | 0 | 0 | 0 | 0 | 0 | NA | NA | NA | NA | NA | NA | NA | NA | NA | NA |
| Olfr212 | 0 | 0 | 0 | 0 | 0 | 0 | 0 | 0 | 0 | 0 | 0 | 0 | 0 | 0 | 0 | NA | NA | NA | NA | NA | NA | NA | NA | NA | NA |
| Olfr213 | 0 | 0 | 0 | 0 | 0 | 0 | 0 | 0 | 0 | 0 | 0 | 0 | 0 | 0 | 0 | NA | NA | NA | NA | NA | NA | NA | NA | NA | NA |
| Olfr214 | 0 | 0 | 0 | 0 | 0 | 0 | 0 | 0 | 0 | 0 | 0 | 0 | 0 | 0 | 0 | NA | NA | NA | NA | NA | NA | NA | NA | NA | NA |
| Olfr215 | 0 | 0 | 0 | 0 | 0 | 0 | 0 | 0 | 0 | 0 | 0 | 0 | 0 | 0 | 0 | NA | NA | NA | NA | NA | NA | NA | NA | NA | NA |
| Olfr218 | 0 | 0 | 0 | 0 | 0 | 0 | 0 | 0 | 0 | 0 | 0 | 0 | 0 | 0 | 0 | NA | NA | NA | NA | NA | NA | NA | NA | NA | NA |
| Olfr220 | 0 | 0 | 0 | 0 | 0 | 0 | 0 | 0 | 0 | 0 | 0 | 0 | 0 | 0 | 0 | NA | NA | NA | NA | NA | NA | NA | NA | NA | NA |
| Olfr221 | 0 | 0 | 0 | 0 | 0 | 0 | 0 | 0 | 0 | 0 | 0 | 0 | 0 | 0 | 0 | NA | NA | NA | NA | NA | NA | NA | NA | NA | NA |
| Olfr222 | 0 | 0 | 0 | 0 | 0 | 0 | 0 | 0 | 0 | 0 | 0 | 0 | 0 | 0 | 0 | NA | NA | NA | NA | NA | NA | NA | NA | NA | NA |
| Olfr223 | 0 | 0 | 0 | 0 | 0 | 0 | 0 | 0 | 0 | 0 | 0 | 0 | 0 | 0 | 0 | NA | NA | NA | NA | NA | NA | NA | NA | NA | NA |
| Olfr224 | 0 | 0 | 0 | 0 | 0 | 0 | 0 | 0 | 0 | 0 | 0 | 0 | 0 | 0 | 0 | NA | NA | NA | NA | NA | NA | NA | NA | NA | NA |
| Olfr225 | 0 | 0 | 0 | 0 | 0 | 0 | 0 | 0 | 0 | 0 | 0 | 0 | 0 | 0 | 0 | NA | NA | NA | NA | NA | NA | NA | NA | NA | NA |
| Olfr228 | 0 | 0 | 0 | 0 | 0 | 0 | 0 | 0 | 0 | 0 | 0 | 0 | 0 | 0 | 0 | NA | NA | NA | NA | NA | NA | NA | NA | NA | NA |
| Olfr229 | 0 | 0 | 0 | 0 | 0 | 0 | 0 | 0 | 0 | 0 | 0 | 0 | 0 | 0 | 0 | NA | NA | NA | NA | NA | NA | NA | NA | NA | NA |
| Olfr23 | 0 | 0 | 0 | 0 | 0 | 0 | 0 | 0 | 0 | 0 | 0 | 0 | 0 | 0 | 0 | NA | NA | NA | NA | NA | NA | NA | NA | NA | NA |
| Olfr231 | 0 | 0 | 0 | 0 | 0 | 0 | 0 | 0 | 0 | 0 | 0 | 0 | 0 | 0 | 0 | NA | NA | NA | NA | NA | NA | NA | NA | NA | NA |
| Olfr235 | 0 | 0 | 0 | 0 | 0 | 0 | 0 | 0 | 0 | 0 | 0 | 0 | 0 | 0 | 0 | NA | NA | NA | NA | NA | NA | NA | NA | NA | NA |
| Olfr237-ps1 | 0 | 0 | 0 | 0 | 0 | 0 | 0 | 0 | 0 | 0 | 0 | 0 | 0 | 0 | 0 | NA | NA | NA | NA | NA | NA | NA | NA | NA | NA |
| Olfr239 | 0 | 0 | 0 | 0 | 0 | 0 | 0 | 0 | 0 | 0 | 0 | 0 | 0 | 0 | 0 | NA | NA | NA | NA | NA | NA | NA | NA | NA | NA |
| Olfr24 | 0 | 0 | 1 | 1 | 0 | 0 | 0 | 0 | 1 | 1 | 0 | 0 | 0.35 | 0 | 0 | 1.075 | 0.105 | 0.97567165 | NA | | 0 | 6.8576 | 6.5143 | 6.5143 | TRUE | 0.5365 |
| Olfr242 | 0 | 0 | 0 | 0 | 0 | 0 | 0 | 0 | 0 | 0 | 0 | 0 | 0 | 0 | 0 | NA | NA | NA | NA | NA | NA | NA | NA | NA | NA |
| Olfr243 | 0 | 0 | 0 | 0 | 0 | 0 | 0 | 0 | 0 | 0 | 0 | 0 | 0 | 0 | 0 | NA | NA | NA | NA | NA | NA | NA | NA | NA | NA |
| Olfr247 | 0 | 0 | 0 | 0 | 0 | 0 | 0 | 0 | 0 | 0 | 0 | 0 | 0 | 0 | 0 | NA | NA | NA | NA | NA | NA | NA | NA | NA | NA |
| Olfr248 | 0 | 0 | 0 | 0 | 0 | 0 | 0 | 0 | 0 | 0 | 0 | 0 | 0 | 0 | 0 | NA | NA | NA | NA | NA | NA | NA | NA | NA | NA |
| Olfr25 | 0 | 0 | 0 | 0 | 0 | 0 | 0 | 0 | 0 | 0 | 0 | 0 | 0 | 0 | 0 | NA | NA | NA | NA | NA | NA | NA | NA | NA | NA |
| Olfr250 | 0 | 0 | 0 | 0 | 0 | 0 | 0 | 0 | 0 | 0 | 0 | 0 | 0 | 0 | 0 | NA | NA | NA | NA | NA | NA | NA | NA | NA | NA |
| Olfr251 | 0 | 0 | 0 | 0 | 0 | 0 | 0 | 0 | 0 | 0 | 0 | 0 | 0 | 0 | 0 | NA | NA | NA | NA | NA | NA | NA | NA | NA | NA |
| Olfr257 | 0 | 0 | 0 | 0 | 0 | 0 | 0 | 0 | 0 | 0 | 0 | 0 | 0 | 0 | 0 | NA | NA | NA | NA | NA | NA | NA | NA | NA | NA |
| Olfr259 | 0 | 0 | 0 | 0 | 0 | 0 | 0 | 0 | 0 | 0 | 0 | 0 | 0 | 0 | 0 | NA | NA | NA | NA | NA | NA | NA | NA | NA | NA |
| Olfr26 | 0 | 1 | 0 | 0 | 0 | 0 | 0 | 1 | 0 | 0 | 0 | 0 | 0.18 | 0 | 0 | 0.552 | -0.857 | 0.8336548 | NA | | 0 | 13.3714 | 10 | 10 | TRUE | 0.5216 |
| Olfr262 | 0 | 0 | 0 | 0 | 0 | 0 | 0 | 0 | 0 | 0 | 0 | 0 | 0 | 0 | 0 | NA | NA | NA | NA | NA | NA | NA | NA | NA | NA |
| Olfr263 | 0 | 0 | 0 | 0 | 0 | 0 | 0 | 0 | 0 | 0 | 0 | 0 | 0 | 0 | 0 | NA | NA | NA | NA | NA | NA | NA | NA | NA | NA |
| Olfr266 | 0 | 0 | 0 | 0 | 0 | 0 | 0 | 0 | 0 | 0 | 0 | 0 | 0 | 0 | 0 | NA | NA | NA | NA | NA | NA | NA | NA | NA | NA |
| Olfr267 | 0 | 0 | 0 | 0 | 0 | 0 | 0 | 0 | 0 | 0 | 0 | 0 | 0 | 0 | 0 | NA | NA | NA | NA | NA | NA | NA | NA | NA | NA |
| Olfr27 | 0 | 0 | 0 | 0 | 0 | 0 | 0 | 0 | 0 | 0 | 0 | 0 | 0 | 0 | 0 | NA | NA | NA | NA | NA | NA | NA | NA | NA | NA |
| Olfr270 | 0 | 0 | 0 | 0 | 0 | 0 | 0 | 0 | 0 | 0 | 0 | 0 | 0 | 0 | 0 | NA | NA | NA | NA | NA | NA | NA | NA | NA | NA |
| Olfr272 | 0 | 0 | 0 | 0 | 0 | 0 | 0 | 0 | 0 | 0 | 0 | 0 | 0 | 0 | 0 | NA | NA | NA | NA | NA | NA | NA | NA | NA | NA |
| Olfr273 | 0 | 0 | 0 | 0 | 0 | 0 | 0 | 0 | 0 | 0 | 0 | 0 | 0 | 0 | 0 | NA | NA | NA | NA | NA | NA | NA | NA | NA | NA |
| Olfr275 | 0 | 0 | 0 | 0 | 0 | 0 | 0 | 0 | 0 | 0 | 0 | 0 | 0 | 0 | 0 | NA | NA | NA | NA | NA | NA | NA | NA | NA | NA |
| Olfr279 | 0 | 0 | 0 | 0 | 0 | 0 | 0 | 0 | 0 | 0 | 0 | 0 | 0 | 0 | 0 | NA | NA | NA | NA | NA | NA | NA | NA | NA | NA |
| Olfr281 | 0 | 0 | 0 | 0 | 0 | 0 | 0 | 0 | 0 | 0 | 0 | 0 | 0 | 0 | 0 | NA | NA | NA | NA | NA | NA | NA | NA | NA | NA |
| Olfr282 | 0 | 0 | 0 | 0 | 0 | 0 | 0 | 0 | 0 | 0 | 0 | 0 | 0 | 0 | 0 | NA | NA | NA | NA | NA | NA | NA | NA | NA | NA |
| Olfr283 | 0 | 0 | 0 | 0 | 0 | 0 | 0 | 0 | 0 | 0 | 0 | 0 | 0 | 0 | 0 | NA | NA | NA | NA | NA | NA | NA | NA | NA | NA |
| Olfr284 | 0 | 0 | 0 | 0 | 0 | 0 | 0 | 0 | 0 | 0 | 0 | 0 | 0 | 0 | 0 | NA | NA | NA | NA | NA | NA | NA | NA | NA | NA |
| Olfr285 | 0 | 0 | 0 | 0 | 0 | 0 | 0 | 0 | 0 | 0 | 0 | 0 | 0 | 0 | 0 | NA | NA | NA | NA | NA | NA | NA | NA | NA | NA |

| | | | | | | | | | | | | | | | | | | | | | | | | | |
|---|---|---|---|---|---|---|---|---|---|---|---|---|---|---|---|---|---|---|---|---|---|---|---|---|---|
| Olfr286 | 0 | 0 | 1 | 0 | 1 | 0 | 0 | 0 | 1 | 0 | 1 | 0 | 0.37 | 0 | 0 | 1.075 | 0.105 | 0.97514588 | NA | 0 | 6.4454 | 6.1578 | 6.1578 | TRUE | 0.5365 |
| Olfr287 | 5 | 2 | 7 | 5 | 2 | 1 | 4 | 2 | 8 | 5 | 2 | 1 | 3.63 | 5 | 3 | 0.611 | -0.711 | 0.5022473 | NA | 0.1456 | 0.6552 | 0.5144 | 0.5144 | TRUE | 0.453 |
| Olfr288 | 0 | 0 | 1 | 1 | 0 | 0 | 0 | 0 | 1 | 1 | 0 | 0 | 0.35 | 0 | 0 | 1.075 | 0.105 | 0.97567165 | NA | 0 | 6.8576 | 6.5143 | 6.5143 | TRUE | 0.5365 |
| Olfr29-ps1 | 0 | 0 | 0 | 0 | 1 | 0 | 0 | 0 | 0 | 0 | 1 | 0 | 0.19 | 0 | 0 | 2.094 | 1.067 | 0.79379329 | NA | 0 | 12.608 | 10 | 10 | TRUE | 0.5075 |
| Olfr290 | 0 | 0 | 0 | 0 | 0 | 0 | 0 | 0 | 0 | 0 | 0 | 0 | 0 | 0 | 0 | NA | NA | NA | NA | NA | NA | NA | NA | NA | NA |
| Olfr291 | 0 | 0 | 0 | 0 | 0 | 0 | 0 | 0 | 0 | 0 | 0 | 0 | 0 | 0 | 0 | NA | NA | NA | NA | NA | NA | NA | NA | NA | NA |
| Olfr292 | 0 | 0 | 0 | 0 | 0 | 0 | 0 | 0 | 0 | 0 | 0 | 0 | 0 | 0 | 0 | NA | NA | NA | NA | NA | NA | NA | NA | NA | NA |
| Olfr293 | 2 | 0 | 0 | 0 | 0 | 2 | 2 | 0 | 0 | 0 | 0 | 2 | 0.59 | 1 | 1 | 1.213 | 0.279 | 0.92253239 | NA | 0 | 4.0336 | 4.2602 | 4.2602 | TRUE | 0.9737 |
| Olfr294 | 0 | 0 | 0 | 0 | 0 | 0 | 0 | 0 | 0 | 0 | 0 | 0 | 0 | 0 | 0 | NA | NA | NA | NA | NA | NA | NA | NA | NA | NA |
| Olfr295 | 0 | 0 | 0 | 0 | 0 | 0 | 0 | 0 | 0 | 0 | 0 | 0 | 0 | 0 | 0 | NA | NA | NA | NA | NA | NA | NA | NA | NA | NA |
| Olfr297 | 0 | 0 | 0 | 0 | 0 | 0 | 0 | 0 | 0 | 0 | 0 | 0 | 0 | 0 | 0 | NA | NA | NA | NA | NA | NA | NA | NA | NA | NA |
| Olfr298 | 0 | 0 | 0 | 0 | 0 | 0 | 0 | 0 | 0 | 0 | 0 | 0 | 0 | 0 | 0 | NA | NA | NA | NA | NA | NA | NA | NA | NA | NA |
| Olfr299 | 0 | 0 | 0 | 0 | 0 | 0 | 0 | 0 | 0 | 0 | 0 | 0 | 0 | 0 | 0 | NA | NA | NA | NA | NA | NA | NA | NA | NA | NA |
| Olfr3 | 0 | 0 | 0 | 0 | 0 | 0 | 0 | 0 | 0 | 0 | 0 | 0 | 0 | 0 | 0 | NA | NA | NA | NA | NA | NA | NA | NA | NA | NA |
| Olfr30 | 0 | 0 | 0 | 0 | 0 | 0 | 0 | 0 | 0 | 0 | 0 | 0 | 0 | 0 | 0 | NA | NA | NA | NA | NA | NA | NA | NA | NA | NA |
| Olfr301 | 0 | 0 | 0 | 0 | 0 | 0 | 0 | 0 | 0 | 0 | 0 | 0 | 0 | 0 | 0 | NA | NA | NA | NA | NA | NA | NA | NA | NA | NA |
| Olfr303 | 0 | 0 | 0 | 0 | 0 | 0 | 0 | 0 | 0 | 0 | 0 | 0 | 0 | 0 | 0 | NA | NA | NA | NA | NA | NA | NA | NA | NA | NA |
| Olfr304 | 0 | 0 | 0 | 0 | 0 | 0 | 0 | 0 | 0 | 0 | 0 | 0 | 0 | 0 | 0 | NA | NA | NA | NA | NA | NA | NA | NA | NA | NA |
| Olfr305 | 0 | 0 | 0 | 0 | 0 | 0 | 0 | 0 | 0 | 0 | 0 | 0 | 0 | 0 | 0 | NA | NA | NA | NA | NA | NA | NA | NA | NA | NA |
| Olfr307 | 3 | 6 | 6 | 2 | 2 | 7 | 2 | 6 | 6 | 2 | 2 | 7 | 4.39 | 5 | 4 | 0.753 | -0.409 | 0.67091856 | NA | 0.1659 | 0.5422 | 0.4312 | 0.4312 | TRUE | 0.9143 |
| Olfr308 | 0 | 0 | 0 | 0 | 0 | 0 | 0 | 0 | 0 | 0 | 0 | 0 | 0 | 0 | 0 | NA | NA | NA | NA | NA | NA | NA | NA | NA | NA |
| Olfr309 | 0 | 0 | 0 | 0 | 0 | 0 | 0 | 0 | 0 | 0 | 0 | 0 | 0 | 0 | 0 | NA | NA | NA | NA | NA | NA | NA | NA | NA | NA |
| Olfr31 | 1 | 0 | 0 | 0 | 1 | 0 | 1 | 0 | 0 | 0 | 1 | 0 | 0.32 | 0 | 0 | 1.075 | 0.105 | 0.97638847 | NA | 0 | 7.4681 | 7.0392 | 7.0392 | TRUE | 0.5075 |
| Olfr310 | 0 | 0 | 0 | 0 | 0 | 0 | 0 | 0 | 0 | 0 | 0 | 0 | 0 | 0 | 0 | NA | NA | NA | NA | NA | NA | NA | NA | NA | NA |
| Olfr311 | 0 | 0 | 0 | 0 | 0 | 0 | 0 | 0 | 0 | 0 | 0 | 0 | 0 | 0 | 0 | NA | NA | NA | NA | NA | NA | NA | NA | NA | NA |
| Olfr312 | 0 | 0 | 0 | 0 | 0 | 0 | 0 | 0 | 0 | 0 | 0 | 0 | 0 | 0 | 0 | NA | NA | NA | NA | NA | NA | NA | NA | NA | NA |
| Olfr313 | 0 | 0 | 0 | 0 | 0 | 0 | 0 | 0 | 0 | 0 | 0 | 0 | 0 | 0 | 0 | NA | NA | NA | NA | NA | NA | NA | NA | NA | NA |
| Olfr314 | 0 | 0 | 0 | 0 | 0 | 0 | 0 | 0 | 0 | 0 | 0 | 0 | 0 | 0 | 0 | NA | NA | NA | NA | NA | NA | NA | NA | NA | NA |
| Olfr315 | 0 | 0 | 0 | 0 | 0 | 0 | 0 | 0 | 0 | 0 | 0 | 0 | 0 | 0 | 0 | NA | NA | NA | NA | NA | NA | NA | NA | NA | NA |
| Olfr316 | 0 | 2 | 1 | 1 | 0 | 0 | 0 | 2 | 1 | 1 | 0 | 0 | 0.7 | 1 | 0 | 0.356 | -1.492 | 0.54563002 | NA | 0 | 3.3868 | 2.9267 | 2.9267 | TRUE | 0.1826 |
| Olfr317 | 0 | 0 | 0 | 0 | 0 | 0 | 0 | 0 | 0 | 0 | 0 | 0 | 0 | 0 | 0 | NA | NA | NA | NA | NA | NA | NA | NA | NA | NA |
| Olfr318 | 0 | 0 | 0 | 0 | 0 | 0 | 0 | 0 | 0 | 0 | 0 | 0 | 0 | 0 | 0 | NA | NA | NA | NA | NA | NA | NA | NA | NA | NA |
| Olfr319 | 0 | 0 | 0 | 0 | 0 | 0 | 0 | 0 | 0 | 0 | 0 | 0 | 0 | 0 | 0 | NA | NA | NA | NA | NA | NA | NA | NA | NA | NA |
| Olfr32 | 0 | 0 | 0 | 0 | 0 | 0 | 0 | 0 | 0 | 0 | 0 | 0 | 0 | 0 | 0 | NA | NA | NA | NA | NA | NA | NA | NA | NA | NA |
| Olfr320 | 0 | 0 | 0 | 0 | 0 | 0 | 0 | 0 | 0 | 0 | 0 | 0 | 0 | 0 | 0 | NA | NA | NA | NA | NA | NA | NA | NA | NA | NA |
| Olfr323 | 2 | 0 | 0 | 0 | 0 | 0 | 2 | 0 | 0 | 0 | 0 | 0 | 0.26 | 1 | 0 | 0.356 | -1.491 | 0.71362796 | NA | 0 | 9.1573 | 10 | 10 | TRUE | 0.968 |
| Olfr324 | 0 | 0 | 0 | 0 | 0 | 0 | 0 | 0 | 0 | 0 | 0 | 0 | 0 | 0 | 0 | NA | NA | NA | NA | NA | NA | NA | NA | NA | NA |
| Olfr325 | 0 | 0 | 0 | 0 | 0 | 0 | 0 | 0 | 0 | 0 | 0 | 0 | 0 | 0 | 0 | NA | NA | NA | NA | NA | NA | NA | NA | NA | NA |
| Olfr328 | 3 | 0 | 1 | 0 | 0 | 0 | 2 | 0 | 1 | 0 | 0 | 0 | 0.57 | 1 | 0 | 0.165 | -2.602 | 0.35448349 | NA | 0 | 4.1739 | 4.2733 | 4.2733 | TRUE | 0.066 |
| Olfr329-ps | 15 | 13 | 5 | 4 | 7 | 3 | 12 | 14 | 5 | 4 | 8 | 3 | 7.62 | 10 | 5 | 0.473 | -1.08 | 0.14893129 | NA | 0.0772 | 0.3135 | 0.2511 | 0.2511 | TRUE | 0.5583 |
| Olfr33 | 0 | 0 | 0 | 0 | 0 | 0 | 0 | 0 | 0 | 0 | 0 | 0 | 0 | 0 | 0 | NA | NA | NA | NA | NA | NA | NA | NA | NA | NA |
| Olfr330 | 0 | 0 | 1 | 0 | 0 | 0 | 0 | 0 | 1 | 0 | 0 | 0 | 0.18 | 0 | 0 | 0.552 | -0.857 | 0.8336548 | NA | 0 | 13.1817 | 10 | 10 | TRUE | 0.5365 |
| Olfr331 | 0 | 0 | 0 | 0 | 0 | 0 | 0 | 0 | 0 | 0 | 0 | 0 | 0 | 0 | 0 | NA | NA | NA | NA | NA | NA | NA | NA | NA | NA |
| Olfr332 | 0 | 0 | 0 | 0 | 0 | 0 | 0 | 0 | 0 | 0 | 0 | 0 | 0 | 0 | 0 | NA | NA | NA | NA | NA | NA | NA | NA | NA | NA |
| Olfr338 | 0 | 0 | 0 | 0 | 0 | 0 | 0 | 0 | 0 | 0 | 0 | 0 | 0 | 0 | 0 | NA | NA | NA | NA | NA | NA | NA | NA | NA | NA |
| Olfr339 | 0 | 0 | 0 | 0 | 0 | 0 | 0 | 0 | 0 | 0 | 0 | 0 | 0 | 0 | 0 | NA | NA | NA | NA | NA | NA | NA | NA | NA | NA |
| Olfr340 | 1 | 1 | 2 | 3 | 1 | 1 | 1 | 1 | 2 | 3 | 1 | 1 | 1.52 | 1 | 2 | 1.308 | 0.387 | 0.79871892 | NA | 0 | 1.5651 | 0.9909 | 0.9909 | TRUE | 0.3467 |
| Olfr341 | 1 | 0 | 0 | 0 | 0 | 0 | 1 | 0 | 0 | 0 | 0 | 0 | 0.13 | 0 | 0 | 0.552 | -0.857 | 0.8336548 | NA | 0 | 18.3121 | 10 | 10 | TRUE | 0.2566 |
| Olfr342 | 0 | 0 | 0 | 0 | 0 | 0 | 0 | 0 | 0 | 0 | 0 | 0 | 0 | 0 | 0 | NA | NA | NA | NA | NA | NA | NA | NA | NA | NA |
| Olfr344 | 0 | 0 | 0 | 0 | 0 | 0 | 0 | 0 | 0 | 0 | 0 | 0 | 0 | 0 | 0 | NA | NA | NA | NA | NA | NA | NA | NA | NA | NA |
| Olfr345 | 0 | 0 | 0 | 0 | 0 | 0 | 0 | 0 | 0 | 0 | 0 | 0 | 0 | 0 | 0 | NA | NA | NA | NA | NA | NA | NA | NA | NA | NA |
| Olfr346 | 0 | 0 | 0 | 0 | 0 | 0 | 0 | 0 | 0 | 0 | 0 | 0 | 0 | 0 | 0 | NA | NA | NA | NA | NA | NA | NA | NA | NA | NA |
| Olfr347 | 0 | 0 | 0 | 0 | 0 | 0 | 0 | 0 | 0 | 0 | 0 | 0 | 0 | 0 | 0 | NA | NA | NA | NA | NA | NA | NA | NA | NA | NA |
| Olfr348 | 0 | 0 | 0 | 0 | 0 | 0 | 0 | 0 | 0 | 0 | 0 | 0 | 0 | 0 | 0 | NA | NA | NA | NA | NA | NA | NA | NA | NA | NA |
| Olfr350 | 0 | 0 | 0 | 0 | 0 | 0 | 0 | 0 | 0 | 0 | 0 | 0 | 0 | 0 | 0 | NA | NA | NA | NA | NA | NA | NA | NA | NA | NA |
| Olfr351 | 0 | 0 | 0 | 0 | 0 | 0 | 0 | 0 | 0 | 0 | 0 | 0 | 0 | 0 | 0 | NA | NA | NA | NA | NA | NA | NA | NA | NA | NA |
| Olfr352 | 0 | 0 | 0 | 0 | 0 | 0 | 0 | 0 | 0 | 0 | 0 | 0 | 0 | 0 | 0 | NA | NA | NA | NA | NA | NA | NA | NA | NA | NA |
| Olfr353 | 0 | 0 | 0 | 0 | 0 | 0 | 0 | 0 | 0 | 0 | 0 | 0 | 0 | 0 | 0 | NA | NA | NA | NA | NA | NA | NA | NA | NA | NA |
| Olfr354 | 0 | 0 | 0 | 0 | 0 | 0 | 0 | 0 | 0 | 0 | 0 | 0 | 0 | 0 | 0 | NA | NA | NA | NA | NA | NA | NA | NA | NA | NA |
| Olfr355 | 0 | 0 | 0 | 0 | 0 | 0 | 0 | 0 | 0 | 0 | 0 | 0 | 0 | 0 | 0 | NA | NA | NA | NA | NA | NA | NA | NA | NA | NA |
| Olfr356 | 0 | 0 | 0 | 0 | 0 | 0 | 0 | 0 | 0 | 0 | 0 | 0 | 0 | 0 | 0 | NA | NA | NA | NA | NA | NA | NA | NA | NA | NA |
| Olfr357 | 0 | 0 | 0 | 0 | 0 | 0 | 0 | 0 | 0 | 0 | 0 | 0 | 0 | 0 | 0 | NA | NA | NA | NA | NA | NA | NA | NA | NA | NA |
| Olfr358 | 0 | 0 | 0 | 0 | 0 | 0 | 0 | 0 | 0 | 0 | 0 | 0 | 0 | 0 | 0 | NA | NA | NA | NA | NA | NA | NA | NA | NA | NA |
| Olfr360 | 0 | 0 | 0 | 0 | 0 | 0 | 0 | 0 | 0 | 0 | 0 | 0 | 0 | 0 | 0 | NA | NA | NA | NA | NA | NA | NA | NA | NA | NA |
| Olfr361 | 0 | 0 | 0 | 0 | 0 | 0 | 0 | 0 | 0 | 0 | 0 | 0 | 0 | 0 | 0 | NA | NA | NA | NA | NA | NA | NA | NA | NA | NA |
| Olfr362 | 0 | 0 | 0 | 0 | 0 | 0 | 0 | 0 | 0 | 0 | 0 | 0 | 0 | 0 | 0 | NA | NA | NA | NA | NA | NA | NA | NA | NA | NA |
| Olfr365 | 0 | 0 | 0 | 0 | 0 | 0 | 0 | 0 | 0 | 0 | 0 | 0 | 0 | 0 | 0 | NA | NA | NA | NA | NA | NA | NA | NA | NA | NA |
| Olfr366 | 0 | 0 | 0 | 0 | 0 | 0 | 0 | 0 | 0 | 0 | 0 | 0 | 0 | 0 | 0 | NA | NA | NA | NA | NA | NA | NA | NA | NA | NA |
| Olfr367-ps | 0 | 0 | 0 | 0 | 0 | 0 | 0 | 0 | 0 | 0 | 0 | 0 | 0 | 0 | 0 | NA | NA | NA | NA | NA | NA | NA | NA | NA | NA |

| | | | | | | | | | | | | | | | | | | | | | | | | | |
|---|---|---|---|---|---|---|---|---|---|---|---|---|---|---|---|---|---|---|---|---|---|---|---|---|---|
| Olfr368 | 0 | 0 | 0 | 0 | 0 | 0 | 0 | 0 | 0 | 0 | 0 | 0 | 0 | 0 | 0 | NA | NA | NA | NA | NA | NA | NA | NA | NA | NA |
| Olfr370 | 0 | 0 | 0 | 0 | 0 | 0 | 0 | 0 | 0 | 0 | 0 | 0 | 0 | 0 | 0 | NA | NA | NA | NA | NA | NA | NA | NA | NA | NA |
| Olfr371 | 0 | 0 | 0 | 0 | 0 | 0 | 0 | 0 | 0 | 0 | 0 | 0 | 0 | 0 | 0 | NA | NA | NA | NA | NA | NA | NA | NA | NA | NA |
| Olfr372 | 0 | 0 | 0 | 0 | 1 | 0 | 0 | 0 | 0 | 0 | 1 | 0 | 0.19 | 0 | 0 | 2.094 | 1.067 | 0.79379329 | NA | 0 | 12.608 | 10 | 10 | TRUE | 0.5075 |
| Olfr373 | 0 | 0 | 0 | 0 | 0 | 0 | 0 | 0 | 0 | 0 | 0 | 0 | 0 | 0 | 0 | NA | NA | NA | NA | NA | NA | NA | NA | NA | NA |
| Olfr374 | 0 | 0 | 0 | 0 | 0 | 0 | 0 | 0 | 0 | 0 | 0 | 0 | 0 | 0 | 0 | NA | NA | NA | NA | NA | NA | NA | NA | NA | NA |
| Olfr376 | 62 | 38 | 22 | 18 | 8 | 15 | 48 | 40 | 24 | 18 | 9 | 15 | 25.68 | 37 | 14 | 0.372 | -1.428 | 0.00115315 | 0.00804002 | 0.0746 | 0.0947 | 0.0893 | 0.0893 | TRUE | 0.28 |
| Olfr378 | 0 | 0 | 0 | 0 | 0 | 0 | 0 | 0 | 0 | 0 | 0 | 0 | 0 | 0 | 0 | NA | NA | NA | NA | NA | NA | NA | NA | NA | NA |
| Olfr38 | 0 | 0 | 0 | 0 | 0 | 0 | 0 | 0 | 0 | 0 | 0 | 0 | 0 | 0 | 0 | NA | NA | NA | NA | NA | NA | NA | NA | NA | NA |
| Olfr380 | 0 | 0 | 0 | 0 | 0 | 0 | 0 | 0 | 0 | 0 | 0 | 0 | 0 | 0 | 0 | NA | NA | NA | NA | NA | NA | NA | NA | NA | NA |
| Olfr381 | 0 | 0 | 0 | 0 | 0 | 0 | 0 | 0 | 0 | 0 | 0 | 0 | 0 | 0 | 0 | NA | NA | NA | NA | NA | NA | NA | NA | NA | NA |
| Olfr382 | 0 | 0 | 0 | 0 | 0 | 0 | 0 | 0 | 0 | 0 | 0 | 0 | 0 | 0 | 0 | NA | NA | NA | NA | NA | NA | NA | NA | NA | NA |
| Olfr384 | 0 | 0 | 0 | 0 | 0 | 0 | 0 | 0 | 0 | 0 | 0 | 0 | 0 | 0 | 0 | NA | NA | NA | NA | NA | NA | NA | NA | NA | NA |
| Olfr385 | 0 | 0 | 0 | 0 | 0 | 0 | 0 | 0 | 0 | 0 | 0 | 0 | 0 | 0 | 0 | NA | NA | NA | NA | NA | NA | NA | NA | NA | NA |
| Olfr389 | 0 | 0 | 0 | 0 | 0 | 0 | 0 | 0 | 0 | 0 | 0 | 0 | 0 | 0 | 0 | NA | NA | NA | NA | NA | NA | NA | NA | NA | NA |
| Olfr39 | 0 | 0 | 0 | 0 | 0 | 0 | 0 | 0 | 0 | 0 | 0 | 0 | 0 | 0 | 0 | NA | NA | NA | NA | NA | NA | NA | NA | NA | NA |
| Olfr390 | 0 | 0 | 0 | 0 | 0 | 0 | 0 | 0 | 0 | 0 | 0 | 0 | 0 | 0 | 0 | NA | NA | NA | NA | NA | NA | NA | NA | NA | NA |
| Olfr391-ps | 0 | 0 | 0 | 0 | 0 | 0 | 0 | 0 | 0 | 0 | 0 | 0 | 0 | 0 | 0 | NA | NA | NA | NA | NA | NA | NA | NA | NA | NA |
| Olfr392 | 0 | 0 | 0 | 0 | 0 | 0 | 0 | 0 | 0 | 0 | 0 | 0 | 0 | 0 | 0 | NA | NA | NA | NA | NA | NA | NA | NA | NA | NA |
| Olfr393 | 0 | 0 | 0 | 0 | 0 | 0 | 0 | 0 | 0 | 0 | 0 | 0 | 0 | 0 | 0 | NA | NA | NA | NA | NA | NA | NA | NA | NA | NA |
| Olfr394 | 0 | 0 | 0 | 0 | 0 | 0 | 0 | 0 | 0 | 0 | 0 | 0 | 0 | 0 | 0 | NA | NA | NA | NA | NA | NA | NA | NA | NA | NA |
| Olfr395 | 0 | 0 | 0 | 0 | 0 | 0 | 0 | 0 | 0 | 0 | 0 | 0 | 0 | 0 | 0 | NA | NA | NA | NA | NA | NA | NA | NA | NA | NA |
| Olfr397 | 0 | 0 | 0 | 0 | 0 | 0 | 0 | 0 | 0 | 0 | 0 | 0 | 0 | 0 | 0 | NA | NA | NA | NA | NA | NA | NA | NA | NA | NA |
| Olfr398 | 0 | 0 | 0 | 0 | 0 | 0 | 0 | 0 | 0 | 0 | 0 | 0 | 0 | 0 | 0 | NA | NA | NA | NA | NA | NA | NA | NA | NA | NA |
| Olfr399 | 0 | 0 | 0 | 0 | 0 | 0 | 0 | 0 | 0 | 0 | 0 | 0 | 0 | 0 | 0 | NA | NA | NA | NA | NA | NA | NA | NA | NA | NA |
| Olfr401 | 0 | 0 | 0 | 0 | 0 | 0 | 0 | 0 | 0 | 0 | 0 | 0 | 0 | 0 | 0 | NA | NA | NA | NA | NA | NA | NA | NA | NA | NA |
| Olfr402 | 0 | 0 | 0 | 0 | 0 | 0 | 0 | 0 | 0 | 0 | 0 | 0 | 0 | 0 | 0 | NA | NA | NA | NA | NA | NA | NA | NA | NA | NA |
| Olfr403 | 0 | 0 | 0 | 0 | 0 | 0 | 0 | 0 | 0 | 0 | 0 | 0 | 0 | 0 | 0 | NA | NA | NA | NA | NA | NA | NA | NA | NA | NA |
| Olfr406 | 0 | 0 | 0 | 0 | 0 | 0 | 0 | 0 | 0 | 0 | 0 | 0 | 0 | 0 | 0 | NA | NA | NA | NA | NA | NA | NA | NA | NA | NA |
| Olfr410 | 0 | 0 | 0 | 0 | 0 | 0 | 0 | 0 | 0 | 0 | 0 | 0 | 0 | 0 | 0 | NA | NA | NA | NA | NA | NA | NA | NA | NA | NA |
| Olfr411 | 0 | 0 | 0 | 0 | 0 | 0 | 0 | 0 | 0 | 0 | 0 | 0 | 0 | 0 | 0 | NA | NA | NA | NA | NA | NA | NA | NA | NA | NA |
| Olfr412 | 0 | 0 | 0 | 0 | 0 | 0 | 0 | 0 | 0 | 0 | 0 | 0 | 0 | 0 | 0 | NA | NA | NA | NA | NA | NA | NA | NA | NA | NA |
| Olfr414 | 0 | 0 | 0 | 0 | 0 | 0 | 0 | 0 | 0 | 0 | 0 | 0 | 0 | 0 | 0 | NA | NA | NA | NA | NA | NA | NA | NA | NA | NA |
| Olfr417 | 0 | 0 | 0 | 0 | 0 | 0 | 0 | 0 | 0 | 0 | 0 | 0 | 0 | 0 | 0 | NA | NA | NA | NA | NA | NA | NA | NA | NA | NA |
| Olfr418-ps1 | 0 | 0 | 0 | 0 | 0 | 0 | 0 | 0 | 0 | 0 | 0 | 0 | 0 | 0 | 0 | NA | NA | NA | NA | NA | NA | NA | NA | NA | NA |
| Olfr419 | 0 | 0 | 0 | 0 | 0 | 0 | 0 | 0 | 0 | 0 | 0 | 0 | 0 | 0 | 0 | NA | NA | NA | NA | NA | NA | NA | NA | NA | NA |
| Olfr420 | 0 | 0 | 0 | 0 | 0 | 0 | 0 | 0 | 0 | 0 | 0 | 0 | 0 | 0 | 0 | NA | NA | NA | NA | NA | NA | NA | NA | NA | NA |
| Olfr421-ps1 | 0 | 0 | 0 | 0 | 0 | 0 | 0 | 0 | 0 | 0 | 0 | 0 | 0 | 0 | 0 | NA | NA | NA | NA | NA | NA | NA | NA | NA | NA |
| Olfr424 | 0 | 0 | 0 | 0 | 0 | 0 | 0 | 0 | 0 | 0 | 0 | 0 | 0 | 0 | 0 | NA | NA | NA | NA | NA | NA | NA | NA | NA | NA |
| Olfr426 | 0 | 0 | 0 | 0 | 0 | 0 | 0 | 0 | 0 | 0 | 0 | 0 | 0 | 0 | 0 | NA | NA | NA | NA | NA | NA | NA | NA | NA | NA |
| Olfr427 | 0 | 0 | 0 | 0 | 0 | 0 | 0 | 0 | 0 | 0 | 0 | 0 | 0 | 0 | 0 | NA | NA | NA | NA | NA | NA | NA | NA | NA | NA |
| Olfr429 | 0 | 0 | 0 | 0 | 0 | 0 | 0 | 0 | 0 | 0 | 0 | 0 | 0 | 0 | 0 | NA | NA | NA | NA | NA | NA | NA | NA | NA | NA |
| Olfr43 | 0 | 0 | 0 | 0 | 0 | 0 | 0 | 0 | 0 | 0 | 0 | 0 | 0 | 0 | 0 | NA | NA | NA | NA | NA | NA | NA | NA | NA | NA |
| Olfr430 | 0 | 0 | 0 | 0 | 0 | 0 | 0 | 0 | 0 | 0 | 0 | 0 | 0 | 0 | 0 | NA | NA | NA | NA | NA | NA | NA | NA | NA | NA |
| Olfr432 | 0 | 0 | 0 | 0 | 0 | 0 | 0 | 0 | 0 | 0 | 0 | 0 | 0 | 0 | 0 | NA | NA | NA | NA | NA | NA | NA | NA | NA | NA |
| Olfr433 | 0 | 0 | 0 | 0 | 0 | 0 | 0 | 0 | 0 | 0 | 0 | 0 | 0 | 0 | 0 | NA | NA | NA | NA | NA | NA | NA | NA | NA | NA |
| Olfr434 | 0 | 0 | 0 | 0 | 0 | 0 | 0 | 0 | 0 | 0 | 0 | 0 | 0 | 0 | 0 | NA | NA | NA | NA | NA | NA | NA | NA | NA | NA |
| Olfr435 | 0 | 0 | 0 | 0 | 0 | 0 | 0 | 0 | 0 | 0 | 0 | 0 | 0 | 0 | 0 | NA | NA | NA | NA | NA | NA | NA | NA | NA | NA |
| Olfr437 | 0 | 0 | 0 | 0 | 0 | 0 | 0 | 0 | 0 | 0 | 0 | 0 | 0 | 0 | 0 | NA | NA | NA | NA | NA | NA | NA | NA | NA | NA |
| Olfr44 | 0 | 0 | 0 | 0 | 0 | 0 | 0 | 0 | 0 | 0 | 0 | 0 | 0 | 0 | 0 | NA | NA | NA | NA | NA | NA | NA | NA | NA | NA |
| Olfr441 | 0 | 0 | 0 | 0 | 0 | 0 | 0 | 0 | 0 | 0 | 0 | 0 | 0 | 0 | 0 | NA | NA | NA | NA | NA | NA | NA | NA | NA | NA |
| Olfr444 | 0 | 0 | 0 | 0 | 0 | 0 | 0 | 0 | 0 | 0 | 0 | 0 | 0 | 0 | 0 | NA | NA | NA | NA | NA | NA | NA | NA | NA | NA |
| Olfr446 | 0 | 0 | 0 | 0 | 0 | 0 | 0 | 0 | 0 | 0 | 0 | 0 | 0 | 0 | 0 | NA | NA | NA | NA | NA | NA | NA | NA | NA | NA |
| Olfr447 | 0 | 0 | 0 | 0 | 0 | 0 | 0 | 0 | 0 | 0 | 0 | 0 | 0 | 0 | 0 | NA | NA | NA | NA | NA | NA | NA | NA | NA | NA |
| Olfr448 | 0 | 0 | 0 | 0 | 0 | 0 | 0 | 0 | 0 | 0 | 0 | 0 | 0 | 0 | 0 | NA | NA | NA | NA | NA | NA | NA | NA | NA | NA |
| Olfr449 | 0 | 0 | 0 | 0 | 0 | 0 | 0 | 0 | 0 | 0 | 0 | 0 | 0 | 0 | 0 | NA | NA | NA | NA | NA | NA | NA | NA | NA | NA |
| Olfr45 | 0 | 0 | 0 | 0 | 0 | 0 | 0 | 0 | 0 | 0 | 0 | 0 | 0 | 0 | 0 | NA | NA | NA | NA | NA | NA | NA | NA | NA | NA |
| Olfr450 | 0 | 0 | 0 | 0 | 0 | 0 | 0 | 0 | 0 | 0 | 0 | 0 | 0 | 0 | 0 | NA | NA | NA | NA | NA | NA | NA | NA | NA | NA |
| Olfr452 | 0 | 0 | 0 | 0 | 0 | 0 | 0 | 0 | 0 | 0 | 0 | 0 | 0 | 0 | 0 | NA | NA | NA | NA | NA | NA | NA | NA | NA | NA |
| Olfr453 | 0 | 0 | 0 | 0 | 0 | 0 | 0 | 0 | 0 | 0 | 0 | 0 | 0 | 0 | 0 | NA | NA | NA | NA | NA | NA | NA | NA | NA | NA |
| Olfr455 | 3 | 1 | 0 | 0 | 2 | 1 | 2 | 1 | 0 | 0 | 2 | 1 | 1.11 | 1 | 1 | 0.907 | -0.141 | 0.94125051 | NA | 0 | 2.145 | 1.7386 | 1.7386 | TRUE | 0.254 |
| Olfr456 | 0 | 0 | 0 | 0 | 0 | 0 | 0 | 0 | 0 | 0 | 0 | 0 | 0 | 0 | 0 | NA | NA | NA | NA | NA | NA | NA | NA | NA | NA |
| Olfr457 | 0 | 0 | 0 | 0 | 0 | 0 | 0 | 0 | 0 | 0 | 0 | 0 | 0 | 0 | 0 | NA | NA | NA | NA | NA | NA | NA | NA | NA | NA |
| Olfr458 | 0 | 0 | 0 | 0 | 0 | 0 | 0 | 0 | 0 | 0 | 0 | 0 | 0 | 0 | 0 | NA | NA | NA | NA | NA | NA | NA | NA | NA | NA |
| Olfr459 | 1 | 0 | 1 | 0 | 0 | 0 | 1 | 0 | 1 | 0 | 0 | 0 | 0.31 | 1 | 0 | 0.307 | -1.706 | 0.62894649 | NA | 0 | 7.6657 | 7.2 | 7.2 | TRUE | 0.2067 |
| Olfr46 | 0 | 0 | 0 | 0 | 0 | 0 | 0 | 0 | 0 | 0 | 0 | 0 | 0 | 0 | 0 | NA | NA | NA | NA | NA | NA | NA | NA | NA | NA |
| Olfr460 | 0 | 0 | 0 | 0 | 0 | 0 | 0 | 0 | 0 | 0 | 0 | 0 | 0 | 0 | 0 | NA | NA | NA | NA | NA | NA | NA | NA | NA | NA |
| Olfr461 | 0 | 0 | 0 | 0 | 0 | 0 | 0 | 0 | 0 | 0 | 0 | 0 | 0 | 0 | 0 | NA | NA | NA | NA | NA | NA | NA | NA | NA | NA |
| Olfr462 | 0 | 0 | 0 | 0 | 0 | 0 | 0 | 0 | 0 | 0 | 0 | 0 | 0 | 0 | 0 | NA | NA | NA | NA | NA | NA | NA | NA | NA | NA |

| | | | | | | | | | | | | | | | | | | | | | | | | | |
|---|---|---|---|---|---|---|---|---|---|---|---|---|---|---|---|---|---|---|---|---|---|---|---|---|---|
| Olfr463 | 0 | 0 | 0 | 0 | 0 | 0 | 0 | 0 | 0 | 0 | 0 | 0 | 0 | 0 | 0 | NA | NA | NA | NA | NA | NA | NA | NA | NA | NA |
| Olfr464 | 1 | 0 | 0 | 0 | 0 | 0 | 1 | 0 | 0 | 0 | 0 | 0 | 0.13 | 0 | 0 | 0.552 | -0.857 | 0.8336548 | NA | 0 | 18.3121 | 10 | 10 | TRUE | 0.2566 |
| Olfr466 | 0 | 0 | 0 | 0 | 0 | 0 | 0 | 0 | 0 | 0 | 0 | 0 | 0 | 0 | 0 | NA | NA | NA | NA | NA | NA | NA | NA | NA | NA |
| Olfr467 | 0 | 0 | 0 | 0 | 0 | 0 | 0 | 0 | 0 | 0 | 0 | 0 | 0 | 0 | 0 | NA | NA | NA | NA | NA | NA | NA | NA | NA | NA |
| Olfr469 | 0 | 0 | 0 | 0 | 0 | 0 | 0 | 0 | 0 | 0 | 0 | 0 | 0 | 0 | 0 | NA | NA | NA | NA | NA | NA | NA | NA | NA | NA |
| Olfr47 | 0 | 0 | 0 | 0 | 0 | 0 | 0 | 0 | 0 | 0 | 0 | 0 | 0 | 0 | 0 | NA | NA | NA | NA | NA | NA | NA | NA | NA | NA |
| Olfr470 | 0 | 0 | 0 | 1 | 0 | 0 | 0 | 0 | 0 | 1 | 0 | 0 | 0.17 | 0 | 0 | 2.094 | 1.067 | 0.79379329 | NA | 0 | 14.2885 | 10 | 10 | TRUE | 0.3906 |
| Olfr472 | 0 | 0 | 0 | 0 | 0 | 0 | 0 | 0 | 0 | 0 | 0 | 0 | 0 | 0 | 0 | NA | NA | NA | NA | NA | NA | NA | NA | NA | NA |
| Olfr473 | 0 | 0 | 0 | 0 | 0 | 0 | 0 | 0 | 0 | 0 | 0 | 0 | 0 | 0 | 0 | NA | NA | NA | NA | NA | NA | NA | NA | NA | NA |
| Olfr474 | 0 | 0 | 0 | 0 | 0 | 0 | 0 | 0 | 0 | 0 | 0 | 0 | 0 | 0 | 0 | NA | NA | NA | NA | NA | NA | NA | NA | NA | NA |
| Olfr476 | 0 | 0 | 0 | 0 | 0 | 0 | 0 | 0 | 0 | 0 | 0 | 0 | 0 | 0 | 0 | NA | NA | NA | NA | NA | NA | NA | NA | NA | NA |
| Olfr477 | 0 | 0 | 0 | 0 | 0 | 0 | 0 | 0 | 0 | 0 | 0 | 0 | 0 | 0 | 0 | NA | NA | NA | NA | NA | NA | NA | NA | NA | NA |
| Olfr478 | 0 | 0 | 0 | 0 | 0 | 0 | 0 | 0 | 0 | 0 | 0 | 0 | 0 | 0 | 0 | NA | NA | NA | NA | NA | NA | NA | NA | NA | NA |
| Olfr479 | 0 | 0 | 0 | 0 | 0 | 0 | 0 | 0 | 0 | 0 | 0 | 0 | 0 | 0 | 0 | NA | NA | NA | NA | NA | NA | NA | NA | NA | NA |
| Olfr48 | 0 | 0 | 0 | 0 | 0 | 0 | 0 | 0 | 0 | 0 | 0 | 0 | 0 | 0 | 0 | NA | NA | NA | NA | NA | NA | NA | NA | NA | NA |
| Olfr480 | 0 | 0 | 0 | 0 | 0 | 0 | 0 | 0 | 0 | 0 | 0 | 0 | 0 | 0 | 0 | NA | NA | NA | NA | NA | NA | NA | NA | NA | NA |
| Olfr481 | 0 | 0 | 0 | 0 | 0 | 0 | 0 | 0 | 0 | 0 | 0 | 0 | 0 | 0 | 0 | NA | NA | NA | NA | NA | NA | NA | NA | NA | NA |
| Olfr482 | 0 | 0 | 0 | 0 | 0 | 0 | 0 | 0 | 0 | 0 | 0 | 0 | 0 | 0 | 0 | NA | NA | NA | NA | NA | NA | NA | NA | NA | NA |
| Olfr483 | 0 | 0 | 0 | 0 | 0 | 0 | 0 | 0 | 0 | 0 | 0 | 0 | 0 | 0 | 0 | NA | NA | NA | NA | NA | NA | NA | NA | NA | NA |
| Olfr484 | 0 | 0 | 0 | 0 | 0 | 0 | 0 | 0 | 0 | 0 | 0 | 0 | 0 | 0 | 0 | NA | NA | NA | NA | NA | NA | NA | NA | NA | NA |
| Olfr485 | 0 | 0 | 0 | 0 | 0 | 0 | 0 | 0 | 0 | 0 | 0 | 0 | 0 | 0 | 0 | NA | NA | NA | NA | NA | NA | NA | NA | NA | NA |
| Olfr486 | 0 | 0 | 0 | 0 | 0 | 0 | 0 | 0 | 0 | 0 | 0 | 0 | 0 | 0 | 0 | NA | NA | NA | NA | NA | NA | NA | NA | NA | NA |
| Olfr487 | 0 | 0 | 0 | 0 | 0 | 0 | 0 | 0 | 0 | 0 | 0 | 0 | 0 | 0 | 0 | NA | NA | NA | NA | NA | NA | NA | NA | NA | NA |
| Olfr488 | 0 | 0 | 0 | 0 | 0 | 0 | 0 | 0 | 0 | 0 | 0 | 0 | 0 | 0 | 0 | NA | NA | NA | NA | NA | NA | NA | NA | NA | NA |
| Olfr49 | 0 | 0 | 0 | 0 | 0 | 0 | 0 | 0 | 0 | 0 | 0 | 0 | 0 | 0 | 0 | NA | NA | NA | NA | NA | NA | NA | NA | NA | NA |
| Olfr490 | 0 | 0 | 0 | 0 | 0 | 0 | 0 | 0 | 0 | 0 | 0 | 0 | 0 | 0 | 0 | NA | NA | NA | NA | NA | NA | NA | NA | NA | NA |
| Olfr491 | 0 | 0 | 0 | 0 | 0 | 0 | 0 | 0 | 0 | 0 | 0 | 0 | 0 | 0 | 0 | NA | NA | NA | NA | NA | NA | NA | NA | NA | NA |
| Olfr492 | 0 | 0 | 0 | 0 | 0 | 0 | 0 | 0 | 0 | 0 | 0 | 0 | 0 | 0 | 0 | NA | NA | NA | NA | NA | NA | NA | NA | NA | NA |
| Olfr493 | 0 | 0 | 0 | 0 | 0 | 0 | 0 | 0 | 0 | 0 | 0 | 0 | 0 | 0 | 0 | NA | NA | NA | NA | NA | NA | NA | NA | NA | NA |
| Olfr494 | 0 | 0 | 0 | 0 | 0 | 0 | 0 | 0 | 0 | 0 | 0 | 0 | 0 | 0 | 0 | NA | NA | NA | NA | NA | NA | NA | NA | NA | NA |
| Olfr495 | 0 | 0 | 0 | 0 | 0 | 0 | 0 | 0 | 0 | 0 | 0 | 0 | 0 | 0 | 0 | NA | NA | NA | NA | NA | NA | NA | NA | NA | NA |
| Olfr497 | 0 | 0 | 0 | 0 | 0 | 0 | 0 | 0 | 0 | 0 | 0 | 0 | 0 | 0 | 0 | NA | NA | NA | NA | NA | NA | NA | NA | NA | NA |
| Olfr498 | 0 | 0 | 0 | 0 | 0 | 0 | 0 | 0 | 0 | 0 | 0 | 0 | 0 | 0 | 0 | NA | NA | NA | NA | NA | NA | NA | NA | NA | NA |
| Olfr5 | 1 | 0 | 0 | 1 | 0 | 0 | 1 | 0 | 0 | 1 | 0 | 0 | 0.3 | 0 | 0 | 1.075 | 0.105 | 0.97698525 | NA | 0 | 8.0272 | 7.5141 | 7.5141 | TRUE | 0.3906 |
| Olfr50 | 0 | 0 | 0 | 0 | 0 | 0 | 0 | 0 | 0 | 0 | 0 | 0 | 0 | 0 | 0 | NA | NA | NA | NA | NA | NA | NA | NA | NA | NA |
| Olfr502 | 0 | 0 | 0 | 0 | 0 | 0 | 0 | 0 | 0 | 0 | 0 | 0 | 0 | 0 | 0 | NA | NA | NA | NA | NA | NA | NA | NA | NA | NA |
| Olfr503 | 7 | 5 | 4 | 8 | 1 | 2 | 5 | 5 | 4 | 8 | 1 | 2 | 4.36 | 5 | 4 | 0.74 | -0.434 | 0.65485534 | NA | 0.1612 | 0.5465 | 0.4454 | 0.4454 | TRUE | 1.6246 |
| Olfr504 | 1 | 0 | 0 | 0 | 0 | 0 | 1 | 0 | 0 | 0 | 0 | 0 | 0.13 | 0 | 0 | 0.552 | -0.857 | 0.8336548 | NA | 0 | 18.3121 | 10 | 10 | TRUE | 0.2566 |
| Olfr506 | 0 | 0 | 0 | 0 | 0 | 0 | 0 | 0 | 0 | 0 | 0 | 0 | 0 | 0 | 0 | NA | NA | NA | NA | NA | NA | NA | NA | NA | NA |
| Olfr507 | 0 | 0 | 0 | 0 | 0 | 0 | 0 | 0 | 0 | 0 | 0 | 0 | 0 | 0 | 0 | NA | NA | NA | NA | NA | NA | NA | NA | NA | NA |
| Olfr508 | 0 | 0 | 0 | 0 | 0 | 0 | 0 | 0 | 0 | 0 | 0 | 0 | 0 | 0 | 0 | NA | NA | NA | NA | NA | NA | NA | NA | NA | NA |
| Olfr509 | 0 | 0 | 0 | 0 | 0 | 0 | 0 | 0 | 0 | 0 | 0 | 0 | 0 | 0 | 0 | NA | NA | NA | NA | NA | NA | NA | NA | NA | NA |
| Olfr51 | 0 | 0 | 0 | 0 | 0 | 0 | 0 | 0 | 0 | 0 | 0 | 0 | 0 | 0 | 0 | NA | NA | NA | NA | NA | NA | NA | NA | NA | NA |
| Olfr510 | 0 | 0 | 0 | 0 | 0 | 0 | 0 | 0 | 0 | 0 | 0 | 0 | 0 | 0 | 0 | NA | NA | NA | NA | NA | NA | NA | NA | NA | NA |
| Olfr512 | 0 | 0 | 0 | 0 | 0 | 0 | 0 | 0 | 0 | 0 | 0 | 0 | 0 | 0 | 0 | NA | NA | NA | NA | NA | NA | NA | NA | NA | NA |
| Olfr513 | 0 | 0 | 0 | 0 | 0 | 0 | 0 | 0 | 0 | 0 | 0 | 0 | 0 | 0 | 0 | NA | NA | NA | NA | NA | NA | NA | NA | NA | NA |
| Olfr514 | 0 | 0 | 0 | 0 | 0 | 0 | 0 | 0 | 0 | 0 | 0 | 0 | 0 | 0 | 0 | NA | NA | NA | NA | NA | NA | NA | NA | NA | NA |
| Olfr516 | 0 | 0 | 0 | 0 | 0 | 0 | 0 | 0 | 0 | 0 | 0 | 0 | 0 | 0 | 0 | NA | NA | NA | NA | NA | NA | NA | NA | NA | NA |
| Olfr517 | 0 | 0 | 0 | 0 | 0 | 0 | 0 | 0 | 0 | 0 | 0 | 0 | 0 | 0 | 0 | NA | NA | NA | NA | NA | NA | NA | NA | NA | NA |
| Olfr518 | 0 | 0 | 0 | 0 | 0 | 0 | 0 | 0 | 0 | 0 | 0 | 0 | 0 | 0 | 0 | NA | NA | NA | NA | NA | NA | NA | NA | NA | NA |
| Olfr519 | 0 | 0 | 0 | 0 | 0 | 0 | 0 | 0 | 0 | 0 | 0 | 0 | 0 | 0 | 0 | NA | NA | NA | NA | NA | NA | NA | NA | NA | NA |
| Olfr52 | 0 | 0 | 0 | 0 | 0 | 0 | 0 | 0 | 0 | 0 | 0 | 0 | 0 | 0 | 0 | NA | NA | NA | NA | NA | NA | NA | NA | NA | NA |
| Olfr520 | 0 | 0 | 0 | 0 | 0 | 0 | 0 | 0 | 0 | 0 | 0 | 0 | 0 | 0 | 0 | NA | NA | NA | NA | NA | NA | NA | NA | NA | NA |
| Olfr521 | 0 | 0 | 0 | 0 | 0 | 0 | 0 | 0 | 0 | 0 | 0 | 0 | 0 | 0 | 0 | NA | NA | NA | NA | NA | NA | NA | NA | NA | NA |
| Olfr522 | 0 | 0 | 0 | 0 | 0 | 0 | 0 | 0 | 0 | 0 | 0 | 0 | 0 | 0 | 0 | NA | NA | NA | NA | NA | NA | NA | NA | NA | NA |
| Olfr523 | 0 | 0 | 0 | 0 | 0 | 0 | 0 | 0 | 0 | 0 | 0 | 0 | 0 | 0 | 0 | NA | NA | NA | NA | NA | NA | NA | NA | NA | NA |
| Olfr524 | 0 | 0 | 0 | 0 | 0 | 0 | 0 | 0 | 0 | 0 | 0 | 0 | 0 | 0 | 0 | NA | NA | NA | NA | NA | NA | NA | NA | NA | NA |
| Olfr525 | 0 | 0 | 0 | 0 | 0 | 0 | 0 | 0 | 0 | 0 | 0 | 0 | 0 | 0 | 0 | NA | NA | NA | NA | NA | NA | NA | NA | NA | NA |
| Olfr527 | 0 | 0 | 0 | 0 | 0 | 0 | 0 | 0 | 0 | 0 | 0 | 0 | 0 | 0 | 0 | NA | NA | NA | NA | NA | NA | NA | NA | NA | NA |
| Olfr53 | 0 | 0 | 0 | 0 | 0 | 0 | 0 | 0 | 0 | 0 | 0 | 0 | 0 | 0 | 0 | NA | NA | NA | NA | NA | NA | NA | NA | NA | NA |
| Olfr530 | 0 | 0 | 0 | 0 | 0 | 0 | 0 | 0 | 0 | 0 | 0 | 0 | 0 | 0 | 0 | NA | NA | NA | NA | NA | NA | NA | NA | NA | NA |
| Olfr531 | 0 | 0 | 0 | 0 | 0 | 0 | 0 | 0 | 0 | 0 | 0 | 0 | 0 | 0 | 0 | NA | NA | NA | NA | NA | NA | NA | NA | NA | NA |
| Olfr532 | 0 | 0 | 0 | 0 | 0 | 0 | 0 | 0 | 0 | 0 | 0 | 0 | 0 | 0 | 0 | NA | NA | NA | NA | NA | NA | NA | NA | NA | NA |
| Olfr533 | 0 | 0 | 0 | 0 | 0 | 0 | 0 | 0 | 0 | 0 | 0 | 0 | 0 | 0 | 0 | NA | NA | NA | NA | NA | NA | NA | NA | NA | NA |
| Olfr535 | 0 | 0 | 0 | 0 | 0 | 0 | 0 | 0 | 0 | 0 | 0 | 0 | 0 | 0 | 0 | NA | NA | NA | NA | NA | NA | NA | NA | NA | NA |
| Olfr536 | 0 | 0 | 0 | 0 | 0 | 0 | 0 | 0 | 0 | 0 | 0 | 0 | 0 | 0 | 0 | NA | NA | NA | NA | NA | NA | NA | NA | NA | NA |
| Olfr538 | 0 | 0 | 0 | 0 | 0 | 0 | 0 | 0 | 0 | 0 | 0 | 0 | 0 | 0 | 0 | NA | NA | NA | NA | NA | NA | NA | NA | NA | NA |
| Olfr539 | 0 | 0 | 0 | 0 | 0 | 0 | 0 | 0 | 0 | 0 | 0 | 0 | 0 | 0 | 0 | NA | NA | NA | NA | NA | NA | NA | NA | NA | NA |
| Olfr54 | 0 | 0 | 0 | 0 | 0 | 0 | 0 | 0 | 0 | 0 | 0 | 0 | 0 | 0 | 0 | NA | NA | NA | NA | NA | NA | NA | NA | NA | NA |

| | | | | | | | | | | | | | | | | | | | | | | | | | |
|---|---|---|---|---|---|---|---|---|---|---|---|---|---|---|---|---|---|---|---|---|---|---|---|---|---|
| Olfr541 | 0 | 0 | 0 | 0 | 0 | 0 | 0 | 0 | 0 | 0 | 0 | 0 | 0 | 0 | 0 | NA | NA | NA | NA | NA | NA | NA | NA | NA | NA |
| Olfr543 | 0 | 0 | 0 | 0 | 0 | 0 | 0 | 0 | 0 | 0 | 0 | 0 | 0 | 0 | 0 | NA | NA | NA | NA | NA | NA | NA | NA | NA | NA |
| Olfr544 | 0 | 0 | 0 | 0 | 0 | 0 | 0 | 0 | 0 | 0 | 0 | 0 | 0 | 0 | 0 | NA | NA | NA | NA | NA | NA | NA | NA | NA | NA |
| Olfr545 | 0 | 0 | 0 | 0 | 0 | 0 | 0 | 0 | 0 | 0 | 0 | 0 | 0 | 0 | 0 | NA | NA | NA | NA | NA | NA | NA | NA | NA | NA |
| Olfr547 | 0 | 0 | 0 | 0 | 0 | 0 | 0 | 0 | 0 | 0 | 0 | 0 | 0 | 0 | 0 | NA | NA | NA | NA | NA | NA | NA | NA | NA | NA |
| Olfr549 | 0 | 0 | 0 | 0 | 0 | 0 | 0 | 0 | 0 | 0 | 0 | 0 | 0 | 0 | 0 | NA | NA | NA | NA | NA | NA | NA | NA | NA | NA |
| Olfr55 | 1 | 0 | 0 | 0 | 0 | 0 | 1 | 0 | 0 | 0 | 0 | 0 | 0.13 | 0 | 0 | 0.552 | -0.857 | 0.8336548 | NA | 0 | 18.3121 | 10 | 10 | TRUE | 0.2566 |
| Olfr550 | 0 | 0 | 0 | 0 | 0 | 0 | 0 | 0 | 0 | 0 | 0 | 0 | 0 | 0 | 0 | NA | NA | NA | NA | NA | NA | NA | NA | NA | NA |
| Olfr551 | 0 | 0 | 0 | 0 | 0 | 0 | 0 | 0 | 0 | 0 | 0 | 0 | 0 | 0 | 0 | NA | NA | NA | NA | NA | NA | NA | NA | NA | NA |
| Olfr552 | 0 | 0 | 0 | 0 | 0 | 0 | 0 | 0 | 0 | 0 | 0 | 0 | 0 | 0 | 0 | NA | NA | NA | NA | NA | NA | NA | NA | NA | NA |
| Olfr553 | 0 | 0 | 0 | 0 | 0 | 0 | 0 | 0 | 0 | 0 | 0 | 0 | 0 | 0 | 0 | NA | NA | NA | NA | NA | NA | NA | NA | NA | NA |
| Olfr554 | 0 | 0 | 0 | 0 | 0 | 0 | 0 | 0 | 0 | 0 | 0 | 0 | 0 | 0 | 0 | NA | NA | NA | NA | NA | NA | NA | NA | NA | NA |
| Olfr555 | 0 | 0 | 0 | 0 | 0 | 0 | 0 | 0 | 0 | 0 | 0 | 0 | 0 | 0 | 0 | NA | NA | NA | NA | NA | NA | NA | NA | NA | NA |
| Olfr556 | 0 | 0 | 0 | 0 | 0 | 0 | 0 | 0 | 0 | 0 | 0 | 0 | 0 | 0 | 0 | NA | NA | NA | NA | NA | NA | NA | NA | NA | NA |
| Olfr557 | 0 | 0 | 1 | 0 | 0 | 0 | 0 | 0 | 1 | 0 | 0 | 0 | 0.18 | 0 | 0 | 0.552 | -0.857 | 0.8336548 | NA | 0 | 13.1817 | 10 | 10 | TRUE | 0.5365 |
| Olfr558 | 1 | 0 | 0 | 0 | 0 | 0 | 1 | 0 | 0 | 0 | 0 | 0 | 0.13 | 0 | 0 | 0.552 | -0.857 | 0.8336548 | NA | 0 | 18.3121 | 10 | 10 | TRUE | 0.2566 |
| Olfr559 | 0 | 0 | 0 | 0 | 0 | 0 | 0 | 0 | 0 | 0 | 0 | 0 | 0 | 0 | 0 | NA | NA | NA | NA | NA | NA | NA | NA | NA | NA |
| Olfr56 | 0 | 0 | 0 | 0 | 0 | 0 | 0 | 0 | 0 | 0 | 0 | 0 | 0 | 0 | 0 | NA | NA | NA | NA | NA | NA | NA | NA | NA | NA |
| Olfr560 | 0 | 0 | 0 | 0 | 0 | 0 | 0 | 0 | 0 | 0 | 0 | 0 | 0 | 0 | 0 | NA | NA | NA | NA | NA | NA | NA | NA | NA | NA |
| Olfr561 | 0 | 0 | 0 | 0 | 0 | 0 | 0 | 0 | 0 | 0 | 0 | 0 | 0 | 0 | 0 | NA | NA | NA | NA | NA | NA | NA | NA | NA | NA |
| Olfr564 | 0 | 0 | 0 | 0 | 0 | 0 | 0 | 0 | 0 | 0 | 0 | 0 | 0 | 0 | 0 | NA | NA | NA | NA | NA | NA | NA | NA | NA | NA |
| Olfr566 | 0 | 0 | 0 | 0 | 0 | 0 | 0 | 0 | 0 | 0 | 0 | 0 | 0 | 0 | 0 | NA | NA | NA | NA | NA | NA | NA | NA | NA | NA |
| Olfr568 | 0 | 0 | 0 | 0 | 0 | 0 | 0 | 0 | 0 | 0 | 0 | 0 | 0 | 0 | 0 | NA | NA | NA | NA | NA | NA | NA | NA | NA | NA |
| Olfr569 | 0 | 0 | 0 | 0 | 0 | 0 | 0 | 0 | 0 | 0 | 0 | 0 | 0 | 0 | 0 | NA | NA | NA | NA | NA | NA | NA | NA | NA | NA |
| Olfr57 | 0 | 0 | 0 | 0 | 0 | 0 | 0 | 0 | 0 | 0 | 0 | 0 | 0 | 0 | 0 | NA | NA | NA | NA | NA | NA | NA | NA | NA | NA |
| Olfr570 | 1 | 0 | 0 | 0 | 0 | 0 | 1 | 0 | 0 | 0 | 0 | 0 | 0.13 | 0 | 0 | 0.552 | -0.857 | 0.8336548 | NA | 0 | 18.3121 | 10 | 10 | TRUE | 0.2566 |
| Olfr571 | 0 | 0 | 0 | 0 | 0 | 0 | 0 | 0 | 0 | 0 | 0 | 0 | 0 | 0 | 0 | NA | NA | NA | NA | NA | NA | NA | NA | NA | NA |
| Olfr572 | 0 | 0 | 0 | 0 | 0 | 0 | 0 | 0 | 0 | 0 | 0 | 0 | 0 | 0 | 0 | NA | NA | NA | NA | NA | NA | NA | NA | NA | NA |
| Olfr574 | 0 | 0 | 0 | 0 | 0 | 0 | 0 | 0 | 0 | 0 | 0 | 0 | 0 | 0 | 0 | NA | NA | NA | NA | NA | NA | NA | NA | NA | NA |
| Olfr575 | 0 | 0 | 0 | 0 | 0 | 0 | 0 | 0 | 0 | 0 | 0 | 0 | 0 | 0 | 0 | NA | NA | NA | NA | NA | NA | NA | NA | NA | NA |
| Olfr576 | 0 | 0 | 0 | 0 | 0 | 0 | 0 | 0 | 0 | 0 | 0 | 0 | 0 | 0 | 0 | NA | NA | NA | NA | NA | NA | NA | NA | NA | NA |
| Olfr577 | 0 | 0 | 0 | 0 | 0 | 0 | 0 | 0 | 0 | 0 | 0 | 0 | 0 | 0 | 0 | NA | NA | NA | NA | NA | NA | NA | NA | NA | NA |
| Olfr578 | 0 | 0 | 0 | 0 | 0 | 0 | 0 | 0 | 0 | 0 | 0 | 0 | 0 | 0 | 0 | NA | NA | NA | NA | NA | NA | NA | NA | NA | NA |
| Olfr58 | 0 | 0 | 0 | 0 | 1 | 1 | 0 | 0 | 0 | 0 | 1 | 1 | 0.35 | 0 | 1 | 3.965 | 1.987 | 0.55529285 | NA | 0 | 6.7252 | 6.4426 | 6.4426 | TRUE | 0.261 |
| Olfr582 | 0 | 0 | 0 | 0 | 0 | 0 | 0 | 0 | 0 | 0 | 0 | 0 | 0 | 0 | 0 | NA | NA | NA | NA | NA | NA | NA | NA | NA | NA |
| Olfr583 | 0 | 0 | 0 | 0 | 0 | 0 | 0 | 0 | 0 | 0 | 0 | 0 | 0 | 0 | 0 | NA | NA | NA | NA | NA | NA | NA | NA | NA | NA |
| Olfr584 | 0 | 0 | 0 | 0 | 0 | 0 | 0 | 0 | 0 | 0 | 0 | 0 | 0 | 0 | 0 | NA | NA | NA | NA | NA | NA | NA | NA | NA | NA |
| Olfr585 | 0 | 0 | 0 | 0 | 0 | 0 | 0 | 0 | 0 | 0 | 0 | 0 | 0 | 0 | 0 | NA | NA | NA | NA | NA | NA | NA | NA | NA | NA |
| Olfr586 | 0 | 0 | 0 | 0 | 0 | 0 | 0 | 0 | 0 | 0 | 0 | 0 | 0 | 0 | 0 | NA | NA | NA | NA | NA | NA | NA | NA | NA | NA |
| Olfr589 | 0 | 0 | 0 | 0 | 0 | 0 | 0 | 0 | 0 | 0 | 0 | 0 | 0 | 0 | 0 | NA | NA | NA | NA | NA | NA | NA | NA | NA | NA |
| Olfr59 | 0 | 0 | 0 | 0 | 0 | 0 | 0 | 0 | 0 | 0 | 0 | 0 | 0 | 0 | 0 | NA | NA | NA | NA | NA | NA | NA | NA | NA | NA |
| Olfr591 | 0 | 0 | 0 | 0 | 0 | 0 | 0 | 0 | 0 | 0 | 0 | 0 | 0 | 0 | 0 | NA | NA | NA | NA | NA | NA | NA | NA | NA | NA |
| Olfr592 | 0 | 0 | 0 | 0 | 0 | 0 | 0 | 0 | 0 | 0 | 0 | 0 | 0 | 0 | 0 | NA | NA | NA | NA | NA | NA | NA | NA | NA | NA |
| Olfr593 | 0 | 0 | 0 | 0 | 0 | 0 | 0 | 0 | 0 | 0 | 0 | 0 | 0 | 0 | 0 | NA | NA | NA | NA | NA | NA | NA | NA | NA | NA |
| Olfr594 | 0 | 0 | 0 | 0 | 0 | 0 | 0 | 0 | 0 | 0 | 0 | 0 | 0 | 0 | 0 | NA | NA | NA | NA | NA | NA | NA | NA | NA | NA |
| Olfr596 | 0 | 0 | 0 | 0 | 0 | 0 | 0 | 0 | 0 | 0 | 0 | 0 | 0 | 0 | 0 | NA | NA | NA | NA | NA | NA | NA | NA | NA | NA |
| Olfr597 | 0 | 0 | 0 | 0 | 0 | 0 | 0 | 0 | 0 | 0 | 0 | 0 | 0 | 0 | 0 | NA | NA | NA | NA | NA | NA | NA | NA | NA | NA |
| Olfr598 | 0 | 0 | 0 | 0 | 0 | 0 | 0 | 0 | 0 | 0 | 0 | 0 | 0 | 0 | 0 | NA | NA | NA | NA | NA | NA | NA | NA | NA | NA |
| Olfr599 | 0 | 0 | 0 | 0 | 0 | 0 | 0 | 0 | 0 | 0 | 0 | 0 | 0 | 0 | 0 | NA | NA | NA | NA | NA | NA | NA | NA | NA | NA |
| Olfr6 | 0 | 0 | 0 | 0 | 0 | 0 | 0 | 0 | 0 | 0 | 0 | 0 | 0 | 0 | 0 | NA | NA | NA | NA | NA | NA | NA | NA | NA | NA |
| Olfr60 | 0 | 0 | 0 | 0 | 0 | 0 | 0 | 0 | 0 | 0 | 0 | 0 | 0 | 0 | 0 | NA | NA | NA | NA | NA | NA | NA | NA | NA | NA |
| Olfr600 | 0 | 0 | 0 | 0 | 0 | 0 | 0 | 0 | 0 | 0 | 0 | 0 | 0 | 0 | 0 | NA | NA | NA | NA | NA | NA | NA | NA | NA | NA |
| Olfr601 | 0 | 0 | 0 | 0 | 0 | 0 | 0 | 0 | 0 | 0 | 0 | 0 | 0 | 0 | 0 | NA | NA | NA | NA | NA | NA | NA | NA | NA | NA |
| Olfr603 | 0 | 0 | 0 | 0 | 0 | 0 | 0 | 0 | 0 | 0 | 0 | 0 | 0 | 0 | 0 | NA | NA | NA | NA | NA | NA | NA | NA | NA | NA |
| Olfr605 | 0 | 0 | 0 | 0 | 0 | 0 | 0 | 0 | 0 | 0 | 0 | 0 | 0 | 0 | 0 | NA | NA | NA | NA | NA | NA | NA | NA | NA | NA |
| Olfr606 | 0 | 0 | 0 | 0 | 0 | 0 | 0 | 0 | 0 | 0 | 0 | 0 | 0 | 0 | 0 | NA | NA | NA | NA | NA | NA | NA | NA | NA | NA |
| Olfr608 | 0 | 0 | 0 | 0 | 0 | 0 | 0 | 0 | 0 | 0 | 0 | 0 | 0 | 0 | 0 | NA | NA | NA | NA | NA | NA | NA | NA | NA | NA |
| Olfr609 | 0 | 0 | 0 | 0 | 0 | 0 | 0 | 0 | 0 | 0 | 0 | 0 | 0 | 0 | 0 | NA | NA | NA | NA | NA | NA | NA | NA | NA | NA |
| Olfr61 | 0 | 0 | 0 | 0 | 0 | 0 | 0 | 0 | 0 | 0 | 0 | 0 | 0 | 0 | 0 | NA | NA | NA | NA | NA | NA | NA | NA | NA | NA |
| Olfr610 | 0 | 0 | 0 | 0 | 0 | 0 | 0 | 0 | 0 | 0 | 0 | 0 | 0 | 0 | 0 | NA | NA | NA | NA | NA | NA | NA | NA | NA | NA |
| Olfr611 | 0 | 0 | 0 | 0 | 0 | 0 | 0 | 0 | 0 | 0 | 0 | 0 | 0 | 0 | 0 | NA | NA | NA | NA | NA | NA | NA | NA | NA | NA |
| Olfr612 | 0 | 0 | 0 | 0 | 0 | 0 | 0 | 0 | 0 | 0 | 0 | 0 | 0 | 0 | 0 | NA | NA | NA | NA | NA | NA | NA | NA | NA | NA |
| Olfr613 | 0 | 0 | 0 | 0 | 0 | 0 | 0 | 0 | 0 | 0 | 0 | 0 | 0 | 0 | 0 | NA | NA | NA | NA | NA | NA | NA | NA | NA | NA |
| Olfr615 | 0 | 0 | 0 | 0 | 0 | 0 | 0 | 0 | 0 | 0 | 0 | 0 | 0 | 0 | 0 | NA | NA | NA | NA | NA | NA | NA | NA | NA | NA |
| Olfr616 | 0 | 0 | 0 | 0 | 0 | 0 | 0 | 0 | 0 | 0 | 0 | 0 | 0 | 0 | 0 | NA | NA | NA | NA | NA | NA | NA | NA | NA | NA |
| Olfr617 | 0 | 0 | 0 | 0 | 0 | 0 | 0 | 0 | 0 | 0 | 0 | 0 | 0 | 0 | 0 | NA | NA | NA | NA | NA | NA | NA | NA | NA | NA |
| Olfr618 | 0 | 0 | 0 | 0 | 0 | 0 | 0 | 0 | 0 | 0 | 0 | 0 | 0 | 0 | 0 | NA | NA | NA | NA | NA | NA | NA | NA | NA | NA |
| Olfr619 | 0 | 0 | 0 | 0 | 0 | 0 | 0 | 0 | 0 | 0 | 0 | 0 | 0 | 0 | 0 | NA | NA | NA | NA | NA | NA | NA | NA | NA | NA |
| Olfr62 | 0 | 0 | 0 | 0 | 0 | 0 | 0 | 0 | 0 | 0 | 0 | 0 | 0 | 0 | 0 | NA | NA | NA | NA | NA | NA | NA | NA | NA | NA |

| | | | | | | | | | | | | | | | | | | | | | | | | | |
|---|---|---|---|---|---|---|---|---|---|---|---|---|---|---|---|---|---|---|---|---|---|---|---|---|---|
| Olfr620 | 0 | 0 | 0 | 0 | 0 | 0 | 0 | 0 | 0 | 0 | 0 | 0 | 0 | 0 | 0 | NA | NA | NA | NA | NA | NA | NA | NA | NA | NA |
| Olfr622 | 0 | 0 | 0 | 0 | 0 | 0 | 0 | 0 | 0 | 0 | 0 | 0 | 0 | 0 | 0 | NA | NA | NA | NA | NA | NA | NA | NA | NA | NA |
| Olfr623 | 0 | 0 | 0 | 0 | 0 | 0 | 0 | 0 | 0 | 0 | 0 | 0 | 0 | 0 | 0 | NA | NA | NA | NA | NA | NA | NA | NA | NA | NA |
| Olfr624 | 0 | 0 | 0 | 0 | 0 | 0 | 0 | 0 | 0 | 0 | 0 | 0 | 0 | 0 | 0 | NA | NA | NA | NA | NA | NA | NA | NA | NA | NA |
| Olfr628 | 0 | 0 | 0 | 0 | 0 | 0 | 0 | 0 | 0 | 0 | 0 | 0 | 0 | 0 | 0 | NA | NA | NA | NA | NA | NA | NA | NA | NA | NA |
| Olfr629 | 0 | 0 | 0 | 0 | 0 | 0 | 0 | 0 | 0 | 0 | 0 | 0 | 0 | 0 | 0 | NA | NA | NA | NA | NA | NA | NA | NA | NA | NA |
| Olfr63 | 0 | 0 | 0 | 0 | 0 | 0 | 0 | 0 | 0 | 0 | 0 | 0 | 0 | 0 | 0 | NA | NA | NA | NA | NA | NA | NA | NA | NA | NA |
| Olfr630 | 0 | 0 | 0 | 0 | 0 | 0 | 0 | 0 | 0 | 0 | 0 | 0 | 0 | 0 | 0 | NA | NA | NA | NA | NA | NA | NA | NA | NA | NA |
| Olfr631 | 0 | 0 | 0 | 0 | 0 | 0 | 0 | 0 | 0 | 0 | 0 | 0 | 0 | 0 | 0 | NA | NA | NA | NA | NA | NA | NA | NA | NA | NA |
| Olfr632 | 0 | 0 | 0 | 0 | 0 | 0 | 0 | 0 | 0 | 0 | 0 | 0 | 0 | 0 | 0 | NA | NA | NA | NA | NA | NA | NA | NA | NA | NA |
| Olfr633 | 0 | 0 | 0 | 0 | 0 | 0 | 0 | 0 | 0 | 0 | 0 | 0 | 0 | 0 | 0 | NA | NA | NA | NA | NA | NA | NA | NA | NA | NA |
| Olfr635 | 0 | 0 | 0 | 0 | 0 | 0 | 0 | 0 | 0 | 0 | 0 | 0 | 0 | 0 | 0 | NA | NA | NA | NA | NA | NA | NA | NA | NA | NA |
| Olfr638 | 0 | 0 | 0 | 1 | 0 | 0 | 0 | 0 | 0 | 1 | 0 | 0 | 0.17 | 0 | 0 | 2.094 | 1.067 | 0.79379329 | NA | | 0 | 14.2885 | 10 | 10 | TRUE | 0.3906 |
| Olfr639 | 0 | 0 | 0 | 0 | 0 | 0 | 0 | 0 | 0 | 0 | 0 | 0 | 0 | 0 | 0 | NA | NA | NA | NA | NA | NA | NA | NA | NA | NA |
| Olfr64 | 0 | 0 | 0 | 0 | 0 | 0 | 0 | 0 | 0 | 0 | 0 | 0 | 0 | 0 | 0 | NA | NA | NA | NA | NA | NA | NA | NA | NA | NA |
| Olfr640 | 0 | 0 | 0 | 0 | 0 | 0 | 0 | 0 | 0 | 0 | 0 | 0 | 0 | 0 | 0 | NA | NA | NA | NA | NA | NA | NA | NA | NA | NA |
| Olfr641 | 0 | 0 | 0 | 0 | 0 | 0 | 0 | 0 | 0 | 0 | 0 | 0 | 0 | 0 | 0 | NA | NA | NA | NA | NA | NA | NA | NA | NA | NA |
| Olfr642 | 0 | 0 | 0 | 0 | 0 | 0 | 0 | 0 | 0 | 0 | 0 | 0 | 0 | 0 | 0 | NA | NA | NA | NA | NA | NA | NA | NA | NA | NA |
| Olfr643 | 0 | 0 | 0 | 0 | 0 | 0 | 0 | 0 | 0 | 0 | 0 | 0 | 0 | 0 | 0 | NA | NA | NA | NA | NA | NA | NA | NA | NA | NA |
| Olfr644 | 0 | 0 | 0 | 0 | 0 | 0 | 0 | 0 | 0 | 0 | 0 | 0 | 0 | 0 | 0 | NA | NA | NA | NA | NA | NA | NA | NA | NA | NA |
| Olfr645 | 0 | 0 | 0 | 0 | 0 | 0 | 0 | 0 | 0 | 0 | 0 | 0 | 0 | 0 | 0 | NA | NA | NA | NA | NA | NA | NA | NA | NA | NA |
| Olfr646 | 0 | 0 | 0 | 0 | 0 | 0 | 0 | 0 | 0 | 0 | 0 | 0 | 0 | 0 | 0 | NA | NA | NA | NA | NA | NA | NA | NA | NA | NA |
| Olfr648 | 0 | 0 | 0 | 0 | 0 | 0 | 0 | 0 | 0 | 0 | 0 | 0 | 0 | 0 | 0 | NA | NA | NA | NA | NA | NA | NA | NA | NA | NA |
| Olfr649 | 0 | 0 | 0 | 0 | 0 | 0 | 0 | 0 | 0 | 0 | 0 | 0 | 0 | 0 | 0 | NA | NA | NA | NA | NA | NA | NA | NA | NA | NA |
| Olfr65 | 0 | 0 | 0 | 0 | 0 | 0 | 0 | 0 | 0 | 0 | 0 | 0 | 0 | 0 | 0 | NA | NA | NA | NA | NA | NA | NA | NA | NA | NA |
| Olfr651 | 0 | 0 | 0 | 0 | 0 | 0 | 0 | 0 | 0 | 0 | 0 | 0 | 0 | 0 | 0 | NA | NA | NA | NA | NA | NA | NA | NA | NA | NA |
| Olfr652 | 0 | 0 | 0 | 0 | 0 | 0 | 0 | 0 | 0 | 0 | 0 | 0 | 0 | 0 | 0 | NA | NA | NA | NA | NA | NA | NA | NA | NA | NA |
| Olfr653 | 0 | 0 | 0 | 0 | 0 | 0 | 0 | 0 | 0 | 0 | 0 | 0 | 0 | 0 | 0 | NA | NA | NA | NA | NA | NA | NA | NA | NA | NA |
| Olfr654 | 0 | 0 | 0 | 0 | 0 | 0 | 0 | 0 | 0 | 0 | 0 | 0 | 0 | 0 | 0 | NA | NA | NA | NA | NA | NA | NA | NA | NA | NA |
| Olfr655 | 0 | 0 | 0 | 0 | 0 | 0 | 0 | 0 | 0 | 0 | 0 | 0 | 0 | 0 | 0 | NA | NA | NA | NA | NA | NA | NA | NA | NA | NA |
| Olfr656 | 0 | 0 | 0 | 0 | 0 | 0 | 0 | 0 | 0 | 0 | 0 | 0 | 0 | 0 | 0 | NA | NA | NA | NA | NA | NA | NA | NA | NA | NA |
| Olfr657 | 0 | 0 | 0 | 0 | 0 | 0 | 0 | 0 | 0 | 0 | 0 | 0 | 0 | 0 | 0 | NA | NA | NA | NA | NA | NA | NA | NA | NA | NA |
| Olfr658 | 6 | 6 | 3 | 8 | 2 | 5 | 5 | 6 | 3 | 8 | 2 | 5 | 4.91 | 5 | 5 | 1.071 | 0.098 | 0.91071731 | NA | | 0 | 0.4856 | 0.3519 | 0.3519 | TRUE | 0.2039 |
| Olfr659 | 0 | 0 | 0 | 0 | 0 | 0 | 0 | 0 | 0 | 0 | 0 | 0 | 0 | 0 | 0 | NA | NA | NA | NA | NA | NA | NA | NA | NA | NA |
| Olfr66 | 0 | 0 | 0 | 0 | 0 | 0 | 0 | 0 | 0 | 0 | 0 | 0 | 0 | 0 | 0 | NA | NA | NA | NA | NA | NA | NA | NA | NA | NA |
| Olfr661 | 0 | 0 | 0 | 0 | 0 | 0 | 0 | 0 | 0 | 0 | 0 | 0 | 0 | 0 | 0 | NA | NA | NA | NA | NA | NA | NA | NA | NA | NA |
| Olfr663 | 0 | 0 | 0 | 0 | 0 | 0 | 0 | 0 | 0 | 0 | 0 | 0 | 0 | 0 | 0 | NA | NA | NA | NA | NA | NA | NA | NA | NA | NA |
| Olfr665 | 0 | 0 | 0 | 0 | 0 | 0 | 0 | 0 | 0 | 0 | 0 | 0 | 0 | 0 | 0 | NA | NA | NA | NA | NA | NA | NA | NA | NA | NA |
| Olfr666 | 0 | 0 | 0 | 0 | 0 | 0 | 0 | 0 | 0 | 0 | 0 | 0 | 0 | 0 | 0 | NA | NA | NA | NA | NA | NA | NA | NA | NA | NA |
| Olfr667 | 0 | 0 | 0 | 0 | 0 | 0 | 0 | 0 | 0 | 0 | 0 | 0 | 0 | 0 | 0 | NA | NA | NA | NA | NA | NA | NA | NA | NA | NA |
| Olfr668 | 0 | 0 | 0 | 0 | 0 | 0 | 0 | 0 | 0 | 0 | 0 | 0 | 0 | 0 | 0 | NA | NA | NA | NA | NA | NA | NA | NA | NA | NA |
| Olfr669 | 0 | 0 | 0 | 0 | 0 | 0 | 0 | 0 | 0 | 0 | 0 | 0 | 0 | 0 | 0 | NA | NA | NA | NA | NA | NA | NA | NA | NA | NA |
| Olfr67 | 0 | 0 | 0 | 0 | 0 | 0 | 0 | 0 | 0 | 0 | 0 | 0 | 0 | 0 | 0 | NA | NA | NA | NA | NA | NA | NA | NA | NA | NA |
| Olfr670 | 0 | 0 | 0 | 0 | 0 | 0 | 0 | 0 | 0 | 0 | 0 | 0 | 0 | 0 | 0 | NA | NA | NA | NA | NA | NA | NA | NA | NA | NA |
| Olfr671 | 0 | 0 | 0 | 0 | 0 | 0 | 0 | 0 | 0 | 0 | 0 | 0 | 0 | 0 | 0 | NA | NA | NA | NA | NA | NA | NA | NA | NA | NA |
| Olfr672 | 0 | 0 | 0 | 0 | 0 | 0 | 0 | 0 | 0 | 0 | 0 | 0 | 0 | 0 | 0 | NA | NA | NA | NA | NA | NA | NA | NA | NA | NA |
| Olfr675 | 0 | 0 | 0 | 0 | 0 | 0 | 0 | 0 | 0 | 0 | 0 | 0 | 0 | 0 | 0 | NA | NA | NA | NA | NA | NA | NA | NA | NA | NA |
| Olfr676 | 0 | 0 | 0 | 0 | 0 | 0 | 0 | 0 | 0 | 0 | 0 | 0 | 0 | 0 | 0 | NA | NA | NA | NA | NA | NA | NA | NA | NA | NA |
| Olfr677 | 0 | 0 | 0 | 0 | 0 | 0 | 0 | 0 | 0 | 0 | 0 | 0 | 0 | 0 | 0 | NA | NA | NA | NA | NA | NA | NA | NA | NA | NA |
| Olfr678 | 0 | 0 | 0 | 0 | 0 | 0 | 0 | 0 | 0 | 0 | 0 | 0 | 0 | 0 | 0 | NA | NA | NA | NA | NA | NA | NA | NA | NA | NA |
| Olfr679 | 0 | 0 | 0 | 0 | 0 | 0 | 0 | 0 | 0 | 0 | 0 | 0 | 0 | 0 | 0 | NA | NA | NA | NA | NA | NA | NA | NA | NA | NA |
| Olfr68 | 0 | 0 | 0 | 0 | 0 | 0 | 0 | 0 | 0 | 0 | 0 | 0 | 0 | 0 | 0 | NA | NA | NA | NA | NA | NA | NA | NA | NA | NA |
| Olfr681 | 0 | 0 | 0 | 0 | 0 | 0 | 0 | 0 | 0 | 0 | 0 | 0 | 0 | 0 | 0 | NA | NA | NA | NA | NA | NA | NA | NA | NA | NA |
| Olfr683 | 0 | 0 | 0 | 0 | 0 | 0 | 0 | 0 | 0 | 0 | 0 | 0 | 0 | 0 | 0 | NA | NA | NA | NA | NA | NA | NA | NA | NA | NA |
| Olfr684 | 0 | 0 | 0 | 0 | 0 | 0 | 0 | 0 | 0 | 0 | 0 | 0 | 0 | 0 | 0 | NA | NA | NA | NA | NA | NA | NA | NA | NA | NA |
| Olfr685 | 0 | 0 | 0 | 0 | 0 | 0 | 0 | 0 | 0 | 0 | 0 | 0 | 0 | 0 | 0 | NA | NA | NA | NA | NA | NA | NA | NA | NA | NA |
| Olfr686 | 0 | 0 | 0 | 0 | 0 | 0 | 0 | 0 | 0 | 0 | 0 | 0 | 0 | 0 | 0 | NA | NA | NA | NA | NA | NA | NA | NA | NA | NA |
| Olfr688 | 0 | 0 | 0 | 0 | 0 | 0 | 0 | 0 | 0 | 0 | 0 | 0 | 0 | 0 | 0 | NA | NA | NA | NA | NA | NA | NA | NA | NA | NA |
| Olfr689 | 0 | 0 | 0 | 0 | 0 | 0 | 0 | 0 | 0 | 0 | 0 | 0 | 0 | 0 | 0 | NA | NA | NA | NA | NA | NA | NA | NA | NA | NA |
| Olfr69 | 0 | 0 | 0 | 0 | 0 | 0 | 0 | 0 | 0 | 0 | 0 | 0 | 0 | 0 | 0 | NA | NA | NA | NA | NA | NA | NA | NA | NA | NA |
| Olfr690 | 1 | 0 | 0 | 0 | 0 | 0 | 1 | 0 | 0 | 0 | 0 | 0 | 0.13 | 0 | 0 | 0.552 | -0.857 | 0.8336548 | NA | | 0 | 18.3121 | 10 | 10 | TRUE | 0.2566 |
| Olfr691 | 0 | 2 | 0 | 1 | 0 | 0 | 0 | 2 | 0 | 1 | 0 | 0 | 0.52 | 1 | 0 | 0.534 | -0.904 | 0.76303124 | NA | | 0 | 4.5562 | 4.7774 | 4.7774 | TRUE | 0.9746 |
| Olfr692 | 0 | 0 | 0 | 0 | 0 | 0 | 0 | 0 | 0 | 0 | 0 | 0 | 0 | 0 | 0 | NA | NA | NA | NA | NA | NA | NA | NA | NA | NA |
| Olfr693 | 0 | 0 | 0 | 0 | 0 | 1 | 0 | 0 | 0 | 0 | 0 | 1 | 0.16 | 0 | 0 | 2.094 | 1.067 | 0.79379329 | NA | | 0 | 14.4078 | 10 | 10 | TRUE | 0.3835 |
| Olfr694 | 0 | 0 | 0 | 0 | 0 | 0 | 0 | 0 | 0 | 0 | 0 | 0 | 0 | 0 | 0 | NA | NA | NA | NA | NA | NA | NA | NA | NA | NA |
| Olfr695 | 0 | 0 | 0 | 0 | 0 | 0 | 0 | 0 | 0 | 0 | 0 | 0 | 0 | 0 | 0 | NA | NA | NA | NA | NA | NA | NA | NA | NA | NA |
| Olfr697 | 1 | 0 | 1 | 0 | 0 | 0 | 1 | 0 | 1 | 0 | 0 | 0 | 0.31 | 1 | 0 | 0.307 | -1.706 | 0.62894649 | NA | | 0 | 7.6657 | 7.2 | 7.2 | TRUE | 0.2067 |
| Olfr698 | 0 | 0 | 0 | 0 | 0 | 0 | 0 | 0 | 0 | 0 | 0 | 0 | 0 | 0 | 0 | NA | NA | NA | NA | NA | NA | NA | NA | NA | NA |
| Olfr699 | 2 | 1 | 1 | 1 | 1 | 0 | 2 | 1 | 1 | 1 | 1 | 0 | 0.97 | 1 | 1 | 0.562 | -0.833 | 0.66403427 | NA | | 0 | 2.446 | 1.5183 | 1.5183 | TRUE | 0.2702 |

| | | | | | | | | | | | | | | | | | | | | | | | | | |
|---|---|---|---|---|---|---|---|---|---|---|---|---|---|---|---|---|---|---|---|---|---|---|---|---|---|
| Olfr70 | 0 | 0 | 0 | 0 | 0 | 0 | 0 | 0 | 0 | 0 | 0 | 0 | 0 | 0 | 0 | NA | NA | NA | NA | NA | NA | NA | NA | NA | NA |
| Olfr700 | 0 | 0 | 0 | 0 | 0 | 0 | 0 | 0 | 0 | 0 | 0 | 0 | 0 | 0 | 0 | NA | NA | NA | NA | NA | NA | NA | NA | NA | NA |
| Olfr701 | 2 | 5 | 3 | 1 | 3 | 1 | 2 | 5 | 3 | 1 | 3 | 1 | 2.58 | 3 | 2 | 0.53 | -0.915 | 0.45912759 | NA | 0 | 0.9213 | 0.6614 | 0.6614 | TRUE | 0.2641 |
| Olfr702 | 0 | 0 | 0 | 0 | 0 | 0 | 0 | 0 | 0 | 0 | 0 | 0 | 0 | 0 | 0 | NA | NA | NA | NA | NA | NA | NA | NA | NA | NA |
| Olfr703 | 0 | 0 | 0 | 0 | 0 | 0 | 0 | 0 | 0 | 0 | 0 | 0 | 0 | 0 | 0 | NA | NA | NA | NA | NA | NA | NA | NA | NA | NA |
| Olfr704 | 0 | 0 | 0 | 0 | 0 | 0 | 0 | 0 | 0 | 0 | 0 | 0 | 0 | 0 | 0 | NA | NA | NA | NA | NA | NA | NA | NA | NA | NA |
| Olfr705 | 0 | 0 | 0 | 1 | 0 | 1 | 0 | 0 | 0 | 1 | 0 | 1 | 0.33 | 0 | 1 | 3.763 | 1.912 | 0.57887588 | NA | 0 | 7.1752 | 6.7756 | 6.7756 | TRUE | 0.2094 |
| Olfr706 | 0 | 0 | 0 | 0 | 0 | 0 | 0 | 0 | 0 | 0 | 0 | 0 | 0 | 0 | 0 | NA | NA | NA | NA | NA | NA | NA | NA | NA | NA |
| Olfr707 | 0 | 0 | 0 | 0 | 0 | 0 | 0 | 0 | 0 | 0 | 0 | 0 | 0 | 0 | 0 | NA | NA | NA | NA | NA | NA | NA | NA | NA | NA |
| Olfr71 | 0 | 0 | 0 | 0 | 0 | 0 | 0 | 0 | 0 | 0 | 0 | 0 | 0 | 0 | 0 | NA | NA | NA | NA | NA | NA | NA | NA | NA | NA |
| Olfr710 | 0 | 0 | 0 | 0 | 0 | 0 | 0 | 0 | 0 | 0 | 0 | 0 | 0 | 0 | 0 | NA | NA | NA | NA | NA | NA | NA | NA | NA | NA |
| Olfr711 | 0 | 0 | 0 | 0 | 0 | 0 | 0 | 0 | 0 | 0 | 0 | 0 | 0 | 0 | 0 | NA | NA | NA | NA | NA | NA | NA | NA | NA | NA |
| Olfr713 | 0 | 0 | 0 | 0 | 0 | 0 | 0 | 0 | 0 | 0 | 0 | 0 | 0 | 0 | 0 | NA | NA | NA | NA | NA | NA | NA | NA | NA | NA |
| Olfr714 | 0 | 0 | 0 | 0 | 0 | 0 | 0 | 0 | 0 | 0 | 0 | 0 | 0 | 0 | 0 | NA | NA | NA | NA | NA | NA | NA | NA | NA | NA |
| Olfr715 | 0 | 0 | 0 | 0 | 0 | 0 | 0 | 0 | 0 | 0 | 0 | 0 | 0 | 0 | 0 | NA | NA | NA | NA | NA | NA | NA | NA | NA | NA |
| Olfr716 | 0 | 0 | 0 | 0 | 0 | 0 | 0 | 0 | 0 | 0 | 0 | 0 | 0 | 0 | 0 | NA | NA | NA | NA | NA | NA | NA | NA | NA | NA |
| Olfr720 | 0 | 0 | 0 | 0 | 0 | 0 | 0 | 0 | 0 | 0 | 0 | 0 | 0 | 0 | 0 | NA | NA | NA | NA | NA | NA | NA | NA | NA | NA |
| Olfr722 | 0 | 0 | 0 | 0 | 0 | 0 | 0 | 0 | 0 | 0 | 0 | 0 | 0 | 0 | 0 | NA | NA | NA | NA | NA | NA | NA | NA | NA | NA |
| Olfr723 | 0 | 0 | 0 | 0 | 0 | 0 | 0 | 0 | 0 | 0 | 0 | 0 | 0 | 0 | 0 | NA | NA | NA | NA | NA | NA | NA | NA | NA | NA |
| Olfr724 | 0 | 0 | 0 | 0 | 0 | 0 | 0 | 0 | 0 | 0 | 0 | 0 | 0 | 0 | 0 | NA | NA | NA | NA | NA | NA | NA | NA | NA | NA |
| Olfr725 | 0 | 0 | 0 | 0 | 0 | 0 | 0 | 0 | 0 | 0 | 0 | 0 | 0 | 0 | 0 | NA | NA | NA | NA | NA | NA | NA | NA | NA | NA |
| Olfr726 | 0 | 0 | 0 | 0 | 0 | 0 | 0 | 0 | 0 | 0 | 0 | 0 | 0 | 0 | 0 | NA | NA | NA | NA | NA | NA | NA | NA | NA | NA |
| Olfr727 | 0 | 0 | 0 | 0 | 0 | 0 | 0 | 0 | 0 | 0 | 0 | 0 | 0 | 0 | 0 | NA | NA | NA | NA | NA | NA | NA | NA | NA | NA |
| Olfr728 | 0 | 0 | 0 | 0 | 0 | 0 | 0 | 0 | 0 | 0 | 0 | 0 | 0 | 0 | 0 | NA | NA | NA | NA | NA | NA | NA | NA | NA | NA |
| Olfr729 | 0 | 0 | 0 | 0 | 0 | 0 | 0 | 0 | 0 | 0 | 0 | 0 | 0 | 0 | 0 | NA | NA | NA | NA | NA | NA | NA | NA | NA | NA |
| Olfr73 | 0 | 0 | 0 | 0 | 0 | 0 | 0 | 0 | 0 | 0 | 0 | 0 | 0 | 0 | 0 | NA | NA | NA | NA | NA | NA | NA | NA | NA | NA |
| Olfr730 | 0 | 0 | 0 | 0 | 0 | 0 | 0 | 0 | 0 | 0 | 0 | 0 | 0 | 0 | 0 | NA | NA | NA | NA | NA | NA | NA | NA | NA | NA |
| Olfr731 | 1 | 0 | 0 | 0 | 1 | 0 | 1 | 0 | 0 | 0 | 1 | 0 | 0.32 | 0 | 0 | 1.075 | 0.105 | 0.97638847 | NA | 0 | 7.4681 | 7.0392 | 7.0392 | TRUE | 0.5075 |
| Olfr732 | 0 | 0 | 0 | 0 | 0 | 0 | 0 | 0 | 0 | 0 | 0 | 0 | 0 | 0 | 0 | NA | NA | NA | NA | NA | NA | NA | NA | NA | NA |
| Olfr733 | 0 | 0 | 0 | 0 | 0 | 0 | 0 | 0 | 0 | 0 | 0 | 0 | 0 | 0 | 0 | NA | NA | NA | NA | NA | NA | NA | NA | NA | NA |
| Olfr734 | 0 | 0 | 0 | 0 | 0 | 0 | 0 | 0 | 0 | 0 | 0 | 0 | 0 | 0 | 0 | NA | NA | NA | NA | NA | NA | NA | NA | NA | NA |
| Olfr735 | 0 | 0 | 0 | 0 | 0 | 0 | 0 | 0 | 0 | 0 | 0 | 0 | 0 | 0 | 0 | NA | NA | NA | NA | NA | NA | NA | NA | NA | NA |
| Olfr736 | 0 | 0 | 0 | 0 | 0 | 0 | 0 | 0 | 0 | 0 | 0 | 0 | 0 | 0 | 0 | NA | NA | NA | NA | NA | NA | NA | NA | NA | NA |
| Olfr738 | 0 | 0 | 0 | 0 | 0 | 0 | 0 | 0 | 0 | 0 | 0 | 0 | 0 | 0 | 0 | NA | NA | NA | NA | NA | NA | NA | NA | NA | NA |
| Olfr739 | 0 | 0 | 0 | 0 | 0 | 0 | 0 | 0 | 0 | 0 | 0 | 0 | 0 | 0 | 0 | NA | NA | NA | NA | NA | NA | NA | NA | NA | NA |
| Olfr74 | 0 | 0 | 0 | 0 | 0 | 0 | 0 | 0 | 0 | 0 | 0 | 0 | 0 | 0 | 0 | NA | NA | NA | NA | NA | NA | NA | NA | NA | NA |
| Olfr740 | 0 | 0 | 0 | 0 | 0 | 0 | 0 | 0 | 0 | 0 | 0 | 0 | 0 | 0 | 0 | NA | NA | NA | NA | NA | NA | NA | NA | NA | NA |
| Olfr741 | 0 | 0 | 0 | 0 | 0 | 0 | 0 | 0 | 0 | 0 | 0 | 0 | 0 | 0 | 0 | NA | NA | NA | NA | NA | NA | NA | NA | NA | NA |
| Olfr742 | 0 | 0 | 0 | 0 | 0 | 0 | 0 | 0 | 0 | 0 | 0 | 0 | 0 | 0 | 0 | NA | NA | NA | NA | NA | NA | NA | NA | NA | NA |
| Olfr743 | 0 | 0 | 0 | 0 | 0 | 0 | 0 | 0 | 0 | 0 | 0 | 0 | 0 | 0 | 0 | NA | NA | NA | NA | NA | NA | NA | NA | NA | NA |
| Olfr744 | 0 | 0 | 0 | 0 | 0 | 0 | 0 | 0 | 0 | 0 | 0 | 0 | 0 | 0 | 0 | NA | NA | NA | NA | NA | NA | NA | NA | NA | NA |
| Olfr745 | 0 | 0 | 0 | 0 | 0 | 0 | 0 | 0 | 0 | 0 | 0 | 0 | 0 | 0 | 0 | NA | NA | NA | NA | NA | NA | NA | NA | NA | NA |
| Olfr746 | 0 | 0 | 0 | 0 | 0 | 0 | 0 | 0 | 0 | 0 | 0 | 0 | 0 | 0 | 0 | NA | NA | NA | NA | NA | NA | NA | NA | NA | NA |
| Olfr747 | 0 | 0 | 0 | 0 | 0 | 0 | 0 | 0 | 0 | 0 | 0 | 0 | 0 | 0 | 0 | NA | NA | NA | NA | NA | NA | NA | NA | NA | NA |
| Olfr748 | 0 | 0 | 0 | 0 | 0 | 0 | 0 | 0 | 0 | 0 | 0 | 0 | 0 | 0 | 0 | NA | NA | NA | NA | NA | NA | NA | NA | NA | NA |
| Olfr749 | 0 | 0 | 0 | 0 | 0 | 0 | 0 | 0 | 0 | 0 | 0 | 0 | 0 | 0 | 0 | NA | NA | NA | NA | NA | NA | NA | NA | NA | NA |
| Olfr75-ps1 | 0 | 0 | 0 | 0 | 0 | 0 | 0 | 0 | 0 | 0 | 0 | 0 | 0 | 0 | 0 | NA | NA | NA | NA | NA | NA | NA | NA | NA | NA |
| Olfr750 | 0 | 0 | 0 | 0 | 0 | 0 | 0 | 0 | 0 | 0 | 0 | 0 | 0 | 0 | 0 | NA | NA | NA | NA | NA | NA | NA | NA | NA | NA |
| Olfr76 | 0 | 0 | 1 | 1 | 0 | 0 | 0 | 0 | 1 | 1 | 0 | 0 | 0.35 | 0 | 0 | 1.075 | 0.105 | 0.97567165 | NA | 0 | 6.8576 | 6.5143 | 6.5143 | TRUE | 0.5365 |
| Olfr761 | 0 | 0 | 0 | 0 | 0 | 0 | 0 | 0 | 0 | 0 | 0 | 0 | 0 | 0 | 0 | NA | NA | NA | NA | NA | NA | NA | NA | NA | NA |
| Olfr763 | 0 | 0 | 0 | 0 | 0 | 0 | 0 | 0 | 0 | 0 | 0 | 0 | 0 | 0 | 0 | NA | NA | NA | NA | NA | NA | NA | NA | NA | NA |
| Olfr765 | 0 | 0 | 0 | 0 | 0 | 0 | 0 | 0 | 0 | 0 | 0 | 0 | 0 | 0 | 0 | NA | NA | NA | NA | NA | NA | NA | NA | NA | NA |
| Olfr767 | 0 | 0 | 0 | 0 | 0 | 0 | 0 | 0 | 0 | 0 | 0 | 0 | 0 | 0 | 0 | NA | NA | NA | NA | NA | NA | NA | NA | NA | NA |
| Olfr768 | 0 | 0 | 0 | 0 | 0 | 0 | 0 | 0 | 0 | 0 | 0 | 0 | 0 | 0 | 0 | NA | NA | NA | NA | NA | NA | NA | NA | NA | NA |
| Olfr769 | 0 | 0 | 0 | 0 | 0 | 0 | 0 | 0 | 0 | 0 | 0 | 0 | 0 | 0 | 0 | NA | NA | NA | NA | NA | NA | NA | NA | NA | NA |
| Olfr77 | 0 | 0 | 0 | 0 | 0 | 0 | 0 | 0 | 0 | 0 | 0 | 0 | 0 | 0 | 0 | NA | NA | NA | NA | NA | NA | NA | NA | NA | NA |
| Olfr770 | 0 | 0 | 0 | 0 | 0 | 0 | 0 | 0 | 0 | 0 | 0 | 0 | 0 | 0 | 0 | NA | NA | NA | NA | NA | NA | NA | NA | NA | NA |
| Olfr771 | 0 | 0 | 0 | 0 | 0 | 0 | 0 | 0 | 0 | 0 | 0 | 0 | 0 | 0 | 0 | NA | NA | NA | NA | NA | NA | NA | NA | NA | NA |
| Olfr772 | 1 | 0 | 1 | 0 | 0 | 0 | 1 | 0 | 1 | 0 | 0 | 0 | 0.31 | 1 | 0 | 0.307 | -1.706 | 0.62894649 | NA | 0 | 7.6657 | 7.2 | 7.2 | TRUE | 0.2067 |
| Olfr773 | 0 | 0 | 0 | 0 | 0 | 1 | 0 | 0 | 0 | 0 | 0 | 1 | 0.16 | 0 | 0 | 2.094 | 1.067 | 0.79379329 | NA | 0 | 14.4078 | 10 | 10 | TRUE | 0.3835 |
| Olfr774 | 0 | 0 | 0 | 0 | 0 | 0 | 0 | 0 | 0 | 0 | 0 | 0 | 0 | 0 | 0 | NA | NA | NA | NA | NA | NA | NA | NA | NA | NA |
| Olfr775 | 0 | 0 | 0 | 0 | 0 | 0 | 0 | 0 | 0 | 0 | 0 | 0 | 0 | 0 | 0 | NA | NA | NA | NA | NA | NA | NA | NA | NA | NA |
| Olfr776 | 0 | 0 | 0 | 0 | 0 | 0 | 0 | 0 | 0 | 0 | 0 | 0 | 0 | 0 | 0 | NA | NA | NA | NA | NA | NA | NA | NA | NA | NA |
| Olfr777 | 0 | 0 | 0 | 0 | 0 | 0 | 0 | 0 | 0 | 0 | 0 | 0 | 0 | 0 | 0 | NA | NA | NA | NA | NA | NA | NA | NA | NA | NA |
| Olfr78 | 0 | 0 | 0 | 0 | 0 | 0 | 0 | 0 | 0 | 0 | 0 | 0 | 0 | 0 | 0 | NA | NA | NA | NA | NA | NA | NA | NA | NA | NA |
| Olfr780 | 0 | 0 | 0 | 0 | 0 | 0 | 0 | 0 | 0 | 0 | 0 | 0 | 0 | 0 | 0 | NA | NA | NA | NA | NA | NA | NA | NA | NA | NA |
| Olfr781 | 0 | 0 | 0 | 0 | 0 | 0 | 0 | 0 | 0 | 0 | 0 | 0 | 0 | 0 | 0 | NA | NA | NA | NA | NA | NA | NA | NA | NA | NA |
| Olfr782 | 0 | 0 | 0 | 0 | 0 | 0 | 0 | 0 | 0 | 0 | 0 | 0 | 0 | 0 | 0 | NA | NA | NA | NA | NA | NA | NA | NA | NA | NA |
| Olfr784 | 0 | 0 | 0 | 0 | 0 | 0 | 0 | 0 | 0 | 0 | 0 | 0 | 0 | 0 | 0 | NA | NA | NA | NA | NA | NA | NA | NA | NA | NA |

| | | | | | | | | | | | | | | | | | | | | | | | | | |
|---|---|---|---|---|---|---|---|---|---|---|---|---|---|---|---|---|---|---|---|---|---|---|---|---|---|
| Olfr786 | 0 | 0 | 0 | 0 | 0 | 0 | 0 | 0 | 0 | 0 | 0 | 0 | 0 | 0 | 0 | NA | NA | NA | NA | NA | NA | NA | NA | NA | NA |
| Olfr787 | 2 | 3 | 0 | 1 | 0 | 3 | 2 | 3 | 0 | 1 | 0 | 3 | 1.45 | 2 | 1 | 0.846 | -0.241 | 0.8887841 | NA | 0.6399 | 1.6367 | 1.4399 | 1.4399 | TRUE | 0.2378 |
| Olfr788 | 0 | 1 | 0 | 0 | 1 | 0 | 0 | 1 | 0 | 0 | 1 | 0 | 0.37 | 0 | 0 | 1.075 | 0.105 | 0.97520514 | NA | 0 | 6.4904 | 6.1969 | 6.1969 | TRUE | 0.5216 |
| Olfr790 | 0 | 0 | 0 | 0 | 0 | 0 | 0 | 0 | 0 | 0 | 0 | 0 | 0 | 0 | 0 | NA | NA | NA | NA | NA | NA | NA | NA | NA | NA |
| Olfr791 | 0 | 0 | 0 | 0 | 0 | 0 | 0 | 0 | 0 | 0 | 0 | 0 | 0 | 0 | 0 | NA | NA | NA | NA | NA | NA | NA | NA | NA | NA |
| Olfr792 | 0 | 0 | 0 | 0 | 0 | 0 | 0 | 0 | 0 | 0 | 0 | 0 | 0 | 0 | 0 | NA | NA | NA | NA | NA | NA | NA | NA | NA | NA |
| Olfr794 | 0 | 0 | 0 | 0 | 0 | 0 | 0 | 0 | 0 | 0 | 0 | 0 | 0 | 0 | 0 | NA | NA | NA | NA | NA | NA | NA | NA | NA | NA |
| Olfr796 | 0 | 0 | 0 | 0 | 0 | 0 | 0 | 0 | 0 | 0 | 0 | 0 | 0 | 0 | 0 | NA | NA | NA | NA | NA | NA | NA | NA | NA | NA |
| Olfr798 | 0 | 0 | 0 | 0 | 0 | 0 | 0 | 0 | 0 | 0 | 0 | 0 | 0 | 0 | 0 | NA | NA | NA | NA | NA | NA | NA | NA | NA | NA |
| Olfr799 | 0 | 0 | 0 | 0 | 0 | 0 | 0 | 0 | 0 | 0 | 0 | 0 | 0 | 0 | 0 | NA | NA | NA | NA | NA | NA | NA | NA | NA | NA |
| Olfr8 | 0 | 0 | 0 | 0 | 0 | 0 | 0 | 0 | 0 | 0 | 0 | 0 | 0 | 0 | 0 | NA | NA | NA | NA | NA | NA | NA | NA | NA | NA |
| Olfr800 | 0 | 0 | 0 | 0 | 0 | 0 | 0 | 0 | 0 | 0 | 0 | 0 | 0 | 0 | 0 | NA | NA | NA | NA | NA | NA | NA | NA | NA | NA |
| Olfr801 | 0 | 0 | 0 | 0 | 0 | 0 | 0 | 0 | 0 | 0 | 0 | 0 | 0 | 0 | 0 | NA | NA | NA | NA | NA | NA | NA | NA | NA | NA |
| Olfr802 | 0 | 0 | 0 | 0 | 0 | 0 | 0 | 0 | 0 | 0 | 0 | 0 | 0 | 0 | 0 | NA | NA | NA | NA | NA | NA | NA | NA | NA | NA |
| Olfr803 | 0 | 0 | 0 | 0 | 0 | 0 | 0 | 0 | 0 | 0 | 0 | 0 | 0 | 0 | 0 | NA | NA | NA | NA | NA | NA | NA | NA | NA | NA |
| Olfr804 | 0 | 0 | 0 | 0 | 0 | 0 | 0 | 0 | 0 | 0 | 0 | 0 | 0 | 0 | 0 | NA | NA | NA | NA | NA | NA | NA | NA | NA | NA |
| Olfr805 | 0 | 0 | 0 | 0 | 0 | 0 | 0 | 0 | 0 | 0 | 0 | 0 | 0 | 0 | 0 | NA | NA | NA | NA | NA | NA | NA | NA | NA | NA |
| Olfr806 | 0 | 0 | 0 | 0 | 0 | 0 | 0 | 0 | 0 | 0 | 0 | 0 | 0 | 0 | 0 | NA | NA | NA | NA | NA | NA | NA | NA | NA | NA |
| Olfr807 | 0 | 0 | 0 | 0 | 0 | 0 | 0 | 0 | 0 | 0 | 0 | 0 | 0 | 0 | 0 | NA | NA | NA | NA | NA | NA | NA | NA | NA | NA |
| Olfr808 | 0 | 0 | 0 | 0 | 0 | 0 | 0 | 0 | 0 | 0 | 0 | 0 | 0 | 0 | 0 | NA | NA | NA | NA | NA | NA | NA | NA | NA | NA |
| Olfr809 | 0 | 0 | 0 | 0 | 0 | 0 | 0 | 0 | 0 | 0 | 0 | 0 | 0 | 0 | 0 | NA | NA | NA | NA | NA | NA | NA | NA | NA | NA |
| Olfr810 | 0 | 0 | 0 | 0 | 0 | 0 | 0 | 0 | 0 | 0 | 0 | 0 | 0 | 0 | 0 | NA | NA | NA | NA | NA | NA | NA | NA | NA | NA |
| Olfr811 | 0 | 0 | 0 | 0 | 0 | 0 | 0 | 0 | 0 | 0 | 0 | 0 | 0 | 0 | 0 | NA | NA | NA | NA | NA | NA | NA | NA | NA | NA |
| Olfr812 | 0 | 0 | 0 | 0 | 0 | 0 | 0 | 0 | 0 | 0 | 0 | 0 | 0 | 0 | 0 | NA | NA | NA | NA | NA | NA | NA | NA | NA | NA |
| Olfr813 | 0 | 0 | 0 | 0 | 0 | 0 | 0 | 0 | 0 | 0 | 0 | 0 | 0 | 0 | 0 | NA | NA | NA | NA | NA | NA | NA | NA | NA | NA |
| Olfr814 | 0 | 0 | 0 | 0 | 0 | 0 | 0 | 0 | 0 | 0 | 0 | 0 | 0 | 0 | 0 | NA | NA | NA | NA | NA | NA | NA | NA | NA | NA |
| Olfr815 | 18 | 13 | 5 | 3 | 3 | 3 | 14 | 14 | 5 | 3 | 3 | 3 | 7.09 | 11 | 3 | 0.278 | -1.845 | 0.02165924 | NA | 0 | 0.3368 | 0.2551 | 0.2551 | TRUE | 0.6901 |
| Olfr816 | 0 | 0 | 0 | 0 | 0 | 0 | 0 | 0 | 0 | 0 | 0 | 0 | 0 | 0 | 0 | NA | NA | NA | NA | NA | NA | NA | NA | NA | NA |
| Olfr818 | 0 | 0 | 0 | 0 | 0 | 0 | 0 | 0 | 0 | 0 | 0 | 0 | 0 | 0 | 0 | NA | NA | NA | NA | NA | NA | NA | NA | NA | NA |
| Olfr819 | 0 | 0 | 1 | 0 | 0 | 0 | 0 | 0 | 1 | 0 | 0 | 0 | 0.18 | 0 | 0 | 0.552 | -0.857 | 0.8336548 | NA | 0 | 13.1817 | 10 | 10 | TRUE | 0.5365 |
| Olfr820 | 0 | 0 | 0 | 0 | 0 | 0 | 0 | 0 | 0 | 0 | 0 | 0 | 0 | 0 | 0 | NA | NA | NA | NA | NA | NA | NA | NA | NA | NA |
| Olfr821 | 0 | 0 | 0 | 0 | 0 | 0 | 0 | 0 | 0 | 0 | 0 | 0 | 0 | 0 | 0 | NA | NA | NA | NA | NA | NA | NA | NA | NA | NA |
| Olfr822 | 0 | 0 | 0 | 0 | 0 | 0 | 0 | 0 | 0 | 0 | 0 | 0 | 0 | 0 | 0 | NA | NA | NA | NA | NA | NA | NA | NA | NA | NA |
| Olfr823 | 0 | 0 | 0 | 0 | 0 | 0 | 0 | 0 | 0 | 0 | 0 | 0 | 0 | 0 | 0 | NA | NA | NA | NA | NA | NA | NA | NA | NA | NA |
| Olfr824 | 0 | 0 | 0 | 0 | 0 | 0 | 0 | 0 | 0 | 0 | 0 | 0 | 0 | 0 | 0 | NA | NA | NA | NA | NA | NA | NA | NA | NA | NA |
| Olfr825 | 0 | 0 | 0 | 0 | 0 | 0 | 0 | 0 | 0 | 0 | 0 | 0 | 0 | 0 | 0 | NA | NA | NA | NA | NA | NA | NA | NA | NA | NA |
| Olfr826 | 0 | 0 | 0 | 0 | 0 | 0 | 0 | 0 | 0 | 0 | 0 | 0 | 0 | 0 | 0 | NA | NA | NA | NA | NA | NA | NA | NA | NA | NA |
| Olfr827 | 0 | 0 | 0 | 0 | 0 | 0 | 0 | 0 | 0 | 0 | 0 | 0 | 0 | 0 | 0 | NA | NA | NA | NA | NA | NA | NA | NA | NA | NA |
| Olfr828 | 0 | 0 | 0 | 0 | 0 | 0 | 0 | 0 | 0 | 0 | 0 | 0 | 0 | 0 | 0 | NA | NA | NA | NA | NA | NA | NA | NA | NA | NA |
| Olfr829 | 0 | 0 | 0 | 0 | 0 | 0 | 0 | 0 | 0 | 0 | 0 | 0 | 0 | 0 | 0 | NA | NA | NA | NA | NA | NA | NA | NA | NA | NA |
| Olfr830 | 0 | 0 | 0 | 0 | 0 | 0 | 0 | 0 | 0 | 0 | 0 | 0 | 0 | 0 | 0 | NA | NA | NA | NA | NA | NA | NA | NA | NA | NA |
| Olfr832 | 0 | 0 | 0 | 0 | 0 | 0 | 0 | 0 | 0 | 0 | 0 | 0 | 0 | 0 | 0 | NA | NA | NA | NA | NA | NA | NA | NA | NA | NA |
| Olfr834 | 0 | 0 | 0 | 0 | 1 | 0 | 0 | 0 | 0 | 0 | 1 | 0 | 0.19 | 0 | 0 | 2.094 | 1.067 | 0.79379329 | NA | 0 | 12.608 | 10 | 10 | TRUE | 0.5075 |
| Olfr835 | 0 | 0 | 0 | 0 | 0 | 0 | 0 | 0 | 0 | 0 | 0 | 0 | 0 | 0 | 0 | NA | NA | NA | NA | NA | NA | NA | NA | NA | NA |
| Olfr836 | 1 | 0 | 1 | 0 | 0 | 0 | 1 | 0 | 1 | 0 | 0 | 0 | 0.31 | 1 | 0 | 0.307 | -1.706 | 0.62894649 | NA | 0 | 7.6657 | 7.2 | 7.2 | TRUE | 0.2067 |
| Olfr837 | 0 | 1 | 0 | 0 | 0 | 0 | 0 | 1 | 0 | 0 | 0 | 0 | 0.18 | 0 | 0 | 0.552 | -0.857 | 0.8336548 | NA | 0 | 13.3714 | 10 | 10 | TRUE | 0.5216 |
| Olfr843 | 0 | 0 | 0 | 0 | 0 | 0 | 0 | 0 | 0 | 0 | 0 | 0 | 0 | 0 | 0 | NA | NA | NA | NA | NA | NA | NA | NA | NA | NA |
| Olfr845 | 0 | 0 | 0 | 0 | 0 | 0 | 0 | 0 | 0 | 0 | 0 | 0 | 0 | 0 | 0 | NA | NA | NA | NA | NA | NA | NA | NA | NA | NA |
| Olfr846 | 2 | 0 | 0 | 0 | 0 | 1 | 2 | 0 | 0 | 0 | 0 | 1 | 0.42 | 1 | 0 | 0.682 | -0.553 | 0.86290097 | NA | 0 | 5.6 | 5.5066 | 5.5066 | TRUE | 0.9727 |
| Olfr847 | 0 | 0 | 0 | 0 | 0 | 0 | 0 | 0 | 0 | 0 | 0 | 0 | 0 | 0 | 0 | NA | NA | NA | NA | NA | NA | NA | NA | NA | NA |
| Olfr849 | 0 | 0 | 0 | 0 | 0 | 0 | 0 | 0 | 0 | 0 | 0 | 0 | 0 | 0 | 0 | NA | NA | NA | NA | NA | NA | NA | NA | NA | NA |
| Olfr850 | 0 | 0 | 0 | 0 | 0 | 0 | 0 | 0 | 0 | 0 | 0 | 0 | 0 | 0 | 0 | NA | NA | NA | NA | NA | NA | NA | NA | NA | NA |
| Olfr851 | 0 | 0 | 0 | 0 | 0 | 0 | 0 | 0 | 0 | 0 | 0 | 0 | 0 | 0 | 0 | NA | NA | NA | NA | NA | NA | NA | NA | NA | NA |
| Olfr853 | 0 | 0 | 0 | 0 | 0 | 0 | 0 | 0 | 0 | 0 | 0 | 0 | 0 | 0 | 0 | NA | NA | NA | NA | NA | NA | NA | NA | NA | NA |
| Olfr854 | 0 | 0 | 0 | 0 | 0 | 0 | 0 | 0 | 0 | 0 | 0 | 0 | 0 | 0 | 0 | NA | NA | NA | NA | NA | NA | NA | NA | NA | NA |
| Olfr855 | 0 | 0 | 0 | 0 | 0 | 0 | 0 | 0 | 0 | 0 | 0 | 0 | 0 | 0 | 0 | NA | NA | NA | NA | NA | NA | NA | NA | NA | NA |
| Olfr856-ps1 | 0 | 0 | 0 | 0 | 0 | 1 | 0 | 0 | 0 | 0 | 0 | 1 | 0.16 | 0 | 0 | 2.094 | 1.067 | 0.79379329 | NA | 0 | 14.4078 | 10 | 10 | TRUE | 0.3835 |
| Olfr857 | 0 | 1 | 1 | 1 | 0 | 1 | 0 | 1 | 1 | 1 | 0 | 1 | 0.69 | 1 | 1 | 0.979 | -0.031 | 0.98943171 | NA | 0 | 3.4496 | 2.4101 | 2.4101 | TRUE | 0.3734 |
| Olfr859 | 2 | 3 | 0 | 0 | 0 | 0 | 2 | 3 | 0 | 0 | 0 | 0 | 0.79 | 2 | 0 | 0.12 | -3.055 | 0.22090216 | NA | 5.3026 | 2.9997 | 3.1827 | 3.1827 | TRUE | 0.0516 |
| Olfr860 | 0 | 0 | 0 | 0 | 0 | 0 | 0 | 0 | 0 | 0 | 0 | 0 | 0 | 0 | 0 | NA | NA | NA | NA | NA | NA | NA | NA | NA | NA |
| Olfr862 | 0 | 0 | 0 | 0 | 0 | 0 | 0 | 0 | 0 | 0 | 0 | 0 | 0 | 0 | 0 | NA | NA | NA | NA | NA | NA | NA | NA | NA | NA |
| Olfr866 | 0 | 0 | 0 | 0 | 0 | 0 | 0 | 0 | 0 | 0 | 0 | 0 | 0 | 0 | 0 | NA | NA | NA | NA | NA | NA | NA | NA | NA | NA |
| Olfr867 | 0 | 0 | 0 | 0 | 0 | 0 | 0 | 0 | 0 | 0 | 0 | 0 | 0 | 0 | 0 | NA | NA | NA | NA | NA | NA | NA | NA | NA | NA |
| Olfr868 | 0 | 0 | 0 | 0 | 0 | 0 | 0 | 0 | 0 | 0 | 0 | 0 | 0 | 0 | 0 | NA | NA | NA | NA | NA | NA | NA | NA | NA | NA |
| Olfr869 | 0 | 0 | 0 | 0 | 0 | 0 | 0 | 0 | 0 | 0 | 0 | 0 | 0 | 0 | 0 | NA | NA | NA | NA | NA | NA | NA | NA | NA | NA |
| Olfr870 | 0 | 0 | 0 | 0 | 0 | 0 | 0 | 0 | 0 | 0 | 0 | 0 | 0 | 0 | 0 | NA | NA | NA | NA | NA | NA | NA | NA | NA | NA |
| Olfr871 | 0 | 0 | 0 | 0 | 0 | 0 | 0 | 0 | 0 | 0 | 0 | 0 | 0 | 0 | 0 | NA | NA | NA | NA | NA | NA | NA | NA | NA | NA |
| Olfr872 | 0 | 0 | 0 | 0 | 0 | 0 | 0 | 0 | 0 | 0 | 0 | 0 | 0 | 0 | 0 | NA | NA | NA | NA | NA | NA | NA | NA | NA | NA |
| Olfr873 | 0 | 0 | 0 | 0 | 0 | 0 | 0 | 0 | 0 | 0 | 0 | 0 | 0 | 0 | 0 | NA | NA | NA | NA | NA | NA | NA | NA | NA | NA |

| | | | | | | | | | | | | | | | | | | | | | | | | | |
|---|---|---|---|---|---|---|---|---|---|---|---|---|---|---|---|---|---|---|---|---|---|---|---|---|---|
| Olfr874 | 0 | 0 | 0 | 0 | 0 | 0 | 0 | 0 | 0 | 0 | 0 | 0 | 0 | 0 | 0 | NA | NA | NA | NA | NA | NA | NA | NA | NA | NA |
| Olfr875 | 0 | 0 | 0 | 0 | 0 | 0 | 0 | 0 | 0 | 0 | 0 | 0 | 0 | 0 | 0 | NA | NA | NA | NA | NA | NA | NA | NA | NA | NA |
| Olfr876 | 0 | 1 | 0 | 0 | 0 | 0 | 0 | 1 | 0 | 0 | 0 | 0 | 0.18 | 0 | 0 | 0.552 | -0.857 | 0.8336548 | NA | 0 | 13.3714 | 10 | 10 | TRUE | 0.5216 |
| Olfr877 | 0 | 0 | 0 | 0 | 0 | 0 | 0 | 0 | 0 | 0 | 0 | 0 | 0 | 0 | 0 | NA | NA | NA | NA | NA | NA | NA | NA | NA | NA |
| Olfr878 | 0 | 0 | 0 | 0 | 0 | 0 | 0 | 0 | 0 | 0 | 0 | 0 | 0 | 0 | 0 | NA | NA | NA | NA | NA | NA | NA | NA | NA | NA |
| Olfr881 | 1 | 2 | 1 | 0 | 1 | 0 | 1 | 2 | 1 | 0 | 1 | 0 | 0.85 | 1 | 0 | 0.282 | -1.824 | 0.39588467 | NA | 0 | 2.7847 | 1.9575 | 1.9575 | TRUE | 0.5075 |
| Olfr883 | 0 | 0 | 0 | 0 | 0 | 1 | 0 | 0 | 0 | 0 | 0 | 1 | 0.16 | 0 | 0 | 2.094 | 1.067 | 0.79379329 | NA | 0 | 14.4078 | 10 | 10 | TRUE | 0.3835 |
| Olfr884 | 0 | 0 | 0 | 0 | 0 | 0 | 0 | 0 | 0 | 0 | 0 | 0 | 0 | 0 | 0 | NA | NA | NA | NA | NA | NA | NA | NA | NA | NA |
| Olfr885 | 0 | 0 | 0 | 0 | 0 | 0 | 0 | 0 | 0 | 0 | 0 | 0 | 0 | 0 | 0 | NA | NA | NA | NA | NA | NA | NA | NA | NA | NA |
| Olfr887 | 0 | 0 | 0 | 0 | 0 | 0 | 0 | 0 | 0 | 0 | 0 | 0 | 0 | 0 | 0 | NA | NA | NA | NA | NA | NA | NA | NA | NA | NA |
| Olfr888 | 0 | 0 | 0 | 0 | 0 | 0 | 0 | 0 | 0 | 0 | 0 | 0 | 0 | 0 | 0 | NA | NA | NA | NA | NA | NA | NA | NA | NA | NA |
| Olfr889 | 0 | 0 | 0 | 0 | 0 | 0 | 0 | 0 | 0 | 0 | 0 | 0 | 0 | 0 | 0 | NA | NA | NA | NA | NA | NA | NA | NA | NA | NA |
| Olfr890 | 0 | 0 | 0 | 0 | 0 | 0 | 0 | 0 | 0 | 0 | 0 | 0 | 0 | 0 | 0 | NA | NA | NA | NA | NA | NA | NA | NA | NA | NA |
| Olfr891 | 0 | 0 | 0 | 0 | 0 | 0 | 0 | 0 | 0 | 0 | 0 | 0 | 0 | 0 | 0 | NA | NA | NA | NA | NA | NA | NA | NA | NA | NA |
| Olfr893 | 0 | 0 | 0 | 0 | 0 | 0 | 0 | 0 | 0 | 0 | 0 | 0 | 0 | 0 | 0 | NA | NA | NA | NA | NA | NA | NA | NA | NA | NA |
| Olfr894 | 0 | 0 | 0 | 0 | 0 | 0 | 0 | 0 | 0 | 0 | 0 | 0 | 0 | 0 | 0 | NA | NA | NA | NA | NA | NA | NA | NA | NA | NA |
| Olfr895 | 0 | 0 | 0 | 0 | 0 | 0 | 0 | 0 | 0 | 0 | 0 | 0 | 0 | 0 | 0 | NA | NA | NA | NA | NA | NA | NA | NA | NA | NA |
| Olfr898 | 0 | 0 | 0 | 0 | 0 | 0 | 0 | 0 | 0 | 0 | 0 | 0 | 0 | 0 | 0 | NA | NA | NA | NA | NA | NA | NA | NA | NA | NA |
| Olfr899 | 0 | 0 | 0 | 0 | 0 | 0 | 0 | 0 | 0 | 0 | 0 | 0 | 0 | 0 | 0 | NA | NA | NA | NA | NA | NA | NA | NA | NA | NA |
| Olfr9 | 0 | 0 | 0 | 0 | 0 | 0 | 0 | 0 | 0 | 0 | 0 | 0 | 0 | 0 | 0 | NA | NA | NA | NA | NA | NA | NA | NA | NA | NA |
| Olfr90 | 0 | 0 | 0 | 0 | 0 | 0 | 0 | 0 | 0 | 0 | 0 | 0 | 0 | 0 | 0 | NA | NA | NA | NA | NA | NA | NA | NA | NA | NA |
| Olfr900 | 0 | 0 | 0 | 0 | 0 | 0 | 0 | 0 | 0 | 0 | 0 | 0 | 0 | 0 | 0 | NA | NA | NA | NA | NA | NA | NA | NA | NA | NA |
| Olfr901 | 4 | 3 | 3 | 0 | 0 | 2 | 3 | 3 | 3 | 0 | 0 | 2 | 1.92 | 3 | 1 | 0.213 | -2.229 | 0.16389317 | NA | 0 | 1.238 | 0.9324 | 0.9324 | TRUE | 0.9733 |
| Olfr902 | 0 | 0 | 0 | 0 | 0 | 0 | 0 | 0 | 0 | 0 | 0 | 0 | 0 | 0 | 0 | NA | NA | NA | NA | NA | NA | NA | NA | NA | NA |
| Olfr904 | 0 | 0 | 0 | 0 | 0 | 0 | 0 | 0 | 0 | 0 | 0 | 0 | 0 | 0 | 0 | NA | NA | NA | NA | NA | NA | NA | NA | NA | NA |
| Olfr905 | 0 | 0 | 0 | 0 | 0 | 0 | 0 | 0 | 0 | 0 | 0 | 0 | 0 | 0 | 0 | NA | NA | NA | NA | NA | NA | NA | NA | NA | NA |
| Olfr906 | 0 | 0 | 0 | 0 | 0 | 0 | 0 | 0 | 0 | 0 | 0 | 0 | 0 | 0 | 0 | NA | NA | NA | NA | NA | NA | NA | NA | NA | NA |
| Olfr907 | 0 | 0 | 0 | 0 | 0 | 0 | 0 | 0 | 0 | 0 | 0 | 0 | 0 | 0 | 0 | NA | NA | NA | NA | NA | NA | NA | NA | NA | NA |
| Olfr908 | 0 | 0 | 0 | 0 | 0 | 0 | 0 | 0 | 0 | 0 | 0 | 0 | 0 | 0 | 0 | NA | NA | NA | NA | NA | NA | NA | NA | NA | NA |
| Olfr91 | 0 | 0 | 0 | 0 | 0 | 0 | 0 | 0 | 0 | 0 | 0 | 0 | 0 | 0 | 0 | NA | NA | NA | NA | NA | NA | NA | NA | NA | NA |
| Olfr910 | 0 | 0 | 0 | 0 | 0 | 0 | 0 | 0 | 0 | 0 | 0 | 0 | 0 | 0 | 0 | NA | NA | NA | NA | NA | NA | NA | NA | NA | NA |
| Olfr911-ps1 | 0 | 0 | 0 | 0 | 0 | 0 | 0 | 0 | 0 | 0 | 0 | 0 | 0 | 0 | 0 | NA | NA | NA | NA | NA | NA | NA | NA | NA | NA |
| Olfr912 | 0 | 0 | 0 | 0 | 0 | 0 | 0 | 0 | 0 | 0 | 0 | 0 | 0 | 0 | 0 | NA | NA | NA | NA | NA | NA | NA | NA | NA | NA |
| Olfr913 | 0 | 0 | 0 | 0 | 0 | 0 | 0 | 0 | 0 | 0 | 0 | 0 | 0 | 0 | 0 | NA | NA | NA | NA | NA | NA | NA | NA | NA | NA |
| Olfr914 | 0 | 0 | 0 | 0 | 0 | 0 | 0 | 0 | 0 | 0 | 0 | 0 | 0 | 0 | 0 | NA | NA | NA | NA | NA | NA | NA | NA | NA | NA |
| Olfr915 | 1 | 0 | 0 | 0 | 0 | 0 | 1 | 0 | 0 | 0 | 0 | 0 | 0.13 | 0 | 0 | 0.552 | -0.857 | 0.8336548 | NA | 0 | 18.3121 | 10 | 10 | TRUE | 0.2566 |
| Olfr916 | 0 | 0 | 0 | 0 | 0 | 0 | 0 | 0 | 0 | 0 | 0 | 0 | 0 | 0 | 0 | NA | NA | NA | NA | NA | NA | NA | NA | NA | NA |
| Olfr917 | 0 | 0 | 0 | 0 | 0 | 0 | 0 | 0 | 0 | 0 | 0 | 0 | 0 | 0 | 0 | NA | NA | NA | NA | NA | NA | NA | NA | NA | NA |
| Olfr918 | 0 | 0 | 0 | 0 | 0 | 0 | 0 | 0 | 0 | 0 | 0 | 0 | 0 | 0 | 0 | NA | NA | NA | NA | NA | NA | NA | NA | NA | NA |
| Olfr919 | 0 | 0 | 0 | 0 | 0 | 0 | 0 | 0 | 0 | 0 | 0 | 0 | 0 | 0 | 0 | NA | NA | NA | NA | NA | NA | NA | NA | NA | NA |
| Olfr92 | 0 | 0 | 0 | 0 | 0 | 0 | 0 | 0 | 0 | 0 | 0 | 0 | 0 | 0 | 0 | NA | NA | NA | NA | NA | NA | NA | NA | NA | NA |
| Olfr920 | 5 | 0 | 4 | 2 | 4 | 1 | 4 | 0 | 4 | 2 | 5 | 1 | 2.61 | 3 | 3 | 0.892 | -0.166 | 0.89728008 | NA | 0.4345 | 0.9088 | 0.802 | 0.802 | TRUE | 0.8326 |
| Olfr921 | 1 | 0 | 0 | 0 | 0 | 0 | 0 | 1 | 0 | 0 | 0 | 0 | 0.13 | 0 | 0 | 0.552 | -0.857 | 0.8336548 | NA | 0 | 18.3121 | 10 | 10 | TRUE | 0.2566 |
| Olfr922 | 0 | 0 | 0 | 0 | 0 | 0 | 0 | 0 | 0 | 0 | 0 | 0 | 0 | 0 | 0 | NA | NA | NA | NA | NA | NA | NA | NA | NA | NA |
| Olfr923 | 0 | 0 | 0 | 0 | 0 | 0 | 0 | 0 | 0 | 0 | 0 | 0 | 0 | 0 | 0 | NA | NA | NA | NA | NA | NA | NA | NA | NA | NA |
| Olfr924 | 0 | 0 | 0 | 0 | 0 | 0 | 0 | 0 | 0 | 0 | 0 | 0 | 0 | 0 | 0 | NA | NA | NA | NA | NA | NA | NA | NA | NA | NA |
| Olfr926 | 0 | 0 | 0 | 0 | 0 | 0 | 0 | 0 | 0 | 0 | 0 | 0 | 0 | 0 | 0 | NA | NA | NA | NA | NA | NA | NA | NA | NA | NA |
| Olfr93 | 0 | 0 | 0 | 0 | 0 | 0 | 0 | 0 | 0 | 0 | 0 | 0 | 0 | 0 | 0 | NA | NA | NA | NA | NA | NA | NA | NA | NA | NA |
| Olfr930 | 0 | 0 | 0 | 0 | 0 | 0 | 0 | 0 | 0 | 0 | 0 | 0 | 0 | 0 | 0 | NA | NA | NA | NA | NA | NA | NA | NA | NA | NA |
| Olfr933 | 0 | 0 | 0 | 0 | 0 | 0 | 0 | 0 | 0 | 0 | 0 | 0 | 0 | 0 | 0 | NA | NA | NA | NA | NA | NA | NA | NA | NA | NA |
| Olfr934 | 0 | 0 | 0 | 0 | 0 | 0 | 0 | 0 | 0 | 0 | 0 | 0 | 0 | 0 | 0 | NA | NA | NA | NA | NA | NA | NA | NA | NA | NA |
| Olfr935 | 0 | 0 | 0 | 0 | 0 | 0 | 0 | 0 | 0 | 0 | 0 | 0 | 0 | 0 | 0 | NA | NA | NA | NA | NA | NA | NA | NA | NA | NA |
| Olfr936 | 0 | 0 | 0 | 0 | 0 | 0 | 0 | 0 | 0 | 0 | 0 | 0 | 0 | 0 | 0 | NA | NA | NA | NA | NA | NA | NA | NA | NA | NA |
| Olfr937 | 0 | 0 | 0 | 0 | 0 | 0 | 0 | 0 | 0 | 0 | 0 | 0 | 0 | 0 | 0 | NA | NA | NA | NA | NA | NA | NA | NA | NA | NA |
| Olfr938 | 0 | 0 | 0 | 0 | 0 | 0 | 0 | 0 | 0 | 0 | 0 | 0 | 0 | 0 | 0 | NA | NA | NA | NA | NA | NA | NA | NA | NA | NA |
| Olfr94 | 0 | 0 | 0 | 0 | 0 | 0 | 0 | 0 | 0 | 0 | 0 | 0 | 0 | 0 | 0 | NA | NA | NA | NA | NA | NA | NA | NA | NA | NA |
| Olfr943 | 0 | 0 | 0 | 0 | 0 | 0 | 0 | 0 | 0 | 0 | 0 | 0 | 0 | 0 | 0 | NA | NA | NA | NA | NA | NA | NA | NA | NA | NA |
| Olfr944 | 0 | 0 | 0 | 0 | 0 | 0 | 0 | 0 | 0 | 0 | 0 | 0 | 0 | 0 | 0 | NA | NA | NA | NA | NA | NA | NA | NA | NA | NA |
| Olfr945 | 0 | 0 | 0 | 0 | 0 | 0 | 0 | 0 | 0 | 0 | 0 | 0 | 0 | 0 | 0 | NA | NA | NA | NA | NA | NA | NA | NA | NA | NA |
| Olfr947-ps1 | 0 | 0 | 0 | 0 | 0 | 0 | 0 | 0 | 0 | 0 | 0 | 0 | 0 | 0 | 0 | NA | NA | NA | NA | NA | NA | NA | NA | NA | NA |
| Olfr948 | 0 | 0 | 0 | 0 | 0 | 0 | 0 | 0 | 0 | 0 | 0 | 0 | 0 | 0 | 0 | NA | NA | NA | NA | NA | NA | NA | NA | NA | NA |
| Olfr95 | 0 | 0 | 0 | 0 | 0 | 0 | 0 | 0 | 0 | 0 | 0 | 0 | 0 | 0 | 0 | NA | NA | NA | NA | NA | NA | NA | NA | NA | NA |
| Olfr951 | 0 | 0 | 0 | 0 | 0 | 0 | 0 | 0 | 0 | 0 | 0 | 0 | 0 | 0 | 0 | NA | NA | NA | NA | NA | NA | NA | NA | NA | NA |
| Olfr952 | 0 | 0 | 0 | 0 | 0 | 0 | 0 | 0 | 0 | 0 | 0 | 0 | 0 | 0 | 0 | NA | NA | NA | NA | NA | NA | NA | NA | NA | NA |
| Olfr954 | 0 | 0 | 0 | 0 | 0 | 0 | 0 | 0 | 0 | 0 | 0 | 0 | 0 | 0 | 0 | NA | NA | NA | NA | NA | NA | NA | NA | NA | NA |
| Olfr955 | 0 | 0 | 0 | 0 | 0 | 0 | 0 | 0 | 0 | 0 | 0 | 0 | 0 | 0 | 0 | NA | NA | NA | NA | NA | NA | NA | NA | NA | NA |
| Olfr957 | 0 | 0 | 0 | 0 | 0 | 0 | 0 | 0 | 0 | 0 | 0 | 0 | 0 | 0 | 0 | NA | NA | NA | NA | NA | NA | NA | NA | NA | NA |
| Olfr958 | 0 | 0 | 0 | 0 | 0 | 0 | 0 | 0 | 0 | 0 | 0 | 0 | 0 | 0 | 0 | NA | NA | NA | NA | NA | NA | NA | NA | NA | NA |
| Olfr959 | 0 | 0 | 0 | 0 | 0 | 0 | 0 | 0 | 0 | 0 | 0 | 0 | 0 | 0 | 0 | NA | NA | NA | NA | NA | NA | NA | NA | NA | NA |

| | | | | | | | | | | | | | | | | | | | | | | | | | |
|---|---|---|---|---|---|---|---|---|---|---|---|---|---|---|---|---|---|---|---|---|---|---|---|---|---|
| Olfr96 | 0 | 0 | 0 | 0 | 0 | 0 | 0 | 0 | 0 | 0 | 0 | 0 | 0 | 0 | 0 | NA | NA | NA | NA | NA | NA | NA | NA | NA | NA |
| Olfr960 | 1 | 2 | 1 | 0 | 0 | 0 | 1 | 2 | 1 | 0 | 0 | 0 | 0.66 | 1 | 0 | 0.145 | -2.79 | 0.24993894 | NA | 0 | 3.5727 | 2.8701 | 2.8701 | TRUE | 0.1613 |
| Olfr961 | 0 | 0 | 0 | 0 | 0 | 0 | 0 | 0 | 0 | 0 | 0 | 0 | 0 | 0 | 0 | NA | NA | NA | NA | NA | NA | NA | NA | NA | NA |
| Olfr963 | 0 | 0 | 0 | 0 | 0 | 0 | 0 | 0 | 0 | 0 | 0 | 0 | 0 | 0 | 0 | NA | NA | NA | NA | NA | NA | NA | NA | NA | NA |
| Olfr965 | 0 | 0 | 0 | 0 | 0 | 0 | 0 | 0 | 0 | 0 | 0 | 0 | 0 | 0 | 0 | NA | NA | NA | NA | NA | NA | NA | NA | NA | NA |
| Olfr967 | 0 | 0 | 0 | 0 | 0 | 0 | 0 | 0 | 0 | 0 | 0 | 0 | 0 | 0 | 0 | NA | NA | NA | NA | NA | NA | NA | NA | NA | NA |
| Olfr968 | 0 | 1 | 0 | 0 | 1 | 0 | 0 | 1 | 0 | 0 | 1 | 0 | 0.37 | 0 | 0 | 1.075 | 0.105 | 0.97520514 | NA | 0 | 6.4904 | 6.1969 | 6.1969 | TRUE | 0.5216 |
| Olfr969 | 0 | 0 | 0 | 0 | 0 | 0 | 0 | 0 | 0 | 0 | 0 | 0 | 0 | 0 | 0 | NA | NA | NA | NA | NA | NA | NA | NA | NA | NA |
| Olfr97 | 0 | 0 | 0 | 0 | 0 | 0 | 0 | 0 | 0 | 0 | 0 | 0 | 0 | 0 | 0 | NA | NA | NA | NA | NA | NA | NA | NA | NA | NA |
| Olfr970 | 0 | 0 | 0 | 0 | 0 | 0 | 0 | 0 | 0 | 0 | 0 | 0 | 0 | 0 | 0 | NA | NA | NA | NA | NA | NA | NA | NA | NA | NA |
| Olfr971 | 0 | 0 | 0 | 0 | 0 | 0 | 0 | 0 | 0 | 0 | 0 | 0 | 0 | 0 | 0 | NA | NA | NA | NA | NA | NA | NA | NA | NA | NA |
| Olfr972 | 0 | 0 | 0 | 0 | 0 | 0 | 0 | 0 | 0 | 0 | 0 | 0 | 0 | 0 | 0 | NA | NA | NA | NA | NA | NA | NA | NA | NA | NA |
| Olfr974 | 0 | 0 | 0 | 0 | 0 | 0 | 0 | 0 | 0 | 0 | 0 | 0 | 0 | 0 | 0 | NA | NA | NA | NA | NA | NA | NA | NA | NA | NA |
| Olfr975 | 2 | 0 | 0 | 0 | 0 | 0 | 2 | 0 | 0 | 0 | 0 | 0 | 0.26 | 1 | 0 | 0.356 | -1.491 | 0.71362796 | NA | 0 | 9.1573 | 10 | 10 | TRUE | 0.968 |
| Olfr976 | 0 | 1 | 1 | 2 | 0 | 0 | 0 | 1 | 1 | 2 | 0 | 0 | 0.69 | 1 | 1 | 0.974 | -0.038 | 0.9877327 | NA | 0 | 3.4427 | 3.0223 | 3.0223 | TRUE | 0.9737 |
| Olfr978 | 0 | 0 | 0 | 0 | 0 | 0 | 0 | 0 | 0 | 0 | 0 | 0 | 0 | 0 | 0 | NA | NA | NA | NA | NA | NA | NA | NA | NA | NA |
| Olfr979 | 1 | 0 | 0 | 0 | 0 | 0 | 1 | 0 | 0 | 0 | 0 | 0 | 0.13 | 0 | 0 | 0.552 | -0.857 | 0.8336548 | NA | 0 | 18.3121 | 10 | 10 | TRUE | 0.2566 |
| Olfr98 | 0 | 0 | 0 | 0 | 0 | 0 | 0 | 0 | 0 | 0 | 0 | 0 | 0 | 0 | 0 | NA | NA | NA | NA | NA | NA | NA | NA | NA | NA |
| Olfr980 | 0 | 0 | 0 | 0 | 0 | 0 | 0 | 0 | 0 | 0 | 0 | 0 | 0 | 0 | 0 | NA | NA | NA | NA | NA | NA | NA | NA | NA | NA |
| Olfr981 | 0 | 0 | 0 | 0 | 0 | 0 | 0 | 0 | 0 | 0 | 0 | 0 | 0 | 0 | 0 | NA | NA | NA | NA | NA | NA | NA | NA | NA | NA |
| Olfr982 | 0 | 0 | 0 | 0 | 0 | 0 | 0 | 0 | 0 | 0 | 0 | 0 | 0 | 0 | 0 | NA | NA | NA | NA | NA | NA | NA | NA | NA | NA |
| Olfr983 | 0 | 0 | 0 | 0 | 0 | 0 | 0 | 0 | 0 | 0 | 0 | 0 | 0 | 0 | 0 | NA | NA | NA | NA | NA | NA | NA | NA | NA | NA |
| Olfr984 | 0 | 1 | 0 | 0 | 0 | 0 | 0 | 1 | 0 | 0 | 0 | 0 | 0.18 | 0 | 0 | 0.552 | -0.857 | 0.8336548 | NA | 0 | 13.3714 | 10 | 10 | TRUE | 0.5216 |
| Olfr985 | 92 | 75 | 77 | 95 | 75 | 100 | 71 | 80 | 83 | 95 | 85 | 99 | 85.37 | 78 | 93 | 1.192 | 0.253 | 0.23182443 | 0.39772736 | 0 | 0.0302 | 0.0205 | 0.0205 | TRUE | 0.0591 |
| Olfr986 | 0 | 0 | 0 | 0 | 0 | 0 | 0 | 0 | 0 | 0 | 0 | 0 | 0 | 0 | 0 | NA | NA | NA | NA | NA | NA | NA | NA | NA | NA |
| Olfr987 | 0 | 0 | 0 | 0 | 0 | 0 | 0 | 0 | 0 | 0 | 0 | 0 | 0 | 0 | 0 | NA | NA | NA | NA | NA | NA | NA | NA | NA | NA |
| Olfr988 | 0 | 0 | 0 | 0 | 0 | 0 | 0 | 0 | 0 | 0 | 0 | 0 | 0 | 0 | 0 | NA | NA | NA | NA | NA | NA | NA | NA | NA | NA |
| Olfr99 | 0 | 0 | 0 | 0 | 0 | 0 | 0 | 0 | 0 | 0 | 0 | 0 | 0 | 0 | 0 | NA | NA | NA | NA | NA | NA | NA | NA | NA | NA |
| Olfr992 | 0 | 0 | 0 | 0 | 0 | 0 | 0 | 0 | 0 | 0 | 0 | 0 | 0 | 0 | 0 | NA | NA | NA | NA | NA | NA | NA | NA | NA | NA |
| Olfr993 | 0 | 0 | 0 | 0 | 0 | 0 | 0 | 0 | 0 | 0 | 0 | 0 | 0 | 0 | 0 | NA | NA | NA | NA | NA | NA | NA | NA | NA | NA |
| Olfr994 | 0 | 0 | 0 | 0 | 0 | 0 | 0 | 0 | 0 | 0 | 0 | 0 | 0 | 0 | 0 | NA | NA | NA | NA | NA | NA | NA | NA | NA | NA |
| Olfr995 | 0 | 0 | 0 | 0 | 0 | 0 | 0 | 0 | 0 | 0 | 0 | 0 | 0 | 0 | 0 | NA | NA | NA | NA | NA | NA | NA | NA | NA | NA |
| Olfr996 | 0 | 0 | 0 | 0 | 0 | 0 | 0 | 0 | 0 | 0 | 0 | 0 | 0 | 0 | 0 | NA | NA | NA | NA | NA | NA | NA | NA | NA | NA |
| Olfr998 | 0 | 0 | 0 | 0 | 0 | 0 | 0 | 0 | 0 | 0 | 0 | 0 | 0 | 0 | 0 | NA | NA | NA | NA | NA | NA | NA | NA | NA | NA |
| Olig1 | 112 | 102 | 85 | 104 | 96 | 91 | 87 | 108 | 92 | 104 | 108 | 90 | 98.14 | 96 | 101 | 1.052 | 0.074 | 0.71737154 | 0.8300325 | 0 | 0.0266 | 0.0196 | 0.0196 | TRUE | 0.1285 |
| Olig2 | 5 | 4 | 3 | 4 | 6 | 8 | 4 | 4 | 3 | 4 | 7 | 8 | 5 | 4 | 6 | 1.635 | 0.71 | 0.40409988 | NA | 0 | 0.4761 | 0.3109 | 0.3109 | TRUE | 0.2477 |
| Olig3 | 22 | 10 | 7 | 13 | 4 | 4 | 17 | 11 | 8 | 13 | 5 | 4 | 9.45 | 12 | 7 | 0.603 | -0.731 | 0.29193975 | NA | 0.1834 | 0.2533 | 0.2338 | 0.2338 | TRUE | 0.998 |
| Olr1 | 8 | 5 | 1 | 0 | 0 | 1 | 6 | 5 | 1 | 0 | 0 | 1 | 2.27 | 4 | 0 | 0.087 | -3.518 | 0.03517032 | NA | 0 | 1.0482 | 0.8963 | 0.8963 | TRUE | 0.6785 |
| Oma1 | 1502 | 1048 | 1034 | 1330 | 1051 | 1127 | 1167 | 1115 | 1116 | 1324 | 1186 | 1113 | 1169.83 | 1133 | 1208 | 1.066 | 0.092 | 0.26793164 | 0.43846427 | 0.0035 | 0.0045 | 0.0041 | 0.0041 | TRUE | 0.0866 |
| Omd | 0 | 0 | 0 | 0 | 0 | 0 | 0 | 0 | 0 | 0 | 0 | 0 | 0 | 0 | 0 | NA | NA | NA | NA | NA | NA | NA | NA | NA | NA |
| Omg | 0 | 0 | 0 | 0 | 0 | 0 | 0 | 0 | 0 | 0 | 0 | 0 | 0 | 0 | 0 | NA | NA | NA | NA | NA | NA | NA | NA | NA | NA |
| Omp | 31 | 16 | 18 | 22 | 15 | 24 | 24 | 17 | 19 | 22 | 17 | 24 | 20.5 | 20 | 21 | 1.029 | 0.041 | 0.92379016 | 0.9577958 | 0 | 0.118 | 0.0816 | 0.0816 | TRUE | 0.1909 |
| Omt2a | 1 | 0 | 0 | 0 | 0 | 0 | 0 | 1 | 0 | 0 | 0 | 0 | 0.13 | 0 | 0 | 0.552 | -0.857 | 0.8336548 | NA | 0 | 18.3121 | 10 | 10 | TRUE | 0.2566 |
| Omt2b | 2 | 1 | 1 | 0 | 1 | 1 | 2 | 1 | 1 | 0 | 1 | 1 | 0.97 | 1 | 1 | 0.56 | -0.836 | 0.66301683 | NA | 0 | 2.4495 | 1.5188 | 1.5188 | TRUE | 0.2653 |
| Onecut1 | 35 | 37 | 25 | 24 | 20 | 14 | 27 | 39 | 27 | 24 | 23 | 14 | 25.63 | 31 | 20 | 0.645 | -0.632 | 0.11631785 | 0.24684301 | 0.0217 | 0.0949 | 0.0753 | 0.0753 | TRUE | 0.4207 |
| Onecut2 | 48 | 43 | 33 | 51 | 47 | 47 | 37 | 46 | 36 | 51 | 53 | 46 | 44.8 | 40 | 50 | 1.268 | 0.342 | 0.23575926 | 0.40225703 | 0 | 0.0553 | 0.0374 | 0.0374 | TRUE | 0.1313 |
| Onecut3 | 8 | 8 | 4 | 14 | 12 | 16 | 6 | 9 | 4 | 14 | 14 | 16 | 10.38 | 6 | 15 | 2.275 | 1.186 | 0.05127929 | NA | 0 | 0.2307 | 0.1551 | 0.1551 | TRUE | 0.1905 |
| Ooep | 2632 | 1888 | 1988 | 1837 | 1494 | 1747 | 2044 | 2008 | 2145 | 1829 | 1685 | 1725 | 1905.99 | 2066 | 1746 | 0.846 | -0.242 | 0.00025252 | 0.00254891 | 9.00E-04 | 0.0037 | 0.0026 | 0.0026 | TRUE | 0.0206 |
| Oog1 | 0 | 0 | 0 | 0 | 0 | 0 | 0 | 0 | 0 | 0 | 0 | 0 | 0 | 0 | 0 | NA | NA | NA | NA | NA | NA | NA | NA | NA | NA |
| Oog2 | 0 | 0 | 0 | 0 | 0 | 0 | 0 | 0 | 0 | 0 | 0 | 0 | 0 | 0 | 0 | NA | NA | NA | NA | NA | NA | NA | NA | NA | NA |
| Oog3 | 0 | 0 | 0 | 0 | 0 | 0 | 0 | 0 | 0 | 0 | 0 | 0 | 0 | 0 | 0 | NA | NA | NA | NA | NA | NA | NA | NA | NA | NA |
| Oog4 | 0 | 6 | 1 | 3 | 3 | 2 | 0 | 6 | 1 | 3 | 3 | 2 | 2.63 | 2 | 3 | 1.151 | 0.203 | 0.87807497 | NA | 0.797 | 0.9021 | 0.8751 | 0.8751 | TRUE | 1.9915 |
| Oosp1 | 0 | 0 | 0 | 0 | 0 | 0 | 0 | 0 | 0 | 0 | 0 | 0 | 0 | 0 | 0 | NA | NA | NA | NA | NA | NA | NA | NA | NA | NA |
| Opa1 | 7573 | 5376 | 5765 | 6748 | 5780 | 6792 | 5882 | 5718 | 6220 | 6717 | 6520 | 6705 | 6293.76 | 5940 | 6647 | 1.119 | 0.163 | 0.00276715 | 0.01551017 | 8.00E-04 | 0.0028 | 0.002 | 0.002 | TRUE | 0.0205 |
| Opa3 | 913 | 691 | 565 | 638 | 592 | 697 | 709 | 735 | 610 | 635 | 668 | 688 | 674.11 | 685 | 664 | 0.969 | -0.046 | 0.63534288 | 0.77174822 | 0.004 | 0.0059 | 0.0053 | 0.0053 | TRUE | 0.1046 |
| Opalin | 0 | 0 | 1 | 0 | 0 | 0 | 0 | 0 | 0 | 1 | 0 | 0 | 0.18 | 0 | 0 | 0.552 | -0.857 | 0.8336548 | NA | 0 | 13.1817 | 10 | 10 | TRUE | 0.5365 |
| Opcml | 0 | 1 | 0 | 2 | 0 | 1 | 0 | 1 | 0 | 2 | 0 | 1 | 0.67 | 0 | 1 | 2.911 | 1.541 | 0.53484694 | NA | 0 | 3.5209 | 2.9288 | 2.9288 | TRUE | 0.2725 |
| Ophn1 | 252 | 168 | 150 | 236 | 184 | 207 | 196 | 179 | 162 | 235 | 208 | 204 | 197.18 | 179 | 216 | 1.203 | 0.266 | 0.07487408 | 0.18004295 | 0 | 0.0144 | 0.011 | 0.011 | TRUE | 0.0807 |
| Oplah | 471 | 398 | 426 | 396 | 399 | 425 | 366 | 423 | 460 | 394 | 450 | 420 | 418.77 | 416 | 421 | 1.014 | 0.021 | 0.8617154 | 0.92348341 | 0.0067 | 0.0081 | 0.0076 | 0.0076 | TRUE | 0.1387 |
| Opn1mw | 0 | 0 | 0 | 0 | 0 | 0 | 0 | 0 | 0 | 0 | 0 | 0 | 0 | 0 | 0 | NA | NA | NA | NA | NA | NA | NA | NA | NA | NA |
| Opn1sw | 1 | 0 | 0 | 1 | 0 | 0 | 1 | 0 | 0 | 1 | 0 | 0 | 0.3 | 0 | 0 | 1.075 | 0.105 | 0.97698525 | NA | 0 | 8.0272 | 7.5141 | 7.5141 | TRUE | 0.3906 |
| Opn3 | 116 | 67 | 72 | 89 | 75 | 87 | 90 | 71 | 78 | 89 | 85 | 86 | 83.02 | 80 | 87 | 1.079 | 0.11 | 0.61094034 | 0.75392458 | 0 | 0.031 | 0.0214 | 0.0214 | TRUE | 0.1374 |
| Opn4 | 37 | 23 | 30 | 33 | 18 | 38 | 29 | 24 | 32 | 33 | 20 | 38 | 29.37 | 28 | 30 | 1.065 | 0.09 | 0.80797796 | 0.88895612 | 0.0176 | 0.0831 | 0.0659 | 0.0659 | TRUE | 0.5031 |
| Opn5 | 0 | 0 | 0 | 0 | 0 | 0 | 0 | 0 | 0 | 0 | 0 | 0 | 0 | 0 | 0 | NA | NA | NA | NA | NA | NA | NA | NA | NA | NA |
| Oprd1 | 45 | 36 | 28 | 38 | 49 | 51 | 35 | 38 | 30 | 38 | 55 | 50 | 41.15 | 34 | 48 | 1.383 | 0.467 | 0.13118834 | 0.26863537 | 0 | 0.06 | 0.0443 | 0.0443 | TRUE | 0.2716 |
| Oprk1 | 113 | 81 | 88 | 112 | 87 | 108 | 88 | 86 | 95 | 111 | 98 | 107 | 97.52 | 90 | 105 | 1.179 | 0.237 | 0.22807378 | 0.39336499 | 0 | 0.0267 | 0.0176 | 0.0176 | TRUE | 0.034 |
| Oprl1 | 235 | 180 | 100 | 155 | 134 | 130 | 183 | 191 | 108 | 154 | 151 | 128 | 152.61 | 161 | 144 | 0.897 | -0.157 | 0.45731553 | 0.62717434 | 0.0443 | 0.018 | 0.0254 | 0.0254 | TRUE | 0.8392 |
| Oprm1 | 0 | 0 | 0 | 0 | 0 | 0 | 0 | 0 | 0 | 0 | 0 | 0 | 0 | 0 | 0 | NA | NA | NA | NA | NA | NA | NA | NA | NA | NA |
| Optc | 15 | 7 | 14 | 12 | 15 | 21 | 12 | 7 | 15 | 12 | 17 | 21 | 13.97 | 11 | 17 | 1.45 | 0.536 | 0.31391613 | NA | 0 | 0.1721 | 0.1303 | 0.1303 | TRUE | 0.241 |

| | | | | | | | | | | | | | | | | | | | | | | | | | |
|---|---|---|---|---|---|---|---|---|---|---|---|---|---|---|---|---|---|---|---|---|---|---|---|---|---|
| Optn | 477 | 384 | 452 | 495 | 490 | 566 | 370 | 408 | 488 | 493 | 553 | 559 | 478.47 | 422 | 535 | 1.27 | 0.345 | 0.00395475 | 0.02042833 | 0.01 | 0.0074 | 0.0082 | 0.0082 | TRUE | 0.2107 |
| Orai1 | 715 | 521 | 536 | 594 | 513 | 675 | 555 | 554 | 578 | 591 | 579 | 666 | 587.36 | 562 | 612 | 1.089 | 0.123 | 0.19749176 | 0.35727582 | 0.0016 | 0.0065 | 0.0049 | 0.0049 | TRUE | 0.0713 |
| Orai2 | 94 | 60 | 66 | 103 | 101 | 99 | 73 | 64 | 71 | 103 | 114 | 98 | 87.04 | 69 | 105 | 1.505 | 0.59 | 0.00507095 | 0.02477916 | 0 | 0.0297 | 0.02 | 0.02 | TRUE | 0.0556 |
| Orai3 | 643 | 459 | 533 | 661 | 626 | 745 | 499 | 488 | 575 | 658 | 706 | 735 | 610.38 | 521 | 700 | 1.345 | 0.428 | 1.73E-05 | 0.0002982 | 0.0036 | 0.0063 | 0.0055 | 0.0055 | TRUE | 0.0935 |
| Oraov1 | 934 | 676 | 551 | 562 | 502 | 615 | 725 | 719 | 595 | 559 | 566 | 607 | 628.63 | 680 | 577 | 0.849 | -0.236 | 0.02149373 | 0.07284496 | 0.0056 | 0.0062 | 0.006 | 0.006 | TRUE | 0.1383 |
| Orc1 | 5794 | 4119 | 4117 | 3975 | 3520 | 4130 | 4500 | 4381 | 4442 | 3957 | 3971 | 4077 | 4221.35 | 4441 | 4002 | 0.901 | -0.151 | 0.00344929 | 0.01842671 | 0 | 0.003 | 0.0017 | 0.0017 | TRUE | 0.0033 |
| Orc2 | 9169 | 6476 | 6319 | 6863 | 5983 | 6832 | 7121 | 6888 | 6818 | 6831 | 6749 | 6744 | 6858.81 | 6942 | 6775 | 0.976 | -0.036 | 0.46582741 | 0.6347065 | 2.00E-04 | 0.0028 | 0.0016 | 0.0016 | TRUE | 0.0063 |
| Orc3 | 5563 | 4043 | 4130 | 4434 | 3943 | 4325 | 4321 | 4300 | 4456 | 4414 | 4448 | 4269 | 4368.07 | 4359 | 4377 | 1.004 | 0.006 | 0.90953297 | 0.95008074 | 2.00E-04 | 0.003 | 0.0018 | 0.0018 | TRUE | 0.0057 |
| Orc4 | 3482 | 2434 | 2594 | 2701 | 2227 | 2635 | 2704 | 2589 | 2799 | 2689 | 2512 | 2601 | 2649.04 | 2697 | 2601 | 0.964 | -0.052 | 0.39593401 | 0.56938171 | 8.00E-04 | 0.0033 | 0.0024 | 0.0024 | TRUE | 0.0146 |
| Orc5 | 4516 | 3324 | 3239 | 3468 | 3118 | 3618 | 3507 | 3536 | 3495 | 3452 | 3517 | 3572 | 3513.16 | 3513 | 3514 | 1 | 0 | 0.99328386 | 0.99603371 | 0 | 0.0031 | 0.0017 | 0.0017 | TRUE | 0.0029 |
| Orc6 | 7008 | 4724 | 4384 | 4596 | 3887 | 4551 | 5443 | 5025 | 4730 | 4575 | 4385 | 4493 | 4775.05 | 5066 | 4484 | 0.885 | -0.177 | 0.00603356 | 0.02832108 | 0.0024 | 0.0029 | 0.0028 | 0.0028 | TRUE | 0.0523 |
| ORF63 | 2 | 6 | 5 | 4 | 2 | 1 | 2 | 6 | 5 | 4 | 2 | 1 | 3.43 | 4 | 2 | 0.553 | -0.854 | 0.43478337 | NA | 0 | 0.6942 | 0.5363 | 0.5363 | TRUE | 0.8029 |
| Orm1 | 0 | 0 | 0 | 0 | 0 | 0 | 0 | 0 | 0 | 0 | 0 | 0 | 0 | 0 | 0 | NA | NA | NA | NA | NA | NA | NA | NA | NA | NA |
| Orm2 | 0 | 0 | 0 | 0 | 0 | 0 | 0 | 0 | 0 | 0 | 0 | 0 | 0 | 0 | 0 | NA | NA | NA | NA | NA | NA | NA | NA | NA | NA |
| Orm3 | 0 | 0 | 0 | 0 | 0 | 0 | 0 | 0 | 0 | 0 | 0 | 0 | 0 | 0 | 0 | NA | NA | NA | NA | NA | NA | NA | NA | NA | NA |
| Ormdl1 | 1368 | 892 | 979 | 1096 | 923 | 1027 | 1062 | 949 | 1056 | 1091 | 1041 | 1014 | 1035.6 | 1022 | 1049 | 1.025 | 0.035 | 0.65928519 | 0.78920965 | 0.0017 | 0.0047 | 0.0037 | 0.0037 | TRUE | 0.0466 |
| Ormdl2 | 3007 | 2054 | 2087 | 2388 | 2100 | 2475 | 2335 | 2185 | 2252 | 2377 | 2369 | 2443 | 2326.89 | 2257 | 2396 | 1.061 | 0.086 | 0.15539498 | 0.3030416 | 3.00E-04 | 0.0034 | 0.0022 | 0.0022 | TRUE | 0.0115 |
| Ormdl3 | 1609 | 1095 | 1081 | 1163 | 1016 | 1133 | 1250 | 1165 | 1166 | 1158 | 1146 | 1118 | 1167.17 | 1194 | 1141 | 0.955 | -0.067 | 0.35533911 | 0.52993133 | 0 | 0.0045 | 0.0029 | 0.0029 | TRUE | 0.0212 |
| Os9 | 1441 | 1186 | 1140 | 1142 | 1084 | 1217 | 1119 | 1262 | 1230 | 1137 | 1223 | 1201 | 1195.29 | 1204 | 1187 | 0.987 | -0.018 | 0.81268661 | 0.8917342 | 0.002 | 0.0044 | 0.0035 | 0.0035 | TRUE | 0.0473 |
| Osbp | 3944 | 2928 | 3055 | 3401 | 3038 | 3484 | 3063 | 3114 | 3296 | 3385 | 3427 | 3439 | 3287.62 | 3158 | 3417 | 1.083 | 0.114 | 0.0445413 | 0.1238294 | 5.00E-04 | 0.0031 | 0.002 | 0.002 | TRUE | 0.0177 |
| Osbp2 | 150 | 101 | 137 | 156 | 159 | 197 | 116 | 107 | 148 | 155 | 179 | 194 | 150.15 | 124 | 176 | 1.426 | 0.512 | 0.00452039 | 0.02274197 | 0.0125 | 0.0182 | 0.0166 | 0.0166 | TRUE | 0.274 |
| Osbpl10 | 1409 | 1128 | 1228 | 1311 | 1242 | 1462 | 1094 | 1200 | 1325 | 1305 | 1401 | 1443 | 1294.74 | 1206 | 1383 | 1.148 | 0.199 | 0.02077053 | 0.07116331 | 0.0047 | 0.0043 | 0.0046 | 0.0046 | TRUE | 0.0886 |
| Osbpl11 | 3397 | 2360 | 2184 | 2123 | 2012 | 2241 | 2638 | 2510 | 2357 | 2113 | 2270 | 2212 | 2350.05 | 2502 | 2198 | 0.878 | -0.188 | 0.00522149 | 0.02534732 | 0.0019 | 0.0034 | 0.0028 | 0.0028 | TRUE | 0.0311 |
| Osbpl1a | 5071 | 3904 | 4829 | 4469 | 4241 | 4790 | 3938 | 4153 | 5210 | 4448 | 4784 | 4728 | 4543.78 | 4434 | 4653 | 1.05 | 0.07 | 0.44597342 | 0.61664874 | 0.0101 | 0.0029 | 0.0059 | 0.0059 | TRUE | 0.2853 |
| Osbpl2 | 1338 | 956 | 1103 | 1131 | 1069 | 1289 | 1039 | 1017 | 1190 | 1126 | 1206 | 1272 | 1141.73 | 1082 | 1201 | 1.111 | 0.152 | 0.0797962 | 0.18886026 | 0.0043 | 0.0045 | 0.0045 | 0.0045 | TRUE | 0.0894 |
| Osbpl3 | 1412 | 977 | 1122 | 1062 | 950 | 1065 | 1097 | 1039 | 1211 | 1057 | 1072 | 1051 | 1087.77 | 1116 | 1060 | 0.951 | -0.073 | 0.3628429 | 0.53754219 | 0.0021 | 0.0046 | 0.0037 | 0.0037 | TRUE | 0.0645 |
| Osbpl5 | 125 | 87 | 121 | 161 | 143 | 164 | 97 | 93 | 131 | 160 | 161 | 162 | 133.94 | 107 | 161 | 1.516 | 0.6 | 0.0011256 | 0.00790192 | 0.0065 | 0.0201 | 0.0167 | 0.0167 | TRUE | 0.3582 |
| Osbpl6 | 153 | 110 | 153 | 283 | 220 | 256 | 119 | 117 | 165 | 282 | 248 | 253 | 197.25 | 134 | 261 | 1.961 | 0.972 | 4.54E-09 | 2.56E-07 | 0.0134 | 0.0144 | 0.0142 | 0.0142 | TRUE | 0.4179 |
| Osbpl7 | 1080 | 910 | 890 | 899 | 840 | 1019 | 839 | 968 | 960 | 895 | 948 | 1006 | 935.9 | 922 | 950 | 1.031 | 0.044 | 0.61787549 | 0.75863709 | 0.004 | 0.005 | 0.0046 | 0.0046 | TRUE | 0.0785 |
| Osbpl8 | 2329 | 1622 | 1596 | 2008 | 1585 | 1889 | 1809 | 1725 | 1722 | 1999 | 1788 | 1865 | 1817.96 | 1752 | 1884 | 1.075 | 0.104 | 0.12970332 | 0.26638397 | 0.0014 | 0.0037 | 0.0029 | 0.0029 | TRUE | 0.0345 |
| Osbpl9 | 3598 | 2559 | 2580 | 2680 | 2373 | 2663 | 2794 | 2722 | 2784 | 2668 | 2677 | 2629 | 2712.27 | 2767 | 2658 | 0.96 | -0.058 | 0.29272377 | 0.46492953 | 0 | 0.0033 | 0.0018 | 0.0018 | TRUE | 0.0024 |
| Oscar | 6 | 5 | 5 | 5 | 6 | 5 | 5 | 5 | 5 | 5 | 7 | 5 | 5.34 | 5 | 6 | 1.085 | 0.118 | 0.88338461 | NA | 0 | 0.4461 | 0.2782 | 0.2782 | TRUE | 0.0707 |
| Oscp1 | 76 | 67 | 47 | 73 | 60 | 57 | 59 | 71 | 51 | 73 | 68 | 56 | 62.94 | 60 | 66 | 1.086 | 0.119 | 0.63969083 | 0.77477341 | 0 | 0.0401 | 0.0308 | 0.0308 | TRUE | 0.201 |
| Osgep | 2464 | 1763 | 1999 | 2351 | 1969 | 2384 | 1914 | 1875 | 2157 | 2340 | 2221 | 2353 | 2143.44 | 1982 | 2305 | 1.164 | 0.219 | 0.00241123 | 0.01396986 | 0.0028 | 0.0035 | 0.0033 | 0.0033 | TRUE | 0.0709 |
| Osgepl1 | 904 | 623 | 626 | 659 | 579 | 649 | 702 | 663 | 675 | 656 | 653 | 641 | 665 | 680 | 650 | 0.955 | -0.067 | 0.43213073 | 0.60400928 | 0 | 0.006 | 0.0037 | 0.0037 | TRUE | 0.0101 |
| Osgin1 | 4 | 3 | 4 | 7 | 13 | 7 | 3 | 3 | 4 | 7 | 15 | 7 | 6.53 | 3 | 10 | 2.69 | 1.427 | 0.07260223 | NA | 0 | 0.3656 | 0.2638 | 0.2638 | TRUE | 0.6814 |
| Osgin2 | 1221 | 884 | 853 | 973 | 775 | 862 | 948 | 940 | 920 | 969 | 874 | 851 | 917.12 | 936 | 898 | 0.959 | -0.061 | 0.46071976 | 0.63028752 | 0.0014 | 0.005 | 0.0038 | 0.0038 | TRUE | 0.0571 |
| Osm | 24 | 15 | 21 | 19 | 19 | 22 | 19 | 16 | 23 | 19 | 21 | 22 | 19.89 | 19 | 21 | 1.085 | 0.117 | 0.78275417 | 0.87214728 | 0 | 0.1216 | 0.0804 | 0.0804 | TRUE | 0.1267 |
| Osmr | 15 | 9 | 12 | 7 | 5 | 14 | 12 | 10 | 13 | 7 | 6 | 14 | 10.1 | 12 | 9 | 0.778 | -0.362 | 0.55957218 | NA | 0 | 0.2371 | 0.1765 | 0.1765 | TRUE | 0.8026 |
| Osr1 | 1 | 0 | 0 | 1 | 0 | 0 | 1 | 0 | 0 | 1 | 0 | 0 | 0.3 | 0 | 0 | 1.075 | 0.105 | 0.97698525 | NA | 0 | 8.0272 | 7.5141 | 7.5141 | TRUE | 0.3906 |
| Osr2 | 21 | 18 | 16 | 8 | 7 | 11 | 16 | 19 | 17 | 8 | 8 | 11 | 13.24 | 17 | 9 | 0.509 | -0.973 | 0.06741739 | NA | 0 | 0.1814 | 0.1186 | 0.1186 | TRUE | 0.1218 |
| Ost4 | 3881 | 2680 | 2534 | 2745 | 2264 | 2640 | 3014 | 2851 | 2734 | 2732 | 2554 | 2606 | 2748.59 | 2866 | 2631 | 0.917 | -0.124 | 0.05124788 | 0.13732119 | 0.0015 | 0.0033 | 0.0026 | 0.0026 | TRUE | 0.0253 |
| Ostc | 5552 | 3967 | 3934 | 4672 | 4010 | 4625 | 4312 | 4220 | 4245 | 4651 | 4524 | 4566 | 4419.37 | 4259 | 4580 | 1.075 | 0.105 | 0.03867215 | 0.11175517 | 0 | 0.003 | 0.0016 | 0.0016 | TRUE | 0.0022 |
| Ostf1 | 2137 | 1490 | 1798 | 2028 | 1702 | 1711 | 1660 | 1585 | 1940 | 2019 | 1920 | 1689 | 1802.06 | 1728 | 1876 | 1.086 | 0.119 | 0.19925303 | 0.35985892 | 0.0091 | 0.0037 | 0.0056 | 0.0056 | TRUE | 0.137 |
| Ostm1 | 1908 | 1390 | 1440 | 1570 | 1453 | 1688 | 1482 | 1479 | 1554 | 1563 | 1639 | 1666 | 1563.73 | 1505 | 1623 | 1.079 | 0.109 | 0.10171994 | 0.22388099 | 0 | 0.0039 | 0.0026 | 0.0026 | TRUE | 0.0128 |
| Ostn | 1 | 0 | 0 | 0 | 0 | 0 | 1 | 0 | 0 | 0 | 0 | 0 | 0.13 | 0 | 0 | 0.552 | -0.857 | 0.8336548 | NA | 0 | 18.3121 | 10 | 10 | TRUE | 0.2566 |
| Otc | 1 | 2 | 1 | 0 | 2 | 0 | 1 | 2 | 1 | 0 | 2 | 0 | 1.04 | 1 | 1 | 0.551 | -0.859 | 0.66722185 | NA | 0 | 2.2816 | 1.7914 | 1.7914 | TRUE | 0.975 |
| Otoa | 1 | 2 | 3 | 0 | 0 | 1 | 1 | 2 | 3 | 0 | 0 | 1 | 1.19 | 2 | 0 | 0.184 | -2.442 | 0.2099393 | NA | 0 | 1.9974 | 1.4928 | 1.4928 | TRUE | 0.2878 |
| Otof | 14 | 18 | 10 | 11 | 15 | 8 | 11 | 19 | 11 | 11 | 17 | 8 | 12.76 | 14 | 12 | 0.876 | -0.191 | 0.73483479 | NA | 0.0496 | 0.1881 | 0.1497 | 0.1497 | TRUE | 0.5143 |
| Otog | 29 | 22 | 17 | 19 | 16 | 30 | 23 | 23 | 18 | 19 | 18 | 30 | 21.81 | 21 | 22 | 1.037 | 0.052 | 0.90067988 | 0.94539307 | 0 | 0.1111 | 0.0816 | 0.0816 | TRUE | 0.5018 |
| Otogl | 1 | 2 | 0 | 1 | 3 | 1 | 1 | 2 | 0 | 1 | 3 | 1 | 1.38 | 1 | 2 | 1.838 | 0.878 | 0.60180657 | NA | 0 | 1.7218 | 1.2469 | 1.2469 | TRUE | 0.4404 |
| Otol1 | 12 | 7 | 4 | 2 | 4 | 3 | 9 | 7 | 4 | 2 | 5 | 3 | 5.09 | 7 | 3 | 0.44 | -1.183 | 0.17945643 | NA | 0 | 0.468 | 0.3265 | 0.3265 | TRUE | 0.2326 |
| Otop1 | 72 | 50 | 69 | 54 | 50 | 74 | 56 | 53 | 74 | 54 | 56 | 73 | 61.13 | 61 | 61 | 1.003 | 0.004 | 0.98907767 | 0.99429134 | 0.0147 | 0.0412 | 0.0338 | 0.0338 | TRUE | 0.2945 |
| Otop2 | 13 | 11 | 12 | 20 | 23 | 24 | 10 | 12 | 13 | 20 | 26 | 24 | 17.38 | 12 | 23 | 2.012 | 1.008 | 0.03026489 | 0.09322909 | 0 | 0.1388 | 0.0919 | 0.0919 | TRUE | 0.0906 |
| Otop3 | 1 | 1 | 0 | 0 | 0 | 0 | 1 | 1 | 0 | 0 | 0 | 0 | 0.31 | 1 | 0 | 0.309 | -1.696 | 0.63183309 | NA | 0 | 7.7294 | 7.2472 | 7.2472 | TRUE | 0.2015 |
| Otor | 0 | 0 | 1 | 0 | 0 | 0 | 0 | 0 | 1 | 0 | 0 | 0 | 0.18 | 0 | 0 | 0.552 | -0.857 | 0.8336548 | NA | 0 | 13.1817 | 10 | 10 | TRUE | 0.5365 |
| Otos | 1 | 3 | 3 | 5 | 7 | 5 | 1 | 3 | 3 | 5 | 8 | 5 | 4.17 | 2 | 6 | 2.555 | 1.353 | 0.17024724 | NA | 0 | 0.5709 | 0.4061 | 0.4061 | TRUE | 0.5456 |
| Otp | 0 | 1 | 1 | 1 | 0 | 0 | 0 | 1 | 1 | 1 | 0 | 0 | 0.52 | 1 | 0 | 0.535 | -0.904 | 0.74017109 | NA | 0 | 4.534 | 3.6534 | 3.6534 | TRUE | 0.3906 |
| Ott | 0 | 0 | 0 | 0 | 0 | 0 | 0 | 0 | 0 | 0 | 0 | 0 | 0 | 0 | 0 | NA | NA | NA | NA | NA | NA | NA | NA | NA | NA |
| OTTMUSG00 | 0 | 0 | 0 | 0 | 0 | 0 | 0 | 0 | 0 | 0 | 0 | 0 | 0 | 0 | 0 | NA | NA | NA | NA | NA | NA | NA | NA | NA | NA |
| Otub1 | 3746 | 2787 | 2909 | 2933 | 2636 | 3011 | 2909 | 2964 | 3139 | 2920 | 2974 | 2972 | 2979.69 | 3004 | 2955 | 0.984 | -0.023 | 0.69110667 | 0.81113382 | 5.00E-04 | 0.0032 | 0.0021 | 0.0021 | TRUE | 0.0185 |
| Otub2 | 352 | 238 | 255 | 284 | 256 | 296 | 273 | 253 | 275 | 283 | 289 | 292 | 277.56 | 267 | 288 | 1.076 | 0.106 | 0.38140118 | 0.55579288 | 0 | 0.011 | 0.007 | 0.007 | TRUE | 0.0227 |
| Otud1 | 579 | 474 | 373 | 396 | 346 | 357 | 450 | 504 | 402 | 394 | 390 | 352 | 415.54 | 452 | 379 | 0.838 | -0.255 | 0.02901341 | 0.09034611 | 0.0058 | 0.0081 | 0.0074 | 0.0074 | TRUE | 0.1125 |
| Otud3 | 909 | 694 | 630 | 741 | 587 | 720 | 706 | 738 | 680 | 738 | 662 | 711 | 705.75 | 708 | 704 | 0.994 | -0.008 | 0.92602797 | 0.95917949 | 7.00E-04 | 0.0058 | 0.0041 | 0.0041 | TRUE | 0.0303 |
| Otud4 | 21782 | 15769 | 14719 | 15649 | 13833 | 15866 | 16917 | 16773 | 15882 | 15577 | 15605 | 15662 | 16069.39 | 16524 | 15615 | 0.945 | -0.082 | 0.08951765 | 0.20399806 | 5.00E-04 | 0.0026 | 0.0016 | 0.0016 | TRUE | 0.0141 |
| Otud5 | 4595 | 3555 | 3234 | 3333 | 3056 | 3441 | 3569 | 3781 | 3489 | 3318 | 3447 | 3397 | 3500.26 | 3613 | 3387 | 0.938 | -0.093 | 0.10791429 | 0.23373525 | 6.00E-04 | 0.0031 | 0.0021 | 0.0021 | TRUE | 0.0198 |
| Otud6a | 249 | 189 | 155 | 115 | 90 | 66 | 193 | 201 | 167 | 114 | 102 | 65 | 140.47 | 187 | 94 | 0.499 | -1.001 | 1.21E-06 | 3.41E-05 | 0.0326 | 0.0193 | 0.0226 | 0.0226 | TRUE | 0.7064 |
| Otud6b | 4594 | 3114 | 2941 | 3051 | 2478 | 2846 | 3568 | 3312 | 3173 | 3037 | 2795 | 2809 | 3115.91 | 3351 | 2880 | 0.859 | -0.219 | 0.00115751 | 0.0080625 | 0.0024 | 0.0032 | 0.0029 | 0.0029 | TRUE | 0.0397 |
| Otud7a | 174 | 110 | 139 | 151 | 140 | 158 | 135 | 117 | 150 | 150 | 158 | 156 | 144.39 | 134 | 155 | 1.154 | 0.207 | 0.2214983 | 0.3854243 | 0 | 0.0188 | 0.0136 | 0.0136 | TRUE | 0.1182 |

| | | | | | | | | | | | | | | | | | | | | | | | | | |
|---|---|---|---|---|---|---|---|---|---|---|---|---|---|---|---|---|---|---|---|---|---|---|---|---|---|
| Otud7b | 3744 | 2657 | 2360 | 2635 | 2357 | 2688 | 2908 | 2826 | 2546 | 2623 | 2659 | 2653 | 2702.62 | 2760 | 2645 | 0.958 | -0.062 | 0.34921961 | 0.52375613 | 0.0018 | 0.0033 | 0.0028 | 0.0028 | TRUE | 0.0552 |
| Otx1 | 37 | 33 | 26 | 27 | 20 | 21 | 29 | 35 | 28 | 27 | 23 | 21 | 27.01 | 31 | 24 | 0.766 | -0.385 | 0.29753517 | 0.4697007 | 0 | 0.0902 | 0.0608 | 0.0608 | TRUE | 0.1035 |
| Otx2 | 2534 | 1916 | 2003 | 1597 | 1502 | 1846 | 1968 | 2038 | 2161 | 1590 | 1694 | 1822 | 1878.94 | 2056 | 1702 | 0.828 | -0.271 | 0.00024292 | 0.00247651 | 0.0027 | 0.0037 | 0.0034 | 0.0034 | TRUE | 0.0467 |
| Otx2os1 | 1 | 2 | 5 | 6 | 3 | 2 | 1 | 2 | 5 | 6 | 3 | 2 | 3.27 | 3 | 4 | 1.409 | 0.495 | 0.65692292 | NA | 0.2201 | 0.7268 | 0.5781 | 0.5781 | TRUE | 0.7389 |
| Ovca2 | 775 | 533 | 383 | 599 | 517 | 511 | 602 | 567 | 413 | 596 | 583 | 504 | 544.34 | 527 | 561 | 1.062 | 0.086 | 0.52063083 | 0.68194433 | 0.0215 | 0.0068 | 0.0112 | 0.0112 | TRUE | 0.4112 |
| Ovch2 | 0 | 0 | 0 | 0 | 0 | 0 | 0 | 0 | 0 | 0 | 0 | 0 | 0 | 0 | 0 | NA | NA | NA | NA | NA | NA | NA | NA | NA | NA |
| Ovgp1 | 147 | 126 | 127 | 116 | 124 | 131 | 114 | 134 | 137 | 115 | 140 | 129 | 128.32 | 128 | 128 | 1.001 | 0.002 | 0.99309371 | 0.99602052 | 0 | 0.0209 | 0.0156 | 0.0156 | TRUE | 0.1077 |
| Ovol1 | 44 | 47 | 40 | 44 | 48 | 47 | 34 | 50 | 43 | 44 | 54 | 46 | 45.28 | 42 | 48 | 1.14 | 0.189 | 0.52256751 | 0.68379176 | 0 | 0.0548 | 0.0406 | 0.0406 | TRUE | 0.2664 |
| Ovol2 | 185 | 164 | 168 | 167 | 127 | 169 | 144 | 174 | 181 | 166 | 143 | 167 | 162.62 | 166 | 159 | 0.959 | -0.06 | 0.717797 | 0.83038049 | 0.0059 | 0.017 | 0.0137 | 0.0137 | TRUE | 0.1684 |
| Oxa1l | 5452 | 4109 | 4581 | 4543 | 4364 | 5061 | 4234 | 4371 | 4943 | 4522 | 4923 | 4996 | 4664.84 | 4516 | 4814 | 1.066 | 0.093 | 0.20104038 | 0.36180686 | 0.0045 | 0.0029 | 0.0036 | 0.0036 | TRUE | 0.083 |
| Oxct1 | 4754 | 3431 | 3500 | 4009 | 3246 | 3876 | 3692 | 3650 | 3776 | 3991 | 3662 | 3826 | 3766.13 | 3706 | 3826 | 1.033 | 0.046 | 0.41927041 | 0.59168956 | 7.00E-04 | 0.0031 | 0.0021 | 0.0021 | TRUE | 0.0172 |
| Oxct2a | 11 | 4 | 7 | 5 | 9 | 2 | 9 | 4 | 8 | 5 | 10 | 2 | 6.24 | 7 | 6 | 0.823 | -0.281 | 0.73118254 | NA | 0.1396 | 0.3821 | 0.3181 | 0.3181 | TRUE | 0.4436 |
| Oxct2b | 19 | 17 | 11 | 8 | 7 | 15 | 15 | 18 | 12 | 8 | 8 | 15 | 12.56 | 15 | 10 | 0.689 | -0.537 | 0.33437306 | NA | 0 | 0.1911 | 0.1405 | 0.1405 | TRUE | 0.5599 |
| Oxgr1 | 3 | 1 | 2 | 1 | 2 | 3 | 2 | 1 | 2 | 1 | 2 | 3 | 1.96 | 2 | 2 | 1.106 | 0.146 | 0.91251659 | NA | 0 | 1.2112 | 0.7625 | 0.7625 | TRUE | 0.1998 |
| Oxld1 | 148 | 130 | 148 | 184 | 157 | 178 | 115 | 138 | 160 | 183 | 177 | 176 | 158.15 | 138 | 179 | 1.306 | 0.385 | 0.02377612 | 0.07842882 | 0.007 | 0.0174 | 0.0145 | 0.0145 | TRUE | 0.241 |
| Oxnad1 | 1214 | 813 | 707 | 725 | 638 | 739 | 943 | 865 | 763 | 722 | 720 | 730 | 790.23 | 857 | 724 | 0.843 | -0.246 | 0.00923599 | 0.03905621 | 0.0047 | 0.0054 | 0.0052 | 0.0052 | TRUE | 0.1084 |
| Oxr1 | 5513 | 4139 | 4452 | 4425 | 3927 | 4615 | 4282 | 4403 | 4804 | 4405 | 4430 | 4556 | 4479.73 | 4496 | 4464 | 0.993 | -0.01 | 0.87215912 | 0.92903341 | 0.0018 | 0.003 | 0.0025 | 0.0025 | TRUE | 0.0434 |
| Oxsm | 993 | 771 | 738 | 796 | 645 | 772 | 771 | 820 | 796 | 792 | 728 | 762 | 778.28 | 796 | 761 | 0.957 | -0.063 | 0.44416406 | 0.61509901 | 0 | 0.0055 | 0.0037 | 0.0037 | TRUE | 0.0166 |
| Oxsr1 | 8913 | 6932 | 7224 | 7692 | 6909 | 8032 | 6922 | 7373 | 7795 | 7657 | 7794 | 7929 | 7578.33 | 7363 | 7793 | 1.059 | 0.082 | 0.15984159 | 0.30934863 | 0.0017 | 0.0027 | 0.0023 | 0.0023 | TRUE | 0.0339 |
| Oxt | 5 | 3 | 3 | 8 | 5 | 6 | 4 | 3 | 3 | 8 | 6 | 6 | 4.97 | 3 | 7 | 1.885 | 0.914 | 0.2821215 | NA | 0 | 0.479 | 0.3023 | 0.3023 | TRUE | 0.0978 |
| Oxtr | 22 | 3 | 9 | 13 | 10 | 18 | 17 | 3 | 10 | 13 | 11 | 18 | 12 | 10 | 14 | 1.371 | 0.456 | 0.46677921 | NA | 0.1937 | 0.2 | 0.1984 | 0.1984 | TRUE | 0.3236 |
| P2rx1 | 43 | 30 | 25 | 33 | 19 | 18 | 33 | 32 | 27 | 33 | 21 | 18 | 27.39 | 31 | 24 | 0.78 | -0.359 | 0.35065437 | 0.52530164 | 0.0142 | 0.089 | 0.0695 | 0.0695 | TRUE | 0.635 |
| P2rx2 | 92 | 56 | 73 | 93 | 78 | 86 | 71 | 60 | 79 | 93 | 88 | 85 | 79.21 | 70 | 89 | 1.265 | 0.339 | 0.12662132 | 0.26194798 | 0 | 0.0323 | 0.0227 | 0.0227 | TRUE | 0.1394 |
| P2rx3 | 60 | 36 | 26 | 21 | 16 | 24 | 47 | 38 | 28 | 21 | 18 | 24 | 29.27 | 38 | 21 | 0.552 | -0.857 | 0.02245655 | 0.07506348 | 0 | 0.0834 | 0.0641 | 0.0641 | TRUE | 0.1502 |
| P2rx4 | 340 | 257 | 287 | 340 | 337 | 334 | 264 | 273 | 310 | 338 | 380 | 330 | 315.9 | 282 | 349 | 1.239 | 0.309 | 0.0127963 | 0.04965993 | 0.0032 | 0.0099 | 0.0079 | 0.0079 | TRUE | 0.0784 |
| P2rx5 | 344 | 271 | 299 | 376 | 331 | 405 | 267 | 288 | 323 | 374 | 373 | 400 | 337.59 | 293 | 382 | 1.311 | 0.391 | 0.00105955 | 0.00754566 | 0.0021 | 0.0094 | 0.0073 | 0.0073 | TRUE | 0.0891 |
| P2rx6 | 8 | 8 | 9 | 11 | 10 | 11 | 6 | 9 | 10 | 11 | 11 | 11 | 9.59 | 8 | 11 | 1.369 | 0.453 | 0.45805193 | NA | 0 | 0.2496 | 0.1614 | 0.1614 | TRUE | 0.172 |
| P2rx7 | 533 | 364 | 556 | 575 | 545 | 640 | 414 | 387 | 600 | 572 | 615 | 632 | 536.67 | 467 | 606 | 1.301 | 0.379 | 0.00818237 | 0.03564482 | 0.0292 | 0.0068 | 0.0129 | 0.0129 | TRUE | 0.7072 |
| P2ry1 | 110 | 89 | 61 | 64 | 56 | 54 | 85 | 95 | 66 | 64 | 63 | 53 | 71.02 | 82 | 60 | 0.731 | -0.452 | 0.06307442 | 0.15908287 | 0 | 0.0358 | 0.0282 | 0.0282 | TRUE | 0.2597 |
| P2ry10 | 0 | 0 | 0 | 0 | 0 | 0 | 0 | 0 | 0 | 0 | 0 | 0 | 0 | 0 | 0 | NA | NA | NA | NA | NA | NA | NA | NA | NA | NA |
| P2ry12 | 0 | 0 | 0 | 0 | 0 | 0 | 0 | 0 | 0 | 0 | 0 | 0 | 0 | 0 | 0 | NA | NA | NA | NA | NA | NA | NA | NA | NA | NA |
| P2ry13 | 0 | 0 | 0 | 0 | 0 | 0 | 0 | 0 | 0 | 0 | 0 | 0 | 0 | 0 | 0 | NA | NA | NA | NA | NA | NA | NA | NA | NA | NA |
| P2ry14 | 0 | 0 | 1 | 0 | 0 | 0 | 0 | 0 | 1 | 0 | 0 | 0 | 0.18 | 0 | 0 | 0.552 | -0.857 | 0.8336548 | NA | 0 | 13.1817 | 10 | 10 | TRUE | 0.5365 |
| P2ry2 | 233 | 153 | 162 | 235 | 225 | 254 | 181 | 163 | 175 | 234 | 254 | 251 | 209.5 | 173 | 246 | 1.421 | 0.507 | 0.00027866 | 0.00273734 | 0 | 0.0137 | 0.0091 | 0.0091 | TRUE | 0.0272 |
| P2ry4 | 6 | 3 | 5 | 4 | 6 | 10 | 5 | 3 | 5 | 4 | 7 | 10 | 5.64 | 4 | 7 | 1.555 | 0.637 | 0.43470273 | NA | 0 | 0.4223 | 0.2944 | 0.2944 | TRUE | 0.1582 |
| P2ry6 | 50 | 26 | 30 | 39 | 30 | 28 | 39 | 28 | 32 | 39 | 34 | 28 | 33.19 | 33 | 34 | 1.008 | 0.011 | 0.97398951 | 0.98592295 | 0 | 0.0738 | 0.0538 | 0.0538 | TRUE | 0.1999 |
| P4ha1 | 4603 | 3578 | 4482 | 5627 | 4497 | 5071 | 3575 | 3806 | 4836 | 5601 | 5073 | 5006 | 4649.49 | 4072 | 5227 | 1.284 | 0.361 | 0.00023888 | 0.00244751 | 0.0152 | 0.0029 | 0.0067 | 0.0067 | TRUE | 0.327 |
| P4ha2 | 3165 | 2544 | 2450 | 2764 | 2427 | 2648 | 2458 | 2706 | 2644 | 2751 | 2738 | 2614 | 2651.81 | 2603 | 2701 | 1.039 | 0.055 | 0.38996916 | 0.56381035 | 0.0013 | 0.0033 | 0.0025 | 0.0025 | TRUE | 0.0295 |
| P4ha3 | 14 | 7 | 4 | 11 | 5 | 7 | 11 | 7 | 4 | 11 | 6 | 7 | 7.69 | 7 | 8 | 1.025 | 0.036 | 0.95919862 | NA | 0 | 0.3107 | 0.2327 | 0.2327 | TRUE | 0.2448 |
| P4hb | 43862 | 32683 | 32611 | 34894 | 31018 | 36199 | 34066 | 34764 | 35187 | 34733 | 34992 | 35734 | 34912.7 | 34672 | 35153 | 1.014 | 0.02 | 0.64973242 | 0.78249288 | 2.00E-04 | 0.0025 | 0.0014 | 0.0014 | TRUE | 0.0029 |
| P4htm | 248 | 210 | 178 | 171 | 156 | 179 | 193 | 223 | 192 | 170 | 176 | 177 | 188.49 | 203 | 174 | 0.861 | -0.216 | 0.14476162 | 0.2882527 | 0 | 0.015 | 0.0104 | 0.0104 | TRUE | 0.0839 |
| Pa2g4 | 29310 | 20838 | 18538 | 19117 | 16801 | 19424 | 22764 | 22165 | 20002 | 19029 | 18953 | 19174 | 20348.01 | 21644 | 19052 | 0.88 | -0.184 | 0.00169851 | 0.01064307 | 0.0022 | 0.0025 | 0.0024 | 0.0024 | TRUE | 0.0538 |
| Pabpc1 | 156489 | 117323 | 114159 | 123392 | 107092 | 126366 | 121538 | 124795 | 123177 | 122824 | 120811 | 124743 | 122981.3 | 123170 | 122793 | 0.997 | -0.004 | 0.91783813 | 0.95477878 | 2.00E-04 | 0.0024 | 0.0013 | 0.0013 | TRUE | 0.0024 |
| Pabpc1l | 44 | 25 | 32 | 43 | 51 | 39 | 34 | 27 | 35 | 43 | 58 | 38 | 39.02 | 32 | 46 | 1.448 | 0.534 | 0.09670694 | 0.21610887 | 0 | 0.0632 | 0.0481 | 0.0481 | TRUE | 0.337 |
| Pabpc2 | 0 | 0 | 0 | 0 | 0 | 0 | 0 | 0 | 0 | 0 | 0 | 0 | 0 | 0 | 0 | NA | NA | NA | NA | NA | NA | NA | NA | NA | NA |
| Pabpc4 | 23509 | 16952 | 15631 | 18561 | 14796 | 17006 | 18258 | 18032 | 16866 | 18476 | 16691 | 16788 | 17518.39 | 17719 | 17318 | 0.977 | -0.033 | 0.58154507 | 0.73220787 | 0.0021 | 0.0026 | 0.0025 | 0.0025 | TRUE | 0.0418 |
| Pabpc4l | 730 | 556 | 512 | 516 | 421 | 454 | 567 | 591 | 552 | 514 | 475 | 448 | 524.59 | 570 | 479 | 0.84 | -0.252 | 0.0104699 | 0.04270726 | 7.00E-04 | 0.0069 | 0.005 | 0.005 | TRUE | 0.0479 |
| Pabpc5 | 6 | 3 | 2 | 3 | 1 | 4 | 5 | 3 | 2 | 3 | 1 | 4 | 3.01 | 3 | 3 | 0.802 | -0.318 | 0.77160602 | NA | 0 | 0.7893 | 0.5309 | 0.5309 | TRUE | 0.2619 |
| Pabpc6 | 77 | 57 | 14 | 27 | 22 | 17 | 60 | 61 | 15 | 27 | 25 | 17 | 34 | 45 | 23 | 0.502 | -0.994 | 0.0318525 | 0.09670089 | 0.2452 | 0.0721 | 0.1213 | 0.1213 | TRUE | 2.5348 |
| Pabpn1 | 15934 | 12151 | 11509 | 11775 | 10092 | 12238 | 12375 | 12925 | 12418 | 11721 | 11385 | 12081 | 12150.78 | 12573 | 11729 | 0.933 | -0.1 | 0.04498758 | 0.12466218 | 6.00E-04 | 0.0026 | 0.0017 | 0.0017 | TRUE | 0.0084 |
| Pabpn1l | 0 | 0 | 0 | 0 | 1 | 1 | 0 | 0 | 0 | 0 | 1 | 1 | 0.35 | 0 | 1 | 3.965 | 1.987 | 0.55529285 | NA | 0 | 6.7252 | 6.4426 | 6.4426 | TRUE | 0.261 |
| Pacrg | 27 | 18 | 10 | 13 | 10 | 10 | 21 | 19 | 11 | 13 | 11 | 10 | 14.17 | 17 | 11 | 0.664 | -0.59 | 0.26048148 | NA | 0 | 0.1697 | 0.1243 | 0.1243 | TRUE | 0.4623 |
| Pacrgl | 657 | 467 | 405 | 395 | 361 | 374 | 510 | 497 | 437 | 393 | 407 | 369 | 435.6 | 481 | 390 | 0.808 | -0.307 | 0.00466457 | 0.02324772 | 0.0025 | 0.0079 | 0.0062 | 0.0062 | TRUE | 0.0736 |
| Pacs1 | 3550 | 2801 | 3171 | 3289 | 2943 | 3458 | 2757 | 2979 | 3421 | 3274 | 3320 | 3414 | 3194.24 | 3052 | 3336 | 1.094 | 0.129 | 0.1022419 | 0.22471837 | 0.0059 | 0.0032 | 0.0042 | 0.0042 | TRUE | 0.1354 |
| Pacs2 | 2613 | 2016 | 2054 | 2076 | 1890 | 2234 | 2029 | 2144 | 2216 | 2066 | 2132 | 2205 | 2132.32 | 2130 | 2134 | 1.003 | 0.004 | 0.94835174 | 0.97180664 | 0.0011 | 0.0035 | 0.0026 | 0.0026 | TRUE | 0.0214 |
| Pacsin1 | 2025 | 1558 | 1644 | 2038 | 2001 | 2126 | 1573 | 1657 | 1774 | 2029 | 2257 | 2099 | 1898.08 | 1668 | 2128 | 1.277 | 0.352 | 1.58E-06 | 4.19E-05 | 0.0025 | 0.0037 | 0.0033 | 0.0033 | TRUE | 0.0371 |
| Pacsin2 | 6809 | 5129 | 4770 | 4854 | 4405 | 5083 | 5288 | 5456 | 5147 | 4832 | 4969 | 5018 | 5118.23 | 5297 | 4940 | 0.933 | -0.101 | 0.05579334 | 0.14581354 | 3.00E-04 | 0.0029 | 0.0018 | 0.0018 | TRUE | 0.0082 |
| Pacsin3 | 833 | 623 | 527 | 650 | 575 | 628 | 647 | 663 | 569 | 647 | 649 | 620 | 632.31 | 626 | 639 | 1.019 | 0.027 | 0.77599401 | 0.86759714 | 0.0019 | 0.0062 | 0.0048 | 0.0048 | TRUE | 0.073 |
| Padi1 | 20 | 19 | 15 | 19 | 9 | 15 | 16 | 20 | 16 | 19 | 10 | 15 | 15.97 | 17 | 15 | 0.852 | -0.231 | 0.63520154 | NA | 0 | 0.1509 | 0.1073 | 0.1073 | TRUE | 0.2272 |
| Padi2 | 639 | 521 | 701 | 798 | 864 | 939 | 496 | 554 | 756 | 794 | 975 | 927 | 750.46 | 602 | 899 | 1.495 | 0.58 | 2.52E-05 | 0.00040171 | 0.029 | 0.0056 | 0.0123 | 0.0123 | TRUE | 0.5849 |
| Padi3 | 482 | 417 | 517 | 609 | 560 | 701 | 374 | 444 | 558 | 606 | 632 | 692 | 550.95 | 459 | 643 | 1.408 | 0.493 | 0.00020253 | 0.00215386 | 0.02 | 0.0067 | 0.0108 | 0.0108 | TRUE | 0.4145 |
| Padi4 | 212 | 180 | 170 | 192 | 180 | 187 | 165 | 191 | 183 | 191 | 203 | 185 | 186.39 | 180 | 193 | 1.075 | 0.104 | 0.48207679 | 0.64862314 | 0 | 0.0151 | 0.0105 | 0.0105 | TRUE | 0.0651 |
| Padi6 | 38 | 22 | 16 | 36 | 28 | 35 | 30 | 23 | 17 | 36 | 32 | 35 | 28.69 | 23 | 34 | 1.438 | 0.524 | 0.15410799 | 0.30132224 | 0 | 0.085 | 0.062 | 0.062 | TRUE | 0.2999 |
| Paf1 | 4059 | 3082 | 2539 | 2640 | 2353 | 2769 | 3152 | 3278 | 2740 | 2628 | 2654 | 2733 | 2864.33 | 3057 | 2672 | 0.874 | -0.195 | 0.00879911 | 0.03771694 | 0.0039 | 0.0033 | 0.0036 | 0.0036 | TRUE | 0.0988 |
| Pafah1b1 | 19260 | 13533 | 13776 | 15214 | 12647 | 14573 | 14958 | 14395 | 14864 | 15144 | 14267 | 14386 | 14669.07 | 14739 | 14599 | 0.99 | -0.014 | 0.78020728 | 0.87046309 | 5.00E-04 | 0.0026 | 0.0017 | 0.0017 | TRUE | 0.0131 |
| Pafah1b2 | 3523 | 2515 | 2558 | 2772 | 2353 | 2682 | 2736 | 2675 | 2760 | 2759 | 2654 | 2648 | 2705.44 | 2724 | 2687 | 0.987 | -0.02 | 0.7305471 | 0.8390285 | 0 | 0.0033 | 0.002 | 0.002 | TRUE | 0.0067 |
| Pafah1b3 | 1451 | 1161 | 1365 | 1272 | 1244 | 1358 | 1127 | 1235 | 1473 | 1266 | 1403 | 1341 | 1307.46 | 1278 | 1337 | 1.047 | 0.066 | 0.49936767 | 0.66394997 | 0.0099 | 0.0042 | 0.0061 | 0.0061 | TRUE | 0.2121 |
| Pafah2 | 1160 | 821 | 743 | 955 | 780 | 796 | 901 | 873 | 802 | 951 | 880 | 786 | 865.37 | 859 | 872 | 1.015 | 0.021 | 0.82361495 | 0.89868745 | 0.0052 | 0.0052 | 0.0052 | 0.0052 | TRUE | 0.0907 |
| Pag1 | 556 | 409 | 389 | 434 | 401 | 436 | 432 | 435 | 420 | 432 | 452 | 430 | 433.56 | 429 | 438 | 1.021 | 0.03 | 0.76267831 | 0.8591534 | 0 | 0.0079 | 0.0049 | 0.0049 | TRUE | 0.0089 |

| | | | | | | | | | | | | | | | | | | | | | | | | | |
|---|---|---|---|---|---|---|---|---|---|---|---|---|---|---|---|---|---|---|---|---|---|---|---|---|---|
| Pagr1a | 1939 | 1346 | 1356 | 1301 | 1198 | 1352 | 1506 | 1432 | 1463 | 1295 | 1351 | 1335 | 1396.98 | 1467 | 1327 | 0.904 | -0.146 | 0.0304061 | 0.09348127 | 0 | 0.0041 | 0.0025 | 0.0025 | TRUE | 0.0068 |
| Pah | 1 | 0 | 0 | 0 | 0 | 0 | 1 | 0 | 0 | 0 | 0 | 0 | 0.13 | 0 | 0 | 0.552 | -0.857 | 0.8336548 | NA | 0 | 18.3121 | 10 | 10 | TRUE | 0.2566 |
| Paics | 31057 | 23020 | 25771 | 26178 | 23528 | 27320 | 24121 | 24486 | 27807 | 26058 | 26542 | 26969 | 25997.01 | 25471 | 26523 | 1.041 | 0.058 | 0.35121765 | 0.52598008 | 0.003 | 0.0025 | 0.0028 | 0.0028 | TRUE | 0.0787 |
| Paip1 | 2295 | 1634 | 1766 | 1688 | 1464 | 1661 | 1782 | 1738 | 1905 | 1680 | 1652 | 1640 | 1732.91 | 1808 | 1657 | 0.917 | -0.126 | 0.05812427 | 0.1502762 | 6.00E-04 | 0.0038 | 0.0026 | 0.0026 | TRUE | 0.0258 |
| Paip2 | 10160 | 7060 | 7710 | 8451 | 7448 | 8492 | 7891 | 7510 | 8319 | 8412 | 8402 | 8383 | 8152.77 | 7907 | 8399 | 1.062 | 0.087 | 0.11164293 | 0.23959739 | 0.0011 | 0.0027 | 0.002 | 0.002 | TRUE | 0.0251 |
| Paip2b | 9204 | 6484 | 6498 | 6985 | 5975 | 6781 | 7148 | 6897 | 7011 | 6953 | 6740 | 6694 | 6907.29 | 7019 | 6796 | 0.968 | -0.047 | 0.34254185 | 0.51674488 | 2.00E-04 | 0.0028 | 0.0016 | 0.0016 | TRUE | 0.005 |
| Pak1 | 1829 | 1355 | 1464 | 1630 | 1457 | 1806 | 1421 | 1441 | 1580 | 1623 | 1644 | 1783 | 1581.73 | 1481 | 1683 | 1.138 | 0.186 | 0.01227561 | 0.04820202 | 0.0021 | 0.0039 | 0.0034 | 0.0034 | TRUE | 0.0408 |
| Pak1ip1 | 6710 | 4784 | 4486 | 4784 | 4216 | 4829 | 5211 | 5089 | 4840 | 4762 | 4756 | 4767 | 4904.24 | 5047 | 4762 | 0.943 | -0.084 | 0.11568763 | 0.24576744 | 5.00E-04 | 0.0029 | 0.0019 | 0.0019 | TRUE | 0.0155 |
| Pak2 | 7766 | 5816 | 5851 | 6310 | 5643 | 6295 | 6032 | 6186 | 6313 | 6281 | 6366 | 6214 | 6232.02 | 6177 | 6287 | 1.018 | 0.026 | 0.60443144 | 0.7493188 | 2.00E-04 | 0.0028 | 0.0016 | 0.0016 | TRUE | 0.0053 |
| Pak3 | 289 | 167 | 172 | 252 | 193 | 244 | 224 | 178 | 186 | 251 | 218 | 241 | 216.18 | 196 | 237 | 1.203 | 0.266 | 0.07306046 | 0.17699206 | 0.0063 | 0.0134 | 0.0113 | 0.0113 | TRUE | 0.1912 |
| Pak4 | 6327 | 4919 | 5237 | 5602 | 5251 | 6183 | 4914 | 5232 | 5651 | 5576 | 5924 | 6104 | 5566.72 | 5266 | 5868 | 1.115 | 0.157 | 0.01905303 | 0.06672996 | 0.0032 | 0.0029 | 0.003 | 0.003 | TRUE | 0.0498 |
| Pak6 | 182 | 183 | 150 | 224 | 209 | 200 | 141 | 195 | 162 | 223 | 236 | 197 | 192.34 | 166 | 219 | 1.323 | 0.403 | 0.01327621 | 0.05102364 | 0.0116 | 0.0147 | 0.0138 | 0.0138 | TRUE | 0.242 |
| Pak7 | 32 | 25 | 27 | 23 | 23 | 37 | 25 | 27 | 29 | 23 | 26 | 37 | 27.66 | 27 | 29 | 1.064 | 0.09 | 0.80949853 | 0.8900125 | 0 | 0.0881 | 0.0639 | 0.0639 | TRUE | 0.4121 |
| Palb2 | 2534 | 1878 | 1798 | 1868 | 1674 | 1922 | 1968 | 1998 | 1940 | 1859 | 1888 | 1897 | 1925.14 | 1969 | 1881 | 0.956 | -0.065 | 0.27815808 | 0.44963632 | 0 | 0.0037 | 0.0021 | 0.0021 | TRUE | 0.0019 |
| Pald1 | 7872 | 6195 | 6940 | 8588 | 7751 | 8869 | 6114 | 6590 | 7488 | 8548 | 8744 | 8755 | 7706.51 | 6731 | 8682 | 1.29 | 0.368 | 4.18E-07 | 1.39E-05 | 0.0053 | 0.0027 | 0.0037 | 0.0037 | TRUE | 0.1182 |
| Palld | 2875 | 2048 | 2019 | 1915 | 1754 | 1971 | 2233 | 2178 | 2178 | 1906 | 1979 | 1946 | 2070.06 | 2196 | 1944 | 0.884 | -0.177 | 0.00293767 | 0.01627141 | 0 | 0.0036 | 0.0021 | 0.0021 | TRUE | 0.0034 |
| Palm | 1733 | 1500 | 1838 | 2232 | 2097 | 2417 | 1346 | 1596 | 1983 | 2222 | 2366 | 2386 | 1983 | 1642 | 2325 | 1.418 | 0.504 | 6.03E-06 | 0.00012358 | 0.0191 | 0.0036 | 0.0084 | 0.0084 | TRUE | 0.3995 |
| Palm2 | 150 | 122 | 144 | 131 | 116 | 155 | 116 | 130 | 155 | 130 | 131 | 153 | 135.98 | 134 | 138 | 1.037 | 0.052 | 0.77509098 | 0.86721285 | 0.0083 | 0.0199 | 0.0165 | 0.0165 | TRUE | 0.1995 |
| Palm3 | 3462 | 2649 | 3010 | 3304 | 3119 | 3751 | 2689 | 2818 | 3248 | 3289 | 3519 | 3703 | 3210.74 | 2918 | 3504 | 1.201 | 0.265 | 0.00095227 | 0.00700018 | 0.0063 | 0.0032 | 0.0043 | 0.0043 | TRUE | 0.1181 |
| Palmd | 5 | 3 | 1 | 2 | 1 | 0 | 4 | 3 | 1 | 2 | 1 | 0 | 1.88 | 3 | 1 | 0.376 | -1.409 | 0.34527086 | NA | 0 | 1.264 | 0.932 | 0.932 | TRUE | 0.3966 |
| Pam | 984 | 678 | 727 | 738 | 645 | 726 | 764 | 721 | 784 | 735 | 728 | 717 | 741.46 | 756 | 727 | 0.96 | -0.059 | 0.47487397 | 0.64299637 | 0 | 0.0056 | 0.0036 | 0.0036 | TRUE | 0.0191 |
| Pam16 | 1023 | 695 | 722 | 873 | 687 | 854 | 795 | 739 | 779 | 869 | 775 | 843 | 799.97 | 771 | 829 | 1.075 | 0.105 | 0.21824495 | 0.38124849 | 0.0011 | 0.0054 | 0.004 | 0.004 | TRUE | 0.0376 |
| Pamr1 | 2 | 4 | 9 | 4 | 7 | 6 | 2 | 4 | 10 | 4 | 8 | 6 | 5.55 | 5 | 6 | 1.172 | 0.229 | 0.79343953 | NA | 0.2247 | 0.4292 | 0.3699 | 0.3699 | TRUE | 0.5776 |
| Pan2 | 4993 | 3841 | 3779 | 3847 | 3473 | 4030 | 3878 | 4086 | 4078 | 3829 | 3918 | 3978 | 3961.07 | 4014 | 3908 | 0.974 | -0.038 | 0.48916003 | 0.65489346 | 4.00E-04 | 0.003 | 0.0019 | 0.0019 | TRUE | 0.0109 |
| Pan3 | 1941 | 1582 | 1503 | 1654 | 1463 | 1733 | 1507 | 1683 | 1622 | 1646 | 1650 | 1711 | 1636.59 | 1604 | 1669 | 1.042 | 0.059 | 0.39603685 | 0.5694723 | 0.0013 | 0.0039 | 0.0029 | 0.0029 | TRUE | 0.0348 |
| Pank1 | 878 | 619 | 594 | 618 | 531 | 683 | 682 | 658 | 641 | 615 | 599 | 674 | 644.94 | 660 | 629 | 0.953 | -0.07 | 0.44329014 | 0.61433424 | 9.00E-04 | 0.0061 | 0.0044 | 0.0044 | TRUE | 0.046 |
| Pank2 | 4434 | 3317 | 3425 | 3775 | 3328 | 3986 | 3444 | 3528 | 3696 | 3758 | 3754 | 3935 | 3685.71 | 3556 | 3816 | 1.074 | 0.102 | 0.07405651 | 0.1786584 | 7.00E-04 | 0.0031 | 0.0021 | 0.0021 | TRUE | 0.0143 |
| Pank3 | 9293 | 6675 | 6276 | 7429 | 6244 | 6975 | 7217 | 7100 | 6772 | 7395 | 7044 | 6885 | 7068.91 | 7030 | 7108 | 1.011 | 0.016 | 0.77418828 | 0.86672461 | 9.00E-04 | 0.0028 | 0.002 | 0.002 | TRUE | 0.0153 |
| Pank4 | 1526 | 1257 | 1403 | 1289 | 1156 | 1433 | 1185 | 1337 | 1514 | 1283 | 1304 | 1415 | 1339.63 | 1345 | 1334 | 0.993 | -0.01 | 0.9145768 | 0.9529197 | 0.0082 | 0.0042 | 0.0056 | 0.0056 | TRUE | 0.1442 |
| Panx1 | 674 | 476 | 496 | 565 | 480 | 568 | 523 | 506 | 535 | 562 | 541 | 561 | 538.26 | 521 | 555 | 1.064 | 0.089 | 0.33156422 | 0.50583321 | 0 | 0.0068 | 0.0043 | 0.0043 | TRUE | 0.0073 |
| Panx2 | 37 | 30 | 33 | 24 | 44 | 54 | 29 | 32 | 36 | 24 | 50 | 53 | 37.18 | 32 | 42 | 1.32 | 0.4 | 0.26718862 | 0.43769936 | 0.0674 | 0.0662 | 0.0665 | 0.0665 | TRUE | 1.1343 |
| Panx3 | 1 | 1 | 0 | 0 | 0 | 0 | 1 | 1 | 0 | 0 | 0 | 0 | 0.31 | 1 | 0 | 0.309 | -1.696 | 0.63183309 | NA | 0 | 7.7294 | 7.2472 | 7.2472 | TRUE | 0.2015 |
| Paox | 1114 | 893 | 949 | 1136 | 1043 | 1228 | 865 | 950 | 1024 | 1131 | 1177 | 1212 | 1059.77 | 946 | 1173 | 1.242 | 0.313 | 0.00020753 | 0.00219073 | 0.0032 | 0.0047 | 0.0042 | 0.0042 | TRUE | 0.0705 |
| Papd4 | 2154 | 1504 | 1544 | 1656 | 1485 | 1625 | 1673 | 1600 | 1666 | 1648 | 1675 | 1604 | 1644.4 | 1646 | 1642 | 0.997 | -0.004 | 0.95134317 | 0.97323401 | 0 | 0.0039 | 0.0024 | 0.0024 | TRUE | 0.0072 |
| Papd5 | 9038 | 6592 | 6192 | 6592 | 5906 | 6914 | 7019 | 7012 | 6681 | 6562 | 6663 | 6825 | 6793.63 | 6904 | 6683 | 0.968 | -0.047 | 0.35573044 | 0.53023831 | 4.00E-04 | 0.0028 | 0.0017 | 0.0017 | TRUE | 0.0097 |
| Papd7 | 3439 | 2641 | 2768 | 2757 | 2563 | 2969 | 2671 | 2809 | 2987 | 2744 | 2891 | 2931 | 2838.88 | 2822 | 2855 | 1.012 | 0.018 | 0.78530683 | 0.87383344 | 0.0018 | 0.0033 | 0.0027 | 0.0027 | TRUE | 0.0314 |
| Papln | 158 | 130 | 128 | 127 | 110 | 120 | 123 | 138 | 138 | 126 | 124 | 118 | 128.01 | 133 | 123 | 0.927 | -0.109 | 0.52817126 | 0.68874916 | 0 | 0.0209 | 0.0139 | 0.0139 | TRUE | 0.0531 |
| Papola | 22396 | 15819 | 15053 | 14552 | 13266 | 15437 | 17394 | 16826 | 16242 | 14485 | 14965 | 15239 | 15858.62 | 16821 | 14896 | 0.885 | -0.175 | 0.00053601 | 0.0045386 | 8.00E-04 | 0.0026 | 0.0018 | 0.0018 | TRUE | 0.0111 |
| Papolb | 71 | 30 | 33 | 20 | 13 | 26 | 55 | 32 | 36 | 20 | 15 | 26 | 30.48 | 41 | 20 | 0.489 | -1.031 | 0.00819626 | 0.03567269 | 0.0529 | 0.0802 | 0.0723 | 0.0723 | TRUE | 0.8005 |
| Papolg | 3038 | 2210 | 2338 | 2502 | 2211 | 2457 | 2359 | 2351 | 2523 | 2490 | 2494 | 2425 | 2440.51 | 2411 | 2470 | 1.025 | 0.035 | 0.55990467 | 0.71465596 | 5.00E-04 | 0.0034 | 0.0022 | 0.0022 | TRUE | 0.0196 |
| Pappa | 5 | 2 | 0 | 2 | 1 | 0 | 4 | 2 | 0 | 2 | 1 | 0 | 1.52 | 2 | 1 | 0.505 | -0.987 | 0.56424988 | NA | 0 | 1.56 | 1.3763 | 1.3763 | TRUE | 0.137 |
| Pappa2 | 26 | 11 | 19 | 5 | 8 | 4 | 20 | 12 | 21 | 5 | 9 | 4 | 11.72 | 18 | 6 | 0.337 | -1.568 | 0.01097916 | NA | 0 | 0.2046 | 0.1588 | 0.1588 | TRUE | 0.4106 |
| Papss1 | 2283 | 1651 | 2009 | 2219 | 1995 | 2293 | 1773 | 1756 | 2168 | 2209 | 2251 | 2264 | 2069.98 | 1899 | 2241 | 1.181 | 0.24 | 0.00455412 | 0.02287143 | 0.0069 | 0.0036 | 0.0047 | 0.0047 | TRUE | 0.1832 |
| Papss2 | 101 | 79 | 44 | 69 | 40 | 37 | 78 | 84 | 47 | 69 | 45 | 37 | 60.05 | 70 | 50 | 0.715 | -0.485 | 0.11807393 | 0.24931757 | 0.0785 | 0.0419 | 0.0521 | 0.0521 | TRUE | 0.8738 |
| Paqr3 | 531 | 402 | 372 | 406 | 353 | 412 | 412 | 428 | 401 | 404 | 398 | 407 | 408.41 | 414 | 403 | 0.974 | -0.038 | 0.7129265 | 0.82750191 | 0 | 0.0082 | 0.0051 | 0.0051 | TRUE | 0.0093 |
| Paqr4 | 178 | 101 | 90 | 127 | 151 | 130 | 138 | 107 | 97 | 126 | 170 | 128 | 127.98 | 114 | 141 | 1.23 | 0.298 | 0.14179646 | 0.28412965 | 0.0241 | 0.0209 | 0.0219 | 0.0219 | TRUE | 0.369 |
| Paqr5 | 159 | 121 | 140 | 177 | 150 | 206 | 123 | 129 | 151 | 176 | 169 | 203 | 158.67 | 134 | 183 | 1.366 | 0.45 | 0.00690094 | 0.03140804 | 0.0042 | 0.0174 | 0.0136 | 0.0136 | TRUE | 0.1169 |
| Paqr6 | 57 | 50 | 69 | 80 | 87 | 86 | 44 | 53 | 74 | 80 | 98 | 85 | 72.43 | 57 | 88 | 1.539 | 0.622 | 0.01448754 | 0.05434917 | 0.0242 | 0.0351 | 0.0322 | 0.0322 | TRUE | 0.5668 |
| Paqr7 | 240 | 200 | 219 | 296 | 249 | 340 | 186 | 213 | 236 | 295 | 281 | 336 | 257.77 | 212 | 304 | 1.441 | 0.527 | 0.00019133 | 0.00206755 | 0.0075 | 0.0116 | 0.0104 | 0.0104 | TRUE | 0.1324 |
| Paqr8 | 422 | 294 | 300 | 273 | 347 | 341 | 328 | 313 | 324 | 272 | 391 | 337 | 327.33 | 322 | 333 | 1.035 | 0.049 | 0.72198241 | 0.83361294 | 0.0135 | 0.0097 | 0.0108 | 0.0108 | TRUE | 0.2242 |
| Paqr9 | 201 | 159 | 110 | 155 | 124 | 134 | 156 | 169 | 119 | 154 | 140 | 132 | 145.06 | 148 | 142 | 0.959 | -0.06 | 0.73960068 | 0.84535086 | 0.012 | 0.0188 | 0.0169 | 0.0169 | TRUE | 0.2962 |
| Pard3 | 7829 | 5702 | 6077 | 6458 | 5739 | 6628 | 6080 | 6065 | 6557 | 6428 | 6474 | 6543 | 6357.99 | 6234 | 6482 | 1.04 | 0.056 | 0.29688127 | 0.46882378 | 8.00E-04 | 0.0028 | 0.002 | 0.002 | TRUE | 0.0248 |
| Pard3b | 20 | 30 | 19 | 13 | 12 | 14 | 16 | 32 | 21 | 13 | 14 | 14 | 18.04 | 23 | 14 | 0.599 | -0.74 | 0.12153217 | 0.25435227 | 0 | 0.1338 | 0.1047 | 0.1047 | TRUE | 0.4303 |
| Pard6a | 809 | 544 | 479 | 513 | 424 | 594 | 628 | 579 | 517 | 511 | 478 | 586 | 549.85 | 575 | 525 | 0.913 | -0.131 | 0.2424812 | 0.41020048 | 0.0084 | 0.0067 | 0.0073 | 0.0073 | TRUE | 0.1223 |
| Pard6b | 1995 | 1436 | 1486 | 1490 | 1394 | 1525 | 1549 | 1527 | 1603 | 1483 | 1573 | 1505 | 1540.23 | 1560 | 1520 | 0.974 | -0.037 | 0.57236316 | 0.72505421 | 0 | 0.004 | 0.0025 | 0.0025 | TRUE | 0.0107 |
| Pard6g | 1087 | 932 | 958 | 887 | 824 | 978 | 844 | 991 | 1034 | 883 | 930 | 965 | 941.2 | 956 | 926 | 0.97 | -0.044 | 0.63911981 | 0.77456433 | 0.0058 | 0.0049 | 0.0052 | 0.0052 | TRUE | 0.1311 |
| Parg | 5694 | 4226 | 3951 | 4559 | 3881 | 4303 | 4422 | 4495 | 4263 | 4538 | 4378 | 4248 | 4390.74 | 4393 | 4388 | 0.999 | -0.002 | 0.9722166 | 0.98494753 | 7.00E-04 | 0.003 | 0.002 | 0.002 | TRUE | 0.011 |
| Park2 | 21 | 15 | 22 | 16 | 16 | 19 | 16 | 16 | 24 | 16 | 18 | 19 | 18.12 | 19 | 18 | 0.946 | -0.08 | 0.85938366 | 0.92152565 | 0 | 0.1332 | 0.0908 | 0.0908 | TRUE | 0.2726 |
| Park7 | 12198 | 8537 | 8363 | 9296 | 8201 | 9449 | 9474 | 9081 | 9024 | 9253 | 9252 | 9328 | 9235.07 | 9193 | 9278 | 1.009 | 0.013 | 0.78693982 | 0.87496934 | 3.00E-04 | 0.0027 | 0.0016 | 0.0016 | TRUE | 0.0089 |
| Parl | 5604 | 4191 | 4197 | 4613 | 4043 | 4650 | 4352 | 4458 | 4529 | 4592 | 4561 | 4590 | 4513.63 | 4446 | 4581 | 1.031 | 0.043 | 0.39237101 | 0.56602329 | 0 | 0.003 | 0.0016 | 0.0016 | TRUE | 0.0043 |
| Parm1 | 624 | 442 | 444 | 305 | 308 | 325 | 485 | 470 | 479 | 304 | 347 | 321 | 400.96 | 478 | 324 | 0.677 | -0.563 | 1.47E-07 | 5.47E-06 | 0 | 0.0083 | 0.0057 | 0.0057 | TRUE | 0.044 |
| Parn | 4056 | 3071 | 3484 | 3706 | 3325 | 3793 | 3150 | 3267 | 3759 | 3689 | 3751 | 3744 | 3560.01 | 3392 | 3728 | 1.1 | 0.137 | 0.05834728 | 0.15069708 | 0.0042 | 0.0031 | 0.0035 | 0.0035 | TRUE | 0.1085 |
| Parp1 | 50727 | 39132 | 42065 | 42971 | 39289 | 44217 | 39398 | 41624 | 45388 | 42773 | 44322 | 43649 | 42858.97 | 42137 | 43581 | 1.034 | 0.049 | 0.41688606 | 0.5895361 | 0.0024 | 0.0025 | 0.0026 | 0.0026 | TRUE | 0.0558 |
| Parp10 | 227 | 167 | 180 | 226 | 189 | 268 | 176 | 178 | 194 | 225 | 213 | 265 | 208.48 | 183 | 234 | 1.285 | 0.362 | 0.01395872 | 0.05292096 | 0.0037 | 0.0138 | 0.0108 | 0.0108 | TRUE | 0.143 |
| Parp11 | 1123 | 849 | 741 | 829 | 752 | 801 | 872 | 903 | 800 | 825 | 848 | 791 | 839.84 | 858 | 821 | 0.956 | -0.064 | 0.44387669 | 0.61487977 | 0.0013 | 0.0052 | 0.0039 | 0.0039 | TRUE | 0.041 |
| Parp12 | 2343 | 1693 | 1418 | 1706 | 1534 | 1604 | 1820 | 1801 | 1530 | 1698 | 1731 | 1583 | 1693.77 | 1717 | 1671 | 0.972 | -0.041 | 0.62016566 | 0.76038736 | 0.0051 | 0.0038 | 0.0043 | 0.0043 | TRUE | 0.1079 |
| Parp14 | 376 | 277 | 251 | 287 | 283 | 327 | 292 | 295 | 271 | 286 | 319 | 323 | 297.54 | 286 | 309 | 1.081 | 0.112 | 0.357156 | 0.53164244 | 0 | 0.0104 | 0.0073 | 0.0073 | TRUE | 0.0512 |
| Parp16 | 462 | 354 | 369 | 341 | 350 | 448 | 359 | 377 | 398 | 339 | 395 | 442 | 385 | 378 | 392 | 1.039 | 0.056 | 0.65109001 | 0.78333581 | 0.0076 | 0.0086 | 0.0083 | 0.0083 | TRUE | 0.1625 |
| Parp2 | 3343 | 2486 | 2514 | 2604 | 2204 | 2537 | 2596 | 2644 | 2713 | 2592 | 2486 | 2504 | 2589.34 | 2651 | 2527 | 0.954 | -0.068 | 0.23834824 | 0.405175 | 0 | 0.0033 | 0.002 | 0.002 | TRUE | 0.0061 |

| | | | | | | | | | | | | | | | | | | | | | | | | | |
|---|---|---|---|---|---|---|---|---|---|---|---|---|---|---|---|---|---|---|---|---|---|---|---|---|---|
| Parp3 | 47 | 29 | 38 | 32 | 32 | 29 | 37 | 31 | 41 | 32 | 36 | 29 | 34.16 | 36 | 32 | 0.889 | -0.169 | 0.60832169 | 0.7520791 | 0 | 0.0718 | 0.0491 | 0.0491 | TRUE | 0.1075 |
| Parp4 | 912 | 621 | 544 | 628 | 541 | 616 | 708 | 661 | 587 | 625 | 610 | 608 | 633.22 | 652 | 614 | 0.941 | -0.088 | 0.36282242 | 0.53754219 | 0.0032 | 0.0062 | 0.0052 | 0.0052 | TRUE | 0.0883 |
| Parp6 | 657 | 552 | 566 | 551 | 506 | 646 | 510 | 587 | 611 | 548 | 571 | 638 | 577.52 | 569 | 586 | 1.031 | 0.044 | 0.67495726 | 0.80032886 | 0.006 | 0.0065 | 0.0063 | 0.0063 | TRUE | 0.1024 |
| Parp8 | 80 | 66 | 40 | 45 | 28 | 30 | 62 | 70 | 43 | 45 | 32 | 30 | 46.92 | 58 | 36 | 0.604 | -0.728 | 0.01995698 | 0.0690499 | 0.0335 | 0.0529 | 0.0477 | 0.0477 | TRUE | 0.4213 |
| Parp9 | 735 | 513 | 448 | 449 | 393 | 488 | 571 | 546 | 483 | 447 | 443 | 482 | 495.32 | 533 | 457 | 0.856 | -0.224 | 0.03218895 | 0.09752338 | 0.0029 | 0.0072 | 0.0058 | 0.0058 | TRUE | 0.0767 |
| Parpbp | 2391 | 1690 | 1565 | 1781 | 1520 | 1649 | 1857 | 1798 | 1689 | 1773 | 1715 | 1628 | 1743.1 | 1781 | 1705 | 0.957 | -0.064 | 0.35950441 | 0.53419275 | 0.0015 | 0.0038 | 0.0029 | 0.0029 | TRUE | 0.0247 |
| Pars2 | 636 | 510 | 488 | 505 | 445 | 508 | 494 | 542 | 527 | 503 | 502 | 501 | 511.52 | 521 | 502 | 0.965 | -0.051 | 0.59138907 | 0.73939153 | 0 | 0.0071 | 0.0046 | 0.0046 | TRUE | 0.0259 |
| Parva | 713 | 458 | 454 | 498 | 451 | 448 | 554 | 487 | 490 | 496 | 509 | 442 | 496.25 | 510 | 482 | 0.942 | -0.085 | 0.41896458 | 0.59160845 | 0.0036 | 0.0072 | 0.006 | 0.006 | TRUE | 0.0686 |
| Parvb | 6129 | 4838 | 5834 | 6775 | 6688 | 7432 | 4760 | 5146 | 6295 | 6744 | 7545 | 7337 | 6304.37 | 5400 | 7209 | 1.335 | 0.417 | 5.41E-06 | 0.00011312 | 0.0123 | 0.0028 | 0.0059 | 0.0059 | TRUE | 0.2556 |
| Parvg | 10 | 15 | 6 | 10 | 17 | 6 | 8 | 16 | 6 | 10 | 19 | 6 | 10.88 | 10 | 12 | 1.159 | 0.213 | 0.74323609 | NA | 0.1891 | 0.2204 | 0.2113 | 0.2113 | TRUE | 0.535 |
| Pask | 4316 | 3234 | 2954 | 3081 | 2803 | 3058 | 3352 | 3440 | 3187 | 3067 | 3162 | 3019 | 3204.5 | 3326 | 3083 | 0.926 | -0.11 | 0.0594187 | 0.15228105 | 7.00E-04 | 0.0032 | 0.0022 | 0.0022 | TRUE | 0.0159 |
| Pate2 | 24 | 11 | 9 | 8 | 5 | 14 | 19 | 12 | 10 | 8 | 6 | 14 | 11.25 | 14 | 9 | 0.681 | -0.554 | 0.36100007 | NA | 0.0704 | 0.2132 | 0.1731 | 0.1731 | TRUE | 0.6648 |
| Pate4 | 0 | 0 | 0 | 0 | 0 | 0 | 0 | 0 | 0 | 0 | 0 | 0 | 0 | 0 | 0 | NA | NA | NA | NA | NA | NA | NA | NA | NA | NA |
| Patl1 | 4587 | 3355 | 3917 | 3855 | 3358 | 4255 | 3563 | 3569 | 4226 | 3837 | 3788 | 4200 | 3863.9 | 3786 | 3942 | 1.042 | 0.059 | 0.45345586 | 0.62319659 | 0.0055 | 0.003 | 0.0042 | 0.0042 | TRUE | 0.1254 |
| Patl2 | 98 | 71 | 88 | 112 | 124 | 121 | 76 | 76 | 95 | 111 | 140 | 119 | 102.9 | 82 | 123 | 1.506 | 0.591 | 0.00374421 | 0.01959917 | 0.0049 | 0.0255 | 0.0198 | 0.0198 | TRUE | 0.1635 |
| Patz1 | 2587 | 1891 | 1800 | 1834 | 1614 | 1833 | 2009 | 2011 | 1942 | 1826 | 1821 | 1809 | 1903.1 | 1987 | 1819 | 0.915 | -0.129 | 0.03357638 | 0.1006728 | 0 | 0.0037 | 0.0021 | 0.0021 | TRUE | 0.0047 |
| Pawr | 3398 | 2478 | 2383 | 2583 | 2262 | 2328 | 2639 | 2636 | 2571 | 2571 | 2552 | 2298 | 2544.52 | 2615 | 2474 | 0.946 | -0.081 | 0.21621665 | 0.37886288 | 0.0014 | 0.0034 | 0.0027 | 0.0027 | TRUE | 0.0471 |
| Pax1 | 8 | 9 | 4 | 7 | 9 | 3 | 6 | 10 | 4 | 7 | 10 | 3 | 6.7 | 7 | 7 | 0.993 | -0.011 | 0.98894852 | NA | 0 | 0.3563 | 0.2791 | 0.2791 | TRUE | 0.2583 |
| Pax2 | 19 | 5 | 8 | 7 | 3 | 2 | 15 | 5 | 9 | 7 | 3 | 2 | 6.84 | 10 | 4 | 0.425 | -1.236 | 0.12618569 | NA | 0.1579 | 0.349 | 0.2951 | 0.2951 | TRUE | 0.3234 |
| Pax3 | 4 | 3 | 3 | 1 | 0 | 2 | 3 | 3 | 3 | 1 | 0 | 2 | 2.08 | 3 | 1 | 0.32 | -1.646 | 0.24292037 | NA | 0 | 1.1397 | 0.7698 | 0.7698 | TRUE | 0.3491 |
| Pax4 | 0 | 0 | 0 | 0 | 0 | 0 | 0 | 0 | 0 | 0 | 0 | 0 | 0 | 0 | 0 | NA | NA | NA | NA | NA | NA | NA | NA | NA | NA |
| Pax5 | 3 | 0 | 2 | 2 | 2 | 0 | 2 | 0 | 2 | 2 | 2 | 0 | 1.46 | 1 | 1 | 0.921 | -0.118 | 0.94409642 | NA | 0 | 1.6304 | 1.3709 | 1.3709 | TRUE | 0.5209 |
| Pax6 | 149 | 103 | 117 | 109 | 99 | 163 | 116 | 110 | 126 | 108 | 112 | 161 | 122.1 | 117 | 127 | 1.086 | 0.119 | 0.55494931 | 0.7102732 | 0.0201 | 0.0218 | 0.0213 | 0.0213 | TRUE | 0.5646 |
| Pax6os1 | 0 | 0 | 0 | 0 | 0 | 0 | 0 | 0 | 0 | 0 | 0 | 0 | 0 | 0 | 0 | NA | NA | NA | NA | NA | NA | NA | NA | NA | NA |
| Pax7 | 6 | 4 | 7 | 11 | 16 | 6 | 5 | 4 | 8 | 11 | 18 | 6 | 8.56 | 6 | 12 | 2.122 | 1.086 | 0.12472307 | NA | 0 | 0.2791 | 0.227 | 0.227 | TRUE | 0.1646 |
| Pax8 | 140 | 124 | 172 | 181 | 208 | 222 | 109 | 132 | 186 | 180 | 235 | 219 | 176.7 | 142 | 211 | 1.495 | 0.58 | 0.00349611 | 0.01859487 | 0.0385 | 0.0158 | 0.0226 | 0.0226 | TRUE | 0.7369 |
| Pax9 | 17 | 10 | 12 | 16 | 6 | 18 | 13 | 11 | 13 | 16 | 7 | 18 | 12.88 | 12 | 14 | 1.105 | 0.144 | 0.79509304 | NA | 0 | 0.1865 | 0.1452 | 0.1452 | TRUE | 0.7232 |
| Paxbp1 | 2185 | 1829 | 1672 | 1484 | 1328 | 1612 | 1697 | 1945 | 1804 | 1477 | 1498 | 1591 | 1668.86 | 1815 | 1522 | 0.839 | -0.253 | 0.00070176 | 0.00557603 | 0.0027 | 0.0038 | 0.0034 | 0.0034 | TRUE | 0.0474 |
| Paxip1 | 5885 | 4325 | 4055 | 3666 | 3300 | 3844 | 4571 | 4600 | 4375 | 3649 | 3723 | 3795 | 4118.81 | 4515 | 3722 | 0.824 | -0.279 | 2.28E-07 | 8.21E-06 | 3.00E-04 | 0.003 | 0.0018 | 0.0018 | TRUE | 0.0089 |
| Pbdc1 | 3968 | 2798 | 2764 | 2830 | 2466 | 3067 | 3082 | 2976 | 2982 | 2817 | 2782 | 3028 | 2944.47 | 3013 | 2876 | 0.954 | -0.068 | 0.26218032 | 0.43251886 | 8.00E-04 | 0.0032 | 0.0023 | 0.0023 | TRUE | 0.0261 |
| Pbk | 3073 | 2207 | 2490 | 2527 | 2176 | 2510 | 2387 | 2348 | 2687 | 2515 | 2455 | 2478 | 2478.13 | 2474 | 2483 | 1.004 | 0.006 | 0.93077782 | 0.96192556 | 0.0024 | 0.0034 | 0.003 | 0.003 | TRUE | 0.0678 |
| Pbld1 | 20 | 18 | 18 | 20 | 19 | 34 | 16 | 19 | 19 | 20 | 21 | 34 | 21.5 | 18 | 25 | 1.396 | 0.481 | 0.26089711 | 0.4308495 | 0 | 0.1127 | 0.0843 | 0.0843 | TRUE | 0.5669 |
| Pbld2 | 38 | 31 | 35 | 33 | 29 | 37 | 30 | 33 | 38 | 33 | 33 | 37 | 33.72 | 34 | 34 | 1.024 | 0.034 | 0.91739074 | 0.95445232 | 0 | 0.0727 | 0.0481 | 0.0481 | TRUE | 0.0878 |
| Pbp2 | 9 | 3 | 6 | 6 | 2 | 1 | 7 | 3 | 6 | 6 | 2 | 1 | 4.31 | 5 | 3 | 0.552 | -0.858 | 0.38041625 | NA | 0.0954 | 0.5521 | 0.436 | 0.436 | TRUE | 0.9378 |
| Pbrm1 | 9912 | 7051 | 7065 | 8022 | 6799 | 7875 | 7698 | 7500 | 7623 | 7985 | 7670 | 7774 | 7708.38 | 7607 | 7810 | 1.027 | 0.038 | 0.43213227 | 0.60400928 | 2.00E-04 | 0.0027 | 0.0015 | 0.0015 | TRUE | 0.0047 |
| Pbsn | 0 | 0 | 0 | 0 | 0 | 0 | 0 | 0 | 0 | 0 | 0 | 0 | 0 | 0 | 0 | NA | NA | NA | NA | NA | NA | NA | NA | NA | NA |
| Pbx1 | 558 | 364 | 445 | 340 | 256 | 375 | 433 | 387 | 480 | 338 | 289 | 370 | 383.02 | 433 | 332 | 0.768 | -0.381 | 0.0029455 | 0.01630355 | 0.0104 | 0.0086 | 0.0091 | 0.0091 | TRUE | 0.1455 |
| Pbx2 | 3089 | 2160 | 2127 | 2404 | 2066 | 2550 | 2399 | 2298 | 2295 | 2393 | 2331 | 2517 | 2372.09 | 2331 | 2414 | 1.035 | 0.05 | 0.41691953 | 0.5895361 | 7.00E-04 | 0.0034 | 0.0023 | 0.0023 | TRUE | 0.0172 |
| Pbx3 | 663 | 478 | 439 | 485 | 352 | 419 | 515 | 508 | 474 | 483 | 397 | 414 | 465.09 | 499 | 431 | 0.864 | -0.211 | 0.05364821 | 0.14156329 | 0.004 | 0.0075 | 0.0064 | 0.0064 | TRUE | 0.1276 |
| Pbx4 | 242 | 194 | 250 | 270 | 243 | 272 | 188 | 206 | 270 | 269 | 274 | 269 | 245.91 | 221 | 271 | 1.227 | 0.295 | 0.05201422 | 0.13879494 | 0.0136 | 0.0121 | 0.0125 | 0.0125 | TRUE | 0.3949 |
| Pbxip1 | 891 | 696 | 560 | 604 | 546 | 597 | 692 | 740 | 604 | 601 | 616 | 589 | 640.51 | 679 | 602 | 0.886 | -0.174 | 0.07698523 | 0.18363649 | 0.0039 | 0.0061 | 0.0054 | 0.0054 | TRUE | 0.1049 |
| Pcbd1 | 228 | 173 | 217 | 268 | 220 | 280 | 177 | 184 | 234 | 267 | 248 | 276 | 231.1 | 198 | 264 | 1.335 | 0.417 | 0.00482712 | 0.02386661 | 0.0081 | 0.0127 | 0.0114 | 0.0114 | TRUE | 0.2619 |
| Pcbd2 | 264 | 175 | 135 | 181 | 136 | 153 | 205 | 186 | 146 | 180 | 153 | 151 | 170.25 | 179 | 161 | 0.9 | -0.153 | 0.37893748 | 0.55369661 | 0.0144 | 0.0163 | 0.0158 | 0.0158 | TRUE | 0.2758 |
| Pcbp1 | 27150 | 18853 | 16005 | 15649 | 13406 | 15899 | 21086 | 20054 | 17269 | 15577 | 15123 | 15695 | 17467.39 | 19470 | 15465 | 0.794 | -0.332 | 3.74E-06 | 8.37E-05 | 0.0052 | 0.0026 | 0.0037 | 0.0037 | TRUE | 0.1195 |
| Pcbp2 | 47576 | 32862 | 33911 | 36151 | 30975 | 36309 | 36950 | 34955 | 36590 | 35985 | 34943 | 35843 | 35877.52 | 36165 | 35590 | 0.984 | -0.023 | 0.62079552 | 0.76065413 | 5.00E-04 | 0.0025 | 0.0015 | 0.0015 | TRUE | 0.0105 |
| Pcbp3 | 1964 | 1468 | 1555 | 1677 | 1612 | 1772 | 1525 | 1561 | 1678 | 1669 | 1819 | 1749 | 1666.95 | 1588 | 1746 | 1.1 | 0.137 | 0.05238946 | 0.13938991 | 0.0015 | 0.0038 | 0.003 | 0.003 | TRUE | 0.029 |
| Pcbp4 | 2889 | 2130 | 2230 | 2216 | 2044 | 2593 | 2244 | 2266 | 2406 | 2206 | 2306 | 2560 | 2331.15 | 2305 | 2357 | 1.023 | 0.033 | 0.65164413 | 0.78367265 | 0.0033 | 0.0034 | 0.0034 | 0.0034 | TRUE | 0.0691 |
| Pcca | 1347 | 965 | 1055 | 1316 | 1152 | 1349 | 1046 | 1026 | 1138 | 1310 | 1300 | 1332 | 1192.02 | 1070 | 1314 | 1.228 | 0.297 | 5.78E-05 | 0.00078569 | 5.00E-04 | 0.0044 | 0.0031 | 0.0031 | TRUE | 0.0358 |
| Pccb | 1611 | 1156 | 1096 | 1240 | 1045 | 1294 | 1251 | 1230 | 1183 | 1234 | 1179 | 1277 | 1225.66 | 1221 | 1230 | 1.007 | 0.01 | 0.88485068 | 0.93609243 | 0 | 0.0044 | 0.0029 | 0.0029 | TRUE | 0.0156 |
| Pcdh1 | 807 | 607 | 690 | 777 | 687 | 834 | 627 | 646 | 745 | 773 | 775 | 823 | 731.44 | 673 | 790 | 1.178 | 0.236 | 0.01155985 | 0.04602367 | 0.0035 | 0.0057 | 0.0049 | 0.0049 | TRUE | 0.1023 |
| Pcdh10 | 18 | 6 | 11 | 7 | 3 | 8 | 14 | 6 | 12 | 7 | 3 | 8 | 8.41 | 11 | 6 | 0.566 | -0.821 | 0.23507874 | NA | 0 | 0.2841 | 0.2156 | 0.2156 | TRUE | 0.4335 |
| Pcdh11x | 0 | 0 | 0 | 0 | 1 | 0 | 0 | 0 | 0 | 0 | 1 | 0 | 0.19 | 0 | 0 | 2.094 | 1.067 | 0.79379329 | NA | 0 | 12.608 | 10 | 10 | TRUE | 0.5075 |
| Pcdh12 | 9 | 1 | 2 | 0 | 0 | 2 | 7 | 1 | 2 | 0 | 0 | 2 | 2.03 | 3 | 1 | 0.194 | -2.363 | 0.15532613 | NA | 0.801 | 1.1694 | 1.0888 | 1.0888 | TRUE | 1.2856 |
| Pcdh15 | 0 | 3 | 3 | 1 | 0 | 1 | 0 | 3 | 3 | 1 | 0 | 1 | 1.4 | 2 | 1 | 0.323 | -1.629 | 0.37812084 | NA | 0 | 1.6932 | 1.4644 | 1.4644 | TRUE | 1.0058 |
| Pcdh17 | 20 | 11 | 5 | 3 | 6 | 3 | 16 | 12 | 5 | 3 | 7 | 3 | 7.56 | 11 | 4 | 0.38 | -1.394 | 0.07028758 | NA | 0 | 0.316 | 0.26 | 0.26 | TRUE | 0.1983 |
| Pcdh18 | 13 | 6 | 8 | 8 | 3 | 4 | 10 | 6 | 9 | 8 | 3 | 4 | 6.73 | 8 | 5 | 0.609 | -0.716 | 0.34146486 | NA | 0 | 0.3544 | 0.25 | 0.25 | TRUE | 0.5064 |
| Pcdh19 | 249 | 165 | 97 | 100 | 104 | 90 | 193 | 176 | 105 | 100 | 117 | 89 | 129.88 | 158 | 102 | 0.642 | -0.64 | 0.00554717 | 0.02656138 | 0.0533 | 0.0207 | 0.0302 | 0.0302 | TRUE | 0.8904 |
| Pcdh20 | 7 | 6 | 1 | 1 | 0 | 4 | 5 | 6 | 1 | 1 | 0 | 4 | 2.97 | 4 | 2 | 0.388 | -1.364 | 0.28372485 | NA | 0.5601 | 0.7995 | 0.7473 | 0.7473 | TRUE | 1.1336 |
| Pcdh7 | 2 | 3 | 0 | 0 | 0 | 3 | 2 | 3 | 0 | 0 | 0 | 3 | 1.28 | 2 | 1 | 0.634 | -0.657 | 0.73702524 | NA | 2.4911 | 1.8478 | 1.9406 | 1.9406 | TRUE | 0.4627 |
| Pcdh8 | 99 | 63 | 43 | 60 | 52 | 39 | 77 | 67 | 46 | 60 | 59 | 38 | 57.86 | 63 | 52 | 0.818 | -0.29 | 0.31858674 | 0.4925266 | 0.0431 | 0.0434 | 0.0433 | 0.0433 | TRUE | 0.4461 |
| Pcdh9 | 19 | 9 | 5 | 4 | 9 | 7 | 15 | 10 | 5 | 4 | 10 | 7 | 8.46 | 10 | 7 | 0.692 | -0.531 | 0.44904133 | NA | 0.1043 | 0.2825 | 0.2318 | 0.2318 | TRUE | 0.2023 |
| Pcdha1 | 0 | 2 | 0 | 2 | 0 | 0 | 0 | 2 | 0 | 2 | 0 | 0 | 0.69 | 1 | 1 | 0.971 | -0.042 | 0.98762268 | NA | 8.094 | 3.4555 | 3.8966 | 3.8966 | TRUE | 0.9747 |
| Pcdha10 | 2 | 4 | 0 | 0 | 0 | 1 | 2 | 4 | 0 | 0 | 0 | 1 | 1.13 | 2 | 0 | 0.192 | -2.38 | 0.26121891 | NA | 0 | 2.0951 | 1.9833 | 1.9833 | TRUE | 0.1845 |
| Pcdha11 | 0 | 0 | 0 | 0 | 0 | 0 | 0 | 0 | 0 | 0 | 0 | 0 | 0 | 0 | 0 | NA | NA | NA | NA | NA | NA | NA | NA | NA | NA |
| Pcdha12 | 0 | 0 | 1 | 2 | 1 | 0 | 0 | 0 | 1 | 2 | 1 | 0 | 0.7 | 0 | 1 | 3.01 | 1.59 | 0.51808674 | NA | 0 | 3.3899 | 2.8624 | 2.8624 | TRUE | 0.3541 |
| Pcdha2 | 0 | 0 | 0 | 0 | 0 | 0 | 0 | 0 | 0 | 0 | 0 | 0 | 0 | 0 | 0 | NA | NA | NA | NA | NA | NA | NA | NA | NA | NA |
| Pcdha3 | 3 | 0 | 0 | 0 | 0 | 0 | 2 | 0 | 0 | 0 | 0 | 0 | 0.39 | 1 | 0 | 0.239 | -2.067 | 0.57190649 | NA | 10 | 6.1057 | 8.0283 | 8.0283 | TRUE | 1.4408 |
| Pcdha4 | 0 | 0 | 0 | 1 | 0 | 0 | 0 | 0 | 0 | 1 | 0 | 0 | 0.17 | 0 | 0 | 2.094 | 1.067 | 0.79379329 | NA | 0 | 14.2885 | 10 | 10 | TRUE | 0.3906 |
| Pcdha4-g | 0 | 0 | 0 | 0 | 0 | 0 | 0 | 0 | 0 | 0 | 0 | 0 | 0 | 0 | 0 | NA | NA | NA | NA | NA | NA | NA | NA | NA | NA |

| | | | | | | | | | | | | | | | | | | | | | | | | | |
|---|---|---|---|---|---|---|---|---|---|---|---|---|---|---|---|---|---|---|---|---|---|---|---|---|---|
| Pcdha5 | 0 | 1 | 0 | 0 | 1 | 0 | 0 | 1 | 0 | 0 | 1 | 0 | 0.37 | 0 | 0 | 1.075 | 0.105 | 0.97520514 | NA | 0 | 6.4904 | 6.1969 | 6.1969 | TRUE | 0.5216 |
| Pcdha6 | 6 | 4 | 2 | 2 | 1 | 1 | 5 | 4 | 2 | 2 | 1 | 1 | 2.53 | 4 | 1 | 0.369 | -1.438 | 0.24714637 | NA | 0 | 0.9393 | 0.6068 | 0.6068 | TRUE | 0.1878 |
| Pcdha7 | 5 | 2 | 2 | 4 | 1 | 1 | 4 | 2 | 2 | 4 | 1 | 1 | 2.38 | 3 | 2 | 0.741 | -0.433 | 0.72758386 | NA | 0 | 0.9993 | 0.6867 | 0.6867 | TRUE | 0.6578 |
| Pcdha8 | 0 | 1 | 0 | 0 | 1 | 0 | 0 | 1 | 0 | 0 | 1 | 0 | 0.37 | 0 | 0 | 1.075 | 0.105 | 0.97520514 | NA | 0 | 6.4904 | 6.1969 | 6.1969 | TRUE | 0.5216 |
| Pcdha9 | 1 | 0 | 0 | 0 | 0 | 0 | 1 | 0 | 0 | 0 | 0 | 0 | 0.13 | 0 | 0 | 0.552 | -0.857 | 0.8336548 | NA | 0 | 18.3121 | 10 | 10 | TRUE | 0.2566 |
| Pcdhac1 | 0 | 1 | 0 | 1 | 0 | 0 | 0 | 1 | 0 | 1 | 0 | 0 | 0.34 | 0 | 0 | 1.075 | 0.105 | 0.97573409 | NA | 0 | 6.9086 | 6.5582 | 6.5582 | TRUE | 0.5216 |
| Pcdhac2 | 41 | 34 | 27 | 27 | 26 | 24 | 32 | 36 | 29 | 27 | 29 | 24 | 29.51 | 32 | 27 | 0.821 | -0.284 | 0.41936011 | 0.59169931 | 0 | 0.0827 | 0.0548 | 0.0548 | TRUE | 0.0661 |
| Pcdhb1 | 1 | 0 | 0 | 0 | 0 | 1 | 0 | 1 | 0 | 0 | 0 | 1 | 0.32 | 0 | 0 | 1.075 | 0.105 | 0.97638847 | NA | 0 | 7.4681 | 7.0392 | 7.0392 | TRUE | 0.5075 |
| Pcdhb10 | 0 | 0 | 0 | 0 | 0 | 0 | 0 | 0 | 0 | 0 | 0 | 0 | 0 | 0 | 0 | NA | NA | NA | NA | NA | NA | NA | NA | NA | NA |
| Pcdhb11 | 3 | 7 | 0 | 3 | 1 | 1 | 2 | 7 | 0 | 3 | 1 | 1 | 2.48 | 3 | 2 | 0.528 | -0.922 | 0.50297426 | NA | 0.7903 | 0.9583 | 0.9143 | 0.9143 | TRUE | 0.3492 |
| Pcdhb12 | 0 | 0 | 0 | 0 | 2 | 0 | 0 | 0 | 0 | 0 | 2 | 0 | 0.38 | 0 | 1 | 4.195 | 2.069 | 0.57837236 | NA | 10 | 6.3052 | 8.2866 | 8.2866 | TRUE | 0.9743 |
| Pcdhb13 | 0 | 0 | 0 | 0 | 0 | 0 | 0 | 0 | 0 | 0 | 0 | 0 | 0 | 0 | 0 | NA | NA | NA | NA | NA | NA | NA | NA | NA | NA |
| Pcdhb14 | 5 | 4 | 2 | 2 | 3 | 3 | 4 | 4 | 2 | 2 | 3 | 3 | 3.11 | 3 | 3 | 0.802 | -0.318 | 0.76395046 | NA | 0 | 0.7656 | 0.4828 | 0.4828 | TRUE | 0.1374 |
| Pcdhb15 | 0 | 0 | 0 | 0 | 0 | 0 | 0 | 0 | 0 | 0 | 0 | 0 | 0 | 0 | 0 | NA | NA | NA | NA | NA | NA | NA | NA | NA | NA |
| Pcdhb16 | 0 | 0 | 1 | 0 | 1 | 1 | 0 | 0 | 1 | 0 | 1 | 1 | 0.53 | 0 | 1 | 2.03 | 1.022 | 0.7058734 | NA | 0 | 4.4543 | 3.5426 | 3.5426 | TRUE | 0.5365 |
| Pcdhb17 | 1 | 0 | 0 | 0 | 1 | 0 | 1 | 0 | 0 | 0 | 1 | 0 | 0.32 | 0 | 0 | 1.075 | 0.105 | 0.97638847 | NA | 0 | 7.4681 | 7.0392 | 7.0392 | TRUE | 0.5075 |
| Pcdhb18 | 9 | 1 | 3 | 2 | 3 | 3 | 7 | 1 | 3 | 2 | 3 | 3 | 3.27 | 4 | 3 | 0.716 | -0.482 | 0.65883537 | NA | 0 | 0.727 | 0.5478 | 0.5478 | TRUE | 0.3499 |
| Pcdhb19 | 1 | 2 | 1 | 0 | 3 | 3 | 1 | 2 | 1 | 0 | 3 | 3 | 1.72 | 1 | 2 | 1.609 | 0.686 | 0.65347997 | NA | 0 | 1.3792 | 1.0661 | 1.0661 | TRUE | 0.7476 |
| Pcdhb2 | 6 | 5 | 3 | 5 | 3 | 2 | 5 | 5 | 3 | 5 | 3 | 2 | 3.93 | 4 | 3 | 0.78 | -0.359 | 0.70750862 | NA | 0 | 0.6062 | 0.3985 | 0.3985 | TRUE | 0.2333 |
| Pcdhb20 | 1 | 1 | 0 | 2 | 2 | 1 | 1 | 1 | 0 | 2 | 2 | 1 | 1.18 | 1 | 2 | 2.782 | 1.476 | 0.4136547 | NA | 0 | 2.0125 | 1.2823 | 1.2823 | TRUE | 0.187 |
| Pcdhb21 | 1 | 1 | 0 | 1 | 2 | 0 | 1 | 1 | 0 | 1 | 2 | 0 | 0.85 | 1 | 1 | 1.716 | 0.779 | 0.7168933 | NA | 0 | 2.7951 | 2.0645 | 2.0645 | TRUE | 0.164 |
| Pcdhb22 | 8 | 7 | 4 | 9 | 4 | 5 | 6 | 7 | 4 | 9 | 5 | 5 | 6.06 | 6 | 6 | 1.026 | 0.037 | 0.96163379 | NA | 0 | 0.3933 | 0.2688 | 0.2688 | TRUE | 0.3955 |
| Pcdhb3 | 5 | 4 | 1 | 1 | 1 | 0 | 4 | 4 | 1 | 1 | 1 | 0 | 1.89 | 3 | 1 | 0.225 | -2.154 | 0.17430075 | NA | 0 | 1.2564 | 0.9115 | 0.9115 | TRUE | 0.3869 |
| Pcdhb4 | 1 | 2 | 0 | 0 | 1 | 0 | 1 | 2 | 0 | 0 | 1 | 0 | 0.67 | 1 | 0 | 0.385 | -1.379 | 0.57963618 | NA | 0 | 3.5292 | 2.9645 | 2.9645 | TRUE | 0.3656 |
| Pcdhb5 | 2 | 0 | 1 | 0 | 0 | 0 | 2 | 0 | 1 | 0 | 0 | 0 | 0.44 | 1 | 0 | 0.214 | -2.223 | 0.47051571 | NA | 0 | 5.4047 | 5.2946 | 5.2946 | TRUE | 0.2788 |
| Pcdhb6 | 1 | 0 | 1 | 0 | 0 | 0 | 1 | 0 | 1 | 0 | 0 | 0 | 0.31 | 1 | 0 | 0.307 | -1.706 | 0.62894649 | NA | 0 | 7.6657 | 7.2 | 7.2 | TRUE | 0.2067 |
| Pcdhb7 | 3 | 4 | 1 | 1 | 1 | 2 | 2 | 4 | 1 | 1 | 1 | 2 | 1.96 | 2 | 1 | 0.538 | -0.894 | 0.51458941 | NA | 0 | 1.2115 | 0.7904 | 0.7904 | TRUE | 0.2134 |
| Pcdhb8 | 4 | 1 | 2 | 1 | 1 | 0 | 3 | 1 | 2 | 1 | 1 | 0 | 1.41 | 2 | 1 | 0.326 | -1.618 | 0.34337779 | NA | 0 | 1.685 | 1.1321 | 1.1321 | TRUE | 0.2424 |
| Pcdhb9 | 0 | 0 | 0 | 0 | 0 | 0 | 0 | 0 | 0 | 0 | 0 | 0 | 0 | 0 | 0 | NA | NA | NA | NA | NA | NA | NA | NA | NA | NA |
| Pcdhga1 | 168 | 107 | 94 | 79 | 70 | 64 | 130 | 114 | 101 | 79 | 79 | 63 | 94.42 | 115 | 74 | 0.635 | -0.655 | 0.00211279 | 0.01257206 | 0 | 0.0275 | 0.0215 | 0.0215 | TRUE | 0.1473 |
| Pcdhga10 | 98 | 67 | 35 | 45 | 39 | 56 | 76 | 71 | 38 | 45 | 44 | 55 | 54.87 | 62 | 48 | 0.774 | -0.369 | 0.2282962 | 0.39365878 | 0.0578 | 0.0456 | 0.0491 | 0.0491 | TRUE | 0.9508 |
| Pcdhga11 | 229 | 153 | 133 | 191 | 169 | 194 | 178 | 163 | 144 | 190 | 191 | 192 | 176.06 | 162 | 191 | 1.178 | 0.236 | 0.12513367 | 0.26021509 | 0 | 0.0159 | 0.0114 | 0.0114 | TRUE | 0.0967 |
| Pcdhga12 | 313 | 182 | 179 | 176 | 163 | 219 | 243 | 194 | 193 | 175 | 184 | 216 | 200.85 | 210 | 192 | 0.91 | -0.136 | 0.39316195 | 0.56676386 | 0.0111 | 0.0142 | 0.0133 | 0.0133 | TRUE | 0.2236 |
| Pcdhga2 | 46 | 49 | 39 | 34 | 30 | 29 | 36 | 52 | 42 | 34 | 34 | 29 | 37.71 | 43 | 32 | 0.745 | -0.425 | 0.18656433 | 0.34318713 | 0 | 0.0653 | 0.0473 | 0.0473 | TRUE | 0.2409 |
| Pcdhga3 | 62 | 51 | 34 | 30 | 24 | 37 | 48 | 54 | 37 | 30 | 27 | 37 | 38.76 | 46 | 31 | 0.673 | -0.572 | 0.07566888 | 0.1812865 | 0 | 0.0636 | 0.0478 | 0.0478 | TRUE | 0.244 |
| Pcdhga4 | 45 | 26 | 28 | 36 | 24 | 32 | 35 | 28 | 30 | 36 | 27 | 32 | 31.22 | 31 | 32 | 1.015 | 0.022 | 0.94901759 | 0.97214279 | 0 | 0.0783 | 0.0534 | 0.0534 | TRUE | 0.1036 |
| Pcdhga5 | 79 | 36 | 35 | 43 | 38 | 40 | 61 | 38 | 38 | 43 | 43 | 39 | 43.76 | 46 | 42 | 0.901 | -0.151 | 0.6286123 | 0.76609445 | 0.0262 | 0.0566 | 0.0474 | 0.0474 | TRUE | 0.7927 |
| Pcdhga6 | 15 | 11 | 3 | 4 | 4 | 10 | 12 | 12 | 3 | 4 | 5 | 10 | 7.49 | 9 | 6 | 0.687 | -0.541 | 0.47914966 | NA | 0.1876 | 0.3188 | 0.2828 | 0.2828 | TRUE | 0.8942 |
| Pcdhga7 | 61 | 52 | 40 | 35 | 25 | 32 | 47 | 55 | 43 | 35 | 28 | 32 | 40.08 | 48 | 32 | 0.651 | -0.62 | 0.04408072 | 0.12283986 | 0 | 0.0616 | 0.0421 | 0.0421 | TRUE | 0.1091 |
| Pcdhga8 | 28 | 18 | 7 | 8 | 12 | 13 | 22 | 19 | 8 | 8 | 14 | 13 | 13.8 | 16 | 12 | 0.699 | -0.516 | 0.36086792 | NA | 0.103 | 0.1742 | 0.1546 | 0.1546 | TRUE | 0.9581 |
| Pcdhga9 | 46 | 33 | 29 | 51 | 46 | 57 | 36 | 35 | 31 | 51 | 52 | 56 | 43.51 | 34 | 53 | 1.553 | 0.635 | 0.0286959 | 0.08966999 | 0 | 0.0569 | 0.0369 | 0.0369 | TRUE | 0.0326 |
| Pcdhgb1 | 8 | 7 | 6 | 6 | 9 | 5 | 6 | 7 | 6 | 6 | 10 | 5 | 6.87 | 6 | 7 | 1.041 | 0.058 | 0.93614804 | NA | 0 | 0.3476 | 0.2332 | 0.2332 | TRUE | 0.3629 |
| Pcdhgb2 | 57 | 27 | 19 | 24 | 28 | 43 | 44 | 29 | 21 | 24 | 32 | 42 | 31.9 | 31 | 33 | 1.035 | 0.05 | 0.89849348 | 0.94384643 | 0.0857 | 0.0767 | 0.0793 | 0.0793 | TRUE | 0.5317 |
| Pcdhgb4 | 54 | 49 | 25 | 40 | 30 | 37 | 42 | 52 | 27 | 40 | 34 | 37 | 38.54 | 40 | 37 | 0.91 | -0.136 | 0.68247447 | 0.80530922 | 0.0269 | 0.0639 | 0.0541 | 0.0541 | TRUE | 0.2704 |
| Pcdhgb5 | 54 | 35 | 22 | 35 | 20 | 21 | 42 | 37 | 24 | 35 | 23 | 21 | 30.17 | 34 | 26 | 0.755 | -0.405 | 0.29149715 | 0.46359951 | 0.049 | 0.081 | 0.0721 | 0.0721 | TRUE | 0.5547 |
| Pcdhgb6 | 31 | 21 | 16 | 22 | 14 | 11 | 24 | 22 | 17 | 22 | 16 | 11 | 18.7 | 21 | 16 | 0.759 | -0.398 | 0.38815451 | 0.56215345 | 0 | 0.1291 | 0.0985 | 0.0985 | TRUE | 0.3213 |
| Pcdhgb7 | 45 | 45 | 36 | 23 | 34 | 33 | 35 | 48 | 39 | 23 | 38 | 33 | 35.91 | 41 | 31 | 0.772 | -0.374 | 0.26955089 | 0.44030753 | 0.0142 | 0.0684 | 0.054 | 0.054 | TRUE | 0.3781 |
| Pcdhgb8 | 87 | 61 | 76 | 81 | 97 | 94 | 68 | 65 | 82 | 81 | 109 | 93 | 82.88 | 72 | 94 | 1.319 | 0.4 | 0.07679287 | 0.18333062 | 0.0071 | 0.031 | 0.0245 | 0.0245 | TRUE | 0.1817 |
| Pcdhgc3 | 396 | 241 | 259 | 349 | 285 | 349 | 308 | 256 | 279 | 347 | 322 | 345 | 309.46 | 281 | 338 | 1.199 | 0.262 | 0.03308976 | 0.09965274 | 0.0017 | 0.0101 | 0.0076 | 0.0076 | TRUE | 0.0822 |
| Pcdhgc4 | 4 | 3 | 4 | 3 | 0 | 3 | 3 | 3 | 4 | 3 | 0 | 3 | 2.76 | 3 | 2 | 0.574 | -0.802 | 0.50133159 | NA | 0 | 0.8611 | 0.6319 | 0.6319 | TRUE | 0.58 |
| Pcdhgc5 | 25 | 7 | 17 | 23 | 19 | 25 | 19 | 7 | 18 | 23 | 21 | 25 | 19.04 | 15 | 23 | 1.512 | 0.596 | 0.20387911 | 0.36525999 | 0.0241 | 0.1269 | 0.1044 | 0.1044 | TRUE | 0.828 |
| Pced1a | 905 | 757 | 719 | 740 | 681 | 814 | 703 | 805 | 776 | 737 | 768 | 804 | 765.38 | 761 | 770 | 1.013 | 0.018 | 0.84003238 | 0.90890647 | 0.0023 | 0.0055 | 0.0044 | 0.0044 | TRUE | 0.0566 |
| Pced1b | 1718 | 1418 | 1474 | 1343 | 1166 | 1523 | 1334 | 1508 | 1590 | 1337 | 1315 | 1503 | 1431.44 | 1477 | 1385 | 0.939 | -0.091 | 0.29687587 | 0.46882378 | 0.0056 | 0.0041 | 0.0048 | 0.0048 | TRUE | 0.0895 |
| Pcf11 | 3526 | 2315 | 1897 | 2264 | 1955 | 1974 | 2738 | 2462 | 2047 | 2254 | 2205 | 1949 | 2275.91 | 2416 | 2136 | 0.883 | -0.179 | 0.0720905 | 0.1754153 | 0.0127 | 0.0035 | 0.0067 | 0.0067 | TRUE | 0.2156 |
| Pcgf1 | 468 | 383 | 389 | 451 | 408 | 496 | 363 | 407 | 420 | 449 | 460 | 490 | 431.57 | 397 | 466 | 1.178 | 0.237 | 0.02754685 | 0.08695386 | 0.0018 | 0.0079 | 0.006 | 0.006 | TRUE | 0.0674 |
| Pcgf2 | 5510 | 4130 | 3942 | 4564 | 4050 | 4867 | 4279 | 4393 | 4253 | 4543 | 4569 | 4804 | 4473.68 | 4308 | 4639 | 1.077 | 0.107 | 0.04666735 | 0.12812468 | 4.00E-04 | 0.003 | 0.0019 | 0.0019 | TRUE | 0.012 |
| Pcgf3 | 5221 | 3721 | 3585 | 3942 | 3509 | 3753 | 4055 | 3958 | 3868 | 3924 | 3959 | 3705 | 3911.38 | 3960 | 3863 | 0.975 | -0.037 | 0.51478557 | 0.67726714 | 7.00E-04 | 0.003 | 0.002 | 0.002 | TRUE | 0.0156 |
| Pcgf5 | 1486 | 1156 | 1282 | 1463 | 1205 | 1384 | 1154 | 1230 | 1383 | 1456 | 1359 | 1366 | 1324.81 | 1256 | 1394 | 1.112 | 0.153 | 0.06616946 | 0.16450862 | 0.0039 | 0.0042 | 0.0042 | 0.0042 | TRUE | 0.0943 |
| Pcgf6 | 4100 | 2888 | 2657 | 2699 | 2390 | 2760 | 3184 | 3072 | 2867 | 2687 | 2696 | 2725 | 2871.73 | 3041 | 2703 | 0.888 | -0.171 | 0.00560631 | 0.02679072 | 0.0012 | 0.0033 | 0.0024 | 0.0024 | TRUE | 0.0303 |
| Pcid2 | 5624 | 3991 | 3894 | 4060 | 3759 | 4141 | 4368 | 4245 | 4202 | 4041 | 4241 | 4088 | 4197.39 | 4272 | 4123 | 0.965 | -0.052 | 0.33444564 | 0.50848117 | 3.00E-04 | 0.003 | 0.0018 | 0.0018 | TRUE | 0.0075 |
| Pcif1 | 2629 | 1883 | 2177 | 2362 | 2157 | 2413 | 2042 | 2003 | 2349 | 2351 | 2433 | 2382 | 2260.03 | 2131 | 2389 | 1.121 | 0.165 | 0.02502988 | 0.08143685 | 0.0034 | 0.0035 | 0.0035 | 0.0035 | TRUE | 0.0953 |
| Pck1 | 5 | 4 | 1 | 0 | 1 | 3 | 4 | 4 | 1 | 0 | 1 | 3 | 2.22 | 3 | 1 | 0.442 | -1.177 | 0.3976196 | NA | 0 | 1.0711 | 0.8609 | 0.8609 | TRUE | 0.6634 |
| Pck2 | 867 | 722 | 645 | 868 | 676 | 813 | 673 | 768 | 696 | 864 | 763 | 803 | 761.08 | 712 | 810 | 1.139 | 0.187 | 0.0405568 | 0.11561195 | 0.003 | 0.0055 | 0.0047 | 0.0047 | TRUE | 0.0544 |
| Pclo | 174 | 124 | 80 | 84 | 70 | 76 | 135 | 132 | 86 | 84 | 79 | 75 | 98.49 | 118 | 79 | 0.67 | -0.578 | 0.0098388 | 0.04089152 | 0.0232 | 0.0265 | 0.0255 | 0.0255 | TRUE | 0.5272 |
| Pcm1 | 18940 | 13771 | 13102 | 14821 | 12724 | 14575 | 14710 | 14648 | 14137 | 14753 | 14354 | 14388 | 14498.24 | 14498 | 14498 | 1 | 0 | 0.99860083 | 0.99936995 | 2.00E-04 | 0.0026 | 0.0015 | 0.0015 | TRUE | 0.0058 |
| Pcmt1 | 2516 | 1793 | 1846 | 2011 | 1764 | 2133 | 1954 | 1907 | 1992 | 2002 | 1990 | 2106 | 1991.73 | 1951 | 2033 | 1.042 | 0.059 | 0.34106877 | 0.51515333 | 0 | 0.0036 | 0.0023 | 0.0023 | TRUE | 0.0121 |
| Pcmtd1 | 3305 | 2395 | 2580 | 2967 | 2316 | 2854 | 2567 | 2548 | 2784 | 2953 | 2613 | 2817 | 2713.59 | 2633 | 2794 | 1.062 | 0.087 | 0.20904041 | 0.37076598 | 0.0026 | 0.0033 | 0.0031 | 0.0031 | TRUE | 0.0389 |
| Pcmtd2 | 1869 | 1380 | 1568 | 1740 | 1483 | 1846 | 1452 | 1468 | 1692 | 1732 | 1673 | 1822 | 1639.76 | 1537 | 1742 | 1.135 | 0.182 | 0.02121653 | 0.07221907 | 0.0038 | 0.0039 | 0.0039 | 0.0039 | TRUE | 0.0922 |
| Pcna | 12399 | 8505 | 8027 | 8146 | 7174 | 8306 | 9630 | 9047 | 8661 | 8109 | 8093 | 8199 | 8623.06 | 9113 | 8134 | 0.892 | -0.164 | 0.00315777 | 0.01718106 | 0.0013 | 0.0027 | 0.0021 | 0.0021 | TRUE | 0.0304 |

| | | | | | | | | | | | | | | | | | | | | | | | |
|---|---|---|---|---|---|---|---|---|---|---|---|---|---|---|---|---|---|---|---|---|---|---|---|
| Pcnp | 2773 | 2017 | 2422 | 2212 | 1985 | 2348 | 2154 | 2145 | 2613 | 2202 | 2239 | 2318 | 2278.57 | 2304 | 2253 | 0.978 | -0.031 | 0.70417228 | 0.82094194 | 0.0064 | 0.0035 | 0.0045 | 0.0045 | TRUE | 0.1654 |
| Pcnt | 9915 | 7153 | 6605 | 6767 | 6363 | 7384 | 7701 | 7609 | 7127 | 6736 | 7178 | 7289 | 7273.16 | 7479 | 7068 | 0.945 | -0.082 | 0.15595074 | 0.30388602 | 0.0015 | 0.0028 | 0.0023 | 0.0023 | TRUE | 0.0207 |
| Pcnx | 3351 | 2487 | 2642 | 2549 | 2299 | 2578 | 2603 | 2645 | 2851 | 2537 | 2594 | 2545 | 2629.06 | 2700 | 2559 | 0.948 | -0.077 | 0.21273718 | 0.37509708 | 9.00E-04 | 0.0033 | 0.0024 | 0.0024 | TRUE | 0.0288 |
| Pcnxl2 | 125 | 94 | 83 | 100 | 95 | 104 | 97 | 100 | 90 | 100 | 107 | 103 | 99.33 | 96 | 103 | 1.078 | 0.108 | 0.57639976 | 0.72784538 | 0 | 0.0263 | 0.017 | 0.017 | TRUE | 0.027 |
| Pcnxl3 | 4275 | 3292 | 2921 | 2990 | 2776 | 3109 | 3320 | 3502 | 3152 | 2976 | 3132 | 3069 | 3191.75 | 3325 | 3059 | 0.92 | -0.12 | 0.05213435 | 0.13891803 | 0.0012 | 0.0032 | 0.0025 | 0.0025 | TRUE | 0.0259 |
| Pcnxl4 | 1752 | 1196 | 1161 | 1211 | 1029 | 1181 | 1361 | 1272 | 1253 | 1205 | 1161 | 1166 | 1236.28 | 1295 | 1177 | 0.908 | -0.139 | 0.05401833 | 0.14221478 | 0 | 0.0043 | 0.0029 | 0.0029 | TRUE | 0.0246 |
| Pcolce | 2848 | 2286 | 2768 | 3239 | 2950 | 3795 | 2212 | 2432 | 2987 | 3224 | 3328 | 3746 | 2988.07 | 2544 | 3433 | 1.351 | 0.434 | 1.59E-05 | 0.00027848 | 0.0151 | 0.0032 | 0.0069 | 0.0069 | TRUE | 0.2813 |
| Pcolce2 | 1878 | 1315 | 1313 | 1531 | 1396 | 1646 | 1459 | 1399 | 1417 | 1524 | 1575 | 1625 | 1499.61 | 1425 | 1575 | 1.105 | 0.144 | 0.03151276 | 0.09601618 | 0 | 0.004 | 0.0025 | 0.0025 | TRUE | 0.0097 |
| Pcp2 | 2 | 1 | 2 | 0 | 0 | 0 | 2 | 1 | 2 | 0 | 0 | 0 | 0.8 | 2 | 0 | 0.12 | -3.062 | 0.17528122 | NA | 0 | 2.9805 | 2.3743 | 2.3743 | TRUE | 0.0718 |
| Pcp4 | 4 | 3 | 3 | 3 | 2 | 2 | 3 | 3 | 3 | 3 | 2 | 2 | 2.79 | 3 | 2 | 0.758 | -0.399 | 0.71799932 | NA | 0 | 0.8513 | 0.5135 | 0.5135 | TRUE | 0.0494 |
| Pcp4l1 | 30 | 23 | 22 | 19 | 23 | 17 | 23 | 24 | 24 | 19 | 26 | 17 | 22.19 | 24 | 21 | 0.859 | -0.22 | 0.59035624 | 0.73882577 | 0 | 0.1092 | 0.0744 | 0.0744 | TRUE | 0.2652 |
| Pcsk1 | 109 | 63 | 55 | 40 | 43 | 36 | 85 | 67 | 59 | 40 | 49 | 36 | 55.81 | 70 | 42 | 0.582 | -0.782 | 0.00485722 | 0.02397862 | 0.0146 | 0.0449 | 0.0361 | 0.0361 | TRUE | 0.3183 |
| Pcsk1n | 1406 | 1271 | 1663 | 1215 | 1246 | 1440 | 1092 | 1352 | 1794 | 1209 | 1406 | 1422 | 1379.14 | 1413 | 1346 | 0.953 | -0.07 | 0.75181788 | 0.8527006 | 0.0343 | 0.0041 | 0.0129 | 0.0343 | TRUE | 0.3773 |
| Pcsk2 | 1 | 4 | 2 | 1 | 2 | 0 | 1 | 4 | 2 | 1 | 2 | 0 | 1.74 | 2 | 1 | 0.451 | -1.15 | 0.46035793 | NA | 0 | 1.3644 | 1.0555 | 1.0555 | TRUE | 0.2483 |
| Pcsk4 | 65 | 63 | 75 | 78 | 71 | 114 | 50 | 67 | 81 | 78 | 80 | 113 | 78.12 | 66 | 90 | 1.374 | 0.459 | 0.07310139 | 0.17700121 | 0.0375 | 0.0328 | 0.0341 | 0.0341 | TRUE | 0.3742 |
| Pcsk5 | 39 | 33 | 26 | 20 | 11 | 13 | 30 | 35 | 28 | 20 | 12 | 13 | 23.1 | 31 | 15 | 0.485 | -1.044 | 0.01217456 | 0.04784465 | 0 | 0.105 | 0.0751 | 0.0751 | TRUE | 0.3726 |
| Pcsk6 | 2612 | 1805 | 1778 | 2044 | 1832 | 1933 | 2029 | 1920 | 1918 | 2035 | 2067 | 1908 | 1979.41 | 1956 | 2003 | 1.023 | 0.033 | 0.61027692 | 0.75351486 | 9.00E-04 | 0.0036 | 0.0026 | 0.0026 | TRUE | 0.0209 |
| Pcsk7 | 662 | 446 | 359 | 390 | 348 | 382 | 514 | 474 | 387 | 388 | 393 | 377 | 422.3 | 458 | 386 | 0.839 | -0.253 | 0.03722476 | 0.10880439 | 0.0087 | 0.008 | 0.0082 | 0.0082 | TRUE | 0.2103 |
| Pcsk9 | 82 | 68 | 76 | 169 | 123 | 140 | 64 | 72 | 82 | 168 | 139 | 138 | 110.53 | 73 | 148 | 2.055 | 1.039 | 2.14E-07 | 7.72E-06 | 0 | 0.0239 | 0.0188 | 0.0188 | TRUE | 0.1447 |
| Pctp | 1273 | 863 | 788 | 846 | 845 | 923 | 989 | 918 | 850 | 842 | 953 | 911 | 910.57 | 919 | 902 | 0.98 | -0.029 | 0.74041022 | 0.84573509 | 0.0038 | 0.005 | 0.0046 | 0.0046 | TRUE | 0.0547 |
| Pcx | 2962 | 2230 | 2530 | 2891 | 2826 | 3138 | 2300 | 2372 | 2730 | 2878 | 3188 | 3098 | 2760.96 | 2467 | 3055 | 1.239 | 0.309 | 7.10E-05 | 0.00092808 | 0.0052 | 0.0033 | 0.004 | 0.004 | TRUE | 0.1042 |
| Pcyox1 | 4083 | 3023 | 2989 | 3283 | 2865 | 3422 | 3171 | 3216 | 3225 | 3268 | 3232 | 3378 | 3248.28 | 3204 | 3293 | 1.028 | 0.04 | 0.46365273 | 0.63284298 | 0 | 0.0032 | 0.0018 | 0.0018 | TRUE | 0.0063 |
| Pcyox1l | 1054 | 845 | 802 | 1043 | 787 | 1038 | 819 | 899 | 865 | 1038 | 888 | 1025 | 922.24 | 861 | 984 | 1.144 | 0.195 | 0.02777531 | 0.08753969 | 0.0037 | 0.005 | 0.0046 | 0.0046 | TRUE | 0.0848 |
| Pcyt1a | 2428 | 1652 | 1440 | 1639 | 1413 | 1542 | 1886 | 1757 | 1554 | 1631 | 1594 | 1522 | 1657.39 | 1732 | 1582 | 0.913 | -0.132 | 0.10640901 | 0.23149224 | 0.0044 | 0.0039 | 0.0042 | 0.0042 | TRUE | 0.0973 |
| Pcyt1b | 2516 | 1898 | 2020 | 2194 | 2020 | 2550 | 1954 | 2019 | 2180 | 2184 | 2279 | 2517 | 2188.74 | 2051 | 2327 | 1.135 | 0.183 | 0.01511302 | 0.05613574 | 0.0036 | 0.0035 | 0.0036 | 0.0036 | TRUE | 0.0627 |
| Pcyt2 | 2422 | 1929 | 1708 | 1716 | 1597 | 1823 | 1881 | 2052 | 1843 | 1708 | 1802 | 1800 | 1847.52 | 1925 | 1770 | 0.919 | -0.121 | 0.07931992 | 0.18798202 | 0.0016 | 0.0037 | 0.0029 | 0.0029 | TRUE | 0.0392 |
| Pdap1 | 10034 | 7145 | 6374 | 6900 | 5809 | 7031 | 7793 | 7600 | 6877 | 6868 | 6553 | 6941 | 7105.43 | 7423 | 6787 | 0.914 | -0.129 | 0.03745607 | 0.10921249 | 0.0024 | 0.0028 | 0.0026 | 0.0026 | TRUE | 0.0503 |
| Pdc | 7 | 1 | 4 | 0 | 0 | 4 | 5 | 1 | 4 | 0 | 0 | 4 | 2.46 | 3 | 1 | 0.369 | -1.44 | 0.32241345 | NA | 1.2075 | 0.9655 | 1.007 | 1.007 | TRUE | 1.8968 |
| Pdcd1 | 16 | 11 | 12 | 6 | 10 | 8 | 12 | 12 | 13 | 6 | 11 | 8 | 10.37 | 12 | 8 | 0.674 | -0.569 | 0.3392423 | NA | 0 | 0.231 | 0.1546 | 0.1546 | TRUE | 0.2485 |
| Pdcd10 | 2454 | 1689 | 1708 | 1861 | 1591 | 1765 | 1906 | 1797 | 1843 | 1852 | 1795 | 1742 | 1822.5 | 1849 | 1796 | 0.971 | -0.042 | 0.51507632 | 0.67752492 | 4.00E-04 | 0.0037 | 0.0024 | 0.0024 | TRUE | 0.0093 |
| Pdcd11 | 13699 | 9610 | 7947 | 8416 | 7734 | 8820 | 10639 | 10222 | 8575 | 8377 | 8725 | 8707 | 9207.49 | 9812 | 8603 | 0.877 | -0.19 | 0.01318261 | 0.05071845 | 0.0063 | 0.0027 | 0.0041 | 0.0041 | TRUE | 0.1483 |
| Pdcd1lg2 | 0 | 0 | 0 | 0 | 0 | 1 | 0 | 0 | 0 | 0 | 0 | 1 | 0.16 | 0 | 0 | 2.094 | 1.067 | 0.79379329 | NA | 0 | 14.4078 | 10 | 10 | TRUE | 0.3835 |
| Pdcd2 | 2513 | 1869 | 1874 | 1912 | 1765 | 1999 | 1952 | 1988 | 2022 | 1903 | 1991 | 1973 | 1971.57 | 1987 | 1956 | 0.984 | -0.023 | 0.71031896 | 0.82554796 | 0 | 0.0036 | 0.0022 | 0.0022 | TRUE | 0.0067 |
| Pdcd2l | 2089 | 1665 | 1687 | 1848 | 1629 | 1865 | 1622 | 1771 | 1820 | 1839 | 1838 | 1841 | 1788.66 | 1738 | 1839 | 1.06 | 0.084 | 0.22250275 | 0.38684246 | 0.0013 | 0.0038 | 0.0028 | 0.0028 | TRUE | 0.0424 |
| Pdcd4 | 2673 | 2150 | 2535 | 2468 | 2274 | 2781 | 2076 | 2287 | 2735 | 2457 | 2565 | 2745 | 2477.57 | 2366 | 2589 | 1.095 | 0.131 | 0.16289882 | 0.3133243 | 0.0108 | 0.0034 | 0.006 | 0.006 | TRUE | 0.2253 |
| Pdcd5 | 2780 | 2060 | 1814 | 1936 | 1653 | 1953 | 2159 | 2191 | 1957 | 1927 | 1865 | 1928 | 2004.56 | 2102 | 1907 | 0.907 | -0.142 | 0.03705819 | 0.108408 | 0.0015 | 0.0036 | 0.0028 | 0.0028 | TRUE | 0.0433 |
| Pdcd6 | 3244 | 2317 | 2237 | 2335 | 1946 | 2574 | 2519 | 2465 | 2414 | 2324 | 2195 | 2541 | 2409.7 | 2466 | 2353 | 0.955 | -0.067 | 0.33412948 | 0.50818557 | 0.0024 | 0.0034 | 0.0031 | 0.0031 | TRUE | 0.0589 |
| Pdcd6ip | 10361 | 7630 | 8261 | 8501 | 7673 | 8958 | 8047 | 8116 | 8914 | 8462 | 8656 | 8843 | 8506.2 | 8359 | 8654 | 1.035 | 0.05 | 0.38514943 | 0.55910628 | 0.0017 | 0.0027 | 0.0023 | 0.0023 | TRUE | 0.041 |
| Pdcd7 | 2535 | 1985 | 1911 | 2010 | 1617 | 1845 | 1969 | 2111 | 2062 | 2001 | 1824 | 1821 | 1964.73 | 2047 | 1882 | 0.92 | -0.12 | 0.07826675 | 0.18616519 | 0.0016 | 0.0036 | 0.0029 | 0.0029 | TRUE | 0.037 |
| Pdcl | 4547 | 3319 | 3333 | 3376 | 2812 | 3066 | 3531 | 3530 | 3596 | 3360 | 3172 | 3027 | 3369.57 | 3552 | 3186 | 0.897 | -0.157 | 0.00912744 | 0.03867728 | 0.001 | 0.0031 | 0.0023 | 0.0023 | TRUE | 0.028 |
| Pdcl2 | 179 | 139 | 184 | 159 | 143 | 160 | 139 | 148 | 199 | 158 | 161 | 158 | 160.49 | 162 | 159 | 0.988 | -0.018 | 0.91953869 | 0.95585372 | 0.0131 | 0.0172 | 0.016 | 0.016 | TRUE | 0.4073 |
| Pdcl3 | 3274 | 2382 | 2461 | 2535 | 2233 | 2715 | 2543 | 2534 | 2655 | 2523 | 2519 | 2680 | 2575.73 | 2577 | 2574 | 0.999 | -0.001 | 0.98396636 | 0.9916677 | 6.00E-04 | 0.0033 | 0.0022 | 0.0022 | TRUE | 0.0158 |
| Pddc1 | 1725 | 1303 | 1352 | 1439 | 1361 | 1642 | 1340 | 1386 | 1459 | 1432 | 1535 | 1621 | 1462.19 | 1395 | 1529 | 1.097 | 0.134 | 0.0740608 | 0.1786584 | 0.0022 | 0.004 | 0.0034 | 0.0034 | TRUE | 0.0377 |
| Pde10a | 166 | 86 | 107 | 120 | 110 | 79 | 129 | 91 | 115 | 119 | 124 | 78 | 109.56 | 112 | 107 | 0.952 | -0.071 | 0.7530918 | 0.85348706 | 0.0362 | 0.0241 | 0.0275 | 0.0275 | TRUE | 0.5733 |
| Pde11a | 32 | 20 | 24 | 25 | 33 | 35 | 25 | 21 | 26 | 25 | 37 | 35 | 28.11 | 24 | 32 | 1.337 | 0.42 | 0.25416706 | 0.42340441 | 0 | 0.0867 | 0.0614 | 0.0614 | TRUE | 0.2773 |
| Pde12 | 2383 | 1695 | 1822 | 1954 | 1727 | 2049 | 1851 | 1803 | 1966 | 1945 | 1948 | 2023 | 1922.6 | 1873 | 1972 | 1.053 | 0.075 | 0.25031348 | 0.41918343 | 7.00E-04 | 0.0037 | 0.0025 | 0.0025 | TRUE | 0.022 |
| Pde1a | 1 | 2 | 0 | 2 | 0 | 1 | 1 | 2 | 0 | 2 | 0 | 1 | 0.98 | 1 | 1 | 1.046 | 0.065 | 0.9739236 | NA | 0 | 2.42 | 1.8677 | 1.8677 | TRUE | 0.2379 |
| Pde1b | 620 | 559 | 663 | 732 | 662 | 759 | 482 | 595 | 715 | 729 | 747 | 749 | 669.36 | 597 | 742 | 1.245 | 0.317 | 0.01104385 | 0.04442717 | 0.017 | 0.006 | 0.0097 | 0.0097 | TRUE | 0.2519 |
| Pde1c | 49 | 33 | 31 | 28 | 32 | 23 | 38 | 35 | 33 | 28 | 36 | 23 | 32.21 | 35 | 29 | 0.807 | -0.309 | 0.37141616 | 0.54612609 | 0 | 0.076 | 0.0543 | 0.0543 | TRUE | 0.2872 |
| Pde2a | 976 | 702 | 714 | 1146 | 978 | 1057 | 758 | 747 | 770 | 1141 | 1103 | 1043 | 927.09 | 758 | 1096 | 1.445 | 0.531 | 1.20E-11 | 1.21E-09 | 0 | 0.005 | 0.0033 | 0.0033 | TRUE | 0.0213 |
| Pde3a | 37 | 34 | 18 | 31 | 35 | 19 | 29 | 36 | 19 | 31 | 39 | 19 | 28.9 | 28 | 30 | 1.051 | 0.071 | 0.85896983 | 0.92137004 | 0.0736 | 0.0844 | 0.0815 | 0.0815 | TRUE | 0.2858 |
| Pde3b | 1632 | 1168 | 1155 | 1316 | 1094 | 1257 | 1268 | 1242 | 1246 | 1310 | 1234 | 1241 | 1256.85 | 1252 | 1262 | 1.008 | 0.011 | 0.87503299 | 0.93063819 | 0 | 0.0043 | 0.0027 | 0.0027 | TRUE | 0.0136 |
| Pde4a | 853 | 685 | 677 | 701 | 640 | 685 | 662 | 729 | 730 | 698 | 722 | 676 | 702.93 | 707 | 699 | 0.989 | -0.016 | 0.85767266 | 0.92032398 | 0 | 0.0058 | 0.0041 | 0.0041 | TRUE | 0.0385 |
| Pde4b | 5 | 13 | 8 | 6 | 6 | 5 | 4 | 14 | 9 | 6 | 7 | 5 | 7.34 | 9 | 6 | 0.681 | -0.554 | 0.45807652 | NA | 0 | 0.3255 | 0.2583 | 0.2583 | TRUE | 0.1591 |
| Pde4c | 291 | 305 | 292 | 398 | 413 | 424 | 226 | 324 | 315 | 396 | 466 | 419 | 357.69 | 288 | 427 | 1.486 | 0.571 | 7.76E-05 | 0.00098796 | 0.0195 | 0.0091 | 0.0122 | 0.0122 | TRUE | 0.4348 |
| Pde4d | 586 | 398 | 378 | 475 | 466 | 527 | 455 | 423 | 408 | 473 | 526 | 520 | 467.51 | 429 | 506 | 1.178 | 0.236 | 0.02195938 | 0.07383331 | 0 | 0.0075 | 0.0055 | 0.0055 | TRUE | 0.0393 |
| Pde4dip | 2447 | 1790 | 1894 | 2005 | 1721 | 1941 | 1900 | 1904 | 2044 | 1996 | 1941 | 1916 | 1950.23 | 1949 | 1951 | 1.001 | 0.002 | 0.97511626 | 0.98653073 | 6.00E-04 | 0.0036 | 0.0025 | 0.0025 | TRUE | 0.0212 |
| Pde5a | 455 | 301 | 300 | 299 | 253 | 290 | 353 | 320 | 324 | 298 | 285 | 286 | 311.09 | 332 | 290 | 0.87 | -0.201 | 0.08716108 | 0.20025613 | 0 | 0.01 | 0.0067 | 0.0067 | TRUE | 0.0373 |
| Pde6a | 21 | 16 | 19 | 11 | 9 | 4 | 16 | 17 | 21 | 11 | 10 | 4 | 13.15 | 18 | 8 | 0.464 | -1.107 | 0.04939445 | NA | 0 | 0.1827 | 0.1392 | 0.1392 | TRUE | 0.6799 |
| Pde6b | 3 | 3 | 3 | 7 | 1 | 4 | 2 | 3 | 3 | 7 | 1 | 4 | 3.47 | 3 | 4 | 1.401 | 0.487 | 0.64312554 | NA | 0 | 0.686 | 0.5009 | 0.5009 | TRUE | 0.1519 |
| Pde6c | 1 | 0 | 0 | 0 | 0 | 0 | 1 | 0 | 0 | 0 | 0 | 0 | 0.13 | 0 | 0 | 0.552 | -0.857 | 0.8336548 | NA | 0 | 18.3121 | 10 | 10 | TRUE | 0.2566 |
| Pde6d | 1592 | 1151 | 1084 | 1309 | 1145 | 1195 | 1236 | 1224 | 1170 | 1303 | 1292 | 1180 | 1234.11 | 1210 | 1258 | 1.039 | 0.055 | 0.46025244 | 0.62988358 | 0.0011 | 0.0043 | 0.0032 | 0.0032 | TRUE | 0.0362 |
| Pde6g | 3 | 1 | 6 | 2 | 3 | 9 | 2 | 1 | 6 | 2 | 3 | 9 | 4.02 | 3 | 5 | 1.47 | 0.555 | 0.59442642 | NA | 0.3754 | 0.5918 | 0.5271 | 0.5271 | TRUE | 1.1587 |
| Pde6h | 1 | 0 | 0 | 1 | 0 | 0 | 1 | 0 | 0 | 1 | 0 | 0 | 0.3 | 0 | 0 | 1.075 | 0.105 | 0.97698525 | NA | 0 | 8.0272 | 7.5141 | 7.5141 | TRUE | 0.3906 |
| Pde7a | 2335 | 1671 | 1664 | 1957 | 1745 | 1948 | 1813 | 1777 | 1795 | 1948 | 1969 | 1923 | 1870.98 | 1795 | 1947 | 1.084 | 0.116 | 0.0542439 | 0.14265097 | 0 | 0.0037 | 0.0021 | 0.0021 | TRUE | 0.0014 |
| Pde7b | 0 | 0 | 0 | 1 | 2 | 0 | 0 | 0 | 0 | 1 | 2 | 0 | 0.54 | 0 | 1 | 6.07 | 2.602 | 0.36554818 | NA | 0 | 4.3758 | 4.4761 | 4.4761 | TRUE | 0.0722 |
| Pde8a | 2169 | 1588 | 1831 | 2101 | 1793 | 2043 | 1685 | 1689 | 1976 | 2091 | 2023 | 2017 | 1913.35 | 1783 | 2044 | 1.147 | 0.198 | 0.01001029 | 0.04139964 | 0.0038 | 0.0037 | 0.0037 | 0.0037 | TRUE | 0.1063 |
| Pde8b | 35 | 31 | 30 | 33 | 30 | 42 | 27 | 33 | 32 | 33 | 34 | 41 | 33.45 | 31 | 36 | 1.176 | 0.234 | 0.48219712 | 0.64866291 | 0 | 0.0733 | 0.0496 | 0.0496 | TRUE | 0.1276 |
| Pde9a | 589 | 382 | 436 | 579 | 489 | 570 | 457 | 406 | 470 | 576 | 552 | 563 | 504.15 | 444 | 564 | 1.266 | 0.341 | 0.00063822 | 0.00520372 | 5.00E-04 | 0.0071 | 0.0052 | 0.0052 | TRUE | 0.0633 |

| | | | | | | | | | | | | | | | | | | | | | | | | | |
|---|---|---|---|---|---|---|---|---|---|---|---|---|---|---|---|---|---|---|---|---|---|---|---|---|---|
| Pdf | 1176 | 847 | 762 | 756 | 671 | 659 | 913 | 901 | 822 | 753 | 757 | 651 | 799.42 | 879 | 720 | 0.818 | -0.29 | 0.00171847 | 0.01074926 | 0.0038 | 0.0054 | 0.0049 | 0.0049 | TRUE | 0.0853 |
| Pdgfa | 747 | 694 | 684 | 924 | 734 | 837 | 580 | 738 | 738 | 920 | 828 | 826 | 771.74 | 685 | 858 | 1.255 | 0.328 | 0.00221214 | 0.01304915 | 0.01 | 0.0055 | 0.007 | 0.007 | TRUE | 0.2238 |
| Pdgfb | 295 | 297 | 358 | 472 | 409 | 500 | 229 | 316 | 386 | 470 | 461 | 494 | 392.68 | 310 | 475 | 1.537 | 0.621 | 7.23E-05 | 0.00093916 | 0.031 | 0.0085 | 0.015 | 0.015 | TRUE | 0.4107 |
| Pdgfc | 82 | 63 | 50 | 51 | 32 | 44 | 64 | 67 | 54 | 51 | 36 | 43 | 52.49 | 62 | 43 | 0.707 | -0.501 | 0.06829753 | 0.1683063 | 0 | 0.0476 | 0.0346 | 0.0346 | TRUE | 0.1686 |
| Pdgfd | 22 | 10 | 14 | 3 | 11 | 8 | 17 | 11 | 15 | 3 | 12 | 8 | 11.02 | 14 | 8 | 0.535 | -0.902 | 0.15245187 | NA | 0.0718 | 0.2175 | 0.185 | 0.185 | TRUE | 0.3803 |
| Pdgfra | 531 | 359 | 318 | 429 | 366 | 384 | 412 | 382 | 343 | 427 | 413 | 379 | 392.73 | 379 | 406 | 1.069 | 0.096 | 0.40372144 | 0.57657779 | 0.0037 | 0.0085 | 0.007 | 0.007 | TRUE | 0.078 |
| Pdgfrb | 1007 | 796 | 909 | 1244 | 1146 | 1287 | 782 | 847 | 981 | 1238 | 1293 | 1270 | 1068.52 | 870 | 1267 | 1.459 | 0.545 | 2.01E-09 | 1.25E-07 | 0.0056 | 0.0046 | 0.005 | 0.005 | TRUE | 0.147 |
| Pdgfrl | 489 | 344 | 248 | 267 | 226 | 228 | 380 | 366 | 268 | 266 | 255 | 225 | 293.18 | 338 | 249 | 0.734 | -0.446 | 0.00309815 | 0.01695748 | 0.0183 | 0.0105 | 0.0129 | 0.0129 | TRUE | 0.3667 |
| Pdha1 | 15289 | 11149 | 10859 | 11566 | 10018 | 11567 | 11874 | 11859 | 11717 | 11513 | 11301 | 11418 | 11613.78 | 11817 | 11411 | 0.966 | -0.05 | 0.25737505 | 0.42738419 | 0 | 0.0026 | 0.0013 | 0.0013 | TRUE | 9.00E-04 |
| Pdha2 | 257 | 194 | 135 | 75 | 73 | 79 | 200 | 206 | 146 | 75 | 82 | 78 | 131.1 | 184 | 78 | 0.425 | -1.235 | 3.70E-10 | 2.73E-08 | 0.0148 | 0.0205 | 0.0188 | 0.0188 | TRUE | 0.3418 |
| Pdhb | 6235 | 4476 | 4370 | 5042 | 4262 | 5044 | 4842 | 4761 | 4715 | 5019 | 4808 | 4979 | 4854.12 | 4773 | 4935 | 1.034 | 0.048 | 0.34677329 | 0.52120448 | 1.00E-04 | 0.0029 | 0.0017 | 0.0017 | TRUE | 0.0061 |
| Pdhx | 3194 | 2120 | 2125 | 2186 | 1913 | 2214 | 2481 | 2255 | 2293 | 2176 | 2158 | 2186 | 2258.01 | 2343 | 2173 | 0.927 | -0.11 | 0.08577584 | 0.19793254 | 0.001 | 0.0035 | 0.0025 | 0.0025 | TRUE | 0.0331 |
| Pdia2 | 10 | 10 | 10 | 18 | 18 | 8 | 8 | 11 | 11 | 18 | 20 | 8 | 12.55 | 10 | 15 | 1.586 | 0.665 | 0.24686567 | NA | 0 | 0.1912 | 0.1541 | 0.1541 | TRUE | 0.8652 |
| Pdia3 | 23000 | 15508 | 15335 | 16846 | 13939 | 16272 | 17863 | 16496 | 16546 | 16768 | 15725 | 16063 | 16576.87 | 16968 | 16185 | 0.954 | -0.068 | 0.21242505 | 0.37479216 | 0.0015 | 0.0026 | 0.0021 | 0.0021 | TRUE | 0.0262 |
| Pdia4 | 9985 | 6927 | 6752 | 7430 | 6343 | 7201 | 7755 | 7368 | 7285 | 7396 | 7156 | 7108 | 7344.72 | 7469 | 7220 | 0.966 | -0.049 | 0.34286555 | 0.51698694 | 6.00E-04 | 0.0027 | 0.0018 | 0.0018 | TRUE | 0.0138 |
| Pdia5 | 1756 | 1262 | 1424 | 1749 | 1495 | 1640 | 1364 | 1342 | 1536 | 1741 | 1687 | 1619 | 1548.18 | 1414 | 1682 | 1.19 | 0.251 | 0.00093322 | 0.00690041 | 0.0025 | 0.004 | 0.0035 | 0.0035 | TRUE | 0.0674 |
| Pdia6 | 22638 | 16489 | 17285 | 20368 | 17308 | 20253 | 17582 | 17539 | 18650 | 20274 | 19525 | 19993 | 18927.3 | 17924 | 19931 | 1.112 | 0.153 | 0.00191519 | 0.0116863 | 7.00E-04 | 0.0026 | 0.0017 | 0.0017 | TRUE | 0.0153 |
| Pdik1l | 671 | 577 | 596 | 592 | 553 | 596 | 521 | 614 | 643 | 589 | 624 | 588 | 596.57 | 593 | 600 | 1.016 | 0.022 | 0.82883015 | 0.90190097 | 0.0052 | 0.0064 | 0.006 | 0.006 | TRUE | 0.1387 |
| Pdilt | 2 | 1 | 1 | 0 | 0 | 0 | 2 | 1 | 1 | 0 | 0 | 0 | 0.62 | 1 | 0 | 0.154 | -2.702 | 0.27412557 | NA | 0 | 3.8499 | 3.0088 | 3.0088 | TRUE | 0.0718 |
| Pdk1 | 21379 | 16562 | 17535 | 21453 | 19517 | 21260 | 16604 | 17617 | 18920 | 21354 | 22017 | 20987 | 19583.24 | 17714 | 21453 | 1.211 | 0.276 | 3.04E-06 | 7.01E-05 | 0.0024 | 0.0025 | 0.0025 | 0.0025 | TRUE | 0.0434 |
| Pdk2 | 582 | 470 | 628 | 734 | 687 | 825 | 452 | 500 | 678 | 731 | 775 | 814 | 658.26 | 543 | 773 | 1.427 | 0.513 | 9.47E-05 | 0.00115758 | 0.0231 | 0.006 | 0.0109 | 0.0109 | TRUE | 0.5439 |
| Pdk3 | 2901 | 2152 | 2589 | 2838 | 2389 | 2889 | 2253 | 2289 | 2794 | 2825 | 2695 | 2852 | 2617.92 | 2445 | 2791 | 1.142 | 0.192 | 0.02300486 | 0.07653827 | 0.0075 | 0.0033 | 0.0047 | 0.0047 | TRUE | 0.1869 |
| Pdk4 | 603 | 451 | 665 | 732 | 674 | 770 | 468 | 480 | 718 | 729 | 760 | 760 | 652.44 | 555 | 750 | 1.353 | 0.436 | 0.00180779 | 0.01118559 | 0.028 | 0.0061 | 0.0125 | 0.0125 | TRUE | 0.7578 |
| Pdlim1 | 3551 | 2560 | 2565 | 2406 | 2187 | 2666 | 2758 | 2723 | 2768 | 2395 | 2467 | 2632 | 2623.73 | 2750 | 2498 | 0.909 | -0.138 | 0.02393843 | 0.07885502 | 8.00E-04 | 0.0033 | 0.0023 | 0.0023 | TRUE | 0.0269 |
| Pdlim2 | 167 | 132 | 132 | 150 | 135 | 157 | 130 | 140 | 142 | 149 | 152 | 155 | 144.85 | 137 | 152 | 1.109 | 0.15 | 0.35683654 | 0.53133292 | 0 | 0.0188 | 0.0121 | 0.0121 | TRUE | 0.0287 |
| Pdlim3 | 186 | 168 | 110 | 108 | 108 | 113 | 144 | 179 | 119 | 108 | 122 | 112 | 130.46 | 147 | 114 | 0.771 | -0.375 | 0.05272572 | 0.14007596 | 0.0158 | 0.0206 | 0.0192 | 0.0192 | TRUE | 0.2058 |
| Pdlim4 | 236 | 170 | 246 | 273 | 253 | 305 | 183 | 181 | 265 | 272 | 285 | 301 | 247.96 | 210 | 286 | 1.368 | 0.452 | 0.00485847 | 0.02397862 | 0.0214 | 0.012 | 0.0144 | 0.0144 | TRUE | 0.5701 |
| Pdlim5 | 4557 | 3254 | 3275 | 3948 | 3435 | 3793 | 3539 | 3461 | 3534 | 3930 | 3875 | 3744 | 3680.55 | 3511 | 3850 | 1.096 | 0.132 | 0.01357861 | 0.05189279 | 0 | 0.0031 | 0.0018 | 0.0018 | TRUE | 0.0071 |
| Pdlim7 | 1680 | 1151 | 1002 | 1145 | 1077 | 1217 | 1305 | 1224 | 1081 | 1140 | 1215 | 1201 | 1194.38 | 1203 | 1185 | 0.983 | -0.024 | 0.77724233 | 0.86830738 | 0.0043 | 0.0044 | 0.0044 | 0.0044 | TRUE | 0.0937 |
| Pdp1 | 972 | 759 | 867 | 975 | 793 | 897 | 755 | 807 | 935 | 971 | 895 | 885 | 874.72 | 832 | 917 | 1.103 | 0.142 | 0.13979682 | 0.28110046 | 0.0058 | 0.0051 | 0.0055 | 0.0055 | TRUE | 0.1377 |
| Pdp2 | 2449 | 1882 | 1899 | 2191 | 1957 | 2273 | 1902 | 2002 | 2049 | 2181 | 2208 | 2244 | 2097.55 | 1984 | 2211 | 1.115 | 0.157 | 0.01145679 | 0.04572163 | 4.00E-04 | 0.0036 | 0.0023 | 0.0023 | TRUE | 0.0165 |
| Pdpk1 | 8992 | 6474 | 6289 | 6852 | 6047 | 7231 | 6984 | 6886 | 6786 | 6820 | 6822 | 7138 | 6906 | 6885 | 6927 | 1.006 | 0.009 | 0.86344982 | 0.92466653 | 2.00E-04 | 0.0028 | 0.0016 | 0.0016 | TRUE | 0.0087 |
| Pdpn | 1179 | 888 | 1179 | 1506 | 1296 | 1445 | 916 | 945 | 1272 | 1499 | 1462 | 1426 | 1253.32 | 1044 | 1462 | 1.403 | 0.488 | 1.04E-05 | 0.00019326 | 0.0171 | 0.0043 | 0.008 | 0.008 | TRUE | 0.4326 |
| Pdpr | 2352 | 1633 | 1641 | 2018 | 1651 | 1910 | 1827 | 1737 | 1771 | 2009 | 1862 | 1885 | 1848.5 | 1778 | 1919 | 1.079 | 0.11 | 0.09324894 | 0.2100007 | 6.00E-04 | 0.0037 | 0.0025 | 0.0025 | TRUE | 0.0204 |
| Pdrg1 | 1289 | 912 | 776 | 935 | 782 | 901 | 1001 | 970 | 837 | 931 | 882 | 889 | 918.47 | 936 | 901 | 0.961 | -0.057 | 0.51870509 | 0.68026262 | 0.0038 | 0.005 | 0.0046 | 0.0046 | TRUE | 0.0999 |
| Pds5a | 10254 | 7174 | 6556 | 6553 | 6061 | 6836 | 7964 | 7631 | 7074 | 6523 | 6837 | 6748 | 7129.51 | 7556 | 6703 | 0.887 | -0.173 | 0.00367058 | 0.01930231 | 0.0019 | 0.0028 | 0.0024 | 0.0024 | TRUE | 0.038 |
| Pds5b | 7294 | 5152 | 4997 | 5399 | 4802 | 5503 | 5665 | 5480 | 5392 | 5374 | 5417 | 5432 | 5460.07 | 5512 | 5408 | 0.981 | -0.028 | 0.57880235 | 0.72988172 | 2.00E-04 | 0.0029 | 0.0016 | 0.0016 | TRUE | 0.0073 |
| Pdss1 | 1003 | 642 | 515 | 572 | 497 | 553 | 779 | 683 | 556 | 569 | 561 | 546 | 615.58 | 673 | 559 | 0.829 | -0.271 | 0.02236559 | 0.07479443 | 0.0132 | 0.0063 | 0.0085 | 0.0085 | TRUE | 0.2257 |
| Pdss2 | 737 | 462 | 438 | 450 | 452 | 542 | 572 | 491 | 473 | 448 | 510 | 535 | 504.88 | 512 | 498 | 0.969 | -0.045 | 0.68919011 | 0.80981538 | 0.0077 | 0.0071 | 0.0073 | 0.0073 | TRUE | 0.1297 |
| Pdx1 | 9 | 3 | 1 | 4 | 0 | 2 | 7 | 3 | 1 | 4 | 0 | 2 | 2.87 | 4 | 2 | 0.526 | -0.927 | 0.46140978 | NA | 0.5418 | 0.8284 | 0.7592 | 0.7592 | TRUE | 0.2927 |
| Pdxdc1 | 6841 | 4826 | 4667 | 5192 | 4545 | 5089 | 5313 | 5133 | 5036 | 5168 | 5127 | 5024 | 5133.52 | 5161 | 5106 | 0.989 | -0.016 | 0.76119309 | 0.85829385 | 3.00E-04 | 0.0029 | 0.0017 | 0.0017 | TRUE | 0.0083 |
| Pdxk | 2330 | 1679 | 1507 | 1714 | 1572 | 1699 | 1810 | 1786 | 1626 | 1706 | 1773 | 1677 | 1729.71 | 1741 | 1719 | 0.987 | -0.019 | 0.78135272 | 0.87115299 | 0.0015 | 0.0038 | 0.0029 | 0.0029 | TRUE | 0.0393 |
| Pdxk-ps | 5 | 2 | 3 | 7 | 5 | 4 | 4 | 2 | 3 | 7 | 6 | 4 | 4.3 | 3 | 6 | 1.767 | 0.822 | 0.3708981 | NA | 0 | 0.5535 | 0.3595 | 0.3595 | TRUE | 0.1428 |
| Pdxp | 1367 | 1027 | 1120 | 1459 | 1209 | 1423 | 1062 | 1092 | 1208 | 1452 | 1364 | 1405 | 1263.91 | 1121 | 1407 | 1.257 | 0.33 | 1.82E-05 | 0.00031011 | 0.0019 | 0.0043 | 0.0035 | 0.0035 | TRUE | 0.0552 |
| Pdyn | 24 | 15 | 10 | 27 | 14 | 23 | 19 | 16 | 11 | 27 | 16 | 23 | 18.46 | 15 | 22 | 1.432 | 0.518 | 0.26394429 | 0.43407614 | 0 | 0.1308 | 0.0996 | 0.0996 | TRUE | 0.2697 |
| Pdzd11 | 1180 | 766 | 731 | 894 | 716 | 788 | 916 | 815 | 789 | 890 | 808 | 778 | 832.58 | 840 | 825 | 0.981 | -0.028 | 0.76316849 | 0.85943643 | 0.0046 | 0.0053 | 0.005 | 0.005 | TRUE | 0.0785 |
| Pdzd2 | 348 | 243 | 228 | 275 | 251 | 269 | 270 | 258 | 246 | 274 | 283 | 266 | 266.2 | 258 | 274 | 1.059 | 0.083 | 0.50467546 | 0.66827185 | 0 | 0.0113 | 0.0074 | 0.0074 | TRUE | 0.0201 |
| Pdzd3 | 37 | 27 | 23 | 33 | 30 | 26 | 29 | 29 | 25 | 33 | 34 | 26 | 29.1 | 28 | 31 | 1.118 | 0.161 | 0.6491999 | 0.7820068 | 0 | 0.0839 | 0.0562 | 0.0562 | TRUE | 0.1464 |
| Pdzd4 | 4337 | 3249 | 3538 | 3994 | 3607 | 4380 | 3368 | 3456 | 3817 | 3976 | 4069 | 4324 | 3835.03 | 3547 | 4123 | 1.163 | 0.218 | 0.00117766 | 0.00816395 | 0.0028 | 0.003 | 0.003 | 0.003 | TRUE | 0.0537 |
| Pdzd7 | 125 | 72 | 71 | 99 | 85 | 98 | 97 | 77 | 77 | 99 | 96 | 97 | 90.24 | 84 | 97 | 1.156 | 0.21 | 0.31706117 | 0.4911405 | 0 | 0.0287 | 0.0206 | 0.0206 | TRUE | 0.2168 |
| Pdzd8 | 4087 | 2911 | 2539 | 2597 | 2185 | 2439 | 3174 | 3096 | 2740 | 2585 | 2465 | 2408 | 2744.63 | 3003 | 2486 | 0.827 | -0.274 | 0.0001342 | 0.00155312 | 0.0031 | 0.0033 | 0.0033 | 0.0033 | TRUE | 0.0711 |
| Pdzd9 | 1 | 0 | 0 | 1 | 2 | 0 | 1 | 0 | 0 | 1 | 2 | 0 | 0.67 | 0 | 1 | 3.106 | 1.635 | 0.51172212 | NA | 0 | 3.5326 | 2.9924 | 2.9924 | TRUE | 0.1121 |
| Pdzk1 | 244 | 207 | 305 | 335 | 375 | 400 | 190 | 220 | 329 | 333 | 423 | 395 | 315.02 | 246 | 384 | 1.564 | 0.645 | 0.00031793 | 0.00303781 | 0.0465 | 0.0099 | 0.0198 | 0.0198 | TRUE | 0.9516 |
| Pdzk1ip1 | 17 | 10 | 10 | 26 | 22 | 22 | 13 | 11 | 11 | 26 | 25 | 22 | 17.84 | 12 | 24 | 2.075 | 1.053 | 0.02158651 | 0.07304355 | 0 | 0.1353 | 0.0885 | 0.0885 | TRUE | 0.0789 |
| Pdzrn3 | 425 | 284 | 269 | 199 | 205 | 205 | 330 | 302 | 290 | 198 | 231 | 202 | 259.02 | 307 | 210 | 0.682 | -0.551 | 3.34E-05 | 0.00050765 | 0 | 0.0116 | 0.0087 | 0.0087 | TRUE | 0.0775 |
| Pdzrn4 | 4 | 4 | 4 | 1 | 2 | 1 | 3 | 4 | 4 | 1 | 2 | 1 | 2.65 | 4 | 1 | 0.36 | -1.474 | 0.22545326 | NA | 0 | 0.8959 | 0.5688 | 0.5688 | TRUE | 0.1539 |
| Pea15a | 1016 | 689 | 663 | 986 | 757 | 926 | 789 | 733 | 715 | 981 | 854 | 914 | 831.15 | 746 | 916 | 1.228 | 0.296 | 0.00076586 | 0.00595238 | 0.0026 | 0.0053 | 0.0044 | 0.0044 | TRUE | 0.0458 |
| Pea15b | 0 | 0 | 0 | 0 | 0 | 0 | 0 | 0 | 0 | 0 | 0 | 0 | 0 | 0 | 0 | NA | NA | NA | NA | NA | NA | NA | NA | NA | NA |
| Pear1 | 138 | 112 | 86 | 123 | 106 | 105 | 107 | 119 | 93 | 122 | 120 | 104 | 110.79 | 106 | 115 | 1.082 | 0.114 | 0.55549227 | 0.7106741 | 0 | 0.0238 | 0.0178 | 0.0178 | TRUE | 0.1136 |
| Pebp1 | 10396 | 7825 | 8218 | 9694 | 8944 | 10080 | 8074 | 8323 | 8867 | 9649 | 10090 | 9951 | 9159.05 | 8421 | 9897 | 1.175 | 0.233 | 2.33E-05 | 0.00037765 | 0.0013 | 0.0027 | 0.0021 | 0.0021 | TRUE | 0.0261 |
| Pebp4 | 0 | 0 | 0 | 0 | 0 | 0 | 0 | 0 | 0 | 0 | 0 | 0 | 0 | 0 | 0 | NA | NA | NA | NA | NA | NA | NA | NA | NA | NA |
| Pecam1 | 8723 | 6314 | 6404 | 6604 | 6075 | 7151 | 6775 | 6716 | 6910 | 6574 | 6853 | 7059 | 6814.46 | 6800 | 6829 | 1.004 | 0.006 | 0.90737876 | 0.94923397 | 6.00E-04 | 0.0028 | 0.0018 | 0.0018 | TRUE | 0.0131 |
| Pecr | 1921 | 1524 | 1560 | 1694 | 1563 | 1722 | 1492 | 1621 | 1683 | 1686 | 1763 | 1700 | 1657.59 | 1599 | 1716 | 1.075 | 0.104 | 0.14317202 | 0.28608383 | 0.0016 | 0.0039 | 0.003 | 0.003 | TRUE | 0.0428 |
| Pef1 | 2673 | 1908 | 2150 | 2512 | 2216 | 2521 | 2076 | 2030 | 2320 | 2500 | 2500 | 2489 | 2319.05 | 2142 | 2496 | 1.166 | 0.222 | 0.00117147 | 0.00812808 | 0.002 | 0.0034 | 0.0029 | 0.0029 | TRUE | 0.0629 |
| Peg10 | 25579 | 15712 | 12531 | 15982 | 12002 | 13551 | 19866 | 16713 | 13521 | 15908 | 13540 | 13377 | 15487.42 | 16700 | 14275 | 0.855 | -0.226 | 0.20443781 | 0.36593984 | 0.0229 | 0.0026 | 0.0087 | 0.0229 | TRUE | 0.1607 |
| Peg12 | 542 | 385 | 280 | 244 | 182 | 209 | 421 | 410 | 302 | 243 | 205 | 206 | 297.85 | 378 | 218 | 0.577 | -0.793 | 1.47E-07 | 5.47E-06 | 0.0182 | 0.0104 | 0.0128 | 0.0128 | TRUE | 0.3421 |
| Peg13 | 458 | 289 | 185 | 247 | 202 | 218 | 356 | 307 | 200 | 246 | 228 | 215 | 258.61 | 288 | 230 | 0.796 | -0.33 | 0.07247672 | 0.17602544 | 0.0413 | 0.0116 | 0.0203 | 0.0203 | TRUE | 0.4864 |
| Peg3 | 2969 | 2232 | 2410 | 1873 | 1926 | 2424 | 2306 | 2374 | 2600 | 1864 | 2173 | 2393 | 2285.06 | 2427 | 2143 | 0.883 | -0.179 | 0.04834583 | 0.13145735 | 0.009 | 0.0035 | 0.0055 | 0.0055 | TRUE | 0.1579 |
| Peg3as | 0 | 0 | 0 | 0 | 0 | 0 | 0 | 0 | 0 | 0 | 0 | 0 | 0 | 0 | 0 | NA | NA | NA | NA | NA | NA | NA | NA | NA | NA |

| | | | | | | | | | | | | | | | | | | | | | | | | | |
|---|---|---|---|---|---|---|---|---|---|---|---|---|---|---|---|---|---|---|---|---|---|---|---|---|---|
| Peli1 | 2673 | 1950 | 1882 | 1908 | 1671 | 1938 | 2076 | 2074 | 2031 | 1899 | 1885 | 1913 | 1979.71 | 2060 | 1899 | 0.922 | -0.118 | 0.04760893 | 0.12998556 | 0 | 0.0036 | 0.002 | 0.002 | TRUE | 0.0019 |
| Peli2 | 681 | 523 | 494 | 526 | 528 | 636 | 529 | 556 | 533 | 524 | 596 | 628 | 560.88 | 539 | 583 | 1.08 | 0.111 | 0.26883796 | 0.4393438 | 0.003 | 0.0067 | 0.0055 | 0.0055 | TRUE | 0.0931 |
| Peli3 | 109 | 78 | 71 | 94 | 91 | 91 | 85 | 83 | 77 | 94 | 103 | 90 | 88.38 | 82 | 96 | 1.168 | 0.224 | 0.27724329 | 0.44851335 | 0 | 0.0292 | 0.0192 | 0.0192 | TRUE | 0.0412 |
| Pelo | 2368 | 1684 | 1367 | 1279 | 1155 | 1284 | 1839 | 1791 | 1475 | 1273 | 1303 | 1268 | 1491.49 | 1702 | 1281 | 0.752 | -0.411 | 3.32E-06 | 7.59E-05 | 0.0067 | 0.004 | 0.0049 | 0.0049 | TRUE | 0.1621 |
| Pelp1 | 19389 | 14122 | 11894 | 11990 | 11210 | 12928 | 15059 | 15021 | 12834 | 11935 | 12646 | 12762 | 13376.06 | 14305 | 12448 | 0.87 | -0.201 | 0.00391302 | 0.02026405 | 0.0045 | 0.0026 | 0.0034 | 0.0034 | TRUE | 0.0987 |
| Pemt | 259 | 207 | 185 | 195 | 157 | 223 | 201 | 220 | 200 | 194 | 177 | 220 | 202.05 | 207 | 197 | 0.955 | -0.067 | 0.64916546 | 0.7820068 | 0.0021 | 0.0142 | 0.0107 | 0.0107 | TRUE | 0.1139 |
| Penk | 115 | 89 | 61 | 86 | 78 | 75 | 89 | 95 | 66 | 86 | 88 | 74 | 82.91 | 83 | 83 | 0.988 | -0.018 | 0.93703049 | 0.96508948 | 0.0086 | 0.031 | 0.025 | 0.025 | TRUE | 0.2956 |
| Peo1 | 3498 | 2426 | 2156 | 2266 | 2035 | 2265 | 2717 | 2581 | 2326 | 2256 | 2296 | 2236 | 2401.79 | 2541 | 2263 | 0.89 | -0.169 | 0.01701796 | 0.06143918 | 0.0025 | 0.0034 | 0.0032 | 0.0032 | TRUE | 0.0659 |
| Pepd | 3071 | 2355 | 2204 | 2370 | 2225 | 2627 | 2385 | 2505 | 2378 | 2359 | 2510 | 2593 | 2455.1 | 2423 | 2487 | 1.027 | 0.038 | 0.54807699 | 0.70484654 | 0.0012 | 0.0034 | 0.0025 | 0.0025 | TRUE | 0.025 |
| Per1 | 2298 | 1730 | 1858 | 1978 | 1767 | 2116 | 1785 | 1840 | 2005 | 1969 | 1993 | 2089 | 1946.8 | 1877 | 2017 | 1.076 | 0.105 | 0.12861974 | 0.26509922 | 0.0018 | 0.0036 | 0.003 | 0.003 | TRUE | 0.0428 |
| Per2 | 42 | 26 | 25 | 35 | 42 | 36 | 33 | 28 | 27 | 35 | 47 | 36 | 34.17 | 29 | 39 | 1.339 | 0.421 | 0.20742761 | 0.36909854 | 0 | 0.0718 | 0.0506 | 0.0506 | TRUE | 0.228 |
| Per3 | 821 | 696 | 770 | 915 | 810 | 952 | 638 | 740 | 831 | 911 | 914 | 940 | 828.85 | 736 | 922 | 1.255 | 0.327 | 0.0011216 | 0.00788539 | 0.0076 | 0.0053 | 0.0061 | 0.0061 | TRUE | 0.1707 |
| Perp | 2274 | 1622 | 1775 | 1780 | 1490 | 1633 | 1766 | 1725 | 1915 | 1772 | 1681 | 1612 | 1745.22 | 1802 | 1688 | 0.937 | -0.094 | 0.19174043 | 0.35000587 | 0.0018 | 0.0038 | 0.0031 | 0.0031 | TRUE | 0.0356 |
| Pes1 | 16608 | 11982 | 11525 | 12040 | 10542 | 12435 | 12899 | 12745 | 12435 | 11985 | 11892 | 12275 | 12371.92 | 12693 | 12051 | 0.949 | -0.075 | 0.10696243 | 0.23241766 | 2.00E-04 | 0.0026 | 0.0015 | 0.0015 | TRUE | 0.0038 |
| Pet100 | 993 | 653 | 529 | 573 | 410 | 558 | 771 | 695 | 571 | 570 | 463 | 551 | 603.39 | 679 | 528 | 0.777 | -0.364 | 0.00325922 | 0.01759878 | 0.016 | 0.0064 | 0.0094 | 0.0094 | TRUE | 0.2265 |
| Pet112 | 1382 | 973 | 941 | 970 | 819 | 955 | 1073 | 1035 | 1015 | 966 | 924 | 943 | 992.64 | 1041 | 944 | 0.906 | -0.142 | 0.05714967 | 0.1485259 | 0 | 0.0048 | 0.003 | 0.003 | TRUE | 0.0092 |
| Pet117 | 1 | 2 | 2 | 0 | 2 | 0 | 1 | 2 | 2 | 0 | 2 | 0 | 1.22 | 2 | 1 | 0.434 | -1.205 | 0.52273097 | NA | 0 | 1.9456 | 1.5507 | 1.5507 | TRUE | 0.9751 |
| Pet2 | 80 | 70 | 36 | 7 | 6 | 6 | 62 | 74 | 39 | 7 | 7 | 6 | 32.52 | 58 | 7 | 0.112 | -3.161 | 1.34E-11 | 1.31E-09 | 0 | 0.0753 | 0.0702 | 0.0702 | TRUE | 0.1452 |
| Pex1 | 1761 | 1234 | 1239 | 1294 | 1191 | 1305 | 1368 | 1313 | 1337 | 1288 | 1344 | 1288 | 1322.83 | 1339 | 1307 | 0.975 | -0.036 | 0.59174812 | 0.73958167 | 0 | 0.0042 | 0.0026 | 0.0026 | TRUE | 0.0072 |
| Pex10 | 967 | 709 | 729 | 786 | 743 | 788 | 751 | 754 | 787 | 782 | 838 | 778 | 781.7 | 764 | 799 | 1.046 | 0.066 | 0.42507192 | 0.59781097 | 0 | 0.0055 | 0.0036 | 0.0036 | TRUE | 0.0208 |
| Pex11a | 287 | 198 | 195 | 206 | 195 | 203 | 223 | 211 | 210 | 205 | 220 | 200 | 211.56 | 215 | 208 | 0.969 | -0.045 | 0.74191647 | 0.84644095 | 0 | 0.0136 | 0.0089 | 0.0089 | TRUE | 0.0244 |
| Pex11b | 1066 | 779 | 773 | 801 | 708 | 848 | 828 | 829 | 834 | 797 | 799 | 837 | 820.62 | 830 | 811 | 0.977 | -0.033 | 0.67199479 | 0.79833942 | 0 | 0.0053 | 0.0032 | 0.0032 | TRUE | 0.0095 |
| Pex11c | 271 | 178 | 225 | 198 | 184 | 229 | 210 | 189 | 243 | 197 | 208 | 226 | 212.22 | 214 | 210 | 0.982 | -0.026 | 0.8633679 | 0.92466653 | 0.0051 | 0.0136 | 0.0112 | 0.0112 | TRUE | 0.1406 |
| Pex12 | 1073 | 774 | 703 | 735 | 667 | 686 | 833 | 823 | 759 | 732 | 752 | 677 | 762.74 | 805 | 720 | 0.894 | -0.162 | 0.06275108 | 0.15849111 | 0.0014 | 0.0055 | 0.0042 | 0.0042 | TRUE | 0.0329 |
| Pex13 | 2475 | 1719 | 1708 | 1831 | 1512 | 1795 | 1922 | 1828 | 1843 | 1823 | 1706 | 1772 | 1815.64 | 1864 | 1767 | 0.947 | -0.078 | 0.22535493 | 0.39009301 | 4.00E-04 | 0.0037 | 0.0024 | 0.0024 | TRUE | 0.0108 |
| Pex14 | 2637 | 1940 | 1893 | 1950 | 1803 | 2077 | 2048 | 2064 | 2043 | 1941 | 2034 | 2050 | 2029.91 | 2052 | 2008 | 0.979 | -0.031 | 0.61383753 | 0.75595208 | 0 | 0.0036 | 0.0022 | 0.0022 | TRUE | 0.0105 |
| Pex16 | 895 | 587 | 568 | 669 | 590 | 635 | 695 | 624 | 613 | 666 | 666 | 627 | 648.45 | 644 | 653 | 1.011 | 0.016 | 0.86245178 | 0.92408891 | 0.0017 | 0.0061 | 0.0046 | 0.0046 | TRUE | 0.0597 |
| Pex19 | 4554 | 3581 | 3918 | 4043 | 3636 | 4108 | 3537 | 3809 | 4227 | 4024 | 4102 | 4055 | 3959.14 | 3858 | 4060 | 1.053 | 0.075 | 0.28991686 | 0.46190856 | 0.0037 | 0.003 | 0.0033 | 0.0033 | TRUE | 0.0854 |
| Pex2 | 468 | 292 | 289 | 249 | 275 | 262 | 363 | 311 | 312 | 248 | 310 | 259 | 300.44 | 329 | 272 | 0.825 | -0.278 | 0.0383966 | 0.11116116 | 0.0079 | 0.0103 | 0.0096 | 0.0096 | TRUE | 0.1618 |
| Pex26 | 1392 | 1034 | 989 | 916 | 940 | 1024 | 1081 | 1100 | 1067 | 912 | 1060 | 1011 | 1038.52 | 1083 | 994 | 0.918 | -0.124 | 0.12694229 | 0.26242207 | 0.0019 | 0.0047 | 0.0038 | 0.0038 | TRUE | 0.0633 |
| Pex3 | 1314 | 931 | 845 | 895 | 764 | 824 | 1021 | 990 | 912 | 891 | 862 | 813 | 914.79 | 974 | 855 | 0.877 | -0.189 | 0.02274713 | 0.075822 | 0.0017 | 0.005 | 0.0039 | 0.0039 | TRUE | 0.0369 |
| Pex5 | 6088 | 4313 | 4519 | 4835 | 4214 | 4834 | 4728 | 4588 | 4876 | 4813 | 4754 | 4772 | 4755.07 | 4731 | 4780 | 1.01 | 0.015 | 0.77572073 | 0.86757979 | 2.00E-04 | 0.0029 | 0.0018 | 0.0018 | TRUE | 0.0086 |
| Pex5l | 18 | 13 | 23 | 17 | 17 | 12 | 14 | 14 | 25 | 17 | 19 | 12 | 16.76 | 18 | 16 | 0.915 | -0.128 | 0.79337292 | NA | 0.0331 | 0.1438 | 0.1125 | 0.1125 | TRUE | 0.6179 |
| Pex6 | 1384 | 1098 | 804 | 938 | 777 | 877 | 1075 | 1168 | 868 | 934 | 877 | 866 | 964.38 | 1037 | 892 | 0.86 | -0.218 | 0.03879226 | 0.11197958 | 0.0112 | 0.0049 | 0.0069 | 0.0069 | TRUE | 0.2386 |
| Pex7 | 2194 | 1564 | 1630 | 1860 | 1591 | 1841 | 1704 | 1664 | 1759 | 1851 | 1795 | 1817 | 1764.99 | 1709 | 1821 | 1.066 | 0.092 | 0.14301814 | 0.28585629 | 0 | 0.0038 | 0.0023 | 0.0023 | TRUE | 0.0076 |
| Pf4 | 1 | 1 | 0 | 0 | 0 | 1 | 1 | 1 | 0 | 0 | 0 | 1 | 0.47 | 1 | 0 | 0.599 | -0.74 | 0.79209856 | NA | 0 | 5.0317 | 3.8925 | 3.8925 | TRUE | 0.3835 |
| Pfas | 15982 | 12188 | 10649 | 11050 | 9844 | 11217 | 12413 | 12964 | 11490 | 10999 | 11105 | 11073 | 11674.01 | 12289 | 11059 | 0.9 | -0.152 | 0.00770492 | 0.03405268 | 0.0017 | 0.0026 | 0.0023 | 0.0023 | TRUE | 0.0393 |
| Pfdn1 | 5660 | 4157 | 4008 | 4475 | 3838 | 4320 | 4396 | 4422 | 4325 | 4454 | 4330 | 4265 | 4365.13 | 4381 | 4350 | 0.993 | -0.01 | 0.84137924 | 0.90973852 | 0 | 0.003 | 0.0017 | 0.0017 | TRUE | 0.0055 |
| Pfdn2 | 4900 | 3640 | 3752 | 3936 | 3388 | 3913 | 3806 | 3872 | 4048 | 3918 | 3822 | 3863 | 3888.08 | 3909 | 3868 | 0.99 | -0.015 | 0.78769367 | 0.87526281 | 3.00E-04 | 0.003 | 0.0019 | 0.0019 | TRUE | 0.0118 |
| Pfdn4 | 2534 | 1849 | 1919 | 2072 | 1851 | 2021 | 1968 | 1967 | 2071 | 2062 | 2088 | 1995 | 2025.17 | 2002 | 2048 | 1.024 | 0.034 | 0.58820148 | 0.73734115 | 0 | 0.0036 | 0.0023 | 0.0023 | TRUE | 0.0107 |
| Pfdn5 | 3469 | 2463 | 2699 | 2939 | 2605 | 3126 | 2694 | 2620 | 2912 | 2925 | 2939 | 3086 | 2862.72 | 2742 | 2983 | 1.088 | 0.122 | 0.05581488 | 0.14581649 | 0.0015 | 0.0033 | 0.0026 | 0.0026 | TRUE | 0.0351 |
| Pfkfb1 | 642 | 403 | 497 | 589 | 500 | 587 | 499 | 429 | 536 | 586 | 564 | 579 | 532.22 | 488 | 576 | 1.182 | 0.241 | 0.02170768 | 0.07328425 | 0.0041 | 0.0069 | 0.006 | 0.006 | TRUE | 0.1255 |
| Pfkfb2 | 908 | 698 | 665 | 793 | 636 | 849 | 705 | 742 | 718 | 789 | 717 | 838 | 751.68 | 722 | 781 | 1.085 | 0.117 | 0.1891316 | 0.34666042 | 0.002 | 0.0056 | 0.0044 | 0.0044 | TRUE | 0.0597 |
| Pfkfb3 | 1894 | 1612 | 1838 | 2060 | 1703 | 1976 | 1471 | 1715 | 1983 | 2051 | 1921 | 1951 | 1848.52 | 1723 | 1974 | 1.147 | 0.198 | 0.0403509 | 0.11517939 | 0.0108 | 0.0037 | 0.0062 | 0.0062 | TRUE | 0.2108 |
| Pfkfb4 | 199 | 147 | 145 | 131 | 126 | 160 | 155 | 156 | 156 | 130 | 142 | 158 | 149.64 | 156 | 143 | 0.922 | -0.118 | 0.47080338 | 0.63967946 | 0 | 0.0183 | 0.0125 | 0.0125 | TRUE | 0.0835 |
| Pfkl | 23247 | 18748 | 23350 | 24956 | 23693 | 27505 | 18055 | 19942 | 25194 | 24841 | 26728 | 27152 | 23652.07 | 21064 | 26240 | 1.246 | 0.317 | 0.00115905 | 0.00806928 | 0.0165 | 0.0025 | 0.0068 | 0.0068 | TRUE | 0.36 |
| Pfkm | 3176 | 2391 | 2768 | 2930 | 2644 | 2881 | 2467 | 2543 | 2987 | 2917 | 2983 | 2844 | 2789.97 | 2666 | 2915 | 1.094 | 0.13 | 0.09576995 | 0.21444386 | 0.0053 | 0.0033 | 0.004 | 0.004 | TRUE | 0.1339 |
| Pfkp | 22067 | 18269 | 21322 | 19967 | 18675 | 21565 | 17139 | 19432 | 23006 | 19875 | 21067 | 21288 | 20301.28 | 19859 | 20743 | 1.045 | 0.063 | 0.47685505 | 0.64457057 | 0.0096 | 0.0025 | 0.0056 | 0.0056 | TRUE | 0.2351 |
| Pfn1 | 28925 | 20196 | 16844 | 13390 | 11720 | 14117 | 22465 | 21482 | 18175 | 13328 | 13221 | 13936 | 17101.17 | 20707 | 13495 | 0.652 | -0.618 | 2.23E-16 | 4.43E-14 | 0.0062 | 0.0026 | 0.004 | 0.004 | TRUE | 0.1399 |
| Pfn2 | 1426 | 996 | 1037 | 1463 | 1183 | 1308 | 1108 | 1059 | 1119 | 1456 | 1335 | 1291 | 1227.98 | 1095 | 1361 | 1.242 | 0.313 | 4.15E-05 | 0.00060373 | 0.0017 | 0.0044 | 0.0034 | 0.0034 | TRUE | 0.0459 |
| Pfn3 | 0 | 0 | 0 | 0 | 0 | 0 | 0 | 0 | 0 | 0 | 0 | 0 | 0 | 0 | 0 | NA | NA | NA | NA | NA | NA | NA | NA | NA | NA |
| Pfn4 | 180 | 125 | 75 | 161 | 107 | 119 | 140 | 133 | 81 | 160 | 121 | 117 | 125.35 | 118 | 133 | 1.122 | 0.166 | 0.46111085 | 0.63056423 | 0.048 | 0.0213 | 0.0288 | 0.0288 | TRUE | 0.7288 |
| Pfpl | 0 | 0 | 0 | 0 | 0 | 0 | 0 | 0 | 0 | 0 | 0 | 0 | 0 | 0 | 0 | NA | NA | NA | NA | NA | NA | NA | NA | NA | NA |
| Pga5 | 84 | 78 | 100 | 69 | 98 | 83 | 65 | 83 | 108 | 69 | 111 | 82 | 86.21 | 85 | 87 | 1.023 | 0.033 | 0.89757182 | 0.94349536 | 0.0492 | 0.0299 | 0.0355 | 0.0355 | TRUE | 0.3065 |
| Pgam1 | 15647 | 11224 | 13045 | 13115 | 11266 | 13471 | 12152 | 11939 | 14075 | 13055 | 12709 | 13298 | 12871.41 | 12722 | 13021 | 1.024 | 0.034 | 0.62201476 | 0.76142749 | 0.0036 | 0.0026 | 0.0033 | 0.0033 | TRUE | 0.1056 |
| Pgam2 | 116 | 73 | 112 | 145 | 132 | 148 | 90 | 78 | 121 | 144 | 149 | 146 | 121.32 | 96 | 146 | 1.526 | 0.61 | 0.00203862 | 0.01226001 | 0.0125 | 0.022 | 0.0196 | 0.0196 | TRUE | 0.4567 |
| Pgam5 | 4924 | 3610 | 3859 | 4381 | 3816 | 4413 | 3824 | 3840 | 4164 | 4361 | 4305 | 4356 | 4141.67 | 3943 | 4341 | 1.101 | 0.139 | 0.01522084 | 0.05643359 | 8.00E-04 | 0.003 | 0.0021 | 0.0021 | TRUE | 0.029 |
| Pgap1 | 2229 | 1602 | 1484 | 1883 | 1561 | 1806 | 1731 | 1704 | 1601 | 1874 | 1761 | 1783 | 1742.42 | 1679 | 1806 | 1.075 | 0.105 | 0.11782162 | 0.24907919 | 8.00E-04 | 0.0038 | 0.0027 | 0.0027 | TRUE | 0.0194 |
| Pgap2 | 967 | 737 | 804 | 881 | 867 | 954 | 751 | 784 | 868 | 877 | 978 | 942 | 866.54 | 801 | 932 | 1.165 | 0.221 | 0.01253036 | 0.04889338 | 0.0031 | 0.0052 | 0.0045 | 0.0045 | TRUE | 0.0622 |
| Pgap3 | 338 | 223 | 238 | 259 | 235 | 248 | 263 | 237 | 257 | 258 | 265 | 245 | 254.04 | 252 | 256 | 1.013 | 0.018 | 0.88496995 | 0.93609243 | 0 | 0.0118 | 0.0078 | 0.0078 | TRUE | 0.0292 |
| Pgbd1 | 289 | 228 | 225 | 265 | 224 | 294 | 224 | 243 | 243 | 264 | 253 | 290 | 252.74 | 237 | 269 | 1.14 | 0.189 | 0.14765772 | 0.2923496 | 0 | 0.0118 | 0.0083 | 0.0083 | TRUE | 0.0543 |
| Pgbd5 | 412 | 286 | 309 | 312 | 290 | 317 | 320 | 304 | 333 | 311 | 327 | 313 | 318.04 | 319 | 317 | 0.992 | -0.011 | 0.92236272 | 0.95711845 | 0 | 0.0099 | 0.0064 | 0.0064 | TRUE | 0.0179 |
| Pgc | 223 | 195 | 241 | 251 | 225 | 275 | 173 | 207 | 260 | 250 | 254 | 271 | 235.96 | 213 | 258 | 1.216 | 0.282 | 0.07563288 | 0.1812865 | 0.0174 | 0.0125 | 0.0139 | 0.0139 | TRUE | 0.3943 |
| Pgd | 10797 | 7891 | 7227 | 7805 | 6885 | 8213 | 8386 | 8394 | 7798 | 7769 | 7767 | 8107 | 8036.76 | 8193 | 7881 | 0.962 | -0.056 | 0.30162836 | 0.47401625 | 0.001 | 0.0027 | 0.002 | 0.002 | TRUE | 0.0215 |
| Pgf | 102 | 70 | 79 | 80 | 83 | 85 | 79 | 74 | 85 | 80 | 94 | 84 | 82.68 | 79 | 86 | 1.075 | 0.105 | 0.62474487 | 0.76346484 | 0 | 0.0311 | 0.0209 | 0.0209 | TRUE | 0.0582 |
| Pggt1b | 1276 | 924 | 757 | 824 | 647 | 786 | 991 | 983 | 817 | 820 | 730 | 776 | 852.78 | 930 | 775 | 0.833 | -0.264 | 0.00655719 | 0.03018856 | 0.0064 | 0.0052 | 0.0056 | 0.0056 | TRUE | 0.133 |
| Pgk1 | 30766 | 23238 | 30164 | 34036 | 30053 | 34976 | 23895 | 24718 | 32547 | 33879 | 33903 | 34527 | 30578.05 | 27053 | 34103 | 1.261 | 0.334 | 0.00044133 | 0.00392039 | 0.0156 | 0.0025 | 0.0065 | 0.0065 | TRUE | 0.3861 |
| Pgk2 | 0 | 0 | 1 | 0 | 0 | 1 | 0 | 0 | 1 | 0 | 0 | 1 | 0.34 | 0 | 0 | 1.075 | 0.105 | 0.97570524 | NA | 0 | 6.885 | 6.5379 | 6.5379 | TRUE | 0.5365 |
| Pgls | 3748 | 2744 | 2832 | 3098 | 2703 | 3093 | 2911 | 2919 | 3056 | 3084 | 3049 | 3053 | 3011.94 | 2962 | 3062 | 1.034 | 0.048 | 0.38314501 | 0.55698653 | 0 | 0.0032 | 0.0019 | 0.0019 | TRUE | 0.0092 |

| | | | | | | | | | | | | | | | | | | | | | | | | | |
|---|---|---|---|---|---|---|---|---|---|---|---|---|---|---|---|---|---|---|---|---|---|---|---|---|---|
| Pglyrp1 | 12 | 13 | 15 | 18 | 9 | 21 | 9 | 14 | 16 | 18 | 10 | 21 | 14.69 | 13 | 16 | 1.262 | 0.336 | 0.52077383 | NA | 0 | 0.1638 | 0.1279 | 0.1279 | TRUE | 0.4672 |
| Pglyrp2 | 0 | 0 | 0 | 0 | 0 | 0 | 0 | 0 | 0 | 0 | 0 | 0 | 0 | 0 | 0 | NA | NA | NA | NA | NA | NA | NA | NA | NA | NA |
| Pglyrp3 | 1 | 0 | 1 | 0 | 0 | 0 | 1 | 0 | 1 | 0 | 0 | 0 | 0.31 | 1 | 0 | 0.307 | -1.706 | 0.62894649 | NA | 0 | 7.6657 | 7.2 | 7.2 | TRUE | 0.2067 |
| Pglyrp4 | 0 | 0 | 0 | 0 | 0 | 0 | 0 | 0 | 0 | 0 | 0 | 0 | 0 | 0 | 0 | NA | NA | NA | NA | NA | NA | NA | NA | NA | NA |
| Pgm1 | 3561 | 2797 | 3275 | 3122 | 2925 | 3441 | 2766 | 2975 | 3534 | 3108 | 3300 | 3397 | 3179.77 | 3092 | 3268 | 1.058 | 0.081 | 0.35029001 | 0.52486583 | 0.0077 | 0.0032 | 0.0051 | 0.0051 | TRUE | 0.1897 |
| Pgm2 | 5275 | 4061 | 5106 | 5841 | 5587 | 6244 | 4097 | 4320 | 5509 | 5814 | 6303 | 6164 | 5367.74 | 4642 | 6094 | 1.313 | 0.393 | 3.56E-05 | 0.00053494 | 0.014 | 0.0029 | 0.0063 | 0.0063 | TRUE | 0.3248 |
| Pgm2l1 | 2405 | 1616 | 1494 | 1551 | 1277 | 1447 | 1868 | 1719 | 1612 | 1544 | 1441 | 1428 | 1601.94 | 1733 | 1471 | 0.848 | -0.238 | 0.00197642 | 0.01201063 | 0.0032 | 0.0039 | 0.0036 | 0.0036 | TRUE | 0.0576 |
| Pgm3 | 3627 | 2689 | 2808 | 3079 | 2819 | 3153 | 2817 | 2860 | 3030 | 3065 | 3180 | 3112 | 3010.74 | 2902 | 3119 | 1.075 | 0.104 | 0.07424467 | 0.17907173 | 6.00E-04 | 0.0032 | 0.0021 | 0.0021 | TRUE | 0.0177 |
| Pgm5 | 27 | 17 | 15 | 21 | 17 | 10 | 21 | 18 | 16 | 21 | 19 | 10 | 17.53 | 18 | 17 | 0.897 | -0.156 | 0.74271981 | 0.84688426 | 0 | 0.1376 | 0.1058 | 0.1058 | TRUE | 0.6422 |
| Pgp | 2887 | 2332 | 2129 | 2332 | 1988 | 2186 | 2242 | 2481 | 2297 | 2321 | 2243 | 2158 | 2290.29 | 2340 | 2241 | 0.958 | -0.062 | 0.3533367 | 0.52780934 | 0.0017 | 0.0035 | 0.0028 | 0.0028 | TRUE | 0.0332 |
| Pgpep1 | 2436 | 1943 | 2294 | 2834 | 2647 | 2882 | 1892 | 2067 | 2475 | 2821 | 2986 | 2845 | 2514.32 | 2145 | 2884 | 1.346 | 0.429 | 1.97E-06 | 5.03E-05 | 0.0096 | 0.0034 | 0.0054 | 0.0054 | TRUE | 0.2194 |
| Pgpep1l | 39 | 37 | 29 | 23 | 26 | 25 | 30 | 39 | 31 | 23 | 29 | 25 | 29.64 | 33 | 26 | 0.762 | -0.391 | 0.27111817 | 0.44203202 | 0 | 0.0824 | 0.0564 | 0.0564 | TRUE | 0.1476 |
| Pgr | 85 | 64 | 51 | 75 | 64 | 45 | 66 | 68 | 55 | 75 | 72 | 44 | 63.4 | 63 | 64 | 1.008 | 0.011 | 0.96690274 | 0.9820879 | 0.028 | 0.0398 | 0.0366 | 0.0366 | TRUE | 0.634 |
| Pgr15l | 101 | 70 | 43 | 73 | 42 | 37 | 78 | 74 | 46 | 73 | 47 | 37 | 59.31 | 66 | 52 | 0.783 | -0.353 | 0.25837058 | 0.42834726 | 0.0808 | 0.0424 | 0.0531 | 0.0531 | TRUE | 0.831 |
| Pgrmc1 | 5536 | 3822 | 3592 | 4284 | 3707 | 4287 | 4300 | 4065 | 3876 | 4264 | 4182 | 4232 | 4153.14 | 4080 | 4226 | 1.035 | 0.05 | 0.39469655 | 0.56811659 | 0.001 | 0.003 | 0.0022 | 0.0022 | TRUE | 0.0274 |
| Pgrmc2 | 3972 | 3096 | 3187 | 3304 | 2885 | 3445 | 3085 | 3293 | 3439 | 3289 | 3255 | 3401 | 3293.49 | 3272 | 3315 | 1.014 | 0.02 | 0.75164663 | 0.8526317 | 0.0015 | 0.0031 | 0.0025 | 0.0025 | TRUE | 0.0313 |
| Pgs1 | 7210 | 5389 | 4908 | 5477 | 4873 | 5610 | 5600 | 5732 | 5296 | 5452 | 5497 | 5538 | 5519.1 | 5543 | 5496 | 0.991 | -0.012 | 0.81745046 | 0.89520392 | 5.00E-04 | 0.0029 | 0.0019 | 0.0019 | TRUE | 0.0182 |
| Phactr1 | 217 | 125 | 165 | 131 | 112 | 131 | 169 | 133 | 178 | 130 | 126 | 129 | 144.27 | 160 | 128 | 0.804 | -0.314 | 0.07192325 | 0.17509776 | 0.0052 | 0.0189 | 0.015 | 0.015 | TRUE | 0.2181 |
| Phactr2 | 363 | 256 | 238 | 242 | 218 | 216 | 282 | 272 | 257 | 241 | 246 | 213 | 251.84 | 270 | 233 | 0.861 | -0.216 | 0.09982466 | 0.22082633 | 0 | 0.0118 | 0.0084 | 0.0084 | TRUE | 0.0639 |
| Phactr3 | 246 | 138 | 158 | 212 | 184 | 207 | 191 | 147 | 170 | 211 | 208 | 204 | 188.54 | 169 | 208 | 1.22 | 0.287 | 0.06125295 | 0.15561473 | 0.0032 | 0.015 | 0.0116 | 0.0116 | TRUE | 0.1454 |
| Phactr4 | 3039 | 2279 | 2522 | 2790 | 2538 | 2923 | 2360 | 2424 | 2721 | 2777 | 2863 | 2885 | 2671.89 | 2502 | 2842 | 1.137 | 0.185 | 0.00760769 | 0.03373138 | 0.0026 | 0.0033 | 0.0031 | 0.0031 | TRUE | 0.0707 |
| Phax | 2676 | 1847 | 1787 | 1657 | 1479 | 1829 | 2078 | 1965 | 1928 | 1649 | 1668 | 1806 | 1849.08 | 1990 | 1708 | 0.858 | -0.222 | 0.00127261 | 0.00857586 | 0.0015 | 0.0037 | 0.0029 | 0.0029 | TRUE | 0.0305 |
| Phb | 13848 | 9552 | 8827 | 10795 | 9556 | 10965 | 10755 | 10160 | 9524 | 10745 | 10780 | 10824 | 10464.9 | 10146 | 10783 | 1.062 | 0.087 | 0.12705763 | 0.26254659 | 0.0017 | 0.0027 | 0.0023 | 0.0023 | TRUE | 0.0351 |
| Phb2 | 26605 | 19468 | 18256 | 19100 | 17342 | 20152 | 20663 | 20708 | 19698 | 19012 | 19564 | 19893 | 19922.96 | 20356 | 19490 | 0.957 | -0.063 | 0.19337436 | 0.352134 | 6.00E-04 | 0.0025 | 0.0016 | 0.0016 | TRUE | 0.0098 |
| Phc1 | 41823 | 30582 | 30777 | 38844 | 33278 | 38581 | 32482 | 32530 | 33208 | 38665 | 37541 | 38085 | 35418.61 | 32740 | 38097 | 1.164 | 0.219 | 4.30E-07 | 1.43E-05 | 1.00E-04 | 0.0025 | 0.0013 | 0.0013 | TRUE | 0.0021 |
| Phc2 | 2304 | 1666 | 1567 | 1429 | 1295 | 1484 | 1789 | 1772 | 1691 | 1422 | 1461 | 1465 | 1600.09 | 1751 | 1449 | 0.827 | -0.273 | 2.70E-05 | 0.00042523 | 0 | 0.0039 | 0.0024 | 0.0024 | TRUE | 0.0107 |
| Phc3 | 1284 | 986 | 1102 | 1315 | 1138 | 1283 | 997 | 1049 | 1189 | 1309 | 1284 | 1267 | 1182.39 | 1078 | 1287 | 1.195 | 0.257 | 0.00188975 | 0.01157726 | 0.0031 | 0.0044 | 0.0041 | 0.0041 | TRUE | 0.0955 |
| Phex | 2 | 3 | 3 | 0 | 0 | 1 | 2 | 3 | 3 | 0 | 0 | 1 | 1.49 | 3 | 0 | 0.141 | -2.828 | 0.11612616 | NA | 0 | 1.588 | 1.1505 | 1.1505 | TRUE | 0.3835 |
| Phf1 | 495 | 421 | 369 | 380 | 377 | 420 | 384 | 448 | 398 | 378 | 425 | 415 | 408.09 | 410 | 406 | 0.991 | -0.013 | 0.90843708 | 0.94963001 | 0.0028 | 0.0082 | 0.0065 | 0.0065 | TRUE | 0.0734 |
| Phf10 | 3353 | 2503 | 2524 | 2561 | 2256 | 2659 | 2604 | 2662 | 2723 | 2549 | 2545 | 2625 | 2618.16 | 2663 | 2573 | 0.967 | -0.049 | 0.39093455 | 0.56463496 | 0 | 0.0033 | 0.002 | 0.002 | TRUE | 0.0047 |
| Phf11a | 48 | 30 | 15 | 18 | 13 | 12 | 37 | 32 | 16 | 18 | 15 | 12 | 21.63 | 28 | 15 | 0.516 | -0.955 | 0.03802725 | 0.11029292 | 0.073 | 0.112 | 0.1007 | 0.1007 | TRUE | 0.576 |
| Phf11b | 38 | 17 | 14 | 20 | 15 | 12 | 30 | 18 | 15 | 20 | 17 | 12 | 18.56 | 21 | 16 | 0.766 | -0.385 | 0.41730138 | 0.58984263 | 0.0541 | 0.1301 | 0.1078 | 0.1078 | TRUE | 0.8592 |
| Phf11c | 74 | 56 | 39 | 37 | 44 | 30 | 57 | 60 | 42 | 37 | 50 | 30 | 45.87 | 53 | 39 | 0.724 | -0.466 | 0.13200956 | 0.26982268 | 0.0242 | 0.0541 | 0.0463 | 0.0463 | TRUE | 0.4232 |
| Phf11d | 3455 | 2572 | 2674 | 2468 | 2270 | 2575 | 2683 | 2736 | 2885 | 2457 | 2561 | 2542 | 2643.96 | 2768 | 2520 | 0.911 | -0.135 | 0.02445958 | 0.08007346 | 6.00E-04 | 0.0033 | 0.0022 | 0.0022 | TRUE | 0.0165 |
| Phf12 | 6373 | 4854 | 4039 | 3940 | 3639 | 4110 | 4950 | 5163 | 4358 | 3922 | 4105 | 4057 | 4425.84 | 4824 | 4028 | 0.835 | -0.26 | 0.00020796 | 0.00219362 | 0.0037 | 0.003 | 0.0033 | 0.0033 | TRUE | 0.0861 |
| Phf13 | 4570 | 3557 | 3701 | 3258 | 2990 | 3328 | 3549 | 3784 | 3993 | 3243 | 3373 | 3285 | 3537.92 | 3775 | 3300 | 0.875 | -0.193 | 0.00200184 | 0.01212386 | 0.0015 | 0.0031 | 0.0025 | 0.0025 | TRUE | 0.0341 |
| Phf14 | 2701 | 1815 | 1693 | 1688 | 1400 | 1694 | 2098 | 1931 | 1827 | 1680 | 1579 | 1672 | 1797.82 | 1952 | 1644 | 0.842 | -0.249 | 0.00069235 | 0.00553646 | 0.0026 | 0.0037 | 0.0033 | 0.0033 | TRUE | 0.0532 |
| Phf15 | 2053 | 1634 | 1456 | 1647 | 1528 | 1646 | 1594 | 1738 | 1571 | 1639 | 1724 | 1625 | 1648.6 | 1634 | 1663 | 1.017 | 0.025 | 0.72384482 | 0.83455196 | 0.0014 | 0.0039 | 0.0029 | 0.0029 | TRUE | 0.0363 |
| Phf16 | 2516 | 1766 | 1892 | 1890 | 1733 | 1889 | 1954 | 1878 | 2041 | 1881 | 1955 | 1865 | 1929.17 | 1958 | 1900 | 0.97 | -0.043 | 0.50257192 | 0.66680359 | 6.00E-04 | 0.0037 | 0.0025 | 0.0025 | TRUE | 0.0162 |
| Phf17 | 19127 | 13153 | 11341 | 11533 | 10005 | 11050 | 14855 | 13991 | 12237 | 11480 | 11287 | 10908 | 12459.55 | 13694 | 11225 | 0.82 | -0.287 | 5.65E-05 | 0.000773 | 0.005 | 0.0026 | 0.0036 | 0.0036 | TRUE | 0.1058 |
| Phf19 | 724 | 620 | 644 | 765 | 712 | 865 | 562 | 659 | 695 | 761 | 803 | 854 | 722.54 | 639 | 806 | 1.265 | 0.339 | 0.00074896 | 0.00586244 | 0.0063 | 0.0057 | 0.0059 | 0.0059 | TRUE | 0.1369 |
| Phf2 | 676 | 469 | 501 | 571 | 476 | 523 | 525 | 499 | 541 | 568 | 537 | 516 | 531.02 | 522 | 540 | 1.036 | 0.052 | 0.58971852 | 0.73845911 | 0 | 0.0069 | 0.0047 | 0.0047 | TRUE | 0.0243 |
| Phf20 | 8100 | 5894 | 6792 | 6696 | 6198 | 7244 | 6291 | 6269 | 7329 | 6665 | 6992 | 7151 | 6782.82 | 6630 | 6936 | 1.046 | 0.066 | 0.35362773 | 0.52792993 | 0.0042 | 0.0028 | 0.0034 | 0.0034 | TRUE | 0.1033 |
| Phf20l1 | 893 | 740 | 613 | 580 | 520 | 596 | 694 | 787 | 661 | 577 | 587 | 588 | 649.07 | 714 | 584 | 0.818 | -0.289 | 0.00241789 | 0.01400279 | 0.0028 | 0.0061 | 0.005 | 0.005 | TRUE | 0.0919 |
| Phf21a | 2577 | 1912 | 1948 | 2451 | 2147 | 2569 | 2001 | 2034 | 2102 | 2440 | 2422 | 2536 | 2255.81 | 2046 | 2466 | 1.206 | 0.27 | 7.02E-06 | 0.00014017 | 0 | 0.0035 | 0.0022 | 0.0022 | TRUE | 0.0075 |
| Phf21b | 282 | 235 | 217 | 200 | 204 | 234 | 219 | 250 | 234 | 199 | 230 | 231 | 227.22 | 234 | 220 | 0.94 | -0.089 | 0.52296666 | 0.68425141 | 0 | 0.0129 | 0.0095 | 0.0095 | TRUE | 0.0777 |
| Phf23 | 5883 | 4229 | 3763 | 3727 | 3221 | 3647 | 4569 | 4498 | 4060 | 3710 | 3634 | 3600 | 4011.88 | 4376 | 3648 | 0.833 | -0.263 | 2.69E-05 | 0.00042471 | 0.0017 | 0.003 | 0.0026 | 0.0026 | TRUE | 0.0481 |
| Phf3 | 12476 | 8790 | 8248 | 8311 | 7549 | 8900 | 9690 | 9350 | 8900 | 8273 | 8516 | 8786 | 8918.9 | 9313 | 8525 | 0.915 | -0.128 | 0.02004749 | 0.06929597 | 0.0012 | 0.0027 | 0.0021 | 0.0021 | TRUE | 0.0184 |
| Phf5a | 5300 | 3342 | 2973 | 3172 | 2805 | 3243 | 4116 | 3555 | 3208 | 3157 | 3164 | 3201 | 3400.34 | 3626 | 3174 | 0.875 | -0.193 | 0.02114006 | 0.07204457 | 0.0076 | 0.0031 | 0.0048 | 0.0048 | TRUE | 0.1718 |
| Phf6 | 2782 | 1733 | 1828 | 2124 | 1778 | 2067 | 2161 | 1843 | 1972 | 2114 | 2006 | 2040 | 2022.81 | 1992 | 2053 | 1.03 | 0.042 | 0.56512748 | 0.71958727 | 0.0031 | 0.0036 | 0.0034 | 0.0034 | TRUE | 0.068 |
| Phf7 | 758 | 494 | 449 | 524 | 456 | 482 | 589 | 525 | 484 | 522 | 514 | 476 | 518.41 | 533 | 504 | 0.943 | -0.085 | 0.42499705 | 0.59781097 | 0.0044 | 0.007 | 0.0062 | 0.0062 | TRUE | 0.1034 |
| Phf8 | 3325 | 2253 | 2477 | 2715 | 2534 | 2846 | 2582 | 2396 | 2673 | 2703 | 2859 | 2809 | 2670.35 | 2550 | 2790 | 1.094 | 0.129 | 0.04486471 | 0.12444233 | 0.0014 | 0.0033 | 0.0026 | 0.0026 | TRUE | 0.0332 |
| Phgdh | 5999 | 4823 | 4952 | 5142 | 4684 | 5366 | 4659 | 5130 | 5343 | 5118 | 5284 | 5297 | 5138.66 | 5044 | 5233 | 1.038 | 0.054 | 0.39774258 | 0.57100066 | 0.0022 | 0.0029 | 0.0027 | 0.0027 | TRUE | 0.0552 |
| Phgr1 | 0 | 0 | 0 | 0 | 0 | 0 | 0 | 0 | 0 | 0 | 0 | 0 | 0 | 0 | 0 | NA | NA | NA | NA | NA | NA | NA | NA | NA | NA |
| Phip | 5377 | 3942 | 3711 | 3878 | 3460 | 3995 | 4176 | 4193 | 4004 | 3860 | 3903 | 3944 | 4013.39 | 4124 | 3902 | 0.946 | -0.08 | 0.13010056 | 0.26686673 | 1.00E-04 | 0.003 | 0.0018 | 0.0018 | TRUE | 0.0078 |
| Phka1 | 1819 | 1307 | 1426 | 1524 | 1399 | 1549 | 1413 | 1390 | 1539 | 1517 | 1578 | 1529 | 1494.32 | 1447 | 1541 | 1.065 | 0.091 | 0.19396641 | 0.35280757 | 0.001 | 0.004 | 0.0029 | 0.0029 | TRUE | 0.0358 |
| Phka2 | 1416 | 1077 | 990 | 1127 | 1016 | 1101 | 1100 | 1146 | 1068 | 1122 | 1146 | 1087 | 1111.39 | 1105 | 1118 | 1.012 | 0.018 | 0.8101348 | 0.89056361 | 0 | 0.0046 | 0.003 | 0.003 | TRUE | 0.0122 |
| Phkb | 3180 | 2189 | 2017 | 2603 | 2267 | 2388 | 2470 | 2328 | 2176 | 2591 | 2557 | 2357 | 2413.38 | 2325 | 2502 | 1.075 | 0.105 | 0.14055462 | 0.28239348 | 0.0026 | 0.0034 | 0.0032 | 0.0032 | TRUE | 0.0377 |
| Phkg1 | 251 | 170 | 162 | 218 | 184 | 211 | 195 | 181 | 175 | 217 | 208 | 208 | 197.24 | 184 | 211 | 1.147 | 0.198 | 0.16337599 | 0.31398857 | 0 | 0.0144 | 0.0094 | 0.0094 | TRUE | 0.0351 |
| Phkg2 | 1519 | 1217 | 1227 | 1460 | 1273 | 1439 | 1180 | 1295 | 1324 | 1453 | 1436 | 1421 | 1351.34 | 1266 | 1437 | 1.136 | 0.184 | 0.01162111 | 0.04619035 | 0.0013 | 0.0042 | 0.0031 | 0.0031 | TRUE | 0.0448 |
| Phlda1 | 328 | 239 | 176 | 177 | 143 | 175 | 255 | 254 | 190 | 176 | 161 | 173 | 201.52 | 233 | 170 | 0.729 | -0.456 | 0.00408576 | 0.02098376 | 0.0105 | 0.0142 | 0.0131 | 0.0131 | TRUE | 0.2781 |
| Phlda2 | 1141 | 812 | 859 | 1541 | 1200 | 1538 | 886 | 864 | 927 | 1534 | 1354 | 1518 | 1180.44 | 892 | 1469 | 1.647 | 0.72 | 1.26E-19 | 3.83E-17 | 0.0022 | 0.0044 | 0.0037 | 0.0037 | TRUE | 0.0554 |
| Phlda3 | 2614 | 1799 | 2018 | 2145 | 1842 | 2066 | 2030 | 1914 | 2177 | 2135 | 2078 | 2039 | 2062.29 | 2040 | 2084 | 1.022 | 0.031 | 0.65344617 | 0.7847833 | 0.0018 | 0.0036 | 0.0029 | 0.0029 | TRUE | 0.0406 |
| Phldb1 | 2156 | 1642 | 1494 | 1660 | 1496 | 1718 | 1674 | 1747 | 1612 | 1652 | 1688 | 1696 | 1678.17 | 1678 | 1679 | 1.001 | 0.001 | 0.99066489 | 0.99533061 | 0 | 0.0038 | 0.0025 | 0.0025 | TRUE | 0.015 |
| Phldb2 | 503 | 326 | 270 | 335 | 277 | 282 | 391 | 347 | 291 | 333 | 312 | 278 | 325.51 | 343 | 308 | 0.895 | -0.159 | 0.24539459 | 0.4137578 | 0.0121 | 0.0097 | 0.0105 | 0.0105 | TRUE | 0.1944 |
| Phldb3 | 128 | 127 | 148 | 207 | 178 | 222 | 99 | 135 | 160 | 206 | 201 | 219 | 170.03 | 131 | 209 | 1.601 | 0.679 | 0.0001961 | 0.00209952 | 0.0214 | 0.0164 | 0.0178 | 0.0178 | TRUE | 0.5206 |
| Phlpp1 | 1192 | 906 | 842 | 778 | 798 | 901 | 926 | 964 | 909 | 774 | 900 | 889 | 893.68 | 933 | 854 | 0.916 | -0.126 | 0.14331549 | 0.28621047 | 0.0026 | 0.0051 | 0.0043 | 0.0043 | TRUE | 0.081 |
| Phlpp2 | 1486 | 1080 | 1224 | 1224 | 988 | 1095 | 1154 | 1149 | 1321 | 1218 | 1115 | 1081 | 1172.91 | 1208 | 1138 | 0.943 | -0.085 | 0.3207849 | 0.49465731 | 0.004 | 0.0044 | 0.0044 | 0.0044 | TRUE | 0.0787 |
| Phospho1 | 1778 | 1156 | 1312 | 1621 | 1273 | 1504 | 1381 | 1230 | 1416 | 1614 | 1436 | 1485 | 1426.74 | 1342 | 1512 | 1.126 | 0.171 | 0.03377977 | 0.10121893 | 0.0035 | 0.0041 | 0.004 | 0.004 | TRUE | 0.0632 |

| | | | | | | | | | | | | | | | | | | | | | | | | | |
|---|---|---|---|---|---|---|---|---|---|---|---|---|---|---|---|---|---|---|---|---|---|---|---|---|---|
| Phospho2 | 930 | 615 | 514 | 575 | 552 | 628 | 722 | 654 | 555 | 572 | 623 | 620 | 624.34 | 644 | 605 | 0.938 | -0.093 | 0.39294142 | 0.5665031 | 0.0085 | 0.0062 | 0.0069 | 0.0069 | TRUE | 0.1705 |
| Phox2a | 173 | 145 | 148 | 177 | 146 | 185 | 134 | 154 | 160 | 176 | 165 | 183 | 161.97 | 149 | 175 | 1.173 | 0.23 | 0.14801978 | 0.29281542 | 0 | 0.0171 | 0.0121 | 0.0121 | TRUE | 0.0911 |
| Phox2b | 1 | 2 | 0 | 0 | 0 | 0 | 1 | 2 | 0 | 0 | 0 | 0 | 0.48 | 1 | 0 | 0.197 | -2.343 | 0.43249452 | NA | 0 | 4.8992 | 4.9172 | 4.9172 | TRUE | 0.1226 |
| Phpt1 | 963 | 706 | 757 | 934 | 790 | 855 | 748 | 751 | 817 | 930 | 891 | 844 | 830.1 | 772 | 888 | 1.152 | 0.204 | 0.01555441 | 0.05738751 | 0.0012 | 0.0053 | 0.0039 | 0.0039 | TRUE | 0.0299 |
| Phrf1 | 9398 | 7043 | 6462 | 6676 | 6106 | 6881 | 7299 | 7492 | 6972 | 6645 | 6888 | 6793 | 7014.85 | 7254 | 6775 | 0.934 | -0.099 | 0.05957445 | 0.15254348 | 6.00E-04 | 0.0028 | 0.0018 | 0.0018 | TRUE | 0.0139 |
| Phtf1 | 1752 | 1338 | 1226 | 1265 | 1080 | 1212 | 1361 | 1423 | 1323 | 1259 | 1218 | 1196 | 1296.79 | 1369 | 1224 | 0.895 | -0.16 | 0.02252254 | 0.07523129 | 0 | 0.0043 | 0.0028 | 0.0028 | TRUE | 0.014 |
| Phtf2 | 4334 | 3097 | 2730 | 3013 | 2658 | 3089 | 3366 | 3294 | 2946 | 2999 | 2998 | 3049 | 3108.81 | 3202 | 3015 | 0.941 | -0.087 | 0.18706373 | 0.34388449 | 0.002 | 0.0032 | 0.0028 | 0.0028 | TRUE | 0.059 |
| Phxr4 | 0 | 0 | 0 | 0 | 0 | 0 | 0 | 0 | 0 | 0 | 0 | 0 | 0 | 0 | 0 | NA | NA | NA | NA | NA | NA | NA | NA | NA | NA |
| Phyh | 2446 | 1893 | 2221 | 2483 | 2360 | 2737 | 1900 | 2014 | 2396 | 2472 | 2662 | 2702 | 2357.57 | 2103 | 2612 | 1.243 | 0.314 | 0.00031654 | 0.00302829 | 0.0081 | 0.0034 | 0.005 | 0.005 | TRUE | 0.1791 |
| Phyhd1 | 52 | 37 | 47 | 52 | 46 | 71 | 40 | 39 | 51 | 52 | 52 | 70 | 50.7 | 43 | 58 | 1.338 | 0.42 | 0.13733025 | 0.27763243 | 0 | 0.0492 | 0.0375 | 0.0375 | TRUE | 0.2942 |
| Phyhip | 586 | 494 | 568 | 694 | 609 | 646 | 455 | 525 | 613 | 691 | 687 | 638 | 601.5 | 531 | 672 | 1.268 | 0.343 | 0.00252201 | 0.01445935 | 0.0103 | 0.0064 | 0.0076 | 0.0076 | TRUE | 0.2113 |
| Phyhipl | 581 | 418 | 400 | 467 | 397 | 450 | 451 | 445 | 432 | 465 | 448 | 444 | 447.4 | 443 | 452 | 1.022 | 0.031 | 0.75478519 | 0.8545742 | 0 | 0.0077 | 0.0048 | 0.0048 | TRUE | 0.0069 |
| Pi15 | 1 | 2 | 0 | 1 | 5 | 2 | 1 | 2 | 0 | 1 | 6 | 2 | 1.92 | 1 | 3 | 2.955 | 1.563 | 0.30421589 | NA | 0 | 1.2374 | 0.9849 | 0.9849 | TRUE | 0.7974 |
| Pi16 | 14 | 5 | 17 | 10 | 19 | 21 | 11 | 5 | 18 | 10 | 21 | 21 | 14.44 | 11 | 17 | 1.51 | 0.594 | 0.29901858 | NA | 0.157 | 0.1665 | 0.1641 | 0.1641 | TRUE | 0.3805 |
| Pi4k2a | 3976 | 3009 | 2810 | 2973 | 2604 | 3110 | 3088 | 3201 | 3032 | 2959 | 2938 | 3070 | 3047.92 | 3107 | 2989 | 0.962 | -0.056 | 0.32966751 | 0.5041123 | 3.00E-04 | 0.0032 | 0.002 | 0.002 | TRUE | 0.0083 |
| Pi4k2b | 3776 | 2821 | 2916 | 3180 | 2750 | 3188 | 2933 | 3001 | 3146 | 3165 | 3102 | 3147 | 3082.39 | 3027 | 3138 | 1.037 | 0.053 | 0.3534083 | 0.52780934 | 4.00E-04 | 0.0032 | 0.002 | 0.002 | TRUE | 0.0145 |
| Pi4ka | 8183 | 6024 | 6387 | 7515 | 6947 | 7773 | 6355 | 6408 | 6892 | 7480 | 7837 | 7673 | 7107.51 | 6552 | 7663 | 1.17 | 0.226 | 4.02E-05 | 0.00059032 | 0.0011 | 0.0028 | 0.002 | 0.002 | TRUE | 0.025 |
| Pi4kb | 2361 | 1619 | 1357 | 1453 | 1343 | 1495 | 1834 | 1722 | 1464 | 1446 | 1515 | 1476 | 1576.19 | 1673 | 1479 | 0.883 | -0.18 | 0.0380112 | 0.11026877 | 0.0057 | 0.0039 | 0.0048 | 0.0048 | TRUE | 0.1429 |
| Pianp | 978 | 847 | 980 | 711 | 719 | 928 | 760 | 901 | 1057 | 708 | 811 | 916 | 858.81 | 906 | 812 | 0.897 | -0.156 | 0.21060419 | 0.37259136 | 0.02 | 0.0052 | 0.01 | 0.01 | TRUE | 0.1897 |
| Pias1 | 2654 | 1848 | 1941 | 2143 | 1883 | 2200 | 2061 | 1966 | 2094 | 2133 | 2124 | 2172 | 2091.73 | 2040 | 2143 | 1.05 | 0.071 | 0.24647003 | 0.41513035 | 0 | 0.0036 | 0.0022 | 0.0022 | TRUE | 0.0121 |
| Pias2 | 10974 | 7805 | 8108 | 8339 | 7369 | 8418 | 8523 | 8302 | 8748 | 8301 | 8313 | 8310 | 8416.18 | 8524 | 8308 | 0.975 | -0.037 | 0.44059462 | 0.61193459 | 2.00E-04 | 0.0027 | 0.0016 | 0.0016 | TRUE | 0.0064 |
| Pias3 | 1003 | 755 | 877 | 1010 | 846 | 1003 | 779 | 803 | 946 | 1005 | 954 | 990 | 913.03 | 843 | 983 | 1.169 | 0.225 | 0.01384644 | 0.0526629 | 0.0045 | 0.005 | 0.0049 | 0.0049 | TRUE | 0.1351 |
| Pias4 | 3312 | 2612 | 2720 | 2802 | 2548 | 2995 | 2572 | 2778 | 2935 | 2789 | 2874 | 2957 | 2817.59 | 2762 | 2873 | 1.041 | 0.058 | 0.38541404 | 0.55931799 | 0.0021 | 0.0033 | 0.0029 | 0.0029 | TRUE | 0.0449 |
| Pibf1 | 800 | 613 | 598 | 525 | 451 | 526 | 621 | 652 | 645 | 523 | 509 | 519 | 578.2 | 639 | 517 | 0.809 | -0.306 | 0.0006404 | 0.00521479 | 0 | 0.0065 | 0.004 | 0.004 | TRUE | 0.0078 |
| Picalm | 12272 | 8850 | 9002 | 9324 | 8106 | 9270 | 9531 | 9414 | 9713 | 9281 | 9144 | 9151 | 9372.38 | 9553 | 9192 | 0.962 | -0.055 | 0.22989592 | 0.39555825 | 0 | 0.0027 | 0.0014 | 0.0014 | TRUE | 0.0026 |
| Pick1 | 2818 | 2039 | 1833 | 1925 | 1713 | 1956 | 2189 | 2169 | 1978 | 1916 | 1932 | 1931 | 2019.12 | 2112 | 1926 | 0.912 | -0.133 | 0.0430112 | 0.12075409 | 0.0011 | 0.0036 | 0.0026 | 0.0026 | TRUE | 0.0367 |
| Pid1 | 534 | 358 | 367 | 343 | 307 | 346 | 415 | 381 | 396 | 341 | 346 | 342 | 370.14 | 397 | 343 | 0.863 | -0.213 | 0.04792913 | 0.13070958 | 0 | 0.0088 | 0.0057 | 0.0057 | TRUE | 0.0185 |
| Piezo1 | 12595 | 8620 | 7644 | 10172 | 8904 | 9982 | 9782 | 9169 | 8248 | 10125 | 10045 | 9854 | 9537.07 | 9066 | 10008 | 1.104 | 0.142 | 0.03222134 | 0.09755127 | 0.0036 | 0.0027 | 0.0031 | 0.0031 | TRUE | 0.076 |
| Piezo2 | 154 | 126 | 95 | 109 | 94 | 103 | 120 | 134 | 103 | 108 | 106 | 102 | 112.06 | 119 | 105 | 0.887 | -0.172 | 0.36538834 | 0.54007193 | 0 | 0.0236 | 0.0172 | 0.0172 | TRUE | 0.1332 |
| Pif1 | 2140 | 1694 | 2020 | 2271 | 2017 | 2344 | 1662 | 1802 | 2180 | 2261 | 2275 | 2314 | 2082.22 | 1881 | 2283 | 1.215 | 0.281 | 0.00210242 | 0.01254367 | 0.0095 | 0.0036 | 0.0055 | 0.0055 | TRUE | 0.2318 |
| Pifo | 30 | 22 | 24 | 29 | 28 | 28 | 23 | 23 | 26 | 29 | 32 | 28 | 26.78 | 24 | 30 | 1.214 | 0.28 | 0.4428079 | 0.61381566 | 0 | 0.0909 | 0.0582 | 0.0582 | TRUE | 0.0266 |
| Piga | 2391 | 1952 | 2001 | 2123 | 1811 | 2146 | 1857 | 2076 | 2159 | 2113 | 2043 | 2118 | 2061.17 | 2031 | 2091 | 1.031 | 0.044 | 0.540211 | 0.69850153 | 0.0028 | 0.0036 | 0.0033 | 0.0033 | TRUE | 0.0699 |
| Pigb | 531 | 429 | 407 | 366 | 315 | 377 | 412 | 456 | 439 | 364 | 355 | 372 | 399.95 | 436 | 364 | 0.836 | -0.258 | 0.01496457 | 0.05574369 | 0 | 0.0084 | 0.0055 | 0.0055 | TRUE | 0.0279 |
| Pigc | 232 | 150 | 162 | 187 | 167 | 205 | 180 | 160 | 175 | 186 | 188 | 202 | 181.91 | 172 | 192 | 1.12 | 0.163 | 0.27215 | 0.44316998 | 0 | 0.0155 | 0.0104 | 0.0104 | TRUE | 0.0386 |
| Pigf | 1388 | 918 | 1011 | 1070 | 875 | 1086 | 1078 | 976 | 1091 | 1065 | 987 | 1072 | 1044.92 | 1048 | 1041 | 0.993 | -0.01 | 0.90321865 | 0.94645925 | 0.0018 | 0.0047 | 0.0037 | 0.0037 | TRUE | 0.0421 |
| Pigg | 1866 | 1384 | 1470 | 1736 | 1484 | 1646 | 1449 | 1472 | 1586 | 1728 | 1674 | 1625 | 1589.08 | 1502 | 1676 | 1.116 | 0.158 | 0.0223738 | 0.07480439 | 0.001 | 0.0039 | 0.0028 | 0.0028 | TRUE | 0.0281 |
| Pigh | 1056 | 815 | 848 | 896 | 694 | 877 | 820 | 867 | 915 | 892 | 783 | 866 | 857.09 | 867 | 847 | 0.978 | -0.032 | 0.71567918 | 0.82931903 | 0.0026 | 0.0052 | 0.0043 | 0.0043 | TRUE | 0.0506 |
| Pigk | 1802 | 1181 | 1264 | 1259 | 1102 | 1295 | 1400 | 1256 | 1364 | 1253 | 1243 | 1278 | 1299.06 | 1340 | 1258 | 0.938 | -0.092 | 0.20801426 | 0.36965657 | 0.001 | 0.0043 | 0.0031 | 0.0031 | TRUE | 0.0354 |
| Pigl | 2134 | 1397 | 927 | 1032 | 903 | 1037 | 1657 | 1486 | 1000 | 1027 | 1019 | 1024 | 1202.2 | 1381 | 1023 | 0.74 | -0.434 | 0.00155623 | 0.00992112 | 0.0327 | 0.0044 | 0.0127 | 0.0127 | TRUE | 0.6849 |
| Pigm | 2182 | 1711 | 1655 | 1861 | 1533 | 1839 | 1695 | 1820 | 1786 | 1852 | 1729 | 1815 | 1782.93 | 1767 | 1799 | 1.019 | 0.028 | 0.67755166 | 0.80217909 | 8.00E-04 | 0.0038 | 0.0026 | 0.0026 | TRUE | 0.016 |
| Pign | 2231 | 1571 | 1513 | 1734 | 1547 | 1628 | 1733 | 1671 | 1633 | 1726 | 1745 | 1607 | 1685.76 | 1679 | 1693 | 1.007 | 0.011 | 0.87398204 | 0.93005063 | 9.00E-04 | 0.0038 | 0.0027 | 0.0027 | TRUE | 0.0238 |
| Pigo | 3133 | 2365 | 2414 | 2611 | 2461 | 2666 | 2433 | 2516 | 2605 | 2599 | 2776 | 2632 | 2593.43 | 2518 | 2669 | 1.06 | 0.084 | 0.16880625 | 0.32046058 | 8.00E-04 | 0.0033 | 0.0023 | 0.0023 | TRUE | 0.0148 |
| Pigp | 1182 | 789 | 893 | 1158 | 966 | 1030 | 918 | 839 | 964 | 1153 | 1090 | 1017 | 996.66 | 907 | 1087 | 1.198 | 0.26 | 0.00243052 | 0.01407021 | 0.0033 | 0.0048 | 0.0043 | 0.0043 | TRUE | 0.049 |
| Pigq | 4387 | 3267 | 3125 | 3202 | 2813 | 3260 | 3407 | 3475 | 3372 | 3187 | 3173 | 3218 | 3305.48 | 3418 | 3193 | 0.934 | -0.098 | 0.06474479 | 0.16223655 | 0 | 0.0031 | 0.0017 | 0.0017 | TRUE | 0.0025 |
| Pigr | 2 | 0 | 0 | 0 | 1 | 0 | 2 | 0 | 0 | 0 | 1 | 0 | 0.45 | 1 | 0 | 0.68 | -0.556 | 0.85993441 | NA | 0 | 5.3057 | 5.2576 | 5.2576 | TRUE | 0.9726 |
| Pigs | 6469 | 4864 | 4910 | 5456 | 5033 | 5747 | 5024 | 5174 | 5298 | 5431 | 5678 | 5673 | 5379.61 | 5165 | 5594 | 1.083 | 0.115 | 0.02900052 | 0.09032564 | 5.00E-04 | 0.0029 | 0.0018 | 0.0018 | TRUE | 0.008 |
| Pigt | 6273 | 4793 | 4764 | 5388 | 4973 | 5612 | 4872 | 5098 | 5140 | 5363 | 5610 | 5540 | 5270.62 | 5037 | 5504 | 1.093 | 0.128 | 0.0154125 | 0.05699634 | 5.00E-04 | 0.0029 | 0.0018 | 0.0018 | TRUE | 0.0102 |
| Pigu | 1810 | 1338 | 1308 | 1528 | 1347 | 1625 | 1406 | 1423 | 1411 | 1521 | 1520 | 1604 | 1480.82 | 1413 | 1548 | 1.096 | 0.132 | 0.04557047 | 0.12588814 | 0 | 0.004 | 0.0025 | 0.0025 | TRUE | 0.0121 |
| Pigv | 1039 | 841 | 867 | 1040 | 961 | 1045 | 807 | 895 | 935 | 1035 | 1084 | 1032 | 964.65 | 879 | 1050 | 1.197 | 0.259 | 0.00188663 | 0.01156823 | 0.0022 | 0.0049 | 0.004 | 0.004 | TRUE | 0.0646 |
| Pigw | 433 | 342 | 355 | 428 | 347 | 418 | 336 | 364 | 383 | 426 | 391 | 413 | 385.54 | 361 | 410 | 1.139 | 0.188 | 0.08775526 | 0.20122712 | 0 | 0.0086 | 0.0061 | 0.0061 | TRUE | 0.0446 |
| Pigx | 1722 | 1225 | 1485 | 1642 | 1485 | 1696 | 1337 | 1303 | 1602 | 1634 | 1675 | 1674 | 1537.77 | 1414 | 1661 | 1.176 | 0.233 | 0.00620608 | 0.02893133 | 0.0058 | 0.004 | 0.0046 | 0.0046 | TRUE | 0.1602 |
| Pigyl | 710 | 515 | 684 | 882 | 781 | 852 | 551 | 548 | 738 | 878 | 881 | 841 | 739.55 | 612 | 867 | 1.418 | 0.503 | 1.04E-05 | 0.00019312 | 0.014 | 0.0056 | 0.008 | 0.008 | TRUE | 0.372 |
| Pigz | 2 | 2 | 2 | 0 | 3 | 4 | 2 | 2 | 2 | 0 | 3 | 4 | 2.2 | 2 | 2 | 1.26 | 0.333 | 0.80434592 | NA | 0 | 1.082 | 0.8411 | 0.8411 | TRUE | 0.8571 |
| Pih1d1 | 1439 | 1126 | 1039 | 1054 | 966 | 1087 | 1118 | 1198 | 1121 | 1049 | 1090 | 1073 | 1108.05 | 1146 | 1071 | 0.935 | -0.097 | 0.18552123 | 0.34182093 | 0 | 0.0046 | 0.003 | 0.003 | TRUE | 0.0187 |
| Pih1d2 | 54 | 36 | 28 | 17 | 29 | 31 | 42 | 38 | 30 | 17 | 33 | 31 | 31.78 | 37 | 27 | 0.721 | -0.472 | 0.19985244 | 0.36048598 | 0.03 | 0.077 | 0.0652 | 0.0652 | TRUE | 0.6658 |
| Pih1d3 | 0 | 0 | 0 | 0 | 0 | 0 | 0 | 0 | 0 | 0 | 0 | 0 | 0 | 0 | 0 | NA | NA | NA | NA | NA | NA | NA | NA | NA | NA |
| Pik3ap1 | 99 | 81 | 75 | 86 | 93 | 76 | 77 | 86 | 81 | 86 | 105 | 75 | 84.92 | 81 | 89 | 1.087 | 0.121 | 0.58262524 | 0.73263262 | 0 | 0.0303 | 0.0231 | 0.0231 | TRUE | 0.2382 |
| Pik3c2a | 5425 | 3792 | 3854 | 4411 | 3802 | 4432 | 4213 | 4034 | 4158 | 4391 | 4289 | 4375 | 4243.35 | 4135 | 4352 | 1.052 | 0.073 | 0.15894058 | 0.30807401 | 0 | 0.003 | 0.0017 | 0.0017 | TRUE | 0.0056 |
| Pik3c2b | 1894 | 1447 | 1457 | 1528 | 1437 | 1534 | 1471 | 1539 | 1572 | 1521 | 1621 | 1514 | 1539.76 | 1527 | 1552 | 1.017 | 0.024 | 0.73002577 | 0.83876674 | 7.00E-04 | 0.004 | 0.0027 | 0.0027 | TRUE | 0.0179 |
| Pik3c2g | 9 | 13 | 14 | 10 | 7 | 3 | 7 | 14 | 15 | 10 | 8 | 3 | 9.46 | 12 | 7 | 0.585 | -0.774 | 0.25313989 | NA | 0.112 | 0.2531 | 0.2152 | 0.2152 | TRUE | 0.6976 |
| Pik3c3 | 3002 | 2229 | 2314 | 2260 | 1966 | 2434 | 2332 | 2371 | 2497 | 2250 | 2218 | 2403 | 2344.91 | 2400 | 2290 | 0.955 | -0.067 | 0.29903526 | 0.47133998 | 0.0011 | 0.0034 | 0.0025 | 0.0025 | TRUE | 0.0226 |
| Pik3ca | 2986 | 2138 | 2087 | 2268 | 2079 | 2288 | 2319 | 2274 | 2252 | 2258 | 2345 | 2259 | 2284.44 | 2282 | 2287 | 1.002 | 0.003 | 0.96283312 | 0.97947032 | 0 | 0.0035 | 0.002 | 0.002 | TRUE | 0.0059 |
| Pik3cb | 4569 | 3430 | 3287 | 3194 | 2886 | 3365 | 3549 | 3648 | 3547 | 3179 | 3256 | 3322 | 3416.74 | 3581 | 3252 | 0.908 | -0.139 | 0.01050259 | 0.04279176 | 0 | 0.0031 | 0.0018 | 0.0018 | TRUE | 0.0047 |
| Pik3cd | 2611 | 2102 | 2285 | 2928 | 2777 | 3139 | 2028 | 2236 | 2465 | 2915 | 3133 | 3099 | 2645.86 | 2243 | 3049 | 1.36 | 0.444 | 1.12E-08 | 5.51E-07 | 0.0049 | 0.0033 | 0.004 | 0.004 | TRUE | 0.0911 |
| Pik3cg | 10 | 4 | 4 | 0 | 3 | 3 | 8 | 4 | 4 | 0 | 3 | 3 | 3.78 | 5 | 2 | 0.379 | -1.398 | 0.19582743 | NA | 0 | 0.6293 | 0.5062 | 0.5062 | TRUE | 0.7549 |
| Pik3ip1 | 219 | 175 | 267 | 299 | 241 | 322 | 170 | 186 | 288 | 298 | 272 | 318 | 255.28 | 215 | 296 | 1.384 | 0.469 | 0.00959742 | 0.04018041 | 0.0393 | 0.0117 | 0.0196 | 0.0196 | TRUE | 0.962 |
| Pik3r1 | 988 | 742 | 630 | 793 | 698 | 721 | 767 | 789 | 680 | 789 | 787 | 712 | 754.14 | 745 | 763 | 1.022 | 0.032 | 0.73175581 | 0.83929626 | 0.0034 | 0.0056 | 0.0048 | 0.0048 | TRUE | 0.068 |
| Pik3r2 | 3468 | 2580 | 1965 | 1780 | 1803 | 1986 | 2693 | 2744 | 2120 | 1772 | 2034 | 1960 | 2220.71 | 2519 | 1922 | 0.762 | -0.391 | 5.71E-05 | 0.00077758 | 0.0112 | 0.0035 | 0.0063 | 0.0063 | TRUE | 0.231 |
| Pik3r3 | 636 | 394 | 286 | 282 | 254 | 307 | 494 | 419 | 309 | 281 | 287 | 303 | 348.66 | 407 | 290 | 0.71 | -0.494 | 0.00152098 | 0.00975698 | 0.0263 | 0.0092 | 0.0145 | 0.0145 | TRUE | 0.2318 |

| | | | | | | | | | | | | | | | | | | | | | | | | | |
|---|---|---|---|---|---|---|---|---|---|---|---|---|---|---|---|---|---|---|---|---|---|---|---|---|---|
| Pik3r4 | 3935 | 2793 | 2467 | 2307 | 2285 | 2430 | 3056 | 2971 | 2662 | 2296 | 2578 | 2399 | 2660.3 | 2896 | 2424 | 0.836 | -0.258 | 0.00048603 | 0.00422556 | 0.0037 | 0.0033 | 0.0036 | 0.0036 | TRUE | 0.0604 |
| Pik3r5 | 689 | 480 | 501 | 697 | 566 | 701 | 535 | 511 | 541 | 694 | 639 | 692 | 601.76 | 529 | 675 | 1.277 | 0.352 | 0.0001058 | 0.00126751 | 0 | 0.0064 | 0.0043 | 0.0043 | TRUE | 0.0254 |
| Pik3r6 | 33 | 31 | 25 | 37 | 27 | 32 | 26 | 33 | 27 | 37 | 30 | 32 | 30.74 | 29 | 33 | 1.161 | 0.216 | 0.53255485 | 0.69217119 | 0 | 0.0795 | 0.0535 | 0.0535 | TRUE | 0.1156 |
| Pikfyve | 6036 | 4451 | 4759 | 5877 | 4982 | 5855 | 4688 | 4734 | 5135 | 5850 | 5620 | 5780 | 5301.21 | 4852 | 5750 | 1.185 | 0.245 | 1.98E-05 | 0.00032868 | 0.0012 | 0.0029 | 0.0022 | 0.0022 | TRUE | 0.0314 |
| Pilra | 0 | 0 | 0 | 1 | 0 | 0 | 0 | 0 | 0 | 1 | 0 | 0 | 0.17 | 0 | 0 | 2.094 | 1.067 | 0.79379329 | NA | 0 | 14.2885 | 10 | 10 | TRUE | 0.3906 |
| Pilrb1 | 0 | 0 | 0 | 0 | 0 | 0 | 0 | 0 | 0 | 0 | 0 | 0 | 0 | 0 | 0 | NA | NA | NA | NA | NA | NA | NA | NA | NA | NA |
| Pilrb2 | 0 | 0 | 0 | 0 | 0 | 0 | 0 | 0 | 0 | 0 | 0 | 0 | 0 | 0 | 0 | NA | NA | NA | NA | NA | NA | NA | NA | NA | NA |
| Pim1 | 6068 | 4586 | 5355 | 6008 | 5127 | 6104 | 4713 | 4878 | 5778 | 5980 | 5784 | 6026 | 5526.43 | 5123 | 5930 | 1.158 | 0.212 | 0.00570247 | 0.02714754 | 0.0062 | 0.0029 | 0.004 | 0.004 | TRUE | 0.152 |
| Pim2 | 887 | 606 | 771 | 783 | 650 | 827 | 689 | 645 | 832 | 779 | 733 | 816 | 749.07 | 722 | 776 | 1.077 | 0.107 | 0.30850681 | 0.48117501 | 0.0083 | 0.0056 | 0.0066 | 0.0066 | TRUE | 0.2053 |
| Pim3 | 10047 | 8323 | 10243 | 13669 | 12498 | 14545 | 7803 | 8853 | 11052 | 13606 | 14099 | 14358 | 11628.59 | 9236 | 14021 | 1.518 | 0.603 | 9.22E-10 | 6.25E-08 | 0.0163 | 0.0026 | 0.0069 | 0.0069 | TRUE | 0.3613 |
| Pin1 | 4784 | 3434 | 2699 | 2837 | 2474 | 2701 | 3716 | 3653 | 2912 | 2824 | 2791 | 2666 | 3093.6 | 3427 | 2760 | 0.805 | -0.313 | 0.0003814 | 0.00350865 | 0.0085 | 0.0032 | 0.0053 | 0.0053 | TRUE | 0.2085 |
| Pin1-ps1 | 1 | 1 | 1 | 0 | 0 | 0 | 1 | 1 | 1 | 0 | 0 | 0 | 0.49 | 1 | 0 | 0.196 | -2.347 | 0.38405676 | NA | 0 | 4.8735 | 3.7415 | 3.7415 | TRUE | 0.0718 |
| Pin4 | 717 | 487 | 415 | 490 | 398 | 461 | 557 | 518 | 448 | 488 | 449 | 455 | 485.74 | 508 | 464 | 0.912 | -0.133 | 0.22804416 | 0.39336499 | 0.0052 | 0.0073 | 0.0066 | 0.0066 | TRUE | 0.1216 |
| Pinc | 0 | 0 | 0 | 0 | 0 | 0 | 0 | 0 | 0 | 0 | 0 | 0 | 0 | 0 | 0 | NA | NA | NA | NA | NA | NA | NA | NA | NA | NA |
| Pink1 | 1400 | 1099 | 1200 | 1279 | 1238 | 1384 | 1087 | 1169 | 1295 | 1273 | 1397 | 1366 | 1264.51 | 1184 | 1345 | 1.138 | 0.186 | 0.0264571 | 0.08469995 | 0.0039 | 0.0043 | 0.0043 | 0.0043 | TRUE | 0.0805 |
| Pinlyp | 2 | 0 | 2 | 0 | 1 | 1 | 2 | 0 | 2 | 0 | 1 | 1 | 0.97 | 1 | 1 | 0.56 | -0.836 | 0.67994454 | NA | 0 | 2.443 | 1.8311 | 1.8311 | TRUE | 0.3957 |
| Pinx1 | 997 | 681 | 567 | 612 | 510 | 619 | 774 | 724 | 612 | 609 | 575 | 611 | 651.01 | 703 | 598 | 0.849 | -0.235 | 0.02327176 | 0.07725745 | 0.0065 | 0.0061 | 0.0062 | 0.0062 | TRUE | 0.1513 |
| Pion | 49 | 43 | 47 | 55 | 41 | 47 | 38 | 46 | 51 | 55 | 46 | 46 | 46.98 | 45 | 49 | 1.103 | 0.142 | 0.61850956 | 0.75908957 | 0 | 0.0529 | 0.0371 | 0.0371 | TRUE | 0.1626 |
| Pip | 0 | 0 | 0 | 0 | 0 | 0 | 0 | 0 | 0 | 0 | 0 | 0 | 0 | 0 | 0 | NA | NA | NA | NA | NA | NA | NA | NA | NA | NA |
| Pip4k2a | 1379 | 1161 | 1114 | 1145 | 1079 | 1252 | 1071 | 1235 | 1202 | 1140 | 1217 | 1236 | 1183.47 | 1169 | 1198 | 1.026 | 0.037 | 0.65382364 | 0.7851047 | 0.003 | 0.0044 | 0.0039 | 0.0039 | TRUE | 0.0677 |
| Pip4k2b | 8604 | 6015 | 5896 | 6490 | 5742 | 6458 | 6682 | 6398 | 6362 | 6460 | 6478 | 6375 | 6459.15 | 6481 | 6438 | 0.993 | -0.01 | 0.84038872 | 0.90907815 | 3.00E-04 | 0.0028 | 0.0016 | 0.0016 | TRUE | 0.0092 |
| Pip4k2c | 3818 | 2951 | 3111 | 2963 | 2629 | 3194 | 2965 | 3139 | 3357 | 2949 | 2966 | 3153 | 3088.18 | 3154 | 3023 | 0.959 | -0.06 | 0.36516545 | 0.53990999 | 0.0021 | 0.0032 | 0.0029 | 0.0029 | TRUE | 0.0385 |
| Pip5k1a | 1779 | 1217 | 1040 | 1089 | 966 | 1001 | 1382 | 1295 | 1122 | 1084 | 1090 | 988 | 1160.04 | 1266 | 1054 | 0.831 | -0.267 | 0.00340082 | 0.01826677 | 0.0063 | 0.0045 | 0.0051 | 0.0051 | TRUE | 0.1176 |
| Pip5k1b | 428 | 304 | 311 | 301 | 307 | 357 | 332 | 323 | 336 | 300 | 346 | 352 | 331.62 | 330 | 333 | 1.006 | 0.009 | 0.93733752 | 0.9651585 | 0 | 0.0096 | 0.007 | 0.007 | TRUE | 0.088 |
| Pip5k1c | 3048 | 2318 | 2000 | 2180 | 2106 | 2345 | 2367 | 2466 | 2158 | 2170 | 2376 | 2315 | 2308.58 | 2330 | 2287 | 0.981 | -0.028 | 0.69808795 | 0.81651154 | 0.0029 | 0.0035 | 0.0032 | 0.0032 | TRUE | 0.0497 |
| Pip5kl1 | 103 | 74 | 92 | 96 | 85 | 86 | 80 | 79 | 99 | 96 | 96 | 85 | 89.05 | 86 | 92 | 1.073 | 0.102 | 0.62838566 | 0.76594885 | 0 | 0.029 | 0.0209 | 0.0209 | TRUE | 0.1636 |
| Pipox | 4776 | 3661 | 4366 | 4601 | 4358 | 5327 | 3709 | 3894 | 4711 | 4580 | 4916 | 5259 | 4511.5 | 4105 | 4918 | 1.199 | 0.261 | 0.00321659 | 0.0174223 | 0.0105 | 0.003 | 0.0055 | 0.0055 | TRUE | 0.2026 |
| Pir | 1594 | 1079 | 1044 | 1255 | 1067 | 1275 | 1238 | 1148 | 1126 | 1249 | 1204 | 1259 | 1203.95 | 1171 | 1237 | 1.056 | 0.078 | 0.28928097 | 0.46130681 | 9.00E-04 | 0.0044 | 0.0031 | 0.0031 | TRUE | 0.0317 |
| Pira1 | 0 | 0 | 0 | 0 | 0 | 0 | 0 | 0 | 0 | 0 | 0 | 0 | 0 | 0 | 0 | NA | NA | NA | NA | NA | NA | NA | NA | NA | NA |
| Pira11 | 0 | 0 | 0 | 0 | 0 | 0 | 0 | 0 | 0 | 0 | 0 | 0 | 0 | 0 | 0 | NA | NA | NA | NA | NA | NA | NA | NA | NA | NA |
| Pira2 | 0 | 0 | 0 | 0 | 0 | 0 | 0 | 0 | 0 | 0 | 0 | 0 | 0 | 0 | 0 | NA | NA | NA | NA | NA | NA | NA | NA | NA | NA |
| Pira4 | 0 | 0 | 0 | 0 | 0 | 0 | 0 | 0 | 0 | 0 | 0 | 0 | 0 | 0 | 0 | NA | NA | NA | NA | NA | NA | NA | NA | NA | NA |
| Pira6 | 0 | 0 | 0 | 0 | 0 | 0 | 0 | 0 | 0 | 0 | 0 | 0 | 0 | 0 | 0 | NA | NA | NA | NA | NA | NA | NA | NA | NA | NA |
| Pira7 | 0 | 0 | 0 | 0 | 0 | 0 | 0 | 0 | 0 | 0 | 0 | 0 | 0 | 0 | 0 | NA | NA | NA | NA | NA | NA | NA | NA | NA | NA |
| Pirt | 88 | 65 | 41 | 53 | 36 | 48 | 68 | 69 | 44 | 53 | 41 | 47 | 53.75 | 60 | 47 | 0.773 | -0.372 | 0.19095644 | 0.34910979 | 0.02 | 0.0465 | 0.0392 | 0.0392 | TRUE | 0.4527 |
| Pisd | 6748 | 5163 | 5589 | 6278 | 5908 | 6825 | 5241 | 5492 | 6030 | 6249 | 6665 | 6737 | 6069.08 | 5588 | 6550 | 1.173 | 0.23 | 0.00050186 | 0.00432372 | 0.0032 | 0.0028 | 0.003 | 0.003 | TRUE | 0.0584 |
| Pisd-ps1 | 196 | 206 | 169 | 203 | 166 | 207 | 152 | 219 | 182 | 202 | 187 | 204 | 191.23 | 184 | 198 | 1.078 | 0.108 | 0.50991877 | 0.67315683 | 0.0129 | 0.0148 | 0.0143 | 0.0143 | TRUE | 0.2588 |
| Pisd-ps2 | 47 | 41 | 44 | 26 | 23 | 32 | 37 | 44 | 47 | 26 | 26 | 32 | 35.17 | 43 | 28 | 0.658 | -0.604 | 0.0664841 | 0.1650274 | 0 | 0.0698 | 0.0481 | 0.0481 | TRUE | 0.139 |
| Pisd-ps3 | 7514 | 7014 | 6385 | 6513 | 5479 | 6399 | 5836 | 7461 | 6889 | 6483 | 6181 | 6317 | 6527.76 | 6729 | 6327 | 0.941 | -0.088 | 0.27860239 | 0.4501033 | 0.0076 | 0.0028 | 0.0046 | 0.0046 | TRUE | 0.1657 |
| Pithd1 | 1823 | 1345 | 1186 | 1375 | 1170 | 1334 | 1416 | 1431 | 1280 | 1369 | 1320 | 1317 | 1355.27 | 1376 | 1335 | 0.97 | -0.044 | 0.55274143 | 0.70844002 | 0.0013 | 0.0042 | 0.0031 | 0.0031 | TRUE | 0.0434 |
| Pitpna | 6602 | 4654 | 4135 | 4371 | 3825 | 4334 | 5127 | 4950 | 4462 | 4351 | 4315 | 4278 | 4580.62 | 4846 | 4315 | 0.89 | -0.168 | 0.00862106 | 0.03705889 | 0.0022 | 0.0029 | 0.0027 | 0.0027 | TRUE | 0.0585 |
| Pitpnb | 10221 | 7239 | 6987 | 7467 | 6672 | 7795 | 7938 | 7700 | 7539 | 7433 | 7527 | 7695 | 7638.56 | 7726 | 7552 | 0.977 | -0.033 | 0.50490353 | 0.66844317 | 3.00E-04 | 0.0027 | 0.0017 | 0.0017 | TRUE | 0.0072 |
| Pitpnc1 | 4030 | 2766 | 3112 | 4483 | 3915 | 4692 | 3130 | 2942 | 3358 | 4462 | 4417 | 4632 | 3823.42 | 3143 | 4504 | 1.433 | 0.519 | 5.24E-16 | 1.01E-13 | 0.0021 | 0.003 | 0.0027 | 0.0027 | TRUE | 0.0424 |
| Pitpnm1 | 378 | 338 | 367 | 486 | 424 | 472 | 294 | 360 | 396 | 484 | 478 | 466 | 412.85 | 350 | 476 | 1.367 | 0.451 | 0.0002438 | 0.00248194 | 0.0089 | 0.0082 | 0.0084 | 0.0084 | TRUE | 0.2424 |
| Pitpnm2 | 1034 | 821 | 794 | 953 | 785 | 917 | 803 | 873 | 857 | 949 | 886 | 905 | 878.74 | 844 | 913 | 1.083 | 0.115 | 0.15254904 | 0.29927984 | 0 | 0.0051 | 0.0035 | 0.0035 | TRUE | 0.0231 |
| Pitpnm3 | 115 | 112 | 175 | 91 | 86 | 94 | 89 | 119 | 189 | 91 | 97 | 93 | 112.94 | 132 | 94 | 0.71 | -0.495 | 0.0498615 | 0.13453809 | 0.0669 | 0.0234 | 0.0366 | 0.0366 | TRUE | 0.4815 |
| Pitrm1 | 11804 | 8625 | 8482 | 8964 | 7800 | 9220 | 9168 | 9174 | 9152 | 8923 | 8799 | 9102 | 9052.92 | 9165 | 8941 | 0.976 | -0.036 | 0.44166983 | 0.61279625 | 0 | 0.0027 | 0.0014 | 0.0014 | TRUE | 0.003 |
| Pitx1 | 52 | 44 | 31 | 34 | 23 | 31 | 40 | 47 | 33 | 34 | 26 | 31 | 35.17 | 40 | 30 | 0.751 | -0.414 | 0.21035324 | 0.37232317 | 0 | 0.0698 | 0.0494 | 0.0494 | TRUE | 0.1481 |
| Pitx2 | 493 | 327 | 337 | 299 | 235 | 297 | 383 | 348 | 364 | 298 | 265 | 293 | 325.04 | 365 | 285 | 0.782 | -0.355 | 0.00248276 | 0.01427439 | 0 | 0.0097 | 0.0068 | 0.0068 | TRUE | 0.0414 |
| Pitx3 | 46 | 29 | 29 | 31 | 37 | 31 | 36 | 31 | 31 | 31 | 42 | 31 | 33.51 | 33 | 35 | 1.047 | 0.066 | 0.84286113 | 0.91072075 | 0 | 0.0732 | 0.0508 | 0.0508 | TRUE | 0.2282 |
| Piwil1 | 240 | 175 | 135 | 171 | 162 | 189 | 186 | 186 | 146 | 170 | 183 | 187 | 176.29 | 173 | 180 | 1.038 | 0.054 | 0.7349505 | 0.84165111 | 0.0045 | 0.0159 | 0.0127 | 0.0127 | TRUE | 0.1914 |
| Piwil2 | 2018 | 1597 | 1552 | 1788 | 1530 | 1638 | 1567 | 1699 | 1675 | 1780 | 1726 | 1617 | 1677.22 | 1647 | 1708 | 1.038 | 0.053 | 0.45020114 | 0.62028903 | 0.0016 | 0.0038 | 0.003 | 0.003 | TRUE | 0.0264 |
| Piwil4 | 435 | 319 | 300 | 375 | 316 | 435 | 338 | 339 | 324 | 373 | 356 | 429 | 360 | 334 | 386 | 1.159 | 0.213 | 0.06813368 | 0.16796046 | 0.0026 | 0.009 | 0.007 | 0.007 | TRUE | 0.11 |
| Pja1 | 2659 | 2011 | 1997 | 1960 | 1825 | 2124 | 2065 | 2139 | 2155 | 1951 | 2059 | 2097 | 2077.57 | 2120 | 2036 | 0.961 | -0.058 | 0.35637082 | 0.53085142 | 5.00E-04 | 0.0036 | 0.0024 | 0.0024 | TRUE | 0.0161 |
| Pja2 | 4163 | 2992 | 3235 | 3706 | 3271 | 3764 | 3233 | 3183 | 3491 | 3689 | 3690 | 3716 | 3500.16 | 3302 | 3698 | 1.12 | 0.164 | 0.00532139 | 0.02569547 | 8.00E-04 | 0.0031 | 0.0022 | 0.0022 | TRUE | 0.0298 |
| Pkd1 | 2014 | 1529 | 1598 | 1987 | 1738 | 1965 | 1564 | 1626 | 1724 | 1978 | 1961 | 1940 | 1798.84 | 1638 | 1960 | 1.197 | 0.26 | 8.39E-05 | 0.00104776 | 7.00E-04 | 0.0037 | 0.0026 | 0.0026 | TRUE | 0.0253 |
| Pkd1l2 | 21 | 33 | 19 | 28 | 30 | 25 | 16 | 35 | 21 | 28 | 34 | 25 | 26.38 | 24 | 29 | 1.211 | 0.277 | 0.49768115 | 0.66213876 | 0.0541 | 0.0923 | 0.0816 | 0.0816 | TRUE | 0.9545 |
| Pkd1l3 | 27 | 16 | 14 | 18 | 18 | 15 | 21 | 17 | 15 | 18 | 20 | 15 | 17.69 | 18 | 18 | 0.989 | -0.016 | 0.9718742 | 0.98489776 | 0 | 0.1364 | 0.0921 | 0.0921 | TRUE | 0.1627 |
| Pkd2 | 848 | 562 | 358 | 363 | 290 | 321 | 659 | 598 | 386 | 361 | 327 | 317 | 441.34 | 548 | 335 | 0.611 | -0.711 | 1.46E-05 | 0.00025853 | 0.0386 | 0.0078 | 0.017 | 0.017 | TRUE | 0.7667 |
| Pkd2l1 | 384 | 405 | 350 | 245 | 281 | 291 | 298 | 431 | 378 | 244 | 317 | 287 | 325.8 | 369 | 283 | 0.768 | -0.381 | 0.01289907 | 0.04991653 | 0.023 | 0.0097 | 0.0138 | 0.0138 | TRUE | 0.2684 |
| Pkd2l2 | 10 | 7 | 0 | 9 | 5 | 1 | 8 | 7 | 0 | 9 | 6 | 1 | 5.13 | 5 | 5 | 1.009 | 0.013 | 0.99028416 | NA | 1.1039 | 0.4641 | 0.5997 | 0.5997 | TRUE | 0.4098 |
| Pkdcc | 119 | 110 | 97 | 138 | 114 | 126 | 92 | 117 | 105 | 137 | 129 | 124 | 117.41 | 105 | 130 | 1.248 | 0.32 | 0.08485553 | 0.19652158 | 0 | 0.0226 | 0.0163 | 0.0163 | TRUE | 0.1182 |
| Pkdrej | 9 | 8 | 4 | 8 | 3 | 4 | 7 | 9 | 4 | 8 | 3 | 4 | 5.85 | 7 | 5 | 0.774 | -0.369 | 0.64545428 | NA | 0 | 0.4074 | 0.2902 | 0.2902 | TRUE | 0.5061 |
| Pkhd1 | 69 | 59 | 29 | 28 | 19 | 16 | 54 | 63 | 31 | 28 | 21 | 16 | 35.46 | 49 | 22 | 0.44 | -1.185 | 0.0017789 | 0.01105954 | 0.0717 | 0.0693 | 0.0699 | 0.0699 | TRUE | 0.3671 |
| Pkhd1l1 | 2 | 6 | 1 | 1 | 0 | 1 | 2 | 6 | 1 | 1 | 0 | 1 | 1.83 | 3 | 1 | 0.228 | -2.132 | 0.19395951 | NA | 0 | 1.2955 | 1.0156 | 1.0156 | TRUE | 1.1845 |
| Pkia | 214 | 172 | 105 | 145 | 107 | 121 | 166 | 183 | 113 | 144 | 121 | 119 | 141.16 | 154 | 128 | 0.83 | -0.268 | 0.18178024 | 0.33742661 | 0.0288 | 0.0192 | 0.022 | 0.022 | TRUE | 0.5396 |
| Pkib | 25 | 11 | 13 | 5 | 4 | 9 | 19 | 12 | 14 | 5 | 5 | 9 | 10.59 | 15 | 6 | 0.406 | -1.3 | 0.03774984 | NA | 0 | 0.2263 | 0.1663 | 0.1663 | TRUE | 0.3833 |
| Pkig | 742 | 532 | 461 | 490 | 393 | 464 | 576 | 566 | 497 | 488 | 443 | 458 | 504.78 | 546 | 463 | 0.846 | -0.24 | 0.0194282 | 0.06766257 | 0.0024 | 0.0071 | 0.0056 | 0.0056 | TRUE | 0.0704 |
| Pklr | 36 | 27 | 26 | 34 | 24 | 30 | 28 | 29 | 28 | 34 | 27 | 30 | 29.21 | 28 | 30 | 1.071 | 0.098 | 0.77848733 | 0.86929058 | 0 | 0.0836 | 0.0541 | 0.0541 | TRUE | 0.0759 |

| | | | | | | | | | | | | | | | | | | | | | | | | | |
|---|---|---|---|---|---|---|---|---|---|---|---|---|---|---|---|---|---|---|---|---|---|---|---|---|---|
| Pkm | 192524 | 148271 | 165549 | 170335 | 157929 | 182783 | 149525 | 157714 | 178626 | 169551 | 178160 | 180435 | 169001.96 | 161955 | 176049 | 1.087 | 0.12 | 0.0771869 | 0.1840254 | 0.0048 | 0.0024 | 0.0033 | 0.0033 | TRUE | 0.0993 |
| Pkmyt1 | 1760 | 1400 | 1209 | 1469 | 1255 | 1480 | 1367 | 1489 | 1305 | 1462 | 1416 | 1461 | 1416.6 | 1387 | 1446 | 1.043 | 0.061 | 0.40729852 | 0.58012135 | 0.0016 | 0.0041 | 0.0032 | 0.0032 | TRUE | 0.0487 |
| Pkn1 | 2843 | 2089 | 2344 | 2882 | 2563 | 2852 | 2208 | 2222 | 2529 | 2869 | 2891 | 2815 | 2589.11 | 2320 | 2858 | 1.233 | 0.302 | 1.28E-05 | 0.00022986 | 0.0026 | 0.0033 | 0.0031 | 0.0031 | TRUE | 0.0747 |
| Pkn2 | 11376 | 8173 | 8199 | 9379 | 7981 | 9174 | 8835 | 8694 | 8847 | 9336 | 9003 | 9056 | 8961.8 | 8792 | 9132 | 1.039 | 0.055 | 0.24443123 | 0.41262024 | 1.00E-04 | 0.0027 | 0.0015 | 0.0015 | TRUE | 0.0047 |
| Pkn3 | 1738 | 1407 | 1399 | 1614 | 1454 | 1654 | 1350 | 1497 | 1510 | 1607 | 1640 | 1633 | 1539.26 | 1452 | 1627 | 1.122 | 0.166 | 0.0201316 | 0.06952099 | 0.0014 | 0.004 | 0.003 | 0.003 | TRUE | 0.0476 |
| Pknox1 | 2107 | 1611 | 1627 | 1674 | 1492 | 1663 | 1636 | 1714 | 1756 | 1666 | 1683 | 1642 | 1682.77 | 1702 | 1664 | 0.978 | -0.032 | 0.623957 | 0.76302339 | 0 | 0.0038 | 0.0024 | 0.0024 | TRUE | 0.0143 |
| Pknox2 | 111 | 57 | 52 | 59 | 44 | 45 | 86 | 61 | 56 | 59 | 50 | 44 | 59.29 | 68 | 51 | 0.747 | -0.421 | 0.12642214 | 0.26172595 | 0.0255 | 0.0424 | 0.0373 | 0.0373 | TRUE | 0.5715 |
| Pkp1 | 19 | 6 | 8 | 15 | 10 | 18 | 15 | 6 | 9 | 15 | 11 | 18 | 12.29 | 10 | 15 | 1.453 | 0.54 | 0.34317213 | NA | 0 | 0.1952 | 0.1506 | 0.1506 | TRUE | 0.7934 |
| Pkp2 | 4954 | 4212 | 4243 | 3130 | 3042 | 3618 | 3848 | 4480 | 4578 | 3116 | 3432 | 3572 | 3837.46 | 4302 | 3373 | 0.784 | -0.35 | 1.20E-05 | 0.0002178 | 0.0063 | 0.003 | 0.0043 | 0.0043 | TRUE | 0.1053 |
| Pkp3 | 550 | 415 | 414 | 483 | 473 | 474 | 427 | 441 | 447 | 481 | 534 | 468 | 466.26 | 438 | 494 | 1.127 | 0.172 | 0.09112599 | 0.20660909 | 0 | 0.0075 | 0.0053 | 0.0053 | TRUE | 0.0554 |
| Pkp4 | 8818 | 6367 | 5841 | 5346 | 4974 | 5812 | 6849 | 6772 | 6302 | 5321 | 5611 | 5737 | 6098.9 | 6641 | 5556 | 0.837 | -0.258 | 1.07E-05 | 0.00019647 | 0.0015 | 0.0028 | 0.0023 | 0.0023 | TRUE | 0.0242 |
| Pla1a | 187 | 136 | 173 | 227 | 211 | 251 | 145 | 145 | 187 | 226 | 238 | 248 | 198.05 | 159 | 237 | 1.498 | 0.583 | 0.00015411 | 0.00174122 | 0.0051 | 0.0144 | 0.0119 | 0.0119 | TRUE | 0.2376 |
| Pla2g10 | 1057 | 823 | 886 | 1152 | 1018 | 1065 | 821 | 875 | 956 | 1147 | 1148 | 1051 | 999.79 | 884 | 1115 | 1.263 | 0.337 | 7.86E-05 | 0.00099608 | 0.0032 | 0.0048 | 0.0042 | 0.0042 | TRUE | 0.0599 |
| Pla2g12a | 2251 | 1704 | 1645 | 1779 | 1548 | 1811 | 1748 | 1813 | 1775 | 1771 | 1746 | 1788 | 1773.43 | 1779 | 1768 | 0.995 | -0.008 | 0.90061242 | 0.94539307 | 0 | 0.0038 | 0.0022 | 0.0022 | TRUE | 0.0034 |
| Pla2g12b | 36 | 32 | 28 | 46 | 35 | 34 | 28 | 34 | 30 | 46 | 39 | 34 | 35.17 | 31 | 40 | 1.293 | 0.371 | 0.2566561 | 0.42655647 | 0 | 0.0698 | 0.0483 | 0.0483 | TRUE | 0.1441 |
| Pla2g15 | 3136 | 2378 | 2669 | 2861 | 2491 | 2904 | 2436 | 2529 | 2880 | 2848 | 2810 | 2867 | 2728.25 | 2615 | 2842 | 1.087 | 0.121 | 0.09443776 | 0.21203251 | 0.0035 | 0.0033 | 0.0034 | 0.0034 | TRUE | 0.0946 |
| Pla2g16 | 704 | 523 | 519 | 711 | 579 | 685 | 547 | 556 | 560 | 708 | 653 | 676 | 616.7 | 554 | 679 | 1.226 | 0.294 | 0.00088986 | 0.00666237 | 0 | 0.0063 | 0.004 | 0.004 | TRUE | 0.0161 |
| Pla2g1b | 832 | 575 | 422 | 514 | 442 | 526 | 646 | 612 | 455 | 512 | 499 | 519 | 540.44 | 571 | 510 | 0.891 | -0.166 | 0.18502529 | 0.34127842 | 0.0154 | 0.0068 | 0.0095 | 0.0095 | TRUE | 0.3615 |
| Pla2g2a | 0 | 0 | 1 | 1 | 0 | 0 | 0 | 0 | 1 | 1 | 0 | 0 | 0.35 | 0 | 0 | 1.075 | 0.105 | 0.97567165 | NA | 0 | 6.8576 | 6.5143 | 6.5143 | TRUE | 0.5365 |
| Pla2g2c | 58 | 34 | 38 | 64 | 58 | 72 | 45 | 36 | 41 | 64 | 65 | 71 | 53.74 | 41 | 67 | 1.631 | 0.706 | 0.00776873 | 0.03425 | 0 | 0.0465 | 0.0311 | 0.0311 | TRUE | 0.0771 |
| Pla2g2d | 7 | 0 | 3 | 2 | 0 | 1 | 5 | 0 | 3 | 2 | 0 | 1 | 1.94 | 3 | 1 | 0.343 | -1.542 | 0.33462027 | NA | 0 | 1.2229 | 1.1656 | 1.1656 | TRUE | 0.1459 |
| Pla2g2e | 15 | 9 | 10 | 9 | 14 | 13 | 12 | 10 | 11 | 9 | 16 | 13 | 11.6 | 11 | 13 | 1.165 | 0.22 | 0.6942352 | NA | 0 | 0.2067 | 0.1399 | 0.1399 | TRUE | 0.2429 |
| Pla2g2f | 14 | 9 | 12 | 16 | 12 | 17 | 11 | 10 | 13 | 16 | 14 | 17 | 13.27 | 11 | 16 | 1.389 | 0.474 | 0.36062331 | NA | 0 | 0.181 | 0.1172 | 0.1172 | TRUE | 0.0674 |
| Pla2g3 | 5 | 1 | 6 | 5 | 5 | 3 | 4 | 1 | 6 | 5 | 6 | 3 | 4.17 | 4 | 5 | 1.185 | 0.245 | 0.79972972 | NA | 0 | 0.5712 | 0.429 | 0.429 | TRUE | 0.2184 |
| Pla2g4a | 425 | 274 | 311 | 171 | 172 | 175 | 330 | 291 | 336 | 170 | 194 | 173 | 249.02 | 319 | 179 | 0.56 | -0.837 | 7.02E-10 | 4.92E-08 | 0 | 0.0119 | 0.0089 | 0.0089 | TRUE | 0.0627 |
| Pla2g4b | 415 | 358 | 383 | 386 | 364 | 420 | 322 | 381 | 413 | 384 | 411 | 415 | 387.64 | 372 | 403 | 1.087 | 0.12 | 0.31821137 | 0.49206915 | 0.0065 | 0.0085 | 0.0079 | 0.0079 | TRUE | 0.1689 |
| Pla2g4c | 51 | 41 | 21 | 28 | 24 | 12 | 40 | 44 | 23 | 28 | 27 | 12 | 28.78 | 36 | 22 | 0.627 | -0.674 | 0.10984993 | 0.23713716 | 0.1096 | 0.0848 | 0.0912 | 0.0912 | TRUE | 0.9954 |
| Pla2g4d | 4 | 5 | 0 | 5 | 5 | 3 | 3 | 5 | 0 | 5 | 6 | 3 | 3.67 | 3 | 5 | 1.597 | 0.675 | 0.52977009 | NA | 0.07 | 0.6487 | 0.5461 | 0.5461 | TRUE | 0.4037 |
| Pla2g4e | 23 | 13 | 21 | 10 | 16 | 20 | 18 | 14 | 23 | 10 | 18 | 20 | 17.02 | 18 | 16 | 0.878 | -0.188 | 0.69813814 | NA | 0 | 0.1417 | 0.1106 | 0.1106 | TRUE | 0.4539 |
| Pla2g4f | 116 | 101 | 71 | 75 | 71 | 66 | 90 | 107 | 77 | 75 | 80 | 65 | 82.34 | 91 | 73 | 0.801 | -0.32 | 0.15782564 | 0.30678661 | 0.0075 | 0.0312 | 0.0246 | 0.0246 | TRUE | 0.2019 |
| Pla2g5 | 362 | 277 | 323 | 314 | 310 | 366 | 281 | 295 | 349 | 313 | 350 | 361 | 324.65 | 308 | 341 | 1.11 | 0.15 | 0.23895851 | 0.405971 | 0.0063 | 0.0097 | 0.0087 | 0.0087 | TRUE | 0.1459 |
| Pla2g6 | 4201 | 3437 | 3597 | 4293 | 3961 | 4648 | 3263 | 3656 | 3881 | 4273 | 4468 | 4588 | 4021.62 | 3600 | 4443 | 1.235 | 0.305 | 2.32E-05 | 0.0003762 | 0.004 | 0.003 | 0.0035 | 0.0035 | TRUE | 0.0831 |
| Pla2g7 | 70 | 46 | 86 | 63 | 68 | 78 | 54 | 49 | 93 | 63 | 77 | 77 | 68.75 | 65 | 72 | 1.109 | 0.149 | 0.5927537 | 0.74044182 | 0.052 | 0.0369 | 0.0411 | 0.0411 | TRUE | 1.138 |
| Pla2r1 | 24 | 21 | 17 | 11 | 9 | 11 | 19 | 22 | 18 | 11 | 10 | 11 | 15.21 | 20 | 11 | 0.54 | -0.888 | 0.07135559 | NA | 0 | 0.1582 | 0.1018 | 0.1018 | TRUE | 0.065 |
| Plaa | 12392 | 8923 | 8767 | 9522 | 8178 | 9442 | 9624 | 9491 | 9460 | 9478 | 9226 | 9321 | 9433.28 | 9525 | 9342 | 0.981 | -0.028 | 0.54050353 | 0.69881655 | 0 | 0.0027 | 0.0014 | 0.0014 | TRUE | 0.002 |
| Plac1 | 16 | 13 | 4 | 2 | 3 | 5 | 12 | 14 | 4 | 2 | 3 | 5 | 6.81 | 10 | 3 | 0.336 | -1.576 | 0.05473364 | NA | 0 | 0.3503 | 0.2872 | 0.2872 | TRUE | 0.8192 |
| Plac1l | 0 | 0 | 0 | 0 | 0 | 0 | 0 | 0 | 0 | 0 | 0 | 0 | 0 | 0 | 0 | NA | NA | NA | NA | NA | NA | NA | NA | NA | NA |
| Plac8 | 470 | 324 | 412 | 559 | 465 | 590 | 365 | 345 | 445 | 556 | 525 | 582 | 469.6 | 385 | 554 | 1.444 | 0.53 | 4.68E-06 | 0.00010036 | 0.0075 | 0.0075 | 0.0075 | 0.0075 | TRUE | 0.2046 |
| Plac8l1 | 1 | 0 | 0 | 0 | 0 | 0 | 1 | 0 | 0 | 0 | 0 | 0 | 0.13 | 0 | 0 | 0.552 | -0.857 | 0.8336548 | NA | 0 | 18.3121 | 10 | 10 | TRUE | 0.2566 |
| Plac9a | 0 | 0 | 0 | 0 | 0 | 0 | 0 | 0 | 0 | 0 | 0 | 0 | 0 | 0 | 0 | NA | NA | NA | NA | NA | NA | NA | NA | NA | NA |
| Plac9b | 0 | 0 | 0 | 0 | 0 | 0 | 0 | 0 | 0 | 0 | 0 | 0 | 0 | 0 | 0 | NA | NA | NA | NA | NA | NA | NA | NA | NA | NA |
| Plag1 | 493 | 383 | 351 | 458 | 401 | 410 | 383 | 407 | 379 | 456 | 452 | 405 | 413.67 | 390 | 438 | 1.123 | 0.168 | 0.11686209 | 0.24771019 | 0 | 0.0082 | 0.0058 | 0.0058 | TRUE | 0.0511 |
| Plagl1 | 1122 | 729 | 520 | 583 | 480 | 536 | 871 | 775 | 561 | 580 | 541 | 529 | 643.14 | 736 | 550 | 0.746 | -0.422 | 0.00201508 | 0.01216793 | 0.0249 | 0.0061 | 0.0119 | 0.0119 | TRUE | 0.4626 |
| Plagl2 | 1368 | 935 | 815 | 915 | 856 | 935 | 1062 | 995 | 879 | 911 | 966 | 923 | 955.97 | 979 | 933 | 0.952 | -0.072 | 0.41923556 | 0.59168956 | 0.0041 | 0.0049 | 0.0046 | 0.0046 | TRUE | 0.0931 |
| Plat | 119 | 61 | 78 | 45 | 72 | 77 | 92 | 65 | 84 | 45 | 81 | 76 | 73.92 | 80 | 67 | 0.832 | -0.266 | 0.32232814 | 0.49614381 | 0.0485 | 0.0345 | 0.0383 | 0.0383 | TRUE | 0.7771 |
| Plau | 174 | 157 | 112 | 114 | 82 | 91 | 135 | 167 | 121 | 113 | 93 | 90 | 119.8 | 141 | 99 | 0.701 | -0.513 | 0.00983478 | 0.04088671 | 0.0139 | 0.0222 | 0.0198 | 0.0198 | TRUE | 0.2582 |
| Plaur | 225 | 163 | 177 | 386 | 315 | 371 | 175 | 173 | 191 | 384 | 355 | 366 | 274.15 | 180 | 368 | 2.055 | 1.039 | 1.91E-16 | 3.95E-14 | 0 | 0.0111 | 0.0074 | 0.0074 | TRUE | 0.0305 |
| Plb1 | 227 | 162 | 182 | 166 | 184 | 204 | 176 | 172 | 196 | 165 | 208 | 201 | 186.53 | 181 | 191 | 1.054 | 0.075 | 0.62817923 | 0.76576254 | 0.0043 | 0.0151 | 0.012 | 0.012 | TRUE | 0.1567 |
| Plbd1 | 151 | 122 | 147 | 120 | 119 | 123 | 117 | 130 | 159 | 119 | 134 | 121 | 130.13 | 135 | 125 | 0.928 | -0.108 | 0.55725985 | 0.7122577 | 0.0075 | 0.0206 | 0.0168 | 0.0168 | TRUE | 0.2315 |
| Plbd2 | 4267 | 3362 | 3243 | 3497 | 3331 | 3587 | 3314 | 3576 | 3499 | 3481 | 3758 | 3541 | 3528.14 | 3463 | 3593 | 1.038 | 0.054 | 0.37703435 | 0.55151527 | 0.0012 | 0.0031 | 0.0024 | 0.0024 | TRUE | 0.0193 |
| Plcb1 | 111 | 80 | 69 | 84 | 71 | 72 | 86 | 85 | 74 | 84 | 80 | 71 | 80.09 | 82 | 78 | 0.953 | -0.07 | 0.74980908 | 0.85153057 | 0 | 0.032 | 0.0217 | 0.0217 | TRUE | 0.0617 |
| Plcb2 | 12 | 10 | 8 | 11 | 12 | 10 | 9 | 11 | 9 | 11 | 14 | 10 | 10.49 | 10 | 12 | 1.199 | 0.262 | 0.65186811 | NA | 0 | 0.2283 | 0.1468 | 0.1468 | TRUE | 0.0871 |
| Plcb3 | 8167 | 6283 | 6341 | 7181 | 6827 | 7889 | 6343 | 6683 | 6842 | 7148 | 7702 | 7788 | 7084.2 | 6623 | 7546 | 1.14 | 0.189 | 0.00122383 | 0.00838152 | 0.0016 | 0.0028 | 0.0023 | 0.0023 | TRUE | 0.026 |
| Plcb4 | 1587 | 1093 | 1093 | 1210 | 1072 | 1219 | 1233 | 1163 | 1179 | 1204 | 1209 | 1203 | 1198.6 | 1192 | 1205 | 1.011 | 0.016 | 0.82057475 | 0.89660385 | 0 | 0.0044 | 0.0027 | 0.0027 | TRUE | 0.0115 |
| Plcd1 | 1879 | 1496 | 1489 | 2014 | 1864 | 2036 | 1459 | 1591 | 1607 | 2005 | 2103 | 2010 | 1795.77 | 1552 | 2039 | 1.315 | 0.395 | 7.02E-09 | 3.68E-07 | 0.0012 | 0.0037 | 0.0028 | 0.0028 | TRUE | 0.0346 |
| Plcd3 | 1571 | 1218 | 1366 | 1465 | 1370 | 1621 | 1220 | 1296 | 1474 | 1458 | 1546 | 1600 | 1432.26 | 1330 | 1535 | 1.155 | 0.208 | 0.01361695 | 0.0519976 | 0.0048 | 0.0041 | 0.0044 | 0.0044 | TRUE | 0.1072 |
| Plcd4 | 60 | 34 | 59 | 44 | 27 | 29 | 47 | 36 | 64 | 44 | 30 | 29 | 41.55 | 49 | 34 | 0.705 | -0.505 | 0.1297498 | 0.26641488 | 0.0432 | 0.0595 | 0.0549 | 0.0549 | TRUE | 0.2618 |
| Plce1 | 1601 | 1070 | 901 | 1259 | 1147 | 1328 | 1243 | 1138 | 972 | 1253 | 1294 | 1311 | 1201.97 | 1118 | 1286 | 1.148 | 0.2 | 0.03029865 | 0.09329138 | 0.0065 | 0.0044 | 0.0053 | 0.0053 | TRUE | 0.1542 |
| Plcg1 | 8965 | 7066 | 7383 | 8230 | 7798 | 9042 | 6963 | 7516 | 7966 | 8192 | 8797 | 8926 | 8059.98 | 7482 | 8638 | 1.155 | 0.208 | 0.00136777 | 0.00902984 | 0.0031 | 0.0027 | 0.0029 | 0.0029 | TRUE | 0.0454 |
| Plcg2 | 4504 | 3253 | 3670 | 5324 | 4806 | 5565 | 3498 | 3460 | 3960 | 5300 | 5422 | 5494 | 4522.14 | 3639 | 5405 | 1.486 | 0.571 | 2.36E-18 | 6.02E-16 | 0.0027 | 0.003 | 0.0028 | 0.0028 | TRUE | 0.0715 |
| Plch1 | 332 | 215 | 183 | 131 | 135 | 122 | 258 | 229 | 197 | 130 | 152 | 120 | 181.19 | 228 | 134 | 0.586 | -0.77 | 4.01E-06 | 8.84E-05 | 0.0109 | 0.0155 | 0.0141 | 0.0141 | TRUE | 0.1318 |
| Plch2 | 72 | 59 | 44 | 60 | 58 | 47 | 56 | 63 | 47 | 60 | 65 | 46 | 56.28 | 55 | 57 | 1.03 | 0.042 | 0.87559285 | 0.93081805 | 0 | 0.0445 | 0.0341 | 0.0341 | TRUE | 0.236 |
| Plcl1 | 14 | 29 | 10 | 11 | 6 | 6 | 11 | 31 | 11 | 11 | 7 | 6 | 12.69 | 18 | 8 | 0.455 | -1.136 | 0.07103323 | NA | 0.1986 | 0.1892 | 0.1921 | 0.1921 | TRUE | 2.14 |
| Plcl2 | 573 | 453 | 430 | 538 | 428 | 477 | 445 | 482 | 464 | 536 | 483 | 471 | 480.01 | 464 | 497 | 1.072 | 0.101 | 0.32350707 | 0.49751307 | 0.0014 | 0.0074 | 0.0054 | 0.0054 | TRUE | 0.0562 |
| Plcxd1 | 19 | 28 | 26 | 23 | 18 | 34 | 15 | 30 | 28 | 23 | 20 | 34 | 24.89 | 24 | 26 | 1.074 | 0.103 | 0.80722367 | 0.8885257 | 0.067 | 0.0976 | 0.0889 | 0.0889 | TRUE | 0.8609 |
| Plcxd2 | 685 | 554 | 471 | 623 | 495 | 594 | 532 | 589 | 508 | 620 | 558 | 586 | 565.74 | 543 | 588 | 1.084 | 0.117 | 0.23751117 | 0.40418468 | 0.0023 | 0.0066 | 0.0052 | 0.0052 | TRUE | 0.062 |
| Plcxd3 | 5 | 0 | 1 | 2 | 3 | 4 | 4 | 0 | 1 | 2 | 3 | 4 | 2.38 | 2 | 3 | 1.775 | 0.828 | 0.52844425 | NA | 0.1799 | 0.9978 | 0.8006 | 0.8006 | TRUE | 1.3683 |
| Plcz1 | 2 | 0 | 0 | 0 | 0 | 0 | 2 | 0 | 0 | 0 | 0 | 0 | 0.26 | 1 | 0 | 0.356 | -1.491 | 0.71362796 | NA | 0 | 9.1573 | 10 | 10 | TRUE | 0.968 |
| Pld1 | 386 | 265 | 310 | 270 | 258 | 285 | 300 | 282 | 334 | 269 | 291 | 281 | 292.88 | 305 | 280 | 0.918 | -0.123 | 0.32253615 | 0.49628693 | 0 | 0.0105 | 0.0077 | 0.0077 | TRUE | 0.0744 |

| | | | | | | | | | | | | | | | | | | | | | | | | | |
|---|---|---|---|---|---|---|---|---|---|---|---|---|---|---|---|---|---|---|---|---|---|---|---|---|---|
| Pld2 | 1079 | 776 | 764 | 1088 | 923 | 1058 | 838 | 825 | 824 | 1083 | 1041 | 1044 | 942.74 | 829 | 1056 | 1.273 | 0.349 | 3.13E-06 | 7.19E-05 | 0 | 0.0049 | 0.003 | 0.003 | TRUE | 0.0059 |
| Pld3 | 3146 | 2291 | 2538 | 3077 | 2773 | 3185 | 2443 | 2437 | 2738 | 3063 | 3128 | 3144 | 2825.65 | 2539 | 3112 | 1.226 | 0.294 | 7.44E-06 | 0.00014731 | 0.0019 | 0.0033 | 0.0027 | 0.0027 | TRUE | 0.0562 |
| Pld4 | 58 | 47 | 55 | 88 | 68 | 72 | 45 | 50 | 59 | 88 | 77 | 71 | 64.96 | 51 | 79 | 1.533 | 0.617 | 0.01292669 | 0.04996189 | 0 | 0.0389 | 0.0283 | 0.0283 | TRUE | 0.1426 |
| Pld5 | 1 | 1 | 3 | 1 | 0 | 3 | 1 | 1 | 3 | 1 | 0 | 3 | 1.51 | 2 | 1 | 0.809 | -0.306 | 0.84993666 | NA | 0 | 1.5765 | 1.2096 | 1.2096 | TRUE | 0.5881 |
| Pld6 | 1093 | 879 | 1010 | 1170 | 1041 | 1229 | 849 | 935 | 1090 | 1165 | 1174 | 1213 | 1070.97 | 958 | 1184 | 1.239 | 0.309 | 0.00109873 | 0.00774748 | 0.0069 | 0.0046 | 0.0055 | 0.0055 | TRUE | 0.1723 |
| Pldi | 2 | 0 | 0 | 0 | 2 | 1 | 2 | 0 | 0 | 0 | 2 | 1 | 0.8 | 1 | 1 | 1.903 | 0.928 | 0.6927597 | NA | 0 | 2.9671 | 2.6342 | 2.6342 | TRUE | 0.4267 |
| Plec | 6756 | 5779 | 5946 | 6718 | 6301 | 7514 | 5247 | 6147 | 6416 | 6687 | 7108 | 7417 | 6503.76 | 5937 | 7071 | 1.192 | 0.253 | 0.00117778 | 0.00816395 | 0.0064 | 0.0028 | 0.0042 | 0.0042 | TRUE | 0.1271 |
| Plek | 5 | 4 | 2 | 1 | 1 | 2 | 4 | 4 | 2 | 1 | 1 | 2 | 2.4 | 3 | 1 | 0.398 | -1.329 | 0.291632 | NA | 0 | 0.9904 | 0.6279 | 0.6279 | TRUE | 0.1388 |
| Plek2 | 406 | 357 | 363 | 381 | 318 | 388 | 315 | 380 | 392 | 379 | 359 | 383 | 367.96 | 362 | 374 | 1.036 | 0.051 | 0.6739798 | 0.79979329 | 0.0051 | 0.0089 | 0.0077 | 0.0077 | TRUE | 0.1584 |
| Plekha1 | 3655 | 2672 | 2742 | 2850 | 2603 | 2996 | 2839 | 2842 | 2959 | 2837 | 2936 | 2958 | 2895.05 | 2880 | 2910 | 1.011 | 0.015 | 0.7856183 | 0.87390788 | 2.00E-04 | 0.0032 | 0.002 | 0.002 | TRUE | 0.0068 |
| Plekha2 | 8413 | 7061 | 7789 | 9272 | 8501 | 9775 | 6534 | 7511 | 8404 | 9229 | 9590 | 9649 | 8486.29 | 7483 | 9489 | 1.269 | 0.343 | 2.29E-05 | 0.00037234 | 0.0079 | 0.0027 | 0.0046 | 0.0046 | TRUE | 0.1513 |
| Plekha3 | 2173 | 1624 | 1570 | 1650 | 1475 | 1599 | 1688 | 1727 | 1694 | 1642 | 1664 | 1578 | 1665.66 | 1703 | 1628 | 0.956 | -0.065 | 0.3086235 | 0.48126096 | 0 | 0.0038 | 0.0023 | 0.0023 | TRUE | 0.0087 |
| Plekha4 | 547 | 381 | 432 | 457 | 446 | 508 | 425 | 405 | 466 | 455 | 503 | 501 | 459.29 | 432 | 486 | 1.126 | 0.171 | 0.1030906 | 0.22620111 | 0.0017 | 0.0076 | 0.0058 | 0.0058 | TRUE | 0.0523 |
| Plekha5 | 5084 | 3877 | 3742 | 3467 | 3263 | 3756 | 3949 | 4124 | 4038 | 3451 | 3681 | 3708 | 3824.97 | 4037 | 3613 | 0.895 | -0.16 | 0.00511387 | 0.0249633 | 6.00E-04 | 0.003 | 0.0021 | 0.0021 | TRUE | 0.0189 |
| Plekha6 | 452 | 362 | 353 | 436 | 417 | 440 | 351 | 385 | 381 | 434 | 470 | 434 | 409.29 | 372 | 446 | 1.2 | 0.263 | 0.01328786 | 0.05105468 | 0 | 0.0082 | 0.0057 | 0.0057 | TRUE | 0.0311 |
| Plekha7 | 4488 | 3539 | 3476 | 3758 | 3579 | 4122 | 3486 | 3764 | 3751 | 3741 | 4037 | 4069 | 3807.97 | 3667 | 3949 | 1.077 | 0.108 | 0.08399942 | 0.19540558 | 0.0016 | 0.003 | 0.0025 | 0.0025 | TRUE | 0.0261 |
| Plekha8 | 4330 | 3010 | 2850 | 3054 | 2618 | 2879 | 3363 | 3202 | 3075 | 3040 | 2953 | 2842 | 3079.18 | 3213 | 2945 | 0.916 | -0.127 | 0.04052155 | 0.11555756 | 0.0013 | 0.0032 | 0.0024 | 0.0024 | TRUE | 0.0206 |
| Plekhb1 | 1083 | 678 | 741 | 1123 | 887 | 1006 | 841 | 721 | 800 | 1118 | 1001 | 993 | 912.23 | 787 | 1037 | 1.316 | 0.396 | 1.10E-05 | 0.00020172 | 0.0042 | 0.005 | 0.0047 | 0.0047 | TRUE | 0.0633 |
| Plekhb2 | 1683 | 1388 | 1415 | 1332 | 1234 | 1394 | 1307 | 1476 | 1527 | 1326 | 1392 | 1376 | 1400.72 | 1437 | 1365 | 0.951 | -0.072 | 0.35707613 | 0.53157892 | 0.0031 | 0.0041 | 0.0037 | 0.0037 | TRUE | 0.0779 |
| Plekhd1 | 810 | 665 | 584 | 715 | 618 | 664 | 629 | 707 | 630 | 712 | 697 | 655 | 671.82 | 655 | 688 | 1.051 | 0.071 | 0.43735704 | 0.60879862 | 0.0017 | 0.006 | 0.0046 | 0.0046 | TRUE | 0.0551 |
| Plekhf1 | 163 | 106 | 125 | 143 | 111 | 129 | 127 | 113 | 135 | 142 | 125 | 127 | 128.19 | 125 | 131 | 1.056 | 0.078 | 0.65846067 | 0.78848679 | 0 | 0.0209 | 0.0147 | 0.0147 | TRUE | 0.0672 |
| Plekhf2 | 5822 | 4078 | 3788 | 3570 | 3084 | 3515 | 4522 | 4338 | 4087 | 3554 | 3479 | 3470 | 3908.19 | 4316 | 3501 | 0.811 | -0.302 | 3.06E-07 | 1.06E-05 | 0.0012 | 0.003 | 0.0023 | 0.0023 | TRUE | 0.026 |
| Plekhg1 | 626 | 460 | 380 | 375 | 280 | 307 | 486 | 489 | 410 | 373 | 316 | 303 | 396.28 | 462 | 331 | 0.716 | -0.483 | 9.42E-05 | 0.00115422 | 0.0084 | 0.0084 | 0.0084 | 0.0084 | TRUE | 0.1462 |
| Plekhg2 | 1548 | 1234 | 1092 | 1210 | 1095 | 1298 | 1202 | 1313 | 1178 | 1204 | 1235 | 1281 | 1235.69 | 1231 | 1240 | 1.008 | 0.012 | 0.87792293 | 0.93183963 | 0.0013 | 0.0043 | 0.0033 | 0.0033 | TRUE | 0.0393 |
| Plekhg3 | 2005 | 1501 | 1490 | 1625 | 1512 | 1708 | 1557 | 1597 | 1608 | 1618 | 1706 | 1686 | 1628.46 | 1587 | 1670 | 1.052 | 0.074 | 0.25375611 | 0.4228898 | 0 | 0.0039 | 0.0024 | 0.0024 | TRUE | 0.0091 |
| Plekhg4 | 331 | 290 | 276 | 344 | 297 | 318 | 257 | 308 | 298 | 342 | 335 | 314 | 309.12 | 288 | 330 | 1.152 | 0.204 | 0.09995388 | 0.22086228 | 0.0028 | 0.0101 | 0.0078 | 0.0078 | TRUE | 0.107 |
| Plekhg5 | 2461 | 2168 | 2370 | 2513 | 2425 | 2701 | 1911 | 2306 | 2557 | 2501 | 2736 | 2666 | 2446.34 | 2258 | 2634 | 1.168 | 0.224 | 0.01828766 | 0.06463398 | 0.0108 | 0.0034 | 0.0061 | 0.0061 | TRUE | 0.2227 |
| Plekhg6 | 268 | 191 | 229 | 376 | 314 | 374 | 208 | 203 | 247 | 374 | 354 | 369 | 292.68 | 219 | 366 | 1.671 | 0.741 | 4.95E-09 | 2.75E-07 | 8.00E-04 | 0.0105 | 0.0079 | 0.0079 | TRUE | 0.1266 |
| Plekhh1 | 2969 | 2165 | 2503 | 2701 | 2343 | 2538 | 2306 | 2303 | 2701 | 2689 | 2643 | 2505 | 2524.44 | 2437 | 2612 | 1.073 | 0.101 | 0.18388 | 0.33995561 | 0.0045 | 0.0034 | 0.0038 | 0.0038 | TRUE | 0.108 |
| Plekhh2 | 1156 | 766 | 653 | 801 | 661 | 778 | 898 | 815 | 705 | 797 | 746 | 768 | 788.03 | 806 | 770 | 0.954 | -0.067 | 0.49889332 | 0.66350443 | 0.0068 | 0.0054 | 0.0059 | 0.0059 | TRUE | 0.1416 |
| Plekhh3 | 251 | 206 | 264 | 265 | 262 | 308 | 195 | 219 | 285 | 264 | 296 | 304 | 260.38 | 233 | 288 | 1.24 | 0.311 | 0.04494829 | 0.12457746 | 0.0179 | 0.0115 | 0.0134 | 0.0134 | TRUE | 0.4135 |
| Plekhj1 | 1892 | 1413 | 1346 | 1385 | 1211 | 1378 | 1469 | 1503 | 1452 | 1379 | 1366 | 1360 | 1421.63 | 1475 | 1368 | 0.928 | -0.108 | 0.09799819 | 0.21817573 | 0 | 0.0041 | 0.0024 | 0.0024 | TRUE | 0.0032 |
| Plekhm1 | 4708 | 3557 | 3614 | 3549 | 3386 | 3912 | 3657 | 3784 | 3899 | 3533 | 3820 | 3862 | 3758.95 | 3780 | 3738 | 0.989 | -0.016 | 0.79617392 | 0.88091917 | 0.0014 | 0.0031 | 0.0024 | 0.0024 | TRUE | 0.0283 |
| Plekhm2 | 803 | 508 | 546 | 591 | 523 | 542 | 624 | 540 | 589 | 588 | 590 | 535 | 577.74 | 584 | 571 | 0.975 | -0.036 | 0.71333304 | 0.82763773 | 0.0025 | 0.0065 | 0.0052 | 0.0052 | TRUE | 0.0505 |
| Plekhm3 | 1589 | 1208 | 1037 | 1189 | 1188 | 1272 | 1234 | 1285 | 1119 | 1184 | 1340 | 1256 | 1236.22 | 1213 | 1260 | 1.038 | 0.054 | 0.51243027 | 0.67504942 | 0.0035 | 0.0043 | 0.0041 | 0.0041 | TRUE | 0.0533 |
| Plekhn1 | 102 | 105 | 147 | 110 | 85 | 123 | 79 | 112 | 159 | 109 | 96 | 121 | 112.72 | 117 | 109 | 0.942 | -0.085 | 0.72572834 | 0.83572689 | 0.0585 | 0.0235 | 0.0338 | 0.0338 | TRUE | 0.2905 |
| Plekho1 | 195 | 146 | 119 | 133 | 113 | 113 | 151 | 155 | 128 | 132 | 127 | 112 | 134.43 | 145 | 124 | 0.852 | -0.232 | 0.18892678 | 0.34647412 | 0 | 0.0201 | 0.0149 | 0.0149 | TRUE | 0.0995 |
| Plekho2 | 994 | 680 | 665 | 741 | 600 | 707 | 772 | 723 | 718 | 738 | 677 | 698 | 720.87 | 738 | 704 | 0.954 | -0.068 | 0.42857394 | 0.60066752 | 0 | 0.0057 | 0.004 | 0.004 | TRUE | 0.0208 |
| Plekhs1 | 47 | 32 | 36 | 29 | 23 | 25 | 37 | 34 | 39 | 29 | 26 | 25 | 31.48 | 37 | 27 | 0.727 | -0.46 | 0.17529249 | 0.32873626 | 0 | 0.0777 | 0.0502 | 0.0502 | TRUE | 0.0397 |
| Plg | 7 | 0 | 2 | 1 | 0 | 0 | 5 | 0 | 2 | 1 | 0 | 0 | 1.43 | 2 | 0 | 0.143 | -2.802 | 0.16024734 | NA | 1.8016 | 1.6579 | 1.6812 | 1.6812 | TRUE | 0.1607 |
| Plgrkt | 437 | 303 | 304 | 377 | 264 | 326 | 339 | 322 | 328 | 375 | 298 | 322 | 330.77 | 330 | 332 | 1.006 | 0.008 | 0.94766384 | 0.97142825 | 0.0042 | 0.0096 | 0.008 | 0.008 | TRUE | 0.1512 |
| Plin1 | 0 | 2 | 0 | 0 | 2 | 3 | 0 | 2 | 0 | 0 | 2 | 3 | 1.22 | 1 | 2 | 2.552 | 1.352 | 0.50518172 | NA | 2.2758 | 1.9385 | 1.9582 | 1.9582 | TRUE | 0.9751 |
| Plin2 | 2232 | 1626 | 1803 | 1815 | 1685 | 1906 | 1733 | 1730 | 1945 | 1807 | 1901 | 1882 | 1832.92 | 1803 | 1863 | 1.034 | 0.048 | 0.49942464 | 0.66396395 | 0.0019 | 0.0037 | 0.0031 | 0.0031 | TRUE | 0.0569 |
| Plin3 | 4666 | 3599 | 3899 | 4066 | 3701 | 4264 | 3624 | 3828 | 4207 | 4047 | 4175 | 4209 | 4015.12 | 3886 | 4144 | 1.067 | 0.093 | 0.16109592 | 0.31078701 | 0.0027 | 0.003 | 0.0029 | 0.0029 | TRUE | 0.0632 |
| Plin4 | 6 | 11 | 13 | 12 | 18 | 16 | 5 | 12 | 14 | 12 | 20 | 16 | 13.07 | 10 | 16 | 1.611 | 0.688 | 0.23200293 | NA | 0.0849 | 0.1837 | 0.1578 | 0.1578 | TRUE | 0.6901 |
| Plin5 | 256 | 206 | 185 | 240 | 260 | 242 | 199 | 219 | 200 | 239 | 293 | 239 | 231.44 | 206 | 257 | 1.248 | 0.319 | 0.02505891 | 0.08146041 | 0.0046 | 0.0127 | 0.0103 | 0.0103 | TRUE | 0.1647 |
| Plk1 | 9913 | 7599 | 7992 | 7610 | 7010 | 8066 | 7699 | 8083 | 8623 | 7575 | 7908 | 7962 | 7975.11 | 8135 | 7815 | 0.961 | -0.058 | 0.3276468 | 0.5019575 | 0.0018 | 0.0027 | 0.0024 | 0.0024 | TRUE | 0.0336 |
| Plk1s1 | 631 | 495 | 472 | 427 | 364 | 482 | 490 | 527 | 509 | 425 | 411 | 476 | 472.89 | 509 | 437 | 0.861 | -0.216 | 0.0365957 | 0.10750303 | 0.0013 | 0.0074 | 0.0056 | 0.0056 | TRUE | 0.0699 |
| Plk2 | 2593 | 1795 | 1851 | 1751 | 1448 | 1578 | 2014 | 1909 | 1997 | 1743 | 1633 | 1558 | 1809.1 | 1973 | 1645 | 0.833 | -0.263 | 0.00013393 | 0.00155121 | 0.0014 | 0.0037 | 0.0029 | 0.0029 | TRUE | 0.0334 |
| Plk3 | 1184 | 709 | 570 | 658 | 545 | 610 | 920 | 754 | 615 | 655 | 615 | 602 | 693.45 | 763 | 624 | 0.816 | -0.293 | 0.02162646 | 0.07312679 | 0.0188 | 0.0058 | 0.0103 | 0.0103 | TRUE | 0.1969 |
| Plk4 | 4513 | 3221 | 3531 | 3439 | 3068 | 3436 | 3505 | 3426 | 3810 | 3423 | 3461 | 3392 | 3502.86 | 3580 | 3425 | 0.957 | -0.064 | 0.29620099 | 0.46812001 | 0.0012 | 0.0031 | 0.0024 | 0.0024 | TRUE | 0.0377 |
| Plk5 | 490 | 441 | 569 | 707 | 627 | 746 | 381 | 469 | 614 | 704 | 707 | 736 | 601.85 | 488 | 716 | 1.472 | 0.558 | 6.85E-05 | 0.00090285 | 0.0269 | 0.0064 | 0.0124 | 0.0124 | TRUE | 0.5608 |
| Pllp | 187 | 125 | 95 | 144 | 113 | 137 | 145 | 133 | 103 | 143 | 127 | 135 | 131.13 | 127 | 135 | 1.062 | 0.087 | 0.63967636 | 0.77477341 | 0.0095 | 0.0205 | 0.0174 | 0.0174 | TRUE | 0.2779 |
| Pln | 0 | 0 | 0 | 0 | 0 | 0 | 0 | 0 | 0 | 0 | 0 | 0 | 0 | 0 | 0 | NA | NA | NA | NA | NA | NA | NA | NA | NA | NA |
| Plod1 | 3218 | 2279 | 2431 | 2732 | 2610 | 2975 | 2499 | 2424 | 2623 | 2719 | 2944 | 2937 | 2691.17 | 2515 | 2867 | 1.14 | 0.189 | 0.0031084 | 0.01698331 | 0.0012 | 0.0033 | 0.0026 | 0.0026 | TRUE | 0.0247 |
| Plod2 | 7819 | 5599 | 6229 | 6395 | 5368 | 6054 | 6073 | 5956 | 6721 | 6366 | 6056 | 5976 | 6191.13 | 6250 | 6133 | 0.981 | -0.027 | 0.66746671 | 0.79551793 | 0.0022 | 0.0028 | 0.0027 | 0.0027 | TRUE | 0.0527 |
| Plod3 | 1979 | 1532 | 1518 | 1575 | 1320 | 1519 | 1537 | 1630 | 1638 | 1568 | 1489 | 1499 | 1560.14 | 1602 | 1519 | 0.949 | -0.075 | 0.26390217 | 0.43407614 | 5.00E-04 | 0.0039 | 0.0026 | 0.0026 | TRUE | 0.0156 |
| Plp1 | 144 | 138 | 157 | 165 | 130 | 203 | 112 | 147 | 169 | 164 | 147 | 200 | 156.55 | 143 | 170 | 1.203 | 0.267 | 0.16898966 | 0.32056364 | 0.0281 | 0.0176 | 0.0207 | 0.0207 | TRUE | 0.4094 |
| Plp2 | 3081 | 2389 | 2921 | 2897 | 2672 | 3123 | 2393 | 2541 | 3152 | 2884 | 3014 | 3083 | 2844.44 | 2695 | 2994 | 1.111 | 0.152 | 0.09990331 | 0.22083019 | 0.0112 | 0.0033 | 0.0058 | 0.0058 | TRUE | 0.2654 |
| Plrg1 | 6278 | 4663 | 4588 | 4587 | 4063 | 4793 | 4876 | 4960 | 4950 | 4566 | 4583 | 4731 | 4777.84 | 4929 | 4627 | 0.939 | -0.091 | 0.07227046 | 0.17570262 | 0 | 0.0029 | 0.0016 | 0.0016 | TRUE | 0.0048 |
| Pls1 | 2171 | 1695 | 1720 | 2256 | 1923 | 2150 | 1686 | 1803 | 1856 | 2246 | 2169 | 2122 | 1980.38 | 1782 | 2179 | 1.224 | 0.292 | 1.09E-05 | 0.00019975 | 0.0011 | 0.0036 | 0.0027 | 0.0027 | TRUE | 0.0276 |
| Pls3 | 1121 | 694 | 669 | 654 | 566 | 667 | 871 | 738 | 722 | 651 | 639 | 658 | 713.1 | 777 | 649 | 0.834 | -0.262 | 0.00688703 | 0.03138912 | 0.0046 | 0.0058 | 0.0054 | 0.0054 | TRUE | 0.1377 |
| Plscr1 | 1620 | 1150 | 1223 | 936 | 984 | 1068 | 1258 | 1223 | 1320 | 932 | 1110 | 1054 | 1149.51 | 1267 | 1032 | 0.814 | -0.297 | 0.00041389 | 0.00373236 | 0.0033 | 0.0045 | 0.0042 | 0.0042 | TRUE | 0.0868 |
| Plscr2 | 11 | 3 | 4 | 3 | 1 | 1 | 9 | 3 | 4 | 3 | 1 | 1 | 3.53 | 5 | 2 | 0.315 | -1.667 | 0.13281069 | NA | 0 | 0.6747 | 0.4951 | 0.4951 | TRUE | 0.7467 |
| Plscr3 | 970 | 809 | 751 | 861 | 777 | 842 | 753 | 861 | 810 | 857 | 877 | 831 | 831.49 | 808 | 855 | 1.059 | 0.083 | 0.32626465 | 0.50035415 | 0.0015 | 0.0053 | 0.004 | 0.004 | TRUE | 0.0442 |
| Plscr4 | 35 | 15 | 18 | 18 | 18 | 24 | 27 | 16 | 19 | 18 | 20 | 24 | 20.75 | 21 | 21 | 0.979 | -0.03 | 0.94418015 | 0.96913228 | 0 | 0.1167 | 0.0856 | 0.0856 | TRUE | 0.4726 |
| Plscr5 | 0 | 0 | 0 | 0 | 0 | 0 | 0 | 0 | 0 | 0 | 0 | 0 | 0 | 0 | 0 | NA | NA | NA | NA | NA | NA | NA | NA | NA | NA |
| Pltp | 2312 | 1945 | 2228 | 3370 | 3156 | 3280 | 1796 | 2069 | 2404 | 3354 | 3560 | 3238 | 2736.86 | 2090 | 3384 | 1.621 | 0.697 | 1.04E-13 | 1.54E-11 | 0.0112 | 0.0033 | 0.0059 | 0.0059 | TRUE | 0.2097 |
| Plvap | 273 | 139 | 109 | 136 | 131 | 152 | 212 | 148 | 118 | 135 | 148 | 150 | 151.78 | 159 | 144 | 0.901 | -0.15 | 0.47183469 | 0.64057116 | 0.0409 | 0.018 | 0.0249 | 0.0249 | TRUE | 0.5355 |

| | | | | | | | | | | | | | | | | | | | | | | | | | |
|---|---|---|---|---|---|---|---|---|---|---|---|---|---|---|---|---|---|---|---|---|---|---|---|---|---|
| Plxdc1 | 982 | 834 | 1011 | 1565 | 1583 | 1780 | 763 | 887 | 1091 | 1558 | 1786 | 1757 | 1306.9 | 914 | 1700 | 1.865 | 0.899 | 2.10E-15 | 3.67E-13 | 0.0179 | 0.0042 | 0.0084 | 0.0084 | TRUE | 0.3428 |
| Plxdc2 | 189 | 141 | 115 | 98 | 71 | 98 | 147 | 150 | 124 | 98 | 80 | 97 | 115.87 | 140 | 92 | 0.652 | -0.616 | 0.00122064 | 0.00836368 | 0 | 0.0229 | 0.0171 | 0.0171 | TRUE | 0.1064 |
| Plxna1 | 12768 | 9354 | 10077 | 10509 | 9647 | 10713 | 9916 | 9950 | 10873 | 10461 | 10883 | 10575 | 10442.99 | 10246 | 10640 | 1.038 | 0.054 | 0.32863547 | 0.50318016 | 0.0015 | 0.0027 | 0.0021 | 0.0021 | TRUE | 0.0348 |
| Plxna2 | 757 | 522 | 516 | 586 | 481 | 559 | 588 | 555 | 557 | 583 | 543 | 552 | 562.95 | 567 | 559 | 0.986 | -0.02 | 0.82803684 | 0.90144951 | 0 | 0.0066 | 0.0043 | 0.0043 | TRUE | 0.0167 |
| Plxna3 | 943 | 864 | 762 | 767 | 670 | 786 | 732 | 919 | 822 | 763 | 756 | 776 | 794.8 | 824 | 765 | 0.93 | -0.105 | 0.27727307 | 0.44851335 | 0.0056 | 0.0054 | 0.0055 | 0.0055 | TRUE | 0.1198 |
| Plxna4 | 35 | 22 | 20 | 33 | 32 | 38 | 27 | 23 | 22 | 33 | 36 | 38 | 29.77 | 24 | 36 | 1.466 | 0.552 | 0.11505714 | 0.24486475 | 0 | 0.082 | 0.0539 | 0.0539 | TRUE | 0.0947 |
| Plxnb1 | 1116 | 829 | 798 | 916 | 887 | 863 | 867 | 882 | 861 | 912 | 1001 | 852 | 895.65 | 870 | 922 | 1.058 | 0.082 | 0.33470202 | 0.50876273 | 0.0022 | 0.0051 | 0.0041 | 0.0041 | TRUE | 0.0661 |
| Plxnb2 | 5714 | 4709 | 5546 | 7185 | 6441 | 7148 | 4438 | 5009 | 5984 | 7152 | 7266 | 7056 | 6150.84 | 5144 | 7158 | 1.392 | 0.477 | 1.12E-07 | 4.34E-06 | 0.0114 | 0.0028 | 0.0057 | 0.0057 | TRUE | 0.2489 |
| Plxnb3 | 46 | 39 | 42 | 72 | 67 | 78 | 36 | 41 | 45 | 72 | 76 | 77 | 57.8 | 41 | 75 | 1.841 | 0.881 | 0.00064082 | 0.00521479 | 0 | 0.0434 | 0.0292 | 0.0292 | TRUE | 0.1102 |
| Plxnc1 | 121 | 96 | 83 | 75 | 83 | 92 | 94 | 102 | 90 | 75 | 94 | 91 | 90.79 | 95 | 87 | 0.906 | -0.142 | 0.49520161 | 0.65996429 | 0 | 0.0285 | 0.0202 | 0.0202 | TRUE | 0.1356 |
| Plxnd1 | 3418 | 2697 | 3110 | 3796 | 3472 | 3819 | 2655 | 2869 | 3356 | 3779 | 3917 | 3770 | 3390.72 | 2960 | 3822 | 1.292 | 0.37 | 6.31E-06 | 0.00012872 | 0.0071 | 0.0031 | 0.0045 | 0.0045 | TRUE | 0.1659 |
| Pm20d1 | 742 | 515 | 554 | 607 | 533 | 606 | 576 | 548 | 598 | 604 | 601 | 598 | 587.59 | 574 | 601 | 1.047 | 0.067 | 0.4567655 | 0.62648012 | 0 | 0.0065 | 0.0041 | 0.0041 | TRUE | 0.0176 |
| Pm20d2 | 585 | 423 | 406 | 518 | 409 | 479 | 454 | 450 | 438 | 516 | 461 | 473 | 465.37 | 447 | 483 | 1.08 | 0.111 | 0.26726669 | 0.43772691 | 0 | 0.0075 | 0.0051 | 0.0051 | TRUE | 0.0401 |
| Pmaip1 | 2747 | 2047 | 2146 | 3049 | 2537 | 2753 | 2133 | 2177 | 2316 | 3035 | 2862 | 2718 | 2540.16 | 2209 | 2872 | 1.301 | 0.379 | 1.74E-08 | 8.25E-07 | 0.0019 | 0.0034 | 0.0029 | 0.0029 | TRUE | 0.0304 |
| Pmch | 0 | 0 | 0 | 0 | 0 | 0 | 0 | 0 | 0 | 0 | 0 | 0 | 0 | 0 | 0 | NA | NA | NA | NA | NA | NA | NA | NA | NA | NA |
| Pmel | 383 | 247 | 267 | 302 | 289 | 350 | 297 | 263 | 288 | 301 | 326 | 346 | 303.4 | 283 | 324 | 1.144 | 0.194 | 0.11350196 | 0.24245757 | 0 | 0.0102 | 0.0076 | 0.0076 | TRUE | 0.0466 |
| Pmepa1 | 1806 | 1406 | 1093 | 1592 | 1339 | 1477 | 1403 | 1496 | 1179 | 1585 | 1511 | 1458 | 1438.46 | 1359 | 1518 | 1.116 | 0.159 | 0.07960743 | 0.18847589 | 0.007 | 0.0041 | 0.0052 | 0.0052 | TRUE | 0.1577 |
| Pmf1 | 3120 | 2186 | 2074 | 2310 | 1815 | 2227 | 2423 | 2325 | 2238 | 2299 | 2048 | 2198 | 2255.25 | 2329 | 2182 | 0.937 | -0.094 | 0.16781778 | 0.31927026 | 0.0018 | 0.0035 | 0.0029 | 0.0029 | TRUE | 0.0346 |
| Pmfbp1 | 53 | 30 | 30 | 39 | 43 | 34 | 41 | 32 | 32 | 39 | 49 | 34 | 37.72 | 35 | 41 | 1.136 | 0.183 | 0.56906867 | 0.72273703 | 0 | 0.0653 | 0.0482 | 0.0482 | TRUE | 0.2233 |
| Pml | 12730 | 9153 | 9119 | 10448 | 9238 | 10096 | 9887 | 9736 | 9839 | 10400 | 10421 | 9966 | 10041.63 | 9821 | 10262 | 1.045 | 0.063 | 0.182301 | 0.33792413 | 2.00E-04 | 0.0027 | 0.0015 | 0.0015 | TRUE | 0.0078 |
| Pmm1 | 10612 | 7865 | 8748 | 8319 | 7783 | 9155 | 8242 | 8366 | 9439 | 8281 | 8780 | 9037 | 8690.83 | 8682 | 8699 | 1.002 | 0.003 | 0.9628902 | 0.97947032 | 0.003 | 0.0027 | 0.0031 | 0.0031 | TRUE | 0.0708 |
| Pmm2 | 652 | 499 | 309 | 311 | 303 | 340 | 506 | 531 | 333 | 310 | 342 | 336 | 392.93 | 457 | 329 | 0.719 | -0.476 | 0.00225204 | 0.01322993 | 0.0299 | 0.0085 | 0.0149 | 0.0149 | TRUE | 0.634 |
| Pmp2 | 0 | 0 | 0 | 0 | 0 | 0 | 0 | 0 | 0 | 0 | 0 | 0 | 0 | 0 | 0 | NA | NA | NA | NA | NA | NA | NA | NA | NA | NA |
| Pmp22 | 144 | 125 | 166 | 192 | 250 | 271 | 112 | 133 | 179 | 191 | 282 | 268 | 194.1 | 141 | 247 | 1.755 | 0.811 | 4.64E-05 | 0.00065662 | 0.044 | 0.0146 | 0.023 | 0.023 | TRUE | 0.5616 |
| Pmpca | 11401 | 7881 | 7680 | 8238 | 7385 | 8104 | 8855 | 8383 | 8287 | 8200 | 8331 | 8000 | 8342.55 | 8508 | 8177 | 0.961 | -0.058 | 0.26639527 | 0.43675013 | 7.00E-04 | 0.0027 | 0.0018 | 0.0018 | TRUE | 0.0157 |
| Pmpcb | 7449 | 5233 | 5572 | 6647 | 5835 | 6527 | 5785 | 5566 | 6012 | 6616 | 6582 | 6443 | 6167.63 | 5788 | 6547 | 1.131 | 0.178 | 0.00079916 | 0.00613778 | 6.00E-04 | 0.0028 | 0.0019 | 0.0019 | TRUE | 0.0138 |
| Pms1 | 2278 | 1606 | 1551 | 1600 | 1370 | 1627 | 1769 | 1708 | 1674 | 1593 | 1546 | 1606 | 1649.21 | 1717 | 1582 | 0.921 | -0.119 | 0.06481703 | 0.16231787 | 0 | 0.0039 | 0.0024 | 0.0024 | TRUE | 0.0089 |
| Pms2 | 1843 | 1224 | 1075 | 1187 | 936 | 1168 | 1431 | 1302 | 1160 | 1182 | 1056 | 1153 | 1213.95 | 1298 | 1130 | 0.87 | -0.201 | 0.02697927 | 0.08578747 | 0.0059 | 0.0044 | 0.0051 | 0.0051 | TRUE | 0.1031 |
| Pmvk | 2282 | 1689 | 1757 | 2159 | 1809 | 2235 | 1772 | 1797 | 1896 | 2149 | 2041 | 2206 | 1976.8 | 1822 | 2132 | 1.171 | 0.228 | 0.00048857 | 0.00424249 | 9.00E-04 | 0.0036 | 0.0026 | 0.0026 | TRUE | 0.0167 |
| Pnck | 211 | 173 | 153 | 229 | 176 | 201 | 164 | 184 | 165 | 228 | 199 | 198 | 189.65 | 171 | 208 | 1.221 | 0.288 | 0.0536589 | 0.14156329 | 0 | 0.0149 | 0.0107 | 0.0107 | TRUE | 0.0753 |
| Pnkd | 2306 | 1825 | 1947 | 2203 | 1945 | 2223 | 1791 | 1941 | 2101 | 2193 | 2194 | 2194 | 2069.08 | 1944 | 2194 | 1.13 | 0.176 | 0.01465796 | 0.0548302 | 0.0024 | 0.0036 | 0.0033 | 0.0033 | TRUE | 0.0599 |
| Pnkp | 1544 | 1164 | 1057 | 1190 | 1098 | 1280 | 1199 | 1238 | 1140 | 1185 | 1239 | 1264 | 1210.75 | 1192 | 1229 | 1.03 | 0.043 | 0.55565195 | 0.71081477 | 5.00E-04 | 0.0044 | 0.003 | 0.003 | TRUE | 0.0168 |
| Pnldc1 | 777 | 595 | 550 | 629 | 554 | 636 | 603 | 633 | 593 | 626 | 625 | 628 | 618.12 | 610 | 626 | 1.027 | 0.039 | 0.65817524 | 0.78827709 | 0 | 0.0063 | 0.0039 | 0.0039 | TRUE | 0.0122 |
| Pnlip | 16 | 21 | 23 | 14 | 11 | 10 | 12 | 22 | 25 | 14 | 12 | 10 | 15.97 | 20 | 12 | 0.615 | -0.702 | 0.16537337 | NA | 0 | 0.1509 | 0.1172 | 0.1172 | TRUE | 0.7174 |
| Pnliprp1 | 0 | 0 | 0 | 0 | 0 | 0 | 0 | 0 | 0 | 0 | 0 | 0 | 0 | 0 | 0 | NA | NA | NA | NA | NA | NA | NA | NA | NA | NA |
| Pnliprp2 | 133 | 101 | 116 | 178 | 122 | 133 | 103 | 107 | 125 | 177 | 138 | 131 | 130.33 | 112 | 149 | 1.333 | 0.414 | 0.02876617 | 0.08983064 | 0.0121 | 0.0206 | 0.0181 | 0.0181 | TRUE | 0.2982 |
| Pnma1 | 34 | 20 | 31 | 39 | 37 | 46 | 26 | 21 | 33 | 39 | 42 | 45 | 34.52 | 27 | 42 | 1.555 | 0.637 | 0.0571544 | 0.1485259 | 0 | 0.0711 | 0.0507 | 0.0507 | TRUE | 0.2432 |
| Pnma2 | 943 | 781 | 757 | 967 | 868 | 1019 | 732 | 831 | 817 | 963 | 979 | 1006 | 887.93 | 793 | 983 | 1.241 | 0.311 | 0.00018253 | 0.0019938 | 0.0015 | 0.0051 | 0.0039 | 0.0039 | TRUE | 0.0569 |
| Pnma3 | 60 | 60 | 44 | 72 | 54 | 70 | 47 | 64 | 47 | 72 | 61 | 69 | 59.93 | 53 | 67 | 1.284 | 0.361 | 0.16159259 | 0.31156656 | 0 | 0.042 | 0.0311 | 0.0311 | TRUE | 0.274 |
| Pnma5 | 3342 | 2686 | 2437 | 1908 | 1641 | 1724 | 2596 | 2857 | 2630 | 1899 | 1851 | 1702 | 2255.74 | 2694 | 1817 | 0.675 | -0.568 | 8.71E-16 | 1.64E-13 | 0.0023 | 0.0035 | 0.0031 | 0.0031 | TRUE | 0.0378 |
| Pnmal1 | 31 | 27 | 35 | 25 | 20 | 28 | 24 | 29 | 38 | 25 | 23 | 28 | 27.61 | 30 | 25 | 0.837 | -0.257 | 0.49129325 | 0.65712656 | 0 | 0.0883 | 0.0638 | 0.0638 | TRUE | 0.31 |
| Pnmal2 | 177 | 122 | 116 | 93 | 93 | 105 | 137 | 130 | 125 | 93 | 105 | 104 | 115.59 | 131 | 101 | 0.766 | -0.386 | 0.0348061 | 0.10340545 | 0 | 0.0229 | 0.0152 | 0.0152 | TRUE | 0.0461 |
| Pnmt | 9 | 4 | 10 | 7 | 3 | 9 | 7 | 4 | 11 | 7 | 3 | 9 | 6.88 | 7 | 6 | 0.882 | -0.181 | 0.81025359 | NA | 0 | 0.347 | 0.264 | 0.264 | TRUE | 0.2395 |
| Pnn | 10088 | 8047 | 8104 | 7963 | 6940 | 8425 | 7835 | 8559 | 8744 | 7926 | 7829 | 8317 | 8201.79 | 8379 | 8024 | 0.958 | -0.062 | 0.29951452 | 0.47178324 | 0.002 | 0.0027 | 0.0025 | 0.0025 | TRUE | 0.0399 |
| Pno1 | 7809 | 5287 | 4669 | 5092 | 4351 | 5016 | 6065 | 5624 | 5038 | 5069 | 4908 | 4952 | 5275.83 | 5576 | 4976 | 0.892 | -0.165 | 0.0201229 | 0.06952099 | 0.0041 | 0.0029 | 0.0034 | 0.0034 | TRUE | 0.0866 |
| Pnoc | 20 | 10 | 15 | 17 | 17 | 20 | 16 | 11 | 16 | 17 | 19 | 20 | 16.37 | 14 | 19 | 1.313 | 0.393 | 0.40509643 | NA | 0 | 0.1472 | 0.0984 | 0.0984 | TRUE | 0.1872 |
| Pnp | 3609 | 2659 | 2342 | 2810 | 2519 | 2891 | 2803 | 2828 | 2527 | 2797 | 2842 | 2854 | 2775.15 | 2719 | 2831 | 1.041 | 0.057 | 0.3725094 | 0.54703348 | 0.0014 | 0.0033 | 0.0026 | 0.0026 | TRUE | 0.0458 |
| Pnp2 | 19 | 16 | 15 | 17 | 8 | 15 | 15 | 17 | 16 | 17 | 9 | 15 | 14.79 | 16 | 14 | 0.857 | -0.223 | 0.65707074 | NA | 0 | 0.1627 | 0.113 | 0.113 | TRUE | 0.3288 |
| Pnpla1 | 20 | 9 | 12 | 17 | 12 | 17 | 16 | 10 | 13 | 17 | 14 | 17 | 14.22 | 13 | 16 | 1.233 | 0.302 | 0.55117015 | NA | 0 | 0.1691 | 0.1145 | 0.1145 | TRUE | 0.2004 |
| Pnpla2 | 2192 | 1651 | 1954 | 1987 | 1928 | 2336 | 1702 | 1756 | 2108 | 1978 | 2175 | 2306 | 2004.29 | 1855 | 2153 | 1.161 | 0.215 | 0.01851653 | 0.0652964 | 0.0092 | 0.0036 | 0.0055 | 0.0055 | TRUE | 0.1703 |
| Pnpla3 | 588 | 406 | 413 | 529 | 441 | 484 | 457 | 432 | 446 | 527 | 497 | 478 | 472.67 | 445 | 501 | 1.125 | 0.17 | 0.0869576 | 0.19988502 | 0 | 0.0074 | 0.005 | 0.005 | TRUE | 0.0245 |
| Pnpla5 | 1 | 4 | 2 | 1 | 2 | 1 | 1 | 4 | 2 | 1 | 2 | 1 | 1.9 | 2 | 1 | 0.596 | -0.747 | 0.59608775 | NA | 0 | 1.2468 | 0.8543 | 0.8543 | TRUE | 0.2339 |
| Pnpla6 | 9660 | 6860 | 6253 | 6854 | 5990 | 6900 | 7503 | 7297 | 6747 | 6822 | 6757 | 6811 | 6989.59 | 7182 | 6797 | 0.946 | -0.08 | 0.15882197 | 0.30805298 | 0.0013 | 0.0028 | 0.0022 | 0.0022 | TRUE | 0.0341 |
| Pnpla7 | 916 | 694 | 646 | 816 | 690 | 745 | 711 | 738 | 697 | 812 | 778 | 735 | 745.45 | 715 | 775 | 1.084 | 0.116 | 0.17193455 | 0.32394063 | 0 | 0.0056 | 0.0038 | 0.0038 | TRUE | 0.0246 |
| Pnpla8 | 3489 | 2464 | 2545 | 2695 | 2339 | 2811 | 2710 | 2621 | 2746 | 2683 | 2639 | 2775 | 2695.47 | 2692 | 2699 | 1.002 | 0.004 | 0.95083204 | 0.97288585 | 2.00E-04 | 0.0033 | 0.0021 | 0.0021 | TRUE | 0.0074 |
| Pnpo | 2997 | 2196 | 2212 | 2401 | 2231 | 2534 | 2328 | 2336 | 2387 | 2390 | 2517 | 2501 | 2409.74 | 2350 | 2469 | 1.051 | 0.072 | 0.22260269 | 0.38687503 | 0 | 0.0034 | 0.0021 | 0.0021 | TRUE | 0.0097 |
| Pnpt1 | 7402 | 5457 | 5932 | 6069 | 5339 | 6336 | 5749 | 5805 | 6401 | 6041 | 6023 | 6255 | 6045.43 | 5985 | 6106 | 1.021 | 0.029 | 0.62211924 | 0.76142749 | 0.0016 | 0.0028 | 0.0024 | 0.0024 | TRUE | 0.0448 |
| Pnrc1 | 2802 | 2198 | 2398 | 2075 | 2050 | 2374 | 2176 | 2338 | 2587 | 2065 | 2313 | 2344 | 2303.86 | 2367 | 2241 | 0.947 | -0.078 | 0.33260533 | 0.50692343 | 0.0055 | 0.0035 | 0.0043 | 0.0043 | TRUE | 0.0801 |
| Pnrc2 | 3445 | 2478 | 2576 | 2507 | 2307 | 2732 | 2676 | 2636 | 2779 | 2495 | 2603 | 2697 | 2647.63 | 2697 | 2598 | 0.964 | -0.054 | 0.3784097 | 0.55319728 | 7.00E-04 | 0.0033 | 0.0023 | 0.0023 | TRUE | 0.0147 |
| Poc1a | 1442 | 1052 | 1002 | 1162 | 1070 | 1244 | 1120 | 1119 | 1081 | 1157 | 1207 | 1228 | 1151.97 | 1107 | 1197 | 1.081 | 0.113 | 0.11385032 | 0.24287638 | 0 | 0.0045 | 0.0028 | 0.0028 | TRUE | 0.0107 |
| Poc1b | 1713 | 1193 | 1282 | 1230 | 1140 | 1383 | 1330 | 1269 | 1383 | 1224 | 1286 | 1365 | 1309.71 | 1327 | 1292 | 0.973 | -0.039 | 0.60160297 | 0.74667517 | 0.0016 | 0.0042 | 0.0033 | 0.0033 | TRUE | 0.0301 |
| Poc5 | 1790 | 1361 | 1195 | 1401 | 1296 | 1381 | 1390 | 1448 | 1289 | 1395 | 1462 | 1363 | 1391.19 | 1376 | 1407 | 1.022 | 0.031 | 0.67265178 | 0.79878796 | 0.0015 | 0.0041 | 0.0032 | 0.0032 | TRUE | 0.0351 |
| Podn | 38 | 28 | 34 | 16 | 22 | 13 | 30 | 30 | 37 | 16 | 25 | 13 | 24.93 | 32 | 18 | 0.555 | -0.848 | 0.03698542 | 0.10832635 | 0.003 | 0.0975 | 0.0749 | 0.0749 | TRUE | 0.5369 |
| Podnl1 | 29 | 17 | 28 | 25 | 15 | 25 | 23 | 18 | 30 | 25 | 17 | 25 | 22.88 | 24 | 22 | 0.944 | -0.083 | 0.84176776 | 0.91008975 | 0 | 0.106 | 0.0798 | 0.0798 | TRUE | 0.323 |
| Podxl | 2838 | 2223 | 2741 | 2671 | 2417 | 2956 | 2204 | 2365 | 2958 | 2659 | 2727 | 2918 | 2638.27 | 2509 | 2768 | 1.104 | 0.143 | 0.14212011 | 0.28461858 | 0.0126 | 0.0033 | 0.0064 | 0.0064 | TRUE | 0.296 |
| Podxl2 | 219 | 155 | 174 | 209 | 201 | 216 | 170 | 165 | 188 | 208 | 227 | 213 | 195.12 | 174 | 216 | 1.24 | 0.31 | 0.03144978 | 0.09588559 | 0 | 0.0146 | 0.0098 | 0.0098 | TRUE | 0.0463 |
| Pof1b | 29 | 32 | 34 | 14 | 15 | 14 | 23 | 34 | 37 | 14 | 17 | 14 | 22.99 | 31 | 15 | 0.482 | -1.054 | 0.01281222 | 0.04969473 | 0 | 0.1055 | 0.0786 | 0.0786 | TRUE | 0.4697 |
| Pofut1 | 2322 | 1540 | 1220 | 1334 | 1227 | 1255 | 1803 | 1638 | 1316 | 1328 | 1384 | 1239 | 1451.46 | 1586 | 1317 | 0.829 | -0.27 | 0.00984481 | 0.0409046 | 0.0127 | 0.0041 | 0.0072 | 0.0072 | TRUE | 0.2647 |
| Pofut2 | 1870 | 1267 | 1176 | 1256 | 1111 | 1310 | 1452 | 1348 | 1269 | 1250 | 1253 | 1293 | 1310.94 | 1356 | 1265 | 0.932 | -0.102 | 0.17926305 | 0.33396857 | 0.0019 | 0.0042 | 0.0034 | 0.0034 | TRUE | 0.0479 |

| | | | | | | | | | | | | | | | | | | | | | | | | | |
|---|---|---|---|---|---|---|---|---|---|---|---|---|---|---|---|---|---|---|---|---|---|---|---|---|---|
| Pogk | 440 | 334 | 266 | 287 | 272 | 266 | 342 | 355 | 287 | 286 | 307 | 263 | 306.52 | 328 | 285 | 0.867 | -0.206 | 0.11325305 | 0.24206728 | 0.0058 | 0.0102 | 0.0089 | 0.0089 | TRUE | 0.1302 |
| Poglut1 | 2824 | 2092 | 1861 | 1844 | 1714 | 1812 | 2193 | 2225 | 2008 | 1836 | 1934 | 1789 | 1997.39 | 2142 | 1853 | 0.864 | -0.21 | 0.00227949 | 0.0133637 | 0.0018 | 0.0036 | 0.0029 | 0.0029 | TRUE | 0.0356 |
| Pogz | 6783 | 4877 | 4787 | 5290 | 4759 | 5262 | 5268 | 5188 | 5165 | 5266 | 5369 | 5194 | 5241.59 | 5207 | 5276 | 1.013 | 0.019 | 0.70514103 | 0.82146717 | 0 | 0.0029 | 0.0016 | 0.0016 | TRUE | 0.0028 |
| Pola1 | 6302 | 4786 | 4596 | 4690 | 4210 | 4904 | 4894 | 5091 | 4959 | 4668 | 4749 | 4841 | 4867.18 | 4981 | 4753 | 0.954 | -0.067 | 0.18898186 | 0.34651928 | 2.00E-04 | 0.0029 | 0.0017 | 0.0017 | TRUE | 0.0045 |
| Pola2 | 4276 | 3000 | 2847 | 2577 | 2413 | 2719 | 3321 | 3191 | 3072 | 2565 | 2722 | 2684 | 2925.88 | 3195 | 2657 | 0.831 | -0.267 | 1.06E-05 | 0.00019639 | 9.00E-04 | 0.0032 | 0.0023 | 0.0023 | TRUE | 0.0149 |
| Polb | 2791 | 2061 | 2024 | 2243 | 2010 | 2282 | 2168 | 2192 | 2184 | 2233 | 2267 | 2253 | 2216.11 | 2181 | 2251 | 1.032 | 0.045 | 0.42849527 | 0.60061615 | 0 | 0.0035 | 0.0019 | 0.0019 | TRUE | 6.00E-04 |
| Pold1 | 10719 | 8304 | 8829 | 9743 | 8808 | 9827 | 8325 | 8833 | 9526 | 9698 | 9936 | 9701 | 9336.59 | 8895 | 9778 | 1.1 | 0.137 | 0.02311725 | 0.07680493 | 0.0022 | 0.0027 | 0.0025 | 0.0025 | TRUE | 0.0471 |
| Pold2 | 8201 | 6384 | 6927 | 7464 | 6972 | 8256 | 6369 | 6791 | 7474 | 7430 | 7865 | 8150 | 7346.48 | 6878 | 7815 | 1.137 | 0.185 | 0.00761751 | 0.03374985 | 0.0042 | 0.0027 | 0.0033 | 0.0033 | TRUE | 0.0701 |
| Pold3 | 5673 | 4208 | 4119 | 4048 | 3639 | 4298 | 4406 | 4476 | 4444 | 4029 | 4105 | 4243 | 4283.95 | 4442 | 4126 | 0.929 | -0.106 | 0.0420618 | 0.11862457 | 1.00E-04 | 0.003 | 0.0017 | 0.0017 | TRUE | 0.0076 |
| Pold4 | 306 | 217 | 271 | 400 | 305 | 403 | 238 | 231 | 292 | 398 | 344 | 398 | 316.82 | 254 | 380 | 1.503 | 0.588 | 8.82E-06 | 0.0001693 | 0.0079 | 0.0099 | 0.0093 | 0.0093 | TRUE | 0.1914 |
| Poldip2 | 7669 | 5587 | 5724 | 6438 | 5620 | 6468 | 5956 | 5943 | 6176 | 6408 | 6340 | 6385 | 6201.4 | 6025 | 6378 | 1.059 | 0.082 | 0.09277559 | 0.20928686 | 1.00E-04 | 0.0028 | 0.0016 | 0.0016 | TRUE | 0.0058 |
| Poldip3 | 12364 | 9043 | 8412 | 8083 | 7033 | 8056 | 9603 | 9619 | 9076 | 8046 | 7934 | 7953 | 8705.05 | 9433 | 7978 | 0.846 | -0.242 | 1.21E-06 | 3.41E-05 | 4.00E-04 | 0.0027 | 0.0017 | 0.0017 | TRUE | 0.0132 |
| Pole | 7594 | 5491 | 4748 | 4469 | 4065 | 4688 | 5898 | 5841 | 5123 | 4448 | 4586 | 4628 | 5087.28 | 5621 | 4554 | 0.81 | -0.304 | 4.53E-06 | 9.78E-05 | 0.0029 | 0.0029 | 0.003 | 0.003 | TRUE | 0.0727 |
| Pole2 | 2430 | 1745 | 1797 | 1773 | 1661 | 1854 | 1887 | 1856 | 1939 | 1765 | 1874 | 1830 | 1858.53 | 1894 | 1823 | 0.962 | -0.055 | 0.37884692 | 0.55366657 | 0 | 0.0037 | 0.0023 | 0.0023 | TRUE | 0.0094 |
| Pole3 | 1064 | 803 | 760 | 739 | 627 | 699 | 826 | 854 | 820 | 736 | 707 | 690 | 772.25 | 833 | 711 | 0.853 | -0.229 | 0.00487839 | 0.02404994 | 0 | 0.0055 | 0.0035 | 0.0035 | TRUE | 0.0111 |
| Pole4 | 3788 | 2550 | 2406 | 2644 | 2221 | 2442 | 2942 | 2712 | 2596 | 2632 | 2506 | 2411 | 2633.07 | 2750 | 2516 | 0.914 | -0.129 | 0.06168424 | 0.15651569 | 0.0025 | 0.0033 | 0.0031 | 0.0031 | TRUE | 0.0462 |
| Polg | 8163 | 6275 | 5912 | 6825 | 6013 | 6695 | 6340 | 6675 | 6379 | 6794 | 6783 | 6609 | 6596.56 | 6465 | 6729 | 1.041 | 0.058 | 0.25223453 | 0.42118886 | 3.00E-04 | 0.0028 | 0.0017 | 0.0017 | TRUE | 0.0098 |
| Polg2 | 512 | 442 | 438 | 580 | 401 | 490 | 398 | 470 | 473 | 577 | 452 | 484 | 475.63 | 447 | 504 | 1.133 | 0.18 | 0.13866841 | 0.27970471 | 0.0109 | 0.0074 | 0.0085 | 0.0085 | TRUE | 0.1868 |
| Polh | 1208 | 932 | 1151 | 1165 | 1064 | 1207 | 938 | 991 | 1242 | 1160 | 1200 | 1191 | 1120.49 | 1057 | 1184 | 1.121 | 0.165 | 0.10267938 | 0.22550651 | 0.0106 | 0.0045 | 0.0065 | 0.0065 | TRUE | 0.2772 |
| Poli | 3053 | 2374 | 2454 | 2478 | 2204 | 2539 | 2371 | 2525 | 2648 | 2467 | 2486 | 2506 | 2500.58 | 2515 | 2486 | 0.99 | -0.015 | 0.81165052 | 0.89135487 | 0.0012 | 0.0034 | 0.0025 | 0.0025 | TRUE | 0.0312 |
| Polk | 1498 | 1119 | 1406 | 1585 | 1351 | 1527 | 1163 | 1190 | 1517 | 1578 | 1524 | 1507 | 1413.32 | 1290 | 1536 | 1.192 | 0.254 | 0.00976422 | 0.04067618 | 0.0109 | 0.0041 | 0.0062 | 0.0062 | TRUE | 0.2814 |
| Poll | 1574 | 1160 | 1170 | 1334 | 1118 | 1263 | 1222 | 1234 | 1262 | 1328 | 1261 | 1247 | 1259.1 | 1239 | 1279 | 1.032 | 0.045 | 0.51517241 | 0.67758896 | 0 | 0.0043 | 0.0027 | 0.0027 | TRUE | 0.0138 |
| Polm | 569 | 452 | 476 | 492 | 442 | 520 | 442 | 481 | 514 | 490 | 499 | 513 | 489.66 | 479 | 501 | 1.048 | 0.068 | 0.50479477 | 0.66836791 | 0.0014 | 0.0073 | 0.0053 | 0.0053 | TRUE | 0.0566 |
| Poln | 80 | 53 | 45 | 62 | 61 | 79 | 62 | 56 | 49 | 62 | 69 | 78 | 62.6 | 56 | 70 | 1.243 | 0.313 | 0.21025077 | 0.37224257 | 0 | 0.0403 | 0.029 | 0.029 | TRUE | 0.1059 |
| Polq | 1494 | 1101 | 935 | 1056 | 986 | 1196 | 1160 | 1171 | 1009 | 1051 | 1112 | 1181 | 1114.07 | 1113 | 1115 | 1 | 0.001 | 0.99461144 | 0.99691164 | 0.0042 | 0.0046 | 0.0044 | 0.0044 | TRUE | 0.079 |
| Polr1a | 11386 | 8109 | 7146 | 7641 | 7040 | 8038 | 8843 | 8625 | 7710 | 7606 | 7942 | 7935 | 8110.23 | 8393 | 7828 | 0.932 | -0.101 | 0.11015609 | 0.23733208 | 0.0027 | 0.0027 | 0.0028 | 0.0028 | TRUE | 0.0616 |
| Polr1b | 7851 | 5826 | 5073 | 5064 | 4517 | 5395 | 6098 | 6197 | 5474 | 5041 | 5096 | 5326 | 5538.39 | 5923 | 5154 | 0.87 | -0.201 | 0.00151991 | 0.00975487 | 0.0023 | 0.0029 | 0.0027 | 0.0027 | TRUE | 0.0532 |
| Polr1c | 8006 | 6041 | 6005 | 5782 | 5355 | 6043 | 6218 | 6426 | 6479 | 5755 | 6041 | 5965 | 6147.46 | 6374 | 5920 | 0.929 | -0.106 | 0.03793763 | 0.11018951 | 4.00E-04 | 0.0028 | 0.0017 | 0.0017 | TRUE | 0.0073 |
| Polr1d | 6135 | 4235 | 4755 | 4844 | 4516 | 5382 | 4765 | 4505 | 5131 | 4822 | 5095 | 5313 | 4938.2 | 4800 | 5077 | 1.058 | 0.081 | 0.22369815 | 0.38816531 | 0.003 | 0.0029 | 0.003 | 0.003 | TRUE | 0.0437 |
| Polr1e | 2593 | 1848 | 1575 | 1711 | 1516 | 1824 | 2014 | 1966 | 1699 | 1703 | 1710 | 1801 | 1815.48 | 1893 | 1738 | 0.917 | -0.124 | 0.11242472 | 0.24079013 | 0.0037 | 0.0037 | 0.0039 | 0.0039 | TRUE | 0.0955 |
| Polr2a | 51257 | 35125 | 29007 | 31112 | 29230 | 32327 | 39809 | 37362 | 31298 | 30969 | 32974 | 31912 | 34054.09 | 36156 | 31952 | 0.884 | -0.178 | 0.02447099 | 0.08009249 | 0.0078 | 0.0025 | 0.0045 | 0.0045 | TRUE | 0.169 |
| Polr2b | 20436 | 14509 | 13671 | 14255 | 13064 | 15070 | 15872 | 15433 | 14751 | 14189 | 14738 | 14876 | 14976.51 | 15352 | 14601 | 0.951 | -0.073 | 0.15750339 | 0.30636854 | 9.00E-04 | 0.0026 | 0.0018 | 0.0018 | TRUE | 0.0143 |
| Polr2c | 6855 | 4908 | 5202 | 5442 | 4740 | 5561 | 5324 | 5221 | 5613 | 5417 | 5347 | 5490 | 5401.87 | 5386 | 5418 | 1.006 | 0.009 | 0.86898352 | 0.92765565 | 5.00E-04 | 0.0029 | 0.0019 | 0.0019 | TRUE | 0.0164 |
| Polr2d | 2759 | 1943 | 1704 | 1839 | 1465 | 1755 | 2143 | 2067 | 1839 | 1831 | 1653 | 1732 | 1877.3 | 2016 | 1739 | 0.862 | -0.214 | 0.00566342 | 0.02700941 | 0.0036 | 0.0037 | 0.0038 | 0.0038 | TRUE | 0.071 |
| Polr2e | 11486 | 8246 | 8396 | 8787 | 7870 | 9279 | 8921 | 8771 | 9059 | 8747 | 8878 | 9160 | 8922.6 | 8917 | 8928 | 1.001 | 0.002 | 0.96971949 | 0.9834812 | 2.00E-04 | 0.0027 | 0.0016 | 0.0016 | TRUE | 0.0063 |
| Polr2f | 3902 | 2712 | 2567 | 2646 | 2503 | 2756 | 3031 | 2885 | 2770 | 2634 | 2824 | 2721 | 2810.51 | 2895 | 2726 | 0.941 | -0.088 | 0.16107348 | 0.31078701 | 0.0013 | 0.0033 | 0.0025 | 0.0025 | TRUE | 0.0209 |
| Polr2g | 4067 | 2696 | 2547 | 3020 | 2543 | 2874 | 3159 | 2868 | 2748 | 3006 | 2869 | 2837 | 2914.42 | 2925 | 2904 | 0.992 | -0.011 | 0.86776608 | 0.92697911 | 0.0028 | 0.0032 | 0.0031 | 0.0031 | TRUE | 0.0606 |
| Polr2h | 3460 | 2503 | 2471 | 2516 | 2272 | 2703 | 2687 | 2662 | 2666 | 2504 | 2563 | 2668 | 2625.26 | 2672 | 2578 | 0.965 | -0.051 | 0.37465959 | 0.54920038 | 0 | 0.0033 | 0.002 | 0.002 | TRUE | 0.0114 |
| Polr2i | 1546 | 1056 | 984 | 1041 | 830 | 1120 | 1201 | 1123 | 1062 | 1036 | 936 | 1106 | 1077.31 | 1129 | 1026 | 0.909 | -0.138 | 0.1138089 | 0.24285914 | 0.0044 | 0.0046 | 0.0045 | 0.0045 | TRUE | 0.0686 |
| Polr2j | 2483 | 1716 | 1510 | 1555 | 1395 | 1583 | 1928 | 1825 | 1629 | 1548 | 1574 | 1563 | 1677.87 | 1794 | 1562 | 0.869 | -0.202 | 0.00813447 | 0.03552275 | 0.0032 | 0.0038 | 0.0036 | 0.0036 | TRUE | 0.0775 |
| Polr2k | 364 | 281 | 304 | 281 | 252 | 305 | 283 | 299 | 328 | 280 | 284 | 301 | 295.78 | 303 | 288 | 0.953 | -0.069 | 0.57393862 | 0.72603789 | 0 | 0.0104 | 0.0074 | 0.0074 | TRUE | 0.057 |
| Polr2l | 1231 | 794 | 624 | 636 | 547 | 636 | 956 | 845 | 673 | 633 | 617 | 628 | 725.32 | 825 | 626 | 0.758 | -0.4 | 0.00066457 | 0.00534537 | 0.0146 | 0.0057 | 0.0086 | 0.0086 | TRUE | 0.2602 |
| Polr2m | 16183 | 11175 | 10534 | 9016 | 8815 | 10144 | 12569 | 11887 | 11366 | 8975 | 9944 | 10014 | 10792.32 | 11941 | 9644 | 0.808 | -0.308 | 1.15E-06 | 3.29E-05 | 0.0029 | 0.0026 | 0.0028 | 0.0028 | TRUE | 0.0452 |
| Polr3a | 3981 | 2802 | 2696 | 2762 | 2482 | 2640 | 3092 | 2980 | 2909 | 2749 | 2800 | 2606 | 2856.1 | 2994 | 2718 | 0.907 | -0.14 | 0.02019798 | 0.06969815 | 8.00E-04 | 0.0033 | 0.0023 | 0.0023 | TRUE | 0.0159 |
| Polr3b | 7797 | 5833 | 5527 | 5996 | 5628 | 6457 | 6056 | 6204 | 5964 | 5968 | 6349 | 6374 | 6152.52 | 6075 | 6230 | 1.026 | 0.037 | 0.49383373 | 0.65904439 | 6.00E-04 | 0.0028 | 0.0019 | 0.0019 | TRUE | 0.0166 |
| Polr3c | 2978 | 2165 | 1932 | 1961 | 1744 | 2173 | 2313 | 2303 | 2085 | 1952 | 1967 | 2145 | 2127.48 | 2234 | 2021 | 0.905 | -0.144 | 0.04307233 | 0.12085305 | 0.0024 | 0.0035 | 0.0032 | 0.0032 | TRUE | 0.0406 |
| Polr3d | 2467 | 1784 | 1685 | 1715 | 1608 | 1830 | 1916 | 1898 | 1818 | 1707 | 1814 | 1806 | 1826.55 | 1877 | 1776 | 0.945 | -0.081 | 0.20980659 | 0.37164031 | 4.00E-04 | 0.0037 | 0.0024 | 0.0024 | TRUE | 0.014 |
| Polr3e | 4981 | 3631 | 3325 | 3492 | 3152 | 3766 | 3869 | 3862 | 3588 | 3476 | 3556 | 3718 | 3677.96 | 3773 | 3583 | 0.95 | -0.075 | 0.21229648 | 0.37461155 | 0.0011 | 0.0031 | 0.0023 | 0.0023 | TRUE | 0.0222 |
| Polr3f | 2289 | 1689 | 1456 | 1600 | 1346 | 1428 | 1778 | 1797 | 1571 | 1593 | 1518 | 1410 | 1611.01 | 1715 | 1507 | 0.878 | -0.188 | 0.01831397 | 0.06468715 | 0.0036 | 0.0039 | 0.0039 | 0.0039 | TRUE | 0.0641 |
| Polr3g | 6603 | 4476 | 4300 | 5249 | 4484 | 5058 | 5128 | 4761 | 4640 | 5225 | 5058 | 4993 | 4967.55 | 4843 | 5092 | 1.051 | 0.072 | 0.22365585 | 0.38816531 | 0.0014 | 0.0029 | 0.0023 | 0.0023 | TRUE | 0.0329 |
| Polr3gl | 310 | 201 | 219 | 283 | 264 | 341 | 241 | 214 | 236 | 282 | 298 | 337 | 267.83 | 230 | 306 | 1.325 | 0.406 | 0.00209485 | 0.01251219 | 0.0029 | 0.0113 | 0.0087 | 0.0087 | TRUE | 0.0925 |
| Polr3h | 1945 | 1307 | 1130 | 1240 | 1097 | 1282 | 1511 | 1390 | 1219 | 1234 | 1238 | 1266 | 1309.58 | 1373 | 1246 | 0.906 | -0.143 | 0.09814153 | 0.21824375 | 0.0052 | 0.0042 | 0.0046 | 0.0046 | TRUE | 0.1149 |
| Polr3k | 1881 | 1356 | 1303 | 1474 | 1191 | 1481 | 1461 | 1442 | 1406 | 1467 | 1344 | 1462 | 1430.32 | 1436 | 1424 | 0.992 | -0.012 | 0.86357557 | 0.92466653 | 6.00E-04 | 0.0041 | 0.0028 | 0.0028 | TRUE | 0.0289 |
| Polrmt | 3860 | 2904 | 2721 | 3048 | 2682 | 3043 | 2998 | 3089 | 2936 | 3034 | 3026 | 3004 | 3014.37 | 3008 | 3021 | 1.005 | 0.007 | 0.9061334 | 0.94880353 | 0 | 0.0032 | 0.0019 | 0.0019 | TRUE | 0.0066 |
| Pom121 | 8350 | 6051 | 5907 | 6195 | 5437 | 6496 | 6485 | 6436 | 6374 | 6166 | 6133 | 6413 | 6334.6 | 6432 | 6237 | 0.97 | -0.044 | 0.37143299 | 0.54612609 | 2.00E-04 | 0.0028 | 0.0016 | 0.0016 | TRUE | 0.0074 |
| Pom121l12 | 0 | 0 | 0 | 0 | 0 | 0 | 0 | 0 | 0 | 0 | 0 | 0 | 0 | 0 | 0 | NA | NA | NA | NA | NA | NA | NA | NA | NA | NA |
| Pom121l2 | 16 | 11 | 11 | 20 | 17 | 28 | 12 | 12 | 12 | 20 | 19 | 28 | 17.12 | 12 | 22 | 1.854 | 0.891 | 0.05750751 | 0.14912554 | 0 | 0.1409 | 0.0953 | 0.0953 | TRUE | 0.2645 |
| Pomc | 68 | 52 | 44 | 44 | 35 | 48 | 53 | 55 | 47 | 44 | 39 | 47 | 47.71 | 52 | 43 | 0.84 | -0.251 | 0.3667554 | 0.5414766 | 0 | 0.0521 | 0.0344 | 0.0344 | TRUE | 0.0479 |
| Pomgnt1 | 1841 | 1225 | 1156 | 1171 | 1031 | 1082 | 1430 | 1303 | 1247 | 1166 | 1163 | 1068 | 1229.49 | 1327 | 1132 | 0.852 | -0.231 | 0.00422833 | 0.02154565 | 0.003 | 0.0044 | 0.0039 | 0.0039 | TRUE | 0.0575 |
| Pomp | 5140 | 3489 | 3647 | 3821 | 3492 | 4046 | 3992 | 3711 | 3935 | 3803 | 3939 | 3994 | 3895.85 | 3879 | 3912 | 1.008 | 0.012 | 0.83808707 | 0.90768951 | 8.00E-04 | 0.003 | 0.0021 | 0.0021 | TRUE | 0.0174 |
| Pomt1 | 1534 | 1251 | 1269 | 1289 | 1178 | 1398 | 1191 | 1331 | 1369 | 1283 | 1329 | 1380 | 1313.89 | 1297 | 1331 | 1.027 | 0.039 | 0.61943938 | 0.75986884 | 0.0027 | 0.0042 | 0.0037 | 0.0037 | TRUE | 0.0636 |
| Pomt2 | 1862 | 1341 | 1328 | 1558 | 1434 | 1527 | 1446 | 1426 | 1433 | 1551 | 1618 | 1507 | 1496.89 | 1435 | 1559 | 1.085 | 0.118 | 0.07445179 | 0.17945519 | 0 | 0.004 | 0.0025 | 0.0025 | TRUE | 0.013 |
| Pon1 | 8 | 2 | 1 | 3 | 4 | 4 | 6 | 2 | 1 | 3 | 5 | 4 | 3.48 | 3 | 4 | 1.167 | 0.223 | 0.83151489 | NA | 0 | 0.6839 | 0.5106 | 0.5106 | TRUE | 1.2899 |
| Pon2 | 196 | 141 | 126 | 149 | 147 | 130 | 152 | 150 | 136 | 148 | 166 | 128 | 146.77 | 146 | 147 | 1.006 | 0.009 | 0.95637448 | 0.97583135 | 0 | 0.0186 | 0.0142 | 0.0142 | TRUE | 0.1364 |
| Pon3 | 131 | 87 | 79 | 89 | 118 | 107 | 102 | 93 | 85 | 89 | 133 | 106 | 101.14 | 93 | 109 | 1.164 | 0.219 | 0.30375518 | 0.47628776 | 0.0154 | 0.0259 | 0.0229 | 0.0229 | TRUE | 0.3046 |
| Pop1 | 1960 | 1213 | 894 | 914 | 757 | 898 | 1522 | 1290 | 965 | 910 | 854 | 886 | 1071.23 | 1259 | 883 | 0.701 | -0.513 | 7.31E-05 | 0.00094583 | 0.0238 | 0.0046 | 0.0111 | 0.0111 | TRUE | 0.2134 |
| Pop4 | 3301 | 2410 | 2126 | 2216 | 1953 | 2219 | 2564 | 2563 | 2294 | 2206 | 2203 | 2190 | 2336.78 | 2474 | 2200 | 0.889 | -0.17 | 0.01033612 | 0.04229436 | 0.0016 | 0.0034 | 0.0027 | 0.0027 | TRUE | 0.0483 |
| Pop5 | 2414 | 1645 | 1664 | 1778 | 1610 | 1774 | 1875 | 1750 | 1795 | 1770 | 1816 | 1751 | 1792.89 | 1807 | 1779 | 0.984 | -0.023 | 0.71436282 | 0.82842903 | 0 | 0.0037 | 0.0024 | 0.0024 | TRUE | 0.0137 |
| Pop7 | 930 | 599 | 535 | 577 | 490 | 540 | 722 | 637 | 577 | 574 | 553 | 533 | 599.48 | 645 | 553 | 0.855 | -0.226 | 0.03072589 | 0.09420102 | 0.0058 | 0.0064 | 0.0062 | 0.0062 | TRUE | 0.133 |

| | | | | | | | | | | | | | | | | | | | | | | | | |
|---|---|---|---|---|---|---|---|---|---|---|---|---|---|---|---|---|---|---|---|---|---|---|---|---|
| Popdc2 | 22 | 16 | 15 | 17 | 20 | 13 | 17 | 17 | 16 | 17 | 23 | 13 | 17.1 | 17 | 18 | 1.034 | 0.049 | 0.91675004 | 0.95406352 | 0 | 0.141 | 0.0978 | 0.0978 | TRUE | 0.2569 |
| Popdc3 | 161 | 129 | 137 | 132 | 143 | 124 | 125 | 137 | 148 | 131 | 161 | 122 | 137.53 | 137 | 138 | 1.013 | 0.018 | 0.91880566 | 0.95538599 | 0.0066 | 0.0197 | 0.0159 | 0.0159 | TRUE | 0.2092 |
| Por | 4229 | 3151 | 3065 | 3390 | 2848 | 3190 | 3284 | 3352 | 3307 | 3374 | 3213 | 3149 | 3279.92 | 3314 | 3245 | 0.979 | -0.03 | 0.59391333 | 0.74083813 | 4.00E-04 | 0.0031 | 0.002 | 0.002 | TRUE | 0.0148 |
| Porcn | 789 | 559 | 497 | 639 | 628 | 668 | 613 | 595 | 536 | 636 | 708 | 659 | 624.6 | 581 | 668 | 1.147 | 0.198 | 0.03796186 | 0.11023749 | 0.0022 | 0.0062 | 0.0049 | 0.0049 | TRUE | 0.0525 |
| Postn | 2 | 8 | 6 | 0 | 1 | 1 | 2 | 9 | 6 | 0 | 1 | 1 | 3.11 | 6 | 1 | 0.128 | -2.968 | 0.03857926 | NA | 0 | 0.7648 | 0.6483 | 0.6483 | TRUE | 0.2999 |
| Pot1a | 3607 | 2592 | 2675 | 2752 | 2463 | 2728 | 2801 | 2757 | 2886 | 2739 | 2779 | 2693 | 2775.93 | 2815 | 2737 | 0.972 | -0.041 | 0.4724318 | 0.64096316 | 0 | 0.0033 | 0.0019 | 0.0019 | TRUE | 0.0058 |
| Pot1b | 514 | 401 | 335 | 324 | 275 | 316 | 399 | 427 | 361 | 323 | 310 | 312 | 355.31 | 396 | 315 | 0.796 | -0.33 | 0.00402722 | 0.02072778 | 0 | 0.0091 | 0.0066 | 0.0066 | TRUE | 0.0624 |
| Poteg | 0 | 2 | 0 | 0 | 0 | 1 | 0 | 2 | 0 | 0 | 0 | 1 | 0.52 | 1 | 0 | 0.534 | -0.904 | 0.76321451 | NA | 0 | 4.5682 | 4.7886 | 4.7886 | TRUE | 0.9746 |
| Pou1f1 | 0 | 0 | 0 | 0 | 0 | 0 | 0 | 0 | 0 | 0 | 0 | 0 | 0 | 0 | 0 | NA | NA | NA | NA | NA | NA | NA | NA | NA | NA |
| Pou2af1 | 1 | 3 | 1 | 1 | 0 | 0 | 1 | 3 | 1 | 1 | 0 | 0 | 1.01 | 2 | 0 | 0.223 | -2.163 | 0.29304645 | NA | 0 | 2.356 | 1.7572 | 1.7572 | TRUE | 0.4465 |
| Pou2f1 | 5132 | 3794 | 3754 | 4069 | 3656 | 4465 | 3986 | 4036 | 4051 | 4050 | 4124 | 4408 | 4109.04 | 4024 | 4194 | 1.042 | 0.06 | 0.28984676 | 0.4618935 | 7.00E-04 | 0.003 | 0.0021 | 0.0021 | TRUE | 0.0243 |
| Pou2f2 | 83 | 60 | 55 | 91 | 96 | 88 | 64 | 64 | 59 | 91 | 108 | 87 | 78.9 | 62 | 95 | 1.518 | 0.602 | 0.00683703 | 0.03120104 | 0 | 0.0325 | 0.0226 | 0.0226 | TRUE | 0.1324 |
| Pou2f3 | 450 | 353 | 413 | 889 | 743 | 862 | 349 | 375 | 446 | 885 | 838 | 851 | 624.1 | 390 | 858 | 2.207 | 1.142 | 5.89E-28 | 4.01E-25 | 0.0057 | 0.0062 | 0.006 | 0.006 | TRUE | 0.1746 |
| Pou3f1 | 407 | 343 | 310 | 379 | 327 | 407 | 316 | 365 | 334 | 377 | 369 | 402 | 360.56 | 338 | 383 | 1.133 | 0.181 | 0.11213735 | 0.24044091 | 0 | 0.009 | 0.0065 | 0.0065 | TRUE | 0.0525 |
| Pou3f2 | 20 | 16 | 10 | 14 | 19 | 24 | 16 | 17 | 11 | 14 | 21 | 24 | 17.07 | 15 | 20 | 1.356 | 0.44 | 0.35618276 | 0.53063584 | 0 | 0.1413 | 0.1042 | 0.1042 | TRUE | 0.3608 |
| Pou3f3 | 75 | 75 | 75 | 64 | 43 | 51 | 58 | 80 | 81 | 64 | 49 | 50 | 63.58 | 73 | 54 | 0.748 | -0.419 | 0.10751072 | 0.23332079 | 0.015 | 0.0397 | 0.0325 | 0.0325 | TRUE | 0.3236 |
| Pou3f4 | 4 | 5 | 1 | 1 | 2 | 1 | 3 | 5 | 1 | 1 | 2 | 1 | 2.29 | 3 | 1 | 0.44 | -1.184 | 0.36790483 | NA | 0 | 1.0372 | 0.7233 | 0.7233 | TRUE | 0.1087 |
| Pou4f1 | 444 | 379 | 265 | 298 | 254 | 245 | 345 | 403 | 286 | 297 | 287 | 242 | 309.82 | 345 | 275 | 0.797 | -0.327 | 0.02589899 | 0.08342897 | 0.0171 | 0.0101 | 0.0122 | 0.0122 | TRUE | 0.1449 |
| Pou4f2 | 294 | 207 | 206 | 343 | 288 | 361 | 228 | 220 | 222 | 341 | 325 | 356 | 282.25 | 223 | 341 | 1.524 | 0.608 | 5.96E-07 | 1.89E-05 | 0 | 0.0108 | 0.007 | 0.007 | TRUE | 0.0186 |
| Pou4f3 | 36 | 23 | 18 | 36 | 25 | 19 | 28 | 24 | 19 | 36 | 28 | 19 | 25.77 | 24 | 28 | 1.145 | 0.195 | 0.62668711 | 0.76482743 | 0.0309 | 0.0944 | 0.0768 | 0.0768 | TRUE | 0.2252 |
| Pou5f1 | 43595 | 31651 | 31531 | 36481 | 32174 | 36506 | 33858 | 33667 | 34022 | 36313 | 36296 | 36037 | 35032.11 | 33849 | 36215 | 1.07 | 0.097 | 0.01878228 | 0.06602438 | 0 | 0.0025 | 0.0012 | 0.0012 | TRUE | 3.00E-04 |
| Pou5f2 | 0 | 2 | 5 | 4 | 1 | 1 | 0 | 2 | 5 | 4 | 1 | 1 | 2.27 | 2 | 2 | 0.841 | -0.25 | 0.86034516 | NA | 0.8833 | 1.0466 | 1.0025 | 1.0025 | TRUE | 0.2914 |
| Pou6f1 | 1399 | 1088 | 909 | 602 | 548 | 525 | 1087 | 1157 | 981 | 599 | 618 | 518 | 826.72 | 1075 | 578 | 0.538 | -0.895 | 8.53E-20 | 2.65E-17 | 0.0063 | 0.0053 | 0.0056 | 0.0056 | TRUE | 0.0986 |
| Pou6f2 | 11 | 10 | 6 | 4 | 12 | 6 | 9 | 11 | 6 | 4 | 14 | 6 | 8.18 | 9 | 8 | 0.903 | -0.147 | 0.8349055 | NA | 0.0849 | 0.2921 | 0.2357 | 0.2357 | TRUE | 1.0714 |
| Pp2d1 | 1 | 0 | 0 | 1 | 1 | 0 | 1 | 0 | 0 | 1 | 1 | 0 | 0.48 | 0 | 1 | 2.037 | 1.027 | 0.71241723 | NA | 0 | 4.9057 | 3.8521 | 3.8521 | TRUE | 0.2664 |
| Ppa1 | 12507 | 8430 | 6974 | 7139 | 6105 | 7275 | 9714 | 8967 | 7525 | 7106 | 6887 | 7182 | 7896.7 | 8735 | 7058 | 0.808 | -0.308 | 0.00019123 | 0.00206755 | 0.0082 | 0.0027 | 0.0048 | 0.0048 | TRUE | 0.1791 |
| Ppa2 | 3611 | 2476 | 2592 | 2556 | 2248 | 2681 | 2805 | 2634 | 2797 | 2544 | 2536 | 2647 | 2660.29 | 2745 | 2576 | 0.938 | -0.092 | 0.12277902 | 0.25618693 | 5.00E-04 | 0.0033 | 0.0022 | 0.0022 | TRUE | 0.0151 |
| Ppan | 5284 | 3998 | 3598 | 3424 | 3218 | 3617 | 4104 | 4253 | 3882 | 3408 | 3630 | 3571 | 3807.95 | 4080 | 3536 | 0.867 | -0.206 | 0.00058777 | 0.00488522 | 0.0012 | 0.003 | 0.0023 | 0.0023 | TRUE | 0.0214 |
| Ppap2a | 1327 | 902 | 957 | 873 | 793 | 940 | 1031 | 959 | 1033 | 869 | 895 | 928 | 952.36 | 1008 | 897 | 0.89 | -0.168 | 0.03136518 | 0.09570933 | 0 | 0.0049 | 0.0033 | 0.0033 | TRUE | 0.0203 |
| Ppap2b | 945 | 677 | 702 | 542 | 530 | 569 | 734 | 720 | 757 | 540 | 598 | 562 | 651.77 | 737 | 567 | 0.768 | -0.381 | 1.68E-05 | 0.00029134 | 0 | 0.0061 | 0.0041 | 0.0041 | TRUE | 0.027 |
| Ppap2c | 1787 | 1369 | 1406 | 1749 | 1587 | 1771 | 1388 | 1456 | 1517 | 1741 | 1790 | 1748 | 1606.77 | 1454 | 1760 | 1.212 | 0.277 | 3.66E-05 | 0.00054771 | 5.00E-04 | 0.0039 | 0.0026 | 0.0026 | TRUE | 0.0198 |
| Ppapdc1a | 1 | 1 | 2 | 1 | 3 | 3 | 1 | 1 | 2 | 1 | 3 | 3 | 1.89 | 1 | 2 | 1.867 | 0.901 | 0.51834845 | NA | 0 | 1.2565 | 0.8204 | 0.8204 | TRUE | 0.2997 |
| Ppapdc1b | 873 | 662 | 775 | 985 | 854 | 942 | 678 | 704 | 836 | 980 | 963 | 930 | 848.69 | 739 | 958 | 1.298 | 0.376 | 7.04E-05 | 0.0009226 | 0.0051 | 0.0052 | 0.0052 | 0.0052 | TRUE | 0.1528 |
| Ppapdc2 | 173 | 155 | 150 | 171 | 143 | 177 | 134 | 165 | 162 | 170 | 161 | 175 | 161.22 | 154 | 169 | 1.104 | 0.142 | 0.37918937 | 0.55384873 | 0 | 0.0171 | 0.0126 | 0.0126 | TRUE | 0.1421 |
| Ppapdc3 | 41 | 29 | 22 | 34 | 27 | 35 | 32 | 31 | 24 | 34 | 30 | 35 | 30.88 | 29 | 33 | 1.14 | 0.189 | 0.58314698 | 0.73299796 | 0 | 0.0792 | 0.0532 | 0.0532 | TRUE | 0.1436 |
| Ppara | 79 | 73 | 84 | 128 | 123 | 116 | 61 | 78 | 91 | 127 | 139 | 115 | 101.72 | 77 | 127 | 1.669 | 0.739 | 0.00049494 | 0.00428216 | 0.0118 | 0.0257 | 0.022 | 0.022 | TRUE | 0.3212 |
| Ppard | 2038 | 1574 | 1662 | 1694 | 1584 | 1916 | 1583 | 1674 | 1793 | 1686 | 1787 | 1891 | 1735.81 | 1683 | 1788 | 1.063 | 0.088 | 0.24163938 | 0.40930928 | 0.0027 | 0.0038 | 0.0035 | 0.0035 | TRUE | 0.0392 |
| Pparg | 14 | 2 | 6 | 9 | 5 | 8 | 11 | 2 | 6 | 9 | 6 | 8 | 7 | 6 | 8 | 1.133 | 0.18 | 0.81493975 | NA | 0.1212 | 0.3412 | 0.2825 | 0.2825 | TRUE | 0.2298 |
| Ppargc1a | 11 | 16 | 5 | 8 | 10 | 9 | 9 | 17 | 5 | 8 | 11 | 9 | 9.85 | 10 | 9 | 0.91 | -0.136 | 0.83325853 | NA | 0.0777 | 0.2431 | 0.1965 | 0.1965 | TRUE | 0.7226 |
| Ppargc1b | 260 | 226 | 266 | 355 | 309 | 371 | 202 | 240 | 287 | 353 | 349 | 366 | 299.59 | 243 | 356 | 1.472 | 0.558 | 6.93E-05 | 0.00091049 | 0.0116 | 0.0103 | 0.0107 | 0.0107 | TRUE | 0.2758 |
| Ppat | 10353 | 7444 | 7191 | 6938 | 6204 | 7284 | 8041 | 7918 | 7759 | 6906 | 6999 | 7190 | 7468.85 | 7906 | 7032 | 0.889 | -0.169 | 0.0005476 | 0.00461763 | 2.00E-04 | 0.0027 | 0.0016 | 0.0016 | TRUE | 0.0048 |
| Ppbp | 0 | 0 | 0 | 1 | 0 | 0 | 0 | 0 | 0 | 1 | 0 | 0 | 0.17 | 0 | 0 | 2.094 | 1.067 | 0.79379329 | NA | 0 | 14.2885 | 10 | 10 | TRUE | 0.3906 |
| Ppcdc | 719 | 560 | 401 | 461 | 389 | 522 | 558 | 596 | 433 | 459 | 439 | 515 | 499.96 | 529 | 471 | 0.89 | -0.168 | 0.1860086 | 0.34228524 | 0.0151 | 0.0072 | 0.0096 | 0.0096 | TRUE | 0.2878 |
| Ppcs | 1089 | 763 | 650 | 758 | 603 | 755 | 846 | 812 | 701 | 755 | 680 | 745 | 756.46 | 786 | 727 | 0.923 | -0.115 | 0.23329181 | 0.39938195 | 0.005 | 0.0056 | 0.0054 | 0.0054 | TRUE | 0.104 |
| Ppdpf | 2660 | 1863 | 1736 | 1782 | 1622 | 1945 | 2066 | 1982 | 1873 | 1774 | 1830 | 1920 | 1907.38 | 1974 | 1841 | 0.932 | -0.101 | 0.13963643 | 0.28083355 | 0.0015 | 0.0037 | 0.0028 | 0.0028 | TRUE | 0.0238 |
| Ppef1 | 32 | 27 | 24 | 35 | 27 | 26 | 25 | 29 | 26 | 35 | 30 | 26 | 28.41 | 27 | 30 | 1.147 | 0.198 | 0.58121234 | 0.7318534 | 0 | 0.0859 | 0.0576 | 0.0576 | TRUE | 0.1259 |
| Ppef2 | 247 | 163 | 182 | 189 | 155 | 201 | 192 | 173 | 196 | 188 | 175 | 198 | 187.17 | 187 | 187 | 1 | 0 | 0.99936083 | 0.99957064 | 0 | 0.0151 | 0.0104 | 0.0104 | TRUE | 0.0427 |
| Ppfia1 | 5918 | 4382 | 4004 | 4032 | 3737 | 4242 | 4596 | 4661 | 4320 | 4013 | 4216 | 4188 | 4332.38 | 4526 | 4139 | 0.914 | -0.129 | 0.0233345 | 0.07740444 | 8.00E-04 | 0.003 | 0.0021 | 0.0021 | TRUE | 0.019 |
| Ppfia2 | 0 | 0 | 0 | 0 | 0 | 1 | 0 | 0 | 0 | 0 | 0 | 1 | 0.16 | 0 | 0 | 2.094 | 1.067 | 0.79379329 | NA | 0 | 14.4078 | 10 | 10 | TRUE | 0.3835 |
| Ppfia3 | 417 | 363 | 349 | 338 | 331 | 362 | 324 | 386 | 377 | 336 | 373 | 357 | 358.96 | 362 | 355 | 0.985 | -0.023 | 0.84888491 | 0.91446428 | 0.0035 | 0.009 | 0.0073 | 0.0073 | TRUE | 0.1058 |
| Ppfia4 | 1233 | 1210 | 1356 | 1307 | 1095 | 1347 | 958 | 1287 | 1463 | 1301 | 1235 | 1330 | 1262.29 | 1236 | 1289 | 1.045 | 0.063 | 0.60728544 | 0.75124899 | 0.0228 | 0.0043 | 0.01 | 0.01 | TRUE | 0.4765 |
| Ppfibp1 | 2194 | 1518 | 1478 | 1753 | 1516 | 1818 | 1704 | 1615 | 1595 | 1745 | 1710 | 1795 | 1693.87 | 1638 | 1750 | 1.068 | 0.095 | 0.14865796 | 0.29388219 | 4.00E-04 | 0.0038 | 0.0025 | 0.0025 | TRUE | 0.0157 |
| Ppfibp2 | 974 | 753 | 935 | 1057 | 1052 | 1296 | 756 | 801 | 1009 | 1052 | 1187 | 1279 | 1014.09 | 855 | 1173 | 1.373 | 0.457 | 5.07E-05 | 0.00070814 | 0.0156 | 0.0048 | 0.0082 | 0.0082 | TRUE | 0.2903 |
| Pphln1 | 2998 | 2178 | 2035 | 1964 | 1670 | 1847 | 2328 | 2317 | 2196 | 1955 | 1884 | 1823 | 2083.84 | 2280 | 1887 | 0.827 | -0.273 | 1.73E-05 | 0.0002982 | 6.00E-04 | 0.0036 | 0.0024 | 0.0024 | TRUE | 0.0125 |
| Ppia | 46827 | 31662 | 32997 | 37161 | 30630 | 36190 | 36369 | 33678 | 35603 | 36990 | 34554 | 35725 | 35486.57 | 35217 | 35756 | 1.015 | 0.022 | 0.67745034 | 0.80217909 | 0.0011 | 0.0025 | 0.002 | 0.002 | TRUE | 0.0179 |
| Ppib | 8266 | 5680 | 5541 | 6192 | 5440 | 6185 | 6420 | 6042 | 5979 | 6164 | 6137 | 6106 | 6141.04 | 6147 | 6136 | 0.998 | -0.003 | 0.95140422 | 0.97323401 | 6.00E-04 | 0.0028 | 0.0018 | 0.0018 | TRUE | 0.0187 |
| Ppic | 1532 | 1032 | 1043 | 1342 | 1235 | 1293 | 1190 | 1098 | 1125 | 1336 | 1393 | 1276 | 1236.4 | 1138 | 1335 | 1.172 | 0.229 | 0.00202782 | 0.01222012 | 0.0011 | 0.0043 | 0.0032 | 0.0032 | TRUE | 0.0202 |
| Ppid | 12486 | 9168 | 9045 | 8590 | 7714 | 9132 | 9697 | 9752 | 9759 | 8550 | 8702 | 9015 | 9246.01 | 9736 | 8756 | 0.899 | -0.153 | 0.0014282 | 0.00931684 | 2.00E-04 | 0.0027 | 0.0016 | 0.0016 | TRUE | 0.0082 |
| Ppie | 3070 | 2213 | 2301 | 2303 | 2077 | 2463 | 2384 | 2354 | 2483 | 2292 | 2343 | 2431 | 2381.31 | 2407 | 2355 | 0.979 | -0.031 | 0.61044077 | 0.75363263 | 4.00E-04 | 0.0034 | 0.0022 | 0.0022 | TRUE | 0.0097 |
| Ppif | 4397 | 3213 | 3047 | 3359 | 2964 | 3622 | 3415 | 3418 | 3288 | 3344 | 3344 | 3575 | 3397.17 | 3374 | 3421 | 1.014 | 0.02 | 0.72813105 | 0.83706082 | 7.00E-04 | 0.0031 | 0.0021 | 0.0021 | TRUE | 0.0191 |
| Ppifos | 0 | 0 | 0 | 0 | 0 | 1 | 0 | 0 | 0 | 0 | 0 | 1 | 0.16 | 0 | 0 | 2.094 | 1.067 | 0.79379329 | NA | 0 | 14.4078 | 10 | 10 | TRUE | 0.3835 |
| Ppig | 10825 | 7977 | 7420 | 8305 | 6957 | 7953 | 8407 | 8485 | 8006 | 8267 | 7848 | 7851 | 8144.05 | 8299 | 7989 | 0.963 | -0.055 | 0.2940301 | 0.46596876 | 7.00E-04 | 0.0027 | 0.0019 | 0.0019 | TRUE | 0.0116 |
| Ppih | 2217 | 1485 | 1290 | 1417 | 1199 | 1503 | 1722 | 1580 | 1392 | 1410 | 1353 | 1484 | 1490.02 | 1565 | 1416 | 0.904 | -0.146 | 0.09386205 | 0.21100507 | 0.0057 | 0.004 | 0.0048 | 0.0048 | TRUE | 0.1115 |
| Ppil1 | 6019 | 4346 | 4115 | 4062 | 3669 | 4263 | 4675 | 4623 | 4440 | 4043 | 4139 | 4208 | 4354.68 | 4579 | 4130 | 0.902 | -0.149 | 0.00528813 | 0.02556944 | 4.00E-04 | 0.003 | 0.0018 | 0.0018 | TRUE | 0.0085 |
| Ppil2 | 4208 | 2941 | 2585 | 2462 | 2443 | 2574 | 3268 | 3128 | 2789 | 2451 | 2756 | 2541 | 2822.21 | 3062 | 2583 | 0.843 | -0.247 | 0.00119524 | 0.00822911 | 0.0045 | 0.0033 | 0.0038 | 0.0038 | TRUE | 0.0733 |
| Ppil3 | 1261 | 826 | 809 | 906 | 746 | 887 | 979 | 879 | 873 | 902 | 842 | 876 | 891.65 | 910 | 873 | 0.958 | -0.062 | 0.45760493 | 0.62745081 | 0.0018 | 0.0051 | 0.0039 | 0.0039 | TRUE | 0.055 |
| Ppil4 | 4143 | 3108 | 2711 | 3073 | 2583 | 2968 | 3218 | 3306 | 2925 | 3059 | 2914 | 2930 | 3058.57 | 3150 | 2968 | 0.942 | -0.086 | 0.18749914 | 0.3444634 | 0.0019 | 0.0032 | 0.0027 | 0.0027 | TRUE | 0.0465 |
| Ppil6 | 21 | 9 | 18 | 25 | 22 | 24 | 16 | 10 | 19 | 25 | 25 | 24 | 19.78 | 15 | 25 | 1.615 | 0.692 | 0.11861475 | 0.25005318 | 0 | 0.1222 | 0.0885 | 0.0885 | TRUE | 0.4238 |
| Ppip5k1 | 1026 | 798 | 816 | 1023 | 908 | 997 | 797 | 849 | 880 | 1018 | 1024 | 984 | 925.49 | 842 | 1009 | 1.2 | 0.263 | 0.00089071 | 0.00666237 | 0 | 0.005 | 0.0034 | 0.0034 | TRUE | 0.0278 |

| | | | | | | | | | | | | | | | | | | | | | | | | | |
|---|---|---|---|---|---|---|---|---|---|---|---|---|---|---|---|---|---|---|---|---|---|---|---|---|---|
| Ppip5k2 | 7233 | 5172 | 4606 | 4917 | 4420 | 5167 | 5618 | 5501 | 4970 | 4894 | 4986 | 5101 | 5178.34 | 5363 | 4994 | 0.931 | -0.103 | 0.09595069 | 0.21470336 | 0.002 | 0.0029 | 0.0026 | 0.0026 | TRUE | 0.0498 |
| Ppl | 827 | 672 | 679 | 699 | 597 | 720 | 642 | 715 | 733 | 696 | 673 | 711 | 694.96 | 697 | 693 | 0.998 | -0.003 | 0.96966827 | 0.9834812 | 0.0016 | 0.0058 | 0.0044 | 0.0044 | TRUE | 0.0584 |
| Ppm1a | 7697 | 5818 | 6091 | 5893 | 5149 | 5904 | 5978 | 6189 | 6572 | 5866 | 5809 | 5828 | 6040.21 | 6246 | 5834 | 0.934 | -0.098 | 0.07668647 | 0.18316839 | 9.00E-04 | 0.0028 | 0.002 | 0.002 | TRUE | 0.0254 |
| Ppm1b | 8988 | 6802 | 7427 | 7946 | 7039 | 7973 | 6981 | 7235 | 8014 | 7909 | 7941 | 7871 | 7658.37 | 7410 | 7907 | 1.067 | 0.094 | 0.1281751 | 0.26437478 | 0.0025 | 0.0027 | 0.0026 | 0.0026 | TRUE | 0.0619 |
| Ppm1d | 2681 | 1708 | 1521 | 1459 | 1263 | 1409 | 2082 | 1817 | 1641 | 1452 | 1425 | 1391 | 1634.69 | 1847 | 1423 | 0.769 | -0.378 | 1.83E-05 | 0.00031033 | 0.0065 | 0.0039 | 0.005 | 0.005 | TRUE | 0.1534 |
| Ppm1e | 229 | 160 | 122 | 161 | 142 | 164 | 178 | 170 | 132 | 160 | 160 | 162 | 160.34 | 160 | 161 | 1.002 | 0.003 | 0.98699651 | 0.99373352 | 0.0062 | 0.0172 | 0.0141 | 0.0141 | TRUE | 0.2424 |
| Ppm1f | 5256 | 3506 | 3496 | 3789 | 3304 | 3611 | 4082 | 3729 | 3772 | 3772 | 3727 | 3565 | 3774.5 | 3861 | 3688 | 0.955 | -0.067 | 0.26881556 | 0.4393438 | 0.0013 | 0.0031 | 0.0024 | 0.0024 | TRUE | 0.0311 |
| Ppm1g | 15562 | 11048 | 9829 | 10759 | 9508 | 11522 | 12086 | 11752 | 10605 | 10709 | 10726 | 11374 | 11208.81 | 11481 | 10936 | 0.952 | -0.07 | 0.26033661 | 0.4305706 | 0.0028 | 0.0026 | 0.0027 | 0.0027 | TRUE | 0.0543 |
| Ppm1h | 572 | 453 | 456 | 470 | 462 | 525 | 444 | 482 | 492 | 468 | 521 | 518 | 487.57 | 473 | 502 | 1.064 | 0.09 | 0.37666263 | 0.55120702 | 0.0014 | 0.0073 | 0.0054 | 0.0054 | TRUE | 0.0428 |
| Ppm1j | 1266 | 1010 | 1344 | 1909 | 1745 | 2056 | 983 | 1074 | 1450 | 1900 | 1969 | 2030 | 1567.68 | 1169 | 1966 | 1.684 | 0.752 | 9.57E-11 | 7.82E-09 | 0.0219 | 0.0039 | 0.009 | 0.009 | TRUE | 0.527 |
| Ppm1k | 658 | 484 | 538 | 692 | 608 | 737 | 511 | 515 | 580 | 689 | 686 | 728 | 618.1 | 535 | 701 | 1.311 | 0.391 | 3.03E-05 | 0.0004674 | 0.0012 | 0.0063 | 0.0047 | 0.0047 | TRUE | 0.0615 |
| Ppm1l | 525 | 394 | 355 | 362 | 331 | 365 | 408 | 419 | 383 | 360 | 373 | 360 | 383.99 | 403 | 364 | 0.904 | -0.146 | 0.16981462 | 0.32137906 | 0 | 0.0086 | 0.0056 | 0.0056 | TRUE | 0.021 |
| Ppm1m | 2410 | 1925 | 1985 | 2504 | 2195 | 2415 | 1872 | 2048 | 2142 | 2492 | 2476 | 2384 | 2235.63 | 2021 | 2451 | 1.214 | 0.28 | 5.09E-05 | 0.00070933 | 0.0022 | 0.0035 | 0.003 | 0.003 | TRUE | 0.0518 |
| Ppm1n | 112 | 84 | 81 | 127 | 115 | 115 | 87 | 89 | 87 | 126 | 130 | 114 | 105.57 | 88 | 123 | 1.401 | 0.487 | 0.01038089 | 0.04242206 | 0 | 0.0249 | 0.0163 | 0.0163 | TRUE | 0.049 |
| Ppme1 | 3741 | 2875 | 3305 | 3330 | 2986 | 3507 | 2905 | 3058 | 3566 | 3315 | 3369 | 3462 | 3279.13 | 3176 | 3382 | 1.065 | 0.091 | 0.24233514 | 0.41011323 | 0.0055 | 0.0031 | 0.0041 | 0.0041 | TRUE | 0.1393 |
| Ppox | 536 | 498 | 473 | 457 | 412 | 481 | 416 | 530 | 510 | 455 | 465 | 475 | 475.14 | 485 | 465 | 0.961 | -0.058 | 0.61047776 | 0.75363263 | 0.0069 | 0.0074 | 0.0072 | 0.0072 | TRUE | 0.1925 |
| Ppp1ca | 16607 | 11626 | 11319 | 11513 | 10115 | 11739 | 12898 | 12366 | 12213 | 11460 | 11411 | 11588 | 11989.41 | 12492 | 11486 | 0.919 | -0.121 | 0.01108924 | 0.04452204 | 3.00E-04 | 0.0026 | 0.0016 | 0.0016 | TRUE | 0.01 |
| Ppp1cb | 15206 | 10751 | 11003 | 10970 | 9283 | 10760 | 11810 | 11436 | 11872 | 10920 | 10472 | 10622 | 11188.53 | 11706 | 10671 | 0.912 | -0.134 | 0.00525095 | 0.02544991 | 3.00E-04 | 0.0026 | 0.0016 | 0.0016 | TRUE | 0.0051 |
| Ppp1cc | 9909 | 7458 | 7275 | 7410 | 6664 | 7536 | 7696 | 7933 | 7850 | 7376 | 7518 | 7439 | 7635.22 | 7826 | 7444 | 0.951 | -0.072 | 0.12675939 | 0.26215772 | 0 | 0.0027 | 0.0015 | 0.0015 | TRUE | 0.0026 |
| Ppp1r10 | 3735 | 2403 | 1577 | 1897 | 1826 | 1683 | 2901 | 2556 | 1702 | 1888 | 2060 | 1661 | 2128 | 2386 | 1870 | 0.783 | -0.352 | 0.12981724 | 0.26651505 | 0.0385 | 0.0035 | 0.0142 | 0.0385 | TRUE | 0.5759 |
| Ppp1r11 | 1393 | 954 | 657 | 879 | 723 | 751 | 1082 | 1015 | 709 | 875 | 816 | 741 | 872.91 | 935 | 811 | 0.866 | -0.208 | 0.1188977 | 0.25043274 | 0.0269 | 0.0051 | 0.0117 | 0.0117 | TRUE | 0.5286 |
| Ppp1r12a | 5623 | 4005 | 3533 | 3888 | 3484 | 3918 | 4367 | 4260 | 3812 | 3870 | 3930 | 3868 | 4017.9 | 4146 | 3889 | 0.938 | -0.093 | 0.15038435 | 0.29610739 | 0.0022 | 0.003 | 0.0028 | 0.0028 | TRUE | 0.0601 |
| Ppp1r12b | 282 | 249 | 276 | 346 | 301 | 337 | 219 | 265 | 298 | 344 | 340 | 333 | 299.72 | 261 | 339 | 1.307 | 0.386 | 0.00430467 | 0.02186838 | 0.0085 | 0.0103 | 0.0098 | 0.0098 | TRUE | 0.2364 |
| Ppp1r12c | 4332 | 3345 | 3214 | 3283 | 2985 | 3413 | 3364 | 3558 | 3468 | 3268 | 3367 | 3369 | 3399.14 | 3463 | 3335 | 0.963 | -0.054 | 0.32817676 | 0.50258549 | 3.00E-04 | 0.0031 | 0.0019 | 0.0019 | TRUE | 0.0078 |
| Ppp1r13b | 1576 | 1165 | 1075 | 1282 | 1149 | 1351 | 1224 | 1239 | 1160 | 1276 | 1296 | 1334 | 1254.84 | 1208 | 1302 | 1.078 | 0.108 | 0.12444543 | 0.25902119 | 0 | 0.0043 | 0.0028 | 0.0028 | TRUE | 0.0139 |
| Ppp1r13l | 1021 | 809 | 966 | 888 | 794 | 985 | 793 | 861 | 1042 | 884 | 896 | 972 | 907.96 | 899 | 917 | 1.023 | 0.032 | 0.75589403 | 0.85511255 | 0.0103 | 0.005 | 0.0068 | 0.0068 | TRUE | 0.2315 |
| Ppp1r14a | 739 | 556 | 525 | 622 | 524 | 518 | 574 | 591 | 566 | 619 | 591 | 511 | 575.57 | 577 | 574 | 0.994 | -0.009 | 0.92911355 | 0.96091317 | 0.0034 | 0.0065 | 0.0055 | 0.0055 | TRUE | 0.1085 |
| Ppp1r14b | 5809 | 4186 | 3684 | 3772 | 3475 | 4070 | 4512 | 4453 | 3975 | 3755 | 3920 | 4018 | 4105.29 | 4313 | 3898 | 0.903 | -0.147 | 0.02634294 | 0.08444673 | 0.0025 | 0.003 | 0.0029 | 0.0029 | TRUE | 0.0568 |
| Ppp1r14c | 240 | 191 | 177 | 166 | 150 | 163 | 186 | 203 | 191 | 165 | 169 | 161 | 179.32 | 193 | 165 | 0.854 | -0.228 | 0.12264417 | 0.25601768 | 0 | 0.0156 | 0.0101 | 0.0101 | TRUE | 0.0202 |
| Ppp1r14d | 66 | 53 | 59 | 58 | 58 | 94 | 51 | 56 | 64 | 58 | 65 | 93 | 64.54 | 57 | 72 | 1.267 | 0.341 | 0.19739452 | 0.3572246 | 0.0246 | 0.0391 | 0.0349 | 0.0349 | TRUE | 0.5927 |
| Ppp1r15a | 1036 | 750 | 599 | 573 | 552 | 552 | 805 | 798 | 646 | 570 | 623 | 545 | 664.45 | 750 | 579 | 0.771 | -0.375 | 0.00048153 | 0.00420177 | 0.0083 | 0.006 | 0.0068 | 0.0068 | TRUE | 0.1684 |
| Ppp1r15b | 6660 | 4894 | 4695 | 4640 | 3981 | 4304 | 5173 | 5206 | 5066 | 4619 | 4491 | 4249 | 4800.41 | 5148 | 4453 | 0.865 | -0.209 | 0.00015737 | 0.0017697 | 7.00E-04 | 0.0029 | 0.002 | 0.002 | TRUE | 0.0197 |
| Ppp1r16a | 1600 | 1301 | 1090 | 1154 | 1043 | 1142 | 1243 | 1384 | 1176 | 1149 | 1177 | 1127 | 1209.21 | 1268 | 1151 | 0.908 | -0.139 | 0.08392813 | 0.19532124 | 0.0028 | 0.0044 | 0.0038 | 0.0038 | TRUE | 0.0754 |
| Ppp1r16b | 428 | 276 | 270 | 327 | 268 | 319 | 332 | 294 | 291 | 325 | 302 | 315 | 310.01 | 306 | 314 | 1.025 | 0.036 | 0.76370781 | 0.85969854 | 0 | 0.0101 | 0.0072 | 0.0072 | TRUE | 0.0708 |
| Ppp1r17 | 0 | 0 | 0 | 0 | 0 | 0 | 0 | 0 | 0 | 0 | 0 | 0 | 0 | 0 | 0 | NA | NA | NA | NA | NA | NA | NA | NA | NA | NA |
| Ppp1r18 | 1106 | 783 | 747 | 763 | 699 | 886 | 859 | 833 | 806 | 759 | 789 | 875 | 820.08 | 833 | 808 | 0.969 | -0.045 | 0.60592342 | 0.75040841 | 0.002 | 0.0053 | 0.0042 | 0.0042 | TRUE | 0.0637 |
| Ppp1r1a | 308 | 266 | 314 | 329 | 317 | 365 | 239 | 283 | 339 | 327 | 358 | 360 | 317.73 | 287 | 348 | 1.219 | 0.286 | 0.04103859 | 0.11654644 | 0.0133 | 0.0099 | 0.0109 | 0.0109 | TRUE | 0.2797 |
| Ppp1r1b | 233 | 179 | 165 | 233 | 232 | 258 | 181 | 190 | 178 | 232 | 262 | 255 | 216.29 | 183 | 250 | 1.362 | 0.445 | 0.00123136 | 0.00840893 | 0 | 0.0134 | 0.009 | 0.009 | TRUE | 0.0424 |
| Ppp1r1c | 0 | 0 | 0 | 0 | 0 | 0 | 0 | 0 | 0 | 0 | 0 | 0 | 0 | 0 | 0 | NA | NA | NA | NA | NA | NA | NA | NA | NA | NA |
| Ppp1r2 | 6386 | 4549 | 4269 | 4786 | 4162 | 4640 | 4960 | 4839 | 4606 | 4764 | 4695 | 4580 | 4740.7 | 4802 | 4680 | 0.974 | -0.038 | 0.49500454 | 0.65992911 | 6.00E-04 | 0.0029 | 0.002 | 0.002 | TRUE | 0.0153 |
| Ppp1r2-ps3 | 2 | 0 | 0 | 0 | 0 | 0 | 2 | 0 | 0 | 0 | 0 | 0 | 0.26 | 1 | 0 | 0.356 | -1.491 | 0.71362796 | NA | 0 | 9.1573 | 10 | 10 | TRUE | 0.968 |
| Ppp1r2-ps7 | 1 | 1 | 0 | 0 | 0 | 0 | 1 | 1 | 0 | 0 | 0 | 0 | 0.31 | 1 | 0 | 0.309 | -1.696 | 0.63183309 | NA | 0 | 7.7294 | 7.2472 | 7.2472 | TRUE | 0.2015 |
| Ppp1r2-ps9 | 0 | 0 | 0 | 0 | 0 | 0 | 0 | 0 | 0 | 0 | 0 | 0 | 0 | 0 | 0 | NA | NA | NA | NA | NA | NA | NA | NA | NA | NA |
| Ppp1r21 | 2194 | 1699 | 1602 | 1606 | 1400 | 1701 | 1704 | 1807 | 1729 | 1599 | 1579 | 1679 | 1682.81 | 1747 | 1619 | 0.928 | -0.108 | 0.09998083 | 0.22086955 | 4.00E-04 | 0.0038 | 0.0025 | 0.0025 | TRUE | 0.0128 |
| Ppp1r26 | 449 | 388 | 316 | 349 | 282 | 331 | 349 | 413 | 341 | 347 | 318 | 327 | 349.11 | 368 | 331 | 0.902 | -0.149 | 0.21464454 | 0.37735725 | 0.0038 | 0.0092 | 0.0076 | 0.0076 | TRUE | 0.1289 |
| Ppp1r27 | 1 | 0 | 0 | 0 | 1 | 3 | 1 | 0 | 0 | 0 | 1 | 3 | 0.81 | 0 | 1 | 3.963 | 1.987 | 0.39811512 | NA | 0 | 2.9247 | 2.6042 | 2.6042 | TRUE | 0.1519 |
| Ppp1r32 | 12 | 5 | 4 | 5 | 7 | 8 | 9 | 5 | 4 | 5 | 8 | 8 | 6.62 | 6 | 7 | 1.073 | 0.102 | 0.89120062 | NA | 0 | 0.3604 | 0.2522 | 0.2522 | TRUE | 0.5472 |
| Ppp1r35 | 1124 | 874 | 871 | 858 | 855 | 930 | 873 | 930 | 940 | 854 | 965 | 918 | 913.18 | 914 | 912 | 0.998 | -0.002 | 0.97708813 | 0.98766058 | 0.0016 | 0.005 | 0.0038 | 0.0038 | TRUE | 0.0372 |
| Ppp1r36 | 53 | 33 | 58 | 47 | 46 | 55 | 41 | 35 | 63 | 47 | 52 | 54 | 48.64 | 46 | 51 | 1.106 | 0.146 | 0.62744584 | 0.76512954 | 0.0254 | 0.0512 | 0.0442 | 0.0442 | TRUE | 0.7044 |
| Ppp1r37 | 1769 | 1398 | 1146 | 1192 | 1038 | 1143 | 1374 | 1487 | 1237 | 1187 | 1171 | 1128 | 1263.88 | 1366 | 1162 | 0.851 | -0.234 | 0.00473883 | 0.02352472 | 0.0038 | 0.0043 | 0.0041 | 0.0041 | TRUE | 0.08 |
| Ppp1r3a | 0 | 0 | 1 | 0 | 0 | 0 | 0 | 0 | 1 | 0 | 0 | 0 | 0.18 | 0 | 0 | 0.552 | -0.857 | 0.8336548 | NA | 0 | 13.1817 | 10 | 10 | TRUE | 0.5365 |
| Ppp1r3b | 957 | 877 | 1121 | 1124 | 1069 | 1263 | 743 | 933 | 1210 | 1119 | 1206 | 1247 | 1076.2 | 962 | 1191 | 1.24 | 0.31 | 0.02127629 | 0.07231489 | 0.0298 | 0.0046 | 0.0121 | 0.0121 | TRUE | 0.3919 |
| Ppp1r3c | 174 | 184 | 195 | 237 | 209 | 221 | 135 | 196 | 210 | 236 | 236 | 218 | 205.18 | 180 | 230 | 1.282 | 0.359 | 0.03765818 | 0.10964193 | 0.0227 | 0.014 | 0.0165 | 0.0165 | TRUE | 0.5672 |
| Ppp1r3d | 36 | 45 | 18 | 41 | 44 | 49 | 28 | 48 | 19 | 41 | 50 | 48 | 39.01 | 32 | 46 | 1.461 | 0.547 | 0.12347937 | 0.25746034 | 0.0683 | 0.0632 | 0.0645 | 0.0645 | TRUE | 0.8756 |
| Ppp1r3e | 12 | 6 | 9 | 10 | 4 | 13 | 9 | 6 | 10 | 10 | 5 | 13 | 8.79 | 8 | 9 | 1.079 | 0.11 | 0.86865397 | NA | 0 | 0.2722 | 0.2043 | 0.2043 | TRUE | 0.3985 |
| Ppp1r3f | 181 | 127 | 124 | 174 | 144 | 157 | 141 | 135 | 134 | 173 | 162 | 155 | 150.01 | 137 | 163 | 1.197 | 0.259 | 0.10613802 | 0.23114898 | 0 | 0.0182 | 0.0119 | 0.0119 | TRUE | 0.0291 |
| Ppp1r3g | 90 | 98 | 83 | 87 | 61 | 62 | 70 | 104 | 90 | 87 | 69 | 61 | 80.05 | 88 | 72 | 0.827 | -0.275 | 0.25875971 | 0.428657 | 0.025 | 0.032 | 0.03 | 0.03 | TRUE | 0.2586 |
| Ppp1r42 | 2 | 2 | 2 | 4 | 2 | 3 | 2 | 2 | 2 | 4 | 2 | 3 | 2.51 | 2 | 3 | 1.598 | 0.676 | 0.56538751 | NA | 0 | 0.9481 | 0.5789 | 0.5789 | TRUE | 0.0913 |
| Ppp1r7 | 4482 | 3369 | 2770 | 3056 | 2676 | 3079 | 3481 | 3584 | 2989 | 3042 | 3019 | 3039 | 3192.26 | 3351 | 3033 | 0.905 | -0.144 | 0.05079654 | 0.13644708 | 0.004 | 0.0032 | 0.0036 | 0.0036 | TRUE | 0.1075 |
| Ppp1r8 | 4846 | 3346 | 3120 | 2760 | 2564 | 2865 | 3764 | 3559 | 3366 | 2747 | 2892 | 2828 | 3192.86 | 3563 | 2822 | 0.792 | -0.337 | 9.27E-08 | 3.67E-06 | 0.0014 | 0.0032 | 0.0026 | 0.0026 | TRUE | 0.0301 |
| Ppp1r9a | 2159 | 1652 | 1553 | 1906 | 1684 | 1958 | 1677 | 1757 | 1676 | 1897 | 1900 | 1933 | 1806.58 | 1703 | 1910 | 1.122 | 0.166 | 0.00776094 | 0.03425 | 0 | 0.0037 | 0.0022 | 0.0022 | TRUE | 0.0091 |
| Ppp1r9b | 2558 | 2021 | 1715 | 1869 | 1683 | 1977 | 1987 | 2150 | 1850 | 1860 | 1899 | 1952 | 1949.58 | 1996 | 1904 | 0.954 | -0.068 | 0.34517089 | 0.51936368 | 0.0023 | 0.0036 | 0.0032 | 0.0032 | TRUE | 0.0538 |
| Ppp2ca | 29123 | 20257 | 18947 | 20093 | 17673 | 19979 | 22619 | 21547 | 20444 | 20001 | 19937 | 19722 | 20711.54 | 21537 | 19887 | 0.923 | -0.115 | 0.02937344 | 0.09119732 | 0.0012 | 0.0025 | 0.002 | 0.002 | TRUE | 0.0241 |
| Ppp2cb | 7474 | 5218 | 5079 | 5073 | 4464 | 5124 | 5805 | 5550 | 5480 | 5050 | 5036 | 5058 | 5329.82 | 5612 | 5048 | 0.899 | -0.153 | 0.00295109 | 0.01632249 | 3.00E-04 | 0.0029 | 0.0017 | 0.0017 | TRUE | 0.0112 |
| Ppp2r1a | 16813 | 11967 | 11633 | 11551 | 10247 | 11761 | 13058 | 12729 | 12552 | 11498 | 11560 | 11610 | 12167.74 | 12780 | 11556 | 0.904 | -0.145 | 0.00147939 | 0.00955487 | 2.00E-04 | 0.0026 | 0.0014 | 0.0014 | TRUE | 0.0045 |
| Ppp2r1b | 6736 | 4898 | 4455 | 4527 | 4016 | 4628 | 5232 | 5210 | 4807 | 4506 | 4530 | 4569 | 4808.93 | 5083 | 4535 | 0.892 | -0.165 | 0.00345744 | 0.01845992 | 8.00E-04 | 0.0029 | 0.0021 | 0.0021 | TRUE | 0.0272 |
| Ppp2r2a | 10311 | 7721 | 7660 | 8091 | 6834 | 8012 | 8008 | 8213 | 8265 | 8054 | 7709 | 7909 | 8026.37 | 8162 | 7891 | 0.967 | -0.048 | 0.31949146 | 0.49335473 | 2.00E-04 | 0.0027 | 0.0016 | 0.0016 | TRUE | 0.0049 |
| Ppp2r2b | 34 | 32 | 29 | 19 | 26 | 27 | 26 | 34 | 31 | 19 | 29 | 27 | 27.77 | 30 | 25 | 0.818 | -0.29 | 0.43576104 | 0.60709497 | 0 | 0.0878 | 0.063 | 0.063 | TRUE | 0.2793 |
| Ppp2r2c | 1786 | 1321 | 1494 | 1636 | 1543 | 1643 | 1387 | 1405 | 1612 | 1628 | 1741 | 1622 | 1565.88 | 1468 | 1664 | 1.134 | 0.181 | 0.02125651 | 0.07230349 | 0.0035 | 0.0039 | 0.0038 | 0.0038 | TRUE | 0.0874 |

| | | | | | | | | | | | | | | | | | | | | | | | | | |
|---|---|---|---|---|---|---|---|---|---|---|---|---|---|---|---|---|---|---|---|---|---|---|---|---|---|
| Ppp2r2d | 5445 | 3715 | 3658 | 3831 | 3220 | 3994 | 4229 | 3952 | 3947 | 3813 | 3632 | 3943 | 3919.34 | 4043 | 3796 | 0.939 | -0.091 | 0.13054637 | 0.26758917 | 0.0013 | 0.003 | 0.0024 | 0.0024 | TRUE | 0.0202 |
| Ppp2r3a | 1237 | 793 | 778 | 843 | 736 | 798 | 961 | 844 | 839 | 839 | 830 | 788 | 850.14 | 881 | 819 | 0.928 | -0.109 | 0.21440326 | 0.37711504 | 0.0027 | 0.0052 | 0.0043 | 0.0043 | TRUE | 0.0775 |
| Ppp2r3c | 1388 | 970 | 904 | 899 | 811 | 978 | 1078 | 1032 | 975 | 895 | 915 | 965 | 976.73 | 1028 | 925 | 0.899 | -0.154 | 0.05272065 | 0.14007596 | 0.0011 | 0.0049 | 0.0035 | 0.0035 | TRUE | 0.0241 |
| Ppp2r3d | 130 | 113 | 97 | 112 | 82 | 78 | 101 | 120 | 105 | 111 | 93 | 77 | 101.14 | 109 | 94 | 0.864 | -0.211 | 0.31511978 | 0.48870278 | 0.0108 | 0.0259 | 0.0217 | 0.0217 | TRUE | 0.2753 |
| Ppp2r4 | 7010 | 4861 | 4458 | 4765 | 4186 | 4722 | 5444 | 5171 | 4810 | 4743 | 4722 | 4661 | 4925.29 | 5142 | 4709 | 0.915 | -0.128 | 0.03556948 | 0.10510536 | 0.0016 | 0.0029 | 0.0024 | 0.0024 | TRUE | 0.0387 |
| Ppp2r5a | 6814 | 4878 | 5035 | 5915 | 4969 | 5809 | 5292 | 5189 | 5433 | 5888 | 5606 | 5734 | 5523.54 | 5305 | 5743 | 1.083 | 0.115 | 0.02713711 | 0.08611693 | 3.00E-04 | 0.0029 | 0.0018 | 0.0018 | TRUE | 0.006 |
| Ppp2r5b | 1258 | 895 | 741 | 845 | 706 | 819 | 977 | 952 | 800 | 841 | 796 | 808 | 862.43 | 910 | 815 | 0.895 | -0.159 | 0.08979128 | 0.20436096 | 0.0051 | 0.0052 | 0.0052 | 0.0052 | TRUE | 0.1308 |
| Ppp2r5c | 10108 | 7630 | 8885 | 10809 | 9602 | 11161 | 7850 | 8116 | 9587 | 10759 | 10832 | 11018 | 9693.7 | 8518 | 10870 | 1.276 | 0.352 | 1.90E-06 | 4.90E-05 | 0.0059 | 0.0027 | 0.0038 | 0.0038 | TRUE | 0.1469 |
| Ppp2r5d | 6030 | 4409 | 4596 | 4885 | 4384 | 4976 | 4683 | 4690 | 4959 | 4863 | 4946 | 4912 | 4842.05 | 4777 | 4907 | 1.027 | 0.039 | 0.46184268 | 0.63114231 | 4.00E-04 | 0.0029 | 0.0018 | 0.0018 | TRUE | 0.0133 |
| Ppp2r5e | 3776 | 2752 | 2732 | 2968 | 2467 | 2994 | 2933 | 2927 | 2948 | 2954 | 2783 | 2956 | 2916.77 | 2936 | 2898 | 0.987 | -0.019 | 0.74531477 | 0.84862455 | 2.00E-04 | 0.0032 | 0.002 | 0.002 | TRUE | 0.0143 |
| Ppp3ca | 1443 | 1062 | 1082 | 1147 | 994 | 1064 | 1121 | 1130 | 1167 | 1142 | 1121 | 1050 | 1121.87 | 1139 | 1104 | 0.97 | -0.044 | 0.54653137 | 0.70368191 | 0 | 0.0045 | 0.003 | 0.003 | TRUE | 0.0224 |
| Ppp3cb | 5366 | 3657 | 3837 | 4263 | 3662 | 4140 | 4168 | 3890 | 4140 | 4243 | 4131 | 4087 | 4109.81 | 4066 | 4154 | 1.021 | 0.031 | 0.58510473 | 0.73448989 | 5.00E-04 | 0.003 | 0.002 | 0.002 | TRUE | 0.0173 |
| Ppp3cc | 731 | 539 | 557 | 717 | 594 | 600 | 568 | 573 | 601 | 714 | 670 | 592 | 619.69 | 581 | 659 | 1.135 | 0.182 | 0.06331405 | 0.15946214 | 0.0035 | 0.0063 | 0.0053 | 0.0053 | TRUE | 0.0935 |
| Ppp3r1 | 7103 | 4963 | 4713 | 4775 | 4033 | 4599 | 5517 | 5279 | 5085 | 4753 | 4550 | 4540 | 4953.93 | 5294 | 4614 | 0.871 | -0.199 | 0.00042693 | 0.00382338 | 9.00E-04 | 0.0029 | 0.0021 | 0.0021 | TRUE | 0.0168 |
| Ppp3r2 | 0 | 0 | 0 | 0 | 0 | 0 | 0 | 0 | 0 | 0 | 0 | 0 | 0 | 0 | 0 | NA | NA | NA | NA | NA | NA | NA | NA | NA | NA |
| Ppp4c | 7881 | 5541 | 5396 | 5198 | 4787 | 5643 | 6121 | 5894 | 5822 | 5174 | 5400 | 5571 | 5663.63 | 5946 | 5382 | 0.905 | -0.144 | 0.00848021 | 0.03660754 | 8.00E-04 | 0.0028 | 0.002 | 0.002 | TRUE | 0.014 |
| Ppp4r1 | 5735 | 4054 | 3731 | 3894 | 3595 | 4124 | 4454 | 4312 | 4026 | 3876 | 4056 | 4071 | 4132.45 | 4264 | 4001 | 0.938 | -0.093 | 0.12413778 | 0.25849377 | 0.0013 | 0.003 | 0.0024 | 0.0024 | TRUE | 0.0289 |
| Ppp4r1l-ps | 283 | 263 | 295 | 306 | 266 | 340 | 220 | 280 | 318 | 305 | 300 | 336 | 293.02 | 273 | 314 | 1.156 | 0.209 | 0.15611913 | 0.30416529 | 0.0159 | 0.0105 | 0.0122 | 0.0122 | TRUE | 0.3468 |
| Ppp4r2 | 7496 | 5573 | 5543 | 5526 | 4843 | 5723 | 5822 | 5928 | 5981 | 5501 | 5463 | 5649 | 5724.01 | 5910 | 5538 | 0.937 | -0.094 | 0.05856337 | 0.15077703 | 0 | 0.0028 | 0.0016 | 0.0016 | TRUE | 0.0038 |
| Ppp4r4 | 208 | 162 | 183 | 202 | 168 | 219 | 162 | 172 | 197 | 201 | 190 | 216 | 189.68 | 177 | 202 | 1.147 | 0.197 | 0.19277756 | 0.35127099 | 0 | 0.0149 | 0.0113 | 0.0113 | TRUE | 0.1059 |
| Ppp5c | 6874 | 5107 | 5244 | 5510 | 4963 | 6059 | 5339 | 5432 | 5658 | 5485 | 5599 | 5981 | 5582.3 | 5476 | 5688 | 1.039 | 0.055 | 0.33698039 | 0.51122815 | 0.0012 | 0.0029 | 0.0022 | 0.0022 | TRUE | 0.0249 |
| Ppp6c | 2317 | 1598 | 1656 | 1712 | 1451 | 1790 | 1800 | 1700 | 1787 | 1704 | 1637 | 1767 | 1732.35 | 1762 | 1703 | 0.966 | -0.049 | 0.45534882 | 0.62501687 | 6.00E-04 | 0.0038 | 0.0026 | 0.0026 | TRUE | 0.0135 |
| Ppp6r1 | 8607 | 6108 | 4912 | 5056 | 4658 | 5297 | 6685 | 6497 | 5300 | 5033 | 5255 | 5229 | 5666.35 | 6161 | 5172 | 0.839 | -0.253 | 0.00202799 | 0.01222012 | 0.0074 | 0.0028 | 0.0047 | 0.0047 | TRUE | 0.1815 |
| Ppp6r2 | 927 | 674 | 775 | 808 | 751 | 791 | 720 | 717 | 836 | 804 | 847 | 781 | 784.24 | 758 | 811 | 1.071 | 0.099 | 0.28088143 | 0.45250657 | 0.0034 | 0.0054 | 0.0048 | 0.0048 | TRUE | 0.0951 |
| Ppp6r3 | 19769 | 14236 | 14780 | 15400 | 14100 | 16011 | 15354 | 15143 | 15947 | 15329 | 15906 | 15805 | 15580.77 | 15481 | 15680 | 1.013 | 0.018 | 0.70054011 | 0.81850931 | 4.00E-04 | 0.0026 | 0.0016 | 0.0016 | TRUE | 0.0084 |
| Pprc1 | 16463 | 12371 | 12428 | 12056 | 11359 | 13601 | 12786 | 13159 | 13410 | 12001 | 12814 | 13426 | 12932.61 | 13118 | 12747 | 0.972 | -0.041 | 0.4682569 | 0.6371664 | 0.0016 | 0.0026 | 0.0023 | 0.0023 | TRUE | 0.0322 |
| Ppt1 | 1806 | 1410 | 1418 | 1409 | 1301 | 1442 | 1403 | 1500 | 1530 | 1403 | 1468 | 1423 | 1454.35 | 1478 | 1431 | 0.969 | -0.045 | 0.51989692 | 0.68148264 | 7.00E-04 | 0.0041 | 0.0028 | 0.0028 | TRUE | 0.0247 |
| Ppt2 | 563 | 401 | 527 | 592 | 500 | 600 | 437 | 427 | 569 | 589 | 564 | 592 | 529.67 | 478 | 582 | 1.221 | 0.288 | 0.01480148 | 0.05526581 | 0.0112 | 0.0069 | 0.0082 | 0.0082 | TRUE | 0.3167 |
| Pptc7 | 2217 | 1516 | 1419 | 1551 | 1544 | 1610 | 1722 | 1613 | 1531 | 1544 | 1742 | 1589 | 1623.41 | 1622 | 1625 | 1.001 | 0.001 | 0.99152278 | 0.99565418 | 0.0031 | 0.0039 | 0.0036 | 0.0036 | TRUE | 0.047 |
| Ppwd1 | 3498 | 2611 | 2575 | 2658 | 2261 | 2738 | 2717 | 2777 | 2778 | 2646 | 2551 | 2703 | 2695.28 | 2757 | 2633 | 0.955 | -0.066 | 0.25079854 | 0.41965154 | 0 | 0.0033 | 0.002 | 0.002 | TRUE | 0.009 |
| Ppy | 2 | 2 | 1 | 0 | 2 | 1 | 2 | 2 | 1 | 0 | 2 | 1 | 1.33 | 2 | 1 | 0.668 | -0.583 | 0.72676016 | NA | 0 | 1.7793 | 1.2116 | 1.2116 | TRUE | 0.3095 |
| Ppyr1 | 54 | 44 | 27 | 35 | 32 | 46 | 42 | 47 | 29 | 35 | 36 | 45 | 39.04 | 39 | 39 | 0.986 | -0.021 | 0.94802101 | 0.97160938 | 0 | 0.0631 | 0.0493 | 0.0493 | TRUE | 0.3505 |
| Pqbp1 | 1890 | 1419 | 1366 | 1310 | 1263 | 1521 | 1468 | 1509 | 1474 | 1304 | 1425 | 1501 | 1446.9 | 1484 | 1410 | 0.95 | -0.073 | 0.32419043 | 0.49809594 | 0.0019 | 0.0041 | 0.0033 | 0.0033 | TRUE | 0.0529 |
| Pqlc1 | 645 | 491 | 496 | 609 | 530 | 683 | 501 | 522 | 535 | 606 | 598 | 674 | 572.79 | 519 | 626 | 1.207 | 0.272 | 0.00441432 | 0.02233411 | 0.0012 | 0.0066 | 0.0048 | 0.0048 | TRUE | 0.0538 |
| Pqlc2 | 553 | 380 | 331 | 370 | 325 | 343 | 429 | 404 | 357 | 368 | 367 | 339 | 377.39 | 397 | 358 | 0.899 | -0.153 | 0.18412276 | 0.34028957 | 0.0031 | 0.0087 | 0.007 | 0.007 | TRUE | 0.0867 |
| Pqlc3 | 895 | 630 | 812 | 785 | 690 | 771 | 695 | 670 | 876 | 781 | 778 | 761 | 760.37 | 747 | 773 | 1.037 | 0.052 | 0.62433173 | 0.76326288 | 0.0095 | 0.0055 | 0.0068 | 0.0068 | TRUE | 0.2649 |
| Pradc1 | 599 | 391 | 328 | 390 | 292 | 347 | 465 | 416 | 354 | 388 | 329 | 343 | 382.53 | 412 | 353 | 0.857 | -0.223 | 0.08251402 | 0.19342383 | 0.0107 | 0.0086 | 0.0093 | 0.0093 | TRUE | 0.1714 |
| Praf2 | 1832 | 1275 | 1428 | 1429 | 1196 | 1474 | 1423 | 1356 | 1541 | 1422 | 1349 | 1455 | 1424.42 | 1440 | 1409 | 0.979 | -0.031 | 0.68378751 | 0.80611802 | 0.002 | 0.0041 | 0.0034 | 0.0034 | TRUE | 0.0438 |
| Pram1 | 18 | 18 | 22 | 14 | 19 | 20 | 14 | 19 | 24 | 14 | 21 | 20 | 18.66 | 19 | 18 | 0.976 | -0.035 | 0.93972081 | 0.96635941 | 0 | 0.1294 | 0.0961 | 0.0961 | TRUE | 0.2537 |
| Prame | 115 | 94 | 69 | 37 | 34 | 35 | 89 | 100 | 74 | 37 | 38 | 35 | 62.25 | 88 | 37 | 0.415 | -1.268 | 1.18E-06 | 3.35E-05 | 0 | 0.0405 | 0.0294 | 0.0294 | TRUE | 0.1485 |
| Pramef12 | 2070 | 1442 | 1643 | 1566 | 1377 | 1532 | 1608 | 1534 | 1773 | 1559 | 1553 | 1512 | 1589.8 | 1638 | 1541 | 0.941 | -0.087 | 0.23735182 | 0.40415765 | 0.002 | 0.0039 | 0.0033 | 0.0033 | TRUE | 0.0608 |
| Pramef17 | 12 | 10 | 5 | 5 | 5 | 5 | 9 | 11 | 5 | 5 | 6 | 5 | 6.82 | 8 | 5 | 0.61 | -0.713 | 0.33262039 | NA | 0 | 0.3501 | 0.2336 | 0.2336 | TRUE | 0.2763 |
| Pramef6 | 22 | 17 | 8 | 15 | 10 | 8 | 17 | 18 | 9 | 15 | 11 | 8 | 12.99 | 15 | 11 | 0.774 | -0.37 | 0.50759722 | NA | 0 | 0.1849 | 0.146 | 0.146 | TRUE | 0.4404 |
| Pramef8 | 2613 | 1888 | 1641 | 1299 | 1201 | 1146 | 2029 | 2008 | 1771 | 1293 | 1355 | 1131 | 1597.9 | 1936 | 1260 | 0.65 | -0.622 | 7.74E-13 | 9.88E-11 | 0.006 | 0.0039 | 0.0048 | 0.0048 | TRUE | 0.0961 |
| Pramel1 | 543 | 385 | 320 | 334 | 308 | 313 | 422 | 410 | 345 | 332 | 347 | 309 | 360.9 | 392 | 329 | 0.838 | -0.255 | 0.03502233 | 0.10389666 | 0.0048 | 0.009 | 0.0077 | 0.0077 | TRUE | 0.1223 |
| Pramel3 | 191 | 138 | 95 | 31 | 18 | 17 | 148 | 147 | 103 | 31 | 20 | 17 | 77.6 | 133 | 23 | 0.171 | -2.549 | 1.79E-18 | 4.64E-16 | 0.0396 | 0.033 | 0.0345 | 0.0345 | TRUE | 0.6046 |
| Pramel4 | 0 | 0 | 0 | 0 | 0 | 0 | 0 | 0 | 0 | 0 | 0 | 0 | 0 | 0 | 0 | NA | NA | NA | NA | NA | NA | NA | NA | NA | NA |
| Pramel5 | 2 | 6 | 1 | 1 | 3 | 0 | 2 | 6 | 1 | 1 | 3 | 0 | 2.23 | 3 | 1 | 0.479 | -1.061 | 0.4611965 | NA | 0.7024 | 1.0641 | 0.9674 | 0.9674 | TRUE | 1.1838 |
| Pramel6 | 176 | 109 | 59 | 24 | 21 | 17 | 137 | 116 | 64 | 24 | 24 | 17 | 63.44 | 106 | 22 | 0.202 | -2.307 | 7.97E-12 | 8.51E-10 | 0.0859 | 0.0398 | 0.0533 | 0.0533 | TRUE | 0.2718 |
| Pramel7 | 357 | 229 | 94 | 52 | 56 | 42 | 277 | 244 | 101 | 52 | 63 | 41 | 129.78 | 207 | 52 | 0.25 | -2 | 7.49E-11 | 6.26E-09 | 0.1456 | 0.0207 | 0.0557 | 0.0557 | TRUE | 0.7176 |
| Prap1 | 0 | 0 | 0 | 0 | 0 | 0 | 0 | 0 | 0 | 0 | 0 | 0 | 0 | 0 | 0 | NA | NA | NA | NA | NA | NA | NA | NA | NA | NA |
| Prb1 | 0 | 0 | 0 | 0 | 0 | 0 | 0 | 0 | 0 | 0 | 0 | 0 | 0 | 0 | 0 | NA | NA | NA | NA | NA | NA | NA | NA | NA | NA |
| Prc1 | 10572 | 7938 | 8717 | 9682 | 8506 | 9399 | 8211 | 8444 | 9406 | 9637 | 9596 | 9278 | 9095.22 | 8687 | 9504 | 1.094 | 0.13 | 0.03713084 | 0.10857428 | 0.0027 | 0.0027 | 0.0027 | 0.0027 | TRUE | 0.0638 |
| Prcc | 6168 | 4372 | 4012 | 4205 | 3692 | 4522 | 4790 | 4650 | 4329 | 4186 | 4165 | 4464 | 4430.72 | 4590 | 4272 | 0.93 | -0.104 | 0.09185125 | 0.20788568 | 0.0018 | 0.003 | 0.0025 | 0.0025 | TRUE | 0.03 |
| Prcp | 1606 | 1062 | 1123 | 1301 | 1101 | 1264 | 1247 | 1130 | 1212 | 1295 | 1242 | 1248 | 1228.91 | 1196 | 1262 | 1.054 | 0.076 | 0.30166612 | 0.47402308 | 8.00E-04 | 0.0044 | 0.0031 | 0.0031 | TRUE | 0.0281 |
| Prdm1 | 620 | 543 | 658 | 567 | 591 | 744 | 482 | 578 | 710 | 564 | 667 | 734 | 622.44 | 590 | 655 | 1.114 | 0.155 | 0.26276155 | 0.43318169 | 0.026 | 0.0062 | 0.0122 | 0.0122 | TRUE | 0.3199 |
| Prdm10 | 748 | 546 | 591 | 527 | 512 | 593 | 581 | 581 | 638 | 525 | 578 | 585 | 581.16 | 600 | 563 | 0.938 | -0.092 | 0.33828143 | 0.51245962 | 0.0015 | 0.0065 | 0.0049 | 0.0049 | TRUE | 0.0414 |
| Prdm11 | 95 | 60 | 42 | 81 | 75 | 65 | 74 | 64 | 45 | 81 | 85 | 64 | 68.72 | 61 | 77 | 1.243 | 0.314 | 0.22379414 | 0.38825359 | 0.024 | 0.0369 | 0.0334 | 0.0334 | TRUE | 0.4165 |
| Prdm12 | 5 | 5 | 1 | 1 | 2 | 4 | 4 | 5 | 1 | 1 | 2 | 4 | 2.91 | 3 | 2 | 0.698 | -0.519 | 0.65331945 | NA | 0 | 0.8159 | 0.6067 | 0.6067 | TRUE | 0.3543 |
| Prdm13 | 26 | 12 | 7 | 9 | 4 | 6 | 20 | 13 | 8 | 9 | 5 | 6 | 9.98 | 14 | 7 | 0.475 | -1.075 | 0.10294548 | NA | 0.0954 | 0.2398 | 0.1979 | 0.1979 | TRUE | 0.1841 |
| Prdm14 | 2506 | 1853 | 2075 | 1851 | 1638 | 1870 | 1946 | 1971 | 2239 | 1842 | 1848 | 1846 | 1948.75 | 2052 | 1845 | 0.9 | -0.152 | 0.03531997 | 0.10462432 | 0.0024 | 0.0036 | 0.0032 | 0.0032 | TRUE | 0.0759 |
| Prdm15 | 1190 | 977 | 1033 | 1132 | 1023 | 1171 | 924 | 1039 | 1115 | 1127 | 1154 | 1156 | 1085.81 | 1026 | 1146 | 1.119 | 0.162 | 0.0574288 | 0.14902504 | 0.0037 | 0.0046 | 0.0043 | 0.0043 | TRUE | 0.0942 |
| Prdm16 | 472 | 420 | 374 | 329 | 289 | 346 | 367 | 447 | 404 | 327 | 326 | 342 | 368.66 | 406 | 332 | 0.82 | -0.286 | 0.01415553 | 0.0534616 | 0.0034 | 0.0089 | 0.0071 | 0.0071 | TRUE | 0.0908 |
| Prdm2 | 7566 | 5619 | 5403 | 5542 | 5040 | 5625 | 5876 | 5977 | 5830 | 5517 | 5686 | 5553 | 5739.62 | 5894 | 5585 | 0.947 | -0.078 | 0.11284213 | 0.24151731 | 0 | 0.0028 | 0.0016 | 0.0016 | TRUE | 0.003 |
| Prdm4 | 4079 | 3099 | 3225 | 3224 | 2895 | 3314 | 3168 | 3296 | 3480 | 3209 | 3266 | 3271 | 3281.76 | 3315 | 3249 | 0.981 | -0.028 | 0.63358688 | 0.77064567 | 9.00E-04 | 0.0031 | 0.0022 | 0.0022 | TRUE | 0.023 |
| Prdm5 | 1913 | 1401 | 1898 | 1986 | 1769 | 2183 | 1486 | 1490 | 2048 | 1977 | 1996 | 2155 | 1858.56 | 1675 | 2043 | 1.221 | 0.288 | 0.00888438 | 0.03800792 | 0.019 | 0.0037 | 0.0082 | 0.0082 | TRUE | 0.4559 |
| Prdm6 | 39 | 23 | 31 | 29 | 16 | 13 | 30 | 24 | 33 | 29 | 18 | 13 | 24.66 | 29 | 20 | 0.678 | -0.561 | 0.17993516 | 0.33482793 | 0.0405 | 0.0985 | 0.0838 | 0.0838 | TRUE | 0.7695 |
| Prdm8 | 11 | 10 | 11 | 7 | 9 | 4 | 9 | 11 | 12 | 7 | 10 | 4 | 8.69 | 11 | 7 | 0.677 | -0.563 | 0.39622229 | NA | 0 | 0.2753 | 0.1962 | 0.1962 | TRUE | 0.268 |
| Prdm9 | 108 | 65 | 54 | 62 | 37 | 56 | 84 | 69 | 58 | 62 | 42 | 55 | 61.67 | 70 | 53 | 0.749 | -0.416 | 0.11950425 | 0.25127395 | 0.0194 | 0.0409 | 0.0349 | 0.0349 | TRUE | 0.2458 |

| | | | | | | | | | | | | | | | | | | | | | | | | | |
|---|---|---|---|---|---|---|---|---|---|---|---|---|---|---|---|---|---|---|---|---|---|---|---|---|---|
| Prdx1 | 55456 | 37178 | 37320 | 40806 | 33814 | 40508 | 43070 | 39546 | 40268 | 40618 | 38146 | 39988 | 40272.61 | 40961 | 39584 | 0.966 | -0.049 | 0.35799562 | 0.53239324 | 0.0015 | 0.0025 | 0.0021 | 0.0021 | TRUE | 0.0249 |
| Prdx2 | 11326 | 8437 | 8598 | 8410 | 7914 | 9207 | 8796 | 8974 | 9277 | 8371 | 8928 | 9089 | 8905.96 | 9016 | 8796 | 0.976 | -0.036 | 0.51485926 | 0.67730174 | 0.0011 | 0.0027 | 0.002 | 0.002 | TRUE | 0.0219 |
| Prdx3 | 3805 | 2744 | 2771 | 3082 | 2701 | 3025 | 2955 | 2919 | 2990 | 3068 | 3047 | 2986 | 2994.13 | 2955 | 3034 | 1.027 | 0.038 | 0.48395654 | 0.65044468 | 0 | 0.0032 | 0.0018 | 0.0018 | TRUE | 0.0023 |
| Prdx4 | 4318 | 2828 | 3111 | 2972 | 2633 | 2924 | 3354 | 3008 | 3357 | 2958 | 2970 | 2886 | 3088.92 | 3240 | 2938 | 0.907 | -0.141 | 0.02721096 | 0.08625556 | 0.0015 | 0.0032 | 0.0026 | 0.0026 | TRUE | 0.0469 |
| Prdx5 | 2141 | 1493 | 1536 | 1842 | 1605 | 1831 | 1663 | 1588 | 1657 | 1834 | 1811 | 1807 | 1726.64 | 1636 | 1817 | 1.11 | 0.151 | 0.01593216 | 0.05838933 | 0 | 0.0038 | 0.0022 | 0.0022 | TRUE | 0.0078 |
| Prdx6 | 12772 | 8880 | 8289 | 7694 | 6939 | 8094 | 9919 | 9446 | 8944 | 7659 | 7828 | 7990 | 8630.89 | 9436 | 7826 | 0.829 | -0.27 | 1.58E-06 | 4.21E-05 | 0.0014 | 0.0027 | 0.0022 | 0.0022 | TRUE | 0.0254 |
| Prdx6b | 0 | 0 | 0 | 0 | 0 | 0 | 0 | 0 | 0 | 0 | 0 | 0 | 0 | 0 | 0 | NA | NA | NA | NA | NA | NA | NA | NA | NA | NA |
| Preb | 4115 | 3036 | 2774 | 2737 | 2477 | 2957 | 3196 | 3229 | 2993 | 2724 | 2794 | 2919 | 2976.03 | 3139 | 2812 | 0.896 | -0.159 | 0.00956824 | 0.04008173 | 9.00E-04 | 0.0032 | 0.0024 | 0.0024 | TRUE | 0.0199 |
| Prelid1 | 10585 | 7847 | 7848 | 7878 | 7097 | 8564 | 8221 | 8347 | 8468 | 7842 | 8006 | 8454 | 8222.92 | 8345 | 8101 | 0.971 | -0.043 | 0.41213756 | 0.58508961 | 7.00E-04 | 0.0027 | 0.0018 | 0.0018 | TRUE | 0.0179 |
| Prelid2 | 2387 | 1882 | 2339 | 2618 | 2348 | 2554 | 1854 | 2002 | 2524 | 2606 | 2649 | 2521 | 2359.24 | 2127 | 2592 | 1.22 | 0.287 | 0.00358561 | 0.01895308 | 0.0134 | 0.0034 | 0.0066 | 0.0066 | TRUE | 0.3221 |
| Prelp | 5 | 3 | 1 | 3 | 1 | 0 | 4 | 3 | 1 | 3 | 1 | 0 | 2.04 | 3 | 1 | 0.499 | -1.002 | 0.48211255 | NA | 0 | 1.1616 | 0.9131 | 0.9131 | TRUE | 0.6241 |
| Prep | 6070 | 4310 | 4349 | 4225 | 3920 | 4650 | 4714 | 4584 | 4693 | 4206 | 4422 | 4590 | 4534.89 | 4664 | 4406 | 0.945 | -0.082 | 0.14392495 | 0.2869865 | 7.00E-04 | 0.0029 | 0.0021 | 0.0021 | TRUE | 0.0194 |
| Prepl | 3398 | 2505 | 2260 | 2621 | 2352 | 2627 | 2639 | 2665 | 2439 | 2609 | 2653 | 2593 | 2599.61 | 2581 | 2618 | 1.014 | 0.02 | 0.74079121 | 0.84589988 | 8.00E-04 | 0.0033 | 0.0023 | 0.0023 | TRUE | 0.0276 |
| Prex1 | 1673 | 1437 | 1785 | 2196 | 2023 | 2345 | 1299 | 1529 | 1926 | 2186 | 2282 | 2315 | 1922.8 | 1585 | 2261 | 1.429 | 0.515 | 4.52E-06 | 9.77E-05 | 0.0197 | 0.0037 | 0.0085 | 0.0085 | TRUE | 0.4276 |
| Prex2 | 2042 | 1522 | 979 | 1010 | 897 | 1005 | 1586 | 1619 | 1056 | 1005 | 1012 | 992 | 1211.76 | 1420 | 1003 | 0.706 | -0.503 | 9.54E-05 | 0.00116392 | 0.0267 | 0.0044 | 0.0111 | 0.0111 | TRUE | 0.5994 |
| Prf1 | 10 | 1 | 2 | 3 | 4 | 8 | 8 | 1 | 2 | 3 | 5 | 8 | 4.4 | 4 | 5 | 1.349 | 0.432 | 0.66283957 | NA | 0.3353 | 0.5414 | 0.4808 | 0.4808 | TRUE | 1.9141 |
| Prg2 | 1 | 0 | 1 | 0 | 0 | 0 | 1 | 0 | 1 | 0 | 0 | 0 | 0.31 | 1 | 0 | 0.307 | -1.706 | 0.62894649 | NA | 0 | 7.6657 | 7.2 | 7.2 | TRUE | 0.2067 |
| Prg3 | 0 | 0 | 0 | 0 | 0 | 0 | 0 | 0 | 0 | 0 | 0 | 0 | 0 | 0 | 0 | NA | NA | NA | NA | NA | NA | NA | NA | NA | NA |
| Prg4 | 0 | 2 | 1 | 1 | 0 | 0 | 0 | 2 | 1 | 1 | 0 | 0 | 0.7 | 1 | 0 | 0.356 | -1.492 | 0.54563002 | NA | 0 | 3.3868 | 2.9267 | 2.9267 | TRUE | 0.1826 |
| Prh1 | 0 | 0 | 0 | 0 | 0 | 0 | 0 | 0 | 0 | 0 | 0 | 0 | 0 | 0 | 0 | NA | NA | NA | NA | NA | NA | NA | NA | NA | NA |
| Prhoxnb | 0 | 1 | 0 | 0 | 1 | 0 | 0 | 1 | 0 | 0 | 1 | 0 | 0.37 | 0 | 0 | 1.075 | 0.105 | 0.97520514 | NA | 0 | 6.4904 | 6.1969 | 6.1969 | TRUE | 0.5216 |
| Prickle1 | 589 | 474 | 398 | 342 | 319 | 328 | 457 | 504 | 429 | 340 | 360 | 324 | 402.53 | 463 | 341 | 0.736 | -0.443 | 7.42E-05 | 0.00095852 | 0.0022 | 0.0083 | 0.0064 | 0.0064 | TRUE | 0.0649 |
| Prickle2 | 675 | 500 | 493 | 537 | 461 | 546 | 524 | 532 | 532 | 535 | 520 | 539 | 530.27 | 529 | 531 | 1.004 | 0.006 | 0.94879396 | 0.97205305 | 0 | 0.0069 | 0.0042 | 0.0042 | TRUE | 0.0038 |
| Prickle3 | 129 | 97 | 134 | 110 | 120 | 156 | 100 | 103 | 145 | 109 | 135 | 154 | 124.47 | 116 | 133 | 1.151 | 0.203 | 0.33162788 | 0.50586524 | 0.0279 | 0.0215 | 0.0233 | 0.0233 | TRUE | 0.4534 |
| Prim1 | 4299 | 3000 | 3044 | 3247 | 2778 | 3284 | 3339 | 3191 | 3284 | 3232 | 3134 | 3242 | 3237.02 | 3271 | 3203 | 0.979 | -0.031 | 0.5753566 | 0.72717057 | 0 | 0.0032 | 0.0019 | 0.0019 | TRUE | 0.0056 |
| Prim2 | 3729 | 2764 | 2835 | 3350 | 2862 | 3319 | 2896 | 2940 | 3059 | 3335 | 3229 | 3276 | 3122.45 | 2965 | 3280 | 1.107 | 0.146 | 0.00906595 | 0.03850805 | 2.00E-04 | 0.0032 | 0.0019 | 0.0019 | TRUE | 0.0092 |
| Prima1 | 74 | 52 | 62 | 63 | 55 | 55 | 57 | 55 | 67 | 63 | 62 | 54 | 59.79 | 60 | 60 | 0.997 | -0.004 | 0.98834613 | 0.99389455 | 0 | 0.0421 | 0.0284 | 0.0284 | TRUE | 0.0841 |
| Prkaa1 | 3687 | 2730 | 2976 | 3182 | 2915 | 3392 | 2864 | 2904 | 3211 | 3167 | 3288 | 3348 | 3130.45 | 2993 | 3268 | 1.092 | 0.128 | 0.04998185 | 0.13477101 | 0.0019 | 0.0032 | 0.0027 | 0.0027 | TRUE | 0.049 |
| Prkaa2 | 4158 | 3076 | 3440 | 4900 | 4178 | 4611 | 3229 | 3272 | 3712 | 4877 | 4713 | 4552 | 4059.24 | 3404 | 4714 | 1.385 | 0.47 | 6.09E-12 | 6.65E-10 | 0.0033 | 0.003 | 0.0031 | 0.0031 | TRUE | 0.0753 |
| Prkab1 | 1644 | 1204 | 1152 | 1176 | 1081 | 1383 | 1277 | 1281 | 1243 | 1171 | 1219 | 1365 | 1259.3 | 1267 | 1252 | 0.988 | -0.017 | 0.8280348 | 0.90144951 | 0.0026 | 0.0043 | 0.0037 | 0.0037 | TRUE | 0.0764 |
| Prkab2 | 1740 | 1335 | 1274 | 1173 | 979 | 1108 | 1351 | 1420 | 1375 | 1168 | 1104 | 1094 | 1251.97 | 1382 | 1122 | 0.812 | -0.3 | 2.28E-05 | 0.0003722 | 0 | 0.0043 | 0.0028 | 0.0028 | TRUE | 0.0154 |
| Prkaca | 8081 | 5699 | 5639 | 6138 | 5141 | 5887 | 6276 | 6062 | 6084 | 6110 | 5800 | 5811 | 6023.88 | 6141 | 5907 | 0.962 | -0.056 | 0.27811269 | 0.44961708 | 4.00E-04 | 0.0028 | 0.0018 | 0.0018 | TRUE | 0.011 |
| Prkacb | 3569 | 2795 | 2829 | 3014 | 2662 | 3095 | 2772 | 2973 | 3052 | 3000 | 3003 | 3055 | 2975.96 | 2932 | 3019 | 1.03 | 0.043 | 0.47578618 | 0.64357368 | 0.001 | 0.0032 | 0.0023 | 0.0023 | TRUE | 0.0287 |
| Prkag1 | 1657 | 1111 | 1128 | 1111 | 1043 | 1248 | 1287 | 1182 | 1217 | 1106 | 1177 | 1232 | 1200.04 | 1229 | 1172 | 0.953 | -0.07 | 0.36181267 | 0.53660525 | 0.0016 | 0.0044 | 0.0034 | 0.0034 | TRUE | 0.0293 |
| Prkag2 | 981 | 685 | 690 | 813 | 702 | 808 | 762 | 729 | 745 | 809 | 792 | 798 | 772.31 | 745 | 800 | 1.073 | 0.101 | 0.20583137 | 0.36769751 | 0 | 0.0055 | 0.0033 | 0.0033 | TRUE | 0.005 |
| Prkag3 | 16 | 10 | 16 | 20 | 21 | 13 | 12 | 11 | 17 | 20 | 24 | 13 | 16.13 | 13 | 19 | 1.399 | 0.484 | 0.32755679 | NA | 0 | 0.1494 | 0.1127 | 0.1127 | TRUE | 0.3734 |
| Prkar1a | 15028 | 10909 | 10864 | 11408 | 10235 | 11539 | 11672 | 11604 | 11722 | 11356 | 11546 | 11391 | 11548.33 | 11666 | 11431 | 0.98 | -0.029 | 0.50748525 | 0.67081167 | 0 | 0.0026 | 0.0013 | 0.0013 | TRUE | 9.00E-04 |
| Prkar1b | 183 | 151 | 227 | 277 | 270 | 294 | 142 | 161 | 245 | 276 | 305 | 290 | 236.37 | 183 | 290 | 1.597 | 0.676 | 0.00020289 | 0.00215604 | 0.0373 | 0.0125 | 0.0194 | 0.0194 | TRUE | 0.9457 |
| Prkar2a | 7267 | 5110 | 5093 | 5095 | 4369 | 4964 | 5644 | 5435 | 5495 | 5072 | 4929 | 4900 | 5245.87 | 5525 | 4967 | 0.899 | -0.154 | 0.00247326 | 0.01423118 | 2.00E-04 | 0.0029 | 0.0017 | 0.0017 | TRUE | 0.0044 |
| Prkar2b | 693 | 488 | 390 | 510 | 407 | 435 | 538 | 519 | 421 | 508 | 459 | 429 | 479.05 | 493 | 465 | 0.943 | -0.085 | 0.47757541 | 0.64514436 | 0.0101 | 0.0074 | 0.0082 | 0.0082 | TRUE | 0.1855 |
| Prkca | 1033 | 719 | 596 | 828 | 780 | 804 | 802 | 765 | 643 | 824 | 880 | 794 | 784.66 | 737 | 833 | 1.128 | 0.173 | 0.08064296 | 0.19021222 | 0.0066 | 0.0054 | 0.0058 | 0.0058 | TRUE | 0.1437 |
| Prkcb | 1634 | 1405 | 1609 | 2306 | 2048 | 2318 | 1269 | 1494 | 1736 | 2295 | 2310 | 2288 | 1898.93 | 1500 | 2298 | 1.534 | 0.618 | 2.01E-10 | 1.53E-08 | 0.0113 | 0.0037 | 0.0062 | 0.0062 | TRUE | 0.2288 |
| Prkcd | 2406 | 1682 | 1791 | 1627 | 1570 | 1675 | 1869 | 1789 | 1932 | 1620 | 1771 | 1653 | 1772.39 | 1863 | 1681 | 0.902 | -0.149 | 0.02963242 | 0.09179371 | 0.0012 | 0.0038 | 0.0028 | 0.0028 | TRUE | 0.0259 |
| Prkcdbp | 38 | 24 | 18 | 16 | 11 | 13 | 30 | 26 | 19 | 16 | 12 | 13 | 19.27 | 25 | 14 | 0.55 | -0.861 | 0.05446853 | 0.14308376 | 0 | 0.1254 | 0.0876 | 0.0876 | TRUE | 0.208 |
| Prkce | 217 | 116 | 131 | 120 | 86 | 90 | 169 | 123 | 141 | 119 | 97 | 89 | 123.1 | 144 | 102 | 0.702 | -0.51 | 0.01052251 | 0.0428607 | 0.0168 | 0.0217 | 0.0203 | 0.0203 | TRUE | 0.2405 |
| Prkcg | 539 | 456 | 434 | 558 | 482 | 592 | 419 | 485 | 468 | 555 | 544 | 584 | 509.25 | 457 | 561 | 1.231 | 0.299 | 0.00304811 | 0.01673133 | 0.0018 | 0.0071 | 0.0054 | 0.0054 | TRUE | 0.0686 |
| Prkch | 435 | 311 | 240 | 174 | 165 | 161 | 338 | 331 | 259 | 173 | 186 | 159 | 240.98 | 309 | 173 | 0.557 | -0.845 | 1.84E-08 | 8.60E-07 | 0.0104 | 0.0123 | 0.0117 | 0.0117 | TRUE | 0.2221 |
| Prkci | 14269 | 10416 | 11561 | 12944 | 11208 | 13455 | 11082 | 11079 | 12474 | 12884 | 12644 | 13282 | 12241.02 | 11545 | 12937 | 1.121 | 0.164 | 0.00709605 | 0.03207587 | 0.0026 | 0.0026 | 0.0026 | 0.0026 | TRUE | 0.0604 |
| Prkcq | 27 | 24 | 8 | 13 | 15 | 15 | 21 | 26 | 9 | 13 | 17 | 15 | 16.63 | 19 | 15 | 0.805 | -0.314 | 0.54001807 | NA | 0.0803 | 0.1449 | 0.1277 | 0.1277 | TRUE | 0.6575 |
| Prkcsh | 7137 | 5533 | 5744 | 6133 | 5565 | 6574 | 5543 | 5885 | 6198 | 6105 | 6278 | 6490 | 6083.06 | 5875 | 6291 | 1.071 | 0.099 | 0.09821541 | 0.21829899 | 0.0018 | 0.0028 | 0.0024 | 0.0024 | TRUE | 0.0303 |
| Prkcz | 2020 | 1365 | 1789 | 1916 | 1711 | 2017 | 1569 | 1452 | 1930 | 1907 | 1930 | 1991 | 1796.59 | 1650 | 1943 | 1.178 | 0.236 | 0.01331875 | 0.05114586 | 0.0108 | 0.0037 | 0.006 | 0.006 | TRUE | 0.2621 |
| Prkd1 | 406 | 313 | 292 | 319 | 295 | 283 | 315 | 333 | 315 | 318 | 333 | 279 | 315.5 | 321 | 310 | 0.965 | -0.052 | 0.66623013 | 0.7948039 | 0 | 0.0099 | 0.0073 | 0.0073 | TRUE | 0.0851 |
| Prkd2 | 442 | 333 | 338 | 358 | 321 | 383 | 343 | 354 | 365 | 356 | 362 | 378 | 359.79 | 354 | 365 | 1.034 | 0.048 | 0.66106248 | 0.79067498 | 0 | 0.009 | 0.0057 | 0.0057 | TRUE | 0.0106 |
| Prkd3 | 4531 | 3512 | 3390 | 3452 | 3111 | 3705 | 3519 | 3736 | 3658 | 3436 | 3510 | 3657 | 3585.92 | 3638 | 3534 | 0.972 | -0.041 | 0.47573931 | 0.64357368 | 7.00E-04 | 0.0031 | 0.0021 | 0.0021 | TRUE | 0.0114 |
| Prkdc | 3223 | 2296 | 1943 | 2703 | 2335 | 2696 | 2503 | 2442 | 2096 | 2691 | 2634 | 2661 | 2504.65 | 2347 | 2662 | 1.133 | 0.181 | 0.01597214 | 0.05850095 | 0.0038 | 0.0034 | 0.0036 | 0.0036 | TRUE | 0.1047 |
| Prkg1 | 19 | 15 | 8 | 13 | 15 | 11 | 15 | 16 | 9 | 13 | 17 | 11 | 13.34 | 13 | 14 | 1.025 | 0.036 | 0.9458243 | NA | 0 | 0.18 | 0.1301 | 0.1301 | TRUE | 0.3381 |
| Prkg2 | 89 | 73 | 52 | 74 | 85 | 58 | 69 | 78 | 56 | 74 | 96 | 57 | 71.61 | 68 | 76 | 1.113 | 0.155 | 0.55055111 | 0.70656613 | 0.0311 | 0.0355 | 0.0343 | 0.0343 | TRUE | 0.2455 |
| Prkra | 1062 | 775 | 1077 | 1221 | 1064 | 1214 | 825 | 824 | 1162 | 1215 | 1200 | 1198 | 1070.89 | 937 | 1204 | 1.287 | 0.364 | 0.00200174 | 0.01212386 | 0.0202 | 0.0046 | 0.0091 | 0.0091 | TRUE | 0.5208 |
| Prkrip1 | 1436 | 1009 | 907 | 1149 | 918 | 1075 | 1115 | 1073 | 979 | 1144 | 1036 | 1061 | 1067.95 | 1056 | 1080 | 1.022 | 0.032 | 0.69897163 | 0.81714392 | 0.0027 | 0.0046 | 0.004 | 0.004 | TRUE | 0.0481 |
| Prkrir | 3212 | 2266 | 2186 | 2450 | 1978 | 2235 | 2495 | 2410 | 2359 | 2439 | 2231 | 2206 | 2356.67 | 2421 | 2292 | 0.946 | -0.079 | 0.22646004 | 0.39149527 | 0.0013 | 0.0034 | 0.0027 | 0.0027 | TRUE | 0.0382 |
| Prkx | 1104 | 758 | 747 | 728 | 662 | 770 | 857 | 806 | 806 | 725 | 747 | 760 | 783.55 | 823 | 744 | 0.903 | -0.148 | 0.06997641 | 0.17149741 | 0 | 0.0055 | 0.0035 | 0.0035 | TRUE | 0.0166 |
| Prl | 0 | 0 | 0 | 0 | 0 | 0 | 0 | 0 | 0 | 0 | 0 | 0 | 0 | 0 | 0 | NA | NA | NA | NA | NA | NA | NA | NA | NA | NA |
| Prl2a1 | 0 | 0 | 0 | 0 | 0 | 0 | 0 | 0 | 0 | 0 | 0 | 0 | 0 | 0 | 0 | NA | NA | NA | NA | NA | NA | NA | NA | NA | NA |
| Prl2b1 | 1 | 0 | 0 | 0 | 0 | 0 | 1 | 0 | 0 | 0 | 0 | 0 | 0.13 | 0 | 0 | 0.552 | -0.857 | 0.8336548 | NA | 0 | 18.3121 | 10 | 10 | TRUE | 0.2566 |
| Prl2c1 | 0 | 0 | 0 | 0 | 0 | 0 | 0 | 0 | 0 | 0 | 0 | 0 | 0 | 0 | 0 | NA | NA | NA | NA | NA | NA | NA | NA | NA | NA |
| Prl2c2 | 0 | 0 | 0 | 0 | 0 | 0 | 0 | 0 | 0 | 0 | 0 | 0 | 0 | 0 | 0 | NA | NA | NA | NA | NA | NA | NA | NA | NA | NA |
| Prl2c3 | 0 | 0 | 0 | 0 | 0 | 0 | 0 | 0 | 0 | 0 | 0 | 0 | 0 | 0 | 0 | NA | NA | NA | NA | NA | NA | NA | NA | NA | NA |
| Prl2c4 | 0 | 0 | 0 | 0 | 0 | 0 | 0 | 0 | 0 | 0 | 0 | 0 | 0 | 0 | 0 | NA | NA | NA | NA | NA | NA | NA | NA | NA | NA |
| Prl2c5 | 0 | 0 | 0 | 0 | 0 | 0 | 0 | 0 | 0 | 0 | 0 | 0 | 0 | 0 | 0 | NA | NA | NA | NA | NA | NA | NA | NA | NA | NA |

| | | | | | | | | | | | | | | | | | | | | | | | | | |
|---|---|---|---|---|---|---|---|---|---|---|---|---|---|---|---|---|---|---|---|---|---|---|---|---|---|
| Prl3a1 | 0 | 0 | 0 | 0 | 0 | 0 | 0 | 0 | 0 | 0 | 0 | 0 | 0 | 0 | 0 | NA | NA | NA | NA | NA | NA | NA | NA | NA | NA |
| Prl3b1 | 0 | 0 | 0 | 0 | 0 | 0 | 0 | 0 | 0 | 0 | 0 | 0 | 0 | 0 | 0 | NA | NA | NA | NA | NA | NA | NA | NA | NA | NA |
| Prl3c1 | 0 | 0 | 0 | 0 | 0 | 0 | 0 | 0 | 0 | 0 | 0 | 0 | 0 | 0 | 0 | NA | NA | NA | NA | NA | NA | NA | NA | NA | NA |
| Prl3d1 | 0 | 0 | 0 | 0 | 0 | 0 | 0 | 0 | 0 | 0 | 0 | 0 | 0 | 0 | 0 | NA | NA | NA | NA | NA | NA | NA | NA | NA | NA |
| Prl3d2 | 0 | 0 | 0 | 0 | 0 | 0 | 0 | 0 | 0 | 0 | 0 | 0 | 0 | 0 | 0 | NA | NA | NA | NA | NA | NA | NA | NA | NA | NA |
| Prl3d3 | 0 | 0 | 0 | 0 | 0 | 0 | 0 | 0 | 0 | 0 | 0 | 0 | 0 | 0 | 0 | NA | NA | NA | NA | NA | NA | NA | NA | NA | NA |
| Prl4a1 | 0 | 0 | 0 | 0 | 0 | 0 | 0 | 0 | 0 | 0 | 0 | 0 | 0 | 0 | 0 | NA | NA | NA | NA | NA | NA | NA | NA | NA | NA |
| Prl5a1 | 0 | 0 | 0 | 0 | 0 | 0 | 0 | 0 | 0 | 0 | 0 | 0 | 0 | 0 | 0 | NA | NA | NA | NA | NA | NA | NA | NA | NA | NA |
| Prl6a1 | 0 | 0 | 0 | 0 | 0 | 0 | 0 | 0 | 0 | 0 | 0 | 0 | 0 | 0 | 0 | NA | NA | NA | NA | NA | NA | NA | NA | NA | NA |
| Prl7a1 | 0 | 0 | 0 | 0 | 0 | 0 | 0 | 0 | 0 | 0 | 0 | 0 | 0 | 0 | 0 | NA | NA | NA | NA | NA | NA | NA | NA | NA | NA |
| Prl7a2 | 0 | 0 | 0 | 0 | 0 | 0 | 0 | 0 | 0 | 0 | 0 | 0 | 0 | 0 | 0 | NA | NA | NA | NA | NA | NA | NA | NA | NA | NA |
| Prl7b1 | 0 | 0 | 0 | 0 | 0 | 0 | 0 | 0 | 0 | 0 | 0 | 0 | 0 | 0 | 0 | NA | NA | NA | NA | NA | NA | NA | NA | NA | NA |
| Prl7c1 | 0 | 0 | 0 | 0 | 0 | 0 | 0 | 0 | 0 | 0 | 0 | 0 | 0 | 0 | 0 | NA | NA | NA | NA | NA | NA | NA | NA | NA | NA |
| Prl7d1 | 0 | 0 | 0 | 0 | 0 | 0 | 0 | 0 | 0 | 0 | 0 | 0 | 0 | 0 | 0 | NA | NA | NA | NA | NA | NA | NA | NA | NA | NA |
| Prl8a1 | 0 | 0 | 0 | 0 | 0 | 0 | 0 | 0 | 0 | 0 | 0 | 0 | 0 | 0 | 0 | NA | NA | NA | NA | NA | NA | NA | NA | NA | NA |
| Prl8a2 | 0 | 0 | 0 | 0 | 0 | 0 | 0 | 0 | 0 | 0 | 0 | 0 | 0 | 0 | 0 | NA | NA | NA | NA | NA | NA | NA | NA | NA | NA |
| Prl8a6 | 0 | 0 | 0 | 0 | 0 | 0 | 0 | 0 | 0 | 0 | 0 | 0 | 0 | 0 | 0 | NA | NA | NA | NA | NA | NA | NA | NA | NA | NA |
| Prl8a8 | 0 | 0 | 0 | 0 | 0 | 0 | 0 | 0 | 0 | 0 | 0 | 0 | 0 | 0 | 0 | NA | NA | NA | NA | NA | NA | NA | NA | NA | NA |
| Prl8a9 | 4 | 1 | 0 | 0 | 0 | 0 | 3 | 1 | 0 | 0 | 0 | 0 | 0.7 | 1 | 0 | 0.134 | -2.897 | 0.27044302 | NA | 6.5513 | 3.4123 | 3.6399 | 3.6399 | TRUE | 0.1458 |
| Prlh | 0 | 0 | 0 | 0 | 0 | 1 | 0 | 0 | 0 | 0 | 0 | 1 | 0.16 | 0 | 0 | 2.094 | 1.067 | 0.79379329 | NA | 0 | 14.4078 | 10 | 10 | TRUE | 0.3835 |
| Prlhr | 3 | 3 | 2 | 0 | 1 | 5 | 2 | 3 | 2 | 0 | 1 | 5 | 2.29 | 2 | 2 | 0.798 | -0.325 | 0.80927078 | NA | 0.316 | 1.0372 | 0.8634 | 0.8634 | TRUE | 1.4468 |
| Prlr | 11 | 19 | 10 | 5 | 11 | 17 | 9 | 20 | 11 | 5 | 12 | 17 | 12.29 | 13 | 11 | 0.872 | -0.198 | 0.74814459 | NA | 0.1796 | 0.1953 | 0.1909 | 0.1909 | TRUE | 0.4604 |
| Prm1 | 0 | 0 | 0 | 0 | 0 | 0 | 0 | 0 | 0 | 0 | 0 | 0 | 0 | 0 | 0 | NA | NA | NA | NA | NA | NA | NA | NA | NA | NA |
| Prm2 | 0 | 0 | 0 | 2 | 0 | 0 | 0 | 0 | 0 | 2 | 0 | 0 | 0.33 | 0 | 1 | 3.77 | 1.915 | 0.62344813 | NA | 0 | 7.1455 | 9.1834 | 9.1834 | TRUE | 0.9739 |
| Prm3 | 0 | 0 | 0 | 0 | 0 | 0 | 0 | 0 | 0 | 0 | 0 | 0 | 0 | 0 | 0 | NA | NA | NA | NA | NA | NA | NA | NA | NA | NA |
| Prmt1 | 15377 | 10484 | 9541 | 9428 | 8550 | 9929 | 11943 | 11152 | 10295 | 9385 | 9645 | 9801 | 10370.06 | 11130 | 9610 | 0.863 | -0.212 | 0.000764 | 0.00594443 | 0.0028 | 0.0027 | 0.0028 | 0.0028 | TRUE | 0.0526 |
| Prmt10 | 3787 | 2785 | 2492 | 2659 | 2376 | 2664 | 2941 | 2962 | 2689 | 2647 | 2680 | 2630 | 2758.22 | 2864 | 2652 | 0.926 | -0.111 | 0.07296977 | 0.17686229 | 0.0011 | 0.0033 | 0.0024 | 0.0024 | TRUE | 0.0342 |
| Prmt2 | 1141 | 819 | 862 | 896 | 804 | 919 | 886 | 871 | 930 | 892 | 907 | 907 | 898.91 | 896 | 902 | 1.007 | 0.01 | 0.892707 | 0.94103261 | 0 | 0.0051 | 0.0031 | 0.0031 | TRUE | 0.0128 |
| Prmt3 | 5352 | 4176 | 4219 | 4420 | 3877 | 4571 | 4157 | 4442 | 4552 | 4400 | 4374 | 4512 | 4406.08 | 4384 | 4429 | 1.011 | 0.016 | 0.78725317 | 0.87518158 | 0.0011 | 0.003 | 0.0022 | 0.0022 | TRUE | 0.0255 |
| Prmt5 | 13841 | 10342 | 10655 | 10466 | 9676 | 11607 | 10750 | 11001 | 11497 | 10418 | 10916 | 11458 | 11006.38 | 11083 | 10931 | 0.986 | -0.02 | 0.7266581 | 0.83617456 | 0.0013 | 0.0026 | 0.0022 | 0.0022 | TRUE | 0.0219 |
| Prmt6 | 1851 | 1240 | 925 | 865 | 785 | 791 | 1438 | 1319 | 998 | 861 | 886 | 781 | 1047.01 | 1252 | 843 | 0.672 | -0.573 | 1.44E-06 | 3.91E-05 | 0.019 | 0.0047 | 0.0092 | 0.0092 | TRUE | 0.3737 |
| Prmt7 | 13821 | 10578 | 11817 | 13258 | 11848 | 13776 | 10734 | 11252 | 12750 | 13197 | 13366 | 13599 | 12483.02 | 11579 | 13387 | 1.156 | 0.21 | 0.00180387 | 0.01117103 | 0.0041 | 0.0026 | 0.0032 | 0.0032 | TRUE | 0.0957 |
| Prmt8 | 759 | 579 | 748 | 872 | 811 | 909 | 589 | 616 | 807 | 868 | 915 | 897 | 782.11 | 671 | 893 | 1.334 | 0.416 | 0.00024928 | 0.0025237 | 0.0141 | 0.0055 | 0.008 | 0.008 | TRUE | 0.3682 |
| Prnd | 3 | 5 | 0 | 4 | 4 | 3 | 2 | 5 | 0 | 4 | 5 | 3 | 3.18 | 2 | 4 | 1.496 | 0.581 | 0.61036225 | NA | 0.0413 | 0.7468 | 0.6125 | 0.6125 | TRUE | 0.3449 |
| Prnp | 785 | 525 | 626 | 794 | 664 | 783 | 610 | 558 | 675 | 790 | 749 | 773 | 692.65 | 614 | 771 | 1.255 | 0.328 | 0.00050529 | 0.00434542 | 0.0027 | 0.0058 | 0.0049 | 0.0049 | TRUE | 0.0842 |
| Prob1 | 117 | 101 | 76 | 69 | 61 | 64 | 91 | 107 | 82 | 69 | 69 | 63 | 80.16 | 93 | 67 | 0.716 | -0.482 | 0.03064455 | 0.09401215 | 0 | 0.032 | 0.0229 | 0.0229 | TRUE | 0.1557 |
| Proc | 32 | 21 | 17 | 15 | 15 | 17 | 25 | 22 | 18 | 15 | 17 | 17 | 19.03 | 22 | 16 | 0.738 | -0.439 | 0.31514542 | 0.48870278 | 0 | 0.127 | 0.0835 | 0.0835 | TRUE | 0.1054 |
| Proca1 | 132 | 121 | 119 | 109 | 81 | 107 | 103 | 129 | 128 | 108 | 91 | 106 | 110.85 | 120 | 102 | 0.855 | -0.226 | 0.24597301 | 0.41458635 | 0 | 0.0238 | 0.0183 | 0.0183 | TRUE | 0.1816 |
| Procr | 260 | 170 | 221 | 209 | 210 | 229 | 202 | 181 | 238 | 208 | 237 | 226 | 215.37 | 207 | 224 | 1.08 | 0.112 | 0.45561775 | 0.62526591 | 0.0065 | 0.0134 | 0.0115 | 0.0115 | TRUE | 0.1811 |
| Prodh | 1157 | 973 | 737 | 696 | 714 | 835 | 899 | 1035 | 795 | 693 | 805 | 824 | 841.89 | 910 | 774 | 0.851 | -0.233 | 0.03254969 | 0.09829553 | 0.0112 | 0.0052 | 0.0073 | 0.0073 | TRUE | 0.1691 |
| Prodh2 | 8 | 12 | 7 | 9 | 6 | 14 | 6 | 13 | 8 | 9 | 7 | 14 | 9.35 | 9 | 10 | 1.132 | 0.178 | 0.7827208 | NA | 0 | 0.256 | 0.1937 | 0.1937 | TRUE | 0.4555 |
| Prok1 | 1 | 0 | 0 | 0 | 0 | 0 | 1 | 0 | 0 | 0 | 0 | 0 | 0.13 | 0 | 0 | 0.552 | -0.857 | 0.8336548 | NA | 0 | 18.3121 | 10 | 10 | TRUE | 0.2566 |
| Prok2 | 63 | 40 | 48 | 75 | 60 | 63 | 49 | 43 | 52 | 75 | 68 | 62 | 57.97 | 48 | 68 | 1.426 | 0.512 | 0.04513189 | 0.12494095 | 0 | 0.0433 | 0.0295 | 0.0295 | TRUE | 0.067 |
| Prokr1 | 60 | 48 | 49 | 61 | 44 | 59 | 47 | 51 | 53 | 61 | 50 | 58 | 53.19 | 50 | 56 | 1.125 | 0.17 | 0.51861577 | 0.68023999 | 0 | 0.047 | 0.0314 | 0.0314 | TRUE | 0.0821 |
| Prokr2 | 39 | 23 | 15 | 29 | 19 | 19 | 30 | 24 | 16 | 29 | 21 | 19 | 23.33 | 23 | 23 | 0.965 | -0.052 | 0.90174938 | 0.94568228 | 0.032 | 0.104 | 0.084 | 0.084 | TRUE | 0.2795 |
| Prol1 | 0 | 0 | 0 | 0 | 0 | 0 | 0 | 0 | 0 | 0 | 0 | 0 | 0 | 0 | 0 | NA | NA | NA | NA | NA | NA | NA | NA | NA | NA |
| Prom1 | 666 | 528 | 691 | 782 | 794 | 851 | 517 | 562 | 746 | 778 | 896 | 840 | 723.11 | 608 | 838 | 1.381 | 0.465 | 0.00020847 | 0.00219585 | 0.0201 | 0.0057 | 0.0099 | 0.0099 | TRUE | 0.4554 |
| Prom2 | 47 | 40 | 60 | 48 | 32 | 46 | 37 | 43 | 65 | 48 | 36 | 45 | 45.51 | 48 | 43 | 0.908 | -0.139 | 0.6589614 | 0.78895421 | 0.037 | 0.0545 | 0.0494 | 0.0494 | TRUE | 0.7304 |
| Prop1 | 1 | 0 | 0 | 0 | 0 | 0 | 1 | 0 | 0 | 0 | 0 | 0 | 0.13 | 0 | 0 | 0.552 | -0.857 | 0.8336548 | NA | 0 | 18.3121 | 10 | 10 | TRUE | 0.2566 |
| Prorsd1 | 705 | 518 | 485 | 543 | 476 | 570 | 548 | 551 | 523 | 541 | 537 | 563 | 543.67 | 541 | 547 | 1.011 | 0.016 | 0.86245013 | 0.92408891 | 0 | 0.0068 | 0.0043 | 0.0043 | TRUE | 0.0088 |
| Pros1 | 2505 | 1611 | 1524 | 1540 | 1447 | 1516 | 1946 | 1714 | 1644 | 1533 | 1632 | 1497 | 1660.89 | 1768 | 1554 | 0.878 | -0.188 | 0.01936496 | 0.06752462 | 0.0041 | 0.0039 | 0.0041 | 0.0041 | TRUE | 0.0957 |
| Prosapip1 | 606 | 399 | 296 | 439 | 404 | 501 | 471 | 424 | 319 | 437 | 456 | 495 | 433.62 | 405 | 463 | 1.139 | 0.188 | 0.16990759 | 0.32144133 | 0.0188 | 0.0079 | 0.0112 | 0.0112 | TRUE | 0.3852 |
| Prosc | 5356 | 3936 | 3728 | 3960 | 3581 | 4090 | 4160 | 4187 | 4022 | 3942 | 4040 | 4037 | 4064.65 | 4123 | 4006 | 0.972 | -0.042 | 0.42705326 | 0.5991237 | 0 | 0.003 | 0.0017 | 0.0017 | TRUE | 0.0054 |
| Proser1 | 2798 | 2025 | 1975 | 2063 | 2028 | 2288 | 2173 | 2154 | 2131 | 2054 | 2288 | 2259 | 2176.33 | 2153 | 2200 | 1.022 | 0.031 | 0.63981797 | 0.77486174 | 0.0013 | 0.0035 | 0.0027 | 0.0027 | TRUE | 0.0413 |
| Proser2 | 813 | 650 | 747 | 724 | 660 | 743 | 631 | 691 | 806 | 721 | 745 | 733 | 721.25 | 709 | 733 | 1.035 | 0.05 | 0.62436644 | 0.76326288 | 0.0065 | 0.0057 | 0.006 | 0.006 | TRUE | 0.1658 |
| Prox1 | 76 | 34 | 45 | 28 | 20 | 34 | 59 | 36 | 49 | 28 | 23 | 34 | 37.96 | 48 | 28 | 0.583 | -0.779 | 0.02112311 | 0.07203775 | 0.0261 | 0.0649 | 0.0539 | 0.0539 | TRUE | 0.1473 |
| Prox2 | 7 | 12 | 6 | 8 | 2 | 8 | 5 | 13 | 6 | 8 | 2 | 8 | 7.13 | 8 | 6 | 0.749 | -0.416 | 0.58622082 | NA | 0.121 | 0.3347 | 0.2775 | 0.2775 | TRUE | 0.668 |
| Proz | 4 | 2 | 2 | 2 | 1 | 1 | 3 | 2 | 2 | 2 | 1 | 1 | 1.92 | 2 | 1 | 0.552 | -0.857 | 0.52698802 | NA | 0 | 1.2392 | 0.7512 | 0.7512 | TRUE | 0.1031 |
| Prp2 | 0 | 0 | 0 | 0 | 0 | 0 | 0 | 0 | 0 | 0 | 0 | 0 | 0 | 0 | 0 | NA | NA | NA | NA | NA | NA | NA | NA | NA | NA |
| Prpf18 | 3701 | 2633 | 2648 | 2650 | 2395 | 2637 | 2874 | 2801 | 2857 | 2638 | 2702 | 2603 | 2745.83 | 2844 | 2648 | 0.931 | -0.104 | 0.06348889 | 0.15984616 | 0 | 0.0033 | 0.0019 | 0.0019 | TRUE | 0.0038 |
| Prpf19 | 12978 | 8967 | 7859 | 7624 | 6912 | 7940 | 10079 | 9538 | 8480 | 7589 | 7797 | 7838 | 8553.62 | 9366 | 7741 | 0.826 | -0.275 | 4.74E-05 | 0.00066804 | 0.0037 | 0.0027 | 0.0032 | 0.0032 | TRUE | 0.0835 |
| Prpf3 | 4368 | 3279 | 3258 | 3407 | 3043 | 3652 | 3392 | 3488 | 3515 | 3391 | 3433 | 3605 | 3470.81 | 3465 | 3476 | 1.004 | 0.005 | 0.92664856 | 0.9596832 | 4.00E-04 | 0.0031 | 0.002 | 0.002 | TRUE | 0.0129 |
| Prpf31 | 5521 | 3903 | 2913 | 3157 | 2830 | 3104 | 4288 | 4152 | 3143 | 3142 | 3193 | 3064 | 3496.96 | 3861 | 3133 | 0.811 | -0.302 | 0.00212284 | 0.0126162 | 0.0129 | 0.0031 | 0.0067 | 0.0067 | TRUE | 0.3204 |
| Prpf38a | 5453 | 3754 | 3632 | 3899 | 3386 | 3851 | 4235 | 3993 | 3919 | 3881 | 3820 | 3802 | 3941.57 | 4049 | 3834 | 0.946 | -0.079 | 0.15825995 | 0.3072965 | 7.00E-04 | 0.003 | 0.002 | 0.002 | TRUE | 0.0201 |
| Prpf38b | 6641 | 5174 | 4978 | 5027 | 4408 | 5451 | 5158 | 5504 | 5371 | 5004 | 4973 | 5381 | 5231.67 | 5344 | 5119 | 0.958 | -0.062 | 0.28789442 | 0.46012244 | 0.0012 | 0.0029 | 0.0022 | 0.0022 | TRUE | 0.0245 |
| Prpf39 | 1849 | 1466 | 1371 | 1165 | 942 | 1093 | 1436 | 1559 | 1479 | 1160 | 1063 | 1079 | 1296 | 1491 | 1101 | 0.738 | -0.437 | 3.40E-09 | 1.96E-07 | 0.0012 | 0.0043 | 0.0032 | 0.0032 | TRUE | 0.0267 |
| Prpf4 | 4602 | 3294 | 3145 | 3341 | 2944 | 3261 | 3574 | 3504 | 3393 | 3326 | 3321 | 3219 | 3389.54 | 3490 | 3289 | 0.942 | -0.086 | 0.11712256 | 0.2481519 | 2.00E-04 | 0.0031 | 0.0019 | 0.0019 | TRUE | 0.0071 |
| Prpf40a | 11577 | 8090 | 7742 | 8263 | 7006 | 8143 | 8991 | 8605 | 8354 | 8225 | 7903 | 8038 | 8352.84 | 8650 | 8055 | 0.931 | -0.103 | 0.04862607 | 0.13203123 | 7.00E-04 | 0.0027 | 0.0018 | 0.0018 | TRUE | 0.0147 |
| Prpf40b | 2298 | 1842 | 1850 | 2128 | 1881 | 2140 | 1785 | 1959 | 1996 | 2118 | 2122 | 2113 | 2015.48 | 1913 | 2118 | 1.108 | 0.148 | 0.02720374 | 0.0862518 | 0.0014 | 0.0036 | 0.0027 | 0.0027 | TRUE | 0.0434 |

| | | | | | | | | | | | | | | | | | | | | | | | | | |
|---|---|---|---|---|---|---|---|---|---|---|---|---|---|---|---|---|---|---|---|---|---|---|---|---|---|
| Prpf4b | 4979 | 3846 | 3473 | 3254 | 2680 | 3286 | 3867 | 4091 | 3747 | 3239 | 3023 | 3244 | 3535.23 | 3902 | 3169 | 0.812 | -0.3 | 1.12E-06 | 3.22E-05 | 0.0014 | 0.0031 | 0.0024 | 0.0024 | TRUE | 0.0216 |
| Prpf6 | 7813 | 5762 | 5496 | 5801 | 5528 | 6523 | 6068 | 6129 | 5930 | 5774 | 6236 | 6439 | 6096.13 | 6042 | 6150 | 1.018 | 0.025 | 0.66306897 | 0.79208064 | 0.0013 | 0.0028 | 0.0023 | 0.0023 | TRUE | 0.035 |
| Prpf8 | 69038 | 48051 | 47554 | 51233 | 46999 | 56123 | 53619 | 51111 | 51310 | 50997 | 53020 | 55402 | 52576.59 | 52013 | 53140 | 1.022 | 0.031 | 0.54838763 | 0.70505571 | 0.0011 | 0.0025 | 0.0019 | 0.0019 | TRUE | 0.017 |
| Prph | 1785 | 1532 | 1626 | 2837 | 2524 | 2940 | 1386 | 1630 | 1754 | 2824 | 2847 | 2902 | 2223.98 | 1590 | 2858 | 1.8 | 0.848 | 3.86E-24 | 1.84E-21 | 0.0063 | 0.0035 | 0.0046 | 0.0046 | TRUE | 0.1562 |
| Prph2 | 42 | 24 | 24 | 32 | 34 | 31 | 33 | 26 | 26 | 32 | 38 | 31 | 30.81 | 28 | 34 | 1.189 | 0.25 | 0.47210805 | 0.64086802 | 0 | 0.0794 | 0.0543 | 0.0543 | TRUE | 0.1602 |
| Prpmp5 | 0 | 0 | 0 | 0 | 0 | 0 | 0 | 0 | 0 | 0 | 0 | 0 | 0 | 0 | 0 | NA | NA | NA | NA | NA | NA | NA | NA | NA | NA |
| Prps1 | 12906 | 10303 | 11810 | 9271 | 9103 | 10522 | 10024 | 10959 | 12743 | 9228 | 10269 | 10387 | 10601.65 | 11242 | 9961 | 0.886 | -0.174 | 0.03868492 | 0.11176946 | 0.0095 | 0.0027 | 0.005 | 0.005 | TRUE | 0.1667 |
| Prps1l1 | 0 | 0 | 0 | 0 | 0 | 0 | 0 | 0 | 0 | 0 | 0 | 0 | 0 | 0 | 0 | NA | NA | NA | NA | NA | NA | NA | NA | NA | NA |
| Prps1l3 | 6814 | 4911 | 4725 | 4985 | 4405 | 4918 | 5292 | 5224 | 5098 | 4962 | 4969 | 4855 | 5066.72 | 5205 | 4929 | 0.947 | -0.079 | 0.11716467 | 0.24816755 | 0 | 0.0029 | 0.0016 | 0.0016 | TRUE | 0.0039 |
| Prps2 | 6125 | 4366 | 4565 | 5286 | 4329 | 5075 | 4757 | 4644 | 4926 | 5262 | 4884 | 5010 | 4913.62 | 4776 | 5052 | 1.058 | 0.081 | 0.14781555 | 0.29256717 | 9.00E-04 | 0.0029 | 0.0021 | 0.0021 | TRUE | 0.0161 |
| Prpsap1 | 9814 | 7268 | 6904 | 7137 | 6721 | 7680 | 7622 | 7731 | 7449 | 7104 | 7582 | 7581 | 7511.64 | 7601 | 7422 | 0.976 | -0.034 | 0.51237871 | 0.67504942 | 7.00E-04 | 0.0027 | 0.0018 | 0.0018 | TRUE | 0.0172 |
| Prpsap2 | 3998 | 2825 | 3182 | 3263 | 2917 | 3417 | 3105 | 3005 | 3433 | 3248 | 3291 | 3373 | 3242.52 | 3181 | 3304 | 1.039 | 0.055 | 0.40270035 | 0.57575233 | 0.0022 | 0.0032 | 0.0028 | 0.0028 | TRUE | 0.0576 |
| Prr11 | 3343 | 2456 | 3006 | 3646 | 3177 | 3604 | 2596 | 2612 | 3243 | 3629 | 3584 | 3558 | 3203.86 | 2817 | 3590 | 1.275 | 0.351 | 3.21E-05 | 0.00049161 | 0.0082 | 0.0032 | 0.0048 | 0.0048 | TRUE | 0.2111 |
| Prr12 | 4663 | 3525 | 3093 | 3075 | 3003 | 3335 | 3622 | 3749 | 3337 | 3061 | 3388 | 3292 | 3408.18 | 3569 | 3247 | 0.909 | -0.137 | 0.04259136 | 0.11979932 | 0.0026 | 0.0031 | 0.003 | 0.003 | TRUE | 0.0388 |
| Prr13 | 1590 | 1396 | 1735 | 1840 | 1857 | 2137 | 1235 | 1485 | 1872 | 1832 | 2095 | 2110 | 1771.3 | 1531 | 2012 | 1.316 | 0.396 | 0.00103536 | 0.00741774 | 0.0242 | 0.0038 | 0.0099 | 0.0099 | TRUE | 0.4576 |
| Prr14 | 1683 | 1087 | 1050 | 821 | 749 | 904 | 1307 | 1156 | 1133 | 817 | 845 | 892 | 1025.14 | 1199 | 851 | 0.709 | -0.495 | 6.73E-09 | 3.55E-07 | 0.0034 | 0.0047 | 0.0043 | 0.0043 | TRUE | 0.0776 |
| Prr14l | 5846 | 4062 | 4303 | 4315 | 3917 | 4330 | 4540 | 4321 | 4643 | 4295 | 4419 | 4274 | 4415.37 | 4501 | 4329 | 0.962 | -0.056 | 0.30485129 | 0.47739853 | 5.00E-04 | 0.003 | 0.002 | 0.002 | TRUE | 0.0148 |
| Prr15 | 23 | 15 | 13 | 22 | 26 | 17 | 18 | 16 | 14 | 22 | 29 | 17 | 19.31 | 16 | 23 | 1.408 | 0.494 | 0.26895029 | 0.43947714 | 0 | 0.1252 | 0.0906 | 0.0906 | TRUE | 0.2243 |
| Prr15l | 183 | 138 | 194 | 203 | 192 | 181 | 142 | 147 | 209 | 202 | 217 | 179 | 182.6 | 166 | 199 | 1.203 | 0.266 | 0.13615457 | 0.27587996 | 0.0219 | 0.0154 | 0.0175 | 0.0175 | TRUE | 0.5369 |
| Prr16 | 19 | 22 | 12 | 14 | 8 | 5 | 15 | 23 | 13 | 14 | 9 | 5 | 13.17 | 17 | 9 | 0.548 | -0.869 | 0.13106352 | NA | 0.074 | 0.1824 | 0.1543 | 0.1543 | TRUE | 0.4229 |
| Prr18 | 33 | 41 | 26 | 38 | 38 | 37 | 26 | 44 | 28 | 38 | 43 | 37 | 35.75 | 33 | 39 | 1.212 | 0.277 | 0.41288988 | 0.58580853 | 0.0163 | 0.0687 | 0.0545 | 0.0545 | TRUE | 0.6026 |
| Prr19 | 87 | 48 | 41 | 91 | 61 | 92 | 68 | 51 | 44 | 91 | 69 | 91 | 68.85 | 54 | 84 | 1.526 | 0.609 | 0.01745416 | 0.06245321 | 0.0204 | 0.0369 | 0.0323 | 0.0323 | TRUE | 0.4365 |
| Prr22 | 20 | 11 | 10 | 8 | 11 | 5 | 16 | 12 | 11 | 8 | 12 | 5 | 10.56 | 13 | 8 | 0.655 | -0.611 | 0.31653611 | NA | 0 | 0.227 | 0.1682 | 0.1682 | TRUE | 0.3925 |
| Prr23a | 120 | 73 | 57 | 77 | 56 | 96 | 93 | 78 | 62 | 77 | 63 | 95 | 77.82 | 78 | 78 | 1.006 | 0.009 | 0.9720965 | 0.98493233 | 0.029 | 0.0329 | 0.0318 | 0.0318 | TRUE | 0.2072 |
| Prr24 | 266 | 220 | 200 | 272 | 186 | 276 | 207 | 234 | 216 | 271 | 210 | 272 | 234.91 | 219 | 251 | 1.151 | 0.203 | 0.16623631 | 0.31730977 | 0.0082 | 0.0125 | 0.0113 | 0.0113 | TRUE | 0.2208 |
| Prr3 | 984 | 790 | 647 | 629 | 595 | 729 | 764 | 840 | 698 | 626 | 671 | 720 | 719.94 | 767 | 672 | 0.876 | -0.191 | 0.05209435 | 0.13886321 | 0.0053 | 0.0057 | 0.0056 | 0.0056 | TRUE | 0.0786 |
| Prr5 | 1483 | 1114 | 992 | 1132 | 1054 | 1183 | 1152 | 1185 | 1070 | 1127 | 1189 | 1168 | 1148.45 | 1136 | 1161 | 1.022 | 0.031 | 0.67735797 | 0.80217727 | 7.00E-04 | 0.0045 | 0.0032 | 0.0032 | TRUE | 0.0293 |
| Prr5l | 164 | 85 | 78 | 102 | 90 | 80 | 127 | 90 | 84 | 102 | 102 | 79 | 97.33 | 100 | 94 | 0.926 | -0.11 | 0.62673696 | 0.76482743 | 0.0272 | 0.0268 | 0.0269 | 0.0269 | TRUE | 0.5763 |
| Prr7 | 325 | 310 | 269 | 284 | 266 | 287 | 252 | 330 | 290 | 283 | 300 | 283 | 289.75 | 291 | 289 | 0.996 | -0.006 | 0.96361587 | 0.97993323 | 0.0066 | 0.0106 | 0.0094 | 0.0094 | TRUE | 0.1629 |
| Prr9 | 0 | 0 | 0 | 0 | 0 | 0 | 0 | 0 | 0 | 0 | 0 | 0 | 0 | 0 | 0 | NA | NA | NA | NA | NA | NA | NA | NA | NA | NA |
| Prrc1 | 9214 | 7461 | 8483 | 7621 | 6936 | 8086 | 7156 | 7936 | 9153 | 7586 | 7825 | 7982 | 7939.66 | 8082 | 7798 | 0.965 | -0.051 | 0.52608412 | 0.6870724 | 0.0069 | 0.0027 | 0.0046 | 0.0046 | TRUE | 0.1641 |
| Prrc2a | 29576 | 22535 | 22831 | 21191 | 19987 | 22755 | 22970 | 23970 | 24634 | 21093 | 22547 | 22463 | 22946.45 | 23858 | 22034 | 0.924 | -0.115 | 0.02981953 | 0.09227334 | 0.0012 | 0.0025 | 0.002 | 0.002 | TRUE | 0.0171 |
| Prrc2b | 20654 | 15681 | 17390 | 19982 | 19011 | 20975 | 16041 | 16680 | 18764 | 19890 | 21446 | 20706 | 18921.07 | 17162 | 20681 | 1.205 | 0.269 | 5.13E-05 | 0.00071516 | 0.0041 | 0.0026 | 0.0031 | 0.0031 | TRUE | 0.0815 |
| Prrc2c | 34938 | 24291 | 22266 | 25322 | 22717 | 25682 | 27135 | 25838 | 24025 | 25205 | 25627 | 25352 | 25530.41 | 25666 | 25395 | 0.989 | -0.015 | 0.78304464 | 0.8723772 | 0.0015 | 0.0025 | 0.0022 | 0.0022 | TRUE | 0.0383 |
| Prrg1 | 197 | 111 | 122 | 150 | 117 | 123 | 153 | 118 | 132 | 149 | 132 | 121 | 134.24 | 134 | 134 | 0.996 | -0.006 | 0.97346121 | 0.98545797 | 0.0076 | 0.0201 | 0.0164 | 0.0164 | TRUE | 0.1691 |
| Prrg2 | 143 | 86 | 103 | 113 | 108 | 113 | 111 | 91 | 111 | 112 | 122 | 112 | 109.92 | 104 | 115 | 1.099 | 0.137 | 0.47096969 | 0.63982223 | 0 | 0.024 | 0.0168 | 0.0168 | TRUE | 0.1125 |
| Prrg3 | 711 | 522 | 627 | 704 | 559 | 624 | 552 | 555 | 677 | 701 | 631 | 616 | 621.89 | 595 | 649 | 1.094 | 0.129 | 0.22869967 | 0.39402174 | 0.0071 | 0.0062 | 0.0067 | 0.0067 | TRUE | 0.1669 |
| Prrg4 | 925 | 715 | 735 | 883 | 670 | 825 | 718 | 761 | 793 | 879 | 756 | 814 | 786.86 | 757 | 816 | 1.08 | 0.111 | 0.21708862 | 0.37997154 | 0.0028 | 0.0054 | 0.0046 | 0.0046 | TRUE | 0.0533 |
| Prrt1 | 1 | 2 | 1 | 2 | 0 | 0 | 1 | 2 | 1 | 2 | 0 | 0 | 1 | 1 | 1 | 0.515 | -0.958 | 0.63579447 | NA | 0 | 2.3829 | 1.8115 | 1.8115 | TRUE | 0.9736 |
| Prrt2 | 52 | 46 | 47 | 51 | 47 | 58 | 40 | 49 | 51 | 51 | 53 | 57 | 50.18 | 47 | 54 | 1.157 | 0.21 | 0.44126881 | 0.61245438 | 0 | 0.0497 | 0.0336 | 0.0336 | TRUE | 0.1311 |
| Prrt3 | 156 | 91 | 44 | 61 | 39 | 44 | 121 | 97 | 47 | 61 | 44 | 43 | 68.93 | 88 | 49 | 0.555 | -0.848 | 0.00852248 | 0.03674523 | 0.1131 | 0.0368 | 0.059 | 0.059 | TRUE | 0.3125 |
| Prrt4 | 151 | 104 | 109 | 126 | 106 | 108 | 117 | 111 | 118 | 125 | 120 | 107 | 116.19 | 115 | 117 | 1.017 | 0.024 | 0.89609219 | 0.94272381 | 0 | 0.0228 | 0.0153 | 0.0153 | TRUE | 0.0649 |
| Prrx1 | 1 | 2 | 1 | 2 | 0 | 0 | 1 | 2 | 1 | 2 | 0 | 0 | 1 | 1 | 1 | 0.515 | -0.958 | 0.63579447 | NA | 0 | 2.3829 | 1.8115 | 1.8115 | TRUE | 0.9736 |
| Prrx2 | 140 | 100 | 116 | 162 | 158 | 170 | 109 | 106 | 125 | 161 | 178 | 168 | 141.26 | 113 | 169 | 1.493 | 0.578 | 0.00061771 | 0.00507427 | 0 | 0.0192 | 0.0132 | 0.0132 | TRUE | 0.0772 |
| Prrxl1 | 2 | 0 | 0 | 1 | 1 | 0 | 2 | 0 | 0 | 1 | 1 | 0 | 0.61 | 1 | 1 | 1.262 | 0.336 | 0.89669251 | NA | 0 | 3.87 | 3.2065 | 3.2065 | TRUE | 0.9721 |
| Prss1 | 0 | 0 | 0 | 0 | 0 | 0 | 0 | 0 | 0 | 0 | 0 | 0 | 0 | 0 | 0 | NA | NA | NA | NA | NA | NA | NA | NA | NA | NA |
| Prss12 | 26 | 12 | 12 | 9 | 12 | 14 | 20 | 13 | 13 | 9 | 14 | 14 | 13.7 | 15 | 12 | 0.781 | -0.357 | 0.49895003 | NA | 0 | 0.1754 | 0.1264 | 0.1264 | TRUE | 0.4444 |
| Prss16 | 276 | 244 | 225 | 222 | 232 | 255 | 214 | 260 | 243 | 221 | 262 | 252 | 241.85 | 239 | 245 | 1.027 | 0.038 | 0.78333239 | 0.87248506 | 0.0048 | 0.0122 | 0.01 | 0.01 | TRUE | 0.0976 |
| Prss2 | 0 | 0 | 0 | 0 | 0 | 0 | 0 | 0 | 0 | 0 | 0 | 0 | 0 | 0 | 0 | NA | NA | NA | NA | NA | NA | NA | NA | NA | NA |
| Prss21 | 0 | 1 | 0 | 2 | 0 | 2 | 0 | 1 | 0 | 2 | 0 | 2 | 0.84 | 0 | 1 | 3.874 | 1.954 | 0.39514891 | NA | 0 | 2.8302 | 2.417 | 2.417 | TRUE | 0.5217 |
| Prss22 | 3 | 3 | 1 | 8 | 5 | 4 | 2 | 3 | 1 | 8 | 6 | 4 | 4.03 | 2 | 6 | 2.648 | 1.405 | 0.15688413 | NA | 0 | 0.5912 | 0.4037 | 0.4037 | TRUE | 0.1773 |
| Prss23 | 54 | 48 | 36 | 35 | 26 | 26 | 42 | 51 | 39 | 35 | 29 | 26 | 36.95 | 44 | 30 | 0.682 | -0.551 | 0.08862015 | 0.20263123 | 0 | 0.0666 | 0.0472 | 0.0472 | TRUE | 0.1463 |
| Prss27 | 3 | 3 | 9 | 10 | 13 | 10 | 2 | 3 | 10 | 10 | 15 | 10 | 8.29 | 5 | 12 | 2.319 | 1.214 | 0.09788799 | NA | 0.1028 | 0.2884 | 0.2448 | 0.2448 | TRUE | 1.2223 |
| Prss28 | 0 | 0 | 0 | 0 | 0 | 0 | 0 | 0 | 0 | 0 | 0 | 0 | 0 | 0 | 0 | NA | NA | NA | NA | NA | NA | NA | NA | NA | NA |
| Prss29 | 0 | 0 | 0 | 0 | 0 | 0 | 0 | 0 | 0 | 0 | 0 | 0 | 0 | 0 | 0 | NA | NA | NA | NA | NA | NA | NA | NA | NA | NA |
| Prss3 | 0 | 0 | 0 | 0 | 0 | 0 | 0 | 0 | 0 | 0 | 0 | 0 | 0 | 0 | 0 | NA | NA | NA | NA | NA | NA | NA | NA | NA | NA |
| Prss30 | 0 | 0 | 0 | 0 | 0 | 0 | 0 | 0 | 0 | 0 | 0 | 0 | 0 | 0 | 0 | NA | NA | NA | NA | NA | NA | NA | NA | NA | NA |
| Prss32 | 10 | 11 | 5 | 9 | 7 | 9 | 8 | 12 | 5 | 9 | 8 | 9 | 8.43 | 8 | 9 | 1.039 | 0.055 | 0.9339217 | NA | 0 | 0.2835 | 0.1933 | 0.1933 | TRUE | 0.349 |
| Prss33 | 23 | 12 | 14 | 29 | 19 | 30 | 18 | 13 | 15 | 29 | 21 | 30 | 20.94 | 15 | 27 | 1.739 | 0.798 | 0.06113154 | 0.15538913 | 0 | 0.1156 | 0.0801 | 0.0801 | TRUE | 0.1682 |
| Prss34 | 0 | 0 | 0 | 2 | 0 | 0 | 0 | 0 | 0 | 2 | 0 | 0 | 0.33 | 0 | 1 | 3.77 | 1.915 | 0.62344813 | NA | 0 | 7.1455 | 9.1834 | 9.1834 | TRUE | 0.9739 |
| Prss35 | 437 | 316 | 318 | 324 | 259 | 274 | 339 | 336 | 343 | 323 | 292 | 270 | 317.3 | 339 | 295 | 0.869 | -0.202 | 0.09035866 | 0.20526006 | 0 | 0.0099 | 0.0071 | 0.0071 | TRUE | 0.0767 |
| Prss36 | 733 | 553 | 571 | 727 | 579 | 743 | 569 | 588 | 616 | 724 | 653 | 733 | 647.32 | 591 | 703 | 1.192 | 0.253 | 0.00567802 | 0.02707004 | 0.0012 | 0.0061 | 0.0045 | 0.0045 | TRUE | 0.045 |
| Prss37 | 0 | 0 | 0 | 0 | 0 | 0 | 0 | 0 | 0 | 0 | 0 | 0 | 0 | 0 | 0 | NA | NA | NA | NA | NA | NA | NA | NA | NA | NA |
| Prss38 | 227 | 163 | 213 | 236 | 239 | 219 | 176 | 173 | 230 | 235 | 270 | 216 | 216.7 | 193 | 240 | 1.245 | 0.317 | 0.04633648 | 0.12749512 | 0.0139 | 0.0134 | 0.0135 | 0.0135 | TRUE | 0.2881 |
| Prss39 | 5 | 2 | 0 | 3 | 1 | 2 | 4 | 2 | 0 | 3 | 1 | 2 | 2.02 | 2 | 2 | 0.986 | -0.02 | 0.988413 | NA | 0 | 1.1777 | 0.917 | 0.917 | TRUE | 0.2314 |
| Prss40 | 0 | 0 | 1 | 0 | 0 | 1 | 0 | 0 | 1 | 0 | 0 | 1 | 0.34 | 0 | 0 | 1.075 | 0.105 | 0.97570524 | NA | 0 | 6.885 | 6.5379 | 6.5379 | TRUE | 0.5365 |
| Prss41 | 128 | 119 | 131 | 146 | 155 | 156 | 99 | 127 | 141 | 145 | 175 | 154 | 140.25 | 122 | 158 | 1.297 | 0.376 | 0.04291591 | 0.12053392 | 0.0131 | 0.0193 | 0.0175 | 0.0175 | TRUE | 0.3083 |
| Prss42 | 129 | 142 | 182 | 96 | 78 | 109 | 100 | 151 | 196 | 96 | 88 | 108 | 123.13 | 149 | 97 | 0.655 | -0.61 | 0.01008918 | 0.04160552 | 0.0554 | 0.0217 | 0.032 | 0.032 | TRUE | 0.1547 |
| Prss43 | 3 | 2 | 1 | 1 | 0 | 0 | 2 | 2 | 1 | 1 | 0 | 0 | 1.09 | 2 | 0 | 0.199 | -2.326 | 0.23818909 | NA | 0 | 2.1795 | 1.5416 | 1.5416 | TRUE | 0.3906 |

| | | | | | | | | | | | | | | | | | | | | | | | | | |
|---|---|---|---|---|---|---|---|---|---|---|---|---|---|---|---|---|---|---|---|---|---|---|---|---|---|
| Prss44 | 13 | 19 | 18 | 32 | 17 | 24 | 10 | 20 | 19 | 32 | 19 | 24 | 20.74 | 16 | 25 | 1.53 | 0.614 | 0.17647196 | 0.33027547 | 0.0495 | 0.1167 | 0.0984 | 0.0984 | TRUE | 0.6647 |
| Prss45 | 9 | 11 | 15 | 10 | 8 | 8 | 7 | 12 | 16 | 10 | 9 | 8 | 10.29 | 12 | 9 | 0.782 | -0.356 | 0.56144354 | NA | 0 | 0.2327 | 0.1704 | 0.1704 | TRUE | 0.183 |
| Prss46 | 0 | 1 | 0 | 1 | 0 | 1 | 0 | 1 | 0 | 1 | 0 | 1 | 0.51 | 0 | 1 | 1.941 | 0.957 | 0.72720315 | NA | 0 | 4.6706 | 3.6514 | 3.6514 | TRUE | 0.5217 |
| Prss48 | 11 | 16 | 14 | 16 | 15 | 12 | 9 | 17 | 15 | 16 | 17 | 12 | 14.23 | 14 | 15 | 1.112 | 0.153 | 0.76903833 | NA | 0 | 0.169 | 0.1258 | 0.1258 | TRUE | 0.5794 |
| Prss50 | 409 | 395 | 444 | 538 | 455 | 523 | 318 | 420 | 479 | 536 | 513 | 516 | 463.66 | 406 | 522 | 1.291 | 0.369 | 0.00662117 | 0.03042866 | 0.0193 | 0.0075 | 0.0111 | 0.0111 | TRUE | 0.4394 |
| Prss51 | 0 | 0 | 0 | 0 | 0 | 0 | 0 | 0 | 0 | 0 | 0 | 0 | 0 | 0 | 0 | NA | NA | NA | NA | NA | NA | NA | NA | NA | NA |
| Prss52 | 0 | 0 | 0 | 0 | 0 | 0 | 0 | 0 | 0 | 0 | 0 | 0 | 0 | 0 | 0 | NA | NA | NA | NA | NA | NA | NA | NA | NA | NA |
| Prss53 | 17 | 14 | 19 | 22 | 16 | 11 | 13 | 15 | 21 | 22 | 18 | 11 | 16.57 | 16 | 17 | 1.052 | 0.073 | 0.88253623 | NA | 0 | 0.1455 | 0.1129 | 0.1129 | TRUE | 0.4595 |
| Prss54 | 1 | 0 | 0 | 0 | 0 | 1 | 1 | 0 | 0 | 0 | 0 | 1 | 0.29 | 0 | 0 | 1.075 | 0.105 | 0.97702352 | NA | 0 | 8.0647 | 7.5458 | 7.5458 | TRUE | 0.3835 |
| Prss55 | 0 | 0 | 0 | 0 | 0 | 0 | 0 | 0 | 0 | 0 | 0 | 0 | 0 | 0 | 0 | NA | NA | NA | NA | NA | NA | NA | NA | NA | NA |
| Prss56 | 24 | 27 | 18 | 15 | 29 | 24 | 19 | 29 | 19 | 15 | 33 | 24 | 23.02 | 22 | 24 | 1.068 | 0.095 | 0.82627809 | 0.90029631 | 0.0574 | 0.1054 | 0.0922 | 0.0922 | TRUE | 0.178 |
| Prss57 | 0 | 1 | 0 | 0 | 1 | 0 | 0 | 1 | 0 | 0 | 1 | 0 | 0.37 | 0 | 0 | 1.075 | 0.105 | 0.97520514 | NA | 0 | 6.4904 | 6.1969 | 6.1969 | TRUE | 0.5216 |
| Prss58 | 0 | 0 | 0 | 0 | 0 | 0 | 0 | 0 | 0 | 0 | 0 | 0 | 0 | 0 | 0 | NA | NA | NA | NA | NA | NA | NA | NA | NA | NA |
| Prss8 | 156 | 133 | 171 | 210 | 169 | 213 | 121 | 141 | 185 | 209 | 191 | 210 | 176.18 | 149 | 203 | 1.373 | 0.457 | 0.00922753 | 0.03903201 | 0.0178 | 0.0159 | 0.0164 | 0.0164 | TRUE | 0.447 |
| Prtg | 1563 | 928 | 895 | 1173 | 916 | 1063 | 1214 | 987 | 966 | 1168 | 1033 | 1049 | 1069.5 | 1056 | 1083 | 1.025 | 0.035 | 0.7262282 | 0.83594908 | 0.0096 | 0.0046 | 0.0063 | 0.0063 | TRUE | 0.2117 |
| Prtn3 | 13 | 10 | 28 | 44 | 48 | 37 | 10 | 11 | 30 | 44 | 54 | 37 | 30.9 | 17 | 45 | 2.681 | 1.423 | 0.0010888 | 0.00769531 | 0.1523 | 0.0791 | 0.0962 | 0.0962 | TRUE | 1.5555 |
| Prune | 3928 | 2845 | 3070 | 3004 | 2727 | 3298 | 3051 | 3026 | 3313 | 2990 | 3076 | 3256 | 3118.59 | 3130 | 3107 | 0.993 | -0.01 | 0.87836704 | 0.93196497 | 0.0019 | 0.0032 | 0.0027 | 0.0027 | TRUE | 0.0313 |
| Prune2 | 1964 | 1402 | 1352 | 1478 | 1550 | 1675 | 1525 | 1491 | 1459 | 1471 | 1749 | 1653 | 1558.11 | 1492 | 1624 | 1.088 | 0.122 | 0.11979872 | 0.25153801 | 0.0034 | 0.0039 | 0.0038 | 0.0038 | TRUE | 0.0824 |
| Prx | 933 | 660 | 647 | 998 | 838 | 917 | 725 | 702 | 698 | 993 | 945 | 905 | 828.12 | 708 | 948 | 1.338 | 0.42 | 2.55E-07 | 9.05E-06 | 0 | 0.0053 | 0.0035 | 0.0035 | TRUE | 0.0214 |
| Psap | 18845 | 13470 | 13838 | 14597 | 12911 | 14747 | 14636 | 14328 | 14931 | 14530 | 14565 | 14558 | 14591.24 | 14632 | 14551 | 0.994 | -0.008 | 0.85943727 | 0.92152565 | 1.00E-04 | 0.0026 | 0.0014 | 0.0014 | TRUE | 0.004 |
| Psapl1 | 0 | 1 | 1 | 0 | 0 | 0 | 0 | 1 | 1 | 0 | 0 | 0 | 0.36 | 1 | 0 | 0.271 | -1.881 | 0.5755355 | NA | 0 | 6.6391 | 6.4428 | 6.4428 | TRUE | 0.3513 |
| Psat1 | 13004 | 9195 | 9139 | 9811 | 8302 | 9842 | 10100 | 9781 | 9861 | 9766 | 9366 | 9716 | 9764.69 | 9914 | 9616 | 0.97 | -0.044 | 0.3585307 | 0.53302261 | 2.00E-04 | 0.0027 | 0.0016 | 0.0016 | TRUE | 0.0063 |
| Psca | 3 | 0 | 2 | 5 | 1 | 4 | 2 | 0 | 2 | 5 | 1 | 4 | 2.42 | 1 | 3 | 2.203 | 1.14 | 0.38498804 | NA | 0 | 0.9803 | 0.7724 | 0.7724 | TRUE | 0.4674 |
| Psd | 122 | 95 | 113 | 146 | 137 | 134 | 95 | 101 | 122 | 145 | 155 | 132 | 124.98 | 106 | 144 | 1.364 | 0.448 | 0.01506036 | 0.05599837 | 0 | 0.0214 | 0.0163 | 0.0163 | TRUE | 0.1676 |
| Psd2 | 154 | 116 | 136 | 173 | 162 | 160 | 120 | 123 | 147 | 172 | 183 | 158 | 150.44 | 130 | 171 | 1.319 | 0.399 | 0.01745614 | 0.06245321 | 0 | 0.0182 | 0.0136 | 0.0136 | TRUE | 0.1268 |
| Psd3 | 392 | 245 | 234 | 269 | 207 | 241 | 304 | 261 | 252 | 268 | 234 | 238 | 259.45 | 272 | 247 | 0.902 | -0.149 | 0.27269708 | 0.44367209 | 0.0048 | 0.0116 | 0.0095 | 0.0095 | TRUE | 0.1262 |
| Psd4 | 475 | 350 | 334 | 252 | 244 | 251 | 369 | 372 | 360 | 251 | 275 | 248 | 312.58 | 367 | 258 | 0.702 | -0.511 | 1.27E-05 | 0.00022872 | 0 | 0.01 | 0.0066 | 0.0066 | TRUE | 0.0368 |
| Psen1 | 3026 | 2294 | 2197 | 2377 | 2153 | 2488 | 2350 | 2440 | 2371 | 2366 | 2429 | 2456 | 2401.95 | 2387 | 2417 | 1.013 | 0.018 | 0.7519643 | 0.852799 | 0 | 0.0034 | 0.002 | 0.002 | TRUE | 0.0046 |
| Psen2 | 309 | 236 | 220 | 284 | 287 | 352 | 240 | 251 | 237 | 283 | 324 | 347 | 280.39 | 243 | 318 | 1.31 | 0.389 | 0.00260533 | 0.01481224 | 0.003 | 0.0109 | 0.0084 | 0.0084 | TRUE | 0.1092 |
| Psenen | 729 | 525 | 573 | 601 | 557 | 647 | 566 | 558 | 618 | 598 | 628 | 639 | 601.36 | 581 | 622 | 1.071 | 0.099 | 0.28235813 | 0.4540667 | 0 | 0.0064 | 0.0044 | 0.0044 | TRUE | 0.0357 |
| Psg-ps1 | 0 | 0 | 0 | 0 | 0 | 0 | 0 | 0 | 0 | 0 | 0 | 0 | 0 | 0 | 0 | NA | NA | NA | NA | NA | NA | NA | NA | NA | NA |
| Psg16 | 9 | 7 | 14 | 8 | 11 | 10 | 7 | 7 | 15 | 8 | 12 | 10 | 9.96 | 10 | 10 | 1.032 | 0.045 | 0.94195051 | NA | 0 | 0.2403 | 0.1801 | 0.1801 | TRUE | 0.6987 |
| Psg17 | 2 | 0 | 1 | 0 | 0 | 0 | 2 | 0 | 1 | 0 | 0 | 0 | 0.44 | 1 | 0 | 0.214 | -2.223 | 0.47051571 | NA | 0 | 5.4047 | 5.2946 | 5.2946 | TRUE | 0.2788 |
| Psg18 | 0 | 0 | 0 | 0 | 0 | 0 | 0 | 0 | 0 | 0 | 0 | 0 | 0 | 0 | 0 | NA | NA | NA | NA | NA | NA | NA | NA | NA | NA |
| Psg19 | 0 | 0 | 0 | 0 | 0 | 0 | 0 | 0 | 0 | 0 | 0 | 0 | 0 | 0 | 0 | NA | NA | NA | NA | NA | NA | NA | NA | NA | NA |
| Psg20 | 0 | 0 | 0 | 0 | 0 | 0 | 0 | 0 | 0 | 0 | 0 | 0 | 0 | 0 | 0 | NA | NA | NA | NA | NA | NA | NA | NA | NA | NA |
| Psg21 | 0 | 0 | 0 | 0 | 0 | 0 | 0 | 0 | 0 | 0 | 0 | 0 | 0 | 0 | 0 | NA | NA | NA | NA | NA | NA | NA | NA | NA | NA |
| Psg22 | 0 | 0 | 0 | 0 | 0 | 0 | 0 | 0 | 0 | 0 | 0 | 0 | 0 | 0 | 0 | NA | NA | NA | NA | NA | NA | NA | NA | NA | NA |
| Psg23 | 0 | 0 | 0 | 0 | 0 | 0 | 0 | 0 | 0 | 0 | 0 | 0 | 0 | 0 | 0 | NA | NA | NA | NA | NA | NA | NA | NA | NA | NA |
| Psg25 | 0 | 0 | 0 | 0 | 0 | 0 | 0 | 0 | 0 | 0 | 0 | 0 | 0 | 0 | 0 | NA | NA | NA | NA | NA | NA | NA | NA | NA | NA |
| Psg26 | 0 | 0 | 0 | 0 | 0 | 0 | 0 | 0 | 0 | 0 | 0 | 0 | 0 | 0 | 0 | NA | NA | NA | NA | NA | NA | NA | NA | NA | NA |
| Psg27 | 0 | 0 | 0 | 0 | 0 | 0 | 0 | 0 | 0 | 0 | 0 | 0 | 0 | 0 | 0 | NA | NA | NA | NA | NA | NA | NA | NA | NA | NA |
| Psg28 | 0 | 0 | 0 | 0 | 0 | 0 | 0 | 0 | 0 | 0 | 0 | 0 | 0 | 0 | 0 | NA | NA | NA | NA | NA | NA | NA | NA | NA | NA |
| Psg29 | 0 | 0 | 0 | 0 | 0 | 0 | 0 | 0 | 0 | 0 | 0 | 0 | 0 | 0 | 0 | NA | NA | NA | NA | NA | NA | NA | NA | NA | NA |
| Psip1 | 15096 | 10955 | 10522 | 10804 | 9487 | 11091 | 11724 | 11653 | 11353 | 10754 | 10702 | 10949 | 11189.23 | 11577 | 10802 | 0.933 | -0.1 | 0.02975158 | 0.09208301 | 1.00E-04 | 0.0026 | 0.0014 | 0.0014 | TRUE | 0.0035 |
| Pskh1 | 1793 | 1330 | 1300 | 1456 | 1337 | 1432 | 1393 | 1415 | 1403 | 1449 | 1508 | 1414 | 1430.19 | 1404 | 1457 | 1.038 | 0.054 | 0.41882698 | 0.59147258 | 0 | 0.0041 | 0.0025 | 0.0025 | TRUE | 0.0112 |
| Psma1 | 12138 | 8036 | 7690 | 8368 | 7207 | 8191 | 9427 | 8548 | 8297 | 8329 | 8130 | 8086 | 8469.64 | 8757 | 8182 | 0.934 | -0.099 | 0.10426849 | 0.2282574 | 0.0022 | 0.0027 | 0.0025 | 0.0025 | TRUE | 0.0551 |
| Psma2 | 11531 | 7680 | 7770 | 8168 | 6820 | 7990 | 8956 | 8169 | 8384 | 8130 | 7694 | 7887 | 8203.33 | 8503 | 7904 | 0.929 | -0.106 | 0.05979918 | 0.15287241 | 0.0013 | 0.0027 | 0.0021 | 0.0021 | TRUE | 0.0268 |
| Psma3 | 9778 | 6699 | 6718 | 7506 | 6225 | 7045 | 7594 | 7126 | 7249 | 7471 | 7022 | 6955 | 7236.15 | 7323 | 7149 | 0.976 | -0.035 | 0.52935907 | 0.68976453 | 0.0011 | 0.0028 | 0.0021 | 0.0021 | TRUE | 0.019 |
| Psma4 | 12947 | 8995 | 8265 | 9124 | 7831 | 9175 | 10055 | 9568 | 8918 | 9082 | 8834 | 9057 | 9252.41 | 9514 | 8991 | 0.945 | -0.082 | 0.15879493 | 0.30804233 | 0.0018 | 0.0027 | 0.0023 | 0.0023 | TRUE | 0.0366 |
| Psma5 | 7844 | 5420 | 5112 | 5366 | 4629 | 5492 | 6092 | 5765 | 5516 | 5341 | 5222 | 5421 | 5559.64 | 5791 | 5328 | 0.92 | -0.121 | 0.03498249 | 0.10384315 | 0.0012 | 0.0029 | 0.0022 | 0.0022 | TRUE | 0.0256 |
| Psma6 | 11461 | 7769 | 7166 | 8299 | 7139 | 8242 | 8901 | 8264 | 7732 | 8261 | 8054 | 8136 | 8224.6 | 8299 | 8150 | 0.982 | -0.027 | 0.66634962 | 0.7948039 | 0.0021 | 0.0027 | 0.0026 | 0.0026 | TRUE | 0.0496 |
| Psma7 | 8955 | 6177 | 6104 | 6175 | 5324 | 6450 | 6955 | 6570 | 6586 | 6147 | 6006 | 6367 | 6438.55 | 6704 | 6173 | 0.921 | -0.119 | 0.02646106 | 0.08469995 | 7.00E-04 | 0.0028 | 0.0019 | 0.0019 | TRUE | 0.0133 |
| Psma8 | 1214 | 899 | 806 | 935 | 699 | 854 | 943 | 956 | 870 | 931 | 789 | 843 | 888.51 | 923 | 854 | 0.926 | -0.112 | 0.20976048 | 0.37160468 | 0.0035 | 0.0051 | 0.0046 | 0.0046 | TRUE | 0.0733 |
| Psmb1 | 11593 | 8067 | 8618 | 8860 | 7859 | 8966 | 9004 | 8581 | 9299 | 8819 | 8866 | 8851 | 8903.19 | 8961 | 8845 | 0.987 | -0.019 | 0.71471993 | 0.82854995 | 6.00E-04 | 0.0027 | 0.0018 | 0.0018 | TRUE | 0.0167 |
| Psmb10 | 1350 | 898 | 873 | 1095 | 969 | 1157 | 1048 | 955 | 942 | 1090 | 1093 | 1142 | 1045.15 | 982 | 1108 | 1.127 | 0.173 | 0.02731607 | 0.08641623 | 0.0013 | 0.0047 | 0.0035 | 0.0035 | TRUE | 0.0441 |
| Psmb11 | 0 | 1 | 0 | 0 | 0 | 0 | 0 | 1 | 0 | 0 | 0 | 0 | 0.18 | 0 | 0 | 0.552 | -0.857 | 0.8336548 | NA | 0 | 13.3714 | 10 | 10 | TRUE | 0.5216 |
| Psmb2 | 7471 | 5232 | 4839 | 4878 | 4488 | 5197 | 5802 | 5565 | 5221 | 4856 | 5063 | 5130 | 5272.93 | 5529 | 5016 | 0.907 | -0.141 | 0.01776397 | 0.06324029 | 0.0015 | 0.0029 | 0.0024 | 0.0024 | TRUE | 0.0289 |
| Psmb3 | 12446 | 8629 | 8357 | 9137 | 7865 | 9180 | 9666 | 9179 | 9017 | 9095 | 8873 | 9062 | 9148.59 | 9287 | 9010 | 0.97 | -0.044 | 0.38721428 | 0.56096226 | 6.00E-04 | 0.0027 | 0.0018 | 0.0018 | TRUE | 0.0157 |
| Psmb4 | 12906 | 9356 | 8782 | 9587 | 8339 | 9403 | 10024 | 9952 | 9476 | 9543 | 9407 | 9282 | 9613.91 | 9817 | 9411 | 0.958 | -0.061 | 0.21537422 | 0.37789635 | 4.00E-04 | 0.0027 | 0.0017 | 0.0017 | TRUE | 0.0112 |
| Psmb5 | 4233 | 2964 | 2753 | 2975 | 2661 | 3035 | 3288 | 3153 | 2970 | 2961 | 3002 | 2996 | 3061.67 | 3137 | 2986 | 0.951 | -0.072 | 0.23556182 | 0.40203197 | 0.001 | 0.0032 | 0.0023 | 0.0023 | TRUE | 0.0261 |
| Psmb6 | 19573 | 13453 | 13106 | 14467 | 12834 | 14968 | 15202 | 14310 | 14141 | 14400 | 14478 | 14776 | 14551.13 | 14551 | 14551 | 1 | 0 | 0.99729516 | 0.99848274 | 6.00E-04 | 0.0026 | 0.0018 | 0.0018 | TRUE | 0.0189 |
| Psmb7 | 14352 | 9721 | 10006 | 10807 | 9376 | 10921 | 11147 | 10340 | 10796 | 10757 | 10577 | 10781 | 10733.03 | 10761 | 10705 | 0.995 | -0.008 | 0.87827705 | 0.93196497 | 5.00E-04 | 0.0026 | 0.0017 | 0.0017 | TRUE | 0.0143 |
| Psmb8 | 8 | 3 | 2 | 4 | 6 | 4 | 6 | 3 | 2 | 4 | 7 | 4 | 4.38 | 4 | 5 | 1.232 | 0.301 | 0.74303713 | NA | 0 | 0.5439 | 0.3805 | 0.3805 | TRUE | 0.611 |
| Psmb9 | 182 | 124 | 109 | 184 | 161 | 183 | 141 | 132 | 118 | 183 | 182 | 181 | 156.05 | 130 | 182 | 1.391 | 0.476 | 0.00301276 | 0.01659952 | 0 | 0.0176 | 0.012 | 0.012 | TRUE | 0.0719 |
| Psmc1 | 11791 | 8036 | 7257 | 7449 | 6433 | 7661 | 9158 | 8548 | 7830 | 7415 | 7257 | 7563 | 7961.67 | 8512 | 7412 | 0.87 | -0.2 | 0.00204231 | 0.01227055 | 0.0031 | 0.0027 | 0.0029 | 0.0029 | TRUE | 0.0598 |
| Psmc2 | 9119 | 6462 | 5775 | 6136 | 5463 | 6489 | 7082 | 6874 | 6231 | 6108 | 6163 | 6406 | 6477.22 | 6729 | 6226 | 0.925 | -0.113 | 0.070125 | 0.17162869 | 0.0023 | 0.0028 | 0.0026 | 0.0026 | TRUE | 0.0509 |
| Psmc3 | 12909 | 8999 | 7266 | 7238 | 6474 | 7547 | 10026 | 9572 | 7840 | 7205 | 7303 | 7450 | 8232.67 | 9146 | 7319 | 0.8 | -0.322 | 8.79E-05 | 0.00108618 | 0.0079 | 0.0027 | 0.0047 | 0.0047 | TRUE | 0.1902 |
| Psmc3ip | 1304 | 948 | 999 | 1015 | 848 | 1014 | 1013 | 1008 | 1078 | 1010 | 957 | 1001 | 1011.16 | 1033 | 989 | 0.959 | -0.061 | 0.41940857 | 0.59170927 | 0 | 0.0048 | 0.0031 | 0.0031 | TRUE | 0.0168 |

| | | | | | | | | | | | | | | | | | | | | | | | | | |
|---|---|---|---|---|---|---|---|---|---|---|---|---|---|---|---|---|---|---|---|---|---|---|---|---|---|
| Psmc4 | 10395 | 7024 | 5554 | 5638 | 5031 | 5882 | 8073 | 7471 | 5993 | 5612 | 5675 | 5806 | 6438.57 | 7179 | 5698 | 0.793 | -0.334 | 0.00024053 | 0.00245562 | 0.011 | 0.0028 | 0.0058 | 0.0058 | TRUE | 0.2546 |
| Psmc5 | 14947 | 10455 | 9135 | 9112 | 8090 | 9510 | 11609 | 11121 | 9857 | 9070 | 9126 | 9388 | 10028.4 | 10862 | 9195 | 0.846 | -0.241 | 0.00027332 | 0.00270534 | 0.0035 | 0.0027 | 0.0031 | 0.0031 | TRUE | 0.08 |
| Psmc6 | 9660 | 6869 | 7096 | 7542 | 6415 | 7415 | 7503 | 7306 | 7657 | 7507 | 7237 | 7320 | 7421.56 | 7489 | 7355 | 0.982 | -0.026 | 0.60031095 | 0.74584879 | 2.00E-04 | 0.0027 | 0.0016 | 0.0016 | TRUE | 0.0054 |
| Psmd1 | 25677 | 17771 | 17413 | 18078 | 15815 | 18178 | 19942 | 18903 | 18788 | 17995 | 17841 | 17944 | 18568.96 | 19211 | 17927 | 0.933 | -0.1 | 0.03576425 | 0.10554997 | 5.00E-04 | 0.0026 | 0.0016 | 0.0016 | TRUE | 0.0136 |
| Psmd10 | 1979 | 1411 | 1433 | 1522 | 1312 | 1599 | 1537 | 1501 | 1546 | 1515 | 1480 | 1578 | 1526.27 | 1528 | 1524 | 0.998 | -0.003 | 0.96295508 | 0.97947032 | 0 | 0.004 | 0.0025 | 0.0025 | TRUE | 0.0116 |
| Psmd11 | 15497 | 11306 | 11241 | 12219 | 10647 | 12598 | 12036 | 12026 | 12129 | 12163 | 12011 | 12436 | 12133.45 | 12064 | 12203 | 1.012 | 0.017 | 0.71247678 | 0.82724863 | 0 | 0.0026 | 0.0014 | 0.0014 | TRUE | 0.0034 |
| Psmd12 | 16731 | 11728 | 10600 | 10965 | 9492 | 11210 | 12994 | 12475 | 11437 | 10915 | 10708 | 11066 | 11599.17 | 12302 | 10896 | 0.886 | -0.175 | 0.00304823 | 0.01673133 | 0.0021 | 0.0026 | 0.0024 | 0.0024 | TRUE | 0.0461 |
| Psmd13 | 11900 | 8231 | 7224 | 8276 | 7383 | 8666 | 9242 | 8755 | 7795 | 8238 | 8329 | 8555 | 8485.58 | 8597 | 8374 | 0.974 | -0.038 | 0.56832584 | 0.72205166 | 0.0033 | 0.0027 | 0.0031 | 0.0031 | TRUE | 0.0813 |
| Psmd14 | 9362 | 6362 | 5760 | 6267 | 5381 | 6101 | 7271 | 6767 | 6215 | 6238 | 6070 | 6023 | 6430.73 | 6751 | 6110 | 0.905 | -0.144 | 0.02728674 | 0.08639995 | 0.003 | 0.0028 | 0.0029 | 0.0029 | TRUE | 0.0588 |
| Psmd2 | 37215 | 25603 | 24383 | 26359 | 23129 | 26305 | 28903 | 27234 | 26309 | 26238 | 26092 | 25967 | 26790.44 | 27482 | 26099 | 0.95 | -0.075 | 0.14750352 | 0.29213979 | 0.0011 | 0.0025 | 0.0019 | 0.0019 | TRUE | 0.0251 |
| Psmd3 | 32133 | 22957 | 21191 | 22545 | 20483 | 23058 | 24956 | 24419 | 22865 | 22441 | 23107 | 22762 | 23425.06 | 24080 | 22770 | 0.946 | -0.081 | 0.11704555 | 0.2480255 | 0.0011 | 0.0025 | 0.0019 | 0.0019 | TRUE | 0.0238 |
| Psmd4 | 11331 | 8351 | 7707 | 8687 | 7365 | 8782 | 8800 | 8883 | 8316 | 8647 | 8308 | 8669 | 8603.94 | 8666 | 8541 | 0.986 | -0.021 | 0.68812737 | 0.8090322 | 7.00E-04 | 0.0027 | 0.0018 | 0.0018 | TRUE | 0.0152 |
| Psmd5 | 6437 | 4618 | 4287 | 4199 | 3932 | 4418 | 4999 | 4912 | 4626 | 4180 | 4436 | 4361 | 4585.62 | 4846 | 4326 | 0.892 | -0.164 | 0.00419722 | 0.02143089 | 9.00E-04 | 0.0029 | 0.0022 | 0.0022 | TRUE | 0.0191 |
| Psmd6 | 6847 | 4946 | 4175 | 4304 | 3753 | 4299 | 5318 | 5261 | 4505 | 4284 | 4234 | 4244 | 4640.88 | 5028 | 4254 | 0.846 | -0.242 | 0.00061259 | 0.00505141 | 0.0038 | 0.0029 | 0.0034 | 0.0034 | TRUE | 0.1003 |
| Psmd7 | 17488 | 12570 | 11483 | 11082 | 9955 | 11618 | 13582 | 13371 | 12390 | 11031 | 11230 | 11469 | 12178.81 | 13114 | 11243 | 0.857 | -0.222 | 4.26E-05 | 0.00061541 | 0.0012 | 0.0026 | 0.002 | 0.002 | TRUE | 0.0284 |
| Psmd8 | 13472 | 9458 | 9451 | 10162 | 9200 | 10769 | 10463 | 10060 | 10198 | 10115 | 10379 | 10631 | 10307.58 | 10240 | 10375 | 1.013 | 0.019 | 0.70195534 | 0.81950718 | 4.00E-04 | 0.0027 | 0.0016 | 0.0016 | TRUE | 0.0059 |
| Psmd9 | 1628 | 1107 | 1032 | 951 | 857 | 936 | 1264 | 1178 | 1114 | 947 | 967 | 924 | 1065.47 | 1185 | 946 | 0.797 | -0.328 | 3.40E-05 | 0.00051521 | 0.0016 | 0.0047 | 0.0035 | 0.0035 | TRUE | 0.0427 |
| Psme1 | 2377 | 1840 | 1990 | 2224 | 2041 | 2299 | 1846 | 1957 | 2147 | 2214 | 2302 | 2269 | 2122.7 | 1983 | 2262 | 1.141 | 0.191 | 0.00757796 | 0.03364767 | 0.0024 | 0.0035 | 0.0032 | 0.0032 | TRUE | 0.0628 |
| Psme2 | 1290 | 1011 | 1036 | 1168 | 1118 | 1341 | 1002 | 1075 | 1118 | 1163 | 1261 | 1324 | 1157.12 | 1065 | 1249 | 1.174 | 0.232 | 0.00446218 | 0.02249664 | 0.0029 | 0.0045 | 0.0039 | 0.0039 | TRUE | 0.0447 |
| Psme3 | 17904 | 12752 | 11729 | 11825 | 10567 | 12184 | 13905 | 13564 | 12655 | 11771 | 11921 | 12027 | 12640.61 | 13375 | 11906 | 0.89 | -0.168 | 0.00162738 | 0.01026937 | 0.0011 | 0.0026 | 0.002 | 0.002 | TRUE | 0.027 |
| Psme4 | 34284 | 24518 | 25783 | 26862 | 23985 | 28032 | 26627 | 26079 | 27820 | 26738 | 27058 | 27672 | 26998.99 | 26842 | 27156 | 1.012 | 0.017 | 0.72712121 | 0.83661541 | 6.00E-04 | 0.0025 | 0.0016 | 0.0016 | TRUE | 0.0124 |
| Psmf1 | 2661 | 1895 | 1723 | 1912 | 1681 | 1911 | 2067 | 2016 | 1859 | 1903 | 1896 | 1886 | 1937.91 | 1981 | 1895 | 0.956 | -0.064 | 0.33047614 | 0.5048627 | 0.001 | 0.0036 | 0.0026 | 0.0026 | TRUE | 0.0343 |
| Psmg1 | 4934 | 3599 | 3780 | 3695 | 3429 | 4099 | 3832 | 3828 | 4079 | 3678 | 3868 | 4046 | 3888.57 | 3913 | 3864 | 0.988 | -0.018 | 0.77098015 | 0.86441946 | 0.0015 | 0.003 | 0.0024 | 0.0024 | TRUE | 0.0218 |
| Psmg2 | 3173 | 2141 | 2191 | 2290 | 1991 | 2303 | 2464 | 2277 | 2364 | 2279 | 2246 | 2273 | 2317.45 | 2368 | 2266 | 0.956 | -0.064 | 0.2894559 | 0.46153426 | 5.00E-04 | 0.0034 | 0.0022 | 0.0022 | TRUE | 0.0157 |
| Psmg3 | 1065 | 776 | 674 | 813 | 697 | 805 | 827 | 825 | 727 | 809 | 786 | 795 | 795 | 793 | 797 | 1.003 | 0.005 | 0.95450376 | 0.97517671 | 0.0014 | 0.0054 | 0.0041 | 0.0041 | TRUE | 0.0611 |
| Psmg4 | 946 | 720 | 669 | 636 | 595 | 663 | 735 | 766 | 722 | 633 | 671 | 654 | 696.87 | 741 | 653 | 0.881 | -0.182 | 0.03106849 | 0.0950882 | 0 | 0.0058 | 0.0037 | 0.0037 | TRUE | 0.0099 |
| Psors1c2 | 92 | 64 | 59 | 71 | 52 | 85 | 71 | 68 | 64 | 71 | 59 | 84 | 69.41 | 68 | 71 | 1.05 | 0.071 | 0.76894711 | 0.86329126 | 0 | 0.0366 | 0.0273 | 0.0273 | TRUE | 0.2011 |
| Pspc1 | 5567 | 4417 | 4690 | 5145 | 4750 | 5218 | 4324 | 4698 | 5060 | 5121 | 5358 | 5151 | 4952.2 | 4694 | 5210 | 1.11 | 0.151 | 0.02386894 | 0.07868054 | 0.0031 | 0.0029 | 0.003 | 0.003 | TRUE | 0.059 |
| Psph | 1573 | 1109 | 975 | 1247 | 987 | 1184 | 1222 | 1180 | 1052 | 1241 | 1113 | 1169 | 1162.8 | 1151 | 1174 | 1.02 | 0.028 | 0.73780677 | 0.84383266 | 0.0036 | 0.0045 | 0.0042 | 0.0042 | TRUE | 0.0669 |
| Pspn | 0 | 0 | 0 | 0 | 0 | 0 | 0 | 0 | 0 | 0 | 0 | 0 | 0 | 0 | 0 | NA | NA | NA | NA | NA | NA | NA | NA | NA | NA |
| Psrc1 | 5792 | 4354 | 4885 | 4978 | 4526 | 5355 | 4498 | 4631 | 5271 | 4955 | 5106 | 5286 | 4957.95 | 4800 | 5116 | 1.066 | 0.092 | 0.18484627 | 0.34114602 | 0.0039 | 0.0029 | 0.0033 | 0.0033 | TRUE | 0.0893 |
| Pstk | 2074 | 1444 | 1454 | 1690 | 1404 | 1611 | 1611 | 1536 | 1569 | 1682 | 1584 | 1590 | 1595.33 | 1572 | 1619 | 1.03 | 0.042 | 0.52565246 | 0.68657138 | 0 | 0.0039 | 0.0025 | 0.0025 | TRUE | 0.0143 |
| Pstpip1 | 233 | 178 | 182 | 197 | 163 | 184 | 181 | 189 | 196 | 196 | 184 | 182 | 188.05 | 189 | 187 | 0.993 | -0.01 | 0.94289155 | 0.96815725 | 0 | 0.015 | 0.0098 | 0.0098 | TRUE | 0.019 |
| Pstpip2 | 594 | 482 | 526 | 585 | 492 | 589 | 461 | 513 | 568 | 582 | 555 | 581 | 543.39 | 514 | 573 | 1.118 | 0.161 | 0.11961303 | 0.25134212 | 0.004 | 0.0068 | 0.0059 | 0.0059 | TRUE | 0.0995 |
| Ptafr | 20 | 18 | 7 | 14 | 10 | 8 | 16 | 19 | 8 | 14 | 11 | 8 | 12.56 | 14 | 11 | 0.781 | -0.358 | 0.53206273 | NA | 0.0506 | 0.1912 | 0.1548 | 0.1548 | TRUE | 0.4581 |
| Ptar1 | 319 | 265 | 238 | 244 | 228 | 290 | 248 | 282 | 257 | 243 | 257 | 286 | 262.13 | 262 | 262 | 1.002 | 0.003 | 0.98357545 | 0.99141354 | 0.0021 | 0.0115 | 0.0087 | 0.0087 | TRUE | 0.0744 |
| Ptbp1 | 42581 | 31398 | 31039 | 30782 | 28081 | 32363 | 33071 | 33398 | 33491 | 30640 | 31678 | 31947 | 32370.87 | 33320 | 31422 | 0.943 | -0.085 | 0.05545487 | 0.14514126 | 2.00E-04 | 0.0025 | 0.0014 | 0.0014 | TRUE | 0.0058 |
| Ptbp2 | 1860 | 1454 | 1523 | 1197 | 1071 | 1196 | 1445 | 1547 | 1643 | 1191 | 1208 | 1181 | 1369.14 | 1545 | 1193 | 0.773 | -0.371 | 5.77E-07 | 1.84E-05 | 0.0017 | 0.0042 | 0.0032 | 0.0032 | TRUE | 0.0404 |
| Ptbp3 | 10235 | 7968 | 7950 | 8369 | 7337 | 8205 | 7949 | 8475 | 8578 | 8330 | 8277 | 8100 | 8284.92 | 8334 | 8236 | 0.988 | -0.017 | 0.74983304 | 0.85153057 | 7.00E-04 | 0.0027 | 0.0019 | 0.0019 | TRUE | 0.0202 |
| Ptcd1 | 1252 | 767 | 693 | 697 | 640 | 752 | 972 | 816 | 748 | 694 | 722 | 742 | 782.35 | 845 | 719 | 0.849 | -0.236 | 0.02378261 | 0.07843213 | 0.0088 | 0.0055 | 0.0066 | 0.0066 | TRUE | 0.2123 |
| Ptcd2 | 3243 | 2362 | 2721 | 2813 | 2486 | 3041 | 2519 | 2512 | 2936 | 2800 | 2804 | 3002 | 2762.25 | 2656 | 2869 | 1.081 | 0.112 | 0.13572833 | 0.27517233 | 0.0044 | 0.0033 | 0.0037 | 0.0037 | TRUE | 0.1024 |
| Ptcd3 | 8986 | 6352 | 5870 | 6146 | 5500 | 6248 | 6979 | 6757 | 6334 | 6118 | 6205 | 6168 | 6426.55 | 6690 | 6164 | 0.921 | -0.119 | 0.03235171 | 0.09790452 | 0.001 | 0.0028 | 0.0021 | 0.0021 | TRUE | 0.0264 |
| Ptch1 | 2168 | 1693 | 2194 | 2360 | 2269 | 2686 | 1684 | 1801 | 2367 | 2349 | 2560 | 2651 | 2235.37 | 1951 | 2520 | 1.293 | 0.371 | 0.00064436 | 0.00523581 | 0.0188 | 0.0035 | 0.0081 | 0.0081 | TRUE | 0.4206 |
| Ptch2 | 779 | 613 | 576 | 624 | 556 | 654 | 605 | 652 | 621 | 621 | 627 | 646 | 628.75 | 626 | 631 | 1.009 | 0.013 | 0.87967384 | 0.93295109 | 0 | 0.0062 | 0.004 | 0.004 | TRUE | 0.0152 |
| Ptchd1 | 90 | 54 | 55 | 95 | 91 | 93 | 70 | 57 | 59 | 95 | 103 | 92 | 79.28 | 62 | 97 | 1.539 | 0.622 | 0.00492632 | 0.02424916 | 0 | 0.0323 | 0.0221 | 0.0221 | TRUE | 0.117 |
| Ptchd2 | 95 | 71 | 67 | 83 | 64 | 76 | 74 | 76 | 72 | 83 | 72 | 75 | 75.24 | 74 | 77 | 1.038 | 0.054 | 0.80700117 | 0.88842854 | 0 | 0.0339 | 0.0219 | 0.0219 | TRUE | 0.0437 |
| Ptchd3 | 108 | 80 | 90 | 122 | 72 | 100 | 84 | 85 | 97 | 121 | 81 | 99 | 94.58 | 89 | 100 | 1.138 | 0.186 | 0.3915343 | 0.56515853 | 0.0128 | 0.0275 | 0.0234 | 0.0234 | TRUE | 0.2357 |
| Ptchd4 | 6 | 4 | 1 | 4 | 3 | 1 | 5 | 4 | 1 | 4 | 3 | 1 | 3.06 | 3 | 3 | 0.82 | -0.286 | 0.79908815 | NA | 0 | 0.7775 | 0.5788 | 0.5788 | TRUE | 0.449 |
| Ptcra | 10 | 7 | 5 | 10 | 7 | 10 | 8 | 7 | 5 | 10 | 8 | 10 | 8.05 | 7 | 9 | 1.34 | 0.422 | 0.52355588 | NA | 0 | 0.2967 | 0.1903 | 0.1903 | TRUE | 0.0836 |
| Ptdss1 | 2888 | 1927 | 1805 | 1940 | 1733 | 1860 | 2243 | 2050 | 1948 | 1931 | 1955 | 1836 | 1993.75 | 2080 | 1907 | 0.916 | -0.127 | 0.0801518 | 0.18942973 | 0.0027 | 0.0036 | 0.0033 | 0.0033 | TRUE | 0.0583 |
| Ptdss2 | 5010 | 3550 | 3779 | 3541 | 3268 | 3690 | 3891 | 3776 | 4078 | 3525 | 3687 | 3643 | 3766.43 | 3915 | 3618 | 0.924 | -0.114 | 0.04655105 | 0.12787895 | 6.00E-04 | 0.0031 | 0.0021 | 0.0021 | TRUE | 0.0158 |
| Pten | 6838 | 5019 | 4823 | 5216 | 4719 | 5442 | 5311 | 5339 | 5204 | 5192 | 5324 | 5372 | 5290.17 | 5285 | 5296 | 1.002 | 0.003 | 0.95215199 | 0.97353948 | 0 | 0.0029 | 0.0016 | 0.0016 | TRUE | 0.0036 |
| Pter | 1163 | 755 | 693 | 782 | 633 | 771 | 903 | 803 | 748 | 778 | 714 | 761 | 784.61 | 818 | 751 | 0.917 | -0.125 | 0.18445708 | 0.34064408 | 0.0045 | 0.0054 | 0.0051 | 0.0051 | TRUE | 0.1028 |
| Ptf1a | 2 | 2 | 0 | 4 | 7 | 7 | 2 | 2 | 0 | 4 | 8 | 7 | 3.74 | 1 | 6 | 4.989 | 2.319 | 0.04191382 | NA | 0 | 0.6353 | 0.4699 | 0.4699 | TRUE | 0.3599 |
| Ptgdr | 2 | 0 | 0 | 1 | 0 | 1 | 2 | 0 | 0 | 1 | 0 | 1 | 0.59 | 1 | 1 | 1.206 | 0.27 | 0.91756405 | NA | 0 | 4.0242 | 3.277 | 3.277 | TRUE | 0.9721 |
| Ptgds | 42 | 22 | 23 | 32 | 36 | 49 | 33 | 23 | 25 | 32 | 41 | 48 | 33.61 | 27 | 40 | 1.482 | 0.568 | 0.10146309 | 0.22355088 | 0 | 0.0729 | 0.056 | 0.056 | TRUE | 0.1645 |
| Ptger1 | 7 | 2 | 2 | 9 | 1 | 7 | 5 | 2 | 2 | 9 | 1 | 7 | 4.45 | 3 | 6 | 1.718 | 0.781 | 0.43036187 | NA | 0.3018 | 0.5347 | 0.4712 | 0.4712 | TRUE | 0.5694 |
| Ptger2 | 30 | 16 | 15 | 35 | 31 | 29 | 23 | 17 | 16 | 35 | 35 | 29 | 25.82 | 19 | 33 | 1.721 | 0.784 | 0.04187727 | 0.11822078 | 0 | 0.0942 | 0.0658 | 0.0658 | TRUE | 0.2787 |
| Ptger3 | 27 | 20 | 5 | 9 | 6 | 11 | 21 | 21 | 5 | 9 | 7 | 11 | 12.37 | 16 | 9 | 0.556 | -0.848 | 0.1735029 | NA | 0.1847 | 0.194 | 0.1914 | 0.1914 | TRUE | 1.4557 |
| Ptger4 | 121 | 76 | 101 | 92 | 88 | 111 | 94 | 81 | 109 | 92 | 99 | 110 | 97.37 | 95 | 100 | 1.059 | 0.083 | 0.6875037 | 0.808565 | 0 | 0.0268 | 0.0205 | 0.0205 | TRUE | 0.1575 |
| Ptges | 433 | 304 | 281 | 396 | 335 | 422 | 336 | 323 | 303 | 394 | 378 | 417 | 358.59 | 321 | 396 | 1.233 | 0.302 | 0.00692002 | 0.03147926 | 0 | 0.009 | 0.0062 | 0.0062 | TRUE | 0.026 |
| Ptges2 | 1277 | 1007 | 939 | 1054 | 930 | 1059 | 992 | 1071 | 1013 | 1049 | 1049 | 1045 | 1036.63 | 1025 | 1048 | 1.023 | 0.032 | 0.66151227 | 0.79103653 | 0 | 0.0047 | 0.003 | 0.003 | TRUE | 0.018 |
| Ptges3 | 22952 | 16567 | 16497 | 17547 | 14751 | 16961 | 17826 | 17622 | 17800 | 17466 | 16641 | 16743 | 17349.69 | 17749 | 16950 | 0.955 | -0.066 | 0.14991999 | 0.29555951 | 3.00E-04 | 0.0026 | 0.0015 | 0.0015 | TRUE | 0.0087 |
| Ptges3l | 82 | 36 | 45 | 56 | 50 | 60 | 64 | 38 | 49 | 56 | 56 | 59 | 53.65 | 50 | 57 | 1.128 | 0.174 | 0.53399127 | 0.69315568 | 0.0154 | 0.0466 | 0.0378 | 0.0378 | TRUE | 0.4009 |
| Ptgfr | 17 | 11 | 5 | 5 | 2 | 2 | 13 | 12 | 5 | 5 | 2 | 2 | 6.58 | 10 | 3 | 0.303 | -1.723 | 0.03808086 | NA | 0 | 0.3624 | 0.282 | 0.282 | TRUE | 0.5372 |
| Ptgfrn | 6225 | 4971 | 5590 | 5697 | 5383 | 6312 | 4835 | 5288 | 6032 | 5671 | 6073 | 6231 | 5688.02 | 5385 | 5992 | 1.113 | 0.155 | 0.05292698 | 0.1404801 | 0.0072 | 0.0028 | 0.0044 | 0.0044 | TRUE | 0.1346 |
| Ptgir | 37 | 27 | 30 | 35 | 38 | 38 | 29 | 29 | 32 | 35 | 43 | 38 | 34.17 | 30 | 39 | 1.283 | 0.359 | 0.27097367 | 0.44196113 | 0 | 0.0718 | 0.0471 | 0.0471 | TRUE | 0.0708 |
| Ptgis | 788 | 551 | 493 | 618 | 562 | 591 | 612 | 586 | 532 | 615 | 634 | 583 | 593.77 | 577 | 611 | 1.057 | 0.08 | 0.40296184 | 0.57589577 | 0.0018 | 0.0064 | 0.0049 | 0.0049 | TRUE | 0.053 |
| Ptgr1 | 2376 | 1645 | 1510 | 1484 | 1262 | 1420 | 1845 | 1750 | 1629 | 1477 | 1424 | 1402 | 1587.83 | 1741 | 1434 | 0.823 | -0.281 | 9.91E-05 | 0.00120243 | 0.0018 | 0.0039 | 0.0031 | 0.0031 | TRUE | 0.0381 |

| | | | | | | | | | | | | | | | | | | | | | | | | | |
|---|---|---|---|---|---|---|---|---|---|---|---|---|---|---|---|---|---|---|---|---|---|---|---|---|---|
| Ptgr2 | 1978 | 1445 | 1421 | 1737 | 1440 | 1628 | 1536 | 1537 | 1533 | 1729 | 1624 | 1607 | 1594.51 | 1535 | 1653 | 1.077 | 0.107 | 0.10432669 | 0.22831747 | 0 | 0.0039 | 0.0025 | 0.0025 | TRUE | 0.0194 |
| Ptgs1 | 167 | 137 | 85 | 167 | 137 | 180 | 130 | 146 | 92 | 166 | 155 | 178 | 144.27 | 123 | 166 | 1.356 | 0.439 | 0.020956 | 0.07163942 | 0.0195 | 0.0189 | 0.019 | 0.019 | TRUE | 0.4576 |
| Ptgs2 | 35 | 17 | 16 | 12 | 8 | 8 | 27 | 18 | 17 | 12 | 9 | 8 | 15.23 | 21 | 10 | 0.458 | -1.127 | 0.02896696 | NA | 0 | 0.158 | 0.1151 | 0.1151 | TRUE | 0.4692 |
| Pth | 0 | 0 | 0 | 0 | 0 | 0 | 0 | 0 | 0 | 0 | 0 | 0 | 0 | 0 | 0 | NA | NA | NA | NA | NA | NA | NA | NA | NA | NA |
| Pth1r | 124 | 96 | 73 | 148 | 112 | 122 | 96 | 102 | 79 | 147 | 126 | 120 | 111.88 | 92 | 131 | 1.42 | 0.506 | 0.00969381 | 0.0404536 | 0.004 | 0.0236 | 0.0184 | 0.0184 | TRUE | 0.1486 |
| Pth2 | 1 | 1 | 5 | 3 | 1 | 2 | 1 | 1 | 5 | 3 | 1 | 2 | 2.22 | 2 | 2 | 0.87 | -0.202 | 0.87943692 | NA | 1.00E-04 | 1.0697 | 0.8158 | 0.8158 | TRUE | 1.1685 |
| Pth2r | 0 | 0 | 0 | 1 | 0 | 2 | 0 | 0 | 0 | 1 | 0 | 2 | 0.49 | 0 | 1 | 5.637 | 2.495 | 0.3989397 | NA | 0 | 4.7909 | 4.7853 | 4.7853 | TRUE | 0.0526 |
| Pthlh | 1 | 0 | 0 | 0 | 0 | 2 | 1 | 0 | 0 | 0 | 0 | 2 | 0.46 | 0 | 1 | 1.93 | 0.948 | 0.7608301 | NA | 0 | 5.1717 | 5.216 | 5.216 | TRUE | 0.9737 |
| Ptk2 | 6799 | 4878 | 4885 | 5596 | 4739 | 5701 | 5280 | 5189 | 5271 | 5570 | 5346 | 5628 | 5380.69 | 5247 | 5515 | 1.051 | 0.072 | 0.1580588 | 0.30702457 | 2.00E-04 | 0.0029 | 0.0017 | 0.0017 | TRUE | 0.0086 |
| Ptk2b | 760 | 596 | 690 | 730 | 670 | 828 | 590 | 634 | 745 | 727 | 756 | 817 | 711.43 | 656 | 767 | 1.171 | 0.227 | 0.02833458 | 0.08879328 | 0.007 | 0.0058 | 0.0063 | 0.0063 | TRUE | 0.1616 |
| Ptk6 | 13 | 15 | 10 | 10 | 18 | 26 | 10 | 16 | 11 | 10 | 20 | 26 | 15.46 | 12 | 19 | 1.527 | 0.611 | 0.24889046 | NA | 0.0821 | 0.1557 | 0.1347 | 0.1347 | TRUE | 0.3556 |
| Ptk7 | 5474 | 3938 | 4289 | 5178 | 4378 | 5012 | 4251 | 4189 | 4628 | 5154 | 4939 | 4948 | 4684.77 | 4356 | 5014 | 1.151 | 0.203 | 0.00064776 | 0.00525355 | 0.0015 | 0.0029 | 0.0023 | 0.0023 | TRUE | 0.0359 |
| Ptma | 76995 | 57652 | 52588 | 52500 | 44923 | 53226 | 59799 | 61324 | 56742 | 52258 | 50678 | 52542 | 55557.16 | 59288 | 51826 | 0.874 | -0.194 | 9.34E-05 | 0.00114591 | 9.00E-04 | 0.0025 | 0.0018 | 0.0018 | TRUE | 0.0173 |
| Ptms | 9209 | 7156 | 7069 | 6993 | 6318 | 7408 | 7152 | 7612 | 7627 | 6961 | 7127 | 7313 | 7298.73 | 7464 | 7134 | 0.956 | -0.065 | 0.22350347 | 0.3879688 | 8.00E-04 | 0.0028 | 0.0019 | 0.0019 | TRUE | 0.0165 |
| Ptn | 9 | 4 | 4 | 8 | 4 | 3 | 7 | 4 | 4 | 8 | 5 | 3 | 5.17 | 5 | 5 | 0.981 | -0.028 | 0.97340864 | NA | 0 | 0.4612 | 0.3248 | 0.3248 | TRUE | 0.4926 |
| Ptov1 | 5664 | 4208 | 4183 | 4002 | 4137 | 4809 | 4399 | 4476 | 4513 | 3984 | 4667 | 4747 | 4464.36 | 4463 | 4466 | 1.001 | 0.001 | 0.98917756 | 0.99429134 | 0.0044 | 0.003 | 0.0035 | 0.0035 | TRUE | 0.1091 |
| Ptp4a1 | 374 | 248 | 245 | 295 | 297 | 267 | 290 | 264 | 264 | 294 | 335 | 264 | 285.15 | 273 | 298 | 1.086 | 0.119 | 0.36737035 | 0.54210453 | 0.0053 | 0.0107 | 0.0091 | 0.0091 | TRUE | 0.1342 |
| Ptp4a2 | 14547 | 10457 | 10177 | 11656 | 9690 | 10775 | 11298 | 11123 | 10981 | 11602 | 10931 | 10637 | 11095.37 | 11134 | 11057 | 0.993 | -0.01 | 0.84775017 | 0.91365505 | 8.00E-04 | 0.0026 | 0.0019 | 0.0019 | TRUE | 0.0228 |
| Ptp4a3 | 4227 | 3131 | 3558 | 4084 | 3465 | 3977 | 3283 | 3330 | 3839 | 4065 | 3909 | 3926 | 3725.4 | 3484 | 3967 | 1.139 | 0.188 | 0.0077326 | 0.03414337 | 0.0037 | 0.0031 | 0.0033 | 0.0033 | TRUE | 0.0959 |
| Ptpdc1 | 678 | 519 | 404 | 502 | 393 | 437 | 527 | 552 | 436 | 500 | 443 | 431 | 481.49 | 505 | 458 | 0.907 | -0.141 | 0.22628032 | 0.39126841 | 0.0085 | 0.0073 | 0.0077 | 0.0077 | TRUE | 0.1608 |
| Ptpla | 372 | 283 | 267 | 309 | 283 | 302 | 289 | 301 | 288 | 308 | 319 | 298 | 300.5 | 293 | 308 | 1.053 | 0.075 | 0.52199284 | 0.68329001 | 0 | 0.0103 | 0.0065 | 0.0065 | TRUE | 0.0105 |
| Ptplad1 | 2462 | 1733 | 1618 | 1717 | 1474 | 1689 | 1912 | 1843 | 1746 | 1709 | 1663 | 1667 | 1756.76 | 1834 | 1680 | 0.915 | -0.128 | 0.05361875 | 0.14156329 | 7.00E-04 | 0.0038 | 0.0026 | 0.0026 | TRUE | 0.0211 |
| Ptplad2 | 156 | 107 | 114 | 127 | 110 | 143 | 121 | 114 | 123 | 126 | 124 | 141 | 124.94 | 119 | 130 | 1.094 | 0.13 | 0.45806153 | 0.62777577 | 0 | 0.0214 | 0.0142 | 0.0142 | TRUE | 0.0527 |
| Ptplb | 2203 | 1407 | 1416 | 1727 | 1475 | 1691 | 1711 | 1497 | 1528 | 1719 | 1664 | 1669 | 1631.29 | 1579 | 1684 | 1.066 | 0.092 | 0.21311332 | 0.37540408 | 0.0023 | 0.0039 | 0.0033 | 0.0033 | TRUE | 0.0672 |
| Ptpmt1 | 1185 | 750 | 757 | 979 | 730 | 842 | 920 | 798 | 817 | 974 | 824 | 831 | 860.68 | 845 | 876 | 1.036 | 0.051 | 0.59868511 | 0.74434641 | 0.0065 | 0.0052 | 0.0056 | 0.0056 | TRUE | 0.1148 |
| Ptpn1 | 3184 | 2278 | 2222 | 2432 | 2170 | 2492 | 2473 | 2423 | 2398 | 2421 | 2448 | 2460 | 2437.04 | 2431 | 2443 | 1.005 | 0.007 | 0.90879935 | 0.94963864 | 0 | 0.0034 | 0.0019 | 0.0019 | TRUE | 0.0028 |
| Ptpn11 | 8212 | 5837 | 5397 | 5648 | 5023 | 5617 | 6378 | 6209 | 5823 | 5622 | 5666 | 5545 | 5873.88 | 6137 | 5611 | 0.914 | -0.13 | 0.01909748 | 0.06680651 | 9.00E-04 | 0.0028 | 0.002 | 0.002 | TRUE | 0.0242 |
| Ptpn12 | 3040 | 2178 | 2048 | 1971 | 1699 | 2096 | 2361 | 2317 | 2210 | 1962 | 1917 | 2069 | 2139.2 | 2296 | 1983 | 0.863 | -0.212 | 0.00096673 | 0.00707864 | 9.00E-04 | 0.0035 | 0.0025 | 0.0025 | TRUE | 0.0177 |
| Ptpn13 | 4144 | 2979 | 3314 | 3452 | 3229 | 3753 | 3218 | 3169 | 3576 | 3436 | 3643 | 3705 | 3457.75 | 3321 | 3595 | 1.083 | 0.115 | 0.08652451 | 0.19924196 | 0.0026 | 0.0031 | 0.0029 | 0.0029 | TRUE | 0.0541 |
| Ptpn14 | 3251 | 2237 | 2228 | 2440 | 2036 | 2175 | 2525 | 2379 | 2404 | 2429 | 2297 | 2147 | 2363.5 | 2436 | 2291 | 0.94 | -0.089 | 0.18666636 | 0.34333062 | 0.0018 | 0.0034 | 0.0029 | 0.0029 | TRUE | 0.037 |
| Ptpn18 | 114 | 101 | 95 | 140 | 104 | 99 | 89 | 107 | 103 | 139 | 117 | 98 | 108.81 | 100 | 118 | 1.192 | 0.253 | 0.21601084 | 0.37873441 | 0.0128 | 0.0242 | 0.0209 | 0.0209 | TRUE | 0.1895 |
| Ptpn2 | 3756 | 2545 | 2529 | 2886 | 2481 | 2921 | 2917 | 2707 | 2729 | 2873 | 2799 | 2883 | 2818 | 2784 | 2852 | 1.024 | 0.034 | 0.57199035 | 0.72498188 | 7.00E-04 | 0.0033 | 0.0022 | 0.0022 | TRUE | 0.0217 |
| Ptpn20 | 200 | 167 | 111 | 104 | 100 | 117 | 155 | 178 | 120 | 104 | 113 | 115 | 130.76 | 151 | 111 | 0.732 | -0.45 | 0.01915221 | 0.06691525 | 0.0142 | 0.0206 | 0.0187 | 0.0187 | TRUE | 0.2583 |
| Ptpn21 | 1285 | 912 | 949 | 933 | 855 | 956 | 998 | 970 | 1024 | 929 | 965 | 944 | 971.5 | 997 | 946 | 0.948 | -0.077 | 0.30226708 | 0.474796 | 0 | 0.0049 | 0.003 | 0.003 | TRUE | 0.0065 |
| Ptpn22 | 12 | 4 | 1 | 3 | 2 | 0 | 9 | 4 | 1 | 3 | 2 | 0 | 3.32 | 5 | 2 | 0.348 | -1.523 | 0.21712837 | NA | 0.6509 | 0.7172 | 0.7014 | 0.7014 | TRUE | 0.1831 |
| Ptpn23 | 7462 | 5582 | 4979 | 5100 | 4807 | 5457 | 5795 | 5938 | 5372 | 5077 | 5423 | 5387 | 5498.57 | 5702 | 5296 | 0.929 | -0.107 | 0.07538871 | 0.18091519 | 0.0017 | 0.0029 | 0.0024 | 0.0024 | TRUE | 0.0308 |
| Ptpn3 | 3760 | 2864 | 3027 | 3064 | 2959 | 3378 | 2920 | 3046 | 3266 | 3050 | 3338 | 3335 | 3159.22 | 3077 | 3241 | 1.053 | 0.075 | 0.26518964 | 0.43561108 | 0.0024 | 0.0032 | 0.003 | 0.003 | TRUE | 0.0348 |
| Ptpn4 | 1097 | 752 | 627 | 756 | 584 | 704 | 852 | 800 | 677 | 753 | 659 | 695 | 739.12 | 776 | 702 | 0.903 | -0.146 | 0.15778346 | 0.30675852 | 0.0079 | 0.0056 | 0.0064 | 0.0064 | TRUE | 0.1473 |
| Ptpn5 | 24 | 18 | 16 | 34 | 23 | 18 | 19 | 19 | 17 | 34 | 26 | 18 | 22.1 | 18 | 26 | 1.407 | 0.493 | 0.24471993 | 0.41288552 | 0 | 0.1097 | 0.0831 | 0.0831 | TRUE | 0.149 |
| Ptpn6 | 2627 | 2172 | 2343 | 2146 | 2010 | 2377 | 2040 | 2310 | 2528 | 2136 | 2267 | 2346 | 2271.46 | 2293 | 2250 | 0.982 | -0.026 | 0.75628477 | 0.85520897 | 0.0065 | 0.0035 | 0.0046 | 0.0046 | TRUE | 0.115 |
| Ptpn7 | 227 | 153 | 134 | 154 | 154 | 179 | 176 | 163 | 145 | 153 | 174 | 177 | 164.56 | 161 | 168 | 1.038 | 0.053 | 0.73993799 | 0.84538261 | 0 | 0.0168 | 0.0126 | 0.0126 | TRUE | 0.084 |
| Ptpn9 | 847 | 585 | 526 | 594 | 461 | 547 | 658 | 622 | 568 | 591 | 520 | 540 | 583.16 | 616 | 550 | 0.893 | -0.164 | 0.10128631 | 0.2232704 | 0.0033 | 0.0065 | 0.0055 | 0.0055 | TRUE | 0.0546 |
| Ptpra | 3384 | 2442 | 2570 | 2710 | 2547 | 2689 | 2628 | 2598 | 2773 | 2698 | 2873 | 2654 | 2704 | 2666 | 2742 | 1.028 | 0.04 | 0.51878753 | 0.68027799 | 0.001 | 0.0033 | 0.0024 | 0.0024 | TRUE | 0.0211 |
| Ptprb | 28 | 16 | 25 | 20 | 16 | 27 | 22 | 17 | 27 | 20 | 18 | 27 | 21.73 | 22 | 22 | 0.986 | -0.02 | 0.96195334 | 0.97911443 | 0 | 0.1115 | 0.0813 | 0.0813 | TRUE | 0.2212 |
| Ptprc | 30 | 18 | 19 | 17 | 15 | 13 | 23 | 19 | 21 | 17 | 17 | 13 | 18.27 | 21 | 16 | 0.737 | -0.44 | 0.32410266 | 0.49809594 | 0 | 0.1321 | 0.0871 | 0.0871 | TRUE | 0.1138 |
| Ptprcap | 32 | 24 | 21 | 13 | 10 | 22 | 25 | 26 | 23 | 13 | 11 | 22 | 19.83 | 25 | 15 | 0.632 | -0.663 | 0.13768566 | 0.2781149 | 0 | 0.1219 | 0.0901 | 0.0901 | TRUE | 0.6331 |
| Ptprd | 132 | 65 | 84 | 61 | 53 | 54 | 103 | 69 | 91 | 61 | 60 | 53 | 72.68 | 88 | 58 | 0.659 | -0.601 | 0.01361365 | 0.0519976 | 0 | 0.035 | 0.0284 | 0.0284 | TRUE | 0.3048 |
| Ptpre | 191 | 133 | 111 | 137 | 106 | 117 | 148 | 141 | 120 | 136 | 120 | 115 | 130.17 | 136 | 124 | 0.905 | -0.145 | 0.42075185 | 0.5929435 | 0 | 0.0206 | 0.0155 | 0.0155 | TRUE | 0.1147 |
| Ptprf | 16950 | 12361 | 12974 | 12988 | 11740 | 13247 | 13164 | 13148 | 13999 | 12928 | 13244 | 13077 | 13260.07 | 13437 | 13083 | 0.974 | -0.038 | 0.43839832 | 0.60989169 | 6.00E-04 | 0.0026 | 0.0017 | 0.0017 | TRUE | 0.0163 |
| Ptprg | 3410 | 2398 | 2450 | 2554 | 2231 | 2682 | 2648 | 2551 | 2644 | 2542 | 2517 | 2648 | 2591.54 | 2614 | 2569 | 0.983 | -0.025 | 0.66416845 | 0.79306262 | 2.00E-04 | 0.0033 | 0.0021 | 0.0021 | TRUE | 0.0088 |
| Ptprh | 1 | 1 | 1 | 1 | 1 | 1 | 1 | 1 | 1 | 1 | 1 | 1 | 1 | 1 | 1 | 1.074 | 0.103 | 0.95418264 | NA | 0 | 2.3607 | 1.3027 | 1.3027 | TRUE | 0.0201 |
| Ptprj | 1046 | 777 | 803 | 985 | 840 | 902 | 812 | 826 | 866 | 980 | 948 | 890 | 887.3 | 835 | 939 | 1.126 | 0.171 | 0.03446155 | 0.10271929 | 7.00E-04 | 0.0051 | 0.0036 | 0.0036 | TRUE | 0.0254 |
| Ptprk | 3143 | 2110 | 2085 | 1912 | 1800 | 2070 | 2441 | 2244 | 2250 | 1903 | 2031 | 2043 | 2152.05 | 2312 | 1992 | 0.861 | -0.216 | 0.00125061 | 0.00849164 | 0.0015 | 0.0035 | 0.0028 | 0.0028 | TRUE | 0.0298 |
| Ptprm | 212 | 143 | 131 | 151 | 105 | 127 | 165 | 152 | 141 | 150 | 118 | 125 | 142.04 | 153 | 131 | 0.859 | -0.219 | 0.20886007 | 0.3706067 | 0.0032 | 0.0191 | 0.0147 | 0.0147 | TRUE | 0.1645 |
| Ptprn | 297 | 232 | 198 | 263 | 203 | 274 | 231 | 247 | 214 | 262 | 229 | 270 | 242.06 | 231 | 254 | 1.103 | 0.141 | 0.29955209 | 0.47179042 | 0.0017 | 0.0122 | 0.0092 | 0.0092 | TRUE | 0.078 |
| Ptprn2 | 194 | 144 | 167 | 228 | 245 | 215 | 151 | 153 | 180 | 227 | 276 | 212 | 199.93 | 161 | 238 | 1.479 | 0.565 | 0.00037072 | 0.0034296 | 0.0098 | 0.0143 | 0.013 | 0.013 | TRUE | 0.2073 |
| Ptpro | 113 | 115 | 104 | 44 | 51 | 44 | 88 | 122 | 112 | 44 | 58 | 43 | 77.84 | 107 | 48 | 0.45 | -1.153 | 2.58E-06 | 6.17E-05 | 0.0163 | 0.0329 | 0.0281 | 0.0281 | TRUE | 0.2823 |
| Ptprq | 36 | 48 | 33 | 10 | 14 | 15 | 28 | 51 | 36 | 10 | 16 | 15 | 25.86 | 38 | 14 | 0.355 | -1.494 | 0.00046713 | 0.00411128 | 0 | 0.0941 | 0.0803 | 0.0803 | TRUE | 0.254 |
| Ptprr | 17 | 14 | 11 | 11 | 12 | 10 | 13 | 15 | 12 | 11 | 14 | 10 | 12.39 | 13 | 12 | 0.857 | -0.223 | 0.67784351 | NA | 0 | 0.1938 | 0.1255 | 0.1255 | TRUE | 0.0869 |
| Ptprs | 4948 | 4077 | 4448 | 6125 | 5314 | 6138 | 3843 | 4337 | 4799 | 6097 | 5995 | 6059 | 5188.27 | 4326 | 6050 | 1.399 | 0.485 | 2.30E-10 | 1.72E-08 | 0.0059 | 0.0029 | 0.004 | 0.004 | TRUE | 0.1179 |
| Ptprt | 77 | 38 | 54 | 28 | 27 | 41 | 60 | 40 | 58 | 28 | 30 | 40 | 42.88 | 53 | 33 | 0.622 | -0.685 | 0.03019732 | 0.09309055 | 0.0189 | 0.0577 | 0.0471 | 0.0471 | TRUE | 0.334 |
| Ptpru | 847 | 584 | 637 | 834 | 698 | 806 | 658 | 621 | 687 | 830 | 787 | 796 | 729.93 | 655 | 804 | 1.227 | 0.296 | 0.00051997 | 0.00444227 | 0 | 0.0057 | 0.0039 | 0.0039 | TRUE | 0.0233 |
| Ptprv | 392 | 307 | 303 | 358 | 314 | 313 | 304 | 327 | 327 | 356 | 354 | 309 | 329.58 | 319 | 340 | 1.065 | 0.091 | 0.44067554 | 0.61198726 | 0 | 0.0096 | 0.007 | 0.007 | TRUE | 0.0733 |
| Ptprz1 | 193 | 132 | 105 | 85 | 83 | 98 | 150 | 140 | 113 | 85 | 94 | 97 | 113.1 | 134 | 92 | 0.679 | -0.559 | 0.00419981 | 0.02143089 | 0 | 0.0234 | 0.0182 | 0.0182 | TRUE | 0.1896 |
| Ptrf | 2968 | 2181 | 2210 | 3168 | 2772 | 3192 | 2305 | 2320 | 2385 | 3153 | 3127 | 3151 | 2740.19 | 2337 | 3144 | 1.346 | 0.429 | 1.08E-14 | 1.75E-12 | 0 | 0.0033 | 0.0018 | 0.0018 | TRUE | 0.0039 |
| Ptrh1 | 426 | 356 | 357 | 491 | 435 | 463 | 331 | 379 | 385 | 489 | 491 | 457 | 421.87 | 365 | 479 | 1.316 | 0.396 | 0.00027651 | 0.00272499 | 0.0019 | 0.008 | 0.0061 | 0.0061 | TRUE | 0.0829 |
| Ptrh2 | 2109 | 1409 | 1224 | 1323 | 1097 | 1315 | 1638 | 1499 | 1321 | 1317 | 1238 | 1298 | 1384.99 | 1486 | 1284 | 0.863 | -0.212 | 0.01462459 | 0.05471971 | 0.0057 | 0.0041 | 0.0047 | 0.0047 | TRUE | 0.113 |
| Ptrhd1 | 425 | 321 | 331 | 364 | 330 | 331 | 330 | 341 | 357 | 362 | 372 | 327 | 348.34 | 343 | 354 | 1.032 | 0.046 | 0.68821246 | 0.80906325 | 0 | 0.0092 | 0.0065 | 0.0065 | TRUE | 0.0518 |
| Pts | 663 | 575 | 536 | 642 | 585 | 568 | 515 | 612 | 578 | 639 | 660 | 561 | 594.1 | 568 | 620 | 1.092 | 0.127 | 0.2247696 | 0.3892273 | 0.0059 | 0.0064 | 0.0063 | 0.0063 | TRUE | 0.0839 |

| | | | | | | | | | | | | | | | | | | | | | | | | | |
|---|---|---|---|---|---|---|---|---|---|---|---|---|---|---|---|---|---|---|---|---|---|---|---|---|---|
| Pttg1 | 7231 | 5341 | 5935 | 4798 | 4473 | 5039 | 5616 | 5681 | 6404 | 4776 | 5046 | 4974 | 5416.2 | 5900 | 4932 | 0.836 | -0.258 | 7.64E-05 | 0.00098107 | 0.0029 | 0.0029 | 0.0029 | 0.0029 | TRUE | 0.0676 |
| Pttg1ip | 4253 | 3162 | 3521 | 3726 | 3412 | 3796 | 3303 | 3363 | 3799 | 3709 | 3849 | 3747 | 3628.47 | 3488 | 3768 | 1.081 | 0.112 | 0.09595782 | 0.21470336 | 0.0028 | 0.0031 | 0.003 | 0.003 | TRUE | 0.0732 |
| Ptx3 | 19 | 4 | 7 | 2 | 2 | 3 | 15 | 4 | 8 | 2 | 2 | 3 | 5.63 | 9 | 2 | 0.267 | -1.903 | 0.03791039 | NA | 0 | 0.4235 | 0.3368 | 0.3368 | TRUE | 0.2782 |
| Ptx4 | 0 | 0 | 0 | 0 | 0 | 0 | 0 | 0 | 0 | 0 | 0 | 0 | 0 | 0 | 0 | NA | NA | NA | NA | NA | NA | NA | NA | NA | NA |
| Puf60 | 10847 | 7866 | 7519 | 6871 | 6273 | 7580 | 8424 | 8367 | 8113 | 6839 | 7077 | 7483 | 7717.15 | 8301 | 7133 | 0.859 | -0.219 | 6.10E-05 | 0.00082366 | 0.001 | 0.0027 | 0.002 | 0.002 | TRUE | 0.0226 |
| Pum1 | 12755 | 9331 | 8931 | 9060 | 8159 | 9288 | 9906 | 9925 | 9636 | 9018 | 9204 | 9169 | 9476.53 | 9822 | 9130 | 0.929 | -0.106 | 0.02304896 | 0.07661367 | 1.00E-04 | 0.0027 | 0.0014 | 0.0014 | TRUE | 0.0033 |
| Pum2 | 15739 | 11654 | 12027 | 12386 | 10675 | 12412 | 12224 | 12396 | 12977 | 12329 | 12043 | 12253 | 12370.19 | 12532 | 12208 | 0.974 | -0.038 | 0.4391694 | 0.61048903 | 4.00E-04 | 0.0026 | 0.0016 | 0.0016 | TRUE | 0.0117 |
| Pura | 1464 | 1013 | 855 | 887 | 768 | 887 | 1137 | 1078 | 923 | 883 | 866 | 876 | 960.33 | 1046 | 875 | 0.836 | -0.259 | 0.0044327 | 0.0223993 | 0.005 | 0.0049 | 0.0049 | 0.0049 | TRUE | 0.1251 |
| Purb | 5382 | 4087 | 3452 | 4090 | 3491 | 3927 | 4180 | 4347 | 3725 | 4071 | 3938 | 3877 | 4022.98 | 4084 | 3962 | 0.97 | -0.044 | 0.51826461 | 0.68007354 | 0.0028 | 0.003 | 0.0031 | 0.0031 | TRUE | 0.0713 |
| Purg | 455 | 360 | 304 | 301 | 259 | 308 | 353 | 383 | 328 | 300 | 292 | 304 | 326.69 | 355 | 299 | 0.842 | -0.248 | 0.03439101 | 0.10257736 | 0 | 0.0097 | 0.0068 | 0.0068 | TRUE | 0.052 |
| Pus1 | 3840 | 3009 | 3016 | 3219 | 2856 | 3289 | 2982 | 3201 | 3254 | 3204 | 3222 | 3247 | 3185.01 | 3146 | 3224 | 1.026 | 0.037 | 0.53616601 | 0.69496878 | 8.00E-04 | 0.0032 | 0.0022 | 0.0022 | TRUE | 0.0258 |
| Pus10 | 2799 | 1975 | 1860 | 1828 | 1597 | 1811 | 2174 | 2101 | 2007 | 1820 | 1802 | 1788 | 1948.41 | 2094 | 1803 | 0.861 | -0.217 | 0.00061952 | 0.00508608 | 5.00E-04 | 0.0036 | 0.0024 | 0.0024 | TRUE | 0.0159 |
| Pus3 | 1045 | 727 | 667 | 809 | 655 | 762 | 812 | 773 | 720 | 805 | 739 | 752 | 766.83 | 768 | 765 | 0.995 | -0.007 | 0.93937185 | 0.966328 | 0.0016 | 0.0055 | 0.0042 | 0.0042 | TRUE | 0.0357 |
| Pus7 | 4277 | 3112 | 3093 | 2937 | 2545 | 2982 | 3322 | 3310 | 3337 | 2923 | 2871 | 2944 | 3117.91 | 3323 | 2913 | 0.877 | -0.19 | 0.00037943 | 0.0034966 | 0 | 0.0032 | 0.0017 | 0.0017 | TRUE | 0.0019 |
| Pus7l | 887 | 643 | 565 | 496 | 484 | 509 | 689 | 684 | 610 | 494 | 546 | 502 | 587.44 | 661 | 514 | 0.776 | -0.365 | 0.00016777 | 0.00186467 | 0.002 | 0.0065 | 0.005 | 0.005 | TRUE | 0.0527 |
| Pusl1 | 991 | 750 | 683 | 654 | 573 | 720 | 770 | 798 | 737 | 651 | 646 | 711 | 718.75 | 768 | 669 | 0.872 | -0.198 | 0.0231025 | 0.07677377 | 6.00E-04 | 0.0057 | 0.0041 | 0.0041 | TRUE | 0.0349 |
| Pvalb | 1 | 1 | 1 | 2 | 2 | 2 | 1 | 1 | 1 | 2 | 2 | 2 | 1.52 | 1 | 2 | 2.153 | 1.106 | 0.46559311 | NA | 0 | 1.5582 | 0.9061 | 0.9061 | TRUE | 0.0205 |
| Pvr | 1343 | 1013 | 1235 | 1521 | 1268 | 1465 | 1043 | 1078 | 1333 | 1514 | 1430 | 1446 | 1307.29 | 1151 | 1463 | 1.273 | 0.349 | 0.00023082 | 0.00238752 | 0.0087 | 0.0042 | 0.0057 | 0.0057 | TRUE | 0.2257 |
| Pvrl1 | 2224 | 1595 | 1468 | 1709 | 1526 | 1718 | 1727 | 1697 | 1584 | 1701 | 1721 | 1696 | 1687.73 | 1669 | 1706 | 1.021 | 0.03 | 0.64527942 | 0.77909941 | 5.00E-04 | 0.0038 | 0.0026 | 0.0026 | TRUE | 0.0237 |
| Pvrl2 | 5495 | 3989 | 4557 | 5193 | 4787 | 5467 | 4268 | 4243 | 4917 | 5169 | 5400 | 5397 | 4898.98 | 4476 | 5322 | 1.189 | 0.25 | 0.00026146 | 0.00262248 | 0.0036 | 0.0029 | 0.0032 | 0.0032 | TRUE | 0.0899 |
| Pvrl3 | 1964 | 1427 | 1574 | 1515 | 1447 | 1568 | 1525 | 1518 | 1698 | 1508 | 1632 | 1548 | 1571.64 | 1580 | 1563 | 0.989 | -0.016 | 0.82966519 | 0.9024661 | 0.0022 | 0.0039 | 0.0033 | 0.0033 | TRUE | 0.0503 |
| Pvrl4 | 259 | 196 | 222 | 232 | 168 | 224 | 201 | 208 | 240 | 231 | 190 | 221 | 215.12 | 216 | 214 | 0.992 | -0.012 | 0.93720297 | 0.96508948 | 0.0047 | 0.0134 | 0.0109 | 0.0109 | TRUE | 0.1041 |
| Pvt1 | 413 | 342 | 399 | 532 | 414 | 474 | 321 | 364 | 431 | 530 | 467 | 468 | 429.93 | 372 | 488 | 1.318 | 0.399 | 0.00144473 | 0.00938188 | 0.0111 | 0.0079 | 0.0089 | 0.0089 | TRUE | 0.2184 |
| Pwp1 | 7851 | 5718 | 5613 | 5981 | 5235 | 6005 | 6098 | 6082 | 6056 | 5953 | 5906 | 5928 | 6003.84 | 6079 | 5929 | 0.975 | -0.036 | 0.44661248 | 0.61723382 | 0 | 0.0028 | 0.0014 | 0.0014 | TRUE | 2.00E-04 |
| Pwp2 | 6545 | 4629 | 4093 | 4580 | 3923 | 4485 | 5083 | 4924 | 4416 | 4559 | 4426 | 4427 | 4639.2 | 4808 | 4471 | 0.93 | -0.105 | 0.10394414 | 0.22765454 | 0.0024 | 0.0029 | 0.0028 | 0.0028 | TRUE | 0.0615 |
| Pwwp2a | 3870 | 3094 | 3125 | 3350 | 2924 | 3779 | 3006 | 3291 | 3372 | 3335 | 3299 | 3730 | 3338.7 | 3223 | 3455 | 1.073 | 0.101 | 0.15917135 | 0.30839584 | 0.0037 | 0.0031 | 0.0034 | 0.0034 | TRUE | 0.0596 |
| Pwwp2b | 356 | 298 | 359 | 549 | 479 | 597 | 276 | 317 | 387 | 546 | 540 | 589 | 442.83 | 327 | 558 | 1.716 | 0.779 | 7.54E-10 | 5.23E-08 | 0.0122 | 0.0078 | 0.0091 | 0.0091 | TRUE | 0.2947 |
| Pxdc1 | 101 | 82 | 139 | 151 | 134 | 120 | 78 | 87 | 150 | 150 | 151 | 118 | 122.6 | 105 | 140 | 1.338 | 0.42 | 0.08076324 | 0.19036098 | 0.0625 | 0.0218 | 0.0333 | 0.0333 | TRUE | 1.3382 |
| Pxdn | 2598 | 2075 | 2078 | 2486 | 2297 | 2577 | 2018 | 2207 | 2242 | 2475 | 2591 | 2544 | 2346.13 | 2156 | 2537 | 1.178 | 0.236 | 0.00032072 | 0.00306016 | 0.0015 | 0.0034 | 0.0027 | 0.0027 | TRUE | 0.0392 |
| Pxk | 1222 | 829 | 946 | 894 | 867 | 998 | 949 | 882 | 1021 | 890 | 978 | 985 | 950.79 | 951 | 951 | 1 | 0.001 | 0.99444833 | 0.99681955 | 0.0031 | 0.0049 | 0.0043 | 0.0043 | TRUE | 0.0478 |
| Pxmp2 | 912 | 716 | 627 | 730 | 649 | 685 | 708 | 762 | 677 | 727 | 732 | 676 | 713.57 | 716 | 712 | 0.994 | -0.008 | 0.92849542 | 0.9605519 | 0.0012 | 0.0057 | 0.0043 | 0.0043 | TRUE | 0.0361 |
| Pxmp4 | 538 | 415 | 420 | 567 | 496 | 539 | 418 | 441 | 453 | 564 | 560 | 532 | 494.74 | 437 | 552 | 1.263 | 0.337 | 0.00050987 | 0.00437956 | 0 | 0.0072 | 0.0047 | 0.0047 | TRUE | 0.0193 |
| Pxn | 4674 | 3578 | 3482 | 3567 | 3397 | 4039 | 3630 | 3806 | 3757 | 3551 | 3832 | 3987 | 3760.48 | 3731 | 3790 | 1.016 | 0.023 | 0.71301953 | 0.82754268 | 0.0018 | 0.0031 | 0.0025 | 0.0025 | TRUE | 0.0374 |
| Pxt1 | 123 | 84 | 92 | 98 | 104 | 93 | 96 | 89 | 99 | 98 | 117 | 92 | 98.47 | 95 | 102 | 1.077 | 0.107 | 0.59656773 | 0.74274039 | 0 | 0.0265 | 0.019 | 0.019 | TRUE | 0.1561 |
| Pycard | 1090 | 950 | 1220 | 1285 | 1290 | 1497 | 847 | 1011 | 1316 | 1279 | 1455 | 1478 | 1230.92 | 1058 | 1404 | 1.329 | 0.411 | 0.00140593 | 0.00920099 | 0.0272 | 0.0044 | 0.0111 | 0.0111 | TRUE | 0.5459 |
| Pycr1 | 1181 | 931 | 1231 | 1529 | 1286 | 1380 | 917 | 990 | 1328 | 1522 | 1451 | 1362 | 1261.79 | 1078 | 1445 | 1.342 | 0.424 | 0.00031452 | 0.00301105 | 0.021 | 0.0043 | 0.0092 | 0.0092 | TRUE | 0.4878 |
| Pycr2 | 5264 | 3573 | 3486 | 3707 | 3515 | 3738 | 4088 | 3801 | 3761 | 3690 | 3965 | 3690 | 3832.58 | 3883 | 3782 | 0.973 | -0.039 | 0.52440463 | 0.6856824 | 0.0015 | 0.003 | 0.0025 | 0.0025 | TRUE | 0.0264 |
| Pycrl | 2084 | 1555 | 1345 | 1666 | 1406 | 1699 | 1619 | 1654 | 1451 | 1658 | 1586 | 1677 | 1607.58 | 1575 | 1640 | 1.042 | 0.059 | 0.42339041 | 0.59579789 | 0.0021 | 0.0039 | 0.0033 | 0.0033 | TRUE | 0.0553 |
| Pydc3 | 0 | 0 | 0 | 0 | 0 | 0 | 0 | 0 | 0 | 0 | 0 | 0 | 0 | 0 | 0 | NA | NA | NA | NA | NA | NA | NA | NA | NA | NA |
| Pydc4 | 0 | 0 | 0 | 0 | 0 | 0 | 0 | 0 | 0 | 0 | 0 | 0 | 0 | 0 | 0 | NA | NA | NA | NA | NA | NA | NA | NA | NA | NA |
| Pygb | 3822 | 2922 | 3350 | 3728 | 3536 | 3913 | 2968 | 3108 | 3615 | 3711 | 3989 | 3863 | 3542.28 | 3230 | 3854 | 1.194 | 0.255 | 0.00099514 | 0.00722005 | 0.0058 | 0.0031 | 0.0041 | 0.0041 | TRUE | 0.131 |
| Pygl | 674 | 647 | 918 | 706 | 727 | 835 | 523 | 688 | 991 | 703 | 820 | 824 | 758.22 | 734 | 782 | 1.067 | 0.093 | 0.7363191 | 0.84267826 | 0.0536 | 0.0056 | 0.019 | 0.0536 | TRUE | 0.4747 |
| Pygm | 257 | 219 | 302 | 534 | 496 | 610 | 200 | 233 | 326 | 532 | 560 | 602 | 408.61 | 253 | 565 | 2.244 | 1.166 | 2.53E-14 | 4.02E-12 | 0.0295 | 0.0082 | 0.014 | 0.014 | TRUE | 0.7038 |
| Pygo1 | 170 | 126 | 126 | 130 | 106 | 140 | 132 | 134 | 136 | 129 | 120 | 138 | 131.53 | 134 | 129 | 0.965 | -0.052 | 0.76227406 | 0.85900698 | 0 | 0.0204 | 0.0134 | 0.0134 | TRUE | 0.0401 |
| Pygo2 | 2447 | 1737 | 1553 | 1772 | 1541 | 1774 | 1900 | 1848 | 1676 | 1764 | 1738 | 1751 | 1779.54 | 1808 | 1751 | 0.968 | -0.047 | 0.5001138 | 0.66457109 | 0.0016 | 0.0038 | 0.003 | 0.003 | TRUE | 0.0488 |
| Pyhin1 | 0 | 0 | 0 | 0 | 0 | 0 | 0 | 0 | 0 | 0 | 0 | 0 | 0 | 0 | 0 | NA | NA | NA | NA | NA | NA | NA | NA | NA | NA |
| Pyroxd1 | 234 | 154 | 130 | 120 | 113 | 125 | 182 | 164 | 140 | 119 | 127 | 123 | 142.69 | 162 | 123 | 0.759 | -0.398 | 0.0216873 | 0.07325247 | 0 | 0.019 | 0.0145 | 0.0145 | TRUE | 0.1416 |
| Pyroxd2 | 11 | 16 | 7 | 15 | 14 | 5 | 9 | 17 | 8 | 15 | 16 | 5 | 11.46 | 11 | 12 | 1.079 | 0.11 | 0.86208452 | NA | 0.172 | 0.2092 | 0.1989 | 0.1989 | TRUE | 1.0622 |
| Pyurf | 615 | 415 | 478 | 383 | 337 | 393 | 478 | 441 | 516 | 381 | 380 | 388 | 430.7 | 478 | 383 | 0.801 | -0.32 | 0.00255171 | 0.01461203 | 0 | 0.0079 | 0.0057 | 0.0057 | TRUE | 0.0516 |
| Pyy | 16 | 12 | 7 | 32 | 15 | 21 | 12 | 13 | 8 | 32 | 17 | 21 | 17.04 | 11 | 23 | 2.115 | 1.08 | 0.02999627 | 0.09261991 | 0 | 0.1415 | 0.1125 | 0.1125 | TRUE | 0.6119 |
| Pzp | 183 | 107 | 126 | 64 | 59 | 79 | 142 | 114 | 136 | 64 | 67 | 78 | 100.02 | 131 | 70 | 0.531 | -0.914 | 9.96E-06 | 0.00018858 | 0 | 0.0261 | 0.0197 | 0.0197 | TRUE | 0.1265 |
| Qars | 10616 | 7806 | 8558 | 8802 | 8126 | 9509 | 8245 | 8303 | 9234 | 8762 | 9167 | 9387 | 8849.58 | 8594 | 9105 | 1.06 | 0.084 | 0.17523698 | 0.32873626 | 0.0025 | 0.0027 | 0.0026 | 0.0026 | TRUE | 0.0516 |
| Qdpr | 7280 | 5391 | 5984 | 7380 | 6185 | 7402 | 5654 | 5734 | 6457 | 7346 | 6977 | 7307 | 6579.23 | 5948 | 7210 | 1.213 | 0.278 | 1.59E-05 | 0.00027848 | 0.0029 | 0.0028 | 0.0028 | 0.0028 | TRUE | 0.0678 |
| Qk | 13452 | 10308 | 10440 | 10945 | 9538 | 10561 | 10448 | 10964 | 11265 | 10895 | 10760 | 10425 | 10792.76 | 10892 | 10693 | 0.982 | -0.026 | 0.61265855 | 0.75508227 | 8.00E-04 | 0.0026 | 0.0019 | 0.0019 | TRUE | 0.0157 |
| Qpct | 340 | 304 | 339 | 402 | 383 | 408 | 264 | 323 | 366 | 400 | 432 | 403 | 364.7 | 318 | 412 | 1.301 | 0.379 | 0.00390078 | 0.02023731 | 0.0113 | 0.0089 | 0.0097 | 0.0097 | TRUE | 0.2667 |
| Qpctl | 393 | 296 | 273 | 259 | 217 | 241 | 305 | 315 | 295 | 258 | 245 | 238 | 275.86 | 305 | 247 | 0.81 | -0.305 | 0.01284606 | 0.04977194 | 0 | 0.011 | 0.0071 | 0.0071 | TRUE | 0.0172 |
| Qprt | 737 | 533 | 609 | 792 | 686 | 839 | 572 | 567 | 657 | 788 | 774 | 828 | 697.82 | 599 | 797 | 1.333 | 0.414 | 7.36E-06 | 0.00014619 | 0.0023 | 0.0058 | 0.0047 | 0.0047 | TRUE | 0.0826 |
| Qrfp | 1 | 2 | 0 | 2 | 2 | 0 | 1 | 2 | 0 | 2 | 2 | 0 | 1.19 | 1 | 1 | 1.458 | 0.544 | 0.76960948 | NA | 0 | 1.991 | 1.6195 | 1.6195 | TRUE | 0.5196 |
| Qrfpr | 3 | 1 | 2 | 1 | 0 | 0 | 2 | 1 | 2 | 1 | 0 | 0 | 1.09 | 2 | 0 | 0.199 | -2.329 | 0.23745412 | NA | 0 | 2.1745 | 1.54 | 1.54 | TRUE | 0.3906 |
| Qrich1 | 7340 | 5084 | 4687 | 4900 | 4226 | 4934 | 5701 | 5408 | 5057 | 4877 | 4767 | 4871 | 5113.52 | 5389 | 4838 | 0.898 | -0.156 | 0.00942783 | 0.03969129 | 0.0016 | 0.0029 | 0.0024 | 0.0024 | TRUE | 0.0352 |
| Qrich2 | 37 | 37 | 38 | 41 | 35 | 51 | 29 | 39 | 41 | 41 | 39 | 50 | 39.96 | 36 | 43 | 1.209 | 0.274 | 0.38268306 | 0.55658084 | 0 | 0.0617 | 0.0458 | 0.0458 | TRUE | 0.2937 |
| Qrsl1 | 1461 | 1093 | 1145 | 1223 | 1161 | 1224 | 1135 | 1163 | 1235 | 1217 | 1310 | 1208 | 1211.36 | 1178 | 1245 | 1.058 | 0.081 | 0.27991201 | 0.45144608 | 0.0011 | 0.0044 | 0.0032 | 0.0032 | TRUE | 0.0243 |
| Qser1 | 5271 | 3927 | 4094 | 4834 | 4380 | 5177 | 4094 | 4177 | 4417 | 4812 | 4941 | 5110 | 4591.93 | 4229 | 4954 | 1.172 | 0.229 | 6.16E-05 | 0.00083113 | 9.00E-04 | 0.0029 | 0.0021 | 0.0021 | TRUE | 0.0183 |
| Qsox1 | 1288 | 896 | 872 | 1137 | 998 | 1100 | 1000 | 953 | 941 | 1132 | 1126 | 1086 | 1039.63 | 965 | 1115 | 1.154 | 0.207 | 0.00529627 | 0.02560015 | 0 | 0.0047 | 0.003 | 0.003 | TRUE | 0.0131 |
| Qsox2 | 1638 | 1219 | 1292 | 1537 | 1377 | 1507 | 1272 | 1297 | 1394 | 1530 | 1553 | 1488 | 1422.3 | 1321 | 1524 | 1.154 | 0.207 | 0.00318545 | 0.01729847 | 7.00E-04 | 0.0041 | 0.0028 | 0.0028 | TRUE | 0.0277 |
| Qtrt1 | 1309 | 1031 | 1061 | 1053 | 950 | 1135 | 1017 | 1097 | 1145 | 1048 | 1072 | 1120 | 1083.06 | 1086 | 1080 | 0.996 | -0.006 | 0.94089052 | 0.96681409 | 0.0016 | 0.0046 | 0.0035 | 0.0035 | TRUE | 0.0393 |
| Qtrtd1 | 2389 | 1746 | 1450 | 1579 | 1353 | 1502 | 1855 | 1857 | 1565 | 1572 | 1526 | 1483 | 1642.99 | 1759 | 1527 | 0.867 | -0.205 | 0.0114779 | 0.04576111 | 0.0046 | 0.0039 | 0.0041 | 0.0041 | TRUE | 0.1113 |
| R3hcc1 | 1686 | 1161 | 881 | 926 | 792 | 875 | 1309 | 1235 | 951 | 922 | 893 | 864 | 1028.99 | 1165 | 893 | 0.766 | -0.386 | 0.00043804 | 0.003896 | 0.0138 | 0.0047 | 0.0077 | 0.0077 | TRUE | 0.3072 |
| R3hcc1l | 1393 | 1030 | 1025 | 1051 | 948 | 1131 | 1082 | 1096 | 1106 | 1046 | 1069 | 1116 | 1085.92 | 1095 | 1077 | 0.985 | -0.022 | 0.75944802 | 0.8571376 | 0 | 0.0046 | 0.0029 | 0.0029 | TRUE | 0.0123 |

| | | | | | | | | | | | | | | | | | | | | | | | | | |
|---|---|---|---|---|---|---|---|---|---|---|---|---|---|---|---|---|---|---|---|---|---|---|---|---|---|
| R3hdm1 | 6345 | 5009 | 5538 | 6253 | 5575 | 6661 | 4928 | 5328 | 5975 | 6224 | 6289 | 6575 | 5886.7 | 5410 | 6363 | 1.177 | 0.235 | 0.00125404 | 0.00849884 | 0.0049 | 0.0028 | 0.0036 | 0.0036 | TRUE | 0.1016 |
| R3hdm2 | 5615 | 4225 | 4611 | 4899 | 4410 | 4994 | 4361 | 4494 | 4975 | 4876 | 4975 | 4930 | 4768.58 | 4610 | 4927 | 1.069 | 0.097 | 0.1255039 | 0.26066038 | 0.0022 | 0.0029 | 0.0026 | 0.0026 | TRUE | 0.0582 |
| R3hdm4 | 2027 | 1481 | 1626 | 1776 | 1682 | 2019 | 1574 | 1575 | 1754 | 1768 | 1897 | 1993 | 1760.4 | 1634 | 1886 | 1.154 | 0.207 | 0.00622316 | 0.02900152 | 0.003 | 0.0038 | 0.0036 | 0.0036 | TRUE | 0.0488 |
| R3hdml | 32 | 21 | 12 | 16 | 18 | 26 | 25 | 22 | 13 | 16 | 20 | 26 | 20.34 | 20 | 21 | 1.021 | 0.03 | 0.94716239 | 0.97107433 | 0.0331 | 0.119 | 0.0953 | 0.0953 | TRUE | 0.4556 |
| R74862 | 71 | 37 | 39 | 68 | 62 | 75 | 55 | 39 | 42 | 68 | 70 | 74 | 58.04 | 45 | 71 | 1.536 | 0.62 | 0.01782736 | 0.06344784 | 0 | 0.0433 | 0.0314 | 0.0314 | TRUE | 0.3135 |
| Rab1 | 19035 | 13338 | 13898 | 15541 | 12990 | 15208 | 14784 | 14187 | 14996 | 15469 | 14654 | 15013 | 14850.53 | 14656 | 15045 | 1.027 | 0.038 | 0.44655558 | 0.61723382 | 7.00E-04 | 0.0026 | 0.0017 | 0.0017 | TRUE | 0.0095 |
| Rab10 | 7814 | 5570 | 5490 | 6077 | 5236 | 6019 | 6069 | 5925 | 5924 | 6049 | 5907 | 5942 | 5969.11 | 5973 | 5966 | 0.999 | -0.002 | 0.97117787 | 0.9844713 | 0 | 0.0028 | 0.0015 | 0.0015 | TRUE | 0.0025 |
| Rab11a | 3371 | 2359 | 2607 | 2892 | 2537 | 2809 | 2618 | 2509 | 2813 | 2879 | 2862 | 2773 | 2742.32 | 2647 | 2838 | 1.072 | 0.101 | 0.11450512 | 0.24390786 | 0.0013 | 0.0033 | 0.0026 | 0.0026 | TRUE | 0.0358 |
| Rab11b | 2875 | 2062 | 1968 | 1974 | 1673 | 2098 | 2233 | 2193 | 2123 | 1965 | 1887 | 2071 | 2078.83 | 2183 | 1974 | 0.904 | -0.145 | 0.02502659 | 0.08143685 | 9.00E-04 | 0.0036 | 0.0025 | 0.0025 | TRUE | 0.0222 |
| Rab11fip1 | 4083 | 3183 | 3393 | 3173 | 3112 | 3621 | 3171 | 3386 | 3661 | 3158 | 3511 | 3574 | 3410.23 | 3406 | 3414 | 1.003 | 0.004 | 0.95400343 | 0.97480491 | 0.0045 | 0.0031 | 0.0036 | 0.0036 | TRUE | 0.0526 |
| Rab11fip2 | 693 | 472 | 569 | 559 | 517 | 575 | 538 | 502 | 614 | 556 | 583 | 568 | 560.25 | 551 | 569 | 1.033 | 0.046 | 0.64981799 | 0.78253 | 0.0035 | 0.0067 | 0.0057 | 0.0057 | TRUE | 0.1106 |
| Rab11fip3 | 2443 | 1864 | 1779 | 1852 | 1723 | 1910 | 1897 | 1983 | 1920 | 1843 | 1944 | 1885 | 1912.05 | 1933 | 1891 | 0.978 | -0.032 | 0.6098483 | 0.75322701 | 0 | 0.0037 | 0.0023 | 0.0023 | TRUE | 0.0071 |
| Rab11fip4 | 2526 | 1997 | 1935 | 2247 | 2022 | 2262 | 1962 | 2124 | 2088 | 2237 | 2281 | 2233 | 2154.08 | 2058 | 2250 | 1.094 | 0.13 | 0.03727474 | 0.10886143 | 5.00E-04 | 0.0035 | 0.0023 | 0.0023 | TRUE | 0.021 |
| Rab11fip5 | 1867 | 1427 | 1519 | 1950 | 1909 | 1948 | 1450 | 1518 | 1639 | 1941 | 2154 | 1923 | 1770.74 | 1536 | 2006 | 1.307 | 0.386 | 4.10E-07 | 1.37E-05 | 0.0031 | 0.0038 | 0.0036 | 0.0036 | TRUE | 0.0496 |
| Rab12 | 1924 | 1431 | 1247 | 1168 | 1045 | 1118 | 1494 | 1522 | 1346 | 1163 | 1179 | 1104 | 1301.18 | 1454 | 1149 | 0.789 | -0.342 | 8.47E-06 | 0.00016373 | 0.002 | 0.0042 | 0.0035 | 0.0035 | TRUE | 0.0501 |
| Rab13 | 234 | 140 | 128 | 161 | 114 | 145 | 182 | 149 | 138 | 160 | 129 | 143 | 150.13 | 156 | 144 | 0.918 | -0.123 | 0.48849842 | 0.65436813 | 0.0106 | 0.0182 | 0.0159 | 0.0159 | TRUE | 0.2324 |
| Rab14 | 8051 | 5812 | 6082 | 6736 | 5652 | 6569 | 6253 | 6182 | 6562 | 6705 | 6376 | 6485 | 6427.19 | 6332 | 6522 | 1.03 | 0.043 | 0.41822985 | 0.59082268 | 6.00E-04 | 0.0028 | 0.0019 | 0.0019 | TRUE | 0.0122 |
| Rab15 | 814 | 633 | 593 | 652 | 611 | 641 | 632 | 673 | 640 | 649 | 689 | 633 | 652.73 | 648 | 657 | 1.013 | 0.019 | 0.82923327 | 0.90220226 | 0 | 0.0061 | 0.0041 | 0.0041 | TRUE | 0.0212 |
| Rab17 | 42 | 48 | 33 | 38 | 33 | 35 | 33 | 51 | 36 | 38 | 37 | 35 | 38.15 | 40 | 37 | 0.924 | -0.114 | 0.72401434 | 0.83461061 | 0 | 0.0646 | 0.0486 | 0.0486 | TRUE | 0.447 |
| Rab18 | 6817 | 4948 | 5614 | 5716 | 4979 | 5686 | 5294 | 5263 | 6057 | 5690 | 5617 | 5613 | 5589.09 | 5538 | 5640 | 1.019 | 0.027 | 0.68540901 | 0.80709763 | 0.0025 | 0.0029 | 0.0029 | 0.0029 | TRUE | 0.0815 |
| Rab19 | 19 | 6 | 14 | 6 | 4 | 10 | 15 | 6 | 15 | 6 | 5 | 10 | 9.43 | 12 | 7 | 0.562 | -0.832 | 0.21130114 | NA | 0 | 0.2537 | 0.2033 | 0.2033 | TRUE | 0.6279 |
| Rab1b | 6064 | 4142 | 3725 | 3711 | 3352 | 3737 | 4710 | 4406 | 4019 | 3694 | 3781 | 3689 | 4049.84 | 4378 | 3721 | 0.849 | -0.235 | 0.00048513 | 0.00422033 | 0.0028 | 0.003 | 0.003 | 0.003 | TRUE | 0.0625 |
| Rab20 | 87 | 77 | 65 | 70 | 54 | 68 | 68 | 82 | 70 | 70 | 61 | 67 | 69.55 | 73 | 66 | 0.904 | -0.146 | 0.53243481 | 0.69209228 | 0 | 0.0365 | 0.0248 | 0.0248 | TRUE | 0.0953 |
| Rab21 | 1850 | 1296 | 1152 | 1425 | 1205 | 1282 | 1437 | 1379 | 1243 | 1418 | 1359 | 1266 | 1350.28 | 1353 | 1348 | 0.995 | -0.007 | 0.93204836 | 0.96225044 | 0.0037 | 0.0042 | 0.004 | 0.004 | TRUE | 0.0599 |
| Rab22a | 1169 | 851 | 955 | 933 | 888 | 1037 | 908 | 905 | 1030 | 929 | 1002 | 1024 | 966.28 | 948 | 985 | 1.04 | 0.056 | 0.51133548 | 0.67440417 | 0.003 | 0.0049 | 0.0042 | 0.0042 | TRUE | 0.0678 |
| Rab23 | 2418 | 1692 | 1667 | 1700 | 1529 | 1782 | 1878 | 1800 | 1799 | 1692 | 1725 | 1759 | 1775.43 | 1826 | 1725 | 0.945 | -0.082 | 0.19213814 | 0.35037387 | 0 | 0.0038 | 0.0023 | 0.0023 | TRUE | 0.008 |
| Rab24 | 1450 | 1155 | 1032 | 1163 | 937 | 1095 | 1126 | 1229 | 1114 | 1158 | 1057 | 1081 | 1127.31 | 1156 | 1099 | 0.951 | -0.073 | 0.3542219 | 0.52859612 | 0.0017 | 0.0045 | 0.0035 | 0.0035 | TRUE | 0.0353 |
| Rab25 | 35 | 38 | 53 | 31 | 31 | 56 | 27 | 40 | 57 | 31 | 35 | 55 | 40.98 | 41 | 40 | 0.981 | -0.027 | 0.94027844 | 0.96671811 | 0.0935 | 0.0603 | 0.07 | 0.07 | TRUE | 0.2483 |
| Rab26 | 154 | 117 | 126 | 119 | 119 | 121 | 120 | 124 | 136 | 118 | 134 | 119 | 125.36 | 127 | 124 | 0.98 | -0.029 | 0.86947662 | 0.92797411 | 0 | 0.0213 | 0.0145 | 0.0145 | TRUE | 0.0501 |
| Rab27a | 2482 | 1944 | 2052 | 1951 | 1743 | 2084 | 1928 | 2068 | 2214 | 1942 | 1966 | 2057 | 2029.18 | 2070 | 1988 | 0.962 | -0.056 | 0.4284313 | 0.60058538 | 0.0024 | 0.0036 | 0.0032 | 0.0032 | TRUE | 0.045 |
| Rab27b | 6 | 4 | 7 | 1 | 6 | 1 | 5 | 4 | 8 | 1 | 7 | 1 | 4.2 | 6 | 3 | 0.521 | -0.941 | 0.3581916 | NA | 0.2323 | 0.5663 | 0.4857 | 0.4857 | TRUE | 1.5157 |
| Rab28 | 1446 | 1026 | 1119 | 1152 | 972 | 1184 | 1123 | 1091 | 1207 | 1147 | 1097 | 1169 | 1138.96 | 1140 | 1138 | 0.998 | -0.003 | 0.96763366 | 0.98269063 | 9.00E-04 | 0.0045 | 0.0033 | 0.0033 | TRUE | 0.0304 |
| Rab2a | 5472 | 3840 | 4115 | 4326 | 3832 | 4228 | 4250 | 4085 | 4440 | 4306 | 4323 | 4174 | 4262.86 | 4258 | 4268 | 1.002 | 0.003 | 0.95554131 | 0.97553943 | 8.00E-04 | 0.003 | 0.0021 | 0.0021 | TRUE | 0.0167 |
| Rab2b | 1111 | 883 | 994 | 1047 | 893 | 963 | 863 | 939 | 1073 | 1042 | 1007 | 951 | 979.14 | 958 | 1000 | 1.045 | 0.064 | 0.49429375 | 0.65952267 | 0.0062 | 0.0048 | 0.0053 | 0.0053 | TRUE | 0.1293 |
| Rab30 | 289 | 202 | 215 | 246 | 220 | 261 | 224 | 215 | 232 | 245 | 248 | 258 | 237 | 224 | 250 | 1.118 | 0.161 | 0.21384416 | 0.37629633 | 0 | 0.0124 | 0.0079 | 0.0079 | TRUE | 0.0124 |
| Rab31 | 2295 | 1677 | 1749 | 1891 | 1687 | 1963 | 1782 | 1784 | 1887 | 1882 | 1903 | 1938 | 1862.76 | 1818 | 1908 | 1.05 | 0.07 | 0.26307244 | 0.43335254 | 0 | 0.0037 | 0.0023 | 0.0023 | TRUE | 0.0132 |
| Rab32 | 124 | 109 | 103 | 81 | 87 | 87 | 96 | 116 | 111 | 81 | 98 | 86 | 98.01 | 108 | 88 | 0.82 | -0.287 | 0.15695716 | 0.30547232 | 0 | 0.0266 | 0.0192 | 0.0192 | TRUE | 0.097 |
| Rab33a | 116 | 92 | 86 | 105 | 92 | 125 | 90 | 98 | 93 | 105 | 104 | 123 | 102.07 | 94 | 111 | 1.184 | 0.244 | 0.21076274 | 0.37273345 | 0 | 0.0256 | 0.0176 | 0.0176 | TRUE | 0.1046 |
| Rab33b | 503 | 316 | 313 | 334 | 300 | 296 | 391 | 336 | 338 | 332 | 338 | 292 | 337.93 | 355 | 321 | 0.902 | -0.149 | 0.22264213 | 0.38689654 | 0.0043 | 0.0094 | 0.0078 | 0.0078 | TRUE | 0.095 |
| Rab34 | 1389 | 996 | 1138 | 1151 | 1051 | 1255 | 1079 | 1059 | 1228 | 1146 | 1186 | 1239 | 1156.05 | 1122 | 1190 | 1.062 | 0.086 | 0.29507063 | 0.4669447 | 0.0028 | 0.0045 | 0.004 | 0.004 | TRUE | 0.0798 |
| Rab35 | 4750 | 3403 | 3131 | 3024 | 2702 | 3229 | 3689 | 3620 | 3378 | 3010 | 3048 | 3188 | 3322.15 | 3562 | 3082 | 0.865 | -0.209 | 0.0005761 | 0.00481254 | 0.0011 | 0.0031 | 0.0024 | 0.0024 | TRUE | 0.0246 |
| Rab36 | 19 | 8 | 6 | 8 | 7 | 7 | 15 | 9 | 6 | 8 | 8 | 7 | 8.75 | 10 | 8 | 0.752 | -0.411 | 0.53314467 | NA | 0 | 0.2732 | 0.1968 | 0.1968 | TRUE | 0.7897 |
| Rab37 | 69 | 49 | 46 | 68 | 57 | 68 | 54 | 52 | 50 | 68 | 64 | 67 | 59.08 | 52 | 66 | 1.28 | 0.357 | 0.14953632 | 0.2950473 | 0 | 0.0425 | 0.027 | 0.027 | TRUE | 0.0104 |
| Rab38 | 117 | 60 | 42 | 45 | 33 | 41 | 91 | 64 | 45 | 45 | 37 | 40 | 53.75 | 67 | 41 | 0.608 | -0.718 | 0.01975557 | 0.06853552 | 0.0531 | 0.0465 | 0.0485 | 0.0485 | TRUE | 0.2165 |
| Rab39 | 209 | 182 | 191 | 205 | 181 | 193 | 162 | 194 | 206 | 204 | 204 | 191 | 193.46 | 187 | 200 | 1.07 | 0.097 | 0.52195187 | 0.68329001 | 0.004 | 0.0147 | 0.0115 | 0.0115 | TRUE | 0.1625 |
| Rab39b | 60 | 39 | 52 | 48 | 39 | 47 | 47 | 41 | 56 | 48 | 44 | 46 | 47.06 | 48 | 46 | 0.96 | -0.058 | 0.83607381 | 0.90647068 | 0 | 0.0528 | 0.0362 | 0.0362 | TRUE | 0.1581 |
| Rab3a | 351 | 287 | 332 | 417 | 342 | 395 | 273 | 305 | 358 | 415 | 386 | 390 | 354.49 | 312 | 397 | 1.277 | 0.353 | 0.005032 | 0.02466476 | 0.0072 | 0.0091 | 0.0085 | 0.0085 | TRUE | 0.1882 |
| Rab3b | 477 | 350 | 328 | 415 | 375 | 388 | 370 | 372 | 354 | 413 | 423 | 383 | 385.97 | 365 | 406 | 1.11 | 0.151 | 0.15834544 | 0.30740152 | 0 | 0.0086 | 0.0057 | 0.0057 | TRUE | 0.0295 |
| Rab3c | 5 | 4 | 2 | 4 | 2 | 4 | 4 | 4 | 2 | 4 | 2 | 4 | 3.41 | 3 | 3 | 0.989 | -0.016 | 0.98775182 | NA | 0 | 0.6967 | 0.4449 | 0.4449 | TRUE | 0.137 |
| Rab3d | 835 | 617 | 680 | 922 | 869 | 976 | 649 | 656 | 734 | 918 | 980 | 963 | 816.68 | 680 | 954 | 1.405 | 0.491 | 1.09E-08 | 5.37E-07 | 0.0014 | 0.0053 | 0.0041 | 0.0041 | TRUE | 0.0561 |
| Rab3gap1 | 4586 | 3342 | 3196 | 3270 | 3001 | 3434 | 3562 | 3555 | 3448 | 3255 | 3385 | 3390 | 3432.56 | 3522 | 3343 | 0.949 | -0.075 | 0.16756945 | 0.31904491 | 0 | 0.0031 | 0.0019 | 0.0019 | TRUE | 0.0066 |
| Rab3gap2 | 6054 | 4436 | 4618 | 4965 | 4736 | 5503 | 4702 | 4719 | 4983 | 4942 | 5343 | 5432 | 5020.06 | 4801 | 5239 | 1.091 | 0.126 | 0.03425248 | 0.10222817 | 0.0015 | 0.0029 | 0.0024 | 0.0024 | TRUE | 0.0301 |
| Rab3il1 | 199 | 163 | 226 | 342 | 319 | 365 | 155 | 173 | 244 | 340 | 360 | 360 | 272.07 | 191 | 353 | 1.865 | 0.899 | 2.01E-08 | 9.31E-07 | 0.0234 | 0.0111 | 0.0145 | 0.0145 | TRUE | 0.6347 |
| Rab3ip | 4735 | 3551 | 3672 | 4002 | 3556 | 4180 | 3677 | 3777 | 3962 | 3984 | 4012 | 4126 | 3923.02 | 3805 | 4041 | 1.062 | 0.087 | 0.12137043 | 0.25410101 | 6.00E-04 | 0.003 | 0.002 | 0.002 | TRUE | 0.0157 |
| Rab40b | 22 | 14 | 17 | 28 | 17 | 17 | 17 | 15 | 18 | 28 | 19 | 17 | 19.03 | 17 | 21 | 1.269 | 0.344 | 0.44076998 | 0.61198726 | 0 | 0.127 | 0.0905 | 0.0905 | TRUE | 0.4224 |
| Rab40c | 2125 | 1511 | 1594 | 1712 | 1577 | 1766 | 1650 | 1607 | 1720 | 1704 | 1779 | 1743 | 1700.67 | 1659 | 1742 | 1.05 | 0.07 | 0.27797872 | 0.44945134 | 0 | 0.0038 | 0.0024 | 0.0024 | TRUE | 0.0119 |
| Rab42 | 10 | 13 | 3 | 18 | 11 | 12 | 8 | 14 | 3 | 18 | 12 | 12 | 11.17 | 8 | 14 | 1.703 | 0.768 | 0.21868218 | NA | 0.0906 | 0.2147 | 0.1857 | 0.1857 | TRUE | 0.4197 |
| Rab43 | 459 | 341 | 219 | 245 | 269 | 210 | 356 | 363 | 236 | 244 | 303 | 207 | 285.02 | 318 | 251 | 0.787 | -0.345 | 0.0549453 | 0.14407141 | 0.042 | 0.0107 | 0.0197 | 0.0197 | TRUE | 0.5647 |
| Rab44 | 62 | 60 | 63 | 61 | 75 | 90 | 48 | 64 | 68 | 61 | 85 | 89 | 69.02 | 60 | 78 | 1.309 | 0.389 | 0.13080495 | 0.26792708 | 0.0233 | 0.0368 | 0.0329 | 0.0329 | TRUE | 0.3564 |
| Rab4a | 1974 | 1268 | 1283 | 1339 | 1202 | 1473 | 1533 | 1349 | 1384 | 1333 | 1356 | 1454 | 1401.52 | 1422 | 1381 | 0.97 | -0.044 | 0.57222091 | 0.72501271 | 0.0028 | 0.0041 | 0.0036 | 0.0036 | TRUE | 0.0581 |
| Rab4b | 1269 | 975 | 1072 | 1261 | 1077 | 1249 | 986 | 1037 | 1157 | 1255 | 1215 | 1233 | 1147.08 | 1060 | 1234 | 1.167 | 0.222 | 0.00594832 | 0.02804936 | 0.0026 | 0.0045 | 0.0038 | 0.0038 | TRUE | 0.0758 |
| Rab5a | 4892 | 3504 | 3391 | 3419 | 2999 | 3392 | 3799 | 3727 | 3659 | 3403 | 3383 | 3348 | 3553.38 | 3728 | 3378 | 0.906 | -0.143 | 0.00703072 | 0.03184097 | 0 | 0.0031 | 0.0017 | 0.0017 | TRUE | 0.0035 |
| Rab5b | 3231 | 2395 | 2821 | 3090 | 2768 | 2959 | 2509 | 2548 | 3044 | 3076 | 3123 | 2921 | 2870.02 | 2700 | 3040 | 1.126 | 0.172 | 0.03219165 | 0.09752338 | 0.0062 | 0.0033 | 0.0043 | 0.0043 | TRUE | 0.1493 |
| Rab5c | 9064 | 6422 | 6461 | 6704 | 6148 | 7066 | 7040 | 6831 | 6971 | 6673 | 6936 | 6975 | 6904.33 | 6947 | 6861 | 0.987 | -0.018 | 0.71397171 | 0.82815031 | 3.00E-04 | 0.0028 | 0.0016 | 0.0016 | TRUE | 0.0071 |
| Rab6a | 8394 | 5916 | 6393 | 7232 | 5806 | 6614 | 6519 | 6293 | 6898 | 7199 | 6550 | 6529 | 6664.59 | 6570 | 6759 | 1.029 | 0.041 | 0.50991728 | 0.67315683 | 0.0022 | 0.0028 | 0.0027 | 0.0027 | TRUE | 0.0396 |
| Rab6b | 1256 | 983 | 1299 | 1296 | 1181 | 1433 | 975 | 1046 | 1402 | 1290 | 1332 | 1415 | 1243.27 | 1141 | 1346 | 1.181 | 0.24 | 0.03905142 | 0.11257806 | 0.0199 | 0.0043 | 0.0089 | 0.0089 | TRUE | 0.476 |
| Rab7 | 2825 | 1936 | 2021 | 2020 | 1776 | 1997 | 2194 | 2059 | 2181 | 2011 | 2004 | 1971 | 2069.93 | 2145 | 1995 | 0.93 | -0.105 | 0.08823315 | 0.20181091 | 0 | 0.0036 | 0.0022 | 0.0022 | TRUE | 0.0145 |
| Rab7l1 | 345 | 241 | 228 | 258 | 234 | 255 | 268 | 256 | 246 | 257 | 264 | 252 | 257.14 | 257 | 258 | 1.001 | 0.001 | 0.991638 | 0.99565418 | 0 | 0.0116 | 0.0075 | 0.0075 | TRUE | 0.0176 |
| Rab8a | 4810 | 3467 | 2969 | 3090 | 2699 | 3194 | 3736 | 3688 | 3204 | 3076 | 3045 | 3153 | 3316.76 | 3543 | 3091 | 0.872 | -0.197 | 0.00506244 | 0.02476299 | 0.0032 | 0.0031 | 0.0033 | 0.0033 | TRUE | 0.0844 |

| | | | | | | | | | | | | | | | | | | | | | | | | | |
|---|---|---|---|---|---|---|---|---|---|---|---|---|---|---|---|---|---|---|---|---|---|---|---|---|---|
| Rab8b | 1825 | 1361 | 1310 | 1699 | 1560 | 1743 | 1417 | 1448 | 1413 | 1691 | 1760 | 1721 | 1575.03 | 1426 | 1724 | 1.209 | 0.273 | 1.90E-05 | 0.00032065 | 0 | 0.0039 | 0.0023 | 0.0023 | TRUE | 0.0039 |
| Rab9 | 2760 | 1948 | 2422 | 2577 | 2355 | 2746 | 2144 | 2072 | 2613 | 2565 | 2657 | 2711 | 2460.25 | 2276 | 2644 | 1.162 | 0.217 | 0.01194762 | 0.04718632 | 0.0081 | 0.0034 | 0.005 | 0.005 | TRUE | 0.2012 |
| Rab9b | 25 | 17 | 15 | 26 | 18 | 23 | 19 | 18 | 16 | 26 | 20 | 23 | 20.43 | 18 | 23 | 1.281 | 0.357 | 0.39449833 | 0.5680601 | 0 | 0.1184 | 0.0773 | 0.0773 | TRUE | 0.0724 |
| Rabac1 | 3155 | 2312 | 2747 | 2932 | 2660 | 2884 | 2450 | 2459 | 2964 | 2919 | 3001 | 2847 | 2773.3 | 2624 | 2922 | 1.114 | 0.156 | 0.05174934 | 0.13822713 | 0.0061 | 0.0033 | 0.0043 | 0.0043 | TRUE | 0.1541 |
| Rabep1 | 10617 | 7658 | 7181 | 8737 | 7546 | 8557 | 8246 | 8146 | 7748 | 8697 | 8513 | 8447 | 8299.38 | 8047 | 8552 | 1.063 | 0.088 | 0.0829759 | 0.19400859 | 5.00E-04 | 0.0027 | 0.0017 | 0.0017 | TRUE | 0.0128 |
| Rabep2 | 1502 | 1141 | 1093 | 1072 | 997 | 1089 | 1167 | 1214 | 1179 | 1067 | 1125 | 1075 | 1137.72 | 1187 | 1089 | 0.918 | -0.124 | 0.08315183 | 0.19415985 | 0 | 0.0045 | 0.0028 | 0.0028 | TRUE | 0.0097 |
| Rabepk | 1491 | 1111 | 1171 | 1375 | 1100 | 1336 | 1158 | 1182 | 1263 | 1369 | 1241 | 1319 | 1255.28 | 1201 | 1310 | 1.092 | 0.126 | 0.09322782 | 0.2100007 | 0.0014 | 0.0043 | 0.0033 | 0.0033 | TRUE | 0.0247 |
| Rabgap1 | 4538 | 3198 | 3307 | 3685 | 3325 | 3719 | 3524 | 3402 | 3568 | 3668 | 3751 | 3671 | 3597.43 | 3498 | 3697 | 1.057 | 0.079 | 0.14003076 | 0.28142009 | 0 | 0.0031 | 0.0018 | 0.0018 | TRUE | 0.0069 |
| Rabgap1l | 965 | 720 | 825 | 1080 | 914 | 947 | 749 | 766 | 890 | 1075 | 1031 | 935 | 907.74 | 802 | 1014 | 1.266 | 0.34 | 0.00031796 | 0.00303781 | 0.0054 | 0.005 | 0.0053 | 0.0053 | TRUE | 0.1085 |
| Rabgef1 | 1422 | 1032 | 1054 | 1027 | 956 | 1075 | 1104 | 1098 | 1137 | 1022 | 1078 | 1061 | 1083.55 | 1113 | 1054 | 0.947 | -0.079 | 0.27533527 | 0.44638918 | 0 | 0.0046 | 0.0028 | 0.0028 | TRUE | 0.0083 |
| Rabggta | 2642 | 2009 | 2023 | 2297 | 2017 | 2413 | 2052 | 2137 | 2183 | 2286 | 2275 | 2382 | 2219.25 | 2124 | 2314 | 1.091 | 0.125 | 0.04134723 | 0.11704813 | 4.00E-04 | 0.0035 | 0.0023 | 0.0023 | TRUE | 0.011 |
| Rabggtb | 6853 | 4934 | 4813 | 5268 | 4545 | 5166 | 5322 | 5248 | 5193 | 5244 | 5127 | 5100 | 5205.75 | 5254 | 5157 | 0.981 | -0.027 | 0.58274662 | 0.73268802 | 0 | 0.0029 | 0.0016 | 0.0016 | TRUE | 0.0027 |
| Rabif | 2691 | 1907 | 1967 | 2163 | 1882 | 2142 | 2090 | 2028 | 2122 | 2153 | 2123 | 2114 | 2105.24 | 2080 | 2130 | 1.024 | 0.034 | 0.56554603 | 0.7199278 | 0 | 0.0036 | 0.0021 | 0.0021 | TRUE | 0.0056 |
| Rabl2 | 342 | 258 | 229 | 185 | 163 | 220 | 266 | 274 | 247 | 184 | 184 | 217 | 228.72 | 262 | 195 | 0.744 | -0.427 | 0.00216097 | 0.01279482 | 0 | 0.0128 | 0.0095 | 0.0095 | TRUE | 0.1085 |
| Rabl3 | 887 | 631 | 649 | 696 | 572 | 648 | 689 | 671 | 700 | 693 | 645 | 640 | 673.02 | 687 | 659 | 0.96 | -0.059 | 0.49621243 | 0.66061075 | 0 | 0.0059 | 0.0039 | 0.0039 | TRUE | 0.0237 |
| Rabl5 | 318 | 231 | 239 | 231 | 201 | 254 | 247 | 246 | 258 | 230 | 227 | 251 | 243 | 250 | 236 | 0.944 | -0.084 | 0.51787297 | 0.6797764 | 0 | 0.0122 | 0.008 | 0.008 | TRUE | 0.0344 |
| Rabl6 | 4625 | 3543 | 3937 | 4043 | 3784 | 4429 | 3592 | 3769 | 4248 | 4024 | 4269 | 4372 | 4045.65 | 3870 | 4222 | 1.092 | 0.126 | 0.08165085 | 0.19200981 | 0.0044 | 0.003 | 0.0035 | 0.0035 | TRUE | 0.0888 |
| Rac1 | 13712 | 9924 | 9963 | 10254 | 9158 | 10593 | 10650 | 10556 | 10750 | 10207 | 10331 | 10457 | 10491.75 | 10652 | 10332 | 0.97 | -0.044 | 0.33140024 | 0.50569314 | 0 | 0.0027 | 0.0014 | 0.0014 | TRUE | 0.0014 |
| Rac2 | 47 | 30 | 49 | 64 | 82 | 69 | 37 | 32 | 53 | 64 | 93 | 68 | 57.6 | 41 | 75 | 1.853 | 0.89 | 0.00200418 | 0.01212773 | 0.0332 | 0.0436 | 0.0408 | 0.0408 | TRUE | 0.5068 |
| Rac3 | 561 | 393 | 483 | 603 | 576 | 693 | 436 | 418 | 521 | 600 | 650 | 684 | 551.5 | 458 | 645 | 1.409 | 0.494 | 6.26E-06 | 0.00012799 | 0.0066 | 0.0067 | 0.0068 | 0.0068 | TRUE | 0.1615 |
| Racgap1 | 8752 | 6434 | 6752 | 7059 | 6452 | 7285 | 6797 | 6844 | 7285 | 7027 | 7279 | 7191 | 7070.48 | 6975 | 7166 | 1.027 | 0.039 | 0.46078559 | 0.63030035 | 7.00E-04 | 0.0028 | 0.0019 | 0.0019 | TRUE | 0.0183 |
| Rad1 | 2160 | 1572 | 1567 | 1598 | 1496 | 1739 | 1678 | 1672 | 1691 | 1591 | 1688 | 1717 | 1672.57 | 1680 | 1665 | 0.991 | -0.013 | 0.83971668 | 0.90890349 | 0 | 0.0038 | 0.0025 | 0.0025 | TRUE | 0.0186 |
| Rad17 | 3169 | 2162 | 2162 | 2329 | 1974 | 2277 | 2461 | 2300 | 2333 | 2318 | 2227 | 2248 | 2314.44 | 2365 | 2264 | 0.957 | -0.063 | 0.30102197 | 0.47337518 | 5.00E-04 | 0.0035 | 0.0023 | 0.0023 | TRUE | 0.016 |
| Rad18 | 3744 | 2709 | 2831 | 3193 | 2702 | 3148 | 2908 | 2882 | 3055 | 3178 | 3048 | 3108 | 3029.66 | 2948 | 3111 | 1.056 | 0.078 | 0.17190934 | 0.32394063 | 3.00E-04 | 0.0032 | 0.002 | 0.002 | TRUE | 0.0119 |
| Rad21 | 29230 | 20070 | 20434 | 20874 | 18544 | 20148 | 22702 | 21348 | 22048 | 20778 | 20920 | 19889 | 21280.79 | 22033 | 20529 | 0.932 | -0.102 | 0.03971706 | 0.1139454 | 8.00E-04 | 0.0025 | 0.0017 | 0.0017 | TRUE | 0.0091 |
| Rad21l | 13 | 13 | 12 | 12 | 6 | 7 | 10 | 14 | 13 | 12 | 7 | 7 | 10.42 | 12 | 9 | 0.702 | -0.511 | 0.39405031 | NA | 0 | 0.23 | 0.1592 | 0.1592 | TRUE | 0.3848 |
| Rad23a | 7314 | 5416 | 4914 | 5058 | 4579 | 5205 | 5680 | 5761 | 5302 | 5035 | 5166 | 5138 | 5347 | 5581 | 5113 | 0.916 | -0.127 | 0.02183223 | 0.07356156 | 8.00E-04 | 0.0029 | 0.002 | 0.002 | TRUE | 0.0231 |
| Rad23b | 23833 | 16591 | 15827 | 16399 | 14489 | 16809 | 18510 | 17648 | 17077 | 16324 | 16345 | 16593 | 17082.77 | 17745 | 16421 | 0.925 | -0.112 | 0.02551298 | 0.08248293 | 8.00E-04 | 0.0026 | 0.0018 | 0.0018 | TRUE | 0.0175 |
| Rad50 | 14326 | 10950 | 11065 | 12048 | 10723 | 12627 | 11126 | 11647 | 11939 | 11993 | 12097 | 12465 | 11877.81 | 11571 | 12185 | 1.053 | 0.075 | 0.14075637 | 0.28263991 | 7.00E-04 | 0.0026 | 0.0018 | 0.0018 | TRUE | 0.0139 |
| Rad51 | 6926 | 4881 | 4404 | 4757 | 4052 | 4594 | 5379 | 5192 | 4752 | 4735 | 4571 | 4535 | 4860.67 | 5108 | 4614 | 0.903 | -0.147 | 0.01794434 | 0.06375302 | 0.0019 | 0.0029 | 0.0026 | 0.0026 | TRUE | 0.045 |
| Rad51ap1 | 2732 | 2008 | 2137 | 2138 | 1837 | 2105 | 2122 | 2136 | 2306 | 2128 | 2072 | 2078 | 2140.33 | 2188 | 2093 | 0.957 | -0.063 | 0.31925712 | 0.493206 | 7.00E-04 | 0.0035 | 0.0024 | 0.0024 | TRUE | 0.0265 |
| Rad51ap2 | 23 | 16 | 14 | 16 | 16 | 6 | 18 | 17 | 15 | 16 | 18 | 6 | 14.98 | 17 | 13 | 0.791 | -0.337 | 0.52441433 | NA | 0.0489 | 0.1606 | 0.1344 | 0.1344 | TRUE | 1.0315 |
| Rad51b | 522 | 391 | 456 | 592 | 510 | 596 | 405 | 416 | 492 | 589 | 575 | 588 | 511.05 | 438 | 584 | 1.338 | 0.42 | 6.04E-05 | 0.00081718 | 0.0033 | 0.0071 | 0.0059 | 0.0059 | TRUE | 0.1327 |
| Rad51c | 2049 | 1552 | 1799 | 1699 | 1453 | 1588 | 1591 | 1651 | 1941 | 1691 | 1639 | 1568 | 1680.21 | 1728 | 1633 | 0.946 | -0.08 | 0.34162956 | 0.51567339 | 0.0057 | 0.0038 | 0.0046 | 0.0046 | TRUE | 0.1397 |
| Rad51d | 2306 | 1766 | 1984 | 1992 | 1701 | 1986 | 1791 | 1878 | 2141 | 1983 | 1919 | 1960 | 1945.4 | 1937 | 1954 | 1.01 | 0.014 | 0.85202935 | 0.91647868 | 0.004 | 0.0036 | 0.0038 | 0.0038 | TRUE | 0.102 |
| Rad52 | 828 | 599 | 550 | 612 | 509 | 616 | 643 | 637 | 593 | 609 | 574 | 608 | 610.86 | 624 | 597 | 0.956 | -0.065 | 0.46880352 | 0.63772806 | 0 | 0.0063 | 0.0042 | 0.0042 | TRUE | 0.0217 |
| Rad54b | 2610 | 2137 | 2244 | 2575 | 2296 | 2726 | 2027 | 2273 | 2421 | 2563 | 2590 | 2691 | 2427.62 | 2240 | 2615 | 1.168 | 0.224 | 0.00266754 | 0.0150522 | 0.0037 | 0.0034 | 0.0036 | 0.0036 | TRUE | 0.0863 |
| Rad54l | 4077 | 3072 | 2914 | 3260 | 2742 | 3315 | 3166 | 3268 | 3144 | 3245 | 3093 | 3272 | 3198.16 | 3193 | 3203 | 1.004 | 0.005 | 0.92498786 | 0.95844943 | 3.00E-04 | 0.0032 | 0.002 | 0.002 | TRUE | 0.0109 |
| Rad54l2 | 7513 | 5101 | 4671 | 5715 | 5279 | 5685 | 5835 | 5426 | 5040 | 5689 | 5955 | 5612 | 5592.8 | 5434 | 5752 | 1.058 | 0.081 | 0.21383223 | 0.37629633 | 0.0029 | 0.0029 | 0.0029 | 0.0029 | TRUE | 0.0515 |
| Rad9a | 2402 | 1844 | 1787 | 1758 | 1509 | 1859 | 1866 | 1961 | 1928 | 1750 | 1702 | 1835 | 1840.41 | 1918 | 1762 | 0.919 | -0.121 | 0.06147599 | 0.15612585 | 5.00E-04 | 0.0037 | 0.0025 | 0.0025 | TRUE | 0.0158 |
| Rad9b | 577 | 417 | 425 | 476 | 374 | 445 | 448 | 444 | 459 | 474 | 422 | 439 | 447.54 | 450 | 445 | 0.99 | -0.015 | 0.88094659 | 0.93352384 | 0 | 0.0077 | 0.0052 | 0.0052 | TRUE | 0.0373 |
| Radil | 659 | 540 | 548 | 669 | 613 | 742 | 512 | 574 | 591 | 666 | 692 | 732 | 627.9 | 559 | 697 | 1.249 | 0.321 | 0.0008182 | 0.00624654 | 0.0026 | 0.0062 | 0.005 | 0.005 | TRUE | 0.0687 |
| Rae1 | 7611 | 5847 | 5691 | 6301 | 5659 | 6461 | 5911 | 6219 | 6141 | 6272 | 6384 | 6378 | 6217.5 | 6090 | 6345 | 1.042 | 0.059 | 0.23611817 | 0.40256661 | 3.00E-04 | 0.0028 | 0.0016 | 0.0016 | TRUE | 0.0082 |
| Raet1a | 10 | 15 | 13 | 19 | 14 | 10 | 8 | 16 | 14 | 19 | 16 | 10 | 13.72 | 13 | 15 | 1.197 | 0.259 | 0.63458194 | NA | 0 | 0.1752 | 0.14 | 0.14 | TRUE | 0.5929 |
| Raet1b | 0 | 0 | 0 | 0 | 0 | 0 | 0 | 0 | 0 | 0 | 0 | 0 | 0 | 0 | 0 | NA | NA | NA | NA | NA | NA | NA | NA | NA | NA |
| Raet1c | 0 | 1 | 0 | 0 | 0 | 0 | 0 | 1 | 0 | 0 | 0 | 0 | 0.18 | 0 | 0 | 0.552 | -0.857 | 0.8336548 | NA | 0 | 13.3714 | 10 | 10 | TRUE | 0.5216 |
| Raet1d | 18 | 4 | 15 | 11 | 17 | 9 | 14 | 4 | 16 | 11 | 19 | 9 | 12.24 | 11 | 13 | 1.119 | 0.163 | 0.79123732 | NA | 0.176 | 0.1961 | 0.1908 | 0.1908 | TRUE | 1.0661 |
| Raet1e | 12 | 9 | 6 | 13 | 7 | 8 | 9 | 10 | 6 | 13 | 8 | 8 | 9.02 | 8 | 10 | 1.131 | 0.177 | 0.78078573 | NA | 0 | 0.2653 | 0.1809 | 0.1809 | TRUE | 0.3247 |
| Raf1 | 11197 | 8277 | 7668 | 7803 | 6942 | 8056 | 8696 | 8804 | 8274 | 7767 | 7831 | 7953 | 8220.83 | 8591 | 7850 | 0.914 | -0.13 | 0.00961328 | 0.04022256 | 4.00E-04 | 0.0027 | 0.0017 | 0.0017 | TRUE | 0.0127 |
| Rag1 | 0 | 0 | 0 | 0 | 0 | 0 | 0 | 0 | 0 | 0 | 0 | 0 | 0 | 0 | 0 | NA | NA | NA | NA | NA | NA | NA | NA | NA | NA |
| Rag2 | 0 | 0 | 0 | 0 | 0 | 0 | 0 | 0 | 0 | 0 | 0 | 0 | 0 | 0 | 0 | NA | NA | NA | NA | NA | NA | NA | NA | NA | NA |
| Rai1 | 925 | 683 | 545 | 643 | 580 | 595 | 718 | 726 | 588 | 640 | 654 | 587 | 652.44 | 677 | 627 | 0.924 | -0.114 | 0.2754619 | 0.44654382 | 0.0069 | 0.0061 | 0.0063 | 0.0063 | TRUE | 0.1538 |
| Rai14 | 6306 | 4548 | 4750 | 5681 | 5161 | 6060 | 4898 | 4838 | 5125 | 5655 | 5822 | 5982 | 5386.6 | 4954 | 5820 | 1.175 | 0.233 | 1.61E-05 | 0.00028081 | 6.00E-04 | 0.0029 | 0.0019 | 0.0019 | TRUE | 0.0111 |
| Rai2 | 6 | 4 | 2 | 10 | 7 | 8 | 5 | 4 | 2 | 10 | 8 | 8 | 6.14 | 4 | 9 | 2.294 | 1.198 | 0.12796632 | NA | 0 | 0.3886 | 0.2575 | 0.2575 | TRUE | 0.183 |
| Rala | 3553 | 2514 | 2548 | 2654 | 2164 | 2629 | 2759 | 2674 | 2749 | 2642 | 2441 | 2595 | 2643.51 | 2727 | 2559 | 0.938 | -0.092 | 0.12738416 | 0.26318325 | 5.00E-04 | 0.0033 | 0.0022 | 0.0022 | TRUE | 0.0195 |
| Ralb | 2445 | 1863 | 1937 | 2193 | 1942 | 2226 | 1899 | 1982 | 2090 | 2183 | 2191 | 2197 | 2090.28 | 1990 | 2190 | 1.101 | 0.139 | 0.02855076 | 0.08931409 | 7.00E-04 | 0.0036 | 0.0024 | 0.0024 | TRUE | 0.0232 |
| Ralbp1 | 8007 | 6139 | 6474 | 6678 | 6041 | 7022 | 6219 | 6530 | 6985 | 6647 | 6815 | 6932 | 6688 | 6578 | 6798 | 1.034 | 0.048 | 0.41711753 | 0.58969939 | 0.0017 | 0.0028 | 0.0024 | 0.0024 | TRUE | 0.0358 |
| Ralgapa1 | 2556 | 1854 | 1893 | 2221 | 1956 | 2288 | 1985 | 1972 | 2043 | 2211 | 2207 | 2259 | 2112.62 | 2000 | 2226 | 1.113 | 0.154 | 0.00905172 | 0.0384746 | 0 | 0.0035 | 0.002 | 0.002 | TRUE | 0.0041 |
| Ralgapa2 | 2874 | 2131 | 2021 | 2235 | 2004 | 2277 | 2232 | 2267 | 2181 | 2225 | 2261 | 2248 | 2235.44 | 2227 | 2245 | 1.008 | 0.011 | 0.84494788 | 0.91201028 | 0 | 0.0035 | 0.002 | 0.002 | TRUE | 0.0038 |
| Ralgapb | 7757 | 5491 | 5650 | 6557 | 5930 | 6661 | 6025 | 5841 | 6096 | 6527 | 6690 | 6575 | 6292.24 | 5987 | 6597 | 1.102 | 0.14 | 0.00464744 | 0.02317722 | 1.00E-04 | 0.0028 | 0.0016 | 0.0016 | TRUE | 0.0055 |
| Ralgds | 2065 | 1557 | 1673 | 1592 | 1438 | 1634 | 1604 | 1656 | 1805 | 1585 | 1622 | 1613 | 1647.5 | 1688 | 1607 | 0.952 | -0.07 | 0.31581531 | 0.48958223 | 0.0014 | 0.0039 | 0.0029 | 0.0029 | TRUE | 0.0437 |
| Ralgps1 | 789 | 531 | 561 | 629 | 578 | 640 | 613 | 565 | 605 | 626 | 652 | 632 | 615.47 | 594 | 637 | 1.07 | 0.098 | 0.27309502 | 0.44411731 | 0 | 0.0063 | 0.0041 | 0.0041 | TRUE | 0.0216 |
| Ralgps2 | 3743 | 2965 | 3298 | 3567 | 3268 | 3661 | 2907 | 3154 | 3559 | 3551 | 3687 | 3614 | 3411.76 | 3207 | 3617 | 1.129 | 0.175 | 0.02115405 | 0.07207505 | 0.005 | 0.0031 | 0.0038 | 0.0038 | TRUE | 0.1119 |
| Raly | 18687 | 13828 | 13072 | 12552 | 11388 | 13501 | 14513 | 14709 | 14105 | 12494 | 12847 | 13328 | 13665.88 | 14442 | 12890 | 0.892 | -0.164 | 0.00097841 | 0.00712034 | 6.00E-04 | 0.0026 | 0.0017 | 0.0017 | TRUE | 0.0108 |
| Ralyl | 1 | 0 | 0 | 0 | 0 | 0 | 1 | 0 | 0 | 0 | 0 | 0 | 0.13 | 0 | 0 | 0.552 | -0.857 | 0.8336548 | NA | 0 | 18.3121 | 10 | 10 | TRUE | 0.2566 |
| Ramp1 | 10 | 13 | 2 | 8 | 5 | 1 | 8 | 14 | 2 | 8 | 6 | 1 | 6.39 | 8 | 5 | 0.612 | -0.708 | 0.42943591 | NA | 0.5144 | 0.3733 | 0.4107 | 0.4107 | TRUE | 0.1518 |
| Ramp2 | 52 | 40 | 50 | 73 | 61 | 70 | 40 | 43 | 54 | 73 | 69 | 69 | 57.91 | 46 | 70 | 1.547 | 0.629 | 0.01480021 | 0.05526581 | 0 | 0.0434 | 0.0301 | 0.0301 | TRUE | 0.1905 |
| Ramp3 | 1033 | 789 | 799 | 836 | 758 | 875 | 802 | 839 | 862 | 832 | 855 | 864 | 842.44 | 834 | 850 | 1.02 | 0.028 | 0.71955229 | 0.83168102 | 0 | 0.0052 | 0.0033 | 0.0033 | TRUE | 0.0144 |
| Ran | 40535 | 27997 | 27598 | 29666 | 25096 | 29657 | 31482 | 29780 | 29778 | 29530 | 28311 | 29276 | 29692.72 | 30347 | 29039 | 0.957 | -0.064 | 0.19077465 | 0.34886669 | 7.00E-04 | 0.0025 | 0.0017 | 0.0017 | TRUE | 0.0132 |

| | | | | | | | | | | | | | | | | | | | | | | | | | |
|---|---|---|---|---|---|---|---|---|---|---|---|---|---|---|---|---|---|---|---|---|---|---|---|---|---|
| Ranbp1 | 15300 | 10948 | 9622 | 9728 | 8463 | 9905 | 11883 | 11645 | 10382 | 9683 | 9547 | 9778 | 10486.38 | 11303 | 9669 | 0.855 | -0.225 | 0.00024712 | 0.00251036 | 0.0025 | 0.0027 | 0.0026 | 0.0026 | TRUE | 0.062 |
| Ranbp10 | 3796 | 2542 | 2034 | 2124 | 2013 | 2230 | 2948 | 2704 | 2195 | 2114 | 2271 | 2201 | 2405.53 | 2616 | 2195 | 0.839 | -0.254 | 0.00778618 | 0.03429517 | 0.0108 | 0.0034 | 0.0061 | 0.0061 | TRUE | 0.2392 |
| Ranbp17 | 7805 | 5661 | 5476 | 5629 | 5113 | 5881 | 6062 | 6022 | 5909 | 5603 | 5768 | 5805 | 5861.41 | 5998 | 5725 | 0.955 | -0.067 | 0.17462985 | 0.32794434 | 1.00E-04 | 0.0028 | 0.0016 | 0.0016 | TRUE | 0.0043 |
| Ranbp2 | 20669 | 14105 | 12727 | 14973 | 13028 | 14772 | 16053 | 15003 | 13732 | 14904 | 14697 | 14582 | 14828.61 | 14929 | 14728 | 0.986 | -0.02 | 0.75169748 | 0.8526317 | 0.0025 | 0.0026 | 0.0028 | 0.0028 | TRUE | 0.0601 |
| Ranbp3 | 5181 | 3867 | 3897 | 3636 | 3204 | 3736 | 4024 | 4113 | 4205 | 3619 | 3614 | 3688 | 3877.28 | 4114 | 3640 | 0.885 | -0.176 | 0.00085741 | 0.00647038 | 0 | 0.003 | 0.0017 | 0.0017 | TRUE | 0.0046 |
| Ranbp3l | 1 | 0 | 0 | 0 | 1 | 0 | 1 | 0 | 0 | 0 | 1 | 0 | 0.32 | 0 | 0 | 1.075 | 0.105 | 0.97638847 | NA | 0 | 7.4681 | 7.0392 | 7.0392 | TRUE | 0.5075 |
| Ranbp6 | 684 | 520 | 412 | 484 | 426 | 437 | 531 | 553 | 445 | 482 | 481 | 431 | 487.1 | 510 | 465 | 0.91 | -0.136 | 0.2267324 | 0.39168596 | 0.0064 | 0.0073 | 0.007 | 0.007 | TRUE | 0.1405 |
| Ranbp9 | 3244 | 2506 | 2425 | 2496 | 2264 | 2514 | 2519 | 2666 | 2617 | 2485 | 2554 | 2482 | 2553.65 | 2601 | 2507 | 0.964 | -0.052 | 0.36806441 | 0.54268049 | 0 | 0.0034 | 0.0021 | 0.0021 | TRUE | 0.0093 |
| Rangap1 | 19405 | 13877 | 13478 | 14165 | 12805 | 15279 | 15071 | 14761 | 14543 | 14100 | 14445 | 15083 | 14667.07 | 14792 | 14543 | 0.983 | -0.025 | 0.62029833 | 0.76038736 | 6.00E-04 | 0.0026 | 0.0017 | 0.0017 | TRUE | 0.0129 |
| Rangrf | 3019 | 2220 | 2447 | 2451 | 2183 | 2708 | 2345 | 2361 | 2640 | 2440 | 2463 | 2673 | 2487 | 2449 | 2525 | 1.032 | 0.045 | 0.52685163 | 0.68794905 | 0.0027 | 0.0034 | 0.0033 | 0.0033 | TRUE | 0.0561 |
| Rap1a | 2623 | 1884 | 2191 | 2354 | 1962 | 2340 | 2037 | 2004 | 2364 | 2343 | 2213 | 2310 | 2211.95 | 2135 | 2289 | 1.073 | 0.101 | 0.18374507 | 0.3397501 | 0.0041 | 0.0035 | 0.0037 | 0.0037 | TRUE | 0.1052 |
| Rap1b | 9486 | 6561 | 6100 | 6716 | 5690 | 6525 | 7367 | 6979 | 6582 | 6685 | 6419 | 6441 | 6745.54 | 6976 | 6515 | 0.934 | -0.099 | 0.09116644 | 0.20666803 | 0.0016 | 0.0028 | 0.0023 | 0.0023 | TRUE | 0.0298 |
| Rap1gap | 1461 | 1090 | 1095 | 1310 | 1172 | 1345 | 1135 | 1159 | 1181 | 1304 | 1322 | 1328 | 1238.24 | 1158 | 1318 | 1.138 | 0.187 | 0.00630733 | 0.02930778 | 0 | 0.0043 | 0.0026 | 0.0026 | TRUE | 0.0041 |
| Rap1gap2 | 1062 | 848 | 950 | 1561 | 1345 | 1503 | 825 | 902 | 1025 | 1554 | 1517 | 1484 | 1217.78 | 917 | 1518 | 1.658 | 0.73 | 6.23E-17 | 1.31E-14 | 0.005 | 0.0044 | 0.0046 | 0.0046 | TRUE | 0.1252 |
| Rap1gds1 | 5145 | 3731 | 3999 | 4340 | 3605 | 4240 | 3996 | 3969 | 4315 | 4320 | 4067 | 4186 | 4141.96 | 4093 | 4191 | 1.024 | 0.035 | 0.56009344 | 0.71470543 | 0.0012 | 0.003 | 0.0023 | 0.0023 | TRUE | 0.027 |
| Rap2a | 1290 | 861 | 944 | 934 | 795 | 864 | 1002 | 916 | 1019 | 930 | 897 | 853 | 935.96 | 979 | 893 | 0.912 | -0.133 | 0.10476221 | 0.22874524 | 0.0015 | 0.005 | 0.0038 | 0.0038 | TRUE | 0.0367 |
| Rap2b | 1760 | 1214 | 1184 | 1189 | 953 | 1070 | 1367 | 1291 | 1278 | 1184 | 1075 | 1056 | 1208.44 | 1312 | 1105 | 0.842 | -0.248 | 0.00126028 | 0.0085164 | 0.0017 | 0.0044 | 0.0034 | 0.0034 | TRUE | 0.0468 |
| Rap2c | 2171 | 1555 | 1513 | 1564 | 1311 | 1532 | 1686 | 1654 | 1633 | 1557 | 1479 | 1512 | 1586.79 | 1658 | 1516 | 0.914 | -0.129 | 0.04575671 | 0.12627933 | 0 | 0.0039 | 0.0024 | 0.0024 | TRUE | 0.0067 |
| Rapgef1 | 8025 | 5511 | 5737 | 6052 | 5751 | 6564 | 6233 | 5862 | 6190 | 6024 | 6488 | 6480 | 6212.73 | 6095 | 6331 | 1.038 | 0.054 | 0.33780907 | 0.51206968 | 0.0012 | 0.0028 | 0.0022 | 0.0022 | TRUE | 0.0219 |
| Rapgef2 | 1631 | 1152 | 1151 | 1168 | 1040 | 1209 | 1267 | 1225 | 1242 | 1163 | 1173 | 1193 | 1210.56 | 1245 | 1176 | 0.945 | -0.082 | 0.2337839 | 0.39993696 | 0 | 0.0044 | 0.0026 | 0.0026 | TRUE | 0.003 |
| Rapgef3 | 299 | 259 | 303 | 557 | 491 | 566 | 232 | 275 | 327 | 554 | 554 | 559 | 416.95 | 278 | 556 | 2.007 | 1.005 | 1.62E-15 | 2.93E-13 | 0.0105 | 0.0081 | 0.0088 | 0.0088 | TRUE | 0.2627 |
| Rapgef4 | 49 | 37 | 40 | 57 | 41 | 52 | 38 | 39 | 43 | 57 | 46 | 51 | 45.82 | 40 | 51 | 1.285 | 0.362 | 0.20232588 | 0.36372875 | 0 | 0.0542 | 0.0359 | 0.0359 | TRUE | 0.0667 |
| Rapgef5 | 1561 | 1151 | 1232 | 1310 | 1132 | 1258 | 1212 | 1224 | 1329 | 1304 | 1277 | 1242 | 1264.8 | 1255 | 1274 | 1.016 | 0.023 | 0.75583119 | 0.85511255 | 8.00E-04 | 0.0043 | 0.003 | 0.003 | TRUE | 0.0313 |
| Rapgef6 | 5397 | 3637 | 3452 | 4079 | 3648 | 3884 | 4192 | 3869 | 3725 | 4060 | 4115 | 3834 | 3965.77 | 3929 | 4003 | 1.018 | 0.026 | 0.68351318 | 0.80598552 | 0.0023 | 0.003 | 0.0027 | 0.0027 | TRUE | 0.0426 |
| Rapgefl1 | 378 | 274 | 271 | 367 | 329 | 361 | 294 | 291 | 292 | 365 | 371 | 356 | 328.38 | 292 | 364 | 1.245 | 0.316 | 0.00470358 | 0.02340003 | 0 | 0.0096 | 0.0059 | 0.0059 | TRUE | 0.0042 |
| Raph1 | 3451 | 2842 | 2734 | 3322 | 3025 | 3724 | 2680 | 3023 | 2950 | 3307 | 3413 | 3676 | 3174.77 | 2884 | 3465 | 1.202 | 0.266 | 0.00013876 | 0.00159555 | 0.003 | 0.0032 | 0.0032 | 0.0032 | TRUE | 0.0476 |
| Rapsn | 75 | 52 | 45 | 57 | 33 | 43 | 58 | 55 | 49 | 57 | 37 | 42 | 49.75 | 54 | 45 | 0.842 | -0.249 | 0.38206526 | 0.55626553 | 0 | 0.0501 | 0.0382 | 0.0382 | TRUE | 0.3728 |
| Rara | 3263 | 2674 | 2894 | 4096 | 3857 | 4679 | 2534 | 2844 | 3123 | 4077 | 4351 | 4619 | 3591.38 | 2834 | 4349 | 1.536 | 0.619 | 3.22E-14 | 5.01E-12 | 0.0069 | 0.0031 | 0.0045 | 0.0045 | TRUE | 0.1056 |
| Rarb | 32 | 17 | 8 | 12 | 8 | 12 | 25 | 18 | 9 | 12 | 9 | 12 | 14.06 | 17 | 11 | 0.63 | -0.667 | 0.22909174 | NA | 0.0874 | 0.1709 | 0.1465 | 0.1465 | TRUE | 0.1934 |
| Rarg | 8645 | 5641 | 4366 | 7216 | 5398 | 6221 | 6714 | 6000 | 4711 | 7183 | 6090 | 6141 | 6139.79 | 5808 | 6471 | 1.114 | 0.155 | 0.1482258 | 0.29310892 | 0.0183 | 0.0028 | 0.0082 | 0.0082 | TRUE | 0.3327 |
| Rarres1 | 3 | 3 | 1 | 3 | 2 | 2 | 2 | 3 | 1 | 3 | 2 | 2 | 2.3 | 2 | 2 | 1.091 | 0.125 | 0.91853452 | NA | 0 | 1.0316 | 0.6446 | 0.6446 | TRUE | 0.1691 |
| Rarres2 | 1 | 2 | 3 | 2 | 0 | 0 | 1 | 2 | 3 | 2 | 0 | 0 | 1.36 | 2 | 1 | 0.336 | -1.572 | 0.39051482 | NA | 0 | 1.7512 | 1.4102 | 1.4102 | TRUE | 0.7002 |
| Rars | 12915 | 9105 | 8667 | 9302 | 8338 | 10046 | 10031 | 9685 | 9352 | 9259 | 9406 | 9917 | 9608.22 | 9689 | 9527 | 0.983 | -0.024 | 0.65136446 | 0.7835341 | 0.001 | 0.0027 | 0.002 | 0.002 | TRUE | 0.0157 |
| Rars2 | 2425 | 1696 | 1525 | 1634 | 1534 | 1741 | 1883 | 1804 | 1645 | 1626 | 1731 | 1719 | 1734.75 | 1777 | 1692 | 0.951 | -0.073 | 0.31908334 | 0.493086 | 0.0024 | 0.0038 | 0.0033 | 0.0033 | TRUE | 0.0506 |
| Rasa1 | 4222 | 3034 | 2998 | 2949 | 2534 | 2953 | 3279 | 3227 | 3235 | 2935 | 2859 | 2915 | 3075.03 | 3247 | 2903 | 0.894 | -0.162 | 0.00269281 | 0.01515884 | 0 | 0.0032 | 0.0018 | 0.0018 | TRUE | 0.0021 |
| Rasa2 | 1641 | 1292 | 1274 | 1392 | 1186 | 1257 | 1274 | 1374 | 1375 | 1386 | 1338 | 1241 | 1331.3 | 1341 | 1322 | 0.986 | -0.02 | 0.7887711 | 0.87598332 | 0.0018 | 0.0042 | 0.0033 | 0.0033 | TRUE | 0.0347 |
| Rasa3 | 2209 | 1524 | 1540 | 1764 | 1481 | 1734 | 1716 | 1621 | 1662 | 1756 | 1671 | 1712 | 1689.45 | 1666 | 1713 | 1.028 | 0.039 | 0.54314687 | 0.7009121 | 0 | 0.0038 | 0.0024 | 0.0024 | TRUE | 0.0085 |
| Rasa4 | 728 | 670 | 730 | 938 | 918 | 994 | 565 | 713 | 788 | 934 | 1036 | 981 | 836.04 | 689 | 984 | 1.432 | 0.518 | 4.92E-06 | 0.00010457 | 0.0141 | 0.0053 | 0.008 | 0.008 | TRUE | 0.3022 |
| Rasal1 | 653 | 566 | 606 | 643 | 606 | 676 | 507 | 602 | 654 | 640 | 684 | 667 | 625.68 | 588 | 664 | 1.132 | 0.179 | 0.09276709 | 0.20928686 | 0.0073 | 0.0062 | 0.0066 | 0.0066 | TRUE | 0.1784 |
| Rasal2 | 2156 | 1597 | 1585 | 1816 | 1567 | 1909 | 1674 | 1699 | 1710 | 1808 | 1768 | 1884 | 1757.21 | 1694 | 1820 | 1.075 | 0.104 | 0.10183112 | 0.22406646 | 0 | 0.0038 | 0.0023 | 0.0023 | TRUE | 0.0117 |
| Rasal3 | 71 | 57 | 41 | 45 | 55 | 76 | 55 | 61 | 44 | 45 | 62 | 75 | 56.98 | 53 | 61 | 1.135 | 0.183 | 0.51658125 | 0.6788173 | 0.0277 | 0.044 | 0.0395 | 0.0395 | TRUE | 0.2499 |
| Rasd1 | 260 | 178 | 229 | 193 | 203 | 241 | 202 | 189 | 247 | 192 | 229 | 238 | 216.23 | 213 | 220 | 1.034 | 0.048 | 0.75920856 | 0.8571376 | 0.0112 | 0.0134 | 0.0128 | 0.0128 | TRUE | 0.2085 |
| Rasd2 | 575 | 427 | 395 | 448 | 439 | 443 | 447 | 454 | 426 | 446 | 495 | 437 | 450.91 | 442 | 459 | 1.037 | 0.053 | 0.60707681 | 0.75124163 | 0 | 0.0077 | 0.0054 | 0.0054 | TRUE | 0.0522 |
| Rasef | 60 | 41 | 42 | 24 | 33 | 27 | 47 | 44 | 45 | 24 | 37 | 27 | 37.22 | 45 | 29 | 0.644 | -0.635 | 0.05011507 | 0.13499712 | 0 | 0.0661 | 0.0474 | 0.0474 | TRUE | 0.3436 |
| Rasgef1a | 212 | 209 | 181 | 261 | 228 | 261 | 165 | 222 | 195 | 260 | 257 | 258 | 226.15 | 194 | 258 | 1.337 | 0.419 | 0.00434364 | 0.02204674 | 0.0065 | 0.0129 | 0.011 | 0.011 | TRUE | 0.2111 |
| Rasgef1b | 186 | 157 | 134 | 123 | 138 | 156 | 144 | 167 | 145 | 122 | 156 | 154 | 148.03 | 152 | 144 | 0.948 | -0.077 | 0.65499665 | 0.7856785 | 0.0052 | 0.0184 | 0.0147 | 0.0147 | TRUE | 0.1823 |
| Rasgef1c | 170 | 118 | 98 | 88 | 88 | 70 | 132 | 126 | 106 | 88 | 99 | 69 | 103.21 | 121 | 85 | 0.701 | -0.513 | 0.01439724 | 0.05406721 | 0.0111 | 0.0254 | 0.0216 | 0.0216 | TRUE | 0.2657 |
| Rasgrf1 | 16 | 10 | 7 | 9 | 7 | 9 | 12 | 11 | 8 | 9 | 8 | 9 | 9.39 | 10 | 9 | 0.835 | -0.26 | 0.67294105 | NA | 0 | 0.2548 | 0.1672 | 0.1672 | TRUE | 0.1691 |
| Rasgrf2 | 58 | 48 | 39 | 41 | 32 | 29 | 45 | 51 | 42 | 41 | 36 | 29 | 40.62 | 46 | 35 | 0.764 | -0.389 | 0.20628434 | 0.36818458 | 0 | 0.0608 | 0.043 | 0.043 | TRUE | 0.1974 |
| Rasgrp1 | 202 | 126 | 227 | 249 | 243 | 290 | 157 | 134 | 245 | 248 | 274 | 286 | 224.02 | 179 | 269 | 1.512 | 0.597 | 0.00193292 | 0.01177631 | 0.0461 | 0.013 | 0.0221 | 0.0221 | TRUE | 1.0965 |
| Rasgrp2 | 961 | 788 | 908 | 909 | 884 | 1088 | 746 | 838 | 980 | 905 | 997 | 1074 | 923.39 | 855 | 992 | 1.163 | 0.217 | 0.04331302 | 0.12124421 | 0.0115 | 0.005 | 0.0073 | 0.0073 | TRUE | 0.1939 |
| Rasgrp3 | 66 | 57 | 43 | 50 | 53 | 48 | 51 | 61 | 46 | 50 | 60 | 47 | 52.54 | 53 | 52 | 0.99 | -0.014 | 0.95750292 | 0.97670491 | 0 | 0.0475 | 0.0339 | 0.0339 | TRUE | 0.131 |
| Rasgrp4 | 675 | 642 | 606 | 711 | 743 | 715 | 524 | 683 | 654 | 708 | 838 | 706 | 685.45 | 620 | 751 | 1.212 | 0.278 | 0.01684796 | 0.06097947 | 0.0133 | 0.0059 | 0.0082 | 0.0082 | TRUE | 0.2265 |
| Rasip1 | 461 | 325 | 299 | 389 | 347 | 403 | 358 | 346 | 323 | 387 | 391 | 398 | 367.14 | 342 | 392 | 1.144 | 0.195 | 0.07381064 | 0.17820532 | 0 | 0.0089 | 0.0058 | 0.0058 | TRUE | 0.0277 |
| Rasl10a | 141 | 113 | 122 | 122 | 113 | 138 | 110 | 120 | 132 | 121 | 127 | 136 | 124.41 | 121 | 128 | 1.07 | 0.097 | 0.58653596 | 0.73576958 | 0 | 0.0215 | 0.015 | 0.015 | TRUE | 0.0718 |
| Rasl10b | 292 | 178 | 170 | 319 | 237 | 234 | 227 | 189 | 183 | 318 | 267 | 231 | 235.91 | 200 | 272 | 1.356 | 0.439 | 0.00524217 | 0.025416 | 0.0159 | 0.0125 | 0.0135 | 0.0135 | TRUE | 0.2234 |
| Rasl11a | 329 | 214 | 179 | 184 | 158 | 182 | 256 | 228 | 193 | 183 | 178 | 180 | 202.89 | 226 | 180 | 0.797 | -0.327 | 0.03175226 | 0.09651182 | 0.0064 | 0.0141 | 0.0117 | 0.0117 | TRUE | 0.1693 |
| Rasl11b | 217 | 174 | 207 | 187 | 167 | 201 | 169 | 185 | 223 | 186 | 188 | 198 | 191.65 | 192 | 191 | 0.997 | -0.004 | 0.97884467 | 0.98866701 | 0.0065 | 0.0148 | 0.0123 | 0.0123 | TRUE | 0.2118 |
| Rasl12 | 54 | 36 | 53 | 38 | 30 | 37 | 42 | 38 | 57 | 38 | 34 | 37 | 40.94 | 46 | 36 | 0.791 | -0.339 | 0.2743272 | 0.44536104 | 0 | 0.0603 | 0.0443 | 0.0443 | TRUE | 0.3492 |
| Rasl2-9 | 23 | 14 | 13 | 25 | 20 | 27 | 18 | 15 | 14 | 25 | 23 | 27 | 20.15 | 16 | 25 | 1.575 | 0.656 | 0.12276021 | 0.25618509 | 0 | 0.1201 | 0.0784 | 0.0784 | TRUE | 0.0956 |
| Rassf1 | 534 | 416 | 474 | 480 | 427 | 549 | 415 | 442 | 511 | 478 | 482 | 542 | 478.35 | 456 | 501 | 1.1 | 0.137 | 0.22242902 | 0.38676132 | 0.0063 | 0.0074 | 0.007 | 0.007 | TRUE | 0.1284 |
| Rassf10 | 184 | 166 | 131 | 126 | 116 | 143 | 143 | 177 | 141 | 125 | 131 | 141 | 143.04 | 154 | 132 | 0.864 | -0.21 | 0.2245322 | 0.3889515 | 0 | 0.019 | 0.0145 | 0.0145 | TRUE | 0.1736 |
| Rassf2 | 705 | 577 | 569 | 659 | 646 | 723 | 548 | 614 | 614 | 656 | 729 | 714 | 645.61 | 592 | 700 | 1.184 | 0.244 | 0.00968699 | 0.04044352 | 0.0023 | 0.0061 | 0.0048 | 0.0048 | TRUE | 0.0535 |
| Rassf3 | 3290 | 2561 | 2607 | 2865 | 2541 | 2806 | 2555 | 2724 | 2813 | 2852 | 2867 | 2770 | 2763.42 | 2697 | 2830 | 1.05 | 0.07 | 0.25734075 | 0.42738419 | 0.0011 | 0.0033 | 0.0024 | 0.0024 | TRUE | 0.0266 |
| Rassf4 | 567 | 460 | 624 | 623 | 618 | 703 | 440 | 489 | 673 | 620 | 697 | 694 | 602.37 | 534 | 670 | 1.258 | 0.331 | 0.01653886 | 0.06004318 | 0.026 | 0.0064 | 0.012 | 0.012 | TRUE | 0.6006 |
| Rassf5 | 396 | 294 | 307 | 356 | 278 | 327 | 308 | 313 | 331 | 354 | 314 | 323 | 323.72 | 317 | 330 | 1.043 | 0.061 | 0.60306996 | 0.7479763 | 0 | 0.0097 | 0.0067 | 0.0067 | TRUE | 0.0467 |
| Rassf6 | 280 | 223 | 291 | 291 | 264 | 316 | 217 | 237 | 314 | 290 | 298 | 312 | 278.01 | 256 | 300 | 1.175 | 0.232 | 0.11903746 | 0.25057473 | 0.0151 | 0.011 | 0.0124 | 0.0124 | TRUE | 0.4264 |
| Rassf7 | 1176 | 899 | 922 | 972 | 883 | 989 | 913 | 956 | 995 | 968 | 996 | 976 | 967.4 | 955 | 980 | 1.027 | 0.039 | 0.61130638 | 0.75404219 | 0 | 0.0049 | 0.0032 | 0.0032 | TRUE | 0.0182 |
| Rassf8 | 218 | 136 | 160 | 153 | 126 | 123 | 169 | 145 | 173 | 152 | 142 | 121 | 150.41 | 162 | 138 | 0.853 | -0.229 | 0.17901907 | 0.333601 | 0.004 | 0.0182 | 0.0142 | 0.0142 | TRUE | 0.1257 |

| | | | | | | | | | | | | | | | | | | | | | | | | | |
|---|---|---|---|---|---|---|---|---|---|---|---|---|---|---|---|---|---|---|---|---|---|---|---|---|---|
| Rassf9 | 10 | 10 | 5 | 1 | 1 | 3 | 8 | 11 | 5 | 1 | 1 | 3 | 4.81 | 8 | 2 | 0.216 | -2.212 | 0.02677446 | NA | 0 | 0.4948 | 0.3557 | 0.3557 | TRUE | 0.2452 |
| Raver1 | 3186 | 2349 | 2007 | 1977 | 1821 | 2094 | 2474 | 2499 | 2166 | 1968 | 2054 | 2067 | 2204.64 | 2380 | 2030 | 0.853 | -0.23 | 0.00151708 | 0.00974866 | 0.0027 | 0.0035 | 0.0033 | 0.0033 | TRUE | 0.0739 |
| Raver1-fdx1l | 0 | 0 | 0 | 0 | 0 | 1 | 0 | 0 | 0 | 0 | 0 | 1 | 0.16 | 0 | 0 | 2.094 | 1.067 | 0.79379329 | NA | 0 | 14.4078 | 10 | 10 | TRUE | 0.3835 |
| Raver2 | 3399 | 2664 | 2867 | 3084 | 2877 | 3121 | 2640 | 2834 | 3093 | 3070 | 3246 | 3081 | 2993.88 | 2856 | 3132 | 1.097 | 0.134 | 0.05749694 | 0.14912554 | 0.0032 | 0.0032 | 0.0033 | 0.0033 | TRUE | 0.0643 |
| Rax | 50 | 32 | 57 | 42 | 49 | 38 | 39 | 34 | 62 | 42 | 55 | 38 | 44.83 | 45 | 45 | 1.003 | 0.004 | 0.9892131 | 0.99429134 | 0.0469 | 0.0553 | 0.053 | 0.053 | TRUE | 0.7871 |
| Rb1 | 2412 | 1693 | 1967 | 2147 | 1820 | 2125 | 1873 | 1801 | 2122 | 2137 | 2053 | 2098 | 2014.08 | 1932 | 2096 | 1.085 | 0.118 | 0.11289421 | 0.2415926 | 0.0033 | 0.0036 | 0.0035 | 0.0035 | TRUE | 0.088 |
| Rb1cc1 | 3392 | 2500 | 2373 | 2564 | 2086 | 2401 | 2634 | 2659 | 2560 | 2552 | 2353 | 2370 | 2521.61 | 2618 | 2425 | 0.926 | -0.11 | 0.07382707 | 0.17821488 | 8.00E-04 | 0.0034 | 0.0023 | 0.0023 | TRUE | 0.0256 |
| Rbak | 555 | 435 | 357 | 429 | 392 | 464 | 431 | 463 | 385 | 427 | 442 | 458 | 434.37 | 426 | 442 | 1.037 | 0.053 | 0.625162 | 0.76371286 | 0.0022 | 0.0079 | 0.0062 | 0.0062 | TRUE | 0.0778 |
| Rbbp4 | 22266 | 15767 | 15687 | 16423 | 14267 | 16082 | 17293 | 16771 | 16926 | 16347 | 16095 | 15875 | 16551.31 | 16997 | 16106 | 0.948 | -0.078 | 0.0841416 | 0.19561416 | 1.00E-04 | 0.0026 | 0.0014 | 0.0014 | TRUE | 0.0029 |
| Rbbp5 | 6523 | 4748 | 4350 | 4708 | 4294 | 4748 | 5066 | 5050 | 4694 | 4686 | 4844 | 4687 | 4837.93 | 4937 | 4739 | 0.96 | -0.059 | 0.28896824 | 0.4610591 | 8.00E-04 | 0.0029 | 0.0021 | 0.0021 | TRUE | 0.0224 |
| Rbbp6 | 7621 | 5602 | 5032 | 5223 | 4685 | 5262 | 5919 | 5959 | 5429 | 5199 | 5285 | 5194 | 5497.62 | 5769 | 5226 | 0.906 | -0.143 | 0.01203399 | 0.04744572 | 0.0011 | 0.0029 | 0.0022 | 0.0022 | TRUE | 0.032 |
| Rbbp7 | 18523 | 13260 | 14199 | 13748 | 12338 | 14272 | 14386 | 14104 | 15321 | 13685 | 13919 | 14089 | 14250.51 | 14604 | 13898 | 0.952 | -0.071 | 0.16730336 | 0.3187081 | 0.001 | 0.0026 | 0.0019 | 0.0019 | TRUE | 0.0224 |
| Rbbp8 | 3151 | 2116 | 1927 | 2108 | 1678 | 2035 | 2447 | 2251 | 2079 | 2098 | 1893 | 2009 | 2129.56 | 2259 | 2000 | 0.885 | -0.177 | 0.02178099 | 0.07351021 | 0.0039 | 0.0035 | 0.0038 | 0.0038 | TRUE | 0.0657 |
| Rbbp8nl | 7 | 2 | 10 | 2 | 4 | 3 | 5 | 2 | 11 | 2 | 5 | 3 | 4.64 | 6 | 3 | 0.514 | -0.961 | 0.31444127 | NA | 0 | 0.5136 | 0.413 | 0.413 | TRUE | 0.2094 |
| Rbbp9 | 1868 | 1285 | 1473 | 1577 | 1321 | 1450 | 1451 | 1367 | 1589 | 1570 | 1490 | 1431 | 1483.06 | 1469 | 1497 | 1.019 | 0.028 | 0.72325045 | 0.83426832 | 0.0032 | 0.004 | 0.0037 | 0.0037 | TRUE | 0.0602 |
| Rbck1 | 2801 | 2053 | 1934 | 1898 | 1629 | 1931 | 2175 | 2184 | 2087 | 1889 | 1838 | 1906 | 2013.18 | 2149 | 1878 | 0.874 | -0.195 | 0.00144136 | 0.00936424 | 0 | 0.0036 | 0.0022 | 0.0022 | TRUE | 0.0075 |
| Rbfa | 2696 | 1969 | 1894 | 2102 | 1815 | 2203 | 2094 | 2094 | 2044 | 2092 | 2048 | 2175 | 2091.07 | 2077 | 2105 | 1.013 | 0.019 | 0.7532645 | 0.85352858 | 0 | 0.0036 | 0.0022 | 0.0022 | TRUE | 0.0102 |
| Rbfox1 | 7 | 3 | 6 | 3 | 6 | 6 | 5 | 3 | 6 | 3 | 7 | 6 | 5.13 | 5 | 5 | 1.031 | 0.044 | 0.95878009 | NA | 0 | 0.4644 | 0.3185 | 0.3185 | TRUE | 0.303 |
| Rbfox2 | 7220 | 5311 | 5565 | 5702 | 5039 | 5895 | 5607 | 5649 | 6005 | 5676 | 5685 | 5819 | 5740.14 | 5754 | 5727 | 0.996 | -0.006 | 0.90330432 | 0.94647963 | 6.00E-04 | 0.0028 | 0.0019 | 0.0019 | TRUE | 0.0176 |
| Rbfox3 | 129 | 101 | 112 | 151 | 123 | 153 | 100 | 107 | 121 | 150 | 139 | 151 | 128.09 | 109 | 147 | 1.346 | 0.428 | 0.01515563 | 0.05623558 | 0 | 0.0209 | 0.0145 | 0.0145 | TRUE | 0.0799 |
| Rbks | 94 | 48 | 57 | 61 | 74 | 65 | 73 | 51 | 62 | 61 | 83 | 64 | 65.65 | 62 | 69 | 1.113 | 0.155 | 0.54703391 | 0.70407525 | 0.0161 | 0.0385 | 0.0322 | 0.0322 | TRUE | 0.2656 |
| Rbl1 | 2232 | 1673 | 1821 | 2084 | 1719 | 1968 | 1733 | 1780 | 1965 | 2074 | 1939 | 1943 | 1905.71 | 1826 | 1985 | 1.088 | 0.122 | 0.08965966 | 0.20415893 | 0.0022 | 0.0037 | 0.0032 | 0.0032 | TRUE | 0.0529 |
| Rbl2 | 749 | 682 | 725 | 817 | 697 | 869 | 582 | 725 | 782 | 813 | 786 | 858 | 757.8 | 696 | 819 | 1.18 | 0.238 | 0.030334 | 0.09336534 | 0.0113 | 0.0056 | 0.0074 | 0.0074 | TRUE | 0.257 |
| Rbm10 | 4264 | 3153 | 2774 | 2876 | 2562 | 2825 | 3312 | 3354 | 2993 | 2863 | 2890 | 2789 | 3033.38 | 3220 | 2847 | 0.884 | -0.178 | 0.00556511 | 0.0266205 | 0.0015 | 0.0032 | 0.0026 | 0.0026 | TRUE | 0.0453 |
| Rbm11 | 113 | 85 | 87 | 75 | 110 | 98 | 88 | 90 | 94 | 75 | 124 | 97 | 94.59 | 91 | 99 | 1.084 | 0.116 | 0.60550149 | 0.75008085 | 0.0215 | 0.0275 | 0.0258 | 0.0258 | TRUE | 0.2214 |
| Rbm12 | 5605 | 3977 | 3512 | 3714 | 3054 | 3385 | 4353 | 4230 | 3789 | 3697 | 3445 | 3342 | 3809.42 | 4124 | 3495 | 0.847 | -0.24 | 0.0006796 | 0.00544481 | 0.0036 | 0.003 | 0.0033 | 0.0033 | TRUE | 0.0611 |
| Rbm12b1 | 278 | 202 | 157 | 135 | 132 | 162 | 216 | 215 | 169 | 134 | 149 | 160 | 173.9 | 200 | 148 | 0.737 | -0.441 | 0.00743275 | 0.03322999 | 0.0077 | 0.0161 | 0.0136 | 0.0136 | TRUE | 0.1877 |
| Rbm12b2 | 573 | 448 | 352 | 369 | 268 | 315 | 445 | 477 | 380 | 367 | 302 | 311 | 380.32 | 434 | 327 | 0.754 | -0.408 | 0.00129812 | 0.00870671 | 0.0095 | 0.0087 | 0.0089 | 0.0089 | TRUE | 0.1342 |
| Rbm14 | 7979 | 5855 | 5717 | 5948 | 5217 | 6358 | 6197 | 6228 | 6169 | 5921 | 5885 | 6276 | 6112.62 | 6198 | 6027 | 0.973 | -0.04 | 0.43991714 | 0.61129066 | 4.00E-04 | 0.0028 | 0.0018 | 0.0018 | TRUE | 0.016 |
| Rbm15 | 1895 | 1447 | 1254 | 1431 | 1182 | 1390 | 1472 | 1539 | 1353 | 1424 | 1333 | 1372 | 1415.66 | 1455 | 1376 | 0.946 | -0.08 | 0.28685798 | 0.4589126 | 0.0019 | 0.0041 | 0.0033 | 0.0033 | TRUE | 0.0436 |
| Rbm15b | 2377 | 1763 | 1420 | 1315 | 1285 | 1398 | 1846 | 1875 | 1532 | 1309 | 1450 | 1380 | 1565.36 | 1751 | 1380 | 0.787 | -0.345 | 8.92E-05 | 0.00109969 | 0.0062 | 0.0039 | 0.0049 | 0.0049 | TRUE | 0.1425 |
| Rbm17 | 8937 | 6678 | 6224 | 6436 | 5557 | 6483 | 6941 | 7103 | 6716 | 6406 | 6269 | 6400 | 6639.15 | 6920 | 6358 | 0.919 | -0.122 | 0.01509417 | 0.05608031 | 2.00E-04 | 0.0028 | 0.0017 | 0.0017 | TRUE | 0.008 |
| Rbm18 | 2389 | 1687 | 1651 | 1907 | 1597 | 1935 | 1855 | 1794 | 1781 | 1898 | 1802 | 1910 | 1840.21 | 1810 | 1870 | 1.033 | 0.046 | 0.4643064 | 0.63356183 | 0 | 0.0037 | 0.0024 | 0.0024 | TRUE | 0.0121 |
| Rbm19 | 6188 | 4520 | 3657 | 3733 | 3443 | 3953 | 4806 | 4808 | 3946 | 3716 | 3884 | 3902 | 4176.97 | 4520 | 3834 | 0.848 | -0.238 | 0.00260215 | 0.01480007 | 0.006 | 0.003 | 0.0043 | 0.0043 | TRUE | 0.1494 |
| Rbm20 | 287 | 215 | 214 | 348 | 296 | 294 | 223 | 229 | 231 | 346 | 334 | 290 | 275.51 | 228 | 323 | 1.422 | 0.508 | 7.42E-05 | 0.00095852 | 0 | 0.011 | 0.0081 | 0.0081 | TRUE | 0.094 |
| Rbm22 | 3602 | 2615 | 2810 | 2667 | 2424 | 2876 | 2798 | 2782 | 3032 | 2655 | 2735 | 2839 | 2806.56 | 2871 | 2743 | 0.956 | -0.065 | 0.30312994 | 0.47569567 | 0.0012 | 0.0033 | 0.0025 | 0.0025 | TRUE | 0.029 |
| Rbm24 | 65 | 45 | 51 | 57 | 57 | 68 | 50 | 48 | 55 | 57 | 64 | 67 | 56.92 | 51 | 63 | 1.227 | 0.295 | 0.2469411 | 0.41568896 | 0 | 0.0441 | 0.0292 | 0.0292 | TRUE | 0.0625 |
| Rbm25 | 13825 | 10455 | 8324 | 8384 | 7417 | 9010 | 10737 | 11121 | 8982 | 8345 | 8367 | 8894 | 9407.75 | 10280 | 8535 | 0.83 | -0.268 | 0.00055349 | 0.00465625 | 0.0067 | 0.0027 | 0.0042 | 0.0042 | TRUE | 0.1487 |
| Rbm26 | 7004 | 5268 | 4952 | 5110 | 4381 | 5199 | 5440 | 5604 | 5343 | 5086 | 4942 | 5132 | 5257.88 | 5462 | 5053 | 0.925 | -0.112 | 0.03009578 | 0.09288684 | 2.00E-04 | 0.0029 | 0.0017 | 0.0017 | TRUE | 0.0062 |
| Rbm27 | 6476 | 4688 | 4007 | 4215 | 3551 | 4139 | 5030 | 4987 | 4324 | 4196 | 4006 | 4086 | 4437.84 | 4780 | 4096 | 0.857 | -0.223 | 0.00123247 | 0.00841248 | 0.0034 | 0.003 | 0.0032 | 0.0032 | TRUE | 0.0844 |
| Rbm28 | 8888 | 6497 | 5698 | 6145 | 5582 | 6363 | 6903 | 6911 | 6148 | 6117 | 6297 | 6281 | 6442.81 | 6654 | 6232 | 0.936 | -0.095 | 0.1212125 | 0.25384766 | 0.0021 | 0.0028 | 0.0025 | 0.0025 | TRUE | 0.0536 |
| Rbm3 | 8540 | 6016 | 5292 | 5030 | 4283 | 4826 | 6633 | 6399 | 5710 | 5007 | 4832 | 4764 | 5557.39 | 6247 | 4868 | 0.779 | -0.36 | 5.67E-08 | 2.34E-06 | 0.003 | 0.0029 | 0.003 | 0.003 | TRUE | 0.0688 |
| Rbm31y | 0 | 0 | 0 | 0 | 0 | 0 | 0 | 0 | 0 | 0 | 0 | 0 | 0 | 0 | 0 | NA | NA | NA | NA | NA | NA | NA | NA | NA | NA |
| Rbm33 | 7121 | 5317 | 5428 | 5325 | 4996 | 6071 | 5531 | 5656 | 5857 | 5300 | 5636 | 5993 | 5662.08 | 5681 | 5643 | 0.994 | -0.009 | 0.87906092 | 0.93262644 | 0.0021 | 0.0028 | 0.0026 | 0.0026 | TRUE | 0.0361 |
| Rbm34 | 4026 | 2846 | 2585 | 2630 | 2392 | 2561 | 3127 | 3027 | 2789 | 2618 | 2698 | 2528 | 2797.95 | 2981 | 2615 | 0.876 | -0.19 | 0.00374259 | 0.01959917 | 0.0017 | 0.0033 | 0.0027 | 0.0027 | TRUE | 0.0382 |
| Rbm38 | 5617 | 4272 | 3908 | 3839 | 3404 | 3997 | 4362 | 4544 | 4217 | 3821 | 3840 | 3946 | 4121.72 | 4374 | 3869 | 0.885 | -0.177 | 0.00144987 | 0.00940247 | 5.00E-04 | 0.003 | 0.002 | 0.002 | TRUE | 0.0138 |
| Rbm39 | 17136 | 13295 | 12372 | 13659 | 11179 | 13819 | 13309 | 14142 | 13349 | 13596 | 12611 | 13641 | 13441.42 | 13600 | 13283 | 0.977 | -0.034 | 0.53925047 | 0.6975414 | 0.0013 | 0.0026 | 0.0021 | 0.0021 | TRUE | 0.0239 |
| Rbm4 | 898 | 656 | 617 | 615 | 493 | 589 | 697 | 698 | 666 | 612 | 556 | 581 | 635.12 | 687 | 583 | 0.849 | -0.236 | 0.00795765 | 0.03492131 | 0 | 0.0062 | 0.0041 | 0.0041 | TRUE | 0.0221 |
| Rbm41 | 1228 | 953 | 878 | 954 | 818 | 876 | 954 | 1014 | 947 | 950 | 923 | 865 | 941.99 | 972 | 913 | 0.939 | -0.09 | 0.25610544 | 0.42612273 | 7.00E-04 | 0.0049 | 0.0035 | 0.0035 | TRUE | 0.0253 |
| Rbm42 | 3836 | 2764 | 2346 | 2400 | 2129 | 2508 | 2979 | 2940 | 2531 | 2389 | 2402 | 2476 | 2619.51 | 2817 | 2422 | 0.859 | -0.219 | 0.00301996 | 0.01662721 | 0.0035 | 0.0033 | 0.0035 | 0.0035 | TRUE | 0.0945 |
| Rbm43 | 479 | 361 | 326 | 251 | 220 | 250 | 372 | 384 | 352 | 250 | 248 | 247 | 308.76 | 369 | 248 | 0.672 | -0.573 | 9.88E-07 | 2.91E-05 | 0 | 0.0101 | 0.0065 | 0.0065 | TRUE | 0.0186 |
| Rbm44 | 1226 | 902 | 1019 | 1121 | 866 | 960 | 952 | 959 | 1099 | 1116 | 977 | 948 | 1008.59 | 1003 | 1014 | 1.011 | 0.016 | 0.86715335 | 0.92677197 | 0.0062 | 0.0048 | 0.0052 | 0.0052 | TRUE | 0.0941 |
| Rbm45 | 1869 | 1438 | 1188 | 1162 | 1096 | 1263 | 1452 | 1530 | 1282 | 1157 | 1236 | 1247 | 1317.14 | 1421 | 1213 | 0.853 | -0.229 | 0.0058366 | 0.02764164 | 0.0041 | 0.0042 | 0.0042 | 0.0042 | TRUE | 0.0861 |
| Rbm46 | 39 | 29 | 34 | 39 | 26 | 24 | 30 | 31 | 37 | 39 | 29 | 24 | 31.61 | 33 | 31 | 0.942 | -0.087 | 0.80599946 | 0.88794129 | 0 | 0.0774 | 0.0577 | 0.0577 | TRUE | 0.381 |
| Rbm47 | 1812 | 1379 | 1449 | 1484 | 1333 | 1550 | 1407 | 1467 | 1563 | 1477 | 1504 | 1530 | 1491.44 | 1479 | 1504 | 1.018 | 0.025 | 0.72056772 | 0.83222698 | 0.001 | 0.004 | 0.0029 | 0.0029 | TRUE | 0.0297 |
| Rbm48 | 811 | 626 | 599 | 607 | 521 | 606 | 630 | 666 | 646 | 604 | 588 | 598 | 622.04 | 647 | 597 | 0.923 | -0.116 | 0.18255831 | 0.33812439 | 0 | 0.0062 | 0.0039 | 0.0039 | TRUE | 0.0074 |
| Rbm4b | 292 | 239 | 226 | 285 | 212 | 252 | 227 | 254 | 244 | 284 | 239 | 249 | 249.41 | 242 | 257 | 1.068 | 0.094 | 0.48215555 | 0.64866291 | 0 | 0.0119 | 0.009 | 0.009 | TRUE | 0.0911 |
| Rbm5 | 5862 | 4274 | 3808 | 4155 | 3845 | 4080 | 4553 | 4546 | 4109 | 4136 | 4338 | 4028 | 4284.8 | 4403 | 4167 | 0.946 | -0.08 | 0.20647109 | 0.36830772 | 0.002 | 0.003 | 0.0027 | 0.0027 | TRUE | 0.0411 |
| Rbm6 | 5316 | 3802 | 3526 | 3761 | 3296 | 3732 | 4129 | 4044 | 3805 | 3744 | 3718 | 3684 | 3853.89 | 3993 | 3715 | 0.93 | -0.104 | 0.06525065 | 0.16310554 | 7.00E-04 | 0.003 | 0.002 | 0.002 | TRUE | 0.0205 |
| Rbm7 | 3297 | 2382 | 2569 | 2607 | 2329 | 2600 | 2561 | 2534 | 2772 | 2595 | 2627 | 2567 | 2609.21 | 2622 | 2596 | 0.99 | -0.014 | 0.82244774 | 0.89813272 | 9.00E-04 | 0.0033 | 0.0024 | 0.0024 | TRUE | 0.0298 |
| Rbm8a | 11871 | 8536 | 7606 | 7715 | 6796 | 7713 | 9220 | 9080 | 8207 | 7680 | 7667 | 7614 | 8244.36 | 8836 | 7654 | 0.866 | -0.208 | 0.00040705 | 0.00369392 | 0.0018 | 0.0027 | 0.0024 | 0.0024 | TRUE | 0.0471 |
| Rbms1 | 1780 | 1174 | 1359 | 1325 | 1122 | 1284 | 1382 | 1249 | 1466 | 1319 | 1266 | 1268 | 1324.95 | 1366 | 1284 | 0.94 | -0.089 | 0.2565262 | 0.42653739 | 0.0027 | 0.0042 | 0.0037 | 0.0037 | TRUE | 0.0657 |
| Rbms2 | 4257 | 2919 | 3038 | 2962 | 2389 | 3039 | 3306 | 3105 | 3278 | 2948 | 2695 | 3000 | 3055.41 | 3230 | 2881 | 0.892 | -0.165 | 0.01064538 | 0.04320114 | 0.0017 | 0.0032 | 0.0027 | 0.0027 | TRUE | 0.0383 |
| Rbms3 | 7 | 10 | 4 | 6 | 2 | 2 | 5 | 11 | 4 | 6 | 2 | 2 | 5.1 | 7 | 3 | 0.506 | -0.983 | 0.27510955 | NA | 0.0954 | 0.4673 | 0.3616 | 0.3616 | TRUE | 0.653 |
| Rbmx | 401 | 390 | 324 | 272 | 226 | 259 | 311 | 415 | 350 | 271 | 255 | 256 | 309.54 | 359 | 261 | 0.729 | -0.457 | 0.00069796 | 0.00556693 | 0.0091 | 0.0101 | 0.0097 | 0.0097 | TRUE | 0.2144 |
| Rbmx2 | 1540 | 1142 | 1121 | 1211 | 1062 | 1164 | 1196 | 1215 | 1210 | 1205 | 1198 | 1149 | 1195.48 | 1207 | 1184 | 0.981 | -0.027 | 0.69583331 | 0.81480793 | 0 | 0.0044 | 0.0026 | 0.0026 | TRUE | 0.0082 |
| Rbmxl1 | 6340 | 4587 | 3879 | 3530 | 3247 | 3785 | 4924 | 4879 | 4185 | 3514 | 3663 | 3736 | 4150.27 | 4663 | 3638 | 0.78 | -0.359 | 6.96E-07 | 2.17E-05 | 0.0041 | 0.003 | 0.0035 | 0.0035 | TRUE | 0.0971 |
| Rbmxl2 | 6182 | 4829 | 4399 | 4672 | 4376 | 4988 | 4801 | 5137 | 4746 | 4651 | 4937 | 4924 | 4865.89 | 4895 | 4837 | 0.988 | -0.017 | 0.76932368 | 0.86344275 | 0.0013 | 0.0029 | 0.0022 | 0.0022 | TRUE | 0.0225 |
| Rbmy | 0 | 0 | 0 | 0 | 0 | 0 | 0 | 0 | 0 | 0 | 0 | 0 | 0 | 0 | 0 | NA | NA | NA | NA | NA | NA | NA | NA | NA | NA |

| | | | | | | | | | | | | | | | | | | | | | | | | | |
|---|---|---|---|---|---|---|---|---|---|---|---|---|---|---|---|---|---|---|---|---|---|---|---|---|---|
| Rbp1 | 375 | 299 | 267 | 404 | 410 | 420 | 291 | 318 | 288 | 402 | 463 | 415 | 362.77 | 299 | 427 | 1.425 | 0.511 | 8.57E-06 | 0.00016506 | 0.0014 | 0.009 | 0.0067 | 0.0067 | TRUE | 0.0617 |
| Rbp2 | 2 | 3 | 0 | 0 | 1 | 0 | 2 | 3 | 0 | 0 | 1 | 0 | 0.98 | 2 | 0 | 0.234 | -2.094 | 0.34068661 | NA | 0 | 2.424 | 2.1736 | 2.1736 | TRUE | 0.2185 |
| Rbp3 | 4 | 2 | 0 | 0 | 0 | 2 | 3 | 2 | 0 | 0 | 0 | 2 | 1.2 | 2 | 1 | 0.378 | -1.403 | 0.48856686 | NA | 0 | 1.9752 | 1.9148 | 1.9148 | TRUE | 0.7374 |
| Rbp4 | 394 | 303 | 275 | 373 | 289 | 307 | 306 | 322 | 297 | 371 | 326 | 303 | 320.9 | 308 | 333 | 1.082 | 0.113 | 0.35693817 | 0.53142889 | 0.0033 | 0.0098 | 0.0078 | 0.0078 | TRUE | 0.1144 |
| Rbp7 | 57 | 32 | 41 | 75 | 45 | 51 | 44 | 34 | 44 | 75 | 51 | 50 | 49.72 | 41 | 59 | 1.432 | 0.518 | 0.07754065 | 0.18483799 | 0.0201 | 0.0501 | 0.0415 | 0.0415 | TRUE | 0.4995 |
| Rbpj | 7581 | 5256 | 5262 | 5588 | 4493 | 5246 | 5888 | 5591 | 5678 | 5562 | 5069 | 5179 | 5494.28 | 5719 | 5270 | 0.921 | -0.118 | 0.04113756 | 0.11666252 | 0.0013 | 0.0029 | 0.0022 | 0.0022 | TRUE | 0.0288 |
| Rbpjl | 2 | 1 | 0 | 0 | 0 | 0 | 2 | 1 | 0 | 0 | 0 | 0 | 0.44 | 1 | 0 | 0.215 | -2.217 | 0.47268385 | NA | 0 | 5.4363 | 5.3181 | 5.3181 | TRUE | 0.2411 |
| Rbpms | 9143 | 7167 | 7554 | 7244 | 6829 | 7846 | 7101 | 7623 | 8151 | 7211 | 7704 | 7745 | 7589.14 | 7625 | 7553 | 0.991 | -0.013 | 0.83695656 | 0.90706168 | 0.003 | 0.0027 | 0.0029 | 0.0029 | TRUE | 0.0446 |
| Rbpms2 | 9373 | 6810 | 7102 | 7271 | 6421 | 7768 | 7280 | 7244 | 7663 | 7238 | 7244 | 7668 | 7389.27 | 7396 | 7383 | 0.999 | -0.002 | 0.96785408 | 0.98270717 | 9.00E-04 | 0.0027 | 0.0019 | 0.0019 | TRUE | 0.014 |
| Rbx1 | 4188 | 2935 | 2800 | 3332 | 2918 | 3292 | 3253 | 3122 | 3021 | 3317 | 3292 | 3250 | 3208.99 | 3132 | 3286 | 1.049 | 0.069 | 0.2295655 | 0.39508485 | 5.00E-04 | 0.0032 | 0.002 | 0.002 | TRUE | 0.0142 |
| Rc3h1 | 2837 | 1953 | 1831 | 2181 | 1842 | 2129 | 2203 | 2077 | 1976 | 2171 | 2078 | 2102 | 2101.16 | 2085 | 2117 | 1.014 | 0.02 | 0.75751573 | 0.85595969 | 0.0013 | 0.0036 | 0.0027 | 0.0027 | TRUE | 0.0305 |
| Rc3h2 | 3570 | 2331 | 2007 | 2218 | 1705 | 1908 | 2773 | 2479 | 2166 | 2208 | 1923 | 1883 | 2238.73 | 2473 | 2005 | 0.81 | -0.303 | 0.00136169 | 0.00899799 | 0.0108 | 0.0035 | 0.006 | 0.006 | TRUE | 0.1426 |
| Rcan1 | 2163 | 1585 | 1450 | 1267 | 1125 | 1184 | 1680 | 1686 | 1565 | 1261 | 1269 | 1169 | 1438.24 | 1644 | 1233 | 0.75 | -0.416 | 7.81E-09 | 4.04E-07 | 0.0012 | 0.0041 | 0.003 | 0.003 | TRUE | 0.0252 |
| Rcan2 | 137 | 85 | 91 | 104 | 61 | 86 | 106 | 90 | 98 | 104 | 69 | 85 | 92.04 | 98 | 86 | 0.872 | -0.197 | 0.36743674 | 0.54214651 | 0.011 | 0.0282 | 0.0236 | 0.0236 | TRUE | 0.2544 |
| Rcan3 | 1833 | 1369 | 1587 | 1666 | 1550 | 1725 | 1424 | 1456 | 1712 | 1658 | 1749 | 1703 | 1616.98 | 1531 | 1703 | 1.114 | 0.155 | 0.05838622 | 0.15071711 | 0.0048 | 0.0039 | 0.0042 | 0.0042 | TRUE | 0.1284 |
| Rcbtb1 | 4340 | 3246 | 3466 | 3799 | 3048 | 3708 | 3371 | 3453 | 3740 | 3782 | 3438 | 3660 | 3573.93 | 3521 | 3627 | 1.031 | 0.044 | 0.50533942 | 0.66877929 | 0.0021 | 0.0031 | 0.0028 | 0.0028 | TRUE | 0.0357 |
| Rcbtb2 | 552 | 378 | 361 | 290 | 264 | 292 | 429 | 402 | 390 | 289 | 298 | 288 | 349.17 | 407 | 292 | 0.715 | -0.483 | 1.57E-05 | 0.00027648 | 0 | 0.0092 | 0.0061 | 0.0061 | TRUE | 0.0275 |
| Rcc1 | 9398 | 6652 | 6178 | 6252 | 5762 | 6553 | 7299 | 7076 | 6666 | 6223 | 6500 | 6469 | 6705.48 | 7014 | 6397 | 0.912 | -0.133 | 0.01718708 | 0.06186224 | 0.0011 | 0.0028 | 0.0021 | 0.0021 | TRUE | 0.0229 |
| Rcc2 | 33511 | 25232 | 22048 | 19977 | 18236 | 20666 | 26027 | 26839 | 23790 | 19885 | 20572 | 20401 | 22918.81 | 25552 | 20286 | 0.794 | -0.333 | 6.49E-09 | 3.47E-07 | 0.002 | 0.0025 | 0.0023 | 0.0023 | TRUE | 0.0444 |
| Rccd1 | 768 | 670 | 461 | 499 | 423 | 509 | 596 | 713 | 497 | 497 | 477 | 502 | 547.15 | 602 | 492 | 0.817 | -0.291 | 0.01907511 | 0.06677457 | 0.0147 | 0.0068 | 0.0092 | 0.0092 | TRUE | 0.1837 |
| Rce1 | 1671 | 1249 | 1131 | 1208 | 1133 | 1265 | 1298 | 1329 | 1220 | 1202 | 1278 | 1249 | 1262.67 | 1282 | 1243 | 0.969 | -0.045 | 0.52867511 | 0.68919677 | 5.00E-04 | 0.0043 | 0.003 | 0.003 | TRUE | 0.0207 |
| Rchy1 | 2757 | 2087 | 2223 | 2344 | 2145 | 2494 | 2141 | 2220 | 2399 | 2333 | 2420 | 2462 | 2329.12 | 2253 | 2405 | 1.068 | 0.095 | 0.15253774 | 0.29927984 | 0.0014 | 0.0034 | 0.0027 | 0.0027 | TRUE | 0.0383 |
| Rcl1 | 5292 | 3810 | 3606 | 3654 | 3333 | 3867 | 4110 | 4053 | 3891 | 3637 | 3760 | 3817 | 3878.01 | 4018 | 3738 | 0.93 | -0.105 | 0.05843318 | 0.15071711 | 5.00E-04 | 0.003 | 0.0019 | 0.0019 | TRUE | 0.0092 |
| Rcn1 | 4882 | 3499 | 3883 | 4166 | 3798 | 4367 | 3792 | 3722 | 4190 | 4147 | 4285 | 4311 | 4074.25 | 3901 | 4248 | 1.089 | 0.123 | 0.05013132 | 0.13501544 | 0.0019 | 0.003 | 0.0026 | 0.0026 | TRUE | 0.0504 |
| Rcn2 | 2254 | 1508 | 1536 | 1688 | 1391 | 1527 | 1751 | 1604 | 1657 | 1680 | 1569 | 1507 | 1628.13 | 1671 | 1585 | 0.948 | -0.076 | 0.29100849 | 0.46307998 | 0.0019 | 0.0039 | 0.0032 | 0.0032 | TRUE | 0.0333 |
| Rcn3 | 1622 | 1255 | 1664 | 2141 | 1814 | 2192 | 1260 | 1335 | 1795 | 2131 | 2046 | 2164 | 1788.58 | 1463 | 2114 | 1.446 | 0.532 | 1.48E-06 | 4.01E-05 | 0.0192 | 0.0038 | 0.0082 | 0.0082 | TRUE | 0.472 |
| Rcor1 | 1866 | 1356 | 1273 | 1484 | 1182 | 1320 | 1449 | 1442 | 1374 | 1477 | 1333 | 1303 | 1396.47 | 1422 | 1371 | 0.964 | -0.052 | 0.48491349 | 0.65090872 | 0.002 | 0.0041 | 0.0033 | 0.0033 | TRUE | 0.0555 |
| Rcor2 | 6049 | 4551 | 4180 | 4087 | 3873 | 4404 | 4698 | 4841 | 4510 | 4068 | 4369 | 4347 | 4472.3 | 4683 | 4261 | 0.91 | -0.136 | 0.01915023 | 0.06691525 | 0.0011 | 0.003 | 0.0022 | 0.0022 | TRUE | 0.0193 |
| Rcor3 | 168 | 129 | 128 | 136 | 103 | 121 | 130 | 137 | 138 | 135 | 116 | 119 | 129.47 | 135 | 123 | 0.916 | -0.126 | 0.46660414 | 0.63552248 | 0 | 0.0207 | 0.0139 | 0.0139 | TRUE | 0.0705 |
| Rcsd1 | 151 | 132 | 110 | 124 | 108 | 109 | 117 | 140 | 119 | 123 | 122 | 108 | 121.54 | 125 | 118 | 0.939 | -0.091 | 0.61864714 | 0.75917079 | 0 | 0.0219 | 0.0158 | 0.0158 | TRUE | 0.1071 |
| Rcvrn | 0 | 1 | 2 | 0 | 1 | 0 | 0 | 1 | 2 | 0 | 1 | 0 | 0.72 | 1 | 0 | 0.355 | -1.496 | 0.54059944 | NA | 0 | 3.2716 | 2.8448 | 2.8448 | TRUE | 0.2584 |
| Rd3 | 1 | 1 | 0 | 0 | 0 | 0 | 1 | 1 | 0 | 0 | 0 | 0 | 0.31 | 1 | 0 | 0.309 | -1.696 | 0.63183309 | NA | 0 | 7.7294 | 7.2472 | 7.2472 | TRUE | 0.2015 |
| Rdh1 | 2 | 1 | 2 | 1 | 1 | 1 | 2 | 1 | 2 | 1 | 1 | 1 | 1.31 | 2 | 1 | 0.651 | -0.619 | 0.69903494 | NA | 0 | 1.8057 | 1.0419 | 1.0419 | TRUE | 0.0615 |
| Rdh10 | 269 | 196 | 164 | 209 | 164 | 192 | 209 | 208 | 177 | 208 | 185 | 190 | 196.16 | 198 | 194 | 0.979 | -0.031 | 0.83553931 | 0.90637245 | 0 | 0.0145 | 0.0107 | 0.0107 | TRUE | 0.0906 |
| Rdh11 | 2712 | 2075 | 2611 | 3311 | 2807 | 3363 | 2106 | 2207 | 2817 | 3296 | 3167 | 3320 | 2818.81 | 2377 | 3261 | 1.373 | 0.457 | 1.81E-06 | 4.70E-05 | 0.0129 | 0.0033 | 0.0063 | 0.0063 | TRUE | 0.3169 |
| Rdh12 | 116 | 71 | 62 | 94 | 67 | 110 | 90 | 76 | 67 | 94 | 76 | 109 | 85.04 | 78 | 93 | 1.191 | 0.252 | 0.27313913 | 0.44413852 | 0.0162 | 0.0303 | 0.0263 | 0.0263 | TRUE | 0.1915 |
| Rdh13 | 1717 | 1224 | 1322 | 1467 | 1230 | 1562 | 1334 | 1302 | 1426 | 1460 | 1388 | 1542 | 1408.61 | 1354 | 1463 | 1.082 | 0.113 | 0.1276681 | 0.26363885 | 0.0017 | 0.0041 | 0.0033 | 0.0033 | TRUE | 0.0267 |
| Rdh14 | 468 | 371 | 380 | 420 | 363 | 406 | 363 | 395 | 410 | 418 | 410 | 401 | 399.41 | 389 | 410 | 1.054 | 0.076 | 0.47876194 | 0.64580449 | 0 | 0.0084 | 0.0057 | 0.0057 | TRUE | 0.0422 |
| Rdh16 | 5 | 4 | 2 | 2 | 3 | 4 | 4 | 4 | 2 | 2 | 3 | 4 | 3.27 | 3 | 3 | 0.899 | -0.153 | 0.88237452 | NA | 0 | 0.7272 | 0.4659 | 0.4659 | TRUE | 0.1382 |
| Rdh18-ps | 1 | 1 | 1 | 1 | 0 | 1 | 1 | 1 | 1 | 1 | 0 | 1 | 0.82 | 1 | 1 | 0.698 | -0.518 | 0.79971927 | NA | 0 | 2.9034 | 1.738 | 1.738 | TRUE | 0.2018 |
| Rdh19 | 0 | 0 | 0 | 0 | 0 | 0 | 0 | 0 | 0 | 0 | 0 | 0 | 0 | 0 | 0 | NA | NA | NA | NA | NA | NA | NA | NA | NA | NA |
| Rdh5 | 18 | 11 | 6 | 11 | 4 | 8 | 14 | 12 | 6 | 11 | 5 | 8 | 9.25 | 11 | 8 | 0.724 | -0.466 | 0.47834367 | NA | 0 | 0.2586 | 0.2001 | 0.2001 | TRUE | 0.2648 |
| Rdh7 | 0 | 1 | 0 | 0 | 0 | 0 | 0 | 1 | 0 | 0 | 0 | 0 | 0.18 | 0 | 0 | 0.552 | -0.857 | 0.8336548 | NA | 0 | 13.3714 | 10 | 10 | TRUE | 0.5216 |
| Rdh8 | 0 | 0 | 0 | 0 | 1 | 0 | 0 | 0 | 0 | 0 | 1 | 0 | 0.19 | 0 | 0 | 2.094 | 1.067 | 0.79379329 | NA | 0 | 12.608 | 10 | 10 | TRUE | 0.5075 |
| Rdh9 | 1 | 2 | 2 | 5 | 3 | 4 | 1 | 2 | 2 | 5 | 3 | 4 | 2.9 | 2 | 4 | 2.514 | 1.33 | 0.24662406 | NA | 0 | 0.821 | 0.5298 | 0.5298 | TRUE | 0.2512 |
| Rdm1 | 3206 | 2555 | 2683 | 2907 | 2582 | 2978 | 2490 | 2718 | 2895 | 2894 | 2913 | 2940 | 2808.13 | 2701 | 2916 | 1.08 | 0.112 | 0.10203937 | 0.22430771 | 0.0023 | 0.0033 | 0.003 | 0.003 | TRUE | 0.0581 |
| Rdx | 15675 | 11352 | 11192 | 11135 | 9924 | 11066 | 12174 | 12075 | 12076 | 11084 | 11195 | 10924 | 11588.01 | 12108 | 11068 | 0.914 | -0.13 | 0.00369663 | 0.01940357 | 0 | 0.0026 | 0.0014 | 0.0014 | TRUE | 0.0016 |
| Rec8 | 1519 | 1051 | 1172 | 1064 | 959 | 1229 | 1180 | 1118 | 1265 | 1059 | 1082 | 1213 | 1152.74 | 1188 | 1118 | 0.942 | -0.086 | 0.30485486 | 0.47739853 | 0.0034 | 0.0045 | 0.0042 | 0.0042 | TRUE | 0.0671 |
| Reck | 821 | 624 | 640 | 749 | 630 | 704 | 638 | 664 | 691 | 746 | 711 | 695 | 690.52 | 664 | 717 | 1.081 | 0.113 | 0.19198803 | 0.35032938 | 0 | 0.0059 | 0.0039 | 0.0039 | TRUE | 0.0152 |
| Recql | 588 | 362 | 311 | 285 | 268 | 275 | 457 | 385 | 336 | 284 | 302 | 271 | 339.13 | 393 | 286 | 0.725 | -0.463 | 0.00059618 | 0.00493984 | 0.0115 | 0.0094 | 0.0101 | 0.0101 | TRUE | 0.2444 |
| Recql4 | 1385 | 1039 | 732 | 690 | 624 | 730 | 1076 | 1105 | 790 | 687 | 704 | 721 | 847.01 | 990 | 704 | 0.71 | -0.494 | 2.46E-05 | 0.00039389 | 0.016 | 0.0052 | 0.0087 | 0.0087 | TRUE | 0.3684 |
| Recql5 | 3027 | 2148 | 2005 | 2113 | 1951 | 2189 | 2351 | 2285 | 2163 | 2103 | 2201 | 2161 | 2210.7 | 2266 | 2155 | 0.95 | -0.074 | 0.24141821 | 0.40912859 | 7.00E-04 | 0.0035 | 0.0024 | 0.0024 | TRUE | 0.019 |
| Reep1 | 1803 | 1095 | 1128 | 1314 | 1092 | 1213 | 1400 | 1165 | 1217 | 1308 | 1232 | 1197 | 1253.24 | 1261 | 1246 | 0.987 | -0.019 | 0.82182888 | 0.89776828 | 0.0051 | 0.0043 | 0.0046 | 0.0046 | TRUE | 0.1162 |
| Reep2 | 585 | 413 | 537 | 657 | 557 | 711 | 454 | 439 | 579 | 654 | 628 | 702 | 576.21 | 491 | 661 | 1.35 | 0.433 | 0.00018848 | 0.00204706 | 0.0112 | 0.0065 | 0.0079 | 0.0079 | TRUE | 0.2816 |
| Reep3 | 1152 | 795 | 688 | 728 | 607 | 681 | 895 | 846 | 742 | 725 | 685 | 672 | 760.72 | 828 | 694 | 0.837 | -0.256 | 0.00657704 | 0.03025609 | 0.0042 | 0.0055 | 0.0051 | 0.0051 | TRUE | 0.095 |
| Reep4 | 2261 | 1684 | 1950 | 2159 | 1947 | 2250 | 1756 | 1791 | 2104 | 2149 | 2196 | 2221 | 2036.31 | 1884 | 2189 | 1.163 | 0.218 | 0.00554435 | 0.02656138 | 0.0046 | 0.0036 | 0.0039 | 0.0039 | TRUE | 0.1252 |
| Reep5 | 2472 | 1815 | 2030 | 2357 | 2156 | 2456 | 1920 | 1931 | 2190 | 2346 | 2432 | 2424 | 2207.27 | 2014 | 2401 | 1.193 | 0.255 | 0.0003007 | 0.00291182 | 0.0024 | 0.0035 | 0.0031 | 0.0031 | TRUE | 0.0704 |
| Reep6 | 1318 | 1106 | 964 | 1039 | 948 | 1150 | 1024 | 1176 | 1040 | 1034 | 1069 | 1135 | 1079.85 | 1080 | 1079 | 1.001 | 0.001 | 0.99174258 | 0.99565418 | 0.0032 | 0.0046 | 0.0041 | 0.0041 | TRUE | 0.0719 |
| Reg1 | 4 | 0 | 4 | 3 | 1 | 0 | 3 | 0 | 4 | 3 | 1 | 0 | 1.92 | 2 | 1 | 0.552 | -0.856 | 0.58476332 | NA | 1.0373 | 1.235 | 1.1991 | 1.1991 | TRUE | 0.5262 |
| Reg2 | 0 | 0 | 0 | 0 | 0 | 0 | 0 | 0 | 0 | 0 | 0 | 0 | 0 | 0 | 0 | NA | NA | NA | NA | NA | NA | NA | NA | NA | NA |
| Reg3a | 0 | 0 | 0 | 0 | 0 | 0 | 0 | 0 | 0 | 0 | 0 | 0 | 0 | 0 | 0 | NA | NA | NA | NA | NA | NA | NA | NA | NA | NA |
| Reg3b | 0 | 0 | 0 | 0 | 0 | 0 | 0 | 0 | 0 | 0 | 0 | 0 | 0 | 0 | 0 | NA | NA | NA | NA | NA | NA | NA | NA | NA | NA |
| Reg3d | 0 | 0 | 0 | 0 | 0 | 0 | 0 | 0 | 0 | 0 | 0 | 0 | 0 | 0 | 0 | NA | NA | NA | NA | NA | NA | NA | NA | NA | NA |
| Reg3g | 0 | 0 | 0 | 0 | 0 | 0 | 0 | 0 | 0 | 0 | 0 | 0 | 0 | 0 | 0 | NA | NA | NA | NA | NA | NA | NA | NA | NA | NA |
| Reg4 | 3 | 0 | 1 | 0 | 0 | 0 | 2 | 0 | 1 | 0 | 0 | 0 | 0.57 | 1 | 0 | 0.165 | -2.602 | 0.35448349 | NA | 0 | 4.1739 | 4.2733 | 4.2733 | TRUE | 0.066 |
| Rel | 98 | 89 | 71 | 70 | 49 | 76 | 76 | 95 | 77 | 70 | 55 | 75 | 74.56 | 83 | 67 | 0.812 | -0.3 | 0.20348783 | 0.36474186 | 0 | 0.0342 | 0.0264 | 0.0264 | TRUE | 0.1851 |
| Rela | 922 | 675 | 579 | 612 | 561 | 598 | 716 | 718 | 625 | 609 | 633 | 590 | 648.53 | 686 | 611 | 0.889 | -0.17 | 0.06976726 | 0.17121967 | 0.0022 | 0.0061 | 0.0048 | 0.0048 | TRUE | 0.0705 |
| Relb | 677 | 482 | 555 | 718 | 647 | 786 | 526 | 513 | 599 | 715 | 730 | 776 | 642.97 | 546 | 740 | 1.358 | 0.441 | 3.76E-06 | 8.40E-05 | 0.0025 | 0.0061 | 0.005 | 0.005 | TRUE | 0.0816 |
| Rell1 | 2514 | 2027 | 2434 | 2230 | 1966 | 2356 | 1953 | 2156 | 2626 | 2220 | 2218 | 2326 | 2249.7 | 2245 | 2255 | 1.005 | 0.007 | 0.93785659 | 0.96554692 | 0.0115 | 0.0035 | 0.0062 | 0.0062 | TRUE | 0.2667 |

| | | | | | | | | | | | | | | | | | | | | | | | | | |
|---|---|---|---|---|---|---|---|---|---|---|---|---|---|---|---|---|---|---|---|---|---|---|---|---|---|
| Rell2 | 128 | 108 | 126 | 115 | 99 | 124 | 99 | 115 | 136 | 114 | 112 | 122 | 116.47 | 117 | 116 | 1.001 | 0.001 | 0.99484436 | 0.99700676 | 0.0061 | 0.0228 | 0.0179 | 0.0179 | TRUE | 0.2058 |
| Reln | 168 | 121 | 116 | 167 | 174 | 152 | 130 | 129 | 125 | 166 | 196 | 150 | 149.49 | 128 | 171 | 1.33 | 0.411 | 0.01554369 | 0.0573773 | 0 | 0.0183 | 0.014 | 0.014 | TRUE | 0.1744 |
| Relt | 619 | 445 | 400 | 477 | 391 | 461 | 481 | 473 | 432 | 475 | 441 | 455 | 459.44 | 462 | 457 | 0.989 | -0.017 | 0.87016503 | 0.92815438 | 0 | 0.0076 | 0.0053 | 0.0053 | TRUE | 0.0371 |
| Rem1 | 22 | 17 | 15 | 28 | 23 | 26 | 17 | 18 | 16 | 28 | 26 | 26 | 21.81 | 17 | 27 | 1.548 | 0.631 | 0.11967696 | 0.25135735 | 0 | 0.1111 | 0.0706 | 0.0706 | TRUE | 0.0133 |
| Rem2 | 115 | 75 | 59 | 78 | 78 | 86 | 89 | 80 | 64 | 78 | 88 | 85 | 80.55 | 78 | 84 | 1.07 | 0.097 | 0.66643426 | 0.79482716 | 0 | 0.0319 | 0.0242 | 0.0242 | TRUE | 0.2185 |
| Ren1 | 0 | 0 | 0 | 0 | 0 | 0 | 0 | 0 | 0 | 0 | 0 | 0 | 0 | 0 | 0 | NA | NA | NA | NA | NA | NA | NA | NA | NA | NA |
| Ren2 | 0 | 1 | 1 | 0 | 0 | 0 | 0 | 1 | 1 | 0 | 0 | 0 | 0.36 | 1 | 0 | 0.271 | -1.881 | 0.5755355 | NA | 0 | 6.6391 | 6.4428 | 6.4428 | TRUE | 0.3513 |
| Renbp | 1237 | 970 | 1149 | 1125 | 1062 | 1184 | 961 | 1032 | 1240 | 1120 | 1198 | 1169 | 1119.82 | 1078 | 1162 | 1.08 | 0.111 | 0.25047525 | 0.41935606 | 0.008 | 0.0045 | 0.0059 | 0.0059 | TRUE | 0.2064 |
| Rep15 | 1 | 0 | 0 | 0 | 0 | 0 | 1 | 0 | 0 | 0 | 0 | 0 | 0.13 | 0 | 0 | 0.552 | -0.857 | 0.8336548 | NA | 0 | 18.3121 | 10 | 10 | TRUE | 0.2566 |
| Repin1 | 543 | 384 | 445 | 565 | 496 | 560 | 422 | 408 | 480 | 562 | 560 | 553 | 497.51 | 437 | 558 | 1.28 | 0.356 | 0.00044972 | 0.00398011 | 0.001 | 0.0072 | 0.0054 | 0.0054 | TRUE | 0.0837 |
| Reps1 | 4737 | 3458 | 3738 | 3778 | 3606 | 3986 | 3679 | 3678 | 4033 | 3761 | 4068 | 3935 | 3858.98 | 3797 | 3921 | 1.033 | 0.047 | 0.45796781 | 0.62770751 | 0.0018 | 0.003 | 0.0026 | 0.0026 | TRUE | 0.0358 |
| Reps2 | 426 | 363 | 435 | 303 | 297 | 344 | 331 | 386 | 469 | 302 | 335 | 340 | 360.43 | 395 | 326 | 0.825 | -0.277 | 0.0456046 | 0.12595683 | 0.0154 | 0.009 | 0.011 | 0.011 | TRUE | 0.2733 |
| Rer1 | 5158 | 3842 | 3929 | 4363 | 3738 | 4391 | 4006 | 4087 | 4239 | 4343 | 4217 | 4335 | 4204.4 | 4111 | 4298 | 1.046 | 0.065 | 0.22599821 | 0.39092247 | 3.00E-04 | 0.003 | 0.0018 | 0.0018 | TRUE | 0.0091 |
| Rere | 6595 | 5031 | 5092 | 5503 | 5230 | 5979 | 5122 | 5351 | 5494 | 5478 | 5900 | 5902 | 5541.26 | 5322 | 5760 | 1.082 | 0.114 | 0.04807081 | 0.13088347 | 0.0013 | 0.0029 | 0.0022 | 0.0022 | TRUE | 0.0225 |
| Rerg | 175 | 126 | 142 | 175 | 163 | 199 | 136 | 134 | 153 | 174 | 184 | 196 | 162.95 | 141 | 185 | 1.313 | 0.392 | 0.0128622 | 0.04982099 | 0 | 0.017 | 0.0117 | 0.0117 | TRUE | 0.0559 |
| Rergl | 0 | 0 | 0 | 0 | 0 | 0 | 0 | 0 | 0 | 0 | 0 | 0 | 0 | 0 | 0 | NA | NA | NA | NA | NA | NA | NA | NA | NA | NA |
| Resp18 | 2 | 3 | 3 | 0 | 1 | 3 | 2 | 3 | 3 | 0 | 1 | 3 | 2.01 | 3 | 1 | 0.522 | -0.939 | 0.50694526 | NA | 0 | 1.1805 | 0.8814 | 0.8814 | TRUE | 0.6114 |
| Rest | 8660 | 6518 | 5989 | 7057 | 5973 | 6498 | 6726 | 6933 | 6462 | 7025 | 6738 | 6415 | 6716.38 | 6707 | 6726 | 1.003 | 0.004 | 0.94508865 | 0.96978621 | 0.0015 | 0.0028 | 0.0022 | 0.0022 | TRUE | 0.0201 |
| Ret | 954 | 765 | 694 | 751 | 723 | 739 | 741 | 814 | 749 | 748 | 816 | 730 | 766.02 | 768 | 765 | 0.995 | -0.007 | 0.93931245 | 0.966328 | 0.0017 | 0.0055 | 0.0042 | 0.0042 | TRUE | 0.0401 |
| Retn | 2 | 2 | 3 | 4 | 2 | 5 | 2 | 2 | 3 | 4 | 2 | 5 | 3.02 | 2 | 4 | 1.652 | 0.724 | 0.50731678 | NA | 0 | 0.7885 | 0.5094 | 0.5094 | TRUE | 0.1638 |
| Retnla | 0 | 0 | 0 | 0 | 0 | 0 | 0 | 0 | 0 | 0 | 0 | 0 | 0 | 0 | 0 | NA | NA | NA | NA | NA | NA | NA | NA | NA | NA |
| Retnlb | 0 | 0 | 0 | 0 | 0 | 0 | 0 | 0 | 0 | 0 | 0 | 0 | 0 | 0 | 0 | NA | NA | NA | NA | NA | NA | NA | NA | NA | NA |
| Retnlg | 0 | 0 | 0 | 0 | 0 | 0 | 0 | 0 | 0 | 0 | 0 | 0 | 0 | 0 | 0 | NA | NA | NA | NA | NA | NA | NA | NA | NA | NA |
| Retsat | 1095 | 803 | 927 | 1020 | 978 | 1038 | 850 | 854 | 1000 | 1015 | 1103 | 1025 | 974.68 | 901 | 1048 | 1.163 | 0.218 | 0.01415743 | 0.0534616 | 0.004 | 0.0049 | 0.0047 | 0.0047 | TRUE | 0.107 |
| Rev1 | 3284 | 2726 | 2869 | 2835 | 2500 | 2861 | 2551 | 2900 | 3096 | 2822 | 2820 | 2824 | 2835.37 | 2849 | 2822 | 0.992 | -0.012 | 0.87196269 | 0.92903189 | 0.0045 | 0.0033 | 0.0037 | 0.0037 | TRUE | 0.1038 |
| Rev3l | 5025 | 3560 | 3435 | 4181 | 3540 | 4461 | 3903 | 3787 | 3706 | 4162 | 3993 | 4404 | 3992.45 | 3799 | 4186 | 1.102 | 0.14 | 0.01875959 | 0.06596084 | 0.0012 | 0.003 | 0.0023 | 0.0023 | TRUE | 0.0252 |
| Rex2 | 1520 | 1141 | 990 | 1089 | 959 | 1042 | 1181 | 1214 | 1068 | 1084 | 1082 | 1029 | 1109.47 | 1154 | 1065 | 0.922 | -0.117 | 0.13699429 | 0.27724912 | 0.0017 | 0.0046 | 0.0036 | 0.0036 | TRUE | 0.0493 |
| Rexo1 | 5653 | 4526 | 4291 | 4724 | 4105 | 4937 | 4390 | 4814 | 4630 | 4702 | 4631 | 4874 | 4673.56 | 4611 | 4736 | 1.027 | 0.039 | 0.50124559 | 0.66551818 | 0.0011 | 0.0029 | 0.0022 | 0.0022 | TRUE | 0.0219 |
| Rexo2 | 2351 | 1697 | 1363 | 1512 | 1236 | 1463 | 1826 | 1805 | 1471 | 1505 | 1394 | 1444 | 1574.21 | 1701 | 1448 | 0.851 | -0.233 | 0.0091096 | 0.03865898 | 0.0066 | 0.0039 | 0.0051 | 0.0051 | TRUE | 0.1667 |
| Rexo4 | 3306 | 2463 | 2324 | 2612 | 2275 | 2653 | 2568 | 2620 | 2508 | 2600 | 2566 | 2619 | 2580.07 | 2565 | 2595 | 1.012 | 0.017 | 0.76581252 | 0.86112488 | 0 | 0.0033 | 0.0019 | 0.0019 | TRUE | 0.0045 |
| Rfc1 | 15424 | 11075 | 11296 | 12638 | 10936 | 12857 | 11979 | 11780 | 12188 | 12580 | 12337 | 12692 | 12259.4 | 11982 | 12536 | 1.046 | 0.065 | 0.15613662 | 0.30416529 | 1.00E-04 | 0.0026 | 0.0014 | 0.0014 | TRUE | 0.0027 |
| Rfc2 | 6018 | 4335 | 4094 | 4782 | 4171 | 4792 | 4674 | 4611 | 4417 | 4760 | 4705 | 4730 | 4649.69 | 4567 | 4732 | 1.036 | 0.051 | 0.33186597 | 0.50601241 | 2.00E-04 | 0.0029 | 0.0018 | 0.0018 | TRUE | 0.01 |
| Rfc3 | 2853 | 2103 | 1990 | 1820 | 1595 | 1830 | 2216 | 2237 | 2147 | 1812 | 1799 | 1806 | 2002.9 | 2200 | 1806 | 0.821 | -0.285 | 1.99E-06 | 5.07E-05 | 0 | 0.0036 | 0.0021 | 0.0021 | TRUE | 0.0052 |
| Rfc4 | 3265 | 2293 | 2570 | 2663 | 2452 | 2669 | 2536 | 2439 | 2773 | 2651 | 2766 | 2635 | 2633.23 | 2583 | 2684 | 1.039 | 0.056 | 0.40666928 | 0.57957164 | 0.002 | 0.0033 | 0.0029 | 0.0029 | TRUE | 0.0495 |
| Rfc5 | 12197 | 8574 | 8227 | 7719 | 7196 | 8175 | 9473 | 9120 | 8877 | 7683 | 8118 | 8070 | 8556.85 | 9157 | 7957 | 0.869 | -0.203 | 0.00012053 | 0.00141907 | 8.00E-04 | 0.0027 | 0.0019 | 0.0019 | TRUE | 0.0113 |
| Rfesd | 409 | 282 | 268 | 288 | 250 | 251 | 318 | 300 | 289 | 287 | 282 | 248 | 287.21 | 302 | 272 | 0.898 | -0.154 | 0.21600056 | 0.37873441 | 0 | 0.0107 | 0.0077 | 0.0077 | TRUE | 0.0703 |
| Rffl | 2687 | 1982 | 1927 | 1857 | 1696 | 1995 | 2087 | 2108 | 2079 | 1848 | 1913 | 1969 | 2000.9 | 2091 | 1910 | 0.913 | -0.131 | 0.0330114 | 0.09943772 | 0 | 0.0036 | 0.0022 | 0.0022 | TRUE | 0.0099 |
| Rfk | 2984 | 2150 | 2104 | 2008 | 1673 | 1865 | 2318 | 2287 | 2270 | 1999 | 1887 | 1841 | 2100.3 | 2292 | 1909 | 0.833 | -0.264 | 2.51E-05 | 0.00040065 | 4.00E-04 | 0.0036 | 0.0023 | 0.0023 | TRUE | 0.0207 |
| Rfng | 1247 | 928 | 909 | 968 | 806 | 1015 | 968 | 987 | 981 | 964 | 909 | 1002 | 968.53 | 979 | 958 | 0.98 | -0.029 | 0.70223005 | 0.81967939 | 0 | 0.0049 | 0.0032 | 0.0032 | TRUE | 0.0233 |
| Rfpl3s | 1 | 0 | 0 | 0 | 0 | 0 | 1 | 0 | 0 | 0 | 0 | 0 | 0.13 | 0 | 0 | 0.552 | -0.857 | 0.8336548 | NA | 0 | 18.3121 | 10 | 10 | TRUE | 0.2566 |
| Rfpl4 | 0 | 0 | 0 | 0 | 0 | 0 | 0 | 0 | 0 | 0 | 0 | 0 | 0 | 0 | 0 | NA | NA | NA | NA | NA | NA | NA | NA | NA | NA |
| Rfpl4b | 12 | 7 | 2 | 4 | 2 | 1 | 9 | 7 | 2 | 4 | 2 | 1 | 4.36 | 6 | 2 | 0.377 | -1.407 | 0.16440027 | NA | 0 | 0.5463 | 0.4477 | 0.4477 | TRUE | 0.4909 |
| Rft1 | 796 | 584 | 495 | 579 | 501 | 566 | 618 | 621 | 534 | 576 | 565 | 559 | 578.96 | 591 | 567 | 0.958 | -0.063 | 0.51855499 | 0.68022269 | 0.0019 | 0.0065 | 0.0051 | 0.0051 | TRUE | 0.081 |
| Rftn1 | 873 | 594 | 622 | 760 | 645 | 712 | 678 | 632 | 671 | 757 | 728 | 703 | 694.66 | 660 | 729 | 1.103 | 0.142 | 0.09969322 | 0.22071177 | 0 | 0.0058 | 0.0039 | 0.0039 | TRUE | 0.0164 |
| Rftn2 | 24 | 12 | 16 | 30 | 16 | 19 | 19 | 13 | 17 | 30 | 18 | 19 | 19.22 | 16 | 22 | 1.365 | 0.449 | 0.32053806 | 0.4944367 | 0 | 0.1257 | 0.0946 | 0.0946 | TRUE | 0.5268 |
| Rfwd2 | 1754 | 1206 | 1271 | 1221 | 1062 | 1267 | 1362 | 1283 | 1371 | 1215 | 1198 | 1251 | 1280.1 | 1339 | 1221 | 0.912 | -0.133 | 0.05877796 | 0.1511267 | 0 | 0.0043 | 0.0028 | 0.0028 | TRUE | 0.0157 |
| Rfwd3 | 15352 | 11011 | 11273 | 12037 | 10490 | 12755 | 11923 | 11712 | 12163 | 11982 | 11834 | 12591 | 12034.26 | 11933 | 12136 | 1.017 | 0.024 | 0.62485215 | 0.76346484 | 6.00E-04 | 0.0026 | 0.0017 | 0.0017 | TRUE | 0.0132 |
| Rfx1 | 3113 | 2531 | 2408 | 2364 | 2162 | 2581 | 2418 | 2692 | 2598 | 2353 | 2439 | 2548 | 2508.01 | 2569 | 2447 | 0.953 | -0.07 | 0.29770521 | 0.46979802 | 0.002 | 0.0034 | 0.0028 | 0.0028 | TRUE | 0.0333 |
| Rfx2 | 3177 | 2589 | 2828 | 2656 | 2552 | 3097 | 2467 | 2754 | 3051 | 2644 | 2879 | 3057 | 2808.77 | 2757 | 2860 | 1.038 | 0.054 | 0.53055484 | 0.69038787 | 0.0079 | 0.0033 | 0.0049 | 0.0049 | TRUE | 0.1055 |
| Rfx3 | 465 | 344 | 401 | 429 | 364 | 465 | 361 | 366 | 433 | 427 | 411 | 459 | 409.4 | 387 | 432 | 1.121 | 0.165 | 0.15005925 | 0.29566776 | 0.0042 | 0.0082 | 0.007 | 0.007 | TRUE | 0.1217 |
| Rfx4 | 29 | 19 | 35 | 25 | 34 | 34 | 23 | 20 | 38 | 25 | 38 | 34 | 29.55 | 27 | 32 | 1.207 | 0.272 | 0.47965734 | 0.64652418 | 0.044 | 0.0826 | 0.0722 | 0.0722 | TRUE | 0.7485 |
| Rfx5 | 1028 | 777 | 826 | 884 | 663 | 829 | 798 | 826 | 891 | 880 | 748 | 818 | 827.06 | 838 | 815 | 0.974 | -0.038 | 0.67455921 | 0.80024529 | 0.0036 | 0.0053 | 0.0047 | 0.0047 | TRUE | 0.0608 |
| Rfx6 | 4 | 3 | 1 | 1 | 1 | 1 | 3 | 3 | 1 | 1 | 1 | 1 | 1.75 | 2 | 1 | 0.417 | -1.262 | 0.38836625 | NA | 0 | 1.3584 | 0.8482 | 0.8482 | TRUE | 0.2351 |
| Rfx7 | 6126 | 4400 | 4039 | 4353 | 3819 | 4464 | 4758 | 4680 | 4358 | 4333 | 4308 | 4407 | 4473.99 | 4599 | 4349 | 0.945 | -0.081 | 0.15441746 | 0.30180347 | 8.00E-04 | 0.003 | 0.0021 | 0.0021 | TRUE | 0.0253 |
| Rfx8 | 23 | 23 | 19 | 27 | 25 | 29 | 18 | 24 | 21 | 27 | 28 | 29 | 24.42 | 21 | 28 | 1.34 | 0.423 | 0.27328332 | 0.44422137 | 0 | 0.0995 | 0.0656 | 0.0656 | TRUE | 0.1168 |
| Rfxank | 599 | 477 | 424 | 424 | 363 | 389 | 465 | 507 | 457 | 422 | 410 | 384 | 440.94 | 476 | 405 | 0.85 | -0.234 | 0.0246151 | 0.08043521 | 0 | 0.0078 | 0.0055 | 0.0055 | TRUE | 0.0356 |
| Rfxap | 1025 | 824 | 838 | 820 | 717 | 837 | 796 | 876 | 904 | 816 | 809 | 826 | 838.01 | 859 | 817 | 0.953 | -0.069 | 0.40860806 | 0.58139857 | 0.0014 | 0.0053 | 0.0039 | 0.0039 | TRUE | 0.0515 |
| Rgag1 | 2 | 2 | 0 | 1 | 1 | 0 | 2 | 2 | 0 | 1 | 1 | 0 | 0.97 | 1 | 1 | 0.564 | -0.825 | 0.68413901 | NA | 0 | 2.4524 | 1.8344 | 1.8344 | TRUE | 0.3849 |
| Rgag4 | 38 | 21 | 29 | 21 | 23 | 27 | 30 | 22 | 31 | 21 | 26 | 27 | 26.11 | 28 | 25 | 0.881 | -0.182 | 0.62980998 | 0.76709621 | 0 | 0.0932 | 0.0643 | 0.0643 | TRUE | 0.1712 |
| Rgcc | 37 | 47 | 90 | 75 | 86 | 91 | 29 | 50 | 97 | 75 | 97 | 90 | 72.89 | 59 | 87 | 1.502 | 0.587 | 0.08897375 | 0.20311482 | 0.1676 | 0.0349 | 0.0714 | 0.0714 | TRUE | 0.916 |
| Rgl1 | 427 | 333 | 314 | 391 | 305 | 394 | 332 | 354 | 339 | 389 | 344 | 389 | 357.81 | 342 | 374 | 1.097 | 0.134 | 0.23438497 | 0.40058166 | 0 | 0.0091 | 0.0064 | 0.0064 | TRUE | 0.0545 |
| Rgl2 | 1868 | 1496 | 1221 | 1210 | 1137 | 1310 | 1451 | 1591 | 1317 | 1204 | 1283 | 1293 | 1356.63 | 1453 | 1260 | 0.867 | -0.206 | 0.01356767 | 0.05188163 | 0.0041 | 0.0042 | 0.0043 | 0.0043 | TRUE | 0.081 |
| Rgl3 | 71 | 68 | 80 | 122 | 143 | 152 | 55 | 72 | 86 | 121 | 161 | 150 | 107.77 | 71 | 144 | 2.04 | 1.029 | 2.77E-06 | 6.50E-05 | 0.0239 | 0.0244 | 0.0243 | 0.0243 | TRUE | 0.4114 |
| Rgma | 123 | 118 | 101 | 108 | 115 | 120 | 96 | 126 | 109 | 108 | 130 | 118 | 114.29 | 110 | 119 | 1.081 | 0.113 | 0.56026632 | 0.71486222 | 0.0061 | 0.0232 | 0.0181 | 0.0181 | TRUE | 0.1508 |
| Rgmb | 359 | 280 | 275 | 278 | 257 | 322 | 279 | 298 | 297 | 277 | 290 | 318 | 292.98 | 291 | 295 | 1.014 | 0.021 | 0.86561707 | 0.92572508 | 0 | 0.0105 | 0.0073 | 0.0073 | TRUE | 0.054 |
| Rgn | 8 | 11 | 9 | 14 | 14 | 10 | 6 | 12 | 10 | 14 | 16 | 10 | 11.2 | 9 | 13 | 1.449 | 0.535 | 0.35855931 | NA | 0 | 0.214 | 0.1526 | 0.1526 | TRUE | 0.3537 |
| Rgp1 | 2177 | 1651 | 1746 | 1998 | 1770 | 1987 | 1691 | 1756 | 1884 | 1989 | 1997 | 1961 | 1879.65 | 1777 | 1982 | 1.117 | 0.159 | 0.0171259 | 0.06171975 | 0.001 | 0.0037 | 0.0027 | 0.0027 | TRUE | 0.0332 |
| Rgr | 0 | 0 | 0 | 0 | 0 | 0 | 0 | 0 | 0 | 0 | 0 | 0 | 0 | 0 | 0 | NA | NA | NA | NA | NA | NA | NA | NA | NA | NA |
| Rgs1 | 0 | 0 | 0 | 0 | 0 | 0 | 0 | 0 | 0 | 0 | 0 | 0 | 0 | 0 | 0 | NA | NA | NA | NA | NA | NA | NA | NA | NA | NA |

| | | | | | | | | | | | | | | | | | | | | | | | | | |
|---|---|---|---|---|---|---|---|---|---|---|---|---|---|---|---|---|---|---|---|---|---|---|---|---|---|
| Rgs10 | 334 | 233 | 236 | 273 | 256 | 346 | 259 | 248 | 255 | 272 | 289 | 342 | 277.33 | 254 | 301 | 1.184 | 0.244 | 0.065135 | 0.16290018 | 0.0044 | 0.011 | 0.0089 | 0.0089 | TRUE | 0.1627 |
| Rgs11 | 268 | 247 | 236 | 274 | 240 | 267 | 208 | 263 | 255 | 273 | 271 | 264 | 255.43 | 242 | 269 | 1.117 | 0.16 | 0.24257404 | 0.41021693 | 0.0046 | 0.0117 | 0.0095 | 0.0095 | TRUE | 0.1805 |
| Rgs12 | 2661 | 2045 | 2162 | 2391 | 2106 | 2582 | 2067 | 2175 | 2333 | 2380 | 2376 | 2549 | 2313.22 | 2192 | 2435 | 1.112 | 0.153 | 0.02626379 | 0.08426833 | 0.002 | 0.0035 | 0.003 | 0.003 | TRUE | 0.0383 |
| Rgs13 | 0 | 0 | 0 | 0 | 0 | 0 | 0 | 0 | 0 | 0 | 0 | 0 | 0 | 0 | 0 | NA | NA | NA | NA | NA | NA | NA | NA | NA | NA |
| Rgs14 | 82 | 66 | 81 | 71 | 85 | 68 | 64 | 70 | 87 | 71 | 96 | 67 | 75.83 | 74 | 78 | 1.058 | 0.081 | 0.74071893 | 0.84588493 | 0.0194 | 0.0337 | 0.0297 | 0.0297 | TRUE | 0.3593 |
| Rgs16 | 174 | 131 | 146 | 136 | 153 | 178 | 135 | 139 | 158 | 135 | 173 | 176 | 152.62 | 144 | 161 | 1.121 | 0.165 | 0.33913139 | 0.51314895 | 0.0072 | 0.018 | 0.0149 | 0.0149 | TRUE | 0.212 |
| Rgs17 | 173 | 135 | 150 | 299 | 239 | 261 | 134 | 144 | 162 | 298 | 270 | 258 | 210.78 | 147 | 275 | 1.883 | 0.913 | 5.44E-10 | 3.90E-08 | 0 | 0.0137 | 0.0104 | 0.0104 | TRUE | 0.0844 |
| Rgs18 | 0 | 0 | 0 | 0 | 0 | 0 | 0 | 0 | 0 | 0 | 0 | 0 | 0 | 0 | 0 | NA | NA | NA | NA | NA | NA | NA | NA | NA | NA |
| Rgs19 | 2070 | 1472 | 1373 | 1639 | 1397 | 1594 | 1608 | 1566 | 1481 | 1631 | 1576 | 1574 | 1572.64 | 1552 | 1594 | 1.026 | 0.038 | 0.57625207 | 0.72772317 | 5.00E-04 | 0.0039 | 0.0026 | 0.0026 | TRUE | 0.0186 |
| Rgs2 | 53 | 42 | 31 | 38 | 33 | 38 | 41 | 45 | 33 | 38 | 37 | 38 | 38.64 | 40 | 38 | 0.942 | -0.086 | 0.78090358 | 0.87083209 | 0 | 0.0638 | 0.0426 | 0.0426 | TRUE | 0.132 |
| Rgs20 | 69 | 38 | 43 | 64 | 54 | 82 | 54 | 40 | 46 | 64 | 61 | 81 | 57.66 | 47 | 69 | 1.458 | 0.544 | 0.04070544 | 0.11594317 | 0 | 0.0435 | 0.0331 | 0.0331 | TRUE | 0.2286 |
| Rgs22 | 182 | 162 | 181 | 251 | 223 | 272 | 141 | 172 | 195 | 250 | 252 | 269 | 213.15 | 169 | 257 | 1.522 | 0.606 | 7.77E-05 | 0.00098867 | 0.0083 | 0.0135 | 0.0121 | 0.0121 | TRUE | 0.2524 |
| Rgs3 | 530 | 497 | 577 | 751 | 761 | 819 | 412 | 529 | 623 | 748 | 858 | 808 | 662.89 | 521 | 805 | 1.55 | 0.632 | 1.34E-06 | 3.72E-05 | 0.0206 | 0.006 | 0.0107 | 0.0107 | TRUE | 0.4125 |
| Rgs4 | 4 | 3 | 3 | 5 | 2 | 1 | 3 | 3 | 3 | 5 | 2 | 1 | 2.96 | 3 | 3 | 0.865 | -0.209 | 0.8505362 | NA | 0 | 0.8033 | 0.5498 | 0.5498 | TRUE | 0.6345 |
| Rgs5 | 31 | 17 | 12 | 4 | 5 | 6 | 24 | 18 | 13 | 4 | 6 | 6 | 11.78 | 18 | 5 | 0.279 | -1.841 | 0.0029583 | NA | 0 | 0.2037 | 0.1503 | 0.1503 | TRUE | 0.1086 |
| Rgs6 | 7 | 3 | 6 | 4 | 4 | 1 | 5 | 3 | 6 | 4 | 5 | 1 | 4.1 | 5 | 3 | 0.621 | -0.687 | 0.47937494 | NA | 0 | 0.5809 | 0.4194 | 0.4194 | TRUE | 0.5164 |
| Rgs7 | 26 | 19 | 15 | 28 | 16 | 15 | 20 | 20 | 16 | 28 | 18 | 15 | 19.55 | 19 | 20 | 1.072 | 0.1 | 0.82360088 | 0.89868745 | 0 | 0.1236 | 0.0933 | 0.0933 | TRUE | 0.6084 |
| Rgs7bp | 19 | 7 | 12 | 7 | 4 | 6 | 15 | 7 | 13 | 7 | 5 | 6 | 8.76 | 12 | 6 | 0.494 | -1.019 | 0.12796296 | NA | 0 | 0.273 | 0.1931 | 0.1931 | TRUE | 0.3712 |
| Rgs8 | 10 | 2 | 3 | 4 | 3 | 3 | 8 | 2 | 3 | 4 | 3 | 3 | 3.91 | 4 | 3 | 0.766 | -0.384 | 0.69543104 | NA | 0 | 0.6086 | 0.4363 | 0.4363 | TRUE | 1.0726 |
| Rgs9 | 339 | 339 | 312 | 354 | 328 | 320 | 263 | 361 | 337 | 352 | 370 | 316 | 333.13 | 320 | 346 | 1.085 | 0.117 | 0.39895897 | 0.57224403 | 0.0138 | 0.0095 | 0.0109 | 0.0109 | TRUE | 0.2941 |
| Rgs9bp | 389 | 311 | 354 | 476 | 417 | 465 | 302 | 331 | 382 | 474 | 470 | 459 | 403.02 | 338 | 468 | 1.387 | 0.472 | 4.41E-05 | 0.0006306 | 0.0042 | 0.0083 | 0.0071 | 0.0071 | TRUE | 0.1436 |
| Rgsl1 | 1 | 0 | 0 | 1 | 1 | 0 | 1 | 0 | 0 | 1 | 1 | 0 | 0.48 | 0 | 1 | 2.037 | 1.027 | 0.71241723 | NA | 0 | 4.9057 | 3.8521 | 3.8521 | TRUE | 0.2664 |
| Rhag | 0 | 0 | 0 | 0 | 0 | 0 | 0 | 0 | 0 | 0 | 0 | 0 | 0 | 0 | 0 | NA | NA | NA | NA | NA | NA | NA | NA | NA | NA |
| Rhbdd1 | 1101 | 751 | 752 | 837 | 796 | 906 | 855 | 799 | 811 | 833 | 898 | 894 | 848.47 | 822 | 875 | 1.064 | 0.089 | 0.27324744 | 0.44421357 | 0 | 0.0052 | 0.0036 | 0.0036 | TRUE | 0.0212 |
| Rhbdd2 | 1324 | 1030 | 983 | 1070 | 961 | 1041 | 1028 | 1096 | 1061 | 1065 | 1084 | 1028 | 1060.23 | 1062 | 1059 | 0.998 | -0.003 | 0.96953498 | 0.9834812 | 0 | 0.0047 | 0.003 | 0.003 | TRUE | 0.0095 |
| Rhbdd3 | 1522 | 1138 | 1064 | 1229 | 1157 | 1273 | 1182 | 1210 | 1148 | 1223 | 1305 | 1257 | 1220.97 | 1180 | 1262 | 1.069 | 0.096 | 0.17651867 | 0.33028533 | 0 | 0.0044 | 0.0028 | 0.0028 | TRUE | 0.0107 |
| Rhbdf1 | 801 | 653 | 752 | 752 | 721 | 776 | 622 | 695 | 811 | 749 | 813 | 766 | 742.67 | 709 | 776 | 1.096 | 0.132 | 0.20767288 | 0.36927948 | 0.008 | 0.0056 | 0.0066 | 0.0066 | TRUE | 0.1864 |
| Rhbdf2 | 4601 | 3705 | 4292 | 4844 | 4531 | 5193 | 3573 | 3941 | 4631 | 4822 | 5111 | 5126 | 4534.14 | 4048 | 5020 | 1.241 | 0.311 | 0.00027683 | 0.00272499 | 0.009 | 0.0029 | 0.005 | 0.005 | TRUE | 0.1926 |
| Rhbdl1 | 42 | 39 | 39 | 41 | 21 | 49 | 33 | 41 | 42 | 41 | 24 | 48 | 38.18 | 39 | 38 | 0.981 | -0.027 | 0.93714348 | 0.96508948 | 0.041 | 0.0645 | 0.0583 | 0.0583 | TRUE | 0.5921 |
| Rhbdl2 | 103 | 54 | 42 | 72 | 55 | 68 | 80 | 57 | 45 | 72 | 62 | 67 | 63.93 | 61 | 67 | 1.089 | 0.123 | 0.64705727 | 0.78045481 | 0.0292 | 0.0395 | 0.0365 | 0.0365 | TRUE | 0.5445 |
| Rhbdl3 | 117 | 85 | 76 | 104 | 99 | 91 | 91 | 90 | 82 | 104 | 112 | 90 | 94.72 | 88 | 102 | 1.155 | 0.208 | 0.3056176 | 0.47828669 | 0 | 0.0274 | 0.0191 | 0.0191 | TRUE | 0.1033 |
| Rhbg | 67 | 33 | 34 | 44 | 37 | 49 | 52 | 35 | 37 | 44 | 42 | 48 | 42.96 | 41 | 45 | 1.072 | 0.101 | 0.74032989 | 0.84571092 | 0 | 0.0576 | 0.0435 | 0.0435 | TRUE | 0.4579 |
| Rhcg | 25 | 12 | 8 | 13 | 11 | 8 | 19 | 13 | 9 | 13 | 12 | 8 | 12.34 | 14 | 11 | 0.801 | -0.32 | 0.57466713 | NA | 0 | 0.1944 | 0.1519 | 0.1519 | TRUE | 0.3474 |
| Rhd | 403 | 269 | 322 | 352 | 263 | 367 | 313 | 286 | 347 | 350 | 297 | 362 | 325.99 | 315 | 336 | 1.068 | 0.095 | 0.45928655 | 0.62915302 | 0.0067 | 0.0097 | 0.0088 | 0.0088 | TRUE | 0.1178 |
| Rheb | 4715 | 3357 | 3278 | 3743 | 3066 | 3614 | 3662 | 3571 | 3537 | 3726 | 3459 | 3568 | 3586.97 | 3590 | 3584 | 0.998 | -0.002 | 0.96564424 | 0.98122801 | 6.00E-04 | 0.0031 | 0.002 | 0.002 | TRUE | 0.0146 |
| Rhebl1 | 1830 | 1358 | 1367 | 1286 | 1202 | 1457 | 1421 | 1444 | 1475 | 1280 | 1356 | 1438 | 1402.52 | 1447 | 1358 | 0.939 | -0.091 | 0.20785756 | 0.36946998 | 0.0011 | 0.0041 | 0.003 | 0.003 | TRUE | 0.0327 |
| Rhno1 | 1885 | 1504 | 1388 | 1275 | 1219 | 1410 | 1464 | 1600 | 1498 | 1269 | 1375 | 1392 | 1432.93 | 1521 | 1345 | 0.885 | -0.176 | 0.01665076 | 0.06041871 | 0.0016 | 0.0041 | 0.0032 | 0.0032 | TRUE | 0.0299 |
| Rho | 24 | 24 | 20 | 18 | 13 | 14 | 19 | 26 | 22 | 18 | 15 | 14 | 18.69 | 22 | 16 | 0.71 | -0.494 | 0.26636133 | 0.43674458 | 0 | 0.1292 | 0.0868 | 0.0868 | TRUE | 0.1202 |
| Rhoa | 22700 | 15843 | 15840 | 16441 | 13543 | 15576 | 17630 | 16852 | 17091 | 16365 | 15278 | 15376 | 16432.09 | 17191 | 15673 | 0.912 | -0.133 | 0.00905269 | 0.0384746 | 9.00E-04 | 0.0026 | 0.0018 | 0.0018 | TRUE | 0.0183 |
| Rhob | 3186 | 1937 | 1168 | 1543 | 1244 | 1051 | 2474 | 2060 | 1260 | 1536 | 1403 | 1037 | 1628.64 | 1931 | 1325 | 0.686 | -0.544 | 0.0804759 | 0.18996566 | 0.0691 | 0.0039 | 0.0223 | 0.0691 | TRUE | 0.3435 |
| Rhobtb1 | 813 | 563 | 483 | 734 | 598 | 671 | 631 | 599 | 521 | 731 | 675 | 662 | 636.51 | 584 | 689 | 1.178 | 0.237 | 0.01789091 | 0.06359483 | 0.0045 | 0.0062 | 0.0056 | 0.0056 | TRUE | 0.1014 |
| Rhobtb2 | 1462 | 1091 | 1198 | 1339 | 1154 | 1268 | 1135 | 1160 | 1293 | 1333 | 1302 | 1252 | 1245.83 | 1196 | 1296 | 1.084 | 0.116 | 0.13398112 | 0.27259675 | 0.0021 | 0.0043 | 0.0035 | 0.0035 | TRUE | 0.0588 |
| Rhobtb3 | 1605 | 1152 | 1291 | 1509 | 1099 | 1224 | 1247 | 1225 | 1393 | 1502 | 1240 | 1208 | 1302.5 | 1288 | 1317 | 1.023 | 0.032 | 0.73399013 | 0.84088818 | 0.0089 | 0.0042 | 0.0058 | 0.0058 | TRUE | 0.1833 |
| Rhoc | 2094 | 1454 | 1441 | 1707 | 1405 | 1667 | 1626 | 1547 | 1555 | 1699 | 1585 | 1646 | 1609.58 | 1576 | 1643 | 1.042 | 0.06 | 0.36846721 | 0.54321707 | 4.00E-04 | 0.0039 | 0.0026 | 0.0026 | TRUE | 0.0114 |
| Rhod | 464 | 309 | 345 | 379 | 351 | 341 | 360 | 329 | 372 | 377 | 396 | 337 | 361.86 | 354 | 370 | 1.044 | 0.063 | 0.59167478 | 0.73955467 | 0.0025 | 0.009 | 0.007 | 0.007 | TRUE | 0.0719 |
| Rhof | 219 | 159 | 148 | 165 | 157 | 169 | 170 | 169 | 160 | 164 | 177 | 167 | 167.85 | 166 | 169 | 1.017 | 0.024 | 0.87291505 | 0.92949227 | 0 | 0.0165 | 0.0106 | 0.0106 | TRUE | 0.0162 |
| Rhog | 517 | 386 | 447 | 483 | 455 | 522 | 402 | 411 | 482 | 481 | 513 | 515 | 467.3 | 432 | 503 | 1.168 | 0.224 | 0.03798074 | 0.11025219 | 0.0035 | 0.0075 | 0.0063 | 0.0063 | TRUE | 0.1199 |
| Rhoh | 40 | 31 | 23 | 17 | 22 | 16 | 31 | 33 | 25 | 17 | 25 | 16 | 24.4 | 30 | 19 | 0.643 | -0.638 | 0.11088977 | 0.23852327 | 0 | 0.0996 | 0.0719 | 0.0719 | TRUE | 0.3185 |
| Rhoj | 4 | 7 | 1 | 1 | 1 | 0 | 3 | 7 | 1 | 1 | 1 | 0 | 2.29 | 4 | 1 | 0.18 | -2.473 | 0.10855196 | NA | 0 | 1.0363 | 0.8432 | 0.8432 | TRUE | 0.2152 |
| Rhoq | 998 | 676 | 771 | 746 | 618 | 713 | 775 | 719 | 832 | 743 | 697 | 704 | 744.94 | 775 | 715 | 0.922 | -0.117 | 0.18691205 | 0.34364985 | 0.0018 | 0.0056 | 0.0044 | 0.0044 | TRUE | 0.046 |
| Rhot1 | 9112 | 6682 | 7279 | 8666 | 7360 | 8733 | 7077 | 7108 | 7854 | 8626 | 8303 | 8621 | 7931.37 | 7346 | 8517 | 1.16 | 0.214 | 0.00027288 | 0.00270292 | 0.0018 | 0.0027 | 0.0024 | 0.0024 | TRUE | 0.0444 |
| Rhot2 | 2609 | 1871 | 1601 | 1612 | 1416 | 1643 | 2026 | 1990 | 1727 | 1605 | 1597 | 1622 | 1761.3 | 1914 | 1608 | 0.839 | -0.253 | 0.00088205 | 0.00663014 | 0.0033 | 0.0038 | 0.0036 | 0.0036 | TRUE | 0.0872 |
| Rhou | 1934 | 1405 | 1751 | 1855 | 1794 | 1935 | 1502 | 1494 | 1889 | 1846 | 2024 | 1910 | 1777.71 | 1628 | 1927 | 1.184 | 0.243 | 0.00911795 | 0.03866878 | 0.0098 | 0.0038 | 0.0057 | 0.0057 | TRUE | 0.2342 |
| Rhov | 205 | 143 | 186 | 232 | 198 | 228 | 159 | 152 | 201 | 231 | 223 | 225 | 198.56 | 171 | 226 | 1.33 | 0.412 | 0.00709024 | 0.03205973 | 0.0046 | 0.0144 | 0.0117 | 0.0117 | TRUE | 0.2406 |
| Rhox1 | 372 | 262 | 215 | 194 | 166 | 162 | 289 | 279 | 232 | 193 | 187 | 160 | 223.31 | 267 | 180 | 0.674 | -0.57 | 0.00013611 | 0.00157145 | 0.0074 | 0.013 | 0.0114 | 0.0114 | TRUE | 0.1396 |
| Rhox10 | 421 | 241 | 204 | 267 | 183 | 218 | 327 | 256 | 220 | 266 | 206 | 215 | 248.47 | 268 | 229 | 0.853 | -0.23 | 0.17092848 | 0.32259143 | 0.0261 | 0.012 | 0.0162 | 0.0162 | TRUE | 0.4387 |
| Rhox11 | 5 | 1 | 0 | 1 | 2 | 2 | 4 | 1 | 0 | 1 | 2 | 2 | 1.7 | 2 | 2 | 1.003 | 0.004 | 0.99772612 | NA | 0 | 1.4003 | 1.0908 | 1.0908 | TRUE | 1.0117 |
| Rhox12 | 3 | 1 | 0 | 1 | 1 | 1 | 2 | 1 | 0 | 1 | 1 | 1 | 1.08 | 1 | 1 | 0.879 | -0.186 | 0.91830054 | NA | 0 | 2.1887 | 1.4564 | 1.4564 | TRUE | 0.2453 |
| Rhox13 | 1910 | 1338 | 1107 | 970 | 741 | 777 | 1483 | 1423 | 1194 | 966 | 836 | 767 | 1111.59 | 1367 | 856 | 0.626 | -0.676 | 1.48E-10 | 1.16E-08 | 0.0117 | 0.0046 | 0.0071 | 0.0071 | TRUE | 0.1501 |
| Rhox2a | 90 | 60 | 46 | 38 | 23 | 36 | 70 | 64 | 50 | 38 | 26 | 36 | 47.11 | 61 | 33 | 0.541 | -0.885 | 0.0031942 | 0.0173394 | 0 | 0.0527 | 0.0415 | 0.0415 | TRUE | 0.2264 |
| Rhox2b | 13 | 9 | 8 | 5 | 8 | 3 | 10 | 10 | 9 | 5 | 9 | 3 | 7.54 | 10 | 6 | 0.591 | -0.759 | 0.28961457 | NA | 0 | 0.3166 | 0.2265 | 0.2265 | TRUE | 0.5425 |
| Rhox2c | 10 | 7 | 4 | 5 | 5 | 3 | 8 | 7 | 4 | 5 | 6 | 3 | 5.52 | 6 | 5 | 0.687 | -0.542 | 0.50605648 | NA | 0 | 0.4319 | 0.2898 | 0.2898 | TRUE | 0.1984 |
| Rhox2d | 13 | 7 | 9 | 5 | 3 | 3 | 10 | 7 | 10 | 5 | 3 | 3 | 6.43 | 9 | 4 | 0.415 | -1.269 | 0.10148831 | NA | 0 | 0.3711 | 0.243 | 0.243 | TRUE | 0.1287 |
| Rhox2e | 0 | 1 | 0 | 0 | 0 | 0 | 0 | 1 | 0 | 0 | 0 | 0 | 0.18 | 0 | 0 | 0.552 | -0.857 | 0.8336548 | NA | 0 | 13.3714 | 10 | 10 | TRUE | 0.5216 |
| Rhox2f | 2 | 3 | 4 | 0 | 1 | 0 | 2 | 3 | 4 | 0 | 1 | 0 | 1.7 | 3 | 0 | 0.124 | -3.008 | 0.08568502 | NA | 0 | 1.3982 | 1.0459 | 1.0459 | TRUE | 0.4672 |
| Rhox2g | 6 | 5 | 2 | 2 | 5 | 2 | 5 | 5 | 2 | 2 | 6 | 2 | 3.62 | 4 | 3 | 0.776 | -0.366 | 0.72012566 | NA | 0 | 0.6565 | 0.4683 | 0.4683 | TRUE | 0.5428 |
| Rhox2h | 6 | 3 | 2 | 1 | 1 | 3 | 5 | 3 | 2 | 1 | 1 | 3 | 2.52 | 3 | 2 | 0.505 | -0.985 | 0.41982434 | NA | 0 | 0.9445 | 0.6278 | 0.6278 | TRUE | 0.3421 |
| Rhox3a | 0 | 0 | 0 | 0 | 0 | 0 | 0 | 0 | 0 | 0 | 0 | 0 | 0 | 0 | 0 | NA | NA | NA | NA | NA | NA | NA | NA | NA | NA |
| Rhox3c | 0 | 1 | 0 | 0 | 0 | 0 | 0 | 1 | 0 | 0 | 0 | 0 | 0.18 | 0 | 0 | 0.552 | -0.857 | 0.8336548 | NA | 0 | 13.3714 | 10 | 10 | TRUE | 0.5216 |

| | | | | | | | | | | | | | | | | | | | | | | | | | |
|---|---|---|---|---|---|---|---|---|---|---|---|---|---|---|---|---|---|---|---|---|---|---|---|---|---|
| Rhox3e | 0 | 0 | 0 | 0 | 0 | 0 | 0 | 0 | 0 | 0 | 0 | 0 | 0 | 0 | 0 | NA | NA | NA | NA | NA | NA | NA | NA | NA | NA |
| Rhox3f | 0 | 0 | 0 | 0 | 0 | 0 | 0 | 0 | 0 | 0 | 0 | 0 | 0 | 0 | 0 | NA | NA | NA | NA | NA | NA | NA | NA | NA | NA |
| Rhox3g-ps | 0 | 0 | 0 | 0 | 0 | 0 | 0 | 0 | 0 | 0 | 0 | 0 | 0 | 0 | 0 | NA | NA | NA | NA | NA | NA | NA | NA | NA | NA |
| Rhox3h | 0 | 0 | 0 | 0 | 0 | 0 | 0 | 0 | 0 | 0 | 0 | 0 | 0 | 0 | 0 | NA | NA | NA | NA | NA | NA | NA | NA | NA | NA |
| Rhox4a | 4 | 3 | 2 | 1 | 0 | 1 | 3 | 3 | 2 | 1 | 0 | 1 | 1.74 | 3 | 1 | 0.24 | -2.059 | 0.19522379 | NA | 0 | 1.3648 | 0.8874 | 0.8874 | TRUE | 0.1975 |
| Rhox4b | 0 | 2 | 2 | 0 | 0 | 0 | 0 | 2 | 2 | 0 | 0 | 0 | 0.71 | 1 | 0 | 0.136 | -2.883 | 0.26779854 | NA | 0 | 3.3208 | 3.5331 | 3.5331 | TRUE | 0.6762 |
| Rhox4c | 0 | 1 | 0 | 0 | 0 | 1 | 0 | 1 | 0 | 0 | 0 | 1 | 0.34 | 0 | 0 | 1.075 | 0.105 | 0.97576789 | NA | 0 | 6.9363 | 6.5821 | 6.5821 | TRUE | 0.5216 |
| Rhox4d | 0 | 2 | 0 | 0 | 0 | 0 | 0 | 2 | 0 | 0 | 0 | 0 | 0.35 | 1 | 0 | 0.272 | -1.878 | 0.62190613 | NA | 10 | 6.6869 | 8.7633 | 8.7633 | TRUE | 0.9743 |
| Rhox4e | 2 | 2 | 5 | 0 | 0 | 0 | 2 | 2 | 5 | 0 | 0 | 0 | 1.51 | 3 | 0 | 0.064 | -3.975 | 0.03358532 | NA | 0 | 1.5693 | 1.3604 | 1.3604 | TRUE | 0.5768 |
| Rhox4f | 1 | 0 | 0 | 0 | 0 | 0 | 1 | 0 | 0 | 0 | 0 | 0 | 0.13 | 0 | 0 | 0.552 | -0.857 | 0.8336548 | NA | 0 | 18.3121 | 10 | 10 | TRUE | 0.2566 |
| Rhox4g | 5 | 6 | 8 | 4 | 2 | 2 | 4 | 6 | 9 | 4 | 2 | 2 | 4.52 | 6 | 3 | 0.441 | -1.18 | 0.20691639 | NA | 0 | 0.527 | 0.3636 | 0.3636 | TRUE | 0.1466 |
| Rhox5 | 3103 | 2627 | 2379 | 1691 | 1692 | 1665 | 2410 | 2794 | 2567 | 1683 | 1909 | 1644 | 2167.8 | 2590 | 1745 | 0.674 | -0.57 | 1.95E-12 | 2.28E-10 | 0.0054 | 0.0035 | 0.0042 | 0.0042 | TRUE | 0.0802 |
| Rhox6 | 249 | 212 | 245 | 124 | 138 | 141 | 193 | 226 | 264 | 123 | 156 | 139 | 183.59 | 228 | 139 | 0.613 | -0.705 | 3.47E-05 | 0.00052521 | 0.0144 | 0.0153 | 0.015 | 0.015 | TRUE | 0.1572 |
| Rhox7 | 23 | 31 | 29 | 22 | 23 | 25 | 18 | 33 | 31 | 22 | 26 | 25 | 25.78 | 27 | 24 | 0.893 | -0.164 | 0.68026209 | 0.80368411 | 0 | 0.0944 | 0.0748 | 0.0748 | TRUE | 0.7207 |
| Rhox8 | 0 | 0 | 0 | 0 | 0 | 0 | 0 | 0 | 0 | 0 | 0 | 0 | 0 | 0 | 0 | NA | NA | NA | NA | NA | NA | NA | NA | NA | NA |
| Rhox9 | 207 | 228 | 245 | 109 | 101 | 96 | 161 | 243 | 264 | 108 | 114 | 95 | 164.14 | 223 | 106 | 0.477 | -1.068 | 1.41E-07 | 5.29E-06 | 0.0352 | 0.0169 | 0.0226 | 0.0226 | TRUE | 0.6927 |
| Rhpn1 | 238 | 214 | 200 | 201 | 166 | 198 | 185 | 228 | 216 | 200 | 187 | 195 | 201.84 | 210 | 194 | 0.932 | -0.102 | 0.48673805 | 0.65262167 | 0 | 0.0142 | 0.0106 | 0.0106 | TRUE | 0.1267 |
| Rhpn2 | 2639 | 1929 | 2231 | 2238 | 1980 | 2285 | 2050 | 2052 | 2407 | 2228 | 2234 | 2256 | 2204.28 | 2170 | 2239 | 1.033 | 0.046 | 0.53845705 | 0.69686405 | 0.0039 | 0.0035 | 0.0037 | 0.0037 | TRUE | 0.1099 |
| Rian | 14512 | 11545 | 10883 | 14815 | 13339 | 15860 | 11271 | 12280 | 11743 | 14747 | 15048 | 15656 | 13457.44 | 11765 | 15150 | 1.288 | 0.365 | 1.44E-11 | 1.38E-09 | 0.0013 | 0.0026 | 0.002 | 0.002 | TRUE | 0.018 |
| Ribc1 | 488 | 333 | 294 | 337 | 291 | 329 | 379 | 354 | 317 | 335 | 328 | 325 | 339.82 | 350 | 329 | 0.939 | -0.091 | 0.43678344 | 0.60817786 | 0 | 0.0094 | 0.007 | 0.007 | TRUE | 0.0758 |
| Ribc2 | 7 | 5 | 1 | 5 | 1 | 1 | 5 | 5 | 1 | 5 | 1 | 1 | 3.15 | 4 | 2 | 0.597 | -0.743 | 0.52217349 | NA | 0.3157 | 0.7538 | 0.6342 | 0.6342 | TRUE | 1.0014 |
| Ric3 | 1349 | 989 | 854 | 911 | 813 | 827 | 1048 | 1052 | 921 | 907 | 917 | 816 | 943.58 | 1007 | 880 | 0.873 | -0.196 | 0.026013 | 0.08370189 | 0.0037 | 0.0049 | 0.0045 | 0.0045 | TRUE | 0.0645 |
| Ric8 | 2245 | 1636 | 1495 | 1286 | 1271 | 1450 | 1744 | 1740 | 1613 | 1280 | 1434 | 1431 | 1540.36 | 1699 | 1382 | 0.813 | -0.299 | 6.37E-05 | 0.00085392 | 0.0023 | 0.004 | 0.0034 | 0.0034 | TRUE | 0.0502 |
| Ric8b | 1421 | 1029 | 1100 | 1320 | 1201 | 1405 | 1104 | 1095 | 1187 | 1314 | 1355 | 1387 | 1240.13 | 1129 | 1352 | 1.199 | 0.262 | 0.00029997 | 0.00290679 | 5.00E-04 | 0.0043 | 0.003 | 0.003 | TRUE | 0.024 |
| Rictor | 4388 | 3183 | 3279 | 3608 | 3383 | 3770 | 3408 | 3386 | 3538 | 3591 | 3816 | 3722 | 3576.84 | 3444 | 3710 | 1.077 | 0.107 | 0.05555595 | 0.1452994 | 4.00E-04 | 0.0031 | 0.002 | 0.002 | TRUE | 0.0095 |
| Rif1 | 30930 | 22176 | 22179 | 24319 | 21197 | 23704 | 24022 | 23588 | 23931 | 24207 | 23912 | 23400 | 23843.39 | 23847 | 23840 | 1 | 0 | 0.99114695 | 0.99553503 | 1.00E-04 | 0.0025 | 0.0013 | 0.0013 | TRUE | 0.0032 |
| Riiad1 | 1 | 2 | 2 | 2 | 1 | 3 | 1 | 2 | 2 | 2 | 1 | 3 | 1.86 | 2 | 2 | 1.24 | 0.311 | 0.82079974 | NA | 0 | 1.2787 | 0.8084 | 0.8084 | TRUE | 0.2441 |
| Rilp | 22 | 12 | 11 | 12 | 6 | 11 | 17 | 13 | 12 | 12 | 7 | 11 | 11.88 | 14 | 10 | 0.707 | -0.5 | 0.37331817 | NA | 0 | 0.2019 | 0.1404 | 0.1404 | TRUE | 0.2279 |
| Rilpl1 | 1126 | 818 | 819 | 900 | 804 | 906 | 875 | 870 | 884 | 896 | 907 | 894 | 887.59 | 876 | 899 | 1.026 | 0.037 | 0.62133645 | 0.76111715 | 0 | 0.0051 | 0.003 | 0.003 | TRUE | 7.00E-04 |
| Rilpl2 | 448 | 335 | 333 | 385 | 369 | 438 | 348 | 356 | 359 | 383 | 416 | 432 | 382.58 | 354 | 410 | 1.159 | 0.213 | 0.0497262 | 0.13427859 | 0 | 0.0086 | 0.0058 | 0.0058 | TRUE | 0.0401 |
| Rimbp2 | 286 | 249 | 325 | 157 | 198 | 230 | 222 | 265 | 351 | 156 | 223 | 227 | 240.72 | 279 | 202 | 0.726 | -0.462 | 0.01451786 | 0.05443434 | 0.0437 | 0.0123 | 0.0215 | 0.0215 | TRUE | 0.3832 |
| Rimbp3 | 187 | 164 | 83 | 88 | 75 | 60 | 145 | 174 | 90 | 88 | 85 | 59 | 106.78 | 136 | 77 | 0.564 | -0.826 | 0.00125863 | 0.0085164 | 0.0681 | 0.0246 | 0.037 | 0.037 | TRUE | 0.2838 |
| Rimkla | 523 | 350 | 311 | 268 | 235 | 259 | 406 | 372 | 336 | 267 | 265 | 256 | 316.93 | 371 | 263 | 0.705 | -0.504 | 4.08E-05 | 0.00059527 | 0 | 0.0099 | 0.0076 | 0.0076 | TRUE | 0.0821 |
| Rimklb | 3120 | 2551 | 2577 | 2697 | 2252 | 2531 | 2423 | 2713 | 2781 | 2685 | 2540 | 2498 | 2606.79 | 2639 | 2574 | 0.976 | -0.034 | 0.62719421 | 0.76496691 | 0.0031 | 0.0033 | 0.0032 | 0.0032 | TRUE | 0.0637 |
| Rims1 | 349 | 270 | 243 | 195 | 190 | 198 | 271 | 287 | 262 | 194 | 214 | 195 | 237.39 | 273 | 201 | 0.735 | -0.443 | 0.00082189 | 0.00626858 | 0 | 0.0124 | 0.0083 | 0.0083 | TRUE | 0.0334 |
| Rims2 | 830 | 572 | 529 | 616 | 488 | 587 | 645 | 608 | 571 | 613 | 551 | 579 | 594.5 | 608 | 581 | 0.955 | -0.067 | 0.48436696 | 0.65044468 | 0.0017 | 0.0064 | 0.0049 | 0.0049 | TRUE | 0.0346 |
| Rims3 | 479 | 355 | 365 | 469 | 347 | 379 | 372 | 378 | 394 | 467 | 391 | 374 | 395.98 | 381 | 411 | 1.079 | 0.11 | 0.34937563 | 0.52388019 | 0.0051 | 0.0084 | 0.0074 | 0.0074 | TRUE | 0.1666 |
| Rims4 | 137 | 114 | 94 | 123 | 115 | 125 | 106 | 121 | 101 | 122 | 130 | 123 | 117.44 | 109 | 125 | 1.142 | 0.191 | 0.29267308 | 0.46492953 | 0 | 0.0226 | 0.0153 | 0.0153 | TRUE | 0.0797 |
| Rin1 | 261 | 270 | 365 | 356 | 272 | 321 | 203 | 287 | 394 | 354 | 307 | 317 | 310.3 | 295 | 326 | 1.112 | 0.153 | 0.41965505 | 0.59194015 | 0.0535 | 0.0101 | 0.0226 | 0.0226 | TRUE | 0.2821 |
| Rin2 | 841 | 609 | 647 | 675 | 606 | 699 | 653 | 648 | 698 | 672 | 684 | 690 | 674.1 | 666 | 682 | 1.024 | 0.034 | 0.69108084 | 0.81113382 | 0 | 0.0059 | 0.0038 | 0.0038 | TRUE | 0.0198 |
| Rin3 | 345 | 224 | 191 | 155 | 144 | 120 | 268 | 238 | 206 | 154 | 162 | 118 | 191.25 | 237 | 145 | 0.608 | -0.718 | 2.46E-05 | 0.00039363 | 0.0163 | 0.0148 | 0.0152 | 0.0152 | TRUE | 0.2523 |
| Ring1 | 400 | 311 | 216 | 202 | 188 | 164 | 311 | 311 | 233 | 201 | 212 | 162 | 241.6 | 292 | 192 | 0.656 | -0.609 | 0.00021981 | 0.00229323 | 0.0223 | 0.0122 | 0.0152 | 0.0152 | TRUE | 0.3352 |
| Rinl | 1757 | 1290 | 1079 | 719 | 738 | 756 | 1365 | 1372 | 1164 | 716 | 833 | 746 | 1032.58 | 1300 | 765 | 0.587 | -0.768 | 2.23E-16 | 4.43E-14 | 0.0063 | 0.0047 | 0.0053 | 0.0053 | TRUE | 0.099 |
| Rint1 | 1215 | 872 | 770 | 894 | 779 | 937 | 944 | 928 | 831 | 890 | 879 | 925 | 899.27 | 901 | 898 | 0.996 | -0.006 | 0.94463454 | 0.96952904 | 0.0016 | 0.0051 | 0.0039 | 0.0039 | TRUE | 0.0536 |
| Riok1 | 2001 | 1392 | 1175 | 1250 | 1097 | 1233 | 1554 | 1481 | 1268 | 1244 | 1238 | 1217 | 1333.58 | 1434 | 1233 | 0.859 | -0.22 | 0.00991242 | 0.04110189 | 0.005 | 0.0042 | 0.0045 | 0.0045 | TRUE | 0.1225 |
| Riok2 | 6543 | 4202 | 3411 | 3740 | 3087 | 3559 | 5082 | 4470 | 3680 | 3723 | 3482 | 3513 | 3991.71 | 4411 | 3573 | 0.81 | -0.305 | 0.00161399 | 0.01019838 | 0.013 | 0.003 | 0.0065 | 0.0065 | TRUE | 0.2139 |
| Riok3 | 6076 | 4416 | 4330 | 5273 | 4345 | 5107 | 4719 | 4697 | 4672 | 5249 | 4902 | 5041 | 4880 | 4696 | 5064 | 1.078 | 0.109 | 0.03960931 | 0.11377328 | 4.00E-04 | 0.0029 | 0.0018 | 0.0018 | TRUE | 0.0124 |
| Ripk1 | 454 | 262 | 212 | 248 | 241 | 267 | 353 | 279 | 229 | 247 | 272 | 264 | 273.72 | 287 | 261 | 0.905 | -0.144 | 0.36314713 | 0.53786674 | 0.0217 | 0.0111 | 0.0144 | 0.0144 | TRUE | 0.3048 |
| Ripk2 | 1015 | 754 | 788 | 798 | 709 | 856 | 788 | 802 | 850 | 794 | 800 | 845 | 813.29 | 813 | 813 | 1 | 0.001 | 0.99338562 | 0.99603371 | 0 | 0.0053 | 0.0036 | 0.0036 | TRUE | 0.0182 |
| Ripk3 | 724 | 539 | 540 | 740 | 669 | 754 | 562 | 573 | 583 | 737 | 755 | 744 | 658.98 | 573 | 745 | 1.302 | 0.38 | 7.10E-06 | 0.00014131 | 0 | 0.006 | 0.0036 | 0.0036 | TRUE | 0.0032 |
| Ripk4 | 1603 | 1178 | 1204 | 1376 | 1237 | 1351 | 1245 | 1253 | 1299 | 1370 | 1395 | 1334 | 1315.98 | 1266 | 1366 | 1.08 | 0.111 | 0.10472927 | 0.22870825 | 0 | 0.0042 | 0.0026 | 0.0026 | TRUE | 0.0063 |
| Ripply1 | 78 | 56 | 106 | 100 | 114 | 150 | 61 | 60 | 114 | 100 | 129 | 148 | 101.79 | 78 | 126 | 1.614 | 0.69 | 0.00926865 | 0.03917117 | 0.0789 | 0.0257 | 0.0404 | 0.0404 | TRUE | 1.0063 |
| Ripply2 | 3 | 1 | 1 | 4 | 5 | 4 | 2 | 1 | 1 | 4 | 6 | 4 | 3.01 | 1 | 5 | 2.927 | 1.549 | 0.17428437 | NA | 0 | 0.7905 | 0.5103 | 0.5103 | TRUE | 0.2286 |
| Ripply3 | 167 | 106 | 87 | 174 | 151 | 167 | 130 | 113 | 94 | 173 | 170 | 165 | 140.79 | 112 | 169 | 1.502 | 0.587 | 0.00090479 | 0.00673899 | 0.004 | 0.0193 | 0.0152 | 0.0152 | TRUE | 0.2089 |
| Rit1 | 537 | 316 | 317 | 347 | 302 | 347 | 417 | 336 | 342 | 345 | 341 | 343 | 353.98 | 365 | 343 | 0.936 | -0.096 | 0.43353776 | 0.60548712 | 0.0055 | 0.0091 | 0.0079 | 0.0079 | TRUE | 0.1886 |
| Rit2 | 0 | 1 | 0 | 1 | 0 | 0 | 0 | 1 | 0 | 1 | 0 | 0 | 0.34 | 0 | 0 | 1.075 | 0.105 | 0.97573409 | NA | 0 | 6.9086 | 6.5582 | 6.5582 | TRUE | 0.5216 |
| Rlbp1 | 279 | 220 | 197 | 213 | 150 | 184 | 217 | 234 | 213 | 212 | 169 | 182 | 204.36 | 221 | 188 | 0.85 | -0.234 | 0.112715 | 0.24131747 | 0.0021 | 0.014 | 0.0107 | 0.0107 | TRUE | 0.1403 |
| Rlf | 6209 | 4801 | 5409 | 5358 | 4613 | 5615 | 4822 | 5107 | 5836 | 5333 | 5204 | 5543 | 5307.58 | 5255 | 5360 | 1.02 | 0.029 | 0.69222914 | 0.81185124 | 0.0052 | 0.0029 | 0.0037 | 0.0037 | TRUE | 0.1138 |
| Rlim | 12752 | 8372 | 9366 | 4917 | 3997 | 5349 | 9904 | 8905 | 10106 | 4894 | 4509 | 5280 | 7266.44 | 9638 | 4894 | 0.508 | -0.977 | 4.93E-41 | 9.07E-38 | 0.0051 | 0.0028 | 0.0037 | 0.0037 | TRUE | 0.0579 |
| Rln1 | 126 | 88 | 66 | 98 | 76 | 79 | 98 | 94 | 71 | 98 | 86 | 78 | 87.32 | 88 | 87 | 0.991 | -0.013 | 0.95149252 | 0.97323401 | 0.0088 | 0.0296 | 0.0238 | 0.0238 | TRUE | 0.2412 |
| Rln3 | 101 | 94 | 85 | 86 | 82 | 103 | 78 | 100 | 92 | 86 | 93 | 102 | 91.65 | 90 | 94 | 1.041 | 0.057 | 0.78382914 | 0.87273272 | 0 | 0.0283 | 0.0207 | 0.0207 | TRUE | 0.1386 |
| Rltpr | 939 | 757 | 854 | 816 | 784 | 842 | 729 | 805 | 921 | 812 | 884 | 831 | 830.64 | 818 | 842 | 1.031 | 0.044 | 0.65553592 | 0.78610411 | 0.0068 | 0.0053 | 0.0058 | 0.0058 | TRUE | 0.1428 |
| Rmdn1 | 167 | 125 | 123 | 158 | 138 | 141 | 130 | 133 | 133 | 157 | 156 | 139 | 141.25 | 132 | 151 | 1.144 | 0.194 | 0.24138111 | 0.40911421 | 0 | 0.0192 | 0.0126 | 0.0126 | TRUE | 0.048 |
| Rmdn2 | 138 | 105 | 68 | 53 | 50 | 63 | 107 | 112 | 73 | 53 | 56 | 62 | 77.26 | 97 | 57 | 0.585 | -0.774 | 0.00152229 | 0.00975698 | 0.0187 | 0.0331 | 0.0289 | 0.0289 | TRUE | 0.4301 |
| Rmdn3 | 2949 | 2169 | 2013 | 2316 | 2099 | 2417 | 2290 | 2307 | 2172 | 2305 | 2368 | 2386 | 2304.78 | 2256 | 2353 | 1.042 | 0.06 | 0.31904647 | 0.493086 | 2.00E-04 | 0.0035 | 0.0022 | 0.0022 | TRUE | 0.0127 |
| Rmi1 | 1200 | 841 | 736 | 835 | 695 | 765 | 932 | 895 | 794 | 831 | 784 | 755 | 831.84 | 874 | 790 | 0.903 | -0.147 | 0.10391307 | 0.22762139 | 0.0035 | 0.0053 | 0.0047 | 0.0047 | TRUE | 0.074 |
| Rmi2 | 382 | 330 | 332 | 448 | 363 | 455 | 297 | 351 | 358 | 446 | 410 | 449 | 385.09 | 335 | 435 | 1.302 | 0.381 | 0.00108472 | 0.00767615 | 0.004 | 0.0086 | 0.0072 | 0.0072 | TRUE | 0.1255 |
| Rmnd1 | 4 | 4 | 9 | 4 | 7 | 8 | 3 | 4 | 10 | 4 | 8 | 8 | 6.14 | 6 | 7 | 1.177 | 0.235 | 0.77052804 | NA | 0.0954 | 0.3883 | 0.3019 | 0.3019 | TRUE | 0.8071 |
| Rmnd5a | 11508 | 8143 | 8456 | 9092 | 7798 | 8684 | 8938 | 8662 | 9124 | 9050 | 8797 | 8572 | 8857.15 | 8908 | 8806 | 0.989 | -0.017 | 0.74473938 | 0.84823969 | 5.00E-04 | 0.0027 | 0.0017 | 0.0017 | TRUE | 0.0072 |

| | | | | | | | | | | | | | | | | | | | | | | | | | |
|---|---|---|---|---|---|---|---|---|---|---|---|---|---|---|---|---|---|---|---|---|---|---|---|---|---|
| Rmnd5b | 3184 | 2460 | 2417 | 2336 | 2367 | 2685 | 2473 | 2617 | 2608 | 2325 | 2670 | 2651 | 2557.24 | 2566 | 2549 | 0.993 | -0.009 | 0.89442283 | 0.94200763 | 0.0031 | 0.0034 | 0.0032 | 0.0032 | TRUE | 0.0717 |
| Rmrp | 8 | 8 | 5 | 7 | 8 | 14 | 6 | 9 | 5 | 7 | 9 | 14 | 8.32 | 7 | 10 | 1.489 | 0.574 | 0.39313505 | NA | 0 | 0.2872 | 0.2014 | 0.2014 | TRUE | 0.4197 |
| Rmst | 1 | 0 | 0 | 0 | 0 | 1 | 1 | 0 | 0 | 0 | 0 | 1 | 0.29 | 0 | 0 | 1.075 | 0.105 | 0.97702352 | NA | 0 | 8.0647 | 7.5458 | 7.5458 | TRUE | 0.3835 |
| Rn4.5s | 0 | 0 | 0 | 0 | 0 | 0 | 0 | 0 | 0 | 0 | 0 | 0 | 0 | 0 | 0 | NA | NA | NA | NA | NA | NA | NA | NA | NA | NA |
| Rn45s | 106700 | 89088 | 75211 | 94271 | 79896 | 101834 | 82869 | 94762 | 81152 | 93837 | 90131 | 100526 | 90546.21 | 86261 | 94831 | 1.099 | 0.137 | 0.04948461 | 0.13380317 | 0.0052 | 0.0025 | 0.0035 | 0.0035 | TRUE | 0.091 |
| Rnase1 | 7 | 7 | 7 | 11 | 5 | 8 | 5 | 7 | 8 | 11 | 6 | 8 | 7.49 | 7 | 8 | 1.214 | 0.28 | 0.68741814 | NA | 0 | 0.319 | 0.2149 | 0.2149 | TRUE | 0.2503 |
| Rnase10 | 1 | 0 | 0 | 0 | 1 | 0 | 1 | 0 | 0 | 0 | 1 | 0 | 0.32 | 0 | 0 | 1.075 | 0.105 | 0.97638847 | NA | 0 | 7.4681 | 7.0392 | 7.0392 | TRUE | 0.5075 |
| Rnase11 | 0 | 0 | 0 | 0 | 0 | 0 | 0 | 0 | 0 | 0 | 0 | 0 | 0 | 0 | 0 | NA | NA | NA | NA | NA | NA | NA | NA | NA | NA |
| Rnase12 | 2 | 0 | 9 | 3 | 1 | 1 | 2 | 0 | 10 | 3 | 1 | 1 | 2.73 | 4 | 2 | 0.462 | -1.115 | 0.42301252 | NA | 1.1911 | 0.8713 | 0.9611 | 0.9611 | TRUE | 1.6078 |
| Rnase13 | 0 | 0 | 0 | 0 | 0 | 0 | 0 | 0 | 0 | 0 | 0 | 0 | 0 | 0 | 0 | NA | NA | NA | NA | NA | NA | NA | NA | NA | NA |
| Rnase4 | 124 | 100 | 110 | 154 | 156 | 178 | 96 | 106 | 119 | 153 | 176 | 176 | 137.73 | 107 | 168 | 1.577 | 0.657 | 0.00017212 | 0.0018997 | 0 | 0.0196 | 0.0145 | 0.0145 | TRUE | 0.088 |
| Rnase6 | 8 | 6 | 6 | 7 | 10 | 3 | 6 | 6 | 6 | 7 | 11 | 3 | 6.71 | 6 | 7 | 1.103 | 0.141 | 0.85293808 | NA | 0 | 0.3555 | 0.2673 | 0.2673 | TRUE | 0.1798 |
| Rnase9 | 0 | 0 | 0 | 0 | 0 | 1 | 0 | 0 | 0 | 0 | 0 | 1 | 0.16 | 0 | 0 | 2.094 | 1.067 | 0.79379329 | NA | 0 | 14.4078 | 10 | 10 | TRUE | 0.3835 |
| Rnaseh1 | 1145 | 852 | 797 | 805 | 728 | 849 | 889 | 906 | 860 | 801 | 821 | 838 | 852.69 | 885 | 820 | 0.926 | -0.11 | 0.15893874 | 0.30807401 | 0 | 0.0052 | 0.0032 | 0.0032 | TRUE | 0.007 |
| Rnaseh2a | 5685 | 4209 | 4242 | 4394 | 3945 | 4675 | 4415 | 4477 | 4577 | 4374 | 4450 | 4615 | 4484.76 | 4490 | 4480 | 0.998 | -0.003 | 0.9560862 | 0.9757358 | 3.00E-04 | 0.003 | 0.0018 | 0.0018 | TRUE | 0.0086 |
| Rnaseh2b | 2034 | 1489 | 1564 | 1389 | 1181 | 1486 | 1580 | 1584 | 1688 | 1383 | 1332 | 1467 | 1505.48 | 1617 | 1394 | 0.863 | -0.213 | 0.00270624 | 0.0152105 | 0.0012 | 0.004 | 0.003 | 0.003 | TRUE | 0.0253 |
| Rnaseh2c | 1098 | 817 | 852 | 866 | 789 | 929 | 853 | 869 | 919 | 862 | 890 | 917 | 885.04 | 880 | 890 | 1.011 | 0.016 | 0.83541137 | 0.90637245 | 0 | 0.0051 | 0.0034 | 0.0034 | TRUE | 0.0175 |
| Rnasek | 2674 | 1786 | 1912 | 2079 | 1732 | 2001 | 2077 | 1900 | 2063 | 2069 | 1954 | 1975 | 2006.36 | 2013 | 1999 | 0.993 | -0.01 | 0.87710206 | 0.93138507 | 0.0012 | 0.0036 | 0.0027 | 0.0027 | TRUE | 0.029 |
| Rnasel | 46 | 36 | 53 | 65 | 63 | 69 | 36 | 38 | 57 | 65 | 71 | 68 | 55.85 | 44 | 68 | 1.565 | 0.646 | 0.01910091 | 0.06680651 | 0.0098 | 0.0449 | 0.0361 | 0.0361 | TRUE | 0.5355 |
| Rnaset2a | 0 | 0 | 0 | 0 | 0 | 0 | 0 | 0 | 0 | 0 | 0 | 0 | 0 | 0 | 0 | NA | NA | NA | NA | NA | NA | NA | NA | NA | NA |
| Rnaset2b | 1 | 0 | 4 | 0 | 1 | 0 | 1 | 0 | 4 | 0 | 1 | 0 | 1.04 | 2 | 0 | 0.221 | -2.176 | 0.3254724 | NA | 2.0129 | 2.2884 | 2.241 | 2.241 | TRUE | 1.0791 |
| Rnd1 | 561 | 390 | 348 | 325 | 278 | 307 | 436 | 415 | 375 | 324 | 314 | 303 | 361.03 | 409 | 314 | 0.765 | -0.386 | 0.00070902 | 0.00562064 | 0 | 0.009 | 0.0065 | 0.0065 | TRUE | 0.057 |
| Rnd2 | 1074 | 732 | 732 | 801 | 728 | 857 | 834 | 779 | 790 | 797 | 821 | 846 | 811.19 | 801 | 821 | 1.025 | 0.035 | 0.66427098 | 0.79308223 | 0 | 0.0053 | 0.0035 | 0.0035 | TRUE | 0.0166 |
| Rnd3 | 884 | 619 | 510 | 545 | 461 | 544 | 687 | 658 | 550 | 542 | 520 | 537 | 582.47 | 632 | 533 | 0.843 | -0.247 | 0.01833554 | 0.06474056 | 0.0055 | 0.0065 | 0.0062 | 0.0062 | TRUE | 0.1467 |
| Rnf10 | 28569 | 20987 | 21099 | 22331 | 19994 | 23015 | 22188 | 22324 | 22766 | 22228 | 22555 | 22719 | 22463.41 | 22426 | 22501 | 1.003 | 0.005 | 0.91102849 | 0.95115633 | 1.00E-04 | 0.0025 | 0.0013 | 0.0013 | TRUE | 0.0021 |
| Rnf103 | 588 | 390 | 357 | 426 | 312 | 358 | 457 | 415 | 385 | 424 | 352 | 353 | 397.69 | 419 | 376 | 0.897 | -0.156 | 0.19541726 | 0.35513018 | 0.007 | 0.0084 | 0.008 | 0.008 | TRUE | 0.1416 |
| Rnf11 | 2057 | 1450 | 1508 | 1647 | 1332 | 1365 | 1598 | 1542 | 1627 | 1639 | 1503 | 1347 | 1542.76 | 1589 | 1496 | 0.942 | -0.087 | 0.28810047 | 0.46024589 | 0.0042 | 0.004 | 0.0041 | 0.0041 | TRUE | 0.0926 |
| Rnf111 | 3964 | 2819 | 2646 | 3019 | 2553 | 3040 | 3079 | 2999 | 2855 | 3005 | 2880 | 3001 | 2969.72 | 2978 | 2962 | 0.995 | -0.008 | 0.89381599 | 0.94181872 | 7.00E-04 | 0.0032 | 0.0022 | 0.0022 | TRUE | 0.0156 |
| Rnf112 | 22 | 20 | 15 | 15 | 14 | 17 | 17 | 21 | 16 | 15 | 16 | 17 | 17.01 | 18 | 16 | 0.873 | -0.196 | 0.66847578 | NA | 0 | 0.1418 | 0.0919 | 0.0919 | TRUE | 0.1053 |
| Rnf113a1 | 259 | 180 | 169 | 159 | 164 | 156 | 201 | 191 | 182 | 158 | 185 | 154 | 178.71 | 191 | 166 | 0.862 | -0.214 | 0.16198971 | 0.31205405 | 0 | 0.0157 | 0.0113 | 0.0113 | TRUE | 0.1051 |
| Rnf113a2 | 837 | 632 | 555 | 624 | 474 | 587 | 650 | 672 | 599 | 621 | 535 | 579 | 609.41 | 640 | 578 | 0.904 | -0.146 | 0.13254303 | 0.27055664 | 0.0027 | 0.0063 | 0.0052 | 0.0052 | TRUE | 0.0498 |
| Rnf114 | 4931 | 3741 | 4178 | 4186 | 3874 | 4477 | 3830 | 3979 | 4508 | 4167 | 4370 | 4419 | 4212.25 | 4106 | 4319 | 1.052 | 0.074 | 0.29648516 | 0.46836117 | 0.0038 | 0.003 | 0.0034 | 0.0034 | TRUE | 0.0891 |
| Rnf115 | 3661 | 2628 | 2920 | 3084 | 2693 | 3250 | 2843 | 2795 | 3151 | 3070 | 3038 | 3208 | 3017.57 | 2930 | 3105 | 1.06 | 0.085 | 0.19988283 | 0.36049531 | 0.0021 | 0.0032 | 0.0028 | 0.0028 | TRUE | 0.0521 |
| Rnf121 | 1060 | 871 | 845 | 886 | 780 | 975 | 823 | 926 | 912 | 882 | 880 | 962 | 897.63 | 887 | 908 | 1.026 | 0.036 | 0.67011079 | 0.79708707 | 0.0024 | 0.0051 | 0.0041 | 0.0041 | TRUE | 0.0498 |
| Rnf122 | 388 | 340 | 340 | 372 | 325 | 379 | 301 | 362 | 367 | 370 | 367 | 374 | 356.82 | 343 | 370 | 1.083 | 0.115 | 0.33343968 | 0.50770783 | 0.0036 | 0.0091 | 0.0073 | 0.0073 | TRUE | 0.1411 |
| Rnf123 | 917 | 709 | 773 | 831 | 649 | 801 | 712 | 754 | 834 | 827 | 732 | 791 | 775.07 | 767 | 783 | 1.024 | 0.034 | 0.71552212 | 0.829237 | 0.0038 | 0.0055 | 0.0049 | 0.0049 | TRUE | 0.0692 |
| Rnf125 | 755 | 604 | 591 | 596 | 561 | 721 | 586 | 642 | 638 | 593 | 633 | 712 | 634.06 | 622 | 646 | 1.04 | 0.056 | 0.56944544 | 0.72308845 | 0.0041 | 0.0062 | 0.0055 | 0.0055 | TRUE | 0.0952 |
| Rnf126 | 5161 | 4050 | 4226 | 4082 | 3849 | 4509 | 4008 | 4308 | 4560 | 4063 | 4342 | 4451 | 4288.74 | 4292 | 4285 | 0.999 | -0.002 | 0.98137886 | 0.99024695 | 0.003 | 0.003 | 0.003 | 0.003 | TRUE | 0.0414 |
| Rnf128 | 772 | 594 | 662 | 874 | 616 | 674 | 600 | 632 | 714 | 870 | 695 | 665 | 695.99 | 649 | 743 | 1.148 | 0.199 | 0.08566073 | 0.19785729 | 0.0131 | 0.0058 | 0.0082 | 0.0082 | TRUE | 0.2647 |
| Rnf13 | 1742 | 1233 | 1376 | 1606 | 1385 | 1644 | 1353 | 1312 | 1485 | 1599 | 1562 | 1623 | 1488.85 | 1383 | 1595 | 1.154 | 0.206 | 0.00438088 | 0.02220421 | 0.0013 | 0.004 | 0.0031 | 0.0031 | TRUE | 0.0483 |
| Rnf130 | 1656 | 1254 | 1375 | 1586 | 1397 | 1469 | 1286 | 1334 | 1484 | 1579 | 1576 | 1450 | 1451.4 | 1368 | 1535 | 1.123 | 0.167 | 0.03338818 | 0.10028062 | 0.0029 | 0.0041 | 0.0038 | 0.0038 | TRUE | 0.0652 |
| Rnf133 | 0 | 0 | 0 | 0 | 0 | 0 | 0 | 0 | 0 | 0 | 0 | 0 | 0 | 0 | 0 | NA | NA | NA | NA | NA | NA | NA | NA | NA | NA |
| Rnf135 | 329 | 243 | 234 | 217 | 207 | 245 | 256 | 258 | 252 | 216 | 234 | 242 | 242.98 | 255 | 231 | 0.902 | -0.149 | 0.24966532 | 0.41849026 | 0 | 0.0122 | 0.0079 | 0.0079 | TRUE | 0.0341 |
| Rnf138 | 4868 | 3774 | 4111 | 4694 | 3902 | 4762 | 3781 | 4014 | 4436 | 4672 | 4402 | 4701 | 4334.33 | 4077 | 4592 | 1.127 | 0.172 | 0.01309918 | 0.05046538 | 0.0036 | 0.003 | 0.0033 | 0.0033 | TRUE | 0.072 |
| Rnf139 | 2099 | 1471 | 1498 | 1622 | 1449 | 1613 | 1630 | 1565 | 1616 | 1615 | 1635 | 1592 | 1608.78 | 1604 | 1614 | 1.006 | 0.008 | 0.89552266 | 0.94268886 | 0 | 0.0039 | 0.0023 | 0.0023 | TRUE | 0.0054 |
| Rnf14 | 4186 | 2873 | 2997 | 3314 | 2940 | 3372 | 3251 | 3056 | 3234 | 3299 | 3317 | 3329 | 3247.48 | 3180 | 3315 | 1.042 | 0.059 | 0.28969894 | 0.46187027 | 3.00E-04 | 0.0032 | 0.002 | 0.002 | TRUE | 0.014 |
| Rnf141 | 729 | 576 | 566 | 650 | 553 | 612 | 566 | 613 | 611 | 647 | 624 | 604 | 610.76 | 597 | 625 | 1.049 | 0.069 | 0.44335393 | 0.61433424 | 0 | 0.0063 | 0.0043 | 0.0043 | TRUE | 0.0249 |
| Rnf144a | 1045 | 733 | 781 | 796 | 714 | 811 | 812 | 780 | 843 | 792 | 805 | 801 | 805.39 | 812 | 799 | 0.985 | -0.021 | 0.7903043 | 0.87673234 | 0 | 0.0054 | 0.0034 | 0.0034 | TRUE | 0.0131 |
| Rnf144b | 222 | 149 | 147 | 170 | 178 | 193 | 172 | 158 | 159 | 169 | 201 | 191 | 175.01 | 163 | 187 | 1.142 | 0.191 | 0.21330975 | 0.37556494 | 0 | 0.016 | 0.0113 | 0.0113 | TRUE | 0.0741 |
| Rnf145 | 9286 | 6957 | 7416 | 6491 | 5912 | 6576 | 7212 | 7400 | 8002 | 6461 | 6669 | 6492 | 7039.32 | 7538 | 6541 | 0.868 | -0.204 | 0.00035924 | 0.00334858 | 0.0014 | 0.0028 | 0.0022 | 0.0022 | TRUE | 0.0353 |
| Rnf146 | 1921 | 1449 | 1355 | 1504 | 1327 | 1503 | 1492 | 1541 | 1462 | 1497 | 1497 | 1484 | 1495.51 | 1498 | 1493 | 0.996 | -0.006 | 0.9317309 | 0.96216772 | 0 | 0.004 | 0.0024 | 0.0024 | TRUE | 0.0073 |
| Rnf148 | 0 | 0 | 0 | 0 | 0 | 0 | 0 | 0 | 0 | 0 | 0 | 0 | 0 | 0 | 0 | NA | NA | NA | NA | NA | NA | NA | NA | NA | NA |
| Rnf149 | 2825 | 1903 | 1616 | 1680 | 1468 | 1586 | 2194 | 2024 | 1744 | 1672 | 1656 | 1566 | 1809.31 | 1987 | 1631 | 0.82 | -0.286 | 0.00095054 | 0.00699502 | 0.0063 | 0.0037 | 0.0048 | 0.0048 | TRUE | 0.1382 |
| Rnf150 | 11 | 9 | 8 | 10 | 4 | 2 | 9 | 10 | 9 | 10 | 5 | 2 | 7.2 | 9 | 6 | 0.617 | -0.697 | 0.35805792 | NA | 0.0491 | 0.3317 | 0.2661 | 0.2661 | TRUE | 0.8633 |
| Rnf151 | 3 | 6 | 1 | 3 | 7 | 4 | 2 | 6 | 1 | 3 | 8 | 4 | 4.1 | 3 | 5 | 1.518 | 0.602 | 0.54730531 | NA | 0.1734 | 0.58 | 0.4672 | 0.4672 | TRUE | 0.895 |
| Rnf152 | 111 | 89 | 118 | 128 | 88 | 128 | 86 | 95 | 127 | 127 | 99 | 126 | 110.21 | 103 | 117 | 1.154 | 0.206 | 0.33091806 | 0.50532178 | 0.0215 | 0.0239 | 0.0232 | 0.0232 | TRUE | 0.4195 |
| Rnf157 | 1144 | 751 | 712 | 862 | 778 | 793 | 888 | 799 | 768 | 858 | 878 | 783 | 829.01 | 818 | 840 | 1.023 | 0.033 | 0.71333265 | 0.82763773 | 0.0036 | 0.0053 | 0.0047 | 0.0047 | TRUE | 0.0694 |
| Rnf165 | 1287 | 926 | 1011 | 1124 | 1030 | 1175 | 1000 | 985 | 1091 | 1119 | 1162 | 1160 | 1086.01 | 1025 | 1147 | 1.119 | 0.163 | 0.03271513 | 0.0987117 | 7.00E-04 | 0.0046 | 0.0033 | 0.0033 | TRUE | 0.0365 |
| Rnf166 | 726 | 636 | 615 | 656 | 664 | 738 | 564 | 677 | 664 | 653 | 749 | 729 | 672.42 | 635 | 710 | 1.121 | 0.165 | 0.10620386 | 0.2312219 | 0.0062 | 0.006 | 0.006 | 0.006 | TRUE | 0.1185 |
| Rnf167 | 5298 | 3971 | 4075 | 4130 | 3934 | 4482 | 4115 | 4224 | 4397 | 4111 | 4438 | 4424 | 4284.82 | 4245 | 4324 | 1.019 | 0.027 | 0.64675862 | 0.78033832 | 0.0011 | 0.003 | 0.0023 | 0.0023 | TRUE | 0.0228 |
| Rnf168 | 3514 | 2626 | 2181 | 2237 | 1962 | 2244 | 2729 | 2793 | 2353 | 2227 | 2213 | 2215 | 2421.82 | 2625 | 2218 | 0.845 | -0.244 | 0.00106827 | 0.00758883 | 0.0035 | 0.0034 | 0.0036 | 0.0036 | TRUE | 0.0983 |
| Rnf169 | 3008 | 2204 | 2281 | 2540 | 2282 | 2693 | 2336 | 2344 | 2461 | 2528 | 2574 | 2658 | 2483.8 | 2380 | 2587 | 1.087 | 0.121 | 0.04327795 | 0.12124102 | 3.00E-04 | 0.0034 | 0.0022 | 0.0022 | TRUE | 0.0105 |
| Rnf17 | 3383 | 2490 | 2594 | 2514 | 1920 | 2043 | 2627 | 2649 | 2799 | 2502 | 2166 | 2017 | 2460.01 | 2692 | 2228 | 0.828 | -0.272 | 0.00086859 | 0.00653749 | 0.0062 | 0.0034 | 0.0044 | 0.0044 | TRUE | 0.1411 |
| Rnf170 | 1278 | 1019 | 1045 | 986 | 862 | 1035 | 993 | 1084 | 1128 | 981 | 972 | 1022 | 1029.93 | 1068 | 992 | 0.93 | -0.104 | 0.19264113 | 0.35111896 | 0.0017 | 0.0047 | 0.0036 | 0.0036 | TRUE | 0.0479 |
| Rnf180 | 57 | 41 | 40 | 35 | 39 | 30 | 44 | 44 | 43 | 35 | 44 | 30 | 39.91 | 44 | 36 | 0.824 | -0.279 | 0.36586825 | 0.5405018 | 0 | 0.0618 | 0.0433 | 0.0433 | TRUE | 0.2411 |
| Rnf181 | 5930 | 4310 | 4505 | 5125 | 4241 | 5035 | 4606 | 4584 | 4861 | 5101 | 4784 | 4970 | 4817.83 | 4684 | 4952 | 1.058 | 0.081 | 0.14676054 | 0.29115071 | 7.00E-04 | 0.0029 | 0.002 | 0.002 | TRUE | 0.0132 |
| Rnf182 | 79 | 68 | 60 | 57 | 44 | 49 | 61 | 72 | 65 | 57 | 50 | 48 | 58.86 | 66 | 52 | 0.782 | -0.354 | 0.1609261 | 0.31065866 | 0 | 0.0427 | 0.0287 | 0.0287 | TRUE | 0.0644 |
| Rnf183 | 0 | 4 | 3 | 2 | 1 | 0 | 0 | 4 | 3 | 2 | 1 | 0 | 1.77 | 2 | 1 | 0.425 | -1.233 | 0.45827716 | NA | 0 | 1.3426 | 1.2937 | 1.2937 | TRUE | 0.9633 |
| Rnf185 | 2121 | 1388 | 1314 | 1296 | 1157 | 1368 | 1647 | 1476 | 1418 | 1290 | 1305 | 1350 | 1414.53 | 1514 | 1315 | 0.868 | -0.205 | 0.00838869 | 0.03632217 | 0.0028 | 0.0041 | 0.0036 | 0.0036 | TRUE | 0.0739 |

| | | | | | | | | | | | | | | | | | | | | | | | | | |
|---|---|---|---|---|---|---|---|---|---|---|---|---|---|---|---|---|---|---|---|---|---|---|---|---|---|
| Rnf186 | 4 | 1 | 2 | 6 | 7 | 8 | 3 | 1 | 2 | 6 | 8 | 8 | 4.68 | 2 | 7 | 3.342 | 1.741 | 0.06342061 | NA | 0 | 0.5086 | 0.3408 | 0.3408 | TRUE | 0.2241 |
| Rnf187 | 14835 | 10490 | 9922 | 10375 | 9056 | 10306 | 11522 | 11158 | 10706 | 10327 | 10216 | 10174 | 10683.76 | 11129 | 10239 | 0.92 | -0.12 | 0.0163242 | 0.0594905 | 6.00E-04 | 0.0026 | 0.0017 | 0.0017 | TRUE | 0.0135 |
| Rnf19a | 2688 | 2036 | 2301 | 2447 | 2175 | 2451 | 2088 | 2166 | 2483 | 2436 | 2454 | 2420 | 2340.83 | 2246 | 2437 | 1.086 | 0.119 | 0.1109021 | 0.23852327 | 0.0037 | 0.0034 | 0.0036 | 0.0036 | TRUE | 0.1029 |
| Rnf19b | 4961 | 3565 | 3276 | 3559 | 3319 | 3638 | 3853 | 3792 | 3535 | 3543 | 3744 | 3591 | 3676.31 | 3727 | 3626 | 0.972 | -0.04 | 0.50032179 | 0.66478566 | 0.001 | 0.0031 | 0.0023 | 0.0023 | TRUE | 0.0244 |
| Rnf2 | 7352 | 5258 | 5002 | 5960 | 4881 | 5541 | 5710 | 5593 | 5397 | 5933 | 5506 | 5470 | 5601.44 | 5567 | 5636 | 1.012 | 0.018 | 0.75627975 | 0.85520897 | 0.001 | 0.0028 | 0.0022 | 0.0022 | TRUE | 0.0259 |
| Rnf20 | 3238 | 2412 | 2155 | 2180 | 1932 | 2350 | 2515 | 2566 | 2325 | 2170 | 2179 | 2320 | 2345.82 | 2469 | 2223 | 0.9 | -0.151 | 0.02223626 | 0.07448392 | 0.0014 | 0.0034 | 0.0027 | 0.0027 | TRUE | 0.0307 |
| Rnf207 | 105 | 128 | 124 | 112 | 123 | 153 | 82 | 136 | 134 | 111 | 139 | 151 | 125.46 | 117 | 134 | 1.15 | 0.201 | 0.36726787 | 0.54206523 | 0.0437 | 0.0213 | 0.0278 | 0.0278 | TRUE | 0.7893 |
| Rnf208 | 483 | 370 | 406 | 476 | 417 | 538 | 375 | 394 | 438 | 474 | 470 | 531 | 447.01 | 402 | 492 | 1.226 | 0.294 | 0.0072372 | 0.03252873 | 0.0034 | 0.0077 | 0.0064 | 0.0064 | TRUE | 0.0687 |
| Rnf214 | 1895 | 1483 | 1627 | 1676 | 1500 | 1816 | 1472 | 1577 | 1756 | 1668 | 1692 | 1793 | 1659.64 | 1602 | 1718 | 1.074 | 0.103 | 0.1962435 | 0.35581737 | 0.0038 | 0.0039 | 0.004 | 0.004 | TRUE | 0.0849 |
| Rnf215 | 380 | 268 | 271 | 282 | 250 | 281 | 295 | 285 | 292 | 281 | 282 | 277 | 285.45 | 291 | 280 | 0.962 | -0.056 | 0.63898711 | 0.77456433 | 0 | 0.0107 | 0.0066 | 0.0066 | TRUE | 0.0034 |
| Rnf216 | 4575 | 3589 | 3833 | 3577 | 3468 | 4094 | 3553 | 3818 | 4136 | 3561 | 3912 | 4041 | 3836.8 | 3836 | 3838 | 1.001 | 0.002 | 0.98271256 | 0.99089635 | 0.0048 | 0.003 | 0.0037 | 0.0037 | TRUE | 0.057 |
| Rnf217 | 299 | 202 | 169 | 148 | 91 | 153 | 232 | 215 | 182 | 147 | 103 | 151 | 171.74 | 210 | 134 | 0.638 | -0.649 | 0.00033903 | 0.003188 | 0.021 | 0.0162 | 0.0175 | 0.0175 | TRUE | 0.4034 |
| Rnf219 | 842 | 591 | 505 | 483 | 379 | 475 | 654 | 629 | 545 | 481 | 428 | 469 | 534.12 | 609 | 459 | 0.753 | -0.409 | 0.00011041 | 0.00131502 | 0.0047 | 0.0069 | 0.0062 | 0.0062 | TRUE | 0.0981 |
| Rnf220 | 8018 | 6167 | 6169 | 6326 | 6085 | 7078 | 6227 | 6560 | 6656 | 6297 | 6865 | 6987 | 6598.63 | 6481 | 6716 | 1.036 | 0.052 | 0.39062612 | 0.56447468 | 0.0019 | 0.0028 | 0.0025 | 0.0025 | TRUE | 0.0365 |
| Rnf222 | 11 | 12 | 6 | 9 | 9 | 11 | 9 | 13 | 6 | 9 | 10 | 11 | 9.63 | 9 | 10 | 1.081 | 0.113 | 0.85468485 | NA | 0 | 0.2487 | 0.1691 | 0.1691 | TRUE | 0.3267 |
| Rnf223 | 6 | 5 | 1 | 1 | 2 | 1 | 5 | 5 | 1 | 1 | 2 | 1 | 2.55 | 4 | 1 | 0.375 | -1.415 | 0.2669664 | NA | 0 | 0.9321 | 0.6699 | 0.6699 | TRUE | 0.5371 |
| Rnf24 | 1272 | 899 | 842 | 1065 | 929 | 1115 | 988 | 956 | 909 | 1060 | 1048 | 1101 | 1010.24 | 951 | 1070 | 1.124 | 0.169 | 0.02675972 | 0.08539328 | 0 | 0.0048 | 0.0032 | 0.0032 | TRUE | 0.0179 |
| Rnf25 | 966 | 696 | 603 | 683 | 560 | 714 | 750 | 740 | 651 | 680 | 632 | 705 | 692.94 | 714 | 672 | 0.941 | -0.087 | 0.35289129 | 0.52775572 | 0.0031 | 0.0058 | 0.0049 | 0.0049 | TRUE | 0.069 |
| Rnf26 | 2272 | 1670 | 1718 | 1823 | 1564 | 1803 | 1765 | 1776 | 1854 | 1815 | 1764 | 1780 | 1792.24 | 1798 | 1786 | 0.994 | -0.009 | 0.88695075 | 0.937038 | 0 | 0.0037 | 0.0023 | 0.0023 | TRUE | 0.0087 |
| Rnf31 | 2938 | 2377 | 2680 | 2736 | 2581 | 2931 | 2282 | 2528 | 2892 | 2723 | 2912 | 2893 | 2705.05 | 2567 | 2843 | 1.108 | 0.148 | 0.07870059 | 0.18691718 | 0.0073 | 0.0033 | 0.0047 | 0.0047 | TRUE | 0.148 |
| Rnf32 | 151 | 113 | 121 | 123 | 122 | 163 | 117 | 120 | 131 | 122 | 138 | 161 | 131.5 | 123 | 140 | 1.146 | 0.197 | 0.27487779 | 0.44590037 | 0 | 0.0204 | 0.0158 | 0.0158 | TRUE | 0.1758 |
| Rnf34 | 3017 | 2253 | 2204 | 2425 | 2149 | 2539 | 2343 | 2396 | 2378 | 2414 | 2424 | 2506 | 2410.38 | 2372 | 2448 | 1.032 | 0.046 | 0.4272652 | 0.5992621 | 0 | 0.0034 | 0.002 | 0.002 | TRUE | 0.0053 |
| Rnf38 | 3872 | 2836 | 3028 | 3257 | 2888 | 3390 | 3007 | 3017 | 3267 | 3242 | 3258 | 3346 | 3189.58 | 3097 | 3282 | 1.06 | 0.084 | 0.15778973 | 0.30675852 | 8.00E-04 | 0.0032 | 0.0023 | 0.0023 | TRUE | 0.0277 |
| Rnf39 | 81 | 67 | 30 | 70 | 63 | 66 | 63 | 71 | 32 | 70 | 71 | 65 | 62.07 | 55 | 69 | 1.23 | 0.299 | 0.30298738 | 0.47552416 | 0.0562 | 0.0406 | 0.0446 | 0.0446 | TRUE | 1.0439 |
| Rnf4 | 16723 | 12087 | 11346 | 11755 | 10396 | 11417 | 12988 | 12857 | 12242 | 11701 | 11728 | 11270 | 12131.02 | 12696 | 11566 | 0.911 | -0.135 | 0.00713564 | 0.03221404 | 6.00E-04 | 0.0026 | 0.0017 | 0.0017 | TRUE | 0.0119 |
| Rnf40 | 6182 | 4564 | 4013 | 3977 | 3614 | 4042 | 4801 | 4855 | 4330 | 3959 | 4077 | 3990 | 4335.28 | 4662 | 4009 | 0.86 | -0.218 | 0.00041046 | 0.00371368 | 0.0017 | 0.003 | 0.0025 | 0.0025 | TRUE | 0.0468 |
| Rnf41 | 1660 | 1231 | 1064 | 1151 | 1023 | 1113 | 1289 | 1309 | 1148 | 1146 | 1154 | 1099 | 1190.86 | 1249 | 1133 | 0.906 | -0.142 | 0.06992027 | 0.17145226 | 0.002 | 0.0044 | 0.0036 | 0.0036 | TRUE | 0.0584 |
| Rnf43 | 56 | 31 | 46 | 44 | 39 | 44 | 43 | 33 | 50 | 44 | 44 | 43 | 42.89 | 42 | 44 | 1.04 | 0.056 | 0.85146512 | 0.91638754 | 0 | 0.0577 | 0.0411 | 0.0411 | TRUE | 0.2504 |
| Rnf44 | 14558 | 11221 | 10411 | 9751 | 9448 | 11052 | 11307 | 11936 | 11233 | 9706 | 10658 | 10910 | 10958.35 | 11492 | 10425 | 0.907 | -0.141 | 0.01959845 | 0.06806799 | 0.0023 | 0.0026 | 0.0025 | 0.0025 | TRUE | 0.0445 |
| Rnf5 | 2019 | 1517 | 1524 | 1847 | 1626 | 1914 | 1568 | 1614 | 1644 | 1839 | 1834 | 1889 | 1731.38 | 1609 | 1854 | 1.153 | 0.206 | 0.00108807 | 0.00769509 | 0 | 0.0038 | 0.0023 | 0.0023 | TRUE | 0.0062 |
| Rnf6 | 3311 | 2420 | 2371 | 2802 | 2307 | 2666 | 2572 | 2574 | 2558 | 2789 | 2603 | 2632 | 2621.22 | 2568 | 2675 | 1.042 | 0.059 | 0.31683788 | 0.49084801 | 3.00E-04 | 0.0033 | 0.0021 | 0.0021 | TRUE | 0.0171 |
| Rnf7 | 4949 | 3361 | 3221 | 3142 | 2716 | 3101 | 3844 | 3575 | 3475 | 3128 | 3064 | 3061 | 3357.8 | 3631 | 3084 | 0.849 | -0.236 | 9.45E-05 | 0.00115637 | 0.0012 | 0.0031 | 0.0023 | 0.0023 | TRUE | 0.0325 |
| Rnf8 | 1573 | 1113 | 990 | 846 | 736 | 835 | 1222 | 1184 | 1068 | 842 | 830 | 824 | 995.07 | 1158 | 832 | 0.718 | -0.478 | 4.06E-09 | 2.31E-07 | 0.0018 | 0.0048 | 0.0037 | 0.0037 | TRUE | 0.0542 |
| Rnft1 | 2778 | 1951 | 2079 | 2462 | 2082 | 2378 | 2158 | 2075 | 2243 | 2451 | 2349 | 2347 | 2270.48 | 2159 | 2382 | 1.104 | 0.142 | 0.02181723 | 0.07356156 | 6.00E-04 | 0.0035 | 0.0023 | 0.0023 | TRUE | 0.0137 |
| Rnft2 | 401 | 326 | 424 | 509 | 417 | 496 | 311 | 347 | 457 | 507 | 470 | 490 | 430.4 | 372 | 489 | 1.319 | 0.4 | 0.00328294 | 0.01770504 | 0.0181 | 0.0079 | 0.011 | 0.011 | TRUE | 0.4597 |
| Rngtt | 3555 | 2491 | 2543 | 2761 | 2347 | 2827 | 2761 | 2650 | 2744 | 2748 | 2648 | 2791 | 2723.53 | 2718 | 2729 | 1.004 | 0.006 | 0.92234066 | 0.95711845 | 2.00E-04 | 0.0033 | 0.002 | 0.002 | TRUE | 0.0081 |
| Rnh1 | 2623 | 1971 | 2130 | 2075 | 1985 | 2248 | 2037 | 2097 | 2298 | 2065 | 2239 | 2219 | 2159.3 | 2144 | 2174 | 1.015 | 0.021 | 0.76435291 | 0.86016031 | 0.0025 | 0.0035 | 0.0031 | 0.0031 | TRUE | 0.0476 |
| Rnls | 133 | 70 | 75 | 93 | 69 | 66 | 103 | 74 | 81 | 93 | 78 | 65 | 82.37 | 86 | 79 | 0.905 | -0.144 | 0.54254981 | 0.70040449 | 0.0197 | 0.0312 | 0.0279 | 0.0279 | TRUE | 0.3146 |
| Rnmt | 6628 | 4857 | 4948 | 4942 | 4455 | 5083 | 5148 | 5166 | 5339 | 4919 | 5026 | 5018 | 5102.59 | 5218 | 4988 | 0.956 | -0.065 | 0.19771847 | 0.35753923 | 0 | 0.0029 | 0.0016 | 0.0016 | TRUE | 0.005 |
| Rnmtl1 | 1145 | 816 | 675 | 906 | 699 | 763 | 889 | 868 | 728 | 902 | 789 | 753 | 821.52 | 828 | 815 | 0.982 | -0.026 | 0.801382 | 0.88476386 | 0.009 | 0.0053 | 0.0065 | 0.0065 | TRUE | 0.1307 |
| Rnpc3 | 1509 | 1186 | 1169 | 1090 | 969 | 1138 | 1172 | 1262 | 1261 | 1085 | 1093 | 1123 | 1166.06 | 1232 | 1100 | 0.895 | -0.161 | 0.02730044 | 0.08640504 | 0 | 0.0045 | 0.003 | 0.003 | TRUE | 0.0226 |
| Rnpep | 2045 | 1558 | 1475 | 1758 | 1648 | 1784 | 1588 | 1657 | 1592 | 1750 | 1859 | 1761 | 1701.19 | 1612 | 1790 | 1.11 | 0.151 | 0.0204133 | 0.07020387 | 0 | 0.0038 | 0.0025 | 0.0025 | TRUE | 0.0135 |
| Rnpepl1 | 1500 | 1118 | 1080 | 1055 | 949 | 999 | 1165 | 1189 | 1165 | 1050 | 1071 | 986 | 1104.4 | 1173 | 1036 | 0.883 | -0.18 | 0.013924 | 0.05291453 | 0 | 0.0046 | 0.003 | 0.003 | TRUE | 0.0211 |
| Rnps1 | 6093 | 4446 | 4984 | 4914 | 4551 | 5324 | 4732 | 4729 | 5378 | 4891 | 5134 | 5256 | 5020 | 4946 | 5094 | 1.03 | 0.043 | 0.52340385 | 0.68476076 | 0.0028 | 0.0029 | 0.003 | 0.003 | TRUE | 0.0705 |
| Rnu11 | 0 | 0 | 0 | 0 | 0 | 0 | 0 | 0 | 0 | 0 | 0 | 0 | 0 | 0 | 0 | NA | NA | NA | NA | NA | NA | NA | NA | NA | NA |
| Rnu12 | 0 | 0 | 0 | 1 | 1 | 0 | 0 | 0 | 0 | 1 | 1 | 0 | 0.35 | 0 | 1 | 3.978 | 1.992 | 0.55384016 | NA | 0 | 6.6991 | 6.4234 | 6.4234 | TRUE | 0.2647 |
| Rnu6 | 0 | 0 | 0 | 0 | 0 | 0 | 0 | 0 | 0 | 0 | 0 | 0 | 0 | 0 | 0 | NA | NA | NA | NA | NA | NA | NA | NA | NA | NA |
| Rnu7 | 0 | 0 | 0 | 0 | 0 | 0 | 0 | 0 | 0 | 0 | 0 | 0 | 0 | 0 | 0 | NA | NA | NA | NA | NA | NA | NA | NA | NA | NA |
| Rnu73b | 0 | 6 | 3 | 1 | 3 | 1 | 0 | 6 | 3 | 1 | 3 | 1 | 2.5 | 3 | 2 | 0.564 | -0.827 | 0.55191807 | NA | 0.9203 | 0.9513 | 0.9433 | 0.9433 | TRUE | 0.1347 |
| Robo1 | 2320 | 1728 | 1812 | 1957 | 1845 | 2048 | 1802 | 1838 | 1955 | 1948 | 2081 | 2022 | 1941.01 | 1865 | 2017 | 1.082 | 0.114 | 0.08452481 | 0.19609882 | 9.00E-04 | 0.0036 | 0.0026 | 0.0026 | TRUE | 0.0213 |
| Robo2 | 15 | 8 | 7 | 3 | 6 | 3 | 12 | 9 | 8 | 3 | 7 | 3 | 6.74 | 10 | 4 | 0.449 | -1.154 | 0.13351625 | NA | 0 | 0.3542 | 0.2507 | 0.2507 | TRUE | 0.4238 |
| Robo3 | 78 | 54 | 51 | 65 | 55 | 51 | 61 | 57 | 55 | 65 | 62 | 50 | 58.36 | 58 | 59 | 1.02 | 0.029 | 0.90875855 | 0.94963864 | 0 | 0.043 | 0.0295 | 0.0295 | TRUE | 0.1466 |
| Robo4 | 1164 | 751 | 618 | 993 | 827 | 846 | 904 | 799 | 667 | 988 | 933 | 835 | 854.36 | 790 | 919 | 1.161 | 0.215 | 0.05651026 | 0.14725635 | 0.0131 | 0.0052 | 0.008 | 0.008 | TRUE | 0.2184 |
| Rock1 | 6103 | 4273 | 3990 | 4219 | 3518 | 4085 | 4740 | 4545 | 4305 | 4200 | 3969 | 4033 | 4298.51 | 4530 | 4067 | 0.897 | -0.156 | 0.00866679 | 0.0372219 | 0.0012 | 0.003 | 0.0023 | 0.0023 | TRUE | 0.0229 |
| Rock2 | 6811 | 4890 | 4700 | 5367 | 4455 | 5087 | 5290 | 5201 | 5071 | 5342 | 5026 | 5022 | 5158.69 | 5187 | 5130 | 0.989 | -0.016 | 0.76412839 | 0.85997536 | 7.00E-04 | 0.0029 | 0.0019 | 0.0019 | TRUE | 0.0161 |
| Rogdi | 2096 | 1642 | 1342 | 1536 | 1378 | 1632 | 1628 | 1747 | 1448 | 1529 | 1555 | 1611 | 1586.16 | 1608 | 1565 | 0.973 | -0.039 | 0.62549009 | 0.76391779 | 0.0041 | 0.0039 | 0.004 | 0.004 | TRUE | 0.0885 |
| Rom1 | 130 | 71 | 104 | 122 | 103 | 105 | 101 | 76 | 112 | 121 | 116 | 104 | 105 | 96 | 114 | 1.18 | 0.239 | 0.24868062 | 0.41777057 | 0.0112 | 0.025 | 0.0214 | 0.0214 | TRUE | 0.3229 |
| Romo1 | 1112 | 768 | 818 | 918 | 766 | 883 | 864 | 817 | 883 | 914 | 864 | 872 | 868.79 | 855 | 883 | 1.034 | 0.048 | 0.54793279 | 0.70472451 | 0 | 0.0052 | 0.0034 | 0.0034 | TRUE | 0.0168 |
| Ropn1 | 2 | 1 | 3 | 3 | 0 | 0 | 2 | 1 | 3 | 3 | 0 | 0 | 1.47 | 2 | 1 | 0.522 | -0.938 | 0.5872164 | NA | 0.5013 | 1.611 | 1.3915 | 1.3915 | TRUE | 1.4414 |
| Ropn1l | 39 | 47 | 52 | 67 | 53 | 59 | 30 | 50 | 56 | 67 | 60 | 58 | 53.52 | 45 | 62 | 1.371 | 0.455 | 0.11998558 | 0.25168093 | 0.0317 | 0.0467 | 0.0426 | 0.0426 | TRUE | 0.7988 |
| Ror1 | 25 | 23 | 19 | 17 | 17 | 19 | 19 | 24 | 21 | 17 | 19 | 19 | 19.87 | 21 | 18 | 0.854 | -0.227 | 0.59236526 | 0.74022353 | 0 | 0.1217 | 0.0788 | 0.0788 | TRUE | 0.0806 |
| Ror2 | 1546 | 1179 | 1364 | 1553 | 1502 | 1690 | 1201 | 1254 | 1472 | 1546 | 1694 | 1668 | 1472.52 | 1309 | 1636 | 1.251 | 0.324 | 0.00021048 | 0.00221205 | 0.0062 | 0.004 | 0.0048 | 0.0048 | TRUE | 0.1411 |
| Rora | 58 | 52 | 53 | 51 | 63 | 51 | 45 | 55 | 57 | 51 | 71 | 50 | 54.95 | 52 | 57 | 1.095 | 0.131 | 0.63399007 | 0.7710347 | 0.0115 | 0.0456 | 0.0359 | 0.0359 | TRUE | 0.3486 |
| Rorb | 2 | 1 | 0 | 0 | 1 | 2 | 2 | 1 | 0 | 0 | 1 | 2 | 0.95 | 1 | 1 | 1.149 | 0.201 | 0.92051796 | NA | 0 | 2.4888 | 1.8818 | 1.8818 | TRUE | 0.2533 |
| Rorc | 5 | 6 | 5 | 10 | 14 | 9 | 4 | 6 | 5 | 10 | 16 | 9 | 8.38 | 5 | 12 | 2.229 | 1.157 | 0.09118294 | NA | 0 | 0.2852 | 0.1993 | 0.1993 | TRUE | 0.3614 |
| Ros1 | 1 | 0 | 0 | 0 | 0 | 0 | 1 | 0 | 0 | 0 | 0 | 0 | 0.13 | 0 | 0 | 0.552 | -0.857 | 0.8336548 | NA | 0 | 18.3121 | 10 | 10 | TRUE | 0.2566 |
| Rp1 | 2 | 0 | 1 | 0 | 0 | 0 | 2 | 0 | 1 | 0 | 0 | 0 | 0.44 | 1 | 0 | 0.214 | -2.223 | 0.47051571 | NA | 0 | 5.4047 | 5.2946 | 5.2946 | TRUE | 0.2788 |
| Rp1l1 | 0 | 0 | 0 | 0 | 0 | 0 | 0 | 0 | 0 | 0 | 0 | 0 | 0 | 0 | 0 | NA | NA | NA | NA | NA | NA | NA | NA | NA | NA |
| Rp2h | 1822 | 1340 | 1524 | 1723 | 1451 | 1622 | 1415 | 1425 | 1644 | 1715 | 1637 | 1601 | 1572.98 | 1495 | 1651 | 1.105 | 0.145 | 0.06565133 | 0.16375242 | 0.0034 | 0.0039 | 0.0038 | 0.0038 | TRUE | 0.0908 |

| | | | | | | | | | | | | | | | | | | | | | | | | | |
|---|---|---|---|---|---|---|---|---|---|---|---|---|---|---|---|---|---|---|---|---|---|---|---|---|---|
| Rp9 | 1899 | 1493 | 1583 | 1661 | 1474 | 1685 | 1475 | 1588 | 1708 | 1653 | 1663 | 1663 | 1625.09 | 1590 | 1660 | 1.045 | 0.063 | 0.3857239 | 0.55954042 | 0.0022 | 0.0039 | 0.0032 | 0.0032 | TRUE | 0.0505 |
| Rpa1 | 38561 | 27656 | 25939 | 27548 | 23990 | 28367 | 29949 | 29417 | 27988 | 27421 | 27063 | 28003 | 28306.84 | 29118 | 27496 | 0.944 | -0.083 | 0.08880766 | 0.20283283 | 7.00E-04 | 0.0025 | 0.0017 | 0.0017 | TRUE | 0.0141 |
| Rpa2 | 6354 | 4631 | 4175 | 4207 | 3706 | 4240 | 4935 | 4926 | 4505 | 4188 | 4181 | 4186 | 4486.59 | 4789 | 4185 | 0.874 | -0.195 | 0.00076112 | 0.00592841 | 0.001 | 0.003 | 0.0022 | 0.0022 | TRUE | 0.0324 |
| Rpa3 | 2263 | 1569 | 1502 | 1723 | 1447 | 1636 | 1758 | 1669 | 1621 | 1715 | 1632 | 1615 | 1668.26 | 1683 | 1654 | 0.982 | -0.025 | 0.70654816 | 0.82263708 | 9.00E-04 | 0.0038 | 0.0027 | 0.0027 | TRUE | 0.0191 |
| Rpain | 1011 | 697 | 812 | 803 | 712 | 880 | 785 | 741 | 876 | 799 | 803 | 869 | 812.32 | 801 | 824 | 1.029 | 0.042 | 0.64610167 | 0.7799604 | 0.0034 | 0.0053 | 0.0047 | 0.0047 | TRUE | 0.0773 |
| Rpap1 | 10891 | 8407 | 8618 | 8107 | 7648 | 8606 | 8459 | 8942 | 9299 | 8070 | 8628 | 8495 | 8648.77 | 8900 | 8398 | 0.944 | -0.084 | 0.14417387 | 0.28743123 | 0.0016 | 0.0027 | 0.0022 | 0.0022 | TRUE | 0.0232 |
| Rpap2 | 2189 | 1591 | 1548 | 1726 | 1519 | 1721 | 1700 | 1692 | 1670 | 1718 | 1714 | 1699 | 1698.87 | 1687 | 1710 | 1.013 | 0.019 | 0.75867502 | 0.85680642 | 0 | 0.0038 | 0.0021 | 0.0021 | TRUE | 0.001 |
| Rpap3 | 3592 | 2492 | 2491 | 2645 | 2300 | 2524 | 2790 | 2651 | 2688 | 2633 | 2595 | 2492 | 2641.21 | 2710 | 2573 | 0.949 | -0.075 | 0.2027148 | 0.36416046 | 4.00E-04 | 0.0033 | 0.0021 | 0.0021 | TRUE | 0.0094 |
| Rpe | 2986 | 2153 | 2026 | 2037 | 1684 | 1960 | 2319 | 2290 | 2186 | 2028 | 1900 | 1935 | 2109.57 | 2265 | 1954 | 0.863 | -0.213 | 0.00071164 | 0.00563513 | 6.00E-04 | 0.0035 | 0.0024 | 0.0024 | TRUE | 0.0132 |
| Rpe65 | 1 | 0 | 0 | 0 | 0 | 0 | 1 | 0 | 0 | 0 | 0 | 0 | 0.13 | 0 | 0 | 0.552 | -0.857 | 0.8336548 | NA | 0 | 18.3121 | 10 | 10 | TRUE | 0.2566 |
| Rpf1 | 3230 | 2277 | 2337 | 2287 | 1992 | 2463 | 2509 | 2422 | 2522 | 2276 | 2247 | 2431 | 2401.21 | 2484 | 2318 | 0.933 | -0.099 | 0.10781089 | 0.2335467 | 7.00E-04 | 0.0034 | 0.0023 | 0.0023 | TRUE | 0.0222 |
| Rpf2 | 3134 | 2154 | 2155 | 2224 | 1920 | 2203 | 2434 | 2291 | 2325 | 2214 | 2166 | 2175 | 2267.48 | 2350 | 2185 | 0.929 | -0.106 | 0.07700998 | 0.18366489 | 0 | 0.0035 | 0.0021 | 0.0021 | TRUE | 0.0122 |
| Rpgr | 1956 | 1560 | 1582 | 1821 | 1674 | 1872 | 1519 | 1659 | 1707 | 1813 | 1888 | 1848 | 1739.08 | 1628 | 1850 | 1.137 | 0.185 | 0.00793781 | 0.03485564 | 0.0015 | 0.0038 | 0.0029 | 0.0029 | TRUE | 0.0433 |
| Rpgrip1 | 149 | 110 | 77 | 95 | 99 | 81 | 116 | 117 | 83 | 95 | 112 | 80 | 100.34 | 105 | 96 | 0.901 | -0.15 | 0.49455066 | 0.65963163 | 0.0208 | 0.026 | 0.0246 | 0.0246 | TRUE | 0.3192 |
| Rpgrip1l | 1044 | 736 | 613 | 616 | 589 | 633 | 811 | 783 | 661 | 613 | 664 | 625 | 692.94 | 752 | 634 | 0.842 | -0.248 | 0.01266629 | 0.04924675 | 0.0054 | 0.0058 | 0.0057 | 0.0057 | TRUE | 0.1282 |
| Rph3a | 67 | 59 | 41 | 59 | 48 | 61 | 52 | 63 | 44 | 59 | 54 | 60 | 55.35 | 53 | 58 | 1.09 | 0.124 | 0.63859721 | 0.77423614 | 0 | 0.0452 | 0.0323 | 0.0323 | TRUE | 0.1976 |
| Rph3al | 168 | 108 | 139 | 150 | 138 | 158 | 130 | 115 | 150 | 149 | 156 | 156 | 142.72 | 132 | 154 | 1.166 | 0.222 | 0.19395245 | 0.35280757 | 0 | 0.019 | 0.014 | 0.014 | TRUE | 0.1387 |
| Rpia | 498 | 374 | 350 | 394 | 337 | 410 | 387 | 398 | 378 | 392 | 380 | 405 | 389.89 | 388 | 392 | 1.013 | 0.019 | 0.85639242 | 0.91929529 | 0 | 0.0085 | 0.0054 | 0.0054 | TRUE | 0.0088 |
| Rpl10 | 41871 | 29487 | 31424 | 36158 | 30965 | 36446 | 32519 | 31365 | 33906 | 35992 | 34932 | 35978 | 34115.3 | 32597 | 35634 | 1.093 | 0.129 | 0.00944679 | 0.03974771 | 9.00E-04 | 0.0025 | 0.0017 | 0.0017 | TRUE | 0.0151 |
| Rpl10a | 33271 | 25274 | 28632 | 30408 | 27584 | 32457 | 25840 | 26884 | 30894 | 30268 | 31118 | 32040 | 29507.21 | 27873 | 31142 | 1.117 | 0.16 | 0.02051554 | 0.07047072 | 0.0049 | 0.0025 | 0.0034 | 0.0034 | TRUE | 0.11 |
| Rpl10l | 5225 | 3624 | 2800 | 1698 | 1550 | 1742 | 4058 | 3855 | 3021 | 1690 | 1749 | 1720 | 2682.06 | 3645 | 1720 | 0.472 | -1.085 | 2.16E-29 | 1.72E-26 | 0.0109 | 0.0033 | 0.0063 | 0.0063 | TRUE | 0.2712 |
| Rpl11 | 34768 | 25190 | 26314 | 29943 | 25771 | 30135 | 27003 | 26794 | 28393 | 29805 | 29072 | 29748 | 28469.2 | 27397 | 29542 | 1.078 | 0.109 | 0.02130952 | 0.07239765 | 5.00E-04 | 0.0025 | 0.0016 | 0.0016 | TRUE | 0.0123 |
| Rpl12 | 60755 | 44222 | 47697 | 50473 | 45390 | 52470 | 47186 | 47038 | 51465 | 50241 | 51205 | 51796 | 49821.71 | 48563 | 51081 | 1.052 | 0.073 | 0.16753932 | 0.31903274 | 0.0014 | 0.0025 | 0.002 | 0.002 | TRUE | 0.0334 |
| Rpl13 | 64917 | 48832 | 51787 | 55653 | 49990 | 58394 | 50418 | 51942 | 55878 | 55397 | 56394 | 57644 | 54612.12 | 52746 | 56478 | 1.071 | 0.099 | 0.06713868 | 0.16625313 | 0.0016 | 0.0025 | 0.0021 | 0.0021 | TRUE | 0.033 |
| Rpl13a | 93118 | 65215 | 68300 | 73376 | 65922 | 76127 | 72321 | 69368 | 73695 | 73038 | 74367 | 75149 | 72989.77 | 71795 | 74185 | 1.033 | 0.047 | 0.30910543 | 0.48184577 | 6.00E-04 | 0.0025 | 0.0015 | 0.0015 | TRUE | 0.0107 |
| Rpl14 | 41330 | 29823 | 31685 | 34343 | 29687 | 34936 | 32099 | 31722 | 34188 | 34185 | 33490 | 34487 | 33361.94 | 32670 | 34054 | 1.042 | 0.06 | 0.22794786 | 0.39324789 | 9.00E-04 | 0.0025 | 0.0018 | 0.0018 | TRUE | 0.0202 |
| Rpl15 | 25646 | 17814 | 18849 | 21007 | 18260 | 21241 | 19918 | 18949 | 20338 | 20910 | 20599 | 20968 | 20280.38 | 19735 | 20826 | 1.055 | 0.078 | 0.11032162 | 0.23761707 | 6.00E-04 | 0.0025 | 0.0017 | 0.0017 | TRUE | 0.0148 |
| Rpl17 | 45742 | 32611 | 36688 | 39864 | 35155 | 41249 | 35526 | 34688 | 39586 | 39681 | 39658 | 40719 | 38309.67 | 36600 | 40019 | 1.093 | 0.129 | 0.03046367 | 0.09359786 | 0.0026 | 0.0025 | 0.0025 | 0.0025 | TRUE | 0.0623 |
| Rpl18 | 41511 | 30581 | 31028 | 33170 | 30153 | 35089 | 32240 | 32529 | 33479 | 33017 | 34016 | 34638 | 33319.79 | 32749 | 33890 | 1.035 | 0.049 | 0.28435428 | 0.45640291 | 4.00E-04 | 0.0025 | 0.0015 | 0.0015 | TRUE | 0.0062 |
| Rpl18a | 50932 | 36899 | 40672 | 43670 | 38905 | 45411 | 39557 | 39249 | 43885 | 43469 | 43889 | 44828 | 42479.34 | 40897 | 44062 | 1.077 | 0.108 | 0.05811576 | 0.1502762 | 0.0021 | 0.0025 | 0.0023 | 0.0023 | TRUE | 0.05 |
| Rpl19 | 68932 | 48554 | 51225 | 56803 | 49458 | 58742 | 53537 | 51646 | 55271 | 56542 | 55794 | 57987 | 55129.5 | 53485 | 56774 | 1.062 | 0.086 | 0.07350261 | 0.17773183 | 7.00E-04 | 0.0025 | 0.0017 | 0.0017 | TRUE | 0.011 |
| Rpl21 | 43231 | 31357 | 31708 | 35447 | 29862 | 35802 | 33576 | 33354 | 34213 | 35284 | 33687 | 35342 | 34242.64 | 33714 | 34771 | 1.031 | 0.045 | 0.33119274 | 0.50557918 | 4.00E-04 | 0.0025 | 0.0015 | 0.0015 | TRUE | 0.0091 |
| Rpl22 | 24635 | 18579 | 21991 | 23340 | 20934 | 24183 | 19133 | 19762 | 23728 | 23233 | 23616 | 23872 | 22224 | 20874 | 23574 | 1.129 | 0.176 | 0.02123582 | 0.07225315 | 0.0071 | 0.0025 | 0.0041 | 0.0041 | TRUE | 0.1748 |
| Rpl22l1 | 11715 | 8633 | 10160 | 10791 | 9213 | 10249 | 9099 | 9183 | 10963 | 10741 | 10393 | 10117 | 10082.64 | 9748 | 10417 | 1.069 | 0.096 | 0.19680943 | 0.35644191 | 0.006 | 0.0027 | 0.0039 | 0.0039 | TRUE | 0.1448 |
| Rpl23 | 60844 | 41934 | 44435 | 51012 | 43046 | 50227 | 47255 | 44605 | 47945 | 50777 | 48560 | 49582 | 48120.67 | 46602 | 49640 | 1.065 | 0.091 | 0.06716391 | 0.16625802 | 9.00E-04 | 0.0025 | 0.0018 | 0.0018 | TRUE | 0.0172 |
| Rpl23a | 44690 | 33591 | 36641 | 39151 | 34039 | 40171 | 34709 | 35730 | 39535 | 38971 | 38400 | 39655 | 37833.31 | 36658 | 39009 | 1.064 | 0.09 | 0.1278281 | 0.26387161 | 0.0025 | 0.0025 | 0.0025 | 0.0025 | TRUE | 0.0577 |
| Rpl24 | 20431 | 14671 | 15859 | 16773 | 14886 | 17652 | 15868 | 15605 | 17112 | 16696 | 16793 | 17425 | 16583.17 | 16195 | 16971 | 1.048 | 0.068 | 0.21223011 | 0.37454069 | 0.0014 | 0.0026 | 0.0021 | 0.0021 | TRUE | 0.0299 |
| Rpl26 | 4 | 1 | 3 | 2 | 3 | 0 | 3 | 1 | 3 | 2 | 3 | 0 | 2.13 | 2 | 2 | 0.709 | -0.497 | 0.7173251 | NA | 0 | 1.1149 | 0.8708 | 0.8708 | TRUE | 0.4044 |
| Rpl27 | 20722 | 14341 | 14839 | 17286 | 14772 | 17245 | 16094 | 15254 | 16011 | 17206 | 16664 | 17023 | 16375.61 | 15786 | 16964 | 1.075 | 0.104 | 0.03038403 | 0.09343351 | 5.00E-04 | 0.0026 | 0.0016 | 0.0016 | TRUE | 0.0106 |
| Rpl27a | 31746 | 23326 | 25307 | 27452 | 24175 | 28463 | 24656 | 24812 | 27306 | 27326 | 27272 | 28097 | 26578.06 | 25591 | 27565 | 1.077 | 0.107 | 0.05314527 | 0.14082154 | 0.0018 | 0.0025 | 0.0022 | 0.0022 | TRUE | 0.042 |
| Rpl28 | 32337 | 23529 | 25181 | 25942 | 22462 | 27480 | 25115 | 25027 | 27170 | 25823 | 25339 | 27127 | 25933.58 | 25771 | 26096 | 1.013 | 0.018 | 0.74067376 | 0.84588493 | 0.0014 | 0.0025 | 0.0021 | 0.0021 | TRUE | 0.0276 |
| Rpl29 | 2380 | 1597 | 1587 | 1906 | 1571 | 2030 | 1848 | 1699 | 1712 | 1897 | 1772 | 2004 | 1822.15 | 1753 | 1891 | 1.078 | 0.109 | 0.13471414 | 0.27358188 | 0.0022 | 0.0037 | 0.0033 | 0.0033 | TRUE | 0.036 |
| Rpl3 | 68903 | 50219 | 56780 | 62862 | 54531 | 61970 | 53514 | 53417 | 61265 | 62573 | 61517 | 61174 | 58909.97 | 56065 | 61755 | 1.102 | 0.139 | 0.02472412 | 0.0807361 | 0.0033 | 0.0025 | 0.0028 | 0.0028 | TRUE | 0.0805 |
| Rpl30 | 184 | 124 | 102 | 136 | 109 | 101 | 143 | 132 | 110 | 135 | 123 | 100 | 123.82 | 128 | 119 | 0.927 | -0.11 | 0.57071796 | 0.72416753 | 0.0126 | 0.0216 | 0.019 | 0.019 | TRUE | 0.2156 |
| Rpl31 | 21311 | 14666 | 15300 | 17285 | 14738 | 17588 | 16551 | 15600 | 16509 | 17205 | 16626 | 17362 | 16642.25 | 16220 | 17064 | 1.052 | 0.073 | 0.14175699 | 0.28409039 | 7.00E-04 | 0.0026 | 0.0017 | 0.0017 | TRUE | 0.0136 |
| Rpl31-ps12 | 0 | 1 | 1 | 1 | 0 | 3 | 0 | 1 | 1 | 1 | 0 | 3 | 1.02 | 1 | 1 | 1.961 | 0.971 | 0.63207857 | NA | 0 | 2.3338 | 1.8722 | 1.8722 | TRUE | 0.3332 |
| Rpl32 | 65126 | 46253 | 50894 | 57298 | 50531 | 59443 | 50581 | 49199 | 54914 | 57034 | 57004 | 58679 | 54568.58 | 51565 | 57572 | 1.117 | 0.159 | 0.00379872 | 0.0198013 | 0.0018 | 0.0025 | 0.0022 | 0.0022 | TRUE | 0.0395 |
| Rpl34 | 280 | 222 | 167 | 172 | 151 | 153 | 217 | 236 | 180 | 171 | 170 | 151 | 187.73 | 211 | 164 | 0.776 | -0.366 | 0.02077194 | 0.07116331 | 0.0066 | 0.0151 | 0.0126 | 0.0126 | TRUE | 0.1714 |
| Rpl34-ps1 | 2 | 0 | 2 | 2 | 0 | 2 | 2 | 0 | 2 | 2 | 0 | 2 | 1.28 | 1 | 1 | 1.076 | 0.106 | 0.95232927 | NA | 0 | 1.8549 | 1.4748 | 1.4748 | TRUE | 0.4009 |
| Rpl35 | 17723 | 13175 | 13476 | 14617 | 12838 | 15059 | 13765 | 14014 | 14540 | 14550 | 14483 | 14866 | 14369.53 | 14106 | 14633 | 1.037 | 0.053 | 0.26556498 | 0.435989 | 4.00E-04 | 0.0026 | 0.0016 | 0.0016 | TRUE | 0.0088 |
| Rpl35a | 18461 | 12749 | 14223 | 16139 | 13724 | 16223 | 14338 | 13561 | 15346 | 16065 | 15482 | 16015 | 15134.46 | 14415 | 15854 | 1.1 | 0.137 | 0.01786018 | 0.06351717 | 0.002 | 0.0026 | 0.0024 | 0.0024 | TRUE | 0.0389 |
| Rpl36 | 27187 | 19879 | 21670 | 24603 | 21574 | 25127 | 21115 | 21145 | 23382 | 24490 | 24338 | 24804 | 23212.25 | 21881 | 24544 | 1.122 | 0.166 | 0.00281156 | 0.01570932 | 0.0017 | 0.0025 | 0.0022 | 0.0022 | TRUE | 0.044 |
| Rpl36a | 17134 | 11851 | 12811 | 14017 | 12100 | 14652 | 13307 | 12606 | 13823 | 13953 | 13650 | 14464 | 13633.72 | 13245 | 14022 | 1.059 | 0.082 | 0.13221259 | 0.27011357 | 0.0014 | 0.0026 | 0.0021 | 0.0021 | TRUE | 0.0217 |
| Rpl36al | 7930 | 5607 | 5772 | 6367 | 5519 | 6754 | 6159 | 5964 | 6228 | 6338 | 6226 | 6667 | 6263.65 | 6117 | 6410 | 1.048 | 0.068 | 0.20451214 | 0.36602704 | 7.00E-04 | 0.0028 | 0.0019 | 0.0019 | TRUE | 0.015 |
| Rpl37 | 21659 | 15688 | 16861 | 18362 | 15544 | 18502 | 16822 | 16687 | 18193 | 18278 | 17535 | 18264 | 17629.79 | 17234 | 18026 | 1.046 | 0.065 | 0.22734309 | 0.39248881 | 0.0014 | 0.0026 | 0.002 | 0.002 | TRUE | 0.0289 |
| Rpl37a | 21256 | 14832 | 15635 | 16404 | 13898 | 17143 | 16509 | 15777 | 16870 | 16329 | 15678 | 16923 | 16347.49 | 16385 | 16310 | 0.995 | -0.007 | 0.90152015 | 0.94568228 | 0.001 | 0.0026 | 0.002 | 0.002 | TRUE | 0.014 |
| Rpl38 | 18640 | 13801 | 14680 | 16166 | 14153 | 16157 | 14477 | 14680 | 15840 | 16092 | 15966 | 15949 | 15500.59 | 14999 | 16002 | 1.067 | 0.094 | 0.07486781 | 0.18004295 | 0.0011 | 0.0026 | 0.0019 | 0.0019 | TRUE | 0.0293 |
| Rpl39 | 13114 | 8460 | 9985 | 11451 | 9769 | 11357 | 10185 | 8999 | 10774 | 11398 | 11020 | 11211 | 10597.91 | 9986 | 11210 | 1.123 | 0.167 | 0.01448744 | 0.05434917 | 0.0041 | 0.0027 | 0.0033 | 0.0033 | TRUE | 0.0909 |
| Rpl39l | 4235 | 3338 | 3080 | 2018 | 2018 | 2387 | 3289 | 3551 | 3323 | 2009 | 2277 | 2356 | 2800.77 | 3388 | 2214 | 0.654 | -0.614 | 5.62E-17 | 1.20E-14 | 0.0037 | 0.0033 | 0.0035 | 0.0035 | TRUE | 0.0802 |
| Rpl3l | 113 | 93 | 83 | 106 | 99 | 94 | 88 | 99 | 90 | 106 | 112 | 93 | 97.7 | 92 | 104 | 1.123 | 0.167 | 0.40228877 | 0.57522142 | 0 | 0.0267 | 0.0183 | 0.0183 | TRUE | 0.0795 |
| Rpl4 | 153276 | 111773 | 118605 | 129788 | 115600 | 132670 | 119043 | 118891 | 127974 | 129191 | 130409 | 130966 | 126078.94 | 121969 | 130189 | 1.067 | 0.094 | 0.05470803 | 0.14362257 | 9.00E-04 | 0.0024 | 0.0017 | 0.0017 | TRUE | 0.0227 |
| Rpl41 | 36649 | 24943 | 25324 | 28800 | 25250 | 29841 | 28464 | 26532 | 27324 | 28667 | 28485 | 29458 | 28154.91 | 27440 | 28870 | 1.052 | 0.073 | 0.13383328 | 0.27241228 | 7.00E-04 | 0.0025 | 0.0017 | 0.0017 | TRUE | 0.0131 |
| Rpl5 | 51241 | 34903 | 37817 | 43836 | 37354 | 43227 | 39797 | 37126 | 40804 | 43634 | 42139 | 42672 | 41028.67 | 39242 | 42815 | 1.091 | 0.126 | 0.01645929 | 0.05984548 | 0.0013 | 0.0025 | 0.002 | 0.002 | TRUE | 0.0272 |
| Rpl6 | 57028 | 41278 | 43592 | 46347 | 41254 | 48107 | 44291 | 43907 | 47035 | 46134 | 46539 | 47489 | 45899.18 | 45078 | 46721 | 1.036 | 0.052 | 0.28793205 | 0.46013113 | 8.00E-04 | 0.0025 | 0.0017 | 0.0017 | TRUE | 0.0176 |
| Rpl7 | 81032 | 56820 | 60369 | 65753 | 57322 | 66025 | 62934 | 60439 | 65138 | 65450 | 64665 | 65177 | 63967.16 | 62837 | 65097 | 1.036 | 0.051 | 0.28464234 | 0.45655853 | 7.00E-04 | 0.0025 | 0.0016 | 0.0016 | TRUE | 0.0136 |
| Rpl7a | 79992 | 56882 | 59342 | 64106 | 57434 | 67607 | 62126 | 60505 | 64030 | 63811 | 64792 | 66739 | 63666.94 | 62220 | 65114 | 1.047 | 0.066 | 0.1646253 | 0.31545641 | 6.00E-04 | 0.0025 | 0.0016 | 0.0016 | TRUE | 0.0079 |
| Rpl7l1 | 13258 | 9296 | 8219 | 8781 | 7917 | 8639 | 10297 | 9888 | 8868 | 8741 | 8931 | 8528 | 9208.84 | 9684 | 8733 | 0.902 | -0.15 | 0.01980417 | 0.06865413 | 0.003 | 0.0027 | 0.0029 | 0.0029 | TRUE | 0.0662 |
| Rpl8 | 91402 | 67345 | 71631 | 78774 | 70028 | 81848 | 70988 | 71634 | 77289 | 78412 | 78999 | 80797 | 76353.05 | 73304 | 79403 | 1.083 | 0.115 | 0.0245372 | 0.0802357 | 0.0012 | 0.0025 | 0.0019 | 0.0019 | TRUE | 0.0277 |
| Rpl9 | 51288 | 34531 | 38704 | 45364 | 38874 | 43449 | 39833 | 36730 | 41761 | 45155 | 43854 | 42891 | 41704.12 | 39441 | 43967 | 1.115 | 0.157 | 0.00744837 | 0.03326859 | 0.0024 | 0.0025 | 0.0024 | 0.0024 | TRUE | 0.0442 |

| | | | | | | | | | | | | | | | | | | | | | | | | | |
|---|---|---|---|---|---|---|---|---|---|---|---|---|---|---|---|---|---|---|---|---|---|---|---|---|---|
| Rplp0 | 52543 | 37314 | 37818 | 37517 | 33538 | 38987 | 40808 | 39690 | 40805 | 37344 | 37834 | 38486 | 39161.43 | 40434 | 37888 | 0.937 | -0.094 | 0.03184269 | 0.09670089 | 2.00E-04 | 0.0025 | 0.0014 | 0.0014 | TRUE | 0.0032 |
| Rplp1 | 64074 | 45607 | 49935 | 56338 | 49152 | 56934 | 49764 | 48512 | 53879 | 56079 | 55449 | 56203 | 53314.09 | 50718 | 55910 | 1.102 | 0.141 | 0.00853228 | 0.03676569 | 0.0015 | 0.0025 | 0.002 | 0.002 | TRUE | 0.0364 |
| Rplp2 | 35704 | 25878 | 27744 | 31096 | 27130 | 31522 | 27730 | 27526 | 29936 | 30953 | 30605 | 31117 | 29644.48 | 28397 | 30892 | 1.088 | 0.122 | 0.01759487 | 0.06283946 | 0.0011 | 0.0025 | 0.0019 | 0.0019 | TRUE | 0.0274 |
| Rplp2-ps1 | 0 | 0 | 0 | 0 | 0 | 0 | 0 | 0 | 0 | 0 | 0 | 0 | 0 | 0 | 0 | NA | NA | NA | NA | NA | NA | NA | NA | NA | NA |
| Rpn1 | 15910 | 11771 | 12602 | 14391 | 12897 | 14964 | 12357 | 12521 | 13597 | 14325 | 14549 | 14772 | 13686.74 | 12825 | 14549 | 1.135 | 0.182 | 0.0008364 | 0.00634877 | 0.0014 | 0.0026 | 0.0021 | 0.0021 | TRUE | 0.0338 |
| Rpn2 | 15897 | 11798 | 12807 | 14280 | 12835 | 14736 | 12347 | 12549 | 13819 | 14214 | 14479 | 14547 | 13659.13 | 12905 | 14413 | 1.117 | 0.16 | 0.00523972 | 0.02541273 | 0.0019 | 0.0026 | 0.0023 | 0.0023 | TRUE | 0.0468 |
| Rpp14 | 1730 | 1134 | 1033 | 1051 | 955 | 998 | 1344 | 1206 | 1115 | 1046 | 1077 | 985 | 1128.85 | 1222 | 1036 | 0.847 | -0.24 | 0.00603185 | 0.02832108 | 0.0048 | 0.0045 | 0.0046 | 0.0046 | TRUE | 0.0947 |
| Rpp21 | 850 | 601 | 608 | 565 | 537 | 629 | 660 | 639 | 656 | 562 | 606 | 621 | 624.1 | 652 | 596 | 0.914 | -0.129 | 0.14876385 | 0.29405085 | 0 | 0.0062 | 0.0041 | 0.0041 | TRUE | 0.0297 |
| Rpp25 | 2195 | 1576 | 1283 | 1161 | 1100 | 1302 | 1705 | 1676 | 1384 | 1156 | 1241 | 1285 | 1407.89 | 1588 | 1227 | 0.772 | -0.373 | 3.95E-05 | 0.00058205 | 0.0073 | 0.0041 | 0.0052 | 0.0052 | TRUE | 0.1505 |
| Rpp25l | 1083 | 765 | 712 | 717 | 571 | 726 | 841 | 814 | 768 | 714 | 644 | 717 | 749.6 | 808 | 692 | 0.856 | -0.224 | 0.01056358 | 0.04296678 | 0.0014 | 0.0056 | 0.0042 | 0.0042 | TRUE | 0.0412 |
| Rpp30 | 2246 | 1662 | 1737 | 1770 | 1536 | 1725 | 1744 | 1768 | 1874 | 1762 | 1733 | 1703 | 1763.98 | 1795 | 1733 | 0.965 | -0.051 | 0.43313252 | 0.60509854 | 4.00E-04 | 0.0038 | 0.0024 | 0.0024 | TRUE | 0.0175 |
| Rpp38 | 649 | 451 | 383 | 363 | 325 | 359 | 504 | 480 | 413 | 361 | 367 | 354 | 413.23 | 466 | 361 | 0.773 | -0.372 | 0.00097163 | 0.00709387 | 0.0036 | 0.0082 | 0.0067 | 0.0067 | TRUE | 0.1101 |
| Rpp40 | 2430 | 1701 | 1676 | 1809 | 1513 | 1841 | 1887 | 1809 | 1808 | 1801 | 1707 | 1817 | 1804.97 | 1835 | 1775 | 0.967 | -0.048 | 0.45262512 | 0.62247387 | 3.00E-04 | 0.0037 | 0.0024 | 0.0024 | TRUE | 0.0134 |
| Rpph1 | 33 | 18 | 17 | 18 | 11 | 27 | 26 | 19 | 18 | 18 | 12 | 27 | 20.02 | 21 | 19 | 0.901 | -0.15 | 0.73998334 | 0.84538261 | 0.0344 | 0.1208 | 0.0975 | 0.0975 | TRUE | 0.3885 |
| Rprd1a | 2799 | 2053 | 2547 | 2494 | 2254 | 2875 | 2174 | 2184 | 2748 | 2483 | 2543 | 2838 | 2494.86 | 2369 | 2621 | 1.107 | 0.147 | 0.1196586 | 0.25135735 | 0.0117 | 0.0034 | 0.006 | 0.006 | TRUE | 0.2367 |
| Rprd1b | 2812 | 1829 | 1594 | 1506 | 1285 | 1467 | 2184 | 1945 | 1720 | 1499 | 1450 | 1448 | 1707.7 | 1950 | 1466 | 0.751 | -0.413 | 2.33E-06 | 5.71E-05 | 0.0064 | 0.0038 | 0.0049 | 0.0049 | TRUE | 0.136 |
| Rprd2 | 3088 | 2346 | 2022 | 2215 | 1942 | 2267 | 2398 | 2495 | 2182 | 2205 | 2191 | 2238 | 2284.82 | 2358 | 2211 | 0.937 | -0.093 | 0.16938298 | 0.32083515 | 0.0017 | 0.0035 | 0.0029 | 0.0029 | TRUE | 0.051 |
| Rprl1 | 0 | 0 | 0 | 0 | 0 | 0 | 0 | 0 | 0 | 0 | 0 | 0 | 0 | 0 | 0 | NA | NA | NA | NA | NA | NA | NA | NA | NA | NA |
| Rprl2 | 0 | 0 | 0 | 0 | 0 | 0 | 0 | 0 | 0 | 0 | 0 | 0 | 0 | 0 | 0 | NA | NA | NA | NA | NA | NA | NA | NA | NA | NA |
| Rprl3 | 2 | 2 | 0 | 0 | 1 | 0 | 2 | 2 | 0 | 0 | 1 | 0 | 0.8 | 1 | 0 | 0.3 | -1.737 | 0.45535154 | NA | 0 | 2.9596 | 2.5115 | 2.5115 | TRUE | 0.5075 |
| Rprm | 119 | 89 | 82 | 94 | 72 | 85 | 92 | 95 | 88 | 94 | 81 | 84 | 89.04 | 92 | 86 | 0.939 | -0.09 | 0.65961483 | 0.78940591 | 0 | 0.029 | 0.019 | 0.019 | TRUE | 0.053 |
| Rprml | 84 | 49 | 55 | 86 | 72 | 81 | 65 | 52 | 59 | 86 | 81 | 80 | 70.58 | 59 | 82 | 1.391 | 0.476 | 0.04010218 | 0.11465304 | 0 | 0.036 | 0.0243 | 0.0243 | TRUE | 0.0881 |
| Rps10 | 32133 | 22854 | 24612 | 27793 | 24764 | 28176 | 24956 | 24310 | 26556 | 27665 | 27936 | 27814 | 26539.59 | 25274 | 27805 | 1.1 | 0.138 | 0.00667637 | 0.03065382 | 0.001 | 0.0025 | 0.0018 | 0.0018 | TRUE | 0.024 |
| Rps11 | 49575 | 33440 | 36330 | 39693 | 34574 | 39525 | 38503 | 35570 | 39200 | 39510 | 39003 | 39017 | 38467.16 | 37758 | 39177 | 1.038 | 0.053 | 0.31073993 | 0.48365521 | 0.0013 | 0.0025 | 0.002 | 0.002 | TRUE | 0.0314 |
| Rps12 | 51781 | 36455 | 39102 | 42674 | 37028 | 44033 | 40216 | 38777 | 42191 | 42478 | 41771 | 43467 | 41483.33 | 40395 | 42572 | 1.054 | 0.076 | 0.1352799 | 0.27453579 | 0.0011 | 0.0025 | 0.0018 | 0.0018 | TRUE | 0.0185 |
| Rps13 | 21784 | 15189 | 16157 | 17348 | 14315 | 17792 | 16919 | 16156 | 17433 | 17268 | 16149 | 17563 | 16914.79 | 16836 | 16993 | 1.009 | 0.013 | 0.80796287 | 0.88895612 | 0.0013 | 0.0026 | 0.0022 | 0.0022 | TRUE | 0.0231 |
| Rps14 | 44944 | 32552 | 34140 | 39801 | 35090 | 38763 | 34906 | 34625 | 36837 | 39618 | 39585 | 38265 | 37306.02 | 35456 | 39156 | 1.104 | 0.143 | 0.00307295 | 0.01684113 | 7.00E-04 | 0.0025 | 0.0017 | 0.0017 | TRUE | 0.0142 |
| Rps15 | 42474 | 30705 | 34083 | 37336 | 33363 | 38489 | 32988 | 32661 | 36775 | 37164 | 37637 | 37995 | 35869.88 | 34141 | 37599 | 1.101 | 0.139 | 0.01590464 | 0.05833332 | 0.0023 | 0.0025 | 0.0024 | 0.0024 | TRUE | 0.0557 |
| Rps15a | 29582 | 20234 | 22508 | 25572 | 22076 | 25140 | 22975 | 21523 | 24286 | 25454 | 24904 | 24817 | 23993.18 | 22928 | 25058 | 1.093 | 0.128 | 0.02269892 | 0.07568289 | 0.0018 | 0.0025 | 0.0022 | 0.0022 | TRUE | 0.0351 |
| Rps15a-ps4 | 512 | 376 | 437 | 466 | 434 | 477 | 398 | 400 | 472 | 464 | 490 | 471 | 448.91 | 423 | 475 | 1.124 | 0.169 | 0.11854964 | 0.25002656 | 0.0027 | 0.0077 | 0.0062 | 0.0062 | TRUE | 0.1128 |
| Rps15a-ps6 | 1130 | 914 | 930 | 994 | 970 | 1113 | 878 | 972 | 1003 | 989 | 1094 | 1099 | 1005.95 | 951 | 1061 | 1.117 | 0.159 | 0.06191826 | 0.1569423 | 0.0033 | 0.0048 | 0.0043 | 0.0043 | TRUE | 0.0572 |
| Rps16 | 36544 | 25856 | 27637 | 29774 | 26412 | 30877 | 28382 | 27503 | 29820 | 29637 | 29795 | 30480 | 29269.63 | 28568 | 29971 | 1.049 | 0.069 | 0.16608861 | 0.31707019 | 9.00E-04 | 0.0025 | 0.0018 | 0.0018 | TRUE | 0.0179 |
| Rps17 | 32077 | 22959 | 24303 | 26467 | 23540 | 28089 | 24913 | 24421 | 26223 | 26345 | 26556 | 27728 | 26030.96 | 25186 | 26876 | 1.067 | 0.094 | 0.06531423 | 0.16323417 | 0.001 | 0.0025 | 0.0018 | 0.0018 | TRUE | 0.0158 |
| Rps18 | 54092 | 38629 | 42289 | 48332 | 42232 | 47817 | 42011 | 41089 | 45629 | 48110 | 47642 | 47203 | 45280.68 | 42910 | 47652 | 1.111 | 0.151 | 0.00497054 | 0.02440449 | 0.0016 | 0.0025 | 0.0021 | 0.0021 | TRUE | 0.0376 |
| Rps19 | 30527 | 22081 | 24138 | 26885 | 23335 | 27096 | 23709 | 23487 | 26045 | 26761 | 26324 | 26748 | 25512.42 | 24414 | 26611 | 1.09 | 0.124 | 0.02351514 | 0.07776535 | 0.0017 | 0.0025 | 0.0021 | 0.0021 | TRUE | 0.0417 |
| Rps19-ps3 | 17 | 16 | 7 | 15 | 9 | 7 | 13 | 17 | 8 | 15 | 10 | 7 | 11.63 | 13 | 11 | 0.845 | -0.242 | 0.68233575 | NA | 0.0536 | 0.2062 | 0.1656 | 0.1656 | TRUE | 0.2714 |
| Rps19bp1 | 1479 | 1131 | 1042 | 1135 | 1033 | 1163 | 1149 | 1203 | 1124 | 1130 | 1165 | 1148 | 1153.2 | 1159 | 1148 | 0.991 | -0.014 | 0.84817835 | 0.91390978 | 0 | 0.0045 | 0.0028 | 0.0028 | TRUE | 0.013 |
| Rps2 | 83437 | 62851 | 64018 | 69468 | 62556 | 69529 | 64802 | 66854 | 69075 | 69148 | 70570 | 68636 | 68180.76 | 66910 | 69451 | 1.038 | 0.054 | 0.24983302 | 0.41862408 | 6.00E-04 | 0.0025 | 0.0016 | 0.0016 | TRUE | 0.0098 |
| Rps20 | 57615 | 41905 | 42921 | 46880 | 41091 | 46912 | 44747 | 44574 | 46311 | 46664 | 46355 | 46309 | 45826.82 | 45211 | 46443 | 1.027 | 0.039 | 0.37176329 | 0.54649756 | 2.00E-04 | 0.0025 | 0.0013 | 0.0013 | TRUE | 0.0055 |
| Rps21 | 21094 | 15031 | 16364 | 18996 | 16270 | 18487 | 16383 | 15988 | 17657 | 18909 | 18354 | 18250 | 17590.01 | 16676 | 18504 | 1.11 | 0.15 | 0.00577245 | 0.027396 | 0.0015 | 0.0026 | 0.0021 | 0.0021 | TRUE | 0.0322 |
| Rps23 | 34684 | 24442 | 26282 | 29099 | 24994 | 28838 | 26938 | 25999 | 28358 | 28965 | 28196 | 28468 | 27820.46 | 27098 | 28543 | 1.053 | 0.075 | 0.13953999 | 0.2807496 | 0.001 | 0.0025 | 0.0018 | 0.0018 | TRUE | 0.0202 |
| Rps24 | 44792 | 32158 | 33742 | 36617 | 31739 | 38554 | 34788 | 34206 | 36407 | 36449 | 35805 | 38059 | 35952.26 | 35134 | 36771 | 1.047 | 0.066 | 0.19109527 | 0.34928325 | 0.001 | 0.0025 | 0.0018 | 0.0018 | TRUE | 0.0123 |
| Rps25 | 29699 | 20538 | 22446 | 24566 | 21330 | 24848 | 23066 | 21846 | 24219 | 24453 | 24062 | 24529 | 23695.87 | 23044 | 24348 | 1.057 | 0.079 | 0.13462793 | 0.27347663 | 0.0013 | 0.0025 | 0.002 | 0.002 | TRUE | 0.0252 |
| Rps26 | 35831 | 26059 | 28088 | 29516 | 26391 | 31799 | 27828 | 27719 | 30307 | 29380 | 29772 | 31391 | 29399.38 | 28618 | 30181 | 1.055 | 0.077 | 0.17035837 | 0.32195322 | 0.0019 | 0.0025 | 0.0022 | 0.0022 | TRUE | 0.0326 |
| Rps27 | 78 | 51 | 52 | 80 | 60 | 88 | 61 | 54 | 56 | 80 | 68 | 87 | 67.52 | 57 | 78 | 1.369 | 0.453 | 0.05773052 | 0.14953649 | 0 | 0.0375 | 0.0261 | 0.0261 | TRUE | 0.1178 |
| Rps27a | 55091 | 38708 | 41204 | 47870 | 40562 | 46924 | 42787 | 41173 | 44459 | 47650 | 45758 | 46321 | 44691.32 | 42806 | 46576 | 1.088 | 0.122 | 0.01418358 | 0.05353207 | 9.00E-04 | 0.0025 | 0.0018 | 0.0018 | TRUE | 0.0139 |
| Rps27l | 11198 | 7717 | 8110 | 9318 | 7795 | 8920 | 8697 | 8208 | 8751 | 9275 | 8794 | 8805 | 8755.04 | 8552 | 8958 | 1.047 | 0.067 | 0.20934026 | 0.37109022 | 9.00E-04 | 0.0027 | 0.0019 | 0.0019 | TRUE | 0.015 |
| Rps28 | 21039 | 14415 | 15419 | 17775 | 14651 | 18033 | 16340 | 15333 | 16637 | 17693 | 16528 | 17801 | 16722.09 | 16103 | 17341 | 1.077 | 0.107 | 0.0555662 | 0.14529962 | 0.0016 | 0.0026 | 0.0022 | 0.0022 | TRUE | 0.0213 |
| Rps29 | 18246 | 12228 | 13441 | 15063 | 12682 | 15512 | 14171 | 13007 | 14503 | 14994 | 14307 | 15313 | 14382.24 | 13894 | 14871 | 1.07 | 0.098 | 0.09414039 | 0.21149773 | 0.0021 | 0.0026 | 0.0024 | 0.0024 | TRUE | 0.038 |
| Rps3 | 80523 | 58918 | 62037 | 69934 | 63448 | 71944 | 62539 | 62670 | 66937 | 69612 | 71576 | 71020 | 67392.42 | 64049 | 70736 | 1.104 | 0.143 | 0.00325779 | 0.01759878 | 8.00E-04 | 0.0025 | 0.0017 | 0.0017 | TRUE | 0.019 |
| Rps3a1 | 16679 | 11755 | 12744 | 13683 | 11756 | 14022 | 12954 | 12504 | 13751 | 13620 | 13262 | 13842 | 13322.02 | 13070 | 13575 | 1.039 | 0.055 | 0.31037905 | 0.48330403 | 0.0013 | 0.0026 | 0.002 | 0.002 | TRUE | 0.0253 |
| Rps4x | 66720 | 46568 | 51287 | 59392 | 51121 | 59489 | 51819 | 49534 | 55338 | 59119 | 57670 | 58725 | 55367.33 | 52230 | 58505 | 1.12 | 0.164 | 0.00247275 | 0.01423118 | 0.0016 | 0.0025 | 0.0021 | 0.0021 | TRUE | 0.0331 |
| Rps4y2 | 12213 | 8895 | 8250 | 8103 | 7256 | 8507 | 9485 | 9461 | 8902 | 8066 | 8186 | 8398 | 8749.58 | 9283 | 8217 | 0.885 | -0.176 | 0.00066383 | 0.00534238 | 7.00E-04 | 0.0027 | 0.0018 | 0.0018 | TRUE | 0.0157 |
| Rps5 | 57279 | 41281 | 42965 | 46133 | 41421 | 47839 | 44486 | 43910 | 46359 | 45921 | 46727 | 47224 | 45771.25 | 44918 | 46624 | 1.038 | 0.054 | 0.24267316 | 0.41028241 | 5.00E-04 | 0.0025 | 0.0015 | 0.0015 | TRUE | 0.0096 |
| Rps6 | 32127 | 22614 | 24153 | 26822 | 23315 | 26827 | 24952 | 24054 | 26061 | 26699 | 26302 | 26482 | 25758.25 | 25022 | 26494 | 1.059 | 0.082 | 0.09405568 | 0.21134065 | 8.00E-04 | 0.0025 | 0.0017 | 0.0017 | TRUE | 0.0161 |
| Rps6ka1 | 8761 | 6873 | 7459 | 7831 | 7019 | 8049 | 6804 | 7311 | 8048 | 7795 | 7918 | 7946 | 7636.99 | 7388 | 7886 | 1.068 | 0.095 | 0.15316465 | 0.3000524 | 0.0034 | 0.0027 | 0.003 | 0.003 | TRUE | 0.0746 |
| Rps6ka2 | 1805 | 1240 | 1359 | 1606 | 1514 | 1833 | 1402 | 1319 | 1466 | 1599 | 1708 | 1809 | 1550.53 | 1396 | 1705 | 1.222 | 0.289 | 0.00013661 | 0.00157335 | 0.0027 | 0.004 | 0.0035 | 0.0035 | TRUE | 0.0366 |
| Rps6ka3 | 1366 | 940 | 919 | 1096 | 871 | 924 | 1061 | 1000 | 992 | 1091 | 983 | 912 | 1006.34 | 1018 | 995 | 0.977 | -0.033 | 0.70472942 | 0.82125541 | 0.0038 | 0.0048 | 0.0044 | 0.0044 | TRUE | 0.0858 |
| Rps6ka4 | 732 | 567 | 596 | 759 | 655 | 722 | 569 | 603 | 643 | 756 | 739 | 713 | 670.31 | 605 | 736 | 1.218 | 0.285 | 0.001474 | 0.00953301 | 9.00E-04 | 0.006 | 0.0043 | 0.0043 | TRUE | 0.0356 |
| Rps6ka5 | 1039 | 705 | 859 | 930 | 888 | 999 | 807 | 750 | 927 | 926 | 1002 | 986 | 899.56 | 828 | 971 | 1.174 | 0.231 | 0.01344321 | 0.05152688 | 0.0051 | 0.0051 | 0.0052 | 0.0052 | TRUE | 0.1258 |
| Rps6ka6 | 6884 | 4751 | 5223 | 5145 | 4338 | 4987 | 5347 | 5054 | 5636 | 5121 | 4894 | 4923 | 5162.27 | 5346 | 4979 | 0.932 | -0.102 | 0.08518313 | 0.19710578 | 0.0015 | 0.0029 | 0.0023 | 0.0023 | TRUE | 0.0274 |
| Rps6kb1 | 6049 | 4430 | 4125 | 4281 | 3921 | 4498 | 4698 | 4712 | 4451 | 4261 | 4423 | 4440 | 4497.63 | 4620 | 4375 | 0.947 | -0.079 | 0.14676749 | 0.29115071 | 4.00E-04 | 0.003 | 0.0019 | 0.0019 | TRUE | 0.0124 |
| Rps6kb2 | 456 | 385 | 292 | 290 | 270 | 295 | 354 | 410 | 315 | 289 | 305 | 291 | 327.2 | 360 | 295 | 0.82 | -0.287 | 0.02441533 | 0.07998445 | 0.0062 | 0.0097 | 0.0086 | 0.0086 | TRUE | 0.1614 |
| Rps6kc1 | 2205 | 1619 | 1442 | 1588 | 1373 | 1567 | 1713 | 1722 | 1556 | 1581 | 1549 | 1547 | 1611.17 | 1664 | 1559 | 0.937 | -0.095 | 0.17250291 | 0.32476081 | 0.0011 | 0.0039 | 0.0028 | 0.0028 | TRUE | 0.0378 |
| Rps6kl1 | 456 | 294 | 285 | 301 | 243 | 289 | 354 | 313 | 308 | 300 | 274 | 285 | 305.57 | 325 | 286 | 0.88 | -0.185 | 0.13008147 | 0.26686673 | 0 | 0.0102 | 0.0075 | 0.0075 | TRUE | 0.0762 |
| Rps7 | 27468 | 19576 | 20500 | 21764 | 19047 | 22652 | 21333 | 20823 | 22119 | 21664 | 21487 | 22361 | 21631.19 | 21425 | 21837 | 1.019 | 0.028 | 0.56911812 | 0.72273703 | 6.00E-04 | 0.0025 | 0.0016 | 0.0016 | TRUE | 0.0098 |
| Rps8 | 59255 | 44431 | 45771 | 50235 | 44481 | 50524 | 46021 | 47261 | 49387 | 50004 | 50179 | 49875 | 48787.68 | 47556 | 50019 | 1.052 | 0.073 | 0.12233142 | 0.2555141 | 6.00E-04 | 0.0025 | 0.0016 | 0.0016 | TRUE | 0.0139 |
| Rps9 | 52396 | 37759 | 39259 | 41016 | 37882 | 43434 | 40694 | 40164 | 42360 | 40827 | 42735 | 42876 | 41609.3 | 41073 | 42146 | 1.026 | 0.037 | 0.44026994 | 0.61166196 | 7.00E-04 | 0.0025 | 0.0017 | 0.0017 | TRUE | 0.0092 |
| Rpsa | 98681 | 71024 | 76188 | 84694 | 74022 | 85814 | 76641 | 75547 | 82206 | 84304 | 83504 | 84712 | 81152.56 | 78131 | 84173 | 1.077 | 0.107 | 0.03194746 | 0.09692742 | 0.0011 | 0.0025 | 0.0018 | 0.0018 | TRUE | 0.0255 |

| | | | | | | | | | | | | | | | | | | | | | | | | | |
|---|---|---|---|---|---|---|---|---|---|---|---|---|---|---|---|---|---|---|---|---|---|---|---|---|---|
| Rptn | 0 | 0 | 0 | 0 | 0 | 0 | 0 | 0 | 0 | 0 | 0 | 0 | 0 | 0 | 0 | NA | NA | NA | NA | NA | NA | NA | NA | NA | NA |
| Rptor | 8691 | 6416 | 5884 | 6138 | 5750 | 6168 | 6750 | 6825 | 6349 | 6110 | 6487 | 6089 | 6434.74 | 6641 | 6229 | 0.938 | -0.093 | 0.09822156 | 0.21829899 | 0.0012 | 0.0028 | 0.0021 | 0.0021 | TRUE | 0.0179 |
| Rpusd1 | 585 | 464 | 397 | 393 | 394 | 418 | 454 | 494 | 428 | 391 | 444 | 413 | 437.43 | 459 | 416 | 0.906 | -0.142 | 0.18857671 | 0.34604274 | 0.002 | 0.0078 | 0.0061 | 0.0061 | TRUE | 0.0487 |
| Rpusd2 | 1574 | 1257 | 1098 | 1236 | 1050 | 1193 | 1222 | 1337 | 1185 | 1230 | 1185 | 1178 | 1222.79 | 1248 | 1198 | 0.96 | -0.059 | 0.43558216 | 0.60705138 | 0.0014 | 0.0044 | 0.0033 | 0.0033 | TRUE | 0.0454 |
| Rpusd3 | 1378 | 964 | 825 | 911 | 799 | 862 | 1070 | 1025 | 890 | 907 | 901 | 851 | 940.81 | 995 | 886 | 0.889 | -0.169 | 0.058225 | 0.15040844 | 0.0042 | 0.0049 | 0.0047 | 0.0047 | TRUE | 0.1003 |
| Rpusd4 | 1792 | 1367 | 1369 | 1493 | 1307 | 1601 | 1392 | 1454 | 1477 | 1486 | 1474 | 1580 | 1477.33 | 1441 | 1513 | 1.051 | 0.072 | 0.29322876 | 0.46521877 | 6.00E-04 | 0.004 | 0.0027 | 0.0027 | TRUE | 0.0181 |
| Rqcd1 | 8316 | 5958 | 5839 | 6054 | 5302 | 6350 | 6459 | 6337 | 6300 | 6026 | 5981 | 6268 | 6228.69 | 6365 | 6092 | 0.957 | -0.063 | 0.20540373 | 0.3671171 | 2.00E-04 | 0.0028 | 0.0016 | 0.0016 | TRUE | 0.0079 |
| Rrad | 65 | 34 | 30 | 32 | 26 | 34 | 50 | 36 | 32 | 32 | 29 | 34 | 35.63 | 39 | 32 | 0.79 | -0.34 | 0.30823709 | 0.48085928 | 0 | 0.0689 | 0.052 | 0.052 | TRUE | 0.4906 |
| Rraga | 4247 | 2918 | 2560 | 2448 | 2276 | 2598 | 3298 | 3104 | 2762 | 2437 | 2568 | 2565 | 2788.91 | 3055 | 2523 | 0.825 | -0.277 | 0.0001921 | 0.00207221 | 0.0038 | 0.0033 | 0.0036 | 0.0036 | TRUE | 0.0848 |
| Rragb | 530 | 419 | 421 | 469 | 458 | 517 | 412 | 446 | 454 | 467 | 517 | 510 | 467.57 | 437 | 498 | 1.141 | 0.19 | 0.06366553 | 0.16020624 | 0 | 0.0075 | 0.0054 | 0.0054 | TRUE | 0.0352 |
| Rragc | 2216 | 1658 | 1751 | 1898 | 1620 | 1926 | 1721 | 1764 | 1889 | 1889 | 1828 | 1901 | 1832.01 | 1791 | 1873 | 1.046 | 0.065 | 0.32457794 | 0.49840917 | 9.00E-04 | 0.0037 | 0.0026 | 0.0026 | TRUE | 0.0274 |
| Rragd | 1361 | 1185 | 1227 | 1116 | 1021 | 1223 | 1057 | 1260 | 1324 | 1111 | 1152 | 1207 | 1185.23 | 1214 | 1157 | 0.955 | -0.067 | 0.46977697 | 0.63880908 | 0.0071 | 0.0044 | 0.0053 | 0.0053 | TRUE | 0.1586 |
| Rras | 1074 | 849 | 1119 | 1310 | 1212 | 1398 | 834 | 903 | 1207 | 1304 | 1367 | 1380 | 1165.98 | 981 | 1350 | 1.378 | 0.463 | 6.99E-05 | 0.00091619 | 0.0196 | 0.0045 | 0.0089 | 0.0089 | TRUE | 0.4808 |
| Rras2 | 4933 | 3663 | 3485 | 3910 | 3365 | 3807 | 3831 | 3896 | 3760 | 3892 | 3796 | 3758 | 3822.33 | 3829 | 3815 | 0.996 | -0.005 | 0.9212118 | 0.95666871 | 0 | 0.003 | 0.0018 | 0.0018 | TRUE | 0.0038 |
| Rrbp1 | 11649 | 8398 | 7503 | 7701 | 6940 | 8173 | 9047 | 8933 | 8096 | 7666 | 7829 | 8068 | 8273.07 | 8692 | 7854 | 0.903 | -0.146 | 0.01387497 | 0.05274332 | 0.0019 | 0.0027 | 0.0024 | 0.0024 | TRUE | 0.0438 |
| Rreb1 | 3191 | 2310 | 2011 | 2191 | 2117 | 2366 | 2478 | 2457 | 2170 | 2181 | 2388 | 2336 | 2335 | 2368 | 2302 | 0.971 | -0.042 | 0.56229166 | 0.71682025 | 0.0034 | 0.0034 | 0.0034 | 0.0034 | TRUE | 0.0644 |
| Rrh | 0 | 1 | 0 | 2 | 2 | 0 | 0 | 1 | 0 | 2 | 2 | 0 | 0.89 | 0 | 1 | 4.079 | 2.028 | 0.37173703 | NA | 0 | 2.6801 | 2.3469 | 2.3469 | TRUE | 0.5217 |
| Rrm1 | 14846 | 10690 | 10971 | 11095 | 9590 | 11140 | 11530 | 11371 | 11838 | 11044 | 10819 | 10997 | 11266.34 | 11580 | 10953 | 0.946 | -0.08 | 0.08432577 | 0.19579476 | 1.00E-04 | 0.0026 | 0.0015 | 0.0015 | TRUE | 0.0046 |
| Rrm2 | 18955 | 13220 | 14296 | 14188 | 11970 | 14005 | 14722 | 14062 | 15425 | 14123 | 13503 | 13825 | 14276.66 | 14736 | 13817 | 0.938 | -0.093 | 0.08269206 | 0.19363001 | 0.0012 | 0.0026 | 0.002 | 0.002 | TRUE | 0.0203 |
| Rrm2b | 1194 | 854 | 984 | 1132 | 913 | 1030 | 927 | 908 | 1062 | 1127 | 1030 | 1017 | 1011.83 | 966 | 1058 | 1.096 | 0.133 | 0.13060829 | 0.2676777 | 0.0039 | 0.0048 | 0.0046 | 0.0046 | TRUE | 0.088 |
| Rrn3 | 16784 | 12129 | 12223 | 14442 | 12741 | 14756 | 13035 | 12901 | 13189 | 14376 | 14373 | 14566 | 13740.09 | 13042 | 14438 | 1.107 | 0.147 | 0.00088573 | 0.0066386 | 0 | 0.0026 | 0.0013 | 0.0013 | TRUE | 0.0012 |
| Rrnad1 | 447 | 397 | 366 | 340 | 304 | 375 | 347 | 422 | 395 | 338 | 343 | 370 | 369.32 | 388 | 350 | 0.906 | -0.143 | 0.2265595 | 0.39151432 | 0.004 | 0.0088 | 0.0073 | 0.0073 | TRUE | 0.1054 |
| Rrp1 | 9306 | 6809 | 6236 | 7181 | 6554 | 7718 | 7228 | 7243 | 6729 | 7148 | 7394 | 7619 | 7226.54 | 7067 | 7387 | 1.045 | 0.064 | 0.25233025 | 0.42118886 | 0.0012 | 0.0028 | 0.0021 | 0.0021 | TRUE | 0.0212 |
| Rrp12 | 8832 | 6340 | 5304 | 5729 | 5080 | 5848 | 6859 | 6744 | 5723 | 5703 | 5731 | 5773 | 6088.75 | 6442 | 5736 | 0.89 | -0.168 | 0.01940805 | 0.06764187 | 0.0045 | 0.0028 | 0.0036 | 0.0036 | TRUE | 0.1158 |
| Rrp15 | 4624 | 3477 | 3072 | 2967 | 2757 | 3234 | 3591 | 3698 | 3315 | 2953 | 3110 | 3192 | 3310.06 | 3535 | 3085 | 0.873 | -0.196 | 0.00240368 | 0.01393747 | 0.0019 | 0.0031 | 0.0027 | 0.0027 | TRUE | 0.0355 |
| Rrp1b | 8629 | 6418 | 5484 | 5895 | 5397 | 6106 | 6702 | 6827 | 5917 | 5868 | 6088 | 6028 | 6238.25 | 6482 | 5995 | 0.925 | -0.113 | 0.08180463 | 0.19224491 | 0.0028 | 0.0028 | 0.0029 | 0.0029 | TRUE | 0.0704 |
| Rrp36 | 1919 | 1408 | 1315 | 1459 | 1289 | 1446 | 1490 | 1498 | 1419 | 1452 | 1454 | 1427 | 1456.8 | 1469 | 1444 | 0.983 | -0.025 | 0.7084529 | 0.82410273 | 0 | 0.0041 | 0.0025 | 0.0025 | TRUE | 0.0104 |
| Rrp7a | 5503 | 3981 | 3188 | 3273 | 3122 | 3492 | 4274 | 4235 | 3440 | 3258 | 3522 | 3447 | 3695.89 | 3983 | 3409 | 0.856 | -0.225 | 0.00712499 | 0.03218631 | 0.0074 | 0.0031 | 0.0048 | 0.0048 | TRUE | 0.1721 |
| Rrp8 | 1895 | 1292 | 1120 | 1077 | 994 | 1063 | 1472 | 1374 | 1208 | 1072 | 1121 | 1049 | 1216.21 | 1351 | 1081 | 0.799 | -0.325 | 0.00017376 | 0.00191338 | 0.0049 | 0.0044 | 0.0045 | 0.0045 | TRUE | 0.1021 |
| Rrp9 | 6758 | 5052 | 4835 | 5188 | 4583 | 5306 | 5249 | 5374 | 5217 | 5164 | 5170 | 5238 | 5235.23 | 5280 | 5191 | 0.983 | -0.024 | 0.62021277 | 0.76038736 | 0 | 0.0029 | 0.0016 | 0.0016 | TRUE | 0.0029 |
| Rrs1 | 3978 | 2948 | 2501 | 2538 | 2245 | 2581 | 3090 | 3136 | 2699 | 2526 | 2533 | 2548 | 2755.1 | 2975 | 2536 | 0.852 | -0.231 | 0.00100912 | 0.0072919 | 0.0027 | 0.0033 | 0.0032 | 0.0032 | TRUE | 0.079 |
| Rs1 | 0 | 0 | 0 | 0 | 0 | 0 | 0 | 0 | 0 | 0 | 0 | 0 | 0 | 0 | 0 | NA | NA | NA | NA | NA | NA | NA | NA | NA | NA |
| Rsad1 | 628 | 415 | 378 | 415 | 356 | 405 | 488 | 441 | 408 | 413 | 402 | 400 | 425.25 | 446 | 405 | 0.906 | -0.142 | 0.19125202 | 0.34937075 | 0.0024 | 0.008 | 0.0062 | 0.0062 | TRUE | 0.0838 |
| Rsad2 | 26 | 29 | 25 | 37 | 28 | 25 | 20 | 31 | 27 | 37 | 32 | 25 | 28.52 | 26 | 31 | 1.203 | 0.267 | 0.47443693 | 0.64280192 | 0 | 0.0855 | 0.0647 | 0.0647 | TRUE | 0.2931 |
| Rsbn1 | 2692 | 2179 | 2461 | 2674 | 2325 | 2759 | 2091 | 2318 | 2655 | 2662 | 2623 | 2724 | 2512.01 | 2355 | 2670 | 1.135 | 0.182 | 0.02936749 | 0.09119732 | 0.0069 | 0.0034 | 0.0047 | 0.0047 | TRUE | 0.1511 |
| Rsbn1l | 2870 | 2296 | 2678 | 2843 | 2414 | 2936 | 2229 | 2442 | 2890 | 2830 | 2723 | 2898 | 2668.7 | 2520 | 2817 | 1.119 | 0.162 | 0.06627836 | 0.16471991 | 0.0089 | 0.0033 | 0.0052 | 0.0052 | TRUE | 0.1988 |
| Rsc1a1 | 0 | 0 | 2 | 1 | 1 | 0 | 0 | 0 | 2 | 1 | 1 | 0 | 0.71 | 1 | 1 | 1.011 | 0.016 | 0.99481353 | NA | 0 | 3.3238 | 2.9707 | 2.9707 | TRUE | 0.9749 |
| Rsf1 | 4568 | 3287 | 3225 | 3353 | 2814 | 3479 | 3548 | 3496 | 3480 | 3338 | 3174 | 3434 | 3411.7 | 3508 | 3315 | 0.945 | -0.081 | 0.15329942 | 0.30015187 | 4.00E-04 | 0.0031 | 0.002 | 0.002 | TRUE | 0.0166 |
| Rsg1 | 48 | 40 | 28 | 37 | 37 | 29 | 37 | 43 | 30 | 37 | 42 | 29 | 36.21 | 37 | 36 | 0.971 | -0.043 | 0.89602003 | 0.94272381 | 0 | 0.0679 | 0.0504 | 0.0504 | TRUE | 0.2216 |
| Rsl1 | 236 | 200 | 175 | 197 | 150 | 191 | 183 | 213 | 189 | 196 | 169 | 189 | 189.78 | 195 | 185 | 0.95 | -0.074 | 0.62021486 | 0.76038736 | 0 | 0.0149 | 0.0108 | 0.0108 | TRUE | 0.0675 |
| Rsl1d1 | 16927 | 12170 | 11983 | 13093 | 11622 | 13254 | 13146 | 12945 | 12930 | 13033 | 13111 | 13084 | 13041.41 | 13007 | 13076 | 1.005 | 0.008 | 0.86420874 | 0.92504572 | 0 | 0.0026 | 0.0013 | 0.0013 | TRUE | 0.0011 |
| Rsl24d1 | 5361 | 3964 | 3669 | 4006 | 3379 | 3897 | 4164 | 4216 | 3959 | 3988 | 3812 | 3847 | 3997.55 | 4113 | 3882 | 0.944 | -0.083 | 0.1342071 | 0.27301767 | 5.00E-04 | 0.003 | 0.002 | 0.002 | TRUE | 0.0129 |
| Rslcan18 | 127 | 94 | 95 | 103 | 122 | 143 | 99 | 100 | 103 | 103 | 138 | 141 | 113.74 | 101 | 127 | 1.266 | 0.34 | 0.08254029 | 0.19342383 | 0.0077 | 0.0233 | 0.0188 | 0.0188 | TRUE | 0.2978 |
| Rsph1 | 170 | 138 | 179 | 160 | 133 | 178 | 132 | 147 | 193 | 159 | 150 | 176 | 159.5 | 157 | 162 | 1.033 | 0.047 | 0.79330109 | 0.87889718 | 0.017 | 0.0173 | 0.0172 | 0.0172 | TRUE | 0.4105 |
| Rsph3a | 118 | 92 | 114 | 126 | 114 | 126 | 92 | 98 | 123 | 125 | 129 | 124 | 115.15 | 104 | 126 | 1.216 | 0.282 | 0.13910688 | 0.28039129 | 0 | 0.023 | 0.0175 | 0.0175 | TRUE | 0.2386 |
| Rsph3b | 197 | 166 | 178 | 191 | 179 | 202 | 153 | 177 | 192 | 190 | 202 | 199 | 185.52 | 174 | 197 | 1.138 | 0.187 | 0.22066767 | 0.38421282 | 0 | 0.0152 | 0.0114 | 0.0114 | TRUE | 0.1309 |
| Rsph4a | 18 | 5 | 4 | 1 | 6 | 6 | 14 | 5 | 4 | 1 | 7 | 6 | 6.22 | 8 | 5 | 0.564 | -0.826 | 0.34361352 | NA | 0.3461 | 0.3837 | 0.3733 | 0.3733 | TRUE | 1.6682 |
| Rsph6a | 11 | 6 | 2 | 9 | 3 | 7 | 9 | 6 | 2 | 9 | 3 | 7 | 6.06 | 6 | 6 | 1.114 | 0.155 | 0.84914219 | NA | 0.0954 | 0.3938 | 0.3178 | 0.3178 | TRUE | 0.4875 |
| Rsph9 | 21 | 27 | 41 | 27 | 26 | 35 | 16 | 29 | 44 | 27 | 29 | 35 | 30 | 30 | 30 | 1.031 | 0.044 | 0.91368006 | 0.95246365 | 0.096 | 0.0814 | 0.0857 | 0.0857 | TRUE | 0.257 |
| Rspo1 | 172 | 131 | 118 | 177 | 155 | 168 | 134 | 139 | 127 | 176 | 175 | 166 | 152.86 | 133 | 172 | 1.291 | 0.368 | 0.0202805 | 0.0699154 | 0 | 0.0179 | 0.0115 | 0.0115 | TRUE | 0.015 |
| Rspo2 | 1 | 0 | 2 | 3 | 3 | 0 | 1 | 0 | 2 | 3 | 3 | 0 | 1.55 | 1 | 2 | 2.18 | 1.124 | 0.51051386 | NA | 0 | 1.5307 | 1.3593 | 1.3593 | TRUE | 0.7566 |
| Rspo3 | 12 | 15 | 5 | 6 | 5 | 6 | 9 | 16 | 5 | 6 | 6 | 6 | 8.03 | 10 | 6 | 0.573 | -0.803 | 0.25633272 | NA | 0 | 0.2974 | 0.2245 | 0.2245 | TRUE | 0.2411 |
| Rspo4 | 100 | 62 | 85 | 134 | 126 | 98 | 78 | 66 | 92 | 133 | 142 | 97 | 101.27 | 79 | 124 | 1.58 | 0.66 | 0.00293826 | 0.01627141 | 0.0234 | 0.0258 | 0.0252 | 0.0252 | TRUE | 0.3842 |
| Rspry1 | 2958 | 2125 | 2108 | 2171 | 1908 | 2166 | 2297 | 2260 | 2275 | 2161 | 2152 | 2138 | 2213.97 | 2277 | 2150 | 0.944 | -0.083 | 0.14887585 | 0.29411535 | 0 | 0.0035 | 0.0019 | 0.0019 | TRUE | 7.00E-04 |
| Rsrc1 | 2029 | 1536 | 1565 | 1538 | 1282 | 1522 | 1576 | 1634 | 1689 | 1531 | 1446 | 1502 | 1562.98 | 1633 | 1493 | 0.915 | -0.128 | 0.0576683 | 0.14948368 | 4.00E-04 | 0.0039 | 0.0026 | 0.0026 | TRUE | 0.0117 |
| Rsrc2 | 7496 | 5784 | 6233 | 6119 | 5211 | 6479 | 5822 | 6152 | 6725 | 6091 | 5879 | 6396 | 6177.45 | 6233 | 6122 | 0.983 | -0.025 | 0.70350233 | 0.82071497 | 0.0028 | 0.0028 | 0.003 | 0.003 | TRUE | 0.0581 |
| Rsu1 | 2958 | 2036 | 1714 | 1906 | 1740 | 1976 | 2297 | 2166 | 1849 | 1897 | 1963 | 1951 | 2020.53 | 2104 | 1937 | 0.92 | -0.121 | 0.1432927 | 0.28620493 | 0.0054 | 0.0036 | 0.0044 | 0.0044 | TRUE | 0.1347 |
| Rtbdn | 22 | 12 | 22 | 15 | 21 | 23 | 17 | 13 | 24 | 15 | 24 | 23 | 19.15 | 18 | 21 | 1.144 | 0.195 | 0.66986243 | 0.79706491 | 0 | 0.1262 | 0.0976 | 0.0976 | TRUE | 0.3393 |
| Rtca | 2767 | 2068 | 1945 | 2026 | 1813 | 2068 | 2149 | 2200 | 2099 | 2017 | 2045 | 2041 | 2091.79 | 2149 | 2034 | 0.947 | -0.079 | 0.18350833 | 0.33940018 | 0 | 0.0036 | 0.0021 | 0.0021 | TRUE | 0.0049 |
| Rtdr1 | 2 | 5 | 2 | 0 | 2 | 1 | 2 | 5 | 2 | 0 | 2 | 1 | 2.05 | 3 | 1 | 0.355 | -1.493 | 0.30693522 | NA | 0 | 1.1611 | 0.8868 | 0.8868 | TRUE | 0.5581 |
| Rtel1 | 5082 | 3647 | 2464 | 2576 | 2213 | 2770 | 3947 | 3879 | 2659 | 2564 | 2496 | 2734 | 3046.66 | 3495 | 2598 | 0.743 | -0.428 | 0.00026807 | 0.00266655 | 0.0221 | 0.0032 | 0.0096 | 0.0096 | TRUE | 0.5308 |
| Rtf1 | 5485 | 4066 | 4097 | 4301 | 3771 | 4442 | 4260 | 4325 | 4421 | 4281 | 4254 | 4385 | 4320.96 | 4335 | 4307 | 0.994 | -0.009 | 0.86074363 | 0.92271872 | 0 | 0.003 | 0.0017 | 0.0017 | TRUE | 0.0036 |
| Rtfdc1 | 3512 | 2626 | 2328 | 2361 | 2091 | 2402 | 2728 | 2793 | 2512 | 2350 | 2359 | 2371 | 2518.82 | 2678 | 2360 | 0.881 | -0.183 | 0.00396796 | 0.02047635 | 0.0012 | 0.0034 | 0.0025 | 0.0025 | TRUE | 0.0349 |
| Rtkn | 471 | 344 | 360 | 463 | 413 | 452 | 366 | 366 | 388 | 461 | 466 | 446 | 415.52 | 373 | 458 | 1.226 | 0.294 | 0.0040908 | 0.02099542 | 0 | 0.0081 | 0.0051 | 0.0051 | TRUE | 0.0137 |
| Rtkn2 | 1896 | 1480 | 1462 | 1622 | 1538 | 1677 | 1473 | 1574 | 1577 | 1615 | 1735 | 1655 | 1604.88 | 1541 | 1668 | 1.083 | 0.115 | 0.09311371 | 0.20988396 | 9.00E-04 | 0.0039 | 0.0028 | 0.0028 | TRUE | 0.0192 |
| Rtl1 | 1 | 0 | 0 | 0 | 0 | 0 | 1 | 0 | 0 | 0 | 0 | 0 | 0.13 | 0 | 0 | 0.552 | -0.857 | 0.8336548 | NA | 0 | 18.3121 | 10 | 10 | TRUE | 0.2566 |
| Rtn1 | 855 | 866 | 892 | 986 | 1044 | 1152 | 664 | 921 | 962 | 981 | 1178 | 1137 | 974.01 | 849 | 1099 | 1.296 | 0.375 | 0.00301049 | 0.01659425 | 0.0232 | 0.0049 | 0.0104 | 0.0104 | TRUE | 0.4465 |
| Rtn2 | 930 | 702 | 662 | 803 | 677 | 786 | 722 | 747 | 714 | 799 | 764 | 776 | 753.7 | 728 | 780 | 1.072 | 0.1 | 0.21889279 | 0.38196003 | 0 | 0.0056 | 0.0034 | 0.0034 | TRUE | 0.0059 |
| Rtn3 | 9315 | 6308 | 5787 | 6549 | 5765 | 6719 | 7235 | 6710 | 6244 | 6519 | 6504 | 6633 | 6640.58 | 6730 | 6552 | 0.973 | -0.039 | 0.53539483 | 0.694347 | 0.0023 | 0.0028 | 0.0027 | 0.0027 | TRUE | 0.0531 |

| | | | | | | | | | | | | | | | | | | | | | | | | | |
|---|---|---|---|---|---|---|---|---|---|---|---|---|---|---|---|---|---|---|---|---|---|---|---|---|---|
| Rtn4 | 36394 | 26677 | 26732 | 28875 | 24941 | 28989 | 28266 | 28376 | 28844 | 28742 | 28136 | 28617 | 28496.68 | 28495 | 28498 | 1 | 0 | 0.99600369 | 0.99753911 | 1.00E-04 | 0.0025 | 0.0013 | 0.0013 | TRUE | 0.0015 |
| Rtn4ip1 | 1682 | 1256 | 1334 | 1654 | 1426 | 1596 | 1306 | 1336 | 1439 | 1646 | 1609 | 1575 | 1485.38 | 1360 | 1610 | 1.184 | 0.244 | 0.00047875 | 0.0041826 | 8.00E-04 | 0.004 | 0.0028 | 0.0028 | TRUE | 0.0304 |
| Rtn4r | 73 | 51 | 85 | 84 | 66 | 93 | 57 | 54 | 92 | 84 | 74 | 92 | 75.42 | 68 | 83 | 1.241 | 0.312 | 0.22386811 | 0.38827168 | 0.0335 | 0.0339 | 0.034 | 0.034 | TRUE | 0.8337 |
| Rtn4rl1 | 218 | 151 | 135 | 146 | 119 | 144 | 169 | 161 | 146 | 145 | 134 | 142 | 149.55 | 159 | 140 | 0.885 | -0.176 | 0.2797747 | 0.45138501 | 0 | 0.0183 | 0.0124 | 0.0124 | TRUE | 0.0517 |
| Rtn4rl2 | 43 | 27 | 32 | 62 | 52 | 56 | 33 | 29 | 35 | 62 | 59 | 55 | 45.38 | 32 | 59 | 1.815 | 0.86 | 0.00281267 | 0.01570932 | 0 | 0.0546 | 0.0359 | 0.0359 | TRUE | 0.0567 |
| Rtp1 | 0 | 0 | 0 | 0 | 0 | 0 | 0 | 0 | 0 | 0 | 0 | 0 | 0 | 0 | 0 | NA | NA | NA | NA | NA | NA | NA | NA | NA | NA |
| Rtp2 | 2 | 0 | 1 | 0 | 1 | 0 | 2 | 0 | 1 | 0 | 1 | 0 | 0.63 | 1 | 0 | 0.414 | -1.271 | 0.61575572 | NA | 0 | 3.784 | 3.0839 | 3.0839 | TRUE | 0.5075 |
| Rtp3 | 48 | 32 | 51 | 67 | 67 | 80 | 37 | 34 | 55 | 67 | 76 | 79 | 57.93 | 42 | 74 | 1.76 | 0.816 | 0.00286601 | 0.01596411 | 0.0104 | 0.0433 | 0.0354 | 0.0354 | TRUE | 0.5125 |
| Rtp4 | 4 | 3 | 6 | 2 | 5 | 5 | 3 | 3 | 6 | 2 | 6 | 5 | 4.22 | 4 | 4 | 0.99 | -0.015 | 0.98727165 | NA | 0 | 0.5636 | 0.3974 | 0.3974 | TRUE | 0.3778 |
| Rttn | 2154 | 1530 | 1590 | 1694 | 1470 | 1683 | 1673 | 1627 | 1716 | 1686 | 1658 | 1661 | 1670.31 | 1672 | 1668 | 0.998 | -0.003 | 0.96476517 | 0.9807531 | 0 | 0.0038 | 0.0023 | 0.0023 | TRUE | 0.0063 |
| Rubie | 4 | 3 | 1 | 1 | 2 | 1 | 3 | 3 | 1 | 1 | 2 | 1 | 1.94 | 2 | 1 | 0.562 | -0.832 | 0.54434651 | NA | 0 | 1.2267 | 0.7903 | 0.7903 | TRUE | 0.2345 |
| Rufy1 | 1083 | 834 | 854 | 948 | 917 | 1000 | 841 | 887 | 921 | 944 | 1034 | 987 | 935.83 | 883 | 988 | 1.12 | 0.163 | 0.04268894 | 0.12001435 | 0.0011 | 0.005 | 0.0036 | 0.0036 | TRUE | 0.0219 |
| Rufy2 | 1906 | 1503 | 1522 | 1525 | 1359 | 1467 | 1480 | 1599 | 1642 | 1518 | 1533 | 1448 | 1536.75 | 1574 | 1500 | 0.954 | -0.068 | 0.33694364 | 0.51122815 | 0.0014 | 0.004 | 0.003 | 0.003 | TRUE | 0.0339 |
| Rufy3 | 691 | 541 | 532 | 562 | 496 | 575 | 537 | 575 | 574 | 559 | 560 | 568 | 562.12 | 562 | 562 | 1.002 | 0.002 | 0.97878962 | 0.98866701 | 0 | 0.0066 | 0.0042 | 0.0042 | TRUE | 0.0196 |
| Rufy4 | 51 | 48 | 28 | 42 | 24 | 40 | 40 | 51 | 30 | 42 | 27 | 39 | 38.21 | 40 | 36 | 0.9 | -0.152 | 0.65084238 | 0.78320543 | 0.0291 | 0.0645 | 0.0551 | 0.0551 | TRUE | 0.2043 |
| Rundc1 | 1145 | 839 | 724 | 731 | 692 | 780 | 889 | 892 | 781 | 728 | 781 | 770 | 806.86 | 854 | 760 | 0.888 | -0.172 | 0.05102744 | 0.13688723 | 0.0023 | 0.0054 | 0.0043 | 0.0043 | TRUE | 0.0648 |
| Rundc3a | 1005 | 948 | 904 | 951 | 883 | 996 | 781 | 1008 | 975 | 947 | 996 | 983 | 948.38 | 921 | 975 | 1.061 | 0.085 | 0.39852928 | 0.57190552 | 0.0089 | 0.0049 | 0.0063 | 0.0063 | TRUE | 0.2221 |
| Rundc3b | 684 | 531 | 432 | 545 | 407 | 497 | 531 | 565 | 466 | 542 | 459 | 491 | 509.07 | 521 | 497 | 0.955 | -0.066 | 0.55038252 | 0.7064132 | 0.0061 | 0.0071 | 0.0068 | 0.0068 | TRUE | 0.0943 |
| Runx1 | 144 | 85 | 92 | 113 | 90 | 106 | 112 | 90 | 99 | 112 | 102 | 105 | 103.36 | 100 | 106 | 1.053 | 0.075 | 0.70138253 | 0.81909147 | 0 | 0.0254 | 0.0177 | 0.0177 | TRUE | 0.1066 |
| Runx1t1 | 126 | 116 | 89 | 89 | 80 | 94 | 98 | 123 | 96 | 89 | 90 | 93 | 98.15 | 106 | 91 | 0.858 | -0.221 | 0.276989 | 0.44815528 | 0 | 0.0266 | 0.0194 | 0.0194 | TRUE | 0.2018 |
| Runx2 | 5 | 8 | 3 | 10 | 3 | 7 | 4 | 9 | 3 | 10 | 3 | 7 | 5.98 | 5 | 7 | 1.316 | 0.397 | 0.62755289 | NA | 0.0954 | 0.3988 | 0.312 | 0.312 | TRUE | 0.258 |
| Runx3 | 231 | 196 | 213 | 277 | 278 | 304 | 179 | 208 | 230 | 276 | 314 | 300 | 251.19 | 206 | 297 | 1.445 | 0.531 | 0.00014731 | 0.00168171 | 0.0056 | 0.0119 | 0.01 | 0.01 | TRUE | 0.1525 |
| Rusc1 | 589 | 381 | 386 | 395 | 360 | 384 | 457 | 405 | 416 | 393 | 406 | 379 | 409.6 | 426 | 393 | 0.919 | -0.122 | 0.25331255 | 0.42242648 | 0 | 0.0082 | 0.0058 | 0.0058 | TRUE | 0.0501 |
| Rusc2 | 1573 | 1184 | 1331 | 1360 | 1358 | 1339 | 1222 | 1259 | 1436 | 1354 | 1532 | 1322 | 1354.12 | 1306 | 1403 | 1.075 | 0.104 | 0.23931276 | 0.40630058 | 0.006 | 0.0042 | 0.0049 | 0.0049 | TRUE | 0.0908 |
| Ruvbl1 | 5317 | 3876 | 4001 | 3675 | 3424 | 4135 | 4129 | 4123 | 4317 | 3658 | 3863 | 4082 | 4028.66 | 4190 | 3868 | 0.923 | -0.115 | 0.05886111 | 0.15125888 | 0.0015 | 0.003 | 0.0024 | 0.0024 | TRUE | 0.0289 |
| Ruvbl2 | 13846 | 10627 | 10804 | 12010 | 10600 | 12583 | 10754 | 11304 | 11657 | 11955 | 11958 | 12421 | 11674.81 | 11238 | 12111 | 1.078 | 0.108 | 0.038824 | 0.11203543 | 9.00E-04 | 0.0026 | 0.0019 | 0.0019 | TRUE | 0.0176 |
| Rwdd1 | 3140 | 2058 | 1990 | 2263 | 1919 | 2149 | 2439 | 2189 | 2147 | 2253 | 2165 | 2121 | 2218.96 | 2258 | 2180 | 0.964 | -0.053 | 0.45469797 | 0.62448334 | 0.0022 | 0.0035 | 0.0031 | 0.0031 | TRUE | 0.0606 |
| Rwdd2a | 541 | 387 | 399 | 470 | 361 | 464 | 420 | 412 | 431 | 468 | 407 | 458 | 432.58 | 421 | 444 | 1.057 | 0.08 | 0.44406216 | 0.61501948 | 0 | 0.0079 | 0.0056 | 0.0056 | TRUE | 0.0598 |
| Rwdd2b | 643 | 471 | 454 | 391 | 361 | 396 | 499 | 501 | 490 | 389 | 407 | 391 | 446.27 | 497 | 396 | 0.796 | -0.329 | 0.00088677 | 0.00664287 | 0 | 0.0077 | 0.0048 | 0.0048 | TRUE | 0.0071 |
| Rwdd3 | 366 | 288 | 250 | 281 | 251 | 263 | 284 | 306 | 270 | 280 | 283 | 260 | 280.47 | 287 | 274 | 0.956 | -0.065 | 0.59799434 | 0.74387581 | 0 | 0.0109 | 0.0075 | 0.0075 | TRUE | 0.0378 |
| Rwdd4a | 2741 | 1902 | 1849 | 2034 | 1690 | 1874 | 2129 | 2023 | 1995 | 2025 | 1906 | 1850 | 1988.01 | 2049 | 1927 | 0.94 | -0.089 | 0.17887118 | 0.33349932 | 0.0012 | 0.0036 | 0.0027 | 0.0027 | TRUE | 0.024 |
| Rxfp1 | 1 | 1 | 0 | 0 | 0 | 0 | 1 | 1 | 0 | 0 | 0 | 0 | 0.31 | 1 | 0 | 0.309 | -1.696 | 0.63183309 | NA | 0 | 7.7294 | 7.2472 | 7.2472 | TRUE | 0.2015 |
| Rxfp2 | 12 | 11 | 3 | 12 | 4 | 8 | 9 | 12 | 3 | 12 | 5 | 8 | 8.1 | 8 | 8 | 1.004 | 0.006 | 0.99321251 | NA | 0.1348 | 0.295 | 0.2533 | 0.2533 | TRUE | 0.3462 |
| Rxfp3 | 1 | 0 | 0 | 0 | 0 | 0 | 1 | 0 | 0 | 0 | 0 | 0 | 0.13 | 0 | 0 | 0.552 | -0.857 | 0.8336548 | NA | 0 | 18.3121 | 10 | 10 | TRUE | 0.2566 |
| Rxfp4 | 1 | 1 | 0 | 2 | 0 | 0 | 1 | 1 | 0 | 2 | 0 | 0 | 0.64 | 1 | 1 | 1.084 | 0.117 | 0.96352112 | NA | 0 | 3.7142 | 3.1343 | 3.1343 | TRUE | 0.9737 |
| Rxra | 1789 | 1353 | 1011 | 1057 | 866 | 924 | 1389 | 1439 | 1091 | 1052 | 977 | 912 | 1143.45 | 1306 | 980 | 0.75 | -0.415 | 8.78E-05 | 0.00108618 | 0.0127 | 0.0045 | 0.0072 | 0.0072 | TRUE | 0.2468 |
| Rxrb | 2326 | 1829 | 1889 | 1895 | 1805 | 2001 | 1807 | 1945 | 2038 | 1886 | 2036 | 1975 | 1948 | 1930 | 1966 | 1.019 | 0.028 | 0.69530966 | 0.8143949 | 0.0021 | 0.0036 | 0.0031 | 0.0031 | TRUE | 0.0392 |
| Rxrg | 58 | 47 | 32 | 38 | 39 | 52 | 45 | 50 | 35 | 38 | 44 | 51 | 43.79 | 43 | 44 | 1.026 | 0.037 | 0.90197015 | 0.94570564 | 0 | 0.0566 | 0.0426 | 0.0426 | TRUE | 0.2181 |
| Rybp | 10723 | 7572 | 7213 | 7724 | 6734 | 7839 | 8328 | 8054 | 7783 | 7688 | 7597 | 7738 | 7864.75 | 8055 | 7674 | 0.953 | -0.07 | 0.16575294 | 0.31676786 | 4.00E-04 | 0.0027 | 0.0017 | 0.0017 | TRUE | 0.0109 |
| Ryk | 4974 | 3603 | 3528 | 3931 | 3448 | 3872 | 3863 | 3832 | 3807 | 3913 | 3890 | 3822 | 3854.52 | 3834 | 3875 | 1.011 | 0.015 | 0.76882398 | 0.86325859 | 0 | 0.003 | 0.0016 | 0.0016 | TRUE | 0.0017 |
| Ryr1 | 1294 | 1007 | 1013 | 1464 | 1369 | 1510 | 1005 | 1071 | 1093 | 1457 | 1544 | 1491 | 1276.9 | 1056 | 1497 | 1.419 | 0.505 | 2.67E-12 | 3.01E-10 | 6.00E-04 | 0.0043 | 0.0029 | 0.0029 | TRUE | 0.0229 |
| Ryr2 | 765 | 554 | 669 | 798 | 696 | 695 | 594 | 589 | 722 | 794 | 785 | 686 | 695.14 | 635 | 755 | 1.19 | 0.251 | 0.01785534 | 0.06351577 | 0.008 | 0.0058 | 0.0067 | 0.0067 | TRUE | 0.1646 |
| Ryr3 | 180 | 148 | 94 | 174 | 161 | 165 | 140 | 157 | 101 | 173 | 182 | 163 | 152.73 | 133 | 173 | 1.295 | 0.373 | 0.04163075 | 0.11771699 | 0.0165 | 0.0179 | 0.0176 | 0.0176 | TRUE | 0.4141 |
| S100a1 | 72 | 54 | 92 | 128 | 103 | 109 | 56 | 57 | 99 | 127 | 116 | 108 | 93.97 | 71 | 117 | 1.664 | 0.734 | 0.00263619 | 0.01492239 | 0.043 | 0.0276 | 0.0317 | 0.0317 | TRUE | 1.0676 |
| S100a10 | 2451 | 1709 | 1869 | 1803 | 1606 | 1779 | 1904 | 1818 | 2017 | 1795 | 1812 | 1756 | 1850.11 | 1913 | 1788 | 0.935 | -0.098 | 0.14220028 | 0.28473922 | 9.00E-04 | 0.0037 | 0.0026 | 0.0026 | TRUE | 0.0265 |
| S100a11 | 1789 | 1357 | 1250 | 1390 | 1176 | 1428 | 1389 | 1443 | 1349 | 1384 | 1327 | 1410 | 1383.59 | 1394 | 1374 | 0.986 | -0.021 | 0.76236644 | 0.85900698 | 0 | 0.0041 | 0.0027 | 0.0027 | TRUE | 0.0112 |
| S100a13 | 53 | 37 | 54 | 69 | 75 | 60 | 41 | 39 | 58 | 69 | 85 | 59 | 58.55 | 46 | 71 | 1.535 | 0.618 | 0.02492559 | 0.08122695 | 0.0206 | 0.0429 | 0.0369 | 0.0369 | TRUE | 0.3832 |
| S100a14 | 17 | 17 | 18 | 13 | 16 | 10 | 13 | 18 | 19 | 13 | 18 | 10 | 15.26 | 17 | 14 | 0.807 | -0.309 | 0.53859908 | NA | 0 | 0.1577 | 0.1144 | 0.1144 | TRUE | 0.3143 |
| S100a16 | 47 | 17 | 24 | 38 | 30 | 35 | 37 | 18 | 26 | 38 | 34 | 35 | 31.12 | 27 | 36 | 1.303 | 0.382 | 0.29724203 | 0.46928978 | 0.026 | 0.0786 | 0.0645 | 0.0645 | TRUE | 0.409 |
| S100a2 | 0 | 0 | 0 | 0 | 0 | 0 | 0 | 0 | 0 | 0 | 0 | 0 | 0 | 0 | 0 | NA | NA | NA | NA | NA | NA | NA | NA | NA | NA |
| S100a3 | 11 | 3 | 1 | 4 | 1 | 2 | 9 | 3 | 1 | 4 | 1 | 2 | 3.32 | 4 | 2 | 0.543 | -0.882 | 0.43980393 | NA | 0.3806 | 0.7171 | 0.6118 | 0.6118 | TRUE | 0.5327 |
| S100a4 | 4 | 0 | 6 | 0 | 1 | 3 | 3 | 0 | 6 | 0 | 1 | 3 | 2.28 | 3 | 1 | 0.429 | -1.22 | 0.41833186 | NA | 1.3818 | 1.0427 | 1.1065 | 1.1065 | TRUE | 0.1294 |
| S100a5 | 0 | 0 | 0 | 0 | 0 | 0 | 0 | 0 | 0 | 0 | 0 | 0 | 0 | 0 | 0 | NA | NA | NA | NA | NA | NA | NA | NA | NA | NA |
| S100a6 | 1001 | 673 | 753 | 445 | 368 | 518 | 777 | 716 | 812 | 443 | 415 | 511 | 612.54 | 768 | 456 | 0.594 | -0.75 | 6.05E-13 | 8.16E-11 | 0.0053 | 0.0063 | 0.0061 | 0.0061 | TRUE | 0.1291 |
| S100a7a | 0 | 0 | 0 | 0 | 0 | 0 | 0 | 0 | 0 | 0 | 0 | 0 | 0 | 0 | 0 | NA | NA | NA | NA | NA | NA | NA | NA | NA | NA |
| S100a8 | 0 | 0 | 0 | 0 | 0 | 0 | 0 | 0 | 0 | 0 | 0 | 0 | 0 | 0 | 0 | NA | NA | NA | NA | NA | NA | NA | NA | NA | NA |
| S100a9 | 0 | 0 | 0 | 0 | 0 | 0 | 0 | 0 | 0 | 0 | 0 | 0 | 0 | 0 | 0 | NA | NA | NA | NA | NA | NA | NA | NA | NA | NA |
| S100b | 0 | 2 | 0 | 3 | 1 | 2 | 0 | 2 | 0 | 3 | 1 | 2 | 1.37 | 1 | 2 | 3.029 | 1.599 | 0.37709759 | NA | 0 | 1.7332 | 1.3935 | 1.3935 | TRUE | 0.97 |
| S100g | 2 | 0 | 0 | 4 | 1 | 1 | 2 | 0 | 0 | 4 | 1 | 1 | 1.28 | 1 | 2 | 3.542 | 1.825 | 0.33556405 | NA | 0 | 1.8613 | 1.4989 | 1.4989 | TRUE | 0.9711 |
| S100pbp | 1965 | 1415 | 1492 | 1552 | 1359 | 1612 | 1526 | 1505 | 1610 | 1545 | 1533 | 1591 | 1551.72 | 1547 | 1556 | 1.007 | 0.009 | 0.88767742 | 0.9375978 | 0 | 0.004 | 0.0025 | 0.0025 | TRUE | 0.0148 |
| S100z | 1 | 0 | 0 | 0 | 0 | 0 | 1 | 0 | 0 | 0 | 0 | 0 | 0.13 | 0 | 0 | 0.552 | -0.857 | 0.8336548 | NA | 0 | 18.3121 | 10 | 10 | TRUE | 0.2566 |
| S1pr1 | 58 | 46 | 29 | 21 | 17 | 24 | 45 | 49 | 31 | 21 | 19 | 24 | 31.51 | 42 | 21 | 0.509 | -0.975 | 0.00689474 | 0.03140424 | 0 | 0.0777 | 0.058 | 0.058 | TRUE | 0.3412 |
| S1pr2 | 2838 | 2201 | 2094 | 2797 | 2307 | 2682 | 2204 | 2341 | 2259 | 2784 | 2603 | 2648 | 2473.16 | 2268 | 2678 | 1.181 | 0.24 | 8.88E-05 | 0.001095 | 7.00E-04 | 0.0034 | 0.0023 | 0.0023 | TRUE | 0.0146 |
| S1pr3 | 162 | 133 | 157 | 216 | 191 | 192 | 126 | 141 | 169 | 215 | 215 | 190 | 176.12 | 145 | 207 | 1.426 | 0.512 | 0.00180745 | 0.01118559 | 0.0076 | 0.0159 | 0.0136 | 0.0136 | TRUE | 0.2106 |
| S1pr4 | 59 | 53 | 66 | 95 | 82 | 81 | 46 | 56 | 71 | 95 | 93 | 80 | 73.41 | 58 | 89 | 1.552 | 0.634 | 0.00942658 | 0.03969129 | 0.0123 | 0.0347 | 0.0287 | 0.0287 | TRUE | 0.3442 |
| S1pr5 | 34 | 23 | 27 | 41 | 29 | 42 | 26 | 24 | 29 | 41 | 33 | 41 | 32.5 | 26 | 38 | 1.441 | 0.527 | 0.11794372 | 0.24916845 | 0 | 0.0754 | 0.0504 | 0.0504 | TRUE | 0.112 |
| Saa1 | 1 | 0 | 0 | 0 | 0 | 0 | 1 | 0 | 0 | 0 | 0 | 0 | 0.13 | 0 | 0 | 0.552 | -0.857 | 0.8336548 | NA | 0 | 18.3121 | 10 | 10 | TRUE | 0.2566 |
| Saa2 | 0 | 0 | 0 | 1 | 0 | 0 | 0 | 0 | 0 | 1 | 0 | 0 | 0.17 | 0 | 0 | 2.094 | 1.067 | 0.79379329 | NA | 0 | 14.2885 | 10 | 10 | TRUE | 0.3906 |
| Saa3 | 13 | 9 | 10 | 9 | 7 | 15 | 10 | 10 | 11 | 9 | 8 | 15 | 10.35 | 10 | 11 | 1.044 | 0.063 | 0.91600921 | NA | 0 | 0.2313 | 0.1588 | 0.1588 | TRUE | 0.4598 |

| | | | | | | | | | | | | | | | | | | | | | | | | | |
|---|---|---|---|---|---|---|---|---|---|---|---|---|---|---|---|---|---|---|---|---|---|---|---|---|---|
| Saa4 | 0 | 0 | 0 | 0 | 0 | 0 | 0 | 0 | 0 | 0 | 0 | 0 | 0 | 0 | 0 | NA | NA | NA | NA | NA | NA | NA | NA | NA | NA |
| Saal1 | 2343 | 1641 | 1700 | 1675 | 1433 | 1702 | 1820 | 1746 | 1834 | 1667 | 1617 | 1680 | 1727.25 | 1800 | 1655 | 0.919 | -0.121 | 0.05640415 | 0.14711397 | 0 | 0.0038 | 0.0023 | 0.0023 | TRUE | 0.0082 |
| Sac3d1 | 1346 | 983 | 891 | 937 | 860 | 966 | 1045 | 1046 | 961 | 933 | 970 | 954 | 984.8 | 1017 | 952 | 0.935 | -0.097 | 0.20832413 | 0.36989141 | 0 | 0.0048 | 0.0033 | 0.0033 | TRUE | 0.027 |
| Sacm1l | 3083 | 2204 | 2153 | 2342 | 2056 | 2330 | 2394 | 2344 | 2323 | 2331 | 2319 | 2300 | 2335.42 | 2354 | 2317 | 0.984 | -0.023 | 0.68310798 | 0.80580546 | 0 | 0.0034 | 0.0019 | 0.0019 | TRUE | 0.0028 |
| Sacs | 343 | 221 | 252 | 338 | 283 | 325 | 266 | 235 | 272 | 336 | 319 | 321 | 291.65 | 258 | 325 | 1.262 | 0.335 | 0.00619149 | 0.02887273 | 0 | 0.0106 | 0.0073 | 0.0073 | TRUE | 0.0631 |
| Sae1 | 19475 | 13824 | 13881 | 13707 | 12031 | 14187 | 15125 | 14704 | 14977 | 13644 | 13572 | 14005 | 14338.03 | 14935 | 13740 | 0.92 | -0.12 | 0.00815194 | 0.03554475 | 1.00E-04 | 0.0026 | 0.0014 | 0.0014 | TRUE | 0.0035 |
| Safb | 9238 | 6772 | 6129 | 5558 | 5064 | 5968 | 7175 | 7203 | 6613 | 5532 | 5713 | 5891 | 6354.61 | 6997 | 5712 | 0.816 | -0.293 | 3.97E-07 | 1.33E-05 | 0.0014 | 0.0028 | 0.0022 | 0.0022 | TRUE | 0.0279 |
| Safb2 | 3955 | 3211 | 2556 | 2447 | 2262 | 2703 | 3072 | 3415 | 2758 | 2436 | 2552 | 2668 | 2816.81 | 3082 | 2552 | 0.828 | -0.272 | 0.00080056 | 0.00614522 | 0.0062 | 0.0033 | 0.0044 | 0.0044 | TRUE | 0.1077 |
| Sag | 5 | 0 | 2 | 0 | 1 | 1 | 4 | 0 | 2 | 0 | 1 | 1 | 1.36 | 2 | 1 | 0.338 | -1.564 | 0.39626055 | NA | 0 | 1.7459 | 1.4712 | 1.4712 | TRUE | 0.1167 |
| Sall1 | 12466 | 9696 | 9066 | 8882 | 8476 | 10252 | 9682 | 10314 | 9782 | 8841 | 9562 | 10120 | 9716.78 | 9926 | 9508 | 0.958 | -0.062 | 0.32127344 | 0.49514355 | 0.0027 | 0.0027 | 0.0027 | 0.0027 | TRUE | 0.0461 |
| Sall2 | 354 | 281 | 202 | 208 | 180 | 225 | 275 | 299 | 218 | 207 | 203 | 222 | 237.33 | 264 | 211 | 0.798 | -0.326 | 0.02879677 | 0.0898673 | 0.01 | 0.0124 | 0.0117 | 0.0117 | TRUE | 0.2475 |
| Sall3 | 158 | 125 | 109 | 71 | 75 | 88 | 123 | 133 | 118 | 71 | 85 | 87 | 102.57 | 125 | 81 | 0.648 | -0.625 | 0.00157251 | 0.00998929 | 0 | 0.0255 | 0.0179 | 0.0179 | TRUE | 0.1134 |
| Sall4 | 11543 | 8219 | 7196 | 7090 | 6510 | 7438 | 8965 | 8742 | 7764 | 7057 | 7344 | 7342 | 7869.27 | 8490 | 7248 | 0.853 | -0.229 | 0.00038308 | 0.00351889 | 0.0029 | 0.0027 | 0.0029 | 0.0029 | TRUE | 0.0681 |
| Samd1 | 6456 | 4719 | 4292 | 4573 | 3942 | 4578 | 5014 | 5020 | 4631 | 4552 | 4447 | 4519 | 4697.13 | 4888 | 4506 | 0.922 | -0.118 | 0.03670088 | 0.10771411 | 8.00E-04 | 0.0029 | 0.0021 | 0.0021 | TRUE | 0.0255 |
| Samd10 | 575 | 425 | 494 | 453 | 440 | 517 | 447 | 452 | 533 | 451 | 496 | 510 | 481.55 | 477 | 486 | 1.02 | 0.028 | 0.79813275 | 0.88215303 | 0.005 | 0.0073 | 0.0066 | 0.0066 | TRUE | 0.1188 |
| Samd11 | 2 | 2 | 3 | 1 | 0 | 1 | 2 | 2 | 3 | 1 | 0 | 1 | 1.48 | 2 | 1 | 0.297 | -1.752 | 0.2947897 | NA | 0 | 1.6002 | 1.0397 | 1.0397 | TRUE | 0.1985 |
| Samd12 | 64 | 37 | 40 | 54 | 40 | 59 | 50 | 39 | 43 | 54 | 45 | 58 | 48.22 | 44 | 52 | 1.185 | 0.245 | 0.38413465 | 0.55814135 | 0 | 0.0516 | 0.0363 | 0.0363 | TRUE | 0.1139 |
| Samd14 | 162 | 130 | 97 | 118 | 111 | 116 | 126 | 138 | 105 | 117 | 125 | 115 | 120.99 | 123 | 119 | 0.967 | -0.049 | 0.79305313 | 0.87869058 | 0 | 0.022 | 0.0165 | 0.0165 | TRUE | 0.1596 |
| Samd3 | 0 | 0 | 0 | 0 | 0 | 0 | 0 | 0 | 0 | 0 | 0 | 0 | 0 | 0 | 0 | NA | NA | NA | NA | NA | NA | NA | NA | NA | NA |
| Samd4 | 133 | 72 | 61 | 64 | 47 | 60 | 103 | 77 | 66 | 64 | 53 | 59 | 70.28 | 82 | 59 | 0.712 | -0.49 | 0.05370273 | 0.14159254 | 0.022 | 0.0362 | 0.0319 | 0.0319 | TRUE | 0.5386 |
| Samd4b | 9251 | 6955 | 6503 | 6675 | 6115 | 6974 | 7185 | 7398 | 7017 | 6644 | 6898 | 6884 | 7004.42 | 7200 | 6809 | 0.946 | -0.081 | 0.1120811 | 0.24035975 | 4.00E-04 | 0.0028 | 0.0017 | 0.0017 | TRUE | 0.007 |
| Samd5 | 12 | 9 | 4 | 9 | 16 | 10 | 9 | 10 | 4 | 9 | 18 | 10 | 10.01 | 8 | 12 | 1.565 | 0.646 | 0.31026603 | NA | 0 | 0.2391 | 0.1877 | 0.1877 | TRUE | 0.6121 |
| Samd7 | 72 | 56 | 55 | 23 | 19 | 30 | 56 | 60 | 59 | 23 | 21 | 30 | 41.46 | 58 | 25 | 0.425 | -1.235 | 7.29E-05 | 0.00094469 | 0 | 0.0596 | 0.0407 | 0.0407 | TRUE | 0.1927 |
| Samd8 | 4076 | 2800 | 2854 | 3512 | 3106 | 3439 | 3166 | 2978 | 3079 | 3496 | 3504 | 3395 | 3269.66 | 3074 | 3465 | 1.126 | 0.172 | 0.00222493 | 0.01310836 | 3.00E-04 | 0.0032 | 0.002 | 0.002 | TRUE | 0.0091 |
| Samd9l | 20 | 9 | 4 | 2 | 2 | 5 | 16 | 10 | 4 | 2 | 2 | 5 | 6.43 | 10 | 3 | 0.31 | -1.691 | 0.0490723 | NA | 0 | 0.3708 | 0.3158 | 0.3158 | TRUE | 0.1013 |
| Samhd1 | 5352 | 3777 | 3810 | 4039 | 3654 | 4289 | 4157 | 4018 | 4111 | 4020 | 4122 | 4234 | 4110.26 | 4095 | 4125 | 1.007 | 0.01 | 0.84490456 | 0.91201028 | 3.00E-04 | 0.003 | 0.0018 | 0.0018 | TRUE | 0.0065 |
| Samm50 | 12247 | 8897 | 9561 | 10903 | 9734 | 11314 | 9512 | 9464 | 10316 | 10853 | 10981 | 11169 | 10382.34 | 9764 | 11001 | 1.127 | 0.172 | 0.00141493 | 0.00925565 | 0.0012 | 0.0027 | 0.002 | 0.002 | TRUE | 0.0298 |
| Samsn1 | 0 | 0 | 0 | 0 | 0 | 0 | 0 | 0 | 0 | 0 | 0 | 0 | 0 | 0 | 0 | NA | NA | NA | NA | NA | NA | NA | NA | NA | NA |
| Samt1 | 0 | 0 | 0 | 0 | 0 | 0 | 0 | 0 | 0 | 0 | 0 | 0 | 0 | 0 | 0 | NA | NA | NA | NA | NA | NA | NA | NA | NA | NA |
| Samt2 | 0 | 0 | 0 | 0 | 0 | 0 | 0 | 0 | 0 | 0 | 0 | 0 | 0 | 0 | 0 | NA | NA | NA | NA | NA | NA | NA | NA | NA | NA |
| Samt3 | 0 | 0 | 0 | 0 | 0 | 0 | 0 | 0 | 0 | 0 | 0 | 0 | 0 | 0 | 0 | NA | NA | NA | NA | NA | NA | NA | NA | NA | NA |
| Samt4 | 0 | 0 | 0 | 0 | 0 | 0 | 0 | 0 | 0 | 0 | 0 | 0 | 0 | 0 | 0 | NA | NA | NA | NA | NA | NA | NA | NA | NA | NA |
| Sap130 | 9845 | 7048 | 7202 | 7503 | 6950 | 8048 | 7646 | 7497 | 7771 | 7468 | 7840 | 7945 | 7694.56 | 7638 | 7751 | 1.015 | 0.021 | 0.67872901 | 0.80267034 | 5.00E-04 | 0.0027 | 0.0018 | 0.0018 | TRUE | 0.0125 |
| Sap18 | 684 | 431 | 475 | 388 | 358 | 455 | 531 | 458 | 513 | 386 | 404 | 449 | 456.91 | 501 | 413 | 0.824 | -0.279 | 0.01078339 | 0.04368679 | 0.0038 | 0.0076 | 0.0064 | 0.0064 | TRUE | 0.0687 |
| Sap25 | 0 | 0 | 0 | 0 | 0 | 0 | 0 | 0 | 0 | 0 | 0 | 0 | 0 | 0 | 0 | NA | NA | NA | NA | NA | NA | NA | NA | NA | NA |
| Sap30 | 4342 | 3429 | 3164 | 3278 | 2917 | 3449 | 3372 | 3647 | 3414 | 3263 | 3291 | 3405 | 3398.64 | 3478 | 3320 | 0.955 | -0.067 | 0.25621624 | 0.42612273 | 9.00E-04 | 0.0031 | 0.0022 | 0.0022 | TRUE | 0.0219 |
| Sap30bp | 3376 | 2360 | 1994 | 2118 | 1807 | 2128 | 2622 | 2510 | 2152 | 2108 | 2038 | 2101 | 2255.2 | 2428 | 2082 | 0.857 | -0.222 | 0.00494336 | 0.02432201 | 0.0046 | 0.0035 | 0.0041 | 0.0041 | TRUE | 0.1193 |
| Sap30l | 2274 | 1725 | 1787 | 1963 | 1689 | 2024 | 1766 | 1835 | 1928 | 1954 | 1905 | 1998 | 1897.75 | 1843 | 1952 | 1.06 | 0.085 | 0.1954539 | 0.35515162 | 8.00E-04 | 0.0037 | 0.0025 | 0.0025 | TRUE | 0.0196 |
| Sapcd1 | 28 | 28 | 43 | 43 | 57 | 52 | 22 | 30 | 46 | 43 | 64 | 51 | 42.73 | 33 | 53 | 1.634 | 0.708 | 0.03866621 | 0.11175517 | 0.0648 | 0.0579 | 0.0598 | 0.0598 | TRUE | 0.733 |
| Sapcd2 | 1529 | 1135 | 1152 | 1224 | 1118 | 1223 | 1188 | 1207 | 1243 | 1218 | 1261 | 1207 | 1220.78 | 1213 | 1229 | 1.014 | 0.02 | 0.77770718 | 0.8686909 | 0 | 0.0044 | 0.0027 | 0.0027 | TRUE | 0.0062 |
| Sar1a | 8144 | 5619 | 5552 | 6051 | 5362 | 6131 | 6325 | 5977 | 5991 | 6023 | 6049 | 6052 | 6069.47 | 6098 | 6041 | 0.99 | -0.014 | 0.78738776 | 0.87519506 | 4.00E-04 | 0.0028 | 0.0017 | 0.0017 | TRUE | 0.0132 |
| Sar1b | 4939 | 3543 | 3304 | 4238 | 3559 | 3836 | 3836 | 3769 | 3565 | 4219 | 4015 | 3787 | 3864.95 | 3723 | 4007 | 1.076 | 0.105 | 0.09385464 | 0.21100507 | 0.0018 | 0.003 | 0.0026 | 0.0026 | TRUE | 0.0283 |
| Sardh | 25 | 11 | 10 | 22 | 9 | 15 | 19 | 12 | 11 | 22 | 10 | 15 | 14.79 | 14 | 16 | 1.108 | 0.148 | 0.77924141 | NA | 0.0629 | 0.1626 | 0.1344 | 0.1344 | TRUE | 0.314 |
| Sarm1 | 1078 | 708 | 655 | 891 | 808 | 856 | 837 | 753 | 707 | 887 | 912 | 845 | 823.41 | 766 | 881 | 1.148 | 0.199 | 0.02697903 | 0.08578747 | 0.0033 | 0.0053 | 0.0046 | 0.0046 | TRUE | 0.083 |
| Sarnp | 5334 | 3864 | 3905 | 3948 | 3405 | 4184 | 4143 | 4110 | 4213 | 3930 | 3841 | 4130 | 4061.25 | 4155 | 3967 | 0.955 | -0.067 | 0.22741127 | 0.39251169 | 4.00E-04 | 0.003 | 0.0019 | 0.0019 | TRUE | 0.0158 |
| Sars | 11136 | 8053 | 7861 | 7993 | 7301 | 8652 | 8649 | 8566 | 8482 | 7956 | 8236 | 8541 | 8405.01 | 8566 | 8244 | 0.962 | -0.055 | 0.27617357 | 0.44722484 | 5.00E-04 | 0.0027 | 0.0017 | 0.0017 | TRUE | 0.0122 |
| Sars2 | 1771 | 1262 | 1080 | 1288 | 1092 | 1235 | 1375 | 1342 | 1165 | 1282 | 1232 | 1219 | 1269.37 | 1294 | 1244 | 0.96 | -0.058 | 0.47540552 | 0.64342464 | 0.0035 | 0.0043 | 0.004 | 0.004 | TRUE | 0.0899 |
| Sart1 | 7467 | 5576 | 4580 | 4571 | 4303 | 4974 | 5799 | 5931 | 4942 | 4550 | 4854 | 4910 | 5164.42 | 5557 | 4771 | 0.858 | -0.22 | 0.00329903 | 0.01778018 | 0.0052 | 0.0029 | 0.0039 | 0.0039 | TRUE | 0.1138 |
| Sart3 | 9440 | 7291 | 6292 | 5931 | 5433 | 6366 | 7332 | 7755 | 6789 | 5904 | 6129 | 6284 | 6698.82 | 7292 | 6106 | 0.837 | -0.256 | 4.52E-05 | 0.00064115 | 0.0025 | 0.0028 | 0.0027 | 0.0027 | TRUE | 0.044 |
| Sash1 | 1790 | 1347 | 1446 | 1522 | 1375 | 1745 | 1390 | 1433 | 1560 | 1515 | 1551 | 1723 | 1528.66 | 1461 | 1596 | 1.094 | 0.129 | 0.10158648 | 0.22369058 | 0.0033 | 0.004 | 0.0038 | 0.0038 | TRUE | 0.0583 |
| Sash3 | 7 | 5 | 11 | 2 | 1 | 6 | 5 | 5 | 12 | 2 | 1 | 6 | 5.28 | 7 | 3 | 0.408 | -1.292 | 0.15998453 | NA | 0.1604 | 0.4515 | 0.3752 | 0.3752 | TRUE | 0.9516 |
| Sass6 | 1097 | 813 | 765 | 847 | 682 | 879 | 852 | 865 | 825 | 843 | 769 | 868 | 837.06 | 847 | 827 | 0.976 | -0.035 | 0.67609365 | 0.8011859 | 8.00E-04 | 0.0053 | 0.0038 | 0.0038 | TRUE | 0.0426 |
| Sat1 | 2207 | 1498 | 1807 | 1861 | 1728 | 2079 | 1714 | 1593 | 1950 | 1852 | 1949 | 2052 | 1851.89 | 1752 | 1951 | 1.114 | 0.156 | 0.0625249 | 0.15817139 | 0.0058 | 0.0037 | 0.0045 | 0.0045 | TRUE | 0.115 |
| Sat2 | 149 | 118 | 106 | 123 | 84 | 109 | 116 | 126 | 114 | 122 | 95 | 108 | 113.4 | 119 | 108 | 0.916 | -0.127 | 0.50144702 | 0.66570278 | 0 | 0.0233 | 0.0169 | 0.0169 | TRUE | 0.13 |
| Satb1 | 1919 | 1388 | 1360 | 1689 | 1501 | 1789 | 1490 | 1476 | 1467 | 1681 | 1693 | 1766 | 1595.79 | 1478 | 1713 | 1.159 | 0.213 | 0.00089979 | 0.00671049 | 0 | 0.0039 | 0.0023 | 0.0023 | TRUE | 0.0088 |
| Satb2 | 471 | 282 | 224 | 195 | 155 | 157 | 366 | 300 | 242 | 194 | 175 | 155 | 238.57 | 303 | 175 | 0.575 | -0.798 | 2.54E-06 | 6.08E-05 | 0.0247 | 0.0124 | 0.0162 | 0.0162 | TRUE | 0.1404 |
| Satl1 | 1 | 0 | 0 | 0 | 0 | 0 | 1 | 0 | 0 | 0 | 0 | 0 | 0.13 | 0 | 0 | 0.552 | -0.857 | 0.8336548 | NA | 0 | 18.3121 | 10 | 10 | TRUE | 0.2566 |
| Sav1 | 3370 | 2347 | 2315 | 2300 | 1967 | 2278 | 2617 | 2496 | 2498 | 2289 | 2219 | 2249 | 2394.8 | 2537 | 2252 | 0.887 | -0.172 | 0.00341411 | 0.01829692 | 0 | 0.0034 | 0.0021 | 0.0021 | TRUE | 0.0096 |
| Saysd1 | 514 | 351 | 327 | 388 | 282 | 399 | 399 | 373 | 353 | 386 | 318 | 394 | 370.6 | 375 | 366 | 0.976 | -0.035 | 0.76943298 | 0.86349761 | 0.0056 | 0.0088 | 0.0079 | 0.0079 | TRUE | 0.1455 |
| Sbds | 2419 | 1670 | 1579 | 1538 | 1411 | 1649 | 1879 | 1776 | 1704 | 1531 | 1592 | 1628 | 1684.89 | 1786 | 1584 | 0.886 | -0.175 | 0.01106265 | 0.04445772 | 0.0012 | 0.0038 | 0.0028 | 0.0028 | TRUE | 0.0256 |
| Sbf1 | 8101 | 6123 | 5747 | 6083 | 5511 | 6351 | 6292 | 6513 | 6201 | 6055 | 6217 | 6269 | 6257.84 | 6335 | 6180 | 0.976 | -0.036 | 0.48152884 | 0.64849796 | 2.00E-04 | 0.0028 | 0.0017 | 0.0017 | TRUE | 0.0073 |
| Sbf2 | 7612 | 5667 | 6508 | 7962 | 7364 | 8555 | 5912 | 6028 | 7022 | 7925 | 8307 | 8445 | 7273.29 | 6321 | 8226 | 1.302 | 0.381 | 1.10E-07 | 4.26E-06 | 0.005 | 0.0028 | 0.0036 | 0.0036 | TRUE | 0.1146 |
| Sbk1 | 2942 | 2201 | 2485 | 2425 | 2086 | 2414 | 2285 | 2341 | 2681 | 2414 | 2353 | 2383 | 2409.58 | 2436 | 2383 | 0.979 | -0.03 | 0.67816902 | 0.80231161 | 0.0035 | 0.0034 | 0.0034 | 0.0034 | TRUE | 0.0935 |
| Sbk2 | 23 | 21 | 18 | 21 | 13 | 13 | 18 | 22 | 19 | 21 | 15 | 13 | 18 | 20 | 16 | 0.816 | -0.294 | 0.51930883 | 0.68083665 | 0 | 0.1341 | 0.0934 | 0.0934 | TRUE | 0.3291 |
| Sbno1 | 15326 | 11548 | 11597 | 12440 | 11025 | 12879 | 11903 | 12283 | 12513 | 12383 | 12437 | 12714 | 12372.21 | 12233 | 12511 | 1.023 | 0.033 | 0.49156924 | 0.65729264 | 3.00E-04 | 0.0026 | 0.0015 | 0.0015 | TRUE | 0.0069 |
| Sbno2 | 3739 | 2939 | 3396 | 3259 | 3299 | 3901 | 2904 | 3126 | 3664 | 3244 | 3722 | 3851 | 3418.48 | 3231 | 3606 | 1.116 | 0.159 | 0.08316285 | 0.19415985 | 0.0108 | 0.0031 | 0.0058 | 0.0058 | TRUE | 0.1665 |
| Sbp | 0 | 2 | 1 | 0 | 1 | 0 | 0 | 2 | 1 | 0 | 1 | 0 | 0.72 | 1 | 0 | 0.356 | -1.491 | 0.54231439 | NA | 0 | 3.2832 | 2.8503 | 2.8503 | TRUE | 0.2627 |
| Sbpl | 0 | 0 | 1 | 0 | 2 | 0 | 0 | 0 | 1 | 0 | 2 | 0 | 0.56 | 0 | 1 | 2.136 | 1.095 | 0.70886258 | NA | 0 | 4.2661 | 4.493 | 4.493 | TRUE | 0.9746 |
| Sbsn | 95 | 63 | 60 | 68 | 55 | 59 | 74 | 67 | 65 | 68 | 62 | 58 | 65.58 | 69 | 63 | 0.912 | -0.133 | 0.57666219 | 0.7280742 | 0 | 0.0386 | 0.0256 | 0.0256 | TRUE | 0.0462 |

| | | | | | | | | | | | | | | | | | | | | | | | | |
|---|---|---|---|---|---|---|---|---|---|---|---|---|---|---|---|---|---|---|---|---|---|---|---|---|
| Sbspon | 4 | 4 | 6 | 2 | 4 | 2 | 3 | 4 | 6 | 2 | 5 | 2 | 3.72 | 4 | 3 | 0.613 | -0.707 | 0.48086639 | NA | 0 | 0.6397 | 0.4336 | 0.4336 | TRUE | 0.2959 |
| Sc4mol | 10759 | 8127 | 9241 | 10696 | 9188 | 10723 | 8356 | 8645 | 9971 | 10647 | 10365 | 10585 | 9761.44 | 8991 | 10532 | 1.172 | 0.229 | 0.00096238 | 0.0070576 | 0.0045 | 0.0027 | 0.0034 | 0.0034 | TRUE | 0.1109 |
| Sc5d | 1478 | 1025 | 1045 | 1205 | 999 | 1147 | 1148 | 1090 | 1128 | 1199 | 1127 | 1132 | 1137.4 | 1122 | 1153 | 1.027 | 0.039 | 0.59279897 | 0.74044182 | 0 | 0.0045 | 0.0029 | 0.0029 | TRUE | 0.0151 |
| Scaf1 | 8289 | 6345 | 5472 | 5329 | 4892 | 5595 | 6438 | 6749 | 5904 | 5304 | 5519 | 5523 | 5906.22 | 6364 | 5449 | 0.856 | -0.224 | 0.00033635 | 0.00316908 | 0.0023 | 0.0028 | 0.0026 | 0.0026 | TRUE | 0.0482 |
| Scaf11 | 10866 | 7747 | 8020 | 8254 | 7253 | 8410 | 8439 | 8240 | 8654 | 8216 | 8182 | 8302 | 8338.86 | 8444 | 8233 | 0.975 | -0.036 | 0.44824319 | 0.61829183 | 1.00E-04 | 0.0027 | 0.0015 | 0.0015 | TRUE | 0.0057 |
| Scaf4 | 4719 | 3466 | 3217 | 3488 | 3147 | 3723 | 3665 | 3687 | 3471 | 3472 | 3550 | 3675 | 3586.7 | 3608 | 3566 | 0.988 | -0.017 | 0.76519712 | 0.86070374 | 7.00E-04 | 0.0031 | 0.0021 | 0.0021 | TRUE | 0.0131 |
| Scaf8 | 5222 | 3815 | 3558 | 3596 | 3100 | 3608 | 4056 | 4058 | 3839 | 3579 | 3497 | 3562 | 3765.16 | 3984 | 3546 | 0.89 | -0.168 | 0.00209025 | 0.01250037 | 3.00E-04 | 0.0031 | 0.0019 | 0.0019 | TRUE | 0.0122 |
| Scai | 1014 | 843 | 883 | 991 | 818 | 953 | 788 | 897 | 953 | 986 | 923 | 941 | 914.49 | 879 | 950 | 1.083 | 0.115 | 0.2012955 | 0.36212921 | 0.0043 | 0.005 | 0.0048 | 0.0048 | TRUE | 0.1035 |
| Scamp1 | 3258 | 2427 | 2380 | 2457 | 2130 | 2464 | 2530 | 2582 | 2568 | 2446 | 2403 | 2432 | 2493.47 | 2560 | 2427 | 0.948 | -0.077 | 0.17236022 | 0.32457769 | 0 | 0.0034 | 0.0019 | 0.0019 | TRUE | 0.0013 |
| Scamp2 | 1792 | 1279 | 1420 | 1407 | 1287 | 1537 | 1392 | 1360 | 1532 | 1401 | 1452 | 1517 | 1442.34 | 1428 | 1457 | 1.02 | 0.029 | 0.69678083 | 0.81551657 | 0.0021 | 0.0041 | 0.0034 | 0.0034 | TRUE | 0.0477 |
| Scamp3 | 3456 | 2647 | 2952 | 2934 | 2601 | 3190 | 2684 | 2816 | 3185 | 2921 | 2934 | 3149 | 2948.1 | 2895 | 3001 | 1.037 | 0.053 | 0.47886207 | 0.6458786 | 0.0045 | 0.0032 | 0.0037 | 0.0037 | TRUE | 0.093 |
| Scamp4 | 3624 | 2383 | 2271 | 2368 | 2240 | 2519 | 2815 | 2535 | 2450 | 2357 | 2527 | 2487 | 2528.41 | 2600 | 2457 | 0.944 | -0.083 | 0.24054662 | 0.40794173 | 0.0027 | 0.0034 | 0.0032 | 0.0032 | TRUE | 0.0647 |
| Scamp5 | 2824 | 1872 | 1835 | 2088 | 1843 | 2148 | 2193 | 1991 | 1980 | 2078 | 2079 | 2120 | 2073.72 | 2055 | 2092 | 1.017 | 0.025 | 0.70822678 | 0.82405449 | 0.0014 | 0.0036 | 0.0027 | 0.0027 | TRUE | 0.0434 |
| Scand1 | 926 | 631 | 753 | 815 | 698 | 817 | 719 | 671 | 812 | 811 | 787 | 807 | 768 | 734 | 802 | 1.093 | 0.128 | 0.16442025 | 0.31523023 | 0.0033 | 0.0055 | 0.0048 | 0.0048 | TRUE | 0.0989 |
| Scap | 3859 | 2730 | 2631 | 3016 | 2716 | 3049 | 2997 | 2904 | 2839 | 3002 | 3064 | 3010 | 2969.28 | 2913 | 3025 | 1.038 | 0.054 | 0.3381402 | 0.51231463 | 0 | 0.0032 | 0.0019 | 0.0019 | TRUE | 0.0079 |
| Scaper | 1644 | 1205 | 1238 | 1409 | 1220 | 1399 | 1277 | 1282 | 1336 | 1403 | 1376 | 1381 | 1342.37 | 1298 | 1387 | 1.069 | 0.096 | 0.15346958 | 0.30040272 | 0 | 0.0042 | 0.0025 | 0.0025 | TRUE | 0.0076 |
| Scara3 | 68 | 46 | 36 | 61 | 45 | 42 | 53 | 49 | 39 | 61 | 51 | 41 | 48.92 | 47 | 51 | 1.083 | 0.115 | 0.69166587 | 0.81149322 | 0 | 0.0509 | 0.0398 | 0.0398 | TRUE | 0.1931 |
| Scara5 | 87 | 57 | 72 | 56 | 48 | 58 | 68 | 61 | 78 | 56 | 54 | 57 | 62.17 | 69 | 56 | 0.813 | -0.299 | 0.22566755 | 0.39058686 | 0 | 0.0405 | 0.0275 | 0.0275 | TRUE | 0.11 |
| Scarb1 | 5148 | 3985 | 4699 | 4929 | 4776 | 5429 | 3998 | 4239 | 5070 | 4906 | 5388 | 5359 | 4826.77 | 4436 | 5218 | 1.177 | 0.235 | 0.00579703 | 0.0274907 | 0.0091 | 0.0029 | 0.005 | 0.005 | TRUE | 0.1902 |
| Scarb2 | 1917 | 1415 | 1815 | 2272 | 2005 | 2200 | 1489 | 1505 | 1958 | 2262 | 2262 | 2172 | 1941.25 | 1651 | 2232 | 1.353 | 0.436 | 8.07E-06 | 0.00015762 | 0.0125 | 0.0036 | 0.0064 | 0.0064 | TRUE | 0.318 |
| Scarf1 | 220 | 149 | 144 | 157 | 150 | 156 | 171 | 158 | 155 | 156 | 169 | 154 | 160.7 | 161 | 160 | 0.986 | -0.02 | 0.8978318 | 0.94349536 | 0 | 0.0172 | 0.0113 | 0.0113 | TRUE | 0.0295 |
| Scarf2 | 108 | 82 | 65 | 92 | 93 | 88 | 84 | 87 | 70 | 92 | 105 | 87 | 87.43 | 80 | 95 | 1.171 | 0.227 | 0.28641717 | 0.45837651 | 0 | 0.0295 | 0.0212 | 0.0212 | TRUE | 0.1084 |
| Scarna10 | 2 | 2 | 4 | 1 | 2 | 1 | 2 | 2 | 4 | 1 | 2 | 1 | 2.04 | 3 | 1 | 0.531 | -0.912 | 0.49873834 | NA | 0 | 1.1647 | 0.7562 | 0.7562 | TRUE | 0.3201 |
| Scarna13 | 5 | 1 | 2 | 2 | 2 | 2 | 4 | 1 | 2 | 2 | 2 | 2 | 2.22 | 2 | 2 | 0.853 | -0.229 | 0.85522692 | NA | 0 | 1.0695 | 0.6834 | 0.6834 | TRUE | 0.4322 |
| Scarna17 | 3 | 8 | 3 | 3 | 1 | 3 | 2 | 9 | 3 | 3 | 1 | 3 | 3.53 | 5 | 2 | 0.515 | -0.956 | 0.37269403 | NA | 0 | 0.6747 | 0.5076 | 0.5076 | TRUE | 0.9264 |
| Scarna2 | 0 | 1 | 0 | 0 | 0 | 0 | 0 | 1 | 0 | 0 | 0 | 0 | 0.18 | 0 | 0 | 0.552 | -0.857 | 0.8336548 | NA | 0 | 13.3714 | 10 | 10 | TRUE | 0.5216 |
| Scarna3a | 0 | 0 | 0 | 1 | 0 | 0 | 0 | 0 | 0 | 1 | 0 | 0 | 0.17 | 0 | 0 | 2.094 | 1.067 | 0.79379329 | NA | 0 | 14.2885 | 10 | 10 | TRUE | 0.3906 |
| Scarna3b | 0 | 0 | 0 | 0 | 0 | 0 | 0 | 0 | 0 | 0 | 0 | 0 | 0 | 0 | 0 | NA | NA | NA | NA | NA | NA | NA | NA | NA | NA |
| Scarna6 | 2 | 3 | 1 | 3 | 0 | 3 | 2 | 3 | 1 | 3 | 0 | 3 | 1.96 | 2 | 2 | 1.044 | 0.062 | 0.96480622 | NA | 0 | 1.2105 | 0.9194 | 0.9194 | TRUE | 0.5869 |
| Scarna8 | 4 | 0 | 1 | 2 | 3 | 1 | 3 | 0 | 1 | 2 | 3 | 1 | 1.76 | 1 | 2 | 1.436 | 0.522 | 0.72734282 | NA | 0 | 1.3506 | 1.0396 | 1.0396 | TRUE | 0.7601 |
| Scarna9 | 1 | 0 | 0 | 0 | 0 | 1 | 1 | 0 | 0 | 0 | 0 | 1 | 0.29 | 0 | 0 | 1.075 | 0.105 | 0.97702352 | NA | 0 | 8.0647 | 7.5458 | 7.5458 | TRUE | 0.3835 |
| Sccpdh | 2722 | 1958 | 2166 | 2572 | 2108 | 2353 | 2114 | 2083 | 2337 | 2560 | 2378 | 2323 | 2299.14 | 2178 | 2420 | 1.112 | 0.153 | 0.03180767 | 0.09662637 | 0.0027 | 0.0035 | 0.0032 | 0.0032 | TRUE | 0.0487 |
| Scd1 | 11183 | 8344 | 9832 | 13954 | 11135 | 12890 | 8685 | 8875 | 10609 | 13890 | 12561 | 12724 | 11224.18 | 9390 | 13058 | 1.391 | 0.476 | 1.65E-09 | 1.05E-07 | 0.0078 | 0.0026 | 0.0044 | 0.0044 | TRUE | 0.1572 |
| Scd2 | 49436 | 35328 | 34349 | 48111 | 37287 | 41223 | 38395 | 37578 | 37062 | 47890 | 42064 | 40693 | 40613.63 | 37678 | 43549 | 1.156 | 0.209 | 0.00137821 | 0.00906524 | 0.004 | 0.0025 | 0.003 | 0.003 | TRUE | 0.0931 |
| Scd3 | 131 | 114 | 118 | 141 | 137 | 165 | 102 | 121 | 127 | 140 | 155 | 163 | 134.68 | 117 | 153 | 1.313 | 0.393 | 0.02657013 | 0.08494003 | 0 | 0.02 | 0.0151 | 0.0151 | TRUE | 0.1444 |
| Scd4 | 1 | 1 | 0 | 1 | 1 | 0 | 1 | 1 | 0 | 1 | 1 | 0 | 0.66 | 1 | 1 | 1.128 | 0.173 | 0.94106052 | NA | 0 | 3.5899 | 2.4471 | 2.4471 | TRUE | 0.2681 |
| Scel | 11 | 5 | 6 | 8 | 6 | 3 | 9 | 5 | 6 | 8 | 7 | 3 | 6.34 | 7 | 6 | 0.859 | -0.219 | 0.77608001 | NA | 0 | 0.3764 | 0.2668 | 0.2668 | TRUE | 0.4615 |
| Scfd1 | 2422 | 1671 | 1576 | 1757 | 1441 | 1728 | 1881 | 1777 | 1700 | 1749 | 1626 | 1706 | 1739.88 | 1786 | 1694 | 0.947 | -0.078 | 0.26227617 | 0.43262704 | 0.0014 | 0.0038 | 0.0029 | 0.0029 | TRUE | 0.0269 |
| Scfd2 | 868 | 628 | 507 | 401 | 410 | 411 | 674 | 668 | 547 | 399 | 463 | 406 | 526.1 | 630 | 423 | 0.669 | -0.579 | 3.61E-07 | 1.23E-05 | 0.0082 | 0.0069 | 0.0073 | 0.0073 | TRUE | 0.1518 |
| Scg2 | 0 | 0 | 0 | 0 | 0 | 0 | 0 | 0 | 0 | 0 | 0 | 0 | 0 | 0 | 0 | NA | NA | NA | NA | NA | NA | NA | NA | NA | NA |
| Scg3 | 21 | 13 | 9 | 11 | 6 | 8 | 16 | 14 | 10 | 11 | 7 | 8 | 10.91 | 13 | 9 | 0.64 | -0.643 | 0.27437069 | NA | 0 | 0.2197 | 0.1533 | 0.1533 | TRUE | 0.2182 |
| Scg5 | 11 | 6 | 14 | 16 | 13 | 18 | 9 | 6 | 15 | 16 | 15 | 18 | 13.07 | 10 | 16 | 1.624 | 0.7 | 0.20169172 | NA | 0 | 0.1838 | 0.136 | 0.136 | TRUE | 0.6211 |
| Scgb1a1 | 0 | 0 | 0 | 0 | 0 | 0 | 0 | 0 | 0 | 0 | 0 | 0 | 0 | 0 | 0 | NA | NA | NA | NA | NA | NA | NA | NA | NA | NA |
| Scgb1b2 | 0 | 0 | 0 | 0 | 0 | 0 | 0 | 0 | 0 | 0 | 0 | 0 | 0 | 0 | 0 | NA | NA | NA | NA | NA | NA | NA | NA | NA | NA |
| Scgb1b20 | 0 | 0 | 0 | 0 | 0 | 0 | 0 | 0 | 0 | 0 | 0 | 0 | 0 | 0 | 0 | NA | NA | NA | NA | NA | NA | NA | NA | NA | NA |
| Scgb1b24 | 0 | 0 | 0 | 0 | 0 | 0 | 0 | 0 | 0 | 0 | 0 | 0 | 0 | 0 | 0 | NA | NA | NA | NA | NA | NA | NA | NA | NA | NA |
| Scgb1b27 | 0 | 0 | 0 | 0 | 0 | 0 | 0 | 0 | 0 | 0 | 0 | 0 | 0 | 0 | 0 | NA | NA | NA | NA | NA | NA | NA | NA | NA | NA |
| Scgb1b29 | 0 | 0 | 0 | 0 | 0 | 0 | 0 | 0 | 0 | 0 | 0 | 0 | 0 | 0 | 0 | NA | NA | NA | NA | NA | NA | NA | NA | NA | NA |
| Scgb1b3 | 0 | 0 | 0 | 0 | 0 | 0 | 0 | 0 | 0 | 0 | 0 | 0 | 0 | 0 | 0 | NA | NA | NA | NA | NA | NA | NA | NA | NA | NA |
| Scgb1b30 | 0 | 0 | 0 | 0 | 0 | 0 | 0 | 0 | 0 | 0 | 0 | 0 | 0 | 0 | 0 | NA | NA | NA | NA | NA | NA | NA | NA | NA | NA |
| Scgb1b7 | 0 | 0 | 0 | 0 | 0 | 0 | 0 | 0 | 0 | 0 | 0 | 0 | 0 | 0 | 0 | NA | NA | NA | NA | NA | NA | NA | NA | NA | NA |
| Scgb1c1 | 1 | 1 | 1 | 0 | 1 | 0 | 1 | 1 | 1 | 0 | 1 | 0 | 0.67 | 1 | 0 | 0.383 | -1.383 | 0.54981825 | NA | 0 | 3.5158 | 2.3538 | 2.3538 | TRUE | 0.5075 |
| Scgb2b19 | 0 | 0 | 0 | 0 | 0 | 0 | 0 | 0 | 0 | 0 | 0 | 0 | 0 | 0 | 0 | NA | NA | NA | NA | NA | NA | NA | NA | NA | NA |
| Scgb2b2 | 0 | 0 | 0 | 0 | 0 | 0 | 0 | 0 | 0 | 0 | 0 | 0 | 0 | 0 | 0 | NA | NA | NA | NA | NA | NA | NA | NA | NA | NA |
| Scgb2b20 | 0 | 0 | 0 | 0 | 0 | 0 | 0 | 0 | 0 | 0 | 0 | 0 | 0 | 0 | 0 | NA | NA | NA | NA | NA | NA | NA | NA | NA | NA |
| Scgb2b23-ps | 0 | 0 | 0 | 0 | 0 | 0 | 0 | 0 | 0 | 0 | 0 | 0 | 0 | 0 | 0 | NA | NA | NA | NA | NA | NA | NA | NA | NA | NA |
| Scgb2b24 | 0 | 0 | 0 | 0 | 0 | 0 | 0 | 0 | 0 | 0 | 0 | 0 | 0 | 0 | 0 | NA | NA | NA | NA | NA | NA | NA | NA | NA | NA |
| Scgb2b26 | 0 | 0 | 0 | 0 | 0 | 0 | 0 | 0 | 0 | 0 | 0 | 0 | 0 | 0 | 0 | NA | NA | NA | NA | NA | NA | NA | NA | NA | NA |
| Scgb2b27 | 0 | 0 | 0 | 0 | 0 | 1 | 0 | 0 | 0 | 0 | 0 | 1 | 0.16 | 0 | 0 | 2.094 | 1.067 | 0.79379329 | NA | 0 | 14.4078 | 10 | 10 | TRUE | 0.3835 |
| Scgb2b3 | 0 | 0 | 0 | 0 | 0 | 0 | 0 | 0 | 0 | 0 | 0 | 0 | 0 | 0 | 0 | NA | NA | NA | NA | NA | NA | NA | NA | NA | NA |
| Scgb2b7 | 0 | 0 | 0 | 0 | 0 | 0 | 0 | 0 | 0 | 0 | 0 | 0 | 0 | 0 | 0 | NA | NA | NA | NA | NA | NA | NA | NA | NA | NA |
| Scgb3a1 | 8 | 4 | 6 | 5 | 7 | 4 | 6 | 4 | 6 | 5 | 8 | 4 | 5.63 | 5 | 6 | 0.982 | -0.026 | 0.97424473 | NA | 0 | 0.4236 | 0.2811 | 0.2811 | TRUE | 0.2524 |
| Scgb3a2 | 0 | 0 | 0 | 0 | 0 | 0 | 0 | 0 | 0 | 0 | 0 | 0 | 0 | 0 | 0 | NA | NA | NA | NA | NA | NA | NA | NA | NA | NA |
| Scgn | 3 | 1 | 2 | 1 | 1 | 2 | 2 | 1 | 2 | 1 | 1 | 2 | 1.61 | 2 | 1 | 0.734 | -0.446 | 0.75921754 | NA | 0 | 1.4761 | 0.8906 | 0.8906 | TRUE | 0.1061 |
| Schip1 | 143 | 106 | 144 | 134 | 133 | 142 | 111 | 113 | 155 | 133 | 150 | 140 | 133.8 | 126 | 141 | 1.121 | 0.165 | 0.38137186 | 0.55579288 | 0.0124 | 0.0201 | 0.0179 | 0.0179 | TRUE | 0.3961 |
| Scimp | 0 | 0 | 1 | 0 | 0 | 0 | 0 | 0 | 1 | 0 | 0 | 0 | 0.18 | 0 | 0 | 0.552 | -0.857 | 0.8336548 | NA | 0 | 13.1817 | 10 | 10 | TRUE | 0.5365 |
| Scin | 31 | 12 | 14 | 9 | 3 | 7 | 24 | 13 | 15 | 9 | 3 | 7 | 11.87 | 17 | 6 | 0.371 | -1.432 | 0.01944407 | NA | 0 | 0.2022 | 0.1633 | 0.1633 | TRUE | 0.699 |
| Sclt1 | 673 | 471 | 438 | 405 | 373 | 402 | 523 | 501 | 473 | 403 | 421 | 397 | 452.84 | 499 | 407 | 0.814 | -0.296 | 0.00344049 | 0.01839004 | 0 | 0.0077 | 0.0052 | 0.0052 | TRUE | 0.0242 |
| Scly | 637 | 456 | 380 | 441 | 423 | 456 | 495 | 485 | 410 | 439 | 477 | 450 | 459.35 | 463 | 455 | 0.981 | -0.028 | 0.79775381 | 0.88189654 | 0.0039 | 0.0076 | 0.0064 | 0.0064 | TRUE | 0.1139 |

| | | | | | | | | | | | | | | | | | | | | | | | | | |
|---|---|---|---|---|---|---|---|---|---|---|---|---|---|---|---|---|---|---|---|---|---|---|---|---|---|
| Scmh1 | 1239 | 830 | 866 | 992 | 884 | 984 | 962 | 883 | 934 | 987 | 997 | 971 | 955.93 | 926 | 985 | 1.062 | 0.087 | 0.2558372 | 0.4257896 | 0 | 0.0049 | 0.0032 | 0.0032 | TRUE | 0.02 |
| Scml2 | 3631 | 2812 | 2511 | 1808 | 1449 | 1471 | 2820 | 2991 | 2709 | 1800 | 1635 | 1452 | 2234.48 | 2840 | 1629 | 0.573 | -0.802 | 3.95E-22 | 1.49E-19 | 0.0062 | 0.0035 | 0.0045 | 0.0045 | TRUE | 0.1099 |
| Scml4 | 20 | 23 | 13 | 18 | 16 | 20 | 16 | 24 | 14 | 18 | 18 | 20 | 18.29 | 18 | 19 | 1.037 | 0.052 | 0.90910712 | 0.94970531 | 0 | 0.132 | 0.0942 | 0.0942 | TRUE | 0.4681 |
| Scn10a | 3 | 3 | 0 | 2 | 2 | 3 | 2 | 3 | 0 | 2 | 2 | 3 | 2.12 | 2 | 2 | 1.291 | 0.368 | 0.78281064 | NA | 0 | 1.1195 | 0.8124 | 0.8124 | TRUE | 0.5545 |
| Scn11a | 31 | 15 | 18 | 14 | 21 | 17 | 24 | 16 | 19 | 14 | 24 | 17 | 18.98 | 20 | 18 | 0.904 | -0.146 | 0.74668046 | 0.84970572 | 0 | 0.1273 | 0.0942 | 0.0942 | TRUE | 0.335 |
| Scn1a | 14 | 5 | 7 | 8 | 13 | 10 | 11 | 5 | 8 | 8 | 15 | 10 | 9.37 | 8 | 11 | 1.341 | 0.424 | 0.50765898 | NA | 0 | 0.2553 | 0.1867 | 0.1867 | TRUE | 0.4033 |
| Scn1b | 474 | 382 | 518 | 551 | 462 | 530 | 368 | 406 | 559 | 548 | 521 | 523 | 487.7 | 444 | 531 | 1.198 | 0.261 | 0.06100362 | 0.15517436 | 0.0231 | 0.0073 | 0.0119 | 0.0119 | TRUE | 0.5823 |
| Scn2a1 | 46 | 34 | 26 | 33 | 20 | 32 | 36 | 36 | 28 | 33 | 23 | 32 | 31.16 | 33 | 29 | 0.871 | -0.199 | 0.5702995 | 0.72384067 | 0 | 0.0785 | 0.0562 | 0.0562 | TRUE | 0.2251 |
| Scn2b | 305 | 208 | 230 | 243 | 220 | 228 | 237 | 221 | 248 | 242 | 248 | 225 | 236.91 | 235 | 238 | 1.012 | 0.017 | 0.89876411 | 0.94399913 | 0 | 0.0124 | 0.0084 | 0.0084 | TRUE | 0.0288 |
| Scn3a | 149 | 80 | 51 | 36 | 46 | 38 | 116 | 85 | 55 | 36 | 52 | 38 | 63.51 | 85 | 42 | 0.484 | -1.046 | 0.0006996 | 0.00557069 | 0.0759 | 0.0397 | 0.0505 | 0.0505 | TRUE | 0.1085 |
| Scn3b | 355 | 283 | 288 | 285 | 250 | 265 | 276 | 301 | 311 | 284 | 282 | 262 | 285.8 | 296 | 276 | 0.934 | -0.099 | 0.42321827 | 0.59567295 | 0 | 0.0107 | 0.0074 | 0.0074 | TRUE | 0.0434 |
| Scn4a | 73 | 52 | 18 | 40 | 35 | 25 | 57 | 55 | 19 | 40 | 39 | 25 | 39.23 | 44 | 35 | 0.785 | -0.35 | 0.38197677 | 0.55626553 | 0.1586 | 0.0628 | 0.0893 | 0.0893 | TRUE | 1.7425 |
| Scn4b | 27 | 20 | 16 | 26 | 25 | 20 | 21 | 21 | 17 | 26 | 28 | 20 | 22.22 | 20 | 25 | 1.234 | 0.304 | 0.45558827 | 0.62526591 | 0 | 0.1091 | 0.0742 | 0.0742 | TRUE | 0.1854 |
| Scn5a | 122 | 94 | 72 | 107 | 102 | 103 | 95 | 100 | 78 | 107 | 115 | 102 | 99.28 | 91 | 108 | 1.183 | 0.243 | 0.22674104 | 0.39168596 | 0 | 0.0263 | 0.0189 | 0.0189 | TRUE | 0.1422 |
| Scn7a | 4 | 4 | 1 | 2 | 0 | 1 | 3 | 4 | 1 | 2 | 0 | 1 | 1.9 | 3 | 1 | 0.359 | -1.477 | 0.32097212 | NA | 0 | 1.2478 | 0.9149 | 0.9149 | TRUE | 0.302 |
| Scn8a | 485 | 374 | 411 | 412 | 405 | 427 | 377 | 398 | 443 | 410 | 457 | 422 | 417.74 | 406 | 430 | 1.059 | 0.083 | 0.4521088 | 0.62194332 | 0.0027 | 0.0081 | 0.0064 | 0.0064 | TRUE | 0.0741 |
| Scn9a | 37 | 29 | 27 | 40 | 31 | 39 | 29 | 31 | 29 | 40 | 35 | 38 | 33.67 | 30 | 38 | 1.28 | 0.356 | 0.27469636 | 0.44572249 | 0 | 0.0728 | 0.0466 | 0.0466 | TRUE | 0.0283 |
| Scnm1 | 907 | 618 | 658 | 627 | 559 | 655 | 704 | 657 | 710 | 624 | 631 | 647 | 662.18 | 690 | 634 | 0.917 | -0.124 | 0.15046777 | 0.29622022 | 0 | 0.006 | 0.0039 | 0.0039 | TRUE | 0.0203 |
| Scnn1a | 45 | 45 | 67 | 81 | 90 | 116 | 35 | 48 | 72 | 81 | 102 | 115 | 75.3 | 52 | 99 | 1.934 | 0.951 | 0.00067495 | 0.00541365 | 0.0632 | 0.0339 | 0.0417 | 0.0417 | TRUE | 0.8741 |
| Scnn1b | 15 | 11 | 12 | 13 | 11 | 6 | 12 | 12 | 13 | 13 | 12 | 6 | 11.26 | 12 | 10 | 0.86 | -0.218 | 0.70622082 | NA | 0 | 0.2129 | 0.1517 | 0.1517 | TRUE | 0.5324 |
| Scnn1g | 0 | 0 | 3 | 3 | 0 | 2 | 0 | 0 | 3 | 3 | 0 | 2 | 1.37 | 1 | 2 | 1.608 | 0.685 | 0.72247407 | NA | 2.9129 | 1.7372 | 1.9061 | 1.9061 | TRUE | 1.0926 |
| Sco1 | 2363 | 1822 | 1837 | 2002 | 1803 | 2136 | 1835 | 1938 | 1982 | 1993 | 2034 | 2109 | 1981.78 | 1918 | 2045 | 1.067 | 0.093 | 0.14788426 | 0.29259547 | 8.00E-04 | 0.0036 | 0.0025 | 0.0025 | TRUE | 0.0181 |
| Sco2 | 140 | 118 | 93 | 108 | 89 | 119 | 109 | 126 | 100 | 108 | 100 | 117 | 109.99 | 112 | 108 | 0.974 | -0.038 | 0.84449439 | 0.91195135 | 0 | 0.024 | 0.0174 | 0.0174 | TRUE | 0.1129 |
| Scoc | 1759 | 1182 | 1234 | 1547 | 1248 | 1383 | 1366 | 1257 | 1331 | 1540 | 1408 | 1365 | 1377.98 | 1318 | 1438 | 1.09 | 0.124 | 0.10198178 | 0.2242156 | 0.0022 | 0.0041 | 0.0034 | 0.0034 | TRUE | 0.047 |
| Scp2 | 3327 | 2458 | 2615 | 2869 | 2411 | 2781 | 2584 | 2615 | 2822 | 2856 | 2720 | 2745 | 2723.5 | 2674 | 2774 | 1.038 | 0.054 | 0.38610203 | 0.55991846 | 0.001 | 0.0033 | 0.0024 | 0.0024 | TRUE | 0.0282 |
| Scp2d1 | 0 | 0 | 0 | 1 | 0 | 0 | 0 | 0 | 0 | 1 | 0 | 0 | 0.17 | 0 | 0 | 2.094 | 1.067 | 0.79379329 | NA | 0 | 14.2885 | 10 | 10 | TRUE | 0.3906 |
| Scpep1 | 14927 | 10945 | 13293 | 14561 | 13337 | 15501 | 11593 | 11642 | 14343 | 14494 | 15046 | 15302 | 13736.61 | 12526 | 14947 | 1.193 | 0.255 | 0.00134219 | 0.00891029 | 0.0082 | 0.0026 | 0.0045 | 0.0045 | TRUE | 0.1965 |
| Scrg1 | 0 | 0 | 0 | 0 | 0 | 0 | 0 | 0 | 0 | 0 | 0 | 0 | 0 | 0 | 0 | NA | NA | NA | NA | NA | NA | NA | NA | NA | NA |
| Scrib | 10044 | 7568 | 7076 | 7614 | 6836 | 7347 | 7801 | 8050 | 7635 | 7579 | 7712 | 7253 | 7671.5 | 7829 | 7515 | 0.96 | -0.059 | 0.2570651 | 0.42708724 | 7.00E-04 | 0.0027 | 0.0018 | 0.0018 | TRUE | 0.0114 |
| Scrn1 | 385 | 311 | 303 | 409 | 373 | 422 | 299 | 331 | 327 | 407 | 421 | 417 | 366.87 | 319 | 415 | 1.303 | 0.382 | 0.00048037 | 0.00419425 | 0 | 0.0089 | 0.0059 | 0.0059 | TRUE | 0.0371 |
| Scrn2 | 774 | 571 | 560 | 640 | 528 | 676 | 601 | 607 | 604 | 637 | 596 | 667 | 618.79 | 604 | 633 | 1.049 | 0.069 | 0.44233736 | 0.61332152 | 0 | 0.0063 | 0.0042 | 0.0042 | TRUE | 0.031 |
| Scrn3 | 999 | 768 | 850 | 1098 | 996 | 1045 | 776 | 817 | 917 | 1093 | 1124 | 1032 | 959.68 | 837 | 1083 | 1.296 | 0.374 | 1.82E-05 | 0.00031011 | 0.0035 | 0.0049 | 0.0044 | 0.0044 | TRUE | 0.0833 |
| Scrt1 | 5 | 2 | 4 | 1 | 7 | 3 | 4 | 2 | 4 | 1 | 8 | 3 | 3.7 | 3 | 4 | 1.127 | 0.172 | 0.8695981 | NA | 0.2017 | 0.6436 | 0.5176 | 0.5176 | TRUE | 0.6264 |
| Scrt2 | 2 | 4 | 4 | 5 | 4 | 6 | 2 | 4 | 4 | 5 | 5 | 6 | 4.26 | 3 | 5 | 1.565 | 0.646 | 0.48696709 | NA | 0 | 0.5593 | 0.375 | 0.375 | TRUE | 0.4551 |
| Sct | 106 | 55 | 66 | 102 | 81 | 75 | 82 | 59 | 71 | 102 | 91 | 74 | 79.83 | 71 | 89 | 1.251 | 0.323 | 0.16953748 | 0.32103858 | 0.0152 | 0.0321 | 0.0273 | 0.0273 | TRUE | 0.212 |
| Sctr | 5 | 2 | 6 | 4 | 0 | 5 | 4 | 2 | 6 | 4 | 0 | 5 | 3.57 | 4 | 3 | 0.728 | -0.458 | 0.67698732 | NA | 0.2449 | 0.6669 | 0.5846 | 0.5846 | TRUE | 0.5986 |
| Scube1 | 141 | 115 | 101 | 102 | 111 | 111 | 110 | 122 | 109 | 102 | 125 | 110 | 112.86 | 114 | 112 | 0.986 | -0.02 | 0.91535166 | 0.95330234 | 0 | 0.0234 | 0.0166 | 0.0166 | TRUE | 0.0995 |
| Scube2 | 99 | 76 | 85 | 134 | 127 | 117 | 77 | 81 | 92 | 133 | 143 | 115 | 106.93 | 83 | 130 | 1.575 | 0.655 | 0.00085485 | 0.00646131 | 0 | 0.0246 | 0.0181 | 0.0181 | TRUE | 0.1078 |
| Scube3 | 649 | 478 | 404 | 492 | 409 | 461 | 504 | 508 | 436 | 490 | 461 | 455 | 475.77 | 483 | 469 | 0.97 | -0.044 | 0.6724108 | 0.7986345 | 0.0022 | 0.0074 | 0.0058 | 0.0058 | TRUE | 0.081 |
| Scx | 22 | 24 | 23 | 25 | 29 | 21 | 17 | 26 | 25 | 25 | 33 | 21 | 24.29 | 23 | 26 | 1.168 | 0.224 | 0.57872488 | 0.72987864 | 0 | 0.1 | 0.0756 | 0.0756 | TRUE | 0.3161 |
| Scyl1 | 2167 | 1585 | 1480 | 1369 | 1236 | 1467 | 1683 | 1686 | 1597 | 1363 | 1394 | 1448 | 1528.51 | 1655 | 1402 | 0.847 | -0.24 | 0.00035249 | 0.00328863 | 0 | 0.004 | 0.0026 | 0.0026 | TRUE | 0.0112 |
| Scyl2 | 3866 | 2677 | 2764 | 2896 | 2586 | 2897 | 3003 | 2847 | 2982 | 2883 | 2917 | 2860 | 2915.35 | 2944 | 2887 | 0.98 | -0.029 | 0.60704852 | 0.75124163 | 0 | 0.0032 | 0.0019 | 0.0019 | TRUE | 0.0099 |
| Scyl3 | 1551 | 1042 | 875 | 1090 | 934 | 983 | 1205 | 1108 | 944 | 1085 | 1054 | 970 | 1061.01 | 1086 | 1036 | 0.953 | -0.07 | 0.47545798 | 0.64342464 | 0.0083 | 0.0047 | 0.0059 | 0.0059 | TRUE | 0.1542 |
| Sdad1 | 10557 | 7843 | 7754 | 7557 | 6948 | 7829 | 8199 | 8342 | 8367 | 7522 | 7838 | 7728 | 7999.48 | 8303 | 7696 | 0.927 | -0.109 | 0.02229146 | 0.07459889 | 1.00E-04 | 0.0027 | 0.0015 | 0.0015 | TRUE | 0.0048 |
| Sdc1 | 2438 | 1911 | 2087 | 2196 | 1917 | 2120 | 1893 | 2033 | 2252 | 2186 | 2163 | 2093 | 2103.22 | 2059 | 2147 | 1.044 | 0.062 | 0.41218867 | 0.58510405 | 0.0032 | 0.0036 | 0.0036 | 0.0036 | TRUE | 0.0807 |
| Sdc2 | 624 | 442 | 451 | 472 | 354 | 408 | 485 | 470 | 487 | 470 | 399 | 403 | 452.22 | 481 | 424 | 0.883 | -0.18 | 0.09080001 | 0.20603343 | 0.0019 | 0.0077 | 0.0059 | 0.0059 | TRUE | 0.1045 |
| Sdc3 | 689 | 566 | 620 | 892 | 778 | 889 | 535 | 602 | 669 | 888 | 878 | 878 | 741.55 | 602 | 881 | 1.468 | 0.553 | 9.18E-09 | 4.60E-07 | 0.0046 | 0.0056 | 0.0053 | 0.0053 | TRUE | 0.1182 |
| Sdc4 | 4484 | 3459 | 4100 | 3715 | 3299 | 3603 | 3483 | 3679 | 4424 | 3698 | 3722 | 3557 | 3760.32 | 3862 | 3659 | 0.948 | -0.077 | 0.35775483 | 0.53214588 | 0.0078 | 0.0031 | 0.0048 | 0.0048 | TRUE | 0.1966 |
| Sdcbp | 3830 | 2498 | 2686 | 2657 | 2379 | 2618 | 2975 | 2657 | 2898 | 2645 | 2684 | 2584 | 2740.46 | 2843 | 2638 | 0.927 | -0.109 | 0.08928697 | 0.2036997 | 0.0016 | 0.0033 | 0.0026 | 0.0026 | TRUE | 0.0396 |
| Sdcbp2 | 176 | 161 | 149 | 244 | 209 | 254 | 137 | 171 | 161 | 243 | 236 | 251 | 199.68 | 156 | 243 | 1.564 | 0.645 | 1.40E-05 | 0.00025009 | 0 | 0.0143 | 0.0107 | 0.0107 | TRUE | 0.1421 |
| Sdccag3 | 3236 | 2294 | 2213 | 2478 | 2098 | 2371 | 2513 | 2440 | 2388 | 2467 | 2367 | 2341 | 2419.18 | 2447 | 2392 | 0.977 | -0.034 | 0.57205314 | 0.72498188 | 3.00E-04 | 0.0034 | 0.0022 | 0.0022 | TRUE | 0.0093 |
| Sdccag8 | 234 | 170 | 157 | 156 | 157 | 181 | 182 | 181 | 169 | 155 | 177 | 179 | 173.84 | 177 | 170 | 0.959 | -0.06 | 0.693751 | 0.81310234 | 0 | 0.0161 | 0.0109 | 0.0109 | TRUE | 0.0651 |
| Sde2 | 5114 | 3732 | 3267 | 3664 | 3321 | 3566 | 3972 | 3970 | 3525 | 3647 | 3746 | 3520 | 3730.06 | 3822 | 3638 | 0.951 | -0.072 | 0.27431258 | 0.44536104 | 0.0023 | 0.0031 | 0.0029 | 0.0029 | TRUE | 0.0557 |
| Sdf2 | 2110 | 1631 | 1913 | 2107 | 1859 | 2184 | 1639 | 1735 | 2064 | 2097 | 2097 | 2156 | 1964.69 | 1813 | 2117 | 1.169 | 0.225 | 0.00851859 | 0.03673996 | 0.0069 | 0.0036 | 0.0048 | 0.0048 | TRUE | 0.177 |
| Sdf2l1 | 2374 | 1716 | 1947 | 2464 | 2037 | 2586 | 1844 | 1825 | 2101 | 2453 | 2298 | 2553 | 2178.88 | 1923 | 2435 | 1.267 | 0.341 | 6.90E-06 | 0.00013845 | 0.0039 | 0.0035 | 0.0037 | 0.0037 | TRUE | 0.0777 |
| Sdf4 | 6771 | 4729 | 4396 | 4994 | 4293 | 4687 | 5259 | 5030 | 4743 | 4971 | 4843 | 4627 | 4912.16 | 5011 | 4814 | 0.96 | -0.058 | 0.33543994 | 0.50961335 | 0.0017 | 0.0029 | 0.0025 | 0.0025 | TRUE | 0.0265 |
| Sdha | 20610 | 14770 | 15708 | 17659 | 15575 | 18078 | 16007 | 15711 | 16949 | 17578 | 17570 | 17846 | 16943.35 | 16222 | 17665 | 1.089 | 0.123 | 0.01357826 | 0.05189279 | 7.00E-04 | 0.0026 | 0.0017 | 0.0017 | TRUE | 0.0187 |
| Sdhaf1 | 547 | 369 | 368 | 410 | 352 | 431 | 425 | 393 | 397 | 408 | 397 | 425 | 407.51 | 405 | 410 | 1.012 | 0.018 | 0.86650459 | 0.92632386 | 0 | 0.0082 | 0.0054 | 0.0054 | TRUE | 0.0232 |
| Sdhaf2 | 1749 | 1220 | 1063 | 1042 | 985 | 1106 | 1358 | 1298 | 1147 | 1037 | 1111 | 1092 | 1173.87 | 1268 | 1080 | 0.851 | -0.233 | 0.00527851 | 0.02554021 | 0.0037 | 0.0044 | 0.0042 | 0.0042 | TRUE | 0.0823 |
| Sdhb | 5447 | 3962 | 4310 | 4711 | 4403 | 4981 | 4230 | 4214 | 4650 | 4689 | 4967 | 4917 | 4611.44 | 4365 | 4858 | 1.113 | 0.155 | 0.01144921 | 0.04571048 | 0.0017 | 0.0029 | 0.0025 | 0.0025 | TRUE | 0.0395 |
| Sdhc | 7299 | 5353 | 6014 | 7451 | 6450 | 7601 | 5669 | 5694 | 6489 | 7417 | 7276 | 7503 | 6674.69 | 5951 | 7399 | 1.244 | 0.315 | 1.03E-06 | 3.01E-05 | 0.003 | 0.0028 | 0.0028 | 0.0028 | TRUE | 0.076 |
| Sdhd | 7371 | 5064 | 5415 | 5980 | 5084 | 5902 | 5725 | 5387 | 5843 | 5952 | 5735 | 5826 | 5744.66 | 5652 | 5838 | 1.033 | 0.047 | 0.39265856 | 0.56631041 | 8.00E-04 | 0.0028 | 0.002 | 0.002 | TRUE | 0.0203 |
| Sdk1 | 85 | 38 | 46 | 41 | 40 | 35 | 66 | 40 | 50 | 41 | 45 | 35 | 46.09 | 52 | 40 | 0.764 | -0.388 | 0.20348272 | 0.36474186 | 0.0235 | 0.0538 | 0.0449 | 0.0449 | TRUE | 0.3799 |
| Sdk2 | 1157 | 861 | 817 | 886 | 802 | 899 | 899 | 916 | 882 | 882 | 905 | 887 | 895.01 | 899 | 891 | 0.992 | -0.012 | 0.87402506 | 0.93005063 | 0 | 0.0051 | 0.003 | 0.003 | TRUE | 0.0032 |
| Sdpr | 13 | 4 | 4 | 4 | 3 | 2 | 10 | 4 | 4 | 4 | 3 | 2 | 4.67 | 6 | 3 | 0.49 | -1.029 | 0.26405953 | NA | 0 | 0.5102 | 0.3693 | 0.3693 | TRUE | 0.9142 |
| Sdr16c5 | 0 | 0 | 0 | 0 | 0 | 0 | 0 | 0 | 0 | 0 | 0 | 0 | 0 | 0 | 0 | NA | NA | NA | NA | NA | NA | NA | NA | NA | NA |
| Sdr16c6 | 0 | 0 | 0 | 0 | 0 | 0 | 0 | 0 | 0 | 0 | 0 | 0 | 0 | 0 | 0 | NA | NA | NA | NA | NA | NA | NA | NA | NA | NA |
| Sdr39u1 | 951 | 756 | 720 | 688 | 683 | 739 | 739 | 804 | 777 | 685 | 770 | 730 | 750.74 | 773 | 728 | 0.942 | -0.086 | 0.32597435 | 0.50001404 | 0.0013 | 0.0056 | 0.0042 | 0.0042 | TRUE | 0.0323 |
| Sdr42e1 | 114 | 73 | 89 | 80 | 59 | 97 | 89 | 78 | 96 | 80 | 67 | 96 | 84.03 | 88 | 81 | 0.925 | -0.113 | 0.61720674 | 0.75833714 | 0.0082 | 0.0306 | 0.0246 | 0.0246 | TRUE | 0.2515 |

| | | | | | | | | | | | | | | | | | | | | | | | | | |
|---|---|---|---|---|---|---|---|---|---|---|---|---|---|---|---|---|---|---|---|---|---|---|---|---|---|
| Sdr9c7 | 4 | 1 | 2 | 1 | 0 | 2 | 3 | 1 | 2 | 1 | 0 | 2 | 1.55 | 2 | 1 | 0.473 | -1.081 | 0.49417044 | NA | 0 | 1.5318 | 1.0639 | 1.0639 | TRUE | 0.3109 |
| Sds | 8 | 2 | 3 | 12 | 2 | 5 | 6 | 2 | 3 | 12 | 2 | 5 | 5.12 | 4 | 6 | 1.627 | 0.702 | 0.44673614 | NA | 0.2837 | 0.4654 | 0.4106 | 0.4106 | TRUE | 0.5451 |
| Sdsl | 4 | 0 | 0 | 5 | 5 | 0 | 3 | 0 | 0 | 5 | 6 | 0 | 2.29 | 1 | 4 | 3.14 | 1.651 | 0.31048864 | NA | 3.4308 | 1.0386 | 1.332 | 1.332 | TRUE | 1.8908 |
| Sebox | 2 | 0 | 3 | 0 | 1 | 0 | 2 | 0 | 3 | 0 | 1 | 0 | 0.99 | 2 | 0 | 0.233 | -2.104 | 0.33755304 | NA | 0 | 2.4052 | 2.1651 | 2.1651 | TRUE | 0.2207 |
| Sec1 | 48 | 32 | 17 | 16 | 17 | 17 | 37 | 34 | 18 | 16 | 19 | 17 | 23.59 | 30 | 17 | 0.573 | -0.803 | 0.05965777 | 0.15268725 | 0 | 0.1029 | 0.0848 | 0.0848 | TRUE | 0.71 |
| Sec11a | 2135 | 1543 | 1630 | 1766 | 1520 | 1728 | 1658 | 1641 | 1759 | 1758 | 1715 | 1706 | 1706.1 | 1686 | 1726 | 1.024 | 0.034 | 0.5953371 | 0.74186165 | 0 | 0.0038 | 0.0024 | 0.0024 | TRUE | 0.0167 |
| Sec11c | 708 | 459 | 493 | 627 | 548 | 633 | 550 | 488 | 532 | 624 | 618 | 625 | 572.87 | 523 | 622 | 1.188 | 0.248 | 0.00765193 | 0.03386037 | 0 | 0.0066 | 0.0045 | 0.0045 | TRUE | 0.0395 |
| Sec13 | 8324 | 5529 | 5620 | 5778 | 5235 | 5966 | 6465 | 5881 | 6064 | 5751 | 5906 | 5889 | 5992.73 | 6137 | 5849 | 0.953 | -0.07 | 0.21245757 | 0.37480327 | 0.001 | 0.0028 | 0.0021 | 0.0021 | TRUE | 0.0271 |
| Sec14l1 | 9326 | 7246 | 7988 | 8182 | 7683 | 8973 | 7243 | 7707 | 8619 | 8144 | 8667 | 8858 | 8206.48 | 7856 | 8556 | 1.089 | 0.124 | 0.08108535 | 0.19099422 | 0.0047 | 0.0027 | 0.0035 | 0.0035 | TRUE | 0.0879 |
| Sec14l2 | 457 | 341 | 328 | 284 | 262 | 299 | 355 | 363 | 354 | 283 | 296 | 295 | 324.16 | 357 | 291 | 0.815 | -0.295 | 0.00894112 | 0.03817284 | 0 | 0.0097 | 0.006 | 0.006 | TRUE | 0.0074 |
| Sec14l3 | 4 | 5 | 4 | 4 | 5 | 4 | 3 | 5 | 4 | 4 | 6 | 4 | 4.39 | 4 | 5 | 1.072 | 0.101 | 0.90994658 | NA | 0 | 0.5429 | 0.3446 | 0.3446 | TRUE | 0.1336 |
| Sec14l4 | 217 | 212 | 228 | 270 | 266 | 293 | 169 | 226 | 246 | 269 | 300 | 289 | 249.69 | 214 | 286 | 1.348 | 0.431 | 0.00541438 | 0.02609162 | 0.0162 | 0.0119 | 0.0132 | 0.0132 | TRUE | 0.4036 |
| Sec14l5 | 115 | 119 | 151 | 152 | 168 | 203 | 89 | 127 | 163 | 151 | 190 | 200 | 153.34 | 126 | 180 | 1.438 | 0.524 | 0.01435084 | 0.05399226 | 0.0475 | 0.0179 | 0.0263 | 0.0263 | TRUE | 0.2981 |
| Sec16a | 7238 | 5196 | 4930 | 5019 | 5146 | 6097 | 5621 | 5527 | 5319 | 4996 | 5805 | 6019 | 5547.93 | 5489 | 5607 | 1.021 | 0.03 | 0.67878796 | 0.80267364 | 0.005 | 0.0029 | 0.0037 | 0.0037 | TRUE | 0.111 |
| Sec16b | 69 | 57 | 56 | 49 | 30 | 39 | 54 | 61 | 60 | 49 | 34 | 38 | 49.29 | 58 | 40 | 0.697 | -0.52 | 0.06505178 | 0.16272053 | 0 | 0.0505 | 0.0362 | 0.0362 | TRUE | 0.2515 |
| Sec22a | 1175 | 773 | 791 | 836 | 744 | 821 | 913 | 822 | 853 | 832 | 839 | 810 | 845.03 | 863 | 827 | 0.957 | -0.063 | 0.44142823 | 0.61261615 | 0 | 0.0052 | 0.0036 | 0.0036 | TRUE | 0.032 |
| Sec22b | 2913 | 1975 | 2100 | 2188 | 1881 | 2148 | 2262 | 2101 | 2266 | 2178 | 2122 | 2120 | 2174.9 | 2210 | 2140 | 0.968 | -0.047 | 0.45666918 | 0.62640812 | 6.00E-04 | 0.0035 | 0.0024 | 0.0024 | TRUE | 0.0221 |
| Sec22c | 1033 | 770 | 776 | 890 | 719 | 798 | 802 | 819 | 837 | 886 | 811 | 788 | 823.9 | 819 | 828 | 1.011 | 0.016 | 0.84550057 | 0.9121221 | 9.00E-04 | 0.0053 | 0.0038 | 0.0038 | TRUE | 0.0447 |
| Sec23a | 6263 | 4642 | 5116 | 5693 | 5028 | 5646 | 4864 | 4938 | 5520 | 5667 | 5672 | 5573 | 5372.39 | 5107 | 5637 | 1.104 | 0.143 | 0.02204457 | 0.07405007 | 0.0022 | 0.0029 | 0.0026 | 0.0026 | TRUE | 0.0606 |
| Sec23b | 6871 | 4601 | 4539 | 4832 | 4221 | 4600 | 5336 | 4894 | 4898 | 4810 | 4762 | 4541 | 4873.4 | 5043 | 4704 | 0.932 | -0.101 | 0.08960553 | 0.20413322 | 0.0014 | 0.0029 | 0.0023 | 0.0023 | TRUE | 0.0322 |
| Sec23ip | 5113 | 3545 | 3173 | 3257 | 2992 | 3394 | 3971 | 3771 | 3424 | 3242 | 3375 | 3350 | 3522.19 | 3722 | 3322 | 0.892 | -0.165 | 0.01447354 | 0.05432518 | 0.0026 | 0.0031 | 0.003 | 0.003 | TRUE | 0.0595 |
| Sec24a | 4130 | 3061 | 3083 | 3948 | 3368 | 3942 | 3208 | 3256 | 3327 | 3930 | 3799 | 3891 | 3568.45 | 3264 | 3873 | 1.187 | 0.248 | 3.66E-06 | 8.21E-05 | 0 | 0.0031 | 0.0018 | 0.0018 | TRUE | 0.0035 |
| Sec24b | 4949 | 3621 | 3455 | 3354 | 3175 | 3795 | 3844 | 3852 | 3728 | 3339 | 3582 | 3746 | 3681.63 | 3808 | 3556 | 0.934 | -0.099 | 0.10768277 | 0.2335169 | 0.0014 | 0.0031 | 0.0025 | 0.0025 | TRUE | 0.0349 |
| Sec24c | 10104 | 7220 | 7455 | 8271 | 7272 | 8512 | 7847 | 7680 | 8044 | 8233 | 8204 | 8403 | 8068.37 | 7857 | 8280 | 1.054 | 0.076 | 0.11795101 | 0.24916845 | 2.00E-04 | 0.0027 | 0.0016 | 0.0016 | TRUE | 0.0052 |
| Sec24d | 3276 | 2329 | 2108 | 2468 | 2056 | 2450 | 2544 | 2477 | 2275 | 2457 | 2319 | 2419 | 2415.12 | 2432 | 2398 | 0.986 | -0.021 | 0.75310843 | 0.85348706 | 0.0017 | 0.0034 | 0.0028 | 0.0028 | TRUE | 0.0385 |
| Sec31a | 13967 | 10229 | 10014 | 10405 | 9941 | 11229 | 10848 | 10880 | 10805 | 10357 | 11214 | 11085 | 10864.9 | 10844 | 10885 | 1.004 | 0.005 | 0.91708037 | 0.95426832 | 6.00E-04 | 0.0026 | 0.0018 | 0.0018 | TRUE | 0.0221 |
| Sec31b | 306 | 289 | 300 | 332 | 324 | 352 | 238 | 307 | 324 | 330 | 366 | 347 | 318.7 | 290 | 348 | 1.206 | 0.27 | 0.04855822 | 0.13192219 | 0.0115 | 0.0099 | 0.0104 | 0.0104 | TRUE | 0.2994 |
| Sec61a1 | 15140 | 11100 | 10796 | 12089 | 10695 | 12349 | 11759 | 11807 | 11649 | 12033 | 12065 | 12190 | 11917.19 | 11738 | 12096 | 1.031 | 0.043 | 0.32674598 | 0.50082363 | 0 | 0.0026 | 0.0013 | 0.0013 | TRUE | 6.00E-04 |
| Sec61a2 | 3603 | 2686 | 2983 | 3617 | 3224 | 3665 | 2798 | 2857 | 3219 | 3600 | 3637 | 3618 | 3288.21 | 2958 | 3618 | 1.224 | 0.292 | 1.29E-05 | 0.00023182 | 0.0025 | 0.0031 | 0.0029 | 0.0029 | TRUE | 0.0716 |
| Sec61b | 4139 | 2686 | 2710 | 2998 | 2628 | 3074 | 3215 | 2857 | 2924 | 2984 | 2965 | 3035 | 2996.52 | 2999 | 2995 | 0.998 | -0.003 | 0.96210878 | 0.97916696 | 0.0018 | 0.0032 | 0.0026 | 0.0026 | TRUE | 0.0493 |
| Sec61g | 1899 | 1165 | 1271 | 1449 | 1170 | 1413 | 1475 | 1239 | 1371 | 1442 | 1320 | 1395 | 1373.75 | 1362 | 1386 | 1.017 | 0.024 | 0.77249415 | 0.86543807 | 0.0041 | 0.0042 | 0.0041 | 0.0041 | TRUE | 0.0743 |
| Sec62 | 6775 | 4851 | 4857 | 5252 | 4510 | 5312 | 5262 | 5160 | 5241 | 5228 | 5088 | 5244 | 5203.63 | 5221 | 5187 | 0.993 | -0.01 | 0.84803462 | 0.91382379 | 0 | 0.0029 | 0.0016 | 0.0016 | TRUE | 0.0033 |
| Sec63 | 4483 | 3102 | 3034 | 3161 | 2820 | 3248 | 3482 | 3300 | 3274 | 3146 | 3181 | 3206 | 3264.83 | 3352 | 3178 | 0.948 | -0.078 | 0.16812081 | 0.31966885 | 4.00E-04 | 0.0032 | 0.002 | 0.002 | TRUE | 0.0144 |
| Secisbp2 | 2202 | 1714 | 1855 | 1850 | 1665 | 2054 | 1710 | 1823 | 2002 | 1841 | 1878 | 2028 | 1880.38 | 1845 | 1916 | 1.04 | 0.056 | 0.47023952 | 0.63931642 | 0.004 | 0.0037 | 0.0038 | 0.0038 | TRUE | 0.0663 |
| Secisbp2l | 3740 | 2697 | 2570 | 2763 | 2503 | 2671 | 2905 | 2869 | 2773 | 2750 | 2824 | 2637 | 2792.85 | 2849 | 2737 | 0.96 | -0.059 | 0.32118752 | 0.49506452 | 5.00E-04 | 0.0033 | 0.0022 | 0.0022 | TRUE | 0.0125 |
| Sectm1a | 1 | 3 | 1 | 1 | 1 | 1 | 1 | 3 | 1 | 1 | 1 | 1 | 1.36 | 2 | 1 | 0.627 | -0.673 | 0.67660429 | NA | 0 | 1.7457 | 1.0863 | 1.0863 | TRUE | 0.4454 |
| Sectm1b | 0 | 0 | 0 | 0 | 0 | 0 | 0 | 0 | 0 | 0 | 0 | 0 | 0 | 0 | 0 | NA | NA | NA | NA | NA | NA | NA | NA | NA | NA |
| Seh1l | 9616 | 7184 | 7895 | 8885 | 7426 | 8838 | 7468 | 7642 | 8519 | 8844 | 8377 | 8724 | 8262.4 | 7876 | 8648 | 1.098 | 0.135 | 0.0314803 | 0.09595819 | 0.0028 | 0.0027 | 0.0027 | 0.0027 | TRUE | 0.062 |
| Sel1l | 5265 | 3850 | 3653 | 4219 | 3749 | 4101 | 4089 | 4095 | 3942 | 4200 | 4229 | 4048 | 4100.5 | 4042 | 4159 | 1.029 | 0.041 | 0.44530197 | 0.61613756 | 3.00E-04 | 0.003 | 0.0018 | 0.0018 | TRUE | 0.0066 |
| Sel1l2 | 1 | 1 | 0 | 1 | 1 | 0 | 1 | 1 | 0 | 1 | 1 | 0 | 0.66 | 1 | 1 | 1.128 | 0.173 | 0.94106052 | NA | 0 | 3.5899 | 2.4471 | 2.4471 | TRUE | 0.2681 |
| Sel1l3 | 87 | 71 | 85 | 76 | 72 | 101 | 68 | 76 | 92 | 76 | 81 | 100 | 81.9 | 79 | 86 | 1.099 | 0.136 | 0.55301338 | 0.708708 | 0.0114 | 0.0314 | 0.0257 | 0.0257 | TRUE | 0.2038 |
| Sele | 1 | 1 | 1 | 1 | 1 | 0 | 1 | 1 | 1 | 1 | 1 | 0 | 0.84 | 1 | 1 | 0.724 | -0.466 | 0.8180547 | NA | 0 | 2.8223 | 1.7212 | 1.7212 | TRUE | 0.2697 |
| Selenbp1 | 336 | 273 | 308 | 405 | 387 | 382 | 261 | 290 | 332 | 403 | 437 | 377 | 350.08 | 294 | 406 | 1.381 | 0.465 | 0.00022367 | 0.00232669 | 0.007 | 0.0092 | 0.0085 | 0.0085 | TRUE | 0.1407 |
| Selenbp2 | 21 | 11 | 24 | 32 | 11 | 22 | 16 | 12 | 26 | 32 | 12 | 22 | 19.98 | 18 | 22 | 1.235 | 0.304 | 0.53444563 | 0.69349354 | 0.1251 | 0.121 | 0.1222 | 0.1222 | TRUE | 0.1703 |
| Selk | 1141 | 785 | 695 | 756 | 592 | 706 | 886 | 835 | 750 | 753 | 668 | 697 | 764.72 | 824 | 706 | 0.856 | -0.224 | 0.01720176 | 0.06189948 | 0.0042 | 0.0055 | 0.0051 | 0.0051 | TRUE | 0.072 |
| Sell | 0 | 1 | 0 | 0 | 0 | 0 | 0 | 1 | 0 | 0 | 0 | 0 | 0.18 | 0 | 0 | 0.552 | -0.857 | 0.8336548 | NA | 0 | 13.3714 | 10 | 10 | TRUE | 0.5216 |
| Selm | 677 | 464 | 480 | 560 | 507 | 578 | 526 | 494 | 518 | 557 | 572 | 571 | 539.54 | 513 | 567 | 1.105 | 0.144 | 0.11860242 | 0.25005318 | 0 | 0.0068 | 0.0043 | 0.0043 | TRUE | 0.0119 |
| Selp | 5 | 5 | 7 | 3 | 6 | 3 | 4 | 5 | 8 | 3 | 7 | 3 | 4.91 | 6 | 4 | 0.759 | -0.397 | 0.64918682 | NA | 0 | 0.4849 | 0.339 | 0.339 | TRUE | 0.4257 |
| Selplg | 41 | 29 | 39 | 50 | 72 | 68 | 32 | 31 | 42 | 50 | 81 | 67 | 50.48 | 35 | 66 | 1.894 | 0.921 | 0.00211127 | 0.01256827 | 0.0267 | 0.0494 | 0.043 | 0.043 | TRUE | 0.1482 |
| Selrc1 | 1466 | 961 | 820 | 904 | 802 | 881 | 1139 | 1022 | 885 | 900 | 905 | 870 | 953.3 | 1015 | 892 | 0.876 | -0.19 | 0.04985823 | 0.13453809 | 0.0073 | 0.0049 | 0.0057 | 0.0057 | TRUE | 0.1497 |
| Sema3a | 27 | 24 | 14 | 2 | 7 | 6 | 21 | 26 | 15 | 2 | 8 | 6 | 12.9 | 21 | 5 | 0.253 | -1.98 | 0.00137486 | NA | 0.0197 | 0.1861 | 0.1531 | 0.1531 | TRUE | 0.4061 |
| Sema3b | 135 | 106 | 95 | 129 | 98 | 127 | 105 | 113 | 103 | 128 | 111 | 125 | 114.07 | 107 | 121 | 1.14 | 0.19 | 0.30263873 | 0.4751857 | 0 | 0.0232 | 0.0156 | 0.0156 | TRUE | 0.0591 |
| Sema3c | 117 | 85 | 64 | 41 | 48 | 44 | 91 | 90 | 69 | 41 | 54 | 43 | 64.79 | 83 | 46 | 0.55 | -0.864 | 0.00075359 | 0.00588091 | 0 | 0.039 | 0.0303 | 0.0303 | TRUE | 0.2034 |
| Sema3d | 33 | 20 | 21 | 17 | 18 | 18 | 26 | 21 | 23 | 17 | 20 | 18 | 20.76 | 23 | 18 | 0.786 | -0.347 | 0.40340743 | 0.57635969 | 0 | 0.1166 | 0.0755 | 0.0755 | TRUE | 0.0605 |
| Sema3e | 341 | 221 | 225 | 165 | 169 | 210 | 265 | 235 | 243 | 164 | 191 | 207 | 217.48 | 248 | 187 | 0.755 | -0.405 | 0.00546897 | 0.02628381 | 0.0037 | 0.0133 | 0.0106 | 0.0106 | TRUE | 0.1295 |
| Sema3f | 239 | 227 | 229 | 291 | 258 | 309 | 186 | 241 | 247 | 290 | 291 | 305 | 259.98 | 225 | 295 | 1.321 | 0.402 | 0.0047793 | 0.02368605 | 0.0087 | 0.0115 | 0.0107 | 0.0107 | TRUE | 0.2797 |
| Sema3g | 32 | 29 | 9 | 33 | 19 | 23 | 25 | 31 | 10 | 33 | 21 | 23 | 23.73 | 22 | 26 | 1.173 | 0.231 | 0.61297275 | 0.75508227 | 0.1227 | 0.1023 | 0.1075 | 0.1075 | TRUE | 0.7936 |
| Sema4a | 658 | 495 | 527 | 674 | 609 | 696 | 511 | 527 | 569 | 671 | 687 | 687 | 608.53 | 536 | 682 | 1.275 | 0.35 | 0.00010851 | 0.00129457 | 0 | 0.0063 | 0.0042 | 0.0042 | TRUE | 0.0338 |
| Sema4b | 3064 | 2434 | 2362 | 2274 | 2130 | 2554 | 2380 | 2589 | 2549 | 2264 | 2403 | 2521 | 2450.81 | 2506 | 2396 | 0.957 | -0.064 | 0.34523713 | 0.51936368 | 0.0018 | 0.0034 | 0.0029 | 0.0029 | TRUE | 0.0286 |
| Sema4c | 309 | 257 | 336 | 352 | 322 | 362 | 240 | 273 | 363 | 350 | 363 | 357 | 324.48 | 292 | 357 | 1.227 | 0.296 | 0.04541483 | 0.12552971 | 0.019 | 0.0097 | 0.0126 | 0.0126 | TRUE | 0.4985 |
| Sema4d | 3097 | 2333 | 2337 | 2548 | 2330 | 2767 | 2405 | 2482 | 2522 | 2536 | 2628 | 2731 | 2550.78 | 2470 | 2632 | 1.066 | 0.093 | 0.12613944 | 0.26129144 | 6.00E-04 | 0.0034 | 0.0022 | 0.0022 | TRUE | 0.0134 |
| Sema4f | 571 | 464 | 441 | 589 | 548 | 563 | 443 | 494 | 476 | 586 | 618 | 556 | 528.85 | 471 | 587 | 1.247 | 0.319 | 0.00117013 | 0.00812808 | 0 | 0.0069 | 0.005 | 0.005 | TRUE | 0.0327 |
| Sema4g | 1366 | 1174 | 1292 | 1694 | 1561 | 1831 | 1061 | 1249 | 1394 | 1686 | 1761 | 1807 | 1493.07 | 1235 | 1751 | 1.421 | 0.507 | 7.83E-08 | 3.13E-06 | 0.0087 | 0.004 | 0.0057 | 0.0057 | TRUE | 0.1877 |
| Sema5a | 299 | 222 | 280 | 249 | 223 | 232 | 232 | 236 | 302 | 248 | 252 | 229 | 249.82 | 257 | 243 | 0.948 | -0.078 | 0.58976497 | 0.73845911 | 0.0087 | 0.0119 | 0.0109 | 0.0109 | TRUE | 0.2582 |
| Sema5b | 1063 | 852 | 868 | 1282 | 1085 | 1215 | 826 | 906 | 937 | 1276 | 1224 | 1199 | 1061.32 | 890 | 1233 | 1.389 | 0.474 | 2.77E-09 | 1.63E-07 | 0.0017 | 0.0047 | 0.0036 | 0.0036 | TRUE | 0.0497 |
| Sema6a | 71 | 46 | 33 | 46 | 43 | 56 | 55 | 49 | 36 | 46 | 49 | 55 | 48.21 | 47 | 50 | 1.064 | 0.09 | 0.75684162 | 0.85561474 | 0 | 0.0516 | 0.0397 | 0.0397 | TRUE | 0.3185 |
| Sema6b | 341 | 263 | 297 | 429 | 377 | 404 | 265 | 280 | 320 | 427 | 425 | 399 | 352.7 | 288 | 417 | 1.45 | 0.536 | 5.54E-06 | 0.00011552 | 0.0022 | 0.0091 | 0.0071 | 0.0071 | TRUE | 0.1046 |
| Sema6c | 868 | 688 | 766 | 1310 | 1143 | 1243 | 674 | 732 | 827 | 1304 | 1289 | 1227 | 1008.82 | 744 | 1273 | 1.715 | 0.778 | 2.52E-18 | 6.32E-16 | 0.0043 | 0.0048 | 0.0047 | 0.0047 | TRUE | 0.1102 |
| Sema6d | 14 | 15 | 10 | 10 | 6 | 12 | 11 | 16 | 11 | 10 | 7 | 12 | 11.03 | 13 | 10 | 0.767 | -0.383 | 0.50984289 | NA | 0 | 0.2173 | 0.1506 | 0.1506 | TRUE | 0.2166 |

| | | | | | | | | | | | | | | | | | | | | | | | | | |
|---|---|---|---|---|---|---|---|---|---|---|---|---|---|---|---|---|---|---|---|---|---|---|---|---|---|
| Sema7a | 671 | 535 | 547 | 877 | 823 | 891 | 521 | 569 | 590 | 873 | 928 | 880 | 726.89 | 560 | 894 | 1.598 | 0.677 | 1.91E-14 | 3.06E-12 | 0.0011 | 0.0057 | 0.0042 | 0.0042 | TRUE | 0.0468 |
| Senp1 | 4102 | 3037 | 2937 | 3017 | 2584 | 2975 | 3186 | 3230 | 3169 | 3003 | 2915 | 2937 | 3073.36 | 3195 | 2952 | 0.924 | -0.114 | 0.03475524 | 0.10331876 | 0 | 0.0032 | 0.0018 | 0.0018 | TRUE | 0.0028 |
| Senp2 | 7082 | 5278 | 5070 | 5581 | 4851 | 5556 | 5500 | 5614 | 5470 | 5555 | 5472 | 5485 | 5516.22 | 5528 | 5504 | 0.996 | -0.006 | 0.89848368 | 0.94384643 | 0 | 0.0029 | 0.0015 | 0.0015 | TRUE | 0.0022 |
| Senp3 | 17890 | 12152 | 10958 | 11810 | 10553 | 11538 | 13894 | 12926 | 11824 | 11756 | 11905 | 11390 | 12282.37 | 12881 | 11684 | 0.907 | -0.141 | 0.02962354 | 0.09178611 | 0.0034 | 0.0026 | 0.003 | 0.003 | TRUE | 0.063 |
| Senp5 | 1753 | 1189 | 1088 | 1446 | 1184 | 1214 | 1361 | 1265 | 1174 | 1439 | 1336 | 1198 | 1295.6 | 1267 | 1324 | 1.044 | 0.063 | 0.48137604 | 0.64841275 | 0.0062 | 0.0043 | 0.0049 | 0.0049 | TRUE | 0.0843 |
| Senp6 | 5436 | 3879 | 3850 | 4123 | 3509 | 3880 | 4222 | 4126 | 4154 | 4104 | 3959 | 3830 | 4065.8 | 4167 | 3964 | 0.951 | -0.072 | 0.18542849 | 0.3417746 | 4.00E-04 | 0.003 | 0.0019 | 0.0019 | TRUE | 0.0117 |
| Senp7 | 330 | 271 | 291 | 402 | 341 | 403 | 256 | 288 | 314 | 400 | 385 | 398 | 340.2 | 286 | 394 | 1.383 | 0.468 | 8.79E-05 | 0.00108618 | 0.0021 | 0.0094 | 0.0072 | 0.0072 | TRUE | 0.1027 |
| Senp8 | 965 | 686 | 611 | 817 | 662 | 746 | 749 | 730 | 659 | 813 | 747 | 736 | 739.15 | 713 | 765 | 1.073 | 0.101 | 0.26276396 | 0.43318169 | 0.0024 | 0.0056 | 0.0045 | 0.0045 | TRUE | 0.05 |
| Sephs1 | 5288 | 3795 | 3549 | 3467 | 3165 | 3670 | 4107 | 4037 | 3829 | 3451 | 3570 | 3623 | 3769.56 | 3991 | 3548 | 0.889 | -0.17 | 0.00282185 | 0.01573174 | 7.00E-04 | 0.0031 | 0.0021 | 0.0021 | TRUE | 0.0152 |
| Sephs2 | 8183 | 5894 | 4742 | 5253 | 4690 | 5258 | 6355 | 6269 | 5117 | 5229 | 5291 | 5190 | 5575.24 | 5914 | 5237 | 0.885 | -0.176 | 0.02692755 | 0.08571836 | 0.0066 | 0.0029 | 0.0044 | 0.0044 | TRUE | 0.1689 |
| Sepn1 | 596 | 451 | 466 | 741 | 661 | 727 | 463 | 480 | 503 | 738 | 746 | 718 | 607.72 | 482 | 734 | 1.525 | 0.608 | 1.13E-11 | 1.16E-09 | 0 | 0.0063 | 0.0041 | 0.0041 | TRUE | 0.0167 |
| Seppl | 483 | 372 | 436 | 353 | 321 | 323 | 375 | 396 | 470 | 351 | 362 | 319 | 378.93 | 414 | 344 | 0.833 | -0.264 | 0.03161465 | 0.09620356 | 0.007 | 0.0087 | 0.0082 | 0.0082 | TRUE | 0.1631 |
| Sepsecs | 1103 | 827 | 785 | 875 | 784 | 869 | 857 | 880 | 847 | 871 | 884 | 858 | 866.1 | 861 | 871 | 1.012 | 0.017 | 0.8299206 | 0.90264098 | 0 | 0.0052 | 0.0031 | 0.0031 | TRUE | 0.0041 |
| Sep-01 | 3624 | 2720 | 2450 | 2689 | 2476 | 3117 | 2815 | 2893 | 2644 | 2677 | 2793 | 3077 | 2816.36 | 2784 | 2849 | 1.023 | 0.033 | 0.63846976 | 0.77423614 | 0.0033 | 0.0033 | 0.0033 | 0.0033 | TRUE | 0.06 |
| Sep-02 | 16354 | 11740 | 12180 | 13947 | 11995 | 13544 | 12701 | 12488 | 13142 | 13883 | 13532 | 13370 | 13185.95 | 12777 | 13595 | 1.064 | 0.09 | 0.062617 | 0.158288 | 4.00E-04 | 0.0026 | 0.0016 | 0.0016 | TRUE | 0.0076 |
| Sep-03 | 1184 | 997 | 1136 | 1454 | 1222 | 1434 | 920 | 1060 | 1226 | 1447 | 1379 | 1416 | 1241.2 | 1069 | 1414 | 1.326 | 0.407 | 3.32E-05 | 0.00050631 | 0.0093 | 0.0043 | 0.0061 | 0.0061 | TRUE | 0.1979 |
| Sep-04 | 104 | 84 | 61 | 38 | 51 | 39 | 81 | 89 | 66 | 38 | 58 | 38 | 61.63 | 79 | 45 | 0.564 | -0.826 | 0.00231343 | 0.0135294 | 0.0184 | 0.0409 | 0.0352 | 0.0352 | TRUE | 0.4683 |
| Sep-05 | 170 | 108 | 101 | 137 | 147 | 169 | 132 | 115 | 109 | 136 | 166 | 167 | 137.49 | 119 | 156 | 1.312 | 0.391 | 0.02768071 | 0.08728003 | 0.0049 | 0.0197 | 0.0154 | 0.0154 | TRUE | 0.1338 |
| Sep-06 | 1590 | 1159 | 1401 | 1950 | 1707 | 1985 | 1235 | 1233 | 1512 | 1941 | 1926 | 1960 | 1634.26 | 1327 | 1942 | 1.466 | 0.552 | 1.19E-10 | 9.52E-09 | 0.0064 | 0.0039 | 0.0047 | 0.0047 | TRUE | 0.1768 |
| Sep-07 | 10076 | 6893 | 6772 | 6604 | 5464 | 6149 | 7826 | 7332 | 7307 | 6574 | 6164 | 6070 | 6878.69 | 7488 | 6269 | 0.837 | -0.257 | 8.15E-06 | 0.00015852 | 0.0015 | 0.0028 | 0.0022 | 0.0022 | TRUE | 0.0221 |
| Sep-08 | 5035 | 3876 | 3587 | 3937 | 3563 | 3973 | 3910 | 4123 | 3870 | 3919 | 4019 | 3922 | 3960.66 | 3968 | 3953 | 0.996 | -0.005 | 0.92477261 | 0.95843484 | 4.00E-04 | 0.003 | 0.0019 | 0.0019 | TRUE | 0.014 |
| Sep-09 | 9250 | 7228 | 7869 | 7794 | 7415 | 9077 | 7184 | 7688 | 8491 | 7758 | 8365 | 8960 | 8074.4 | 7788 | 8361 | 1.074 | 0.103 | 0.17108317 | 0.3228108 | 0.006 | 0.0027 | 0.0039 | 0.0039 | TRUE | 0.0761 |
| Sep-10 | 2634 | 1916 | 2008 | 2313 | 1978 | 2053 | 2046 | 2038 | 2167 | 2302 | 2231 | 2027 | 2135.12 | 2084 | 2187 | 1.05 | 0.07 | 0.31806645 | 0.49189826 | 0.002 | 0.0035 | 0.0031 | 0.0031 | TRUE | 0.0502 |
| Sep-11 | 19191 | 13649 | 11966 | 10698 | 9774 | 11393 | 14905 | 14518 | 12911 | 10649 | 11026 | 11247 | 12542.64 | 14111 | 10974 | 0.778 | -0.363 | 1.27E-08 | 6.25E-07 | 0.0031 | 0.0026 | 0.0029 | 0.0029 | TRUE | 0.0675 |
| Sep-12 | 9 | 4 | 6 | 7 | 2 | 2 | 7 | 4 | 6 | 7 | 2 | 2 | 4.82 | 6 | 4 | 0.633 | -0.661 | 0.46415777 | NA | 0 | 0.4942 | 0.3679 | 0.3679 | TRUE | 0.9285 |
| Sep-14 | 0 | 2 | 1 | 2 | 1 | 4 | 0 | 2 | 1 | 2 | 1 | 4 | 1.71 | 1 | 2 | 2.336 | 1.224 | 0.43043665 | NA | 0 | 1.3866 | 1.0439 | 1.0439 | TRUE | 0.4849 |
| Sep-15 | 9916 | 6864 | 6683 | 7818 | 6623 | 7497 | 7701 | 7301 | 7211 | 7782 | 7471 | 7401 | 7477.92 | 7404 | 7551 | 1.02 | 0.028 | 0.59861512 | 0.74432414 | 8.00E-04 | 0.0027 | 0.0019 | 0.0019 | TRUE | 0.0152 |
| Sepw1 | 2692 | 1630 | 1658 | 1963 | 1693 | 1867 | 2091 | 1734 | 1789 | 1954 | 1910 | 1843 | 1886.73 | 1871 | 1902 | 1.015 | 0.022 | 0.78751542 | 0.87523421 | 0.0051 | 0.0037 | 0.0042 | 0.0042 | TRUE | 0.1303 |
| Serac1 | 1752 | 1420 | 1380 | 1437 | 1266 | 1515 | 1361 | 1510 | 1489 | 1430 | 1428 | 1496 | 1452.38 | 1453 | 1451 | 1 | 0 | 0.99891065 | 0.99939385 | 0.0014 | 0.0041 | 0.0031 | 0.0031 | TRUE | 0.0391 |
| Serbp1 | 72390 | 53871 | 53346 | 58429 | 50585 | 59310 | 56222 | 57302 | 57560 | 58160 | 57065 | 58548 | 57476.24 | 57028 | 57924 | 1.016 | 0.023 | 0.59666586 | 0.74274039 | 1.00E-04 | 0.0025 | 0.0013 | 0.0013 | TRUE | 0.0021 |
| Serf1 | 189 | 147 | 131 | 133 | 109 | 144 | 147 | 156 | 141 | 132 | 123 | 142 | 140.33 | 148 | 132 | 0.896 | -0.159 | 0.34126279 | 0.51528301 | 0 | 0.0193 | 0.013 | 0.013 | TRUE | 0.0419 |
| Serf2 | 4310 | 3070 | 3242 | 3646 | 3207 | 3590 | 3347 | 3266 | 3498 | 3629 | 3618 | 3544 | 3483.66 | 3370 | 3597 | 1.067 | 0.094 | 0.09261856 | 0.20919675 | 4.00E-04 | 0.0031 | 0.002 | 0.002 | TRUE | 0.0131 |
| Sergef | 441 | 339 | 339 | 337 | 273 | 346 | 343 | 361 | 366 | 335 | 308 | 342 | 342.31 | 357 | 328 | 0.923 | -0.115 | 0.30957568 | 0.4823158 | 0 | 0.0094 | 0.0063 | 0.0063 | TRUE | 0.0322 |
| Serhl | 397 | 261 | 328 | 381 | 346 | 421 | 308 | 278 | 354 | 379 | 390 | 416 | 354.17 | 313 | 395 | 1.262 | 0.336 | 0.00595466 | 0.0280613 | 0.0048 | 0.0091 | 0.0079 | 0.0079 | TRUE | 0.1377 |
| Serinc1 | 5695 | 4048 | 4045 | 4229 | 3733 | 4022 | 4423 | 4306 | 4365 | 4210 | 4211 | 3970 | 4247.41 | 4365 | 4130 | 0.946 | -0.08 | 0.14059598 | 0.28243687 | 4.00E-04 | 0.003 | 0.0019 | 0.0019 | TRUE | 0.0141 |
| Serinc2 | 677 | 522 | 608 | 764 | 716 | 769 | 526 | 555 | 656 | 760 | 808 | 759 | 677.4 | 579 | 776 | 1.342 | 0.425 | 2.54E-05 | 0.00040444 | 0.0053 | 0.0059 | 0.0058 | 0.0058 | TRUE | 0.1565 |
| Serinc3 | 8643 | 6440 | 7064 | 7483 | 6584 | 7456 | 6713 | 6850 | 7622 | 7449 | 7427 | 7360 | 7236.84 | 7062 | 7412 | 1.05 | 0.07 | 0.24894189 | 0.41806209 | 0.0022 | 0.0028 | 0.0025 | 0.0025 | TRUE | 0.0586 |
| Serinc4 | 6 | 2 | 3 | 5 | 6 | 4 | 5 | 2 | 3 | 5 | 7 | 4 | 4.29 | 3 | 5 | 1.531 | 0.614 | 0.50304166 | NA | 0 | 0.5553 | 0.3655 | 0.3655 | TRUE | 0.2299 |
| Serinc5 | 1920 | 1424 | 1798 | 1706 | 1590 | 1800 | 1491 | 1515 | 1940 | 1698 | 1794 | 1777 | 1702.44 | 1649 | 1756 | 1.066 | 0.092 | 0.34101653 | 0.51512887 | 0.011 | 0.0038 | 0.0062 | 0.0062 | TRUE | 0.2861 |
| Serp1 | 4879 | 3403 | 3380 | 3753 | 3121 | 3506 | 3789 | 3620 | 3647 | 3736 | 3521 | 3461 | 3628.92 | 3685 | 3573 | 0.969 | -0.045 | 0.43579335 | 0.60709497 | 7.00E-04 | 0.0031 | 0.0022 | 0.0022 | TRUE | 0.0196 |
| Serp2 | 165 | 129 | 140 | 115 | 118 | 126 | 128 | 137 | 151 | 114 | 133 | 124 | 131.4 | 139 | 124 | 0.895 | -0.161 | 0.35877337 | 0.5333279 | 0 | 0.0205 | 0.0144 | 0.0144 | TRUE | 0.0606 |
| Serpina10 | 0 | 0 | 0 | 0 | 0 | 0 | 0 | 0 | 0 | 0 | 0 | 0 | 0 | 0 | 0 | NA | NA | NA | NA | NA | NA | NA | NA | NA | NA |
| Serpina11 | 0 | 0 | 0 | 0 | 0 | 0 | 0 | 0 | 0 | 0 | 0 | 0 | 0 | 0 | 0 | NA | NA | NA | NA | NA | NA | NA | NA | NA | NA |
| Serpina12 | 12 | 6 | 5 | 3 | 5 | 6 | 9 | 6 | 5 | 3 | 6 | 6 | 5.94 | 7 | 5 | 0.68 | -0.556 | 0.48034392 | NA | 0 | 0.4014 | 0.273 | 0.273 | TRUE | 0.2779 |
| Serpina1a | 0 | 0 | 0 | 0 | 0 | 0 | 0 | 0 | 0 | 0 | 0 | 0 | 0 | 0 | 0 | NA | NA | NA | NA | NA | NA | NA | NA | NA | NA |
| Serpina1b | 1 | 0 | 1 | 0 | 0 | 0 | 1 | 0 | 1 | 0 | 0 | 0 | 0.31 | 1 | 0 | 0.307 | -1.706 | 0.62894649 | NA | 0 | 7.6657 | 7.2 | 7.2 | TRUE | 0.2067 |
| Serpina1c | 0 | 0 | 0 | 0 | 0 | 0 | 0 | 0 | 0 | 0 | 0 | 0 | 0 | 0 | 0 | NA | NA | NA | NA | NA | NA | NA | NA | NA | NA |
| Serpina1d | 0 | 0 | 0 | 0 | 0 | 0 | 0 | 0 | 0 | 0 | 0 | 0 | 0 | 0 | 0 | NA | NA | NA | NA | NA | NA | NA | NA | NA | NA |
| Serpina1e | 0 | 0 | 0 | 0 | 0 | 0 | 0 | 0 | 0 | 0 | 0 | 0 | 0 | 0 | 0 | NA | NA | NA | NA | NA | NA | NA | NA | NA | NA |
| Serpina1f | 0 | 0 | 0 | 0 | 0 | 0 | 0 | 0 | 0 | 0 | 0 | 0 | 0 | 0 | 0 | NA | NA | NA | NA | NA | NA | NA | NA | NA | NA |
| Serpina3a | 0 | 0 | 0 | 0 | 0 | 0 | 0 | 0 | 0 | 0 | 0 | 0 | 0 | 0 | 0 | NA | NA | NA | NA | NA | NA | NA | NA | NA | NA |
| Serpina3b | 0 | 0 | 0 | 0 | 0 | 0 | 0 | 0 | 0 | 0 | 0 | 0 | 0 | 0 | 0 | NA | NA | NA | NA | NA | NA | NA | NA | NA | NA |
| Serpina3c | 0 | 0 | 0 | 0 | 0 | 0 | 0 | 0 | 0 | 0 | 0 | 0 | 0 | 0 | 0 | NA | NA | NA | NA | NA | NA | NA | NA | NA | NA |
| Serpina3f | 0 | 0 | 0 | 0 | 0 | 0 | 0 | 0 | 0 | 0 | 0 | 0 | 0 | 0 | 0 | NA | NA | NA | NA | NA | NA | NA | NA | NA | NA |
| Serpina3g | 0 | 0 | 1 | 0 | 0 | 0 | 0 | 0 | 1 | 0 | 0 | 0 | 0.18 | 0 | 0 | 0.552 | -0.857 | 0.8336548 | NA | 0 | 13.1817 | 10 | 10 | TRUE | 0.5365 |
| Serpina3h | 1 | 5 | 3 | 5 | 1 | 3 | 1 | 5 | 3 | 5 | 1 | 3 | 3.07 | 3 | 3 | 1.007 | 0.01 | 0.99306357 | NA | 0 | 0.7753 | 0.6207 | 0.6207 | TRUE | 0.2745 |
| Serpina3i | 11 | 7 | 9 | 9 | 8 | 10 | 9 | 7 | 10 | 9 | 9 | 10 | 8.93 | 9 | 9 | 1.085 | 0.117 | 0.85073131 | NA | 0 | 0.268 | 0.1688 | 0.1688 | TRUE | 0.037 |
| Serpina3j | 0 | 0 | 0 | 0 | 0 | 1 | 0 | 0 | 0 | 0 | 0 | 1 | 0.16 | 0 | 0 | 2.094 | 1.067 | 0.79379329 | NA | 0 | 14.4078 | 10 | 10 | TRUE | 0.3835 |
| Serpina3k | 6 | 6 | 3 | 3 | 0 | 1 | 5 | 6 | 3 | 3 | 0 | 1 | 3.04 | 5 | 1 | 0.285 | -1.813 | 0.13846181 | NA | 0 | 0.7815 | 0.5951 | 0.5951 | TRUE | 0.6662 |
| Serpina3m | 65 | 67 | 59 | 18 | 21 | 20 | 50 | 71 | 64 | 18 | 24 | 20 | 41.13 | 62 | 21 | 0.331 | -1.594 | 1.39E-06 | 3.80E-05 | 0 | 0.0601 | 0.0453 | 0.0453 | TRUE | 0.2306 |
| Serpina3n | 2 | 2 | 2 | 2 | 1 | 1 | 2 | 2 | 2 | 2 | 1 | 1 | 1.66 | 2 | 1 | 0.711 | -0.492 | 0.73096965 | NA | 0 | 1.4324 | 0.8486 | 0.8486 | TRUE | 0.103 |
| Serpina4-ps1 | 0 | 0 | 0 | 0 | 0 | 0 | 0 | 0 | 0 | 0 | 0 | 0 | 0 | 0 | 0 | NA | NA | NA | NA | NA | NA | NA | NA | NA | NA |
| Serpina5 | 0 | 0 | 0 | 0 | 0 | 0 | 0 | 0 | 0 | 0 | 0 | 0 | 0 | 0 | 0 | NA | NA | NA | NA | NA | NA | NA | NA | NA | NA |
| Serpina6 | 0 | 0 | 0 | 0 | 0 | 0 | 0 | 0 | 0 | 0 | 0 | 0 | 0 | 0 | 0 | NA | NA | NA | NA | NA | NA | NA | NA | NA | NA |
| Serpina7 | 9 | 13 | 9 | 3 | 2 | 6 | 7 | 14 | 10 | 3 | 2 | 6 | 6.95 | 10 | 4 | 0.372 | -1.425 | 0.06843475 | NA | 0 | 0.3435 | 0.2552 | 0.2552 | TRUE | 0.3034 |
| Serpina9 | 3 | 1 | 0 | 1 | 3 | 1 | 2 | 1 | 0 | 1 | 3 | 1 | 1.46 | 1 | 2 | 1.492 | 0.578 | 0.7221879 | NA | 0 | 1.6257 | 1.2012 | 1.2012 | TRUE | 0.3955 |
| Serpinb10 | 2 | 0 | 0 | 0 | 0 | 0 | 2 | 0 | 0 | 0 | 0 | 0 | 0.26 | 1 | 0 | 0.356 | -1.491 | 0.71362796 | NA | 0 | 9.1573 | 10 | 10 | TRUE | 0.968 |
| Serpinb11 | 0 | 0 | 0 | 0 | 0 | 0 | 0 | 0 | 0 | 0 | 0 | 0 | 0 | 0 | 0 | NA | NA | NA | NA | NA | NA | NA | NA | NA | NA |
| Serpinb12 | 2 | 0 | 0 | 0 | 0 | 0 | 2 | 0 | 0 | 0 | 0 | 0 | 0.26 | 1 | 0 | 0.356 | -1.491 | 0.71362796 | NA | 0 | 9.1573 | 10 | 10 | TRUE | 0.968 |

| | | | | | | | | | | | | | | | | | | | | | | | | | |
|---|---|---|---|---|---|---|---|---|---|---|---|---|---|---|---|---|---|---|---|---|---|---|---|---|---|
| Serpinb13 | 0 | 0 | 0 | 0 | 0 | 0 | 0 | 0 | 0 | 0 | 0 | 0 | 0 | 0 | 0 | NA | NA | NA | NA | NA | NA | NA | NA | NA | NA |
| Serpinb1a | 252 | 188 | 230 | 192 | 187 | 176 | 196 | 200 | 248 | 191 | 211 | 174 | 203.28 | 215 | 192 | 0.895 | -0.159 | 0.30606887 | 0.47857664 | 0.0087 | 0.0141 | 0.0125 | 0.0125 | TRUE | 0.1991 |
| Serpinb1b | 0 | 0 | 0 | 0 | 0 | 0 | 0 | 0 | 0 | 0 | 0 | 0 | 0 | 0 | 0 | NA | NA | NA | NA | NA | NA | NA | NA | NA | NA |
| Serpinb1c | 5 | 2 | 9 | 8 | 3 | 9 | 4 | 2 | 10 | 8 | 3 | 9 | 5.99 | 5 | 7 | 1.31 | 0.389 | 0.64300386 | NA | 0.1807 | 0.3979 | 0.3394 | 0.3394 | TRUE | 1.0641 |
| Serpinb2 | 1 | 1 | 0 | 0 | 0 | 0 | 1 | 1 | 0 | 0 | 0 | 0 | 0.31 | 1 | 0 | 0.309 | -1.696 | 0.63183309 | NA | 0 | 7.7294 | 7.2472 | 7.2472 | TRUE | 0.2015 |
| Serpinb3a | 0 | 0 | 0 | 0 | 0 | 0 | 0 | 0 | 0 | 0 | 0 | 0 | 0 | 0 | 0 | NA | NA | NA | NA | NA | NA | NA | NA | NA | NA |
| Serpinb3b | 0 | 0 | 0 | 0 | 0 | 0 | 0 | 0 | 0 | 0 | 0 | 0 | 0 | 0 | 0 | NA | NA | NA | NA | NA | NA | NA | NA | NA | NA |
| Serpinb3c | 0 | 0 | 0 | 0 | 0 | 0 | 0 | 0 | 0 | 0 | 0 | 0 | 0 | 0 | 0 | NA | NA | NA | NA | NA | NA | NA | NA | NA | NA |
| Serpinb3d | 0 | 0 | 0 | 0 | 0 | 0 | 0 | 0 | 0 | 0 | 0 | 0 | 0 | 0 | 0 | NA | NA | NA | NA | NA | NA | NA | NA | NA | NA |
| Serpinb5 | 42 | 61 | 80 | 75 | 66 | 61 | 33 | 65 | 86 | 75 | 74 | 60 | 65.53 | 61 | 70 | 1.15 | 0.202 | 0.52474122 | 0.68594533 | 0.1041 | 0.0386 | 0.0573 | 0.0573 | TRUE | 0.4055 |
| Serpinb6a | 3655 | 2539 | 2514 | 2145 | 1822 | 2125 | 2839 | 2701 | 2713 | 2135 | 2055 | 2098 | 2423.37 | 2751 | 2096 | 0.762 | -0.393 | 3.37E-11 | 3.05E-09 | 0 | 0.0034 | 0.0021 | 0.0021 | TRUE | 0.0098 |
| Serpinb6b | 72 | 49 | 42 | 45 | 29 | 34 | 56 | 52 | 45 | 45 | 33 | 34 | 44.07 | 51 | 37 | 0.723 | -0.467 | 0.11634001 | 0.24684301 | 0 | 0.0562 | 0.0405 | 0.0405 | TRUE | 0.2496 |
| Serpinb6c | 121 | 96 | 113 | 110 | 114 | 103 | 94 | 102 | 122 | 109 | 129 | 102 | 109.63 | 106 | 113 | 1.071 | 0.098 | 0.62036565 | 0.76038815 | 0.0073 | 0.024 | 0.0193 | 0.0193 | TRUE | 0.1664 |
| Serpinb6d | 0 | 1 | 0 | 0 | 0 | 0 | 0 | 1 | 0 | 0 | 0 | 0 | 0.18 | 0 | 0 | 0.552 | -0.857 | 0.8336548 | NA | 0 | 13.3714 | 10 | 10 | TRUE | 0.5216 |
| Serpinb6e | 0 | 0 | 0 | 0 | 0 | 0 | 0 | 0 | 0 | 0 | 0 | 0 | 0 | 0 | 0 | NA | NA | NA | NA | NA | NA | NA | NA | NA | NA |
| Serpinb7 | 0 | 0 | 0 | 0 | 0 | 0 | 0 | 0 | 0 | 0 | 0 | 0 | 0 | 0 | 0 | NA | NA | NA | NA | NA | NA | NA | NA | NA | NA |
| Serpinb8 | 15 | 17 | 16 | 15 | 10 | 6 | 12 | 18 | 17 | 15 | 11 | 6 | 13.19 | 16 | 11 | 0.688 | -0.539 | 0.33276101 | NA | 0 | 0.1821 | 0.1434 | 0.1434 | TRUE | 0.4493 |
| Serpinb9 | 52 | 30 | 25 | 27 | 22 | 21 | 40 | 32 | 27 | 27 | 25 | 21 | 28.62 | 33 | 24 | 0.724 | -0.467 | 0.20476356 | 0.36629355 | 0 | 0.0852 | 0.0616 | 0.0616 | TRUE | 0.3024 |
| Serpinb9b | 143 | 94 | 60 | 63 | 62 | 49 | 111 | 100 | 65 | 63 | 70 | 48 | 76.14 | 92 | 60 | 0.652 | -0.618 | 0.01910293 | 0.06680651 | 0.0431 | 0.0336 | 0.0363 | 0.0363 | TRUE | 0.6184 |
| Serpinb9c | 0 | 0 | 0 | 0 | 0 | 0 | 0 | 0 | 0 | 0 | 0 | 0 | 0 | 0 | 0 | NA | NA | NA | NA | NA | NA | NA | NA | NA | NA |
| Serpinb9d | 0 | 0 | 0 | 0 | 0 | 0 | 0 | 0 | 0 | 0 | 0 | 0 | 0 | 0 | 0 | NA | NA | NA | NA | NA | NA | NA | NA | NA | NA |
| Serpinb9e | 0 | 2 | 0 | 0 | 0 | 0 | 0 | 2 | 0 | 0 | 0 | 0 | 0.35 | 1 | 0 | 0.272 | -1.878 | 0.62190613 | NA | 10 | 6.6869 | 8.7633 | 8.7633 | TRUE | 0.9743 |
| Serpinb9f | 0 | 0 | 0 | 0 | 0 | 0 | 0 | 0 | 0 | 0 | 0 | 0 | 0 | 0 | 0 | NA | NA | NA | NA | NA | NA | NA | NA | NA | NA |
| Serpinb9g | 0 | 0 | 0 | 0 | 0 | 0 | 0 | 0 | 0 | 0 | 0 | 0 | 0 | 0 | 0 | NA | NA | NA | NA | NA | NA | NA | NA | NA | NA |
| Serpinc1 | 3 | 0 | 3 | 1 | 2 | 1 | 2 | 0 | 3 | 1 | 2 | 1 | 1.63 | 2 | 1 | 0.748 | -0.419 | 0.78548503 | NA | 0 | 1.4526 | 1.0789 | 1.0789 | TRUE | 0.5779 |
| Serpind1 | 0 | 0 | 0 | 0 | 0 | 0 | 0 | 0 | 0 | 0 | 0 | 0 | 0 | 0 | 0 | NA | NA | NA | NA | NA | NA | NA | NA | NA | NA |
| Serpine1 | 1042 | 825 | 735 | 566 | 463 | 505 | 809 | 878 | 793 | 563 | 522 | 499 | 677.35 | 827 | 528 | 0.639 | -0.646 | 2.67E-12 | 3.01E-10 | 0.0018 | 0.0059 | 0.0046 | 0.0046 | TRUE | 0.0409 |
| Serpine2 | 2400 | 1498 | 1771 | 1442 | 1363 | 1507 | 1864 | 1593 | 1911 | 1435 | 1538 | 1488 | 1638.15 | 1789 | 1487 | 0.83 | -0.268 | 0.00102892 | 0.00738274 | 0.0048 | 0.0039 | 0.0042 | 0.0042 | TRUE | 0.1092 |
| Serpine3 | 6 | 4 | 6 | 3 | 2 | 6 | 5 | 4 | 6 | 3 | 2 | 6 | 4.43 | 5 | 4 | 0.734 | -0.446 | 0.62337003 | NA | 0 | 0.538 | 0.3628 | 0.3628 | TRUE | 0.432 |
| Serpinf1 | 24 | 9 | 12 | 24 | 11 | 10 | 19 | 10 | 13 | 24 | 12 | 10 | 14.56 | 14 | 15 | 1.111 | 0.152 | 0.78132314 | NA | 0.1041 | 0.1653 | 0.1475 | 0.1475 | TRUE | 1.0972 |
| Serpinf2 | 11 | 9 | 5 | 7 | 5 | 8 | 9 | 10 | 5 | 7 | 6 | 8 | 7.34 | 8 | 7 | 0.872 | -0.198 | 0.77639112 | NA | 0 | 0.3255 | 0.2148 | 0.2148 | TRUE | 0.1926 |
| Serping1 | 331 | 241 | 251 | 213 | 173 | 167 | 257 | 256 | 271 | 212 | 195 | 165 | 226.05 | 261 | 191 | 0.729 | -0.455 | 0.00143412 | 0.00934219 | 0.0026 | 0.0129 | 0.0101 | 0.0101 | TRUE | 0.1564 |
| Serpinh1 | 2851 | 1769 | 1870 | 2341 | 1989 | 2248 | 2214 | 1882 | 2018 | 2330 | 2244 | 2219 | 2151.13 | 2038 | 2264 | 1.11 | 0.151 | 0.04042313 | 0.11532292 | 0.003 | 0.0035 | 0.0034 | 0.0034 | TRUE | 0.0712 |
| Serpini1 | 379 | 215 | 234 | 312 | 256 | 261 | 294 | 229 | 252 | 311 | 289 | 258 | 272.09 | 258 | 286 | 1.101 | 0.139 | 0.32453972 | 0.49840401 | 0.0093 | 0.0111 | 0.0106 | 0.0106 | TRUE | 0.1756 |
| Serpini2 | 0 | 0 | 0 | 0 | 0 | 0 | 0 | 0 | 0 | 0 | 0 | 0 | 0 | 0 | 0 | NA | NA | NA | NA | NA | NA | NA | NA | NA | NA |
| Sertad1 | 504 | 355 | 332 | 366 | 290 | 356 | 391 | 378 | 358 | 364 | 327 | 351 | 361.69 | 376 | 347 | 0.925 | -0.113 | 0.31012305 | 0.48301065 | 0 | 0.009 | 0.0062 | 0.0062 | TRUE | 0.0292 |
| Sertad2 | 2369 | 1696 | 1785 | 2054 | 1755 | 1991 | 1840 | 1804 | 1926 | 2045 | 1980 | 1965 | 1926.62 | 1857 | 1997 | 1.076 | 0.105 | 0.09448166 | 0.21206649 | 2.00E-04 | 0.0037 | 0.0023 | 0.0023 | TRUE | 0.0125 |
| Sertad3 | 603 | 442 | 454 | 489 | 443 | 511 | 468 | 470 | 490 | 487 | 500 | 504 | 486.55 | 476 | 497 | 1.044 | 0.063 | 0.51213798 | 0.67496433 | 0 | 0.0073 | 0.0045 | 0.0045 | TRUE | 0.0072 |
| Sertad4 | 273 | 236 | 252 | 342 | 288 | 381 | 212 | 251 | 272 | 340 | 325 | 376 | 296.06 | 245 | 347 | 1.424 | 0.51 | 0.00014334 | 0.00164037 | 0.0073 | 0.0104 | 0.0095 | 0.0095 | TRUE | 0.1681 |
| Sertm1 | 134 | 82 | 90 | 100 | 60 | 60 | 104 | 87 | 97 | 100 | 68 | 59 | 85.81 | 96 | 76 | 0.784 | -0.35 | 0.1435991 | 0.28656173 | 0.0278 | 0.03 | 0.0294 | 0.0294 | TRUE | 0.7271 |
| Sesn1 | 2650 | 1834 | 1653 | 2146 | 1882 | 2050 | 2058 | 1951 | 1784 | 2136 | 2123 | 2024 | 2012.57 | 1931 | 2094 | 1.084 | 0.116 | 0.10650407 | 0.23166096 | 0.0023 | 0.0036 | 0.0032 | 0.0032 | TRUE | 0.0534 |
| Sesn2 | 3102 | 2231 | 2045 | 1954 | 1829 | 1936 | 2409 | 2373 | 2207 | 1945 | 2063 | 1911 | 2151.38 | 2330 | 1973 | 0.846 | -0.241 | 0.00029618 | 0.00287978 | 0.0014 | 0.0035 | 0.0027 | 0.0027 | TRUE | 0.0256 |
| Sesn3 | 986 | 676 | 710 | 702 | 612 | 707 | 766 | 719 | 766 | 699 | 690 | 698 | 723 | 750 | 696 | 0.927 | -0.11 | 0.18641754 | 0.34296124 | 0 | 0.0057 | 0.0036 | 0.0036 | TRUE | 0.0153 |
| Sestd1 | 2122 | 1635 | 1752 | 1860 | 1592 | 1927 | 1648 | 1739 | 1890 | 1851 | 1796 | 1902 | 1804.54 | 1759 | 1850 | 1.053 | 0.074 | 0.30377382 | 0.47628776 | 0.0023 | 0.0037 | 0.0032 | 0.0032 | TRUE | 0.0511 |
| Set | 54069 | 39734 | 35584 | 36332 | 30385 | 36443 | 41993 | 42265 | 38395 | 36165 | 34277 | 35975 | 38178.27 | 40884 | 35472 | 0.868 | -0.205 | 0.00022869 | 0.00237372 | 0.0018 | 0.0025 | 0.0022 | 0.0022 | TRUE | 0.0347 |
| Setbp1 | 210 | 169 | 147 | 148 | 122 | 131 | 163 | 180 | 159 | 147 | 138 | 129 | 152.62 | 167 | 138 | 0.827 | -0.275 | 0.09011053 | 0.2048918 | 0 | 0.018 | 0.0123 | 0.0123 | TRUE | 0.0441 |
| Setd1a | 6510 | 4871 | 4583 | 4156 | 4014 | 4880 | 5056 | 5181 | 4945 | 4137 | 4528 | 4817 | 4777.45 | 5061 | 4494 | 0.888 | -0.171 | 0.00953289 | 0.0399688 | 0.0028 | 0.0029 | 0.0029 | 0.0029 | TRUE | 0.0592 |
| Setd1b | 1126 | 739 | 449 | 505 | 430 | 489 | 875 | 786 | 484 | 503 | 485 | 483 | 602.59 | 715 | 490 | 0.684 | -0.547 | 0.00081652 | 0.00624424 | 0.045 | 0.0064 | 0.0175 | 0.0175 | TRUE | 0.8825 |
| Setd2 | 8127 | 5705 | 5600 | 5490 | 4888 | 5702 | 6312 | 6068 | 6042 | 5465 | 5514 | 5629 | 5838.37 | 6141 | 5536 | 0.901 | -0.15 | 0.00301377 | 0.01659952 | 3.00E-04 | 0.0028 | 0.0017 | 0.0017 | TRUE | 0.0074 |
| Setd3 | 7032 | 5196 | 5560 | 5993 | 5351 | 6245 | 5461 | 5527 | 5999 | 5965 | 6036 | 6165 | 5859.04 | 5662 | 6055 | 1.07 | 0.097 | 0.08813491 | 0.20174764 | 0.0012 | 0.0028 | 0.0022 | 0.0022 | TRUE | 0.0328 |
| Setd4 | 777 | 570 | 521 | 564 | 521 | 592 | 603 | 606 | 562 | 561 | 588 | 584 | 584.24 | 590 | 578 | 0.977 | -0.033 | 0.71715736 | 0.82994953 | 0 | 0.0065 | 0.0042 | 0.0042 | TRUE | 0.0201 |
| Setd5 | 22581 | 16494 | 15433 | 15419 | 14230 | 16136 | 17538 | 17544 | 16652 | 15348 | 16053 | 15929 | 16510.65 | 17245 | 15777 | 0.915 | -0.128 | 0.00906219 | 0.03850355 | 6.00E-04 | 0.0026 | 0.0017 | 0.0017 | TRUE | 0.011 |
| Setd6 | 4934 | 3890 | 4465 | 4868 | 4482 | 5190 | 3832 | 4138 | 4818 | 4846 | 5056 | 5123 | 4635.43 | 4263 | 5008 | 1.176 | 0.233 | 0.00360808 | 0.01904368 | 0.0071 | 0.0029 | 0.0044 | 0.0044 | TRUE | 0.1578 |
| Setd7 | 1952 | 1233 | 1186 | 1250 | 1092 | 1198 | 1516 | 1312 | 1280 | 1244 | 1232 | 1183 | 1294.33 | 1369 | 1220 | 0.889 | -0.169 | 0.04169029 | 0.11783258 | 0.0041 | 0.0043 | 0.0042 | 0.0042 | TRUE | 0.1093 |
| Setd8 | 7510 | 5467 | 5655 | 5844 | 5150 | 6141 | 5833 | 5815 | 6102 | 5817 | 5810 | 6062 | 5906.42 | 5917 | 5896 | 0.997 | -0.005 | 0.92888214 | 0.96081289 | 5.00E-04 | 0.0028 | 0.0018 | 0.0018 | TRUE | 0.0091 |
| Setdb1 | 7550 | 5721 | 5106 | 5046 | 4550 | 5174 | 5864 | 6085 | 5509 | 5023 | 5133 | 5108 | 5453.61 | 5819 | 5088 | 0.874 | -0.194 | 0.00061289 | 0.00505141 | 0.001 | 0.0029 | 0.0021 | 0.0021 | TRUE | 0.0262 |
| Setdb2 | 1114 | 786 | 776 | 757 | 665 | 814 | 865 | 836 | 837 | 754 | 750 | 804 | 807.63 | 846 | 769 | 0.909 | -0.138 | 0.08654173 | 0.19924951 | 0 | 0.0054 | 0.0034 | 0.0034 | TRUE | 0.0184 |
| Setmar | 456 | 335 | 337 | 333 | 304 | 351 | 354 | 356 | 364 | 331 | 343 | 346 | 349.17 | 358 | 340 | 0.951 | -0.073 | 0.5039234 | 0.66805555 | 0 | 0.0092 | 0.0057 | 0.0057 | TRUE | 0.006 |
| Setx | 19405 | 13734 | 13569 | 15349 | 13765 | 15850 | 15071 | 14609 | 14641 | 15278 | 15528 | 15646 | 15128.95 | 14774 | 15484 | 1.048 | 0.068 | 0.13443548 | 0.27321014 | 1.00E-04 | 0.0026 | 0.0014 | 0.0014 | TRUE | 0.0038 |
| Sez6 | 83 | 65 | 67 | 106 | 88 | 75 | 64 | 69 | 72 | 106 | 99 | 74 | 80.79 | 68 | 93 | 1.356 | 0.439 | 0.05494241 | 0.14407141 | 0 | 0.0318 | 0.0251 | 0.0251 | TRUE | 0.3129 |
| Sez6l | 99 | 69 | 78 | 105 | 99 | 117 | 77 | 73 | 84 | 105 | 112 | 115 | 94.36 | 78 | 111 | 1.416 | 0.502 | 0.01239618 | 0.04852877 | 0 | 0.0275 | 0.0181 | 0.0181 | TRUE | 0.0385 |
| Sez6l2 | 62 | 31 | 26 | 21 | 31 | 23 | 48 | 33 | 28 | 21 | 35 | 23 | 31.29 | 36 | 26 | 0.709 | -0.495 | 0.19033071 | 0.34836686 | 0.05 | 0.0782 | 0.0701 | 0.0701 | TRUE | 0.664 |
| Sf1 | 22564 | 16827 | 15228 | 14648 | 13795 | 15756 | 17525 | 17899 | 16431 | 14581 | 15562 | 15554 | 16258.41 | 17285 | 15232 | 0.881 | -0.182 | 0.00100143 | 0.00724482 | 0.0016 | 0.0026 | 0.0022 | 0.0022 | TRUE | 0.0227 |
| Sf3a1 | 11778 | 8854 | 7952 | 7774 | 7102 | 8020 | 9147 | 9418 | 8580 | 7738 | 8012 | 7917 | 8468.75 | 9048 | 7889 | 0.872 | -0.198 | 0.00028988 | 0.00283207 | 0.0011 | 0.0027 | 0.002 | 0.002 | TRUE | 0.0248 |
| Sf3a2 | 4863 | 3459 | 3096 | 2871 | 2646 | 3077 | 3777 | 3679 | 3341 | 2858 | 2985 | 3037 | 3279.49 | 3599 | 2960 | 0.822 | -0.283 | 1.73E-05 | 0.0002982 | 0.002 | 0.0031 | 0.0028 | 0.0028 | TRUE | 0.0476 |
| Sf3a3 | 6540 | 4727 | 4095 | 3905 | 3621 | 4282 | 5079 | 5028 | 4418 | 3887 | 4085 | 4227 | 4454.13 | 4842 | 4066 | 0.84 | -0.252 | 0.00026581 | 0.00265306 | 0.0034 | 0.003 | 0.0032 | 0.0032 | TRUE | 0.0709 |
| Sf3b1 | 32093 | 23941 | 26318 | 27392 | 23581 | 27337 | 24925 | 25466 | 28397 | 27266 | 26602 | 26986 | 26606.92 | 26263 | 26951 | 1.026 | 0.037 | 0.52890916 | 0.68931326 | 0.0022 | 0.0025 | 0.0025 | 0.0025 | TRUE | 0.0618 |
| Sf3b2 | 23194 | 17639 | 17287 | 16955 | 15527 | 18277 | 18014 | 18762 | 18653 | 16877 | 17516 | 18042 | 17977.33 | 18476 | 17478 | 0.946 | -0.08 | 0.10672198 | 0.23203852 | 7.00E-04 | 0.0026 | 0.0017 | 0.0017 | TRUE | 0.0111 |
| Sf3b3 | 35519 | 25060 | 21435 | 20576 | 18709 | 20927 | 27586 | 26656 | 23128 | 20481 | 21106 | 20658 | 23269.25 | 25790 | 20748 | 0.804 | -0.314 | 3.50E-06 | 7.87E-05 | 0.0042 | 0.0025 | 0.0033 | 0.0033 | TRUE | 0.0997 |

| | | | | | | | | | | | | | | | | | | | | | | | | | |
|---|---|---|---|---|---|---|---|---|---|---|---|---|---|---|---|---|---|---|---|---|---|---|---|---|---|
| Sf3b4 | 4870 | 3396 | 2647 | 2621 | 2414 | 2756 | 3782 | 3612 | 2856 | 2609 | 2723 | 2721 | 3050.58 | 3417 | 2684 | 0.785 | -0.349 | 0.00016609 | 0.00184881 | 0.0102 | 0.0032 | 0.0059 | 0.0059 | TRUE | 0.2494 |
| Sf3b5 | 4370 | 3239 | 3255 | 3248 | 2920 | 3441 | 3394 | 3445 | 3512 | 3233 | 3294 | 3397 | 3379.22 | 3450 | 3308 | 0.959 | -0.06 | 0.27067835 | 0.44170151 | 2.00E-04 | 0.0031 | 0.0019 | 0.0019 | TRUE | 0.0068 |
| Sfi1 | 290 | 288 | 247 | 195 | 194 | 252 | 225 | 306 | 267 | 194 | 219 | 249 | 243.3 | 266 | 221 | 0.832 | -0.265 | 0.08866761 | 0.20270733 | 0.0159 | 0.0122 | 0.0133 | 0.0133 | TRUE | 0.1736 |
| Sfmbt1 | 2382 | 1609 | 1478 | 1498 | 1200 | 1479 | 1850 | 1711 | 1595 | 1491 | 1354 | 1460 | 1576.84 | 1719 | 1435 | 0.834 | -0.261 | 0.0008337 | 0.00633834 | 0.0035 | 0.0039 | 0.0038 | 0.0038 | TRUE | 0.0554 |
| Sfmbt2 | 1687 | 1332 | 1205 | 656 | 577 | 637 | 1310 | 1417 | 1300 | 653 | 651 | 629 | 993.33 | 1342 | 644 | 0.48 | -1.059 | 5.08E-41 | 9.07E-38 | 0 | 0.0048 | 0.0033 | 0.0033 | TRUE | 0.0277 |
| Sfn | 2435 | 1708 | 1524 | 1165 | 998 | 1157 | 1891 | 1817 | 1644 | 1160 | 1126 | 1142 | 1463.32 | 1784 | 1143 | 0.64 | -0.644 | 1.31E-17 | 2.97E-15 | 0.0023 | 0.004 | 0.0034 | 0.0034 | TRUE | 0.0561 |
| Sfpi1 | 30 | 24 | 24 | 17 | 14 | 24 | 23 | 26 | 26 | 17 | 16 | 24 | 21.86 | 25 | 19 | 0.759 | -0.398 | 0.33425425 | 0.50829832 | 0 | 0.1109 | 0.0755 | 0.0755 | TRUE | 0.2772 |
| Sfpq | 31502 | 24120 | 19215 | 18150 | 16470 | 18138 | 24466 | 25656 | 20733 | 18066 | 18580 | 17905 | 20901.1 | 23618 | 18184 | 0.77 | -0.377 | 3.88E-07 | 1.31E-05 | 0.006 | 0.0025 | 0.0039 | 0.0039 | TRUE | 0.1396 |
| Sfr1 | 10760 | 7366 | 7173 | 7897 | 6687 | 7813 | 8357 | 7835 | 7740 | 7861 | 7544 | 7713 | 7841.42 | 7977 | 7706 | 0.966 | -0.05 | 0.34845786 | 0.52310768 | 9.00E-04 | 0.0027 | 0.0019 | 0.0019 | TRUE | 0.0214 |
| Sfrp1 | 3446 | 2797 | 3210 | 3634 | 3388 | 3918 | 2676 | 2975 | 3464 | 3617 | 3822 | 3868 | 3403.67 | 3038 | 3769 | 1.241 | 0.312 | 0.0003043 | 0.00294076 | 0.0087 | 0.0031 | 0.0051 | 0.0051 | TRUE | 0.1818 |
| Sfrp2 | 1136 | 915 | 924 | 1114 | 936 | 1083 | 882 | 973 | 997 | 1109 | 1056 | 1069 | 1014.4 | 951 | 1078 | 1.136 | 0.184 | 0.02159933 | 0.07306965 | 0.0015 | 0.0048 | 0.0036 | 0.0036 | TRUE | 0.0501 |
| Sfrp4 | 308 | 233 | 208 | 235 | 220 | 278 | 239 | 248 | 224 | 234 | 248 | 274 | 244.67 | 237 | 252 | 1.063 | 0.088 | 0.50669232 | 0.67002033 | 0 | 0.0121 | 0.0087 | 0.0087 | TRUE | 0.0681 |
| Sfrp5 | 231 | 136 | 172 | 197 | 187 | 224 | 179 | 145 | 186 | 196 | 211 | 221 | 189.64 | 170 | 209 | 1.23 | 0.299 | 0.05326947 | 0.14099638 | 0.0041 | 0.0149 | 0.0119 | 0.0119 | TRUE | 0.1711 |
| Sfrs18 | 4549 | 3780 | 3517 | 3452 | 2884 | 3474 | 3533 | 4021 | 3795 | 3436 | 3253 | 3429 | 3577.92 | 3783 | 3373 | 0.892 | -0.164 | 0.01177653 | 0.04667831 | 0.0021 | 0.0031 | 0.0028 | 0.0028 | TRUE | 0.0414 |
| Sfswap | 2901 | 2240 | 1873 | 1995 | 1765 | 2073 | 2253 | 2383 | 2021 | 1986 | 1991 | 2046 | 2113.33 | 2219 | 2008 | 0.905 | -0.145 | 0.04842213 | 0.13162753 | 0.0028 | 0.0035 | 0.0034 | 0.0034 | TRUE | 0.0722 |
| Sft2d1 | 1070 | 714 | 705 | 782 | 724 | 751 | 831 | 759 | 761 | 778 | 817 | 741 | 781.28 | 784 | 779 | 0.992 | -0.012 | 0.88990613 | 0.93898039 | 0.0014 | 0.0055 | 0.0041 | 0.0041 | TRUE | 0.0348 |
| Sft2d2 | 2088 | 1554 | 1724 | 2060 | 1812 | 1991 | 1622 | 1653 | 1860 | 2051 | 2044 | 1965 | 1865.81 | 1712 | 2020 | 1.181 | 0.24 | 0.00092623 | 0.0068594 | 0.0023 | 0.0037 | 0.0032 | 0.0032 | TRUE | 0.0687 |
| Sft2d3 | 266 | 226 | 221 | 239 | 189 | 231 | 207 | 240 | 238 | 238 | 213 | 228 | 227.43 | 228 | 226 | 0.995 | -0.008 | 0.95593312 | 0.97573035 | 0 | 0.0128 | 0.0094 | 0.0094 | TRUE | 0.085 |
| Sfta2 | 1 | 2 | 0 | 1 | 1 | 0 | 1 | 2 | 0 | 1 | 1 | 0 | 0.84 | 1 | 1 | 0.726 | -0.461 | 0.83029328 | NA | 0 | 2.8309 | 2.0932 | 2.0932 | TRUE | 0.3091 |
| Sftpa1 | 0 | 0 | 0 | 0 | 0 | 0 | 0 | 0 | 0 | 0 | 0 | 0 | 0 | 0 | 0 | NA | NA | NA | NA | NA | NA | NA | NA | NA | NA |
| Sftpb | 2 | 0 | 0 | 4 | 0 | 1 | 2 | 0 | 0 | 4 | 0 | 1 | 1.09 | 1 | 2 | 2.951 | 1.561 | 0.46599016 | NA | 1.9403 | 2.1828 | 2.1442 | 2.1442 | TRUE | 0.6426 |
| Sftpc | 0 | 0 | 0 | 0 | 0 | 0 | 0 | 0 | 0 | 0 | 0 | 0 | 0 | 0 | 0 | NA | NA | NA | NA | NA | NA | NA | NA | NA | NA |
| Sftpd | 8 | 2 | 3 | 3 | 1 | 5 | 6 | 2 | 3 | 3 | 1 | 5 | 3.44 | 4 | 3 | 0.775 | -0.369 | 0.72696858 | NA | 0 | 0.6918 | 0.5109 | 0.5109 | TRUE | 0.2851 |
| Sfxn1 | 4473 | 3297 | 3648 | 4356 | 3804 | 4645 | 3474 | 3507 | 3936 | 4336 | 4291 | 4585 | 4021.62 | 3639 | 4404 | 1.211 | 0.276 | 3.29E-05 | 0.00050283 | 0.0028 | 0.003 | 0.0029 | 0.0029 | TRUE | 0.0616 |
| Sfxn2 | 866 | 651 | 632 | 584 | 514 | 585 | 673 | 692 | 682 | 581 | 580 | 577 | 630.94 | 682 | 579 | 0.85 | -0.235 | 0.00614431 | 0.02872773 | 0 | 0.0062 | 0.0037 | 0.0037 | TRUE | 0.002 |
| Sfxn3 | 502 | 441 | 480 | 634 | 556 | 660 | 390 | 469 | 518 | 631 | 627 | 652 | 547.79 | 459 | 637 | 1.392 | 0.477 | 2.22E-05 | 0.0003635 | 0.0084 | 0.0068 | 0.0073 | 0.0073 | TRUE | 0.2146 |
| Sfxn4 | 46 | 40 | 17 | 29 | 27 | 21 | 36 | 43 | 18 | 29 | 30 | 21 | 29.45 | 32 | 27 | 0.824 | -0.28 | 0.48175295 | 0.64849796 | 0.07 | 0.0829 | 0.0794 | 0.0794 | TRUE | 0.6058 |
| Sfxn5 | 232 | 165 | 189 | 180 | 173 | 206 | 180 | 176 | 204 | 179 | 195 | 203 | 189.55 | 187 | 192 | 1.033 | 0.048 | 0.74930137 | 0.8514003 | 0 | 0.0149 | 0.0106 | 0.0106 | TRUE | 0.0681 |
| Sgca | 5 | 2 | 0 | 6 | 2 | 5 | 4 | 2 | 0 | 6 | 2 | 5 | 3.2 | 2 | 4 | 2.11 | 1.077 | 0.35032466 | NA | 0.0954 | 0.744 | 0.6112 | 0.6112 | TRUE | 0.489 |
| Sgcb | 1432 | 1203 | 1222 | 988 | 950 | 1115 | 1112 | 1280 | 1319 | 983 | 1072 | 1101 | 1144.36 | 1237 | 1052 | 0.852 | -0.232 | 0.00854925 | 0.03682775 | 0.0051 | 0.0045 | 0.0047 | 0.0047 | TRUE | 0.0967 |
| Sgcd | 31 | 20 | 39 | 37 | 27 | 22 | 24 | 21 | 42 | 37 | 30 | 22 | 29.41 | 29 | 30 | 1.024 | 0.034 | 0.93258294 | 0.96255113 | 0.0685 | 0.083 | 0.079 | 0.079 | TRUE | 0.7256 |
| Sgce | 580 | 329 | 462 | 663 | 507 | 588 | 450 | 350 | 498 | 660 | 572 | 580 | 518.54 | 433 | 604 | 1.395 | 0.48 | 0.00017079 | 0.00188831 | 0.0156 | 0.007 | 0.0098 | 0.0098 | TRUE | 0.315 |
| Sgcg | 3 | 3 | 2 | 1 | 2 | 0 | 2 | 3 | 2 | 1 | 2 | 0 | 1.82 | 2 | 1 | 0.414 | -1.273 | 0.39036136 | NA | 0 | 1.3034 | 0.9028 | 0.9028 | TRUE | 0.3962 |
| Sgcz | 17 | 8 | 2 | 1 | 1 | 0 | 13 | 9 | 2 | 1 | 1 | 0 | 4.33 | 8 | 1 | 0.086 | -3.539 | 0.00889904 | NA | 0 | 0.5495 | 0.5137 | 0.5137 | TRUE | 0.1121 |
| Sgip1 | 10 | 10 | 9 | 5 | 8 | 6 | 8 | 11 | 10 | 5 | 9 | 6 | 8.01 | 10 | 7 | 0.708 | -0.498 | 0.46150431 | NA | 0 | 0.2984 | 0.1989 | 0.1989 | TRUE | 0.223 |
| Sgk1 | 14489 | 9938 | 9605 | 8972 | 8078 | 8881 | 11253 | 10571 | 10364 | 8931 | 9113 | 8767 | 9833.01 | 10729 | 8937 | 0.833 | -0.264 | 7.04E-07 | 2.18E-05 | 0.001 | 0.0027 | 0.0019 | 0.0019 | TRUE | 0.0225 |
| Sgk2 | 134 | 75 | 67 | 53 | 62 | 65 | 104 | 80 | 72 | 53 | 70 | 64 | 73.83 | 85 | 62 | 0.724 | -0.467 | 0.0575525 | 0.14921056 | 0.0177 | 0.0345 | 0.0296 | 0.0296 | TRUE | 0.3825 |
| Sgk3 | 1205 | 909 | 1073 | 1227 | 981 | 1103 | 936 | 967 | 1158 | 1221 | 1107 | 1089 | 1079.56 | 1020 | 1139 | 1.118 | 0.161 | 0.09202141 | 0.20816554 | 0.0075 | 0.0046 | 0.0056 | 0.0056 | TRUE | 0.1644 |
| Sgms1 | 1780 | 1285 | 1428 | 1394 | 1360 | 1576 | 1382 | 1367 | 1541 | 1388 | 1534 | 1556 | 1461.28 | 1430 | 1493 | 1.044 | 0.062 | 0.43118865 | 0.60332642 | 0.0031 | 0.004 | 0.0038 | 0.0038 | TRUE | 0.0542 |
| Sgms2 | 410 | 321 | 330 | 399 | 321 | 365 | 318 | 341 | 356 | 397 | 362 | 360 | 355.92 | 338 | 373 | 1.105 | 0.143 | 0.20564569 | 0.36745766 | 0 | 0.0091 | 0.0064 | 0.0064 | TRUE | 0.0367 |
| Sgol1 | 3760 | 2675 | 2827 | 2973 | 2465 | 2861 | 2920 | 2845 | 3050 | 2959 | 2781 | 2824 | 2896.71 | 2938 | 2855 | 0.972 | -0.041 | 0.49014411 | 0.65595785 | 6.00E-04 | 0.0032 | 0.0022 | 0.0022 | TRUE | 0.0131 |
| Sgol2 | 10163 | 7560 | 8132 | 7937 | 7129 | 8156 | 7893 | 8041 | 8774 | 7900 | 8042 | 8051 | 8117.16 | 8236 | 7998 | 0.971 | -0.042 | 0.4605483 | 0.63015669 | 0.0015 | 0.0027 | 0.0022 | 0.0022 | TRUE | 0.0397 |
| Sgpl1 | 6183 | 4433 | 4938 | 5498 | 4926 | 5489 | 4802 | 4715 | 5328 | 5473 | 5557 | 5418 | 5215.62 | 4948 | 5483 | 1.108 | 0.148 | 0.01617033 | 0.05905022 | 0.002 | 0.0029 | 0.0025 | 0.0025 | TRUE | 0.0544 |
| Sgpp1 | 1843 | 1347 | 1421 | 1366 | 1126 | 1264 | 1431 | 1433 | 1533 | 1360 | 1270 | 1248 | 1379.19 | 1466 | 1293 | 0.882 | -0.181 | 0.01240844 | 0.0485343 | 0.0011 | 0.0041 | 0.003 | 0.003 | TRUE | 0.0251 |
| Sgpp2 | 175 | 178 | 233 | 206 | 208 | 173 | 136 | 189 | 251 | 205 | 235 | 171 | 197.86 | 192 | 204 | 1.063 | 0.088 | 0.67158877 | 0.79792338 | 0.0538 | 0.0144 | 0.0259 | 0.0259 | TRUE | 0.2397 |
| Sgsh | 375 | 275 | 237 | 301 | 260 | 293 | 291 | 293 | 256 | 300 | 293 | 289 | 286.94 | 280 | 294 | 1.049 | 0.069 | 0.57114647 | 0.72441179 | 0 | 0.0107 | 0.0074 | 0.0074 | TRUE | 0.0609 |
| Sgsm1 | 1473 | 1146 | 1260 | 1427 | 1446 | 1614 | 1144 | 1219 | 1360 | 1420 | 1631 | 1593 | 1394.58 | 1241 | 1548 | 1.249 | 0.321 | 0.00022203 | 0.00231301 | 0.0054 | 0.0041 | 0.0047 | 0.0047 | TRUE | 0.0836 |
| Sgsm2 | 3552 | 2731 | 2718 | 3168 | 2851 | 3165 | 2759 | 2905 | 2933 | 3153 | 3216 | 3124 | 3015.05 | 2866 | 3164 | 1.105 | 0.144 | 0.01172695 | 0.04650756 | 4.00E-04 | 0.0032 | 0.002 | 0.002 | TRUE | 0.0133 |
| Sgsm3 | 2707 | 2099 | 1882 | 2214 | 1868 | 2317 | 2102 | 2233 | 2031 | 2204 | 2107 | 2287 | 2160.68 | 2122 | 2199 | 1.037 | 0.052 | 0.4329048 | 0.60483953 | 0.0013 | 0.0035 | 0.0027 | 0.0027 | TRUE | 0.0247 |
| Sgta | 6433 | 4542 | 4270 | 4489 | 4039 | 4779 | 4996 | 4831 | 4607 | 4468 | 4556 | 4718 | 4696.19 | 4811 | 4581 | 0.952 | -0.071 | 0.20922613 | 0.37093389 | 9.00E-04 | 0.0029 | 0.0021 | 0.0021 | TRUE | 0.0168 |
| Sgtb | 913 | 730 | 868 | 993 | 890 | 1074 | 709 | 776 | 937 | 988 | 1004 | 1060 | 912.47 | 807 | 1017 | 1.263 | 0.337 | 0.00106936 | 0.00759282 | 0.0095 | 0.005 | 0.0065 | 0.0065 | TRUE | 0.231 |
| Sh2b1 | 3099 | 2295 | 2194 | 2216 | 2021 | 2304 | 2407 | 2441 | 2367 | 2206 | 2280 | 2274 | 2329.24 | 2405 | 2253 | 0.937 | -0.094 | 0.10439335 | 0.22842838 | 0 | 0.0034 | 0.002 | 0.002 | TRUE | 0.0042 |
| Sh2b2 | 331 | 307 | 312 | 367 | 348 | 443 | 257 | 327 | 337 | 365 | 393 | 437 | 352.58 | 307 | 398 | 1.304 | 0.383 | 0.0041905 | 0.02140629 | 0.012 | 0.0091 | 0.01 | 0.01 | TRUE | 0.2447 |
| Sh2b3 | 1332 | 1128 | 1100 | 1277 | 1143 | 1339 | 1035 | 1200 | 1187 | 1271 | 1289 | 1322 | 1217.26 | 1141 | 1294 | 1.137 | 0.185 | 0.02127814 | 0.07231489 | 0.0028 | 0.0044 | 0.0038 | 0.0038 | TRUE | 0.0828 |
| Sh2d1a | 9 | 1 | 1 | 2 | 2 | 3 | 7 | 1 | 1 | 2 | 2 | 3 | 2.72 | 3 | 2 | 0.759 | -0.398 | 0.74423365 | NA | 0.3615 | 0.8726 | 0.7027 | 0.7027 | TRUE | 2.1683 |
| Sh2d1b1 | 0 | 0 | 0 | 0 | 0 | 0 | 0 | 0 | 0 | 0 | 0 | 0 | 0 | 0 | 0 | NA | NA | NA | NA | NA | NA | NA | NA | NA | NA |
| Sh2d1b2 | 0 | 0 | 0 | 0 | 0 | 0 | 0 | 0 | 0 | 0 | 0 | 0 | 0 | 0 | 0 | NA | NA | NA | NA | NA | NA | NA | NA | NA | NA |
| Sh2d2a | 5 | 3 | 3 | 5 | 4 | 3 | 4 | 3 | 3 | 5 | 5 | 3 | 3.79 | 3 | 4 | 1.199 | 0.261 | 0.78402088 | NA | 0 | 0.6272 | 0.3905 | 0.3905 | TRUE | 0.1137 |
| Sh2d3c | 511 | 311 | 340 | 429 | 336 | 390 | 397 | 331 | 367 | 427 | 379 | 385 | 380.93 | 365 | 397 | 1.086 | 0.119 | 0.30601677 | 0.47857664 | 0.0038 | 0.0086 | 0.0071 | 0.0071 | TRUE | 0.0756 |
| Sh2d4a | 207 | 164 | 145 | 131 | 121 | 136 | 161 | 174 | 156 | 130 | 137 | 134 | 148.8 | 164 | 134 | 0.816 | -0.293 | 0.06901736 | 0.16981668 | 0 | 0.0184 | 0.0119 | 0.0119 | TRUE | 0.032 |
| Sh2d4b | 21 | 23 | 20 | 13 | 10 | 17 | 16 | 24 | 22 | 13 | 11 | 17 | 17.23 | 21 | 14 | 0.665 | -0.589 | 0.21037445 | 0.37232317 | 0 | 0.14 | 0.0984 | 0.0984 | TRUE | 0.2428 |
| Sh2d5 | 287 | 277 | 324 | 417 | 327 | 385 | 223 | 295 | 350 | 415 | 369 | 380 | 338.53 | 289 | 388 | 1.349 | 0.432 | 0.00452729 | 0.02276066 | 0.0238 | 0.0094 | 0.0137 | 0.0137 | TRUE | 0.3854 |
| Sh2d7 | 5 | 4 | 0 | 2 | 1 | 3 | 4 | 4 | 0 | 2 | 1 | 3 | 2.37 | 3 | 2 | 0.742 | -0.431 | 0.7434633 | NA | 0 | 1.0026 | 0.8189 | 0.8189 | TRUE | 0.8114 |
| Sh3bgr | 19 | 11 | 11 | 10 | 6 | 13 | 15 | 12 | 12 | 10 | 7 | 13 | 11.31 | 13 | 10 | 0.771 | -0.374 | 0.51197825 | NA | 0 | 0.2119 | 0.1452 | 0.1452 | TRUE | 0.2369 |
| Sh3bgrl | 2151 | 1433 | 1351 | 1442 | 1161 | 1246 | 1671 | 1524 | 1458 | 1435 | 1310 | 1230 | 1437.94 | 1551 | 1325 | 0.854 | -0.228 | 0.0063178 | 0.02932781 | 0.0045 | 0.0041 | 0.0043 | 0.0043 | TRUE | 0.0645 |
| Sh3bgrl2 | 5140 | 3644 | 3356 | 3531 | 3051 | 3333 | 3992 | 3876 | 3621 | 3515 | 3442 | 3290 | 3622.66 | 3830 | 3416 | 0.891 | -0.166 | 0.00715397 | 0.03226622 | 0.0014 | 0.0031 | 0.0025 | 0.0025 | TRUE | 0.0275 |
| Sh3bgrl3 | 2173 | 1529 | 1529 | 1704 | 1496 | 1709 | 1688 | 1626 | 1650 | 1696 | 1688 | 1687 | 1672.45 | 1655 | 1690 | 1.021 | 0.03 | 0.62886496 | 0.76633702 | 0 | 0.0038 | 0.0022 | 0.0022 | TRUE | 0.0039 |
| Sh3bp1 | 952 | 820 | 823 | 759 | 779 | 883 | 739 | 872 | 888 | 756 | 879 | 872 | 834.26 | 833 | 836 | 1.004 | 0.006 | 0.95313308 | 0.97419416 | 0.0075 | 0.0053 | 0.006 | 0.006 | TRUE | 0.1205 |
| Sh3bp2 | 300 | 270 | 220 | 224 | 212 | 226 | 233 | 287 | 237 | 223 | 239 | 223 | 240.47 | 252 | 228 | 0.906 | -0.143 | 0.30534026 | 0.47795733 | 0.004 | 0.0123 | 0.0098 | 0.0098 | TRUE | 0.1567 |

| | | | | | | | | | | | | | | | | | | | | | | | | | |
|---|---|---|---|---|---|---|---|---|---|---|---|---|---|---|---|---|---|---|---|---|---|---|---|---|---|
| Sh3bp4 | 2456 | 1829 | 1813 | 1617 | 1425 | 1660 | 1907 | 1945 | 1956 | 1610 | 1608 | 1639 | 1777.49 | 1936 | 1619 | 0.836 | -0.258 | 2.60E-05 | 0.00041188 | 0 | 0.0038 | 0.0021 | 0.0021 | TRUE | 0.0022 |
| Sh3bp5 | 529 | 405 | 423 | 484 | 441 | 488 | 411 | 431 | 456 | 482 | 497 | 482 | 459.84 | 433 | 487 | 1.127 | 0.173 | 0.08437345 | 0.19583463 | 0 | 0.0076 | 0.005 | 0.005 | TRUE | 0.0264 |
| Sh3bp5l | 3424 | 2434 | 2201 | 2208 | 1949 | 2277 | 2659 | 2589 | 2375 | 2198 | 2199 | 2248 | 2377.9 | 2541 | 2215 | 0.871 | -0.199 | 0.00228528 | 0.01338122 | 0.0015 | 0.0034 | 0.0026 | 0.0026 | TRUE | 0.0393 |
| Sh3d19 | 3193 | 2300 | 2469 | 2711 | 2619 | 2920 | 2480 | 2446 | 2664 | 2699 | 2955 | 2882 | 2687.65 | 2530 | 2845 | 1.125 | 0.169 | 0.00945346 | 0.03976408 | 0.0016 | 0.0033 | 0.0027 | 0.0027 | TRUE | 0.0256 |
| Sh3d21 | 167 | 117 | 108 | 95 | 98 | 119 | 130 | 124 | 117 | 95 | 111 | 117 | 115.55 | 124 | 108 | 0.868 | -0.204 | 0.27402343 | 0.4450701 | 0 | 0.0229 | 0.0162 | 0.0162 | TRUE | 0.1134 |
| Sh3gl1 | 4367 | 3129 | 2708 | 2849 | 2544 | 2744 | 3392 | 3328 | 2922 | 2836 | 2870 | 2709 | 3009.4 | 3214 | 2805 | 0.872 | -0.197 | 0.00535283 | 0.0258124 | 0.0031 | 0.0032 | 0.0033 | 0.0033 | TRUE | 0.0761 |
| Sh3gl2 | 1620 | 1283 | 1496 | 1597 | 1404 | 1595 | 1258 | 1365 | 1614 | 1590 | 1584 | 1575 | 1497.51 | 1412 | 1583 | 1.122 | 0.166 | 0.06572489 | 0.1637818 | 0.0074 | 0.004 | 0.0052 | 0.0052 | TRUE | 0.1871 |
| Sh3gl3 | 882 | 593 | 598 | 587 | 565 | 650 | 685 | 631 | 645 | 584 | 637 | 642 | 637.39 | 654 | 621 | 0.949 | -0.076 | 0.40348705 | 0.57641583 | 0 | 0.0061 | 0.0044 | 0.0044 | TRUE | 0.0321 |
| Sh3glb1 | 4499 | 3281 | 3114 | 3354 | 2958 | 3392 | 3494 | 3490 | 3360 | 3339 | 3337 | 3348 | 3394.67 | 3448 | 3341 | 0.969 | -0.046 | 0.39405679 | 0.56765306 | 0 | 0.0031 | 0.0018 | 0.0018 | TRUE | 0.006 |
| Sh3glb2 | 1356 | 1066 | 908 | 871 | 875 | 1022 | 1053 | 1134 | 980 | 867 | 987 | 1009 | 1004.95 | 1056 | 954 | 0.904 | -0.146 | 0.10451075 | 0.22844046 | 0.0049 | 0.0048 | 0.0048 | 0.0048 | TRUE | 0.0773 |
| Sh3kbp1 | 485 | 357 | 301 | 349 | 301 | 347 | 377 | 380 | 325 | 347 | 340 | 343 | 351.78 | 361 | 343 | 0.951 | -0.073 | 0.527777 | 0.68871694 | 0 | 0.0092 | 0.0067 | 0.0067 | TRUE | 0.082 |
| Sh3pxd2a | 2803 | 2073 | 2202 | 2658 | 2523 | 2604 | 2177 | 2205 | 2376 | 2646 | 2846 | 2571 | 2470.08 | 2253 | 2688 | 1.193 | 0.255 | 0.00016401 | 0.00182994 | 0.0019 | 0.0034 | 0.0029 | 0.0029 | TRUE | 0.032 |
| Sh3pxd2b | 1953 | 1301 | 1330 | 1438 | 1369 | 1535 | 1517 | 1384 | 1435 | 1431 | 1544 | 1515 | 1471.13 | 1445 | 1497 | 1.035 | 0.049 | 0.49164735 | 0.65729264 | 0.0012 | 0.004 | 0.003 | 0.003 | TRUE | 0.0235 |
| Sh3rf1 | 2171 | 1484 | 1436 | 1543 | 1351 | 1411 | 1686 | 1579 | 1549 | 1536 | 1524 | 1393 | 1544.49 | 1605 | 1484 | 0.924 | -0.114 | 0.11766706 | 0.24886287 | 0.0019 | 0.004 | 0.0032 | 0.0032 | TRUE | 0.0353 |
| Sh3rf2 | 49 | 23 | 35 | 11 | 17 | 26 | 38 | 24 | 38 | 11 | 19 | 26 | 26.01 | 33 | 19 | 0.554 | -0.851 | 0.04304802 | 0.12081002 | 0.0602 | 0.0935 | 0.0852 | 0.0852 | TRUE | 0.4605 |
| Sh3rf3 | 128 | 104 | 85 | 113 | 115 | 113 | 99 | 111 | 92 | 112 | 130 | 112 | 109.25 | 101 | 118 | 1.171 | 0.228 | 0.23280457 | 0.39888224 | 0 | 0.0241 | 0.017 | 0.017 | TRUE | 0.0739 |
| Sh3tc1 | 584 | 416 | 463 | 486 | 450 | 462 | 454 | 442 | 500 | 484 | 508 | 456 | 473.85 | 465 | 483 | 1.037 | 0.053 | 0.60742059 | 0.75128613 | 0.0012 | 0.0074 | 0.0055 | 0.0055 | TRUE | 0.0465 |
| Sh3tc2 | 175 | 133 | 105 | 130 | 103 | 127 | 136 | 141 | 113 | 129 | 116 | 125 | 126.94 | 130 | 123 | 0.949 | -0.076 | 0.67225699 | 0.79851817 | 0 | 0.0211 | 0.0152 | 0.0152 | TRUE | 0.1247 |
| Sh3yl1 | 352 | 245 | 214 | 187 | 184 | 197 | 273 | 261 | 231 | 186 | 208 | 194 | 225.51 | 255 | 196 | 0.767 | -0.384 | 0.00594032 | 0.02803995 | 0 | 0.0129 | 0.0095 | 0.0095 | TRUE | 0.0739 |
| Shank1 | 77 | 76 | 40 | 76 | 71 | 61 | 60 | 81 | 43 | 76 | 80 | 60 | 66.63 | 61 | 72 | 1.174 | 0.231 | 0.39717337 | 0.5705899 | 0.04 | 0.038 | 0.0385 | 0.0385 | TRUE | 0.2796 |
| Shank2 | 93 | 64 | 47 | 70 | 60 | 56 | 72 | 68 | 51 | 70 | 68 | 55 | 63.94 | 64 | 64 | 1.003 | 0.004 | 0.9875376 | 0.99373352 | 0 | 0.0395 | 0.0311 | 0.0311 | TRUE | 0.2634 |
| Shank3 | 617 | 501 | 467 | 587 | 570 | 637 | 479 | 533 | 504 | 584 | 643 | 629 | 562.02 | 505 | 619 | 1.226 | 0.294 | 0.00214974 | 0.01275006 | 0 | 0.0066 | 0.0048 | 0.0048 | TRUE | 0.0281 |
| Sharpin | 1332 | 1049 | 983 | 971 | 968 | 1052 | 1035 | 1116 | 1061 | 967 | 1092 | 1038 | 1051.33 | 1071 | 1032 | 0.965 | -0.052 | 0.5143798 | 0.6770449 | 0.0016 | 0.0047 | 0.0036 | 0.0036 | TRUE | 0.0373 |
| Shb | 810 | 667 | 702 | 656 | 609 | 707 | 629 | 709 | 757 | 653 | 687 | 698 | 688.99 | 698 | 679 | 0.974 | -0.037 | 0.69594211 | 0.81484692 | 0.0038 | 0.0059 | 0.0052 | 0.0052 | TRUE | 0.0948 |
| Shbg | 6 | 7 | 2 | 1 | 0 | 1 | 5 | 7 | 2 | 1 | 0 | 1 | 2.71 | 5 | 1 | 0.143 | -2.805 | 0.05311531 | NA | 0 | 0.8777 | 0.6542 | 0.6542 | TRUE | 0.1097 |
| Shc1 | 4642 | 3288 | 2960 | 3516 | 3118 | 3527 | 3605 | 3497 | 3194 | 3500 | 3517 | 3482 | 3465.9 | 3432 | 3500 | 1.019 | 0.027 | 0.66328419 | 0.79227151 | 0.0017 | 0.0031 | 0.0025 | 0.0025 | TRUE | 0.0445 |
| Shc2 | 448 | 347 | 274 | 341 | 306 | 321 | 348 | 369 | 296 | 339 | 345 | 317 | 335.7 | 338 | 334 | 0.988 | -0.018 | 0.88296658 | 0.93490194 | 0.0041 | 0.0095 | 0.0079 | 0.0079 | TRUE | 0.1279 |
| Shc3 | 25 | 15 | 11 | 8 | 3 | 12 | 19 | 16 | 12 | 8 | 3 | 12 | 11.74 | 16 | 8 | 0.493 | -1.02 | 0.09168572 | NA | 0.0396 | 0.2043 | 0.1669 | 0.1669 | TRUE | 0.3955 |
| Shc4 | 30 | 19 | 12 | 18 | 13 | 23 | 23 | 20 | 13 | 18 | 15 | 23 | 18.62 | 19 | 19 | 0.974 | -0.038 | 0.93364718 | 0.96302394 | 0 | 0.1297 | 0.0979 | 0.0979 | TRUE | 0.3626 |
| Shcbp1 | 4306 | 3158 | 2943 | 2995 | 2572 | 2888 | 3344 | 3359 | 3175 | 2981 | 2901 | 2851 | 3102.08 | 3293 | 2911 | 0.884 | -0.178 | 0.00188924 | 0.01157726 | 4.00E-04 | 0.0032 | 0.002 | 0.002 | TRUE | 0.0116 |
| Shcbp1l | 141 | 130 | 97 | 165 | 113 | 128 | 110 | 138 | 105 | 164 | 127 | 126 | 128.42 | 118 | 139 | 1.19 | 0.25 | 0.19591366 | 0.35547397 | 0.015 | 0.0209 | 0.0192 | 0.0192 | TRUE | 0.2561 |
| Shd | 13 | 9 | 8 | 17 | 13 | 14 | 10 | 10 | 9 | 17 | 15 | 14 | 12.28 | 10 | 15 | 1.599 | 0.678 | 0.20926181 | NA | 0 | 0.1954 | 0.1248 | 0.1248 | TRUE | 0.0504 |
| She | 304 | 254 | 222 | 307 | 237 | 299 | 236 | 270 | 240 | 306 | 267 | 295 | 268.99 | 249 | 289 | 1.167 | 0.223 | 0.08484169 | 0.19652158 | 0 | 0.0112 | 0.0083 | 0.0083 | TRUE | 0.0623 |
| Shf | 290 | 245 | 207 | 256 | 232 | 316 | 225 | 261 | 223 | 255 | 262 | 312 | 256.28 | 236 | 276 | 1.171 | 0.227 | 0.10269844 | 0.22551373 | 0.0063 | 0.0117 | 0.01 | 0.01 | TRUE | 0.1465 |
| Shfm1 | 4005 | 2782 | 3017 | 3339 | 2748 | 3228 | 3111 | 2959 | 3255 | 3324 | 3100 | 3187 | 3155.87 | 3108 | 3204 | 1.031 | 0.044 | 0.48186377 | 0.64851967 | 0.0013 | 0.0032 | 0.0025 | 0.0025 | TRUE | 0.0209 |
| Shh | 5 | 0 | 1 | 1 | 1 | 4 | 4 | 0 | 1 | 1 | 1 | 4 | 1.84 | 2 | 2 | 1.176 | 0.234 | 0.87622338 | NA | 0.6029 | 1.2912 | 1.1001 | 1.1001 | TRUE | 1.0969 |
| Shisa2 | 432 | 353 | 323 | 398 | 376 | 409 | 336 | 375 | 349 | 396 | 424 | 404 | 380.6 | 353 | 408 | 1.157 | 0.21 | 0.05386153 | 0.14190652 | 0 | 0.0087 | 0.0059 | 0.0059 | TRUE | 0.0345 |
| Shisa3 | 18 | 22 | 7 | 22 | 14 | 22 | 14 | 23 | 8 | 22 | 16 | 22 | 17.39 | 15 | 20 | 1.328 | 0.409 | 0.41601997 | 0.58878834 | 0.0782 | 0.1387 | 0.1237 | 0.1237 | TRUE | 0.3826 |
| Shisa4 | 206 | 163 | 186 | 277 | 246 | 219 | 160 | 173 | 201 | 276 | 278 | 216 | 217.25 | 178 | 257 | 1.444 | 0.53 | 0.00072457 | 0.00571221 | 0.0121 | 0.0133 | 0.013 | 0.013 | TRUE | 0.2137 |
| Shisa5 | 1900 | 1406 | 1455 | 1740 | 1538 | 1598 | 1476 | 1496 | 1570 | 1732 | 1735 | 1577 | 1597.6 | 1514 | 1681 | 1.111 | 0.152 | 0.03153663 | 0.09606843 | 0.0014 | 0.0039 | 0.003 | 0.003 | TRUE | 0.0357 |
| Shisa6 | 33 | 22 | 12 | 22 | 26 | 10 | 26 | 23 | 13 | 22 | 29 | 10 | 20.51 | 21 | 20 | 0.973 | -0.039 | 0.9360146 | 0.96464483 | 0.1403 | 0.118 | 0.124 | 0.124 | TRUE | 0.3781 |
| Shisa7 | 505 | 334 | 295 | 352 | 290 | 299 | 392 | 355 | 318 | 350 | 327 | 295 | 339.75 | 355 | 324 | 0.91 | -0.136 | 0.28083473 | 0.45248233 | 0.0065 | 0.0094 | 0.0085 | 0.0085 | TRUE | 0.1003 |
| Shisa9 | 3 | 2 | 3 | 2 | 3 | 2 | 2 | 2 | 3 | 2 | 3 | 2 | 2.51 | 2 | 2 | 0.953 | -0.07 | 0.95205348 | NA | 0 | 0.9477 | 0.5794 | 0.5794 | TRUE | 0.1058 |
| Shkbp1 | 2047 | 1447 | 1435 | 1566 | 1408 | 1672 | 1590 | 1539 | 1548 | 1559 | 1588 | 1651 | 1579.17 | 1559 | 1599 | 1.025 | 0.036 | 0.5787245 | 0.72987864 | 0 | 0.0039 | 0.0024 | 0.0024 | TRUE | 0.0096 |
| Shmt1 | 13097 | 9379 | 9042 | 8752 | 7902 | 9453 | 10172 | 9976 | 9756 | 8712 | 8914 | 9332 | 9477 | 9968 | 8986 | 0.901 | -0.15 | 0.00354264 | 0.01874677 | 7.00E-04 | 0.0027 | 0.0018 | 0.0018 | TRUE | 0.0139 |
| Shmt2 | 10071 | 7769 | 8022 | 9205 | 8029 | 9471 | 7822 | 8264 | 8656 | 9163 | 9058 | 9349 | 8718.45 | 8247 | 9190 | 1.115 | 0.157 | 0.00464104 | 0.02315519 | 0.0012 | 0.0027 | 0.0021 | 0.0021 | TRUE | 0.0252 |
| Shoc2 | 4176 | 3113 | 3140 | 3075 | 2758 | 3105 | 3243 | 3311 | 3388 | 3061 | 3111 | 3065 | 3196.65 | 3314 | 3079 | 0.929 | -0.106 | 0.05169852 | 0.138143 | 0 | 0.0032 | 0.0018 | 0.0018 | TRUE | 0.0046 |
| Shox2 | 10 | 3 | 5 | 2 | 2 | 1 | 8 | 3 | 5 | 2 | 2 | 1 | 3.6 | 5 | 2 | 0.314 | -1.672 | 0.12169166 | NA | 0 | 0.6612 | 0.4544 | 0.4544 | TRUE | 0.1114 |
| Shpk | 220 | 150 | 162 | 225 | 166 | 217 | 171 | 160 | 175 | 224 | 187 | 214 | 188.44 | 169 | 208 | 1.239 | 0.309 | 0.03828802 | 0.11089174 | 0 | 0.015 | 0.0107 | 0.0107 | TRUE | 0.0829 |
| Shprh | 3214 | 2295 | 2266 | 2526 | 2183 | 2516 | 2496 | 2441 | 2445 | 2514 | 2463 | 2484 | 2473.84 | 2461 | 2487 | 1.01 | 0.015 | 0.7906882 | 0.8770061 | 0 | 0.0034 | 0.0019 | 0.0019 | TRUE | 0.002 |
| Shq1 | 3365 | 2475 | 2357 | 2280 | 2087 | 2425 | 2613 | 2633 | 2543 | 2270 | 2354 | 2394 | 2467.83 | 2596 | 2339 | 0.901 | -0.151 | 0.01011375 | 0.04165893 | 0 | 0.0034 | 0.0021 | 0.0021 | TRUE | 0.0083 |
| Shroom1 | 59 | 49 | 24 | 24 | 28 | 22 | 46 | 52 | 26 | 24 | 32 | 22 | 33.51 | 41 | 26 | 0.619 | -0.692 | 0.06261164 | 0.158288 | 0.0535 | 0.0732 | 0.0677 | 0.0677 | TRUE | 0.7283 |
| Shroom2 | 2704 | 1925 | 1861 | 2460 | 2072 | 2401 | 2100 | 2048 | 2008 | 2449 | 2337 | 2370 | 2218.66 | 2052 | 2385 | 1.162 | 0.217 | 0.00029962 | 0.00290664 | 0 | 0.0035 | 0.0021 | 0.0021 | TRUE | 0.0065 |
| Shroom3 | 2870 | 2131 | 2030 | 2039 | 1821 | 2160 | 2229 | 2267 | 2190 | 2030 | 2054 | 2132 | 2150.37 | 2229 | 2072 | 0.93 | -0.105 | 0.0801515 | 0.18942973 | 0 | 0.0035 | 0.0021 | 0.0021 | TRUE | 0.0079 |
| Shroom4 | 479 | 315 | 298 | 338 | 292 | 274 | 372 | 335 | 322 | 336 | 329 | 270 | 327.49 | 343 | 312 | 0.908 | -0.14 | 0.2761949 | 0.44722484 | 0.0069 | 0.0097 | 0.0088 | 0.0088 | TRUE | 0.1571 |
| Siae | 1294 | 907 | 1093 | 1128 | 993 | 1081 | 1005 | 965 | 1179 | 1123 | 1120 | 1067 | 1076.54 | 1050 | 1103 | 1.052 | 0.073 | 0.41303563 | 0.585899 | 0.0046 | 0.0046 | 0.0048 | 0.0048 | TRUE | 0.1364 |
| Siah1a | 1412 | 1054 | 937 | 1077 | 965 | 1061 | 1097 | 1121 | 1011 | 1072 | 1089 | 1047 | 1072.8 | 1076 | 1069 | 0.993 | -0.01 | 0.89270034 | 0.94103261 | 6.00E-04 | 0.0046 | 0.0032 | 0.0032 | TRUE | 0.0325 |
| Siah1b | 2196 | 1438 | 1332 | 1354 | 1288 | 1373 | 1706 | 1530 | 1437 | 1348 | 1453 | 1355 | 1471.41 | 1558 | 1385 | 0.888 | -0.171 | 0.03491198 | 0.10367691 | 0.0037 | 0.004 | 0.0041 | 0.0041 | TRUE | 0.0857 |
| Siah2 | 834 | 673 | 623 | 723 | 647 | 705 | 648 | 716 | 672 | 720 | 730 | 696 | 696.88 | 679 | 715 | 1.055 | 0.077 | 0.37273708 | 0.54725538 | 0 | 0.0058 | 0.004 | 0.004 | TRUE | 0.0271 |
| Siah3 | 0 | 0 | 0 | 0 | 0 | 1 | 0 | 0 | 0 | 0 | 0 | 1 | 0.16 | 0 | 0 | 2.094 | 1.067 | 0.79379329 | NA | 0 | 14.4078 | 10 | 10 | TRUE | 0.3835 |
| Sidt1 | 10 | 4 | 7 | 5 | 3 | 6 | 8 | 4 | 8 | 5 | 3 | 6 | 5.64 | 7 | 5 | 0.729 | -0.456 | 0.56970532 | NA | 0 | 0.4224 | 0.2817 | 0.2817 | TRUE | 0.2152 |
| Sidt2 | 3335 | 2305 | 2394 | 2454 | 2090 | 2376 | 2590 | 2452 | 2583 | 2443 | 2358 | 2345 | 2461.83 | 2542 | 2382 | 0.937 | -0.094 | 0.11499919 | 0.24477786 | 3.00E-04 | 0.0034 | 0.0022 | 0.0022 | TRUE | 0.0114 |
| Sigirr | 697 | 503 | 599 | 595 | 609 | 753 | 541 | 535 | 646 | 592 | 687 | 743 | 624.21 | 574 | 674 | 1.175 | 0.233 | 0.03950822 | 0.11352856 | 0.0106 | 0.0062 | 0.0076 | 0.0076 | TRUE | 0.1384 |
| Siglec1 | 36 | 14 | 21 | 18 | 17 | 16 | 28 | 15 | 23 | 18 | 19 | 16 | 19.73 | 22 | 18 | 0.799 | -0.324 | 0.46572772 | 0.63463117 | 0 | 0.1225 | 0.0911 | 0.0911 | TRUE | 0.2556 |
| Siglec15 | 1 | 3 | 2 | 2 | 2 | 1 | 1 | 3 | 2 | 2 | 2 | 1 | 1.89 | 2 | 2 | 0.87 | -0.2 | 0.88384194 | NA | 0 | 1.2542 | 0.8129 | 0.8129 | TRUE | 0.3482 |
| Siglec5 | 223 | 183 | 181 | 166 | 188 | 164 | 173 | 195 | 195 | 165 | 212 | 162 | 183.73 | 188 | 180 | 0.957 | -0.063 | 0.69672447 | 0.81551657 | 0.0081 | 0.0153 | 0.0133 | 0.0133 | TRUE | 0.2567 |
| Siglece | 7 | 3 | 5 | 11 | 6 | 3 | 5 | 3 | 5 | 11 | 7 | 3 | 5.78 | 4 | 7 | 1.465 | 0.551 | 0.50539485 | NA | 0 | 0.4122 | 0.3155 | 0.3155 | TRUE | 0.1559 |
| Siglecg | 44 | 22 | 40 | 36 | 34 | 30 | 34 | 23 | 43 | 36 | 38 | 30 | 34.09 | 33 | 35 | 1.029 | 0.041 | 0.90704755 | 0.94914932 | 0.0189 | 0.072 | 0.0582 | 0.0582 | TRUE | 0.2242 |
| Siglech | 0 | 0 | 0 | 1 | 0 | 0 | 0 | 0 | 0 | 1 | 0 | 0 | 0.17 | 0 | 0 | 2.094 | 1.067 | 0.79379329 | NA | 0 | 14.2885 | 10 | 10 | TRUE | 0.3906 |

| | | | | | | | | | | | | | | | | | | | | | | | | | |
|---|---|---|---|---|---|---|---|---|---|---|---|---|---|---|---|---|---|---|---|---|---|---|---|---|---|
| Sigmar1 | 5150 | 3550 | 3232 | 3333 | 3045 | 3519 | 4000 | 3776 | 3487 | 3318 | 3435 | 3474 | 3581.62 | 3754 | 3409 | 0.907 | -0.14 | 0.03257567 | 0.09835322 | 0.0022 | 0.0031 | 0.0028 | 0.0028 | TRUE | 0.0469 |
| Sik1 | 10583 | 7973 | 7575 | 8056 | 7248 | 7522 | 8219 | 8481 | 8173 | 8019 | 8176 | 7425 | 8082.38 | 8291 | 7873 | 0.95 | -0.075 | 0.18186756 | 0.33745782 | 0.0013 | 0.0027 | 0.0021 | 0.0021 | TRUE | 0.0304 |
| Sik2 | 1143 | 813 | 801 | 826 | 711 | 858 | 888 | 865 | 864 | 822 | 802 | 847 | 848 | 872 | 824 | 0.944 | -0.083 | 0.28876517 | 0.46093761 | 0 | 0.0052 | 0.0032 | 0.0032 | TRUE | 0.0073 |
| Sik3 | 2698 | 1956 | 1991 | 1981 | 1940 | 2262 | 2095 | 2081 | 2148 | 1972 | 2189 | 2233 | 2119.6 | 2108 | 2131 | 1.011 | 0.016 | 0.81967784 | 0.8964401 | 0.0019 | 0.0035 | 0.0029 | 0.0029 | TRUE | 0.0521 |
| Sike1 | 2363 | 1773 | 1784 | 1865 | 1713 | 1853 | 1835 | 1886 | 1925 | 1856 | 1932 | 1829 | 1877.36 | 1882 | 1872 | 0.995 | -0.007 | 0.91288589 | 0.95219135 | 0 | 0.0037 | 0.0023 | 0.0023 | TRUE | 0.0093 |
| Sil1 | 1285 | 1016 | 1134 | 1466 | 1286 | 1539 | 998 | 1081 | 1224 | 1459 | 1451 | 1519 | 1288.59 | 1101 | 1476 | 1.344 | 0.426 | 5.49E-07 | 1.77E-05 | 0.0045 | 0.0043 | 0.0044 | 0.0044 | TRUE | 0.1134 |
| Sim1 | 17 | 4 | 5 | 10 | 7 | 5 | 13 | 4 | 5 | 10 | 8 | 5 | 7.61 | 7 | 8 | 0.972 | -0.041 | 0.9561144 | NA | 0.1389 | 0.314 | 0.2613 | 0.2613 | TRUE | 1.4699 |
| Sim2 | 9 | 7 | 6 | 2 | 5 | 11 | 7 | 7 | 6 | 2 | 6 | 11 | 6.57 | 7 | 6 | 0.886 | -0.175 | 0.82455022 | NA | 0.1007 | 0.3634 | 0.2947 | 0.2947 | TRUE | 0.3499 |
| Simc1 | 1841 | 1379 | 1382 | 1600 | 1478 | 1740 | 1430 | 1467 | 1491 | 1593 | 1667 | 1718 | 1560.91 | 1463 | 1659 | 1.135 | 0.183 | 0.00637535 | 0.02951827 | 0 | 0.0039 | 0.0026 | 0.0026 | TRUE | 0.0151 |
| Sin3a | 16609 | 12302 | 12839 | 13506 | 12074 | 13767 | 12900 | 13085 | 13853 | 13444 | 13621 | 13590 | 13415.48 | 13279 | 13552 | 1.021 | 0.029 | 0.55388226 | 0.70963062 | 6.00E-04 | 0.0026 | 0.0017 | 0.0017 | TRUE | 0.0174 |
| Sin3b | 7154 | 5385 | 5425 | 5199 | 4524 | 5286 | 5556 | 5728 | 5854 | 5175 | 5104 | 5218 | 5439.07 | 5713 | 5166 | 0.905 | -0.145 | 0.00450828 | 0.02269703 | 3.00E-04 | 0.0029 | 0.0017 | 0.0017 | TRUE | 0.0071 |
| Sipa1 | 1443 | 1123 | 1220 | 1246 | 1205 | 1458 | 1121 | 1195 | 1316 | 1240 | 1359 | 1439 | 1278.42 | 1211 | 1346 | 1.113 | 0.155 | 0.0744923 | 0.17945702 | 0.0049 | 0.0043 | 0.0047 | 0.0047 | TRUE | 0.0698 |
| Sipa1l1 | 4925 | 3828 | 4199 | 3581 | 3588 | 3927 | 3825 | 4072 | 4531 | 3565 | 4048 | 3877 | 3986.04 | 4143 | 3830 | 0.925 | -0.113 | 0.13910159 | 0.28039129 | 0.0054 | 0.003 | 0.0039 | 0.0039 | TRUE | 0.0816 |
| Sipa1l2 | 7065 | 5308 | 5950 | 6245 | 5819 | 6538 | 5487 | 5646 | 6420 | 6216 | 6564 | 6454 | 6131.31 | 5851 | 6411 | 1.096 | 0.132 | 0.05203491 | 0.13879494 | 0.0038 | 0.0028 | 0.0032 | 0.0032 | TRUE | 0.0879 |
| Sipa1l3 | 3464 | 2622 | 2337 | 2455 | 2243 | 2500 | 2690 | 2789 | 2522 | 2444 | 2530 | 2468 | 2573.81 | 2667 | 2481 | 0.93 | -0.105 | 0.09400049 | 0.21124984 | 0.001 | 0.0033 | 0.0024 | 0.0024 | TRUE | 0.0269 |
| Sirpa | 696 | 549 | 633 | 506 | 522 | 575 | 541 | 584 | 683 | 504 | 589 | 568 | 577.95 | 603 | 554 | 0.92 | -0.121 | 0.2811927 | 0.45275857 | 0.0089 | 0.0065 | 0.0073 | 0.0073 | TRUE | 0.1591 |
| Sirpb1a | 0 | 0 | 0 | 0 | 0 | 0 | 0 | 0 | 0 | 0 | 0 | 0 | 0 | 0 | 0 | NA | NA | NA | NA | NA | NA | NA | NA | NA | NA |
| Sirpb1b | 0 | 0 | 0 | 0 | 0 | 0 | 0 | 0 | 0 | 0 | 0 | 0 | 0 | 0 | 0 | NA | NA | NA | NA | NA | NA | NA | NA | NA | NA |
| Sirt1 | 7489 | 5308 | 4389 | 4953 | 4211 | 4511 | 5816 | 5646 | 4736 | 4930 | 4750 | 4453 | 5055.31 | 5399 | 4711 | 0.872 | -0.197 | 0.01392833 | 0.05291453 | 0.0068 | 0.0029 | 0.0044 | 0.0044 | TRUE | 0.1404 |
| Sirt2 | 1237 | 879 | 848 | 902 | 795 | 958 | 961 | 935 | 915 | 898 | 897 | 946 | 925.18 | 937 | 914 | 0.975 | -0.037 | 0.62779906 | 0.76543728 | 0 | 0.005 | 0.0032 | 0.0032 | TRUE | 0.0115 |
| Sirt3 | 882 | 657 | 807 | 875 | 795 | 969 | 685 | 699 | 871 | 871 | 897 | 957 | 829.83 | 752 | 908 | 1.21 | 0.276 | 0.00748848 | 0.03340357 | 0.0088 | 0.0053 | 0.0064 | 0.0064 | TRUE | 0.2247 |
| Sirt4 | 411 | 341 | 359 | 386 | 291 | 352 | 319 | 363 | 387 | 384 | 328 | 347 | 354.88 | 356 | 353 | 0.995 | -0.007 | 0.95177648 | 0.97336443 | 0.0055 | 0.0091 | 0.008 | 0.008 | TRUE | 0.1026 |
| Sirt5 | 428 | 327 | 334 | 368 | 336 | 399 | 332 | 348 | 360 | 366 | 379 | 394 | 363.31 | 347 | 380 | 1.096 | 0.133 | 0.22493477 | 0.38946004 | 0 | 0.009 | 0.0059 | 0.0059 | TRUE | 0.0165 |
| Sirt6 | 1112 | 891 | 785 | 702 | 654 | 762 | 864 | 948 | 847 | 699 | 738 | 752 | 807.86 | 886 | 730 | 0.824 | -0.28 | 0.0010522 | 0.007512 | 0.0014 | 0.0054 | 0.004 | 0.004 | TRUE | 0.0429 |
| Sirt7 | 991 | 824 | 726 | 731 | 705 | 766 | 770 | 876 | 783 | 728 | 795 | 756 | 784.77 | 810 | 760 | 0.939 | -0.091 | 0.302464 | 0.47501571 | 0.0023 | 0.0054 | 0.0044 | 0.0044 | TRUE | 0.0606 |
| Sis | 4 | 0 | 6 | 3 | 7 | 3 | 3 | 0 | 6 | 3 | 8 | 3 | 3.9 | 3 | 5 | 1.436 | 0.522 | 0.63078951 | NA | 0.5046 | 0.6095 | 0.5869 | 0.5869 | TRUE | 0.2694 |
| Sit1 | 22 | 15 | 16 | 13 | 9 | 19 | 17 | 16 | 17 | 13 | 10 | 19 | 15.36 | 17 | 14 | 0.836 | -0.259 | 0.59858612 | NA | 0 | 0.1567 | 0.1091 | 0.1091 | TRUE | 0.4028 |
| Siva1 | 3654 | 2640 | 2801 | 3011 | 2656 | 2992 | 2838 | 2808 | 3022 | 2997 | 2996 | 2954 | 2935.88 | 2889 | 2982 | 1.032 | 0.046 | 0.42954064 | 0.60160944 | 5.00E-04 | 0.0032 | 0.0021 | 0.0021 | TRUE | 0.0193 |
| Six1 | 1004 | 752 | 718 | 843 | 740 | 815 | 780 | 800 | 775 | 839 | 835 | 805 | 805.47 | 785 | 826 | 1.053 | 0.074 | 0.34877364 | 0.52344945 | 0 | 0.0054 | 0.0033 | 0.0033 | TRUE | 0.0063 |
| Six2 | 36 | 27 | 24 | 24 | 22 | 21 | 28 | 29 | 26 | 24 | 25 | 21 | 25.34 | 28 | 23 | 0.839 | -0.253 | 0.50041965 | 0.66485388 | 0 | 0.096 | 0.0617 | 0.0617 | TRUE | 0.0502 |
| Six3 | 12 | 9 | 6 | 3 | 0 | 5 | 9 | 10 | 6 | 3 | 0 | 5 | 5.55 | 8 | 3 | 0.319 | -1.65 | 0.07098616 | NA | 0 | 0.4296 | 0.3535 | 0.3535 | TRUE | 0.7004 |
| Six3os1 | 7 | 5 | 5 | 7 | 3 | 1 | 5 | 5 | 5 | 7 | 3 | 1 | 4.58 | 5 | 4 | 0.703 | -0.509 | 0.58386083 | NA | 0 | 0.5197 | 0.3964 | 0.3964 | TRUE | 0.4546 |
| Six4 | 1471 | 1190 | 1213 | 1353 | 1161 | 1314 | 1142 | 1266 | 1309 | 1347 | 1310 | 1297 | 1278.45 | 1239 | 1318 | 1.065 | 0.091 | 0.23411508 | 0.40031186 | 0.002 | 0.0043 | 0.0035 | 0.0035 | TRUE | 0.0583 |
| Six5 | 326 | 279 | 277 | 389 | 347 | 357 | 253 | 297 | 299 | 387 | 391 | 352 | 329.99 | 283 | 377 | 1.336 | 0.418 | 0.00059529 | 0.00493531 | 0.0031 | 0.0096 | 0.0076 | 0.0076 | TRUE | 0.104 |
| Six6 | 10 | 5 | 5 | 7 | 5 | 9 | 8 | 5 | 5 | 7 | 6 | 9 | 6.66 | 6 | 7 | 1.155 | 0.207 | 0.7761301 | NA | 0 | 0.3582 | 0.2339 | 0.2339 | TRUE | 0.1643 |
| Ska1 | 1631 | 1117 | 1075 | 1021 | 906 | 1147 | 1267 | 1188 | 1160 | 1016 | 1022 | 1132 | 1130.9 | 1205 | 1057 | 0.877 | -0.19 | 0.01672733 | 0.06065038 | 0.0021 | 0.0045 | 0.0037 | 0.0037 | TRUE | 0.0471 |
| Ska2 | 2134 | 1483 | 1701 | 1792 | 1553 | 1813 | 1657 | 1577 | 1835 | 1784 | 1752 | 1790 | 1732.6 | 1690 | 1775 | 1.051 | 0.071 | 0.32905622 | 0.50355466 | 0.0023 | 0.0038 | 0.0033 | 0.0033 | TRUE | 0.0666 |
| Ska3 | 3492 | 2397 | 2324 | 2440 | 2275 | 2557 | 2712 | 2550 | 2508 | 2429 | 2566 | 2524 | 2548.11 | 2590 | 2506 | 0.967 | -0.048 | 0.43539943 | 0.60701951 | 9.00E-04 | 0.0034 | 0.0024 | 0.0024 | TRUE | 0.0213 |
| Skap1 | 168 | 130 | 148 | 232 | 210 | 229 | 130 | 138 | 160 | 231 | 237 | 226 | 187.06 | 143 | 231 | 1.625 | 0.7 | 3.34E-06 | 7.61E-05 | 0 | 0.0151 | 0.0107 | 0.0107 | TRUE | 0.1079 |
| Skap2 | 3017 | 2203 | 2273 | 2588 | 2273 | 2747 | 2343 | 2343 | 2453 | 2576 | 2564 | 2712 | 2498.5 | 2380 | 2617 | 1.1 | 0.138 | 0.02147545 | 0.07282821 | 4.00E-04 | 0.0034 | 0.0022 | 0.0022 | TRUE | 0.0122 |
| Ski | 9719 | 7156 | 6639 | 7902 | 7242 | 7790 | 7548 | 7612 | 7163 | 7866 | 8170 | 7690 | 7674.8 | 7441 | 7909 | 1.063 | 0.088 | 0.09976464 | 0.22082633 | 8.00E-04 | 0.0027 | 0.0019 | 0.0019 | TRUE | 0.0129 |
| Skida1 | 241 | 227 | 171 | 140 | 133 | 132 | 187 | 241 | 185 | 139 | 150 | 130 | 172.14 | 204 | 140 | 0.685 | -0.546 | 0.00115418 | 0.00804322 | 0.0097 | 0.0162 | 0.0142 | 0.0142 | TRUE | 0.2669 |
| Skil | 10798 | 8272 | 9426 | 10998 | 9711 | 11586 | 8386 | 8799 | 10171 | 10947 | 10955 | 11437 | 10115.89 | 9119 | 11113 | 1.219 | 0.286 | 7.70E-05 | 0.00098558 | 0.0054 | 0.0027 | 0.0037 | 0.0037 | TRUE | 0.1242 |
| Skint1 | 0 | 0 | 0 | 0 | 0 | 0 | 0 | 0 | 0 | 0 | 0 | 0 | 0 | 0 | 0 | NA | NA | NA | NA | NA | NA | NA | NA | NA | NA |
| Skint10 | 0 | 0 | 0 | 0 | 0 | 0 | 0 | 0 | 0 | 0 | 0 | 0 | 0 | 0 | 0 | NA | NA | NA | NA | NA | NA | NA | NA | NA | NA |
| Skint11 | 0 | 0 | 0 | 0 | 0 | 0 | 0 | 0 | 0 | 0 | 0 | 0 | 0 | 0 | 0 | NA | NA | NA | NA | NA | NA | NA | NA | NA | NA |
| Skint2 | 3 | 0 | 0 | 0 | 0 | 1 | 2 | 0 | 0 | 0 | 0 | 1 | 0.55 | 1 | 0 | 0.458 | -1.128 | 0.69846567 | NA | 0 | 4.2894 | 4.5119 | 4.5119 | TRUE | 1.4399 |
| Skint3 | 0 | 0 | 0 | 0 | 0 | 0 | 0 | 0 | 0 | 0 | 0 | 0 | 0 | 0 | 0 | NA | NA | NA | NA | NA | NA | NA | NA | NA | NA |
| Skint4 | 0 | 0 | 0 | 0 | 0 | 0 | 0 | 0 | 0 | 0 | 0 | 0 | 0 | 0 | 0 | NA | NA | NA | NA | NA | NA | NA | NA | NA | NA |
| Skint5 | 1 | 0 | 0 | 0 | 0 | 0 | 1 | 0 | 0 | 0 | 0 | 0 | 0.13 | 0 | 0 | 0.552 | -0.857 | 0.8336548 | NA | 0 | 18.3121 | 10 | 10 | TRUE | 0.2566 |
| Skint6 | 0 | 0 | 0 | 0 | 0 | 0 | 0 | 0 | 0 | 0 | 0 | 0 | 0 | 0 | 0 | NA | NA | NA | NA | NA | NA | NA | NA | NA | NA |
| Skint7 | 0 | 0 | 0 | 0 | 0 | 0 | 0 | 0 | 0 | 0 | 0 | 0 | 0 | 0 | 0 | NA | NA | NA | NA | NA | NA | NA | NA | NA | NA |
| Skint8 | 0 | 0 | 0 | 0 | 0 | 0 | 0 | 0 | 0 | 0 | 0 | 0 | 0 | 0 | 0 | NA | NA | NA | NA | NA | NA | NA | NA | NA | NA |
| Skint9 | 0 | 0 | 0 | 0 | 0 | 0 | 0 | 0 | 0 | 0 | 0 | 0 | 0 | 0 | 0 | NA | NA | NA | NA | NA | NA | NA | NA | NA | NA |
| Skiv2l | 2043 | 1643 | 1311 | 1463 | 1248 | 1466 | 1587 | 1748 | 1415 | 1456 | 1408 | 1447 | 1510.04 | 1583 | 1437 | 0.908 | -0.139 | 0.09509538 | 0.21310728 | 0.0047 | 0.004 | 0.0044 | 0.0044 | TRUE | 0.1017 |
| Skiv2l2 | 8103 | 5725 | 5994 | 5830 | 5336 | 6295 | 6293 | 6090 | 6467 | 5803 | 6020 | 6214 | 6147.87 | 6283 | 6012 | 0.957 | -0.064 | 0.24183971 | 0.4094545 | 8.00E-04 | 0.0028 | 0.002 | 0.002 | TRUE | 0.0114 |
| Skor1 | 7 | 4 | 0 | 11 | 3 | 8 | 5 | 4 | 0 | 11 | 3 | 8 | 5.32 | 3 | 7 | 2.239 | 1.163 | 0.22157879 | NA | 0.3927 | 0.4479 | 0.438 | 0.438 | TRUE | 0.4241 |
| Skor2 | 0 | 0 | 0 | 0 | 1 | 1 | 0 | 0 | 0 | 0 | 1 | 1 | 0.35 | 0 | 1 | 3.965 | 1.987 | 0.55529285 | NA | 0 | 6.7252 | 6.4426 | 6.4426 | TRUE | 0.261 |
| Skp1a | 23970 | 17250 | 17051 | 14611 | 12306 | 14663 | 18616 | 18349 | 18398 | 14544 | 13882 | 14475 | 16377.31 | 18454 | 14300 | 0.775 | -0.368 | 1.58E-15 | 2.90E-13 | 3.00E-04 | 0.0026 | 0.0015 | 0.0015 | TRUE | 0.008 |
| Skp2 | 3474 | 2400 | 2300 | 2250 | 1972 | 2394 | 2698 | 2553 | 2482 | 2240 | 2225 | 2363 | 2426.69 | 2578 | 2276 | 0.883 | -0.18 | 0.00450752 | 0.02269703 | 0.0011 | 0.0034 | 0.0025 | 0.0025 | TRUE | 0.0209 |
| Sla | 0 | 0 | 0 | 0 | 0 | 0 | 0 | 0 | 0 | 0 | 0 | 0 | 0 | 0 | 0 | NA | NA | NA | NA | NA | NA | NA | NA | NA | NA |
| Sla2 | 4 | 3 | 4 | 4 | 3 | 1 | 3 | 3 | 4 | 4 | 3 | 1 | 3.16 | 3 | 3 | 0.787 | -0.345 | 0.74720264 | NA | 0 | 0.7522 | 0.5029 | 0.5029 | TRUE | 0.4085 |
| Slain1 | 1598 | 1254 | 1366 | 1175 | 1099 | 1222 | 1241 | 1334 | 1474 | 1170 | 1240 | 1206 | 1277.42 | 1350 | 1205 | 0.894 | -0.162 | 0.04760447 | 0.12998556 | 0.0035 | 0.0043 | 0.004 | 0.004 | TRUE | 0.0777 |
| Slain2 | 2165 | 1612 | 1695 | 1704 | 1556 | 1854 | 1681 | 1715 | 1829 | 1696 | 1755 | 1830 | 1751.12 | 1742 | 1760 | 1.012 | 0.017 | 0.80878998 | 0.88964408 | 0.0012 | 0.0038 | 0.0028 | 0.0028 | TRUE | 0.0229 |
| Slamf1 | 1 | 0 | 0 | 0 | 0 | 0 | 1 | 0 | 0 | 0 | 0 | 0 | 0.13 | 0 | 0 | 0.552 | -0.857 | 0.8336548 | NA | 0 | 18.3121 | 10 | 10 | TRUE | 0.2566 |
| Slamf6 | 12 | 6 | 3 | 1 | 5 | 1 | 9 | 6 | 3 | 1 | 6 | 1 | 4.43 | 6 | 3 | 0.388 | -1.367 | 0.18053635 | NA | 0.2515 | 0.5378 | 0.4665 | 0.4665 | TRUE | 0.4991 |
| Slamf7 | 5 | 2 | 5 | 1 | 0 | 0 | 4 | 2 | 5 | 1 | 0 | 0 | 2.07 | 4 | 0 | 0.098 | -3.358 | 0.04345528 | NA | 0 | 1.1491 | 0.8657 | 0.8657 | TRUE | 0.3233 |
| Slamf8 | 0 | 1 | 1 | 0 | 1 | 0 | 0 | 1 | 1 | 0 | 1 | 0 | 0.55 | 1 | 0 | 0.535 | -0.902 | 0.73748201 | NA | 0 | 4.3501 | 3.5264 | 3.5264 | TRUE | 0.5075 |
| Slamf9 | 86 | 74 | 38 | 53 | 47 | 44 | 67 | 79 | 41 | 53 | 53 | 43 | 55.95 | 62 | 50 | 0.797 | -0.327 | 0.26502659 | 0.43545523 | 0.0411 | 0.0448 | 0.0438 | 0.0438 | TRUE | 0.4496 |

| | | | | | | | | | | | | | | | | | | | | | | | | | |
|---|---|---|---|---|---|---|---|---|---|---|---|---|---|---|---|---|---|---|---|---|---|---|---|---|---|
| Slbp | 7229 | 5059 | 4539 | 4146 | 3579 | 4117 | 5614 | 5381 | 4898 | 4127 | 4037 | 4064 | 4686.95 | 5298 | 4076 | 0.769 | -0.379 | 2.47E-09 | 1.48E-07 | 0.0021 | 0.0029 | 0.0027 | 0.0027 | TRUE | 0.053 |
| Slc10a1 | 2 | 0 | 0 | 0 | 1 | 0 | 2 | 0 | 0 | 0 | 1 | 0 | 0.45 | 1 | 0 | 0.68 | -0.556 | 0.85993441 | NA | 0 | 5.3057 | 5.2576 | 5.2576 | TRUE | 0.9726 |
| Slc10a2 | 0 | 0 | 0 | 0 | 0 | 0 | 0 | 0 | 0 | 0 | 0 | 0 | 0 | 0 | 0 | NA | NA | NA | NA | NA | NA | NA | NA | NA | NA |
| Slc10a3 | 149 | 109 | 52 | 62 | 54 | 66 | 116 | 116 | 56 | 62 | 61 | 65 | 79.26 | 96 | 63 | 0.65 | -0.622 | 0.02562181 | 0.0827412 | 0.0679 | 0.0323 | 0.0426 | 0.0426 | TRUE | 1.2243 |
| Slc10a4 | 18 | 12 | 12 | 5 | 5 | 6 | 14 | 13 | 13 | 5 | 6 | 6 | 9.37 | 13 | 6 | 0.416 | -1.266 | 0.04705869 | NA | 0 | 0.2553 | 0.1626 | 0.1626 | TRUE | 0.0167 |
| Slc10a5 | 2 | 2 | 0 | 1 | 0 | 2 | 2 | 2 | 0 | 1 | 0 | 2 | 1.11 | 1 | 1 | 0.812 | -0.3 | 0.87369046 | NA | 0 | 2.1407 | 1.662 | 1.662 | TRUE | 0.2238 |
| Slc10a6 | 69 | 45 | 50 | 51 | 55 | 38 | 54 | 48 | 54 | 51 | 62 | 38 | 50.95 | 52 | 50 | 0.963 | -0.055 | 0.84722832 | 0.91329922 | 0.0115 | 0.0489 | 0.0388 | 0.0388 | TRUE | 0.2392 |
| Slc10a7 | 1913 | 1328 | 1158 | 1117 | 991 | 1179 | 1486 | 1413 | 1249 | 1112 | 1118 | 1164 | 1256.91 | 1383 | 1131 | 0.817 | -0.291 | 0.00039423 | 0.00360716 | 0.0036 | 0.0043 | 0.0041 | 0.0041 | TRUE | 0.0843 |
| Slc11a1 | 179 | 127 | 175 | 189 | 224 | 202 | 139 | 135 | 189 | 188 | 253 | 199 | 183.86 | 154 | 213 | 1.385 | 0.47 | 0.00917887 | 0.03884917 | 0.0235 | 0.0153 | 0.0178 | 0.0178 | TRUE | 0.3885 |
| Slc11a2 | 3534 | 2392 | 2420 | 2342 | 2025 | 2470 | 2745 | 2544 | 2611 | 2331 | 2284 | 2438 | 2492.35 | 2633 | 2351 | 0.893 | -0.164 | 0.00861336 | 0.03703693 | 0.001 | 0.0034 | 0.0024 | 0.0024 | TRUE | 0.0171 |
| Slc12a1 | 1 | 3 | 4 | 3 | 0 | 2 | 1 | 3 | 4 | 3 | 0 | 2 | 2.21 | 3 | 2 | 0.623 | -0.682 | 0.61937974 | NA | 0 | 1.0761 | 0.8736 | 0.8736 | TRUE | 0.5892 |
| Slc12a2 | 5813 | 3938 | 3458 | 3432 | 3219 | 3490 | 4515 | 4189 | 3731 | 3416 | 3631 | 3445 | 3821.23 | 4145 | 3497 | 0.843 | -0.246 | 0.00097833 | 0.00712034 | 0.0047 | 0.003 | 0.0037 | 0.0037 | TRUE | 0.0925 |
| Slc12a3 | 77 | 60 | 48 | 56 | 43 | 62 | 60 | 64 | 52 | 56 | 49 | 61 | 56.81 | 59 | 55 | 0.944 | -0.083 | 0.74876368 | 0.85099231 | 0 | 0.0441 | 0.0306 | 0.0306 | TRUE | 0.0868 |
| Slc12a4 | 3496 | 2505 | 2600 | 3140 | 2892 | 3234 | 2715 | 2665 | 2805 | 3126 | 3262 | 3192 | 2960.93 | 2728 | 3193 | 1.17 | 0.227 | 6.28E-05 | 0.00084318 | 2.00E-04 | 0.0032 | 0.002 | 0.002 | TRUE | 0.0072 |
| Slc12a5 | 135 | 134 | 104 | 166 | 138 | 140 | 105 | 143 | 112 | 165 | 156 | 138 | 136.45 | 120 | 153 | 1.282 | 0.358 | 0.05165697 | 0.13808361 | 0.0097 | 0.0198 | 0.0169 | 0.0169 | TRUE | 0.2707 |
| Slc12a6 | 4003 | 2823 | 3030 | 3190 | 2945 | 3301 | 3109 | 3003 | 3269 | 3175 | 3322 | 3259 | 3189.55 | 3127 | 3252 | 1.04 | 0.057 | 0.33983465 | 0.51388666 | 7.00E-04 | 0.0032 | 0.0022 | 0.0022 | TRUE | 0.0188 |
| Slc12a7 | 9488 | 7429 | 8699 | 8779 | 8638 | 9235 | 7369 | 7902 | 9386 | 8739 | 9745 | 9116 | 8709.46 | 8219 | 9200 | 1.12 | 0.163 | 0.05290415 | 0.14047167 | 0.0094 | 0.0027 | 0.005 | 0.005 | TRUE | 0.1882 |
| Slc12a8 | 540 | 502 | 670 | 685 | 673 | 777 | 419 | 534 | 723 | 682 | 759 | 767 | 647.4 | 559 | 736 | 1.321 | 0.401 | 0.00843642 | 0.03647361 | 0.0373 | 0.0061 | 0.0152 | 0.0152 | TRUE | 0.5041 |
| Slc12a9 | 841 | 681 | 647 | 701 | 586 | 723 | 653 | 724 | 698 | 698 | 661 | 714 | 691.37 | 692 | 691 | 1 | 0.001 | 0.9940491 | 0.99662906 | 0 | 0.0059 | 0.0042 | 0.0042 | TRUE | 0.0302 |
| Slc13a1 | 0 | 0 | 0 | 0 | 0 | 0 | 0 | 0 | 0 | 0 | 0 | 0 | 0 | 0 | 0 | NA | NA | NA | NA | NA | NA | NA | NA | NA | NA |
| Slc13a2 | 11 | 10 | 10 | 13 | 10 | 10 | 9 | 11 | 11 | 13 | 11 | 10 | 10.68 | 10 | 11 | 1.145 | 0.195 | 0.73476185 | NA | 0 | 0.2244 | 0.1443 | 0.1443 | TRUE | 0.0731 |
| Slc13a3 | 17 | 15 | 13 | 17 | 21 | 25 | 13 | 16 | 14 | 17 | 24 | 25 | 18.08 | 14 | 22 | 1.516 | 0.6 | 0.18550508 | 0.34182093 | 0 | 0.1335 | 0.0905 | 0.0905 | TRUE | 0.2205 |
| Slc13a4 | 191 | 131 | 128 | 128 | 120 | 144 | 148 | 139 | 138 | 127 | 135 | 142 | 138.45 | 142 | 135 | 0.949 | -0.075 | 0.6532047 | 0.78460939 | 0 | 0.0195 | 0.0128 | 0.0128 | TRUE | 0.0256 |
| Slc13a5 | 515 | 390 | 417 | 641 | 539 | 604 | 400 | 415 | 450 | 638 | 608 | 596 | 517.85 | 422 | 614 | 1.459 | 0.545 | 2.56E-08 | 1.16E-06 | 0 | 0.007 | 0.0049 | 0.0049 | TRUE | 0.0392 |
| Slc14a1 | 2 | 0 | 0 | 0 | 0 | 0 | 2 | 0 | 0 | 0 | 0 | 0 | 0.26 | 1 | 0 | 0.356 | -1.491 | 0.71362796 | NA | 0 | 9.1573 | 10 | 10 | TRUE | 0.968 |
| Slc14a2 | 2 | 4 | 1 | 1 | 2 | 2 | 2 | 4 | 1 | 1 | 2 | 2 | 2.02 | 2 | 2 | 0.763 | -0.39 | 0.77085011 | NA | 0 | 1.1764 | 0.7811 | 0.7811 | TRUE | 0.5252 |
| Slc15a1 | 1264 | 983 | 1222 | 1383 | 1307 | 1449 | 982 | 1046 | 1319 | 1377 | 1474 | 1430 | 1271.21 | 1116 | 1427 | 1.281 | 0.358 | 0.00048716 | 0.00423281 | 0.0123 | 0.0043 | 0.0068 | 0.0068 | TRUE | 0.3021 |
| Slc15a2 | 313 | 239 | 329 | 224 | 236 | 280 | 243 | 254 | 355 | 223 | 266 | 276 | 269.65 | 284 | 255 | 0.901 | -0.151 | 0.35382378 | 0.52811229 | 0.0241 | 0.0112 | 0.0154 | 0.0154 | TRUE | 0.5267 |
| Slc15a3 | 3 | 3 | 3 | 3 | 4 | 3 | 2 | 3 | 3 | 3 | 5 | 3 | 3.2 | 3 | 4 | 1.199 | 0.262 | 0.8003035 | NA | 0 | 0.7424 | 0.4575 | 0.4575 | TRUE | 0.0867 |
| Slc15a4 | 1751 | 1310 | 1225 | 1242 | 1094 | 1350 | 1360 | 1393 | 1322 | 1236 | 1234 | 1333 | 1313.04 | 1358 | 1268 | 0.933 | -0.099 | 0.1627166 | 0.31312974 | 5.00E-04 | 0.0042 | 0.0029 | 0.0029 | TRUE | 0.0244 |
| Slc15a5 | 0 | 0 | 1 | 2 | 1 | 1 | 0 | 0 | 1 | 2 | 1 | 1 | 0.86 | 0 | 1 | 3.977 | 1.992 | 0.35112459 | NA | 0 | 2.745 | 1.9107 | 1.9107 | TRUE | 0.5365 |
| Slc16a1 | 13302 | 9623 | 9358 | 9851 | 8647 | 9555 | 10331 | 10236 | 10097 | 9806 | 9755 | 9432 | 9942.8 | 10221 | 9664 | 0.945 | -0.081 | 0.08512449 | 0.19700643 | 1.00E-04 | 0.0027 | 0.0015 | 0.0015 | TRUE | 0.0054 |
| Slc16a10 | 1143 | 914 | 815 | 1035 | 836 | 1022 | 888 | 972 | 879 | 1030 | 943 | 1009 | 953.59 | 913 | 994 | 1.09 | 0.124 | 0.1277594 | 0.26376789 | 0.0015 | 0.0049 | 0.0037 | 0.0037 | TRUE | 0.0373 |
| Slc16a11 | 30 | 29 | 43 | 61 | 44 | 43 | 23 | 31 | 46 | 61 | 50 | 42 | 42.22 | 33 | 51 | 1.537 | 0.621 | 0.06390949 | 0.16059394 | 0.0495 | 0.0586 | 0.0561 | 0.0561 | TRUE | 0.7509 |
| Slc16a12 | 10 | 7 | 6 | 6 | 2 | 5 | 8 | 7 | 6 | 6 | 2 | 5 | 5.81 | 7 | 4 | 0.611 | -0.71 | 0.37302957 | NA | 0 | 0.4105 | 0.2748 | 0.2748 | TRUE | 0.2829 |
| Slc16a13 | 393 | 254 | 196 | 255 | 206 | 228 | 305 | 270 | 211 | 254 | 232 | 225 | 249.7 | 262 | 237 | 0.901 | -0.151 | 0.32520502 | 0.49915758 | 0.0154 | 0.0119 | 0.013 | 0.013 | TRUE | 0.3136 |
| Slc16a14 | 140 | 103 | 77 | 73 | 68 | 76 | 109 | 110 | 83 | 73 | 77 | 75 | 87.63 | 101 | 75 | 0.742 | -0.43 | 0.04613563 | 0.12710159 | 0 | 0.0295 | 0.0218 | 0.0218 | TRUE | 0.2121 |
| Slc16a2 | 77 | 55 | 39 | 45 | 46 | 39 | 60 | 59 | 42 | 45 | 52 | 38 | 49.26 | 54 | 45 | 0.837 | -0.256 | 0.37305931 | 0.54755974 | 0 | 0.0505 | 0.0391 | 0.0391 | TRUE | 0.2715 |
| Slc16a3 | 15116 | 11977 | 14334 | 14614 | 13313 | 14911 | 11740 | 12740 | 15466 | 14547 | 15018 | 14719 | 14038.44 | 13315 | 14761 | 1.109 | 0.149 | 0.0823573 | 0.19319429 | 0.0105 | 0.0026 | 0.0052 | 0.0052 | TRUE | 0.244 |
| Slc16a4 | 16 | 7 | 5 | 9 | 7 | 11 | 12 | 7 | 5 | 9 | 8 | 11 | 8.83 | 8 | 9 | 1.079 | 0.11 | 0.86578702 | NA | 0 | 0.2708 | 0.1945 | 0.1945 | TRUE | 0.6735 |
| Slc16a5 | 24 | 16 | 7 | 10 | 11 | 6 | 19 | 17 | 8 | 10 | 12 | 6 | 11.92 | 15 | 9 | 0.645 | -0.633 | 0.28953818 | NA | 0.0806 | 0.2013 | 0.1686 | 0.1686 | TRUE | 0.7109 |
| Slc16a6 | 2460 | 1842 | 1579 | 1657 | 1446 | 1654 | 1911 | 1959 | 1704 | 1649 | 1631 | 1633 | 1747.83 | 1858 | 1638 | 0.881 | -0.183 | 0.0115895 | 0.04611602 | 0.0022 | 0.0038 | 0.0032 | 0.0032 | TRUE | 0.0623 |
| Slc16a7 | 34 | 31 | 23 | 28 | 24 | 28 | 26 | 33 | 25 | 28 | 27 | 28 | 27.8 | 28 | 28 | 0.983 | -0.025 | 0.94588012 | 0.97029019 | 0 | 0.0877 | 0.0579 | 0.0579 | TRUE | 0.1405 |
| Slc16a8 | 1 | 1 | 3 | 2 | 2 | 1 | 1 | 1 | 3 | 2 | 2 | 1 | 1.72 | 2 | 2 | 1.05 | 0.071 | 0.96076943 | NA | 0 | 1.3815 | 0.9007 | 0.9007 | TRUE | 0.4552 |
| Slc16a9 | 275 | 197 | 205 | 254 | 207 | 206 | 214 | 210 | 221 | 253 | 234 | 203 | 222.34 | 215 | 230 | 1.07 | 0.098 | 0.48707747 | 0.6530155 | 0 | 0.0131 | 0.0098 | 0.0098 | TRUE | 0.1157 |
| Slc17a1 | 0 | 0 | 0 | 0 | 0 | 0 | 0 | 0 | 0 | 0 | 0 | 0 | 0 | 0 | 0 | NA | NA | NA | NA | NA | NA | NA | NA | NA | NA |
| Slc17a2 | 0 | 0 | 0 | 0 | 0 | 0 | 0 | 0 | 0 | 0 | 0 | 0 | 0 | 0 | 0 | NA | NA | NA | NA | NA | NA | NA | NA | NA | NA |
| Slc17a3 | 0 | 0 | 0 | 0 | 0 | 0 | 0 | 0 | 0 | 0 | 0 | 0 | 0 | 0 | 0 | NA | NA | NA | NA | NA | NA | NA | NA | NA | NA |
| Slc17a4 | 0 | 0 | 0 | 0 | 0 | 0 | 0 | 0 | 0 | 0 | 0 | 0 | 0 | 0 | 0 | NA | NA | NA | NA | NA | NA | NA | NA | NA | NA |
| Slc17a5 | 691 | 512 | 581 | 648 | 632 | 678 | 537 | 545 | 627 | 645 | 713 | 669 | 622.57 | 570 | 676 | 1.188 | 0.249 | 0.01106851 | 0.04446382 | 0.0033 | 0.0062 | 0.0053 | 0.0053 | TRUE | 0.0894 |
| Slc17a6 | 0 | 0 | 0 | 0 | 0 | 1 | 0 | 0 | 0 | 0 | 0 | 1 | 0.16 | 0 | 0 | 2.094 | 1.067 | 0.79379329 | NA | 0 | 14.4078 | 10 | 10 | TRUE | 0.3835 |
| Slc17a7 | 885 | 780 | 713 | 972 | 950 | 947 | 687 | 830 | 769 | 968 | 1072 | 935 | 876.73 | 762 | 992 | 1.303 | 0.381 | 6.24E-05 | 0.00083965 | 0.0059 | 0.0051 | 0.0054 | 0.0054 | TRUE | 0.0919 |
| Slc17a8 | 1 | 0 | 0 | 1 | 0 | 0 | 1 | 0 | 0 | 1 | 0 | 0 | 0.3 | 0 | 0 | 1.075 | 0.105 | 0.97698525 | NA | 0 | 8.0272 | 7.5141 | 7.5141 | TRUE | 0.3906 |
| Slc17a9 | 812 | 905 | 973 | 1003 | 1060 | 1335 | 631 | 963 | 1050 | 998 | 1196 | 1318 | 1025.86 | 881 | 1171 | 1.33 | 0.411 | 0.09821456 | 0.21829899 | 0.0435 | 0.0047 | 0.0157 | 0.0435 | TRUE | 0.7457 |
| Slc18a1 | 88 | 44 | 20 | 21 | 17 | 23 | 68 | 47 | 22 | 21 | 19 | 23 | 33.25 | 46 | 21 | 0.455 | -1.135 | 0.00737069 | 0.03305593 | 0.1406 | 0.0737 | 0.094 | 0.094 | TRUE | 0.1195 |
| Slc18a2 | 29 | 21 | 15 | 14 | 28 | 18 | 23 | 22 | 16 | 14 | 32 | 18 | 20.72 | 20 | 21 | 1.026 | 0.036 | 0.93609626 | 0.96464483 | 0.0567 | 0.1168 | 0.1003 | 0.1003 | TRUE | 0.9729 |
| Slc18a3 | 46 | 45 | 19 | 26 | 18 | 20 | 36 | 48 | 21 | 26 | 20 | 20 | 28.34 | 35 | 22 | 0.633 | -0.659 | 0.10305765 | 0.22616353 | 0.066 | 0.0861 | 0.0804 | 0.0804 | TRUE | 0.1607 |
| Slc18b1 | 192 | 140 | 129 | 139 | 100 | 130 | 149 | 149 | 139 | 138 | 113 | 128 | 136.12 | 146 | 126 | 0.869 | -0.203 | 0.23640261 | 0.40282576 | 0 | 0.0198 | 0.0138 | 0.0138 | TRUE | 0.0866 |
| Slc19a1 | 1286 | 954 | 991 | 1109 | 1086 | 1141 | 999 | 1015 | 1069 | 1104 | 1225 | 1126 | 1089.7 | 1028 | 1152 | 1.121 | 0.165 | 0.03381593 | 0.1012848 | 0.0012 | 0.0046 | 0.0034 | 0.0034 | TRUE | 0.0365 |
| Slc19a2 | 5450 | 4096 | 4775 | 5061 | 4368 | 4836 | 4233 | 4357 | 5152 | 5038 | 4928 | 4774 | 4746.83 | 4581 | 4913 | 1.073 | 0.102 | 0.18422296 | 0.34036954 | 0.006 | 0.0029 | 0.004 | 0.004 | TRUE | 0.1446 |
| Slc19a3 | 270 | 161 | 245 | 305 | 247 | 284 | 210 | 171 | 264 | 304 | 279 | 280 | 251.32 | 215 | 288 | 1.338 | 0.42 | 0.0075597 | 0.03361879 | 0.0187 | 0.0119 | 0.0138 | 0.0138 | TRUE | 0.3852 |
| Slc1a1 | 1035 | 760 | 855 | 1067 | 960 | 972 | 804 | 808 | 923 | 1062 | 1083 | 960 | 939.89 | 845 | 1035 | 1.226 | 0.294 | 0.00097791 | 0.00712034 | 0.004 | 0.0049 | 0.0046 | 0.0046 | TRUE | 0.0752 |
| Slc1a2 | 11 | 14 | 14 | 13 | 9 | 8 | 9 | 15 | 15 | 13 | 10 | 8 | 11.59 | 13 | 10 | 0.814 | -0.297 | 0.60480855 | NA | 0 | 0.2069 | 0.1497 | 0.1497 | TRUE | 0.4561 |
| Slc1a3 | 1136 | 824 | 1007 | 1006 | 864 | 1148 | 882 | 876 | 1087 | 1001 | 975 | 1133 | 992.44 | 948 | 1036 | 1.094 | 0.13 | 0.20128812 | 0.36212921 | 0.0098 | 0.0048 | 0.0065 | 0.0065 | TRUE | 0.1907 |
| Slc1a4 | 728 | 472 | 609 | 489 | 460 | 586 | 565 | 502 | 657 | 487 | 519 | 578 | 551.45 | 575 | 528 | 0.919 | -0.122 | 0.29925964 | 0.47153765 | 0.0104 | 0.0067 | 0.0081 | 0.0081 | TRUE | 0.1776 |
| Slc1a5 | 2223 | 1677 | 1994 | 2259 | 2042 | 2407 | 1727 | 1784 | 2152 | 2249 | 2304 | 2376 | 2098.35 | 1888 | 2310 | 1.225 | 0.293 | 0.00060354 | 0.0049921 | 0.0071 | 0.0036 | 0.0048 | 0.0048 | TRUE | 0.1799 |
| Slc1a6 | 151 | 138 | 156 | 197 | 185 | 232 | 117 | 147 | 168 | 196 | 209 | 229 | 177.7 | 144 | 211 | 1.475 | 0.561 | 0.00097506 | 0.00711048 | 0.0132 | 0.0158 | 0.015 | 0.015 | TRUE | 0.3086 |
| Slc1a7 | 13 | 9 | 7 | 7 | 7 | 6 | 10 | 10 | 8 | 7 | 8 | 6 | 8 | 9 | 7 | 0.758 | -0.399 | 0.54796769 | NA | 0 | 0.2986 | 0.1907 | 0.1907 | TRUE | 0.0641 |
| Slc20a1 | 7065 | 5347 | 5295 | 5676 | 4840 | 5947 | 5487 | 5688 | 5713 | 5650 | 5460 | 5871 | 5644.73 | 5629 | 5660 | 1.006 | 0.008 | 0.87630211 | 0.93096567 | 7.00E-04 | 0.0028 | 0.0019 | 0.0019 | TRUE | 0.0129 |

| | | | | | | | | | | | | | | | | | | | | | | | | | |
|---|---|---|---|---|---|---|---|---|---|---|---|---|---|---|---|---|---|---|---|---|---|---|---|---|---|
| Slc20a2 | 7801 | 5466 | 5651 | 6212 | 5770 | 6569 | 6059 | 5814 | 6097 | 6183 | 6509 | 6485 | 6191.23 | 5990 | 6392 | 1.067 | 0.094 | 0.07432207 | 0.17919794 | 5.00E-04 | 0.0028 | 0.0018 | 0.0018 | TRUE | 0.01 |
| Slc22a1 | 13 | 3 | 3 | 6 | 1 | 5 | 10 | 3 | 3 | 6 | 1 | 5 | 4.76 | 5 | 4 | 0.719 | -0.476 | 0.61629076 | NA | 0.2795 | 0.5003 | 0.4366 | 0.4366 | TRUE | 1.4778 |
| Slc22a12 | 0 | 0 | 0 | 0 | 0 | 0 | 0 | 0 | 0 | 0 | 0 | 0 | 0 | 0 | 0 | NA | NA | NA | NA | NA | NA | NA | NA | NA | NA |
| Slc22a13 | 3 | 1 | 3 | 4 | 0 | 2 | 2 | 1 | 3 | 4 | 0 | 2 | 2.1 | 2 | 2 | 0.911 | -0.134 | 0.92208817 | NA | 0 | 1.1322 | 0.884 | 0.884 | TRUE | 0.2012 |
| Slc22a13b-ps | 20 | 13 | 7 | 10 | 10 | 16 | 16 | 14 | 8 | 10 | 11 | 16 | 12.32 | 13 | 12 | 0.995 | -0.008 | 0.98913333 | NA | 0 | 0.1947 | 0.1451 | 0.1451 | TRUE | 0.4205 |
| Slc22a14 | 21 | 15 | 14 | 12 | 13 | 19 | 16 | 16 | 15 | 12 | 15 | 19 | 15.46 | 16 | 15 | 0.957 | -0.063 | 0.89602025 | NA | 0 | 0.1558 | 0.1038 | 0.1038 | TRUE | 0.212 |
| Slc22a15 | 36 | 25 | 24 | 27 | 30 | 36 | 28 | 27 | 26 | 27 | 34 | 36 | 29.45 | 27 | 32 | 1.193 | 0.255 | 0.46820949 | 0.63716255 | 0 | 0.0829 | 0.0551 | 0.0551 | TRUE | 0.1421 |
| Slc22a16 | 4 | 3 | 5 | 2 | 6 | 2 | 3 | 3 | 5 | 2 | 7 | 2 | 3.74 | 4 | 4 | 0.911 | -0.134 | 0.89472041 | NA | 0 | 0.6365 | 0.4637 | 0.4637 | TRUE | 0.8249 |
| Slc22a17 | 240 | 257 | 218 | 211 | 201 | 226 | 186 | 273 | 235 | 210 | 227 | 223 | 225.81 | 231 | 220 | 0.954 | -0.069 | 0.66631975 | 0.7948039 | 0.0156 | 0.0129 | 0.0138 | 0.0138 | TRUE | 0.2653 |
| Slc22a18 | 83 | 79 | 131 | 207 | 205 | 239 | 64 | 84 | 141 | 206 | 231 | 236 | 160.51 | 96 | 224 | 2.343 | 1.228 | 1.02E-07 | 4.01E-06 | 0.0715 | 0.0172 | 0.031 | 0.031 | TRUE | 1.5693 |
| Slc22a19 | 0 | 0 | 0 | 0 | 0 | 0 | 0 | 0 | 0 | 0 | 0 | 0 | 0 | 0 | 0 | NA | NA | NA | NA | NA | NA | NA | NA | NA | NA |
| Slc22a2 | 1 | 0 | 0 | 0 | 1 | 0 | 1 | 0 | 0 | 0 | 1 | 0 | 0.32 | 0 | 0 | 1.075 | 0.105 | 0.97638847 | NA | 0 | 7.4681 | 7.0392 | 7.0392 | TRUE | 0.5075 |
| Slc22a20 | 7 | 10 | 4 | 4 | 3 | 4 | 5 | 11 | 4 | 4 | 3 | 4 | 5.28 | 7 | 4 | 0.559 | -0.838 | 0.32417648 | NA | 0 | 0.451 | 0.3124 | 0.3124 | TRUE | 0.5891 |
| Slc22a21 | 150 | 99 | 82 | 107 | 85 | 85 | 116 | 105 | 88 | 107 | 96 | 84 | 99.43 | 103 | 96 | 0.919 | -0.122 | 0.55491268 | 0.7102732 | 0.0072 | 0.0263 | 0.0209 | 0.0209 | TRUE | 0.1523 |
| Slc22a22 | 0 | 0 | 0 | 0 | 0 | 0 | 0 | 0 | 0 | 0 | 0 | 0 | 0 | 0 | 0 | NA | NA | NA | NA | NA | NA | NA | NA | NA | NA |
| Slc22a23 | 449 | 292 | 282 | 302 | 248 | 283 | 349 | 311 | 304 | 301 | 280 | 279 | 303.89 | 321 | 287 | 0.89 | -0.168 | 0.16782354 | 0.31927026 | 0 | 0.0102 | 0.0074 | 0.0074 | TRUE | 0.0685 |
| Slc22a26 | 0 | 0 | 0 | 0 | 0 | 0 | 0 | 0 | 0 | 0 | 0 | 0 | 0 | 0 | 0 | NA | NA | NA | NA | NA | NA | NA | NA | NA | NA |
| Slc22a27 | 0 | 1 | 0 | 0 | 0 | 1 | 0 | 1 | 0 | 0 | 0 | 1 | 0.34 | 0 | 0 | 1.075 | 0.105 | 0.97576789 | NA | 0 | 6.9363 | 6.5821 | 6.5821 | TRUE | 0.5216 |
| Slc22a28 | 0 | 0 | 1 | 0 | 0 | 0 | 0 | 0 | 1 | 0 | 0 | 0 | 0.18 | 0 | 0 | 0.552 | -0.857 | 0.8336548 | NA | 0 | 13.1817 | 10 | 10 | TRUE | 0.5365 |
| Slc22a29 | 0 | 1 | 0 | 0 | 0 | 0 | 0 | 1 | 0 | 0 | 0 | 0 | 0.18 | 0 | 0 | 0.552 | -0.857 | 0.8336548 | NA | 0 | 13.3714 | 10 | 10 | TRUE | 0.5216 |
| Slc22a3 | 3 | 2 | 0 | 0 | 2 | 0 | 2 | 2 | 0 | 0 | 2 | 0 | 1.12 | 1 | 1 | 0.481 | -1.056 | 0.60992644 | NA | 1.491 | 2.1206 | 2.0517 | 2.0517 | TRUE | 0.9749 |
| Slc22a30 | 0 | 0 | 1 | 1 | 0 | 0 | 0 | 0 | 1 | 1 | 0 | 0 | 0.35 | 0 | 0 | 1.075 | 0.105 | 0.97567165 | NA | 0 | 6.8576 | 6.5143 | 6.5143 | TRUE | 0.5365 |
| Slc22a4 | 4 | 0 | 1 | 0 | 3 | 1 | 3 | 0 | 1 | 0 | 3 | 1 | 1.43 | 1 | 1 | 0.988 | -0.018 | 0.99200896 | NA | 1.1147 | 1.6642 | 1.5422 | 1.5422 | TRUE | 0.5556 |
| Slc22a5 | 799 | 660 | 695 | 717 | 630 | 736 | 621 | 702 | 750 | 714 | 711 | 727 | 703.91 | 691 | 717 | 1.04 | 0.057 | 0.54513322 | 0.70230023 | 0.0035 | 0.0058 | 0.005 | 0.005 | TRUE | 0.099 |
| Slc22a6 | 2 | 1 | 1 | 1 | 1 | 1 | 2 | 1 | 1 | 1 | 1 | 1 | 1.13 | 1 | 1 | 0.831 | -0.267 | 0.87456621 | NA | 0 | 2.0916 | 1.174 | 1.174 | TRUE | 0.0408 |
| Slc22a7 | 259 | 219 | 263 | 347 | 325 | 393 | 201 | 233 | 284 | 345 | 367 | 388 | 302.98 | 239 | 367 | 1.539 | 0.622 | 1.04E-05 | 0.00019326 | 0.0126 | 0.0102 | 0.0109 | 0.0109 | TRUE | 0.2904 |
| Slc22a8 | 0 | 0 | 0 | 0 | 0 | 0 | 0 | 0 | 0 | 0 | 0 | 0 | 0 | 0 | 0 | NA | NA | NA | NA | NA | NA | NA | NA | NA | NA |
| Slc23a1 | 33 | 28 | 28 | 21 | 12 | 28 | 26 | 30 | 30 | 21 | 14 | 28 | 24.62 | 29 | 21 | 0.732 | -0.45 | 0.26526819 | 0.43561108 | 0 | 0.0987 | 0.0759 | 0.0759 | TRUE | 0.2618 |
| Slc23a2 | 3696 | 2680 | 2369 | 2434 | 2204 | 2645 | 2871 | 2851 | 2556 | 2423 | 2486 | 2611 | 2632.92 | 2759 | 2507 | 0.908 | -0.139 | 0.04199047 | 0.11844677 | 0.0022 | 0.0033 | 0.003 | 0.003 | TRUE | 0.0497 |
| Slc23a3 | 5 | 1 | 1 | 2 | 1 | 3 | 4 | 1 | 1 | 2 | 1 | 3 | 2.02 | 2 | 2 | 0.981 | -0.027 | 0.98385392 | NA | 0 | 1.177 | 0.8021 | 0.8021 | TRUE | 0.7933 |
| Slc24a1 | 395 | 282 | 305 | 395 | 336 | 356 | 307 | 300 | 329 | 393 | 379 | 351 | 343.25 | 312 | 374 | 1.201 | 0.265 | 0.02064025 | 0.070814 | 0 | 0.0093 | 0.0065 | 0.0065 | TRUE | 0.0343 |
| Slc24a2 | 60 | 23 | 21 | 8 | 11 | 12 | 47 | 24 | 23 | 8 | 12 | 12 | 20.99 | 31 | 11 | 0.339 | -1.561 | 0.0013321 | 0.00885981 | 0.0883 | 0.1153 | 0.1069 | 0.1069 | TRUE | 1.4351 |
| Slc24a3 | 10 | 6 | 2 | 6 | 3 | 4 | 8 | 6 | 2 | 6 | 3 | 4 | 4.94 | 5 | 4 | 0.807 | -0.31 | 0.7241932 | NA | 0 | 0.4826 | 0.3532 | 0.3532 | TRUE | 0.5482 |
| Slc24a4 | 2 | 1 | 2 | 1 | 1 | 1 | 2 | 1 | 2 | 1 | 1 | 1 | 1.31 | 2 | 1 | 0.651 | -0.619 | 0.69903494 | NA | 0 | 1.8057 | 1.0419 | 1.0419 | TRUE | 0.0615 |
| Slc24a5 | 31 | 20 | 21 | 28 | 18 | 25 | 24 | 21 | 23 | 28 | 20 | 25 | 23.48 | 23 | 24 | 1.071 | 0.1 | 0.79951685 | 0.88311394 | 0 | 0.1034 | 0.0684 | 0.0684 | TRUE | 0.1121 |
| Slc24a6 | 2040 | 1648 | 1644 | 1918 | 1660 | 2080 | 1584 | 1753 | 1774 | 1909 | 1873 | 2053 | 1824.38 | 1704 | 1945 | 1.143 | 0.193 | 0.00833979 | 0.03615429 | 0.0026 | 0.0037 | 0.0033 | 0.0033 | TRUE | 0.047 |
| Slc25a1 | 6208 | 4840 | 5365 | 6409 | 5778 | 6714 | 4821 | 5148 | 5789 | 6380 | 6518 | 6628 | 5880.66 | 5253 | 6509 | 1.24 | 0.31 | 1.07E-05 | 0.00019764 | 0.0043 | 0.0028 | 0.0034 | 0.0034 | TRUE | 0.0969 |
| Slc25a10 | 676 | 476 | 304 | 278 | 308 | 266 | 525 | 506 | 328 | 277 | 347 | 263 | 374.35 | 453 | 296 | 0.65 | -0.621 | 0.00026809 | 0.00266655 | 0.0408 | 0.0088 | 0.0181 | 0.0181 | TRUE | 0.6653 |
| Slc25a11 | 5648 | 4020 | 3728 | 3808 | 3329 | 3878 | 4387 | 4276 | 4022 | 3790 | 3755 | 3828 | 4009.86 | 4228 | 3791 | 0.896 | -0.158 | 0.0055458 | 0.02656138 | 8.00E-04 | 0.003 | 0.0021 | 0.0021 | TRUE | 0.0219 |
| Slc25a12 | 2308 | 1853 | 1774 | 1754 | 1656 | 1823 | 1793 | 1971 | 1914 | 1746 | 1868 | 1800 | 1848.55 | 1893 | 1805 | 0.954 | -0.068 | 0.31717877 | 0.49116318 | 0.0013 | 0.0037 | 0.0028 | 0.0028 | TRUE | 0.0268 |
| Slc25a13 | 11032 | 8088 | 8517 | 9806 | 8505 | 9665 | 8568 | 8603 | 9190 | 9761 | 9595 | 9541 | 9209.53 | 8787 | 9632 | 1.096 | 0.133 | 0.01002012 | 0.04141691 | 7.00E-04 | 0.0027 | 0.0018 | 0.0018 | TRUE | 0.0195 |
| Slc25a14 | 323 | 214 | 217 | 212 | 204 | 196 | 251 | 228 | 234 | 211 | 230 | 193 | 224.55 | 238 | 211 | 0.888 | -0.172 | 0.21509833 | 0.37765193 | 0 | 0.013 | 0.0093 | 0.0093 | TRUE | 0.0622 |
| Slc25a15 | 5796 | 4160 | 3908 | 4604 | 3826 | 4377 | 4502 | 4425 | 4217 | 4583 | 4316 | 4321 | 4393.81 | 4381 | 4407 | 1.006 | 0.008 | 0.8869454 | 0.937038 | 9.00E-04 | 0.003 | 0.0021 | 0.0021 | TRUE | 0.015 |
| Slc25a16 | 1497 | 970 | 920 | 1038 | 882 | 979 | 1163 | 1032 | 993 | 1033 | 995 | 966 | 1030.29 | 1063 | 998 | 0.938 | -0.092 | 0.27604658 | 0.44713664 | 0.0033 | 0.0047 | 0.0042 | 0.0042 | TRUE | 0.0847 |
| Slc25a17 | 3638 | 2583 | 2893 | 3371 | 2753 | 3178 | 2825 | 2748 | 3122 | 3355 | 3106 | 3137 | 3048.81 | 2898 | 3199 | 1.104 | 0.143 | 0.03635128 | 0.10695117 | 0.0027 | 0.0032 | 0.003 | 0.003 | TRUE | 0.0544 |
| Slc25a18 | 8 | 3 | 2 | 2 | 3 | 4 | 6 | 3 | 2 | 2 | 3 | 4 | 3.48 | 4 | 3 | 0.788 | -0.344 | 0.73706272 | NA | 0 | 0.6833 | 0.4678 | 0.4678 | TRUE | 0.6075 |
| Slc25a19 | 1339 | 1023 | 1261 | 1272 | 1157 | 1443 | 1040 | 1088 | 1361 | 1266 | 1305 | 1424 | 1247.42 | 1163 | 1332 | 1.147 | 0.198 | 0.05279041 | 0.14022177 | 0.0118 | 0.0043 | 0.0067 | 0.0067 | TRUE | 0.263 |
| Slc25a2 | 16 | 8 | 8 | 6 | 6 | 8 | 12 | 9 | 9 | 6 | 7 | 8 | 8.37 | 10 | 7 | 0.692 | -0.531 | 0.41850367 | NA | 0 | 0.2857 | 0.1872 | 0.1872 | TRUE | 0.2273 |
| Slc25a20 | 3683 | 2549 | 2353 | 2049 | 1982 | 1958 | 2860 | 2711 | 2539 | 2040 | 2236 | 1933 | 2386.49 | 2703 | 2070 | 0.765 | -0.387 | 2.69E-07 | 9.46E-06 | 0.0039 | 0.0034 | 0.0036 | 0.0036 | TRUE | 0.0593 |
| Slc25a21 | 2 | 0 | 1 | 1 | 1 | 1 | 2 | 0 | 1 | 1 | 1 | 1 | 0.96 | 1 | 1 | 1.143 | 0.193 | 0.91871967 | NA | 0 | 2.4786 | 1.5559 | 1.5559 | TRUE | 0.2848 |
| Slc25a22 | 1009 | 746 | 713 | 763 | 619 | 814 | 784 | 794 | 769 | 759 | 698 | 804 | 767.97 | 782 | 754 | 0.964 | -0.052 | 0.54517202 | 0.70230023 | 0.0011 | 0.0055 | 0.0041 | 0.0041 | TRUE | 0.0478 |
| Slc25a23 | 1026 | 861 | 980 | 980 | 974 | 1159 | 797 | 916 | 1057 | 975 | 1099 | 1144 | 998.08 | 923 | 1073 | 1.164 | 0.219 | 0.04108663 | 0.11661065 | 0.0118 | 0.0048 | 0.0073 | 0.0073 | TRUE | 0.1921 |
| Slc25a24 | 617 | 399 | 373 | 363 | 267 | 266 | 479 | 424 | 402 | 361 | 301 | 263 | 371.86 | 435 | 308 | 0.707 | -0.5 | 0.00020924 | 0.00220066 | 0.0139 | 0.0088 | 0.0103 | 0.0103 | TRUE | 0.2603 |
| Slc25a25 | 1670 | 1214 | 1155 | 1451 | 1239 | 1394 | 1297 | 1291 | 1246 | 1444 | 1398 | 1376 | 1342.12 | 1278 | 1406 | 1.1 | 0.137 | 0.04347627 | 0.12162973 | 0 | 0.0042 | 0.0026 | 0.0026 | TRUE | 0.0069 |
| Slc25a26 | 1133 | 789 | 730 | 781 | 667 | 774 | 880 | 839 | 788 | 777 | 752 | 764 | 800.13 | 836 | 764 | 0.914 | -0.13 | 0.11903625 | 0.25057473 | 0 | 0.0054 | 0.0037 | 0.0037 | TRUE | 0.0296 |
| Slc25a27 | 84 | 87 | 81 | 102 | 78 | 82 | 65 | 93 | 87 | 102 | 88 | 81 | 85.94 | 82 | 90 | 1.111 | 0.152 | 0.50258693 | 0.66680359 | 0.0134 | 0.03 | 0.0252 | 0.0252 | TRUE | 0.3332 |
| Slc25a28 | 1004 | 865 | 765 | 690 | 624 | 679 | 780 | 920 | 825 | 687 | 704 | 670 | 764.39 | 842 | 687 | 0.817 | -0.291 | 0.00133624 | 0.00887497 | 0.0029 | 0.0055 | 0.0046 | 0.0046 | TRUE | 0.0784 |
| Slc25a29 | 84 | 78 | 65 | 82 | 91 | 84 | 65 | 83 | 70 | 82 | 103 | 83 | 80.92 | 73 | 89 | 1.226 | 0.294 | 0.19199119 | 0.35032938 | 0 | 0.0317 | 0.024 | 0.024 | TRUE | 0.1617 |
| Slc25a3 | 31333 | 22026 | 20958 | 23108 | 20005 | 23061 | 24335 | 23429 | 22614 | 23002 | 22568 | 22765 | 23118.57 | 23459 | 22778 | 0.971 | -0.043 | 0.38031767 | 0.55490817 | 7.00E-04 | 0.0025 | 0.0017 | 0.0017 | TRUE | 0.0131 |
| Slc25a30 | 1740 | 1461 | 1990 | 2147 | 1868 | 2139 | 1351 | 1554 | 2147 | 2137 | 2107 | 2112 | 1901.43 | 1684 | 2119 | 1.258 | 0.332 | 0.10353185 | 0.22692543 | 0.0294 | 0.0037 | 0.0109 | 0.0294 | TRUE | 0.697 |
| Slc25a31 | 368 | 273 | 225 | 222 | 150 | 139 | 286 | 290 | 243 | 221 | 169 | 137 | 224.4 | 273 | 176 | 0.644 | -0.636 | 0.00024544 | 0.00249509 | 0.0265 | 0.013 | 0.0169 | 0.0169 | TRUE | 0.5521 |
| Slc25a32 | 1440 | 1031 | 1007 | 1192 | 951 | 1144 | 1118 | 1097 | 1087 | 1187 | 1073 | 1129 | 1115.04 | 1101 | 1130 | 1.026 | 0.038 | 0.61254394 | 0.75500953 | 4.00E-04 | 0.0046 | 0.0031 | 0.0031 | TRUE | 0.0233 |
| Slc25a33 | 1371 | 1056 | 1085 | 1487 | 1279 | 1379 | 1065 | 1123 | 1171 | 1480 | 1443 | 1361 | 1273.84 | 1120 | 1428 | 1.277 | 0.353 | 2.24E-06 | 5.52E-05 | 0.0013 | 0.0043 | 0.0032 | 0.0032 | TRUE | 0.0231 |
| Slc25a34 | 42 | 19 | 25 | 28 | 27 | 26 | 33 | 20 | 27 | 28 | 30 | 26 | 27.3 | 27 | 28 | 1.042 | 0.06 | 0.87284181 | 0.92948353 | 0 | 0.0892 | 0.0641 | 0.0641 | TRUE | 0.2558 |
| Slc25a35 | 210 | 131 | 179 | 246 | 169 | 187 | 163 | 139 | 193 | 245 | 191 | 185 | 185.95 | 165 | 207 | 1.253 | 0.325 | 0.06218274 | 0.15744515 | 0.0197 | 0.0152 | 0.0165 | 0.0165 | TRUE | 0.2879 |
| Slc25a36 | 5288 | 4116 | 5026 | 7252 | 5760 | 7009 | 4107 | 4378 | 5423 | 7219 | 6498 | 6919 | 5757.26 | 4636 | 6879 | 1.484 | 0.57 | 5.19E-10 | 3.74E-08 | 0.0123 | 0.0028 | 0.0059 | 0.0059 | TRUE | 0.2681 |
| Slc25a37 | 1325 | 1182 | 976 | 1140 | 1031 | 1251 | 1029 | 1257 | 1053 | 1135 | 1163 | 1235 | 1145.37 | 1113 | 1178 | 1.059 | 0.083 | 0.3618903 | 0.53662186 | 0.0057 | 0.0045 | 0.0051 | 0.0051 | TRUE | 0.1521 |
| Slc25a38 | 976 | 766 | 728 | 768 | 659 | 817 | 758 | 815 | 786 | 764 | 743 | 807 | 778.78 | 786 | 771 | 0.983 | -0.025 | 0.76232598 | 0.85900698 | 0 | 0.0055 | 0.0037 | 0.0037 | TRUE | 0.0188 |
| Slc25a39 | 14557 | 10536 | 11295 | 11367 | 10363 | 11370 | 11306 | 11207 | 12187 | 11315 | 11691 | 11224 | 11488.2 | 11567 | 11410 | 0.986 | -0.02 | 0.71552367 | 0.829237 | 0.001 | 0.0026 | 0.002 | 0.002 | TRUE | 0.0268 |

| | | | | | | | | | | | | | | | | | | | | | | | | |
|---|---|---|---|---|---|---|---|---|---|---|---|---|---|---|---|---|---|---|---|---|---|---|---|---|
| Slc25a4 | 19520 | 13541 | 13608 | 11316 | 9457 | 10789 | 15160 | 14403 | 14683 | 11264 | 10668 | 10650 | 12804.91 | 14749 | 10861 | 0.736 | -0.442 | 2.81E-18 | 6.92E-16 | 7.00E-04 | 0.0026 | 0.0018 | 0.0018 | TRUE | 0.0129 |
| Slc25a40 | 1750 | 1237 | 1233 | 1325 | 1184 | 1241 | 1359 | 1316 | 1330 | 1319 | 1336 | 1225 | 1314.16 | 1335 | 1293 | 0.968 | -0.047 | 0.50650533 | 0.6698881 | 4.00E-04 | 0.0042 | 0.0028 | 0.0028 | TRUE | 0.0258 |
| Slc25a41 | 62 | 46 | 30 | 37 | 35 | 41 | 48 | 49 | 32 | 37 | 39 | 40 | 41.04 | 43 | 39 | 0.898 | -0.155 | 0.61593179 | 0.75749645 | 0 | 0.0602 | 0.0445 | 0.0445 | TRUE | 0.3433 |
| Slc25a42 | 708 | 567 | 612 | 950 | 741 | 846 | 550 | 603 | 660 | 946 | 836 | 835 | 738.34 | 604 | 872 | 1.447 | 0.533 | 5.98E-08 | 2.45E-06 | 0.0054 | 0.0056 | 0.0056 | 0.0056 | TRUE | 0.0777 |
| Slc25a43 | 68 | 63 | 53 | 62 | 53 | 68 | 53 | 67 | 57 | 62 | 60 | 67 | 60.94 | 59 | 63 | 1.071 | 0.099 | 0.69233243 | 0.81190576 | 0 | 0.0413 | 0.0283 | 0.0283 | TRUE | 0.118 |
| Slc25a44 | 871 | 549 | 542 | 596 | 555 | 657 | 676 | 584 | 585 | 593 | 626 | 649 | 618.86 | 615 | 623 | 1.01 | 0.014 | 0.88496867 | 0.93609243 | 0.0034 | 0.0063 | 0.0053 | 0.0053 | TRUE | 0.0944 |
| Slc25a45 | 15 | 8 | 7 | 4 | 3 | 11 | 12 | 9 | 8 | 4 | 3 | 11 | 7.66 | 10 | 6 | 0.658 | -0.604 | 0.40792492 | NA | 0.0675 | 0.312 | 0.2478 | 0.2478 | TRUE | 1.1563 |
| Slc25a46 | 3284 | 2269 | 2381 | 2465 | 2043 | 2372 | 2551 | 2414 | 2569 | 2454 | 2305 | 2342 | 2438.84 | 2511 | 2367 | 0.942 | -0.086 | 0.16496418 | 0.31576711 | 7.00E-04 | 0.0034 | 0.0023 | 0.0023 | TRUE | 0.0138 |
| Slc25a47 | 33 | 16 | 16 | 17 | 15 | 13 | 26 | 17 | 17 | 17 | 17 | 13 | 17.76 | 20 | 16 | 0.77 | -0.377 | 0.4143431 | 0.58693815 | 0 | 0.1358 | 0.0963 | 0.0963 | TRUE | 0.4023 |
| Slc25a48 | 12 | 13 | 15 | 11 | 17 | 15 | 9 | 14 | 16 | 11 | 19 | 15 | 14.04 | 13 | 15 | 1.151 | 0.203 | 0.69913337 | NA | 0 | 0.1712 | 0.1263 | 0.1263 | TRUE | 0.2422 |
| Slc25a5 | 37570 | 25821 | 25981 | 28178 | 24033 | 28997 | 29179 | 27465 | 28033 | 28048 | 27112 | 28625 | 28077.06 | 28226 | 27928 | 0.989 | -0.015 | 0.75562568 | 0.85511255 | 7.00E-04 | 0.0025 | 0.0017 | 0.0017 | TRUE | 0.0107 |
| Slc25a51 | 2393 | 1692 | 1795 | 1721 | 1536 | 1813 | 1859 | 1800 | 1937 | 1713 | 1733 | 1790 | 1805.11 | 1865 | 1745 | 0.936 | -0.096 | 0.1387085 | 0.27974611 | 3.00E-04 | 0.0037 | 0.0024 | 0.0024 | TRUE | 0.0132 |
| Slc25a53 | 79 | 57 | 54 | 80 | 45 | 63 | 61 | 61 | 58 | 80 | 51 | 62 | 62.14 | 60 | 64 | 1.071 | 0.099 | 0.70261104 | 0.81985628 | 0.0093 | 0.0406 | 0.032 | 0.032 | TRUE | 0.316 |
| Slc25a54 | 0 | 0 | 0 | 0 | 0 | 0 | 0 | 0 | 0 | 0 | 0 | 0 | 0 | 0 | 0 | NA | NA | NA | NA | NA | NA | NA | NA | NA | NA |
| Slc26a1 | 5 | 2 | 0 | 1 | 0 | 1 | 4 | 2 | 0 | 1 | 0 | 1 | 1.33 | 2 | 1 | 0.329 | -1.603 | 0.38712656 | NA | 0 | 1.7815 | 1.4752 | 1.4752 | TRUE | 0.107 |
| Slc26a10 | 132 | 112 | 135 | 116 | 136 | 148 | 103 | 119 | 146 | 115 | 153 | 146 | 130.38 | 123 | 138 | 1.134 | 0.181 | 0.35678852 | 0.53131677 | 0.0193 | 0.0206 | 0.0202 | 0.0202 | TRUE | 0.2756 |
| Slc26a11 | 435 | 306 | 339 | 405 | 360 | 413 | 338 | 325 | 366 | 403 | 406 | 408 | 374.34 | 343 | 406 | 1.183 | 0.243 | 0.02515523 | 0.08164025 | 0 | 0.0088 | 0.0058 | 0.0058 | TRUE | 0.0362 |
| Slc26a2 | 2345 | 1632 | 1521 | 1568 | 1378 | 1590 | 1821 | 1736 | 1641 | 1561 | 1555 | 1570 | 1647.21 | 1733 | 1562 | 0.9 | -0.151 | 0.02606091 | 0.0838183 | 9.00E-04 | 0.0039 | 0.0027 | 0.0027 | TRUE | 0.0257 |
| Slc26a3 | 2 | 2 | 1 | 0 | 0 | 1 | 2 | 2 | 1 | 0 | 0 | 1 | 0.96 | 2 | 0 | 0.234 | -2.097 | 0.30586543 | NA | 0 | 2.4769 | 1.7179 | 1.7179 | TRUE | 0.3835 |
| Slc26a4 | 27 | 6 | 9 | 10 | 7 | 11 | 21 | 6 | 10 | 10 | 8 | 11 | 10.96 | 12 | 10 | 0.761 | -0.394 | 0.53101601 | NA | 0.1339 | 0.2186 | 0.1925 | 0.1925 | TRUE | 1.1712 |
| Slc26a5 | 9 | 4 | 6 | 6 | 4 | 3 | 7 | 4 | 6 | 6 | 5 | 3 | 5.19 | 6 | 5 | 0.754 | -0.408 | 0.62432282 | NA | 0 | 0.4587 | 0.3035 | 0.3035 | TRUE | 0.1723 |
| Slc26a6 | 309 | 209 | 259 | 275 | 233 | 253 | 240 | 222 | 279 | 274 | 263 | 250 | 254.68 | 247 | 262 | 1.061 | 0.086 | 0.52818862 | 0.68874916 | 0.0035 | 0.0117 | 0.0094 | 0.0094 | TRUE | 0.1367 |
| Slc26a7 | 4 | 0 | 3 | 0 | 2 | 2 | 3 | 0 | 3 | 0 | 2 | 2 | 1.76 | 2 | 1 | 0.653 | -0.615 | 0.69737944 | NA | 0 | 1.3473 | 1.2081 | 1.2081 | TRUE | 0.6624 |
| Slc26a9 | 27 | 24 | 21 | 16 | 25 | 21 | 21 | 26 | 23 | 16 | 28 | 21 | 22.34 | 23 | 22 | 0.936 | -0.095 | 0.81877052 | 0.89590869 | 0 | 0.1085 | 0.0788 | 0.0788 | TRUE | 0.3326 |
| Slc27a1 | 1216 | 1114 | 1203 | 1123 | 972 | 1085 | 944 | 1185 | 1298 | 1118 | 1097 | 1071 | 1118.8 | 1142 | 1095 | 0.96 | -0.058 | 0.58239507 | 0.73253195 | 0.0126 | 0.0045 | 0.0072 | 0.0072 | TRUE | 0.284 |
| Slc27a2 | 512 | 460 | 411 | 390 | 395 | 526 | 398 | 489 | 443 | 388 | 446 | 519 | 447.24 | 443 | 451 | 1.019 | 0.027 | 0.83112655 | 0.9033735 | 0.0139 | 0.0077 | 0.0096 | 0.0096 | TRUE | 0.2066 |
| Slc27a3 | 1202 | 929 | 861 | 987 | 851 | 991 | 934 | 988 | 929 | 982 | 960 | 978 | 961.91 | 950 | 973 | 1.025 | 0.036 | 0.63439717 | 0.77136867 | 0 | 0.0049 | 0.0031 | 0.0031 | TRUE | 0.0141 |
| Slc27a4 | 2463 | 1710 | 1570 | 1697 | 1431 | 1542 | 1913 | 1819 | 1694 | 1689 | 1614 | 1522 | 1708.59 | 1809 | 1608 | 0.889 | -0.17 | 0.02194939 | 0.07381708 | 0.0027 | 0.0038 | 0.0034 | 0.0034 | TRUE | 0.0369 |
| Slc27a5 | 37 | 21 | 31 | 41 | 24 | 26 | 29 | 22 | 33 | 41 | 27 | 26 | 29.68 | 28 | 31 | 1.108 | 0.148 | 0.68891633 | 0.8096268 | 0.0197 | 0.0823 | 0.0652 | 0.0652 | TRUE | 0.5033 |
| Slc27a6 | 9 | 3 | 3 | 3 | 9 | 2 | 7 | 3 | 3 | 3 | 10 | 2 | 4.76 | 4 | 5 | 1.089 | 0.123 | 0.89640794 | NA | 0.2787 | 0.5008 | 0.4329 | 0.4329 | TRUE | 1.4477 |
| Slc28a1 | 285 | 212 | 241 | 268 | 251 | 325 | 221 | 226 | 260 | 267 | 283 | 321 | 262.94 | 236 | 290 | 1.235 | 0.304 | 0.02503687 | 0.08144104 | 0.0048 | 0.0114 | 0.0094 | 0.0094 | TRUE | 0.0977 |
| Slc28a2 | 6 | 2 | 2 | 1 | 0 | 1 | 5 | 2 | 2 | 1 | 0 | 1 | 1.82 | 3 | 1 | 0.224 | -2.157 | 0.17362659 | NA | 0 | 1.3037 | 0.8901 | 0.8901 | TRUE | 0.4099 |
| Slc28a3 | 73 | 52 | 57 | 37 | 53 | 46 | 57 | 55 | 62 | 37 | 60 | 45 | 52.59 | 58 | 47 | 0.816 | -0.294 | 0.29329884 | 0.46521877 | 0.0061 | 0.0475 | 0.037 | 0.037 | TRUE | 0.3954 |
| Slc29a1 | 15542 | 11964 | 11616 | 13445 | 12027 | 14301 | 12071 | 12726 | 12534 | 13383 | 13568 | 14117 | 13066.41 | 12444 | 13689 | 1.1 | 0.138 | 0.00575274 | 0.02732598 | 6.00E-04 | 0.0026 | 0.0017 | 0.0017 | TRUE | 0.0092 |
| Slc29a2 | 929 | 669 | 737 | 718 | 680 | 801 | 722 | 712 | 795 | 715 | 767 | 791 | 750.14 | 743 | 758 | 1.02 | 0.029 | 0.74137938 | 0.8460229 | 0.0018 | 0.0056 | 0.0043 | 0.0043 | TRUE | 0.0438 |
| Slc29a3 | 1276 | 969 | 915 | 1056 | 895 | 1133 | 991 | 1031 | 987 | 1051 | 1010 | 1118 | 1031.37 | 1003 | 1060 | 1.057 | 0.08 | 0.29524562 | 0.46705464 | 6.00E-04 | 0.0047 | 0.0033 | 0.0033 | TRUE | 0.0282 |
| Slc29a4 | 1564 | 1209 | 1172 | 1717 | 1563 | 1664 | 1215 | 1286 | 1265 | 1709 | 1763 | 1643 | 1480.04 | 1255 | 1705 | 1.359 | 0.443 | 9.67E-11 | 7.86E-09 | 5.00E-04 | 0.004 | 0.0027 | 0.0027 | TRUE | 0.0124 |
| Slc2a1 | 61481 | 43709 | 44289 | 49603 | 42305 | 46587 | 47750 | 46493 | 47787 | 49375 | 47724 | 45989 | 47519.6 | 47343 | 47696 | 1.007 | 0.011 | 0.82416508 | 0.89901206 | 6.00E-04 | 0.0025 | 0.0016 | 0.0016 | TRUE | 0.012 |
| Slc2a10 | 3 | 1 | 4 | 7 | 8 | 7 | 2 | 1 | 4 | 7 | 9 | 7 | 5.1 | 2 | 8 | 2.985 | 1.578 | 0.078831 | NA | 0 | 0.467 | 0.3242 | 0.3242 | TRUE | 0.342 |
| Slc2a12 | 57 | 42 | 45 | 43 | 42 | 40 | 44 | 45 | 49 | 43 | 47 | 39 | 44.53 | 46 | 43 | 0.943 | -0.085 | 0.7659726 | 0.86116938 | 0 | 0.0557 | 0.0363 | 0.0363 | TRUE | 0.0509 |
| Slc2a13 | 5 | 7 | 2 | 3 | 1 | 3 | 4 | 7 | 2 | 3 | 1 | 3 | 3.43 | 4 | 2 | 0.532 | -0.911 | 0.39388729 | NA | 0 | 0.694 | 0.497 | 0.497 | TRUE | 0.3348 |
| Slc2a2 | 3 | 4 | 0 | 1 | 0 | 1 | 2 | 4 | 0 | 1 | 0 | 1 | 1.43 | 2 | 1 | 0.307 | -1.704 | 0.3488457 | NA | 0 | 1.6623 | 1.3973 | 1.3973 | TRUE | 0.1017 |
| Slc2a3 | 165404 | 125212 | 139259 | 138901 | 124750 | 136391 | 128462 | 133186 | 150259 | 138262 | 140731 | 134639 | 137589.96 | 137302 | 137877 | 1.004 | 0.006 | 0.92456896 | 0.95836863 | 0.003 | 0.0024 | 0.0029 | 0.0029 | TRUE | 0.0835 |
| Slc2a4 | 141 | 127 | 175 | 248 | 218 | 230 | 110 | 135 | 189 | 247 | 246 | 227 | 192.21 | 145 | 240 | 1.672 | 0.741 | 6.89E-05 | 0.00090777 | 0.032 | 0.0148 | 0.0195 | 0.0195 | TRUE | 0.743 |
| Slc2a4rg-ps | 183 | 166 | 130 | 101 | 96 | 129 | 142 | 177 | 140 | 101 | 108 | 127 | 132.52 | 153 | 112 | 0.734 | -0.446 | 0.01623372 | 0.05922116 | 0.0087 | 0.0203 | 0.017 | 0.017 | TRUE | 0.185 |
| Slc2a5 | 493 | 355 | 411 | 472 | 412 | 462 | 383 | 378 | 443 | 470 | 465 | 456 | 432.44 | 401 | 464 | 1.157 | 0.21 | 0.04998872 | 0.13477101 | 0.0014 | 0.0079 | 0.006 | 0.006 | TRUE | 0.0937 |
| Slc2a6 | 50 | 36 | 35 | 49 | 41 | 47 | 39 | 38 | 38 | 49 | 46 | 46 | 42.72 | 38 | 47 | 1.231 | 0.299 | 0.30026221 | 0.47244031 | 0 | 0.0579 | 0.0366 | 0.0366 | TRUE | 0.0075 |
| Slc2a7 | 75 | 44 | 39 | 50 | 37 | 58 | 58 | 47 | 42 | 50 | 42 | 57 | 49.32 | 49 | 50 | 1.007 | 0.01 | 0.97251619 | 0.98512927 | 0 | 0.0505 | 0.0386 | 0.0386 | TRUE | 0.2502 |
| Slc2a8 | 1332 | 1198 | 1214 | 1319 | 1253 | 1349 | 1035 | 1274 | 1310 | 1313 | 1414 | 1332 | 1279.47 | 1206 | 1353 | 1.123 | 0.168 | 0.07503985 | 0.18035052 | 0.0082 | 0.0043 | 0.0056 | 0.0056 | TRUE | 0.1925 |
| Slc2a9 | 58 | 41 | 37 | 35 | 21 | 19 | 45 | 44 | 40 | 35 | 24 | 19 | 34.31 | 43 | 26 | 0.601 | -0.735 | 0.03475451 | 0.10331876 | 0.0056 | 0.0715 | 0.056 | 0.056 | TRUE | 0.6069 |
| Slc30a1 | 2628 | 2113 | 1777 | 2164 | 1834 | 2032 | 2041 | 2248 | 1917 | 2154 | 2069 | 2006 | 2072.48 | 2069 | 2076 | 1.004 | 0.005 | 0.9410423 | 0.96681409 | 0.0033 | 0.0036 | 0.0035 | 0.0035 | TRUE | 0.0679 |
| Slc30a10 | 86 | 86 | 96 | 117 | 91 | 90 | 67 | 91 | 104 | 116 | 103 | 89 | 94.97 | 87 | 103 | 1.185 | 0.245 | 0.28055208 | 0.45212886 | 0.024 | 0.0274 | 0.0264 | 0.0264 | TRUE | 0.4546 |
| Slc30a2 | 1850 | 1374 | 1234 | 1265 | 1232 | 1349 | 1437 | 1462 | 1331 | 1259 | 1390 | 1332 | 1368.41 | 1410 | 1327 | 0.94 | -0.089 | 0.23317251 | 0.39931812 | 0.0016 | 0.0042 | 0.0033 | 0.0033 | TRUE | 0.0278 |
| Slc30a3 | 147 | 125 | 165 | 253 | 209 | 255 | 114 | 133 | 178 | 252 | 236 | 252 | 194.08 | 142 | 247 | 1.751 | 0.808 | 2.65E-06 | 6.29E-05 | 0.0192 | 0.0146 | 0.0158 | 0.0158 | TRUE | 0.5134 |
| Slc30a4 | 1114 | 842 | 689 | 724 | 693 | 748 | 865 | 896 | 743 | 721 | 782 | 738 | 790.85 | 835 | 747 | 0.894 | -0.162 | 0.08408871 | 0.19555483 | 0.0044 | 0.0054 | 0.0051 | 0.0051 | TRUE | 0.1056 |
| Slc30a5 | 4303 | 3075 | 2997 | 3272 | 2722 | 3173 | 3342 | 3271 | 3234 | 3257 | 3071 | 3132 | 3217.74 | 3282 | 3153 | 0.961 | -0.058 | 0.30173068 | 0.4740724 | 3.00E-04 | 0.0032 | 0.002 | 0.002 | TRUE | 0.0101 |
| Slc30a6 | 936 | 634 | 632 | 778 | 659 | 714 | 727 | 674 | 682 | 774 | 743 | 705 | 717.65 | 694 | 741 | 1.066 | 0.092 | 0.29125056 | 0.46341359 | 0 | 0.0057 | 0.004 | 0.004 | TRUE | 0.022 |
| Slc30a7 | 1945 | 1372 | 1501 | 1500 | 1383 | 1649 | 1511 | 1459 | 1620 | 1493 | 1560 | 1628 | 1545.1 | 1530 | 1560 | 1.02 | 0.029 | 0.68931121 | 0.80989109 | 0.0017 | 0.004 | 0.0031 | 0.0031 | TRUE | 0.0308 |
| Slc30a8 | 0 | 0 | 1 | 0 | 0 | 0 | 0 | 0 | 1 | 0 | 0 | 0 | 0.18 | 0 | 0 | 0.552 | -0.857 | 0.8336548 | NA | 0 | 13.1817 | 10 | 10 | TRUE | 0.5365 |
| Slc30a9 | 4694 | 3430 | 3436 | 3979 | 3561 | 3950 | 3646 | 3648 | 3707 | 3961 | 4017 | 3899 | 3813.1 | 3667 | 3959 | 1.08 | 0.11 | 0.03312819 | 0.09970366 | 0 | 0.003 | 0.0017 | 0.0017 | TRUE | 0.0021 |
| Slc31a1 | 4556 | 3440 | 4121 | 4577 | 3958 | 4626 | 3538 | 3659 | 4447 | 4556 | 4465 | 4567 | 4205.27 | 3881 | 4529 | 1.168 | 0.223 | 0.00647803 | 0.02990647 | 0.0078 | 0.003 | 0.0046 | 0.0046 | TRUE | 0.1967 |
| Slc31a2 | 684 | 432 | 525 | 565 | 570 | 748 | 531 | 460 | 566 | 562 | 643 | 738 | 583.5 | 519 | 648 | 1.248 | 0.32 | 0.00755848 | 0.03361879 | 0.0132 | 0.0065 | 0.0086 | 0.0086 | TRUE | 0.1783 |
| Slc32a1 | 107 | 76 | 87 | 151 | 127 | 150 | 83 | 81 | 94 | 150 | 143 | 148 | 116.58 | 86 | 147 | 1.717 | 0.779 | 2.05E-05 | 0.00033825 | 0 | 0.0228 | 0.015 | 0.015 | TRUE | 0.0573 |
| Slc33a1 | 1180 | 831 | 724 | 783 | 655 | 665 | 916 | 884 | 781 | 779 | 739 | 656 | 792.72 | 860 | 725 | 0.841 | -0.249 | 0.01072627 | 0.04348001 | 0.006 | 0.0054 | 0.0056 | 0.0056 | TRUE | 0.0821 |
| Slc34a1 | 0 | 0 | 0 | 0 | 1 | 2 | 0 | 0 | 0 | 0 | 1 | 2 | 0.52 | 0 | 1 | 5.84 | 2.546 | 0.38239357 | NA | 0 | 4.5861 | 4.6218 | 4.6218 | TRUE | 0.0847 |
| Slc34a2 | 89 | 50 | 23 | 31 | 23 | 20 | 69 | 53 | 25 | 31 | 26 | 20 | 37.28 | 49 | 26 | 0.516 | -0.955 | 0.01647914 | 0.05987196 | 0.1281 | 0.066 | 0.0842 | 0.0842 | TRUE | 0.2425 |
| Slc34a3 | 63 | 46 | 32 | 68 | 47 | 65 | 49 | 49 | 35 | 68 | 53 | 64 | 52.88 | 44 | 62 | 1.393 | 0.478 | 0.0842906 | 0.1957695 | 0 | 0.0472 | 0.0359 | 0.0359 | TRUE | 0.2617 |
| Slc35a1 | 1437 | 1047 | 925 | 1132 | 911 | 1044 | 1116 | 1114 | 998 | 1127 | 1028 | 1031 | 1068.81 | 1076 | 1062 | 0.986 | -0.02 | 0.80689709 | 0.88842854 | 0.0024 | 0.0046 | 0.0039 | 0.0039 | TRUE | 0.0469 |
| Slc35a2 | 974 | 692 | 717 | 838 | 707 | 872 | 756 | 736 | 774 | 834 | 798 | 861 | 793.11 | 755 | 831 | 1.1 | 0.138 | 0.08971476 | 0.20422136 | 0 | 0.0054 | 0.0035 | 0.0035 | TRUE | 0.0142 |

| | | | | | | | | | | | | | | | | | | | | | | | | | |
|---|---|---|---|---|---|---|---|---|---|---|---|---|---|---|---|---|---|---|---|---|---|---|---|---|---|
| Slc35a3 | 2250 | 1572 | 1770 | 2228 | 1909 | 1974 | 1747 | 1672 | 1910 | 2218 | 2154 | 1949 | 1941.56 | 1776 | 2107 | 1.186 | 0.246 | 0.00144756 | 0.00939174 | 0.0038 | 0.0036 | 0.0038 | 0.0038 | TRUE | 0.0525 |
| Slc35a4 | 3139 | 2199 | 1822 | 2089 | 1853 | 2197 | 2438 | 2339 | 1966 | 2079 | 2090 | 2169 | 2180.24 | 2248 | 2113 | 0.939 | -0.09 | 0.27497751 | 0.44596092 | 0.0057 | 0.0035 | 0.0045 | 0.0045 | TRUE | 0.1443 |
| Slc35a5 | 809 | 627 | 582 | 611 | 541 | 639 | 628 | 667 | 628 | 608 | 610 | 631 | 628.75 | 641 | 616 | 0.962 | -0.056 | 0.52528403 | 0.68634104 | 0 | 0.0062 | 0.0039 | 0.0039 | TRUE | 0.0143 |
| Slc35b1 | 3603 | 2515 | 2017 | 2307 | 1994 | 2239 | 2798 | 2675 | 2176 | 2296 | 2249 | 2210 | 2400.98 | 2550 | 2252 | 0.883 | -0.18 | 0.04112398 | 0.11664716 | 0.0079 | 0.0034 | 0.0052 | 0.0052 | TRUE | 0.1978 |
| Slc35b2 | 1582 | 1117 | 1231 | 1457 | 1335 | 1433 | 1229 | 1188 | 1328 | 1450 | 1506 | 1415 | 1352.66 | 1248 | 1457 | 1.167 | 0.223 | 0.00237833 | 0.01382972 | 0.0013 | 0.0042 | 0.0031 | 0.0031 | TRUE | 0.0364 |
| Slc35b3 | 338 | 228 | 214 | 229 | 195 | 228 | 263 | 243 | 231 | 228 | 220 | 225 | 234.82 | 246 | 224 | 0.913 | -0.132 | 0.31914399 | 0.493086 | 0 | 0.0125 | 0.0084 | 0.0084 | TRUE | 0.0453 |
| Slc35b4 | 1918 | 1436 | 1288 | 1415 | 1261 | 1362 | 1490 | 1527 | 1390 | 1408 | 1423 | 1345 | 1430.39 | 1469 | 1392 | 0.947 | -0.079 | 0.26783503 | 0.43840213 | 9.00E-04 | 0.0041 | 0.0029 | 0.0029 | TRUE | 0.026 |
| Slc35c1 | 763 | 545 | 618 | 651 | 636 | 721 | 593 | 580 | 667 | 648 | 717 | 712 | 652.72 | 613 | 692 | 1.13 | 0.176 | 0.06387886 | 0.16059394 | 0.0028 | 0.0061 | 0.005 | 0.005 | TRUE | 0.067 |
| Slc35c2 | 1837 | 1330 | 1328 | 1347 | 1336 | 1532 | 1427 | 1415 | 1433 | 1341 | 1507 | 1512 | 1439.1 | 1425 | 1453 | 1.02 | 0.028 | 0.69953516 | 0.81766896 | 0.0016 | 0.0041 | 0.0032 | 0.0032 | TRUE | 0.0558 |
| Slc35d1 | 445 | 360 | 292 | 301 | 251 | 306 | 346 | 383 | 315 | 300 | 283 | 302 | 321.41 | 348 | 295 | 0.848 | -0.237 | 0.05125646 | 0.13732119 | 0.0022 | 0.0098 | 0.0075 | 0.0075 | TRUE | 0.084 |
| Slc35d2 | 815 | 548 | 659 | 818 | 712 | 750 | 633 | 583 | 711 | 814 | 803 | 740 | 714.12 | 642 | 786 | 1.224 | 0.291 | 0.00266721 | 0.0150522 | 0.0042 | 0.0057 | 0.0054 | 0.0054 | TRUE | 0.099 |
| Slc35d3 | 63 | 28 | 37 | 48 | 33 | 44 | 49 | 30 | 40 | 48 | 37 | 43 | 41.18 | 40 | 43 | 1.076 | 0.105 | 0.73791623 | 0.84383266 | 0.0142 | 0.06 | 0.0474 | 0.0474 | TRUE | 0.2403 |
| Slc35e1 | 6284 | 4496 | 4045 | 4348 | 4045 | 4378 | 4881 | 4782 | 4365 | 4328 | 4563 | 4322 | 4540.05 | 4676 | 4404 | 0.941 | -0.087 | 0.16205572 | 0.31212269 | 0.0019 | 0.0029 | 0.0026 | 0.0026 | TRUE | 0.041 |
| Slc35e2 | 2255 | 1715 | 1423 | 1588 | 1410 | 1494 | 1751 | 1824 | 1535 | 1581 | 1591 | 1475 | 1626.19 | 1703 | 1549 | 0.909 | -0.138 | 0.08388218 | 0.19526515 | 0.0038 | 0.0039 | 0.004 | 0.004 | TRUE | 0.0882 |
| Slc35e3 | 987 | 659 | 594 | 574 | 532 | 573 | 767 | 701 | 641 | 571 | 600 | 566 | 640.93 | 703 | 579 | 0.822 | -0.283 | 0.00348602 | 0.01855705 | 0.0033 | 0.0061 | 0.0052 | 0.0052 | TRUE | 0.0779 |
| Slc35e4 | 233 | 188 | 191 | 230 | 194 | 202 | 181 | 200 | 206 | 229 | 219 | 199 | 205.7 | 196 | 216 | 1.105 | 0.144 | 0.31457887 | 0.48814199 | 0 | 0.0139 | 0.0099 | 0.0099 | TRUE | 0.0518 |
| Slc35f1 | 85 | 50 | 58 | 65 | 56 | 52 | 66 | 53 | 63 | 65 | 63 | 51 | 60.16 | 61 | 60 | 0.982 | -0.027 | 0.91651052 | 0.95388891 | 0 | 0.0418 | 0.0298 | 0.0298 | TRUE | 0.1342 |
| Slc35f2 | 4201 | 3123 | 3599 | 4079 | 3548 | 4366 | 3263 | 3322 | 3883 | 4060 | 4003 | 4310 | 3806.76 | 3489 | 4124 | 1.183 | 0.242 | 0.00131213 | 0.00876107 | 0.0052 | 0.003 | 0.0038 | 0.0038 | TRUE | 0.118 |
| Slc35f3 | 200 | 151 | 141 | 134 | 127 | 145 | 155 | 161 | 152 | 133 | 143 | 143 | 147.98 | 156 | 140 | 0.897 | -0.157 | 0.32655207 | 0.5006795 | 0 | 0.0184 | 0.0118 | 0.0118 | TRUE | 0.0177 |
| Slc35f4 | 10 | 9 | 6 | 6 | 3 | 8 | 8 | 10 | 6 | 6 | 3 | 8 | 6.84 | 8 | 6 | 0.731 | -0.453 | 0.53689175 | NA | 0 | 0.3487 | 0.2376 | 0.2376 | TRUE | 0.2569 |
| Slc35f5 | 1410 | 1096 | 1276 | 1422 | 1218 | 1451 | 1095 | 1166 | 1377 | 1415 | 1374 | 1432 | 1309.92 | 1213 | 1407 | 1.162 | 0.217 | 0.01537033 | 0.05687241 | 0.0064 | 0.0042 | 0.005 | 0.005 | TRUE | 0.1671 |
| Slc35f6 | 1168 | 841 | 776 | 835 | 832 | 864 | 907 | 895 | 837 | 831 | 939 | 853 | 876.94 | 880 | 874 | 0.992 | -0.011 | 0.89553133 | 0.94268886 | 0.0017 | 0.0051 | 0.004 | 0.004 | TRUE | 0.0483 |
| Slc35g1 | 4526 | 3305 | 3236 | 3169 | 2615 | 3160 | 3515 | 3515 | 3492 | 3154 | 2950 | 3119 | 3291.01 | 3507 | 3074 | 0.877 | -0.19 | 0.00072228 | 0.0057036 | 2.00E-04 | 0.0031 | 0.002 | 0.002 | TRUE | 0.0151 |
| Slc35g2 | 82 | 51 | 52 | 81 | 53 | 89 | 64 | 54 | 56 | 81 | 60 | 88 | 67.05 | 58 | 76 | 1.311 | 0.39 | 0.11765302 | 0.24886287 | 0 | 0.0378 | 0.0298 | 0.0298 | TRUE | 0.304 |
| Slc35g3 | 1 | 1 | 2 | 2 | 2 | 4 | 1 | 1 | 2 | 2 | 2 | 4 | 2.03 | 1 | 3 | 2.105 | 1.074 | 0.42429721 | NA | 0 | 1.1686 | 0.7423 | 0.7423 | TRUE | 0.1883 |
| Slc36a1 | 1921 | 1494 | 1480 | 1522 | 1446 | 1549 | 1492 | 1589 | 1597 | 1515 | 1631 | 1529 | 1558.89 | 1559 | 1558 | 1 | 0 | 0.99750219 | 0.99855014 | 9.00E-04 | 0.0039 | 0.0028 | 0.0028 | TRUE | 0.0198 |
| Slc36a2 | 22 | 20 | 16 | 24 | 20 | 19 | 17 | 21 | 17 | 24 | 23 | 19 | 20.14 | 18 | 22 | 1.175 | 0.233 | 0.58241748 | 0.73253195 | 0 | 0.1201 | 0.0792 | 0.0792 | TRUE | 0.0847 |
| Slc36a3 | 15 | 8 | 10 | 16 | 18 | 16 | 12 | 9 | 11 | 16 | 20 | 16 | 13.83 | 11 | 17 | 1.667 | 0.737 | 0.15223461 | NA | 0 | 0.1738 | 0.1147 | 0.1147 | TRUE | 0.1015 |
| Slc36a4 | 298 | 222 | 184 | 204 | 169 | 171 | 231 | 236 | 199 | 203 | 191 | 169 | 204.77 | 222 | 188 | 0.843 | -0.246 | 0.097881 | 0.21801669 | 0.0036 | 0.014 | 0.011 | 0.011 | TRUE | 0.09 |
| Slc37a1 | 585 | 458 | 533 | 601 | 555 | 655 | 454 | 487 | 575 | 598 | 626 | 647 | 564.59 | 505 | 624 | 1.237 | 0.306 | 0.00430274 | 0.02186838 | 0.0063 | 0.0066 | 0.0065 | 0.0065 | TRUE | 0.1668 |
| Slc37a2 | 335 | 223 | 150 | 184 | 165 | 154 | 260 | 237 | 162 | 183 | 186 | 152 | 196.76 | 220 | 174 | 0.787 | -0.345 | 0.06202677 | 0.15713374 | 0.0314 | 0.0145 | 0.0195 | 0.0195 | TRUE | 0.568 |
| Slc37a3 | 2455 | 1632 | 1416 | 1429 | 1279 | 1385 | 1907 | 1736 | 1528 | 1422 | 1443 | 1367 | 1567.16 | 1724 | 1411 | 0.817 | -0.291 | 0.0007266 | 0.00572511 | 0.0055 | 0.0039 | 0.0047 | 0.0047 | TRUE | 0.1183 |
| Slc37a4 | 583 | 432 | 537 | 551 | 431 | 525 | 453 | 460 | 579 | 548 | 486 | 518 | 507.44 | 497 | 517 | 1.044 | 0.062 | 0.59837936 | 0.74416177 | 0.0096 | 0.0071 | 0.0079 | 0.0079 | TRUE | 0.2388 |
| Slc38a1 | 5805 | 4111 | 3670 | 3585 | 3585 | 4045 | 4508 | 4373 | 3960 | 3569 | 4044 | 3993 | 4074.5 | 4280 | 3869 | 0.903 | -0.147 | 0.04254939 | 0.11971622 | 0.0041 | 0.003 | 0.0035 | 0.0035 | TRUE | 0.0562 |
| Slc38a10 | 10189 | 7355 | 6961 | 7858 | 6800 | 7670 | 7913 | 7823 | 7511 | 7822 | 7671 | 7571 | 7718.68 | 7749 | 7688 | 0.992 | -0.012 | 0.81533827 | 0.89389387 | 3.00E-04 | 0.0027 | 0.0017 | 0.0017 | TRUE | 0.0088 |
| Slc38a11 | 9 | 3 | 3 | 3 | 0 | 4 | 7 | 3 | 3 | 3 | 0 | 4 | 3.39 | 4 | 2 | 0.518 | -0.948 | 0.39679729 | NA | 0.1179 | 0.7011 | 0.5755 | 0.5755 | TRUE | 0.6703 |
| Slc38a2 | 16365 | 12198 | 12042 | 11655 | 11417 | 13749 | 12710 | 12975 | 12993 | 11601 | 12880 | 13572 | 12788.57 | 12893 | 12684 | 0.984 | -0.023 | 0.71209447 | 0.82700628 | 0.0025 | 0.0026 | 0.0028 | 0.0028 | TRUE | 0.0683 |
| Slc38a3 | 333 | 282 | 345 | 459 | 377 | 523 | 259 | 300 | 372 | 457 | 425 | 516 | 388.22 | 310 | 466 | 1.509 | 0.594 | 2.41E-05 | 0.00038623 | 0.0181 | 0.0085 | 0.0116 | 0.0116 | TRUE | 0.3432 |
| Slc38a4 | 25534 | 18385 | 21423 | 23712 | 22197 | 24846 | 19831 | 19556 | 23115 | 23603 | 25041 | 24527 | 22612.09 | 20834 | 24390 | 1.171 | 0.227 | 0.0010021 | 0.00724482 | 0.0049 | 0.0025 | 0.0034 | 0.0034 | TRUE | 0.1121 |
| Slc38a5 | 102 | 65 | 86 | 153 | 148 | 129 | 79 | 69 | 93 | 152 | 167 | 127 | 114.63 | 80 | 149 | 1.851 | 0.888 | 9.51E-06 | 0.00018101 | 0.0099 | 0.0231 | 0.0195 | 0.0195 | TRUE | 0.1694 |
| Slc38a6 | 286 | 216 | 167 | 180 | 163 | 179 | 222 | 230 | 180 | 179 | 184 | 177 | 195.3 | 211 | 180 | 0.852 | -0.231 | 0.12767845 | 0.26363885 | 0.0032 | 0.0146 | 0.0114 | 0.0114 | TRUE | 0.1668 |
| Slc38a7 | 1713 | 1266 | 1218 | 1136 | 1016 | 1092 | 1330 | 1347 | 1314 | 1131 | 1146 | 1078 | 1224.36 | 1330 | 1118 | 0.84 | -0.251 | 0.00032576 | 0.00309787 | 0 | 0.0044 | 0.0027 | 0.0027 | TRUE | 0.012 |
| Slc38a8 | 168 | 107 | 103 | 147 | 176 | 210 | 130 | 114 | 111 | 146 | 199 | 207 | 151.27 | 118 | 184 | 1.547 | 0.63 | 0.00061465 | 0.00506061 | 0.016 | 0.0181 | 0.0175 | 0.0175 | TRUE | 0.3518 |
| Slc38a9 | 1681 | 1331 | 1154 | 1453 | 1284 | 1480 | 1306 | 1416 | 1245 | 1446 | 1448 | 1461 | 1387.05 | 1322 | 1452 | 1.098 | 0.135 | 0.06350011 | 0.15984626 | 0.0013 | 0.0041 | 0.0031 | 0.0031 | TRUE | 0.0447 |
| Slc39a1 | 2181 | 1584 | 1550 | 1547 | 1448 | 1638 | 1694 | 1685 | 1672 | 1540 | 1633 | 1617 | 1640.26 | 1684 | 1597 | 0.948 | -0.077 | 0.2304057 | 0.39614588 | 0 | 0.0039 | 0.0024 | 0.0024 | TRUE | 0.0118 |
| Slc39a10 | 6985 | 4763 | 4954 | 5283 | 4383 | 5447 | 5425 | 5066 | 5345 | 5259 | 4944 | 5377 | 5236.14 | 5279 | 5193 | 0.984 | -0.024 | 0.68517557 | 0.8070222 | 0.0011 | 0.0029 | 0.0022 | 0.0022 | TRUE | 0.0213 |
| Slc39a11 | 2060 | 1531 | 1773 | 2043 | 2032 | 2097 | 1600 | 1629 | 1913 | 2034 | 2292 | 2070 | 1922.91 | 1714 | 2132 | 1.245 | 0.316 | 0.00019152 | 0.00206755 | 0.0065 | 0.0037 | 0.0046 | 0.0046 | TRUE | 0.1236 |
| Slc39a12 | 0 | 0 | 0 | 0 | 1 | 0 | 0 | 0 | 0 | 0 | 1 | 0 | 0.19 | 0 | 0 | 2.094 | 1.067 | 0.79379329 | NA | 0 | 12.608 | 10 | 10 | TRUE | 0.5075 |
| Slc39a13 | 354 | 264 | 287 | 351 | 304 | 343 | 275 | 281 | 310 | 349 | 343 | 339 | 316.06 | 289 | 344 | 1.193 | 0.255 | 0.02897804 | 0.09027532 | 0 | 0.0099 | 0.0066 | 0.0066 | TRUE | 0.0451 |
| Slc39a14 | 3602 | 2779 | 2733 | 2603 | 2464 | 3209 | 2798 | 2956 | 2949 | 2591 | 2780 | 3168 | 2873.47 | 2901 | 2846 | 0.982 | -0.027 | 0.73026307 | 0.8388371 | 0.0054 | 0.0033 | 0.004 | 0.004 | TRUE | 0.1194 |
| Slc39a2 | 27 | 15 | 16 | 17 | 11 | 11 | 21 | 16 | 17 | 17 | 12 | 11 | 15.73 | 18 | 13 | 0.738 | -0.438 | 0.36759991 | NA | 0 | 0.1531 | 0.1049 | 0.1049 | TRUE | 0.2325 |
| Slc39a3 | 237 | 159 | 142 | 190 | 170 | 173 | 184 | 169 | 153 | 189 | 192 | 171 | 176.35 | 169 | 184 | 1.085 | 0.118 | 0.44499645 | 0.61583407 | 0 | 0.0159 | 0.0115 | 0.0115 | TRUE | 0.073 |
| Slc39a4 | 1119 | 928 | 1164 | 1223 | 1247 | 1435 | 869 | 987 | 1256 | 1217 | 1407 | 1417 | 1192.14 | 1037 | 1347 | 1.3 | 0.379 | 0.00143646 | 0.00934844 | 0.0205 | 0.0044 | 0.0093 | 0.0093 | TRUE | 0.4048 |
| Slc39a5 | 7 | 5 | 7 | 2 | 9 | 3 | 5 | 5 | 8 | 2 | 10 | 3 | 5.57 | 6 | 5 | 0.815 | -0.296 | 0.73229017 | NA | 0.1571 | 0.428 | 0.3539 | 0.3539 | TRUE | 1.4468 |
| Slc39a6 | 2378 | 1820 | 1882 | 1905 | 1694 | 1894 | 1847 | 1936 | 2031 | 1896 | 1911 | 1870 | 1915.06 | 1938 | 1892 | 0.977 | -0.033 | 0.6093481 | 0.75282326 | 8.00E-04 | 0.0037 | 0.0025 | 0.0025 | TRUE | 0.0212 |
| Slc39a7 | 6201 | 4313 | 3878 | 3638 | 3275 | 3934 | 4816 | 4588 | 4184 | 3621 | 3695 | 3883 | 4131.22 | 4529 | 3733 | 0.824 | -0.28 | 3.03E-05 | 0.00046756 | 0.0028 | 0.003 | 0.003 | 0.003 | TRUE | 0.0539 |
| Slc39a8 | 102 | 63 | 74 | 98 | 77 | 93 | 79 | 67 | 80 | 98 | 87 | 92 | 83.72 | 75 | 92 | 1.22 | 0.287 | 0.17780142 | 0.33219813 | 0 | 0.0307 | 0.0208 | 0.0208 | TRUE | 0.0813 |
| Slc39a9 | 2732 | 1892 | 1803 | 1997 | 1826 | 2019 | 2122 | 2012 | 1945 | 1988 | 2060 | 1993 | 2020.09 | 2026 | 2014 | 0.993 | -0.011 | 0.86790644 | 0.92704068 | 7.00E-04 | 0.0036 | 0.0025 | 0.0025 | TRUE | 0.0211 |
| Slc3a1 | 3 | 0 | 3 | 6 | 1 | 1 | 2 | 0 | 3 | 6 | 1 | 1 | 2.28 | 2 | 3 | 1.451 | 0.537 | 0.69940037 | NA | 0.6562 | 1.0439 | 0.9437 | 0.9437 | TRUE | 1.3599 |
| Slc3a2 | 12662 | 9381 | 9314 | 10325 | 9114 | 10525 | 9834 | 9978 | 10050 | 10277 | 10282 | 10390 | 10135.17 | 9954 | 10316 | 1.036 | 0.052 | 0.25214669 | 0.42111856 | 0 | 0.0027 | 0.0014 | 0.0014 | TRUE | 0.0014 |
| Slc40a1 | 368 | 255 | 278 | 169 | 117 | 147 | 286 | 271 | 300 | 168 | 132 | 145 | 217.06 | 286 | 148 | 0.521 | -0.942 | 1.23E-10 | 9.74E-09 | 0.001 | 0.0133 | 0.0103 | 0.0103 | TRUE | 0.1426 |
| Slc41a1 | 2836 | 2084 | 2125 | 2480 | 2192 | 2406 | 2203 | 2217 | 2293 | 2469 | 2473 | 2375 | 2338.11 | 2238 | 2439 | 1.09 | 0.125 | 0.03474672 | 0.10331876 | 0 | 0.0034 | 0.0021 | 0.0021 | TRUE | 0.0064 |
| Slc41a2 | 200 | 144 | 162 | 211 | 169 | 227 | 155 | 153 | 175 | 210 | 191 | 224 | 184.68 | 161 | 208 | 1.296 | 0.374 | 0.0134016 | 0.05140878 | 0 | 0.0153 | 0.011 | 0.011 | TRUE | 0.0573 |
| Slc41a3 | 282 | 219 | 275 | 340 | 280 | 323 | 219 | 233 | 297 | 338 | 316 | 319 | 286.97 | 250 | 324 | 1.305 | 0.384 | 0.00582249 | 0.027584 | 0.0098 | 0.0107 | 0.0104 | 0.0104 | TRUE | 0.2974 |
| Slc43a1 | 473 | 333 | 364 | 511 | 497 | 528 | 367 | 354 | 393 | 509 | 561 | 521 | 450.81 | 371 | 530 | 1.427 | 0.513 | 6.66E-07 | 2.09E-05 | 0 | 0.0077 | 0.0054 | 0.0054 | TRUE | 0.0288 |
| Slc43a2 | 2518 | 1827 | 1834 | 2083 | 1838 | 2107 | 1956 | 1943 | 1979 | 2073 | 2073 | 2080 | 2017.44 | 1959 | 2075 | 1.059 | 0.083 | 0.15637042 | 0.30453773 | 0 | 0.0036 | 0.002 | 0.002 | TRUE | 9.00E-04 |
| Slc43a3 | 681 | 426 | 332 | 349 | 258 | 284 | 529 | 453 | 358 | 347 | 291 | 280 | 376.51 | 447 | 306 | 0.684 | -0.548 | 0.00023392 | 0.00241225 | 0.0234 | 0.0087 | 0.0132 | 0.0132 | TRUE | 0.1771 |
| Slc44a1 | 4064 | 2965 | 3222 | 3446 | 3168 | 3485 | 3156 | 3154 | 3477 | 3430 | 3574 | 3440 | 3371.81 | 3262 | 3481 | 1.067 | 0.094 | 0.12967788 | 0.26638397 | 0.0014 | 0.0031 | 0.0025 | 0.0025 | TRUE | 0.0397 |
| Slc44a2 | 2847 | 2353 | 2498 | 3181 | 2750 | 3220 | 2211 | 2503 | 2695 | 3166 | 3102 | 3179 | 2809.43 | 2470 | 3149 | 1.276 | 0.352 | 3.45E-06 | 7.81E-05 | 0.0044 | 0.0033 | 0.0038 | 0.0038 | TRUE | 0.1041 |

| | | | | | | | | | | | | | | | | | | | | | | | | | |
|---|---|---|---|---|---|---|---|---|---|---|---|---|---|---|---|---|---|---|---|---|---|---|---|---|---|
| Slc44a3 | 165 | 121 | 168 | 198 | 198 | 202 | 128 | 129 | 181 | 197 | 223 | 199 | 176.33 | 146 | 206 | 1.419 | 0.505 | 0.00352799 | 0.01870386 | 0.0155 | 0.0159 | 0.0158 | 0.0158 | TRUE | 0.4488 |
| Slc44a4 | 412 | 284 | 364 | 442 | 412 | 431 | 320 | 302 | 393 | 440 | 465 | 425 | 390.84 | 338 | 443 | 1.312 | 0.392 | 0.00132306 | 0.0088161 | 0.0072 | 0.0085 | 0.0081 | 0.0081 | TRUE | 0.2175 |
| Slc44a5 | 11 | 6 | 6 | 14 | 7 | 12 | 9 | 6 | 6 | 14 | 8 | 12 | 9.18 | 7 | 11 | 1.566 | 0.647 | 0.30690421 | NA | 0 | 0.2606 | 0.1765 | 0.1765 | TRUE | 0.2264 |
| Slc45a1 | 22 | 9 | 20 | 22 | 17 | 22 | 17 | 10 | 22 | 22 | 19 | 22 | 18.51 | 16 | 21 | 1.301 | 0.38 | 0.41112654 | 0.58400234 | 0 | 0.1305 | 0.099 | 0.099 | TRUE | 0.4332 |
| Slc45a2 | 3 | 1 | 0 | 2 | 1 | 0 | 2 | 1 | 0 | 2 | 1 | 0 | 1.09 | 1 | 1 | 0.885 | -0.177 | 0.92674766 | NA | 0 | 2.186 | 1.7483 | 1.7483 | TRUE | 0.2457 |
| Slc45a3 | 565 | 393 | 420 | 553 | 443 | 451 | 439 | 418 | 453 | 550 | 500 | 445 | 467.57 | 437 | 498 | 1.141 | 0.191 | 0.08231353 | 0.19312333 | 0.0045 | 0.0075 | 0.0065 | 0.0065 | TRUE | 0.1042 |
| Slc45a4 | 4519 | 3462 | 3758 | 3763 | 3543 | 3800 | 3510 | 3682 | 4055 | 3746 | 3997 | 3751 | 3790.13 | 3749 | 3831 | 1.022 | 0.032 | 0.64090752 | 0.77572116 | 0.0032 | 0.0031 | 0.0031 | 0.0031 | TRUE | 0.0617 |
| Slc46a1 | 1189 | 784 | 618 | 645 | 584 | 605 | 923 | 834 | 667 | 642 | 659 | 597 | 720.38 | 808 | 633 | 0.781 | -0.356 | 0.00214558 | 0.01273909 | 0.0137 | 0.0057 | 0.0083 | 0.0083 | TRUE | 0.2745 |
| Slc46a2 | 0 | 1 | 1 | 2 | 0 | 2 | 0 | 1 | 1 | 2 | 0 | 2 | 1.02 | 1 | 1 | 1.967 | 0.976 | 0.62585884 | NA | 0 | 2.3307 | 1.7931 | 1.7931 | TRUE | 0.4032 |
| Slc46a3 | 82 | 45 | 65 | 112 | 115 | 139 | 64 | 48 | 70 | 111 | 130 | 137 | 93.35 | 61 | 126 | 2.078 | 1.056 | 1.57E-06 | 4.19E-05 | 0.0073 | 0.0278 | 0.0228 | 0.0228 | TRUE | 0.2678 |
| Slc47a1 | 412 | 240 | 254 | 372 | 345 | 392 | 320 | 255 | 274 | 370 | 389 | 387 | 332.63 | 283 | 382 | 1.344 | 0.427 | 0.00053556 | 0.0045386 | 0.004 | 0.0096 | 0.0079 | 0.0079 | TRUE | 0.1571 |
| Slc47a2 | 440 | 451 | 439 | 413 | 452 | 458 | 342 | 480 | 474 | 411 | 510 | 452 | 444.71 | 432 | 458 | 1.063 | 0.088 | 0.53046972 | 0.69038787 | 0.0215 | 0.0078 | 0.012 | 0.012 | TRUE | 0.4052 |
| Slc48a1 | 1041 | 735 | 854 | 1084 | 924 | 1051 | 809 | 782 | 921 | 1079 | 1042 | 1037 | 945.11 | 837 | 1053 | 1.259 | 0.332 | 0.00010179 | 0.00122673 | 0.0025 | 0.0049 | 0.0042 | 0.0042 | TRUE | 0.0892 |
| Slc4a1 | 114 | 82 | 97 | 123 | 95 | 99 | 89 | 87 | 105 | 122 | 107 | 98 | 101.29 | 94 | 109 | 1.17 | 0.226 | 0.25983292 | 0.42983702 | 0 | 0.0258 | 0.0192 | 0.0192 | TRUE | 0.1165 |
| Slc4a10 | 3 | 3 | 3 | 2 | 0 | 1 | 2 | 3 | 3 | 2 | 0 | 1 | 1.96 | 3 | 1 | 0.35 | -1.515 | 0.29358004 | NA | 0 | 1.2141 | 0.8227 | 0.8227 | TRUE | 0.3539 |
| Slc4a11 | 1144 | 864 | 960 | 918 | 821 | 1027 | 888 | 919 | 1036 | 914 | 926 | 1014 | 949.52 | 948 | 951 | 1.005 | 0.007 | 0.93312609 | 0.9627836 | 0.0039 | 0.0049 | 0.0046 | 0.0046 | TRUE | 0.0778 |
| Slc4a1ap | 2603 | 1876 | 1809 | 1965 | 1708 | 1913 | 2022 | 1995 | 1952 | 1956 | 1927 | 1888 | 1956.7 | 1990 | 1924 | 0.967 | -0.049 | 0.41742062 | 0.58995283 | 0 | 0.0036 | 0.0021 | 0.0021 | TRUE | 0.0033 |
| Slc4a2 | 941 | 679 | 497 | 402 | 393 | 490 | 731 | 722 | 536 | 400 | 443 | 484 | 552.76 | 663 | 442 | 0.666 | -0.586 | 5.70E-06 | 0.00011773 | 0.0176 | 0.0067 | 0.0101 | 0.0101 | TRUE | 0.3245 |
| Slc4a3 | 344 | 266 | 237 | 259 | 202 | 272 | 267 | 283 | 256 | 258 | 228 | 269 | 260 | 269 | 252 | 0.937 | -0.093 | 0.47296685 | 0.64149887 | 0 | 0.0115 | 0.0084 | 0.0084 | TRUE | 0.0715 |
| Slc4a4 | 434 | 279 | 226 | 241 | 195 | 231 | 337 | 297 | 244 | 240 | 220 | 228 | 260.93 | 293 | 229 | 0.781 | -0.356 | 0.01457552 | 0.05459327 | 0.0112 | 0.0115 | 0.0114 | 0.0114 | TRUE | 0.2113 |
| Slc4a5 | 1470 | 1077 | 918 | 702 | 670 | 766 | 1142 | 1146 | 991 | 699 | 756 | 756 | 914.76 | 1093 | 737 | 0.674 | -0.57 | 1.12E-10 | 9.01E-09 | 0.0034 | 0.005 | 0.0045 | 0.0045 | TRUE | 0.0782 |
| Slc4a7 | 8134 | 5935 | 6278 | 7162 | 6286 | 7206 | 6317 | 6313 | 6774 | 7129 | 7091 | 7113 | 6789.66 | 6468 | 7111 | 1.1 | 0.137 | 0.00912002 | 0.03866878 | 6.00E-04 | 0.0028 | 0.0018 | 0.0018 | TRUE | 0.0207 |
| Slc4a8 | 1448 | 1170 | 1175 | 1175 | 1079 | 1288 | 1125 | 1245 | 1268 | 1170 | 1217 | 1271 | 1215.87 | 1213 | 1219 | 1.007 | 0.01 | 0.893794 | 0.94181872 | 0.0023 | 0.0044 | 0.0036 | 0.0036 | TRUE | 0.0502 |
| Slc4a9 | 40 | 37 | 42 | 35 | 38 | 49 | 31 | 39 | 45 | 35 | 43 | 48 | 40.3 | 38 | 42 | 1.097 | 0.134 | 0.67104043 | 0.79747076 | 0 | 0.0612 | 0.0466 | 0.0466 | TRUE | 0.2568 |
| Slc50a1 | 494 | 393 | 378 | 350 | 307 | 420 | 384 | 418 | 408 | 348 | 346 | 415 | 386.48 | 403 | 370 | 0.919 | -0.122 | 0.29288503 | 0.46503064 | 0.0037 | 0.0086 | 0.0071 | 0.0071 | TRUE | 0.1306 |
| Slc51a | 56 | 59 | 36 | 35 | 29 | 29 | 43 | 63 | 39 | 35 | 33 | 29 | 40.21 | 48 | 32 | 0.664 | -0.59 | 0.06768071 | 0.16730549 | 0.0176 | 0.0614 | 0.0489 | 0.0489 | TRUE | 0.5178 |
| Slc51b | 0 | 0 | 0 | 0 | 0 | 0 | 0 | 0 | 0 | 0 | 0 | 0 | 0 | 0 | 0 | NA | NA | NA | NA | NA | NA | NA | NA | NA | NA |
| Slc52a2 | 1216 | 833 | 816 | 820 | 700 | 840 | 944 | 886 | 880 | 816 | 790 | 829 | 857.67 | 903 | 812 | 0.898 | -0.156 | 0.0503114 | 0.13539839 | 0 | 0.0052 | 0.0034 | 0.0034 | TRUE | 0.0196 |
| Slc52a3 | 633 | 526 | 556 | 642 | 582 | 642 | 492 | 559 | 600 | 639 | 657 | 634 | 596.73 | 550 | 643 | 1.172 | 0.228 | 0.02217204 | 0.07435592 | 0.0036 | 0.0064 | 0.0055 | 0.0055 | TRUE | 0.1088 |
| Slc5a1 | 111 | 72 | 55 | 40 | 39 | 34 | 86 | 77 | 59 | 40 | 44 | 34 | 56.59 | 74 | 39 | 0.525 | -0.931 | 0.00075378 | 0.00588091 | 0 | 0.0443 | 0.0352 | 0.0352 | TRUE | 0.2589 |
| Slc5a10 | 7 | 2 | 8 | 9 | 10 | 8 | 5 | 2 | 9 | 9 | 11 | 8 | 7.39 | 5 | 9 | 1.734 | 0.794 | 0.2789949 | NA | 0 | 0.3232 | 0.243 | 0.243 | TRUE | 0.2851 |
| Slc5a11 | 143 | 108 | 111 | 111 | 116 | 170 | 111 | 115 | 120 | 110 | 131 | 168 | 125.81 | 115 | 136 | 1.186 | 0.246 | 0.21101977 | 0.37307385 | 0.0164 | 0.0213 | 0.0198 | 0.0198 | TRUE | 0.3847 |
| Slc5a12 | 1 | 0 | 0 | 0 | 0 | 0 | 1 | 0 | 0 | 0 | 0 | 0 | 0.13 | 0 | 0 | 0.552 | -0.857 | 0.8336548 | NA | 0 | 18.3121 | 10 | 10 | TRUE | 0.2566 |
| Slc5a2 | 58 | 40 | 41 | 75 | 56 | 54 | 45 | 43 | 44 | 75 | 63 | 53 | 53.83 | 44 | 64 | 1.448 | 0.534 | 0.04756105 | 0.12992931 | 0 | 0.0465 | 0.0333 | 0.0333 | TRUE | 0.1731 |
| Slc5a3 | 1208 | 979 | 1100 | 1317 | 1061 | 1268 | 938 | 1041 | 1187 | 1311 | 1197 | 1252 | 1154.34 | 1055 | 1253 | 1.19 | 0.251 | 0.00710202 | 0.03209267 | 0.0067 | 0.0045 | 0.0054 | 0.0054 | TRUE | 0.1415 |
| Slc5a4a | 9 | 12 | 8 | 10 | 6 | 7 | 7 | 13 | 9 | 10 | 7 | 7 | 8.67 | 10 | 8 | 0.842 | -0.248 | 0.70435951 | NA | 0 | 0.2758 | 0.19 | 0.19 | TRUE | 0.2982 |
| Slc5a4b | 420 | 297 | 279 | 186 | 119 | 154 | 326 | 316 | 301 | 185 | 134 | 152 | 235.76 | 314 | 157 | 0.501 | -0.999 | 1.31E-11 | 1.30E-09 | 0.0058 | 0.0125 | 0.0109 | 0.0109 | TRUE | 0.257 |
| Slc5a5 | 176 | 140 | 198 | 227 | 203 | 252 | 137 | 149 | 214 | 226 | 229 | 249 | 200.5 | 167 | 235 | 1.417 | 0.503 | 0.00413322 | 0.02118942 | 0.0246 | 0.0142 | 0.017 | 0.017 | TRUE | 0.6413 |
| Slc5a6 | 1054 | 672 | 539 | 553 | 472 | 549 | 819 | 715 | 582 | 550 | 532 | 542 | 623.31 | 705 | 541 | 0.767 | -0.384 | 0.00130293 | 0.00871753 | 0.0136 | 0.0062 | 0.0086 | 0.0086 | TRUE | 0.2017 |
| Slc5a7 | 6 | 0 | 0 | 1 | 1 | 0 | 5 | 0 | 0 | 1 | 1 | 0 | 1.13 | 2 | 1 | 0.432 | -1.21 | 0.57241325 | NA | 3.1014 | 2.0988 | 2.2754 | 2.2754 | TRUE | 2.7689 |
| Slc5a8 | 4 | 6 | 5 | 3 | 1 | 3 | 3 | 6 | 5 | 3 | 1 | 3 | 3.66 | 5 | 2 | 0.487 | -1.039 | 0.30941408 | NA | 0 | 0.65 | 0.4379 | 0.4379 | TRUE | 0.2883 |
| Slc5a9 | 3 | 3 | 8 | 1 | 0 | 3 | 2 | 3 | 9 | 1 | 0 | 3 | 3.02 | 5 | 1 | 0.29 | -1.786 | 0.15971188 | NA | 0.346 | 0.7877 | 0.6751 | 0.6751 | TRUE | 0.9547 |
| Slc6a1 | 140 | 109 | 82 | 158 | 105 | 136 | 109 | 116 | 88 | 157 | 118 | 134 | 120.52 | 104 | 136 | 1.309 | 0.389 | 0.04583977 | 0.12643531 | 0.0105 | 0.0221 | 0.0189 | 0.0189 | TRUE | 0.1802 |
| Slc6a11 | 41 | 26 | 20 | 10 | 14 | 12 | 32 | 28 | 22 | 10 | 16 | 12 | 19.78 | 27 | 13 | 0.459 | -1.125 | 0.0129132 | 0.04995084 | 0 | 0.1223 | 0.0881 | 0.0881 | TRUE | 0.1872 |
| Slc6a12 | 58 | 44 | 42 | 42 | 45 | 44 | 45 | 47 | 45 | 42 | 51 | 43 | 45.53 | 46 | 45 | 0.99 | -0.014 | 0.95947563 | 0.97761443 | 0 | 0.0545 | 0.0355 | 0.0355 | TRUE | 0.0803 |
| Slc6a13 | 110 | 67 | 82 | 73 | 58 | 69 | 85 | 71 | 88 | 73 | 65 | 68 | 75.23 | 81 | 69 | 0.841 | -0.251 | 0.26676683 | 0.43719529 | 0 | 0.0339 | 0.0233 | 0.0233 | TRUE | 0.1106 |
| Slc6a14 | 70 | 36 | 29 | 12 | 6 | 8 | 54 | 38 | 31 | 12 | 7 | 8 | 25.09 | 41 | 9 | 0.214 | -2.222 | 1.35E-06 | 3.73E-05 | 0 | 0.0969 | 0.0828 | 0.0828 | TRUE | 0.2799 |
| Slc6a15 | 576 | 383 | 424 | 550 | 467 | 561 | 447 | 407 | 457 | 547 | 527 | 554 | 490.05 | 437 | 543 | 1.24 | 0.311 | 0.00165856 | 0.01043387 | 0 | 0.0073 | 0.005 | 0.005 | TRUE | 0.0399 |
| Slc6a17 | 68 | 42 | 35 | 43 | 45 | 36 | 53 | 45 | 38 | 43 | 51 | 36 | 44.06 | 45 | 43 | 0.946 | -0.08 | 0.79075301 | 0.8770061 | 0 | 0.0562 | 0.0429 | 0.0429 | TRUE | 0.2036 |
| Slc6a18 | 0 | 0 | 0 | 0 | 0 | 0 | 0 | 0 | 0 | 0 | 0 | 0 | 0 | 0 | 0 | NA | NA | NA | NA | NA | NA | NA | NA | NA | NA |
| Slc6a19 | 18 | 11 | 7 | 10 | 9 | 13 | 14 | 12 | 8 | 10 | 10 | 13 | 11.03 | 11 | 11 | 0.982 | -0.026 | 0.96440259 | NA | 0 | 0.2173 | 0.1496 | 0.1496 | TRUE | 0.2677 |
| Slc6a2 | 41 | 37 | 46 | 33 | 41 | 34 | 32 | 39 | 50 | 33 | 46 | 34 | 38.92 | 40 | 38 | 0.936 | -0.096 | 0.77000741 | 0.86393877 | 0.02 | 0.0633 | 0.0511 | 0.0511 | TRUE | 0.2626 |
| Slc6a20a | 4 | 0 | 2 | 3 | 3 | 1 | 3 | 0 | 2 | 3 | 3 | 1 | 2.1 | 2 | 2 | 1.35 | 0.432 | 0.75249889 | NA | 0 | 1.129 | 0.8759 | 0.8759 | TRUE | 0.5535 |
| Slc6a20b | 0 | 0 | 0 | 0 | 0 | 0 | 0 | 0 | 0 | 0 | 0 | 0 | 0 | 0 | 0 | NA | NA | NA | NA | NA | NA | NA | NA | NA | NA |
| Slc6a3 | 25 | 12 | 15 | 18 | 16 | 17 | 19 | 13 | 16 | 18 | 18 | 17 | 16.85 | 16 | 18 | 1.081 | 0.113 | 0.80737025 | NA | 0 | 0.1431 | 0.0952 | 0.0952 | TRUE | 0.1902 |
| Slc6a4 | 7 | 1 | 2 | 4 | 3 | 5 | 5 | 1 | 2 | 4 | 3 | 5 | 3.49 | 3 | 4 | 1.374 | 0.459 | 0.65879435 | NA | 0 | 0.6809 | 0.4899 | 0.4899 | TRUE | 0.9868 |
| Slc6a5 | 2 | 1 | 4 | 3 | 2 | 0 | 2 | 1 | 4 | 3 | 2 | 0 | 2.03 | 2 | 2 | 0.761 | -0.395 | 0.78068351 | NA | 0 | 1.1704 | 0.9378 | 0.9378 | TRUE | 0.6397 |
| Slc6a6 | 12294 | 9267 | 10048 | 10154 | 9418 | 11184 | 9548 | 9857 | 10842 | 10107 | 10624 | 11040 | 10336.54 | 10082 | 10590 | 1.051 | 0.071 | 0.26363724 | 0.43397745 | 0.0031 | 0.0027 | 0.0028 | 0.0028 | TRUE | 0.0529 |
| Slc6a7 | 67 | 84 | 83 | 133 | 113 | 127 | 52 | 89 | 90 | 132 | 127 | 125 | 102.7 | 77 | 128 | 1.686 | 0.754 | 0.00096965 | 0.00708925 | 0.0321 | 0.0255 | 0.0273 | 0.0273 | TRUE | 0.8549 |
| Slc6a8 | 4703 | 3377 | 3262 | 3893 | 3278 | 3789 | 3653 | 3592 | 3520 | 3875 | 3698 | 3740 | 3679.62 | 3588 | 3771 | 1.051 | 0.072 | 0.18671325 | 0.34337269 | 2.00E-04 | 0.0031 | 0.0018 | 0.0018 | TRUE | 0.0071 |
| Slc6a9 | 108 | 84 | 69 | 54 | 68 | 85 | 84 | 89 | 74 | 54 | 77 | 84 | 77.01 | 82 | 72 | 0.864 | -0.211 | 0.38003159 | 0.55466062 | 0.0149 | 0.0332 | 0.0283 | 0.0283 | TRUE | 0.4359 |
| Slc7a1 | 7680 | 5482 | 5065 | 6371 | 5661 | 6300 | 5965 | 5831 | 5465 | 6342 | 6386 | 6219 | 6034.65 | 5754 | 6316 | 1.097 | 0.134 | 0.01474259 | 0.0550891 | 9.00E-04 | 0.0028 | 0.002 | 0.002 | TRUE | 0.0233 |
| Slc7a10 | 4 | 3 | 3 | 3 | 4 | 4 | 3 | 3 | 3 | 3 | 5 | 4 | 3.5 | 3 | 4 | 1.198 | 0.261 | 0.79173845 | NA | 0 | 0.6802 | 0.4156 | 0.4156 | TRUE | 0.0593 |
| Slc7a11 | 17 | 20 | 9 | 7 | 5 | 11 | 13 | 21 | 10 | 7 | 6 | 11 | 11.28 | 15 | 8 | 0.535 | -0.901 | 0.13798433 | NA | 0 | 0.2126 | 0.1674 | 0.1674 | TRUE | 0.397 |
| Slc7a12 | 0 | 0 | 0 | 0 | 0 | 0 | 0 | 0 | 0 | 0 | 0 | 0 | 0 | 0 | 0 | NA | NA | NA | NA | NA | NA | NA | NA | NA | NA |
| Slc7a13 | 3 | 4 | 1 | 2 | 5 | 0 | 2 | 4 | 1 | 2 | 6 | 0 | 2.55 | 2 | 3 | 0.98 | -0.029 | 0.98225469 | NA | 0.5703 | 0.9322 | 0.8446 | 0.8446 | TRUE | 0.4296 |
| Slc7a14 | 51 | 33 | 24 | 23 | 10 | 35 | 40 | 35 | 26 | 23 | 11 | 35 | 28.22 | 34 | 23 | 0.685 | -0.546 | 0.19675116 | 0.35644191 | 0.109 | 0.0864 | 0.092 | 0.092 | TRUE | 0.2709 |
| Slc7a15 | 4 | 3 | 6 | 6 | 3 | 2 | 3 | 3 | 6 | 6 | 3 | 2 | 4.02 | 4 | 4 | 0.898 | -0.155 | 0.8720732 | NA | 0 | 0.5924 | 0.4196 | 0.4196 | TRUE | 0.4281 |
| Slc7a2 | 872 | 556 | 510 | 580 | 480 | 502 | 677 | 591 | 550 | 577 | 541 | 496 | 572.22 | 606 | 538 | 0.885 | -0.176 | 0.10577542 | 0.23060527 | 0.0071 | 0.0066 | 0.0067 | 0.0067 | TRUE | 0.1286 |

| | | | | | | | | | | | | | | | | | | | | | | | | | |
|---|---|---|---|---|---|---|---|---|---|---|---|---|---|---|---|---|---|---|---|---|---|---|---|---|---|
| Slc7a3 | 3698 | 2650 | 2230 | 1979 | 1821 | 2349 | 2872 | 2819 | 2406 | 1970 | 2054 | 2319 | 2406.67 | 2699 | 2114 | 0.783 | -0.353 | 4.47E-05 | 0.00063513 | 0.0075 | 0.0034 | 0.005 | 0.005 | TRUE | 0.1085 |
| Slc7a4 | 288 | 231 | 236 | 379 | 264 | 327 | 224 | 246 | 255 | 377 | 298 | 323 | 286.98 | 242 | 333 | 1.383 | 0.468 | 0.00049242 | 0.00426298 | 0.0066 | 0.0107 | 0.0094 | 0.0094 | TRUE | 0.1573 |
| Slc7a5 | 22321 | 16721 | 14600 | 15125 | 14448 | 15961 | 17336 | 17786 | 15753 | 15055 | 16299 | 15756 | 16330.86 | 16958 | 15703 | 0.926 | -0.111 | 0.06991179 | 0.17145226 | 0.0027 | 0.0026 | 0.0026 | 0.0026 | TRUE | 0.0471 |
| Slc7a6 | 6491 | 4349 | 4195 | 4842 | 4037 | 4597 | 5041 | 4626 | 4526 | 4820 | 4554 | 4538 | 4684.24 | 4731 | 4637 | 0.98 | -0.03 | 0.63401461 | 0.7710347 | 0.0017 | 0.0029 | 0.0026 | 0.0026 | TRUE | 0.0407 |
| Slc7a6os | 3721 | 2821 | 2798 | 2724 | 2352 | 2863 | 2890 | 3001 | 3019 | 2711 | 2653 | 2826 | 2850.1 | 2970 | 2730 | 0.92 | -0.121 | 0.03915728 | 0.11274678 | 5.00E-04 | 0.0033 | 0.0021 | 0.0021 | TRUE | 0.0115 |
| Slc7a7 | 2850 | 2027 | 1839 | 3218 | 2992 | 3591 | 2213 | 2156 | 1984 | 3203 | 3375 | 3545 | 2746.2 | 2118 | 3374 | 1.592 | 0.671 | 9.96E-23 | 4.07E-20 | 0.0023 | 0.0033 | 0.003 | 0.003 | TRUE | 0.0365 |
| Slc7a8 | 115 | 77 | 76 | 91 | 99 | 103 | 89 | 82 | 82 | 91 | 112 | 102 | 92.86 | 84 | 102 | 1.196 | 0.258 | 0.20664011 | 0.36835958 | 0 | 0.0279 | 0.0192 | 0.0192 | TRUE | 0.0846 |
| Slc7a9 | 5 | 4 | 2 | 1 | 0 | 1 | 4 | 4 | 2 | 1 | 0 | 1 | 2.05 | 3 | 1 | 0.197 | -2.342 | 0.12545242 | NA | 0 | 1.1606 | 0.7825 | 0.7825 | TRUE | 0.1968 |
| Slc8a1 | 43 | 22 | 35 | 27 | 29 | 19 | 33 | 23 | 38 | 27 | 33 | 19 | 28.82 | 31 | 26 | 0.824 | -0.28 | 0.46291122 | 0.63217944 | 0.0261 | 0.0847 | 0.0694 | 0.0694 | TRUE | 0.3468 |
| Slc8a2 | 348 | 272 | 278 | 350 | 299 | 343 | 270 | 289 | 300 | 348 | 337 | 339 | 313.97 | 286 | 341 | 1.194 | 0.256 | 0.02752 | 0.08688831 | 0 | 0.01 | 0.0065 | 0.0065 | TRUE | 0.0303 |
| Slc8a3 | 53 | 39 | 35 | 44 | 35 | 39 | 41 | 41 | 38 | 44 | 39 | 38 | 40.37 | 40 | 40 | 1.011 | 0.015 | 0.95913693 | 0.97739513 | 0 | 0.0611 | 0.0393 | 0.0393 | TRUE | 0.0369 |
| Slc9a1 | 3350 | 2438 | 2190 | 2334 | 2208 | 2469 | 2602 | 2593 | 2363 | 2323 | 2491 | 2437 | 2468.24 | 2519 | 2417 | 0.959 | -0.061 | 0.35771221 | 0.53214291 | 0.0015 | 0.0034 | 0.0027 | 0.0027 | TRUE | 0.0352 |
| Slc9a2 | 8 | 5 | 8 | 12 | 2 | 2 | 6 | 5 | 9 | 12 | 2 | 2 | 6.06 | 7 | 5 | 0.81 | -0.303 | 0.72694226 | NA | 0.3324 | 0.3938 | 0.3764 | 0.3764 | TRUE | 2.478 |
| Slc9a3 | 0 | 0 | 0 | 0 | 0 | 0 | 0 | 0 | 0 | 0 | 0 | 0 | 0 | 0 | 0 | NA | NA | NA | NA | NA | NA | NA | NA | NA | NA |
| Slc9a3r1 | 5756 | 4691 | 4039 | 3894 | 3829 | 4442 | 4470 | 4990 | 4358 | 3876 | 4320 | 4385 | 4399.8 | 4606 | 4194 | 0.91 | -0.135 | 0.06298979 | 0.15892551 | 0.0045 | 0.003 | 0.0036 | 0.0036 | TRUE | 0.0642 |
| Slc9a3r2 | 765 | 596 | 607 | 642 | 611 | 737 | 594 | 634 | 655 | 639 | 689 | 728 | 656.48 | 628 | 685 | 1.093 | 0.129 | 0.16570598 | 0.31676786 | 0.002 | 0.006 | 0.0047 | 0.0047 | TRUE | 0.042 |
| Slc9a4 | 1 | 1 | 0 | 0 | 0 | 0 | 1 | 1 | 0 | 0 | 0 | 0 | 0.31 | 1 | 0 | 0.309 | -1.696 | 0.63183309 | NA | 0 | 7.7294 | 7.2472 | 7.2472 | TRUE | 0.2015 |
| Slc9a5 | 1466 | 1203 | 1386 | 1982 | 1728 | 1964 | 1139 | 1280 | 1495 | 1973 | 1949 | 1939 | 1629.12 | 1305 | 1954 | 1.5 | 0.585 | 2.09E-10 | 1.58E-08 | 0.0086 | 0.0039 | 0.0055 | 0.0055 | TRUE | 0.1965 |
| Slc9a6 | 1766 | 1245 | 1137 | 1372 | 1217 | 1420 | 1372 | 1324 | 1227 | 1366 | 1373 | 1402 | 1343.84 | 1308 | 1380 | 1.054 | 0.076 | 0.28975084 | 0.46187814 | 0.001 | 0.0042 | 0.003 | 0.003 | TRUE | 0.0346 |
| Slc9a7 | 112 | 66 | 57 | 74 | 68 | 64 | 87 | 70 | 62 | 74 | 77 | 63 | 72.04 | 73 | 71 | 0.969 | -0.046 | 0.84937961 | 0.91479443 | 0 | 0.0353 | 0.0277 | 0.0277 | TRUE | 0.2914 |
| Slc9a8 | 1290 | 969 | 837 | 998 | 905 | 1007 | 1002 | 1031 | 903 | 993 | 1021 | 994 | 990.69 | 979 | 1003 | 1.024 | 0.034 | 0.67050243 | 0.79723222 | 0.0013 | 0.0048 | 0.0036 | 0.0036 | TRUE | 0.0525 |
| Slc9a9 | 18 | 11 | 9 | 4 | 13 | 7 | 14 | 12 | 10 | 4 | 15 | 7 | 10.16 | 12 | 9 | 0.709 | -0.495 | 0.44206957 | NA | 0.0815 | 0.2357 | 0.1968 | 0.1968 | TRUE | 0.9417 |
| Slc9b1 | 1568 | 1130 | 1045 | 1158 | 1002 | 1093 | 1218 | 1202 | 1128 | 1153 | 1130 | 1079 | 1151.55 | 1183 | 1121 | 0.947 | -0.079 | 0.28879146 | 0.46093761 | 6.00E-04 | 0.0045 | 0.0031 | 0.0031 | TRUE | 0.0194 |
| Slc9b2 | 183 | 131 | 145 | 188 | 179 | 195 | 142 | 139 | 156 | 187 | 202 | 192 | 169.91 | 146 | 194 | 1.329 | 0.41 | 0.00722942 | 0.03250409 | 0 | 0.0164 | 0.0108 | 0.0108 | TRUE | 0.0387 |
| Slc9c1 | 0 | 0 | 0 | 0 | 0 | 0 | 0 | 0 | 0 | 0 | 0 | 0 | 0 | 0 | 0 | NA | NA | NA | NA | NA | NA | NA | NA | NA | NA |
| Slco1a1 | 0 | 0 | 0 | 0 | 0 | 0 | 0 | 0 | 0 | 0 | 0 | 0 | 0 | 0 | 0 | NA | NA | NA | NA | NA | NA | NA | NA | NA | NA |
| Slco1a4 | 0 | 2 | 0 | 2 | 0 | 0 | 0 | 2 | 0 | 2 | 0 | 0 | 0.69 | 1 | 1 | 0.971 | -0.042 | 0.98762268 | NA | 8.094 | 3.4555 | 3.8966 | 3.8966 | TRUE | 0.9747 |
| Slco1a5 | 6 | 2 | 0 | 1 | 1 | 0 | 5 | 2 | 0 | 1 | 1 | 0 | 1.49 | 2 | 1 | 0.301 | -1.734 | 0.33745826 | NA | 0 | 1.5983 | 1.4039 | 1.4039 | TRUE | 0.1089 |
| Slco1a6 | 0 | 0 | 0 | 0 | 0 | 0 | 0 | 0 | 0 | 0 | 0 | 0 | 0 | 0 | 0 | NA | NA | NA | NA | NA | NA | NA | NA | NA | NA |
| Slco1b2 | 1 | 1 | 0 | 0 | 0 | 0 | 1 | 1 | 0 | 0 | 0 | 0 | 0.31 | 1 | 0 | 0.309 | -1.696 | 0.63183309 | NA | 0 | 7.7294 | 7.2472 | 7.2472 | TRUE | 0.2015 |
| Slco1c1 | 0 | 0 | 0 | 0 | 0 | 0 | 0 | 0 | 0 | 0 | 0 | 0 | 0 | 0 | 0 | NA | NA | NA | NA | NA | NA | NA | NA | NA | NA |
| Slco2a1 | 86 | 69 | 59 | 55 | 42 | 55 | 67 | 73 | 64 | 55 | 47 | 54 | 60.04 | 68 | 52 | 0.769 | -0.379 | 0.12818283 | 0.26437478 | 0 | 0.0419 | 0.0277 | 0.0277 | TRUE | 0.0487 |
| Slco2b1 | 8 | 3 | 7 | 2 | 6 | 3 | 6 | 3 | 8 | 2 | 7 | 3 | 4.78 | 6 | 4 | 0.68 | -0.556 | 0.53946517 | NA | 0 | 0.4983 | 0.3716 | 0.3716 | TRUE | 0.5984 |
| Slco3a1 | 288 | 178 | 165 | 193 | 154 | 198 | 224 | 189 | 178 | 192 | 174 | 195 | 192.06 | 197 | 187 | 0.947 | -0.079 | 0.60637851 | 0.7508545 | 0.0049 | 0.0148 | 0.0118 | 0.0118 | TRUE | 0.1644 |
| Slco4a1 | 778 | 677 | 1063 | 1493 | 1403 | 1777 | 604 | 720 | 1147 | 1486 | 1583 | 1754 | 1215.73 | 824 | 1608 | 1.953 | 0.965 | 0.00112345 | 0.0078945 | 0.0624 | 0.0044 | 0.0186 | 0.0624 | TRUE | 1.3968 |
| Slco4c1 | 482 | 284 | 384 | 332 | 336 | 371 | 374 | 302 | 414 | 330 | 379 | 366 | 361.09 | 363 | 358 | 0.985 | -0.021 | 0.87143399 | 0.92866769 | 0.0121 | 0.009 | 0.0099 | 0.0099 | TRUE | 0.2411 |
| Slco5a1 | 412 | 298 | 329 | 299 | 281 | 382 | 320 | 317 | 355 | 298 | 317 | 377 | 330.61 | 331 | 331 | 1.001 | 0.002 | 0.98881073 | 0.99416655 | 0.0067 | 0.0096 | 0.0087 | 0.0087 | TRUE | 0.1756 |
| Slco6b1 | 0 | 0 | 0 | 0 | 0 | 0 | 0 | 0 | 0 | 0 | 0 | 0 | 0 | 0 | 0 | NA | NA | NA | NA | NA | NA | NA | NA | NA | NA |
| Slco6c1 | 0 | 0 | 0 | 0 | 0 | 0 | 0 | 0 | 0 | 0 | 0 | 0 | 0 | 0 | 0 | NA | NA | NA | NA | NA | NA | NA | NA | NA | NA |
| Slco6d1 | 0 | 0 | 0 | 0 | 0 | 0 | 0 | 0 | 0 | 0 | 0 | 0 | 0 | 0 | 0 | NA | NA | NA | NA | NA | NA | NA | NA | NA | NA |
| Slfn1 | 0 | 0 | 0 | 0 | 0 | 0 | 0 | 0 | 0 | 0 | 0 | 0 | 0 | 0 | 0 | NA | NA | NA | NA | NA | NA | NA | NA | NA | NA |
| Slfn10-ps | 143 | 99 | 80 | 81 | 66 | 70 | 111 | 105 | 86 | 81 | 74 | 69 | 87.81 | 101 | 75 | 0.738 | -0.438 | 0.04143615 | 0.11723019 | 0 | 0.0294 | 0.0216 | 0.0216 | TRUE | 0.15 |
| Slfn14-ps | 0 | 0 | 0 | 0 | 0 | 1 | 0 | 0 | 0 | 0 | 0 | 1 | 0.16 | 0 | 0 | 2.094 | 1.067 | 0.79379329 | NA | 0 | 14.4078 | 10 | 10 | TRUE | 0.3835 |
| Slfn2 | 0 | 0 | 0 | 0 | 0 | 0 | 0 | 0 | 0 | 0 | 0 | 0 | 0 | 0 | 0 | NA | NA | NA | NA | NA | NA | NA | NA | NA | NA |
| Slfn3 | 196 | 134 | 117 | 69 | 50 | 44 | 152 | 143 | 126 | 69 | 56 | 43 | 98.25 | 140 | 56 | 0.399 | -1.324 | 2.60E-09 | 1.54E-07 | 0.0109 | 0.0265 | 0.023 | 0.023 | TRUE | 0.3485 |
| Slfn4 | 30 | 21 | 14 | 10 | 9 | 8 | 23 | 22 | 15 | 10 | 10 | 8 | 14.79 | 20 | 9 | 0.458 | -1.127 | 0.02899368 | NA | 0 | 0.1627 | 0.1127 | 0.1127 | TRUE | 0.2502 |
| Slfn5 | 79 | 57 | 66 | 101 | 101 | 124 | 61 | 61 | 71 | 101 | 114 | 122 | 88.35 | 64 | 112 | 1.747 | 0.805 | 0.00016086 | 0.00180042 | 0 | 0.0293 | 0.0207 | 0.0207 | TRUE | 0.086 |
| Slfn8 | 126 | 85 | 79 | 69 | 59 | 90 | 98 | 90 | 85 | 69 | 67 | 89 | 82.93 | 91 | 75 | 0.819 | -0.288 | 0.19344329 | 0.35221464 | 0 | 0.031 | 0.0232 | 0.0232 | TRUE | 0.2559 |
| Slfn9 | 1360 | 1032 | 958 | 651 | 503 | 674 | 1056 | 1098 | 1034 | 648 | 567 | 665 | 844.74 | 1063 | 627 | 0.591 | -0.759 | 4.35E-18 | 1.04E-15 | 0.002 | 0.0052 | 0.0042 | 0.0042 | TRUE | 0.0787 |
| Slfnl1 | 3 | 1 | 0 | 0 | 0 | 0 | 2 | 1 | 0 | 0 | 0 | 0 | 0.57 | 1 | 0 | 0.165 | -2.597 | 0.35609012 | NA | 0 | 4.1927 | 4.2873 | 4.2873 | TRUE | 0.0685 |
| Slirp | 2502 | 1700 | 1609 | 1891 | 1579 | 1841 | 1943 | 1808 | 1736 | 1882 | 1781 | 1817 | 1828.08 | 1829 | 1827 | 0.998 | -0.003 | 0.96596138 | 0.98144468 | 0.0016 | 0.0037 | 0.0029 | 0.0029 | TRUE | 0.0371 |
| Slit1 | 37 | 34 | 36 | 38 | 37 | 36 | 29 | 36 | 39 | 38 | 42 | 36 | 36.47 | 35 | 39 | 1.115 | 0.158 | 0.62228067 | 0.76149466 | 0 | 0.0674 | 0.0463 | 0.0463 | TRUE | 0.1871 |
| Slit2 | 451 | 353 | 378 | 210 | 232 | 247 | 350 | 375 | 408 | 209 | 262 | 244 | 308.03 | 378 | 238 | 0.631 | -0.665 | 3.26E-07 | 1.12E-05 | 0.0054 | 0.0101 | 0.0088 | 0.0088 | TRUE | 0.128 |
| Slit3 | 485 | 308 | 352 | 356 | 303 | 352 | 377 | 328 | 380 | 354 | 342 | 347 | 354.63 | 362 | 348 | 0.962 | -0.056 | 0.62208024 | 0.76142749 | 0 | 0.0091 | 0.0066 | 0.0066 | TRUE | 0.0737 |
| Slitrk1 | 0 | 0 | 0 | 0 | 0 | 0 | 0 | 0 | 0 | 0 | 0 | 0 | 0 | 0 | 0 | NA | NA | NA | NA | NA | NA | NA | NA | NA | NA |
| Slitrk2 | 1 | 2 | 1 | 1 | 1 | 1 | 1 | 2 | 1 | 1 | 1 | 1 | 1.18 | 1 | 1 | 0.792 | -0.336 | 0.84173106 | NA | 0 | 2.0071 | 1.1654 | 1.1654 | TRUE | 0.1607 |
| Slitrk3 | 1 | 0 | 0 | 0 | 0 | 0 | 1 | 0 | 0 | 0 | 0 | 0 | 0.13 | 0 | 0 | 0.552 | -0.857 | 0.8336548 | NA | 0 | 18.3121 | 10 | 10 | TRUE | 0.2566 |
| Slitrk4 | 80 | 43 | 47 | 49 | 46 | 48 | 62 | 46 | 51 | 49 | 52 | 47 | 51.11 | 53 | 49 | 0.927 | -0.109 | 0.68905778 | 0.80972645 | 0 | 0.0488 | 0.0343 | 0.0343 | TRUE | 0.224 |
| Slitrk5 | 65 | 51 | 31 | 38 | 37 | 33 | 50 | 54 | 33 | 38 | 42 | 33 | 41.72 | 46 | 38 | 0.807 | -0.309 | 0.32661423 | 0.5006795 | 2.00E-04 | 0.0592 | 0.0472 | 0.0472 | TRUE | 0.4217 |
| Slitrk6 | 0 | 0 | 1 | 0 | 0 | 0 | 0 | 0 | 1 | 0 | 0 | 0 | 0.18 | 0 | 0 | 0.552 | -0.857 | 0.8336548 | NA | 0 | 13.1817 | 10 | 10 | TRUE | 0.5365 |
| Slk | 6594 | 5173 | 5243 | 5814 | 5389 | 6061 | 5121 | 5502 | 5657 | 5787 | 6079 | 5983 | 5688.44 | 5427 | 5950 | 1.097 | 0.133 | 0.02193922 | 0.07380025 | 0.0013 | 0.0028 | 0.0023 | 0.0023 | TRUE | 0.0301 |
| Slmap | 6020 | 4314 | 4220 | 4308 | 3794 | 4310 | 4675 | 4589 | 4553 | 4288 | 4280 | 4255 | 4440.07 | 4606 | 4274 | 0.928 | -0.108 | 0.03139034 | 0.09576565 | 0 | 0.003 | 0.0016 | 0.0016 | TRUE | 0.0022 |
| Slmo1 | 280 | 204 | 185 | 190 | 163 | 195 | 217 | 217 | 200 | 189 | 184 | 192 | 199.93 | 211 | 188 | 0.891 | -0.166 | 0.23742458 | 0.40417188 | 0 | 0.0143 | 0.0092 | 0.0092 | TRUE | 0.0247 |
| Slmo2 | 3683 | 2552 | 2131 | 2168 | 1762 | 2241 | 2860 | 2715 | 2299 | 2158 | 1988 | 2212 | 2372.04 | 2625 | 2119 | 0.807 | -0.309 | 0.00031125 | 0.00299262 | 0.007 | 0.0034 | 0.0049 | 0.0049 | TRUE | 0.1418 |
| Sln | 0 | 0 | 0 | 0 | 0 | 0 | 0 | 0 | 0 | 0 | 0 | 0 | 0 | 0 | 0 | NA | NA | NA | NA | NA | NA | NA | NA | NA | NA |
| Slpi | 2 | 2 | 1 | 1 | 0 | 1 | 2 | 2 | 1 | 1 | 0 | 1 | 1.12 | 2 | 1 | 0.426 | -1.23 | 0.49943099 | NA | 0 | 2.1116 | 1.3217 | 1.3217 | TRUE | 0.2 |
| Sltm | 5897 | 4224 | 3671 | 3810 | 3339 | 4243 | 4580 | 4493 | 3961 | 3792 | 3767 | 4188 | 4130.27 | 4345 | 3916 | 0.901 | -0.15 | 0.03982487 | 0.11418592 | 0.0044 | 0.003 | 0.0036 | 0.0036 | TRUE | 0.0722 |
| Slu7 | 4227 | 3065 | 2843 | 2955 | 2636 | 3133 | 3283 | 3260 | 3068 | 2941 | 2974 | 3093 | 3103.09 | 3204 | 3003 | 0.937 | -0.094 | 0.11120941 | 0.23898904 | 7.00E-04 | 0.0032 | 0.0022 | 0.0022 | TRUE | 0.0166 |
| Slurp1 | 0 | 0 | 0 | 0 | 0 | 0 | 0 | 0 | 0 | 0 | 0 | 0 | 0 | 0 | 0 | NA | NA | NA | NA | NA | NA | NA | NA | NA | NA |

| | | | | | | | | | | | | | | | | | | | | | | | | | |
|---|---|---|---|---|---|---|---|---|---|---|---|---|---|---|---|---|---|---|---|---|---|---|---|---|---|
| Slx | 0 | 0 | 0 | 0 | 0 | 0 | 0 | 0 | 0 | 0 | 0 | 0 | 0 | 0 | 0 | NA | NA | NA | NA | NA | NA | NA | NA | NA | NA |
| Slx1b | 994 | 689 | 754 | 754 | 739 | 959 | 772 | 733 | 814 | 751 | 834 | 947 | 808.22 | 773 | 844 | 1.092 | 0.127 | 0.20241374 | 0.36377463 | 0.007 | 0.0054 | 0.0059 | 0.0059 | TRUE | 0.1378 |
| Slx4 | 2438 | 1745 | 1590 | 1627 | 1393 | 1600 | 1893 | 1856 | 1716 | 1620 | 1571 | 1579 | 1705.94 | 1822 | 1590 | 0.872 | -0.197 | 0.00368056 | 0.0193283 | 0.001 | 0.0038 | 0.0027 | 0.0027 | TRUE | 0.0309 |
| Slx4ip | 654 | 504 | 532 | 478 | 449 | 530 | 508 | 536 | 574 | 476 | 507 | 523 | 520.59 | 539 | 502 | 0.932 | -0.102 | 0.30423737 | 0.47675308 | 0.0014 | 0.007 | 0.0051 | 0.0051 | TRUE | 0.0367 |
| Slxl1 | 0 | 0 | 0 | 0 | 0 | 0 | 0 | 0 | 0 | 0 | 0 | 0 | 0 | 0 | 0 | NA | NA | NA | NA | NA | NA | NA | NA | NA | NA |
| Sly | 0 | 0 | 0 | 0 | 0 | 0 | 0 | 0 | 0 | 0 | 0 | 0 | 0 | 0 | 0 | NA | NA | NA | NA | NA | NA | NA | NA | NA | NA |
| Smad1 | 1195 | 848 | 788 | 755 | 704 | 709 | 928 | 902 | 850 | 752 | 794 | 700 | 820.99 | 893 | 749 | 0.836 | -0.258 | 0.00273911 | 0.01535898 | 0.0017 | 0.0053 | 0.0041 | 0.0041 | TRUE | 0.0385 |
| Smad2 | 5377 | 3985 | 4221 | 4877 | 4264 | 4846 | 4176 | 4239 | 4554 | 4855 | 4810 | 4784 | 4569.64 | 4323 | 4816 | 1.114 | 0.156 | 0.00554246 | 0.02656138 | 8.00E-04 | 0.0029 | 0.0021 | 0.0021 | TRUE | 0.0265 |
| Smad3 | 1096 | 831 | 838 | 845 | 760 | 906 | 851 | 884 | 904 | 841 | 857 | 894 | 872.03 | 880 | 864 | 0.983 | -0.024 | 0.75828089 | 0.85642898 | 0 | 0.0051 | 0.0033 | 0.0033 | TRUE | 0.0112 |
| Smad4 | 6473 | 4707 | 4978 | 4853 | 4376 | 5028 | 5027 | 5007 | 5371 | 4831 | 4937 | 4963 | 5022.66 | 5135 | 4910 | 0.956 | -0.064 | 0.2379936 | 0.40471655 | 6.00E-04 | 0.0029 | 0.0019 | 0.0019 | TRUE | 0.0195 |
| Smad5 | 4108 | 2975 | 2998 | 3290 | 2879 | 3258 | 3191 | 3164 | 3235 | 3275 | 3248 | 3216 | 3221.44 | 3197 | 3246 | 1.016 | 0.022 | 0.67495086 | 0.80032886 | 0 | 0.0032 | 0.0017 | 0.0017 | TRUE | 0.0013 |
| Smad6 | 373 | 274 | 313 | 254 | 214 | 292 | 290 | 291 | 338 | 253 | 241 | 288 | 283.56 | 306 | 261 | 0.854 | -0.228 | 0.08319894 | 0.19416945 | 0.0045 | 0.0108 | 0.0089 | 0.0089 | TRUE | 0.0953 |
| Smad7 | 4856 | 3781 | 3265 | 4396 | 3995 | 4115 | 3771 | 4022 | 3523 | 4376 | 4507 | 4062 | 4043.47 | 3772 | 4315 | 1.144 | 0.194 | 0.00467918 | 0.02328673 | 0.0032 | 0.003 | 0.0031 | 0.0031 | TRUE | 0.0402 |
| Smad9 | 163 | 122 | 113 | 115 | 97 | 102 | 127 | 130 | 122 | 114 | 109 | 101 | 117.15 | 126 | 108 | 0.858 | -0.222 | 0.21879797 | 0.38193028 | 0 | 0.0227 | 0.0148 | 0.0148 | TRUE | 0.0379 |
| Smagp | 1155 | 935 | 1117 | 1380 | 1317 | 1554 | 897 | 995 | 1205 | 1374 | 1486 | 1534 | 1248.37 | 1032 | 1465 | 1.421 | 0.507 | 8.33E-07 | 2.51E-05 | 0.012 | 0.0043 | 0.0068 | 0.0068 | TRUE | 0.2556 |
| Smap1 | 5451 | 3999 | 4081 | 4180 | 3741 | 4317 | 4234 | 4254 | 4403 | 4161 | 4220 | 4262 | 4255.53 | 4297 | 4214 | 0.981 | -0.028 | 0.59278574 | 0.74044182 | 0 | 0.003 | 0.0017 | 0.0017 | TRUE | 0.0057 |
| Smap2 | 2244 | 1665 | 1533 | 1586 | 1400 | 1643 | 1743 | 1771 | 1654 | 1579 | 1579 | 1622 | 1657.98 | 1723 | 1593 | 0.925 | -0.113 | 0.08287916 | 0.19387755 | 0 | 0.0039 | 0.0024 | 0.0024 | TRUE | 0.0142 |
| Smarca1 | 829 | 595 | 570 | 870 | 709 | 780 | 644 | 633 | 615 | 866 | 800 | 770 | 721.26 | 631 | 812 | 1.287 | 0.364 | 3.03E-05 | 0.0004674 | 0 | 0.0057 | 0.0041 | 0.0041 | TRUE | 0.0409 |
| Smarca2 | 214 | 167 | 143 | 152 | 115 | 139 | 166 | 178 | 154 | 151 | 130 | 137 | 152.73 | 166 | 139 | 0.84 | -0.251 | 0.12395615 | 0.25819081 | 0 | 0.0179 | 0.0125 | 0.0125 | TRUE | 0.0579 |
| Smarca4 | 19912 | 14693 | 14942 | 16471 | 14878 | 17453 | 15465 | 15629 | 16122 | 16395 | 16784 | 17229 | 16270.63 | 15739 | 16803 | 1.068 | 0.094 | 0.04798027 | 0.13077845 | 5.00E-04 | 0.0026 | 0.0016 | 0.0016 | TRUE | 0.006 |
| Smarca5 | 32788 | 23258 | 21203 | 21259 | 18144 | 20838 | 25465 | 24739 | 22878 | 21161 | 20468 | 20570 | 22546.99 | 24361 | 20733 | 0.851 | -0.233 | 2.39E-05 | 0.0003854 | 0.0016 | 0.0025 | 0.0021 | 0.0021 | TRUE | 0.0347 |
| Smarca5-ps | 120 | 97 | 39 | 51 | 47 | 43 | 93 | 103 | 42 | 51 | 53 | 42 | 64.12 | 79 | 49 | 0.61 | -0.712 | 0.02512271 | 0.08157836 | 0.0979 | 0.0394 | 0.0562 | 0.0562 | TRUE | 1.5031 |
| Smarcad1 | 7997 | 5304 | 4370 | 4528 | 3893 | 4364 | 6211 | 5642 | 4715 | 4507 | 4392 | 4308 | 4962.46 | 5523 | 4402 | 0.797 | -0.328 | 0.00016984 | 0.00188033 | 0.0093 | 0.0029 | 0.0053 | 0.0053 | TRUE | 0.199 |
| Smarcal1 | 2778 | 2037 | 1840 | 1950 | 1747 | 2092 | 2158 | 2167 | 1985 | 1941 | 1971 | 2065 | 2047.76 | 2103 | 1992 | 0.947 | -0.079 | 0.23580579 | 0.40228839 | 0.0012 | 0.0036 | 0.0027 | 0.0027 | TRUE | 0.0285 |
| Smarcb1 | 4726 | 3330 | 2963 | 2805 | 2565 | 2840 | 3670 | 3542 | 3197 | 2792 | 2894 | 2804 | 3149.8 | 3470 | 2830 | 0.815 | -0.295 | 1.01E-05 | 0.00019108 | 0.0022 | 0.0032 | 0.0029 | 0.0029 | TRUE | 0.0571 |
| Smarcc1 | 24687 | 18397 | 18293 | 19248 | 17920 | 20444 | 19173 | 19569 | 19738 | 19159 | 20216 | 20181 | 19672.74 | 19493 | 19852 | 1.018 | 0.026 | 0.58028014 | 0.73107457 | 5.00E-04 | 0.0025 | 0.0016 | 0.0016 | TRUE | 0.0114 |
| Smarcc2 | 2419 | 1827 | 1746 | 1739 | 1507 | 1808 | 1879 | 1943 | 1884 | 1731 | 1700 | 1785 | 1820.31 | 1902 | 1739 | 0.914 | -0.129 | 0.03879682 | 0.11197958 | 0 | 0.0037 | 0.0023 | 0.0023 | TRUE | 0.0065 |
| Smarcd1 | 9286 | 6891 | 6930 | 7062 | 6636 | 7709 | 7212 | 7330 | 7477 | 7030 | 7486 | 7610 | 7357.48 | 7340 | 7375 | 1.005 | 0.007 | 0.89521982 | 0.94249977 | 9.00E-04 | 0.0027 | 0.0019 | 0.0019 | TRUE | 0.0206 |
| Smarcd2 | 6779 | 5188 | 5289 | 5214 | 4481 | 5226 | 5265 | 5518 | 5707 | 5190 | 5055 | 5159 | 5315.68 | 5497 | 5135 | 0.935 | -0.098 | 0.0728575 | 0.17665007 | 6.00E-04 | 0.0029 | 0.0019 | 0.0019 | TRUE | 0.0169 |
| Smarcd3 | 271 | 219 | 267 | 398 | 353 | 438 | 210 | 233 | 288 | 396 | 398 | 432 | 326.38 | 244 | 409 | 1.685 | 0.752 | 2.21E-08 | 1.01E-06 | 0.0097 | 0.0097 | 0.0098 | 0.0098 | TRUE | 0.2753 |
| Smarce1 | 5108 | 3928 | 3742 | 3959 | 3543 | 4030 | 3967 | 4178 | 4038 | 3941 | 3997 | 3978 | 4016.47 | 4061 | 3972 | 0.978 | -0.032 | 0.54977224 | 0.70613719 | 2.00E-04 | 0.003 | 0.0018 | 0.0018 | TRUE | 0.0077 |
| Smc1a | 12485 | 9107 | 9486 | 9408 | 8306 | 9880 | 9697 | 9687 | 10235 | 9365 | 9370 | 9753 | 9684.45 | 9873 | 9496 | 0.962 | -0.056 | 0.27116328 | 0.44203202 | 6.00E-04 | 0.0027 | 0.0018 | 0.0018 | TRUE | 0.0125 |
| Smc1b | 2119 | 1571 | 1800 | 2051 | 1561 | 1685 | 1646 | 1671 | 1942 | 2042 | 1761 | 1663 | 1787.48 | 1753 | 1822 | 1.04 | 0.057 | 0.54168849 | 0.69973309 | 0.0094 | 0.0038 | 0.0057 | 0.0057 | TRUE | 0.1351 |
| Smc2 | 16852 | 12115 | 11336 | 11692 | 10334 | 11368 | 13088 | 12887 | 12231 | 11638 | 11658 | 11222 | 12120.71 | 12735 | 11506 | 0.903 | -0.147 | 0.00390989 | 0.02026251 | 7.00E-04 | 0.0026 | 0.0018 | 0.0018 | TRUE | 0.0146 |
| Smc3 | 19759 | 14853 | 15614 | 15839 | 14515 | 17186 | 15346 | 15799 | 16847 | 15766 | 16374 | 16965 | 16183.01 | 15997 | 16368 | 1.023 | 0.033 | 0.55446855 | 0.70999991 | 0.0016 | 0.0026 | 0.0022 | 0.0022 | TRUE | 0.0264 |
| Smc4 | 16989 | 12374 | 14027 | 14983 | 12900 | 14684 | 13195 | 13162 | 15135 | 14914 | 14553 | 14495 | 14242.28 | 13831 | 14654 | 1.06 | 0.084 | 0.19074259 | 0.34885269 | 0.0034 | 0.0026 | 0.0029 | 0.0029 | TRUE | 0.083 |
| Smc5 | 7689 | 5865 | 5919 | 5989 | 5189 | 6234 | 5972 | 6239 | 6387 | 5961 | 5854 | 6154 | 6094.32 | 6199 | 5990 | 0.967 | -0.049 | 0.36086435 | 0.5358232 | 7.00E-04 | 0.0028 | 0.0019 | 0.0019 | TRUE | 0.0128 |
| Smc6 | 9029 | 6235 | 6340 | 6220 | 5391 | 6045 | 7012 | 6632 | 6841 | 6191 | 6082 | 5967 | 6454.28 | 6828 | 6080 | 0.89 | -0.168 | 0.00102568 | 0.00736317 | 4.00E-04 | 0.0028 | 0.0017 | 0.0017 | TRUE | 0.0077 |
| Smchd1 | 8120 | 5770 | 6116 | 6827 | 5833 | 6743 | 6306 | 6137 | 6599 | 6796 | 6580 | 6656 | 6512.54 | 6347 | 6677 | 1.052 | 0.073 | 0.16386192 | 0.31462634 | 6.00E-04 | 0.0028 | 0.0018 | 0.0018 | TRUE | 0.0145 |
| Smcp | 0 | 0 | 0 | 0 | 0 | 0 | 0 | 0 | 0 | 0 | 0 | 0 | 0 | 0 | 0 | NA | NA | NA | NA | NA | NA | NA | NA | NA | NA |
| Smcr7 | 1225 | 849 | 759 | 804 | 701 | 817 | 951 | 903 | 819 | 800 | 791 | 807 | 845.17 | 891 | 799 | 0.896 | -0.159 | 0.06294399 | 0.15883801 | 0.0019 | 0.0052 | 0.0041 | 0.0041 | TRUE | 0.059 |
| Smcr7l | 4502 | 3162 | 2954 | 3189 | 2825 | 3104 | 3497 | 3363 | 3187 | 3174 | 3187 | 3064 | 3245.43 | 3349 | 3142 | 0.938 | -0.093 | 0.1209686 | 0.25340821 | 0.001 | 0.0032 | 0.0023 | 0.0023 | TRUE | 0.0216 |
| Smcr8 | 4730 | 3167 | 2841 | 2863 | 2657 | 2959 | 3674 | 3369 | 3065 | 2850 | 2997 | 2921 | 3145.98 | 3369 | 2923 | 0.867 | -0.206 | 0.00487965 | 0.02404994 | 0.0039 | 0.0032 | 0.0035 | 0.0035 | TRUE | 0.0771 |
| Smek1 | 6015 | 4308 | 3750 | 3513 | 3212 | 3740 | 4672 | 4582 | 4046 | 3497 | 3623 | 3692 | 4018.74 | 4433 | 3604 | 0.813 | -0.299 | 1.05E-05 | 0.00019326 | 0.0029 | 0.003 | 0.0031 | 0.0031 | TRUE | 0.0706 |
| Smek2 | 8070 | 5474 | 5306 | 5618 | 4846 | 5412 | 6268 | 5823 | 5725 | 5592 | 5467 | 5342 | 5702.8 | 5939 | 5467 | 0.92 | -0.12 | 0.03599072 | 0.1060869 | 0.0012 | 0.0028 | 0.0022 | 0.0022 | TRUE | 0.0291 |
| Smg1 | 14484 | 10373 | 10155 | 12129 | 11025 | 12601 | 11249 | 11034 | 10957 | 12073 | 12437 | 12439 | 11698.26 | 11080 | 12316 | 1.111 | 0.152 | 0.00093804 | 0.00692893 | 1.00E-04 | 0.0026 | 0.0014 | 0.0014 | TRUE | 0.0037 |
| Smg5 | 13558 | 9288 | 8583 | 8694 | 7942 | 9132 | 10530 | 9880 | 9261 | 8654 | 8959 | 9015 | 9383.09 | 9890 | 8876 | 0.897 | -0.156 | 0.00908327 | 0.03857015 | 0.0021 | 0.0027 | 0.0025 | 0.0025 | TRUE | 0.0394 |
| Smg6 | 8039 | 5978 | 5846 | 6402 | 5580 | 6689 | 6244 | 6359 | 6308 | 6373 | 6295 | 6603 | 6363.42 | 6304 | 6424 | 1.019 | 0.027 | 0.57952135 | 0.73041403 | 2.00E-04 | 0.0028 | 0.0016 | 0.0016 | TRUE | 0.0073 |
| Smg7 | 7833 | 6095 | 5747 | 6073 | 5317 | 6359 | 6084 | 6483 | 6201 | 6045 | 5998 | 6277 | 6181.37 | 6256 | 6107 | 0.976 | -0.034 | 0.51709204 | 0.67923873 | 6.00E-04 | 0.0028 | 0.0019 | 0.0019 | TRUE | 0.0123 |
| Smg8 | 3801 | 2587 | 2370 | 2646 | 2263 | 2489 | 2952 | 2752 | 2557 | 2634 | 2553 | 2457 | 2650.8 | 2754 | 2548 | 0.925 | -0.113 | 0.10551951 | 0.23018779 | 0.0026 | 0.0033 | 0.0031 | 0.0031 | TRUE | 0.0492 |
| Smg9 | 885 | 699 | 794 | 786 | 769 | 930 | 687 | 744 | 857 | 782 | 868 | 918 | 809.25 | 763 | 856 | 1.124 | 0.169 | 0.09889055 | 0.21930839 | 0.0077 | 0.0054 | 0.0063 | 0.0063 | TRUE | 0.1374 |
| Smgc | 0 | 0 | 0 | 0 | 0 | 0 | 0 | 0 | 0 | 0 | 0 | 0 | 0 | 0 | 0 | NA | NA | NA | NA | NA | NA | NA | NA | NA | NA |
| Smim1 | 121 | 74 | 76 | 77 | 89 | 102 | 94 | 79 | 82 | 77 | 100 | 101 | 88.74 | 85 | 93 | 1.085 | 0.118 | 0.58569333 | 0.7350351 | 0 | 0.0291 | 0.0225 | 0.0225 | TRUE | 0.2222 |
| Smim11 | 413 | 278 | 262 | 258 | 232 | 244 | 321 | 296 | 283 | 257 | 262 | 241 | 276.43 | 300 | 253 | 0.842 | -0.248 | 0.04781823 | 0.13045732 | 0 | 0.011 | 0.0076 | 0.0076 | TRUE | 0.0458 |
| Smim12 | 1393 | 1003 | 1007 | 1062 | 927 | 1108 | 1082 | 1067 | 1087 | 1057 | 1046 | 1094 | 1071.99 | 1079 | 1066 | 0.988 | -0.017 | 0.81116342 | 0.89108899 | 0 | 0.0046 | 0.0028 | 0.0028 | TRUE | 0.0065 |
| Smim13 | 1110 | 805 | 777 | 934 | 772 | 847 | 862 | 856 | 838 | 930 | 871 | 836 | 865.58 | 852 | 879 | 1.031 | 0.044 | 0.58482213 | 0.734344 | 0 | 0.0052 | 0.0035 | 0.0035 | TRUE | 0.031 |
| Smim14 | 1894 | 1363 | 1358 | 1821 | 1487 | 1806 | 1471 | 1450 | 1465 | 1813 | 1677 | 1783 | 1609.83 | 1462 | 1758 | 1.202 | 0.266 | 5.59E-05 | 0.00076745 | 0 | 0.0039 | 0.0025 | 0.0025 | TRUE | 0.0189 |
| Smim15 | 2665 | 1863 | 1875 | 2103 | 1740 | 1985 | 2070 | 1982 | 2023 | 2093 | 1963 | 1960 | 2015.05 | 2025 | 2005 | 0.99 | -0.014 | 0.82011901 | 0.89651557 | 5.00E-04 | 0.0036 | 0.0024 | 0.0024 | TRUE | 0.018 |
| Smim18 | 0 | 0 | 0 | 0 | 0 | 0 | 0 | 0 | 0 | 0 | 0 | 0 | 0 | 0 | 0 | NA | NA | NA | NA | NA | NA | NA | NA | NA | NA |
| Smim3 | 367 | 282 | 244 | 287 | 210 | 257 | 285 | 300 | 263 | 286 | 237 | 254 | 270.76 | 283 | 259 | 0.916 | -0.127 | 0.33317395 | 0.50741133 | 0.0025 | 0.0112 | 0.0087 | 0.0087 | TRUE | 0.0927 |
| Smim4 | 136 | 93 | 102 | 114 | 102 | 146 | 106 | 99 | 110 | 113 | 115 | 144 | 114.55 | 105 | 124 | 1.186 | 0.246 | 0.19656588 | 0.3563115 | 0 | 0.0231 | 0.0173 | 0.0173 | TRUE | 0.2037 |
| Smim5 | 0 | 0 | 0 | 1 | 0 | 0 | 0 | 0 | 0 | 1 | 0 | 0 | 0.17 | 0 | 0 | 2.094 | 1.067 | 0.79379329 | NA | 0 | 14.2885 | 10 | 10 | TRUE | 0.3906 |
| Smim6 | 0 | 0 | 0 | 0 | 0 | 0 | 0 | 0 | 0 | 0 | 0 | 0 | 0 | 0 | 0 | NA | NA | NA | NA | NA | NA | NA | NA | NA | NA |
| Smim7 | 3388 | 2343 | 2516 | 2960 | 2536 | 2793 | 2631 | 2492 | 2715 | 2946 | 2861 | 2757 | 2733.78 | 2613 | 2855 | 1.093 | 0.128 | 0.03994111 | 0.11436973 | 9.00E-04 | 0.0033 | 0.0024 | 0.0024 | TRUE | 0.0193 |
| Smim8 | 686 | 437 | 380 | 464 | 410 | 387 | 533 | 465 | 410 | 462 | 463 | 382 | 452.34 | 469 | 436 | 0.925 | -0.112 | 0.37135776 | 0.54612609 | 0.0124 | 0.0077 | 0.0092 | 0.0092 | TRUE | 0.1705 |
| Smim9 | 0 | 1 | 1 | 0 | 1 | 0 | 0 | 1 | 1 | 0 | 1 | 0 | 0.55 | 1 | 0 | 0.535 | -0.902 | 0.73748201 | NA | 0 | 4.3501 | 3.5264 | 3.5264 | TRUE | 0.5075 |
| Smlr1 | 0 | 0 | 0 | 0 | 0 | 0 | 0 | 0 | 0 | 0 | 0 | 0 | 0 | 0 | 0 | NA | NA | NA | NA | NA | NA | NA | NA | NA | NA |
| Smn1 | 4082 | 2941 | 2695 | 2701 | 2483 | 2952 | 3170 | 3128 | 2908 | 2689 | 2801 | 2914 | 2935.04 | 3069 | 2801 | 0.912 | -0.132 | 0.03739605 | 0.10905974 | 0.0014 | 0.0032 | 0.0026 | 0.0026 | TRUE | 0.0253 |
| Smndc1 | 5869 | 4261 | 3842 | 4074 | 3388 | 4022 | 4558 | 4532 | 4145 | 4055 | 3822 | 3970 | 4180.61 | 4412 | 3949 | 0.895 | -0.16 | 0.00848658 | 0.036624 | 0.0015 | 0.003 | 0.0024 | 0.0024 | TRUE | 0.0337 |

| | | | | | | | | | | | | | | | | | | | | | | | | | |
|---|---|---|---|---|---|---|---|---|---|---|---|---|---|---|---|---|---|---|---|---|---|---|---|---|---|
| Smo | 3095 | 2144 | 2135 | 2588 | 2209 | 2439 | 2404 | 2281 | 2304 | 2576 | 2492 | 2408 | 2410.62 | 2330 | 2492 | 1.069 | 0.097 | 0.11400965 | 0.24303355 | 6.00E-04 | 0.0034 | 0.0023 | 0.0023 | TRUE | 0.0108 |
| Smoc1 | 1958 | 1473 | 1388 | 1257 | 1156 | 1318 | 1521 | 1567 | 1498 | 1251 | 1304 | 1301 | 1406.92 | 1529 | 1285 | 0.841 | -0.25 | 0.00019303 | 0.00207725 | 0 | 0.0041 | 0.0025 | 0.0025 | TRUE | 0.0066 |
| Smoc2 | 10 | 8 | 9 | 7 | 9 | 15 | 8 | 9 | 10 | 7 | 10 | 15 | 9.65 | 9 | 11 | 1.235 | 0.305 | 0.62379657 | NA | 0 | 0.248 | 0.1741 | 0.1741 | TRUE | 0.2791 |
| Smok2a | 0 | 0 | 0 | 0 | 0 | 0 | 0 | 0 | 0 | 0 | 0 | 0 | 0 | 0 | 0 | NA | NA | NA | NA | NA | NA | NA | NA | NA | NA |
| Smok2b | 0 | 0 | 0 | 0 | 0 | 0 | 0 | 0 | 0 | 0 | 0 | 0 | 0 | 0 | 0 | NA | NA | NA | NA | NA | NA | NA | NA | NA | NA |
| Smok3a | 0 | 0 | 0 | 0 | 0 | 0 | 0 | 0 | 0 | 0 | 0 | 0 | 0 | 0 | 0 | NA | NA | NA | NA | NA | NA | NA | NA | NA | NA |
| Smok3b | 0 | 0 | 0 | 0 | 0 | 0 | 0 | 0 | 0 | 0 | 0 | 0 | 0 | 0 | 0 | NA | NA | NA | NA | NA | NA | NA | NA | NA | NA |
| Smok4a | 2 | 0 | 0 | 0 | 0 | 1 | 2 | 0 | 0 | 0 | 0 | 1 | 0.42 | 1 | 0 | 0.682 | -0.553 | 0.86290097 | NA | 0 | 5.6 | 5.5066 | 5.5066 | TRUE | 0.9727 |
| Smox | 670 | 529 | 502 | 531 | 492 | 493 | 520 | 563 | 542 | 529 | 555 | 487 | 532.49 | 542 | 524 | 0.967 | -0.049 | 0.61808266 | 0.75876109 | 0.0011 | 0.0069 | 0.005 | 0.005 | TRUE | 0.0444 |
| Smpd1 | 709 | 483 | 570 | 676 | 618 | 676 | 551 | 514 | 615 | 673 | 697 | 667 | 619.47 | 560 | 679 | 1.213 | 0.279 | 0.00367141 | 0.01930231 | 0.0023 | 0.0063 | 0.005 | 0.005 | TRUE | 0.0832 |
| Smpd2 | 798 | 594 | 570 | 577 | 490 | 606 | 620 | 632 | 615 | 574 | 553 | 598 | 598.66 | 622 | 575 | 0.925 | -0.113 | 0.20636866 | 0.3682718 | 0 | 0.0064 | 0.0041 | 0.0041 | TRUE | 0.0145 |
| Smpd3 | 85 | 69 | 67 | 84 | 91 | 100 | 66 | 73 | 72 | 84 | 103 | 99 | 82.78 | 70 | 95 | 1.348 | 0.431 | 0.04640203 | 0.1275922 | 0 | 0.0311 | 0.0214 | 0.0214 | TRUE | 0.1077 |
| Smpd4 | 8397 | 5964 | 5778 | 6277 | 5441 | 6588 | 6522 | 6344 | 6234 | 6248 | 6138 | 6503 | 6331.56 | 6367 | 6296 | 0.989 | -0.016 | 0.75695274 | 0.85564566 | 4.00E-04 | 0.0028 | 0.0018 | 0.0018 | TRUE | 0.0101 |
| Smpd5 | 3 | 3 | 2 | 2 | 3 | 2 | 2 | 3 | 2 | 2 | 3 | 2 | 2.5 | 2 | 2 | 0.954 | -0.069 | 0.95315019 | NA | 0 | 0.9487 | 0.5793 | 0.5793 | TRUE | 0.1058 |
| Smpdl3a | 77 | 53 | 81 | 54 | 33 | 37 | 60 | 56 | 87 | 54 | 37 | 37 | 55.18 | 68 | 43 | 0.63 | -0.668 | 0.02302455 | 0.07656816 | 0.0364 | 0.0454 | 0.0429 | 0.0429 | TRUE | 0.5414 |
| Smpdl3b | 628 | 370 | 311 | 270 | 296 | 273 | 488 | 394 | 336 | 269 | 334 | 269 | 348.17 | 406 | 291 | 0.713 | -0.487 | 0.0012794 | 0.008615 | 0.0232 | 0.0092 | 0.0135 | 0.0135 | TRUE | 0.2828 |
| Smpx | 0 | 0 | 0 | 0 | 0 | 0 | 0 | 0 | 0 | 0 | 0 | 0 | 0 | 0 | 0 | NA | NA | NA | NA | NA | NA | NA | NA | NA | NA |
| Smr2 | 0 | 0 | 0 | 0 | 0 | 0 | 0 | 0 | 0 | 0 | 0 | 0 | 0 | 0 | 0 | NA | NA | NA | NA | NA | NA | NA | NA | NA | NA |
| Smr3a | 0 | 3 | 0 | 1 | 0 | 1 | 0 | 3 | 0 | 1 | 0 | 1 | 0.86 | 1 | 1 | 0.647 | -0.627 | 0.7874777 | NA | 2.2711 | 2.7511 | 2.6648 | 2.6648 | TRUE | 1.4437 |
| Sms | 2958 | 2172 | 2041 | 1816 | 1597 | 1954 | 2297 | 2310 | 2202 | 1808 | 1802 | 1929 | 2058 | 2270 | 1846 | 0.813 | -0.298 | 2.64E-06 | 6.29E-05 | 5.00E-04 | 0.0036 | 0.0024 | 0.0024 | TRUE | 0.0188 |
| Smtn | 1375 | 1085 | 1120 | 1259 | 1179 | 1209 | 1068 | 1154 | 1208 | 1253 | 1330 | 1193 | 1201.2 | 1143 | 1259 | 1.102 | 0.14 | 0.08007961 | 0.18934292 | 0.0026 | 0.0044 | 0.0038 | 0.0038 | TRUE | 0.0419 |
| Smtnl1 | 149 | 127 | 95 | 78 | 90 | 80 | 116 | 135 | 103 | 78 | 102 | 79 | 101.91 | 118 | 86 | 0.729 | -0.456 | 0.02915765 | 0.09071622 | 0.0099 | 0.0257 | 0.0214 | 0.0214 | TRUE | 0.2225 |
| Smtnl2 | 7817 | 6542 | 6765 | 7237 | 6677 | 7714 | 6071 | 6959 | 7299 | 7204 | 7532 | 7615 | 7113.35 | 6776 | 7450 | 1.1 | 0.137 | 0.05472389 | 0.14362257 | 0.0046 | 0.0028 | 0.0035 | 0.0035 | TRUE | 0.1021 |
| Smu1 | 9279 | 6751 | 6670 | 6993 | 6323 | 7366 | 7207 | 7181 | 7197 | 6961 | 7133 | 7271 | 7158.27 | 7195 | 7122 | 0.99 | -0.015 | 0.75921771 | 0.8571376 | 1.00E-04 | 0.0028 | 0.0015 | 0.0015 | TRUE | 0.0048 |
| Smug1 | 1261 | 1036 | 1186 | 1343 | 1139 | 1396 | 979 | 1102 | 1280 | 1337 | 1285 | 1378 | 1226.8 | 1120 | 1333 | 1.192 | 0.254 | 0.00830198 | 0.0360122 | 0.0085 | 0.0044 | 0.0058 | 0.0058 | TRUE | 0.185 |
| Smurf1 | 3525 | 2520 | 2548 | 2526 | 2251 | 2545 | 2738 | 2680 | 2749 | 2514 | 2539 | 2512 | 2622.25 | 2722 | 2522 | 0.926 | -0.111 | 0.04661747 | 0.12803679 | 0 | 0.0033 | 0.0018 | 0.0018 | TRUE | 0.0022 |
| Smurf2 | 7490 | 5541 | 5697 | 6607 | 5703 | 6412 | 5817 | 5894 | 6147 | 6577 | 6434 | 6330 | 6199.65 | 5953 | 6447 | 1.083 | 0.115 | 0.02524211 | 0.08184787 | 4.00E-04 | 0.0028 | 0.0018 | 0.0018 | TRUE | 0.0099 |
| Smyd1 | 174 | 109 | 75 | 48 | 38 | 52 | 135 | 116 | 81 | 48 | 43 | 51 | 79 | 111 | 47 | 0.426 | -1.23 | 1.54E-06 | 4.13E-05 | 0.0309 | 0.0324 | 0.032 | 0.032 | TRUE | 0.2259 |
| Smyd2 | 2712 | 1865 | 1695 | 1796 | 1655 | 1791 | 2106 | 1984 | 1829 | 1788 | 1867 | 1768 | 1890.28 | 1973 | 1808 | 0.915 | -0.128 | 0.07610386 | 0.18200333 | 0.0025 | 0.0037 | 0.0032 | 0.0032 | TRUE | 0.0491 |
| Smyd3 | 1195 | 875 | 1052 | 1236 | 1184 | 1278 | 928 | 931 | 1135 | 1230 | 1336 | 1262 | 1136.92 | 998 | 1276 | 1.28 | 0.356 | 0.00011053 | 0.00131536 | 0.0066 | 0.0045 | 0.0052 | 0.0052 | TRUE | 0.1698 |
| Smyd4 | 2276 | 1597 | 1571 | 1665 | 1475 | 1816 | 1768 | 1699 | 1695 | 1657 | 1664 | 1793 | 1712.57 | 1721 | 1705 | 0.991 | -0.014 | 0.83689866 | 0.90706168 | 7.00E-04 | 0.0038 | 0.0026 | 0.0026 | TRUE | 0.0249 |
| Smyd5 | 6932 | 5067 | 4677 | 4875 | 4293 | 5066 | 5384 | 5390 | 5046 | 4853 | 4843 | 5001 | 5086.06 | 5273 | 4899 | 0.929 | -0.106 | 0.05000285 | 0.13477101 | 6.00E-04 | 0.0029 | 0.0019 | 0.0019 | TRUE | 0.0171 |
| Snai1 | 116 | 75 | 45 | 97 | 64 | 70 | 90 | 80 | 49 | 97 | 72 | 69 | 76.05 | 73 | 79 | 1.083 | 0.115 | 0.66861139 | 0.79617284 | 0.0518 | 0.0336 | 0.0387 | 0.0387 | TRUE | 0.7371 |
| Snai2 | 75 | 49 | 44 | 34 | 35 | 26 | 58 | 52 | 47 | 34 | 39 | 26 | 42.81 | 52 | 33 | 0.623 | -0.683 | 0.02558251 | 0.08265164 | 0 | 0.0578 | 0.0426 | 0.0426 | TRUE | 0.268 |
| Snai3 | 300 | 266 | 263 | 356 | 316 | 317 | 233 | 283 | 284 | 354 | 356 | 313 | 303.91 | 267 | 341 | 1.284 | 0.361 | 0.00554441 | 0.02656138 | 0.0058 | 0.0102 | 0.0089 | 0.0089 | TRUE | 0.1481 |
| Snap23 | 2599 | 1743 | 1875 | 2146 | 1863 | 2080 | 2019 | 1854 | 2023 | 2136 | 2102 | 2053 | 2031.12 | 1965 | 2097 | 1.067 | 0.093 | 0.15269364 | 0.29941695 | 9.00E-04 | 0.0036 | 0.0026 | 0.0026 | TRUE | 0.0291 |
| Snap25 | 15 | 14 | 16 | 13 | 9 | 12 | 12 | 15 | 17 | 13 | 10 | 12 | 13.12 | 15 | 12 | 0.806 | -0.312 | 0.55295181 | NA | 0 | 0.183 | 0.1214 | 0.1214 | TRUE | 0.1784 |
| Snap29 | 1800 | 1208 | 1163 | 1366 | 1176 | 1286 | 1398 | 1285 | 1255 | 1360 | 1327 | 1269 | 1315.61 | 1313 | 1319 | 1.003 | 0.005 | 0.94983104 | 0.97250044 | 0.0016 | 0.0042 | 0.0033 | 0.0033 | TRUE | 0.0405 |
| Snap47 | 6506 | 4607 | 4545 | 4975 | 4396 | 4920 | 5053 | 4900 | 4904 | 4952 | 4959 | 4857 | 4937.57 | 4952 | 4923 | 0.994 | -0.009 | 0.8566561 | 0.91944023 | 0 | 0.0029 | 0.0016 | 0.0016 | TRUE | 0.0039 |
| Snap91 | 372 | 311 | 323 | 354 | 263 | 263 | 289 | 331 | 349 | 352 | 297 | 260 | 312.82 | 323 | 303 | 0.941 | -0.088 | 0.5310731 | 0.69099935 | 0.0133 | 0.01 | 0.011 | 0.011 | TRUE | 0.2351 |
| Snapc1 | 1630 | 1203 | 1257 | 1291 | 1032 | 1342 | 1266 | 1280 | 1356 | 1285 | 1164 | 1325 | 1279.31 | 1301 | 1258 | 0.968 | -0.046 | 0.54695868 | 0.70406727 | 0.0021 | 0.0043 | 0.0035 | 0.0035 | TRUE | 0.0501 |
| Snapc2 | 2060 | 1501 | 1409 | 1441 | 1338 | 1485 | 1600 | 1597 | 1520 | 1434 | 1509 | 1466 | 1521.08 | 1572 | 1470 | 0.934 | -0.098 | 0.13981272 | 0.28110046 | 0 | 0.004 | 0.0025 | 0.0025 | TRUE | 0.0099 |
| Snapc3 | 1418 | 1054 | 1052 | 1261 | 1044 | 1248 | 1101 | 1121 | 1135 | 1255 | 1178 | 1232 | 1170.41 | 1119 | 1222 | 1.092 | 0.127 | 0.07243899 | 0.17599363 | 0 | 0.0045 | 0.0028 | 0.0028 | TRUE | 0.0116 |
| Snapc4 | 2305 | 1992 | 1590 | 1549 | 1424 | 1811 | 1790 | 2119 | 1716 | 1542 | 1606 | 1788 | 1760.11 | 1875 | 1645 | 0.878 | -0.188 | 0.04019515 | 0.11483794 | 0.0087 | 0.0038 | 0.0055 | 0.0055 | TRUE | 0.1549 |
| Snapc5 | 925 | 608 | 536 | 552 | 435 | 581 | 718 | 647 | 578 | 549 | 491 | 574 | 592.87 | 648 | 538 | 0.829 | -0.27 | 0.0128321 | 0.04974485 | 0.0075 | 0.0064 | 0.0068 | 0.0068 | TRUE | 0.1115 |
| Snapin | 2366 | 1636 | 1743 | 1973 | 1674 | 1851 | 1838 | 1740 | 1881 | 1964 | 1888 | 1827 | 1856.34 | 1820 | 1893 | 1.04 | 0.057 | 0.38913402 | 0.56322963 | 9.00E-04 | 0.0037 | 0.0026 | 0.0026 | TRUE | 0.0171 |
| Snca | 99 | 54 | 46 | 58 | 38 | 30 | 77 | 57 | 50 | 58 | 43 | 30 | 52.36 | 61 | 44 | 0.703 | -0.508 | 0.10450775 | 0.22844046 | 0.0585 | 0.0477 | 0.0507 | 0.0507 | TRUE | 0.3118 |
| Sncaip | 47 | 25 | 19 | 22 | 15 | 10 | 37 | 27 | 21 | 22 | 17 | 10 | 22.05 | 28 | 16 | 0.577 | -0.794 | 0.07892717 | 0.18729269 | 0.0654 | 0.1099 | 0.0979 | 0.0979 | TRUE | 0.3477 |
| Sncb | 120 | 100 | 135 | 146 | 159 | 177 | 93 | 106 | 146 | 145 | 179 | 175 | 140.78 | 115 | 166 | 1.454 | 0.54 | 0.0063144 | 0.02932781 | 0.025 | 0.0193 | 0.0209 | 0.0209 | TRUE | 0.5314 |
| Sncg | 268 | 174 | 190 | 246 | 189 | 211 | 208 | 185 | 205 | 245 | 213 | 208 | 210.77 | 199 | 222 | 1.113 | 0.154 | 0.28201789 | 0.45367268 | 0 | 0.0137 | 0.0101 | 0.0101 | TRUE | 0.0897 |
| Snd1 | 19188 | 14090 | 14373 | 15553 | 14175 | 16523 | 14903 | 14987 | 15508 | 15481 | 15991 | 16311 | 15530.21 | 15133 | 15928 | 1.053 | 0.074 | 0.12474475 | 0.25945527 | 5.00E-04 | 0.0026 | 0.0016 | 0.0016 | TRUE | 0.0074 |
| Sned1 | 249 | 206 | 148 | 207 | 170 | 201 | 193 | 219 | 160 | 206 | 192 | 198 | 194.74 | 191 | 199 | 1.042 | 0.059 | 0.70537875 | 0.82167713 | 0.0069 | 0.0146 | 0.0125 | 0.0125 | TRUE | 0.2052 |
| Snf8 | 4152 | 2817 | 2513 | 2731 | 2326 | 2759 | 3225 | 2996 | 2712 | 2718 | 2624 | 2724 | 2833.09 | 2978 | 2689 | 0.902 | -0.148 | 0.04078812 | 0.11608614 | 0.0034 | 0.0033 | 0.0034 | 0.0034 | TRUE | 0.0739 |
| Snhg1 | 3202 | 2344 | 3045 | 3311 | 2762 | 3510 | 2487 | 2493 | 3286 | 3296 | 3116 | 3465 | 3023.69 | 2755 | 3292 | 1.196 | 0.258 | 0.00940024 | 0.03962184 | 0.0148 | 0.0032 | 0.0068 | 0.0068 | TRUE | 0.3425 |
| Snhg10 | 48 | 37 | 52 | 35 | 39 | 51 | 37 | 39 | 56 | 35 | 44 | 50 | 43.65 | 44 | 43 | 0.978 | -0.031 | 0.91954024 | 0.95585372 | 0.0208 | 0.0567 | 0.0467 | 0.0467 | TRUE | 0.4094 |
| Snhg11 | 2042 | 2028 | 1905 | 1822 | 1524 | 2059 | 1586 | 2157 | 2055 | 1814 | 1719 | 2033 | 1893.99 | 1933 | 1855 | 0.961 | -0.057 | 0.59354239 | 0.74083813 | 0.0164 | 0.0037 | 0.0078 | 0.0078 | TRUE | 0.3034 |
| Snhg12 | 950 | 700 | 875 | 966 | 811 | 1075 | 738 | 745 | 944 | 962 | 915 | 1061 | 894.03 | 809 | 979 | 1.213 | 0.278 | 0.00950114 | 0.03986267 | 0.0117 | 0.0051 | 0.0072 | 0.0072 | TRUE | 0.2506 |
| Snhg3 | 1095 | 768 | 907 | 963 | 721 | 910 | 850 | 817 | 979 | 959 | 813 | 898 | 886.04 | 882 | 890 | 1.01 | 0.015 | 0.87653566 | 0.93106882 | 0.0067 | 0.0051 | 0.0056 | 0.0056 | TRUE | 0.1068 |
| Snhg4 | 1079 | 947 | 1032 | 995 | 852 | 1081 | 838 | 1007 | 1114 | 990 | 961 | 1067 | 996.25 | 986 | 1006 | 1.023 | 0.032 | 0.75735575 | 0.85587269 | 0.0108 | 0.0048 | 0.0068 | 0.0068 | TRUE | 0.2139 |
| Snhg5 | 1957 | 1574 | 1230 | 1282 | 1089 | 1189 | 1520 | 1674 | 1327 | 1276 | 1229 | 1174 | 1366.61 | 1507 | 1226 | 0.813 | -0.298 | 0.00094181 | 0.00694598 | 0.0064 | 0.0042 | 0.0051 | 0.0051 | TRUE | 0.1284 |
| Snhg6 | 1111 | 914 | 1219 | 1362 | 1209 | 1379 | 863 | 972 | 1315 | 1356 | 1364 | 1361 | 1205.21 | 1050 | 1360 | 1.297 | 0.376 | 0.00225603 | 0.01324792 | 0.024 | 0.0044 | 0.0101 | 0.0101 | TRUE | 0.5812 |
| Snhg7 | 520 | 453 | 660 | 762 | 671 | 835 | 404 | 482 | 712 | 758 | 757 | 824 | 656.26 | 533 | 780 | 1.468 | 0.554 | 0.00042548 | 0.00381514 | 0.0441 | 0.006 | 0.0162 | 0.0162 | TRUE | 1.0119 |
| Snhg8 | 1347 | 1047 | 1097 | 1266 | 1118 | 1201 | 1046 | 1114 | 1184 | 1260 | 1261 | 1186 | 1175.08 | 1115 | 1236 | 1.11 | 0.15 | 0.05215423 | 0.1389451 | 0.0017 | 0.0044 | 0.0035 | 0.0035 | TRUE | 0.0361 |
| Snhg9 | 12 | 7 | 4 | 4 | 2 | 2 | 9 | 7 | 4 | 4 | 2 | 2 | 4.88 | 7 | 3 | 0.387 | -1.37 | 0.13190757 | NA | 0 | 0.4879 | 0.3391 | 0.3391 | TRUE | 0.218 |
| Snip1 | 2226 | 1673 | 1472 | 1641 | 1404 | 1661 | 1729 | 1780 | 1588 | 1633 | 1584 | 1640 | 1658.94 | 1699 | 1619 | 0.953 | -0.07 | 0.31668667 | 0.49072014 | 0.0013 | 0.0039 | 0.0029 | 0.0029 | TRUE | 0.0381 |
| Snn | 1637 | 1159 | 1278 | 1347 | 1172 | 1367 | 1271 | 1233 | 1379 | 1341 | 1322 | 1349 | 1315.92 | 1294 | 1337 | 1.034 | 0.048 | 0.50835664 | 0.67162045 | 9.00E-04 | 0.0042 | 0.003 | 0.003 | TRUE | 0.038 |
| Snora15 | 2 | 3 | 7 | 3 | 4 | 2 | 2 | 3 | 8 | 3 | 5 | 2 | 3.63 | 4 | 3 | 0.782 | -0.355 | 0.73366319 | NA | 0 | 0.6556 | 0.4999 | 0.4999 | TRUE | 0.6657 |
| Snora16a | 54 | 43 | 45 | 45 | 27 | 41 | 42 | 46 | 49 | 45 | 30 | 40 | 41.99 | 46 | 38 | 0.855 | -0.226 | 0.45428214 | 0.62403216 | 0 | 0.0589 | 0.0419 | 0.0419 | TRUE | 0.2303 |
| Snora17 | 36 | 23 | 16 | 26 | 17 | 22 | 28 | 24 | 17 | 26 | 19 | 22 | 22.74 | 23 | 22 | 0.953 | -0.07 | 0.86506596 | 0.92551298 | 0 | 0.1066 | 0.0771 | 0.0771 | TRUE | 0.2774 |

| | | | | | | | | | | | | | | | | | | | | | | | | | |
|---|---|---|---|---|---|---|---|---|---|---|---|---|---|---|---|---|---|---|---|---|---|---|---|---|---|
| Snora19 | 0 | 0 | 0 | 0 | 1 | 1 | 0 | 0 | 0 | 0 | 1 | 1 | 0.35 | 0 | 1 | 3.965 | 1.987 | 0.55529285 | NA | 0 | 6.7252 | 6.4426 | 6.4426 | TRUE | 0.261 |
| Snora20 | 0 | 0 | 1 | 0 | 0 | 0 | 0 | 0 | 1 | 0 | 0 | 0 | 0.18 | 0 | 0 | 0.552 | -0.857 | 0.8336548 | NA | 0 | 13.1817 | 10 | 10 | TRUE | 0.5365 |
| Snora21 | 4 | 1 | 4 | 2 | 5 | 4 | 3 | 1 | 4 | 2 | 6 | 4 | 3.34 | 3 | 4 | 1.351 | 0.434 | 0.68214268 | NA | 0 | 0.7111 | 0.5053 | 0.5053 | TRUE | 0.2263 |
| Snora23 | 5 | 9 | 7 | 8 | 7 | 12 | 4 | 10 | 8 | 8 | 8 | 12 | 8.12 | 7 | 9 | 1.347 | 0.429 | 0.53174587 | NA | 0 | 0.2943 | 0.2136 | 0.2136 | TRUE | 0.5479 |
| Snora24 | 0 | 0 | 0 | 0 | 0 | 0 | 0 | 0 | 0 | 0 | 0 | 0 | 0 | 0 | 0 | NA | NA | NA | NA | NA | NA | NA | NA | NA | NA |
| Snora26 | 0 | 0 | 0 | 0 | 0 | 0 | 0 | 0 | 0 | 0 | 0 | 0 | 0 | 0 | 0 | NA | NA | NA | NA | NA | NA | NA | NA | NA | NA |
| Snora28 | 29 | 23 | 22 | 24 | 17 | 27 | 23 | 24 | 24 | 24 | 19 | 27 | 23.41 | 24 | 23 | 0.991 | -0.014 | 0.97204253 | 0.98493233 | 0 | 0.1037 | 0.0684 | 0.0684 | TRUE | 0.1245 |
| Snora2b | 2 | 4 | 0 | 5 | 0 | 5 | 2 | 4 | 0 | 5 | 0 | 5 | 2.62 | 2 | 3 | 1.734 | 0.794 | 0.56707087 | NA | 1.3355 | 0.907 | 0.9801 | 0.9801 | TRUE | 0.578 |
| Snora3 | 24 | 12 | 11 | 12 | 8 | 9 | 19 | 13 | 12 | 12 | 9 | 9 | 12.19 | 15 | 10 | 0.684 | -0.548 | 0.32228264 | NA | 0 | 0.1969 | 0.1357 | 0.1357 | TRUE | 0.3595 |
| Snora30 | 1 | 4 | 4 | 9 | 3 | 5 | 1 | 4 | 4 | 9 | 3 | 5 | 4.44 | 3 | 6 | 1.923 | 0.943 | 0.33200627 | NA | 0 | 0.5365 | 0.4318 | 0.4318 | TRUE | 0.8285 |
| Snora31 | 19 | 19 | 11 | 11 | 9 | 20 | 15 | 20 | 12 | 11 | 10 | 20 | 14.61 | 16 | 14 | 0.877 | -0.189 | 0.71829401 | NA | 0.0331 | 0.1646 | 0.129 | 0.129 | TRUE | 0.6816 |
| Snora33 | 22 | 15 | 19 | 19 | 3 | 19 | 17 | 16 | 21 | 19 | 3 | 19 | 15.77 | 18 | 14 | 0.776 | -0.367 | 0.51522687 | NA | 0.2059 | 0.1528 | 0.1642 | 0.1642 | TRUE | 1.6653 |
| Snora34 | 13 | 5 | 5 | 13 | 3 | 12 | 10 | 5 | 5 | 13 | 3 | 12 | 8.16 | 7 | 9 | 1.342 | 0.425 | 0.55649563 | NA | 0.1453 | 0.2928 | 0.253 | 0.253 | TRUE | 0.923 |
| Snora35 | 0 | 0 | 0 | 0 | 0 | 0 | 0 | 0 | 0 | 0 | 0 | 0 | 0 | 0 | 0 | NA | NA | NA | NA | NA | NA | NA | NA | NA | NA |
| Snora36b | 0 | 0 | 0 | 0 | 0 | 0 | 0 | 0 | 0 | 0 | 0 | 0 | 0 | 0 | 0 | NA | NA | NA | NA | NA | NA | NA | NA | NA | NA |
| Snora41 | 8 | 8 | 7 | 7 | 4 | 7 | 6 | 9 | 8 | 7 | 5 | 7 | 6.78 | 8 | 6 | 0.835 | -0.261 | 0.71834474 | NA | 0 | 0.3521 | 0.2268 | 0.2268 | TRUE | 0.1116 |
| Snora43 | 29 | 21 | 22 | 39 | 28 | 32 | 23 | 22 | 24 | 39 | 32 | 32 | 28.43 | 23 | 34 | 1.489 | 0.575 | 0.10868688 | 0.2351236 | 0 | 0.0858 | 0.0562 | 0.0562 | TRUE | 0.1102 |
| Snora44 | 25 | 12 | 15 | 9 | 9 | 13 | 19 | 13 | 16 | 9 | 10 | 13 | 13.39 | 16 | 11 | 0.657 | -0.607 | 0.25051641 | NA | 0 | 0.1795 | 0.1225 | 0.1225 | TRUE | 0.1797 |
| Snora47 | 1 | 0 | 0 | 0 | 0 | 0 | 1 | 0 | 0 | 0 | 0 | 0 | 0.13 | 0 | 0 | 0.552 | -0.857 | 0.8336548 | NA | 0 | 18.3121 | 10 | 10 | TRUE | 0.2566 |
| Snora52 | 23 | 21 | 14 | 13 | 8 | 11 | 18 | 22 | 15 | 13 | 9 | 11 | 14.69 | 18 | 11 | 0.596 | -0.747 | 0.14021771 | NA | 0 | 0.1638 | 0.1114 | 0.1114 | TRUE | 0.1628 |
| Snora5c | 26 | 15 | 17 | 11 | 6 | 5 | 20 | 16 | 18 | 11 | 7 | 5 | 12.86 | 18 | 8 | 0.415 | -1.268 | 0.02444409 | NA | 0 | 0.1868 | 0.1341 | 0.1341 | TRUE | 0.4552 |
| Snora61 | 90 | 72 | 72 | 65 | 36 | 58 | 70 | 77 | 78 | 65 | 41 | 57 | 64.46 | 75 | 54 | 0.73 | -0.455 | 0.07475336 | 0.17993429 | 0.0046 | 0.0392 | 0.031 | 0.031 | TRUE | 0.3716 |
| Snora62 | 4 | 3 | 2 | 2 | 2 | 3 | 3 | 3 | 2 | 2 | 2 | 3 | 2.61 | 3 | 2 | 0.85 | -0.235 | 0.83698996 | NA | 0 | 0.9103 | 0.5517 | 0.5517 | TRUE | 0.0462 |
| Snora64 | 48 | 26 | 27 | 29 | 23 | 28 | 37 | 28 | 29 | 29 | 26 | 28 | 29.42 | 31 | 28 | 0.871 | -0.199 | 0.5745004 | 0.72653815 | 0 | 0.083 | 0.0562 | 0.0562 | TRUE | 0.2196 |
| Snora65 | 175 | 114 | 177 | 267 | 170 | 229 | 136 | 121 | 191 | 266 | 192 | 226 | 188.63 | 149 | 228 | 1.531 | 0.615 | 0.00124051 | 0.00844673 | 0.0352 | 0.015 | 0.0206 | 0.0206 | TRUE | 0.4068 |
| Snora68 | 52 | 32 | 33 | 33 | 36 | 23 | 40 | 34 | 36 | 33 | 41 | 23 | 34.37 | 37 | 32 | 0.867 | -0.206 | 0.54916569 | 0.70563439 | 0 | 0.0714 | 0.0555 | 0.0555 | TRUE | 0.3052 |
| Snora69 | 2 | 3 | 5 | 3 | 5 | 7 | 2 | 3 | 5 | 3 | 6 | 7 | 4.28 | 3 | 5 | 1.57 | 0.65 | 0.49335228 | NA | 0 | 0.5563 | 0.4036 | 0.4036 | TRUE | 0.4159 |
| Snora70 | 31 | 22 | 15 | 17 | 14 | 15 | 24 | 23 | 16 | 17 | 16 | 15 | 18.53 | 21 | 16 | 0.743 | -0.429 | 0.33658535 | 0.51075638 | 0 | 0.1303 | 0.0884 | 0.0884 | TRUE | 0.223 |
| Snora74a | 9 | 16 | 5 | 3 | 6 | 10 | 7 | 17 | 5 | 3 | 7 | 10 | 8.17 | 10 | 7 | 0.672 | -0.573 | 0.44229415 | NA | 0.2213 | 0.2925 | 0.2717 | 0.2717 | TRUE | 0.7106 |
| Snora75 | 7 | 6 | 4 | 7 | 3 | 12 | 5 | 6 | 4 | 7 | 3 | 12 | 6.39 | 5 | 7 | 1.386 | 0.471 | 0.54672779 | NA | 0 | 0.3734 | 0.2806 | 0.2806 | TRUE | 0.2167 |
| Snora78 | 11 | 10 | 3 | 8 | 4 | 10 | 9 | 11 | 3 | 8 | 5 | 10 | 7.46 | 8 | 8 | 0.997 | -0.004 | 0.99551196 | NA | 0 | 0.3201 | 0.2529 | 0.2529 | TRUE | 0.6137 |
| Snora7a | 43 | 30 | 38 | 37 | 43 | 32 | 33 | 32 | 41 | 37 | 49 | 32 | 37.21 | 35 | 39 | 1.098 | 0.135 | 0.67825269 | 0.80231161 | 0 | 0.0661 | 0.0497 | 0.0497 | TRUE | 0.3176 |
| Snora81 | 44 | 40 | 49 | 48 | 31 | 32 | 34 | 43 | 53 | 48 | 35 | 32 | 40.66 | 43 | 38 | 0.889 | -0.169 | 0.60147438 | 0.74667517 | 0.0257 | 0.0607 | 0.0507 | 0.0507 | TRUE | 0.2786 |
| Snord100 | 33 | 13 | 9 | 16 | 9 | 16 | 26 | 14 | 10 | 16 | 10 | 16 | 15.17 | 17 | 14 | 0.841 | -0.251 | 0.63990119 | NA | 0.0988 | 0.1586 | 0.1409 | 0.1409 | TRUE | 0.9331 |
| Snord104 | 135 | 94 | 61 | 70 | 58 | 72 | 105 | 100 | 66 | 70 | 65 | 71 | 79.47 | 90 | 69 | 0.759 | -0.398 | 0.09812733 | 0.21824375 | 0.0207 | 0.0322 | 0.0289 | 0.0289 | TRUE | 0.5104 |
| Snord11 | 1 | 0 | 0 | 0 | 0 | 1 | 1 | 0 | 0 | 0 | 0 | 1 | 0.29 | 0 | 0 | 1.075 | 0.105 | 0.97702352 | NA | 0 | 8.0647 | 7.5458 | 7.5458 | TRUE | 0.3835 |
| Snord110 | 1 | 0 | 0 | 1 | 0 | 0 | 1 | 0 | 0 | 1 | 0 | 0 | 0.3 | 0 | 0 | 1.075 | 0.105 | 0.97698525 | NA | 0 | 8.0272 | 7.5141 | 7.5141 | TRUE | 0.3906 |
| Snord111 | 1 | 1 | 1 | 4 | 1 | 3 | 1 | 1 | 1 | 4 | 1 | 3 | 1.83 | 1 | 3 | 2.82 | 1.496 | 0.30305892 | NA | 0 | 1.2963 | 0.8288 | 0.8288 | TRUE | 0.2407 |
| Snord116 | 0 | 0 | 0 | 0 | 0 | 0 | 0 | 0 | 0 | 0 | 0 | 0 | 0 | 0 | 0 | NA | NA | NA | NA | NA | NA | NA | NA | NA | NA |
| Snord116l1 | 0 | 0 | 0 | 0 | 0 | 0 | 0 | 0 | 0 | 0 | 0 | 0 | 0 | 0 | 0 | NA | NA | NA | NA | NA | NA | NA | NA | NA | NA |
| Snord116l2 | 0 | 0 | 0 | 0 | 0 | 0 | 0 | 0 | 0 | 0 | 0 | 0 | 0 | 0 | 0 | NA | NA | NA | NA | NA | NA | NA | NA | NA | NA |
| Snord118 | 0 | 0 | 0 | 0 | 0 | 0 | 0 | 0 | 0 | 0 | 0 | 0 | 0 | 0 | 0 | NA | NA | NA | NA | NA | NA | NA | NA | NA | NA |
| Snord12 | 69 | 45 | 32 | 37 | 24 | 41 | 54 | 48 | 35 | 37 | 27 | 40 | 40.06 | 46 | 35 | 0.766 | -0.385 | 0.2347692 | 0.40095067 | 0.0195 | 0.0616 | 0.0502 | 0.0502 | TRUE | 0.3246 |
| Snord123 | 0 | 0 | 0 | 2 | 0 | 0 | 0 | 0 | 0 | 2 | 0 | 0 | 0.33 | 0 | 1 | 3.77 | 1.915 | 0.62344813 | NA | 0 | 7.1455 | 9.1834 | 9.1834 | TRUE | 0.9739 |
| Snord14a | 1 | 1 | 0 | 0 | 2 | 2 | 1 | 1 | 0 | 0 | 2 | 2 | 1.01 | 1 | 1 | 2.248 | 1.169 | 0.56154412 | NA | 0 | 2.3448 | 1.7784 | 1.7784 | TRUE | 0.5111 |
| Snord14c | 0 | 0 | 0 | 0 | 0 | 0 | 0 | 0 | 0 | 0 | 0 | 0 | 0 | 0 | 0 | NA | NA | NA | NA | NA | NA | NA | NA | NA | NA |
| Snord14d | 0 | 0 | 0 | 0 | 0 | 0 | 0 | 0 | 0 | 0 | 0 | 0 | 0 | 0 | 0 | NA | NA | NA | NA | NA | NA | NA | NA | NA | NA |
| Snord15a | 4 | 1 | 0 | 3 | 0 | 0 | 3 | 1 | 0 | 3 | 0 | 0 | 1.19 | 1 | 1 | 0.703 | -0.508 | 0.80218094 | NA | 2.7772 | 1.9895 | 2.1142 | 2.1142 | TRUE | 0.8361 |
| Snord15b | 16 | 13 | 9 | 13 | 12 | 23 | 12 | 14 | 10 | 13 | 14 | 23 | 14.19 | 12 | 17 | 1.369 | 0.453 | 0.38427872 | NA | 0 | 0.1694 | 0.1236 | 0.1236 | TRUE | 0.5619 |
| Snord16a | 9 | 9 | 5 | 6 | 4 | 7 | 7 | 10 | 5 | 6 | 5 | 7 | 6.56 | 7 | 6 | 0.796 | -0.328 | 0.657275 | NA | 0 | 0.3638 | 0.2398 | 0.2398 | TRUE | 0.1806 |
| Snord17 | 15 | 14 | 6 | 5 | 5 | 8 | 12 | 15 | 6 | 5 | 6 | 8 | 8.59 | 11 | 6 | 0.561 | -0.835 | 0.21733141 | NA | 0 | 0.2784 | 0.2027 | 0.2027 | TRUE | 0.2199 |
| Snord19 | 6 | 11 | 12 | 6 | 7 | 10 | 5 | 12 | 13 | 6 | 8 | 10 | 8.84 | 10 | 8 | 0.825 | -0.278 | 0.68072697 | NA | 0 | 0.2705 | 0.2126 | 0.2126 | TRUE | 0.938 |
| Snord1a | 0 | 0 | 0 | 0 | 0 | 0 | 0 | 0 | 0 | 0 | 0 | 0 | 0 | 0 | 0 | NA | NA | NA | NA | NA | NA | NA | NA | NA | NA |
| Snord1b | 10 | 3 | 6 | 10 | 7 | 11 | 8 | 3 | 6 | 10 | 8 | 11 | 7.69 | 6 | 10 | 1.627 | 0.702 | 0.31436119 | NA | 0 | 0.3106 | 0.2154 | 0.2154 | TRUE | 0.3288 |
| Snord1c | 7 | 3 | 4 | 3 | 1 | 1 | 5 | 3 | 4 | 3 | 1 | 1 | 3.01 | 4 | 2 | 0.394 | -1.345 | 0.23919322 | NA | 0 | 0.7905 | 0.5273 | 0.5273 | TRUE | 0.3483 |
| Snord2 | 29 | 24 | 21 | 20 | 14 | 28 | 23 | 26 | 23 | 20 | 16 | 28 | 22.34 | 24 | 21 | 0.901 | -0.15 | 0.71587524 | 0.82941668 | 0 | 0.1085 | 0.0779 | 0.0779 | TRUE | 0.4136 |
| Snord22 | 115 | 105 | 94 | 93 | 61 | 71 | 89 | 112 | 101 | 93 | 69 | 70 | 88.98 | 101 | 77 | 0.769 | -0.379 | 0.08572842 | 0.19785729 | 0.009 | 0.0291 | 0.0235 | 0.0235 | TRUE | 0.2858 |
| Snord23 | 1 | 0 | 0 | 1 | 0 | 0 | 1 | 0 | 0 | 1 | 0 | 0 | 0.3 | 0 | 0 | 1.075 | 0.105 | 0.97698525 | NA | 0 | 8.0272 | 7.5141 | 7.5141 | TRUE | 0.3906 |
| Snord32a | 19 | 21 | 14 | 10 | 15 | 12 | 15 | 22 | 15 | 10 | 17 | 12 | 15.15 | 17 | 13 | 0.741 | -0.432 | 0.39219481 | NA | 0 | 0.1588 | 0.1153 | 0.1153 | TRUE | 0.2918 |
| Snord33 | 13 | 6 | 1 | 7 | 5 | 5 | 10 | 6 | 1 | 7 | 6 | 5 | 5.85 | 6 | 6 | 0.976 | -0.035 | 0.96700246 | NA | 0.2075 | 0.4075 | 0.3563 | 0.3563 | TRUE | 0.2827 |
| Snord34 | 16 | 11 | 7 | 4 | 2 | 10 | 12 | 12 | 8 | 4 | 2 | 10 | 7.96 | 11 | 5 | 0.512 | -0.967 | 0.18808082 | NA | 0.0764 | 0.3 | 0.2478 | 0.2478 | TRUE | 1.1772 |
| Snord35a | 11 | 3 | 15 | 5 | 6 | 4 | 9 | 3 | 16 | 5 | 7 | 4 | 7.27 | 9 | 5 | 0.561 | -0.834 | 0.2866562 | NA | 0.1971 | 0.3285 | 0.29 | 0.29 | TRUE | 0.1807 |
| Snord35b | 9 | 5 | 7 | 8 | 2 | 7 | 7 | 5 | 8 | 8 | 2 | 7 | 6.17 | 7 | 6 | 0.87 | -0.201 | 0.79719286 | NA | 0 | 0.3869 | 0.2802 | 0.2802 | TRUE | 0.5512 |
| Snord37 | 11 | 7 | 1 | 9 | 4 | 9 | 9 | 7 | 1 | 9 | 5 | 9 | 6.57 | 6 | 8 | 1.292 | 0.37 | 0.64910657 | NA | 0.1889 | 0.3631 | 0.3235 | 0.3235 | TRUE | 1.0171 |
| Snord38a | 1 | 1 | 3 | 1 | 0 | 1 | 1 | 1 | 3 | 1 | 0 | 1 | 1.18 | 2 | 1 | 0.407 | -1.298 | 0.47882381 | NA | 0 | 2.0167 | 1.3656 | 1.3656 | TRUE | 0.4567 |
| Snord42a | 3 | 2 | 1 | 1 | 4 | 1 | 2 | 2 | 1 | 1 | 5 | 1 | 2.01 | 2 | 2 | 1.14 | 0.189 | 0.88920939 | NA | 0 | 1.1844 | 0.8228 | 0.8228 | TRUE | 0.7749 |
| Snord42b | 37 | 20 | 34 | 21 | 27 | 30 | 29 | 21 | 37 | 21 | 30 | 30 | 27.95 | 29 | 27 | 0.933 | -0.1 | 0.79248995 | 0.87847517 | 0.0176 | 0.0872 | 0.0686 | 0.0686 | TRUE | 0.173 |
| Snord43 | 4 | 3 | 4 | 2 | 3 | 2 | 3 | 3 | 4 | 2 | 3 | 2 | 2.99 | 3 | 2 | 0.691 | -0.534 | 0.62108993 | NA | 0 | 0.7941 | 0.4918 | 0.4918 | TRUE | 0.1055 |
| Snord45b | 13 | 2 | 0 | 1 | 1 | 1 | 10 | 2 | 0 | 1 | 1 | 1 | 2.56 | 4 | 1 | 0.247 | -2.019 | 0.17413678 | NA | 0 | 0.9298 | 0.9752 | 0.9752 | TRUE | 0.6652 |

| | | | | | | | | | | | | | | | | | | | | | | | | | |
|---|---|---|---|---|---|---|---|---|---|---|---|---|---|---|---|---|---|---|---|---|---|---|---|---|---|
| Snord45c | 0 | 0 | 0 | 0 | 0 | 0 | 0 | 0 | 0 | 0 | 0 | 0 | 0 | 0 | 0 | NA | NA | NA | NA | NA | NA | NA | NA | NA | NA |
| Snord47 | 43 | 23 | 17 | 29 | 11 | 27 | 33 | 24 | 18 | 29 | 12 | 27 | 24.02 | 25 | 23 | 0.889 | -0.17 | 0.6980348 | 0.81651154 | 0.0869 | 0.1011 | 0.0973 | 0.0973 | TRUE | 0.524 |
| Snord49a | 28 | 29 | 17 | 11 | 15 | 18 | 22 | 31 | 18 | 11 | 17 | 18 | 19.43 | 24 | 15 | 0.644 | -0.636 | 0.16283068 | 0.31327756 | 0 | 0.1244 | 0.095 | 0.095 | TRUE | 0.3926 |
| Snord49b | 28 | 24 | 18 | 22 | 15 | 12 | 22 | 26 | 19 | 22 | 17 | 12 | 19.56 | 22 | 17 | 0.76 | -0.396 | 0.37298959 | 0.54751363 | 0 | 0.1236 | 0.0902 | 0.0902 | TRUE | 0.2487 |
| Snord4a | 15 | 5 | 5 | 7 | 5 | 5 | 12 | 5 | 5 | 7 | 6 | 5 | 6.65 | 7 | 6 | 0.77 | -0.377 | 0.61638527 | NA | 0 | 0.3588 | 0.257 | 0.257 | TRUE | 0.8595 |
| Snord52 | 15 | 14 | 13 | 14 | 12 | 7 | 12 | 15 | 14 | 14 | 14 | 7 | 12.49 | 14 | 12 | 0.849 | -0.237 | 0.66771078 | NA | 0 | 0.1922 | 0.1375 | 0.1375 | TRUE | 0.4805 |
| Snord53 | 10 | 10 | 2 | 2 | 6 | 10 | 8 | 11 | 2 | 2 | 7 | 10 | 6.53 | 7 | 6 | 0.901 | -0.15 | 0.85823674 | NA | 0.329 | 0.3653 | 0.3556 | 0.3556 | TRUE | 0.3793 |
| Snord55 | 13 | 13 | 10 | 8 | 3 | 3 | 10 | 14 | 11 | 8 | 3 | 3 | 8.17 | 12 | 5 | 0.416 | -1.265 | 0.07507407 | NA | 0 | 0.2925 | 0.2144 | 0.2144 | TRUE | 0.6868 |
| Snord57 | 8 | 10 | 11 | 9 | 5 | 5 | 6 | 11 | 12 | 9 | 6 | 5 | 8.04 | 10 | 7 | 0.69 | -0.535 | 0.43724094 | NA | 0 | 0.2971 | 0.2119 | 0.2119 | TRUE | 0.4137 |
| Snord58b | 5 | 5 | 2 | 5 | 2 | 5 | 4 | 5 | 2 | 5 | 2 | 5 | 3.92 | 4 | 4 | 1.077 | 0.107 | 0.9110649 | NA | 0 | 0.6068 | 0.4101 | 0.4101 | TRUE | 0.2226 |
| Snord61 | 0 | 0 | 0 | 0 | 0 | 0 | 0 | 0 | 0 | 0 | 0 | 0 | 0 | 0 | 0 | NA | NA | NA | NA | NA | NA | NA | NA | NA | NA |
| Snord64 | 2 | 1 | 4 | 3 | 0 | 3 | 2 | 1 | 4 | 3 | 0 | 3 | 2.15 | 2 | 2 | 0.882 | -0.181 | 0.89521932 | NA | 0 | 1.1064 | 0.8875 | 0.8875 | TRUE | 0.5862 |
| Snord65 | 9 | 15 | 8 | 19 | 12 | 9 | 7 | 16 | 9 | 19 | 14 | 9 | 12.15 | 11 | 14 | 1.325 | 0.406 | 0.49033824 | NA | 0.0853 | 0.1975 | 0.1655 | 0.1655 | TRUE | 0.3127 |
| Snord66 | 0 | 1 | 0 | 1 | 0 | 3 | 0 | 1 | 0 | 1 | 0 | 3 | 0.84 | 0 | 1 | 3.866 | 1.951 | 0.40188466 | NA | 0 | 2.8349 | 2.5155 | 2.5155 | TRUE | 0.3413 |
| Snord67 | 1 | 0 | 1 | 1 | 0 | 0 | 1 | 0 | 1 | 1 | 0 | 0 | 0.48 | 1 | 0 | 0.596 | -0.748 | 0.78941713 | NA | 0 | 4.9902 | 3.8695 | 3.8695 | TRUE | 0.3906 |
| Snord68 | 11 | 11 | 11 | 5 | 9 | 7 | 9 | 12 | 12 | 5 | 10 | 7 | 9.03 | 11 | 7 | 0.686 | -0.543 | 0.3991865 | NA | 0 | 0.265 | 0.1826 | 0.1826 | TRUE | 0.2741 |
| Snord69 | 9 | 5 | 7 | 3 | 4 | 3 | 7 | 5 | 8 | 3 | 5 | 3 | 5.05 | 7 | 4 | 0.522 | -0.937 | 0.27183343 | NA | 0 | 0.4714 | 0.3013 | 0.3013 | TRUE | 0.086 |
| Snord7 | 0 | 0 | 0 | 0 | 0 | 0 | 0 | 0 | 0 | 0 | 0 | 0 | 0 | 0 | 0 | NA | NA | NA | NA | NA | NA | NA | NA | NA | NA |
| Snord70 | 2 | 0 | 0 | 7 | 1 | 1 | 2 | 0 | 0 | 7 | 1 | 1 | 1.77 | 1 | 3 | 5.239 | 2.389 | 0.17783329 | NA | 1.0448 | 1.3394 | 1.2687 | 1.2687 | TRUE | 1.7272 |
| Snord71 | 1 | 1 | 2 | 1 | 2 | 0 | 1 | 1 | 2 | 1 | 2 | 0 | 1.21 | 1 | 1 | 0.81 | -0.304 | 0.86231033 | NA | 0 | 1.9639 | 1.3587 | 1.3587 | TRUE | 0.2723 |
| Snord72 | 15 | 5 | 4 | 11 | 4 | 7 | 12 | 5 | 4 | 11 | 5 | 7 | 7.28 | 7 | 8 | 1.034 | 0.048 | 0.94903792 | NA | 0.1178 | 0.3282 | 0.2663 | 0.2663 | TRUE | 0.8272 |
| Snord73a | 5 | 4 | 3 | 5 | 1 | 2 | 4 | 4 | 3 | 5 | 1 | 2 | 3.24 | 4 | 3 | 0.717 | -0.481 | 0.65249642 | NA | 0 | 0.7334 | 0.5031 | 0.5031 | TRUE | 0.6573 |
| Snord8 | 1 | 2 | 3 | 1 | 2 | 1 | 1 | 2 | 3 | 1 | 2 | 1 | 1.73 | 2 | 1 | 0.699 | -0.518 | 0.72005363 | NA | 0 | 1.3724 | 0.8912 | 0.8912 | TRUE | 0.2499 |
| Snord82 | 0 | 0 | 0 | 0 | 0 | 0 | 0 | 0 | 0 | 0 | 0 | 0 | 0 | 0 | 0 | NA | NA | NA | NA | NA | NA | NA | NA | NA | NA |
| Snord83b | 10 | 4 | 9 | 9 | 5 | 5 | 8 | 4 | 10 | 9 | 6 | 5 | 6.88 | 7 | 7 | 0.899 | -0.154 | 0.83475604 | NA | 0 | 0.347 | 0.2464 | 0.2464 | TRUE | 0.3214 |
| Snord85 | 0 | 0 | 0 | 0 | 1 | 0 | 0 | 0 | 0 | 0 | 1 | 0 | 0.19 | 0 | 0 | 2.094 | 1.067 | 0.79379329 | NA | 0 | 12.608 | 10 | 10 | TRUE | 0.5075 |
| Snord87 | 5 | 1 | 2 | 3 | 3 | 1 | 4 | 1 | 2 | 3 | 3 | 1 | 2.41 | 2 | 2 | 1.004 | 0.006 | 0.99628753 | NA | 0 | 0.9857 | 0.6791 | 0.6791 | TRUE | 0.4382 |
| Snord88a | 5 | 3 | 1 | 3 | 2 | 4 | 4 | 3 | 1 | 3 | 2 | 4 | 2.89 | 3 | 3 | 1.112 | 0.153 | 0.89028095 | NA | 0 | 0.8223 | 0.544 | 0.544 | TRUE | 0.2972 |
| Snord88c | 3 | 5 | 7 | 3 | 2 | 4 | 2 | 5 | 8 | 3 | 2 | 4 | 4.07 | 5 | 3 | 0.619 | -0.691 | 0.47590097 | NA | 0 | 0.5854 | 0.4126 | 0.4126 | TRUE | 0.2299 |
| Snord89 | 5 | 3 | 4 | 0 | 1 | 5 | 4 | 3 | 4 | 0 | 1 | 5 | 2.91 | 4 | 2 | 0.537 | -0.896 | 0.46244124 | NA | 0.1799 | 0.8172 | 0.6916 | 0.6916 | TRUE | 1.5002 |
| Snord90 | 0 | 0 | 0 | 0 | 0 | 0 | 0 | 0 | 0 | 0 | 0 | 0 | 0 | 0 | 0 | NA | NA | NA | NA | NA | NA | NA | NA | NA | NA |
| Snord91a | 0 | 2 | 0 | 0 | 0 | 1 | 0 | 2 | 0 | 0 | 0 | 1 | 0.52 | 1 | 0 | 0.534 | -0.904 | 0.76321451 | NA | 0 | 4.5682 | 4.7886 | 4.7886 | TRUE | 0.9746 |
| Snord92 | 0 | 0 | 0 | 0 | 0 | 0 | 0 | 0 | 0 | 0 | 0 | 0 | 0 | 0 | 0 | NA | NA | NA | NA | NA | NA | NA | NA | NA | NA |
| Snord93 | 0 | 0 | 0 | 0 | 0 | 0 | 0 | 0 | 0 | 0 | 0 | 0 | 0 | 0 | 0 | NA | NA | NA | NA | NA | NA | NA | NA | NA | NA |
| Snord95 | 9 | 11 | 8 | 8 | 6 | 6 | 7 | 12 | 9 | 8 | 7 | 6 | 8 | 9 | 7 | 0.762 | -0.391 | 0.56063448 | NA | 0 | 0.2988 | 0.1974 | 0.1974 | TRUE | 0.1937 |
| Snord96a | 148 | 89 | 65 | 101 | 52 | 87 | 115 | 95 | 70 | 101 | 59 | 86 | 87.47 | 93 | 82 | 0.874 | -0.194 | 0.45021188 | 0.62028903 | 0.0535 | 0.0295 | 0.0362 | 0.0362 | TRUE | 0.2585 |
| Snord98 | 0 | 0 | 0 | 0 | 0 | 0 | 0 | 0 | 0 | 0 | 0 | 0 | 0 | 0 | 0 | NA | NA | NA | NA | NA | NA | NA | NA | NA | NA |
| Snord99 | 26 | 27 | 20 | 22 | 21 | 13 | 20 | 29 | 22 | 22 | 24 | 13 | 21.49 | 24 | 20 | 0.829 | -0.27 | 0.53175568 | 0.69157269 | 0 | 0.1127 | 0.087 | 0.087 | TRUE | 0.4901 |
| Snph | 40 | 27 | 32 | 63 | 39 | 57 | 31 | 29 | 35 | 63 | 44 | 56 | 42.88 | 32 | 54 | 1.734 | 0.794 | 0.00900022 | 0.03835427 | 0 | 0.0577 | 0.0417 | 0.0417 | TRUE | 0.216 |
| Snrk | 743 | 530 | 477 | 527 | 467 | 512 | 577 | 564 | 515 | 525 | 527 | 505 | 535.39 | 552 | 519 | 0.939 | -0.091 | 0.34271778 | 0.51687931 | 0 | 0.0069 | 0.0047 | 0.0047 | TRUE | 0.0396 |
| Snrnp200 | 24727 | 17965 | 18427 | 19203 | 17714 | 20122 | 19204 | 19109 | 19883 | 19115 | 19983 | 19864 | 19526.25 | 19399 | 19654 | 1.013 | 0.019 | 0.68972117 | 0.81023956 | 4.00E-04 | 0.0025 | 0.0016 | 0.0016 | TRUE | 0.0071 |
| Snrnp25 | 2655 | 1942 | 1823 | 1997 | 1822 | 2017 | 2062 | 2066 | 1967 | 1988 | 2055 | 1991 | 2021.5 | 2032 | 2011 | 0.99 | -0.015 | 0.80577769 | 0.88783384 | 0 | 0.0036 | 0.0022 | 0.0022 | TRUE | 0.0091 |
| Snrnp27 | 2742 | 1892 | 1886 | 2341 | 1887 | 2205 | 2130 | 2012 | 2035 | 2330 | 2129 | 2177 | 2135.45 | 2059 | 2212 | 1.074 | 0.103 | 0.11485424 | 0.24455542 | 0.0011 | 0.0035 | 0.0026 | 0.0026 | TRUE | 0.0266 |
| Snrnp35 | 443 | 301 | 265 | 263 | 222 | 298 | 344 | 320 | 286 | 262 | 250 | 294 | 292.76 | 317 | 269 | 0.847 | -0.239 | 0.06616451 | 0.16450862 | 0.0046 | 0.0105 | 0.0087 | 0.0087 | TRUE | 0.0805 |
| Snrnp40 | 8731 | 6302 | 5872 | 5822 | 5302 | 6101 | 6781 | 6703 | 6336 | 5795 | 5981 | 6023 | 6269.87 | 6607 | 5933 | 0.898 | -0.156 | 0.00349845 | 0.01859552 | 6.00E-04 | 0.0028 | 0.0019 | 0.0019 | TRUE | 0.0156 |
| Snrnp48 | 1611 | 1374 | 1472 | 1661 | 1425 | 1704 | 1251 | 1462 | 1588 | 1653 | 1608 | 1682 | 1540.67 | 1434 | 1648 | 1.151 | 0.203 | 0.02151735 | 0.07287856 | 0.0063 | 0.004 | 0.005 | 0.005 | TRUE | 0.1539 |
| Snrnp70 | 23967 | 17210 | 16255 | 16140 | 14017 | 16940 | 18614 | 18306 | 17539 | 16066 | 15813 | 16722 | 17176.66 | 18153 | 16200 | 0.892 | -0.164 | 0.00108923 | 0.00769531 | 8.00E-04 | 0.0026 | 0.0018 | 0.0018 | TRUE | 0.0107 |
| Snrpa | 9361 | 6921 | 7215 | 7360 | 6287 | 7537 | 7270 | 7362 | 7785 | 7326 | 7092 | 7440 | 7379.28 | 7472 | 7286 | 0.975 | -0.036 | 0.49809445 | 0.66262695 | 8.00E-04 | 0.0027 | 0.0019 | 0.0019 | TRUE | 0.0163 |
| Snrpa1 | 7438 | 5056 | 5099 | 5327 | 4440 | 5465 | 5777 | 5378 | 5502 | 5302 | 5009 | 5395 | 5393.77 | 5552 | 5235 | 0.943 | -0.085 | 0.13772193 | 0.27814887 | 0.0011 | 0.0029 | 0.0022 | 0.0022 | TRUE | 0.0173 |
| Snrpb | 12894 | 9195 | 8463 | 7970 | 7235 | 8395 | 10014 | 9781 | 9132 | 7933 | 8162 | 8287 | 8884.78 | 9642 | 8127 | 0.843 | -0.247 | 7.62E-06 | 0.00015018 | 0.0012 | 0.0027 | 0.0021 | 0.0021 | TRUE | 0.0261 |
| Snrpb2 | 5102 | 3506 | 3278 | 3555 | 3014 | 3560 | 3963 | 3729 | 3537 | 3539 | 3400 | 3514 | 3613.62 | 3743 | 3484 | 0.93 | -0.104 | 0.09303988 | 0.20975063 | 0.0014 | 0.0031 | 0.0025 | 0.0025 | TRUE | 0.0327 |
| Snrpc | 5524 | 3711 | 3805 | 4320 | 3830 | 4480 | 4290 | 3947 | 4106 | 4300 | 4321 | 4422 | 4231.06 | 4114 | 4348 | 1.056 | 0.079 | 0.16009571 | 0.30951548 | 8.00E-04 | 0.003 | 0.002 | 0.002 | TRUE | 0.0174 |
| Snrpd1 | 6468 | 4391 | 4352 | 4521 | 3990 | 4539 | 5023 | 4671 | 4696 | 4500 | 4501 | 4481 | 4645.31 | 4797 | 4494 | 0.936 | -0.095 | 0.08432887 | 0.19579476 | 7.00E-04 | 0.0029 | 0.002 | 0.002 | TRUE | 0.0213 |
| Snrpd2 | 9268 | 6300 | 6882 | 7492 | 6433 | 7858 | 7198 | 6701 | 7426 | 7458 | 7257 | 7757 | 7299.43 | 7108 | 7491 | 1.054 | 0.076 | 0.19756556 | 0.35735315 | 0.0017 | 0.0028 | 0.0023 | 0.0023 | TRUE | 0.0304 |
| Snrpd3 | 7889 | 5444 | 4995 | 5209 | 4517 | 5235 | 6127 | 5791 | 5390 | 5185 | 5096 | 5168 | 5459.29 | 5769 | 5150 | 0.892 | -0.164 | 0.00688238 | 0.0313852 | 0.0018 | 0.0029 | 0.0025 | 0.0025 | TRUE | 0.0403 |
| Snrpe | 5552 | 3913 | 3966 | 4346 | 3695 | 4282 | 4312 | 4162 | 4279 | 4326 | 4168 | 4227 | 4245.81 | 4251 | 4240 | 0.997 | -0.004 | 0.94278948 | 0.968122 | 0 | 0.003 | 0.0017 | 0.0017 | TRUE | 0.0041 |
| Snrpf | 4324 | 3148 | 3114 | 3067 | 2746 | 3357 | 3358 | 3348 | 3360 | 3053 | 3098 | 3314 | 3255.21 | 3355 | 3155 | 0.94 | -0.089 | 0.12602388 | 0.2612035 | 6.00E-04 | 0.0032 | 0.0021 | 0.0021 | TRUE | 0.0238 |
| Snrpg | 3959 | 2733 | 2680 | 2839 | 2460 | 2914 | 3075 | 2907 | 2892 | 2826 | 2775 | 2877 | 2891.86 | 2958 | 2826 | 0.955 | -0.066 | 0.25269615 | 0.42159287 | 5.00E-04 | 0.0032 | 0.0021 | 0.0021 | TRUE | 0.0149 |
| Snrpn | 0 | 0 | 0 | 0 | 0 | 0 | 0 | 0 | 0 | 0 | 0 | 0 | 0 | 0 | 0 | NA | NA | NA | NA | NA | NA | NA | NA | NA | NA |
| Snta1 | 1038 | 742 | 727 | 933 | 823 | 935 | 806 | 789 | 784 | 929 | 928 | 923 | 860 | 793 | 927 | 1.168 | 0.224 | 0.00347945 | 0.01853312 | 0 | 0.0052 | 0.003 | 0.003 | TRUE | 0.0026 |
| Sntb1 | 173 | 105 | 64 | 83 | 70 | 65 | 134 | 112 | 69 | 83 | 79 | 64 | 90.14 | 105 | 75 | 0.712 | -0.491 | 0.05650777 | 0.14725635 | 0.0545 | 0.0287 | 0.0363 | 0.0363 | TRUE | 0.3358 |
| Sntb2 | 5774 | 4338 | 4297 | 5152 | 4411 | 5188 | 4484 | 4614 | 4636 | 5128 | 4976 | 5121 | 4826.81 | 4578 | 5075 | 1.109 | 0.149 | 0.00346188 | 0.01847672 | 0 | 0.0029 | 0.0017 | 0.0017 | TRUE | 0.004 |
| Sntg1 | 0 | 2 | 5 | 0 | 1 | 1 | 0 | 2 | 5 | 0 | 1 | 1 | 1.61 | 2 | 1 | 0.284 | -1.814 | 0.31506214 | NA | 0 | 1.4779 | 1.4086 | 1.4086 | TRUE | 0.1522 |
| Sntg2 | 18 | 23 | 27 | 25 | 23 | 31 | 14 | 24 | 29 | 25 | 26 | 31 | 24.84 | 22 | 27 | 1.223 | 0.291 | 0.47783909 | 0.64541242 | 0.0333 | 0.0979 | 0.0801 | 0.0801 | TRUE | 0.7949 |
| Sntn | 0 | 0 | 0 | 0 | 0 | 0 | 0 | 0 | 0 | 0 | 0 | 0 | 0 | 0 | 0 | NA | NA | NA | NA | NA | NA | NA | NA | NA | NA |
| Snupn | 1410 | 977 | 1018 | 1184 | 974 | 1101 | 1095 | 1039 | 1098 | 1179 | 1099 | 1087 | 1099.48 | 1077 | 1122 | 1.04 | 0.057 | 0.44559533 | 0.61646566 | 6.00E-04 | 0.0046 | 0.0032 | 0.0032 | TRUE | 0.0241 |
| Snurf | 0 | 0 | 0 | 0 | 0 | 0 | 0 | 0 | 0 | 0 | 0 | 0 | 0 | 0 | 0 | NA | NA | NA | NA | NA | NA | NA | NA | NA | NA |
| Snw1 | 6833 | 4866 | 4650 | 4630 | 4247 | 4909 | 5307 | 5176 | 5017 | 4609 | 4791 | 4846 | 4957.63 | 5167 | 4749 | 0.919 | -0.122 | 0.02298902 | 0.07650338 | 6.00E-04 | 0.0029 | 0.0019 | 0.0019 | TRUE | 0.0081 |
| Snx1 | 2864 | 1973 | 1855 | 1826 | 1713 | 2011 | 2224 | 2099 | 2002 | 1818 | 1932 | 1985 | 2009.96 | 2108 | 1912 | 0.906 | -0.142 | 0.04064346 | 0.11581278 | 0.002 | 0.0036 | 0.003 | 0.003 | TRUE | 0.0289 |

| | | | | | | | | | | | | | | | | | | | | | | | | | |
|---|---|---|---|---|---|---|---|---|---|---|---|---|---|---|---|---|---|---|---|---|---|---|---|---|---|
| Snx10 | 3035 | 2312 | 2770 | 2762 | 2673 | 3254 | 2357 | 2459 | 2989 | 2749 | 3015 | 3212 | 2797.02 | 2602 | 2992 | 1.151 | 0.203 | 0.02867414 | 0.08962839 | 0.011 | 0.0033 | 0.0058 | 0.0058 | TRUE | 0.2046 |
| Snx11 | 1548 | 1240 | 1027 | 1159 | 988 | 1110 | 1202 | 1319 | 1108 | 1154 | 1115 | 1096 | 1165.56 | 1210 | 1122 | 0.927 | -0.109 | 0.18432374 | 0.34051173 | 0.0032 | 0.0045 | 0.004 | 0.004 | TRUE | 0.0724 |
| Snx12 | 1927 | 1436 | 1250 | 1519 | 1351 | 1560 | 1497 | 1527 | 1349 | 1512 | 1524 | 1540 | 1491.48 | 1458 | 1525 | 1.046 | 0.065 | 0.37027011 | 0.5449938 | 0.0015 | 0.004 | 0.0031 | 0.0031 | TRUE | 0.05 |
| Snx13 | 2569 | 1921 | 2014 | 2222 | 1934 | 2169 | 1995 | 2043 | 2173 | 2212 | 2182 | 2141 | 2124.39 | 2070 | 2178 | 1.053 | 0.074 | 0.2416258 | 0.40930928 | 7.00E-04 | 0.0035 | 0.0024 | 0.0024 | TRUE | 0.0225 |
| Snx14 | 2515 | 1810 | 1776 | 2213 | 1837 | 2049 | 1953 | 1925 | 1916 | 2203 | 2072 | 2023 | 2015.45 | 1931 | 2099 | 1.087 | 0.12 | 0.05918825 | 0.15188109 | 6.00E-04 | 0.0036 | 0.0024 | 0.0024 | TRUE | 0.0228 |
| Snx15 | 599 | 432 | 438 | 523 | 481 | 544 | 465 | 460 | 473 | 521 | 543 | 537 | 499.59 | 466 | 534 | 1.145 | 0.195 | 0.03812574 | 0.1105337 | 0 | 0.0072 | 0.0044 | 0.0044 | TRUE | 0.0052 |
| Snx16 | 1282 | 890 | 857 | 935 | 809 | 883 | 996 | 947 | 925 | 931 | 913 | 872 | 930.34 | 956 | 905 | 0.946 | -0.08 | 0.30528677 | 0.47792594 | 0 | 0.005 | 0.0033 | 0.0033 | TRUE | 0.0168 |
| Snx17 | 3026 | 2148 | 1986 | 2125 | 2057 | 2340 | 2350 | 2285 | 2143 | 2115 | 2321 | 2310 | 2253.92 | 2259 | 2249 | 0.995 | -0.008 | 0.90784653 | 0.94923397 | 0.002 | 0.0035 | 0.0029 | 0.0029 | TRUE | 0.0328 |
| Snx18 | 4417 | 3403 | 3463 | 3132 | 3114 | 3413 | 3430 | 3620 | 3737 | 3118 | 3513 | 3369 | 3464.41 | 3596 | 3333 | 0.927 | -0.109 | 0.09873252 | 0.2190599 | 0.0023 | 0.0031 | 0.0029 | 0.0029 | TRUE | 0.039 |
| Snx19 | 1756 | 1214 | 1053 | 1008 | 1030 | 1061 | 1364 | 1291 | 1136 | 1003 | 1162 | 1047 | 1167.33 | 1264 | 1071 | 0.846 | -0.241 | 0.00799276 | 0.03499042 | 0.0059 | 0.0045 | 0.0051 | 0.0051 | TRUE | 0.0927 |
| Snx2 | 8805 | 6441 | 6709 | 7404 | 6414 | 7467 | 6838 | 6851 | 7239 | 7370 | 7236 | 7371 | 7150.88 | 6976 | 7326 | 1.05 | 0.071 | 0.16245743 | 0.31272783 | 4.00E-04 | 0.0028 | 0.0017 | 0.0017 | TRUE | 0.0131 |
| Snx20 | 93 | 65 | 43 | 48 | 42 | 35 | 72 | 69 | 46 | 48 | 47 | 35 | 52.91 | 62 | 43 | 0.686 | -0.543 | 0.06182549 | 0.15676276 | 0.0253 | 0.0472 | 0.0412 | 0.0412 | TRUE | 0.424 |
| Snx21 | 288 | 263 | 347 | 362 | 372 | 414 | 224 | 280 | 374 | 360 | 420 | 409 | 344.42 | 293 | 396 | 1.359 | 0.443 | 0.00674719 | 0.03089959 | 0.0326 | 0.0093 | 0.0163 | 0.0163 | TRUE | 0.5347 |
| Snx22 | 128 | 91 | 92 | 105 | 74 | 87 | 99 | 97 | 99 | 105 | 83 | 86 | 94.89 | 98 | 91 | 0.928 | -0.108 | 0.59386663 | 0.74083813 | 0 | 0.0274 | 0.0191 | 0.0191 | TRUE | 0.156 |
| Snx24 | 512 | 348 | 346 | 387 | 344 | 429 | 398 | 370 | 373 | 385 | 388 | 423 | 389.65 | 380 | 399 | 1.048 | 0.067 | 0.5331335 | 0.69263607 | 0 | 0.0085 | 0.0058 | 0.0058 | TRUE | 0.034 |
| Snx25 | 2362 | 1695 | 1477 | 1602 | 1366 | 1571 | 1834 | 1803 | 1594 | 1595 | 1541 | 1551 | 1652.92 | 1744 | 1562 | 0.895 | -0.16 | 0.03142537 | 0.09583164 | 0.0025 | 0.0039 | 0.0034 | 0.0034 | TRUE | 0.0674 |
| Snx27 | 8999 | 6559 | 6939 | 8059 | 6800 | 8029 | 6989 | 6977 | 7487 | 8022 | 7671 | 7926 | 7511.98 | 7151 | 7873 | 1.101 | 0.139 | 0.00968315 | 0.04044352 | 9.00E-04 | 0.0027 | 0.0019 | 0.0019 | TRUE | 0.0205 |
| Snx29 | 697 | 522 | 537 | 646 | 651 | 670 | 541 | 555 | 579 | 643 | 734 | 661 | 619.14 | 558 | 679 | 1.217 | 0.283 | 0.00255324 | 0.01461494 | 0.0015 | 0.0063 | 0.0047 | 0.0047 | TRUE | 0.0573 |
| Snx3 | 9136 | 6573 | 6894 | 6846 | 6023 | 7122 | 7096 | 6992 | 7439 | 6815 | 6795 | 7031 | 7027.55 | 7176 | 6880 | 0.959 | -0.06 | 0.24155245 | 0.40930755 | 5.00E-04 | 0.0028 | 0.0018 | 0.0018 | TRUE | 0.0124 |
| Snx30 | 2586 | 2016 | 1998 | 2267 | 2143 | 2439 | 2008 | 2144 | 2156 | 2257 | 2418 | 2408 | 2231.74 | 2103 | 2361 | 1.123 | 0.168 | 0.00941957 | 0.0396799 | 0.0011 | 0.0035 | 0.0026 | 0.0026 | TRUE | 0.0194 |
| Snx31 | 140 | 115 | 107 | 92 | 101 | 110 | 109 | 122 | 115 | 92 | 114 | 109 | 110.1 | 115 | 105 | 0.907 | -0.141 | 0.45998563 | 0.62988358 | 0 | 0.024 | 0.017 | 0.017 | TRUE | 0.1202 |
| Snx32 | 99 | 76 | 58 | 79 | 68 | 85 | 77 | 81 | 63 | 79 | 77 | 84 | 76.59 | 74 | 80 | 1.084 | 0.117 | 0.60212762 | 0.74706685 | 0 | 0.0334 | 0.0233 | 0.0233 | TRUE | 0.1413 |
| Snx33 | 94 | 67 | 44 | 54 | 54 | 39 | 73 | 71 | 47 | 54 | 61 | 38 | 57.49 | 64 | 51 | 0.793 | -0.334 | 0.24234355 | 0.41011323 | 0.0347 | 0.0437 | 0.0412 | 0.0412 | TRUE | 0.4213 |
| Snx4 | 6839 | 4823 | 4815 | 5250 | 4647 | 5158 | 5312 | 5130 | 5195 | 5226 | 5242 | 5092 | 5199.49 | 5212 | 5187 | 0.995 | -0.008 | 0.88154082 | 0.93382702 | 0 | 0.0029 | 0.0016 | 0.0016 | TRUE | 0.0035 |
| Snx5 | 39149 | 26266 | 21698 | 23084 | 20225 | 21954 | 30405 | 27939 | 23412 | 22978 | 22816 | 21672 | 24870.3 | 27252 | 22489 | 0.825 | -0.277 | 0.00078888 | 0.00608167 | 0.0089 | 0.0025 | 0.0049 | 0.0049 | TRUE | 0.1859 |
| Snx6 | 5938 | 4282 | 4332 | 4140 | 3752 | 4240 | 4612 | 4555 | 4674 | 4121 | 4233 | 4186 | 4396.64 | 4614 | 4180 | 0.906 | -0.143 | 0.00498722 | 0.02447044 | 0 | 0.003 | 0.0016 | 0.0016 | TRUE | 0.0018 |
| Snx7 | 734 | 502 | 423 | 421 | 370 | 385 | 570 | 534 | 456 | 419 | 417 | 380 | 462.83 | 520 | 405 | 0.778 | -0.363 | 0.00139911 | 0.00916485 | 0.0061 | 0.0075 | 0.0071 | 0.0071 | TRUE | 0.1317 |
| Snx8 | 2060 | 1633 | 1680 | 1978 | 1815 | 2096 | 1600 | 1737 | 1813 | 1969 | 2048 | 2069 | 1872.52 | 1717 | 2029 | 1.183 | 0.242 | 0.00053116 | 0.00451094 | 0.0018 | 0.0037 | 0.003 | 0.003 | TRUE | 0.0443 |
| Snx9 | 858 | 584 | 556 | 623 | 516 | 578 | 666 | 621 | 600 | 620 | 582 | 571 | 610.05 | 629 | 591 | 0.938 | -0.092 | 0.31803115 | 0.49189826 | 0 | 0.0063 | 0.0045 | 0.0045 | TRUE | 0.0334 |
| Soat1 | 1964 | 1536 | 1931 | 2111 | 1994 | 2424 | 1525 | 1634 | 2084 | 2101 | 2249 | 2393 | 1997.72 | 1748 | 2248 | 1.288 | 0.365 | 0.00047324 | 0.00414927 | 0.0157 | 0.0036 | 0.0073 | 0.0073 | TRUE | 0.34 |
| Soat2 | 117 | 81 | 94 | 141 | 138 | 136 | 91 | 86 | 101 | 140 | 156 | 134 | 118.12 | 93 | 143 | 1.545 | 0.628 | 0.00065413 | 0.00528815 | 0 | 0.0225 | 0.0156 | 0.0156 | TRUE | 0.0584 |
| Sobp | 656 | 535 | 345 | 417 | 433 | 454 | 509 | 569 | 372 | 415 | 488 | 448 | 467.09 | 483 | 450 | 0.93 | -0.104 | 0.46436673 | 0.63356183 | 0.0241 | 0.0075 | 0.0125 | 0.0125 | TRUE | 0.4594 |
| Socs1 | 278 | 241 | 213 | 269 | 248 | 304 | 216 | 256 | 230 | 268 | 280 | 300 | 258.29 | 234 | 283 | 1.211 | 0.276 | 0.03683079 | 0.10798411 | 0 | 0.0116 | 0.0087 | 0.0087 | TRUE | 0.0761 |
| Socs2 | 2763 | 2024 | 1774 | 1783 | 1512 | 1795 | 2146 | 2153 | 1914 | 1775 | 1706 | 1772 | 1910.89 | 2071 | 1751 | 0.845 | -0.243 | 0.00054379 | 0.00459635 | 0.002 | 0.0037 | 0.003 | 0.003 | TRUE | 0.0522 |
| Socs3 | 20841 | 17741 | 22349 | 20219 | 20603 | 25676 | 16186 | 18871 | 24114 | 20126 | 23242 | 25346 | 21314.34 | 19724 | 22905 | 1.161 | 0.216 | 0.26561083 | 0.43601419 | 0.027 | 0.0025 | 0.0097 | 0.027 | TRUE | 0.4639 |
| Socs4 | 703 | 533 | 562 | 612 | 518 | 604 | 546 | 567 | 606 | 609 | 584 | 596 | 584.85 | 573 | 596 | 1.043 | 0.06 | 0.51130604 | 0.67440417 | 0 | 0.0065 | 0.0044 | 0.0044 | TRUE | 0.0299 |
| Socs5 | 854 | 588 | 602 | 676 | 560 | 633 | 663 | 625 | 650 | 673 | 632 | 625 | 644.63 | 646 | 643 | 0.995 | -0.007 | 0.93364743 | 0.96302394 | 0 | 0.0061 | 0.004 | 0.004 | TRUE | 0.0195 |
| Socs6 | 2036 | 1481 | 1613 | 1629 | 1410 | 1602 | 1581 | 1575 | 1740 | 1622 | 1591 | 1581 | 1615.09 | 1632 | 1598 | 0.979 | -0.03 | 0.66627802 | 0.7948039 | 0.0011 | 0.0039 | 0.0028 | 0.0028 | TRUE | 0.0397 |
| Socs7 | 3087 | 2382 | 2333 | 2547 | 2273 | 2606 | 2398 | 2534 | 2517 | 2535 | 2564 | 2573 | 2520.09 | 2483 | 2557 | 1.031 | 0.043 | 0.45514395 | 0.62481957 | 0 | 0.0034 | 0.002 | 0.002 | TRUE | 0.0114 |
| Sod1 | 25300 | 17599 | 18192 | 20456 | 17900 | 20956 | 19649 | 18720 | 19629 | 20362 | 20193 | 20687 | 19873.34 | 19333 | 20414 | 1.056 | 0.078 | 0.09263959 | 0.2092112 | 4.00E-04 | 0.0025 | 0.0015 | 0.0015 | TRUE | 0.0094 |
| Sod2 | 6176 | 4918 | 5059 | 5515 | 5114 | 5908 | 4797 | 5231 | 5459 | 5490 | 5769 | 5832 | 5429.55 | 5162 | 5697 | 1.104 | 0.143 | 0.0244156 | 0.07998445 | 0.0024 | 0.0029 | 0.0027 | 0.0027 | TRUE | 0.0475 |
| Sod3 | 2 | 4 | 0 | 0 | 0 | 1 | 2 | 4 | 0 | 0 | 0 | 1 | 1.13 | 2 | 0 | 0.192 | -2.38 | 0.26121891 | NA | 0 | 2.0951 | 1.9833 | 1.9833 | TRUE | 0.1845 |
| Soga2 | 6199 | 4569 | 4383 | 5275 | 4518 | 5045 | 4815 | 4860 | 4729 | 5251 | 5097 | 4980 | 4955.23 | 4801 | 5109 | 1.064 | 0.09 | 0.08345763 | 0.19449859 | 3.00E-04 | 0.0029 | 0.0017 | 0.0017 | TRUE | 0.0072 |
| Soga3 | 161 | 151 | 129 | 288 | 228 | 319 | 125 | 161 | 139 | 287 | 257 | 315 | 213.94 | 142 | 286 | 2.032 | 1.023 | 3.08E-11 | 2.82E-09 | 0.0077 | 0.0135 | 0.0119 | 0.0119 | TRUE | 0.1421 |
| Sohlh1 | 14 | 8 | 10 | 4 | 8 | 6 | 11 | 9 | 11 | 4 | 9 | 6 | 8.18 | 10 | 6 | 0.621 | -0.688 | 0.30976758 | NA | 0 | 0.292 | 0.2002 | 0.2002 | TRUE | 0.3077 |
| Sohlh2 | 11088 | 8594 | 8511 | 7434 | 6774 | 7506 | 8612 | 9141 | 9183 | 7400 | 7642 | 7410 | 8231.23 | 8979 | 7484 | 0.834 | -0.263 | 3.96E-07 | 1.33E-05 | 6.00E-04 | 0.0027 | 0.0018 | 0.0018 | TRUE | 0.0158 |
| Solh | 1317 | 945 | 937 | 994 | 895 | 986 | 1023 | 1005 | 1011 | 989 | 1010 | 973 | 1001.91 | 1013 | 991 | 0.978 | -0.033 | 0.65391718 | 0.78515106 | 0 | 0.0048 | 0.0028 | 0.0028 | TRUE | 0.0032 |
| Son | 19288 | 13575 | 12265 | 13222 | 11358 | 13199 | 14980 | 14440 | 13234 | 13161 | 12813 | 13029 | 13609.53 | 14218 | 13001 | 0.914 | -0.129 | 0.02612721 | 0.08392982 | 0.002 | 0.0026 | 0.0024 | 0.0024 | TRUE | 0.0447 |
| Sorbs1 | 2513 | 1923 | 2108 | 1944 | 1763 | 2249 | 1952 | 2045 | 2275 | 1935 | 1989 | 2220 | 2069.29 | 2091 | 2048 | 0.98 | -0.028 | 0.72431869 | 0.83482679 | 0.0053 | 0.0036 | 0.0042 | 0.0042 | TRUE | 0.0713 |
| Sorbs2 | 181 | 119 | 108 | 103 | 104 | 112 | 141 | 127 | 117 | 103 | 117 | 111 | 119.02 | 128 | 110 | 0.857 | -0.222 | 0.22589277 | 0.3909066 | 0 | 0.0223 | 0.0158 | 0.0158 | TRUE | 0.0848 |
| Sorbs3 | 594 | 406 | 474 | 557 | 513 | 541 | 461 | 432 | 511 | 554 | 579 | 534 | 511.97 | 468 | 556 | 1.187 | 0.247 | 0.01526839 | 0.05658052 | 0.002 | 0.0071 | 0.0055 | 0.0055 | TRUE | 0.072 |
| Sorcs1 | 7 | 10 | 7 | 8 | 4 | 8 | 5 | 11 | 8 | 8 | 5 | 8 | 7.33 | 8 | 7 | 0.877 | -0.189 | 0.79026239 | NA | 0 | 0.3256 | 0.2268 | 0.2268 | TRUE | 0.2834 |
| Sorcs2 | 278 | 174 | 188 | 221 | 206 | 200 | 216 | 185 | 203 | 220 | 232 | 197 | 208.94 | 201 | 216 | 1.072 | 0.101 | 0.48491039 | 0.65090872 | 0 | 0.0138 | 0.0103 | 0.0103 | TRUE | 0.0665 |
| Sorcs3 | 16 | 14 | 20 | 10 | 17 | 18 | 12 | 15 | 22 | 10 | 19 | 18 | 15.97 | 16 | 16 | 0.964 | -0.052 | 0.9170638 | NA | 0 | 0.1509 | 0.1188 | 0.1188 | TRUE | 0.4844 |
| Sord | 3336 | 2387 | 2362 | 2488 | 2251 | 2614 | 2591 | 2539 | 2549 | 2477 | 2539 | 2580 | 2545.81 | 2560 | 2532 | 0.989 | -0.016 | 0.78045437 | 0.87053897 | 0 | 0.0034 | 0.0019 | 0.0019 | TRUE | 0.0045 |
| Sorl1 | 4990 | 3741 | 4457 | 4997 | 4738 | 5121 | 3876 | 3979 | 4809 | 4974 | 5345 | 5055 | 4673 | 4221 | 5125 | 1.214 | 0.28 | 0.00058173 | 0.00484819 | 0.0077 | 0.0029 | 0.0046 | 0.0046 | TRUE | 0.18 |
| Sort1 | 2586 | 1889 | 1755 | 2334 | 2101 | 2422 | 2008 | 2009 | 1894 | 2323 | 2370 | 2391 | 2165.94 | 1970 | 2361 | 1.198 | 0.261 | 1.88E-05 | 0.00031743 | 0 | 0.0035 | 0.0022 | 0.0022 | TRUE | 0.0138 |
| Sos1 | 4535 | 3312 | 3165 | 3289 | 2952 | 3451 | 3522 | 3523 | 3415 | 3274 | 3330 | 3407 | 3411.8 | 3487 | 3337 | 0.957 | -0.064 | 0.24095679 | 0.40854038 | 0 | 0.0031 | 0.0018 | 0.0018 | TRUE | 0.0041 |
| Sos2 | 1574 | 1158 | 1249 | 1293 | 1053 | 1303 | 1222 | 1232 | 1348 | 1287 | 1188 | 1286 | 1260.51 | 1267 | 1254 | 0.99 | -0.014 | 0.85578688 | 0.91887185 | 0.0017 | 0.0043 | 0.0034 | 0.0034 | TRUE | 0.0363 |
| Sost | 34 | 17 | 9 | 21 | 13 | 12 | 26 | 18 | 10 | 21 | 15 | 12 | 16.94 | 18 | 16 | 0.863 | -0.212 | 0.67964757 | NA | 0.1028 | 0.1424 | 0.1311 | 0.1311 | TRUE | 0.1727 |
| Sostdc1 | 1 | 3 | 7 | 0 | 1 | 1 | 1 | 3 | 8 | 0 | 1 | 1 | 2.27 | 4 | 1 | 0.184 | -2.439 | 0.11995732 | NA | 0 | 1.0453 | 0.9002 | 0.9002 | TRUE | 0.1677 |
| Sowaha | 142 | 123 | 164 | 204 | 200 | 222 | 110 | 131 | 177 | 203 | 226 | 219 | 177.65 | 139 | 216 | 1.559 | 0.641 | 0.00041175 | 0.00372002 | 0.0226 | 0.0158 | 0.0178 | 0.0178 | TRUE | 0.5689 |
| Sowahb | 391 | 286 | 323 | 348 | 322 | 306 | 304 | 304 | 349 | 346 | 363 | 302 | 328.02 | 319 | 337 | 1.058 | 0.082 | 0.51094927 | 0.67419755 | 0.0045 | 0.0097 | 0.0081 | 0.0081 | TRUE | 0.0959 |
| Sowahc | 555 | 422 | 397 | 406 | 319 | 386 | 431 | 449 | 428 | 404 | 360 | 381 | 408.89 | 436 | 382 | 0.876 | -0.191 | 0.07007781 | 0.17162425 | 0 | 0.0082 | 0.0055 | 0.0055 | TRUE | 0.0304 |
| Sowahd | 0 | 0 | 1 | 0 | 0 | 0 | 0 | 0 | 1 | 0 | 0 | 0 | 0.18 | 0 | 0 | 0.552 | -0.857 | 0.8336548 | NA | 0 | 13.1817 | 10 | 10 | TRUE | 0.5365 |
| Sox1 | 233 | 185 | 135 | 251 | 198 | 213 | 181 | 197 | 146 | 250 | 223 | 210 | 201.15 | 175 | 228 | 1.304 | 0.383 | 0.01496223 | 0.05574369 | 0.0092 | 0.0142 | 0.0128 | 0.0128 | TRUE | 0.2098 |
| Sox10 | 42 | 39 | 32 | 49 | 38 | 42 | 33 | 41 | 35 | 49 | 43 | 41 | 40.29 | 36 | 44 | 1.231 | 0.299 | 0.32407135 | 0.49809594 | 0 | 0.0613 | 0.0414 | 0.0414 | TRUE | 0.1126 |
| Sox11 | 852 | 671 | 662 | 883 | 739 | 956 | 662 | 714 | 714 | 879 | 834 | 944 | 791.01 | 697 | 886 | 1.274 | 0.349 | 6.07E-05 | 0.00082077 | 0.0018 | 0.0054 | 0.0042 | 0.0042 | TRUE | 0.0396 |
| Sox12 | 3486 | 2866 | 2827 | 3030 | 2674 | 3072 | 2707 | 3049 | 3050 | 3016 | 3017 | 3033 | 2978.57 | 2935 | 3022 | 1.03 | 0.043 | 0.5115864 | 0.67467286 | 0.0021 | 0.0032 | 0.0028 | 0.0028 | TRUE | 0.0575 |

| | | | | | | | | | | | | | | | | | | | | | | | | | |
|---|---|---|---|---|---|---|---|---|---|---|---|---|---|---|---|---|---|---|---|---|---|---|---|---|---|
| Sox13 | 1761 | 1363 | 1389 | 1306 | 1331 | 1605 | 1368 | 1450 | 1499 | 1300 | 1502 | 1584 | 1450.35 | 1439 | 1462 | 1.017 | 0.024 | 0.78011532 | 0.87046309 | 0.0055 | 0.0041 | 0.0046 | 0.0046 | TRUE | 0.1141 |
| Sox14 | 0 | 0 | 4 | 1 | 1 | 3 | 0 | 0 | 4 | 1 | 1 | 3 | 1.57 | 1 | 2 | 1.227 | 0.295 | 0.86507613 | NA | 1.5324 | 1.5151 | 1.5193 | 1.5193 | TRUE | 1.9021 |
| Sox15 | 1456 | 1156 | 1430 | 1313 | 1311 | 1488 | 1131 | 1230 | 1543 | 1307 | 1479 | 1469 | 1359.7 | 1301 | 1418 | 1.091 | 0.126 | 0.24113713 | 0.40878154 | 0.0135 | 0.0042 | 0.0076 | 0.0076 | TRUE | 0.3161 |
| Sox17 | 88 | 55 | 39 | 77 | 63 | 59 | 68 | 59 | 42 | 77 | 71 | 58 | 62.48 | 56 | 69 | 1.209 | 0.274 | 0.30417584 | 0.47670891 | 0.0218 | 0.0404 | 0.0353 | 0.0353 | TRUE | 0.3938 |
| Sox18 | 68 | 51 | 39 | 79 | 76 | 82 | 53 | 54 | 42 | 79 | 86 | 81 | 65.74 | 50 | 82 | 1.639 | 0.713 | 0.00328025 | 0.01769898 | 0 | 0.0385 | 0.0263 | 0.0263 | TRUE | 0.1346 |
| Sox2 | 38188 | 29022 | 29122 | 28732 | 26574 | 31879 | 29659 | 30870 | 31422 | 28600 | 29978 | 31469 | 30333.21 | 30650 | 30016 | 0.979 | -0.03 | 0.57803047 | 0.72933171 | 0.0013 | 0.0025 | 0.0021 | 0.0021 | TRUE | 0.022 |
| Sox21 | 299 | 219 | 198 | 127 | 102 | 147 | 232 | 233 | 214 | 126 | 115 | 145 | 177.57 | 226 | 129 | 0.57 | -0.81 | 2.46E-07 | 8.78E-06 | 0 | 0.0158 | 0.0116 | 0.0116 | TRUE | 0.1255 |
| Sox2ot | 58 | 42 | 38 | 29 | 29 | 32 | 45 | 45 | 41 | 29 | 33 | 32 | 37.32 | 44 | 31 | 0.711 | -0.492 | 0.11486046 | 0.24455542 | 0 | 0.0659 | 0.0424 | 0.0424 | TRUE | 0.0261 |
| Sox3 | 459 | 362 | 254 | 432 | 313 | 323 | 356 | 385 | 274 | 430 | 353 | 319 | 352.93 | 338 | 367 | 1.084 | 0.116 | 0.43939654 | 0.61068599 | 0.0239 | 0.0091 | 0.0135 | 0.0135 | TRUE | 0.3058 |
| Sox30 | 30 | 24 | 23 | 25 | 21 | 22 | 23 | 26 | 25 | 25 | 24 | 22 | 23.99 | 25 | 24 | 0.956 | -0.065 | 0.86616952 | 0.9261042 | 0 | 0.1012 | 0.0644 | 0.0644 | TRUE | 0.0252 |
| Sox4 | 1610 | 1239 | 1159 | 1470 | 1285 | 1494 | 1250 | 1318 | 1251 | 1463 | 1450 | 1475 | 1367.76 | 1273 | 1463 | 1.149 | 0.201 | 0.00282209 | 0.01573174 | 0 | 0.0042 | 0.0025 | 0.0025 | TRUE | 0.0112 |
| Sox5 | 25 | 10 | 9 | 2 | 12 | 10 | 19 | 11 | 10 | 2 | 14 | 10 | 10.86 | 13 | 9 | 0.626 | -0.675 | 0.31869145 | NA | 0.271 | 0.2207 | 0.2329 | 0.2329 | TRUE | 0.866 |
| Sox6 | 43 | 28 | 15 | 21 | 26 | 10 | 33 | 30 | 16 | 21 | 29 | 10 | 23.25 | 26 | 20 | 0.747 | -0.42 | 0.37291022 | 0.54745334 | 0.148 | 0.1044 | 0.116 | 0.116 | TRUE | 0.3572 |
| Sox7 | 69 | 56 | 25 | 57 | 34 | 39 | 54 | 60 | 27 | 57 | 38 | 38 | 45.62 | 47 | 44 | 0.951 | -0.073 | 0.83078366 | 0.9033735 | 0.0823 | 0.0544 | 0.062 | 0.062 | TRUE | 1.0165 |
| Sox8 | 114 | 89 | 62 | 73 | 68 | 71 | 89 | 95 | 67 | 73 | 77 | 70 | 78.26 | 84 | 73 | 0.875 | -0.193 | 0.39900539 | 0.57224403 | 0 | 0.0327 | 0.0249 | 0.0249 | TRUE | 0.2624 |
| Sox9 | 16 | 10 | 12 | 10 | 8 | 7 | 12 | 11 | 13 | 10 | 9 | 7 | 10.32 | 12 | 9 | 0.717 | -0.479 | 0.41589638 | NA | 0 | 0.2322 | 0.1499 | 0.1499 | TRUE | 0.0996 |
| Sp1 | 15732 | 11580 | 12556 | 15263 | 12469 | 15084 | 12218 | 12317 | 13548 | 15193 | 14066 | 14890 | 13705.5 | 12694 | 14716 | 1.159 | 0.213 | 0.00035983 | 0.00335052 | 0.0024 | 0.0026 | 0.0025 | 0.0025 | TRUE | 0.0422 |
| Sp100 | 5 | 6 | 0 | 3 | 1 | 5 | 4 | 6 | 0 | 3 | 1 | 5 | 3.22 | 3 | 3 | 0.885 | -0.176 | 0.88177133 | NA | 0.5709 | 0.7386 | 0.7025 | 0.7025 | TRUE | 0.2854 |
| Sp110 | 153 | 139 | 64 | 45 | 46 | 44 | 119 | 148 | 69 | 45 | 52 | 43 | 79.31 | 112 | 47 | 0.416 | -1.265 | 9.01E-06 | 0.00017248 | 0.0684 | 0.0323 | 0.0431 | 0.0431 | TRUE | 0.197 |
| Sp140 | 102 | 55 | 43 | 25 | 45 | 34 | 79 | 59 | 46 | 25 | 51 | 34 | 48.89 | 61 | 37 | 0.586 | -0.771 | 0.02033809 | 0.0700311 | 0.0748 | 0.0509 | 0.0574 | 0.0574 | TRUE | 0.4218 |
| Sp2 | 3516 | 2487 | 2603 | 2664 | 2571 | 2966 | 2731 | 2645 | 2809 | 2652 | 2900 | 2928 | 2777.45 | 2728 | 2827 | 1.036 | 0.051 | 0.42652655 | 0.59879619 | 0.0014 | 0.0033 | 0.0026 | 0.0026 | TRUE | 0.0358 |
| Sp3 | 7458 | 5384 | 5586 | 5771 | 4878 | 5745 | 5792 | 5727 | 6027 | 5744 | 5503 | 5671 | 5744.16 | 5849 | 5639 | 0.964 | -0.052 | 0.31380912 | 0.48721201 | 3.00E-04 | 0.0028 | 0.0018 | 0.0018 | TRUE | 0.0086 |
| Sp4 | 1591 | 1265 | 1292 | 1298 | 1023 | 1250 | 1236 | 1346 | 1394 | 1292 | 1154 | 1234 | 1275.88 | 1325 | 1227 | 0.927 | -0.109 | 0.16927515 | 0.32079683 | 0.0028 | 0.0043 | 0.0037 | 0.0037 | TRUE | 0.0436 |
| Sp5 | 82 | 59 | 78 | 88 | 81 | 108 | 64 | 63 | 84 | 88 | 91 | 107 | 82.7 | 70 | 95 | 1.362 | 0.446 | 0.04807516 | 0.13088347 | 0.0061 | 0.0311 | 0.0243 | 0.0243 | TRUE | 0.2579 |
| Sp6 | 236 | 215 | 263 | 215 | 197 | 252 | 183 | 229 | 284 | 214 | 222 | 249 | 230.13 | 232 | 228 | 0.989 | -0.015 | 0.92734951 | 0.95992226 | 0.0231 | 0.0127 | 0.0159 | 0.0159 | TRUE | 0.2361 |
| Sp7 | 7 | 7 | 3 | 11 | 11 | 12 | 5 | 7 | 3 | 11 | 12 | 12 | 8.55 | 5 | 12 | 2.179 | 1.124 | 0.09291292 | NA | 0 | 0.2795 | 0.1887 | 0.1887 | TRUE | 0.2214 |
| Sp8 | 19 | 21 | 6 | 19 | 16 | 13 | 15 | 22 | 6 | 19 | 18 | 13 | 15.56 | 14 | 17 | 1.14 | 0.189 | 0.7236851 | NA | 0.1009 | 0.1547 | 0.1411 | 0.1411 | TRUE | 0.2174 |
| Sp9 | 18 | 19 | 8 | 15 | 8 | 19 | 14 | 20 | 9 | 15 | 9 | 19 | 14.26 | 14 | 14 | 1.003 | 0.004 | 0.99371107 | NA | 0.0654 | 0.1687 | 0.1415 | 0.1415 | TRUE | 0.2089 |
| Spa17 | 82 | 43 | 39 | 52 | 33 | 41 | 64 | 46 | 42 | 52 | 37 | 40 | 46.83 | 51 | 43 | 0.849 | -0.236 | 0.4347224 | 0.60648973 | 0.0228 | 0.053 | 0.0442 | 0.0442 | TRUE | 0.4834 |
| Spaca1 | 44 | 34 | 27 | 13 | 15 | 15 | 34 | 36 | 29 | 13 | 17 | 15 | 24.02 | 33 | 15 | 0.447 | -1.161 | 0.00396977 | 0.02047635 | 0 | 0.1011 | 0.0676 | 0.0676 | TRUE | 0.0724 |
| Spaca3 | 0 | 1 | 1 | 0 | 0 | 0 | 0 | 1 | 1 | 0 | 0 | 0 | 0.36 | 1 | 0 | 0.271 | -1.881 | 0.5755355 | NA | 0 | 6.6391 | 6.4428 | 6.4428 | TRUE | 0.3513 |
| Spaca4 | 0 | 0 | 0 | 0 | 0 | 0 | 0 | 0 | 0 | 0 | 0 | 0 | 0 | 0 | 0 | NA | NA | NA | NA | NA | NA | NA | NA | NA | NA |
| Spaca5 | 0 | 0 | 0 | 0 | 0 | 0 | 0 | 0 | 0 | 0 | 0 | 0 | 0 | 0 | 0 | NA | NA | NA | NA | NA | NA | NA | NA | NA | NA |
| Spaca7 | 0 | 1 | 0 | 0 | 0 | 1 | 0 | 1 | 0 | 0 | 0 | 1 | 0.34 | 0 | 0 | 1.075 | 0.105 | 0.97576789 | NA | 0 | 6.9363 | 6.5821 | 6.5821 | TRUE | 0.5216 |
| Spag1 | 1438 | 1215 | 1245 | 1335 | 1222 | 1403 | 1117 | 1292 | 1343 | 1329 | 1379 | 1385 | 1307.49 | 1251 | 1364 | 1.092 | 0.127 | 0.12651936 | 0.26185756 | 0.0043 | 0.0042 | 0.0042 | 0.0042 | TRUE | 0.1097 |
| Spag11a | 0 | 0 | 0 | 0 | 0 | 0 | 0 | 0 | 0 | 0 | 0 | 0 | 0 | 0 | 0 | NA | NA | NA | NA | NA | NA | NA | NA | NA | NA |
| Spag11b | 1 | 1 | 0 | 0 | 0 | 1 | 1 | 1 | 0 | 0 | 0 | 1 | 0.47 | 1 | 0 | 0.599 | -0.74 | 0.79209856 | NA | 0 | 5.0317 | 3.8925 | 3.8925 | TRUE | 0.3835 |
| Spag16 | 710 | 587 | 586 | 565 | 530 | 555 | 551 | 624 | 632 | 562 | 598 | 548 | 586.04 | 602 | 569 | 0.946 | -0.08 | 0.41256423 | 0.58546273 | 0.0024 | 0.0065 | 0.0051 | 0.0051 | TRUE | 0.0693 |
| Spag17 | 3 | 0 | 0 | 0 | 2 | 2 | 2 | 0 | 0 | 0 | 2 | 2 | 1.09 | 1 | 1 | 1.679 | 0.748 | 0.71689736 | NA | 1.8421 | 2.17 | 2.1184 | 2.1184 | TRUE | 1.4382 |
| Spag4 | 117 | 108 | 104 | 136 | 140 | 144 | 91 | 115 | 112 | 135 | 158 | 142 | 125.57 | 106 | 145 | 1.375 | 0.46 | 0.01245037 | 0.04863436 | 0 | 0.0213 | 0.0162 | 0.0162 | TRUE | 0.1753 |
| Spag5 | 10017 | 7361 | 7328 | 8297 | 7538 | 8468 | 7780 | 7830 | 7907 | 8259 | 8504 | 8359 | 8106.35 | 7839 | 8374 | 1.068 | 0.095 | 0.04120034 | 0.11677105 | 0 | 0.0027 | 0.0014 | 0.0014 | TRUE | 0.0022 |
| Spag6 | 29 | 22 | 24 | 25 | 28 | 19 | 23 | 23 | 26 | 25 | 32 | 19 | 24.51 | 24 | 25 | 1.045 | 0.064 | 0.87074188 | 0.92839601 | 0 | 0.0991 | 0.0709 | 0.0709 | TRUE | 0.1903 |
| Spag7 | 5031 | 3789 | 4130 | 4555 | 3899 | 4654 | 3907 | 4030 | 4456 | 4534 | 4398 | 4594 | 4320.11 | 4131 | 4509 | 1.092 | 0.127 | 0.04813525 | 0.13099716 | 0.0023 | 0.003 | 0.0027 | 0.0027 | TRUE | 0.0573 |
| Spag8 | 62 | 46 | 39 | 37 | 44 | 57 | 48 | 49 | 42 | 37 | 50 | 56 | 46.98 | 46 | 48 | 1.024 | 0.034 | 0.90773913 | 0.94923397 | 0 | 0.0529 | 0.0397 | 0.0397 | TRUE | 0.3219 |
| Spag9 | 6621 | 5016 | 5820 | 6268 | 5652 | 6375 | 5142 | 5335 | 6280 | 6239 | 6376 | 6293 | 5944.29 | 5586 | 6303 | 1.129 | 0.175 | 0.01955284 | 0.06798072 | 0.0057 | 0.0028 | 0.0039 | 0.0039 | TRUE | 0.1436 |
| Spam1 | 0 | 0 | 0 | 0 | 0 | 0 | 0 | 0 | 0 | 0 | 0 | 0 | 0 | 0 | 0 | NA | NA | NA | NA | NA | NA | NA | NA | NA | NA |
| Sparc | 17251 | 11444 | 11160 | 12582 | 10433 | 11650 | 13398 | 12173 | 12042 | 12524 | 11770 | 11500 | 12234.41 | 12538 | 11931 | 0.951 | -0.072 | 0.24244896 | 0.41019448 | 0.0026 | 0.0026 | 0.0026 | 0.0026 | TRUE | 0.0443 |
| Sparcl1 | 2 | 7 | 6 | 2 | 1 | 1 | 2 | 7 | 6 | 2 | 1 | 1 | 3.26 | 5 | 1 | 0.271 | -1.882 | 0.11605595 | NA | 0 | 0.7287 | 0.5612 | 0.5612 | TRUE | 1.035 |
| Spast | 4438 | 3142 | 3071 | 3351 | 2913 | 3229 | 3447 | 3342 | 3314 | 3336 | 3286 | 3188 | 3318.63 | 3368 | 3270 | 0.971 | -0.043 | 0.43608756 | 0.60732725 | 2.00E-04 | 0.0031 | 0.0019 | 0.0019 | TRUE | 0.0059 |
| Spata1 | 45 | 33 | 38 | 36 | 26 | 30 | 35 | 35 | 41 | 36 | 29 | 30 | 34.31 | 37 | 32 | 0.856 | -0.224 | 0.49367889 | 0.65889928 | 0 | 0.0715 | 0.0476 | 0.0476 | TRUE | 0.0957 |
| Spata13 | 2098 | 1679 | 1672 | 1569 | 1659 | 1961 | 1629 | 1786 | 1804 | 1562 | 1872 | 1936 | 1764.76 | 1740 | 1790 | 1.029 | 0.041 | 0.63908551 | 0.77456433 | 0.0075 | 0.0038 | 0.0051 | 0.0051 | TRUE | 0.1508 |
| Spata16 | 23 | 13 | 13 | 10 | 9 | 7 | 18 | 14 | 14 | 10 | 10 | 7 | 12.12 | 15 | 9 | 0.586 | -0.771 | 0.16370016 | NA | 0 | 0.1979 | 0.1319 | 0.1319 | TRUE | 0.1389 |
| Spata17 | 31 | 15 | 9 | 13 | 14 | 9 | 24 | 16 | 10 | 13 | 16 | 9 | 14.56 | 17 | 13 | 0.743 | -0.429 | 0.42696455 | NA | 0.0793 | 0.1652 | 0.1403 | 0.1403 | TRUE | 0.2239 |
| Spata18 | 16 | 16 | 23 | 13 | 12 | 9 | 12 | 17 | 25 | 13 | 14 | 9 | 14.94 | 18 | 12 | 0.657 | -0.607 | 0.24322957 | NA | 0 | 0.1611 | 0.124 | 0.124 | TRUE | 0.2807 |
| Spata19 | 0 | 0 | 0 | 0 | 1 | 0 | 0 | 0 | 0 | 0 | 1 | 0 | 0.19 | 0 | 0 | 2.094 | 1.067 | 0.79379329 | NA | 0 | 12.608 | 10 | 10 | TRUE | 0.5075 |
| Spata2 | 2311 | 1619 | 1450 | 1577 | 1377 | 1531 | 1795 | 1722 | 1565 | 1570 | 1553 | 1511 | 1619.33 | 1694 | 1545 | 0.911 | -0.134 | 0.06600775 | 0.16424106 | 0.0021 | 0.0039 | 0.0032 | 0.0032 | TRUE | 0.0532 |
| Spata20 | 25 | 13 | 6 | 25 | 12 | 13 | 19 | 14 | 6 | 25 | 14 | 13 | 15.16 | 13 | 17 | 1.273 | 0.348 | 0.52700207 | NA | 0.1347 | 0.1587 | 0.1521 | 0.1521 | TRUE | 0.3222 |
| Spata21 | 50 | 55 | 58 | 62 | 59 | 80 | 39 | 59 | 63 | 62 | 67 | 79 | 61.19 | 54 | 69 | 1.309 | 0.389 | 0.15104755 | 0.29712669 | 0.0234 | 0.0412 | 0.0362 | 0.0362 | TRUE | 0.5512 |
| Spata22 | 63 | 61 | 43 | 33 | 26 | 31 | 49 | 65 | 46 | 33 | 29 | 31 | 42.17 | 53 | 31 | 0.581 | -0.783 | 0.01080933 | 0.04374283 | 0 | 0.0586 | 0.0423 | 0.0423 | TRUE | 0.28 |
| Spata24 | 375 | 281 | 279 | 279 | 294 | 301 | 291 | 299 | 301 | 278 | 332 | 297 | 299.62 | 297 | 302 | 1.017 | 0.024 | 0.84705766 | 0.91321597 | 0 | 0.0103 | 0.0075 | 0.0075 | TRUE | 0.0796 |
| Spata25 | 0 | 0 | 0 | 2 | 1 | 1 | 0 | 0 | 0 | 2 | 1 | 1 | 0.68 | 0 | 1 | 7.743 | 2.953 | 0.21812153 | NA | 0 | 3.4657 | 2.7652 | 2.7652 | TRUE | 0.1019 |
| Spata2l | 29 | 21 | 17 | 25 | 13 | 32 | 23 | 22 | 18 | 25 | 15 | 32 | 22.39 | 21 | 24 | 1.129 | 0.175 | 0.68196515 | 0.80494909 | 0.0268 | 0.1083 | 0.0868 | 0.0868 | TRUE | 0.2827 |
| Spata3 | 1 | 1 | 0 | 1 | 2 | 1 | 1 | 1 | 0 | 1 | 2 | 1 | 1.01 | 1 | 1 | 2.246 | 1.167 | 0.54025274 | NA | 0 | 2.3416 | 1.4771 | 1.4771 | TRUE | 0.1884 |
| Spata31 | 0 | 0 | 0 | 0 | 0 | 0 | 0 | 0 | 0 | 0 | 0 | 0 | 0 | 0 | 0 | NA | NA | NA | NA | NA | NA | NA | NA | NA | NA |
| Spata31d1a | 234 | 187 | 125 | 174 | 132 | 166 | 182 | 199 | 135 | 173 | 149 | 164 | 166.92 | 172 | 162 | 0.942 | -0.086 | 0.6247901 | 0.76346484 | 0.016 | 0.0166 | 0.0164 | 0.0164 | TRUE | 0.3588 |
| Spata31d1b | 150 | 109 | 89 | 181 | 132 | 180 | 116 | 116 | 96 | 180 | 149 | 178 | 139.21 | 109 | 169 | 1.541 | 0.624 | 0.00041052 | 0.00371368 | 0 | 0.0195 | 0.015 | 0.015 | TRUE | 0.1096 |
| Spata31d1c | 3 | 4 | 2 | 5 | 4 | 2 | 2 | 4 | 2 | 5 | 5 | 2 | 3.37 | 3 | 4 | 1.318 | 0.398 | 0.70095923 | NA | 0 | 0.7062 | 0.471 | 0.471 | TRUE | 0.301 |
| Spata31d1d | 4 | 5 | 4 | 3 | 0 | 5 | 3 | 5 | 4 | 3 | 0 | 5 | 3.44 | 4 | 3 | 0.638 | -0.648 | 0.55448678 | NA | 0.0165 | 0.6906 | 0.5594 | 0.5594 | TRUE | 0.4461 |
| Spata32 | 5 | 4 | 6 | 5 | 3 | 3 | 4 | 4 | 6 | 5 | 3 | 3 | 4.32 | 5 | 4 | 0.782 | -0.354 | 0.69609641 | NA | 0 | 0.5507 | 0.3546 | 0.3546 | TRUE | 0.1517 |

| | | | | | | | | | | | | | | | | | | | | | | | | | |
|---|---|---|---|---|---|---|---|---|---|---|---|---|---|---|---|---|---|---|---|---|---|---|---|---|---|
| Spata4 | 2 | 3 | 3 | 9 | 0 | 3 | 2 | 3 | 3 | 9 | 0 | 3 | 3.32 | 3 | 4 | 1.533 | 0.616 | 0.60469847 | NA | 0.6822 | 0.717 | 0.7084 | 0.7084 | TRUE | 0.3685 |
| Spata5 | 4552 | 3276 | 3078 | 3252 | 2817 | 3429 | 3535 | 3485 | 3321 | 3237 | 3178 | 3385 | 3356.83 | 3447 | 3267 | 0.948 | -0.078 | 0.18236095 | 0.33796172 | 8.00E-04 | 0.0031 | 0.0022 | 0.0022 | TRUE | 0.0123 |
| Spata5l1 | 343 | 297 | 268 | 342 | 317 | 347 | 266 | 316 | 289 | 340 | 358 | 343 | 318.68 | 290 | 347 | 1.197 | 0.259 | 0.03036154 | 0.09340453 | 0 | 0.0099 | 0.0072 | 0.0072 | TRUE | 0.066 |
| Spata6 | 1162 | 802 | 912 | 925 | 810 | 1002 | 902 | 853 | 984 | 921 | 914 | 989 | 927.21 | 913 | 941 | 1.031 | 0.044 | 0.59983233 | 0.74551335 | 0.0024 | 0.005 | 0.0041 | 0.0041 | TRUE | 0.0528 |
| Spata7 | 729 | 586 | 568 | 550 | 516 | 543 | 566 | 623 | 613 | 547 | 582 | 536 | 577.99 | 601 | 555 | 0.925 | -0.113 | 0.23041555 | 0.39614588 | 0 | 0.0065 | 0.0046 | 0.0046 | TRUE | 0.0319 |
| Spata9 | 1 | 0 | 0 | 0 | 0 | 0 | 1 | 0 | 0 | 0 | 0 | 0 | 0.13 | 0 | 0 | 0.552 | -0.857 | 0.8336548 | NA | 0 | 18.3121 | 10 | 10 | TRUE | 0.2566 |
| Spatc1 | 5 | 7 | 14 | 12 | 13 | 19 | 4 | 7 | 15 | 12 | 15 | 19 | 11.97 | 9 | 15 | 1.758 | 0.814 | 0.18511204 | NA | 0.1279 | 0.2005 | 0.1816 | 0.1816 | TRUE | 0.8812 |
| Spatc1l | 1 | 0 | 0 | 0 | 1 | 0 | 1 | 0 | 0 | 0 | 1 | 0 | 0.32 | 0 | 0 | 1.075 | 0.105 | 0.97638847 | NA | 0 | 7.4681 | 7.0392 | 7.0392 | TRUE | 0.5075 |
| Spats1 | 184 | 131 | 130 | 139 | 128 | 143 | 143 | 139 | 140 | 138 | 144 | 141 | 141.07 | 141 | 141 | 1.002 | 0.003 | 0.98294179 | 0.99091459 | 0 | 0.0192 | 0.012 | 0.012 | TRUE | 0.0036 |
| Spats2 | 3022 | 2236 | 2341 | 2365 | 2102 | 2491 | 2347 | 2378 | 2526 | 2354 | 2371 | 2459 | 2405.96 | 2417 | 2395 | 0.991 | -0.013 | 0.83748533 | 0.9073128 | 6.00E-04 | 0.0034 | 0.0023 | 0.0023 | TRUE | 0.0186 |
| Spats2l | 158 | 126 | 129 | 96 | 82 | 91 | 123 | 134 | 139 | 96 | 93 | 90 | 112.3 | 132 | 93 | 0.704 | -0.507 | 0.00617685 | 0.02883271 | 0 | 0.0235 | 0.0155 | 0.0155 | TRUE | 0.0431 |
| Spc24 | 2996 | 2342 | 2424 | 2706 | 2403 | 2802 | 2327 | 2491 | 2615 | 2694 | 2711 | 2766 | 2600.65 | 2478 | 2724 | 1.1 | 0.138 | 0.03240652 | 0.09797329 | 0.0015 | 0.0033 | 0.0026 | 0.0026 | TRUE | 0.0355 |
| Spc25 | 4914 | 3455 | 3599 | 3982 | 3279 | 3827 | 3816 | 3675 | 3883 | 3964 | 3699 | 3778 | 3802.57 | 3791 | 3814 | 1.006 | 0.008 | 0.88252966 | 0.93457779 | 7.00E-04 | 0.003 | 0.0021 | 0.0021 | TRUE | 0.0145 |
| Spcs1 | 1710 | 1197 | 1320 | 1313 | 1079 | 1248 | 1328 | 1273 | 1424 | 1307 | 1217 | 1232 | 1296.96 | 1342 | 1252 | 0.934 | -0.099 | 0.18543777 | 0.3417746 | 0.0015 | 0.0043 | 0.0033 | 0.0033 | TRUE | 0.0336 |
| Spcs2 | 7235 | 5198 | 5157 | 5312 | 4734 | 5405 | 5619 | 5529 | 5564 | 5288 | 5340 | 5336 | 5446.02 | 5571 | 5321 | 0.955 | -0.066 | 0.16937995 | 0.32083515 | 0 | 0.0029 | 0.0015 | 0.0015 | TRUE | 7.00E-04 |
| Spcs3 | 14455 | 10555 | 11616 | 12942 | 10907 | 12674 | 11227 | 11227 | 12534 | 12882 | 12304 | 12511 | 12114.21 | 11663 | 12566 | 1.078 | 0.108 | 0.06900853 | 0.16981668 | 0.0022 | 0.0026 | 0.0025 | 0.0025 | TRUE | 0.052 |
| Spdef | 32 | 35 | 34 | 44 | 49 | 49 | 25 | 37 | 37 | 44 | 55 | 48 | 41.04 | 33 | 49 | 1.506 | 0.59 | 0.05988044 | 0.15299806 | 0 | 0.0602 | 0.0456 | 0.0456 | TRUE | 0.3903 |
| Spdl1 | 4854 | 3428 | 3048 | 3178 | 2826 | 3215 | 3770 | 3646 | 3289 | 3163 | 3188 | 3174 | 3371.68 | 3568 | 3175 | 0.889 | -0.169 | 0.00961593 | 0.04022256 | 0.002 | 0.0031 | 0.0028 | 0.0028 | TRUE | 0.0567 |
| Spdya | 47 | 23 | 33 | 40 | 20 | 31 | 37 | 24 | 36 | 40 | 23 | 31 | 31.59 | 32 | 31 | 0.962 | -0.055 | 0.87968809 | 0.93295109 | 0.0271 | 0.0774 | 0.0641 | 0.0641 | TRUE | 0.2275 |
| Spdyb | 4 | 6 | 2 | 3 | 2 | 4 | 3 | 6 | 2 | 3 | 2 | 4 | 3.47 | 4 | 3 | 0.798 | -0.325 | 0.75093367 | NA | 0 | 0.6849 | 0.4629 | 0.4629 | TRUE | 0.4749 |
| Specc1 | 2640 | 1900 | 1759 | 2128 | 1719 | 2072 | 2050 | 2021 | 1898 | 2118 | 1939 | 2045 | 2012.02 | 1990 | 2034 | 1.022 | 0.032 | 0.63609906 | 0.77255774 | 0.0013 | 0.0036 | 0.0027 | 0.0027 | TRUE | 0.0199 |
| Specc1l | 5623 | 4113 | 3882 | 3797 | 3500 | 3964 | 4367 | 4375 | 4189 | 3780 | 3948 | 3913 | 4095.28 | 4310 | 3880 | 0.9 | -0.152 | 0.00482909 | 0.02386661 | 3.00E-04 | 0.003 | 0.0018 | 0.0018 | TRUE | 0.0073 |
| Speer1-ps1 | 0 | 0 | 0 | 0 | 0 | 0 | 0 | 0 | 0 | 0 | 0 | 0 | 0 | 0 | 0 | NA | NA | NA | NA | NA | NA | NA | NA | NA | NA |
| Speer2 | 0 | 0 | 0 | 0 | 0 | 0 | 0 | 0 | 0 | 0 | 0 | 0 | 0 | 0 | 0 | NA | NA | NA | NA | NA | NA | NA | NA | NA | NA |
| Speer3 | 0 | 0 | 0 | 0 | 0 | 0 | 0 | 0 | 0 | 0 | 0 | 0 | 0 | 0 | 0 | NA | NA | NA | NA | NA | NA | NA | NA | NA | NA |
| Speer4a | 2 | 0 | 1 | 0 | 0 | 1 | 2 | 0 | 1 | 0 | 0 | 1 | 0.6 | 1 | 0 | 0.414 | -1.271 | 0.61985863 | NA | 0 | 3.9313 | 3.1876 | 3.1876 | TRUE | 0.3835 |
| Speer4b | 10 | 4 | 6 | 1 | 3 | 1 | 8 | 4 | 6 | 1 | 3 | 1 | 3.98 | 6 | 2 | 0.282 | -1.828 | 0.08270059 | NA | 0 | 0.5984 | 0.4257 | 0.4257 | TRUE | 0.4327 |
| Speer4d | 0 | 1 | 0 | 0 | 0 | 0 | 0 | 1 | 0 | 0 | 0 | 0 | 0.18 | 0 | 0 | 0.552 | -0.857 | 0.8336548 | NA | 0 | 13.3714 | 10 | 10 | TRUE | 0.5216 |
| Speer4e | 0 | 0 | 0 | 0 | 1 | 0 | 0 | 0 | 0 | 0 | 1 | 0 | 0.19 | 0 | 0 | 2.094 | 1.067 | 0.79379329 | NA | 0 | 12.608 | 10 | 10 | TRUE | 0.5075 |
| Speer4f | 1 | 2 | 0 | 0 | 0 | 0 | 1 | 2 | 0 | 0 | 0 | 0 | 0.48 | 1 | 0 | 0.197 | -2.343 | 0.43249452 | NA | 0 | 4.8992 | 4.9172 | 4.9172 | TRUE | 0.1226 |
| Speer5-ps1 | 20 | 22 | 25 | 6 | 5 | 7 | 16 | 23 | 27 | 6 | 6 | 7 | 14.07 | 22 | 6 | 0.284 | -1.816 | 0.00131946 | NA | 0 | 0.1709 | 0.1246 | 0.1246 | TRUE | 0.3586 |
| Speer6-ps1 | 0 | 0 | 0 | 0 | 0 | 0 | 0 | 0 | 0 | 0 | 0 | 0 | 0 | 0 | 0 | NA | NA | NA | NA | NA | NA | NA | NA | NA | NA |
| Speer7-ps1 | 15 | 14 | 4 | 2 | 1 | 4 | 12 | 15 | 4 | 2 | 1 | 4 | 6.32 | 10 | 2 | 0.231 | -2.112 | 0.01896667 | NA | 0 | 0.3774 | 0.3206 | 0.3206 | TRUE | 0.25 |
| Speer8-ps1 | 9 | 13 | 2 | 5 | 2 | 5 | 7 | 14 | 2 | 5 | 2 | 5 | 5.86 | 8 | 4 | 0.535 | -0.904 | 0.30388991 | NA | 0.2672 | 0.407 | 0.3669 | 0.3669 | TRUE | 0.1717 |
| Speer9-ps1 | 4 | 2 | 1 | 2 | 0 | 1 | 3 | 2 | 1 | 2 | 0 | 1 | 1.55 | 2 | 1 | 0.474 | -1.077 | 0.49588817 | NA | 0 | 1.533 | 1.0642 | 1.0642 | TRUE | 0.3114 |
| Spef1 | 88 | 48 | 69 | 47 | 38 | 61 | 68 | 51 | 74 | 47 | 43 | 60 | 57.29 | 64 | 50 | 0.774 | -0.37 | 0.17664686 | 0.33042972 | 0.0158 | 0.0438 | 0.0362 | 0.0362 | TRUE | 0.2796 |
| Spef2 | 21 | 17 | 11 | 14 | 12 | 20 | 16 | 18 | 12 | 14 | 14 | 20 | 15.58 | 15 | 16 | 1.021 | 0.029 | 0.95211828 | NA | 0 | 0.1546 | 0.1077 | 0.1077 | TRUE | 0.2394 |
| Speg | 404 | 321 | 331 | 473 | 368 | 491 | 314 | 341 | 357 | 471 | 415 | 485 | 397.17 | 337 | 457 | 1.358 | 0.442 | 0.00010129 | 0.00122375 | 0.0031 | 0.0084 | 0.0067 | 0.0067 | TRUE | 0.072 |
| Spem1 | 3 | 2 | 3 | 6 | 3 | 1 | 2 | 2 | 3 | 6 | 3 | 1 | 3.01 | 2 | 3 | 1.35 | 0.433 | 0.6995914 | NA | 0 | 0.7908 | 0.5664 | 0.5664 | TRUE | 0.1651 |
| Spen | 6141 | 4165 | 3591 | 3815 | 3568 | 4067 | 4769 | 4430 | 3875 | 3797 | 4025 | 4015 | 4151.94 | 4358 | 3946 | 0.905 | -0.144 | 0.06118539 | 0.15549836 | 0.0055 | 0.003 | 0.004 | 0.004 | TRUE | 0.1143 |
| Spert | 0 | 0 | 0 | 0 | 0 | 0 | 0 | 0 | 0 | 0 | 0 | 0 | 0 | 0 | 0 | NA | NA | NA | NA | NA | NA | NA | NA | NA | NA |
| Spesp1 | 257 | 222 | 254 | 198 | 181 | 213 | 200 | 236 | 274 | 197 | 204 | 210 | 220.22 | 237 | 204 | 0.865 | -0.209 | 0.17025945 | 0.32193655 | 0.0098 | 0.0132 | 0.0121 | 0.0121 | TRUE | 0.1771 |
| Spg11 | 2196 | 1769 | 2086 | 2216 | 2018 | 2170 | 1706 | 1882 | 2251 | 2206 | 2277 | 2142 | 2077.07 | 1946 | 2208 | 1.136 | 0.184 | 0.04793812 | 0.13070958 | 0.01 | 0.0036 | 0.0057 | 0.0057 | TRUE | 0.2263 |
| Spg20 | 2570 | 1800 | 1731 | 1680 | 1461 | 1544 | 1996 | 1915 | 1868 | 1672 | 1648 | 1524 | 1770.5 | 1926 | 1615 | 0.838 | -0.255 | 0.00019249 | 0.00207325 | 0.0012 | 0.0038 | 0.0028 | 0.0028 | TRUE | 0.0295 |
| Spg21 | 4403 | 3355 | 3664 | 3711 | 3257 | 3855 | 3420 | 3569 | 3953 | 3694 | 3674 | 3805 | 3685.89 | 3647 | 3724 | 1.022 | 0.031 | 0.64065354 | 0.77554511 | 0.0027 | 0.0031 | 0.0029 | 0.0029 | TRUE | 0.0653 |
| Spg7 | 6004 | 4298 | 4159 | 4945 | 4265 | 4702 | 4663 | 4572 | 4488 | 4922 | 4811 | 4642 | 4682.92 | 4574 | 4792 | 1.047 | 0.067 | 0.21103352 | 0.37307385 | 4.00E-04 | 0.0029 | 0.0018 | 0.0018 | TRUE | 0.0092 |
| Sphk1 | 235 | 154 | 184 | 257 | 232 | 254 | 183 | 164 | 199 | 256 | 262 | 251 | 218.86 | 182 | 256 | 1.41 | 0.495 | 0.00037699 | 0.00348081 | 0 | 0.0133 | 0.0093 | 0.0093 | TRUE | 0.0732 |
| Sphk2 | 1458 | 1062 | 782 | 843 | 773 | 848 | 1132 | 1130 | 844 | 839 | 872 | 837 | 942.34 | 1035 | 849 | 0.819 | -0.287 | 0.00865122 | 0.03716619 | 0.013 | 0.0049 | 0.0076 | 0.0076 | TRUE | 0.3082 |
| Sphkap | 6 | 2 | 4 | 9 | 1 | 5 | 5 | 2 | 4 | 9 | 1 | 5 | 4.35 | 4 | 5 | 1.354 | 0.437 | 0.65356802 | NA | 0.195 | 0.5467 | 0.4523 | 0.4523 | TRUE | 0.1496 |
| Spib | 9 | 8 | 7 | 15 | 11 | 13 | 7 | 9 | 8 | 15 | 12 | 13 | 10.54 | 8 | 13 | 1.752 | 0.809 | 0.16662276 | NA | 0 | 0.2273 | 0.145 | 0.145 | TRUE | 0.0437 |
| Spic | 103 | 81 | 94 | 80 | 54 | 75 | 80 | 86 | 101 | 80 | 61 | 74 | 80.36 | 89 | 72 | 0.807 | -0.31 | 0.17054887 | 0.32218543 | 0 | 0.0319 | 0.0242 | 0.0242 | TRUE | 0.1425 |
| Spice1 | 2889 | 2357 | 2333 | 2372 | 2196 | 2541 | 2244 | 2507 | 2517 | 2361 | 2477 | 2508 | 2435.82 | 2423 | 2449 | 1.012 | 0.017 | 0.80668478 | 0.88831923 | 0.0023 | 0.0034 | 0.003 | 0.003 | TRUE | 0.0521 |
| Spin1 | 7899 | 5735 | 6039 | 6614 | 5792 | 6693 | 6135 | 6100 | 6516 | 6584 | 6534 | 6607 | 6412.61 | 6250 | 6575 | 1.052 | 0.073 | 0.15794736 | 0.30687686 | 5.00E-04 | 0.0028 | 0.0018 | 0.0018 | TRUE | 0.0167 |
| Spin2 | 72 | 44 | 37 | 27 | 25 | 23 | 56 | 47 | 40 | 27 | 28 | 23 | 36.74 | 48 | 26 | 0.541 | -0.886 | 0.00716049 | 0.03228547 | 0 | 0.0669 | 0.048 | 0.048 | TRUE | 0.2013 |
| Spin4 | 236 | 216 | 194 | 231 | 205 | 217 | 183 | 230 | 209 | 230 | 231 | 214 | 216.3 | 207 | 225 | 1.089 | 0.123 | 0.39438402 | 0.56800996 | 0.0032 | 0.0134 | 0.0103 | 0.0103 | TRUE | 0.125 |
| Spink10 | 9 | 8 | 3 | 7 | 9 | 8 | 7 | 9 | 3 | 7 | 10 | 8 | 7.29 | 6 | 8 | 1.323 | 0.404 | 0.57235072 | NA | 0 | 0.3274 | 0.2299 | 0.2299 | TRUE | 0.3799 |
| Spink11 | 0 | 0 | 0 | 0 | 0 | 0 | 0 | 0 | 0 | 0 | 0 | 0 | 0 | 0 | 0 | NA | NA | NA | NA | NA | NA | NA | NA | NA | NA |
| Spink12 | 0 | 0 | 0 | 0 | 0 | 0 | 0 | 0 | 0 | 0 | 0 | 0 | 0 | 0 | 0 | NA | NA | NA | NA | NA | NA | NA | NA | NA | NA |
| Spink13 | 0 | 0 | 0 | 0 | 0 | 0 | 0 | 0 | 0 | 0 | 0 | 0 | 0 | 0 | 0 | NA | NA | NA | NA | NA | NA | NA | NA | NA | NA |
| Spink14 | 0 | 0 | 0 | 0 | 0 | 0 | 0 | 0 | 0 | 0 | 0 | 0 | 0 | 0 | 0 | NA | NA | NA | NA | NA | NA | NA | NA | NA | NA |
| Spink2 | 116 | 84 | 90 | 78 | 93 | 80 | 90 | 89 | 97 | 78 | 105 | 79 | 89.68 | 92 | 87 | 0.944 | -0.084 | 0.6957677 | 0.81479785 | 0 | 0.0289 | 0.0219 | 0.0219 | TRUE | 0.2858 |
| Spink3 | 4 | 2 | 2 | 2 | 0 | 1 | 3 | 2 | 2 | 2 | 0 | 1 | 1.73 | 2 | 1 | 0.408 | -1.292 | 0.39038235 | NA | 0 | 1.3737 | 0.9261 | 0.9261 | TRUE | 0.3388 |
| Spink4 | 0 | 0 | 0 | 0 | 0 | 0 | 0 | 0 | 0 | 0 | 0 | 0 | 0 | 0 | 0 | NA | NA | NA | NA | NA | NA | NA | NA | NA | NA |
| Spink5 | 1 | 2 | 2 | 1 | 1 | 2 | 1 | 2 | 2 | 1 | 1 | 2 | 1.53 | 2 | 1 | 0.831 | -0.268 | 0.85824055 | NA | 0 | 1.5549 | 0.9496 | 0.9496 | TRUE | 0.2415 |
| Spink6 | 0 | 0 | 0 | 0 | 0 | 0 | 0 | 0 | 0 | 0 | 0 | 0 | 0 | 0 | 0 | NA | NA | NA | NA | NA | NA | NA | NA | NA | NA |
| Spink7 | 0 | 1 | 0 | 0 | 0 | 0 | 0 | 1 | 0 | 0 | 0 | 0 | 0.18 | 0 | 0 | 0.552 | -0.857 | 0.8336548 | NA | 0 | 13.3714 | 10 | 10 | TRUE | 0.5216 |
| Spink8 | 0 | 0 | 0 | 0 | 0 | 0 | 0 | 0 | 0 | 0 | 0 | 0 | 0 | 0 | 0 | NA | NA | NA | NA | NA | NA | NA | NA | NA | NA |
| Spinkl | 0 | 0 | 0 | 0 | 0 | 0 | 0 | 0 | 0 | 0 | 0 | 0 | 0 | 0 | 0 | NA | NA | NA | NA | NA | NA | NA | NA | NA | NA |
| Spint1 | 3011 | 2212 | 2245 | 2303 | 2114 | 2362 | 2339 | 2353 | 2422 | 2292 | 2385 | 2332 | 2353.77 | 2371 | 2336 | 0.985 | -0.021 | 0.71506051 | 0.82883463 | 0 | 0.0034 | 0.002 | 0.002 | TRUE | 0.0042 |

| | | | | | | | | | | | | | | | | | | | | | | | | | |
|---|---|---|---|---|---|---|---|---|---|---|---|---|---|---|---|---|---|---|---|---|---|---|---|---|---|
| Spint2 | 8561 | 6442 | 7189 | 8234 | 7548 | 8528 | 6649 | 6852 | 7757 | 8196 | 8515 | 8418 | 7731.27 | 7086 | 8376 | 1.182 | 0.242 | 0.00024856 | 0.00252137 | 0.0035 | 0.0027 | 0.003 | 0.003 | TRUE | 0.0835 |
| Spint3 | 0 | 0 | 0 | 0 | 0 | 0 | 0 | 0 | 0 | 0 | 0 | 0 | 0 | 0 | 0 | NA | NA | NA | NA | NA | NA | NA | NA | NA | NA |
| Spint4 | 0 | 0 | 0 | 0 | 0 | 0 | 0 | 0 | 0 | 0 | 0 | 0 | 0 | 0 | 0 | NA | NA | NA | NA | NA | NA | NA | NA | NA | NA |
| Spint5 | 12 | 5 | 2 | 1 | 1 | 2 | 9 | 5 | 2 | 1 | 1 | 2 | 3.48 | 5 | 1 | 0.241 | -2.053 | 0.07852441 | NA | 0 | 0.683 | 0.5194 | 0.5194 | TRUE | 0.1099 |
| Spire1 | 4037 | 3072 | 3328 | 3932 | 3331 | 3904 | 3135 | 3268 | 3591 | 3914 | 3758 | 3854 | 3586.56 | 3331 | 3842 | 1.154 | 0.207 | 0.00156501 | 0.0099616 | 0.0023 | 0.0031 | 0.0028 | 0.0028 | TRUE | 0.0562 |
| Spire2 | 2058 | 1646 | 1638 | 1725 | 1651 | 1970 | 1598 | 1751 | 1767 | 1717 | 1862 | 1945 | 1773.47 | 1705 | 1841 | 1.081 | 0.112 | 0.13569151 | 0.27513672 | 0.003 | 0.0038 | 0.0035 | 0.0035 | TRUE | 0.0426 |
| Spn | 948 | 703 | 742 | 1014 | 927 | 1057 | 736 | 748 | 801 | 1009 | 1046 | 1043 | 897.19 | 762 | 1033 | 1.357 | 0.441 | 2.19E-08 | 1.01E-06 | 0 | 0.0051 | 0.0033 | 0.0033 | TRUE | 0.0233 |
| Spn-ps | 0 | 0 | 0 | 0 | 0 | 0 | 0 | 0 | 0 | 0 | 0 | 0 | 0 | 0 | 0 | NA | NA | NA | NA | NA | NA | NA | NA | NA | NA |
| Spns1 | 3358 | 2453 | 2063 | 2070 | 1972 | 2202 | 2608 | 2609 | 2226 | 2060 | 2225 | 2174 | 2317 | 2481 | 2153 | 0.867 | -0.206 | 0.00744613 | 0.03326859 | 0.0041 | 0.0034 | 0.0038 | 0.0038 | TRUE | 0.0969 |
| Spns2 | 306 | 235 | 220 | 309 | 254 | 295 | 238 | 250 | 237 | 308 | 287 | 291 | 268.39 | 242 | 295 | 1.222 | 0.29 | 0.01888598 | 0.06633997 | 0 | 0.0113 | 0.0072 | 0.0072 | TRUE | 0.0155 |
| Spns3 | 29 | 21 | 16 | 30 | 17 | 22 | 23 | 22 | 17 | 30 | 19 | 22 | 22.15 | 21 | 24 | 1.138 | 0.186 | 0.65206188 | 0.78392575 | 0 | 0.1094 | 0.078 | 0.078 | TRUE | 0.3241 |
| Spo11 | 3 | 1 | 1 | 0 | 1 | 1 | 2 | 1 | 1 | 0 | 1 | 1 | 1.1 | 1 | 1 | 0.459 | -1.125 | 0.54242351 | NA | 0 | 2.161 | 1.3949 | 1.3949 | TRUE | 0.2656 |
| Spock1 | 207 | 184 | 179 | 224 | 184 | 207 | 161 | 196 | 193 | 223 | 208 | 204 | 197.42 | 183 | 212 | 1.16 | 0.214 | 0.15003391 | 0.29566175 | 0 | 0.0144 | 0.0109 | 0.0109 | TRUE | 0.137 |
| Spock2 | 902 | 692 | 739 | 857 | 819 | 902 | 701 | 736 | 797 | 853 | 924 | 890 | 816.9 | 745 | 889 | 1.195 | 0.257 | 0.00276969 | 0.01551224 | 0.0017 | 0.0053 | 0.0041 | 0.0041 | TRUE | 0.045 |
| Spock3 | 65 | 53 | 59 | 52 | 51 | 42 | 50 | 56 | 64 | 52 | 58 | 41 | 53.55 | 57 | 50 | 0.885 | -0.176 | 0.51830699 | 0.68007354 | 0 | 0.0467 | 0.0343 | 0.0343 | TRUE | 0.1946 |
| Spon1 | 18 | 30 | 22 | 15 | 16 | 18 | 14 | 32 | 24 | 15 | 18 | 18 | 20.06 | 23 | 17 | 0.737 | -0.44 | 0.33863371 | 0.51283023 | 0.0556 | 0.1206 | 0.1009 | 0.1009 | TRUE | 0.2002 |
| Spon2 | 119 | 80 | 76 | 106 | 68 | 103 | 92 | 85 | 82 | 106 | 77 | 102 | 90.57 | 86 | 95 | 1.094 | 0.13 | 0.54411641 | 0.70164704 | 0 | 0.0286 | 0.022 | 0.022 | TRUE | 0.2504 |
| Spop | 5033 | 3439 | 3366 | 3808 | 3190 | 3989 | 3909 | 3658 | 3632 | 3790 | 3599 | 3938 | 3754.29 | 3733 | 3776 | 1.011 | 0.016 | 0.79426616 | 0.87961466 | 0.0016 | 0.0031 | 0.0025 | 0.0025 | TRUE | 0.0211 |
| Spopl | 843 | 537 | 544 | 638 | 495 | 592 | 655 | 571 | 587 | 635 | 558 | 584 | 598.46 | 604 | 592 | 0.979 | -0.03 | 0.7604989 | 0.85778178 | 0.0033 | 0.0064 | 0.0054 | 0.0054 | TRUE | 0.066 |
| Spp1 | 11754 | 7920 | 7394 | 4992 | 4761 | 5530 | 9129 | 8424 | 7978 | 4969 | 5371 | 5459 | 6888.37 | 8510 | 5266 | 0.619 | -0.693 | 2.18E-25 | 1.15E-22 | 0.0033 | 0.0028 | 0.003 | 0.003 | TRUE | 0.0497 |
| Spp2 | 0 | 0 | 0 | 0 | 0 | 1 | 0 | 0 | 0 | 0 | 0 | 1 | 0.16 | 0 | 0 | 2.094 | 1.067 | 0.79379329 | NA | 0 | 14.4078 | 10 | 10 | TRUE | 0.3835 |
| Sppl2a | 1972 | 1393 | 1461 | 1724 | 1402 | 1563 | 1532 | 1482 | 1576 | 1716 | 1582 | 1543 | 1571.71 | 1530 | 1614 | 1.055 | 0.077 | 0.27845616 | 0.44996878 | 0.0014 | 0.0039 | 0.003 | 0.003 | TRUE | 0.0376 |
| Sppl2b | 564 | 394 | 294 | 338 | 264 | 286 | 438 | 419 | 317 | 336 | 298 | 282 | 348.49 | 391 | 305 | 0.779 | -0.361 | 0.0104227 | 0.04253902 | 0.0162 | 0.0092 | 0.0113 | 0.0113 | TRUE | 0.3091 |
| Sppl2c | 0 | 0 | 0 | 0 | 0 | 0 | 0 | 0 | 0 | 0 | 0 | 0 | 0 | 0 | 0 | NA | NA | NA | NA | NA | NA | NA | NA | NA | NA |
| Sppl3 | 4421 | 3223 | 3089 | 3203 | 2804 | 3284 | 3434 | 3428 | 3333 | 3188 | 3163 | 3242 | 3298.03 | 3398 | 3198 | 0.941 | -0.088 | 0.10147611 | 0.22355088 | 0 | 0.0031 | 0.0018 | 0.0018 | TRUE | 0.0034 |
| Spr | 3066 | 2020 | 1854 | 1879 | 1770 | 1989 | 2381 | 2149 | 2000 | 1870 | 1997 | 1963 | 2060.15 | 2177 | 1943 | 0.892 | -0.165 | 0.03124634 | 0.09540782 | 0.0037 | 0.0036 | 0.0038 | 0.0038 | TRUE | 0.0835 |
| Spred1 | 3368 | 2443 | 2333 | 2663 | 2272 | 2604 | 2616 | 2599 | 2517 | 2651 | 2563 | 2571 | 2586 | 2577 | 2595 | 1.007 | 0.01 | 0.8675737 | 0.92684288 | 0 | 0.0033 | 0.002 | 0.002 | TRUE | 0.0049 |
| Spred2 | 3612 | 2526 | 2326 | 2582 | 2391 | 2590 | 2805 | 2687 | 2510 | 2570 | 2697 | 2557 | 2637.67 | 2667 | 2608 | 0.977 | -0.034 | 0.60479666 | 0.74959753 | 0.0016 | 0.0033 | 0.0027 | 0.0027 | TRUE | 0.0322 |
| Spred3 | 172 | 139 | 77 | 148 | 100 | 142 | 134 | 148 | 83 | 147 | 113 | 140 | 127.47 | 122 | 133 | 1.097 | 0.133 | 0.54510475 | 0.70230023 | 0.0435 | 0.021 | 0.0272 | 0.0272 | TRUE | 0.7381 |
| Sprn | 1 | 4 | 2 | 1 | 1 | 2 | 1 | 4 | 2 | 1 | 1 | 2 | 1.88 | 2 | 1 | 0.585 | -0.773 | 0.5842578 | NA | 0 | 1.2623 | 0.8531 | 0.8531 | TRUE | 0.2292 |
| Sprr1a | 0 | 0 | 1 | 0 | 0 | 0 | 0 | 0 | 1 | 0 | 0 | 0 | 0.18 | 0 | 0 | 0.552 | -0.857 | 0.8336548 | NA | 0 | 13.1817 | 10 | 10 | TRUE | 0.5365 |
| Sprr1b | 0 | 0 | 0 | 0 | 0 | 0 | 0 | 0 | 0 | 0 | 0 | 0 | 0 | 0 | 0 | NA | NA | NA | NA | NA | NA | NA | NA | NA | NA |
| Sprr2a1 | 0 | 0 | 0 | 0 | 0 | 0 | 0 | 0 | 0 | 0 | 0 | 0 | 0 | 0 | 0 | NA | NA | NA | NA | NA | NA | NA | NA | NA | NA |
| Sprr2a2 | 0 | 0 | 0 | 0 | 0 | 0 | 0 | 0 | 0 | 0 | 0 | 0 | 0 | 0 | 0 | NA | NA | NA | NA | NA | NA | NA | NA | NA | NA |
| Sprr2b | 0 | 0 | 0 | 0 | 0 | 0 | 0 | 0 | 0 | 0 | 0 | 0 | 0 | 0 | 0 | NA | NA | NA | NA | NA | NA | NA | NA | NA | NA |
| Sprr2d | 0 | 0 | 0 | 0 | 0 | 0 | 0 | 0 | 0 | 0 | 0 | 0 | 0 | 0 | 0 | NA | NA | NA | NA | NA | NA | NA | NA | NA | NA |
| Sprr2e | 3 | 2 | 1 | 0 | 1 | 0 | 2 | 2 | 1 | 0 | 1 | 0 | 1.11 | 2 | 0 | 0.199 | -2.326 | 0.23599045 | NA | 0 | 2.1362 | 1.5158 | 1.5158 | TRUE | 0.5075 |
| Sprr2f | 0 | 0 | 0 | 0 | 0 | 0 | 0 | 0 | 0 | 0 | 0 | 0 | 0 | 0 | 0 | NA | NA | NA | NA | NA | NA | NA | NA | NA | NA |
| Sprr2g | 0 | 0 | 0 | 0 | 0 | 0 | 0 | 0 | 0 | 0 | 0 | 0 | 0 | 0 | 0 | NA | NA | NA | NA | NA | NA | NA | NA | NA | NA |
| Sprr2h | 0 | 0 | 0 | 0 | 0 | 0 | 0 | 0 | 0 | 0 | 0 | 0 | 0 | 0 | 0 | NA | NA | NA | NA | NA | NA | NA | NA | NA | NA |
| Sprr2i | 0 | 0 | 0 | 0 | 0 | 0 | 0 | 0 | 0 | 0 | 0 | 0 | 0 | 0 | 0 | NA | NA | NA | NA | NA | NA | NA | NA | NA | NA |
| Sprr2j-ps | 0 | 0 | 0 | 0 | 0 | 0 | 0 | 0 | 0 | 0 | 0 | 0 | 0 | 0 | 0 | NA | NA | NA | NA | NA | NA | NA | NA | NA | NA |
| Sprr2k | 2 | 0 | 0 | 1 | 0 | 0 | 2 | 0 | 0 | 1 | 0 | 0 | 0.42 | 1 | 0 | 0.681 | -0.553 | 0.8627148 | NA | 0 | 5.5819 | 5.4906 | 5.4906 | TRUE | 0.9727 |
| Sprr3 | 0 | 0 | 0 | 0 | 0 | 0 | 0 | 0 | 0 | 0 | 0 | 0 | 0 | 0 | 0 | NA | NA | NA | NA | NA | NA | NA | NA | NA | NA |
| Sprr4 | 0 | 0 | 0 | 0 | 0 | 0 | 0 | 0 | 0 | 0 | 0 | 0 | 0 | 0 | 0 | NA | NA | NA | NA | NA | NA | NA | NA | NA | NA |
| Sprtn | 1751 | 1216 | 1021 | 1062 | 1023 | 1086 | 1360 | 1293 | 1102 | 1057 | 1154 | 1072 | 1173.04 | 1252 | 1094 | 0.873 | -0.196 | 0.03068318 | 0.09411044 | 0.0063 | 0.0044 | 0.0051 | 0.0051 | TRUE | 0.1304 |
| Spry1 | 983 | 788 | 823 | 1015 | 925 | 1072 | 763 | 838 | 888 | 1010 | 1043 | 1058 | 933.62 | 830 | 1037 | 1.252 | 0.325 | 0.00010804 | 0.00129003 | 0.0021 | 0.005 | 0.004 | 0.004 | TRUE | 0.0616 |
| Spry2 | 5162 | 3738 | 3510 | 3375 | 3283 | 3918 | 4009 | 3976 | 3787 | 3359 | 3704 | 3868 | 3783.86 | 3924 | 3644 | 0.928 | -0.107 | 0.10754689 | 0.23336386 | 0.0026 | 0.0031 | 0.0029 | 0.0029 | TRUE | 0.0569 |
| Spry3 | 18 | 12 | 10 | 5 | 3 | 4 | 14 | 13 | 11 | 5 | 3 | 4 | 8.31 | 13 | 4 | 0.328 | -1.608 | 0.02155275 | NA | 0 | 0.2877 | 0.1886 | 0.1886 | TRUE | 0.0602 |
| Spry4 | 11079 | 8648 | 6365 | 13351 | 12181 | 12264 | 8605 | 9199 | 6868 | 13290 | 13741 | 12106 | 10634.77 | 8224 | 13046 | 1.586 | 0.666 | 9.87E-13 | 1.23E-10 | 0.0128 | 0.0026 | 0.0062 | 0.0062 | TRUE | 0.2533 |
| Spryd3 | 1468 | 1122 | 1108 | 1272 | 1114 | 1325 | 1140 | 1193 | 1196 | 1266 | 1257 | 1308 | 1226.66 | 1176 | 1277 | 1.086 | 0.12 | 0.08669867 | 0.19951442 | 0 | 0.0044 | 0.0027 | 0.0027 | TRUE | 0.0092 |
| Spryd4 | 1537 | 1121 | 1037 | 1052 | 958 | 1089 | 1194 | 1192 | 1119 | 1047 | 1081 | 1075 | 1117.99 | 1168 | 1068 | 0.913 | -0.131 | 0.0711432 | 0.17352385 | 0 | 0.0045 | 0.0029 | 0.0029 | TRUE | 0.016 |
| Spryd7 | 1630 | 1201 | 1353 | 1467 | 1229 | 1484 | 1266 | 1277 | 1460 | 1460 | 1386 | 1465 | 1385.82 | 1334 | 1437 | 1.078 | 0.109 | 0.16565864 | 0.31676786 | 0.0027 | 0.0041 | 0.0037 | 0.0037 | TRUE | 0.08 |
| Spsb1 | 985 | 782 | 801 | 962 | 858 | 953 | 765 | 832 | 864 | 958 | 968 | 941 | 887.89 | 820 | 956 | 1.166 | 0.222 | 0.00651813 | 0.03004687 | 0.001 | 0.0051 | 0.0037 | 0.0037 | TRUE | 0.0439 |
| Spsb2 | 202 | 151 | 136 | 107 | 97 | 135 | 157 | 161 | 147 | 107 | 109 | 133 | 135.57 | 155 | 116 | 0.753 | -0.41 | 0.01924381 | 0.06716771 | 0 | 0.0199 | 0.0145 | 0.0145 | TRUE | 0.1654 |
| Spsb3 | 1120 | 839 | 708 | 775 | 704 | 767 | 870 | 892 | 764 | 771 | 794 | 757 | 808.16 | 842 | 774 | 0.918 | -0.123 | 0.16475616 | 0.31552065 | 0.0024 | 0.0054 | 0.0044 | 0.0044 | TRUE | 0.0758 |
| Spsb4 | 119 | 69 | 61 | 58 | 43 | 64 | 92 | 73 | 66 | 58 | 49 | 63 | 66.84 | 77 | 57 | 0.729 | -0.457 | 0.06996198 | 0.17149144 | 0.0121 | 0.0379 | 0.0304 | 0.0304 | TRUE | 0.3048 |
| Spt1 | 0 | 0 | 0 | 0 | 0 | 0 | 0 | 0 | 0 | 0 | 0 | 0 | 0 | 0 | 0 | NA | NA | NA | NA | NA | NA | NA | NA | NA | NA |
| Spta1 | 117 | 71 | 25 | 59 | 68 | 78 | 91 | 76 | 27 | 59 | 77 | 77 | 67.63 | 65 | 71 | 1.09 | 0.124 | 0.72053944 | 0.83222698 | 0.1661 | 0.0375 | 0.0718 | 0.0718 | TRUE | 1.1545 |
| Sptan1 | 30334 | 20677 | 18858 | 19399 | 17593 | 19722 | 23559 | 21994 | 20348 | 19310 | 19847 | 19469 | 20754.29 | 21967 | 19542 | 0.889 | -0.169 | 0.00554078 | 0.02656138 | 0.0027 | 0.0025 | 0.0026 | 0.0026 | TRUE | 0.0509 |
| Sptb | 1970 | 1525 | 1708 | 2079 | 1889 | 2150 | 1530 | 1622 | 1843 | 2069 | 2131 | 2122 | 1886.31 | 1665 | 2107 | 1.267 | 0.342 | 1.14E-05 | 0.00020853 | 0.004 | 0.0037 | 0.0038 | 0.0038 | TRUE | 0.1047 |
| Sptbn1 | 27995 | 19320 | 18652 | 21604 | 19380 | 21803 | 21743 | 20550 | 20125 | 21505 | 21863 | 21523 | 21218.08 | 20806 | 21630 | 1.039 | 0.056 | 0.26105855 | 0.4310165 | 8.00E-04 | 0.0025 | 0.0017 | 0.0017 | TRUE | 0.0191 |
| Sptbn2 | 2223 | 1736 | 1734 | 1566 | 1541 | 1658 | 1727 | 1847 | 1871 | 1559 | 1738 | 1637 | 1729.66 | 1815 | 1645 | 0.907 | -0.141 | 0.04676477 | 0.12829344 | 0.0018 | 0.0038 | 0.0031 | 0.0031 | TRUE | 0.0295 |
| Sptbn4 | 1411 | 1231 | 1395 | 1873 | 1614 | 1946 | 1096 | 1309 | 1505 | 1864 | 1821 | 1921 | 1586.1 | 1303 | 1869 | 1.436 | 0.522 | 1.89E-07 | 6.91E-06 | 0.0117 | 0.0039 | 0.0066 | 0.0066 | TRUE | 0.2399 |
| Sptlc1 | 3171 | 2192 | 2028 | 2227 | 2118 | 2217 | 2463 | 2332 | 2188 | 2217 | 2389 | 2189 | 2296.2 | 2328 | 2265 | 0.972 | -0.041 | 0.55835061 | 0.71311815 | 0.0025 | 0.0035 | 0.0031 | 0.0031 | TRUE | 0.0331 |
| Sptlc2 | 4476 | 3053 | 3348 | 3219 | 2865 | 3374 | 3476 | 3247 | 3612 | 3204 | 3232 | 3331 | 3350.51 | 3445 | 3256 | 0.945 | -0.082 | 0.18217464 | 0.33776393 | 0.0012 | 0.0031 | 0.0024 | 0.0024 | TRUE | 0.0302 |
| Sptlc3 | 0 | 0 | 0 | 0 | 0 | 0 | 0 | 0 | 0 | 0 | 0 | 0 | 0 | 0 | 0 | NA | NA | NA | NA | NA | NA | NA | NA | NA | NA |
| Sptssa | 1606 | 1118 | 1164 | 1137 | 944 | 1164 | 1247 | 1189 | 1256 | 1132 | 1065 | 1149 | 1173.03 | 1231 | 1115 | 0.906 | -0.142 | 0.05162222 | 0.13801653 | 0 | 0.0044 | 0.003 | 0.003 | TRUE | 0.0182 |
| Sptssb | 0 | 0 | 0 | 0 | 0 | 0 | 0 | 0 | 0 | 0 | 0 | 0 | 0 | 0 | 0 | NA | NA | NA | NA | NA | NA | NA | NA | NA | NA |

| | | | | | | | | | | | | | | | | | | | | | | | | | |
|---|---|---|---|---|---|---|---|---|---|---|---|---|---|---|---|---|---|---|---|---|---|---|---|---|---|
| Spty2d1 | 1687 | 1205 | 1175 | 1362 | 1190 | 1337 | 1310 | 1282 | 1268 | 1356 | 1342 | 1320 | 1312.96 | 1287 | 1339 | 1.04 | 0.057 | 0.39344648 | 0.56700248 | 0 | 0.0042 | 0.0025 | 0.0025 | TRUE | 0.0033 |
| Spz1 | 57 | 34 | 15 | 21 | 15 | 13 | 44 | 36 | 16 | 21 | 17 | 13 | 24.55 | 32 | 17 | 0.52 | -0.945 | 0.03855053 | 0.11149387 | 0.1188 | 0.099 | 0.1047 | 0.1047 | TRUE | 0.4185 |
| Sqle | 8580 | 6414 | 6424 | 6667 | 6071 | 7121 | 6664 | 6822 | 6931 | 6636 | 6849 | 7030 | 6822.04 | 6806 | 6838 | 1.005 | 0.007 | 0.89003819 | 0.93898109 | 5.00E-04 | 0.0028 | 0.0017 | 0.0017 | TRUE | 0.0082 |
| Sqrdl | 41 | 40 | 33 | 27 | 22 | 35 | 32 | 43 | 36 | 27 | 25 | 35 | 32.71 | 37 | 29 | 0.79 | -0.341 | 0.32012356 | 0.49401058 | 0 | 0.0749 | 0.0536 | 0.0536 | TRUE | 0.2073 |
| Sqstm1 | 14468 | 10323 | 9973 | 11022 | 10115 | 11961 | 11237 | 10980 | 10761 | 10971 | 11411 | 11807 | 11194.55 | 10993 | 11396 | 1.037 | 0.052 | 0.3127659 | 0.48606752 | 8.00E-04 | 0.0026 | 0.0018 | 0.0018 | TRUE | 0.0131 |
| Sra1 | 2535 | 1790 | 1876 | 1852 | 1629 | 1907 | 1969 | 1904 | 2024 | 1843 | 1838 | 1883 | 1910.11 | 1966 | 1855 | 0.944 | -0.084 | 0.17652345 | 0.33028533 | 0 | 0.0037 | 0.0022 | 0.0022 | TRUE | 0.0088 |
| Srbd1 | 3979 | 3024 | 3144 | 3541 | 3039 | 3720 | 3090 | 3217 | 3392 | 3525 | 3428 | 3672 | 3387.41 | 3233 | 3542 | 1.096 | 0.133 | 0.03118003 | 0.09532751 | 0.0013 | 0.0031 | 0.0024 | 0.0024 | TRUE | 0.0225 |
| Src | 4332 | 3409 | 3725 | 3938 | 3545 | 3975 | 3364 | 3626 | 4019 | 3920 | 3999 | 3924 | 3808.8 | 3670 | 3948 | 1.076 | 0.106 | 0.13354768 | 0.27206343 | 0.0037 | 0.003 | 0.0033 | 0.0033 | TRUE | 0.0842 |
| Srcin1 | 678 | 595 | 595 | 740 | 652 | 749 | 527 | 633 | 642 | 737 | 736 | 739 | 668.83 | 601 | 737 | 1.231 | 0.3 | 0.00234906 | 0.01369855 | 0.0045 | 0.006 | 0.0055 | 0.0055 | TRUE | 0.1449 |
| Srcrb4d | 366 | 298 | 254 | 328 | 312 | 353 | 284 | 317 | 274 | 326 | 352 | 348 | 317.04 | 292 | 342 | 1.174 | 0.231 | 0.0522965 | 0.13924625 | 0 | 0.0099 | 0.007 | 0.007 | TRUE | 0.0614 |
| Srd5a1 | 491 | 316 | 217 | 259 | 246 | 284 | 381 | 336 | 234 | 258 | 278 | 280 | 294.55 | 317 | 272 | 0.854 | -0.228 | 0.16415143 | 0.31507508 | 0.0285 | 0.0105 | 0.0159 | 0.0159 | TRUE | 0.5264 |
| Srd5a2 | 2 | 2 | 0 | 3 | 1 | 1 | 2 | 2 | 0 | 3 | 1 | 1 | 1.46 | 1 | 2 | 1.374 | 0.459 | 0.77410506 | NA | 0 | 1.6217 | 1.1496 | 1.1496 | TRUE | 0.3768 |
| Srd5a3 | 805 | 584 | 501 | 522 | 518 | 535 | 625 | 621 | 541 | 520 | 584 | 528 | 569.84 | 596 | 544 | 0.911 | -0.134 | 0.18584299 | 0.34221256 | 0.0035 | 0.0066 | 0.0056 | 0.0056 | TRUE | 0.0747 |
| Srebf1 | 9546 | 7109 | 7322 | 8345 | 7668 | 8816 | 7414 | 7562 | 7900 | 8307 | 8650 | 8703 | 8089.29 | 7625 | 8553 | 1.122 | 0.166 | 0.00143263 | 0.00933721 | 7.00E-04 | 0.0027 | 0.0018 | 0.0018 | TRUE | 0.0121 |
| Srebf2 | 18338 | 13363 | 15185 | 16208 | 14662 | 16611 | 14242 | 14214 | 16384 | 16133 | 16540 | 16398 | 15652.03 | 14947 | 16357 | 1.094 | 0.13 | 0.04233822 | 0.1192735 | 0.0035 | 0.0026 | 0.0029 | 0.0029 | TRUE | 0.0864 |
| Srek1 | 3780 | 2906 | 2738 | 2658 | 2343 | 2836 | 2936 | 3091 | 2954 | 2646 | 2643 | 2800 | 2844.93 | 2994 | 2696 | 0.901 | -0.15 | 0.0110333 | 0.04440975 | 6.00E-04 | 0.0033 | 0.0022 | 0.0022 | TRUE | 0.0138 |
| Srek1ip1 | 620 | 426 | 522 | 516 | 439 | 520 | 482 | 453 | 563 | 514 | 495 | 513 | 503.35 | 499 | 507 | 1.017 | 0.025 | 0.81534278 | 0.89389387 | 0.0044 | 0.0071 | 0.0063 | 0.0063 | TRUE | 0.1408 |
| Srf | 2925 | 2319 | 1800 | 1791 | 1691 | 2002 | 2272 | 2467 | 1942 | 1783 | 1908 | 1976 | 2057.88 | 2227 | 1889 | 0.848 | -0.238 | 0.00718994 | 0.03239781 | 0.0077 | 0.0036 | 0.0051 | 0.0051 | TRUE | 0.1492 |
| Srfbp1 | 3466 | 2434 | 2325 | 2583 | 2180 | 2504 | 2692 | 2589 | 2509 | 2571 | 2459 | 2472 | 2548.63 | 2597 | 2501 | 0.963 | -0.055 | 0.36284944 | 0.53754219 | 6.00E-04 | 0.0034 | 0.0022 | 0.0022 | TRUE | 0.0129 |
| Srgap1 | 754 | 528 | 532 | 554 | 583 | 563 | 586 | 562 | 574 | 551 | 658 | 556 | 581.03 | 574 | 588 | 1.024 | 0.034 | 0.73723167 | 0.843385 | 0.0034 | 0.0065 | 0.0055 | 0.0055 | TRUE | 0.1221 |
| Srgap2 | 4701 | 3423 | 3597 | 3715 | 3245 | 3851 | 3651 | 3641 | 3881 | 3698 | 3661 | 3802 | 3722.22 | 3724 | 3720 | 0.999 | -0.001 | 0.98301322 | 0.99091671 | 6.00E-04 | 0.0031 | 0.002 | 0.002 | TRUE | 0.0163 |
| Srgap3 | 1183 | 844 | 895 | 814 | 729 | 833 | 919 | 898 | 966 | 810 | 822 | 822 | 872.86 | 928 | 818 | 0.883 | -0.18 | 0.02096803 | 0.07164621 | 0 | 0.0051 | 0.0032 | 0.0032 | TRUE | 0.0149 |
| Srgn | 61 | 30 | 31 | 58 | 34 | 37 | 47 | 32 | 33 | 58 | 38 | 37 | 40.89 | 37 | 44 | 1.168 | 0.224 | 0.49611614 | 0.66061075 | 0.0367 | 0.0604 | 0.0535 | 0.0535 | TRUE | 0.5682 |
| Sri | 1804 | 1285 | 1201 | 1385 | 1184 | 1273 | 1401 | 1367 | 1296 | 1379 | 1336 | 1257 | 1339.12 | 1355 | 1324 | 0.976 | -0.034 | 0.63663792 | 0.77290969 | 0.0012 | 0.0042 | 0.0031 | 0.0031 | TRUE | 0.0239 |
| Srl | 3912 | 2766 | 2732 | 2948 | 2752 | 3369 | 3038 | 2942 | 2948 | 2934 | 3105 | 3326 | 3048.82 | 2976 | 3122 | 1.049 | 0.069 | 0.28664904 | 0.45864488 | 0.0016 | 0.0032 | 0.0027 | 0.0027 | TRUE | 0.0402 |
| Srm | 9637 | 7314 | 6878 | 7819 | 7180 | 8341 | 7485 | 7780 | 7421 | 7783 | 8100 | 8234 | 7800.4 | 7562 | 8039 | 1.063 | 0.088 | 0.08570025 | 0.19785729 | 5.00E-04 | 0.0027 | 0.0018 | 0.0018 | TRUE | 0.0095 |
| Srms | 93 | 51 | 47 | 83 | 63 | 65 | 72 | 54 | 51 | 83 | 71 | 64 | 65.84 | 59 | 73 | 1.22 | 0.287 | 0.25860781 | 0.42850468 | 0.0131 | 0.0384 | 0.0312 | 0.0312 | TRUE | 0.3704 |
| Srp14 | 4417 | 3066 | 2891 | 3363 | 2844 | 3339 | 3430 | 3261 | 3119 | 3348 | 3208 | 3296 | 3277.18 | 3270 | 3284 | 1.004 | 0.005 | 0.92953855 | 0.96100503 | 0.0011 | 0.0031 | 0.0023 | 0.0023 | TRUE | 0.0228 |
| Srp19 | 2772 | 1947 | 1802 | 1906 | 1603 | 1962 | 2153 | 2071 | 1944 | 1897 | 1808 | 1937 | 1968.44 | 2056 | 1881 | 0.914 | -0.129 | 0.05650185 | 0.14725635 | 0.0015 | 0.0036 | 0.0028 | 0.0028 | TRUE | 0.0271 |
| Srp54a | 436 | 286 | 272 | 358 | 306 | 307 | 339 | 304 | 293 | 356 | 345 | 303 | 323.49 | 312 | 335 | 1.07 | 0.097 | 0.43033978 | 0.60255157 | 0.0037 | 0.0098 | 0.0079 | 0.0079 | TRUE | 0.0803 |
| Srp54b | 0 | 0 | 0 | 0 | 0 | 0 | 0 | 0 | 0 | 0 | 0 | 0 | 0 | 0 | 0 | NA | NA | NA | NA | NA | NA | NA | NA | NA | NA |
| Srp54c | 0 | 0 | 0 | 0 | 0 | 0 | 0 | 0 | 0 | 0 | 0 | 0 | 0 | 0 | 0 | NA | NA | NA | NA | NA | NA | NA | NA | NA | NA |
| Srp68 | 13520 | 10293 | 10736 | 12139 | 10848 | 12459 | 10500 | 10949 | 11584 | 12083 | 12238 | 12299 | 11608.79 | 11011 | 12207 | 1.109 | 0.149 | 0.00543554 | 0.02614512 | 0.0011 | 0.0026 | 0.002 | 0.002 | TRUE | 0.0253 |
| Srp72 | 16486 | 12040 | 11562 | 13295 | 11837 | 13383 | 12804 | 12807 | 12475 | 13234 | 13353 | 13211 | 12980.72 | 12695 | 13266 | 1.045 | 0.063 | 0.1566035 | 0.30486705 | 0 | 0.0026 | 0.0014 | 0.0014 | TRUE | 0.0028 |
| Srp9 | 3588 | 2570 | 2472 | 2790 | 2443 | 2775 | 2787 | 2734 | 2667 | 2777 | 2756 | 2739 | 2743.34 | 2729 | 2757 | 1.01 | 0.014 | 0.79715354 | 0.88160891 | 0 | 0.0033 | 0.0019 | 0.0019 | TRUE | 0.0048 |
| Srpk1 | 9372 | 7070 | 6329 | 6645 | 5953 | 6652 | 7279 | 7520 | 6829 | 6614 | 6716 | 6567 | 6920.77 | 7209 | 6632 | 0.92 | -0.121 | 0.02896909 | 0.09026709 | 0.001 | 0.0028 | 0.0021 | 0.0021 | TRUE | 0.0257 |
| Srpk2 | 4896 | 3496 | 3760 | 4626 | 3921 | 4675 | 3803 | 3719 | 4057 | 4605 | 4423 | 4615 | 4203.52 | 3860 | 4548 | 1.179 | 0.237 | 4.21E-05 | 0.00060853 | 0.001 | 0.003 | 0.0022 | 0.0022 | TRUE | 0.024 |
| Srpk3 | 4 | 6 | 3 | 3 | 6 | 4 | 3 | 6 | 3 | 3 | 7 | 4 | 4.4 | 4 | 5 | 1.079 | 0.11 | 0.90410655 | NA | 0 | 0.5405 | 0.3726 | 0.3726 | TRUE | 0.3205 |
| Srpr | 8408 | 6237 | 6366 | 6473 | 5959 | 6953 | 6530 | 6634 | 6869 | 6443 | 6722 | 6864 | 6677.08 | 6678 | 6676 | 1 | 0 | 0.99904388 | 0.99939385 | 7.00E-04 | 0.0028 | 0.0019 | 0.0019 | TRUE | 0.0115 |
| Srprb | 3286 | 2443 | 2367 | 2733 | 2481 | 2827 | 2552 | 2599 | 2554 | 2720 | 2799 | 2791 | 2669.1 | 2568 | 2770 | 1.079 | 0.109 | 0.0500318 | 0.13482362 | 0 | 0.0033 | 0.0019 | 0.0019 | TRUE | 0.003 |
| Srpx | 4 | 1 | 0 | 2 | 3 | 3 | 3 | 1 | 0 | 2 | 3 | 3 | 2.08 | 1 | 3 | 1.884 | 0.914 | 0.50617884 | NA | 0 | 1.1394 | 0.8477 | 0.8477 | TRUE | 0.9617 |
| Srpx2 | 1 | 0 | 0 | 0 | 0 | 0 | 1 | 0 | 0 | 0 | 0 | 0 | 0.13 | 0 | 0 | 0.552 | -0.857 | 0.8336548 | NA | 0 | 18.3121 | 10 | 10 | TRUE | 0.2566 |
| Srr | 400 | 229 | 249 | 333 | 268 | 264 | 311 | 244 | 269 | 331 | 302 | 261 | 286.22 | 275 | 298 | 1.083 | 0.115 | 0.41699802 | 0.58958875 | 0.0117 | 0.0107 | 0.011 | 0.011 | TRUE | 0.1607 |
| Srrd | 572 | 360 | 283 | 326 | 259 | 293 | 444 | 383 | 305 | 325 | 292 | 289 | 339.74 | 377 | 302 | 0.797 | -0.326 | 0.02284944 | 0.07607432 | 0.0172 | 0.0094 | 0.0118 | 0.0118 | TRUE | 0.2041 |
| Srrm1 | 20529 | 15683 | 15762 | 15045 | 13596 | 15974 | 15944 | 16682 | 17007 | 14976 | 15338 | 15769 | 15952.53 | 16544 | 15361 | 0.929 | -0.107 | 0.03395545 | 0.10161751 | 8.00E-04 | 0.0026 | 0.0018 | 0.0018 | TRUE | 0.0124 |
| Srrm2 | 37901 | 27741 | 27063 | 30257 | 27557 | 31849 | 29436 | 29508 | 29201 | 30118 | 31087 | 31440 | 30131.57 | 29382 | 30882 | 1.051 | 0.072 | 0.10444806 | 0.22844046 | 2.00E-04 | 0.0025 | 0.0014 | 0.0014 | TRUE | 0.0057 |
| Srrm3 | 39 | 31 | 30 | 44 | 32 | 45 | 30 | 33 | 32 | 44 | 36 | 44 | 36.66 | 32 | 41 | 1.305 | 0.384 | 0.22297809 | 0.387292 | 0 | 0.0671 | 0.0441 | 0.0441 | TRUE | 0.0895 |
| Srrm4 | 56 | 39 | 39 | 38 | 36 | 23 | 43 | 41 | 42 | 38 | 41 | 23 | 38.03 | 42 | 34 | 0.793 | -0.336 | 0.30733558 | 0.47997678 | 0.0075 | 0.0647 | 0.051 | 0.051 | TRUE | 0.5884 |
| Srrt | 11506 | 8428 | 6489 | 5758 | 5314 | 6384 | 8936 | 8965 | 7002 | 5732 | 5995 | 6302 | 7155.13 | 8301 | 6010 | 0.724 | -0.466 | 1.35E-07 | 5.07E-06 | 0.0101 | 0.0028 | 0.0055 | 0.0055 | TRUE | 0.2283 |
| Srsf1 | 28709 | 22479 | 18994 | 17302 | 15618 | 18177 | 22297 | 23911 | 20494 | 17222 | 17619 | 17944 | 19914.44 | 22234 | 17595 | 0.791 | -0.338 | 7.44E-08 | 3.01E-06 | 0.0031 | 0.0025 | 0.0028 | 0.0028 | TRUE | 0.0571 |
| Srsf10 | 10049 | 7574 | 6511 | 6184 | 5354 | 6350 | 7805 | 8056 | 7025 | 6156 | 6040 | 6268 | 6891.69 | 7629 | 6155 | 0.807 | -0.31 | 7.91E-07 | 2.41E-05 | 0.0024 | 0.0028 | 0.0027 | 0.0027 | TRUE | 0.058 |
| Srsf11 | 10678 | 8209 | 7915 | 8209 | 7242 | 8674 | 8293 | 8732 | 8540 | 8171 | 8170 | 8563 | 8411.45 | 8522 | 8301 | 0.974 | -0.038 | 0.46099775 | 0.6304699 | 5.00E-04 | 0.0027 | 0.0017 | 0.0017 | TRUE | 0.0093 |
| Srsf12 | 252 | 215 | 221 | 337 | 226 | 329 | 196 | 229 | 238 | 335 | 255 | 325 | 263.01 | 221 | 305 | 1.387 | 0.472 | 0.00135205 | 0.00895498 | 0.0125 | 0.0114 | 0.0118 | 0.0118 | TRUE | 0.2263 |
| Srsf2 | 27544 | 20597 | 15641 | 13748 | 12962 | 14458 | 21392 | 21909 | 16877 | 13685 | 14622 | 14272 | 17126.17 | 20059 | 14193 | 0.708 | -0.499 | 7.82E-09 | 4.04E-07 | 0.01 | 0.0026 | 0.0053 | 0.0053 | TRUE | 0.2354 |
| Srsf3 | 21140 | 15392 | 12442 | 12613 | 10809 | 11888 | 16419 | 16372 | 13425 | 12555 | 12194 | 11735 | 13783.26 | 15405 | 12161 | 0.789 | -0.341 | 9.65E-06 | 0.00018309 | 0.0067 | 0.0026 | 0.0042 | 0.0042 | TRUE | 0.1544 |
| Srsf4 | 5664 | 4181 | 3996 | 4039 | 3648 | 4514 | 4399 | 4447 | 4312 | 4020 | 4115 | 4456 | 4291.61 | 4386 | 4197 | 0.957 | -0.063 | 0.28572032 | 0.45767125 | 0.0013 | 0.003 | 0.0023 | 0.0023 | TRUE | 0.0356 |
| Srsf5 | 11633 | 8888 | 6779 | 7125 | 5971 | 6444 | 9035 | 9454 | 7314 | 7092 | 6736 | 6361 | 7665.46 | 8601 | 6730 | 0.782 | -0.354 | 5.25E-05 | 0.00072881 | 0.01 | 0.0027 | 0.0054 | 0.0054 | TRUE | 0.2085 |
| Srsf6 | 19221 | 14172 | 11306 | 9923 | 9304 | 10749 | 14928 | 15075 | 12199 | 9877 | 10496 | 10611 | 12197.65 | 14067 | 10328 | 0.734 | -0.446 | 1.76E-08 | 8.34E-07 | 0.0072 | 0.0026 | 0.0044 | 0.0044 | TRUE | 0.1647 |
| Srsf7 | 16652 | 12761 | 12196 | 12215 | 10472 | 12124 | 12933 | 13574 | 13159 | 12159 | 11814 | 11968 | 12601.09 | 13222 | 11980 | 0.906 | -0.142 | 0.0026556 | 0.01500852 | 3.00E-04 | 0.0026 | 0.0015 | 0.0015 | TRUE | 0.0066 |
| Srsf9 | 4851 | 3537 | 3145 | 3014 | 2596 | 3082 | 3768 | 3762 | 3393 | 3000 | 2929 | 3042 | 3315.73 | 3641 | 2990 | 0.821 | -0.284 | 6.39E-06 | 0.00012998 | 0.0015 | 0.0031 | 0.0026 | 0.0026 | TRUE | 0.0426 |
| Srxn1 | 1678 | 1125 | 934 | 1263 | 1033 | 1176 | 1303 | 1197 | 1008 | 1257 | 1165 | 1161 | 1181.85 | 1169 | 1194 | 1.02 | 0.029 | 0.76466919 | 0.86038075 | 0.0086 | 0.0044 | 0.0058 | 0.0058 | TRUE | 0.1732 |
| Sry | 0 | 0 | 0 | 0 | 0 | 0 | 0 | 0 | 0 | 0 | 0 | 0 | 0 | 0 | 0 | NA | NA | NA | NA | NA | NA | NA | NA | NA | NA |
| Ss18 | 6853 | 4951 | 4887 | 4885 | 4180 | 4727 | 5322 | 5266 | 5273 | 4863 | 4715 | 4666 | 5017.68 | 5287 | 4748 | 0.898 | -0.155 | 0.00203054 | 0.01223032 | 0 | 0.0029 | 0.0016 | 0.0016 | TRUE | 0.0055 |
| Ss18l1 | 644 | 517 | 614 | 656 | 567 | 678 | 500 | 550 | 663 | 653 | 640 | 669 | 612.42 | 571 | 654 | 1.149 | 0.2 | 0.07005549 | 0.17161517 | 0.0084 | 0.0063 | 0.0071 | 0.0071 | TRUE | 0.2292 |
| Ssb | 18357 | 13088 | 12925 | 13887 | 11863 | 13730 | 14257 | 13922 | 13946 | 13823 | 13383 | 13554 | 13814.01 | 14042 | 13587 | 0.968 | -0.048 | 0.29459935 | 0.46650881 | 1.00E-04 | 0.0026 | 0.0014 | 0.0014 | TRUE | 0.0028 |
| Ssbp1 | 3741 | 2569 | 2416 | 2535 | 2157 | 2504 | 2905 | 2733 | 2607 | 2523 | 2433 | 2472 | 2612.24 | 2748 | 2476 | 0.9 | -0.151 | 0.01746133 | 0.06245616 | 0.0014 | 0.0033 | 0.0025 | 0.0025 | TRUE | 0.0311 |
| Ssbp2 | 152 | 112 | 113 | 112 | 118 | 82 | 118 | 119 | 122 | 111 | 133 | 81 | 114.11 | 120 | 108 | 0.904 | -0.145 | 0.48420843 | 0.65044468 | 0.0199 | 0.0232 | 0.0223 | 0.0223 | TRUE | 0.332 |
| Ssbp3 | 3895 | 2895 | 2803 | 2917 | 2596 | 3031 | 3025 | 3079 | 3024 | 2904 | 2929 | 2992 | 2992.18 | 3043 | 2942 | 0.967 | -0.049 | 0.36967179 | 0.54443265 | 0 | 0.0032 | 0.0018 | 0.0018 | TRUE | 0.0028 |
| Ssbp4 | 4801 | 3716 | 3635 | 4170 | 3492 | 4032 | 3729 | 3953 | 3922 | 4151 | 3939 | 3980 | 3945.65 | 3868 | 4023 | 1.041 | 0.058 | 0.30482017 | 0.47739853 | 7.00E-04 | 0.003 | 0.002 | 0.002 | TRUE | 0.0124 |

| | | | | | | | | | | | | | | | | | | | | | | | | | |
|---|---|---|---|---|---|---|---|---|---|---|---|---|---|---|---|---|---|---|---|---|---|---|---|---|---|
| Ssc5d | 58 | 56 | 44 | 64 | 58 | 60 | 45 | 60 | 47 | 64 | 65 | 59 | 56.74 | 51 | 63 | 1.243 | 0.314 | 0.2268282 | 0.39178918 | 0 | 0.0442 | 0.0308 | 0.0308 | TRUE | 0.1845 |
| Ssfa2 | 2496 | 1803 | 2081 | 2076 | 1830 | 2172 | 1939 | 1918 | 2245 | 2066 | 2064 | 2144 | 2062.79 | 2034 | 2091 | 1.029 | 0.041 | 0.58304646 | 0.73293606 | 0.0037 | 0.0036 | 0.0036 | 0.0036 | TRUE | 0.0987 |
| Ssh1 | 1409 | 1004 | 957 | 945 | 894 | 948 | 1094 | 1068 | 1033 | 941 | 1009 | 936 | 1013.31 | 1065 | 962 | 0.902 | -0.149 | 0.05084793 | 0.13650817 | 0 | 0.0048 | 0.0032 | 0.0032 | TRUE | 0.0212 |
| Ssh2 | 4234 | 2780 | 2087 | 2943 | 2524 | 2763 | 3288 | 2957 | 2252 | 2929 | 2847 | 2728 | 2833.6 | 2832 | 2835 | 1 | 0 | 0.99867229 | 0.99937086 | 0.0187 | 0.0033 | 0.0081 | 0.0081 | TRUE | 0.3888 |
| Ssh3 | 229 | 226 | 235 | 247 | 263 | 263 | 178 | 240 | 254 | 246 | 297 | 260 | 245.66 | 224 | 268 | 1.199 | 0.262 | 0.09827192 | 0.21837696 | 0.0184 | 0.0121 | 0.014 | 0.014 | TRUE | 0.387 |
| Ssna1 | 1758 | 1252 | 1215 | 1228 | 1075 | 1214 | 1365 | 1332 | 1311 | 1222 | 1213 | 1198 | 1273.59 | 1336 | 1211 | 0.906 | -0.142 | 0.0360282 | 0.10613172 | 0 | 0.0043 | 0.0025 | 0.0025 | TRUE | 0.0047 |
| Sspn | 5 | 7 | 15 | 7 | 11 | 12 | 4 | 7 | 16 | 7 | 12 | 12 | 9.79 | 9 | 10 | 1.155 | 0.208 | 0.75837179 | NA | 0.1849 | 0.2445 | 0.2272 | 0.2272 | TRUE | 0.7769 |
| Sspo | 9 | 12 | 6 | 13 | 14 | 13 | 7 | 13 | 6 | 13 | 16 | 13 | 11.3 | 9 | 14 | 1.595 | 0.673 | 0.24593812 | NA | 0 | 0.2122 | 0.1497 | 0.1497 | TRUE | 0.4723 |
| Ssr1 | 13088 | 9274 | 9734 | 10102 | 8934 | 9869 | 10165 | 9865 | 10503 | 10056 | 10078 | 9742 | 10068.11 | 10178 | 9959 | 0.978 | -0.031 | 0.53034622 | 0.69033619 | 5.00E-04 | 0.0027 | 0.0017 | 0.0017 | TRUE | 0.0095 |
| Ssr2 | 8319 | 5683 | 6054 | 6684 | 5987 | 7121 | 6461 | 6045 | 6532 | 6653 | 6754 | 7030 | 6579.15 | 6346 | 6812 | 1.073 | 0.102 | 0.06561194 | 0.163692 | 0.001 | 0.0028 | 0.0021 | 0.0021 | TRUE | 0.0208 |
| Ssr3 | 17983 | 12731 | 13295 | 15802 | 13515 | 15431 | 13967 | 13542 | 14345 | 15729 | 15246 | 15233 | 14677 | 13951 | 15403 | 1.104 | 0.143 | 0.00310195 | 0.01696107 | 5.00E-04 | 0.0026 | 0.0016 | 0.0016 | TRUE | 0.008 |
| Ssr4 | 1916 | 1426 | 1476 | 1621 | 1429 | 1544 | 1488 | 1517 | 1593 | 1614 | 1612 | 1524 | 1557.88 | 1533 | 1583 | 1.034 | 0.048 | 0.48170386 | 0.64849796 | 5.00E-04 | 0.0039 | 0.0027 | 0.0027 | TRUE | 0.014 |
| Ssrp1 | 21552 | 15752 | 15152 | 13913 | 12448 | 14841 | 16739 | 16755 | 16349 | 13849 | 14043 | 14650 | 15397.43 | 16614 | 14181 | 0.854 | -0.229 | 1.72E-06 | 4.50E-05 | 4.00E-04 | 0.0026 | 0.0016 | 0.0016 | TRUE | 0.0103 |
| Sssca1 | 1341 | 859 | 672 | 566 | 526 | 627 | 1041 | 914 | 725 | 563 | 593 | 619 | 742.67 | 893 | 592 | 0.661 | -0.597 | 8.53E-07 | 2.57E-05 | 0.0167 | 0.0056 | 0.0092 | 0.0092 | TRUE | 0.2193 |
| Sst | 0 | 1 | 1 | 1 | 1 | 1 | 0 | 1 | 1 | 1 | 1 | 1 | 0.88 | 1 | 1 | 1.524 | 0.607 | 0.7620868 | NA | 0 | 2.7093 | 1.693 | 1.693 | TRUE | 0.3841 |
| Sstr1 | 0 | 0 | 0 | 0 | 1 | 0 | 0 | 0 | 0 | 0 | 1 | 0 | 0.19 | 0 | 0 | 2.094 | 1.067 | 0.79379329 | NA | 0 | 12.608 | 10 | 10 | TRUE | 0.5075 |
| Sstr2 | 31 | 38 | 31 | 44 | 49 | 86 | 24 | 40 | 33 | 44 | 55 | 85 | 46.99 | 32 | 61 | 1.898 | 0.924 | 0.00617512 | 0.02883271 | 0.0729 | 0.0529 | 0.0587 | 0.0587 | TRUE | 0.4815 |
| Sstr3 | 0 | 1 | 0 | 0 | 1 | 1 | 0 | 1 | 0 | 0 | 1 | 1 | 0.53 | 0 | 1 | 2.03 | 1.022 | 0.70625413 | NA | 0 | 4.4757 | 3.5573 | 3.5573 | TRUE | 0.5217 |
| Sstr4 | 8 | 0 | 2 | 1 | 1 | 0 | 6 | 0 | 2 | 1 | 1 | 0 | 1.75 | 3 | 1 | 0.244 | -2.037 | 0.24473051 | NA | 0 | 1.3574 | 1.2991 | 1.2991 | TRUE | 0.1995 |
| Sstr5 | 30 | 15 | 20 | 37 | 45 | 25 | 23 | 16 | 22 | 37 | 51 | 25 | 28.85 | 20 | 38 | 1.827 | 0.87 | 0.02830362 | 0.08871573 | 0.0538 | 0.0846 | 0.0757 | 0.0757 | TRUE | 0.1314 |
| Ssty1 | 0 | 0 | 0 | 0 | 0 | 0 | 0 | 0 | 0 | 0 | 0 | 0 | 0 | 0 | 0 | NA | NA | NA | NA | NA | NA | NA | NA | NA | NA |
| Ssty2 | 0 | 0 | 0 | 0 | 0 | 0 | 0 | 0 | 0 | 0 | 0 | 0 | 0 | 0 | 0 | NA | NA | NA | NA | NA | NA | NA | NA | NA | NA |
| Ssu2 | 26 | 6 | 20 | 27 | 25 | 23 | 20 | 6 | 22 | 27 | 28 | 23 | 20.99 | 16 | 26 | 1.6 | 0.678 | 0.15013101 | 0.29569004 | 0.0904 | 0.1153 | 0.11 | 0.11 | TRUE | 1.2153 |
| Ssu72 | 5169 | 3484 | 3380 | 3335 | 3034 | 3481 | 4015 | 3706 | 3647 | 3320 | 3423 | 3436 | 3591 | 3789 | 3393 | 0.895 | -0.16 | 0.0079666 | 0.0349456 | 0.0011 | 0.0031 | 0.0023 | 0.0023 | TRUE | 0.0336 |
| Ssx2ip | 2532 | 1907 | 1937 | 1905 | 1664 | 1921 | 1966 | 2028 | 2090 | 1896 | 1877 | 1896 | 1959.11 | 2028 | 1890 | 0.932 | -0.101 | 0.1000535 | 0.22099593 | 0 | 0.0036 | 0.0022 | 0.0022 | TRUE | 0.0089 |
| Ssx9 | 0 | 0 | 0 | 0 | 0 | 0 | 0 | 0 | 0 | 0 | 0 | 0 | 0 | 0 | 0 | NA | NA | NA | NA | NA | NA | NA | NA | NA | NA |
| Ssxb1 | 0 | 0 | 0 | 0 | 0 | 0 | 0 | 0 | 0 | 0 | 0 | 0 | 0 | 0 | 0 | NA | NA | NA | NA | NA | NA | NA | NA | NA | NA |
| Ssxb10 | 0 | 0 | 0 | 0 | 0 | 0 | 0 | 0 | 0 | 0 | 0 | 0 | 0 | 0 | 0 | NA | NA | NA | NA | NA | NA | NA | NA | NA | NA |
| Ssxb2 | 0 | 0 | 0 | 0 | 0 | 0 | 0 | 0 | 0 | 0 | 0 | 0 | 0 | 0 | 0 | NA | NA | NA | NA | NA | NA | NA | NA | NA | NA |
| Ssxb3 | 0 | 0 | 0 | 0 | 0 | 0 | 0 | 0 | 0 | 0 | 0 | 0 | 0 | 0 | 0 | NA | NA | NA | NA | NA | NA | NA | NA | NA | NA |
| Ssxb5 | 0 | 0 | 0 | 0 | 0 | 0 | 0 | 0 | 0 | 0 | 0 | 0 | 0 | 0 | 0 | NA | NA | NA | NA | NA | NA | NA | NA | NA | NA |
| Ssxb6 | 0 | 0 | 1 | 0 | 0 | 0 | 0 | 0 | 1 | 0 | 0 | 0 | 0.18 | 0 | 0 | 0.552 | -0.857 | 0.8336548 | NA | 0 | 13.1817 | 10 | 10 | TRUE | 0.5365 |
| Ssxb8 | 0 | 0 | 0 | 0 | 0 | 0 | 0 | 0 | 0 | 0 | 0 | 0 | 0 | 0 | 0 | NA | NA | NA | NA | NA | NA | NA | NA | NA | NA |
| Ssxb9 | 1 | 0 | 0 | 0 | 0 | 0 | 1 | 0 | 0 | 0 | 0 | 0 | 0.13 | 0 | 0 | 0.552 | -0.857 | 0.8336548 | NA | 0 | 18.3121 | 10 | 10 | TRUE | 0.2566 |
| St13 | 21319 | 15615 | 15997 | 16356 | 14338 | 16191 | 16558 | 16609 | 17261 | 16281 | 16175 | 15983 | 16477.7 | 16809 | 16146 | 0.961 | -0.058 | 0.20537137 | 0.36710517 | 2.00E-04 | 0.0026 | 0.0014 | 0.0014 | TRUE | 0.0067 |
| St14 | 4012 | 3130 | 4035 | 3643 | 3273 | 3857 | 3116 | 3329 | 4354 | 3626 | 3692 | 3807 | 3654.17 | 3600 | 3708 | 1.031 | 0.044 | 0.66823471 | 0.79609449 | 0.0161 | 0.0031 | 0.0072 | 0.0072 | TRUE | 0.4076 |
| St18 | 1 | 1 | 0 | 2 | 1 | 0 | 1 | 1 | 0 | 2 | 1 | 0 | 0.83 | 1 | 1 | 1.668 | 0.738 | 0.73249436 | NA | 0 | 2.8699 | 2.0884 | 2.0884 | TRUE | 0.1969 |
| St3gal1 | 360 | 302 | 279 | 348 | 330 | 347 | 280 | 321 | 301 | 346 | 372 | 343 | 327.18 | 301 | 354 | 1.179 | 0.237 | 0.04378221 | 0.12222248 | 0 | 0.0097 | 0.0069 | 0.0069 | TRUE | 0.0462 |
| St3gal2 | 1941 | 1301 | 1329 | 1489 | 1235 | 1399 | 1507 | 1384 | 1434 | 1482 | 1393 | 1381 | 1430.29 | 1442 | 1419 | 0.983 | -0.024 | 0.73385483 | 0.84080056 | 0.0011 | 0.0041 | 0.003 | 0.003 | TRUE | 0.0199 |
| St3gal3 | 587 | 489 | 438 | 411 | 407 | 470 | 456 | 520 | 473 | 409 | 459 | 464 | 463.47 | 483 | 444 | 0.921 | -0.119 | 0.26202132 | 0.43235636 | 0.0027 | 0.0075 | 0.006 | 0.006 | TRUE | 0.0558 |
| St3gal4 | 674 | 589 | 660 | 791 | 774 | 821 | 523 | 627 | 712 | 787 | 873 | 810 | 722.18 | 621 | 823 | 1.33 | 0.412 | 0.00023622 | 0.00242655 | 0.0109 | 0.0057 | 0.0076 | 0.0076 | TRUE | 0.232 |
| St3gal5 | 915 | 618 | 659 | 764 | 640 | 729 | 711 | 657 | 711 | 760 | 722 | 720 | 713.53 | 693 | 734 | 1.059 | 0.082 | 0.33579345 | 0.51004206 | 0 | 0.0057 | 0.0039 | 0.0039 | TRUE | 0.0233 |
| St3gal6 | 96 | 63 | 74 | 89 | 43 | 69 | 75 | 67 | 80 | 89 | 49 | 68 | 71.1 | 74 | 69 | 0.93 | -0.105 | 0.68685526 | 0.80798786 | 0.0305 | 0.0358 | 0.0344 | 0.0344 | TRUE | 0.2081 |
| St5 | 1899 | 1401 | 1332 | 1427 | 1238 | 1399 | 1475 | 1490 | 1437 | 1420 | 1397 | 1381 | 1433.4 | 1467 | 1399 | 0.953 | -0.069 | 0.29449219 | 0.46649416 | 0 | 0.0041 | 0.0024 | 0.0024 | TRUE | 0.0038 |
| St6gal1 | 981 | 741 | 730 | 606 | 534 | 631 | 762 | 788 | 788 | 603 | 602 | 623 | 694.38 | 779 | 609 | 0.783 | -0.353 | 2.36E-05 | 0.00038187 | 0 | 0.0058 | 0.0036 | 0.0036 | TRUE | 0.0048 |
| St6gal2 | 16 | 13 | 7 | 11 | 5 | 5 | 12 | 14 | 8 | 11 | 6 | 5 | 9.22 | 11 | 7 | 0.637 | -0.652 | 0.32043394 | NA | 0 | 0.2594 | 0.1954 | 0.1954 | TRUE | 0.5898 |
| St6galnac1 | 4 | 1 | 3 | 0 | 1 | 0 | 3 | 1 | 3 | 0 | 1 | 0 | 1.42 | 2 | 0 | 0.149 | -2.744 | 0.13600608 | NA | 0 | 1.6685 | 1.2462 | 1.2462 | TRUE | 0.5075 |
| St6galnac2 | 508 | 474 | 357 | 290 | 274 | 305 | 395 | 504 | 385 | 289 | 309 | 301 | 363.8 | 428 | 300 | 0.701 | -0.513 | 7.76E-05 | 0.00098796 | 0.01 | 0.0089 | 0.0093 | 0.0093 | TRUE | 0.2747 |
| St6galnac3 | 318 | 242 | 306 | 371 | 363 | 368 | 247 | 257 | 330 | 369 | 410 | 363 | 329.44 | 278 | 381 | 1.372 | 0.456 | 0.00073722 | 0.00578963 | 0.0108 | 0.0096 | 0.0101 | 0.0101 | TRUE | 0.2931 |
| St6galnac4 | 755 | 548 | 569 | 662 | 630 | 666 | 586 | 583 | 614 | 659 | 711 | 657 | 635.05 | 594 | 676 | 1.137 | 0.185 | 0.03718755 | 0.10871786 | 0 | 0.0062 | 0.0041 | 0.0041 | TRUE | 0.0236 |
| St6galnac5 | 4 | 5 | 8 | 4 | 5 | 2 | 3 | 5 | 9 | 4 | 6 | 2 | 4.78 | 6 | 4 | 0.685 | -0.547 | 0.54568996 | NA | 0 | 0.4987 | 0.3682 | 0.3682 | TRUE | 0.2438 |
| St6galnac6 | 1662 | 1329 | 1757 | 2205 | 2030 | 2350 | 1291 | 1414 | 1896 | 2195 | 2290 | 2320 | 1900.83 | 1534 | 2268 | 1.481 | 0.566 | 6.36E-07 | 2.00E-05 | 0.0213 | 0.0037 | 0.0088 | 0.0088 | TRUE | 0.513 |
| St7 | 222 | 163 | 171 | 176 | 143 | 184 | 172 | 173 | 185 | 175 | 161 | 182 | 174.74 | 177 | 173 | 0.979 | -0.031 | 0.83866604 | 0.9080766 | 0 | 0.016 | 0.0105 | 0.0105 | TRUE | 0.0339 |
| St7l | 349 | 265 | 335 | 329 | 348 | 380 | 271 | 282 | 361 | 327 | 393 | 375 | 334.93 | 305 | 365 | 1.2 | 0.263 | 0.05667064 | 0.14759355 | 0.0136 | 0.0095 | 0.0107 | 0.0107 | TRUE | 0.2929 |
| St8sia1 | 656 | 496 | 507 | 369 | 388 | 464 | 509 | 528 | 547 | 367 | 438 | 458 | 474.53 | 528 | 421 | 0.798 | -0.326 | 0.00306681 | 0.01681392 | 0.0047 | 0.0074 | 0.0066 | 0.0066 | TRUE | 0.1464 |
| St8sia2 | 10 | 5 | 4 | 7 | 1 | 4 | 8 | 5 | 4 | 7 | 1 | 4 | 4.91 | 6 | 4 | 0.693 | -0.529 | 0.55687497 | NA | 0 | 0.4854 | 0.3747 | 0.3747 | TRUE | 0.2765 |
| St8sia3 | 26 | 26 | 13 | 25 | 17 | 18 | 20 | 28 | 14 | 25 | 19 | 18 | 20.62 | 21 | 21 | 1 | 0.001 | 0.99899617 | 0.99939385 | 0 | 0.1174 | 0.0909 | 0.0909 | TRUE | 0.22 |
| St8sia4 | 17 | 13 | 8 | 7 | 9 | 3 | 13 | 14 | 9 | 7 | 10 | 3 | 9.29 | 12 | 7 | 0.555 | -0.848 | 0.20266258 | NA | 0 | 0.2575 | 0.201 | 0.201 | TRUE | 0.4253 |
| St8sia5 | 162 | 106 | 145 | 111 | 107 | 109 | 126 | 113 | 156 | 110 | 121 | 108 | 122.3 | 132 | 113 | 0.858 | -0.221 | 0.24644214 | 0.41513035 | 0.0079 | 0.0218 | 0.0179 | 0.0179 | TRUE | 0.2632 |
| St8sia6 | 740 | 500 | 517 | 419 | 377 | 426 | 575 | 532 | 558 | 417 | 425 | 421 | 487.88 | 555 | 421 | 0.758 | -0.4 | 3.56E-05 | 0.00053494 | 0 | 0.0073 | 0.0046 | 0.0046 | TRUE | 0.0152 |
| Stab1 | 49 | 29 | 23 | 33 | 15 | 29 | 38 | 31 | 25 | 33 | 17 | 29 | 28.69 | 31 | 26 | 0.835 | -0.26 | 0.50049661 | 0.66489432 | 0.0359 | 0.085 | 0.0722 | 0.0722 | TRUE | 0.4547 |
| Stab2 | 73 | 71 | 73 | 31 | 30 | 35 | 57 | 76 | 79 | 31 | 34 | 35 | 51.71 | 71 | 33 | 0.473 | -1.08 | 0.00015194 | 0.00172595 | 0 | 0.0483 | 0.0361 | 0.0361 | TRUE | 0.2949 |
| Stac | 3 | 2 | 4 | 3 | 0 | 0 | 2 | 2 | 4 | 3 | 0 | 0 | 1.96 | 3 | 1 | 0.349 | -1.519 | 0.32463401 | NA | 0.1179 | 1.2117 | 1.0379 | 1.0379 | TRUE | 1.4412 |
| Stac2 | 2144 | 1657 | 2141 | 2730 | 2462 | 2795 | 1665 | 1763 | 2310 | 2717 | 2777 | 2759 | 2331.96 | 1913 | 2751 | 1.44 | 0.526 | 3.14E-07 | 1.08E-05 | 0.0159 | 0.0034 | 0.0072 | 0.0072 | TRUE | 0.3978 |
| Stac3 | 21 | 8 | 14 | 10 | 4 | 18 | 16 | 9 | 15 | 10 | 5 | 18 | 12.03 | 13 | 11 | 0.81 | -0.305 | 0.61897279 | NA | 0.1516 | 0.1995 | 0.187 | 0.187 | TRUE | 0.3717 |
| Stag1 | 7641 | 5294 | 5265 | 5590 | 4965 | 5639 | 5934 | 5631 | 5681 | 5564 | 5601 | 5567 | 5663.06 | 5749 | 5577 | 0.97 | -0.044 | 0.38388118 | 0.55785229 | 3.00E-04 | 0.0028 | 0.0017 | 0.0017 | TRUE | 0.0099 |
| Stag2 | 5585 | 3908 | 4162 | 4898 | 4059 | 4623 | 4338 | 4157 | 4491 | 4875 | 4579 | 4564 | 4500.56 | 4329 | 4673 | 1.08 | 0.111 | 0.0579375 | 0.14996391 | 0.0011 | 0.003 | 0.0022 | 0.0022 | TRUE | 0.0176 |
| Stag3 | 3637 | 3135 | 3243 | 3935 | 3316 | 3485 | 2825 | 3335 | 3499 | 3917 | 3741 | 3440 | 3459.41 | 3220 | 3699 | 1.15 | 0.201 | 0.01720793 | 0.061905 | 0.0077 | 0.0031 | 0.0049 | 0.0049 | TRUE | 0.142 |
| Stam | 4218 | 3032 | 3428 | 3460 | 2817 | 3546 | 3276 | 3225 | 3699 | 3444 | 3178 | 3500 | 3387.04 | 3400 | 3374 | 0.993 | -0.01 | 0.88642328 | 0.93690919 | 0.0039 | 0.0031 | 0.0034 | 0.0034 | TRUE | 0.0712 |

| | | | | | | | | | | | | | | | | | | | | | | | | | |
|---|---|---|---|---|---|---|---|---|---|---|---|---|---|---|---|---|---|---|---|---|---|---|---|---|---|
| Stam2 | 4136 | 2895 | 2951 | 2974 | 2708 | 3078 | 3212 | 3079 | 3184 | 2960 | 3055 | 3038 | 3088.24 | 3158 | 3018 | 0.955 | -0.066 | 0.23177282 | 0.39772736 | 0 | 0.0032 | 0.0019 | 0.0019 | TRUE | 0.0058 |
| Stambp | 240 | 209 | 198 | 213 | 180 | 213 | 186 | 222 | 214 | 212 | 203 | 210 | 207.95 | 207 | 208 | 1.009 | 0.012 | 0.93087507 | 0.96192556 | 0 | 0.0138 | 0.0098 | 0.0098 | TRUE | 0.0949 |
| Stambpl1 | 394 | 324 | 331 | 456 | 417 | 492 | 306 | 345 | 357 | 454 | 470 | 486 | 402.96 | 336 | 470 | 1.403 | 0.489 | 8.53E-06 | 0.00016458 | 0 | 0.0083 | 0.0061 | 0.0061 | TRUE | 0.0754 |
| Stap1 | 1 | 0 | 2 | 0 | 0 | 2 | 1 | 0 | 2 | 0 | 0 | 2 | 0.82 | 1 | 1 | 0.69 | -0.536 | 0.81780122 | NA | 0 | 2.8993 | 2.6298 | 2.6298 | TRUE | 0.7053 |
| Stap2 | 281 | 220 | 209 | 271 | 212 | 267 | 218 | 234 | 226 | 270 | 239 | 264 | 241.71 | 226 | 258 | 1.142 | 0.192 | 0.14319612 | 0.286092 | 0 | 0.0122 | 0.0082 | 0.0082 | TRUE | 0.0415 |
| Star | 113 | 101 | 72 | 98 | 80 | 100 | 88 | 107 | 78 | 98 | 90 | 99 | 93.23 | 91 | 96 | 1.052 | 0.073 | 0.72839182 | 0.8372403 | 0 | 0.0278 | 0.021 | 0.021 | TRUE | 0.2255 |
| Stard10 | 524 | 380 | 372 | 365 | 299 | 354 | 407 | 404 | 401 | 363 | 337 | 349 | 377.11 | 404 | 350 | 0.866 | -0.207 | 0.05137048 | 0.13752357 | 0 | 0.0087 | 0.0055 | 0.0055 | TRUE | 0.0126 |
| Stard13 | 352 | 238 | 276 | 284 | 205 | 281 | 273 | 253 | 298 | 283 | 231 | 277 | 269.28 | 275 | 264 | 0.962 | -0.056 | 0.67648046 | 0.80153653 | 0.0048 | 0.0112 | 0.0094 | 0.0094 | TRUE | 0.125 |
| Stard3 | 1424 | 1019 | 1078 | 1033 | 1011 | 1107 | 1106 | 1084 | 1163 | 1028 | 1141 | 1093 | 1102.43 | 1118 | 1087 | 0.973 | -0.04 | 0.60042068 | 0.74592028 | 0.001 | 0.0046 | 0.0033 | 0.0033 | TRUE | 0.027 |
| Stard3nl | 2991 | 2156 | 2132 | 2315 | 1974 | 2191 | 2323 | 2293 | 2300 | 2304 | 2227 | 2163 | 2268.46 | 2305 | 2231 | 0.968 | -0.047 | 0.4264948 | 0.59879619 | 0 | 0.0035 | 0.0021 | 0.0021 | TRUE | 0.0101 |
| Stard4 | 4979 | 3738 | 4030 | 4559 | 3711 | 4158 | 3867 | 3976 | 4348 | 4538 | 4186 | 4105 | 4170.06 | 4064 | 4276 | 1.053 | 0.074 | 0.27093429 | 0.44196113 | 0.003 | 0.003 | 0.003 | 0.003 | TRUE | 0.0455 |
| Stard5 | 930 | 658 | 646 | 696 | 645 | 706 | 722 | 700 | 697 | 693 | 728 | 697 | 706.1 | 706 | 706 | 0.998 | -0.003 | 0.97341436 | 0.98545797 | 0 | 0.0058 | 0.0036 | 0.0036 | TRUE | 0.0084 |
| Stard6 | 17 | 6 | 5 | 8 | 3 | 5 | 13 | 6 | 5 | 8 | 3 | 5 | 6.88 | 8 | 5 | 0.644 | -0.634 | 0.40765318 | NA | 0 | 0.347 | 0.2711 | 0.2711 | TRUE | 1.0007 |
| Stard7 | 5310 | 3911 | 3904 | 4783 | 4155 | 4726 | 4124 | 4160 | 4212 | 4761 | 4687 | 4665 | 4435.01 | 4165 | 4704 | 1.13 | 0.176 | 0.00047348 | 0.00414927 | 0 | 0.003 | 0.0016 | 0.0016 | TRUE | 0.0014 |
| Stard8 | 495 | 379 | 427 | 517 | 472 | 497 | 384 | 403 | 461 | 515 | 532 | 491 | 464.34 | 416 | 513 | 1.234 | 0.303 | 0.00467014 | 0.02324983 | 0.003 | 0.0075 | 0.0061 | 0.0061 | TRUE | 0.0997 |
| Stat1 | 920 | 663 | 656 | 810 | 732 | 886 | 715 | 705 | 708 | 806 | 826 | 875 | 772.37 | 709 | 836 | 1.178 | 0.236 | 0.00387293 | 0.02011472 | 0 | 0.0055 | 0.0035 | 0.0035 | TRUE | 0.0203 |
| Stat2 | 4326 | 3420 | 3496 | 3861 | 3702 | 4150 | 3360 | 3638 | 3772 | 3843 | 4176 | 4097 | 3814.33 | 3590 | 4039 | 1.126 | 0.171 | 0.00892147 | 0.03812097 | 0.0022 | 0.003 | 0.0028 | 0.0028 | TRUE | 0.039 |
| Stat3 | 8270 | 6614 | 7335 | 7942 | 7528 | 9057 | 6423 | 7035 | 7914 | 7905 | 8492 | 8941 | 7785.18 | 7124 | 8446 | 1.186 | 0.246 | 0.00185583 | 0.01142352 | 0.0072 | 0.0027 | 0.0044 | 0.0044 | TRUE | 0.1149 |
| Stat4 | 594 | 413 | 458 | 454 | 439 | 453 | 461 | 439 | 494 | 452 | 495 | 447 | 464.86 | 465 | 465 | 0.999 | -0.001 | 0.99025903 | 0.99506274 | 9.00E-04 | 0.0075 | 0.0055 | 0.0055 | TRUE | 0.0371 |
| Stat5a | 670 | 567 | 596 | 536 | 626 | 677 | 520 | 603 | 643 | 534 | 706 | 668 | 612.43 | 589 | 636 | 1.081 | 0.113 | 0.35303102 | 0.52775572 | 0.0149 | 0.0063 | 0.009 | 0.009 | TRUE | 0.2354 |
| Stat5b | 1580 | 1244 | 1369 | 1615 | 1471 | 1676 | 1227 | 1323 | 1477 | 1608 | 1659 | 1654 | 1491.49 | 1342 | 1640 | 1.224 | 0.291 | 0.00027658 | 0.00272499 | 0.0035 | 0.004 | 0.0039 | 0.0039 | TRUE | 0.0921 |
| Stat6 | 2265 | 1748 | 1811 | 2067 | 1982 | 2347 | 1759 | 1859 | 1954 | 2057 | 2236 | 2317 | 2030.46 | 1857 | 2203 | 1.187 | 0.247 | 0.00063684 | 0.00519538 | 0.0024 | 0.0036 | 0.0033 | 0.0033 | TRUE | 0.041 |
| Stau1 | 4831 | 3429 | 3327 | 3474 | 3071 | 3476 | 3752 | 3647 | 3590 | 3458 | 3464 | 3431 | 3557.17 | 3663 | 3451 | 0.942 | -0.086 | 0.10461894 | 0.22857208 | 0 | 0.0031 | 0.0018 | 0.0018 | TRUE | 0.0056 |
| Stau2 | 616 | 455 | 435 | 473 | 379 | 446 | 478 | 484 | 469 | 471 | 428 | 440 | 461.73 | 477 | 446 | 0.935 | -0.096 | 0.33169105 | 0.50590761 | 0 | 0.0076 | 0.0049 | 0.0049 | TRUE | 0.0271 |
| Stbd1 | 167 | 121 | 147 | 198 | 180 | 229 | 130 | 129 | 159 | 197 | 203 | 226 | 173.87 | 139 | 209 | 1.506 | 0.591 | 0.00021808 | 0.00228017 | 0.003 | 0.0161 | 0.0125 | 0.0125 | TRUE | 0.1503 |
| Stc1 | 360 | 330 | 354 | 361 | 342 | 381 | 280 | 351 | 382 | 359 | 386 | 376 | 355.64 | 338 | 374 | 1.112 | 0.153 | 0.24545097 | 0.41380404 | 0.0108 | 0.0091 | 0.0096 | 0.0096 | TRUE | 0.2757 |
| Stc2 | 519 | 473 | 513 | 478 | 506 | 502 | 403 | 503 | 554 | 476 | 571 | 496 | 500.32 | 487 | 514 | 1.059 | 0.082 | 0.5201047 | 0.68169248 | 0.0157 | 0.0072 | 0.0098 | 0.0098 | TRUE | 0.2762 |
| Steap1 | 1018 | 646 | 586 | 653 | 564 | 644 | 791 | 687 | 632 | 650 | 636 | 636 | 672.01 | 703 | 641 | 0.909 | -0.138 | 0.16888554 | 0.32052596 | 0.0054 | 0.006 | 0.0058 | 0.0058 | TRUE | 0.1453 |
| Steap2 | 2195 | 1477 | 1442 | 1540 | 1256 | 1460 | 1705 | 1571 | 1556 | 1533 | 1417 | 1441 | 1537.13 | 1611 | 1464 | 0.908 | -0.139 | 0.05322443 | 0.14094286 | 0.0016 | 0.004 | 0.0031 | 0.0031 | TRUE | 0.0327 |
| Steap3 | 4101 | 2595 | 2292 | 2649 | 2385 | 2681 | 3185 | 2760 | 2473 | 2637 | 2691 | 2647 | 2732.05 | 2806 | 2658 | 0.946 | -0.08 | 0.35158996 | 0.52609837 | 0.0074 | 0.0033 | 0.0049 | 0.0049 | TRUE | 0.1717 |
| Steap4 | 61 | 35 | 15 | 15 | 17 | 20 | 47 | 37 | 16 | 15 | 19 | 20 | 25.77 | 33 | 18 | 0.529 | -0.919 | 0.04047289 | 0.11544184 | 0.1203 | 0.0944 | 0.102 | 0.102 | TRUE | 0.3158 |
| Stfa1 | 0 | 0 | 0 | 0 | 0 | 0 | 0 | 0 | 0 | 0 | 0 | 0 | 0 | 0 | 0 | NA | NA | NA | NA | NA | NA | NA | NA | NA | NA |
| Stfa2 | 0 | 0 | 0 | 0 | 0 | 0 | 0 | 0 | 0 | 0 | 0 | 0 | 0 | 0 | 0 | NA | NA | NA | NA | NA | NA | NA | NA | NA | NA |
| Stfa2l1 | 0 | 0 | 0 | 1 | 0 | 0 | 0 | 0 | 0 | 1 | 0 | 0 | 0.17 | 0 | 0 | 2.094 | 1.067 | 0.79379329 | NA | 0 | 14.2885 | 10 | 10 | TRUE | 0.3906 |
| Stfa3 | 0 | 0 | 0 | 0 | 0 | 0 | 0 | 0 | 0 | 0 | 0 | 0 | 0 | 0 | 0 | NA | NA | NA | NA | NA | NA | NA | NA | NA | NA |
| Stil | 1653 | 1189 | 1310 | 1414 | 1306 | 1402 | 1284 | 1265 | 1413 | 1407 | 1473 | 1384 | 1371.13 | 1321 | 1421 | 1.077 | 0.107 | 0.14987 | 0.29554249 | 0.0016 | 0.0042 | 0.0032 | 0.0032 | TRUE | 0.0443 |
| Stim1 | 2988 | 2144 | 2284 | 2838 | 2474 | 2782 | 2321 | 2281 | 2464 | 2825 | 2791 | 2746 | 2571.29 | 2355 | 2787 | 1.184 | 0.243 | 5.03E-05 | 0.00070259 | 4.00E-04 | 0.0033 | 0.0022 | 0.0022 | TRUE | 0.0195 |
| Stim2 | 1532 | 1049 | 948 | 969 | 821 | 920 | 1190 | 1116 | 1023 | 965 | 926 | 908 | 1021.24 | 1110 | 933 | 0.84 | -0.252 | 0.00239553 | 0.01390485 | 0.0026 | 0.0047 | 0.004 | 0.004 | TRUE | 0.0554 |
| Stip1 | 37136 | 26839 | 25992 | 26844 | 23717 | 27488 | 28842 | 28548 | 28045 | 26720 | 26755 | 27135 | 27674.34 | 28478 | 26870 | 0.944 | -0.084 | 0.05151038 | 0.13779485 | 1.00E-04 | 0.0025 | 0.0013 | 0.0013 | TRUE | 0.0022 |
| Stk10 | 2618 | 2034 | 2075 | 2102 | 1915 | 2198 | 2033 | 2164 | 2239 | 2092 | 2160 | 2170 | 2143.02 | 2145 | 2141 | 0.999 | -0.002 | 0.97678899 | 0.98763759 | 0.001 | 0.0035 | 0.0025 | 0.0025 | TRUE | 0.0261 |
| Stk11 | 3620 | 2688 | 2663 | 2691 | 2371 | 2933 | 2812 | 2859 | 2873 | 2679 | 2675 | 2895 | 2798.79 | 2848 | 2750 | 0.966 | -0.05 | 0.40357451 | 0.57648315 | 7.00E-04 | 0.0033 | 0.0022 | 0.0022 | TRUE | 0.0263 |
| Stk11ip | 2096 | 1609 | 1377 | 1325 | 1183 | 1363 | 1628 | 1711 | 1486 | 1319 | 1335 | 1345 | 1470.68 | 1608 | 1333 | 0.829 | -0.271 | 0.00025376 | 0.00255964 | 0.0019 | 0.004 | 0.0033 | 0.0033 | TRUE | 0.0521 |
| Stk16 | 1493 | 1056 | 865 | 872 | 808 | 976 | 1160 | 1123 | 933 | 868 | 912 | 963 | 993.18 | 1072 | 914 | 0.852 | -0.231 | 0.0157343 | 0.05785704 | 0.0071 | 0.0048 | 0.0056 | 0.0056 | TRUE | 0.1509 |
| Stk17b | 1594 | 1111 | 1187 | 1255 | 1131 | 1226 | 1238 | 1182 | 1281 | 1249 | 1276 | 1210 | 1239.31 | 1234 | 1245 | 1.009 | 0.013 | 0.85474409 | 0.91842259 | 0 | 0.0043 | 0.0029 | 0.0029 | TRUE | 0.0155 |
| Stk19 | 582 | 428 | 314 | 347 | 319 | 330 | 452 | 455 | 339 | 345 | 360 | 326 | 379.52 | 415 | 344 | 0.825 | -0.277 | 0.03541983 | 0.10481483 | 0.0124 | 0.0087 | 0.0098 | 0.0098 | TRUE | 0.2914 |
| Stk24 | 5491 | 4202 | 4426 | 4496 | 4126 | 4621 | 4265 | 4470 | 4776 | 4475 | 4655 | 4562 | 4533.56 | 4504 | 4564 | 1.014 | 0.02 | 0.74214267 | 0.84658231 | 0.0016 | 0.0029 | 0.0024 | 0.0024 | TRUE | 0.034 |
| Stk25 | 4117 | 3204 | 3006 | 3130 | 2762 | 3107 | 3197 | 3408 | 3243 | 3116 | 3116 | 3067 | 3191.25 | 3283 | 3100 | 0.944 | -0.083 | 0.14269705 | 0.28539162 | 3.00E-04 | 0.0032 | 0.002 | 0.002 | TRUE | 0.0134 |
| Stk3 | 1273 | 802 | 796 | 953 | 782 | 867 | 989 | 853 | 859 | 949 | 882 | 856 | 897.88 | 900 | 896 | 0.993 | -0.01 | 0.9109687 | 0.95115633 | 0.004 | 0.0051 | 0.0047 | 0.0047 | TRUE | 0.0917 |
| Stk30 | 624 | 516 | 571 | 469 | 450 | 499 | 485 | 549 | 616 | 467 | 508 | 493 | 519.45 | 550 | 489 | 0.891 | -0.166 | 0.13296751 | 0.27111335 | 0.0066 | 0.007 | 0.0069 | 0.0069 | TRUE | 0.1334 |
| Stk31 | 1449 | 1038 | 872 | 1049 | 798 | 893 | 1125 | 1104 | 941 | 1044 | 900 | 882 | 999.38 | 1057 | 942 | 0.891 | -0.167 | 0.08768799 | 0.20111112 | 0.0075 | 0.0048 | 0.0059 | 0.0059 | TRUE | 0.1084 |
| Stk32a | 260 | 201 | 181 | 190 | 177 | 193 | 202 | 214 | 195 | 189 | 200 | 191 | 198.39 | 204 | 193 | 0.948 | -0.077 | 0.583257 | 0.73307178 | 0 | 0.0144 | 0.0093 | 0.0093 | TRUE | 0.0194 |
| Stk32b | 39 | 39 | 34 | 57 | 46 | 54 | 30 | 41 | 37 | 57 | 52 | 53 | 45.07 | 36 | 54 | 1.504 | 0.589 | 0.04251616 | 0.11966986 | 0 | 0.055 | 0.0378 | 0.0378 | TRUE | 0.1769 |
| Stk32c | 210 | 142 | 133 | 218 | 149 | 182 | 163 | 151 | 144 | 217 | 168 | 180 | 170.4 | 153 | 188 | 1.233 | 0.302 | 0.06417789 | 0.16107015 | 0.0059 | 0.0163 | 0.0133 | 0.0133 | TRUE | 0.1938 |
| Stk33 | 23 | 23 | 23 | 15 | 16 | 19 | 18 | 24 | 25 | 15 | 18 | 19 | 19.81 | 22 | 17 | 0.776 | -0.367 | 0.39659216 | 0.56998409 | 0 | 0.122 | 0.083 | 0.083 | TRUE | 0.2218 |
| Stk35 | 3425 | 2375 | 2043 | 1707 | 1547 | 1737 | 2660 | 2526 | 2204 | 1699 | 1745 | 1715 | 2091.62 | 2463 | 1720 | 0.697 | -0.52 | 2.52E-11 | 2.34E-09 | 0.0044 | 0.0036 | 0.0039 | 0.0039 | TRUE | 0.1019 |
| Stk36 | 254 | 197 | 177 | 204 | 174 | 206 | 197 | 210 | 191 | 203 | 196 | 203 | 200.08 | 199 | 201 | 1.009 | 0.013 | 0.92765369 | 0.96016758 | 0 | 0.0143 | 0.0091 | 0.0091 | TRUE | 0.0208 |
| Stk38 | 6136 | 4856 | 4683 | 4716 | 4528 | 5440 | 4766 | 5165 | 5053 | 4694 | 5108 | 5370 | 5026.04 | 4995 | 5057 | 1.013 | 0.018 | 0.77942756 | 0.8699327 | 0.003 | 0.0029 | 0.0029 | 0.0029 | TRUE | 0.0483 |
| Stk38l | 817 | 623 | 525 | 533 | 548 | 562 | 635 | 663 | 566 | 531 | 618 | 555 | 594.53 | 621 | 568 | 0.913 | -0.132 | 0.19611132 | 0.35566794 | 0.0045 | 0.0064 | 0.0058 | 0.0058 | TRUE | 0.0689 |
| Stk39 | 426 | 316 | 404 | 460 | 409 | 468 | 331 | 336 | 436 | 458 | 461 | 462 | 414.03 | 368 | 460 | 1.256 | 0.328 | 0.00776747 | 0.03425 | 0.0093 | 0.0082 | 0.0085 | 0.0085 | TRUE | 0.2954 |
| Stk4 | 7149 | 5196 | 4775 | 4760 | 4324 | 5043 | 5552 | 5527 | 5152 | 4738 | 4878 | 4978 | 5137.61 | 5410 | 4865 | 0.899 | -0.154 | 0.00619015 | 0.02887273 | 9.00E-04 | 0.0029 | 0.0021 | 0.0021 | TRUE | 0.0211 |
| Stk40 | 3613 | 2661 | 2487 | 2558 | 2399 | 2858 | 2806 | 2830 | 2683 | 2546 | 2706 | 2821 | 2732.3 | 2773 | 2691 | 0.97 | -0.044 | 0.49164443 | 0.65729264 | 0.0012 | 0.0033 | 0.0025 | 0.0025 | TRUE | 0.0273 |
| Stmn1 | 6375 | 4645 | 4764 | 4906 | 4556 | 5524 | 4951 | 4941 | 5140 | 4883 | 5140 | 5453 | 5084.74 | 5011 | 5159 | 1.03 | 0.042 | 0.47805191 | 0.64546861 | 0.0015 | 0.0029 | 0.0023 | 0.0023 | TRUE | 0.0306 |
| Stmn1-rs1 | 1 | 0 | 2 | 0 | 2 | 0 | 1 | 0 | 2 | 0 | 2 | 0 | 0.87 | 1 | 1 | 0.751 | -0.414 | 0.85669157 | NA | 0 | 2.7419 | 2.5715 | 2.5715 | TRUE | 0.7602 |
| Stmn2 | 16437 | 11906 | 13265 | 15839 | 14190 | 16204 | 12766 | 12664 | 14313 | 15766 | 16008 | 15996 | 14585.46 | 13248 | 15923 | 1.202 | 0.266 | 7.91E-06 | 0.00015493 | 0.0024 | 0.0026 | 0.0025 | 0.0025 | TRUE | 0.0603 |
| Stmn3 | 117 | 86 | 75 | 110 | 95 | 98 | 91 | 91 | 81 | 109 | 107 | 97 | 96.11 | 88 | 104 | 1.188 | 0.249 | 0.21129056 | 0.37334356 | 0 | 0.0271 | 0.0181 | 0.0181 | TRUE | 0.0424 |
| Stmn4 | 1 | 0 | 1 | 0 | 0 | 0 | 1 | 0 | 1 | 0 | 0 | 0 | 0.31 | 1 | 0 | 0.307 | -1.706 | 0.62894649 | NA | 0 | 7.6657 | 7.2 | 7.2 | TRUE | 0.2067 |
| Stmnd1 | 2 | 0 | 0 | 0 | 1 | 0 | 2 | 0 | 0 | 0 | 1 | 0 | 0.45 | 1 | 0 | 0.68 | -0.556 | 0.85993441 | NA | 0 | 5.3057 | 5.2576 | 5.2576 | TRUE | 0.9726 |
| Stom | 1247 | 1016 | 1234 | 1590 | 1337 | 1601 | 968 | 1081 | 1331 | 1583 | 1508 | 1580 | 1342.01 | 1127 | 1557 | 1.384 | 0.469 | 5.60E-06 | 0.00011652 | 0.0128 | 0.0042 | 0.0069 | 0.0069 | TRUE | 0.3009 |
| Stoml1 | 2388 | 1934 | 1952 | 2247 | 1911 | 2210 | 1855 | 2057 | 2106 | 2237 | 2156 | 2182 | 2098.68 | 2006 | 2192 | 1.094 | 0.129 | 0.06173264 | 0.15657854 | 0.002 | 0.0036 | 0.003 | 0.003 | TRUE | 0.0545 |

| | | | | | | | | | | | | | | | | | | | | | | | | | |
|---|---|---|---|---|---|---|---|---|---|---|---|---|---|---|---|---|---|---|---|---|---|---|---|---|---|
| Stoml2 | 6387 | 4619 | 4431 | 4527 | 3791 | 4683 | 4961 | 4913 | 4781 | 4506 | 4277 | 4623 | 4676.72 | 4885 | 4469 | 0.915 | -0.128 | 0.02029446 | 0.06992978 | 6.00E-04 | 0.0029 | 0.002 | 0.002 | TRUE | 0.017 |
| Stoml3 | 1 | 2 | 1 | 0 | 0 | 0 | 1 | 2 | 1 | 0 | 0 | 0 | 0.66 | 1 | 0 | 0.145 | -2.79 | 0.24993894 | NA | 0 | 3.5727 | 2.8701 | 2.8701 | TRUE | 0.1613 |
| Ston1 | 670 | 499 | 532 | 567 | 498 | 583 | 520 | 531 | 574 | 564 | 562 | 576 | 554.48 | 542 | 567 | 1.049 | 0.068 | 0.46269138 | 0.63199999 | 0 | 0.0067 | 0.0044 | 0.0044 | TRUE | 0.0311 |
| Ston2 | 4304 | 2954 | 3081 | 2912 | 2661 | 3159 | 3343 | 3142 | 3324 | 2899 | 3002 | 3118 | 3138.02 | 3270 | 3006 | 0.919 | -0.122 | 0.04246944 | 0.11958544 | 9.00E-04 | 0.0032 | 0.0023 | 0.0023 | TRUE | 0.014 |
| Stox1 | 72 | 56 | 36 | 44 | 37 | 29 | 56 | 60 | 39 | 44 | 42 | 29 | 44.75 | 52 | 38 | 0.736 | -0.442 | 0.1576958 | 0.30665933 | 0.0253 | 0.0554 | 0.0474 | 0.0474 | TRUE | 0.3573 |
| Stox2 | 2107 | 1452 | 1365 | 1349 | 1140 | 1372 | 1636 | 1544 | 1473 | 1343 | 1286 | 1354 | 1439.49 | 1551 | 1328 | 0.855 | -0.225 | 0.00165282 | 0.01040236 | 0.0012 | 0.0041 | 0.003 | 0.003 | TRUE | 0.0288 |
| Stpg1 | 123 | 62 | 53 | 77 | 57 | 60 | 96 | 66 | 57 | 77 | 64 | 59 | 69.81 | 73 | 67 | 0.908 | -0.14 | 0.59614085 | 0.74235543 | 0.0339 | 0.0364 | 0.0356 | 0.0356 | TRUE | 0.7423 |
| Stpg2 | 1 | 0 | 2 | 1 | 0 | 0 | 1 | 0 | 2 | 1 | 0 | 0 | 0.66 | 1 | 0 | 0.382 | -1.39 | 0.57951664 | NA | 0 | 3.6208 | 3.037 | 3.037 | TRUE | 0.2643 |
| Stra13 | 1898 | 1269 | 1457 | 1726 | 1442 | 1761 | 1474 | 1350 | 1572 | 1718 | 1627 | 1738 | 1579.86 | 1465 | 1694 | 1.157 | 0.21 | 0.00564223 | 0.02693891 | 0.0026 | 0.0039 | 0.0035 | 0.0035 | TRUE | 0.0556 |
| Stra6 | 110 | 62 | 54 | 93 | 71 | 67 | 85 | 66 | 58 | 93 | 80 | 66 | 74.74 | 70 | 80 | 1.131 | 0.177 | 0.47319574 | 0.64174843 | 0.0217 | 0.0341 | 0.0305 | 0.0305 | TRUE | 0.3573 |
| Stra8 | 3620 | 2194 | 1639 | 1224 | 987 | 1180 | 2812 | 2334 | 1768 | 1218 | 1113 | 1165 | 1735.06 | 2305 | 1165 | 0.505 | -0.985 | 7.31E-15 | 1.21E-12 | 0.0257 | 0.0038 | 0.0109 | 0.0109 | TRUE | 0.131 |
| Strada | 2687 | 1972 | 1818 | 2054 | 1844 | 2146 | 2087 | 2098 | 1962 | 2045 | 2080 | 2118 | 2064.88 | 2049 | 2081 | 1.015 | 0.022 | 0.72246647 | 0.83390238 | 3.00E-04 | 0.0036 | 0.0023 | 0.0023 | TRUE | 0.0164 |
| Stradb | 935 | 635 | 702 | 863 | 729 | 789 | 726 | 675 | 757 | 859 | 822 | 779 | 769.89 | 719 | 820 | 1.139 | 0.188 | 0.0308305 | 0.09450145 | 0.0014 | 0.0055 | 0.0042 | 0.0042 | TRUE | 0.0328 |
| Strap | 21358 | 15014 | 14896 | 15744 | 13629 | 15898 | 16588 | 15970 | 16073 | 15672 | 15375 | 15694 | 15895.16 | 16210 | 15580 | 0.961 | -0.057 | 0.20804409 | 0.36966361 | 2.00E-04 | 0.0026 | 0.0014 | 0.0014 | TRUE | 0.0051 |
| Strbp | 8732 | 6329 | 7081 | 8103 | 6720 | 8174 | 6782 | 6732 | 7640 | 8066 | 7581 | 8069 | 7478.29 | 7051 | 7905 | 1.121 | 0.165 | 0.01037613 | 0.04242206 | 0.0031 | 0.0027 | 0.0029 | 0.0029 | TRUE | 0.0648 |
| Strc | 3 | 5 | 2 | 5 | 3 | 1 | 2 | 5 | 2 | 5 | 3 | 1 | 3.19 | 3 | 3 | 0.961 | -0.058 | 0.95791719 | NA | 0 | 0.7448 | 0.5473 | 0.5473 | TRUE | 0.3537 |
| Strip1 | 4636 | 3344 | 3146 | 3243 | 2861 | 3353 | 3601 | 3557 | 3395 | 3228 | 3228 | 3310 | 3386.26 | 3518 | 3255 | 0.925 | -0.112 | 0.04394754 | 0.12255312 | 3.00E-04 | 0.0031 | 0.0019 | 0.0019 | TRUE | 0.0112 |
| Strip2 | 283 | 208 | 171 | 244 | 216 | 199 | 220 | 221 | 185 | 243 | 244 | 196 | 218.09 | 209 | 228 | 1.089 | 0.123 | 0.41336808 | 0.5860164 | 0.0077 | 0.0133 | 0.0117 | 0.0117 | TRUE | 0.1614 |
| Strn | 1827 | 1401 | 1285 | 1397 | 1248 | 1480 | 1419 | 1490 | 1387 | 1391 | 1408 | 1461 | 1425.85 | 1432 | 1420 | 0.992 | -0.012 | 0.86250554 | 0.92408891 | 0 | 0.0041 | 0.0027 | 0.0027 | TRUE | 0.0148 |
| Strn3 | 6879 | 4903 | 4728 | 5153 | 4488 | 5159 | 5343 | 5215 | 5101 | 5129 | 5063 | 5093 | 5157.38 | 5220 | 5095 | 0.976 | -0.035 | 0.48491402 | 0.65090872 | 0 | 0.0029 | 0.0016 | 0.0016 | TRUE | 0.0053 |
| Strn4 | 5790 | 4443 | 4285 | 4592 | 4194 | 4609 | 4497 | 4726 | 4623 | 4571 | 4731 | 4550 | 4616.37 | 4615 | 4617 | 1.001 | 0.001 | 0.98789837 | 0.99380852 | 3.00E-04 | 0.0029 | 0.0018 | 0.0018 | TRUE | 0.0063 |
| Stt3a | 13613 | 9887 | 11074 | 13099 | 11273 | 13206 | 10573 | 10517 | 11949 | 13039 | 12717 | 13036 | 11971.71 | 11013 | 12931 | 1.174 | 0.232 | 0.00015882 | 0.00178321 | 0.0027 | 0.0026 | 0.0026 | 0.0026 | TRUE | 0.0673 |
| Stt3b | 11086 | 8413 | 8952 | 9488 | 8246 | 9157 | 8610 | 8949 | 9659 | 9444 | 9302 | 9039 | 9167.34 | 9073 | 9262 | 1.021 | 0.03 | 0.60516639 | 0.74984457 | 0.0016 | 0.0027 | 0.0023 | 0.0023 | TRUE | 0.039 |
| Stub1 | 4041 | 2908 | 2624 | 2717 | 2418 | 2796 | 3138 | 3093 | 2831 | 2704 | 2728 | 2760 | 2875.88 | 3021 | 2731 | 0.904 | -0.146 | 0.01892229 | 0.06641852 | 0.0013 | 0.0033 | 0.0025 | 0.0025 | TRUE | 0.0362 |
| Stx11 | 120 | 115 | 123 | 101 | 116 | 124 | 93 | 122 | 133 | 101 | 131 | 122 | 117.01 | 116 | 118 | 1.021 | 0.029 | 0.88500784 | 0.93609243 | 0.018 | 0.0227 | 0.0213 | 0.0213 | TRUE | 0.3338 |
| Stx12 | 3749 | 2749 | 3141 | 3753 | 3172 | 3585 | 2912 | 2924 | 3389 | 3736 | 3578 | 3539 | 3346.32 | 3075 | 3618 | 1.177 | 0.235 | 0.00104129 | 0.00745281 | 0.0039 | 0.0031 | 0.0034 | 0.0034 | TRUE | 0.0962 |
| Stx16 | 2741 | 2108 | 2025 | 2385 | 2017 | 2343 | 2129 | 2242 | 2185 | 2374 | 2275 | 2313 | 2253.06 | 2185 | 2321 | 1.063 | 0.088 | 0.14361167 | 0.28656173 | 0 | 0.0035 | 0.0021 | 0.0021 | TRUE | 0.0064 |
| Stx17 | 1154 | 844 | 771 | 910 | 811 | 879 | 896 | 898 | 832 | 906 | 915 | 868 | 885.72 | 875 | 896 | 1.023 | 0.033 | 0.68100035 | 0.80422523 | 0 | 0.0051 | 0.0034 | 0.0034 | TRUE | 0.0217 |
| Stx18 | 603 | 497 | 407 | 417 | 389 | 439 | 468 | 529 | 439 | 415 | 439 | 433 | 453.9 | 479 | 429 | 0.897 | -0.158 | 0.14223155 | 0.28474364 | 0.0027 | 0.0076 | 0.0061 | 0.0061 | TRUE | 0.093 |
| Stx19 | 0 | 0 | 0 | 0 | 0 | 0 | 0 | 0 | 0 | 0 | 0 | 0 | 0 | 0 | 0 | NA | NA | NA | NA | NA | NA | NA | NA | NA | NA |
| Stx1a | 375 | 314 | 318 | 383 | 293 | 336 | 291 | 334 | 343 | 381 | 331 | 332 | 335.3 | 323 | 348 | 1.081 | 0.113 | 0.36093458 | 0.53587181 | 0.0046 | 0.0095 | 0.008 | 0.008 | TRUE | 0.0897 |
| Stx1b | 87 | 82 | 71 | 97 | 76 | 97 | 68 | 87 | 77 | 97 | 86 | 96 | 84.91 | 77 | 93 | 1.209 | 0.273 | 0.20491454 | 0.36644055 | 0 | 0.0303 | 0.0216 | 0.0216 | TRUE | 0.1254 |
| Stx2 | 438 | 356 | 372 | 380 | 335 | 408 | 340 | 379 | 401 | 378 | 378 | 403 | 379.86 | 373 | 386 | 1.038 | 0.053 | 0.63663218 | 0.77290969 | 0.0021 | 0.0087 | 0.0065 | 0.0065 | TRUE | 0.0751 |
| Stx3 | 1043 | 802 | 1028 | 1003 | 891 | 1140 | 810 | 853 | 1109 | 998 | 1005 | 1125 | 983.54 | 924 | 1043 | 1.131 | 0.177 | 0.11960245 | 0.25134212 | 0.0161 | 0.0048 | 0.0083 | 0.0083 | TRUE | 0.3629 |
| Stx4a | 974 | 685 | 675 | 608 | 559 | 695 | 756 | 729 | 728 | 605 | 631 | 686 | 689.22 | 738 | 641 | 0.868 | -0.204 | 0.02113598 | 0.07204457 | 7.00E-04 | 0.0059 | 0.0042 | 0.0042 | TRUE | 0.0463 |
| Stx5a | 1451 | 992 | 887 | 896 | 810 | 866 | 1127 | 1055 | 957 | 892 | 914 | 855 | 966.62 | 1046 | 887 | 0.846 | -0.241 | 0.00488257 | 0.02405605 | 0.0031 | 0.0049 | 0.0042 | 0.0042 | TRUE | 0.066 |
| Stx6 | 1816 | 1374 | 1342 | 1449 | 1354 | 1491 | 1410 | 1462 | 1448 | 1442 | 1527 | 1472 | 1460.26 | 1440 | 1480 | 1.028 | 0.04 | 0.54460865 | 0.70202845 | 0 | 0.004 | 0.0025 | 0.0025 | TRUE | 0.0091 |
| Stx7 | 2615 | 1848 | 2112 | 2225 | 1889 | 2118 | 2031 | 1966 | 2279 | 2215 | 2131 | 2091 | 2118.67 | 2092 | 2146 | 1.026 | 0.037 | 0.60870578 | 0.75233722 | 0.003 | 0.0035 | 0.0033 | 0.0033 | TRUE | 0.0727 |
| Stx8 | 1346 | 935 | 928 | 996 | 861 | 942 | 1045 | 995 | 1001 | 991 | 971 | 930 | 988.97 | 1014 | 964 | 0.95 | -0.074 | 0.33083636 | 0.50530499 | 0 | 0.0048 | 0.0031 | 0.0031 | TRUE | 0.0118 |
| Stxbp1 | 1859 | 1384 | 1481 | 1773 | 1626 | 1920 | 1444 | 1472 | 1598 | 1765 | 1834 | 1895 | 1668.07 | 1505 | 1831 | 1.218 | 0.285 | 5.23E-05 | 0.00072684 | 0.0015 | 0.0038 | 0.003 | 0.003 | TRUE | 0.035 |
| Stxbp2 | 4820 | 3844 | 4435 | 3923 | 3978 | 4528 | 3743 | 4089 | 4785 | 3905 | 4488 | 4470 | 4246.67 | 4206 | 4288 | 1.02 | 0.028 | 0.75338253 | 0.85359465 | 0.0106 | 0.003 | 0.0056 | 0.0056 | TRUE | 0.1766 |
| Stxbp3a | 1828 | 1332 | 1416 | 1545 | 1293 | 1498 | 1420 | 1417 | 1528 | 1538 | 1459 | 1479 | 1473.28 | 1455 | 1492 | 1.026 | 0.037 | 0.59062335 | 0.73901229 | 6.00E-04 | 0.004 | 0.0028 | 0.0028 | TRUE | 0.0227 |
| Stxbp3b | 8 | 9 | 11 | 16 | 12 | 13 | 6 | 10 | 12 | 16 | 14 | 13 | 11.66 | 9 | 14 | 1.554 | 0.636 | 0.2623469 | NA | 0 | 0.2057 | 0.1419 | 0.1419 | TRUE | 0.3564 |
| Stxbp4 | 1631 | 1146 | 999 | 929 | 953 | 1111 | 1267 | 1219 | 1078 | 925 | 1075 | 1097 | 1110.03 | 1188 | 1032 | 0.868 | -0.204 | 0.02798149 | 0.08801484 | 0.0068 | 0.0046 | 0.0053 | 0.0053 | TRUE | 0.1 |
| Stxbp5 | 927 | 690 | 828 | 802 | 711 | 908 | 720 | 734 | 893 | 798 | 802 | 896 | 807.34 | 782 | 832 | 1.065 | 0.092 | 0.37065234 | 0.54531486 | 0.0082 | 0.0054 | 0.0063 | 0.0063 | TRUE | 0.1798 |
| Stxbp5l | 39 | 35 | 20 | 29 | 23 | 20 | 30 | 37 | 22 | 29 | 26 | 20 | 27.28 | 30 | 25 | 0.835 | -0.26 | 0.4993584 | 0.66394997 | 0 | 0.0893 | 0.0696 | 0.0696 | TRUE | 0.1993 |
| Stxbp6 | 259 | 156 | 191 | 179 | 122 | 186 | 201 | 166 | 206 | 178 | 138 | 184 | 178.77 | 191 | 167 | 0.872 | -0.197 | 0.2399564 | 0.407134 | 0.0121 | 0.0157 | 0.0147 | 0.0147 | TRUE | 0.2335 |
| Styk1 | 41 | 32 | 24 | 48 | 49 | 44 | 32 | 34 | 26 | 48 | 55 | 43 | 39.71 | 31 | 49 | 1.59 | 0.669 | 0.03122871 | 0.09539999 | 0 | 0.0621 | 0.0431 | 0.0431 | TRUE | 0.1092 |
| Styx | 828 | 590 | 648 | 740 | 605 | 690 | 643 | 628 | 699 | 737 | 683 | 681 | 678.34 | 657 | 700 | 1.067 | 0.094 | 0.29649002 | 0.46836117 | 0.001 | 0.0059 | 0.0043 | 0.0043 | TRUE | 0.0365 |
| Styxl1 | 11 | 3 | 4 | 6 | 3 | 7 | 9 | 3 | 4 | 6 | 3 | 7 | 5.39 | 5 | 5 | 0.998 | -0.003 | 0.99682061 | NA | 0 | 0.4425 | 0.322 | 0.322 | TRUE | 0.7553 |
| Sub1 | 13706 | 10464 | 9421 | 9627 | 8818 | 9725 | 10645 | 11130 | 10165 | 9583 | 9948 | 9600 | 10178.48 | 10647 | 9710 | 0.912 | -0.133 | 0.01355419 | 0.05185495 | 0.0011 | 0.0027 | 0.002 | 0.002 | TRUE | 0.0191 |
| Sucla2 | 6053 | 4433 | 4467 | 4679 | 3955 | 4544 | 4701 | 4715 | 4820 | 4657 | 4462 | 4486 | 4640.17 | 4745 | 4535 | 0.956 | -0.065 | 0.20723522 | 0.36907176 | 1.00E-04 | 0.0029 | 0.0017 | 0.0017 | TRUE | 0.0068 |
| Suclg1 | 6406 | 4382 | 4386 | 4917 | 4239 | 4813 | 4975 | 4661 | 4732 | 4894 | 4782 | 4751 | 4799.4 | 4789 | 4809 | 1.004 | 0.005 | 0.92048926 | 0.95642287 | 5.00E-04 | 0.0029 | 0.0019 | 0.0019 | TRUE | 0.0143 |
| Suclg2 | 11617 | 8528 | 9601 | 9758 | 8401 | 9603 | 9022 | 9071 | 10359 | 9713 | 9477 | 9480 | 9520.48 | 9484 | 9557 | 1.008 | 0.011 | 0.86055329 | 0.92258387 | 0.0025 | 0.0027 | 0.0028 | 0.0028 | TRUE | 0.0793 |
| Sucnr1 | 0 | 0 | 0 | 0 | 0 | 0 | 0 | 0 | 0 | 0 | 0 | 0 | 0 | 0 | 0 | NA | NA | NA | NA | NA | NA | NA | NA | NA | NA |
| Suco | 4739 | 3424 | 3477 | 4212 | 3405 | 3904 | 3681 | 3642 | 3752 | 4193 | 3841 | 3854 | 3826.99 | 3692 | 3963 | 1.074 | 0.102 | 0.08193397 | 0.19242231 | 0.001 | 0.003 | 0.0022 | 0.0022 | TRUE | 0.0315 |
| Suds3 | 5335 | 3923 | 3573 | 3551 | 3425 | 3771 | 4143 | 4173 | 3855 | 3535 | 3864 | 3723 | 3882.09 | 4057 | 3707 | 0.913 | -0.131 | 0.03397037 | 0.10164087 | 0.0015 | 0.003 | 0.0025 | 0.0025 | TRUE | 0.0228 |
| Sufu | 1064 | 767 | 677 | 691 | 621 | 690 | 826 | 816 | 730 | 688 | 701 | 681 | 740.37 | 791 | 690 | 0.871 | -0.199 | 0.02227932 | 0.07457708 | 0.0011 | 0.0056 | 0.0041 | 0.0041 | TRUE | 0.0516 |
| Sugp1 | 4194 | 3020 | 2385 | 2211 | 2121 | 2298 | 3257 | 3212 | 2573 | 2201 | 2393 | 2268 | 2650.84 | 3014 | 2287 | 0.758 | -0.399 | 7.10E-06 | 0.00014131 | 0.0084 | 0.0033 | 0.0053 | 0.0053 | TRUE | 0.1975 |
| Sugp2 | 3271 | 2991 | 3340 | 3322 | 2922 | 3435 | 2540 | 3181 | 3604 | 3307 | 3296 | 3391 | 3219.95 | 3108 | 3331 | 1.073 | 0.101 | 0.31805236 | 0.49189826 | 0.0152 | 0.0032 | 0.0071 | 0.0071 | TRUE | 0.3143 |
| Sugt1 | 5997 | 4231 | 4408 | 4282 | 3743 | 4537 | 4658 | 4500 | 4756 | 4262 | 4222 | 4479 | 4479.63 | 4638 | 4321 | 0.932 | -0.102 | 0.06481857 | 0.16231787 | 6.00E-04 | 0.003 | 0.002 | 0.002 | TRUE | 0.0124 |
| Sulf1 | 1612 | 1119 | 1140 | 1214 | 1134 | 1324 | 1252 | 1190 | 1230 | 1208 | 1279 | 1307 | 1244.49 | 1224 | 1265 | 1.033 | 0.046 | 0.51633893 | 0.67856126 | 0 | 0.0043 | 0.0029 | 0.0029 | TRUE | 0.0186 |
| Sulf2 | 11504 | 8274 | 7868 | 7340 | 6713 | 7457 | 8935 | 8801 | 8490 | 7306 | 7573 | 7361 | 8077.59 | 8742 | 7413 | 0.848 | -0.238 | 1.72E-06 | 4.50E-05 | 3.00E-04 | 0.0027 | 0.0017 | 0.0017 | TRUE | 0.0077 |
| Sult1a1 | 2 | 0 | 1 | 1 | 1 | 1 | 2 | 0 | 1 | 1 | 1 | 1 | 0.96 | 1 | 1 | 1.143 | 0.193 | 0.91871967 | NA | 0 | 2.4786 | 1.5559 | 1.5559 | TRUE | 0.2848 |
| Sult1b1 | 0 | 0 | 0 | 0 | 0 | 0 | 0 | 0 | 0 | 0 | 0 | 0 | 0 | 0 | 0 | NA | NA | NA | NA | NA | NA | NA | NA | NA | NA |
| Sult1c1 | 0 | 2 | 0 | 0 | 0 | 0 | 0 | 2 | 0 | 0 | 0 | 0 | 0.35 | 1 | 0 | 0.272 | -1.878 | 0.62190613 | NA | 10 | 6.6869 | 8.7633 | 8.7633 | TRUE | 0.9743 |
| Sult1c2 | 0 | 0 | 0 | 0 | 0 | 0 | 0 | 0 | 0 | 0 | 0 | 0 | 0 | 0 | 0 | NA | NA | NA | NA | NA | NA | NA | NA | NA | NA |
| Sult1d1 | 0 | 0 | 0 | 0 | 0 | 0 | 0 | 0 | 0 | 0 | 0 | 0 | 0 | 0 | 0 | NA | NA | NA | NA | NA | NA | NA | NA | NA | NA |
| Sult1e1 | 0 | 0 | 0 | 0 | 0 | 0 | 0 | 0 | 0 | 0 | 0 | 0 | 0 | 0 | 0 | NA | NA | NA | NA | NA | NA | NA | NA | NA | NA |

| | | | | | | | | | | | | | | | | | | | | | | | | | |
|---|---|---|---|---|---|---|---|---|---|---|---|---|---|---|---|---|---|---|---|---|---|---|---|---|---|
| Sult2a1 | 0 | 0 | 0 | 0 | 0 | 0 | 0 | 0 | 0 | 0 | 0 | 0 | 0 | 0 | 0 | NA | NA | NA | NA | NA | NA | NA | NA | NA | NA |
| Sult2a2 | 0 | 0 | 0 | 0 | 0 | 0 | 0 | 0 | 0 | 0 | 0 | 0 | 0 | 0 | 0 | NA | NA | NA | NA | NA | NA | NA | NA | NA | NA |
| Sult2a3 | 2 | 2 | 1 | 0 | 1 | 1 | 2 | 2 | 1 | 0 | 1 | 1 | 1.15 | 2 | 1 | 0.439 | -1.188 | 0.5118568 | NA | 0 | 2.0709 | 1.3166 | 1.3166 | TRUE | 0.2658 |
| Sult2a4 | 5 | 2 | 3 | 5 | 2 | 3 | 4 | 2 | 3 | 5 | 2 | 3 | 3.24 | 3 | 3 | 1.097 | 0.133 | 0.89803664 | NA | 0 | 0.7338 | 0.4732 | 0.4732 | TRUE | 0.2434 |
| Sult2a5 | 5 | 2 | 1 | 2 | 3 | 3 | 4 | 2 | 1 | 2 | 3 | 3 | 2.57 | 2 | 3 | 1.141 | 0.19 | 0.87139042 | NA | 0 | 0.9243 | 0.6053 | 0.6053 | TRUE | 0.442 |
| Sult2a6 | 0 | 0 | 0 | 0 | 0 | 0 | 0 | 0 | 0 | 0 | 0 | 0 | 0 | 0 | 0 | NA | NA | NA | NA | NA | NA | NA | NA | NA | NA |
| Sult2a7 | 1 | 0 | 0 | 0 | 0 | 0 | 1 | 0 | 0 | 0 | 0 | 0 | 0.13 | 0 | 0 | 0.552 | -0.857 | 0.8336548 | NA | 0 | 18.3121 | 10 | 10 | TRUE | 0.2566 |
| Sult2b1 | 123 | 83 | 125 | 131 | 117 | 126 | 96 | 88 | 135 | 130 | 132 | 124 | 117.58 | 106 | 129 | 1.218 | 0.284 | 0.15995758 | 0.30945258 | 0.0162 | 0.0226 | 0.0209 | 0.0209 | TRUE | 0.5243 |
| Sult3a1 | 0 | 0 | 0 | 0 | 0 | 0 | 0 | 0 | 0 | 0 | 0 | 0 | 0 | 0 | 0 | NA | NA | NA | NA | NA | NA | NA | NA | NA | NA |
| Sult4a1 | 2792 | 2024 | 2044 | 2115 | 1976 | 2286 | 2168 | 2153 | 2205 | 2105 | 2229 | 2257 | 2186.3 | 2175 | 2197 | 1.01 | 0.014 | 0.81785052 | 0.89536894 | 3.00E-04 | 0.0035 | 0.0022 | 0.0022 | TRUE | 0.0163 |
| Sult5a1 | 190 | 134 | 159 | 203 | 172 | 209 | 148 | 143 | 172 | 202 | 194 | 206 | 177.35 | 154 | 201 | 1.308 | 0.387 | 0.01125841 | 0.04506203 | 0 | 0.0158 | 0.0111 | 0.0111 | TRUE | 0.1002 |
| Sult6b1 | 485 | 423 | 392 | 486 | 391 | 434 | 377 | 450 | 423 | 484 | 441 | 428 | 433.81 | 417 | 451 | 1.086 | 0.119 | 0.28640689 | 0.45837651 | 0.0041 | 0.0079 | 0.0067 | 0.0067 | TRUE | 0.0869 |
| Sumf1 | 1745 | 1233 | 1172 | 1208 | 1036 | 1152 | 1355 | 1312 | 1265 | 1202 | 1169 | 1137 | 1239.96 | 1311 | 1169 | 0.892 | -0.165 | 0.02024729 | 0.06985146 | 0 | 0.0043 | 0.0028 | 0.0028 | TRUE | 0.0112 |
| Sumf2 | 2001 | 1445 | 1413 | 1567 | 1452 | 1658 | 1554 | 1537 | 1525 | 1560 | 1638 | 1637 | 1575.04 | 1539 | 1612 | 1.047 | 0.066 | 0.30460327 | 0.4771695 | 0 | 0.0039 | 0.0024 | 0.0024 | TRUE | 0.0096 |
| Sumo1 | 6211 | 4277 | 4027 | 4102 | 3518 | 4018 | 4824 | 4549 | 4345 | 4083 | 3969 | 3966 | 4289.42 | 4573 | 4006 | 0.876 | -0.192 | 0.00121146 | 0.00831274 | 0.0012 | 0.003 | 0.0023 | 0.0023 | TRUE | 0.0286 |
| Sumo2 | 3549 | 2353 | 2366 | 2605 | 2078 | 2585 | 2756 | 2503 | 2553 | 2593 | 2344 | 2552 | 2550.19 | 2604 | 2496 | 0.958 | -0.061 | 0.36941288 | 0.54416348 | 0.0022 | 0.0034 | 0.003 | 0.003 | TRUE | 0.034 |
| Sumo3 | 7936 | 5516 | 5682 | 5774 | 5143 | 6310 | 6164 | 5867 | 6131 | 5747 | 5802 | 6229 | 5989.98 | 6054 | 5926 | 0.979 | -0.031 | 0.58355641 | 0.73325466 | 0.001 | 0.0028 | 0.0021 | 0.0021 | TRUE | 0.0245 |
| Sun1 | 12105 | 9420 | 10739 | 11307 | 10070 | 11506 | 9401 | 10020 | 11587 | 11255 | 11360 | 11358 | 10830.31 | 10336 | 11324 | 1.096 | 0.132 | 0.07246161 | 0.17601865 | 0.0058 | 0.0026 | 0.0038 | 0.0038 | TRUE | 0.1368 |
| Sun2 | 2253 | 1851 | 1774 | 2208 | 1964 | 2320 | 1750 | 1969 | 1914 | 2198 | 2216 | 2290 | 2056.08 | 1878 | 2235 | 1.191 | 0.253 | 0.00020387 | 0.00216164 | 0.0017 | 0.0036 | 0.0028 | 0.0028 | TRUE | 0.0445 |
| Sun3 | 13 | 5 | 8 | 7 | 3 | 7 | 10 | 5 | 9 | 7 | 3 | 7 | 6.88 | 8 | 6 | 0.717 | -0.48 | 0.51555526 | NA | 0 | 0.3467 | 0.2434 | 0.2434 | TRUE | 0.2573 |
| Sun5 | 0 | 0 | 0 | 0 | 0 | 1 | 0 | 0 | 0 | 0 | 0 | 1 | 0.16 | 0 | 0 | 2.094 | 1.067 | 0.79379329 | NA | 0 | 14.4078 | 10 | 10 | TRUE | 0.3835 |
| Suox | 494 | 358 | 328 | 445 | 384 | 447 | 384 | 381 | 354 | 443 | 433 | 441 | 405.96 | 373 | 439 | 1.177 | 0.235 | 0.02350934 | 0.07776418 | 0 | 0.0083 | 0.0053 | 0.0053 | TRUE | 0.0217 |
| Supt16 | 31629 | 22839 | 21615 | 22930 | 20081 | 23293 | 24565 | 24294 | 23322 | 22824 | 22653 | 22994 | 23442.1 | 24060 | 22824 | 0.949 | -0.076 | 0.09693795 | 0.21645588 | 3.00E-04 | 0.0025 | 0.0015 | 0.0015 | TRUE | 0.0088 |
| Supt20 | 4290 | 3140 | 3095 | 3521 | 3146 | 3688 | 3332 | 3340 | 3339 | 3505 | 3549 | 3641 | 3450.96 | 3337 | 3565 | 1.068 | 0.095 | 0.07228106 | 0.17570262 | 0 | 0.0031 | 0.0017 | 0.0017 | TRUE | 0.0043 |
| Supt3 | 976 | 780 | 796 | 868 | 749 | 919 | 758 | 830 | 859 | 864 | 845 | 907 | 843.79 | 816 | 872 | 1.071 | 0.1 | 0.24078567 | 0.40829869 | 0.0017 | 0.0052 | 0.004 | 0.004 | TRUE | 0.0478 |
| Supt4a | 2814 | 1986 | 1814 | 1787 | 1507 | 1711 | 2186 | 2112 | 1957 | 1779 | 1700 | 1689 | 1903.86 | 2085 | 1723 | 0.826 | -0.276 | 5.49E-05 | 0.00075576 | 0.0015 | 0.0037 | 0.0029 | 0.0029 | TRUE | 0.0345 |
| Supt5 | 17495 | 12802 | 11857 | 11351 | 10479 | 12177 | 13588 | 13617 | 12794 | 11299 | 11821 | 12021 | 12523.22 | 13333 | 11714 | 0.878 | -0.187 | 0.00038068 | 0.00350586 | 0.001 | 0.0026 | 0.0019 | 0.0019 | TRUE | 0.0152 |
| Supt6 | 17908 | 12221 | 9843 | 10469 | 9420 | 10767 | 13908 | 12999 | 10621 | 10421 | 10627 | 10629 | 11534.08 | 12509 | 10559 | 0.844 | -0.245 | 0.00378891 | 0.01979239 | 0.0091 | 0.0026 | 0.0051 | 0.0051 | TRUE | 0.213 |
| Supt7l | 1557 | 1138 | 1014 | 1170 | 945 | 1175 | 1209 | 1210 | 1094 | 1165 | 1066 | 1160 | 1150.74 | 1171 | 1130 | 0.965 | -0.052 | 0.51099014 | 0.67419755 | 0.0019 | 0.0045 | 0.0036 | 0.0036 | TRUE | 0.0389 |
| Supv3l1 | 3882 | 2934 | 2667 | 2959 | 2632 | 3120 | 3015 | 3121 | 2878 | 2945 | 2969 | 3080 | 3001.33 | 3005 | 2998 | 0.998 | -0.003 | 0.95897336 | 0.97739513 | 7.00E-04 | 0.0032 | 0.0022 | 0.0022 | TRUE | 0.0162 |
| Surf1 | 1146 | 887 | 963 | 1007 | 914 | 1095 | 890 | 943 | 1039 | 1002 | 1031 | 1081 | 997.83 | 957 | 1038 | 1.086 | 0.119 | 0.15897294 | 0.30809495 | 0.0029 | 0.0048 | 0.0041 | 0.0041 | TRUE | 0.0657 |
| Surf2 | 1135 | 753 | 755 | 648 | 625 | 723 | 882 | 801 | 815 | 645 | 705 | 714 | 760.15 | 833 | 688 | 0.825 | -0.277 | 0.00158818 | 0.01006643 | 0.0017 | 0.0055 | 0.0042 | 0.0042 | TRUE | 0.0357 |
| Surf4 | 8995 | 6140 | 6383 | 6839 | 5992 | 6778 | 6986 | 6531 | 6887 | 6808 | 6760 | 6691 | 6777.06 | 6801 | 6753 | 0.993 | -0.011 | 0.83599712 | 0.90645629 | 5.00E-04 | 0.0028 | 0.0018 | 0.0018 | TRUE | 0.0147 |
| Surf6 | 3857 | 2788 | 2610 | 2728 | 2450 | 2725 | 2996 | 2966 | 2816 | 2715 | 2764 | 2690 | 2824.43 | 2926 | 2723 | 0.93 | -0.104 | 0.07079932 | 0.17295071 | 3.00E-04 | 0.0033 | 0.002 | 0.002 | TRUE | 0.0129 |
| Susd1 | 733 | 621 | 671 | 953 | 856 | 1012 | 569 | 661 | 724 | 949 | 966 | 999 | 811.19 | 651 | 971 | 1.495 | 0.58 | 2.90E-09 | 1.69E-07 | 0.0062 | 0.0053 | 0.0056 | 0.0056 | TRUE | 0.1513 |
| Susd2 | 1499 | 1322 | 1707 | 1791 | 1785 | 1970 | 1164 | 1406 | 1842 | 1783 | 2014 | 1945 | 1692.23 | 1471 | 1914 | 1.303 | 0.381 | 0.00259774 | 0.01480007 | 0.0283 | 0.0038 | 0.011 | 0.011 | TRUE | 0.5703 |
| Susd3 | 57 | 40 | 62 | 62 | 52 | 55 | 44 | 43 | 67 | 62 | 59 | 54 | 54.73 | 51 | 58 | 1.142 | 0.192 | 0.49064261 | 0.65644059 | 0.0148 | 0.0457 | 0.0374 | 0.0374 | TRUE | 0.5547 |
| Susd4 | 546 | 435 | 506 | 552 | 524 | 662 | 424 | 463 | 546 | 549 | 591 | 653 | 537.8 | 478 | 598 | 1.255 | 0.328 | 0.00487833 | 0.02404994 | 0.0103 | 0.0068 | 0.0079 | 0.0079 | TRUE | 0.1808 |
| Susd5 | 749 | 608 | 745 | 910 | 856 | 916 | 582 | 647 | 804 | 906 | 966 | 904 | 801.33 | 678 | 925 | 1.369 | 0.453 | 5.32E-05 | 0.00073717 | 0.013 | 0.0054 | 0.0078 | 0.0078 | TRUE | 0.3125 |
| Suv39h1 | 1901 | 1334 | 1420 | 1694 | 1392 | 1612 | 1476 | 1419 | 1532 | 1686 | 1570 | 1591 | 1545.9 | 1476 | 1616 | 1.095 | 0.131 | 0.05706869 | 0.14844092 | 7.00E-04 | 0.004 | 0.0028 | 0.0028 | TRUE | 0.0175 |
| Suv39h2 | 2546 | 1874 | 1860 | 1959 | 1729 | 1933 | 1977 | 1993 | 2007 | 1950 | 1950 | 1908 | 1964.38 | 1992 | 1936 | 0.972 | -0.041 | 0.48801907 | 0.65397117 | 0 | 0.0036 | 0.002 | 0.002 | TRUE | 0.002 |
| Suv420h1 | 2977 | 2266 | 2079 | 2219 | 1870 | 2182 | 2312 | 2410 | 2243 | 2209 | 2110 | 2154 | 2239.66 | 2322 | 2158 | 0.929 | -0.106 | 0.08626994 | 0.1988159 | 4.00E-04 | 0.0035 | 0.0023 | 0.0023 | TRUE | 0.0131 |
| Suv420h2 | 6102 | 4789 | 4831 | 5274 | 4780 | 5207 | 4739 | 5094 | 5213 | 5250 | 5392 | 5140 | 5137.99 | 5015 | 5261 | 1.049 | 0.069 | 0.23158676 | 0.3975582 | 0.0012 | 0.0029 | 0.0022 | 0.0022 | TRUE | 0.0288 |
| Suz12 | 38051 | 27405 | 28977 | 32199 | 27614 | 31140 | 29553 | 29150 | 31266 | 32051 | 31151 | 30740 | 30651.86 | 29990 | 31314 | 1.044 | 0.062 | 0.20972377 | 0.37158571 | 9.00E-04 | 0.0025 | 0.0018 | 0.0018 | TRUE | 0.0169 |
| Sv2a | 807 | 562 | 677 | 915 | 824 | 977 | 627 | 598 | 730 | 911 | 930 | 964 | 793.3 | 652 | 935 | 1.436 | 0.522 | 2.45E-08 | 1.11E-06 | 0.0042 | 0.0054 | 0.005 | 0.005 | TRUE | 0.1275 |
| Sv2b | 20 | 15 | 19 | 11 | 13 | 11 | 16 | 16 | 21 | 11 | 15 | 11 | 14.74 | 18 | 12 | 0.702 | -0.511 | 0.30547656 | NA | 0 | 0.1632 | 0.1087 | 0.1087 | TRUE | 0.1193 |
| Sv2c | 39 | 36 | 47 | 58 | 45 | 42 | 30 | 38 | 51 | 58 | 51 | 41 | 44.88 | 40 | 50 | 1.268 | 0.342 | 0.2701801 | 0.44113367 | 0.0252 | 0.0552 | 0.0469 | 0.0469 | TRUE | 0.4076 |
| Sva | 3 | 2 | 1 | 1 | 0 | 0 | 2 | 2 | 1 | 1 | 0 | 0 | 1.09 | 2 | 0 | 0.199 | -2.326 | 0.23818909 | NA | 0 | 2.1795 | 1.5416 | 1.5416 | TRUE | 0.3906 |
| Sval1 | 0 | 0 | 0 | 0 | 0 | 0 | 0 | 0 | 0 | 0 | 0 | 0 | 0 | 0 | 0 | NA | NA | NA | NA | NA | NA | NA | NA | NA | NA |
| Sval2 | 0 | 0 | 0 | 0 | 0 | 0 | 0 | 0 | 0 | 0 | 0 | 0 | 0 | 0 | 0 | NA | NA | NA | NA | NA | NA | NA | NA | NA | NA |
| Sval3 | 0 | 0 | 0 | 0 | 0 | 0 | 0 | 0 | 0 | 0 | 0 | 0 | 0 | 0 | 0 | NA | NA | NA | NA | NA | NA | NA | NA | NA | NA |
| Svep1 | 5 | 2 | 2 | 2 | 4 | 3 | 4 | 2 | 2 | 2 | 5 | 3 | 2.94 | 3 | 3 | 1.13 | 0.176 | 0.87173403 | NA | 0 | 0.8089 | 0.5215 | 0.5215 | TRUE | 0.2226 |
| Svil | 1280 | 942 | 854 | 873 | 765 | 845 | 994 | 1002 | 921 | 869 | 863 | 834 | 913.95 | 972 | 855 | 0.879 | -0.186 | 0.01799269 | 0.06387663 | 0 | 0.005 | 0.0034 | 0.0034 | TRUE | 0.0244 |
| Svip | 1058 | 849 | 958 | 1113 | 935 | 1086 | 822 | 903 | 1034 | 1108 | 1055 | 1072 | 998.86 | 920 | 1078 | 1.175 | 0.233 | 0.0121406 | 0.04774374 | 0.0056 | 0.0048 | 0.0052 | 0.0052 | TRUE | 0.14 |
| Svop | 58 | 37 | 49 | 77 | 70 | 62 | 45 | 39 | 53 | 77 | 79 | 61 | 59.01 | 46 | 72 | 1.579 | 0.659 | 0.01184943 | 0.04688923 | 0 | 0.0426 | 0.0317 | 0.0317 | TRUE | 0.1663 |
| Svopl | 7 | 5 | 5 | 5 | 10 | 5 | 5 | 5 | 5 | 5 | 5 | 11 | 5 | 6.22 | 5 | 7 | 1.3 | 0.378 | 0.62507085 | NA | 0 | 0.3832 | 0.2681 | 0.2681 | TRUE | 0.6556 |
| Svs1 | 0 | 0 | 0 | 0 | 0 | 0 | 0 | 0 | 0 | 0 | 0 | 0 | 0 | 0 | 0 | NA | NA | NA | NA | NA | NA | NA | NA | NA | NA |
| Svs2 | 0 | 0 | 0 | 0 | 0 | 0 | 0 | 0 | 0 | 0 | 0 | 0 | 0 | 0 | 0 | NA | NA | NA | NA | NA | NA | NA | NA | NA | NA |
| Svs3a | 0 | 0 | 0 | 0 | 0 | 0 | 0 | 0 | 0 | 0 | 0 | 0 | 0 | 0 | 0 | NA | NA | NA | NA | NA | NA | NA | NA | NA | NA |
| Svs3b | 0 | 0 | 0 | 0 | 0 | 0 | 0 | 0 | 0 | 0 | 0 | 0 | 0 | 0 | 0 | NA | NA | NA | NA | NA | NA | NA | NA | NA | NA |
| Svs4 | 0 | 0 | 0 | 0 | 0 | 0 | 0 | 0 | 0 | 0 | 0 | 0 | 0 | 0 | 0 | NA | NA | NA | NA | NA | NA | NA | NA | NA | NA |
| Svs5 | 6 | 3 | 2 | 0 | 1 | 1 | 5 | 3 | 2 | 0 | 1 | 1 | 2.02 | 3 | 1 | 0.206 | -2.281 | 0.13708048 | NA | 0 | 1.1753 | 0.8073 | 0.8073 | TRUE | 0.2675 |
| Svs6 | 0 | 0 | 0 | 0 | 0 | 0 | 0 | 0 | 0 | 0 | 0 | 0 | 0 | 0 | 0 | NA | NA | NA | NA | NA | NA | NA | NA | NA | NA |
| Swap70 | 4289 | 3314 | 3076 | 3348 | 3008 | 3356 | 3331 | 3525 | 3319 | 3333 | 3393 | 3313 | 3368.99 | 3392 | 3346 | 0.987 | -0.019 | 0.73085718 | 0.83911492 | 3.00E-04 | 0.0031 | 0.002 | 0.002 | TRUE | 0.0142 |
| Swi5 | 7548 | 5127 | 5487 | 5834 | 4983 | 5757 | 5862 | 5454 | 5920 | 5807 | 5621 | 5683 | 5724.62 | 5745 | 5704 | 0.993 | -0.011 | 0.84784749 | 0.91369103 | 9.00E-04 | 0.0028 | 0.002 | 0.002 | TRUE | 0.0239 |
| Swsap1 | 711 | 480 | 437 | 553 | 524 | 640 | 552 | 511 | 472 | 550 | 591 | 632 | 551.28 | 512 | 591 | 1.153 | 0.206 | 0.04491542 | 0.12451691 | 0.0038 | 0.0067 | 0.0058 | 0.0058 | TRUE | 0.06 |
| Swt1 | 3062 | 2180 | 2200 | 2536 | 2064 | 2373 | 2378 | 2319 | 2374 | 2524 | 2328 | 2343 | 2377.67 | 2357 | 2398 | 1.018 | 0.025 | 0.68333197 | 0.80591334 | 7.00E-04 | 0.0034 | 0.0023 | 0.0023 | TRUE | 0.0257 |
| Syap1 | 913 | 688 | 790 | 714 | 716 | 790 | 709 | 732 | 852 | 711 | 808 | 780 | 765.27 | 764 | 766 | 1.003 | 0.005 | 0.9625935 | 0.97941456 | 0.0058 | 0.0055 | 0.0056 | 0.0056 | TRUE | 0.1184 |
| Sybu | 30 | 25 | 19 | 21 | 19 | 20 | 23 | 27 | 21 | 21 | 21 | 20 | 22.08 | 24 | 21 | 0.881 | -0.182 | 0.65081721 | 0.78320543 | 0 | 0.1098 | 0.0712 | 0.0712 | TRUE | 0.0727 |

| | | | | | | | | | | | | | | | | | | | | | | | | | |
|---|---|---|---|---|---|---|---|---|---|---|---|---|---|---|---|---|---|---|---|---|---|---|---|---|---|
| Syce1 | 1971 | 1524 | 1130 | 1264 | 1084 | 1174 | 1531 | 1621 | 1219 | 1258 | 1223 | 1159 | 1335.18 | 1457 | 1213 | 0.832 | -0.265 | 0.00826398 | 0.03589665 | 0.0102 | 0.0042 | 0.0065 | 0.0065 | TRUE | 0.2415 |
| Syce1l | 1 | 1 | 0 | 1 | 3 | 2 | 1 | 1 | 0 | 1 | 3 | 2 | 1.37 | 1 | 2 | 3.364 | 1.75 | 0.31395414 | NA | 0 | 1.7378 | 1.1617 | 1.1617 | TRUE | 0.1609 |
| Syce2 | 2709 | 2180 | 2423 | 2726 | 2320 | 2772 | 2104 | 2319 | 2614 | 2713 | 2617 | 2736 | 2517.37 | 2346 | 2689 | 1.148 | 0.199 | 0.01342908 | 0.05148656 | 0.0056 | 0.0034 | 0.0042 | 0.0042 | TRUE | 0.1215 |
| Syce3 | 58 | 56 | 74 | 91 | 94 | 84 | 45 | 60 | 80 | 91 | 106 | 83 | 77.33 | 62 | 93 | 1.527 | 0.611 | 0.01621882 | 0.05918191 | 0.0331 | 0.0331 | 0.0331 | 0.0331 | TRUE | 0.4391 |
| Sycn | 0 | 0 | 0 | 0 | 0 | 0 | 0 | 0 | 0 | 0 | 0 | 0 | 0 | 0 | 0 | NA | NA | NA | NA | NA | NA | NA | NA | NA | NA |
| Sycp1 | 3728 | 2778 | 1903 | 1653 | 1233 | 1314 | 2895 | 2955 | 2053 | 1645 | 1391 | 1297 | 2039.52 | 2634 | 1444 | 0.548 | -0.867 | 2.82E-12 | 3.15E-10 | 0.0253 | 0.0036 | 0.0106 | 0.0106 | TRUE | 0.4495 |
| Sycp1-ps1 | 1 | 2 | 1 | 0 | 1 | 1 | 1 | 2 | 1 | 0 | 1 | 1 | 1.02 | 1 | 1 | 0.532 | -0.91 | 0.63226269 | NA | 0 | 2.3343 | 1.4964 | 1.4964 | TRUE | 0.2654 |
| Sycp2 | 147 | 106 | 85 | 87 | 71 | 73 | 114 | 113 | 92 | 87 | 80 | 72 | 92.9 | 106 | 80 | 0.747 | -0.42 | 0.04457919 | 0.1238294 | 0 | 0.0279 | 0.0205 | 0.0205 | TRUE | 0.1343 |
| Sycp3 | 660 | 507 | 646 | 725 | 627 | 666 | 513 | 539 | 697 | 722 | 707 | 657 | 639.22 | 583 | 695 | 1.195 | 0.258 | 0.02861777 | 0.08950411 | 0.0134 | 0.0061 | 0.0084 | 0.0084 | TRUE | 0.3395 |
| Syde1 | 243 | 183 | 133 | 157 | 164 | 144 | 189 | 195 | 144 | 156 | 185 | 142 | 168.39 | 176 | 161 | 0.914 | -0.13 | 0.46020239 | 0.62988358 | 0.0159 | 0.0165 | 0.0163 | 0.0163 | TRUE | 0.262 |
| Syde2 | 3477 | 2682 | 2653 | 2575 | 2402 | 2786 | 2700 | 2853 | 2863 | 2563 | 2710 | 2750 | 2739.81 | 2805 | 2674 | 0.954 | -0.068 | 0.26283983 | 0.43322121 | 9.00E-04 | 0.0033 | 0.0023 | 0.0023 | TRUE | 0.0162 |
| Syf2 | 2381 | 1760 | 1766 | 1792 | 1656 | 1922 | 1849 | 1872 | 1905 | 1784 | 1868 | 1897 | 1862.67 | 1875 | 1850 | 0.986 | -0.02 | 0.75309476 | 0.85348706 | 0 | 0.0037 | 0.0023 | 0.0023 | TRUE | 0.0119 |
| Syk | 1724 | 1299 | 1332 | 1464 | 1423 | 1604 | 1339 | 1382 | 1437 | 1457 | 1605 | 1583 | 1467.31 | 1386 | 1548 | 1.118 | 0.161 | 0.02541444 | 0.08229464 | 0.0013 | 0.004 | 0.003 | 0.003 | TRUE | 0.0323 |
| Sympk | 14566 | 11163 | 11374 | 11904 | 11211 | 12776 | 11313 | 11874 | 12272 | 11849 | 12647 | 12612 | 12094.58 | 11820 | 12369 | 1.047 | 0.066 | 0.23206729 | 0.39790521 | 0.0014 | 0.0026 | 0.0021 | 0.0021 | TRUE | 0.0174 |
| Syn1 | 877 | 638 | 654 | 655 | 580 | 700 | 681 | 679 | 706 | 652 | 654 | 691 | 677.12 | 689 | 666 | 0.968 | -0.048 | 0.57619716 | 0.72771813 | 0 | 0.0059 | 0.0037 | 0.0037 | TRUE | 0.0132 |
| Syn2 | 31 | 29 | 20 | 24 | 21 | 17 | 24 | 31 | 22 | 24 | 24 | 17 | 23.48 | 26 | 22 | 0.841 | -0.25 | 0.53493767 | 0.694006 | 0 | 0.1034 | 0.0736 | 0.0736 | TRUE | 0.2114 |
| Syn3 | 92 | 60 | 75 | 80 | 59 | 61 | 71 | 64 | 81 | 80 | 67 | 60 | 70.43 | 72 | 69 | 0.956 | -0.066 | 0.78360902 | 0.87262359 | 0 | 0.0361 | 0.0269 | 0.0269 | TRUE | 0.1745 |
| Syna | 55 | 38 | 33 | 42 | 35 | 42 | 43 | 40 | 36 | 42 | 39 | 41 | 40.25 | 40 | 41 | 1.031 | 0.044 | 0.88308119 | 0.93495403 | 0 | 0.0613 | 0.0397 | 0.0397 | TRUE | 0.0547 |
| Synb | 0 | 0 | 0 | 0 | 0 | 0 | 0 | 0 | 0 | 0 | 0 | 0 | 0 | 0 | 0 | NA | NA | NA | NA | NA | NA | NA | NA | NA | NA |
| Sync | 136 | 79 | 66 | 62 | 80 | 70 | 106 | 84 | 71 | 62 | 90 | 69 | 80.32 | 87 | 74 | 0.84 | -0.252 | 0.30327576 | 0.47586926 | 0.0272 | 0.0319 | 0.0306 | 0.0306 | TRUE | 0.3684 |
| Syncrip | 15593 | 11077 | 9960 | 11059 | 9193 | 10724 | 12110 | 11782 | 10747 | 11008 | 10371 | 10586 | 11100.78 | 11546 | 10655 | 0.923 | -0.116 | 0.05291974 | 0.1404801 | 0.0022 | 0.0026 | 0.0025 | 0.0025 | TRUE | 0.0448 |
| Syndig1 | 97 | 62 | 55 | 62 | 60 | 63 | 75 | 66 | 59 | 62 | 68 | 62 | 65.37 | 67 | 64 | 0.95 | -0.075 | 0.75633934 | 0.85520897 | 0 | 0.0387 | 0.0265 | 0.0265 | TRUE | 0.1236 |
| Syndig1l | 13 | 13 | 15 | 8 | 9 | 13 | 10 | 14 | 16 | 8 | 10 | 13 | 11.84 | 13 | 10 | 0.779 | -0.36 | 0.52114169 | NA | 0 | 0.2025 | 0.1407 | 0.1407 | TRUE | 0.2484 |
| Syne1 | 797 | 591 | 775 | 850 | 848 | 925 | 619 | 629 | 836 | 846 | 957 | 913 | 799.95 | 695 | 905 | 1.305 | 0.384 | 0.00091976 | 0.00682914 | 0.0156 | 0.0054 | 0.0084 | 0.0084 | TRUE | 0.3707 |
| Syne2 | 2867 | 1944 | 1976 | 2266 | 2109 | 2367 | 2227 | 2068 | 2132 | 2256 | 2379 | 2337 | 2232.99 | 2142 | 2324 | 1.084 | 0.116 | 0.06265432 | 0.15833066 | 7.00E-04 | 0.0035 | 0.0024 | 0.0024 | TRUE | 0.0149 |
| Syne3 | 344 | 254 | 312 | 374 | 328 | 380 | 267 | 270 | 337 | 372 | 370 | 375 | 331.9 | 291 | 372 | 1.282 | 0.358 | 0.00418918 | 0.02140629 | 0.005 | 0.0096 | 0.0082 | 0.0082 | TRUE | 0.2023 |
| Syne4 | 523 | 414 | 510 | 511 | 498 | 641 | 406 | 440 | 550 | 509 | 562 | 633 | 516.68 | 465 | 568 | 1.222 | 0.29 | 0.02436798 | 0.07988338 | 0.0162 | 0.007 | 0.01 | 0.01 | TRUE | 0.2908 |
| Syngr1 | 1419 | 1009 | 1212 | 1382 | 1335 | 1493 | 1102 | 1073 | 1308 | 1376 | 1506 | 1474 | 1306.43 | 1161 | 1452 | 1.251 | 0.323 | 0.00024932 | 0.0025237 | 0.006 | 0.0042 | 0.0048 | 0.0048 | TRUE | 0.1437 |
| Syngr2 | 6860 | 5281 | 5491 | 6616 | 6171 | 7004 | 5328 | 5617 | 5925 | 6586 | 6962 | 6914 | 6221.84 | 5623 | 6821 | 1.213 | 0.279 | 2.26E-06 | 5.54E-05 | 0.0016 | 0.0028 | 0.0023 | 0.0023 | TRUE | 0.0268 |
| Syngr3 | 494 | 399 | 501 | 432 | 473 | 581 | 384 | 424 | 541 | 430 | 534 | 574 | 480.97 | 450 | 513 | 1.142 | 0.192 | 0.17896739 | 0.333601 | 0.0237 | 0.0074 | 0.0126 | 0.0126 | TRUE | 0.3607 |
| Syngr4 | 451 | 342 | 444 | 567 | 463 | 509 | 350 | 364 | 479 | 564 | 522 | 502 | 463.72 | 398 | 529 | 1.336 | 0.417 | 0.00107122 | 0.00760225 | 0.0143 | 0.0075 | 0.0095 | 0.0095 | TRUE | 0.3622 |
| Synj1 | 2301 | 1679 | 1468 | 1617 | 1382 | 1652 | 1787 | 1786 | 1584 | 1610 | 1559 | 1631 | 1659.39 | 1719 | 1600 | 0.93 | -0.104 | 0.15000752 | 0.29565051 | 0.002 | 0.0039 | 0.0032 | 0.0032 | TRUE | 0.0559 |
| Synj2 | 458 | 398 | 412 | 516 | 473 | 591 | 356 | 423 | 445 | 514 | 534 | 583 | 475.7 | 408 | 544 | 1.337 | 0.419 | 0.00023512 | 0.00242117 | 0.0068 | 0.0074 | 0.0072 | 0.0072 | TRUE | 0.1548 |
| Synj2bp | 1599 | 1159 | 1210 | 1509 | 1231 | 1415 | 1242 | 1233 | 1306 | 1502 | 1389 | 1397 | 1344.64 | 1260 | 1429 | 1.135 | 0.182 | 0.01049505 | 0.04278544 | 7.00E-04 | 0.0042 | 0.0029 | 0.0029 | TRUE | 0.0241 |
| Synm | 1297 | 1035 | 1008 | 967 | 904 | 1003 | 1007 | 1101 | 1088 | 963 | 1020 | 990 | 1028.06 | 1065 | 991 | 0.931 | -0.103 | 0.18054151 | 0.33560668 | 8.00E-04 | 0.0047 | 0.0033 | 0.0033 | TRUE | 0.0286 |
| Synpo | 248 | 198 | 250 | 302 | 311 | 326 | 193 | 211 | 270 | 301 | 351 | 322 | 274.37 | 225 | 325 | 1.451 | 0.537 | 0.00027513 | 0.0027174 | 0.0135 | 0.0111 | 0.012 | 0.012 | TRUE | 0.3391 |
| Synpo2 | 142 | 114 | 74 | 71 | 63 | 76 | 110 | 121 | 80 | 71 | 71 | 75 | 88.03 | 104 | 72 | 0.695 | -0.525 | 0.0199756 | 0.06909758 | 0.0137 | 0.0294 | 0.0249 | 0.0249 | TRUE | 0.378 |
| Synpo2l | 9 | 4 | 0 | 4 | 1 | 1 | 7 | 4 | 0 | 4 | 1 | 1 | 2.89 | 4 | 2 | 0.534 | -0.905 | 0.48339915 | NA | 0.8181 | 0.8224 | 0.8216 | 0.8216 | TRUE | 0.2018 |
| Synpr | 77 | 45 | 50 | 66 | 61 | 68 | 60 | 48 | 54 | 66 | 69 | 67 | 60.54 | 54 | 67 | 1.242 | 0.312 | 0.20793119 | 0.3695549 | 0 | 0.0416 | 0.0277 | 0.0277 | TRUE | 0.0899 |
| Synrg | 3031 | 2162 | 2327 | 2563 | 2361 | 2542 | 2354 | 2300 | 2511 | 2551 | 2663 | 2509 | 2481.43 | 2388 | 2574 | 1.078 | 0.109 | 0.0859373 | 0.19817712 | 9.00E-04 | 0.0034 | 0.0025 | 0.0025 | TRUE | 0.0239 |
| Syp | 59 | 60 | 63 | 87 | 66 | 98 | 46 | 64 | 68 | 87 | 74 | 97 | 72.57 | 59 | 86 | 1.466 | 0.552 | 0.02533399 | 0.08208995 | 0.0155 | 0.0351 | 0.0297 | 0.0297 | TRUE | 0.3935 |
| Sypl | 4565 | 3380 | 3801 | 4153 | 3609 | 4210 | 3545 | 3595 | 4101 | 4134 | 4071 | 4156 | 3933.85 | 3747 | 4120 | 1.1 | 0.138 | 0.04178725 | 0.11801244 | 0.003 | 0.003 | 0.003 | 0.003 | TRUE | 0.0825 |
| Sypl2 | 16 | 7 | 9 | 12 | 6 | 8 | 12 | 7 | 10 | 12 | 7 | 8 | 9.37 | 10 | 9 | 0.894 | -0.161 | 0.79799626 | NA | 0 | 0.2555 | 0.1781 | 0.1781 | TRUE | 0.2972 |
| Sys1 | 1973 | 1401 | 1275 | 1273 | 1203 | 1452 | 1532 | 1490 | 1376 | 1267 | 1357 | 1433 | 1409.31 | 1466 | 1352 | 0.922 | -0.117 | 0.13015611 | 0.26694235 | 0.0028 | 0.0041 | 0.0036 | 0.0036 | TRUE | 0.0372 |
| Syt1 | 51 | 44 | 33 | 42 | 39 | 56 | 40 | 47 | 36 | 42 | 44 | 55 | 43.85 | 41 | 47 | 1.158 | 0.212 | 0.47488069 | 0.64299637 | 0 | 0.0565 | 0.0407 | 0.0407 | TRUE | 0.1929 |
| Syt10 | 4 | 0 | 1 | 0 | 0 | 1 | 3 | 0 | 1 | 0 | 0 | 1 | 0.86 | 1 | 0 | 0.259 | -1.95 | 0.39665209 | NA | 0 | 2.7515 | 2.4763 | 2.4763 | TRUE | 0.2206 |
| Syt11 | 686 | 460 | 525 | 572 | 478 | 641 | 533 | 489 | 566 | 569 | 539 | 633 | 554.99 | 529 | 580 | 1.097 | 0.134 | 0.19559684 | 0.35527584 | 0.0041 | 0.0067 | 0.0059 | 0.0059 | TRUE | 0.0733 |
| Syt12 | 94 | 77 | 60 | 61 | 45 | 58 | 73 | 82 | 65 | 61 | 51 | 57 | 64.73 | 73 | 56 | 0.769 | -0.378 | 0.12155732 | 0.25435227 | 0 | 0.039 | 0.0272 | 0.0272 | TRUE | 0.0907 |
| Syt13 | 158 | 112 | 150 | 229 | 219 | 236 | 123 | 119 | 162 | 228 | 247 | 233 | 185.28 | 135 | 236 | 1.759 | 0.815 | 4.68E-07 | 1.53E-05 | 0.0066 | 0.0152 | 0.013 | 0.013 | TRUE | 0.3088 |
| Syt14 | 207 | 159 | 152 | 173 | 189 | 168 | 161 | 169 | 164 | 172 | 213 | 166 | 174.19 | 165 | 184 | 1.115 | 0.156 | 0.32661692 | 0.5006795 | 0.004 | 0.016 | 0.0126 | 0.0126 | TRUE | 0.2041 |
| Syt15 | 14 | 16 | 10 | 12 | 11 | 9 | 11 | 17 | 11 | 12 | 12 | 9 | 11.99 | 13 | 11 | 0.862 | -0.214 | 0.6996601 | NA | 0 | 0.2002 | 0.1378 | 0.1378 | TRUE | 0.3044 |
| Syt16 | 22 | 21 | 19 | 35 | 27 | 29 | 17 | 22 | 21 | 35 | 30 | 29 | 25.64 | 20 | 31 | 1.578 | 0.658 | 0.08327664 | 0.19423514 | 0 | 0.0949 | 0.0633 | 0.0633 | TRUE | 0.1098 |
| Syt17 | 249 | 146 | 160 | 200 | 174 | 207 | 193 | 155 | 173 | 199 | 196 | 204 | 186.84 | 174 | 200 | 1.146 | 0.197 | 0.19207227 | 0.35037387 | 0 | 0.0151 | 0.0111 | 0.0111 | TRUE | 0.1143 |
| Syt2 | 18 | 8 | 7 | 19 | 10 | 7 | 14 | 9 | 8 | 19 | 11 | 7 | 11.19 | 10 | 12 | 1.218 | 0.285 | 0.64189056 | NA | 0.1019 | 0.2142 | 0.1814 | 0.1814 | TRUE | 0.3545 |
| Syt3 | 320 | 249 | 257 | 310 | 245 | 299 | 249 | 265 | 277 | 309 | 276 | 295 | 278.47 | 264 | 293 | 1.116 | 0.159 | 0.20237512 | 0.36375096 | 0 | 0.0109 | 0.0076 | 0.0076 | TRUE | 0.0304 |
| Syt4 | 35 | 19 | 12 | 18 | 13 | 21 | 27 | 20 | 13 | 18 | 15 | 21 | 18.94 | 20 | 18 | 0.875 | -0.192 | 0.67797132 | 0.80231161 | 0 | 0.1275 | 0.1011 | 0.1011 | TRUE | 0.184 |
| Syt5 | 124 | 121 | 162 | 176 | 185 | 178 | 96 | 129 | 175 | 175 | 209 | 176 | 159.9 | 133 | 187 | 1.408 | 0.494 | 0.0166593 | 0.06043435 | 0.0417 | 0.0172 | 0.0242 | 0.0242 | TRUE | 0.4275 |
| Syt6 | 13 | 10 | 5 | 7 | 10 | 12 | 10 | 11 | 5 | 7 | 11 | 12 | 9.37 | 9 | 10 | 1.142 | 0.192 | 0.76241108 | NA | 0 | 0.2554 | 0.1826 | 0.1826 | TRUE | 0.3127 |
| Syt7 | 596 | 498 | 530 | 692 | 620 | 748 | 463 | 530 | 572 | 689 | 699 | 738 | 615.18 | 522 | 709 | 1.363 | 0.447 | 1.02E-05 | 0.00019211 | 0.0046 | 0.0063 | 0.0057 | 0.0057 | TRUE | 0.1205 |
| Syt8 | 1 | 1 | 0 | 0 | 0 | 0 | 1 | 1 | 0 | 0 | 0 | 0 | 0.31 | 1 | 0 | 0.309 | -1.696 | 0.63183309 | NA | 0 | 7.7294 | 7.2472 | 7.2472 | TRUE | 0.2015 |
| Syt9 | 2680 | 1956 | 1876 | 2156 | 1959 | 2167 | 2081 | 2081 | 2024 | 2146 | 2210 | 2139 | 2113.57 | 2062 | 2165 | 1.05 | 0.07 | 0.2391951 | 0.40620822 | 0 | 0.0035 | 0.0021 | 0.0021 | TRUE | 0.0039 |
| Sytl1 | 107 | 92 | 89 | 163 | 118 | 148 | 83 | 98 | 96 | 162 | 133 | 146 | 119.74 | 92 | 147 | 1.602 | 0.68 | 0.00026278 | 0.00263015 | 0 | 0.0222 | 0.0163 | 0.0163 | TRUE | 0.0845 |
| Sytl2 | 21 | 15 | 15 | 9 | 14 | 11 | 16 | 16 | 16 | 9 | 16 | 11 | 14.01 | 16 | 12 | 0.73 | -0.453 | 0.37725048 | NA | 0 | 0.1716 | 0.1163 | 0.1163 | TRUE | 0.2942 |
| Sytl3 | 78 | 75 | 91 | 139 | 136 | 150 | 61 | 80 | 98 | 138 | 153 | 148 | 113.07 | 80 | 146 | 1.86 | 0.895 | 1.80E-05 | 0.00030868 | 0.017 | 0.0234 | 0.0218 | 0.0218 | TRUE | 0.4222 |
| Sytl4 | 75 | 34 | 27 | 23 | 24 | 25 | 58 | 36 | 29 | 23 | 27 | 25 | 33.03 | 41 | 25 | 0.597 | -0.745 | 0.04571785 | 0.12619646 | 0.0535 | 0.0742 | 0.0677 | 0.0677 | TRUE | 0.8202 |
| Sytl5 | 57 | 38 | 34 | 28 | 28 | 25 | 44 | 40 | 37 | 28 | 32 | 25 | 34.25 | 40 | 28 | 0.689 | -0.537 | 0.10343757 | 0.22678834 | 0 | 0.0716 | 0.048 | 0.048 | TRUE | 0.0721 |
| Syvn1 | 2004 | 1499 | 1460 | 1425 | 1394 | 1686 | 1556 | 1594 | 1575 | 1418 | 1573 | 1664 | 1563.6 | 1575 | 1552 | 0.985 | -0.022 | 0.77512541 | 0.86721285 | 0.0027 | 0.0039 | 0.0035 | 0.0035 | TRUE | 0.0689 |
| Szrd1 | 5142 | 3868 | 3346 | 3358 | 2968 | 3414 | 3994 | 4114 | 3610 | 3343 | 3348 | 3370 | 3629.85 | 3906 | 3354 | 0.858 | -0.22 | 0.00057493 | 0.00480553 | 0.0019 | 0.0031 | 0.0027 | 0.0027 | TRUE | 0.0527 |
| Szt2 | 2420 | 1930 | 1696 | 1783 | 1671 | 1880 | 1880 | 2053 | 1830 | 1775 | 1885 | 1856 | 1879.68 | 1921 | 1839 | 0.957 | -0.063 | 0.36599546 | 0.54063385 | 0.0015 | 0.0037 | 0.003 | 0.003 | TRUE | 0.0429 |

| | | | | | | | | | | | | | | | | | | | | | | | | | |
|---|---|---|---|---|---|---|---|---|---|---|---|---|---|---|---|---|---|---|---|---|---|---|---|---|---|
| T | 60 | 50 | 48 | 87 | 90 | 95 | 47 | 53 | 52 | 87 | 102 | 94 | 72.25 | 51 | 94 | 1.865 | 0.899 | 0.00010657 | 0.00127463 | 0 | 0.0352 | 0.0237 | 0.0237 | TRUE | 0.0456 |
| T2 | 9 | 1 | 6 | 5 | 5 | 13 | 7 | 1 | 6 | 5 | 6 | 13 | 6.33 | 5 | 8 | 1.595 | 0.673 | 0.42227135 | NA | 0.2449 | 0.3769 | 0.3432 | 0.3432 | TRUE | 0.938 |
| Taar1 | 0 | 0 | 0 | 0 | 0 | 0 | 0 | 0 | 0 | 0 | 0 | 0 | 0 | 0 | 0 | NA | NA | NA | NA | NA | NA | NA | NA | NA | NA |
| Taar2 | 0 | 0 | 2 | 1 | 1 | 0 | 0 | 0 | 2 | 1 | 1 | 0 | 0.71 | 1 | 1 | 1.011 | 0.016 | 0.99481353 | NA | 0 | 3.3238 | 2.9707 | 2.9707 | TRUE | 0.9749 |
| Taar3 | 0 | 0 | 0 | 0 | 0 | 0 | 0 | 0 | 0 | 0 | 0 | 0 | 0 | 0 | 0 | NA | NA | NA | NA | NA | NA | NA | NA | NA | NA |
| Taar4 | 0 | 0 | 0 | 0 | 0 | 0 | 0 | 0 | 0 | 0 | 0 | 0 | 0 | 0 | 0 | NA | NA | NA | NA | NA | NA | NA | NA | NA | NA |
| Taar5 | 0 | 0 | 0 | 0 | 0 | 0 | 0 | 0 | 0 | 0 | 0 | 0 | 0 | 0 | 0 | NA | NA | NA | NA | NA | NA | NA | NA | NA | NA |
| Taar6 | 0 | 0 | 0 | 0 | 0 | 0 | 0 | 0 | 0 | 0 | 0 | 0 | 0 | 0 | 0 | NA | NA | NA | NA | NA | NA | NA | NA | NA | NA |
| Taar7a | 0 | 0 | 0 | 0 | 0 | 0 | 0 | 0 | 0 | 0 | 0 | 0 | 0 | 0 | 0 | NA | NA | NA | NA | NA | NA | NA | NA | NA | NA |
| Taar7b | 0 | 0 | 0 | 0 | 0 | 0 | 0 | 0 | 0 | 0 | 0 | 0 | 0 | 0 | 0 | NA | NA | NA | NA | NA | NA | NA | NA | NA | NA |
| Taar7d | 0 | 0 | 0 | 0 | 0 | 0 | 0 | 0 | 0 | 0 | 0 | 0 | 0 | 0 | 0 | NA | NA | NA | NA | NA | NA | NA | NA | NA | NA |
| Taar7e | 0 | 0 | 0 | 0 | 0 | 0 | 0 | 0 | 0 | 0 | 0 | 0 | 0 | 0 | 0 | NA | NA | NA | NA | NA | NA | NA | NA | NA | NA |
| Taar7f | 0 | 0 | 0 | 0 | 0 | 0 | 0 | 0 | 0 | 0 | 0 | 0 | 0 | 0 | 0 | NA | NA | NA | NA | NA | NA | NA | NA | NA | NA |
| Taar8a | 0 | 0 | 0 | 0 | 0 | 0 | 0 | 0 | 0 | 0 | 0 | 0 | 0 | 0 | 0 | NA | NA | NA | NA | NA | NA | NA | NA | NA | NA |
| Taar8b | 0 | 0 | 0 | 0 | 0 | 0 | 0 | 0 | 0 | 0 | 0 | 0 | 0 | 0 | 0 | NA | NA | NA | NA | NA | NA | NA | NA | NA | NA |
| Taar8c | 0 | 0 | 0 | 0 | 0 | 0 | 0 | 0 | 0 | 0 | 0 | 0 | 0 | 0 | 0 | NA | NA | NA | NA | NA | NA | NA | NA | NA | NA |
| Taar9 | 0 | 0 | 0 | 0 | 0 | 0 | 0 | 0 | 0 | 0 | 0 | 0 | 0 | 0 | 0 | NA | NA | NA | NA | NA | NA | NA | NA | NA | NA |
| Tab1 | 1072 | 876 | 803 | 888 | 908 | 907 | 833 | 932 | 866 | 884 | 1024 | 895 | 905.73 | 877 | 934 | 1.066 | 0.092 | 0.30351436 | 0.47603761 | 0.0039 | 0.005 | 0.0047 | 0.0047 | TRUE | 0.0827 |
| Tab2 | 6794 | 4912 | 5148 | 5230 | 4434 | 5295 | 5277 | 5225 | 5555 | 5206 | 5002 | 5227 | 5248.5 | 5352 | 5145 | 0.962 | -0.057 | 0.29432023 | 0.46637683 | 6.00E-04 | 0.0029 | 0.0019 | 0.0019 | TRUE | 0.0132 |
| Tab3 | 3996 | 2853 | 2811 | 2859 | 2452 | 2709 | 3104 | 3035 | 3033 | 2846 | 2766 | 2674 | 2909.57 | 3057 | 2762 | 0.903 | -0.147 | 0.01002027 | 0.04141691 | 2.00E-04 | 0.0032 | 0.002 | 0.002 | TRUE | 0.0095 |
| Tac1 | 2 | 1 | 1 | 1 | 0 | 0 | 2 | 1 | 1 | 1 | 0 | 0 | 0.78 | 1 | 0 | 0.299 | -1.743 | 0.425346 | NA | 0 | 3.0336 | 2.0559 | 2.0559 | TRUE | 0.3906 |
| Tac2 | 4 | 5 | 4 | 7 | 8 | 5 | 3 | 5 | 4 | 7 | 9 | 5 | 5.61 | 4 | 7 | 1.656 | 0.728 | 0.36826228 | NA | 0 | 0.4248 | 0.2819 | 0.2819 | TRUE | 0.1401 |
| Tac4 | 0 | 0 | 0 | 0 | 0 | 1 | 0 | 0 | 0 | 0 | 0 | 1 | 0.16 | 0 | 0 | 2.094 | 1.067 | 0.79379329 | NA | 0 | 14.4078 | 10 | 10 | TRUE | 0.3835 |
| Tacc1 | 7827 | 5533 | 5921 | 7661 | 6802 | 7561 | 6079 | 5885 | 6389 | 7626 | 7673 | 7464 | 6852.66 | 6118 | 7588 | 1.24 | 0.311 | 5.58E-09 | 3.08E-07 | 7.00E-04 | 0.0028 | 0.0019 | 0.0019 | TRUE | 0.0181 |
| Tacc2 | 3802 | 3064 | 3308 | 3310 | 3034 | 3510 | 2953 | 3259 | 3569 | 3295 | 3423 | 3465 | 3327.27 | 3260 | 3394 | 1.042 | 0.059 | 0.42689392 | 0.59901775 | 0.0046 | 0.0031 | 0.0037 | 0.0037 | TRUE | 0.0843 |
| Tacc3 | 7250 | 5211 | 4910 | 4944 | 4325 | 4992 | 5631 | 5543 | 5298 | 4921 | 4879 | 4928 | 5199.94 | 5491 | 4909 | 0.894 | -0.162 | 0.00186716 | 0.01148133 | 3.00E-04 | 0.0029 | 0.0018 | 0.0018 | TRUE | 0.0114 |
| Taco1 | 1167 | 786 | 635 | 795 | 624 | 762 | 906 | 836 | 685 | 791 | 704 | 752 | 779.18 | 809 | 749 | 0.925 | -0.113 | 0.2950077 | 0.4669447 | 0.0106 | 0.0055 | 0.0071 | 0.0071 | TRUE | 0.2107 |
| Tacr1 | 8 | 4 | 6 | 5 | 10 | 8 | 6 | 4 | 6 | 5 | 11 | 8 | 6.85 | 5 | 8 | 1.412 | 0.497 | 0.49976992 | NA | 0 | 0.3484 | 0.2419 | 0.2419 | TRUE | 0.1617 |
| Tacr2 | 35 | 23 | 21 | 31 | 26 | 35 | 27 | 24 | 23 | 31 | 29 | 35 | 28.17 | 25 | 32 | 1.271 | 0.346 | 0.33322741 | 0.50743868 | 0 | 0.0865 | 0.0564 | 0.0564 | TRUE | 0.0538 |
| Tacr3 | 38 | 31 | 11 | 15 | 11 | 11 | 30 | 33 | 12 | 15 | 12 | 11 | 18.76 | 25 | 13 | 0.511 | -0.968 | 0.05362666 | 0.14156329 | 0.1027 | 0.1288 | 0.1213 | 0.1213 | TRUE | 1.1693 |
| Tacstd2 | 12 | 15 | 10 | 9 | 4 | 2 | 9 | 16 | 11 | 9 | 5 | 2 | 8.59 | 12 | 5 | 0.431 | -1.213 | 0.09140294 | NA | 0.0753 | 0.2785 | 0.2311 | 0.2311 | TRUE | 0.8293 |
| Tada1 | 489 | 363 | 359 | 310 | 281 | 332 | 380 | 386 | 387 | 309 | 317 | 328 | 351.09 | 384 | 318 | 0.827 | -0.274 | 0.01214568 | 0.0477443 | 0 | 0.0092 | 0.0057 | 0.0057 | TRUE | 0.0088 |
| Tada2a | 6481 | 4727 | 4520 | 4706 | 4215 | 4935 | 5034 | 5028 | 4877 | 4684 | 4755 | 4872 | 4874.92 | 4980 | 4770 | 0.958 | -0.062 | 0.22602928 | 0.39092891 | 1.00E-04 | 0.0029 | 0.0017 | 0.0017 | TRUE | 0.0042 |
| Tada2b | 983 | 735 | 606 | 634 | 561 | 621 | 763 | 782 | 654 | 631 | 633 | 613 | 679.35 | 733 | 626 | 0.853 | -0.23 | 0.01583419 | 0.05814954 | 0.0033 | 0.0059 | 0.0051 | 0.0051 | TRUE | 0.1024 |
| Tada3 | 5809 | 4357 | 3957 | 4385 | 3980 | 4857 | 4512 | 4634 | 4270 | 4365 | 4490 | 4795 | 4510.82 | 4472 | 4550 | 1.017 | 0.025 | 0.68491666 | 0.80694783 | 0.0015 | 0.003 | 0.0025 | 0.0025 | TRUE | 0.0272 |
| Taf1 | 9780 | 6885 | 6346 | 7228 | 6037 | 7107 | 7596 | 7323 | 6847 | 7195 | 6810 | 7016 | 7131.21 | 7255 | 7007 | 0.966 | -0.051 | 0.38233811 | 0.55640702 | 0.0015 | 0.0028 | 0.0023 | 0.0023 | TRUE | 0.0295 |
| Taf10 | 4519 | 3283 | 3110 | 3301 | 2973 | 3339 | 3510 | 3492 | 3356 | 3286 | 3354 | 3296 | 3382.21 | 3453 | 3312 | 0.959 | -0.06 | 0.26564307 | 0.43601704 | 0 | 0.0031 | 0.0018 | 0.0018 | TRUE | 0.0072 |
| Taf11 | 1819 | 1265 | 1143 | 1117 | 1048 | 1120 | 1413 | 1346 | 1233 | 1112 | 1182 | 1106 | 1231.89 | 1331 | 1133 | 0.85 | -0.234 | 0.00291843 | 0.01619296 | 0.0023 | 0.0044 | 0.0036 | 0.0036 | TRUE | 0.0486 |
| Taf12 | 1114 | 780 | 625 | 732 | 599 | 691 | 865 | 830 | 674 | 729 | 676 | 682 | 742.62 | 790 | 696 | 0.879 | -0.185 | 0.07545802 | 0.18096474 | 0.0082 | 0.0056 | 0.0065 | 0.0065 | TRUE | 0.1904 |
| Taf13 | 1113 | 789 | 679 | 785 | 574 | 726 | 864 | 839 | 733 | 781 | 648 | 717 | 763.65 | 812 | 715 | 0.88 | -0.184 | 0.06700451 | 0.16603598 | 0.0068 | 0.0055 | 0.0059 | 0.0059 | TRUE | 0.0855 |
| Taf15 | 14653 | 11056 | 10128 | 10340 | 9143 | 10703 | 11380 | 11760 | 10928 | 10292 | 10314 | 10566 | 10873.45 | 11356 | 10391 | 0.915 | -0.128 | 0.01124749 | 0.04504745 | 6.00E-04 | 0.0026 | 0.0017 | 0.0017 | TRUE | 0.0132 |
| Taf1a | 649 | 527 | 445 | 474 | 404 | 441 | 504 | 561 | 480 | 472 | 456 | 435 | 484.61 | 515 | 454 | 0.883 | -0.18 | 0.08122472 | 0.19119646 | 0.0019 | 0.0073 | 0.0056 | 0.0056 | TRUE | 0.0675 |
| Taf1b | 1936 | 1410 | 1388 | 1476 | 1296 | 1535 | 1504 | 1500 | 1498 | 1469 | 1462 | 1515 | 1491.26 | 1501 | 1482 | 0.988 | -0.017 | 0.78668469 | 0.87477829 | 0 | 0.004 | 0.0023 | 0.0023 | TRUE | 0.0046 |
| Taf1c | 2135 | 1593 | 1277 | 1348 | 1267 | 1376 | 1658 | 1694 | 1378 | 1342 | 1429 | 1358 | 1476.65 | 1577 | 1376 | 0.872 | -0.197 | 0.02338958 | 0.0774756 | 0.0056 | 0.004 | 0.0048 | 0.0048 | TRUE | 0.1444 |
| Taf1d | 3867 | 2826 | 2374 | 2445 | 2084 | 2394 | 3003 | 3006 | 2562 | 2434 | 2351 | 2363 | 2619.8 | 2857 | 2383 | 0.834 | -0.263 | 0.00039952 | 0.00364644 | 0.0036 | 0.0033 | 0.0036 | 0.0036 | TRUE | 0.0983 |
| Taf2 | 7992 | 5629 | 5145 | 5794 | 4870 | 5640 | 6207 | 5987 | 5551 | 5767 | 5494 | 5568 | 5762.45 | 5915 | 5610 | 0.948 | -0.077 | 0.19703883 | 0.35676159 | 0.0016 | 0.0028 | 0.0024 | 0.0024 | TRUE | 0.0352 |
| Taf3 | 2626 | 1824 | 1782 | 1879 | 1671 | 1860 | 2040 | 1940 | 1923 | 1870 | 1885 | 1836 | 1915.66 | 1968 | 1864 | 0.947 | -0.079 | 0.20408546 | 0.36544644 | 0 | 0.0037 | 0.0023 | 0.0023 | TRUE | 0.0129 |
| Taf4a | 3665 | 2892 | 2672 | 2524 | 2463 | 2753 | 2846 | 3076 | 2883 | 2512 | 2779 | 2718 | 2802.37 | 2935 | 2670 | 0.91 | -0.137 | 0.03673022 | 0.10777768 | 0.0017 | 0.0033 | 0.0027 | 0.0027 | TRUE | 0.0323 |
| Taf4b | 3724 | 2587 | 2459 | 2596 | 2388 | 2594 | 2892 | 2752 | 2653 | 2584 | 2694 | 2561 | 2689.32 | 2766 | 2613 | 0.944 | -0.083 | 0.17719259 | 0.33123381 | 0.001 | 0.0033 | 0.0024 | 0.0024 | TRUE | 0.02 |
| Taf5 | 2567 | 1892 | 1977 | 2273 | 2002 | 2364 | 1994 | 2012 | 2133 | 2263 | 2258 | 2334 | 2165.66 | 2046 | 2285 | 1.117 | 0.16 | 0.00958618 | 0.04014511 | 4.00E-04 | 0.0035 | 0.0023 | 0.0023 | TRUE | 0.0164 |
| Taf5l | 7071 | 5334 | 5586 | 5389 | 5181 | 5793 | 5492 | 5674 | 6027 | 5364 | 5845 | 5719 | 5686.7 | 5731 | 5643 | 0.985 | -0.022 | 0.71431497 | 0.82842903 | 0.0019 | 0.0028 | 0.0025 | 0.0025 | TRUE | 0.0249 |
| Taf6 | 3575 | 2668 | 2722 | 3031 | 2834 | 3223 | 2777 | 2838 | 2937 | 3017 | 3197 | 3182 | 2991.2 | 2851 | 3132 | 1.099 | 0.136 | 0.02009126 | 0.06943046 | 6.00E-04 | 0.0032 | 0.0021 | 0.0021 | TRUE | 0.0126 |
| Taf6l | 1505 | 1115 | 791 | 851 | 643 | 773 | 1169 | 1186 | 853 | 847 | 725 | 763 | 923.98 | 1069 | 778 | 0.728 | -0.459 | 0.00013233 | 0.00153642 | 0.0187 | 0.005 | 0.0093 | 0.0093 | TRUE | 0.3684 |
| Taf7 | 7829 | 5637 | 4909 | 4596 | 4058 | 4641 | 6080 | 5996 | 5297 | 4575 | 4578 | 4581 | 5184.55 | 5791 | 4578 | 0.79 | -0.34 | 1.42E-07 | 5.34E-06 | 0.0025 | 0.0029 | 0.0028 | 0.0028 | TRUE | 0.0676 |
| Taf7l | 549 | 360 | 290 | 302 | 235 | 214 | 426 | 383 | 313 | 301 | 265 | 211 | 316.53 | 374 | 259 | 0.691 | -0.534 | 0.00060091 | 0.00497328 | 0.0243 | 0.0099 | 0.0142 | 0.0142 | TRUE | 0.2961 |
| Taf8 | 2852 | 2186 | 2204 | 2366 | 2176 | 2562 | 2215 | 2325 | 2378 | 2355 | 2455 | 2529 | 2376.22 | 2306 | 2446 | 1.061 | 0.086 | 0.1712607 | 0.32306046 | 9.00E-04 | 0.0034 | 0.0024 | 0.0024 | TRUE | 0.015 |
| Taf9 | 6955 | 4707 | 4824 | 5186 | 4611 | 5271 | 5402 | 5007 | 5205 | 5162 | 5202 | 5203 | 5196.77 | 5205 | 5189 | 0.997 | -0.005 | 0.92842408 | 0.96054762 | 6.00E-04 | 0.0029 | 0.0019 | 0.0019 | TRUE | 0.0136 |
| Taf9b | 365 | 323 | 457 | 298 | 213 | 232 | 283 | 344 | 493 | 297 | 240 | 229 | 314.35 | 373 | 255 | 0.686 | -0.544 | 0.00309729 | 0.01695748 | 0.0469 | 0.01 | 0.021 | 0.021 | TRUE | 0.5185 |
| Tagap | 159 | 124 | 140 | 154 | 127 | 148 | 123 | 132 | 151 | 153 | 143 | 146 | 141.52 | 135 | 147 | 1.093 | 0.129 | 0.44677933 | 0.61740471 | 0 | 0.0192 | 0.0135 | 0.0135 | TRUE | 0.1018 |
| Tagap1 | 210 | 163 | 175 | 170 | 171 | 180 | 163 | 173 | 189 | 169 | 193 | 178 | 177.52 | 175 | 180 | 1.029 | 0.041 | 0.78609139 | 0.87423002 | 0 | 0.0158 | 0.0111 | 0.0111 | TRUE | 0.0495 |
| Tagln | 87 | 47 | 41 | 73 | 43 | 63 | 68 | 50 | 44 | 73 | 49 | 62 | 57.53 | 54 | 61 | 1.127 | 0.172 | 0.53865662 | 0.69705922 | 0.0282 | 0.0436 | 0.0393 | 0.0393 | TRUE | 0.389 |
| Tagln2 | 7909 | 6272 | 7751 | 9874 | 9314 | 10555 | 6143 | 6671 | 8363 | 9829 | 10507 | 10419 | 8655.41 | 7059 | 10252 | 1.453 | 0.539 | 9.96E-09 | 4.96E-07 | 0.014 | 0.0027 | 0.0062 | 0.0062 | TRUE | 0.3185 |
| Tagln3 | 294 | 234 | 274 | 302 | 249 | 323 | 228 | 249 | 296 | 301 | 281 | 319 | 278.87 | 258 | 300 | 1.17 | 0.226 | 0.09696578 | 0.21648422 | 0.0071 | 0.0109 | 0.0098 | 0.0098 | TRUE | 0.1829 |
| Tal1 | 37 | 33 | 44 | 47 | 54 | 50 | 29 | 35 | 47 | 47 | 61 | 49 | 44.73 | 37 | 52 | 1.421 | 0.507 | 0.09984323 | 0.22082633 | 0.0179 | 0.0554 | 0.0453 | 0.0453 | TRUE | 0.4218 |
| Tal2 | 43 | 19 | 30 | 39 | 48 | 65 | 33 | 20 | 32 | 39 | 54 | 64 | 40.52 | 28 | 52 | 1.816 | 0.861 | 0.01062214 | 0.04315307 | 0.0396 | 0.0609 | 0.0552 | 0.0552 | TRUE | 0.2276 |
| Taldo1 | 12852 | 9411 | 9901 | 11583 | 10268 | 11929 | 9982 | 10010 | 10683 | 11530 | 11583 | 11776 | 10927.32 | 10225 | 11630 | 1.138 | 0.186 | 0.00024537 | 0.00249509 | 7.00E-04 | 0.0026 | 0.0018 | 0.0018 | TRUE | 0.0187 |
| Tamm41 | 2224 | 1595 | 1694 | 1598 | 1440 | 1847 | 1727 | 1697 | 1828 | 1591 | 1624 | 1823 | 1715.01 | 1751 | 1679 | 0.96 | -0.059 | 0.43060704 | 0.60268963 | 0.0027 | 0.0038 | 0.0035 | 0.0035 | TRUE | 0.0683 |
| Tanc1 | 5991 | 4488 | 4830 | 4839 | 4552 | 5272 | 4653 | 4774 | 5212 | 4817 | 5135 | 5204 | 4965.74 | 4880 | 5052 | 1.036 | 0.051 | 0.42646171 | 0.59879619 | 0.0023 | 0.0029 | 0.0027 | 0.0027 | TRUE | 0.043 |
| Tanc2 | 1785 | 1219 | 1120 | 1322 | 1008 | 1161 | 1386 | 1297 | 1208 | 1316 | 1137 | 1146 | 1248.43 | 1297 | 1200 | 0.924 | -0.114 | 0.18787089 | 0.34501332 | 0.0047 | 0.0043 | 0.0046 | 0.0046 | TRUE | 0.0866 |

| | | | | | | | | | | | | | | | | | | | | | | | | | |
|---|---|---|---|---|---|---|---|---|---|---|---|---|---|---|---|---|---|---|---|---|---|---|---|---|---|
| Tango2 | 631 | 435 | 384 | 389 | 341 | 387 | 490 | 463 | 414 | 387 | 385 | 382 | 420.17 | 456 | 385 | 0.842 | -0.248 | 0.02209564 | 0.07416933 | 0 | 0.0081 | 0.006 | 0.006 | TRUE | 0.0718 |
| Tango6 | 4979 | 3645 | 3599 | 3762 | 3393 | 3798 | 3867 | 3877 | 3883 | 3745 | 3828 | 3749 | 3824.83 | 3876 | 3774 | 0.974 | -0.039 | 0.45250107 | 0.62236314 | 0 | 0.003 | 0.0016 | 0.0016 | TRUE | 0.0019 |
| Tank | 633 | 460 | 457 | 484 | 388 | 491 | 492 | 489 | 493 | 482 | 438 | 485 | 479.7 | 491 | 468 | 0.953 | -0.069 | 0.48413785 | 0.65044468 | 0 | 0.0074 | 0.0049 | 0.0049 | TRUE | 0.0361 |
| Taok1 | 10949 | 7740 | 6934 | 9204 | 7391 | 8477 | 8504 | 8233 | 7482 | 9162 | 8338 | 8368 | 8347.64 | 8073 | 8623 | 1.068 | 0.095 | 0.15145924 | 0.29764978 | 0.0034 | 0.0027 | 0.003 | 0.003 | TRUE | 0.0499 |
| Taok2 | 5207 | 3871 | 3425 | 3127 | 2918 | 3338 | 4044 | 4118 | 3696 | 3113 | 3292 | 3295 | 3592.78 | 3953 | 3233 | 0.818 | -0.29 | 4.74E-06 | 0.00010159 | 0.0017 | 0.0031 | 0.0026 | 0.0026 | TRUE | 0.0388 |
| Taok3 | 1278 | 912 | 1007 | 1161 | 1131 | 1231 | 993 | 970 | 1087 | 1156 | 1276 | 1215 | 1115.99 | 1017 | 1216 | 1.196 | 0.258 | 0.00118681 | 0.00819866 | 0.002 | 0.0046 | 0.0037 | 0.0037 | TRUE | 0.0423 |
| Tap1 | 318 | 277 | 297 | 347 | 317 | 318 | 247 | 295 | 320 | 345 | 358 | 314 | 313.17 | 287 | 339 | 1.184 | 0.243 | 0.06572813 | 0.1637818 | 0.0079 | 0.01 | 0.0094 | 0.0094 | TRUE | 0.1843 |
| Tap2 | 159 | 115 | 132 | 204 | 166 | 210 | 123 | 122 | 142 | 203 | 187 | 207 | 164.31 | 129 | 199 | 1.544 | 0.626 | 7.54E-05 | 0.00097141 | 0 | 0.0169 | 0.0117 | 0.0117 | TRUE | 0.075 |
| Tapbp | 1195 | 792 | 923 | 1027 | 910 | 1079 | 928 | 842 | 996 | 1022 | 1027 | 1065 | 980.07 | 922 | 1038 | 1.126 | 0.171 | 0.0419758 | 0.11842876 | 0.0025 | 0.0048 | 0.0041 | 0.0041 | TRUE | 0.0655 |
| Tapbpl | 85 | 75 | 60 | 49 | 55 | 77 | 66 | 80 | 65 | 49 | 62 | 76 | 66.23 | 70 | 62 | 0.889 | -0.169 | 0.50726573 | 0.67064556 | 0.0149 | 0.0382 | 0.0319 | 0.0319 | TRUE | 0.2272 |
| Tapt1 | 951 | 732 | 738 | 829 | 705 | 844 | 739 | 779 | 796 | 825 | 795 | 833 | 794.53 | 771 | 818 | 1.062 | 0.087 | 0.28526881 | 0.45727025 | 0 | 0.0054 | 0.0035 | 0.0035 | TRUE | 0.0172 |
| Tarbp2 | 3369 | 2549 | 2291 | 2429 | 2230 | 2595 | 2617 | 2711 | 2472 | 2418 | 2516 | 2562 | 2549.17 | 2600 | 2499 | 0.961 | -0.058 | 0.35962394 | 0.53425891 | 0.0011 | 0.0034 | 0.0025 | 0.0025 | TRUE | 0.0219 |
| Tardbp | 28669 | 21402 | 18738 | 19694 | 15844 | 19126 | 22266 | 22765 | 20218 | 19603 | 17874 | 18880 | 20267.76 | 21750 | 18786 | 0.864 | -0.211 | 0.00065847 | 0.00532025 | 0.0029 | 0.0025 | 0.0027 | 0.0027 | TRUE | 0.0463 |
| Tarm1 | 12 | 13 | 9 | 31 | 20 | 17 | 9 | 14 | 10 | 31 | 23 | 17 | 17.18 | 11 | 24 | 2.151 | 1.105 | 0.0244573 | 0.08007346 | 0 | 0.1404 | 0.1075 | 0.1075 | TRUE | 0.1806 |
| Tars | 10408 | 7619 | 7303 | 7197 | 6195 | 7631 | 8083 | 8104 | 7880 | 7164 | 6989 | 7533 | 7625.5 | 8022 | 7229 | 0.901 | -0.15 | 0.00401457 | 0.02067758 | 7.00E-04 | 0.0027 | 0.0018 | 0.0018 | TRUE | 0.0166 |
| Tars2 | 4910 | 3734 | 3695 | 3913 | 3393 | 4059 | 3813 | 3972 | 3987 | 3895 | 3828 | 4007 | 3916.93 | 3924 | 3910 | 0.997 | -0.005 | 0.93302767 | 0.9627836 | 3.00E-04 | 0.003 | 0.0019 | 0.0019 | TRUE | 0.0076 |
| Tarsl2 | 2395 | 1609 | 1721 | 2121 | 1891 | 2107 | 1860 | 1711 | 1857 | 2111 | 2133 | 2080 | 1958.82 | 1809 | 2108 | 1.164 | 0.22 | 0.00066328 | 0.00534099 | 6.00E-04 | 0.0036 | 0.0025 | 0.0025 | TRUE | 0.0266 |
| Tas1r1 | 31 | 22 | 20 | 11 | 13 | 27 | 24 | 23 | 22 | 11 | 15 | 27 | 20.22 | 23 | 18 | 0.758 | -0.4 | 0.38205479 | 0.55626553 | 0.0439 | 0.1196 | 0.1001 | 0.1001 | TRUE | 1.1105 |
| Tas1r2 | 5 | 2 | 4 | 2 | 3 | 1 | 4 | 2 | 4 | 2 | 3 | 1 | 2.78 | 3 | 2 | 0.606 | -0.722 | 0.52965317 | NA | 0 | 0.8545 | 0.5648 | 0.5648 | TRUE | 0.2266 |
| Tas1r3 | 3 | 1 | 1 | 5 | 1 | 0 | 2 | 1 | 1 | 5 | 1 | 0 | 1.76 | 1 | 2 | 1.343 | 0.426 | 0.78196793 | NA | 0.5945 | 1.3468 | 1.1347 | 1.1347 | TRUE | 1.0462 |
| Tas2r102 | 0 | 0 | 0 | 0 | 0 | 0 | 0 | 0 | 0 | 0 | 0 | 0 | 0 | 0 | 0 | NA | NA | NA | NA | NA | NA | NA | NA | NA | NA |
| Tas2r103 | 0 | 0 | 0 | 0 | 0 | 0 | 0 | 0 | 0 | 0 | 0 | 0 | 0 | 0 | 0 | NA | NA | NA | NA | NA | NA | NA | NA | NA | NA |
| Tas2r104 | 0 | 0 | 0 | 0 | 0 | 0 | 0 | 0 | 0 | 0 | 0 | 0 | 0 | 0 | 0 | NA | NA | NA | NA | NA | NA | NA | NA | NA | NA |
| Tas2r105 | 0 | 0 | 0 | 0 | 0 | 0 | 0 | 0 | 0 | 0 | 0 | 0 | 0 | 0 | 0 | NA | NA | NA | NA | NA | NA | NA | NA | NA | NA |
| Tas2r106 | 0 | 0 | 0 | 0 | 0 | 0 | 0 | 0 | 0 | 0 | 0 | 0 | 0 | 0 | 0 | NA | NA | NA | NA | NA | NA | NA | NA | NA | NA |
| Tas2r107 | 1 | 0 | 0 | 0 | 0 | 0 | 1 | 0 | 0 | 0 | 0 | 0 | 0.13 | 0 | 0 | 0.552 | -0.857 | 0.8336548 | NA | 0 | 18.3121 | 10 | 10 | TRUE | 0.2566 |
| Tas2r108 | 0 | 0 | 0 | 1 | 0 | 0 | 0 | 0 | 0 | 1 | 0 | 0 | 0.17 | 0 | 0 | 2.094 | 1.067 | 0.79379329 | NA | 0 | 14.2885 | 10 | 10 | TRUE | 0.3906 |
| Tas2r109 | 0 | 0 | 0 | 0 | 0 | 0 | 0 | 0 | 0 | 0 | 0 | 0 | 0 | 0 | 0 | NA | NA | NA | NA | NA | NA | NA | NA | NA | NA |
| Tas2r110 | 0 | 0 | 0 | 0 | 0 | 0 | 0 | 0 | 0 | 0 | 0 | 0 | 0 | 0 | 0 | NA | NA | NA | NA | NA | NA | NA | NA | NA | NA |
| Tas2r113 | 0 | 0 | 0 | 0 | 0 | 0 | 0 | 0 | 0 | 0 | 0 | 0 | 0 | 0 | 0 | NA | NA | NA | NA | NA | NA | NA | NA | NA | NA |
| Tas2r114 | 0 | 0 | 0 | 0 | 0 | 0 | 0 | 0 | 0 | 0 | 0 | 0 | 0 | 0 | 0 | NA | NA | NA | NA | NA | NA | NA | NA | NA | NA |
| Tas2r115 | 0 | 0 | 0 | 0 | 0 | 0 | 0 | 0 | 0 | 0 | 0 | 0 | 0 | 0 | 0 | NA | NA | NA | NA | NA | NA | NA | NA | NA | NA |
| Tas2r116 | 0 | 0 | 0 | 0 | 0 | 0 | 0 | 0 | 0 | 0 | 0 | 0 | 0 | 0 | 0 | NA | NA | NA | NA | NA | NA | NA | NA | NA | NA |
| Tas2r117 | 0 | 0 | 0 | 0 | 0 | 0 | 0 | 0 | 0 | 0 | 0 | 0 | 0 | 0 | 0 | NA | NA | NA | NA | NA | NA | NA | NA | NA | NA |
| Tas2r118 | 0 | 0 | 0 | 0 | 0 | 0 | 0 | 0 | 0 | 0 | 0 | 0 | 0 | 0 | 0 | NA | NA | NA | NA | NA | NA | NA | NA | NA | NA |
| Tas2r119 | 0 | 0 | 0 | 0 | 0 | 0 | 0 | 0 | 0 | 0 | 0 | 0 | 0 | 0 | 0 | NA | NA | NA | NA | NA | NA | NA | NA | NA | NA |
| Tas2r120 | 0 | 0 | 0 | 0 | 0 | 0 | 0 | 0 | 0 | 0 | 0 | 0 | 0 | 0 | 0 | NA | NA | NA | NA | NA | NA | NA | NA | NA | NA |
| Tas2r121 | 0 | 0 | 0 | 0 | 0 | 0 | 0 | 0 | 0 | 0 | 0 | 0 | 0 | 0 | 0 | NA | NA | NA | NA | NA | NA | NA | NA | NA | NA |
| Tas2r122 | 0 | 0 | 0 | 0 | 0 | 0 | 0 | 0 | 0 | 0 | 0 | 0 | 0 | 0 | 0 | NA | NA | NA | NA | NA | NA | NA | NA | NA | NA |
| Tas2r123 | 0 | 0 | 0 | 0 | 0 | 0 | 0 | 0 | 0 | 0 | 0 | 0 | 0 | 0 | 0 | NA | NA | NA | NA | NA | NA | NA | NA | NA | NA |
| Tas2r124 | 0 | 0 | 0 | 0 | 0 | 0 | 0 | 0 | 0 | 0 | 0 | 0 | 0 | 0 | 0 | NA | NA | NA | NA | NA | NA | NA | NA | NA | NA |
| Tas2r125 | 0 | 0 | 0 | 0 | 0 | 0 | 0 | 0 | 0 | 0 | 0 | 0 | 0 | 0 | 0 | NA | NA | NA | NA | NA | NA | NA | NA | NA | NA |
| Tas2r126 | 0 | 0 | 0 | 0 | 0 | 0 | 0 | 0 | 0 | 0 | 0 | 0 | 0 | 0 | 0 | NA | NA | NA | NA | NA | NA | NA | NA | NA | NA |
| Tas2r129 | 0 | 0 | 0 | 0 | 0 | 0 | 0 | 0 | 0 | 0 | 0 | 0 | 0 | 0 | 0 | NA | NA | NA | NA | NA | NA | NA | NA | NA | NA |
| Tas2r130 | 0 | 0 | 0 | 0 | 0 | 0 | 0 | 0 | 0 | 0 | 0 | 0 | 0 | 0 | 0 | NA | NA | NA | NA | NA | NA | NA | NA | NA | NA |
| Tas2r131 | 0 | 0 | 0 | 0 | 0 | 0 | 0 | 0 | 0 | 0 | 0 | 0 | 0 | 0 | 0 | NA | NA | NA | NA | NA | NA | NA | NA | NA | NA |
| Tas2r134 | 0 | 0 | 0 | 0 | 0 | 0 | 0 | 0 | 0 | 0 | 0 | 0 | 0 | 0 | 0 | NA | NA | NA | NA | NA | NA | NA | NA | NA | NA |
| Tas2r135 | 0 | 0 | 0 | 0 | 0 | 0 | 0 | 0 | 0 | 0 | 0 | 0 | 0 | 0 | 0 | NA | NA | NA | NA | NA | NA | NA | NA | NA | NA |
| Tas2r136 | 0 | 0 | 0 | 0 | 0 | 0 | 0 | 0 | 0 | 0 | 0 | 0 | 0 | 0 | 0 | NA | NA | NA | NA | NA | NA | NA | NA | NA | NA |
| Tas2r137 | 0 | 0 | 0 | 1 | 0 | 0 | 0 | 0 | 0 | 1 | 0 | 0 | 0.17 | 0 | 0 | 2.094 | 1.067 | 0.79379329 | NA | 0 | 14.2885 | 10 | 10 | TRUE | 0.3906 |
| Tas2r138 | 0 | 0 | 0 | 0 | 0 | 0 | 0 | 0 | 0 | 0 | 0 | 0 | 0 | 0 | 0 | NA | NA | NA | NA | NA | NA | NA | NA | NA | NA |
| Tas2r139 | 0 | 0 | 0 | 1 | 0 | 0 | 0 | 0 | 0 | 1 | 0 | 0 | 0.17 | 0 | 0 | 2.094 | 1.067 | 0.79379329 | NA | 0 | 14.2885 | 10 | 10 | TRUE | 0.3906 |
| Tas2r140 | 0 | 0 | 0 | 0 | 0 | 0 | 0 | 0 | 0 | 0 | 0 | 0 | 0 | 0 | 0 | NA | NA | NA | NA | NA | NA | NA | NA | NA | NA |
| Tas2r143 | 0 | 0 | 0 | 0 | 0 | 0 | 0 | 0 | 0 | 0 | 0 | 0 | 0 | 0 | 0 | NA | NA | NA | NA | NA | NA | NA | NA | NA | NA |
| Tas2r144 | 0 | 0 | 0 | 0 | 0 | 0 | 0 | 0 | 0 | 0 | 0 | 0 | 0 | 0 | 0 | NA | NA | NA | NA | NA | NA | NA | NA | NA | NA |
| Tasp1 | 932 | 651 | 628 | 639 | 578 | 662 | 724 | 692 | 678 | 636 | 652 | 653 | 672.59 | 698 | 647 | 0.926 | -0.111 | 0.1943832 | 0.35347572 | 0 | 0.006 | 0.0037 | 0.0037 | TRUE | 0.0132 |
| Tat | 2 | 3 | 3 | 3 | 5 | 2 | 2 | 3 | 3 | 3 | 6 | 2 | 3.1 | 3 | 4 | 1.34 | 0.422 | 0.69717651 | NA | 0 | 0.7677 | 0.5167 | 0.5167 | TRUE | 0.3634 |
| Tatdn1 | 521 | 324 | 390 | 432 | 371 | 413 | 405 | 345 | 421 | 430 | 419 | 408 | 404.39 | 390 | 419 | 1.073 | 0.101 | 0.3695081 | 0.54424766 | 0.0029 | 0.0083 | 0.0067 | 0.0067 | TRUE | 0.1144 |
| Tatdn2 | 2961 | 1983 | 1612 | 1624 | 1509 | 1748 | 2300 | 2109 | 1739 | 1617 | 1702 | 1726 | 1865.45 | 2049 | 1682 | 0.82 | -0.287 | 0.00239727 | 0.01390589 | 0.0094 | 0.0037 | 0.0059 | 0.0059 | TRUE | 0.2107 |
| Tatdn3 | 350 | 259 | 245 | 251 | 215 | 281 | 272 | 275 | 264 | 250 | 243 | 277 | 263.58 | 270 | 257 | 0.949 | -0.075 | 0.55076281 | 0.70677437 | 0 | 0.0114 | 0.0077 | 0.0077 | TRUE | 0.0567 |
| Tax1bp1 | 9150 | 6429 | 6130 | 6054 | 5351 | 6217 | 7106 | 6838 | 6614 | 6026 | 6036 | 6137 | 6459.81 | 6853 | 6066 | 0.885 | -0.176 | 0.00071374 | 0.00564863 | 5.00E-04 | 0.0028 | 0.0018 | 0.0018 | TRUE | 0.013 |
| Tax1bp3 | 439 | 329 | 297 | 248 | 205 | 224 | 341 | 350 | 320 | 247 | 231 | 221 | 285.1 | 337 | 233 | 0.691 | -0.533 | 1.51E-05 | 0.00026641 | 0 | 0.0107 | 0.0073 | 0.0073 | TRUE | 0.0304 |
| Taz | 790 | 690 | 684 | 707 | 658 | 690 | 614 | 734 | 738 | 704 | 742 | 681 | 702.12 | 695 | 709 | 1.022 | 0.031 | 0.75026772 | 0.85168585 | 0.0052 | 0.0058 | 0.0056 | 0.0056 | TRUE | 0.1315 |
| Tbata | 0 | 1 | 1 | 2 | 1 | 0 | 0 | 1 | 1 | 2 | 1 | 0 | 0.88 | 1 | 1 | 1.518 | 0.602 | 0.77693369 | NA | 0 | 2.7051 | 2.052 | 2.052 | TRUE | 0.2258 |
| Tbc1d1 | 6054 | 4453 | 4709 | 5284 | 4666 | 5343 | 4702 | 4737 | 5081 | 5260 | 5264 | 5274 | 5052.87 | 4840 | 5266 | 1.088 | 0.122 | 0.0254694 | 0.08241661 | 6.00E-04 | 0.0029 | 0.002 | 0.002 | TRUE | 0.023 |
| Tbc1d10a | 1831 | 1331 | 1333 | 1554 | 1412 | 1540 | 1422 | 1416 | 1438 | 1547 | 1593 | 1520 | 1489.35 | 1425 | 1553 | 1.09 | 0.124 | 0.05692764 | 0.14812794 | 0 | 0.004 | 0.0024 | 0.0024 | TRUE | 0.0059 |
| Tbc1d10b | 6937 | 4815 | 4540 | 4426 | 3838 | 4399 | 5388 | 5122 | 4899 | 4406 | 4330 | 4342 | 4747.62 | 5136 | 4359 | 0.848 | -0.237 | 2.95E-05 | 0.00045785 | 9.00E-04 | 0.0029 | 0.0021 | 0.0021 | TRUE | 0.0228 |
| Tbc1d10c | 91 | 78 | 73 | 73 | 78 | 111 | 71 | 83 | 79 | 73 | 88 | 110 | 83.77 | 78 | 90 | 1.167 | 0.222 | 0.33228135 | 0.50659171 | 0.0142 | 0.0307 | 0.0259 | 0.0259 | TRUE | 0.2673 |
| Tbc1d12 | 708 | 478 | 322 | 404 | 356 | 338 | 550 | 508 | 347 | 402 | 402 | 334 | 423.86 | 468 | 379 | 0.807 | -0.309 | 0.04709087 | 0.12898999 | 0.0319 | 0.008 | 0.0151 | 0.0151 | TRUE | 0.5845 |

| | | | | | | | | | | | | | | | | | | | | | | | | | |
|---|---|---|---|---|---|---|---|---|---|---|---|---|---|---|---|---|---|---|---|---|---|---|---|---|---|
| Tbc1d13 | 4235 | 2965 | 2925 | 3017 | 2758 | 3292 | 3289 | 3154 | 3156 | 3003 | 3111 | 3250 | 3160.53 | 3200 | 3121 | 0.975 | -0.036 | 0.53927324 | 0.6975414 | 8.00E-04 | 0.0032 | 0.0022 | 0.0022 | TRUE | 0.0159 |
| Tbc1d14 | 4447 | 3301 | 3574 | 3690 | 3402 | 3886 | 3454 | 3511 | 3856 | 3673 | 3838 | 3836 | 3694.71 | 3607 | 3782 | 1.049 | 0.069 | 0.27194292 | 0.4429977 | 0.0017 | 0.0031 | 0.0026 | 0.0026 | TRUE | 0.0441 |
| Tbc1d15 | 5509 | 4181 | 4945 | 4919 | 4649 | 5441 | 4279 | 4447 | 5336 | 4896 | 5245 | 5371 | 4928.92 | 4687 | 5171 | 1.104 | 0.142 | 0.08523442 | 0.19719256 | 0.0082 | 0.0029 | 0.0047 | 0.0047 | TRUE | 0.1779 |
| Tbc1d16 | 3120 | 2457 | 2395 | 3258 | 2778 | 3089 | 2423 | 2613 | 2584 | 3243 | 3134 | 3049 | 2841.17 | 2540 | 3142 | 1.238 | 0.308 | 4.84E-07 | 1.58E-05 | 0.001 | 0.0033 | 0.0023 | 0.0023 | TRUE | 0.0204 |
| Tbc1d17 | 1668 | 1251 | 1253 | 1189 | 1164 | 1372 | 1295 | 1331 | 1352 | 1184 | 1313 | 1354 | 1304.86 | 1326 | 1284 | 0.968 | -0.046 | 0.54170199 | 0.69973309 | 0.0019 | 0.0042 | 0.0034 | 0.0034 | TRUE | 0.0566 |
| Tbc1d19 | 793 | 619 | 570 | 581 | 498 | 623 | 616 | 658 | 615 | 578 | 562 | 615 | 607.41 | 630 | 585 | 0.93 | -0.105 | 0.250185 | 0.41906648 | 0 | 0.0063 | 0.0043 | 0.0043 | TRUE | 0.0237 |
| Tbc1d2 | 338 | 271 | 287 | 312 | 309 | 296 | 263 | 288 | 310 | 311 | 349 | 292 | 301.96 | 287 | 317 | 1.107 | 0.147 | 0.25073292 | 0.41959087 | 0.0043 | 0.0103 | 0.0085 | 0.0085 | TRUE | 0.083 |
| Tbc1d20 | 2256 | 1685 | 1794 | 1801 | 1662 | 1802 | 1752 | 1792 | 1936 | 1793 | 1875 | 1779 | 1821.11 | 1827 | 1816 | 0.994 | -0.008 | 0.90367069 | 0.9467247 | 0.0012 | 0.0037 | 0.0028 | 0.0028 | TRUE | 0.0325 |
| Tbc1d21 | 1 | 0 | 1 | 3 | 1 | 0 | 1 | 0 | 1 | 3 | 1 | 0 | 0.99 | 1 | 1 | 2.197 | 1.136 | 0.5788429 | NA | 0 | 2.3844 | 1.8693 | 1.8693 | TRUE | 0.2259 |
| Tbc1d22a | 1466 | 1152 | 1257 | 1235 | 1174 | 1393 | 1139 | 1225 | 1356 | 1229 | 1324 | 1375 | 1274.84 | 1240 | 1309 | 1.057 | 0.08 | 0.34516835 | 0.51936368 | 0.0043 | 0.0043 | 0.0045 | 0.0045 | TRUE | 0.0803 |
| Tbc1d22b | 2455 | 1916 | 2078 | 2050 | 2004 | 2230 | 1907 | 2038 | 2242 | 2041 | 2261 | 2201 | 2114.92 | 2062 | 2168 | 1.052 | 0.073 | 0.3454708 | 0.51955115 | 0.004 | 0.0035 | 0.0038 | 0.0038 | TRUE | 0.0702 |
| Tbc1d23 | 3210 | 2328 | 2297 | 2492 | 2246 | 2647 | 2493 | 2476 | 2478 | 2481 | 2534 | 2613 | 2512.51 | 2482 | 2543 | 1.024 | 0.034 | 0.55015826 | 0.70637909 | 0 | 0.0034 | 0.002 | 0.002 | TRUE | 0.0073 |
| Tbc1d24 | 1514 | 1229 | 1206 | 1481 | 1371 | 1440 | 1176 | 1307 | 1301 | 1474 | 1547 | 1422 | 1371.12 | 1261 | 1481 | 1.175 | 0.233 | 0.00204697 | 0.01227751 | 0.002 | 0.0042 | 0.0034 | 0.0034 | TRUE | 0.0443 |
| Tbc1d25 | 993 | 695 | 574 | 541 | 533 | 579 | 771 | 739 | 619 | 539 | 601 | 572 | 640.2 | 710 | 571 | 0.802 | -0.319 | 0.00235224 | 0.0137115 | 0.0067 | 0.0061 | 0.0063 | 0.0063 | TRUE | 0.1446 |
| Tbc1d2b | 1361 | 1044 | 937 | 902 | 857 | 839 | 1057 | 1110 | 1011 | 898 | 967 | 828 | 978.57 | 1059 | 898 | 0.847 | -0.24 | 0.00474017 | 0.02352472 | 0.0029 | 0.0048 | 0.0042 | 0.0042 | TRUE | 0.0547 |
| Tbc1d30 | 87 | 68 | 76 | 91 | 87 | 120 | 68 | 72 | 82 | 91 | 98 | 118 | 88.18 | 74 | 102 | 1.39 | 0.475 | 0.02846684 | 0.0890997 | 0 | 0.0293 | 0.0223 | 0.0223 | TRUE | 0.1896 |
| Tbc1d4 | 363 | 320 | 272 | 347 | 311 | 306 | 282 | 340 | 293 | 345 | 351 | 302 | 319.02 | 305 | 333 | 1.092 | 0.126 | 0.31871413 | 0.4926334 | 0.0052 | 0.0099 | 0.0085 | 0.0085 | TRUE | 0.1126 |
| Tbc1d5 | 1665 | 1170 | 1116 | 1111 | 973 | 1108 | 1293 | 1245 | 1204 | 1106 | 1098 | 1094 | 1173.18 | 1247 | 1099 | 0.88 | -0.184 | 0.00986214 | 0.04095282 | 0 | 0.0044 | 0.0028 | 0.0028 | TRUE | 0.013 |
| Tbc1d7 | 759 | 554 | 522 | 542 | 478 | 555 | 589 | 589 | 563 | 540 | 539 | 548 | 561.43 | 580 | 542 | 0.933 | -0.099 | 0.27008631 | 0.44103092 | 0 | 0.0066 | 0.0041 | 0.0041 | TRUE | 0.0078 |
| Tbc1d8 | 1255 | 949 | 1017 | 865 | 794 | 888 | 975 | 1009 | 1097 | 861 | 896 | 877 | 952.47 | 1027 | 878 | 0.855 | -0.225 | 0.00505609 | 0.02474042 | 0.0012 | 0.0049 | 0.0036 | 0.0036 | TRUE | 0.0422 |
| Tbc1d8b | 247 | 173 | 165 | 233 | 170 | 195 | 192 | 184 | 178 | 232 | 192 | 192 | 195.01 | 185 | 205 | 1.112 | 0.153 | 0.30488307 | 0.47739853 | 0 | 0.0146 | 0.0109 | 0.0109 | TRUE | 0.1409 |
| Tbc1d9 | 931 | 595 | 643 | 676 | 533 | 712 | 723 | 633 | 694 | 673 | 601 | 703 | 671.13 | 683 | 659 | 0.964 | -0.053 | 0.58779332 | 0.73702342 | 0.0039 | 0.006 | 0.0053 | 0.0053 | TRUE | 0.0675 |
| Tbc1d9b | 3602 | 2579 | 2647 | 2936 | 2614 | 2807 | 2798 | 2743 | 2856 | 2922 | 2949 | 2771 | 2839.86 | 2799 | 2881 | 1.029 | 0.041 | 0.4775955 | 0.64514436 | 4.00E-04 | 0.0033 | 0.0021 | 0.0021 | TRUE | 0.0136 |
| Tbca | 4189 | 2979 | 3181 | 3558 | 3076 | 3513 | 3253 | 3169 | 3432 | 3542 | 3470 | 3468 | 3388.99 | 3285 | 3493 | 1.064 | 0.089 | 0.12038229 | 0.25236492 | 5.00E-04 | 0.0031 | 0.0021 | 0.0021 | TRUE | 0.0184 |
| Tbcb | 3713 | 2587 | 2452 | 2669 | 2431 | 2810 | 2884 | 2752 | 2646 | 2657 | 2742 | 2774 | 2742.37 | 2761 | 2724 | 0.986 | -0.02 | 0.74367249 | 0.84754073 | 9.00E-04 | 0.0033 | 0.0023 | 0.0023 | TRUE | 0.0191 |
| Tbcc | 1101 | 774 | 566 | 581 | 581 | 581 | 855 | 823 | 611 | 578 | 655 | 574 | 682.73 | 763 | 602 | 0.788 | -0.344 | 0.0055567 | 0.0265892 | 0.0178 | 0.0059 | 0.0096 | 0.0096 | TRUE | 0.356 |
| Tbccd1 | 1377 | 1015 | 925 | 894 | 789 | 861 | 1069 | 1080 | 998 | 890 | 890 | 850 | 962.84 | 1049 | 877 | 0.835 | -0.26 | 0.00078323 | 0.00606152 | 0 | 0.0049 | 0.0033 | 0.0033 | TRUE | 0.021 |
| Tbcd | 9557 | 6860 | 6494 | 7310 | 6339 | 7097 | 7423 | 7297 | 7007 | 7276 | 7151 | 7006 | 7193.27 | 7242 | 7144 | 0.986 | -0.02 | 0.69623211 | 0.81507458 | 4.00E-04 | 0.0028 | 0.0017 | 0.0017 | TRUE | 0.0098 |
| Tbce | 1046 | 770 | 740 | 750 | 650 | 740 | 812 | 819 | 798 | 747 | 733 | 730 | 773.36 | 810 | 737 | 0.91 | -0.137 | 0.08571297 | 0.19785729 | 0 | 0.0055 | 0.0033 | 0.0033 | TRUE | 0.0017 |
| Tbcel | 2105 | 1643 | 1936 | 2041 | 1920 | 2103 | 1635 | 1748 | 2089 | 2032 | 2166 | 2076 | 1957.5 | 1824 | 2091 | 1.148 | 0.199 | 0.02550766 | 0.08248293 | 0.0082 | 0.0036 | 0.0052 | 0.0052 | TRUE | 0.1944 |
| Tbck | 704 | 516 | 489 | 541 | 528 | 550 | 547 | 549 | 528 | 539 | 596 | 543 | 550.06 | 541 | 559 | 1.032 | 0.045 | 0.62954961 | 0.76684443 | 0 | 0.0067 | 0.0046 | 0.0046 | TRUE | 0.0374 |
| Tbk1 | 2609 | 1913 | 1928 | 1871 | 1632 | 2004 | 2026 | 2035 | 2080 | 1862 | 1841 | 1978 | 1970.52 | 2047 | 1894 | 0.926 | -0.112 | 0.07612229 | 0.18200333 | 3.00E-04 | 0.0036 | 0.0023 | 0.0023 | TRUE | 0.0185 |
| Tbkbp1 | 1392 | 1094 | 1181 | 1361 | 1209 | 1422 | 1081 | 1164 | 1274 | 1355 | 1364 | 1404 | 1273.57 | 1173 | 1374 | 1.173 | 0.231 | 0.00370628 | 0.0194471 | 0.0028 | 0.0043 | 0.0038 | 0.0038 | TRUE | 0.0681 |
| Tbl1x | 3126 | 2398 | 2486 | 2509 | 2219 | 2487 | 2428 | 2551 | 2682 | 2497 | 2503 | 2455 | 2519.45 | 2554 | 2485 | 0.974 | -0.038 | 0.53976988 | 0.69799429 | 0.001 | 0.0034 | 0.0024 | 0.0024 | TRUE | 0.0236 |
| Tbl1xr1 | 3942 | 2941 | 3233 | 3629 | 2964 | 3922 | 3062 | 3128 | 3488 | 3612 | 3344 | 3872 | 3417.65 | 3226 | 3609 | 1.119 | 0.163 | 0.02998011 | 0.09260983 | 0.0048 | 0.0031 | 0.0038 | 0.0038 | TRUE | 0.0611 |
| Tbl2 | 4780 | 4215 | 4994 | 5255 | 4754 | 5898 | 3712 | 4483 | 5388 | 5231 | 5363 | 5822 | 5000.07 | 4528 | 5472 | 1.209 | 0.274 | 0.00941561 | 0.03967493 | 0.0185 | 0.0029 | 0.0078 | 0.0078 | TRUE | 0.279 |
| Tbl3 | 7524 | 5353 | 5197 | 5718 | 4951 | 5578 | 5844 | 5694 | 5608 | 5692 | 5585 | 5506 | 5654.71 | 5715 | 5594 | 0.979 | -0.031 | 0.53705602 | 0.69586997 | 2.00E-04 | 0.0028 | 0.0016 | 0.0016 | TRUE | 0.0048 |
| Tbp | 2139 | 1467 | 1470 | 1605 | 1372 | 1525 | 1661 | 1560 | 1586 | 1598 | 1548 | 1505 | 1576.43 | 1602 | 1550 | 0.967 | -0.049 | 0.46465459 | 0.63377308 | 5.00E-04 | 0.0039 | 0.0026 | 0.0026 | TRUE | 0.0129 |
| Tbpl1 | 2316 | 1662 | 1595 | 1515 | 1290 | 1514 | 1799 | 1768 | 1721 | 1508 | 1455 | 1495 | 1624.24 | 1763 | 1486 | 0.843 | -0.247 | 0.00011852 | 0.0013989 | 0 | 0.0039 | 0.0023 | 0.0023 | TRUE | 0.0051 |
| Tbpl2 | 0 | 0 | 0 | 0 | 0 | 0 | 0 | 0 | 0 | 0 | 0 | 0 | 0 | 0 | 0 | NA | NA | NA | NA | NA | NA | NA | NA | NA | NA |
| Tbr1 | 4 | 4 | 0 | 4 | 5 | 3 | 3 | 4 | 0 | 4 | 6 | 3 | 3.32 | 2 | 4 | 1.68 | 0.748 | 0.50086606 | NA | 0 | 0.7154 | 0.572 | 0.572 | TRUE | 0.7216 |
| Tbrg1 | 12345 | 8909 | 9063 | 8953 | 8085 | 9188 | 9588 | 9476 | 9779 | 8912 | 9121 | 9070 | 9324.27 | 9614 | 9034 | 0.94 | -0.09 | 0.05367183 | 0.14156329 | 1.00E-04 | 0.0027 | 0.0015 | 0.0015 | TRUE | 0.0027 |
| Tbrg3 | 34 | 20 | 19 | 51 | 36 | 52 | 26 | 21 | 21 | 51 | 41 | 51 | 35.15 | 23 | 48 | 2.081 | 1.057 | 0.00157948 | 0.01002017 | 0 | 0.0699 | 0.0488 | 0.0488 | TRUE | 0.1437 |
| Tbrg4 | 6201 | 4402 | 4037 | 4035 | 3804 | 4320 | 4816 | 4682 | 4356 | 4016 | 4291 | 4265 | 4404.42 | 4618 | 4191 | 0.907 | -0.141 | 0.02164999 | 0.07318906 | 0.0016 | 0.003 | 0.0025 | 0.0025 | TRUE | 0.0299 |
| Tbx1 | 85 | 63 | 68 | 120 | 93 | 98 | 66 | 67 | 73 | 119 | 105 | 97 | 87.92 | 69 | 107 | 1.559 | 0.641 | 0.00249982 | 0.01435514 | 0 | 0.0294 | 0.0205 | 0.0205 | TRUE | 0.1046 |
| Tbx10 | 2 | 2 | 1 | 0 | 4 | 2 | 2 | 2 | 1 | 0 | 5 | 2 | 1.87 | 2 | 2 | 1.343 | 0.425 | 0.77133381 | NA | 0 | 1.2669 | 0.9939 | 0.9939 | TRUE | 0.1962 |
| Tbx15 | 33 | 28 | 27 | 22 | 18 | 25 | 26 | 30 | 29 | 22 | 20 | 25 | 25.24 | 28 | 22 | 0.795 | -0.331 | 0.38145898 | 0.55579288 | 0 | 0.0963 | 0.0626 | 0.0626 | TRUE | 0.0512 |
| Tbx18 | 26 | 16 | 28 | 29 | 15 | 15 | 20 | 17 | 30 | 29 | 17 | 15 | 21.34 | 22 | 20 | 0.904 | -0.146 | 0.74633933 | 0.84945278 | 0.0621 | 0.1135 | 0.0992 | 0.0992 | TRUE | 0.7845 |
| Tbx19 | 1 | 1 | 1 | 1 | 2 | 1 | 1 | 1 | 1 | 1 | 2 | 1 | 1.19 | 1 | 1 | 1.452 | 0.538 | 0.74839225 | NA | 0 | 1.989 | 1.1562 | 1.1562 | TRUE | 0.1556 |
| Tbx2 | 45 | 29 | 29 | 47 | 28 | 29 | 35 | 31 | 31 | 47 | 32 | 29 | 34.01 | 32 | 36 | 1.1 | 0.138 | 0.68673296 | 0.80798786 | 0 | 0.0721 | 0.0552 | 0.0552 | TRUE | 0.5448 |
| Tbx20 | 97 | 44 | 56 | 52 | 53 | 42 | 75 | 47 | 60 | 52 | 60 | 41 | 55.93 | 61 | 51 | 0.83 | -0.268 | 0.34484207 | 0.51910603 | 0.0283 | 0.0448 | 0.0401 | 0.0401 | TRUE | 0.1918 |
| Tbx21 | 93 | 67 | 68 | 73 | 56 | 74 | 72 | 71 | 73 | 73 | 63 | 73 | 70.96 | 72 | 70 | 0.965 | -0.052 | 0.82020448 | 0.89651557 | 0 | 0.0358 | 0.0233 | 0.0233 | TRUE | 0.0553 |
| Tbx22 | 3 | 0 | 0 | 0 | 0 | 0 | 2 | 0 | 0 | 0 | 0 | 0 | 0.39 | 1 | 0 | 0.239 | -2.067 | 0.57190649 | NA | 10 | 6.1057 | 8.0283 | 8.0283 | TRUE | 1.4408 |
| Tbx3 | 7560 | 5735 | 4519 | 3489 | 3259 | 3657 | 5872 | 6100 | 4876 | 3473 | 3676 | 3610 | 4601.2 | 5616 | 3586 | 0.638 | -0.647 | 1.38E-15 | 2.55E-13 | 0.0068 | 0.0029 | 0.0045 | 0.0045 | TRUE | 0.1611 |
| Tbx4 | 152 | 93 | 114 | 198 | 124 | 135 | 118 | 99 | 123 | 197 | 140 | 133 | 135.04 | 113 | 157 | 1.383 | 0.468 | 0.01901895 | 0.06664327 | 0.0238 | 0.02 | 0.0211 | 0.0211 | TRUE | 0.5409 |
| Tbx5 | 20 | 14 | 20 | 29 | 8 | 12 | 16 | 15 | 22 | 29 | 9 | 12 | 16.96 | 18 | 17 | 0.965 | -0.052 | 0.92260269 | NA | 0.1479 | 0.1422 | 0.1438 | 0.1438 | TRUE | 2.061 |
| Tbx6 | 36 | 34 | 51 | 55 | 45 | 52 | 28 | 36 | 55 | 55 | 51 | 51 | 46 | 40 | 52 | 1.331 | 0.413 | 0.18905393 | 0.34660695 | 0.0355 | 0.054 | 0.0489 | 0.0489 | TRUE | 0.7899 |
| Tbxa2r | 44 | 34 | 43 | 59 | 44 | 60 | 34 | 36 | 46 | 59 | 50 | 59 | 47.39 | 39 | 56 | 1.446 | 0.532 | 0.06310607 | 0.15913463 | 0 | 0.0524 | 0.0373 | 0.0373 | TRUE | 0.1994 |
| Tbxas1 | 0 | 0 | 0 | 1 | 1 | 1 | 0 | 0 | 0 | 1 | 1 | 1 | 0.52 | 0 | 1 | 5.852 | 2.549 | 0.3348364 | NA | 0 | 4.5739 | 3.5345 | 3.5345 | TRUE | 0.0847 |
| Tc2n | 96 | 63 | 64 | 74 | 92 | 70 | 75 | 67 | 69 | 74 | 104 | 69 | 76.2 | 70 | 82 | 1.164 | 0.219 | 0.3598824 | 0.53453176 | 0.0141 | 0.0335 | 0.028 | 0.028 | TRUE | 0.4723 |
| Tcaim | 988 | 760 | 738 | 942 | 833 | 923 | 767 | 808 | 796 | 938 | 940 | 911 | 860.09 | 790 | 930 | 1.176 | 0.234 | 0.00257663 | 0.01470171 | 0 | 0.0052 | 0.0032 | 0.0032 | TRUE | 0.0085 |
| Tcam1 | 86 | 56 | 50 | 63 | 42 | 52 | 67 | 60 | 54 | 63 | 47 | 51 | 56.96 | 60 | 54 | 0.893 | -0.163 | 0.53414375 | 0.69329064 | 0 | 0.044 | 0.0318 | 0.0318 | TRUE | 0.1762 |
| Tcap | 1 | 0 | 0 | 2 | 0 | 0 | 1 | 0 | 0 | 2 | 0 | 0 | 0.46 | 0 | 1 | 1.942 | 0.958 | 0.7580687 | NA | 0 | 5.1409 | 5.1921 | 5.1921 | TRUE | 0.9738 |
| Tcea1 | 8292 | 6012 | 6184 | 5886 | 5249 | 6085 | 6440 | 6395 | 6672 | 5859 | 5921 | 6007 | 6215.76 | 6502 | 5929 | 0.912 | -0.133 | 0.00735763 | 0.03301836 | 2.00E-04 | 0.0028 | 0.0016 | 0.0016 | TRUE | 0.0063 |
| Tcea2 | 344 | 280 | 286 | 337 | 273 | 328 | 267 | 298 | 309 | 335 | 308 | 324 | 306.8 | 291 | 322 | 1.111 | 0.152 | 0.21008402 | 0.37203951 | 0 | 0.0102 | 0.0073 | 0.0073 | TRUE | 0.0641 |
| Tcea3 | 3300 | 2755 | 3245 | 3489 | 3397 | 4063 | 2563 | 2930 | 3501 | 3473 | 3832 | 4011 | 3385.11 | 2998 | 3772 | 1.259 | 0.332 | 0.00086294 | 0.00650527 | 0.0146 | 0.0031 | 0.0069 | 0.0069 | TRUE | 0.2615 |
| Tceal1 | 110 | 71 | 79 | 62 | 48 | 61 | 85 | 76 | 85 | 62 | 54 | 60 | 70.38 | 82 | 59 | 0.715 | -0.483 | 0.03657884 | 0.10748795 | 0 | 0.0361 | 0.0239 | 0.0239 | TRUE | 0.0435 |
| Tceal3 | 91 | 72 | 61 | 71 | 66 | 73 | 71 | 77 | 66 | 71 | 74 | 72 | 71.71 | 71 | 72 | 1.019 | 0.027 | 0.90515573 | 0.94807203 | 0 | 0.0355 | 0.023 | 0.023 | TRUE | 0.0389 |
| Tceal5 | 68 | 38 | 35 | 39 | 44 | 32 | 53 | 40 | 38 | 39 | 50 | 32 | 41.84 | 44 | 40 | 0.907 | -0.141 | 0.65434534 | 0.78533529 | 0.0189 | 0.0591 | 0.0479 | 0.0479 | TRUE | 0.3103 |

| | | | | | | | | | | | | | | | | | | | | | | | | | |
|---|---|---|---|---|---|---|---|---|---|---|---|---|---|---|---|---|---|---|---|---|---|---|---|---|---|
| Tceal6 | 11 | 11 | 6 | 13 | 4 | 10 | 9 | 12 | 6 | 13 | 5 | 10 | 9.01 | 9 | 9 | 1.031 | 0.045 | 0.94608909 | NA | 0 | 0.2656 | 0.2047 | 0.2047 | TRUE | 0.3662 |
| Tceal7 | 47 | 21 | 29 | 43 | 30 | 34 | 37 | 22 | 31 | 43 | 34 | 34 | 33.39 | 30 | 37 | 1.214 | 0.28 | 0.41722097 | 0.58978729 | 0 | 0.0734 | 0.0559 | 0.0559 | TRUE | 0.3199 |
| Tceal8 | 2271 | 1528 | 1366 | 1411 | 1136 | 1290 | 1764 | 1625 | 1474 | 1405 | 1282 | 1273 | 1470.41 | 1621 | 1320 | 0.813 | -0.298 | 0.00037934 | 0.0034966 | 0.0045 | 0.004 | 0.0044 | 0.0044 | TRUE | 0.0756 |
| Tceanc | 627 | 462 | 609 | 715 | 545 | 660 | 487 | 491 | 657 | 712 | 615 | 652 | 602.26 | 545 | 660 | 1.212 | 0.278 | 0.02416621 | 0.07940406 | 0.0161 | 0.0064 | 0.0093 | 0.0093 | TRUE | 0.3715 |
| Tceanc2 | 327 | 236 | 256 | 317 | 271 | 301 | 254 | 251 | 276 | 316 | 306 | 297 | 283.27 | 260 | 306 | 1.177 | 0.235 | 0.05333327 | 0.14101745 | 0 | 0.0108 | 0.0071 | 0.0071 | TRUE | 0.03 |
| Tceb1 | 4129 | 2699 | 2517 | 2634 | 2166 | 2468 | 3207 | 2871 | 2716 | 2622 | 2443 | 2436 | 2715.86 | 2931 | 2500 | 0.852 | -0.23 | 0.00202217 | 0.01219531 | 0.004 | 0.0033 | 0.0036 | 0.0036 | TRUE | 0.0836 |
| Tceb2 | 7774 | 5821 | 5850 | 6085 | 5497 | 6419 | 6038 | 6192 | 6312 | 6057 | 6201 | 6337 | 6189.38 | 6181 | 6198 | 1.003 | 0.004 | 0.93061216 | 0.96190625 | 4.00E-04 | 0.0028 | 0.0017 | 0.0017 | TRUE | 0.0051 |
| Tceb3 | 2875 | 2053 | 1898 | 2234 | 1925 | 2132 | 2233 | 2184 | 2048 | 2224 | 2172 | 2105 | 2160.75 | 2155 | 2167 | 1.005 | 0.007 | 0.91348811 | 0.95246365 | 9.00E-04 | 0.0035 | 0.0025 | 0.0025 | TRUE | 0.0225 |
| Tcerg1 | 11741 | 8752 | 8479 | 8524 | 7616 | 9339 | 9119 | 9309 | 9149 | 8485 | 8592 | 9219 | 8978.72 | 9192 | 8765 | 0.954 | -0.069 | 0.19703025 | 0.35676159 | 9.00E-04 | 0.0027 | 0.0019 | 0.0019 | TRUE | 0.0251 |
| Tcerg1l | 120 | 79 | 55 | 54 | 59 | 70 | 93 | 84 | 59 | 54 | 67 | 69 | 71 | 79 | 63 | 0.796 | -0.33 | 0.19350436 | 0.3522542 | 0.0224 | 0.0358 | 0.032 | 0.032 | TRUE | 0.416 |
| Tcf12 | 2130 | 1571 | 1523 | 1698 | 1451 | 1743 | 1654 | 1671 | 1643 | 1690 | 1637 | 1721 | 1669.39 | 1656 | 1683 | 1.016 | 0.023 | 0.71388979 | 0.82814923 | 0 | 0.0038 | 0.0023 | 0.0023 | TRUE | 0.0067 |
| Tcf15 | 2637 | 1963 | 1482 | 2565 | 1959 | 2214 | 2048 | 2088 | 1599 | 2553 | 2210 | 2186 | 2113.97 | 1912 | 2316 | 1.211 | 0.276 | 0.00632119 | 0.02933403 | 0.0133 | 0.0035 | 0.0069 | 0.0069 | TRUE | 0.2449 |
| Tcf19 | 943 | 666 | 663 | 653 | 610 | 736 | 732 | 708 | 715 | 650 | 688 | 727 | 703.48 | 718 | 688 | 0.957 | -0.063 | 0.46488502 | 0.6339059 | 0 | 0.0058 | 0.004 | 0.004 | TRUE | 0.0288 |
| Tcf20 | 13634 | 9852 | 9653 | 10455 | 9227 | 10639 | 10589 | 10479 | 10416 | 10407 | 10409 | 10502 | 10467.02 | 10495 | 10439 | 0.995 | -0.008 | 0.86358017 | 0.92466653 | 0 | 0.0027 | 0.0013 | 0.0013 | TRUE | 8.00E-04 |
| Tcf21 | 1 | 2 | 0 | 2 | 0 | 0 | 1 | 2 | 0 | 2 | 0 | 0 | 0.82 | 1 | 1 | 0.698 | -0.518 | 0.82391604 | NA | 0 | 2.9076 | 2.6343 | 2.6343 | TRUE | 0.7035 |
| Tcf23 | 17 | 7 | 8 | 12 | 7 | 6 | 13 | 7 | 9 | 12 | 8 | 6 | 9.17 | 10 | 9 | 0.87 | -0.201 | 0.75502088 | NA | 0 | 0.2608 | 0.1901 | 0.1901 | TRUE | 0.4114 |
| Tcf25 | 20372 | 15072 | 14906 | 15635 | 14038 | 16215 | 15822 | 16032 | 16083 | 15563 | 15836 | 16007 | 15890.58 | 15979 | 15802 | 0.989 | -0.016 | 0.7171837 | 0.82994953 | 1.00E-04 | 0.0026 | 0.0013 | 0.0013 | TRUE | 0.0021 |
| Tcf3 | 7162 | 5608 | 5736 | 5521 | 5231 | 6139 | 5562 | 5965 | 6189 | 5496 | 5901 | 6060 | 5862.25 | 5905 | 5819 | 0.986 | -0.021 | 0.74215884 | 0.84658231 | 0.0026 | 0.0028 | 0.0027 | 0.0027 | TRUE | 0.0319 |
| Tcf4 | 3540 | 2820 | 2547 | 2507 | 2388 | 2703 | 2749 | 3000 | 2748 | 2495 | 2694 | 2668 | 2725.8 | 2832 | 2619 | 0.925 | -0.113 | 0.08424829 | 0.19572816 | 0.0016 | 0.0033 | 0.0027 | 0.0027 | TRUE | 0.032 |
| Tcf7 | 5002 | 3810 | 4469 | 4754 | 4511 | 5472 | 3885 | 4053 | 4822 | 4732 | 5089 | 5402 | 4663.7 | 4253 | 5074 | 1.194 | 0.255 | 0.00252029 | 0.01445935 | 0.0088 | 0.0029 | 0.0049 | 0.0049 | TRUE | 0.1662 |
| Tcf7l1 | 3969 | 2887 | 2559 | 1712 | 1492 | 1988 | 3083 | 3071 | 2761 | 1704 | 1683 | 1962 | 2377.38 | 2972 | 1783 | 0.6 | -0.737 | 8.83E-21 | 3.01E-18 | 0.005 | 0.0034 | 0.004 | 0.004 | TRUE | 0.0938 |
| Tcf7l2 | 291 | 175 | 181 | 154 | 132 | 156 | 226 | 186 | 195 | 153 | 149 | 154 | 177.27 | 202 | 152 | 0.749 | -0.418 | 0.00696561 | 0.03161619 | 0 | 0.0158 | 0.0115 | 0.0115 | TRUE | 0.1218 |
| Tcfl5 | 4493 | 3136 | 2693 | 2925 | 2514 | 2851 | 3490 | 3336 | 2906 | 2912 | 2836 | 2814 | 3048.82 | 3244 | 2854 | 0.879 | -0.185 | 0.01236981 | 0.04845209 | 0.004 | 0.0032 | 0.0036 | 0.0036 | TRUE | 0.1003 |
| Tchh | 143 | 101 | 46 | 73 | 63 | 78 | 111 | 107 | 50 | 73 | 71 | 77 | 81.48 | 89 | 74 | 0.819 | -0.288 | 0.30759995 | 0.48024087 | 0.0789 | 0.0315 | 0.0448 | 0.0448 | TRUE | 1.3831 |
| Tchhl1 | 1 | 0 | 0 | 0 | 0 | 0 | 1 | 0 | 0 | 0 | 0 | 0 | 0.13 | 0 | 0 | 0.552 | -0.857 | 0.8336548 | NA | 0 | 18.3121 | 10 | 10 | TRUE | 0.2566 |
| Tchp | 702 | 563 | 518 | 502 | 423 | 502 | 545 | 599 | 559 | 500 | 477 | 496 | 529.24 | 568 | 491 | 0.866 | -0.208 | 0.02876594 | 0.08983064 | 0 | 0.0069 | 0.0046 | 0.0046 | TRUE | 0.0267 |
| Tcirg1 | 672 | 549 | 524 | 561 | 501 | 531 | 522 | 584 | 565 | 558 | 565 | 524 | 553.17 | 557 | 549 | 0.987 | -0.018 | 0.8470872 | 0.91321597 | 0 | 0.0067 | 0.0048 | 0.0048 | TRUE | 0.0383 |
| Tcl1 | 3080 | 2523 | 2995 | 3482 | 3354 | 3717 | 2392 | 2684 | 3232 | 3466 | 3784 | 3669 | 3204.38 | 2769 | 3640 | 1.315 | 0.395 | 3.09E-05 | 0.00047627 | 0.0123 | 0.0032 | 0.0062 | 0.0062 | TRUE | 0.2587 |
| Tcl1b1 | 0 | 0 | 0 | 0 | 0 | 0 | 0 | 0 | 0 | 0 | 0 | 0 | 0 | 0 | 0 | NA | NA | NA | NA | NA | NA | NA | NA | NA | NA |
| Tcl1b2 | 1 | 1 | 0 | 0 | 0 | 0 | 1 | 1 | 0 | 0 | 0 | 0 | 0.31 | 1 | 0 | 0.309 | -1.696 | 0.63183309 | NA | 0 | 7.7294 | 7.2472 | 7.2472 | TRUE | 0.2015 |
| Tcl1b3 | 0 | 0 | 0 | 0 | 0 | 0 | 0 | 0 | 0 | 0 | 0 | 0 | 0 | 0 | 0 | NA | NA | NA | NA | NA | NA | NA | NA | NA | NA |
| Tcl1b4 | 0 | 0 | 0 | 0 | 0 | 0 | 0 | 0 | 0 | 0 | 0 | 0 | 0 | 0 | 0 | NA | NA | NA | NA | NA | NA | NA | NA | NA | NA |
| Tcl1b5 | 0 | 0 | 0 | 0 | 0 | 0 | 0 | 0 | 0 | 0 | 0 | 0 | 0 | 0 | 0 | NA | NA | NA | NA | NA | NA | NA | NA | NA | NA |
| Tcn2 | 1402 | 1117 | 1231 | 1341 | 1264 | 1429 | 1089 | 1188 | 1328 | 1335 | 1426 | 1411 | 1296.11 | 1202 | 1391 | 1.159 | 0.212 | 0.0126726 | 0.04924675 | 0.0045 | 0.0043 | 0.0045 | 0.0045 | TRUE | 0.1014 |
| Tcof1 | 16613 | 12478 | 10943 | 10960 | 10152 | 12240 | 12903 | 13273 | 11807 | 10910 | 11453 | 12083 | 12071.26 | 12661 | 11482 | 0.907 | -0.141 | 0.02567781 | 0.08285449 | 0.003 | 0.0026 | 0.0028 | 0.0028 | TRUE | 0.0423 |
| Tcp1 | 48006 | 34067 | 34063 | 36571 | 31159 | 35897 | 37284 | 36237 | 36754 | 36403 | 35151 | 35436 | 36210.62 | 36758 | 35663 | 0.97 | -0.044 | 0.32208252 | 0.49587577 | 2.00E-04 | 0.0025 | 0.0014 | 0.0014 | TRUE | 0.004 |
| Tcp10a | 0 | 0 | 1 | 0 | 0 | 0 | 0 | 0 | 1 | 0 | 0 | 0 | 0.18 | 0 | 0 | 0.552 | -0.857 | 0.8336548 | NA | 0 | 13.1817 | 10 | 10 | TRUE | 0.5365 |
| Tcp10b | 2 | 1 | 3 | 3 | 4 | 3 | 2 | 1 | 3 | 3 | 5 | 3 | 2.72 | 2 | 4 | 1.8 | 0.848 | 0.46336511 | NA | 0 | 0.8741 | 0.5641 | 0.5641 | TRUE | 0.2583 |
| Tcp10c | 1 | 1 | 0 | 0 | 0 | 1 | 1 | 1 | 0 | 0 | 0 | 1 | 0.47 | 1 | 0 | 0.599 | -0.74 | 0.79209856 | NA | 0 | 5.0317 | 3.8925 | 3.8925 | TRUE | 0.3835 |
| Tcp11 | 848 | 706 | 741 | 754 | 757 | 832 | 659 | 751 | 800 | 751 | 854 | 821 | 772.49 | 737 | 809 | 1.1 | 0.137 | 0.16026478 | 0.30963294 | 0.0058 | 0.0055 | 0.0056 | 0.0056 | TRUE | 0.1062 |
| Tcp11l1 | 263 | 198 | 197 | 245 | 177 | 229 | 204 | 211 | 213 | 244 | 200 | 226 | 216.17 | 209 | 223 | 1.07 | 0.097 | 0.49043377 | 0.65622261 | 0 | 0.0134 | 0.0097 | 0.0097 | TRUE | 0.0896 |
| Tcp11l2 | 475 | 347 | 382 | 418 | 347 | 447 | 369 | 369 | 412 | 416 | 391 | 441 | 399.83 | 383 | 416 | 1.088 | 0.122 | 0.26812756 | 0.43873465 | 0.0011 | 0.0084 | 0.0062 | 0.0062 | TRUE | 0.0479 |
| Tcstv1 | 61 | 35 | 24 | 17 | 26 | 14 | 47 | 37 | 26 | 17 | 29 | 14 | 28.43 | 37 | 20 | 0.536 | -0.901 | 0.02926258 | 0.09096348 | 0.0791 | 0.0858 | 0.084 | 0.084 | TRUE | 0.2827 |
| Tcstv3 | 138 | 105 | 43 | 63 | 68 | 40 | 107 | 112 | 46 | 63 | 77 | 39 | 74.03 | 88 | 60 | 0.67 | -0.577 | 0.08122346 | 0.19119646 | 0.1436 | 0.0344 | 0.0648 | 0.0648 | TRUE | 1.1541 |
| Tcta | 292 | 222 | 208 | 235 | 204 | 228 | 227 | 236 | 224 | 234 | 230 | 225 | 229.41 | 229 | 230 | 1.003 | 0.004 | 0.97494871 | 0.98647473 | 0 | 0.0128 | 0.008 | 0.008 | TRUE | 0.0075 |
| Tcte1 | 17 | 27 | 10 | 23 | 20 | 26 | 13 | 29 | 11 | 23 | 23 | 26 | 20.64 | 18 | 24 | 1.361 | 0.445 | 0.33990471 | 0.51393822 | 0.0781 | 0.1173 | 0.107 | 0.107 | TRUE | 1.4566 |
| Tcte2 | 26 | 22 | 13 | 16 | 16 | 10 | 20 | 23 | 14 | 16 | 18 | 10 | 16.91 | 19 | 15 | 0.756 | -0.403 | 0.40436768 | NA | 0 | 0.1426 | 0.1079 | 0.1079 | TRUE | 0.3737 |
| Tcte3 | 0 | 0 | 0 | 0 | 0 | 0 | 0 | 0 | 0 | 0 | 0 | 0 | 0 | 0 | 0 | NA | NA | NA | NA | NA | NA | NA | NA | NA | NA |
| Tctex1d1 | 0 | 2 | 1 | 0 | 2 | 0 | 0 | 2 | 1 | 0 | 2 | 0 | 0.91 | 1 | 1 | 0.701 | -0.513 | 0.82081793 | NA | 0 | 2.6057 | 2.5168 | 2.5168 | TRUE | 0.4908 |
| Tctex1d2 | 763 | 579 | 666 | 857 | 768 | 784 | 593 | 616 | 719 | 853 | 866 | 774 | 736.74 | 643 | 831 | 1.296 | 0.374 | 0.00015274 | 0.00172865 | 0.0052 | 0.0056 | 0.0056 | 0.0056 | TRUE | 0.125 |
| Tctex1d4 | 5 | 2 | 4 | 1 | 0 | 3 | 4 | 2 | 4 | 1 | 0 | 3 | 2.38 | 3 | 1 | 0.391 | -1.356 | 0.30683218 | NA | 0 | 0.998 | 0.7452 | 0.7452 | TRUE | 0.728 |
| Tctn1 | 544 | 427 | 454 | 442 | 386 | 393 | 423 | 454 | 490 | 440 | 435 | 388 | 438.32 | 456 | 421 | 0.926 | -0.111 | 0.30927189 | 0.48202942 | 0.0031 | 0.0078 | 0.0063 | 0.0063 | TRUE | 0.0558 |
| Tctn2 | 993 | 701 | 702 | 764 | 655 | 832 | 771 | 746 | 757 | 760 | 739 | 821 | 765.84 | 758 | 773 | 1.02 | 0.029 | 0.72760236 | 0.83665491 | 0 | 0.0055 | 0.0038 | 0.0038 | TRUE | 0.0348 |
| Tctn3 | 706 | 526 | 464 | 557 | 498 | 518 | 548 | 559 | 501 | 554 | 562 | 511 | 539.34 | 536 | 542 | 1.011 | 0.015 | 0.87471395 | 0.93043734 | 9.00E-04 | 0.0068 | 0.005 | 0.005 | TRUE | 0.0376 |
| Tdg | 41 | 37 | 24 | 25 | 32 | 27 | 32 | 39 | 26 | 25 | 36 | 27 | 30.79 | 32 | 29 | 0.899 | -0.153 | 0.66967396 | 0.79697335 | 0 | 0.0794 | 0.06 | 0.06 | TRUE | 0.2466 |
| Tdgf1 | 80521 | 59249 | 66219 | 74528 | 65715 | 73415 | 62537 | 63022 | 71450 | 74185 | 74133 | 72472 | 69633.29 | 65670 | 73597 | 1.121 | 0.164 | 0.00680197 | 0.03110062 | 0.003 | 0.0025 | 0.0026 | 0.0026 | TRUE | 0.0726 |
| Tdh | 10969 | 9322 | 10584 | 11084 | 11690 | 14025 | 8519 | 9916 | 11420 | 11033 | 13188 | 13845 | 11320.04 | 9952 | 12689 | 1.275 | 0.351 | 0.00055535 | 0.00466916 | 0.0172 | 0.0026 | 0.0073 | 0.0073 | TRUE | 0.2036 |
| Tdo2 | 0 | 1 | 0 | 0 | 0 | 0 | 0 | 1 | 0 | 0 | 0 | 0 | 0.18 | 0 | 0 | 0.552 | -0.857 | 0.8336548 | NA | 0 | 13.3714 | 10 | 10 | TRUE | 0.5216 |
| Tdp1 | 3114 | 2202 | 2222 | 2236 | 2156 | 2513 | 2419 | 2342 | 2398 | 2226 | 2432 | 2481 | 2382.82 | 2386 | 2380 | 0.997 | -0.004 | 0.946055 | 0.97029019 | 0.0014 | 0.0034 | 0.0026 | 0.0026 | TRUE | 0.0391 |
| Tdp2 | 4157 | 3018 | 3010 | 3104 | 2621 | 3197 | 3229 | 3210 | 3248 | 3090 | 2957 | 3156 | 3148.16 | 3229 | 3068 | 0.95 | -0.074 | 0.18936816 | 0.3470495 | 2.00E-04 | 0.0032 | 0.0019 | 0.0019 | TRUE | 0.0119 |
| Tdpoz1 | 3 | 7 | 2 | 1 | 0 | 0 | 2 | 7 | 2 | 1 | 0 | 0 | 2.15 | 4 | 0 | 0.094 | -3.411 | 0.04201177 | NA | 0 | 1.1023 | 0.9053 | 0.9053 | TRUE | 0.9159 |
| Tdpoz2 | 4 | 3 | 1 | 0 | 0 | 0 | 3 | 3 | 1 | 0 | 0 | 0 | 1.23 | 2 | 0 | 0.077 | -3.699 | 0.06124435 | NA | 0 | 1.9302 | 1.6193 | 1.6193 | TRUE | 0.2395 |
| Tdpoz3 | 108 | 86 | 22 | 18 | 9 | 23 | 84 | 91 | 24 | 18 | 10 | 23 | 41.64 | 66 | 17 | 0.256 | -1.967 | 2.46E-05 | 0.00039363 | 0.2746 | 0.0593 | 0.119 | 0.119 | TRUE | 2.5221 |
| Tdpoz4 | 51 | 29 | 14 | 7 | 5 | 4 | 40 | 31 | 15 | 7 | 6 | 4 | 17.02 | 29 | 6 | 0.192 | -2.383 | 4.55E-05 | NA | 0 | 0.1417 | 0.1354 | 0.1354 | TRUE | 0.1556 |
| Tdpoz5 | 0 | 0 | 0 | 0 | 0 | 0 | 0 | 0 | 0 | 0 | 0 | 0 | 0 | 0 | 0 | NA | NA | NA | NA | NA | NA | NA | NA | NA | NA |
| Tdrd1 | 305 | 280 | 260 | 268 | 266 | 303 | 237 | 298 | 281 | 267 | 300 | 299 | 280.2 | 272 | 289 | 1.066 | 0.092 | 0.49445303 | 0.65958273 | 0.006 | 0.0109 | 0.0094 | 0.0094 | TRUE | 0.1535 |
| Tdrd12 | 1282 | 1054 | 1124 | 596 | 502 | 643 | 996 | 1121 | 1213 | 593 | 566 | 635 | 853.98 | 1110 | 598 | 0.54 | -0.889 | 3.01E-20 | 9.77E-18 | 0.0058 | 0.0052 | 0.0054 | 0.0054 | TRUE | 0.1006 |
| Tdrd3 | 1041 | 759 | 818 | 779 | 644 | 780 | 809 | 807 | 883 | 775 | 726 | 770 | 795.06 | 833 | 757 | 0.91 | -0.135 | 0.10674129 | 0.23203852 | 7.00E-04 | 0.0054 | 0.0038 | 0.0038 | TRUE | 0.0317 |
| Tdrd5 | 972 | 748 | 744 | 829 | 740 | 966 | 755 | 796 | 803 | 825 | 835 | 954 | 827.82 | 785 | 871 | 1.112 | 0.153 | 0.08317957 | 0.19416716 | 0.0028 | 0.0053 | 0.0044 | 0.0044 | TRUE | 0.0824 |

| | | | | | | | | | | | | | | | | | | | | | | | | |
|---|---|---|---|---|---|---|---|---|---|---|---|---|---|---|---|---|---|---|---|---|---|---|---|---|
| Tdrd6 | 32 | 19 | 13 | 29 | 12 | 8 | 25 | 20 | 14 | 29 | 14 | 8 | 18.23 | 20 | 17 | 0.847 | -0.24 | 0.6505393 | 0.78306874 | 0.1819 | 0.1324 | 0.1463 | 0.1463 | TRUE | 0.9919 |
| Tdrd7 | 624 | 440 | 467 | 520 | 459 | 538 | 485 | 468 | 504 | 518 | 518 | 531 | 503.84 | 486 | 522 | 1.076 | 0.105 | 0.26664459 | 0.43705861 | 0 | 0.0071 | 0.0045 | 0.0045 | TRUE | 0.0119 |
| Tdrd9 | 116 | 54 | 64 | 110 | 88 | 101 | 90 | 57 | 69 | 109 | 99 | 100 | 87.51 | 72 | 103 | 1.412 | 0.498 | 0.02812062 | 0.08833583 | 0.0146 | 0.0295 | 0.0254 | 0.0254 | TRUE | 0.4797 |
| Tdrkh | 771 | 529 | 498 | 792 | 582 | 624 | 599 | 563 | 537 | 788 | 657 | 616 | 626.62 | 566 | 687 | 1.212 | 0.277 | 0.01094915 | 0.04413998 | 0.0085 | 0.0062 | 0.007 | 0.007 | TRUE | 0.199 |
| Tead1 | 14218 | 10059 | 9850 | 8204 | 7827 | 9257 | 11043 | 10700 | 10628 | 8166 | 8830 | 9138 | 9750.7 | 10790 | 8711 | 0.807 | -0.309 | 7.50E-08 | 3.02E-06 | 0.0017 | 0.0027 | 0.0023 | 0.0023 | TRUE | 0.0367 |
| Tead2 | 6574 | 4581 | 4912 | 5277 | 4866 | 5767 | 5106 | 4873 | 5300 | 5253 | 5489 | 5693 | 5285.58 | 5093 | 5478 | 1.076 | 0.105 | 0.07317806 | 0.17706687 | 0.0014 | 0.0029 | 0.0023 | 0.0023 | TRUE | 0.0172 |
| Tead3 | 229 | 233 | 219 | 260 | 219 | 251 | 178 | 248 | 236 | 259 | 247 | 248 | 235.94 | 221 | 251 | 1.145 | 0.195 | 0.19898041 | 0.3594574 | 0.012 | 0.0125 | 0.0123 | 0.0123 | TRUE | 0.3457 |
| Tead4 | 1236 | 994 | 948 | 1288 | 1137 | 1172 | 960 | 1057 | 1023 | 1282 | 1283 | 1157 | 1126.97 | 1013 | 1241 | 1.225 | 0.293 | 0.00023602 | 0.00242655 | 0.0021 | 0.0045 | 0.0037 | 0.0037 | TRUE | 0.0421 |
| Tec | 204 | 169 | 182 | 202 | 189 | 195 | 158 | 180 | 196 | 201 | 213 | 192 | 190.23 | 178 | 202 | 1.139 | 0.187 | 0.21515212 | 0.37765193 | 0 | 0.0149 | 0.0112 | 0.0112 | TRUE | 0.1117 |
| Tecpr1 | 1128 | 920 | 958 | 1046 | 945 | 1105 | 876 | 979 | 1034 | 1041 | 1066 | 1091 | 1014.4 | 963 | 1066 | 1.109 | 0.15 | 0.07534536 | 0.18084152 | 0.0029 | 0.0048 | 0.0041 | 0.0041 | TRUE | 0.0778 |
| Tecpr2 | 3079 | 2059 | 1975 | 2184 | 1843 | 2158 | 2391 | 2190 | 2131 | 2174 | 2079 | 2130 | 2182.63 | 2237 | 2128 | 0.95 | -0.074 | 0.27413103 | 0.44519428 | 0.0017 | 0.0035 | 0.0028 | 0.0028 | TRUE | 0.045 |
| Tecr | 14477 | 10713 | 11594 | 13289 | 12208 | 14059 | 11244 | 11395 | 12510 | 13228 | 13772 | 13878 | 12671.15 | 11716 | 13626 | 1.163 | 0.218 | 0.00016301 | 0.00182165 | 0.002 | 0.0026 | 0.0023 | 0.0023 | TRUE | 0.0428 |
| Tecrl | 0 | 0 | 0 | 0 | 0 | 0 | 0 | 0 | 0 | 0 | 0 | 0 | 0 | 0 | 0 | NA | NA | NA | NA | NA | NA | NA | NA | NA | NA |
| Tecta | 8 | 7 | 4 | 2 | 2 | 3 | 6 | 7 | 4 | 2 | 2 | 3 | 4.2 | 6 | 2 | 0.401 | -1.318 | 0.16883248 | NA | 0 | 0.5671 | 0.3658 | 0.3658 | TRUE | 0.1256 |
| Tectb | 0 | 0 | 0 | 0 | 0 | 0 | 0 | 0 | 0 | 0 | 0 | 0 | 0 | 0 | 0 | NA | NA | NA | NA | NA | NA | NA | NA | NA | NA |
| Teddm1 | 0 | 0 | 0 | 0 | 0 | 0 | 0 | 0 | 0 | 0 | 0 | 0 | 0 | 0 | 0 | NA | NA | NA | NA | NA | NA | NA | NA | NA | NA |
| Tef | 1490 | 1135 | 1387 | 1532 | 1412 | 1486 | 1157 | 1207 | 1497 | 1525 | 1593 | 1467 | 1407.64 | 1287 | 1528 | 1.189 | 0.249 | 0.00973834 | 0.04060386 | 0.0099 | 0.0041 | 0.006 | 0.006 | TRUE | 0.2416 |
| Tefm | 1730 | 1165 | 1124 | 1353 | 1212 | 1422 | 1344 | 1239 | 1213 | 1347 | 1367 | 1404 | 1318.89 | 1265 | 1373 | 1.084 | 0.116 | 0.1114342 | 0.2393036 | 0.0011 | 0.0042 | 0.003 | 0.003 | TRUE | 0.0368 |
| Tek | 623 | 424 | 375 | 397 | 368 | 382 | 484 | 451 | 405 | 395 | 415 | 377 | 421.15 | 447 | 396 | 0.884 | -0.178 | 0.10858263 | 0.2349692 | 0.0031 | 0.0081 | 0.0065 | 0.0065 | TRUE | 0.0767 |
| Tekt1 | 608 | 477 | 471 | 458 | 426 | 501 | 472 | 507 | 508 | 456 | 481 | 495 | 486.47 | 496 | 477 | 0.963 | -0.054 | 0.58207898 | 0.73253195 | 0 | 0.0073 | 0.0049 | 0.0049 | TRUE | 0.0219 |
| Tekt2 | 54 | 28 | 45 | 46 | 32 | 51 | 42 | 30 | 49 | 46 | 36 | 50 | 42.08 | 40 | 44 | 1.101 | 0.139 | 0.65883904 | 0.78887378 | 0.0147 | 0.0587 | 0.0474 | 0.0474 | TRUE | 0.3515 |
| Tekt3 | 3 | 0 | 0 | 1 | 2 | 0 | 2 | 0 | 0 | 1 | 2 | 0 | 0.93 | 1 | 1 | 1.289 | 0.366 | 0.86815116 | NA | 1.8421 | 2.5502 | 2.4435 | 2.4435 | TRUE | 0.6393 |
| Tekt4 | 1 | 0 | 0 | 1 | 0 | 1 | 1 | 0 | 0 | 1 | 0 | 1 | 0.46 | 0 | 1 | 1.94 | 0.956 | 0.73503911 | NA | 0 | 5.1562 | 3.9771 | 3.9771 | TRUE | 0.2566 |
| Tekt5 | 10 | 13 | 7 | 5 | 18 | 7 | 8 | 14 | 8 | 5 | 20 | 7 | 10.22 | 10 | 11 | 1.098 | 0.135 | 0.84243554 | NA | 0.2378 | 0.2343 | 0.2353 | 0.2353 | TRUE | 2.0337 |
| Telo2 | 1692 | 1301 | 1075 | 1143 | 1049 | 1182 | 1314 | 1384 | 1160 | 1138 | 1183 | 1167 | 1224.3 | 1286 | 1163 | 0.904 | -0.146 | 0.07467267 | 0.1798309 | 0.0034 | 0.0044 | 0.004 | 0.004 | TRUE | 0.0859 |
| Ten1 | 717 | 446 | 483 | 528 | 485 | 547 | 557 | 474 | 521 | 526 | 547 | 540 | 527.52 | 517 | 538 | 1.037 | 0.052 | 0.60291417 | 0.74784798 | 0.0017 | 0.0069 | 0.0052 | 0.0052 | TRUE | 0.0613 |
| Tenc1 | 399 | 369 | 413 | 443 | 444 | 572 | 310 | 393 | 446 | 441 | 501 | 565 | 442.42 | 383 | 502 | 1.317 | 0.397 | 0.00496161 | 0.02436988 | 0.022 | 0.0078 | 0.0121 | 0.0121 | TRUE | 0.3364 |
| Tenm1 | 7 | 5 | 3 | 8 | 9 | 2 | 5 | 5 | 3 | 8 | 10 | 2 | 5.68 | 4 | 7 | 1.415 | 0.501 | 0.55559415 | NA | 0 | 0.4197 | 0.3403 | 0.3403 | TRUE | 0.6927 |
| Tenm2 | 5 | 3 | 0 | 3 | 4 | 2 | 4 | 3 | 0 | 3 | 5 | 2 | 2.76 | 2 | 3 | 1.297 | 0.375 | 0.75627887 | NA | 0 | 0.8618 | 0.6869 | 0.6869 | TRUE | 0.7102 |
| Tenm3 | 4621 | 3170 | 2796 | 2885 | 2744 | 3114 | 3589 | 3372 | 3017 | 2872 | 3096 | 3074 | 3169.82 | 3326 | 3014 | 0.905 | -0.143 | 0.05314246 | 0.14082154 | 0.0041 | 0.0032 | 0.0036 | 0.0036 | TRUE | 0.0799 |
| Tenm4 | 808 | 542 | 372 | 419 | 382 | 439 | 628 | 577 | 401 | 417 | 431 | 433 | 481.14 | 535 | 427 | 0.796 | -0.329 | 0.02224325 | 0.07448987 | 0.0252 | 0.0074 | 0.0128 | 0.0128 | TRUE | 0.5501 |
| Tep1 | 2184 | 1595 | 1767 | 2096 | 1880 | 2081 | 1696 | 1697 | 1907 | 2086 | 2121 | 2054 | 1926.81 | 1767 | 2087 | 1.182 | 0.242 | 0.00052559 | 0.00447689 | 0.0017 | 0.0037 | 0.003 | 0.003 | TRUE | 0.0572 |
| Tepp | 48 | 38 | 32 | 44 | 46 | 38 | 37 | 40 | 35 | 44 | 52 | 38 | 40.9 | 37 | 45 | 1.184 | 0.243 | 0.4232888 | 0.59571355 | 0 | 0.0604 | 0.0419 | 0.0419 | TRUE | 0.159 |
| Terc | 58 | 54 | 26 | 27 | 24 | 11 | 45 | 57 | 28 | 27 | 27 | 11 | 32.56 | 43 | 22 | 0.494 | -1.018 | 0.01425432 | 0.05371396 | 0.1299 | 0.0752 | 0.0891 | 0.0891 | TRUE | 0.306 |
| Terf1 | 6443 | 4812 | 5563 | 5931 | 5009 | 5786 | 5004 | 5118 | 6002 | 5904 | 5651 | 5712 | 5565.16 | 5375 | 5756 | 1.071 | 0.099 | 0.17568896 | 0.32921205 | 0.0052 | 0.0029 | 0.0037 | 0.0037 | TRUE | 0.1267 |
| Terf2 | 2718 | 2161 | 1973 | 1917 | 1852 | 2154 | 2111 | 2299 | 2129 | 1908 | 2089 | 2126 | 2110.37 | 2180 | 2041 | 0.937 | -0.094 | 0.17996497 | 0.33483979 | 0.0021 | 0.0035 | 0.0031 | 0.0031 | TRUE | 0.0397 |
| Terf2ip | 2372 | 1622 | 1492 | 1499 | 1274 | 1518 | 1842 | 1725 | 1610 | 1492 | 1437 | 1499 | 1600.87 | 1726 | 1476 | 0.854 | -0.227 | 0.00188012 | 0.01154106 | 0.0021 | 0.0039 | 0.0032 | 0.0032 | TRUE | 0.0434 |
| Tert | 1231 | 987 | 1152 | 1120 | 1028 | 1182 | 956 | 1050 | 1243 | 1115 | 1160 | 1167 | 1115.05 | 1083 | 1147 | 1.061 | 0.085 | 0.37773004 | 0.55231667 | 0.0084 | 0.0046 | 0.0058 | 0.0058 | TRUE | 0.199 |
| Tes | 1601 | 1171 | 1468 | 1490 | 1438 | 1571 | 1243 | 1246 | 1584 | 1483 | 1622 | 1551 | 1454.86 | 1358 | 1552 | 1.144 | 0.194 | 0.0450786 | 0.12484178 | 0.0104 | 0.0041 | 0.0061 | 0.0061 | TRUE | 0.253 |
| Tesc | 432 | 316 | 293 | 458 | 453 | 439 | 336 | 336 | 316 | 456 | 511 | 433 | 398.01 | 329 | 467 | 1.415 | 0.501 | 6.96E-06 | 0.00013936 | 0.002 | 0.0084 | 0.0064 | 0.0064 | TRUE | 0.0779 |
| Tescl | 20 | 15 | 11 | 11 | 11 | 18 | 16 | 16 | 12 | 11 | 12 | 18 | 14.08 | 15 | 14 | 0.947 | -0.078 | 0.87896315 | NA | 0 | 0.1707 | 0.1178 | 0.1178 | TRUE | 0.3 |
| Tesk1 | 583 | 390 | 407 | 342 | 308 | 380 | 453 | 415 | 439 | 340 | 347 | 375 | 394.96 | 436 | 354 | 0.813 | -0.299 | 0.00548018 | 0.02632882 | 0 | 0.0084 | 0.0058 | 0.0058 | TRUE | 0.0308 |
| Tesk2 | 1577 | 1077 | 1077 | 1038 | 955 | 1044 | 1225 | 1146 | 1162 | 1033 | 1077 | 1031 | 1112.27 | 1178 | 1047 | 0.888 | -0.171 | 0.01945823 | 0.06770121 | 0 | 0.0046 | 0.003 | 0.003 | TRUE | 0.0155 |
| Tespa1 | 0 | 0 | 0 | 0 | 0 | 0 | 0 | 0 | 0 | 0 | 0 | 0 | 0 | 0 | 0 | NA | NA | NA | NA | NA | NA | NA | NA | NA | NA |
| Tet1 | 44442 | 30885 | 29903 | 45783 | 38206 | 45277 | 34516 | 32852 | 32265 | 45572 | 43100 | 44695 | 38833.56 | 33211 | 44456 | 1.339 | 0.421 | 5.43E-17 | 1.18E-14 | 0.001 | 0.0025 | 0.0018 | 0.0018 | TRUE | 0.0145 |
| Tet2 | 5424 | 3617 | 3233 | 4171 | 3501 | 4017 | 4213 | 3847 | 3488 | 4152 | 3949 | 3965 | 3935.84 | 3849 | 4022 | 1.044 | 0.062 | 0.39395915 | 0.56756961 | 0.004 | 0.003 | 0.0036 | 0.0036 | TRUE | 0.084 |
| Tet3 | 1175 | 871 | 785 | 839 | 766 | 832 | 913 | 926 | 847 | 835 | 864 | 821 | 867.77 | 895 | 840 | 0.938 | -0.093 | 0.2472049 | 0.41592698 | 0 | 0.0052 | 0.0035 | 0.0035 | TRUE | 0.0256 |
| Tex10 | 9634 | 6853 | 6135 | 6524 | 5531 | 6471 | 7482 | 7289 | 6620 | 6494 | 6240 | 6388 | 6752.13 | 7130 | 6374 | 0.894 | -0.162 | 0.00756232 | 0.03361999 | 0.002 | 0.0028 | 0.0025 | 0.0025 | TRUE | 0.0477 |
| Tex101 | 126 | 79 | 68 | 82 | 80 | 78 | 98 | 84 | 73 | 82 | 90 | 77 | 84.02 | 85 | 83 | 0.968 | -0.046 | 0.83320712 | 0.90494144 | 0 | 0.0306 | 0.0228 | 0.0228 | TRUE | 0.1819 |
| Tex11 | 830 | 587 | 570 | 673 | 579 | 651 | 645 | 624 | 615 | 670 | 653 | 643 | 641.62 | 628 | 655 | 1.043 | 0.06 | 0.4852981 | 0.65117966 | 0 | 0.0061 | 0.0038 | 0.0038 | TRUE | 0.0067 |
| Tex12 | 126 | 63 | 50 | 86 | 53 | 64 | 98 | 67 | 54 | 86 | 60 | 63 | 71.23 | 73 | 70 | 0.946 | -0.08 | 0.77286492 | 0.86564989 | 0.0531 | 0.0357 | 0.0408 | 0.0408 | TRUE | 0.6935 |
| Tex13 | 421 | 423 | 302 | 274 | 234 | 236 | 327 | 450 | 326 | 273 | 264 | 233 | 312.08 | 368 | 257 | 0.699 | -0.517 | 0.00053272 | 0.00451877 | 0.0186 | 0.01 | 0.0127 | 0.0127 | TRUE | 0.433 |
| Tex13a | 0 | 0 | 0 | 0 | 0 | 0 | 0 | 0 | 0 | 0 | 0 | 0 | 0 | 0 | 0 | NA | NA | NA | NA | NA | NA | NA | NA | NA | NA |
| Tex14 | 4084 | 3096 | 3066 | 3477 | 3150 | 3548 | 3172 | 3293 | 3308 | 3461 | 3554 | 3502 | 3381.7 | 3258 | 3506 | 1.076 | 0.106 | 0.05025701 | 0.13530296 | 0 | 0.0031 | 0.0018 | 0.0018 | TRUE | 0.0067 |
| Tex15 | 2328 | 1706 | 1319 | 1515 | 1342 | 1392 | 1808 | 1815 | 1423 | 1508 | 1514 | 1374 | 1573.66 | 1682 | 1465 | 0.87 | -0.2 | 0.03765548 | 0.10964193 | 0.0094 | 0.0039 | 0.006 | 0.006 | TRUE | 0.216 |
| Tex16 | 56 | 45 | 24 | 48 | 32 | 24 | 43 | 48 | 26 | 48 | 36 | 24 | 37.47 | 39 | 36 | 0.914 | -0.13 | 0.72227703 | 0.83375105 | 0.0792 | 0.0657 | 0.0694 | 0.0694 | TRUE | 0.2227 |
| Tex19.1 | 11733 | 7286 | 4183 | 4421 | 3837 | 3839 | 9113 | 7750 | 4513 | 4401 | 4329 | 3790 | 5649.14 | 7125 | 4173 | 0.586 | -0.772 | 0.00653455 | 0.03009939 | 0.0579 | 0.0028 | 0.0187 | 0.0579 | TRUE | 0.4246 |
| Tex19.2 | 506 | 379 | 221 | 243 | 183 | 194 | 393 | 403 | 238 | 242 | 206 | 192 | 279.07 | 345 | 213 | 0.617 | -0.696 | 0.00016899 | 0.00187524 | 0.0447 | 0.0109 | 0.0208 | 0.0208 | TRUE | 0.8131 |
| Tex2 | 5419 | 3708 | 3903 | 4251 | 3749 | 4346 | 4209 | 3944 | 4211 | 4231 | 4229 | 4290 | 4185.84 | 4121 | 4250 | 1.031 | 0.044 | 0.42011435 | 0.59229576 | 5.00E-04 | 0.003 | 0.0019 | 0.0019 | TRUE | 0.017 |
| Tex21 | 227 | 145 | 148 | 149 | 165 | 162 | 176 | 154 | 160 | 148 | 186 | 160 | 164.1 | 163 | 165 | 1.004 | 0.006 | 0.97133689 | 0.98452324 | 0.0032 | 0.0169 | 0.0129 | 0.0129 | TRUE | 0.1309 |
| Tex22 | 1 | 0 | 2 | 0 | 0 | 0 | 1 | 0 | 2 | 0 | 0 | 0 | 0.49 | 1 | 0 | 0.195 | -2.355 | 0.4284796 | NA | 0 | 4.8481 | 4.8763 | 4.8763 | TRUE | 0.127 |
| Tex24 | 0 | 0 | 1 | 0 | 1 | 3 | 0 | 0 | 1 | 0 | 1 | 3 | 0.86 | 0 | 1 | 3.963 | 1.987 | 0.39022271 | NA | 0 | 2.7537 | 2.4741 | 2.4741 | TRUE | 0.311 |
| Tex26 | 32 | 36 | 23 | 40 | 37 | 39 | 25 | 38 | 25 | 40 | 42 | 38 | 34.67 | 29 | 40 | 1.372 | 0.456 | 0.17516366 | 0.3286877 | 0 | 0.0708 | 0.0521 | 0.0521 | TRUE | 0.4466 |
| Tex261 | 1853 | 1337 | 1275 | 1261 | 1144 | 1346 | 1439 | 1422 | 1376 | 1255 | 1291 | 1329 | 1351.91 | 1412 | 1292 | 0.914 | -0.13 | 0.05768852 | 0.14950898 | 0 | 0.0042 | 0.0026 | 0.0026 | TRUE | 0.0078 |
| Tex264 | 1813 | 1347 | 1490 | 1361 | 1251 | 1328 | 1408 | 1433 | 1608 | 1355 | 1411 | 1311 | 1420.92 | 1483 | 1359 | 0.917 | -0.125 | 0.10450108 | 0.22844046 | 0.0026 | 0.0041 | 0.0036 | 0.0036 | TRUE | 0.0645 |
| Tex28 | 0 | 0 | 0 | 0 | 0 | 0 | 0 | 0 | 0 | 0 | 0 | 0 | 0 | 0 | 0 | NA | NA | NA | NA | NA | NA | NA | NA | NA | NA |
| Tex29 | 0 | 0 | 0 | 0 | 0 | 0 | 0 | 0 | 0 | 0 | 0 | 0 | 0 | 0 | 0 | NA | NA | NA | NA | NA | NA | NA | NA | NA | NA |
| Tex30 | 1873 | 1315 | 1318 | 1467 | 1230 | 1495 | 1455 | 1399 | 1422 | 1460 | 1388 | 1476 | 1433.19 | 1425 | 1441 | 1.011 | 0.016 | 0.81214945 | 0.89148699 | 0 | 0.0041 | 0.0026 | 0.0026 | TRUE | 0.0125 |
| Tex33 | 0 | 0 | 0 | 0 | 0 | 0 | 0 | 0 | 0 | 0 | 0 | 0 | 0 | 0 | 0 | NA | NA | NA | NA | NA | NA | NA | NA | NA | NA |

| | | | | | | | | | | | | | | | | | | | | | | | | | |
|---|---|---|---|---|---|---|---|---|---|---|---|---|---|---|---|---|---|---|---|---|---|---|---|---|---|
| Tex35 | 17 | 5 | 7 | 17 | 2 | 12 | 13 | 5 | 8 | 17 | 2 | 12 | 9.52 | 9 | 10 | 1.182 | 0.242 | 0.74010666 | NA | 0.3508 | 0.2515 | 0.2781 | 0.2781 | TRUE | 0.3771 |
| Tex36 | 0 | 0 | 2 | 1 | 0 | 1 | 0 | 0 | 2 | 1 | 0 | 1 | 0.69 | 1 | 1 | 0.966 | -0.049 | 0.98423749 | NA | 0 | 3.4369 | 3.0198 | 3.0198 | TRUE | 0.9749 |
| Tex37 | 0 | 0 | 0 | 0 | 0 | 0 | 0 | 0 | 0 | 0 | 0 | 0 | 0 | 0 | 0 | NA | NA | NA | NA | NA | NA | NA | NA | NA | NA |
| Tex38 | 1 | 1 | 1 | 0 | 0 | 1 | 1 | 1 | 1 | 0 | 0 | 1 | 0.65 | 1 | 0 | 0.383 | -1.383 | 0.55329037 | NA | 0 | 3.6426 | 2.4227 | 2.4227 | TRUE | 0.3835 |
| Tex40 | 219 | 151 | 139 | 200 | 151 | 182 | 170 | 161 | 150 | 199 | 170 | 180 | 171.63 | 160 | 183 | 1.141 | 0.19 | 0.2198772 | 0.38306982 | 0 | 0.0162 | 0.0115 | 0.0115 | TRUE | 0.0636 |
| Tex9 | 986 | 766 | 616 | 527 | 444 | 561 | 766 | 815 | 665 | 525 | 501 | 554 | 637.41 | 749 | 527 | 0.703 | -0.508 | 6.06E-07 | 1.92E-05 | 0.0052 | 0.0061 | 0.0058 | 0.0058 | TRUE | 0.1097 |
| Tfam | 6950 | 5007 | 5105 | 5310 | 4731 | 5277 | 5398 | 5326 | 5508 | 5286 | 5337 | 5209 | 5343.96 | 5411 | 5277 | 0.975 | -0.036 | 0.46791801 | 0.63694781 | 0 | 0.0029 | 0.0016 | 0.0016 | TRUE | 0.003 |
| Tfap2a | 150 | 102 | 106 | 117 | 85 | 100 | 116 | 108 | 114 | 116 | 96 | 99 | 108.41 | 113 | 104 | 0.917 | -0.125 | 0.50876978 | 0.67201243 | 0 | 0.0243 | 0.0167 | 0.0167 | TRUE | 0.1157 |
| Tfap2b | 0 | 1 | 0 | 0 | 0 | 0 | 0 | 1 | 0 | 0 | 0 | 0 | 0.18 | 0 | 0 | 0.552 | -0.857 | 0.8336548 | NA | 0 | 13.3714 | 10 | 10 | TRUE | 0.5216 |
| Tfap2c | 4239 | 3024 | 3147 | 2706 | 2432 | 2933 | 3292 | 3217 | 3396 | 2694 | 2744 | 2895 | 3039.47 | 3302 | 2778 | 0.841 | -0.249 | 2.62E-05 | 0.00041452 | 6.00E-04 | 0.0032 | 0.0022 | 0.0022 | TRUE | 0.0169 |
| Tfap2d | 0 | 0 | 0 | 0 | 0 | 0 | 0 | 0 | 0 | 0 | 0 | 0 | 0 | 0 | 0 | NA | NA | NA | NA | NA | NA | NA | NA | NA | NA |
| Tfap2e | 49 | 25 | 25 | 48 | 37 | 37 | 38 | 27 | 27 | 48 | 42 | 37 | 36.28 | 31 | 42 | 1.362 | 0.446 | 0.17660338 | 0.33039163 | 0 | 0.0678 | 0.0505 | 0.0505 | TRUE | 0.3692 |
| Tfap4 | 1045 | 923 | 833 | 828 | 793 | 876 | 812 | 982 | 899 | 824 | 895 | 865 | 879.29 | 898 | 861 | 0.961 | -0.057 | 0.53022997 | 0.69033619 | 0.0044 | 0.0051 | 0.0049 | 0.0049 | TRUE | 0.0874 |
| Tfb1m | 1666 | 1256 | 1163 | 1416 | 1295 | 1373 | 1294 | 1336 | 1255 | 1409 | 1461 | 1355 | 1351.75 | 1295 | 1408 | 1.087 | 0.121 | 0.08488262 | 0.1965377 | 4.00E-04 | 0.0042 | 0.0028 | 0.0028 | TRUE | 0.0132 |
| Tfb2m | 655 | 498 | 512 | 480 | 411 | 499 | 509 | 530 | 552 | 478 | 464 | 493 | 504.15 | 530 | 478 | 0.903 | -0.147 | 0.12562057 | 0.26077638 | 0 | 0.0071 | 0.0047 | 0.0047 | TRUE | 0.0159 |
| Tfcp2 | 1406 | 1068 | 977 | 989 | 1000 | 1162 | 1092 | 1136 | 1054 | 984 | 1128 | 1147 | 1090.3 | 1094 | 1086 | 0.993 | -0.01 | 0.90149275 | 0.94568228 | 0.0032 | 0.0046 | 0.0041 | 0.0041 | TRUE | 0.0817 |
| Tfcp2l1 | 10356 | 7970 | 9037 | 10000 | 9333 | 10316 | 8043 | 8478 | 9751 | 9954 | 10529 | 10183 | 9489.59 | 8757 | 10222 | 1.168 | 0.223 | 0.00201692 | 0.0121739 | 0.0054 | 0.0027 | 0.0037 | 0.0037 | TRUE | 0.1202 |
| Tfdp1 | 11830 | 8756 | 7994 | 7360 | 6669 | 7833 | 9188 | 9314 | 8625 | 7326 | 7523 | 7732 | 8284.8 | 9042 | 7527 | 0.832 | -0.265 | 1.04E-06 | 3.02E-05 | 0.001 | 0.0027 | 0.002 | 0.002 | TRUE | 0.0197 |
| Tfdp2 | 5424 | 4156 | 4380 | 4553 | 4135 | 4816 | 4213 | 4421 | 4726 | 4532 | 4665 | 4754 | 4551.69 | 4453 | 4650 | 1.045 | 0.063 | 0.30003287 | 0.47223541 | 0.0016 | 0.0029 | 0.0025 | 0.0025 | TRUE | 0.0349 |
| Tfe3 | 3190 | 2463 | 2835 | 2679 | 2556 | 3196 | 2478 | 2620 | 3059 | 2667 | 2883 | 3155 | 2810.23 | 2719 | 2902 | 1.068 | 0.095 | 0.28554459 | 0.45754359 | 0.0086 | 0.0033 | 0.0053 | 0.0053 | TRUE | 0.1449 |
| Tfeb | 1256 | 1034 | 1071 | 1162 | 1047 | 1336 | 975 | 1100 | 1156 | 1157 | 1181 | 1319 | 1147.93 | 1077 | 1219 | 1.134 | 0.181 | 0.04194378 | 0.11836178 | 0.0055 | 0.0045 | 0.0048 | 0.0048 | TRUE | 0.0847 |
| Tfec | 6 | 3 | 0 | 4 | 2 | 2 | 5 | 3 | 0 | 4 | 2 | 2 | 2.68 | 3 | 3 | 1.02 | 0.028 | 0.98176276 | NA | 0 | 0.8877 | 0.7267 | 0.7267 | TRUE | 0.5424 |
| Tff1 | 0 | 0 | 0 | 0 | 0 | 0 | 0 | 0 | 0 | 0 | 0 | 0 | 0 | 0 | 0 | NA | NA | NA | NA | NA | NA | NA | NA | NA | NA |
| Tff2 | 1 | 2 | 1 | 0 | 2 | 1 | 1 | 2 | 1 | 0 | 2 | 1 | 1.2 | 1 | 1 | 0.811 | -0.303 | 0.86289797 | NA | 0 | 1.9703 | 1.3597 | 1.3597 | TRUE | 0.2764 |
| Tff3 | 1 | 0 | 0 | 0 | 1 | 1 | 1 | 0 | 0 | 0 | 1 | 1 | 0.48 | 0 | 1 | 2.031 | 1.022 | 0.71376068 | NA | 0 | 4.9197 | 3.859 | 3.859 | TRUE | 0.2623 |
| Tfg | 5082 | 3522 | 3449 | 3295 | 2914 | 3481 | 3947 | 3746 | 3721 | 3280 | 3287 | 3436 | 3569.69 | 3805 | 3334 | 0.876 | -0.191 | 0.00079674 | 0.00613233 | 6.00E-04 | 0.0031 | 0.0021 | 0.0021 | TRUE | 0.0133 |
| Tfip11 | 3330 | 2436 | 2056 | 2134 | 1893 | 2106 | 2586 | 2591 | 2218 | 2124 | 2136 | 2079 | 2289.07 | 2465 | 2113 | 0.857 | -0.223 | 0.00254695 | 0.01459647 | 0.0032 | 0.0035 | 0.0035 | 0.0035 | TRUE | 0.0918 |
| Tfpi | 6203 | 4442 | 4870 | 4021 | 3816 | 4546 | 4818 | 4725 | 5255 | 4002 | 4305 | 4488 | 4598.69 | 4933 | 4265 | 0.865 | -0.21 | 0.00168233 | 0.01055093 | 0.003 | 0.0029 | 0.003 | 0.003 | TRUE | 0.0395 |
| Tfpi2 | 15 | 3 | 1 | 1 | 4 | 1 | 12 | 3 | 1 | 1 | 5 | 1 | 3.74 | 5 | 2 | 0.39 | -1.36 | 0.24789541 | NA | 0.7278 | 0.6368 | 0.6652 | 0.6652 | TRUE | 1.0071 |
| Tfpt | 859 | 603 | 475 | 489 | 423 | 538 | 667 | 641 | 513 | 487 | 477 | 531 | 552.68 | 607 | 498 | 0.82 | -0.287 | 0.01271008 | 0.04936552 | 0.0098 | 0.0067 | 0.0077 | 0.0077 | TRUE | 0.2137 |
| Tfr2 | 445 | 338 | 375 | 524 | 425 | 433 | 346 | 360 | 405 | 522 | 479 | 427 | 423.04 | 370 | 476 | 1.289 | 0.367 | 0.00170001 | 0.01064777 | 0.0062 | 0.008 | 0.0074 | 0.0074 | TRUE | 0.0951 |
| Tfrc | 36581 | 28702 | 36821 | 46088 | 38293 | 43649 | 28411 | 30530 | 39730 | 45876 | 43198 | 43088 | 38472.19 | 32890 | 44054 | 1.34 | 0.422 | 1.85E-05 | 0.00031323 | 0.0173 | 0.0025 | 0.007 | 0.007 | TRUE | 0.405 |
| Tg | 28 | 17 | 11 | 10 | 7 | 2 | 22 | 18 | 12 | 10 | 8 | 2 | 11.92 | 17 | 7 | 0.379 | -1.399 | 0.02741843 | NA | 0.1139 | 0.2012 | 0.1834 | 0.1834 | TRUE | 0.741 |
| Tgds | 1170 | 878 | 883 | 1000 | 929 | 996 | 909 | 934 | 953 | 995 | 1048 | 983 | 970.33 | 932 | 1009 | 1.083 | 0.115 | 0.12928536 | 0.2660728 | 0 | 0.0049 | 0.0031 | 0.0031 | TRUE | 0.0134 |
| Tgfa | 359 | 246 | 291 | 432 | 393 | 421 | 279 | 262 | 314 | 430 | 443 | 416 | 357.24 | 285 | 430 | 1.509 | 0.594 | 2.84E-07 | 9.87E-06 | 5.00E-04 | 0.0091 | 0.0067 | 0.0067 | TRUE | 0.0848 |
| Tgfb1 | 989 | 820 | 808 | 1237 | 1053 | 1361 | 768 | 872 | 872 | 1231 | 1188 | 1344 | 1045.81 | 837 | 1254 | 1.501 | 0.586 | 1.09E-11 | 1.13E-09 | 0.0038 | 0.0047 | 0.0044 | 0.0044 | TRUE | 0.0657 |
| Tgfb1i1 | 60 | 48 | 38 | 70 | 48 | 63 | 47 | 51 | 41 | 70 | 54 | 62 | 54.11 | 46 | 62 | 1.343 | 0.426 | 0.11013002 | 0.23731167 | 0 | 0.0462 | 0.0324 | 0.0324 | TRUE | 0.1018 |
| Tgfb2 | 17 | 16 | 9 | 3 | 10 | 6 | 13 | 17 | 10 | 3 | 11 | 6 | 10.02 | 13 | 7 | 0.499 | -1.002 | 0.12768949 | NA | 0.0609 | 0.2389 | 0.1979 | 0.1979 | TRUE | 0.5321 |
| Tgfb3 | 7 | 10 | 10 | 8 | 10 | 13 | 5 | 11 | 11 | 8 | 11 | 13 | 9.82 | 9 | 11 | 1.213 | 0.278 | 0.6549607 | NA | 0 | 0.2437 | 0.176 | 0.176 | TRUE | 0.5041 |
| Tgfbi | 813 | 739 | 1020 | 1251 | 1282 | 1456 | 631 | 786 | 1101 | 1245 | 1446 | 1437 | 1107.8 | 839 | 1376 | 1.641 | 0.714 | 0.00363616 | 0.01916359 | 0.0425 | 0.0046 | 0.0149 | 0.0425 | TRUE | 0.8797 |
| Tgfbr1 | 3072 | 2246 | 2157 | 2177 | 1890 | 2104 | 2386 | 2389 | 2327 | 2167 | 2132 | 2077 | 2246.4 | 2367 | 2125 | 0.898 | -0.156 | 0.00797751 | 0.03497625 | 0 | 0.0035 | 0.002 | 0.002 | TRUE | 0.0049 |
| Tgfbr2 | 200 | 123 | 154 | 152 | 137 | 151 | 155 | 131 | 166 | 151 | 155 | 149 | 151.21 | 151 | 152 | 1.005 | 0.007 | 0.96838404 | 0.98296379 | 0 | 0.0181 | 0.0132 | 0.0132 | TRUE | 0.1322 |
| Tgfbr3 | 325 | 212 | 216 | 280 | 197 | 252 | 252 | 226 | 233 | 279 | 222 | 249 | 243.45 | 237 | 250 | 1.054 | 0.076 | 0.58520259 | 0.73454823 | 0.004 | 0.0122 | 0.0097 | 0.0097 | TRUE | 0.1129 |
| Tgfbrap1 | 726 | 533 | 474 | 433 | 440 | 514 | 564 | 567 | 511 | 431 | 496 | 507 | 512.84 | 547 | 478 | 0.873 | -0.197 | 0.05989728 | 0.15301374 | 0.0033 | 0.007 | 0.0059 | 0.0059 | TRUE | 0.0882 |
| Tgif1 | 8728 | 6438 | 6530 | 7655 | 6742 | 7163 | 6779 | 6848 | 7046 | 7620 | 7606 | 7071 | 7161.49 | 6891 | 7432 | 1.079 | 0.109 | 0.04375486 | 0.12217 | 9.00E-04 | 0.0028 | 0.002 | 0.002 | TRUE | 0.0221 |
| Tgif2 | 3001 | 2165 | 2295 | 2469 | 2198 | 2713 | 2331 | 2303 | 2476 | 2458 | 2480 | 2678 | 2454.21 | 2370 | 2539 | 1.072 | 0.1 | 0.1256536 | 0.26077638 | 0.0013 | 0.0034 | 0.0026 | 0.0026 | TRUE | 0.0283 |
| Tgif2lx1 | 0 | 0 | 0 | 0 | 0 | 0 | 0 | 0 | 0 | 0 | 0 | 0 | 0 | 0 | 0 | NA | NA | NA | NA | NA | NA | NA | NA | NA | NA |
| Tgif2lx2 | 0 | 0 | 0 | 0 | 0 | 0 | 0 | 0 | 0 | 0 | 0 | 0 | 0 | 0 | 0 | NA | NA | NA | NA | NA | NA | NA | NA | NA | NA |
| Tgm1 | 519 | 361 | 354 | 459 | 407 | 504 | 403 | 384 | 382 | 457 | 459 | 498 | 430.43 | 390 | 471 | 1.208 | 0.273 | 0.00795197 | 0.0349071 | 0 | 0.0079 | 0.0053 | 0.0053 | TRUE | 0.0282 |
| Tgm2 | 1004 | 741 | 1177 | 1238 | 1177 | 1290 | 780 | 788 | 1270 | 1232 | 1328 | 1273 | 1111.91 | 946 | 1278 | 1.351 | 0.434 | 0.06742371 | 0.16678558 | 0.0398 | 0.0046 | 0.0145 | 0.0398 | TRUE | 1.0674 |
| Tgm3 | 89 | 64 | 115 | 102 | 113 | 166 | 69 | 68 | 124 | 102 | 127 | 164 | 109.03 | 87 | 131 | 1.512 | 0.596 | 0.02220889 | 0.07446202 | 0.0785 | 0.0242 | 0.0394 | 0.0394 | TRUE | 0.5607 |
| Tgm4 | 0 | 0 | 0 | 1 | 1 | 0 | 0 | 0 | 0 | 1 | 1 | 0 | 0.35 | 0 | 1 | 3.978 | 1.992 | 0.55384016 | NA | 0 | 6.6991 | 6.4234 | 6.4234 | TRUE | 0.2647 |
| Tgm5 | 7 | 3 | 4 | 4 | 2 | 0 | 5 | 3 | 4 | 4 | 2 | 0 | 3.2 | 4 | 2 | 0.477 | -1.068 | 0.35260438 | NA | 0 | 0.7438 | 0.5899 | 0.5899 | TRUE | 0.5169 |
| Tgm6 | 0 | 4 | 3 | 1 | 2 | 1 | 0 | 4 | 3 | 1 | 2 | 1 | 1.96 | 2 | 1 | 0.576 | -0.795 | 0.59341324 | NA | 0 | 1.2147 | 1.0284 | 1.0284 | TRUE | 1.1644 |
| Tgm7 | 0 | 0 | 1 | 0 | 0 | 0 | 0 | 0 | 1 | 0 | 0 | 0 | 0.18 | 0 | 0 | 0.552 | -0.857 | 0.8336548 | NA | 0 | 13.1817 | 10 | 10 | TRUE | 0.5365 |
| Tgoln1 | 2371 | 1661 | 1708 | 1916 | 1706 | 1901 | 1841 | 1767 | 1843 | 1907 | 1925 | 1877 | 1859.91 | 1817 | 1903 | 1.047 | 0.066 | 0.28375664 | 0.45564922 | 0 | 0.0037 | 0.0022 | 0.0022 | TRUE | 0.0069 |
| Tgoln2 | 0 | 0 | 0 | 0 | 0 | 0 | 0 | 0 | 0 | 0 | 0 | 0 | 0 | 0 | 0 | NA | NA | NA | NA | NA | NA | NA | NA | NA | NA |
| Tgs1 | 3871 | 2689 | 2732 | 2926 | 2577 | 2936 | 3006 | 2860 | 2948 | 2913 | 2907 | 2898 | 2922.07 | 2938 | 2906 | 0.989 | -0.016 | 0.76782483 | 0.86270893 | 0 | 0.0032 | 0.0019 | 0.0019 | TRUE | 0.0065 |
| Tgtp1 | 1 | 2 | 2 | 1 | 2 | 0 | 1 | 2 | 2 | 1 | 2 | 0 | 1.39 | 2 | 1 | 0.64 | -0.643 | 0.69840412 | NA | 0 | 1.7129 | 1.2046 | 1.2046 | TRUE | 0.3161 |
| Tgtp2 | 0 | 0 | 0 | 0 | 0 | 0 | 0 | 0 | 0 | 0 | 0 | 0 | 0 | 0 | 0 | NA | NA | NA | NA | NA | NA | NA | NA | NA | NA |
| Th | 0 | 0 | 1 | 3 | 1 | 0 | 0 | 0 | 1 | 3 | 1 | 0 | 0.87 | 0 | 1 | 3.982 | 1.993 | 0.38811729 | NA | 0 | 2.7406 | 2.4679 | 2.4679 | TRUE | 0.3126 |
| Tha1 | 59 | 41 | 37 | 33 | 43 | 29 | 46 | 44 | 40 | 33 | 49 | 29 | 39.89 | 43 | 37 | 0.844 | -0.245 | 0.4446588 | 0.61554559 | 0.0095 | 0.0618 | 0.0485 | 0.0485 | TRUE | 0.5402 |
| Thada | 4961 | 3216 | 2497 | 2630 | 2252 | 2516 | 3853 | 3421 | 2694 | 2618 | 2540 | 2484 | 2935.02 | 3323 | 2547 | 0.766 | -0.384 | 0.00021629 | 0.00226316 | 0.0154 | 0.0032 | 0.0074 | 0.0074 | TRUE | 0.3025 |
| Thap1 | 1354 | 943 | 818 | 891 | 770 | 858 | 1052 | 1003 | 883 | 887 | 869 | 847 | 923.3 | 979 | 868 | 0.885 | -0.177 | 0.04357455 | 0.12185698 | 0.0034 | 0.005 | 0.0044 | 0.0044 | TRUE | 0.0873 |
| Thap11 | 3806 | 2788 | 2564 | 2677 | 2441 | 2693 | 2956 | 2966 | 2767 | 2665 | 2754 | 2658 | 2794.14 | 2896 | 2692 | 0.929 | -0.106 | 0.07470695 | 0.17988317 | 6.00E-04 | 0.0033 | 0.0022 | 0.0022 | TRUE | 0.0183 |
| Thap2 | 240 | 157 | 150 | 187 | 166 | 192 | 186 | 167 | 162 | 186 | 187 | 190 | 179.7 | 172 | 188 | 1.089 | 0.123 | 0.40864108 | 0.58139857 | 0 | 0.0156 | 0.0105 | 0.0105 | TRUE | 0.0653 |
| Thap3 | 318 | 275 | 334 | 379 | 387 | 432 | 247 | 293 | 360 | 377 | 437 | 426 | 356.69 | 300 | 413 | 1.383 | 0.468 | 0.0009824 | 0.00714578 | 0.0178 | 0.0091 | 0.0117 | 0.0117 | TRUE | 0.349 |
| Thap4 | 2991 | 2217 | 2275 | 2260 | 2030 | 2439 | 2323 | 2358 | 2455 | 2250 | 2290 | 2408 | 2347.2 | 2379 | 2316 | 0.974 | -0.038 | 0.53829534 | 0.69678096 | 6.00E-04 | 0.0034 | 0.0023 | 0.0023 | TRUE | 0.0148 |
| Thap6 | 665 | 438 | 296 | 370 | 288 | 317 | 516 | 466 | 319 | 368 | 325 | 313 | 384.65 | 434 | 335 | 0.771 | -0.375 | 0.01819219 | 0.06437756 | 0.0317 | 0.0086 | 0.0155 | 0.0155 | TRUE | 0.6068 |

| | | | | | | | | | | | | | | | | | | | | | | | | |
|---|---|---|---|---|---|---|---|---|---|---|---|---|---|---|---|---|---|---|---|---|---|---|---|---|
| Thap7 | 799 | 527 | 493 | 586 | 469 | 557 | 621 | 561 | 532 | 583 | 529 | 550 | 562.55 | 571 | 554 | 0.969 | -0.046 | 0.64492742 | 0.77893761 | 0.0028 | 0.0066 | 0.0054 | 0.0054 | TRUE | 0.0713 |
| Thbd | 36 | 32 | 35 | 18 | 20 | 25 | 28 | 34 | 38 | 18 | 23 | 25 | 27.49 | 33 | 22 | 0.656 | -0.607 | 0.10375312 | 0.22730585 | 0 | 0.0887 | 0.062 | 0.062 | TRUE | 0.1614 |
| Thbs1 | 189 | 117 | 90 | 54 | 59 | 50 | 147 | 124 | 97 | 54 | 67 | 49 | 89.67 | 123 | 57 | 0.457 | -1.13 | 1.92E-06 | 4.94E-05 | 0.024 | 0.0289 | 0.0274 | 0.0274 | TRUE | 0.1301 |
| Thbs2 | 22 | 11 | 13 | 24 | 13 | 16 | 17 | 12 | 14 | 24 | 15 | 16 | 16.19 | 14 | 18 | 1.263 | 0.337 | 0.4872241 | NA | 0 | 0.1488 | 0.1072 | 0.1072 | TRUE | 0.4038 |
| Thbs3 | 187 | 116 | 129 | 215 | 166 | 171 | 145 | 123 | 139 | 214 | 187 | 169 | 162.98 | 136 | 190 | 1.395 | 0.48 | 0.00379009 | 0.01979239 | 0.0053 | 0.017 | 0.0135 | 0.0135 | TRUE | 0.1349 |
| Thbs4 | 110 | 89 | 54 | 64 | 60 | 60 | 85 | 95 | 58 | 64 | 68 | 59 | 71.5 | 79 | 64 | 0.797 | -0.328 | 0.19110019 | 0.34928325 | 0.0191 | 0.0356 | 0.031 | 0.031 | TRUE | 0.4741 |
| Theg | 0 | 2 | 4 | 5 | 4 | 0 | 0 | 2 | 4 | 5 | 5 | 0 | 2.66 | 2 | 3 | 1.516 | 0.6 | 0.67070126 | NA | 1.67 | 0.895 | 1.0378 | 1.0378 | TRUE | 0.3264 |
| Them4 | 400 | 261 | 217 | 264 | 223 | 221 | 311 | 278 | 234 | 263 | 252 | 218 | 259.16 | 274 | 244 | 0.887 | -0.172 | 0.23698771 | 0.40363029 | 0.0112 | 0.0116 | 0.0114 | 0.0114 | TRUE | 0.1793 |
| Them5 | 21 | 17 | 13 | 18 | 26 | 22 | 16 | 18 | 14 | 18 | 29 | 22 | 19.56 | 16 | 23 | 1.418 | 0.503 | 0.25336865 | 0.42242648 | 0 | 0.1236 | 0.0876 | 0.0876 | TRUE | 0.3126 |
| Them6 | 850 | 585 | 593 | 704 | 588 | 692 | 660 | 622 | 640 | 701 | 663 | 683 | 661.58 | 641 | 682 | 1.064 | 0.09 | 0.29465166 | 0.46653995 | 0 | 0.006 | 0.0038 | 0.0038 | TRUE | 0.0088 |
| Themis | 29 | 29 | 28 | 26 | 25 | 38 | 23 | 31 | 30 | 26 | 28 | 38 | 29.2 | 28 | 31 | 1.105 | 0.145 | 0.69169677 | 0.81149322 | 0 | 0.0836 | 0.0614 | 0.0614 | TRUE | 0.2764 |
| Themis2 | 154 | 82 | 92 | 77 | 62 | 95 | 120 | 87 | 99 | 77 | 70 | 94 | 91.08 | 102 | 80 | 0.783 | -0.353 | 0.11201754 | 0.24029217 | 0.0143 | 0.0284 | 0.0244 | 0.0244 | TRUE | 0.2453 |
| Themis3 | 31 | 17 | 13 | 11 | 4 | 10 | 24 | 18 | 14 | 11 | 5 | 10 | 13.59 | 19 | 9 | 0.451 | -1.148 | 0.04013605 | NA | 0 | 0.1769 | 0.139 | 0.139 | TRUE | 0.3184 |
| Thg1l | 3026 | 2191 | 2189 | 2045 | 2023 | 2396 | 2350 | 2331 | 2362 | 2036 | 2282 | 2365 | 2287.6 | 2348 | 2228 | 0.949 | -0.076 | 0.28166714 | 0.45315943 | 0.0023 | 0.0035 | 0.0031 | 0.0031 | TRUE | 0.0695 |
| Thnsl1 | 1449 | 1024 | 1046 | 1310 | 1004 | 1134 | 1125 | 1089 | 1129 | 1304 | 1133 | 1119 | 1149.87 | 1114 | 1185 | 1.064 | 0.089 | 0.27658475 | 0.44775466 | 0.003 | 0.0045 | 0.004 | 0.004 | TRUE | 0.0927 |
| Thnsl2 | 644 | 410 | 449 | 516 | 421 | 475 | 500 | 436 | 484 | 514 | 475 | 469 | 479.7 | 473 | 486 | 1.025 | 0.035 | 0.73451692 | 0.8412894 | 0.0017 | 0.0074 | 0.0056 | 0.0056 | TRUE | 0.0546 |
| Thoc1 | 3088 | 2077 | 1771 | 1838 | 1608 | 1826 | 2398 | 2209 | 1911 | 1830 | 1814 | 1803 | 1994.09 | 2173 | 1816 | 0.835 | -0.261 | 0.00182422 | 0.01126286 | 0.0057 | 0.0036 | 0.0045 | 0.0045 | TRUE | 0.1338 |
| Thoc2 | 6432 | 4817 | 4236 | 4846 | 4098 | 4664 | 4995 | 5124 | 4571 | 4824 | 4623 | 4604 | 4790.1 | 4897 | 4684 | 0.956 | -0.064 | 0.29394659 | 0.46590435 | 0.0018 | 0.0029 | 0.0025 | 0.0025 | TRUE | 0.0409 |
| Thoc3 | 5959 | 4274 | 3877 | 4053 | 3645 | 4152 | 4628 | 4546 | 4183 | 4034 | 4112 | 4099 | 4267.08 | 4452 | 4082 | 0.916 | -0.126 | 0.03254838 | 0.09829553 | 0.0011 | 0.003 | 0.0023 | 0.0023 | TRUE | 0.0338 |
| Thoc5 | 3955 | 2915 | 2619 | 2820 | 2538 | 3055 | 3072 | 3101 | 2826 | 2807 | 2863 | 3016 | 2947.35 | 3000 | 2895 | 0.965 | -0.051 | 0.42167268 | 0.59396547 | 0.0015 | 0.0032 | 0.0026 | 0.0026 | TRUE | 0.0306 |
| Thoc6 | 3225 | 2321 | 2239 | 2540 | 2342 | 2715 | 2505 | 2469 | 2416 | 2528 | 2642 | 2680 | 2539.98 | 2463 | 2617 | 1.062 | 0.087 | 0.13947219 | 0.28069219 | 3.00E-04 | 0.0034 | 0.0021 | 0.0021 | TRUE | 0.0107 |
| Thoc7 | 2555 | 1793 | 1884 | 2003 | 1739 | 1967 | 1984 | 1907 | 2033 | 1994 | 1962 | 1942 | 1970.28 | 1975 | 1966 | 0.995 | -0.007 | 0.91387027 | 0.95259246 | 0 | 0.0036 | 0.0022 | 0.0022 | TRUE | 0.0105 |
| Thop1 | 9606 | 7110 | 6435 | 6971 | 6266 | 7273 | 7461 | 7563 | 6943 | 6939 | 7069 | 7180 | 7192.32 | 7322 | 7063 | 0.964 | -0.052 | 0.33814988 | 0.51231463 | 0.001 | 0.0028 | 0.002 | 0.002 | TRUE | 0.0248 |
| Thpo | 160 | 121 | 151 | 131 | 115 | 157 | 124 | 129 | 163 | 130 | 130 | 155 | 138.5 | 139 | 138 | 1.002 | 0.003 | 0.98470263 | 0.99216018 | 0.0095 | 0.0195 | 0.0166 | 0.0166 | TRUE | 0.2358 |
| Thra | 595 | 467 | 534 | 717 | 672 | 709 | 462 | 497 | 576 | 714 | 758 | 700 | 617.79 | 512 | 724 | 1.418 | 0.504 | 9.90E-07 | 2.91E-05 | 0.0049 | 0.0063 | 0.006 | 0.006 | TRUE | 0.1402 |
| Thrap3 | 14722 | 10563 | 8174 | 7230 | 6820 | 7793 | 11434 | 11236 | 8820 | 7197 | 7694 | 7693 | 9012.11 | 10497 | 7528 | 0.717 | -0.48 | 5.14E-08 | 2.15E-06 | 0.0102 | 0.0027 | 0.0055 | 0.0055 | TRUE | 0.2382 |
| Thrb | 84 | 67 | 57 | 55 | 55 | 67 | 65 | 71 | 62 | 55 | 62 | 66 | 63.49 | 66 | 61 | 0.924 | -0.114 | 0.6385929 | 0.77423614 | 0 | 0.0398 | 0.0267 | 0.0267 | TRUE | 0.0714 |
| Thrsp | 132 | 62 | 76 | 119 | 79 | 130 | 103 | 66 | 82 | 118 | 89 | 128 | 97.73 | 84 | 112 | 1.335 | 0.417 | 0.07052655 | 0.17246125 | 0.0311 | 0.0267 | 0.0279 | 0.0279 | TRUE | 0.3776 |
| Thsd1 | 801 | 652 | 590 | 756 | 660 | 782 | 622 | 694 | 637 | 753 | 745 | 772 | 703.54 | 651 | 757 | 1.164 | 0.219 | 0.01164554 | 0.04626172 | 0 | 0.0058 | 0.004 | 0.004 | TRUE | 0.0384 |
| Thsd4 | 142 | 94 | 87 | 61 | 78 | 74 | 110 | 100 | 94 | 61 | 88 | 73 | 87.65 | 101 | 74 | 0.725 | -0.464 | 0.0354379 | 0.10482488 | 0.0057 | 0.0295 | 0.0233 | 0.0233 | TRUE | 0.2389 |
| Thsd7a | 11 | 7 | 6 | 2 | 5 | 0 | 9 | 7 | 6 | 2 | 6 | 0 | 5.02 | 7 | 3 | 0.329 | -1.602 | 0.10099088 | NA | 0 | 0.475 | 0.4141 | 0.4141 | TRUE | 0.9727 |
| Thsd7b | 9 | 9 | 5 | 4 | 6 | 5 | 7 | 10 | 5 | 4 | 7 | 5 | 6.27 | 7 | 5 | 0.711 | -0.492 | 0.51890105 | NA | 0 | 0.3802 | 0.2529 | 0.2529 | TRUE | 0.181 |
| Thtpa | 695 | 515 | 558 | 602 | 482 | 559 | 540 | 548 | 602 | 599 | 544 | 552 | 564.07 | 563 | 565 | 1.005 | 0.007 | 0.94484081 | 0.96967112 | 0.0014 | 0.0066 | 0.0049 | 0.0049 | TRUE | 0.0417 |
| Thumpd1 | 8331 | 6556 | 6177 | 6212 | 5345 | 6307 | 6470 | 6974 | 6665 | 6183 | 6030 | 6226 | 6424.65 | 6703 | 6146 | 0.917 | -0.125 | 0.01894137 | 0.06645269 | 6.00E-04 | 0.0028 | 0.0019 | 0.0019 | TRUE | 0.0152 |
| Thumpd2 | 731 | 539 | 617 | 723 | 590 | 718 | 568 | 573 | 666 | 720 | 666 | 709 | 650.14 | 602 | 698 | 1.161 | 0.216 | 0.02519377 | 0.08170968 | 0.0032 | 0.0061 | 0.0051 | 0.0051 | TRUE | 0.0975 |
| Thumpd3 | 6279 | 4558 | 4520 | 5074 | 4539 | 5354 | 4877 | 4848 | 4877 | 5051 | 5120 | 5285 | 5009.72 | 4867 | 5152 | 1.059 | 0.082 | 0.10448236 | 0.22844046 | 0 | 0.0029 | 0.0016 | 0.0016 | TRUE | 0.0063 |
| Thy1 | 479 | 355 | 458 | 620 | 585 | 698 | 372 | 378 | 494 | 617 | 660 | 689 | 534.99 | 415 | 655 | 1.585 | 0.664 | 3.05E-08 | 1.34E-06 | 0.0124 | 0.0069 | 0.0084 | 0.0084 | TRUE | 0.3184 |
| Thyn1 | 2751 | 1977 | 1945 | 2376 | 1919 | 2340 | 2137 | 2103 | 2099 | 2365 | 2165 | 2310 | 2196.33 | 2113 | 2280 | 1.079 | 0.11 | 0.07878051 | 0.1870759 | 6.00E-04 | 0.0035 | 0.0024 | 0.0024 | TRUE | 0.0233 |
| Tia1 | 1588 | 1225 | 1033 | 1190 | 901 | 1229 | 1233 | 1303 | 1115 | 1185 | 1016 | 1213 | 1177.52 | 1217 | 1138 | 0.936 | -0.096 | 0.2931878 | 0.46521877 | 0.0062 | 0.0044 | 0.0052 | 0.0052 | TRUE | 0.103 |
| Tial1 | 4844 | 3740 | 4167 | 4379 | 3794 | 4444 | 3762 | 3978 | 4496 | 4359 | 4280 | 4387 | 4210.38 | 4079 | 4342 | 1.065 | 0.091 | 0.1997749 | 0.3604826 | 0.004 | 0.003 | 0.0034 | 0.0034 | TRUE | 0.0972 |
| Tiam1 | 3936 | 2771 | 3081 | 3206 | 2932 | 3214 | 3057 | 2947 | 3324 | 3191 | 3308 | 3173 | 3166.72 | 3109 | 3224 | 1.037 | 0.052 | 0.41499845 | 0.58774352 | 0.0017 | 0.0032 | 0.0026 | 0.0026 | TRUE | 0.0436 |
| Tiam2 | 138 | 102 | 79 | 116 | 90 | 120 | 107 | 108 | 85 | 115 | 102 | 118 | 106.06 | 100 | 112 | 1.113 | 0.155 | 0.43128473 | 0.60334273 | 0 | 0.0248 | 0.0185 | 0.0185 | TRUE | 0.159 |
| Ticam1 | 255 | 151 | 140 | 207 | 183 | 190 | 198 | 161 | 151 | 206 | 206 | 188 | 184.96 | 170 | 200 | 1.17 | 0.227 | 0.15323423 | 0.30014758 | 0.007 | 0.0152 | 0.0128 | 0.0128 | TRUE | 0.243 |
| Ticam2 | 82 | 65 | 69 | 100 | 98 | 126 | 64 | 69 | 74 | 100 | 111 | 124 | 90.29 | 69 | 112 | 1.62 | 0.696 | 0.00099908 | 0.00723765 | 0 | 0.0287 | 0.0206 | 0.0206 | TRUE | 0.1052 |
| Ticrr | 7165 | 4848 | 3923 | 3916 | 3658 | 4275 | 5565 | 5157 | 4233 | 3898 | 4127 | 4220 | 4533.18 | 4985 | 4082 | 0.818 | -0.289 | 0.00118354 | 0.00818977 | 0.0099 | 0.0029 | 0.0055 | 0.0055 | TRUE | 0.2116 |
| Tie1 | 58 | 35 | 48 | 81 | 59 | 86 | 45 | 37 | 52 | 81 | 67 | 85 | 61.02 | 45 | 78 | 1.734 | 0.794 | 0.00215912 | 0.01278919 | 0 | 0.0413 | 0.0309 | 0.0309 | TRUE | 0.151 |
| Tifa | 4 | 6 | 3 | 3 | 4 | 3 | 3 | 6 | 3 | 3 | 5 | 3 | 3.86 | 4 | 4 | 0.825 | -0.277 | 0.77365857 | NA | 0 | 0.6157 | 0.4035 | 0.4035 | TRUE | 0.3209 |
| Tifab | 50 | 32 | 35 | 18 | 21 | 16 | 39 | 34 | 38 | 18 | 24 | 16 | 28.01 | 37 | 19 | 0.515 | -0.956 | 0.01027225 | 0.04212948 | 0 | 0.0871 | 0.0598 | 0.0598 | TRUE | 0.2076 |
| Tigd2 | 1052 | 806 | 738 | 705 | 633 | 667 | 817 | 857 | 796 | 702 | 714 | 658 | 757.49 | 823 | 691 | 0.839 | -0.253 | 0.00270069 | 0.0151972 | 0 | 0.0056 | 0.0038 | 0.0038 | TRUE | 0.0208 |
| Tigd3 | 175 | 146 | 111 | 105 | 111 | 99 | 136 | 155 | 120 | 105 | 125 | 98 | 123.07 | 137 | 109 | 0.795 | -0.331 | 0.08210663 | 0.19270116 | 0.0076 | 0.0217 | 0.0178 | 0.0178 | TRUE | 0.1572 |
| Tigd4 | 0 | 0 | 0 | 0 | 0 | 0 | 0 | 0 | 0 | 0 | 0 | 0 | 0 | 0 | 0 | NA | NA | NA | NA | NA | NA | NA | NA | NA | NA |
| Tigd5 | 58 | 32 | 15 | 31 | 18 | 20 | 45 | 34 | 16 | 31 | 20 | 20 | 27.7 | 32 | 24 | 0.738 | -0.439 | 0.30975311 | 0.48253964 | 0.1215 | 0.088 | 0.0976 | 0.0976 | TRUE | 0.2733 |
| Tigit | 2 | 0 | 0 | 1 | 2 | 3 | 2 | 0 | 0 | 1 | 2 | 3 | 1.29 | 1 | 2 | 3.579 | 1.839 | 0.32080763 | NA | 0 | 1.8334 | 1.3923 | 1.3923 | TRUE | 0.971 |
| Timd2 | 38 | 18 | 13 | 18 | 18 | 8 | 30 | 19 | 14 | 18 | 20 | 8 | 18.13 | 21 | 15 | 0.724 | -0.466 | 0.35836179 | 0.53282691 | 0.1186 | 0.1331 | 0.1291 | 0.1291 | TRUE | 0.527 |
| Timd4 | 2 | 3 | 1 | 3 | 0 | 2 | 2 | 3 | 1 | 3 | 0 | 2 | 1.8 | 2 | 2 | 0.871 | -0.199 | 0.89158905 | NA | 0 | 1.3211 | 0.976 | 0.976 | TRUE | 0.4604 |
| Timeless | 5961 | 4530 | 4160 | 4490 | 4113 | 4789 | 4630 | 4819 | 4489 | 4469 | 4640 | 4727 | 4628.91 | 4646 | 4612 | 0.993 | -0.01 | 0.85237882 | 0.9167369 | 8.00E-04 | 0.0029 | 0.002 | 0.002 | TRUE | 0.0127 |
| Timm10 | 952 | 608 | 544 | 633 | 487 | 582 | 739 | 647 | 587 | 630 | 549 | 575 | 621.18 | 658 | 585 | 0.887 | -0.172 | 0.11010935 | 0.23730291 | 0.0079 | 0.0062 | 0.0068 | 0.0068 | TRUE | 0.1449 |
| Timm10b | 1058 | 814 | 767 | 860 | 723 | 890 | 822 | 866 | 828 | 856 | 816 | 879 | 844.23 | 839 | 850 | 1.015 | 0.021 | 0.78899122 | 0.8761597 | 0 | 0.0052 | 0.0034 | 0.0034 | TRUE | 0.0146 |
| Timm13 | 3735 | 2519 | 2349 | 2610 | 2215 | 2519 | 2901 | 2679 | 2535 | 2598 | 2499 | 2487 | 2616.36 | 2705 | 2528 | 0.934 | -0.099 | 0.14481347 | 0.28828933 | 0.002 | 0.0033 | 0.0029 | 0.0029 | TRUE | 0.0498 |
| Timm17a | 5314 | 3516 | 3400 | 3763 | 3227 | 3773 | 4127 | 3740 | 3669 | 3746 | 3640 | 3725 | 3774.38 | 3845 | 3704 | 0.963 | -0.055 | 0.38272197 | 0.55658084 | 0.0018 | 0.0031 | 0.0026 | 0.0026 | TRUE | 0.051 |
| Timm17b | 2595 | 1861 | 1929 | 2080 | 1981 | 2202 | 2015 | 1980 | 2081 | 2070 | 2235 | 2174 | 2092.54 | 2025 | 2160 | 1.066 | 0.092 | 0.14350568 | 0.28647768 | 6.00E-04 | 0.0036 | 0.0024 | 0.0024 | TRUE | 0.0158 |
| Timm21 | 570 | 400 | 345 | 606 | 452 | 478 | 443 | 425 | 372 | 603 | 510 | 472 | 470.9 | 413 | 528 | 1.276 | 0.352 | 0.00349495 | 0.01859487 | 0.0101 | 0.0075 | 0.0083 | 0.0083 | TRUE | 0.182 |
| Timm22 | 1748 | 1182 | 988 | 1035 | 900 | 1002 | 1358 | 1257 | 1066 | 1030 | 1015 | 989 | 1119.26 | 1227 | 1011 | 0.823 | -0.281 | 0.00267936 | 0.01510692 | 0.007 | 0.0045 | 0.0054 | 0.0054 | TRUE | 0.1564 |
| Timm23 | 3978 | 2810 | 2681 | 2823 | 2546 | 2712 | 3090 | 2989 | 2893 | 2810 | 2872 | 2677 | 2888.43 | 2991 | 2786 | 0.931 | -0.103 | 0.08817169 | 0.2017995 | 8.00E-04 | 0.0032 | 0.0023 | 0.0023 | TRUE | 0.0143 |
| Timm44 | 5763 | 4227 | 4388 | 4359 | 4070 | 4637 | 4476 | 4496 | 4735 | 4339 | 4591 | 4577 | 4535.74 | 4569 | 4502 | 0.986 | -0.021 | 0.70772018 | 0.82353566 | 6.00E-04 | 0.0029 | 0.002 | 0.002 | TRUE | 0.0124 |
| Timm50 | 5518 | 4214 | 4285 | 4492 | 3995 | 4727 | 4286 | 4482 | 4623 | 4471 | 4507 | 4666 | 4505.97 | 4464 | 4548 | 1.019 | 0.028 | 0.62059431 | 0.7606032 | 7.00E-04 | 0.003 | 0.002 | 0.002 | TRUE | 0.0152 |
| Timm8a1 | 653 | 496 | 429 | 453 | 392 | 442 | 507 | 528 | 463 | 451 | 442 | 436 | 471.18 | 499 | 443 | 0.887 | -0.173 | 0.08693091 | 0.19988502 | 0 | 0.0075 | 0.0052 | 0.0052 | TRUE | 0.0449 |
| Timm8a2 | 126 | 106 | 111 | 99 | 68 | 79 | 98 | 113 | 120 | 99 | 77 | 78 | 97.27 | 110 | 85 | 0.77 | -0.377 | 0.07067372 | 0.17279156 | 0 | 0.0268 | 0.0208 | 0.0208 | TRUE | 0.2051 |
| Timm8b | 1886 | 1303 | 1379 | 1520 | 1277 | 1545 | 1465 | 1386 | 1488 | 1513 | 1441 | 1525 | 1469.57 | 1446 | 1493 | 1.032 | 0.046 | 0.50268872 | 0.66683001 | 4.00E-04 | 0.004 | 0.0027 | 0.0027 | TRUE | 0.0155 |

| | | | | | | | | | | | | | | | | | | | | | | | | | |
|---|---|---|---|---|---|---|---|---|---|---|---|---|---|---|---|---|---|---|---|---|---|---|---|---|---|
| Timm9 | 951 | 690 | 731 | 770 | 651 | 783 | 739 | 734 | 789 | 766 | 734 | 773 | 755.85 | 754 | 758 | 1.007 | 0.009 | 0.91028905 | 0.95073156 | 0 | 0.0056 | 0.0036 | 0.0036 | TRUE | 0.0189 |
| Timmdc1 | 2092 | 1556 | 1561 | 1877 | 1617 | 1832 | 1625 | 1655 | 1684 | 1868 | 1824 | 1808 | 1744.19 | 1655 | 1833 | 1.109 | 0.149 | 0.01697192 | 0.06131944 | 0 | 0.0038 | 0.0022 | 0.0022 | TRUE | 0.0034 |
| Timp1 | 699 | 515 | 686 | 775 | 717 | 799 | 543 | 548 | 740 | 771 | 809 | 789 | 699.98 | 610 | 790 | 1.296 | 0.374 | 0.00143963 | 0.00935725 | 0.0151 | 0.0058 | 0.0085 | 0.0085 | TRUE | 0.4014 |
| Timp2 | 1175 | 841 | 792 | 1062 | 960 | 1041 | 913 | 895 | 855 | 1057 | 1083 | 1028 | 971.57 | 888 | 1056 | 1.189 | 0.25 | 0.0010204 | 0.00734 | 0 | 0.0049 | 0.0031 | 0.0031 | TRUE | 0.0121 |
| Timp3 | 689 | 530 | 569 | 658 | 623 | 639 | 535 | 564 | 614 | 655 | 703 | 631 | 616.9 | 571 | 663 | 1.162 | 0.217 | 0.02352499 | 0.07776887 | 0.0023 | 0.0063 | 0.005 | 0.005 | TRUE | 0.0504 |
| Timp4 | 1 | 2 | 6 | 5 | 5 | 11 | 1 | 2 | 6 | 5 | 6 | 11 | 5.14 | 3 | 7 | 2.383 | 1.253 | 0.1845829 | NA | 0.255 | 0.4633 | 0.4104 | 0.4104 | TRUE | 1.1255 |
| Tinag | 0 | 0 | 0 | 0 | 0 | 0 | 0 | 0 | 0 | 0 | 0 | 0 | 0 | 0 | 0 | NA | NA | NA | NA | NA | NA | NA | NA | NA | NA |
| Tinagl1 | 630 | 448 | 485 | 592 | 527 | 546 | 489 | 477 | 523 | 589 | 595 | 539 | 535.32 | 496 | 574 | 1.157 | 0.21 | 0.03001602 | 0.09266069 | 0 | 0.0069 | 0.0049 | 0.0049 | TRUE | 0.0343 |
| Tinf2 | 3405 | 2382 | 2276 | 2348 | 2140 | 2472 | 2645 | 2534 | 2456 | 2337 | 2414 | 2440 | 2470.93 | 2545 | 2397 | 0.941 | -0.087 | 0.15212972 | 0.29872099 | 6.00E-04 | 0.0034 | 0.0023 | 0.0023 | TRUE | 0.0147 |
| Tiparp | 3566 | 3016 | 3051 | 3970 | 3633 | 3886 | 2770 | 3208 | 3292 | 3952 | 4098 | 3836 | 3525.98 | 3090 | 3962 | 1.283 | 0.36 | 1.08E-06 | 3.14E-05 | 0.0042 | 0.0031 | 0.0036 | 0.0036 | TRUE | 0.1019 |
| Tipin | 8056 | 5480 | 5298 | 5450 | 4760 | 5625 | 6257 | 5829 | 5716 | 5425 | 5370 | 5553 | 5691.62 | 5934 | 5449 | 0.918 | -0.123 | 0.02846976 | 0.0890997 | 0.001 | 0.0028 | 0.0021 | 0.0021 | TRUE | 0.028 |
| Tiprl | 3265 | 2359 | 2568 | 2859 | 2479 | 2797 | 2536 | 2509 | 2771 | 2846 | 2797 | 2761 | 2703.23 | 2605 | 2801 | 1.076 | 0.105 | 0.09425752 | 0.21169434 | 0.0011 | 0.0033 | 0.0025 | 0.0025 | TRUE | 0.0369 |
| Tirap | 870 | 625 | 599 | 633 | 590 | 658 | 676 | 665 | 646 | 630 | 666 | 650 | 655.34 | 662 | 649 | 0.978 | -0.032 | 0.7099453 | 0.82524789 | 0 | 0.006 | 0.0038 | 0.0038 | TRUE | 0.0072 |
| Tjap1 | 4554 | 3639 | 3666 | 3674 | 3393 | 3897 | 3537 | 3871 | 3956 | 3657 | 3828 | 3847 | 3782.49 | 3788 | 3777 | 0.998 | -0.003 | 0.95950496 | 0.97761443 | 0.0019 | 0.0031 | 0.0026 | 0.0026 | TRUE | 0.0417 |
| Tjp1 | 17397 | 12850 | 12202 | 12936 | 11652 | 13575 | 13512 | 13668 | 13166 | 12876 | 13145 | 13401 | 13294.59 | 13449 | 13141 | 0.977 | -0.033 | 0.47580236 | 0.64357368 | 3.00E-04 | 0.0026 | 0.0015 | 0.0015 | TRUE | 0.0041 |
| Tjp2 | 11249 | 7910 | 7323 | 6303 | 5899 | 6944 | 8737 | 8414 | 7901 | 6274 | 6655 | 6855 | 7472.55 | 8351 | 6595 | 0.79 | -0.341 | 1.78E-08 | 8.39E-07 | 0.0021 | 0.0027 | 0.0025 | 0.0025 | TRUE | 0.027 |
| Tjp3 | 846 | 826 | 1273 | 1378 | 1379 | 1615 | 657 | 879 | 1374 | 1372 | 1556 | 1594 | 1238.46 | 970 | 1507 | 1.555 | 0.637 | 0.04723661 | 0.12926535 | 0.0734 | 0.0043 | 0.0216 | 0.0734 | TRUE | 0.9916 |
| Tk1 | 4469 | 3246 | 3198 | 3429 | 3068 | 3603 | 3471 | 3453 | 3451 | 3413 | 3461 | 3557 | 3467.53 | 3458 | 3477 | 1.005 | 0.008 | 0.88275609 | 0.93474832 | 0 | 0.0031 | 0.0018 | 0.0018 | TRUE | 0.0049 |
| Tk2 | 1569 | 1200 | 1193 | 1221 | 1191 | 1269 | 1219 | 1276 | 1287 | 1215 | 1344 | 1253 | 1265.65 | 1261 | 1271 | 1.008 | 0.011 | 0.87626414 | 0.93096567 | 0.001 | 0.0043 | 0.0031 | 0.0031 | TRUE | 0.0298 |
| Tkt | 36090 | 27228 | 28449 | 31534 | 28817 | 32731 | 28030 | 28962 | 30696 | 31389 | 32509 | 32311 | 30649.32 | 29229 | 32070 | 1.097 | 0.134 | 0.00997582 | 0.04129292 | 0.0012 | 0.0025 | 0.0019 | 0.0019 | TRUE | 0.0236 |
| Tktl1 | 314 | 170 | 143 | 113 | 83 | 83 | 244 | 181 | 154 | 112 | 94 | 82 | 144.51 | 193 | 96 | 0.495 | -1.014 | 1.32E-06 | 3.67E-05 | 0.0352 | 0.0188 | 0.0238 | 0.0238 | TRUE | 0.4058 |
| Tktl2 | 1921 | 1312 | 1098 | 572 | 556 | 536 | 1492 | 1396 | 1185 | 569 | 627 | 529 | 966.33 | 1358 | 575 | 0.423 | -1.241 | 3.08E-33 | 3.39E-30 | 0.0096 | 0.0049 | 0.0064 | 0.0064 | TRUE | 0.1476 |
| Tlcd1 | 715 | 518 | 360 | 359 | 313 | 336 | 555 | 551 | 388 | 357 | 353 | 332 | 422.81 | 498 | 347 | 0.696 | -0.523 | 0.00015718 | 0.00176891 | 0.0187 | 0.008 | 0.0114 | 0.0114 | TRUE | 0.4237 |
| Tlcd2 | 293 | 222 | 238 | 276 | 253 | 269 | 228 | 236 | 257 | 275 | 285 | 266 | 257.7 | 240 | 275 | 1.148 | 0.199 | 0.11958381 | 0.25134212 | 0 | 0.0116 | 0.0078 | 0.0078 | TRUE | 0.0397 |
| Tle1 | 3115 | 2320 | 2203 | 2175 | 1817 | 2293 | 2419 | 2468 | 2377 | 2165 | 2050 | 2264 | 2290.39 | 2421 | 2160 | 0.892 | -0.165 | 0.00953051 | 0.0399688 | 7.00E-04 | 0.0035 | 0.0025 | 0.0025 | TRUE | 0.0235 |
| Tle2 | 84 | 78 | 60 | 85 | 68 | 87 | 65 | 83 | 65 | 85 | 77 | 86 | 76.69 | 71 | 83 | 1.165 | 0.221 | 0.32914922 | 0.50362565 | 0 | 0.0333 | 0.0238 | 0.0238 | TRUE | 0.1881 |
| Tle3 | 6808 | 5406 | 5330 | 5915 | 5584 | 6069 | 5287 | 5750 | 5751 | 5888 | 6299 | 5991 | 5827.83 | 5596 | 6059 | 1.083 | 0.115 | 0.05054584 | 0.13591994 | 0.0015 | 0.0028 | 0.0023 | 0.0023 | TRUE | 0.0289 |
| Tle4 | 4756 | 3081 | 2741 | 2357 | 2138 | 2432 | 3694 | 3277 | 2958 | 2346 | 2412 | 2401 | 2847.89 | 3310 | 2386 | 0.72 | -0.473 | 4.12E-09 | 2.34E-07 | 0.0058 | 0.0033 | 0.0043 | 0.0043 | TRUE | 0.1271 |
| Tle6 | 939 | 734 | 697 | 645 | 531 | 685 | 729 | 781 | 752 | 642 | 599 | 676 | 696.56 | 754 | 639 | 0.849 | -0.236 | 0.00760033 | 0.03371555 | 8.00E-04 | 0.0058 | 0.0042 | 0.0042 | TRUE | 0.0344 |
| Tlk1 | 4994 | 3694 | 3602 | 3788 | 3299 | 3859 | 3879 | 3929 | 3887 | 3771 | 3722 | 3809 | 3832.67 | 3898 | 3767 | 0.967 | -0.049 | 0.3394531 | 0.513527 | 0 | 0.003 | 0.0016 | 0.0016 | TRUE | 0.0013 |
| Tlk2 | 7867 | 5574 | 5643 | 6005 | 5048 | 5941 | 6110 | 5929 | 6089 | 5977 | 5695 | 5865 | 5944.07 | 6043 | 5846 | 0.967 | -0.048 | 0.34419798 | 0.51840061 | 2.00E-04 | 0.0028 | 0.0017 | 0.0017 | TRUE | 0.0062 |
| Tll1 | 19 | 12 | 8 | 6 | 11 | 7 | 15 | 13 | 9 | 6 | 12 | 7 | 10.24 | 12 | 8 | 0.688 | -0.539 | 0.38260804 | NA | 0 | 0.2339 | 0.1727 | 0.1727 | TRUE | 0.4679 |
| Tll2 | 3 | 4 | 4 | 5 | 3 | 3 | 2 | 4 | 4 | 5 | 3 | 3 | 3.7 | 3 | 4 | 1.056 | 0.079 | 0.93555331 | NA | 0 | 0.6423 | 0.4122 | 0.4122 | TRUE | 0.2097 |
| Tln1 | 10219 | 7000 | 5897 | 6395 | 5558 | 6141 | 7937 | 7446 | 6363 | 6366 | 6270 | 6062 | 6740.5 | 7249 | 6233 | 0.86 | -0.218 | 0.00476975 | 0.02364692 | 0.0063 | 0.0028 | 0.0042 | 0.0042 | TRUE | 0.1391 |
| Tln2 | 5133 | 3533 | 3576 | 4394 | 3961 | 4330 | 3987 | 3758 | 3858 | 4374 | 4468 | 4274 | 4119.94 | 3868 | 4372 | 1.13 | 0.176 | 0.00125855 | 0.0085164 | 5.00E-04 | 0.003 | 0.0019 | 0.0019 | TRUE | 0.009 |
| Tlr1 | 2 | 0 | 0 | 0 | 0 | 2 | 2 | 0 | 0 | 0 | 0 | 2 | 0.59 | 1 | 1 | 1.213 | 0.279 | 0.92253239 | NA | 0 | 4.0336 | 4.2602 | 4.2602 | TRUE | 0.9737 |
| Tlr11 | 0 | 0 | 0 | 0 | 0 | 0 | 0 | 0 | 0 | 0 | 0 | 0 | 0 | 0 | 0 | NA | NA | NA | NA | NA | NA | NA | NA | NA | NA |
| Tlr12 | 0 | 0 | 1 | 0 | 0 | 0 | 0 | 0 | 1 | 0 | 0 | 0 | 0.18 | 0 | 0 | 0.552 | -0.857 | 0.8336548 | NA | 0 | 13.1817 | 10 | 10 | TRUE | 0.5365 |
| Tlr13 | 1 | 0 | 0 | 0 | 0 | 0 | 1 | 0 | 0 | 0 | 0 | 0 | 0.13 | 0 | 0 | 0.552 | -0.857 | 0.8336548 | NA | 0 | 18.3121 | 10 | 10 | TRUE | 0.2566 |
| Tlr2 | 344 | 255 | 217 | 233 | 182 | 163 | 267 | 271 | 234 | 232 | 205 | 161 | 228.45 | 257 | 199 | 0.773 | -0.371 | 0.01824812 | 0.06452756 | 0.0147 | 0.0128 | 0.0133 | 0.0133 | TRUE | 0.3177 |
| Tlr3 | 131 | 99 | 84 | 75 | 74 | 76 | 102 | 105 | 91 | 75 | 83 | 75 | 88.47 | 99 | 78 | 0.781 | -0.356 | 0.08666883 | 0.19947787 | 0 | 0.0292 | 0.0195 | 0.0195 | TRUE | 0.0524 |
| Tlr4 | 0 | 0 | 2 | 0 | 0 | 1 | 0 | 0 | 2 | 0 | 0 | 1 | 0.52 | 1 | 0 | 0.529 | -0.92 | 0.75853134 | NA | 0 | 4.5238 | 4.7579 | 4.7579 | TRUE | 0.9746 |
| Tlr5 | 36 | 14 | 5 | 16 | 20 | 14 | 28 | 15 | 5 | 16 | 23 | 14 | 16.76 | 16 | 18 | 1.059 | 0.083 | 0.88316764 | NA | 0.244 | 0.1438 | 0.1716 | 0.1716 | TRUE | 0.3218 |
| Tlr6 | 33 | 10 | 22 | 19 | 21 | 34 | 26 | 11 | 24 | 19 | 24 | 34 | 22.7 | 20 | 26 | 1.259 | 0.333 | 0.46022602 | 0.62988358 | 0.0887 | 0.1069 | 0.1021 | 0.1021 | TRUE | 0.7953 |
| Tlr7 | 6 | 2 | 2 | 1 | 2 | 4 | 5 | 2 | 2 | 1 | 2 | 4 | 2.69 | 3 | 2 | 0.79 | -0.339 | 0.77128499 | NA | 0 | 0.8832 | 0.6088 | 0.6088 | TRUE | 0.369 |
| Tlr8 | 0 | 0 | 0 | 0 | 0 | 0 | 0 | 0 | 0 | 0 | 0 | 0 | 0 | 0 | 0 | NA | NA | NA | NA | NA | NA | NA | NA | NA | NA |
| Tlr9 | 0 | 0 | 0 | 0 | 0 | 1 | 0 | 0 | 0 | 0 | 0 | 1 | 0.16 | 0 | 0 | 2.094 | 1.067 | 0.79379329 | NA | 0 | 14.4078 | 10 | 10 | TRUE | 0.3835 |
| Tlx1 | 24 | 21 | 23 | 22 | 30 | 16 | 19 | 22 | 25 | 22 | 34 | 16 | 22.89 | 22 | 24 | 1.086 | 0.12 | 0.77955517 | 0.87000719 | 0.0395 | 0.106 | 0.0875 | 0.0875 | TRUE | 0.4352 |
| Tlx2 | 10 | 7 | 3 | 14 | 9 | 8 | 8 | 7 | 3 | 14 | 10 | 8 | 8.41 | 6 | 11 | 1.713 | 0.776 | 0.25409168 | NA | 0 | 0.2844 | 0.208 | 0.208 | TRUE | 0.3679 |
| Tlx3 | 135 | 127 | 96 | 104 | 99 | 119 | 105 | 135 | 104 | 104 | 112 | 117 | 112.7 | 115 | 111 | 0.971 | -0.043 | 0.82619386 | 0.90029631 | 0.0049 | 0.0235 | 0.0182 | 0.0182 | TRUE | 0.2388 |
| Tm2d1 | 772 | 556 | 684 | 708 | 571 | 675 | 600 | 591 | 738 | 705 | 644 | 666 | 657.37 | 643 | 672 | 1.047 | 0.066 | 0.53244573 | 0.69209228 | 0.0073 | 0.006 | 0.0064 | 0.0064 | TRUE | 0.1926 |
| Tm2d2 | 1355 | 888 | 916 | 1004 | 914 | 1047 | 1052 | 945 | 988 | 999 | 1031 | 1034 | 1008.22 | 995 | 1021 | 1.025 | 0.036 | 0.64747931 | 0.78051363 | 8.00E-04 | 0.0048 | 0.0034 | 0.0034 | TRUE | 0.0318 |
| Tm2d3 | 1171 | 900 | 1000 | 1102 | 1015 | 1146 | 909 | 957 | 1079 | 1097 | 1145 | 1131 | 1053.17 | 982 | 1124 | 1.147 | 0.197 | 0.01894537 | 0.06645269 | 0.0031 | 0.0047 | 0.0041 | 0.0041 | TRUE | 0.0884 |
| Tm4sf1 | 2 | 3 | 2 | 0 | 1 | 1 | 2 | 3 | 2 | 0 | 1 | 1 | 1.5 | 2 | 1 | 0.305 | -1.713 | 0.30336182 | NA | 0 | 1.5794 | 1.0395 | 1.0395 | TRUE | 0.2666 |
| Tm4sf19 | 0 | 3 | 2 | 4 | 1 | 1 | 0 | 3 | 2 | 4 | 1 | 1 | 1.91 | 2 | 2 | 1.189 | 0.25 | 0.86621098 | NA | 0 | 1.2448 | 1.0519 | 1.0519 | TRUE | 0.8244 |
| Tm4sf20 | 0 | 0 | 0 | 0 | 0 | 0 | 0 | 0 | 0 | 0 | 0 | 0 | 0 | 0 | 0 | NA | NA | NA | NA | NA | NA | NA | NA | NA | NA |
| Tm4sf4 | 0 | 0 | 0 | 0 | 0 | 0 | 0 | 0 | 0 | 0 | 0 | 0 | 0 | 0 | 0 | NA | NA | NA | NA | NA | NA | NA | NA | NA | NA |
| Tm4sf5 | 293 | 225 | 302 | 390 | 413 | 513 | 228 | 239 | 326 | 388 | 466 | 506 | 358.88 | 264 | 453 | 1.722 | 0.784 | 2.58E-07 | 9.12E-06 | 0.0249 | 0.009 | 0.0137 | 0.0137 | TRUE | 0.4561 |
| Tm6sf1 | 599 | 423 | 469 | 577 | 545 | 582 | 465 | 450 | 506 | 574 | 615 | 575 | 530.82 | 474 | 588 | 1.241 | 0.312 | 0.00131972 | 0.00879791 | 0 | 0.0069 | 0.0049 | 0.0049 | TRUE | 0.0394 |
| Tm6sf2 | 46 | 34 | 37 | 40 | 29 | 48 | 36 | 36 | 40 | 40 | 33 | 47 | 38.62 | 37 | 40 | 1.077 | 0.108 | 0.7308133 | 0.83911492 | 0 | 0.0638 | 0.0447 | 0.0447 | TRUE | 0.1898 |
| Tm7sf2 | 761 | 601 | 654 | 982 | 885 | 941 | 591 | 639 | 706 | 977 | 998 | 929 | 806.79 | 645 | 968 | 1.503 | 0.588 | 9.06E-11 | 7.44E-09 | 0.0032 | 0.0054 | 0.0046 | 0.0046 | TRUE | 0.0784 |
| Tm7sf3 | 3966 | 2784 | 3099 | 3855 | 3322 | 3755 | 3080 | 2961 | 3344 | 3837 | 3748 | 3707 | 3446.15 | 3128 | 3764 | 1.203 | 0.267 | 2.21E-05 | 0.00036191 | 0.0017 | 0.0031 | 0.0026 | 0.0026 | TRUE | 0.0433 |
| Tm9sf1 | 1740 | 1241 | 1107 | 1100 | 989 | 1067 | 1351 | 1320 | 1194 | 1095 | 1116 | 1053 | 1188.3 | 1288 | 1088 | 0.843 | -0.246 | 0.00150789 | 0.00970822 | 0.0018 | 0.0044 | 0.0035 | 0.0035 | TRUE | 0.0483 |
| Tm9sf2 | 8107 | 5651 | 5751 | 6222 | 5366 | 6082 | 6296 | 6011 | 6205 | 6193 | 6053 | 6004 | 6127.2 | 6171 | 6083 | 0.986 | -0.021 | 0.67922912 | 0.8028453 | 3.00E-04 | 0.0028 | 0.0017 | 0.0017 | TRUE | 0.0063 |
| Tm9sf3 | 9002 | 6401 | 6974 | 7696 | 6679 | 7483 | 6991 | 6809 | 7525 | 7661 | 7535 | 7387 | 7317.85 | 7108 | 7528 | 1.059 | 0.083 | 0.14224233 | 0.28474364 | 0.0013 | 0.0028 | 0.0022 | 0.0022 | TRUE | 0.0318 |
| Tm9sf4 | 4322 | 3084 | 3289 | 3528 | 3229 | 3701 | 3357 | 3280 | 3549 | 3512 | 3643 | 3653 | 3498.97 | 3395 | 3603 | 1.061 | 0.086 | 0.1391628 | 0.2804249 | 6.00E-04 | 0.0031 | 0.0021 | 0.0021 | TRUE | 0.0187 |
| Tma16 | 2277 | 1595 | 1605 | 1591 | 1399 | 1621 | 1768 | 1697 | 1732 | 1584 | 1578 | 1600 | 1659.81 | 1732 | 1587 | 0.916 | -0.127 | 0.04418853 | 0.12306832 | 0 | 0.0039 | 0.0022 | 0.0022 | TRUE | 0.0042 |
| Tma7 | 2516 | 1725 | 1550 | 1606 | 1289 | 1583 | 1954 | 1835 | 1672 | 1599 | 1454 | 1563 | 1679.46 | 1820 | 1539 | 0.845 | -0.244 | 0.00180872 | 0.0111865 | 0.0033 | 0.0038 | 0.0038 | 0.0038 | TRUE | 0.0607 |
| Tmbim1 | 888 | 764 | 795 | 1034 | 846 | 1098 | 690 | 813 | 858 | 1029 | 954 | 1084 | 904.61 | 787 | 1022 | 1.303 | 0.382 | 9.48E-05 | 0.00115758 | 0.0073 | 0.005 | 0.0058 | 0.0058 | TRUE | 0.1448 |

| | | | | | | | | | | | | | | | | | | | | | | | | | |
|---|---|---|---|---|---|---|---|---|---|---|---|---|---|---|---|---|---|---|---|---|---|---|---|---|---|
| Tmbim4 | 2099 | 1458 | 1420 | 1432 | 1291 | 1394 | 1630 | 1551 | 1532 | 1425 | 1456 | 1376 | 1495.19 | 1571 | 1419 | 0.902 | -0.148 | 0.02878097 | 0.08983762 | 0 | 0.004 | 0.0026 | 0.0026 | TRUE | 0.0136 |
| Tmbim6 | 13710 | 9910 | 9770 | 9632 | 8529 | 9910 | 10648 | 10541 | 10542 | 9588 | 9622 | 9783 | 10120.47 | 10577 | 9664 | 0.914 | -0.13 | 0.00387532 | 0.02011984 | 0 | 0.0027 | 0.0014 | 0.0014 | TRUE | 0.0014 |
| Tmc1 | 5 | 5 | 2 | 4 | 7 | 4 | 4 | 5 | 2 | 4 | 8 | 4 | 4.53 | 4 | 5 | 1.38 | 0.465 | 0.60728206 | NA | 0 | 0.5255 | 0.363 | 0.363 | TRUE | 0.3567 |
| Tmc2 | 5 | 2 | 3 | 2 | 2 | 4 | 4 | 2 | 3 | 2 | 2 | 4 | 2.91 | 3 | 3 | 0.881 | -0.183 | 0.8668835 | NA | 0 | 0.8177 | 0.5181 | 0.5181 | TRUE | 0.189 |
| Tmc3 | 273 | 234 | 199 | 219 | 184 | 219 | 212 | 249 | 215 | 218 | 208 | 216 | 219.57 | 225 | 214 | 0.952 | -0.071 | 0.60956387 | 0.75302475 | 0 | 0.0132 | 0.0094 | 0.0094 | TRUE | 0.0906 |
| Tmc4 | 131 | 121 | 112 | 128 | 97 | 134 | 102 | 129 | 121 | 127 | 109 | 132 | 120.07 | 117 | 123 | 1.057 | 0.08 | 0.66983684 | 0.79706491 | 0 | 0.0222 | 0.0171 | 0.0171 | TRUE | 0.1491 |
| Tmc5 | 60 | 33 | 33 | 57 | 30 | 32 | 47 | 35 | 36 | 57 | 34 | 32 | 39.91 | 39 | 41 | 1.037 | 0.053 | 0.87554177 | 0.93081805 | 0.0447 | 0.0618 | 0.0569 | 0.0569 | TRUE | 0.9097 |
| Tmc6 | 2372 | 1964 | 2045 | 2152 | 2146 | 2371 | 1842 | 2089 | 2207 | 2142 | 2421 | 2341 | 2173.57 | 2046 | 2301 | 1.126 | 0.171 | 0.0362104 | 0.10660253 | 0.0054 | 0.0035 | 0.0043 | 0.0043 | TRUE | 0.0941 |
| Tmc7 | 1316 | 951 | 1086 | 1694 | 1232 | 1450 | 1022 | 1012 | 1172 | 1686 | 1390 | 1431 | 1285.47 | 1069 | 1502 | 1.408 | 0.493 | 1.61E-07 | 5.93E-06 | 0.008 | 0.0043 | 0.0056 | 0.0056 | TRUE | 0.1383 |
| Tmc8 | 657 | 548 | 614 | 908 | 843 | 1003 | 510 | 583 | 663 | 904 | 951 | 990 | 766.77 | 585 | 948 | 1.625 | 0.7 | 1.03E-11 | 1.08E-09 | 0.0074 | 0.0055 | 0.0063 | 0.0063 | TRUE | 0.1565 |
| Tmcc1 | 1636 | 1114 | 1104 | 1022 | 901 | 1062 | 1271 | 1185 | 1191 | 1017 | 1016 | 1048 | 1121.47 | 1216 | 1027 | 0.844 | -0.244 | 0.00085549 | 0.00646278 | 0 | 0.0045 | 0.003 | 0.003 | TRUE | 0.0197 |
| Tmcc2 | 503 | 386 | 404 | 412 | 348 | 382 | 391 | 411 | 436 | 410 | 393 | 377 | 402.82 | 413 | 393 | 0.955 | -0.066 | 0.53613094 | 0.69496878 | 0 | 0.0083 | 0.0058 | 0.0058 | TRUE | 0.0285 |
| Tmcc3 | 399 | 283 | 246 | 255 | 271 | 244 | 310 | 301 | 265 | 254 | 306 | 241 | 279.46 | 292 | 267 | 0.91 | -0.136 | 0.32041811 | 0.49435839 | 0.0073 | 0.0109 | 0.0099 | 0.0099 | TRUE | 0.1766 |
| Tmco1 | 2210 | 1459 | 1524 | 1805 | 1531 | 1719 | 1716 | 1552 | 1644 | 1797 | 1727 | 1697 | 1688.91 | 1637 | 1740 | 1.062 | 0.087 | 0.20818131 | 0.36975192 | 0.0012 | 0.0038 | 0.0028 | 0.0028 | TRUE | 0.0252 |
| Tmco2 | 0 | 0 | 0 | 0 | 0 | 0 | 0 | 0 | 0 | 0 | 0 | 0 | 0 | 0 | 0 | NA | NA | NA | NA | NA | NA | NA | NA | NA | NA |
| Tmco3 | 1156 | 803 | 790 | 786 | 670 | 867 | 898 | 854 | 852 | 782 | 756 | 856 | 833.07 | 868 | 798 | 0.919 | -0.122 | 0.14888845 | 0.29411535 | 0.0013 | 0.0053 | 0.0039 | 0.0039 | TRUE | 0.048 |
| Tmco4 | 456 | 335 | 302 | 404 | 287 | 359 | 354 | 356 | 326 | 402 | 324 | 354 | 352.77 | 345 | 360 | 1.043 | 0.06 | 0.61602251 | 0.75753331 | 0.0042 | 0.0091 | 0.0076 | 0.0076 | TRUE | 0.1195 |
| Tmco5 | 0 | 0 | 0 | 0 | 0 | 0 | 0 | 0 | 0 | 0 | 0 | 0 | 0 | 0 | 0 | NA | NA | NA | NA | NA | NA | NA | NA | NA | NA |
| Tmco5b | 1 | 0 | 0 | 0 | 0 | 0 | 1 | 0 | 0 | 0 | 0 | 0 | 0.13 | 0 | 0 | 0.552 | -0.857 | 0.8336548 | NA | 0 | 18.3121 | 10 | 10 | TRUE | 0.2566 |
| Tmco6 | 1663 | 1300 | 1367 | 1629 | 1509 | 1685 | 1292 | 1383 | 1475 | 1622 | 1702 | 1663 | 1522.75 | 1383 | 1662 | 1.203 | 0.267 | 0.00026488 | 0.00264561 | 0.0018 | 0.004 | 0.0032 | 0.0032 | TRUE | 0.0421 |
| Tmed1 | 1565 | 1151 | 1131 | 1377 | 1251 | 1354 | 1215 | 1224 | 1220 | 1371 | 1411 | 1337 | 1296.44 | 1220 | 1373 | 1.125 | 0.17 | 0.01232422 | 0.04832652 | 0 | 0.0043 | 0.0025 | 0.0025 | TRUE | 0.0071 |
| Tmed10 | 7549 | 5461 | 5598 | 6618 | 5943 | 6675 | 5863 | 5809 | 6040 | 6588 | 6704 | 6589 | 6265.52 | 5904 | 6627 | 1.123 | 0.167 | 0.00066054 | 0.0053279 | 1.00E-04 | 0.0028 | 0.0016 | 0.0016 | TRUE | 0.0049 |
| Tmed11 | 0 | 0 | 0 | 0 | 0 | 0 | 0 | 0 | 0 | 0 | 0 | 0 | 0 | 0 | 0 | NA | NA | NA | NA | NA | NA | NA | NA | NA | NA |
| Tmed2 | 1550 | 1253 | 1327 | 1351 | 1198 | 1374 | 1204 | 1333 | 1432 | 1345 | 1351 | 1356 | 1336.84 | 1323 | 1351 | 1.023 | 0.032 | 0.68361873 | 0.80598552 | 0.0032 | 0.0042 | 0.0038 | 0.0038 | TRUE | 0.0774 |
| Tmed3 | 1359 | 1021 | 1071 | 1253 | 1061 | 1094 | 1055 | 1086 | 1156 | 1247 | 1197 | 1080 | 1136.87 | 1099 | 1175 | 1.069 | 0.097 | 0.23745727 | 0.40417188 | 0.003 | 0.0045 | 0.004 | 0.004 | TRUE | 0.0605 |
| Tmed4 | 1602 | 1119 | 1167 | 1315 | 1112 | 1223 | 1244 | 1190 | 1259 | 1309 | 1254 | 1207 | 1244.06 | 1231 | 1257 | 1.021 | 0.029 | 0.68201084 | 0.80494909 | 5.00E-04 | 0.0043 | 0.0029 | 0.0029 | TRUE | 0.016 |
| Tmed5 | 1763 | 1275 | 1329 | 1578 | 1212 | 1368 | 1369 | 1356 | 1434 | 1571 | 1367 | 1350 | 1407.98 | 1386 | 1429 | 1.032 | 0.045 | 0.57278943 | 0.72540132 | 0.0033 | 0.0041 | 0.0038 | 0.0038 | TRUE | 0.0906 |
| Tmed6 | 11 | 10 | 20 | 21 | 9 | 12 | 9 | 11 | 22 | 21 | 10 | 12 | 13.94 | 14 | 14 | 1.07 | 0.098 | 0.86350171 | NA | 0.13 | 0.1724 | 0.16 | 0.16 | TRUE | 1.0758 |
| Tmed7 | 6752 | 4891 | 5040 | 5978 | 5243 | 5841 | 5244 | 5202 | 5438 | 5950 | 5915 | 5766 | 5585.95 | 5295 | 5877 | 1.11 | 0.151 | 0.00303408 | 0.01667284 | 2.00E-04 | 0.0029 | 0.0017 | 0.0017 | TRUE | 0.0067 |
| Tmed8 | 2895 | 2097 | 1930 | 2109 | 1830 | 2047 | 2248 | 2231 | 2082 | 2099 | 2064 | 2021 | 2124.31 | 2187 | 2061 | 0.942 | -0.086 | 0.17195393 | 0.32394063 | 6.00E-04 | 0.0035 | 0.0024 | 0.0024 | TRUE | 0.0209 |
| Tmed9 | 6099 | 4164 | 4156 | 4466 | 3725 | 4304 | 4737 | 4429 | 4484 | 4445 | 4202 | 4249 | 4424.44 | 4550 | 4299 | 0.944 | -0.082 | 0.145328 | 0.28917905 | 8.00E-04 | 0.003 | 0.0021 | 0.0021 | TRUE | 0.016 |
| Tmeff1 | 6409 | 4863 | 5685 | 6254 | 6217 | 7245 | 4978 | 5173 | 6134 | 6225 | 7013 | 7152 | 6112.49 | 5428 | 6797 | 1.252 | 0.325 | 0.00011383 | 0.00135247 | 0.009 | 0.0028 | 0.0049 | 0.0049 | TRUE | 0.1574 |
| Tmeff2 | 78 | 53 | 46 | 44 | 46 | 47 | 61 | 56 | 50 | 44 | 52 | 46 | 51.45 | 56 | 47 | 0.849 | -0.237 | 0.37935705 | 0.55390237 | 0 | 0.0485 | 0.0327 | 0.0327 | TRUE | 0.0697 |
| Tmem100 | 6 | 1 | 1 | 0 | 1 | 3 | 5 | 1 | 1 | 0 | 1 | 3 | 1.82 | 2 | 1 | 0.586 | -0.771 | 0.61324366 | NA | 0.5284 | 1.308 | 1.0916 | 1.0916 | TRUE | 0.7304 |
| Tmem101 | 1986 | 1453 | 1428 | 1730 | 1437 | 1687 | 1542 | 1546 | 1541 | 1722 | 1621 | 1665 | 1606.21 | 1543 | 1669 | 1.082 | 0.114 | 0.07603812 | 0.18189343 | 0 | 0.0039 | 0.0024 | 0.0024 | TRUE | 0.0092 |
| Tmem102 | 233 | 203 | 167 | 189 | 157 | 188 | 181 | 216 | 180 | 188 | 177 | 186 | 187.98 | 192 | 184 | 0.957 | -0.064 | 0.67089646 | 0.79743227 | 0 | 0.015 | 0.0109 | 0.0109 | TRUE | 0.1201 |
| Tmem104 | 691 | 571 | 470 | 525 | 486 | 567 | 537 | 607 | 507 | 523 | 548 | 560 | 546.95 | 550 | 544 | 0.988 | -0.017 | 0.86292624 | 0.92443447 | 0.003 | 0.0068 | 0.0056 | 0.0056 | TRUE | 0.0925 |
| Tmem106a | 186 | 141 | 117 | 147 | 110 | 145 | 144 | 150 | 126 | 146 | 124 | 143 | 139.04 | 140 | 138 | 0.983 | -0.024 | 0.88804028 | 0.93791175 | 0 | 0.0195 | 0.0141 | 0.0141 | TRUE | 0.075 |
| Tmem106b | 2268 | 1696 | 2123 | 2369 | 2017 | 2298 | 1761 | 1804 | 2291 | 2358 | 2275 | 2268 | 2126.36 | 1952 | 2300 | 1.18 | 0.238 | 0.01110163 | 0.04455329 | 0.0109 | 0.0035 | 0.0059 | 0.0059 | TRUE | 0.2766 |
| Tmem106c | 975 | 766 | 970 | 1136 | 961 | 1095 | 757 | 815 | 1047 | 1131 | 1084 | 1081 | 985.74 | 873 | 1099 | 1.261 | 0.334 | 0.0023595 | 0.0137482 | 0.0142 | 0.0048 | 0.0077 | 0.0077 | TRUE | 0.358 |
| Tmem107 | 954 | 672 | 761 | 775 | 749 | 848 | 741 | 715 | 821 | 771 | 845 | 837 | 788.39 | 759 | 818 | 1.078 | 0.108 | 0.22417164 | 0.38856237 | 0.0024 | 0.0054 | 0.0044 | 0.0044 | TRUE | 0.0588 |
| Tmem108 | 509 | 421 | 279 | 306 | 329 | 345 | 395 | 448 | 301 | 305 | 371 | 341 | 360.08 | 381 | 339 | 0.887 | -0.173 | 0.23921182 | 0.40620822 | 0.0218 | 0.009 | 0.0128 | 0.0128 | TRUE | 0.3718 |
| Tmem109 | 1107 | 734 | 750 | 806 | 733 | 809 | 860 | 781 | 809 | 802 | 827 | 799 | 812.92 | 817 | 809 | 0.989 | -0.015 | 0.85203727 | 0.91647868 | 0 | 0.0053 | 0.0036 | 0.0036 | TRUE | 0.0269 |
| Tmem11 | 4396 | 3183 | 3238 | 3425 | 3081 | 3536 | 3414 | 3386 | 3494 | 3409 | 3476 | 3491 | 3444.86 | 3431 | 3459 | 1.008 | 0.012 | 0.82868875 | 0.90181577 | 0 | 0.0031 | 0.0017 | 0.0017 | TRUE | 0.003 |
| Tmem110 | 1676 | 1269 | 1307 | 1322 | 1291 | 1322 | 1302 | 1350 | 1410 | 1316 | 1456 | 1305 | 1356.51 | 1354 | 1359 | 1.004 | 0.006 | 0.93954619 | 0.966328 | 0.0019 | 0.0042 | 0.0034 | 0.0034 | TRUE | 0.0463 |
| Tmem115 | 962 | 664 | 567 | 483 | 413 | 506 | 747 | 706 | 612 | 481 | 466 | 500 | 585.23 | 688 | 482 | 0.699 | -0.516 | 5.23E-07 | 1.69E-05 | 0.0046 | 0.0065 | 0.0058 | 0.0058 | TRUE | 0.1101 |
| Tmem116 | 84 | 54 | 62 | 68 | 54 | 56 | 65 | 57 | 67 | 68 | 61 | 55 | 62.24 | 63 | 61 | 0.969 | -0.046 | 0.85304961 | 0.91687758 | 0 | 0.0405 | 0.0275 | 0.0275 | TRUE | 0.0747 |
| Tmem117 | 41 | 45 | 48 | 52 | 41 | 45 | 32 | 48 | 52 | 52 | 46 | 44 | 45.66 | 44 | 47 | 1.094 | 0.13 | 0.666635 | 0.79491786 | 0 | 0.0543 | 0.0435 | 0.0435 | TRUE | 0.5292 |
| Tmem119 | 28 | 25 | 17 | 19 | 16 | 19 | 22 | 27 | 18 | 19 | 18 | 19 | 20.4 | 22 | 19 | 0.836 | -0.258 | 0.54147714 | 0.69969557 | 0 | 0.1186 | 0.0788 | 0.0788 | TRUE | 0.1544 |
| Tmem120a | 1275 | 821 | 992 | 918 | 882 | 990 | 990 | 873 | 1070 | 914 | 995 | 977 | 969.99 | 978 | 962 | 0.983 | -0.024 | 0.78967195 | 0.87639639 | 0.0051 | 0.0049 | 0.0049 | 0.0049 | TRUE | 0.1015 |
| Tmem120b | 597 | 405 | 344 | 433 | 385 | 418 | 464 | 431 | 371 | 431 | 434 | 413 | 423.93 | 422 | 426 | 1.007 | 0.01 | 0.93096735 | 0.96192556 | 0.0045 | 0.008 | 0.0069 | 0.0069 | TRUE | 0.1251 |
| Tmem121 | 748 | 592 | 600 | 713 | 603 | 631 | 581 | 630 | 647 | 710 | 680 | 623 | 645.15 | 619 | 671 | 1.085 | 0.117 | 0.2131537 | 0.37542894 | 0.0024 | 0.0061 | 0.0049 | 0.0049 | TRUE | 0.0472 |
| Tmem123 | 2257 | 1713 | 1893 | 1993 | 1777 | 1892 | 1753 | 1822 | 2043 | 1984 | 2005 | 1868 | 1912.28 | 1873 | 1952 | 1.043 | 0.061 | 0.41810584 | 0.59080442 | 0.0031 | 0.0037 | 0.0036 | 0.0036 | TRUE | 0.0756 |
| Tmem125 | 95 | 71 | 71 | 84 | 79 | 99 | 74 | 76 | 77 | 84 | 89 | 98 | 82.73 | 76 | 90 | 1.199 | 0.261 | 0.21865686 | 0.3817814 | 0 | 0.0311 | 0.0203 | 0.0203 | TRUE | 0.0534 |
| Tmem126a | 1254 | 860 | 766 | 833 | 759 | 888 | 974 | 915 | 827 | 829 | 856 | 877 | 879.53 | 905 | 854 | 0.942 | -0.086 | 0.32241854 | 0.49621278 | 0.0028 | 0.0051 | 0.0043 | 0.0043 | TRUE | 0.0679 |
| Tmem126b | 644 | 481 | 450 | 448 | 397 | 469 | 500 | 512 | 486 | 446 | 448 | 463 | 475.69 | 499 | 452 | 0.906 | -0.142 | 0.14101692 | 0.28296432 | 0 | 0.0074 | 0.0046 | 0.0046 | TRUE | 0.0062 |
| Tmem127 | 1205 | 839 | 722 | 825 | 713 | 750 | 936 | 892 | 779 | 821 | 804 | 740 | 828.87 | 869 | 788 | 0.906 | -0.143 | 0.1284912 | 0.26493433 | 0.0048 | 0.0053 | 0.0051 | 0.0051 | TRUE | 0.0963 |
| Tmem128 | 1197 | 846 | 845 | 890 | 744 | 830 | 930 | 900 | 912 | 886 | 839 | 819 | 880.97 | 914 | 848 | 0.928 | -0.108 | 0.1686969 | 0.32029553 | 0 | 0.0051 | 0.0033 | 0.0033 | TRUE | 0.0183 |
| Tmem129 | 446 | 330 | 336 | 409 | 284 | 359 | 346 | 351 | 363 | 407 | 320 | 354 | 356.97 | 353 | 360 | 1.023 | 0.032 | 0.78978246 | 0.87642552 | 0.0045 | 0.0091 | 0.0077 | 0.0077 | TRUE | 0.1456 |
| Tmem130 | 194 | 147 | 200 | 242 | 209 | 236 | 151 | 156 | 216 | 241 | 236 | 233 | 205.41 | 174 | 237 | 1.363 | 0.447 | 0.00589378 | 0.02787553 | 0.0141 | 0.014 | 0.014 | 0.014 | TRUE | 0.4515 |
| Tmem131 | 15454 | 11147 | 10609 | 12642 | 11683 | 13521 | 12002 | 11857 | 11447 | 12584 | 13180 | 13347 | 12402.86 | 11769 | 13037 | 1.108 | 0.147 | 0.00332048 | 0.01787554 | 7.00E-04 | 0.0026 | 0.0017 | 0.0017 | TRUE | 0.0113 |
| Tmem132a | 1496 | 1140 | 1070 | 1277 | 995 | 1151 | 1162 | 1213 | 1155 | 1271 | 1122 | 1136 | 1176.47 | 1177 | 1176 | 1.001 | 0.001 | 0.98800999 | 0.99383851 | 0.0019 | 0.0044 | 0.0035 | 0.0035 | TRUE | 0.0596 |
| Tmem132b | 58 | 53 | 53 | 40 | 47 | 47 | 45 | 56 | 57 | 40 | 53 | 46 | 49.64 | 53 | 46 | 0.88 | -0.184 | 0.51245845 | 0.67504942 | 0 | 0.0502 | 0.0363 | 0.0363 | TRUE | 0.1622 |
| Tmem132c | 187 | 115 | 132 | 176 | 160 | 164 | 145 | 122 | 142 | 175 | 180 | 162 | 154.59 | 136 | 172 | 1.259 | 0.332 | 0.04110886 | 0.1166342 | 0 | 0.0178 | 0.0125 | 0.0125 | TRUE | 0.0836 |
| Tmem132d | 58 | 41 | 48 | 65 | 71 | 75 | 45 | 44 | 52 | 65 | 80 | 74 | 59.88 | 47 | 73 | 1.559 | 0.641 | 0.0115378 | 0.04596149 | 0 | 0.042 | 0.0289 | 0.0289 | TRUE | 0.0897 |
| Tmem132e | 148 | 94 | 74 | 112 | 91 | 94 | 115 | 100 | 80 | 111 | 103 | 93 | 100.29 | 98 | 102 | 1.035 | 0.05 | 0.81190436 | 0.89135487 | 0.0113 | 0.0261 | 0.0219 | 0.0219 | TRUE | 0.2531 |
| Tmem134 | 1145 | 906 | 888 | 998 | 858 | 988 | 889 | 964 | 958 | 993 | 968 | 975 | 957.96 | 937 | 979 | 1.046 | 0.065 | 0.3984505 | 0.57184988 | 0 | 0.0049 | 0.0032 | 0.0032 | TRUE | 0.0251 |
| Tmem135 | 2165 | 1514 | 1580 | 1749 | 1478 | 1677 | 1681 | 1610 | 1705 | 1741 | 1667 | 1655 | 1676.74 | 1665 | 1688 | 1.013 | 0.019 | 0.76885761 | 0.86325859 | 0 | 0.0038 | 0.0025 | 0.0025 | TRUE | 0.0098 |
| Tmem136 | 552 | 437 | 499 | 613 | 526 | 493 | 429 | 465 | 538 | 610 | 593 | 487 | 520.37 | 477 | 563 | 1.182 | 0.242 | 0.04617312 | 0.12710947 | 0.0123 | 0.007 | 0.0086 | 0.0086 | TRUE | 0.1685 |

| | | | | | | | | | | | | | | | | | | | | | | | | | |
|---|---|---|---|---|---|---|---|---|---|---|---|---|---|---|---|---|---|---|---|---|---|---|---|---|---|
| Tmem138 | 626 | 481 | 528 | 553 | 496 | 637 | 486 | 512 | 570 | 550 | 560 | 629 | 551.06 | 523 | 580 | 1.112 | 0.153 | 0.14071116 | 0.28258882 | 0.0043 | 0.0067 | 0.0059 | 0.0059 | TRUE | 0.072 |
| Tmem139 | 15 | 6 | 10 | 3 | 8 | 5 | 12 | 6 | 11 | 3 | 9 | 5 | 7.63 | 10 | 6 | 0.578 | -0.791 | 0.27613144 | NA | 0 | 0.3131 | 0.2377 | 0.2377 | TRUE | 0.5443 |
| Tmem140 | 14 | 8 | 5 | 5 | 3 | 11 | 11 | 9 | 5 | 5 | 3 | 11 | 7.33 | 8 | 6 | 0.774 | -0.37 | 0.61847 | NA | 0.0812 | 0.3256 | 0.2589 | 0.2589 | TRUE | 0.7894 |
| Tmem141 | 292 | 206 | 173 | 275 | 232 | 262 | 227 | 219 | 187 | 274 | 262 | 259 | 237.78 | 211 | 265 | 1.252 | 0.324 | 0.01721611 | 0.061905 | 0 | 0.0124 | 0.0091 | 0.0091 | TRUE | 0.1059 |
| Tmem143 | 861 | 641 | 690 | 841 | 718 | 822 | 669 | 682 | 745 | 837 | 810 | 811 | 758.93 | 699 | 819 | 1.175 | 0.233 | 0.00598911 | 0.02819578 | 0 | 0.0055 | 0.0038 | 0.0038 | TRUE | 0.0387 |
| Tmem144 | 348 | 311 | 297 | 261 | 230 | 274 | 270 | 331 | 320 | 260 | 259 | 270 | 285.21 | 307 | 263 | 0.86 | -0.217 | 0.0924458 | 0.20890558 | 0.0037 | 0.0107 | 0.0085 | 0.0085 | TRUE | 0.1354 |
| Tmem145 | 29 | 32 | 31 | 57 | 65 | 90 | 23 | 34 | 33 | 57 | 73 | 89 | 51.49 | 30 | 73 | 2.46 | 1.299 | 2.11E-05 | 0.00034856 | 0.0319 | 0.0485 | 0.0439 | 0.0439 | TRUE | 0.1354 |
| Tmem147 | 1891 | 1308 | 1414 | 1595 | 1333 | 1471 | 1469 | 1391 | 1526 | 1588 | 1504 | 1452 | 1488.2 | 1462 | 1515 | 1.036 | 0.051 | 0.47805004 | 0.64546861 | 0.0014 | 0.004 | 0.0031 | 0.0031 | TRUE | 0.0218 |
| Tmem14a | 184 | 158 | 148 | 146 | 137 | 170 | 143 | 168 | 160 | 145 | 155 | 168 | 156.39 | 157 | 156 | 0.997 | -0.004 | 0.97817155 | 0.98826649 | 0 | 0.0176 | 0.0126 | 0.0126 | TRUE | 0.0713 |
| Tmem14c | 4318 | 2924 | 3136 | 3253 | 2791 | 3325 | 3354 | 3110 | 3384 | 3238 | 3149 | 3282 | 3252.73 | 3283 | 3223 | 0.982 | -0.027 | 0.65630144 | 0.7866921 | 8.00E-04 | 0.0032 | 0.0023 | 0.0023 | TRUE | 0.0253 |
| Tmem150a | 714 | 520 | 529 | 604 | 572 | 588 | 555 | 553 | 571 | 601 | 645 | 580 | 584.23 | 560 | 609 | 1.088 | 0.122 | 0.18490432 | 0.34118739 | 0 | 0.0065 | 0.0044 | 0.0044 | TRUE | 0.0311 |
| Tmem150b | 0 | 0 | 0 | 0 | 0 | 1 | 0 | 0 | 0 | 0 | 0 | 1 | 0.16 | 0 | 0 | 2.094 | 1.067 | 0.79379329 | NA | 0 | 14.4078 | 10 | 10 | TRUE | 0.3835 |
| Tmem150c | 52 | 35 | 28 | 41 | 37 | 34 | 40 | 37 | 30 | 41 | 42 | 34 | 37.32 | 36 | 39 | 1.07 | 0.098 | 0.75812591 | 0.85640931 | 0 | 0.0659 | 0.0458 | 0.0458 | TRUE | 0.1321 |
| Tmem151a | 57 | 47 | 65 | 93 | 89 | 92 | 44 | 50 | 70 | 93 | 100 | 91 | 74.7 | 55 | 95 | 1.739 | 0.798 | 0.00104418 | 0.00746594 | 0.0106 | 0.0342 | 0.0284 | 0.0284 | TRUE | 0.4841 |
| Tmem151b | 352 | 271 | 255 | 474 | 426 | 431 | 273 | 288 | 275 | 472 | 481 | 425 | 369.11 | 279 | 459 | 1.647 | 0.72 | 1.33E-10 | 1.05E-08 | 0 | 0.0088 | 0.0062 | 0.0062 | TRUE | 0.0488 |
| Tmem154 | 4 | 5 | 4 | 6 | 9 | 5 | 3 | 5 | 4 | 6 | 10 | 5 | 5.63 | 4 | 7 | 1.663 | 0.733 | 0.36785727 | NA | 0 | 0.4231 | 0.289 | 0.289 | TRUE | 0.3641 |
| Tmem158 | 409 | 323 | 313 | 373 | 306 | 364 | 318 | 344 | 338 | 371 | 345 | 359 | 345.79 | 333 | 358 | 1.079 | 0.11 | 0.32477272 | 0.49865469 | 0 | 0.0093 | 0.0061 | 0.0061 | TRUE | 0.0201 |
| Tmem159 | 427 | 324 | 332 | 286 | 254 | 371 | 332 | 345 | 358 | 285 | 287 | 366 | 328.66 | 345 | 313 | 0.908 | -0.139 | 0.2853741 | 0.45732168 | 0.0081 | 0.0096 | 0.0092 | 0.0092 | TRUE | 0.2599 |
| Tmem160 | 1724 | 1277 | 1226 | 1679 | 1486 | 1600 | 1339 | 1358 | 1323 | 1671 | 1676 | 1579 | 1491.2 | 1340 | 1642 | 1.225 | 0.293 | 1.03E-05 | 0.00019285 | 0 | 0.004 | 0.0025 | 0.0025 | TRUE | 0.0137 |
| Tmem161a | 2852 | 2090 | 2303 | 2900 | 2533 | 2907 | 2215 | 2223 | 2485 | 2887 | 2857 | 2870 | 2589.48 | 2308 | 2871 | 1.245 | 0.316 | 1.44E-06 | 3.91E-05 | 0.0016 | 0.0033 | 0.0027 | 0.0027 | TRUE | 0.0539 |
| Tmem161b | 1537 | 1121 | 1140 | 1275 | 1172 | 1298 | 1194 | 1192 | 1230 | 1269 | 1322 | 1281 | 1248.13 | 1205 | 1291 | 1.071 | 0.099 | 0.14991856 | 0.29555951 | 0 | 0.0043 | 0.0026 | 0.0026 | TRUE | 0.0053 |
| Tmem163 | 60 | 42 | 69 | 54 | 69 | 66 | 47 | 45 | 74 | 54 | 78 | 65 | 60.41 | 55 | 66 | 1.191 | 0.252 | 0.37479189 | 0.54920038 | 0.0407 | 0.0417 | 0.0414 | 0.0414 | TRUE | 0.5831 |
| Tmem164 | 1412 | 989 | 970 | 972 | 801 | 984 | 1097 | 1052 | 1047 | 968 | 904 | 971 | 1006.29 | 1065 | 948 | 0.889 | -0.169 | 0.02597307 | 0.08362988 | 0 | 0.0048 | 0.0032 | 0.0032 | TRUE | 0.0191 |
| Tmem165 | 4507 | 3332 | 3391 | 3691 | 3374 | 3678 | 3500 | 3544 | 3659 | 3674 | 3806 | 3631 | 3635.74 | 3568 | 3704 | 1.038 | 0.054 | 0.32434086 | 0.49820569 | 3.00E-04 | 0.0031 | 0.0019 | 0.0019 | TRUE | 0.0071 |
| Tmem167 | 4059 | 2836 | 2802 | 3191 | 2567 | 2982 | 3152 | 3017 | 3023 | 3176 | 2896 | 2944 | 3034.71 | 3064 | 3005 | 0.981 | -0.028 | 0.64693908 | 0.78037811 | 0.0012 | 0.0032 | 0.0024 | 0.0024 | TRUE | 0.0302 |
| Tmem167b | 2069 | 1439 | 1437 | 1589 | 1337 | 1469 | 1607 | 1531 | 1551 | 1582 | 1508 | 1450 | 1538.03 | 1563 | 1513 | 0.968 | -0.047 | 0.49188757 | 0.65736783 | 7.00E-04 | 0.004 | 0.0027 | 0.0027 | TRUE | 0.0191 |
| Tmem168 | 2456 | 1700 | 1774 | 1799 | 1632 | 1820 | 1907 | 1808 | 1914 | 1791 | 1841 | 1797 | 1843.05 | 1876 | 1810 | 0.964 | -0.053 | 0.39729302 | 0.57070444 | 0 | 0.0037 | 0.0023 | 0.0023 | TRUE | 0.012 |
| Tmem169 | 36 | 30 | 23 | 26 | 26 | 25 | 28 | 32 | 25 | 26 | 29 | 25 | 27.43 | 28 | 27 | 0.942 | -0.086 | 0.81188761 | 0.89135487 | 0 | 0.0888 | 0.0586 | 0.0586 | TRUE | 0.0765 |
| Tmem17 | 220 | 147 | 140 | 170 | 148 | 165 | 171 | 156 | 151 | 169 | 167 | 163 | 162.89 | 159 | 166 | 1.041 | 0.057 | 0.7111232 | 0.8262229 | 0 | 0.017 | 0.0112 | 0.0112 | TRUE | 0.0459 |
| Tmem170 | 2711 | 2045 | 2202 | 2475 | 2111 | 2536 | 2106 | 2175 | 2376 | 2464 | 2381 | 2503 | 2334.19 | 2219 | 2449 | 1.105 | 0.144 | 0.03240871 | 0.09797329 | 0.0017 | 0.0034 | 0.0028 | 0.0028 | TRUE | 0.046 |
| Tmem170b | 1088 | 862 | 1009 | 1171 | 906 | 1082 | 845 | 917 | 1089 | 1166 | 1022 | 1068 | 1017.73 | 950 | 1085 | 1.144 | 0.195 | 0.05451605 | 0.14318227 | 0.0095 | 0.0048 | 0.0064 | 0.0064 | TRUE | 0.1927 |
| Tmem171 | 0 | 1 | 0 | 4 | 1 | 2 | 0 | 1 | 0 | 4 | 1 | 2 | 1.36 | 0 | 2 | 6.885 | 2.784 | 0.13799269 | NA | 0 | 1.7477 | 1.3197 | 1.3197 | TRUE | 0.519 |
| Tmem173 | 136 | 91 | 85 | 148 | 102 | 117 | 106 | 97 | 92 | 147 | 115 | 115 | 112 | 98 | 126 | 1.282 | 0.359 | 0.06542289 | 0.16336292 | 0 | 0.0236 | 0.0183 | 0.0183 | TRUE | 0.2273 |
| Tmem174 | 0 | 1 | 2 | 4 | 6 | 2 | 0 | 1 | 2 | 4 | 7 | 2 | 2.66 | 1 | 4 | 4.186 | 2.066 | 0.12638698 | NA | 0 | 0.8942 | 0.7158 | 0.7158 | TRUE | 0.3067 |
| Tmem175 | 824 | 577 | 567 | 671 | 575 | 616 | 640 | 614 | 612 | 668 | 649 | 608 | 631.69 | 622 | 642 | 1.031 | 0.044 | 0.62457487 | 0.76345237 | 0 | 0.0062 | 0.0041 | 0.0041 | TRUE | 0.0251 |
| Tmem176a | 18 | 13 | 14 | 26 | 18 | 20 | 14 | 14 | 15 | 26 | 20 | 20 | 18.14 | 14 | 22 | 1.54 | 0.623 | 0.16431809 | 0.31522329 | 0 | 0.1331 | 0.0872 | 0.0872 | TRUE | 0.1408 |
| Tmem176b | 9 | 10 | 8 | 17 | 20 | 12 | 7 | 11 | 9 | 17 | 23 | 12 | 12.93 | 9 | 17 | 1.965 | 0.974 | 0.07915414 | NA | 0 | 0.1857 | 0.136 | 0.136 | TRUE | 0.1234 |
| Tmem177 | 1332 | 1050 | 1085 | 1301 | 1129 | 1307 | 1035 | 1117 | 1171 | 1295 | 1274 | 1290 | 1196.82 | 1108 | 1286 | 1.163 | 0.218 | 0.00374486 | 0.01959917 | 0.0012 | 0.0044 | 0.0032 | 0.0032 | TRUE | 0.0417 |
| Tmem178 | 24 | 30 | 20 | 19 | 16 | 26 | 19 | 32 | 22 | 19 | 18 | 26 | 22.46 | 24 | 21 | 0.877 | -0.19 | 0.65276889 | 0.78431896 | 0 | 0.108 | 0.0832 | 0.0832 | TRUE | 0.4725 |
| Tmem178b | 29 | 22 | 16 | 27 | 31 | 28 | 23 | 23 | 17 | 27 | 35 | 28 | 25.45 | 21 | 30 | 1.408 | 0.494 | 0.19781938 | 0.35763121 | 0 | 0.0956 | 0.0656 | 0.0656 | TRUE | 0.1394 |
| Tmem179 | 136 | 105 | 79 | 158 | 134 | 142 | 106 | 112 | 85 | 157 | 151 | 140 | 125.19 | 101 | 149 | 1.479 | 0.565 | 0.00204495 | 0.01227055 | 0 | 0.0214 | 0.016 | 0.016 | TRUE | 0.1666 |
| Tmem179b | 965 | 708 | 663 | 692 | 690 | 806 | 749 | 753 | 715 | 689 | 778 | 796 | 746.8 | 739 | 754 | 1.02 | 0.028 | 0.75384814 | 0.85391912 | 0.002 | 0.0056 | 0.0044 | 0.0044 | TRUE | 0.0692 |
| Tmem18 | 1867 | 1341 | 1213 | 1133 | 1072 | 1183 | 1450 | 1426 | 1309 | 1128 | 1209 | 1168 | 1281.69 | 1395 | 1168 | 0.837 | -0.257 | 0.00055616 | 0.00467322 | 0.0014 | 0.0043 | 0.0032 | 0.0032 | TRUE | 0.0346 |
| Tmem180 | 2161 | 1622 | 2282 | 3009 | 2825 | 3452 | 1678 | 1725 | 2462 | 2995 | 3187 | 3408 | 2575.94 | 1955 | 3197 | 1.635 | 0.71 | 0.00025708 | 0.00258761 | 0.0267 | 0.0033 | 0.0098 | 0.0267 | TRUE | 0.6208 |
| Tmem181a | 893 | 753 | 764 | 801 | 690 | 809 | 694 | 801 | 824 | 797 | 778 | 799 | 782.2 | 773 | 791 | 1.026 | 0.038 | 0.68031722 | 0.80368411 | 0.0033 | 0.0055 | 0.0047 | 0.0047 | TRUE | 0.1011 |
| Tmem181b-[illegible] | 409 | 471 | 353 | 314 | 266 | 313 | 318 | 501 | 381 | 313 | 300 | 309 | 353.52 | 400 | 307 | 0.771 | -0.376 | 0.01436377 | 0.05399997 | 0.0249 | 0.0091 | 0.0141 | 0.0141 | TRUE | 0.3692 |
| Tmem181c-[illegible] | 20 | 10 | 13 | 11 | 9 | 11 | 16 | 11 | 14 | 11 | 10 | 11 | 12.03 | 14 | 11 | 0.791 | -0.338 | 0.53554703 | NA | 0 | 0.1995 | 0.1298 | 0.1298 | TRUE | 0.1305 |
| Tmem182 | 0 | 0 | 0 | 0 | 0 | 0 | 0 | 0 | 0 | 0 | 0 | 0 | 0 | 0 | 0 | NA | NA | NA | NA | NA | NA | NA | NA | NA | NA |
| Tmem183a | 6884 | 4832 | 4671 | 4876 | 4342 | 4806 | 5347 | 5140 | 5040 | 4854 | 4898 | 4744 | 5003.71 | 5176 | 4832 | 0.933 | -0.1 | 0.05898023 | 0.15151051 | 4.00E-04 | 0.0029 | 0.0018 | 0.0018 | TRUE | 0.0104 |
| Tmem184a | 71 | 56 | 86 | 59 | 69 | 79 | 55 | 60 | 93 | 59 | 78 | 78 | 70.34 | 69 | 72 | 1.039 | 0.056 | 0.83505049 | 0.90624396 | 0.0399 | 0.0361 | 0.0372 | 0.0372 | TRUE | 0.7749 |
| Tmem184b | 5405 | 4048 | 4539 | 5131 | 4644 | 5256 | 4198 | 4306 | 4898 | 5107 | 5239 | 5188 | 4822.66 | 4467 | 5178 | 1.16 | 0.214 | 0.00153589 | 0.00982212 | 0.0033 | 0.0029 | 0.0031 | 0.0031 | TRUE | 0.0861 |
| Tmem184c | 3220 | 2315 | 2177 | 2304 | 1996 | 2238 | 2501 | 2462 | 2349 | 2293 | 2252 | 2209 | 2344.43 | 2437 | 2251 | 0.923 | -0.115 | 0.05553281 | 0.14526546 | 3.00E-04 | 0.0034 | 0.0022 | 0.0022 | TRUE | 0.0121 |
| Tmem185b | 621 | 472 | 367 | 409 | 370 | 351 | 482 | 502 | 396 | 407 | 417 | 346 | 425.23 | 460 | 390 | 0.847 | -0.24 | 0.0530736 | 0.14074549 | 0.0103 | 0.008 | 0.0087 | 0.0087 | TRUE | 0.167 |
| Tmem186 | 1117 | 802 | 883 | 1046 | 864 | 1051 | 868 | 853 | 953 | 1041 | 975 | 1037 | 954.45 | 891 | 1018 | 1.143 | 0.193 | 0.01700949 | 0.06143918 | 0.0013 | 0.0049 | 0.0037 | 0.0037 | TRUE | 0.0423 |
| Tmem189 | 2096 | 1518 | 1482 | 1366 | 1277 | 1480 | 1628 | 1615 | 1599 | 1360 | 1441 | 1461 | 1517.15 | 1614 | 1421 | 0.88 | -0.185 | 0.00551056 | 0.0264392 | 0 | 0.004 | 0.0025 | 0.0025 | TRUE | 0.017 |
| Tmem19 | 2904 | 2166 | 2133 | 2430 | 2155 | 2387 | 2255 | 2304 | 2301 | 2419 | 2431 | 2356 | 2344.51 | 2287 | 2402 | 1.05 | 0.071 | 0.21674962 | 0.37953812 | 0 | 0.0034 | 0.002 | 0.002 | TRUE | 0.0034 |
| Tmem190 | 0 | 0 | 0 | 0 | 0 | 1 | 0 | 0 | 0 | 0 | 0 | 1 | 0.16 | 0 | 0 | 2.094 | 1.067 | 0.79379329 | NA | 0 | 14.4078 | 10 | 10 | TRUE | 0.3835 |
| Tmem191c | 467 | 480 | 421 | 462 | 383 | 473 | 363 | 511 | 454 | 460 | 432 | 467 | 447.73 | 443 | 453 | 1.028 | 0.039 | 0.76032331 | 0.85765141 | 0.0137 | 0.0077 | 0.0096 | 0.0096 | TRUE | 0.3056 |
| Tmem192 | 1687 | 1249 | 1312 | 1243 | 1082 | 1295 | 1310 | 1329 | 1416 | 1237 | 1221 | 1278 | 1298.44 | 1352 | 1245 | 0.922 | -0.117 | 0.09963498 | 0.220617 | 0 | 0.0043 | 0.0028 | 0.0028 | TRUE | 0.0204 |
| Tmem194 | 4114 | 3346 | 3657 | 4412 | 4038 | 4628 | 3195 | 3559 | 3946 | 4392 | 4555 | 4569 | 4035.95 | 3567 | 4505 | 1.264 | 0.338 | 9.37E-06 | 0.00017851 | 0.0054 | 0.003 | 0.0039 | 0.0039 | TRUE | 0.1052 |
| Tmem194b | 943 | 800 | 942 | 1013 | 888 | 1036 | 732 | 851 | 1016 | 1008 | 1002 | 1023 | 938.76 | 866 | 1011 | 1.169 | 0.225 | 0.03726 | 0.10886143 | 0.012 | 0.005 | 0.0074 | 0.0074 | TRUE | 0.2718 |
| Tmem196 | 0 | 0 | 0 | 0 | 0 | 0 | 0 | 0 | 0 | 0 | 0 | 0 | 0 | 0 | 0 | NA | NA | NA | NA | NA | NA | NA | NA | NA | NA |
| Tmem198 | 19 | 25 | 11 | 30 | 19 | 30 | 15 | 27 | 12 | 30 | 21 | 30 | 22.35 | 18 | 27 | 1.533 | 0.617 | 0.15999822 | 0.30945258 | 0.0444 | 0.1084 | 0.0923 | 0.0923 | TRUE | 0.896 |
| Tmem198b | 641 | 459 | 470 | 505 | 461 | 554 | 498 | 488 | 507 | 503 | 520 | 547 | 510.47 | 498 | 523 | 1.051 | 0.072 | 0.44777356 | 0.61788255 | 0 | 0.0071 | 0.0046 | 0.0046 | TRUE | 0.0188 |
| Tmem199 | 2733 | 2063 | 2409 | 2531 | 2265 | 2666 | 2123 | 2194 | 2599 | 2519 | 2555 | 2632 | 2437.09 | 2305 | 2569 | 1.115 | 0.157 | 0.05108159 | 0.13700678 | 0.0058 | 0.0034 | 0.0043 | 0.0043 | TRUE | 0.1496 |
| Tmem2 | 6228 | 4820 | 5023 | 5431 | 5111 | 5975 | 4837 | 5127 | 5420 | 5406 | 5766 | 5898 | 5408.96 | 5128 | 5690 | 1.11 | 0.151 | 0.01732387 | 0.06216672 | 0.0023 | 0.0029 | 0.0027 | 0.0027 | TRUE | 0.0305 |
| Tmem200a | 16 | 3 | 10 | 12 | 7 | 4 | 12 | 3 | 11 | 12 | 8 | 4 | 8.37 | 9 | 8 | 0.89 | -0.168 | 0.81806732 | NA | 0.2115 | 0.2857 | 0.2655 | 0.2655 | TRUE | 0.3213 |
| Tmem200b | 45 | 31 | 25 | 49 | 29 | 26 | 35 | 33 | 27 | 49 | 33 | 26 | 33.68 | 32 | 36 | 1.126 | 0.171 | 0.63198055 | 0.76915003 | 0.0356 | 0.0728 | 0.0623 | 0.0623 | TRUE | 0.583 |
| Tmem200c | 31 | 13 | 12 | 16 | 7 | 10 | 24 | 14 | 13 | 16 | 8 | 10 | 14.09 | 17 | 11 | 0.657 | -0.605 | 0.26714024 | NA | 0.0656 | 0.1706 | 0.1402 | 0.1402 | TRUE | 0.8033 |

| | | | | | | | | | | | | | | | | | | | | | | | | | |
|---|---|---|---|---|---|---|---|---|---|---|---|---|---|---|---|---|---|---|---|---|---|---|---|---|---|
| Tmem201 | 6048 | 4892 | 4872 | 4996 | 4755 | 5491 | 4697 | 5204 | 5257 | 4973 | 5364 | 5420 | 5152.54 | 5053 | 5252 | 1.04 | 0.057 | 0.38550064 | 0.55936317 | 0.0026 | 0.0029 | 0.0029 | 0.0029 | TRUE | 0.0468 |
| Tmem202 | 1 | 0 | 0 | 1 | 1 | 0 | 1 | 0 | 0 | 1 | 1 | 0 | 0.48 | 0 | 1 | 2.037 | 1.027 | 0.71241723 | NA | 0 | 4.9057 | 3.8521 | 3.8521 | TRUE | 0.2664 |
| Tmem203 | 261 | 179 | 187 | 216 | 208 | 192 | 203 | 190 | 202 | 215 | 235 | 190 | 205.68 | 198 | 213 | 1.072 | 0.101 | 0.48865476 | 0.65445488 | 0 | 0.0139 | 0.0103 | 0.0103 | TRUE | 0.1031 |
| Tmem204 | 0 | 0 | 1 | 0 | 0 | 0 | 0 | 0 | 1 | 0 | 0 | 0 | 0.18 | 0 | 0 | 0.552 | -0.857 | 0.8336548 | NA | 0 | 13.1817 | 10 | 10 | TRUE | 0.5365 |
| Tmem205 | 519 | 398 | 448 | 689 | 668 | 675 | 403 | 423 | 483 | 686 | 754 | 666 | 569.26 | 436 | 702 | 1.61 | 0.687 | 3.74E-11 | 3.34E-09 | 0.0045 | 0.0066 | 0.0059 | 0.0059 | TRUE | 0.0999 |
| Tmem206 | 788 | 671 | 628 | 831 | 712 | 832 | 612 | 714 | 678 | 827 | 803 | 821 | 742.51 | 668 | 817 | 1.227 | 0.295 | 0.00093459 | 0.00690694 | 0.0019 | 0.0056 | 0.0044 | 0.0044 | TRUE | 0.0672 |
| Tmem207 | 0 | 0 | 0 | 0 | 0 | 0 | 0 | 0 | 0 | 0 | 0 | 0 | 0 | 0 | 0 | NA | NA | NA | NA | NA | NA | NA | NA | NA | NA |
| Tmem208 | 924 | 685 | 655 | 741 | 637 | 749 | 718 | 729 | 707 | 738 | 719 | 739 | 724.76 | 718 | 732 | 1.02 | 0.029 | 0.72639784 | 0.83598639 | 0 | 0.0057 | 0.0034 | 0.0034 | TRUE | 0.0029 |
| Tmem209 | 7447 | 5553 | 6648 | 7141 | 6170 | 7585 | 5784 | 5907 | 7173 | 7108 | 6960 | 7488 | 6736.61 | 6288 | 7185 | 1.143 | 0.193 | 0.01683685 | 0.06097947 | 0.008 | 0.0028 | 0.0045 | 0.0045 | TRUE | 0.1846 |
| Tmem210 | 0 | 0 | 0 | 0 | 0 | 0 | 0 | 0 | 0 | 0 | 0 | 0 | 0 | 0 | 0 | NA | NA | NA | NA | NA | NA | NA | NA | NA | NA |
| Tmem211 | 12 | 14 | 23 | 27 | 29 | 35 | 9 | 15 | 25 | 27 | 33 | 35 | 23.86 | 16 | 32 | 1.956 | 0.968 | 0.02715687 | 0.08616053 | 0.0604 | 0.1018 | 0.0917 | 0.0917 | TRUE | 0.7514 |
| Tmem212 | 0 | 0 | 0 | 0 | 0 | 0 | 0 | 0 | 0 | 0 | 0 | 0 | 0 | 0 | 0 | NA | NA | NA | NA | NA | NA | NA | NA | NA | NA |
| Tmem213 | 36 | 44 | 53 | 61 | 66 | 65 | 28 | 47 | 57 | 61 | 74 | 64 | 55.21 | 44 | 66 | 1.528 | 0.612 | 0.04103785 | 0.11654644 | 0.0466 | 0.0454 | 0.0457 | 0.0457 | TRUE | 0.7578 |
| Tmem214 | 3034 | 2312 | 2179 | 2207 | 2132 | 2405 | 2356 | 2459 | 2351 | 2197 | 2405 | 2374 | 2357.14 | 2389 | 2325 | 0.973 | -0.039 | 0.54128469 | 0.69951013 | 0.0011 | 0.0034 | 0.0025 | 0.0025 | TRUE | 0.0285 |
| Tmem215 | 73 | 52 | 57 | 40 | 33 | 42 | 57 | 55 | 62 | 40 | 37 | 41 | 48.67 | 58 | 39 | 0.684 | -0.547 | 0.04635384 | 0.12750873 | 0 | 0.0511 | 0.033 | 0.033 | TRUE | 0.0244 |
| Tmem216 | 427 | 317 | 278 | 253 | 237 | 258 | 332 | 337 | 300 | 252 | 267 | 255 | 290.44 | 323 | 258 | 0.798 | -0.326 | 0.00738294 | 0.03306935 | 0 | 0.0106 | 0.0072 | 0.0072 | TRUE | 0.042 |
| Tmem217 | 4 | 4 | 7 | 9 | 3 | 2 | 3 | 4 | 8 | 9 | 3 | 2 | 4.87 | 5 | 5 | 0.977 | -0.034 | 0.97048469 | NA | 0.1931 | 0.4889 | 0.4005 | 0.4005 | TRUE | 1.1779 |
| Tmem218 | 358 | 266 | 243 | 291 | 250 | 284 | 278 | 283 | 262 | 290 | 282 | 280 | 279.2 | 274 | 284 | 1.035 | 0.049 | 0.68451488 | 0.80664285 | 0 | 0.0109 | 0.0069 | 0.0069 | TRUE | 0.016 |
| Tmem219 | 272 | 166 | 165 | 223 | 163 | 200 | 211 | 177 | 178 | 222 | 184 | 197 | 194.86 | 189 | 201 | 1.063 | 0.089 | 0.56369004 | 0.71807681 | 0.0053 | 0.0146 | 0.0118 | 0.0118 | TRUE | 0.1292 |
| Tmem220 | 168 | 127 | 125 | 151 | 127 | 178 | 130 | 135 | 135 | 150 | 143 | 176 | 144.95 | 133 | 156 | 1.174 | 0.232 | 0.16634199 | 0.31738434 | 0 | 0.0188 | 0.0133 | 0.0133 | TRUE | 0.1241 |
| Tmem221 | 28 | 29 | 27 | 31 | 34 | 52 | 22 | 31 | 29 | 31 | 38 | 51 | 33.71 | 27 | 40 | 1.488 | 0.573 | 0.10459548 | 0.22856085 | 0.0258 | 0.0727 | 0.0592 | 0.0592 | TRUE | 0.3079 |
| Tmem222 | 1722 | 1263 | 1451 | 1432 | 1309 | 1414 | 1337 | 1343 | 1566 | 1425 | 1477 | 1396 | 1424.07 | 1415 | 1433 | 1.013 | 0.018 | 0.81978039 | 0.8964401 | 0.0038 | 0.0041 | 0.004 | 0.004 | TRUE | 0.1021 |
| Tmem223 | 584 | 387 | 464 | 464 | 421 | 533 | 454 | 412 | 501 | 462 | 475 | 526 | 471.47 | 456 | 488 | 1.072 | 0.1 | 0.36521508 | 0.53992751 | 0.0048 | 0.0075 | 0.0066 | 0.0066 | TRUE | 0.0836 |
| Tmem225 | 1 | 0 | 0 | 0 | 0 | 0 | 1 | 0 | 0 | 0 | 0 | 0 | 0.13 | 0 | 0 | 0.552 | -0.857 | 0.8336548 | NA | 0 | 18.3121 | 10 | 10 | TRUE | 0.2566 |
| Tmem229a | 45 | 30 | 32 | 26 | 12 | 26 | 35 | 32 | 35 | 26 | 14 | 26 | 27.75 | 34 | 22 | 0.646 | -0.631 | 0.0983753 | 0.21850476 | 0 | 0.0878 | 0.0672 | 0.0672 | TRUE | 0.5503 |
| Tmem229b | 663 | 419 | 439 | 548 | 464 | 524 | 515 | 446 | 474 | 545 | 523 | 517 | 503.41 | 478 | 528 | 1.104 | 0.142 | 0.15548138 | 0.30313674 | 0.0014 | 0.0071 | 0.0052 | 0.0052 | TRUE | 0.0564 |
| Tmem230 | 2201 | 1550 | 1847 | 2002 | 1853 | 2035 | 1709 | 1649 | 1993 | 1993 | 2090 | 2009 | 1907.18 | 1784 | 2031 | 1.139 | 0.188 | 0.0189565 | 0.06647333 | 0.0048 | 0.0037 | 0.0041 | 0.0041 | TRUE | 0.1251 |
| Tmem231 | 3500 | 2707 | 2590 | 2386 | 2144 | 2414 | 2718 | 2879 | 2795 | 2375 | 2419 | 2383 | 2594.83 | 2797 | 2392 | 0.855 | -0.225 | 9.64E-05 | 0.00117512 | 0 | 0.0033 | 0.002 | 0.002 | TRUE | 0.008 |
| Tmem232 | 1 | 1 | 2 | 1 | 1 | 1 | 1 | 1 | 2 | 1 | 1 | 1 | 1.18 | 1 | 1 | 0.791 | -0.339 | 0.84019023 | NA | 0 | 2.0028 | 1.1649 | 1.1649 | TRUE | 0.1671 |
| Tmem233 | 5 | 2 | 3 | 2 | 1 | 2 | 4 | 2 | 3 | 2 | 1 | 2 | 2.39 | 3 | 2 | 0.55 | -0.863 | 0.4793951 | NA | 0 | 0.994 | 0.6148 | 0.6148 | TRUE | 0.0921 |
| Tmem234 | 2129 | 1632 | 1316 | 1479 | 1321 | 1597 | 1654 | 1736 | 1420 | 1472 | 1490 | 1576 | 1558.05 | 1603 | 1513 | 0.943 | -0.084 | 0.31756485 | 0.49149479 | 0.0049 | 0.0039 | 0.0044 | 0.0044 | TRUE | 0.118 |
| Tmem235 | 34 | 31 | 40 | 39 | 47 | 46 | 26 | 33 | 43 | 39 | 53 | 45 | 39.97 | 34 | 46 | 1.348 | 0.431 | 0.18186788 | 0.33745782 | 0.0164 | 0.0617 | 0.0494 | 0.0494 | TRUE | 0.3612 |
| Tmem236 | 8 | 0 | 2 | 2 | 1 | 0 | 6 | 0 | 2 | 2 | 1 | 0 | 1.92 | 3 | 1 | 0.362 | -1.467 | 0.36765836 | NA | 1.2423 | 1.24 | 1.2417 | 1.2417 | TRUE | 0.2391 |
| Tmem237 | 1558 | 1176 | 1245 | 1451 | 1266 | 1461 | 1210 | 1251 | 1343 | 1444 | 1428 | 1442 | 1353.17 | 1268 | 1438 | 1.135 | 0.183 | 0.01004524 | 0.04149611 | 7.00E-04 | 0.0042 | 0.0029 | 0.0029 | TRUE | 0.032 |
| Tmem238 | 197 | 155 | 140 | 166 | 159 | 174 | 153 | 165 | 151 | 165 | 179 | 172 | 164.22 | 156 | 172 | 1.101 | 0.139 | 0.36566678 | 0.54031585 | 0 | 0.0169 | 0.011 | 0.011 | TRUE | 0.023 |
| Tmem239 | 0 | 0 | 0 | 0 | 0 | 0 | 0 | 0 | 0 | 0 | 0 | 0 | 0 | 0 | 0 | NA | NA | NA | NA | NA | NA | NA | NA | NA | NA |
| Tmem240 | 4 | 4 | 9 | 10 | 6 | 4 | 3 | 4 | 10 | 10 | 7 | 4 | 6.29 | 6 | 7 | 1.233 | 0.302 | 0.70747995 | NA | 0.1272 | 0.3792 | 0.3057 | 0.3057 | TRUE | 0.4237 |
| Tmem241 | 126 | 82 | 80 | 105 | 73 | 78 | 98 | 87 | 86 | 105 | 82 | 77 | 89.21 | 90 | 88 | 0.97 | -0.043 | 0.84021512 | 0.90902996 | 0 | 0.029 | 0.0223 | 0.0223 | TRUE | 0.2635 |
| Tmem242 | 2465 | 1702 | 1616 | 1689 | 1613 | 1845 | 1914 | 1810 | 1744 | 1681 | 1820 | 1821 | 1798.44 | 1823 | 1774 | 0.972 | -0.041 | 0.56114675 | 0.71579389 | 0.0016 | 0.0037 | 0.0029 | 0.0029 | TRUE | 0.0255 |
| Tmem243 | 687 | 535 | 531 | 562 | 500 | 612 | 534 | 569 | 573 | 559 | 564 | 604 | 567.2 | 559 | 576 | 1.033 | 0.046 | 0.61778588 | 0.75859223 | 0 | 0.0066 | 0.0045 | 0.0045 | TRUE | 0.022 |
| Tmem245 | 1338 | 1137 | 1125 | 1519 | 1342 | 1561 | 1039 | 1209 | 1214 | 1512 | 1514 | 1541 | 1338.22 | 1154 | 1522 | 1.321 | 0.402 | 4.66E-07 | 1.53E-05 | 0.0032 | 0.0042 | 0.0038 | 0.0038 | TRUE | 0.0953 |
| Tmem246 | 508 | 305 | 325 | 478 | 332 | 443 | 395 | 324 | 351 | 476 | 375 | 437 | 392.88 | 357 | 429 | 1.202 | 0.266 | 0.03460414 | 0.10299813 | 0.0097 | 0.0085 | 0.0088 | 0.0088 | TRUE | 0.1397 |
| Tmem247 | 0 | 0 | 2 | 0 | 0 | 0 | 0 | 0 | 2 | 0 | 0 | 0 | 0.36 | 1 | 0 | 0.269 | -1.896 | 0.61680529 | NA | 10 | 6.5921 | 8.6623 | 8.6623 | TRUE | 0.9744 |
| Tmem248 | 6634 | 4832 | 4859 | 5111 | 4461 | 5217 | 5152 | 5140 | 5243 | 5087 | 5032 | 5150 | 5134.14 | 5178 | 5090 | 0.983 | -0.025 | 0.61669952 | 0.75810494 | 0 | 0.0029 | 0.0016 | 0.0016 | TRUE | 0.0014 |
| Tmem25 | 110 | 95 | 98 | 98 | 94 | 91 | 85 | 101 | 106 | 98 | 106 | 90 | 97.61 | 97 | 98 | 1.007 | 0.01 | 0.9590289 | 0.97739513 | 0 | 0.0267 | 0.0193 | 0.0193 | TRUE | 0.1276 |
| Tmem251 | 349 | 222 | 186 | 192 | 166 | 200 | 271 | 236 | 201 | 191 | 187 | 197 | 213.95 | 236 | 192 | 0.81 | -0.303 | 0.04577459 | 0.12630428 | 0.0083 | 0.0135 | 0.0119 | 0.0119 | TRUE | 0.1635 |
| Tmem252 | 43 | 34 | 20 | 39 | 28 | 36 | 33 | 36 | 22 | 39 | 32 | 36 | 32.85 | 30 | 36 | 1.159 | 0.213 | 0.53837451 | 0.69682033 | 0 | 0.0746 | 0.0562 | 0.0562 | TRUE | 0.3927 |
| Tmem253 | 47 | 43 | 28 | 27 | 28 | 23 | 37 | 46 | 30 | 27 | 32 | 23 | 32.27 | 38 | 27 | 0.72 | -0.474 | 0.17607181 | 0.32969924 | 0 | 0.0759 | 0.0563 | 0.0563 | TRUE | 0.2296 |
| Tmem254a | 0 | 0 | 0 | 0 | 0 | 0 | 0 | 0 | 0 | 0 | 0 | 0 | 0 | 0 | 0 | NA | NA | NA | NA | NA | NA | NA | NA | NA | NA |
| Tmem254b | 0 | 0 | 0 | 0 | 0 | 0 | 0 | 0 | 0 | 0 | 0 | 0 | 0 | 0 | 0 | NA | NA | NA | NA | NA | NA | NA | NA | NA | NA |
| Tmem254c | 0 | 0 | 0 | 0 | 0 | 0 | 0 | 0 | 0 | 0 | 0 | 0 | 0 | 0 | 0 | NA | NA | NA | NA | NA | NA | NA | NA | NA | NA |
| Tmem255a | 4 | 0 | 1 | 1 | 2 | 2 | 3 | 0 | 1 | 1 | 2 | 2 | 1.57 | 1 | 2 | 1.187 | 0.247 | 0.87366987 | NA | 0 | 1.5134 | 1.1157 | 1.1157 | TRUE | 0.6463 |
| Tmem255b | 0 | 2 | 2 | 2 | 2 | 2 | 0 | 2 | 2 | 2 | 2 | 2 | 1.75 | 1 | 2 | 1.518 | 0.602 | 0.68385382 | NA | 0 | 1.3559 | 0.971 | 0.971 | TRUE | 0.7347 |
| Tmem256 | 1961 | 1373 | 1573 | 1910 | 1577 | 1883 | 1523 | 1460 | 1697 | 1901 | 1779 | 1859 | 1703.29 | 1560 | 1846 | 1.184 | 0.244 | 0.00118087 | 0.00818141 | 0.0028 | 0.0038 | 0.0035 | 0.0035 | TRUE | 0.0695 |
| Tmem258 | 4070 | 2849 | 3019 | 3770 | 3287 | 3944 | 3161 | 3030 | 3257 | 3753 | 3708 | 3893 | 3467.16 | 3149 | 3785 | 1.202 | 0.265 | 4.08E-06 | 8.95E-05 | 5.00E-04 | 0.0031 | 0.0021 | 0.0021 | TRUE | 0.013 |
| Tmem259 | 6281 | 4837 | 4515 | 4707 | 4180 | 4888 | 4878 | 5145 | 4872 | 4685 | 4715 | 4825 | 4853.49 | 4965 | 4742 | 0.955 | -0.066 | 0.21293771 | 0.3752326 | 4.00E-04 | 0.0029 | 0.0018 | 0.0018 | TRUE | 0.0122 |
| Tmem26 | 15 | 16 | 8 | 16 | 2 | 12 | 12 | 17 | 9 | 16 | 2 | 12 | 11.22 | 13 | 10 | 0.814 | -0.296 | 0.65417527 | NA | 0.2617 | 0.2136 | 0.2249 | 0.2249 | TRUE | 0.7335 |
| Tmem27 | 0 | 0 | 0 | 0 | 0 | 0 | 0 | 0 | 0 | 0 | 0 | 0 | 0 | 0 | 0 | NA | NA | NA | NA | NA | NA | NA | NA | NA | NA |
| Tmem28 | 122 | 102 | 78 | 125 | 103 | 122 | 95 | 108 | 84 | 124 | 116 | 120 | 108.08 | 96 | 120 | 1.257 | 0.33 | 0.08535597 | 0.19737791 | 0 | 0.0244 | 0.0172 | 0.0172 | TRUE | 0.1202 |
| Tmem29 | 613 | 446 | 407 | 426 | 366 | 479 | 476 | 474 | 439 | 424 | 413 | 473 | 449.9 | 463 | 437 | 0.942 | -0.086 | 0.41527377 | 0.58799094 | 0.0013 | 0.0077 | 0.0057 | 0.0057 | TRUE | 0.0617 |
| Tmem30a | 4427 | 2946 | 3332 | 3827 | 3082 | 3608 | 3438 | 3134 | 3595 | 3809 | 3477 | 3562 | 3502.49 | 3389 | 3616 | 1.067 | 0.094 | 0.17574253 | 0.32921205 | 0.0031 | 0.0031 | 0.0032 | 0.0032 | TRUE | 0.0521 |
| Tmem30b | 415 | 345 | 312 | 348 | 296 | 301 | 322 | 367 | 337 | 346 | 334 | 297 | 333.9 | 342 | 326 | 0.954 | -0.068 | 0.57051408 | 0.72399518 | 0.0025 | 0.0095 | 0.0074 | 0.0074 | TRUE | 0.0689 |
| Tmem30c | 0 | 0 | 0 | 0 | 0 | 1 | 0 | 0 | 0 | 0 | 0 | 1 | 0.16 | 0 | 0 | 2.094 | 1.067 | 0.79379329 | NA | 0 | 14.4078 | 10 | 10 | TRUE | 0.3835 |
| Tmem33 | 2703 | 1944 | 2038 | 2029 | 1673 | 2008 | 2099 | 2068 | 2199 | 2020 | 1887 | 1982 | 2042.55 | 2122 | 1963 | 0.926 | -0.112 | 0.07953081 | 0.1883569 | 6.00E-04 | 0.0036 | 0.0024 | 0.0024 | TRUE | 0.0135 |
| Tmem35 | 95 | 70 | 57 | 46 | 42 | 36 | 74 | 74 | 62 | 46 | 47 | 36 | 56.41 | 70 | 43 | 0.611 | -0.71 | 0.0075823 | 0.03365647 | 0 | 0.0444 | 0.032 | 0.032 | TRUE | 0.1789 |
| Tmem37 | 257 | 180 | 208 | 234 | 190 | 225 | 200 | 191 | 224 | 233 | 214 | 222 | 214.14 | 205 | 223 | 1.089 | 0.123 | 0.37782003 | 0.55239174 | 0 | 0.0135 | 0.0094 | 0.0094 | TRUE | 0.0695 |
| Tmem38a | 704 | 497 | 512 | 594 | 489 | 600 | 547 | 529 | 552 | 591 | 552 | 592 | 560.51 | 543 | 578 | 1.066 | 0.093 | 0.31250646 | 0.48572717 | 0 | 0.0067 | 0.0043 | 0.0043 | TRUE | 0.0186 |
| Tmem38b | 1851 | 1286 | 1510 | 1827 | 1524 | 1817 | 1438 | 1368 | 1629 | 1819 | 1719 | 1794 | 1627.71 | 1478 | 1777 | 1.203 | 0.267 | 0.00069529 | 0.00555182 | 0.0037 | 0.0039 | 0.0038 | 0.0038 | TRUE | 0.094 |
| Tmem39a | 3724 | 2636 | 2776 | 3657 | 3057 | 3488 | 2892 | 2804 | 2995 | 3640 | 3449 | 3443 | 3203.9 | 2897 | 3511 | 1.212 | 0.277 | 2.25E-06 | 5.53E-05 | 6.00E-04 | 0.0032 | 0.0022 | 0.0022 | TRUE | 0.0127 |

| | | | | | | | | | | | | | | | | | | | | | | | | | |
|---|---|---|---|---|---|---|---|---|---|---|---|---|---|---|---|---|---|---|---|---|---|---|---|---|---|
| Tmem39b | 994 | 708 | 551 | 600 | 511 | 580 | 772 | 753 | 595 | 597 | 576 | 573 | 644.31 | 707 | 582 | 0.823 | -0.282 | 0.00989604 | 0.04106972 | 0.0089 | 0.0061 | 0.007 | 0.007 | TRUE | 0.2228 |
| Tmem40 | 387 | 265 | 274 | 220 | 204 | 229 | 301 | 282 | 296 | 219 | 230 | 226 | 258.88 | 293 | 225 | 0.768 | -0.381 | 0.00236922 | 0.01378799 | 0 | 0.0116 | 0.0074 | 0.0074 | TRUE | 0.0115 |
| Tmem41a | 1430 | 994 | 921 | 1020 | 879 | 1008 | 1111 | 1057 | 994 | 1015 | 992 | 995 | 1027.27 | 1054 | 1001 | 0.948 | -0.076 | 0.32251662 | 0.49628693 | 8.00E-04 | 0.0047 | 0.0033 | 0.0033 | TRUE | 0.0295 |
| Tmem41b | 2842 | 1955 | 1817 | 1795 | 1614 | 1853 | 2207 | 2080 | 1961 | 1787 | 1821 | 1829 | 1947.33 | 2083 | 1812 | 0.869 | -0.202 | 0.00288331 | 0.01604171 | 0.0015 | 0.0036 | 0.0028 | 0.0028 | TRUE | 0.0343 |
| Tmem42 | 48 | 37 | 34 | 31 | 22 | 27 | 37 | 39 | 37 | 31 | 25 | 27 | 32.61 | 38 | 28 | 0.728 | -0.458 | 0.17046531 | 0.32211273 | 0 | 0.0751 | 0.0488 | 0.0488 | TRUE | 0.0764 |
| Tmem43 | 2112 | 1571 | 1742 | 1969 | 1646 | 1945 | 1640 | 1671 | 1880 | 1960 | 1857 | 1920 | 1821.3 | 1730 | 1912 | 1.106 | 0.146 | 0.04608106 | 0.12698058 | 0.0024 | 0.0037 | 0.0033 | 0.0033 | TRUE | 0.0679 |
| Tmem44 | 466 | 390 | 394 | 388 | 349 | 377 | 362 | 415 | 425 | 386 | 394 | 372 | 392.33 | 401 | 384 | 0.961 | -0.058 | 0.60644122 | 0.7508545 | 0.0022 | 0.0085 | 0.0064 | 0.0064 | TRUE | 0.0887 |
| Tmem45a | 434 | 317 | 340 | 421 | 387 | 436 | 337 | 337 | 367 | 419 | 437 | 430 | 387.86 | 347 | 429 | 1.236 | 0.306 | 0.00402378 | 0.02071752 | 0 | 0.0085 | 0.0056 | 0.0056 | TRUE | 0.0274 |
| Tmem45b | 47 | 26 | 17 | 22 | 28 | 34 | 37 | 28 | 18 | 22 | 32 | 34 | 28.26 | 28 | 29 | 1.042 | 0.06 | 0.87835878 | 0.93196497 | 0.0471 | 0.0863 | 0.0754 | 0.0754 | TRUE | 0.2173 |
| Tmem47 | 1648 | 1172 | 1209 | 1495 | 1179 | 1464 | 1280 | 1247 | 1305 | 1488 | 1330 | 1445 | 1349.07 | 1277 | 1421 | 1.113 | 0.155 | 0.0335413 | 0.10058872 | 0.0011 | 0.0042 | 0.0031 | 0.0031 | TRUE | 0.0371 |
| Tmem48 | 5348 | 3815 | 3968 | 4396 | 3869 | 4521 | 4154 | 4058 | 4281 | 4376 | 4365 | 4463 | 4282.72 | 4164 | 4401 | 1.057 | 0.08 | 0.12934977 | 0.26608271 | 2.00E-04 | 0.003 | 0.0018 | 0.0018 | TRUE | 0.0072 |
| Tmem5 | 1155 | 833 | 755 | 805 | 673 | 814 | 897 | 886 | 815 | 801 | 759 | 804 | 826.96 | 866 | 788 | 0.91 | -0.137 | 0.09851337 | 0.21870948 | 7.00E-04 | 0.0053 | 0.0037 | 0.0037 | TRUE | 0.0312 |
| Tmem50a | 2815 | 2129 | 2393 | 2759 | 2590 | 2994 | 2186 | 2265 | 2582 | 2746 | 2922 | 2956 | 2609.42 | 2344 | 2875 | 1.227 | 0.295 | 8.35E-05 | 0.00104432 | 0.0042 | 0.0033 | 0.0037 | 0.0037 | TRUE | 0.0947 |
| Tmem50b | 1733 | 1231 | 1422 | 1390 | 1207 | 1364 | 1346 | 1309 | 1534 | 1384 | 1362 | 1346 | 1380.23 | 1396 | 1364 | 0.977 | -0.033 | 0.67297656 | 0.79910725 | 0.003 | 0.0041 | 0.0037 | 0.0037 | TRUE | 0.0877 |
| Tmem51 | 738 | 527 | 543 | 560 | 533 | 611 | 573 | 561 | 586 | 557 | 601 | 603 | 580.25 | 573 | 587 | 1.024 | 0.035 | 0.70351919 | 0.82071497 | 0 | 0.0065 | 0.0042 | 0.0042 | TRUE | 0.0236 |
| Tmem51as1 | 35 | 43 | 32 | 47 | 34 | 44 | 27 | 46 | 35 | 47 | 38 | 43 | 39.34 | 36 | 43 | 1.209 | 0.274 | 0.39580822 | 0.56928589 | 0.0142 | 0.0627 | 0.0492 | 0.0492 | TRUE | 0.3841 |
| Tmem52 | 20 | 21 | 12 | 22 | 21 | 15 | 16 | 22 | 13 | 22 | 24 | 15 | 18.54 | 17 | 20 | 1.188 | 0.249 | 0.58837967 | 0.73749983 | 0 | 0.1303 | 0.0982 | 0.0982 | TRUE | 0.3518 |
| Tmem52b | 0 | 0 | 0 | 1 | 0 | 0 | 0 | 0 | 0 | 1 | 0 | 0 | 0.17 | 0 | 0 | 2.094 | 1.067 | 0.79379329 | NA | 0 | 14.2885 | 10 | 10 | TRUE | 0.3906 |
| Tmem53 | 100 | 81 | 89 | 84 | 82 | 96 | 78 | 86 | 96 | 84 | 93 | 95 | 88.46 | 87 | 91 | 1.046 | 0.065 | 0.75563357 | 0.85511255 | 0 | 0.0292 | 0.0203 | 0.0203 | TRUE | 0.0884 |
| Tmem54 | 53 | 33 | 26 | 39 | 38 | 38 | 41 | 35 | 28 | 39 | 43 | 38 | 37.25 | 35 | 40 | 1.133 | 0.18 | 0.57278658 | 0.72540132 | 0 | 0.066 | 0.0466 | 0.0466 | TRUE | 0.216 |
| Tmem55a | 1602 | 1193 | 1421 | 1478 | 1284 | 1522 | 1244 | 1269 | 1533 | 1471 | 1448 | 1502 | 1411.43 | 1349 | 1474 | 1.094 | 0.13 | 0.13862898 | 0.27970413 | 0.0062 | 0.0041 | 0.0048 | 0.0048 | TRUE | 0.1701 |
| Tmem55b | 4358 | 3269 | 3854 | 4451 | 4014 | 4481 | 3385 | 3477 | 4158 | 4431 | 4528 | 4423 | 4067.08 | 3673 | 4461 | 1.215 | 0.281 | 0.00032817 | 0.00311459 | 0.0063 | 0.003 | 0.0042 | 0.0042 | TRUE | 0.1615 |
| Tmem56 | 93 | 77 | 62 | 80 | 71 | 59 | 72 | 82 | 67 | 80 | 80 | 58 | 73.17 | 74 | 73 | 0.985 | -0.022 | 0.92698341 | 0.95969289 | 0 | 0.0348 | 0.0271 | 0.0271 | TRUE | 0.2806 |
| Tmem57 | 4819 | 3576 | 3273 | 3624 | 3086 | 3751 | 3743 | 3804 | 3532 | 3607 | 3481 | 3703 | 3644.91 | 3693 | 3597 | 0.974 | -0.038 | 0.51834613 | 0.68007354 | 8.00E-04 | 0.0031 | 0.0022 | 0.0022 | TRUE | 0.0174 |
| Tmem59 | 4220 | 2938 | 3359 | 3554 | 3130 | 3405 | 3277 | 3125 | 3624 | 3538 | 3531 | 3361 | 3409.47 | 3342 | 3477 | 1.04 | 0.057 | 0.40329949 | 0.57626308 | 0.0028 | 0.0031 | 0.0031 | 0.0031 | TRUE | 0.0653 |
| Tmem59l | 346 | 248 | 253 | 396 | 373 | 422 | 269 | 264 | 273 | 394 | 421 | 417 | 339.51 | 269 | 411 | 1.528 | 0.612 | 4.53E-08 | 1.94E-06 | 0 | 0.0094 | 0.006 | 0.006 | TRUE | 0.0141 |
| Tmem60 | 1134 | 804 | 981 | 1134 | 946 | 1129 | 881 | 855 | 1058 | 1129 | 1067 | 1114 | 1017.48 | 931 | 1103 | 1.186 | 0.246 | 0.00778367 | 0.03429517 | 0.006 | 0.0048 | 0.0052 | 0.0052 | TRUE | 0.1661 |
| Tmem62 | 150 | 89 | 79 | 97 | 66 | 99 | 116 | 95 | 85 | 97 | 74 | 98 | 94.19 | 99 | 90 | 0.904 | -0.146 | 0.50418557 | 0.66824225 | 0.014 | 0.0276 | 0.0237 | 0.0237 | TRUE | 0.2637 |
| Tmem63a | 521 | 416 | 486 | 531 | 489 | 593 | 405 | 442 | 524 | 529 | 552 | 585 | 506.18 | 457 | 555 | 1.218 | 0.284 | 0.01274341 | 0.04946809 | 0.008 | 0.0071 | 0.0074 | 0.0074 | TRUE | 0.19 |
| Tmem63b | 4764 | 3640 | 3824 | 3918 | 3520 | 3894 | 3700 | 3872 | 4126 | 3900 | 3971 | 3844 | 3902.13 | 3899 | 3905 | 1.002 | 0.003 | 0.96393086 | 0.98011409 | 0.0014 | 0.003 | 0.0024 | 0.0024 | TRUE | 0.0314 |
| Tmem63c | 274 | 216 | 231 | 278 | 293 | 307 | 213 | 230 | 249 | 277 | 331 | 303 | 267.02 | 231 | 304 | 1.318 | 0.398 | 0.00276859 | 0.01551212 | 0.0035 | 0.0113 | 0.009 | 0.009 | TRUE | 0.0672 |
| Tmem64 | 15142 | 9728 | 8124 | 9190 | 7744 | 8099 | 11760 | 10348 | 8766 | 9148 | 8736 | 7995 | 9458.69 | 10291 | 8626 | 0.838 | -0.255 | 0.00602149 | 0.02829228 | 0.0127 | 0.0027 | 0.0061 | 0.0061 | TRUE | 0.181 |
| Tmem65 | 2035 | 1495 | 1650 | 1726 | 1542 | 1780 | 1580 | 1590 | 1780 | 1718 | 1740 | 1757 | 1694.3 | 1650 | 1738 | 1.054 | 0.076 | 0.28504418 | 0.45710047 | 0.0015 | 0.0038 | 0.003 | 0.003 | TRUE | 0.0564 |
| Tmem66 | 4028 | 3089 | 3509 | 4273 | 3745 | 4426 | 3128 | 3286 | 3786 | 4253 | 4225 | 4369 | 3841.25 | 3400 | 4282 | 1.26 | 0.334 | 6.91E-06 | 0.00013845 | 0.0049 | 0.003 | 0.0037 | 0.0037 | TRUE | 0.1195 |
| Tmem67 | 434 | 331 | 270 | 287 | 271 | 296 | 337 | 352 | 291 | 286 | 306 | 292 | 310.68 | 327 | 295 | 0.9 | -0.152 | 0.21569015 | 0.37831136 | 0.002 | 0.0101 | 0.0077 | 0.0077 | TRUE | 0.0976 |
| Tmem68 | 747 | 492 | 553 | 654 | 568 | 618 | 580 | 523 | 597 | 651 | 641 | 610 | 600.33 | 567 | 634 | 1.118 | 0.161 | 0.08688937 | 0.19985675 | 0.001 | 0.0064 | 0.0047 | 0.0047 | TRUE | 0.0505 |
| Tmem69 | 954 | 673 | 621 | 686 | 625 | 688 | 741 | 716 | 670 | 683 | 705 | 679 | 698.99 | 709 | 689 | 0.97 | -0.043 | 0.61519991 | 0.75699128 | 0 | 0.0058 | 0.0039 | 0.0039 | TRUE | 0.0268 |
| Tmem70 | 1878 | 1393 | 1493 | 1577 | 1239 | 1464 | 1459 | 1482 | 1611 | 1570 | 1398 | 1445 | 1493.98 | 1517 | 1471 | 0.971 | -0.043 | 0.5697313 | 0.72332292 | 0.0026 | 0.004 | 0.0035 | 0.0035 | TRUE | 0.0417 |
| Tmem71 | 0 | 0 | 0 | 0 | 0 | 0 | 0 | 0 | 0 | 0 | 0 | 0 | 0 | 0 | 0 | NA | NA | NA | NA | NA | NA | NA | NA | NA | NA |
| Tmem72 | 2 | 1 | 0 | 1 | 1 | 1 | 2 | 1 | 0 | 1 | 1 | 1 | 0.95 | 1 | 1 | 1.147 | 0.198 | 0.91694087 | NA | 0 | 2.4852 | 1.5573 | 1.5573 | TRUE | 0.2767 |
| Tmem74 | 47 | 48 | 38 | 46 | 40 | 40 | 37 | 51 | 41 | 46 | 45 | 39 | 43.16 | 43 | 43 | 1.019 | 0.027 | 0.92706078 | 0.95969289 | 0 | 0.0573 | 0.0407 | 0.0407 | TRUE | 0.2107 |
| Tmem74b | 69 | 51 | 67 | 85 | 65 | 89 | 54 | 54 | 72 | 85 | 73 | 88 | 70.99 | 60 | 82 | 1.373 | 0.457 | 0.05674829 | 0.14774196 | 0 | 0.0358 | 0.0271 | 0.0271 | TRUE | 0.2604 |
| Tmem79 | 341 | 219 | 170 | 250 | 202 | 245 | 265 | 233 | 183 | 249 | 228 | 242 | 233.3 | 227 | 240 | 1.051 | 0.071 | 0.643575 | 0.77809422 | 0.0136 | 0.0126 | 0.0129 | 0.0129 | TRUE | 0.3039 |
| Tmem8 | 2298 | 1677 | 1554 | 1359 | 1371 | 1575 | 1785 | 1784 | 1677 | 1353 | 1547 | 1555 | 1616.58 | 1749 | 1485 | 0.849 | -0.237 | 0.00191078 | 0.01166628 | 0.0027 | 0.0039 | 0.0036 | 0.0036 | TRUE | 0.0734 |
| Tmem80 | 33 | 14 | 16 | 14 | 16 | 9 | 26 | 15 | 17 | 14 | 18 | 9 | 16.44 | 19 | 14 | 0.696 | -0.523 | 0.29589556 | NA | 0.0403 | 0.1466 | 0.1171 | 0.1171 | TRUE | 0.3997 |
| Tmem81 | 378 | 304 | 270 | 313 | 263 | 312 | 294 | 323 | 291 | 312 | 297 | 308 | 304.08 | 303 | 306 | 1.01 | 0.015 | 0.90214337 | 0.94574851 | 0 | 0.0102 | 0.0068 | 0.0068 | TRUE | 0.0386 |
| Tmem82 | 153 | 135 | 120 | 131 | 122 | 135 | 119 | 144 | 129 | 130 | 138 | 133 | 132.2 | 131 | 134 | 1.027 | 0.038 | 0.82552273 | 0.89994633 | 0 | 0.0204 | 0.014 | 0.014 | TRUE | 0.0768 |
| Tmem86a | 287 | 236 | 272 | 326 | 291 | 337 | 223 | 251 | 293 | 325 | 328 | 333 | 292.14 | 256 | 329 | 1.289 | 0.367 | 0.00563289 | 0.02690873 | 0.0059 | 0.0105 | 0.0092 | 0.0092 | TRUE | 0.1833 |
| Tmem86b | 80 | 45 | 43 | 67 | 40 | 53 | 62 | 48 | 46 | 67 | 45 | 52 | 53.42 | 52 | 55 | 1.045 | 0.064 | 0.82042837 | 0.89651557 | 0.0159 | 0.0468 | 0.0381 | 0.0381 | TRUE | 0.3089 |
| Tmem87a | 4501 | 3470 | 3438 | 3686 | 3168 | 3710 | 3496 | 3691 | 3710 | 3669 | 3574 | 3662 | 3633.59 | 3632 | 3635 | 1.001 | 0.002 | 0.97136687 | 0.98452324 | 4.00E-04 | 0.0031 | 0.0019 | 0.0019 | TRUE | 0.0135 |
| Tmem87b | 1778 | 1436 | 1667 | 1817 | 1651 | 1827 | 1381 | 1527 | 1799 | 1809 | 1862 | 1804 | 1696.95 | 1569 | 1825 | 1.164 | 0.22 | 0.01619289 | 0.0591024 | 0.0084 | 0.0038 | 0.0054 | 0.0054 | TRUE | 0.197 |
| Tmem88 | 271 | 207 | 151 | 185 | 179 | 241 | 210 | 220 | 163 | 184 | 202 | 238 | 202.93 | 198 | 208 | 1.05 | 0.07 | 0.6707894 | 0.79737134 | 0.0158 | 0.0141 | 0.0146 | 0.0146 | TRUE | 0.2474 |
| Tmem88b | 8 | 9 | 8 | 2 | 7 | 7 | 6 | 10 | 9 | 2 | 8 | 7 | 6.87 | 8 | 6 | 0.688 | -0.539 | 0.47917627 | NA | 0 | 0.3474 | 0.2645 | 0.2645 | TRUE | 0.7289 |
| Tmem89 | 0 | 0 | 0 | 0 | 1 | 0 | 0 | 0 | 0 | 0 | 1 | 0 | 0.19 | 0 | 0 | 2.094 | 1.067 | 0.79379329 | NA | 0 | 12.608 | 10 | 10 | TRUE | 0.5075 |
| Tmem8b | 454 | 401 | 493 | 667 | 516 | 641 | 353 | 427 | 532 | 664 | 582 | 633 | 531.65 | 437 | 626 | 1.438 | 0.525 | 9.91E-05 | 0.00120243 | 0.0205 | 0.0069 | 0.0111 | 0.0111 | TRUE | 0.4169 |
| Tmem8c | 0 | 0 | 0 | 1 | 0 | 1 | 0 | 0 | 0 | 1 | 0 | 1 | 0.33 | 0 | 1 | 3.763 | 1.912 | 0.57887588 | NA | 0 | 7.1752 | 6.7756 | 6.7756 | TRUE | 0.2094 |
| Tmem9 | 1864 | 1448 | 1578 | 1635 | 1468 | 1709 | 1448 | 1540 | 1703 | 1627 | 1656 | 1687 | 1610.19 | 1564 | 1657 | 1.061 | 0.085 | 0.26205785 | 0.43236672 | 0.0026 | 0.0039 | 0.0036 | 0.0036 | TRUE | 0.0727 |
| Tmem91 | 48 | 26 | 44 | 54 | 45 | 62 | 37 | 28 | 47 | 54 | 51 | 61 | 46.36 | 37 | 55 | 1.477 | 0.563 | 0.06034 | 0.1538697 | 0.0069 | 0.0536 | 0.0424 | 0.0424 | TRUE | 0.2595 |
| Tmem92 | 560 | 430 | 192 | 214 | 211 | 172 | 435 | 457 | 207 | 213 | 238 | 170 | 286.72 | 366 | 207 | 0.564 | -0.826 | 0.02790876 | 0.08786341 | 0.0978 | 0.0107 | 0.0345 | 0.0978 | TRUE | 1.6433 |
| Tmem95 | 9 | 4 | 2 | 6 | 7 | 9 | 7 | 4 | 2 | 6 | 8 | 9 | 6.03 | 4 | 8 | 1.654 | 0.726 | 0.36163156 | NA | 0 | 0.3957 | 0.2829 | 0.2829 | TRUE | 0.4935 |
| Tmem97 | 6771 | 4938 | 5396 | 5872 | 5177 | 6014 | 5259 | 5252 | 5822 | 5845 | 5840 | 5937 | 5659.23 | 5444 | 5874 | 1.079 | 0.11 | 0.06242698 | 0.15800759 | 0.0016 | 0.0028 | 0.0023 | 0.0023 | TRUE | 0.0446 |
| Tmem98 | 801 | 639 | 724 | 718 | 726 | 782 | 622 | 680 | 781 | 715 | 819 | 772 | 731.44 | 694 | 769 | 1.109 | 0.149 | 0.14959412 | 0.2950811 | 0.0076 | 0.0057 | 0.0063 | 0.0063 | TRUE | 0.1406 |
| Tmem9b | 706 | 461 | 487 | 625 | 473 | 533 | 548 | 490 | 525 | 622 | 534 | 526 | 541 | 521 | 561 | 1.074 | 0.104 | 0.32062958 | 0.49452451 | 0.0043 | 0.0068 | 0.006 | 0.006 | TRUE | 0.1091 |
| Tmf1 | 4742 | 3509 | 3643 | 3804 | 3254 | 3491 | 3683 | 3732 | 3931 | 3786 | 3671 | 3446 | 3708.28 | 3782 | 3634 | 0.961 | -0.057 | 0.34960318 | 0.52416639 | 0.0013 | 0.0031 | 0.0024 | 0.0024 | TRUE | 0.0252 |
| Tmie | 51 | 44 | 56 | 92 | 60 | 79 | 40 | 47 | 60 | 92 | 68 | 78 | 64.01 | 49 | 79 | 1.631 | 0.706 | 0.00727697 | 0.0326869 | 0.017 | 0.0395 | 0.0334 | 0.0334 | TRUE | 0.3302 |
| Tmigd1 | 2 | 1 | 2 | 0 | 0 | 1 | 2 | 1 | 2 | 0 | 0 | 1 | 0.96 | 2 | 0 | 0.233 | -2.1 | 0.30489818 | NA | 0 | 2.4703 | 1.7158 | 1.7158 | TRUE | 0.3835 |
| Tmlhe | 227 | 133 | 141 | 163 | 115 | 132 | 176 | 141 | 152 | 162 | 130 | 130 | 148.7 | 156 | 141 | 0.896 | -0.158 | 0.36885051 | 0.54353597 | 0.0087 | 0.0184 | 0.0155 | 0.0155 | TRUE | 0.1876 |
| Tmod1 | 17 | 15 | 8 | 11 | 10 | 19 | 13 | 16 | 9 | 11 | 11 | 19 | 13.13 | 13 | 14 | 1.085 | 0.118 | 0.82823339 | NA | 0 | 0.1829 | 0.1373 | 0.1373 | TRUE | 0.4708 |
| Tmod2 | 788 | 650 | 619 | 860 | 709 | 816 | 612 | 691 | 668 | 856 | 800 | 806 | 738.78 | 657 | 821 | 1.251 | 0.323 | 0.00023723 | 0.00243309 | 0.0014 | 0.0056 | 0.0042 | 0.0042 | TRUE | 0.0454 |

| | | | | | | | | | | | | | | | | | | | | | | | | | |
|---|---|---|---|---|---|---|---|---|---|---|---|---|---|---|---|---|---|---|---|---|---|---|---|---|---|
| Tmod3 | 5487 | 3823 | 3885 | 4365 | 3808 | 4245 | 4262 | 4066 | 4192 | 4345 | 4296 | 4190 | 4225.18 | 4173 | 4277 | 1.025 | 0.035 | 0.50824565 | 0.67156838 | 2.00E-04 | 0.003 | 0.0018 | 0.0018 | TRUE | 0.0061 |
| Tmod4 | 29 | 21 | 9 | 23 | 22 | 13 | 23 | 22 | 10 | 23 | 25 | 13 | 19.19 | 18 | 20 | 1.097 | 0.133 | 0.78336254 | 0.87248506 | 0.0931 | 0.126 | 0.1173 | 0.1173 | TRUE | 0.7834 |
| Tmpo | 16647 | 12005 | 11573 | 12190 | 10510 | 11955 | 12929 | 12770 | 12487 | 12134 | 11856 | 11801 | 12329.58 | 12729 | 11930 | 0.937 | -0.094 | 0.04272156 | 0.12008244 | 1.00E-04 | 0.0026 | 0.0015 | 0.0015 | TRUE | 0.0034 |
| Tmppe | 2053 | 1608 | 1341 | 1323 | 1210 | 1325 | 1594 | 1710 | 1447 | 1317 | 1365 | 1308 | 1456.95 | 1584 | 1330 | 0.839 | -0.253 | 0.00122943 | 0.00840378 | 0.0031 | 0.0041 | 0.0037 | 0.0037 | TRUE | 0.0671 |
| Tmprss11a | 0 | 0 | 0 | 0 | 0 | 0 | 0 | 0 | 0 | 0 | 0 | 0 | 0 | 0 | 0 | NA | NA | NA | NA | NA | NA | NA | NA | NA | NA |
| Tmprss11bnl | 3 | 0 | 0 | 0 | 0 | 0 | 2 | 0 | 0 | 0 | 0 | 0 | 0.39 | 1 | 0 | 0.239 | -2.067 | 0.57190649 | NA | 10 | 6.1057 | 8.0283 | 8.0283 | TRUE | 1.4408 |
| Tmprss11c | 0 | 0 | 0 | 0 | 0 | 0 | 0 | 0 | 0 | 0 | 0 | 0 | 0 | 0 | 0 | NA | NA | NA | NA | NA | NA | NA | NA | NA | NA |
| Tmprss11d | 35 | 32 | 11 | 21 | 5 | 15 | 27 | 34 | 12 | 21 | 6 | 15 | 19.07 | 24 | 14 | 0.569 | -0.814 | 0.13387532 | 0.27245907 | 0.2383 | 0.1267 | 0.1556 | 0.1556 | TRUE | 0.428 |
| Tmprss11e | 7 | 4 | 1 | 6 | 3 | 4 | 5 | 4 | 1 | 6 | 3 | 4 | 4.01 | 3 | 4 | 1.214 | 0.279 | 0.77346616 | NA | 0 | 0.5931 | 0.4303 | 0.4303 | TRUE | 0.5127 |
| Tmprss11f | 0 | 1 | 2 | 0 | 0 | 0 | 0 | 1 | 2 | 0 | 0 | 0 | 0.54 | 1 | 0 | 0.18 | -2.47 | 0.39300405 | NA | 0 | 4.4164 | 4.5678 | 4.5678 | TRUE | 0.0584 |
| Tmprss11g | 0 | 0 | 0 | 0 | 0 | 0 | 0 | 0 | 0 | 0 | 0 | 0 | 0 | 0 | 0 | NA | NA | NA | NA | NA | NA | NA | NA | NA | NA |
| Tmprss12 | 41 | 25 | 30 | 42 | 17 | 28 | 32 | 27 | 32 | 42 | 19 | 28 | 29.9 | 30 | 30 | 0.979 | -0.03 | 0.93640307 | 0.96471383 | 0.0419 | 0.0817 | 0.071 | 0.071 | TRUE | 0.3927 |
| Tmprss13 | 11 | 13 | 24 | 17 | 14 | 17 | 9 | 14 | 26 | 17 | 16 | 17 | 16.29 | 16 | 17 | 1.043 | 0.06 | 0.90753831 | 0.96054762 | 0.0977 | 0.1479 | 0.1333 | 0.1333 | TRUE | 0.6378 |
| Tmprss15 | 0 | 0 | 0 | 1 | 0 | 0 | 0 | 0 | 0 | 1 | 0 | 0 | 0.17 | 0 | 0 | 2.094 | 1.067 | 0.79379329 | NA | 0 | 14.2885 | 10 | 10 | TRUE | 0.3906 |
| Tmprss2 | 31 | 22 | 20 | 23 | 15 | 23 | 24 | 23 | 22 | 23 | 17 | 23 | 21.93 | 23 | 21 | 0.907 | -0.141 | 0.72737609 | 0.83661541 | 0 | 0.1105 | 0.0728 | 0.0728 | TRUE | 0.1353 |
| Tmprss3 | 42 | 14 | 13 | 23 | 17 | 19 | 33 | 15 | 14 | 23 | 19 | 19 | 20.39 | 21 | 20 | 0.971 | -0.042 | 0.92831732 | 0.96054762 | 0.0872 | 0.1186 | 0.1092 | 0.1092 | TRUE | 1.7334 |
| Tmprss4 | 5 | 2 | 3 | 2 | 1 | 1 | 4 | 2 | 3 | 2 | 1 | 1 | 2.23 | 3 | 1 | 0.442 | -1.178 | 0.36073579 | NA | 0 | 1.0673 | 0.6658 | 0.6658 | TRUE | 0.1032 |
| Tmprss5 | 9 | 6 | 13 | 7 | 3 | 4 | 7 | 6 | 14 | 7 | 3 | 4 | 6.95 | 9 | 5 | 0.529 | -0.919 | 0.22827157 | NA | 0 | 0.3434 | 0.2581 | 0.2581 | TRUE | 0.6626 |
| Tmprss6 | 17 | 11 | 13 | 12 | 19 | 15 | 13 | 12 | 14 | 12 | 21 | 15 | 14.52 | 13 | 16 | 1.231 | 0.3 | 0.55484508 | NA | 0 | 0.1657 | 0.1159 | 0.1159 | TRUE | 0.3711 |
| Tmprss7 | 3 | 2 | 3 | 2 | 0 | 3 | 2 | 2 | 3 | 2 | 0 | 3 | 2.11 | 2 | 2 | 0.658 | -0.603 | 0.65303076 | NA | 0 | 1.1269 | 0.8019 | 0.8019 | TRUE | 0.4894 |
| Tmprss9 | 5 | 4 | 2 | 3 | 5 | 7 | 4 | 4 | 2 | 3 | 6 | 7 | 4.31 | 3 | 5 | 1.498 | 0.583 | 0.52736987 | NA | 0 | 0.5529 | 0.3729 | 0.3729 | TRUE | 0.2965 |
| Tmsb10 | 18536 | 13964 | 14439 | 13220 | 12514 | 15400 | 14396 | 14853 | 15580 | 13159 | 14117 | 15202 | 14551.24 | 14943 | 14159 | 0.948 | -0.078 | 0.22596168 | 0.3909066 | 0.0033 | 0.0026 | 0.0029 | 0.0029 | TRUE | 0.0509 |
| Tmsb15a | 15 | 10 | 11 | 13 | 3 | 11 | 12 | 11 | 12 | 13 | 3 | 11 | 10.22 | 12 | 9 | 0.803 | -0.316 | 0.61596235 | NA | 0.0074 | 0.2343 | 0.1868 | 0.1868 | TRUE | 0.8544 |
| Tmsb15b1 | 89 | 58 | 58 | 64 | 67 | 63 | 69 | 62 | 63 | 64 | 76 | 62 | 65.81 | 65 | 67 | 1.037 | 0.052 | 0.82743769 | 0.90114042 | 0 | 0.0384 | 0.026 | 0.026 | TRUE | 0.1009 |
| Tmsb15b2 | 52 | 35 | 32 | 43 | 39 | 37 | 40 | 37 | 35 | 43 | 44 | 37 | 39.24 | 37 | 41 | 1.095 | 0.131 | 0.66845066 | 0.79617284 | 0 | 0.0628 | 0.0415 | 0.0415 | TRUE | 0.074 |
| Tmsb15l | 22 | 14 | 27 | 22 | 31 | 29 | 17 | 15 | 29 | 22 | 35 | 29 | 24.43 | 20 | 29 | 1.404 | 0.49 | 0.2424429 | 0.41019448 | 0.0436 | 0.0994 | 0.0846 | 0.0846 | TRUE | 0.446 |
| Tmsb4x | 10505 | 7079 | 6578 | 6473 | 5502 | 6598 | 8159 | 7530 | 7098 | 6443 | 6207 | 6513 | 6991.59 | 7596 | 6388 | 0.841 | -0.25 | 7.30E-05 | 0.00094469 | 0.0026 | 0.0028 | 0.0027 | 0.0027 | TRUE | 0.0519 |
| Tmtc1 | 284 | 186 | 224 | 268 | 223 | 247 | 221 | 198 | 242 | 267 | 252 | 244 | 237.05 | 220 | 254 | 1.155 | 0.208 | 0.12843689 | 0.26486056 | 0 | 0.0124 | 0.0092 | 0.0092 | TRUE | 0.0798 |
| Tmtc2 | 160 | 141 | 139 | 147 | 135 | 153 | 124 | 150 | 150 | 146 | 152 | 151 | 145.65 | 141 | 150 | 1.064 | 0.09 | 0.59102416 | 0.73920352 | 0 | 0.0187 | 0.0133 | 0.0133 | TRUE | 0.1309 |
| Tmtc3 | 4545 | 3112 | 2956 | 3129 | 2597 | 3010 | 3530 | 3310 | 3189 | 3115 | 2930 | 2971 | 3174.2 | 3343 | 3005 | 0.898 | -0.154 | 0.01393797 | 0.0529194 | 0.0014 | 0.0032 | 0.0025 | 0.0025 | TRUE | 0.0296 |
| Tmtc4 | 1498 | 1098 | 1315 | 1358 | 1192 | 1297 | 1163 | 1168 | 1419 | 1352 | 1345 | 1280 | 1287.84 | 1250 | 1326 | 1.061 | 0.086 | 0.3339308 | 0.50810999 | 0.0058 | 0.0043 | 0.0049 | 0.0049 | TRUE | 0.1652 |
| Tmub1 | 198 | 114 | 160 | 158 | 131 | 170 | 154 | 121 | 173 | 157 | 148 | 168 | 153.42 | 149 | 158 | 1.056 | 0.079 | 0.65188526 | 0.78383074 | 0.0099 | 0.0179 | 0.0157 | 0.0157 | TRUE | 0.2656 |
| Tmub2 | 1235 | 948 | 902 | 1082 | 955 | 1002 | 959 | 1008 | 973 | 1077 | 1077 | 989 | 1014.05 | 980 | 1048 | 1.069 | 0.096 | 0.21074966 | 0.37273345 | 6.00E-04 | 0.0048 | 0.0033 | 0.0033 | TRUE | 0.0291 |
| Tmx1 | 4571 | 3296 | 3336 | 3431 | 2860 | 3252 | 3550 | 3506 | 3600 | 3415 | 3226 | 3210 | 3417.89 | 3552 | 3284 | 0.925 | -0.113 | 0.04384129 | 0.12236352 | 4.00E-04 | 0.0031 | 0.002 | 0.002 | TRUE | 0.015 |
| Tmx2 | 4645 | 3406 | 3902 | 4594 | 3987 | 4525 | 3608 | 3623 | 4210 | 4573 | 4498 | 4467 | 4163.03 | 3814 | 4513 | 1.184 | 0.243 | 0.00049023 | 0.00425074 | 0.0038 | 0.003 | 0.0033 | 0.0033 | TRUE | 0.1 |
| Tmx3 | 4596 | 3180 | 3043 | 3072 | 2585 | 2851 | 3570 | 3383 | 3283 | 3058 | 2916 | 2814 | 3170.63 | 3412 | 2929 | 0.858 | -0.221 | 0.00041276 | 0.00372572 | 0.0013 | 0.0032 | 0.0025 | 0.0025 | TRUE | 0.0203 |
| Tmx4 | 1791 | 1350 | 1362 | 1393 | 1157 | 1269 | 1391 | 1436 | 1470 | 1387 | 1305 | 1253 | 1373.51 | 1432 | 1315 | 0.919 | -0.122 | 0.08823108 | 0.20181091 | 0.001 | 0.0042 | 0.003 | 0.003 | TRUE | 0.0278 |
| Tnc | 7 | 5 | 2 | 3 | 3 | 8 | 5 | 5 | 2 | 3 | 3 | 8 | 4.53 | 4 | 5 | 1.099 | 0.136 | 0.88238151 | NA | 0 | 0.5256 | 0.3869 | 0.3869 | TRUE | 0.6695 |
| Tnf | 0 | 0 | 1 | 0 | 0 | 0 | 0 | 0 | 1 | 0 | 0 | 0 | 0.18 | 0 | 0 | 0.552 | -0.857 | 0.8336548 | NA | 0 | 13.1817 | 10 | 10 | TRUE | 0.5365 |
| Tnfaip1 | 4926 | 3332 | 3213 | 3432 | 2872 | 3416 | 3826 | 3544 | 3467 | 3416 | 3240 | 3372 | 3477.51 | 3612 | 3343 | 0.925 | -0.113 | 0.0677848 | 0.16750067 | 0.0013 | 0.0031 | 0.0024 | 0.0024 | TRUE | 0.0332 |
| Tnfaip2 | 57 | 47 | 55 | 72 | 59 | 86 | 44 | 50 | 59 | 72 | 67 | 85 | 62.79 | 51 | 75 | 1.463 | 0.549 | 0.03056032 | 0.093794 | 0 | 0.0402 | 0.03 | 0.03 | TRUE | 0.154 |
| Tnfaip3 | 223 | 190 | 232 | 215 | 204 | 250 | 173 | 202 | 250 | 214 | 230 | 247 | 219.43 | 208 | 230 | 1.11 | 0.15 | 0.3489107 | 0.52351256 | 0.0154 | 0.0132 | 0.0139 | 0.0139 | TRUE | 0.3325 |
| Tnfaip6 | 36 | 14 | 27 | 16 | 27 | 19 | 28 | 15 | 29 | 16 | 30 | 19 | 22.85 | 24 | 22 | 0.896 | -0.159 | 0.71917176 | 0.83150918 | 0.0722 | 0.1061 | 0.097 | 0.097 | TRUE | 0.6476 |
| Tnfaip8 | 758 | 585 | 602 | 722 | 586 | 654 | 589 | 622 | 650 | 719 | 661 | 646 | 647.64 | 620 | 675 | 1.09 | 0.125 | 0.17447868 | 0.32770351 | 0.0014 | 0.0061 | 0.0045 | 0.0045 | TRUE | 0.0382 |
| Tnfaip8l1 | 122 | 95 | 110 | 137 | 105 | 125 | 95 | 101 | 119 | 136 | 118 | 123 | 115.45 | 105 | 126 | 1.208 | 0.273 | 0.14698564 | 0.29136488 | 0 | 0.023 | 0.0168 | 0.0168 | TRUE | 0.128 |
| Tnfaip8l2 | 42 | 22 | 27 | 30 | 29 | 28 | 33 | 23 | 29 | 30 | 33 | 28 | 29.23 | 28 | 30 | 1.052 | 0.073 | 0.83711338 | 0.90711611 | 0 | 0.0835 | 0.0568 | 0.0568 | TRUE | 0.1465 |
| Tnfaip8l3 | 26 | 23 | 34 | 39 | 43 | 50 | 20 | 24 | 37 | 39 | 49 | 49 | 36.34 | 27 | 46 | 1.698 | 0.764 | 0.02619533 | 0.08411813 | 0.021 | 0.0676 | 0.0557 | 0.0557 | TRUE | 0.5791 |
| Tnfrsf10b | 3044 | 2365 | 2803 | 3181 | 2647 | 3140 | 2364 | 2516 | 3024 | 3166 | 2986 | 3100 | 2859.38 | 2635 | 3084 | 1.172 | 0.228 | 0.00829448 | 0.0359906 | 0.0085 | 0.0033 | 0.005 | 0.005 | TRUE | 0.2023 |
| Tnfrsf11a | 142 | 102 | 101 | 127 | 101 | 107 | 110 | 108 | 109 | 126 | 114 | 106 | 112.29 | 109 | 115 | 1.055 | 0.078 | 0.67461614 | 0.80024529 | 0 | 0.0235 | 0.0158 | 0.0158 | TRUE | 0.0732 |
| Tnfrsf11b | 1 | 1 | 1 | 0 | 0 | 0 | 1 | 1 | 1 | 0 | 0 | 0 | 0.49 | 1 | 0 | 0.196 | -2.347 | 0.38405676 | NA | 0 | 4.8735 | 3.7415 | 3.7415 | TRUE | 0.0718 |
| Tnfrsf12a | 383 | 269 | 301 | 338 | 282 | 357 | 297 | 286 | 325 | 336 | 318 | 352 | 319.23 | 303 | 335 | 1.11 | 0.151 | 0.20259772 | 0.36405949 | 0 | 0.0099 | 0.0069 | 0.0069 | TRUE | 0.0431 |
| Tnfrsf13b | 12 | 14 | 8 | 10 | 10 | 8 | 9 | 15 | 9 | 10 | 11 | 8 | 10.33 | 11 | 10 | 0.889 | -0.17 | 0.77625408 | NA | 0 | 0.2319 | 0.1596 | 0.1596 | TRUE | 0.3424 |
| Tnfrsf13c | 50 | 36 | 38 | 54 | 35 | 45 | 39 | 38 | 41 | 54 | 39 | 44 | 42.63 | 39 | 46 | 1.168 | 0.225 | 0.44822932 | 0.61829183 | 0 | 0.058 | 0.0397 | 0.0397 | TRUE | 0.1787 |
| Tnfrsf14 | 0 | 1 | 3 | 5 | 6 | 4 | 0 | 1 | 3 | 5 | 7 | 4 | 3.33 | 1 | 5 | 3.875 | 1.954 | 0.10269186 | NA | 0 | 0.7136 | 0.5769 | 0.5769 | TRUE | 0.7746 |
| Tnfrsf17 | 5 | 2 | 2 | 4 | 4 | 3 | 4 | 2 | 2 | 4 | 5 | 3 | 3.27 | 3 | 4 | 1.371 | 0.456 | 0.65920148 | NA | 0 | 0.7271 | 0.4597 | 0.4597 | TRUE | 0.2237 |
| Tnfrsf18 | 2 | 2 | 5 | 2 | 2 | 1 | 2 | 2 | 5 | 2 | 2 | 1 | 2.38 | 3 | 2 | 0.583 | -0.778 | 0.53598501 | NA | 0 | 0.9962 | 0.6773 | 0.6773 | TRUE | 0.5744 |
| Tnfrsf19 | 1254 | 889 | 841 | 966 | 762 | 995 | 974 | 946 | 907 | 962 | 860 | 982 | 938.39 | 942 | 935 | 0.992 | -0.012 | 0.88517717 | 0.93620226 | 0.002 | 0.005 | 0.0039 | 0.0039 | TRUE | 0.0572 |
| Tnfrsf1a | 595 | 448 | 484 | 539 | 432 | 591 | 462 | 477 | 522 | 537 | 487 | 583 | 511.36 | 487 | 536 | 1.103 | 0.141 | 0.18249279 | 0.33809067 | 0.0041 | 0.0071 | 0.0061 | 0.0061 | TRUE | 0.0711 |
| Tnfrsf1b | 1681 | 1250 | 1595 | 1905 | 1773 | 1929 | 1306 | 1330 | 1721 | 1896 | 2000 | 1904 | 1692.79 | 1452 | 1933 | 1.333 | 0.414 | 2.84E-05 | 0.00044328 | 0.0124 | 0.0038 | 0.0065 | 0.0065 | TRUE | 0.3134 |
| Tnfrsf21 | 1347 | 1040 | 907 | 1096 | 870 | 1049 | 1046 | 1106 | 979 | 1091 | 981 | 1036 | 1039.83 | 1044 | 1036 | 0.993 | -0.01 | 0.90081641 | 0.94546695 | 0.0022 | 0.0047 | 0.0038 | 0.0038 | TRUE | 0.0341 |
| Tnfrsf22 | 901 | 653 | 596 | 842 | 792 | 871 | 700 | 695 | 643 | 838 | 893 | 860 | 771.47 | 679 | 864 | 1.27 | 0.345 | 3.71E-05 | 0.00055308 | 0 | 0.0055 | 0.0037 | 0.0037 | TRUE | 0.0247 |
| Tnfrsf23 | 99 | 75 | 47 | 85 | 61 | 74 | 77 | 80 | 51 | 85 | 69 | 73 | 72.31 | 69 | 76 | 1.089 | 0.123 | 0.62104547 | 0.76089515 | 0.0184 | 0.0352 | 0.0308 | 0.0308 | TRUE | 0.4564 |
| Tnfrsf25 | 101 | 102 | 90 | 89 | 87 | 92 | 78 | 108 | 97 | 89 | 98 | 91 | 93.6 | 94 | 93 | 0.982 | -0.026 | 0.90067545 | 0.94539307 | 0 | 0.0277 | 0.0214 | 0.0214 | TRUE | 0.2473 |
| Tnfrsf26 | 11 | 6 | 9 | 15 | 7 | 11 | 9 | 6 | 10 | 15 | 8 | 11 | 9.72 | 8 | 11 | 1.371 | 0.456 | 0.46115264 | NA | 0 | 0.2462 | 0.171 | 0.171 | TRUE | 0.2314 |
| Tnfrsf4 | 18 | 13 | 14 | 15 | 8 | 10 | 14 | 14 | 15 | 15 | 9 | 10 | 12.79 | 14 | 11 | 0.792 | -0.337 | 0.52790641 | NA | 0 | 0.1877 | 0.126 | 0.126 | TRUE | 0.3095 |
| Tnfrsf8 | 64 | 59 | 49 | 69 | 76 | 76 | 50 | 63 | 53 | 69 | 86 | 75 | 65.8 | 55 | 77 | 1.391 | 0.476 | 0.0517439 | 0.13822713 | 0 | 0.0384 | 0.0276 | 0.0276 | TRUE | 0.1198 |
| Tnfrsf9 | 48 | 38 | 37 | 63 | 38 | 58 | 37 | 40 | 40 | 63 | 43 | 57 | 46.74 | 39 | 54 | 1.391 | 0.476 | 0.09988402 | 0.22082633 | 0 | 0.0531 | 0.0386 | 0.0386 | TRUE | 0.2634 |
| Tnfsf10 | 0 | 1 | 0 | 0 | 0 | 0 | 0 | 1 | 0 | 0 | 0 | 0 | 0.18 | 0 | 0 | 0.552 | -0.857 | 0.8336548 | NA | 0 | 13.3714 | 10 | 10 | TRUE | 0.5216 |
| Tnfsf11 | 46 | 35 | 18 | 25 | 23 | 24 | 36 | 37 | 19 | 25 | 26 | 24 | 27.82 | 31 | 25 | 0.802 | -0.319 | 0.40760857 | 0.58038944 | 0 | 0.0876 | 0.0704 | 0.0704 | TRUE | 0.6497 |

| | | | | | | | | | | | | | | | | | | | | | | | | | |
|---|---|---|---|---|---|---|---|---|---|---|---|---|---|---|---|---|---|---|---|---|---|---|---|---|---|
| Tnfsf12 | 435 | 332 | 396 | 489 | 478 | 540 | 338 | 353 | 427 | 487 | 539 | 533 | 446.22 | 373 | 520 | 1.397 | 0.483 | 3.33E-05 | 0.00050654 | 0.0068 | 0.0077 | 0.0075 | 0.0075 | TRUE | 0.1839 |
| Tnfsf13 | 23 | 17 | 9 | 27 | 24 | 24 | 18 | 18 | 10 | 27 | 27 | 24 | 20.55 | 15 | 26 | 1.685 | 0.753 | 0.08393361 | 0.19532124 | 0 | 0.1178 | 0.0854 | 0.0854 | TRUE | 0.4162 |
| Tnfsf13b | 13 | 15 | 8 | 13 | 17 | 5 | 10 | 16 | 9 | 13 | 19 | 5 | 11.96 | 12 | 12 | 1.064 | 0.089 | 0.88443281 | NA | 0.1559 | 0.2007 | 0.1885 | 0.1885 | TRUE | 0.2495 |
| Tnfsf14 | 0 | 0 | 0 | 0 | 0 | 0 | 0 | 0 | 0 | 0 | 0 | 0 | 0 | 0 | 0 | NA | NA | NA | NA | NA | NA | NA | NA | NA | NA |
| Tnfsf15 | 1 | 0 | 0 | 1 | 2 | 1 | 1 | 0 | 0 | 1 | 2 | 1 | 0.84 | 0 | 1 | 4.069 | 2.025 | 0.34772519 | NA | 0 | 2.8378 | 1.9813 | 1.9813 | TRUE | 0.2566 |
| Tnfsf18 | 11 | 3 | 6 | 4 | 4 | 5 | 9 | 3 | 6 | 4 | 5 | 5 | 5.27 | 6 | 5 | 0.727 | -0.459 | 0.58285838 | NA | 0 | 0.4519 | 0.3096 | 0.3096 | TRUE | 0.2737 |
| Tnfsf4 | 1 | 5 | 3 | 5 | 2 | 4 | 1 | 5 | 3 | 5 | 2 | 4 | 3.42 | 3 | 4 | 1.241 | 0.312 | 0.77076346 | NA | 0 | 0.6956 | 0.5281 | 0.5281 | TRUE | 0.3361 |
| Tnfsf8 | 0 | 0 | 0 | 0 | 0 | 0 | 0 | 0 | 0 | 0 | 0 | 0 | 0 | 0 | 0 | NA | NA | NA | NA | NA | NA | NA | NA | NA | NA |
| Tnfsf9 | 65 | 44 | 48 | 89 | 60 | 92 | 50 | 47 | 52 | 89 | 68 | 91 | 66.03 | 50 | 83 | 1.661 | 0.732 | 0.00303781 | 0.01668691 | 0 | 0.0383 | 0.0279 | 0.0279 | TRUE | 0.215 |
| Tnik | 1022 | 655 | 715 | 727 | 605 | 729 | 794 | 697 | 771 | 724 | 683 | 720 | 731.29 | 754 | 709 | 0.939 | -0.091 | 0.30570004 | 0.47836334 | 0.0015 | 0.0057 | 0.0043 | 0.0043 | TRUE | 0.0514 |
| Tnip1 | 3963 | 3032 | 3456 | 3295 | 3223 | 3901 | 3078 | 3225 | 3729 | 3280 | 3636 | 3851 | 3466.43 | 3344 | 3589 | 1.074 | 0.103 | 0.22450786 | 0.3889515 | 0.0079 | 0.0031 | 0.0049 | 0.0049 | TRUE | 0.1229 |
| Tnip2 | 860 | 723 | 674 | 616 | 541 | 624 | 668 | 769 | 727 | 613 | 610 | 616 | 667.28 | 721 | 613 | 0.851 | -0.232 | 0.01064027 | 0.04319267 | 0.0015 | 0.006 | 0.0044 | 0.0044 | TRUE | 0.0528 |
| Tnip3 | 0 | 3 | 2 | 1 | 1 | 3 | 0 | 3 | 2 | 1 | 1 | 3 | 1.74 | 2 | 2 | 0.991 | -0.013 | 0.99315971 | NA | 0 | 1.3653 | 1.0916 | 1.0916 | TRUE | 0.7244 |
| Tnk1 | 919 | 810 | 945 | 862 | 817 | 980 | 714 | 862 | 1020 | 858 | 922 | 967 | 890.35 | 865 | 916 | 1.061 | 0.085 | 0.46653569 | 0.63548982 | 0.0166 | 0.0051 | 0.0087 | 0.0087 | TRUE | 0.2131 |
| Tnk2 | 7487 | 5840 | 5558 | 5836 | 5446 | 6458 | 5815 | 6212 | 5997 | 5809 | 6144 | 6375 | 6058.61 | 6008 | 6109 | 1.017 | 0.024 | 0.67318152 | 0.79921783 | 0.0012 | 0.0028 | 0.0023 | 0.0023 | TRUE | 0.0227 |
| Tnks | 8771 | 6785 | 6956 | 7729 | 7124 | 8042 | 6812 | 7217 | 7505 | 7693 | 8037 | 7939 | 7533.9 | 7178 | 7890 | 1.099 | 0.137 | 0.01456735 | 0.05459327 | 0.0012 | 0.0027 | 0.0021 | 0.0021 | TRUE | 0.0246 |
| Tnks1bp1 | 1371 | 1058 | 928 | 886 | 835 | 979 | 1065 | 1125 | 1001 | 882 | 942 | 966 | 996.97 | 1064 | 930 | 0.874 | -0.194 | 0.01701448 | 0.06143918 | 0.0017 | 0.0048 | 0.0038 | 0.0038 | TRUE | 0.0303 |
| Tnks2 | 32342 | 23683 | 26502 | 28142 | 25023 | 28869 | 25119 | 25191 | 28595 | 28013 | 28229 | 28498 | 27274.1 | 26302 | 28247 | 1.074 | 0.103 | 0.0898061 | 0.20436214 | 0.0028 | 0.0025 | 0.0026 | 0.0026 | TRUE | 0.0711 |
| Tnmd | 0 | 0 | 0 | 0 | 0 | 0 | 0 | 0 | 0 | 0 | 0 | 0 | 0 | 0 | 0 | NA | NA | NA | NA | NA | NA | NA | NA | NA | NA |
| Tnn | 7 | 10 | 9 | 6 | 8 | 6 | 5 | 11 | 10 | 6 | 9 | 6 | 7.78 | 9 | 7 | 0.818 | -0.29 | 0.67541399 | NA | 0 | 0.3069 | 0.2136 | 0.2136 | TRUE | 0.4244 |
| Tnnc1 | 16 | 11 | 11 | 16 | 16 | 22 | 12 | 12 | 12 | 16 | 18 | 22 | 15.28 | 12 | 19 | 1.546 | 0.628 | 0.19663602 | NA | 0 | 0.1575 | 0.1026 | 0.1026 | TRUE | 0.1191 |
| Tnnc2 | 8 | 4 | 3 | 8 | 7 | 6 | 6 | 4 | 3 | 8 | 8 | 6 | 5.91 | 4 | 7 | 1.56 | 0.642 | 0.41117646 | NA | 0 | 0.4031 | 0.2647 | 0.2647 | TRUE | 0.2477 |
| Tnni1 | 63 | 38 | 59 | 66 | 47 | 71 | 49 | 40 | 64 | 66 | 53 | 70 | 56.97 | 51 | 63 | 1.239 | 0.309 | 0.25908487 | 0.42899676 | 0.0145 | 0.044 | 0.0363 | 0.0363 | TRUE | 0.3558 |
| Tnni2 | 6 | 3 | 8 | 4 | 3 | 5 | 5 | 3 | 9 | 4 | 3 | 5 | 4.8 | 6 | 4 | 0.754 | -0.408 | 0.64192351 | NA | 0 | 0.4965 | 0.3411 | 0.3411 | TRUE | 0.4931 |
| Tnni3 | 68 | 70 | 78 | 103 | 90 | 95 | 53 | 74 | 84 | 103 | 102 | 94 | 84.88 | 70 | 100 | 1.421 | 0.507 | 0.0282628 | 0.08861178 | 0.0158 | 0.0303 | 0.0264 | 0.0264 | TRUE | 0.5028 |
| Tnni3k | 0 | 0 | 0 | 0 | 0 | 0 | 0 | 0 | 0 | 0 | 0 | 0 | 0 | 0 | 0 | NA | NA | NA | NA | NA | NA | NA | NA | NA | NA |
| Tnnt1 | 344 | 266 | 271 | 390 | 287 | 372 | 267 | 283 | 292 | 388 | 324 | 367 | 320.29 | 281 | 360 | 1.285 | 0.361 | 0.00305096 | 0.01673988 | 0.0023 | 0.0098 | 0.0076 | 0.0076 | TRUE | 0.0846 |
| Tnnt2 | 14 | 17 | 2 | 12 | 11 | 13 | 11 | 18 | 2 | 12 | 12 | 13 | 11.38 | 10 | 12 | 1.192 | 0.253 | 0.69839723 | NA | 0.2518 | 0.2106 | 0.2201 | 0.2201 | TRUE | 0.2641 |
| Tnnt3 | 3 | 5 | 2 | 0 | 0 | 2 | 2 | 5 | 2 | 0 | 0 | 2 | 1.96 | 3 | 1 | 0.208 | -2.262 | 0.16199287 | NA | 0 | 1.2095 | 0.9729 | 0.9729 | TRUE | 0.9733 |
| Tnp1 | 5 | 3 | 0 | 0 | 1 | 1 | 4 | 3 | 0 | 0 | 1 | 1 | 1.53 | 2 | 1 | 0.29 | -1.783 | 0.31581897 | NA | 0 | 1.5499 | 1.3331 | 1.3331 | TRUE | 0.7263 |
| Tnp2 | 3 | 2 | 2 | 0 | 0 | 2 | 2 | 2 | 2 | 0 | 0 | 2 | 1.43 | 2 | 1 | 0.306 | -1.709 | 0.33012839 | NA | 0 | 1.658 | 1.2355 | 1.2355 | TRUE | 0.9734 |
| Tnpo1 | 22731 | 16080 | 16062 | 18211 | 15938 | 18081 | 17654 | 17104 | 17331 | 18127 | 17980 | 17849 | 17674.12 | 17363 | 17985 | 1.036 | 0.051 | 0.25086347 | 0.41967867 | 1.00E-04 | 0.0026 | 0.0013 | 0.0013 | TRUE | 0.0027 |
| Tnpo2 | 18920 | 13459 | 13924 | 13719 | 11930 | 13318 | 14694 | 14316 | 15024 | 13656 | 13458 | 13147 | 14049.25 | 14678 | 13420 | 0.914 | -0.129 | 0.00656365 | 0.0302042 | 4.00E-04 | 0.0026 | 0.0016 | 0.0016 | TRUE | 0.0056 |
| Tnpo3 | 15273 | 10963 | 11126 | 12517 | 11254 | 12980 | 11862 | 11661 | 12005 | 12459 | 12696 | 12813 | 12249.38 | 11843 | 12656 | 1.069 | 0.096 | 0.03572441 | 0.10547593 | 1.00E-04 | 0.0026 | 0.0014 | 0.0014 | TRUE | 0.0023 |
| Tnr | 2 | 1 | 0 | 0 | 0 | 1 | 2 | 1 | 0 | 0 | 0 | 1 | 0.6 | 1 | 0 | 0.416 | -1.265 | 0.62184281 | NA | 0 | 3.948 | 3.1959 | 3.1959 | TRUE | 0.3835 |
| Tnrc18 | 2242 | 1864 | 1819 | 2426 | 2224 | 2513 | 1741 | 1983 | 1963 | 2415 | 2509 | 2481 | 2181.85 | 1896 | 2468 | 1.304 | 0.382 | 4.49E-08 | 1.93E-06 | 0.0023 | 0.0035 | 0.0031 | 0.0031 | TRUE | 0.0635 |
| Tnrc6a | 6372 | 4741 | 4618 | 4875 | 4454 | 4900 | 4949 | 5043 | 4983 | 4853 | 5025 | 4837 | 4948.13 | 4992 | 4905 | 0.983 | -0.025 | 0.61583741 | 0.75749645 | 0 | 0.0029 | 0.0016 | 0.0016 | TRUE | 0.0055 |
| Tnrc6b | 3526 | 2333 | 2307 | 2837 | 2431 | 2873 | 2738 | 2482 | 2489 | 2824 | 2742 | 2836 | 2685.3 | 2570 | 2801 | 1.089 | 0.123 | 0.05366458 | 0.14156329 | 0.0014 | 0.0033 | 0.0026 | 0.0026 | TRUE | 0.0411 |
| Tnrc6c | 5592 | 4404 | 4039 | 4726 | 4624 | 5217 | 4343 | 4684 | 4358 | 4704 | 5216 | 5150 | 4742.7 | 4462 | 5023 | 1.126 | 0.171 | 0.00672496 | 0.03080763 | 0.0022 | 0.0029 | 0.0027 | 0.0027 | TRUE | 0.0378 |
| Tns1 | 4123 | 3139 | 2965 | 3464 | 3382 | 3712 | 3202 | 3339 | 3199 | 3448 | 3815 | 3664 | 3444.65 | 3247 | 3642 | 1.122 | 0.166 | 0.00640549 | 0.02961942 | 0.0011 | 0.0031 | 0.0024 | 0.0024 | TRUE | 0.0266 |
| Tns3 | 10963 | 7654 | 6806 | 7084 | 6606 | 8083 | 8514 | 8141 | 7344 | 7051 | 7452 | 7979 | 7747.07 | 8000 | 7494 | 0.937 | -0.094 | 0.18102056 | 0.33640968 | 0.0045 | 0.0027 | 0.0035 | 0.0035 | TRUE | 0.0627 |
| Tns4 | 54 | 50 | 61 | 71 | 73 | 87 | 42 | 53 | 66 | 71 | 82 | 86 | 66.64 | 54 | 80 | 1.497 | 0.582 | 0.02264146 | 0.07554025 | 0.0135 | 0.038 | 0.0314 | 0.0314 | TRUE | 0.3387 |
| Tnxb | 39 | 26 | 29 | 46 | 45 | 33 | 30 | 28 | 31 | 46 | 51 | 33 | 36.39 | 30 | 43 | 1.442 | 0.528 | 0.10864658 | 0.23507199 | 0 | 0.0675 | 0.0499 | 0.0499 | TRUE | 0.357 |
| Tob1 | 2373 | 1950 | 2067 | 2163 | 1900 | 2267 | 1843 | 2074 | 2230 | 2153 | 2143 | 2238 | 2113.63 | 2049 | 2178 | 1.064 | 0.09 | 0.24823824 | 0.4171745 | 0.0041 | 0.0035 | 0.0039 | 0.0039 | TRUE | 0.0963 |
| Tob2 | 1331 | 980 | 849 | 805 | 764 | 826 | 1034 | 1042 | 916 | 801 | 862 | 815 | 911.79 | 997 | 826 | 0.827 | -0.273 | 0.00125036 | 0.00849164 | 0.0022 | 0.005 | 0.0041 | 0.0041 | TRUE | 0.0592 |
| Toe1 | 1331 | 903 | 709 | 658 | 672 | 700 | 1034 | 961 | 765 | 655 | 758 | 691 | 810.55 | 920 | 701 | 0.761 | -0.394 | 0.00051337 | 0.00439381 | 0.0136 | 0.0054 | 0.008 | 0.008 | TRUE | 0.2551 |
| Tollip | 4546 | 3368 | 3808 | 4353 | 3691 | 4214 | 3531 | 3582 | 4109 | 4333 | 4164 | 4160 | 3979.78 | 3741 | 4219 | 1.128 | 0.174 | 0.01208704 | 0.04761853 | 0.0036 | 0.003 | 0.0032 | 0.0032 | TRUE | 0.0896 |
| Tom1 | 1270 | 890 | 806 | 865 | 801 | 910 | 986 | 947 | 870 | 861 | 904 | 898 | 910.94 | 934 | 888 | 0.949 | -0.076 | 0.35544892 | 0.53000655 | 0.0014 | 0.005 | 0.0038 | 0.0038 | TRUE | 0.0429 |
| Tom1l1 | 2140 | 1637 | 1652 | 1670 | 1488 | 1667 | 1662 | 1741 | 1782 | 1662 | 1679 | 1646 | 1695.39 | 1728 | 1662 | 0.962 | -0.056 | 0.38936337 | 0.5633074 | 0 | 0.0038 | 0.0024 | 0.0024 | TRUE | 0.0143 |
| Tom1l2 | 1307 | 945 | 1005 | 1133 | 1044 | 1106 | 1015 | 1005 | 1084 | 1128 | 1178 | 1092 | 1083.66 | 1035 | 1133 | 1.094 | 0.13 | 0.08561474 | 0.19785729 | 6.00E-04 | 0.0046 | 0.0032 | 0.0032 | TRUE | 0.0203 |
| Tomm20 | 20097 | 14742 | 15379 | 17237 | 15106 | 16577 | 15608 | 15681 | 16594 | 17158 | 17041 | 16364 | 16407.68 | 15961 | 16854 | 1.056 | 0.079 | 0.11964161 | 0.25135735 | 8.00E-04 | 0.0026 | 0.0018 | 0.0018 | TRUE | 0.0147 |
| Tomm20l | 23 | 14 | 12 | 11 | 11 | 10 | 18 | 15 | 13 | 11 | 12 | 10 | 13.16 | 15 | 11 | 0.721 | -0.472 | 0.36739621 | NA | 0 | 0.1826 | 0.1196 | 0.1196 | TRUE | 0.1303 |
| Tomm22 | 6883 | 4946 | 4855 | 5341 | 4665 | 5287 | 5346 | 5261 | 5239 | 5316 | 5263 | 5219 | 5273.89 | 5282 | 5266 | 0.997 | -0.004 | 0.92681868 | 0.95969289 | 0 | 0.0029 | 0.0015 | 0.0015 | TRUE | 0.0014 |
| Tomm34 | 5410 | 4182 | 4643 | 4081 | 4010 | 4785 | 4202 | 4448 | 5010 | 4062 | 4524 | 4724 | 4494.88 | 4553 | 4437 | 0.975 | -0.037 | 0.64270071 | 0.7773516 | 0.0068 | 0.003 | 0.0044 | 0.0044 | TRUE | 0.0935 |
| Tomm40 | 11147 | 7872 | 6602 | 7107 | 6206 | 6941 | 8657 | 8373 | 7124 | 7074 | 7001 | 6852 | 7513.57 | 8051 | 6976 | 0.866 | -0.207 | 0.00462262 | 0.0230937 | 0.0051 | 0.0027 | 0.0037 | 0.0037 | TRUE | 0.1237 |
| Tomm40l | 1693 | 1294 | 1175 | 1375 | 1240 | 1490 | 1315 | 1376 | 1268 | 1369 | 1399 | 1471 | 1366.25 | 1320 | 1413 | 1.071 | 0.099 | 0.16659774 | 0.31770268 | 7.00E-04 | 0.0042 | 0.0029 | 0.0029 | TRUE | 0.0164 |
| Tomm5 | 3070 | 2076 | 2026 | 2206 | 1921 | 2264 | 2384 | 2208 | 2186 | 2196 | 2167 | 2235 | 2229.41 | 2259 | 2199 | 0.973 | -0.04 | 0.52837502 | 0.68886838 | 9.00E-04 | 0.0035 | 0.0025 | 0.0025 | TRUE | 0.0292 |
| Tomm6 | 4433 | 3307 | 3219 | 3430 | 3020 | 3712 | 3443 | 3518 | 3473 | 3414 | 3407 | 3664 | 3486.54 | 3478 | 3495 | 1.005 | 0.008 | 0.89577024 | 0.94272381 | 6.00E-04 | 0.0031 | 0.0021 | 0.0021 | TRUE | 0.0219 |
| Tomm7 | 5002 | 3713 | 3903 | 4422 | 3804 | 4274 | 3885 | 3949 | 4211 | 4402 | 4291 | 4219 | 4159.61 | 4015 | 4304 | 1.072 | 0.101 | 0.07810702 | 0.18587806 | 8.00E-04 | 0.003 | 0.0021 | 0.0021 | TRUE | 0.0221 |
| Tomm70a | 24755 | 17918 | 16961 | 19086 | 16027 | 18346 | 19226 | 19059 | 18301 | 18998 | 18080 | 18110 | 18629.12 | 18862 | 18396 | 0.975 | -0.036 | 0.46160745 | 0.63095143 | 7.00E-04 | 0.0026 | 0.0017 | 0.0017 | TRUE | 0.01 |
| Tomt | 1 | 0 | 0 | 0 | 0 | 0 | 1 | 0 | 0 | 0 | 0 | 0 | 0.13 | 0 | 0 | 0.552 | -0.857 | 0.8336548 | NA | 0 | 18.3121 | 10 | 10 | TRUE | 0.2566 |
| Tonsl | 3344 | 2642 | 2498 | 2450 | 2230 | 2666 | 2597 | 2810 | 2695 | 2439 | 2516 | 2632 | 2614.81 | 2701 | 2529 | 0.937 | -0.094 | 0.13394978 | 0.27257179 | 0.0012 | 0.0033 | 0.0025 | 0.0025 | TRUE | 0.0156 |
| Top1 | 11488 | 8091 | 7551 | 7928 | 6600 | 7853 | 8922 | 8606 | 8147 | 7892 | 7445 | 7752 | 8127.52 | 8558 | 7696 | 0.899 | -0.153 | 0.00613071 | 0.02867352 | 0.0013 | 0.0027 | 0.0021 | 0.0021 | TRUE | 0.0215 |
| Top1mt | 1150 | 919 | 977 | 1036 | 966 | 1154 | 893 | 978 | 1054 | 1031 | 1090 | 1139 | 1030.84 | 975 | 1087 | 1.116 | 0.159 | 0.06569136 | 0.1637818 | 0.0038 | 0.0047 | 0.0044 | 0.0044 | TRUE | 0.0673 |
| Top2a | 82905 | 61722 | 68699 | 79577 | 70984 | 80446 | 64389 | 65653 | 74126 | 79211 | 80077 | 79413 | 73811.35 | 68056 | 79567 | 1.169 | 0.225 | 0.00021566 | 0.00225987 | 0.003 | 0.0025 | 0.0027 | 0.0027 | TRUE | 0.0745 |
| Top2b | 5799 | 4042 | 4357 | 4987 | 4205 | 5077 | 4504 | 4299 | 4701 | 4964 | 4744 | 5012 | 4703.99 | 4501 | 4907 | 1.09 | 0.125 | 0.03140399 | 0.09578685 | 0.0011 | 0.0029 | 0.0022 | 0.0022 | TRUE | 0.0185 |
| Top3a | 2599 | 1836 | 1628 | 1611 | 1505 | 1676 | 2019 | 1953 | 1757 | 1604 | 1698 | 1654 | 1780.65 | 1910 | 1652 | 0.864 | -0.21 | 0.00404504 | 0.02079702 | 0.0025 | 0.0038 | 0.0033 | 0.0033 | TRUE | 0.0585 |
| Top3b | 5353 | 3830 | 3600 | 3059 | 2795 | 2992 | 4157 | 4074 | 3884 | 3045 | 3153 | 2954 | 3544.55 | 4038 | 3051 | 0.755 | -0.406 | 3.81E-12 | 4.23E-10 | 7.00E-04 | 0.0031 | 0.0022 | 0.0022 | TRUE | 0.0135 |
| Topaz1 | 152 | 104 | 84 | 63 | 70 | 65 | 118 | 111 | 91 | 63 | 79 | 64 | 87.53 | 107 | 69 | 0.641 | -0.642 | 0.00362621 | 0.01911821 | 0 | 0.0295 | 0.023 | 0.023 | TRUE | 0.1608 |

| | | | | | | | | | | | | | | | | | | | | | | | | | |
|---|---|---|---|---|---|---|---|---|---|---|---|---|---|---|---|---|---|---|---|---|---|---|---|---|---|
| Topbp1 | 17248 | 12899 | 13235 | 13171 | 11556 | 13318 | 13396 | 13720 | 14280 | 13110 | 13036 | 13147 | 13448.4 | 13799 | 13098 | 0.949 | -0.075 | 0.11932045 | 0.25102254 | 4.00E-04 | 0.0026 | 0.0016 | 0.0016 | TRUE | 0.0114 |
| Topors | 4875 | 3399 | 3122 | 3095 | 2689 | 3120 | 3786 | 3615 | 3369 | 3081 | 3033 | 3080 | 3327.41 | 3590 | 3065 | 0.853 | -0.229 | 0.00023035 | 0.00238582 | 0.0015 | 0.0031 | 0.0025 | 0.0025 | TRUE | 0.0353 |
| Tor1a | 1462 | 1028 | 960 | 1098 | 968 | 1100 | 1135 | 1093 | 1036 | 1093 | 1092 | 1086 | 1089.27 | 1088 | 1090 | 1.001 | 0.001 | 0.987841 | 0.99380852 | 0 | 0.0046 | 0.003 | 0.003 | TRUE | 0.021 |
| Tor1aip1 | 1112 | 754 | 606 | 714 | 609 | 671 | 864 | 802 | 654 | 711 | 687 | 662 | 729.94 | 773 | 687 | 0.886 | -0.174 | 0.10539474 | 0.22995069 | 0.0095 | 0.0057 | 0.0069 | 0.0069 | TRUE | 0.2129 |
| Tor1aip2 | 10328 | 7472 | 7289 | 7811 | 6847 | 7653 | 8021 | 7948 | 7865 | 7775 | 7724 | 7555 | 7814.64 | 7945 | 7685 | 0.967 | -0.048 | 0.30594146 | 0.47853155 | 0 | 0.0027 | 0.0015 | 0.0015 | TRUE | 0.0027 |
| Tor1b | 1644 | 1287 | 1036 | 1048 | 952 | 1037 | 1277 | 1369 | 1118 | 1043 | 1074 | 1024 | 1150.74 | 1255 | 1047 | 0.834 | -0.262 | 0.0025672 | 0.01467083 | 0.0047 | 0.0045 | 0.0046 | 0.0046 | TRUE | 0.1061 |
| Tor2a | 1949 | 1377 | 1517 | 1661 | 1431 | 1657 | 1514 | 1465 | 1637 | 1653 | 1614 | 1636 | 1586.44 | 1539 | 1634 | 1.063 | 0.088 | 0.20492264 | 0.36644055 | 0.001 | 0.0039 | 0.0028 | 0.0028 | TRUE | 0.0366 |
| Tor3a | 981 | 717 | 797 | 798 | 679 | 827 | 762 | 763 | 860 | 794 | 766 | 816 | 793.53 | 795 | 792 | 0.998 | -0.003 | 0.97489894 | 0.98647473 | 0.0017 | 0.0054 | 0.0042 | 0.0042 | TRUE | 0.0595 |
| Tor4a | 354 | 317 | 266 | 292 | 268 | 257 | 275 | 337 | 287 | 291 | 302 | 254 | 290.97 | 300 | 282 | 0.943 | -0.085 | 0.52447474 | 0.6856824 | 0.0067 | 0.0106 | 0.0094 | 0.0094 | TRUE | 0.133 |
| Tox | 2 | 1 | 2 | 1 | 1 | 0 | 2 | 1 | 2 | 1 | 1 | 0 | 1.15 | 2 | 1 | 0.439 | -1.188 | 0.51151502 | NA | 0 | 2.0638 | 1.3155 | 1.3155 | TRUE | 0.2709 |
| Tox2 | 70 | 45 | 48 | 64 | 57 | 64 | 54 | 48 | 52 | 64 | 64 | 63 | 57.53 | 51 | 64 | 1.238 | 0.308 | 0.21971964 | 0.38284198 | 0 | 0.0436 | 0.0279 | 0.0279 | TRUE | 0.0279 |
| Tox3 | 117 | 117 | 89 | 73 | 60 | 75 | 91 | 124 | 96 | 73 | 68 | 74 | 87.62 | 104 | 72 | 0.692 | -0.532 | 0.01571349 | 0.05781029 | 0 | 0.0295 | 0.023 | 0.023 | TRUE | 0.2917 |
| Tox4 | 7236 | 5398 | 5120 | 5579 | 4836 | 5600 | 5620 | 5742 | 5524 | 5553 | 5456 | 5528 | 5570.5 | 5629 | 5512 | 0.979 | -0.03 | 0.54315416 | 0.7009121 | 0 | 0.0029 | 0.0016 | 0.0016 | TRUE | 0.0037 |
| Tpbg | 457 | 369 | 363 | 282 | 256 | 312 | 355 | 393 | 392 | 281 | 289 | 308 | 336.1 | 380 | 293 | 0.772 | -0.373 | 0.00126377 | 0.00852839 | 0 | 0.0095 | 0.0066 | 0.0066 | TRUE | 0.0403 |
| Tpbpa | 0 | 0 | 0 | 0 | 0 | 0 | 0 | 0 | 0 | 0 | 0 | 0 | 0 | 0 | 0 | NA | NA | NA | NA | NA | NA | NA | NA | NA | NA |
| Tpbpb | 0 | 0 | 0 | 0 | 0 | 0 | 0 | 0 | 0 | 0 | 0 | 0 | 0 | 0 | 0 | NA | NA | NA | NA | NA | NA | NA | NA | NA | NA |
| Tpcn1 | 2110 | 1484 | 1589 | 1700 | 1574 | 1732 | 1639 | 1579 | 1715 | 1692 | 1776 | 1710 | 1684.89 | 1644 | 1726 | 1.05 | 0.07 | 0.29253709 | 0.46479694 | 5.00E-04 | 0.0038 | 0.0026 | 0.0026 | TRUE | 0.0164 |
| Tpcn2 | 337 | 244 | 183 | 190 | 170 | 195 | 262 | 260 | 197 | 189 | 192 | 192 | 215.35 | 240 | 191 | 0.796 | -0.33 | 0.02995033 | 0.09254167 | 0.0083 | 0.0134 | 0.0119 | 0.0119 | TRUE | 0.2522 |
| Tpd52 | 11552 | 9125 | 10771 | 10994 | 9986 | 11256 | 8972 | 9706 | 11622 | 10943 | 11265 | 11111 | 10603.33 | 10100 | 11106 | 1.1 | 0.137 | 0.09753375 | 0.21737874 | 0.0091 | 0.0027 | 0.0049 | 0.0049 | TRUE | 0.2121 |
| Tpd52l1 | 1154 | 797 | 986 | 1090 | 930 | 938 | 896 | 848 | 1064 | 1085 | 1049 | 926 | 978 | 936 | 1020 | 1.09 | 0.125 | 0.21722122 | 0.380002 | 0.0091 | 0.0048 | 0.0063 | 0.0063 | TRUE | 0.1669 |
| Tpd52l2 | 5229 | 3753 | 3516 | 3936 | 3576 | 4066 | 4061 | 3992 | 3794 | 3918 | 4034 | 4014 | 3968.78 | 3949 | 3989 | 1.01 | 0.014 | 0.80232834 | 0.88553506 | 5.00E-04 | 0.003 | 0.0019 | 0.0019 | TRUE | 0.0143 |
| Tpgs1 | 601 | 482 | 430 | 492 | 484 | 523 | 467 | 513 | 464 | 490 | 546 | 516 | 499.24 | 481 | 517 | 1.075 | 0.105 | 0.29506882 | 0.4669447 | 9.00E-04 | 0.0072 | 0.0052 | 0.0052 | TRUE | 0.0371 |
| Tpgs2 | 1778 | 1222 | 1246 | 1423 | 1195 | 1370 | 1381 | 1300 | 1344 | 1416 | 1348 | 1352 | 1357.01 | 1342 | 1372 | 1.022 | 0.032 | 0.64374699 | 0.77809422 | 0 | 0.0042 | 0.0027 | 0.0027 | TRUE | 0.0096 |
| Tph1 | 17 | 12 | 5 | 10 | 2 | 12 | 13 | 13 | 5 | 10 | 2 | 12 | 9.24 | 10 | 8 | 0.769 | -0.379 | 0.59182634 | NA | 0.2194 | 0.259 | 0.2495 | 0.2495 | TRUE | 1.0272 |
| Tph2 | 53 | 33 | 38 | 29 | 26 | 13 | 41 | 35 | 41 | 29 | 29 | 13 | 31.38 | 39 | 24 | 0.602 | -0.732 | 0.05401785 | 0.14221478 | 0.04 | 0.0779 | 0.0696 | 0.0696 | TRUE | 0.9862 |
| Tpi1 | 35814 | 29175 | 36879 | 32799 | 30955 | 36526 | 27815 | 31033 | 39792 | 32648 | 34920 | 36057 | 33710.96 | 32880 | 34542 | 1.051 | 0.071 | 0.48475718 | 0.65088158 | 0.0153 | 0.0025 | 0.0075 | 0.0075 | TRUE | 0.4139 |
| Tpk1 | 695 | 522 | 458 | 353 | 354 | 433 | 540 | 555 | 494 | 351 | 399 | 427 | 461.23 | 530 | 392 | 0.741 | -0.433 | 8.69E-05 | 0.0010781 | 0.0041 | 0.0076 | 0.0065 | 0.0065 | TRUE | 0.0997 |
| Tpm1 | 3614 | 2570 | 2502 | 2664 | 2274 | 2651 | 2807 | 2734 | 2700 | 2652 | 2565 | 2617 | 2679.02 | 2747 | 2611 | 0.95 | -0.073 | 0.19570245 | 0.3553322 | 0 | 0.0033 | 0.0019 | 0.0019 | TRUE | 0.0046 |
| Tpm2 | 504 | 399 | 390 | 480 | 440 | 615 | 391 | 424 | 421 | 478 | 496 | 607 | 469.65 | 412 | 527 | 1.281 | 0.357 | 0.0021089 | 0.01255935 | 0.0078 | 0.0075 | 0.0076 | 0.0076 | TRUE | 0.2086 |
| Tpm3 | 8892 | 6234 | 5195 | 5311 | 4501 | 5161 | 6906 | 6631 | 5605 | 5287 | 5078 | 5095 | 5766.88 | 6381 | 5153 | 0.807 | -0.309 | 5.48E-05 | 0.00075576 | 0.0058 | 0.0028 | 0.004 | 0.004 | TRUE | 0.1374 |
| Tpm4 | 5328 | 3896 | 3948 | 4001 | 3352 | 3860 | 4138 | 4144 | 4260 | 3983 | 3781 | 3810 | 4019.4 | 4181 | 3858 | 0.923 | -0.116 | 0.03180594 | 0.09662637 | 3.00E-04 | 0.003 | 0.0018 | 0.0018 | TRUE | 0.0097 |
| Tpmt | 991 | 696 | 731 | 740 | 637 | 771 | 770 | 740 | 789 | 737 | 719 | 761 | 752.5 | 766 | 739 | 0.964 | -0.052 | 0.52564675 | 0.68657138 | 0 | 0.0056 | 0.0036 | 0.0036 | TRUE | 0.0099 |
| Tpo | 28 | 19 | 30 | 26 | 14 | 21 | 22 | 20 | 32 | 26 | 16 | 21 | 22.79 | 25 | 21 | 0.847 | -0.24 | 0.56670247 | 0.720758 | 0 | 0.1064 | 0.0823 | 0.0823 | TRUE | 0.3561 |
| Tpp1 | 2663 | 1870 | 2035 | 2047 | 1915 | 2087 | 2068 | 1989 | 2196 | 2038 | 2160 | 2060 | 2085.2 | 2084 | 2086 | 1.001 | 0.001 | 0.98754571 | 0.99373352 | 0.0012 | 0.0036 | 0.0027 | 0.0027 | TRUE | 0.0257 |
| Tpp2 | 8872 | 6428 | 6364 | 7026 | 6000 | 6854 | 6891 | 6837 | 6867 | 6994 | 6769 | 6766 | 6853.81 | 6865 | 6843 | 0.997 | -0.005 | 0.92281137 | 0.95719768 | 0 | 0.0028 | 0.0015 | 0.0015 | TRUE | 0.0046 |
| Tppp | 92 | 77 | 59 | 83 | 82 | 101 | 71 | 82 | 64 | 83 | 93 | 100 | 81.97 | 72 | 92 | 1.267 | 0.341 | 0.12154886 | 0.25435227 | 0 | 0.0313 | 0.0226 | 0.0226 | TRUE | 0.1119 |
| Tppp2 | 0 | 0 | 0 | 0 | 1 | 0 | 0 | 0 | 0 | 0 | 1 | 0 | 0.19 | 0 | 0 | 2.094 | 1.067 | 0.79379329 | NA | 0 | 12.608 | 10 | 10 | TRUE | 0.5075 |
| Tppp3 | 561 | 408 | 435 | 708 | 689 | 757 | 436 | 434 | 469 | 705 | 777 | 747 | 594.72 | 446 | 743 | 1.666 | 0.736 | 3.27E-15 | 5.64E-13 | 0 | 0.0064 | 0.0045 | 0.0045 | TRUE | 0.0242 |
| Tpr | 23355 | 15906 | 14074 | 13934 | 12236 | 14383 | 18139 | 16919 | 15186 | 13870 | 13803 | 14198 | 15352.53 | 16748 | 13957 | 0.833 | -0.263 | 8.38E-05 | 0.00104742 | 0.0039 | 0.0026 | 0.0032 | 0.0032 | TRUE | 0.0814 |
| Tpra1 | 565 | 483 | 517 | 524 | 447 | 565 | 439 | 514 | 558 | 522 | 504 | 558 | 515.67 | 504 | 528 | 1.052 | 0.073 | 0.51174619 | 0.67469683 | 0.007 | 0.007 | 0.007 | 0.007 | TRUE | 0.1563 |
| Tprg | 0 | 1 | 0 | 0 | 0 | 0 | 0 | 1 | 0 | 0 | 0 | 0 | 0.18 | 0 | 0 | 0.552 | -0.857 | 0.8336548 | NA | 0 | 13.3714 | 10 | 10 | TRUE | 0.5216 |
| Tprgl | 4189 | 3051 | 3085 | 3435 | 3128 | 3563 | 3253 | 3245 | 3329 | 3419 | 3529 | 3517 | 3382.09 | 3276 | 3488 | 1.065 | 0.091 | 0.09018741 | 0.20499458 | 0 | 0.0031 | 0.0018 | 0.0018 | TRUE | 0.0037 |
| Tprkb | 2600 | 1804 | 1748 | 1857 | 1618 | 1987 | 2019 | 1919 | 1886 | 1848 | 1825 | 1961 | 1909.91 | 1941 | 1878 | 0.967 | -0.048 | 0.46347131 | 0.63270253 | 9.00E-04 | 0.0037 | 0.0026 | 0.0026 | TRUE | 0.0182 |
| Tprn | 1612 | 1282 | 1246 | 1086 | 1111 | 1200 | 1252 | 1364 | 1344 | 1081 | 1253 | 1185 | 1246.49 | 1320 | 1173 | 0.889 | -0.17 | 0.0344537 | 0.10271929 | 0.003 | 0.0043 | 0.0038 | 0.0038 | TRUE | 0.0566 |
| Tpsab1 | 0 | 0 | 0 | 0 | 0 | 0 | 0 | 0 | 0 | 0 | 0 | 0 | 0 | 0 | 0 | NA | NA | NA | NA | NA | NA | NA | NA | NA | NA |
| Tpsb2 | 0 | 1 | 0 | 0 | 0 | 0 | 0 | 1 | 0 | 0 | 0 | 0 | 0.18 | 0 | 0 | 0.552 | -0.857 | 0.8336548 | NA | 0 | 13.3714 | 10 | 10 | TRUE | 0.5216 |
| Tpsg1 | 0 | 0 | 1 | 2 | 2 | 0 | 0 | 0 | 1 | 2 | 2 | 0 | 0.89 | 0 | 1 | 4.079 | 2.028 | 0.37136419 | NA | 0 | 2.6724 | 2.3411 | 2.3411 | TRUE | 0.5365 |
| Tpst1 | 4349 | 3078 | 3077 | 2979 | 2689 | 3204 | 3378 | 3274 | 3320 | 2965 | 3033 | 3163 | 3188.9 | 3324 | 3054 | 0.919 | -0.122 | 0.03045779 | 0.09359786 | 3.00E-04 | 0.0032 | 0.002 | 0.002 | TRUE | 0.012 |
| Tpst2 | 1641 | 1115 | 1126 | 1222 | 1159 | 1265 | 1274 | 1186 | 1215 | 1216 | 1307 | 1249 | 1241.34 | 1225 | 1257 | 1.025 | 0.036 | 0.62031181 | 0.76038736 | 6.00E-04 | 0.0043 | 0.003 | 0.003 | TRUE | 0.0156 |
| Tpt1 | 60213 | 41384 | 41084 | 46573 | 39881 | 46969 | 46765 | 44020 | 44329 | 46359 | 44990 | 46366 | 45471.34 | 45038 | 45905 | 1.019 | 0.027 | 0.56530111 | 0.71974423 | 6.00E-04 | 0.0025 | 0.0016 | 0.0016 | TRUE | 0.0138 |
| Tpte | 8 | 10 | 9 | 11 | 12 | 15 | 6 | 11 | 10 | 11 | 14 | 15 | 10.98 | 9 | 13 | 1.499 | 0.584 | 0.31475514 | NA | 0 | 0.2184 | 0.1484 | 0.1484 | TRUE | 0.2901 |
| Tpx2 | 15474 | 11130 | 11458 | 12948 | 11061 | 12738 | 12018 | 11839 | 12363 | 12888 | 12478 | 12574 | 12360.11 | 12073 | 12647 | 1.048 | 0.067 | 0.15497655 | 0.30248256 | 3.00E-04 | 0.0026 | 0.0015 | 0.0015 | TRUE | 0.0053 |
| Tra2a | 7992 | 6167 | 6578 | 5830 | 5208 | 6287 | 6207 | 6560 | 7098 | 5803 | 5875 | 6206 | 6291.5 | 6622 | 5961 | 0.901 | -0.151 | 0.0184281 | 0.06503526 | 0.0027 | 0.0028 | 0.0028 | 0.0028 | TRUE | 0.0482 |
| Tra2b | 11401 | 7985 | 7132 | 6761 | 6138 | 7281 | 8855 | 8494 | 7695 | 6730 | 6924 | 7187 | 7647.54 | 8348 | 6947 | 0.832 | -0.265 | 3.66E-05 | 0.00054785 | 0.0029 | 0.0027 | 0.0028 | 0.0028 | TRUE | 0.0569 |
| Trabd | 2209 | 1573 | 1546 | 1621 | 1437 | 1648 | 1716 | 1673 | 1668 | 1614 | 1621 | 1627 | 1653.07 | 1686 | 1621 | 0.961 | -0.057 | 0.35755873 | 0.53202029 | 0 | 0.0039 | 0.0022 | 0.0022 | TRUE | 0.0031 |
| Trabd2b | 59 | 62 | 31 | 16 | 25 | 18 | 46 | 66 | 33 | 16 | 28 | 18 | 34.52 | 48 | 21 | 0.424 | -1.238 | 0.00129669 | 0.00870121 | 0.0728 | 0.0711 | 0.0716 | 0.0716 | TRUE | 0.1862 |
| Tradd | 723 | 569 | 580 | 725 | 660 | 773 | 562 | 605 | 626 | 722 | 745 | 763 | 670.31 | 598 | 743 | 1.246 | 0.317 | 0.00034503 | 0.00322745 | 0 | 0.006 | 0.0042 | 0.0042 | TRUE | 0.035 |
| Traf1 | 60 | 49 | 42 | 42 | 54 | 57 | 47 | 52 | 45 | 42 | 61 | 56 | 50.5 | 48 | 53 | 1.103 | 0.141 | 0.61407923 | 0.75613092 | 0 | 0.0494 | 0.0364 | 0.0364 | TRUE | 0.2875 |
| Traf2 | 878 | 598 | 625 | 566 | 554 | 579 | 682 | 636 | 674 | 563 | 625 | 572 | 625.38 | 664 | 587 | 0.882 | -0.181 | 0.04717982 | 0.12915939 | 0 | 0.0062 | 0.0044 | 0.0044 | TRUE | 0.0373 |
| Traf3 | 2041 | 1452 | 1269 | 1370 | 1233 | 1343 | 1585 | 1544 | 1369 | 1364 | 1391 | 1326 | 1429.88 | 1499 | 1360 | 0.906 | -0.142 | 0.06401203 | 0.16082333 | 0.0026 | 0.0041 | 0.0036 | 0.0036 | TRUE | 0.0686 |
| Traf3ip1 | 1302 | 992 | 942 | 854 | 800 | 858 | 1011 | 1055 | 1016 | 850 | 902 | 847 | 947.05 | 1027 | 866 | 0.843 | -0.246 | 0.00129959 | 0.00870843 | 0 | 0.0049 | 0.0032 | 0.0032 | TRUE | 0.0153 |
| Traf3ip2 | 729 | 558 | 544 | 550 | 542 | 666 | 566 | 594 | 587 | 547 | 611 | 657 | 593.84 | 582 | 605 | 1.041 | 0.057 | 0.55978241 | 0.71456372 | 0.0029 | 0.0064 | 0.0053 | 0.0053 | TRUE | 0.0841 |
| Traf3ip3 | 2 | 2 | 0 | 2 | 2 | 0 | 2 | 2 | 0 | 2 | 2 | 0 | 1.32 | 1 | 1 | 1.131 | 0.177 | 0.91971667 | NA | 0 | 1.7962 | 1.4631 | 1.4631 | TRUE | 0.5204 |
| Traf4 | 6570 | 4757 | 5244 | 5769 | 5202 | 6049 | 5103 | 5060 | 5658 | 5742 | 5868 | 5971 | 5567.16 | 5274 | 5860 | 1.112 | 0.153 | 0.01199692 | 0.0473418 | 0.0019 | 0.0029 | 0.0025 | 0.0025 | TRUE | 0.0492 |
| Traf5 | 82 | 81 | 60 | 65 | 66 | 67 | 64 | 86 | 65 | 65 | 74 | 66 | 69.98 | 72 | 68 | 0.959 | -0.061 | 0.80103969 | 0.88445425 | 0 | 0.0363 | 0.0274 | 0.0274 | TRUE | 0.2791 |
| Traf6 | 2039 | 1410 | 1243 | 1340 | 1184 | 1242 | 1584 | 1500 | 1341 | 1334 | 1336 | 1226 | 1386.69 | 1475 | 1299 | 0.879 | -0.186 | 0.02361318 | 0.07799936 | 0.0041 | 0.0041 | 0.0041 | 0.0041 | TRUE | 0.075 |
| Traf7 | 9213 | 6906 | 6384 | 6829 | 6398 | 7358 | 7155 | 7346 | 6888 | 6798 | 7218 | 7263 | 7111.35 | 7130 | 7093 | 0.995 | -0.008 | 0.88924182 | 0.93844044 | 0.001 | 0.0028 | 0.002 | 0.002 | TRUE | 0.0162 |
| Trafd1 | 1121 | 831 | 774 | 674 | 583 | 617 | 871 | 884 | 835 | 671 | 658 | 609 | 754.56 | 863 | 646 | 0.748 | -0.419 | 7.53E-07 | 2.30E-05 | 0 | 0.0056 | 0.0038 | 0.0038 | TRUE | 0.0299 |
| Traip | 1996 | 1511 | 1380 | 1401 | 1394 | 1558 | 1550 | 1607 | 1489 | 1395 | 1573 | 1538 | 1525.26 | 1549 | 1502 | 0.969 | -0.045 | 0.541212 | 0.69947944 | 0.002 | 0.004 | 0.0033 | 0.0033 | TRUE | 0.0472 |

| | | | | | | | | | | | | | | | | | | | | | | | | | |
|---|---|---|---|---|---|---|---|---|---|---|---|---|---|---|---|---|---|---|---|---|---|---|---|---|---|
| Trak1 | 3253 | 2505 | 2824 | 2725 | 2684 | 3249 | 2526 | 2665 | 3047 | 2712 | 3028 | 3207 | 2864.27 | 2746 | 2982 | 1.087 | 0.12 | 0.16007305 | 0.30951354 | 0.0078 | 0.0033 | 0.0049 | 0.0049 | TRUE | 0.1112 |
| Trak2 | 4765 | 3262 | 2735 | 3260 | 2879 | 3250 | 3701 | 3470 | 2951 | 3245 | 3248 | 3208 | 3303.77 | 3374 | 3234 | 0.958 | -0.062 | 0.43811917 | 0.60962202 | 0.0057 | 0.0031 | 0.0043 | 0.0043 | TRUE | 0.1454 |
| Tram1 | 5786 | 4178 | 4158 | 4243 | 3734 | 4265 | 4494 | 4444 | 4486 | 4223 | 4212 | 4210 | 4345.05 | 4475 | 4215 | 0.942 | -0.086 | 0.0832213 | 0.19416945 | 0 | 0.003 | 0.0016 | 0.0016 | TRUE | 4.00E-04 |
| Tram1l1 | 189 | 126 | 105 | 116 | 84 | 91 | 147 | 134 | 113 | 115 | 95 | 90 | 115.69 | 131 | 100 | 0.759 | -0.397 | 0.04316579 | 0.12099979 | 0.0092 | 0.0229 | 0.019 | 0.019 | TRUE | 0.1822 |
| Tram2 | 508 | 384 | 427 | 444 | 349 | 423 | 395 | 408 | 461 | 442 | 394 | 418 | 419.49 | 421 | 418 | 0.994 | -0.008 | 0.93924001 | 0.966328 | 0.0027 | 0.0081 | 0.0064 | 0.0064 | TRUE | 0.0761 |
| Trank1 | 585 | 444 | 425 | 665 | 498 | 549 | 454 | 472 | 459 | 662 | 562 | 542 | 525.15 | 462 | 589 | 1.276 | 0.351 | 0.00089903 | 0.00671008 | 0.0045 | 0.0069 | 0.0061 | 0.0061 | TRUE | 0.1413 |
| Trap1 | 13077 | 9633 | 9444 | 11405 | 10101 | 11239 | 10156 | 10246 | 10190 | 11353 | 11395 | 11095 | 10739.16 | 10197 | 11281 | 1.106 | 0.146 | 0.00128715 | 0.00865561 | 0 | 0.0026 | 0.0014 | 0.0014 | TRUE | 0.0026 |
| Trap1a | 7528 | 5285 | 5194 | 5901 | 5044 | 5852 | 5847 | 5622 | 5604 | 5874 | 5690 | 5777 | 5735.56 | 5691 | 5780 | 1.016 | 0.022 | 0.66080777 | 0.79043647 | 3.00E-04 | 0.0028 | 0.0017 | 0.0017 | TRUE | 0.0071 |
| Trappc1 | 2073 | 1430 | 1416 | 1414 | 1156 | 1308 | 1610 | 1521 | 1528 | 1407 | 1304 | 1291 | 1443.62 | 1553 | 1334 | 0.859 | -0.22 | 0.00192946 | 0.01176528 | 0.001 | 0.0041 | 0.0029 | 0.0029 | TRUE | 0.028 |
| Trappc10 | 3116 | 2285 | 2368 | 2862 | 2462 | 2821 | 2420 | 2431 | 2555 | 2849 | 2777 | 2785 | 2636.1 | 2469 | 2804 | 1.136 | 0.184 | 0.00146526 | 0.00948073 | 0 | 0.0033 | 0.002 | 0.002 | TRUE | 0.0112 |
| Trappc11 | 8165 | 5729 | 5732 | 5822 | 5305 | 6102 | 6341 | 6094 | 6185 | 5795 | 5985 | 6024 | 6070.58 | 6207 | 5935 | 0.956 | -0.065 | 0.19667235 | 0.3563689 | 3.00E-04 | 0.0028 | 0.0017 | 0.0017 | TRUE | 0.0052 |
| Trappc12 | 2172 | 1631 | 1624 | 1671 | 1521 | 1880 | 1687 | 1735 | 1752 | 1663 | 1716 | 1856 | 1734.84 | 1725 | 1745 | 1.012 | 0.018 | 0.79735775 | 0.88168299 | 0.0013 | 0.0038 | 0.0028 | 0.0028 | TRUE | 0.0377 |
| Trappc13 | 3419 | 2257 | 2057 | 2151 | 1940 | 2080 | 2655 | 2401 | 2219 | 2141 | 2189 | 2053 | 2276.42 | 2425 | 2128 | 0.876 | -0.19 | 0.01258391 | 0.04905665 | 0.0039 | 0.0035 | 0.0038 | 0.0038 | TRUE | 0.0852 |
| Trappc2 | 574 | 374 | 380 | 444 | 334 | 411 | 446 | 398 | 410 | 442 | 377 | 406 | 413.02 | 418 | 408 | 0.976 | -0.035 | 0.75019677 | 0.85168585 | 0.0026 | 0.0082 | 0.0064 | 0.0064 | TRUE | 0.0605 |
| Trappc2l | 1165 | 807 | 899 | 974 | 802 | 942 | 905 | 858 | 970 | 970 | 905 | 930 | 922.9 | 911 | 935 | 1.027 | 0.038 | 0.64450045 | 0.77868512 | 0.0013 | 0.005 | 0.0037 | 0.0037 | TRUE | 0.0365 |
| Trappc3 | 2141 | 1460 | 1355 | 1538 | 1461 | 1546 | 1663 | 1553 | 1462 | 1531 | 1648 | 1526 | 1563.84 | 1559 | 1568 | 1.004 | 0.006 | 0.93245739 | 0.96249104 | 0.0024 | 0.0039 | 0.0034 | 0.0034 | TRUE | 0.042 |
| Trappc4 | 2320 | 1638 | 1732 | 1925 | 1699 | 1957 | 1802 | 1742 | 1869 | 1916 | 1917 | 1932 | 1862.94 | 1804 | 1922 | 1.065 | 0.091 | 0.14652878 | 0.29090266 | 0 | 0.0037 | 0.0023 | 0.0023 | TRUE | 0.0114 |
| Trappc5 | 1253 | 924 | 830 | 924 | 829 | 965 | 973 | 983 | 896 | 920 | 935 | 953 | 943.19 | 951 | 936 | 0.984 | -0.023 | 0.76573394 | 0.86110426 | 0 | 0.0049 | 0.0034 | 0.0034 | TRUE | 0.0295 |
| Trappc6a | 631 | 450 | 562 | 650 | 588 | 691 | 490 | 479 | 606 | 647 | 663 | 682 | 594.6 | 525 | 664 | 1.267 | 0.342 | 0.00143167 | 0.00933524 | 0.0069 | 0.0064 | 0.0066 | 0.0066 | TRUE | 0.2089 |
| Trappc6b | 2049 | 1429 | 1365 | 1547 | 1304 | 1488 | 1591 | 1520 | 1473 | 1540 | 1471 | 1469 | 1510.67 | 1528 | 1493 | 0.977 | -0.034 | 0.61548844 | 0.7572023 | 6.00E-04 | 0.004 | 0.0027 | 0.0027 | TRUE | 0.0165 |
| Trappc8 | 7299 | 5325 | 5194 | 5602 | 5004 | 6037 | 5669 | 5664 | 5604 | 5576 | 5645 | 5959 | 5686.32 | 5646 | 5727 | 1.014 | 0.021 | 0.69337321 | 0.81291342 | 4.00E-04 | 0.0028 | 0.0018 | 0.0018 | TRUE | 0.0155 |
| Trappc9 | 1870 | 1387 | 1712 | 1957 | 1768 | 1992 | 1452 | 1475 | 1847 | 1948 | 1994 | 1966 | 1780.64 | 1591 | 1969 | 1.239 | 0.309 | 0.00069894 | 0.00557069 | 0.0089 | 0.0038 | 0.0054 | 0.0054 | TRUE | 0.2357 |
| Trat1 | 0 | 0 | 0 | 0 | 0 | 0 | 0 | 0 | 0 | 0 | 0 | 0 | 0 | 0 | 0 | NA | NA | NA | NA | NA | NA | NA | NA | NA | NA |
| Trcg1 | 1 | 0 | 0 | 0 | 0 | 0 | 1 | 0 | 0 | 0 | 0 | 0 | 0.13 | 0 | 0 | 0.552 | -0.857 | 0.8336548 | NA | 0 | 18.3121 | 10 | 10 | TRUE | 0.2566 |
| Trdmt1 | 742 | 550 | 493 | 536 | 456 | 467 | 576 | 585 | 532 | 534 | 514 | 461 | 533.7 | 564 | 503 | 0.89 | -0.168 | 0.09504354 | 0.21302451 | 0.0021 | 0.0069 | 0.0054 | 0.0054 | TRUE | 0.0635 |
| Trdn | 0 | 0 | 0 | 0 | 0 | 0 | 0 | 0 | 0 | 0 | 0 | 0 | 0 | 0 | 0 | NA | NA | NA | NA | NA | NA | NA | NA | NA | NA |
| Treh | 1 | 1 | 0 | 1 | 0 | 1 | 1 | 1 | 0 | 1 | 0 | 1 | 0.64 | 1 | 1 | 1.082 | 0.113 | 0.96183421 | NA | 0 | 3.7222 | 2.4899 | 2.4899 | TRUE | 0.2041 |
| Trem1 | 0 | 0 | 0 | 0 | 0 | 1 | 0 | 0 | 0 | 0 | 0 | 1 | 0.16 | 0 | 0 | 2.094 | 1.067 | 0.79379329 | NA | 0 | 14.4078 | 10 | 10 | TRUE | 0.3835 |
| Trem2 | 0 | 1 | 0 | 2 | 1 | 1 | 0 | 1 | 0 | 2 | 1 | 1 | 0.86 | 0 | 1 | 3.977 | 1.992 | 0.3514665 | NA | 0 | 2.7531 | 1.9153 | 1.9153 | TRUE | 0.5217 |
| Trem3 | 0 | 0 | 0 | 0 | 0 | 0 | 0 | 0 | 0 | 0 | 0 | 0 | 0 | 0 | 0 | NA | NA | NA | NA | NA | NA | NA | NA | NA | NA |
| Treml1 | 2 | 1 | 0 | 2 | 1 | 2 | 2 | 1 | 0 | 2 | 1 | 2 | 1.29 | 1 | 2 | 1.88 | 0.911 | 0.58799782 | NA | 0 | 1.8468 | 1.2019 | 1.2019 | TRUE | 0.2741 |
| Treml2 | 296 | 233 | 182 | 246 | 240 | 282 | 230 | 248 | 196 | 245 | 271 | 278 | 244.68 | 225 | 265 | 1.177 | 0.235 | 0.0923215 | 0.2087378 | 0.0042 | 0.0121 | 0.0099 | 0.0099 | TRUE | 0.1258 |
| Treml4 | 7 | 6 | 3 | 3 | 8 | 6 | 5 | 6 | 3 | 3 | 9 | 6 | 5.5 | 5 | 6 | 1.177 | 0.236 | 0.77655115 | NA | 0 | 0.4335 | 0.3138 | 0.3138 | TRUE | 0.1675 |
| Trerf1 | 1130 | 831 | 832 | 1027 | 967 | 1152 | 878 | 884 | 898 | 1022 | 1091 | 1137 | 984.94 | 887 | 1083 | 1.223 | 0.29 | 0.00019551 | 0.00209639 | 0 | 0.0048 | 0.0033 | 0.0033 | TRUE | 0.0297 |
| Trex1 | 259 | 222 | 225 | 192 | 166 | 245 | 201 | 236 | 243 | 191 | 187 | 242 | 216.72 | 227 | 207 | 0.916 | -0.127 | 0.4134464 | 0.5860164 | 0.011 | 0.0134 | 0.0127 | 0.0127 | TRUE | 0.2439 |
| Trex2 | 5 | 5 | 8 | 3 | 2 | 4 | 4 | 5 | 9 | 3 | 2 | 4 | 4.5 | 6 | 3 | 0.525 | -0.931 | 0.31073798 | NA | 0 | 0.5286 | 0.3575 | 0.3575 | TRUE | 0.3442 |
| Trf | 730 | 477 | 536 | 842 | 752 | 810 | 567 | 507 | 578 | 838 | 848 | 800 | 689.79 | 551 | 829 | 1.503 | 0.588 | 6.47E-11 | 5.44E-09 | 9.00E-04 | 0.0059 | 0.0043 | 0.0043 | TRUE | 0.0537 |
| Trh | 3623 | 2613 | 2442 | 2501 | 2832 | 3700 | 2814 | 2779 | 2635 | 2489 | 3195 | 3652 | 2927.48 | 2743 | 3112 | 1.135 | 0.182 | 0.09241108 | 0.20886015 | 0.0178 | 0.0032 | 0.0081 | 0.0081 | TRUE | 0.3021 |
| Trhde | 0 | 2 | 0 | 2 | 0 | 0 | 0 | 2 | 0 | 2 | 0 | 0 | 0.69 | 1 | 1 | 0.971 | -0.042 | 0.98762268 | NA | 8.094 | 3.4555 | 3.8966 | 3.8966 | TRUE | 0.9747 |
| Trhr | 1 | 0 | 0 | 1 | 1 | 0 | 1 | 0 | 0 | 1 | 1 | 0 | 0.48 | 0 | 1 | 2.037 | 1.027 | 0.71241723 | NA | 0 | 4.9057 | 3.8521 | 3.8521 | TRUE | 0.2664 |
| Trhr2 | 1 | 0 | 0 | 0 | 1 | 0 | 1 | 0 | 0 | 0 | 1 | 0 | 0.32 | 0 | 0 | 1.075 | 0.105 | 0.97638847 | NA | 0 | 7.4681 | 7.0392 | 7.0392 | TRUE | 0.5075 |
| Triap1 | 1588 | 1074 | 956 | 1142 | 1029 | 1127 | 1233 | 1142 | 1032 | 1137 | 1161 | 1113 | 1136.22 | 1136 | 1137 | 0.999 | -0.001 | 0.9876475 | 0.99376597 | 0.0034 | 0.0045 | 0.0041 | 0.0041 | TRUE | 0.0766 |
| Trib1 | 470 | 332 | 274 | 283 | 277 | 318 | 365 | 353 | 296 | 282 | 312 | 314 | 320.32 | 338 | 303 | 0.893 | -0.163 | 0.19854099 | 0.35879963 | 0.0051 | 0.0098 | 0.0084 | 0.0084 | TRUE | 0.1325 |
| Trib2 | 227 | 182 | 217 | 266 | 247 | 300 | 176 | 194 | 234 | 265 | 279 | 296 | 240.6 | 201 | 280 | 1.396 | 0.481 | 0.00090049 | 0.00671049 | 0.0074 | 0.0123 | 0.0109 | 0.0109 | TRUE | 0.217 |
| Trib3 | 596 | 432 | 356 | 366 | 348 | 390 | 463 | 460 | 384 | 364 | 393 | 385 | 408.07 | 436 | 381 | 0.872 | -0.197 | 0.08293025 | 0.19393358 | 0.0038 | 0.0082 | 0.0069 | 0.0069 | TRUE | 0.1194 |
| Tril | 35 | 14 | 9 | 10 | 16 | 18 | 27 | 15 | 10 | 10 | 18 | 18 | 16.26 | 17 | 15 | 0.867 | -0.206 | 0.69916646 | NA | 0.1289 | 0.1482 | 0.1425 | 0.1425 | TRUE | 0.6672 |
| Trim10 | 6 | 6 | 5 | 16 | 7 | 6 | 5 | 6 | 5 | 16 | 8 | 6 | 7.7 | 5 | 10 | 1.826 | 0.869 | 0.22944287 | NA | 0 | 0.3103 | 0.2358 | 0.2358 | TRUE | 0.9951 |
| Trim11 | 1563 | 1243 | 1310 | 1194 | 1060 | 1223 | 1214 | 1322 | 1413 | 1189 | 1196 | 1207 | 1256.86 | 1316 | 1197 | 0.911 | -0.135 | 0.08358432 | 0.19473035 | 0.0023 | 0.0043 | 0.0036 | 0.0036 | TRUE | 0.0582 |
| Trim12a | 0 | 0 | 0 | 0 | 0 | 0 | 0 | 0 | 0 | 0 | 0 | 0 | 0 | 0 | 0 | NA | NA | NA | NA | NA | NA | NA | NA | NA | NA |
| Trim12c | 670 | 473 | 556 | 501 | 433 | 500 | 520 | 503 | 600 | 499 | 488 | 494 | 517.36 | 541 | 494 | 0.913 | -0.131 | 0.20043992 | 0.36118102 | 0.0026 | 0.007 | 0.0056 | 0.0056 | TRUE | 0.102 |
| Trim13 | 1103 | 853 | 1078 | 1016 | 895 | 1101 | 857 | 907 | 1163 | 1011 | 1010 | 1087 | 1005.83 | 976 | 1036 | 1.064 | 0.089 | 0.41503501 | 0.58774352 | 0.0134 | 0.0048 | 0.0076 | 0.0076 | TRUE | 0.3344 |
| Trim14 | 82 | 66 | 107 | 131 | 135 | 134 | 64 | 70 | 115 | 130 | 152 | 132 | 110.72 | 83 | 138 | 1.675 | 0.744 | 0.00132684 | 0.008833 | 0.0437 | 0.0238 | 0.0291 | 0.0291 | TRUE | 1.0568 |
| Trim15 | 1 | 2 | 1 | 0 | 1 | 2 | 1 | 2 | 1 | 0 | 1 | 2 | 1.18 | 1 | 1 | 0.79 | -0.34 | 0.84691257 | NA | 0 | 2.0094 | 1.3651 | 1.3651 | TRUE | 0.3331 |
| Trim16 | 273 | 190 | 168 | 227 | 179 | 191 | 212 | 202 | 181 | 226 | 202 | 189 | 201.97 | 198 | 206 | 1.033 | 0.047 | 0.75052031 | 0.85183728 | 0 | 0.0142 | 0.0108 | 0.0108 | TRUE | 0.0849 |
| Trim17 | 30 | 23 | 30 | 45 | 34 | 35 | 23 | 24 | 32 | 45 | 38 | 35 | 32.97 | 26 | 39 | 1.478 | 0.563 | 0.09917736 | 0.21977395 | 0 | 0.0743 | 0.0528 | 0.0528 | TRUE | 0.2031 |
| Trim2 | 1417 | 1138 | 1410 | 1306 | 1394 | 1896 | 1101 | 1210 | 1521 | 1300 | 1573 | 1872 | 1429.43 | 1277 | 1582 | 1.239 | 0.309 | 0.01831588 | 0.06468715 | 0.0299 | 0.0041 | 0.0117 | 0.0117 | TRUE | 0.1835 |
| Trim21 | 451 | 341 | 360 | 373 | 337 | 389 | 350 | 363 | 388 | 371 | 380 | 384 | 372.81 | 367 | 378 | 1.032 | 0.046 | 0.67063828 | 0.79732432 | 0 | 0.0088 | 0.0058 | 0.0058 | TRUE | 0.029 |
| Trim23 | 2719 | 1897 | 2001 | 2084 | 1838 | 2058 | 2112 | 2018 | 2159 | 2074 | 2073 | 2032 | 2078.01 | 2096 | 2060 | 0.982 | -0.025 | 0.67764013 | 0.80217909 | 0 | 0.0036 | 0.0022 | 0.0022 | TRUE | 0.0126 |
| Trim24 | 10195 | 7792 | 7156 | 7812 | 6889 | 7668 | 7918 | 8288 | 7721 | 7776 | 7772 | 7569 | 7840.77 | 7976 | 7706 | 0.966 | -0.05 | 0.33495225 | 0.50903483 | 6.00E-04 | 0.0027 | 0.0018 | 0.0018 | TRUE | 0.0142 |
| Trim25 | 14203 | 10481 | 10218 | 11941 | 10644 | 11456 | 11031 | 11149 | 11025 | 11886 | 12008 | 11309 | 11401.16 | 11068 | 11734 | 1.06 | 0.084 | 0.08290658 | 0.19390995 | 4.00E-04 | 0.0026 | 0.0016 | 0.0016 | TRUE | 0.0123 |
| Trim26 | 2443 | 1858 | 1789 | 1891 | 1824 | 2077 | 1897 | 1976 | 1930 | 1882 | 2058 | 2050 | 1965.72 | 1934 | 1997 | 1.032 | 0.046 | 0.48668293 | 0.65260897 | 0.001 | 0.0036 | 0.0026 | 0.0026 | TRUE | 0.0306 |
| Trim27 | 9185 | 6717 | 6611 | 6492 | 6036 | 7048 | 7134 | 7145 | 7133 | 6462 | 6809 | 6957 | 6940.07 | 7137 | 6743 | 0.945 | -0.082 | 0.11307572 | 0.24187238 | 5.00E-04 | 0.0028 | 0.0018 | 0.0018 | TRUE | 0.0163 |
| Trim28 | 117317 | 81958 | 67092 | 66331 | 61518 | 69930 | 91115 | 87178 | 72392 | 66026 | 69399 | 69032 | 75856.78 | 83562 | 68152 | 0.816 | -0.294 | 0.00015502 | 0.00174873 | 0.0073 | 0.0025 | 0.0043 | 0.0043 | TRUE | 0.1674 |
| Trim29 | 102 | 97 | 139 | 222 | 230 | 288 | 79 | 103 | 150 | 221 | 259 | 284 | 182.85 | 111 | 255 | 2.32 | 1.214 | 4.80E-09 | 2.68E-07 | 0.0509 | 0.0154 | 0.0245 | 0.0245 | TRUE | 0.9441 |
| Trim3 | 1045 | 860 | 803 | 877 | 723 | 925 | 812 | 915 | 866 | 873 | 816 | 913 | 865.75 | 864 | 867 | 1.005 | 0.008 | 0.92896023 | 0.96082412 | 0.0023 | 0.0052 | 0.0042 | 0.0042 | TRUE | 0.0357 |
| Trim30a | 17 | 11 | 7 | 9 | 3 | 5 | 13 | 12 | 8 | 9 | 3 | 5 | 8.29 | 11 | 6 | 0.533 | -0.909 | 0.19112526 | NA | 0 | 0.2883 | 0.2146 | 0.2146 | TRUE | 0.5428 |
| Trim30b | 0 | 0 | 0 | 0 | 0 | 0 | 0 | 0 | 0 | 0 | 0 | 0 | 0 | 0 | 0 | NA | NA | NA | NA | NA | NA | NA | NA | NA | NA |
| Trim30d | 12 | 3 | 8 | 7 | 6 | 4 | 9 | 3 | 9 | 7 | 7 | 4 | 6.47 | 7 | 6 | 0.825 | -0.277 | 0.7200658 | NA | 0 | 0.3687 | 0.2742 | 0.2742 | TRUE | 0.5655 |
| Trim30e-ps1 | 0 | 0 | 0 | 0 | 0 | 0 | 0 | 0 | 0 | 0 | 0 | 0 | 0 | 0 | 0 | NA | NA | NA | NA | NA | NA | NA | NA | NA | NA |

| | | | | | | | | | | | | | | | | | | | | | | | | | |
|---|---|---|---|---|---|---|---|---|---|---|---|---|---|---|---|---|---|---|---|---|---|---|---|---|---|
| Trim31 | 3 | 3 | 1 | 0 | 1 | 1 | 2 | 3 | 1 | 0 | 1 | 1 | 1.45 | 2 | 1 | 0.315 | -1.665 | 0.32491817 | NA | 0 | 1.6341 | 1.0987 | 1.0987 | TRUE | 0.2665 |
| Trim32 | 1461 | 1080 | 955 | 861 | 811 | 899 | 1135 | 1149 | 1030 | 857 | 915 | 887 | 995.55 | 1105 | 886 | 0.802 | -0.319 | 6.68E-05 | 0.00088584 | 0.0013 | 0.0048 | 0.0036 | 0.0036 | TRUE | 0.0402 |
| Trim33 | 7404 | 5393 | 5367 | 5510 | 4877 | 5837 | 5750 | 5736 | 5791 | 5485 | 5502 | 5762 | 5671.04 | 5759 | 5583 | 0.969 | -0.045 | 0.37600493 | 0.55058278 | 2.00E-04 | 0.0028 | 0.0017 | 0.0017 | TRUE | 0.0097 |
| Trim34a | 57 | 55 | 71 | 66 | 50 | 57 | 44 | 59 | 77 | 66 | 56 | 56 | 59.62 | 60 | 59 | 1.003 | 0.005 | 0.98565981 | 0.99281435 | 0.027 | 0.0422 | 0.0378 | 0.0378 | TRUE | 0.254 |
| Trim34b | 52 | 40 | 54 | 53 | 41 | 56 | 40 | 43 | 58 | 53 | 46 | 55 | 49.25 | 47 | 51 | 1.1 | 0.138 | 0.62696866 | 0.76493923 | 0 | 0.0506 | 0.0379 | 0.0379 | TRUE | 0.3292 |
| Trim35 | 4189 | 2778 | 2447 | 3106 | 2883 | 3073 | 3253 | 2955 | 2640 | 3092 | 3252 | 3034 | 3037.7 | 2949 | 3126 | 1.059 | 0.082 | 0.29536446 | 0.46709939 | 0.0053 | 0.0032 | 0.0041 | 0.0041 | TRUE | 0.1019 |
| Trim36 | 345 | 211 | 267 | 299 | 323 | 283 | 268 | 224 | 288 | 298 | 364 | 279 | 286.97 | 260 | 314 | 1.204 | 0.267 | 0.06546014 | 0.16342738 | 0.014 | 0.0107 | 0.0117 | 0.0117 | TRUE | 0.22 |
| Trim37 | 13939 | 10513 | 11305 | 11671 | 10148 | 11864 | 10826 | 11183 | 12198 | 11617 | 11448 | 11712 | 11497.21 | 11402 | 11592 | 1.017 | 0.024 | 0.67590896 | 0.80105861 | 0.0016 | 0.0026 | 0.0023 | 0.0023 | TRUE | 0.0455 |
| Trim38 | 14 | 11 | 4 | 4 | 7 | 6 | 11 | 12 | 4 | 4 | 8 | 6 | 7.45 | 9 | 6 | 0.653 | -0.614 | 0.40116878 | NA | 0 | 0.3206 | 0.2443 | 0.2443 | TRUE | 0.5983 |
| Trim39 | 989 | 808 | 571 | 548 | 517 | 610 | 768 | 859 | 616 | 545 | 583 | 602 | 662.43 | 748 | 577 | 0.771 | -0.375 | 0.00147805 | 0.0095528 | 0.0139 | 0.006 | 0.0085 | 0.0085 | TRUE | 0.2741 |
| Trim40 | 41 | 32 | 22 | 60 | 40 | 46 | 32 | 34 | 24 | 60 | 45 | 45 | 39.98 | 30 | 50 | 1.673 | 0.743 | 0.0192753 | 0.06726118 | 0 | 0.0617 | 0.0462 | 0.0462 | TRUE | 0.2342 |
| Trim41 | 7694 | 5598 | 4901 | 5173 | 4664 | 5313 | 5976 | 5955 | 5288 | 5149 | 5261 | 5245 | 5478.95 | 5740 | 5218 | 0.909 | -0.138 | 0.02729474 | 0.08640504 | 0.0021 | 0.0029 | 0.0026 | 0.0026 | TRUE | 0.0573 |
| Trim42 | 0 | 0 | 0 | 0 | 1 | 0 | 0 | 0 | 0 | 0 | 1 | 0 | 0.19 | 0 | 0 | 2.094 | 1.067 | 0.79379329 | NA | 0 | 12.608 | 10 | 10 | TRUE | 0.5075 |
| Trim43a | 141 | 98 | 33 | 8 | 18 | 8 | 110 | 104 | 36 | 8 | 20 | 8 | 47.59 | 83 | 12 | 0.143 | -2.81 | 1.70E-09 | 1.07E-07 | 0.2736 | 0.0522 | 0.1074 | 0.1074 | TRUE | 2.2768 |
| Trim43b | 28 | 13 | 5 | 1 | 0 | 0 | 22 | 14 | 5 | 1 | 0 | 0 | 6.99 | 14 | 0 | 0.027 | -5.218 | 0.00016082 | NA | 0 | 0.3413 | 0.3428 | 0.3428 | TRUE | 0.2064 |
| Trim43c | 7 | 3 | 1 | 0 | 0 | 3 | 5 | 3 | 1 | 0 | 0 | 3 | 2.11 | 3 | 1 | 0.307 | -1.702 | 0.27088085 | NA | 0.7718 | 1.125 | 1.0624 | 1.0624 | TRUE | 0.576 |
| Trim44 | 3264 | 2332 | 2414 | 3076 | 2360 | 2815 | 2535 | 2481 | 2605 | 3062 | 2662 | 2779 | 2687.2 | 2540 | 2834 | 1.116 | 0.159 | 0.02051508 | 0.07047072 | 0.0024 | 0.0033 | 0.003 | 0.003 | TRUE | 0.06 |
| Trim45 | 997 | 710 | 758 | 813 | 728 | 853 | 774 | 755 | 818 | 809 | 821 | 842 | 803.33 | 782 | 824 | 1.054 | 0.075 | 0.35096273 | 0.52570847 | 0 | 0.0054 | 0.0035 | 0.0035 | TRUE | 0.0178 |
| Trim46 | 105 | 109 | 107 | 122 | 128 | 107 | 82 | 116 | 115 | 121 | 144 | 106 | 114.07 | 104 | 124 | 1.192 | 0.254 | 0.22908794 | 0.39449868 | 0.0234 | 0.0232 | 0.0233 | 0.0233 | TRUE | 0.4038 |
| Trim47 | 176 | 161 | 200 | 302 | 285 | 283 | 137 | 171 | 216 | 301 | 322 | 279 | 237.54 | 175 | 301 | 1.731 | 0.792 | 1.74E-06 | 4.51E-05 | 0.0226 | 0.0124 | 0.0152 | 0.0152 | TRUE | 0.4543 |
| Trim50 | 7 | 4 | 9 | 12 | 14 | 9 | 5 | 4 | 10 | 12 | 16 | 9 | 9.34 | 6 | 12 | 1.896 | 0.923 | 0.15746043 | NA | 0 | 0.2562 | 0.1893 | 0.1893 | TRUE | 0.3197 |
| Trim52 | 312 | 331 | 353 | 347 | 268 | 295 | 242 | 352 | 381 | 345 | 302 | 291 | 319.04 | 325 | 313 | 0.967 | -0.048 | 0.76518939 | 0.86070374 | 0.0292 | 0.0099 | 0.0157 | 0.0157 | TRUE | 0.5972 |
| Trim54 | 79 | 53 | 38 | 77 | 67 | 71 | 61 | 56 | 41 | 77 | 76 | 70 | 63.51 | 53 | 74 | 1.391 | 0.476 | 0.05985562 | 0.15298429 | 0 | 0.0397 | 0.0301 | 0.0301 | TRUE | 0.3027 |
| Trim55 | 0 | 0 | 0 | 0 | 0 | 0 | 0 | 0 | 0 | 0 | 0 | 0 | 0 | 0 | 0 | NA | NA | NA | NA | NA | NA | NA | NA | NA | NA |
| Trim56 | 212 | 155 | 199 | 246 | 236 | 277 | 165 | 165 | 215 | 245 | 266 | 273 | 221.46 | 182 | 261 | 1.446 | 0.532 | 0.00040371 | 0.00367293 | 0.0078 | 0.0131 | 0.0116 | 0.0116 | TRUE | 0.2671 |
| Trim58 | 2 | 0 | 1 | 0 | 0 | 0 | 2 | 0 | 1 | 0 | 0 | 0 | 0.44 | 1 | 0 | 0.214 | -2.223 | 0.47051571 | NA | 0 | 5.4047 | 5.2946 | 5.2946 | TRUE | 0.2788 |
| Trim59 | 7569 | 5441 | 5566 | 5700 | 4699 | 5521 | 5879 | 5788 | 6006 | 5674 | 5301 | 5450 | 5682.75 | 5891 | 5475 | 0.93 | -0.105 | 0.04671159 | 0.12819678 | 5.00E-04 | 0.0028 | 0.0018 | 0.0018 | TRUE | 0.0123 |
| Trim6 | 8402 | 6224 | 5553 | 5928 | 5106 | 5668 | 6525 | 6620 | 5992 | 5901 | 5760 | 5595 | 6065.59 | 6379 | 5752 | 0.902 | -0.15 | 0.01083897 | 0.04382499 | 0.0015 | 0.0028 | 0.0023 | 0.0023 | TRUE | 0.0342 |
| Trim60 | 53 | 40 | 44 | 33 | 30 | 27 | 41 | 43 | 47 | 33 | 34 | 27 | 37.42 | 44 | 31 | 0.712 | -0.49 | 0.11989143 | 0.25155727 | 0 | 0.0658 | 0.0439 | 0.0439 | TRUE | 0.1095 |
| Trim61 | 10 | 6 | 4 | 6 | 8 | 8 | 8 | 6 | 4 | 6 | 9 | 8 | 6.89 | 6 | 8 | 1.222 | 0.29 | 0.68787882 | NA | 0 | 0.3463 | 0.2292 | 0.2292 | TRUE | 0.1651 |
| Trim62 | 192 | 153 | 144 | 161 | 187 | 210 | 149 | 163 | 155 | 160 | 211 | 207 | 174.29 | 156 | 193 | 1.239 | 0.309 | 0.05883918 | 0.15122971 | 0.0072 | 0.016 | 0.0134 | 0.0134 | TRUE | 0.2398 |
| Trim63 | 257 | 218 | 248 | 317 | 297 | 395 | 200 | 232 | 268 | 316 | 335 | 390 | 289.93 | 233 | 347 | 1.495 | 0.58 | 5.66E-05 | 0.00077318 | 0.0132 | 0.0106 | 0.0114 | 0.0114 | TRUE | 0.1892 |
| Trim65 | 38 | 33 | 37 | 68 | 60 | 65 | 30 | 35 | 40 | 68 | 68 | 64 | 50.68 | 35 | 67 | 1.923 | 0.943 | 0.00066302 | 0.00534099 | 0 | 0.0492 | 0.0336 | 0.0336 | TRUE | 0.1562 |
| Trim66 | 150 | 96 | 90 | 82 | 76 | 78 | 116 | 102 | 97 | 82 | 86 | 77 | 93.35 | 105 | 82 | 0.77 | -0.376 | 0.06525134 | 0.16310554 | 0 | 0.0278 | 0.0191 | 0.0191 | TRUE | 0.096 |
| Trim67 | 63 | 45 | 33 | 46 | 50 | 48 | 49 | 48 | 36 | 46 | 56 | 47 | 47 | 44 | 50 | 1.123 | 0.167 | 0.56195996 | 0.71665003 | 0 | 0.0529 | 0.0383 | 0.0383 | TRUE | 0.2078 |
| Trim68 | 491 | 416 | 330 | 330 | 333 | 315 | 381 | 442 | 356 | 328 | 376 | 311 | 365.83 | 393 | 338 | 0.86 | -0.218 | 0.08310698 | 0.19410862 | 0.0081 | 0.0089 | 0.0086 | 0.0086 | TRUE | 0.133 |
| Trim69 | 0 | 0 | 1 | 2 | 0 | 0 | 0 | 0 | 1 | 2 | 0 | 0 | 0.51 | 0 | 1 | 1.943 | 0.958 | 0.74995199 | NA | 0 | 4.6348 | 4.7457 | 4.7457 | TRUE | 0.9738 |
| Trim7 | 72 | 52 | 60 | 75 | 63 | 67 | 56 | 55 | 65 | 75 | 71 | 66 | 64.64 | 59 | 71 | 1.206 | 0.271 | 0.25955573 | 0.42956694 | 0 | 0.0391 | 0.0259 | 0.0259 | TRUE | 0.066 |
| Trim71 | 7432 | 5294 | 4013 | 5447 | 4607 | 4930 | 5772 | 5631 | 4330 | 5422 | 5197 | 4867 | 5203.18 | 5244 | 5162 | 0.984 | -0.023 | 0.80695749 | 0.88842854 | 0.0135 | 0.0029 | 0.0064 | 0.0064 | TRUE | 0.2826 |
| Trim72 | 22 | 16 | 19 | 31 | 33 | 31 | 17 | 17 | 21 | 31 | 37 | 31 | 25.55 | 18 | 33 | 1.81 | 0.856 | 0.02524831 | 0.08184941 | 0 | 0.0952 | 0.0631 | 0.0631 | TRUE | 0.0853 |
| Trim75 | 1 | 1 | 2 | 0 | 0 | 1 | 1 | 1 | 2 | 0 | 0 | 1 | 0.83 | 1 | 0 | 0.282 | -1.828 | 0.39771905 | NA | 0 | 2.8548 | 1.9998 | 1.9998 | TRUE | 0.3835 |
| Trim8 | 1442 | 1203 | 869 | 1072 | 921 | 1067 | 1120 | 1280 | 938 | 1067 | 1039 | 1053 | 1082.76 | 1113 | 1053 | 0.947 | -0.079 | 0.4421842 | 0.61324847 | 0.0111 | 0.0046 | 0.0067 | 0.0067 | TRUE | 0.2035 |
| Trim9 | 2 | 1 | 1 | 2 | 0 | 3 | 2 | 1 | 1 | 2 | 0 | 3 | 1.44 | 1 | 2 | 1.346 | 0.428 | 0.78916252 | NA | 0 | 1.6467 | 1.1528 | 1.1528 | TRUE | 0.3902 |
| Triml1 | 5414 | 4501 | 4457 | 5511 | 5076 | 5981 | 4205 | 4788 | 4809 | 5486 | 5726 | 5904 | 5152.94 | 4601 | 5705 | 1.241 | 0.311 | 3.81E-06 | 8.50E-05 | 0.0032 | 0.0029 | 0.0031 | 0.0031 | TRUE | 0.07 |
| Triml2 | 9496 | 7063 | 6674 | 7140 | 6513 | 7539 | 7375 | 7513 | 7201 | 7107 | 7347 | 7442 | 7330.97 | 7363 | 7299 | 0.991 | -0.013 | 0.79983827 | 0.88333244 | 3.00E-04 | 0.0027 | 0.0017 | 0.0017 | TRUE | 0.0065 |
| Trio | 1960 | 1399 | 1329 | 1432 | 1277 | 1384 | 1522 | 1488 | 1434 | 1425 | 1441 | 1366 | 1446.09 | 1481 | 1411 | 0.952 | -0.072 | 0.28982432 | 0.4618935 | 0 | 0.0041 | 0.0026 | 0.0026 | TRUE | 0.0094 |
| Triobp | 2881 | 2271 | 2611 | 2777 | 2548 | 2882 | 2238 | 2416 | 2817 | 2764 | 2874 | 2845 | 2659.01 | 2490 | 2828 | 1.137 | 0.185 | 0.02550063 | 0.08248032 | 0.0067 | 0.0033 | 0.0045 | 0.0045 | TRUE | 0.1596 |
| Trip10 | 3683 | 2945 | 2709 | 2490 | 2315 | 2609 | 2860 | 3133 | 2923 | 2479 | 2612 | 2575 | 2763.59 | 2972 | 2555 | 0.86 | -0.218 | 0.00048427 | 0.00421793 | 0.0012 | 0.0033 | 0.0024 | 0.0024 | TRUE | 0.027 |
| Trip11 | 1564 | 1127 | 1014 | 1191 | 1030 | 1165 | 1215 | 1199 | 1094 | 1186 | 1162 | 1150 | 1167.51 | 1169 | 1166 | 0.996 | -0.005 | 0.94263694 | 0.9680349 | 9.00E-04 | 0.0045 | 0.0032 | 0.0032 | TRUE | 0.037 |
| Trip12 | 54837 | 37430 | 34845 | 36821 | 31641 | 35266 | 42590 | 39814 | 37597 | 36652 | 35694 | 34813 | 37859.96 | 40000 | 35720 | 0.893 | -0.163 | 0.00478594 | 0.02371074 | 0.0022 | 0.0025 | 0.0024 | 0.0024 | TRUE | 0.0393 |
| Trip13 | 4542 | 3021 | 2598 | 2508 | 2250 | 2456 | 3528 | 3213 | 2803 | 2496 | 2538 | 2424 | 2833.89 | 3181 | 2486 | 0.781 | -0.357 | 1.34E-05 | 0.00024071 | 0.0062 | 0.0033 | 0.0045 | 0.0045 | TRUE | 0.1309 |
| Trip4 | 2766 | 2004 | 2014 | 2243 | 1983 | 2313 | 2148 | 2132 | 2173 | 2233 | 2237 | 2283 | 2200.99 | 2151 | 2251 | 1.047 | 0.066 | 0.25660234 | 0.42653739 | 0 | 0.0035 | 0.002 | 0.002 | TRUE | 0.0019 |
| Trip6 | 1029 | 690 | 676 | 683 | 619 | 721 | 799 | 734 | 729 | 680 | 698 | 712 | 725.4 | 754 | 697 | 0.922 | -0.117 | 0.17301283 | 0.32542077 | 0 | 0.0057 | 0.0039 | 0.0039 | TRUE | 0.0341 |
| Triqk | 103 | 89 | 88 | 94 | 105 | 85 | 80 | 95 | 95 | 94 | 118 | 84 | 94.26 | 90 | 99 | 1.099 | 0.136 | 0.5282107 | 0.68874916 | 0.0103 | 0.0276 | 0.0227 | 0.0227 | TRUE | 0.2875 |
| Trit1 | 2050 | 1531 | 1786 | 1940 | 1700 | 2042 | 1592 | 1629 | 1927 | 1931 | 1918 | 2016 | 1835.39 | 1716 | 1955 | 1.14 | 0.19 | 0.0205719 | 0.07061338 | 0.0053 | 0.0037 | 0.0043 | 0.0043 | TRUE | 0.1385 |
| Trmt1 | 6986 | 5321 | 4947 | 5261 | 4878 | 5472 | 5426 | 5660 | 5338 | 5237 | 5503 | 5402 | 5427.46 | 5475 | 5381 | 0.983 | -0.025 | 0.63891704 | 0.7745582 | 5.00E-04 | 0.0029 | 0.0019 | 0.0019 | TRUE | 0.0106 |
| Trmt10a | 2736 | 1890 | 1770 | 1869 | 1629 | 1930 | 2125 | 2010 | 1910 | 1860 | 1838 | 1905 | 1941.4 | 2015 | 1868 | 0.926 | -0.111 | 0.09713178 | 0.21675325 | 0.0012 | 0.0036 | 0.0027 | 0.0027 | TRUE | 0.0284 |
| Trmt10b | 1248 | 856 | 770 | 937 | 831 | 956 | 969 | 911 | 831 | 933 | 937 | 944 | 920.75 | 904 | 938 | 1.036 | 0.052 | 0.53774102 | 0.69625259 | 0.002 | 0.005 | 0.004 | 0.004 | TRUE | 0.0584 |
| Trmt10c | 2196 | 1611 | 1451 | 1608 | 1303 | 1583 | 1706 | 1714 | 1566 | 1601 | 1470 | 1563 | 1602.99 | 1662 | 1545 | 0.929 | -0.106 | 0.13861227 | 0.27970413 | 0.0015 | 0.0039 | 0.003 | 0.003 | TRUE | 0.0302 |
| Trmt11 | 864 | 603 | 539 | 479 | 411 | 502 | 671 | 641 | 582 | 477 | 464 | 496 | 555 | 631 | 479 | 0.757 | -0.401 | 4.21E-05 | 0.00060834 | 0.0018 | 0.0067 | 0.0051 | 0.0051 | TRUE | 0.055 |
| Trmt112 | 1758 | 1285 | 1262 | 1247 | 1247 | 1409 | 1365 | 1367 | 1362 | 1241 | 1407 | 1391 | 1355.47 | 1365 | 1346 | 0.986 | -0.02 | 0.78754546 | 0.87523421 | 0.0016 | 0.0042 | 0.0032 | 0.0032 | TRUE | 0.0564 |
| Trmt12 | 230 | 150 | 90 | 88 | 96 | 86 | 179 | 160 | 97 | 88 | 108 | 85 | 119.35 | 145 | 94 | 0.641 | -0.642 | 0.00608716 | 0.0285352 | 0.0505 | 0.0223 | 0.0306 | 0.0306 | TRUE | 0.7846 |
| Trmt13 | 638 | 486 | 495 | 523 | 405 | 590 | 496 | 517 | 534 | 521 | 457 | 582 | 517.74 | 516 | 520 | 1.011 | 0.015 | 0.88720153 | 0.93716439 | 0.0059 | 0.007 | 0.0067 | 0.0067 | TRUE | 0.1289 |
| Trmt1l | 1720 | 1210 | 1107 | 1185 | 1038 | 1267 | 1336 | 1287 | 1194 | 1180 | 1171 | 1251 | 1236.43 | 1272 | 1201 | 0.943 | -0.085 | 0.26081123 | 0.43080723 | 0.0015 | 0.0043 | 0.0033 | 0.0033 | TRUE | 0.0341 |
| Trmt2a | 2965 | 2058 | 1701 | 1812 | 1611 | 1824 | 2303 | 2189 | 1835 | 1804 | 1817 | 1801 | 1958.14 | 2109 | 1807 | 0.856 | -0.224 | 0.00815073 | 0.03554475 | 0.0059 | 0.0036 | 0.0047 | 0.0047 | TRUE | 0.1547 |
| Trmt2b | 746 | 615 | 667 | 619 | 553 | 664 | 579 | 654 | 720 | 616 | 624 | 655 | 641.45 | 651 | 632 | 0.973 | -0.04 | 0.69464716 | 0.81381901 | 0.0051 | 0.0061 | 0.0058 | 0.0058 | TRUE | 0.1155 |
| Trmt44 | 783 | 528 | 473 | 544 | 511 | 564 | 608 | 562 | 510 | 541 | 576 | 557 | 559.14 | 560 | 558 | 0.994 | -0.008 | 0.93321234 | 0.9627836 | 0.0027 | 0.0067 | 0.0054 | 0.0054 | TRUE | 0.0698 |
| Trmt5 | 1749 | 1262 | 1084 | 1229 | 1037 | 1196 | 1358 | 1342 | 1170 | 1223 | 1170 | 1181 | 1240.7 | 1290 | 1191 | 0.923 | -0.116 | 0.14735145 | 0.29194543 | 0.0029 | 0.0043 | 0.0038 | 0.0038 | TRUE | 0.0787 |
| Trmt6 | 3714 | 2353 | 1989 | 1875 | 1711 | 1845 | 2885 | 2503 | 2146 | 1866 | 1930 | 1821 | 2191.89 | 2511 | 1872 | 0.745 | -0.425 | 7.98E-06 | 0.00015602 | 0.0105 | 0.0035 | 0.0061 | 0.0061 | TRUE | 0.1554 |

| | | | | | | | | | | | | | | | | | | | | | | | | | |
|---|---|---|---|---|---|---|---|---|---|---|---|---|---|---|---|---|---|---|---|---|---|---|---|---|---|
| Trmt61a | 1437 | 1095 | 867 | 932 | 894 | 921 | 1116 | 1165 | 935 | 928 | 1009 | 909 | 1010.28 | 1072 | 949 | 0.884 | -0.178 | 0.06103304 | 0.15519395 | 0.0071 | 0.0048 | 0.0055 | 0.0055 | TRUE | 0.1453 |
| Trmt61b | 317 | 208 | 166 | 212 | 151 | 196 | 246 | 221 | 179 | 211 | 170 | 193 | 203.57 | 215 | 191 | 0.887 | -0.173 | 0.28564951 | 0.4576604 | 0.0138 | 0.0141 | 0.014 | 0.014 | TRUE | 0.2336 |
| Trmu | 217 | 211 | 125 | 133 | 130 | 153 | 169 | 224 | 135 | 132 | 147 | 151 | 159.65 | 176 | 143 | 0.815 | -0.295 | 0.12813354 | 0.2643494 | 0.029 | 0.0173 | 0.0207 | 0.0207 | TRUE | 0.3042 |
| Trnau1ap | 1876 | 1318 | 1320 | 1478 | 1282 | 1466 | 1457 | 1402 | 1424 | 1471 | 1446 | 1447 | 1441.3 | 1428 | 1455 | 1.019 | 0.027 | 0.68423923 | 0.80651751 | 0 | 0.0041 | 0.0024 | 0.0024 | TRUE | 0.0041 |
| Trnp1 | 86 | 69 | 63 | 77 | 66 | 83 | 67 | 73 | 68 | 77 | 74 | 82 | 73.53 | 69 | 78 | 1.122 | 0.166 | 0.45769979 | 0.62749036 | 0 | 0.0347 | 0.0224 | 0.0224 | TRUE | 0.0221 |
| Trnt1 | 8403 | 5970 | 5129 | 5435 | 4752 | 5237 | 6526 | 6350 | 5534 | 5410 | 5361 | 5170 | 5725.18 | 6137 | 5314 | 0.866 | -0.208 | 0.00262462 | 0.01488051 | 0.0037 | 0.0028 | 0.0033 | 0.0033 | TRUE | 0.0896 |
| Tro | 3713 | 2806 | 2621 | 2836 | 2462 | 2708 | 2884 | 2985 | 2828 | 2823 | 2777 | 2673 | 2828.34 | 2899 | 2758 | 0.951 | -0.072 | 0.21719124 | 0.380002 | 4.00E-04 | 0.0033 | 0.0021 | 0.0021 | TRUE | 0.0088 |
| Troap | 1301 | 1022 | 933 | 1036 | 920 | 1099 | 1010 | 1087 | 1007 | 1031 | 1038 | 1085 | 1043.03 | 1035 | 1051 | 1.017 | 0.024 | 0.7523784 | 0.85313324 | 0 | 0.0047 | 0.0032 | 0.0032 | TRUE | 0.0229 |
| Trove2 | 2522 | 1839 | 1866 | 2253 | 1906 | 2305 | 1959 | 1956 | 2013 | 2243 | 2150 | 2275 | 2099.41 | 1976 | 2223 | 1.125 | 0.17 | 0.00505565 | 0.02474042 | 0 | 0.0036 | 0.0022 | 0.0022 | TRUE | 0.0097 |
| Trp53 | 24965 | 17572 | 15369 | 14387 | 13675 | 16502 | 19389 | 18691 | 16583 | 14321 | 15427 | 16290 | 16783.51 | 18221 | 15346 | 0.842 | -0.248 | 0.00052217 | 0.00445764 | 0.0052 | 0.0026 | 0.0036 | 0.0036 | TRUE | 0.0756 |
| Trp53bp1 | 6756 | 5159 | 5021 | 5074 | 4805 | 5456 | 5247 | 5488 | 5418 | 5051 | 5421 | 5386 | 5334.9 | 5384 | 5286 | 0.982 | -0.026 | 0.63087775 | 0.76820034 | 7.00E-04 | 0.0029 | 0.002 | 0.002 | TRUE | 0.0185 |
| Trp53bp2 | 3097 | 2168 | 1988 | 2162 | 1921 | 2209 | 2405 | 2306 | 2145 | 2152 | 2167 | 2181 | 2226.03 | 2285 | 2167 | 0.947 | -0.078 | 0.23189338 | 0.39777528 | 0.0013 | 0.0035 | 0.0026 | 0.0026 | TRUE | 0.0347 |
| Trp53i11 | 3204 | 2492 | 3336 | 4627 | 4304 | 4964 | 2488 | 2651 | 3600 | 4606 | 4855 | 4900 | 3849.99 | 2913 | 4787 | 1.644 | 0.718 | 4.70E-11 | 4.07E-09 | 0.0214 | 0.003 | 0.0083 | 0.0083 | TRUE | 0.5151 |
| Trp53i13 | 1103 | 799 | 745 | 904 | 790 | 894 | 857 | 850 | 804 | 900 | 891 | 883 | 863.99 | 837 | 891 | 1.064 | 0.09 | 0.24903462 | 0.41806702 | 0 | 0.0052 | 0.0032 | 0.0032 | TRUE | 0.0137 |
| Trp53inp1 | 14589 | 10401 | 11159 | 12901 | 10449 | 12300 | 11331 | 11063 | 12040 | 12842 | 11788 | 12142 | 11867.62 | 11478 | 12257 | 1.068 | 0.095 | 0.09745112 | 0.21729623 | 0.0018 | 0.0026 | 0.0023 | 0.0023 | TRUE | 0.0223 |
| Trp53inp2 | 3666 | 2572 | 2411 | 3189 | 2630 | 3096 | 2847 | 2736 | 2601 | 3174 | 2967 | 3056 | 2896.99 | 2728 | 3066 | 1.123 | 0.168 | 0.00705802 | 0.03194436 | 0.0011 | 0.0032 | 0.0024 | 0.0024 | TRUE | 0.0199 |
| Trp53rk | 264 | 175 | 130 | 151 | 155 | 158 | 205 | 186 | 140 | 150 | 175 | 156 | 168.76 | 177 | 160 | 0.901 | -0.151 | 0.39055086 | 0.564423 | 0.0162 | 0.0165 | 0.0164 | 0.0164 | TRUE | 0.3445 |
| Trp53tg5 | 0 | 0 | 0 | 1 | 0 | 0 | 0 | 0 | 0 | 1 | 0 | 0 | 0.17 | 0 | 0 | 2.094 | 1.067 | 0.79379329 | NA | 0 | 14.2885 | 10 | 10 | TRUE | 0.3906 |
| Trp63 | 38 | 33 | 14 | 19 | 9 | 17 | 30 | 35 | 15 | 19 | 10 | 17 | 20.93 | 27 | 15 | 0.576 | -0.795 | 0.09057162 | 0.2056131 | 0.0841 | 0.1157 | 0.1072 | 0.1072 | TRUE | 0.4819 |
| Trp73 | 175 | 100 | 111 | 144 | 118 | 169 | 136 | 106 | 120 | 143 | 133 | 167 | 134.22 | 121 | 148 | 1.221 | 0.288 | 0.11369124 | 0.24275305 | 0.0072 | 0.0201 | 0.0164 | 0.0164 | TRUE | 0.1366 |
| Trpa1 | 675 | 463 | 332 | 274 | 243 | 297 | 524 | 492 | 358 | 273 | 274 | 293 | 369.17 | 458 | 280 | 0.61 | -0.714 | 7.08E-07 | 2.19E-05 | 0.019 | 0.0088 | 0.012 | 0.012 | TRUE | 0.4154 |
| Trpc1 | 379 | 255 | 268 | 342 | 282 | 332 | 294 | 271 | 289 | 340 | 318 | 328 | 306.84 | 285 | 329 | 1.153 | 0.205 | 0.07921514 | 0.18776485 | 0 | 0.0102 | 0.0066 | 0.0066 | TRUE | 0.0193 |
| Trpc2 | 662 | 459 | 464 | 453 | 413 | 486 | 514 | 488 | 501 | 451 | 466 | 480 | 483.27 | 501 | 466 | 0.928 | -0.107 | 0.26751297 | 0.43808007 | 0 | 0.0073 | 0.0046 | 0.0046 | TRUE | 0.009 |
| Trpc3 | 48 | 36 | 17 | 23 | 17 | 25 | 37 | 38 | 18 | 23 | 19 | 25 | 26.78 | 31 | 22 | 0.708 | -0.498 | 0.21842304 | 0.38146635 | 0.049 | 0.0909 | 0.0794 | 0.0794 | TRUE | 0.827 |
| Trpc4 | 0 | 1 | 3 | 3 | 0 | 1 | 0 | 1 | 3 | 3 | 0 | 1 | 1.38 | 1 | 1 | 0.968 | -0.047 | 0.97934616 | NA | 1.1676 | 1.7211 | 1.5961 | 1.5961 | TRUE | 0.4926 |
| Trpc4ap | 6690 | 5010 | 5253 | 5331 | 4796 | 5616 | 5196 | 5329 | 5668 | 5306 | 5410 | 5544 | 5408.93 | 5398 | 5420 | 1.005 | 0.007 | 0.90800205 | 0.94931412 | 0.0011 | 0.0029 | 0.0021 | 0.0021 | TRUE | 0.0233 |
| Trpc5 | 0 | 0 | 0 | 0 | 0 | 0 | 0 | 0 | 0 | 0 | 0 | 0 | 0 | 0 | 0 | NA | NA | NA | NA | NA | NA | NA | NA | NA | NA |
| Trpc6 | 31 | 11 | 4 | 7 | 5 | 9 | 24 | 12 | 4 | 7 | 6 | 9 | 10.26 | 13 | 7 | 0.528 | -0.922 | 0.18802207 | NA | 0.2733 | 0.2333 | 0.2456 | 0.2456 | TRUE | 0.2389 |
| Trpc7 | 12 | 12 | 11 | 9 | 8 | 13 | 9 | 13 | 12 | 9 | 9 | 13 | 10.79 | 11 | 10 | 0.916 | -0.127 | 0.82640524 | NA | 0 | 0.222 | 0.1471 | 0.1471 | TRUE | 0.1741 |
| Trpd52l3 | 10 | 7 | 1 | 1 | 3 | 1 | 8 | 7 | 1 | 1 | 3 | 1 | 3.61 | 5 | 2 | 0.319 | -1.646 | 0.15331517 | NA | 0 | 0.659 | 0.5749 | 0.5749 | TRUE | 0.9714 |
| Trpm1 | 17 | 22 | 19 | 19 | 12 | 25 | 13 | 23 | 21 | 19 | 14 | 25 | 19.04 | 19 | 19 | 1.016 | 0.023 | 0.96106541 | 0.97845108 | 0.0331 | 0.1269 | 0.1014 | 0.1014 | TRUE | 0.2546 |
| Trpm2 | 28 | 11 | 12 | 24 | 21 | 26 | 22 | 12 | 13 | 24 | 24 | 26 | 19.94 | 16 | 25 | 1.553 | 0.635 | 0.15275759 | 0.29950126 | 0 | 0.1213 | 0.0901 | 0.0901 | TRUE | 0.7265 |
| Trpm3 | 22 | 10 | 9 | 11 | 17 | 8 | 17 | 11 | 10 | 11 | 19 | 8 | 12.58 | 13 | 13 | 0.995 | -0.007 | 0.99046573 | NA | 0.0747 | 0.1909 | 0.1578 | 0.1578 | TRUE | 0.7467 |
| Trpm4 | 705 | 599 | 742 | 811 | 780 | 828 | 548 | 637 | 801 | 807 | 880 | 817 | 748.31 | 662 | 835 | 1.264 | 0.338 | 0.00570378 | 0.02714754 | 0.0179 | 0.0056 | 0.0094 | 0.0094 | TRUE | 0.3959 |
| Trpm5 | 1 | 1 | 5 | 1 | 0 | 0 | 1 | 1 | 5 | 1 | 0 | 0 | 1.37 | 2 | 0 | 0.156 | -2.678 | 0.16499353 | NA | 0.7906 | 1.7301 | 1.4766 | 1.4766 | TRUE | 1.1718 |
| Trpm6 | 108 | 90 | 49 | 87 | 69 | 70 | 84 | 96 | 53 | 87 | 78 | 69 | 77.67 | 78 | 78 | 1.002 | 0.003 | 0.99196218 | 0.99565418 | 0.0361 | 0.0329 | 0.0338 | 0.0338 | TRUE | 0.6691 |
| Trpm7 | 8847 | 6567 | 6741 | 6762 | 5742 | 6744 | 6871 | 6985 | 7273 | 6731 | 6478 | 6657 | 6832.61 | 7043 | 6622 | 0.94 | -0.089 | 0.08364346 | 0.19480462 | 4.00E-04 | 0.0028 | 0.0017 | 0.0017 | TRUE | 0.0099 |
| Trpm8 | 1 | 0 | 0 | 0 | 0 | 0 | 1 | 0 | 0 | 0 | 0 | 0 | 0.13 | 0 | 0 | 0.552 | -0.857 | 0.8336548 | NA | 0 | 18.3121 | 10 | 10 | TRUE | 0.2566 |
| Trps1 | 1982 | 1326 | 1409 | 1617 | 1464 | 1561 | 1539 | 1410 | 1520 | 1610 | 1652 | 1541 | 1545.36 | 1490 | 1601 | 1.073 | 0.102 | 0.14463937 | 0.28812968 | 0.001 | 0.004 | 0.0029 | 0.0029 | TRUE | 0.0259 |
| Trpt1 | 175 | 118 | 118 | 131 | 125 | 138 | 136 | 126 | 127 | 130 | 141 | 136 | 132.73 | 130 | 136 | 1.046 | 0.065 | 0.70100788 | 0.81872084 | 0 | 0.0203 | 0.0131 | 0.0131 | TRUE | 0.0209 |
| Trpv1 | 7 | 3 | 4 | 1 | 2 | 3 | 5 | 3 | 4 | 1 | 2 | 3 | 3.19 | 4 | 2 | 0.475 | -1.073 | 0.32300982 | NA | 0 | 0.7447 | 0.4877 | 0.4877 | TRUE | 0.2014 |
| Trpv2 | 391 | 288 | 229 | 293 | 235 | 278 | 304 | 306 | 247 | 292 | 265 | 274 | 281.38 | 286 | 277 | 0.968 | -0.046 | 0.72556296 | 0.83572689 | 0.0045 | 0.0108 | 0.009 | 0.009 | TRUE | 0.1508 |
| Trpv3 | 43 | 15 | 16 | 18 | 9 | 11 | 33 | 16 | 17 | 18 | 10 | 11 | 17.59 | 22 | 13 | 0.579 | -0.789 | 0.1204366 | 0.25244176 | 0.0967 | 0.1372 | 0.1249 | 0.1249 | TRUE | 1.3113 |
| Trpv4 | 113 | 114 | 109 | 136 | 104 | 133 | 88 | 121 | 118 | 135 | 117 | 131 | 118.44 | 109 | 128 | 1.185 | 0.244 | 0.21043031 | 0.37232983 | 0.0102 | 0.0224 | 0.0189 | 0.0189 | TRUE | 0.3233 |
| Trpv5 | 6 | 1 | 0 | 3 | 0 | 4 | 5 | 1 | 0 | 3 | 0 | 4 | 2.11 | 2 | 2 | 1.174 | 0.232 | 0.87826988 | NA | 1.4288 | 1.1258 | 1.1858 | 1.1858 | TRUE | 1.7463 |
| Trpv6 | 2 | 0 | 2 | 0 | 0 | 0 | 2 | 0 | 2 | 0 | 0 | 0 | 0.62 | 1 | 0 | 0.153 | -2.705 | 0.31809589 | NA | 0 | 3.8341 | 3.9014 | 3.9014 | TRUE | 0.3054 |
| Trrap | 12190 | 8410 | 7393 | 7583 | 7337 | 8197 | 9467 | 8946 | 7977 | 7548 | 8277 | 8092 | 8384.46 | 8797 | 7972 | 0.906 | -0.142 | 0.04482439 | 0.12440291 | 0.0046 | 0.0027 | 0.0035 | 0.0035 | TRUE | 0.081 |
| Trub1 | 2326 | 1750 | 1620 | 1836 | 1592 | 1947 | 1807 | 1861 | 1748 | 1828 | 1796 | 1922 | 1826.9 | 1805 | 1849 | 1.024 | 0.034 | 0.59744437 | 0.74351523 | 5.00E-04 | 0.0037 | 0.0025 | 0.0025 | TRUE | 0.0147 |
| Trub2 | 2373 | 1784 | 1398 | 1521 | 1414 | 1468 | 1843 | 1898 | 1508 | 1514 | 1595 | 1449 | 1634.56 | 1750 | 1519 | 0.868 | -0.205 | 0.0241882 | 0.07945803 | 0.0075 | 0.0039 | 0.0053 | 0.0053 | TRUE | 0.1732 |
| Try10 | 0 | 0 | 0 | 0 | 0 | 0 | 0 | 0 | 0 | 0 | 0 | 0 | 0 | 0 | 0 | NA | NA | NA | NA | NA | NA | NA | NA | NA | NA |
| Try4 | 0 | 0 | 0 | 0 | 0 | 0 | 0 | 0 | 0 | 0 | 0 | 0 | 0 | 0 | 0 | NA | NA | NA | NA | NA | NA | NA | NA | NA | NA |
| Try5 | 0 | 0 | 0 | 0 | 0 | 0 | 0 | 0 | 0 | 0 | 0 | 0 | 0 | 0 | 0 | NA | NA | NA | NA | NA | NA | NA | NA | NA | NA |
| Tsacc | 224 | 184 | 185 | 194 | 205 | 218 | 174 | 196 | 200 | 193 | 231 | 215 | 201.48 | 190 | 213 | 1.125 | 0.17 | 0.24741925 | 0.41623862 | 0 | 0.0142 | 0.0107 | 0.0107 | TRUE | 0.0747 |
| Tsc1 | 3779 | 2929 | 3040 | 3125 | 2927 | 3447 | 2935 | 3116 | 3280 | 3111 | 3302 | 3403 | 3190.99 | 3110 | 3272 | 1.053 | 0.074 | 0.26314047 | 0.43335254 | 0.0021 | 0.0032 | 0.0028 | 0.0028 | TRUE | 0.0302 |
| Tsc2 | 7352 | 5443 | 5247 | 6088 | 5566 | 6392 | 5710 | 5790 | 5661 | 6060 | 6279 | 6310 | 5968.33 | 5720 | 6216 | 1.087 | 0.12 | 0.01566022 | 0.05765882 | 1.00E-04 | 0.0028 | 0.0016 | 0.0016 | TRUE | 0.0059 |
| Tsc22d1 | 6972 | 4705 | 4572 | 5051 | 4382 | 4592 | 5415 | 5005 | 4933 | 5028 | 4943 | 4533 | 4976.13 | 5118 | 4835 | 0.944 | -0.083 | 0.1991068 | 0.35964028 | 0.0025 | 0.0029 | 0.0028 | 0.0028 | TRUE | 0.0365 |
| Tsc22d2 | 2026 | 1425 | 1301 | 1310 | 1212 | 1353 | 1574 | 1516 | 1404 | 1304 | 1367 | 1336 | 1416.65 | 1498 | 1336 | 0.891 | -0.167 | 0.0213587 | 0.07251363 | 0.0013 | 0.0041 | 0.0031 | 0.0031 | TRUE | 0.0358 |
| Tsc22d3 | 403 | 310 | 319 | 484 | 401 | 434 | 313 | 330 | 344 | 482 | 452 | 428 | 391.58 | 329 | 454 | 1.383 | 0.468 | 1.90E-05 | 0.00032065 | 0 | 0.0085 | 0.006 | 0.006 | TRUE | 0.0335 |
| Tsc22d4 | 2026 | 1548 | 1560 | 1802 | 1618 | 1817 | 1574 | 1647 | 1683 | 1794 | 1825 | 1794 | 1719.33 | 1635 | 1804 | 1.105 | 0.144 | 0.02505992 | 0.08146041 | 0 | 0.0038 | 0.0024 | 0.0024 | TRUE | 0.0134 |
| Tsen15 | 1101 | 751 | 773 | 966 | 836 | 1003 | 855 | 799 | 834 | 962 | 943 | 990 | 897.13 | 829 | 965 | 1.163 | 0.218 | 0.00516882 | 0.02518381 | 0 | 0.0051 | 0.0032 | 0.0032 | TRUE | 0.0122 |
| Tsen2 | 1873 | 1435 | 1350 | 1465 | 1307 | 1580 | 1455 | 1526 | 1457 | 1458 | 1474 | 1560 | 1488.35 | 1479 | 1497 | 1.013 | 0.018 | 0.78714238 | 0.87512648 | 4.00E-04 | 0.004 | 0.0027 | 0.0027 | TRUE | 0.0161 |
| Tsen34 | 2866 | 2074 | 1916 | 1769 | 1596 | 1906 | 2226 | 2206 | 2067 | 1761 | 1800 | 1882 | 1990.36 | 2166 | 1814 | 0.837 | -0.256 | 8.57E-05 | 0.00106551 | 9.00E-04 | 0.0036 | 0.0026 | 0.0026 | TRUE | 0.0191 |
| Tsen54 | 3190 | 2508 | 2267 | 2381 | 2085 | 2418 | 2478 | 2668 | 2446 | 2370 | 2352 | 2387 | 2450.07 | 2531 | 2370 | 0.937 | -0.094 | 0.12662072 | 0.26194798 | 7.00E-04 | 0.0034 | 0.0023 | 0.0023 | TRUE | 0.0269 |
| Tsfm | 3290 | 2476 | 2489 | 2809 | 2433 | 2833 | 2555 | 2634 | 2686 | 2796 | 2745 | 2797 | 2701.98 | 2625 | 2779 | 1.059 | 0.083 | 0.14244458 | 0.28506866 | 0 | 0.0033 | 0.0019 | 0.0019 | TRUE | 0.0068 |
| Tsg101 | 3727 | 2675 | 2758 | 2953 | 2582 | 3060 | 2895 | 2845 | 2976 | 2939 | 2913 | 3021 | 2931.45 | 2905 | 2958 | 1.018 | 0.026 | 0.64274592 | 0.7773516 | 0 | 0.0032 | 0.0019 | 0.0019 | TRUE | 0.0053 |
| Tsga10 | 100 | 74 | 94 | 85 | 66 | 93 | 78 | 79 | 101 | 85 | 74 | 92 | 84.78 | 86 | 84 | 0.978 | -0.032 | 0.8854327 | 0.93634542 | 0 | 0.0304 | 0.0234 | 0.0234 | TRUE | 0.2253 |
| Tsga13 | 0 | 0 | 0 | 0 | 0 | 0 | 0 | 0 | 0 | 0 | 0 | 0 | 0 | 0 | 0 | NA | NA | NA | NA | NA | NA | NA | NA | NA | NA |
| Tsga14 | 1322 | 989 | 808 | 965 | 812 | 944 | 1027 | 1052 | 872 | 961 | 916 | 932 | 959.83 | 984 | 936 | 0.951 | -0.072 | 0.41819307 | 0.59082268 | 0.0043 | 0.0049 | 0.0047 | 0.0047 | TRUE | 0.1148 |
| Tsga8 | 7 | 9 | 1 | 1 | 6 | 2 | 5 | 10 | 1 | 1 | 7 | 2 | 4.3 | 5 | 3 | 0.594 | -0.751 | 0.47861496 | NA | 0.5694 | 0.553 | 0.5576 | 0.5576 | TRUE | 0.2206 |

| | | | | | | | | | | | | | | | | | | | | | | | | | |
|---|---|---|---|---|---|---|---|---|---|---|---|---|---|---|---|---|---|---|---|---|---|---|---|---|---|
| Tshb | 1 | 1 | 0 | 1 | 1 | 1 | 1 | 1 | 0 | 1 | 1 | 1 | 0.83 | 1 | 1 | 1.662 | 0.733 | 0.71902164 | NA | 0 | 2.8746 | 1.72 | 1.72 | TRUE | 0.1898 |
| Tshr | 16 | 16 | 7 | 9 | 4 | 3 | 12 | 17 | 8 | 9 | 5 | 3 | 8.91 | 12 | 6 | 0.445 | -1.168 | 0.09565035 | NA | 0.0808 | 0.2686 | 0.2206 | 0.2206 | TRUE | 0.2531 |
| Tshz1 | 116 | 88 | 71 | 90 | 66 | 89 | 90 | 94 | 77 | 90 | 74 | 88 | 85.37 | 87 | 84 | 0.968 | -0.047 | 0.82649286 | 0.90038586 | 0 | 0.0302 | 0.0214 | 0.0214 | TRUE | 0.0929 |
| Tshz2 | 10 | 8 | 6 | 5 | 4 | 3 | 8 | 9 | 6 | 5 | 5 | 3 | 5.87 | 8 | 4 | 0.546 | -0.874 | 0.26839715 | NA | 0 | 0.4064 | 0.2611 | 0.2611 | TRUE | 0.1145 |
| Tshz3 | 147 | 111 | 104 | 100 | 81 | 119 | 114 | 118 | 112 | 100 | 91 | 117 | 108.81 | 115 | 103 | 0.897 | -0.157 | 0.41341941 | 0.5860164 | 0 | 0.0242 | 0.0172 | 0.0172 | TRUE | 0.1549 |
| Tsix | 215 | 192 | 134 | 114 | 90 | 124 | 167 | 204 | 145 | 113 | 102 | 122 | 142.2 | 172 | 112 | 0.655 | -0.609 | 0.00101576 | 0.00732135 | 0.0132 | 0.0191 | 0.0174 | 0.0174 | TRUE | 0.2553 |
| Tsks | 96 | 57 | 77 | 117 | 103 | 122 | 75 | 61 | 83 | 116 | 116 | 120 | 95.23 | 73 | 117 | 1.617 | 0.693 | 0.00078677 | 0.00607768 | 0 | 0.0273 | 0.0197 | 0.0197 | TRUE | 0.1775 |
| Tsku | 490 | 396 | 369 | 356 | 306 | 402 | 381 | 421 | 398 | 354 | 345 | 397 | 382.72 | 400 | 365 | 0.916 | -0.127 | 0.25669212 | 0.42656674 | 0.0014 | 0.0086 | 0.0064 | 0.0064 | TRUE | 0.0653 |
| Tslp | 1 | 1 | 1 | 3 | 1 | 3 | 1 | 1 | 1 | 3 | 1 | 3 | 1.67 | 1 | 2 | 2.47 | 1.305 | 0.38067152 | NA | 0 | 1.4252 | 0.8786 | 0.8786 | TRUE | 0.1897 |
| Tsn | 8181 | 5799 | 5253 | 5363 | 4754 | 5235 | 6354 | 6168 | 5668 | 5338 | 5363 | 5168 | 5676.53 | 6063 | 5290 | 0.872 | -0.197 | 0.00096905 | 0.00708925 | 0.0016 | 0.0028 | 0.0024 | 0.0024 | TRUE | 0.0395 |
| Tsnax | 8509 | 6241 | 6361 | 6357 | 5687 | 6519 | 6609 | 6638 | 6863 | 6328 | 6416 | 6435 | 6548.18 | 6703 | 6393 | 0.954 | -0.068 | 0.16092426 | 0.31065866 | 1.00E-04 | 0.0028 | 0.0016 | 0.0016 | TRUE | 0.0053 |
| Tsnaxip1 | 17 | 14 | 19 | 18 | 10 | 15 | 13 | 15 | 21 | 18 | 11 | 15 | 15.43 | 16 | 15 | 0.916 | -0.127 | 0.79646214 | NA | 0 | 0.156 | 0.1105 | 0.1105 | TRUE | 0.244 |
| Tspan1 | 75 | 55 | 40 | 51 | 55 | 57 | 58 | 59 | 43 | 51 | 62 | 56 | 54.83 | 53 | 56 | 1.052 | 0.073 | 0.78507089 | 0.87370694 | 0 | 0.0457 | 0.0333 | 0.0333 | TRUE | 0.215 |
| Tspan10 | 0 | 0 | 0 | 0 | 0 | 1 | 0 | 0 | 0 | 0 | 0 | 1 | 0.16 | 0 | 0 | 2.094 | 1.067 | 0.79379329 | NA | 0 | 14.4078 | 10 | 10 | TRUE | 0.3835 |
| Tspan11 | 33 | 27 | 27 | 38 | 55 | 59 | 26 | 29 | 29 | 38 | 62 | 58 | 40.27 | 28 | 53 | 1.897 | 0.923 | 0.00456629 | 0.02290783 | 0 | 0.0613 | 0.0492 | 0.0492 | TRUE | 0.5133 |
| Tspan12 | 164 | 99 | 103 | 112 | 79 | 123 | 127 | 105 | 111 | 111 | 89 | 121 | 110.97 | 114 | 107 | 0.936 | -0.096 | 0.62850368 | 0.76602738 | 0.0076 | 0.0238 | 0.0193 | 0.0193 | TRUE | 0.2069 |
| Tspan13 | 3265 | 2400 | 2574 | 3001 | 2525 | 2949 | 2536 | 2553 | 2777 | 2987 | 2848 | 2911 | 2768.79 | 2622 | 2915 | 1.113 | 0.154 | 0.01374479 | 0.05241 | 0.0011 | 0.0033 | 0.0025 | 0.0025 | TRUE | 0.0322 |
| Tspan14 | 3151 | 2552 | 2747 | 2855 | 2835 | 3218 | 2447 | 2715 | 2964 | 2842 | 3198 | 3177 | 2890.41 | 2709 | 3072 | 1.135 | 0.183 | 0.02422041 | 0.07952731 | 0.0061 | 0.0032 | 0.0044 | 0.0044 | TRUE | 0.0881 |
| Tspan15 | 522 | 410 | 471 | 644 | 609 | 684 | 405 | 436 | 508 | 641 | 687 | 675 | 558.83 | 450 | 668 | 1.488 | 0.573 | 5.75E-08 | 2.36E-06 | 0.0047 | 0.0067 | 0.0062 | 0.0062 | TRUE | 0.147 |
| Tspan17 | 531 | 385 | 448 | 598 | 511 | 671 | 412 | 410 | 483 | 595 | 576 | 662 | 523.23 | 435 | 611 | 1.408 | 0.494 | 4.24E-06 | 9.25E-05 | 0.005 | 0.007 | 0.0063 | 0.0063 | TRUE | 0.1057 |
| Tspan18 | 129 | 88 | 115 | 119 | 93 | 109 | 100 | 94 | 124 | 118 | 105 | 108 | 108.14 | 106 | 110 | 1.044 | 0.062 | 0.75120545 | 0.85220887 | 0 | 0.0243 | 0.0185 | 0.0185 | TRUE | 0.2088 |
| Tspan2 | 1074 | 827 | 815 | 800 | 666 | 763 | 834 | 880 | 879 | 796 | 751 | 753 | 815.67 | 864 | 767 | 0.888 | -0.171 | 0.0341607 | 0.1020608 | 0 | 0.0053 | 0.0035 | 0.0035 | TRUE | 0.0135 |
| Tspan3 | 4449 | 3205 | 3659 | 4536 | 4014 | 4416 | 3455 | 3409 | 3948 | 4515 | 4528 | 4359 | 4035.85 | 3604 | 4467 | 1.24 | 0.31 | 5.44E-06 | 0.00011377 | 0.0033 | 0.003 | 0.0031 | 0.0031 | TRUE | 0.0839 |
| Tspan31 | 3176 | 2296 | 2381 | 2771 | 2468 | 2736 | 2467 | 2442 | 2569 | 2758 | 2784 | 2701 | 2620.21 | 2493 | 2748 | 1.102 | 0.141 | 0.01451224 | 0.05442758 | 0 | 0.0033 | 0.002 | 0.002 | TRUE | 0.0085 |
| Tspan32 | 12 | 6 | 5 | 9 | 13 | 8 | 9 | 6 | 5 | 9 | 15 | 8 | 8.77 | 7 | 11 | 1.464 | 0.55 | 0.40260445 | NA | 0 | 0.2727 | 0.1945 | 0.1945 | TRUE | 0.3899 |
| Tspan33 | 305 | 255 | 254 | 298 | 249 | 312 | 237 | 271 | 274 | 297 | 281 | 308 | 277.95 | 261 | 295 | 1.136 | 0.184 | 0.14568965 | 0.28966556 | 0 | 0.011 | 0.008 | 0.008 | TRUE | 0.0783 |
| Tspan4 | 1554 | 1126 | 1243 | 1495 | 1282 | 1427 | 1207 | 1198 | 1341 | 1488 | 1446 | 1409 | 1348.14 | 1249 | 1448 | 1.16 | 0.214 | 0.00396838 | 0.02047635 | 0.0016 | 0.0042 | 0.0032 | 0.0032 | TRUE | 0.0493 |
| Tspan5 | 1038 | 691 | 778 | 920 | 844 | 979 | 806 | 735 | 839 | 916 | 952 | 966 | 869.16 | 793 | 945 | 1.19 | 0.251 | 0.00263819 | 0.01492779 | 0.0012 | 0.0052 | 0.0039 | 0.0039 | TRUE | 0.0475 |
| Tspan6 | 485 | 354 | 342 | 324 | 263 | 252 | 377 | 377 | 369 | 323 | 297 | 249 | 331.7 | 374 | 290 | 0.773 | -0.372 | 0.00298393 | 0.01647329 | 0.0048 | 0.0096 | 0.0082 | 0.0082 | TRUE | 0.1738 |
| Tspan7 | 1456 | 997 | 1044 | 1111 | 912 | 1072 | 1131 | 1060 | 1126 | 1106 | 1029 | 1058 | 1085.12 | 1106 | 1064 | 0.962 | -0.056 | 0.45774724 | 0.62749036 | 0 | 0.0046 | 0.0031 | 0.0031 | TRUE | 0.0151 |
| Tspan8 | 233 | 147 | 132 | 159 | 128 | 137 | 181 | 156 | 142 | 158 | 144 | 135 | 152.94 | 160 | 146 | 0.909 | -0.138 | 0.41680438 | 0.58950096 | 0.0052 | 0.0179 | 0.0141 | 0.0141 | TRUE | 0.1525 |
| Tspan9 | 4601 | 3270 | 3450 | 3282 | 2991 | 3347 | 3573 | 3478 | 3723 | 3267 | 3374 | 3304 | 3453.21 | 3591 | 3315 | 0.923 | -0.116 | 0.03970821 | 0.11394288 | 4.00E-04 | 0.0031 | 0.002 | 0.002 | TRUE | 0.0121 |
| Tspo | 719 | 502 | 670 | 832 | 757 | 812 | 558 | 534 | 723 | 828 | 854 | 802 | 716.5 | 605 | 828 | 1.37 | 0.454 | 5.68E-05 | 0.00077504 | 0.0128 | 0.0057 | 0.0077 | 0.0077 | TRUE | 0.3336 |
| Tspo2 | 22 | 13 | 28 | 31 | 53 | 47 | 17 | 14 | 30 | 31 | 60 | 46 | 33.03 | 20 | 46 | 2.252 | 1.171 | 0.00336218 | 0.01807961 | 0.0949 | 0.0742 | 0.0797 | 0.0797 | TRUE | 0.2339 |
| Tspy-ps | 0 | 0 | 0 | 0 | 0 | 0 | 0 | 0 | 0 | 0 | 0 | 0 | 0 | 0 | 0 | NA | NA | NA | NA | NA | NA | NA | NA | NA | NA |
| Tspyl1 | 3194 | 2383 | 2229 | 2413 | 2031 | 2389 | 2481 | 2535 | 2405 | 2402 | 2291 | 2358 | 2411.98 | 2474 | 2350 | 0.95 | -0.073 | 0.21557642 | 0.37815828 | 2.00E-04 | 0.0034 | 0.0021 | 0.0021 | TRUE | 0.0069 |
| Tspyl2 | 1032 | 838 | 701 | 686 | 609 | 726 | 802 | 891 | 756 | 683 | 687 | 717 | 755.96 | 816 | 696 | 0.852 | -0.231 | 0.01079235 | 0.04371068 | 0.0026 | 0.0056 | 0.0046 | 0.0046 | TRUE | 0.0741 |
| Tspyl3 | 182 | 162 | 126 | 144 | 114 | 141 | 141 | 172 | 136 | 143 | 129 | 139 | 143.46 | 150 | 137 | 0.917 | -0.125 | 0.46672269 | 0.63562339 | 0 | 0.0189 | 0.0144 | 0.0144 | TRUE | 0.1723 |
| Tspyl4 | 412 | 269 | 187 | 278 | 250 | 261 | 320 | 286 | 202 | 277 | 282 | 258 | 270.71 | 269 | 272 | 1.006 | 0.009 | 0.95534548 | 0.97553943 | 0.0248 | 0.0112 | 0.0152 | 0.0152 | TRUE | 0.5255 |
| Tspyl5 | 159 | 100 | 103 | 106 | 86 | 105 | 123 | 106 | 111 | 106 | 97 | 104 | 107.86 | 113 | 102 | 0.896 | -0.159 | 0.40033334 | 0.57345805 | 0 | 0.0244 | 0.0163 | 0.0163 | TRUE | 0.0638 |
| Tsr1 | 16626 | 11840 | 11176 | 11555 | 10482 | 12154 | 12913 | 12594 | 12059 | 11502 | 11825 | 11998 | 12148.34 | 12522 | 11775 | 0.94 | -0.089 | 0.07893732 | 0.18729269 | 7.00E-04 | 0.0026 | 0.0018 | 0.0018 | TRUE | 0.0128 |
| Tsr2 | 1818 | 1285 | 1107 | 1276 | 1091 | 1199 | 1412 | 1367 | 1194 | 1270 | 1231 | 1184 | 1276.29 | 1324 | 1228 | 0.926 | -0.11 | 0.17958783 | 0.33446994 | 0.0038 | 0.0043 | 0.0041 | 0.0041 | TRUE | 0.0873 |
| Tsr3 | 1373 | 1084 | 930 | 933 | 845 | 937 | 1066 | 1153 | 1003 | 929 | 953 | 925 | 1004.96 | 1074 | 936 | 0.871 | -0.2 | 0.01320699 | 0.05077996 | 0.0016 | 0.0048 | 0.0037 | 0.0037 | TRUE | 0.0474 |
| Tssc1 | 1885 | 1440 | 1374 | 1215 | 1097 | 1373 | 1464 | 1532 | 1483 | 1209 | 1238 | 1355 | 1380.09 | 1493 | 1267 | 0.85 | -0.235 | 0.001304 | 0.00871753 | 0.0013 | 0.0041 | 0.0031 | 0.0031 | TRUE | 0.0448 |
| Tssc4 | 1137 | 837 | 722 | 674 | 625 | 688 | 883 | 890 | 779 | 671 | 705 | 679 | 767.92 | 851 | 685 | 0.804 | -0.314 | 0.00036649 | 0.00339488 | 0.0019 | 0.0055 | 0.0043 | 0.0043 | TRUE | 0.0629 |
| Tssk1 | 1 | 1 | 0 | 0 | 0 | 0 | 1 | 1 | 0 | 0 | 0 | 0 | 0.31 | 1 | 0 | 0.309 | -1.696 | 0.63183309 | NA | 0 | 7.7294 | 7.2472 | 7.2472 | TRUE | 0.2015 |
| Tssk2 | 0 | 0 | 0 | 0 | 0 | 0 | 0 | 0 | 0 | 0 | 0 | 0 | 0 | 0 | 0 | NA | NA | NA | NA | NA | NA | NA | NA | NA | NA |
| Tssk3 | 5 | 1 | 3 | 3 | 4 | 6 | 4 | 1 | 3 | 3 | 5 | 6 | 3.6 | 3 | 5 | 1.61 | 0.687 | 0.49849776 | NA | 0 | 0.6606 | 0.4539 | 0.4539 | TRUE | 0.31 |
| Tssk4 | 0 | 0 | 1 | 1 | 0 | 0 | 0 | 0 | 1 | 1 | 0 | 0 | 0.35 | 0 | 0 | 1.075 | 0.105 | 0.97567165 | NA | 0 | 6.8576 | 6.5143 | 6.5143 | TRUE | 0.5365 |
| Tssk5 | 4 | 3 | 3 | 1 | 0 | 3 | 3 | 3 | 3 | 1 | 0 | 3 | 2.25 | 3 | 1 | 0.425 | -1.235 | 0.35673984 | NA | 0 | 1.0564 | 0.7593 | 0.7593 | TRUE | 0.7281 |
| Tssk6 | 3 | 5 | 0 | 0 | 1 | 0 | 2 | 5 | 0 | 0 | 1 | 0 | 1.46 | 2 | 0 | 0.146 | -2.781 | 0.15926534 | NA | 1.491 | 1.6227 | 1.6222 | 1.6222 | TRUE | 0.2076 |
| Tst | 1507 | 1158 | 1153 | 1294 | 1138 | 1468 | 1170 | 1232 | 1244 | 1288 | 1284 | 1449 | 1277.87 | 1215 | 1340 | 1.104 | 0.143 | 0.06565907 | 0.16375242 | 0.0022 | 0.0043 | 0.0035 | 0.0035 | TRUE | 0.0611 |
| Tsta3 | 2599 | 1818 | 1842 | 2071 | 1865 | 2052 | 2019 | 1934 | 1988 | 2061 | 2104 | 2026 | 2021.81 | 1980 | 2064 | 1.042 | 0.059 | 0.33008567 | 0.50448598 | 0 | 0.0036 | 0.0021 | 0.0021 | TRUE | 0.005 |
| Tstd1 | 34 | 34 | 29 | 43 | 30 | 54 | 26 | 36 | 31 | 43 | 34 | 53 | 37.3 | 31 | 43 | 1.397 | 0.483 | 0.14427309 | 0.28752025 | 0 | 0.066 | 0.0513 | 0.0513 | TRUE | 0.175 |
| Tstd2 | 2478 | 1812 | 1824 | 1866 | 1638 | 1893 | 1925 | 1927 | 1968 | 1857 | 1848 | 1869 | 1899 | 1940 | 1858 | 0.958 | -0.062 | 0.30101409 | 0.47337518 | 0 | 0.0037 | 0.0021 | 0.0021 | TRUE | 0.0019 |
| Tstd3 | 1087 | 781 | 775 | 984 | 791 | 965 | 844 | 831 | 836 | 979 | 892 | 953 | 889.27 | 837 | 941 | 1.125 | 0.17 | 0.03104407 | 0.09503382 | 0 | 0.0051 | 0.0034 | 0.0034 | TRUE | 0.0242 |
| Tsx | 87 | 80 | 86 | 82 | 81 | 84 | 68 | 85 | 93 | 82 | 91 | 83 | 83.56 | 82 | 85 | 1.048 | 0.068 | 0.75960438 | 0.8571787 | 0 | 0.0308 | 0.0233 | 0.0233 | TRUE | 0.2487 |
| Ttbk1 | 46 | 47 | 52 | 51 | 39 | 51 | 36 | 50 | 56 | 51 | 44 | 50 | 47.82 | 47 | 48 | 1.034 | 0.048 | 0.87051552 | 0.9283204 | 0 | 0.052 | 0.0405 | 0.0405 | TRUE | 0.4308 |
| Ttbk2 | 1858 | 1342 | 1356 | 1650 | 1421 | 1623 | 1443 | 1427 | 1463 | 1642 | 1603 | 1602 | 1530.2 | 1444 | 1616 | 1.119 | 0.162 | 0.0113776 | 0.04548814 | 0 | 0.004 | 0.0023 | 0.0023 | TRUE | 0.0025 |
| Ttc1 | 4524 | 3322 | 3444 | 3757 | 3409 | 4136 | 3514 | 3534 | 3716 | 3740 | 3846 | 4083 | 3738.58 | 3588 | 3890 | 1.084 | 0.117 | 0.05048024 | 0.13580164 | 0.0011 | 0.0031 | 0.0023 | 0.0023 | TRUE | 0.0232 |
| Ttc12 | 451 | 396 | 468 | 534 | 494 | 533 | 350 | 421 | 505 | 532 | 557 | 526 | 481.91 | 425 | 538 | 1.269 | 0.344 | 0.00707002 | 0.03197843 | 0.0149 | 0.0073 | 0.0097 | 0.0097 | TRUE | 0.2936 |
| Ttc13 | 3699 | 2730 | 2528 | 2554 | 2557 | 2882 | 2873 | 2904 | 2728 | 2542 | 2885 | 2845 | 2796.03 | 2835 | 2757 | 0.972 | -0.041 | 0.55204526 | 0.7078682 | 0.0025 | 0.0033 | 0.003 | 0.003 | TRUE | 0.0567 |
| Ttc14 | 2867 | 2565 | 2415 | 2497 | 2065 | 2478 | 2227 | 2728 | 2606 | 2486 | 2330 | 2446 | 2470.34 | 2520 | 2421 | 0.961 | -0.057 | 0.48158767 | 0.64849796 | 0.0058 | 0.0034 | 0.0043 | 0.0043 | TRUE | 0.1287 |
| Ttc16 | 76 | 64 | 44 | 73 | 88 | 65 | 59 | 68 | 47 | 73 | 99 | 64 | 68.45 | 58 | 79 | 1.347 | 0.43 | 0.09943761 | 0.22028236 | 0.0266 | 0.0371 | 0.0341 | 0.0341 | TRUE | 0.4629 |
| Ttc17 | 592 | 405 | 429 | 370 | 345 | 361 | 460 | 431 | 463 | 368 | 389 | 356 | 411.22 | 451 | 371 | 0.822 | -0.283 | 0.00695882 | 0.03160549 | 0 | 0.0082 | 0.0055 | 0.0055 | TRUE | 0.0197 |
| Ttc18 | 30 | 23 | 25 | 17 | 27 | 28 | 23 | 24 | 27 | 17 | 30 | 28 | 24.96 | 25 | 25 | 1.003 | 0.004 | 0.9912897 | 0.99553848 | 0 | 0.0974 | 0.0726 | 0.0726 | TRUE | 0.4978 |
| Ttc19 | 1886 | 1411 | 1409 | 1683 | 1421 | 1608 | 1465 | 1501 | 1520 | 1675 | 1603 | 1587 | 1558.6 | 1495 | 1622 | 1.085 | 0.118 | 0.07139808 | 0.17399705 | 0 | 0.0039 | 0.0024 | 0.0024 | TRUE | 0.0101 |
| Ttc21a | 63 | 55 | 36 | 49 | 35 | 44 | 49 | 59 | 39 | 49 | 39 | 43 | 46.33 | 49 | 44 | 0.901 | -0.15 | 0.60859296 | 0.75228048 | 0 | 0.0536 | 0.0401 | 0.0401 | TRUE | 0.1648 |
| Ttc21b | 1706 | 1154 | 1332 | 1537 | 1406 | 1488 | 1325 | 1227 | 1437 | 1530 | 1586 | 1469 | 1429.1 | 1330 | 1528 | 1.149 | 0.2 | 0.01034551 | 0.04232067 | 0.0028 | 0.0041 | 0.0037 | 0.0037 | TRUE | 0.0579 |

| | | | | | | | | | | | | | | | | | | | | | | | | | |
|---|---|---|---|---|---|---|---|---|---|---|---|---|---|---|---|---|---|---|---|---|---|---|---|---|---|
| Ttc22 | 4 | 5 | 2 | 3 | 4 | 3 | 3 | 5 | 2 | 3 | 5 | 3 | 3.51 | 3 | 4 | 0.987 | -0.019 | 0.98529665 | NA | 0 | 0.6782 | 0.4409 | 0.4409 | TRUE | 0.2676 |
| Ttc23 | 527 | 394 | 396 | 434 | 407 | 480 | 409 | 419 | 427 | 432 | 459 | 474 | 436.77 | 418 | 455 | 1.088 | 0.121 | 0.23190803 | 0.39777528 | 0 | 0.0079 | 0.0051 | 0.0051 | TRUE | 0.0232 |
| Ttc23l | 0 | 0 | 0 | 0 | 0 | 0 | 0 | 0 | 0 | 0 | 0 | 0 | 0 | 0 | 0 | NA | NA | NA | NA | NA | NA | NA | NA | NA | NA |
| Ttc24 | 5 | 1 | 4 | 0 | 1 | 2 | 4 | 1 | 4 | 0 | 1 | 2 | 2.06 | 3 | 1 | 0.333 | -1.588 | 0.27570003 | NA | 0 | 1.1524 | 0.8736 | 0.8736 | TRUE | 0.4032 |
| Ttc25 | 122 | 119 | 129 | 130 | 154 | 129 | 95 | 127 | 139 | 129 | 174 | 127 | 131.83 | 120 | 143 | 1.198 | 0.261 | 0.20233732 | 0.36372875 | 0.0278 | 0.0204 | 0.0226 | 0.0226 | TRUE | 0.385 |
| Ttc26 | 922 | 674 | 633 | 706 | 649 | 687 | 716 | 717 | 683 | 703 | 732 | 678 | 704.85 | 705 | 704 | 0.998 | -0.003 | 0.96820518 | 0.98285203 | 0 | 0.0058 | 0.0038 | 0.0038 | TRUE | 0.0137 |
| Ttc27 | 4897 | 3644 | 3584 | 3685 | 3374 | 3968 | 3803 | 3876 | 3867 | 3668 | 3806 | 3917 | 3822.96 | 3849 | 3797 | 0.987 | -0.019 | 0.72368092 | 0.83449548 | 3.00E-04 | 0.003 | 0.0019 | 0.0019 | TRUE | 0.0109 |
| Ttc28 | 664 | 458 | 430 | 515 | 456 | 492 | 516 | 487 | 464 | 513 | 514 | 486 | 496.59 | 489 | 504 | 1.029 | 0.042 | 0.67118076 | 0.79757121 | 0 | 0.0072 | 0.0049 | 0.0049 | TRUE | 0.0285 |
| Ttc29 | 108 | 90 | 68 | 55 | 44 | 35 | 84 | 96 | 73 | 55 | 50 | 35 | 65.32 | 84 | 47 | 0.548 | -0.867 | 0.00094593 | 0.00697275 | 0.014 | 0.0387 | 0.0326 | 0.0326 | TRUE | 0.4061 |
| Ttc3 | 7144 | 5216 | 4947 | 5709 | 5021 | 5895 | 5548 | 5548 | 5338 | 5683 | 5664 | 5819 | 5600.1 | 5478 | 5722 | 1.044 | 0.063 | 0.21295262 | 0.3752326 | 2.00E-04 | 0.0028 | 0.0016 | 0.0016 | TRUE | 0.0061 |
| Ttc30a1 | 305 | 236 | 143 | 180 | 156 | 168 | 237 | 251 | 154 | 179 | 176 | 166 | 193.87 | 214 | 174 | 0.809 | -0.305 | 0.09440719 | 0.21199716 | 0.0288 | 0.0147 | 0.0188 | 0.0188 | TRUE | 0.6295 |
| Ttc30a2 | 6 | 2 | 4 | 3 | 3 | 1 | 5 | 2 | 4 | 3 | 3 | 1 | 3.08 | 4 | 2 | 0.652 | -0.617 | 0.57332731 | NA | 0 | 0.7727 | 0.5237 | 0.5237 | TRUE | 0.31 |
| Ttc30b | 143 | 135 | 72 | 89 | 74 | 76 | 111 | 144 | 78 | 89 | 83 | 75 | 96.57 | 111 | 82 | 0.743 | -0.429 | 0.07359781 | 0.17784168 | 0.0399 | 0.027 | 0.0307 | 0.0307 | TRUE | 0.1382 |
| Ttc32 | 215 | 153 | 168 | 160 | 124 | 147 | 167 | 163 | 181 | 159 | 140 | 145 | 159.21 | 170 | 148 | 0.871 | -0.2 | 0.20735974 | 0.36908856 | 0 | 0.0173 | 0.0117 | 0.0117 | TRUE | 0.046 |
| Ttc33 | 1582 | 1196 | 1195 | 1182 | 1002 | 1257 | 1229 | 1272 | 1289 | 1177 | 1130 | 1241 | 1223 | 1263 | 1183 | 0.937 | -0.094 | 0.19713044 | 0.35688225 | 6.00E-04 | 0.0044 | 0.003 | 0.003 | TRUE | 0.0223 |
| Ttc34 | 9 | 9 | 8 | 7 | 12 | 1 | 7 | 10 | 9 | 7 | 14 | 1 | 7.78 | 9 | 7 | 0.846 | -0.241 | 0.75954026 | NA | 0.3431 | 0.307 | 0.3155 | 0.3155 | TRUE | 0.2397 |
| Ttc36 | 2 | 0 | 0 | 0 | 2 | 2 | 2 | 0 | 0 | 0 | 2 | 2 | 0.96 | 1 | 1 | 2.479 | 1.31 | 0.54958602 | NA | 0 | 2.4611 | 2.2311 | 2.2311 | TRUE | 0.9717 |
| Ttc37 | 5418 | 3732 | 3939 | 4492 | 3739 | 4398 | 4208 | 3970 | 4250 | 4471 | 4218 | 4342 | 4243.1 | 4143 | 4344 | 1.049 | 0.068 | 0.22853764 | 0.39383754 | 7.00E-04 | 0.003 | 0.0021 | 0.0021 | TRUE | 0.016 |
| Ttc38 | 405 | 325 | 351 | 360 | 333 | 388 | 315 | 346 | 379 | 358 | 376 | 383 | 359.33 | 347 | 372 | 1.078 | 0.108 | 0.35105634 | 0.5257936 | 0.0024 | 0.009 | 0.007 | 0.007 | TRUE | 0.0795 |
| Ttc39a | 234 | 174 | 163 | 270 | 264 | 276 | 182 | 185 | 176 | 269 | 298 | 272 | 230.29 | 181 | 280 | 1.544 | 0.627 | 2.81E-06 | 6.56E-05 | 0 | 0.0127 | 0.0084 | 0.0084 | TRUE | 0.0349 |
| Ttc39b | 5597 | 3768 | 3813 | 4801 | 4180 | 4592 | 4347 | 4008 | 4114 | 4779 | 4715 | 4533 | 4416.09 | 4156 | 4676 | 1.124 | 0.169 | 0.00321265 | 0.01741311 | 9.00E-04 | 0.003 | 0.0021 | 0.0021 | TRUE | 0.02 |
| Ttc39c | 811 | 543 | 539 | 588 | 560 | 586 | 630 | 578 | 582 | 585 | 632 | 578 | 597.42 | 597 | 598 | 1.001 | 0.002 | 0.98251576 | 0.99089635 | 0 | 0.0064 | 0.0045 | 0.0045 | TRUE | 0.0302 |
| Ttc39d | 0 | 0 | 0 | 1 | 0 | 0 | 0 | 0 | 0 | 1 | 0 | 0 | 0.17 | 0 | 0 | 2.094 | 1.067 | 0.79379329 | NA | 0 | 14.2885 | 10 | 10 | TRUE | 0.3906 |
| Ttc4 | 3081 | 2210 | 2184 | 2458 | 2246 | 2413 | 2393 | 2351 | 2357 | 2447 | 2534 | 2382 | 2410.43 | 2367 | 2454 | 1.036 | 0.052 | 0.37924493 | 0.55385183 | 0 | 0.0034 | 0.0021 | 0.0021 | TRUE | 0.0096 |
| Ttc5 | 3731 | 2880 | 2841 | 2918 | 2641 | 3001 | 2898 | 3063 | 3065 | 2905 | 2979 | 2962 | 2978.81 | 3009 | 2949 | 0.98 | -0.028 | 0.61830135 | 0.75896438 | 3.00E-04 | 0.0032 | 0.002 | 0.002 | TRUE | 0.0131 |
| Ttc7 | 1288 | 989 | 1038 | 1351 | 1185 | 1379 | 1000 | 1052 | 1120 | 1345 | 1337 | 1361 | 1202.53 | 1057 | 1348 | 1.276 | 0.352 | 1.98E-06 | 5.05E-05 | 8.00E-04 | 0.0044 | 0.0031 | 0.0031 | TRUE | 0.0319 |
| Ttc7b | 4439 | 3054 | 3298 | 3381 | 3014 | 3452 | 3448 | 3249 | 3559 | 3365 | 3400 | 3408 | 3404.63 | 3419 | 3391 | 0.992 | -0.012 | 0.84190227 | 0.91013957 | 8.00E-04 | 0.0031 | 0.0022 | 0.0022 | TRUE | 0.0225 |
| Ttc8 | 410 | 237 | 208 | 210 | 178 | 220 | 318 | 252 | 224 | 209 | 201 | 217 | 236.99 | 265 | 209 | 0.785 | -0.348 | 0.02453131 | 0.08023478 | 0.0142 | 0.0124 | 0.013 | 0.013 | TRUE | 0.3679 |
| Ttc9 | 117 | 104 | 98 | 127 | 101 | 125 | 91 | 111 | 106 | 126 | 114 | 123 | 111.83 | 103 | 121 | 1.19 | 0.251 | 0.18196941 | 0.33758977 | 0 | 0.0236 | 0.0165 | 0.0165 | TRUE | 0.1087 |
| Ttc9b | 73 | 66 | 60 | 40 | 38 | 51 | 57 | 70 | 65 | 40 | 43 | 50 | 54.11 | 64 | 44 | 0.697 | -0.52 | 0.05203961 | 0.13879494 | 0 | 0.0462 | 0.0324 | 0.0324 | TRUE | 0.1134 |
| Ttc9c | 3798 | 2743 | 3519 | 3796 | 3298 | 3820 | 2950 | 2918 | 3797 | 3779 | 3720 | 3771 | 3489.06 | 3222 | 3757 | 1.167 | 0.222 | 0.01540225 | 0.05697318 | 0.0116 | 0.0031 | 0.0058 | 0.0058 | TRUE | 0.2953 |
| Ttf1 | 2092 | 1576 | 1501 | 1445 | 1325 | 1556 | 1625 | 1676 | 1620 | 1438 | 1495 | 1536 | 1564.97 | 1640 | 1490 | 0.908 | -0.139 | 0.03533349 | 0.10462432 | 0 | 0.0039 | 0.0025 | 0.0025 | TRUE | 0.0111 |
| Ttf2 | 6579 | 5154 | 5665 | 6111 | 5318 | 6094 | 5110 | 5482 | 6112 | 6083 | 5999 | 6016 | 5800.37 | 5568 | 6033 | 1.084 | 0.116 | 0.09268009 | 0.20921947 | 0.0039 | 0.0028 | 0.0033 | 0.0033 | TRUE | 0.0891 |
| Tti1 | 4941 | 3631 | 3719 | 3855 | 3550 | 4074 | 3837 | 3862 | 4013 | 3837 | 4005 | 4022 | 3929.36 | 3904 | 3955 | 1.013 | 0.019 | 0.73164189 | 0.83929626 | 4.00E-04 | 0.003 | 0.0019 | 0.0019 | TRUE | 0.0082 |
| Tti2 | 1316 | 1032 | 1037 | 995 | 948 | 1064 | 1022 | 1098 | 1119 | 990 | 1069 | 1050 | 1058.15 | 1080 | 1036 | 0.961 | -0.057 | 0.46123795 | 0.63067771 | 0.0011 | 0.0047 | 0.0034 | 0.0034 | TRUE | 0.0273 |
| Ttk | 7203 | 5383 | 5579 | 6336 | 5286 | 6543 | 5594 | 5726 | 6020 | 6307 | 5963 | 6459 | 6011.46 | 5780 | 6243 | 1.081 | 0.112 | 0.05145902 | 0.137709 | 0.0013 | 0.0028 | 0.0022 | 0.0022 | TRUE | 0.0186 |
| Ttl | 1204 | 782 | 749 | 850 | 662 | 766 | 935 | 832 | 808 | 846 | 747 | 756 | 820.69 | 858 | 783 | 0.911 | -0.134 | 0.14782901 | 0.29256717 | 0.0044 | 0.0053 | 0.005 | 0.005 | TRUE | 0.0758 |
| Ttll1 | 1561 | 1052 | 1130 | 1368 | 1158 | 1376 | 1212 | 1119 | 1219 | 1362 | 1306 | 1358 | 1262.83 | 1183 | 1342 | 1.134 | 0.181 | 0.01202919 | 0.04744293 | 5.00E-04 | 0.0043 | 0.003 | 0.003 | TRUE | 0.0266 |
| Ttll10 | 3 | 1 | 4 | 7 | 3 | 3 | 2 | 1 | 4 | 7 | 3 | 3 | 3.5 | 2 | 4 | 1.742 | 0.801 | 0.44510735 | NA | 0 | 0.6788 | 0.4872 | 0.4872 | TRUE | 0.4665 |
| Ttll11 | 17 | 6 | 11 | 5 | 21 | 12 | 13 | 6 | 12 | 5 | 24 | 12 | 11.99 | 10 | 14 | 1.267 | 0.342 | 0.59510212 | NA | 0.2465 | 0.2 | 0.2133 | 0.2133 | TRUE | 0.3163 |
| Ttll12 | 6060 | 4398 | 4044 | 3966 | 3624 | 4278 | 4707 | 4678 | 4363 | 3948 | 4088 | 4223 | 4334.52 | 4583 | 4086 | 0.891 | -0.166 | 0.00466761 | 0.02324772 | 0.0011 | 0.003 | 0.0022 | 0.0022 | TRUE | 0.0211 |
| Ttll13 | 77 | 40 | 33 | 38 | 55 | 58 | 60 | 43 | 36 | 38 | 62 | 57 | 49.18 | 46 | 52 | 1.126 | 0.171 | 0.58396187 | 0.73357067 | 0.0489 | 0.0506 | 0.0501 | 0.0501 | TRUE | 0.6212 |
| Ttll2 | 2 | 3 | 5 | 0 | 1 | 0 | 2 | 3 | 5 | 0 | 1 | 0 | 1.88 | 3 | 0 | 0.111 | -3.169 | 0.0650686 | NA | 0 | 1.2645 | 0.9817 | 0.9817 | TRUE | 0.3852 |
| Ttll3 | 268 | 232 | 183 | 271 | 230 | 230 | 208 | 247 | 197 | 270 | 259 | 227 | 234.77 | 217 | 252 | 1.16 | 0.214 | 0.13748058 | 0.27777918 | 0.0063 | 0.0125 | 0.0107 | 0.0107 | TRUE | 0.1462 |
| Ttll4 | 3443 | 2458 | 2372 | 2281 | 1994 | 2290 | 2674 | 2615 | 2559 | 2271 | 2249 | 2261 | 2438.08 | 2616 | 2260 | 0.864 | -0.211 | 0.00022777 | 0.00236595 | 0 | 0.0034 | 0.002 | 0.002 | TRUE | 0.0047 |
| Ttll5 | 1798 | 1311 | 1107 | 1192 | 996 | 1235 | 1396 | 1394 | 1194 | 1187 | 1124 | 1219 | 1252.44 | 1328 | 1177 | 0.885 | -0.176 | 0.03480021 | 0.10340545 | 0.004 | 0.0043 | 0.0042 | 0.0042 | TRUE | 0.0918 |
| Ttll6 | 256 | 200 | 195 | 184 | 155 | 184 | 199 | 213 | 210 | 183 | 175 | 182 | 193.6 | 207 | 180 | 0.869 | -0.202 | 0.15494283 | 0.3024758 | 0 | 0.0147 | 0.0093 | 0.0093 | TRUE | 0.0155 |
| Ttll7 | 299 | 218 | 186 | 194 | 167 | 175 | 232 | 232 | 201 | 193 | 188 | 173 | 203.17 | 222 | 185 | 0.832 | -0.265 | 0.0668115 | 0.16561513 | 0 | 0.0141 | 0.0101 | 0.0101 | TRUE | 0.0717 |
| Ttll8 | 12 | 14 | 18 | 15 | 14 | 9 | 9 | 15 | 19 | 15 | 16 | 9 | 13.87 | 14 | 13 | 0.916 | -0.126 | 0.81533475 | NA | 0 | 0.1733 | 0.1368 | 0.1368 | TRUE | 0.2656 |
| Ttll9 | 56 | 39 | 53 | 72 | 47 | 79 | 43 | 41 | 57 | 72 | 53 | 78 | 57.47 | 47 | 68 | 1.435 | 0.521 | 0.05740837 | 0.14901083 | 0.0161 | 0.0437 | 0.0362 | 0.0362 | TRUE | 0.297 |
| Ttn | 373 | 238 | 123 | 146 | 111 | 126 | 290 | 253 | 133 | 145 | 125 | 124 | 178.42 | 225 | 131 | 0.583 | -0.779 | 0.00092052 | 0.00683127 | 0.0781 | 0.0157 | 0.0337 | 0.0337 | TRUE | 0.7353 |
| Ttpa | 369 | 289 | 301 | 271 | 263 | 342 | 287 | 307 | 325 | 270 | 297 | 338 | 303.8 | 306 | 302 | 0.986 | -0.02 | 0.87521097 | 0.93071867 | 0.0052 | 0.0102 | 0.0087 | 0.0087 | TRUE | 0.1282 |
| Ttpal | 938 | 579 | 464 | 497 | 466 | 461 | 729 | 616 | 501 | 495 | 526 | 455 | 553.42 | 615 | 492 | 0.797 | -0.327 | 0.01229025 | 0.04824624 | 0.0186 | 0.0067 | 0.0104 | 0.0104 | TRUE | 0.1553 |
| Ttr | 0 | 0 | 1 | 1 | 0 | 0 | 0 | 0 | 1 | 1 | 0 | 0 | 0.35 | 0 | 0 | 1.075 | 0.105 | 0.97567165 | NA | 0 | 6.8576 | 6.5143 | 6.5143 | TRUE | 0.5365 |
| Ttyh1 | 590 | 461 | 447 | 508 | 482 | 547 | 458 | 490 | 482 | 506 | 544 | 540 | 503.38 | 477 | 530 | 1.112 | 0.153 | 0.11299276 | 0.24176729 | 0 | 0.0071 | 0.0047 | 0.0047 | TRUE | 0.0188 |
| Ttyh2 | 2323 | 1987 | 2533 | 2040 | 2072 | 2439 | 1804 | 2114 | 2733 | 2031 | 2337 | 2408 | 2237.75 | 2217 | 2259 | 1.02 | 0.028 | 0.81801673 | 0.89536894 | 0.0257 | 0.0035 | 0.0102 | 0.0102 | TRUE | 0.5018 |
| Ttyh3 | 1391 | 987 | 810 | 939 | 845 | 903 | 1080 | 1050 | 874 | 935 | 953 | 891 | 963.92 | 1001 | 926 | 0.924 | -0.114 | 0.22128996 | 0.38520245 | 0.006 | 0.0049 | 0.0053 | 0.0053 | TRUE | 0.1454 |
| Tub | 2050 | 1641 | 1863 | 2224 | 2059 | 2381 | 1592 | 1746 | 2010 | 2214 | 2323 | 2350 | 2039.13 | 1783 | 2296 | 1.289 | 0.367 | 1.72E-05 | 0.00029644 | 0.0068 | 0.0036 | 0.0047 | 0.0047 | TRUE | 0.1502 |
| Tuba1a | 12891 | 9078 | 8432 | 10174 | 9342 | 10682 | 10012 | 9656 | 9098 | 10127 | 10539 | 10545 | 9996.14 | 9589 | 10404 | 1.085 | 0.117 | 0.03294048 | 0.0992735 | 0.0013 | 0.0027 | 0.0021 | 0.0021 | TRUE | 0.0244 |
| Tuba1b | 25039 | 17137 | 15550 | 16627 | 14357 | 16388 | 19447 | 18228 | 16778 | 16550 | 16196 | 16177 | 17229.6 | 18151 | 16308 | 0.898 | -0.155 | 0.01158391 | 0.0461066 | 0.0027 | 0.0026 | 0.0026 | 0.0026 | TRUE | 0.0535 |
| Tuba1c | 10369 | 7364 | 6674 | 6528 | 6258 | 6624 | 8053 | 7833 | 7201 | 6498 | 7060 | 6539 | 7197.32 | 7696 | 6699 | 0.87 | -0.201 | 0.00138366 | 0.0090927 | 0.0025 | 0.0028 | 0.0027 | 0.0027 | TRUE | 0.0385 |
| Tuba3a | 3212 | 2637 | 2544 | 2897 | 2599 | 2852 | 2495 | 2805 | 2745 | 2884 | 2932 | 2815 | 2779.25 | 2682 | 2877 | 1.074 | 0.103 | 0.11571321 | 0.24578524 | 0.0018 | 0.0033 | 0.0027 | 0.0027 | TRUE | 0.0464 |
| Tuba3b | 608 | 449 | 337 | 324 | 284 | 254 | 472 | 478 | 364 | 323 | 320 | 251 | 367.84 | 438 | 298 | 0.679 | -0.559 | 8.52E-05 | 0.00106186 | 0.0184 | 0.0089 | 0.0117 | 0.0117 | TRUE | 0.2482 |
| Tuba4a | 4939 | 3621 | 3231 | 3587 | 3341 | 4137 | 3836 | 3852 | 3486 | 3570 | 3769 | 4084 | 3766.18 | 3725 | 3808 | 1.022 | 0.032 | 0.65162559 | 0.78367265 | 0.0036 | 0.0031 | 0.0033 | 0.0033 | TRUE | 0.0493 |
| Tuba8 | 70 | 48 | 49 | 45 | 31 | 42 | 54 | 51 | 53 | 45 | 35 | 41 | 46.59 | 53 | 40 | 0.767 | -0.383 | 0.17413708 | 0.32714791 | 0 | 0.0533 | 0.0353 | 0.0353 | TRUE | 0.0959 |
| Tubal3 | 0 | 0 | 0 | 0 | 0 | 0 | 0 | 0 | 0 | 0 | 0 | 0 | 0 | 0 | 0 | NA | NA | NA | NA | NA | NA | NA | NA | NA | NA |
| Tubb1 | 20 | 9 | 14 | 25 | 16 | 15 | 16 | 10 | 15 | 25 | 18 | 15 | 16.33 | 14 | 19 | 1.428 | 0.514 | 0.29298463 | NA | 0 | 0.1476 | 0.1095 | 0.1095 | TRUE | 0.358 |
| Tubb2a | 3037 | 2210 | 1912 | 2108 | 1953 | 2090 | 2359 | 2351 | 2063 | 2098 | 2203 | 2063 | 2189.52 | 2258 | 2121 | 0.939 | -0.091 | 0.20962257 | 0.37145244 | 0.0027 | 0.0035 | 0.0033 | 0.0033 | TRUE | 0.0678 |
| Tubb2a-ps2 | 7 | 10 | 11 | 13 | 9 | 9 | 5 | 11 | 12 | 13 | 10 | 9 | 9.99 | 9 | 11 | 1.164 | 0.22 | 0.7221661 | NA | 0 | 0.2397 | 0.1739 | 0.1739 | TRUE | 0.5811 |

| | | | | | | | | | | | | | | | | | | | | | | | | | |
|---|---|---|---|---|---|---|---|---|---|---|---|---|---|---|---|---|---|---|---|---|---|---|---|---|---|
| Tubb2b | 17243 | 13483 | 15592 | 17241 | 15905 | 18139 | 13392 | 14342 | 16824 | 17162 | 17942 | 17906 | 16261.23 | 14853 | 17670 | 1.19 | 0.251 | 0.00119078 | 0.00821106 | 0.0073 | 0.0026 | 0.0043 | 0.0043 | TRUE | 0.1647 |
| Tubb3 | 2939 | 2141 | 1920 | 1940 | 1725 | 2153 | 2283 | 2277 | 2072 | 1931 | 1946 | 2125 | 2105.67 | 2211 | 2001 | 0.905 | -0.144 | 0.04190526 | 0.11827645 | 0.0022 | 0.0036 | 0.0031 | 0.0031 | TRUE | 0.0361 |
| Tubb4a | 137 | 117 | 101 | 123 | 124 | 157 | 106 | 124 | 109 | 122 | 140 | 155 | 126.19 | 113 | 139 | 1.231 | 0.299 | 0.10143041 | 0.22351909 | 0 | 0.0212 | 0.0161 | 0.0161 | TRUE | 0.1171 |
| Tubb4b | 36957 | 24930 | 22416 | 23172 | 20115 | 23119 | 28703 | 26518 | 24187 | 23065 | 22692 | 22822 | 24664.42 | 26469 | 22860 | 0.864 | -0.212 | 0.00108451 | 0.00767615 | 0.0036 | 0.0025 | 0.003 | 0.003 | TRUE | 0.0696 |
| Tubb5 | 113018 | 83280 | 84175 | 87634 | 78819 | 89480 | 87776 | 88584 | 90824 | 87231 | 88916 | 88331 | 88610.27 | 89061 | 88159 | 0.99 | -0.015 | 0.73129442 | 0.83929626 | 2.00E-04 | 0.0025 | 0.0013 | 0.0013 | TRUE | 0.0037 |
| Tubb6 | 8806 | 6089 | 5476 | 5667 | 4906 | 5625 | 6839 | 6477 | 5909 | 5641 | 5534 | 5553 | 5992.12 | 6408 | 5576 | 0.87 | -0.201 | 0.00155695 | 0.00992131 | 0.0025 | 0.0028 | 0.0027 | 0.0027 | TRUE | 0.0566 |
| Tubd1 | 718 | 469 | 427 | 513 | 458 | 498 | 558 | 499 | 461 | 511 | 517 | 492 | 506.03 | 506 | 507 | 0.998 | -0.003 | 0.98015782 | 0.98964367 | 0.0033 | 0.0071 | 0.0059 | 0.0059 | TRUE | 0.0993 |
| Tube1 | 1127 | 813 | 1114 | 1024 | 902 | 1018 | 875 | 865 | 1202 | 1019 | 1018 | 1005 | 997.3 | 981 | 1014 | 1.035 | 0.05 | 0.66648034 | 0.79482716 | 0.0169 | 0.0048 | 0.0086 | 0.0086 | TRUE | 0.4603 |
| Tubg1 | 6148 | 4133 | 4120 | 3889 | 3544 | 4090 | 4775 | 4396 | 4445 | 3871 | 3998 | 4037 | 4253.85 | 4539 | 3969 | 0.874 | -0.194 | 0.00078469 | 0.00606902 | 9.00E-04 | 0.003 | 0.0022 | 0.0022 | TRUE | 0.0257 |
| Tubg2 | 104 | 80 | 74 | 90 | 80 | 96 | 81 | 85 | 80 | 90 | 90 | 95 | 86.72 | 82 | 92 | 1.118 | 0.161 | 0.43214153 | 0.60400928 | 0 | 0.0298 | 0.0189 | 0.0189 | TRUE | 0.0102 |
| Tubgcp2 | 3652 | 2350 | 2174 | 2087 | 1934 | 2111 | 2836 | 2500 | 2346 | 2077 | 2182 | 2084 | 2337.46 | 2561 | 2114 | 0.825 | -0.278 | 0.00036472 | 0.00338721 | 0.0045 | 0.0034 | 0.0039 | 0.0039 | TRUE | 0.1096 |
| Tubgcp3 | 3508 | 2512 | 2417 | 2178 | 2042 | 2292 | 2725 | 2672 | 2608 | 2168 | 2304 | 2263 | 2456.42 | 2668 | 2245 | 0.841 | -0.25 | 2.75E-05 | 0.00043257 | 3.00E-04 | 0.0034 | 0.0022 | 0.0022 | TRUE | 0.0109 |
| Tubgcp4 | 2547 | 1930 | 1885 | 1906 | 1571 | 1825 | 1978 | 2053 | 2034 | 1897 | 1772 | 1802 | 1922.67 | 2022 | 1824 | 0.903 | -0.148 | 0.01929996 | 0.0673308 | 3.00E-04 | 0.0037 | 0.0024 | 0.0024 | TRUE | 0.0151 |
| Tubgcp5 | 1962 | 1409 | 1456 | 1354 | 1232 | 1403 | 1524 | 1499 | 1571 | 1348 | 1390 | 1385 | 1452.69 | 1531 | 1374 | 0.897 | -0.156 | 0.01799449 | 0.06387663 | 0 | 0.0041 | 0.0024 | 0.0024 | TRUE | 0.006 |
| Tubgcp6 | 3368 | 2599 | 2253 | 2121 | 2102 | 2169 | 2616 | 2765 | 2431 | 2111 | 2371 | 2141 | 2405.82 | 2604 | 2208 | 0.848 | -0.239 | 0.00120341 | 0.00826939 | 0.0035 | 0.0034 | 0.0035 | 0.0035 | TRUE | 0.0502 |
| Tufm | 8625 | 6066 | 5669 | 6087 | 5329 | 6422 | 6699 | 6452 | 6117 | 6059 | 6012 | 6340 | 6279.66 | 6423 | 6137 | 0.955 | -0.066 | 0.24481073 | 0.41296823 | 0.0012 | 0.0028 | 0.0022 | 0.0022 | TRUE | 0.0211 |
| Tuft1 | 607 | 513 | 497 | 557 | 465 | 521 | 471 | 546 | 536 | 554 | 525 | 514 | 524.45 | 518 | 531 | 1.028 | 0.04 | 0.69018296 | 0.81040066 | 0.0024 | 0.0069 | 0.0054 | 0.0054 | TRUE | 0.0767 |
| Tug1 | 3445 | 2675 | 2164 | 2264 | 1880 | 2338 | 2676 | 2845 | 2335 | 2254 | 2121 | 2308 | 2423.05 | 2619 | 2228 | 0.851 | -0.233 | 0.00347817 | 0.01853312 | 0.0052 | 0.0034 | 0.0042 | 0.0042 | TRUE | 0.1074 |
| Tulp1 | 78 | 72 | 83 | 140 | 113 | 144 | 61 | 77 | 90 | 139 | 127 | 142 | 105.95 | 76 | 136 | 1.82 | 0.864 | 2.75E-05 | 0.00043257 | 0.0074 | 0.0248 | 0.0204 | 0.0204 | TRUE | 0.3219 |
| Tulp2 | 245 | 195 | 205 | 297 | 252 | 273 | 190 | 207 | 221 | 296 | 284 | 269 | 244.72 | 206 | 283 | 1.376 | 0.461 | 0.00051907 | 0.00443992 | 0 | 0.0121 | 0.0085 | 0.0085 | TRUE | 0.0557 |
| Tulp3 | 6027 | 4574 | 4865 | 5653 | 5077 | 5699 | 4681 | 4865 | 5249 | 5627 | 5727 | 5626 | 5295.95 | 4932 | 5660 | 1.148 | 0.199 | 0.00075114 | 0.00587313 | 0.0015 | 0.0029 | 0.0023 | 0.0023 | TRUE | 0.0385 |
| Tulp4 | 1336 | 915 | 827 | 906 | 749 | 916 | 1038 | 973 | 892 | 902 | 845 | 904 | 925.7 | 968 | 884 | 0.912 | -0.133 | 0.12022155 | 0.25206493 | 0.0026 | 0.005 | 0.0042 | 0.0042 | TRUE | 0.0549 |
| Tusc1 | 98 | 70 | 62 | 47 | 46 | 57 | 76 | 74 | 67 | 47 | 52 | 56 | 62.07 | 72 | 52 | 0.711 | -0.492 | 0.04616065 | 0.12710159 | 0 | 0.0406 | 0.0271 | 0.0271 | TRUE | 0.058 |
| Tusc2 | 1184 | 927 | 746 | 782 | 674 | 839 | 920 | 986 | 805 | 778 | 760 | 828 | 846.25 | 904 | 789 | 0.873 | -0.196 | 0.0370588 | 0.108408 | 0.005 | 0.0052 | 0.0052 | 0.0052 | TRUE | 0.105 |
| Tusc3 | 2627 | 1839 | 2014 | 1821 | 1686 | 1831 | 2040 | 1956 | 2173 | 1813 | 1902 | 1807 | 1948.6 | 2056 | 1841 | 0.895 | -0.16 | 0.01731531 | 0.0621516 | 0.0013 | 0.0036 | 0.0027 | 0.0027 | TRUE | 0.029 |
| Tusc5 | 8 | 5 | 9 | 7 | 6 | 9 | 6 | 5 | 10 | 7 | 7 | 9 | 7.31 | 7 | 8 | 1.073 | 0.101 | 0.88485417 | NA | 0 | 0.3266 | 0.2167 | 0.2167 | TRUE | 0.2555 |
| Tut1 | 2614 | 1829 | 1544 | 1575 | 1353 | 1505 | 2030 | 1945 | 1666 | 1568 | 1526 | 1486 | 1703.56 | 1880 | 1527 | 0.811 | -0.302 | 0.00024924 | 0.0025237 | 0.0052 | 0.0038 | 0.0043 | 0.0043 | TRUE | 0.1192 |
| Tvp23a | 136 | 90 | 74 | 105 | 86 | 74 | 106 | 96 | 80 | 105 | 97 | 73 | 92.63 | 94 | 92 | 0.972 | -0.042 | 0.85148675 | 0.91638754 | 0.0162 | 0.028 | 0.0247 | 0.0247 | TRUE | 0.3076 |
| Tvp23b | 2877 | 2149 | 2410 | 2728 | 2387 | 2608 | 2234 | 2286 | 2600 | 2715 | 2693 | 2574 | 2517.23 | 2373 | 2661 | 1.122 | 0.166 | 0.02192429 | 0.07378477 | 0.0033 | 0.0034 | 0.0034 | 0.0034 | TRUE | 0.084 |
| Twf1 | 5983 | 4382 | 4349 | 4209 | 3816 | 4287 | 4647 | 4661 | 4693 | 4190 | 4305 | 4232 | 4454.46 | 4667 | 4242 | 0.909 | -0.138 | 0.00608351 | 0.02852742 | 0 | 0.003 | 0.0016 | 0.0016 | TRUE | 0.002 |
| Twf2 | 4558 | 3174 | 3631 | 4025 | 3885 | 4767 | 3540 | 3376 | 3918 | 4006 | 4383 | 4706 | 3988.15 | 3611 | 4365 | 1.209 | 0.274 | 0.00040812 | 0.00369895 | 0.0058 | 0.003 | 0.0041 | 0.0041 | TRUE | 0.0664 |
| Twist1 | 166 | 129 | 137 | 191 | 132 | 177 | 129 | 137 | 148 | 190 | 149 | 175 | 154.62 | 138 | 171 | 1.246 | 0.317 | 0.05813175 | 0.1502762 | 0 | 0.0178 | 0.0137 | 0.0137 | TRUE | 0.1329 |
| Twist2 | 6 | 2 | 4 | 1 | 3 | 1 | 5 | 2 | 4 | 1 | 3 | 1 | 2.75 | 4 | 2 | 0.47 | -1.088 | 0.36109467 | NA | 0 | 0.8658 | 0.5979 | 0.5979 | TRUE | 0.4345 |
| Twistnb | 3448 | 2459 | 2341 | 2857 | 2383 | 2922 | 2678 | 2616 | 2526 | 2844 | 2688 | 2884 | 2706.01 | 2607 | 2805 | 1.076 | 0.106 | 0.08017498 | 0.18944304 | 7.00E-04 | 0.0033 | 0.0023 | 0.0023 | TRUE | 0.016 |
| Twsg1 | 4959 | 3739 | 4455 | 5213 | 4492 | 5173 | 3851 | 3977 | 4807 | 5189 | 5067 | 5107 | 4666.41 | 4212 | 5121 | 1.216 | 0.283 | 0.00043512 | 0.00387728 | 0.0073 | 0.0029 | 0.0044 | 0.0044 | TRUE | 0.1853 |
| Txk | 1 | 0 | 1 | 0 | 0 | 1 | 1 | 0 | 1 | 0 | 0 | 1 | 0.47 | 1 | 0 | 0.596 | -0.748 | 0.78959815 | NA | 0 | 5.0047 | 3.8792 | 3.8792 | TRUE | 0.3835 |
| Txlna | 8428 | 6141 | 5642 | 5623 | 4951 | 5815 | 6546 | 6532 | 6088 | 5597 | 5585 | 5740 | 6014.69 | 6389 | 5641 | 0.883 | -0.18 | 0.00088275 | 0.00663014 | 7.00E-04 | 0.0028 | 0.0019 | 0.0019 | TRUE | 0.0205 |
| Txlnb | 189 | 195 | 180 | 145 | 142 | 167 | 147 | 207 | 194 | 144 | 160 | 165 | 169.63 | 183 | 156 | 0.86 | -0.218 | 0.21069481 | 0.37270556 | 0.0144 | 0.0164 | 0.0158 | 0.0158 | TRUE | 0.3498 |
| Txlng | 2979 | 2140 | 2142 | 2180 | 2050 | 2299 | 2314 | 2276 | 2311 | 2170 | 2313 | 2269 | 2275.53 | 2300 | 2251 | 0.978 | -0.032 | 0.59289231 | 0.74048805 | 0 | 0.0035 | 0.0021 | 0.0021 | TRUE | 0.012 |
| Txn1 | 17067 | 12507 | 12732 | 13306 | 11790 | 13722 | 13255 | 13304 | 13738 | 13245 | 13300 | 13546 | 13397.89 | 13432 | 13364 | 0.995 | -0.007 | 0.87419464 | 0.93016184 | 1.00E-04 | 0.0026 | 0.0014 | 0.0014 | TRUE | 0.0048 |
| Txn2 | 4643 | 3183 | 3336 | 3626 | 3118 | 3919 | 3606 | 3386 | 3600 | 3609 | 3517 | 3869 | 3597.78 | 3531 | 3665 | 1.038 | 0.054 | 0.38210768 | 0.55627063 | 0.0014 | 0.0031 | 0.0025 | 0.0025 | TRUE | 0.0288 |
| Txndc11 | 1581 | 1214 | 1101 | 1267 | 1087 | 1278 | 1228 | 1291 | 1188 | 1261 | 1226 | 1262 | 1242.7 | 1236 | 1250 | 1.012 | 0.017 | 0.8152687 | 0.89389387 | 0 | 0.0043 | 0.0028 | 0.0028 | TRUE | 0.0179 |
| Txndc12 | 1843 | 1322 | 1483 | 1684 | 1468 | 1626 | 1431 | 1406 | 1600 | 1676 | 1656 | 1605 | 1562.52 | 1479 | 1646 | 1.113 | 0.155 | 0.03500088 | 0.10387616 | 0.0018 | 0.0039 | 0.0032 | 0.0032 | TRUE | 0.0602 |
| Txndc15 | 949 | 648 | 598 | 644 | 518 | 658 | 737 | 689 | 645 | 641 | 584 | 650 | 657.75 | 690 | 625 | 0.904 | -0.145 | 0.12227726 | 0.25547564 | 0.0024 | 0.006 | 0.0048 | 0.0048 | TRUE | 0.0431 |
| Txndc16 | 866 | 587 | 717 | 903 | 772 | 952 | 673 | 624 | 774 | 899 | 871 | 940 | 796.69 | 690 | 903 | 1.31 | 0.389 | 4.42E-05 | 0.0006306 | 0.0049 | 0.0054 | 0.0053 | 0.0053 | TRUE | 0.1274 |
| Txndc17 | 6610 | 4669 | 4445 | 5003 | 4069 | 4707 | 5134 | 4966 | 4796 | 4980 | 4590 | 4647 | 4852.16 | 4965 | 4739 | 0.954 | -0.068 | 0.24788194 | 0.41672273 | 0.0013 | 0.0029 | 0.0023 | 0.0023 | TRUE | 0.0242 |
| Txndc2 | 2 | 1 | 3 | 1 | 3 | 3 | 2 | 1 | 3 | 1 | 3 | 3 | 2.2 | 2 | 2 | 1.259 | 0.333 | 0.79510202 | NA | 0 | 1.0801 | 0.7203 | 0.7203 | TRUE | 0.3001 |
| Txndc5 | 9909 | 7370 | 7592 | 8324 | 7174 | 8339 | 7696 | 7839 | 8192 | 8286 | 8093 | 8232 | 8056.26 | 7909 | 8204 | 1.037 | 0.053 | 0.29065617 | 0.4627254 | 4.00E-04 | 0.0027 | 0.0017 | 0.0017 | TRUE | 0.0119 |
| Txndc8 | 0 | 0 | 0 | 0 | 0 | 0 | 0 | 0 | 0 | 0 | 0 | 0 | 0 | 0 | 0 | NA | NA | NA | NA | NA | NA | NA | NA | NA | NA |
| Txndc9 | 3781 | 2493 | 2181 | 2122 | 1823 | 2031 | 2937 | 2652 | 2353 | 2112 | 2057 | 2005 | 2352.55 | 2647 | 2058 | 0.777 | -0.365 | 8.95E-06 | 0.00017141 | 0.0058 | 0.0034 | 0.0044 | 0.0044 | TRUE | 0.1141 |
| Txnip | 24788 | 19118 | 31005 | 39565 | 32348 | 38031 | 19252 | 20336 | 33454 | 39383 | 36492 | 37542 | 31076.47 | 24347 | 37806 | 1.553 | 0.635 | 0.01973631 | 0.06848534 | 0.0534 | 0.0025 | 0.0146 | 0.0534 | TRUE | 1.3102 |
| Txnl1 | 8470 | 6221 | 6275 | 6877 | 5738 | 6525 | 6578 | 6617 | 6771 | 6845 | 6473 | 6441 | 6620.96 | 6655 | 6586 | 0.99 | -0.015 | 0.77529181 | 0.86721285 | 4.00E-04 | 0.0028 | 0.0018 | 0.0018 | TRUE | 0.0145 |
| Txnl4a | 11201 | 7735 | 7151 | 7110 | 6433 | 7324 | 8699 | 8228 | 7716 | 7077 | 7257 | 7230 | 7701.19 | 8214 | 7188 | 0.875 | -0.193 | 0.00096224 | 0.0070576 | 0.0017 | 0.0027 | 0.0023 | 0.0023 | TRUE | 0.0344 |
| Txnl4b | 444 | 319 | 263 | 251 | 218 | 275 | 345 | 339 | 284 | 250 | 246 | 271 | 289.19 | 323 | 256 | 0.792 | -0.337 | 0.00932151 | 0.0393364 | 0.004 | 0.0106 | 0.0086 | 0.0086 | TRUE | 0.1219 |
| Txnrd1 | 19596 | 13359 | 13063 | 15179 | 13403 | 14734 | 15219 | 14210 | 14095 | 15109 | 15120 | 14545 | 14716.32 | 14508 | 14925 | 1.029 | 0.041 | 0.43852566 | 0.60989659 | 0.0011 | 0.0026 | 0.0019 | 0.0019 | TRUE | 0.0227 |
| Txnrd2 | 452 | 345 | 364 | 335 | 286 | 392 | 351 | 367 | 393 | 333 | 323 | 387 | 358.97 | 370 | 348 | 0.941 | -0.087 | 0.46234585 | 0.63170914 | 0.0036 | 0.009 | 0.0074 | 0.0074 | TRUE | 0.1127 |
| Txnrd3 | 1111 | 880 | 938 | 1035 | 890 | 1020 | 863 | 936 | 1012 | 1030 | 1004 | 1007 | 975.36 | 937 | 1014 | 1.084 | 0.116 | 0.16353819 | 0.31417234 | 0.0024 | 0.0049 | 0.004 | 0.004 | TRUE | 0.0601 |
| Tyk2 | 3036 | 2266 | 2292 | 2387 | 2243 | 2408 | 2358 | 2410 | 2473 | 2376 | 2530 | 2377 | 2420.79 | 2414 | 2428 | 1.006 | 0.008 | 0.88919417 | 0.93844044 | 5.00E-04 | 0.0034 | 0.0023 | 0.0023 | TRUE | 0.0163 |
| Tymp | 10 | 23 | 20 | 36 | 20 | 25 | 8 | 24 | 22 | 36 | 23 | 25 | 22.81 | 18 | 28 | 1.577 | 0.657 | 0.164327 | 0.31522329 | 0.1377 | 0.1063 | 0.1147 | 0.1147 | TRUE | 1.5911 |
| Tyms | 3400 | 2192 | 2059 | 1848 | 1664 | 2082 | 2641 | 2332 | 2222 | 1839 | 1877 | 2055 | 2160.97 | 2398 | 1924 | 0.802 | -0.319 | 7.67E-05 | 0.00098351 | 0.0052 | 0.0035 | 0.0042 | 0.0042 | TRUE | 0.0968 |
| Tyms-ps | 0 | 1 | 2 | 1 | 0 | 1 | 0 | 1 | 2 | 1 | 0 | 1 | 0.87 | 1 | 1 | 0.648 | -0.625 | 0.77012475 | NA | 0 | 2.7349 | 2.0642 | 2.0642 | TRUE | 0.1602 |
| Tyr | 1 | 1 | 0 | 0 | 0 | 0 | 1 | 1 | 0 | 0 | 0 | 0 | 0.31 | 1 | 0 | 0.309 | -1.696 | 0.63183309 | NA | 0 | 7.7294 | 7.2472 | 7.2472 | TRUE | 0.2015 |
| Tyro3 | 3882 | 2829 | 3087 | 3280 | 3043 | 3419 | 3015 | 3009 | 3331 | 3265 | 3433 | 3375 | 3237.97 | 3118 | 3358 | 1.077 | 0.107 | 0.09059504 | 0.20562233 | 0.0016 | 0.0032 | 0.0026 | 0.0026 | TRUE | 0.0427 |
| Tyrobp | 1 | 0 | 0 | 2 | 2 | 1 | 1 | 0 | 0 | 2 | 2 | 1 | 1 | 0 | 2 | 5.042 | 2.334 | 0.24985589 | NA | 0 | 2.3682 | 1.6672 | 1.6672 | TRUE | 0.2566 |
| Tyrp1 | 6 | 2 | 1 | 2 | 1 | 0 | 5 | 2 | 1 | 2 | 1 | 0 | 1.83 | 3 | 1 | 0.387 | -1.371 | 0.36677007 | NA | 0 | 1.2969 | 0.9824 | 0.9824 | TRUE | 0.5547 |
| Tysnd1 | 1471 | 1083 | 792 | 1154 | 896 | 988 | 1142 | 1152 | 855 | 1149 | 1011 | 975 | 1047.3 | 1050 | 1045 | 0.995 | -0.008 | 0.94592422 | 0.97029019 | 0.0166 | 0.0047 | 0.0084 | 0.0084 | TRUE | 0.3116 |
| Tyw1 | 3152 | 2377 | 2404 | 2333 | 2267 | 2589 | 2448 | 2528 | 2594 | 2322 | 2557 | 2556 | 2500.95 | 2523 | 2478 | 0.982 | -0.026 | 0.69415556 | 0.81337641 | 0.0015 | 0.0034 | 0.0027 | 0.0027 | TRUE | 0.0371 |
| Tyw3 | 373 | 243 | 207 | 228 | 205 | 243 | 290 | 258 | 223 | 227 | 231 | 240 | 244.94 | 257 | 233 | 0.902 | -0.149 | 0.28688058 | 0.4589126 | 0.0056 | 0.0121 | 0.0101 | 0.0101 | TRUE | 0.146 |
| Tyw5 | 537 | 390 | 382 | 364 | 313 | 336 | 417 | 415 | 412 | 362 | 353 | 332 | 381.86 | 415 | 349 | 0.841 | -0.249 | 0.01919741 | 0.06703849 | 0 | 0.0086 | 0.0055 | 0.0055 | TRUE | 0.0222 |

| | | | | | | | | | | | | | | | | | | | | | | | | | |
|---|---|---|---|---|---|---|---|---|---|---|---|---|---|---|---|---|---|---|---|---|---|---|---|---|---|
| U05342 | 0 | 0 | 1 | 0 | 0 | 0 | 0 | 0 | 1 | 0 | 0 | 0 | 0.18 | 0 | 0 | 0.552 | -0.857 | 0.8336548 | NA | 0 | 13.1817 | 10 | 10 | TRUE | 0.5365 |
| U2af1 | 9755 | 7247 | 7004 | 7366 | 6595 | 7743 | 7576 | 7709 | 7557 | 7332 | 7440 | 7644 | 7542.93 | 7614 | 7472 | 0.981 | -0.027 | 0.57349206 | 0.7257767 | 1.00E-04 | 0.0027 | 0.0015 | 0.0015 | TRUE | 0.005 |
| U2af1l4 | 897 | 697 | 654 | 694 | 610 | 680 | 697 | 741 | 706 | 691 | 688 | 671 | 698.99 | 715 | 683 | 0.957 | -0.064 | 0.44925776 | 0.61944307 | 0 | 0.0058 | 0.0036 | 0.0036 | TRUE | 0.0125 |
| U2af2 | 19483 | 13879 | 12138 | 12866 | 11101 | 13140 | 15132 | 14763 | 13097 | 12807 | 12523 | 12971 | 13548.74 | 14331 | 12767 | 0.891 | -0.167 | 0.00798519 | 0.03498395 | 0.0029 | 0.0026 | 0.0028 | 0.0028 | TRUE | 0.0692 |
| U2surp | 12696 | 8807 | 7458 | 7557 | 6753 | 7146 | 9860 | 9368 | 8047 | 7522 | 7618 | 7054 | 8245 | 9092 | 7398 | 0.813 | -0.298 | 8.53E-05 | 0.00106186 | 0.006 | 0.0027 | 0.004 | 0.004 | TRUE | 0.1232 |
| U90926 | 0 | 0 | 0 | 0 | 0 | 0 | 0 | 0 | 0 | 0 | 0 | 0 | 0 | 0 | 0 | NA | NA | NA | NA | NA | NA | NA | NA | NA | NA |
| Uaca | 4595 | 3344 | 3595 | 3841 | 3559 | 4023 | 3569 | 3557 | 3879 | 3823 | 4015 | 3971 | 3802.38 | 3668 | 3936 | 1.073 | 0.102 | 0.08779228 | 0.20125342 | 0.0012 | 0.003 | 0.0023 | 0.0023 | TRUE | 0.0304 |
| Uap1 | 3221 | 2293 | 2425 | 2459 | 2185 | 2480 | 2502 | 2439 | 2617 | 2448 | 2465 | 2448 | 2486.32 | 2519 | 2454 | 0.974 | -0.038 | 0.52130423 | 0.68258888 | 2.00E-04 | 0.0034 | 0.0021 | 0.0021 | TRUE | 0.0135 |
| Uap1l1 | 954 | 818 | 1032 | 1026 | 998 | 1005 | 741 | 870 | 1114 | 1021 | 1126 | 992 | 977.29 | 908 | 1046 | 1.154 | 0.207 | 0.09810962 | 0.21824375 | 0.022 | 0.0049 | 0.0102 | 0.0102 | TRUE | 0.465 |
| Uba1 | 42248 | 30647 | 30879 | 32303 | 29086 | 33485 | 32812 | 32599 | 33318 | 32154 | 32812 | 33055 | 32791.75 | 32910 | 32674 | 0.993 | -0.01 | 0.80991101 | 0.89039751 | 1.00E-04 | 0.0025 | 0.0013 | 0.0013 | TRUE | 0.0024 |
| Uba1y | 3099 | 2377 | 1947 | 1017 | 996 | 1050 | 2407 | 2528 | 2101 | 1012 | 1124 | 1037 | 1701.41 | 2345 | 1058 | 0.451 | -1.15 | 2.61E-42 | 6.21E-39 | 0.0056 | 0.0038 | 0.0044 | 0.0044 | TRUE | 0.0994 |
| Uba2 | 21807 | 15820 | 16001 | 17468 | 15279 | 17603 | 16937 | 16828 | 17265 | 17388 | 17236 | 17377 | 17171.64 | 17010 | 17334 | 1.019 | 0.027 | 0.53226134 | 0.69204142 | 0 | 0.0026 | 0.0013 | 0.0013 | TRUE | 0.0021 |
| Uba3 | 10253 | 7427 | 8022 | 8477 | 7624 | 8885 | 7963 | 7900 | 8656 | 8438 | 8601 | 8771 | 8388.04 | 8173 | 8603 | 1.053 | 0.074 | 0.18307944 | 0.33887003 | 0.0013 | 0.0027 | 0.0021 | 0.0021 | TRUE | 0.0324 |
| Uba5 | 2554 | 1736 | 1473 | 1533 | 1336 | 1657 | 1984 | 1847 | 1589 | 1526 | 1507 | 1636 | 1681.38 | 1807 | 1556 | 0.861 | -0.216 | 0.01297494 | 0.05009498 | 0.0062 | 0.0038 | 0.0049 | 0.0049 | TRUE | 0.1326 |
| Uba52 | 25426 | 18899 | 20473 | 21644 | 18759 | 22941 | 19747 | 20103 | 22090 | 21544 | 21162 | 22646 | 21215.49 | 20647 | 21784 | 1.055 | 0.078 | 0.19050817 | 0.34864703 | 0.0024 | 0.0025 | 0.0025 | 0.0025 | TRUE | 0.0457 |
| Uba6 | 6084 | 4410 | 4175 | 4660 | 3955 | 4756 | 4725 | 4691 | 4505 | 4639 | 4462 | 4695 | 4619.32 | 4640 | 4599 | 0.991 | -0.013 | 0.80645729 | 0.88818896 | 4.00E-04 | 0.0029 | 0.0019 | 0.0019 | TRUE | 0.0082 |
| Uba7 | 0 | 0 | 2 | 1 | 1 | 2 | 0 | 0 | 2 | 1 | 1 | 2 | 1.04 | 1 | 1 | 2.004 | 1.003 | 0.61459692 | NA | 0 | 2.2756 | 1.7771 | 1.7771 | TRUE | 0.9753 |
| Ubac1 | 5531 | 3791 | 3827 | 4049 | 3431 | 4104 | 4296 | 4032 | 4129 | 4030 | 3871 | 4051 | 4068.27 | 4152 | 3984 | 0.959 | -0.06 | 0.27980788 | 0.45138759 | 6.00E-04 | 0.003 | 0.002 | 0.002 | TRUE | 0.0113 |
| Ubac2 | 3370 | 2281 | 2388 | 2525 | 2265 | 2605 | 2617 | 2426 | 2577 | 2513 | 2555 | 2572 | 2543.39 | 2540 | 2547 | 1.002 | 0.003 | 0.95890913 | 0.97739513 | 5.00E-04 | 0.0034 | 0.0022 | 0.0022 | TRUE | 0.0184 |
| Ubald1 | 1178 | 875 | 832 | 768 | 699 | 843 | 915 | 931 | 898 | 764 | 789 | 832 | 854.76 | 915 | 795 | 0.87 | -0.202 | 0.01127974 | 0.04512213 | 0 | 0.0052 | 0.0034 | 0.0034 | TRUE | 0.0202 |
| Ubald2 | 438 | 367 | 441 | 510 | 528 | 544 | 340 | 390 | 476 | 508 | 596 | 537 | 474.45 | 402 | 547 | 1.363 | 0.447 | 0.00052239 | 0.00445764 | 0.0147 | 0.0074 | 0.0098 | 0.0098 | TRUE | 0.2944 |
| Ubap1 | 3378 | 2474 | 2491 | 2853 | 2356 | 2912 | 2624 | 2632 | 2688 | 2840 | 2658 | 2875 | 2719.19 | 2648 | 2791 | 1.055 | 0.077 | 0.1995249 | 0.36017505 | 6.00E-04 | 0.0033 | 0.0022 | 0.0022 | TRUE | 0.0208 |
| Ubap2 | 18242 | 13588 | 13200 | 13946 | 12755 | 14960 | 14168 | 14453 | 14243 | 13882 | 14389 | 14768 | 14317.08 | 14288 | 14346 | 1.004 | 0.006 | 0.90192624 | 0.94570564 | 4.00E-04 | 0.0026 | 0.0016 | 0.0016 | TRUE | 0.0098 |
| Ubap2l | 15843 | 11603 | 11324 | 11737 | 10397 | 11688 | 12305 | 12342 | 12219 | 11683 | 11729 | 11538 | 11969.13 | 12289 | 11650 | 0.948 | -0.077 | 0.08159872 | 0.19195037 | 0 | 0.0026 | 0.0013 | 0.0013 | TRUE | 9.00E-04 |
| Ubash3a | 1678 | 1283 | 1388 | 1562 | 1383 | 1620 | 1303 | 1365 | 1498 | 1555 | 1560 | 1599 | 1479.96 | 1389 | 1571 | 1.133 | 0.18 | 0.0149471 | 0.05570766 | 0.002 | 0.004 | 0.0033 | 0.0033 | TRUE | 0.0564 |
| Ubash3b | 270 | 186 | 180 | 219 | 179 | 200 | 210 | 198 | 194 | 218 | 202 | 197 | 203.19 | 201 | 206 | 1.024 | 0.035 | 0.80389925 | 0.88644746 | 0 | 0.0141 | 0.0093 | 0.0093 | TRUE | 0.03 |
| Ubb | 3748 | 2868 | 2896 | 2780 | 2775 | 3054 | 2911 | 3051 | 3125 | 2767 | 3130 | 3015 | 2999.8 | 3029 | 2971 | 0.981 | -0.028 | 0.67777454 | 0.80227176 | 0.0023 | 0.0032 | 0.0029 | 0.0029 | TRUE | 0.0438 |
| Ubc | 535 | 380 | 396 | 477 | 430 | 514 | 416 | 404 | 427 | 475 | 485 | 507 | 452.38 | 416 | 489 | 1.177 | 0.235 | 0.01804402 | 0.0640276 | 0 | 0.0077 | 0.0049 | 0.0049 | TRUE | 0.0128 |
| Ubd | 0 | 0 | 0 | 0 | 0 | 0 | 0 | 0 | 0 | 0 | 0 | 0 | 0 | 0 | 0 | NA | NA | NA | NA | NA | NA | NA | NA | NA | NA |
| Ube2a | 2815 | 1980 | 1792 | 1723 | 1421 | 1619 | 2186 | 2106 | 1934 | 1715 | 1603 | 1598 | 1857.04 | 2075 | 1639 | 0.789 | -0.342 | 2.01E-06 | 5.10E-05 | 0.0023 | 0.0037 | 0.0032 | 0.0032 | TRUE | 0.0428 |
| Ube2b | 4993 | 3896 | 3891 | 4254 | 3644 | 4092 | 3878 | 4144 | 4198 | 4234 | 4111 | 4039 | 4100.83 | 4073 | 4128 | 1.014 | 0.02 | 0.72852722 | 0.83731421 | 0.001 | 0.003 | 0.0021 | 0.0021 | TRUE | 0.022 |
| Ube2c | 9180 | 6667 | 7553 | 8470 | 7509 | 8649 | 7130 | 7092 | 8150 | 8431 | 8471 | 8538 | 7968.47 | 7457 | 8480 | 1.137 | 0.186 | 0.00379794 | 0.0198013 | 0.0031 | 0.0027 | 0.0028 | 0.0028 | TRUE | 0.0802 |
| Ube2cbp | 611 | 411 | 424 | 366 | 358 | 441 | 475 | 437 | 457 | 364 | 404 | 435 | 428.79 | 456 | 401 | 0.878 | -0.188 | 0.08564623 | 0.19785729 | 0.0024 | 0.008 | 0.0062 | 0.0062 | TRUE | 0.0762 |
| Ube2d1 | 1692 | 1227 | 1178 | 1280 | 1151 | 1428 | 1314 | 1305 | 1271 | 1274 | 1298 | 1410 | 1312.09 | 1297 | 1327 | 1.024 | 0.034 | 0.64232194 | 0.77710407 | 9.00E-04 | 0.0042 | 0.003 | 0.003 | TRUE | 0.0358 |
| Ube2d2a | 6251 | 4348 | 4287 | 4495 | 3834 | 4337 | 4855 | 4625 | 4626 | 4474 | 4325 | 4281 | 4531.03 | 4702 | 4360 | 0.927 | -0.109 | 0.04259569 | 0.11979932 | 5.00E-04 | 0.0029 | 0.0019 | 0.0019 | TRUE | 0.0101 |
| Ube2d2b | 1 | 3 | 2 | 5 | 0 | 1 | 1 | 3 | 2 | 5 | 0 | 1 | 2.01 | 2 | 2 | 1.007 | 0.01 | 0.99432711 | NA | 0.6413 | 1.1786 | 1.0333 | 1.0333 | TRUE | 1.5577 |
| Ube2d3 | 23390 | 16457 | 15820 | 19059 | 15905 | 18192 | 18166 | 17505 | 17070 | 18971 | 17942 | 17958 | 17935.48 | 17580 | 18290 | 1.04 | 0.057 | 0.2639245 | 0.43407614 | 9.00E-04 | 0.0026 | 0.0018 | 0.0018 | TRUE | 0.013 |
| Ube2dnl1 | 75 | 58 | 34 | 8 | 21 | 24 | 58 | 62 | 37 | 8 | 24 | 24 | 35.33 | 52 | 19 | 0.351 | -1.511 | 0.00020693 | 0.00218602 | 0.1279 | 0.0695 | 0.082 | 0.082 | TRUE | 1.3183 |
| Ube2dnl2 | 7 | 5 | 0 | 1 | 1 | 0 | 5 | 5 | 0 | 1 | 1 | 0 | 2.15 | 3 | 1 | 0.192 | -2.38 | 0.14783715 | NA | 0 | 1.1066 | 1.0728 | 1.0728 | TRUE | 1.0626 |
| Ube2e1 | 3107 | 2157 | 2292 | 2588 | 2221 | 2591 | 2413 | 2294 | 2473 | 2576 | 2506 | 2558 | 2469.97 | 2393 | 2547 | 1.064 | 0.089 | 0.13626184 | 0.27605818 | 3.00E-04 | 0.0034 | 0.0022 | 0.0022 | TRUE | 0.0155 |
| Ube2e2 | 502 | 359 | 348 | 349 | 330 | 386 | 390 | 382 | 375 | 347 | 372 | 381 | 374.66 | 382 | 367 | 0.959 | -0.061 | 0.57177111 | 0.72494672 | 0 | 0.0088 | 0.0057 | 0.0057 | TRUE | 0.0253 |
| Ube2e3 | 2873 | 1955 | 1768 | 1712 | 1570 | 1664 | 2231 | 2080 | 1908 | 1704 | 1771 | 1643 | 1889.4 | 2073 | 1706 | 0.822 | -0.283 | 0.00017608 | 0.00193449 | 0.0033 | 0.0037 | 0.0036 | 0.0036 | TRUE | 0.0585 |
| Ube2f | 1967 | 1280 | 1331 | 1545 | 1309 | 1504 | 1528 | 1362 | 1436 | 1538 | 1477 | 1485 | 1470.77 | 1442 | 1500 | 1.039 | 0.055 | 0.43904471 | 0.61048068 | 0.0013 | 0.004 | 0.003 | 0.003 | TRUE | 0.034 |
| Ube2g1 | 6784 | 4828 | 5179 | 5256 | 4415 | 5135 | 5269 | 5135 | 5588 | 5232 | 4981 | 5069 | 5212.31 | 5331 | 5094 | 0.956 | -0.065 | 0.24698701 | 0.41568896 | 0.001 | 0.0029 | 0.0021 | 0.0021 | TRUE | 0.0215 |
| Ube2g2 | 2194 | 1563 | 1559 | 1695 | 1467 | 1652 | 1704 | 1663 | 1682 | 1687 | 1655 | 1631 | 1670.26 | 1683 | 1658 | 0.985 | -0.022 | 0.72322853 | 0.83426832 | 0 | 0.0038 | 0.0022 | 0.0022 | TRUE | 0.003 |
| Ube2h | 4292 | 3016 | 3202 | 3335 | 2976 | 3470 | 3333 | 3208 | 3455 | 3320 | 3357 | 3425 | 3349.79 | 3332 | 3367 | 1.011 | 0.015 | 0.7886355 | 0.87590079 | 5.00E-04 | 0.0031 | 0.002 | 0.002 | TRUE | 0.0126 |
| Ube2i | 6460 | 4558 | 4490 | 4168 | 3816 | 4543 | 5017 | 4848 | 4845 | 4149 | 4305 | 4485 | 4608.08 | 4903 | 4313 | 0.879 | -0.185 | 0.00083323 | 0.00633834 | 6.00E-04 | 0.0029 | 0.002 | 0.002 | TRUE | 0.0149 |
| Ube2j1 | 711 | 521 | 484 | 473 | 402 | 463 | 552 | 554 | 522 | 471 | 453 | 457 | 501.67 | 543 | 460 | 0.848 | -0.238 | 0.01234881 | 0.04839635 | 0 | 0.0072 | 0.0045 | 0.0045 | TRUE | 0.0125 |
| Ube2j2 | 991 | 768 | 651 | 709 | 631 | 746 | 770 | 817 | 702 | 706 | 712 | 736 | 740.5 | 763 | 718 | 0.941 | -0.088 | 0.32170557 | 0.49543559 | 0.0017 | 0.0056 | 0.0043 | 0.0043 | TRUE | 0.0546 |
| Ube2k | 7275 | 5399 | 5691 | 5834 | 5019 | 5912 | 5650 | 5743 | 6141 | 5807 | 5662 | 5836 | 5806.46 | 5845 | 5768 | 0.987 | -0.018 | 0.7380948 | 0.84389961 | 8.00E-04 | 0.0028 | 0.002 | 0.002 | TRUE | 0.0238 |
| Ube2l3 | 7636 | 5352 | 4890 | 4739 | 4327 | 4996 | 5931 | 5693 | 5276 | 4717 | 4881 | 4932 | 5238.33 | 5633 | 4843 | 0.859 | -0.218 | 0.0003128 | 0.00300055 | 0.0017 | 0.0029 | 0.0025 | 0.0025 | TRUE | 0.0373 |
| Ube2l6 | 171 | 122 | 97 | 99 | 74 | 89 | 133 | 130 | 105 | 99 | 83 | 88 | 106.19 | 123 | 90 | 0.734 | -0.447 | 0.02429728 | 0.07968816 | 0 | 0.0247 | 0.0187 | 0.0187 | TRUE | 0.1562 |
| Ube2m | 3852 | 2747 | 2247 | 2115 | 1952 | 2124 | 2992 | 2922 | 2424 | 2105 | 2202 | 2097 | 2457.03 | 2779 | 2135 | 0.767 | -0.382 | 3.86E-06 | 8.58E-05 | 0.0059 | 0.0034 | 0.0045 | 0.0045 | TRUE | 0.1502 |
| Ube2n | 4938 | 3444 | 3399 | 3352 | 2861 | 3403 | 3835 | 3663 | 3667 | 3337 | 3228 | 3359 | 3514.89 | 3722 | 3308 | 0.889 | -0.171 | 0.00203526 | 0.01225357 | 3.00E-04 | 0.0031 | 0.0019 | 0.0019 | TRUE | 0.0088 |
| Ube2o | 14777 | 11575 | 11511 | 11884 | 11096 | 12863 | 11477 | 12312 | 12420 | 11829 | 12517 | 12698 | 12208.94 | 12070 | 12348 | 1.023 | 0.033 | 0.55206092 | 0.7078682 | 0.0014 | 0.0026 | 0.0022 | 0.0022 | TRUE | 0.0228 |
| Ube2q1 | 4081 | 2875 | 2765 | 2912 | 2497 | 2864 | 3170 | 3058 | 2983 | 2899 | 2817 | 2827 | 2958.96 | 3070 | 2848 | 0.927 | -0.109 | 0.05524397 | 0.1447218 | 3.00E-04 | 0.0032 | 0.002 | 0.002 | TRUE | 0.01 |
| Ube2q2 | 4146 | 3214 | 3185 | 3358 | 2998 | 3324 | 3220 | 3419 | 3437 | 3343 | 3382 | 3281 | 3346.87 | 3359 | 3335 | 0.994 | -0.009 | 0.86963301 | 0.92801725 | 5.00E-04 | 0.0031 | 0.002 | 0.002 | TRUE | 0.0163 |
| Ube2ql1 | 87 | 61 | 54 | 66 | 54 | 57 | 68 | 65 | 58 | 66 | 61 | 56 | 62.27 | 64 | 61 | 0.956 | -0.064 | 0.79256086 | 0.87848564 | 0 | 0.0405 | 0.0269 | 0.0269 | TRUE | 0.0446 |
| Ube2r2 | 4980 | 3589 | 3796 | 4026 | 3344 | 4063 | 3868 | 3818 | 4096 | 4007 | 3772 | 4011 | 3928.64 | 3927 | 3930 | 1.001 | 0.002 | 0.97701861 | 0.98766016 | 0.001 | 0.003 | 0.0022 | 0.0022 | TRUE | 0.017 |
| Ube2s | 10743 | 7653 | 7344 | 8183 | 7409 | 8477 | 8344 | 8140 | 7924 | 8145 | 8358 | 8368 | 8213.29 | 8136 | 8290 | 1.019 | 0.027 | 0.58483438 | 0.734344 | 3.00E-04 | 0.0027 | 0.0016 | 0.0016 | TRUE | 0.0063 |
| Ube2t | 2731 | 2090 | 2480 | 2939 | 2669 | 3027 | 2121 | 2223 | 2676 | 2925 | 3011 | 2988 | 2657.43 | 2340 | 2975 | 1.272 | 0.347 | 3.68E-05 | 0.00054961 | 0.0075 | 0.0033 | 0.0047 | 0.0047 | TRUE | 0.19 |
| Ube2u | 25 | 15 | 19 | 16 | 16 | 13 | 19 | 16 | 21 | 16 | 18 | 13 | 17.11 | 19 | 16 | 0.834 | -0.262 | 0.56993781 | 0.72352084 | 0 | 0.1409 | 0.0936 | 0.0936 | TRUE | 0.1156 |
| Ube2v1 | 2828 | 2076 | 1830 | 1891 | 1729 | 2011 | 2196 | 2208 | 1975 | 1882 | 1950 | 1985 | 2032.85 | 2126 | 1939 | 0.912 | -0.134 | 0.05246935 | 0.13956948 | 0.0018 | 0.0036 | 0.0029 | 0.0029 | TRUE | 0.0464 |
| Ube2v2 | 1250 | 873 | 996 | 878 | 829 | 892 | 971 | 929 | 1075 | 874 | 935 | 881 | 943.97 | 992 | 897 | 0.904 | -0.145 | 0.08628563 | 0.19882 | 0.0024 | 0.0049 | 0.0041 | 0.0041 | TRUE | 0.0625 |
| Ube2w | 1167 | 809 | 805 | 855 | 705 | 795 | 906 | 861 | 869 | 851 | 795 | 785 | 844.44 | 879 | 810 | 0.922 | -0.117 | 0.14632608 | 0.29067946 | 0 | 0.0052 | 0.0035 | 0.0035 | TRUE | 0.0232 |
| Ube2z | 21251 | 15561 | 14903 | 14760 | 13129 | 14764 | 16505 | 16552 | 16080 | 14692 | 14811 | 14574 | 15535.72 | 16379 | 14692 | 0.897 | -0.157 | 0.00042292 | 0.00379696 | 1.00E-04 | 0.0026 | 0.0014 | 0.0014 | TRUE | 0.0031 |
| Ube3a | 6359 | 4456 | 4077 | 4753 | 4052 | 4769 | 4939 | 4740 | 4399 | 4731 | 4571 | 4708 | 4681.26 | 4693 | 4670 | 0.995 | -0.007 | 0.90161305 | 0.94568228 | 0.0017 | 0.0029 | 0.0024 | 0.0024 | TRUE | 0.0363 |
| Ube3b | 6086 | 4608 | 4633 | 4662 | 4295 | 4881 | 4727 | 4901 | 4999 | 4641 | 4845 | 4818 | 4821.87 | 4876 | 4768 | 0.978 | -0.032 | 0.55132272 | 0.70717541 | 5.00E-04 | 0.0029 | 0.0019 | 0.0019 | TRUE | 0.0089 |
| Ube3c | 10701 | 7588 | 7161 | 8221 | 7272 | 8353 | 8311 | 8071 | 7727 | 8183 | 8204 | 8246 | 8123.56 | 8036 | 8211 | 1.021 | 0.031 | 0.54665848 | 0.70378217 | 5.00E-04 | 0.0027 | 0.0017 | 0.0017 | TRUE | 0.0138 |

| | | | | | | | | | | | | | | | | | | | | | | | | | |
|---|---|---|---|---|---|---|---|---|---|---|---|---|---|---|---|---|---|---|---|---|---|---|---|---|---|
| Ube4a | 3735 | 2380 | 1579 | 1934 | 1686 | 1851 | 2901 | 2532 | 1704 | 1925 | 1902 | 1827 | 2131.74 | 2379 | 1885 | 0.792 | -0.336 | 0.11802572 | 0.24928944 | 0.0329 | 0.0035 | 0.0129 | 0.0329 | TRUE | 0.4929 |
| Ube4b | 13737 | 10282 | 11310 | 11042 | 10571 | 12441 | 10669 | 10937 | 12203 | 10991 | 11925 | 12281 | 11501.12 | 11270 | 11732 | 1.041 | 0.058 | 0.3900606 | 0.56385036 | 0.0038 | 0.0026 | 0.0032 | 0.0032 | TRUE | 0.0641 |
| Ubfd1 | 6268 | 4722 | 4048 | 4165 | 3595 | 4110 | 4868 | 5023 | 4368 | 4146 | 4056 | 4057 | 4419.53 | 4753 | 4086 | 0.86 | -0.218 | 0.00073294 | 0.00576549 | 0.0023 | 0.003 | 0.0028 | 0.0028 | TRUE | 0.0606 |
| Ubiad1 | 1207 | 850 | 766 | 871 | 799 | 827 | 937 | 904 | 827 | 867 | 901 | 816 | 875.47 | 889 | 861 | 0.967 | -0.048 | 0.57307662 | 0.72548197 | 0.0022 | 0.0051 | 0.0041 | 0.0041 | TRUE | 0.0448 |
| Ubl3 | 3080 | 2401 | 2526 | 2625 | 2454 | 2854 | 2392 | 2554 | 2726 | 2613 | 2768 | 2817 | 2645.03 | 2557 | 2733 | 1.069 | 0.097 | 0.15865393 | 0.30789419 | 0.0023 | 0.0033 | 0.003 | 0.003 | TRUE | 0.0402 |
| Ubl4 | 1148 | 746 | 861 | 858 | 783 | 848 | 892 | 794 | 929 | 854 | 883 | 837 | 864.76 | 872 | 858 | 0.984 | -0.023 | 0.78969489 | 0.87639639 | 0.0024 | 0.0052 | 0.0042 | 0.0042 | TRUE | 0.0704 |
| Ubl4b | 0 | 0 | 0 | 0 | 0 | 0 | 0 | 0 | 0 | 0 | 0 | 0 | 0 | 0 | 0 | NA | NA | NA | NA | NA | NA | NA | NA | NA | NA |
| Ubl5 | 2645 | 1761 | 1928 | 2078 | 1840 | 2116 | 2054 | 1873 | 2080 | 2068 | 2076 | 2089 | 2040.11 | 2002 | 2078 | 1.037 | 0.053 | 0.42461681 | 0.59737271 | 0.0011 | 0.0036 | 0.0026 | 0.0026 | TRUE | 0.0379 |
| Ubl7 | 2546 | 1821 | 1820 | 1857 | 1758 | 2102 | 1977 | 1937 | 1964 | 1848 | 1983 | 2075 | 1964.13 | 1959 | 1969 | 1.005 | 0.007 | 0.91966307 | 0.95591188 | 0.0012 | 0.0036 | 0.0027 | 0.0027 | TRUE | 0.0351 |
| Ublcp1 | 2597 | 1762 | 1573 | 1838 | 1569 | 1737 | 2017 | 1874 | 1697 | 1830 | 1770 | 1715 | 1817.11 | 1863 | 1772 | 0.95 | -0.074 | 0.33720745 | 0.51142892 | 0.0034 | 0.0037 | 0.0037 | 0.0037 | TRUE | 0.0726 |
| Ubn1 | 3181 | 2426 | 2407 | 2600 | 2407 | 2739 | 2471 | 2581 | 2597 | 2588 | 2715 | 2704 | 2609.23 | 2550 | 2669 | 1.047 | 0.067 | 0.25794439 | 0.42795116 | 4.00E-04 | 0.0033 | 0.0021 | 0.0021 | TRUE | 0.0092 |
| Ubn2 | 3610 | 2768 | 2503 | 3099 | 2607 | 3202 | 2804 | 2944 | 2701 | 3085 | 2941 | 3161 | 2939.22 | 2816 | 3062 | 1.088 | 0.121 | 0.05087069 | 0.13654361 | 0.0011 | 0.0032 | 0.0024 | 0.0024 | TRUE | 0.0188 |
| Ubox5 | 1613 | 1122 | 1007 | 1097 | 1087 | 1122 | 1253 | 1193 | 1087 | 1092 | 1226 | 1108 | 1159.76 | 1178 | 1142 | 0.968 | -0.047 | 0.57759183 | 0.72902862 | 0.0037 | 0.0045 | 0.0042 | 0.0042 | TRUE | 0.0542 |
| Ubp1 | 6072 | 4721 | 4374 | 4234 | 3761 | 4402 | 4716 | 5022 | 4720 | 4215 | 4243 | 4345 | 4543.3 | 4819 | 4268 | 0.886 | -0.175 | 0.00134517 | 0.00891771 | 6.00E-04 | 0.0029 | 0.0019 | 0.0019 | TRUE | 0.0163 |
| Ubqln1 | 20322 | 14593 | 14974 | 15765 | 13670 | 15773 | 15783 | 15522 | 16157 | 15692 | 15421 | 15570 | 15691.08 | 15821 | 15561 | 0.984 | -0.024 | 0.59853232 | 0.74428593 | 1.00E-04 | 0.0026 | 0.0014 | 0.0014 | TRUE | 0.0042 |
| Ubqln2 | 3699 | 2728 | 2233 | 2143 | 1949 | 2324 | 2873 | 2902 | 2409 | 2133 | 2199 | 2294 | 2468.32 | 2728 | 2209 | 0.809 | -0.305 | 0.00013601 | 0.00157145 | 0.0052 | 0.0034 | 0.0042 | 0.0042 | TRUE | 0.1253 |
| Ubqln3 | 0 | 0 | 0 | 0 | 0 | 0 | 0 | 0 | 0 | 0 | 0 | 0 | 0 | 0 | 0 | NA | NA | NA | NA | NA | NA | NA | NA | NA | NA |
| Ubqln4 | 12598 | 9499 | 10512 | 10174 | 9435 | 11346 | 9784 | 10104 | 11342 | 10127 | 10644 | 11200 | 10533.63 | 10410 | 10657 | 1.024 | 0.034 | 0.61798057 | 0.75870094 | 0.0035 | 0.0027 | 0.0033 | 0.0033 | TRUE | 0.0748 |
| Ubqlnl | 0 | 0 | 0 | 0 | 0 | 0 | 0 | 0 | 0 | 0 | 0 | 0 | 0 | 0 | 0 | NA | NA | NA | NA | NA | NA | NA | NA | NA | NA |
| Ubr1 | 4110 | 2698 | 2713 | 3188 | 2864 | 3270 | 3192 | 2870 | 2927 | 3173 | 3231 | 3228 | 3103.57 | 2996 | 3211 | 1.071 | 0.099 | 0.11312699 | 0.24190963 | 0.0012 | 0.0032 | 0.0025 | 0.0025 | TRUE | 0.0406 |
| Ubr2 | 5204 | 3717 | 3525 | 3835 | 3452 | 3849 | 4042 | 3954 | 3803 | 3817 | 3894 | 3800 | 3885 | 3933 | 3837 | 0.975 | -0.036 | 0.50462509 | 0.66826706 | 3.00E-04 | 0.003 | 0.0019 | 0.0019 | TRUE | 0.01 |
| Ubr3 | 10405 | 7725 | 7783 | 9103 | 7888 | 8881 | 8081 | 8217 | 8398 | 9061 | 8898 | 8767 | 8570.4 | 8232 | 8909 | 1.082 | 0.114 | 0.01722601 | 0.06190876 | 2.00E-04 | 0.0027 | 0.0015 | 0.0015 | TRUE | 0.0038 |
| Ubr4 | 29575 | 18996 | 16479 | 21036 | 19679 | 21859 | 22970 | 20206 | 17781 | 20939 | 22200 | 21578 | 20945.59 | 20319 | 21572 | 1.062 | 0.086 | 0.2874644 | 0.45954509 | 0.0075 | 0.0025 | 0.0047 | 0.0047 | TRUE | 0.1597 |
| Ubr5 | 43809 | 31162 | 29340 | 29844 | 28642 | 33925 | 34025 | 33147 | 31658 | 29707 | 32311 | 33489 | 32389.31 | 32943 | 31836 | 0.966 | -0.049 | 0.40508461 | 0.57788945 | 0.0023 | 0.0025 | 0.0025 | 0.0025 | TRUE | 0.0419 |
| Ubr7 | 12025 | 8886 | 10049 | 12817 | 11240 | 13254 | 9339 | 9452 | 10843 | 12758 | 12680 | 13084 | 11359.28 | 9878 | 12841 | 1.3 | 0.379 | 6.76E-09 | 3.55E-07 | 0.0036 | 0.0026 | 0.003 | 0.003 | TRUE | 0.0889 |
| Ubtd1 | 879 | 633 | 776 | 826 | 685 | 805 | 683 | 673 | 837 | 822 | 773 | 795 | 763.82 | 731 | 797 | 1.091 | 0.126 | 0.2077876 | 0.36939156 | 0.0064 | 0.0055 | 0.0059 | 0.0059 | TRUE | 0.1871 |
| Ubtd2 | 3131 | 2209 | 2195 | 2389 | 1990 | 2327 | 2432 | 2350 | 2368 | 2378 | 2245 | 2297 | 2344.97 | 2383 | 2307 | 0.968 | -0.047 | 0.42609861 | 0.59860697 | 0 | 0.0034 | 0.0021 | 0.0021 | TRUE | 0.0089 |
| Ubtf | 16482 | 12177 | 10584 | 11483 | 10434 | 11847 | 12801 | 12953 | 11420 | 11430 | 11771 | 11695 | 12011.51 | 12391 | 11632 | 0.939 | -0.091 | 0.12963598 | 0.26637249 | 0.0023 | 0.0026 | 0.0025 | 0.0025 | TRUE | 0.0572 |
| Ubtfl1 | 152 | 115 | 54 | 65 | 71 | 45 | 118 | 122 | 58 | 65 | 80 | 44 | 81.31 | 99 | 63 | 0.63 | -0.667 | 0.02682937 | 0.08550106 | 0.106 | 0.0316 | 0.0524 | 0.0524 | TRUE | 0.8776 |
| Ubxn1 | 8109 | 5970 | 5831 | 5614 | 5049 | 5871 | 6298 | 6350 | 6292 | 5588 | 5696 | 5796 | 6003.21 | 6313 | 5693 | 0.902 | -0.149 | 0.00216499 | 0.01281334 | 0 | 0.0028 | 0.0015 | 0.0015 | TRUE | 0.0032 |
| Ubxn10 | 184 | 193 | 183 | 180 | 197 | 202 | 143 | 205 | 197 | 179 | 222 | 199 | 191.08 | 182 | 200 | 1.106 | 0.145 | 0.40163197 | 0.57479982 | 0.0199 | 0.0148 | 0.0163 | 0.0163 | TRUE | 0.4134 |
| Ubxn11 | 118 | 104 | 69 | 74 | 78 | 64 | 92 | 111 | 74 | 74 | 88 | 63 | 83.59 | 92 | 75 | 0.81 | -0.303 | 0.19767934 | 0.35751371 | 0.02 | 0.0308 | 0.0278 | 0.0278 | TRUE | 0.1643 |
| Ubxn2a | 4562 | 3027 | 2678 | 3383 | 2841 | 3285 | 3543 | 3220 | 2890 | 3367 | 3205 | 3243 | 3244.6 | 3218 | 3272 | 1.016 | 0.023 | 0.76292651 | 0.85929934 | 0.0052 | 0.0032 | 0.0039 | 0.0039 | TRUE | 0.0967 |
| Ubxn2b | 1581 | 1345 | 1280 | 1249 | 1071 | 1306 | 1228 | 1431 | 1381 | 1243 | 1208 | 1289 | 1296.72 | 1347 | 1247 | 0.928 | -0.109 | 0.17483212 | 0.32816264 | 0.0031 | 0.0043 | 0.0038 | 0.0038 | TRUE | 0.0743 |
| Ubxn4 | 5004 | 3735 | 3559 | 3854 | 3367 | 3828 | 3886 | 3973 | 3840 | 3836 | 3798 | 3779 | 3852.14 | 3900 | 3804 | 0.976 | -0.036 | 0.49231471 | 0.65756719 | 0 | 0.003 | 0.0017 | 0.0017 | TRUE | 0.0032 |
| Ubxn6 | 1988 | 1453 | 1441 | 1589 | 1457 | 1676 | 1544 | 1546 | 1555 | 1582 | 1644 | 1654 | 1587.36 | 1548 | 1627 | 1.051 | 0.071 | 0.2640095 | 0.43412627 | 0 | 0.0039 | 0.0023 | 0.0023 | TRUE | 0.0071 |
| Ubxn7 | 9057 | 6658 | 7110 | 8824 | 7348 | 8908 | 7034 | 7082 | 7672 | 8783 | 8289 | 8794 | 7942.35 | 7263 | 8622 | 1.188 | 0.248 | 1.55E-05 | 0.00027329 | 0.0016 | 0.0027 | 0.0022 | 0.0022 | TRUE | 0.0295 |
| Ubxn8 | 1386 | 1007 | 827 | 919 | 804 | 907 | 1076 | 1071 | 892 | 915 | 907 | 895 | 959.5 | 1013 | 906 | 0.893 | -0.164 | 0.06946562 | 0.17071391 | 0.0046 | 0.0049 | 0.0048 | 0.0048 | TRUE | 0.1276 |
| Uchl1 | 5522 | 4079 | 3682 | 3962 | 3567 | 4125 | 4289 | 4339 | 3973 | 3944 | 4024 | 4072 | 4106.68 | 4200 | 4013 | 0.955 | -0.066 | 0.25443801 | 0.42375699 | 9.00E-04 | 0.003 | 0.0022 | 0.0022 | TRUE | 0.0269 |
| Uchl3 | 4979 | 3544 | 3474 | 3764 | 3165 | 3721 | 3867 | 3770 | 3748 | 3747 | 3570 | 3673 | 3729.24 | 3795 | 3663 | 0.965 | -0.051 | 0.34289766 | 0.51698694 | 2.00E-04 | 0.0031 | 0.0018 | 0.0018 | TRUE | 0.0059 |
| Uchl4 | 26 | 14 | 14 | 13 | 18 | 20 | 20 | 15 | 15 | 13 | 20 | 20 | 17.2 | 17 | 18 | 1.045 | 0.064 | 0.89162192 | 0.94021713 | 0 | 0.1402 | 0.0992 | 0.0992 | TRUE | 0.2826 |
| Uchl5 | 4734 | 3314 | 3261 | 3542 | 3138 | 3648 | 3677 | 3525 | 3519 | 3526 | 3540 | 3601 | 3564.53 | 3574 | 3556 | 0.995 | -0.008 | 0.88601805 | 0.93667574 | 0 | 0.0031 | 0.0018 | 0.0018 | TRUE | 0.008 |
| Uck1 | 1855 | 1362 | 1320 | 1310 | 1132 | 1351 | 1441 | 1449 | 1424 | 1304 | 1277 | 1334 | 1371.39 | 1438 | 1305 | 0.908 | -0.14 | 0.03520909 | 0.10436408 | 0 | 0.0042 | 0.0024 | 0.0024 | TRUE | 0.0045 |
| Uck2 | 8087 | 6432 | 7278 | 7497 | 6977 | 8019 | 6281 | 6842 | 7853 | 7463 | 7871 | 7916 | 7370.77 | 6992 | 7750 | 1.109 | 0.149 | 0.05545074 | 0.14514126 | 0.0068 | 0.0027 | 0.0042 | 0.0042 | TRUE | 0.1416 |
| Uckl1 | 787 | 599 | 580 | 716 | 634 | 686 | 611 | 637 | 626 | 713 | 715 | 677 | 663.22 | 625 | 702 | 1.124 | 0.168 | 0.04933007 | 0.13346104 | 0 | 0.006 | 0.0038 | 0.0038 | TRUE | 0.0112 |
| Ucma | 9 | 8 | 7 | 10 | 2 | 5 | 7 | 9 | 8 | 10 | 2 | 5 | 6.7 | 8 | 6 | 0.753 | -0.409 | 0.59493542 | NA | 0.0109 | 0.3562 | 0.2729 | 0.2729 | TRUE | 0.581 |
| Ucn | 0 | 0 | 0 | 2 | 0 | 2 | 0 | 0 | 0 | 2 | 0 | 2 | 0.66 | 0 | 1 | 7.523 | 2.911 | 0.27166297 | NA | 0 | 3.5888 | 3.6663 | 3.6663 | TRUE | 0.41 |
| Ucn2 | 5 | 2 | 3 | 0 | 2 | 1 | 4 | 2 | 3 | 0 | 2 | 1 | 2.08 | 3 | 1 | 0.34 | -1.558 | 0.27282141 | NA | 0 | 1.1409 | 0.8091 | 0.8091 | TRUE | 0.3994 |
| Ucn3 | 0 | 0 | 0 | 0 | 0 | 0 | 0 | 0 | 0 | 0 | 0 | 0 | 0 | 0 | 0 | NA | NA | NA | NA | NA | NA | NA | NA | NA | NA |
| Ucp1 | 18 | 14 | 15 | 41 | 36 | 47 | 14 | 15 | 16 | 41 | 41 | 46 | 28.81 | 15 | 43 | 2.85 | 1.511 | 5.02E-05 | 0.00070171 | 0 | 0.0847 | 0.0558 | 0.0558 | TRUE | 0.0476 |
| Ucp2 | 10168 | 7767 | 7297 | 8712 | 7664 | 9011 | 7897 | 8262 | 7873 | 8672 | 8646 | 8895 | 8374.18 | 8011 | 8738 | 1.091 | 0.125 | 0.01091604 | 0.04404938 | 3.00E-04 | 0.0027 | 0.0016 | 0.0016 | TRUE | 0.0091 |
| Ucp3 | 10 | 8 | 6 | 9 | 6 | 9 | 8 | 9 | 6 | 9 | 7 | 9 | 7.89 | 8 | 8 | 1.084 | 0.116 | 0.86165183 | NA | 0 | 0.3027 | 0.1925 | 0.1925 | TRUE | 0.0622 |
| Uevld | 2566 | 1982 | 2229 | 2566 | 2137 | 2542 | 1993 | 2108 | 2405 | 2554 | 2411 | 2509 | 2330.08 | 2169 | 2491 | 1.15 | 0.202 | 0.00928177 | 0.03920621 | 0.0046 | 0.0034 | 0.0039 | 0.0039 | TRUE | 0.1095 |
| Ufc1 | 2339 | 1628 | 1614 | 1864 | 1563 | 1846 | 1817 | 1732 | 1741 | 1855 | 1763 | 1822 | 1788.45 | 1763 | 1813 | 1.028 | 0.04 | 0.52812084 | 0.68874916 | 0 | 0.0038 | 0.0024 | 0.0024 | TRUE | 0.0088 |
| Ufd1l | 2176 | 1418 | 1453 | 1467 | 1238 | 1501 | 1690 | 1508 | 1568 | 1460 | 1397 | 1482 | 1517.44 | 1589 | 1446 | 0.91 | -0.137 | 0.05845971 | 0.15071711 | 0.0016 | 0.004 | 0.0031 | 0.0031 | TRUE | 0.0389 |
| Ufl1 | 3377 | 2356 | 2426 | 2542 | 2185 | 2552 | 2623 | 2506 | 2618 | 2530 | 2465 | 2519 | 2543.48 | 2582 | 2505 | 0.97 | -0.044 | 0.4449479 | 0.61582651 | 0 | 0.0034 | 0.002 | 0.002 | TRUE | 0.0079 |
| Ufm1 | 2464 | 1732 | 1568 | 1659 | 1396 | 1713 | 1914 | 1842 | 1692 | 1651 | 1575 | 1691 | 1727.51 | 1816 | 1639 | 0.902 | -0.149 | 0.0390998 | 0.11267206 | 0.0021 | 0.0038 | 0.0032 | 0.0032 | TRUE | 0.0427 |
| Ufsp1 | 167 | 125 | 112 | 126 | 67 | 103 | 130 | 133 | 121 | 125 | 76 | 102 | 114.36 | 128 | 101 | 0.792 | -0.337 | 0.10523268 | 0.22966724 | 0.0201 | 0.0232 | 0.0223 | 0.0223 | TRUE | 0.2523 |
| Ufsp2 | 2935 | 2064 | 2266 | 2472 | 2171 | 2540 | 2279 | 2195 | 2445 | 2461 | 2449 | 2507 | 2389.51 | 2306 | 2472 | 1.072 | 0.1 | 0.1152631 | 0.24515708 | 0.001 | 0.0034 | 0.0025 | 0.0025 | TRUE | 0.0325 |
| Ugcg | 4077 | 2931 | 3132 | 3248 | 2831 | 3187 | 3166 | 3118 | 3379 | 3233 | 3194 | 3146 | 3206.05 | 3221 | 3191 | 0.991 | -0.013 | 0.82024222 | 0.89651557 | 7.00E-04 | 0.0032 | 0.0021 | 0.0021 | TRUE | 0.0221 |
| Ugdh | 2624 | 1898 | 1928 | 1883 | 1561 | 1879 | 2038 | 2019 | 2080 | 1874 | 1761 | 1855 | 1937.88 | 2046 | 1830 | 0.895 | -0.16 | 0.01014998 | 0.04174403 | 0 | 0.0036 | 0.0023 | 0.0023 | TRUE | 0.0129 |
| Uggt1 | 11422 | 8188 | 8020 | 9366 | 8186 | 9477 | 8871 | 8709 | 8654 | 9323 | 9235 | 9355 | 9024.46 | 8745 | 9304 | 1.064 | 0.089 | 0.05141892 | 0.13762746 | 0 | 0.0027 | 0.0014 | 0.0014 | TRUE | 0.002 |
| Uggt2 | 435 | 294 | 340 | 353 | 307 | 309 | 338 | 313 | 367 | 351 | 346 | 305 | 336.69 | 339 | 334 | 0.985 | -0.021 | 0.85917131 | 0.92142036 | 0.0029 | 0.0095 | 0.0075 | 0.0075 | TRUE | 0.0678 |
| Ugp2 | 2960 | 2064 | 2147 | 2060 | 1698 | 1927 | 2299 | 2195 | 2317 | 2051 | 1916 | 1902 | 2113.21 | 2270 | 1956 | 0.862 | -0.215 | 0.00078818 | 0.00608167 | 8.00E-04 | 0.0035 | 0.0025 | 0.0025 | TRUE | 0.0217 |
| Ugt1a1 | 0 | 0 | 0 | 0 | 1 | 0 | 0 | 0 | 0 | 0 | 1 | 0 | 0.19 | 0 | 0 | 2.094 | 1.067 | 0.79379329 | NA | 0 | 12.608 | 10 | 10 | TRUE | 0.5075 |
| Ugt1a10 | 0 | 0 | 0 | 0 | 0 | 0 | 0 | 0 | 0 | 0 | 0 | 0 | 0 | 0 | 0 | NA | NA | NA | NA | NA | NA | NA | NA | NA | NA |
| Ugt1a2 | 0 | 0 | 0 | 0 | 0 | 0 | 0 | 0 | 0 | 0 | 0 | 0 | 0 | 0 | 0 | NA | NA | NA | NA | NA | NA | NA | NA | NA | NA |
| Ugt1a5 | 0 | 0 | 0 | 0 | 0 | 0 | 0 | 0 | 0 | 0 | 0 | 0 | 0 | 0 | 0 | NA | NA | NA | NA | NA | NA | NA | NA | NA | NA |
| Ugt1a6a | 0 | 1 | 0 | 0 | 0 | 0 | 0 | 1 | 0 | 0 | 0 | 0 | 0.18 | 0 | 0 | 0.552 | -0.857 | 0.8336548 | NA | 0 | 13.3714 | 10 | 10 | TRUE | 0.5216 |

| | | | | | | | | | | | | | | | | | | | | | | | | | |
|---|---|---|---|---|---|---|---|---|---|---|---|---|---|---|---|---|---|---|---|---|---|---|---|---|---|
| Ugt1a6b | 0 | 0 | 0 | 0 | 0 | 0 | 0 | 0 | 0 | 0 | 0 | 0 | 0 | 0 | 0 | NA | NA | NA | NA | NA | NA | NA | NA | NA | NA |
| Ugt1a7c | 0 | 0 | 0 | 0 | 0 | 0 | 0 | 0 | 0 | 0 | 0 | 0 | 0 | 0 | 0 | NA | NA | NA | NA | NA | NA | NA | NA | NA | NA |
| Ugt1a9 | 0 | 0 | 0 | 0 | 0 | 0 | 0 | 0 | 0 | 0 | 0 | 0 | 0 | 0 | 0 | NA | NA | NA | NA | NA | NA | NA | NA | NA | NA |
| Ugt2a1 | 0 | 0 | 0 | 0 | 0 | 0 | 0 | 0 | 0 | 0 | 0 | 0 | 0 | 0 | 0 | NA | NA | NA | NA | NA | NA | NA | NA | NA | NA |
| Ugt2a2 | 0 | 0 | 0 | 0 | 0 | 0 | 0 | 0 | 0 | 0 | 0 | 0 | 0 | 0 | 0 | NA | NA | NA | NA | NA | NA | NA | NA | NA | NA |
| Ugt2a3 | 0 | 0 | 0 | 0 | 0 | 0 | 0 | 0 | 0 | 0 | 0 | 0 | 0 | 0 | 0 | NA | NA | NA | NA | NA | NA | NA | NA | NA | NA |
| Ugt2b1 | 0 | 0 | 0 | 0 | 0 | 0 | 0 | 0 | 0 | 0 | 0 | 0 | 0 | 0 | 0 | NA | NA | NA | NA | NA | NA | NA | NA | NA | NA |
| Ugt2b34 | 1 | 0 | 0 | 0 | 1 | 0 | 1 | 0 | 0 | 0 | 1 | 0 | 0.32 | 0 | 0 | 1.075 | 0.105 | 0.97638847 | NA | 0 | 7.4681 | 7.0392 | 7.0392 | TRUE | 0.5075 |
| Ugt2b35 | 0 | 0 | 0 | 0 | 0 | 0 | 0 | 0 | 0 | 0 | 0 | 0 | 0 | 0 | 0 | NA | NA | NA | NA | NA | NA | NA | NA | NA | NA |
| Ugt2b36 | 0 | 0 | 0 | 0 | 0 | 0 | 0 | 0 | 0 | 0 | 0 | 0 | 0 | 0 | 0 | NA | NA | NA | NA | NA | NA | NA | NA | NA | NA |
| Ugt2b37 | 0 | 0 | 0 | 0 | 0 | 0 | 0 | 0 | 0 | 0 | 0 | 0 | 0 | 0 | 0 | NA | NA | NA | NA | NA | NA | NA | NA | NA | NA |
| Ugt2b38 | 0 | 0 | 0 | 0 | 0 | 0 | 0 | 0 | 0 | 0 | 0 | 0 | 0 | 0 | 0 | NA | NA | NA | NA | NA | NA | NA | NA | NA | NA |
| Ugt2b5 | 0 | 0 | 0 | 0 | 0 | 0 | 0 | 0 | 0 | 0 | 0 | 0 | 0 | 0 | 0 | NA | NA | NA | NA | NA | NA | NA | NA | NA | NA |
| Ugt3a1 | 3 | 2 | 0 | 1 | 2 | 3 | 2 | 2 | 0 | 1 | 2 | 3 | 1.78 | 1 | 2 | 1.359 | 0.442 | 0.76193542 | NA | 0 | 1.3351 | 0.9674 | 0.9674 | TRUE | 0.458 |
| Ugt3a2 | 1 | 0 | 2 | 0 | 0 | 0 | 1 | 0 | 2 | 0 | 0 | 0 | 0.49 | 1 | 0 | 0.195 | -2.355 | 0.4284796 | NA | 0 | 4.8481 | 4.8763 | 4.8763 | TRUE | 0.127 |
| Ugt8a | 655 | 480 | 396 | 490 | 369 | 352 | 509 | 511 | 427 | 488 | 416 | 347 | 449.68 | 482 | 417 | 0.864 | -0.21 | 0.11328692 | 0.24207056 | 0.0169 | 0.0077 | 0.0105 | 0.0105 | TRUE | 0.2277 |
| Uhmk1 | 988 | 760 | 804 | 920 | 719 | 901 | 767 | 808 | 868 | 916 | 811 | 889 | 843.26 | 814 | 872 | 1.073 | 0.102 | 0.24778181 | 0.41665236 | 0.0027 | 0.0052 | 0.0044 | 0.0044 | TRUE | 0.0435 |
| Uhrf1 | 19294 | 14398 | 13465 | 13564 | 12365 | 14400 | 14985 | 15315 | 14529 | 13502 | 13949 | 14215 | 14415.67 | 14943 | 13889 | 0.929 | -0.106 | 0.03175646 | 0.09651182 | 6.00E-04 | 0.0026 | 0.0017 | 0.0017 | TRUE | 0.0073 |
| Uhrf1bp1 | 6889 | 5252 | 5291 | 5693 | 5256 | 5875 | 5350 | 5586 | 5709 | 5667 | 5929 | 5800 | 5673.58 | 5548 | 5799 | 1.045 | 0.064 | 0.23087884 | 0.39662875 | 5.00E-04 | 0.0028 | 0.0019 | 0.0019 | TRUE | 0.0121 |
| Uhrf1bp1l | 3621 | 2742 | 2776 | 2415 | 2325 | 2574 | 2812 | 2917 | 2995 | 2404 | 2623 | 2541 | 2715.31 | 2908 | 2523 | 0.868 | -0.205 | 0.00098894 | 0.00718601 | 0.0011 | 0.0033 | 0.0024 | 0.0024 | TRUE | 0.0206 |
| Uhrf2 | 6684 | 4871 | 4987 | 5090 | 4642 | 5315 | 5191 | 5181 | 5381 | 5067 | 5237 | 5247 | 5217.22 | 5251 | 5184 | 0.987 | -0.019 | 0.71624547 | 0.82949644 | 2.00E-04 | 0.0029 | 0.0017 | 0.0017 | TRUE | 0.0056 |
| Uimc1 | 3210 | 2256 | 2013 | 2224 | 1962 | 2280 | 2493 | 2400 | 2172 | 2214 | 2213 | 2251 | 2290.43 | 2355 | 2226 | 0.945 | -0.082 | 0.23061094 | 0.39635909 | 0.0022 | 0.0035 | 0.003 | 0.003 | TRUE | 0.0554 |
| Ulbp1 | 185 | 106 | 148 | 158 | 121 | 111 | 144 | 113 | 160 | 157 | 137 | 110 | 136.58 | 139 | 135 | 0.968 | -0.046 | 0.81441555 | 0.89319411 | 0.0238 | 0.0198 | 0.0209 | 0.0209 | TRUE | 0.2774 |
| Ulk1 | 20803 | 17090 | 18813 | 17656 | 17311 | 19384 | 16157 | 18178 | 20299 | 17575 | 19529 | 19135 | 18478.78 | 18211 | 18746 | 1.029 | 0.042 | 0.59838041 | 0.74416177 | 0.0065 | 0.0026 | 0.0045 | 0.0045 | TRUE | 0.123 |
| Ulk2 | 1529 | 1103 | 1281 | 1562 | 1255 | 1344 | 1188 | 1173 | 1382 | 1555 | 1416 | 1327 | 1340.04 | 1248 | 1433 | 1.149 | 0.2 | 0.02537596 | 0.08220728 | 0.0065 | 0.0042 | 0.005 | 0.005 | TRUE | 0.1051 |
| Ulk3 | 379 | 314 | 371 | 341 | 285 | 424 | 294 | 334 | 400 | 339 | 322 | 419 | 351.36 | 343 | 360 | 1.053 | 0.075 | 0.60391592 | 0.74889556 | 0.0192 | 0.0092 | 0.0122 | 0.0122 | TRUE | 0.2431 |
| Ulk4 | 256 | 252 | 255 | 240 | 233 | 271 | 199 | 268 | 275 | 239 | 263 | 268 | 251.88 | 247 | 257 | 1.042 | 0.059 | 0.6980768 | 0.81651154 | 0.0142 | 0.0118 | 0.0125 | 0.0125 | TRUE | 0.3543 |
| Umod | 1 | 3 | 1 | 2 | 0 | 0 | 1 | 3 | 1 | 2 | 0 | 0 | 1.17 | 2 | 1 | 0.408 | -1.295 | 0.50272732 | NA | 0 | 2.0231 | 1.6328 | 1.6328 | TRUE | 0.9735 |
| Umodl1 | 25 | 14 | 10 | 16 | 16 | 21 | 19 | 15 | 11 | 16 | 18 | 21 | 16.63 | 15 | 18 | 1.199 | 0.262 | 0.58223983 | NA | 0 | 0.1449 | 0.1032 | 0.1032 | TRUE | 0.2921 |
| Umps | 11601 | 7953 | 6643 | 6934 | 6241 | 6990 | 9010 | 8460 | 7168 | 6902 | 7040 | 6900 | 7580.01 | 8213 | 6947 | 0.846 | -0.242 | 0.00189134 | 0.01157726 | 0.0065 | 0.0027 | 0.0042 | 0.0042 | TRUE | 0.1509 |
| Unc119 | 2121 | 1571 | 1698 | 1745 | 1576 | 1778 | 1647 | 1671 | 1832 | 1737 | 1778 | 1755 | 1736.75 | 1717 | 1757 | 1.024 | 0.034 | 0.61924667 | 0.75973328 | 0.0012 | 0.0038 | 0.0028 | 0.0028 | TRUE | 0.0411 |
| Unc119b | 7653 | 5551 | 5561 | 5634 | 5063 | 6069 | 5944 | 5905 | 6000 | 5608 | 5712 | 5991 | 5859.88 | 5950 | 5770 | 0.97 | -0.044 | 0.39618706 | 0.56958266 | 4.00E-04 | 0.0028 | 0.0018 | 0.0018 | TRUE | 0.0138 |
| Unc13a | 511 | 488 | 545 | 773 | 684 | 731 | 397 | 519 | 588 | 769 | 772 | 722 | 627.78 | 501 | 754 | 1.51 | 0.594 | 2.66E-06 | 6.32E-05 | 0.0184 | 0.0062 | 0.0099 | 0.0099 | TRUE | 0.4077 |
| Unc13b | 3069 | 2264 | 1913 | 2303 | 2177 | 2619 | 2384 | 2408 | 2064 | 2292 | 2456 | 2585 | 2364.92 | 2285 | 2444 | 1.069 | 0.097 | 0.21780884 | 0.3807193 | 0.0047 | 0.0034 | 0.004 | 0.004 | TRUE | 0.0857 |
| Unc13c | 8 | 10 | 6 | 2 | 5 | 3 | 6 | 11 | 6 | 2 | 6 | 3 | 5.65 | 8 | 4 | 0.451 | -1.149 | 0.17264347 | NA | 0 | 0.4217 | 0.3003 | 0.3003 | TRUE | 0.3612 |
| Unc13d | 287 | 226 | 237 | 316 | 286 | 342 | 223 | 240 | 256 | 315 | 323 | 338 | 282.3 | 240 | 325 | 1.359 | 0.443 | 0.00035208 | 0.0032869 | 0 | 0.0108 | 0.0075 | 0.0075 | TRUE | 0.0458 |
| Unc45a | 2548 | 1787 | 1501 | 1588 | 1457 | 1593 | 1979 | 1901 | 1620 | 1581 | 1644 | 1573 | 1716.03 | 1833 | 1599 | 0.871 | -0.199 | 0.01615926 | 0.05902551 | 0.0047 | 0.0038 | 0.0043 | 0.0043 | TRUE | 0.1241 |
| Unc45b | 85 | 60 | 51 | 47 | 43 | 59 | 66 | 64 | 55 | 47 | 49 | 58 | 56.4 | 62 | 51 | 0.829 | -0.27 | 0.29865144 | 0.47094274 | 0 | 0.0444 | 0.0308 | 0.0308 | TRUE | 0.1231 |
| Unc50 | 2092 | 1503 | 1575 | 1826 | 1597 | 1746 | 1625 | 1599 | 1699 | 1818 | 1802 | 1724 | 1710.94 | 1641 | 1781 | 1.085 | 0.118 | 0.06969732 | 0.1711068 | 3.00E-04 | 0.0038 | 0.0025 | 0.0025 | TRUE | 0.0114 |
| Unc5a | 330 | 283 | 403 | 600 | 544 | 593 | 256 | 301 | 435 | 597 | 614 | 585 | 464.74 | 331 | 599 | 1.818 | 0.862 | 2.85E-08 | 1.27E-06 | 0.0362 | 0.0075 | 0.015 | 0.015 | TRUE | 0.856 |
| Unc5b | 1727 | 1310 | 1459 | 1582 | 1431 | 1588 | 1341 | 1393 | 1574 | 1575 | 1614 | 1568 | 1510.93 | 1436 | 1586 | 1.105 | 0.144 | 0.06209453 | 0.15727752 | 0.0028 | 0.004 | 0.0036 | 0.0036 | TRUE | 0.0841 |
| Unc5c | 10 | 3 | 1 | 5 | 3 | 2 | 8 | 3 | 1 | 5 | 3 | 2 | 3.73 | 4 | 3 | 0.833 | -0.264 | 0.80087111 | NA | 0.2504 | 0.6381 | 0.5208 | 0.5208 | TRUE | 0.5727 |
| Unc5cl | 72 | 54 | 54 | 37 | 56 | 71 | 56 | 57 | 58 | 37 | 63 | 70 | 56.95 | 57 | 57 | 0.99 | -0.014 | 0.96163725 | 0.97889618 | 0.0348 | 0.044 | 0.0415 | 0.0415 | TRUE | 0.8153 |
| Unc5d | 61 | 47 | 26 | 27 | 24 | 28 | 47 | 50 | 28 | 27 | 27 | 28 | 34.5 | 42 | 27 | 0.648 | -0.626 | 0.07317124 | 0.17706687 | 0 | 0.0711 | 0.0575 | 0.0575 | TRUE | 0.5896 |
| Unc79 | 15 | 20 | 12 | 20 | 9 | 14 | 12 | 21 | 13 | 20 | 10 | 14 | 14.96 | 15 | 15 | 0.969 | -0.046 | 0.9293635 | NA | 0.0448 | 0.1609 | 0.1284 | 0.1284 | TRUE | 0.4518 |
| Unc80 | 61 | 51 | 33 | 47 | 43 | 53 | 47 | 54 | 36 | 47 | 49 | 52 | 47.47 | 46 | 49 | 1.074 | 0.103 | 0.71949275 | 0.83168102 | 0 | 0.0524 | 0.0384 | 0.0384 | TRUE | 0.2706 |
| Unc93a | 6 | 6 | 3 | 9 | 6 | 5 | 5 | 6 | 3 | 9 | 7 | 5 | 5.82 | 5 | 7 | 1.447 | 0.534 | 0.50053062 | NA | 0 | 0.4094 | 0.2755 | 0.2755 | TRUE | 0.1889 |
| Unc93b1 | 209 | 159 | 152 | 251 | 191 | 260 | 162 | 169 | 164 | 250 | 215 | 257 | 202.9 | 165 | 241 | 1.46 | 0.546 | 0.0001621 | 0.00181294 | 0 | 0.0141 | 0.01 | 0.01 | TRUE | 0.0893 |
| Uncx | 14 | 4 | 7 | 3 | 4 | 4 | 11 | 4 | 8 | 3 | 5 | 4 | 5.69 | 8 | 4 | 0.496 | -1.011 | 0.22257739 | NA | 0 | 0.4191 | 0.2958 | 0.2958 | TRUE | 0.122 |
| Ung | 6922 | 5256 | 5102 | 5191 | 4578 | 5432 | 5376 | 5591 | 5505 | 5167 | 5164 | 5362 | 5360.93 | 5491 | 5231 | 0.953 | -0.069 | 0.17483793 | 0.32816264 | 3.00E-04 | 0.0029 | 0.0017 | 0.0017 | TRUE | 0.0059 |
| Unk | 3470 | 2743 | 2775 | 2991 | 2764 | 3243 | 2695 | 2918 | 2994 | 2977 | 3118 | 3201 | 2983.93 | 2869 | 3099 | 1.081 | 0.112 | 0.08307116 | 0.19410862 | 0.0016 | 0.0032 | 0.0027 | 0.0027 | TRUE | 0.0351 |
| Unkl | 631 | 475 | 506 | 471 | 417 | 520 | 490 | 505 | 546 | 469 | 470 | 513 | 498.98 | 514 | 484 | 0.944 | -0.083 | 0.4065064 | 0.57939728 | 0 | 0.0072 | 0.0052 | 0.0052 | TRUE | 0.0345 |
| Uox | 10 | 5 | 2 | 4 | 1 | 0 | 8 | 5 | 2 | 4 | 1 | 0 | 3.39 | 5 | 2 | 0.333 | -1.587 | 0.18429124 | NA | 0.3794 | 0.7011 | 0.634 | 0.634 | TRUE | 0.5068 |
| Upb1 | 74 | 42 | 52 | 55 | 46 | 50 | 57 | 45 | 56 | 55 | 52 | 49 | 52.38 | 53 | 52 | 0.983 | -0.025 | 0.92454502 | 0.95836863 | 0 | 0.0477 | 0.0326 | 0.0326 | TRUE | 0.1411 |
| Upf1 | 17194 | 13301 | 13074 | 12873 | 12017 | 13708 | 13354 | 14148 | 14107 | 12814 | 13556 | 13532 | 13585.13 | 13870 | 13301 | 0.959 | -0.06 | 0.24473281 | 0.41288552 | 9.00E-04 | 0.0026 | 0.0019 | 0.0019 | TRUE | 0.013 |
| Upf2 | 4964 | 3601 | 3815 | 4066 | 3516 | 4406 | 3855 | 3830 | 4116 | 4047 | 3966 | 4349 | 4027.52 | 3934 | 4121 | 1.048 | 0.068 | 0.27344897 | 0.44438955 | 0.0016 | 0.003 | 0.0025 | 0.0025 | TRUE | 0.0287 |
| Upf3a | 3397 | 2682 | 2363 | 2459 | 2414 | 2600 | 2638 | 2853 | 2550 | 2448 | 2723 | 2567 | 2629.72 | 2680 | 2579 | 0.962 | -0.056 | 0.42196669 | 0.59426248 | 0.0026 | 0.0033 | 0.0031 | 0.0031 | TRUE | 0.0379 |
| Upf3b | 2371 | 1762 | 1727 | 1917 | 1644 | 2032 | 1841 | 1874 | 1863 | 1908 | 1855 | 2006 | 1891.29 | 1859 | 1923 | 1.034 | 0.049 | 0.44072852 | 0.61198726 | 3.00E-04 | 0.0037 | 0.0024 | 0.0024 | TRUE | 0.0173 |
| Upk1a | 65 | 51 | 65 | 81 | 73 | 92 | 50 | 54 | 70 | 81 | 82 | 91 | 71.44 | 58 | 85 | 1.46 | 0.546 | 0.02182777 | 0.07356156 | 0 | 0.0356 | 0.0265 | 0.0265 | TRUE | 0.2588 |
| Upk1b | 1 | 6 | 3 | 8 | 2 | 3 | 1 | 6 | 3 | 8 | 2 | 3 | 3.93 | 3 | 4 | 1.312 | 0.391 | 0.71020232 | NA | 0.3819 | 0.6056 | 0.5387 | 0.5387 | TRUE | 0.3309 |
| Upk2 | 79 | 55 | 49 | 66 | 45 | 60 | 61 | 59 | 53 | 66 | 51 | 59 | 58.07 | 58 | 59 | 1.017 | 0.024 | 0.92568548 | 0.95896373 | 0 | 0.0432 | 0.0298 | 0.0298 | TRUE | 0.1081 |
| Upk3a | 4 | 4 | 6 | 9 | 9 | 6 | 3 | 4 | 6 | 9 | 10 | 6 | 6.48 | 4 | 8 | 1.834 | 0.875 | 0.25273317 | NA | 0 | 0.3683 | 0.253 | 0.253 | TRUE | 0.2218 |
| Upk3b | 66 | 46 | 47 | 85 | 63 | 79 | 51 | 49 | 51 | 85 | 71 | 78 | 64.09 | 50 | 78 | 1.549 | 0.631 | 0.00902621 | 0.03843061 | 0 | 0.0394 | 0.0259 | 0.0259 | TRUE | 0.0526 |
| Upk3bl | 55 | 34 | 49 | 58 | 45 | 55 | 43 | 36 | 53 | 58 | 51 | 54 | 49.09 | 44 | 54 | 1.239 | 0.309 | 0.27217142 | 0.44316998 | 0 | 0.0507 | 0.0364 | 0.0364 | TRUE | 0.2253 |
| Upp1 | 27485 | 20419 | 24083 | 21840 | 20663 | 23895 | 21346 | 21719 | 25985 | 21740 | 23310 | 23588 | 22948.13 | 23017 | 22879 | 0.994 | -0.009 | 0.91066806 | 0.95105791 | 0.0058 | 0.0025 | 0.0042 | 0.0042 | TRUE | 0.1556 |
| Upp2 | 0 | 1 | 1 | 0 | 0 | 0 | 0 | 1 | 1 | 0 | 0 | 0 | 0.36 | 1 | 0 | 0.271 | -1.881 | 0.5755355 | NA | 0 | 6.6391 | 6.4428 | 6.4428 | TRUE | 0.3513 |
| Uprt | 98 | 68 | 76 | 96 | 58 | 79 | 76 | 72 | 82 | 96 | 65 | 78 | 78.24 | 77 | 80 | 1.04 | 0.057 | 0.80585477 | 0.88785032 | 0.0061 | 0.0327 | 0.0255 | 0.0255 | TRUE | 0.2837 |
| Uqcc | 1820 | 1262 | 1285 | 1388 | 1139 | 1420 | 1414 | 1342 | 1387 | 1382 | 1285 | 1402 | 1368.45 | 1381 | 1356 | 0.982 | -0.026 | 0.71309536 | 0.82756349 | 6.00E-04 | 0.0042 | 0.0029 | 0.0029 | TRUE | 0.0248 |
| Uqcr10 | 4710 | 3081 | 3307 | 3571 | 2964 | 3645 | 3658 | 3277 | 3568 | 3555 | 3344 | 3598 | 3499.99 | 3501 | 3499 | 0.999 | -0.001 | 0.98423934 | 0.99180294 | 0.0021 | 0.0031 | 0.0027 | 0.0027 | TRUE | 0.0379 |

| | | | | | | | | | | | | | | | | | | | | | | | | | |
|---|---|---|---|---|---|---|---|---|---|---|---|---|---|---|---|---|---|---|---|---|---|---|---|---|---|
| Uqcr11 | 3238 | 2118 | 2267 | 2516 | 2120 | 2404 | 2515 | 2253 | 2446 | 2504 | 2392 | 2373 | 2413.82 | 2405 | 2423 | 1.007 | 0.01 | 0.87611842 | 0.93096567 | 0.0016 | 0.0034 | 0.0027 | 0.0027 | TRUE | 0.0366 |
| Uqcrb | 5652 | 3800 | 4128 | 4714 | 3902 | 4481 | 4390 | 4042 | 4454 | 4692 | 4402 | 4423 | 4400.56 | 4295 | 4506 | 1.049 | 0.069 | 0.25972326 | 0.42970535 | 0.0016 | 0.003 | 0.0025 | 0.0025 | TRUE | 0.032 |
| Uqcrc1 | 24277 | 17851 | 18723 | 21276 | 18936 | 21452 | 18855 | 18988 | 20202 | 21178 | 21362 | 21176 | 20293.51 | 19348 | 21239 | 1.098 | 0.135 | 0.00590519 | 0.02790178 | 7.00E-04 | 0.0025 | 0.0017 | 0.0017 | TRUE | 0.0182 |
| Uqcrc2 | 11947 | 8341 | 8940 | 10101 | 9009 | 9991 | 9279 | 8872 | 9646 | 10055 | 10163 | 9863 | 9646.23 | 9266 | 10027 | 1.082 | 0.114 | 0.02949377 | 0.09150302 | 8.00E-04 | 0.0027 | 0.0019 | 0.0019 | TRUE | 0.0167 |
| Uqcrfs1 | 9124 | 6603 | 6687 | 7295 | 6297 | 7359 | 7086 | 7024 | 7215 | 7261 | 7104 | 7264 | 7159.09 | 7108 | 7210 | 1.014 | 0.021 | 0.6651575 | 0.7937289 | 0 | 0.0028 | 0.0015 | 0.0015 | TRUE | 0.0021 |
| Uqcrh | 4817 | 3220 | 3336 | 3480 | 3034 | 3586 | 3741 | 3425 | 3600 | 3464 | 3423 | 3540 | 3532.06 | 3589 | 3476 | 0.968 | -0.047 | 0.42316145 | 0.59565162 | 9.00E-04 | 0.0031 | 0.0022 | 0.0022 | TRUE | 0.0193 |
| Uqcrq | 6701 | 4651 | 4470 | 5098 | 4174 | 4867 | 5204 | 4947 | 4823 | 5075 | 4709 | 4804 | 4927.07 | 4991 | 4863 | 0.974 | -0.038 | 0.51262761 | 0.67504942 | 0.0012 | 0.0029 | 0.0022 | 0.0022 | TRUE | 0.0177 |
| Urb1 | 8801 | 6364 | 5781 | 6462 | 5777 | 6883 | 6835 | 6769 | 6238 | 6432 | 6517 | 6795 | 6597.7 | 6614 | 6581 | 0.995 | -0.007 | 0.89724213 | 0.94337809 | 0.0015 | 0.0028 | 0.0022 | 0.0022 | TRUE | 0.0301 |
| Urb2 | 13478 | 9720 | 9856 | 10211 | 9398 | 10694 | 10468 | 10339 | 10635 | 10164 | 10602 | 10557 | 10460.66 | 10481 | 10441 | 0.996 | -0.006 | 0.90785897 | 0.94923397 | 2.00E-04 | 0.0027 | 0.0015 | 0.0015 | TRUE | 0.0066 |
| Urgcp | 757 | 514 | 544 | 348 | 319 | 299 | 588 | 547 | 587 | 346 | 360 | 295 | 453.84 | 574 | 334 | 0.581 | -0.784 | 8.96E-13 | 1.13E-10 | 0.0025 | 0.0076 | 0.0063 | 0.0063 | TRUE | 0.1186 |
| Uri1 | 12392 | 9218 | 8896 | 9772 | 8608 | 9557 | 9624 | 9805 | 9599 | 9727 | 9711 | 9434 | 9650.02 | 9676 | 9624 | 0.995 | -0.008 | 0.86726757 | 0.92678292 | 1.00E-04 | 0.0027 | 0.0015 | 0.0015 | TRUE | 0.0036 |
| Urm1 | 1916 | 1379 | 1416 | 1351 | 1191 | 1612 | 1488 | 1467 | 1528 | 1345 | 1344 | 1591 | 1460.4 | 1494 | 1427 | 0.955 | -0.066 | 0.41910439 | 0.59168417 | 0.004 | 0.004 | 0.0041 | 0.0041 | TRUE | 0.1237 |
| Uroc1 | 15 | 11 | 6 | 0 | 0 | 3 | 12 | 12 | 6 | 0 | 0 | 3 | 5.46 | 10 | 1 | 0.102 | -3.289 | 0.00393615 | NA | 0.0032 | 0.4362 | 0.383 | 0.383 | TRUE | 1.4404 |
| Urod | 2379 | 1751 | 1750 | 1685 | 1530 | 1922 | 1848 | 1863 | 1888 | 1677 | 1726 | 1897 | 1816.5 | 1866 | 1767 | 0.947 | -0.078 | 0.25963292 | 0.42960562 | 0.0016 | 0.0037 | 0.0029 | 0.0029 | TRUE | 0.0509 |
| Uros | 430 | 314 | 151 | 193 | 177 | 165 | 334 | 334 | 163 | 192 | 200 | 163 | 230.93 | 277 | 185 | 0.666 | -0.587 | 0.00835574 | 0.03620144 | 0.0782 | 0.0127 | 0.0312 | 0.0312 | TRUE | 1.426 |
| Usb1 | 3052 | 2109 | 2176 | 2377 | 2082 | 2393 | 2370 | 2243 | 2348 | 2366 | 2349 | 2362 | 2339.77 | 2320 | 2359 | 1.016 | 0.023 | 0.69199785 | 0.81166791 | 0 | 0.0034 | 0.0021 | 0.0021 | TRUE | 0.0101 |
| Use1 | 2259 | 1733 | 1502 | 1636 | 1480 | 1692 | 1754 | 1843 | 1621 | 1628 | 1670 | 1670 | 1697.8 | 1739 | 1656 | 0.952 | -0.071 | 0.31203035 | 0.48529581 | 0.0016 | 0.0038 | 0.003 | 0.003 | TRUE | 0.0418 |
| Usf1 | 3261 | 2283 | 2161 | 2187 | 2049 | 2458 | 2533 | 2428 | 2332 | 2177 | 2311 | 2426 | 2367.94 | 2431 | 2305 | 0.948 | -0.078 | 0.24971125 | 0.41849354 | 0.0017 | 0.0034 | 0.0028 | 0.0028 | TRUE | 0.0289 |
| Usf2 | 1581 | 1268 | 1386 | 1419 | 1228 | 1430 | 1228 | 1349 | 1495 | 1412 | 1385 | 1412 | 1380.26 | 1357 | 1403 | 1.035 | 0.05 | 0.544562 | 0.70202845 | 0.0043 | 0.0041 | 0.0042 | 0.0042 | TRUE | 0.0951 |
| Ush1c | 80 | 64 | 75 | 89 | 79 | 59 | 62 | 68 | 81 | 89 | 89 | 58 | 74.51 | 70 | 79 | 1.12 | 0.163 | 0.51235278 | 0.67504942 | 0.0237 | 0.0342 | 0.0313 | 0.0313 | TRUE | 0.4887 |
| Ush1g | 13 | 16 | 20 | 36 | 24 | 35 | 10 | 17 | 22 | 36 | 27 | 35 | 24.36 | 16 | 33 | 2.041 | 1.029 | 0.01410388 | 0.05334393 | 0.0217 | 0.0997 | 0.0806 | 0.0806 | TRUE | 0.6815 |
| Ush2a | 8 | 7 | 7 | 3 | 3 | 5 | 6 | 7 | 8 | 3 | 3 | 5 | 5.42 | 7 | 4 | 0.537 | -0.898 | 0.27420491 | NA | 0 | 0.4397 | 0.2806 | 0.2806 | TRUE | 0.1224 |
| Ushbp1 | 147 | 128 | 157 | 113 | 94 | 130 | 114 | 136 | 169 | 112 | 106 | 128 | 127.76 | 140 | 115 | 0.832 | -0.266 | 0.177825 | 0.33219875 | 0.0183 | 0.021 | 0.0202 | 0.0202 | TRUE | 0.2763 |
| Usmg5 | 3385 | 2133 | 2153 | 2557 | 2053 | 2419 | 2629 | 2269 | 2323 | 2545 | 2316 | 2388 | 2411.68 | 2407 | 2416 | 1.003 | 0.005 | 0.95158086 | 0.97323401 | 0.004 | 0.0034 | 0.0036 | 0.0036 | TRUE | 0.0806 |
| Uso1 | 9655 | 6602 | 5919 | 6739 | 5571 | 6502 | 7499 | 7022 | 6387 | 6708 | 6285 | 6418 | 6719.8 | 6969 | 6470 | 0.928 | -0.108 | 0.11069744 | 0.238314 | 0.0035 | 0.0028 | 0.0031 | 0.0031 | TRUE | 0.0652 |
| Usp1 | 8878 | 6477 | 5736 | 6565 | 5566 | 6423 | 6895 | 6890 | 6189 | 6535 | 6279 | 6340 | 6521.35 | 6658 | 6385 | 0.959 | -0.061 | 0.31230768 | 0.48561942 | 0.0018 | 0.0028 | 0.0024 | 0.0024 | TRUE | 0.046 |
| Usp10 | 18906 | 13998 | 13745 | 12536 | 12001 | 13506 | 14683 | 14889 | 14831 | 12478 | 13538 | 13333 | 13958.82 | 14801 | 13116 | 0.886 | -0.174 | 0.00065175 | 0.00528085 | 8.00E-04 | 0.0026 | 0.0018 | 0.0018 | TRUE | 0.0222 |
| Usp11 | 2527 | 2048 | 2319 | 3069 | 2753 | 3168 | 1963 | 2178 | 2502 | 3055 | 3106 | 3127 | 2655.18 | 2214 | 3096 | 1.4 | 0.485 | 6.21E-09 | 3.36E-07 | 0.007 | 0.0033 | 0.0046 | 0.0046 | TRUE | 0.1563 |
| Usp12 | 4274 | 3218 | 2965 | 3164 | 2840 | 3036 | 3319 | 3423 | 3199 | 3149 | 3204 | 2997 | 3215.31 | 3314 | 3117 | 0.94 | -0.089 | 0.13364462 | 0.27218333 | 7.00E-04 | 0.0032 | 0.0022 | 0.0022 | TRUE | 0.0138 |
| Usp13 | 450 | 327 | 313 | 354 | 270 | 310 | 349 | 348 | 338 | 352 | 305 | 306 | 333 | 345 | 321 | 0.931 | -0.104 | 0.37471036 | 0.54920038 | 0 | 0.0095 | 0.0068 | 0.0068 | TRUE | 0.084 |
| Usp14 | 11830 | 8322 | 8402 | 9830 | 8234 | 9611 | 9188 | 8852 | 9066 | 9785 | 9289 | 9488 | 9277.78 | 9035 | 9521 | 1.054 | 0.075 | 0.12515816 | 0.26021509 | 4.00E-04 | 0.0027 | 0.0016 | 0.0016 | TRUE | 0.0072 |
| Usp15 | 10401 | 7497 | 7381 | 8148 | 7171 | 8558 | 8078 | 7974 | 7964 | 8111 | 8090 | 8448 | 8110.79 | 8005 | 8216 | 1.026 | 0.037 | 0.43750721 | 0.60894835 | 1.00E-04 | 0.0027 | 0.0016 | 0.0016 | TRUE | 0.0075 |
| Usp16 | 3253 | 2220 | 1944 | 1951 | 1770 | 1976 | 2526 | 2361 | 2098 | 1942 | 1997 | 1951 | 2145.8 | 2328 | 1963 | 0.842 | -0.248 | 0.00122451 | 0.00838216 | 0.0037 | 0.0035 | 0.0038 | 0.0038 | TRUE | 0.0906 |
| Usp17la | 1012 | 714 | 313 | 337 | 320 | 268 | 786 | 759 | 338 | 335 | 361 | 265 | 474.03 | 628 | 320 | 0.51 | -0.971 | 0.01405156 | 0.0532447 | 0.1103 | 0.0074 | 0.0344 | 0.1103 | TRUE | 1.9144 |
| Usp17lb | 859 | 612 | 271 | 339 | 302 | 216 | 667 | 651 | 292 | 337 | 341 | 213 | 416.98 | 537 | 297 | 0.553 | -0.854 | 0.04330807 | 0.12124421 | 0.1262 | 0.0081 | 0.0386 | 0.1262 | TRUE | 1.8466 |
| Usp17lc | 437 | 369 | 137 | 121 | 138 | 122 | 339 | 393 | 148 | 120 | 156 | 120 | 212.71 | 293 | 132 | 0.45 | -1.151 | 0.00737771 | 0.03305908 | 0.1274 | 0.0136 | 0.0448 | 0.1274 | TRUE | 0.7279 |
| Usp17ld | 171 | 126 | 63 | 81 | 74 | 56 | 133 | 134 | 68 | 81 | 83 | 55 | 92.37 | 112 | 73 | 0.652 | -0.617 | 0.0248939 | 0.08117921 | 0.0813 | 0.0281 | 0.0432 | 0.0432 | TRUE | 1.1247 |
| Usp17le | 1427 | 1013 | 493 | 629 | 575 | 442 | 1108 | 1078 | 532 | 626 | 649 | 436 | 738.14 | 906 | 570 | 0.629 | -0.668 | 0.07195723 | 0.17515066 | 0.0979 | 0.0056 | 0.0293 | 0.0979 | TRUE | 1.5489 |
| Usp18 | 24 | 10 | 14 | 15 | 19 | 17 | 19 | 11 | 15 | 15 | 21 | 17 | 16.25 | 15 | 18 | 1.182 | 0.241 | 0.61781972 | NA | 0 | 0.1482 | 0.1064 | 0.1064 | TRUE | 0.2922 |
| Usp19 | 8055 | 6165 | 5605 | 5276 | 5033 | 5671 | 6256 | 6558 | 6048 | 5252 | 5678 | 5598 | 5898.16 | 6287 | 5509 | 0.876 | -0.191 | 0.00105313 | 0.00751495 | 0.0014 | 0.0028 | 0.0023 | 0.0023 | TRUE | 0.0204 |
| Usp2 | 816 | 650 | 806 | 837 | 796 | 937 | 634 | 691 | 870 | 833 | 898 | 925 | 808.48 | 732 | 885 | 1.213 | 0.278 | 0.01411774 | 0.05338223 | 0.0137 | 0.0054 | 0.008 | 0.008 | TRUE | 0.3201 |
| Usp20 | 789 | 544 | 462 | 571 | 549 | 581 | 613 | 579 | 498 | 568 | 619 | 574 | 575.19 | 563 | 587 | 1.039 | 0.056 | 0.59070509 | 0.73901229 | 0.0049 | 0.0065 | 0.006 | 0.006 | TRUE | 0.1164 |
| Usp21 | 2562 | 1921 | 2119 | 1965 | 1734 | 2154 | 1990 | 2043 | 2286 | 1956 | 1956 | 2126 | 2059.66 | 2106 | 2013 | 0.956 | -0.064 | 0.38867279 | 0.56275488 | 0.0033 | 0.0036 | 0.0035 | 0.0035 | TRUE | 0.067 |
| Usp22 | 25018 | 17582 | 16565 | 19224 | 16372 | 18306 | 19430 | 18702 | 17873 | 19136 | 18469 | 18071 | 18613.56 | 18668 | 18559 | 0.994 | -0.009 | 0.86971436 | 0.92801725 | 0.001 | 0.0026 | 0.002 | 0.002 | TRUE | 0.017 |
| Usp24 | 9435 | 6938 | 6854 | 7051 | 6309 | 7420 | 7328 | 7380 | 7395 | 7019 | 7117 | 7325 | 7260.58 | 7368 | 7154 | 0.971 | -0.042 | 0.37743054 | 0.55199168 | 1.00E-04 | 0.0028 | 0.0015 | 0.0015 | TRUE | 0.0054 |
| Usp25 | 1291 | 953 | 1117 | 1080 | 885 | 1082 | 1003 | 1014 | 1205 | 1075 | 998 | 1068 | 1060.52 | 1074 | 1047 | 0.977 | -0.034 | 0.70395381 | 0.82094194 | 0.0053 | 0.0047 | 0.0049 | 0.0049 | TRUE | 0.135 |
| Usp26 | 3369 | 2737 | 2522 | 1297 | 1168 | 1154 | 2617 | 2911 | 2721 | 1291 | 1318 | 1139 | 1999.49 | 2750 | 1249 | 0.454 | -1.138 | 1.61E-49 | 5.75E-46 | 0.0036 | 0.0036 | 0.0037 | 0.0037 | TRUE | 0.072 |
| Usp27x | 313 | 267 | 223 | 235 | 184 | 207 | 243 | 284 | 241 | 234 | 208 | 204 | 235.59 | 256 | 215 | 0.843 | -0.247 | 0.07660661 | 0.18303883 | 0.0031 | 0.0125 | 0.0097 | 0.0097 | TRUE | 0.0996 |
| Usp28 | 13975 | 11361 | 15814 | 17609 | 16943 | 19822 | 10854 | 12085 | 17063 | 17528 | 19113 | 19567 | 16035.05 | 13334 | 18736 | 1.405 | 0.491 | 0.01985018 | 0.06874693 | 0.0319 | 0.0026 | 0.0105 | 0.0319 | TRUE | 0.7318 |
| Usp29 | 515 | 347 | 342 | 285 | 252 | 300 | 400 | 369 | 369 | 284 | 284 | 296 | 333.7 | 379 | 288 | 0.758 | -0.399 | 0.00044518 | 0.00394477 | 0 | 0.0095 | 0.0063 | 0.0063 | TRUE | 0.0278 |
| Usp3 | 1374 | 1068 | 1254 | 1478 | 1190 | 1377 | 1067 | 1136 | 1353 | 1471 | 1342 | 1359 | 1288.19 | 1185 | 1391 | 1.175 | 0.233 | 0.01302227 | 0.05025036 | 0.008 | 0.0043 | 0.0056 | 0.0056 | TRUE | 0.1822 |
| Usp30 | 1118 | 885 | 1067 | 1076 | 950 | 1097 | 868 | 941 | 1151 | 1071 | 1072 | 1083 | 1031.1 | 987 | 1075 | 1.091 | 0.126 | 0.21253463 | 0.3748753 | 0.0094 | 0.0047 | 0.0064 | 0.0064 | TRUE | 0.2515 |
| Usp31 | 3236 | 2400 | 2359 | 2549 | 2165 | 2834 | 2513 | 2553 | 2545 | 2537 | 2442 | 2798 | 2564.78 | 2537 | 2592 | 1.022 | 0.032 | 0.63738474 | 0.7734222 | 0.0018 | 0.0034 | 0.0029 | 0.0029 | TRUE | 0.0583 |
| Usp32 | 7556 | 5520 | 5694 | 6209 | 5451 | 6413 | 5868 | 5872 | 6144 | 6180 | 6149 | 6331 | 6090.68 | 5961 | 6220 | 1.044 | 0.062 | 0.22402882 | 0.38840902 | 2.00E-04 | 0.0028 | 0.0017 | 0.0017 | TRUE | 0.0087 |
| Usp33 | 5020 | 3668 | 3614 | 3841 | 3404 | 3841 | 3899 | 3902 | 3899 | 3823 | 3840 | 3792 | 3859.16 | 3900 | 3818 | 0.979 | -0.031 | 0.54770657 | 0.70456035 | 0 | 0.003 | 0.0016 | 0.0016 | TRUE | 5.00E-04 |
| Usp34 | 14527 | 10007 | 9717 | 10978 | 9701 | 11098 | 11283 | 10644 | 10485 | 10927 | 10944 | 10955 | 10873.01 | 10804 | 10942 | 1.013 | 0.018 | 0.72060737 | 0.83222698 | 6.00E-04 | 0.0026 | 0.0017 | 0.0017 | TRUE | 0.0185 |
| Usp35 | 396 | 380 | 387 | 426 | 341 | 416 | 308 | 404 | 418 | 424 | 385 | 411 | 391.45 | 377 | 407 | 1.085 | 0.117 | 0.37069482 | 0.54532123 | 0.0126 | 0.0085 | 0.0098 | 0.0098 | TRUE | 0.3138 |
| Usp36 | 6041 | 4544 | 4090 | 3830 | 3614 | 4124 | 4692 | 4833 | 4413 | 3812 | 4077 | 4071 | 4316.44 | 4646 | 3987 | 0.858 | -0.221 | 0.00025 | 0.00252884 | 0.0014 | 0.003 | 0.0024 | 0.0024 | TRUE | 0.0231 |
| Usp37 | 7894 | 5601 | 5252 | 5983 | 4978 | 5727 | 6131 | 5958 | 5667 | 5955 | 5616 | 5653 | 5830.02 | 5919 | 5741 | 0.97 | -0.044 | 0.43332277 | 0.60530516 | 0.0011 | 0.0028 | 0.0021 | 0.0021 | TRUE | 0.0168 |
| Usp38 | 6226 | 4311 | 3830 | 4040 | 3812 | 4074 | 4835 | 4586 | 4133 | 4021 | 4300 | 4022 | 4316.18 | 4518 | 4114 | 0.91 | -0.136 | 0.05095241 | 0.13671163 | 0.0035 | 0.003 | 0.0033 | 0.0033 | TRUE | 0.0675 |
| Usp39 | 7292 | 5218 | 5206 | 5680 | 4882 | 5700 | 5663 | 5550 | 5617 | 5654 | 5507 | 5627 | 5603.16 | 5610 | 5596 | 0.997 | -0.004 | 0.93940587 | 0.966328 | 0 | 0.0028 | 0.0015 | 0.0015 | TRUE | 0.0023 |
| Usp4 | 7086 | 5062 | 5073 | 5020 | 4488 | 5086 | 5503 | 5384 | 5474 | 4997 | 5063 | 5021 | 5240.33 | 5454 | 5027 | 0.922 | -0.118 | 0.01582494 | 0.05813054 | 0 | 0.0029 | 0.0015 | 0.0015 | TRUE | 0.0015 |
| Usp40 | 1477 | 1053 | 955 | 1008 | 894 | 992 | 1147 | 1120 | 1030 | 1003 | 1009 | 979 | 1048.13 | 1099 | 997 | 0.906 | -0.142 | 0.06454187 | 0.16178481 | 8.00E-04 | 0.0047 | 0.0033 | 0.0033 | TRUE | 0.0352 |
| Usp42 | 2343 | 1838 | 1681 | 1642 | 1512 | 1653 | 1820 | 1955 | 1814 | 1634 | 1706 | 1632 | 1760.08 | 1863 | 1657 | 0.89 | -0.169 | 0.01093163 | 0.04409985 | 7.00E-04 | 0.0038 | 0.0026 | 0.0026 | TRUE | 0.0223 |
| Usp43 | 1186 | 992 | 1165 | 1050 | 1065 | 1175 | 921 | 1055 | 1257 | 1045 | 1201 | 1160 | 1106.64 | 1078 | 1135 | 1.055 | 0.077 | 0.47680995 | 0.64457056 | 0.0139 | 0.0046 | 0.0076 | 0.0076 | TRUE | 0.2528 |
| Usp44 | 1908 | 1417 | 1539 | 1900 | 1677 | 2047 | 1482 | 1507 | 1661 | 1891 | 1892 | 2021 | 1742.24 | 1550 | 1935 | 1.249 | 0.321 | 7.66E-06 | 0.00015076 | 0.0018 | 0.0038 | 0.0031 | 0.0031 | TRUE | 0.0464 |
| Usp45 | 4390 | 3173 | 3191 | 3850 | 3200 | 3631 | 3410 | 3375 | 3443 | 3832 | 3610 | 3584 | 3542.37 | 3409 | 3675 | 1.078 | 0.109 | 0.05271661 | 0.14007596 | 5.00E-04 | 0.0031 | 0.002 | 0.002 | TRUE | 0.017 |
| Usp46 | 1057 | 828 | 951 | 983 | 909 | 1097 | 821 | 881 | 1026 | 978 | 1025 | 1083 | 969.1 | 909 | 1029 | 1.134 | 0.181 | 0.05817844 | 0.15036969 | 0.0065 | 0.0049 | 0.0055 | 0.0055 | TRUE | 0.1494 |
| Usp47 | 7206 | 4874 | 4953 | 4863 | 4269 | 4543 | 5597 | 5184 | 5344 | 4841 | 4816 | 4485 | 5044.4 | 5375 | 4714 | 0.877 | -0.19 | 0.00124301 | 0.00845615 | 0.0013 | 0.0029 | 0.0023 | 0.0023 | TRUE | 0.0221 |

| | | | | | | | | | | | | | | | | | | | | | | | | | |
|---|---|---|---|---|---|---|---|---|---|---|---|---|---|---|---|---|---|---|---|---|---|---|---|---|---|
| Usp48 | 15780 | 12184 | 12025 | 11927 | 10262 | 11212 | 12256 | 12960 | 12975 | 11872 | 11577 | 11068 | 12117.87 | 12730 | 11506 | 0.904 | -0.146 | 0.00584799 | 0.02767725 | 0.001 | 0.0026 | 0.0019 | 0.0019 | TRUE | 0.0136 |
| Usp49 | 1943 | 1471 | 1580 | 1605 | 1435 | 1732 | 1509 | 1565 | 1705 | 1598 | 1619 | 1710 | 1617.46 | 1593 | 1642 | 1.032 | 0.045 | 0.53345438 | 0.6928378 | 0.002 | 0.0039 | 0.0032 | 0.0032 | TRUE | 0.0452 |
| Usp5 | 16202 | 12053 | 11576 | 12262 | 10976 | 12516 | 12583 | 12821 | 12490 | 12206 | 12382 | 12355 | 12472.88 | 12631 | 12314 | 0.975 | -0.037 | 0.41245587 | 0.58536707 | 0 | 0.0026 | 0.0014 | 0.0014 | TRUE | 0.0021 |
| Usp50 | 54 | 44 | 41 | 36 | 27 | 25 | 42 | 47 | 44 | 36 | 30 | 25 | 37.33 | 44 | 30 | 0.685 | -0.546 | 0.08783302 | 0.20131453 | 0 | 0.0659 | 0.0456 | 0.0456 | TRUE | 0.186 |
| Usp51 | 213 | 133 | 157 | 161 | 131 | 167 | 165 | 141 | 169 | 160 | 148 | 165 | 158.2 | 158 | 158 | 0.993 | -0.011 | 0.9462935 | 0.97046517 | 0 | 0.0174 | 0.0124 | 0.0124 | TRUE | 0.0913 |
| Usp53 | 988 | 774 | 654 | 667 | 693 | 654 | 767 | 823 | 706 | 664 | 782 | 646 | 731.27 | 765 | 697 | 0.91 | -0.137 | 0.17805682 | 0.33250145 | 0.0065 | 0.0057 | 0.006 | 0.006 | TRUE | 0.1309 |
| Usp54 | 3098 | 2171 | 2211 | 1907 | 1754 | 1987 | 2406 | 2309 | 2386 | 1898 | 1979 | 1961 | 2156.57 | 2367 | 1946 | 0.822 | -0.283 | 2.44E-06 | 5.89E-05 | 0 | 0.0035 | 0.0021 | 0.0021 | TRUE | 0.0056 |
| Usp6nl | 2140 | 1527 | 1476 | 1296 | 1275 | 1453 | 1662 | 1624 | 1593 | 1290 | 1438 | 1434 | 1506.93 | 1626 | 1387 | 0.853 | -0.23 | 0.00135973 | 0.00899334 | 0.0014 | 0.004 | 0.003 | 0.003 | TRUE | 0.0459 |
| Usp7 | 26046 | 18962 | 18413 | 17586 | 16498 | 19005 | 20229 | 20170 | 19867 | 17505 | 18611 | 18761 | 19190.56 | 20089 | 18292 | 0.911 | -0.135 | 0.00594498 | 0.02804346 | 7.00E-04 | 0.0025 | 0.0017 | 0.0017 | TRUE | 0.0174 |
| Usp8 | 7850 | 5843 | 5552 | 6317 | 5532 | 6593 | 6097 | 6215 | 5991 | 6288 | 6241 | 6508 | 6223.23 | 6101 | 6346 | 1.04 | 0.057 | 0.25739335 | 0.42738419 | 2.00E-04 | 0.0028 | 0.0017 | 0.0017 | TRUE | 0.0062 |
| Usp9x | 30756 | 20820 | 19715 | 22905 | 20214 | 22186 | 23887 | 22146 | 21272 | 22800 | 22803 | 21901 | 22468.21 | 22435 | 22501 | 1.003 | 0.004 | 0.94310935 | 0.96831132 | 0.0016 | 0.0025 | 0.0023 | 0.0023 | TRUE | 0.0394 |
| Usp9y | 204 | 153 | 126 | 28 | 28 | 20 | 158 | 163 | 136 | 28 | 32 | 20 | 89.39 | 152 | 27 | 0.172 | -2.539 | 9.21E-24 | 3.99E-21 | 0 | 0.0289 | 0.0231 | 0.0231 | TRUE | 0.3152 |
| Uspl1 | 4933 | 3723 | 3421 | 3704 | 3093 | 3649 | 3831 | 3960 | 3691 | 3687 | 3489 | 3602 | 3710.15 | 3827 | 3593 | 0.939 | -0.091 | 0.11071188 | 0.238314 | 6.00E-04 | 0.0031 | 0.0021 | 0.0021 | TRUE | 0.0115 |
| Ust | 170 | 138 | 88 | 63 | 67 | 76 | 132 | 147 | 95 | 63 | 76 | 75 | 97.85 | 125 | 71 | 0.569 | -0.813 | 0.00029469 | 0.00286922 | 0.0216 | 0.0266 | 0.0252 | 0.0252 | TRUE | 0.4184 |
| Utf1 | 17387 | 13330 | 14474 | 16333 | 14660 | 17949 | 13504 | 14179 | 15617 | 16258 | 16538 | 17718 | 15635.72 | 14433 | 16838 | 1.167 | 0.223 | 0.00069788 | 0.00556693 | 0.0038 | 0.0026 | 0.003 | 0.003 | TRUE | 0.0629 |
| Utp11l | 2656 | 1872 | 1922 | 2229 | 1872 | 2271 | 2063 | 1991 | 2074 | 2219 | 2112 | 2242 | 2116.7 | 2043 | 2191 | 1.073 | 0.101 | 0.09982976 | 0.22082633 | 2.00E-04 | 0.0035 | 0.0023 | 0.0023 | TRUE | 0.0118 |
| Utp14a | 4746 | 3459 | 3488 | 3810 | 3277 | 4075 | 3686 | 3679 | 3764 | 3792 | 3697 | 4023 | 3773.46 | 3710 | 3837 | 1.035 | 0.049 | 0.38938481 | 0.5633074 | 6.00E-04 | 0.0031 | 0.0021 | 0.0021 | TRUE | 0.0218 |
| Utp14b | 929 | 646 | 467 | 584 | 461 | 436 | 722 | 687 | 504 | 581 | 520 | 430 | 574.05 | 638 | 510 | 0.8 | -0.323 | 0.02359587 | 0.07796019 | 0.0277 | 0.0066 | 0.0129 | 0.0129 | TRUE | 0.3905 |
| Utp15 | 5120 | 3607 | 3391 | 3442 | 3077 | 3347 | 3976 | 3837 | 3659 | 3426 | 3471 | 3304 | 3612.24 | 3824 | 3400 | 0.889 | -0.17 | 0.00367153 | 0.01930231 | 0.001 | 0.0031 | 0.0022 | 0.0022 | TRUE | 0.0173 |
| Utp18 | 8683 | 6276 | 6318 | 6810 | 5961 | 6731 | 6744 | 6676 | 6817 | 6779 | 6725 | 6645 | 6730.72 | 6746 | 6716 | 0.996 | -0.006 | 0.89355249 | 0.9417388 | 0 | 0.0028 | 0.0015 | 0.0015 | TRUE | 0.0011 |
| Utp20 | 13597 | 8948 | 7658 | 8000 | 7069 | 8096 | 10560 | 9518 | 8263 | 7963 | 7975 | 7992 | 8711.79 | 9447 | 7977 | 0.844 | -0.244 | 0.0020438 | 0.01227055 | 0.0072 | 0.0027 | 0.0044 | 0.0044 | TRUE | 0.1467 |
| Utp23 | 1210 | 812 | 738 | 773 | 727 | 798 | 940 | 864 | 796 | 769 | 820 | 788 | 829.51 | 867 | 792 | 0.913 | -0.132 | 0.13719052 | 0.27754623 | 0.003 | 0.0053 | 0.0045 | 0.0045 | TRUE | 0.0678 |
| Utp3 | 4101 | 3040 | 2556 | 2458 | 2200 | 2534 | 3185 | 3234 | 2758 | 2447 | 2482 | 2501 | 2767.76 | 3059 | 2477 | 0.809 | -0.305 | 2.28E-05 | 0.0003722 | 0.0032 | 0.0033 | 0.0034 | 0.0034 | TRUE | 0.0889 |
| Utp6 | 11332 | 7740 | 7728 | 8291 | 7096 | 8220 | 8801 | 8233 | 8338 | 8253 | 8005 | 8114 | 8290.79 | 8457 | 8124 | 0.96 | -0.058 | 0.25576417 | 0.42571763 | 6.00E-04 | 0.0027 | 0.0018 | 0.0018 | TRUE | 0.0156 |
| Utrn | 4874 | 3416 | 4094 | 3986 | 3829 | 4471 | 3785 | 3634 | 4417 | 3968 | 4320 | 4414 | 4089.52 | 3945 | 4234 | 1.073 | 0.102 | 0.20056377 | 0.36126754 | 0.0067 | 0.003 | 0.0043 | 0.0043 | TRUE | 0.1323 |
| Uts2 | 1 | 0 | 0 | 1 | 0 | 0 | 1 | 0 | 0 | 1 | 0 | 0 | 0.3 | 0 | 0 | 1.075 | 0.105 | 0.97698525 | NA | 0 | 8.0272 | 7.5141 | 7.5141 | TRUE | 0.3906 |
| Uts2d | 0 | 0 | 0 | 0 | 0 | 0 | 0 | 0 | 0 | 0 | 0 | 0 | 0 | 0 | 0 | NA | NA | NA | NA | NA | NA | NA | NA | NA | NA |
| Uts2r | 26 | 13 | 10 | 22 | 5 | 9 | 20 | 14 | 11 | 22 | 6 | 9 | 13.54 | 15 | 12 | 0.811 | -0.303 | 0.61237844 | NA | 0.1976 | 0.1775 | 0.1831 | 0.1831 | TRUE | 1.931 |
| Uty | 903 | 675 | 626 | 519 | 461 | 515 | 701 | 718 | 675 | 517 | 520 | 508 | 606.64 | 698 | 515 | 0.737 | -0.44 | 6.66E-07 | 2.09E-05 | 0 | 0.0063 | 0.0039 | 0.0039 | TRUE | 0.0092 |
| Uvrag | 2078 | 1486 | 1434 | 1721 | 1576 | 1760 | 1614 | 1581 | 1547 | 1713 | 1778 | 1737 | 1661.7 | 1581 | 1743 | 1.102 | 0.14 | 0.02750282 | 0.08685324 | 0 | 0.0039 | 0.0023 | 0.0023 | TRUE | 0.0043 |
| Uvssa | 324 | 176 | 184 | 182 | 137 | 168 | 252 | 187 | 199 | 181 | 155 | 166 | 189.82 | 213 | 167 | 0.784 | -0.351 | 0.0323785 | 0.09794411 | 0.0121 | 0.0149 | 0.014 | 0.014 | TRUE | 0.3043 |
| Uxs1 | 1109 | 747 | 763 | 816 | 681 | 847 | 861 | 795 | 823 | 812 | 768 | 836 | 815.96 | 826 | 805 | 0.974 | -0.038 | 0.64941674 | 0.7821786 | 0 | 0.0053 | 0.0037 | 0.0037 | TRUE | 0.019 |
| Uxt | 890 | 635 | 668 | 687 | 643 | 697 | 691 | 675 | 721 | 684 | 725 | 688 | 697.45 | 696 | 699 | 1.004 | 0.006 | 0.93955504 | 0.966328 | 0 | 0.0058 | 0.0038 | 0.0038 | TRUE | 0.0124 |
| V1ra8 | 0 | 0 | 0 | 0 | 0 | 0 | 0 | 0 | 0 | 0 | 0 | 0 | 0 | 0 | 0 | NA | NA | NA | NA | NA | NA | NA | NA | NA | NA |
| V1rd18 | 0 | 0 | 0 | 0 | 0 | 0 | 0 | 0 | 0 | 0 | 0 | 0 | 0 | 0 | 0 | NA | NA | NA | NA | NA | NA | NA | NA | NA | NA |
| V1rd19 | 0 | 0 | 0 | 0 | 0 | 0 | 0 | 0 | 0 | 0 | 0 | 0 | 0 | 0 | 0 | NA | NA | NA | NA | NA | NA | NA | NA | NA | NA |
| Vac14 | 6469 | 4792 | 4306 | 4444 | 4012 | 4677 | 5024 | 5097 | 4646 | 4424 | 4526 | 4617 | 4722.32 | 4922 | 4522 | 0.918 | -0.123 | 0.03486064 | 0.10354595 | 0.0011 | 0.0029 | 0.0022 | 0.0022 | TRUE | 0.0291 |
| Vamp1 | 207 | 172 | 117 | 148 | 134 | 133 | 161 | 183 | 126 | 147 | 151 | 131 | 149.96 | 157 | 143 | 0.913 | -0.131 | 0.46546085 | 0.63450949 | 0.0128 | 0.0182 | 0.0167 | 0.0167 | TRUE | 0.2861 |
| Vamp2 | 1480 | 1261 | 1338 | 1249 | 1125 | 1325 | 1149 | 1341 | 1444 | 1243 | 1269 | 1308 | 1292.47 | 1311 | 1273 | 0.973 | -0.04 | 0.65501609 | 0.7856785 | 0.0064 | 0.0043 | 0.005 | 0.005 | TRUE | 0.1452 |
| Vamp3 | 4982 | 3476 | 3352 | 3629 | 3000 | 3301 | 3869 | 3697 | 3617 | 3612 | 3384 | 3259 | 3573.11 | 3728 | 3418 | 0.917 | -0.125 | 0.04431892 | 0.12331133 | 0.0016 | 0.0031 | 0.0025 | 0.0025 | TRUE | 0.0302 |
| Vamp4 | 576 | 427 | 512 | 548 | 471 | 491 | 447 | 454 | 552 | 545 | 531 | 485 | 502.58 | 484 | 520 | 1.076 | 0.105 | 0.34806456 | 0.52268195 | 0.0069 | 0.0071 | 0.0071 | 0.0071 | TRUE | 0.1704 |
| Vamp5 | 31 | 24 | 24 | 34 | 33 | 23 | 24 | 26 | 26 | 34 | 37 | 23 | 28.21 | 25 | 31 | 1.24 | 0.311 | 0.40063675 | 0.57372017 | 0 | 0.0864 | 0.0628 | 0.0628 | TRUE | 0.3983 |
| Vamp7 | 1911 | 1387 | 1454 | 1496 | 1326 | 1442 | 1484 | 1475 | 1569 | 1489 | 1496 | 1423 | 1489.47 | 1509 | 1469 | 0.974 | -0.038 | 0.56958192 | 0.72319751 | 0 | 0.004 | 0.0026 | 0.0026 | TRUE | 0.0139 |
| Vamp8 | 628 | 395 | 471 | 475 | 415 | 485 | 488 | 420 | 508 | 473 | 468 | 479 | 472.64 | 472 | 473 | 1.002 | 0.003 | 0.97896494 | 0.98871862 | 0.0026 | 0.0074 | 0.0059 | 0.0059 | TRUE | 0.1033 |
| Vangl1 | 2312 | 1831 | 1911 | 1807 | 1702 | 1896 | 1796 | 1948 | 2062 | 1799 | 1920 | 1872 | 1899.26 | 1935 | 1864 | 0.964 | -0.053 | 0.46129403 | 0.63069405 | 0.0025 | 0.0037 | 0.0032 | 0.0032 | TRUE | 0.0496 |
| Vangl2 | 1185 | 1009 | 969 | 1148 | 1051 | 1198 | 920 | 1073 | 1046 | 1143 | 1186 | 1183 | 1091.68 | 1013 | 1171 | 1.157 | 0.211 | 0.01025237 | 0.04206885 | 0.0027 | 0.0046 | 0.0039 | 0.0039 | TRUE | 0.0805 |
| Vapa | 14138 | 10649 | 11183 | 12304 | 10816 | 12368 | 10980 | 11327 | 12066 | 12247 | 12202 | 12209 | 11838.67 | 11458 | 12219 | 1.067 | 0.093 | 0.07867776 | 0.18691718 | 0.001 | 0.0026 | 0.0019 | 0.0019 | TRUE | 0.0263 |
| Vapb | 6066 | 4521 | 4962 | 4845 | 4551 | 5168 | 4711 | 4809 | 5354 | 4823 | 5134 | 5102 | 4988.74 | 4958 | 5020 | 1.013 | 0.018 | 0.78015549 | 0.87046309 | 0.0028 | 0.0029 | 0.0029 | 0.0029 | TRUE | 0.0592 |
| Vars | 19105 | 14552 | 13448 | 13926 | 12052 | 14062 | 14838 | 15479 | 14510 | 13862 | 13596 | 13881 | 14361.05 | 14942 | 13780 | 0.922 | -0.117 | 0.01615944 | 0.05902551 | 5.00E-04 | 0.0026 | 0.0016 | 0.0016 | TRUE | 0.012 |
| Vars2 | 3915 | 2897 | 2879 | 2857 | 2660 | 2940 | 3041 | 3082 | 3106 | 2844 | 3001 | 2902 | 2995.9 | 3076 | 2916 | 0.948 | -0.077 | 0.16573558 | 0.31676786 | 0 | 0.0032 | 0.0019 | 0.0019 | TRUE | 0.0078 |
| Vash1 | 1947 | 1496 | 1370 | 1746 | 1632 | 1751 | 1512 | 1591 | 1478 | 1738 | 1841 | 1729 | 1648.2 | 1527 | 1769 | 1.158 | 0.212 | 0.00167706 | 0.01053172 | 6.00E-04 | 0.0039 | 0.0027 | 0.0027 | TRUE | 0.0158 |
| Vash2 | 782 | 586 | 503 | 779 | 595 | 763 | 607 | 623 | 543 | 775 | 671 | 753 | 662.21 | 591 | 733 | 1.24 | 0.311 | 0.00139913 | 0.00916485 | 0.0038 | 0.006 | 0.0053 | 0.0053 | TRUE | 0.0634 |
| Vasn | 218 | 120 | 93 | 133 | 98 | 122 | 169 | 128 | 100 | 132 | 111 | 120 | 126.78 | 132 | 121 | 0.909 | -0.137 | 0.51630719 | 0.67856126 | 0.0317 | 0.0211 | 0.0243 | 0.0243 | TRUE | 0.3247 |
| Vasp | 5566 | 3866 | 3576 | 3535 | 3159 | 3776 | 4323 | 4112 | 3858 | 3519 | 3564 | 3727 | 3850.58 | 4098 | 3603 | 0.879 | -0.186 | 0.00293258 | 0.0162564 | 0.0017 | 0.003 | 0.0026 | 0.0026 | TRUE | 0.0317 |
| Vat1 | 15532 | 11764 | 13092 | 17297 | 15358 | 17631 | 12063 | 12513 | 14126 | 17217 | 17325 | 17405 | 15108.29 | 12901 | 17316 | 1.342 | 0.425 | 3.74E-11 | 3.34E-09 | 0.0034 | 0.0026 | 0.0029 | 0.0029 | TRUE | 0.0843 |
| Vat1l | 158 | 95 | 57 | 59 | 60 | 44 | 123 | 101 | 62 | 59 | 68 | 43 | 75.85 | 95 | 57 | 0.59 | -0.76 | 0.00778585 | 0.03429517 | 0.0714 | 0.0337 | 0.0445 | 0.0445 | TRUE | 0.2787 |
| Vaultrc5 | 0 | 2 | 0 | 1 | 0 | 0 | 0 | 2 | 0 | 1 | 0 | 0 | 0.52 | 1 | 0 | 0.534 | -0.904 | 0.76303124 | NA | 0 | 4.5562 | 4.7774 | 4.7774 | TRUE | 0.9746 |
| Vav1 | 90 | 58 | 61 | 120 | 119 | 118 | 70 | 62 | 66 | 119 | 134 | 116 | 94.6 | 66 | 123 | 1.868 | 0.901 | 1.03E-05 | 0.00019285 | 0 | 0.0275 | 0.0186 | 0.0186 | TRUE | 0.0576 |
| Vav2 | 1894 | 1572 | 1569 | 1568 | 1486 | 1577 | 1471 | 1672 | 1693 | 1561 | 1676 | 1557 | 1604.99 | 1612 | 1598 | 0.992 | -0.011 | 0.88431598 | 0.93584543 | 0.0033 | 0.0039 | 0.0037 | 0.0037 | TRUE | 0.0731 |
| Vav3 | 279 | 196 | 164 | 218 | 185 | 201 | 217 | 208 | 177 | 217 | 209 | 198 | 204.37 | 201 | 208 | 1.034 | 0.048 | 0.74397392 | 0.84763805 | 0 | 0.014 | 0.0105 | 0.0105 | TRUE | 0.1123 |
| Vax1 | 11 | 10 | 4 | 8 | 7 | 4 | 9 | 11 | 4 | 8 | 8 | 4 | 7.22 | 8 | 7 | 0.837 | -0.257 | 0.72486688 | NA | 0 | 0.3308 | 0.2445 | 0.2445 | TRUE | 0.3991 |
| Vax2 | 18 | 9 | 24 | 22 | 19 | 21 | 14 | 10 | 26 | 22 | 21 | 21 | 18.92 | 17 | 21 | 1.304 | 0.383 | 0.41990895 | 0.59218142 | 0.0523 | 0.1277 | 0.1086 | 0.1086 | TRUE | 0.9459 |
| Vax2os | 68 | 50 | 78 | 70 | 59 | 87 | 53 | 53 | 84 | 70 | 67 | 86 | 68.71 | 63 | 74 | 1.176 | 0.234 | 0.37384723 | 0.54846343 | 0.0312 | 0.0369 | 0.0353 | 0.0353 | TRUE | 0.6895 |
| Vbp1 | 7820 | 5810 | 6941 | 8093 | 6841 | 7810 | 6073 | 6180 | 7489 | 8056 | 7717 | 7710 | 7204.26 | 6581 | 7828 | 1.19 | 0.251 | 0.00142032 | 0.00928664 | 0.0073 | 0.0028 | 0.0043 | 0.0043 | TRUE | 0.1774 |
| Vcam1 | 1 | 2 | 1 | 0 | 1 | 0 | 1 | 2 | 1 | 0 | 1 | 0 | 0.85 | 1 | 0 | 0.282 | -1.824 | 0.39588467 | NA | 0 | 2.7847 | 1.9575 | 1.9575 | TRUE | 0.5075 |
| Vcan | 1135 | 828 | 1171 | 897 | 900 | 1151 | 882 | 881 | 1263 | 893 | 1015 | 1136 | 1011.69 | 1009 | 1015 | 1.007 | 0.01 | 0.93795275 | 0.96554692 | 0.0285 | 0.0048 | 0.0119 | 0.0119 | TRUE | 0.5797 |
| Vcl | 9753 | 6502 | 6192 | 6691 | 5781 | 6682 | 7575 | 6916 | 6681 | 6660 | 6522 | 6596 | 6824.98 | 7057 | 6593 | 0.934 | -0.099 | 0.10169985 | 0.22387123 | 0.002 | 0.0028 | 0.0025 | 0.0025 | TRUE | 0.0507 |
| Vcp | 31558 | 21260 | 20638 | 22117 | 18837 | 21675 | 24510 | 22614 | 22268 | 22015 | 21250 | 21397 | 22342.31 | 23131 | 21554 | 0.932 | -0.102 | 0.05928011 | 0.15208951 | 0.0015 | 0.0025 | 0.0021 | 0.0021 | TRUE | 0.0334 |
| Vcpip1 | 3002 | 2121 | 2017 | 2199 | 1948 | 2148 | 2332 | 2256 | 2176 | 2189 | 2198 | 2120 | 2211.79 | 2255 | 2169 | 0.961 | -0.057 | 0.3530176 | 0.52775572 | 4.00E-04 | 0.0035 | 0.0023 | 0.0023 | TRUE | 0.0111 |

| | | | | | | | | | | | | | | | | | | | | | | | | | |
|---|---|---|---|---|---|---|---|---|---|---|---|---|---|---|---|---|---|---|---|---|---|---|---|---|---|
| Vdac1 | 46606 | 35063 | 38457 | 39044 | 34080 | 39688 | 36197 | 37296 | 41495 | 38864 | 38446 | 39178 | 38579.34 | 38329 | 38829 | 1.013 | 0.019 | 0.75300878 | 0.85348706 | 0.0022 | 0.0025 | 0.0025 | 0.0025 | TRUE | 0.0639 |
| Vdac2 | 17502 | 12503 | 12678 | 14357 | 12658 | 14547 | 13593 | 13299 | 13679 | 14291 | 14280 | 14360 | 13917.07 | 13524 | 14310 | 1.058 | 0.082 | 0.06608089 | 0.16437421 | 0 | 0.0026 | 0.0013 | 0.0013 | TRUE | 0.0026 |
| Vdac3 | 7927 | 5540 | 5776 | 6300 | 5138 | 6253 | 6157 | 5893 | 6232 | 6271 | 5796 | 6173 | 6086.92 | 6094 | 6080 | 0.998 | -0.003 | 0.95473288 | 0.97527137 | 0.0011 | 0.0028 | 0.0021 | 0.0021 | TRUE | 0.0202 |
| Vdr | 71 | 64 | 45 | 50 | 49 | 39 | 55 | 68 | 49 | 50 | 55 | 38 | 52.55 | 57 | 48 | 0.834 | -0.261 | 0.35212062 | 0.52672528 | 0.0093 | 0.0475 | 0.0376 | 0.0376 | TRUE | 0.2391 |
| Vegfa | 2291 | 2248 | 2514 | 2411 | 2277 | 2519 | 1779 | 2391 | 2713 | 2400 | 2569 | 2487 | 2389.72 | 2294 | 2485 | 1.084 | 0.117 | 0.31658419 | 0.49061454 | 0.0228 | 0.0034 | 0.0093 | 0.0093 | TRUE | 0.4735 |
| Vegfb | 2238 | 1854 | 1996 | 2175 | 2117 | 2502 | 1738 | 1972 | 2154 | 2165 | 2388 | 2470 | 2147.83 | 1955 | 2341 | 1.199 | 0.262 | 0.00257579 | 0.01470171 | 0.0072 | 0.0035 | 0.005 | 0.005 | TRUE | 0.1162 |
| Vegfc | 615 | 395 | 340 | 496 | 385 | 461 | 478 | 420 | 367 | 494 | 434 | 455 | 441.3 | 422 | 461 | 1.091 | 0.125 | 0.29457376 | 0.46650881 | 0.0086 | 0.0078 | 0.0081 | 0.0081 | TRUE | 0.1648 |
| Veph1 | 81 | 77 | 36 | 39 | 31 | 37 | 63 | 82 | 39 | 39 | 35 | 37 | 49 | 61 | 37 | 0.6 | -0.736 | 0.02110285 | 0.07198641 | 0.0529 | 0.0508 | 0.0514 | 0.0514 | TRUE | 0.1571 |
| Vezf1 | 3849 | 3012 | 3270 | 3293 | 2762 | 3336 | 2989 | 3204 | 3528 | 3278 | 3116 | 3293 | 3234.72 | 3240 | 3229 | 0.997 | -0.004 | 0.95534402 | 0.97553943 | 0.0037 | 0.0032 | 0.0034 | 0.0034 | TRUE | 0.0732 |
| Vezt | 859 | 675 | 619 | 693 | 667 | 704 | 667 | 718 | 668 | 690 | 752 | 695 | 698.37 | 684 | 712 | 1.041 | 0.058 | 0.50795062 | 0.67130266 | 0 | 0.0058 | 0.0041 | 0.0041 | TRUE | 0.0279 |
| Vgf | 47 | 39 | 17 | 25 | 19 | 20 | 37 | 41 | 18 | 25 | 21 | 20 | 27.07 | 32 | 22 | 0.683 | -0.55 | 0.17676091 | 0.33055654 | 0.0574 | 0.09 | 0.081 | 0.081 | TRUE | 0.8934 |
| Vgll1 | 11 | 10 | 0 | 2 | 1 | 2 | 9 | 11 | 0 | 2 | 1 | 2 | 4.05 | 7 | 2 | 0.265 | -1.913 | 0.10917074 | NA | 0 | 0.5883 | 0.6503 | 0.6503 | TRUE | 0.7706 |
| Vgll2 | 11 | 2 | 3 | 4 | 2 | 1 | 9 | 2 | 3 | 4 | 2 | 1 | 3.52 | 5 | 2 | 0.508 | -0.978 | 0.36907047 | NA | 0 | 0.6753 | 0.5383 | 0.5383 | TRUE | 1.3198 |
| Vgll3 | 76 | 45 | 40 | 25 | 17 | 22 | 59 | 48 | 43 | 25 | 19 | 22 | 35.97 | 50 | 22 | 0.437 | -1.194 | 0.00041509 | 0.00373742 | 0 | 0.0683 | 0.0492 | 0.0492 | TRUE | 0.2296 |
| Vgll4 | 224 | 203 | 152 | 132 | 110 | 136 | 174 | 216 | 164 | 131 | 124 | 134 | 157.27 | 185 | 130 | 0.705 | -0.504 | 0.00303137 | 0.01666437 | 0.0066 | 0.0175 | 0.0142 | 0.0142 | TRUE | 0.2278 |
| Vhl | 2878 | 2038 | 2026 | 2215 | 1940 | 2103 | 2235 | 2168 | 2186 | 2205 | 2189 | 2076 | 2176.39 | 2196 | 2157 | 0.981 | -0.027 | 0.65600728 | 0.7864714 | 0 | 0.0035 | 0.0022 | 0.0022 | TRUE | 0.013 |
| Vil1 | 217 | 164 | 179 | 223 | 197 | 200 | 169 | 174 | 193 | 222 | 222 | 197 | 196.29 | 179 | 214 | 1.199 | 0.261 | 0.07362261 | 0.17787152 | 0 | 0.0145 | 0.0103 | 0.0103 | TRUE | 0.052 |
| Vill | 1877 | 1545 | 1554 | 1840 | 1667 | 1949 | 1458 | 1643 | 1677 | 1832 | 1881 | 1924 | 1735.66 | 1593 | 1879 | 1.181 | 0.24 | 0.001142 | 0.00798561 | 0.0026 | 0.0038 | 0.0034 | 0.0034 | TRUE | 0.0687 |
| Vim | 10880 | 8047 | 9086 | 10761 | 10030 | 12284 | 8450 | 8559 | 9804 | 10711 | 11315 | 12126 | 10160.97 | 8938 | 11384 | 1.274 | 0.349 | 1.28E-06 | 3.58E-05 | 0.0053 | 0.0027 | 0.0036 | 0.0036 | TRUE | 0.0875 |
| Vimp | 1471 | 940 | 942 | 1063 | 900 | 1038 | 1142 | 1000 | 1016 | 1058 | 1015 | 1025 | 1042.8 | 1053 | 1033 | 0.979 | -0.03 | 0.70991876 | 0.82524789 | 0.0022 | 0.0047 | 0.0038 | 0.0038 | TRUE | 0.069 |
| Vip | 0 | 0 | 0 | 0 | 0 | 0 | 0 | 0 | 0 | 0 | 0 | 0 | 0 | 0 | 0 | NA | NA | NA | NA | NA | NA | NA | NA | NA | NA |
| Vipas39 | 2011 | 1350 | 1187 | 1339 | 1187 | 1342 | 1562 | 1436 | 1281 | 1333 | 1339 | 1325 | 1379.21 | 1426 | 1332 | 0.933 | -0.1 | 0.22530183 | 0.39004833 | 0.0043 | 0.0041 | 0.0042 | 0.0042 | TRUE | 0.0951 |
| Vipr1 | 136 | 124 | 97 | 131 | 91 | 96 | 106 | 132 | 105 | 130 | 103 | 95 | 111.67 | 114 | 109 | 0.961 | -0.058 | 0.77618659 | 0.86766476 | 0.0141 | 0.0237 | 0.0209 | 0.0209 | TRUE | 0.29 |
| Vipr2 | 14 | 16 | 9 | 10 | 11 | 7 | 11 | 17 | 10 | 10 | 12 | 7 | 11.15 | 13 | 10 | 0.778 | -0.362 | 0.53617478 | NA | 0 | 0.2151 | 0.1554 | 0.1554 | TRUE | 0.3706 |
| Vit | 1 | 2 | 0 | 1 | 0 | 1 | 1 | 2 | 0 | 1 | 0 | 1 | 0.81 | 1 | 1 | 0.699 | -0.518 | 0.81129103 | NA | 0 | 2.9125 | 2.1155 | 2.1155 | TRUE | 0.297 |
| Vkorc1 | 1045 | 781 | 868 | 1049 | 959 | 1055 | 812 | 831 | 937 | 1044 | 1082 | 1041 | 957.73 | 860 | 1056 | 1.23 | 0.298 | 0.00033428 | 0.00315368 | 0.002 | 0.0049 | 0.0039 | 0.0039 | TRUE | 0.0716 |
| Vkorc1l1 | 646 | 518 | 493 | 546 | 479 | 526 | 502 | 551 | 532 | 543 | 540 | 519 | 531.29 | 528 | 534 | 1.013 | 0.019 | 0.84434604 | 0.91195135 | 0 | 0.0069 | 0.0046 | 0.0046 | TRUE | 0.0242 |
| Vldlr | 283 | 202 | 255 | 243 | 231 | 283 | 220 | 215 | 275 | 242 | 261 | 279 | 248.61 | 237 | 261 | 1.104 | 0.143 | 0.31914493 | 0.493086 | 0.0079 | 0.012 | 0.0108 | 0.0108 | TRUE | 0.2166 |
| Vma21 | 1373 | 1042 | 1025 | 1065 | 925 | 1040 | 1066 | 1108 | 1106 | 1060 | 1043 | 1027 | 1068.49 | 1093 | 1043 | 0.955 | -0.067 | 0.35330738 | 0.52780934 | 0 | 0.0046 | 0.0028 | 0.0028 | TRUE | 0.006 |
| Vmac | 941 | 741 | 816 | 1005 | 864 | 1003 | 731 | 788 | 880 | 1000 | 975 | 990 | 894.11 | 800 | 988 | 1.238 | 0.308 | 0.00047246 | 0.00414791 | 0.0033 | 0.0051 | 0.0045 | 0.0045 | TRUE | 0.0917 |
| Vmn1r-ps10 | 0 | 0 | 0 | 0 | 0 | 0 | 0 | 0 | 0 | 0 | 0 | 0 | 0 | 0 | 0 | NA | NA | NA | NA | NA | NA | NA | NA | NA | NA |
| Vmn1r-ps79 | 0 | 0 | 0 | 0 | 0 | 0 | 0 | 0 | 0 | 0 | 0 | 0 | 0 | 0 | 0 | NA | NA | NA | NA | NA | NA | NA | NA | NA | NA |
| Vmn1r1 | 0 | 1 | 0 | 0 | 1 | 0 | 0 | 1 | 0 | 0 | 1 | 0 | 0.37 | 0 | 0 | 1.075 | 0.105 | 0.97520514 | NA | 0 | 6.4904 | 6.1969 | 6.1969 | TRUE | 0.5216 |
| Vmn1r10 | 0 | 0 | 0 | 0 | 0 | 0 | 0 | 0 | 0 | 0 | 0 | 0 | 0 | 0 | 0 | NA | NA | NA | NA | NA | NA | NA | NA | NA | NA |
| Vmn1r100 | 0 | 0 | 0 | 0 | 0 | 0 | 0 | 0 | 0 | 0 | 0 | 0 | 0 | 0 | 0 | NA | NA | NA | NA | NA | NA | NA | NA | NA | NA |
| Vmn1r101 | 0 | 0 | 0 | 0 | 0 | 0 | 0 | 0 | 0 | 0 | 0 | 0 | 0 | 0 | 0 | NA | NA | NA | NA | NA | NA | NA | NA | NA | NA |
| Vmn1r103 | 0 | 0 | 0 | 0 | 0 | 0 | 0 | 0 | 0 | 0 | 0 | 0 | 0 | 0 | 0 | NA | NA | NA | NA | NA | NA | NA | NA | NA | NA |
| Vmn1r104 | 0 | 0 | 0 | 0 | 0 | 0 | 0 | 0 | 0 | 0 | 0 | 0 | 0 | 0 | 0 | NA | NA | NA | NA | NA | NA | NA | NA | NA | NA |
| Vmn1r107 | 0 | 0 | 0 | 0 | 0 | 0 | 0 | 0 | 0 | 0 | 0 | 0 | 0 | 0 | 0 | NA | NA | NA | NA | NA | NA | NA | NA | NA | NA |
| Vmn1r11 | 0 | 0 | 0 | 0 | 0 | 0 | 0 | 0 | 0 | 0 | 0 | 0 | 0 | 0 | 0 | NA | NA | NA | NA | NA | NA | NA | NA | NA | NA |
| Vmn1r111 | 0 | 0 | 0 | 0 | 0 | 0 | 0 | 0 | 0 | 0 | 0 | 0 | 0 | 0 | 0 | NA | NA | NA | NA | NA | NA | NA | NA | NA | NA |
| Vmn1r112 | 0 | 0 | 0 | 0 | 0 | 0 | 0 | 0 | 0 | 0 | 0 | 0 | 0 | 0 | 0 | NA | NA | NA | NA | NA | NA | NA | NA | NA | NA |
| Vmn1r113 | 0 | 0 | 0 | 0 | 0 | 0 | 0 | 0 | 0 | 0 | 0 | 0 | 0 | 0 | 0 | NA | NA | NA | NA | NA | NA | NA | NA | NA | NA |
| Vmn1r114 | 0 | 0 | 0 | 0 | 0 | 0 | 0 | 0 | 0 | 0 | 0 | 0 | 0 | 0 | 0 | NA | NA | NA | NA | NA | NA | NA | NA | NA | NA |
| Vmn1r115 | 0 | 0 | 0 | 0 | 0 | 0 | 0 | 0 | 0 | 0 | 0 | 0 | 0 | 0 | 0 | NA | NA | NA | NA | NA | NA | NA | NA | NA | NA |
| Vmn1r116 | 0 | 0 | 0 | 0 | 0 | 0 | 0 | 0 | 0 | 0 | 0 | 0 | 0 | 0 | 0 | NA | NA | NA | NA | NA | NA | NA | NA | NA | NA |
| Vmn1r117 | 0 | 0 | 0 | 0 | 0 | 0 | 0 | 0 | 0 | 0 | 0 | 0 | 0 | 0 | 0 | NA | NA | NA | NA | NA | NA | NA | NA | NA | NA |
| Vmn1r118 | 0 | 0 | 0 | 0 | 0 | 0 | 0 | 0 | 0 | 0 | 0 | 0 | 0 | 0 | 0 | NA | NA | NA | NA | NA | NA | NA | NA | NA | NA |
| Vmn1r119 | 0 | 0 | 0 | 0 | 0 | 0 | 0 | 0 | 0 | 0 | 0 | 0 | 0 | 0 | 0 | NA | NA | NA | NA | NA | NA | NA | NA | NA | NA |
| Vmn1r12 | 0 | 0 | 0 | 0 | 0 | 0 | 0 | 0 | 0 | 0 | 0 | 0 | 0 | 0 | 0 | NA | NA | NA | NA | NA | NA | NA | NA | NA | NA |
| Vmn1r120 | 0 | 0 | 0 | 0 | 0 | 0 | 0 | 0 | 0 | 0 | 0 | 0 | 0 | 0 | 0 | NA | NA | NA | NA | NA | NA | NA | NA | NA | NA |
| Vmn1r121 | 0 | 0 | 0 | 0 | 0 | 0 | 0 | 0 | 0 | 0 | 0 | 0 | 0 | 0 | 0 | NA | NA | NA | NA | NA | NA | NA | NA | NA | NA |
| Vmn1r122 | 0 | 0 | 0 | 0 | 0 | 0 | 0 | 0 | 0 | 0 | 0 | 0 | 0 | 0 | 0 | NA | NA | NA | NA | NA | NA | NA | NA | NA | NA |
| Vmn1r123 | 0 | 0 | 0 | 0 | 0 | 0 | 0 | 0 | 0 | 0 | 0 | 0 | 0 | 0 | 0 | NA | NA | NA | NA | NA | NA | NA | NA | NA | NA |
| Vmn1r124 | 0 | 0 | 0 | 0 | 0 | 0 | 0 | 0 | 0 | 0 | 0 | 0 | 0 | 0 | 0 | NA | NA | NA | NA | NA | NA | NA | NA | NA | NA |
| Vmn1r125 | 0 | 0 | 0 | 0 | 0 | 0 | 0 | 0 | 0 | 0 | 0 | 0 | 0 | 0 | 0 | NA | NA | NA | NA | NA | NA | NA | NA | NA | NA |
| Vmn1r126 | 0 | 0 | 0 | 0 | 0 | 0 | 0 | 0 | 0 | 0 | 0 | 0 | 0 | 0 | 0 | NA | NA | NA | NA | NA | NA | NA | NA | NA | NA |
| Vmn1r127 | 0 | 0 | 0 | 0 | 0 | 0 | 0 | 0 | 0 | 0 | 0 | 0 | 0 | 0 | 0 | NA | NA | NA | NA | NA | NA | NA | NA | NA | NA |
| Vmn1r128 | 0 | 0 | 0 | 0 | 0 | 0 | 0 | 0 | 0 | 0 | 0 | 0 | 0 | 0 | 0 | NA | NA | NA | NA | NA | NA | NA | NA | NA | NA |
| Vmn1r129 | 0 | 0 | 0 | 0 | 0 | 0 | 0 | 0 | 0 | 0 | 0 | 0 | 0 | 0 | 0 | NA | NA | NA | NA | NA | NA | NA | NA | NA | NA |
| Vmn1r13 | 0 | 0 | 0 | 0 | 0 | 0 | 0 | 0 | 0 | 0 | 0 | 0 | 0 | 0 | 0 | NA | NA | NA | NA | NA | NA | NA | NA | NA | NA |
| Vmn1r130 | 0 | 0 | 0 | 0 | 0 | 0 | 0 | 0 | 0 | 0 | 0 | 0 | 0 | 0 | 0 | NA | NA | NA | NA | NA | NA | NA | NA | NA | NA |
| Vmn1r132 | 0 | 0 | 0 | 0 | 0 | 0 | 0 | 0 | 0 | 0 | 0 | 0 | 0 | 0 | 0 | NA | NA | NA | NA | NA | NA | NA | NA | NA | NA |
| Vmn1r135 | 0 | 0 | 0 | 0 | 0 | 0 | 0 | 0 | 0 | 0 | 0 | 0 | 0 | 0 | 0 | NA | NA | NA | NA | NA | NA | NA | NA | NA | NA |
| Vmn1r137 | 0 | 0 | 0 | 0 | 0 | 0 | 0 | 0 | 0 | 0 | 0 | 0 | 0 | 0 | 0 | NA | NA | NA | NA | NA | NA | NA | NA | NA | NA |
| Vmn1r139 | 0 | 0 | 0 | 0 | 0 | 0 | 0 | 0 | 0 | 0 | 0 | 0 | 0 | 0 | 0 | NA | NA | NA | NA | NA | NA | NA | NA | NA | NA |
| Vmn1r14 | 3 | 4 | 2 | 1 | 0 | 0 | 2 | 4 | 2 | 1 | 0 | 0 | 1.62 | 3 | 0 | 0.128 | -2.968 | 0.09168078 | NA | 0 | 1.4627 | 1.0648 | 1.0648 | TRUE | 0.3906 |
| Vmn1r142 | 0 | 0 | 0 | 0 | 0 | 0 | 0 | 0 | 0 | 0 | 0 | 0 | 0 | 0 | 0 | NA | NA | NA | NA | NA | NA | NA | NA | NA | NA |
| Vmn1r143 | 0 | 0 | 0 | 0 | 0 | 0 | 0 | 0 | 0 | 0 | 0 | 0 | 0 | 0 | 0 | NA | NA | NA | NA | NA | NA | NA | NA | NA | NA |

| | | | | | | | | | | | | | | | | | | | | | | | | | |
|---|---|---|---|---|---|---|---|---|---|---|---|---|---|---|---|---|---|---|---|---|---|---|---|---|---|
| Vmn1r148 | 0 | 0 | 0 | 0 | 0 | 0 | 0 | 0 | 0 | 0 | 0 | 0 | 0 | 0 | 0 | NA | NA | NA | NA | NA | NA | NA | NA | NA | NA |
| Vmn1r149 | 0 | 0 | 0 | 0 | 0 | 0 | 0 | 0 | 0 | 0 | 0 | 0 | 0 | 0 | 0 | NA | NA | NA | NA | NA | NA | NA | NA | NA | NA |
| Vmn1r15 | 0 | 1 | 0 | 0 | 1 | 0 | 0 | 1 | 0 | 0 | 1 | 0 | 0.37 | 0 | 0 | 1.075 | 0.105 | 0.97520514 | NA | 0 | 6.4904 | 6.1969 | 6.1969 | TRUE | 0.5216 |
| Vmn1r151 | 0 | 0 | 0 | 0 | 0 | 0 | 0 | 0 | 0 | 0 | 0 | 0 | 0 | 0 | 0 | NA | NA | NA | NA | NA | NA | NA | NA | NA | NA |
| Vmn1r152 | 0 | 0 | 0 | 0 | 0 | 0 | 0 | 0 | 0 | 0 | 0 | 0 | 0 | 0 | 0 | NA | NA | NA | NA | NA | NA | NA | NA | NA | NA |
| Vmn1r155 | 0 | 0 | 0 | 0 | 0 | 0 | 0 | 0 | 0 | 0 | 0 | 0 | 0 | 0 | 0 | NA | NA | NA | NA | NA | NA | NA | NA | NA | NA |
| Vmn1r157 | 0 | 0 | 0 | 0 | 0 | 0 | 0 | 0 | 0 | 0 | 0 | 0 | 0 | 0 | 0 | NA | NA | NA | NA | NA | NA | NA | NA | NA | NA |
| Vmn1r158 | 0 | 0 | 0 | 0 | 0 | 0 | 0 | 0 | 0 | 0 | 0 | 0 | 0 | 0 | 0 | NA | NA | NA | NA | NA | NA | NA | NA | NA | NA |
| Vmn1r159 | 0 | 0 | 0 | 0 | 0 | 0 | 0 | 0 | 0 | 0 | 0 | 0 | 0 | 0 | 0 | NA | NA | NA | NA | NA | NA | NA | NA | NA | NA |
| Vmn1r16 | 0 | 0 | 0 | 0 | 0 | 0 | 0 | 0 | 0 | 0 | 0 | 0 | 0 | 0 | 0 | NA | NA | NA | NA | NA | NA | NA | NA | NA | NA |
| Vmn1r160 | 0 | 0 | 0 | 0 | 0 | 0 | 0 | 0 | 0 | 0 | 0 | 0 | 0 | 0 | 0 | NA | NA | NA | NA | NA | NA | NA | NA | NA | NA |
| Vmn1r163 | 0 | 0 | 0 | 0 | 0 | 0 | 0 | 0 | 0 | 0 | 0 | 0 | 0 | 0 | 0 | NA | NA | NA | NA | NA | NA | NA | NA | NA | NA |
| Vmn1r165 | 0 | 0 | 0 | 0 | 0 | 0 | 0 | 0 | 0 | 0 | 0 | 0 | 0 | 0 | 0 | NA | NA | NA | NA | NA | NA | NA | NA | NA | NA |
| Vmn1r167 | 0 | 0 | 0 | 0 | 0 | 0 | 0 | 0 | 0 | 0 | 0 | 0 | 0 | 0 | 0 | NA | NA | NA | NA | NA | NA | NA | NA | NA | NA |
| Vmn1r168 | 0 | 0 | 0 | 0 | 0 | 0 | 0 | 0 | 0 | 0 | 0 | 0 | 0 | 0 | 0 | NA | NA | NA | NA | NA | NA | NA | NA | NA | NA |
| Vmn1r169 | 0 | 0 | 0 | 0 | 0 | 0 | 0 | 0 | 0 | 0 | 0 | 0 | 0 | 0 | 0 | NA | NA | NA | NA | NA | NA | NA | NA | NA | NA |
| Vmn1r17 | 0 | 0 | 0 | 0 | 0 | 0 | 0 | 0 | 0 | 0 | 0 | 0 | 0 | 0 | 0 | NA | NA | NA | NA | NA | NA | NA | NA | NA | NA |
| Vmn1r170 | 0 | 0 | 0 | 0 | 0 | 0 | 0 | 0 | 0 | 0 | 0 | 0 | 0 | 0 | 0 | NA | NA | NA | NA | NA | NA | NA | NA | NA | NA |
| Vmn1r171 | 0 | 0 | 0 | 0 | 0 | 0 | 0 | 0 | 0 | 0 | 0 | 0 | 0 | 0 | 0 | NA | NA | NA | NA | NA | NA | NA | NA | NA | NA |
| Vmn1r172 | 0 | 0 | 0 | 0 | 0 | 0 | 0 | 0 | 0 | 0 | 0 | 0 | 0 | 0 | 0 | NA | NA | NA | NA | NA | NA | NA | NA | NA | NA |
| Vmn1r173 | 0 | 0 | 0 | 0 | 0 | 0 | 0 | 0 | 0 | 0 | 0 | 0 | 0 | 0 | 0 | NA | NA | NA | NA | NA | NA | NA | NA | NA | NA |
| Vmn1r174 | 0 | 0 | 0 | 0 | 0 | 0 | 0 | 0 | 0 | 0 | 0 | 0 | 0 | 0 | 0 | NA | NA | NA | NA | NA | NA | NA | NA | NA | NA |
| Vmn1r175 | 0 | 0 | 0 | 0 | 0 | 0 | 0 | 0 | 0 | 0 | 0 | 0 | 0 | 0 | 0 | NA | NA | NA | NA | NA | NA | NA | NA | NA | NA |
| Vmn1r176 | 0 | 0 | 0 | 0 | 0 | 0 | 0 | 0 | 0 | 0 | 0 | 0 | 0 | 0 | 0 | NA | NA | NA | NA | NA | NA | NA | NA | NA | NA |
| Vmn1r177 | 0 | 0 | 0 | 0 | 0 | 0 | 0 | 0 | 0 | 0 | 0 | 0 | 0 | 0 | 0 | NA | NA | NA | NA | NA | NA | NA | NA | NA | NA |
| Vmn1r178 | 1 | 0 | 0 | 1 | 0 | 0 | 1 | 0 | 0 | 1 | 0 | 0 | 0.3 | 0 | 0 | 1.075 | 0.105 | 0.97698525 | NA | 0 | 8.0272 | 7.5141 | 7.5141 | TRUE | 0.3906 |
| Vmn1r179 | 0 | 0 | 0 | 0 | 0 | 0 | 0 | 0 | 0 | 0 | 0 | 0 | 0 | 0 | 0 | NA | NA | NA | NA | NA | NA | NA | NA | NA | NA |
| Vmn1r18 | 0 | 0 | 0 | 0 | 0 | 0 | 0 | 0 | 0 | 0 | 0 | 0 | 0 | 0 | 0 | NA | NA | NA | NA | NA | NA | NA | NA | NA | NA |
| Vmn1r180 | 0 | 0 | 0 | 0 | 0 | 0 | 0 | 0 | 0 | 0 | 0 | 0 | 0 | 0 | 0 | NA | NA | NA | NA | NA | NA | NA | NA | NA | NA |
| Vmn1r181 | 0 | 0 | 0 | 0 | 0 | 0 | 0 | 0 | 0 | 0 | 0 | 0 | 0 | 0 | 0 | NA | NA | NA | NA | NA | NA | NA | NA | NA | NA |
| Vmn1r183 | 0 | 0 | 0 | 0 | 0 | 0 | 0 | 0 | 0 | 0 | 0 | 0 | 0 | 0 | 0 | NA | NA | NA | NA | NA | NA | NA | NA | NA | NA |
| Vmn1r184 | 0 | 0 | 0 | 0 | 0 | 0 | 0 | 0 | 0 | 0 | 0 | 0 | 0 | 0 | 0 | NA | NA | NA | NA | NA | NA | NA | NA | NA | NA |
| Vmn1r185 | 0 | 0 | 0 | 0 | 0 | 0 | 0 | 0 | 0 | 0 | 0 | 0 | 0 | 0 | 0 | NA | NA | NA | NA | NA | NA | NA | NA | NA | NA |
| Vmn1r186 | 0 | 0 | 0 | 0 | 0 | 0 | 0 | 0 | 0 | 0 | 0 | 0 | 0 | 0 | 0 | NA | NA | NA | NA | NA | NA | NA | NA | NA | NA |
| Vmn1r187 | 0 | 0 | 0 | 0 | 0 | 0 | 0 | 0 | 0 | 0 | 0 | 0 | 0 | 0 | 0 | NA | NA | NA | NA | NA | NA | NA | NA | NA | NA |
| Vmn1r188 | 0 | 0 | 0 | 0 | 0 | 0 | 0 | 0 | 0 | 0 | 0 | 0 | 0 | 0 | 0 | NA | NA | NA | NA | NA | NA | NA | NA | NA | NA |
| Vmn1r189 | 0 | 0 | 0 | 0 | 0 | 0 | 0 | 0 | 0 | 0 | 0 | 0 | 0 | 0 | 0 | NA | NA | NA | NA | NA | NA | NA | NA | NA | NA |
| Vmn1r19 | 0 | 0 | 0 | 0 | 0 | 0 | 0 | 0 | 0 | 0 | 0 | 0 | 0 | 0 | 0 | NA | NA | NA | NA | NA | NA | NA | NA | NA | NA |
| Vmn1r191 | 0 | 0 | 0 | 0 | 0 | 0 | 0 | 0 | 0 | 0 | 0 | 0 | 0 | 0 | 0 | NA | NA | NA | NA | NA | NA | NA | NA | NA | NA |
| Vmn1r192 | 0 | 0 | 0 | 0 | 0 | 0 | 0 | 0 | 0 | 0 | 0 | 0 | 0 | 0 | 0 | NA | NA | NA | NA | NA | NA | NA | NA | NA | NA |
| Vmn1r193 | 0 | 0 | 0 | 0 | 0 | 0 | 0 | 0 | 0 | 0 | 0 | 0 | 0 | 0 | 0 | NA | NA | NA | NA | NA | NA | NA | NA | NA | NA |
| Vmn1r194 | 0 | 0 | 0 | 0 | 0 | 0 | 0 | 0 | 0 | 0 | 0 | 0 | 0 | 0 | 0 | NA | NA | NA | NA | NA | NA | NA | NA | NA | NA |
| Vmn1r195 | 0 | 0 | 0 | 0 | 0 | 0 | 0 | 0 | 0 | 0 | 0 | 0 | 0 | 0 | 0 | NA | NA | NA | NA | NA | NA | NA | NA | NA | NA |
| Vmn1r196 | 0 | 0 | 0 | 0 | 0 | 0 | 0 | 0 | 0 | 0 | 0 | 0 | 0 | 0 | 0 | NA | NA | NA | NA | NA | NA | NA | NA | NA | NA |
| Vmn1r197 | 0 | 0 | 0 | 0 | 0 | 0 | 0 | 0 | 0 | 0 | 0 | 0 | 0 | 0 | 0 | NA | NA | NA | NA | NA | NA | NA | NA | NA | NA |
| Vmn1r198 | 0 | 0 | 0 | 0 | 0 | 0 | 0 | 0 | 0 | 0 | 0 | 0 | 0 | 0 | 0 | NA | NA | NA | NA | NA | NA | NA | NA | NA | NA |
| Vmn1r199 | 0 | 0 | 0 | 0 | 0 | 0 | 0 | 0 | 0 | 0 | 0 | 0 | 0 | 0 | 0 | NA | NA | NA | NA | NA | NA | NA | NA | NA | NA |
| Vmn1r2 | 0 | 0 | 0 | 0 | 0 | 0 | 0 | 0 | 0 | 0 | 0 | 0 | 0 | 0 | 0 | NA | NA | NA | NA | NA | NA | NA | NA | NA | NA |
| Vmn1r20 | 0 | 0 | 0 | 0 | 0 | 0 | 0 | 0 | 0 | 0 | 0 | 0 | 0 | 0 | 0 | NA | NA | NA | NA | NA | NA | NA | NA | NA | NA |
| Vmn1r200 | 0 | 0 | 0 | 0 | 0 | 0 | 0 | 0 | 0 | 0 | 0 | 0 | 0 | 0 | 0 | NA | NA | NA | NA | NA | NA | NA | NA | NA | NA |
| Vmn1r201 | 0 | 0 | 0 | 0 | 0 | 0 | 0 | 0 | 0 | 0 | 0 | 0 | 0 | 0 | 0 | NA | NA | NA | NA | NA | NA | NA | NA | NA | NA |
| Vmn1r202 | 0 | 0 | 0 | 0 | 0 | 0 | 0 | 0 | 0 | 0 | 0 | 0 | 0 | 0 | 0 | NA | NA | NA | NA | NA | NA | NA | NA | NA | NA |
| Vmn1r203 | 0 | 0 | 0 | 0 | 0 | 0 | 0 | 0 | 0 | 0 | 0 | 0 | 0 | 0 | 0 | NA | NA | NA | NA | NA | NA | NA | NA | NA | NA |
| Vmn1r204 | 0 | 0 | 0 | 0 | 0 | 0 | 0 | 0 | 0 | 0 | 0 | 0 | 0 | 0 | 0 | NA | NA | NA | NA | NA | NA | NA | NA | NA | NA |
| Vmn1r205 | 0 | 0 | 0 | 0 | 0 | 0 | 0 | 0 | 0 | 0 | 0 | 0 | 0 | 0 | 0 | NA | NA | NA | NA | NA | NA | NA | NA | NA | NA |
| Vmn1r206 | 0 | 0 | 0 | 0 | 0 | 0 | 0 | 0 | 0 | 0 | 0 | 0 | 0 | 0 | 0 | NA | NA | NA | NA | NA | NA | NA | NA | NA | NA |
| Vmn1r207-p | 0 | 0 | 0 | 0 | 0 | 0 | 0 | 0 | 0 | 0 | 0 | 0 | 0 | 0 | 0 | NA | NA | NA | NA | NA | NA | NA | NA | NA | NA |
| Vmn1r208 | 0 | 0 | 0 | 0 | 0 | 0 | 0 | 0 | 0 | 0 | 0 | 0 | 0 | 0 | 0 | NA | NA | NA | NA | NA | NA | NA | NA | NA | NA |
| Vmn1r209 | 0 | 0 | 0 | 0 | 0 | 0 | 0 | 0 | 0 | 0 | 0 | 0 | 0 | 0 | 0 | NA | NA | NA | NA | NA | NA | NA | NA | NA | NA |
| Vmn1r21 | 0 | 0 | 0 | 0 | 0 | 0 | 0 | 0 | 0 | 0 | 0 | 0 | 0 | 0 | 0 | NA | NA | NA | NA | NA | NA | NA | NA | NA | NA |
| Vmn1r210 | 0 | 1 | 0 | 0 | 1 | 0 | 0 | 1 | 0 | 0 | 1 | 0 | 0.37 | 0 | 0 | 1.075 | 0.105 | 0.97520514 | NA | 0 | 6.4904 | 6.1969 | 6.1969 | TRUE | 0.5216 |
| Vmn1r211 | 0 | 0 | 0 | 0 | 0 | 0 | 0 | 0 | 0 | 0 | 0 | 0 | 0 | 0 | 0 | NA | NA | NA | NA | NA | NA | NA | NA | NA | NA |
| Vmn1r212 | 0 | 0 | 0 | 0 | 0 | 0 | 0 | 0 | 0 | 0 | 0 | 0 | 0 | 0 | 0 | NA | NA | NA | NA | NA | NA | NA | NA | NA | NA |
| Vmn1r213 | 0 | 0 | 0 | 0 | 0 | 0 | 0 | 0 | 0 | 0 | 0 | 0 | 0 | 0 | 0 | NA | NA | NA | NA | NA | NA | NA | NA | NA | NA |
| Vmn1r214 | 0 | 0 | 0 | 0 | 0 | 0 | 0 | 0 | 0 | 0 | 0 | 0 | 0 | 0 | 0 | NA | NA | NA | NA | NA | NA | NA | NA | NA | NA |
| Vmn1r215 | 0 | 0 | 0 | 0 | 0 | 0 | 0 | 0 | 0 | 0 | 0 | 0 | 0 | 0 | 0 | NA | NA | NA | NA | NA | NA | NA | NA | NA | NA |
| Vmn1r216 | 0 | 0 | 0 | 0 | 0 | 0 | 0 | 0 | 0 | 0 | 0 | 0 | 0 | 0 | 0 | NA | NA | NA | NA | NA | NA | NA | NA | NA | NA |
| Vmn1r217 | 0 | 0 | 0 | 0 | 0 | 0 | 0 | 0 | 0 | 0 | 0 | 0 | 0 | 0 | 0 | NA | NA | NA | NA | NA | NA | NA | NA | NA | NA |
| Vmn1r218 | 0 | 0 | 0 | 0 | 0 | 0 | 0 | 0 | 0 | 0 | 0 | 0 | 0 | 0 | 0 | NA | NA | NA | NA | NA | NA | NA | NA | NA | NA |

| | | | | | | | | | | | | | | | | | | | | | | | | | |
|---|---|---|---|---|---|---|---|---|---|---|---|---|---|---|---|---|---|---|---|---|---|---|---|---|---|
| Vmn1r219 | 0 | 0 | 0 | 0 | 0 | 0 | 0 | 0 | 0 | 0 | 0 | 0 | 0 | 0 | 0 | NA | NA | NA | NA | NA | NA | NA | NA | NA | NA |
| Vmn1r22 | 0 | 0 | 0 | 0 | 0 | 0 | 0 | 0 | 0 | 0 | 0 | 0 | 0 | 0 | 0 | NA | NA | NA | NA | NA | NA | NA | NA | NA | NA |
| Vmn1r220 | 0 | 0 | 0 | 0 | 0 | 0 | 0 | 0 | 0 | 0 | 0 | 0 | 0 | 0 | 0 | NA | NA | NA | NA | NA | NA | NA | NA | NA | NA |
| Vmn1r221 | 0 | 0 | 0 | 0 | 0 | 0 | 0 | 0 | 0 | 0 | 0 | 0 | 0 | 0 | 0 | NA | NA | NA | NA | NA | NA | NA | NA | NA | NA |
| Vmn1r222 | 0 | 0 | 0 | 0 | 0 | 0 | 0 | 0 | 0 | 0 | 0 | 0 | 0 | 0 | 0 | NA | NA | NA | NA | NA | NA | NA | NA | NA | NA |
| Vmn1r223 | 0 | 0 | 0 | 0 | 0 | 0 | 0 | 0 | 0 | 0 | 0 | 0 | 0 | 0 | 0 | NA | NA | NA | NA | NA | NA | NA | NA | NA | NA |
| Vmn1r224 | 0 | 0 | 0 | 0 | 0 | 0 | 0 | 0 | 0 | 0 | 0 | 0 | 0 | 0 | 0 | NA | NA | NA | NA | NA | NA | NA | NA | NA | NA |
| Vmn1r225 | 0 | 0 | 0 | 0 | 0 | 0 | 0 | 0 | 0 | 0 | 0 | 0 | 0 | 0 | 0 | NA | NA | NA | NA | NA | NA | NA | NA | NA | NA |
| Vmn1r226 | 0 | 0 | 0 | 0 | 0 | 0 | 0 | 0 | 0 | 0 | 0 | 0 | 0 | 0 | 0 | NA | NA | NA | NA | NA | NA | NA | NA | NA | NA |
| Vmn1r227 | 1 | 1 | 0 | 1 | 0 | 0 | 1 | 1 | 0 | 1 | 0 | 0 | 0.47 | 1 | 0 | 0.599 | -0.74 | 0.79191656 | NA | 0 | 5.0171 | 3.8827 | 3.8827 | TRUE | 0.3906 |
| Vmn1r228 | 0 | 0 | 0 | 0 | 0 | 0 | 0 | 0 | 0 | 0 | 0 | 0 | 0 | 0 | 0 | NA | NA | NA | NA | NA | NA | NA | NA | NA | NA |
| Vmn1r229 | 0 | 0 | 0 | 0 | 0 | 0 | 0 | 0 | 0 | 0 | 0 | 0 | 0 | 0 | 0 | NA | NA | NA | NA | NA | NA | NA | NA | NA | NA |
| Vmn1r23 | 0 | 0 | 0 | 0 | 0 | 0 | 0 | 0 | 0 | 0 | 0 | 0 | 0 | 0 | 0 | NA | NA | NA | NA | NA | NA | NA | NA | NA | NA |
| Vmn1r230 | 0 | 0 | 0 | 0 | 0 | 0 | 0 | 0 | 0 | 0 | 0 | 0 | 0 | 0 | 0 | NA | NA | NA | NA | NA | NA | NA | NA | NA | NA |
| Vmn1r231 | 0 | 0 | 0 | 0 | 0 | 0 | 0 | 0 | 0 | 0 | 0 | 0 | 0 | 0 | 0 | NA | NA | NA | NA | NA | NA | NA | NA | NA | NA |
| Vmn1r232 | 0 | 0 | 0 | 0 | 0 | 0 | 0 | 0 | 0 | 0 | 0 | 0 | 0 | 0 | 0 | NA | NA | NA | NA | NA | NA | NA | NA | NA | NA |
| Vmn1r233 | 0 | 0 | 0 | 0 | 0 | 0 | 0 | 0 | 0 | 0 | 0 | 0 | 0 | 0 | 0 | NA | NA | NA | NA | NA | NA | NA | NA | NA | NA |
| Vmn1r234 | 0 | 0 | 0 | 0 | 0 | 0 | 0 | 0 | 0 | 0 | 0 | 0 | 0 | 0 | 0 | NA | NA | NA | NA | NA | NA | NA | NA | NA | NA |
| Vmn1r235 | 0 | 0 | 0 | 0 | 0 | 0 | 0 | 0 | 0 | 0 | 0 | 0 | 0 | 0 | 0 | NA | NA | NA | NA | NA | NA | NA | NA | NA | NA |
| Vmn1r236 | 0 | 0 | 0 | 0 | 0 | 0 | 0 | 0 | 0 | 0 | 0 | 0 | 0 | 0 | 0 | NA | NA | NA | NA | NA | NA | NA | NA | NA | NA |
| Vmn1r237 | 0 | 0 | 0 | 0 | 0 | 0 | 0 | 0 | 0 | 0 | 0 | 0 | 0 | 0 | 0 | NA | NA | NA | NA | NA | NA | NA | NA | NA | NA |
| Vmn1r238 | 0 | 0 | 0 | 0 | 0 | 0 | 0 | 0 | 0 | 0 | 0 | 0 | 0 | 0 | 0 | NA | NA | NA | NA | NA | NA | NA | NA | NA | NA |
| Vmn1r24 | 0 | 0 | 0 | 0 | 0 | 0 | 0 | 0 | 0 | 0 | 0 | 0 | 0 | 0 | 0 | NA | NA | NA | NA | NA | NA | NA | NA | NA | NA |
| Vmn1r25 | 0 | 0 | 0 | 0 | 0 | 0 | 0 | 0 | 0 | 0 | 0 | 0 | 0 | 0 | 0 | NA | NA | NA | NA | NA | NA | NA | NA | NA | NA |
| Vmn1r26 | 0 | 0 | 0 | 0 | 0 | 0 | 0 | 0 | 0 | 0 | 0 | 0 | 0 | 0 | 0 | NA | NA | NA | NA | NA | NA | NA | NA | NA | NA |
| Vmn1r27 | 0 | 0 | 0 | 0 | 0 | 0 | 0 | 0 | 0 | 0 | 0 | 0 | 0 | 0 | 0 | NA | NA | NA | NA | NA | NA | NA | NA | NA | NA |
| Vmn1r28 | 0 | 0 | 0 | 0 | 0 | 0 | 0 | 0 | 0 | 0 | 0 | 0 | 0 | 0 | 0 | NA | NA | NA | NA | NA | NA | NA | NA | NA | NA |
| Vmn1r29 | 1 | 2 | 0 | 0 | 1 | 2 | 1 | 2 | 0 | 0 | 1 | 2 | 1 | 1 | 1 | 1.074 | 0.102 | 0.95894372 | NA | 0 | 2.3699 | 1.8538 | 1.8538 | TRUE | 0.3275 |
| Vmn1r3 | 0 | 0 | 0 | 0 | 0 | 0 | 0 | 0 | 0 | 0 | 0 | 0 | 0 | 0 | 0 | NA | NA | NA | NA | NA | NA | NA | NA | NA | NA |
| Vmn1r30 | 0 | 0 | 0 | 0 | 0 | 0 | 0 | 0 | 0 | 0 | 0 | 0 | 0 | 0 | 0 | NA | NA | NA | NA | NA | NA | NA | NA | NA | NA |
| Vmn1r31 | 0 | 0 | 0 | 0 | 0 | 0 | 0 | 0 | 0 | 0 | 0 | 0 | 0 | 0 | 0 | NA | NA | NA | NA | NA | NA | NA | NA | NA | NA |
| Vmn1r32 | 4 | 1 | 0 | 0 | 1 | 1 | 3 | 1 | 0 | 0 | 1 | 1 | 1.05 | 1 | 1 | 0.485 | -1.045 | 0.60183738 | NA | 0 | 2.2648 | 1.813 | 1.813 | TRUE | 0.3335 |
| Vmn1r33 | 0 | 0 | 0 | 0 | 0 | 0 | 0 | 0 | 0 | 0 | 0 | 0 | 0 | 0 | 0 | NA | NA | NA | NA | NA | NA | NA | NA | NA | NA |
| Vmn1r34 | 0 | 0 | 0 | 0 | 0 | 0 | 0 | 0 | 0 | 0 | 0 | 0 | 0 | 0 | 0 | NA | NA | NA | NA | NA | NA | NA | NA | NA | NA |
| Vmn1r35 | 0 | 0 | 0 | 0 | 0 | 0 | 0 | 0 | 0 | 0 | 0 | 0 | 0 | 0 | 0 | NA | NA | NA | NA | NA | NA | NA | NA | NA | NA |
| Vmn1r36 | 0 | 0 | 0 | 0 | 0 | 0 | 0 | 0 | 0 | 0 | 0 | 0 | 0 | 0 | 0 | NA | NA | NA | NA | NA | NA | NA | NA | NA | NA |
| Vmn1r37 | 0 | 0 | 0 | 0 | 0 | 0 | 0 | 0 | 0 | 0 | 0 | 0 | 0 | 0 | 0 | NA | NA | NA | NA | NA | NA | NA | NA | NA | NA |
| Vmn1r38 | 0 | 0 | 0 | 0 | 0 | 0 | 0 | 0 | 0 | 0 | 0 | 0 | 0 | 0 | 0 | NA | NA | NA | NA | NA | NA | NA | NA | NA | NA |
| Vmn1r39 | 0 | 0 | 0 | 0 | 0 | 0 | 0 | 0 | 0 | 0 | 0 | 0 | 0 | 0 | 0 | NA | NA | NA | NA | NA | NA | NA | NA | NA | NA |
| Vmn1r4 | 0 | 0 | 0 | 0 | 0 | 0 | 0 | 0 | 0 | 0 | 0 | 0 | 0 | 0 | 0 | NA | NA | NA | NA | NA | NA | NA | NA | NA | NA |
| Vmn1r40 | 0 | 0 | 0 | 0 | 0 | 0 | 0 | 0 | 0 | 0 | 0 | 0 | 0 | 0 | 0 | NA | NA | NA | NA | NA | NA | NA | NA | NA | NA |
| Vmn1r41 | 0 | 0 | 0 | 0 | 0 | 0 | 0 | 0 | 0 | 0 | 0 | 0 | 0 | 0 | 0 | NA | NA | NA | NA | NA | NA | NA | NA | NA | NA |
| Vmn1r42 | 0 | 0 | 0 | 0 | 0 | 0 | 0 | 0 | 0 | 0 | 0 | 0 | 0 | 0 | 0 | NA | NA | NA | NA | NA | NA | NA | NA | NA | NA |
| Vmn1r43 | 0 | 1 | 0 | 0 | 0 | 0 | 0 | 1 | 0 | 0 | 0 | 0 | 0.18 | 0 | 0 | 0.552 | -0.857 | 0.8336548 | NA | 0 | 13.3714 | 10 | 10 | TRUE | 0.5216 |
| Vmn1r44 | 0 | 0 | 0 | 0 | 0 | 0 | 0 | 0 | 0 | 0 | 0 | 0 | 0 | 0 | 0 | NA | NA | NA | NA | NA | NA | NA | NA | NA | NA |
| Vmn1r45 | 0 | 0 | 0 | 0 | 0 | 0 | 0 | 0 | 0 | 0 | 0 | 0 | 0 | 0 | 0 | NA | NA | NA | NA | NA | NA | NA | NA | NA | NA |
| Vmn1r46 | 0 | 0 | 0 | 0 | 0 | 0 | 0 | 0 | 0 | 0 | 0 | 0 | 0 | 0 | 0 | NA | NA | NA | NA | NA | NA | NA | NA | NA | NA |
| Vmn1r47 | 0 | 0 | 0 | 0 | 0 | 0 | 0 | 0 | 0 | 0 | 0 | 0 | 0 | 0 | 0 | NA | NA | NA | NA | NA | NA | NA | NA | NA | NA |
| Vmn1r48 | 0 | 0 | 0 | 0 | 0 | 0 | 0 | 0 | 0 | 0 | 0 | 0 | 0 | 0 | 0 | NA | NA | NA | NA | NA | NA | NA | NA | NA | NA |
| Vmn1r49 | 0 | 0 | 0 | 0 | 0 | 0 | 0 | 0 | 0 | 0 | 0 | 0 | 0 | 0 | 0 | NA | NA | NA | NA | NA | NA | NA | NA | NA | NA |
| Vmn1r5 | 0 | 0 | 0 | 0 | 0 | 0 | 0 | 0 | 0 | 0 | 0 | 0 | 0 | 0 | 0 | NA | NA | NA | NA | NA | NA | NA | NA | NA | NA |
| Vmn1r50 | 0 | 0 | 0 | 0 | 0 | 0 | 0 | 0 | 0 | 0 | 0 | 0 | 0 | 0 | 0 | NA | NA | NA | NA | NA | NA | NA | NA | NA | NA |
| Vmn1r51 | 0 | 0 | 0 | 0 | 0 | 0 | 0 | 0 | 0 | 0 | 0 | 0 | 0 | 0 | 0 | NA | NA | NA | NA | NA | NA | NA | NA | NA | NA |
| Vmn1r52 | 0 | 0 | 0 | 0 | 0 | 0 | 0 | 0 | 0 | 0 | 0 | 0 | 0 | 0 | 0 | NA | NA | NA | NA | NA | NA | NA | NA | NA | NA |
| Vmn1r53 | 83 | 46 | 53 | 41 | 36 | 35 | 64 | 49 | 57 | 41 | 41 | 35 | 47.76 | 57 | 39 | 0.676 | -0.564 | 0.04740371 | 0.12959856 | 0 | 0.0521 | 0.0365 | 0.0365 | TRUE | 0.1325 |
| Vmn1r54 | 0 | 0 | 0 | 0 | 0 | 0 | 0 | 0 | 0 | 0 | 0 | 0 | 0 | 0 | 0 | NA | NA | NA | NA | NA | NA | NA | NA | NA | NA |
| Vmn1r55 | 0 | 0 | 0 | 0 | 0 | 0 | 0 | 0 | 0 | 0 | 0 | 0 | 0 | 0 | 0 | NA | NA | NA | NA | NA | NA | NA | NA | NA | NA |
| Vmn1r56 | 0 | 0 | 0 | 0 | 0 | 0 | 0 | 0 | 0 | 0 | 0 | 0 | 0 | 0 | 0 | NA | NA | NA | NA | NA | NA | NA | NA | NA | NA |
| Vmn1r57 | 0 | 0 | 0 | 0 | 0 | 0 | 0 | 0 | 0 | 0 | 0 | 0 | 0 | 0 | 0 | NA | NA | NA | NA | NA | NA | NA | NA | NA | NA |
| Vmn1r58 | 0 | 0 | 0 | 0 | 0 | 0 | 0 | 0 | 0 | 0 | 0 | 0 | 0 | 0 | 0 | NA | NA | NA | NA | NA | NA | NA | NA | NA | NA |
| Vmn1r59 | 0 | 0 | 0 | 0 | 0 | 0 | 0 | 0 | 0 | 0 | 0 | 0 | 0 | 0 | 0 | NA | NA | NA | NA | NA | NA | NA | NA | NA | NA |
| Vmn1r6 | 0 | 0 | 0 | 0 | 0 | 0 | 0 | 0 | 0 | 0 | 0 | 0 | 0 | 0 | 0 | NA | NA | NA | NA | NA | NA | NA | NA | NA | NA |
| Vmn1r60 | 0 | 0 | 0 | 0 | 0 | 0 | 0 | 0 | 0 | 0 | 0 | 0 | 0 | 0 | 0 | NA | NA | NA | NA | NA | NA | NA | NA | NA | NA |
| Vmn1r61 | 0 | 0 | 0 | 0 | 0 | 0 | 0 | 0 | 0 | 0 | 0 | 0 | 0 | 0 | 0 | NA | NA | NA | NA | NA | NA | NA | NA | NA | NA |
| Vmn1r62 | 0 | 0 | 0 | 0 | 0 | 0 | 0 | 0 | 0 | 0 | 0 | 0 | 0 | 0 | 0 | NA | NA | NA | NA | NA | NA | NA | NA | NA | NA |
| Vmn1r63 | 0 | 0 | 0 | 0 | 0 | 0 | 0 | 0 | 0 | 0 | 0 | 0 | 0 | 0 | 0 | NA | NA | NA | NA | NA | NA | NA | NA | NA | NA |
| Vmn1r64 | 0 | 0 | 0 | 0 | 0 | 0 | 0 | 0 | 0 | 0 | 0 | 0 | 0 | 0 | 0 | NA | NA | NA | NA | NA | NA | NA | NA | NA | NA |
| Vmn1r65 | 0 | 0 | 0 | 0 | 0 | 0 | 0 | 0 | 0 | 0 | 0 | 0 | 0 | 0 | 0 | NA | NA | NA | NA | NA | NA | NA | NA | NA | NA |
| Vmn1r66 | 0 | 0 | 0 | 0 | 0 | 0 | 0 | 0 | 0 | 0 | 0 | 0 | 0 | 0 | 0 | NA | NA | NA | NA | NA | NA | NA | NA | NA | NA |

| | | | | | | | | | | | | | | | | | | | | | | | | | | |
|---|---|---|---|---|---|---|---|---|---|---|---|---|---|---|---|---|---|---|---|---|---|---|---|---|---|---|
| Vmn1r67 | 0 | 0 | 0 | 0 | 0 | 0 | 0 | 0 | 0 | 0 | 0 | 0 | 0 | 0 | 0 | NA | NA | NA | NA | NA | NA | NA | NA | NA | NA | |
| Vmn1r68 | 0 | 0 | 0 | 0 | 0 | 0 | 0 | 0 | 0 | 0 | 0 | 0 | 0 | 0 | 0 | NA | NA | NA | NA | NA | NA | NA | NA | NA | NA | |
| Vmn1r69 | 0 | 0 | 0 | 0 | 0 | 0 | 0 | 0 | 0 | 0 | 0 | 0 | 0 | 0 | 0 | NA | NA | NA | NA | NA | NA | NA | NA | NA | NA | |
| Vmn1r7 | 0 | 0 | 0 | 0 | 0 | 0 | 0 | 0 | 0 | 0 | 0 | 0 | 0 | 0 | 0 | NA | NA | NA | NA | NA | NA | NA | NA | NA | NA | |
| Vmn1r70 | 0 | 0 | 0 | 0 | 0 | 0 | 0 | 0 | 0 | 0 | 0 | 0 | 0 | 0 | 0 | NA | NA | NA | NA | NA | NA | NA | NA | NA | NA | |
| Vmn1r71 | 0 | 0 | 0 | 0 | 0 | 0 | 0 | 0 | 0 | 0 | 0 | 0 | 0 | 0 | 0 | NA | NA | NA | NA | NA | NA | NA | NA | NA | NA | |
| Vmn1r72 | 0 | 0 | 0 | 0 | 0 | 0 | 0 | 0 | 0 | 0 | 0 | 0 | 0 | 0 | 0 | NA | NA | NA | NA | NA | NA | NA | NA | NA | NA | |
| Vmn1r73 | 0 | 0 | 0 | 0 | 0 | 0 | 0 | 0 | 0 | 0 | 0 | 0 | 0 | 0 | 0 | NA | NA | NA | NA | NA | NA | NA | NA | NA | NA | |
| Vmn1r74 | 0 | 0 | 0 | 0 | 0 | 0 | 0 | 0 | 0 | 0 | 0 | 0 | 0 | 0 | 0 | NA | NA | NA | NA | NA | NA | NA | NA | NA | NA | |
| Vmn1r75 | 0 | 0 | 0 | 0 | 0 | 0 | 0 | 0 | 0 | 0 | 0 | 0 | 0 | 0 | 0 | NA | NA | NA | NA | NA | NA | NA | NA | NA | NA | |
| Vmn1r76 | 0 | 0 | 0 | 0 | 0 | 0 | 0 | 0 | 0 | 0 | 0 | 0 | 0 | 0 | 0 | NA | NA | NA | NA | NA | NA | NA | NA | NA | NA | |
| Vmn1r77 | 0 | 0 | 0 | 0 | 0 | 0 | 0 | 0 | 0 | 0 | 0 | 0 | 0 | 0 | 0 | NA | NA | NA | NA | NA | NA | NA | NA | NA | NA | |
| Vmn1r78 | 0 | 0 | 0 | 0 | 0 | 0 | 0 | 0 | 0 | 0 | 0 | 0 | 0 | 0 | 0 | NA | NA | NA | NA | NA | NA | NA | NA | NA | NA | |
| Vmn1r79 | 0 | 0 | 0 | 0 | 0 | 0 | 0 | 0 | 0 | 0 | 0 | 0 | 0 | 0 | 0 | NA | NA | NA | NA | NA | NA | NA | NA | NA | NA | |
| Vmn1r8 | 0 | 0 | 0 | 0 | 0 | 0 | 0 | 0 | 0 | 0 | 0 | 0 | 0 | 0 | 0 | NA | NA | NA | NA | NA | NA | NA | NA | NA | NA | |
| Vmn1r80 | 0 | 0 | 0 | 0 | 0 | 0 | 0 | 0 | 0 | 0 | 0 | 0 | 0 | 0 | 0 | NA | NA | NA | NA | NA | NA | NA | NA | NA | NA | |
| Vmn1r81 | 0 | 0 | 0 | 0 | 0 | 0 | 0 | 0 | 0 | 0 | 0 | 0 | 0 | 0 | 0 | NA | NA | NA | NA | NA | NA | NA | NA | NA | NA | |
| Vmn1r82 | 0 | 0 | 0 | 0 | 0 | 0 | 0 | 0 | 0 | 0 | 0 | 0 | 0 | 0 | 0 | NA | NA | NA | NA | NA | NA | NA | NA | NA | NA | |
| Vmn1r83 | 0 | 0 | 0 | 0 | 0 | 0 | 0 | 0 | 0 | 0 | 0 | 0 | 0 | 0 | 0 | NA | NA | NA | NA | NA | NA | NA | NA | NA | NA | |
| Vmn1r84 | 0 | 0 | 0 | 0 | 0 | 0 | 0 | 0 | 0 | 0 | 0 | 0 | 0 | 0 | 0 | NA | NA | NA | NA | NA | NA | NA | NA | NA | NA | |
| Vmn1r85 | 0 | 0 | 0 | 0 | 0 | 0 | 0 | 0 | 0 | 0 | 0 | 0 | 0 | 0 | 0 | NA | NA | NA | NA | NA | NA | NA | NA | NA | NA | |
| Vmn1r86 | 0 | 0 | 0 | 0 | 0 | 0 | 0 | 0 | 0 | 0 | 0 | 0 | 0 | 0 | 0 | NA | NA | NA | NA | NA | NA | NA | NA | NA | NA | |
| Vmn1r87 | 0 | 0 | 0 | 0 | 0 | 0 | 0 | 0 | 0 | 0 | 0 | 0 | 0 | 0 | 0 | NA | NA | NA | NA | NA | NA | NA | NA | NA | NA | |
| Vmn1r88 | 0 | 0 | 0 | 0 | 0 | 0 | 0 | 0 | 0 | 0 | 0 | 0 | 0 | 0 | 0 | NA | NA | NA | NA | NA | NA | NA | NA | NA | NA | |
| Vmn1r89 | 0 | 0 | 0 | 0 | 0 | 0 | 0 | 0 | 0 | 0 | 0 | 0 | 0 | 0 | 0 | NA | NA | NA | NA | NA | NA | NA | NA | NA | NA | |
| Vmn1r9 | 0 | 0 | 0 | 0 | 0 | 0 | 0 | 0 | 0 | 0 | 0 | 0 | 0 | 0 | 0 | NA | NA | NA | NA | NA | NA | NA | NA | NA | NA | |
| Vmn1r90 | 0 | 0 | 0 | 0 | 0 | 0 | 0 | 0 | 0 | 0 | 0 | 0 | 0 | 0 | 0 | NA | NA | NA | NA | NA | NA | NA | NA | NA | NA | |
| Vmn1r91 | 0 | 0 | 0 | 0 | 0 | 0 | 0 | 0 | 0 | 0 | 0 | 0 | 0 | 0 | 0 | NA | NA | NA | NA | NA | NA | NA | NA | NA | NA | |
| Vmn1r93 | 0 | 0 | 0 | 0 | 0 | 0 | 0 | 0 | 0 | 0 | 0 | 0 | 0 | 0 | 0 | NA | NA | NA | NA | NA | NA | NA | NA | NA | NA | |
| Vmn1r94 | 0 | 0 | 0 | 0 | 0 | 0 | 0 | 0 | 0 | 0 | 0 | 0 | 0 | 0 | 0 | NA | NA | NA | NA | NA | NA | NA | NA | NA | NA | |
| Vmn1r95 | 0 | 0 | 0 | 0 | 0 | 0 | 0 | 0 | 0 | 0 | 0 | 0 | 0 | 0 | 0 | NA | NA | NA | NA | NA | NA | NA | NA | NA | NA | |
| Vmn2r-ps11 | 0 | 0 | 0 | 0 | 0 | 0 | 0 | 0 | 0 | 0 | 0 | 0 | 0 | 0 | 0 | NA | NA | NA | NA | NA | NA | NA | NA | NA | NA | |
| Vmn2r-ps12 | 0 | 0 | 0 | 1 | 0 | 0 | 0 | 0 | 0 | 1 | 0 | 0 | 0.17 | 0 | 0 | 2.094 | 1.067 | 0.79379329 | NA | 0 | 14.2885 | 10 | 10 | TRUE | | 0.3906 |
| Vmn2r-ps15 | 0 | 0 | 0 | 0 | 0 | 0 | 0 | 0 | 0 | 0 | 0 | 0 | 0 | 0 | 0 | NA | NA | NA | NA | NA | NA | NA | NA | NA | NA | |
| Vmn2r-ps54 | 1 | 1 | 2 | 1 | 2 | 3 | 1 | 1 | 2 | 1 | 2 | 3 | 1.7 | 1 | 2 | 1.586 | 0.665 | 0.64486667 | NA | 0 | 1.395 | 0.8824 | 0.8824 | TRUE | | 0.1994 |
| Vmn2r-ps60 | 0 | 0 | 0 | 0 | 0 | 0 | 0 | 0 | 0 | 0 | 0 | 0 | 0 | 0 | 0 | NA | NA | NA | NA | NA | NA | NA | NA | NA | NA | |
| Vmn2r1 | 5 | 14 | 6 | 5 | 6 | 9 | 4 | 15 | 6 | 5 | 7 | 9 | 7.65 | 8 | 7 | 0.831 | -0.267 | 0.72068841 | NA | 0.1685 | 0.3124 | 0.2681 | 0.2681 | TRUE | | 0.9726 |
| Vmn2r10 | 0 | 0 | 0 | 0 | 0 | 0 | 0 | 0 | 0 | 0 | 0 | 0 | 0 | 0 | 0 | NA | NA | NA | NA | NA | NA | NA | NA | NA | NA | |
| Vmn2r100 | 0 | 0 | 0 | 0 | 0 | 0 | 0 | 0 | 0 | 0 | 0 | 0 | 0 | 0 | 0 | NA | NA | NA | NA | NA | NA | NA | NA | NA | NA | |
| Vmn2r101 | 0 | 0 | 0 | 0 | 0 | 0 | 0 | 0 | 0 | 0 | 0 | 0 | 0 | 0 | 0 | NA | NA | NA | NA | NA | NA | NA | NA | NA | NA | |
| Vmn2r102 | 0 | 0 | 0 | 0 | 0 | 0 | 0 | 0 | 0 | 0 | 0 | 0 | 0 | 0 | 0 | NA | NA | NA | NA | NA | NA | NA | NA | NA | NA | |
| Vmn2r103 | 0 | 0 | 0 | 0 | 0 | 0 | 0 | 0 | 0 | 0 | 0 | 0 | 0 | 0 | 0 | NA | NA | NA | NA | NA | NA | NA | NA | NA | NA | |
| Vmn2r104 | 0 | 0 | 0 | 0 | 0 | 0 | 0 | 0 | 0 | 0 | 0 | 0 | 0 | 0 | 0 | NA | NA | NA | NA | NA | NA | NA | NA | NA | NA | |
| Vmn2r105 | 0 | 0 | 0 | 0 | 0 | 0 | 0 | 0 | 0 | 0 | 0 | 0 | 0 | 0 | 0 | NA | NA | NA | NA | NA | NA | NA | NA | NA | NA | |
| Vmn2r106 | 0 | 0 | 0 | 0 | 0 | 0 | 0 | 0 | 0 | 0 | 0 | 0 | 0 | 0 | 0 | NA | NA | NA | NA | NA | NA | NA | NA | NA | NA | |
| Vmn2r107 | 0 | 0 | 0 | 0 | 0 | 0 | 0 | 0 | 0 | 0 | 0 | 0 | 0 | 0 | 0 | NA | NA | NA | NA | NA | NA | NA | NA | NA | NA | |
| Vmn2r108 | 0 | 0 | 0 | 0 | 0 | 0 | 0 | 0 | 0 | 0 | 0 | 0 | 0 | 0 | 0 | NA | NA | NA | NA | NA | NA | NA | NA | NA | NA | |
| Vmn2r109 | 0 | 0 | 0 | 0 | 0 | 0 | 0 | 0 | 0 | 0 | 0 | 0 | 0 | 0 | 0 | NA | NA | NA | NA | NA | NA | NA | NA | NA | NA | |
| Vmn2r11 | 0 | 0 | 0 | 0 | 0 | 0 | 0 | 0 | 0 | 0 | 0 | 0 | 0 | 0 | 0 | NA | NA | NA | NA | NA | NA | NA | NA | NA | NA | |
| Vmn2r110 | 0 | 0 | 0 | 0 | 0 | 0 | 0 | 0 | 0 | 0 | 0 | 0 | 0 | 0 | 0 | NA | NA | NA | NA | NA | NA | NA | NA | NA | NA | |
| Vmn2r111 | 1 | 2 | 0 | 2 | 0 | 0 | 1 | 2 | 0 | 2 | 0 | 0 | 0.82 | 1 | 1 | 0.698 | -0.518 | 0.82391604 | NA | 0 | 2.9076 | 2.6343 | 2.6343 | TRUE | | 0.7035 |
| Vmn2r112 | 0 | 0 | 0 | 0 | 0 | 0 | 0 | 0 | 0 | 0 | 0 | 0 | 0 | 0 | 0 | NA | NA | NA | NA | NA | NA | NA | NA | NA | NA | |
| Vmn2r113 | 0 | 1 | 0 | 0 | 1 | 1 | 0 | 1 | 0 | 0 | 1 | 1 | 0.53 | 0 | 1 | 2.03 | 1.022 | 0.70625413 | NA | 0 | 4.4757 | 3.5573 | 3.5573 | TRUE | | 0.5217 |
| Vmn2r114 | 0 | 0 | 0 | 0 | 0 | 0 | 0 | 0 | 0 | 0 | 0 | 0 | 0 | 0 | 0 | NA | NA | NA | NA | NA | NA | NA | NA | NA | NA | |
| Vmn2r115 | 0 | 0 | 0 | 0 | 0 | 0 | 0 | 0 | 0 | 0 | 0 | 0 | 0 | 0 | 0 | NA | NA | NA | NA | NA | NA | NA | NA | NA | NA | |
| Vmn2r116 | 0 | 0 | 0 | 0 | 0 | 0 | 0 | 0 | 0 | 0 | 0 | 0 | 0 | 0 | 0 | NA | NA | NA | NA | NA | NA | NA | NA | NA | NA | |
| Vmn2r117 | 0 | 0 | 0 | 0 | 0 | 0 | 0 | 0 | 0 | 0 | 0 | 0 | 0 | 0 | 0 | NA | NA | NA | NA | NA | NA | NA | NA | NA | NA | |
| Vmn2r118 | 0 | 0 | 0 | 0 | 0 | 0 | 0 | 0 | 0 | 0 | 0 | 0 | 0 | 0 | 0 | NA | NA | NA | NA | NA | NA | NA | NA | NA | NA | |
| Vmn2r12 | 0 | 0 | 0 | 0 | 0 | 0 | 0 | 0 | 0 | 0 | 0 | 0 | 0 | 0 | 0 | NA | NA | NA | NA | NA | NA | NA | NA | NA | NA | |
| Vmn2r120 | 0 | 0 | 0 | 0 | 0 | 0 | 0 | 0 | 0 | 0 | 0 | 0 | 0 | 0 | 0 | NA | NA | NA | NA | NA | NA | NA | NA | NA | NA | |
| Vmn2r121 | 0 | 0 | 0 | 0 | 0 | 0 | 0 | 0 | 0 | 0 | 0 | 0 | 0 | 0 | 0 | NA | NA | NA | NA | NA | NA | NA | NA | NA | NA | |
| Vmn2r122 | 0 | 0 | 0 | 0 | 0 | 0 | 0 | 0 | 0 | 0 | 0 | 0 | 0 | 0 | 0 | NA | NA | NA | NA | NA | NA | NA | NA | NA | NA | |
| Vmn2r123 | 0 | 0 | 0 | 0 | 0 | 0 | 0 | 0 | 0 | 0 | 0 | 0 | 0 | 0 | 0 | NA | NA | NA | NA | NA | NA | NA | NA | NA | NA | |
| Vmn2r124 | 0 | 0 | 0 | 0 | 0 | 0 | 0 | 0 | 0 | 0 | 0 | 0 | 0 | 0 | 0 | NA | NA | NA | NA | NA | NA | NA | NA | NA | NA | |
| Vmn2r13 | 0 | 0 | 0 | 0 | 0 | 0 | 0 | 0 | 0 | 0 | 0 | 0 | 0 | 0 | 0 | NA | NA | NA | NA | NA | NA | NA | NA | NA | NA | |
| Vmn2r14 | 0 | 0 | 0 | 0 | 0 | 0 | 0 | 0 | 0 | 0 | 0 | 0 | 0 | 0 | 0 | NA | NA | NA | NA | NA | NA | NA | NA | NA | NA | |
| Vmn2r15 | 0 | 0 | 0 | 0 | 0 | 0 | 0 | 0 | 0 | 0 | 0 | 0 | 0 | 0 | 0 | NA | NA | NA | NA | NA | NA | NA | NA | NA | NA | |
| Vmn2r16 | 0 | 0 | 0 | 0 | 0 | 0 | 0 | 0 | 0 | 0 | 0 | 0 | 0 | 0 | 0 | NA | NA | NA | NA | NA | NA | NA | NA | NA | NA | |
| Vmn2r17 | 0 | 0 | 0 | 0 | 0 | 0 | 0 | 0 | 0 | 0 | 0 | 0 | 0 | 0 | 0 | NA | NA | NA | NA | NA | NA | NA | NA | NA | NA | |

| | | | | | | | | | | | | | | | | | | | | | | | | | |
|---|---|---|---|---|---|---|---|---|---|---|---|---|---|---|---|---|---|---|---|---|---|---|---|---|---|
| Vmn2r18 | 4 | 2 | 0 | 1 | 1 | 2 | 3 | 2 | 0 | 1 | 1 | 2 | 1.56 | 2 | 1 | 0.764 | -0.388 | 0.80391439 | NA | 0 | 1.5263 | 1.1091 | 1.1091 | TRUE | 0.4538 |
| Vmn2r19 | 0 | 0 | 0 | 0 | 0 | 0 | 0 | 0 | 0 | 0 | 0 | 0 | 0 | 0 | 0 | NA | NA | NA | NA | NA | NA | NA | NA | NA | NA |
| Vmn2r2 | 0 | 0 | 0 | 0 | 0 | 0 | 0 | 0 | 0 | 0 | 0 | 0 | 0 | 0 | 0 | NA | NA | NA | NA | NA | NA | NA | NA | NA | NA |
| Vmn2r20 | 0 | 0 | 0 | 0 | 0 | 0 | 0 | 0 | 0 | 0 | 0 | 0 | 0 | 0 | 0 | NA | NA | NA | NA | NA | NA | NA | NA | NA | NA |
| Vmn2r21 | 0 | 0 | 0 | 0 | 0 | 0 | 0 | 0 | 0 | 0 | 0 | 0 | 0 | 0 | 0 | NA | NA | NA | NA | NA | NA | NA | NA | NA | NA |
| Vmn2r22 | 0 | 0 | 0 | 0 | 0 | 0 | 0 | 0 | 0 | 0 | 0 | 0 | 0 | 0 | 0 | NA | NA | NA | NA | NA | NA | NA | NA | NA | NA |
| Vmn2r23 | 0 | 0 | 0 | 0 | 0 | 0 | 0 | 0 | 0 | 0 | 0 | 0 | 0 | 0 | 0 | NA | NA | NA | NA | NA | NA | NA | NA | NA | NA |
| Vmn2r24 | 0 | 0 | 0 | 0 | 0 | 0 | 0 | 0 | 0 | 0 | 0 | 0 | 0 | 0 | 0 | NA | NA | NA | NA | NA | NA | NA | NA | NA | NA |
| Vmn2r25 | 0 | 0 | 0 | 0 | 0 | 0 | 0 | 0 | 0 | 0 | 0 | 0 | 0 | 0 | 0 | NA | NA | NA | NA | NA | NA | NA | NA | NA | NA |
| Vmn2r26 | 0 | 0 | 0 | 0 | 0 | 0 | 0 | 0 | 0 | 0 | 0 | 0 | 0 | 0 | 0 | NA | NA | NA | NA | NA | NA | NA | NA | NA | NA |
| Vmn2r27 | 4 | 2 | 2 | 1 | 4 | 2 | 3 | 2 | 2 | 1 | 5 | 2 | 2.48 | 2 | 3 | 0.988 | -0.017 | 0.98857953 | NA | 0 | 0.9585 | 0.6408 | 0.6408 | TRUE | 0.4887 |
| Vmn2r28 | 0 | 1 | 0 | 0 | 0 | 0 | 0 | 1 | 0 | 0 | 0 | 0 | 0.18 | 0 | 0 | 0.552 | -0.857 | 0.8336548 | NA | 0 | 13.3714 | 10 | 10 | TRUE | 0.5216 |
| Vmn2r29 | 1 | 1 | 1 | 1 | 0 | 2 | 1 | 1 | 1 | 1 | 0 | 2 | 0.98 | 1 | 1 | 1.044 | 0.062 | 0.97363744 | NA | 0 | 2.4171 | 1.5475 | 1.5475 | TRUE | 0.1802 |
| Vmn2r3 | 0 | 2 | 1 | 0 | 0 | 0 | 0 | 2 | 1 | 0 | 0 | 0 | 0.53 | 1 | 0 | 0.181 | -2.465 | 0.39476468 | NA | 0 | 4.4375 | 4.5834 | 4.5834 | TRUE | 0.0598 |
| Vmn2r30 | 0 | 0 | 0 | 0 | 0 | 0 | 0 | 0 | 0 | 0 | 0 | 0 | 0 | 0 | 0 | NA | NA | NA | NA | NA | NA | NA | NA | NA | NA |
| Vmn2r31 | 0 | 0 | 0 | 0 | 0 | 0 | 0 | 0 | 0 | 0 | 0 | 0 | 0 | 0 | 0 | NA | NA | NA | NA | NA | NA | NA | NA | NA | NA |
| Vmn2r32 | 0 | 0 | 0 | 0 | 0 | 0 | 0 | 0 | 0 | 0 | 0 | 0 | 0 | 0 | 0 | NA | NA | NA | NA | NA | NA | NA | NA | NA | NA |
| Vmn2r33 | 0 | 0 | 0 | 0 | 0 | 0 | 0 | 0 | 0 | 0 | 0 | 0 | 0 | 0 | 0 | NA | NA | NA | NA | NA | NA | NA | NA | NA | NA |
| Vmn2r34 | 0 | 0 | 0 | 0 | 0 | 0 | 0 | 0 | 0 | 0 | 0 | 0 | 0 | 0 | 0 | NA | NA | NA | NA | NA | NA | NA | NA | NA | NA |
| Vmn2r35 | 0 | 0 | 0 | 0 | 0 | 0 | 0 | 0 | 0 | 0 | 0 | 0 | 0 | 0 | 0 | NA | NA | NA | NA | NA | NA | NA | NA | NA | NA |
| Vmn2r36 | 0 | 0 | 0 | 0 | 0 | 0 | 0 | 0 | 0 | 0 | 0 | 0 | 0 | 0 | 0 | NA | NA | NA | NA | NA | NA | NA | NA | NA | NA |
| Vmn2r37 | 0 | 0 | 0 | 0 | 0 | 0 | 0 | 0 | 0 | 0 | 0 | 0 | 0 | 0 | 0 | NA | NA | NA | NA | NA | NA | NA | NA | NA | NA |
| Vmn2r38 | 0 | 0 | 0 | 0 | 0 | 0 | 0 | 0 | 0 | 0 | 0 | 0 | 0 | 0 | 0 | NA | NA | NA | NA | NA | NA | NA | NA | NA | NA |
| Vmn2r39 | 0 | 0 | 0 | 0 | 0 | 0 | 0 | 0 | 0 | 0 | 0 | 0 | 0 | 0 | 0 | NA | NA | NA | NA | NA | NA | NA | NA | NA | NA |
| Vmn2r4 | 0 | 0 | 0 | 0 | 0 | 0 | 0 | 0 | 0 | 0 | 0 | 0 | 0 | 0 | 0 | NA | NA | NA | NA | NA | NA | NA | NA | NA | NA |
| Vmn2r40 | 0 | 0 | 0 | 0 | 0 | 0 | 0 | 0 | 0 | 0 | 0 | 0 | 0 | 0 | 0 | NA | NA | NA | NA | NA | NA | NA | NA | NA | NA |
| Vmn2r41 | 0 | 0 | 0 | 0 | 0 | 0 | 0 | 0 | 0 | 0 | 0 | 0 | 0 | 0 | 0 | NA | NA | NA | NA | NA | NA | NA | NA | NA | NA |
| Vmn2r42 | 0 | 0 | 0 | 0 | 0 | 0 | 0 | 0 | 0 | 0 | 0 | 0 | 0 | 0 | 0 | NA | NA | NA | NA | NA | NA | NA | NA | NA | NA |
| Vmn2r43 | 0 | 0 | 0 | 0 | 0 | 0 | 0 | 0 | 0 | 0 | 0 | 0 | 0 | 0 | 0 | NA | NA | NA | NA | NA | NA | NA | NA | NA | NA |
| Vmn2r44 | 0 | 0 | 0 | 0 | 0 | 0 | 0 | 0 | 0 | 0 | 0 | 0 | 0 | 0 | 0 | NA | NA | NA | NA | NA | NA | NA | NA | NA | NA |
| Vmn2r45 | 0 | 0 | 0 | 0 | 0 | 0 | 0 | 0 | 0 | 0 | 0 | 0 | 0 | 0 | 0 | NA | NA | NA | NA | NA | NA | NA | NA | NA | NA |
| Vmn2r46 | 0 | 0 | 0 | 0 | 0 | 0 | 0 | 0 | 0 | 0 | 0 | 0 | 0 | 0 | 0 | NA | NA | NA | NA | NA | NA | NA | NA | NA | NA |
| Vmn2r47 | 0 | 0 | 0 | 0 | 0 | 0 | 0 | 0 | 0 | 0 | 0 | 0 | 0 | 0 | 0 | NA | NA | NA | NA | NA | NA | NA | NA | NA | NA |
| Vmn2r48 | 0 | 0 | 0 | 0 | 0 | 0 | 0 | 0 | 0 | 0 | 0 | 0 | 0 | 0 | 0 | NA | NA | NA | NA | NA | NA | NA | NA | NA | NA |
| Vmn2r49 | 0 | 0 | 0 | 0 | 0 | 0 | 0 | 0 | 0 | 0 | 0 | 0 | 0 | 0 | 0 | NA | NA | NA | NA | NA | NA | NA | NA | NA | NA |
| Vmn2r5 | 0 | 0 | 0 | 0 | 0 | 0 | 0 | 0 | 0 | 0 | 0 | 0 | 0 | 0 | 0 | NA | NA | NA | NA | NA | NA | NA | NA | NA | NA |
| Vmn2r50 | 0 | 0 | 0 | 0 | 0 | 0 | 0 | 0 | 0 | 0 | 0 | 0 | 0 | 0 | 0 | NA | NA | NA | NA | NA | NA | NA | NA | NA | NA |
| Vmn2r51 | 0 | 0 | 0 | 0 | 0 | 0 | 0 | 0 | 0 | 0 | 0 | 0 | 0 | 0 | 0 | NA | NA | NA | NA | NA | NA | NA | NA | NA | NA |
| Vmn2r52 | 0 | 0 | 0 | 0 | 0 | 0 | 0 | 0 | 0 | 0 | 0 | 0 | 0 | 0 | 0 | NA | NA | NA | NA | NA | NA | NA | NA | NA | NA |
| Vmn2r53 | 0 | 0 | 0 | 0 | 0 | 0 | 0 | 0 | 0 | 0 | 0 | 0 | 0 | 0 | 0 | NA | NA | NA | NA | NA | NA | NA | NA | NA | NA |
| Vmn2r54 | 0 | 1 | 0 | 0 | 0 | 0 | 0 | 1 | 0 | 0 | 0 | 0 | 0.18 | 0 | 0 | 0.552 | -0.857 | 0.8336548 | NA | 0 | 13.3714 | 10 | 10 | TRUE | 0.5216 |
| Vmn2r55 | 0 | 0 | 0 | 0 | 0 | 0 | 0 | 0 | 0 | 0 | 0 | 0 | 0 | 0 | 0 | NA | NA | NA | NA | NA | NA | NA | NA | NA | NA |
| Vmn2r56 | 0 | 0 | 0 | 0 | 0 | 0 | 0 | 0 | 0 | 0 | 0 | 0 | 0 | 0 | 0 | NA | NA | NA | NA | NA | NA | NA | NA | NA | NA |
| Vmn2r57 | 15 | 8 | 11 | 25 | 13 | 16 | 12 | 9 | 12 | 25 | 15 | 16 | 14.56 | 11 | 19 | 1.726 | 0.787 | 0.12764377 | NA | 0 | 0.1652 | 0.1195 | 0.1195 | TRUE | 0.4889 |
| Vmn2r58 | 0 | 0 | 0 | 0 | 0 | 0 | 0 | 0 | 0 | 0 | 0 | 0 | 0 | 0 | 0 | NA | NA | NA | NA | NA | NA | NA | NA | NA | NA |
| Vmn2r59 | 0 | 0 | 0 | 0 | 0 | 0 | 0 | 0 | 0 | 0 | 0 | 0 | 0 | 0 | 0 | NA | NA | NA | NA | NA | NA | NA | NA | NA | NA |
| Vmn2r6 | 0 | 0 | 0 | 0 | 0 | 0 | 0 | 0 | 0 | 0 | 0 | 0 | 0 | 0 | 0 | NA | NA | NA | NA | NA | NA | NA | NA | NA | NA |
| Vmn2r60 | 0 | 0 | 0 | 0 | 0 | 0 | 0 | 0 | 0 | 0 | 0 | 0 | 0 | 0 | 0 | NA | NA | NA | NA | NA | NA | NA | NA | NA | NA |
| Vmn2r61 | 0 | 0 | 1 | 0 | 0 | 0 | 0 | 0 | 1 | 0 | 0 | 0 | 0.18 | 0 | 0 | 0.552 | -0.857 | 0.8336548 | NA | 0 | 13.1817 | 10 | 10 | TRUE | 0.5365 |
| Vmn2r62 | 0 | 0 | 0 | 0 | 1 | 0 | 0 | 0 | 0 | 0 | 1 | 0 | 0.19 | 0 | 0 | 2.094 | 1.067 | 0.79379329 | NA | 0 | 12.608 | 10 | 10 | TRUE | 0.5075 |
| Vmn2r63 | 0 | 0 | 0 | 0 | 0 | 0 | 0 | 0 | 0 | 0 | 0 | 0 | 0 | 0 | 0 | NA | NA | NA | NA | NA | NA | NA | NA | NA | NA |
| Vmn2r65 | 0 | 0 | 0 | 0 | 0 | 0 | 0 | 0 | 0 | 0 | 0 | 0 | 0 | 0 | 0 | NA | NA | NA | NA | NA | NA | NA | NA | NA | NA |
| Vmn2r66 | 0 | 0 | 0 | 0 | 0 | 0 | 0 | 0 | 0 | 0 | 0 | 0 | 0 | 0 | 0 | NA | NA | NA | NA | NA | NA | NA | NA | NA | NA |
| Vmn2r67 | 0 | 0 | 0 | 0 | 0 | 0 | 0 | 0 | 0 | 0 | 0 | 0 | 0 | 0 | 0 | NA | NA | NA | NA | NA | NA | NA | NA | NA | NA |
| Vmn2r68 | 0 | 0 | 0 | 0 | 0 | 0 | 0 | 0 | 0 | 0 | 0 | 0 | 0 | 0 | 0 | NA | NA | NA | NA | NA | NA | NA | NA | NA | NA |
| Vmn2r69 | 0 | 0 | 0 | 0 | 0 | 0 | 0 | 0 | 0 | 0 | 0 | 0 | 0 | 0 | 0 | NA | NA | NA | NA | NA | NA | NA | NA | NA | NA |
| Vmn2r7 | 0 | 0 | 0 | 0 | 0 | 0 | 0 | 0 | 0 | 0 | 0 | 0 | 0 | 0 | 0 | NA | NA | NA | NA | NA | NA | NA | NA | NA | NA |
| Vmn2r70 | 0 | 0 | 0 | 0 | 0 | 0 | 0 | 0 | 0 | 0 | 0 | 0 | 0 | 0 | 0 | NA | NA | NA | NA | NA | NA | NA | NA | NA | NA |
| Vmn2r71 | 0 | 0 | 0 | 0 | 0 | 0 | 0 | 0 | 0 | 0 | 0 | 0 | 0 | 0 | 0 | NA | NA | NA | NA | NA | NA | NA | NA | NA | NA |
| Vmn2r72 | 0 | 0 | 0 | 0 | 0 | 0 | 0 | 0 | 0 | 0 | 0 | 0 | 0 | 0 | 0 | NA | NA | NA | NA | NA | NA | NA | NA | NA | NA |
| Vmn2r73 | 0 | 0 | 0 | 0 | 0 | 0 | 0 | 0 | 0 | 0 | 0 | 0 | 0 | 0 | 0 | NA | NA | NA | NA | NA | NA | NA | NA | NA | NA |
| Vmn2r74 | 0 | 0 | 0 | 0 | 0 | 0 | 0 | 0 | 0 | 0 | 0 | 0 | 0 | 0 | 0 | NA | NA | NA | NA | NA | NA | NA | NA | NA | NA |
| Vmn2r75 | 0 | 0 | 0 | 0 | 0 | 0 | 0 | 0 | 0 | 0 | 0 | 0 | 0 | 0 | 0 | NA | NA | NA | NA | NA | NA | NA | NA | NA | NA |
| Vmn2r76 | 0 | 0 | 0 | 0 | 0 | 0 | 0 | 0 | 0 | 0 | 0 | 0 | 0 | 0 | 0 | NA | NA | NA | NA | NA | NA | NA | NA | NA | NA |
| Vmn2r77 | 0 | 0 | 0 | 0 | 0 | 0 | 0 | 0 | 0 | 0 | 0 | 0 | 0 | 0 | 0 | NA | NA | NA | NA | NA | NA | NA | NA | NA | NA |
| Vmn2r78 | 0 | 0 | 0 | 0 | 0 | 0 | 0 | 0 | 0 | 0 | 0 | 0 | 0 | 0 | 0 | NA | NA | NA | NA | NA | NA | NA | NA | NA | NA |
| Vmn2r79 | 0 | 0 | 0 | 0 | 0 | 0 | 0 | 0 | 0 | 0 | 0 | 0 | 0 | 0 | 0 | NA | NA | NA | NA | NA | NA | NA | NA | NA | NA |
| Vmn2r8 | 0 | 0 | 0 | 0 | 0 | 0 | 0 | 0 | 0 | 0 | 0 | 0 | 0 | 0 | 0 | NA | NA | NA | NA | NA | NA | NA | NA | NA | NA |
| Vmn2r80 | 0 | 0 | 0 | 0 | 0 | 0 | 0 | 0 | 0 | 0 | 0 | 0 | 0 | 0 | 0 | NA | NA | NA | NA | NA | NA | NA | NA | NA | NA |

| | | | | | | | | | | | | | | | | | | | | | | | | | |
|---|---|---|---|---|---|---|---|---|---|---|---|---|---|---|---|---|---|---|---|---|---|---|---|---|---|
| Vmn2r81 | 0 | 0 | 0 | 0 | 0 | 0 | 0 | 0 | 0 | 0 | 0 | 0 | 0 | 0 | 0 | NA | NA | NA | NA | NA | NA | NA | NA | NA | NA |
| Vmn2r82 | 0 | 0 | 0 | 0 | 0 | 0 | 0 | 0 | 0 | 0 | 0 | 0 | 0 | 0 | 0 | NA | NA | NA | NA | NA | NA | NA | NA | NA | NA |
| Vmn2r83 | 0 | 0 | 0 | 0 | 0 | 0 | 0 | 0 | 0 | 0 | 0 | 0 | 0 | 0 | 0 | NA | NA | NA | NA | NA | NA | NA | NA | NA | NA |
| Vmn2r84 | 1 | 2 | 0 | 0 | 0 | 0 | 1 | 2 | 0 | 0 | 0 | 0 | 0.48 | 1 | 0 | 0.197 | -2.343 | 0.43249452 | NA | 0 | 4.8992 | 4.9172 | 4.9172 | TRUE | 0.1226 |
| Vmn2r85 | 0 | 0 | 0 | 0 | 0 | 1 | 0 | 0 | 0 | 0 | 0 | 1 | 0.16 | 0 | 0 | 2.094 | 1.067 | 0.79379329 | NA | 0 | 14.4078 | 10 | 10 | TRUE | 0.3835 |
| Vmn2r86 | 0 | 0 | 0 | 0 | 0 | 0 | 0 | 0 | 0 | 0 | 0 | 0 | 0 | 0 | 0 | NA | NA | NA | NA | NA | NA | NA | NA | NA | NA |
| Vmn2r87 | 0 | 0 | 0 | 0 | 0 | 1 | 0 | 0 | 0 | 0 | 0 | 1 | 0.16 | 0 | 0 | 2.094 | 1.067 | 0.79379329 | NA | 0 | 14.4078 | 10 | 10 | TRUE | 0.3835 |
| Vmn2r88 | 0 | 0 | 0 | 0 | 0 | 0 | 0 | 0 | 0 | 0 | 0 | 0 | 0 | 0 | 0 | NA | NA | NA | NA | NA | NA | NA | NA | NA | NA |
| Vmn2r89 | 1 | 0 | 1 | 0 | 0 | 0 | 1 | 0 | 1 | 0 | 0 | 0 | 0.31 | 1 | 0 | 0.307 | -1.706 | 0.62894649 | NA | 0 | 7.6657 | 7.2 | 7.2 | TRUE | 0.2067 |
| Vmn2r9 | 0 | 0 | 0 | 0 | 0 | 0 | 0 | 0 | 0 | 0 | 0 | 0 | 0 | 0 | 0 | NA | NA | NA | NA | NA | NA | NA | NA | NA | NA |
| Vmn2r90 | 3 | 5 | 3 | 2 | 0 | 3 | 2 | 5 | 3 | 2 | 0 | 3 | 2.64 | 3 | 2 | 0.469 | -1.091 | 0.3804242 | NA | 0 | 0.9003 | 0.6742 | 0.6742 | TRUE | 0.4867 |
| Vmn2r91 | 0 | 0 | 0 | 0 | 0 | 0 | 0 | 0 | 0 | 0 | 0 | 0 | 0 | 0 | 0 | NA | NA | NA | NA | NA | NA | NA | NA | NA | NA |
| Vmn2r92 | 0 | 0 | 0 | 0 | 0 | 0 | 0 | 0 | 0 | 0 | 0 | 0 | 0 | 0 | 0 | NA | NA | NA | NA | NA | NA | NA | NA | NA | NA |
| Vmn2r93 | 0 | 0 | 0 | 0 | 0 | 0 | 0 | 0 | 0 | 0 | 0 | 0 | 0 | 0 | 0 | NA | NA | NA | NA | NA | NA | NA | NA | NA | NA |
| Vmn2r94 | 4 | 4 | 0 | 2 | 2 | 1 | 3 | 4 | 0 | 2 | 2 | 1 | 2.1 | 2 | 2 | 0.7 | -0.514 | 0.71093656 | NA | 0 | 1.1314 | 0.8922 | 0.8922 | TRUE | 0.6151 |
| Vmn2r95 | 0 | 1 | 0 | 0 | 0 | 0 | 0 | 1 | 0 | 0 | 0 | 0 | 0.18 | 0 | 0 | 0.552 | -0.857 | 0.8336548 | NA | 0 | 13.3714 | 10 | 10 | TRUE | 0.5216 |
| Vmn2r96 | 0 | 0 | 0 | 0 | 0 | 0 | 0 | 0 | 0 | 0 | 0 | 0 | 0 | 0 | 0 | NA | NA | NA | NA | NA | NA | NA | NA | NA | NA |
| Vmn2r97 | 0 | 0 | 0 | 0 | 0 | 0 | 0 | 0 | 0 | 0 | 0 | 0 | 0 | 0 | 0 | NA | NA | NA | NA | NA | NA | NA | NA | NA | NA |
| Vmn2r98 | 0 | 0 | 0 | 0 | 0 | 0 | 0 | 0 | 0 | 0 | 0 | 0 | 0 | 0 | 0 | NA | NA | NA | NA | NA | NA | NA | NA | NA | NA |
| Vmn2r99 | 0 | 0 | 0 | 0 | 0 | 0 | 0 | 0 | 0 | 0 | 0 | 0 | 0 | 0 | 0 | NA | NA | NA | NA | NA | NA | NA | NA | NA | NA |
| Vmo1 | 1 | 2 | 3 | 7 | 3 | 4 | 1 | 2 | 3 | 7 | 3 | 4 | 3.41 | 2 | 5 | 2.418 | 1.274 | 0.23875042 | NA | 0 | 0.6981 | 0.4913 | 0.4913 | TRUE | 0.3946 |
| Vmp1 | 5953 | 3961 | 4068 | 4652 | 3910 | 4418 | 4623 | 4213 | 4389 | 4631 | 4411 | 4361 | 4438.13 | 4408 | 4468 | 1.013 | 0.019 | 0.75382093 | 0.85391912 | 0.0014 | 0.003 | 0.0023 | 0.0023 | TRUE | 0.0225 |
| Vnn1 | 83 | 46 | 31 | 10 | 16 | 8 | 64 | 49 | 33 | 10 | 18 | 8 | 30.46 | 49 | 12 | 0.24 | -2.058 | 2.17E-06 | 5.37E-05 | 0.0891 | 0.0802 | 0.0825 | 0.0825 | TRUE | 0.1608 |
| Vnn3 | 0 | 1 | 0 | 0 | 0 | 0 | 0 | 0 | 1 | 0 | 0 | 0 | 0.18 | 0 | 0 | 0.552 | -0.857 | 0.8336548 | NA | 0 | 13.3714 | 10 | 10 | TRUE | 0.5216 |
| Vopp1 | 581 | 415 | 480 | 470 | 350 | 421 | 451 | 441 | 518 | 468 | 395 | 416 | 448.14 | 470 | 426 | 0.908 | -0.139 | 0.21714346 | 0.380002 | 0.0053 | 0.0077 | 0.007 | 0.007 | TRUE | 0.0888 |
| Vprbp | 9750 | 7021 | 6985 | 7624 | 6850 | 7820 | 7572 | 7468 | 7537 | 7589 | 7728 | 7720 | 7602.22 | 7526 | 7679 | 1.02 | 0.029 | 0.53359441 | 0.6929552 | 0 | 0.0027 | 0.0014 | 0.0014 | TRUE | 0.0013 |
| Vpreb1 | 21 | 14 | 21 | 9 | 12 | 16 | 16 | 15 | 23 | 9 | 14 | 16 | 15.36 | 18 | 13 | 0.713 | -0.488 | 0.33008788 | NA | 0 | 0.1567 | 0.1136 | 0.1136 | TRUE | 0.2906 |
| Vpreb2 | 1 | 0 | 1 | 0 | 0 | 2 | 1 | 0 | 1 | 0 | 0 | 2 | 0.64 | 1 | 1 | 1.073 | 0.101 | 0.96834165 | NA | 0 | 3.7154 | 3.1348 | 3.1348 | TRUE | 0.9736 |
| Vpreb3 | 8 | 8 | 11 | 18 | 3 | 13 | 6 | 9 | 12 | 18 | 3 | 13 | 10.12 | 9 | 11 | 1.308 | 0.387 | 0.56766946 | NA | 0.2159 | 0.2366 | 0.231 | 0.231 | TRUE | 0.3451 |
| Vps11 | 4084 | 2938 | 2866 | 3015 | 2784 | 3204 | 3172 | 3125 | 3092 | 3001 | 3141 | 3163 | 3115.66 | 3130 | 3102 | 0.991 | -0.013 | 0.80930479 | 0.8899364 | 0 | 0.0032 | 0.0019 | 0.0019 | TRUE | 0.0098 |
| Vps13a | 4369 | 3201 | 3175 | 4010 | 3418 | 3934 | 3393 | 3405 | 3426 | 3992 | 3856 | 3883 | 3659.13 | 3408 | 3910 | 1.148 | 0.199 | 0.00015171 | 0.00172505 | 0 | 0.0031 | 0.0017 | 0.0017 | TRUE | 0.004 |
| Vps13b | 4615 | 3308 | 3311 | 3760 | 3521 | 3854 | 3584 | 3519 | 3573 | 3743 | 3972 | 3804 | 3699.12 | 3559 | 3840 | 1.079 | 0.109 | 0.04458309 | 0.1238294 | 2.00E-04 | 0.0031 | 0.0019 | 0.0019 | TRUE | 0.0109 |
| Vps13c | 6152 | 4254 | 4266 | 5624 | 4722 | 5311 | 4778 | 4525 | 4603 | 5598 | 5327 | 5243 | 5012.28 | 4635 | 5389 | 1.162 | 0.217 | 8.31E-05 | 0.00104018 | 7.00E-04 | 0.0029 | 0.002 | 0.002 | TRUE | 0.0141 |
| Vps13d | 4525 | 3214 | 3245 | 4097 | 3862 | 4455 | 3514 | 3419 | 3501 | 4078 | 4357 | 4398 | 3877.84 | 3478 | 4278 | 1.23 | 0.298 | 1.46E-07 | 5.45E-06 | 7.00E-04 | 0.003 | 0.0021 | 0.0021 | TRUE | 0.0204 |
| Vps16 | 5158 | 3967 | 4051 | 4244 | 3838 | 4564 | 4006 | 4220 | 4371 | 4224 | 4330 | 4505 | 4276.02 | 4199 | 4353 | 1.037 | 0.053 | 0.37096274 | 0.5456592 | 0.0011 | 0.003 | 0.0023 | 0.0023 | TRUE | 0.0201 |
| Vps18 | 1944 | 1325 | 1194 | 1281 | 1071 | 1153 | 1510 | 1409 | 1288 | 1275 | 1208 | 1138 | 1304.84 | 1402 | 1207 | 0.86 | -0.218 | 0.00859065 | 0.03698379 | 0.0041 | 0.0042 | 0.0042 | 0.0042 | TRUE | 0.0606 |
| Vps25 | 4983 | 3441 | 3295 | 3476 | 3182 | 3597 | 3870 | 3660 | 3555 | 3460 | 3590 | 3551 | 3614.32 | 3695 | 3534 | 0.956 | -0.065 | 0.26048735 | 0.43057079 | 9.00E-04 | 0.0031 | 0.0022 | 0.0022 | TRUE | 0.0213 |
| Vps26a | 4301 | 3243 | 3304 | 3711 | 3222 | 3676 | 3340 | 3450 | 3565 | 3694 | 3635 | 3629 | 3552.06 | 3452 | 3653 | 1.059 | 0.082 | 0.13606508 | 0.27573772 | 3.00E-04 | 0.0031 | 0.0019 | 0.0019 | TRUE | 0.01 |
| Vps26b | 4579 | 3311 | 3456 | 3362 | 3023 | 3411 | 3556 | 3522 | 3729 | 3347 | 3410 | 3367 | 3488.52 | 3602 | 3375 | 0.937 | -0.094 | 0.0868467 | 0.19979074 | 2.00E-04 | 0.0031 | 0.0019 | 0.0019 | TRUE | 0.0113 |
| Vps28 | 1822 | 1329 | 1518 | 1572 | 1523 | 1668 | 1415 | 1414 | 1638 | 1565 | 1718 | 1647 | 1566.01 | 1489 | 1643 | 1.104 | 0.143 | 0.07405409 | 0.1786584 | 0.0039 | 0.0039 | 0.004 | 0.004 | TRUE | 0.091 |
| Vps29 | 3835 | 2665 | 2595 | 2733 | 2414 | 2755 | 2978 | 2835 | 2800 | 2720 | 2723 | 2720 | 2796.08 | 2871 | 2721 | 0.947 | -0.078 | 0.17270363 | 0.32505324 | 3.00E-04 | 0.0033 | 0.002 | 0.002 | TRUE | 0.0134 |
| Vps33a | 2894 | 1991 | 1898 | 1834 | 1711 | 1954 | 2248 | 2118 | 2048 | 1826 | 1930 | 1929 | 2016.34 | 2138 | 1895 | 0.886 | -0.175 | 0.00819162 | 0.03566794 | 0.0012 | 0.0036 | 0.0027 | 0.0027 | TRUE | 0.0252 |
| Vps33b | 1416 | 1124 | 1011 | 992 | 954 | 1069 | 1100 | 1196 | 1091 | 987 | 1076 | 1055 | 1084.18 | 1129 | 1039 | 0.921 | -0.118 | 0.12960992 | 0.26635725 | 0.0013 | 0.0046 | 0.0035 | 0.0035 | TRUE | 0.0315 |
| Vps35 | 25394 | 18351 | 19009 | 19912 | 17547 | 19854 | 19722 | 19520 | 20511 | 19820 | 19795 | 19599 | 19827.82 | 19918 | 19738 | 0.991 | -0.013 | 0.77636358 | 0.86766476 | 3.00E-04 | 0.0025 | 0.0015 | 0.0015 | TRUE | 0.0083 |
| Vps36 | 4305 | 3184 | 3168 | 3609 | 3049 | 3553 | 3344 | 3387 | 3418 | 3592 | 3440 | 3507 | 3447.98 | 3383 | 3513 | 1.039 | 0.055 | 0.30631779 | 0.47877557 | 0 | 0.0031 | 0.0018 | 0.0018 | TRUE | 0.0047 |
| Vps37a | 2776 | 2031 | 2028 | 2192 | 1872 | 2116 | 2156 | 2160 | 2188 | 2182 | 2112 | 2089 | 2147.85 | 2168 | 2128 | 0.981 | -0.027 | 0.6467969 | 0.78033832 | 0 | 0.0035 | 0.0021 | 0.0021 | TRUE | 0.0061 |
| Vps37b | 1300 | 976 | 852 | 841 | 783 | 821 | 1010 | 1038 | 919 | 837 | 883 | 810 | 916.33 | 989 | 843 | 0.852 | -0.231 | 0.00568851 | 0.02710195 | 0.0018 | 0.005 | 0.0039 | 0.0039 | TRUE | 0.044 |
| Vps37c | 898 | 662 | 648 | 665 | 696 | 753 | 697 | 704 | 699 | 662 | 785 | 743 | 715.2 | 700 | 730 | 1.042 | 0.059 | 0.51567001 | 0.67800192 | 0.0023 | 0.0057 | 0.0046 | 0.0046 | TRUE | 0.0794 |
| Vps37d | 155 | 115 | 172 | 187 | 167 | 194 | 120 | 122 | 186 | 186 | 188 | 192 | 165.72 | 143 | 189 | 1.328 | 0.409 | 0.02763225 | 0.0871657 | 0.0246 | 0.0167 | 0.0187 | 0.0187 | TRUE | 0.6951 |
| Vps39 | 3731 | 2759 | 3005 | 3269 | 2907 | 3364 | 2898 | 2935 | 3242 | 3254 | 3279 | 3321 | 3154.82 | 3025 | 3285 | 1.086 | 0.12 | 0.05847103 | 0.15071711 | 0.0015 | 0.0032 | 0.0026 | 0.0026 | TRUE | 0.0476 |
| Vps41 | 7896 | 5805 | 6083 | 6601 | 5758 | 6484 | 6132 | 6175 | 6564 | 6571 | 6496 | 6401 | 6389.61 | 6290 | 6489 | 1.032 | 0.045 | 0.39070969 | 0.5645332 | 6.00E-04 | 0.0028 | 0.0018 | 0.0018 | TRUE | 0.0175 |
| Vps45 | 988 | 617 | 637 | 847 | 701 | 778 | 767 | 656 | 687 | 843 | 791 | 768 | 752.14 | 703 | 801 | 1.136 | 0.183 | 0.04652221 | 0.12783363 | 0.0032 | 0.0056 | 0.0048 | 0.0048 | TRUE | 0.0779 |
| Vps4a | 3399 | 2383 | 2465 | 2722 | 2433 | 2815 | 2640 | 2535 | 2660 | 2709 | 2745 | 2779 | 2677.89 | 2612 | 2744 | 1.051 | 0.071 | 0.21016028 | 0.37212846 | 0 | 0.0033 | 0.002 | 0.002 | TRUE | 0.0078 |
| Vps4b | 6326 | 4590 | 4479 | 4208 | 3770 | 4400 | 4913 | 4882 | 4833 | 4189 | 4253 | 4343 | 4568.89 | 4876 | 4262 | 0.874 | -0.194 | 0.00012526 | 0.00146625 | 0 | 0.0029 | 0.0016 | 0.0016 | TRUE | 0.0035 |
| Vps51 | 1646 | 1190 | 1211 | 1199 | 1167 | 1281 | 1278 | 1266 | 1307 | 1193 | 1316 | 1265 | 1270.89 | 1284 | 1258 | 0.98 | -0.029 | 0.68270019 | 0.80543407 | 4.00E-04 | 0.0043 | 0.0029 | 0.0029 | TRUE | 0.0243 |
| Vps52 | 2121 | 1585 | 1229 | 1328 | 1214 | 1409 | 1647 | 1686 | 1326 | 1322 | 1370 | 1391 | 1456.94 | 1553 | 1361 | 0.876 | -0.192 | 0.03782518 | 0.10992992 | 0.0074 | 0.0041 | 0.0055 | 0.0055 | TRUE | 0.1941 |
| Vps53 | 3664 | 2598 | 2141 | 2437 | 2301 | 2694 | 2846 | 2763 | 2310 | 2426 | 2596 | 2659 | 2600.03 | 2640 | 2560 | 0.969 | -0.045 | 0.58898216 | 0.73786678 | 0.007 | 0.0033 | 0.0046 | 0.0046 | TRUE | 0.1436 |
| Vps54 | 7490 | 5400 | 5567 | 5094 | 4422 | 5064 | 5817 | 5744 | 6007 | 5071 | 4988 | 4999 | 5437.63 | 5856 | 5019 | 0.857 | -0.222 | 9.62E-06 | 0.00018279 | 1.00E-04 | 0.0029 | 0.0016 | 0.0016 | TRUE | 0.0061 |
| Vps72 | 2808 | 2119 | 2007 | 2095 | 1966 | 2256 | 2181 | 2254 | 2166 | 2085 | 2218 | 2227 | 2188.43 | 2200 | 2177 | 0.989 | -0.015 | 0.80327182 | 0.88613128 | 4.00E-04 | 0.0035 | 0.0023 | 0.0023 | TRUE | 0.0165 |
| Vps8 | 1564 | 1146 | 1125 | 1277 | 1126 | 1249 | 1215 | 1219 | 1214 | 1271 | 1270 | 1233 | 1236.98 | 1216 | 1258 | 1.035 | 0.049 | 0.46961246 | 0.63864614 | 0 | 0.0043 | 0.0025 | 0.0025 | TRUE | 0.0037 |
| Vps9d1 | 794 | 598 | 545 | 506 | 511 | 558 | 617 | 636 | 588 | 504 | 576 | 551 | 578.63 | 614 | 544 | 0.885 | -0.176 | 0.06533867 | 0.16323816 | 0.0011 | 0.0065 | 0.0048 | 0.0048 | TRUE | 0.0487 |
| Vrk1 | 4317 | 3064 | 3062 | 3167 | 2842 | 3294 | 3353 | 3259 | 3304 | 3152 | 3206 | 3252 | 3254.34 | 3305 | 3203 | 0.969 | -0.045 | 0.39823235 | 0.57159419 | 0 | 0.0032 | 0.0018 | 0.0018 | TRUE | 0.0024 |
| Vrk2 | 784 | 555 | 587 | 633 | 464 | 623 | 609 | 590 | 633 | 630 | 523 | 615 | 600.19 | 611 | 589 | 0.966 | -0.049 | 0.61957072 | 0.75986994 | 0.0035 | 0.0064 | 0.0055 | 0.0055 | TRUE | 0.1097 |
| Vrk3 | 2171 | 1678 | 1854 | 1896 | 1776 | 2123 | 1686 | 1785 | 2000 | 1887 | 2004 | 2096 | 1909.66 | 1824 | 1996 | 1.095 | 0.131 | 0.09938816 | 0.22020695 | 0.0045 | 0.0037 | 0.004 | 0.004 | TRUE | 0.0863 |
| Vrtn | 60 | 24 | 19 | 66 | 53 | 45 | 47 | 26 | 21 | 66 | 60 | 44 | 43.76 | 31 | 57 | 1.802 | 0.849 | 0.01537103 | 0.05687241 | 0.084 | 0.0566 | 0.064 | 0.064 | TRUE | 1.5494 |
| Vsig1 | 11 | 4 | 14 | 8 | 11 | 13 | 9 | 4 | 15 | 8 | 12 | 13 | 10.18 | 9 | 11 | 1.193 | 0.255 | 0.69071124 | NA | 0.0835 | 0.2351 | 0.1959 | 0.1959 | TRUE | 0.3627 |
| Vsig10 | 1138 | 822 | 970 | 1107 | 1024 | 1214 | 884 | 874 | 1047 | 1102 | 1155 | 1198 | 1043.38 | 935 | 1152 | 1.233 | 0.303 | 0.00070183 | 0.00557603 | 0.0048 | 0.0047 | 0.0048 | 0.0048 | TRUE | 0.1275 |
| Vsig2 | 149 | 92 | 127 | 116 | 109 | 120 | 116 | 98 | 137 | 115 | 123 | 118 | 117.92 | 117 | 119 | 1.018 | 0.026 | 0.89249999 | 0.94095319 | 0.0046 | 0.0225 | 0.0177 | 0.0177 | TRUE | 0.2126 |
| Vsig4 | 0 | 0 | 0 | 0 | 0 | 0 | 0 | 0 | 0 | 0 | 0 | 0 | 0 | 0 | 0 | NA | NA | NA | NA | NA | NA | NA | NA | NA | NA |

| | | | | | | | | | | | | | | | | | | | | | | | | | |
|---|---|---|---|---|---|---|---|---|---|---|---|---|---|---|---|---|---|---|---|---|---|---|---|---|---|
| Vsig8 | 6 | 2 | 6 | 9 | 15 | 17 | 5 | 2 | 6 | 9 | 17 | 17 | 9.32 | 4 | 14 | 3.205 | 1.68 | 0.01515533 | NA | 0 | 0.2567 | 0.1998 | 0.1998 | TRUE | 0.4729 |
| Vsnl1 | 15 | 8 | 7 | 6 | 10 | 7 | 12 | 9 | 8 | 6 | 11 | 7 | 8.65 | 10 | 8 | 0.858 | -0.222 | 0.73486703 | NA | 0 | 0.2765 | 0.1922 | 0.1922 | TRUE | 0.3252 |
| Vstm2a | 17 | 9 | 6 | 1 | 7 | 8 | 13 | 10 | 6 | 1 | 8 | 8 | 7.67 | 10 | 6 | 0.565 | -0.824 | 0.2862812 | NA | 0.2028 | 0.3113 | 0.2879 | 0.2879 | TRUE | 0.6833 |
| Vstm2b | 45 | 35 | 22 | 32 | 25 | 23 | 35 | 37 | 24 | 32 | 28 | 23 | 29.78 | 32 | 28 | 0.859 | -0.219 | 0.54838135 | 0.70505571 | 0 | 0.082 | 0.0619 | 0.0619 | TRUE | 0.3202 |
| Vstm2l | 107 | 88 | 113 | 163 | 182 | 172 | 83 | 94 | 122 | 162 | 205 | 170 | 139.33 | 100 | 179 | 1.808 | 0.854 | 1.01E-05 | 0.00019108 | 0.0186 | 0.0194 | 0.0192 | 0.0192 | TRUE | 0.3685 |
| Vstm4 | 10 | 4 | 4 | 6 | 9 | 8 | 8 | 4 | 4 | 6 | 10 | 8 | 6.73 | 5 | 8 | 1.438 | 0.524 | 0.47969601 | NA | 0 | 0.3548 | 0.2439 | 0.2439 | TRUE | 0.3967 |
| Vstm5 | 100 | 85 | 90 | 120 | 84 | 109 | 78 | 90 | 97 | 119 | 95 | 108 | 97.83 | 88 | 107 | 1.221 | 0.288 | 0.16080604 | 0.31055273 | 0 | 0.0267 | 0.0201 | 0.0201 | TRUE | 0.1226 |
| Vsx1 | 10 | 5 | 7 | 17 | 10 | 19 | 8 | 5 | 8 | 17 | 11 | 19 | 11.27 | 7 | 16 | 2.27 | 1.182 | 0.04477077 | NA | 0 | 0.2128 | 0.1477 | 0.1477 | TRUE | 0.2528 |
| Vsx2 | 16 | 9 | 9 | 18 | 9 | 12 | 12 | 10 | 10 | 18 | 10 | 12 | 11.94 | 11 | 13 | 1.254 | 0.326 | 0.55823578 | NA | 0 | 0.201 | 0.14 | 0.14 | TRUE | 0.3937 |
| Vta1 | 2525 | 1872 | 1972 | 2050 | 1866 | 2103 | 1961 | 1991 | 2128 | 2041 | 2105 | 2076 | 2050.27 | 2027 | 2074 | 1.024 | 0.034 | 0.59341716 | 0.74082553 | 6.00E-04 | 0.0036 | 0.0024 | 0.0024 | TRUE | 0.0227 |
| Vtcn1 | 7 | 5 | 3 | 1 | 2 | 2 | 5 | 5 | 3 | 1 | 2 | 2 | 3.2 | 4 | 2 | 0.369 | -1.438 | 0.19364971 | NA | 0 | 0.7424 | 0.4804 | 0.4804 | TRUE | 0.1245 |
| Vti1a | 748 | 533 | 571 | 491 | 506 | 536 | 581 | 567 | 616 | 489 | 571 | 529 | 558.78 | 588 | 530 | 0.9 | -0.152 | 0.12241388 | 0.25564898 | 0.0018 | 0.0067 | 0.0052 | 0.0052 | TRUE | 0.0532 |
| Vti1b | 2832 | 1885 | 1843 | 2171 | 1798 | 2119 | 2199 | 2005 | 1989 | 2161 | 2028 | 2092 | 2079.04 | 2064 | 2094 | 1.013 | 0.019 | 0.7765917 | 0.867784 | 0.0017 | 0.0036 | 0.0028 | 0.0028 | TRUE | 0.0409 |
| Vtn | 5 | 1 | 3 | 8 | 2 | 2 | 4 | 1 | 3 | 8 | 2 | 2 | 3.4 | 3 | 4 | 1.474 | 0.56 | 0.60798359 | NA | 0.2284 | 0.7003 | 0.5567 | 0.5567 | TRUE | 1.2196 |
| Vwa1 | 115 | 82 | 59 | 100 | 79 | 94 | 89 | 87 | 64 | 100 | 89 | 93 | 86.94 | 80 | 94 | 1.167 | 0.223 | 0.30798222 | 0.48060171 | 1.00E-04 | 0.0297 | 0.0231 | 0.0231 | TRUE | 0.2821 |
| Vwa2 | 252 | 171 | 207 | 212 | 201 | 224 | 196 | 182 | 223 | 211 | 227 | 221 | 209.98 | 200 | 220 | 1.097 | 0.134 | 0.35156752 | 0.52609837 | 0 | 0.0137 | 0.01 | 0.01 | TRUE | 0.1034 |
| Vwa3a | 3 | 1 | 3 | 1 | 4 | 0 | 2 | 1 | 3 | 1 | 5 | 0 | 2.02 | 2 | 2 | 0.81 | -0.304 | 0.83294344 | NA | 0.4097 | 1.1739 | 0.9855 | 0.9855 | TRUE | 1.2095 |
| Vwa5a | 1977 | 1550 | 1990 | 1845 | 1760 | 1952 | 1535 | 1649 | 2147 | 1837 | 1985 | 1927 | 1846.71 | 1777 | 1916 | 1.079 | 0.11 | 0.30330714 | 0.47586926 | 0.0162 | 0.0037 | 0.0077 | 0.0077 | TRUE | 0.3991 |
| Vwa5b1 | 4 | 5 | 1 | 4 | 3 | 3 | 3 | 5 | 1 | 4 | 3 | 3 | 3.31 | 3 | 3 | 1.086 | 0.119 | 0.90961379 | NA | 0 | 0.7195 | 0.4941 | 0.4941 | TRUE | 0.1997 |
| Vwa5b2 | 50 | 39 | 32 | 35 | 33 | 42 | 39 | 41 | 35 | 35 | 37 | 41 | 38.06 | 38 | 38 | 0.988 | -0.017 | 0.95572261 | 0.97558515 | 0 | 0.0647 | 0.0426 | 0.0426 | TRUE | 0.0534 |
| Vwa7 | 9 | 15 | 11 | 20 | 16 | 22 | 7 | 16 | 12 | 20 | 18 | 22 | 15.75 | 12 | 20 | 1.747 | 0.805 | 0.11038787 | NA | 0 | 0.1529 | 0.1149 | 0.1149 | TRUE | 0.4725 |
| Vwa8 | 1808 | 1343 | 1162 | 1361 | 1171 | 1387 | 1404 | 1429 | 1254 | 1355 | 1321 | 1369 | 1355.24 | 1362 | 1348 | 0.989 | -0.015 | 0.83777241 | 0.90748625 | 0.0018 | 0.0042 | 0.0033 | 0.0033 | TRUE | 0.0568 |
| Vwa9 | 1962 | 1319 | 1222 | 1235 | 1055 | 1218 | 1524 | 1403 | 1319 | 1229 | 1190 | 1202 | 1311.19 | 1415 | 1207 | 0.852 | -0.231 | 0.00268916 | 0.01514425 | 0.0022 | 0.0042 | 0.0035 | 0.0035 | TRUE | 0.0562 |
| Vwc2 | 24 | 15 | 7 | 9 | 16 | 4 | 19 | 16 | 8 | 9 | 18 | 4 | 12.18 | 14 | 10 | 0.72 | -0.473 | 0.45929427 | NA | 0.2454 | 0.1969 | 0.21 | 0.21 | TRUE | 0.5601 |
| Vwc2l | 0 | 0 | 0 | 0 | 0 | 0 | 0 | 0 | 0 | 0 | 0 | 0 | 0 | 0 | 0 | NA | NA | NA | NA | NA | NA | NA | NA | NA | NA |
| Vwce | 55 | 31 | 40 | 52 | 47 | 57 | 43 | 33 | 43 | 52 | 53 | 56 | 46.65 | 40 | 54 | 1.351 | 0.434 | 0.12548661 | 0.26066038 | 0 | 0.0532 | 0.0361 | 0.0361 | TRUE | 0.1503 |
| Vwde | 9 | 2 | 2 | 1 | 0 | 1 | 7 | 2 | 2 | 1 | 0 | 1 | 2.21 | 4 | 1 | 0.177 | -2.498 | 0.10394854 | NA | 0 | 1.075 | 0.8137 | 0.8137 | TRUE | 1.1509 |
| Vwf | 783 | 675 | 920 | 933 | 1075 | 1174 | 608 | 718 | 993 | 929 | 1213 | 1159 | 936.52 | 773 | 1100 | 1.424 | 0.51 | 0.0339323 | 0.1015695 | 0.0406 | 0.005 | 0.0149 | 0.0406 | TRUE | 0.7319 |
| Wac | 4964 | 3667 | 4062 | 3721 | 3411 | 3769 | 3855 | 3901 | 4383 | 3704 | 3848 | 3721 | 3901.86 | 4046 | 3758 | 0.929 | -0.106 | 0.10417433 | 0.2280888 | 0.0024 | 0.003 | 0.0028 | 0.0028 | TRUE | 0.064 |
| Wap | 0 | 0 | 0 | 0 | 0 | 0 | 0 | 0 | 0 | 0 | 0 | 0 | 0 | 0 | 0 | NA | NA | NA | NA | NA | NA | NA | NA | NA | NA |
| Wapal | 14835 | 10597 | 10585 | 11396 | 9897 | 11494 | 11522 | 11272 | 11421 | 11344 | 11165 | 11346 | 11344.92 | 11405 | 11285 | 0.989 | -0.015 | 0.73343752 | 0.84073346 | 0 | 0.0026 | 0.0014 | 0.0014 | TRUE | 0.0013 |
| Wars | 6871 | 4803 | 4217 | 4250 | 3898 | 4356 | 5336 | 5109 | 4550 | 4230 | 4397 | 4300 | 4653.87 | 4998 | 4309 | 0.862 | -0.215 | 0.00155157 | 0.00989583 | 0.0031 | 0.0029 | 0.0031 | 0.0031 | TRUE | 0.0747 |
| Wars2 | 219 | 207 | 152 | 134 | 112 | 131 | 170 | 220 | 164 | 133 | 126 | 129 | 157.22 | 185 | 129 | 0.704 | -0.507 | 0.00334877 | 0.0180143 | 0.0091 | 0.0175 | 0.0149 | 0.0149 | TRUE | 0.2934 |
| Was | 112 | 93 | 69 | 75 | 94 | 72 | 87 | 99 | 74 | 75 | 106 | 71 | 85.36 | 87 | 84 | 0.963 | -0.054 | 0.81811071 | 0.89536894 | 0.0216 | 0.0302 | 0.0278 | 0.0278 | TRUE | 0.4773 |
| Wasf1 | 4292 | 3026 | 2907 | 2847 | 2603 | 3103 | 3333 | 3219 | 3137 | 2834 | 2936 | 3063 | 3087.04 | 3230 | 2944 | 0.911 | -0.134 | 0.02558066 | 0.08265164 | 9.00E-04 | 0.0032 | 0.0023 | 0.0023 | TRUE | 0.0153 |
| Wasf2 | 9379 | 6386 | 6332 | 7071 | 6366 | 7049 | 7284 | 6793 | 6832 | 7038 | 7182 | 6958 | 7014.6 | 6970 | 7059 | 1.013 | 0.018 | 0.73483049 | 0.8415811 | 6.00E-04 | 0.0028 | 0.0019 | 0.0019 | TRUE | 0.0193 |
| Wasf3 | 2965 | 2374 | 2800 | 3022 | 2797 | 3280 | 2303 | 2525 | 3021 | 3008 | 3155 | 3238 | 2875.07 | 2616 | 3134 | 1.199 | 0.261 | 0.00381724 | 0.0198722 | 0.01 | 0.0033 | 0.0055 | 0.0055 | TRUE | 0.2217 |
| Wash | 1810 | 1408 | 1443 | 1444 | 1384 | 1561 | 1406 | 1498 | 1557 | 1437 | 1561 | 1541 | 1500 | 1487 | 1513 | 1.019 | 0.026 | 0.71608643 | 0.82948564 | 0.0017 | 0.004 | 0.0031 | 0.0031 | TRUE | 0.0285 |
| Wasl | 5218 | 3931 | 3792 | 3930 | 3583 | 4088 | 4053 | 4181 | 4092 | 3912 | 4042 | 4035 | 4052.48 | 4109 | 3996 | 0.973 | -0.04 | 0.44720233 | 0.61781006 | 0 | 0.003 | 0.0017 | 0.0017 | TRUE | 0.0042 |
| Wbp1 | 1018 | 730 | 692 | 880 | 786 | 871 | 791 | 776 | 747 | 876 | 887 | 860 | 822.71 | 771 | 874 | 1.132 | 0.179 | 0.02316633 | 0.07695011 | 0 | 0.0053 | 0.0033 | 0.0033 | TRUE | 0.0091 |
| Wbp11 | 18001 | 13222 | 13036 | 12519 | 11515 | 13429 | 13981 | 14064 | 14066 | 12461 | 12990 | 13256 | 13469.74 | 14037 | 12902 | 0.919 | -0.122 | 0.01112438 | 0.04462553 | 4.00E-04 | 0.0026 | 0.0016 | 0.0016 | TRUE | 0.011 |
| Wbp1l | 2879 | 2196 | 1897 | 1932 | 1826 | 1983 | 2236 | 2336 | 2047 | 1923 | 2060 | 1958 | 2093.21 | 2206 | 1980 | 0.897 | -0.157 | 0.02659944 | 0.08500085 | 0.0021 | 0.0036 | 0.0031 | 0.0031 | TRUE | 0.0473 |
| Wbp2 | 9126 | 6521 | 7709 | 8186 | 7555 | 8769 | 7088 | 6936 | 8318 | 8148 | 8523 | 8656 | 7944.92 | 7447 | 8442 | 1.134 | 0.181 | 0.0130444 | 0.05030859 | 0.0055 | 0.0027 | 0.0037 | 0.0037 | TRUE | 0.1272 |
| Wbp2nl | 151 | 110 | 94 | 129 | 108 | 99 | 117 | 117 | 101 | 128 | 122 | 98 | 113.95 | 112 | 116 | 1.034 | 0.048 | 0.80330253 | 0.88613128 | 0 | 0.0232 | 0.018 | 0.018 | TRUE | 0.1959 |
| Wbp4 | 2126 | 1511 | 1491 | 1574 | 1317 | 1492 | 1651 | 1607 | 1609 | 1567 | 1486 | 1473 | 1565.41 | 1622 | 1509 | 0.93 | -0.105 | 0.10870604 | 0.23512944 | 0 | 0.0039 | 0.0025 | 0.0025 | TRUE | 0.0139 |
| Wbp5 | 5323 | 3805 | 4128 | 4230 | 3653 | 4215 | 4134 | 4047 | 4454 | 4211 | 4121 | 4161 | 4187.99 | 4212 | 4164 | 0.989 | -0.016 | 0.78194375 | 0.87149433 | 9.00E-04 | 0.003 | 0.0022 | 0.0022 | TRUE | 0.0304 |
| Wbp7 | 7598 | 5635 | 5591 | 5489 | 5381 | 6058 | 5901 | 5994 | 6033 | 5464 | 6070 | 5980 | 5906.97 | 5976 | 5838 | 0.977 | -0.034 | 0.56196834 | 0.71665003 | 0.0013 | 0.0028 | 0.0023 | 0.0023 | TRUE | 0.0385 |
| Wbscr16 | 2020 | 1547 | 1479 | 1780 | 1496 | 1680 | 1569 | 1646 | 1596 | 1772 | 1688 | 1658 | 1654.68 | 1604 | 1706 | 1.064 | 0.09 | 0.17000918 | 0.3215484 | 0 | 0.0039 | 0.0025 | 0.0025 | TRUE | 0.0139 |
| Wbscr17 | 204 | 163 | 128 | 203 | 163 | 178 | 158 | 173 | 138 | 202 | 184 | 176 | 171.93 | 156 | 187 | 1.195 | 0.257 | 0.10625319 | 0.23129406 | 0.0021 | 0.0162 | 0.0123 | 0.0123 | TRUE | 0.1048 |
| Wbscr22 | 3208 | 2181 | 1843 | 2194 | 1858 | 2125 | 2492 | 2320 | 1989 | 2184 | 2096 | 2098 | 2196.27 | 2267 | 2126 | 0.937 | -0.094 | 0.26012521 | 0.43027076 | 0.006 | 0.0035 | 0.0045 | 0.0045 | TRUE | 0.1386 |
| Wbscr25 | 3 | 4 | 1 | 0 | 1 | 2 | 2 | 4 | 1 | 0 | 1 | 2 | 1.79 | 2 | 1 | 0.405 | -1.303 | 0.39110521 | NA | 0 | 1.3233 | 0.9735 | 0.9735 | TRUE | 0.2791 |
| Wbscr27 | 515 | 413 | 426 | 451 | 430 | 429 | 400 | 439 | 460 | 449 | 485 | 423 | 442.74 | 433 | 452 | 1.046 | 0.065 | 0.54515384 | 0.70230023 | 0.0027 | 0.0078 | 0.0062 | 0.0062 | TRUE | 0.0552 |
| Wbscr28 | 0 | 0 | 0 | 0 | 0 | 0 | 0 | 0 | 0 | 0 | 0 | 0 | 0 | 0 | 0 | NA | NA | NA | NA | NA | NA | NA | NA | NA | NA |
| Wdfy1 | 3994 | 3267 | 3859 | 3981 | 3683 | 4431 | 3102 | 3475 | 4164 | 3963 | 4155 | 4374 | 3872.07 | 3580 | 4164 | 1.164 | 0.219 | 0.01942607 | 0.06766257 | 0.0122 | 0.003 | 0.0061 | 0.0061 | TRUE | 0.247 |
| Wdfy2 | 612 | 476 | 429 | 491 | 470 | 476 | 475 | 506 | 463 | 489 | 530 | 470 | 488.89 | 481 | 496 | 1.03 | 0.043 | 0.67050504 | 0.79723222 | 0 | 0.0073 | 0.0052 | 0.0052 | TRUE | 0.0405 |
| Wdfy3 | 2859 | 2100 | 1826 | 2171 | 1988 | 2277 | 2220 | 2234 | 1970 | 2161 | 2243 | 2248 | 2179.31 | 2141 | 2217 | 1.035 | 0.049 | 0.47867222 | 0.64580449 | 0.002 | 0.0035 | 0.003 | 0.003 | TRUE | 0.0582 |
| Wdfy4 | 182 | 153 | 130 | 131 | 165 | 136 | 141 | 163 | 140 | 130 | 186 | 134 | 149.19 | 148 | 150 | 1.013 | 0.018 | 0.92132826 | 0.95666871 | 0.0165 | 0.0183 | 0.0178 | 0.0178 | TRUE | 0.4413 |
| Wdhd1 | 7024 | 5226 | 5145 | 4963 | 4511 | 5205 | 5455 | 5559 | 5551 | 4940 | 5089 | 5138 | 5288.78 | 5522 | 5056 | 0.916 | -0.127 | 0.01115212 | 0.04472426 | 0 | 0.0029 | 0.0016 | 0.0016 | TRUE | 0.0049 |
| Wdpcp | 95 | 59 | 45 | 52 | 45 | 73 | 74 | 63 | 49 | 52 | 51 | 72 | 59.95 | 62 | 58 | 0.939 | -0.09 | 0.74147022 | 0.8460229 | 0.026 | 0.042 | 0.0375 | 0.0375 | TRUE | 0.3106 |
| Wdr1 | 6754 | 4845 | 4331 | 3505 | 3205 | 3862 | 5246 | 5154 | 4673 | 3489 | 3616 | 3812 | 4331.51 | 5024 | 3639 | 0.724 | -0.466 | 1.27E-12 | 1.54E-10 | 0.0025 | 0.003 | 0.0029 | 0.0029 | TRUE | 0.0452 |
| Wdr11 | 5008 | 3712 | 3770 | 4174 | 3728 | 4138 | 3890 | 3948 | 4068 | 4155 | 4206 | 4085 | 4058.49 | 3969 | 4149 | 1.046 | 0.064 | 0.22338975 | 0.38781849 | 0 | 0.003 | 0.0018 | 0.0018 | TRUE | 0.0058 |
| Wdr12 | 5954 | 4118 | 4063 | 4334 | 3798 | 4214 | 4624 | 4380 | 4384 | 4314 | 4285 | 4160 | 4357.82 | 4463 | 4253 | 0.953 | -0.07 | 0.19579574 | 0.3554113 | 5.00E-04 | 0.003 | 0.0019 | 0.0019 | TRUE | 0.0125 |
| Wdr13 | 565 | 430 | 543 | 543 | 418 | 534 | 439 | 457 | 586 | 541 | 472 | 527 | 503.55 | 494 | 513 | 1.041 | 0.059 | 0.63496772 | 0.77160293 | 0.0131 | 0.0071 | 0.009 | 0.009 | TRUE | 0.3036 |
| Wdr16 | 40 | 33 | 19 | 41 | 23 | 29 | 31 | 35 | 21 | 41 | 26 | 29 | 30.34 | 29 | 32 | 1.1 | 0.137 | 0.71196361 | 0.82692149 | 0.0292 | 0.0805 | 0.0669 | 0.0669 | TRUE | 0.4248 |
| Wdr17 | 91 | 54 | 57 | 71 | 49 | 72 | 71 | 57 | 62 | 71 | 55 | 71 | 64.44 | 63 | 66 | 1.037 | 0.053 | 0.83117606 | 0.9033735 | 0 | 0.0392 | 0.0285 | 0.0285 | TRUE | 0.1587 |
| Wdr18 | 8084 | 6039 | 5215 | 5890 | 5162 | 5727 | 6279 | 6424 | 5627 | 5863 | 5823 | 5653 | 5944.78 | 6110 | 5780 | 0.946 | -0.08 | 0.20162645 | 0.36258768 | 0.0023 | 0.0028 | 0.0027 | 0.0027 | TRUE | 0.0578 |
| Wdr19 | 798 | 617 | 730 | 727 | 717 | 816 | 620 | 656 | 788 | 724 | 809 | 806 | 733.63 | 688 | 780 | 1.135 | 0.182 | 0.08269142 | 0.19363001 | 0.0081 | 0.0057 | 0.0066 | 0.0066 | TRUE | 0.1879 |
| Wdr20a | 1113 | 762 | 792 | 811 | 733 | 819 | 864 | 811 | 855 | 807 | 827 | 808 | 828.69 | 843 | 814 | 0.965 | -0.052 | 0.51458256 | 0.67718705 | 0 | 0.0053 | 0.0033 | 0.0033 | TRUE | 0.0134 |
| Wdr20b | 175 | 165 | 134 | 106 | 68 | 91 | 136 | 176 | 145 | 106 | 77 | 90 | 121.34 | 152 | 91 | 0.6 | -0.738 | 0.0001915 | 0.00206755 | 0.0125 | 0.022 | 0.0193 | 0.0193 | TRUE | 0.1958 |

| | | | | | | | | | | | | | | | | | | | | | | | | | |
|---|---|---|---|---|---|---|---|---|---|---|---|---|---|---|---|---|---|---|---|---|---|---|---|---|---|
| Wdr24 | 1996 | 1587 | 1457 | 1499 | 1440 | 1601 | 1550 | 1688 | 1572 | 1492 | 1624 | 1580 | 1584.56 | 1603 | 1565 | 0.977 | -0.034 | 0.63025223 | 0.76756946 | 0.0013 | 0.0039 | 0.003 | 0.003 | TRUE | 0.0254 |
| Wdr25 | 417 | 298 | 301 | 351 | 341 | 355 | 324 | 317 | 325 | 349 | 385 | 350 | 341.69 | 322 | 361 | 1.122 | 0.166 | 0.13925642 | 0.28048626 | 0 | 0.0094 | 0.0061 | 0.0061 | TRUE | 0.0348 |
| Wdr26 | 10507 | 7707 | 7670 | 7832 | 7204 | 8202 | 8160 | 8198 | 8276 | 7796 | 8127 | 8097 | 8108.92 | 8211 | 8007 | 0.975 | -0.036 | 0.44659659 | 0.61723382 | 2.00E-04 | 0.0027 | 0.0015 | 0.0015 | TRUE | 0.0065 |
| Wdr27 | 9 | 13 | 7 | 13 | 19 | 14 | 7 | 14 | 8 | 13 | 21 | 14 | 12.76 | 10 | 16 | 1.71 | 0.774 | 0.16958048 | NA | 0 | 0.1882 | 0.1447 | 0.1447 | TRUE | 0.5137 |
| Wdr3 | 7045 | 4911 | 4526 | 4298 | 3912 | 4497 | 5472 | 5224 | 4884 | 4278 | 4413 | 4439 | 4784.91 | 5193 | 4377 | 0.842 | -0.247 | 3.92E-05 | 0.00058071 | 0.0015 | 0.0029 | 0.0024 | 0.0024 | TRUE | 0.033 |
| Wdr31 | 1074 | 790 | 804 | 789 | 778 | 890 | 834 | 840 | 868 | 785 | 878 | 879 | 847.26 | 847 | 847 | 1 | 0 | 0.99874158 | 0.99937086 | 0.001 | 0.0052 | 0.0038 | 0.0038 | TRUE | 0.049 |
| Wdr33 | 8586 | 5877 | 5104 | 5556 | 4864 | 5347 | 6668 | 6251 | 5507 | 5530 | 5487 | 5278 | 5787.12 | 6142 | 5432 | 0.884 | -0.178 | 0.01397467 | 0.05296736 | 0.0046 | 0.0028 | 0.0036 | 0.0036 | TRUE | 0.0995 |
| Wdr34 | 2038 | 1531 | 1640 | 1661 | 1724 | 2011 | 1583 | 1629 | 1770 | 1653 | 1945 | 1985 | 1760.71 | 1661 | 1861 | 1.121 | 0.165 | 0.05054205 | 0.13591994 | 0.0056 | 0.0038 | 0.0046 | 0.0046 | TRUE | 0.1159 |
| Wdr35 | 3808 | 2757 | 2989 | 3019 | 2567 | 3095 | 2958 | 2933 | 3225 | 3005 | 2896 | 3055 | 3011.9 | 3039 | 2985 | 0.983 | -0.025 | 0.69397229 | 0.81322833 | 0.0012 | 0.0032 | 0.0025 | 0.0025 | TRUE | 0.0346 |
| Wdr36 | 11772 | 8658 | 9294 | 9640 | 8608 | 9919 | 9143 | 9209 | 10028 | 9596 | 9711 | 9792 | 9579.72 | 9460 | 9700 | 1.025 | 0.036 | 0.50837904 | 0.67162045 | 0.0012 | 0.0027 | 0.0021 | 0.0021 | TRUE | 0.0336 |
| Wdr37 | 2297 | 1705 | 1742 | 1929 | 1654 | 1892 | 1784 | 1814 | 1880 | 1920 | 1866 | 1868 | 1855.15 | 1826 | 1885 | 1.033 | 0.047 | 0.45415634 | 0.62391932 | 0 | 0.0037 | 0.0022 | 0.0022 | TRUE | 0.008 |
| Wdr38 | 1 | 0 | 0 | 0 | 0 | 1 | 1 | 0 | 0 | 0 | 0 | 1 | 0.29 | 0 | 0 | 1.075 | 0.105 | 0.97702352 | NA | 0 | 8.0647 | 7.5458 | 7.5458 | TRUE | 0.3835 |
| Wdr4 | 2379 | 1759 | 1597 | 1628 | 1430 | 1724 | 1848 | 1871 | 1723 | 1621 | 1613 | 1702 | 1729.57 | 1814 | 1645 | 0.907 | -0.141 | 0.03574263 | 0.10550794 | 8.00E-04 | 0.0038 | 0.0027 | 0.0027 | TRUE | 0.0226 |
| Wdr41 | 2258 | 1639 | 1644 | 1689 | 1637 | 1757 | 1754 | 1743 | 1774 | 1681 | 1847 | 1734 | 1755.55 | 1757 | 1754 | 0.998 | -0.003 | 0.96492248 | 0.9807735 | 5.00E-04 | 0.0038 | 0.0026 | 0.0026 | TRUE | 0.0253 |
| Wdr43 | 13549 | 10202 | 10413 | 10895 | 9792 | 11478 | 10523 | 10852 | 11236 | 10845 | 11046 | 11331 | 10972 | 10870 | 11074 | 1.019 | 0.027 | 0.59266214 | 0.74044182 | 6.00E-04 | 0.0026 | 0.0018 | 0.0018 | TRUE | 0.0106 |
| Wdr44 | 1065 | 688 | 577 | 680 | 544 | 653 | 827 | 732 | 623 | 677 | 614 | 645 | 686.12 | 727 | 645 | 0.885 | -0.176 | 0.11006775 | 0.23724903 | 0.01 | 0.0059 | 0.0072 | 0.0072 | TRUE | 0.1856 |
| Wdr45 | 521 | 339 | 252 | 204 | 185 | 206 | 405 | 361 | 272 | 203 | 209 | 203 | 275.37 | 346 | 205 | 0.591 | -0.759 | 9.47E-07 | 2.81E-05 | 0.0183 | 0.011 | 0.0133 | 0.0133 | TRUE | 0.3257 |
| Wdr45b | 9213 | 6474 | 6458 | 6671 | 6052 | 6877 | 7155 | 6886 | 6968 | 6640 | 6827 | 6789 | 6877.67 | 7003 | 6752 | 0.964 | -0.053 | 0.2775382 | 0.44878985 | 2.00E-04 | 0.0028 | 0.0016 | 0.0016 | TRUE | 0.0045 |
| Wdr46 | 4755 | 3394 | 2551 | 2511 | 2357 | 2670 | 3693 | 3610 | 2753 | 2499 | 2659 | 2636 | 2974.96 | 3352 | 2598 | 0.775 | -0.368 | 0.00015799 | 0.00177522 | 0.0121 | 0.0032 | 0.0065 | 0.0065 | TRUE | 0.2959 |
| Wdr47 | 1787 | 1357 | 1194 | 1335 | 1121 | 1273 | 1388 | 1443 | 1288 | 1329 | 1265 | 1257 | 1328.29 | 1373 | 1284 | 0.934 | -0.098 | 0.18528261 | 0.34166487 | 0.0013 | 0.0042 | 0.0032 | 0.0032 | TRUE | 0.034 |
| Wdr48 | 6123 | 4426 | 4753 | 5012 | 4474 | 5406 | 4755 | 4708 | 5128 | 4989 | 5047 | 5337 | 4994.07 | 4864 | 5124 | 1.054 | 0.076 | 0.20291388 | 0.36436932 | 0.0015 | 0.0029 | 0.0023 | 0.0023 | TRUE | 0.0273 |
| Wdr5 | 14062 | 9885 | 10026 | 10763 | 9611 | 10762 | 10921 | 10515 | 10818 | 10713 | 10842 | 10624 | 10738.89 | 10751 | 10726 | 0.998 | -0.004 | 0.93980479 | 0.96637625 | 1.00E-04 | 0.0026 | 0.0015 | 0.0015 | TRUE | 0.0045 |
| Wdr52 | 32 | 23 | 22 | 21 | 18 | 39 | 25 | 24 | 24 | 21 | 20 | 38 | 25.46 | 24 | 26 | 1.094 | 0.13 | 0.74749267 | 0.85008854 | 0.0347 | 0.0955 | 0.0782 | 0.0782 | TRUE | 0.989 |
| Wdr53 | 651 | 441 | 408 | 487 | 390 | 516 | 506 | 469 | 440 | 485 | 440 | 509 | 474.83 | 472 | 478 | 1.013 | 0.018 | 0.86580323 | 0.92585108 | 0.0031 | 0.0074 | 0.006 | 0.006 | TRUE | 0.0548 |
| Wdr54 | 103 | 71 | 69 | 75 | 79 | 67 | 80 | 76 | 74 | 75 | 89 | 66 | 76.65 | 77 | 77 | 0.996 | -0.006 | 0.97742935 | 0.98786577 | 0 | 0.0333 | 0.0239 | 0.0239 | TRUE | 0.177 |
| Wdr55 | 3157 | 2310 | 1913 | 2124 | 1912 | 2233 | 2452 | 2457 | 2064 | 2114 | 2157 | 2204 | 2241.44 | 2324 | 2158 | 0.928 | -0.108 | 0.16630079 | 0.31738434 | 0.0043 | 0.0035 | 0.0039 | 0.0039 | TRUE | 0.1148 |
| Wdr59 | 5885 | 4310 | 4647 | 5271 | 4663 | 5447 | 4571 | 4584 | 5014 | 5247 | 5260 | 5377 | 5008.89 | 4723 | 5295 | 1.121 | 0.165 | 0.00430621 | 0.02186838 | 0.0012 | 0.0029 | 0.0022 | 0.0022 | TRUE | 0.0351 |
| Wdr5b | 44 | 48 | 19 | 31 | 20 | 22 | 34 | 51 | 21 | 31 | 23 | 22 | 30.14 | 35 | 25 | 0.712 | -0.491 | 0.22330675 | 0.38773143 | 0.0865 | 0.081 | 0.0826 | 0.0826 | TRUE | 0.1729 |
| Wdr6 | 10167 | 6680 | 4893 | 4798 | 4318 | 4769 | 7896 | 7105 | 5280 | 4776 | 4871 | 4708 | 5772.67 | 6760 | 4785 | 0.708 | -0.499 | 4.98E-06 | 0.00010571 | 0.0196 | 0.0028 | 0.0084 | 0.0084 | TRUE | 0.4474 |
| Wdr60 | 698 | 528 | 470 | 453 | 438 | 476 | 542 | 562 | 507 | 451 | 494 | 470 | 504.3 | 537 | 472 | 0.878 | -0.188 | 0.05534392 | 0.14490394 | 0 | 0.0071 | 0.005 | 0.005 | TRUE | 0.0262 |
| Wdr61 | 4952 | 3594 | 3519 | 3918 | 3576 | 3939 | 3846 | 3823 | 3797 | 3900 | 4034 | 3888 | 3881.39 | 3822 | 3941 | 1.031 | 0.044 | 0.40056993 | 0.57368196 | 0 | 0.003 | 0.0017 | 0.0017 | TRUE | 0.0052 |
| Wdr62 | 2895 | 2275 | 2092 | 1940 | 1913 | 2382 | 2248 | 2420 | 2257 | 1931 | 2158 | 2351 | 2227.68 | 2308 | 2147 | 0.93 | -0.104 | 0.18982695 | 0.34766743 | 0.005 | 0.0035 | 0.0041 | 0.0041 | TRUE | 0.0944 |
| Wdr63 | 44 | 44 | 28 | 14 | 21 | 20 | 34 | 47 | 30 | 14 | 24 | 20 | 28.09 | 37 | 19 | 0.515 | -0.958 | 0.01387095 | 0.05274206 | 0.0217 | 0.0868 | 0.069 | 0.069 | TRUE | 0.3638 |
| Wdr64 | 84 | 80 | 74 | 35 | 35 | 58 | 65 | 85 | 80 | 35 | 39 | 57 | 60.29 | 77 | 44 | 0.575 | -0.798 | 0.00381048 | 0.01984809 | 0.0228 | 0.0417 | 0.0367 | 0.0367 | TRUE | 0.5714 |
| Wdr65 | 22 | 20 | 13 | 16 | 17 | 29 | 17 | 21 | 14 | 16 | 19 | 29 | 19.35 | 17 | 21 | 1.219 | 0.286 | 0.52743093 | 0.68851679 | 0 | 0.1249 | 0.0955 | 0.0955 | TRUE | 0.5343 |
| Wdr67 | 2964 | 2265 | 2440 | 2456 | 2169 | 2527 | 2302 | 2409 | 2633 | 2445 | 2447 | 2495 | 2455.02 | 2448 | 2462 | 1.007 | 0.009 | 0.88897188 | 0.93834097 | 0.002 | 0.0034 | 0.0029 | 0.0029 | TRUE | 0.0525 |
| Wdr69 | 11 | 8 | 5 | 8 | 8 | 13 | 9 | 9 | 5 | 8 | 9 | 13 | 8.71 | 8 | 10 | 1.322 | 0.403 | 0.53384381 | NA | 0 | 0.2745 | 0.186 | 0.186 | TRUE | 0.2326 |
| Wdr7 | 2038 | 1423 | 1343 | 1334 | 1201 | 1442 | 1583 | 1514 | 1449 | 1328 | 1355 | 1423 | 1441.96 | 1515 | 1369 | 0.903 | -0.148 | 0.03727199 | 0.10886143 | 0.001 | 0.0041 | 0.0029 | 0.0029 | TRUE | 0.0191 |
| Wdr70 | 2165 | 1536 | 1507 | 1607 | 1356 | 1721 | 1681 | 1634 | 1626 | 1600 | 1530 | 1699 | 1628.26 | 1647 | 1610 | 0.977 | -0.033 | 0.62669363 | 0.76482743 | 9.00E-04 | 0.0039 | 0.0028 | 0.0028 | TRUE | 0.0286 |
| Wdr72 | 19 | 10 | 13 | 18 | 17 | 17 | 15 | 11 | 14 | 18 | 19 | 17 | 15.55 | 13 | 18 | 1.358 | 0.442 | 0.35779992 | NA | 0 | 0.1548 | 0.1009 | 0.1009 | TRUE | 0.1085 |
| Wdr73 | 2348 | 1719 | 1663 | 1589 | 1454 | 1621 | 1824 | 1828 | 1794 | 1582 | 1640 | 1600 | 1711.43 | 1815 | 1607 | 0.885 | -0.176 | 0.00466809 | 0.02324772 | 0 | 0.0038 | 0.0022 | 0.0022 | TRUE | 0.0038 |
| Wdr74 | 3693 | 2738 | 2422 | 2487 | 2389 | 2759 | 2868 | 2912 | 2613 | 2476 | 2695 | 2724 | 2714.67 | 2798 | 2632 | 0.94 | -0.089 | 0.19615126 | 0.35569526 | 0.0024 | 0.0033 | 0.0031 | 0.0031 | TRUE | 0.0399 |
| Wdr75 | 8802 | 6191 | 6118 | 6453 | 5456 | 6445 | 6836 | 6585 | 6601 | 6423 | 6155 | 6362 | 6493.86 | 6674 | 6313 | 0.946 | -0.08 | 0.11085539 | 0.23852327 | 3.00E-04 | 0.0028 | 0.0017 | 0.0017 | TRUE | 0.0058 |
| Wdr76 | 5124 | 3986 | 4529 | 4636 | 4119 | 4728 | 3980 | 4240 | 4887 | 4615 | 4647 | 4667 | 4505.8 | 4369 | 4643 | 1.063 | 0.089 | 0.23940708 | 0.40634667 | 0.0052 | 0.003 | 0.0039 | 0.0039 | TRUE | 0.1307 |
| Wdr77 | 16951 | 11859 | 11293 | 11211 | 10126 | 11718 | 13165 | 12614 | 12185 | 11159 | 11423 | 11567 | 12019.08 | 12655 | 11383 | 0.899 | -0.153 | 0.00287175 | 0.01598985 | 8.00E-04 | 0.0026 | 0.0018 | 0.0018 | TRUE | 0.0153 |
| Wdr78 | 43 | 31 | 22 | 27 | 27 | 26 | 33 | 33 | 24 | 27 | 30 | 26 | 28.85 | 30 | 28 | 0.915 | -0.127 | 0.72300706 | 0.83418953 | 0 | 0.0846 | 0.0584 | 0.0584 | TRUE | 0.2076 |
| Wdr8 | 1225 | 912 | 843 | 841 | 720 | 920 | 951 | 970 | 910 | 837 | 812 | 908 | 898.1 | 944 | 852 | 0.904 | -0.146 | 0.07358289 | 0.17783684 | 0.001 | 0.0051 | 0.0037 | 0.0037 | TRUE | 0.0391 |
| Wdr81 | 1916 | 1440 | 1196 | 1249 | 1148 | 1275 | 1488 | 1532 | 1290 | 1243 | 1295 | 1259 | 1351.2 | 1437 | 1266 | 0.88 | -0.184 | 0.0230111 | 0.07654122 | 0.0036 | 0.0042 | 0.004 | 0.004 | TRUE | 0.0933 |
| Wdr82 | 11181 | 8219 | 7510 | 8067 | 7246 | 8443 | 8684 | 8742 | 8103 | 8030 | 8174 | 8335 | 8344.69 | 8510 | 8180 | 0.961 | -0.057 | 0.28164919 | 0.45315943 | 9.00E-04 | 0.0027 | 0.0019 | 0.0019 | TRUE | 0.0212 |
| Wdr83 | 935 | 656 | 528 | 619 | 517 | 563 | 726 | 698 | 570 | 616 | 583 | 556 | 624.8 | 665 | 585 | 0.879 | -0.186 | 0.0848619 | 0.19652158 | 0.0081 | 0.0062 | 0.0068 | 0.0068 | TRUE | 0.1803 |
| Wdr85 | 1168 | 869 | 799 | 890 | 768 | 824 | 907 | 924 | 862 | 886 | 866 | 813 | 876.55 | 898 | 855 | 0.952 | -0.071 | 0.37894496 | 0.55369661 | 0 | 0.0051 | 0.0035 | 0.0035 | TRUE | 0.0222 |
| Wdr86 | 323 | 206 | 206 | 340 | 255 | 309 | 251 | 219 | 222 | 338 | 288 | 305 | 270.56 | 231 | 310 | 1.342 | 0.425 | 0.00122814 | 0.00839892 | 0.0029 | 0.0112 | 0.0087 | 0.0087 | TRUE | 0.071 |
| Wdr89 | 1186 | 896 | 839 | 951 | 856 | 984 | 921 | 953 | 905 | 947 | 966 | 971 | 943.85 | 926 | 961 | 1.038 | 0.053 | 0.4784335 | 0.64578809 | 0 | 0.0049 | 0.003 | 0.003 | TRUE | 0.0072 |
| Wdr90 | 1471 | 1249 | 1204 | 1027 | 997 | 1219 | 1142 | 1329 | 1299 | 1022 | 1125 | 1203 | 1186.74 | 1257 | 1117 | 0.89 | -0.169 | 0.05963224 | 0.15266408 | 0.0059 | 0.0044 | 0.0049 | 0.0049 | TRUE | 0.0786 |
| Wdr91 | 2400 | 1707 | 1687 | 1877 | 1605 | 1816 | 1864 | 1816 | 1820 | 1868 | 1811 | 1793 | 1828.6 | 1833 | 1824 | 0.995 | -0.008 | 0.89882349 | 0.94399913 | 0 | 0.0037 | 0.0022 | 0.0022 | TRUE | 0.0056 |
| Wdr92 | 3356 | 2451 | 2349 | 2461 | 2247 | 2545 | 2606 | 2607 | 2535 | 2450 | 2535 | 2512 | 2540.82 | 2583 | 2499 | 0.967 | -0.048 | 0.39946207 | 0.57284151 | 0 | 0.0034 | 0.0019 | 0.0019 | TRUE | 0.0036 |
| Wdr93 | 102 | 79 | 96 | 140 | 112 | 124 | 79 | 84 | 104 | 139 | 126 | 122 | 109.16 | 89 | 129 | 1.461 | 0.547 | 0.00515389 | 0.02512433 | 0 | 0.0241 | 0.0182 | 0.0182 | TRUE | 0.1905 |
| Wdr95 | 50 | 38 | 63 | 30 | 28 | 27 | 39 | 40 | 68 | 30 | 32 | 27 | 39.22 | 49 | 30 | 0.601 | -0.734 | 0.0310266 | 0.09501418 | 0.04 | 0.0629 | 0.0559 | 0.0559 | TRUE | 0.8817 |
| Wdr96 | 186 | 159 | 138 | 170 | 148 | 139 | 144 | 169 | 149 | 169 | 167 | 137 | 155.98 | 154 | 158 | 1.025 | 0.035 | 0.83329474 | 0.90496784 | 0.004 | 0.0176 | 0.0137 | 0.0137 | TRUE | 0.1405 |
| Wdsub1 | 1114 | 809 | 786 | 718 | 707 | 827 | 865 | 861 | 848 | 715 | 798 | 816 | 817.07 | 858 | 776 | 0.904 | -0.145 | 0.08699416 | 0.19993688 | 0.0012 | 0.0053 | 0.0039 | 0.0039 | TRUE | 0.0579 |
| Wdtc1 | 16062 | 11214 | 10356 | 10644 | 9205 | 10433 | 12475 | 11928 | 11174 | 10595 | 10384 | 10299 | 11142.52 | 11859 | 10426 | 0.879 | -0.186 | 0.00089077 | 0.00666237 | 0.0015 | 0.0026 | 0.0022 | 0.0022 | TRUE | 0.0312 |
| Wdyhv1 | 690 | 558 | 470 | 504 | 488 | 582 | 536 | 594 | 507 | 502 | 551 | 575 | 543.88 | 546 | 543 | 0.994 | -0.008 | 0.93504842 | 0.96412113 | 0.0036 | 0.0068 | 0.0058 | 0.0058 | TRUE | 0.0667 |
| Wee1 | 4715 | 3468 | 3711 | 3767 | 3363 | 4061 | 3662 | 3689 | 4004 | 3750 | 3794 | 4009 | 3817.88 | 3785 | 3851 | 1.018 | 0.025 | 0.67927015 | 0.8028453 | 0.0016 | 0.003 | 0.0025 | 0.0025 | TRUE | 0.0309 |
| Wee2 | 5 | 1 | 3 | 1 | 3 | 7 | 4 | 1 | 3 | 1 | 3 | 7 | 3.25 | 3 | 4 | 1.361 | 0.444 | 0.69075947 | NA | 0 | 0.7327 | 0.5878 | 0.5878 | TRUE | 0.2529 |
| Wfdc1 | 33 | 20 | 26 | 34 | 20 | 30 | 26 | 21 | 28 | 34 | 23 | 30 | 26.83 | 25 | 29 | 1.15 | 0.202 | 0.59004238 | 0.73868977 | 0 | 0.0908 | 0.0642 | 0.0642 | TRUE | 0.2069 |
| Wfdc10 | 77 | 41 | 57 | 72 | 64 | 78 | 60 | 44 | 62 | 72 | 72 | 77 | 64.3 | 55 | 74 | 1.335 | 0.417 | 0.09282734 | 0.20933751 | 0 | 0.0393 | 0.0286 | 0.0286 | TRUE | 0.2569 |
| Wfdc11 | 0 | 0 | 0 | 0 | 0 | 0 | 0 | 0 | 0 | 0 | 0 | 0 | 0 | 0 | 0 | NA | NA | NA | NA | NA | NA | NA | NA | NA | NA |
| Wfdc12 | 2 | 3 | 0 | 0 | 0 | 2 | 2 | 3 | 0 | 0 | 0 | 2 | 1.12 | 2 | 1 | 0.424 | -1.237 | 0.551706 | NA | 1.491 | 2.119 | 2.0351 | 2.0351 | TRUE | 0.3265 |
| Wfdc13 | 3 | 2 | 1 | 1 | 0 | 3 | 2 | 2 | 1 | 1 | 0 | 3 | 1.58 | 2 | 1 | 0.721 | -0.471 | 0.76077144 | NA | 0 | 1.5004 | 1.0809 | 1.0809 | TRUE | 0.6211 |

| | | | | | | | | | | | | | | | | | | | | | | | | | |
|---|---|---|---|---|---|---|---|---|---|---|---|---|---|---|---|---|---|---|---|---|---|---|---|---|---|
| Wfdc15a | 171 | 126 | 100 | 83 | 50 | 62 | 133 | 134 | 108 | 83 | 56 | 61 | 95.83 | 125 | 67 | 0.535 | -0.903 | 4.42E-05 | 0.0006306 | 0.0134 | 0.0272 | 0.0236 | 0.0236 | TRUE | 0.3886 |
| Wfdc15b | 0 | 0 | 0 | 0 | 0 | 0 | 0 | 0 | 0 | 0 | 0 | 0 | 0 | 0 | 0 | NA | NA | NA | NA | NA | NA | NA | NA | NA | NA |
| Wfdc16 | 1 | 1 | 0 | 1 | 0 | 0 | 1 | 1 | 0 | 1 | 0 | 0 | 0.47 | 1 | 0 | 0.599 | -0.74 | 0.79191656 | NA | 0 | 5.0171 | 3.8827 | 3.8827 | TRUE | 0.3906 |
| Wfdc17 | 0 | 0 | 0 | 0 | 0 | 0 | 0 | 0 | 0 | 0 | 0 | 0 | 0 | 0 | 0 | NA | NA | NA | NA | NA | NA | NA | NA | NA | NA |
| Wfdc18 | 0 | 0 | 0 | 0 | 0 | 0 | 0 | 0 | 0 | 0 | 0 | 0 | 0 | 0 | 0 | NA | NA | NA | NA | NA | NA | NA | NA | NA | NA |
| Wfdc2 | 2154 | 1692 | 2063 | 2052 | 1764 | 1951 | 1673 | 1800 | 2226 | 2043 | 1990 | 1926 | 1942.85 | 1900 | 1986 | 1.047 | 0.066 | 0.49446785 | 0.65958273 | 0.0111 | 0.0036 | 0.0062 | 0.0062 | TRUE | 0.272 |
| Wfdc3 | 2 | 0 | 1 | 1 | 0 | 2 | 2 | 0 | 1 | 1 | 0 | 2 | 0.93 | 1 | 1 | 1.115 | 0.156 | 0.93833771 | NA | 0 | 2.5409 | 1.8967 | 1.8967 | TRUE | 0.1672 |
| Wfdc5 | 1 | 1 | 0 | 1 | 1 | 1 | 1 | 1 | 0 | 1 | 1 | 1 | 0.83 | 1 | 1 | 1.662 | 0.733 | 0.71902164 | NA | 0 | 2.8746 | 1.72 | 1.72 | TRUE | 0.1898 |
| Wfdc6a | 0 | 1 | 0 | 1 | 0 | 0 | 0 | 1 | 0 | 1 | 0 | 0 | 0.34 | 0 | 0 | 1.075 | 0.105 | 0.97573409 | NA | 0 | 6.9086 | 6.5582 | 6.5582 | TRUE | 0.5216 |
| Wfdc6b | 0 | 0 | 0 | 0 | 0 | 0 | 0 | 0 | 0 | 0 | 0 | 0 | 0 | 0 | 0 | NA | NA | NA | NA | NA | NA | NA | NA | NA | NA |
| Wfdc8 | 5 | 2 | 8 | 1 | 5 | 4 | 4 | 2 | 9 | 1 | 6 | 4 | 4.2 | 5 | 4 | 0.721 | -0.471 | 0.63763034 | NA | 0.2325 | 0.5661 | 0.4741 | 0.4741 | TRUE | 0.6291 |
| Wfdc9 | 2 | 0 | 1 | 8 | 4 | 5 | 2 | 0 | 1 | 8 | 5 | 5 | 3.34 | 1 | 6 | 6.289 | 2.653 | 0.03331148 | NA | 0 | 0.7119 | 0.5124 | 0.5124 | TRUE | 0.2746 |
| Wfikkn1 | 8 | 6 | 3 | 4 | 1 | 5 | 6 | 6 | 3 | 4 | 1 | 5 | 4.31 | 5 | 3 | 0.638 | -0.647 | 0.49257811 | NA | 0 | 0.5519 | 0.3976 | 0.3976 | TRUE | 0.4155 |
| Wfikkn2 | 115 | 90 | 56 | 66 | 51 | 85 | 89 | 96 | 60 | 66 | 58 | 84 | 75.43 | 82 | 69 | 0.843 | -0.246 | 0.33615209 | 0.51047831 | 0.0329 | 0.0338 | 0.0336 | 0.0336 | TRUE | 0.4624 |
| Wfs1 | 1363 | 1158 | 1168 | 1460 | 1257 | 1468 | 1059 | 1232 | 1260 | 1453 | 1418 | 1449 | 1311.84 | 1184 | 1440 | 1.219 | 0.286 | 0.00048509 | 0.00422033 | 0.0038 | 0.0042 | 0.0041 | 0.0041 | TRUE | 0.1067 |
| Whamm | 867 | 735 | 761 | 871 | 795 | 876 | 673 | 782 | 821 | 867 | 897 | 865 | 817.48 | 759 | 876 | 1.157 | 0.211 | 0.02330368 | 0.07733445 | 0.0043 | 0.0053 | 0.005 | 0.005 | TRUE | 0.1212 |
| Whrn | 706 | 667 | 765 | 719 | 678 | 838 | 548 | 709 | 825 | 716 | 765 | 827 | 731.83 | 694 | 769 | 1.111 | 0.151 | 0.24331811 | 0.41108106 | 0.0222 | 0.0057 | 0.0108 | 0.0108 | TRUE | 0.3349 |
| Whsc1 | 6236 | 4356 | 4338 | 4493 | 3732 | 4222 | 4843 | 4633 | 4681 | 4472 | 4210 | 4168 | 4501.25 | 4719 | 4283 | 0.908 | -0.14 | 0.01226493 | 0.04817331 | 7.00E-04 | 0.003 | 0.002 | 0.002 | TRUE | 0.0182 |
| Whsc1l1 | 3263 | 2372 | 2355 | 2504 | 2295 | 2652 | 2534 | 2523 | 2541 | 2492 | 2589 | 2618 | 2549.62 | 2533 | 2566 | 1.013 | 0.019 | 0.73981134 | 0.84538261 | 0 | 0.0034 | 0.002 | 0.002 | TRUE | 0.0078 |
| Wibg | 2506 | 1844 | 1533 | 1522 | 1441 | 1617 | 1946 | 1961 | 1654 | 1515 | 1626 | 1596 | 1716.44 | 1854 | 1579 | 0.851 | -0.233 | 0.00391284 | 0.02026405 | 0.0041 | 0.0038 | 0.0041 | 0.0041 | TRUE | 0.1058 |
| Wif1 | 123 | 112 | 95 | 99 | 88 | 130 | 96 | 119 | 103 | 99 | 99 | 128 | 107.22 | 106 | 109 | 1.033 | 0.047 | 0.81803263 | 0.89536894 | 0.0094 | 0.0245 | 0.0202 | 0.0202 | TRUE | 0.252 |
| Wipf1 | 1048 | 730 | 761 | 763 | 651 | 844 | 814 | 776 | 821 | 759 | 734 | 833 | 789.76 | 804 | 775 | 0.965 | -0.051 | 0.55244179 | 0.70822949 | 0.0012 | 0.0054 | 0.004 | 0.004 | TRUE | 0.0502 |
| Wipf2 | 3817 | 2853 | 2975 | 3100 | 2841 | 3261 | 2965 | 3035 | 3210 | 3086 | 3205 | 3219 | 3119.83 | 3070 | 3170 | 1.033 | 0.047 | 0.42827488 | 0.600425 | 8.00E-04 | 0.0032 | 0.0022 | 0.0022 | TRUE | 0.0193 |
| Wipf3 | 420 | 310 | 358 | 427 | 414 | 473 | 326 | 330 | 386 | 425 | 467 | 467 | 400.2 | 347 | 453 | 1.306 | 0.385 | 0.0007065 | 0.00560381 | 0.0032 | 0.0083 | 0.0068 | 0.0068 | TRUE | 0.1061 |
| Wipi1 | 1559 | 1309 | 1559 | 1747 | 1611 | 1816 | 1211 | 1392 | 1682 | 1739 | 1817 | 1793 | 1605.72 | 1428 | 1783 | 1.25 | 0.322 | 0.00168169 | 0.01055093 | 0.0132 | 0.0039 | 0.0069 | 0.0069 | TRUE | 0.2901 |
| Wipi2 | 4866 | 3405 | 3267 | 3385 | 2994 | 3451 | 3779 | 3622 | 3525 | 3369 | 3378 | 3407 | 3513.3 | 3642 | 3385 | 0.929 | -0.106 | 0.0560204 | 0.1462732 | 4.00E-04 | 0.0031 | 0.002 | 0.002 | TRUE | 0.0135 |
| Wisp1 | 45 | 30 | 17 | 36 | 17 | 26 | 35 | 32 | 18 | 36 | 19 | 26 | 27.65 | 28 | 27 | 0.944 | -0.084 | 0.83544172 | 0.90637245 | 0.0633 | 0.0882 | 0.0814 | 0.0814 | TRUE | 0.397 |
| Wisp2 | 23 | 14 | 14 | 18 | 12 | 9 | 18 | 15 | 15 | 18 | 14 | 9 | 14.7 | 16 | 14 | 0.839 | -0.253 | 0.61801266 | NA | 0 | 0.1637 | 0.1176 | 0.1176 | TRUE | 0.2398 |
| Wisp3 | 21 | 7 | 9 | 13 | 16 | 12 | 16 | 7 | 10 | 13 | 18 | 12 | 12.72 | 11 | 14 | 1.253 | 0.325 | 0.55975914 | NA | 0 | 0.1888 | 0.1449 | 0.1449 | TRUE | 0.7691 |
| Wiz | 3327 | 2596 | 2836 | 2800 | 2603 | 3172 | 2584 | 2761 | 3060 | 2787 | 2936 | 3131 | 2876.69 | 2802 | 2951 | 1.054 | 0.076 | 0.3214365 | 0.49524911 | 0.0047 | 0.0033 | 0.0039 | 0.0039 | TRUE | 0.0787 |
| Wls | 376 | 239 | 195 | 221 | 165 | 196 | 292 | 254 | 210 | 220 | 186 | 193 | 226.04 | 252 | 200 | 0.79 | -0.34 | 0.02972321 | 0.09201512 | 0.0136 | 0.0129 | 0.0131 | 0.0131 | TRUE | 0.1789 |
| Wnk1 | 40493 | 29463 | 29090 | 31750 | 28793 | 31865 | 31449 | 31339 | 31388 | 31604 | 32481 | 31456 | 31619.59 | 31392 | 31847 | 1.014 | 0.021 | 0.63033249 | 0.7676018 | 1.00E-04 | 0.0025 | 0.0013 | 0.0013 | TRUE | 0.0037 |
| Wnk2 | 943 | 798 | 888 | 1111 | 1105 | 1210 | 732 | 849 | 958 | 1106 | 1247 | 1194 | 1014.38 | 846 | 1182 | 1.399 | 0.485 | 1.70E-06 | 4.47E-05 | 0.0092 | 0.0048 | 0.0064 | 0.0064 | TRUE | 0.1722 |
| Wnk3 | 5887 | 3885 | 3427 | 3993 | 3314 | 3801 | 4572 | 4132 | 3698 | 3975 | 3739 | 3752 | 3977.94 | 4134 | 3822 | 0.924 | -0.114 | 0.14278862 | 0.28548421 | 0.0058 | 0.003 | 0.0041 | 0.0041 | TRUE | 0.1058 |
| Wnk4 | 56 | 40 | 39 | 68 | 63 | 62 | 43 | 43 | 42 | 68 | 71 | 61 | 54.68 | 43 | 67 | 1.558 | 0.639 | 0.01394281 | 0.0529194 | 0 | 0.0458 | 0.0297 | 0.0297 | TRUE | 0.0462 |
| Wnt1 | 0 | 1 | 0 | 2 | 0 | 3 | 0 | 1 | 0 | 2 | 0 | 3 | 1 | 0 | 2 | 4.836 | 2.274 | 0.29661342 | NA | 0 | 2.3662 | 2.1098 | 2.1098 | TRUE | 0.4076 |
| Wnt10a | 16 | 10 | 10 | 13 | 16 | 13 | 12 | 11 | 11 | 13 | 18 | 13 | 12.95 | 11 | 15 | 1.284 | 0.361 | 0.49376154 | NA | 0 | 0.1855 | 0.122 | 0.122 | TRUE | 0.1728 |
| Wnt10b | 4 | 6 | 4 | 3 | 1 | 5 | 3 | 6 | 4 | 3 | 1 | 5 | 3.81 | 4 | 3 | 0.671 | -0.576 | 0.56393088 | NA | 0 | 0.6246 | 0.4433 | 0.4433 | TRUE | 0.2719 |
| Wnt11 | 115 | 67 | 81 | 104 | 68 | 82 | 89 | 71 | 87 | 104 | 77 | 81 | 84.86 | 82 | 87 | 1.052 | 0.073 | 0.74314756 | 0.84710169 | 0.0087 | 0.0304 | 0.0244 | 0.0244 | TRUE | 0.2623 |
| Wnt16 | 14 | 16 | 29 | 20 | 23 | 19 | 11 | 17 | 31 | 20 | 26 | 19 | 20.63 | 20 | 22 | 1.105 | 0.144 | 0.76203667 | 0.85895197 | 0.1075 | 0.1173 | 0.1145 | 0.1145 | TRUE | 0.6929 |
| Wnt2 | 8 | 7 | 4 | 2 | 3 | 3 | 6 | 7 | 4 | 2 | 3 | 3 | 4.39 | 6 | 3 | 0.461 | -1.117 | 0.22800295 | NA | 0 | 0.5429 | 0.3516 | 0.3516 | TRUE | 0.1254 |
| Wnt2b | 46 | 23 | 37 | 32 | 20 | 17 | 36 | 24 | 40 | 32 | 23 | 17 | 28.55 | 33 | 24 | 0.71 | -0.494 | 0.20663239 | 0.36835958 | 0.0416 | 0.0854 | 0.0738 | 0.0738 | TRUE | 0.4975 |
| Wnt3 | 46 | 21 | 35 | 42 | 23 | 31 | 36 | 22 | 38 | 42 | 26 | 31 | 32.36 | 32 | 33 | 1.026 | 0.036 | 0.92010233 | 0.95615985 | 0.0334 | 0.0757 | 0.0644 | 0.0644 | TRUE | 0.4426 |
| Wnt3a | 71 | 46 | 55 | 73 | 55 | 70 | 55 | 49 | 59 | 73 | 62 | 69 | 61.2 | 54 | 68 | 1.248 | 0.32 | 0.1958979 | 0.35547397 | 0 | 0.0412 | 0.0277 | 0.0277 | TRUE | 0.0607 |
| Wnt4 | 72 | 48 | 41 | 43 | 48 | 51 | 56 | 51 | 44 | 43 | 54 | 50 | 49.75 | 50 | 49 | 0.968 | -0.046 | 0.86718312 | 0.92677197 | 0 | 0.0501 | 0.0349 | 0.0349 | TRUE | 0.103 |
| Wnt5a | 53 | 47 | 70 | 42 | 57 | 69 | 41 | 50 | 76 | 42 | 64 | 68 | 56.82 | 56 | 58 | 1.052 | 0.073 | 0.81230383 | 0.89151249 | 0.0649 | 0.0441 | 0.0501 | 0.0501 | TRUE | 0.8105 |
| Wnt5b | 185 | 125 | 153 | 203 | 170 | 189 | 144 | 133 | 165 | 202 | 192 | 187 | 170.36 | 147 | 194 | 1.316 | 0.396 | 0.01169207 | 0.0463821 | 0 | 0.0163 | 0.0118 | 0.0118 | TRUE | 0.1095 |
| Wnt6 | 84 | 83 | 69 | 137 | 127 | 169 | 65 | 88 | 74 | 136 | 143 | 167 | 112.41 | 76 | 149 | 1.971 | 0.979 | 1.03E-06 | 3.01E-05 | 0.0073 | 0.0235 | 0.0191 | 0.0191 | TRUE | 0.182 |
| Wnt7a | 38 | 25 | 22 | 34 | 24 | 28 | 30 | 27 | 24 | 34 | 27 | 28 | 28.07 | 27 | 30 | 1.106 | 0.145 | 0.68791904 | 0.80889723 | 0 | 0.0869 | 0.058 | 0.058 | TRUE | 0.1103 |
| Wnt7b | 135 | 111 | 110 | 129 | 125 | 118 | 105 | 118 | 119 | 128 | 141 | 116 | 121.25 | 114 | 128 | 1.131 | 0.178 | 0.32749431 | 0.5019168 | 0 | 0.022 | 0.0155 | 0.0155 | TRUE | 0.0704 |
| Wnt8a | 38 | 19 | 11 | 14 | 12 | 15 | 30 | 20 | 12 | 14 | 14 | 15 | 17.31 | 21 | 14 | 0.678 | -0.561 | 0.25615001 | 0.42612273 | 0.0611 | 0.1393 | 0.1159 | 0.1159 | TRUE | 0.1633 |
| Wnt8b | 21 | 10 | 8 | 10 | 15 | 6 | 16 | 11 | 9 | 10 | 17 | 6 | 11.4 | 12 | 11 | 0.903 | -0.147 | 0.80828342 | NA | 0.0941 | 0.2104 | 0.1779 | 0.1779 | TRUE | 0.409 |
| Wnt9a | 239 | 180 | 180 | 234 | 211 | 218 | 186 | 191 | 194 | 233 | 238 | 215 | 209.58 | 190 | 229 | 1.201 | 0.265 | 0.05517467 | 0.14459323 | 0 | 0.0137 | 0.0089 | 0.0089 | TRUE | 0.0299 |
| Wnt9b | 7 | 2 | 6 | 3 | 5 | 2 | 5 | 2 | 6 | 3 | 6 | 2 | 4.11 | 4 | 4 | 0.743 | -0.428 | 0.65783843 | NA | 0 | 0.5796 | 0.4213 | 0.4213 | TRUE | 0.396 |
| Wrap53 | 2845 | 1928 | 1494 | 1249 | 1111 | 1265 | 2210 | 2051 | 1612 | 1243 | 1253 | 1249 | 1602.95 | 1958 | 1248 | 0.637 | -0.65 | 1.99E-10 | 1.52E-08 | 0.0128 | 0.0039 | 0.0069 | 0.0069 | TRUE | 0.2862 |
| Wrb | 3126 | 2168 | 2043 | 2038 | 1865 | 2079 | 2428 | 2306 | 2204 | 2029 | 2104 | 2052 | 2187.19 | 2313 | 2062 | 0.891 | -0.167 | 0.009153 | 0.0387741 | 0.001 | 0.0035 | 0.0025 | 0.0025 | TRUE | 0.0236 |
| Wrn | 2338 | 1599 | 1583 | 1805 | 1551 | 1753 | 1816 | 1701 | 1708 | 1797 | 1750 | 1730 | 1750.26 | 1742 | 1759 | 1.009 | 0.013 | 0.838573 | 0.9080766 | 4.00E-04 | 0.0038 | 0.0025 | 0.0025 | TRUE | 0.0175 |
| Wrnip1 | 3555 | 2400 | 2353 | 2309 | 2078 | 2304 | 2761 | 2553 | 2539 | 2298 | 2344 | 2274 | 2461.62 | 2618 | 2305 | 0.88 | -0.184 | 0.00313021 | 0.01705684 | 0.001 | 0.0034 | 0.0024 | 0.0024 | TRUE | 0.0287 |
| Wsb1 | 7708 | 6470 | 6144 | 5997 | 5039 | 5634 | 5986 | 6882 | 6629 | 5969 | 5685 | 5562 | 6118.9 | 6499 | 5739 | 0.883 | -0.179 | 0.00632619 | 0.02933817 | 0.0029 | 0.0028 | 0.0029 | 0.0029 | TRUE | 0.0588 |
| Wsb2 | 4090 | 2991 | 3029 | 3782 | 3559 | 4025 | 3177 | 3181 | 3268 | 3765 | 4015 | 3973 | 3563.18 | 3209 | 3918 | 1.221 | 0.288 | 2.74E-07 | 9.62E-06 | 5.00E-04 | 0.0031 | 0.002 | 0.002 | TRUE | 0.0143 |
| Wscd1 | 662 | 471 | 438 | 562 | 495 | 556 | 514 | 501 | 473 | 559 | 558 | 549 | 525.74 | 496 | 555 | 1.119 | 0.162 | 0.08424235 | 0.19572816 | 0 | 0.0069 | 0.0044 | 0.0044 | TRUE | 0.0193 |
| Wscd2 | 319 | 248 | 236 | 306 | 274 | 266 | 248 | 264 | 255 | 305 | 309 | 263 | 273.74 | 256 | 292 | 1.144 | 0.194 | 0.12882254 | 0.26538781 | 0 | 0.0111 | 0.0081 | 0.0081 | TRUE | 0.0895 |
| Wt1 | 91 | 59 | 38 | 34 | 36 | 45 | 71 | 63 | 41 | 34 | 41 | 44 | 48.89 | 58 | 40 | 0.677 | -0.563 | 0.06390724 | 0.16059394 | 0.0307 | 0.0509 | 0.0451 | 0.0451 | TRUE | 0.541 |
| Wtap | 5073 | 3625 | 3486 | 3316 | 2967 | 3579 | 3940 | 3856 | 3761 | 3301 | 3347 | 3533 | 3623.01 | 3852 | 3394 | 0.881 | -0.183 | 0.00130085 | 0.00871279 | 6.00E-04 | 0.0031 | 0.0021 | 0.0021 | TRUE | 0.0158 |
| Wtip | 3664 | 2704 | 2370 | 3050 | 2623 | 2945 | 2846 | 2876 | 2557 | 3036 | 2959 | 2907 | 2863.54 | 2760 | 2967 | 1.075 | 0.104 | 0.11164336 | 0.23959739 | 0.0017 | 0.0033 | 0.0027 | 0.0027 | TRUE | 0.0492 |
| Wwc1 | 4261 | 3380 | 3498 | 3354 | 3022 | 3547 | 3309 | 3595 | 3774 | 3339 | 3409 | 3501 | 3488.01 | 3559 | 3416 | 0.96 | -0.058 | 0.37035785 | 0.5449938 | 0.002 | 0.0031 | 0.0028 | 0.0028 | TRUE | 0.047 |
| Wwc2 | 11784 | 8583 | 8352 | 8206 | 7655 | 8262 | 9152 | 9130 | 9012 | 8168 | 8636 | 8156 | 8708.87 | 9098 | 8320 | 0.914 | -0.129 | 0.00937512 | 0.03952761 | 4.00E-04 | 0.0027 | 0.0017 | 0.0017 | TRUE | 0.0134 |
| Wwox | 747 | 505 | 540 | 525 | 487 | 491 | 580 | 537 | 583 | 523 | 549 | 485 | 542.77 | 567 | 519 | 0.914 | -0.129 | 0.18477903 | 0.3410883 | 9.00E-04 | 0.0068 | 0.005 | 0.005 | TRUE | 0.0391 |
| Wwp1 | 4465 | 3405 | 3743 | 4040 | 3373 | 3834 | 3468 | 3622 | 4039 | 4021 | 3805 | 3785 | 3789.93 | 3710 | 3870 | 1.044 | 0.062 | 0.36995676 | 0.54468396 | 0.0032 | 0.0031 | 0.0032 | 0.0032 | TRUE | 0.073 |

| | | | | | | | | | | | | | | | | | | | | | | | | | |
|---|---|---|---|---|---|---|---|---|---|---|---|---|---|---|---|---|---|---|---|---|---|---|---|---|---|
| Wwp2 | 7838 | 5883 | 5913 | 5283 | 5173 | 6017 | 6087 | 6258 | 6380 | 5259 | 5836 | 5940 | 5959.88 | 6242 | 5678 | 0.91 | -0.136 | 0.02722371 | 0.08625771 | 0.0021 | 0.0028 | 0.0026 | 0.0026 | TRUE | 0.0511 |
| Wwtr1 | 3263 | 2267 | 2601 | 3277 | 2640 | 3101 | 2534 | 2411 | 2806 | 3262 | 2978 | 3061 | 2842.23 | 2584 | 3100 | 1.2 | 0.263 | 0.00027288 | 0.00270292 | 0.0036 | 0.0033 | 0.0034 | 0.0034 | TRUE | 0.0677 |
| Xab2 | 8953 | 6783 | 6305 | 6673 | 5860 | 6805 | 6953 | 7215 | 6803 | 6642 | 6611 | 6718 | 6823.67 | 6990 | 6657 | 0.952 | -0.07 | 0.1598579 | 0.30934863 | 3.00E-04 | 0.0028 | 0.0017 | 0.0017 | TRUE | 0.0095 |
| Xaf1 | 781 | 615 | 713 | 828 | 863 | 891 | 607 | 654 | 769 | 824 | 974 | 880 | 784.56 | 677 | 893 | 1.321 | 0.401 | 0.00015236 | 0.00172833 | 0.0093 | 0.0054 | 0.0068 | 0.0068 | TRUE | 0.1678 |
| Xbp1 | 4682 | 3613 | 3277 | 3574 | 3247 | 3546 | 3636 | 3843 | 3536 | 3558 | 3663 | 3500 | 3622.7 | 3672 | 3574 | 0.973 | -0.039 | 0.50147777 | 0.66570278 | 8.00E-04 | 0.0031 | 0.0022 | 0.0022 | TRUE | 0.0199 |
| Xcl1 | 0 | 0 | 0 | 0 | 0 | 0 | 0 | 0 | 0 | 0 | 0 | 0 | 0 | 0 | 0 | NA | NA | NA | NA | NA | NA | NA | NA | NA | NA |
| Xcr1 | 0 | 0 | 0 | 0 | 0 | 0 | 0 | 0 | 0 | 0 | 0 | 0 | 0 | 0 | 0 | NA | NA | NA | NA | NA | NA | NA | NA | NA | NA |
| Xdh | 1 | 1 | 0 | 0 | 1 | 0 | 1 | 1 | 0 | 0 | 1 | 0 | 0.49 | 1 | 0 | 0.599 | -0.74 | 0.78905321 | NA | 0 | 4.7929 | 3.7323 | 3.7323 | TRUE | 0.5075 |
| Xiap | 4451 | 3129 | 3463 | 3643 | 3062 | 3531 | 3457 | 3328 | 3737 | 3626 | 3454 | 3486 | 3514.64 | 3507 | 3522 | 1.004 | 0.006 | 0.91850443 | 0.95533284 | 0.0018 | 0.0031 | 0.0026 | 0.0026 | TRUE | 0.0391 |
| Xirp1 | 5 | 5 | 3 | 4 | 3 | 4 | 4 | 5 | 3 | 4 | 3 | 4 | 3.96 | 4 | 4 | 0.913 | -0.131 | 0.88836376 | NA | 0 | 0.6011 | 0.3732 | 0.3732 | TRUE | 0.0953 |
| Xirp2 | 26 | 22 | 11 | 1 | 6 | 2 | 20 | 23 | 12 | 1 | 7 | 2 | 10.87 | 18 | 3 | 0.17 | -2.555 | 0.00055143 | NA | 0.1211 | 0.2205 | 0.204 | 0.204 | TRUE | 0.9987 |
| Xist | 206 | 169 | 102 | 3 | 4 | 1 | 160 | 180 | 110 | 3 | 5 | 1 | 76.38 | 150 | 3 | 0.018 | -5.765 | 1.18E-24 | 5.82E-22 | 0 | 0.0335 | 0.0376 | 0.0376 | TRUE | 0.3501 |
| Xk | 817 | 617 | 678 | 771 | 679 | 722 | 635 | 656 | 732 | 767 | 766 | 713 | 711.42 | 674 | 749 | 1.112 | 0.153 | 0.09338548 | 0.21019822 | 0.0023 | 0.0058 | 0.0046 | 0.0046 | TRUE | 0.0645 |
| Xkr4 | 2 | 0 | 3 | 3 | 1 | 1 | 2 | 0 | 3 | 3 | 1 | 1 | 1.65 | 2 | 2 | 1.073 | 0.102 | 0.94722717 | NA | 0 | 1.44 | 1.0891 | 1.0891 | TRUE | 0.196 |
| Xkr5 | 593 | 496 | 448 | 536 | 443 | 478 | 461 | 528 | 483 | 534 | 500 | 472 | 496.11 | 491 | 502 | 1.024 | 0.035 | 0.73565581 | 0.84225637 | 0.0024 | 0.0072 | 0.0057 | 0.0057 | TRUE | 0.0505 |
| Xkr6 | 31 | 19 | 17 | 10 | 9 | 15 | 24 | 20 | 18 | 10 | 10 | 15 | 16.26 | 21 | 12 | 0.556 | -0.846 | 0.08050987 | NA | 0 | 0.1482 | 0.1016 | 0.1016 | TRUE | 0.2271 |
| Xkr7 | 31 | 21 | 21 | 33 | 26 | 35 | 24 | 22 | 23 | 33 | 29 | 35 | 27.63 | 23 | 32 | 1.4 | 0.485 | 0.1783726 | 0.33291716 | 0 | 0.0882 | 0.0567 | 0.0567 | TRUE | 0.0395 |
| Xkr8 | 458 | 336 | 378 | 399 | 352 | 418 | 356 | 357 | 408 | 397 | 397 | 413 | 387.98 | 374 | 402 | 1.079 | 0.109 | 0.32078233 | 0.49465731 | 0 | 0.0085 | 0.0061 | 0.0061 | TRUE | 0.0711 |
| Xkr9 | 4 | 1 | 1 | 2 | 3 | 2 | 3 | 1 | 1 | 2 | 3 | 2 | 2.1 | 2 | 2 | 1.345 | 0.428 | 0.74139344 | NA | 0 | 1.1311 | 0.7308 | 0.7308 | TRUE | 0.4893 |
| Xkrx | 82 | 53 | 26 | 27 | 28 | 17 | 64 | 56 | 28 | 27 | 32 | 17 | 37.23 | 49 | 25 | 0.503 | -0.992 | 0.0105304 | 0.0428684 | 0.1084 | 0.0661 | 0.0779 | 0.0779 | TRUE | 0.9245 |
| Xlr | 133 | 116 | 72 | 71 | 34 | 39 | 103 | 123 | 78 | 71 | 38 | 38 | 75.32 | 101 | 49 | 0.486 | -1.042 | 0.00027757 | 0.00272853 | 0.0718 | 0.0339 | 0.0439 | 0.0439 | TRUE | 0.4745 |
| Xlr3a | 3477 | 2692 | 3130 | 3924 | 3262 | 3863 | 2700 | 2863 | 3377 | 3906 | 3680 | 3813 | 3390.05 | 2980 | 3800 | 1.276 | 0.351 | 1.58E-05 | 0.00027733 | 0.007 | 0.0031 | 0.0045 | 0.0045 | TRUE | 0.1642 |
| Xlr3b | 1328 | 1007 | 1002 | 1196 | 1019 | 1148 | 1031 | 1071 | 1081 | 1190 | 1150 | 1133 | 1109.5 | 1061 | 1158 | 1.092 | 0.127 | 0.07892428 | 0.18729269 | 0 | 0.0046 | 0.0028 | 0.0028 | TRUE | 0.0076 |
| Xlr3c | 1808 | 1450 | 1378 | 1364 | 1172 | 1236 | 1404 | 1542 | 1487 | 1358 | 1322 | 1220 | 1388.9 | 1478 | 1300 | 0.88 | -0.184 | 0.01437249 | 0.0540027 | 0.002 | 0.0041 | 0.0033 | 0.0033 | TRUE | 0.0352 |
| Xlr4a | 544 | 444 | 428 | 474 | 364 | 455 | 423 | 472 | 462 | 472 | 411 | 449 | 448.03 | 452 | 444 | 0.984 | -0.023 | 0.82945142 | 0.90237092 | 0.002 | 0.0077 | 0.0059 | 0.0059 | TRUE | 0.0481 |
| Xlr4b | 495 | 320 | 372 | 406 | 320 | 341 | 384 | 340 | 401 | 404 | 361 | 337 | 371.33 | 375 | 367 | 0.978 | -0.032 | 0.78798122 | 0.87535675 | 0.0051 | 0.0088 | 0.0077 | 0.0077 | TRUE | 0.0902 |
| Xlr4c | 263 | 223 | 221 | 224 | 194 | 166 | 204 | 237 | 238 | 223 | 219 | 164 | 214.27 | 226 | 202 | 0.892 | -0.165 | 0.29805105 | 0.4702554 | 0.0131 | 0.0135 | 0.0134 | 0.0134 | TRUE | 0.3013 |
| Xlr5a | 810 | 604 | 572 | 665 | 531 | 597 | 629 | 642 | 617 | 662 | 599 | 589 | 623.17 | 629 | 617 | 0.98 | -0.029 | 0.74854766 | 0.85088211 | 0 | 0.0062 | 0.0044 | 0.0044 | TRUE | 0.049 |
| Xlr5b | 635 | 482 | 511 | 527 | 465 | 535 | 493 | 513 | 551 | 525 | 525 | 528 | 522.42 | 519 | 526 | 1.014 | 0.021 | 0.82853683 | 0.9017191 | 0 | 0.007 | 0.0047 | 0.0047 | TRUE | 0.0341 |
| Xlr5c | 272 | 187 | 192 | 195 | 176 | 201 | 211 | 199 | 207 | 194 | 199 | 198 | 201.4 | 206 | 197 | 0.956 | -0.064 | 0.64372964 | 0.77809422 | 0 | 0.0142 | 0.0089 | 0.0089 | TRUE | 0.0091 |
| Xpa | 295 | 253 | 168 | 201 | 158 | 170 | 229 | 269 | 181 | 200 | 178 | 168 | 204.27 | 226 | 182 | 0.803 | -0.316 | 0.06018513 | 0.15358436 | 0.0185 | 0.014 | 0.0153 | 0.0153 | TRUE | 0.1831 |
| Xpc | 2493 | 1845 | 2077 | 2052 | 1941 | 2151 | 1936 | 1963 | 2241 | 2043 | 2190 | 2123 | 2082.56 | 2047 | 2119 | 1.036 | 0.051 | 0.49676889 | 0.66123279 | 0.0034 | 0.0036 | 0.0035 | 0.0035 | TRUE | 0.0827 |
| Xpnpep1 | 12877 | 9687 | 10333 | 9983 | 9405 | 10988 | 10001 | 10304 | 11149 | 9937 | 10610 | 10847 | 10474.65 | 10485 | 10465 | 0.998 | -0.003 | 0.96663626 | 0.98188701 | 0.002 | 0.0027 | 0.0026 | 0.0026 | TRUE | 0.0375 |
| Xpnpep2 | 65 | 46 | 40 | 46 | 37 | 40 | 50 | 49 | 43 | 46 | 42 | 39 | 44.93 | 47 | 42 | 0.889 | -0.17 | 0.55101714 | 0.70697379 | 0 | 0.0552 | 0.0362 | 0.0362 | TRUE | 0.0488 |
| Xpnpep3 | 1161 | 775 | 726 | 842 | 677 | 805 | 902 | 824 | 783 | 838 | 764 | 795 | 817.65 | 836 | 799 | 0.954 | -0.068 | 0.43912068 | 0.61048068 | 0.0026 | 0.0053 | 0.0044 | 0.0044 | TRUE | 0.0579 |
| Xpo1 | 33854 | 24206 | 23162 | 24893 | 20713 | 23504 | 26293 | 25748 | 24992 | 24778 | 23366 | 23202 | 24729.86 | 25678 | 23782 | 0.926 | -0.111 | 0.02817578 | 0.08847022 | 9.00E-04 | 0.0025 | 0.0018 | 0.0018 | TRUE | 0.0164 |
| Xpo4 | 4597 | 3309 | 3203 | 3768 | 3239 | 3734 | 3570 | 3520 | 3456 | 3751 | 3654 | 3686 | 3606.11 | 3515 | 3697 | 1.051 | 0.072 | 0.17035098 | 0.32195322 | 0 | 0.0031 | 0.0017 | 0.0017 | TRUE | 0.0026 |
| Xpo5 | 7005 | 4674 | 3770 | 3512 | 3103 | 3423 | 5440 | 4972 | 4068 | 3496 | 3501 | 3379 | 4142.56 | 4827 | 3459 | 0.716 | -0.482 | 1.13E-07 | 4.34E-06 | 0.0103 | 0.003 | 0.0057 | 0.0057 | TRUE | 0.2299 |
| Xpo6 | 9078 | 6686 | 6979 | 7084 | 6692 | 7928 | 7050 | 7112 | 7530 | 7051 | 7549 | 7826 | 7353.24 | 7231 | 7475 | 1.034 | 0.048 | 0.41584756 | 0.58860262 | 0.0018 | 0.0027 | 0.0024 | 0.0024 | TRUE | 0.0302 |
| Xpo7 | 10482 | 7358 | 7863 | 8522 | 7451 | 8718 | 8141 | 7827 | 8484 | 8483 | 8406 | 8606 | 8324.32 | 8151 | 8498 | 1.043 | 0.06 | 0.24689954 | 0.41568896 | 7.00E-04 | 0.0027 | 0.0018 | 0.0018 | TRUE | 0.0155 |
| Xpot | 11557 | 7895 | 7562 | 7560 | 6695 | 7390 | 8976 | 8398 | 8159 | 7525 | 7553 | 7295 | 7984.32 | 8511 | 7458 | 0.876 | -0.191 | 0.00058225 | 0.00484972 | 0.0012 | 0.0027 | 0.0021 | 0.0021 | TRUE | 0.0282 |
| Xpr1 | 3811 | 2711 | 3033 | 3469 | 2839 | 3256 | 2960 | 2884 | 3273 | 3453 | 3203 | 3214 | 3164.33 | 3039 | 3290 | 1.083 | 0.115 | 0.09131499 | 0.20697196 | 0.0027 | 0.0032 | 0.003 | 0.003 | TRUE | 0.0543 |
| Xrcc1 | 3815 | 2877 | 2888 | 2876 | 2657 | 3280 | 2963 | 3060 | 3116 | 2863 | 2997 | 3238 | 3039.55 | 3046 | 3033 | 0.996 | -0.006 | 0.92651103 | 0.9596103 | 0.002 | 0.0032 | 0.0027 | 0.0027 | TRUE | 0.0429 |
| Xrcc2 | 1178 | 881 | 904 | 973 | 745 | 800 | 915 | 937 | 975 | 969 | 840 | 790 | 904.35 | 942 | 866 | 0.92 | -0.121 | 0.19024984 | 0.34831415 | 0.0045 | 0.005 | 0.005 | 0.005 | TRUE | 0.1286 |
| Xrcc3 | 865 | 680 | 658 | 667 | 591 | 710 | 672 | 723 | 710 | 664 | 667 | 701 | 689.44 | 702 | 677 | 0.966 | -0.049 | 0.56541174 | 0.71982096 | 0 | 0.0059 | 0.0039 | 0.0039 | TRUE | 0.0175 |
| Xrcc4 | 425 | 339 | 337 | 387 | 320 | 377 | 330 | 361 | 364 | 385 | 361 | 372 | 362.11 | 352 | 373 | 1.063 | 0.089 | 0.42244586 | 0.59472469 | 0 | 0.009 | 0.006 | 0.006 | TRUE | 0.0351 |
| Xrcc5 | 8287 | 6036 | 6252 | 6621 | 5844 | 6637 | 6436 | 6420 | 6746 | 6591 | 6593 | 6552 | 6556.22 | 6534 | 6579 | 1.007 | 0.01 | 0.84220073 | 0.91032497 | 2.00E-04 | 0.0028 | 0.0016 | 0.0016 | TRUE | 0.0097 |
| Xrcc6 | 8112 | 5813 | 5713 | 6205 | 5512 | 6398 | 6300 | 6183 | 6164 | 6176 | 6218 | 6316 | 6226.35 | 6216 | 6237 | 1.003 | 0.005 | 0.9225817 | 0.95719768 | 0 | 0.0028 | 0.0015 | 0.0015 | TRUE | 0.0018 |
| Xrcc6bp1 | 772 | 587 | 677 | 694 | 648 | 679 | 600 | 624 | 730 | 691 | 731 | 670 | 674.42 | 651 | 697 | 1.072 | 0.1 | 0.31368151 | 0.48713345 | 0.005 | 0.0059 | 0.0056 | 0.0056 | TRUE | 0.1306 |
| Xrn1 | 3213 | 2361 | 2302 | 2759 | 2402 | 2742 | 2495 | 2511 | 2484 | 2746 | 2710 | 2707 | 2608.9 | 2497 | 2721 | 1.09 | 0.124 | 0.02479219 | 0.08092138 | 0 | 0.0033 | 0.0018 | 0.0018 | TRUE | 8.00E-04 |
| Xrn2 | 12640 | 9383 | 9890 | 8790 | 7940 | 9263 | 9817 | 9981 | 10671 | 8750 | 8957 | 9144 | 9553.24 | 10156 | 8950 | 0.881 | -0.182 | 0.00073564 | 0.00578355 | 0.0011 | 0.0027 | 0.002 | 0.002 | TRUE | 0.024 |
| Xrra1 | 62 | 42 | 42 | 48 | 31 | 42 | 48 | 45 | 45 | 48 | 35 | 41 | 43.73 | 46 | 41 | 0.9 | -0.152 | 0.60247403 | 0.74743176 | 0 | 0.0566 | 0.0385 | 0.0385 | TRUE | 0.1379 |
| Xylb | 1376 | 994 | 923 | 957 | 848 | 1028 | 1069 | 1057 | 996 | 953 | 957 | 1015 | 1007.65 | 1041 | 975 | 0.936 | -0.095 | 0.21427076 | 0.37693193 | 0 | 0.0048 | 0.0032 | 0.0032 | TRUE | 0.0165 |
| Xylt1 | 196 | 142 | 114 | 112 | 109 | 122 | 152 | 151 | 123 | 111 | 123 | 120 | 130.19 | 142 | 118 | 0.83 | -0.269 | 0.13128859 | 0.2686689 | 0 | 0.0206 | 0.0151 | 0.0151 | TRUE | 0.1369 |
| Xylt2 | 3800 | 2893 | 2497 | 2771 | 2626 | 2767 | 2951 | 3077 | 2694 | 2758 | 2962 | 2731 | 2862.48 | 2907 | 2817 | 0.969 | -0.046 | 0.5062099 | 0.66968328 | 0.0026 | 0.0033 | 0.0031 | 0.0031 | TRUE | 0.0492 |
| Yae1d1 | 1646 | 1172 | 1301 | 1407 | 1226 | 1427 | 1278 | 1247 | 1404 | 1401 | 1383 | 1409 | 1353.51 | 1310 | 1398 | 1.068 | 0.095 | 0.19550972 | 0.35520789 | 0.0012 | 0.0042 | 0.0031 | 0.0031 | TRUE | 0.0462 |
| Yaf2 | 1210 | 834 | 833 | 860 | 724 | 805 | 940 | 887 | 899 | 856 | 817 | 795 | 865.52 | 909 | 823 | 0.905 | -0.145 | 0.06927114 | 0.17041176 | 0 | 0.0052 | 0.0034 | 0.0034 | TRUE | 0.0154 |
| Yap1 | 10278 | 7189 | 6928 | 6782 | 5909 | 6962 | 7982 | 7647 | 7475 | 6751 | 6666 | 6873 | 7232.32 | 7701 | 6763 | 0.878 | -0.188 | 0.00025914 | 0.00260109 | 5.00E-04 | 0.0028 | 0.0018 | 0.0018 | TRUE | 0.0126 |
| Yars | 7891 | 5584 | 4844 | 4944 | 4440 | 5124 | 6129 | 5940 | 5227 | 4921 | 5009 | 5058 | 5380.51 | 5765 | 4996 | 0.866 | -0.207 | 0.00215027 | 0.01275006 | 0.0032 | 0.0029 | 0.0031 | 0.0031 | TRUE | 0.081 |
| Yars2 | 1835 | 1485 | 1386 | 1392 | 1156 | 1418 | 1425 | 1580 | 1495 | 1386 | 1304 | 1400 | 1431.62 | 1500 | 1363 | 0.91 | -0.136 | 0.06045103 | 0.15407036 | 0.0014 | 0.0041 | 0.0031 | 0.0031 | TRUE | 0.0259 |
| Ybey | 264 | 215 | 207 | 210 | 203 | 243 | 205 | 229 | 223 | 209 | 229 | 240 | 222.5 | 219 | 226 | 1.034 | 0.048 | 0.72803543 | 0.83701821 | 0 | 0.0131 | 0.0092 | 0.0092 | TRUE | 0.0485 |
| Ybx1 | 70148 | 51825 | 49589 | 55071 | 48011 | 55820 | 54481 | 55126 | 53506 | 54818 | 54161 | 55103 | 54532.44 | 54371 | 54694 | 1.006 | 0.009 | 0.84045048 | 0.90907815 | 1.00E-04 | 0.0025 | 0.0013 | 0.0013 | TRUE | 0.0024 |
| Ybx2 | 549 | 569 | 612 | 812 | 694 | 798 | 426 | 605 | 660 | 808 | 783 | 788 | 678.48 | 564 | 793 | 1.411 | 0.497 | 0.00020131 | 0.00214459 | 0.024 | 0.0059 | 0.0113 | 0.0113 | TRUE | 0.5576 |
| Ydjc | 2028 | 1511 | 1474 | 1675 | 1339 | 1655 | 1575 | 1607 | 1590 | 1667 | 1511 | 1634 | 1597.38 | 1591 | 1604 | 1.009 | 0.013 | 0.85271998 | 0.9167369 | 7.00E-04 | 0.0039 | 0.0027 | 0.0027 | TRUE | 0.0306 |
| Yeats2 | 4220 | 3027 | 3008 | 3104 | 2990 | 3306 | 3277 | 3220 | 3246 | 3090 | 3373 | 3264 | 3244.86 | 3248 | 3242 | 0.998 | -0.003 | 0.95978349 | 0.97773647 | 7.00E-04 | 0.0032 | 0.0021 | 0.0021 | TRUE | 0.0206 |
| Yeats4 | 5018 | 3561 | 3547 | 3878 | 3462 | 3953 | 3897 | 3788 | 3827 | 3860 | 3905 | 3902 | 3863.35 | 3837 | 3889 | 1.013 | 0.019 | 0.71193158 | 0.82692149 | 0 | 0.003 | 0.0017 | 0.0017 | TRUE | 0.0023 |
| Yes1 | 5307 | 3835 | 4108 | 3648 | 3113 | 3774 | 4122 | 4079 | 4432 | 3631 | 3512 | 3726 | 3917 | 4211 | 3623 | 0.861 | -0.217 | 0.00026313 | 0.00263185 | 0.0011 | 0.003 | 0.0023 | 0.0023 | TRUE | 0.0254 |
| Yif1a | 1384 | 1042 | 1093 | 1185 | 1080 | 1128 | 1075 | 1108 | 1179 | 1180 | 1218 | 1114 | 1145.67 | 1121 | 1171 | 1.045 | 0.063 | 0.4093168 | 0.58206796 | 0.0013 | 0.0045 | 0.0034 | 0.0034 | TRUE | 0.0247 |
| Yif1b | 3181 | 2448 | 2653 | 3020 | 2568 | 3048 | 2471 | 2604 | 2863 | 3006 | 2897 | 3009 | 2808.16 | 2646 | 2971 | 1.124 | 0.168 | 0.01421227 | 0.05361201 | 0.0025 | 0.0033 | 0.003 | 0.003 | TRUE | 0.0621 |

| | | | | | | | | | | | | | | | | | | | | | | | | | |
|---|---|---|---|---|---|---|---|---|---|---|---|---|---|---|---|---|---|---|---|---|---|---|---|---|---|
| Yipf1 | 738 | 566 | 563 | 598 | 562 | 619 | 573 | 602 | 607 | 595 | 634 | 611 | 603.83 | 594 | 613 | 1.033 | 0.047 | 0.59999751 | 0.7456538 | 0 | 0.0064 | 0.0041 | 0.0041 | TRUE | 0.0121 |
| Yipf2 | 664 | 514 | 530 | 554 | 481 | 572 | 516 | 547 | 572 | 551 | 543 | 565 | 548.84 | 545 | 553 | 1.017 | 0.024 | 0.79750795 | 0.88178085 | 0 | 0.0067 | 0.0045 | 0.0045 | TRUE | 0.0273 |
| Yipf3 | 2058 | 1426 | 1381 | 1371 | 1322 | 1467 | 1598 | 1517 | 1490 | 1365 | 1491 | 1448 | 1484.91 | 1535 | 1435 | 0.934 | -0.099 | 0.16026169 | 0.30963294 | 0.001 | 0.004 | 0.0029 | 0.0029 | TRUE | 0.022 |
| Yipf4 | 2301 | 1744 | 1779 | 2151 | 1911 | 2238 | 1787 | 1855 | 1920 | 2141 | 2156 | 2209 | 2011.31 | 1854 | 2169 | 1.171 | 0.227 | 0.000269 | 0.00267369 | 3.00E-04 | 0.0036 | 0.0023 | 0.0023 | TRUE | 0.0125 |
| Yipf5 | 2894 | 2048 | 1838 | 1996 | 1654 | 1884 | 2248 | 2178 | 1983 | 1987 | 1866 | 1860 | 2020.29 | 2136 | 1904 | 0.891 | -0.167 | 0.01858075 | 0.06546085 | 0.0023 | 0.0036 | 0.0031 | 0.0031 | TRUE | 0.0472 |
| Yipf6 | 1499 | 1092 | 1114 | 1186 | 999 | 1143 | 1164 | 1162 | 1202 | 1181 | 1127 | 1128 | 1160.6 | 1176 | 1145 | 0.974 | -0.037 | 0.59611674 | 0.74235543 | 0 | 0.0045 | 0.0027 | 0.0027 | TRUE | 0.0088 |
| Yipf7 | 8 | 6 | 5 | 6 | 3 | 4 | 6 | 6 | 5 | 6 | 3 | 4 | 5.22 | 6 | 4 | 0.742 | -0.431 | 0.60011937 | NA | 0 | 0.4568 | 0.2911 | 0.2911 | TRUE | 0.1712 |
| Ykt6 | 4520 | 3089 | 2738 | 2986 | 2539 | 2975 | 3510 | 3286 | 2954 | 2972 | 2864 | 2937 | 3087.3 | 3250 | 2924 | 0.899 | -0.153 | 0.03247258 | 0.0981456 | 0.0034 | 0.0032 | 0.0034 | 0.0034 | TRUE | 0.0767 |
| Ylpm1 | 6577 | 4791 | 3983 | 4201 | 3698 | 4437 | 5108 | 5096 | 4298 | 4182 | 4172 | 4380 | 4539.2 | 4834 | 4245 | 0.878 | -0.188 | 0.01089869 | 0.04399182 | 0.0047 | 0.0029 | 0.0037 | 0.0037 | TRUE | 0.1141 |
| Yme1l1 | 20013 | 14157 | 14168 | 15712 | 13238 | 15193 | 15543 | 15059 | 15287 | 15640 | 14934 | 14998 | 15243.4 | 15296 | 15191 | 0.993 | -0.01 | 0.83098442 | 0.9033735 | 3.00E-04 | 0.0026 | 0.0015 | 0.0015 | TRUE | 0.0082 |
| Yod1 | 301 | 224 | 228 | 233 | 224 | 212 | 234 | 238 | 246 | 232 | 253 | 209 | 235.32 | 239 | 231 | 0.966 | -0.05 | 0.71070769 | 0.8259326 | 0 | 0.0125 | 0.009 | 0.009 | TRUE | 0.0772 |
| Ypel1 | 427 | 305 | 334 | 352 | 313 | 334 | 332 | 324 | 360 | 350 | 353 | 330 | 341.6 | 339 | 344 | 1.017 | 0.024 | 0.83128353 | 0.90340142 | 0 | 0.0094 | 0.0063 | 0.0063 | TRUE | 0.0337 |
| Ypel2 | 1955 | 1648 | 2004 | 2091 | 1995 | 2417 | 1518 | 1753 | 2162 | 2081 | 2251 | 2386 | 2025.25 | 1811 | 2239 | 1.238 | 0.308 | 0.00471522 | 0.02344978 | 0.0178 | 0.0036 | 0.008 | 0.008 | TRUE | 0.3469 |
| Ypel3 | 853 | 654 | 746 | 781 | 668 | 771 | 662 | 696 | 805 | 777 | 754 | 761 | 742.52 | 721 | 764 | 1.062 | 0.086 | 0.36060131 | 0.53548827 | 0.004 | 0.0056 | 0.0051 | 0.0051 | TRUE | 0.121 |
| Ypel4 | 0 | 2 | 0 | 0 | 1 | 0 | 0 | 2 | 0 | 0 | 1 | 0 | 0.54 | 1 | 0 | 0.535 | -0.903 | 0.76014553 | NA | 0 | 4.3706 | 4.6038 | 4.6038 | TRUE | 0.9746 |
| Ypel5 | 1789 | 1271 | 1314 | 1454 | 1196 | 1403 | 1389 | 1352 | 1418 | 1447 | 1349 | 1385 | 1390.11 | 1386 | 1394 | 1.006 | 0.008 | 0.90591857 | 0.94866248 | 0 | 0.0041 | 0.0027 | 0.0027 | TRUE | 0.0136 |
| Yrdc | 2893 | 1972 | 1850 | 2082 | 1789 | 2038 | 2247 | 2098 | 1996 | 2072 | 2018 | 2012 | 2073.84 | 2114 | 2034 | 0.962 | -0.057 | 0.40015537 | 0.573415 | 0.0015 | 0.0036 | 0.0028 | 0.0028 | TRUE | 0.038 |
| Ythdc1 | 6679 | 4917 | 4338 | 4678 | 4089 | 4622 | 5187 | 5230 | 4681 | 4656 | 4613 | 4563 | 4821.67 | 5033 | 4611 | 0.916 | -0.127 | 0.03619234 | 0.10657131 | 0.0016 | 0.0029 | 0.0024 | 0.0024 | TRUE | 0.0452 |
| Ythdc2 | 3234 | 2236 | 2178 | 2110 | 1929 | 2203 | 2512 | 2378 | 2350 | 2100 | 2176 | 2175 | 2281.88 | 2413 | 2150 | 0.89 | -0.168 | 0.00623654 | 0.02904491 | 5.00E-04 | 0.0035 | 0.0023 | 0.0023 | TRUE | 0.0159 |
| Ythdf1 | 10246 | 7316 | 6589 | 6872 | 6262 | 7276 | 7958 | 7782 | 7109 | 6840 | 7064 | 7183 | 7322.69 | 7616 | 7029 | 0.923 | -0.116 | 0.05076592 | 0.13639046 | 0.0018 | 0.0028 | 0.0024 | 0.0024 | TRUE | 0.0412 |
| Ythdf2 | 5988 | 4395 | 3950 | 4010 | 3457 | 3827 | 4651 | 4675 | 4262 | 3992 | 3900 | 3778 | 4209.47 | 4529 | 3890 | 0.859 | -0.22 | 0.00025661 | 0.00258477 | 0.0013 | 0.003 | 0.0024 | 0.0024 | TRUE | 0.0321 |
| Ythdf3 | 7708 | 5373 | 5502 | 5611 | 4798 | 5520 | 5986 | 5715 | 5937 | 5585 | 5413 | 5449 | 5680.87 | 5879 | 5482 | 0.932 | -0.101 | 0.04713095 | 0.12907504 | 3.00E-04 | 0.0028 | 0.0017 | 0.0017 | TRUE | 0.0072 |
| Ywhab | 14850 | 10879 | 9736 | 9733 | 8921 | 10048 | 11533 | 11572 | 10505 | 9688 | 10064 | 9919 | 10546.87 | 11203 | 9890 | 0.883 | -0.18 | 0.00137791 | 0.00906524 | 0.0015 | 0.0027 | 0.0022 | 0.0022 | TRUE | 0.0362 |
| Ywhae | 104133 | 75538 | 80326 | 87038 | 76699 | 89042 | 80876 | 80349 | 86671 | 86638 | 86524 | 87898 | 84825.95 | 82632 | 87020 | 1.053 | 0.075 | 0.12864536 | 0.26509922 | 9.00E-04 | 0.0025 | 0.0017 | 0.0017 | TRUE | 0.0224 |
| Ywhag | 33024 | 23563 | 19392 | 20531 | 17724 | 19908 | 25648 | 25064 | 20924 | 20437 | 19995 | 19652 | 21953.19 | 23879 | 20028 | 0.839 | -0.254 | 0.00061773 | 0.00507427 | 0.0059 | 0.0025 | 0.0039 | 0.0039 | TRUE | 0.1432 |
| Ywhah | 10144 | 6927 | 6715 | 7153 | 6145 | 6901 | 7878 | 7368 | 7245 | 7120 | 6932 | 6812 | 7226.11 | 7497 | 6955 | 0.927 | -0.109 | 0.04800913 | 0.13077845 | 0.001 | 0.0028 | 0.002 | 0.002 | TRUE | 0.0244 |
| Ywhaq | 11449 | 8125 | 8127 | 9065 | 7900 | 9080 | 8892 | 8642 | 8769 | 9023 | 8912 | 8963 | 8867.01 | 8768 | 8966 | 1.023 | 0.032 | 0.48520617 | 0.65111743 | 0 | 0.0027 | 0.0014 | 0.0014 | TRUE | 0.0019 |
| Ywhaz | 20461 | 14523 | 14404 | 14150 | 12325 | 14510 | 15891 | 15448 | 15542 | 14085 | 13904 | 14324 | 14865.55 | 15627 | 14104 | 0.903 | -0.148 | 0.00106606 | 0.00758278 | 1.00E-04 | 0.0026 | 0.0014 | 0.0014 | TRUE | 0.0027 |
| Yy1 | 7260 | 5511 | 5017 | 4914 | 4340 | 4779 | 5639 | 5862 | 5413 | 4891 | 4896 | 4718 | 5236.46 | 5638 | 4835 | 0.858 | -0.222 | 5.66E-05 | 0.00077314 | 7.00E-04 | 0.0029 | 0.002 | 0.002 | TRUE | 0.0146 |
| Yy2 | 534 | 385 | 293 | 264 | 212 | 264 | 415 | 410 | 316 | 263 | 239 | 261 | 317.16 | 380 | 254 | 0.668 | -0.583 | 1.78E-05 | 0.00030491 | 0.0101 | 0.0099 | 0.01 | 0.01 | TRUE | 0.2419 |
| Zadh2 | 345 | 286 | 308 | 423 | 413 | 404 | 268 | 304 | 332 | 421 | 466 | 399 | 365.04 | 301 | 429 | 1.425 | 0.511 | 3.22E-05 | 0.00049329 | 0.0061 | 0.0089 | 0.0081 | 0.0081 | TRUE | 0.1162 |
| Zan | 79 | 62 | 60 | 85 | 75 | 106 | 61 | 66 | 65 | 85 | 85 | 105 | 77.65 | 64 | 92 | 1.43 | 0.516 | 0.02140538 | 0.07264297 | 0 | 0.0329 | 0.023 | 0.023 | TRUE | 0.1601 |
| Zap70 | 448 | 382 | 478 | 382 | 342 | 399 | 348 | 406 | 516 | 380 | 386 | 394 | 404.99 | 423 | 387 | 0.916 | -0.126 | 0.36229828 | 0.53696431 | 0.018 | 0.0083 | 0.0113 | 0.0113 | TRUE | 0.4191 |
| Zar1 | 9 | 10 | 3 | 6 | 8 | 0 | 7 | 11 | 3 | 6 | 9 | 0 | 5.98 | 7 | 5 | 0.71 | -0.494 | 0.60006598 | NA | 0.7415 | 0.399 | 0.4651 | 0.4651 | TRUE | 0.4877 |
| Zar1l | 4 | 2 | 2 | 4 | 4 | 1 | 3 | 2 | 2 | 4 | 5 | 1 | 2.81 | 2 | 3 | 1.259 | 0.333 | 0.77017712 | NA | 0 | 0.8452 | 0.5764 | 0.5764 | TRUE | 0.5142 |
| Zbbx | 52 | 31 | 29 | 20 | 20 | 19 | 40 | 33 | 31 | 20 | 23 | 19 | 27.65 | 35 | 21 | 0.581 | -0.784 | 0.03414196 | 0.10204885 | 0 | 0.0882 | 0.0595 | 0.0595 | TRUE | 0.1583 |
| Zbed3 | 603 | 434 | 395 | 433 | 344 | 423 | 468 | 462 | 426 | 431 | 388 | 418 | 432.13 | 452 | 412 | 0.912 | -0.134 | 0.20155268 | 0.36250062 | 0 | 0.0079 | 0.0056 | 0.0056 | TRUE | 0.0292 |
| Zbed4 | 1703 | 1176 | 1216 | 1266 | 1117 | 1335 | 1323 | 1251 | 1312 | 1260 | 1260 | 1318 | 1287.29 | 1295 | 1279 | 0.988 | -0.018 | 0.79431795 | 0.87961466 | 0 | 0.0043 | 0.0027 | 0.0027 | TRUE | 0.0106 |
| Zbed6 | 3437 | 2647 | 2579 | 2904 | 2520 | 3100 | 2669 | 2816 | 2783 | 2891 | 2843 | 3060 | 2843.55 | 2756 | 2931 | 1.064 | 0.09 | 0.13588916 | 0.27542027 | 8.00E-04 | 0.0033 | 0.0023 | 0.0023 | TRUE | 0.0181 |
| Zbp1 | 43 | 43 | 44 | 54 | 57 | 67 | 33 | 46 | 47 | 54 | 64 | 66 | 51.8 | 42 | 61 | 1.467 | 0.552 | 0.04803524 | 0.13082463 | 0 | 0.0482 | 0.0363 | 0.0363 | TRUE | 0.3069 |
| Zbtb1 | 924 | 701 | 667 | 673 | 622 | 728 | 718 | 746 | 720 | 670 | 702 | 719 | 712.2 | 728 | 697 | 0.958 | -0.062 | 0.45838473 | 0.6281242 | 0 | 0.0058 | 0.0037 | 0.0037 | TRUE | 0.0137 |
| Zbtb10 | 3267 | 2273 | 2217 | 1825 | 1607 | 1666 | 2537 | 2418 | 2392 | 1817 | 1813 | 1645 | 2103.55 | 2449 | 1758 | 0.717 | -0.479 | 1.27E-12 | 1.54E-10 | 0.0013 | 0.0036 | 0.0028 | 0.0028 | TRUE | 0.0388 |
| Zbtb11 | 3825 | 2906 | 2843 | 3092 | 2669 | 3260 | 2971 | 3091 | 3068 | 3078 | 3011 | 3218 | 3072.69 | 3043 | 3102 | 1.02 | 0.029 | 0.6213269 | 0.76111715 | 5.00E-04 | 0.0032 | 0.0021 | 0.0021 | TRUE | 0.013 |
| Zbtb12 | 598 | 502 | 494 | 695 | 603 | 748 | 464 | 534 | 533 | 692 | 680 | 738 | 606.98 | 510 | 703 | 1.382 | 0.467 | 1.44E-06 | 3.92E-05 | 0.0025 | 0.0063 | 0.0051 | 0.0051 | TRUE | 0.078 |
| Zbtb14 | 1249 | 970 | 906 | 855 | 791 | 963 | 970 | 1032 | 978 | 851 | 892 | 951 | 945.57 | 993 | 898 | 0.905 | -0.144 | 0.07233024 | 0.17575928 | 0.001 | 0.0049 | 0.0036 | 0.0036 | TRUE | 0.0319 |
| Zbtb16 | 31 | 29 | 22 | 19 | 17 | 18 | 24 | 31 | 24 | 19 | 19 | 18 | 22.42 | 26 | 19 | 0.712 | -0.491 | 0.22427386 | 0.38869241 | 0 | 0.1081 | 0.0713 | 0.0713 | TRUE | 0.1403 |
| Zbtb17 | 2654 | 1991 | 2010 | 2006 | 1904 | 2165 | 2061 | 2118 | 2169 | 1997 | 2148 | 2137 | 2104.95 | 2116 | 2094 | 0.99 | -0.015 | 0.81692511 | 0.89487832 | 7.00E-04 | 0.0036 | 0.0024 | 0.0024 | TRUE | 0.0201 |
| Zbtb18 | 1627 | 1251 | 1242 | 1125 | 1055 | 1186 | 1264 | 1331 | 1340 | 1120 | 1190 | 1171 | 1235.86 | 1312 | 1160 | 0.885 | -0.176 | 0.01360064 | 0.05196308 | 0 | 0.0043 | 0.0028 | 0.0028 | TRUE | 0.0128 |
| Zbtb2 | 494 | 328 | 279 | 234 | 231 | 256 | 384 | 349 | 301 | 233 | 261 | 253 | 296.64 | 345 | 249 | 0.719 | -0.475 | 0.00034034 | 0.00319401 | 0.0065 | 0.0104 | 0.0092 | 0.0092 | TRUE | 0.1367 |
| Zbtb20 | 98 | 64 | 66 | 78 | 97 | 87 | 76 | 68 | 71 | 78 | 109 | 86 | 81.39 | 72 | 91 | 1.261 | 0.334 | 0.13915619 | 0.2804249 | 0 | 0.0315 | 0.0244 | 0.0244 | TRUE | 0.2872 |
| Zbtb21 | 2366 | 1775 | 1493 | 1713 | 1456 | 1595 | 1838 | 1888 | 1611 | 1705 | 1643 | 1575 | 1709.78 | 1779 | 1641 | 0.922 | -0.117 | 0.13190685 | 0.26968168 | 0.0034 | 0.0038 | 0.0038 | 0.0038 | TRUE | 0.0808 |
| Zbtb22 | 1252 | 869 | 743 | 809 | 710 | 770 | 972 | 924 | 802 | 805 | 801 | 760 | 844.13 | 899 | 789 | 0.875 | -0.192 | 0.03763366 | 0.10964084 | 0.0044 | 0.0052 | 0.0049 | 0.0049 | TRUE | 0.106 |
| Zbtb24 | 1022 | 801 | 727 | 796 | 765 | 784 | 794 | 852 | 784 | 792 | 863 | 774 | 809.91 | 810 | 810 | 0.999 | -0.001 | 0.9912707 | 0.99553848 | 0.0013 | 0.0054 | 0.004 | 0.004 | TRUE | 0.0384 |
| Zbtb25 | 828 | 560 | 632 | 626 | 571 | 686 | 643 | 596 | 682 | 623 | 644 | 677 | 644.19 | 640 | 648 | 1.012 | 0.018 | 0.84688658 | 0.91313744 | 0.0015 | 0.0061 | 0.0046 | 0.0046 | TRUE | 0.0416 |
| Zbtb26 | 632 | 513 | 560 | 605 | 567 | 652 | 491 | 546 | 604 | 602 | 640 | 644 | 587.71 | 547 | 629 | 1.152 | 0.204 | 0.0456601 | 0.12608575 | 0.0044 | 0.0065 | 0.0058 | 0.0058 | TRUE | 0.1002 |
| Zbtb3 | 357 | 274 | 251 | 300 | 275 | 282 | 277 | 291 | 271 | 299 | 310 | 278 | 287.79 | 280 | 296 | 1.056 | 0.079 | 0.51470448 | 0.67722278 | 0 | 0.0107 | 0.0071 | 0.0071 | TRUE | 0.0301 |
| Zbtb32 | 434 | 310 | 308 | 464 | 347 | 421 | 337 | 330 | 332 | 462 | 391 | 416 | 378.01 | 333 | 423 | 1.271 | 0.346 | 0.00204838 | 0.01228083 | 0 | 0.0087 | 0.0064 | 0.0064 | TRUE | 0.075 |
| Zbtb33 | 1140 | 790 | 742 | 828 | 702 | 824 | 885 | 840 | 801 | 824 | 792 | 813 | 825.97 | 842 | 810 | 0.961 | -0.058 | 0.47896511 | 0.64589568 | 0 | 0.0053 | 0.0036 | 0.0036 | TRUE | 0.0253 |
| Zbtb34 | 2324 | 1794 | 1748 | 1940 | 1628 | 1926 | 1805 | 1908 | 1886 | 1931 | 1837 | 1901 | 1878.03 | 1866 | 1890 | 1.013 | 0.019 | 0.76357525 | 0.85969854 | 0 | 0.0037 | 0.0024 | 0.0024 | TRUE | 0.0105 |
| Zbtb37 | 232 | 173 | 166 | 176 | 162 | 176 | 180 | 184 | 179 | 175 | 183 | 174 | 179.17 | 181 | 177 | 0.978 | -0.032 | 0.82822173 | 0.90158212 | 0 | 0.0157 | 0.0098 | 0.0098 | TRUE | 0.0076 |
| Zbtb38 | 970 | 733 | 702 | 645 | 652 | 744 | 753 | 780 | 757 | 642 | 736 | 734 | 733.75 | 763 | 704 | 0.922 | -0.117 | 0.18782163 | 0.34496717 | 0.0016 | 0.0057 | 0.0043 | 0.0043 | TRUE | 0.0709 |
| Zbtb39 | 1189 | 893 | 899 | 856 | 751 | 905 | 923 | 950 | 970 | 852 | 847 | 893 | 906 | 948 | 864 | 0.913 | -0.132 | 0.0873735 | 0.20058862 | 0 | 0.005 | 0.0032 | 0.0032 | TRUE | 0.0105 |
| Zbtb4 | 134 | 124 | 74 | 157 | 125 | 135 | 104 | 132 | 80 | 156 | 141 | 133 | 124.4 | 105 | 143 | 1.363 | 0.447 | 0.02824694 | 0.08859635 | 0.0226 | 0.0215 | 0.0218 | 0.0218 | TRUE | 0.285 |
| Zbtb40 | 1722 | 1366 | 1238 | 1282 | 1127 | 1373 | 1337 | 1453 | 1336 | 1276 | 1271 | 1355 | 1338.17 | 1375 | 1301 | 0.947 | -0.079 | 0.27586597 | 0.44699607 | 0.0011 | 0.0042 | 0.0031 | 0.0031 | TRUE | 0.0288 |
| Zbtb41 | 1049 | 755 | 814 | 905 | 732 | 819 | 815 | 803 | 878 | 901 | 826 | 808 | 838.53 | 832 | 845 | 1.016 | 0.023 | 0.78308259 | 0.8723772 | 0.0016 | 0.0053 | 0.004 | 0.004 | TRUE | 0.0402 |
| Zbtb42 | 242 | 187 | 165 | 166 | 177 | 195 | 188 | 199 | 178 | 165 | 200 | 192 | 187.05 | 188 | 186 | 0.986 | -0.021 | 0.89046752 | 0.93936469 | 0 | 0.0151 | 0.011 | 0.011 | TRUE | 0.1023 |
| Zbtb43 | 2789 | 2131 | 1959 | 1984 | 1826 | 2049 | 2166 | 2267 | 2114 | 1975 | 2060 | 2023 | 2100.67 | 2182 | 2019 | 0.925 | -0.112 | 0.0711104 | 0.17351974 | 3.00E-04 | 0.0036 | 0.0023 | 0.0023 | TRUE | 0.0135 |
| Zbtb44 | 6546 | 4802 | 5397 | 5197 | 4545 | 5176 | 5084 | 5108 | 5823 | 5173 | 5127 | 5110 | 5237.5 | 5338 | 5137 | 0.963 | -0.055 | 0.3973828 | 0.57074956 | 0.0028 | 0.0029 | 0.0029 | 0.0029 | TRUE | 0.0765 |
| Zbtb45 | 1666 | 1197 | 1007 | 969 | 826 | 1002 | 1294 | 1273 | 1087 | 965 | 932 | 989 | 1089.86 | 1218 | 962 | 0.789 | -0.342 | 7.94E-05 | 0.00100431 | 0.0042 | 0.0046 | 0.0045 | 0.0045 | TRUE | 0.1049 |

| | | | | | | | | | | | | | | | | | | | | | | | | | |
|---|---|---|---|---|---|---|---|---|---|---|---|---|---|---|---|---|---|---|---|---|---|---|---|---|---|
| Zbtb46 | 442 | 359 | 320 | 376 | 375 | 389 | 343 | 382 | 345 | 374 | 423 | 384 | 375.29 | 357 | 394 | 1.104 | 0.142 | 0.20500669 | 0.36654499 | 0 | 0.0087 | 0.0064 | 0.0064 | TRUE | 0.0472 |
| Zbtb48 | 784 | 583 | 532 | 545 | 530 | 521 | 609 | 620 | 574 | 542 | 598 | 514 | 576.29 | 601 | 551 | 0.916 | -0.126 | 0.19409338 | 0.35299359 | 0.0017 | 0.0065 | 0.005 | 0.005 | TRUE | 0.0616 |
| Zbtb49 | 264 | 230 | 177 | 202 | 202 | 225 | 205 | 245 | 191 | 201 | 228 | 222 | 215.29 | 214 | 217 | 1.017 | 0.024 | 0.87213846 | 0.92903341 | 0.0054 | 0.0134 | 0.0111 | 0.0111 | TRUE | 0.1697 |
| Zbtb5 | 773 | 646 | 648 | 673 | 562 | 643 | 600 | 687 | 699 | 670 | 634 | 635 | 654.22 | 662 | 646 | 0.978 | -0.032 | 0.73668688 | 0.84296418 | 0.0027 | 0.006 | 0.0049 | 0.0049 | TRUE | 0.0836 |
| Zbtb6 | 770 | 547 | 462 | 602 | 480 | 567 | 598 | 582 | 498 | 599 | 541 | 560 | 563.13 | 559 | 567 | 1.012 | 0.017 | 0.86869452 | 0.92755496 | 0.0042 | 0.0066 | 0.0059 | 0.0059 | TRUE | 0.1038 |
| Zbtb7a | 1251 | 1186 | 1268 | 1584 | 1538 | 1596 | 972 | 1262 | 1368 | 1577 | 1735 | 1575 | 1414.75 | 1201 | 1629 | 1.359 | 0.443 | 5.42E-05 | 0.00074875 | 0.0151 | 0.0041 | 0.0079 | 0.0079 | TRUE | 0.3427 |
| Zbtb7b | 1022 | 787 | 760 | 826 | 756 | 841 | 794 | 837 | 820 | 822 | 853 | 830 | 826.02 | 817 | 835 | 1.023 | 0.032 | 0.68129573 | 0.80444113 | 0 | 0.0053 | 0.0033 | 0.0033 | TRUE | 0.0078 |
| Zbtb7c | 401 | 274 | 229 | 349 | 321 | 397 | 311 | 291 | 247 | 347 | 362 | 392 | 325.23 | 283 | 367 | 1.292 | 0.37 | 0.00349703 | 0.01859487 | 0.0054 | 0.0097 | 0.0084 | 0.0084 | TRUE | 0.1369 |
| Zbtb8a | 817 | 636 | 626 | 663 | 593 | 722 | 635 | 677 | 675 | 660 | 669 | 713 | 671.35 | 662 | 681 | 1.029 | 0.042 | 0.63493176 | 0.77160293 | 0 | 0.006 | 0.004 | 0.004 | TRUE | 0.0205 |
| Zbtb8b | 435 | 349 | 388 | 489 | 460 | 504 | 338 | 371 | 419 | 487 | 519 | 498 | 438.49 | 376 | 501 | 1.337 | 0.419 | 0.00016831 | 0.0018692 | 0.0038 | 0.0078 | 0.0066 | 0.0066 | TRUE | 0.1126 |
| Zbtb8os | 1583 | 1064 | 915 | 988 | 764 | 889 | 1229 | 1132 | 987 | 983 | 862 | 878 | 1011.9 | 1116 | 908 | 0.812 | -0.3 | 0.00200306 | 0.0121261 | 0.0078 | 0.0048 | 0.0058 | 0.0058 | TRUE | 0.1211 |
| Zbtb9 | 1450 | 1035 | 1012 | 1004 | 859 | 1045 | 1126 | 1101 | 1092 | 999 | 969 | 1032 | 1053.17 | 1106 | 1000 | 0.904 | -0.146 | 0.04607156 | 0.12698058 | 0 | 0.0047 | 0.0029 | 0.0029 | TRUE | 0.0091 |
| Zbtbd6 | 162 | 94 | 57 | 78 | 67 | 52 | 126 | 100 | 62 | 78 | 76 | 51 | 81.98 | 96 | 68 | 0.707 | -0.501 | 0.07446643 | 0.17945519 | 0.076 | 0.0313 | 0.0442 | 0.0442 | TRUE | 0.2337 |
| Zc2hc1a | 1680 | 1173 | 1464 | 1635 | 1448 | 1679 | 1305 | 1248 | 1580 | 1627 | 1633 | 1657 | 1508.42 | 1378 | 1639 | 1.191 | 0.252 | 0.00472653 | 0.02348965 | 0.0073 | 0.004 | 0.0051 | 0.0051 | TRUE | 0.1952 |
| Zc2hc1b | 0 | 0 | 0 | 0 | 0 | 0 | 0 | 0 | 0 | 0 | 0 | 0 | 0 | 0 | 0 | NA | NA | NA | NA | NA | NA | NA | NA | NA | NA |
| Zc2hc1c | 89 | 79 | 82 | 73 | 59 | 75 | 69 | 84 | 88 | 73 | 67 | 74 | 75.81 | 80 | 71 | 0.888 | -0.171 | 0.45029311 | 0.62034115 | 0 | 0.0337 | 0.0238 | 0.0238 | TRUE | 0.164 |
| Zc3h10 | 1430 | 1195 | 1359 | 1340 | 1254 | 1516 | 1111 | 1271 | 1466 | 1334 | 1415 | 1497 | 1348.85 | 1283 | 1415 | 1.105 | 0.144 | 0.14627753 | 0.2906234 | 0.0102 | 0.0042 | 0.0063 | 0.0063 | TRUE | 0.1883 |
| Zc3h11a | 10504 | 7339 | 6828 | 7448 | 6577 | 7710 | 8158 | 7806 | 7367 | 7414 | 7420 | 7611 | 7629.33 | 7777 | 7482 | 0.962 | -0.056 | 0.31291108 | 0.48624027 | 0.0012 | 0.0027 | 0.0021 | 0.0021 | TRUE | 0.0259 |
| Zc3h12a | 363 | 299 | 269 | 333 | 306 | 308 | 282 | 318 | 290 | 331 | 345 | 304 | 311.82 | 297 | 327 | 1.103 | 0.141 | 0.24344919 | 0.41122859 | 0 | 0.01 | 0.0073 | 0.0073 | TRUE | 0.0437 |
| Zc3h12b | 42 | 14 | 20 | 24 | 16 | 23 | 33 | 15 | 22 | 24 | 18 | 23 | 22.29 | 23 | 22 | 0.926 | -0.112 | 0.79718476 | 0.88160891 | 0.0488 | 0.1088 | 0.0914 | 0.0914 | TRUE | 0.3923 |
| Zc3h12c | 250 | 166 | 166 | 180 | 139 | 156 | 194 | 177 | 179 | 179 | 157 | 154 | 173.3 | 183 | 163 | 0.89 | -0.168 | 0.27394732 | 0.44499705 | 0 | 0.0161 | 0.0113 | 0.0113 | TRUE | 0.0778 |
| Zc3h12d | 18 | 4 | 5 | 5 | 4 | 6 | 14 | 4 | 5 | 5 | 5 | 6 | 6.51 | 8 | 5 | 0.639 | -0.647 | 0.41488577 | NA | 0 | 0.3667 | 0.2928 | 0.2928 | TRUE | 1.6728 |
| Zc3h13 | 5849 | 4339 | 3622 | 4221 | 3679 | 4203 | 4543 | 4615 | 3908 | 4202 | 4150 | 4149 | 4261.17 | 4355 | 4167 | 0.956 | -0.064 | 0.36160228 | 0.53649194 | 0.0037 | 0.003 | 0.0033 | 0.0033 | TRUE | 0.0974 |
| Zc3h14 | 11435 | 8383 | 8875 | 8980 | 7924 | 9213 | 8881 | 8917 | 9576 | 8939 | 8939 | 9095 | 9057.74 | 9125 | 8991 | 0.985 | -0.021 | 0.68720377 | 0.80827876 | 7.00E-04 | 0.0027 | 0.0019 | 0.0019 | TRUE | 0.0228 |
| Zc3h15 | 9482 | 6653 | 6109 | 6346 | 5220 | 6253 | 7364 | 7077 | 6592 | 6317 | 5889 | 6173 | 6568.45 | 7011 | 6126 | 0.874 | -0.195 | 0.00127962 | 0.008615 | 0.002 | 0.0028 | 0.0025 | 0.0025 | TRUE | 0.0333 |
| Zc3h18 | 13274 | 9951 | 8131 | 8603 | 7898 | 9283 | 10309 | 10585 | 8773 | 8563 | 8910 | 9164 | 9384.05 | 9889 | 8879 | 0.898 | -0.156 | 0.03346923 | 0.10041473 | 0.0053 | 0.0027 | 0.0038 | 0.0038 | TRUE | 0.1186 |
| Zc3h3 | 797 | 644 | 564 | 538 | 468 | 614 | 619 | 685 | 609 | 536 | 528 | 606 | 597.03 | 638 | 557 | 0.874 | -0.194 | 0.0498715 | 0.13453809 | 0.0032 | 0.0064 | 0.0054 | 0.0054 | TRUE | 0.0719 |
| Zc3h4 | 5807 | 3937 | 3245 | 3512 | 3229 | 3505 | 4510 | 4188 | 3501 | 3496 | 3643 | 3460 | 3799.6 | 4066 | 3533 | 0.868 | -0.204 | 0.01657601 | 0.06016274 | 0.008 | 0.0031 | 0.0049 | 0.0049 | TRUE | 0.1792 |
| Zc3h6 | 1234 | 921 | 1303 | 1641 | 1490 | 1578 | 958 | 980 | 1406 | 1633 | 1681 | 1558 | 1369.34 | 1115 | 1624 | 1.459 | 0.545 | 8.17E-06 | 0.00015867 | 0.0252 | 0.0042 | 0.01 | 0.01 | TRUE | 0.6214 |
| Zc3h7a | 2122 | 1570 | 1699 | 1827 | 1585 | 1811 | 1648 | 1670 | 1833 | 1819 | 1788 | 1788 | 1757.61 | 1717 | 1798 | 1.048 | 0.068 | 0.32209548 | 0.49587577 | 0.0012 | 0.0038 | 0.0028 | 0.0028 | TRUE | 0.0416 |
| Zc3h7b | 6085 | 4545 | 4748 | 4865 | 4478 | 5023 | 4726 | 4834 | 5123 | 4843 | 5052 | 4958 | 4922.7 | 4894 | 4951 | 1.012 | 0.017 | 0.76250753 | 0.85909823 | 9.00E-04 | 0.0029 | 0.002 | 0.002 | TRUE | 0.0202 |
| Zc3h8 | 991 | 743 | 657 | 815 | 615 | 773 | 770 | 790 | 709 | 811 | 694 | 763 | 756.16 | 756 | 756 | 1 | 0 | 0.99989697 | 0.99993023 | 0.0032 | 0.0056 | 0.0048 | 0.0048 | TRUE | 0.06 |
| Zc3hav1 | 3637 | 2779 | 2956 | 2810 | 2607 | 2762 | 2825 | 2956 | 3189 | 2797 | 2941 | 2727 | 2905.79 | 2990 | 2822 | 0.944 | -0.083 | 0.21385679 | 0.37629633 | 0.0021 | 0.0032 | 0.0029 | 0.0029 | TRUE | 0.0412 |
| Zc3hav1l | 1851 | 1365 | 1418 | 1501 | 1289 | 1575 | 1438 | 1452 | 1530 | 1494 | 1454 | 1555 | 1487.09 | 1473 | 1501 | 1.02 | 0.028 | 0.68042305 | 0.8037427 | 4.00E-04 | 0.004 | 0.0027 | 0.0027 | TRUE | 0.0135 |
| Zc3hc1 | 7930 | 5620 | 5662 | 6075 | 5501 | 6416 | 6159 | 5978 | 6109 | 6047 | 6206 | 6334 | 6138.73 | 6082 | 6196 | 1.019 | 0.026 | 0.5966473 | 0.74274039 | 2.00E-04 | 0.0028 | 0.0016 | 0.0016 | TRUE | 0.0054 |
| Zc4h2 | 350 | 272 | 274 | 327 | 257 | 307 | 272 | 289 | 296 | 325 | 290 | 303 | 295.88 | 286 | 306 | 1.075 | 0.104 | 0.3907851 | 0.5645332 | 0 | 0.0104 | 0.0071 | 0.0071 | TRUE | 0.0346 |
| Zcchc10 | 3090 | 2211 | 1911 | 1978 | 1733 | 2105 | 2400 | 2352 | 2062 | 1969 | 1955 | 2078 | 2135.92 | 2271 | 2001 | 0.88 | -0.184 | 0.01338835 | 0.05137171 | 0.0031 | 0.0035 | 0.0035 | 0.0035 | TRUE | 0.0778 |
| Zcchc11 | 6807 | 4933 | 4778 | 5322 | 4648 | 5300 | 5287 | 5247 | 5155 | 5298 | 5243 | 5232 | 5243.69 | 5230 | 5258 | 1.005 | 0.007 | 0.87823329 | 0.93196497 | 0 | 0.0029 | 0.0015 | 0.0015 | TRUE | 0.0019 |
| Zcchc12 | 104 | 83 | 60 | 51 | 59 | 65 | 81 | 88 | 65 | 51 | 67 | 64 | 69.21 | 78 | 61 | 0.774 | -0.37 | 0.13224071 | 0.2701324 | 0 | 0.0367 | 0.0287 | 0.0287 | TRUE | 0.1891 |
| Zcchc13 | 6 | 3 | 10 | 17 | 15 | 16 | 5 | 3 | 11 | 17 | 17 | 16 | 11.38 | 6 | 17 | 2.701 | 1.434 | 0.02011323 | NA | 0 | 0.2107 | 0.1643 | 0.1643 | TRUE | 0.9002 |
| Zcchc14 | 5509 | 4068 | 4016 | 4251 | 3954 | 4602 | 4279 | 4327 | 4333 | 4231 | 4461 | 4543 | 4362.29 | 4313 | 4412 | 1.023 | 0.033 | 0.54719168 | 0.7040944 | 5.00E-04 | 0.003 | 0.0019 | 0.0019 | TRUE | 0.0157 |
| Zcchc16 | 1 | 0 | 0 | 0 | 0 | 0 | 1 | 0 | 0 | 0 | 0 | 0 | 0.13 | 0 | 0 | 0.552 | -0.857 | 0.8336548 | NA | 0 | 18.3121 | 10 | 10 | TRUE | 0.2566 |
| Zcchc17 | 2428 | 1607 | 1211 | 1379 | 1181 | 1227 | 1886 | 1709 | 1307 | 1373 | 1332 | 1211 | 1469.65 | 1634 | 1305 | 0.798 | -0.325 | 0.00441819 | 0.02234581 | 0.0175 | 0.004 | 0.0087 | 0.0087 | TRUE | 0.3679 |
| Zcchc18 | 149 | 115 | 132 | 149 | 139 | 162 | 116 | 122 | 142 | 148 | 157 | 160 | 140.92 | 127 | 155 | 1.226 | 0.294 | 0.08335874 | 0.1943632 | 0 | 0.0192 | 0.0137 | 0.0137 | TRUE | 0.1147 |
| Zcchc2 | 2427 | 1809 | 1927 | 1855 | 1607 | 1856 | 1885 | 1924 | 2079 | 1846 | 1813 | 1832 | 1896.64 | 1963 | 1830 | 0.933 | -0.1 | 0.13129693 | 0.2686689 | 9.00E-04 | 0.0037 | 0.0026 | 0.0026 | TRUE | 0.0322 |
| Zcchc24 | 1792 | 1316 | 1449 | 1672 | 1523 | 1657 | 1392 | 1400 | 1563 | 1664 | 1718 | 1636 | 1562.19 | 1452 | 1673 | 1.153 | 0.205 | 0.00466572 | 0.02324772 | 0.0016 | 0.0039 | 0.0032 | 0.0032 | TRUE | 0.0537 |
| Zcchc3 | 1857 | 1465 | 1096 | 1007 | 905 | 1104 | 1442 | 1558 | 1183 | 1002 | 1021 | 1090 | 1216.04 | 1394 | 1038 | 0.744 | -0.427 | 1.81E-05 | 0.00030897 | 0.0092 | 0.0044 | 0.0063 | 0.0063 | TRUE | 0.2086 |
| Zcchc4 | 900 | 614 | 638 | 654 | 571 | 672 | 699 | 653 | 688 | 651 | 644 | 663 | 666.5 | 680 | 653 | 0.959 | -0.06 | 0.48314384 | 0.64950855 | 0 | 0.006 | 0.0038 | 0.0038 | TRUE | 0.014 |
| Zcchc5 | 0 | 0 | 0 | 0 | 0 | 0 | 0 | 0 | 0 | 0 | 0 | 0 | 0 | 0 | 0 | NA | NA | NA | NA | NA | NA | NA | NA | NA | NA |
| Zcchc6 | 3267 | 2307 | 2325 | 2586 | 2194 | 2706 | 2537 | 2454 | 2509 | 2574 | 2475 | 2671 | 2536.72 | 2500 | 2573 | 1.029 | 0.042 | 0.48513286 | 0.65111743 | 5.00E-04 | 0.0034 | 0.0022 | 0.0022 | TRUE | 0.0134 |
| Zcchc7 | 1562 | 1082 | 829 | 959 | 770 | 956 | 1213 | 1151 | 894 | 955 | 869 | 944 | 1004.25 | 1086 | 923 | 0.848 | -0.237 | 0.02895384 | 0.09023927 | 0.0132 | 0.0048 | 0.0075 | 0.0075 | TRUE | 0.2818 |
| Zcchc8 | 4716 | 3417 | 3411 | 3371 | 3027 | 3466 | 3663 | 3635 | 3680 | 3355 | 3415 | 3421 | 3528.25 | 3659 | 3397 | 0.928 | -0.107 | 0.03948393 | 0.11349754 | 0 | 0.0031 | 0.0017 | 0.0017 | TRUE | 0.0014 |
| Zcchc9 | 1071 | 781 | 892 | 916 | 782 | 951 | 832 | 831 | 962 | 912 | 882 | 939 | 892.96 | 875 | 911 | 1.043 | 0.06 | 0.49095188 | 0.65673142 | 0.0029 | 0.0051 | 0.0044 | 0.0044 | TRUE | 0.089 |
| Zcrb1 | 1468 | 1128 | 1026 | 992 | 830 | 966 | 1140 | 1200 | 1107 | 987 | 936 | 954 | 1054.06 | 1149 | 959 | 0.835 | -0.26 | 0.00054079 | 0.00457368 | 0 | 0.0047 | 0.0031 | 0.0031 | TRUE | 0.0173 |
| Zcwpw1 | 438 | 348 | 341 | 495 | 337 | 354 | 340 | 370 | 368 | 493 | 380 | 349 | 383.44 | 359 | 407 | 1.135 | 0.183 | 0.17732324 | 0.33143469 | 0.0152 | 0.0086 | 0.0107 | 0.0107 | TRUE | 0.3908 |
| Zdbf2 | 6966 | 4595 | 2976 | 3679 | 3467 | 3929 | 5410 | 4888 | 3211 | 3662 | 3911 | 3879 | 4160.11 | 4503 | 3817 | 0.848 | -0.239 | 0.26535226 | 0.43568982 | 0.0328 | 0.003 | 0.0124 | 0.0328 | TRUE | 0.7665 |
| Zdhhc1 | 234 | 221 | 207 | 304 | 275 | 300 | 182 | 235 | 223 | 303 | 310 | 296 | 258.19 | 213 | 303 | 1.427 | 0.513 | 0.00019988 | 0.00213357 | 0.005 | 0.0116 | 0.0097 | 0.0097 | TRUE | 0.2037 |
| Zdhhc11 | 0 | 0 | 0 | 0 | 0 | 0 | 0 | 0 | 0 | 0 | 0 | 0 | 0 | 0 | 0 | NA | NA | NA | NA | NA | NA | NA | NA | NA | NA |
| Zdhhc12 | 2074 | 1500 | 1616 | 1822 | 1635 | 1820 | 1611 | 1596 | 1744 | 1814 | 1844 | 1797 | 1734.11 | 1650 | 1818 | 1.102 | 0.141 | 0.03450912 | 0.10282237 | 6.00E-04 | 0.0038 | 0.0026 | 0.0026 | TRUE | 0.0291 |
| Zdhhc13 | 3079 | 2111 | 2365 | 2662 | 2326 | 2710 | 2391 | 2245 | 2552 | 2650 | 2624 | 2675 | 2522.92 | 2396 | 2650 | 1.106 | 0.145 | 0.02645812 | 0.08469995 | 0.0015 | 0.0034 | 0.0027 | 0.0027 | TRUE | 0.0381 |
| Zdhhc14 | 131 | 78 | 91 | 150 | 114 | 123 | 102 | 83 | 98 | 149 | 129 | 121 | 113.71 | 94 | 133 | 1.408 | 0.494 | 0.01014172 | 0.04173787 | 0 | 0.0233 | 0.0176 | 0.0176 | TRUE | 0.119 |
| Zdhhc15 | 703 | 438 | 373 | 468 | 357 | 414 | 546 | 466 | 402 | 466 | 403 | 409 | 448.6 | 471 | 426 | 0.901 | -0.151 | 0.23516655 | 0.40153707 | 0.013 | 0.0077 | 0.0094 | 0.0094 | TRUE | 0.2319 |
| Zdhhc16 | 1718 | 1223 | 1211 | 1326 | 1128 | 1241 | 1334 | 1301 | 1307 | 1320 | 1273 | 1225 | 1293.22 | 1314 | 1273 | 0.968 | -0.047 | 0.50103848 | 0.66530499 | 0 | 0.0043 | 0.0027 | 0.0027 | TRUE | 0.013 |
| Zdhhc17 | 998 | 706 | 718 | 798 | 689 | 785 | 775 | 751 | 775 | 794 | 777 | 775 | 774.55 | 767 | 782 | 1.02 | 0.028 | 0.72480383 | 0.83525124 | 0 | 0.0055 | 0.0033 | 0.0033 | TRUE | 0.0038 |
| Zdhhc18 | 9032 | 6605 | 6450 | 8191 | 7262 | 8104 | 7015 | 7026 | 6959 | 8153 | 8192 | 8000 | 7557.57 | 7000 | 8115 | 1.159 | 0.213 | 4.80E-06 | 0.00010249 | 0 | 0.0027 | 0.0014 | 0.0014 | TRUE | 0.0019 |
| Zdhhc19 | 4 | 1 | 2 | 5 | 0 | 0 | 3 | 1 | 2 | 5 | 0 | 0 | 1.88 | 2 | 2 | 0.787 | -0.346 | 0.8273988 | NA | 1.3644 | 1.2603 | 1.282 | 1.282 | TRUE | 2.3419 |
| Zdhhc2 | 1214 | 888 | 776 | 678 | 576 | 646 | 943 | 945 | 837 | 675 | 650 | 638 | 781.18 | 908 | 654 | 0.72 | -0.475 | 5.39E-08 | 2.23E-06 | 0.0017 | 0.0055 | 0.0042 | 0.0042 | TRUE | 0.0542 |
| Zdhhc20 | 2921 | 2165 | 2381 | 2873 | 2436 | 3129 | 2269 | 2303 | 2569 | 2860 | 2748 | 3089 | 2639.54 | 2380 | 2899 | 1.219 | 0.286 | 9.36E-05 | 0.00114711 | 0.0036 | 0.0033 | 0.0035 | 0.0035 | TRUE | 0.0579 |
| Zdhhc21 | 3006 | 2314 | 2464 | 2869 | 2394 | 2836 | 2335 | 2461 | 2659 | 2856 | 2701 | 2800 | 2635.11 | 2485 | 2786 | 1.122 | 0.166 | 0.01354961 | 0.0518513 | 0.002 | 0.0033 | 0.0029 | 0.0029 | TRUE | 0.0452 |

| | | | | | | | | | | | | | | | | | | | | | | | | | |
|---|---|---|---|---|---|---|---|---|---|---|---|---|---|---|---|---|---|---|---|---|---|---|---|---|---|
| Zdhhc22 | 0 | 2 | 1 | 2 | 3 | 3 | 0 | 2 | 1 | 2 | 3 | 3 | 1.92 | 1 | 3 | 2.742 | 1.455 | 0.32083429 | NA | 0 | 1.2344 | 0.8856 | 0.8856 | TRUE | 0.5466 |
| Zdhhc23 | 160 | 124 | 114 | 117 | 99 | 123 | 124 | 132 | 123 | 116 | 112 | 121 | 121.46 | 126 | 116 | 0.923 | -0.115 | 0.51204803 | 0.67490801 | 0 | 0.0219 | 0.0141 | 0.0141 | TRUE | 0.0141 |
| Zdhhc24 | 664 | 525 | 582 | 722 | 612 | 735 | 516 | 558 | 628 | 719 | 690 | 726 | 639.46 | 567 | 712 | 1.257 | 0.331 | 0.00072992 | 0.00574491 | 0.0036 | 0.0061 | 0.0053 | 0.0053 | TRUE | 0.1015 |
| Zdhhc25 | 0 | 1 | 1 | 0 | 0 | 0 | 0 | 1 | 1 | 0 | 0 | 0 | 0.36 | 1 | 0 | 0.271 | -1.881 | 0.5755355 | NA | 0 | 6.6391 | 6.4428 | 6.4428 | TRUE | 0.3513 |
| Zdhhc3 | 4002 | 2700 | 2411 | 2358 | 2127 | 2429 | 3108 | 2872 | 2601 | 2347 | 2399 | 2398 | 2621 | 2860 | 2381 | 0.832 | -0.266 | 0.00029447 | 0.00286907 | 0.0034 | 0.0033 | 0.0035 | 0.0035 | TRUE | 0.0759 |
| Zdhhc4 | 808 | 555 | 464 | 506 | 467 | 455 | 628 | 590 | 501 | 504 | 527 | 449 | 533.03 | 573 | 493 | 0.859 | -0.22 | 0.05179266 | 0.13831697 | 0.0082 | 0.0069 | 0.0073 | 0.0073 | TRUE | 0.1401 |
| Zdhhc5 | 5413 | 3695 | 3066 | 3121 | 2781 | 3112 | 4204 | 3930 | 3308 | 3107 | 3137 | 3072 | 3459.75 | 3814 | 3105 | 0.814 | -0.297 | 0.00031436 | 0.00301105 | 0.0069 | 0.0031 | 0.0046 | 0.0046 | TRUE | 0.1632 |
| Zdhhc6 | 3094 | 2166 | 1968 | 2061 | 1777 | 2007 | 2403 | 2304 | 2123 | 2052 | 2005 | 1981 | 2144.63 | 2277 | 2013 | 0.883 | -0.179 | 0.0081437 | 0.03554133 | 0.0017 | 0.0035 | 0.0028 | 0.0028 | TRUE | 0.0417 |
| Zdhhc7 | 2237 | 1561 | 1621 | 1762 | 1629 | 1744 | 1737 | 1660 | 1749 | 1754 | 1838 | 1722 | 1743.34 | 1715 | 1771 | 1.032 | 0.045 | 0.48856127 | 0.65439098 | 3.00E-04 | 0.0038 | 0.0025 | 0.0025 | TRUE | 0.0129 |
| Zdhhc8 | 2215 | 1656 | 1525 | 1795 | 1573 | 1849 | 1720 | 1761 | 1645 | 1787 | 1775 | 1825 | 1752.29 | 1709 | 1796 | 1.051 | 0.071 | 0.26598625 | 0.43637988 | 0 | 0.0038 | 0.0024 | 0.0024 | TRUE | 0.0124 |
| Zdhhc9 | 1398 | 1002 | 1051 | 963 | 893 | 1069 | 1086 | 1066 | 1134 | 959 | 1007 | 1055 | 1051.14 | 1095 | 1007 | 0.92 | -0.121 | 0.1154419 | 0.24530321 | 7.00E-04 | 0.0047 | 0.0033 | 0.0033 | TRUE | 0.0217 |
| Zeb1 | 1177 | 799 | 726 | 817 | 720 | 826 | 914 | 850 | 783 | 813 | 812 | 815 | 831.37 | 849 | 813 | 0.957 | -0.064 | 0.45641247 | 0.62611609 | 0.002 | 0.0053 | 0.0041 | 0.0041 | TRUE | 0.0559 |
| Zeb2 | 128 | 69 | 52 | 93 | 62 | 96 | 99 | 73 | 56 | 93 | 70 | 95 | 81.03 | 76 | 86 | 1.117 | 0.159 | 0.53241882 | 0.69209228 | 0.0419 | 0.0317 | 0.0346 | 0.0346 | TRUE | 0.3935 |
| Zer1 | 1106 | 823 | 969 | 1011 | 995 | 1102 | 859 | 875 | 1046 | 1006 | 1122 | 1088 | 999.43 | 927 | 1072 | 1.158 | 0.212 | 0.02407817 | 0.07916939 | 0.0062 | 0.0048 | 0.0054 | 0.0054 | TRUE | 0.148 |
| Zf12 | 1 | 0 | 0 | 0 | 0 | 0 | 1 | 0 | 0 | 0 | 0 | 0 | 0.13 | 0 | 0 | 0.552 | -0.857 | 0.8336548 | NA | 0 | 18.3121 | 10 | 10 | TRUE | 0.2566 |
| Zfa-ps | 6 | 4 | 2 | 1 | 2 | 0 | 5 | 4 | 2 | 1 | 2 | 0 | 2.39 | 4 | 1 | 0.283 | -1.819 | 0.18371487 | NA | 0 | 0.9952 | 0.728 | 0.728 | TRUE | 0.4053 |
| Zfand1 | 1586 | 1188 | 1263 | 1429 | 1169 | 1459 | 1232 | 1264 | 1363 | 1422 | 1319 | 1440 | 1339.94 | 1286 | 1394 | 1.085 | 0.118 | 0.11620204 | 0.24667693 | 0.0017 | 0.0042 | 0.0033 | 0.0033 | TRUE | 0.0323 |
| Zfand2a | 835 | 525 | 429 | 482 | 372 | 451 | 649 | 558 | 463 | 480 | 420 | 445 | 502.41 | 557 | 448 | 0.804 | -0.316 | 0.01284138 | 0.04976731 | 0.0148 | 0.0071 | 0.0096 | 0.0096 | TRUE | 0.1613 |
| Zfand2b | 739 | 583 | 554 | 580 | 506 | 692 | 574 | 620 | 598 | 577 | 571 | 683 | 603.85 | 597 | 610 | 1.024 | 0.034 | 0.73867803 | 0.84443135 | 0.0043 | 0.0064 | 0.0057 | 0.0057 | TRUE | 0.1289 |
| Zfand3 | 3096 | 2333 | 2203 | 2118 | 1853 | 2244 | 2405 | 2482 | 2377 | 2108 | 2090 | 2215 | 2279.49 | 2421 | 2138 | 0.883 | -0.179 | 0.0031248 | 0.01704687 | 3.00E-04 | 0.0035 | 0.0022 | 0.0022 | TRUE | 0.0122 |
| Zfand4 | 231 | 186 | 123 | 190 | 180 | 197 | 179 | 198 | 133 | 189 | 203 | 194 | 182.77 | 170 | 195 | 1.148 | 0.199 | 0.23871975 | 0.4056618 | 0.0141 | 0.0154 | 0.015 | 0.015 | TRUE | 0.3707 |
| Zfand5 | 5051 | 3444 | 3109 | 3078 | 2604 | 2891 | 3923 | 3663 | 3355 | 3064 | 2938 | 2854 | 3299.35 | 3647 | 2952 | 0.809 | -0.306 | 1.41E-05 | 0.00025103 | 0.0032 | 0.0031 | 0.0033 | 0.0033 | TRUE | 0.0596 |
| Zfand6 | 3262 | 2373 | 2514 | 2713 | 2367 | 2778 | 2533 | 2524 | 2713 | 2701 | 2670 | 2742 | 2647.2 | 2590 | 2704 | 1.045 | 0.063 | 0.29209196 | 0.46428718 | 5.00E-04 | 0.0033 | 0.0022 | 0.0022 | TRUE | 0.0204 |
| Zfat | 811 | 553 | 514 | 472 | 390 | 487 | 630 | 588 | 555 | 470 | 440 | 481 | 527.2 | 591 | 464 | 0.784 | -0.352 | 0.00038264 | 0.00351713 | 0.0015 | 0.0069 | 0.0051 | 0.0051 | TRUE | 0.0413 |
| Zfc3h1 | 4874 | 3895 | 3654 | 3847 | 3398 | 4000 | 3785 | 4143 | 3943 | 3829 | 3833 | 3949 | 3913.72 | 3957 | 3870 | 0.979 | -0.031 | 0.58951391 | 0.7382741 | 8.00E-04 | 0.003 | 0.0022 | 0.0022 | TRUE | 0.0206 |
| Zfhx2 | 1509 | 840 | 519 | 962 | 726 | 701 | 1172 | 893 | 560 | 958 | 819 | 692 | 849.01 | 875 | 823 | 0.94 | -0.09 | 0.78594286 | 0.87416062 | 0.0773 | 0.0052 | 0.0242 | 0.0773 | TRUE | 0.2212 |
| Zfhx2as | 6 | 6 | 4 | 5 | 6 | 5 | 5 | 6 | 4 | 5 | 7 | 5 | 5.34 | 5 | 6 | 1.086 | 0.119 | 0.88317218 | NA | 0 | 0.4463 | 0.2827 | 0.2827 | TRUE | 0.0878 |
| Zfhx3 | 2334 | 1439 | 1396 | 1632 | 1540 | 1867 | 1813 | 1531 | 1506 | 1624 | 1737 | 1843 | 1675.74 | 1617 | 1735 | 1.072 | 0.1 | 0.25209304 | 0.42107817 | 0.0066 | 0.0038 | 0.0049 | 0.0049 | TRUE | 0.1391 |
| Zfhx4 | 51 | 29 | 17 | 21 | 26 | 17 | 40 | 31 | 18 | 21 | 29 | 17 | 25.97 | 30 | 22 | 0.744 | -0.427 | 0.31015997 | 0.48301553 | 0.0738 | 0.0937 | 0.0881 | 0.0881 | TRUE | 0.2292 |
| Zfml | 10433 | 7398 | 6763 | 7216 | 6279 | 7418 | 8103 | 7869 | 7297 | 7183 | 7083 | 7323 | 7476.35 | 7756 | 7196 | 0.928 | -0.108 | 0.05645022 | 0.14720726 | 0.0014 | 0.0027 | 0.0022 | 0.0022 | TRUE | 0.0326 |
| Zfp1 | 1713 | 1255 | 1243 | 1359 | 1156 | 1490 | 1330 | 1335 | 1341 | 1353 | 1304 | 1471 | 1355.7 | 1335 | 1376 | 1.031 | 0.044 | 0.54780118 | 0.70461863 | 0.0012 | 0.0042 | 0.0031 | 0.0031 | TRUE | 0.0441 |
| Zfp101 | 500 | 323 | 312 | 348 | 296 | 307 | 388 | 344 | 337 | 346 | 334 | 303 | 341.99 | 356 | 328 | 0.918 | -0.124 | 0.29937209 | 0.47166284 | 0.003 | 0.0094 | 0.0073 | 0.0073 | TRUE | 0.0761 |
| Zfp105 | 906 | 609 | 551 | 549 | 521 | 607 | 704 | 648 | 595 | 546 | 588 | 599 | 613.23 | 649 | 578 | 0.889 | -0.17 | 0.08363868 | 0.19480462 | 0.0034 | 0.0063 | 0.0053 | 0.0053 | TRUE | 0.0681 |
| Zfp106 | 46888 | 33452 | 36061 | 37996 | 34057 | 38765 | 36416 | 35582 | 38910 | 37821 | 38420 | 38267 | 37569.32 | 36969 | 38169 | 1.032 | 0.046 | 0.36533657 | 0.54005125 | 0.0011 | 0.0025 | 0.0018 | 0.0018 | TRUE | 0.0257 |
| Zfp108 | 60 | 46 | 29 | 47 | 48 | 47 | 47 | 49 | 31 | 47 | 54 | 46 | 45.69 | 42 | 49 | 1.155 | 0.208 | 0.48287835 | 0.64927378 | 0 | 0.0543 | 0.0414 | 0.0414 | TRUE | 0.3656 |
| Zfp109 | 255 | 166 | 165 | 201 | 196 | 192 | 198 | 177 | 178 | 200 | 221 | 190 | 193.9 | 184 | 204 | 1.101 | 0.139 | 0.34696037 | 0.52135246 | 0 | 0.0146 | 0.0105 | 0.0105 | TRUE | 0.0595 |
| Zfp11 | 177 | 137 | 122 | 157 | 136 | 137 | 137 | 146 | 132 | 156 | 153 | 135 | 143.3 | 138 | 148 | 1.072 | 0.1 | 0.5456273 | 0.70277112 | 0 | 0.019 | 0.0129 | 0.0129 | TRUE | 0.0638 |
| Zfp110 | 4744 | 3398 | 3289 | 3960 | 3434 | 3928 | 3684 | 3614 | 3549 | 3942 | 3874 | 3878 | 3756.82 | 3616 | 3898 | 1.078 | 0.108 | 0.03963045 | 0.11380845 | 0 | 0.0031 | 0.0017 | 0.0017 | TRUE | 0.0034 |
| Zfp111 | 1232 | 889 | 701 | 753 | 669 | 713 | 957 | 946 | 756 | 750 | 755 | 704 | 811.15 | 886 | 736 | 0.829 | -0.27 | 0.00810832 | 0.0354194 | 0.0081 | 0.0053 | 0.0063 | 0.0063 | TRUE | 0.192 |
| Zfp112 | 26 | 21 | 16 | 18 | 11 | 12 | 20 | 22 | 17 | 18 | 12 | 12 | 16.99 | 20 | 14 | 0.706 | -0.502 | 0.28376396 | NA | 0 | 0.1419 | 0.097 | 0.097 | TRUE | 0.2595 |
| Zfp113 | 414 | 329 | 285 | 341 | 290 | 313 | 322 | 350 | 308 | 339 | 327 | 309 | 325.76 | 327 | 325 | 0.997 | -0.005 | 0.96782829 | 0.98270717 | 0 | 0.0097 | 0.0068 | 0.0068 | TRUE | 0.0433 |
| Zfp114 | 37 | 32 | 33 | 26 | 24 | 31 | 29 | 34 | 36 | 26 | 27 | 31 | 30.32 | 33 | 28 | 0.854 | -0.228 | 0.51088504 | 0.67418335 | 0 | 0.0806 | 0.0533 | 0.0533 | TRUE | 0.0976 |
| Zfp119a | 95 | 75 | 63 | 71 | 56 | 69 | 74 | 80 | 68 | 71 | 63 | 68 | 70.58 | 74 | 67 | 0.912 | -0.132 | 0.56476608 | 0.71920734 | 0 | 0.036 | 0.0237 | 0.0237 | TRUE | 0.0414 |
| Zfp119b | 324 | 208 | 218 | 263 | 241 | 257 | 252 | 221 | 235 | 262 | 272 | 254 | 249.24 | 236 | 263 | 1.109 | 0.149 | 0.25007934 | 0.41893859 | 0 | 0.0119 | 0.0081 | 0.0081 | TRUE | 0.0404 |
| Zfp12 | 708 | 577 | 559 | 541 | 473 | 599 | 550 | 614 | 603 | 539 | 534 | 591 | 571.7 | 589 | 555 | 0.944 | -0.084 | 0.38688988 | 0.56071448 | 0.0018 | 0.0066 | 0.005 | 0.005 | TRUE | 0.0421 |
| Zfp120 | 778 | 576 | 613 | 630 | 585 | 705 | 604 | 613 | 661 | 627 | 660 | 696 | 643.56 | 626 | 661 | 1.057 | 0.08 | 0.38224452 | 0.55640702 | 0.001 | 0.0061 | 0.0044 | 0.0044 | TRUE | 0.028 |
| Zfp128 | 90 | 74 | 93 | 103 | 87 | 86 | 70 | 79 | 100 | 103 | 98 | 85 | 89.09 | 83 | 95 | 1.153 | 0.205 | 0.35332781 | 0.52780934 | 0.0112 | 0.029 | 0.024 | 0.024 | TRUE | 0.3063 |
| Zfp13 | 993 | 735 | 733 | 666 | 587 | 769 | 771 | 782 | 791 | 663 | 662 | 759 | 738.03 | 781 | 695 | 0.89 | -0.168 | 0.05903243 | 0.15156286 | 0.0017 | 0.0056 | 0.0044 | 0.0044 | TRUE | 0.0786 |
| Zfp131 | 1849 | 1377 | 1070 | 1064 | 921 | 1051 | 1436 | 1465 | 1155 | 1059 | 1039 | 1037 | 1198.47 | 1352 | 1045 | 0.773 | -0.372 | 8.01E-05 | 0.0010116 | 0.0078 | 0.0044 | 0.0056 | 0.0056 | TRUE | 0.1926 |
| Zfp133-ps | 22 | 12 | 18 | 12 | 12 | 16 | 17 | 13 | 19 | 12 | 14 | 16 | 15.09 | 16 | 14 | 0.837 | -0.256 | 0.60226692 | NA | 0 | 0.1595 | 0.1076 | 0.1076 | TRUE | 0.1673 |
| Zfp14 | 185 | 133 | 145 | 161 | 164 | 181 | 144 | 141 | 156 | 160 | 185 | 179 | 160.92 | 147 | 175 | 1.187 | 0.247 | 0.11826508 | 0.24968431 | 0 | 0.0172 | 0.0118 | 0.0118 | TRUE | 0.0561 |
| Zfp142 | 2005 | 1468 | 1243 | 1230 | 1137 | 1272 | 1557 | 1561 | 1341 | 1224 | 1283 | 1256 | 1370.42 | 1486 | 1254 | 0.843 | -0.247 | 0.00200158 | 0.01212386 | 0.0033 | 0.0042 | 0.0039 | 0.0039 | TRUE | 0.0866 |
| Zfp143 | 3252 | 2427 | 2597 | 2691 | 2355 | 2906 | 2526 | 2582 | 2802 | 2679 | 2657 | 2869 | 2685.56 | 2637 | 2735 | 1.038 | 0.054 | 0.4191424 | 0.59168417 | 0.0019 | 0.0033 | 0.0028 | 0.0028 | TRUE | 0.0364 |
| Zfp146 | 2189 | 1563 | 1456 | 1674 | 1473 | 1624 | 1700 | 1663 | 1571 | 1666 | 1662 | 1603 | 1644.13 | 1645 | 1644 | 0.999 | -0.002 | 0.97767219 | 0.98804134 | 5.00E-04 | 0.0039 | 0.0026 | 0.0026 | TRUE | 0.0182 |
| Zfp148 | 2693 | 1897 | 1895 | 2162 | 1870 | 2098 | 2092 | 2018 | 2045 | 2152 | 2110 | 2071 | 2081.12 | 2052 | 2111 | 1.029 | 0.041 | 0.49625535 | 0.66061075 | 0 | 0.0036 | 0.0021 | 0.0021 | TRUE | 0.0037 |
| Zfp157 | 1319 | 997 | 1091 | 1215 | 1042 | 1229 | 1024 | 1060 | 1177 | 1209 | 1175 | 1213 | 1143.37 | 1087 | 1199 | 1.105 | 0.143 | 0.0682081 | 0.16811491 | 0.0019 | 0.0045 | 0.0036 | 0.0036 | TRUE | 0.0617 |
| Zfp160 | 399 | 303 | 189 | 224 | 168 | 205 | 310 | 322 | 204 | 223 | 190 | 202 | 241.83 | 279 | 205 | 0.734 | -0.445 | 0.01019928 | 0.04190237 | 0.0297 | 0.0122 | 0.0174 | 0.0174 | TRUE | 0.5999 |
| Zfp169 | 201 | 175 | 129 | 128 | 153 | 161 | 156 | 186 | 139 | 127 | 173 | 159 | 156.73 | 160 | 153 | 0.952 | -0.07 | 0.69598063 | 0.81484692 | 0.0156 | 0.0175 | 0.017 | 0.017 | TRUE | 0.2269 |
| Zfp174 | 281 | 198 | 163 | 231 | 195 | 226 | 218 | 211 | 176 | 230 | 220 | 223 | 212.96 | 202 | 224 | 1.11 | 0.151 | 0.29438973 | 0.46643525 | 0 | 0.0136 | 0.0102 | 0.0102 | TRUE | 0.1301 |
| Zfp180 | 1560 | 1260 | 1013 | 1121 | 1135 | 1211 | 1212 | 1340 | 1093 | 1116 | 1280 | 1195 | 1206.09 | 1215 | 1197 | 0.985 | -0.022 | 0.8101867 | 0.89056361 | 0.0066 | 0.0044 | 0.0051 | 0.0051 | TRUE | 0.095 |
| Zfp182 | 420 | 303 | 260 | 240 | 183 | 223 | 326 | 322 | 281 | 239 | 206 | 220 | 265.75 | 310 | 222 | 0.716 | -0.482 | 0.00026188 | 0.00262482 | 0.0021 | 0.0113 | 0.0087 | 0.0087 | TRUE | 0.0741 |
| Zfp184 | 820 | 650 | 670 | 690 | 606 | 706 | 637 | 691 | 723 | 687 | 684 | 697 | 686.43 | 684 | 689 | 1.01 | 0.014 | 0.8730763 | 0.92959473 | 0 | 0.0059 | 0.0042 | 0.0042 | TRUE | 0.0452 |
| Zfp185 | 1968 | 1387 | 1316 | 1557 | 1324 | 1481 | 1528 | 1475 | 1420 | 1550 | 1494 | 1462 | 1488.19 | 1474 | 1502 | 1.018 | 0.025 | 0.71088731 | 0.82607417 | 5.00E-04 | 0.004 | 0.0027 | 0.0027 | TRUE | 0.0128 |
| Zfp189 | 361 | 274 | 317 | 307 | 254 | 318 | 280 | 291 | 342 | 306 | 287 | 314 | 303.32 | 304 | 302 | 0.995 | -0.008 | 0.95001522 | 0.97253743 | 0.0036 | 0.0102 | 0.0082 | 0.0082 | TRUE | 0.1275 |
| Zfp191 | 2726 | 1809 | 1736 | 1833 | 1555 | 1878 | 2117 | 1924 | 1873 | 1825 | 1754 | 1854 | 1891.19 | 1971 | 1811 | 0.918 | -0.124 | 0.07990131 | 0.18907773 | 0.0021 | 0.0037 | 0.0031 | 0.0031 | TRUE | 0.052 |
| Zfp2 | 950 | 768 | 837 | 969 | 803 | 1012 | 738 | 817 | 903 | 965 | 906 | 999 | 887.88 | 819 | 957 | 1.17 | 0.227 | 0.01502979 | 0.05594292 | 0.0052 | 0.0051 | 0.0051 | 0.0051 | TRUE | 0.0951 |
| Zfp202 | 79 | 64 | 36 | 33 | 31 | 32 | 61 | 68 | 39 | 33 | 35 | 32 | 44.61 | 56 | 33 | 0.588 | -0.765 | 0.01494442 | 0.05570766 | 0.0256 | 0.0556 | 0.047 | 0.047 | TRUE | 0.5718 |
| Zfp207 | 15065 | 11531 | 11148 | 11073 | 9306 | 11341 | 11700 | 12265 | 12029 | 11022 | 10498 | 11195 | 11451.64 | 11998 | 10905 | 0.909 | -0.138 | 0.00682459 | 0.03116419 | 7.00E-04 | 0.0026 | 0.0018 | 0.0018 | TRUE | 0.013 |
| Zfp212 | 1435 | 1058 | 958 | 1047 | 950 | 1038 | 1115 | 1125 | 1034 | 1042 | 1072 | 1025 | 1068.68 | 1091 | 1046 | 0.958 | -0.062 | 0.41017326 | 0.58305384 | 0 | 0.0046 | 0.0031 | 0.0031 | TRUE | 0.0247 |

| | | | | | | | | | | | | | | | | | | | | | | | | | |
|---|---|---|---|---|---|---|---|---|---|---|---|---|---|---|---|---|---|---|---|---|---|---|---|---|---|
| Zfp213 | 442 | 296 | 241 | 261 | 263 | 247 | 343 | 315 | 260 | 260 | 297 | 244 | 286.41 | 306 | 267 | 0.868 | -0.204 | 0.14959474 | 0.2950811 | 0.0116 | 0.0107 | 0.011 | 0.011 | TRUE | 0.1915 |
| Zfp217 | 6907 | 5172 | 5027 | 4898 | 4626 | 5150 | 5364 | 5501 | 5424 | 4875 | 5219 | 5084 | 5244.63 | 5430 | 5059 | 0.932 | -0.102 | 0.05348525 | 0.14135812 | 4.00E-04 | 0.0029 | 0.0018 | 0.0018 | TRUE | 0.0123 |
| Zfp219 | 5218 | 4154 | 3887 | 3965 | 3596 | 4185 | 4053 | 4419 | 4194 | 3947 | 4057 | 4131 | 4133.31 | 4222 | 4045 | 0.958 | -0.061 | 0.28897288 | 0.4610591 | 9.00E-04 | 0.003 | 0.0022 | 0.0022 | TRUE | 0.0202 |
| Zfp229 | 3938 | 2655 | 2524 | 2842 | 2547 | 2663 | 3058 | 2824 | 2723 | 2829 | 2873 | 2629 | 2822.82 | 2868 | 2777 | 0.967 | -0.048 | 0.48072744 | 0.64760012 | 0.0023 | 0.0033 | 0.003 | 0.003 | TRUE | 0.0416 |
| Zfp235 | 258 | 204 | 131 | 172 | 151 | 169 | 200 | 217 | 141 | 171 | 170 | 167 | 177.85 | 186 | 169 | 0.908 | -0.139 | 0.42753619 | 0.59950699 | 0.0187 | 0.0158 | 0.0166 | 0.0166 | TRUE | 0.4573 |
| Zfp236 | 2048 | 1527 | 1646 | 1717 | 1594 | 1821 | 1591 | 1624 | 1776 | 1709 | 1798 | 1798 | 1715.96 | 1664 | 1768 | 1.064 | 0.089 | 0.20510626 | 0.36667714 | 0.0016 | 0.0038 | 0.003 | 0.003 | TRUE | 0.0416 |
| Zfp239 | 660 | 515 | 402 | 537 | 443 | 540 | 513 | 548 | 434 | 535 | 500 | 533 | 510.25 | 498 | 523 | 1.049 | 0.068 | 0.52945693 | 0.68977559 | 0.0055 | 0.0071 | 0.0066 | 0.0066 | TRUE | 0.1424 |
| Zfp248 | 902 | 627 | 605 | 618 | 584 | 656 | 701 | 667 | 653 | 615 | 659 | 648 | 656.97 | 674 | 641 | 0.95 | -0.074 | 0.39510004 | 0.56864011 | 0 | 0.006 | 0.004 | 0.004 | TRUE | 0.0156 |
| Zfp251 | 781 | 500 | 410 | 440 | 402 | 450 | 607 | 532 | 442 | 438 | 453 | 444 | 486.08 | 527 | 445 | 0.842 | -0.247 | 0.04105262 | 0.1165604 | 0.011 | 0.0073 | 0.0085 | 0.0085 | TRUE | 0.1983 |
| Zfp259 | 5835 | 4369 | 4214 | 4072 | 3801 | 4515 | 4532 | 4647 | 4547 | 4053 | 4288 | 4457 | 4420.68 | 4575 | 4266 | 0.932 | -0.101 | 0.0781423 | 0.18592588 | 9.00E-04 | 0.003 | 0.0021 | 0.0021 | TRUE | 0.0234 |
| Zfp26 | 1346 | 987 | 842 | 878 | 698 | 849 | 1045 | 1050 | 909 | 874 | 787 | 838 | 917.2 | 1001 | 833 | 0.832 | -0.266 | 0.00265214 | 0.01499488 | 0.0036 | 0.005 | 0.0045 | 0.0045 | TRUE | 0.0766 |
| Zfp260 | 2522 | 1903 | 1887 | 2349 | 1924 | 2275 | 1959 | 2024 | 2036 | 2338 | 2170 | 2246 | 2128.9 | 2006 | 2251 | 1.123 | 0.167 | 0.00724425 | 0.03255017 | 5.00E-04 | 0.0035 | 0.0023 | 0.0023 | TRUE | 0.0137 |
| Zfp263 | 1628 | 1286 | 1411 | 1383 | 1290 | 1428 | 1264 | 1368 | 1522 | 1377 | 1455 | 1410 | 1399.38 | 1385 | 1414 | 1.022 | 0.031 | 0.70053335 | 0.81850931 | 0.0041 | 0.0041 | 0.0041 | 0.0041 | TRUE | 0.0905 |
| Zfp266 | 3346 | 2552 | 2698 | 3221 | 2807 | 3348 | 2599 | 2715 | 2911 | 3206 | 3167 | 3305 | 2983.68 | 2742 | 3226 | 1.178 | 0.236 | 0.00019659 | 0.00210314 | 0.0014 | 0.0032 | 0.0026 | 0.0026 | TRUE | 0.0354 |
| Zfp27 | 340 | 220 | 198 | 251 | 208 | 232 | 264 | 234 | 214 | 250 | 235 | 229 | 237.54 | 237 | 238 | 0.999 | -0.001 | 0.99244482 | 0.99592003 | 0.0032 | 0.0124 | 0.0096 | 0.0096 | TRUE | 0.1167 |
| Zfp273 | 264 | 205 | 200 | 219 | 159 | 206 | 205 | 218 | 216 | 218 | 179 | 203 | 206.6 | 213 | 200 | 0.943 | -0.085 | 0.55340635 | 0.70914802 | 0 | 0.0139 | 0.01 | 0.01 | TRUE | 0.0863 |
| Zfp275 | 1246 | 960 | 1055 | 1060 | 853 | 1014 | 968 | 1021 | 1138 | 1055 | 962 | 1001 | 1024.26 | 1042 | 1006 | 0.967 | -0.049 | 0.57170363 | 0.72492548 | 0.0036 | 0.0047 | 0.0044 | 0.0044 | TRUE | 0.0768 |
| Zfp276 | 531 | 413 | 352 | 263 | 258 | 302 | 412 | 439 | 380 | 262 | 291 | 298 | 347.08 | 410 | 284 | 0.691 | -0.534 | 6.27E-06 | 0.00012799 | 0.0019 | 0.0093 | 0.0071 | 0.0071 | TRUE | 0.0522 |
| Zfp277 | 3078 | 2248 | 2499 | 2844 | 2557 | 2949 | 2391 | 2391 | 2696 | 2831 | 2885 | 2911 | 2684.12 | 2493 | 2876 | 1.154 | 0.207 | 0.00198484 | 0.01204642 | 0.002 | 0.0033 | 0.0029 | 0.0029 | TRUE | 0.0612 |
| Zfp28 | 275 | 225 | 264 | 240 | 266 | 287 | 214 | 239 | 285 | 239 | 300 | 283 | 260.01 | 246 | 274 | 1.117 | 0.16 | 0.28332136 | 0.45520596 | 0.0137 | 0.0115 | 0.0122 | 0.0122 | TRUE | 0.2107 |
| Zfp280b | 6876 | 4828 | 4481 | 4950 | 4391 | 4792 | 5340 | 5135 | 4835 | 4927 | 4954 | 4730 | 4986.99 | 5103 | 4870 | 0.954 | -0.068 | 0.24403032 | 0.41199213 | 0.0013 | 0.0029 | 0.0023 | 0.0023 | TRUE | 0.0257 |
| Zfp280c | 1069 | 819 | 940 | 1156 | 922 | 1240 | 830 | 871 | 1014 | 1151 | 1040 | 1224 | 1021.75 | 905 | 1138 | 1.26 | 0.333 | 0.00057954 | 0.00483558 | 0.0075 | 0.0047 | 0.0058 | 0.0058 | TRUE | 0.1309 |
| Zfp280d | 2073 | 1498 | 1468 | 1428 | 1328 | 1588 | 1610 | 1593 | 1584 | 1421 | 1498 | 1568 | 1545.75 | 1596 | 1496 | 0.937 | -0.094 | 0.16921009 | 0.3207586 | 6.00E-04 | 0.004 | 0.0027 | 0.0027 | TRUE | 0.0231 |
| Zfp281 | 8194 | 5848 | 4994 | 4978 | 4150 | 4798 | 6364 | 6220 | 5388 | 4955 | 4682 | 4736 | 5390.99 | 5991 | 4791 | 0.8 | -0.323 | 4.97E-06 | 0.00010553 | 0.004 | 0.0029 | 0.0034 | 0.0034 | TRUE | 0.094 |
| Zfp282 | 730 | 528 | 528 | 556 | 473 | 517 | 567 | 562 | 570 | 553 | 534 | 510 | 549.28 | 566 | 532 | 0.94 | -0.089 | 0.33608346 | 0.51042832 | 0 | 0.0067 | 0.0043 | 0.0043 | TRUE | 0.0159 |
| Zfp286 | 536 | 399 | 362 | 391 | 289 | 351 | 416 | 424 | 391 | 389 | 326 | 346 | 382.17 | 410 | 354 | 0.863 | -0.213 | 0.06055846 | 0.15420667 | 0.0019 | 0.0086 | 0.0066 | 0.0066 | TRUE | 0.0873 |
| Zfp287 | 282 | 199 | 181 | 192 | 196 | 252 | 219 | 212 | 195 | 191 | 221 | 249 | 214.5 | 209 | 220 | 1.054 | 0.076 | 0.6076016 | 0.75144498 | 0.0059 | 0.0135 | 0.0113 | 0.0113 | TRUE | 0.1517 |
| Zfp292 | 6347 | 5149 | 5428 | 5881 | 5349 | 6428 | 4929 | 5477 | 5857 | 5854 | 6034 | 6345 | 5749.46 | 5421 | 6078 | 1.122 | 0.166 | 0.01940047 | 0.06763192 | 0.0043 | 0.0028 | 0.0034 | 0.0034 | TRUE | 0.0776 |
| Zfp296 | 2280 | 1745 | 1724 | 1723 | 1591 | 1713 | 1771 | 1856 | 1860 | 1715 | 1795 | 1691 | 1781.33 | 1829 | 1734 | 0.948 | -0.077 | 0.23348704 | 0.39962044 | 3.00E-04 | 0.0038 | 0.0024 | 0.0024 | TRUE | 0.0113 |
| Zfp3 | 237 | 189 | 155 | 172 | 163 | 160 | 184 | 201 | 167 | 171 | 184 | 158 | 177.56 | 184 | 171 | 0.928 | -0.108 | 0.48381526 | 0.65034999 | 0 | 0.0158 | 0.0116 | 0.0116 | TRUE | 0.0649 |
| Zfp30 | 184 | 165 | 187 | 190 | 151 | 186 | 143 | 176 | 202 | 189 | 170 | 184 | 177.21 | 174 | 181 | 1.05 | 0.071 | 0.67214618 | 0.79845291 | 0.0114 | 0.0158 | 0.0145 | 0.0145 | TRUE | 0.2783 |
| Zfp300 | 32 | 10 | 3 | 10 | 8 | 4 | 25 | 11 | 3 | 10 | 9 | 4 | 10.28 | 13 | 8 | 0.579 | -0.788 | 0.28858351 | NA | 0.4324 | 0.2331 | 0.2928 | 0.2928 | TRUE | 0.3153 |
| Zfp316 | 291 | 216 | 217 | 260 | 265 | 248 | 226 | 230 | 234 | 259 | 299 | 245 | 248.75 | 230 | 268 | 1.162 | 0.217 | 0.10576001 | 0.23060527 | 0 | 0.012 | 0.0089 | 0.0089 | TRUE | 0.1143 |
| Zfp317 | 3910 | 2882 | 2801 | 2811 | 2596 | 2831 | 3037 | 3066 | 3022 | 2798 | 2929 | 2795 | 2940.97 | 3042 | 2841 | 0.934 | -0.099 | 0.0758626 | 0.18165592 | 0 | 0.0032 | 0.0019 | 0.0019 | TRUE | 0.0088 |
| Zfp318 | 4884 | 3779 | 3636 | 3616 | 3356 | 3863 | 3793 | 4020 | 3923 | 3599 | 3786 | 3813 | 3822.46 | 3912 | 3733 | 0.954 | -0.067 | 0.2344928 | 0.40061384 | 7.00E-04 | 0.003 | 0.002 | 0.002 | TRUE | 0.012 |
| Zfp319 | 752 | 563 | 415 | 364 | 403 | 356 | 584 | 599 | 448 | 362 | 455 | 351 | 466.51 | 544 | 389 | 0.715 | -0.485 | 0.00041004 | 0.00371368 | 0.0202 | 0.0075 | 0.0113 | 0.0113 | TRUE | 0.272 |
| Zfp322a | 3102 | 2323 | 2575 | 2715 | 2342 | 2784 | 2409 | 2471 | 2778 | 2703 | 2642 | 2748 | 2625.22 | 2553 | 2698 | 1.058 | 0.081 | 0.24607202 | 0.41470433 | 0.0026 | 0.0033 | 0.0031 | 0.0031 | TRUE | 0.0719 |
| Zfp324 | 183 | 149 | 79 | 93 | 75 | 100 | 142 | 158 | 85 | 93 | 85 | 99 | 110.29 | 128 | 92 | 0.714 | -0.486 | 0.03767239 | 0.10964193 | 0.0452 | 0.0239 | 0.03 | 0.03 | TRUE | 0.8516 |
| Zfp326 | 1346 | 1065 | 1033 | 1184 | 1020 | 1114 | 1045 | 1133 | 1115 | 1179 | 1151 | 1100 | 1120.29 | 1098 | 1143 | 1.042 | 0.06 | 0.42525043 | 0.59794435 | 8.00E-04 | 0.0045 | 0.0032 | 0.0032 | TRUE | 0.0218 |
| Zfp329 | 809 | 684 | 536 | 626 | 485 | 596 | 628 | 728 | 578 | 623 | 547 | 588 | 615.47 | 645 | 586 | 0.91 | -0.136 | 0.20273297 | 0.36416046 | 0.0072 | 0.0063 | 0.0066 | 0.0066 | TRUE | 0.1443 |
| Zfp330 | 6126 | 4560 | 5339 | 5598 | 5213 | 5589 | 4758 | 4850 | 5761 | 5572 | 5881 | 5517 | 5389.87 | 5123 | 5657 | 1.104 | 0.143 | 0.0609392 | 0.1550381 | 0.0062 | 0.0029 | 0.004 | 0.004 | TRUE | 0.144 |
| Zfp334 | 1034 | 714 | 694 | 795 | 671 | 736 | 803 | 759 | 749 | 791 | 757 | 727 | 764.37 | 770 | 758 | 0.983 | -0.025 | 0.77010543 | 0.86394025 | 0 | 0.0055 | 0.0038 | 0.0038 | TRUE | 0.0176 |
| Zfp335 | 2036 | 1516 | 1470 | 1495 | 1459 | 1764 | 1581 | 1613 | 1586 | 1488 | 1646 | 1741 | 1609.22 | 1593 | 1625 | 1.02 | 0.029 | 0.70259876 | 0.81985628 | 0.0026 | 0.0039 | 0.0034 | 0.0034 | TRUE | 0.0664 |
| Zfp341 | 625 | 514 | 640 | 638 | 607 | 675 | 485 | 547 | 691 | 635 | 685 | 666 | 618.14 | 574 | 662 | 1.156 | 0.209 | 0.0871787 | 0.20026441 | 0.0151 | 0.0063 | 0.0091 | 0.0091 | TRUE | 0.3659 |
| Zfp345 | 507 | 400 | 235 | 265 | 224 | 240 | 394 | 425 | 254 | 264 | 253 | 237 | 304.37 | 358 | 251 | 0.701 | -0.512 | 0.00268717 | 0.01513904 | 0.035 | 0.0102 | 0.0176 | 0.0176 | TRUE | 0.7219 |
| Zfp346 | 2980 | 2227 | 2178 | 2329 | 2162 | 2424 | 2314 | 2369 | 2350 | 2318 | 2439 | 2393 | 2363.9 | 2344 | 2383 | 1.017 | 0.024 | 0.68320039 | 0.80582466 | 0 | 0.0034 | 0.002 | 0.002 | TRUE | 0.0069 |
| Zfp35 | 1853 | 1309 | 1152 | 1236 | 1127 | 1288 | 1439 | 1392 | 1243 | 1230 | 1271 | 1271 | 1307.94 | 1358 | 1257 | 0.925 | -0.113 | 0.14701767 | 0.29136488 | 0.0024 | 0.0042 | 0.0036 | 0.0036 | TRUE | 0.0652 |
| Zfp352 | 194 | 150 | 64 | 43 | 36 | 33 | 151 | 160 | 69 | 43 | 41 | 33 | 82.55 | 127 | 39 | 0.305 | -1.715 | 2.92E-08 | 1.29E-06 | 0.1007 | 0.0311 | 0.0516 | 0.0516 | TRUE | 1.5574 |
| Zfp354a | 523 | 390 | 350 | 428 | 382 | 375 | 406 | 415 | 378 | 426 | 431 | 370 | 404.3 | 400 | 409 | 1.022 | 0.032 | 0.77262128 | 0.86551265 | 0.0021 | 0.0083 | 0.0064 | 0.0064 | TRUE | 0.0809 |
| Zfp354b | 77 | 55 | 44 | 50 | 49 | 66 | 60 | 59 | 47 | 50 | 55 | 65 | 56 | 55 | 57 | 1.024 | 0.034 | 0.89718474 | 0.94337809 | 0 | 0.0448 | 0.0324 | 0.0324 | TRUE | 0.1477 |
| Zfp354c | 1483 | 1114 | 1122 | 1223 | 1095 | 1274 | 1152 | 1185 | 1211 | 1217 | 1235 | 1258 | 1209.61 | 1183 | 1237 | 1.047 | 0.066 | 0.34480453 | 0.51910603 | 0 | 0.0044 | 0.0027 | 0.0027 | TRUE | 0.0065 |
| Zfp358 | 269 | 195 | 175 | 217 | 177 | 217 | 209 | 207 | 189 | 216 | 200 | 214 | 205.84 | 202 | 210 | 1.04 | 0.057 | 0.68436109 | 0.80659463 | 0 | 0.0139 | 0.0093 | 0.0093 | TRUE | 0.0326 |
| Zfp36 | 700 | 513 | 766 | 631 | 598 | 787 | 544 | 546 | 827 | 628 | 675 | 777 | 665.91 | 639 | 693 | 1.087 | 0.121 | 0.41464331 | 0.58730521 | 0.0343 | 0.006 | 0.0143 | 0.0143 | TRUE | 0.7729 |
| Zfp362 | 591 | 466 | 481 | 563 | 486 | 560 | 459 | 496 | 519 | 560 | 548 | 553 | 522.53 | 491 | 554 | 1.13 | 0.176 | 0.06783997 | 0.16755428 | 0 | 0.007 | 0.0048 | 0.0048 | TRUE | 0.0413 |
| Zfp365 | 1679 | 1235 | 1208 | 1401 | 1149 | 1302 | 1304 | 1314 | 1303 | 1395 | 1296 | 1285 | 1316.18 | 1307 | 1325 | 1.014 | 0.02 | 0.7709181 | 0.86441946 | 0 | 0.0042 | 0.0028 | 0.0028 | TRUE | 0.0253 |
| Zfp366 | 3 | 1 | 3 | 2 | 2 | 0 | 2 | 1 | 3 | 2 | 2 | 0 | 1.81 | 2 | 1 | 0.632 | -0.662 | 0.65104018 | NA | 0 | 1.3097 | 0.9537 | 0.9537 | TRUE | 0.524 |
| Zfp367 | 908 | 751 | 824 | 977 | 775 | 969 | 705 | 799 | 889 | 973 | 874 | 957 | 866.08 | 798 | 935 | 1.174 | 0.232 | 0.01855362 | 0.0653814 | 0.0072 | 0.0052 | 0.0058 | 0.0058 | TRUE | 0.1279 |
| Zfp369 | 777 | 572 | 528 | 646 | 523 | 612 | 603 | 608 | 570 | 643 | 590 | 604 | 603.13 | 594 | 612 | 1.031 | 0.044 | 0.62634967 | 0.76470623 | 0 | 0.0064 | 0.0043 | 0.0043 | TRUE | 0.0227 |
| Zfp36l1 | 4119 | 3316 | 3123 | 2705 | 2429 | 2984 | 3199 | 3527 | 3370 | 2693 | 2740 | 2946 | 3079.05 | 3365 | 2793 | 0.83 | -0.268 | 3.95E-05 | 0.00058202 | 0.0019 | 0.0032 | 0.0027 | 0.0027 | TRUE | 0.028 |
| Zfp36l2 | 2075 | 1827 | 1841 | 1797 | 1693 | 1958 | 1612 | 1943 | 1986 | 1789 | 1910 | 1933 | 1862.13 | 1847 | 1877 | 1.018 | 0.025 | 0.7716256 | 0.86486091 | 0.0069 | 0.0037 | 0.0048 | 0.0048 | TRUE | 0.1543 |
| Zfp36l3 | 233 | 163 | 160 | 222 | 194 | 228 | 181 | 173 | 173 | 221 | 219 | 225 | 198.65 | 176 | 222 | 1.26 | 0.334 | 0.0168991 | 0.06111811 | 0 | 0.0144 | 0.009 | 0.009 | TRUE | 0.0083 |
| Zfp37 | 592 | 427 | 451 | 478 | 385 | 496 | 460 | 454 | 487 | 476 | 434 | 490 | 466.73 | 467 | 467 | 1.001 | 0.001 | 0.99103806 | 0.99553503 | 0 | 0.0075 | 0.0053 | 0.0053 | TRUE | 0.0411 |
| Zfp382 | 376 | 258 | 258 | 275 | 260 | 304 | 292 | 274 | 278 | 274 | 293 | 300 | 285.33 | 281 | 289 | 1.025 | 0.036 | 0.76902429 | 0.8633101 | 0 | 0.0107 | 0.007 | 0.007 | TRUE | 0.0247 |
| Zfp383 | 693 | 560 | 719 | 753 | 645 | 835 | 538 | 596 | 776 | 750 | 728 | 824 | 701.85 | 637 | 767 | 1.208 | 0.273 | 0.02891685 | 0.09016329 | 0.019 | 0.0058 | 0.0098 | 0.0098 | TRUE | 0.4281 |
| Zfp384 | 3642 | 2650 | 2504 | 2504 | 2310 | 2728 | 2829 | 2819 | 2702 | 2492 | 2606 | 2693 | 2690.08 | 2783 | 2597 | 0.933 | -0.1 | 0.0975146 | 0.21736995 | 7.00E-04 | 0.0033 | 0.0023 | 0.0023 | TRUE | 0.0153 |
| Zfp385a | 2646 | 2019 | 2227 | 2351 | 2141 | 2584 | 2055 | 2148 | 2403 | 2340 | 2415 | 2551 | 2318.63 | 2202 | 2435 | 1.107 | 0.147 | 0.04945354 | 0.13374445 | 0.0037 | 0.0034 | 0.0036 | 0.0036 | TRUE | 0.0768 |
| Zfp385b | 44 | 56 | 37 | 30 | 35 | 30 | 34 | 60 | 40 | 30 | 39 | 30 | 38.77 | 45 | 33 | 0.743 | -0.429 | 0.20414647 | 0.36550989 | 0.036 | 0.0636 | 0.0554 | 0.0554 | TRUE | 0.6628 |
| Zfp385c | 159 | 98 | 138 | 205 | 195 | 260 | 123 | 104 | 149 | 204 | 220 | 257 | 176.22 | 125 | 227 | 1.81 | 0.856 | 8.28E-07 | 2.50E-05 | 0.0148 | 0.0159 | 0.0156 | 0.0156 | TRUE | 0.2506 |
| Zfp386 | 201 | 115 | 87 | 84 | 82 | 81 | 156 | 122 | 94 | 84 | 93 | 80 | 104.73 | 124 | 86 | 0.683 | -0.551 | 0.01383143 | 0.05264516 | 0.0285 | 0.0251 | 0.0261 | 0.0261 | TRUE | 0.1751 |

| | | | | | | | | | | | | | | | | | | | | | | | | |
|---|---|---|---|---|---|---|---|---|---|---|---|---|---|---|---|---|---|---|---|---|---|---|---|---|
| Zfp389 | 5 | 4 | 4 | 4 | 2 | 5 | 4 | 4 | 4 | 4 | 2 | 5 | 3.94 | 4 | 4 | 0.906 | -0.142 | 0.87972842 | NA | 0 | 0.6043 | 0.3828 | 0.3828 | TRUE | 0.1626 |
| Zfp39 | 681 | 536 | 559 | 574 | 553 | 530 | 529 | 570 | 603 | 571 | 624 | 523 | 570.1 | 567 | 573 | 1.01 | 0.015 | 0.88661538 | 0.93690919 | 0.0043 | 0.0066 | 0.0058 | 0.0058 | TRUE | 0.0697 |
| Zfp395 | 2419 | 1809 | 2146 | 2148 | 1914 | 2230 | 1879 | 1924 | 2316 | 2138 | 2159 | 2201 | 2102.85 | 2040 | 2166 | 1.063 | 0.088 | 0.28976851 | 0.46187814 | 0.0059 | 0.0036 | 0.0045 | 0.0045 | TRUE | 0.1682 |
| Zfp397 | 1876 | 1504 | 1565 | 1689 | 1561 | 1766 | 1457 | 1600 | 1689 | 1681 | 1761 | 1743 | 1655.15 | 1582 | 1728 | 1.094 | 0.13 | 0.08072583 | 0.19033559 | 0.0025 | 0.0039 | 0.0034 | 0.0034 | TRUE | 0.0596 |
| Zfp398 | 360 | 247 | 221 | 247 | 215 | 262 | 280 | 263 | 238 | 246 | 243 | 259 | 254.64 | 260 | 249 | 0.955 | -0.067 | 0.60836071 | 0.7520791 | 0 | 0.0117 | 0.0083 | 0.0083 | TRUE | 0.0587 |
| Zfp40 | 301 | 227 | 170 | 206 | 173 | 190 | 234 | 241 | 183 | 205 | 195 | 188 | 207.74 | 219 | 196 | 0.891 | -0.167 | 0.27197263 | 0.4429977 | 0.0069 | 0.0138 | 0.0118 | 0.0118 | TRUE | 0.2171 |
| Zfp407 | 2169 | 1495 | 1309 | 1311 | 1314 | 1404 | 1685 | 1590 | 1412 | 1305 | 1482 | 1386 | 1476.74 | 1562 | 1391 | 0.889 | -0.17 | 0.04557093 | 0.12588814 | 0.0049 | 0.004 | 0.0045 | 0.0045 | TRUE | 0.0843 |
| Zfp408 | 489 | 381 | 306 | 308 | 269 | 317 | 380 | 405 | 330 | 307 | 303 | 313 | 339.7 | 372 | 308 | 0.827 | -0.274 | 0.02227973 | 0.07457708 | 0.0027 | 0.0094 | 0.0074 | 0.0074 | TRUE | 0.1043 |
| Zfp41 | 1582 | 1217 | 1117 | 1094 | 1008 | 1199 | 1229 | 1295 | 1205 | 1089 | 1137 | 1184 | 1189.68 | 1243 | 1137 | 0.915 | -0.129 | 0.08163975 | 0.19200981 | 7.00E-04 | 0.0044 | 0.0031 | 0.0031 | TRUE | 0.0164 |
| Zfp410 | 1234 | 872 | 799 | 835 | 676 | 786 | 958 | 928 | 862 | 831 | 763 | 776 | 852.95 | 916 | 790 | 0.862 | -0.215 | 0.01053382 | 0.04287012 | 0.0014 | 0.0052 | 0.0039 | 0.0039 | TRUE | 0.0311 |
| Zfp414 | 1830 | 1435 | 1524 | 1571 | 1448 | 1774 | 1421 | 1526 | 1644 | 1564 | 1633 | 1751 | 1590.09 | 1530 | 1649 | 1.079 | 0.11 | 0.16472101 | 0.31551292 | 0.0034 | 0.0039 | 0.0038 | 0.0038 | TRUE | 0.0508 |
| Zfp418 | 339 | 251 | 224 | 229 | 203 | 239 | 263 | 267 | 242 | 228 | 229 | 236 | 244.14 | 257 | 231 | 0.897 | -0.157 | 0.22413488 | 0.38854576 | 0 | 0.0121 | 0.0079 | 0.0079 | TRUE | 0.0299 |
| Zfp42 | 50192 | 33356 | 25934 | 27971 | 25022 | 26985 | 38982 | 35480 | 27983 | 27842 | 28227 | 26638 | 30858.83 | 34148 | 27569 | 0.807 | -0.309 | 0.00113363 | 0.00794658 | 0.0139 | 0.0025 | 0.0065 | 0.0065 | TRUE | 0.3053 |
| Zfp420 | 396 | 291 | 288 | 341 | 293 | 282 | 308 | 310 | 311 | 339 | 331 | 278 | 312.7 | 310 | 316 | 1.022 | 0.031 | 0.80247664 | 0.88561013 | 0.0024 | 0.01 | 0.0077 | 0.0077 | TRUE | 0.1262 |
| Zfp422 | 1117 | 826 | 809 | 692 | 651 | 755 | 868 | 879 | 873 | 689 | 734 | 745 | 797.92 | 873 | 723 | 0.828 | -0.272 | 0.00075159 | 0.00587317 | 0 | 0.0054 | 0.0034 | 0.0034 | TRUE | 0.0204 |
| Zfp423 | 1584 | 1093 | 1084 | 1626 | 1372 | 1628 | 1230 | 1163 | 1170 | 1619 | 1548 | 1607 | 1389.3 | 1188 | 1591 | 1.339 | 0.421 | 6.99E-10 | 4.92E-08 | 0 | 0.0041 | 0.0026 | 0.0026 | TRUE | 0.0125 |
| Zfp426 | 813 | 494 | 381 | 462 | 376 | 450 | 631 | 525 | 411 | 460 | 424 | 444 | 482.71 | 522 | 443 | 0.845 | -0.243 | 0.08043739 | 0.18990611 | 0.0219 | 0.0073 | 0.0119 | 0.0119 | TRUE | 0.1724 |
| Zfp428 | 2220 | 1592 | 1286 | 1857 | 1632 | 1718 | 1724 | 1693 | 1388 | 1848 | 1841 | 1696 | 1698.43 | 1602 | 1795 | 1.12 | 0.163 | 0.06653004 | 0.1650892 | 0.007 | 0.0038 | 0.0051 | 0.0051 | TRUE | 0.1628 |
| Zfp429 | 31 | 16 | 14 | 16 | 24 | 31 | 24 | 17 | 15 | 16 | 27 | 31 | 21.63 | 19 | 25 | 1.294 | 0.372 | 0.39754314 | 0.57083425 | 0.0436 | 0.112 | 0.0928 | 0.0928 | TRUE | 0.5843 |
| Zfp433 | 64 | 36 | 35 | 43 | 45 | 35 | 50 | 38 | 38 | 43 | 51 | 35 | 42.31 | 42 | 43 | 1.009 | 0.013 | 0.96650186 | 0.98188155 | 0 | 0.0584 | 0.0446 | 0.0446 | TRUE | 0.204 |
| Zfp438 | 359 | 221 | 171 | 203 | 185 | 207 | 279 | 235 | 185 | 202 | 209 | 204 | 218.92 | 233 | 205 | 0.876 | -0.19 | 0.2432988 | 0.41108106 | 0.0175 | 0.0133 | 0.0146 | 0.0146 | TRUE | 0.1952 |
| Zfp442 | 148 | 157 | 104 | 119 | 113 | 98 | 115 | 167 | 112 | 118 | 127 | 97 | 122.8 | 131 | 114 | 0.871 | -0.2 | 0.34076362 | 0.51480123 | 0.0279 | 0.0217 | 0.0235 | 0.0235 | TRUE | 0.5615 |
| Zfp444 | 1288 | 1013 | 1040 | 1136 | 956 | 1201 | 1000 | 1078 | 1122 | 1131 | 1078 | 1186 | 1099.14 | 1067 | 1132 | 1.063 | 0.088 | 0.27028559 | 0.44122953 | 0.002 | 0.0046 | 0.0037 | 0.0037 | TRUE | 0.0371 |
| Zfp445 | 6836 | 4947 | 4480 | 4574 | 4032 | 4698 | 5309 | 5262 | 4834 | 4553 | 4549 | 4638 | 4857.38 | 5135 | 4580 | 0.892 | -0.166 | 0.00409292 | 0.0209979 | 0.0011 | 0.0029 | 0.0022 | 0.0022 | TRUE | 0.0318 |
| Zfp446 | 883 | 612 | 505 | 514 | 454 | 494 | 686 | 651 | 545 | 512 | 512 | 488 | 565.52 | 627 | 504 | 0.802 | -0.319 | 0.00282192 | 0.01573174 | 0.0061 | 0.0066 | 0.0064 | 0.0064 | TRUE | 0.1522 |
| Zfp449 | 588 | 438 | 447 | 507 | 397 | 547 | 457 | 466 | 482 | 505 | 448 | 540 | 482.9 | 468 | 498 | 1.064 | 0.09 | 0.39113412 | 0.56486611 | 0.0026 | 0.0073 | 0.0059 | 0.0059 | TRUE | 0.0862 |
| Zfp451 | 4596 | 3154 | 3271 | 3632 | 3173 | 3653 | 3570 | 3355 | 3529 | 3615 | 3579 | 3606 | 3542.43 | 3485 | 3600 | 1.033 | 0.047 | 0.39575913 | 0.56928589 | 3.00E-04 | 0.0031 | 0.0019 | 0.0019 | TRUE | 0.0128 |
| Zfp454 | 347 | 246 | 205 | 221 | 224 | 258 | 270 | 262 | 221 | 220 | 253 | 255 | 246.62 | 251 | 243 | 0.964 | -0.053 | 0.70451826 | 0.82107628 | 0.0047 | 0.012 | 0.0099 | 0.0099 | TRUE | 0.1145 |
| Zfp455 | 90 | 57 | 61 | 64 | 62 | 74 | 70 | 61 | 66 | 64 | 70 | 73 | 67.17 | 66 | 69 | 1.05 | 0.07 | 0.76539135 | 0.86085447 | 0 | 0.0377 | 0.0249 | 0.0249 | TRUE | 0.0401 |
| Zfp456 | 291 | 197 | 221 | 251 | 191 | 251 | 226 | 210 | 238 | 250 | 215 | 248 | 231.18 | 225 | 238 | 1.059 | 0.083 | 0.54508534 | 0.70230023 | 0 | 0.0127 | 0.0092 | 0.0092 | TRUE | 0.0711 |
| Zfp457 | 16 | 6 | 8 | 8 | 9 | 5 | 12 | 6 | 9 | 8 | 10 | 5 | 8.42 | 9 | 8 | 0.825 | -0.278 | 0.67968025 | NA | 0 | 0.2841 | 0.2058 | 0.2058 | TRUE | 0.407 |
| Zfp458 | 43 | 23 | 11 | 31 | 22 | 17 | 33 | 24 | 12 | 31 | 25 | 17 | 23.7 | 23 | 24 | 1.025 | 0.036 | 0.93709164 | 0.96508948 | 0.1228 | 0.1024 | 0.1081 | 0.1081 | TRUE | 0.2954 |
| Zfp459 | 2384 | 2071 | 2349 | 2209 | 2129 | 2520 | 1852 | 2203 | 2535 | 2199 | 2402 | 2488 | 2279.53 | 2197 | 2363 | 1.077 | 0.107 | 0.2889966 | 0.4610591 | 0.0138 | 0.0035 | 0.0069 | 0.0069 | TRUE | 0.2174 |
| Zfp46 | 303 | 249 | 194 | 212 | 236 | 238 | 235 | 265 | 209 | 211 | 266 | 235 | 236.95 | 236 | 237 | 1.002 | 0.003 | 0.98171155 | 0.99051272 | 0.0089 | 0.0124 | 0.0114 | 0.0114 | TRUE | 0.1209 |
| Zfp462 | 9975 | 7300 | 7434 | 7976 | 7390 | 8906 | 7747 | 7765 | 8021 | 7939 | 8337 | 8792 | 8100.15 | 7844 | 8356 | 1.065 | 0.091 | 0.10328908 | 0.22649751 | 0.0013 | 0.0027 | 0.0021 | 0.0021 | TRUE | 0.0255 |
| Zfp467 | 31 | 28 | 21 | 62 | 49 | 37 | 24 | 30 | 23 | 62 | 55 | 37 | 38.34 | 26 | 51 | 2.008 | 1.006 | 0.00295693 | 0.01633051 | 0 | 0.0642 | 0.0535 | 0.0535 | TRUE | 0.5287 |
| Zfp472 | 790 | 597 | 616 | 654 | 570 | 647 | 614 | 635 | 665 | 651 | 643 | 639 | 640.99 | 638 | 644 | 1.011 | 0.016 | 0.85177202 | 0.91640026 | 0 | 0.0061 | 0.0039 | 0.0039 | TRUE | 0.0159 |
| Zfp473 | 1864 | 1407 | 1518 | 1621 | 1500 | 1774 | 1448 | 1497 | 1638 | 1614 | 1692 | 1751 | 1606.52 | 1528 | 1686 | 1.105 | 0.144 | 0.05208911 | 0.13886321 | 0.0023 | 0.0039 | 0.0033 | 0.0033 | TRUE | 0.0478 |
| Zfp474 | 1 | 0 | 2 | 1 | 1 | 1 | 1 | 0 | 2 | 1 | 1 | 1 | 1.01 | 1 | 1 | 1.07 | 0.097 | 0.95858839 | NA | 0 | 2.3547 | 1.5349 | 1.5349 | TRUE | 0.3886 |
| Zfp488 | 6 | 11 | 10 | 5 | 3 | 9 | 5 | 12 | 11 | 5 | 3 | 9 | 7.4 | 9 | 6 | 0.65 | -0.622 | 0.40489714 | NA | 0 | 0.3227 | 0.2584 | 0.2584 | TRUE | 0.7598 |
| Zfp493 | 455 | 341 | 225 | 294 | 251 | 308 | 353 | 363 | 243 | 293 | 283 | 304 | 306.45 | 320 | 293 | 0.916 | -0.127 | 0.40420759 | 0.57715675 | 0.021 | 0.0102 | 0.0134 | 0.0134 | TRUE | 0.4864 |
| Zfp503 | 41 | 24 | 29 | 22 | 20 | 21 | 32 | 26 | 31 | 22 | 23 | 21 | 25.64 | 30 | 22 | 0.733 | -0.449 | 0.23319866 | 0.39931812 | 0 | 0.0949 | 0.0619 | 0.0619 | TRUE | 0.0886 |
| Zfp507 | 3572 | 2625 | 2492 | 2431 | 2367 | 2570 | 2774 | 2792 | 2689 | 2420 | 2670 | 2537 | 2647.04 | 2752 | 2542 | 0.924 | -0.115 | 0.06438563 | 0.1614498 | 8.00E-04 | 0.0033 | 0.0024 | 0.0024 | TRUE | 0.0232 |
| Zfp51 | 783 | 538 | 522 | 656 | 565 | 624 | 608 | 572 | 563 | 653 | 637 | 616 | 608.33 | 581 | 635 | 1.092 | 0.127 | 0.15835705 | 0.30740152 | 0 | 0.0063 | 0.0042 | 0.0042 | TRUE | 0.0205 |
| Zfp511 | 1110 | 778 | 798 | 847 | 785 | 924 | 862 | 828 | 861 | 843 | 886 | 912 | 865.24 | 850 | 880 | 1.035 | 0.05 | 0.53164813 | 0.69157269 | 0 | 0.0052 | 0.0034 | 0.0034 | TRUE | 0.0166 |
| Zfp512 | 3410 | 2529 | 2554 | 3029 | 2722 | 2971 | 2648 | 2690 | 2756 | 3015 | 3071 | 2933 | 2852.13 | 2698 | 3006 | 1.114 | 0.156 | 0.00578781 | 0.02745607 | 0 | 0.0033 | 0.002 | 0.002 | TRUE | 0.0056 |
| Zfp513 | 1203 | 882 | 842 | 776 | 771 | 871 | 934 | 938 | 909 | 772 | 870 | 860 | 880.5 | 927 | 834 | 0.899 | -0.153 | 0.06195768 | 0.15701438 | 9.00E-04 | 0.0051 | 0.0037 | 0.0037 | TRUE | 0.0501 |
| Zfp516 | 6474 | 5228 | 5990 | 6207 | 6114 | 6666 | 5028 | 5561 | 6463 | 6178 | 6897 | 6580 | 6118.04 | 5684 | 6552 | 1.153 | 0.205 | 0.0161857 | 0.05909123 | 0.0094 | 0.0028 | 0.0051 | 0.0051 | TRUE | 0.1752 |
| Zfp518a | 8038 | 5454 | 4617 | 4744 | 4080 | 4478 | 6243 | 5801 | 4982 | 4722 | 4603 | 4420 | 5128.52 | 5675 | 4582 | 0.807 | -0.309 | 9.94E-05 | 0.00120467 | 0.0066 | 0.0029 | 0.0044 | 0.0044 | TRUE | 0.139 |
| Zfp518b | 5935 | 4096 | 3493 | 3213 | 2989 | 3406 | 4609 | 4357 | 3769 | 3198 | 3372 | 3362 | 3777.93 | 4245 | 3311 | 0.779 | -0.359 | 2.84E-06 | 6.60E-05 | 0.0052 | 0.0031 | 0.004 | 0.004 | TRUE | 0.1167 |
| Zfp52 | 291 | 231 | 207 | 198 | 201 | 208 | 226 | 246 | 223 | 197 | 227 | 205 | 220.71 | 232 | 210 | 0.905 | -0.144 | 0.29504996 | 0.4669447 | 0 | 0.0132 | 0.0091 | 0.0091 | TRUE | 0.0529 |
| Zfp521 | 119 | 77 | 112 | 73 | 81 | 85 | 92 | 82 | 121 | 73 | 91 | 84 | 90.52 | 98 | 83 | 0.841 | -0.251 | 0.26338772 | 0.43370975 | 0.0158 | 0.0286 | 0.025 | 0.025 | TRUE | 0.3676 |
| Zfp523 | 1329 | 988 | 896 | 1087 | 950 | 1087 | 1032 | 1051 | 967 | 1082 | 1072 | 1073 | 1046.1 | 1017 | 1076 | 1.058 | 0.081 | 0.27741717 | 0.44869567 | 0 | 0.0047 | 0.003 | 0.003 | TRUE | 0.0212 |
| Zfp524 | 231 | 193 | 200 | 205 | 168 | 208 | 179 | 205 | 216 | 204 | 190 | 205 | 199.9 | 200 | 200 | 1.002 | 0.002 | 0.98752671 | 0.99373352 | 0 | 0.0143 | 0.0104 | 0.0104 | TRUE | 0.0986 |
| Zfp526 | 544 | 451 | 370 | 360 | 345 | 326 | 423 | 480 | 399 | 358 | 389 | 322 | 395.13 | 434 | 356 | 0.821 | -0.284 | 0.01750199 | 0.06256671 | 0.0062 | 0.0084 | 0.0077 | 0.0077 | TRUE | 0.0957 |
| Zfp53 | 653 | 466 | 435 | 454 | 405 | 442 | 507 | 496 | 469 | 452 | 457 | 436 | 469.55 | 491 | 448 | 0.912 | -0.132 | 0.17853821 | 0.33318275 | 0 | 0.0075 | 0.0048 | 0.0048 | TRUE | 0.0165 |
| Zfp532 | 3703 | 2469 | 2419 | 3005 | 2607 | 3235 | 2876 | 2626 | 2610 | 2991 | 2941 | 3193 | 2872.98 | 2704 | 3042 | 1.124 | 0.169 | 0.01073849 | 0.04351721 | 0.002 | 0.0033 | 0.0028 | 0.0028 | TRUE | 0.0384 |
| Zfp534 | 2016 | 1379 | 1137 | 1221 | 1051 | 1204 | 1566 | 1467 | 1227 | 1215 | 1186 | 1189 | 1308.16 | 1420 | 1197 | 0.842 | -0.249 | 0.00670663 | 0.03074338 | 0.0073 | 0.0042 | 0.0053 | 0.0053 | TRUE | 0.1684 |
| Zfp536 | 74 | 54 | 47 | 44 | 32 | 44 | 57 | 57 | 51 | 44 | 36 | 43 | 48.16 | 55 | 41 | 0.745 | -0.424 | 0.12675213 | 0.26215772 | 0 | 0.0516 | 0.0344 | 0.0344 | TRUE | 0.0793 |
| Zfp54 | 269 | 180 | 146 | 194 | 173 | 172 | 209 | 191 | 158 | 193 | 195 | 170 | 186 | 186 | 186 | 0.996 | -0.006 | 0.96966225 | 0.9834812 | 0.0079 | 0.0152 | 0.0131 | 0.0131 | TRUE | 0.1873 |
| Zfp541 | 7 | 2 | 3 | 4 | 3 | 1 | 5 | 2 | 3 | 4 | 3 | 1 | 3.19 | 3 | 3 | 0.757 | -0.402 | 0.71041142 | NA | 0 | 0.7448 | 0.5278 | 0.5278 | TRUE | 0.4081 |
| Zfp551 | 160 | 114 | 109 | 125 | 97 | 116 | 124 | 121 | 118 | 124 | 109 | 115 | 118.58 | 121 | 116 | 0.959 | -0.06 | 0.73686085 | 0.84309575 | 0 | 0.0224 | 0.0146 | 0.0146 | TRUE | 0.0396 |
| Zfp553 | 2138 | 1574 | 1456 | 1566 | 1515 | 1659 | 1660 | 1674 | 1571 | 1559 | 1709 | 1638 | 1635.22 | 1635 | 1635 | 0.999 | -0.001 | 0.98836446 | 0.99389455 | 0.001 | 0.0039 | 0.0028 | 0.0028 | TRUE | 0.0202 |
| Zfp558 | 224 | 200 | 176 | 191 | 175 | 173 | 174 | 213 | 190 | 190 | 197 | 171 | 189.15 | 192 | 186 | 0.97 | -0.044 | 0.77513645 | 0.86721285 | 0 | 0.015 | 0.0115 | 0.0115 | TRUE | 0.094 |
| Zfp560 | 453 | 286 | 218 | 217 | 204 | 180 | 352 | 304 | 235 | 216 | 230 | 178 | 252.51 | 297 | 208 | 0.697 | -0.521 | 0.00178424 | 0.01108305 | 0.0255 | 0.0118 | 0.0159 | 0.0159 | TRUE | 0.2225 |
| Zfp563 | 342 | 229 | 249 | 265 | 264 | 274 | 266 | 244 | 269 | 264 | 298 | 270 | 268.33 | 260 | 277 | 1.068 | 0.095 | 0.45516137 | 0.62481957 | 0 | 0.0113 | 0.0078 | 0.0078 | TRUE | 0.0451 |
| Zfp566 | 577 | 411 | 368 | 322 | 295 | 327 | 448 | 437 | 397 | 321 | 333 | 323 | 376.41 | 427 | 326 | 0.76 | -0.396 | 0.00031134 | 0.00299262 | 0 | 0.0087 | 0.006 | 0.006 | TRUE | 0.0435 |
| Zfp568 | 15416 | 10994 | 10001 | 10126 | 9205 | 10013 | 11973 | 11694 | 10791 | 10079 | 10384 | 9884 | 10801.02 | 11486 | 10116 | 0.881 | -0.184 | 0.00124104 | 0.00844673 | 0.0016 | 0.0026 | 0.0022 | 0.0022 | TRUE | 0.0341 |
| Zfp57 | 11709 | 9189 | 7338 | 8361 | 6372 | 7278 | 9094 | 9774 | 7918 | 8323 | 7188 | 7185 | 8246.84 | 8929 | 7565 | 0.847 | -0.239 | 0.0044619 | 0.02249664 | 0.0091 | 0.0027 | 0.005 | 0.005 | TRUE | 0.1194 |

| | | | | | | | | | | | | | | | | | | | | | | | | | |
|---|---|---|---|---|---|---|---|---|---|---|---|---|---|---|---|---|---|---|---|---|---|---|---|---|---|
| Zfp572 | 0 | 0 | 0 | 2 | 1 | 0 | 0 | 0 | 0 | 2 | 1 | 0 | 0.52 | 0 | 1 | 5.866 | 2.552 | 0.38036449 | NA | 0 | 4.5618 | 4.6025 | 4.6025 | TRUE | 0.0821 |
| Zfp574 | 1786 | 1341 | 1275 | 1172 | 1017 | 1255 | 1387 | 1426 | 1376 | 1167 | 1147 | 1239 | 1290.33 | 1396 | 1184 | 0.848 | -0.237 | 0.00074755 | 0.00585466 | 0 | 0.0043 | 0.0028 | 0.0028 | TRUE | 0.0197 |
| Zfp575 | 92 | 72 | 75 | 87 | 75 | 99 | 71 | 77 | 81 | 87 | 85 | 98 | 82.98 | 76 | 90 | 1.179 | 0.237 | 0.26617873 | 0.436634 | 0 | 0.031 | 0.0207 | 0.0207 | TRUE | 0.0605 |
| Zfp579 | 134 | 120 | 89 | 101 | 69 | 119 | 104 | 128 | 96 | 101 | 78 | 117 | 103.93 | 109 | 99 | 0.906 | -0.142 | 0.51113426 | 0.67432545 | 0.0206 | 0.0252 | 0.024 | 0.024 | TRUE | 0.2882 |
| Zfp58 | 403 | 249 | 278 | 329 | 276 | 325 | 313 | 265 | 300 | 327 | 311 | 321 | 306.25 | 293 | 320 | 1.091 | 0.126 | 0.29900813 | 0.47133998 | 0 | 0.0102 | 0.0073 | 0.0073 | TRUE | 0.0758 |
| Zfp580 | 385 | 334 | 346 | 385 | 315 | 379 | 299 | 355 | 373 | 383 | 355 | 374 | 356.72 | 342 | 371 | 1.087 | 0.121 | 0.31979175 | 0.49360512 | 0.005 | 0.0091 | 0.0078 | 0.0078 | TRUE | 0.1522 |
| Zfp583 | 343 | 225 | 245 | 280 | 210 | 273 | 266 | 239 | 264 | 279 | 237 | 269 | 259.2 | 256 | 262 | 1.02 | 0.028 | 0.83039788 | 0.90305691 | 0 | 0.0116 | 0.0086 | 0.0086 | TRUE | 0.0737 |
| Zfp59 | 2535 | 1796 | 1330 | 1469 | 1390 | 1530 | 1969 | 1910 | 1435 | 1462 | 1568 | 1510 | 1642.49 | 1771 | 1513 | 0.854 | -0.228 | 0.02992245 | 0.0925117 | 0.0134 | 0.0039 | 0.0074 | 0.0074 | TRUE | 0.3303 |
| Zfp592 | 2141 | 1456 | 1369 | 1398 | 1186 | 1462 | 1663 | 1549 | 1477 | 1392 | 1338 | 1443 | 1476.9 | 1563 | 1391 | 0.889 | -0.169 | 0.02191135 | 0.07375857 | 0.0019 | 0.004 | 0.0032 | 0.0032 | TRUE | 0.039 |
| Zfp593 | 1857 | 1204 | 1185 | 1300 | 1121 | 1205 | 1442 | 1281 | 1279 | 1294 | 1265 | 1190 | 1291.61 | 1334 | 1250 | 0.935 | -0.097 | 0.22137888 | 0.3852634 | 0.0028 | 0.0043 | 0.0037 | 0.0037 | TRUE | 0.0629 |
| Zfp595 | 246 | 170 | 193 | 220 | 192 | 208 | 191 | 181 | 208 | 219 | 217 | 205 | 203.51 | 193 | 214 | 1.105 | 0.144 | 0.30770105 | 0.48028142 | 0 | 0.0141 | 0.0095 | 0.0095 | TRUE | 0.0457 |
| Zfp597 | 737 | 515 | 485 | 503 | 430 | 450 | 572 | 548 | 523 | 501 | 485 | 444 | 512.25 | 548 | 477 | 0.869 | -0.203 | 0.04075633 | 0.11606498 | 0 | 0.0071 | 0.0051 | 0.0051 | TRUE | 0.0422 |
| Zfp598 | 7479 | 5660 | 5517 | 5401 | 5019 | 5782 | 5809 | 6020 | 5953 | 5376 | 5662 | 5708 | 5754.62 | 5927 | 5582 | 0.942 | -0.086 | 0.09987271 | 0.22082633 | 4.00E-04 | 0.0028 | 0.0018 | 0.0018 | TRUE | 0.0127 |
| Zfp599 | 1119 | 738 | 541 | 696 | 646 | 687 | 869 | 785 | 584 | 693 | 729 | 678 | 722.92 | 746 | 700 | 0.936 | -0.095 | 0.45109529 | 0.62102725 | 0.0199 | 0.0057 | 0.0101 | 0.0101 | TRUE | 0.4233 |
| Zfp60 | 1753 | 1367 | 1361 | 1368 | 1205 | 1506 | 1361 | 1454 | 1469 | 1362 | 1359 | 1487 | 1415.3 | 1428 | 1403 | 0.983 | -0.024 | 0.73982444 | 0.84538261 | 0.0015 | 0.0041 | 0.0031 | 0.0031 | TRUE | 0.0334 |
| Zfp600 | 2200 | 1667 | 1345 | 1363 | 1200 | 1419 | 1709 | 1773 | 1451 | 1357 | 1354 | 1401 | 1507.38 | 1644 | 1371 | 0.833 | -0.264 | 0.00166938 | 0.01048809 | 0.0051 | 0.004 | 0.0044 | 0.0044 | TRUE | 0.1249 |
| Zfp605 | 513 | 353 | 322 | 388 | 333 | 380 | 398 | 375 | 347 | 386 | 376 | 375 | 376.39 | 373 | 379 | 1.012 | 0.017 | 0.87631793 | 0.93096567 | 0 | 0.0087 | 0.006 | 0.006 | TRUE | 0.0429 |
| Zfp606 | 1031 | 764 | 817 | 894 | 775 | 921 | 801 | 813 | 882 | 890 | 874 | 909 | 861.38 | 832 | 891 | 1.073 | 0.101 | 0.20910714 | 0.37081493 | 0 | 0.0052 | 0.0035 | 0.0035 | TRUE | 0.032 |
| Zfp607 | 358 | 235 | 221 | 201 | 187 | 187 | 278 | 250 | 238 | 200 | 211 | 185 | 227.02 | 255 | 199 | 0.774 | -0.369 | 0.00804599 | 0.03520089 | 0 | 0.0129 | 0.0095 | 0.0095 | TRUE | 0.0715 |
| Zfp608 | 60 | 38 | 48 | 47 | 54 | 48 | 47 | 40 | 52 | 47 | 61 | 47 | 48.98 | 46 | 52 | 1.114 | 0.156 | 0.57927271 | 0.73024739 | 0 | 0.0508 | 0.0365 | 0.0365 | TRUE | 0.188 |
| Zfp609 | 7789 | 5632 | 5645 | 6261 | 5621 | 6622 | 6049 | 5991 | 6091 | 6232 | 6341 | 6537 | 6206.86 | 6044 | 6370 | 1.054 | 0.076 | 0.12560059 | 0.26077638 | 2.00E-04 | 0.0028 | 0.0016 | 0.0016 | TRUE | 0.0065 |
| Zfp61 | 495 | 486 | 416 | 430 | 369 | 426 | 384 | 517 | 449 | 428 | 416 | 421 | 435.85 | 450 | 422 | 0.94 | -0.09 | 0.45975302 | 0.62973166 | 0.0093 | 0.0079 | 0.0083 | 0.0083 | TRUE | 0.1904 |
| Zfp612 | 440 | 293 | 193 | 248 | 190 | 193 | 342 | 312 | 208 | 247 | 214 | 191 | 252.23 | 287 | 217 | 0.754 | -0.407 | 0.02400487 | 0.07896468 | 0.0374 | 0.0118 | 0.0194 | 0.0194 | TRUE | 0.6371 |
| Zfp616 | 0 | 0 | 0 | 0 | 0 | 0 | 0 | 0 | 0 | 0 | 0 | 0 | 0 | 0 | 0 | NA | NA | NA | NA | NA | NA | NA | NA | NA | NA |
| Zfp617 | 588 | 378 | 312 | 389 | 292 | 321 | 457 | 402 | 337 | 387 | 329 | 317 | 371.48 | 399 | 344 | 0.862 | -0.214 | 0.11749237 | 0.24870412 | 0.0149 | 0.0088 | 0.0107 | 0.0107 | TRUE | 0.2027 |
| Zfp618 | 217 | 170 | 100 | 105 | 109 | 112 | 169 | 181 | 108 | 105 | 123 | 111 | 132.55 | 153 | 113 | 0.737 | -0.441 | 0.03550457 | 0.10495694 | 0.0329 | 0.0203 | 0.0239 | 0.0239 | TRUE | 0.6569 |
| Zfp619 | 258 | 155 | 164 | 179 | 172 | 155 | 200 | 165 | 177 | 178 | 194 | 153 | 177.9 | 181 | 175 | 0.964 | -0.053 | 0.74334258 | 0.84725642 | 0.0067 | 0.0157 | 0.0131 | 0.0131 | TRUE | 0.1316 |
| Zfp62 | 2519 | 1670 | 1184 | 1563 | 1325 | 1447 | 1956 | 1776 | 1278 | 1556 | 1495 | 1428 | 1581.54 | 1670 | 1493 | 0.893 | -0.163 | 0.18065751 | 0.33577865 | 0.0221 | 0.0039 | 0.0101 | 0.0101 | TRUE | 0.507 |
| Zfp622 | 4930 | 3668 | 3394 | 3481 | 3228 | 3455 | 3829 | 3902 | 3662 | 3465 | 3642 | 3411 | 3651.62 | 3798 | 3506 | 0.923 | -0.116 | 0.04535247 | 0.12540585 | 7.00E-04 | 0.0031 | 0.0021 | 0.0021 | TRUE | 0.0138 |
| Zfp623 | 507 | 386 | 365 | 384 | 307 | 397 | 394 | 411 | 394 | 382 | 346 | 392 | 386.44 | 400 | 373 | 0.936 | -0.095 | 0.38143698 | 0.55579288 | 0 | 0.0086 | 0.0058 | 0.0058 | TRUE | 0.0447 |
| Zfp628 | 427 | 262 | 170 | 240 | 197 | 218 | 332 | 279 | 183 | 239 | 222 | 215 | 245.01 | 265 | 225 | 0.849 | -0.237 | 0.20275582 | 0.36416046 | 0.0415 | 0.0121 | 0.0207 | 0.0207 | TRUE | 0.3649 |
| Zfp629 | 98 | 68 | 72 | 136 | 112 | 119 | 76 | 72 | 78 | 135 | 126 | 117 | 100.89 | 75 | 126 | 1.676 | 0.745 | 0.00014263 | 0.00163477 | 0 | 0.0259 | 0.0172 | 0.0172 | TRUE | 0.0406 |
| Zfp637 | 501 | 312 | 309 | 406 | 277 | 349 | 389 | 332 | 333 | 404 | 312 | 345 | 352.59 | 351 | 354 | 1.005 | 0.007 | 0.95563866 | 0.97556913 | 0.01 | 0.0091 | 0.0094 | 0.0094 | TRUE | 0.178 |
| Zfp639 | 1971 | 1408 | 1407 | 1363 | 1315 | 1519 | 1531 | 1498 | 1518 | 1357 | 1483 | 1499 | 1481.05 | 1516 | 1446 | 0.954 | -0.068 | 0.33313231 | 0.50740197 | 8.00E-04 | 0.004 | 0.0029 | 0.0029 | TRUE | 0.0359 |
| Zfp64 | 3656 | 2871 | 2607 | 2758 | 2532 | 2933 | 2839 | 3054 | 2813 | 2745 | 2856 | 2895 | 2867.2 | 2902 | 2832 | 0.976 | -0.035 | 0.57080189 | 0.72416753 | 9.00E-04 | 0.0033 | 0.0024 | 0.0024 | TRUE | 0.0251 |
| Zfp641 | 15 | 18 | 18 | 17 | 9 | 12 | 12 | 19 | 19 | 17 | 10 | 12 | 14.86 | 17 | 13 | 0.786 | -0.347 | 0.49790434 | NA | 0 | 0.162 | 0.1205 | 0.1205 | TRUE | 0.4353 |
| Zfp644 | 3264 | 2163 | 2041 | 2194 | 1910 | 2197 | 2535 | 2301 | 2202 | 2184 | 2155 | 2169 | 2257.56 | 2346 | 2169 | 0.924 | -0.114 | 0.09905316 | 0.21960088 | 0.0023 | 0.0035 | 0.003 | 0.003 | TRUE | 0.0617 |
| Zfp646 | 3960 | 2922 | 2874 | 3056 | 2838 | 3194 | 3076 | 3108 | 3101 | 3042 | 3202 | 3153 | 3113.52 | 3095 | 3132 | 1.012 | 0.017 | 0.75474734 | 0.8545742 | 0 | 0.0032 | 0.0019 | 0.0019 | TRUE | 0.0077 |
| Zfp647 | 381 | 255 | 243 | 329 | 288 | 317 | 296 | 271 | 262 | 327 | 325 | 313 | 299.11 | 276 | 322 | 1.161 | 0.215 | 0.07228233 | 0.17570262 | 0 | 0.0104 | 0.007 | 0.007 | TRUE | 0.0462 |
| Zfp648 | 1 | 0 | 0 | 0 | 0 | 0 | 1 | 0 | 0 | 0 | 0 | 0 | 0.13 | 0 | 0 | 0.552 | -0.857 | 0.8336548 | NA | 0 | 18.3121 | 10 | 10 | TRUE | 0.2566 |
| Zfp651 | 123 | 105 | 90 | 118 | 109 | 99 | 96 | 112 | 97 | 117 | 123 | 98 | 107.08 | 102 | 113 | 1.112 | 0.153 | 0.43417404 | 0.60584297 | 0 | 0.0246 | 0.0182 | 0.0182 | TRUE | 0.1378 |
| Zfp652 | 2211 | 1613 | 1847 | 2165 | 1958 | 2286 | 1717 | 1716 | 1993 | 2155 | 2209 | 2257 | 2007.72 | 1809 | 2207 | 1.221 | 0.288 | 0.00012129 | 0.00142566 | 0.0034 | 0.0036 | 0.0035 | 0.0035 | TRUE | 0.0946 |
| Zfp653 | 518 | 461 | 406 | 419 | 375 | 458 | 402 | 490 | 438 | 417 | 423 | 452 | 437.16 | 443 | 431 | 0.974 | -0.039 | 0.72785922 | 0.83688294 | 0.004 | 0.0078 | 0.0066 | 0.0066 | TRUE | 0.0984 |
| Zfp654 | 1495 | 1104 | 1177 | 1151 | 1039 | 1265 | 1161 | 1174 | 1270 | 1146 | 1172 | 1249 | 1195.32 | 1202 | 1189 | 0.99 | -0.014 | 0.85016915 | 0.9153648 | 0.0014 | 0.0044 | 0.0033 | 0.0033 | TRUE | 0.029 |
| Zfp655 | 4042 | 3067 | 3665 | 4351 | 3789 | 4422 | 3139 | 3262 | 3955 | 4331 | 4274 | 4365 | 3887.77 | 3452 | 4323 | 1.253 | 0.326 | 7.73E-05 | 0.00098723 | 0.0078 | 0.003 | 0.0046 | 0.0046 | TRUE | 0.1963 |
| Zfp658 | 324 | 203 | 155 | 207 | 203 | 206 | 252 | 216 | 167 | 206 | 229 | 203 | 212.2 | 212 | 213 | 1 | 0 | 0.99904424 | 0.99939385 | 0.0183 | 0.0136 | 0.015 | 0.015 | TRUE | 0.2787 |
| Zfp661 | 524 | 430 | 421 | 460 | 397 | 461 | 407 | 457 | 454 | 458 | 448 | 455 | 446.57 | 439 | 454 | 1.035 | 0.049 | 0.63350825 | 0.77062871 | 0 | 0.0077 | 0.0054 | 0.0054 | TRUE | 0.0525 |
| Zfp663 | 18 | 13 | 14 | 14 | 10 | 10 | 14 | 14 | 15 | 14 | 11 | 10 | 13 | 14 | 12 | 0.819 | -0.288 | 0.58143861 | NA | 0 | 0.1847 | 0.1193 | 0.1193 | TRUE | 0.113 |
| Zfp664 | 7707 | 5704 | 5878 | 6298 | 5528 | 6361 | 5986 | 6067 | 6342 | 6269 | 6236 | 6279 | 6196.63 | 6132 | 6261 | 1.021 | 0.031 | 0.54600041 | 0.70317212 | 3.00E-04 | 0.0028 | 0.0017 | 0.0017 | TRUE | 0.0109 |
| Zfp667 | 660 | 481 | 473 | 528 | 472 | 532 | 513 | 512 | 510 | 526 | 532 | 525 | 519.63 | 512 | 528 | 1.031 | 0.045 | 0.62720543 | 0.76496691 | 0 | 0.007 | 0.0042 | 0.0042 | TRUE | 7.00E-04 |
| Zfp668 | 670 | 484 | 419 | 459 | 406 | 474 | 520 | 515 | 452 | 457 | 458 | 468 | 478.35 | 496 | 461 | 0.928 | -0.107 | 0.2933899 | 0.46526371 | 0 | 0.0074 | 0.0054 | 0.0054 | TRUE | 0.067 |
| Zfp672 | 162 | 110 | 97 | 145 | 115 | 129 | 126 | 117 | 105 | 144 | 130 | 127 | 124.82 | 116 | 134 | 1.152 | 0.204 | 0.25474312 | 0.42421571 | 0 | 0.0214 | 0.0151 | 0.0151 | TRUE | 0.0694 |
| Zfp677 | 160 | 113 | 118 | 114 | 85 | 109 | 124 | 120 | 127 | 113 | 96 | 108 | 114.79 | 124 | 106 | 0.854 | -0.228 | 0.21275533 | 0.37509708 | 0 | 0.0231 | 0.0153 | 0.0153 | TRUE | 0.0609 |
| Zfp68 | 951 | 709 | 831 | 926 | 833 | 951 | 739 | 754 | 897 | 922 | 940 | 939 | 864.94 | 797 | 934 | 1.174 | 0.231 | 0.0121422 | 0.04774374 | 0.0043 | 0.0052 | 0.005 | 0.005 | TRUE | 0.1412 |
| Zfp687 | 3968 | 2998 | 2889 | 2858 | 2501 | 3144 | 3082 | 3189 | 3117 | 2845 | 2821 | 3104 | 3026.29 | 3129 | 2923 | 0.935 | -0.098 | 0.11306731 | 0.24187238 | 0.0011 | 0.0032 | 0.0024 | 0.0024 | TRUE | 0.0356 |
| Zfp688 | 541 | 418 | 474 | 466 | 353 | 424 | 420 | 445 | 511 | 464 | 398 | 419 | 442.81 | 459 | 427 | 0.933 | -0.1 | 0.38461511 | 0.55866908 | 0.0061 | 0.0078 | 0.0072 | 0.0072 | TRUE | 0.1156 |
| Zfp689 | 497 | 396 | 378 | 381 | 391 | 446 | 386 | 421 | 408 | 379 | 441 | 440 | 412.61 | 405 | 420 | 1.038 | 0.054 | 0.62329432 | 0.7624087 | 0.0023 | 0.0082 | 0.0063 | 0.0063 | TRUE | 0.0851 |
| Zfp69 | 326 | 219 | 228 | 223 | 179 | 228 | 253 | 233 | 246 | 222 | 202 | 225 | 230.19 | 244 | 216 | 0.886 | -0.174 | 0.19265752 | 0.35111896 | 0 | 0.0127 | 0.0085 | 0.0085 | TRUE | 0.0355 |
| Zfp691 | 216 | 165 | 207 | 173 | 168 | 214 | 168 | 176 | 223 | 172 | 190 | 211 | 189.93 | 189 | 191 | 1.015 | 0.021 | 0.89788175 | 0.94349536 | 0.0124 | 0.0149 | 0.0141 | 0.0141 | TRUE | 0.2685 |
| Zfp692 | 1456 | 1337 | 1342 | 1398 | 1331 | 1539 | 1131 | 1422 | 1448 | 1392 | 1502 | 1519 | 1402.21 | 1334 | 1471 | 1.105 | 0.144 | 0.13738289 | 0.27766029 | 0.0097 | 0.0041 | 0.006 | 0.006 | TRUE | 0.2192 |
| Zfp697 | 103 | 80 | 73 | 115 | 93 | 94 | 80 | 85 | 79 | 114 | 105 | 93 | 92.67 | 81 | 104 | 1.281 | 0.357 | 0.08111224 | 0.19102608 | 0 | 0.028 | 0.0192 | 0.0192 | TRUE | 0.0914 |
| Zfp7 | 2879 | 2250 | 2207 | 2230 | 2039 | 2390 | 2236 | 2393 | 2381 | 2220 | 2300 | 2359 | 2314.98 | 2337 | 2293 | 0.982 | -0.026 | 0.67394106 | 0.79979329 | 8.00E-04 | 0.0035 | 0.0024 | 0.0024 | TRUE | 0.0179 |
| Zfp703 | 135 | 83 | 76 | 74 | 64 | 82 | 105 | 88 | 82 | 74 | 72 | 81 | 83.66 | 92 | 76 | 0.821 | -0.285 | 0.19131917 | 0.34937075 | 0 | 0.0308 | 0.0221 | 0.0221 | TRUE | 0.1678 |
| Zfp704 | 3579 | 2796 | 2619 | 2554 | 2502 | 2777 | 2780 | 2974 | 2826 | 2542 | 2823 | 2741 | 2780.95 | 2860 | 2702 | 0.945 | -0.082 | 0.20608653 | 0.3679234 | 0.0015 | 0.0033 | 0.0027 | 0.0027 | TRUE | 0.0326 |
| Zfp706 | 11757 | 8716 | 9101 | 9520 | 8092 | 9483 | 9131 | 9271 | 9820 | 9476 | 9129 | 9361 | 9364.69 | 9407 | 9322 | 0.991 | -0.013 | 0.8049578 | 0.88713562 | 7.00E-04 | 0.0027 | 0.0018 | 0.0018 | TRUE | 0.0179 |
| Zfp707 | 486 | 317 | 273 | 287 | 268 | 288 | 377 | 337 | 295 | 286 | 302 | 284 | 313.59 | 336 | 291 | 0.861 | -0.216 | 0.09280514 | 0.20932048 | 0.0058 | 0.01 | 0.0087 | 0.0087 | TRUE | 0.1378 |
| Zfp708 | 263 | 201 | 232 | 224 | 215 | 281 | 204 | 214 | 250 | 223 | 243 | 277 | 235.22 | 223 | 248 | 1.115 | 0.157 | 0.28363667 | 0.45550774 | 0.0079 | 0.0125 | 0.0111 | 0.0111 | TRUE | 0.1256 |
| Zfp709 | 187 | 118 | 98 | 88 | 68 | 92 | 145 | 126 | 106 | 88 | 77 | 91 | 105.27 | 126 | 85 | 0.676 | -0.566 | 0.00571618 | 0.02718848 | 0.0093 | 0.0249 | 0.0203 | 0.0203 | TRUE | 0.1422 |
| Zfp71-rs1 | 932 | 672 | 430 | 541 | 515 | 578 | 724 | 715 | 464 | 539 | 581 | 571 | 598.78 | 634 | 564 | 0.887 | -0.173 | 0.22655871 | 0.39151432 | 0.0291 | 0.0064 | 0.0131 | 0.0131 | TRUE | 0.6388 |

| | | | | | | | | | | | | | | | | | | | | | | | | | |
|---|---|---|---|---|---|---|---|---|---|---|---|---|---|---|---|---|---|---|---|---|---|---|---|---|---|
| Zfp710 | 11508 | 9297 | 10693 | 12117 | 11606 | 13244 | 8938 | 9889 | 11538 | 12061 | 13093 | 13074 | 11432.07 | 10122 | 12743 | 1.259 | 0.333 | 7.21E-05 | 0.00093739 | 0.0094 | 0.0026 | 0.005 | 0.005 | TRUE | 0.183 |
| Zfp711 | 196 | 127 | 90 | 111 | 89 | 92 | 152 | 135 | 97 | 110 | 100 | 91 | 114.36 | 128 | 100 | 0.781 | -0.357 | 0.09083533 | 0.20608086 | 0.0232 | 0.0232 | 0.0232 | 0.0232 | TRUE | 0.4438 |
| Zfp712 | 275 | 216 | 236 | 272 | 202 | 269 | 214 | 230 | 255 | 271 | 228 | 266 | 243.69 | 233 | 255 | 1.099 | 0.136 | 0.32582027 | 0.49988719 | 0.0041 | 0.0122 | 0.0098 | 0.0098 | TRUE | 0.0911 |
| Zfp715 | 1656 | 1291 | 1230 | 1248 | 1108 | 1166 | 1286 | 1373 | 1327 | 1242 | 1250 | 1151 | 1271.62 | 1329 | 1214 | 0.914 | -0.129 | 0.07527269 | 0.18072513 | 8.00E-04 | 0.0043 | 0.003 | 0.003 | TRUE | 0.0253 |
| Zfp719 | 757 | 550 | 352 | 378 | 351 | 363 | 588 | 585 | 380 | 376 | 396 | 358 | 447.22 | 518 | 377 | 0.727 | -0.461 | 0.00224632 | 0.01321803 | 0.0289 | 0.0077 | 0.0141 | 0.0141 | TRUE | 0.6213 |
| Zfp72 | 289 | 214 | 214 | 282 | 231 | 246 | 224 | 228 | 231 | 281 | 261 | 243 | 244.52 | 228 | 262 | 1.149 | 0.2 | 0.12642244 | 0.26172595 | 0 | 0.0121 | 0.0082 | 0.0082 | TRUE | 0.048 |
| Zfp735 | 0 | 0 | 0 | 1 | 0 | 0 | 0 | 0 | 0 | 1 | 0 | 0 | 0.17 | 0 | 0 | 2.094 | 1.067 | 0.79379329 | NA | 0 | 14.2885 | 10 | 10 | TRUE | 0.3906 |
| Zfp738 | 323 | 274 | 249 | 338 | 251 | 312 | 251 | 291 | 269 | 336 | 283 | 308 | 289.76 | 270 | 309 | 1.147 | 0.198 | 0.12156163 | 0.25435227 | 0.0032 | 0.0106 | 0.0084 | 0.0084 | TRUE | 0.0672 |
| Zfp74 | 733 | 574 | 538 | 534 | 419 | 576 | 569 | 611 | 580 | 532 | 473 | 569 | 555.53 | 587 | 525 | 0.895 | -0.16 | 0.11222886 | 0.24056495 | 0.0028 | 0.0067 | 0.0055 | 0.0055 | TRUE | 0.084 |
| Zfp740 | 6677 | 4939 | 4561 | 5827 | 5013 | 5666 | 5186 | 5254 | 4921 | 5800 | 5655 | 5593 | 5401.53 | 5120 | 5683 | 1.11 | 0.15 | 0.0048282 | 0.02386661 | 5.00E-04 | 0.0029 | 0.0019 | 0.0019 | TRUE | 0.0139 |
| Zfp746 | 1246 | 957 | 992 | 1097 | 950 | 1088 | 968 | 1018 | 1070 | 1092 | 1072 | 1074 | 1048.95 | 1019 | 1079 | 1.061 | 0.085 | 0.26065638 | 0.43077006 | 0 | 0.0047 | 0.0032 | 0.0032 | TRUE | 0.0242 |
| Zfp747 | 401 | 270 | 261 | 280 | 218 | 300 | 311 | 287 | 282 | 279 | 246 | 296 | 283.51 | 293 | 274 | 0.932 | -0.102 | 0.42676176 | 0.59899397 | 0.0021 | 0.0108 | 0.0082 | 0.0082 | TRUE | 0.0844 |
| Zfp748 | 1215 | 823 | 829 | 940 | 816 | 913 | 944 | 875 | 894 | 936 | 921 | 901 | 911.84 | 904 | 919 | 1.015 | 0.022 | 0.78143312 | 0.87115299 | 0 | 0.005 | 0.0032 | 0.0032 | TRUE | 0.018 |
| Zfp750 | 463 | 403 | 429 | 329 | 318 | 338 | 360 | 429 | 463 | 327 | 359 | 334 | 378.5 | 417 | 340 | 0.817 | -0.291 | 0.01844402 | 0.06507539 | 0.0075 | 0.0087 | 0.0083 | 0.0083 | TRUE | 0.1787 |
| Zfp758 | 160 | 102 | 68 | 91 | 68 | 67 | 124 | 108 | 73 | 91 | 77 | 66 | 89.93 | 102 | 78 | 0.758 | -0.399 | 0.10134305 | 0.22336101 | 0.0377 | 0.0288 | 0.0313 | 0.0313 | TRUE | 0.4652 |
| Zfp759 | 261 | 183 | 120 | 146 | 99 | 142 | 203 | 195 | 129 | 145 | 112 | 140 | 154 | 176 | 132 | 0.753 | -0.41 | 0.03969059 | 0.11391518 | 0.0322 | 0.0178 | 0.022 | 0.022 | TRUE | 0.5469 |
| Zfp760 | 1085 | 818 | 700 | 854 | 666 | 809 | 843 | 870 | 755 | 850 | 751 | 799 | 811.34 | 823 | 800 | 0.972 | -0.04 | 0.65509407 | 0.78570609 | 0.0032 | 0.0053 | 0.0047 | 0.0047 | TRUE | 0.059 |
| Zfp763 | 688 | 467 | 575 | 619 | 551 | 570 | 534 | 497 | 620 | 616 | 622 | 563 | 575.32 | 550 | 600 | 1.091 | 0.125 | 0.23814875 | 0.4049322 | 0.006 | 0.0065 | 0.0064 | 0.0064 | TRUE | 0.1393 |
| Zfp764 | 418 | 279 | 229 | 241 | 263 | 260 | 325 | 297 | 247 | 240 | 297 | 257 | 276.96 | 290 | 265 | 0.909 | -0.137 | 0.34028262 | 0.51440079 | 0.0119 | 0.011 | 0.0112 | 0.0112 | TRUE | 0.181 |
| Zfp768 | 122 | 74 | 73 | 76 | 80 | 90 | 95 | 79 | 79 | 76 | 90 | 89 | 84.5 | 84 | 85 | 1.004 | 0.006 | 0.97637366 | 0.98728943 | 0 | 0.0305 | 0.0219 | 0.0219 | TRUE | 0.1305 |
| Zfp770 | 2221 | 1610 | 1361 | 1478 | 1211 | 1335 | 1725 | 1713 | 1469 | 1471 | 1366 | 1318 | 1510.2 | 1636 | 1385 | 0.846 | -0.241 | 0.00393566 | 0.02034444 | 0.0045 | 0.004 | 0.0043 | 0.0043 | TRUE | 0.0947 |
| Zfp771 | 708 | 568 | 412 | 565 | 537 | 617 | 550 | 604 | 445 | 562 | 606 | 609 | 562.64 | 533 | 592 | 1.111 | 0.152 | 0.19131061 | 0.34937075 | 0.0108 | 0.0066 | 0.0079 | 0.0079 | TRUE | 0.237 |
| Zfp772 | 383 | 250 | 282 | 313 | 242 | 331 | 297 | 266 | 304 | 312 | 273 | 327 | 296.49 | 289 | 304 | 1.051 | 0.072 | 0.57304299 | 0.72548197 | 0.003 | 0.0104 | 0.0083 | 0.0083 | TRUE | 0.0856 |
| Zfp773 | 221 | 157 | 173 | 171 | 152 | 174 | 172 | 167 | 187 | 170 | 171 | 172 | 173.13 | 175 | 171 | 0.978 | -0.032 | 0.8321018 | 0.90401574 | 0 | 0.0161 | 0.0104 | 0.0104 | TRUE | 0.0336 |
| Zfp775 | 896 | 751 | 814 | 967 | 843 | 1063 | 696 | 799 | 878 | 963 | 951 | 1049 | 889.32 | 791 | 988 | 1.252 | 0.324 | 0.00093943 | 0.00693559 | 0.0071 | 0.0051 | 0.0058 | 0.0058 | TRUE | 0.1376 |
| Zfp777 | 1528 | 1177 | 1135 | 1102 | 944 | 1094 | 1187 | 1252 | 1225 | 1097 | 1065 | 1080 | 1150.86 | 1221 | 1081 | 0.886 | -0.175 | 0.01337423 | 0.05133132 | 0 | 0.0045 | 0.0027 | 0.0027 | TRUE | 0.0077 |
| Zfp78 | 94 | 65 | 59 | 72 | 71 | 85 | 73 | 69 | 64 | 72 | 80 | 84 | 73.58 | 69 | 79 | 1.142 | 0.191 | 0.39682377 | 0.57025962 | 0 | 0.0346 | 0.0232 | 0.0232 | TRUE | 0.0561 |
| Zfp780b | 296 | 180 | 110 | 107 | 97 | 118 | 230 | 191 | 119 | 107 | 109 | 116 | 145.41 | 180 | 111 | 0.613 | -0.707 | 0.00151344 | 0.00973686 | 0.051 | 0.0187 | 0.0284 | 0.0284 | TRUE | 0.3022 |
| Zfp781 | 37 | 39 | 50 | 33 | 33 | 39 | 29 | 41 | 54 | 33 | 37 | 38 | 38.79 | 41 | 36 | 0.883 | -0.18 | 0.59072381 | 0.73901229 | 0.0328 | 0.0635 | 0.0544 | 0.0544 | TRUE | 0.1862 |
| Zfp783 | 68 | 40 | 28 | 27 | 36 | 33 | 53 | 43 | 30 | 27 | 41 | 33 | 37.61 | 42 | 34 | 0.788 | -0.344 | 0.31326908 | 0.48674366 | 0.0347 | 0.0655 | 0.0568 | 0.0568 | TRUE | 0.1878 |
| Zfp784 | 42 | 29 | 30 | 21 | 25 | 28 | 33 | 31 | 32 | 21 | 28 | 28 | 28.76 | 32 | 26 | 0.799 | -0.324 | 0.36384976 | 0.53841025 | 0 | 0.0848 | 0.0564 | 0.0564 | TRUE | 0.1616 |
| Zfp786 | 139 | 100 | 64 | 95 | 83 | 76 | 108 | 106 | 69 | 95 | 94 | 75 | 91.1 | 94 | 88 | 0.924 | -0.114 | 0.62388003 | 0.76299455 | 0.0266 | 0.0284 | 0.0279 | 0.0279 | TRUE | 0.5087 |
| Zfp787 | 290 | 185 | 198 | 227 | 212 | 257 | 225 | 197 | 214 | 226 | 239 | 254 | 225.74 | 212 | 240 | 1.128 | 0.174 | 0.20271398 | 0.36416046 | 0 | 0.0129 | 0.0091 | 0.0091 | TRUE | 0.0419 |
| Zfp788 | 714 | 544 | 528 | 523 | 467 | 566 | 555 | 579 | 570 | 521 | 527 | 559 | 551.51 | 568 | 536 | 0.944 | -0.083 | 0.36751468 | 0.54220555 | 0 | 0.0067 | 0.0043 | 0.0043 | TRUE | 0.0173 |
| Zfp790 | 589 | 410 | 422 | 427 | 354 | 379 | 457 | 436 | 455 | 425 | 399 | 374 | 424.57 | 449 | 399 | 0.888 | -0.171 | 0.10169964 | 0.22387123 | 0 | 0.008 | 0.0055 | 0.0055 | TRUE | 0.037 |
| Zfp791 | 173 | 125 | 130 | 148 | 133 | 163 | 134 | 133 | 140 | 147 | 150 | 161 | 144.31 | 136 | 153 | 1.125 | 0.17 | 0.29527081 | 0.46705464 | 0 | 0.0188 | 0.0121 | 0.0121 | TRUE | 0.0235 |
| Zfp799 | 1163 | 750 | 855 | 1045 | 893 | 1104 | 903 | 798 | 923 | 1040 | 1007 | 1090 | 960.16 | 875 | 1046 | 1.195 | 0.258 | 0.00233323 | 0.01362292 | 0.0025 | 0.0049 | 0.0041 | 0.0041 | TRUE | 0.0682 |
| Zfp800 | 823 | 608 | 564 | 698 | 600 | 594 | 639 | 647 | 609 | 695 | 677 | 586 | 642.08 | 632 | 653 | 1.033 | 0.047 | 0.62892088 | 0.76633983 | 0.003 | 0.0061 | 0.0051 | 0.0051 | TRUE | 0.0946 |
| Zfp804a | 0 | 0 | 0 | 0 | 0 | 0 | 0 | 0 | 0 | 0 | 0 | 0 | 0 | 0 | 0 | NA | NA | NA | NA | NA | NA | NA | NA | NA | NA |
| Zfp804b | 0 | 0 | 0 | 1 | 1 | 1 | 0 | 0 | 0 | 1 | 1 | 1 | 0.52 | 0 | 1 | 5.852 | 2.549 | 0.3348364 | NA | 0 | 4.5739 | 3.5345 | 3.5345 | TRUE | 0.0847 |
| Zfp808 | 1229 | 835 | 722 | 818 | 648 | 818 | 955 | 888 | 779 | 814 | 731 | 807 | 829.08 | 874 | 784 | 0.896 | -0.158 | 0.10183556 | 0.22406646 | 0.0059 | 0.0053 | 0.0055 | 0.0055 | TRUE | 0.1061 |
| Zfp809 | 2055 | 1449 | 1253 | 1331 | 1181 | 1314 | 1596 | 1541 | 1352 | 1325 | 1332 | 1297 | 1407.26 | 1496 | 1318 | 0.88 | -0.185 | 0.01960175 | 0.06806799 | 0.0032 | 0.0041 | 0.0038 | 0.0038 | TRUE | 0.0847 |
| Zfp81 | 699 | 491 | 538 | 595 | 496 | 574 | 543 | 522 | 580 | 592 | 560 | 567 | 560.68 | 548 | 573 | 1.045 | 0.063 | 0.49843132 | 0.66301339 | 0 | 0.0067 | 0.0045 | 0.0045 | TRUE | 0.0288 |
| Zfp810 | 390 | 258 | 274 | 345 | 283 | 308 | 303 | 274 | 296 | 343 | 319 | 304 | 306.61 | 291 | 322 | 1.106 | 0.145 | 0.22593817 | 0.3909066 | 0 | 0.0102 | 0.0071 | 0.0071 | TRUE | 0.0384 |
| Zfp811 | 541 | 407 | 383 | 374 | 392 | 488 | 420 | 433 | 413 | 372 | 442 | 482 | 427.1 | 422 | 432 | 1.024 | 0.034 | 0.77406931 | 0.8666593 | 0.0064 | 0.008 | 0.0075 | 0.0075 | TRUE | 0.1727 |
| Zfp819 | 425 | 310 | 267 | 273 | 270 | 305 | 330 | 330 | 288 | 272 | 305 | 301 | 304.22 | 316 | 293 | 0.924 | -0.114 | 0.35340464 | 0.52780934 | 0 | 0.0102 | 0.0076 | 0.0076 | TRUE | 0.0649 |
| Zfp82 | 517 | 367 | 387 | 344 | 322 | 399 | 402 | 390 | 418 | 342 | 363 | 394 | 384.84 | 403 | 366 | 0.909 | -0.137 | 0.21222496 | 0.37454069 | 0 | 0.0086 | 0.0061 | 0.0061 | TRUE | 0.0502 |
| Zfp820 | 770 | 588 | 491 | 553 | 387 | 503 | 598 | 625 | 530 | 550 | 437 | 497 | 539.47 | 584 | 495 | 0.847 | -0.24 | 0.03250655 | 0.09820674 | 0.0081 | 0.0068 | 0.0072 | 0.0072 | TRUE | 0.1191 |
| Zfp821 | 933 | 706 | 615 | 626 | 545 | 641 | 725 | 751 | 664 | 623 | 615 | 633 | 668.31 | 713 | 624 | 0.874 | -0.194 | 0.02947644 | 0.09148004 | 0 | 0.006 | 0.0042 | 0.0042 | TRUE | 0.0417 |
| Zfp825 | 551 | 391 | 411 | 473 | 380 | 450 | 428 | 416 | 443 | 471 | 429 | 444 | 438.5 | 429 | 448 | 1.045 | 0.063 | 0.536957 | 0.69580475 | 0 | 0.0078 | 0.0052 | 0.0052 | TRUE | 0.0233 |
| Zfp827 | 626 | 474 | 366 | 372 | 362 | 395 | 486 | 504 | 395 | 370 | 408 | 390 | 425.65 | 462 | 389 | 0.842 | -0.248 | 0.03662285 | 0.10755095 | 0.0073 | 0.008 | 0.0078 | 0.0078 | TRUE | 0.1798 |
| Zfp828 | 7006 | 5046 | 4840 | 4941 | 4472 | 5100 | 5441 | 5367 | 5222 | 4918 | 5045 | 5034 | 5171.43 | 5343 | 4999 | 0.935 | -0.096 | 0.05673224 | 0.14772708 | 1.00E-04 | 0.0029 | 0.0017 | 0.0017 | TRUE | 0.0048 |
| Zfp830 | 2015 | 1484 | 1326 | 1268 | 1094 | 1348 | 1565 | 1579 | 1431 | 1262 | 1234 | 1331 | 1400.2 | 1525 | 1276 | 0.836 | -0.258 | 0.00045239 | 0.00399873 | 0.0015 | 0.0041 | 0.0032 | 0.0032 | TRUE | 0.0342 |
| Zfp831 | 49 | 30 | 35 | 37 | 29 | 39 | 38 | 32 | 38 | 37 | 33 | 38 | 35.96 | 36 | 36 | 1.002 | 0.003 | 0.99181013 | 0.99565418 | 0 | 0.0683 | 0.0449 | 0.0449 | TRUE | 0.0643 |
| Zfp839 | 1030 | 740 | 789 | 852 | 794 | 872 | 800 | 787 | 851 | 848 | 896 | 861 | 840.5 | 813 | 868 | 1.068 | 0.095 | 0.2360594 | 0.40252888 | 0 | 0.0052 | 0.0035 | 0.0035 | TRUE | 0.0196 |
| Zfp84 | 665 | 434 | 391 | 416 | 400 | 433 | 516 | 462 | 422 | 414 | 451 | 427 | 448.79 | 467 | 431 | 0.92 | -0.12 | 0.27993889 | 0.45144608 | 0.0043 | 0.0077 | 0.0066 | 0.0066 | TRUE | 0.1069 |
| Zfp846 | 523 | 419 | 394 | 362 | 307 | 393 | 406 | 446 | 425 | 360 | 346 | 388 | 395.27 | 426 | 365 | 0.859 | -0.219 | 0.04285468 | 0.12040926 | 0 | 0.0084 | 0.0059 | 0.0059 | TRUE | 0.0353 |
| Zfp85-rs1 | 299 | 206 | 211 | 232 | 189 | 210 | 232 | 219 | 228 | 231 | 213 | 207 | 221.74 | 226 | 217 | 0.959 | -0.061 | 0.6530261 | 0.78454209 | 0 | 0.0131 | 0.0086 | 0.0086 | TRUE | 0.0346 |
| Zfp862 | 383 | 279 | 209 | 259 | 212 | 240 | 297 | 297 | 226 | 258 | 239 | 237 | 258.94 | 273 | 245 | 0.893 | -0.163 | 0.25637957 | 0.42634476 | 0.0094 | 0.0116 | 0.0109 | 0.0109 | TRUE | 0.2523 |
| Zfp865 | 739 | 497 | 443 | 425 | 418 | 481 | 574 | 529 | 478 | 423 | 472 | 475 | 491.67 | 527 | 457 | 0.864 | -0.21 | 0.05121964 | 0.13729975 | 0.0045 | 0.0072 | 0.0063 | 0.0063 | TRUE | 0.0761 |
| Zfp866 | 1900 | 1465 | 1275 | 1277 | 1188 | 1418 | 1476 | 1558 | 1376 | 1271 | 1340 | 1400 | 1403.46 | 1470 | 1337 | 0.91 | -0.137 | 0.07309909 | 0.17700121 | 0.0024 | 0.0041 | 0.0035 | 0.0035 | TRUE | 0.0366 |
| Zfp867 | 767 | 514 | 494 | 482 | 493 | 535 | 596 | 547 | 533 | 480 | 556 | 528 | 539.92 | 559 | 521 | 0.931 | -0.103 | 0.30906216 | 0.48183087 | 0.0027 | 0.0068 | 0.0055 | 0.0055 | TRUE | 0.0571 |
| Zfp868 | 1418 | 1022 | 839 | 888 | 762 | 815 | 1101 | 1087 | 905 | 884 | 860 | 805 | 940.29 | 1031 | 850 | 0.823 | -0.282 | 0.00278526 | 0.01558109 | 0.006 | 0.0049 | 0.0053 | 0.0053 | TRUE | 0.1339 |
| Zfp869 | 1344 | 1001 | 957 | 798 | 730 | 833 | 1044 | 1065 | 1033 | 794 | 824 | 822 | 930.22 | 1047 | 813 | 0.777 | -0.364 | 1.31E-06 | 3.65E-05 | 0 | 0.005 | 0.003 | 0.003 | TRUE | 0.005 |
| Zfp87 | 661 | 493 | 489 | 579 | 480 | 504 | 513 | 524 | 528 | 576 | 541 | 498 | 530.12 | 522 | 538 | 1.032 | 0.046 | 0.63954527 | 0.77477341 | 0 | 0.0069 | 0.005 | 0.005 | TRUE | 0.053 |
| Zfp870 | 1090 | 794 | 815 | 826 | 742 | 876 | 847 | 845 | 879 | 822 | 837 | 865 | 849.08 | 857 | 841 | 0.982 | -0.026 | 0.74240341 | 0.84665858 | 0 | 0.0052 | 0.0032 | 0.0032 | TRUE | 0.0072 |
| Zfp871 | 3190 | 2464 | 2547 | 2587 | 2313 | 2885 | 2478 | 2621 | 2748 | 2575 | 2609 | 2848 | 2646.5 | 2616 | 2677 | 1.024 | 0.035 | 0.61152402 | 0.75404219 | 0.0025 | 0.0033 | 0.003 | 0.003 | TRUE | 0.0379 |
| Zfp872 | 25 | 19 | 21 | 39 | 30 | 47 | 19 | 20 | 23 | 39 | 34 | 46 | 30.22 | 21 | 40 | 1.922 | 0.943 | 0.00826781 | 0.03589665 | 0 | 0.0808 | 0.0555 | 0.0555 | TRUE | 0.1655 |
| Zfp873 | 427 | 335 | 302 | 343 | 249 | 300 | 332 | 356 | 326 | 341 | 281 | 296 | 322.05 | 338 | 306 | 0.907 | -0.14 | 0.25267333 | 0.42159287 | 0.0028 | 0.0098 | 0.0077 | 0.0077 | TRUE | 0.1155 |

| | | | | | | | | | | | | | | | | | | | | | | | | | |
|---|---|---|---|---|---|---|---|---|---|---|---|---|---|---|---|---|---|---|---|---|---|---|---|---|---|
| Zfp874a | 520 | 388 | 405 | 494 | 395 | 443 | 404 | 413 | 437 | 492 | 446 | 437 | 438.03 | 418 | 458 | 1.098 | 0.135 | 0.196636 | 0.35635711 | 0 | 0.0078 | 0.0056 | 0.0056 | TRUE | 0.0482 |
| Zfp874b | 779 | 608 | 593 | 682 | 530 | 658 | 605 | 647 | 640 | 679 | 598 | 650 | 636.31 | 631 | 642 | 1.02 | 0.029 | 0.75293564 | 0.85348706 | 0.0011 | 0.0062 | 0.0045 | 0.0045 | TRUE | 0.0415 |
| Zfp879 | 71 | 47 | 61 | 55 | 49 | 53 | 55 | 50 | 66 | 55 | 55 | 52 | 55.55 | 57 | 54 | 0.951 | -0.073 | 0.77913665 | 0.86967591 | 0 | 0.0451 | 0.0308 | 0.0308 | TRUE | 0.1441 |
| Zfp882 | 349 | 240 | 218 | 167 | 200 | 199 | 271 | 255 | 235 | 166 | 226 | 196 | 224.98 | 254 | 196 | 0.77 | -0.378 | 0.01190839 | 0.04708345 | 0.0085 | 0.013 | 0.0117 | 0.0117 | TRUE | 0.1934 |
| Zfp9 | 247 | 203 | 192 | 210 | 208 | 194 | 192 | 216 | 207 | 209 | 235 | 192 | 208.35 | 205 | 212 | 1.033 | 0.048 | 0.74387628 | 0.84759436 | 0 | 0.0138 | 0.0104 | 0.0104 | TRUE | 0.0946 |
| Zfp90 | 655 | 446 | 474 | 441 | 459 | 498 | 509 | 474 | 511 | 439 | 518 | 492 | 490.49 | 498 | 483 | 0.968 | -0.047 | 0.64812919 | 0.78115444 | 0.0021 | 0.0073 | 0.0057 | 0.0057 | TRUE | 0.0738 |
| Zfp91 | 2567 | 1812 | 1816 | 1864 | 1675 | 1833 | 1994 | 1927 | 1959 | 1855 | 1890 | 1809 | 1905.83 | 1960 | 1851 | 0.944 | -0.083 | 0.17535863 | 0.32879456 | 0 | 0.0037 | 0.0022 | 0.0022 | TRUE | 0.0048 |
| Zfp92 | 50 | 18 | 23 | 29 | 26 | 30 | 39 | 19 | 25 | 29 | 29 | 30 | 28.43 | 28 | 29 | 1.046 | 0.065 | 0.86537098 | 0.92566588 | 0.0357 | 0.0858 | 0.0713 | 0.0713 | TRUE | 0.877 |
| Zfp93 | 186 | 118 | 122 | 114 | 111 | 121 | 144 | 126 | 132 | 113 | 125 | 119 | 126.63 | 134 | 119 | 0.889 | -0.17 | 0.33124256 | 0.50560123 | 0 | 0.0211 | 0.0142 | 0.0142 | TRUE | 0.0546 |
| Zfp930 | 2447 | 1860 | 1797 | 2188 | 1795 | 2245 | 1900 | 1978 | 1939 | 2178 | 2025 | 2216 | 2039.49 | 1939 | 2140 | 1.104 | 0.143 | 0.02716561 | 0.08616913 | 8.00E-04 | 0.0036 | 0.0025 | 0.0025 | TRUE | 0.0262 |
| Zfp931 | 101 | 66 | 71 | 90 | 74 | 82 | 78 | 70 | 77 | 90 | 83 | 81 | 79.88 | 75 | 85 | 1.126 | 0.171 | 0.42679275 | 0.59899397 | 0 | 0.0321 | 0.0209 | 0.0209 | TRUE | 0.0285 |
| Zfp932 | 729 | 510 | 444 | 510 | 437 | 479 | 566 | 542 | 479 | 508 | 493 | 473 | 510.2 | 529 | 491 | 0.926 | -0.111 | 0.28124761 | 0.45279028 | 0.0026 | 0.0071 | 0.0056 | 0.0056 | TRUE | 0.0786 |
| Zfp933 | 289 | 194 | 148 | 170 | 144 | 144 | 224 | 206 | 160 | 169 | 162 | 142 | 177.39 | 197 | 158 | 0.799 | -0.324 | 0.05844985 | 0.15071711 | 0.0143 | 0.0158 | 0.0153 | 0.0153 | TRUE | 0.2871 |
| Zfp934 | 276 | 201 | 161 | 216 | 182 | 261 | 214 | 214 | 174 | 215 | 205 | 258 | 213.31 | 201 | 226 | 1.125 | 0.17 | 0.26815988 | 0.43873732 | 0.0095 | 0.0135 | 0.0124 | 0.0124 | TRUE | 0.167 |
| Zfp935 | 461 | 286 | 231 | 219 | 208 | 237 | 358 | 304 | 249 | 218 | 235 | 234 | 266.35 | 304 | 229 | 0.75 | -0.415 | 0.00565463 | 0.0269765 | 0.0146 | 0.0113 | 0.0123 | 0.0123 | TRUE | 0.1502 |
| Zfp936 | 1792 | 1219 | 673 | 710 | 732 | 712 | 1392 | 1297 | 726 | 707 | 826 | 703 | 941.65 | 1138 | 745 | 0.654 | -0.612 | 0.03777405 | 0.10987067 | 0.0614 | 0.0049 | 0.0203 | 0.0614 | TRUE | 1.1979 |
| Zfp937 | 522 | 393 | 411 | 481 | 396 | 467 | 405 | 418 | 443 | 479 | 447 | 461 | 442.24 | 422 | 462 | 1.096 | 0.133 | 0.19168187 | 0.34994366 | 0 | 0.0078 | 0.0052 | 0.0052 | TRUE | 0.0218 |
| Zfp938 | 526 | 364 | 389 | 482 | 410 | 467 | 409 | 387 | 420 | 480 | 463 | 461 | 436.46 | 405 | 468 | 1.154 | 0.207 | 0.03994502 | 0.11436973 | 0 | 0.0079 | 0.005 | 0.005 | TRUE | 0.0164 |
| Zfp939 | 149 | 113 | 88 | 116 | 107 | 127 | 116 | 120 | 95 | 115 | 121 | 125 | 115.4 | 110 | 120 | 1.091 | 0.125 | 0.50270016 | 0.66683001 | 0 | 0.023 | 0.0164 | 0.0164 | TRUE | 0.1381 |
| Zfp94 | 404 | 242 | 212 | 256 | 226 | 258 | 314 | 257 | 229 | 255 | 255 | 255 | 260.73 | 267 | 255 | 0.951 | -0.072 | 0.61593953 | 0.75749645 | 0.01 | 0.0115 | 0.011 | 0.011 | TRUE | 0.2842 |
| Zfp940 | 738 | 519 | 556 | 620 | 503 | 622 | 573 | 552 | 600 | 617 | 567 | 614 | 587.29 | 575 | 599 | 1.043 | 0.061 | 0.5064583 | 0.66988788 | 0 | 0.0065 | 0.0044 | 0.0044 | TRUE | 0.025 |
| Zfp941 | 935 | 690 | 656 | 708 | 612 | 627 | 726 | 734 | 708 | 705 | 690 | 619 | 697 | 723 | 671 | 0.929 | -0.107 | 0.22908449 | 0.39449868 | 0.001 | 0.0058 | 0.0043 | 0.0043 | TRUE | 0.056 |
| Zfp942 | 458 | 309 | 264 | 295 | 248 | 280 | 356 | 329 | 285 | 294 | 280 | 276 | 303.18 | 323 | 283 | 0.874 | -0.194 | 0.12755554 | 0.26346117 | 0.0041 | 0.0102 | 0.0083 | 0.0083 | TRUE | 0.1191 |
| Zfp943 | 161 | 107 | 100 | 113 | 122 | 113 | 125 | 114 | 108 | 112 | 138 | 112 | 118.07 | 116 | 121 | 1.038 | 0.053 | 0.77453646 | 0.86691069 | 0 | 0.0225 | 0.0167 | 0.0167 | TRUE | 0.1485 |
| Zfp944 | 506 | 400 | 349 | 334 | 298 | 287 | 393 | 425 | 377 | 332 | 336 | 283 | 357.83 | 398 | 317 | 0.796 | -0.329 | 0.00558112 | 0.02668816 | 0.0031 | 0.009 | 0.0073 | 0.0073 | TRUE | 0.1014 |
| Zfp945 | 1490 | 1114 | 1093 | 1207 | 1026 | 1198 | 1157 | 1185 | 1179 | 1201 | 1157 | 1183 | 1177.17 | 1174 | 1180 | 1.006 | 0.009 | 0.89718357 | 0.94337809 | 0 | 0.0044 | 0.0026 | 0.0026 | TRUE | 0.0034 |
| Zfp946 | 293 | 169 | 152 | 136 | 128 | 154 | 228 | 180 | 164 | 135 | 144 | 152 | 167.19 | 191 | 144 | 0.752 | -0.411 | 0.01718275 | 0.06186222 | 0.0126 | 0.0166 | 0.0153 | 0.0153 | TRUE | 0.3364 |
| Zfp947 | 122 | 89 | 55 | 70 | 56 | 65 | 95 | 95 | 59 | 70 | 63 | 64 | 74.3 | 83 | 66 | 0.789 | -0.342 | 0.16976969 | 0.32136735 | 0.0226 | 0.0343 | 0.031 | 0.031 | TRUE | 0.5503 |
| Zfp948 | 560 | 377 | 394 | 442 | 353 | 409 | 435 | 401 | 425 | 440 | 398 | 404 | 417.17 | 420 | 414 | 0.984 | -0.023 | 0.82833581 | 0.90163764 | 0 | 0.0081 | 0.0056 | 0.0056 | TRUE | 0.0351 |
| Zfp949 | 551 | 402 | 399 | 426 | 371 | 498 | 428 | 428 | 431 | 424 | 419 | 492 | 436.7 | 429 | 445 | 1.038 | 0.054 | 0.61283585 | 0.75508227 | 0.0018 | 0.0079 | 0.006 | 0.006 | TRUE | 0.0997 |
| Zfp951 | 1497 | 1190 | 908 | 756 | 746 | 879 | 1163 | 1266 | 980 | 753 | 842 | 868 | 978.33 | 1136 | 821 | 0.722 | -0.47 | 5.03E-06 | 0.00010634 | 0.0101 | 0.0048 | 0.0066 | 0.0066 | TRUE | 0.1698 |
| Zfp952 | 629 | 438 | 495 | 503 | 415 | 525 | 489 | 466 | 534 | 501 | 468 | 518 | 495.94 | 496 | 496 | 1 | 0 | 0.99714785 | 0.99840519 | 0.0015 | 0.0072 | 0.0054 | 0.0054 | TRUE | 0.0497 |
| Zfp953 | 451 | 319 | 257 | 275 | 256 | 225 | 350 | 339 | 277 | 274 | 289 | 222 | 291.92 | 322 | 262 | 0.809 | -0.306 | 0.03318748 | 0.09981562 | 0.0134 | 0.0105 | 0.0114 | 0.0114 | TRUE | 0.1968 |
| Zfp954 | 562 | 383 | 403 | 405 | 370 | 439 | 436 | 407 | 435 | 403 | 417 | 433 | 422.1 | 426 | 418 | 0.98 | -0.029 | 0.77648734 | 0.86773522 | 0 | 0.008 | 0.0053 | 0.0053 | TRUE | 0.0168 |
| Zfp955a | 1409 | 1047 | 1029 | 1157 | 962 | 1024 | 1094 | 1114 | 1110 | 1152 | 1085 | 1011 | 1094.34 | 1106 | 1083 | 0.979 | -0.031 | 0.69159412 | 0.81149322 | 0.0013 | 0.0046 | 0.0034 | 0.0034 | TRUE | 0.041 |
| Zfp955b | 1381 | 934 | 665 | 836 | 694 | 753 | 1073 | 993 | 718 | 832 | 783 | 743 | 856.99 | 928 | 786 | 0.846 | -0.241 | 0.0574129 | 0.14901083 | 0.0223 | 0.0052 | 0.0104 | 0.0104 | TRUE | 0.4636 |
| Zfp956 | 439 | 348 | 266 | 278 | 273 | 323 | 341 | 370 | 287 | 277 | 308 | 319 | 316.95 | 333 | 301 | 0.904 | -0.145 | 0.26855429 | 0.43913127 | 0.0076 | 0.0099 | 0.0092 | 0.0092 | TRUE | 0.1566 |
| Zfp957 | 0 | 0 | 0 | 0 | 0 | 0 | 0 | 0 | 0 | 0 | 0 | 0 | 0 | 0 | 0 | NA | NA | NA | NA | NA | NA | NA | NA | NA | NA |
| Zfp958 | 2202 | 1549 | 1485 | 1480 | 1404 | 1522 | 1710 | 1648 | 1602 | 1473 | 1584 | 1502 | 1586.61 | 1653 | 1520 | 0.918 | -0.123 | 0.06969157 | 0.1711068 | 6.00E-04 | 0.0039 | 0.0027 | 0.0027 | TRUE | 0.0161 |
| Zfp959 | 369 | 238 | 167 | 240 | 184 | 220 | 287 | 253 | 180 | 239 | 208 | 217 | 230.6 | 240 | 221 | 0.919 | -0.122 | 0.47269963 | 0.64119729 | 0.026 | 0.0127 | 0.0166 | 0.0166 | TRUE | 0.5134 |
| Zfp960 | 863 | 617 | 603 | 774 | 693 | 765 | 670 | 656 | 651 | 770 | 782 | 755 | 714.1 | 659 | 769 | 1.166 | 0.222 | 0.00699352 | 0.03171572 | 0 | 0.0057 | 0.0035 | 0.0035 | TRUE | 0.003 |
| Zfp961 | 6440 | 4370 | 4130 | 5069 | 4243 | 4829 | 5002 | 4648 | 4456 | 5046 | 4787 | 4767 | 4784.24 | 4702 | 4867 | 1.034 | 0.049 | 0.43001735 | 0.60221811 | 0.0019 | 0.0029 | 0.0026 | 0.0026 | TRUE | 0.0384 |
| Zfp963 | 208 | 114 | 92 | 124 | 89 | 104 | 162 | 121 | 99 | 123 | 100 | 103 | 118.09 | 127 | 109 | 0.85 | -0.235 | 0.27421604 | 0.44528174 | 0.0301 | 0.0225 | 0.0248 | 0.0248 | TRUE | 0.4088 |
| Zfp964 | 104 | 60 | 36 | 59 | 40 | 37 | 81 | 64 | 39 | 59 | 45 | 37 | 53.97 | 61 | 47 | 0.76 | -0.397 | 0.2152316 | 0.37769249 | 0.076 | 0.0463 | 0.0549 | 0.0549 | TRUE | 0.2434 |
| Zfp97 | 1059 | 812 | 811 | 963 | 789 | 947 | 822 | 864 | 875 | 959 | 890 | 935 | 890.79 | 854 | 928 | 1.088 | 0.122 | 0.12273211 | 0.25616384 | 0 | 0.0051 | 0.0034 | 0.0034 | TRUE | 0.0147 |
| Zfpl1 | 1145 | 816 | 822 | 801 | 736 | 806 | 889 | 868 | 887 | 797 | 830 | 796 | 844.57 | 881 | 808 | 0.916 | -0.127 | 0.10316836 | 0.22633697 | 0 | 0.0052 | 0.0032 | 0.0032 | TRUE | 0.0069 |
| Zfpm1 | 2290 | 1964 | 1920 | 2100 | 1858 | 1994 | 1779 | 2089 | 2072 | 2090 | 2096 | 1968 | 2015.67 | 1980 | 2051 | 1.037 | 0.053 | 0.49508673 | 0.65996429 | 0.0042 | 0.0036 | 0.0038 | 0.0038 | TRUE | 0.0984 |
| Zfpm2 | 23 | 9 | 3 | 6 | 7 | 12 | 18 | 10 | 3 | 6 | 8 | 12 | 9.4 | 10 | 9 | 0.822 | -0.283 | 0.68972144 | NA | 0.2541 | 0.2546 | 0.2545 | 0.2545 | TRUE | 0.2364 |
| Zfr | 9651 | 6783 | 6727 | 7200 | 6546 | 7326 | 7496 | 7215 | 7258 | 7167 | 7385 | 7232 | 7292.04 | 7323 | 7261 | 0.991 | -0.013 | 0.79614223 | 0.88091917 | 2.00E-04 | 0.0028 | 0.0016 | 0.0016 | TRUE | 0.0053 |
| Zfr2 | 52 | 45 | 71 | 99 | 79 | 80 | 40 | 48 | 77 | 99 | 89 | 79 | 71.92 | 55 | 89 | 1.632 | 0.707 | 0.00880053 | 0.03771694 | 0.0445 | 0.0354 | 0.0378 | 0.0378 | TRUE | 0.9669 |
| Zfx | 2944 | 2162 | 2121 | 2304 | 1868 | 2148 | 2286 | 2300 | 2289 | 2293 | 2107 | 2120 | 2232.64 | 2292 | 2173 | 0.949 | -0.076 | 0.2259233 | 0.3909066 | 6.00E-04 | 0.0035 | 0.0024 | 0.0024 | TRUE | 0.0283 |
| Zfy1 | 276 | 208 | 243 | 143 | 128 | 129 | 214 | 221 | 262 | 142 | 144 | 127 | 185.31 | 232 | 138 | 0.594 | -0.751 | 1.67E-06 | 4.41E-05 | 0 | 0.0152 | 0.0119 | 0.0119 | TRUE | 0.1334 |
| Zfy2 | 66 | 83 | 62 | 26 | 19 | 21 | 51 | 88 | 67 | 26 | 21 | 21 | 45.75 | 69 | 23 | 0.332 | -1.591 | 1.43E-06 | 3.90E-05 | 0.0331 | 0.0542 | 0.0487 | 0.0487 | TRUE | 0.1305 |
| Zfyve1 | 2330 | 1661 | 1708 | 1846 | 1639 | 1930 | 1810 | 1767 | 1843 | 1838 | 1849 | 1905 | 1835.17 | 1807 | 1864 | 1.032 | 0.045 | 0.46498745 | 0.63398508 | 0 | 0.0037 | 0.0022 | 0.0022 | TRUE | 0.0046 |
| Zfyve16 | 1805 | 1197 | 1225 | 1390 | 1122 | 1239 | 1402 | 1273 | 1322 | 1384 | 1266 | 1223 | 1311.55 | 1332 | 1291 | 0.968 | -0.047 | 0.54893872 | 0.70553666 | 0.0025 | 0.0042 | 0.0036 | 0.0036 | TRUE | 0.048 |
| Zfyve19 | 634 | 457 | 367 | 415 | 387 | 452 | 492 | 486 | 396 | 413 | 437 | 446 | 445.06 | 458 | 432 | 0.941 | -0.088 | 0.44152102 | 0.61268543 | 0.0058 | 0.0078 | 0.0071 | 0.0071 | TRUE | 0.1587 |
| Zfyve20 | 5935 | 4106 | 3877 | 4378 | 3827 | 4419 | 4609 | 4367 | 4183 | 4358 | 4317 | 4362 | 4366.26 | 4386 | 4346 | 0.99 | -0.014 | 0.80419332 | 0.88656654 | 0.001 | 0.003 | 0.0021 | 0.0021 | TRUE | 0.0245 |
| Zfyve21 | 1991 | 1580 | 1820 | 2004 | 1857 | 2037 | 1546 | 1681 | 1964 | 1995 | 2095 | 2011 | 1881.87 | 1730 | 2034 | 1.176 | 0.234 | 0.00699419 | 0.03171572 | 0.0071 | 0.0037 | 0.0049 | 0.0049 | TRUE | 0.1676 |
| Zfyve26 | 2531 | 1839 | 1848 | 1667 | 1548 | 1872 | 1966 | 1956 | 1994 | 1659 | 1746 | 1848 | 1861.57 | 1972 | 1751 | 0.888 | -0.171 | 0.01008566 | 0.04160552 | 9.00E-04 | 0.0037 | 0.0026 | 0.0026 | TRUE | 0.0287 |
| Zfyve27 | 1963 | 1655 | 1603 | 1687 | 1572 | 1713 | 1525 | 1760 | 1730 | 1679 | 1773 | 1691 | 1693.04 | 1672 | 1714 | 1.027 | 0.038 | 0.61310106 | 0.75517524 | 0.003 | 0.0038 | 0.0035 | 0.0035 | TRUE | 0.0739 |
| Zfyve28 | 1066 | 854 | 762 | 997 | 886 | 949 | 828 | 908 | 822 | 992 | 999 | 937 | 914.54 | 853 | 976 | 1.145 | 0.196 | 0.01602428 | 0.05864021 | 0.001 | 0.005 | 0.0037 | 0.0037 | TRUE | 0.0377 |
| Zfyve9 | 1095 | 705 | 705 | 792 | 667 | 810 | 850 | 750 | 761 | 788 | 752 | 800 | 783.57 | 787 | 780 | 0.99 | -0.015 | 0.8651633 | 0.92551298 | 0.0019 | 0.0055 | 0.0042 | 0.0042 | TRUE | 0.062 |
| Zg16 | 1 | 0 | 0 | 1 | 1 | 0 | 1 | 0 | 0 | 1 | 1 | 0 | 0.48 | 0 | 1 | 2.037 | 1.027 | 0.71241723 | NA | 0 | 4.9057 | 3.8521 | 3.8521 | TRUE | 0.2664 |
| Zglp1 | 378 | 281 | 289 | 367 | 368 | 387 | 294 | 299 | 312 | 365 | 415 | 382 | 344.46 | 302 | 387 | 1.286 | 0.362 | 0.0014784 | 0.0095528 | 0 | 0.0093 | 0.0064 | 0.0064 | TRUE | 0.0434 |
| Zgpat | 964 | 749 | 647 | 576 | 522 | 611 | 749 | 797 | 698 | 573 | 589 | 603 | 668.15 | 748 | 588 | 0.787 | -0.346 | 0.00012859 | 0.00150214 | 0.0011 | 0.006 | 0.0044 | 0.0044 | TRUE | 0.0381 |
| Zhx1 | 608 | 431 | 364 | 466 | 440 | 524 | 472 | 458 | 393 | 464 | 496 | 517 | 466.82 | 441 | 492 | 1.114 | 0.156 | 0.15242454 | 0.29913551 | 0.004 | 0.0075 | 0.0064 | 0.0064 | TRUE | 0.1036 |
| Zhx2 | 45 | 31 | 34 | 41 | 29 | 48 | 35 | 33 | 37 | 41 | 33 | 47 | 37.59 | 35 | 40 | 1.159 | 0.213 | 0.5007958 | 0.66510634 | 0 | 0.0655 | 0.0457 | 0.0457 | TRUE | 0.1887 |
| Zhx3 | 97 | 90 | 55 | 79 | 77 | 81 | 75 | 96 | 59 | 79 | 87 | 80 | 79.31 | 77 | 82 | 1.065 | 0.09 | 0.70166278 | 0.81928737 | 0.0144 | 0.0323 | 0.0275 | 0.0275 | TRUE | 0.2552 |
| Zic1 | 6 | 11 | 5 | 6 | 4 | 3 | 5 | 12 | 5 | 6 | 5 | 3 | 5.87 | 7 | 5 | 0.625 | -0.679 | 0.40766606 | NA | 0 | 0.4064 | 0.3019 | 0.3019 | TRUE | 0.7471 |

| | | | | | | | | | | | | | | | | | | | | | | | | |
|---|---|---|---|---|---|---|---|---|---|---|---|---|---|---|---|---|---|---|---|---|---|---|---|---|
| Zic2 | 1872 | 1224 | 918 | 599 | 504 | 592 | 1454 | 1302 | 991 | 596 | 569 | 584 | 915.93 | 1249 | 583 | 0.466 | -1.1 | 1.78E-19 | 5.08E-17 | 0.0188 | 0.005 | 0.0094 | 0.0094 | TRUE | 0.285 |
| Zic3 | 10329 | 6929 | 5995 | 8465 | 6651 | 7847 | 8022 | 7370 | 6469 | 8426 | 7503 | 7746 | 7589.37 | 7287 | 7892 | 1.083 | 0.115 | 0.15054794 | 0.29626625 | 0.0074 | 0.0027 | 0.0045 | 0.0045 | TRUE | 0.1176 |
| Zic4 | 0 | 0 | 0 | 0 | 0 | 0 | 0 | 0 | 0 | 0 | 0 | 0 | 0 | 0 | 0 | NA | NA | NA | NA | NA | NA | NA | NA | NA | NA |
| Zic5 | 773 | 540 | 440 | 461 | 403 | 440 | 600 | 574 | 475 | 459 | 455 | 434 | 499.56 | 550 | 449 | 0.815 | -0.295 | 0.008141 | 0.03554042 | 0.0063 | 0.0072 | 0.0069 | 0.0069 | TRUE | 0.1634 |
| Zik1 | 1451 | 1093 | 1083 | 1142 | 957 | 1155 | 1127 | 1163 | 1169 | 1137 | 1080 | 1140 | 1135.77 | 1153 | 1119 | 0.972 | -0.042 | 0.56047075 | 0.71505922 | 0 | 0.0045 | 0.0028 | 0.0028 | TRUE | 0.011 |
| Zim1 | 94 | 73 | 68 | 88 | 79 | 67 | 73 | 78 | 73 | 88 | 89 | 66 | 77.81 | 75 | 81 | 1.083 | 0.115 | 0.61151493 | 0.75404219 | 0 | 0.0329 | 0.0242 | 0.0242 | TRUE | 0.2448 |
| Zim3 | 98 | 79 | 58 | 51 | 44 | 52 | 76 | 84 | 63 | 51 | 50 | 51 | 62.41 | 74 | 51 | 0.681 | -0.555 | 0.02668592 | 0.08525298 | 0 | 0.0404 | 0.0284 | 0.0284 | TRUE | 0.1599 |
| Zkscan1 | 1341 | 983 | 909 | 1158 | 991 | 992 | 1041 | 1046 | 981 | 1153 | 1118 | 979 | 1052.96 | 1023 | 1083 | 1.059 | 0.082 | 0.33275056 | 0.50702065 | 0.0034 | 0.0047 | 0.0042 | 0.0042 | TRUE | 0.0856 |
| Zkscan14 | 262 | 196 | 167 | 190 | 178 | 207 | 203 | 208 | 180 | 189 | 201 | 204 | 197.74 | 197 | 198 | 1.002 | 0.003 | 0.98241287 | 0.99089635 | 0 | 0.0144 | 0.0099 | 0.0099 | TRUE | 0.0595 |
| Zkscan16 | 213 | 191 | 158 | 208 | 183 | 223 | 165 | 203 | 170 | 207 | 206 | 220 | 195.45 | 179 | 211 | 1.179 | 0.237 | 0.11091919 | 0.23852327 | 0 | 0.0146 | 0.0108 | 0.0108 | TRUE | 0.1365 |
| Zkscan17 | 6218 | 4944 | 5906 | 5975 | 5554 | 6340 | 4829 | 5259 | 6373 | 5948 | 6265 | 6259 | 5822.03 | 5487 | 6157 | 1.123 | 0.167 | 0.05897424 | 0.15151051 | 0.0107 | 0.0028 | 0.0055 | 0.0055 | TRUE | 0.2428 |
| Zkscan2 | 402 | 295 | 278 | 347 | 342 | 305 | 312 | 314 | 300 | 345 | 386 | 301 | 326.38 | 309 | 344 | 1.113 | 0.154 | 0.21833488 | 0.38135897 | 0.0049 | 0.0097 | 0.0082 | 0.0082 | TRUE | 0.1374 |
| Zkscan3 | 895 | 642 | 645 | 670 | 599 | 662 | 695 | 683 | 696 | 667 | 676 | 653 | 678.35 | 691 | 665 | 0.962 | -0.056 | 0.50494514 | 0.66844317 | 0 | 0.0059 | 0.0035 | 0.0035 | TRUE | 0.0029 |
| Zkscan4 | 37 | 35 | 28 | 33 | 36 | 24 | 29 | 37 | 30 | 33 | 41 | 24 | 32.22 | 32 | 33 | 1.01 | 0.014 | 0.96884266 | 0.98315003 | 0 | 0.076 | 0.0588 | 0.0588 | TRUE | 0.2316 |
| Zkscan5 | 1841 | 1412 | 1361 | 1671 | 1384 | 1670 | 1430 | 1502 | 1469 | 1663 | 1561 | 1649 | 1545.57 | 1467 | 1624 | 1.108 | 0.149 | 0.02610501 | 0.08390352 | 0 | 0.004 | 0.0026 | 0.0026 | TRUE | 0.0136 |
| Zkscan6 | 787 | 569 | 563 | 569 | 525 | 575 | 611 | 605 | 607 | 566 | 592 | 568 | 591.7 | 608 | 575 | 0.946 | -0.08 | 0.36323206 | 0.53786674 | 0 | 0.0064 | 0.0039 | 0.0039 | TRUE | 0.0074 |
| Zkscan7 | 166 | 126 | 122 | 113 | 91 | 101 | 129 | 134 | 132 | 112 | 103 | 100 | 118.24 | 132 | 105 | 0.799 | -0.324 | 0.07049114 | 0.17240415 | 0 | 0.0225 | 0.0146 | 0.0146 | TRUE | 0.0399 |
| Zkscan8 | 1521 | 1142 | 1313 | 1346 | 1246 | 1382 | 1181 | 1215 | 1417 | 1340 | 1406 | 1364 | 1320.4 | 1271 | 1370 | 1.079 | 0.11 | 0.18972219 | 0.34752008 | 0.0042 | 0.0042 | 0.0043 | 0.0043 | TRUE | 0.1195 |
| Zmat1 | 67 | 52 | 46 | 67 | 52 | 58 | 52 | 55 | 50 | 67 | 59 | 57 | 56.6 | 52 | 61 | 1.164 | 0.219 | 0.38985533 | 0.56375983 | 0 | 0.0443 | 0.0291 | 0.0291 | TRUE | 0.062 |
| Zmat2 | 5846 | 3954 | 3635 | 3869 | 3478 | 3958 | 4540 | 4206 | 3922 | 3851 | 3924 | 3907 | 4058.37 | 4223 | 3894 | 0.922 | -0.118 | 0.07087899 | 0.17307769 | 0.0024 | 0.003 | 0.0028 | 0.0028 | TRUE | 0.0535 |
| Zmat3 | 8461 | 6064 | 6394 | 7666 | 6378 | 7458 | 6571 | 6450 | 6899 | 7631 | 7195 | 7362 | 7018.09 | 6640 | 7396 | 1.114 | 0.156 | 0.00379202 | 0.01979508 | 8.00E-04 | 0.0028 | 0.0019 | 0.0019 | TRUE | 0.014 |
| Zmat4 | 228 | 210 | 232 | 218 | 219 | 281 | 177 | 223 | 250 | 217 | 247 | 277 | 232.04 | 217 | 247 | 1.145 | 0.195 | 0.22777639 | 0.39299951 | 0.0188 | 0.0126 | 0.0145 | 0.0145 | TRUE | 0.3071 |
| Zmat5 | 675 | 457 | 449 | 455 | 441 | 474 | 524 | 486 | 484 | 453 | 497 | 468 | 485.52 | 498 | 473 | 0.947 | -0.079 | 0.42679316 | 0.59899397 | 0 | 0.0073 | 0.005 | 0.005 | TRUE | 0.0259 |
| Zmiz1 | 2095 | 1264 | 905 | 1076 | 1025 | 1110 | 1627 | 1345 | 976 | 1071 | 1156 | 1096 | 1211.86 | 1316 | 1108 | 0.84 | -0.251 | 0.06609484 | 0.16438029 | 0.0303 | 0.0044 | 0.0126 | 0.0126 | TRUE | 0.2241 |
| Zmiz2 | 8852 | 6539 | 6427 | 6584 | 6080 | 6818 | 6875 | 6955 | 6935 | 6554 | 6859 | 6730 | 6818.02 | 6922 | 6714 | 0.97 | -0.044 | 0.36667045 | 0.54146304 | 1.00E-04 | 0.0028 | 0.0016 | 0.0016 | TRUE | 0.0053 |
| Zmpste24 | 3345 | 2354 | 2216 | 2471 | 2209 | 2402 | 2598 | 2504 | 2391 | 2460 | 2492 | 2371 | 2469.27 | 2498 | 2441 | 0.977 | -0.034 | 0.58078378 | 0.73144265 | 8.00E-04 | 0.0034 | 0.0024 | 0.0024 | TRUE | 0.0168 |
| Zmym1 | 5712 | 3993 | 3305 | 3761 | 3240 | 3677 | 4436 | 4247 | 3566 | 3744 | 3655 | 3630 | 3879.69 | 4083 | 3676 | 0.9 | -0.152 | 0.05490767 | 0.14402558 | 0.006 | 0.003 | 0.0043 | 0.0043 | TRUE | 0.1487 |
| Zmym2 | 11655 | 8450 | 8292 | 9130 | 7907 | 9083 | 9052 | 8988 | 8947 | 9088 | 8920 | 8966 | 8993.55 | 8996 | 8991 | 1 | -0.001 | 0.9873898 | 0.99373352 | 0 | 0.0027 | 0.0014 | 0.0014 | TRUE | 0.0011 |
| Zmym3 | 5689 | 4074 | 3394 | 3543 | 3226 | 3861 | 4418 | 4333 | 3662 | 3527 | 3639 | 3811 | 3898.55 | 4138 | 3659 | 0.884 | -0.178 | 0.02100718 | 0.07172853 | 0.0054 | 0.003 | 0.004 | 0.004 | TRUE | 0.1225 |
| Zmym4 | 6296 | 4240 | 3989 | 4216 | 3763 | 4430 | 4890 | 4510 | 4304 | 4197 | 4245 | 4373 | 4419.79 | 4568 | 4272 | 0.935 | -0.098 | 0.12130418 | 0.25399952 | 0.002 | 0.003 | 0.0026 | 0.0026 | TRUE | 0.047 |
| Zmym5 | 2948 | 2032 | 1728 | 1928 | 1520 | 1755 | 2290 | 2161 | 1864 | 1919 | 1715 | 1732 | 1946.97 | 2105 | 1789 | 0.849 | -0.236 | 0.00616152 | 0.02877995 | 0.0066 | 0.0036 | 0.0048 | 0.0048 | TRUE | 0.1202 |
| Zmym6 | 1277 | 1031 | 1174 | 1442 | 1138 | 1265 | 992 | 1097 | 1267 | 1435 | 1284 | 1249 | 1220.51 | 1119 | 1323 | 1.185 | 0.244 | 0.01265713 | 0.0492331 | 0.0093 | 0.0044 | 0.0061 | 0.0061 | TRUE | 0.1607 |
| Zmynd10 | 15 | 16 | 51 | 64 | 35 | 64 | 12 | 17 | 55 | 64 | 39 | 63 | 41.68 | 28 | 55 | 2.019 | 1.013 | 0.02884676 | 0.08998402 | 0.3163 | 0.0593 | 0.128 | 0.128 | TRUE | 4.511 |
| Zmynd11 | 1967 | 1312 | 1268 | 1280 | 1147 | 1222 | 1528 | 1396 | 1368 | 1274 | 1294 | 1206 | 1344.29 | 1431 | 1258 | 0.878 | -0.187 | 0.01265876 | 0.0492331 | 0.0018 | 0.0042 | 0.0033 | 0.0033 | TRUE | 0.0441 |
| Zmynd12 | 8 | 3 | 1 | 9 | 4 | 2 | 6 | 3 | 1 | 9 | 5 | 2 | 4.32 | 3 | 5 | 1.432 | 0.518 | 0.60319649 | NA | 0.3116 | 0.5509 | 0.4814 | 0.4814 | TRUE | 0.334 |
| Zmynd15 | 65 | 52 | 56 | 66 | 65 | 77 | 50 | 55 | 60 | 66 | 73 | 76 | 63.54 | 55 | 72 | 1.298 | 0.377 | 0.12166314 | 0.2545274 | 0 | 0.0397 | 0.0267 | 0.0267 | TRUE | 0.0597 |
| Zmynd19 | 4417 | 3322 | 3271 | 3184 | 2927 | 3427 | 3430 | 3534 | 3529 | 3169 | 3302 | 3383 | 3391.29 | 3498 | 3285 | 0.939 | -0.09 | 0.10776957 | 0.23352795 | 4.00E-04 | 0.0031 | 0.002 | 0.002 | TRUE | 0.0116 |
| Zmynd8 | 6860 | 5423 | 5979 | 7006 | 6151 | 7613 | 5328 | 5768 | 6451 | 6974 | 6939 | 7515 | 6495.91 | 5849 | 7143 | 1.222 | 0.289 | 9.65E-05 | 0.00117512 | 0.0055 | 0.0028 | 0.0038 | 0.0038 | TRUE | 0.0988 |
| Znf41-ps | 114 | 117 | 54 | 77 | 69 | 64 | 89 | 124 | 58 | 77 | 78 | 63 | 81.49 | 90 | 73 | 0.802 | -0.319 | 0.24050494 | 0.40791944 | 0.0636 | 0.0315 | 0.0406 | 0.0406 | TRUE | 0.1817 |
| Znf512b | 3722 | 2645 | 2272 | 2061 | 1930 | 2227 | 2891 | 2813 | 2451 | 2052 | 2177 | 2198 | 2430.47 | 2718 | 2142 | 0.788 | -0.345 | 5.04E-06 | 0.0001065 | 0.0038 | 0.0034 | 0.0037 | 0.0037 | TRUE | 0.0888 |
| Znfx1 | 1943 | 1539 | 1664 | 1771 | 1773 | 2189 | 1509 | 1637 | 1795 | 1763 | 2000 | 2161 | 1810.89 | 1647 | 1975 | 1.2 | 0.263 | 0.00376654 | 0.01969097 | 0.0081 | 0.0037 | 0.0054 | 0.0054 | TRUE | 0.1072 |
| Znhit1 | 1195 | 889 | 751 | 797 | 778 | 882 | 928 | 946 | 810 | 793 | 878 | 871 | 870.95 | 895 | 847 | 0.946 | -0.08 | 0.3735499 | 0.54816722 | 0.0038 | 0.0051 | 0.0047 | 0.0047 | TRUE | 0.0788 |
| Znhit2 | 613 | 478 | 359 | 383 | 329 | 376 | 476 | 508 | 387 | 381 | 371 | 371 | 415.91 | 457 | 374 | 0.818 | -0.289 | 0.01602523 | 0.05864021 | 0.0076 | 0.0081 | 0.008 | 0.008 | TRUE | 0.1999 |
| Znhit3 | 2646 | 1601 | 1318 | 1566 | 1322 | 1402 | 2055 | 1703 | 1422 | 1559 | 1491 | 1384 | 1602.37 | 1727 | 1478 | 0.855 | -0.226 | 0.04363359 | 0.12199821 | 0.0177 | 0.0039 | 0.0084 | 0.0084 | TRUE | 0.2191 |
| Znhit6 | 2842 | 2186 | 1994 | 1915 | 1799 | 2114 | 2207 | 2325 | 2152 | 1906 | 2029 | 2087 | 2117.75 | 2228 | 2007 | 0.901 | -0.15 | 0.02371657 | 0.07828658 | 0.0014 | 0.0035 | 0.0027 | 0.0027 | TRUE | 0.0238 |
| Znrd1 | 1577 | 1085 | 979 | 1193 | 969 | 1110 | 1225 | 1154 | 1056 | 1188 | 1093 | 1096 | 1135.27 | 1145 | 1126 | 0.982 | -0.026 | 0.74780281 | 0.85030594 | 0.0031 | 0.0045 | 0.004 | 0.004 | TRUE | 0.0546 |
| Znrd1as | 1080 | 727 | 829 | 885 | 819 | 857 | 839 | 773 | 894 | 881 | 924 | 846 | 859.57 | 835 | 884 | 1.057 | 0.08 | 0.3541783 | 0.52858625 | 0.0023 | 0.0052 | 0.0042 | 0.0042 | TRUE | 0.0484 |
| Znrf1 | 2071 | 1661 | 1474 | 1672 | 1481 | 1823 | 1608 | 1767 | 1590 | 1664 | 1671 | 1800 | 1683.38 | 1655 | 1712 | 1.035 | 0.049 | 0.49707275 | 0.66151404 | 0.0018 | 0.0038 | 0.0032 | 0.0032 | TRUE | 0.0412 |
| Znrf2 | 3215 | 2300 | 2293 | 2227 | 1865 | 2234 | 2497 | 2446 | 2474 | 2217 | 2104 | 2205 | 2323.92 | 2472 | 2175 | 0.88 | -0.185 | 0.00172598 | 0.01078679 | 0 | 0.0034 | 0.0021 | 0.0021 | TRUE | 0.0098 |
| Znrf3 | 1048 | 726 | 697 | 719 | 627 | 769 | 814 | 772 | 752 | 716 | 707 | 759 | 753.39 | 779 | 727 | 0.932 | -0.101 | 0.23170322 | 0.3976626 | 0 | 0.0056 | 0.0038 | 0.0038 | TRUE | 0.0188 |
| Znrf4 | 0 | 0 | 0 | 0 | 0 | 0 | 0 | 0 | 0 | 0 | 0 | 0 | 0 | 0 | 0 | NA | NA | NA | NA | NA | NA | NA | NA | NA | NA |
| Zp1 | 1 | 1 | 1 | 1 | 0 | 1 | 1 | 1 | 1 | 1 | 0 | 1 | 0.82 | 1 | 1 | 0.698 | -0.518 | 0.79971927 | NA | 0 | 2.9034 | 1.738 | 1.738 | TRUE | 0.2018 |
| Zp2 | 7 | 5 | 9 | 2 | 2 | 5 | 5 | 5 | 10 | 2 | 2 | 5 | 4.94 | 7 | 3 | 0.455 | -1.137 | 0.2050023 | NA | 0 | 0.4821 | 0.3411 | 0.3411 | TRUE | 0.3956 |
| Zp3 | 154 | 152 | 115 | 130 | 119 | 147 | 120 | 162 | 124 | 129 | 134 | 145 | 135.69 | 135 | 136 | 1.012 | 0.017 | 0.9245794 | 0.95836863 | 0.0088 | 0.0199 | 0.0167 | 0.0167 | TRUE | 0.2972 |
| Zp3r | 7 | 2 | 2 | 2 | 3 | 5 | 5 | 2 | 2 | 2 | 3 | 5 | 3.34 | 3 | 3 | 1.034 | 0.048 | 0.9632163 | NA | 0 | 0.7123 | 0.497 | 0.497 | TRUE | 0.6088 |
| Zp4-ps | 0 | 0 | 0 | 0 | 0 | 0 | 0 | 0 | 0 | 0 | 0 | 0 | 0 | 0 | 0 | NA | NA | NA | NA | NA | NA | NA | NA | NA | NA |
| Zpbp | 41 | 26 | 26 | 42 | 25 | 24 | 32 | 28 | 28 | 42 | 28 | 24 | 30.21 | 29 | 31 | 1.068 | 0.095 | 0.794888 | 0.87990506 | 0 | 0.0809 | 0.063 | 0.063 | TRUE | 0.6089 |
| Zpbp2 | 14 | 15 | 6 | 16 | 16 | 9 | 11 | 16 | 6 | 16 | 18 | 9 | 12.69 | 11 | 14 | 1.282 | 0.358 | 0.53087292 | NA | 0.0626 | 0.1891 | 0.156 | 0.156 | TRUE | 0.2551 |
| Zpld1 | 0 | 0 | 0 | 0 | 0 | 0 | 0 | 0 | 0 | 0 | 0 | 0 | 0 | 0 | 0 | NA | NA | NA | NA | NA | NA | NA | NA | NA | NA |
| Zranb1 | 1670 | 1222 | 1332 | 1377 | 1269 | 1390 | 1297 | 1300 | 1437 | 1371 | 1432 | 1372 | 1368.07 | 1345 | 1392 | 1.035 | 0.05 | 0.49352186 | 0.65875121 | 0.0013 | 0.0042 | 0.0031 | 0.0031 | TRUE | 0.0427 |
| Zranb2 | 8847 | 6900 | 6946 | 7186 | 6183 | 7220 | 6871 | 7339 | 7495 | 7153 | 6975 | 7127 | 7160.08 | 7235 | 7085 | 0.98 | -0.03 | 0.58454199 | 0.73423488 | 9.00E-04 | 0.0028 | 0.002 | 0.002 | TRUE | 0.024 |
| Zranb3 | 2680 | 1974 | 2027 | 2214 | 2000 | 2349 | 2081 | 2100 | 2187 | 2204 | 2256 | 2319 | 2191.19 | 2123 | 2260 | 1.065 | 0.091 | 0.13561067 | 0.27501181 | 0 | 0.0035 | 0.0022 | 0.0022 | TRUE | 0.0084 |
| Zrsr1 | 1490 | 1031 | 857 | 962 | 813 | 898 | 1157 | 1097 | 925 | 958 | 917 | 886 | 989.96 | 1060 | 920 | 0.867 | -0.205 | 0.02995157 | 0.09254167 | 0.0066 | 0.0048 | 0.0054 | 0.0054 | TRUE | 0.1462 |
| Zrsr2 | 3139 | 2418 | 2436 | 2519 | 2086 | 2445 | 2438 | 2572 | 2628 | 2507 | 2353 | 2414 | 2485.43 | 2546 | 2425 | 0.953 | -0.069 | 0.26512666 | 0.43556958 | 9.00E-04 | 0.0034 | 0.0024 | 0.0024 | TRUE | 0.0173 |
| Zscan10 | 3151 | 2596 | 2512 | 2286 | 2214 | 2739 | 2447 | 2761 | 2710 | 2275 | 2498 | 2704 | 2565.99 | 2639 | 2492 | 0.945 | -0.082 | 0.29963444 | 0.47186812 | 0.0051 | 0.0034 | 0.0041 | 0.0041 | TRUE | 0.0708 |
| Zscan12 | 930 | 649 | 796 | 853 | 712 | 832 | 722 | 690 | 859 | 849 | 803 | 821 | 790.85 | 757 | 824 | 1.09 | 0.125 | 0.19862436 | 0.35890492 | 0.0055 | 0.0054 | 0.0055 | 0.0055 | TRUE | 0.1593 |
| Zscan18 | 1009 | 883 | 774 | 782 | 710 | 760 | 784 | 939 | 835 | 778 | 801 | 750 | 814.6 | 853 | 776 | 0.912 | -0.133 | 0.14565269 | 0.28966556 | 0.0038 | 0.0053 | 0.0048 | 0.0048 | TRUE | 0.0934 |
| Zscan2 | 229 | 182 | 197 | 185 | 195 | 197 | 178 | 194 | 213 | 184 | 220 | 194 | 197.1 | 195 | 199 | 1.026 | 0.038 | 0.80171495 | 0.88499473 | 0.0032 | 0.0145 | 0.0112 | 0.0112 | TRUE | 0.0841 |
| Zscan20 | 933 | 693 | 613 | 576 | 547 | 605 | 725 | 737 | 661 | 573 | 617 | 597 | 651.8 | 708 | 596 | 0.841 | -0.25 | 0.00574444 | 0.02729565 | 0 | 0.0061 | 0.0043 | 0.0043 | TRUE | 0.0373 |

| | | | | | | | | | | | | | | | | | | | | | | | | | |
|---|---|---|---|---|---|---|---|---|---|---|---|---|---|---|---|---|---|---|---|---|---|---|---|---|---|
| Zscan21 | 1493 | 1007 | 948 | 991 | 933 | 1036 | 1160 | 1071 | 1023 | 986 | 1053 | 1023 | 1052.54 | 1085 | 1021 | 0.939 | -0.09 | 0.25839293 | 0.42834726 | 0.0018 | 0.0047 | 0.0036 | 0.0036 | TRUE | 0.0457 |
| Zscan22 | 412 | 248 | 184 | 212 | 173 | 148 | 320 | 264 | 199 | 211 | 195 | 146 | 222.43 | 261 | 184 | 0.703 | -0.508 | 0.00761475 | 0.03374809 | 0.0417 | 0.0131 | 0.0215 | 0.0215 | TRUE | 0.1629 |
| Zscan25 | 611 | 462 | 449 | 458 | 395 | 416 | 475 | 491 | 484 | 456 | 446 | 411 | 460.43 | 483 | 438 | 0.905 | -0.144 | 0.14963824 | 0.29512618 | 0 | 0.0076 | 0.005 | 0.005 | TRUE | 0.0338 |
| Zscan26 | 2220 | 1724 | 1780 | 1596 | 1526 | 1684 | 1724 | 1834 | 1921 | 1589 | 1721 | 1662 | 1741.85 | 1826 | 1657 | 0.908 | -0.139 | 0.04922138 | 0.13324266 | 0.0018 | 0.0038 | 0.003 | 0.003 | TRUE | 0.0299 |
| Zscan29 | 630 | 448 | 357 | 416 | 341 | 349 | 489 | 477 | 385 | 414 | 385 | 345 | 415.72 | 450 | 381 | 0.845 | -0.244 | 0.05067241 | 0.13619043 | 0.0102 | 0.0081 | 0.0088 | 0.0088 | TRUE | 0.1811 |
| Zscan4a | 1962 | 1242 | 442 | 571 | 551 | 457 | 1524 | 1321 | 477 | 568 | 622 | 451 | 827.15 | 1107 | 547 | 0.494 | -1.018 | 0.04126835 | 0.11689429 | 0.1778 | 0.0053 | 0.0459 | 0.1778 | TRUE | 0.9931 |
| Zscan4b | 413 | 325 | 101 | 133 | 135 | 109 | 321 | 346 | 109 | 132 | 152 | 108 | 194.62 | 259 | 131 | 0.505 | -0.984 | 0.05310637 | 0.14079936 | 0.1808 | 0.0146 | 0.0587 | 0.1808 | TRUE | 2.8297 |
| Zscan4c | 1713 | 1160 | 506 | 534 | 567 | 477 | 1330 | 1234 | 546 | 532 | 640 | 471 | 792.05 | 1037 | 548 | 0.528 | -0.922 | 0.02260881 | 0.07546655 | 0.1165 | 0.0054 | 0.0333 | 0.1165 | TRUE | 2.0452 |
| Zscan4d | 1907 | 1346 | 454 | 550 | 638 | 475 | 1481 | 1432 | 490 | 547 | 720 | 469 | 856.46 | 1134 | 579 | 0.51 | -0.971 | 0.05571743 | 0.14564178 | 0.1844 | 0.0052 | 0.047 | 0.1844 | TRUE | 2.9532 |
| Zscan4e | 891 | 594 | 251 | 282 | 299 | 276 | 692 | 632 | 271 | 281 | 337 | 272 | 414.19 | 532 | 297 | 0.558 | -0.842 | 0.04139833 | 0.11714636 | 0.12 | 0.0081 | 0.0375 | 0.12 | TRUE | 2.1096 |
| Zscan4f | 1061 | 831 | 314 | 390 | 376 | 337 | 824 | 884 | 339 | 388 | 424 | 333 | 531.97 | 682 | 382 | 0.559 | -0.838 | 0.04830702 | 0.1314396 | 0.1279 | 0.0069 | 0.0377 | 0.1279 | TRUE | 2.2811 |
| Zscan5b | 197 | 157 | 154 | 114 | 102 | 106 | 153 | 167 | 166 | 113 | 115 | 105 | 136.56 | 162 | 111 | 0.686 | -0.543 | 0.0013288 | 0.00884195 | 0 | 0.0198 | 0.013 | 0.013 | TRUE | 0.028 |
| Zswim1 | 3829 | 3097 | 3863 | 4126 | 3796 | 4776 | 2974 | 3294 | 4168 | 4107 | 4282 | 4715 | 3923.36 | 3479 | 4368 | 1.256 | 0.329 | 0.00160163 | 0.01013376 | 0.018 | 0.003 | 0.0076 | 0.0076 | TRUE | 0.3649 |
| Zswim2 | 5 | 2 | 1 | 0 | 0 | 1 | 4 | 2 | 1 | 0 | 0 | 1 | 1.35 | 2 | 0 | 0.154 | -2.695 | 0.15032806 | NA | 0 | 1.7631 | 1.3278 | 1.3278 | TRUE | 0.3283 |
| Zswim3 | 1080 | 811 | 735 | 785 | 687 | 863 | 839 | 863 | 793 | 781 | 775 | 852 | 817.13 | 832 | 803 | 0.966 | -0.051 | 0.54824675 | 0.70500143 | 0.001 | 0.0053 | 0.0039 | 0.0039 | TRUE | 0.0344 |
| Zswim4 | 3077 | 2283 | 1960 | 2251 | 2052 | 2383 | 2390 | 2428 | 2115 | 2241 | 2315 | 2352 | 2306.82 | 2311 | 2303 | 0.996 | -0.006 | 0.93319551 | 0.9627836 | 0.0026 | 0.0035 | 0.0032 | 0.0032 | TRUE | 0.0657 |
| Zswim5 | 1115 | 883 | 768 | 825 | 756 | 859 | 866 | 939 | 829 | 821 | 853 | 848 | 859.32 | 878 | 841 | 0.958 | -0.062 | 0.45050643 | 0.62051541 | 0.001 | 0.0052 | 0.0038 | 0.0038 | TRUE | 0.0428 |
| Zswim6 | 1494 | 1083 | 1085 | 1139 | 1053 | 1120 | 1160 | 1152 | 1171 | 1134 | 1188 | 1106 | 1151.71 | 1161 | 1143 | 0.984 | -0.024 | 0.73824236 | 0.84400081 | 0 | 0.0045 | 0.0028 | 0.0028 | TRUE | 0.0142 |
| Zswim7 | 1686 | 1187 | 1336 | 1396 | 1235 | 1314 | 1309 | 1263 | 1442 | 1390 | 1393 | 1297 | 1348.91 | 1338 | 1360 | 1.017 | 0.024 | 0.75579242 | 0.85511255 | 0.0024 | 0.0042 | 0.0035 | 0.0035 | TRUE | 0.0538 |
| Zswim8 | 1471 | 1184 | 1090 | 1005 | 896 | 1075 | 1142 | 1259 | 1176 | 1000 | 1011 | 1061 | 1108.39 | 1192 | 1024 | 0.86 | -0.218 | 0.00415018 | 0.02124586 | 0.001 | 0.0046 | 0.0033 | 0.0033 | TRUE | 0.0285 |
| Zufsp | 744 | 466 | 441 | 452 | 410 | 430 | 578 | 496 | 476 | 450 | 463 | 424 | 481.04 | 517 | 446 | 0.86 | -0.217 | 0.0466674 | 0.12812468 | 0.0048 | 0.0074 | 0.0065 | 0.0065 | TRUE | 0.1328 |
| Zw10 | 3393 | 2385 | 2086 | 2179 | 1955 | 2169 | 2635 | 2537 | 2251 | 2169 | 2205 | 2141 | 2323.07 | 2474 | 2172 | 0.877 | -0.189 | 0.00859791 | 0.03698427 | 0.0027 | 0.0034 | 0.0033 | 0.0033 | TRUE | 0.075 |
| Zwilch | 7737 | 5758 | 5880 | 6232 | 5413 | 6079 | 6009 | 6125 | 6344 | 6203 | 6106 | 6001 | 6131.47 | 6159 | 6103 | 0.991 | -0.013 | 0.80144816 | 0.88476856 | 4.00E-04 | 0.0028 | 0.0017 | 0.0017 | TRUE | 0.0084 |
| Zwint | 6373 | 4468 | 3882 | 4339 | 3732 | 4090 | 4950 | 4753 | 4189 | 4319 | 4210 | 4037 | 4409.57 | 4631 | 4189 | 0.904 | -0.145 | 0.04077328 | 0.11608368 | 0.0039 | 0.003 | 0.0034 | 0.0034 | TRUE | 0.0844 |
| Zxda | 45 | 54 | 45 | 53 | 29 | 42 | 35 | 57 | 49 | 53 | 33 | 41 | 44.65 | 47 | 42 | 0.91 | -0.136 | 0.67035808 | 0.79719012 | 0.0378 | 0.0555 | 0.0505 | 0.0505 | TRUE | 0.3534 |
| Zxdb | 102 | 78 | 103 | 100 | 96 | 86 | 79 | 83 | 111 | 100 | 108 | 85 | 94.34 | 91 | 98 | 1.074 | 0.103 | 0.63682897 | 0.77307597 | 0.0142 | 0.0275 | 0.0238 | 0.0238 | TRUE | 0.3421 |
| Zxdc | 1683 | 1265 | 1308 | 1397 | 1204 | 1334 | 1307 | 1346 | 1411 | 1391 | 1358 | 1317 | 1354.95 | 1355 | 1355 | 1.001 | 0.002 | 0.98124397 | 0.99018074 | 5.00E-04 | 0.0042 | 0.0028 | 0.0028 | TRUE | 0.016 |
| Zyg11a | 2764 | 2044 | 2060 | 2692 | 2423 | 2553 | 2147 | 2174 | 2223 | 2680 | 2733 | 2520 | 2412.8 | 2181 | 2644 | 1.212 | 0.278 | 6.26E-06 | 0.00012799 | 7.00E-04 | 0.0034 | 0.0023 | 0.0023 | TRUE | 0.0206 |
| Zyg11b | 4688 | 3415 | 3529 | 4324 | 3585 | 4250 | 3641 | 3632 | 3808 | 4304 | 4044 | 4195 | 3937.5 | 3694 | 4181 | 1.132 | 0.179 | 0.00131915 | 0.00879791 | 5.00E-04 | 0.003 | 0.002 | 0.002 | TRUE | 0.0099 |
| Zyx | 1700 | 1163 | 1007 | 1068 | 989 | 1194 | 1320 | 1237 | 1087 | 1063 | 1116 | 1179 | 1166.9 | 1215 | 1119 | 0.92 | -0.12 | 0.17552844 | 0.32906978 | 0.0054 | 0.0045 | 0.0048 | 0.0048 | TRUE | 0.1011 |
| Zzef1 | 3713 | 2413 | 2013 | 2233 | 2056 | 2310 | 2884 | 2567 | 2172 | 2223 | 2319 | 2280 | 2407.48 | 2541 | 2274 | 0.894 | -0.161 | 0.07832575 | 0.18624355 | 0.0095 | 0.0034 | 0.0056 | 0.0056 | TRUE | 0.1948 |
| Zzz3 | 5525 | 3955 | 3736 | 3777 | 3353 | 3959 | 4291 | 4207 | 4031 | 3760 | 3783 | 3908 | 3996.55 | 4176 | 3817 | 0.914 | -0.13 | 0.01815023 | 0.06426088 | 5.00E-04 | 0.003 | 0.0019 | 0.0019 | TRUE | 0.0112 |

**Table S4 RNF12 WT vs Control RNA-Seq analysis**

| Id | control1 | control2 | control3 | RNF12WT1 | RNF12WT2 | RNF12WT3 | norm.control1 | norm.control2 | norm.control3 | norm.RNF12WT1 | norm.RNF12WT2 | norm.RNF12WT3 | baseMean | Control | Rnf12WT | FoldChange | log2FoldChange | pvalue | padj | dispGeneEst | dispFit | dispMAP | dispersion | betaConv | maxCooks |
|---|---|---|---|---|---|---|---|---|---|---|---|---|---|---|---|---|---|---|---|---|---|---|---|---|---|
| 0610005C13 | 29 | 5 | 3 | 26 | 18 | 2 | 31 | 5 | 3 | 20 | 19 | 2 | 13.47 | 13 | 14 | 1.091 | 0.126 | 0.89866616 | 0.95078955 | 1.0212 | 0.2515 | 0.6266 | 0.6266 | TRUE | 5.8557 |
| 0610007N19 | 44 | 66 | 68 | 56 | 47 | 55 | 47 | 66 | 68 | 44 | 51 | 60 | 55.95 | 60 | 52 | 0.85 | -0.234 | 0.41461489 | 0.62619037 | 0.0151 | 0.0658 | 0.0414 | 0.0414 | TRUE | 0.3252 |
| 0610007P14 | 1845 | 2169 | 2161 | 2426 | 1873 | 1923 | 1960 | 2166 | 2170 | 1907 | 2015 | 2105 | 2053.86 | 2099 | 2009 | 0.957 | -0.064 | 0.45270251 | 0.65809138 | 0.0021 | 0.0085 | 0.0047 | 0.0047 | TRUE | 0.0404 |
| 0610008F07 | 0 | 0 | 0 | 0 | 0 | 1 | 0 | 0 | 0 | 0 | 0 | 1 | 0.18 | 0 | 0 | 1.859 | 0.894 | 0.82650734 | NA | 0 | 18.0643 | 10 | 10 | TRUE | 0.5383 |
| 0610009B14 | 30 | 34 | 27 | 23 | 16 | 20 | 32 | 34 | 27 | 18 | 17 | 22 | 25.02 | 31 | 19 | 0.614 | -0.704 | 0.0652989 | 0.19695312 | 0 | 0.1386 | 0.0631 | 0.0631 | TRUE | 0.0815 |
| 0610009B22 | 1473 | 1430 | 1422 | 1812 | 1293 | 1261 | 1565 | 1428 | 1428 | 1424 | 1391 | 1380 | 1436.1 | 1474 | 1398 | 0.95 | -0.075 | 0.35383147 | 0.57222912 | 8.00E-04 | 0.0092 | 0.004 | 0.004 | TRUE | 0.035 |
| 0610009D07 | 2631 | 2682 | 2645 | 3618 | 2593 | 2496 | 2795 | 2679 | 2656 | 2844 | 2790 | 2732 | 2749.22 | 2710 | 2789 | 1.03 | 0.042 | 0.52521948 | 0.7148075 | 2.00E-04 | 0.0081 | 0.0028 | 0.0028 | TRUE | 0.0091 |
| 0610009L18 | 11 | 19 | 16 | 29 | 19 | 11 | 12 | 19 | 16 | 23 | 20 | 12 | 17 | 16 | 18 | 1.191 | 0.252 | 0.60956782 | 0.77891178 | 0 | 0.2007 | 0.116 | 0.116 | TRUE | 0.437 |
| 0610009O20 | 1929 | 2737 | 2903 | 2613 | 2007 | 2432 | 2049 | 2733 | 2915 | 2054 | 2159 | 2662 | 2428.85 | 2566 | 2292 | 0.893 | -0.163 | 0.29616322 | 0.51688461 | 0.0247 | 0.0083 | 0.0172 | 0.0172 | TRUE | 0.3761 |
| 0610010B08 | 0 | 0 | 0 | 0 | 0 | 0 | 0 | 0 | 0 | 0 | 0 | 0 | 0 | 0 | 0 | NA | NA | NA | NA | NA | NA | NA | NA | NA | NA |
| 0610010F05 | 890 | 1040 | 1166 | 1486 | 1046 | 1062 | 945 | 1039 | 1171 | 1168 | 1125 | 1163 | 1101.82 | 1052 | 1152 | 1.095 | 0.131 | 0.21027403 | 0.42076148 | 0.0046 | 0.0099 | 0.007 | 0.007 | TRUE | 0.1176 |
| 0610010K14 | 2046 | 2673 | 2812 | 3359 | 2392 | 2683 | 2173 | 2670 | 2824 | 2641 | 2574 | 2937 | 2636.25 | 2556 | 2717 | 1.063 | 0.088 | 0.46025623 | 0.66439331 | 0.0113 | 0.0082 | 0.0099 | 0.0099 | TRUE | 0.2072 |
| 0610011F06 | 673 | 845 | 844 | 824 | 636 | 694 | 715 | 844 | 847 | 648 | 684 | 760 | 749.66 | 802 | 697 | 0.868 | -0.204 | 0.08306261 | 0.23068521 | 0.0066 | 0.0113 | 0.0086 | 0.0086 | TRUE | 0.1067 |
| 0610012G03 | 558 | 782 | 814 | 955 | 752 | 793 | 593 | 781 | 817 | 751 | 809 | 868 | 769.82 | 730 | 809 | 1.107 | 0.147 | 0.30640733 | 0.52689232 | 0.0156 | 0.0112 | 0.0135 | 0.0135 | TRUE | 0.3199 |
| 0610012H03 | 10 | 18 | 13 | 17 | 10 | 6 | 11 | 18 | 13 | 13 | 11 | 7 | 12.06 | 14 | 10 | 0.745 | -0.424 | 0.45891858 | 0.66355554 | 0 | 0.2801 | 0.1528 | 0.1528 | TRUE | 0.3189 |
| 0610030E20 | 628 | 840 | 923 | 1064 | 848 | 838 | 667 | 839 | 927 | 836 | 912 | 917 | 849.82 | 811 | 888 | 1.095 | 0.131 | 0.33960281 | 0.55980513 | 0.0138 | 0.0108 | 0.0124 | 0.0124 | TRUE | 0.2845 |
| 0610031J06 | 2430 | 2714 | 2758 | 3083 | 2316 | 2458 | 2581 | 2710 | 2769 | 2424 | 2492 | 2691 | 2611.18 | 2687 | 2536 | 0.943 | -0.084 | 0.2808817 | 0.50220609 | 0.0016 | 0.0082 | 0.004 | 0.004 | TRUE | 0.0346 |
| 0610031O16 | 0 | 0 | 0 | 0 | 0 | 0 | 0 | 0 | 0 | 0 | 0 | 0 | 0 | 0 | 0 | NA | NA | NA | NA | NA | NA | NA | NA | NA | NA |
| 0610037L13 | 822 | 914 | 903 | 1143 | 806 | 876 | 873 | 913 | 907 | 899 | 867 | 959 | 902.88 | 898 | 908 | 1.012 | 0.017 | 0.85131833 | 0.92587032 | 0 | 0.0106 | 0.0045 | 0.0045 | TRUE | 0.0274 |
| 0610038B21 | 6 | 14 | 4 | 17 | 12 | 12 | 6 | 14 | 4 | 13 | 13 | 13 | 10.63 | 8 | 13 | 1.615 | 0.691 | 0.28390497 | 0.50542537 | 0.0441 | 0.3168 | 0.1998 | 0.1998 | TRUE | 1.2075 |
| 0610038L08 | 0 | 0 | 0 | 0 | 0 | 0 | 0 | 0 | 0 | 0 | 0 | 0 | 0 | 0 | 0 | NA | NA | NA | NA | NA | NA | NA | NA | NA | NA |
| 0610039K10 | 2 | 1 | 0 | 2 | 1 | 0 | 2 | 1 | 0 | 2 | 1 | 0 | 0.96 | 1 | 1 | 0.879 | -0.185 | 0.92762908 | NA | 0 | 3.4319 | 1.9663 | 1.9663 | TRUE | 0.1856 |
| 0610040B10 | 5 | 4 | 5 | 11 | 6 | 5 | 5 | 4 | 5 | 9 | 6 | 5 | 5.82 | 5 | 7 | 1.452 | 0.539 | 0.48305553 | 0.68129391 | 0 | 0.5732 | 0.2472 | 0.2472 | TRUE | 0.1751 |
| 0610040F04 | 82 | 97 | 152 | 79 | 77 | 79 | 87 | 97 | 153 | 62 | 83 | 86 | 94.67 | 112 | 77 | 0.684 | -0.548 | 0.05399323 | 0.17278444 | 0.0529 | 0.0417 | 0.0473 | 0.0473 | TRUE | 0.9953 |
| 0610040J01 | 101 | 104 | 99 | 169 | 130 | 106 | 107 | 104 | 99 | 133 | 140 | 116 | 116.55 | 103 | 130 | 1.253 | 0.326 | 0.0806617 | 0.2258631 | 0 | 0.0352 | 0.0164 | 0.0164 | TRUE | 0.0801 |
| 0610043K17 | 6 | 4 | 2 | 6 | 6 | 4 | 6 | 4 | 2 | 5 | 6 | 4 | 4.65 | 4 | 5 | 1.258 | 0.331 | 0.70785574 | 0.84081193 | 0 | 0.7147 | 0.3448 | 0.3448 | TRUE | 0.2657 |
| 1100001G20 | 0 | 1 | 0 | 0 | 0 | 1 | 0 | 1 | 0 | 0 | 0 | 1 | 0.35 | 0 | 0 | 0.954 | -0.067 | 0.98567705 | NA | 0 | 9.4494 | 8.1588 | 8.1588 | TRUE | 0.5383 |
| 1110001A16 | 1221 | 1689 | 1556 | 1541 | 1051 | 1222 | 1297 | 1687 | 1562 | 1211 | 1131 | 1338 | 1371 | 1515 | 1227 | 0.809 | -0.305 | 0.01510724 | 0.07112887 | 0.0111 | 0.0093 | 0.0106 | 0.0106 | TRUE | 0.191 |
| 1110001J03 | 359 | 541 | 536 | 586 | 408 | 456 | 381 | 540 | 538 | 461 | 439 | 499 | 476.44 | 486 | 466 | 0.958 | -0.063 | 0.69610542 | 0.83364698 | 0.0188 | 0.0138 | 0.0164 | 0.0164 | TRUE | 0.4139 |
| 1110002L01 | 1055 | 674 | 735 | 1224 | 824 | 724 | 1121 | 673 | 738 | 962 | 887 | 793 | 862.18 | 844 | 881 | 1.044 | 0.062 | 0.74627415 | 0.86526766 | 0.0414 | 0.0107 | 0.0254 | 0.0254 | TRUE | 0.9758 |
| 1110004E09 | 1453 | 1743 | 1812 | 1956 | 1504 | 1459 | 1543 | 1741 | 1819 | 1538 | 1618 | 1597 | 1642.75 | 1701 | 1584 | 0.931 | -0.103 | 0.26226998 | 0.48166304 | 0.0033 | 0.0089 | 0.0055 | 0.0055 | TRUE | 0.0791 |
| 1110004F10 | 3494 | 3509 | 3392 | 5058 | 3354 | 3212 | 3711 | 3504 | 3406 | 3976 | 3609 | 3516 | 3620.43 | 3540 | 3700 | 1.046 | 0.064 | 0.43667588 | 0.64496661 | 0.0027 | 0.0078 | 0.0047 | 0.0047 | TRUE | 0.0524 |
| 1110006O24 | 10 | 6 | 6 | 14 | 11 | 10 | 11 | 6 | 6 | 11 | 12 | 11 | 9.4 | 8 | 11 | 1.497 | 0.582 | 0.34777373 | 0.56662263 | 0 | 0.3572 | 0.1654 | 0.1654 | TRUE | 0.34 |
| 1110007C09 | 862 | 1018 | 1093 | 844 | 584 | 659 | 916 | 1017 | 1097 | 663 | 628 | 721 | 840.5 | 1010 | 671 | 0.664 | -0.59 | 1.13E-07 | 3.97E-06 | 0.0053 | 0.0108 | 0.0077 | 0.0077 | TRUE | 0.0793 |
| 1110008F13 | 4280 | 3855 | 3586 | 5360 | 3645 | 3352 | 4546 | 3850 | 3601 | 4214 | 3922 | 3669 | 3966.93 | 3999 | 3935 | 0.984 | -0.023 | 0.83729131 | 0.91728754 | 0.0094 | 0.0077 | 0.0087 | 0.0087 | TRUE | 0.1742 |
| 1110008J03 | 1204 | 1090 | 1098 | 1187 | 847 | 917 | 1279 | 1089 | 1103 | 933 | 911 | 1004 | 1053.03 | 1157 | 949 | 0.821 | -0.285 | 0.00591408 | 0.03552049 | 0.0041 | 0.01 | 0.0068 | 0.0068 | TRUE | 0.1019 |
| 1110008L16 | 1437 | 1469 | 1513 | 1989 | 1474 | 1437 | 1526 | 1467 | 1519 | 1564 | 1586 | 1573 | 1539.2 | 1504 | 1574 | 1.046 | 0.066 | 0.35317201 | 0.57165733 | 0 | 0.009 | 0.0029 | 0.0029 | TRUE | 0.0056 |
| 1110008P14 | 889 | 807 | 848 | 911 | 642 | 645 | 944 | 806 | 851 | 716 | 691 | 706 | 785.78 | 867 | 704 | 0.813 | -0.299 | 0.0027861 | 0.01999197 | 0.0023 | 0.0111 | 0.0059 | 0.0059 | TRUE | 0.0714 |
| 1110012L19 | 549 | 584 | 526 | 694 | 516 | 504 | 583 | 583 | 528 | 546 | 555 | 552 | 557.83 | 565 | 551 | 0.975 | -0.036 | 0.71592486 | 0.84571767 | 0 | 0.0128 | 0.0054 | 0.0054 | TRUE | 0.038 |
| 1110015O18 | 0 | 0 | 0 | 0 | 0 | 0 | 0 | 0 | 0 | 0 | 0 | 0 | 0 | 0 | 0 | NA | NA | NA | NA | NA | NA | NA | NA | NA | NA |
| 1110017D15 | 7 | 5 | 1 | 5 | 3 | 2 | 7 | 5 | 1 | 4 | 3 | 2 | 3.8 | 4 | 3 | 0.706 | -0.502 | 0.63078966 | NA | 0 | 0.8746 | 0.5173 | 0.5173 | TRUE | 0.2825 |
| 1110018G07 | 1992 | 2047 | 2013 | 2751 | 1998 | 1899 | 2116 | 2044 | 2021 | 2163 | 2150 | 2079 | 2095.43 | 2060 | 2131 | 1.034 | 0.049 | 0.47720526 | 0.67672014 | 0 | 0.0085 | 0.0029 | 0.0029 | TRUE | 0.0066 |
| 1110019D14 | 256 | 258 | 256 | 346 | 228 | 209 | 272 | 258 | 257 | 272 | 245 | 229 | 255.46 | 262 | 249 | 0.951 | -0.072 | 0.60122166 | 0.77281728 | 0 | 0.0198 | 0.0097 | 0.0097 | TRUE | 0.0803 |
| 1110020A21 | 7 | 9 | 7 | 9 | 9 | 3 | 7 | 9 | 7 | 7 | 10 | 3 | 7.25 | 8 | 7 | 0.857 | -0.222 | 0.75740604 | 0.8719099 | 0 | 0.4614 | 0.235 | 0.235 | TRUE | 0.3087 |
| 1110025L11 | 0 | 0 | 0 | 0 | 0 | 0 | 0 | 0 | 0 | 0 | 0 | 0 | 0 | 0 | 0 | NA | NA | NA | NA | NA | NA | NA | NA | NA | NA |
| 1110028F11 | 1 | 0 | 0 | 0 | 0 | 0 | 1 | 0 | 0 | 0 | 0 | 0 | 0.18 | 0 | 0 | 0.49 | -1.029 | 0.80087 | NA | 0 | 18.6165 | 10 | 10 | TRUE | 0.4614 |
| 1110028F18 | 3 | 2 | 1 | 6 | 1 | 4 | 3 | 2 | 1 | 5 | 1 | 4 | 2.73 | 2 | 3 | 1.671 | 0.741 | 0.53311934 | NA | 0 | 1.2152 | 0.6264 | 0.6264 | TRUE | 0.4704 |
| 1110032A03 | 113 | 135 | 131 | 244 | 155 | 113 | 120 | 135 | 132 | 192 | 167 | 124 | 144.78 | 129 | 161 | 1.253 | 0.325 | 0.12209073 | 0.29718298 | 0.0203 | 0.0297 | 0.0249 | 0.0249 | TRUE | 0.3216 |
| 1110032F04 | 57 | 98 | 131 | 143 | 121 | 106 | 61 | 98 | 132 | 112 | 130 | 116 | 108.1 | 97 | 119 | 1.234 | 0.304 | 0.28613491 | 0.50736877 | 0.0623 | 0.0374 | 0.049 | 0.049 | TRUE | 0.2597 |
| 1110034G24 | 246 | 291 | 281 | 304 | 192 | 208 | 261 | 291 | 282 | 239 | 207 | 228 | 251.22 | 278 | 225 | 0.809 | -0.307 | 0.02604034 | 0.10447933 | 0 | 0.02 | 0.0097 | 0.0097 | TRUE | 0.0522 |
| 1110035M17 | 4 | 4 | 2 | 4 | 1 | 1 | 4 | 4 | 2 | 3 | 1 | 1 | 2.59 | 3 | 2 | 0.539 | -0.891 | 0.45219389 | NA | 0 | 1.2767 | 0.5967 | 0.5967 | TRUE | 0.4915 |
| 1110036E04 | 1 | 0 | 0 | 0 | 3 | 1 | 1 | 0 | 0 | 0 | 3 | 1 | 0.9 | 0 | 1 | 3.864 | 1.95 | 0.42601043 | NA | 0 | 3.6778 | 2.9783 | 2.9783 | TRUE | 0.2815 |
| 1110037F02 | 6162 | 5067 | 5084 | 7537 | 5165 | 4544 | 6545 | 5060 | 5105 | 5925 | 5557 | 4974 | 5527.8 | 5570 | 5485 | 0.985 | -0.022 | 0.8628872 | 0.93221844 | 0.0122 | 0.0075 | 0.0113 | 0.0113 | TRUE | 0.2857 |
| 1110038B12 | 2001 | 2854 | 3452 | 2250 | 1706 | 2594 | 2125 | 2850 | 3466 | 1769 | 1836 | 2840 | 2480.96 | 2814 | 2148 | 0.763 | -0.39 | 0.07053999 | 0.20679493 | 0.0668 | 0.0082 | 0.0331 | 0.0331 | TRUE | 0.3087 |
| 1110038F14 | 1032 | 976 | 853 | 1693 | 1165 | 982 | 1096 | 975 | 857 | 1331 | 1253 | 1075 | 1097.79 | 976 | 1220 | 1.251 | 0.323 | 0.01322238 | 0.06450461 | 0.0119 | 0.0099 | 0.0113 | 0.0113 | TRUE | 0.1384 |
| 1110046J04 | 6 | 8 | 5 | 8 | 9 | 9 | 6 | 8 | 5 | 6 | 10 | 10 | 7.53 | 6 | 9 | 1.318 | 0.399 | 0.5613505 | 0.74256648 | 0 | 0.4441 | 0.2029 | 0.2029 | TRUE | 0.2261 |
| 1110051M2 | 467 | 544 | 574 | 634 | 455 | 490 | 496 | 543 | 576 | 498 | 490 | 536 | 523.34 | 538 | 508 | 0.943 | -0.085 | 0.4425245 | 0.64943673 | 0.002 | 0.0132 | 0.0069 | 0.0069 | TRUE | 0.0552 |
| 1110054M0 | 71 | 76 | 95 | 60 | 53 | 48 | 75 | 76 | 95 | 47 | 57 | 53 | 67.24 | 82 | 52 | 0.633 | -0.661 | 0.00797769 | 0.04421089 | 0 | 0.0559 | 0.0291 | 0.0291 | TRUE | 0.1845 |
| 1110057K04 | 1820 | 1358 | 1355 | 2433 | 1792 | 1514 | 1933 | 1356 | 1361 | 1913 | 1928 | 1657 | 1691.33 | 1550 | 1833 | 1.183 | 0.242 | 0.11735845 | 0.28984926 | 0.0255 | 0.0089 | 0.0166 | 0.0166 | TRUE | 0.5626 |
| 1110058L19 | 634 | 807 | 809 | 877 | 667 | 675 | 673 | 806 | 812 | 689 | 718 | 739 | 739.61 | 764 | 715 | 0.936 | -0.096 | 0.38789599 | 0.6017358 | 0.0047 | 0.0114 | 0.0076 | 0.0076 | TRUE | 0.1263 |
| 1110059E24 | 1423 | 1629 | 1612 | 2200 | 1407 | 1490 | 1511 | 1627 | 1619 | 1729 | 1514 | 1631 | 1605.22 | 1586 | 1625 | 1.025 | 0.036 | 0.68258453 | 0.82407557 | 0.0025 | 0.009 | 0.005 | 0.005 | TRUE | 0.0429 |
| 1110059G10 | 328 | 363 | 364 | 525 | 363 | 332 | 348 | 363 | 365 | 413 | 391 | 363 | 373.85 | 359 | 389 | 1.086 | 0.118 | 0.30767223 | 0.5282036 | 0 | 0.0157 | 0.007 | 0.007 | TRUE | 0.0372 |
| 1110059M1 | 0 | 0 | 0 | 0 | 0 | 0 | 0 | 0 | 0 | 0 | 0 | 0 | 0 | 0 | 0 | NA | NA | NA | NA | NA | NA | NA | NA | NA | NA |
| 1110065P20 | 389 | 439 | 448 | 510 | 449 | 394 | 413 | 438 | 450 | 401 | 483 | 431 | 436.13 | 434 | 438 | 1.009 | 0.012 | 0.91871749 | 0.96017575 | 0.0033 | 0.0145 | 0.0083 | 0.0083 | TRUE | 0.0915 |
| 1190002F15 | 56 | 108 | 119 | 89 | 73 | 92 | 59 | 108 | 119 | 70 | 79 | 101 | 89.34 | 95 | 83 | 0.865 | -0.209 | 0.49455888 | 0.69052608 | 0.0695 | 0.0438 | 0.0561 | 0.0561 | TRUE | 1.0388 |
| 1190002N15 | 1708 | 1718 | 1770 | 2413 | 1933 | 1800 | 1814 | 1716 | 1777 | 1897 | 2080 | 1970 | 1875.72 | 1769 | 1982 | 1.12 | 0.164 | 0.03350507 | 0.12416399 | 0.001 | 0.0087 | 0.0037 | 0.0037 | TRUE | 0.0223 |

| | | | | | | | | | | | | | | | | | | | | | | | | | |
|---|---|---|---|---|---|---|---|---|---|---|---|---|---|---|---|---|---|---|---|---|---|---|---|---|---|
| 1190003J15F | 354 | 402 | 426 | 508 | 378 | 382 | 376 | 401 | 428 | 399 | 407 | 418 | 404.91 | 402 | 408 | 1.015 | 0.021 | 0.85106382 | 0.92587032 | 0 | 0.015 | 0.0067 | 0.0067 | TRUE | 0.0369 |
| 1190003K10I | 0 | 0 | 0 | 0 | 0 | 0 | 0 | 0 | 0 | 0 | 0 | 0 | 0 | 0 | 0 | NA | NA | NA | NA | NA | NA | NA | NA | NA | NA |
| 1190005I06R | 431 | 558 | 524 | 460 | 336 | 340 | 458 | 557 | 526 | 362 | 362 | 372 | 439.42 | 514 | 365 | 0.71 | -0.494 | 4.53E-05 | 0.00071376 | 0.0033 | 0.0144 | 0.0082 | 0.0082 | TRUE | 0.1048 |
| 1190007I07R | 303 | 522 | 579 | 462 | 378 | 450 | 322 | 521 | 581 | 363 | 407 | 493 | 447.84 | 475 | 421 | 0.885 | -0.176 | 0.42951448 | 0.63926605 | 0.0554 | 0.0143 | 0.0336 | 0.0336 | TRUE | 0.9189 |
| 1200011I18R | 2439 | 2356 | 2426 | 3679 | 2636 | 2692 | 2591 | 2353 | 2436 | 2892 | 2836 | 2947 | 2675.78 | 2460 | 2892 | 1.176 | 0.234 | 0.00132833 | 0.01110842 | 8.00E-04 | 0.0081 | 0.0034 | 0.0034 | TRUE | 0.0261 |
| 1200014J11F | 3921 | 4059 | 4043 | 5894 | 4169 | 4071 | 4165 | 4054 | 4060 | 4633 | 4486 | 4456 | 4308.92 | 4093 | 4525 | 1.106 | 0.145 | 0.01575956 | 0.07356015 | 0 | 0.0077 | 0.0024 | 0.0024 | TRUE | 0.0054 |
| 1300002E11I | 308 | 301 | 281 | 459 | 388 | 259 | 327 | 301 | 282 | 361 | 417 | 284 | 328.62 | 303 | 354 | 1.168 | 0.224 | 0.18609902 | 0.38984643 | 0.0182 | 0.0169 | 0.0176 | 0.0176 | TRUE | 0.2382 |
| 1300002K09I | 18 | 14 | 13 | 21 | 15 | 12 | 19 | 14 | 13 | 17 | 16 | 13 | 15.32 | 15 | 15 | 0.997 | -0.004 | 0.99362571 | 0.9964688 | 0 | 0.2219 | 0.1023 | 0.1023 | TRUE | 0.2 |
| 1300015D01 | 2 | 1 | 1 | 4 | 2 | 3 | 2 | 1 | 1 | 3 | 2 | 3 | 2.12 | 1 | 3 | 2.098 | 1.069 | 0.40341539 | NA | 0 | 1.5624 | 0.6357 | 0.6357 | TRUE | 0.1347 |
| 1300017J02F | 50 | 78 | 73 | 84 | 70 | 50 | 53 | 78 | 73 | 66 | 75 | 55 | 66.73 | 68 | 65 | 0.959 | -0.06 | 0.82235926 | 0.90923689 | 0.0154 | 0.0563 | 0.037 | 0.037 | TRUE | 0.3226 |
| 1300018J18F | 344 | 508 | 504 | 351 | 247 | 343 | 365 | 507 | 506 | 276 | 266 | 375 | 382.66 | 459 | 306 | 0.664 | -0.591 | 0.00209745 | 0.0159888 | 0.0313 | 0.0155 | 0.0238 | 0.0238 | TRUE | 0.4423 |
| 1500002O10 | 2 | 6 | 1 | 5 | 0 | 1 | 2 | 6 | 1 | 4 | 0 | 1 | 2.36 | 3 | 2 | 0.576 | -0.795 | 0.58442044 | NA | 0.7011 | 1.4042 | 1.0772 | 1.0772 | TRUE | 1.362 |
| 1500004A13 | 3 | 3 | 4 | 9 | 7 | 5 | 3 | 3 | 4 | 7 | 8 | 5 | 5.05 | 3 | 7 | 1.973 | 0.98 | 0.23926644 | 0.45521235 | 0 | 0.6597 | 0.2768 | 0.2768 | TRUE | 0.0579 |
| 1500009C09I | 17 | 12 | 16 | 15 | 7 | 15 | 18 | 12 | 16 | 12 | 8 | 16 | 13.64 | 15 | 12 | 0.775 | -0.367 | 0.49671554 | 0.69206691 | 0 | 0.2484 | 0.1362 | 0.1362 | TRUE | 0.228 |
| 1500009L16F | 2299 | 2509 | 2378 | 2197 | 1681 | 1809 | 2442 | 2506 | 2388 | 1727 | 1809 | 1980 | 2141.91 | 2445 | 1839 | 0.751 | -0.412 | 7.65E-07 | 2.18E-05 | 0.002 | 0.0084 | 0.0045 | 0.0045 | TRUE | 0.0545 |
| 1500011B03 | 963 | 920 | 836 | 865 | 687 | 739 | 1023 | 919 | 839 | 680 | 739 | 809 | 834.87 | 927 | 743 | 0.8 | -0.321 | 0.00692446 | 0.03981499 | 0.0075 | 0.0108 | 0.009 | 0.009 | TRUE | 0.097 |
| 1500011K16I | 336 | 656 | 641 | 604 | 445 | 484 | 357 | 655 | 644 | 475 | 479 | 530 | 523.18 | 552 | 495 | 0.895 | -0.159 | 0.46513449 | 0.66788976 | 0.0542 | 0.0132 | 0.0324 | 0.0324 | TRUE | 1.1135 |
| 1500012F01I | 2985 | 5368 | 6137 | 3413 | 2743 | 4293 | 3171 | 5361 | 6162 | 2683 | 2951 | 4699 | 4171.26 | 4898 | 3444 | 0.703 | -0.508 | 0.1734883 | 0.37263482 | 0.1001 | 0.0077 | 0.0465 | 0.1001 | TRUE | 0.6606 |
| 1500012K07I | 0 | 0 | 0 | 0 | 0 | 0 | 0 | 0 | 0 | 0 | 0 | 0 | 0 | 0 | 0 | NA | NA | NA | NA | NA | NA | NA | NA | NA | NA |
| 1500015A07 | 227 | 223 | 175 | 287 | 240 | 183 | 241 | 223 | 176 | 226 | 258 | 200 | 220.62 | 213 | 228 | 1.07 | 0.098 | 0.58744283 | 0.76212456 | 0.016 | 0.0218 | 0.0189 | 0.0189 | TRUE | 0.2596 |
| 1500015L24F | 0 | 0 | 0 | 0 | 0 | 0 | 0 | 0 | 0 | 0 | 0 | 0 | 0 | 0 | 0 | NA | NA | NA | NA | NA | NA | NA | NA | NA | NA |
| 1500015O10 | 17 | 11 | 10 | 24 | 11 | 11 | 18 | 11 | 10 | 19 | 12 | 12 | 13.64 | 13 | 14 | 1.109 | 0.149 | 0.78397373 | 0.88846976 | 0 | 0.2485 | 0.1397 | 0.1397 | TRUE | 0.4549 |
| 1500016L03F | 0 | 0 | 0 | 1 | 0 | 0 | 0 | 0 | 0 | 1 | 0 | 0 | 0.13 | 0 | 0 | 1.859 | 0.894 | 0.82650734 | NA | 0 | 25.1515 | 10 | 10 | TRUE | 0.2563 |
| 1500017E21I | 2 | 1 | 0 | 2 | 2 | 0 | 2 | 1 | 0 | 2 | 2 | 0 | 1.14 | 1 | 1 | 1.213 | 0.279 | 0.8846965 | NA | 0 | 2.8937 | 1.7759 | 1.7759 | TRUE | 0.2368 |
| 1500032L24F | 1858 | 2043 | 2209 | 2604 | 1882 | 1778 | 1974 | 2040 | 2218 | 2047 | 2025 | 1946 | 2041.71 | 2077 | 2006 | 0.966 | -0.05 | 0.53382077 | 0.72173954 | 0.0015 | 0.0085 | 0.0042 | 0.0042 | TRUE | 0.0426 |
| 1600002D24 | 1 | 0 | 0 | 0 | 0 | 0 | 1 | 0 | 0 | 0 | 0 | 0 | 0.18 | 0 | 0 | 0.49 | -1.029 | 0.80087 | NA | 0 | 18.6165 | 10 | 10 | TRUE | 0.4614 |
| 1600002H07 | 1094 | 1229 | 1275 | 1538 | 1154 | 1117 | 1162 | 1227 | 1280 | 1209 | 1242 | 1223 | 1223.85 | 1223 | 1225 | 1.001 | 0.001 | 0.9908912 | 0.99518186 | 4.00E-04 | 0.0096 | 0.0039 | 0.0039 | TRUE | 0.0228 |
| 1600002K03I | 63 | 60 | 43 | 71 | 64 | 63 | 67 | 60 | 43 | 56 | 69 | 69 | 60.61 | 57 | 65 | 1.136 | 0.183 | 0.50789417 | 0.7020602 | 0.0139 | 0.0613 | 0.0387 | 0.0387 | TRUE | 0.3698 |
| 1600010M07 | 0 | 0 | 0 | 1 | 0 | 0 | 0 | 0 | 0 | 1 | 0 | 0 | 0.13 | 0 | 0 | 1.859 | 0.894 | 0.82650734 | NA | 0 | 25.1515 | 10 | 10 | TRUE | 0.2563 |
| 1600012H06 | 1015 | 954 | 928 | 1274 | 896 | 861 | 1078 | 953 | 932 | 1002 | 964 | 943 | 978.46 | 988 | 970 | 0.982 | -0.026 | 0.79045123 | 0.89222048 | 0.0026 | 0.0103 | 0.0057 | 0.0057 | TRUE | 0.0763 |
| 1600014C10I | 304 | 394 | 368 | 477 | 321 | 328 | 323 | 393 | 370 | 375 | 345 | 359 | 360.88 | 362 | 360 | 0.994 | -0.008 | 0.95022657 | 0.97481103 | 0.003 | 0.016 | 0.009 | 0.009 | TRUE | 0.1007 |
| 1600014C23I | 5 | 6 | 15 | 21 | 22 | 12 | 5 | 6 | 15 | 17 | 24 | 13 | 13.28 | 9 | 18 | 2.014 | 1.01 | 0.1003267 | 0.26184886 | 0.103 | 0.255 | 0.1868 | 0.1868 | TRUE | 1.1229 |
| 1600014K23I | 6 | 7 | 11 | 27 | 15 | 17 | 6 | 7 | 11 | 21 | 16 | 19 | 13.4 | 8 | 19 | 2.298 | 1.201 | 0.02693606 | 0.10717343 | 0 | 0.2528 | 0.1234 | 0.1234 | TRUE | 0.2944 |
| 1600015I10R | 5 | 0 | 1 | 1 | 1 | 0 | 5 | 0 | 1 | 1 | 1 | 0 | 1.36 | 2 | 1 | 0.301 | -1.731 | 0.38696204 | NA | 1.1625 | 2.4242 | 1.8666 | 1.8666 | TRUE | 0.9913 |
| 1600016N20 | 209 | 279 | 267 | 245 | 243 | 277 | 222 | 279 | 268 | 193 | 261 | 303 | 254.33 | 256 | 252 | 0.981 | -0.027 | 0.89178164 | 0.94753967 | 0.0294 | 0.0199 | 0.0247 | 0.0247 | TRUE | 0.402 |
| 1600019K03I | 0 | 0 | 0 | 0 | 0 | 0 | 0 | 0 | 0 | 0 | 0 | 0 | 0 | 0 | 0 | NA | NA | NA | NA | NA | NA | NA | NA | NA | NA |
| 1600020E01I | 83 | 99 | 101 | 133 | 134 | 122 | 88 | 99 | 101 | 105 | 144 | 134 | 111.79 | 96 | 128 | 1.318 | 0.399 | 0.05844167 | 0.18266785 | 0.0102 | 0.0364 | 0.0228 | 0.0228 | TRUE | 0.2781 |
| 1600023N17 | 3 | 6 | 7 | 5 | 14 | 8 | 3 | 6 | 7 | 4 | 15 | 9 | 7.33 | 5 | 9 | 1.682 | 0.75 | 0.34832031 | 0.56702733 | 0.1757 | 0.4566 | 0.3129 | 0.3129 | TRUE | 0.1756 |
| 1600025M17 | 0 | 0 | 0 | 1 | 0 | 1 | 0 | 0 | 0 | 1 | 0 | 1 | 0.31 | 0 | 1 | 3.348 | 1.743 | 0.65064889 | NA | 0 | 10.5167 | 8.8968 | 8.8968 | TRUE | 0.2082 |
| 1600029D21 | 248 | 242 | 276 | 312 | 218 | 245 | 263 | 242 | 277 | 245 | 235 | 268 | 255.04 | 261 | 249 | 0.956 | -0.065 | 0.63627815 | 0.79538551 | 0 | 0.0198 | 0.0098 | 0.0098 | TRUE | 0.0462 |
| 1600029I14R | 0 | 0 | 0 | 0 | 0 | 0 | 0 | 0 | 0 | 0 | 0 | 0 | 0 | 0 | 0 | NA | NA | NA | NA | NA | NA | NA | NA | NA | NA |
| 1600029O15 | 6 | 13 | 7 | 11 | 9 | 6 | 6 | 13 | 7 | 9 | 10 | 7 | 8.55 | 9 | 9 | 0.944 | -0.084 | 0.89907677 | 0.95105559 | 0 | 0.3923 | 0.1985 | 0.1985 | TRUE | 0.5562 |
| 1700001C02I | 1 | 4 | 7 | 1 | 2 | 5 | 1 | 4 | 7 | 1 | 2 | 5 | 3.42 | 4 | 3 | 0.675 | -0.568 | 0.63012124 | NA | 0.4191 | 0.9713 | 0.6958 | 0.6958 | TRUE | 0.2179 |
| 1700001C19I | 32 | 24 | 27 | 42 | 24 | 19 | 34 | 24 | 27 | 33 | 26 | 21 | 27.45 | 28 | 27 | 0.946 | -0.08 | 0.83454098 | 0.91571799 | 0 | 0.1269 | 0.0703 | 0.0703 | TRUE | 0.3378 |
| 1700001D01 | 0 | 0 | 0 | 0 | 0 | 0 | 0 | 0 | 0 | 0 | 0 | 0 | 0 | 0 | 0 | NA | NA | NA | NA | NA | NA | NA | NA | NA | NA |
| 1700001F09I | 2 | 2 | 0 | 2 | 0 | 0 | 2 | 2 | 0 | 2 | 0 | 0 | 0.95 | 1 | 1 | 0.414 | -1.273 | 0.58519735 | NA | 0 | 3.4784 | 2.6849 | 2.6849 | TRUE | 0.9719 |
| 1700001G11 | 17 | 38 | 32 | 45 | 35 | 31 | 18 | 38 | 32 | 35 | 38 | 34 | 32.52 | 29 | 36 | 1.21 | 0.275 | 0.46524543 | 0.66796058 | 0.025 | 0.1082 | 0.0715 | 0.0715 | TRUE | 0.7194 |
| 1700001G17 | 5 | 3 | 2 | 7 | 7 | 6 | 5 | 3 | 2 | 6 | 8 | 7 | 4.99 | 3 | 7 | 1.9 | 0.926 | 0.28185315 | 0.50334119 | 0 | 0.6676 | 0.309 | 0.309 | TRUE | 0.3142 |
| 1700001J03F | 0 | 2 | 3 | 5 | 5 | 3 | 0 | 2 | 3 | 4 | 5 | 3 | 2.93 | 2 | 4 | 2.487 | 1.314 | 0.2730401 | NA | 0 | 1.1298 | 0.6164 | 0.6164 | TRUE | 0.5382 |
| 1700001J11F | 0 | 0 | 0 | 0 | 0 | 0 | 0 | 0 | 0 | 0 | 0 | 0 | 0 | 0 | 0 | NA | NA | NA | NA | NA | NA | NA | NA | NA | NA |
| 1700001K19I | 1 | 0 | 0 | 1 | 0 | 0 | 1 | 0 | 0 | 1 | 0 | 0 | 0.31 | 0 | 0 | 0.954 | -0.067 | 0.98625928 | NA | 0 | 10.7014 | 9.0384 | 9.0384 | TRUE | 0.4614 |
| 1700001K23I | 1 | 0 | 1 | 1 | 1 | 0 | 1 | 0 | 1 | 1 | 1 | 0 | 0.65 | 1 | 1 | 0.917 | -0.125 | 0.95730863 | NA | 0 | 5.0387 | 2.3854 | 2.3854 | TRUE | 0.2549 |
| 1700001L05F | 1911 | 2045 | 2232 | 3132 | 2716 | 2572 | 2030 | 2042 | 2241 | 2462 | 2922 | 2815 | 2418.86 | 2104 | 2733 | 1.298 | 0.376 | 0.0001152 | 0.00154332 | 0.0048 | 0.0083 | 0.0064 | 0.0064 | TRUE | 0.0927 |
| 1700001L19F | 20 | 15 | 11 | 32 | 30 | 12 | 21 | 15 | 11 | 25 | 32 | 13 | 19.64 | 16 | 23 | 1.5 | 0.585 | 0.25408574 | 0.47207448 | 0.0931 | 0.1746 | 0.136 | 0.136 | TRUE | 0.2643 |
| 1700001O22 | 22 | 33 | 25 | 43 | 38 | 29 | 23 | 33 | 25 | 34 | 41 | 32 | 31.31 | 27 | 36 | 1.304 | 0.383 | 0.27592352 | 0.49661826 | 0 | 0.1121 | 0.0563 | 0.0563 | TRUE | 0.2252 |
| 1700001P01I | 15 | 8 | 21 | 8 | 11 | 10 | 16 | 8 | 21 | 6 | 12 | 11 | 12.35 | 15 | 10 | 0.638 | -0.649 | 0.28911645 | 0.50996392 | 0.092 | 0.2737 | 0.1848 | 0.1848 | TRUE | 0.2429 |
| 1700003C15I | 0 | 0 | 0 | 0 | 0 | 0 | 0 | 0 | 0 | 0 | 0 | 0 | 0 | 0 | 0 | NA | NA | NA | NA | NA | NA | NA | NA | NA | NA |
| 1700003D09 | 0 | 0 | 1 | 0 | 2 | 0 | 0 | 0 | 1 | 0 | 2 | 0 | 0.53 | 0 | 1 | 1.927 | 0.947 | 0.77736506 | NA | 0 | 6.2701 | 6.3814 | 6.3814 | TRUE | 0.9744 |
| 1700003E16I | 5 | 6 | 1 | 12 | 9 | 4 | 5 | 6 | 1 | 9 | 10 | 4 | 5.97 | 4 | 8 | 1.924 | 0.944 | 0.27995371 | 0.50109666 | 0.0772 | 0.559 | 0.3635 | 0.3635 | TRUE | 0.7594 |
| 1700003F12I | 9 | 23 | 23 | 15 | 3 | 11 | 10 | 23 | 23 | 12 | 3 | 12 | 13.78 | 19 | 9 | 0.49 | -1.029 | 0.11402428 | 0.28478082 | 0.1957 | 0.246 | 0.224 | 0.224 | TRUE | 0.9281 |
| 1700003F17I | 0 | 0 | 0 | 0 | 0 | 0 | 0 | 0 | 0 | 0 | 0 | 0 | 0 | 0 | 0 | NA | NA | NA | NA | NA | NA | NA | NA | NA | NA |
| 1700003G13 | 0 | 0 | 0 | 0 | 0 | 0 | 0 | 0 | 0 | 0 | 0 | 0 | 0 | 0 | 0 | NA | NA | NA | NA | NA | NA | NA | NA | NA | NA |
| 1700003G18 | 1 | 3 | 3 | 2 | 5 | 1 | 1 | 3 | 3 | 2 | 5 | 1 | 2.52 | 2 | 3 | 1.118 | 0.161 | 0.89636998 | NA | 0 | 1.3145 | 0.7028 | 0.7028 | TRUE | 0.838 |
| 1700003H04 | 0 | 0 | 0 | 0 | 0 | 0 | 0 | 0 | 0 | 0 | 0 | 0 | 0 | 0 | 0 | NA | NA | NA | NA | NA | NA | NA | NA | NA | NA |
| 1700003L19F | 0 | 0 | 0 | 0 | 0 | 0 | 0 | 0 | 0 | 0 | 0 | 0 | 0 | 0 | 0 | NA | NA | NA | NA | NA | NA | NA | NA | NA | NA |
| 1700003M02 | 13 | 10 | 10 | 26 | 17 | 13 | 14 | 10 | 10 | 20 | 18 | 14 | 14.47 | 11 | 17 | 1.577 | 0.657 | 0.19240783 | 0.39803992 | 0 | 0.2346 | 0.1104 | 0.1104 | TRUE | 0.1384 |
| 1700003M07 | 32 | 45 | 30 | 67 | 34 | 30 | 34 | 45 | 30 | 53 | 37 | 33 | 38.52 | 36 | 41 | 1.129 | 0.175 | 0.61833209 | 0.78432795 | 0.0318 | 0.0924 | 0.0624 | 0.0624 | TRUE | 0.5703 |
| 1700003P14I | 0 | 0 | 0 | 0 | 0 | 0 | 0 | 0 | 0 | 0 | 0 | 0 | 0 | 0 | 0 | NA | NA | NA | NA | NA | NA | NA | NA | NA | NA |
| 1700006A11 | 0 | 0 | 0 | 0 | 0 | 0 | 0 | 0 | 0 | 0 | 0 | 0 | 0 | 0 | 0 | NA | NA | NA | NA | NA | NA | NA | NA | NA | NA |

| | | | | | | | | | | | | | | | | | | | | | | | | | |
|---|---|---|---|---|---|---|---|---|---|---|---|---|---|---|---|---|---|---|---|---|---|---|---|---|---|
| 1700006E09I | 0 | 1 | 1 | 1 | 0 | 1 | 0 | 1 | 1 | 1 | 0 | 1 | 0.65 | 1 | 1 | 0.94 | -0.089 | 0.96942491 | NA | 0 | 5.0967 | 2.3884 | 2.3884 | TRUE | 0.2246 |
| 1700006F04I | 0 | 0 | 0 | 0 | 0 | 0 | 0 | 0 | 0 | 0 | 0 | 0 | 0 | 0 | 0 | NA | NA | NA | NA | NA | NA | NA | NA | NA | NA |
| 1700006H21 | 0 | 0 | 0 | 0 | 2 | 0 | 0 | 0 | 0 | 0 | 2 | 0 | 0.36 | 0 | 1 | 3.775 | 1.916 | 0.63423067 | NA | 10 | 9.1927 | 10 | 10 | TRUE | 0.9743 |
| 1700007B14 | 1 | 0 | 1 | 2 | 0 | 2 | 1 | 0 | 1 | 2 | 0 | 2 | 0.97 | 1 | 1 | 1.841 | 0.881 | 0.6673686 | NA | 0 | 3.3987 | 1.894 | 1.894 | TRUE | 0.3972 |
| 1700007F19I | 0 | 0 | 0 | 0 | 0 | 0 | 0 | 0 | 0 | 0 | 0 | 0 | 0 | 0 | 0 | NA | NA | NA | NA | NA | NA | NA | NA | NA | NA |
| 1700007G11 | 1 | 2 | 4 | 6 | 8 | 4 | 1 | 2 | 4 | 5 | 9 | 4 | 4.13 | 2 | 6 | 2.476 | 1.308 | 0.18884213 | NA | 0 | 0.8046 | 0.415 | 0.415 | TRUE | 0.3966 |
| 1700007J10F | 0 | 0 | 1 | 0 | 0 | 0 | 0 | 0 | 1 | 0 | 0 | 0 | 0.17 | 0 | 0 | 0.49 | -1.029 | 0.80087 | NA | 0 | 19.6928 | 10 | 10 | TRUE | 0.4098 |
| 1700007K09I | 5 | 2 | 1 | 0 | 0 | 1 | 5 | 2 | 1 | 0 | 0 | 1 | 1.57 | 3 | 0 | 0.127 | -2.983 | 0.11099196 | NA | 0 | 2.1082 | 1.3391 | 1.3391 | TRUE | 0.5564 |
| 1700007K13I | 223 | 305 | 298 | 354 | 282 | 325 | 237 | 305 | 299 | 278 | 303 | 356 | 296.36 | 280 | 312 | 1.112 | 0.154 | 0.34749119 | 0.56641496 | 0.0138 | 0.018 | 0.0159 | 0.0159 | TRUE | 0.2033 |
| 1700007L15F | 40 | 82 | 77 | 141 | 93 | 71 | 42 | 82 | 77 | 111 | 100 | 78 | 81.72 | 67 | 96 | 1.432 | 0.518 | 0.08395451 | 0.2321814 | 0.058 | 0.0472 | 0.0522 | 0.0522 | TRUE | 0.8998 |
| 1700007P06I | 0 | 0 | 0 | 1 | 0 | 0 | 0 | 0 | 0 | 1 | 0 | 0 | 0.13 | 0 | 0 | 1.859 | 0.894 | 0.82650734 | NA | 0 | 25.1515 | 10 | 10 | TRUE | 0.2563 |
| 1700008A04 | 0 | 0 | 0 | 0 | 0 | 0 | 0 | 0 | 0 | 0 | 0 | 0 | 0 | 0 | 0 | NA | NA | NA | NA | NA | NA | NA | NA | NA | NA |
| 1700008F21I | 0 | 0 | 0 | 0 | 0 | 0 | 0 | 0 | 0 | 0 | 0 | 0 | 0 | 0 | 0 | NA | NA | NA | NA | NA | NA | NA | NA | NA | NA |
| 1700008I05R | 69 | 59 | 48 | 96 | 76 | 50 | 73 | 59 | 48 | 75 | 82 | 55 | 65.4 | 60 | 71 | 1.179 | 0.237 | 0.4021249 | 0.61492282 | 0.026 | 0.0573 | 0.0424 | 0.0424 | TRUE | 0.3293 |
| 1700008J07F | 29 | 40 | 47 | 67 | 54 | 56 | 31 | 40 | 47 | 53 | 58 | 61 | 48.34 | 39 | 57 | 1.453 | 0.539 | 0.06572339 | 0.19770918 | 0 | 0.0751 | 0.0403 | 0.0403 | TRUE | 0.2574 |
| 1700008K24I | 0 | 0 | 0 | 1 | 1 | 0 | 0 | 0 | 0 | 1 | 1 | 0 | 0.31 | 0 | 1 | 3.32 | 1.731 | 0.65383052 | NA | 0 | 10.6224 | 8.9541 | 8.9541 | TRUE | 0.202 |
| 1700008O03 | 46 | 53 | 70 | 118 | 75 | 55 | 49 | 53 | 70 | 93 | 81 | 60 | 67.62 | 57 | 78 | 1.363 | 0.447 | 0.112635 | 0.28247403 | 0.0279 | 0.0556 | 0.0421 | 0.0421 | TRUE | 0.2952 |
| 1700008P02I | 0 | 0 | 0 | 0 | 0 | 0 | 0 | 0 | 0 | 0 | 0 | 0 | 0 | 0 | 0 | NA | NA | NA | NA | NA | NA | NA | NA | NA | NA |
| 1700009C05I | 3 | 3 | 2 | 0 | 3 | 2 | 3 | 3 | 2 | 0 | 3 | 2 | 2.27 | 3 | 2 | 0.635 | -0.656 | 0.62843678 | NA | 0 | 1.4595 | 0.8559 | 0.8559 | TRUE | 0.8967 |
| 1700009J07F | 0 | 0 | 0 | 0 | 0 | 0 | 0 | 0 | 0 | 0 | 0 | 0 | 0 | 0 | 0 | NA | NA | NA | NA | NA | NA | NA | NA | NA | NA |
| 1700009N14 | 0 | 0 | 0 | 0 | 0 | 0 | 0 | 0 | 0 | 0 | 0 | 0 | 0 | 0 | 0 | NA | NA | NA | NA | NA | NA | NA | NA | NA | NA |
| 1700009P17I | 37 | 46 | 50 | 37 | 34 | 36 | 39 | 46 | 50 | 29 | 37 | 39 | 40.09 | 45 | 35 | 0.77 | -0.378 | 0.22623098 | 0.43904737 | 0 | 0.0891 | 0.0448 | 0.0448 | TRUE | 0.1887 |
| 1700010B08 | 0 | 0 | 0 | 1 | 0 | 0 | 0 | 0 | 0 | 1 | 0 | 0 | 0.13 | 0 | 0 | 1.859 | 0.894 | 0.82650734 | NA | 0 | 25.1515 | 10 | 10 | TRUE | 0.2563 |
| 1700010D01 | 1 | 2 | 0 | 2 | 6 | 1 | 1 | 2 | 0 | 2 | 6 | 1 | 2.03 | 1 | 3 | 2.949 | 1.56 | 0.31414923 | NA | 0 | 1.6295 | 1.0681 | 1.0681 | TRUE | 1.1837 |
| 1700010I02R | 0 | 0 | 0 | 1 | 0 | 0 | 0 | 0 | 0 | 1 | 0 | 0 | 0.13 | 0 | 0 | 1.859 | 0.894 | 0.82650734 | NA | 0 | 25.1515 | 10 | 10 | TRUE | 0.2563 |
| 1700010I14R | 24 | 25 | 25 | 26 | 22 | 28 | 25 | 25 | 25 | 20 | 24 | 31 | 25.05 | 25 | 25 | 0.982 | -0.026 | 0.94581346 | 0.97280132 | 0 | 0.1384 | 0.0658 | 0.0658 | TRUE | 0.2335 |
| 1700010J16F | 0 | 0 | 0 | 0 | 0 | 0 | 0 | 0 | 0 | 0 | 0 | 0 | 0 | 0 | 0 | NA | NA | NA | NA | NA | NA | NA | NA | NA | NA |
| 1700010K23I | 0 | 0 | 0 | 0 | 0 | 0 | 0 | 0 | 0 | 0 | 0 | 0 | 0 | 0 | 0 | NA | NA | NA | NA | NA | NA | NA | NA | NA | NA |
| 1700011A15 | 0 | 1 | 1 | 0 | 0 | 1 | 0 | 1 | 1 | 0 | 0 | 1 | 0.52 | 1 | 0 | 0.52 | -0.943 | 0.73639299 | NA | 0 | 6.3885 | 3.8941 | 3.8941 | TRUE | 0.5383 |
| 1700011B04 | 0 | 0 | 0 | 0 | 0 | 0 | 0 | 0 | 0 | 0 | 0 | 0 | 0 | 0 | 0 | NA | NA | NA | NA | NA | NA | NA | NA | NA | NA |
| 1700011E24I | 2 | 2 | 5 | 0 | 4 | 4 | 2 | 2 | 5 | 0 | 4 | 4 | 2.97 | 3 | 3 | 0.92 | -0.121 | 0.92577223 | NA | 0.5356 | 1.1159 | 0.8657 | 0.8657 | TRUE | 1.3204 |
| 1700011H14 | 1 | 0 | 0 | 0 | 0 | 0 | 1 | 0 | 0 | 0 | 0 | 0 | 0.18 | 0 | 0 | 0.49 | -1.029 | 0.80087 | NA | 0 | 18.6165 | 10 | 10 | TRUE | 0.4614 |
| 1700011I03R | 0 | 0 | 0 | 0 | 0 | 0 | 0 | 0 | 0 | 0 | 0 | 0 | 0 | 0 | 0 | NA | NA | NA | NA | NA | NA | NA | NA | NA | NA |
| 1700011J10F | 159 | 228 | 235 | 330 | 185 | 209 | 169 | 228 | 236 | 259 | 199 | 229 | 219.97 | 211 | 229 | 1.088 | 0.122 | 0.51694696 | 0.7086118 | 0.02 | 0.0219 | 0.0209 | 0.0209 | TRUE | 0.3269 |
| 1700011L22F | 0 | 0 | 0 | 1 | 0 | 0 | 0 | 0 | 0 | 1 | 0 | 0 | 0.13 | 0 | 0 | 1.859 | 0.894 | 0.82650734 | NA | 0 | 25.1515 | 10 | 10 | TRUE | 0.2563 |
| 1700011M02 | 0 | 0 | 0 | 1 | 0 | 0 | 0 | 0 | 0 | 1 | 0 | 0 | 0.13 | 0 | 0 | 1.859 | 0.894 | 0.82650734 | NA | 0 | 25.1515 | 10 | 10 | TRUE | 0.2563 |
| 1700012A03 | 0 | 0 | 0 | 0 | 0 | 0 | 0 | 0 | 0 | 0 | 0 | 0 | 0 | 0 | 0 | NA | NA | NA | NA | NA | NA | NA | NA | NA | NA |
| 1700012B07 | 16 | 10 | 8 | 32 | 21 | 12 | 17 | 10 | 8 | 25 | 23 | 13 | 15.98 | 12 | 20 | 1.756 | 0.813 | 0.13141213 | 0.31223744 | 0.059 | 0.213 | 0.1413 | 0.1413 | TRUE | 0.5889 |
| 1700012B09 | 0 | 1 | 0 | 0 | 1 | 0 | 0 | 1 | 0 | 0 | 1 | 0 | 0.35 | 0 | 0 | 0.954 | -0.067 | 0.98571955 | NA | 0 | 9.5346 | 8.2194 | 8.2194 | TRUE | 0.5204 |
| 1700012B15 | 284 | 251 | 207 | 338 | 239 | 186 | 302 | 251 | 208 | 266 | 257 | 204 | 247.77 | 254 | 242 | 0.958 | -0.062 | 0.74183566 | 0.86183206 | 0.0231 | 0.0202 | 0.0217 | 0.0217 | TRUE | 0.2215 |
| 1700012D01 | 7 | 6 | 10 | 7 | 7 | 15 | 7 | 6 | 10 | 6 | 8 | 16 | 8.82 | 8 | 10 | 1.237 | 0.307 | 0.66383409 | 0.81170547 | 0.1098 | 0.3804 | 0.2429 | 0.2429 | TRUE | 1.1116 |
| 1700012D14 | 14 | 26 | 20 | 18 | 24 | 33 | 15 | 26 | 20 | 14 | 26 | 36 | 22.84 | 20 | 25 | 1.233 | 0.303 | 0.5363091 | 0.72314843 | 0.1044 | 0.1512 | 0.128 | 0.128 | TRUE | 0.195 |
| 1700012I11R | 0 | 0 | 0 | 0 | 0 | 0 | 0 | 0 | 0 | 0 | 0 | 0 | 0 | 0 | 0 | NA | NA | NA | NA | NA | NA | NA | NA | NA | NA |
| 1700012L04F | 0 | 0 | 0 | 0 | 0 | 0 | 0 | 0 | 0 | 0 | 0 | 0 | 0 | 0 | 0 | NA | NA | NA | NA | NA | NA | NA | NA | NA | NA |
| 1700012P22I | 0 | 0 | 0 | 0 | 0 | 0 | 0 | 0 | 0 | 0 | 0 | 0 | 0 | 0 | 0 | NA | NA | NA | NA | NA | NA | NA | NA | NA | NA |
| 1700013D24 | 0 | 0 | 0 | 0 | 0 | 0 | 0 | 0 | 0 | 0 | 0 | 0 | 0 | 0 | 0 | NA | NA | NA | NA | NA | NA | NA | NA | NA | NA |
| 1700013F07I | 2 | 5 | 3 | 10 | 3 | 4 | 2 | 5 | 3 | 8 | 3 | 4 | 4.27 | 3 | 5 | 1.549 | 0.631 | 0.50137146 | NA | 0 | 0.7791 | 0.3906 | 0.3906 | TRUE | 0.5614 |
| 1700013G23 | 3 | 9 | 13 | 13 | 9 | 19 | 3 | 9 | 13 | 10 | 10 | 21 | 10.99 | 8 | 14 | 1.597 | 0.675 | 0.32850917 | 0.54950119 | 0.1783 | 0.3067 | 0.2472 | 0.2472 | TRUE | 0.451 |
| 1700013G24 | 1 | 2 | 3 | 3 | 3 | 2 | 1 | 2 | 3 | 2 | 3 | 2 | 2.31 | 2 | 2 | 1.271 | 0.346 | 0.77237314 | NA | 0 | 1.4344 | 0.5918 | 0.5918 | TRUE | 0.1659 |
| 1700013H16 | 70 | 94 | 68 | 1723 | 1288 | 943 | 74 | 94 | 68 | 1354 | 1386 | 1032 | 668.18 | 79 | 1257 | 15.947 | 3.995 | 3.24E-109 | 2.56E-105 | 0.022 | 0.0118 | 0.0165 | 0.0165 | TRUE | 0.2925 |
| 1700015E13I | 0 | 0 | 0 | 0 | 0 | 0 | 0 | 0 | 0 | 0 | 0 | 0 | 0 | 0 | 0 | NA | NA | NA | NA | NA | NA | NA | NA | NA | NA |
| 1700015F17I | 1 | 0 | 2 | 0 | 0 | 1 | 1 | 0 | 2 | 0 | 0 | 1 | 0.69 | 1 | 0 | 0.341 | -1.551 | 0.54644411 | NA | 0 | 4.7528 | 3.2643 | 3.2643 | TRUE | 0.3117 |
| 1700015G11 | 0 | 0 | 0 | 0 | 0 | 0 | 0 | 0 | 0 | 0 | 0 | 0 | 0 | 0 | 0 | NA | NA | NA | NA | NA | NA | NA | NA | NA | NA |
| 1700016C15I | 0 | 0 | 0 | 0 | 0 | 0 | 0 | 0 | 0 | 0 | 0 | 0 | 0 | 0 | 0 | NA | NA | NA | NA | NA | NA | NA | NA | NA | NA |
| 1700016D06 | 5 | 0 | 0 | 1 | 0 | 2 | 5 | 0 | 0 | 1 | 0 | 2 | 1.38 | 2 | 1 | 0.56 | -0.837 | 0.70933884 | NA | 3.7952 | 2.3924 | 2.8436 | 2.8436 | TRUE | 2.3447 |
| 1700016G22 | 2 | 0 | 0 | 3 | 1 | 2 | 2 | 0 | 0 | 2 | 1 | 2 | 1.29 | 1 | 2 | 2.716 | 1.442 | 0.44085401 | NA | 0 | 2.5581 | 1.5265 | 1.5265 | TRUE | 0.9745 |
| 1700016H13 | 21 | 12 | 9 | 30 | 17 | 20 | 22 | 12 | 9 | 24 | 18 | 22 | 17.85 | 14 | 21 | 1.482 | 0.567 | 0.25863194 | 0.47748367 | 0.0383 | 0.1915 | 0.1233 | 0.1233 | TRUE | 0.9698 |
| 1700016K19I | 80 | 60 | 76 | 129 | 117 | 111 | 85 | 60 | 76 | 101 | 126 | 122 | 95 | 74 | 116 | 1.573 | 0.654 | 0.00413114 | 0.02711456 | 0.0098 | 0.0416 | 0.0263 | 0.0263 | TRUE | 0.2472 |
| 1700016L04F | 2 | 2 | 2 | 1 | 0 | 1 | 2 | 2 | 2 | 1 | 0 | 1 | 1.34 | 2 | 1 | 0.311 | -1.685 | 0.31776282 | NA | 0 | 2.4744 | 1.0343 | 1.0343 | TRUE | 0.1918 |
| 1700016L21F | 0 | 0 | 0 | 0 | 0 | 0 | 0 | 0 | 0 | 0 | 0 | 0 | 0 | 0 | 0 | NA | NA | NA | NA | NA | NA | NA | NA | NA | NA |
| 1700016P04I | 0 | 0 | 0 | 0 | 0 | 0 | 0 | 0 | 0 | 0 | 0 | 0 | 0 | 0 | 0 | NA | NA | NA | NA | NA | NA | NA | NA | NA | NA |
| 1700017B05 | 2242 | 1749 | 1576 | 2633 | 1872 | 1655 | 2381 | 1747 | 1582 | 2070 | 2014 | 1812 | 1934.38 | 1903 | 1965 | 1.033 | 0.046 | 0.76529084 | 0.87700158 | 0.0251 | 0.0086 | 0.0169 | 0.0169 | TRUE | 0.5821 |
| 1700017D01 | 0 | 0 | 0 | 0 | 0 | 0 | 0 | 0 | 0 | 0 | 0 | 0 | 0 | 0 | 0 | NA | NA | NA | NA | NA | NA | NA | NA | NA | NA |
| 1700017G19 | 0 | 0 | 0 | 0 | 0 | 0 | 0 | 0 | 0 | 0 | 0 | 0 | 0 | 0 | 0 | NA | NA | NA | NA | NA | NA | NA | NA | NA | NA |
| 1700017J07F | 3 | 3 | 4 | 1 | 2 | 4 | 3 | 3 | 4 | 1 | 2 | 4 | 2.92 | 3 | 2 | 0.694 | -0.526 | 0.64001539 | NA | 0 | 1.1354 | 0.5561 | 0.5561 | TRUE | 0.4571 |
| 1700017N19 | 0 | 0 | 0 | 0 | 0 | 0 | 0 | 0 | 0 | 0 | 0 | 0 | 0 | 0 | 0 | NA | NA | NA | NA | NA | NA | NA | NA | NA | NA |
| 1700018A04 | 0 | 0 | 0 | 0 | 0 | 0 | 0 | 0 | 0 | 0 | 0 | 0 | 0 | 0 | 0 | NA | NA | NA | NA | NA | NA | NA | NA | NA | NA |
| 1700018B08 | 76 | 88 | 93 | 172 | 126 | 107 | 81 | 88 | 93 | 135 | 136 | 117 | 108.32 | 87 | 129 | 1.482 | 0.568 | 0.00350311 | 0.02386655 | 0 | 0.0373 | 0.0176 | 0.0176 | TRUE | 0.066 |
| 1700018B24 | 237 | 347 | 284 | 754 | 526 | 495 | 252 | 347 | 285 | 593 | 566 | 542 | 430.66 | 295 | 567 | 1.925 | 0.945 | 4.75E-11 | 3.71E-09 | 0.0105 | 0.0146 | 0.0123 | 0.0123 | TRUE | 0.2704 |

| | | | | | | | | | | | | | | | | | | | | | | | | | |
|---|---|---|---|---|---|---|---|---|---|---|---|---|---|---|---|---|---|---|---|---|---|---|---|---|---|
| 1700018C11I | 0 | 0 | 0 | 0 | 0 | 0 | 0 | 0 | 0 | 0 | 0 | 0 | 0 | 0 | 0 | NA | NA | NA | NA | NA | NA | NA | NA | NA | NA |
| 1700018F24I | 1 | 0 | 0 | 0 | 0 | 0 | 1 | 0 | 0 | 0 | 0 | 0 | 0.18 | 0 | 0 | 0.49 | -1.029 | 0.80087 | NA | 0 | 18.6165 | 10 | 10 | TRUE | 0.4614 |
| 1700018G05 | 0 | 0 | 0 | 1 | 0 | 0 | 0 | 0 | 0 | 1 | 0 | 0 | 0.13 | 0 | 0 | 1.859 | 0.894 | 0.82650734 | NA | 0 | 25.1515 | 10 | 10 | TRUE | 0.2563 |
| 1700018L02F | 1 | 1 | 1 | 6 | 1 | 1 | 1 | 1 | 1 | 5 | 1 | 1 | 1.66 | 1 | 2 | 2.323 | 1.216 | 0.4220366 | NA | 0 | 1.993 | 0.9474 | 0.9474 | TRUE | 1.1183 |
| 1700019A02I | 27 | 44 | 35 | 114 | 105 | 96 | 29 | 44 | 35 | 90 | 113 | 105 | 69.24 | 36 | 103 | 2.841 | 1.507 | 2.16E-08 | 9.51E-07 | 0.0075 | 0.0545 | 0.0332 | 0.0332 | TRUE | 0.2758 |
| 1700019B03I | 5 | 3 | 9 | 2 | 2 | 5 | 5 | 3 | 9 | 2 | 2 | 5 | 4.42 | 6 | 3 | 0.518 | -0.949 | 0.33386119 | NA | 3.00E-04 | 0.7517 | 0.4442 | 0.4442 | TRUE | 0.3337 |
| 1700019B21I | 81 | 95 | 70 | 202 | 144 | 153 | 86 | 95 | 70 | 159 | 155 | 167 | 122.07 | 84 | 160 | 1.915 | 0.938 | 1.41E-06 | 3.71E-05 | 0 | 0.0339 | 0.0181 | 0.0181 | TRUE | 0.1883 |
| 1700019D03I | 428 | 731 | 844 | 641 | 473 | 535 | 455 | 730 | 847 | 504 | 509 | 586 | 605.1 | 677 | 533 | 0.786 | -0.347 | 0.10115092 | 0.26335947 | 0.0512 | 0.0123 | 0.0307 | 0.0307 | TRUE | 0.5273 |
| 1700019E08I | 4 | 9 | 5 | 34 | 24 | 32 | 4 | 9 | 5 | 27 | 26 | 35 | 17.64 | 6 | 29 | 4.767 | 2.253 | 2.79E-05 | 0.00047345 | 0 | 0.1937 | 0.108 | 0.108 | TRUE | 0.4203 |
| 1700019G17 | 310 | 317 | 302 | 453 | 345 | 330 | 329 | 317 | 303 | 356 | 371 | 361 | 339.61 | 316 | 363 | 1.147 | 0.197 | 0.08751152 | 0.23954038 | 0 | 0.0166 | 0.0066 | 0.0066 | TRUE | 0.0149 |
| 1700019G24 | 2 | 2 | 2 | 7 | 2 | 5 | 2 | 2 | 2 | 6 | 2 | 5 | 3.21 | 2 | 4 | 2.16 | 1.111 | 0.29911793 | NA | 0 | 1.0333 | 0.4639 | 0.4639 | TRUE | 0.3255 |
| 1700019L03F | 1 | 0 | 3 | 3 | 1 | 2 | 1 | 0 | 3 | 2 | 1 | 2 | 1.62 | 1 | 2 | 1.39 | 0.475 | 0.7594719 | NA | 0 | 2.0451 | 1.1013 | 1.1013 | TRUE | 0.5616 |
| 1700019M22 | 0 | 0 | 0 | 0 | 0 | 0 | 0 | 0 | 0 | 0 | 0 | 0 | 0 | 0 | 0 | NA | NA | NA | NA | NA | NA | NA | NA | NA | NA |
| 1700019N19 | 28 | 40 | 43 | 41 | 23 | 23 | 30 | 40 | 43 | 32 | 25 | 25 | 32.5 | 38 | 27 | 0.732 | -0.45 | 0.19972279 | 0.40732488 | 0 | 0.1083 | 0.0574 | 0.0574 | TRUE | 0.2376 |
| 1700019O17 | 3 | 0 | 2 | 1 | 1 | 2 | 3 | 0 | 2 | 1 | 1 | 2 | 1.54 | 2 | 1 | 0.77 | -0.376 | 0.81420823 | NA | 0 | 2.1447 | 1.1819 | 1.1819 | TRUE | 0.3387 |
| 1700020A23I | 0 | 0 | 0 | 0 | 0 | 0 | 0 | 0 | 0 | 0 | 0 | 0 | 0 | 0 | 0 | NA | NA | NA | NA | NA | NA | NA | NA | NA | NA |
| 1700020D05I | 14 | 12 | 10 | 15 | 10 | 10 | 15 | 12 | 10 | 12 | 11 | 11 | 11.73 | 12 | 11 | 0.912 | -0.134 | 0.80492493 | 0.89931073 | 0 | 0.2877 | 0.1258 | 0.1258 | TRUE | 0.1279 |
| 1700020G17 | 1 | 1 | 0 | 0 | 0 | 0 | 1 | 1 | 0 | 0 | 0 | 0 | 0.34 | 1 | 0 | 0.261 | -1.936 | 0.60454921 | NA | 0 | 9.5983 | 8.3269 | 8.3269 | TRUE | 0.2507 |
| 1700020I14R | 769 | 1019 | 1079 | 1426 | 1134 | 996 | 817 | 1018 | 1083 | 1121 | 1220 | 1090 | 1058.22 | 973 | 1144 | 1.175 | 0.233 | 0.06635292 | 0.19873115 | 0.0105 | 0.01 | 0.0107 | 0.0107 | TRUE | 0.2333 |
| 1700020L24F | 3 | 2 | 2 | 6 | 4 | 2 | 3 | 2 | 2 | 5 | 4 | 2 | 3.07 | 2 | 4 | 1.58 | 0.66 | 0.53327504 | NA | 0 | 1.0811 | 0.4651 | 0.4651 | TRUE | 0.1848 |
| 1700020M21 | 0 | 2 | 1 | 0 | 1 | 1 | 0 | 2 | 1 | 0 | 1 | 1 | 0.86 | 1 | 1 | 0.688 | -0.539 | 0.8028361 | NA | 0 | 3.8287 | 2.1472 | 2.1472 | TRUE | 0.1651 |
| 1700020N01 | 55 | 32 | 25 | 97 | 78 | 59 | 58 | 32 | 25 | 76 | 84 | 65 | 56.71 | 38 | 75 | 1.953 | 0.965 | 0.00598626 | 0.03587208 | 0.0742 | 0.065 | 0.0691 | 0.0691 | TRUE | 1.4766 |
| 1700020N15 | 6 | 5 | 3 | 5 | 7 | 3 | 6 | 5 | 3 | 4 | 8 | 3 | 4.85 | 5 | 5 | 1.021 | 0.03 | 0.97267104 | 0.98548003 | 0 | 0.6856 | 0.3366 | 0.3366 | TRUE | 0.3948 |
| 1700020N18 | 0 | 2 | 1 | 1 | 1 | 0 | 0 | 2 | 1 | 1 | 1 | 0 | 0.81 | 1 | 1 | 0.625 | -0.678 | 0.7553135 | NA | 0 | 4.0711 | 2.1302 | 2.1302 | TRUE | 0.1304 |
| 1700021F05I | 1093 | 1300 | 1226 | 1459 | 990 | 1112 | 1161 | 1298 | 1231 | 1147 | 1065 | 1217 | 1186.62 | 1230 | 1143 | 0.929 | -0.106 | 0.26260353 | 0.48199498 | 0.0028 | 0.0097 | 0.0056 | 0.0056 | TRUE | 0.0415 |
| 1700021F07I | 6 | 8 | 5 | 5 | 1 | 3 | 6 | 8 | 5 | 4 | 1 | 3 | 4.61 | 6 | 3 | 0.437 | -1.195 | 0.19122779 | 0.3962054 | 0 | 0.7212 | 0.3509 | 0.3509 | TRUE | 0.3089 |
| 1700021K19I | 2177 | 2045 | 2179 | 2496 | 1747 | 1824 | 2312 | 2042 | 2188 | 1962 | 1880 | 1997 | 2063.52 | 2181 | 1946 | 0.893 | -0.164 | 0.04534302 | 0.15363055 | 0.0019 | 0.0085 | 0.0044 | 0.0044 | TRUE | 0.0374 |
| 1700021N21 | 1 | 0 | 0 | 0 | 0 | 0 | 1 | 0 | 0 | 0 | 0 | 0 | 0.18 | 0 | 0 | 0.49 | -1.029 | 0.80087 | NA | 0 | 18.6165 | 10 | 10 | TRUE | 0.4614 |
| 1700022A21I | 0 | 0 | 2 | 0 | 2 | 2 | 0 | 0 | 2 | 0 | 2 | 2 | 1.06 | 1 | 1 | 2.103 | 1.073 | 0.64149152 | NA | 0 | 3.1201 | 2.7207 | 2.7207 | TRUE | 0.9739 |
| 1700022A22I | 0 | 0 | 0 | 0 | 0 | 0 | 0 | 0 | 0 | 0 | 0 | 0 | 0 | 0 | 0 | NA | NA | NA | NA | NA | NA | NA | NA | NA | NA |
| 1700022E09I | 0 | 0 | 0 | 0 | 0 | 0 | 0 | 0 | 0 | 0 | 0 | 0 | 0 | 0 | 0 | NA | NA | NA | NA | NA | NA | NA | NA | NA | NA |
| 1700022H16I | 0 | 0 | 0 | 0 | 0 | 0 | 0 | 0 | 0 | 0 | 0 | 0 | 0 | 0 | 0 | NA | NA | NA | NA | NA | NA | NA | NA | NA | NA |
| 1700022I11R | 0 | 0 | 0 | 0 | 0 | 0 | 0 | 0 | 0 | 0 | 0 | 0 | 0 | 0 | 0 | NA | NA | NA | NA | NA | NA | NA | NA | NA | NA |
| 1700022P22I | 5 | 6 | 0 | 9 | 6 | 3 | 5 | 6 | 0 | 7 | 6 | 3 | 4.69 | 4 | 5 | 1.501 | 0.586 | 0.58056047 | 0.75708116 | 0.3578 | 0.7099 | 0.5866 | 0.5866 | TRUE | 1.2345 |
| 1700023A16I | 0 | 0 | 0 | 2 | 1 | 1 | 0 | 0 | 0 | 2 | 1 | 1 | 0.62 | 0 | 1 | 6.676 | 2.739 | 0.27800695 | NA | 0 | 5.2881 | 3.2034 | 3.2034 | TRUE | 0.07 |
| 1700023C21I | 0 | 0 | 0 | 0 | 0 | 0 | 0 | 0 | 0 | 0 | 0 | 0 | 0 | 0 | 0 | NA | NA | NA | NA | NA | NA | NA | NA | NA | NA |
| 1700023E05I | 13 | 5 | 2 | 3 | 4 | 2 | 14 | 5 | 2 | 2 | 4 | 2 | 4.94 | 7 | 3 | 0.424 | -1.238 | 0.22231491 | 0.43444058 | 0.3465 | 0.6733 | 0.5009 | 0.5009 | TRUE | 0.5289 |
| 1700023F02I | 0 | 0 | 0 | 0 | 0 | 0 | 0 | 0 | 0 | 0 | 0 | 0 | 0 | 0 | 0 | NA | NA | NA | NA | NA | NA | NA | NA | NA | NA |
| 1700023F06I | 73 | 71 | 96 | 112 | 88 | 82 | 78 | 71 | 96 | 88 | 95 | 90 | 86.22 | 82 | 91 | 1.112 | 0.153 | 0.49212996 | 0.6883982 | 0 | 0.0451 | 0.0239 | 0.0239 | TRUE | 0.2372 |
| 1700023L04F | 9 | 14 | 13 | 13 | 9 | 11 | 10 | 14 | 13 | 10 | 10 | 12 | 11.42 | 12 | 11 | 0.87 | -0.201 | 0.71438024 | 0.8451249 | 0 | 0.2953 | 0.1306 | 0.1306 | TRUE | 0.1357 |
| 1700024B18I | 0 | 0 | 0 | 0 | 0 | 0 | 0 | 0 | 0 | 0 | 0 | 0 | 0 | 0 | 0 | NA | NA | NA | NA | NA | NA | NA | NA | NA | NA |
| 1700024F13I | 4 | 1 | 0 | 2 | 3 | 0 | 4 | 1 | 0 | 2 | 3 | 0 | 1.67 | 2 | 2 | 0.922 | -0.116 | 0.94771317 | NA | 1.2105 | 1.9742 | 1.674 | 1.674 | TRUE | 0.4241 |
| 1700024G13 | 2 | 0 | 0 | 0 | 0 | 0 | 2 | 0 | 0 | 0 | 0 | 0 | 0.35 | 1 | 0 | 0.254 | -1.975 | 0.62446185 | NA | 10 | 9.3117 | 10 | 10 | TRUE | 0.9741 |
| 1700024P03I | 0 | 0 | 0 | 0 | 0 | 0 | 0 | 0 | 0 | 0 | 0 | 0 | 0 | 0 | 0 | NA | NA | NA | NA | NA | NA | NA | NA | NA | NA |
| 1700024P04I | 0 | 0 | 0 | 0 | 0 | 0 | 0 | 0 | 0 | 0 | 0 | 0 | 0 | 0 | 0 | NA | NA | NA | NA | NA | NA | NA | NA | NA | NA |
| 1700024P16I | 24 | 10 | 9 | 10 | 10 | 8 | 25 | 10 | 9 | 8 | 11 | 9 | 11.98 | 15 | 9 | 0.614 | -0.703 | 0.27016539 | 0.4903393 | 0.1318 | 0.2818 | 0.2046 | 0.2046 | TRUE | 1.7233 |
| 1700025B11I | 0 | 1 | 0 | 1 | 1 | 1 | 0 | 1 | 0 | 1 | 1 | 1 | 0.66 | 0 | 1 | 2.676 | 1.42 | 0.53620179 | NA | 0 | 5.0043 | 2.2992 | 2.2992 | TRUE | 0.405 |
| 1700025C18I | 0 | 0 | 0 | 0 | 0 | 0 | 0 | 0 | 0 | 0 | 0 | 0 | 0 | 0 | 0 | NA | NA | NA | NA | NA | NA | NA | NA | NA | NA |
| 1700025E21I | 5 | 6 | 5 | 5 | 2 | 5 | 5 | 6 | 5 | 4 | 2 | 5 | 4.65 | 5 | 4 | 0.708 | -0.499 | 0.56353553 | 0.74445785 | 0 | 0.7159 | 0.3171 | 0.3171 | TRUE | 0.2095 |
| 1700025F22I | 0 | 0 | 0 | 0 | 0 | 0 | 0 | 0 | 0 | 0 | 0 | 0 | 0 | 0 | 0 | NA | NA | NA | NA | NA | NA | NA | NA | NA | NA |
| 1700025F24I | 2 | 1 | 0 | 11 | 12 | 5 | 2 | 1 | 0 | 9 | 13 | 5 | 5.03 | 1 | 9 | 8.773 | 3.133 | 0.00673222 | 0.03902025 | 0 | 0.6624 | 0.4117 | 0.4117 | TRUE | 0.2459 |
| 1700025G04 | 589 | 587 | 574 | 1000 | 758 | 707 | 626 | 586 | 576 | 786 | 816 | 774 | 693.97 | 596 | 792 | 1.329 | 0.41 | 9.86E-06 | 0.00019646 | 0 | 0.0117 | 0.0047 | 0.0047 | TRUE | 0.0218 |
| 1700025K24I | 0 | 0 | 0 | 0 | 0 | 0 | 0 | 0 | 0 | 0 | 0 | 0 | 0 | 0 | 0 | NA | NA | NA | NA | NA | NA | NA | NA | NA | NA |
| 1700025M24 | 0 | 0 | 0 | 0 | 0 | 0 | 0 | 0 | 0 | 0 | 0 | 0 | 0 | 0 | 0 | NA | NA | NA | NA | NA | NA | NA | NA | NA | NA |
| 1700025N23 | 0 | 0 | 0 | 0 | 0 | 0 | 0 | 0 | 0 | 0 | 0 | 0 | 0 | 0 | 0 | NA | NA | NA | NA | NA | NA | NA | NA | NA | NA |
| 1700026D08I | 8 | 6 | 12 | 8 | 4 | 8 | 8 | 6 | 12 | 6 | 4 | 9 | 7.65 | 9 | 6 | 0.728 | -0.459 | 0.51501108 | 0.70743379 | 0 | 0.4377 | 0.2239 | 0.2239 | TRUE | 0.2466 |
| 1700026D11I | 0 | 0 | 0 | 0 | 0 | 0 | 0 | 0 | 0 | 0 | 0 | 0 | 0 | 0 | 0 | NA | NA | NA | NA | NA | NA | NA | NA | NA | NA |
| 1700026F02I | 0 | 0 | 0 | 0 | 0 | 0 | 0 | 0 | 0 | 0 | 0 | 0 | 0 | 0 | 0 | NA | NA | NA | NA | NA | NA | NA | NA | NA | NA |
| 1700026L06F | 7 | 12 | 13 | 18 | 29 | 20 | 7 | 12 | 13 | 14 | 31 | 22 | 16.62 | 11 | 22 | 2.047 | 1.034 | 0.05403959 | 0.17281409 | 0.0714 | 0.2051 | 0.1383 | 0.1383 | TRUE | 0.136 |
| 1700027A15I | 0 | 0 | 0 | 0 | 0 | 0 | 0 | 0 | 0 | 0 | 0 | 0 | 0 | 0 | 0 | NA | NA | NA | NA | NA | NA | NA | NA | NA | NA |
| 1700027F09I | 0 | 0 | 0 | 0 | 0 | 0 | 0 | 0 | 0 | 0 | 0 | 0 | 0 | 0 | 0 | NA | NA | NA | NA | NA | NA | NA | NA | NA | NA |
| 1700027H10I | 0 | 0 | 0 | 0 | 0 | 0 | 0 | 0 | 0 | 0 | 0 | 0 | 0 | 0 | 0 | NA | NA | NA | NA | NA | NA | NA | NA | NA | NA |
| 1700027I24R | 1 | 1 | 2 | 1 | 1 | 0 | 1 | 1 | 2 | 1 | 1 | 0 | 0.99 | 1 | 1 | 0.465 | -1.104 | 0.55512019 | NA | 0 | 3.3396 | 1.3765 | 1.3765 | TRUE | 0.1869 |
| 1700027J07F | 1 | 0 | 0 | 0 | 0 | 0 | 1 | 0 | 0 | 0 | 0 | 0 | 0.18 | 0 | 0 | 0.49 | -1.029 | 0.80087 | NA | 0 | 18.6165 | 10 | 10 | TRUE | 0.4614 |
| 1700028B04I | 56 | 53 | 54 | 50 | 48 | 42 | 59 | 53 | 54 | 39 | 52 | 46 | 50.59 | 55 | 46 | 0.818 | -0.291 | 0.28870893 | 0.5097411 | 0 | 0.072 | 0.0341 | 0.0341 | TRUE | 0.1382 |
| 1700028D13I | 0 | 0 | 0 | 0 | 1 | 1 | 0 | 0 | 0 | 0 | 1 | 1 | 0.36 | 0 | 1 | 3.794 | 1.924 | 0.60247382 | NA | 0 | 9.1135 | 8.1425 | 8.1425 | TRUE | 0.3482 |
| 1700028E10I | 6 | 24 | 10 | 22 | 14 | 13 | 6 | 24 | 10 | 17 | 15 | 14 | 14.5 | 13 | 15 | 1.153 | 0.206 | 0.73282009 | 0.8565538 | 0.1474 | 0.2342 | 0.1928 | 0.1928 | TRUE | 1.682 |
| 1700028I16R | 69 | 85 | 68 | 119 | 85 | 58 | 73 | 85 | 68 | 94 | 91 | 63 | 79.16 | 75 | 83 | 1.101 | 0.139 | 0.5780001 | 0.75534159 | 0.0153 | 0.0485 | 0.0325 | 0.0325 | TRUE | 0.3711 |
| 1700028J19F | 13 | 36 | 28 | 36 | 31 | 25 | 14 | 36 | 28 | 28 | 33 | 27 | 27.82 | 26 | 29 | 1.139 | 0.187 | 0.66355227 | 0.81147138 | 0.0644 | 0.1253 | 0.0977 | 0.0977 | TRUE | 0.4844 |

| | | | | | | | | | | | | | | | | | | | | | | | | | |
|---|---|---|---|---|---|---|---|---|---|---|---|---|---|---|---|---|---|---|---|---|---|---|---|---|---|
| 1700028K03I | 20 | 33 | 40 | 33 | 41 | 26 | 21 | 33 | 40 | 26 | 44 | 28 | 32.15 | 31 | 33 | 1.037 | 0.052 | 0.89717326 | 0.95016592 | 0.059 | 0.1094 | 0.0852 | 0.0852 | TRUE | 0.4201 |
| 1700028M03 | 0 | 0 | 0 | 0 | 0 | 0 | 0 | 0 | 0 | 0 | 0 | 0 | 0 | 0 | 0 | NA | NA | NA | NA | NA | NA | NA | NA | NA | NA |
| 1700028P14I | 2 | 0 | 0 | 2 | 1 | 1 | 2 | 0 | 0 | 2 | 1 | 1 | 0.98 | 1 | 1 | 1.807 | 0.853 | 0.67681949 | NA | 0 | 3.3759 | 1.8982 | 1.8982 | TRUE | 0.9744 |
| 1700028P15I | 1 | 0 | 1 | 1 | 3 | 2 | 1 | 0 | 1 | 1 | 3 | 2 | 1.38 | 1 | 2 | 2.966 | 1.569 | 0.36347209 | NA | 0 | 2.3972 | 1.1594 | 1.1594 | TRUE | 0.2032 |
| 1700029B22I | 0 | 0 | 0 | 0 | 0 | 0 | 0 | 0 | 0 | 0 | 0 | 0 | 0 | 0 | 0 | NA | NA | NA | NA | NA | NA | NA | NA | NA | NA |
| 1700029F12I | 0 | 0 | 0 | 0 | 0 | 0 | 0 | 0 | 0 | 0 | 0 | 0 | 0 | 0 | 0 | NA | NA | NA | NA | NA | NA | NA | NA | NA | NA |
| 1700029H14I | 2 | 1 | 2 | 0 | 0 | 1 | 2 | 1 | 2 | 0 | 0 | 1 | 1.04 | 2 | 0 | 0.204 | -2.29 | 0.25432785 | NA | 0 | 3.1818 | 1.6101 | 1.6101 | TRUE | 0.5383 |
| 1700029I15R | 3 | 1 | 2 | 6 | 3 | 1 | 3 | 1 | 2 | 5 | 3 | 1 | 2.54 | 2 | 3 | 1.496 | 0.581 | 0.63129667 | NA | 0 | 1.3046 | 0.6498 | 0.6498 | TRUE | 0.218 |
| 1700029J03F | 0 | 0 | 0 | 0 | 0 | 0 | 0 | 0 | 0 | 0 | 0 | 0 | 0 | 0 | 0 | NA | NA | NA | NA | NA | NA | NA | NA | NA | NA |
| 1700029J07F | 38 | 65 | 81 | 46 | 32 | 56 | 40 | 65 | 81 | 36 | 34 | 61 | 53.08 | 62 | 44 | 0.703 | -0.509 | 0.1707782 | 0.36937928 | 0.0921 | 0.069 | 0.08 | 0.08 | TRUE | 0.2845 |
| 1700029M20 | 109 | 73 | 81 | 45 | 37 | 35 | 116 | 73 | 81 | 35 | 40 | 38 | 63.92 | 90 | 38 | 0.42 | -1.252 | 1.38E-05 | 0.00026151 | 0.025 | 0.0584 | 0.0412 | 0.0412 | TRUE | 0.5898 |
| 1700029N11 | 0 | 0 | 0 | 0 | 0 | 0 | 0 | 0 | 0 | 0 | 0 | 0 | 0 | 0 | 0 | NA | NA | NA | NA | NA | NA | NA | NA | NA | NA |
| 1700029P11I | 224 | 202 | 186 | 478 | 401 | 362 | 238 | 202 | 187 | 376 | 431 | 396 | 304.99 | 209 | 401 | 1.921 | 0.942 | 9.16E-11 | 6.75E-09 | 0.0055 | 0.0177 | 0.0115 | 0.0115 | TRUE | 0.16 |
| 1700030A11I | 1 | 0 | 0 | 1 | 0 | 0 | 1 | 0 | 0 | 1 | 0 | 0 | 0.31 | 0 | 0 | 0.954 | -0.067 | 0.98625928 | NA | 0 | 10.7014 | 9.0384 | 9.0384 | TRUE | 0.4614 |
| 1700030C10I | 203 | 134 | 154 | 254 | 220 | 168 | 216 | 134 | 155 | 200 | 237 | 184 | 187.39 | 168 | 207 | 1.231 | 0.3 | 0.16585245 | 0.36290577 | 0.0325 | 0.0245 | 0.0284 | 0.0284 | TRUE | 0.6463 |
| 1700030F04I | 0 | 0 | 0 | 0 | 0 | 0 | 0 | 0 | 0 | 0 | 0 | 0 | 0 | 0 | 0 | NA | NA | NA | NA | NA | NA | NA | NA | NA | NA |
| 1700030F18I | 0 | 1 | 0 | 0 | 1 | 0 | 0 | 1 | 0 | 0 | 1 | 0 | 0.35 | 0 | 0 | 0.954 | -0.067 | 0.98571955 | NA | 0 | 9.5346 | 8.2194 | 8.2194 | TRUE | 0.5204 |
| 1700030J22F | 326 | 308 | 322 | 509 | 442 | 364 | 346 | 308 | 323 | 400 | 476 | 398 | 375.22 | 326 | 425 | 1.303 | 0.382 | 0.00353196 | 0.02398037 | 0.0046 | 0.0157 | 0.0096 | 0.0096 | TRUE | 0.1242 |
| 1700030K09I | 498 | 635 | 664 | 892 | 596 | 666 | 529 | 634 | 667 | 701 | 641 | 729 | 650.23 | 610 | 690 | 1.132 | 0.179 | 0.15010057 | 0.34072916 | 0.0075 | 0.012 | 0.0096 | 0.0096 | TRUE | 0.1573 |
| 1700030L20F | 7 | 2 | 0 | 11 | 6 | 3 | 7 | 2 | 0 | 9 | 6 | 3 | 4.64 | 3 | 6 | 1.982 | 0.987 | 0.3839213 | 0.59804511 | 0.6293 | 0.7174 | 0.6833 | 0.6833 | TRUE | 1.2072 |
| 1700030M09 | 2 | 10 | 3 | 8 | 7 | 5 | 2 | 10 | 3 | 6 | 8 | 5 | 5.74 | 5 | 6 | 1.27 | 0.344 | 0.69211422 | 0.83110773 | 0.1303 | 0.5812 | 0.3678 | 0.3678 | TRUE | 1.5181 |
| 1700030N03 | 0 | 0 | 0 | 0 | 0 | 0 | 0 | 0 | 0 | 0 | 0 | 0 | 0 | 0 | 0 | NA | NA | NA | NA | NA | NA | NA | NA | NA | NA |
| 1700030O20 | 8 | 10 | 14 | 4 | 1 | 3 | 8 | 10 | 14 | 3 | 1 | 3 | 6.67 | 11 | 2 | 0.234 | -2.096 | 0.01150068 | 0.05841826 | 0 | 0.5005 | 0.2574 | 0.2574 | TRUE | 0.2479 |
| 1700031A10I | 1 | 0 | 1 | 2 | 1 | 0 | 1 | 0 | 1 | 2 | 1 | 0 | 0.79 | 1 | 1 | 1.325 | 0.406 | 0.85251519 | NA | 0 | 4.1997 | 2.1499 | 2.1499 | TRUE | 0.2789 |
| 1700031F05I | 1 | 0 | 2 | 1 | 0 | 2 | 1 | 0 | 2 | 1 | 0 | 2 | 1.01 | 1 | 1 | 0.961 | -0.058 | 0.97727667 | NA | 0 | 3.2764 | 1.9788 | 1.9788 | TRUE | 0.273 |
| 1700031M16 | 0 | 0 | 0 | 0 | 0 | 0 | 0 | 0 | 0 | 0 | 0 | 0 | 0 | 0 | 0 | NA | NA | NA | NA | NA | NA | NA | NA | NA | NA |
| 1700031P21I | 0 | 0 | 0 | 0 | 0 | 0 | 0 | 0 | 0 | 0 | 0 | 0 | 0 | 0 | 0 | NA | NA | NA | NA | NA | NA | NA | NA | NA | NA |
| 1700034E13I | 0 | 0 | 0 | 0 | 0 | 0 | 0 | 0 | 0 | 0 | 0 | 0 | 0 | 0 | 0 | NA | NA | NA | NA | NA | NA | NA | NA | NA | NA |
| 1700034F02I | 20 | 28 | 32 | 30 | 18 | 16 | 21 | 28 | 32 | 24 | 19 | 18 | 23.63 | 27 | 20 | 0.748 | -0.419 | 0.29802887 | 0.51872997 | 0 | 0.1463 | 0.0742 | 0.0742 | TRUE | 0.2185 |
| 1700034G24 | 0 | 0 | 0 | 0 | 0 | 0 | 0 | 0 | 0 | 0 | 0 | 0 | 0 | 0 | 0 | NA | NA | NA | NA | NA | NA | NA | NA | NA | NA |
| 1700034H15I | 16 | 21 | 19 | 31 | 25 | 21 | 17 | 21 | 19 | 24 | 27 | 23 | 21.88 | 19 | 25 | 1.3 | 0.378 | 0.34293811 | 0.56306682 | 0 | 0.1574 | 0.0681 | 0.0681 | TRUE | 0.0434 |
| 1700034I23R | 0 | 0 | 0 | 0 | 0 | 0 | 0 | 0 | 0 | 0 | 0 | 0 | 0 | 0 | 0 | NA | NA | NA | NA | NA | NA | NA | NA | NA | NA |
| 1700034J05F | 0 | 0 | 0 | 0 | 0 | 0 | 0 | 0 | 0 | 0 | 0 | 0 | 0 | 0 | 0 | NA | NA | NA | NA | NA | NA | NA | NA | NA | NA |
| 1700034K08I | 0 | 0 | 0 | 0 | 0 | 0 | 0 | 0 | 0 | 0 | 0 | 0 | 0 | 0 | 0 | NA | NA | NA | NA | NA | NA | NA | NA | NA | NA |
| 1700034O15 | 0 | 0 | 0 | 0 | 0 | 0 | 0 | 0 | 0 | 0 | 0 | 0 | 0 | 0 | 0 | NA | NA | NA | NA | NA | NA | NA | NA | NA | NA |
| 1700034P13I | 3 | 7 | 8 | 19 | 12 | 5 | 3 | 7 | 8 | 15 | 13 | 5 | 8.59 | 6 | 11 | 1.841 | 0.88 | 0.22460101 | 0.43701051 | 0 | 0.3905 | 0.2513 | 0.2513 | TRUE | 0.6737 |
| 1700036G14 | 0 | 0 | 0 | 0 | 0 | 0 | 0 | 0 | 0 | 0 | 0 | 0 | 0 | 0 | 0 | NA | NA | NA | NA | NA | NA | NA | NA | NA | NA |
| 1700037C18I | 20 | 34 | 38 | 55 | 41 | 27 | 21 | 34 | 38 | 43 | 44 | 30 | 35.04 | 31 | 39 | 1.254 | 0.327 | 0.37673294 | 0.59206639 | 0.0322 | 0.1009 | 0.0694 | 0.0694 | TRUE | 0.5013 |
| 1700037H04I | 5630 | 5567 | 5326 | 6182 | 4633 | 4672 | 5980 | 5560 | 5348 | 4860 | 4985 | 5114 | 5307.77 | 5629 | 4986 | 0.886 | -0.175 | 0.01734238 | 0.07832875 | 0.0017 | 0.0075 | 0.0037 | 0.0037 | TRUE | 0.0362 |
| 1700039E15I | 0 | 0 | 0 | 0 | 0 | 0 | 0 | 0 | 0 | 0 | 0 | 0 | 0 | 0 | 0 | NA | NA | NA | NA | NA | NA | NA | NA | NA | NA |
| 1700039E22I | 0 | 0 | 0 | 0 | 0 | 0 | 0 | 0 | 0 | 0 | 0 | 0 | 0 | 0 | 0 | NA | NA | NA | NA | NA | NA | NA | NA | NA | NA |
| 1700040L02F | 4 | 2 | 8 | 1 | 2 | 6 | 4 | 2 | 8 | 1 | 2 | 7 | 3.96 | 5 | 3 | 0.647 | -0.628 | 0.5716731 | NA | 0.4114 | 0.838 | 0.6237 | 0.6237 | TRUE | 0.585 |
| 1700040N02 | 2 | 2 | 0 | 2 | 1 | 2 | 2 | 2 | 0 | 2 | 1 | 2 | 1.49 | 1 | 2 | 1.176 | 0.234 | 0.88082424 | NA | 0 | 2.2132 | 1.0809 | 1.0809 | TRUE | 0.4915 |
| 1700041C23I | 0 | 0 | 0 | 0 | 0 | 0 | 0 | 0 | 0 | 0 | 0 | 0 | 0 | 0 | 0 | NA | NA | NA | NA | NA | NA | NA | NA | NA | NA |
| 1700041M19 | 0 | 0 | 0 | 0 | 0 | 0 | 0 | 0 | 0 | 0 | 0 | 0 | 0 | 0 | 0 | NA | NA | NA | NA | NA | NA | NA | NA | NA | NA |
| 1700042B14I | 0 | 0 | 0 | 0 | 0 | 0 | 0 | 0 | 0 | 0 | 0 | 0 | 0 | 0 | 0 | NA | NA | NA | NA | NA | NA | NA | NA | NA | NA |
| 1700042G07 | 0 | 1 | 0 | 0 | 1 | 1 | 0 | 1 | 0 | 0 | 1 | 1 | 0.53 | 0 | 1 | 1.924 | 0.944 | 0.7364225 | NA | 0 | 6.2438 | 3.9727 | 3.9727 | TRUE | 0.405 |
| 1700042G15 | 9 | 5 | 7 | 7 | 7 | 4 | 10 | 5 | 7 | 6 | 8 | 4 | 6.5 | 7 | 6 | 0.808 | -0.308 | 0.6774125 | 0.82064429 | 0 | 0.5138 | 0.2406 | 0.2406 | TRUE | 0.2069 |
| 1700042O10 | 50 | 27 | 19 | 58 | 43 | 35 | 53 | 27 | 19 | 46 | 46 | 38 | 38.22 | 33 | 43 | 1.318 | 0.398 | 0.34128345 | 0.56159399 | 0.1068 | 0.0931 | 0.0995 | 0.0995 | TRUE | 1.4803 |
| 1700044C05I | 0 | 1 | 1 | 3 | 3 | 3 | 0 | 1 | 1 | 2 | 3 | 3 | 1.81 | 1 | 3 | 4.346 | 2.12 | 0.17160889 | NA | 0 | 1.8249 | 0.8109 | 0.8109 | TRUE | 0.2212 |
| 1700044K03I | 0 | 0 | 0 | 0 | 0 | 0 | 0 | 0 | 0 | 0 | 0 | 0 | 0 | 0 | 0 | NA | NA | NA | NA | NA | NA | NA | NA | NA | NA |
| 1700045H11I | 0 | 0 | 0 | 0 | 0 | 0 | 0 | 0 | 0 | 0 | 0 | 0 | 0 | 0 | 0 | NA | NA | NA | NA | NA | NA | NA | NA | NA | NA |
| 1700045I19R | 1 | 1 | 0 | 1 | 1 | 2 | 1 | 1 | 0 | 1 | 1 | 2 | 1.02 | 1 | 1 | 1.944 | 0.959 | 0.60783444 | NA | 0 | 3.2408 | 1.4013 | 1.4013 | TRUE | 0.2529 |
| 1700046C09I | 0 | 0 | 0 | 0 | 0 | 0 | 0 | 0 | 0 | 0 | 0 | 0 | 0 | 0 | 0 | NA | NA | NA | NA | NA | NA | NA | NA | NA | NA |
| 1700047E10I | 0 | 0 | 0 | 0 | 0 | 0 | 0 | 0 | 0 | 0 | 0 | 0 | 0 | 0 | 0 | NA | NA | NA | NA | NA | NA | NA | NA | NA | NA |
| 1700047G03 | 0 | 1 | 0 | 0 | 1 | 0 | 0 | 1 | 0 | 0 | 1 | 0 | 0.35 | 0 | 0 | 0.954 | -0.067 | 0.98571955 | NA | 0 | 9.5346 | 8.2194 | 8.2194 | TRUE | 0.5204 |
| 1700047I17R | 0 | 0 | 0 | 0 | 0 | 0 | 0 | 0 | 0 | 0 | 0 | 0 | 0 | 0 | 0 | NA | NA | NA | NA | NA | NA | NA | NA | NA | NA |
| 1700047L14F | 0 | 0 | 0 | 0 | 0 | 0 | 0 | 0 | 0 | 0 | 0 | 0 | 0 | 0 | 0 | NA | NA | NA | NA | NA | NA | NA | NA | NA | NA |
| 1700047M11 | 1 | 0 | 0 | 1 | 1 | 2 | 1 | 0 | 0 | 1 | 1 | 2 | 0.85 | 0 | 1 | 3.644 | 1.866 | 0.38437313 | NA | 0 | 3.8725 | 1.9476 | 1.9476 | TRUE | 0.4614 |
| 1700048M11 | 0 | 0 | 0 | 0 | 0 | 0 | 0 | 0 | 0 | 0 | 0 | 0 | 0 | 0 | 0 | NA | NA | NA | NA | NA | NA | NA | NA | NA | NA |
| 1700048O20 | 107 | 127 | 94 | 125 | 100 | 93 | 114 | 127 | 94 | 98 | 108 | 102 | 107.09 | 112 | 103 | 0.918 | -0.124 | 0.54218856 | 0.72790245 | 0 | 0.0377 | 0.0206 | 0.0206 | TRUE | 0.1844 |
| 1700049E15I | 0 | 0 | 0 | 0 | 0 | 0 | 0 | 0 | 0 | 0 | 0 | 0 | 0 | 0 | 0 | NA | NA | NA | NA | NA | NA | NA | NA | NA | NA |
| 1700049E22I | 28 | 57 | 40 | 60 | 44 | 39 | 30 | 57 | 40 | 47 | 47 | 43 | 44.01 | 42 | 46 | 1.081 | 0.112 | 0.73520235 | 0.85798087 | 0.0279 | 0.0818 | 0.0564 | 0.0564 | TRUE | 0.3901 |
| 1700049G17 | 214 | 213 | 206 | 255 | 171 | 182 | 227 | 213 | 207 | 200 | 184 | 199 | 205.09 | 216 | 194 | 0.903 | -0.146 | 0.30679107 | 0.52732223 | 0 | 0.023 | 0.0099 | 0.0099 | TRUE | 0.0242 |
| 1700049L16F | 0 | 0 | 0 | 0 | 0 | 0 | 0 | 0 | 0 | 0 | 0 | 0 | 0 | 0 | 0 | NA | NA | NA | NA | NA | NA | NA | NA | NA | NA |
| 1700051A21I | 1 | 2 | 0 | 3 | 0 | 1 | 1 | 2 | 0 | 2 | 0 | 1 | 1.09 | 1 | 1 | 1.165 | 0.22 | 0.91107747 | NA | 0 | 3.042 | 1.8904 | 1.8904 | TRUE | 0.2272 |
| 1700052I22R | 0 | 0 | 0 | 0 | 0 | 0 | 0 | 0 | 0 | 0 | 0 | 0 | 0 | 0 | 0 | NA | NA | NA | NA | NA | NA | NA | NA | NA | NA |
| 1700052K11I | 49 | 52 | 40 | 166 | 114 | 51 | 52 | 52 | 40 | 130 | 123 | 56 | 75.52 | 48 | 103 | 2.151 | 1.105 | 0.00169706 | 0.01351141 | 0.099 | 0.0505 | 0.074 | 0.074 | TRUE | 1.5273 |
| 1700052N19 | 2575 | 2923 | 2991 | 2846 | 2126 | 2378 | 2735 | 2919 | 3003 | 2237 | 2287 | 2603 | 2630.91 | 2886 | 2376 | 0.823 | -0.281 | 0.00218713 | 0.01652082 | 0.004 | 0.0082 | 0.0057 | 0.0057 | TRUE | 0.0843 |

| | | | | | | | | | | | | | | | | | | | | | | | | | |
|---|---|---|---|---|---|---|---|---|---|---|---|---|---|---|---|---|---|---|---|---|---|---|---|---|---|
| 1700054A03 | 0 | 1 | 0 | 1 | 0 | 0 | 0 | 1 | 0 | 1 | 0 | 0 | 0.3 | 0 | 0 | 0.954 | -0.067 | 0.98642001 | NA | 0 | 11.0817 | 9.3014 | 9.3014 | TRUE | 0.405 |
| 1700054K19I | 0 | 0 | 0 | 1 | 0 | 0 | 0 | 0 | 0 | 1 | 0 | 0 | 0.13 | 0 | 0 | 1.859 | 0.894 | 0.82650734 | NA | 0 | 25.1515 | 10 | 10 | TRUE | 0.2563 |
| 1700054M17 | 0 | 0 | 0 | 0 | 0 | 0 | 0 | 0 | 0 | 0 | 0 | 0 | 0 | 0 | 0 | NA | NA | NA | NA | NA | NA | NA | NA | NA | NA |
| 1700054O13 | 0 | 0 | 0 | 1 | 0 | 0 | 0 | 0 | 0 | 1 | 0 | 0 | 0.13 | 0 | 0 | 1.859 | 0.894 | 0.82650734 | NA | 0 | 25.1515 | 10 | 10 | TRUE | 0.2563 |
| 1700055C04I | 0 | 0 | 0 | 1 | 2 | 0 | 0 | 0 | 0 | 1 | 2 | 0 | 0.49 | 0 | 1 | 5.204 | 2.38 | 0.47267317 | NA | 0 | 6.7348 | 6.4099 | 6.4099 | TRUE | 0.1222 |
| 1700055N04 | 1 | 1 | 2 | 0 | 2 | 2 | 1 | 1 | 2 | 0 | 2 | 2 | 1.4 | 1 | 1 | 1.025 | 0.035 | 0.98301921 | NA | 0 | 2.3572 | 1.2378 | 1.2378 | TRUE | 0.7215 |
| 1700056E22I | 21 | 23 | 30 | 38 | 34 | 21 | 22 | 23 | 30 | 30 | 37 | 23 | 27.47 | 25 | 30 | 1.186 | 0.246 | 0.51991113 | 0.71100654 | 0 | 0.1268 | 0.0687 | 0.0687 | TRUE | 0.1502 |
| 1700057G04 | 0 | 0 | 0 | 1 | 0 | 0 | 0 | 0 | 0 | 1 | 0 | 0 | 0.13 | 0 | 0 | 1.859 | 0.894 | 0.82650734 | NA | 0 | 25.1515 | 10 | 10 | TRUE | 0.2563 |
| 1700057H15I | 0 | 0 | 0 | 0 | 0 | 1 | 0 | 0 | 0 | 0 | 0 | 1 | 0.18 | 0 | 0 | 1.859 | 0.894 | 0.82650734 | NA | 0 | 18.0643 | 10 | 10 | TRUE | 0.5383 |
| 1700058G18 | 0 | 0 | 0 | 0 | 0 | 0 | 0 | 0 | 0 | 0 | 0 | 0 | 0 | 0 | 0 | NA | NA | NA | NA | NA | NA | NA | NA | NA | NA |
| 1700060C16I | 0 | 1 | 0 | 0 | 0 | 0 | 0 | 1 | 0 | 0 | 0 | 0 | 0.17 | 0 | 0 | 0.49 | -1.029 | 0.80087 | NA | 0 | 19.7991 | 10 | 10 | TRUE | 0.405 |
| 1700060C20I | 3 | 1 | 1 | 0 | 1 | 0 | 3 | 1 | 1 | 0 | 1 | 0 | 1.04 | 2 | 0 | 0.203 | -2.302 | 0.2619563 | NA | 0 | 3.1619 | 1.7394 | 1.7394 | TRUE | 0.5204 |
| 1700061F12I | 0 | 0 | 0 | 0 | 0 | 0 | 0 | 0 | 0 | 0 | 0 | 0 | 0 | 0 | 0 | NA | NA | NA | NA | NA | NA | NA | NA | NA | NA |
| 1700061G19 | 390 | 279 | 263 | 494 | 389 | 369 | 414 | 279 | 264 | 388 | 419 | 404 | 361.3 | 319 | 404 | 1.266 | 0.34 | 0.06795309 | 0.20177545 | 0.0285 | 0.016 | 0.0222 | 0.0222 | TRUE | 0.769 |
| 1700061I17R | 0 | 0 | 0 | 0 | 1 | 0 | 0 | 0 | 0 | 0 | 1 | 0 | 0.18 | 0 | 0 | 1.859 | 0.894 | 0.82650734 | NA | 0 | 18.3785 | 10 | 10 | TRUE | 0.5204 |
| 1700063A18I | 0 | 0 | 0 | 0 | 0 | 0 | 0 | 0 | 0 | 0 | 0 | 0 | 0 | 0 | 0 | NA | NA | NA | NA | NA | NA | NA | NA | NA | NA |
| 1700063D05I | 24 | 24 | 14 | 22 | 12 | 16 | 25 | 24 | 14 | 17 | 13 | 18 | 18.54 | 21 | 16 | 0.755 | -0.406 | 0.38389409 | 0.59804511 | 0 | 0.1846 | 0.1022 | 0.1022 | TRUE | 0.4872 |
| 1700063O14 | 0 | 0 | 0 | 0 | 0 | 0 | 0 | 0 | 0 | 0 | 0 | 0 | 0 | 0 | 0 | NA | NA | NA | NA | NA | NA | NA | NA | NA | NA |
| 1700064J06F | 0 | 0 | 0 | 0 | 0 | 0 | 0 | 0 | 0 | 0 | 0 | 0 | 0 | 0 | 0 | NA | NA | NA | NA | NA | NA | NA | NA | NA | NA |
| 1700064M15 | 0 | 3 | 1 | 0 | 0 | 2 | 0 | 3 | 1 | 0 | 0 | 2 | 1.03 | 1 | 1 | 0.525 | -0.93 | 0.69249688 | NA | 2.276 | 3.2005 | 2.9093 | 2.9093 | TRUE | 0.6178 |
| 1700065D16I | 0 | 1 | 2 | 1 | 2 | 1 | 0 | 1 | 2 | 1 | 2 | 1 | 1.17 | 1 | 1 | 1.311 | 0.39 | 0.82325362 | NA | 0 | 2.8148 | 1.3246 | 1.3246 | TRUE | 0.2291 |
| 1700065I16R | 0 | 0 | 0 | 0 | 0 | 0 | 0 | 0 | 0 | 0 | 0 | 0 | 0 | 0 | 0 | NA | NA | NA | NA | NA | NA | NA | NA | NA | NA |
| 1700065I17R | 0 | 0 | 0 | 0 | 0 | 1 | 0 | 0 | 0 | 0 | 0 | 1 | 0.18 | 0 | 0 | 1.859 | 0.894 | 0.82650734 | NA | 0 | 18.0643 | 10 | 10 | TRUE | 0.5383 |
| 1700065J11F | 0 | 1 | 1 | 0 | 0 | 0 | 0 | 1 | 1 | 0 | 0 | 0 | 0.33 | 1 | 0 | 0.268 | -1.901 | 0.61401143 | NA | 0 | 9.8764 | 8.4772 | 8.4772 | TRUE | 0.2278 |
| 1700065J18F | 0 | 0 | 0 | 0 | 0 | 0 | 0 | 0 | 0 | 0 | 0 | 0 | 0 | 0 | 0 | NA | NA | NA | NA | NA | NA | NA | NA | NA | NA |
| 1700065O20 | 0 | 0 | 0 | 0 | 0 | 0 | 0 | 0 | 0 | 0 | 0 | 0 | 0 | 0 | 0 | NA | NA | NA | NA | NA | NA | NA | NA | NA | NA |
| 1700066B17 | 0 | 1 | 1 | 9 | 2 | 2 | 0 | 1 | 1 | 7 | 2 | 2 | 2.24 | 1 | 4 | 5.745 | 2.522 | 0.10315105 | NA | 0 | 1.4799 | 0.8447 | 0.8447 | TRUE | 1.149 |
| 1700066B19 | 11 | 9 | 8 | 12 | 4 | 4 | 12 | 9 | 8 | 9 | 4 | 4 | 7.8 | 10 | 6 | 0.648 | -0.627 | 0.3723805 | 0.58896764 | 0 | 0.4291 | 0.2235 | 0.2235 | TRUE | 0.7352 |
| 1700066J24F | 0 | 0 | 0 | 0 | 0 | 0 | 0 | 0 | 0 | 0 | 0 | 0 | 0 | 0 | 0 | NA | NA | NA | NA | NA | NA | NA | NA | NA | NA |
| 1700066M21 | 378 | 522 | 539 | 625 | 499 | 528 | 402 | 521 | 541 | 491 | 537 | 578 | 511.71 | 488 | 535 | 1.095 | 0.132 | 0.37051857 | 0.58743656 | 0.0138 | 0.0133 | 0.0136 | 0.0136 | TRUE | 0.2777 |
| 1700066N21 | 0 | 0 | 0 | 0 | 0 | 0 | 0 | 0 | 0 | 0 | 0 | 0 | 0 | 0 | 0 | NA | NA | NA | NA | NA | NA | NA | NA | NA | NA |
| 1700066O22 | 0 | 0 | 0 | 0 | 0 | 0 | 0 | 0 | 0 | 0 | 0 | 0 | 0 | 0 | 0 | NA | NA | NA | NA | NA | NA | NA | NA | NA | NA |
| 1700067G17 | 0 | 0 | 0 | 0 | 2 | 0 | 0 | 0 | 0 | 0 | 2 | 0 | 0.36 | 0 | 1 | 3.775 | 1.916 | 0.63423067 | NA | 10 | 9.1927 | 10 | 10 | TRUE | 0.9743 |
| 1700067K01I | 60 | 99 | 100 | 78 | 63 | 106 | 64 | 99 | 100 | 61 | 68 | 116 | 84.69 | 88 | 82 | 0.927 | -0.109 | 0.73284899 | 0.8565538 | 0.0795 | 0.0458 | 0.062 | 0.062 | TRUE | 1.2297 |
| 1700067P10I | 0 | 0 | 0 | 0 | 0 | 0 | 0 | 0 | 0 | 0 | 0 | 0 | 0 | 0 | 0 | NA | NA | NA | NA | NA | NA | NA | NA | NA | NA |
| 1700069L16F | 2 | 0 | 3 | 2 | 3 | 2 | 2 | 0 | 3 | 2 | 3 | 2 | 2.02 | 2 | 2 | 1.35 | 0.433 | 0.75638819 | NA | 0 | 1.637 | 0.8901 | 0.8901 | TRUE | 0.6111 |
| 1700069P05I | 0 | 0 | 0 | 0 | 0 | 0 | 0 | 0 | 0 | 0 | 0 | 0 | 0 | 0 | 0 | NA | NA | NA | NA | NA | NA | NA | NA | NA | NA |
| 1700071K01I | 0 | 0 | 0 | 0 | 0 | 0 | 0 | 0 | 0 | 0 | 0 | 0 | 0 | 0 | 0 | NA | NA | NA | NA | NA | NA | NA | NA | NA | NA |
| 1700071M16 | 0 | 0 | 0 | 0 | 0 | 0 | 0 | 0 | 0 | 0 | 0 | 0 | 0 | 0 | 0 | NA | NA | NA | NA | NA | NA | NA | NA | NA | NA |
| 1700072B07 | 0 | 0 | 0 | 0 | 0 | 0 | 0 | 0 | 0 | 0 | 0 | 0 | 0 | 0 | 0 | NA | NA | NA | NA | NA | NA | NA | NA | NA | NA |
| 1700072O05 | 0 | 0 | 0 | 0 | 0 | 0 | 0 | 0 | 0 | 0 | 0 | 0 | 0 | 0 | 0 | NA | NA | NA | NA | NA | NA | NA | NA | NA | NA |
| 1700073E17I | 0 | 0 | 0 | 0 | 0 | 0 | 0 | 0 | 0 | 0 | 0 | 0 | 0 | 0 | 0 | NA | NA | NA | NA | NA | NA | NA | NA | NA | NA |
| 1700074H08 | 0 | 0 | 0 | 0 | 0 | 0 | 0 | 0 | 0 | 0 | 0 | 0 | 0 | 0 | 0 | NA | NA | NA | NA | NA | NA | NA | NA | NA | NA |
| 1700074P13I | 0 | 0 | 0 | 0 | 0 | 0 | 0 | 0 | 0 | 0 | 0 | 0 | 0 | 0 | 0 | NA | NA | NA | NA | NA | NA | NA | NA | NA | NA |
| 1700080E11I | 0 | 0 | 0 | 0 | 1 | 0 | 0 | 0 | 0 | 0 | 1 | 0 | 0.18 | 0 | 0 | 1.859 | 0.894 | 0.82650734 | NA | 0 | 18.3785 | 10 | 10 | TRUE | 0.5204 |
| 1700080N15 | 0 | 0 | 0 | 0 | 0 | 0 | 0 | 0 | 0 | 0 | 0 | 0 | 0 | 0 | 0 | NA | NA | NA | NA | NA | NA | NA | NA | NA | NA |
| 1700080O16 | 3 | 2 | 3 | 62 | 52 | 29 | 3 | 2 | 3 | 49 | 56 | 32 | 24.1 | 3 | 46 | 16.72 | 4.063 | 2.57E-10 | 1.80E-08 | 0 | 0.1436 | 0.0995 | 0.0995 | TRUE | 0.2005 |
| 1700081H04 | 0 | 0 | 0 | 0 | 0 | 0 | 0 | 0 | 0 | 0 | 0 | 0 | 0 | 0 | 0 | NA | NA | NA | NA | NA | NA | NA | NA | NA | NA |
| 1700081H22 | 0 | 0 | 0 | 0 | 0 | 0 | 0 | 0 | 0 | 0 | 0 | 0 | 0 | 0 | 0 | NA | NA | NA | NA | NA | NA | NA | NA | NA | NA |
| 1700084C01I | 3 | 1 | 1 | 7 | 6 | 2 | 3 | 1 | 1 | 6 | 6 | 2 | 3.22 | 2 | 5 | 2.765 | 1.467 | 0.20009531 | NA | 0 | 1.0291 | 0.5442 | 0.5442 | TRUE | 0.402 |
| 1700084E18I | 9 | 10 | 16 | 10 | 10 | 10 | 10 | 10 | 16 | 8 | 11 | 11 | 10.86 | 12 | 10 | 0.822 | -0.283 | 0.62785124 | 0.79061634 | 0 | 0.3102 | 0.1518 | 0.1518 | TRUE | 0.3773 |
| 1700084F23I | 0 | 0 | 0 | 0 | 0 | 0 | 0 | 0 | 0 | 0 | 0 | 0 | 0 | 0 | 0 | NA | NA | NA | NA | NA | NA | NA | NA | NA | NA |
| 1700084J12F | 1 | 0 | 0 | 3 | 0 | 0 | 1 | 0 | 0 | 2 | 0 | 0 | 0.57 | 0 | 1 | 2.223 | 1.153 | 0.72182708 | NA | 0 | 5.7857 | 5.9519 | 5.9519 | TRUE | 1.4404 |
| 1700085B03 | 0 | 0 | 0 | 0 | 0 | 0 | 0 | 0 | 0 | 0 | 0 | 0 | 0 | 0 | 0 | NA | NA | NA | NA | NA | NA | NA | NA | NA | NA |
| 1700085C21I | 0 | 0 | 0 | 0 | 0 | 0 | 0 | 0 | 0 | 0 | 0 | 0 | 0 | 0 | 0 | NA | NA | NA | NA | NA | NA | NA | NA | NA | NA |
| 1700086L19F | 1 | 0 | 0 | 3 | 7 | 6 | 1 | 0 | 0 | 2 | 8 | 7 | 2.92 | 0 | 6 | 14.82 | 3.889 | 0.01450535 | NA | 0 | 1.1351 | 0.734 | 0.734 | TRUE | 0.7024 |
| 1700086O06 | 27 | 29 | 27 | 30 | 24 | 24 | 29 | 29 | 27 | 24 | 26 | 26 | 26.74 | 28 | 25 | 0.89 | -0.167 | 0.63936508 | 0.79678145 | 0 | 0.1301 | 0.0545 | 0.0545 | TRUE | 0.0244 |
| 1700088E04I | 9 | 8 | 11 | 15 | 11 | 8 | 10 | 8 | 11 | 12 | 12 | 9 | 10.16 | 10 | 11 | 1.137 | 0.186 | 0.74973161 | 0.86736352 | 0 | 0.3311 | 0.1454 | 0.1454 | TRUE | 0.0898 |
| 1700091E21I | 0 | 0 | 0 | 0 | 0 | 1 | 0 | 0 | 0 | 0 | 0 | 1 | 0.18 | 0 | 0 | 1.859 | 0.894 | 0.82650734 | NA | 0 | 18.0643 | 10 | 10 | TRUE | 0.5383 |
| 1700091H14 | 0 | 0 | 0 | 1 | 0 | 1 | 0 | 0 | 0 | 1 | 0 | 1 | 0.31 | 0 | 1 | 3.348 | 1.743 | 0.65064889 | NA | 0 | 10.5167 | 8.8968 | 8.8968 | TRUE | 0.2082 |
| 1700092C02I | 0 | 0 | 0 | 0 | 0 | 0 | 0 | 0 | 0 | 0 | 0 | 0 | 0 | 0 | 0 | NA | NA | NA | NA | NA | NA | NA | NA | NA | NA |
| 1700092C10I | 0 | 1 | 0 | 0 | 1 | 0 | 0 | 1 | 0 | 0 | 1 | 0 | 0.35 | 0 | 0 | 0.954 | -0.067 | 0.98571955 | NA | 0 | 9.5346 | 8.2194 | 8.2194 | TRUE | 0.5204 |
| 1700092E19I | 0 | 0 | 0 | 0 | 0 | 0 | 0 | 0 | 0 | 0 | 0 | 0 | 0 | 0 | 0 | NA | NA | NA | NA | NA | NA | NA | NA | NA | NA |
| 1700092K14I | 0 | 0 | 0 | 0 | 0 | 0 | 0 | 0 | 0 | 0 | 0 | 0 | 0 | 0 | 0 | NA | NA | NA | NA | NA | NA | NA | NA | NA | NA |
| 1700092M07 | 8 | 2 | 4 | 12 | 3 | 7 | 8 | 2 | 4 | 9 | 3 | 8 | 5.81 | 5 | 7 | 1.423 | 0.509 | 0.56305492 | 0.74413459 | 0.1671 | 0.5743 | 0.381 | 0.381 | TRUE | 0.7252 |
| 1700093K21I | 2 | 0 | 2 | 3 | 1 | 1 | 2 | 0 | 2 | 2 | 1 | 1 | 1.44 | 1 | 1 | 1.124 | 0.169 | 0.91580851 | NA | 0 | 2.289 | 1.152 | 1.152 | TRUE | 0.4966 |
| 1700094D03 | 164 | 154 | 130 | 160 | 135 | 128 | 174 | 154 | 131 | 126 | 145 | 140 | 144.95 | 153 | 137 | 0.895 | -0.16 | 0.38983675 | 0.6037429 | 0.0066 | 0.0296 | 0.018 | 0.018 | TRUE | 0.1717 |
| 1700094J05F | 0 | 0 | 1 | 0 | 0 | 0 | 0 | 0 | 1 | 0 | 0 | 0 | 0.17 | 0 | 0 | 0.49 | -1.029 | 0.80087 | NA | 0 | 19.6928 | 10 | 10 | TRUE | 0.4098 |
| 1700094M24 | 0 | 0 | 0 | 0 | 0 | 0 | 0 | 0 | 0 | 0 | 0 | 0 | 0 | 0 | 0 | NA | NA | NA | NA | NA | NA | NA | NA | NA | NA |

| | | | | | | | | | | | | | | | | | | | | | | | | | |
|---|---|---|---|---|---|---|---|---|---|---|---|---|---|---|---|---|---|---|---|---|---|---|---|---|---|
| 1700095A21 | 0 | 0 | 0 | 0 | 0 | 0 | 0 | 0 | 0 | 0 | 0 | 0 | 0 | 0 | 0 | NA | NA | NA | NA | NA | NA | NA | NA | NA | NA |
| 1700095B10 | 5 | 3 | 1 | 3 | 2 | 1 | 5 | 3 | 1 | 2 | 2 | 1 | 2.49 | 3 | 2 | 0.613 | -0.707 | 0.56417711 | NA | 0 | 1.3321 | 0.6607 | 0.6607 | TRUE | 0.1508 |
| 1700095B22 | 0 | 0 | 0 | 1 | 0 | 0 | 0 | 0 | 0 | 1 | 0 | 0 | 0.13 | 0 | 0 | 1.859 | 0.894 | 0.82650734 | NA | 0 | 25.1515 | 10 | 10 | TRUE | 0.2563 |
| 1700096J18F | 0 | 0 | 0 | 0 | 0 | 0 | 0 | 0 | 0 | 0 | 0 | 0 | 0 | 0 | 0 | NA | NA | NA | NA | NA | NA | NA | NA | NA | NA |
| 1700096K18 | 28 | 36 | 51 | 70 | 37 | 43 | 30 | 36 | 51 | 55 | 40 | 47 | 43.14 | 39 | 47 | 1.218 | 0.284 | 0.39326203 | 0.60672295 | 0.0268 | 0.0833 | 0.0564 | 0.0564 | TRUE | 0.5626 |
| 1700097N02 | 356 | 433 | 439 | 139 | 86 | 121 | 378 | 432 | 441 | 109 | 93 | 132 | 264.27 | 417 | 111 | 0.267 | -1.907 | 1.13E-29 | 6.15E-27 | 0.009 | 0.0194 | 0.0149 | 0.0149 | TRUE | 0.2498 |
| 1700100L14F | 0 | 1 | 0 | 0 | 0 | 0 | 0 | 1 | 0 | 0 | 0 | 0 | 0.17 | 0 | 0 | 0.49 | -1.029 | 0.80087 | NA | 0 | 19.7991 | 10 | 10 | TRUE | 0.405 |
| 1700101E01 | 21 | 40 | 39 | 79 | 61 | 52 | 22 | 40 | 39 | 62 | 66 | 57 | 47.68 | 34 | 62 | 1.817 | 0.861 | 0.00640631 | 0.03778622 | 0.0078 | 0.076 | 0.0489 | 0.0489 | TRUE | 0.5961 |
| 1700101I11R | 3 | 0 | 1 | 0 | 1 | 0 | 3 | 0 | 1 | 0 | 1 | 0 | 0.88 | 1 | 0 | 0.251 | -1.993 | 0.41455294 | NA | 0 | 3.7601 | 2.9308 | 2.9308 | TRUE | 0.3522 |
| 1700101O22 | 0 | 0 | 0 | 0 | 0 | 0 | 0 | 0 | 0 | 0 | 0 | 0 | 0 | 0 | 0 | NA | NA | NA | NA | NA | NA | NA | NA | NA | NA |
| 1700102H20 | 6 | 11 | 2 | 3 | 2 | 4 | 6 | 11 | 2 | 2 | 2 | 4 | 4.71 | 6 | 3 | 0.454 | -1.139 | 0.24765141 | 0.46433167 | 0 | 0.7065 | 0.4537 | 0.4537 | TRUE | 0.11 |
| 1700102P08 | 58 | 56 | 56 | 65 | 38 | 43 | 62 | 56 | 56 | 51 | 41 | 47 | 52.14 | 58 | 46 | 0.804 | -0.315 | 0.236979 | 0.45233363 | 0 | 0.0701 | 0.0318 | 0.0318 | TRUE | 0.0812 |
| 1700104A03 | 0 | 0 | 0 | 0 | 0 | 0 | 0 | 0 | 0 | 0 | 0 | 0 | 0 | 0 | 0 | NA | NA | NA | NA | NA | NA | NA | NA | NA | NA |
| 1700105P06 | 1 | 2 | 6 | 1 | 7 | 2 | 1 | 2 | 6 | 1 | 8 | 2 | 3.27 | 3 | 4 | 1.13 | 0.176 | 0.88762556 | NA | 0.6182 | 1.0159 | 0.8055 | 0.8055 | TRUE | 1.325 |
| 1700106J16F | 9 | 6 | 6 | 13 | 10 | 5 | 10 | 6 | 6 | 10 | 11 | 5 | 8 | 7 | 9 | 1.237 | 0.307 | 0.6524649 | 0.80502827 | 0 | 0.4185 | 0.2085 | 0.2085 | TRUE | 0.3114 |
| 1700108F19 | 6 | 20 | 11 | 31 | 20 | 21 | 6 | 20 | 11 | 24 | 22 | 23 | 17.71 | 12 | 23 | 1.838 | 0.878 | 0.09162162 | 0.24703325 | 0.0451 | 0.1929 | 0.133 | 0.133 | TRUE | 0.8205 |
| 1700108J01F | 0 | 0 | 0 | 0 | 0 | 0 | 0 | 0 | 0 | 0 | 0 | 0 | 0 | 0 | 0 | NA | NA | NA | NA | NA | NA | NA | NA | NA | NA |
| 1700109F18 | 0 | 0 | 0 | 0 | 0 | 0 | 0 | 0 | 0 | 0 | 0 | 0 | 0 | 0 | 0 | NA | NA | NA | NA | NA | NA | NA | NA | NA | NA |
| 1700109G14 | 4 | 0 | 1 | 1 | 1 | 1 | 4 | 0 | 1 | 1 | 1 | 1 | 1.37 | 2 | 1 | 0.563 | -0.83 | 0.63468466 | NA | 0 | 2.4147 | 1.4041 | 1.4041 | TRUE | 0.7833 |
| 1700109G15 | 1 | 1 | 0 | 1 | 1 | 1 | 1 | 1 | 0 | 1 | 1 | 1 | 0.84 | 1 | 1 | 1.429 | 0.515 | 0.79427198 | NA | 0 | 3.9464 | 1.567 | 1.567 | TRUE | 0.2527 |
| 1700109H08 | 19 | 16 | 14 | 16 | 11 | 20 | 20 | 16 | 14 | 13 | 12 | 22 | 16.09 | 17 | 16 | 0.916 | -0.126 | 0.80131338 | 0.89741637 | 0 | 0.2117 | 0.1181 | 0.1181 | TRUE | 0.5716 |
| 1700109I08R | 0 | 0 | 0 | 0 | 0 | 0 | 0 | 0 | 0 | 0 | 0 | 0 | 0 | 0 | 0 | NA | NA | NA | NA | NA | NA | NA | NA | NA | NA |
| 1700110C19 | 0 | 1 | 2 | 4 | 0 | 1 | 0 | 1 | 2 | 3 | 0 | 1 | 1.21 | 1 | 1 | 1.447 | 0.533 | 0.78222137 | NA | 0 | 2.7348 | 1.8427 | 1.8427 | TRUE | 0.4018 |
| 1700110I01R | 0 | 0 | 0 | 0 | 0 | 0 | 0 | 0 | 0 | 0 | 0 | 0 | 0 | 0 | 0 | NA | NA | NA | NA | NA | NA | NA | NA | NA | NA |
| 1700110K17 | 2 | 1 | 0 | 0 | 0 | 0 | 2 | 1 | 0 | 0 | 0 | 0 | 0.52 | 1 | 0 | 0.173 | -2.534 | 0.43568356 | NA | 0 | 6.3362 | 6.1226 | 6.1226 | TRUE | 0.0614 |
| 1700110M21 | 0 | 0 | 0 | 0 | 0 | 0 | 0 | 0 | 0 | 0 | 0 | 0 | 0 | 0 | 0 | NA | NA | NA | NA | NA | NA | NA | NA | NA | NA |
| 1700111N16 | 0 | 0 | 0 | 0 | 0 | 0 | 0 | 0 | 0 | 0 | 0 | 0 | 0 | 0 | 0 | NA | NA | NA | NA | NA | NA | NA | NA | NA | NA |
| 1700112E06 | 1 | 1 | 2 | 4 | 1 | 3 | 1 | 1 | 2 | 3 | 1 | 3 | 1.93 | 1 | 2 | 1.856 | 0.893 | 0.50784717 | NA | 0 | 1.7148 | 0.7395 | 0.7395 | TRUE | 0.2473 |
| 1700112H15 | 0 | 0 | 1 | 4 | 2 | 1 | 0 | 0 | 1 | 3 | 2 | 1 | 1.23 | 0 | 2 | 5.894 | 2.559 | 0.18276267 | NA | 0 | 2.6798 | 1.432 | 1.432 | TRUE | 0.3504 |
| 1700112J05F | 0 | 0 | 0 | 0 | 0 | 0 | 0 | 0 | 0 | 0 | 0 | 0 | 0 | 0 | 0 | NA | NA | NA | NA | NA | NA | NA | NA | NA | NA |
| 1700113A16 | 26 | 38 | 35 | 53 | 28 | 27 | 28 | 38 | 35 | 42 | 30 | 30 | 33.68 | 34 | 34 | 1.013 | 0.019 | 0.95611498 | 0.97741618 | 0 | 0.1047 | 0.0573 | 0.0573 | TRUE | 0.3407 |
| 1700113H08 | 0 | 0 | 0 | 0 | 0 | 0 | 0 | 0 | 0 | 0 | 0 | 0 | 0 | 0 | 0 | NA | NA | NA | NA | NA | NA | NA | NA | NA | NA |
| 1700119H24 | 0 | 0 | 0 | 0 | 0 | 0 | 0 | 0 | 0 | 0 | 0 | 0 | 0 | 0 | 0 | NA | NA | NA | NA | NA | NA | NA | NA | NA | NA |
| 1700120C14 | 1 | 2 | 1 | 1 | 0 | 0 | 1 | 2 | 1 | 1 | 0 | 0 | 0.81 | 1 | 0 | 0.257 | -1.958 | 0.36424053 | NA | 0 | 4.0827 | 1.9787 | 1.9787 | TRUE | 0.2562 |
| 1700120E14 | 0 | 0 | 0 | 0 | 0 | 0 | 0 | 0 | 0 | 0 | 0 | 0 | 0 | 0 | 0 | NA | NA | NA | NA | NA | NA | NA | NA | NA | NA |
| 1700120G07 | 0 | 0 | 0 | 0 | 0 | 0 | 0 | 0 | 0 | 0 | 0 | 0 | 0 | 0 | 0 | NA | NA | NA | NA | NA | NA | NA | NA | NA | NA |
| 1700120K04 | 0 | 0 | 0 | 0 | 0 | 0 | 0 | 0 | 0 | 0 | 0 | 0 | 0 | 0 | 0 | NA | NA | NA | NA | NA | NA | NA | NA | NA | NA |
| 1700121L16F | 0 | 0 | 0 | 0 | 0 | 0 | 0 | 0 | 0 | 0 | 0 | 0 | 0 | 0 | 0 | NA | NA | NA | NA | NA | NA | NA | NA | NA | NA |
| 1700121N20 | 7 | 2 | 4 | 2 | 0 | 1 | 7 | 2 | 4 | 2 | 0 | 1 | 2.69 | 4 | 1 | 0.208 | -2.263 | 0.09873361 | NA | 0 | 1.2334 | 0.7172 | 0.7172 | TRUE | 0.1719 |
| 1700122O11 | 1 | 1 | 0 | 1 | 2 | 0 | 1 | 1 | 0 | 1 | 2 | 0 | 0.83 | 1 | 1 | 1.425 | 0.511 | 0.81397997 | NA | 0 | 3.9611 | 2.1459 | 2.1459 | TRUE | 0.2989 |
| 1700123I01R | 25 | 25 | 33 | 55 | 38 | 43 | 27 | 25 | 33 | 43 | 41 | 47 | 35.98 | 28 | 44 | 1.548 | 0.631 | 0.05053872 | 0.16534682 | 0 | 0.0985 | 0.0457 | 0.0457 | TRUE | 0.1521 |
| 1700123K08 | 0 | 0 | 0 | 0 | 0 | 0 | 0 | 0 | 0 | 0 | 0 | 0 | 0 | 0 | 0 | NA | NA | NA | NA | NA | NA | NA | NA | NA | NA |
| 1700123L14F | 0 | 0 | 0 | 0 | 0 | 0 | 0 | 0 | 0 | 0 | 0 | 0 | 0 | 0 | 0 | NA | NA | NA | NA | NA | NA | NA | NA | NA | NA |
| 1700123M08 | 25 | 27 | 29 | 24 | 28 | 23 | 27 | 27 | 29 | 19 | 30 | 25 | 26.13 | 28 | 25 | 0.889 | -0.17 | 0.65678113 | 0.80670515 | 0 | 0.133 | 0.0665 | 0.0665 | TRUE | 0.3215 |
| 1700123O12 | 0 | 0 | 0 | 0 | 0 | 0 | 0 | 0 | 0 | 0 | 0 | 0 | 0 | 0 | 0 | NA | NA | NA | NA | NA | NA | NA | NA | NA | NA |
| 1700123O20 | 986 | 826 | 724 | 1395 | 946 | 794 | 1047 | 825 | 727 | 1097 | 1018 | 869 | 930.47 | 866 | 995 | 1.149 | 0.2 | 0.20482244 | 0.41330025 | 0.0231 | 0.0104 | 0.0169 | 0.0169 | TRUE | 0.3957 |
| 1700123O21 | 0 | 0 | 0 | 0 | 0 | 0 | 0 | 0 | 0 | 0 | 0 | 0 | 0 | 0 | 0 | NA | NA | NA | NA | NA | NA | NA | NA | NA | NA |
| 1700125G02 | 0 | 0 | 0 | 0 | 0 | 0 | 0 | 0 | 0 | 0 | 0 | 0 | 0 | 0 | 0 | NA | NA | NA | NA | NA | NA | NA | NA | NA | NA |
| 1700125G22 | 2 | 2 | 2 | 3 | 4 | 1 | 2 | 2 | 2 | 2 | 4 | 1 | 2.31 | 2 | 2 | 1.263 | 0.337 | 0.78121645 | NA | 0 | 1.4303 | 0.6216 | 0.6216 | TRUE | 0.2621 |
| 1700125H03 | 3 | 0 | 1 | 2 | 0 | 1 | 3 | 0 | 1 | 2 | 0 | 1 | 1.14 | 1 | 1 | 0.659 | -0.601 | 0.75967187 | NA | 0 | 2.8894 | 1.8829 | 1.8829 | TRUE | 0.4216 |
| 1700125H20 | 6 | 27 | 21 | 21 | 8 | 17 | 6 | 27 | 21 | 17 | 9 | 19 | 16.36 | 18 | 15 | 0.803 | -0.316 | 0.61270296 | 0.78070375 | 0.2303 | 0.2083 | 0.2186 | 0.2186 | TRUE | 0.5972 |
| 1700126H18 | 0 | 0 | 0 | 0 | 0 | 0 | 0 | 0 | 0 | 0 | 0 | 0 | 0 | 0 | 0 | NA | NA | NA | NA | NA | NA | NA | NA | NA | NA |
| 1700128A07 | 0 | 0 | 0 | 2 | 0 | 0 | 0 | 0 | 0 | 2 | 0 | 0 | 0.26 | 0 | 1 | 2.886 | 1.529 | 0.70676491 | NA | 0 | 12.5792 | 10 | 10 | TRUE | 0.9669 |
| 1700128F08 | 0 | 0 | 0 | 0 | 0 | 1 | 0 | 0 | 0 | 0 | 0 | 1 | 0.18 | 0 | 0 | 1.859 | 0.894 | 0.82650734 | NA | 0 | 18.0643 | 10 | 10 | TRUE | 0.5383 |
| 1700129C05 | 0 | 0 | 0 | 0 | 0 | 0 | 0 | 0 | 0 | 0 | 0 | 0 | 0 | 0 | 0 | NA | NA | NA | NA | NA | NA | NA | NA | NA | NA |
| 1810006J02F | 0 | 0 | 0 | 0 | 0 | 0 | 0 | 0 | 0 | 0 | 0 | 0 | 0 | 0 | 0 | NA | NA | NA | NA | NA | NA | NA | NA | NA | NA |
| 1810007C17 | 0 | 0 | 0 | 0 | 0 | 0 | 0 | 0 | 0 | 0 | 0 | 0 | 0 | 0 | 0 | NA | NA | NA | NA | NA | NA | NA | NA | NA | NA |
| 1810007D17 | 2 | 0 | 0 | 0 | 1 | 1 | 2 | 0 | 0 | 0 | 1 | 1 | 0.72 | 1 | 1 | 1.003 | 0.004 | 0.99874028 | NA | 0 | 4.6092 | 3.4914 | 3.4914 | TRUE | 0.9743 |
| 1810007I06R | 2 | 1 | 2 | 1 | 1 | 2 | 2 | 1 | 2 | 1 | 1 | 2 | 1.53 | 2 | 1 | 0.774 | -0.369 | 0.80016733 | NA | 0 | 2.1595 | 0.8657 | 0.8657 | TRUE | 0.1675 |
| 1810008I18R | 0 | 0 | 0 | 0 | 0 | 0 | 0 | 0 | 0 | 0 | 0 | 0 | 0 | 0 | 0 | NA | NA | NA | NA | NA | NA | NA | NA | NA | NA |
| 1810009A15 | 1487 | 1673 | 1601 | 2978 | 1959 | 1695 | 1579 | 1671 | 1608 | 2341 | 2108 | 1855 | 1860.36 | 1619 | 2101 | 1.299 | 0.377 | 0.00037893 | 0.00415028 | 0.0063 | 0.0087 | 0.0076 | 0.0076 | TRUE | 0.1265 |
| 1810009J06F | 0 | 0 | 1 | 0 | 0 | 0 | 0 | 0 | 1 | 0 | 0 | 0 | 0.17 | 0 | 0 | 0.49 | -1.029 | 0.80087 | NA | 0 | 19.6928 | 10 | 10 | TRUE | 0.4098 |
| 1810010D01 | 1 | 2 | 0 | 1 | 1 | 1 | 1 | 2 | 0 | 1 | 1 | 1 | 1 | 1 | 1 | 0.958 | -0.063 | 0.97296134 | NA | 0 | 3.2924 | 1.457 | 1.457 | TRUE | 0.2158 |
| 1810010H24 | 101 | 169 | 170 | 125 | 101 | 115 | 107 | 169 | 171 | 98 | 109 | 126 | 129.93 | 149 | 111 | 0.742 | -0.43 | 0.06983087 | 0.20540329 | 0.0332 | 0.0323 | 0.0327 | 0.0327 | TRUE | 0.6167 |
| 1810011H11 | 0 | 0 | 0 | 2 | 0 | 0 | 0 | 0 | 0 | 2 | 0 | 0 | 0.26 | 0 | 1 | 2.886 | 1.529 | 0.70676491 | NA | 0 | 12.5792 | 10 | 10 | TRUE | 0.9669 |
| 1810011O10 | 30 | 18 | 5 | 46 | 33 | 22 | 32 | 18 | 5 | 36 | 36 | 24 | 25.1 | 18 | 32 | 1.754 | 0.811 | 0.16419539 | 0.36032972 | 0.2848 | 0.1381 | 0.2014 | 0.2014 | TRUE | 0.3668 |
| 1810012K16 | 1 | 9 | 4 | 9 | 4 | 7 | 1 | 9 | 4 | 7 | 4 | 8 | 5.52 | 5 | 6 | 1.35 | 0.433 | 0.63524741 | 0.79491607 | 0.1772 | 0.6039 | 0.4149 | 0.4149 | TRUE | 0.5084 |
| 1810013A23 | 0 | 0 | 0 | 0 | 0 | 0 | 0 | 0 | 0 | 0 | 0 | 0 | 0 | 0 | 0 | NA | NA | NA | NA | NA | NA | NA | NA | NA | NA |
| 1810013D10 | 727 | 750 | 683 | 991 | 632 | 603 | 772 | 749 | 686 | 779 | 680 | 660 | 721.03 | 736 | 706 | 0.962 | -0.056 | 0.614839 | 0.78208006 | 0.0046 | 0.0115 | 0.0076 | 0.0076 | TRUE | 0.0993 |

| | | | | | | | | | | | | | | | | | | | | | | | | | |
|---|---|---|---|---|---|---|---|---|---|---|---|---|---|---|---|---|---|---|---|---|---|---|---|---|---|
| 1810013L24I | 3009 | 2950 | 2917 | 3848 | 2862 | 2599 | 3196 | 2946 | 2929 | 3025 | 3079 | 2845 | 3003.43 | 3024 | 2983 | 0.987 | -0.019 | 0.80280289 | 0.89812791 | 0.0017 | 0.008 | 0.0039 | 0.0039 | TRUE | 0.0301 |
| 1810014B01 | 143 | 171 | 165 | 218 | 152 | 145 | 152 | 171 | 166 | 171 | 164 | 159 | 163.67 | 163 | 165 | 1.012 | 0.017 | 0.91375094 | 0.95745946 | 0 | 0.027 | 0.0117 | 0.0117 | TRUE | 0.0356 |
| 1810018F18I | 0 | 0 | 0 | 0 | 0 | 0 | 0 | 0 | 0 | 0 | 0 | 0 | 0 | 0 | 0 | NA | NA | NA | NA | NA | NA | NA | NA | NA | NA |
| 1810019D21 | 37 | 43 | 34 | 63 | 47 | 42 | 39 | 43 | 34 | 50 | 51 | 46 | 43.74 | 39 | 49 | 1.256 | 0.329 | 0.25395511 | 0.47188737 | 0 | 0.0822 | 0.0367 | 0.0367 | TRUE | 0.0831 |
| 1810019J16F | 276 | 353 | 383 | 370 | 311 | 326 | 293 | 353 | 385 | 291 | 335 | 357 | 335.44 | 344 | 328 | 0.951 | -0.072 | 0.63879604 | 0.79655189 | 0.0119 | 0.0167 | 0.0142 | 0.0142 | TRUE | 0.1849 |
| 1810020O05 | 6 | 9 | 5 | 5 | 2 | 7 | 6 | 9 | 5 | 4 | 2 | 8 | 5.69 | 7 | 5 | 0.669 | -0.58 | 0.48599859 | 0.6828659 | 0 | 0.5861 | 0.317 | 0.317 | TRUE | 0.5711 |
| 1810021B22 | 3 | 1 | 4 | 4 | 1 | 2 | 3 | 1 | 4 | 3 | 1 | 2 | 2.44 | 3 | 2 | 0.798 | -0.325 | 0.78894163 | NA | 0 | 1.3597 | 0.6522 | 0.6522 | TRUE | 0.3776 |
| 1810022K09I | 1201 | 1216 | 1177 | 1644 | 1193 | 1157 | 1276 | 1214 | 1182 | 1292 | 1284 | 1267 | 1252.41 | 1224 | 1281 | 1.047 | 0.066 | 0.3957437 | 0.60912166 | 0 | 0.0095 | 0.0035 | 0.0035 | TRUE | 0.0162 |
| 1810024B03 | 3 | 1 | 2 | 6 | 4 | 0 | 3 | 1 | 2 | 5 | 4 | 0 | 2.54 | 2 | 3 | 1.487 | 0.572 | 0.67126709 | NA | 0 | 1.3062 | 0.9009 | 0.9009 | TRUE | 0.8874 |
| 1810026B05 | 197 | 238 | 241 | 325 | 273 | 256 | 209 | 238 | 242 | 255 | 294 | 280 | 253.06 | 230 | 276 | 1.201 | 0.264 | 0.06131378 | 0.18883831 | 0 | 0.0199 | 0.0104 | 0.0104 | TRUE | 0.0654 |
| 1810026J23F | 1763 | 1700 | 1507 | 2104 | 1503 | 1359 | 1873 | 1698 | 1513 | 1654 | 1617 | 1488 | 1640.4 | 1695 | 1586 | 0.937 | -0.095 | 0.37465521 | 0.59076389 | 0.0062 | 0.0089 | 0.0076 | 0.0076 | TRUE | 0.1058 |
| 1810030O07 | 1853 | 1913 | 1959 | 2639 | 1895 | 1821 | 1968 | 1911 | 1967 | 2075 | 2039 | 1993 | 1992.11 | 1949 | 2036 | 1.045 | 0.064 | 0.34732372 | 0.56625901 | 0 | 0.0086 | 0.0028 | 0.0028 | TRUE | 0.004 |
| 1810032O08 | 1545 | 2118 | 2082 | 1819 | 1379 | 1614 | 1641 | 2115 | 2091 | 1430 | 1484 | 1767 | 1754.55 | 1949 | 1560 | 0.8 | -0.322 | 0.01703576 | 0.07747634 | 0.0152 | 0.0088 | 0.0125 | 0.0125 | TRUE | 0.2304 |
| 1810033B17 | 2 | 3 | 0 | 4 | 2 | 3 | 2 | 3 | 0 | 3 | 2 | 3 | 2.28 | 2 | 3 | 1.683 | 0.751 | 0.56812589 | NA | 0 | 1.4496 | 0.7767 | 0.7767 | TRUE | 0.6049 |
| 1810034E14I | 2 | 6 | 5 | 5 | 5 | 2 | 2 | 6 | 5 | 4 | 5 | 2 | 4.11 | 4 | 4 | 0.874 | -0.195 | 0.83689584 | NA | 0 | 0.8092 | 0.4009 | 0.4009 | TRUE | 0.3454 |
| 1810035I16R | 0 | 0 | 0 | 0 | 0 | 0 | 0 | 0 | 0 | 0 | 0 | 0 | 0 | 0 | 0 | NA | NA | NA | NA | NA | NA | NA | NA | NA | NA |
| 1810037I17R | 1194 | 1498 | 1546 | 1843 | 1283 | 1375 | 1268 | 1496 | 1552 | 1449 | 1380 | 1505 | 1441.84 | 1439 | 1445 | 1.004 | 0.006 | 0.95724929 | 0.97783263 | 0.0058 | 0.0092 | 0.0072 | 0.0072 | TRUE | 0.129 |
| 1810041L15F | 47 | 57 | 57 | 75 | 37 | 68 | 50 | 57 | 57 | 59 | 40 | 74 | 56.21 | 55 | 58 | 1.055 | 0.078 | 0.7985025 | 0.89619481 | 0.0298 | 0.0655 | 0.0486 | 0.0486 | TRUE | 0.2214 |
| 1810043G02 | 245 | 289 | 284 | 350 | 282 | 276 | 260 | 289 | 285 | 275 | 303 | 302 | 285.78 | 278 | 293 | 1.054 | 0.075 | 0.5595956 | 0.74136428 | 0 | 0.0184 | 0.0085 | 0.0085 | TRUE | 0.0368 |
| 1810043H04 | 673 | 750 | 765 | 983 | 783 | 726 | 715 | 749 | 768 | 773 | 842 | 795 | 773.66 | 744 | 803 | 1.079 | 0.109 | 0.23386612 | 0.44850881 | 0 | 0.0112 | 0.0048 | 0.0048 | TRUE | 0.0214 |
| 1810044D09 | 32 | 40 | 43 | 77 | 39 | 46 | 34 | 40 | 43 | 61 | 42 | 50 | 44.99 | 39 | 51 | 1.312 | 0.391 | 0.19698098 | 0.40371603 | 0 | 0.0801 | 0.0437 | 0.0437 | TRUE | 0.218 |
| 1810046K07I | 0 | 0 | 0 | 0 | 0 | 0 | 0 | 0 | 0 | 0 | 0 | 0 | 0 | 0 | 0 | NA | NA | NA | NA | NA | NA | NA | NA | NA | NA |
| 1810053B23 | 0 | 0 | 0 | 0 | 0 | 0 | 0 | 0 | 0 | 0 | 0 | 0 | 0 | 0 | 0 | NA | NA | NA | NA | NA | NA | NA | NA | NA | NA |
| 1810055G02 | 1374 | 1226 | 1112 | 1761 | 1228 | 1106 | 1459 | 1224 | 1117 | 1384 | 1321 | 1211 | 1286.12 | 1267 | 1305 | 1.031 | 0.044 | 0.71990086 | 0.8481969 | 0.0108 | 0.0095 | 0.0102 | 0.0102 | TRUE | 0.2118 |
| 1810058I24R | 2215 | 2178 | 2053 | 2636 | 1818 | 1985 | 2353 | 2175 | 2061 | 2072 | 1956 | 2173 | 2131.75 | 2196 | 2067 | 0.941 | -0.087 | 0.3221185 | 0.54330584 | 0.0029 | 0.0084 | 0.0052 | 0.0052 | TRUE | 0.0467 |
| 1810062G17 | 16 | 11 | 6 | 9 | 13 | 9 | 17 | 11 | 6 | 7 | 14 | 10 | 10.82 | 11 | 10 | 0.902 | -0.148 | 0.81776981 | 0.90711028 | 0.0904 | 0.3114 | 0.2042 | 0.2042 | TRUE | 0.2157 |
| 1810062O18 | 1 | 1 | 0 | 1 | 0 | 2 | 1 | 1 | 0 | 1 | 0 | 2 | 0.84 | 1 | 1 | 1.436 | 0.522 | 0.80963223 | NA | 0 | 3.9317 | 2.1449 | 2.1449 | TRUE | 0.3069 |
| 1810064F22I | 0 | 1 | 0 | 0 | 0 | 0 | 0 | 1 | 0 | 0 | 0 | 0 | 0.17 | 0 | 0 | 0.49 | -1.029 | 0.80087 | NA | 0 | 19.7991 | 10 | 10 | TRUE | 0.405 |
| 1810065E05I | 0 | 0 | 0 | 1 | 0 | 0 | 0 | 0 | 0 | 1 | 0 | 0 | 0.13 | 0 | 0 | 1.859 | 0.894 | 0.82650734 | NA | 0 | 25.1515 | 10 | 10 | TRUE | 0.2563 |
| 2010001E11I | 0 | 1 | 0 | 1 | 1 | 1 | 0 | 1 | 0 | 1 | 1 | 1 | 0.66 | 0 | 1 | 2.676 | 1.42 | 0.53620179 | NA | 0 | 5.0043 | 2.2992 | 2.2992 | TRUE | 0.405 |
| 2010001M06 | 54 | 75 | 97 | 99 | 81 | 85 | 57 | 75 | 97 | 78 | 87 | 93 | 81.28 | 76 | 86 | 1.119 | 0.163 | 0.53159684 | 0.71978876 | 0.0244 | 0.0474 | 0.0364 | 0.0364 | TRUE | 0.2893 |
| 2010002M12 | 4 | 0 | 2 | 3 | 0 | 2 | 4 | 0 | 2 | 2 | 0 | 2 | 1.8 | 2 | 1 | 0.743 | -0.429 | 0.79804482 | NA | 0.7906 | 1.8364 | 1.4634 | 1.4634 | TRUE | 0.1619 |
| 2010003K11I | 1 | 0 | 0 | 1 | 0 | 0 | 1 | 0 | 0 | 1 | 0 | 0 | 0.31 | 0 | 0 | 0.954 | -0.067 | 0.98625928 | NA | 0 | 10.7014 | 9.0384 | 9.0384 | TRUE | 0.4614 |
| 2010003O02 | 51 | 36 | 34 | 62 | 41 | 24 | 54 | 36 | 34 | 49 | 44 | 26 | 40.56 | 41 | 40 | 0.966 | -0.05 | 0.89266065 | 0.94801992 | 0.0565 | 0.0881 | 0.073 | 0.073 | TRUE | 0.6165 |
| 2010005H15 | 1 | 0 | 0 | 0 | 0 | 0 | 1 | 0 | 0 | 0 | 0 | 0 | 0.18 | 0 | 0 | 0.49 | -1.029 | 0.80087 | NA | 0 | 18.6165 | 10 | 10 | TRUE | 0.4614 |
| 2010009K17I | 0 | 0 | 0 | 0 | 0 | 0 | 0 | 0 | 0 | 0 | 0 | 0 | 0 | 0 | 0 | NA | NA | NA | NA | NA | NA | NA | NA | NA | NA |
| 2010010A06 | 5 | 6 | 6 | 1 | 8 | 3 | 5 | 6 | 6 | 1 | 9 | 3 | 5 | 6 | 4 | 0.711 | -0.491 | 0.61379654 | 0.78152895 | 0.2563 | 0.6657 | 0.4758 | 0.4758 | TRUE | 0.7373 |
| 2010012O05 | 220 | 253 | 218 | 301 | 200 | 220 | 234 | 253 | 219 | 237 | 215 | 241 | 232.98 | 235 | 231 | 0.983 | -0.025 | 0.85884461 | 0.9304418 | 0 | 0.021 | 0.0101 | 0.0101 | TRUE | 0.048 |
| 2010015L04F | 86 | 91 | 69 | 96 | 98 | 69 | 91 | 91 | 69 | 75 | 105 | 76 | 84.66 | 84 | 85 | 1.017 | 0.024 | 0.92188815 | 0.96126008 | 0.0193 | 0.0458 | 0.0328 | 0.0328 | TRUE | 0.3812 |
| 2010016I18R | 0 | 0 | 0 | 0 | 0 | 1 | 0 | 0 | 0 | 0 | 0 | 1 | 0.18 | 0 | 0 | 1.859 | 0.894 | 0.82650734 | NA | 0 | 18.0643 | 10 | 10 | TRUE | 0.5383 |
| 2010106C02I | 0 | 0 | 0 | 0 | 0 | 0 | 0 | 0 | 0 | 0 | 0 | 0 | 0 | 0 | 0 | NA | NA | NA | NA | NA | NA | NA | NA | NA | NA |
| 2010106E10I | 0 | 0 | 0 | 0 | 0 | 0 | 0 | 0 | 0 | 0 | 0 | 0 | 0 | 0 | 0 | NA | NA | NA | NA | NA | NA | NA | NA | NA | NA |
| 2010107E04I | 2845 | 3192 | 3079 | 3788 | 2626 | 2695 | 3022 | 3188 | 3092 | 2978 | 2825 | 2950 | 3009.13 | 3101 | 2918 | 0.941 | -0.087 | 0.19117857 | 0.39617243 | 4.00E-04 | 0.008 | 0.0029 | 0.0029 | TRUE | 0.0092 |
| 2010107G12 | 19 | 7 | 10 | 35 | 28 | 16 | 20 | 7 | 10 | 28 | 30 | 18 | 18.73 | 12 | 25 | 2.034 | 1.024 | 0.05595728 | 0.17743431 | 0.0999 | 0.1828 | 0.146 | 0.146 | TRUE | 1.1638 |
| 2010107G23 | 547 | 877 | 855 | 826 | 585 | 752 | 581 | 876 | 859 | 649 | 629 | 823 | 736.22 | 772 | 700 | 0.907 | -0.141 | 0.44240678 | 0.64932438 | 0.0347 | 0.0114 | 0.0228 | 0.0228 | TRUE | 0.5523 |
| 2010109A12 | 1 | 1 | 0 | 2 | 0 | 2 | 1 | 1 | 0 | 2 | 0 | 2 | 0.97 | 1 | 1 | 1.844 | 0.883 | 0.66663732 | NA | 0 | 3.4018 | 1.8936 | 1.8936 | TRUE | 0.3972 |
| 2010109I03R | 0 | 0 | 0 | 0 | 0 | 0 | 0 | 0 | 0 | 0 | 0 | 0 | 0 | 0 | 0 | NA | NA | NA | NA | NA | NA | NA | NA | NA | NA |
| 2010111I01R | 226 | 156 | 178 | 280 | 182 | 155 | 240 | 156 | 179 | 220 | 196 | 170 | 193.36 | 192 | 195 | 1.022 | 0.032 | 0.87817836 | 0.93976544 | 0.028 | 0.0239 | 0.0259 | 0.0259 | TRUE | 0.5259 |
| 2010204K13I | 575 | 549 | 509 | 556 | 434 | 426 | 611 | 548 | 511 | 437 | 467 | 466 | 506.75 | 557 | 457 | 0.82 | -0.286 | 0.01285334 | 0.06328543 | 0.003 | 0.0134 | 0.0076 | 0.0076 | TRUE | 0.0837 |
| 2010300C02I | 302 | 363 | 352 | 408 | 331 | 284 | 321 | 363 | 353 | 321 | 356 | 311 | 337.42 | 346 | 329 | 0.952 | -0.071 | 0.57277759 | 0.7512479 | 0.0014 | 0.0167 | 0.0085 | 0.0085 | TRUE | 0.0559 |
| 2010308F09I | 4 | 3 | 1 | 0 | 4 | 0 | 4 | 3 | 1 | 0 | 4 | 0 | 2.09 | 3 | 1 | 0.507 | -0.98 | 0.56195431 | NA | 1.4166 | 1.5816 | 1.521 | 1.521 | TRUE | 1.5067 |
| 2010310C07I | 0 | 1 | 0 | 0 | 1 | 0 | 0 | 1 | 0 | 0 | 1 | 0 | 0.35 | 0 | 0 | 0.954 | -0.067 | 0.98571955 | NA | 0 | 9.5346 | 8.2194 | 8.2194 | TRUE | 0.5204 |
| 2010315B03 | 25 | 24 | 22 | 28 | 22 | 20 | 27 | 24 | 22 | 22 | 24 | 22 | 23.36 | 24 | 23 | 0.93 | -0.104 | 0.78603722 | 0.89016867 | 0 | 0.1479 | 0.0629 | 0.0629 | TRUE | 0.0415 |
| 2010320M18 | 226 | 239 | 239 | 281 | 252 | 221 | 240 | 239 | 240 | 221 | 271 | 242 | 242.11 | 240 | 245 | 1.018 | 0.026 | 0.85346259 | 0.92686167 | 0 | 0.0205 | 0.0106 | 0.0106 | TRUE | 0.0991 |
| 2200002D01 | 57 | 52 | 49 | 158 | 84 | 84 | 61 | 52 | 49 | 124 | 90 | 92 | 78.04 | 54 | 102 | 1.908 | 0.932 | 0.00021639 | 0.00258943 | 0.015 | 0.0491 | 0.0315 | 0.0315 | TRUE | 0.3795 |
| 2200002J24F | 4 | 7 | 5 | 2 | 4 | 1 | 4 | 7 | 5 | 2 | 4 | 1 | 3.87 | 5 | 2 | 0.422 | -1.246 | 0.21783693 | NA | 0 | 0.8578 | 0.4299 | 0.4299 | TRUE | 0.5167 |
| 2210010C04I | 0 | 0 | 0 | 0 | 0 | 0 | 0 | 0 | 0 | 0 | 0 | 0 | 0 | 0 | 0 | NA | NA | NA | NA | NA | NA | NA | NA | NA | NA |
| 2210013O21 | 381 | 358 | 379 | 463 | 326 | 331 | 405 | 358 | 381 | 364 | 351 | 362 | 369.97 | 381 | 359 | 0.943 | -0.084 | 0.46471731 | 0.66783795 | 0 | 0.0158 | 0.0069 | 0.0069 | TRUE | 0.0338 |
| 2210015D19 | 370 | 383 | 325 | 515 | 342 | 332 | 393 | 383 | 326 | 405 | 368 | 363 | 373.02 | 367 | 379 | 1.033 | 0.047 | 0.71677294 | 0.84630504 | 0.004 | 0.0157 | 0.0094 | 0.0094 | TRUE | 0.1094 |
| 2210016F16I | 777 | 810 | 812 | 1654 | 1183 | 1010 | 825 | 809 | 815 | 1300 | 1273 | 1106 | 1021.38 | 816 | 1226 | 1.503 | 0.588 | 2.39E-09 | 1.33E-07 | 0.0032 | 0.0101 | 0.006 | 0.006 | TRUE | 0.0877 |
| 2210016L21F | 1516 | 1248 | 1350 | 1853 | 1384 | 1206 | 1610 | 1246 | 1356 | 1457 | 1489 | 1320 | 1413.02 | 1404 | 1422 | 1.013 | 0.019 | 0.87459318 | 0.93811777 | 0.0099 | 0.0092 | 0.0096 | 0.0096 | TRUE | 0.198 |
| 2210018M11 | 3369 | 2778 | 2945 | 4180 | 3082 | 2843 | 3578 | 2774 | 2957 | 3286 | 3316 | 3112 | 3170.69 | 3103 | 3238 | 1.044 | 0.062 | 0.58051052 | 0.75708116 | 0.0091 | 0.0079 | 0.0086 | 0.0086 | TRUE | 0.2176 |
| 2210019I11R | 0 | 0 | 0 | 0 | 1 | 0 | 0 | 0 | 0 | 0 | 1 | 0 | 0.18 | 0 | 0 | 1.859 | 0.894 | 0.82650734 | NA | 0 | 18.3785 | 10 | 10 | TRUE | 0.5204 |
| 2210039B01 | 8 | 13 | 14 | 4 | 5 | 9 | 8 | 13 | 14 | 3 | 5 | 10 | 8.99 | 12 | 6 | 0.505 | -0.987 | 0.15520157 | 0.34749626 | 7.00E-04 | 0.3735 | 0.2228 | 0.2228 | TRUE | 0.6364 |
| 2210404J11F | 207 | 239 | 207 | 362 | 227 | 185 | 220 | 239 | 208 | 285 | 244 | 203 | 232.96 | 222 | 244 | 1.101 | 0.139 | 0.41781843 | 0.62868081 | 0.0131 | 0.021 | 0.0169 | 0.0169 | TRUE | 0.1803 |
| 2210404O07 | 40 | 45 | 58 | 36 | 26 | 35 | 42 | 45 | 58 | 28 | 28 | 38 | 40.04 | 48 | 31 | 0.646 | -0.63 | 0.05170812 | 0.16770316 | 0 | 0.0892 | 0.0496 | 0.0496 | TRUE | 0.2463 |
| 2210404O09 | 114 | 86 | 102 | 211 | 124 | 107 | 121 | 86 | 102 | 166 | 133 | 117 | 120.97 | 103 | 139 | 1.353 | 0.436 | 0.05374022 | 0.17235682 | 0.0225 | 0.0341 | 0.0283 | 0.0283 | TRUE | 0.3224 |
| 2210407C18I | 0 | 0 | 0 | 0 | 0 | 1 | 0 | 0 | 0 | 0 | 0 | 1 | 0.18 | 0 | 0 | 1.859 | 0.894 | 0.82650734 | NA | 0 | 18.0643 | 10 | 10 | TRUE | 0.5383 |
| 2210408F21I | 0 | 1 | 1 | 2 | 1 | 2 | 0 | 1 | 1 | 2 | 1 | 2 | 1.14 | 1 | 2 | 2.386 | 1.254 | 0.48062816 | NA | 0 | 2.8966 | 1.2192 | 1.2192 | TRUE | 0.2224 |

| | | | | | | | | | | | | | | | | | | | | | | | | | |
|---|---|---|---|---|---|---|---|---|---|---|---|---|---|---|---|---|---|---|---|---|---|---|---|---|---|
| 2210408I21R | 66 | 67 | 79 | 148 | 108 | 77 | 70 | 67 | 79 | 116 | 116 | 84 | 88.86 | 72 | 105 | 1.469 | 0.554 | 0.0171911 | 0.07784612 | 0.0103 | 0.044 | 0.0273 | 0.0273 | TRUE | 0.2933 |
| 2210409D07 | 0 | 0 | 0 | 0 | 0 | 0 | 0 | 0 | 0 | 0 | 0 | 0 | 0 | 0 | 0 | NA | NA | NA | NA | NA | NA | NA | NA | NA | NA |
| 2210409E12 | 14 | 17 | 17 | 119 | 61 | 32 | 15 | 17 | 17 | 94 | 66 | 35 | 40.52 | 16 | 65 | 3.992 | 1.997 | 5.99E-06 | 0.00012897 | 0.1143 | 0.0882 | 0.1014 | 0.1014 | TRUE | 0.081 |
| 2210414B05 | 0 | 0 | 0 | 0 | 0 | 0 | 0 | 0 | 0 | 0 | 0 | 0 | 0 | 0 | 0 | NA | NA | NA | NA | NA | NA | NA | NA | NA | NA |
| 2210416O15 | 4 | 2 | 3 | 12 | 6 | 8 | 4 | 2 | 3 | 9 | 6 | 9 | 5.65 | 3 | 8 | 2.685 | 1.425 | 0.08558514 | 0.23565796 | 0 | 0.5899 | 0.2698 | 0.2698 | TRUE | 0.1363 |
| 2210417A02 | 9 | 18 | 16 | 27 | 19 | 17 | 10 | 18 | 16 | 21 | 20 | 19 | 17.31 | 15 | 20 | 1.381 | 0.466 | 0.32065603 | 0.54182417 | 0 | 0.1972 | 0.0994 | 0.0994 | TRUE | 0.3808 |
| 2210417K05 | 11 | 21 | 19 | 6 | 12 | 16 | 12 | 21 | 19 | 5 | 13 | 18 | 14.48 | 17 | 12 | 0.665 | -0.588 | 0.33455052 | 0.5548783 | 0.1517 | 0.2344 | 0.1957 | 0.1957 | TRUE | 0.7191 |
| 2210420H20 | 9 | 8 | 6 | 6 | 10 | 5 | 10 | 8 | 6 | 5 | 11 | 5 | 7.42 | 8 | 7 | 0.879 | -0.186 | 0.79517443 | 0.8946193 | 0 | 0.4509 | 0.2332 | 0.2332 | TRUE | 0.5569 |
| 2300002M23 | 6 | 17 | 12 | 7 | 11 | 12 | 6 | 17 | 12 | 6 | 12 | 13 | 10.98 | 12 | 10 | 0.845 | -0.242 | 0.71055953 | 0.84224748 | 0.1167 | 0.307 | 0.2148 | 0.2148 | TRUE | 0.217 |
| 2300003K06 | 0 | 0 | 0 | 0 | 0 | 0 | 0 | 0 | 0 | 0 | 0 | 0 | 0 | 0 | 0 | NA | NA | NA | NA | NA | NA | NA | NA | NA | NA |
| 2300005B03 | 0 | 0 | 1 | 0 | 0 | 0 | 0 | 0 | 1 | 0 | 0 | 0 | 0.17 | 0 | 0 | 0.49 | -1.029 | 0.80087 | NA | 0 | 19.6928 | 10 | 10 | TRUE | 0.4098 |
| 2300009A05 | 93 | 147 | 153 | 117 | 103 | 105 | 99 | 147 | 154 | 92 | 111 | 115 | 119.49 | 133 | 106 | 0.793 | -0.335 | 0.14964243 | 0.34028031 | 0.0266 | 0.0345 | 0.0306 | 0.0306 | TRUE | 0.5139 |
| 2310001H17 | 6 | 7 | 6 | 4 | 7 | 12 | 6 | 7 | 6 | 3 | 8 | 13 | 7.2 | 6 | 8 | 1.204 | 0.267 | 0.7300419 | 0.85500295 | 0 | 0.4645 | 0.2911 | 0.2911 | TRUE | 0.2184 |
| 2310001K24 | 0 | 0 | 0 | 1 | 0 | 0 | 0 | 0 | 0 | 1 | 0 | 0 | 0.13 | 0 | 0 | 1.859 | 0.894 | 0.82650734 | NA | 0 | 25.1515 | 10 | 10 | TRUE | 0.2563 |
| 2310002D06 | 0 | 0 | 0 | 0 | 0 | 0 | 0 | 0 | 0 | 0 | 0 | 0 | 0 | 0 | 0 | NA | NA | NA | NA | NA | NA | NA | NA | NA | NA |
| 2310002F09 | 9 | 8 | 5 | 5 | 6 | 8 | 10 | 8 | 5 | 4 | 6 | 9 | 6.95 | 8 | 6 | 0.836 | -0.259 | 0.72553298 | 0.85186527 | 0 | 0.4808 | 0.245 | 0.245 | TRUE | 0.301 |
| 2310002J15F | 12 | 15 | 19 | 8 | 5 | 9 | 13 | 15 | 19 | 6 | 5 | 10 | 11.39 | 16 | 7 | 0.456 | -1.133 | 0.05380759 | 0.1723792 | 0 | 0.2962 | 0.1476 | 0.1476 | TRUE | 0.2543 |
| 2310002L09F | 0 | 0 | 0 | 0 | 0 | 0 | 0 | 0 | 0 | 0 | 0 | 0 | 0 | 0 | 0 | NA | NA | NA | NA | NA | NA | NA | NA | NA | NA |
| 2310003H01 | 1583 | 1820 | 1734 | 2445 | 1834 | 1552 | 1681 | 1818 | 1741 | 1922 | 1973 | 1699 | 1805.74 | 1747 | 1865 | 1.068 | 0.095 | 0.29900119 | 0.51932012 | 0.003 | 0.0087 | 0.0054 | 0.0054 | TRUE | 0.0721 |
| 2310005A03 | 0 | 0 | 0 | 0 | 0 | 0 | 0 | 0 | 0 | 0 | 0 | 0 | 0 | 0 | 0 | NA | NA | NA | NA | NA | NA | NA | NA | NA | NA |
| 2310005E17 | 1 | 1 | 0 | 0 | 0 | 2 | 1 | 1 | 0 | 0 | 0 | 2 | 0.71 | 1 | 1 | 1.031 | 0.044 | 0.9865942 | NA | 0 | 4.6577 | 3.4964 | 3.4964 | TRUE | 0.9748 |
| 2310005G13 | 7 | 16 | 15 | 48 | 31 | 11 | 7 | 16 | 15 | 38 | 33 | 12 | 20.27 | 13 | 28 | 2.169 | 1.117 | 0.05025302 | 0.16477974 | 0.1861 | 0.1695 | 0.1773 | 0.1773 | TRUE | 1.3698 |
| 2310007B03 | 20 | 28 | 36 | 35 | 23 | 39 | 21 | 28 | 36 | 28 | 25 | 43 | 30.05 | 28 | 32 | 1.106 | 0.146 | 0.71570007 | 0.84567137 | 0.0433 | 0.1165 | 0.0816 | 0.0816 | TRUE | 0.4278 |
| 2310007L24F | 2 | 0 | 2 | 1 | 0 | 2 | 2 | 0 | 2 | 1 | 0 | 2 | 1.18 | 1 | 1 | 0.715 | -0.484 | 0.80082038 | NA | 0 | 2.7878 | 1.7781 | 1.7781 | TRUE | 0.4861 |
| 2310008H04 | 1867 | 2445 | 2709 | 2826 | 2025 | 2393 | 1983 | 2442 | 2720 | 2222 | 2179 | 2620 | 2360.82 | 2382 | 2340 | 0.982 | -0.026 | 0.84988458 | 0.92526408 | 0.0173 | 0.0083 | 0.0131 | 0.0131 | TRUE | 0.2593 |
| 2310008H09 | 10068 | 8424 | 8355 | 12599 | 8807 | 7914 | 10694 | 8413 | 8389 | 9904 | 9476 | 8663 | 9256.61 | 9165 | 9348 | 1.02 | 0.029 | 0.80900662 | 0.90174222 | 0.01 | 0.0073 | 0.01 | 0.01 | TRUE | 0.2599 |
| 2310008N11 | 0 | 2 | 0 | 4 | 1 | 1 | 0 | 2 | 0 | 3 | 1 | 1 | 1.22 | 1 | 2 | 2.685 | 1.425 | 0.46282576 | NA | 0 | 2.71 | 1.684 | 1.684 | TRUE | 0.9737 |
| 2310009A05 | 109 | 94 | 100 | 173 | 133 | 118 | 116 | 94 | 100 | 136 | 143 | 129 | 119.72 | 103 | 136 | 1.318 | 0.398 | 0.0330127 | 0.12282999 | 0 | 0.0344 | 0.0166 | 0.0166 | TRUE | 0.1064 |
| 2310009B15 | 121 | 91 | 74 | 142 | 117 | 86 | 129 | 91 | 74 | 112 | 126 | 94 | 104.23 | 98 | 111 | 1.131 | 0.178 | 0.49142461 | 0.68773282 | 0.0382 | 0.0385 | 0.0384 | 0.0384 | TRUE | 0.6894 |
| 2310010J17F | 24 | 27 | 31 | 24 | 20 | 16 | 25 | 27 | 31 | 19 | 22 | 18 | 23.58 | 28 | 20 | 0.692 | -0.532 | 0.17038923 | 0.36901257 | 0 | 0.1466 | 0.0647 | 0.0647 | TRUE | 0.0679 |
| 2310010M2C | 0 | 2 | 0 | 0 | 2 | 0 | 0 | 2 | 0 | 0 | 2 | 0 | 0.69 | 1 | 1 | 1.052 | 0.073 | 0.98136341 | NA | 8.094 | 4.7708 | 5.501 | 5.501 | TRUE | 0.9745 |
| 2310011J03F | 391 | 354 | 379 | 757 | 553 | 413 | 415 | 354 | 381 | 595 | 595 | 452 | 465.27 | 383 | 547 | 1.431 | 0.517 | 0.00050118 | 0.00517876 | 0.0134 | 0.014 | 0.0137 | 0.0137 | TRUE | 0.2665 |
| 2310014F07 | 1 | 1 | 5 | 3 | 0 | 1 | 1 | 1 | 5 | 2 | 0 | 1 | 1.76 | 2 | 1 | 0.508 | -0.978 | 0.53799747 | NA | 0.4192 | 1.8833 | 1.194 | 1.194 | TRUE | 0.735 |
| 2310014L17F | 32 | 45 | 44 | 72 | 52 | 54 | 34 | 45 | 44 | 57 | 56 | 59 | 49.13 | 41 | 57 | 1.392 | 0.477 | 0.08577468 | 0.23605624 | 0 | 0.074 | 0.0345 | 0.0345 | TRUE | 0.1648 |
| 2310015A10 | 190 | 202 | 200 | 252 | 203 | 211 | 202 | 202 | 201 | 198 | 218 | 231 | 208.64 | 202 | 216 | 1.068 | 0.096 | 0.50888049 | 0.70257778 | 0 | 0.0227 | 0.0103 | 0.0103 | TRUE | 0.0619 |
| 2310015B20 | 1 | 0 | 2 | 0 | 0 | 2 | 1 | 0 | 2 | 0 | 0 | 2 | 0.88 | 1 | 1 | 0.688 | -0.539 | 0.82569208 | NA | 0 | 3.765 | 3.1169 | 3.1169 | TRUE | 0.5573 |
| 2310015D24 | 1 | 0 | 0 | 0 | 0 | 0 | 1 | 0 | 0 | 0 | 0 | 0 | 0.18 | 0 | 0 | 0.49 | -1.029 | 0.80087 | NA | 0 | 18.6165 | 10 | 10 | TRUE | 0.4614 |
| 2310016D03 | 0 | 0 | 0 | 0 | 0 | 0 | 0 | 0 | 0 | 0 | 0 | 0 | 0 | 0 | 0 | NA | NA | NA | NA | NA | NA | NA | NA | NA | NA |
| 2310020H05 | 0 | 0 | 0 | 0 | 0 | 0 | 0 | 0 | 0 | 0 | 0 | 0 | 0 | 0 | 0 | NA | NA | NA | NA | NA | NA | NA | NA | NA | NA |
| 2310022A10 | 2229 | 2591 | 2646 | 2916 | 2141 | 2178 | 2368 | 2588 | 2657 | 2292 | 2304 | 2384 | 2432.03 | 2538 | 2327 | 0.917 | -0.125 | 0.10893906 | 0.2765969 | 0.0016 | 0.0083 | 0.004 | 0.004 | TRUE | 0.0414 |
| 2310022B05 | 2549 | 3139 | 3146 | 3924 | 3040 | 3289 | 2707 | 3135 | 3159 | 3085 | 3271 | 3600 | 3159.55 | 3000 | 3319 | 1.106 | 0.145 | 0.15209158 | 0.3431743 | 0.0062 | 0.0079 | 0.0071 | 0.0071 | TRUE | 0.0884 |
| 2310030A07 | 3 | 3 | 3 | 2 | 1 | 2 | 3 | 3 | 3 | 2 | 1 | 2 | 2.34 | 3 | 2 | 0.525 | -0.93 | 0.43619542 | NA | 0 | 1.4155 | 0.5578 | 0.5578 | TRUE | 0.0601 |
| 2310030G06 | 129 | 178 | 158 | 162 | 101 | 109 | 137 | 178 | 159 | 127 | 109 | 119 | 138.13 | 158 | 118 | 0.752 | -0.412 | 0.02748335 | 0.10855682 | 0.0049 | 0.0308 | 0.0178 | 0.0178 | TRUE | 0.1377 |
| 2310033P09 | 791 | 664 | 657 | 1204 | 914 | 795 | 840 | 663 | 660 | 946 | 983 | 870 | 827.2 | 721 | 933 | 1.295 | 0.373 | 0.00345747 | 0.02362743 | 0.0103 | 0.0109 | 0.0105 | 0.0105 | TRUE | 0.2461 |
| 2310034C09 | 0 | 0 | 0 | 0 | 0 | 0 | 0 | 0 | 0 | 0 | 0 | 0 | 0 | 0 | 0 | NA | NA | NA | NA | NA | NA | NA | NA | NA | NA |
| 2310034G01 | 10 | 14 | 20 | 16 | 16 | 15 | 11 | 14 | 20 | 13 | 17 | 16 | 15.15 | 15 | 15 | 1.025 | 0.036 | 0.94336016 | 0.9717062 | 0 | 0.2244 | 0.1181 | 0.1181 | TRUE | 0.4228 |
| 2310034O05 | 0 | 0 | 0 | 1 | 0 | 0 | 0 | 0 | 0 | 1 | 0 | 0 | 0.13 | 0 | 0 | 1.859 | 0.894 | 0.82650734 | NA | 0 | 25.1515 | 10 | 10 | TRUE | 0.2563 |
| 2310035C23 | 2183 | 1691 | 1757 | 2973 | 2227 | 1912 | 2319 | 1689 | 1764 | 2337 | 2396 | 2093 | 2099.67 | 1924 | 2275 | 1.183 | 0.242 | 0.07657943 | 0.21848569 | 0.0178 | 0.0085 | 0.013 | 0.013 | TRUE | 0.3889 |
| 2310036O22 | 2528 | 2494 | 2405 | 4520 | 3345 | 2597 | 2685 | 2491 | 2415 | 3553 | 3599 | 2843 | 2930.99 | 2530 | 3332 | 1.317 | 0.397 | 0.00047888 | 0.00501184 | 0.0091 | 0.008 | 0.009 | 0.009 | TRUE | 0.1993 |
| 2310039H08 | 122 | 220 | 235 | 228 | 212 | 215 | 130 | 220 | 236 | 179 | 228 | 235 | 204.66 | 195 | 214 | 1.095 | 0.131 | 0.58952081 | 0.76400346 | 0.0539 | 0.023 | 0.0377 | 0.0377 | TRUE | 0.9243 |
| 2310039L15F | 35 | 29 | 29 | 44 | 27 | 31 | 37 | 29 | 29 | 35 | 29 | 34 | 32.14 | 32 | 33 | 1.028 | 0.04 | 0.90583599 | 0.953297 | 0 | 0.1094 | 0.0509 | 0.0509 | TRUE | 0.147 |
| 2310040G24 | 31 | 32 | 42 | 56 | 47 | 38 | 33 | 32 | 42 | 44 | 51 | 42 | 40.54 | 36 | 46 | 1.27 | 0.345 | 0.2610717 | 0.48052549 | 0 | 0.0882 | 0.0429 | 0.0429 | TRUE | 0.1831 |
| 2310042D19 | 1 | 0 | 0 | 0 | 0 | 0 | 1 | 0 | 0 | 0 | 0 | 0 | 0.18 | 0 | 0 | 0.49 | -1.029 | 0.80087 | NA | 0 | 18.6165 | 10 | 10 | TRUE | 0.4614 |
| 2310042E22 | 24 | 12 | 11 | 9 | 5 | 9 | 25 | 12 | 11 | 7 | 5 | 10 | 11.8 | 16 | 7 | 0.46 | -1.121 | 0.07587004 | 0.21720036 | 0.0954 | 0.286 | 0.1899 | 0.1899 | TRUE | 1.1708 |
| 2310043L19F | 0 | 0 | 0 | 1 | 0 | 0 | 0 | 0 | 0 | 1 | 0 | 0 | 0.13 | 0 | 0 | 1.859 | 0.894 | 0.82650734 | NA | 0 | 25.1515 | 10 | 10 | TRUE | 0.2563 |
| 2310043O21 | 0 | 0 | 0 | 0 | 0 | 0 | 0 | 0 | 0 | 0 | 0 | 0 | 0 | 0 | 0 | NA | NA | NA | NA | NA | NA | NA | NA | NA | NA |
| 2310044G17 | 1166 | 1282 | 1310 | 1620 | 1194 | 1190 | 1238 | 1280 | 1315 | 1274 | 1285 | 1303 | 1282.51 | 1278 | 1287 | 1.007 | 0.01 | 0.89968196 | 0.95140627 | 0 | 0.0095 | 0.0033 | 0.0033 | TRUE | 0.0087 |
| 2310045N01 | 298 | 278 | 327 | 350 | 268 | 251 | 317 | 278 | 328 | 275 | 288 | 275 | 293.46 | 308 | 279 | 0.908 | -0.138 | 0.29061163 | 0.51144015 | 0 | 0.0181 | 0.009 | 0.009 | TRUE | 0.0825 |
| 2310047M1C | 532 | 533 | 499 | 662 | 543 | 483 | 565 | 532 | 501 | 520 | 584 | 529 | 538.63 | 533 | 544 | 1.021 | 0.03 | 0.78224071 | 0.88739843 | 0.0019 | 0.013 | 0.0068 | 0.0068 | TRUE | 0.047 |
| 2310050B05 | 0 | 0 | 0 | 0 | 0 | 0 | 0 | 0 | 0 | 0 | 0 | 0 | 0 | 0 | 0 | NA | NA | NA | NA | NA | NA | NA | NA | NA | NA |
| 2310050C09 | 0 | 0 | 0 | 0 | 0 | 0 | 0 | 0 | 0 | 0 | 0 | 0 | 0 | 0 | 0 | NA | NA | NA | NA | NA | NA | NA | NA | NA | NA |
| 2310057J18F | 0 | 0 | 0 | 0 | 0 | 0 | 0 | 0 | 0 | 0 | 0 | 0 | 0 | 0 | 0 | NA | NA | NA | NA | NA | NA | NA | NA | NA | NA |
| 2310057M21 | 581 | 623 | 606 | 836 | 585 | 612 | 617 | 622 | 608 | 657 | 629 | 670 | 634.06 | 616 | 652 | 1.059 | 0.083 | 0.36698645 | 0.58453273 | 0 | 0.0121 | 0.0045 | 0.0045 | TRUE | 0.0105 |
| 2310057N15 | 0 | 0 | 0 | 0 | 0 | 0 | 0 | 0 | 0 | 0 | 0 | 0 | 0 | 0 | 0 | NA | NA | NA | NA | NA | NA | NA | NA | NA | NA |
| 2310061I04R | 2022 | 2309 | 2352 | 2321 | 1926 | 1951 | 2148 | 2306 | 2362 | 1825 | 2072 | 2136 | 2141.32 | 2272 | 2011 | 0.884 | -0.177 | 0.05898944 | 0.18387917 | 0.0038 | 0.0084 | 0.0059 | 0.0059 | TRUE | 0.081 |
| 2310061J03F | 49 | 48 | 53 | 78 | 60 | 53 | 52 | 48 | 53 | 61 | 65 | 58 | 56.18 | 51 | 61 | 1.201 | 0.264 | 0.29571059 | 0.5165093 | 0 | 0.0655 | 0.0279 | 0.0279 | TRUE | 0.0239 |
| 2310061N02 | 0 | 0 | 0 | 0 | 0 | 0 | 0 | 0 | 0 | 0 | 0 | 0 | 0 | 0 | 0 | NA | NA | NA | NA | NA | NA | NA | NA | NA | NA |
| 2310065F04 | 0 | 0 | 0 | 0 | 0 | 0 | 0 | 0 | 0 | 0 | 0 | 0 | 0 | 0 | 0 | NA | NA | NA | NA | NA | NA | NA | NA | NA | NA |
| 2310067B10 | 4449 | 5238 | 5524 | 5147 | 3792 | 4370 | 4726 | 5231 | 5547 | 4046 | 4080 | 4784 | 4735.55 | 5168 | 4303 | 0.833 | -0.264 | 0.01090135 | 0.05615157 | 0.0076 | 0.0076 | 0.0076 | 0.0076 | TRUE | 0.1158 |

| | | | | | | | | | | | | | | | | | | | | | | | | | |
|---|---|---|---|---|---|---|---|---|---|---|---|---|---|---|---|---|---|---|---|---|---|---|---|---|---|
| 2310068J16I | 16 | 22 | 23 | 27 | 23 | 18 | 17 | 22 | 23 | 21 | 25 | 20 | 21.29 | 21 | 22 | 1.056 | 0.078 | 0.84841215 | 0.92417684 | 0 | 0.1616 | 0.0739 | 0.0739 | TRUE | 0.1284 |
| 2310069B03 | 14 | 11 | 12 | 23 | 15 | 11 | 15 | 11 | 12 | 18 | 16 | 12 | 14.03 | 13 | 15 | 1.231 | 0.299 | 0.55419831 | 0.7374912 | 0 | 0.2418 | 0.1126 | 0.1126 | TRUE | 0.1574 |
| 2310069G16 | 2 | 0 | 1 | 5 | 3 | 2 | 2 | 0 | 1 | 4 | 3 | 2 | 2.08 | 1 | 3 | 3.047 | 1.608 | 0.25728436 | NA | 0 | 1.5913 | 0.8 | 0.8 | TRUE | 0.3809 |
| 2310079G19 | 0 | 0 | 0 | 0 | 0 | 0 | 0 | 0 | 0 | 0 | 0 | 0 | 0 | 0 | 0 | NA | NA | NA | NA | NA | NA | NA | NA | NA | NA |
| 2310081J21I | 0 | 0 | 0 | 0 | 0 | 0 | 0 | 0 | 0 | 0 | 0 | 0 | 0 | 0 | 0 | NA | NA | NA | NA | NA | NA | NA | NA | NA | NA |
| 2410002F23I | 2678 | 2671 | 2424 | 3472 | 2706 | 2538 | 2845 | 2668 | 2434 | 2729 | 2911 | 2778 | 2727.53 | 2649 | 2806 | 1.059 | 0.083 | 0.33738381 | 0.55742902 | 0.003 | 0.0081 | 0.0051 | 0.0051 | TRUE | 0.061 |
| 2410003L11I | 1070 | 631 | 622 | 1011 | 673 | 564 | 1137 | 630 | 625 | 795 | 724 | 617 | 754.59 | 797 | 712 | 0.894 | -0.162 | 0.48282381 | 0.68129391 | 0.0688 | 0.0113 | 0.0369 | 0.0369 | TRUE | 1.644 |
| 2410004B18 | 567 | 425 | 407 | 798 | 542 | 494 | 602 | 424 | 409 | 627 | 583 | 541 | 531.1 | 478 | 584 | 1.221 | 0.289 | 0.08749731 | 0.23954038 | 0.0251 | 0.0131 | 0.0187 | 0.0187 | TRUE | 0.5928 |
| 2410004I01I | 35 | 66 | 60 | 60 | 42 | 48 | 37 | 66 | 60 | 47 | 45 | 53 | 51.37 | 54 | 48 | 0.885 | -0.177 | 0.56834782 | 0.74749682 | 0.0256 | 0.071 | 0.0495 | 0.0495 | TRUE | 0.6254 |
| 2410004N09 | 840 | 1509 | 1576 | 907 | 801 | 1147 | 892 | 1507 | 1582 | 713 | 862 | 1256 | 1135.36 | 1327 | 944 | 0.71 | -0.493 | 0.04582885 | 0.15485794 | 0.0835 | 0.0098 | 0.0431 | 0.0431 | TRUE | 0.9967 |
| 2410004P03I | 110 | 138 | 128 | 177 | 109 | 71 | 117 | 138 | 129 | 139 | 117 | 78 | 119.55 | 128 | 111 | 0.876 | -0.19 | 0.4382103 | 0.64586981 | 0.0358 | 0.0345 | 0.0351 | 0.0351 | TRUE | 0.4816 |
| 2410006H16 | 4949 | 10901 | 11560 | 7009 | 5225 | 7883 | 5257 | 10887 | 11607 | 5510 | 5622 | 8629 | 7918.66 | 9250 | 6587 | 0.712 | -0.49 | 0.20284776 | 0.41048433 | 0.1065 | 0.0073 | 0.0517 | 0.1065 | TRUE | 1.7423 |
| 2410007B07 | 1129 | 1313 | 1349 | 1461 | 1115 | 1176 | 1199 | 1311 | 1355 | 1149 | 1200 | 1287 | 1250.09 | 1288 | 1212 | 0.94 | -0.09 | 0.33827664 | 0.5583911 | 0.0028 | 0.0095 | 0.0055 | 0.0055 | TRUE | 0.0437 |
| 2410012E07I | 0 | 0 | 0 | 1 | 0 | 0 | 0 | 0 | 0 | 1 | 0 | 0 | 0.13 | 0 | 0 | 1.859 | 0.894 | 0.82650734 | NA | 0 | 25.1515 | 10 | 10 | TRUE | 0.2563 |
| 2410012M07 | 483 | 428 | 479 | 720 | 525 | 454 | 513 | 427 | 481 | 566 | 565 | 497 | 508.22 | 474 | 543 | 1.147 | 0.197 | 0.10480494 | 0.26978678 | 0.0048 | 0.0134 | 0.0087 | 0.0087 | TRUE | 0.0855 |
| 2410015M20 | 1393 | 1501 | 1452 | 2024 | 1419 | 1313 | 1480 | 1499 | 1458 | 1591 | 1527 | 1437 | 1498.63 | 1479 | 1518 | 1.027 | 0.039 | 0.6210435 | 0.78565991 | 8.00E-04 | 0.0091 | 0.0038 | 0.0038 | TRUE | 0.0261 |
| 2410016O06 | 1280 | 1421 | 1377 | 2038 | 1425 | 1211 | 1360 | 1419 | 1383 | 1602 | 1533 | 1326 | 1437.06 | 1387 | 1487 | 1.073 | 0.101 | 0.3052414 | 0.526077 | 0.0044 | 0.0092 | 0.0063 | 0.0063 | TRUE | 0.1075 |
| 2410017I17R | 0 | 0 | 0 | 0 | 1 | 0 | 0 | 0 | 0 | 0 | 1 | 0 | 0.18 | 0 | 0 | 1.859 | 0.894 | 0.82650734 | NA | 0 | 18.3785 | 10 | 10 | TRUE | 0.5204 |
| 2410018L13I | 16 | 17 | 22 | 19 | 10 | 28 | 17 | 17 | 22 | 15 | 11 | 31 | 18.73 | 19 | 19 | 0.998 | -0.003 | 0.99554416 | 0.99782172 | 0.1 | 0.1827 | 0.1428 | 0.1428 | TRUE | 1.3658 |
| 2410018M08 | 687 | 658 | 765 | 645 | 581 | 562 | 730 | 657 | 768 | 507 | 625 | 615 | 650.4 | 718 | 582 | 0.809 | -0.305 | 0.01569656 | 0.07335292 | 0.0083 | 0.012 | 0.0099 | 0.0099 | TRUE | 0.1572 |
| 2410021H03 | 33 | 62 | 56 | 30 | 31 | 32 | 35 | 62 | 56 | 24 | 33 | 35 | 40.86 | 51 | 31 | 0.595 | -0.75 | 0.03561042 | 0.12953124 | 0.0427 | 0.0875 | 0.0657 | 0.0657 | TRUE | 0.5997 |
| 2410057H14 | 1 | 0 | 0 | 2 | 0 | 3 | 1 | 0 | 0 | 2 | 0 | 3 | 0.99 | 0 | 2 | 4.416 | 2.143 | 0.35587542 | NA | 0 | 3.3468 | 2.5787 | 2.5787 | TRUE | 0.192 |
| 2410066E13I | 221 | 295 | 314 | 187 | 177 | 210 | 235 | 295 | 315 | 147 | 190 | 230 | 235.33 | 282 | 189 | 0.669 | -0.58 | 0.00483072 | 0.03053712 | 0.0315 | 0.0209 | 0.0262 | 0.0262 | TRUE | 0.3384 |
| 2410076I21R | 1447 | 1866 | 1890 | 1934 | 1411 | 1568 | 1537 | 1864 | 1898 | 1520 | 1518 | 1716 | 1675.54 | 1766 | 1585 | 0.897 | -0.157 | 0.16706395 | 0.3645851 | 0.008 | 0.0089 | 0.0087 | 0.0087 | TRUE | 0.155 |
| 2410088K16I | 139 | 61 | 42 | 99 | 81 | 71 | 148 | 61 | 42 | 78 | 87 | 78 | 82.24 | 84 | 81 | 0.969 | -0.045 | 0.91405949 | 0.95770315 | 0.1957 | 0.047 | 0.1155 | 0.1155 | TRUE | 2.0649 |
| 2410089E03I | 3397 | 2361 | 2821 | 3558 | 2610 | 2275 | 3608 | 2358 | 2833 | 2797 | 2808 | 2490 | 2815.72 | 2933 | 2698 | 0.92 | -0.12 | 0.43277145 | 0.64199587 | 0.0235 | 0.0081 | 0.0165 | 0.0165 | TRUE | 0.3427 |
| 2410114N07 | 17 | 9 | 12 | 9 | 16 | 16 | 18 | 9 | 12 | 7 | 17 | 18 | 13.48 | 13 | 14 | 1.056 | 0.079 | 0.89456813 | 0.9490474 | 0.1068 | 0.2512 | 0.1802 | 0.1802 | TRUE | 1.0194 |
| 2410124H12 | 33 | 38 | 48 | 34 | 26 | 25 | 35 | 38 | 48 | 27 | 28 | 27 | 33.88 | 40 | 27 | 0.676 | -0.565 | 0.08998549 | 0.24406725 | 0 | 0.1041 | 0.0496 | 0.0496 | TRUE | 0.2158 |
| 2410127L17I | 746 | 1046 | 1001 | 1044 | 893 | 912 | 792 | 1045 | 1005 | 821 | 961 | 998 | 937 | 947 | 927 | 0.977 | -0.034 | 0.80893576 | 0.90174222 | 0.015 | 0.0104 | 0.0129 | 0.0129 | TRUE | 0.2432 |
| 2410131K14I | 588 | 578 | 507 | 667 | 492 | 424 | 625 | 577 | 509 | 524 | 529 | 464 | 538.12 | 570 | 506 | 0.888 | -0.171 | 0.16584264 | 0.36290577 | 0.006 | 0.013 | 0.0091 | 0.0091 | TRUE | 0.1035 |
| 2410137F16I | 85 | 150 | 179 | 132 | 120 | 112 | 90 | 150 | 180 | 104 | 129 | 123 | 129.22 | 140 | 119 | 0.844 | -0.244 | 0.36419172 | 0.58164551 | 0.0573 | 0.0324 | 0.0444 | 0.0444 | TRUE | 0.4288 |
| 2410137M14 | 51 | 48 | 38 | 138 | 72 | 52 | 54 | 48 | 38 | 108 | 77 | 57 | 63.86 | 47 | 81 | 1.744 | 0.802 | 0.01233029 | 0.0613868 | 0.0555 | 0.0585 | 0.0571 | 0.0571 | TRUE | 0.216 |
| 2410141K09I | 257 | 265 | 224 | 372 | 267 | 245 | 273 | 265 | 225 | 292 | 287 | 268 | 268.41 | 254 | 282 | 1.113 | 0.155 | 0.26777416 | 0.48754524 | 0.0021 | 0.0192 | 0.0104 | 0.0104 | TRUE | 0.1135 |
| 2500004C02I | 16 | 31 | 25 | 52 | 59 | 24 | 17 | 31 | 25 | 41 | 63 | 26 | 33.95 | 24 | 43 | 1.784 | 0.835 | 0.05631913 | 0.1781519 | 0.1078 | 0.1039 | 0.1057 | 0.1057 | TRUE | 0.2048 |
| 2510002D24 | 223 | 300 | 310 | 335 | 272 | 296 | 237 | 300 | 311 | 263 | 293 | 324 | 287.96 | 283 | 293 | 1.035 | 0.05 | 0.75529193 | 0.87077942 | 0.0123 | 0.0183 | 0.0152 | 0.0152 | TRUE | 0.2223 |
| 2510003D18 | 0 | 0 | 0 | 0 | 0 | 0 | 0 | 0 | 0 | 0 | 0 | 0 | 0 | 0 | 0 | NA | NA | NA | NA | NA | NA | NA | NA | NA | NA |
| 2510003E04I | 1337 | 1494 | 1581 | 2033 | 1519 | 1581 | 1420 | 1492 | 1587 | 1598 | 1634 | 1731 | 1577.15 | 1500 | 1654 | 1.102 | 0.141 | 0.09571577 | 0.25362095 | 0.0017 | 0.009 | 0.0045 | 0.0045 | TRUE | 0.0314 |
| 2510009E07I | 97 | 84 | 76 | 141 | 75 | 72 | 103 | 84 | 76 | 111 | 81 | 79 | 88.93 | 88 | 90 | 1.034 | 0.048 | 0.84423963 | 0.92161735 | 0.0212 | 0.0439 | 0.0325 | 0.0325 | TRUE | 0.4235 |
| 2510039O18 | 2060 | 2593 | 2644 | 3534 | 2702 | 2542 | 2188 | 2590 | 2655 | 2778 | 2907 | 2783 | 2650.09 | 2478 | 2823 | 1.139 | 0.188 | 0.05351234 | 0.17185385 | 0.005 | 0.0081 | 0.0064 | 0.0064 | TRUE | 0.1263 |
| 2510049J12I | 1 | 0 | 0 | 1 | 0 | 1 | 1 | 0 | 0 | 1 | 0 | 1 | 0.49 | 0 | 1 | 1.724 | 0.785 | 0.78200058 | NA | 0 | 6.7235 | 4.0295 | 4.0295 | TRUE | 0.4614 |
| 2600006L11I | 0 | 0 | 0 | 0 | 1 | 0 | 0 | 0 | 0 | 0 | 1 | 0 | 0.18 | 0 | 0 | 1.859 | 0.894 | 0.82650734 | NA | 0 | 18.3785 | 10 | 10 | TRUE | 0.5204 |
| 2610001J05I | 1355 | 1260 | 1203 | 1608 | 1115 | 1159 | 1439 | 1258 | 1208 | 1264 | 1200 | 1269 | 1273 | 1302 | 1244 | 0.956 | -0.065 | 0.51037125 | 0.70388033 | 0.0037 | 0.0095 | 0.0062 | 0.0062 | TRUE | 0.1019 |
| 2610002J02I | 653 | 726 | 761 | 988 | 686 | 647 | 694 | 725 | 764 | 777 | 738 | 708 | 734.3 | 728 | 741 | 1.019 | 0.028 | 0.77361594 | 0.88192776 | 0.001 | 0.0114 | 0.0053 | 0.0053 | TRUE | 0.0228 |
| 2610002M06 | 475 | 723 | 725 | 965 | 737 | 745 | 505 | 722 | 728 | 759 | 793 | 816 | 720.28 | 652 | 789 | 1.21 | 0.275 | 0.07342716 | 0.21257217 | 0.0196 | 0.0115 | 0.0156 | 0.0156 | TRUE | 0.4565 |
| 2610005L07I | 536 | 708 | 657 | 834 | 605 | 670 | 569 | 707 | 660 | 656 | 651 | 733 | 662.68 | 645 | 680 | 1.053 | 0.074 | 0.53907211 | 0.7252619 | 0.0066 | 0.0119 | 0.009 | 0.009 | TRUE | 0.1242 |
| 2610008E11I | 78 | 63 | 94 | 187 | 122 | 96 | 83 | 63 | 94 | 147 | 131 | 105 | 103.92 | 80 | 128 | 1.602 | 0.68 | 0.00474706 | 0.03014373 | 0.0241 | 0.0386 | 0.0315 | 0.0315 | TRUE | 0.3308 |
| 2610015P09I | 894 | 1044 | 1107 | 1249 | 1045 | 982 | 950 | 1043 | 1112 | 982 | 1124 | 1075 | 1047.49 | 1035 | 1060 | 1.024 | 0.034 | 0.74413346 | 0.86342055 | 0.0046 | 0.01 | 0.0068 | 0.0068 | TRUE | 0.0614 |
| 2610016A17 | 2 | 3 | 2 | 2 | 5 | 3 | 2 | 3 | 2 | 2 | 5 | 3 | 2.89 | 2 | 3 | 1.406 | 0.492 | 0.65797286 | NA | 0 | 1.1452 | 0.5283 | 0.5283 | TRUE | 0.2027 |
| 2610017I09R | 1 | 1 | 1 | 3 | 2 | 2 | 1 | 1 | 1 | 2 | 2 | 2 | 1.63 | 1 | 2 | 2.192 | 1.132 | 0.42884344 | NA | 0 | 2.0312 | 0.7567 | 0.7567 | TRUE | 0.0032 |
| 2610018G03 | 459 | 528 | 452 | 761 | 522 | 518 | 488 | 527 | 454 | 598 | 562 | 567 | 532.6 | 490 | 576 | 1.177 | 0.235 | 0.03009592 | 0.11554841 | 0.0014 | 0.0131 | 0.0065 | 0.0065 | TRUE | 0.0535 |
| 2610019E17I | 138 | 192 | 171 | 212 | 177 | 143 | 147 | 192 | 172 | 167 | 190 | 157 | 170.61 | 170 | 171 | 1.006 | 0.008 | 0.96282068 | 0.98130897 | 0.0076 | 0.0262 | 0.0169 | 0.0169 | TRUE | 0.1523 |
| 2610019F03I | 1709 | 1803 | 1753 | 2286 | 1661 | 1534 | 1815 | 1801 | 1760 | 1797 | 1787 | 1679 | 1773.26 | 1792 | 1754 | 0.979 | -0.03 | 0.68084314 | 0.82311098 | 0 | 0.0088 | 0.0033 | 0.0033 | TRUE | 0.0167 |
| 2610020C07I | 33 | 16 | 16 | 43 | 37 | 19 | 35 | 16 | 16 | 34 | 40 | 21 | 26.92 | 22 | 32 | 1.414 | 0.5 | 0.29034702 | 0.51138358 | 0.1152 | 0.1293 | 0.1226 | 0.1226 | TRUE | 1.0716 |
| 2610020H08 | 174 | 287 | 298 | 395 | 277 | 351 | 185 | 287 | 299 | 311 | 298 | 384 | 293.91 | 257 | 331 | 1.286 | 0.363 | 0.08081837 | 0.22612512 | 0.0381 | 0.0181 | 0.0277 | 0.0277 | TRUE | 0.6646 |
| 2610027K06I | 0 | 0 | 2 | 3 | 1 | 1 | 0 | 0 | 2 | 2 | 1 | 1 | 1.09 | 1 | 1 | 2.273 | 1.184 | 0.55074087 | NA | 0 | 3.0306 | 1.7632 | 1.7632 | TRUE | 0.9738 |
| 2610028E06I | 8 | 6 | 4 | 11 | 9 | 5 | 8 | 6 | 4 | 9 | 10 | 5 | 7.05 | 6 | 8 | 1.295 | 0.373 | 0.60586089 | 0.77641012 | 0 | 0.4741 | 0.2322 | 0.2322 | TRUE | 0.2485 |
| 2610028H24 | 1 | 2 | 4 | 5 | 1 | 2 | 1 | 2 | 4 | 4 | 1 | 2 | 2.38 | 2 | 2 | 1.038 | 0.053 | 0.96572457 | NA | 0 | 1.3919 | 0.6886 | 0.6886 | TRUE | 0.4371 |
| 2610034B18 | 150 | 256 | 266 | 333 | 226 | 250 | 159 | 256 | 267 | 262 | 243 | 274 | 243.45 | 227 | 260 | 1.141 | 0.19 | 0.36258525 | 0.58035501 | 0.0344 | 0.0204 | 0.0272 | 0.0272 | TRUE | 0.7463 |
| 2610034M16 | 7 | 2 | 2 | 17 | 6 | 7 | 7 | 2 | 2 | 13 | 6 | 8 | 6.49 | 4 | 9 | 2.449 | 1.292 | 0.13326055 | 0.3151575 | 0.144 | 0.5147 | 0.3458 | 0.3458 | TRUE | 1.0531 |
| 2610035D17 | 124 | 102 | 108 | 96 | 97 | 94 | 132 | 102 | 108 | 75 | 104 | 103 | 104.13 | 114 | 94 | 0.823 | -0.282 | 0.21426589 | 0.42604661 | 0.0162 | 0.0385 | 0.0274 | 0.0274 | TRUE | 0.3292 |
| 2610035F20I | 3 | 1 | 0 | 4 | 3 | 1 | 3 | 1 | 0 | 3 | 3 | 1 | 1.94 | 1 | 2 | 1.818 | 0.862 | 0.55926978 | NA | 0 | 1.7033 | 1.007 | 1.007 | TRUE | 0.7668 |
| 2610037D02 | 1 | 4 | 8 | 4 | 11 | 3 | 1 | 4 | 8 | 3 | 12 | 3 | 5.23 | 4 | 6 | 1.375 | 0.459 | 0.65131913 | 0.80449923 | 0.4549 | 0.6373 | 0.5459 | 0.5459 | TRUE | 0.5306 |
| 2610044O15 | 369 | 342 | 336 | 455 | 352 | 326 | 392 | 342 | 337 | 358 | 379 | 357 | 360.69 | 357 | 365 | 1.021 | 0.03 | 0.80673501 | 0.90035554 | 0.0011 | 0.016 | 0.008 | 0.008 | TRUE | 0.083 |
| 2610100L16I | 0 | 0 | 0 | 0 | 0 | 0 | 0 | 0 | 0 | 0 | 0 | 0 | 0 | 0 | 0 | NA | NA | NA | NA | NA | NA | NA | NA | NA | NA |
| 2610203C20I | 29 | 6 | 10 | 8 | 9 | 4 | 31 | 6 | 10 | 6 | 10 | 4 | 11.2 | 16 | 7 | 0.435 | -1.2 | 0.12115427 | 0.29597972 | 0.3514 | 0.3011 | 0.327 | 0.327 | TRUE | 1.4453 |
| 2610203C22I | 0 | 0 | 0 | 0 | 0 | 0 | 0 | 0 | 0 | 0 | 0 | 0 | 0 | 0 | 0 | NA | NA | NA | NA | NA | NA | NA | NA | NA | NA |
| 2610204G22 | 9 | 13 | 10 | 19 | 10 | 12 | 10 | 13 | 10 | 15 | 11 | 13 | 11.9 | 11 | 13 | 1.197 | 0.26 | 0.63147453 | 0.79271062 | 0 | 0.2837 | 0.1271 | 0.1271 | TRUE | 0.1091 |
| 2610206C17I | 2 | 3 | 4 | 7 | 4 | 3 | 2 | 3 | 4 | 6 | 4 | 3 | 3.7 | 3 | 4 | 1.444 | 0.53 | 0.58010027 | NA | 0 | 0.8962 | 0.3829 | 0.3829 | TRUE | 0.1234 |
| 2610301B20 | 955 | 1190 | 1180 | 1146 | 934 | 908 | 1014 | 1188 | 1185 | 901 | 1005 | 994 | 1047.91 | 1129 | 967 | 0.855 | -0.226 | 0.03217672 | 0.12083124 | 0.0049 | 0.01 | 0.0071 | 0.0071 | TRUE | 0.0943 |

| | | | | | | | | | | | | | | | | | | | | | | | | | |
|---|---|---|---|---|---|---|---|---|---|---|---|---|---|---|---|---|---|---|---|---|---|---|---|---|---|
| 2610301G19 | 2732 | 2616 | 2609 | 3540 | 2826 | 2573 | 2902 | 2613 | 2620 | 2783 | 3041 | 2817 | 2795.7 | 2712 | 2880 | 1.062 | 0.087 | 0.29633221 | 0.51705948 | 0.0025 | 0.0081 | 0.0046 | 0.0046 | TRUE | 0.0457 |
| 2610305D13 | 6311 | 5684 | 5570 | 8606 | 6234 | 5666 | 6703 | 5677 | 5593 | 6765 | 6707 | 6202 | 6274.67 | 5991 | 6558 | 1.095 | 0.131 | 0.17805238 | 0.37877557 | 0.0062 | 0.0074 | 0.0066 | 0.0066 | TRUE | 0.1319 |
| 2610306M01 | 29 | 50 | 42 | 76 | 67 | 40 | 31 | 50 | 42 | 60 | 72 | 44 | 49.76 | 41 | 59 | 1.428 | 0.514 | 0.11471967 | 0.28592898 | 0.0361 | 0.0731 | 0.0556 | 0.0556 | TRUE | 0.183 |
| 2610307P16I | 28 | 32 | 32 | 23 | 9 | 15 | 30 | 32 | 32 | 18 | 10 | 16 | 23 | 31 | 15 | 0.476 | -1.072 | 0.01058755 | 0.05499872 | 0 | 0.1501 | 0.0776 | 0.0776 | TRUE | 0.3724 |
| 2610316D01 | 3 | 12 | 7 | 6 | 3 | 2 | 3 | 12 | 7 | 5 | 3 | 2 | 5.39 | 7 | 3 | 0.464 | -1.109 | 0.21625634 | 0.42827661 | 0 | 0.6182 | 0.3693 | 0.3693 | TRUE | 0.1453 |
| 2610318N02 | 950 | 862 | 877 | 778 | 638 | 574 | 1009 | 861 | 881 | 612 | 686 | 628 | 779.49 | 917 | 642 | 0.7 | -0.515 | 2.44E-06 | 5.99E-05 | 0.0044 | 0.0111 | 0.0073 | 0.0073 | TRUE | 0.0917 |
| 2610507B11 | 40145 | 40002 | 41773 | 54177 | 38516 | 37911 | 42641 | 39950 | 41945 | 42590 | 41441 | 41500 | 41677.68 | 41512 | 41844 | 1.008 | 0.012 | 0.84381371 | 0.92139504 | 5.00E-04 | 0.007 | 0.0024 | 0.0024 | TRUE | 0.0133 |
| 2610507I01R | 7 | 16 | 17 | 10 | 12 | 12 | 7 | 16 | 17 | 8 | 13 | 13 | 12.4 | 13 | 11 | 0.826 | -0.276 | 0.63763446 | 0.79603827 | 0 | 0.2726 | 0.1656 | 0.1656 | TRUE | 0.6267 |
| 2610524H06 | 572 | 668 | 621 | 783 | 544 | 696 | 608 | 667 | 624 | 616 | 585 | 762 | 643.5 | 633 | 654 | 1.033 | 0.047 | 0.71938997 | 0.8479218 | 0.0095 | 0.012 | 0.0107 | 0.0107 | TRUE | 0.2398 |
| 2610528A11 | 9 | 13 | 14 | 15 | 9 | 15 | 10 | 13 | 14 | 12 | 10 | 16 | 12.42 | 12 | 13 | 1.03 | 0.043 | 0.93663484 | 0.96825513 | 0 | 0.2722 | 0.1326 | 0.1326 | TRUE | 0.2508 |
| 2610528J11F | 20 | 74 | 67 | 22 | 15 | 31 | 21 | 74 | 67 | 17 | 16 | 34 | 38.3 | 54 | 22 | 0.412 | -1.279 | 0.01470689 | 0.06982452 | 0.2498 | 0.0929 | 0.1668 | 0.1668 | TRUE | 2.3083 |
| 2700029M09 | 3770 | 5031 | 5075 | 5777 | 4237 | 4649 | 4004 | 5025 | 5096 | 4541 | 4559 | 5089 | 4719 | 4708 | 4730 | 1.004 | 0.006 | 0.95615775 | 0.97741618 | 0.0107 | 0.0076 | 0.0094 | 0.0094 | TRUE | 0.2081 |
| 2700038G22 | 405 | 356 | 379 | 489 | 413 | 380 | 430 | 356 | 381 | 384 | 444 | 416 | 401.84 | 389 | 415 | 1.066 | 0.092 | 0.47421544 | 0.67490502 | 0.0049 | 0.0151 | 0.0095 | 0.0095 | TRUE | 0.0989 |
| 2700046A07 | 3 | 2 | 2 | 6 | 6 | 9 | 3 | 2 | 2 | 5 | 6 | 10 | 4.7 | 2 | 7 | 2.903 | 1.537 | 0.09771642 | 0.25690672 | 0 | 0.7074 | 0.3368 | 0.3368 | TRUE | 0.2769 |
| 2700046G09 | 10 | 14 | 25 | 28 | 18 | 15 | 11 | 14 | 25 | 22 | 19 | 16 | 17.92 | 17 | 19 | 1.165 | 0.221 | 0.65769937 | 0.8072556 | 0.0465 | 0.1908 | 0.1231 | 0.1231 | TRUE | 0.9888 |
| 2700049A03 | 2791 | 2915 | 3079 | 3557 | 2588 | 2746 | 2965 | 2911 | 3092 | 2796 | 2785 | 3006 | 2925.68 | 2989 | 2862 | 0.957 | -0.063 | 0.3849546 | 0.59900394 | 9.00E-04 | 0.008 | 0.0034 | 0.0034 | TRUE | 0.0232 |
| 2700050L05F | 4400 | 4304 | 4405 | 5443 | 3987 | 4137 | 4674 | 4298 | 4423 | 4279 | 4290 | 4529 | 4415.39 | 4465 | 4366 | 0.978 | -0.033 | 0.64292941 | 0.79907039 | 0.0011 | 0.0076 | 0.0033 | 0.0033 | TRUE | 0.0203 |
| 2700054A10 | 45 | 52 | 72 | 37 | 27 | 30 | 48 | 52 | 72 | 29 | 29 | 33 | 43.83 | 57 | 30 | 0.527 | -0.923 | 0.00361013 | 0.02441637 | 0 | 0.0821 | 0.0474 | 0.0474 | TRUE | 0.4442 |
| 2700060E02I | 6300 | 7085 | 7036 | 7876 | 5718 | 5627 | 6692 | 7076 | 7065 | 6191 | 6152 | 6160 | 6555.97 | 6944 | 6168 | 0.888 | -0.171 | 0.00422338 | 0.02755469 | 3.00E-04 | 0.0074 | 0.0024 | 0.0024 | TRUE | 0.0123 |
| 2700062C07I | 536 | 642 | 612 | 798 | 612 | 608 | 569 | 641 | 615 | 627 | 658 | 666 | 629.39 | 608 | 650 | 1.068 | 0.095 | 0.33973013 | 0.55983947 | 0 | 0.0121 | 0.0055 | 0.0055 | TRUE | 0.0366 |
| 2700069I18R | 0 | 0 | 0 | 0 | 0 | 0 | 0 | 0 | 0 | 0 | 0 | 0 | 0 | 0 | 0 | NA | NA | NA | NA | NA | NA | NA | NA | NA | NA |
| 2700070H01 | 15 | 13 | 13 | 13 | 8 | 14 | 16 | 13 | 13 | 10 | 9 | 15 | 12.69 | 14 | 11 | 0.81 | -0.303 | 0.57211547 | 0.7508626 | 0 | 0.2666 | 0.128 | 0.128 | TRUE | 0.3117 |
| 2700081L22F | 1 | 0 | 0 | 2 | 2 | 1 | 1 | 0 | 0 | 2 | 2 | 1 | 0.98 | 0 | 2 | 4.39 | 2.134 | 0.29289438 | NA | 0 | 3.368 | 1.662 | 1.662 | TRUE | 0.4614 |
| 2700081O15 | 711 | 644 | 498 | 896 | 774 | 534 | 755 | 643 | 500 | 704 | 833 | 585 | 670.02 | 633 | 707 | 1.118 | 0.161 | 0.38350281 | 0.5978656 | 0.0348 | 0.0118 | 0.023 | 0.023 | TRUE | 0.2528 |
| 2700086A05 | 0 | 0 | 2 | 0 | 0 | 0 | 0 | 0 | 2 | 0 | 0 | 0 | 0.33 | 1 | 0 | 0.267 | -1.904 | 0.63742563 | NA | 0.0046 | 9.8498 | 10 | 10 | TRUE | 0.974 |
| 2700089E24I | 356 | 359 | 362 | 722 | 530 | 436 | 378 | 359 | 363 | 568 | 570 | 477 | 452.54 | 367 | 538 | 1.47 | 0.556 | 4.56E-06 | 0.00010192 | 0.0036 | 0.0142 | 0.0083 | 0.0083 | TRUE | 0.1122 |
| 2700089I24R | 0 | 0 | 0 | 0 | 0 | 0 | 0 | 0 | 0 | 0 | 0 | 0 | 0 | 0 | 0 | NA | NA | NA | NA | NA | NA | NA | NA | NA | NA |
| 2700094K13I | 4642 | 5268 | 5038 | 7230 | 4896 | 4888 | 4931 | 5261 | 5059 | 5684 | 5268 | 5351 | 5258.77 | 5084 | 5434 | 1.069 | 0.097 | 0.16161151 | 0.35669419 | 0.0011 | 0.0075 | 0.0032 | 0.0032 | TRUE | 0.0199 |
| 2700097O09 | 649 | 793 | 787 | 1068 | 837 | 794 | 689 | 792 | 790 | 840 | 901 | 869 | 813.48 | 757 | 870 | 1.148 | 0.199 | 0.04669088 | 0.15657569 | 0.0023 | 0.011 | 0.006 | 0.006 | TRUE | 0.0721 |
| 2700099C18I | 314 | 350 | 307 | 442 | 389 | 399 | 334 | 350 | 308 | 347 | 419 | 437 | 365.68 | 331 | 401 | 1.21 | 0.275 | 0.04743421 | 0.15806925 | 0.0072 | 0.0159 | 0.0111 | 0.0111 | TRUE | 0.1656 |
| 2810001G20 | 154 | 89 | 91 | 178 | 116 | 101 | 164 | 89 | 91 | 140 | 125 | 111 | 119.86 | 115 | 125 | 1.095 | 0.131 | 0.63587569 | 0.79536224 | 0.0599 | 0.0344 | 0.0467 | 0.0467 | TRUE | 1.3838 |
| 2810002D19 | 98 | 62 | 48 | 112 | 79 | 81 | 104 | 62 | 48 | 88 | 85 | 89 | 79.32 | 71 | 87 | 1.224 | 0.292 | 0.34213837 | 0.56239774 | 0.0627 | 0.0484 | 0.0552 | 0.0552 | TRUE | 1.2348 |
| 2810004N23 | 5112 | 5541 | 5457 | 7721 | 5529 | 4844 | 5430 | 5534 | 5479 | 6070 | 5949 | 5303 | 5627.35 | 5481 | 5774 | 1.054 | 0.075 | 0.33655584 | 0.55668597 | 0.0023 | 0.0075 | 0.0043 | 0.0043 | TRUE | 0.0619 |
| 2810006K23I | 436 | 571 | 676 | 720 | 507 | 485 | 463 | 570 | 679 | 566 | 546 | 531 | 559.09 | 571 | 548 | 0.959 | -0.06 | 0.69308453 | 0.83159723 | 0.0167 | 0.0128 | 0.0149 | 0.0149 | TRUE | 0.1793 |
| 2810007J24F | 0 | 0 | 0 | 0 | 0 | 0 | 0 | 0 | 0 | 0 | 0 | 0 | 0 | 0 | 0 | NA | NA | NA | NA | NA | NA | NA | NA | NA | NA |
| 2810008D09 | 296 | 377 | 386 | 467 | 478 | 527 | 314 | 377 | 388 | 367 | 514 | 577 | 422.8 | 360 | 486 | 1.349 | 0.432 | 0.02112244 | 0.09040097 | 0.0309 | 0.0147 | 0.0228 | 0.0228 | TRUE | 0.5217 |
| 2810011L19F | 304 | 352 | 379 | 298 | 227 | 246 | 323 | 352 | 381 | 234 | 244 | 269 | 300.47 | 352 | 249 | 0.707 | -0.5 | 0.00022491 | 0.00266912 | 0.0027 | 0.0179 | 0.0098 | 0.0098 | TRUE | 0.059 |
| 2810013P06I | 243 | 326 | 314 | 396 | 399 | 271 | 258 | 326 | 315 | 311 | 429 | 297 | 322.71 | 300 | 346 | 1.152 | 0.204 | 0.26683382 | 0.48648172 | 0.0248 | 0.0171 | 0.0213 | 0.0213 | TRUE | 0.5022 |
| 2810021J22F | 613 | 687 | 676 | 896 | 693 | 651 | 651 | 686 | 679 | 704 | 746 | 713 | 696.44 | 672 | 721 | 1.072 | 0.1 | 0.267321 | 0.48702174 | 0 | 0.0116 | 0.0045 | 0.0045 | TRUE | 0.0105 |
| 2810025M15 | 3293 | 3654 | 3728 | 4399 | 3187 | 3281 | 3498 | 3649 | 3743 | 3458 | 3429 | 3592 | 3561.51 | 3630 | 3493 | 0.962 | -0.056 | 0.40369976 | 0.61649426 | 6.00E-04 | 0.0078 | 0.0029 | 0.0029 | TRUE | 0.0123 |
| 2810029C07I | 29 | 26 | 20 | 19 | 24 | 22 | 31 | 26 | 20 | 15 | 26 | 24 | 23.62 | 26 | 22 | 0.835 | -0.26 | 0.54151526 | 0.72712232 | 0 | 0.1464 | 0.0874 | 0.0874 | TRUE | 0.4909 |
| 2810032G03 | 9 | 17 | 15 | 29 | 21 | 17 | 10 | 17 | 15 | 23 | 23 | 19 | 17.6 | 14 | 22 | 1.538 | 0.621 | 0.18192716 | 0.38376383 | 0 | 0.1941 | 0.0965 | 0.0965 | TRUE | 0.304 |
| 2810047C21I | 23 | 19 | 10 | 53 | 26 | 29 | 24 | 19 | 10 | 42 | 28 | 32 | 25.81 | 18 | 34 | 1.912 | 0.935 | 0.03502062 | 0.12812461 | 0.0528 | 0.1346 | 0.0986 | 0.0986 | TRUE | 0.6597 |
| 2810049E08I | 2 | 0 | 0 | 1 | 1 | 2 | 2 | 0 | 0 | 1 | 1 | 2 | 1.03 | 1 | 1 | 1.915 | 0.938 | 0.64606439 | NA | 0 | 3.2076 | 1.903 | 1.903 | TRUE | 0.9744 |
| 2810055G20 | 0 | 0 | 0 | 0 | 0 | 0 | 0 | 0 | 0 | 0 | 0 | 0 | 0 | 0 | 0 | NA | NA | NA | NA | NA | NA | NA | NA | NA | NA |
| 2810403A07 | 3658 | 4073 | 4263 | 4159 | 3298 | 3816 | 3885 | 4068 | 4281 | 3269 | 3548 | 4177 | 3871.47 | 4078 | 3665 | 0.898 | -0.155 | 0.15695351 | 0.34988986 | 0.0088 | 0.0078 | 0.0083 | 0.0083 | TRUE | 0.1815 |
| 2810403D21 | 9 | 5 | 5 | 12 | 4 | 5 | 10 | 5 | 5 | 9 | 4 | 5 | 6.46 | 7 | 6 | 1.002 | 0.003 | 0.9970823 | 0.99828918 | 0 | 0.5166 | 0.2737 | 0.2737 | TRUE | 0.543 |
| 2810404M03 | 0 | 0 | 0 | 0 | 0 | 0 | 0 | 0 | 0 | 0 | 0 | 0 | 0 | 0 | 0 | NA | NA | NA | NA | NA | NA | NA | NA | NA | NA |
| 2810405F15I | 0 | 0 | 0 | 0 | 0 | 0 | 0 | 0 | 0 | 0 | 0 | 0 | 0 | 0 | 0 | NA | NA | NA | NA | NA | NA | NA | NA | NA | NA |
| 2810407C02I | 5728 | 6311 | 6484 | 8156 | 5671 | 5654 | 6084 | 6303 | 6511 | 6412 | 6102 | 6189 | 6266.68 | 6299 | 6234 | 0.99 | -0.015 | 0.8173668 | 0.90708637 | 6.00E-04 | 0.0074 | 0.0028 | 0.0028 | TRUE | 0.0109 |
| 2810408A11 | 350 | 559 | 643 | 441 | 407 | 455 | 372 | 558 | 646 | 347 | 438 | 498 | 476.39 | 525 | 428 | 0.813 | -0.299 | 0.17070756 | 0.3693278 | 0.0528 | 0.0138 | 0.0323 | 0.0323 | TRUE | 0.4658 |
| 2810408I11R | 438 | 410 | 373 | 592 | 426 | 396 | 465 | 409 | 375 | 465 | 458 | 433 | 434.41 | 416 | 452 | 1.088 | 0.121 | 0.33033345 | 0.55109763 | 0.0041 | 0.0145 | 0.0089 | 0.0089 | TRUE | 0.1202 |
| 2810408M09 | 799 | 839 | 718 | 936 | 667 | 585 | 849 | 838 | 721 | 736 | 718 | 640 | 750.23 | 803 | 698 | 0.871 | -0.2 | 0.07823842 | 0.22134105 | 0.0054 | 0.0113 | 0.0079 | 0.0079 | TRUE | 0.0941 |
| 2810410L24F | 42 | 80 | 107 | 97 | 63 | 117 | 45 | 80 | 107 | 76 | 68 | 128 | 84.01 | 77 | 91 | 1.17 | 0.226 | 0.54325398 | 0.72861181 | 0.1351 | 0.0461 | 0.0875 | 0.0875 | TRUE | 0.3012 |
| 2810417H13 | 1774 | 2131 | 2238 | 2910 | 1887 | 1895 | 1884 | 2128 | 2247 | 2288 | 2030 | 2074 | 2108.67 | 2086 | 2131 | 1.022 | 0.031 | 0.75909376 | 0.87295942 | 0.0056 | 0.0085 | 0.0068 | 0.0068 | TRUE | 0.0867 |
| 2810428I15R | 2076 | 2498 | 2587 | 2613 | 1994 | 1965 | 2205 | 2495 | 2598 | 2054 | 2145 | 2151 | 2274.67 | 2433 | 2117 | 0.87 | -0.201 | 0.0251785 | 0.10230464 | 0.0032 | 0.0084 | 0.0054 | 0.0054 | TRUE | 0.0808 |
| 2810429I04R | 14 | 15 | 12 | 72 | 51 | 34 | 15 | 15 | 12 | 57 | 55 | 37 | 31.77 | 14 | 50 | 3.564 | 1.834 | 2.74E-06 | 6.61E-05 | 0 | 0.1106 | 0.0638 | 0.0638 | TRUE | 0.3637 |
| 2810433D01 | 2 | 0 | 0 | 5 | 3 | 2 | 2 | 0 | 0 | 4 | 3 | 2 | 1.91 | 1 | 3 | 4.532 | 2.18 | 0.18452293 | NA | 0 | 1.7299 | 1.0513 | 1.0513 | TRUE | 0.9746 |
| 2810442I21R | 3 | 11 | 10 | 21 | 11 | 9 | 3 | 11 | 10 | 17 | 12 | 10 | 10.4 | 8 | 13 | 1.584 | 0.663 | 0.31241187 | 0.53274517 | 0.0576 | 0.3236 | 0.2095 | 0.2095 | TRUE | 0.7843 |
| 2810442N19 | 0 | 0 | 0 | 0 | 0 | 0 | 0 | 0 | 0 | 0 | 0 | 0 | 0 | 0 | 0 | NA | NA | NA | NA | NA | NA | NA | NA | NA | NA |
| 2810454H06 | 16 | 16 | 10 | 30 | 17 | 14 | 17 | 16 | 10 | 24 | 18 | 15 | 16.7 | 14 | 19 | 1.342 | 0.425 | 0.38169343 | 0.59592841 | 0 | 0.2041 | 0.1087 | 0.1087 | TRUE | 0.3082 |
| 2810459M11 | 52 | 48 | 32 | 93 | 56 | 36 | 55 | 48 | 32 | 73 | 60 | 39 | 51.35 | 45 | 57 | 1.286 | 0.363 | 0.29758233 | 0.5184073 | 0.0636 | 0.0711 | 0.0675 | 0.0675 | TRUE | 0.2876 |
| 2810468N07 | 67 | 105 | 101 | 70 | 55 | 72 | 71 | 105 | 101 | 55 | 59 | 79 | 78.41 | 92 | 64 | 0.692 | -0.532 | 0.04673637 | 0.15663537 | 0.0271 | 0.0489 | 0.0383 | 0.0383 | TRUE | 0.3828 |
| 2810471M01 | 0 | 0 | 0 | 0 | 0 | 0 | 0 | 0 | 0 | 0 | 0 | 0 | 0 | 0 | 0 | NA | NA | NA | NA | NA | NA | NA | NA | NA | NA |
| 2810474O19 | 2612 | 2268 | 2224 | 3399 | 2415 | 2117 | 2774 | 2265 | 2233 | 2672 | 2598 | 2317 | 2476.74 | 2424 | 2529 | 1.044 | 0.062 | 0.59208767 | 0.76555069 | 0.0099 | 0.0082 | 0.0092 | 0.0092 | TRUE | 0.1932 |
| 2900002K06I | 7 | 13 | 8 | 22 | 9 | 10 | 7 | 13 | 8 | 17 | 10 | 11 | 11.06 | 9 | 13 | 1.347 | 0.429 | 0.47318904 | 0.67411593 | 0 | 0.3047 | 0.1662 | 0.1662 | TRUE | 0.5216 |
| 2900005J15F | 23 | 15 | 19 | 12 | 30 | 12 | 24 | 15 | 19 | 9 | 32 | 13 | 18.89 | 19 | 18 | 0.929 | -0.107 | 0.85203179 | 0.92621941 | 0.1837 | 0.1813 | 0.1826 | 0.1826 | TRUE | 2.1564 |
| 2900008C10I | 4 | 5 | 2 | 5 | 12 | 9 | 4 | 5 | 2 | 4 | 13 | 10 | 6.32 | 4 | 9 | 2.34 | 1.227 | 0.15012983 | 0.34074653 | 0 | 0.5278 | 0.3316 | 0.3316 | TRUE | 0.31 |
| 2900009J06F | 48 | 34 | 40 | 53 | 34 | 26 | 51 | 34 | 40 | 42 | 37 | 28 | 38.64 | 42 | 36 | 0.859 | -0.218 | 0.5134485 | 0.70639477 | 0 | 0.0922 | 0.0546 | 0.0546 | TRUE | 0.2811 |

| | | | | | | | | | | | | | | | | | | | | | | | | | |
|---|---|---|---|---|---|---|---|---|---|---|---|---|---|---|---|---|---|---|---|---|---|---|---|---|---|
| 2900011O08 | 546 | 369 | 355 | 652 | 384 | 341 | 580 | 369 | 356 | 513 | 413 | 373 | 433.99 | 435 | 433 | 0.997 | -0.004 | 0.98499841 | 0.99199132 | 0.0492 | 0.0145 | 0.0311 | 0.0311 | TRUE | 0.9796 |
| 2900026A02 | 1077 | 908 | 834 | 1196 | 855 | 820 | 1144 | 907 | 837 | 940 | 920 | 898 | 940.99 | 963 | 919 | 0.955 | -0.066 | 0.61863569 | 0.78452377 | 0.0126 | 0.0104 | 0.0116 | 0.0116 | TRUE | 0.3224 |
| 2900041M22 | 309 | 207 | 227 | 148 | 102 | 110 | 328 | 207 | 228 | 116 | 110 | 120 | 184.9 | 254 | 115 | 0.455 | -1.137 | 1.24E-07 | 4.30E-06 | 0.0293 | 0.0247 | 0.0271 | 0.0271 | TRUE | 0.7147 |
| 2900052N01 | 9 | 4 | 5 | 10 | 3 | 2 | 10 | 4 | 5 | 8 | 3 | 2 | 5.31 | 6 | 4 | 0.738 | -0.438 | 0.62449494 | 0.78834114 | 0.1419 | 0.6275 | 0.3859 | 0.3859 | TRUE | 1.0777 |
| 2900055J20F | 0 | 0 | 0 | 0 | 0 | 0 | 0 | 0 | 0 | 0 | 0 | 0 | 0 | 0 | 0 | NA | NA | NA | NA | NA | NA | NA | NA | NA | NA |
| 2900056M2C | 247 | 307 | 337 | 311 | 274 | 252 | 262 | 307 | 338 | 244 | 295 | 276 | 287.08 | 302 | 272 | 0.896 | -0.158 | 0.30325526 | 0.52410059 | 0.0092 | 0.0184 | 0.0136 | 0.0136 | TRUE | 0.15 |
| 2900057B20 | 0 | 0 | 0 | 0 | 0 | 1 | 0 | 0 | 0 | 0 | 0 | 1 | 0.18 | 0 | 0 | 1.859 | 0.894 | 0.82650734 | NA | 0 | 18.0643 | 10 | 10 | TRUE | 0.5383 |
| 2900060B14 | 2 | 0 | 3 | 2 | 4 | 1 | 2 | 0 | 3 | 2 | 4 | 1 | 2.02 | 2 | 2 | 1.348 | 0.431 | 0.76640847 | NA | 0 | 1.6395 | 1.0057 | 1.0057 | TRUE | 0.6104 |
| 2900076A07 | 80 | 67 | 71 | 90 | 63 | 54 | 85 | 67 | 71 | 71 | 68 | 59 | 70.14 | 74 | 66 | 0.889 | -0.17 | 0.47846464 | 0.67795745 | 0 | 0.0539 | 0.0271 | 0.0271 | TRUE | 0.1382 |
| 2900079G21 | 0 | 0 | 0 | 0 | 0 | 0 | 0 | 0 | 0 | 0 | 0 | 0 | 0 | 0 | 0 | NA | NA | NA | NA | NA | NA | NA | NA | NA | NA |
| 2900092C05I | 0 | 0 | 0 | 0 | 0 | 0 | 0 | 0 | 0 | 0 | 0 | 0 | 0 | 0 | 0 | NA | NA | NA | NA | NA | NA | NA | NA | NA | NA |
| 2900092D14 | 1 | 1 | 1 | 2 | 5 | 2 | 1 | 1 | 1 | 2 | 5 | 2 | 2.03 | 1 | 3 | 2.94 | 1.556 | 0.26520452 | NA | 0 | 1.6263 | 0.7426 | 0.7426 | TRUE | 0.5562 |
| 2900093L17F | 0 | 0 | 0 | 0 | 0 | 1 | 0 | 0 | 0 | 0 | 0 | 1 | 0.18 | 0 | 0 | 1.859 | 0.894 | 0.82650734 | NA | 0 | 18.0643 | 10 | 10 | TRUE | 0.5383 |
| 2900097C17I | 2023 | 2251 | 2409 | 2930 | 2184 | 2271 | 2149 | 2248 | 2419 | 2303 | 2350 | 2486 | 2325.82 | 2272 | 2380 | 1.047 | 0.066 | 0.41867044 | 0.62930126 | 0.002 | 0.0083 | 0.0044 | 0.0044 | TRUE | 0.0388 |
| 3000002C10I | 25 | 24 | 17 | 41 | 26 | 19 | 27 | 24 | 17 | 32 | 28 | 21 | 24.77 | 23 | 27 | 1.208 | 0.272 | 0.50177961 | 0.69709047 | 0 | 0.1399 | 0.0779 | 0.0779 | TRUE | 0.2617 |
| 3010001F23I | 1 | 3 | 2 | 1 | 2 | 0 | 1 | 3 | 2 | 1 | 2 | 0 | 1.5 | 2 | 1 | 0.479 | -1.061 | 0.50778004 | NA | 0 | 2.2021 | 1.0952 | 1.0952 | TRUE | 0.4013 |
| 3010026O09 | 830 | 867 | 915 | 1457 | 1005 | 966 | 882 | 866 | 919 | 1145 | 1081 | 1057 | 991.73 | 889 | 1094 | 1.233 | 0.302 | 0.0004076 | 0.00439275 | 0 | 0.0102 | 0.0042 | 0.0042 | TRUE | 0.0204 |
| 3100003L05F | 1 | 1 | 0 | 1 | 0 | 1 | 1 | 1 | 0 | 1 | 0 | 1 | 0.66 | 1 | 1 | 0.924 | -0.114 | 0.96080322 | NA | 0 | 5.0217 | 2.3843 | 2.3843 | TRUE | 0.2521 |
| 3110001D03 | 1099 | 1017 | 997 | 1530 | 1091 | 1027 | 1167 | 1016 | 1001 | 1203 | 1174 | 1124 | 1114.16 | 1061 | 1167 | 1.1 | 0.138 | 0.15794166 | 0.35154612 | 0.003 | 0.0099 | 0.006 | 0.006 | TRUE | 0.0907 |
| 3110001I22R | 218 | 149 | 146 | 278 | 197 | 139 | 232 | 149 | 147 | 219 | 212 | 152 | 184.94 | 176 | 194 | 1.109 | 0.149 | 0.53560706 | 0.72271961 | 0.0482 | 0.0247 | 0.0361 | 0.0361 | TRUE | 0.8216 |
| 3110002H16 | 1725 | 1597 | 1490 | 1969 | 1474 | 1493 | 1832 | 1595 | 1496 | 1548 | 1586 | 1634 | 1615.24 | 1641 | 1589 | 0.968 | -0.046 | 0.64645577 | 0.80150149 | 0.0052 | 0.0089 | 0.0067 | 0.0067 | TRUE | 0.1248 |
| 3110007F17I | 2 | 1 | 1 | 11 | 4 | 1 | 2 | 1 | 1 | 9 | 4 | 1 | 3.03 | 1 | 5 | 3.493 | 1.805 | 0.1553458 | NA | 0 | 1.0946 | 0.6866 | 0.6866 | TRUE | 0.1272 |
| 3110009E18I | 65 | 61 | 65 | 82 | 49 | 52 | 69 | 61 | 65 | 64 | 53 | 57 | 61.56 | 65 | 58 | 0.896 | -0.158 | 0.52126007 | 0.71159696 | 0 | 0.0604 | 0.0277 | 0.0277 | TRUE | 0.0922 |
| 3110009F21I | 0 | 0 | 0 | 0 | 0 | 0 | 0 | 0 | 0 | 0 | 0 | 0 | 0 | 0 | 0 | NA | NA | NA | NA | NA | NA | NA | NA | NA | NA |
| 3110015C05I | 0 | 0 | 0 | 0 | 0 | 0 | 0 | 0 | 0 | 0 | 0 | 0 | 0 | 0 | 0 | NA | NA | NA | NA | NA | NA | NA | NA | NA | NA |
| 3110021A11 | 19 | 12 | 16 | 14 | 20 | 11 | 20 | 12 | 16 | 11 | 22 | 12 | 15.47 | 16 | 15 | 0.918 | -0.124 | 0.81343554 | 0.90450545 | 0.0397 | 0.2199 | 0.1333 | 0.1333 | TRUE | 0.4965 |
| 3110021N24 | 1 | 2 | 7 | 1 | 1 | 1 | 1 | 2 | 7 | 1 | 1 | 1 | 2.17 | 3 | 1 | 0.289 | -1.792 | 0.21287797 | NA | 0 | 1.5222 | 0.8408 | 0.8408 | TRUE | 1.3156 |
| 3110035E14I | 1 | 1 | 1 | 1 | 0 | 1 | 1 | 1 | 1 | 1 | 0 | 1 | 0.82 | 1 | 1 | 0.621 | -0.688 | 0.72801999 | NA | 0 | 4.0036 | 1.5539 | 1.5539 | TRUE | 0.1955 |
| 3110039I08R | 0 | 0 | 0 | 0 | 0 | 0 | 0 | 0 | 0 | 0 | 0 | 0 | 0 | 0 | 0 | NA | NA | NA | NA | NA | NA | NA | NA | NA | NA |
| 3110039M2C | 0 | 0 | 0 | 0 | 0 | 0 | 0 | 0 | 0 | 0 | 0 | 0 | 0 | 0 | 0 | NA | NA | NA | NA | NA | NA | NA | NA | NA | NA |
| 3110040N11 | 553 | 851 | 789 | 811 | 588 | 636 | 587 | 850 | 792 | 638 | 633 | 696 | 699.32 | 743 | 656 | 0.881 | -0.182 | 0.23009678 | 0.44377649 | 0.0184 | 0.0116 | 0.0152 | 0.0152 | TRUE | 0.3962 |
| 3110043O21 | 1098 | 1272 | 1279 | 1535 | 1188 | 1181 | 1166 | 1270 | 1284 | 1207 | 1278 | 1293 | 1249.76 | 1240 | 1259 | 1.014 | 0.021 | 0.80865394 | 0.90160787 | 0.0013 | 0.0095 | 0.0044 | 0.0044 | TRUE | 0.0325 |
| 3110045C21I | 16 | 5 | 13 | 9 | 7 | 11 | 17 | 5 | 13 | 7 | 8 | 12 | 10.28 | 12 | 9 | 0.757 | -0.402 | 0.54906362 | 0.73382183 | 0.1154 | 0.3273 | 0.2253 | 0.2253 | TRUE | 0.43 |
| 3110047P20I | 2 | 0 | 0 | 7 | 2 | 4 | 2 | 0 | 0 | 6 | 2 | 4 | 2.36 | 1 | 4 | 5.849 | 2.548 | 0.10528432 | NA | 0 | 1.4031 | 0.9234 | 0.9234 | TRUE | 0.9746 |
| 3110052M02 | 80 | 85 | 94 | 102 | 107 | 103 | 85 | 85 | 94 | 80 | 115 | 113 | 95.38 | 88 | 103 | 1.159 | 0.212 | 0.3547063 | 0.57316787 | 0.014 | 0.0414 | 0.0274 | 0.0274 | TRUE | 0.4028 |
| 3110056K07I | 80 | 87 | 104 | 81 | 92 | 77 | 85 | 87 | 104 | 64 | 99 | 84 | 87.21 | 92 | 82 | 0.888 | -0.171 | 0.48802746 | 0.6847406 | 0.0202 | 0.0447 | 0.0325 | 0.0325 | TRUE | 0.3672 |
| 3110057O12 | 204 | 216 | 242 | 338 | 233 | 275 | 217 | 216 | 243 | 266 | 251 | 301 | 248.81 | 225 | 273 | 1.209 | 0.274 | 0.05822557 | 0.18235677 | 0.0026 | 0.0201 | 0.011 | 0.011 | TRUE | 0.0899 |
| 3110062M04 | 216 | 243 | 209 | 305 | 237 | 188 | 229 | 243 | 210 | 240 | 255 | 206 | 230.42 | 227 | 234 | 1.028 | 0.04 | 0.79210543 | 0.89305706 | 0.0038 | 0.0212 | 0.0123 | 0.0123 | TRUE | 0.1131 |
| 3110070M22 | 2 | 1 | 1 | 2 | 2 | 4 | 2 | 1 | 1 | 2 | 2 | 4 | 2.04 | 1 | 3 | 1.943 | 0.958 | 0.47052576 | NA | 0 | 1.6232 | 0.7134 | 0.7134 | TRUE | 0.3215 |
| 3110079O15 | 5 | 10 | 6 | 6 | 5 | 9 | 5 | 10 | 6 | 5 | 5 | 10 | 6.88 | 7 | 7 | 0.922 | -0.117 | 0.87540742 | 0.93854434 | 0 | 0.4859 | 0.2523 | 0.2523 | TRUE | 0.4162 |
| 3110082I17R | 1410 | 1422 | 1453 | 1955 | 1475 | 1509 | 1498 | 1420 | 1459 | 1537 | 1587 | 1652 | 1525.42 | 1459 | 1592 | 1.091 | 0.125 | 0.10045135 | 0.26204412 | 4.00E-04 | 0.0091 | 0.0035 | 0.0035 | TRUE | 0.013 |
| 3110082J24F | 2 | 8 | 6 | 5 | 4 | 9 | 2 | 8 | 6 | 4 | 4 | 10 | 5.7 | 5 | 6 | 1.102 | 0.141 | 0.87118497 | 0.93669162 | 0.1307 | 0.5845 | 0.3658 | 0.3658 | TRUE | 0.5967 |
| 3110099E03I | 0 | 0 | 0 | 0 | 0 | 0 | 0 | 0 | 0 | 0 | 0 | 0 | 0 | 0 | 0 | NA | NA | NA | NA | NA | NA | NA | NA | NA | NA |
| 3200001D21 | 0 | 0 | 0 | 0 | 0 | 0 | 0 | 0 | 0 | 0 | 0 | 0 | 0 | 0 | 0 | NA | NA | NA | NA | NA | NA | NA | NA | NA | NA |
| 3230401D17 | 563 | 520 | 552 | 783 | 516 | 480 | 598 | 519 | 554 | 616 | 555 | 525 | 561.29 | 557 | 565 | 1.017 | 0.024 | 0.8368994 | 0.91714685 | 0.0041 | 0.0128 | 0.0079 | 0.0079 | TRUE | 0.0736 |
| 3300002I08R | 15 | 29 | 20 | 27 | 19 | 18 | 16 | 29 | 20 | 21 | 20 | 20 | 21.06 | 22 | 20 | 0.944 | -0.083 | 0.8470537 | 0.92352627 | 0 | 0.1633 | 0.0864 | 0.0864 | TRUE | 0.4958 |
| 3300005D01 | 0 | 0 | 0 | 0 | 0 | 1 | 0 | 0 | 0 | 0 | 0 | 1 | 0.18 | 0 | 0 | 1.859 | 0.894 | 0.82650734 | NA | 0 | 18.0643 | 10 | 10 | TRUE | 0.5383 |
| 3425401B19 | 11 | 12 | 5 | 18 | 14 | 9 | 12 | 12 | 5 | 14 | 15 | 10 | 11.29 | 10 | 13 | 1.368 | 0.452 | 0.45126386 | 0.65683457 | 0 | 0.2986 | 0.1685 | 0.1685 | TRUE | 0.5888 |
| 3632451O06 | 1459 | 2189 | 2330 | 1693 | 1406 | 1864 | 1550 | 2186 | 2340 | 1331 | 1513 | 2040 | 1826.6 | 2025 | 1628 | 0.803 | -0.316 | 0.10305607 | 0.26724768 | 0.0462 | 0.0087 | 0.0265 | 0.0265 | TRUE | 0.5912 |
| 3632454L22F | 4 | 7 | 5 | 7 | 9 | 10 | 4 | 7 | 5 | 6 | 10 | 11 | 7.07 | 5 | 9 | 1.583 | 0.663 | 0.36330983 | 0.58083215 | 0 | 0.4732 | 0.2322 | 0.2322 | TRUE | 0.4243 |
| 3830403N18 | 15 | 17 | 12 | 108 | 61 | 49 | 16 | 17 | 12 | 85 | 66 | 54 | 41.52 | 15 | 68 | 4.567 | 2.191 | 3.17E-09 | 1.70E-07 | 0 | 0.0862 | 0.0574 | 0.0574 | TRUE | 0.1501 |
| 3830406C13I | 406 | 569 | 564 | 728 | 510 | 533 | 431 | 568 | 566 | 572 | 549 | 583 | 545.05 | 522 | 568 | 1.088 | 0.122 | 0.37150474 | 0.58813674 | 0.0103 | 0.0129 | 0.0115 | 0.0115 | TRUE | 0.2676 |
| 3830408C21I | 42 | 47 | 62 | 51 | 52 | 60 | 45 | 47 | 62 | 40 | 56 | 66 | 52.59 | 51 | 54 | 1.043 | 0.061 | 0.84435635 | 0.92161735 | 0.0305 | 0.0696 | 0.0502 | 0.0502 | TRUE | 0.4022 |
| 3830417A13 | 9 | 5 | 6 | 77 | 57 | 41 | 10 | 5 | 6 | 61 | 61 | 45 | 31.22 | 7 | 56 | 8.164 | 3.029 | 3.92E-11 | 3.10E-09 | 0 | 0.1124 | 0.0678 | 0.0678 | TRUE | 0.2923 |
| 3930402G23 | 1 | 0 | 1 | 1 | 2 | 0 | 1 | 0 | 1 | 1 | 2 | 0 | 0.83 | 1 | 1 | 1.422 | 0.508 | 0.81476789 | NA | 0 | 3.9569 | 2.146 | 2.146 | TRUE | 0.2993 |
| 4430402I18R | 11 | 23 | 32 | 16 | 8 | 20 | 12 | 23 | 32 | 13 | 9 | 22 | 18.31 | 22 | 15 | 0.641 | -0.642 | 0.25850903 | 0.4773685 | 0.1626 | 0.1868 | 0.1752 | 0.1752 | TRUE | 0.1865 |
| 4632415K11I | 760 | 972 | 941 | 998 | 685 | 851 | 807 | 971 | 945 | 785 | 737 | 932 | 862.66 | 908 | 818 | 0.9 | -0.152 | 0.24096924 | 0.45755776 | 0.0106 | 0.0107 | 0.0109 | 0.0109 | TRUE | 0.1733 |
| 4632415L05F | 254 | 339 | 387 | 372 | 299 | 290 | 270 | 339 | 389 | 292 | 322 | 317 | 321.42 | 333 | 310 | 0.933 | -0.1 | 0.53782143 | 0.72419528 | 0.0147 | 0.0172 | 0.0159 | 0.0159 | TRUE | 0.2661 |
| 4632427E13I | 6 | 15 | 6 | 15 | 7 | 5 | 6 | 15 | 6 | 12 | 8 | 5 | 8.7 | 9 | 8 | 0.917 | -0.125 | 0.86024248 | 0.9309173 | 0.113 | 0.3858 | 0.2499 | 0.2499 | TRUE | 1.0375 |
| 4632428C04I | 5 | 11 | 11 | 11 | 9 | 6 | 5 | 11 | 11 | 9 | 10 | 7 | 8.71 | 9 | 9 | 0.91 | -0.136 | 0.83460856 | 0.91571799 | 0 | 0.3853 | 0.1927 | 0.1927 | TRUE | 0.4095 |
| 4632428N05 | 258 | 278 | 274 | 319 | 252 | 246 | 274 | 278 | 275 | 251 | 271 | 269 | 269.67 | 276 | 264 | 0.955 | -0.066 | 0.59973857 | 0.77185449 | 0 | 0.0191 | 0.0077 | 0.0077 | TRUE | 0.0226 |
| 4632434I11R | 2655 | 2863 | 2717 | 3138 | 2415 | 2486 | 2820 | 2859 | 2728 | 2467 | 2598 | 2721 | 2699.02 | 2802 | 2595 | 0.926 | -0.111 | 0.13114285 | 0.31177432 | 0.0012 | 0.0081 | 0.0036 | 0.0036 | TRUE | 0.0233 |
| 4732415M23 | 15 | 17 | 24 | 23 | 14 | 16 | 16 | 17 | 24 | 18 | 15 | 18 | 17.94 | 19 | 17 | 0.89 | -0.168 | 0.70952434 | 0.84171221 | 0 | 0.1905 | 0.0908 | 0.0908 | TRUE | 0.2913 |
| 4732416N19 | 0 | 2 | 1 | 3 | 5 | 2 | 0 | 2 | 1 | 2 | 5 | 2 | 2.15 | 1 | 3 | 3.247 | 1.699 | 0.23448211 | NA | 0 | 1.5358 | 0.818 | 0.818 | TRUE | 0.3941 |
| 4732456N10 | 2 | 0 | 0 | 1 | 0 | 1 | 2 | 0 | 0 | 1 | 0 | 1 | 0.67 | 1 | 1 | 0.902 | -0.15 | 0.95485009 | NA | 0 | 4.9422 | 3.5197 | 3.5197 | TRUE | 0.9743 |
| 4732471J01F | 22 | 28 | 23 | 38 | 20 | 29 | 23 | 28 | 23 | 30 | 22 | 32 | 26.26 | 25 | 28 | 1.119 | 0.163 | 0.66749576 | 0.81402174 | 0 | 0.1324 | 0.065 | 0.065 | TRUE | 0.2261 |
| 4732490B19 | 5 | 2 | 3 | 3 | 0 | 0 | 5 | 2 | 3 | 2 | 0 | 0 | 2.11 | 3 | 1 | 0.254 | -1.976 | 0.2085302 | NA | 0.1296 | 1.5659 | 1.0722 | 1.0722 | TRUE | 1.4363 |
| 4732491K20I | 14 | 21 | 11 | 30 | 20 | 6 | 15 | 21 | 11 | 24 | 22 | 7 | 16.43 | 16 | 18 | 1.112 | 0.153 | 0.79593763 | 0.8952099 | 0.1729 | 0.2075 | 0.1913 | 0.1913 | TRUE | 1.223 |
| 4831440E17I | 50 | 56 | 78 | 71 | 77 | 84 | 53 | 56 | 78 | 56 | 83 | 92 | 69.66 | 62 | 77 | 1.223 | 0.29 | 0.32781967 | 0.54900652 | 0.0432 | 0.0542 | 0.0487 | 0.0487 | TRUE | 0.5905 |

| | | | | | | | | | | | | | | | | | | | | | | | | | |
|---|---|---|---|---|---|---|---|---|---|---|---|---|---|---|---|---|---|---|---|---|---|---|---|---|---|
| 4833403I15R | 10 | 10 | 6 | 38 | 12 | 20 | 11 | 10 | 6 | 30 | 13 | 22 | 15.22 | 9 | 22 | 2.454 | 1.295 | 0.02214572 | 0.09359118 | 0 | 0.2234 | 0.1508 | 0.1508 | TRUE | 0.1089 |
| 4833412C05I | 0 | 3 | 1 | 1 | 1 | 0 | 0 | 3 | 1 | 1 | 1 | 0 | 0.98 | 1 | 1 | 0.47 | -1.091 | 0.60223219 | NA | 0 | 3.3787 | 1.9999 | 1.9999 | TRUE | 0.3082 |
| 4833417C18I | 97 | 112 | 133 | 118 | 119 | 96 | 103 | 112 | 134 | 93 | 128 | 105 | 112.39 | 116 | 109 | 0.931 | -0.103 | 0.63540171 | 0.79504608 | 0.0139 | 0.0362 | 0.025 | 0.025 | TRUE | 0.2397 |
| 4833418N02 | 5 | 4 | 9 | 8 | 4 | 5 | 5 | 4 | 9 | 6 | 4 | 5 | 5.73 | 6 | 5 | 0.881 | -0.182 | 0.81779109 | 0.90711028 | 0 | 0.5814 | 0.2758 | 0.2758 | TRUE | 0.4234 |
| 4833419F23I | 1 | 2 | 3 | 4 | 3 | 3 | 1 | 2 | 3 | 3 | 3 | 3 | 2.62 | 2 | 3 | 1.582 | 0.662 | 0.55923142 | NA | 0 | 1.2637 | 0.5221 | 0.5221 | TRUE | 0.1658 |
| 4833420G17 | 1334 | 1159 | 1140 | 1447 | 1122 | 1091 | 1417 | 1158 | 1145 | 1138 | 1207 | 1194 | 1209.69 | 1240 | 1180 | 0.951 | -0.072 | 0.52051037 | 0.71109881 | 0.0067 | 0.0096 | 0.0082 | 0.0082 | TRUE | 0.187 |
| 4833422C13I | 1 | 0 | 0 | 0 | 0 | 1 | 1 | 0 | 0 | 0 | 0 | 1 | 0.36 | 0 | 0 | 0.954 | -0.067 | 0.98553519 | NA | 0 | 9.1716 | 7.9605 | 7.9605 | TRUE | 0.5383 |
| 4833423E24I | 0 | 1 | 0 | 4 | 1 | 4 | 0 | 1 | 0 | 3 | 1 | 4 | 1.6 | 0 | 3 | 7.845 | 2.972 | 0.10254641 | NA | 0 | 2.0664 | 1.217 | 1.217 | TRUE | 0.338 |
| 4833424O15 | 0 | 2 | 0 | 0 | 1 | 0 | 0 | 2 | 0 | 0 | 1 | 0 | 0.51 | 1 | 0 | 0.522 | -0.937 | 0.77976772 | NA | 0 | 6.4385 | 6.3352 | 6.3352 | TRUE | 0.9739 |
| 4833427F10I | 0 | 0 | 0 | 0 | 0 | 0 | 0 | 0 | 0 | 0 | 0 | 0 | 0 | 0 | 0 | NA | NA | NA | NA | NA | NA | NA | NA | NA | NA |
| 4833427G06 | 2 | 4 | 0 | 5 | 7 | 3 | 2 | 4 | 0 | 4 | 8 | 3 | 3.48 | 2 | 5 | 2.398 | 1.262 | 0.28000528 | NA | 0.1046 | 0.9542 | 0.633 | 0.633 | TRUE | 0.479 |
| 4833428L15F | 2 | 1 | 1 | 2 | 0 | 5 | 2 | 1 | 1 | 2 | 0 | 5 | 1.86 | 1 | 2 | 1.695 | 0.762 | 0.62542769 | NA | 0 | 1.7761 | 1.1705 | 1.1705 | TRUE | 0.4678 |
| 4833439L19F | 8331 | 7237 | 7005 | 9161 | 6290 | 6045 | 8849 | 7228 | 7034 | 7202 | 6768 | 6617 | 7282.82 | 7704 | 6862 | 0.891 | -0.167 | 0.12177264 | 0.29675497 | 0.0091 | 0.0074 | 0.0082 | 0.0082 | TRUE | 0.2065 |
| 4921501E09I | 0 | 0 | 0 | 0 | 0 | 0 | 0 | 0 | 0 | 0 | 0 | 0 | 0 | 0 | 0 | NA | NA | NA | NA | NA | NA | NA | NA | NA | NA |
| 4921504E06I | 1 | 1 | 1 | 1 | 1 | 1 | 1 | 1 | 1 | 1 | 1 | 1 | 1 | 1 | 1 | 0.955 | -0.066 | 0.96933682 | NA | 0 | 3.2895 | 1.1311 | 1.1311 | TRUE | 0.0204 |
| 4921506M07 | 6 | 10 | 8 | 25 | 12 | 10 | 6 | 10 | 8 | 20 | 13 | 11 | 11.32 | 8 | 15 | 1.8 | 0.848 | 0.15069706 | 0.3416207 | 0 | 0.298 | 0.155 | 0.155 | TRUE | 0.5246 |
| 4921507L20F | 0 | 0 | 0 | 0 | 0 | 0 | 0 | 0 | 0 | 0 | 0 | 0 | 0 | 0 | 0 | NA | NA | NA | NA | NA | NA | NA | NA | NA | NA |
| 4921507P07I | 58 | 37 | 55 | 67 | 45 | 34 | 62 | 37 | 55 | 53 | 48 | 37 | 48.68 | 51 | 46 | 0.904 | -0.145 | 0.64901161 | 0.8032811 | 0.0283 | 0.0746 | 0.0525 | 0.0525 | TRUE | 0.4941 |
| 4921508A21 | 0 | 0 | 0 | 0 | 0 | 0 | 0 | 0 | 0 | 0 | 0 | 0 | 0 | 0 | 0 | NA | NA | NA | NA | NA | NA | NA | NA | NA | NA |
| 4921508D12 | 0 | 0 | 0 | 0 | 0 | 0 | 0 | 0 | 0 | 0 | 0 | 0 | 0 | 0 | 0 | NA | NA | NA | NA | NA | NA | NA | NA | NA | NA |
| 4921509C19I | 11 | 6 | 6 | 14 | 7 | 9 | 12 | 6 | 6 | 11 | 8 | 10 | 8.68 | 8 | 10 | 1.209 | 0.274 | 0.67623301 | 0.81979608 | 0 | 0.3864 | 0.1941 | 0.1941 | TRUE | 0.4876 |
| 4921509O07 | 0 | 0 | 0 | 0 | 1 | 0 | 0 | 0 | 0 | 0 | 1 | 0 | 0.18 | 0 | 0 | 1.859 | 0.894 | 0.82650734 | NA | 0 | 18.3785 | 10 | 10 | TRUE | 0.5204 |
| 4921511C10I | 0 | 2 | 1 | 0 | 0 | 0 | 0 | 2 | 1 | 0 | 0 | 0 | 0.5 | 1 | 0 | 0.179 | -2.485 | 0.44922046 | NA | 0 | 6.5925 | 6.2606 | 6.2606 | TRUE | 0.0526 |
| 4921511C20I | 0 | 0 | 1 | 0 | 0 | 0 | 0 | 0 | 1 | 0 | 0 | 0 | 0.17 | 0 | 0 | 0.49 | -1.029 | 0.80087 | NA | 0 | 19.6928 | 10 | 10 | TRUE | 0.4098 |
| 4921511H03 | 0 | 0 | 0 | 0 | 0 | 0 | 0 | 0 | 0 | 0 | 0 | 0 | 0 | 0 | 0 | NA | NA | NA | NA | NA | NA | NA | NA | NA | NA |
| 4921511I17R | 0 | 0 | 0 | 1 | 0 | 0 | 0 | 0 | 0 | 1 | 0 | 0 | 0.13 | 0 | 0 | 1.859 | 0.894 | 0.82650734 | NA | 0 | 25.1515 | 10 | 10 | TRUE | 0.2563 |
| 4921511M17 | 0 | 0 | 0 | 0 | 0 | 0 | 0 | 0 | 0 | 0 | 0 | 0 | 0 | 0 | 0 | NA | NA | NA | NA | NA | NA | NA | NA | NA | NA |
| 4921513I03R | 3 | 0 | 0 | 2 | 0 | 0 | 3 | 0 | 0 | 2 | 0 | 0 | 0.79 | 1 | 1 | 0.523 | -0.934 | 0.754563 | NA | 8.9049 | 4.1606 | 5.0595 | 5.0595 | TRUE | 1.4426 |
| 4921515E04I | 0 | 0 | 0 | 0 | 0 | 0 | 0 | 0 | 0 | 0 | 0 | 0 | 0 | 0 | 0 | NA | NA | NA | NA | NA | NA | NA | NA | NA | NA |
| 4921517D22 | 0 | 0 | 0 | 0 | 0 | 2 | 0 | 0 | 0 | 0 | 0 | 2 | 0.36 | 0 | 1 | 3.832 | 1.938 | 0.63022034 | NA | 10 | 9.0356 | 10 | 10 | TRUE | 0.9744 |
| 4921524J17F | 2056 | 2185 | 2119 | 2050 | 1668 | 1672 | 2184 | 2182 | 2128 | 1612 | 1795 | 1830 | 1955.03 | 2165 | 1746 | 0.806 | -0.312 | 0.00016013 | 0.00201401 | 0.0019 | 0.0086 | 0.0044 | 0.0044 | TRUE | 0.056 |
| 4921524L21F | 0 | 0 | 0 | 0 | 0 | 0 | 0 | 0 | 0 | 0 | 0 | 0 | 0 | 0 | 0 | NA | NA | NA | NA | NA | NA | NA | NA | NA | NA |
| 4921525B02 | 0 | 0 | 0 | 0 | 1 | 0 | 0 | 0 | 0 | 0 | 1 | 0 | 0.18 | 0 | 0 | 1.859 | 0.894 | 0.82650734 | NA | 0 | 18.3785 | 10 | 10 | TRUE | 0.5204 |
| 4921525O09 | 92 | 186 | 155 | 238 | 199 | 207 | 98 | 186 | 156 | 187 | 214 | 227 | 177.82 | 147 | 209 | 1.426 | 0.512 | 0.03306127 | 0.12295269 | 0.0476 | 0.0254 | 0.0358 | 0.0358 | TRUE | 0.6827 |
| 4921530D09 | 0 | 0 | 0 | 0 | 0 | 0 | 0 | 0 | 0 | 0 | 0 | 0 | 0 | 0 | 0 | NA | NA | NA | NA | NA | NA | NA | NA | NA | NA |
| 4921530L18F | 7 | 9 | 5 | 16 | 13 | 8 | 7 | 9 | 5 | 13 | 14 | 9 | 9.46 | 7 | 12 | 1.652 | 0.724 | 0.24872973 | 0.46563387 | 0 | 0.3551 | 0.1712 | 0.1712 | TRUE | 0.1868 |
| 4921530L21F | 0 | 0 | 0 | 0 | 0 | 0 | 0 | 0 | 0 | 0 | 0 | 0 | 0 | 0 | 0 | NA | NA | NA | NA | NA | NA | NA | NA | NA | NA |
| 4921531C22I | 12 | 10 | 12 | 22 | 22 | 12 | 13 | 10 | 12 | 17 | 24 | 13 | 14.81 | 12 | 18 | 1.556 | 0.637 | 0.21133824 | 0.42189873 | 0 | 0.2293 | 0.1162 | 0.1162 | TRUE | 0.2268 |
| 4921531P14I | 0 | 0 | 0 | 0 | 0 | 0 | 0 | 0 | 0 | 0 | 0 | 0 | 0 | 0 | 0 | NA | NA | NA | NA | NA | NA | NA | NA | NA | NA |
| 4921533I20R | 2 | 0 | 0 | 2 | 0 | 0 | 2 | 0 | 0 | 2 | 0 | 0 | 0.62 | 1 | 1 | 0.783 | -0.353 | 0.91186553 | NA | 0 | 5.3542 | 5.6997 | 5.6997 | TRUE | 0.9742 |
| 4921536K21I | 50 | 89 | 84 | 58 | 46 | 50 | 53 | 89 | 84 | 46 | 49 | 55 | 62.69 | 75 | 50 | 0.659 | -0.601 | 0.03788063 | 0.13526222 | 0.0279 | 0.0595 | 0.0439 | 0.0439 | TRUE | 0.6008 |
| 4921539E11I | 1 | 0 | 0 | 2 | 2 | 1 | 1 | 0 | 0 | 2 | 2 | 1 | 0.98 | 0 | 2 | 4.39 | 2.134 | 0.29289438 | NA | 0 | 3.368 | 1.662 | 1.662 | TRUE | 0.4614 |
| 4922501C03I | 987 | 963 | 962 | 1489 | 1218 | 979 | 1048 | 962 | 966 | 1171 | 1310 | 1072 | 1088.13 | 992 | 1184 | 1.194 | 0.256 | 0.01662898 | 0.07637515 | 0.005 | 0.0099 | 0.0073 | 0.0073 | TRUE | 0.1024 |
| 4922501L14F | 6 | 10 | 4 | 4 | 4 | 10 | 6 | 10 | 4 | 3 | 4 | 11 | 6.46 | 7 | 6 | 0.886 | -0.174 | 0.83440146 | 0.91571799 | 0.1693 | 0.5167 | 0.3426 | 0.3426 | TRUE | 0.7797 |
| 4922502D21 | 0 | 0 | 0 | 1 | 0 | 0 | 0 | 0 | 0 | 1 | 0 | 0 | 0.13 | 0 | 0 | 1.859 | 0.894 | 0.82650734 | NA | 0 | 25.1515 | 10 | 10 | TRUE | 0.2563 |
| 4922502H24 | 0 | 0 | 0 | 0 | 0 | 0 | 0 | 0 | 0 | 0 | 0 | 0 | 0 | 0 | 0 | NA | NA | NA | NA | NA | NA | NA | NA | NA | NA |
| 4922502N22 | 0 | 0 | 0 | 2 | 1 | 1 | 0 | 0 | 0 | 2 | 1 | 1 | 0.62 | 0 | 1 | 6.676 | 2.739 | 0.27800695 | NA | 0 | 5.2881 | 3.2034 | 3.2034 | TRUE | 0.07 |
| 4922505E12I | 0 | 1 | 0 | 3 | 1 | 1 | 0 | 1 | 0 | 2 | 1 | 1 | 0.92 | 0 | 1 | 4.189 | 2.067 | 0.3213176 | NA | 0 | 3.5829 | 1.8098 | 1.8098 | TRUE | 0.405 |
| 4930401C15I | 0 | 0 | 0 | 0 | 0 | 0 | 0 | 0 | 0 | 0 | 0 | 0 | 0 | 0 | 0 | NA | NA | NA | NA | NA | NA | NA | NA | NA | NA |
| 4930401O10 | 0 | 0 | 0 | 0 | 0 | 0 | 0 | 0 | 0 | 0 | 0 | 0 | 0 | 0 | 0 | NA | NA | NA | NA | NA | NA | NA | NA | NA | NA |
| 4930401O12 | 0 | 0 | 0 | 0 | 0 | 0 | 0 | 0 | 0 | 0 | 0 | 0 | 0 | 0 | 0 | NA | NA | NA | NA | NA | NA | NA | NA | NA | NA |
| 4930402F06I | 0 | 0 | 1 | 0 | 0 | 0 | 0 | 0 | 1 | 0 | 0 | 0 | 0.17 | 0 | 0 | 0.49 | -1.029 | 0.80087 | NA | 0 | 19.6928 | 10 | 10 | TRUE | 0.4098 |
| 4930402F11I | 0 | 0 | 0 | 0 | 0 | 0 | 0 | 0 | 0 | 0 | 0 | 0 | 0 | 0 | 0 | NA | NA | NA | NA | NA | NA | NA | NA | NA | NA |
| 4930402H24 | 1986 | 2545 | 2796 | 2479 | 1884 | 2004 | 2109 | 2542 | 2807 | 1949 | 2027 | 2194 | 2271.37 | 2486 | 2057 | 0.827 | -0.274 | 0.02418966 | 0.09932812 | 0.0112 | 0.0084 | 0.0102 | 0.0102 | TRUE | 0.2126 |
| 4930402K13I | 0 | 0 | 0 | 5 | 0 | 0 | 0 | 0 | 0 | 4 | 0 | 0 | 0.66 | 0 | 1 | 7.081 | 2.824 | 0.45988427 | NA | 10 | 5.0358 | 9.153 | 9.153 | TRUE | 2.3415 |
| 4930404A05 | 0 | 0 | 0 | 0 | 0 | 0 | 0 | 0 | 0 | 0 | 0 | 0 | 0 | 0 | 0 | NA | NA | NA | NA | NA | NA | NA | NA | NA | NA |
| 4930404A10 | 3 | 2 | 8 | 15 | 19 | 13 | 3 | 2 | 8 | 12 | 20 | 14 | 9.95 | 4 | 15 | 3.49 | 1.803 | 0.01234282 | 0.06139101 | 0.0645 | 0.3381 | 0.2268 | 0.2268 | TRUE | 0.9579 |
| 4930404H11 | 0 | 0 | 0 | 0 | 0 | 0 | 0 | 0 | 0 | 0 | 0 | 0 | 0 | 0 | 0 | NA | NA | NA | NA | NA | NA | NA | NA | NA | NA |
| 4930404I05R | 0 | 0 | 0 | 1 | 0 | 0 | 0 | 0 | 0 | 1 | 0 | 0 | 0.13 | 0 | 0 | 1.859 | 0.894 | 0.82650734 | NA | 0 | 25.1515 | 10 | 10 | TRUE | 0.2563 |
| 4930404N11 | 1 | 2 | 0 | 0 | 3 | 1 | 1 | 2 | 0 | 0 | 3 | 1 | 1.23 | 1 | 1 | 1.375 | 0.459 | 0.8150902 | NA | 0 | 2.6846 | 1.9283 | 1.9283 | TRUE | 0.3847 |
| 4930405A10 | 0 | 0 | 0 | 0 | 0 | 0 | 0 | 0 | 0 | 0 | 0 | 0 | 0 | 0 | 0 | NA | NA | NA | NA | NA | NA | NA | NA | NA | NA |
| 4930405A21 | 11 | 12 | 14 | 11 | 15 | 11 | 12 | 12 | 14 | 9 | 16 | 12 | 12.43 | 13 | 12 | 0.965 | -0.052 | 0.92397679 | 0.96274037 | 0 | 0.272 | 0.1332 | 0.1332 | TRUE | 0.2659 |
| 4930405D11 | 0 | 0 | 0 | 0 | 0 | 0 | 0 | 0 | 0 | 0 | 0 | 0 | 0 | 0 | 0 | NA | NA | NA | NA | NA | NA | NA | NA | NA | NA |
| 4930405J17F | 0 | 0 | 0 | 0 | 0 | 0 | 0 | 0 | 0 | 0 | 0 | 0 | 0 | 0 | 0 | NA | NA | NA | NA | NA | NA | NA | NA | NA | NA |
| 4930405P13I | 0 | 0 | 0 | 0 | 0 | 1 | 0 | 0 | 0 | 0 | 0 | 1 | 0.18 | 0 | 0 | 1.859 | 0.894 | 0.82650734 | NA | 0 | 18.0643 | 10 | 10 | TRUE | 0.5383 |
| 4930406D18 | 0 | 0 | 0 | 0 | 0 | 0 | 0 | 0 | 0 | 0 | 0 | 0 | 0 | 0 | 0 | NA | NA | NA | NA | NA | NA | NA | NA | NA | NA |
| 4930407I10R | 0 | 0 | 0 | 1 | 0 | 0 | 0 | 0 | 0 | 1 | 0 | 0 | 0.13 | 0 | 0 | 1.859 | 0.894 | 0.82650734 | NA | 0 | 25.1515 | 10 | 10 | TRUE | 0.2563 |
| 4930408F14I | 0 | 5 | 5 | 3 | 5 | 2 | 0 | 5 | 5 | 2 | 5 | 2 | 3.32 | 3 | 3 | 0.978 | -0.032 | 0.9788569 | NA | 0.3729 | 0.9981 | 0.7476 | 0.7476 | TRUE | 1.0284 |
| 4930412B13 | 0 | 0 | 0 | 0 | 0 | 0 | 0 | 0 | 0 | 0 | 0 | 0 | 0 | 0 | 0 | NA | NA | NA | NA | NA | NA | NA | NA | NA | NA |

| | | | | | | | | | | | | | | | | | | | | | | | | | |
|---|---|---|---|---|---|---|---|---|---|---|---|---|---|---|---|---|---|---|---|---|---|---|---|---|---|
| 4930412C18I | 31 | 27 | 15 | 32 | 31 | 14 | 33 | 27 | 15 | 25 | 33 | 15 | 24.8 | 25 | 24 | 0.988 | -0.018 | 0.96969581 | 0.98448447 | 0.0984 | 0.1398 | 0.1203 | 0.1203 | TRUE | 0.2665 |
| 4930412O13 | 1 | 1 | 0 | 4 | 4 | 0 | 1 | 1 | 0 | 3 | 4 | 0 | 1.58 | 1 | 2 | 3.656 | 1.87 | 0.30821441 | NA | 0 | 2.0856 | 1.485 | 1.485 | TRUE | 0.3981 |
| 4930413E15I | 0 | 0 | 0 | 0 | 0 | 0 | 0 | 0 | 0 | 0 | 0 | 0 | 0 | 0 | 0 | NA | NA | NA | NA | NA | NA | NA | NA | NA | NA |
| 4930413F20I | 1 | 0 | 0 | 1 | 0 | 0 | 1 | 0 | 0 | 1 | 0 | 0 | 0.31 | 0 | 0 | 0.954 | -0.067 | 0.98625928 | NA | 0 | 10.7014 | 9.0384 | 9.0384 | TRUE | 0.4614 |
| 4930413G21 | 2 | 2 | 7 | 4 | 6 | 6 | 2 | 2 | 7 | 3 | 6 | 7 | 4.55 | 4 | 5 | 1.427 | 0.513 | 0.59045888 | NA | 0.0954 | 0.7305 | 0.4252 | 0.4252 | TRUE | 0.9636 |
| 4930413M19 | 0 | 0 | 0 | 0 | 0 | 0 | 0 | 0 | 0 | 0 | 0 | 0 | 0 | 0 | 0 | NA | NA | NA | NA | NA | NA | NA | NA | NA | NA |
| 4930414L22F | 15 | 52 | 61 | 48 | 40 | 31 | 16 | 52 | 61 | 38 | 43 | 34 | 40.64 | 43 | 38 | 0.887 | -0.174 | 0.71154684 | 0.84297429 | 0.1878 | 0.088 | 0.134 | 0.134 | TRUE | 1.3732 |
| 4930414N06 | 0 | 0 | 0 | 0 | 0 | 0 | 0 | 0 | 0 | 0 | 0 | 0 | 0 | 0 | 0 | NA | NA | NA | NA | NA | NA | NA | NA | NA | NA |
| 4930415F15I | 1 | 0 | 0 | 4 | 3 | 1 | 1 | 0 | 0 | 3 | 3 | 1 | 1.42 | 0 | 2 | 6.85 | 2.776 | 0.13485814 | NA | 0 | 2.3245 | 1.2887 | 1.2887 | TRUE | 0.4614 |
| 4930415L06F | 0 | 0 | 0 | 0 | 0 | 0 | 0 | 0 | 0 | 0 | 0 | 0 | 0 | 0 | 0 | NA | NA | NA | NA | NA | NA | NA | NA | NA | NA |
| 4930415O20 | 1 | 3 | 2 | 3 | 1 | 1 | 1 | 3 | 2 | 2 | 1 | 1 | 1.77 | 2 | 1 | 0.764 | -0.389 | 0.77617093 | NA | 0 | 1.8725 | 0.7843 | 0.7843 | TRUE | 0.225 |
| 4930417O13 | 1 | 1 | 1 | 3 | 1 | 3 | 1 | 1 | 1 | 2 | 1 | 3 | 1.63 | 1 | 2 | 2.195 | 1.134 | 0.43744803 | NA | 0 | 2.0274 | 0.8193 | 0.8193 | TRUE | 0.1809 |
| 4930417O22 | 28 | 15 | 13 | 41 | 49 | 18 | 30 | 15 | 13 | 32 | 53 | 20 | 27.07 | 19 | 35 | 1.815 | 0.86 | 0.09479547 | 0.25228627 | 0.1739 | 0.1286 | 0.1503 | 0.1503 | TRUE | 0.2429 |
| 4930419G24 | 0 | 0 | 0 | 0 | 0 | 0 | 0 | 0 | 0 | 0 | 0 | 0 | 0 | 0 | 0 | NA | NA | NA | NA | NA | NA | NA | NA | NA | NA |
| 4930422G04 | 1323 | 1276 | 1502 | 1894 | 1416 | 1261 | 1405 | 1274 | 1508 | 1489 | 1524 | 1380 | 1430.1 | 1396 | 1464 | 1.049 | 0.069 | 0.4787842 | 0.67817611 | 0.0038 | 0.0092 | 0.0062 | 0.0062 | TRUE | 0.0701 |
| 4930423M02 | 0 | 0 | 0 | 0 | 0 | 0 | 0 | 0 | 0 | 0 | 0 | 0 | 0 | 0 | 0 | NA | NA | NA | NA | NA | NA | NA | NA | NA | NA |
| 4930425K10I | 0 | 0 | 0 | 0 | 0 | 0 | 0 | 0 | 0 | 0 | 0 | 0 | 0 | 0 | 0 | NA | NA | NA | NA | NA | NA | NA | NA | NA | NA |
| 4930425O10 | 0 | 1 | 0 | 0 | 1 | 1 | 0 | 1 | 0 | 0 | 1 | 1 | 0.53 | 0 | 1 | 1.924 | 0.944 | 0.7364225 | NA | 0 | 6.2438 | 3.9727 | 3.9727 | TRUE | 0.405 |
| 4930426D05I | 1 | 3 | 4 | 2 | 3 | 0 | 1 | 3 | 4 | 2 | 3 | 0 | 2.15 | 3 | 2 | 0.593 | -0.753 | 0.58999521 | NA | 0 | 1.5422 | 0.9131 | 0.9131 | TRUE | 0.2742 |
| 4930426L09F | 0 | 3 | 5 | 1 | 1 | 1 | 0 | 3 | 5 | 1 | 1 | 1 | 1.83 | 3 | 1 | 0.363 | -1.461 | 0.35757098 | NA | 0 | 1.8082 | 1.1265 | 1.1265 | TRUE | 0.1565 |
| 4930427A07I | 2431 | 2645 | 2680 | 2755 | 2124 | 2159 | 2582 | 2642 | 2691 | 2166 | 2285 | 2363 | 2454.86 | 2638 | 2271 | 0.86 | -0.217 | 0.00265913 | 0.01928269 | 8.00E-04 | 0.0082 | 0.0033 | 0.0033 | TRUE | 0.0206 |
| 4930428D18I | 0 | 0 | 0 | 0 | 0 | 0 | 0 | 0 | 0 | 0 | 0 | 0 | 0 | 0 | 0 | NA | NA | NA | NA | NA | NA | NA | NA | NA | NA |
| 4930428E07I | 0 | 0 | 0 | 0 | 0 | 0 | 0 | 0 | 0 | 0 | 0 | 0 | 0 | 0 | 0 | NA | NA | NA | NA | NA | NA | NA | NA | NA | NA |
| 4930428G15 | 0 | 0 | 0 | 0 | 0 | 0 | 0 | 0 | 0 | 0 | 0 | 0 | 0 | 0 | 0 | NA | NA | NA | NA | NA | NA | NA | NA | NA | NA |
| 4930428O21 | 0 | 0 | 0 | 0 | 0 | 0 | 0 | 0 | 0 | 0 | 0 | 0 | 0 | 0 | 0 | NA | NA | NA | NA | NA | NA | NA | NA | NA | NA |
| 4930429B21I | 13 | 15 | 10 | 44 | 38 | 16 | 14 | 15 | 10 | 35 | 41 | 18 | 21.97 | 13 | 31 | 2.403 | 1.265 | 0.00969619 | 0.05143904 | 0.0751 | 0.1569 | 0.1171 | 0.1171 | TRUE | 0.4334 |
| 4930429D17I | 0 | 0 | 0 | 0 | 1 | 1 | 0 | 0 | 0 | 0 | 1 | 1 | 0.36 | 0 | 1 | 3.794 | 1.924 | 0.60247382 | NA | 0 | 9.1135 | 8.1425 | 8.1425 | TRUE | 0.3482 |
| 4930429F11I | 0 | 0 | 0 | 0 | 0 | 0 | 0 | 0 | 0 | 0 | 0 | 0 | 0 | 0 | 0 | NA | NA | NA | NA | NA | NA | NA | NA | NA | NA |
| 4930429F24I | 0 | 0 | 0 | 0 | 1 | 1 | 0 | 0 | 0 | 0 | 1 | 1 | 0.36 | 0 | 1 | 3.794 | 1.924 | 0.60247382 | NA | 0 | 9.1135 | 8.1425 | 8.1425 | TRUE | 0.3482 |
| 4930430A15I | 0 | 0 | 0 | 0 | 0 | 0 | 0 | 0 | 0 | 0 | 0 | 0 | 0 | 0 | 0 | NA | NA | NA | NA | NA | NA | NA | NA | NA | NA |
| 4930430D24I | 0 | 0 | 0 | 0 | 0 | 0 | 0 | 0 | 0 | 0 | 0 | 0 | 0 | 0 | 0 | NA | NA | NA | NA | NA | NA | NA | NA | NA | NA |
| 4930430F08I | 1281 | 1450 | 1434 | 1707 | 1196 | 1364 | 1361 | 1448 | 1440 | 1342 | 1287 | 1493 | 1395.08 | 1416 | 1374 | 0.97 | -0.044 | 0.6285213 | 0.79095491 | 0.0027 | 0.0093 | 0.0053 | 0.0053 | TRUE | 0.0679 |
| 4930430F21I | 0 | 0 | 0 | 0 | 0 | 0 | 0 | 0 | 0 | 0 | 0 | 0 | 0 | 0 | 0 | NA | NA | NA | NA | NA | NA | NA | NA | NA | NA |
| 4930430J02F | 0 | 0 | 0 | 0 | 0 | 0 | 0 | 0 | 0 | 0 | 0 | 0 | 0 | 0 | 0 | NA | NA | NA | NA | NA | NA | NA | NA | NA | NA |
| 4930431F12I | 1 | 0 | 1 | 1 | 0 | 0 | 1 | 0 | 1 | 1 | 0 | 0 | 0.48 | 1 | 0 | 0.509 | -0.975 | 0.7336918 | NA | 0 | 6.9367 | 4.1806 | 4.1806 | TRUE | 0.2562 |
| 4930431P03I | 0 | 0 | 0 | 0 | 0 | 0 | 0 | 0 | 0 | 0 | 0 | 0 | 0 | 0 | 0 | NA | NA | NA | NA | NA | NA | NA | NA | NA | NA |
| 4930432J09F | 0 | 0 | 0 | 0 | 0 | 0 | 0 | 0 | 0 | 0 | 0 | 0 | 0 | 0 | 0 | NA | NA | NA | NA | NA | NA | NA | NA | NA | NA |
| 4930432K09I | 0 | 0 | 0 | 0 | 0 | 0 | 0 | 0 | 0 | 0 | 0 | 0 | 0 | 0 | 0 | NA | NA | NA | NA | NA | NA | NA | NA | NA | NA |
| 4930432K21I | 130 | 142 | 143 | 198 | 151 | 121 | 138 | 142 | 144 | 156 | 162 | 132 | 145.68 | 141 | 150 | 1.065 | 0.091 | 0.59372567 | 0.76728255 | 0 | 0.0295 | 0.0142 | 0.0142 | TRUE | 0.1051 |
| 4930432M17 | 0 | 0 | 0 | 0 | 0 | 0 | 0 | 0 | 0 | 0 | 0 | 0 | 0 | 0 | 0 | NA | NA | NA | NA | NA | NA | NA | NA | NA | NA |
| 4930433B08I | 0 | 0 | 0 | 0 | 0 | 0 | 0 | 0 | 0 | 0 | 0 | 0 | 0 | 0 | 0 | NA | NA | NA | NA | NA | NA | NA | NA | NA | NA |
| 4930433I11R | 0 | 0 | 0 | 0 | 0 | 0 | 0 | 0 | 0 | 0 | 0 | 0 | 0 | 0 | 0 | NA | NA | NA | NA | NA | NA | NA | NA | NA | NA |
| 4930433N12 | 54 | 70 | 85 | 95 | 67 | 84 | 57 | 70 | 85 | 75 | 72 | 92 | 75.22 | 71 | 80 | 1.119 | 0.163 | 0.52095635 | 0.71144892 | 0.0136 | 0.0507 | 0.0329 | 0.0329 | TRUE | 0.2707 |
| 4930434J06F | 0 | 0 | 0 | 0 | 0 | 0 | 0 | 0 | 0 | 0 | 0 | 0 | 0 | 0 | 0 | NA | NA | NA | NA | NA | NA | NA | NA | NA | NA |
| 4930435E12I | 1 | 0 | 0 | 0 | 0 | 0 | 1 | 0 | 0 | 0 | 0 | 0 | 0.18 | 0 | 0 | 0.49 | -1.029 | 0.80087 | NA | 0 | 18.6165 | 10 | 10 | TRUE | 0.4614 |
| 4930438E09I | 0 | 0 | 0 | 0 | 0 | 0 | 0 | 0 | 0 | 0 | 0 | 0 | 0 | 0 | 0 | NA | NA | NA | NA | NA | NA | NA | NA | NA | NA |
| 4930440C22I | 0 | 0 | 0 | 0 | 0 | 0 | 0 | 0 | 0 | 0 | 0 | 0 | 0 | 0 | 0 | NA | NA | NA | NA | NA | NA | NA | NA | NA | NA |
| 4930441J16F | 0 | 0 | 0 | 0 | 0 | 0 | 0 | 0 | 0 | 0 | 0 | 0 | 0 | 0 | 0 | NA | NA | NA | NA | NA | NA | NA | NA | NA | NA |
| 4930441O14 | 10 | 10 | 9 | 9 | 7 | 4 | 11 | 10 | 9 | 7 | 8 | 4 | 8.11 | 10 | 6 | 0.645 | -0.633 | 0.33456506 | 0.5548783 | 0 | 0.4134 | 0.1818 | 0.1818 | TRUE | 0.1554 |
| 4930442J19F | 0 | 0 | 0 | 2 | 3 | 1 | 0 | 0 | 0 | 2 | 3 | 1 | 0.98 | 0 | 2 | 10.427 | 3.382 | 0.12480123 | NA | 0 | 3.3602 | 2.2486 | 2.2486 | TRUE | 0.2581 |
| 4930442L01F | 0 | 0 | 0 | 0 | 0 | 1 | 0 | 0 | 0 | 0 | 0 | 1 | 0.18 | 0 | 0 | 1.859 | 0.894 | 0.82650734 | NA | 0 | 18.0643 | 10 | 10 | TRUE | 0.5383 |
| 4930443O20 | 0 | 0 | 0 | 0 | 0 | 0 | 0 | 0 | 0 | 0 | 0 | 0 | 0 | 0 | 0 | NA | NA | NA | NA | NA | NA | NA | NA | NA | NA |
| 4930444A02I | 973 | 1241 | 1208 | 1369 | 1019 | 1043 | 1033 | 1239 | 1213 | 1076 | 1096 | 1142 | 1133.36 | 1162 | 1105 | 0.95 | -0.074 | 0.47041015 | 0.67209973 | 0.0043 | 0.0098 | 0.0066 | 0.0066 | TRUE | 0.1114 |
| 4930444F02I | 4 | 3 | 5 | 1 | 0 | 1 | 4 | 3 | 5 | 1 | 0 | 1 | 2.36 | 4 | 1 | 0.156 | -2.682 | 0.06730741 | NA | 0 | 1.4042 | 0.6617 | 0.6617 | TRUE | 0.1881 |
| 4930444G20 | 0 | 0 | 0 | 0 | 0 | 0 | 0 | 0 | 0 | 0 | 0 | 0 | 0 | 0 | 0 | NA | NA | NA | NA | NA | NA | NA | NA | NA | NA |
| 4930444M15 | 80 | 100 | 79 | 123 | 91 | 82 | 85 | 100 | 79 | 97 | 98 | 90 | 91.42 | 88 | 95 | 1.077 | 0.107 | 0.61009437 | 0.77891178 | 0 | 0.0429 | 0.0207 | 0.0207 | TRUE | 0.1329 |
| 4930444P10I | 1 | 2 | 0 | 1 | 1 | 3 | 1 | 2 | 0 | 1 | 1 | 3 | 1.37 | 1 | 2 | 1.649 | 0.722 | 0.67017936 | NA | 0 | 2.4158 | 1.279 | 1.279 | TRUE | 0.3619 |
| 4930447A16I | 0 | 0 | 0 | 0 | 0 | 0 | 0 | 0 | 0 | 0 | 0 | 0 | 0 | 0 | 0 | NA | NA | NA | NA | NA | NA | NA | NA | NA | NA |
| 4930447C04I | 881 | 1218 | 1238 | 1274 | 1016 | 1030 | 936 | 1216 | 1243 | 1002 | 1093 | 1127 | 1102.91 | 1132 | 1074 | 0.948 | -0.077 | 0.56013317 | 0.74190289 | 0.013 | 0.0099 | 0.0115 | 0.0115 | TRUE | 0.2738 |
| 4930447J18F | 0 | 0 | 0 | 0 | 0 | 0 | 0 | 0 | 0 | 0 | 0 | 0 | 0 | 0 | 0 | NA | NA | NA | NA | NA | NA | NA | NA | NA | NA |
| 4930447K03I | 0 | 6 | 6 | 2 | 0 | 2 | 0 | 6 | 6 | 2 | 0 | 2 | 2.63 | 4 | 1 | 0.315 | -1.668 | 0.28140035 | NA | 0 | 1.2597 | 1.2154 | 1.2154 | TRUE | 1.2157 |
| 4930447N08 | 1 | 0 | 0 | 0 | 1 | 0 | 1 | 0 | 0 | 0 | 1 | 0 | 0.36 | 0 | 0 | 0.954 | -0.067 | 0.98557668 | NA | 0 | 9.2518 | 8.0179 | 8.0179 | TRUE | 0.5204 |
| 4930448C13I | 0 | 0 | 0 | 0 | 0 | 0 | 0 | 0 | 0 | 0 | 0 | 0 | 0 | 0 | 0 | NA | NA | NA | NA | NA | NA | NA | NA | NA | NA |
| 4930448F12I | 0 | 0 | 0 | 0 | 0 | 0 | 0 | 0 | 0 | 0 | 0 | 0 | 0 | 0 | 0 | NA | NA | NA | NA | NA | NA | NA | NA | NA | NA |
| 4930448H16I | 0 | 0 | 0 | 0 | 0 | 0 | 0 | 0 | 0 | 0 | 0 | 0 | 0 | 0 | 0 | NA | NA | NA | NA | NA | NA | NA | NA | NA | NA |
| 4930448I06R | 0 | 0 | 0 | 0 | 0 | 0 | 0 | 0 | 0 | 0 | 0 | 0 | 0 | 0 | 0 | NA | NA | NA | NA | NA | NA | NA | NA | NA | NA |
| 4930448I18R | 0 | 0 | 0 | 0 | 0 | 0 | 0 | 0 | 0 | 0 | 0 | 0 | 0 | 0 | 0 | NA | NA | NA | NA | NA | NA | NA | NA | NA | NA |
| 4930448K20I | 0 | 0 | 0 | 0 | 0 | 2 | 0 | 0 | 0 | 0 | 0 | 2 | 0.36 | 0 | 1 | 3.832 | 1.938 | 0.63022034 | NA | 10 | 9.0356 | 10 | 10 | TRUE | 0.9744 |
| 4930449E01I | 2 | 0 | 0 | 2 | 0 | 1 | 2 | 0 | 0 | 2 | 0 | 1 | 0.8 | 1 | 1 | 1.296 | 0.374 | 0.87990755 | NA | 0 | 4.1325 | 3.1495 | 3.1495 | TRUE | 0.9743 |

| | | | | | | | | | | | | | | | | | | | | | | | | | |
|---|---|---|---|---|---|---|---|---|---|---|---|---|---|---|---|---|---|---|---|---|---|---|---|---|---|
| 4930449E18 | 0 | 0 | 0 | 0 | 0 | 0 | 0 | 0 | 0 | 0 | 0 | 0 | 0 | 0 | 0 | NA | NA | NA | NA | NA | NA | NA | NA | NA | NA |
| 4930449I24R | 0 | 0 | 0 | 0 | 0 | 0 | 0 | 0 | 0 | 0 | 0 | 0 | 0 | 0 | 0 | NA | NA | NA | NA | NA | NA | NA | NA | NA | NA |
| 4930451C15 | 19 | 23 | 24 | 19 | 14 | 13 | 20 | 23 | 24 | 15 | 15 | 14 | 18.58 | 22 | 15 | 0.658 | -0.605 | 0.16101579 | 0.35572784 | 0 | 0.1842 | 0.0782 | 0.0782 | TRUE | 0.042 |
| 4930451G09 | 27 | 27 | 17 | 52 | 38 | 28 | 29 | 27 | 17 | 41 | 41 | 31 | 30.85 | 24 | 38 | 1.552 | 0.635 | 0.0885484 | 0.24141492 | 0 | 0.1137 | 0.0659 | 0.0659 | TRUE | 0.4054 |
| 4930451I11R | 0 | 0 | 0 | 2 | 0 | 1 | 0 | 0 | 0 | 2 | 0 | 1 | 0.44 | 0 | 1 | 4.782 | 2.258 | 0.50653358 | NA | 0 | 7.4188 | 6.7775 | 6.7775 | TRUE | 0.2804 |
| 4930452A19 | 0 | 0 | 0 | 0 | 0 | 0 | 0 | 0 | 0 | 0 | 0 | 0 | 0 | 0 | 0 | NA | NA | NA | NA | NA | NA | NA | NA | NA | NA |
| 4930452B06 | 1421 | 1336 | 1369 | 1842 | 1351 | 1445 | 1509 | 1334 | 1375 | 1448 | 1454 | 1582 | 1450.28 | 1406 | 1495 | 1.063 | 0.088 | 0.33196705 | 0.55247275 | 0.0024 | 0.0092 | 0.0052 | 0.0052 | TRUE | 0.0494 |
| 4930452G13 | 0 | 0 | 0 | 1 | 1 | 0 | 0 | 0 | 0 | 1 | 1 | 0 | 0.31 | 0 | 1 | 3.32 | 1.731 | 0.65383052 | NA | 0 | 10.6224 | 8.9541 | 8.9541 | TRUE | 0.202 |
| 4930452N14 | 1 | 0 | 0 | 0 | 0 | 0 | 1 | 0 | 0 | 0 | 0 | 0 | 0.18 | 0 | 0 | 0.49 | -1.029 | 0.80087 | NA | 0 | 18.6165 | 10 | 10 | TRUE | 0.4614 |
| 4930453H23 | 1 | 0 | 0 | 2 | 4 | 0 | 1 | 0 | 0 | 2 | 4 | 0 | 1.16 | 0 | 2 | 5.341 | 2.417 | 0.28284963 | NA | 0 | 2.8559 | 2.3991 | 2.3991 | TRUE | 0.2338 |
| 4930453L07 | 0 | 0 | 0 | 0 | 0 | 0 | 0 | 0 | 0 | 0 | 0 | 0 | 0 | 0 | 0 | NA | NA | NA | NA | NA | NA | NA | NA | NA | NA |
| 4930453N24 | 1403 | 1159 | 1259 | 2147 | 1415 | 1210 | 1490 | 1158 | 1264 | 1688 | 1522 | 1325 | 1407.78 | 1304 | 1512 | 1.16 | 0.214 | 0.11038215 | 0.27886389 | 0.0147 | 0.0092 | 0.0123 | 0.0123 | TRUE | 0.1871 |
| 4930455B14 | 1 | 4 | 2 | 5 | 2 | 9 | 1 | 4 | 2 | 4 | 2 | 10 | 3.83 | 2 | 5 | 2.226 | 1.154 | 0.28941821 | NA | 0.2348 | 0.8663 | 0.547 | 0.547 | TRUE | 0.6935 |
| 4930455C13 | 0 | 1 | 0 | 0 | 0 | 0 | 0 | 1 | 0 | 0 | 0 | 0 | 0.17 | 0 | 0 | 0.49 | -1.029 | 0.80087 | NA | 0 | 19.7991 | 10 | 10 | TRUE | 0.405 |
| 4930455D15 | 0 | 0 | 1 | 0 | 0 | 0 | 0 | 0 | 1 | 0 | 0 | 0 | 0.17 | 0 | 0 | 0.49 | -1.029 | 0.80087 | NA | 0 | 19.6928 | 10 | 10 | TRUE | 0.4098 |
| 4930455F16 | 0 | 0 | 0 | 0 | 0 | 0 | 0 | 0 | 0 | 0 | 0 | 0 | 0 | 0 | 0 | NA | NA | NA | NA | NA | NA | NA | NA | NA | NA |
| 4930455F23 | 463 | 513 | 553 | 812 | 584 | 545 | 492 | 512 | 555 | 638 | 628 | 597 | 570.44 | 520 | 621 | 1.195 | 0.257 | 0.01243614 | 0.06171896 | 0 | 0.0127 | 0.0059 | 0.0059 | TRUE | 0.0417 |
| 4930455H04 | 0 | 0 | 0 | 0 | 0 | 0 | 0 | 0 | 0 | 0 | 0 | 0 | 0 | 0 | 0 | NA | NA | NA | NA | NA | NA | NA | NA | NA | NA |
| 4930455J16 | 0 | 0 | 0 | 0 | 0 | 0 | 0 | 0 | 0 | 0 | 0 | 0 | 0 | 0 | 0 | NA | NA | NA | NA | NA | NA | NA | NA | NA | NA |
| 4930456L15 | 0 | 0 | 0 | 1 | 0 | 0 | 0 | 0 | 0 | 1 | 0 | 0 | 0.13 | 0 | 0 | 1.859 | 0.894 | 0.82650734 | NA | 0 | 25.1515 | 10 | 10 | TRUE | 0.2563 |
| 4930459L07 | 0 | 0 | 0 | 0 | 0 | 0 | 0 | 0 | 0 | 0 | 0 | 0 | 0 | 0 | 0 | NA | NA | NA | NA | NA | NA | NA | NA | NA | NA |
| 4930461G14 | 624 | 522 | 575 | 608 | 416 | 457 | 663 | 521 | 577 | 478 | 448 | 500 | 531.22 | 587 | 475 | 0.81 | -0.304 | 0.01682717 | 0.07690471 | 0.0071 | 0.0131 | 0.0098 | 0.0098 | TRUE | 0.1481 |
| 4930465A12 | 0 | 0 | 0 | 0 | 0 | 0 | 0 | 0 | 0 | 0 | 0 | 0 | 0 | 0 | 0 | NA | NA | NA | NA | NA | NA | NA | NA | NA | NA |
| 4930465K10 | 1 | 1 | 0 | 1 | 3 | 2 | 1 | 1 | 0 | 1 | 3 | 2 | 1.38 | 1 | 2 | 2.97 | 1.57 | 0.36300414 | NA | 0 | 2.3988 | 1.1589 | 1.1589 | TRUE | 0.2008 |
| 4930465M20 | 0 | 0 | 1 | 4 | 1 | 0 | 0 | 0 | 1 | 3 | 1 | 0 | 0.87 | 0 | 1 | 3.94 | 1.978 | 0.41505635 | NA | 0 | 3.7903 | 2.9051 | 2.9051 | TRUE | 0.2422 |
| 4930467D21 | 8 | 17 | 12 | 20 | 12 | 14 | 8 | 17 | 12 | 16 | 13 | 15 | 13.58 | 12 | 15 | 1.171 | 0.228 | 0.66513242 | 0.81258471 | 0 | 0.2495 | 0.1262 | 0.1262 | TRUE | 0.3282 |
| 4930467E23 | 0 | 0 | 1 | 0 | 0 | 0 | 0 | 0 | 1 | 0 | 0 | 0 | 0.17 | 0 | 0 | 0.49 | -1.029 | 0.80087 | NA | 0 | 19.6928 | 10 | 10 | TRUE | 0.4098 |
| 4930467K11 | 0 | 1 | 2 | 1 | 2 | 0 | 0 | 1 | 2 | 1 | 2 | 0 | 0.99 | 1 | 1 | 0.966 | -0.05 | 0.98057163 | NA | 0 | 3.3319 | 1.9761 | 1.9761 | TRUE | 0.2365 |
| 4930468A15 | 0 | 0 | 0 | 0 | 0 | 0 | 0 | 0 | 0 | 0 | 0 | 0 | 0 | 0 | 0 | NA | NA | NA | NA | NA | NA | NA | NA | NA | NA |
| 4930469G21 | 0 | 0 | 0 | 0 | 2 | 0 | 0 | 0 | 0 | 0 | 2 | 0 | 0.36 | 0 | 1 | 3.775 | 1.916 | 0.63423067 | NA | 10 | 9.1927 | 10 | 10 | TRUE | 0.9743 |
| 4930470H14 | 0 | 0 | 0 | 0 | 0 | 0 | 0 | 0 | 0 | 0 | 0 | 0 | 0 | 0 | 0 | NA | NA | NA | NA | NA | NA | NA | NA | NA | NA |
| 4930470P17 | 2 | 1 | 0 | 4 | 3 | 4 | 2 | 1 | 0 | 3 | 3 | 4 | 2.31 | 1 | 3 | 3.469 | 1.794 | 0.18909136 | NA | 0 | 1.4316 | 0.7121 | 0.7121 | TRUE | 0.3784 |
| 4930471C04 | 0 | 0 | 0 | 0 | 0 | 0 | 0 | 0 | 0 | 0 | 0 | 0 | 0 | 0 | 0 | NA | NA | NA | NA | NA | NA | NA | NA | NA | NA |
| 4930471G03 | 0 | 0 | 0 | 0 | 0 | 0 | 0 | 0 | 0 | 0 | 0 | 0 | 0 | 0 | 0 | NA | NA | NA | NA | NA | NA | NA | NA | NA | NA |
| 4930471I20R | 0 | 0 | 0 | 0 | 0 | 0 | 0 | 0 | 0 | 0 | 0 | 0 | 0 | 0 | 0 | NA | NA | NA | NA | NA | NA | NA | NA | NA | NA |
| 4930471M0 | 0 | 1 | 0 | 1 | 0 | 0 | 0 | 1 | 0 | 1 | 0 | 0 | 0.3 | 0 | 0 | 0.954 | -0.067 | 0.98642001 | NA | 0 | 11.0817 | 9.3014 | 9.3014 | TRUE | 0.405 |
| 4930473A02 | 3 | 0 | 2 | 1 | 0 | 1 | 3 | 0 | 2 | 1 | 0 | 1 | 1.18 | 2 | 1 | 0.367 | -1.444 | 0.46429227 | NA | 0 | 2.8006 | 1.7412 | 1.7412 | TRUE | 0.2145 |
| 4930473O22 | 0 | 0 | 0 | 0 | 0 | 0 | 0 | 0 | 0 | 0 | 0 | 0 | 0 | 0 | 0 | NA | NA | NA | NA | NA | NA | NA | NA | NA | NA |
| 4930474G06 | 0 | 0 | 0 | 0 | 0 | 0 | 0 | 0 | 0 | 0 | 0 | 0 | 0 | 0 | 0 | NA | NA | NA | NA | NA | NA | NA | NA | NA | NA |
| 4930474H20 | 0 | 0 | 0 | 0 | 1 | 0 | 0 | 0 | 0 | 0 | 1 | 0 | 0.18 | 0 | 0 | 1.859 | 0.894 | 0.82650734 | NA | 0 | 18.3785 | 10 | 10 | TRUE | 0.5204 |
| 4930474M22 | 0 | 0 | 0 | 0 | 0 | 0 | 0 | 0 | 0 | 0 | 0 | 0 | 0 | 0 | 0 | NA | NA | NA | NA | NA | NA | NA | NA | NA | NA |
| 4930474N05 | 0 | 0 | 0 | 0 | 0 | 0 | 0 | 0 | 0 | 0 | 0 | 0 | 0 | 0 | 0 | NA | NA | NA | NA | NA | NA | NA | NA | NA | NA |
| 4930474N09 | 0 | 1 | 1 | 0 | 0 | 1 | 0 | 1 | 1 | 0 | 0 | 1 | 0.52 | 1 | 0 | 0.52 | -0.943 | 0.73639299 | NA | 0 | 6.3885 | 3.8941 | 3.8941 | TRUE | 0.5383 |
| 4930478L05 | 0 | 0 | 0 | 0 | 0 | 0 | 0 | 0 | 0 | 0 | 0 | 0 | 0 | 0 | 0 | NA | NA | NA | NA | NA | NA | NA | NA | NA | NA |
| 4930478P22 | 0 | 0 | 0 | 0 | 0 | 0 | 0 | 0 | 0 | 0 | 0 | 0 | 0 | 0 | 0 | NA | NA | NA | NA | NA | NA | NA | NA | NA | NA |
| 4930479D17 | 0 | 0 | 0 | 2 | 1 | 1 | 0 | 0 | 0 | 2 | 1 | 1 | 0.62 | 0 | 1 | 6.676 | 2.739 | 0.27800695 | NA | 0 | 5.2881 | 3.2034 | 3.2034 | TRUE | 0.07 |
| 4930479M11 | 10 | 12 | 9 | 13 | 6 | 10 | 11 | 12 | 9 | 10 | 6 | 11 | 9.88 | 11 | 9 | 0.876 | -0.19 | 0.75015155 | 0.86769716 | 0 | 0.3404 | 0.1558 | 0.1558 | TRUE | 0.2017 |
| 4930480E11 | 0 | 0 | 0 | 0 | 0 | 0 | 0 | 0 | 0 | 0 | 0 | 0 | 0 | 0 | 0 | NA | NA | NA | NA | NA | NA | NA | NA | NA | NA |
| 4930480G23 | 0 | 0 | 0 | 0 | 1 | 0 | 0 | 0 | 0 | 0 | 1 | 0 | 0.18 | 0 | 0 | 1.859 | 0.894 | 0.82650734 | NA | 0 | 18.3785 | 10 | 10 | TRUE | 0.5204 |
| 4930480K15 | 0 | 0 | 0 | 0 | 0 | 0 | 0 | 0 | 0 | 0 | 0 | 0 | 0 | 0 | 0 | NA | NA | NA | NA | NA | NA | NA | NA | NA | NA |
| 4930480M12 | 0 | 0 | 0 | 0 | 0 | 0 | 0 | 0 | 0 | 0 | 0 | 0 | 0 | 0 | 0 | NA | NA | NA | NA | NA | NA | NA | NA | NA | NA |
| 4930481A15 | 5 | 3 | 2 | 8 | 9 | 4 | 5 | 3 | 2 | 6 | 10 | 4 | 5.11 | 3 | 7 | 1.98 | 0.985 | 0.26195835 | 0.48137098 | 0 | 0.6515 | 0.3343 | 0.3343 | TRUE | 0.2756 |
| 4930483J18 | 2 | 2 | 0 | 1 | 2 | 1 | 2 | 2 | 0 | 1 | 2 | 1 | 1.36 | 1 | 1 | 0.966 | -0.049 | 0.97621505 | NA | 0 | 2.4309 | 1.2183 | 1.2183 | TRUE | 0.4911 |
| 4930483K19 | 0 | 0 | 0 | 0 | 0 | 0 | 0 | 0 | 0 | 0 | 0 | 0 | 0 | 0 | 0 | NA | NA | NA | NA | NA | NA | NA | NA | NA | NA |
| 4930483O08 | 0 | 0 | 0 | 0 | 0 | 1 | 0 | 0 | 0 | 0 | 0 | 1 | 0.18 | 0 | 0 | 1.859 | 0.894 | 0.82650734 | NA | 0 | 18.0643 | 10 | 10 | TRUE | 0.5383 |
| 4930486F22 | 0 | 1 | 0 | 0 | 0 | 0 | 0 | 1 | 0 | 0 | 0 | 0 | 0.17 | 0 | 0 | 0.49 | -1.029 | 0.80087 | NA | 0 | 19.7991 | 10 | 10 | TRUE | 0.405 |
| 4930486I03R | 0 | 0 | 0 | 0 | 0 | 0 | 0 | 0 | 0 | 0 | 0 | 0 | 0 | 0 | 0 | NA | NA | NA | NA | NA | NA | NA | NA | NA | NA |
| 4930486L24 | 37 | 21 | 26 | 119 | 87 | 90 | 39 | 21 | 26 | 94 | 94 | 99 | 62.01 | 29 | 96 | 3.317 | 1.73 | 3.76E-09 | 1.96E-07 | 0 | 0.06 | 0.0391 | 0.0391 | TRUE | 0.641 |
| 4930487D11 | 0 | 0 | 0 | 0 | 0 | 0 | 0 | 0 | 0 | 0 | 0 | 0 | 0 | 0 | 0 | NA | NA | NA | NA | NA | NA | NA | NA | NA | NA |
| 4930487H11 | 1 | 0 | 0 | 0 | 0 | 1 | 1 | 0 | 0 | 0 | 0 | 1 | 0.36 | 0 | 0 | 0.954 | -0.067 | 0.98553519 | NA | 0 | 9.1716 | 7.9605 | 7.9605 | TRUE | 0.5383 |
| 4930488B22 | 0 | 0 | 0 | 0 | 0 | 0 | 0 | 0 | 0 | 0 | 0 | 0 | 0 | 0 | 0 | NA | NA | NA | NA | NA | NA | NA | NA | NA | NA |
| 4930488L21 | 0 | 0 | 0 | 0 | 0 | 0 | 0 | 0 | 0 | 0 | 0 | 0 | 0 | 0 | 0 | NA | NA | NA | NA | NA | NA | NA | NA | NA | NA |
| 4930500F04 | 1 | 1 | 0 | 0 | 0 | 0 | 1 | 1 | 0 | 0 | 0 | 0 | 0.34 | 1 | 0 | 0.261 | -1.936 | 0.60454921 | NA | 0 | 9.5983 | 8.3269 | 8.3269 | TRUE | 0.2507 |
| 4930500J02 | 404 | 632 | 638 | 1157 | 912 | 960 | 429 | 631 | 641 | 910 | 981 | 1051 | 773.77 | 567 | 981 | 1.728 | 0.789 | 1.40E-06 | 3.69E-05 | 0.0247 | 0.0112 | 0.0179 | 0.0179 | TRUE | 0.5271 |
| 4930500L23 | 0 | 0 | 0 | 0 | 0 | 0 | 0 | 0 | 0 | 0 | 0 | 0 | 0 | 0 | 0 | NA | NA | NA | NA | NA | NA | NA | NA | NA | NA |
| 4930502A04 | 0 | 0 | 0 | 0 | 0 | 0 | 0 | 0 | 0 | 0 | 0 | 0 | 0 | 0 | 0 | NA | NA | NA | NA | NA | NA | NA | NA | NA | NA |
| 4930502E09 | 2 | 6 | 12 | 3 | 6 | 8 | 2 | 6 | 12 | 2 | 6 | 9 | 6.29 | 7 | 6 | 0.853 | -0.23 | 0.80011219 | 0.89686328 | 0.3323 | 0.5307 | 0.432 | 0.432 | TRUE | 0.3075 |
| 4930502E18 | 121 | 113 | 110 | 378 | 294 | 230 | 129 | 113 | 110 | 297 | 316 | 252 | 202.85 | 117 | 288 | 2.463 | 1.3 | 1.02E-14 | 1.35E-12 | 0.006 | 0.0231 | 0.0143 | 0.0143 | TRUE | 0.1337 |
| 4930503B20 | 6 | 1 | 3 | 5 | 6 | 1 | 6 | 1 | 3 | 4 | 6 | 1 | 3.64 | 3 | 4 | 1.114 | 0.156 | 0.88901076 | NA | 0.3037 | 0.9109 | 0.6221 | 0.6221 | TRUE | 0.2615 |

| | | | | | | | | | | | | | | | | | | | | | | | | | |
|---|---|---|---|---|---|---|---|---|---|---|---|---|---|---|---|---|---|---|---|---|---|---|---|---|---|
| 4930503E14I | 1 | 0 | 0 | 3 | 4 | 0 | 1 | 0 | 0 | 2 | 4 | 0 | 1.29 | 0 | 2 | 6.082 | 2.605 | 0.23055126 | NA | | 0 | 2.5659 | 2.1804 | 2.1804 | TRUE | 0.2002 |
| 4930503E24I | 0 | 0 | 0 | 0 | 0 | 0 | 0 | 0 | 0 | 0 | 0 | 0 | 0 | 0 | 0 | NA | NA | NA | NA | NA | NA | NA | NA | NA | NA | |
| 4930503H13 | 0 | 0 | 0 | 0 | 0 | 0 | 0 | 0 | 0 | 0 | 0 | 0 | 0 | 0 | 0 | NA | NA | NA | NA | NA | NA | NA | NA | NA | NA | |
| 4930503L19F | 870 | 961 | 948 | 1299 | 906 | 868 | 924 | 960 | 952 | 1021 | 975 | 950 | 963.65 | 945 | 982 | 1.04 | 0.056 | 0.49944249 | 0.6951908 | 0 | 0.0103 | 0.0039 | 0.0039 | TRUE | 0.0152 |
| 4930503O07 | 0 | 0 | 0 | 0 | 0 | 0 | 0 | 0 | 0 | 0 | 0 | 0 | 0 | 0 | 0 | NA | NA | NA | NA | NA | NA | NA | NA | NA | NA | |
| 4930504O13 | 4 | 6 | 6 | 16 | 10 | 3 | 4 | 6 | 6 | 13 | 11 | 3 | 7.15 | 5 | 9 | 1.655 | 0.727 | 0.35053247 | 0.56931295 | 0 | 0.4678 | 0.2879 | 0.2879 | TRUE | 0.8765 |
| 4930505A04 | 9 | 6 | 3 | 11 | 15 | 5 | 10 | 6 | 3 | 9 | 16 | 5 | 8.14 | 6 | 10 | 1.631 | 0.706 | 0.35480894 | 0.57324796 | 0.154 | 0.4118 | 0.288 | 0.288 | TRUE | 0.4438 |
| 4930505G20 | 0 | 0 | 0 | 0 | 0 | 0 | 0 | 0 | 0 | 0 | 0 | 0 | 0 | 0 | 0 | NA | NA | NA | NA | NA | NA | NA | NA | NA | NA | |
| 4930506C21I | 1 | 3 | 0 | 0 | 0 | 0 | 1 | 3 | 0 | 0 | 0 | 0 | 0.68 | 1 | 0 | 0.132 | -2.917 | 0.33163516 | NA | 6.5513 | 4.8776 | 5.1281 | 5.1281 | TRUE | 0.1294 |
| 4930506M07 | 470 | 622 | 591 | 857 | 677 | 629 | 499 | 621 | 593 | 674 | 728 | 689 | 634.08 | 571 | 697 | 1.219 | 0.285 | 0.01526412 | 0.07165051 | 0.0053 | 0.0121 | 0.0084 | 0.0084 | TRUE | 0.1414 |
| 4930507D05 | 3 | 5 | 6 | 4 | 5 | 1 | 3 | 5 | 6 | 3 | 5 | 1 | 3.97 | 5 | 3 | 0.676 | -0.564 | 0.56451904 | NA | 0 | 0.8366 | 0.4316 | 0.4316 | TRUE | 0.2524 |
| 4930509E16I | 26 | 31 | 30 | 23 | 12 | 13 | 28 | 31 | 30 | 18 | 13 | 14 | 22.32 | 30 | 15 | 0.515 | -0.958 | 0.01873597 | 0.08301738 | 0 | 0.1545 | 0.0704 | 0.0704 | TRUE | 0.1761 |
| 4930509J09F | 0 | 0 | 0 | 0 | 0 | 0 | 0 | 0 | 0 | 0 | 0 | 0 | 0 | 0 | 0 | NA | NA | NA | NA | NA | NA | NA | NA | NA | NA | |
| 4930509K18I | 0 | 0 | 0 | 0 | 0 | 0 | 0 | 0 | 0 | 0 | 0 | 0 | 0 | 0 | 0 | NA | NA | NA | NA | NA | NA | NA | NA | NA | NA | |
| 4930511A02 | 1 | 0 | 0 | 1 | 1 | 1 | 1 | 0 | 0 | 1 | 1 | 1 | 0.67 | 0 | 1 | 2.676 | 1.42 | 0.53486561 | NA | 0 | 4.9253 | 2.274 | 2.274 | TRUE | 0.4614 |
| 4930511E03I | 0 | 0 | 0 | 1 | 0 | 0 | 0 | 0 | 0 | 1 | 0 | 0 | 0.13 | 0 | 0 | 1.859 | 0.894 | 0.82650734 | NA | 0 | 25.1515 | 10 | 10 | TRUE | 0.2563 |
| 4930511M06 | 16 | 15 | 7 | 24 | 9 | 15 | 17 | 15 | 7 | 19 | 10 | 16 | 14 | 13 | 15 | 1.164 | 0.219 | 0.70129684 | 0.83650923 | 0.0691 | 0.2423 | 0.1621 | 0.1621 | TRUE | 0.6789 |
| 4930511M11 | 0 | 0 | 0 | 0 | 0 | 0 | 0 | 0 | 0 | 0 | 0 | 0 | 0 | 0 | 0 | NA | NA | NA | NA | NA | NA | NA | NA | NA | NA | |
| 4930512B01 | 0 | 0 | 0 | 0 | 0 | 0 | 0 | 0 | 0 | 0 | 0 | 0 | 0 | 0 | 0 | NA | NA | NA | NA | NA | NA | NA | NA | NA | NA | |
| 4930513D17 | 2 | 0 | 0 | 3 | 0 | 0 | 2 | 0 | 0 | 2 | 0 | 0 | 0.75 | 1 | 1 | 1.163 | 0.218 | 0.94242245 | NA | 8.7098 | 4.4164 | 5.2331 | 5.2331 | TRUE | 1.4402 |
| 4930513N10 | 87 | 70 | 63 | 221 | 146 | 108 | 92 | 70 | 63 | 174 | 157 | 118 | 112.44 | 75 | 150 | 1.999 | 0.999 | 4.12E-05 | 0.00065762 | 0.0291 | 0.0362 | 0.0327 | 0.0327 | TRUE | 0.3598 |
| 4930513O06 | 0 | 0 | 0 | 1 | 0 | 0 | 0 | 0 | 0 | 1 | 0 | 0 | 0.13 | 0 | 0 | 1.859 | 0.894 | 0.82650734 | NA | 0 | 25.1515 | 10 | 10 | TRUE | 0.2563 |
| 4930515B02 | 0 | 0 | 0 | 0 | 0 | 0 | 0 | 0 | 0 | 0 | 0 | 0 | 0 | 0 | 0 | NA | NA | NA | NA | NA | NA | NA | NA | NA | NA | |
| 4930515G01 | 0 | 0 | 0 | 0 | 1 | 0 | 0 | 0 | 0 | 0 | 1 | 0 | 0.18 | 0 | 0 | 1.859 | 0.894 | 0.82650734 | NA | 0 | 18.3785 | 10 | 10 | TRUE | 0.5204 |
| 4930515G16 | 0 | 0 | 0 | 0 | 0 | 0 | 0 | 0 | 0 | 0 | 0 | 0 | 0 | 0 | 0 | NA | NA | NA | NA | NA | NA | NA | NA | NA | NA | |
| 4930515L03F | 0 | 0 | 0 | 0 | 0 | 0 | 0 | 0 | 0 | 0 | 0 | 0 | 0 | 0 | 0 | NA | NA | NA | NA | NA | NA | NA | NA | NA | NA | |
| 4930515L19F | 0 | 0 | 0 | 0 | 0 | 0 | 0 | 0 | 0 | 0 | 0 | 0 | 0 | 0 | 0 | NA | NA | NA | NA | NA | NA | NA | NA | NA | NA | |
| 4930517E11I | 0 | 0 | 0 | 0 | 0 | 0 | 0 | 0 | 0 | 0 | 0 | 0 | 0 | 0 | 0 | NA | NA | NA | NA | NA | NA | NA | NA | NA | NA | |
| 4930518P08I | 2 | 3 | 3 | 9 | 4 | 5 | 2 | 3 | 3 | 7 | 4 | 5 | 4.16 | 3 | 5 | 2.086 | 1.061 | 0.25101768 | NA | 0 | 0.798 | 0.3412 | 0.3412 | TRUE | 0.1489 |
| 4930519D14 | 0 | 0 | 0 | 0 | 0 | 0 | 0 | 0 | 0 | 0 | 0 | 0 | 0 | 0 | 0 | NA | NA | NA | NA | NA | NA | NA | NA | NA | NA | |
| 4930519F09I | 7 | 1 | 0 | 4 | 1 | 1 | 7 | 1 | 0 | 3 | 1 | 1 | 2.29 | 3 | 2 | 0.65 | -0.622 | 0.68950885 | NA | 1.1328 | 1.4446 | 1.2925 | 1.2925 | TRUE | 2.4254 |
| 4930519F16I | 6 | 4 | 4 | 13 | 18 | 10 | 6 | 4 | 4 | 10 | 19 | 11 | 9.15 | 5 | 13 | 2.809 | 1.49 | 0.0305483 | 0.11680189 | 0 | 0.3668 | 0.1988 | 0.1988 | TRUE | 0.5807 |
| 4930519F24I | 0 | 0 | 0 | 0 | 0 | 0 | 0 | 0 | 0 | 0 | 0 | 0 | 0 | 0 | 0 | NA | NA | NA | NA | NA | NA | NA | NA | NA | NA | |
| 4930519G04 | 0 | 0 | 0 | 0 | 0 | 0 | 0 | 0 | 0 | 0 | 0 | 0 | 0 | 0 | 0 | NA | NA | NA | NA | NA | NA | NA | NA | NA | NA | |
| 4930519H02 | 0 | 0 | 0 | 0 | 0 | 0 | 0 | 0 | 0 | 0 | 0 | 0 | 0 | 0 | 0 | NA | NA | NA | NA | NA | NA | NA | NA | NA | NA | |
| 4930520O04 | 13 | 16 | 24 | 28 | 20 | 16 | 14 | 16 | 24 | 22 | 22 | 18 | 19.16 | 18 | 21 | 1.135 | 0.182 | 0.68554342 | 0.82631951 | 0 | 0.1789 | 0.0935 | 0.0935 | TRUE | 0.457 |
| 4930520P13I | 0 | 0 | 0 | 0 | 0 | 0 | 0 | 0 | 0 | 0 | 0 | 0 | 0 | 0 | 0 | NA | NA | NA | NA | NA | NA | NA | NA | NA | NA | |
| 4930521E06I | 0 | 0 | 0 | 0 | 0 | 0 | 0 | 0 | 0 | 0 | 0 | 0 | 0 | 0 | 0 | NA | NA | NA | NA | NA | NA | NA | NA | NA | NA | |
| 4930522H14 | 0 | 0 | 0 | 0 | 0 | 0 | 0 | 0 | 0 | 0 | 0 | 0 | 0 | 0 | 0 | NA | NA | NA | NA | NA | NA | NA | NA | NA | NA | |
| 4930522O17 | 0 | 0 | 0 | 0 | 0 | 0 | 0 | 0 | 0 | 0 | 0 | 0 | 0 | 0 | 0 | NA | NA | NA | NA | NA | NA | NA | NA | NA | NA | |
| 4930523C07I | 18 | 19 | 14 | 29 | 21 | 12 | 19 | 19 | 14 | 23 | 23 | 13 | 18.45 | 17 | 20 | 1.13 | 0.176 | 0.70305265 | 0.83762068 | 0 | 0.1855 | 0.1001 | 0.1001 | TRUE | 0.3975 |
| 4930523O13 | 0 | 0 | 0 | 0 | 0 | 0 | 0 | 0 | 0 | 0 | 0 | 0 | 0 | 0 | 0 | NA | NA | NA | NA | NA | NA | NA | NA | NA | NA | |
| 4930524B15 | 58 | 67 | 72 | 230 | 156 | 122 | 62 | 67 | 72 | 181 | 168 | 134 | 113.84 | 67 | 161 | 2.407 | 1.267 | 3.45E-09 | 1.81E-07 | 0 | 0.0358 | 0.0225 | 0.0225 | TRUE | 0.2247 |
| 4930524C18I | 0 | 0 | 0 | 1 | 0 | 0 | 0 | 0 | 0 | 1 | 0 | 0 | 0.13 | 0 | 0 | 1.859 | 0.894 | 0.82650734 | NA | 0 | 25.1515 | 10 | 10 | TRUE | 0.2563 |
| 4930524L23F | 4 | 2 | 4 | 16 | 7 | 4 | 4 | 2 | 4 | 13 | 8 | 4 | 5.79 | 3 | 8 | 2.429 | 1.28 | 0.13717222 | 0.32118176 | 0 | 0.5757 | 0.3259 | 0.3259 | TRUE | 0.213 |
| 4930524N10 | 0 | 0 | 0 | 0 | 0 | 0 | 0 | 0 | 0 | 0 | 0 | 0 | 0 | 0 | 0 | NA | NA | NA | NA | NA | NA | NA | NA | NA | NA | |
| 4930524O05 | 0 | 0 | 0 | 0 | 0 | 0 | 0 | 0 | 0 | 0 | 0 | 0 | 0 | 0 | 0 | NA | NA | NA | NA | NA | NA | NA | NA | NA | NA | |
| 4930524O08 | 0 | 0 | 0 | 0 | 0 | 0 | 0 | 0 | 0 | 0 | 0 | 0 | 0 | 0 | 0 | NA | NA | NA | NA | NA | NA | NA | NA | NA | NA | |
| 4930525D18 | 2 | 0 | 0 | 0 | 0 | 1 | 2 | 0 | 0 | 0 | 0 | 1 | 0.54 | 1 | 0 | 0.497 | -1.009 | 0.76081378 | NA | 0 | 6.1475 | 6.1929 | 6.1929 | TRUE | 0.9742 |
| 4930525G20 | 5 | 4 | 5 | 5 | 3 | 3 | 5 | 4 | 5 | 4 | 3 | 3 | 4.13 | 5 | 3 | 0.734 | -0.446 | 0.61662953 | NA | 0 | 0.805 | 0.3249 | 0.3249 | TRUE | 0.0405 |
| 4930525M21 | 0 | 0 | 0 | 0 | 0 | 0 | 0 | 0 | 0 | 0 | 0 | 0 | 0 | 0 | 0 | NA | NA | NA | NA | NA | NA | NA | NA | NA | NA | |
| 4930526D03 | 2 | 1 | 1 | 2 | 2 | 1 | 2 | 1 | 1 | 2 | 2 | 1 | 1.49 | 1 | 2 | 1.173 | 0.231 | 0.87447545 | NA | 0 | 2.2165 | 0.8574 | 0.8574 | TRUE | 0.1349 |
| 4930526I15R | 83 | 93 | 100 | 106 | 97 | 72 | 88 | 93 | 100 | 83 | 104 | 79 | 91.33 | 94 | 89 | 0.945 | -0.082 | 0.70629144 | 0.8396577 | 0 | 0.043 | 0.023 | 0.023 | TRUE | 0.2116 |
| 4930526L06F | 30 | 25 | 18 | 30 | 16 | 22 | 32 | 25 | 18 | 24 | 17 | 24 | 23.3 | 25 | 22 | 0.87 | -0.201 | 0.63301706 | 0.79357325 | 0 | 0.1483 | 0.0841 | 0.0841 | TRUE | 0.1629 |
| 4930527F14I | 0 | 0 | 0 | 0 | 1 | 0 | 0 | 0 | 0 | 0 | 1 | 0 | 0.18 | 0 | 0 | 1.859 | 0.894 | 0.82650734 | NA | 0 | 18.3785 | 10 | 10 | TRUE | 0.5204 |
| 4930527G23 | 0 | 0 | 0 | 0 | 0 | 0 | 0 | 0 | 0 | 0 | 0 | 0 | 0 | 0 | 0 | NA | NA | NA | NA | NA | NA | NA | NA | NA | NA | |
| 4930528A17 | 18 | 21 | 11 | 41 | 18 | 14 | 19 | 21 | 11 | 32 | 19 | 15 | 19.68 | 17 | 22 | 1.325 | 0.406 | 0.41812626 | 0.62897231 | 0.083 | 0.1743 | 0.1294 | 0.1294 | TRUE | 1.0188 |
| 4930528D03 | 1 | 5 | 8 | 6 | 9 | 5 | 1 | 5 | 8 | 5 | 10 | 5 | 5.66 | 5 | 7 | 1.393 | 0.478 | 0.5947733 | 0.7677918 | 0.1615 | 0.5889 | 0.3987 | 0.3987 | TRUE | 0.3442 |
| 4930528P14I | 0 | 0 | 0 | 0 | 0 | 0 | 0 | 0 | 0 | 0 | 0 | 0 | 0 | 0 | 0 | NA | NA | NA | NA | NA | NA | NA | NA | NA | NA | |
| 4930529C04I | 0 | 0 | 0 | 0 | 1 | 0 | 0 | 0 | 0 | 0 | 1 | 0 | 0.18 | 0 | 0 | 1.859 | 0.894 | 0.82650734 | NA | 0 | 18.3785 | 10 | 10 | TRUE | 0.5204 |
| 4930529F24I | 0 | 0 | 0 | 0 | 0 | 0 | 0 | 0 | 0 | 0 | 0 | 0 | 0 | 0 | 0 | NA | NA | NA | NA | NA | NA | NA | NA | NA | NA | |
| 4930529K09I | 0 | 0 | 0 | 0 | 0 | 0 | 0 | 0 | 0 | 0 | 0 | 0 | 0 | 0 | 0 | NA | NA | NA | NA | NA | NA | NA | NA | NA | NA | |
| 4930529L06F | 0 | 0 | 0 | 0 | 0 | 0 | 0 | 0 | 0 | 0 | 0 | 0 | 0 | 0 | 0 | NA | NA | NA | NA | NA | NA | NA | NA | NA | NA | |
| 4930529M08 | 9 | 18 | 18 | 12 | 15 | 17 | 10 | 18 | 18 | 9 | 16 | 19 | 14.97 | 15 | 15 | 0.955 | -0.067 | 0.90112549 | 0.95196343 | 0.0506 | 0.227 | 0.1428 | 0.1428 | TRUE | 0.5631 |
| 4930533B01 | 0 | 0 | 0 | 0 | 0 | 0 | 0 | 0 | 0 | 0 | 0 | 0 | 0 | 0 | 0 | NA | NA | NA | NA | NA | NA | NA | NA | NA | NA | |
| 4930533P14I | 0 | 0 | 0 | 0 | 0 | 0 | 0 | 0 | 0 | 0 | 0 | 0 | 0 | 0 | 0 | NA | NA | NA | NA | NA | NA | NA | NA | NA | NA | |
| 4930538K18I | 18 | 20 | 8 | 10 | 11 | 18 | 19 | 20 | 8 | 8 | 12 | 20 | 14.42 | 16 | 13 | 0.826 | -0.275 | 0.64770085 | 0.80218563 | 0.1436 | 0.2353 | 0.1908 | 0.1908 | TRUE | 0.6027 |
| 4930539C22I | 0 | 0 | 0 | 0 | 0 | 0 | 0 | 0 | 0 | 0 | 0 | 0 | 0 | 0 | 0 | NA | NA | NA | NA | NA | NA | NA | NA | NA | NA | |
| 4930539E08I | 364 | 403 | 340 | 474 | 373 | 393 | 387 | 402 | 341 | 373 | 401 | 430 | 389.11 | 377 | 401 | 1.063 | 0.089 | 0.48401181 | 0.68159234 | 0.0037 | 0.0154 | 0.009 | 0.009 | TRUE | 0.0782 |

| | | | | | | | | | | | | | | | | | | | | | | | | | | |
|---|---|---|---|---|---|---|---|---|---|---|---|---|---|---|---|---|---|---|---|---|---|---|---|---|---|---|
| 4930539J05F | 3 | 2 | 2 | 4 | 1 | 2 | 3 | 2 | 2 | 3 | 1 | 2 | 2.27 | 2 | 2 | 0.912 | -0.133 | 0.91229569 | NA | | 0 | 1.4601 | 0.6137 | 0.6137 | TRUE | 0.2186 |
| 4930539M17 | 8 | 3 | 2 | 7 | 5 | 5 | 8 | 3 | 2 | 6 | 5 | 5 | 4.98 | 4 | 5 | 1.22 | 0.287 | 0.74843256 | 0.86655986 | | 0 | 0.6689 | 0.3717 | 0.3717 | TRUE | 1.0633 |
| 4930539N22 | 0 | 0 | 0 | 0 | 0 | 0 | 0 | 0 | 0 | 0 | 0 | 0 | 0 | 0 | 0 | NA | NA | NA | NA | NA | NA | NA | NA | NA | NA | |
| 4930540M03 | 0 | 0 | 0 | 0 | 0 | 0 | 0 | 0 | 0 | 0 | 0 | 0 | 0 | 0 | 0 | NA | NA | NA | NA | NA | NA | NA | NA | NA | NA | |
| 4930542C21I | 0 | 0 | 0 | 0 | 0 | 0 | 0 | 0 | 0 | 0 | 0 | 0 | 0 | 0 | 0 | NA | NA | NA | NA | NA | NA | NA | NA | NA | NA | |
| 4930542D17I | 0 | 2 | 0 | 0 | 1 | 0 | 0 | 2 | 0 | 0 | 1 | 0 | 0.51 | 1 | 0 | 0.522 | -0.937 | 0.77976772 | NA | | 0 | 6.4385 | 6.3352 | 6.3352 | TRUE | 0.9739 |
| 4930542H20I | 0 | 0 | 0 | 0 | 0 | 0 | 0 | 0 | 0 | 0 | 0 | 0 | 0 | 0 | 0 | NA | NA | NA | NA | NA | NA | NA | NA | NA | NA | |
| 4930543E12I | 3 | 1 | 3 | 6 | 3 | 1 | 3 | 1 | 3 | 5 | 3 | 1 | 2.71 | 2 | 3 | 1.286 | 0.363 | 0.75763163 | NA | | 0 | 1.2243 | 0.6229 | 0.6229 | TRUE | 0.2441 |
| 4930544D05I | 0 | 0 | 0 | 1 | 0 | 0 | 0 | 0 | 0 | 1 | 0 | 0 | 0.13 | 0 | 0 | 1.859 | 0.894 | 0.82650734 | NA | | 0 | 25.1515 | 10 | 10 | TRUE | 0.2563 |
| 4930544G11 | 0 | 0 | 0 | 0 | 0 | 0 | 0 | 0 | 0 | 0 | 0 | 0 | 0 | 0 | 0 | NA | NA | NA | NA | NA | NA | NA | NA | NA | NA | |
| 4930544M13 | 0 | 0 | 0 | 0 | 0 | 0 | 0 | 0 | 0 | 0 | 0 | 0 | 0 | 0 | 0 | NA | NA | NA | NA | NA | NA | NA | NA | NA | NA | |
| 4930545E07I | 0 | 0 | 0 | 0 | 0 | 0 | 0 | 0 | 0 | 0 | 0 | 0 | 0 | 0 | 0 | NA | NA | NA | NA | NA | NA | NA | NA | NA | NA | |
| 4930545H06I | 0 | 1 | 0 | 0 | 0 | 1 | 0 | 1 | 0 | 0 | 0 | 1 | 0.35 | 0 | 0 | 0.954 | -0.067 | 0.98567705 | NA | | 0 | 9.4494 | 8.1588 | 8.1588 | TRUE | 0.5383 |
| 4930545L23F | 0 | 0 | 0 | 0 | 0 | 0 | 0 | 0 | 0 | 0 | 0 | 0 | 0 | 0 | 0 | NA | NA | NA | NA | NA | NA | NA | NA | NA | NA | |
| 4930546C10I | 0 | 0 | 0 | 0 | 0 | 0 | 0 | 0 | 0 | 0 | 0 | 0 | 0 | 0 | 0 | NA | NA | NA | NA | NA | NA | NA | NA | NA | NA | |
| 4930546K05I | 1 | 2 | 3 | 3 | 2 | 2 | 1 | 2 | 3 | 2 | 2 | 2 | 2.13 | 2 | 2 | 1.102 | 0.14 | 0.90978489 | NA | | 0 | 1.5546 | 0.6261 | 0.6261 | TRUE | 0.1659 |
| 4930547C10I | 1 | 1 | 1 | 8 | 1 | 0 | 1 | 1 | 1 | 6 | 1 | 0 | 1.74 | 1 | 2 | 2.491 | 1.317 | 0.43502191 | NA | | 0 | 1.9021 | 1.3571 | 1.3571 | TRUE | 1.7285 |
| 4930547E08I | 2 | 2 | 3 | 2 | 3 | 5 | 2 | 2 | 3 | 2 | 3 | 5 | 2.9 | 2 | 3 | 1.409 | 0.494 | 0.65644057 | NA | | 0 | 1.1424 | 0.53 | 0.53 | TRUE | 0.2214 |
| 4930547E14I | 0 | 0 | 0 | 0 | 0 | 0 | 0 | 0 | 0 | 0 | 0 | 0 | 0 | 0 | 0 | NA | NA | NA | NA | NA | NA | NA | NA | NA | NA | |
| 4930548G14 | 0 | 0 | 0 | 0 | 0 | 0 | 0 | 0 | 0 | 0 | 0 | 0 | 0 | 0 | 0 | NA | NA | NA | NA | NA | NA | NA | NA | NA | NA | |
| 4930548H24I | 1 | 2 | 2 | 0 | 0 | 1 | 1 | 2 | 2 | 0 | 0 | 1 | 1.03 | 2 | 0 | 0.206 | -2.277 | 0.2572845 | NA | | 0 | 3.2145 | 1.6104 | 1.6104 | TRUE | 0.5383 |
| 4930548J01F | 0 | 0 | 0 | 0 | 0 | 0 | 0 | 0 | 0 | 0 | 0 | 0 | 0 | 0 | 0 | NA | NA | NA | NA | NA | NA | NA | NA | NA | NA | |
| 4930548K13I | 0 | 0 | 0 | 0 | 0 | 0 | 0 | 0 | 0 | 0 | 0 | 0 | 0 | 0 | 0 | NA | NA | NA | NA | NA | NA | NA | NA | NA | NA | |
| 4930549C01I | 1 | 1 | 0 | 1 | 0 | 0 | 1 | 1 | 0 | 1 | 0 | 0 | 0.47 | 1 | 0 | 0.51 | -0.973 | 0.73452857 | NA | | 0 | 6.9499 | 4.1835 | 4.1835 | TRUE | 0.2562 |
| 4930549G23 | 1 | 0 | 1 | 1 | 1 | 2 | 1 | 0 | 1 | 1 | 1 | 2 | 1.02 | 1 | 1 | 1.941 | 0.957 | 0.60849107 | NA | | 0 | 3.238 | 1.4016 | 1.4016 | TRUE | 0.256 |
| 4930550C14I | 21 | 19 | 21 | 24 | 16 | 13 | 22 | 19 | 21 | 19 | 17 | 14 | 18.78 | 21 | 17 | 0.812 | -0.301 | 0.48506151 | 0.68237017 | | 0 | 0.1823 | 0.0802 | 0.0802 | TRUE | 0.0793 |
| 4930550L24F | 13 | 6 | 7 | 161 | 134 | 83 | 14 | 6 | 7 | 127 | 144 | 91 | 64.74 | 9 | 121 | 13.588 | 3.764 | 8.37E-21 | 2.36E-18 | | 0.0504 | 0.0578 | 0.0551 | 0.0551 | TRUE | 0.4298 |
| 4930552N02 | 0 | 0 | 0 | 0 | 0 | 0 | 0 | 0 | 0 | 0 | 0 | 0 | 0 | 0 | 0 | NA | NA | NA | NA | NA | NA | NA | NA | NA | NA | |
| 4930552P12I | 18 | 17 | 24 | 23 | 15 | 28 | 19 | 17 | 24 | 18 | 16 | 31 | 20.84 | 20 | 22 | 1.071 | 0.099 | 0.82764363 | 0.91271118 | | 0.031 | 0.165 | 0.1006 | 0.1006 | TRUE | 0.6969 |
| 4930553E22I | 0 | 0 | 0 | 0 | 0 | 0 | 0 | 0 | 0 | 0 | 0 | 0 | 0 | 0 | 0 | NA | NA | NA | NA | NA | NA | NA | NA | NA | NA | |
| 4930554C24I | 0 | 0 | 0 | 0 | 0 | 0 | 0 | 0 | 0 | 0 | 0 | 0 | 0 | 0 | 0 | NA | NA | NA | NA | NA | NA | NA | NA | NA | NA | |
| 4930555B11I | 0 | 0 | 0 | 0 | 0 | 0 | 0 | 0 | 0 | 0 | 0 | 0 | 0 | 0 | 0 | NA | NA | NA | NA | NA | NA | NA | NA | NA | NA | |
| 4930555G01 | 1 | 3 | 0 | 1 | 2 | 3 | 1 | 3 | 0 | 1 | 2 | 3 | 1.71 | 1 | 2 | 1.499 | 0.584 | 0.7080666 | NA | | 0 | 1.9297 | 1.1353 | 1.1353 | TRUE | 0.5351 |
| 4930556C24I | 0 | 0 | 1 | 0 | 0 | 0 | 0 | 0 | 1 | 0 | 0 | 0 | 0.17 | 0 | 0 | 0.49 | -1.029 | 0.80087 | NA | | 0 | 19.6928 | 10 | 10 | TRUE | 0.4098 |
| 4930556G01 | 0 | 0 | 1 | 2 | 0 | 1 | 0 | 0 | 1 | 2 | 0 | 1 | 0.61 | 0 | 1 | 2.474 | 1.307 | 0.62050174 | NA | | 0 | 5.3914 | 3.4771 | 3.4771 | TRUE | 0.4098 |
| 4930556J02F | 0 | 0 | 0 | 0 | 0 | 0 | 0 | 0 | 0 | 0 | 0 | 0 | 0 | 0 | 0 | NA | NA | NA | NA | NA | NA | NA | NA | NA | NA | |
| 4930556M19 | 22 | 31 | 23 | 18 | 17 | 19 | 23 | 31 | 23 | 14 | 18 | 21 | 21.78 | 26 | 18 | 0.681 | -0.554 | 0.18555041 | 0.3890589 | | 0 | 0.1582 | 0.0788 | 0.0788 | TRUE | 0.2011 |
| 4930556N09 | 0 | 0 | 0 | 0 | 0 | 0 | 0 | 0 | 0 | 0 | 0 | 0 | 0 | 0 | 0 | NA | NA | NA | NA | NA | NA | NA | NA | NA | NA | |
| 4930557A04I | 0 | 0 | 0 | 0 | 0 | 0 | 0 | 0 | 0 | 0 | 0 | 0 | 0 | 0 | 0 | NA | NA | NA | NA | NA | NA | NA | NA | NA | NA | |
| 4930557J02F | 0 | 0 | 0 | 0 | 0 | 0 | 0 | 0 | 0 | 0 | 0 | 0 | 0 | 0 | 0 | NA | NA | NA | NA | NA | NA | NA | NA | NA | NA | |
| 4930558C23I | 3 | 2 | 0 | 8 | 4 | 7 | 3 | 2 | 0 | 6 | 4 | 8 | 3.91 | 2 | 6 | 3.553 | 1.829 | 0.09758844 | NA | | 0 | 0.8502 | 0.4999 | 0.4999 | TRUE | 0.6101 |
| 4930558G05 | 0 | 0 | 0 | 0 | 0 | 0 | 0 | 0 | 0 | 0 | 0 | 0 | 0 | 0 | 0 | NA | NA | NA | NA | NA | NA | NA | NA | NA | NA | |
| 4930558J18F | 135 | 173 | 171 | 204 | 148 | 167 | 143 | 173 | 172 | 160 | 159 | 183 | 165.05 | 163 | 167 | 1.028 | 0.039 | 0.81550928 | 0.90592684 | | 0 | 0.0269 | 0.0144 | 0.0144 | TRUE | 0.1112 |
| 4930558K02I | 1 | 2 | 1 | 1 | 2 | 1 | 1 | 2 | 1 | 1 | 2 | 1 | 1.35 | 1 | 1 | 0.972 | -0.041 | 0.97884658 | NA | | 0 | 2.4483 | 0.9574 | 0.9574 | TRUE | 0.16 |
| 4930562C15I | 8 | 7 | 4 | 11 | 3 | 5 | 8 | 7 | 4 | 9 | 3 | 5 | 6.14 | 6 | 6 | 0.908 | -0.14 | 0.86026627 | 0.9309173 | | 0 | 0.5433 | 0.2919 | 0.2919 | TRUE | 0.3931 |
| 4930562F07I | 0 | 0 | 1 | 2 | 0 | 1 | 0 | 0 | 1 | 2 | 0 | 1 | 0.61 | 0 | 1 | 2.474 | 1.307 | 0.62050174 | NA | | 0 | 5.3914 | 3.4771 | 3.4771 | TRUE | 0.4098 |
| 4930563D23I | 0 | 1 | 0 | 0 | 1 | 0 | 0 | 1 | 0 | 0 | 1 | 0 | 0.35 | 0 | 0 | 0.954 | -0.067 | 0.98571955 | NA | | 0 | 9.5346 | 8.2194 | 8.2194 | TRUE | 0.5204 |
| 4930563E18I | 0 | 0 | 0 | 3 | 0 | 1 | 0 | 0 | 0 | 2 | 0 | 1 | 0.58 | 0 | 1 | 6.21 | 2.635 | 0.40034089 | NA | | 0 | 5.7314 | 5.6518 | 5.6518 | TRUE | 0.0653 |
| 4930563E22I | 76 | 87 | 75 | 101 | 108 | 87 | 81 | 87 | 75 | 79 | 116 | 95 | 88.96 | 81 | 97 | 1.191 | 0.253 | 0.27886707 | 0.49999026 | | 0.0123 | 0.0439 | 0.0278 | 0.0278 | TRUE | 0.2308 |
| 4930563F08I | 0 | 0 | 0 | 0 | 0 | 0 | 0 | 0 | 0 | 0 | 0 | 0 | 0 | 0 | 0 | NA | NA | NA | NA | NA | NA | NA | NA | NA | NA | |
| 4930563M2C | 0 | 0 | 0 | 1 | 1 | 0 | 0 | 0 | 0 | 1 | 1 | 0 | 0.31 | 0 | 1 | 3.32 | 1.731 | 0.65383052 | NA | | 0 | 10.6224 | 8.9541 | 8.9541 | TRUE | 0.202 |
| 4930564B18I | 0 | 0 | 0 | 0 | 0 | 0 | 0 | 0 | 0 | 0 | 0 | 0 | 0 | 0 | 0 | NA | NA | NA | NA | NA | NA | NA | NA | NA | NA | |
| 4930564C03I | 0 | 0 | 0 | 0 | 0 | 0 | 0 | 0 | 0 | 0 | 0 | 0 | 0 | 0 | 0 | NA | NA | NA | NA | NA | NA | NA | NA | NA | NA | |
| 4930564D02I | 0 | 0 | 0 | 0 | 0 | 0 | 0 | 0 | 0 | 0 | 0 | 0 | 0 | 0 | 0 | NA | NA | NA | NA | NA | NA | NA | NA | NA | NA | |
| 4930564G21 | 0 | 0 | 0 | 0 | 1 | 2 | 0 | 0 | 0 | 0 | 1 | 2 | 0.54 | 0 | 1 | 5.713 | 2.514 | 0.43612448 | NA | | 0 | 6.0606 | 6.0547 | 6.0547 | TRUE | 0.0576 |
| 4930564K09I | 0 | 0 | 0 | 0 | 0 | 0 | 0 | 0 | 0 | 0 | 0 | 0 | 0 | 0 | 0 | NA | NA | NA | NA | NA | NA | NA | NA | NA | NA | |
| 4930565D16I | 0 | 0 | 0 | 0 | 0 | 0 | 0 | 0 | 0 | 0 | 0 | 0 | 0 | 0 | 0 | NA | NA | NA | NA | NA | NA | NA | NA | NA | NA | |
| 4930565N06 | 0 | 0 | 0 | 0 | 0 | 0 | 0 | 0 | 0 | 0 | 0 | 0 | 0 | 0 | 0 | NA | NA | NA | NA | NA | NA | NA | NA | NA | NA | |
| 4930567H12I | 0 | 0 | 0 | 0 | 0 | 0 | 0 | 0 | 0 | 0 | 0 | 0 | 0 | 0 | 0 | NA | NA | NA | NA | NA | NA | NA | NA | NA | NA | |
| 4930567H17I | 0 | 0 | 0 | 0 | 0 | 0 | 0 | 0 | 0 | 0 | 0 | 0 | 0 | 0 | 0 | NA | NA | NA | NA | NA | NA | NA | NA | NA | NA | |
| 4930567J20F | 0 | 0 | 0 | 0 | 0 | 0 | 0 | 0 | 0 | 0 | 0 | 0 | 0 | 0 | 0 | NA | NA | NA | NA | NA | NA | NA | NA | NA | NA | |
| 4930567K20I | 0 | 0 | 0 | 0 | 0 | 0 | 0 | 0 | 0 | 0 | 0 | 0 | 0 | 0 | 0 | NA | NA | NA | NA | NA | NA | NA | NA | NA | NA | |
| 4930568D16I | 0 | 0 | 0 | 0 | 0 | 0 | 0 | 0 | 0 | 0 | 0 | 0 | 0 | 0 | 0 | NA | NA | NA | NA | NA | NA | NA | NA | NA | NA | |
| 4930568E12I | 0 | 0 | 0 | 23 | 13 | 8 | 0 | 0 | 0 | 18 | 14 | 9 | 6.8 | 0 | 14 | 72.876 | 6.187 | 8.03E-06 | 0.00016393 | | 0.0625 | 0.4911 | 0.3485 | 0.3485 | TRUE | 0.0698 |
| 4930568G15 | 0 | 0 | 0 | 0 | 0 | 0 | 0 | 0 | 0 | 0 | 0 | 0 | 0 | 0 | 0 | NA | NA | NA | NA | NA | NA | NA | NA | NA | NA | |
| 4930568K20I | 0 | 0 | 0 | 0 | 0 | 0 | 0 | 0 | 0 | 0 | 0 | 0 | 0 | 0 | 0 | NA | NA | NA | NA | NA | NA | NA | NA | NA | NA | |
| 4930570G19 | 0 | 0 | 0 | 0 | 0 | 0 | 0 | 0 | 0 | 0 | 0 | 0 | 0 | 0 | 0 | NA | NA | NA | NA | NA | NA | NA | NA | NA | NA | |
| 4930571K23I | 0 | 0 | 0 | 0 | 0 | 0 | 0 | 0 | 0 | 0 | 0 | 0 | 0 | 0 | 0 | NA | NA | NA | NA | NA | NA | NA | NA | NA | NA | |
| 4930572K03I | 0 | 0 | 0 | 0 | 0 | 0 | 0 | 0 | 0 | 0 | 0 | 0 | 0 | 0 | 0 | NA | NA | NA | NA | NA | NA | NA | NA | NA | NA | |

| | | | | | | | | | | | | | | | | | | | | | | | | |
|---|---|---|---|---|---|---|---|---|---|---|---|---|---|---|---|---|---|---|---|---|---|---|---|---|
| 4930572O03 | 0 | 0 | 0 | 1 | 0 | 0 | 0 | 0 | 0 | 1 | 0 | 0 | 0.13 | 0 | 0 | 1.859 | 0.894 | 0.82650734 | NA | 0 | 25.1515 | 10 | 10 | TRUE | 0.2563 |
| 4930572O13 | 0 | 0 | 0 | 0 | 0 | 0 | 0 | 0 | 0 | 0 | 0 | 0 | 0 | 0 | 0 | NA | NA | NA | NA | NA | NA | NA | NA | NA | NA |
| 4930573O16 | 0 | 0 | 0 | 0 | 0 | 0 | 0 | 0 | 0 | 0 | 0 | 0 | 0 | 0 | 0 | NA | NA | NA | NA | NA | NA | NA | NA | NA | NA |
| 4930577N17 | 7 | 2 | 11 | 11 | 3 | 1 | 7 | 2 | 11 | 9 | 3 | 1 | 5.57 | 7 | 4 | 0.653 | -0.615 | 0.54349732 | 0.7286671 | 0.5081 | 0.5979 | 0.554 | 0.554 | TRUE | 0.4389 |
| 4930578C19I | 0 | 0 | 0 | 0 | 0 | 0 | 0 | 0 | 0 | 0 | 0 | 0 | 0 | 0 | 0 | NA | NA | NA | NA | NA | NA | NA | NA | NA | NA |
| 4930578E11I | 0 | 0 | 0 | 0 | 0 | 0 | 0 | 0 | 0 | 0 | 0 | 0 | 0 | 0 | 0 | NA | NA | NA | NA | NA | NA | NA | NA | NA | NA |
| 4930578I06R | 0 | 0 | 0 | 0 | 0 | 0 | 0 | 0 | 0 | 0 | 0 | 0 | 0 | 0 | 0 | NA | NA | NA | NA | NA | NA | NA | NA | NA | NA |
| 4930578M01 | 1 | 0 | 0 | 3 | 1 | 0 | 1 | 0 | 0 | 2 | 1 | 0 | 0.75 | 0 | 1 | 3.204 | 1.68 | 0.5025945 | NA | 0 | 4.4029 | 3.1058 | 3.1058 | TRUE | 0.2352 |
| 4930578N18 | 0 | 0 | 0 | 0 | 0 | 0 | 0 | 0 | 0 | 0 | 0 | 0 | 0 | 0 | 0 | NA | NA | NA | NA | NA | NA | NA | NA | NA | NA |
| 4930579F01I | 1 | 0 | 0 | 0 | 0 | 1 | 1 | 0 | 0 | 0 | 0 | 1 | 0.36 | 0 | 0 | 0.954 | -0.067 | 0.98553519 | NA | 0 | 9.1716 | 7.9605 | 7.9605 | TRUE | 0.5383 |
| 4930579G18 | 6 | 4 | 5 | 5 | 4 | 5 | 6 | 4 | 5 | 4 | 4 | 5 | 4.85 | 5 | 4 | 0.887 | -0.174 | 0.83423184 | 0.91571799 | 0 | 0.6863 | 0.2885 | 0.2885 | TRUE | 0.088 |
| 4930579G24 | 794 | 762 | 725 | 1172 | 855 | 796 | 843 | 761 | 728 | 921 | 920 | 871 | 840.83 | 777 | 904 | 1.164 | 0.219 | 0.02527528 | 0.1025577 | 0.002 | 0.0108 | 0.0057 | 0.0057 | TRUE | 0.0647 |
| 4930579K19I | 4 | 11 | 15 | 18 | 13 | 13 | 4 | 11 | 15 | 14 | 14 | 14 | 12.11 | 10 | 14 | 1.393 | 0.478 | 0.42609186 | 0.63621327 | 0.0214 | 0.2789 | 0.175 | 0.175 | TRUE | 0.4856 |
| 4930581F22I | 10 | 8 | 6 | 11 | 13 | 11 | 11 | 8 | 6 | 9 | 14 | 12 | 9.89 | 8 | 12 | 1.398 | 0.484 | 0.43011852 | 0.6399839 | 0 | 0.3402 | 0.1659 | 0.1659 | TRUE | 0.2392 |
| 4930583K01I | 0 | 0 | 0 | 2 | 0 | 1 | 0 | 0 | 0 | 2 | 0 | 1 | 0.44 | 0 | 1 | 4.782 | 2.258 | 0.50653358 | NA | 0 | 7.4188 | 6.7775 | 6.7775 | TRUE | 0.2804 |
| 4930583P06I | 0 | 0 | 0 | 0 | 0 | 0 | 0 | 0 | 0 | 0 | 0 | 0 | 0 | 0 | 0 | NA | NA | NA | NA | NA | NA | NA | NA | NA | NA |
| 4930584F24I | 0 | 0 | 0 | 0 | 0 | 0 | 0 | 0 | 0 | 0 | 0 | 0 | 0 | 0 | 0 | NA | NA | NA | NA | NA | NA | NA | NA | NA | NA |
| 4930590J08R | 20 | 19 | 27 | 21 | 19 | 14 | 21 | 19 | 27 | 17 | 20 | 15 | 19.93 | 22 | 17 | 0.775 | -0.368 | 0.39162079 | 0.60534847 | 0 | 0.1722 | 0.0823 | 0.0823 | TRUE | 0.1932 |
| 4930590L20R | 2 | 0 | 0 | 4 | 3 | 3 | 2 | 0 | 0 | 3 | 3 | 3 | 1.96 | 1 | 3 | 4.652 | 2.218 | 0.16947309 | NA | 0 | 1.6848 | 0.9873 | 0.9873 | TRUE | 0.9746 |
| 4930591A17 | 115 | 94 | 81 | 239 | 160 | 130 | 122 | 94 | 81 | 188 | 172 | 142 | 133.28 | 99 | 167 | 1.695 | 0.762 | 0.0005486 | 0.0055895 | 0.0219 | 0.0316 | 0.0269 | 0.0269 | TRUE | 0.3956 |
| 4930592A05 | 0 | 0 | 0 | 0 | 0 | 0 | 0 | 0 | 0 | 0 | 0 | 0 | 0 | 0 | 0 | NA | NA | NA | NA | NA | NA | NA | NA | NA | NA |
| 4930592I03R | 1 | 0 | 0 | 0 | 0 | 0 | 1 | 0 | 0 | 0 | 0 | 0 | 0.18 | 0 | 0 | 0.49 | -1.029 | 0.80087 | NA | 0 | 18.6165 | 10 | 10 | TRUE | 0.4614 |
| 4930593A02 | 0 | 0 | 0 | 0 | 0 | 0 | 0 | 0 | 0 | 0 | 0 | 0 | 0 | 0 | 0 | NA | NA | NA | NA | NA | NA | NA | NA | NA | NA |
| 4930593C16I | 0 | 0 | 0 | 0 | 0 | 0 | 0 | 0 | 0 | 0 | 0 | 0 | 0 | 0 | 0 | NA | NA | NA | NA | NA | NA | NA | NA | NA | NA |
| 4930594C11I | 3 | 4 | 6 | 4 | 0 | 3 | 3 | 4 | 6 | 3 | 0 | 3 | 3.27 | 4 | 2 | 0.495 | -1.013 | 0.37554035 | NA | 0 | 1.0136 | 0.6026 | 0.6026 | TRUE | 0.6476 |
| 4930595M18 | 0 | 0 | 0 | 0 | 0 | 0 | 0 | 0 | 0 | 0 | 0 | 0 | 0 | 0 | 0 | NA | NA | NA | NA | NA | NA | NA | NA | NA | NA |
| 4930596D02 | 0 | 0 | 0 | 0 | 0 | 0 | 0 | 0 | 0 | 0 | 0 | 0 | 0 | 0 | 0 | NA | NA | NA | NA | NA | NA | NA | NA | NA | NA |
| 4930596M17 | 0 | 0 | 0 | 0 | 0 | 0 | 0 | 0 | 0 | 0 | 0 | 0 | 0 | 0 | 0 | NA | NA | NA | NA | NA | NA | NA | NA | NA | NA |
| 4930597G03 | 0 | 0 | 0 | 0 | 0 | 0 | 0 | 0 | 0 | 0 | 0 | 0 | 0 | 0 | 0 | NA | NA | NA | NA | NA | NA | NA | NA | NA | NA |
| 4930598F16I | 0 | 0 | 0 | 0 | 0 | 0 | 0 | 0 | 0 | 0 | 0 | 0 | 0 | 0 | 0 | NA | NA | NA | NA | NA | NA | NA | NA | NA | NA |
| 4930599N23 | 1 | 0 | 2 | 4 | 4 | 2 | 1 | 0 | 2 | 3 | 4 | 2 | 2.12 | 1 | 3 | 3.143 | 1.652 | 0.24220218 | NA | 0 | 1.5624 | 0.7862 | 0.7862 | TRUE | 0.378 |
| 4931402G19 | 0 | 0 | 0 | 0 | 0 | 0 | 0 | 0 | 0 | 0 | 0 | 0 | 0 | 0 | 0 | NA | NA | NA | NA | NA | NA | NA | NA | NA | NA |
| 4931403E22I | 1 | 0 | 0 | 0 | 0 | 0 | 1 | 0 | 0 | 0 | 0 | 0 | 0.18 | 0 | 0 | 0.49 | -1.029 | 0.80087 | NA | 0 | 18.6165 | 10 | 10 | TRUE | 0.4614 |
| 4931403G20 | 0 | 0 | 0 | 0 | 0 | 0 | 0 | 0 | 0 | 0 | 0 | 0 | 0 | 0 | 0 | NA | NA | NA | NA | NA | NA | NA | NA | NA | NA |
| 4931406B18 | 3 | 2 | 0 | 2 | 1 | 3 | 3 | 2 | 0 | 2 | 1 | 3 | 1.85 | 2 | 2 | 1.142 | 0.191 | 0.89695719 | NA | 0 | 1.7851 | 1.0205 | 1.0205 | TRUE | 0.429 |
| 4931406C07I | 1007 | 1086 | 1076 | 1006 | 686 | 753 | 1070 | 1085 | 1080 | 791 | 738 | 824 | 931.31 | 1078 | 784 | 0.728 | -0.459 | 1.33E-07 | 4.58E-06 | 0 | 0.0104 | 0.0044 | 0.0044 | TRUE | 0.0304 |
| 4931406H21 | 0 | 0 | 0 | 3 | 1 | 0 | 0 | 0 | 0 | 0 | 2 | 1 | 0 | 0.57 | 0 | 1 | 6.182 | 2.628 | 0.40213576 | NA | 0 | 5.7626 | 5.6688 | 5.6688 | TRUE | 0.0684 |
| 4931406P16I | 4773 | 4412 | 4238 | 5795 | 4281 | 3923 | 5070 | 4406 | 4255 | 4556 | 4606 | 4294 | 4531.25 | 4577 | 4485 | 0.98 | -0.029 | 0.7531241 | 0.86975715 | 0.0041 | 0.0076 | 0.0059 | 0.0059 | TRUE | 0.1078 |
| 4931408C20I | 1 | 0 | 0 | 0 | 0 | 0 | 1 | 0 | 0 | 0 | 0 | 0 | 0.18 | 0 | 0 | 0.49 | -1.029 | 0.80087 | NA | 0 | 18.6165 | 10 | 10 | TRUE | 0.4614 |
| 4931408D14 | 3 | 3 | 10 | 7 | 6 | 4 | 3 | 3 | 10 | 6 | 6 | 4 | 5.43 | 5 | 5 | 1.005 | 0.008 | 0.99292042 | 0.99616826 | 0.0954 | 0.614 | 0.3545 | 0.3545 | TRUE | 1.2238 |
| 4931409K22I | 10 | 11 | 11 | 14 | 9 | 13 | 11 | 11 | 11 | 11 | 10 | 14 | 11.26 | 11 | 12 | 1.066 | 0.093 | 0.86650868 | 0.93389495 | 0 | 0.2994 | 0.1298 | 0.1298 | TRUE | 0.1301 |
| 4931414P19I | 362 | 369 | 360 | 442 | 343 | 377 | 385 | 369 | 361 | 347 | 369 | 413 | 373.95 | 372 | 376 | 1.011 | 0.016 | 0.89476014 | 0.94909927 | 0.0019 | 0.0157 | 0.0081 | 0.0081 | TRUE | 0.0804 |
| 4931417E11I | 0 | 0 | 0 | 0 | 0 | 0 | 0 | 0 | 0 | 0 | 0 | 0 | 0 | 0 | 0 | NA | NA | NA | NA | NA | NA | NA | NA | NA | NA |
| 4931417G12 | 8 | 14 | 13 | 36 | 26 | 16 | 8 | 14 | 13 | 28 | 28 | 18 | 18.22 | 12 | 25 | 2.081 | 1.057 | 0.02842772 | 0.11092579 | 0 | 0.1877 | 0.105 | 0.105 | TRUE | 0.3544 |
| 4931419H13 | 1 | 0 | 0 | 0 | 0 | 0 | 1 | 0 | 0 | 0 | 0 | 0 | 0.18 | 0 | 0 | 0.49 | -1.029 | 0.80087 | NA | 0 | 18.6165 | 10 | 10 | TRUE | 0.4614 |
| 4931420L22I | 0 | 0 | 0 | 0 | 1 | 1 | 0 | 0 | 0 | 0 | 1 | 1 | 0.36 | 0 | 1 | 3.794 | 1.924 | 0.60247382 | NA | 0 | 9.1135 | 8.1425 | 8.1425 | TRUE | 0.3482 |
| 4931423N10 | 4 | 3 | 1 | 3 | 2 | 3 | 4 | 3 | 1 | 2 | 2 | 3 | 2.67 | 3 | 2 | 0.944 | -0.082 | 0.94245769 | NA | 0 | 1.239 | 0.5642 | 0.5642 | TRUE | 0.3463 |
| 4931428F04I | 435 | 644 | 718 | 942 | 780 | 744 | 462 | 643 | 721 | 741 | 839 | 814 | 703.39 | 609 | 798 | 1.31 | 0.39 | 0.01955376 | 0.08547172 | 0.0256 | 0.0116 | 0.0186 | 0.0186 | TRUE | 0.5194 |
| 4931428L18I | 1 | 0 | 0 | 3 | 1 | 3 | 1 | 0 | 0 | 2 | 1 | 3 | 1.3 | 0 | 2 | 6.124 | 2.615 | 0.16830101 | NA | 0 | 2.5475 | 1.3768 | 1.3768 | TRUE | 0.4282 |
| 4931429I11R | 10 | 9 | 15 | 9 | 3 | 8 | 11 | 9 | 15 | 7 | 3 | 9 | 8.96 | 12 | 6 | 0.552 | -0.858 | 0.20154567 | 0.40908409 | 0 | 0.3748 | 0.2044 | 0.2044 | TRUE | 0.4033 |
| 4931429L15I | 0 | 1 | 3 | 1 | 3 | 0 | 0 | 1 | 3 | 1 | 3 | 0 | 1.34 | 1 | 1 | 0.985 | -0.021 | 0.99104873 | NA | 1.0082 | 2.4701 | 1.8674 | 1.8674 | TRUE | 0.6102 |
| 4931429P17I | 0 | 0 | 0 | 0 | 0 | 0 | 0 | 0 | 0 | 0 | 0 | 0 | 0 | 0 | 0 | NA | NA | NA | NA | NA | NA | NA | NA | NA | NA |
| 4931430N09 | 0 | 0 | 0 | 0 | 0 | 1 | 0 | 0 | 0 | 0 | 0 | 1 | 0.18 | 0 | 0 | 1.859 | 0.894 | 0.82650734 | NA | 0 | 18.0643 | 10 | 10 | TRUE | 0.5383 |
| 4931431B13 | 0 | 0 | 0 | 0 | 0 | 0 | 0 | 0 | 0 | 0 | 0 | 0 | 0 | 0 | 0 | NA | NA | NA | NA | NA | NA | NA | NA | NA | NA |
| 4931431C16I | 1 | 0 | 0 | 0 | 0 | 0 | 1 | 0 | 0 | 0 | 0 | 0 | 0.18 | 0 | 0 | 0.49 | -1.029 | 0.80087 | NA | 0 | 18.6165 | 10 | 10 | TRUE | 0.4614 |
| 4931431F19I | 0 | 0 | 0 | 0 | 0 | 0 | 0 | 0 | 0 | 0 | 0 | 0 | 0 | 0 | 0 | NA | NA | NA | NA | NA | NA | NA | NA | NA | NA |
| 4931440F15I | 8 | 9 | 4 | 12 | 2 | 4 | 8 | 9 | 4 | 9 | 2 | 4 | 6.24 | 7 | 5 | 0.763 | -0.39 | 0.64421134 | 0.79991349 | 0.1583 | 0.5345 | 0.3532 | 0.3532 | TRUE | 0.9363 |
| 4931440J10R | 0 | 0 | 0 | 0 | 0 | 0 | 0 | 0 | 0 | 0 | 0 | 0 | 0 | 0 | 0 | NA | NA | NA | NA | NA | NA | NA | NA | NA | NA |
| 4931440L10R | 31 | 14 | 25 | 27 | 27 | 21 | 33 | 14 | 25 | 21 | 29 | 23 | 24.21 | 24 | 24 | 1.016 | 0.023 | 0.95905701 | 0.97898705 | 0.0542 | 0.143 | 0.1012 | 0.1012 | TRUE | 0.3083 |
| 4931440P22I | 13 | 15 | 20 | 25 | 13 | 23 | 14 | 15 | 20 | 20 | 14 | 25 | 17.95 | 16 | 20 | 1.202 | 0.265 | 0.57304444 | 0.75135563 | 0 | 0.1905 | 0.1032 | 0.1032 | TRUE | 0.1499 |
| 4932411E22I | 0 | 0 | 0 | 0 | 1 | 0 | 0 | 0 | 0 | 0 | 1 | 0 | 0.18 | 0 | 0 | 1.859 | 0.894 | 0.82650734 | NA | 0 | 18.3785 | 10 | 10 | TRUE | 0.5204 |
| 4932411N23 | 3 | 4 | 1 | 6 | 3 | 0 | 3 | 4 | 1 | 5 | 3 | 0 | 2.69 | 3 | 3 | 0.996 | -0.006 | 0.99609264 | NA | 0.3393 | 1.2324 | 0.8712 | 0.8712 | TRUE | 0.5338 |
| 4932412D23 | 56 | 33 | 18 | 51 | 54 | 34 | 59 | 33 | 18 | 40 | 58 | 37 | 40.99 | 37 | 45 | 1.226 | 0.294 | 0.51561634 | 0.70792965 | 0.1627 | 0.0873 | 0.1227 | 0.1227 | TRUE | 0.5165 |
| 4932413F04I | 2 | 3 | 2 | 1 | 1 | 2 | 2 | 3 | 2 | 1 | 1 | 2 | 1.86 | 2 | 1 | 0.554 | -0.851 | 0.52669385 | NA | 0 | 1.7749 | 0.7193 | 0.7193 | TRUE | 0.1672 |
| 4932414J04R | 0 | 0 | 0 | 0 | 0 | 0 | 0 | 0 | 0 | 0 | 0 | 0 | 0 | 0 | 0 | NA | NA | NA | NA | NA | NA | NA | NA | NA | NA |
| 4932414N04 | 0 | 0 | 0 | 1 | 0 | 0 | 0 | 0 | 0 | 1 | 0 | 0 | 0.13 | 0 | 0 | 1.859 | 0.894 | 0.82650734 | NA | 0 | 25.1515 | 10 | 10 | TRUE | 0.2563 |
| 4932415G12 | 41 | 90 | 56 | 86 | 75 | 42 | 44 | 90 | 56 | 68 | 81 | 46 | 63.99 | 63 | 65 | 1.024 | 0.034 | 0.92179822 | 0.96126008 | 0.0896 | 0.0584 | 0.0733 | 0.0733 | TRUE | 0.7801 |
| 4932415M13 | 0 | 0 | 0 | 9 | 4 | 2 | 0 | 0 | 0 | 7 | 4 | 2 | 2.26 | 0 | 4 | 24.315 | 4.604 | 0.00816096 | NA | 0 | 1.4637 | 1.0753 | 1.0753 | TRUE | 0.08 |
| 4932416H05 | 34 | 49 | 47 | 79 | 55 | 52 | 36 | 49 | 47 | 62 | 59 | 57 | 51.74 | 44 | 59 | 1.348 | 0.431 | 0.11512226 | 0.28648781 | 0 | 0.0706 | 0.034 | 0.034 | TRUE | 0.1877 |

| | | | | | | | | | | | | | | | | | | | | | | | | | |
|---|---|---|---|---|---|---|---|---|---|---|---|---|---|---|---|---|---|---|---|---|---|---|---|---|---|
| 4932416K20I | 0 | 0 | 0 | 0 | 0 | 0 | 0 | 0 | 0 | 0 | 0 | 0 | 0 | 0 | 0 | NA | NA | NA | NA | NA | NA | NA | NA | NA | NA |
| 4932418E24I | 0 | 1 | 0 | 0 | 0 | 0 | 0 | 1 | 0 | 0 | 0 | 0 | 0.17 | 0 | 0 | 0.49 | -1.029 | 0.80087 | NA | | 0 | 19.7991 | 10 | 10 | TRUE | 0.405 |
| 4932429P05I | 0 | 0 | 0 | 0 | 0 | 0 | 0 | 0 | 0 | 0 | 0 | 0 | 0 | 0 | 0 | NA | NA | NA | NA | NA | NA | NA | NA | NA | NA |
| 4932435O22 | 1 | 0 | 2 | 1 | 0 | 0 | 1 | 0 | 2 | 1 | 0 | 0 | 0.64 | 1 | 0 | 0.341 | -1.551 | 0.55438633 | NA | | 0 | 5.1324 | 3.4582 | 3.4582 | TRUE | 0.1148 |
| 4932438A13 | 3678 | 3254 | 4294 | 5083 | 3567 | 3520 | 3907 | 3250 | 4312 | 3996 | 3838 | 3853 | 3859.17 | 3823 | 3896 | 1.019 | 0.027 | 0.80976417 | 0.90213132 | 0.0101 | 0.0078 | 0.0091 | 0.0091 | TRUE | 0.2094 |
| 4932438H23 | 0 | 0 | 0 | 0 | 1 | 1 | 0 | 0 | 0 | 0 | 1 | 1 | 0.36 | 0 | 1 | 3.794 | 1.924 | 0.60247382 | NA | | 0 | 9.1135 | 8.1425 | 8.1425 | TRUE | 0.3482 |
| 4932441J04F | 0 | 0 | 0 | 0 | 0 | 0 | 0 | 0 | 0 | 0 | 0 | 0 | 0 | 0 | 0 | NA | NA | NA | NA | NA | NA | NA | NA | NA | NA |
| 4932443I19R | 1 | 0 | 0 | 0 | 0 | 1 | 1 | 0 | 0 | 0 | 0 | 1 | 0.36 | 0 | 0 | 0.954 | -0.067 | 0.98553519 | NA | | 0 | 9.1716 | 7.9605 | 7.9605 | TRUE | 0.5383 |
| 4932702P03I | 0 | 1 | 0 | 0 | 0 | 0 | 0 | 1 | 0 | 0 | 0 | 0 | 0.17 | 0 | 0 | 0.49 | -1.029 | 0.80087 | NA | | 0 | 19.7991 | 10 | 10 | TRUE | 0.405 |
| 4933400A11 | 0 | 0 | 0 | 2 | 0 | 0 | 0 | 0 | 0 | 2 | 0 | 0 | 0.26 | 0 | 1 | 2.886 | 1.529 | 0.70676491 | NA | | 0 | 12.5792 | 10 | 10 | TRUE | 0.9669 |
| 4933400B14 | 0 | 0 | 0 | 0 | 0 | 0 | 0 | 0 | 0 | 0 | 0 | 0 | 0 | 0 | 0 | NA | NA | NA | NA | NA | NA | NA | NA | NA | NA |
| 4933400C23I | 0 | 0 | 0 | 0 | 0 | 0 | 0 | 0 | 0 | 0 | 0 | 0 | 0 | 0 | 0 | NA | NA | NA | NA | NA | NA | NA | NA | NA | NA |
| 4933400F21I | 0 | 0 | 0 | 0 | 0 | 0 | 0 | 0 | 0 | 0 | 0 | 0 | 0 | 0 | 0 | NA | NA | NA | NA | NA | NA | NA | NA | NA | NA |
| 4933400L20F | 0 | 0 | 0 | 0 | 0 | 0 | 0 | 0 | 0 | 0 | 0 | 0 | 0 | 0 | 0 | NA | NA | NA | NA | NA | NA | NA | NA | NA | NA |
| 4933401B06 | 0 | 0 | 0 | 0 | 0 | 0 | 0 | 0 | 0 | 0 | 0 | 0 | 0 | 0 | 0 | NA | NA | NA | NA | NA | NA | NA | NA | NA | NA |
| 4933401D09 | 1 | 1 | 1 | 2 | 3 | 0 | 1 | 1 | 1 | 2 | 3 | 0 | 1.31 | 1 | 2 | 1.569 | 0.65 | 0.70132179 | NA | | 0 | 2.5201 | 1.2622 | 1.2622 | TRUE | 0.1275 |
| 4933401H06 | 0 | 0 | 0 | 0 | 0 | 0 | 0 | 0 | 0 | 0 | 0 | 0 | 0 | 0 | 0 | NA | NA | NA | NA | NA | NA | NA | NA | NA | NA |
| 4933401P06I | 0 | 0 | 0 | 0 | 0 | 0 | 0 | 0 | 0 | 0 | 0 | 0 | 0 | 0 | 0 | NA | NA | NA | NA | NA | NA | NA | NA | NA | NA |
| 4933402C06I | 0 | 0 | 0 | 0 | 0 | 0 | 0 | 0 | 0 | 0 | 0 | 0 | 0 | 0 | 0 | NA | NA | NA | NA | NA | NA | NA | NA | NA | NA |
| 4933402D24 | 0 | 0 | 0 | 0 | 0 | 0 | 0 | 0 | 0 | 0 | 0 | 0 | 0 | 0 | 0 | NA | NA | NA | NA | NA | NA | NA | NA | NA | NA |
| 4933402E13I | 78 | 49 | 41 | 140 | 118 | 91 | 83 | 49 | 41 | 110 | 127 | 100 | 84.93 | 58 | 112 | 1.95 | 0.964 | 0.00114606 | 0.00987744 | 0.0554 | 0.0457 | 0.05 | 0.05 | TRUE | 1.2179 |
| 4933402J07F | 3 | 1 | 2 | 2 | 0 | 0 | 3 | 1 | 2 | 2 | 0 | 0 | 1.29 | 2 | 1 | 0.284 | -1.818 | 0.33398583 | NA | | 0 | 2.5523 | 1.4688 | 1.4688 | TRUE | 0.7418 |
| 4933402J10F | 0 | 0 | 0 | 0 | 0 | 0 | 0 | 0 | 0 | 0 | 0 | 0 | 0 | 0 | 0 | NA | NA | NA | NA | NA | NA | NA | NA | NA | NA |
| 4933402J15F | 0 | 0 | 0 | 1 | 0 | 0 | 0 | 0 | 0 | 1 | 0 | 0 | 0.13 | 0 | 0 | 1.859 | 0.894 | 0.82650734 | NA | | 0 | 25.1515 | 10 | 10 | TRUE | 0.2563 |
| 4933402N03 | 0 | 0 | 0 | 0 | 0 | 0 | 0 | 0 | 0 | 0 | 0 | 0 | 0 | 0 | 0 | NA | NA | NA | NA | NA | NA | NA | NA | NA | NA |
| 4933402N22 | 0 | 0 | 0 | 1 | 0 | 0 | 0 | 0 | 0 | 1 | 0 | 0 | 0.13 | 0 | 0 | 1.859 | 0.894 | 0.82650734 | NA | | 0 | 25.1515 | 10 | 10 | TRUE | 0.2563 |
| 4933402P03I | 0 | 0 | 0 | 0 | 0 | 0 | 0 | 0 | 0 | 0 | 0 | 0 | 0 | 0 | 0 | NA | NA | NA | NA | NA | NA | NA | NA | NA | NA |
| 4933403G14 | 571 | 554 | 518 | 823 | 676 | 548 | 607 | 553 | 520 | 647 | 727 | 600 | 609.02 | 560 | 658 | 1.175 | 0.233 | 0.05311028 | 0.17112251 | 0.006 | 0.0123 | 0.0088 | 0.0088 | TRUE | 0.0975 |
| 4933403O08 | 3 | 1 | 9 | 9 | 4 | 2 | 3 | 1 | 9 | 7 | 4 | 2 | 4.47 | 4 | 4 | 1.042 | 0.059 | 0.95495071 | NA | 0.3918 | 0.7447 | 0.5692 | 0.5692 | TRUE | 0.8432 |
| 4933404G15 | 0 | 0 | 0 | 0 | 0 | 0 | 0 | 0 | 0 | 0 | 0 | 0 | 0 | 0 | 0 | NA | NA | NA | NA | NA | NA | NA | NA | NA | NA |
| 4933404K08I | 0 | 0 | 0 | 0 | 0 | 0 | 0 | 0 | 0 | 0 | 0 | 0 | 0 | 0 | 0 | NA | NA | NA | NA | NA | NA | NA | NA | NA | NA |
| 4933404O12 | 2090 | 2249 | 2298 | 2187 | 1631 | 1868 | 2220 | 2246 | 2307 | 1719 | 1755 | 2045 | 2048.73 | 2258 | 1840 | 0.814 | -0.296 | 0.00165677 | 0.01324406 | 0.0042 | 0.0085 | 0.0059 | 0.0059 | TRUE | 0.1141 |
| 4933405D12 | 4 | 4 | 7 | 3 | 4 | 4 | 4 | 4 | 7 | 2 | 4 | 4 | 4.39 | 5 | 3 | 0.71 | -0.494 | 0.58103762 | NA | | 0 | 0.7581 | 0.3425 | 0.3425 | TRUE | 0.2305 |
| 4933405E24I | 0 | 0 | 0 | 0 | 0 | 0 | 0 | 0 | 0 | 0 | 0 | 0 | 0 | 0 | 0 | NA | NA | NA | NA | NA | NA | NA | NA | NA | NA |
| 4933405L10F | 16 | 12 | 13 | 26 | 12 | 19 | 17 | 12 | 13 | 20 | 13 | 21 | 16.03 | 14 | 18 | 1.296 | 0.374 | 0.44281214 | 0.64973793 | 0 | 0.2124 | 0.1076 | 0.1076 | TRUE | 0.2928 |
| 4933405O20 | 0 | 0 | 0 | 0 | 0 | 0 | 0 | 0 | 0 | 0 | 0 | 0 | 0 | 0 | 0 | NA | NA | NA | NA | NA | NA | NA | NA | NA | NA |
| 4933406C10I | 3 | 3 | 3 | 1 | 3 | 5 | 3 | 3 | 3 | 1 | 3 | 5 | 3.11 | 3 | 3 | 1.002 | 0.002 | 0.9982854 | NA | | 0 | 1.065 | 0.5612 | 0.5612 | TRUE | 0.2425 |
| 4933406D12 | 1 | 0 | 0 | 0 | 0 | 0 | 1 | 0 | 0 | 0 | 0 | 0 | 0.18 | 0 | 0 | 0.49 | -1.029 | 0.80087 | NA | | 0 | 18.6165 | 10 | 10 | TRUE | 0.4614 |
| 4933406F09I | 0 | 0 | 0 | 0 | 0 | 0 | 0 | 0 | 0 | 0 | 0 | 0 | 0 | 0 | 0 | NA | NA | NA | NA | NA | NA | NA | NA | NA | NA |
| 4933406G16 | 0 | 0 | 0 | 0 | 0 | 0 | 0 | 0 | 0 | 0 | 0 | 0 | 0 | 0 | 0 | NA | NA | NA | NA | NA | NA | NA | NA | NA | NA |
| 4933406I18R | 0 | 0 | 0 | 0 | 0 | 0 | 0 | 0 | 0 | 0 | 0 | 0 | 0 | 0 | 0 | NA | NA | NA | NA | NA | NA | NA | NA | NA | NA |
| 4933406J08F | 52 | 92 | 94 | 181 | 138 | 141 | 55 | 92 | 94 | 142 | 148 | 154 | 114.44 | 80 | 148 | 1.838 | 0.878 | 0.00021282 | 0.00255644 | 0.0245 | 0.0357 | 0.0309 | 0.0309 | TRUE | 0.6844 |
| 4933406J10F | 1 | 0 | 1 | 0 | 0 | 1 | 1 | 0 | 1 | 0 | 0 | 1 | 0.53 | 1 | 0 | 0.509 | -0.975 | 0.72674596 | NA | | 0 | 6.2603 | 3.8683 | 3.8683 | TRUE | 0.5383 |
| 4933406K04I | 0 | 0 | 0 | 0 | 0 | 0 | 0 | 0 | 0 | 0 | 0 | 0 | 0 | 0 | 0 | NA | NA | NA | NA | NA | NA | NA | NA | NA | NA |
| 4933406M09 | 4 | 1 | 3 | 1 | 1 | 2 | 4 | 1 | 3 | 1 | 1 | 2 | 2.05 | 3 | 1 | 0.481 | -1.056 | 0.4346043 | NA | | 0 | 1.6125 | 0.7611 | 0.7611 | TRUE | 0.2492 |
| 4933407E24I | 0 | 0 | 0 | 0 | 0 | 0 | 0 | 0 | 0 | 0 | 0 | 0 | 0 | 0 | 0 | NA | NA | NA | NA | NA | NA | NA | NA | NA | NA |
| 4933407G14 | 0 | 1 | 0 | 0 | 0 | 0 | 0 | 1 | 0 | 0 | 0 | 0 | 0.17 | 0 | 0 | 0.49 | -1.029 | 0.80087 | NA | | 0 | 19.7991 | 10 | 10 | TRUE | 0.405 |
| 4933407I05R | 0 | 0 | 0 | 0 | 0 | 0 | 0 | 0 | 0 | 0 | 0 | 0 | 0 | 0 | 0 | NA | NA | NA | NA | NA | NA | NA | NA | NA | NA |
| 4933407K13I | 41 | 44 | 43 | 60 | 38 | 51 | 44 | 44 | 43 | 47 | 41 | 56 | 45.76 | 44 | 48 | 1.1 | 0.138 | 0.63097662 | 0.79265149 | 0 | 0.0789 | 0.0372 | 0.0372 | TRUE | 0.15 |
| 4933407L21F | 0 | 0 | 0 | 0 | 0 | 0 | 0 | 0 | 0 | 0 | 0 | 0 | 0 | 0 | 0 | NA | NA | NA | NA | NA | NA | NA | NA | NA | NA |
| 4933408B17 | 14 | 15 | 9 | 4 | 7 | 4 | 15 | 15 | 9 | 3 | 8 | 4 | 8.99 | 13 | 5 | 0.38 | -1.396 | 0.04272358 | 0.1475652 | 0 | 0.3733 | 0.2034 | 0.2034 | TRUE | 0.369 |
| 4933408J17F | 0 | 0 | 0 | 1 | 0 | 0 | 0 | 0 | 0 | 1 | 0 | 0 | 0.13 | 0 | 0 | 1.859 | 0.894 | 0.82650734 | NA | | 0 | 25.1515 | 10 | 10 | TRUE | 0.2563 |
| 4933408N05 | 6 | 2 | 3 | 22 | 11 | 11 | 6 | 2 | 3 | 17 | 12 | 12 | 8.76 | 4 | 14 | 3.669 | 1.875 | 0.01036164 | 0.05416765 | 0 | 0.383 | 0.2146 | 0.2146 | TRUE | 0.5343 |
| 4933409G03 | 0 | 0 | 0 | 0 | 0 | 0 | 0 | 0 | 0 | 0 | 0 | 0 | 0 | 0 | 0 | NA | NA | NA | NA | NA | NA | NA | NA | NA | NA |
| 4933409K07I | 0 | 0 | 0 | 0 | 0 | 0 | 0 | 0 | 0 | 0 | 0 | 0 | 0 | 0 | 0 | NA | NA | NA | NA | NA | NA | NA | NA | NA | NA |
| 4933411E08I | 1 | 1 | 0 | 1 | 2 | 2 | 1 | 1 | 0 | 1 | 2 | 2 | 1.2 | 1 | 2 | 2.455 | 1.296 | 0.46556115 | NA | | 0 | 2.7568 | 1.2338 | 1.2338 | TRUE | 0.2531 |
| 4933411G06 | 0 | 0 | 0 | 0 | 0 | 0 | 0 | 0 | 0 | 0 | 0 | 0 | 0 | 0 | 0 | NA | NA | NA | NA | NA | NA | NA | NA | NA | NA |
| 4933411G11 | 3 | 1 | 3 | 7 | 3 | 3 | 3 | 1 | 3 | 6 | 3 | 3 | 3.2 | 2 | 4 | 1.697 | 0.763 | 0.47221606 | NA | | 0 | 1.0358 | 0.4778 | 0.4778 | TRUE | 0.2837 |
| 4933411K16I | 3 | 0 | 0 | 8 | 8 | 3 | 3 | 0 | 0 | 6 | 9 | 3 | 3.56 | 1 | 6 | 5.841 | 2.546 | 0.06160816 | NA | 0.2357 | 0.932 | 0.7649 | 0.7649 | TRUE | 0.9548 |
| 4933411K20I | 1634 | 1963 | 2120 | 2770 | 2006 | 1868 | 1736 | 1960 | 2129 | 2178 | 2158 | 2045 | 2034.25 | 1942 | 2127 | 1.095 | 0.132 | 0.18425185 | 0.38709544 | 0.005 | 0.0085 | 0.0066 | 0.0066 | TRUE | 0.1038 |
| 4933412E12I | 90 | 65 | 47 | 69 | 28 | 34 | 96 | 65 | 47 | 54 | 30 | 37 | 54.88 | 69 | 40 | 0.591 | -0.759 | 0.0388763 | 0.13788103 | 0.0896 | 0.0669 | 0.078 | 0.078 | TRUE | 0.2579 |
| 4933412E24I | 0 | 0 | 0 | 0 | 2 | 0 | 0 | 0 | 0 | 0 | 2 | 0 | 0.36 | 0 | 1 | 3.775 | 1.916 | 0.63423067 | NA | 10 | 9.1927 | 10 | 10 | TRUE | 0.9743 |
| 4933412O06 | 0 | 0 | 0 | 0 | 0 | 0 | 0 | 0 | 0 | 0 | 0 | 0 | 0 | 0 | 0 | NA | NA | NA | NA | NA | NA | NA | NA | NA | NA |
| 4933413G19 | 2 | 0 | 2 | 2 | 1 | 0 | 2 | 0 | 2 | 2 | 1 | 0 | 1.13 | 1 | 1 | 0.664 | -0.59 | 0.75862464 | NA | | 0 | 2.922 | 1.7522 | 1.7522 | TRUE | 0.4951 |
| 4933413J09F | 7 | 8 | 6 | 17 | 14 | 8 | 7 | 8 | 6 | 13 | 15 | 9 | 9.77 | 7 | 12 | 1.74 | 0.799 | 0.19373587 | 0.39947733 | 0 | 0.344 | 0.1614 | 0.1614 | TRUE | 0.2376 |
| 4933413L06F | 0 | 0 | 0 | 0 | 0 | 0 | 0 | 0 | 0 | 0 | 0 | 0 | 0 | 0 | 0 | NA | NA | NA | NA | NA | NA | NA | NA | NA | NA |
| 4933415F23I | 0 | 0 | 0 | 1 | 1 | 1 | 0 | 0 | 0 | 1 | 1 | 1 | 0.49 | 0 | 1 | 5.224 | 2.385 | 0.38465 | NA | | 0 | 6.6923 | 3.9249 | 3.9249 | TRUE | 0.07 |
| 4933416C03I | 48 | 66 | 52 | 169 | 140 | 121 | 51 | 66 | 52 | 133 | 151 | 132 | 97.51 | 56 | 139 | 2.455 | 1.296 | 2.46E-09 | 1.36E-07 | 0 | 0.0407 | 0.0214 | 0.0214 | TRUE | 0.1876 |
| 4933416E03I | 0 | 0 | 0 | 0 | 0 | 0 | 0 | 0 | 0 | 0 | 0 | 0 | 0 | 0 | 0 | NA | NA | NA | NA | NA | NA | NA | NA | NA | NA |

| | | | | | | | | | | | | | | | | | | | | | | | | | |
|---|---|---|---|---|---|---|---|---|---|---|---|---|---|---|---|---|---|---|---|---|---|---|---|---|---|
| 4933416I08R | 2 | 2 | 0 | 4 | 2 | 3 | 2 | 2 | 0 | 3 | 2 | 3 | 2.12 | 1 | 3 | 2.097 | 1.068 | 0.43064619 | NA | 0 | 1.5631 | 0.7807 | 0.7807 | TRUE | 0.4924 |
| 4933416M06 | 0 | 0 | 0 | 1 | 0 | 0 | 0 | 0 | 0 | 1 | 0 | 0 | 0.13 | 0 | 0 | 1.859 | 0.894 | 0.82650734 | NA | 0 | 25.1515 | 10 | 10 | TRUE | 0.2563 |
| 4933416M07 | 27 | 20 | 21 | 49 | 24 | 23 | 29 | 20 | 21 | 39 | 26 | 25 | 26.54 | 23 | 30 | 1.298 | 0.376 | 0.345606 | 0.56509411 | 0 | 0.131 | 0.0762 | 0.0762 | TRUE | 0.5008 |
| 4933417A18 | 0 | 0 | 0 | 0 | 0 | 0 | 0 | 0 | 0 | 0 | 0 | 0 | 0 | 0 | 0 | NA | NA | NA | NA | NA | NA | NA | NA | NA | NA |
| 4933417D19 | 0 | 2 | 2 | 5 | 2 | 0 | 0 | 2 | 2 | 4 | 2 | 0 | 1.68 | 1 | 2 | 1.545 | 0.628 | 0.71171996 | NA | 0 | 1.9663 | 1.4655 | 1.4655 | TRUE | 0.1602 |
| 4933417E11I | 1 | 0 | 1 | 1 | 0 | 0 | 1 | 0 | 1 | 1 | 0 | 0 | 0.48 | 1 | 0 | 0.509 | -0.975 | 0.7336918 | NA | 0 | 6.9367 | 4.1806 | 4.1806 | TRUE | 0.2562 |
| 4933417G07 | 0 | 0 | 0 | 0 | 0 | 0 | 0 | 0 | 0 | 0 | 0 | 0 | 0 | 0 | 0 | NA | NA | NA | NA | NA | NA | NA | NA | NA | NA |
| 4933417O13 | 0 | 0 | 0 | 0 | 0 | 0 | 0 | 0 | 0 | 0 | 0 | 0 | 0 | 0 | 0 | NA | NA | NA | NA | NA | NA | NA | NA | NA | NA |
| 4933421I07R | 1 | 0 | 2 | 1 | 1 | 0 | 1 | 0 | 2 | 1 | 1 | 0 | 0.82 | 1 | 1 | 0.615 | -0.7 | 0.74728665 | NA | 0 | 4.0144 | 2.1281 | 2.1281 | TRUE | 0.1545 |
| 4933421O10 | 2 | 9 | 7 | 15 | 11 | 9 | 2 | 9 | 7 | 12 | 12 | 10 | 8.6 | 6 | 11 | 1.839 | 0.879 | 0.20978597 | 0.42010465 | 0 | 0.3898 | 0.2259 | 0.2259 | TRUE | 0.715 |
| 4933422A05 | 3 | 0 | 0 | 2 | 0 | 0 | 3 | 0 | 0 | 2 | 0 | 0 | 0.79 | 1 | 1 | 0.523 | -0.934 | 0.754563 | NA | 8.9049 | 4.1606 | 5.0595 | 5.0595 | TRUE | 1.4426 |
| 4933422H20 | 31 | 14 | 23 | 23 | 17 | 14 | 33 | 14 | 23 | 18 | 18 | 15 | 20.28 | 23 | 17 | 0.742 | -0.431 | 0.3620414 | 0.58006064 | 0.0495 | 0.1693 | 0.111 | 0.111 | TRUE | 0.1544 |
| 4933424G05 | 2 | 0 | 0 | 0 | 0 | 2 | 2 | 0 | 0 | 0 | 0 | 2 | 0.72 | 1 | 1 | 1.016 | 0.022 | 0.99423081 | NA | 9.003 | 4.5892 | 5.4427 | 5.4427 | TRUE | 0.9746 |
| 4933424G06 | 4 | 1 | 0 | 1 | 0 | 2 | 4 | 1 | 0 | 1 | 0 | 2 | 1.37 | 2 | 1 | 0.565 | -0.823 | 0.66804686 | NA | 1.1043 | 2.4108 | 1.8639 | 1.8639 | TRUE | 0.7965 |
| 4933425B07 | 0 | 0 | 0 | 1 | 0 | 0 | 0 | 0 | 0 | 1 | 0 | 0 | 0.13 | 0 | 0 | 1.859 | 0.894 | 0.82650734 | NA | 0 | 25.1515 | 10 | 10 | TRUE | 0.2563 |
| 4933425H06 | 0 | 0 | 0 | 0 | 0 | 0 | 0 | 0 | 0 | 0 | 0 | 0 | 0 | 0 | 0 | NA | NA | NA | NA | NA | NA | NA | NA | NA | NA |
| 4933425L06F | 1 | 2 | 0 | 26 | 16 | 15 | 1 | 2 | 0 | 20 | 17 | 16 | 9.52 | 1 | 18 | 17.713 | 4.147 | 3.48E-05 | 0.00056705 | 0 | 0.3529 | 0.1959 | 0.1959 | TRUE | 0.3767 |
| 4933426D04 | 0 | 0 | 1 | 0 | 0 | 0 | 0 | 0 | 1 | 0 | 0 | 0 | 0.17 | 0 | 0 | 0.49 | -1.029 | 0.80087 | NA | 0 | 19.6928 | 10 | 10 | TRUE | 0.4098 |
| 4933426M11 | 3475 | 4167 | 4005 | 4112 | 2964 | 3229 | 3691 | 4162 | 4021 | 3233 | 3189 | 3535 | 3638.4 | 3958 | 3319 | 0.838 | -0.254 | 0.0026749 | 0.01937034 | 0.0031 | 0.0078 | 0.0049 | 0.0049 | TRUE | 0.0423 |
| 4933427D06 | 283 | 287 | 219 | 689 | 483 | 375 | 301 | 287 | 220 | 542 | 520 | 410 | 379.82 | 269 | 491 | 1.826 | 0.869 | 4.36E-07 | 1.33E-05 | 0.0212 | 0.0156 | 0.0184 | 0.0184 | TRUE | 0.2868 |
| 4933427D14 | 658 | 802 | 868 | 969 | 888 | 815 | 699 | 801 | 872 | 762 | 955 | 892 | 830.13 | 791 | 870 | 1.099 | 0.136 | 0.30178605 | 0.52237237 | 0.0116 | 0.0109 | 0.0113 | 0.0113 | TRUE | 0.1444 |
| 4933427E11I | 0 | 3 | 3 | 3 | 1 | 0 | 0 | 3 | 3 | 2 | 1 | 0 | 1.57 | 2 | 1 | 0.592 | -0.757 | 0.66423058 | NA | 0 | 2.1002 | 1.5297 | 1.5297 | TRUE | 0.4347 |
| 4933427E13I | 0 | 0 | 0 | 0 | 0 | 0 | 0 | 0 | 0 | 0 | 0 | 0 | 0 | 0 | 0 | NA | NA | NA | NA | NA | NA | NA | NA | NA | NA |
| 4933427G17 | 2 | 10 | 7 | 8 | 5 | 0 | 2 | 10 | 7 | 6 | 5 | 0 | 5.13 | 6 | 4 | 0.619 | -0.691 | 0.52619788 | 0.71544767 | 0.6596 | 0.6485 | 0.6528 | 0.6528 | TRUE | 0.4643 |
| 4933427I22R | 0 | 0 | 1 | 0 | 0 | 0 | 0 | 0 | 1 | 0 | 0 | 0 | 0.17 | 0 | 0 | 0.49 | -1.029 | 0.80087 | NA | 0 | 19.6928 | 10 | 10 | TRUE | 0.4098 |
| 4933428C19I | 1 | 0 | 0 | 2 | 3 | 2 | 1 | 0 | 0 | 2 | 3 | 2 | 1.34 | 0 | 2 | 6.316 | 2.659 | 0.15211161 | NA | 0 | 2.4619 | 1.2722 | 1.2722 | TRUE | 0.4614 |
| 4933429K18I | 0 | 0 | 0 | 0 | 0 | 0 | 0 | 0 | 0 | 0 | 0 | 0 | 0 | 0 | 0 | NA | NA | NA | NA | NA | NA | NA | NA | NA | NA |
| 4933429O19 | 0 | 0 | 0 | 1 | 0 | 0 | 0 | 0 | 0 | 1 | 0 | 0 | 0.13 | 0 | 0 | 1.859 | 0.894 | 0.82650734 | NA | 0 | 25.1515 | 10 | 10 | TRUE | 0.2563 |
| 4933430H16 | 0 | 0 | 0 | 0 | 1 | 0 | 0 | 0 | 0 | 0 | 1 | 0 | 0.18 | 0 | 0 | 1.859 | 0.894 | 0.82650734 | NA | 0 | 18.3785 | 10 | 10 | TRUE | 0.5204 |
| 4933430I17R | 24 | 42 | 45 | 38 | 29 | 27 | 25 | 42 | 45 | 30 | 31 | 30 | 33.88 | 37 | 30 | 0.803 | -0.317 | 0.37521698 | 0.59102546 | 0 | 0.1042 | 0.0621 | 0.0621 | TRUE | 0.5571 |
| 4933430M04 | 0 | 0 | 0 | 0 | 0 | 0 | 0 | 0 | 0 | 0 | 0 | 0 | 0 | 0 | 0 | NA | NA | NA | NA | NA | NA | NA | NA | NA | NA |
| 4933430N04 | 0 | 0 | 0 | 0 | 0 | 0 | 0 | 0 | 0 | 0 | 0 | 0 | 0 | 0 | 0 | NA | NA | NA | NA | NA | NA | NA | NA | NA | NA |
| 4933431E20I | 446 | 461 | 433 | 434 | 273 | 317 | 474 | 460 | 435 | 341 | 294 | 347 | 391.8 | 456 | 327 | 0.718 | -0.478 | 9.19E-05 | 0.00129127 | 0.0016 | 0.0153 | 0.0081 | 0.0081 | TRUE | 0.0882 |
| 4933431G14 | 0 | 0 | 0 | 0 | 0 | 0 | 0 | 0 | 0 | 0 | 0 | 0 | 0 | 0 | 0 | NA | NA | NA | NA | NA | NA | NA | NA | NA | NA |
| 4933432G23 | 5 | 10 | 1 | 3 | 2 | 9 | 5 | 10 | 1 | 2 | 2 | 10 | 5.11 | 5 | 5 | 0.868 | -0.204 | 0.84922176 | 0.92493099 | 0.6121 | 0.6515 | 0.6331 | 0.6331 | TRUE | 0.2824 |
| 4933432I03R | 0 | 0 | 0 | 0 | 1 | 1 | 0 | 0 | 0 | 0 | 1 | 1 | 0.36 | 0 | 1 | 3.794 | 1.924 | 0.60247382 | NA | 0 | 9.1135 | 8.1425 | 8.1425 | TRUE | 0.3482 |
| 4933432I09R | 0 | 0 | 0 | 0 | 0 | 0 | 0 | 0 | 0 | 0 | 0 | 0 | 0 | 0 | 0 | NA | NA | NA | NA | NA | NA | NA | NA | NA | NA |
| 4933433C11I | 14 | 10 | 16 | 25 | 21 | 19 | 15 | 10 | 16 | 20 | 23 | 21 | 17.33 | 14 | 21 | 1.539 | 0.622 | 0.177388 | 0.3779353 | 0 | 0.197 | 0.0923 | 0.0923 | TRUE | 0.241 |
| 4933433F19I | 0 | 0 | 0 | 0 | 0 | 0 | 0 | 0 | 0 | 0 | 0 | 0 | 0 | 0 | 0 | NA | NA | NA | NA | NA | NA | NA | NA | NA | NA |
| 4933433G15 | 1 | 14 | 12 | 5 | 10 | 6 | 1 | 14 | 12 | 4 | 11 | 7 | 8.06 | 9 | 7 | 0.774 | -0.37 | 0.68031526 | 0.82277546 | 0.5017 | 0.4157 | 0.4553 | 0.4553 | TRUE | 1.3596 |
| 4933433G19 | 4 | 5 | 6 | 9 | 10 | 2 | 4 | 5 | 6 | 7 | 11 | 2 | 5.88 | 5 | 7 | 1.314 | 0.394 | 0.63997243 | 0.79734933 | 0 | 0.567 | 0.3382 | 0.3382 | TRUE | 0.1997 |
| 4933433H22 | 1 | 0 | 2 | 0 | 0 | 0 | 1 | 0 | 2 | 0 | 0 | 0 | 0.51 | 1 | 0 | 0.175 | -2.513 | 0.44147732 | NA | 0 | 6.4448 | 6.181 | 6.181 | TRUE | 0.063 |
| 4933434E20I | 709 | 981 | 996 | 1242 | 997 | 1031 | 753 | 980 | 1000 | 976 | 1073 | 1129 | 985.1 | 911 | 1059 | 1.162 | 0.216 | 0.10762375 | 0.27453369 | 0.0135 | 0.0102 | 0.012 | 0.012 | TRUE | 0.2725 |
| 4933434I20R | 11 | 21 | 14 | 10 | 6 | 4 | 12 | 21 | 14 | 8 | 6 | 4 | 10.9 | 16 | 6 | 0.405 | -1.303 | 0.03470944 | 0.12737024 | 0 | 0.3091 | 0.1646 | 0.1646 | TRUE | 0.4341 |
| 4933436C20I | 93 | 166 | 120 | 101 | 84 | 126 | 99 | 166 | 120 | 79 | 90 | 138 | 115.46 | 128 | 102 | 0.795 | -0.33 | 0.25243152 | 0.47049751 | 0.0683 | 0.0354 | 0.0512 | 0.0512 | TRUE | 0.5977 |
| 4933436E23I | 1 | 0 | 0 | 0 | 0 | 0 | 1 | 0 | 0 | 0 | 0 | 0 | 0.18 | 0 | 0 | 0.49 | -1.029 | 0.80087 | NA | 0 | 18.6165 | 10 | 10 | TRUE | 0.4614 |
| 4933436H12 | 0 | 0 | 0 | 0 | 0 | 0 | 0 | 0 | 0 | 0 | 0 | 0 | 0 | 0 | 0 | NA | NA | NA | NA | NA | NA | NA | NA | NA | NA |
| 4933436I01R | 0 | 1 | 0 | 0 | 0 | 0 | 0 | 1 | 0 | 0 | 0 | 0 | 0.17 | 0 | 0 | 0.49 | -1.029 | 0.80087 | NA | 0 | 19.7991 | 10 | 10 | TRUE | 0.405 |
| 4933438B17 | 0 | 0 | 0 | 0 | 0 | 0 | 0 | 0 | 0 | 0 | 0 | 0 | 0 | 0 | 0 | NA | NA | NA | NA | NA | NA | NA | NA | NA | NA |
| 4933438K21I | 1 | 2 | 0 | 4 | 2 | 1 | 1 | 2 | 0 | 3 | 2 | 1 | 1.58 | 1 | 2 | 2.12 | 1.084 | 0.49042591 | NA | 0 | 2.0985 | 1.0543 | 1.0543 | TRUE | 0.3288 |
| 4933439C10I | 96 | 148 | 135 | 95 | 88 | 130 | 102 | 148 | 136 | 75 | 95 | 142 | 116.17 | 129 | 104 | 0.804 | -0.314 | 0.26427143 | 0.48381628 | 0.0624 | 0.0353 | 0.0483 | 0.0483 | TRUE | 0.8208 |
| 4933439K11I | 0 | 2 | 1 | 1 | 1 | 1 | 0 | 2 | 1 | 1 | 1 | 1 | 0.99 | 1 | 1 | 0.968 | -0.047 | 0.97991429 | NA | 0 | 3.3244 | 1.4549 | 1.4549 | TRUE | 0.1811 |
| 4933440J02F | 0 | 0 | 0 | 0 | 1 | 0 | 0 | 0 | 0 | 0 | 1 | 0 | 0.18 | 0 | 0 | 1.859 | 0.894 | 0.82650734 | NA | 0 | 18.3785 | 10 | 10 | TRUE | 0.5204 |
| 4933440M02 | 5 | 14 | 18 | 7 | 3 | 11 | 5 | 14 | 18 | 6 | 3 | 12 | 9.69 | 12 | 7 | 0.55 | -0.862 | 0.25362628 | 0.47155412 | 0.2461 | 0.3469 | 0.2991 | 0.2991 | TRUE | 0.4707 |
| 5031410I06R | 1 | 1 | 2 | 1 | 2 | 0 | 1 | 1 | 2 | 1 | 2 | 0 | 1.17 | 1 | 1 | 0.717 | -0.481 | 0.78281428 | NA | 0 | 2.8279 | 1.3158 | 1.3158 | TRUE | 0.4306 |
| 5031414D18 | 14 | 22 | 19 | 26 | 13 | 14 | 15 | 22 | 19 | 20 | 14 | 15 | 17.61 | 19 | 16 | 0.897 | -0.158 | 0.7320076 | 0.85627965 | 0 | 0.194 | 0.096 | 0.096 | TRUE | 0.2473 |
| 5031425E22I | 112 | 164 | 158 | 164 | 134 | 148 | 119 | 164 | 159 | 129 | 144 | 162 | 146.08 | 147 | 145 | 0.982 | -0.026 | 0.89583135 | 0.94959689 | 0.0142 | 0.0295 | 0.0218 | 0.0218 | TRUE | 0.2879 |
| 5031425F14I | 0 | 0 | 1 | 0 | 0 | 0 | 0 | 0 | 1 | 0 | 0 | 0 | 0.17 | 0 | 0 | 0.49 | -1.029 | 0.80087 | NA | 0 | 19.6928 | 10 | 10 | TRUE | 0.4098 |
| 5031426D15 | 0 | 0 | 0 | 0 | 1 | 0 | 0 | 0 | 0 | 0 | 1 | 0 | 0.18 | 0 | 0 | 1.859 | 0.894 | 0.82650734 | NA | 0 | 18.3785 | 10 | 10 | TRUE | 0.5204 |
| 5031434C07I | 11 | 4 | 8 | 8 | 11 | 12 | 12 | 4 | 8 | 6 | 12 | 13 | 9.16 | 8 | 10 | 1.307 | 0.386 | 0.57586435 | 0.75380936 | 0.0817 | 0.3665 | 0.2309 | 0.2309 | TRUE | 0.2709 |
| 5031434O11 | 6 | 6 | 8 | 10 | 9 | 6 | 6 | 6 | 8 | 8 | 10 | 7 | 7.42 | 7 | 8 | 1.181 | 0.24 | 0.7227571 | 0.85005881 | 0 | 0.451 | 0.1941 | 0.1941 | TRUE | 0.0843 |
| 5031439G07 | 2926 | 3328 | 3271 | 4090 | 2955 | 3039 | 3108 | 3324 | 3284 | 3215 | 3179 | 3327 | 3239.56 | 3239 | 3240 | 1 | 0.001 | 0.99361683 | 0.9964688 | 6.00E-04 | 0.0079 | 0.003 | 0.003 | TRUE | 0.0151 |
| 5033403H07 | 0 | 0 | 0 | 0 | 0 | 0 | 0 | 0 | 0 | 0 | 0 | 0 | 0 | 0 | 0 | NA | NA | NA | NA | NA | NA | NA | NA | NA | NA |
| 5033404E19I | 16 | 22 | 22 | 8 | 12 | 10 | 17 | 22 | 22 | 6 | 13 | 11 | 15.2 | 20 | 10 | 0.484 | -1.045 | 0.04381514 | 0.14996797 | 0 | 0.2236 | 0.1193 | 0.1193 | TRUE | 0.4995 |
| 5033406O09 | 46 | 54 | 39 | 48 | 36 | 29 | 49 | 54 | 39 | 38 | 39 | 32 | 41.69 | 47 | 36 | 0.764 | -0.388 | 0.204415 | 0.41280838 | 0 | 0.0859 | 0.0432 | 0.0432 | TRUE | 0.1839 |
| 5033411D12 | 5 | 7 | 14 | 15 | 5 | 6 | 5 | 7 | 14 | 12 | 5 | 7 | 8.35 | 9 | 8 | 0.913 | -0.131 | 0.85641452 | 0.92891615 | 0.1118 | 0.4015 | 0.2574 | 0.2574 | TRUE | 0.8752 |
| 5330411J11F | 40 | 33 | 30 | 74 | 46 | 42 | 42 | 33 | 30 | 58 | 49 | 46 | 43.2 | 35 | 51 | 1.464 | 0.55 | 0.07301369 | 0.21164712 | 0 | 0.0832 | 0.0438 | 0.0438 | TRUE | 0.2267 |
| 5330413P13I | 11 | 9 | 14 | 7 | 1 | 11 | 12 | 9 | 14 | 6 | 1 | 12 | 8.89 | 12 | 6 | 0.533 | -0.907 | 0.25016769 | 0.46733938 | 0.2432 | 0.3774 | 0.3252 | 0.3252 | TRUE | 0.5514 |
| 5330417C22I | 32 | 52 | 60 | 48 | 40 | 48 | 34 | 52 | 60 | 38 | 43 | 53 | 46.58 | 49 | 45 | 0.906 | -0.142 | 0.66586156 | 0.81297171 | 0.0329 | 0.0776 | 0.0562 | 0.0562 | TRUE | 0.5179 |

| | | | | | | | | | | | | | | | | | | | | | | | | | |
|---|---|---|---|---|---|---|---|---|---|---|---|---|---|---|---|---|---|---|---|---|---|---|---|---|---|
| 5330426P16I | 65 | 74 | 69 | 107 | 86 | 72 | 69 | 74 | 69 | 84 | 93 | 79 | 77.95 | 71 | 85 | 1.203 | 0.267 | 0.22302152 | 0.43505698 | 0 | 0.0492 | 0.0216 | 0.0216 | TRUE | 0.0503 |
| 5330434G04 | 0 | 0 | 0 | 0 | 0 | 0 | 0 | 0 | 0 | 0 | 0 | 0 | 0 | 0 | 0 | NA | NA | NA | NA | NA | NA | NA | NA | NA | NA |
| 5330439B14 | 0 | 0 | 0 | 0 | 0 | 0 | 0 | 0 | 0 | 0 | 0 | 0 | 0 | 0 | 0 | NA | NA | NA | NA | NA | NA | NA | NA | NA | NA |
| 5430401F13I | 0 | 0 | 0 | 0 | 1 | 1 | 0 | 0 | 0 | 0 | 1 | 1 | 0.36 | 0 | 1 | 3.794 | 1.924 | 0.60247382 | NA | 0 | 9.1135 | 8.1425 | 8.1425 | TRUE | 0.3482 |
| 5430402E10I | 0 | 0 | 0 | 0 | 0 | 0 | 0 | 0 | 0 | 0 | 0 | 0 | 0 | 0 | 0 | NA | NA | NA | NA | NA | NA | NA | NA | NA | NA |
| 5430402O13 | 19 | 6 | 3 | 36 | 27 | 7 | 20 | 6 | 3 | 28 | 29 | 8 | 15.7 | 10 | 22 | 2.247 | 1.168 | 0.13461941 | 0.3173718 | 0.5314 | 0.2167 | 0.3637 | 0.3637 | TRUE | 2.893 |
| 5430403N17 | 0 | 0 | 0 | 0 | 0 | 0 | 0 | 0 | 0 | 0 | 0 | 0 | 0 | 0 | 0 | NA | NA | NA | NA | NA | NA | NA | NA | NA | NA |
| 5430405H02 | 9 | 12 | 12 | 30 | 11 | 19 | 10 | 12 | 12 | 24 | 12 | 21 | 14.97 | 11 | 19 | 1.684 | 0.752 | 0.149929 | 0.34065871 | 0 | 0.227 | 0.1251 | 0.1251 | TRUE | 0.5088 |
| 5430416N02 | 384 | 572 | 578 | 424 | 343 | 367 | 408 | 571 | 580 | 333 | 369 | 402 | 443.94 | 520 | 368 | 0.707 | -0.501 | 0.00278029 | 0.01997754 | 0.0212 | 0.0143 | 0.0179 | 0.0179 | TRUE | 0.4118 |
| 5430416O09 | 2 | 3 | 2 | 3 | 4 | 1 | 2 | 3 | 2 | 2 | 4 | 1 | 2.48 | 2 | 2 | 1.085 | 0.117 | 0.92067445 | NA | 0 | 1.3348 | 0.5932 | 0.5932 | TRUE | 0.2856 |
| 5430417L22I | 179 | 351 | 379 | 402 | 351 | 386 | 190 | 351 | 381 | 316 | 378 | 423 | 339.57 | 307 | 372 | 1.21 | 0.275 | 0.26946631 | 0.48946478 | 0.0714 | 0.0166 | 0.0416 | 0.0416 | TRUE | 1.2465 |
| 5430419D17 | 1 | 2 | 1 | 2 | 1 | 2 | 1 | 2 | 1 | 2 | 1 | 2 | 1.48 | 1 | 2 | 1.185 | 0.245 | 0.86647192 | NA | 0 | 2.2276 | 0.8551 | 0.8551 | TRUE | 0.1096 |
| 5430421F17I | 7 | 2 | 1 | 18 | 10 | 8 | 7 | 2 | 1 | 14 | 11 | 9 | 7.35 | 3 | 11 | 3.279 | 1.713 | 0.04418704 | 0.15075017 | 0.1046 | 0.4551 | 0.3305 | 0.3305 | TRUE | 1.3943 |
| 5430421N21 | 1 | 0 | 1 | 0 | 1 | 0 | 1 | 0 | 1 | 0 | 1 | 0 | 0.52 | 1 | 0 | 0.509 | -0.975 | 0.72714826 | NA | 0 | 6.2976 | 3.8857 | 3.8857 | TRUE | 0.5204 |
| 5430425J12I | 0 | 0 | 0 | 0 | 0 | 0 | 0 | 0 | 0 | 0 | 0 | 0 | 0 | 0 | 0 | NA | NA | NA | NA | NA | NA | NA | NA | NA | NA |
| 5430427M07 | 0 | 0 | 0 | 0 | 0 | 0 | 0 | 0 | 0 | 0 | 0 | 0 | 0 | 0 | 0 | NA | NA | NA | NA | NA | NA | NA | NA | NA | NA |
| 5430427O19 | 0 | 0 | 0 | 0 | 0 | 0 | 0 | 0 | 0 | 0 | 0 | 0 | 0 | 0 | 0 | NA | NA | NA | NA | NA | NA | NA | NA | NA | NA |
| 5430428K19I | 0 | 0 | 0 | 0 | 0 | 0 | 0 | 0 | 0 | 0 | 0 | 0 | 0 | 0 | 0 | NA | NA | NA | NA | NA | NA | NA | NA | NA | NA |
| 5430434I15R | 0 | 0 | 0 | 0 | 0 | 0 | 0 | 0 | 0 | 0 | 0 | 0 | 0 | 0 | 0 | NA | NA | NA | NA | NA | NA | NA | NA | NA | NA |
| 5430435G22 | 2 | 1 | 2 | 5 | 1 | 1 | 2 | 1 | 2 | 4 | 1 | 1 | 1.87 | 2 | 2 | 1.234 | 0.304 | 0.82442671 | NA | 0 | 1.7667 | 0.8178 | 0.8178 | TRUE | 0.7923 |
| 5430437J10I | 0 | 0 | 0 | 0 | 0 | 0 | 0 | 0 | 0 | 0 | 0 | 0 | 0 | 0 | 0 | NA | NA | NA | NA | NA | NA | NA | NA | NA | NA |
| 5430440P10I | 8 | 5 | 3 | 7 | 3 | 3 | 8 | 5 | 3 | 6 | 3 | 3 | 4.75 | 5 | 4 | 0.742 | -0.43 | 0.62774232 | 0.79060543 | 0 | 0.7 | 0.353 | 0.353 | TRUE | 0.327 |
| 5530400C23I | 0 | 0 | 0 | 0 | 0 | 0 | 0 | 0 | 0 | 0 | 0 | 0 | 0 | 0 | 0 | NA | NA | NA | NA | NA | NA | NA | NA | NA | NA |
| 5530401A14 | 0 | 0 | 0 | 0 | 0 | 0 | 0 | 0 | 0 | 0 | 0 | 0 | 0 | 0 | 0 | NA | NA | NA | NA | NA | NA | NA | NA | NA | NA |
| 5530601H04 | 123 | 131 | 117 | 184 | 124 | 101 | 131 | 131 | 117 | 145 | 133 | 111 | 127.93 | 126 | 130 | 1.03 | 0.042 | 0.8249319 | 0.91128174 | 0 | 0.0327 | 0.0181 | 0.0181 | TRUE | 0.1623 |
| 5730403I07R | 3 | 5 | 2 | 3 | 6 | 3 | 3 | 5 | 2 | 2 | 6 | 3 | 3.71 | 3 | 4 | 1.169 | 0.225 | 0.82156456 | NA | 0 | 0.8938 | 0.4454 | 0.4454 | TRUE | 0.4396 |
| 5730405O15 | 48 | 62 | 93 | 57 | 47 | 42 | 51 | 62 | 93 | 45 | 51 | 46 | 57.94 | 69 | 47 | 0.684 | -0.549 | 0.07569332 | 0.21690316 | 0.0384 | 0.0638 | 0.051 | 0.051 | TRUE | 0.6866 |
| 5730408K05I | 114 | 122 | 102 | 140 | 127 | 102 | 121 | 122 | 102 | 110 | 137 | 112 | 117.28 | 115 | 120 | 1.036 | 0.05 | 0.7980895 | 0.89608114 | 0 | 0.035 | 0.0195 | 0.0195 | TRUE | 0.1551 |
| 5730409E04I | 146 | 156 | 180 | 223 | 211 | 176 | 155 | 156 | 181 | 175 | 227 | 193 | 181.1 | 164 | 198 | 1.207 | 0.271 | 0.11838723 | 0.29119031 | 0.0078 | 0.0251 | 0.0161 | 0.0161 | TRUE | 0.1724 |
| 5730412P04I | 0 | 0 | 0 | 0 | 0 | 0 | 0 | 0 | 0 | 0 | 0 | 0 | 0 | 0 | 0 | NA | NA | NA | NA | NA | NA | NA | NA | NA | NA |
| 5730416F02I | 0 | 0 | 0 | 2 | 0 | 0 | 0 | 0 | 0 | 2 | 0 | 0 | 0.26 | 0 | 1 | 2.886 | 1.529 | 0.70676491 | NA | 0 | 12.5792 | 10 | 10 | TRUE | 0.9669 |
| 5730420D15 | 1 | 7 | 8 | 3 | 4 | 4 | 1 | 7 | 8 | 2 | 4 | 4 | 4.52 | 5 | 3 | 0.674 | -0.569 | 0.56505922 | NA | 0 | 0.7356 | 0.4761 | 0.4761 | TRUE | 0.9961 |
| 5730422E09I | 0 | 2 | 2 | 1 | 2 | 1 | 0 | 2 | 2 | 1 | 2 | 1 | 1.34 | 1 | 1 | 0.983 | -0.024 | 0.988347 | NA | 0 | 2.466 | 1.2122 | 1.2122 | TRUE | 0.4385 |
| 5730435O14 | 0 | 0 | 0 | 1 | 0 | 1 | 0 | 0 | 0 | 1 | 0 | 1 | 0.31 | 0 | 1 | 3.348 | 1.743 | 0.65064889 | NA | 0 | 10.5167 | 8.8968 | 8.8968 | TRUE | 0.2082 |
| 5730455P16I | 1560 | 2068 | 2144 | 2212 | 1544 | 1608 | 1657 | 2065 | 2153 | 1739 | 1661 | 1760 | 1839.25 | 1958 | 1720 | 0.878 | -0.187 | 0.10659563 | 0.27266637 | 0.0088 | 0.0087 | 0.0092 | 0.0092 | TRUE | 0.2185 |
| 5730457N03 | 0 | 0 | 0 | 3 | 5 | 1 | 0 | 0 | 0 | 2 | 5 | 1 | 1.47 | 0 | 3 | 15.632 | 3.966 | 0.04718259 | NA | 0.5179 | 2.2448 | 1.7113 | 1.7113 | TRUE | 0.2115 |
| 5730460C07I | 0 | 0 | 0 | 0 | 0 | 0 | 0 | 0 | 0 | 0 | 0 | 0 | 0 | 0 | 0 | NA | NA | NA | NA | NA | NA | NA | NA | NA | NA |
| 5730480H06 | 58 | 48 | 41 | 62 | 38 | 35 | 62 | 48 | 41 | 49 | 41 | 38 | 46.44 | 50 | 43 | 0.854 | -0.227 | 0.4540745 | 0.65921051 | 0 | 0.0778 | 0.0447 | 0.0447 | TRUE | 0.3108 |
| 5730488B01 | 0 | 0 | 0 | 0 | 0 | 2 | 0 | 0 | 0 | 0 | 0 | 2 | 0.36 | 0 | 1 | 3.832 | 1.938 | 0.63022034 | NA | 10 | 9.0356 | 10 | 10 | TRUE | 0.9744 |
| 5730507C01I | 61 | 50 | 57 | 115 | 64 | 73 | 65 | 50 | 57 | 90 | 69 | 80 | 68.52 | 57 | 80 | 1.398 | 0.483 | 0.05379994 | 0.1723792 | 0 | 0.055 | 0.0302 | 0.0302 | TRUE | 0.1424 |
| 5730508B09 | 89 | 102 | 116 | 178 | 119 | 110 | 95 | 102 | 116 | 140 | 128 | 120 | 116.88 | 104 | 129 | 1.244 | 0.315 | 0.10034714 | 0.2618589 | 0 | 0.0351 | 0.0178 | 0.0178 | TRUE | 0.1032 |
| 5730522E02I | 2 | 3 | 2 | 6 | 1 | 4 | 2 | 3 | 2 | 5 | 1 | 4 | 2.88 | 2 | 3 | 1.444 | 0.53 | 0.63716877 | NA | 0 | 1.1495 | 0.5526 | 0.5526 | TRUE | 0.4687 |
| 5730559C18I | 737 | 834 | 851 | 717 | 525 | 575 | 783 | 833 | 854 | 564 | 565 | 629 | 704.7 | 823 | 586 | 0.711 | -0.492 | 8.35E-07 | 2.36E-05 | 0.0014 | 0.0116 | 0.0057 | 0.0057 | TRUE | 0.0481 |
| 5730577I03R | 0 | 5 | 2 | 9 | 3 | 4 | 0 | 5 | 2 | 7 | 3 | 4 | 3.61 | 2 | 5 | 2.102 | 1.072 | 0.35410924 | NA | 0.2049 | 0.9185 | 0.6476 | 0.6476 | TRUE | 0.6179 |
| 5830403L16I | 1 | 3 | 4 | 1 | 1 | 1 | 1 | 3 | 4 | 1 | 1 | 1 | 1.84 | 3 | 1 | 0.36 | -1.473 | 0.30335077 | NA | 0 | 1.7988 | 0.7862 | 0.7862 | TRUE | 0.2819 |
| 5830403M04 | 1 | 0 | 0 | 1 | 2 | 1 | 1 | 0 | 0 | 1 | 2 | 1 | 0.85 | 0 | 1 | 3.632 | 1.861 | 0.38570833 | NA | 0 | 3.8867 | 1.9484 | 1.9484 | TRUE | 0.4614 |
| 5830411N06 | 1 | 0 | 0 | 0 | 0 | 0 | 1 | 0 | 0 | 0 | 0 | 0 | 0.18 | 0 | 0 | 0.49 | -1.029 | 0.80087 | NA | 0 | 18.6165 | 10 | 10 | TRUE | 0.4614 |
| 5830415F09I | 337 | 428 | 423 | 739 | 563 | 506 | 358 | 427 | 425 | 581 | 606 | 554 | 491.79 | 403 | 580 | 1.438 | 0.524 | 9.72E-06 | 0.00019474 | 0.003 | 0.0136 | 0.008 | 0.008 | TRUE | 0.1104 |
| 5830416I19R | 2 | 1 | 2 | 2 | 1 | 0 | 2 | 1 | 2 | 2 | 1 | 0 | 1.3 | 2 | 1 | 0.539 | -0.892 | 0.58964573 | NA | 0 | 2.5478 | 1.1441 | 1.1441 | TRUE | 0.2725 |
| 5830416P10I | 29 | 23 | 25 | 15 | 23 | 11 | 31 | 23 | 25 | 12 | 25 | 12 | 21.24 | 26 | 16 | 0.611 | -0.712 | 0.12785381 | 0.30739486 | 0.0417 | 0.162 | 0.1075 | 0.1075 | TRUE | 0.9622 |
| 5830417I10R | 1402 | 1351 | 1426 | 1997 | 1446 | 1426 | 1489 | 1349 | 1432 | 1570 | 1556 | 1561 | 1492.83 | 1423 | 1562 | 1.098 | 0.135 | 0.08369225 | 0.23178124 | 5.00E-04 | 0.0091 | 0.0037 | 0.0037 | TRUE | 0.0251 |
| 5830418K08I | 3463 | 3851 | 4301 | 4690 | 3479 | 3545 | 3678 | 3846 | 4319 | 3687 | 3743 | 3881 | 3858.94 | 3948 | 3770 | 0.955 | -0.067 | 0.44130243 | 0.6484276 | 0.0035 | 0.0078 | 0.0051 | 0.0051 | TRUE | 0.0823 |
| 5830418P13I | 31 | 29 | 41 | 85 | 45 | 34 | 33 | 29 | 41 | 67 | 48 | 37 | 42.59 | 34 | 51 | 1.491 | 0.576 | 0.09774385 | 0.25693601 | 0.0412 | 0.0843 | 0.0627 | 0.0627 | TRUE | 0.2885 |
| 5830428M24 | 0 | 2 | 0 | 3 | 1 | 2 | 0 | 2 | 0 | 2 | 1 | 2 | 1.27 | 1 | 2 | 2.802 | 1.487 | 0.4276653 | NA | 0 | 2.6006 | 1.5131 | 1.5131 | TRUE | 0.9737 |
| 5830432E09I | 0 | 0 | 0 | 0 | 0 | 0 | 0 | 0 | 0 | 0 | 0 | 0 | 0 | 0 | 0 | NA | NA | NA | NA | NA | NA | NA | NA | NA | NA |
| 5830454E08I | 22 | 16 | 12 | 15 | 12 | 16 | 23 | 16 | 12 | 12 | 13 | 18 | 15.6 | 17 | 14 | 0.818 | -0.29 | 0.56887193 | 0.7478913 | 0 | 0.2181 | 0.122 | 0.122 | TRUE | 0.3863 |
| 5830473C10I | 6 | 2 | 2 | 3 | 4 | 2 | 6 | 2 | 2 | 2 | 4 | 2 | 3.21 | 3 | 3 | 0.852 | -0.232 | 0.83047195 | NA | 0 | 1.0348 | 0.5289 | 0.5289 | TRUE | 0.7606 |
| 5930403L14I | 9 | 17 | 22 | 52 | 33 | 26 | 10 | 17 | 22 | 41 | 36 | 28 | 25.58 | 16 | 35 | 2.158 | 1.11 | 0.01057615 | 0.05499872 | 0.0281 | 0.1357 | 0.0905 | 0.0905 | TRUE | 0.5306 |
| 5930412G12 | 56 | 114 | 107 | 62 | 52 | 63 | 59 | 114 | 107 | 49 | 56 | 69 | 75.74 | 93 | 58 | 0.615 | -0.701 | 0.02452405 | 0.1003355 | 0.0619 | 0.0504 | 0.056 | 0.056 | TRUE | 0.9618 |
| 5930438M14 | 2 | 0 | 0 | 5 | 1 | 1 | 2 | 0 | 0 | 4 | 1 | 1 | 1.37 | 1 | 2 | 2.989 | 1.58 | 0.40703439 | NA | 0.5179 | 2.41 | 1.6459 | 1.6459 | TRUE | 0.9745 |
| 6030407O03 | 4 | 2 | 4 | 6 | 8 | 3 | 4 | 2 | 4 | 5 | 9 | 3 | 4.48 | 3 | 6 | 1.616 | 0.692 | 0.45222513 | NA | 0 | 0.7425 | 0.3723 | 0.3723 | TRUE | 0.4719 |
| 6030408B16 | 3 | 0 | 4 | 3 | 5 | 3 | 3 | 0 | 4 | 2 | 5 | 3 | 3.04 | 2 | 3 | 1.52 | 0.604 | 0.61768424 | NA | 0 | 1.0915 | 0.7064 | 0.7064 | TRUE | 0.8341 |
| 6030419C18I | 109 | 149 | 106 | 272 | 199 | 141 | 116 | 149 | 106 | 214 | 214 | 154 | 158.88 | 124 | 194 | 1.572 | 0.652 | 0.00245037 | 0.01805949 | 0.0258 | 0.0276 | 0.0268 | 0.0268 | TRUE | 0.3372 |
| 6030440G07 | 1 | 5 | 0 | 1 | 2 | 0 | 1 | 5 | 0 | 1 | 2 | 0 | 1.5 | 2 | 1 | 0.481 | -1.054 | 0.57724948 | NA | 1.2983 | 2.2047 | 1.8245 | 1.8245 | TRUE | 0.9587 |
| 6030458C11I | 3086 | 3333 | 3261 | 3948 | 2834 | 2756 | 3278 | 3329 | 3274 | 3104 | 3049 | 3017 | 3175.11 | 3294 | 3057 | 0.928 | -0.108 | 0.0755032 | 0.21651571 | 0 | 0.0079 | 0.0023 | 0.0023 | TRUE | 0.0023 |
| 6030466F02I | 0 | 0 | 0 | 3 | 0 | 2 | 0 | 0 | 0 | 2 | 0 | 2 | 0.76 | 0 | 1 | 8.123 | 3.022 | 0.28653847 | NA | 0 | 4.3534 | 4.4726 | 4.4726 | TRUE | 0.4916 |
| 6030468B19 | 0 | 0 | 0 | 1 | 1 | 0 | 0 | 0 | 0 | 1 | 1 | 0 | 0.31 | 0 | 1 | 3.32 | 1.731 | 0.65383052 | NA | 0 | 10.6224 | 8.9541 | 8.9541 | TRUE | 0.202 |
| 6030498E09I | 0 | 0 | 0 | 0 | 0 | 0 | 0 | 0 | 0 | 0 | 0 | 0 | 0 | 0 | 0 | NA | NA | NA | NA | NA | NA | NA | NA | NA | NA |
| 6230400D17 | 25 | 23 | 36 | 32 | 23 | 28 | 27 | 23 | 36 | 25 | 25 | 31 | 27.7 | 29 | 27 | 0.937 | -0.093 | 0.80476321 | 0.89931073 | 0 | 0.1258 | 0.0662 | 0.0662 | TRUE | 0.3556 |

| | | | | | | | | | | | | | | | | | | | | | | | | | |
|---|---|---|---|---|---|---|---|---|---|---|---|---|---|---|---|---|---|---|---|---|---|---|---|---|---|
| 6330403A02 | 1 | 3 | 1 | 3 | 3 | 2 | 1 | 3 | 1 | 2 | 3 | 2 | 2.14 | 2 | 2 | 1.525 | 0.609 | 0.62984907 | NA | 0 | 1.5466 | 0.6601 | 0.6601 | TRUE | 0.3594 |
| 6330403K07 | 174 | 182 | 182 | 234 | 156 | 157 | 185 | 182 | 183 | 184 | 168 | 172 | 178.83 | 183 | 175 | 0.955 | -0.066 | 0.65716501 | 0.80696343 | 0 | 0.0253 | 0.0104 | 0.0104 | TRUE | 0.0262 |
| 6330403M23 | 93 | 93 | 129 | 131 | 108 | 104 | 99 | 93 | 130 | 103 | 116 | 114 | 109.04 | 107 | 111 | 1.034 | 0.049 | 0.81838975 | 0.90726388 | 0.0092 | 0.0371 | 0.0232 | 0.0232 | TRUE | 0.3364 |
| 6330407A03 | 0 | 0 | 1 | 0 | 0 | 0 | 0 | 0 | 1 | 0 | 0 | 0 | 0.17 | 0 | 0 | 0.49 | -1.029 | 0.80087 | NA | 0 | 19.6928 | 10 | 10 | TRUE | 0.4098 |
| 6330408A02 | 181 | 191 | 237 | 298 | 187 | 191 | 192 | 191 | 238 | 234 | 201 | 209 | 210.92 | 207 | 215 | 1.04 | 0.057 | 0.72850645 | 0.85395791 | 0.0068 | 0.0225 | 0.0144 | 0.0144 | TRUE | 0.188 |
| 6330409D20 | 0 | 0 | 1 | 1 | 0 | 0 | 0 | 0 | 1 | 1 | 0 | 0 | 0.3 | 0 | 0 | 0.954 | -0.067 | 0.98640617 | NA | 0 | 11.0483 | 9.2783 | 9.2783 | TRUE | 0.4098 |
| 6330410L21 | 98 | 41 | 31 | 160 | 115 | 60 | 104 | 41 | 31 | 126 | 124 | 66 | 81.89 | 59 | 105 | 1.795 | 0.844 | 0.0620089 | 0.19038434 | 0.2468 | 0.0471 | 0.134 | 0.134 | TRUE | 3.8474 |
| 6330415B21 | 0 | 0 | 0 | 0 | 0 | 0 | 0 | 0 | 0 | 0 | 0 | 0 | 0 | 0 | 0 | NA | NA | NA | NA | NA | NA | NA | NA | NA | NA |
| 6330416G13 | 387 | 662 | 598 | 705 | 577 | 582 | 411 | 661 | 600 | 554 | 621 | 637 | 580.8 | 557 | 604 | 1.082 | 0.114 | 0.52449827 | 0.71418343 | 0.0305 | 0.0126 | 0.0214 | 0.0214 | TRUE | 0.6154 |
| 6330418K02 | 14 | 15 | 16 | 36 | 20 | 14 | 15 | 15 | 16 | 28 | 22 | 15 | 18.51 | 15 | 22 | 1.433 | 0.519 | 0.26118153 | 0.48061843 | 0 | 0.1849 | 0.0981 | 0.0981 | TRUE | 0.171 |
| 6330549D23 | 10 | 15 | 8 | 15 | 7 | 7 | 11 | 15 | 8 | 12 | 8 | 8 | 10.1 | 11 | 9 | 0.812 | -0.3 | 0.62360546 | 0.78778577 | 0 | 0.333 | 0.1708 | 0.1708 | TRUE | 0.3321 |
| 6430411K18 | 32 | 15 | 20 | 31 | 29 | 18 | 34 | 15 | 20 | 24 | 31 | 20 | 24.05 | 23 | 25 | 1.093 | 0.128 | 0.77859855 | 0.88542689 | 0.0672 | 0.1439 | 0.1074 | 0.1074 | TRUE | 0.8784 |
| 6430531B16 | 26 | 47 | 56 | 62 | 55 | 53 | 28 | 47 | 56 | 49 | 59 | 58 | 49.45 | 44 | 55 | 1.263 | 0.336 | 0.30864469 | 0.52923941 | 0.0397 | 0.0735 | 0.0581 | 0.0581 | TRUE | 0.6517 |
| 6430548M08 | 198 | 151 | 161 | 220 | 159 | 166 | 210 | 151 | 162 | 173 | 171 | 182 | 174.75 | 174 | 175 | 1.006 | 0.009 | 0.96007106 | 0.97957308 | 0.0103 | 0.0258 | 0.018 | 0.018 | TRUE | 0.3455 |
| 6430550D23 | 12 | 18 | 18 | 14 | 6 | 16 | 13 | 18 | 18 | 11 | 6 | 18 | 13.96 | 16 | 12 | 0.714 | -0.487 | 0.38088679 | 0.59526263 | 0.0266 | 0.2429 | 0.149 | 0.149 | TRUE | 0.3709 |
| 6430562O15 | 0 | 0 | 1 | 1 | 0 | 2 | 0 | 0 | 1 | 1 | 0 | 2 | 0.66 | 0 | 1 | 2.693 | 1.429 | 0.58329919 | NA | 0 | 4.974 | 3.3955 | 3.3955 | TRUE | 0.2781 |
| 6430571L13 | 18 | 26 | 9 | 21 | 16 | 17 | 19 | 26 | 9 | 17 | 17 | 19 | 17.74 | 18 | 18 | 0.965 | -0.051 | 0.9211228 | 0.96116102 | 0.0671 | 0.1926 | 0.1338 | 0.1338 | TRUE | 0.3112 |
| 6430573F11 | 67 | 70 | 62 | 54 | 49 | 36 | 71 | 70 | 62 | 42 | 53 | 39 | 56.32 | 68 | 45 | 0.661 | -0.598 | 0.02388591 | 0.09844049 | 0 | 0.0654 | 0.032 | 0.032 | TRUE | 0.1705 |
| 6430584L05 | 1 | 0 | 0 | 1 | 0 | 0 | 1 | 0 | 0 | 1 | 0 | 0 | 0.31 | 0 | 0 | 0.954 | -0.067 | 0.98625928 | NA | 0 | 10.7014 | 9.0384 | 9.0384 | TRUE | 0.4614 |
| 6430706D22 | 0 | 0 | 0 | 0 | 0 | 0 | 0 | 0 | 0 | 0 | 0 | 0 | 0 | 0 | 0 | NA | NA | NA | NA | NA | NA | NA | NA | NA | NA |
| 6530402F18 | 7 | 22 | 11 | 14 | 10 | 13 | 7 | 22 | 11 | 11 | 11 | 14 | 12.74 | 13 | 12 | 0.885 | -0.177 | 0.76752731 | 0.87860653 | 0.0891 | 0.2655 | 0.1791 | 0.1791 | TRUE | 0.9324 |
| 6530411M01 | 0 | 2 | 0 | 1 | 1 | 0 | 0 | 2 | 0 | 1 | 1 | 0 | 0.64 | 1 | 1 | 0.935 | -0.096 | 0.97106576 | NA | 0 | 5.1285 | 3.5446 | 3.5446 | TRUE | 0.9738 |
| 6720401G13 | 2700 | 3345 | 3081 | 2987 | 2822 | 2501 | 2868 | 3341 | 3094 | 2348 | 3036 | 2738 | 2904.06 | 3101 | 2707 | 0.873 | -0.196 | 0.09450009 | 0.25193271 | 0.0105 | 0.008 | 0.0096 | 0.0096 | TRUE | 0.1653 |
| 6720456H20 | 68 | 53 | 69 | 142 | 104 | 67 | 72 | 53 | 69 | 112 | 112 | 73 | 81.89 | 65 | 99 | 1.532 | 0.616 | 0.02083162 | 0.08940307 | 0.0291 | 0.0471 | 0.0383 | 0.0383 | TRUE | 0.478 |
| 6720468P15 | 41 | 34 | 38 | 29 | 25 | 34 | 44 | 34 | 38 | 23 | 27 | 37 | 33.76 | 39 | 29 | 0.746 | -0.424 | 0.22891845 | 0.44224019 | 0 | 0.1045 | 0.059 | 0.059 | TRUE | 0.3862 |
| 6720483E21 | 7 | 10 | 18 | 3 | 5 | 8 | 7 | 10 | 18 | 2 | 5 | 9 | 8.67 | 12 | 5 | 0.453 | -1.144 | 0.13013503 | 0.31070082 | 0.1571 | 0.3871 | 0.2775 | 0.2775 | TRUE | 0.4979 |
| 6720489N17 | 156 | 128 | 130 | 262 | 184 | 150 | 166 | 128 | 131 | 206 | 198 | 164 | 165.37 | 142 | 189 | 1.344 | 0.426 | 0.02259437 | 0.09482663 | 0.0111 | 0.0268 | 0.019 | 0.019 | TRUE | 0.2319 |
| 6820408C15 | 7 | 5 | 6 | 12 | 7 | 13 | 7 | 5 | 6 | 9 | 8 | 14 | 8.27 | 6 | 10 | 1.687 | 0.755 | 0.2617114 | 0.48125362 | 0 | 0.405 | 0.1955 | 0.1955 | TRUE | 0.3337 |
| 6820431F20 | 156 | 205 | 198 | 218 | 185 | 200 | 166 | 205 | 199 | 171 | 199 | 219 | 193.1 | 190 | 196 | 1.031 | 0.045 | 0.79625971 | 0.89536272 | 0.009 | 0.024 | 0.0163 | 0.0163 | TRUE | 0.1464 |
| 7420426K07 | 7 | 0 | 0 | 3 | 3 | 1 | 7 | 0 | 0 | 2 | 3 | 1 | 2.35 | 2 | 2 | 0.907 | -0.14 | 0.93435863 | NA | 2.0478 | 1.4072 | 1.6596 | 1.6596 | TRUE | 3.2034 |
| 7420461P10 | 0 | 0 | 0 | 0 | 0 | 1 | 0 | 0 | 0 | 0 | 0 | 1 | 0.18 | 0 | 0 | 1.859 | 0.894 | 0.82650734 | NA | 0 | 18.0643 | 10 | 10 | TRUE | 0.5383 |
| 7420700N18 | 0 | 0 | 0 | 0 | 0 | 0 | 0 | 0 | 0 | 0 | 0 | 0 | 0 | 0 | 0 | NA | NA | NA | NA | NA | NA | NA | NA | NA | NA |
| 7420701I03 | 0 | 0 | 0 | 0 | 0 | 0 | 0 | 0 | 0 | 0 | 0 | 0 | 0 | 0 | 0 | NA | NA | NA | NA | NA | NA | NA | NA | NA | NA |
| 7530416G11 | 0 | 0 | 0 | 0 | 0 | 0 | 0 | 0 | 0 | 0 | 0 | 0 | 0 | 0 | 0 | NA | NA | NA | NA | NA | NA | NA | NA | NA | NA |
| 7530420F21 | 0 | 0 | 0 | 0 | 0 | 0 | 0 | 0 | 0 | 0 | 0 | 0 | 0 | 0 | 0 | NA | NA | NA | NA | NA | NA | NA | NA | NA | NA |
| 7630403G23 | 0 | 0 | 0 | 0 | 0 | 0 | 0 | 0 | 0 | 0 | 0 | 0 | 0 | 0 | 0 | NA | NA | NA | NA | NA | NA | NA | NA | NA | NA |
| 8030411F24 | 0 | 0 | 0 | 1 | 0 | 0 | 0 | 0 | 0 | 1 | 0 | 0 | 0.13 | 0 | 0 | 1.859 | 0.894 | 0.82650734 | NA | 0 | 25.1515 | 10 | 10 | TRUE | 0.2563 |
| 8030423F21 | 0 | 1 | 1 | 1 | 1 | 0 | 0 | 1 | 1 | 1 | 1 | 0 | 0.64 | 1 | 1 | 0.935 | -0.097 | 0.96674856 | NA | 0 | 5.1213 | 2.3899 | 2.3899 | TRUE | 0.2246 |
| 8030423J24 | 0 | 0 | 0 | 0 | 0 | 0 | 0 | 0 | 0 | 0 | 0 | 0 | 0 | 0 | 0 | NA | NA | NA | NA | NA | NA | NA | NA | NA | NA |
| 8030442B05 | 0 | 0 | 0 | 0 | 0 | 0 | 0 | 0 | 0 | 0 | 0 | 0 | 0 | 0 | 0 | NA | NA | NA | NA | NA | NA | NA | NA | NA | NA |
| 8030443G20 | 0 | 0 | 0 | 0 | 0 | 0 | 0 | 0 | 0 | 0 | 0 | 0 | 0 | 0 | 0 | NA | NA | NA | NA | NA | NA | NA | NA | NA | NA |
| 8030462N17 | 2694 | 2490 | 2570 | 3450 | 2493 | 2456 | 2862 | 2487 | 2581 | 2712 | 2682 | 2688 | 2668.63 | 2643 | 2694 | 1.02 | 0.028 | 0.73152637 | 0.85590756 | 0.0023 | 0.0081 | 0.0044 | 0.0044 | TRUE | 0.0633 |
| 8430403D17 | 0 | 0 | 0 | 0 | 0 | 0 | 0 | 0 | 0 | 0 | 0 | 0 | 0 | 0 | 0 | NA | NA | NA | NA | NA | NA | NA | NA | NA | NA |
| 8430408G22 | 2 | 0 | 0 | 1 | 0 | 0 | 2 | 0 | 0 | 1 | 0 | 0 | 0.49 | 1 | 0 | 0.497 | -1.01 | 0.76786142 | NA | 0 | 6.7985 | 6.7083 | 6.7083 | TRUE | 0.9742 |
| 8430410A17 | 1561 | 1529 | 1453 | 2174 | 1686 | 1705 | 1658 | 1527 | 1459 | 1709 | 1814 | 1866 | 1672.25 | 1548 | 1796 | 1.16 | 0.214 | 0.01469044 | 0.06978845 | 0.0022 | 0.0089 | 0.005 | 0.005 | TRUE | 0.0464 |
| 8430419L09 | 4013 | 5406 | 5551 | 4170 | 3372 | 3749 | 4263 | 5399 | 5574 | 3278 | 3628 | 4104 | 4374.23 | 5079 | 3670 | 0.722 | -0.469 | 0.00038829 | 0.00423523 | 0.0159 | 0.0077 | 0.0124 | 0.0124 | TRUE | 0.2406 |
| 8430422H06 | 0 | 0 | 0 | 0 | 0 | 0 | 0 | 0 | 0 | 0 | 0 | 0 | 0 | 0 | 0 | NA | NA | NA | NA | NA | NA | NA | NA | NA | NA |
| 8430423G03 | 3 | 0 | 0 | 0 | 1 | 0 | 3 | 0 | 0 | 0 | 1 | 0 | 0.71 | 1 | 0 | 0.331 | -1.596 | 0.60418455 | NA | 8.2582 | 4.6443 | 5.349 | 5.349 | TRUE | 1.4426 |
| 8430427H17 | 854 | 998 | 963 | 1166 | 886 | 1034 | 907 | 997 | 967 | 917 | 953 | 1132 | 978.76 | 957 | 1001 | 1.044 | 0.063 | 0.57814328 | 0.7554661 | 0.0066 | 0.0103 | 0.0082 | 0.0082 | TRUE | 0.156 |
| 8430429K09 | 33 | 31 | 35 | 66 | 38 | 17 | 35 | 31 | 35 | 52 | 41 | 19 | 35.42 | 34 | 37 | 1.111 | 0.152 | 0.71607808 | 0.84573786 | 0.0962 | 0.0999 | 0.098 | 0.098 | TRUE | 0.4581 |
| 8430431K14 | 0 | 0 | 0 | 0 | 0 | 0 | 0 | 0 | 0 | 0 | 0 | 0 | 0 | 0 | 0 | NA | NA | NA | NA | NA | NA | NA | NA | NA | NA |
| 8430436N08 | 0 | 0 | 0 | 0 | 0 | 0 | 0 | 0 | 0 | 0 | 0 | 0 | 0 | 0 | 0 | NA | NA | NA | NA | NA | NA | NA | NA | NA | NA |
| 8430437L04 | 0 | 0 | 0 | 0 | 0 | 0 | 0 | 0 | 0 | 0 | 0 | 0 | 0 | 0 | 0 | NA | NA | NA | NA | NA | NA | NA | NA | NA | NA |
| 9030025P20 | 67 | 52 | 59 | 82 | 74 | 48 | 71 | 52 | 59 | 64 | 80 | 53 | 63.16 | 61 | 66 | 1.079 | 0.11 | 0.68918862 | 0.82917062 | 0.0165 | 0.0591 | 0.0388 | 0.0388 | TRUE | 0.2243 |
| 9030204H09 | 0 | 0 | 0 | 0 | 0 | 0 | 0 | 0 | 0 | 0 | 0 | 0 | 0 | 0 | 0 | NA | NA | NA | NA | NA | NA | NA | NA | NA | NA |
| 9030404E10 | 4 | 1 | 7 | 11 | 1 | 3 | 4 | 1 | 7 | 9 | 1 | 3 | 4.21 | 4 | 4 | 1.087 | 0.121 | 0.91392598 | NA | 0.5354 | 0.7887 | 0.6622 | 0.6622 | TRUE | 0.4487 |
| 9030612E09 | 1 | 8 | 9 | 5 | 5 | 7 | 1 | 8 | 9 | 4 | 5 | 8 | 5.84 | 6 | 6 | 0.925 | -0.113 | 0.90042491 | 0.95169273 | 0.2209 | 0.5707 | 0.4144 | 0.4144 | TRUE | 1.1575 |
| 9030617O03 | 208 | 327 | 296 | 385 | 255 | 306 | 221 | 327 | 297 | 303 | 274 | 335 | 292.79 | 282 | 304 | 1.079 | 0.109 | 0.53993673 | 0.72592976 | 0.0208 | 0.0182 | 0.0195 | 0.0195 | TRUE | 0.3944 |
| 9030619P08 | 0 | 0 | 0 | 2 | 2 | 1 | 0 | 0 | 0 | 2 | 2 | 1 | 0.8 | 0 | 2 | 8.55 | 3.096 | 0.18206245 | NA | 0 | 4.1089 | 2.5737 | 2.5737 | TRUE | 0.07 |
| 9030624G23 | 406 | 529 | 467 | 506 | 399 | 413 | 431 | 528 | 469 | 398 | 429 | 452 | 451.27 | 476 | 426 | 0.894 | -0.162 | 0.19981439 | 0.40732488 | 0.0052 | 0.0142 | 0.0093 | 0.0093 | TRUE | 0.107 |
| 9030624J02 | 3004 | 3943 | 4124 | 3873 | 2939 | 3723 | 3191 | 3938 | 4141 | 3045 | 3162 | 4075 | 3591.98 | 3757 | 3427 | 0.912 | -0.133 | 0.3622825 | 0.58015226 | 0.0199 | 0.0078 | 0.015 | 0.015 | TRUE | 0.332 |
| 9130008F23 | 14 | 6 | 9 | 11 | 7 | 9 | 15 | 6 | 9 | 9 | 8 | 10 | 9.32 | 10 | 9 | 0.873 | -0.195 | 0.76174742 | 0.87437467 | 0 | 0.3603 | 0.1919 | 0.1919 | TRUE | 0.4151 |
| 9130011E15 | 1340 | 1252 | 1301 | 1611 | 1155 | 1161 | 1423 | 1250 | 1306 | 1266 | 1243 | 1271 | 1293.35 | 1326 | 1260 | 0.95 | -0.074 | 0.38960254 | 0.60362205 | 0.0015 | 0.0095 | 0.0046 | 0.0046 | TRUE | 0.0485 |
| 9130015A21 | 2 | 0 | 0 | 0 | 1 | 0 | 2 | 0 | 0 | 0 | 1 | 0 | 0.53 | 1 | 0 | 0.497 | -1.009 | 0.76131406 | NA | 0 | 6.1834 | 6.228 | 6.228 | TRUE | 0.9742 |
| 9130015L21 | 0 | 0 | 0 | 0 | 0 | 0 | 0 | 0 | 0 | 0 | 0 | 0 | 0 | 0 | 0 | NA | NA | NA | NA | NA | NA | NA | NA | NA | NA |
| 9130017N09 | 215 | 387 | 410 | 137 | 138 | 190 | 228 | 387 | 412 | 108 | 148 | 208 | 248.45 | 342 | 155 | 0.45 | -1.151 | 5.77E-05 | 0.00087748 | 0.0956 | 0.0202 | 0.0543 | 0.0543 | TRUE | 0.768 |
| 9130019O22 | 137 | 134 | 101 | 170 | 118 | 76 | 146 | 134 | 101 | 134 | 127 | 83 | 120.76 | 127 | 115 | 0.906 | -0.142 | 0.57033447 | 0.7491102 | 0.0392 | 0.0342 | 0.0366 | 0.0366 | TRUE | 0.5549 |
| 9130019P16 | 0 | 0 | 1 | 0 | 0 | 0 | 0 | 0 | 1 | 0 | 0 | 0 | 0.17 | 0 | 0 | 0.49 | -1.029 | 0.80087 | NA | 0 | 19.6928 | 10 | 10 | TRUE | 0.4098 |
| 9130023H24 | 43 | 39 | 19 | 105 | 54 | 30 | 46 | 39 | 19 | 83 | 58 | 33 | 46.2 | 35 | 58 | 1.684 | 0.752 | 0.09196281 | 0.24745015 | 0.1678 | 0.0782 | 0.1203 | 0.1203 | TRUE | 0.1307 |

| | | | | | | | | | | | | | | | | | | | | | | | | | |
|---|---|---|---|---|---|---|---|---|---|---|---|---|---|---|---|---|---|---|---|---|---|---|---|---|---|
| 9130024F11 | 2 | 0 | 1 | 6 | 7 | 3 | 2 | 0 | 1 | 5 | 8 | 3 | 3.11 | 1 | 5 | 5.023 | 2.329 | 0.06984162 | NA | 0 | 1.0662 | 0.5979 | 0.5979 | TRUE | 0.3564 |
| 9130204L05 | 0 | 0 | 0 | 0 | 1 | 0 | 0 | 0 | 0 | 0 | 1 | 0 | 0.18 | 0 | 0 | 1.859 | 0.894 | 0.82650734 | NA | 0 | 18.3785 | 10 | 10 | TRUE | 0.5204 |
| 9130206I24R | 2 | 2 | 5 | 23 | 12 | 11 | 2 | 2 | 5 | 18 | 13 | 12 | 8.7 | 3 | 14 | 4.724 | 2.24 | 0.00297334 | 0.02107661 | 0 | 0.3858 | 0.2088 | 0.2088 | TRUE | 0.4288 |
| 9130209A04 | 0 | 0 | 0 | 0 | 0 | 0 | 0 | 0 | 0 | 0 | 0 | 0 | 0 | 0 | 0 | NA | NA | NA | NA | NA | NA | NA | NA | NA | NA |
| 9130221F21 | 0 | 0 | 0 | 0 | 0 | 0 | 0 | 0 | 0 | 0 | 0 | 0 | 0 | 0 | 0 | NA | NA | NA | NA | NA | NA | NA | NA | NA | NA |
| 9130221H12 | 169 | 175 | 144 | 231 | 178 | 178 | 180 | 175 | 145 | 182 | 192 | 195 | 177.81 | 167 | 190 | 1.138 | 0.186 | 0.25113248 | 0.46874101 | 0 | 0.0254 | 0.0133 | 0.0133 | TRUE | 0.1394 |
| 9130227L01 | 4 | 1 | 0 | 11 | 7 | 6 | 4 | 1 | 0 | 9 | 8 | 7 | 4.67 | 2 | 8 | 4.416 | 2.143 | 0.04743481 | 0.15806925 | 0 | 0.713 | 0.481 | 0.481 | TRUE | 1.176 |
| 9130230L23 | 1 | 4 | 6 | 12 | 5 | 8 | 1 | 4 | 6 | 9 | 5 | 9 | 5.78 | 4 | 8 | 2.125 | 1.088 | 0.20627808 | 0.41518859 | 0 | 0.5773 | 0.3343 | 0.3343 | TRUE | 0.5062 |
| 9130401M01 | 3487 | 3639 | 3701 | 4610 | 3369 | 3287 | 3704 | 3634 | 3716 | 3624 | 3625 | 3598 | 3650.22 | 3685 | 3616 | 0.981 | -0.027 | 0.64110421 | 0.79813687 | 0 | 0.0078 | 0.0022 | 0.0022 | TRUE | 0.0018 |
| 9130409I23R | 1 | 0 | 0 | 1 | 2 | 1 | 1 | 0 | 0 | 1 | 2 | 1 | 0.85 | 0 | 1 | 3.632 | 1.861 | 0.38570833 | NA | 0 | 3.8867 | 1.9484 | 1.9484 | TRUE | 0.4614 |
| 9230009I02R | 0 | 0 | 0 | 0 | 0 | 0 | 0 | 0 | 0 | 0 | 0 | 0 | 0 | 0 | 0 | NA | NA | NA | NA | NA | NA | NA | NA | NA | NA |
| 9230102K24 | 0 | 0 | 0 | 1 | 0 | 0 | 0 | 0 | 0 | 1 | 0 | 0 | 0.13 | 0 | 0 | 1.859 | 0.894 | 0.82650734 | NA | 0 | 25.1515 | 10 | 10 | TRUE | 0.2563 |
| 9230102O04 | 0 | 0 | 0 | 3 | 0 | 1 | 0 | 0 | 0 | 2 | 0 | 1 | 0.58 | 0 | 1 | 6.21 | 2.635 | 0.40034089 | NA | 0 | 5.7314 | 5.6518 | 5.6518 | TRUE | 0.0653 |
| 9230104L09 | 0 | 0 | 0 | 0 | 1 | 0 | 0 | 0 | 0 | 0 | 1 | 0 | 0.18 | 0 | 0 | 1.859 | 0.894 | 0.82650734 | NA | 0 | 18.3785 | 10 | 10 | TRUE | 0.5204 |
| 9230105E05 | 13 | 8 | 13 | 7 | 7 | 8 | 14 | 8 | 13 | 6 | 8 | 9 | 9.44 | 12 | 8 | 0.619 | -0.692 | 0.27158427 | 0.49163634 | 0 | 0.3559 | 0.1737 | 0.1737 | TRUE | 0.2926 |
| 9230110C19 | 348 | 435 | 405 | 436 | 313 | 322 | 370 | 434 | 407 | 343 | 337 | 352 | 373.79 | 404 | 344 | 0.852 | -0.231 | 0.0537749 | 0.1723792 | 0 | 0.0157 | 0.0077 | 0.0077 | TRUE | 0.0616 |
| 9230110F15 | 0 | 0 | 0 | 0 | 0 | 0 | 0 | 0 | 0 | 0 | 0 | 0 | 0 | 0 | 0 | NA | NA | NA | NA | NA | NA | NA | NA | NA | NA |
| 9230112D13 | 0 | 1 | 0 | 1 | 0 | 0 | 0 | 1 | 0 | 1 | 0 | 0 | 0.3 | 0 | 0 | 0.954 | -0.067 | 0.98642001 | NA | 0 | 11.0817 | 9.3014 | 9.3014 | TRUE | 0.405 |
| 9230112J17 | 3 | 0 | 4 | 2 | 1 | 2 | 3 | 0 | 4 | 2 | 1 | 2 | 2.01 | 2 | 2 | 0.672 | -0.573 | 0.69164797 | NA | 0 | 1.6486 | 0.9896 | 0.9896 | TRUE | 0.8318 |
| 9230114K14 | 91 | 82 | 67 | 99 | 82 | 65 | 97 | 82 | 67 | 78 | 88 | 71 | 80.51 | 82 | 79 | 0.966 | -0.05 | 0.83329179 | 0.91529202 | 0.0087 | 0.0478 | 0.029 | 0.029 | TRUE | 0.1792 |
| 9230115E21 | 0 | 0 | 0 | 3 | 0 | 0 | 0 | 0 | 0 | 2 | 0 | 0 | 0.39 | 0 | 1 | 4.278 | 2.097 | 0.60136958 | NA | 10 | 8.3884 | 10 | 10 | TRUE | 1.4411 |
| 9230116N13 | 37 | 52 | 55 | 46 | 33 | 46 | 39 | 52 | 55 | 36 | 36 | 50 | 44.75 | 49 | 41 | 0.829 | -0.271 | 0.38587351 | 0.59978289 | 0.0115 | 0.0805 | 0.048 | 0.048 | TRUE | 0.3067 |
| 9330020H09 | 21 | 31 | 14 | 26 | 36 | 22 | 22 | 31 | 14 | 20 | 39 | 24 | 25.1 | 22 | 28 | 1.229 | 0.297 | 0.52227309 | 0.71226036 | 0.0905 | 0.1382 | 0.1151 | 0.1151 | TRUE | 0.2669 |
| 9330111N05 | 0 | 0 | 1 | 0 | 0 | 0 | 0 | 0 | 1 | 0 | 0 | 0 | 0.17 | 0 | 0 | 0.49 | -1.029 | 0.80087 | NA | 0 | 19.6928 | 10 | 10 | TRUE | 0.4098 |
| 9330117O12 | 0 | 0 | 0 | 0 | 0 | 0 | 0 | 0 | 0 | 0 | 0 | 0 | 0 | 0 | 0 | NA | NA | NA | NA | NA | NA | NA | NA | NA | NA |
| 9330133O14 | 71 | 72 | 47 | 102 | 89 | 50 | 75 | 72 | 47 | 80 | 96 | 55 | 70.87 | 65 | 77 | 1.188 | 0.249 | 0.4170607 | 0.62807995 | 0.0534 | 0.0534 | 0.0534 | 0.0534 | TRUE | 0.304 |
| 9330151L19 | 84 | 148 | 163 | 128 | 137 | 157 | 89 | 148 | 164 | 101 | 147 | 172 | 136.76 | 134 | 140 | 1.043 | 0.061 | 0.83174867 | 0.91463474 | 0.0765 | 0.031 | 0.0526 | 0.0526 | TRUE | 0.6233 |
| 9330158H04 | 0 | 0 | 0 | 0 | 0 | 0 | 0 | 0 | 0 | 0 | 0 | 0 | 0 | 0 | 0 | NA | NA | NA | NA | NA | NA | NA | NA | NA | NA |
| 9330159F19 | 97 | 58 | 60 | 36 | 31 | 34 | 103 | 58 | 60 | 28 | 33 | 37 | 53.35 | 74 | 33 | 0.445 | -1.167 | 0.0005731 | 0.00578738 | 0.0533 | 0.0687 | 0.0609 | 0.0609 | TRUE | 1.083 |
| 9330159M07 | 34 | 51 | 36 | 69 | 58 | 43 | 36 | 51 | 36 | 54 | 62 | 47 | 47.82 | 41 | 54 | 1.328 | 0.409 | 0.17295468 | 0.37179245 | 0 | 0.0758 | 0.0435 | 0.0435 | TRUE | 0.3389 |
| 9330162B11 | 0 | 0 | 0 | 0 | 0 | 0 | 0 | 0 | 0 | 0 | 0 | 0 | 0 | 0 | 0 | NA | NA | NA | NA | NA | NA | NA | NA | NA | NA |
| 9330175E14 | 1 | 0 | 0 | 1 | 1 | 0 | 1 | 0 | 0 | 1 | 1 | 0 | 0.49 | 0 | 1 | 1.712 | 0.776 | 0.78488768 | NA | 0 | 6.7665 | 4.0387 | 4.0387 | TRUE | 0.4614 |
| 9330175M2 | 0 | 0 | 0 | 0 | 0 | 0 | 0 | 0 | 0 | 0 | 0 | 0 | 0 | 0 | 0 | NA | NA | NA | NA | NA | NA | NA | NA | NA | NA |
| 9330178D15 | 0 | 0 | 0 | 0 | 0 | 0 | 0 | 0 | 0 | 0 | 0 | 0 | 0 | 0 | 0 | NA | NA | NA | NA | NA | NA | NA | NA | NA | NA |
| 9330179D12 | 0 | 0 | 0 | 1 | 0 | 0 | 0 | 0 | 0 | 1 | 0 | 0 | 0.13 | 0 | 0 | 1.859 | 0.894 | 0.82650734 | NA | 0 | 25.1515 | 10 | 10 | TRUE | 0.2563 |
| 9330182L06 | 347 | 443 | 447 | 437 | 311 | 308 | 369 | 442 | 449 | 344 | 335 | 337 | 379.19 | 420 | 339 | 0.806 | -0.312 | 0.01445286 | 0.06886723 | 0.0036 | 0.0156 | 0.009 | 0.009 | TRUE | 0.1307 |
| 9330182O14 | 2 | 4 | 4 | 3 | 0 | 2 | 2 | 4 | 4 | 2 | 0 | 2 | 2.45 | 3 | 1 | 0.459 | -1.123 | 0.38007139 | NA | 0 | 1.3531 | 0.7143 | 0.7143 | TRUE | 0.4672 |
| 9430007A20 | 0 | 0 | 0 | 0 | 0 | 0 | 0 | 0 | 0 | 0 | 0 | 0 | 0 | 0 | 0 | NA | NA | NA | NA | NA | NA | NA | NA | NA | NA |
| 9430008C03 | 989 | 1127 | 1048 | 1194 | 1040 | 932 | 1050 | 1126 | 1052 | 939 | 1119 | 1020 | 1051.03 | 1076 | 1026 | 0.952 | -0.071 | 0.48306574 | 0.68129391 | 0.0038 | 0.01 | 0.0063 | 0.0063 | TRUE | 0.0753 |
| 9430014N10 | 0 | 0 | 0 | 0 | 0 | 0 | 0 | 0 | 0 | 0 | 0 | 0 | 0 | 0 | 0 | NA | NA | NA | NA | NA | NA | NA | NA | NA | NA |
| 9430015G10 | 1291 | 1781 | 1710 | 1705 | 1485 | 1303 | 1371 | 1779 | 1717 | 1340 | 1598 | 1426 | 1538.57 | 1622 | 1455 | 0.896 | -0.158 | 0.21741332 | 0.4294633 | 0.0122 | 0.009 | 0.0112 | 0.0112 | TRUE | 0.2204 |
| 9430016H08 | 631 | 629 | 708 | 1051 | 758 | 722 | 670 | 628 | 711 | 826 | 816 | 790 | 740.24 | 670 | 811 | 1.211 | 0.276 | 0.0036789 | 0.02478579 | 6.00E-04 | 0.0114 | 0.0051 | 0.0051 | TRUE | 0.0352 |
| 9430018G01 | 17 | 21 | 17 | 76 | 62 | 40 | 18 | 21 | 17 | 60 | 67 | 44 | 37.72 | 19 | 57 | 3.038 | 1.603 | 5.56E-06 | 0.000121 | 0 | 0.0942 | 0.0539 | 0.0539 | TRUE | 0.3006 |
| 9430019J16 | 0 | 0 | 0 | 0 | 0 | 0 | 0 | 0 | 0 | 0 | 0 | 0 | 0 | 0 | 0 | NA | NA | NA | NA | NA | NA | NA | NA | NA | NA |
| 9430020K01 | 1635 | 1663 | 1676 | 2257 | 1699 | 1665 | 1737 | 1661 | 1683 | 1774 | 1828 | 1823 | 1750.88 | 1694 | 1808 | 1.068 | 0.094 | 0.17613301 | 0.37637393 | 0 | 0.0088 | 0.0029 | 0.0029 | TRUE | 0.006 |
| 9430021M05 | 1 | 0 | 2 | 0 | 0 | 1 | 1 | 0 | 2 | 0 | 0 | 1 | 0.69 | 1 | 0 | 0.341 | -1.551 | 0.54644411 | NA | 0 | 4.7528 | 3.2643 | 3.2643 | TRUE | 0.3117 |
| 9430023L20 | 2553 | 2190 | 2007 | 3141 | 2206 | 1946 | 2712 | 2187 | 2015 | 2469 | 2374 | 2130 | 2314.51 | 2305 | 2324 | 1.009 | 0.013 | 0.92145827 | 0.96126008 | 0.0144 | 0.0083 | 0.0116 | 0.0116 | TRUE | 0.2887 |
| 9430037G07 | 21 | 16 | 23 | 38 | 29 | 12 | 22 | 16 | 23 | 30 | 31 | 13 | 22.6 | 20 | 25 | 1.217 | 0.284 | 0.5461086 | 0.73111152 | 0.0708 | 0.1527 | 0.1144 | 0.1144 | TRUE | 0.9393 |
| 9430038I01R | 532 | 517 | 415 | 676 | 483 | 318 | 565 | 516 | 417 | 531 | 520 | 348 | 482.89 | 499 | 466 | 0.935 | -0.097 | 0.61616072 | 0.78295898 | 0.0361 | 0.0137 | 0.0246 | 0.0246 | TRUE | 0.5638 |
| 9430041J12 | 3 | 6 | 3 | 7 | 12 | 5 | 3 | 6 | 3 | 6 | 13 | 5 | 6.01 | 4 | 8 | 1.939 | 0.955 | 0.25249736 | 0.47053511 | 0 | 0.5548 | 0.3147 | 0.3147 | TRUE | 0.813 |
| 9430060I03R | 1 | 4 | 7 | 10 | 9 | 5 | 1 | 4 | 7 | 8 | 10 | 5 | 5.85 | 4 | 8 | 1.896 | 0.923 | 0.28573142 | 0.50703825 | 0.0223 | 0.57 | 0.3486 | 0.3486 | TRUE | 0.3483 |
| 9430069I07R | 16 | 30 | 37 | 3 | 4 | 6 | 17 | 30 | 37 | 2 | 4 | 7 | 16.22 | 28 | 4 | 0.152 | -2.715 | 2.01E-05 | 0.00035865 | 0 | 0.21 | 0.1602 | 0.1602 | TRUE | 0.1853 |
| 9430076C15 | 1 | 1 | 1 | 4 | 1 | 3 | 1 | 1 | 1 | 3 | 1 | 3 | 1.76 | 1 | 2 | 2.469 | 1.304 | 0.36259163 | NA | 0 | 1.8771 | 0.7846 | 0.7846 | TRUE | 0.2478 |
| 9430076G02 | 1 | 0 | 1 | 4 | 0 | 1 | 1 | 0 | 1 | 3 | 0 | 1 | 1.05 | 1 | 1 | 2.129 | 1.09 | 0.59527377 | NA | 0 | 3.1418 | 1.9522 | 1.9522 | TRUE | 0.3095 |
| 9430083A17 | 4 | 5 | 5 | 9 | 21 | 5 | 4 | 5 | 5 | 7 | 23 | 5 | 8.23 | 5 | 12 | 2.443 | 1.289 | 0.11163279 | 0.28112285 | 0.2507 | 0.407 | 0.3234 | 0.3234 | TRUE | 2.4217 |
| 9430091E24 | 13 | 9 | 6 | 17 | 11 | 5 | 14 | 9 | 6 | 13 | 12 | 5 | 9.92 | 10 | 10 | 1.078 | 0.109 | 0.86884546 | 0.93544206 | 0.0692 | 0.3392 | 0.2128 | 0.2128 | TRUE | 0.4125 |
| 9530002B09 | 0 | 0 | 0 | 0 | 0 | 0 | 0 | 0 | 0 | 0 | 0 | 0 | 0 | 0 | 0 | NA | NA | NA | NA | NA | NA | NA | NA | NA | NA |
| 9530003J23 | 0 | 0 | 0 | 0 | 0 | 0 | 0 | 0 | 0 | 0 | 0 | 0 | 0 | 0 | 0 | NA | NA | NA | NA | NA | NA | NA | NA | NA | NA |
| 9530026F06 | 0 | 0 | 0 | 0 | 0 | 0 | 0 | 0 | 0 | 0 | 0 | 0 | 0 | 0 | 0 | NA | NA | NA | NA | NA | NA | NA | NA | NA | NA |
| 9530026P05 | 5 | 4 | 7 | 8 | 8 | 9 | 5 | 4 | 7 | 6 | 9 | 10 | 6.85 | 5 | 8 | 1.502 | 0.587 | 0.41684281 | 0.62807995 | 0 | 0.488 | 0.2225 | 0.2225 | TRUE | 0.1698 |
| 9530027J09 | 9 | 6 | 3 | 5 | 4 | 1 | 10 | 6 | 3 | 4 | 4 | 1 | 4.65 | 6 | 3 | 0.511 | -0.97 | 0.30995722 | 0.53027935 | 0 | 0.7156 | 0.4195 | 0.4195 | TRUE | 0.1491 |
| 9530036O11 | 0 | 0 | 0 | 0 | 0 | 1 | 0 | 0 | 0 | 0 | 0 | 1 | 0.18 | 0 | 0 | 1.859 | 0.894 | 0.82650734 | NA | 0 | 18.0643 | 10 | 10 | TRUE | 0.5383 |
| 9530048J24 | 0 | 0 | 0 | 0 | 0 | 0 | 0 | 0 | 0 | 0 | 0 | 0 | 0 | 0 | 0 | NA | NA | NA | NA | NA | NA | NA | NA | NA | NA |
| 9530051G07 | 17 | 13 | 26 | 9 | 11 | 12 | 18 | 13 | 26 | 7 | 12 | 13 | 14.87 | 19 | 11 | 0.553 | -0.855 | 0.11765698 | 0.29020869 | 0 | 0.2285 | 0.1422 | 0.1422 | TRUE | 0.2327 |
| 9530052E02 | 3 | 0 | 0 | 0 | 0 | 1 | 3 | 0 | 0 | 0 | 0 | 1 | 0.71 | 1 | 0 | 0.331 | -1.596 | 0.60371867 | NA | 8.2583 | 4.624 | 5.3312 | 5.3312 | TRUE | 1.4426 |
| 9530053A07 | 40 | 46 | 35 | 51 | 31 | 31 | 42 | 46 | 35 | 40 | 33 | 34 | 38.49 | 41 | 36 | 0.873 | -0.195 | 0.5310056 | 0.71913278 | 0 | 0.0925 | 0.0442 | 0.0442 | TRUE | 0.1269 |
| 9530059O14 | 0 | 0 | 1 | 2 | 0 | 0 | 0 | 0 | 1 | 2 | 0 | 0 | 0.43 | 0 | 1 | 1.493 | 0.579 | 0.86952142 | NA | 0 | 7.6793 | 7.0425 | 7.0425 | TRUE | 0.9718 |
| 9530068E07 | 1773 | 2212 | 2143 | 2524 | 1941 | 1992 | 1883 | 2209 | 2152 | 1984 | 2088 | 2181 | 2082.88 | 2081 | 2084 | 1.001 | 0.001 | 0.98886138 | 0.99415615 | 0.0042 | 0.0085 | 0.0059 | 0.0059 | TRUE | 0.0836 |
| 9530077C05 | 174 | 141 | 138 | 242 | 165 | 143 | 185 | 141 | 139 | 190 | 178 | 157 | 164.75 | 155 | 175 | 1.133 | 0.18 | 0.33765043 | 0.55771127 | 0.0118 | 0.0269 | 0.0193 | 0.0193 | TRUE | 0.3 |

| | | | | | | | | | | | | | | | | | | | | | | | | | |
|---|---|---|---|---|---|---|---|---|---|---|---|---|---|---|---|---|---|---|---|---|---|---|---|---|---|
| 9530080O11 | 6 | 8 | 2 | 9 | 5 | 5 | 6 | 8 | 2 | 7 | 5 | 5 | 5.72 | 5 | 6 | 1.103 | 0.142 | 0.8635358 | 0.9324082 | 0 | 0.5832 | 0.3154 | 0.3154 | TRUE | 0.676 |
| 9530082P21I | 44 | 36 | 32 | 40 | 32 | 24 | 47 | 36 | 32 | 31 | 34 | 26 | 34.49 | 38 | 30 | 0.805 | -0.314 | 0.35543781 | 0.57387109 | 0 | 0.1024 | 0.0537 | 0.0537 | TRUE | 0.2676 |
| 9530091C08I | 1 | 2 | 1 | 0 | 0 | 1 | 1 | 2 | 1 | 0 | 0 | 1 | 0.86 | 1 | 0 | 0.257 | -1.958 | 0.35814405 | NA | 0 | 3.8389 | 1.8948 | 1.8948 | TRUE | 0.5383 |
| 9630013A20 | 0 | 0 | 0 | 0 | 0 | 0 | 0 | 0 | 0 | 0 | 0 | 0 | 0 | 0 | 0 | NA | NA | NA | NA | NA | NA | NA | NA | NA | NA |
| 9630028B13 | 0 | 0 | 0 | 0 | 1 | 1 | 0 | 0 | 0 | 0 | 1 | 1 | 0.36 | 0 | 1 | 3.794 | 1.924 | 0.60247382 | NA | 0 | 9.1135 | 8.1425 | 8.1425 | TRUE | 0.3482 |
| 9630028H03 | 0 | 0 | 0 | 0 | 0 | 0 | 0 | 0 | 0 | 0 | 0 | 0 | 0 | 0 | 0 | NA | NA | NA | NA | NA | NA | NA | NA | NA | NA |
| 9630033F20I | 3132 | 2617 | 2358 | 3696 | 2632 | 2108 | 3327 | 2614 | 2368 | 2906 | 2832 | 2308 | 2725.49 | 2770 | 2682 | 0.969 | -0.046 | 0.75762292 | 0.8719099 | 0.0229 | 0.0081 | 0.0156 | 0.0156 | TRUE | 0.3759 |
| 9830001H06 | 554 | 629 | 600 | 1118 | 760 | 706 | 588 | 628 | 602 | 879 | 818 | 773 | 714.75 | 606 | 823 | 1.359 | 0.443 | 7.94E-06 | 0.00016271 | 0.0016 | 0.0115 | 0.0056 | 0.0056 | TRUE | 0.0434 |
| 9830107B12 | 0 | 0 | 0 | 0 | 0 | 0 | 0 | 0 | 0 | 0 | 0 | 0 | 0 | 0 | 0 | NA | NA | NA | NA | NA | NA | NA | NA | NA | NA |
| 9830132P13I | 1 | 0 | 0 | 0 | 0 | 0 | 1 | 0 | 0 | 0 | 0 | 0 | 0.18 | 0 | 0 | 0.49 | -1.029 | 0.80087 | NA | 0 | 18.6165 | 10 | 10 | TRUE | 0.4614 |
| 9830147E19I | 1062 | 897 | 903 | 1223 | 923 | 848 | 1128 | 896 | 907 | 961 | 993 | 928 | 968.89 | 977 | 961 | 0.984 | -0.023 | 0.8445219 | 0.92165786 | 0.0081 | 0.0103 | 0.0091 | 0.0091 | TRUE | 0.2178 |
| 9830166K06I | 0 | 0 | 0 | 2 | 1 | 2 | 0 | 0 | 0 | 2 | 1 | 2 | 0.81 | 0 | 2 | 8.576 | 3.1 | 0.18121756 | NA | 0 | 4.093 | 2.5705 | 2.5705 | TRUE | 0.07 |
| 9930012K11I | 208 | 212 | 195 | 262 | 199 | 158 | 221 | 212 | 196 | 206 | 214 | 173 | 203.58 | 210 | 198 | 0.945 | -0.081 | 0.6031585 | 0.77412127 | 0.0026 | 0.0231 | 0.0127 | 0.0127 | TRUE | 0.1232 |
| 9930013L23I | 18 | 8 | 8 | 37 | 19 | 16 | 19 | 8 | 8 | 29 | 20 | 18 | 17.03 | 12 | 22 | 1.93 | 0.949 | 0.08194173 | 0.22845159 | 0.0885 | 0.2003 | 0.1489 | 0.1489 | TRUE | 1.137 |
| 9930014A18 | 1 | 3 | 1 | 14 | 6 | 5 | 1 | 3 | 1 | 11 | 6 | 5 | 4.67 | 2 | 7 | 4.559 | 2.189 | 0.03168098 | 0.11971732 | 0 | 0.7129 | 0.3836 | 0.3836 | TRUE | 0.5257 |
| 9930021J03I | 2607 | 2496 | 2686 | 3159 | 2327 | 2389 | 2769 | 2493 | 2697 | 2483 | 2504 | 2615 | 2593.52 | 2653 | 2534 | 0.955 | -0.066 | 0.38714268 | 0.60099965 | 0.0013 | 0.0082 | 0.0039 | 0.0039 | TRUE | 0.0339 |
| 9930104L06I | 820 | 768 | 695 | 1064 | 789 | 639 | 871 | 767 | 698 | 836 | 849 | 699 | 786.78 | 779 | 795 | 1.022 | 0.031 | 0.80926607 | 0.90182956 | 0.0105 | 0.0111 | 0.0108 | 0.0108 | TRUE | 0.1289 |
| 9930111J21I | 3 | 2 | 0 | 3 | 0 | 1 | 3 | 2 | 0 | 2 | 0 | 1 | 1.44 | 2 | 1 | 0.693 | -0.529 | 0.76905053 | NA | 0 | 2.2955 | 1.634 | 1.634 | TRUE | 0.3039 |
| 9930111J21I | 0 | 0 | 0 | 0 | 0 | 0 | 0 | 0 | 0 | 0 | 0 | 0 | 0 | 0 | 0 | NA | NA | NA | NA | NA | NA | NA | NA | NA | NA |
| a | 3 | 5 | 2 | 2 | 4 | 8 | 3 | 5 | 2 | 2 | 4 | 9 | 4.14 | 3 | 5 | 1.405 | 0.49 | 0.63205307 | NA | 0 | 0.8033 | 0.5021 | 0.5021 | TRUE | 0.2787 |
| A030009H04 | 229 | 219 | 219 | 402 | 276 | 230 | 243 | 219 | 220 | 316 | 297 | 252 | 257.77 | 227 | 288 | 1.272 | 0.347 | 0.01971622 | 0.08601559 | 0.0052 | 0.0197 | 0.012 | 0.012 | TRUE | 0.1347 |
| A130010J15I | 574 | 804 | 812 | 925 | 699 | 675 | 610 | 803 | 815 | 727 | 752 | 739 | 741.02 | 743 | 739 | 0.995 | -0.007 | 0.9569899 | 0.97770215 | 0.0115 | 0.0113 | 0.0114 | 0.0114 | TRUE | 0.2887 |
| A130049A11 | 30 | 38 | 47 | 29 | 40 | 48 | 32 | 38 | 47 | 23 | 43 | 53 | 39.23 | 39 | 40 | 1.001 | 0.001 | 0.99779667 | 0.99860587 | 0.0819 | 0.0909 | 0.0865 | 0.0865 | TRUE | 0.6237 |
| A130077B15 | 0 | 0 | 0 | 0 | 0 | 0 | 0 | 0 | 0 | 0 | 0 | 0 | 0 | 0 | 0 | NA | NA | NA | NA | NA | NA | NA | NA | NA | NA |
| A1bg | 0 | 0 | 0 | 0 | 0 | 1 | 0 | 0 | 0 | 0 | 0 | 1 | 0.18 | 0 | 0 | 1.859 | 0.894 | 0.82650734 | NA | 0 | 18.0643 | 10 | 10 | TRUE | 0.5383 |
| A1cf | 0 | 0 | 0 | 0 | 1 | 0 | 0 | 0 | 0 | 0 | 1 | 0 | 0.18 | 0 | 0 | 1.859 | 0.894 | 0.82650734 | NA | 0 | 18.3785 | 10 | 10 | TRUE | 0.5204 |
| A230001M10 | 0 | 1 | 0 | 0 | 1 | 0 | 0 | 1 | 0 | 0 | 1 | 0 | 0.35 | 0 | 0 | 0.954 | -0.067 | 0.98571955 | NA | 0 | 9.5346 | 8.2194 | 8.2194 | TRUE | 0.5204 |
| A230020J21I | 2 | 0 | 3 | 6 | 1 | 0 | 2 | 0 | 3 | 5 | 1 | 0 | 1.82 | 2 | 2 | 1.169 | 0.226 | 0.89667165 | NA | 1.3492 | 1.8155 | 1.6309 | 1.6309 | TRUE | 1.3724 |
| A230028O05 | 5 | 1 | 1 | 10 | 1 | 3 | 5 | 1 | 1 | 8 | 1 | 3 | 3.26 | 2 | 4 | 1.722 | 0.784 | 0.52752102 | NA | 0.5544 | 1.0188 | 0.7806 | 0.7806 | TRUE | 0.4877 |
| A230046K03 | 9062 | 8475 | 8472 | 9262 | 6722 | 7062 | 9625 | 8464 | 8507 | 7281 | 7232 | 7730 | 8140.05 | 8865 | 7414 | 0.836 | -0.258 | 0.00164482 | 0.01316192 | 0.0033 | 0.0073 | 0.0047 | 0.0047 | TRUE | 0.0686 |
| A230050P20 | 744 | 641 | 646 | 660 | 589 | 601 | 790 | 640 | 649 | 519 | 634 | 658 | 648.26 | 693 | 604 | 0.87 | -0.202 | 0.15398768 | 0.34586923 | 0.0136 | 0.012 | 0.0129 | 0.0129 | TRUE | 0.1833 |
| A230056J06I | 2 | 4 | 3 | 2 | 5 | 1 | 2 | 4 | 3 | 2 | 5 | 1 | 2.86 | 3 | 3 | 0.866 | -0.207 | 0.85689633 | NA | 0 | 1.1576 | 0.6004 | 0.6004 | TRUE | 0.8356 |
| A230056P14 | 13 | 17 | 17 | 19 | 13 | 9 | 14 | 17 | 17 | 15 | 14 | 10 | 14.44 | 16 | 13 | 0.815 | -0.295 | 0.5523448 | 0.73624731 | 0 | 0.2351 | 0.1077 | 0.1077 | TRUE | 0.1625 |
| A230057D06 | 2 | 0 | 0 | 2 | 1 | 1 | 2 | 0 | 0 | 2 | 1 | 1 | 0.98 | 1 | 1 | 1.807 | 0.853 | 0.67681949 | NA | 0 | 3.3759 | 1.8982 | 1.8982 | TRUE | 0.9744 |
| A230065H16 | 1 | 3 | 1 | 13 | 2 | 2 | 1 | 3 | 1 | 10 | 2 | 2 | 3.27 | 2 | 5 | 2.938 | 1.555 | 0.20808976 | NA | 0.434 | 1.0142 | 0.6991 | 0.6991 | TRUE | 2.3032 |
| A230070E04 | 15 | 25 | 24 | 39 | 21 | 19 | 16 | 25 | 24 | 31 | 23 | 21 | 23.18 | 22 | 25 | 1.147 | 0.197 | 0.63646758 | 0.79543785 | 0 | 0.1491 | 0.0824 | 0.0824 | TRUE | 0.3222 |
| A230072C01 | 67 | 107 | 96 | 105 | 95 | 89 | 71 | 107 | 96 | 83 | 102 | 97 | 92.77 | 91 | 94 | 1.024 | 0.034 | 0.88678642 | 0.94438861 | 0.016 | 0.0424 | 0.0296 | 0.0296 | TRUE | 0.3536 |
| A230072E10 | 0 | 0 | 0 | 0 | 0 | 1 | 0 | 0 | 0 | 0 | 0 | 1 | 0.18 | 0 | 0 | 1.859 | 0.894 | 0.82650734 | NA | 0 | 18.0643 | 10 | 10 | TRUE | 0.5383 |
| A230073K19 | 6 | 5 | 2 | 9 | 1 | 9 | 6 | 5 | 2 | 7 | 1 | 10 | 5.23 | 4 | 6 | 1.354 | 0.437 | 0.65526065 | 0.80609504 | 0.3301 | 0.6369 | 0.4954 | 0.4954 | TRUE | 0.4289 |
| A230077H06 | 7 | 2 | 4 | 6 | 6 | 3 | 7 | 2 | 4 | 5 | 6 | 3 | 4.65 | 4 | 5 | 1.08 | 0.111 | 0.90252255 | 0.95205059 | 0 | 0.7152 | 0.3795 | 0.3795 | TRUE | 0.3973 |
| A230108P19 | 0 | 0 | 0 | 0 | 0 | 0 | 0 | 0 | 0 | 0 | 0 | 0 | 0 | 0 | 0 | NA | NA | NA | NA | NA | NA | NA | NA | NA | NA |
| A2m | 8111 | 14904 | 16490 | 11379 | 10307 | 14205 | 8615 | 14885 | 16558 | 8945 | 11090 | 15550 | 12607.06 | 13353 | 11862 | 0.888 | -0.171 | 0.60085616 | 0.77266232 | 0.0768 | 0.0072 | 0.0432 | 0.0768 | TRUE | 0.7997 |
| A330009N23 | 31 | 39 | 30 | 46 | 26 | 17 | 33 | 39 | 30 | 36 | 28 | 19 | 30.79 | 34 | 28 | 0.82 | -0.286 | 0.45320304 | 0.6586729 | 0.0257 | 0.1139 | 0.0725 | 0.0725 | TRUE | 0.2705 |
| A330021E22 | 67 | 99 | 114 | 136 | 127 | 107 | 71 | 99 | 114 | 107 | 137 | 117 | 107.53 | 95 | 120 | 1.263 | 0.337 | 0.15626402 | 0.34905281 | 0.0246 | 0.0375 | 0.0313 | 0.0313 | TRUE | 0.4505 |
| A330023F24 | 2 | 10 | 2 | 13 | 17 | 9 | 2 | 10 | 2 | 10 | 18 | 10 | 8.75 | 5 | 13 | 2.694 | 1.43 | 0.07784366 | 0.22065963 | 0.2512 | 0.3835 | 0.3273 | 0.3273 | TRUE | 1.878 |
| A330032B11 | 0 | 1 | 0 | 0 | 0 | 0 | 0 | 1 | 0 | 0 | 0 | 0 | 0.17 | 0 | 0 | 0.49 | -1.029 | 0.80087 | NA | 0 | 19.7991 | 10 | 10 | TRUE | 0.405 |
| A330035P11 | 10 | 9 | 12 | 18 | 14 | 15 | 11 | 9 | 12 | 14 | 15 | 16 | 12.88 | 11 | 15 | 1.438 | 0.524 | 0.31218156 | 0.53255705 | 0 | 0.2626 | 0.1129 | 0.1129 | TRUE | 0.0624 |
| A330040F15 | 0 | 0 | 0 | 2 | 0 | 0 | 0 | 0 | 0 | 2 | 0 | 0 | 0.26 | 0 | 1 | 2.886 | 1.529 | 0.70676491 | NA | 0 | 12.5792 | 10 | 10 | TRUE | 0.9669 |
| A330041J22I | 1 | 3 | 7 | 1 | 2 | 1 | 1 | 3 | 7 | 1 | 2 | 1 | 2.52 | 4 | 1 | 0.355 | -1.495 | 0.2569403 | NA | 0 | 1.3143 | 0.745 | 0.745 | TRUE | 0.282 |
| A330048O09 | 9 | 17 | 14 | 17 | 14 | 9 | 10 | 17 | 14 | 13 | 15 | 10 | 13.15 | 14 | 13 | 0.943 | -0.085 | 0.874162 | 0.93791982 | 0 | 0.2575 | 0.1297 | 0.1297 | TRUE | 0.2681 |
| A330049N07 | 0 | 0 | 0 | 0 | 0 | 0 | 0 | 0 | 0 | 0 | 0 | 0 | 0 | 0 | 0 | NA | NA | NA | NA | NA | NA | NA | NA | NA | NA |
| A330050B17 | 4 | 2 | 2 | 6 | 1 | 1 | 4 | 2 | 2 | 5 | 1 | 1 | 2.52 | 3 | 2 | 0.872 | -0.198 | 0.87233445 | NA | 0 | 1.3124 | 0.7008 | 0.7008 | TRUE | 1.1242 |
| A330050F15 | 9 | 39 | 32 | 7 | 7 | 6 | 10 | 39 | 32 | 6 | 8 | 7 | 16.71 | 27 | 7 | 0.241 | -2.053 | 0.00162455 | 0.01305937 | 0.2192 | 0.2041 | 0.2117 | 0.2117 | TRUE | 0.4828 |
| A330069E16 | 56 | 45 | 52 | 54 | 51 | 54 | 59 | 45 | 52 | 42 | 55 | 59 | 52.18 | 52 | 52 | 0.993 | -0.01 | 0.97264811 | 0.98548003 | 0 | 0.07 | 0.0394 | 0.0394 | TRUE | 0.254 |
| A330070K13 | 0 | 0 | 0 | 0 | 0 | 0 | 0 | 0 | 0 | 0 | 0 | 0 | 0 | 0 | 0 | NA | NA | NA | NA | NA | NA | NA | NA | NA | NA |
| A330076C08 | 0 | 0 | 0 | 0 | 0 | 0 | 0 | 0 | 0 | 0 | 0 | 0 | 0 | 0 | 0 | NA | NA | NA | NA | NA | NA | NA | NA | NA | NA |
| A330076H08 | 1 | 2 | 6 | 3 | 3 | 5 | 1 | 2 | 6 | 2 | 3 | 5 | 3.36 | 3 | 3 | 1.197 | 0.259 | 0.81328995 | NA | 0.1456 | 0.9882 | 0.5645 | 0.5645 | TRUE | 0.9874 |
| A330093E20 | 0 | 0 | 0 | 1 | 0 | 0 | 0 | 0 | 0 | 1 | 0 | 0 | 0.13 | 0 | 0 | 1.859 | 0.894 | 0.82650734 | NA | 0 | 25.1515 | 10 | 10 | TRUE | 0.2563 |
| A330102I10I | 14 | 18 | 11 | 26 | 23 | 18 | 15 | 18 | 11 | 20 | 25 | 20 | 18.13 | 15 | 22 | 1.476 | 0.562 | 0.21630783 | 0.42832481 | 0 | 0.1886 | 0.0911 | 0.0911 | TRUE | 0.2105 |
| A3galt2 | 0 | 0 | 0 | 0 | 0 | 0 | 0 | 0 | 0 | 0 | 0 | 0 | 0 | 0 | 0 | NA | NA | NA | NA | NA | NA | NA | NA | NA | NA |
| A430005L14I | 1674 | 1835 | 1673 | 2629 | 1940 | 1792 | 1778 | 1833 | 1680 | 2067 | 2087 | 1962 | 1901.04 | 1764 | 2039 | 1.156 | 0.209 | 0.00651758 | 0.0381713 | 9.00E-04 | 0.0086 | 0.0037 | 0.0037 | TRUE | 0.0209 |
| A430033K04 | 175 | 240 | 246 | 292 | 205 | 219 | 186 | 240 | 247 | 230 | 221 | 240 | 227.07 | 224 | 230 | 1.025 | 0.035 | 0.82701166 | 0.91239703 | 0.0075 | 0.0214 | 0.0144 | 0.0144 | TRUE | 0.2421 |
| A430035B10 | 0 | 1 | 0 | 1 | 1 | 3 | 0 | 1 | 0 | 1 | 1 | 3 | 1.02 | 0 | 2 | 4.619 | 2.208 | 0.28640721 | NA | 0 | 3.2238 | 1.8007 | 1.8007 | TRUE | 0.4588 |
| A430071A18 | 0 | 0 | 1 | 1 | 0 | 0 | 0 | 0 | 1 | 1 | 0 | 0 | 0.3 | 0 | 0 | 0.954 | -0.067 | 0.98640617 | NA | 0 | 11.0483 | 9.2783 | 9.2783 | TRUE | 0.4098 |
| A430078G23 | 24 | 20 | 28 | 26 | 21 | 9 | 25 | 20 | 28 | 20 | 23 | 10 | 21.08 | 24 | 18 | 0.724 | -0.467 | 0.31199646 | 0.53255705 | 0.0284 | 0.1632 | 0.1051 | 0.1051 | TRUE | 0.674 |
| A430088P11 | 0 | 0 | 0 | 0 | 0 | 0 | 0 | 0 | 0 | 0 | 0 | 0 | 0 | 0 | 0 | NA | NA | NA | NA | NA | NA | NA | NA | NA | NA |
| A430089I19I | 0 | 0 | 0 | 0 | 0 | 1 | 0 | 0 | 0 | 0 | 0 | 1 | 0.18 | 0 | 0 | 1.859 | 0.894 | 0.82650734 | NA | 0 | 18.0643 | 10 | 10 | TRUE | 0.5383 |
| A430090L17I | 0 | 0 | 0 | 0 | 1 | 0 | 0 | 0 | 0 | 0 | 1 | 0 | 0.18 | 0 | 0 | 1.859 | 0.894 | 0.82650734 | NA | 0 | 18.3785 | 10 | 10 | TRUE | 0.5204 |
| A430092G05 | 0 | 0 | 0 | 0 | 0 | 0 | 0 | 0 | 0 | 0 | 0 | 0 | 0 | 0 | 0 | NA | NA | NA | NA | NA | NA | NA | NA | NA | NA |

| | | | | | | | | | | | | | | | | | | | | | | | | | |
|---|---|---|---|---|---|---|---|---|---|---|---|---|---|---|---|---|---|---|---|---|---|---|---|---|---|
| A430093F15 | 0 | 0 | 0 | 2 | 0 | 1 | 0 | 0 | 0 | 2 | 0 | 1 | 0.44 | 0 | 1 | 4.782 | 2.258 | 0.50653358 | NA | 0 | 7.4188 | 6.7775 | 6.7775 | TRUE | 0.2804 |
| A430105I19F | 63 | 88 | 76 | 61 | 57 | 73 | 67 | 88 | 76 | 48 | 61 | 80 | 70.05 | 77 | 63 | 0.812 | -0.3 | 0.27885525 | 0.49999026 | 0.0272 | 0.0539 | 0.0409 | 0.0409 | TRUE | 0.3464 |
| A430107P09 | 1 | 5 | 1 | 3 | 5 | 6 | 1 | 5 | 1 | 2 | 5 | 7 | 3.56 | 2 | 5 | 1.987 | 0.991 | 0.37391914 | NA | 0.1824 | 0.9321 | 0.5744 | 0.5744 | TRUE | 1.0206 |
| A4galt | 310 | 416 | 449 | 566 | 366 | 407 | 329 | 415 | 451 | 445 | 394 | 446 | 413.31 | 398 | 428 | 1.074 | 0.103 | 0.48956986 | 0.68569518 | 0.0125 | 0.0149 | 0.0137 | 0.0137 | TRUE | 0.2634 |
| A4gnt | 0 | 0 | 1 | 0 | 0 | 0 | 0 | 0 | 1 | 0 | 0 | 0 | 0.17 | 0 | 0 | 0.49 | -1.029 | 0.80087 | NA | 0 | 19.6928 | 10 | 10 | TRUE | 0.4098 |
| A530006G24 | 0 | 0 | 0 | 0 | 0 | 0 | 0 | 0 | 0 | 0 | 0 | 0 | 0 | 0 | 0 | NA | NA | NA | NA | NA | NA | NA | NA | NA | NA |
| A530013C23 | 0 | 0 | 0 | 1 | 0 | 0 | 0 | 0 | 0 | 1 | 0 | 0 | 0.13 | 0 | 0 | 1.859 | 0.894 | 0.82650734 | NA | 0 | 25.1515 | 10 | 10 | TRUE | 0.2563 |
| A530016L24 | 0 | 0 | 0 | 1 | 1 | 0 | 0 | 0 | 0 | 1 | 1 | 0 | 0.31 | 0 | 1 | 3.32 | 1.731 | 0.65383052 | NA | 0 | 10.6224 | 8.9541 | 8.9541 | TRUE | 0.202 |
| A530032D15 | 2 | 1 | 0 | 4 | 2 | 4 | 2 | 1 | 0 | 3 | 2 | 4 | 2.13 | 1 | 3 | 3.128 | 1.645 | 0.24439059 | NA | 0 | 1.5514 | 0.7915 | 0.7915 | TRUE | 0.3786 |
| A530046M15 | 0 | 0 | 0 | 1 | 1 | 0 | 0 | 0 | 0 | 1 | 1 | 0 | 0.31 | 0 | 1 | 3.32 | 1.731 | 0.65383052 | NA | 0 | 10.6224 | 8.9541 | 8.9541 | TRUE | 0.202 |
| A530050N04 | 1 | 0 | 0 | 0 | 0 | 0 | 1 | 0 | 0 | 0 | 0 | 0 | 0.18 | 0 | 0 | 0.49 | -1.029 | 0.80087 | NA | 0 | 18.6165 | 10 | 10 | TRUE | 0.4614 |
| A530053G22 | 0 | 0 | 0 | 0 | 0 | 0 | 0 | 0 | 0 | 0 | 0 | 0 | 0 | 0 | 0 | NA | NA | NA | NA | NA | NA | NA | NA | NA | NA |
| A530054K11 | 799 | 822 | 894 | 966 | 791 | 817 | 849 | 821 | 898 | 759 | 851 | 894 | 845.35 | 856 | 835 | 0.974 | -0.038 | 0.71551858 | 0.84564699 | 0.0035 | 0.0108 | 0.0065 | 0.0065 | TRUE | 0.0775 |
| A530058N18 | 0 | 0 | 0 | 0 | 0 | 0 | 0 | 0 | 0 | 0 | 0 | 0 | 0 | 0 | 0 | NA | NA | NA | NA | NA | NA | NA | NA | NA | NA |
| A530064D06 | 1 | 1 | 0 | 3 | 3 | 1 | 1 | 1 | 0 | 2 | 3 | 1 | 1.46 | 1 | 2 | 3.268 | 1.709 | 0.30761936 | NA | 0 | 2.2681 | 1.0545 | 1.0545 | TRUE | 0.2533 |
| A530065N20 | 0 | 0 | 0 | 0 | 0 | 0 | 0 | 0 | 0 | 0 | 0 | 0 | 0 | 0 | 0 | NA | NA | NA | NA | NA | NA | NA | NA | NA | NA |
| A530072M11 | 14 | 9 | 13 | 18 | 13 | 9 | 15 | 9 | 13 | 14 | 14 | 10 | 12.48 | 12 | 13 | 1.036 | 0.051 | 0.92471984 | 0.96292 | 0 | 0.2708 | 0.1307 | 0.1307 | TRUE | 0.2263 |
| A530088E08 | 0 | 5 | 1 | 0 | 2 | 1 | 0 | 5 | 1 | 0 | 2 | 1 | 1.54 | 2 | 1 | 0.52 | -0.942 | 0.61842699 | NA | 1.4564 | 2.1452 | 1.858 | 1.858 | TRUE | 1.0341 |
| A530099J19 | 0 | 0 | 0 | 0 | 0 | 0 | 0 | 0 | 0 | 0 | 0 | 0 | 0 | 0 | 0 | NA | NA | NA | NA | NA | NA | NA | NA | NA | NA |
| A630001G21 | 28 | 25 | 20 | 36 | 18 | 19 | 30 | 25 | 20 | 28 | 19 | 21 | 23.88 | 25 | 23 | 0.925 | -0.113 | 0.78085568 | 0.88649531 | 0 | 0.1449 | 0.0767 | 0.0767 | TRUE | 0.308 |
| A630007B06 | 1036 | 918 | 990 | 1240 | 864 | 909 | 1100 | 917 | 994 | 975 | 930 | 995 | 985.12 | 1004 | 967 | 0.963 | -0.054 | 0.59523904 | 0.76817352 | 0.0037 | 0.0102 | 0.0065 | 0.0065 | TRUE | 0.0843 |
| A630010A05 | 0 | 0 | 0 | 1 | 0 | 0 | 0 | 0 | 0 | 1 | 0 | 0 | 0.13 | 0 | 0 | 1.859 | 0.894 | 0.82650734 | NA | 0 | 25.1515 | 10 | 10 | TRUE | 0.2563 |
| A630012P03 | 0 | 0 | 0 | 0 | 0 | 0 | 0 | 0 | 0 | 0 | 0 | 0 | 0 | 0 | 0 | NA | NA | NA | NA | NA | NA | NA | NA | NA | NA |
| A630019I02 | 9 | 8 | 5 | 10 | 19 | 7 | 10 | 8 | 5 | 8 | 20 | 8 | 9.76 | 8 | 12 | 1.583 | 0.663 | 0.34094365 | 0.56119425 | 0.1389 | 0.3446 | 0.2398 | 0.2398 | TRUE | 1.4167 |
| A630020A06 | 46 | 50 | 53 | 91 | 44 | 41 | 49 | 50 | 53 | 72 | 47 | 45 | 52.63 | 51 | 55 | 1.087 | 0.12 | 0.68708738 | 0.8274849 | 0.0218 | 0.0695 | 0.0453 | 0.0453 | TRUE | 0.6926 |
| A630023A22 | 0 | 0 | 0 | 0 | 0 | 0 | 0 | 0 | 0 | 0 | 0 | 0 | 0 | 0 | 0 | NA | NA | NA | NA | NA | NA | NA | NA | NA | NA |
| A630033H20 | 0 | 0 | 0 | 0 | 0 | 0 | 0 | 0 | 0 | 0 | 0 | 0 | 0 | 0 | 0 | NA | NA | NA | NA | NA | NA | NA | NA | NA | NA |
| A630066F11 | 6 | 10 | 8 | 21 | 18 | 8 | 6 | 10 | 8 | 17 | 19 | 9 | 11.5 | 8 | 15 | 1.833 | 0.874 | 0.14559747 | 0.33399418 | 0 | 0.2933 | 0.1647 | 0.1647 | TRUE | 0.5307 |
| A630072M18 | 104 | 74 | 88 | 164 | 149 | 76 | 110 | 74 | 88 | 129 | 160 | 83 | 107.53 | 91 | 124 | 1.368 | 0.452 | 0.11512552 | 0.28648781 | 0.0628 | 0.0375 | 0.0497 | 0.0497 | TRUE | 0.2289 |
| A630073D07 | 0 | 0 | 0 | 0 | 0 | 0 | 0 | 0 | 0 | 0 | 0 | 0 | 0 | 0 | 0 | NA | NA | NA | NA | NA | NA | NA | NA | NA | NA |
| A630075F10 | 2 | 3 | 0 | 2 | 0 | 1 | 2 | 3 | 0 | 2 | 0 | 1 | 1.3 | 2 | 1 | 0.537 | -0.896 | 0.62933876 | NA | 0 | 2.5452 | 1.6535 | 1.6535 | TRUE | 0.5123 |
| A630076J17 | 0 | 0 | 0 | 0 | 0 | 0 | 0 | 0 | 0 | 0 | 0 | 0 | 0 | 0 | 0 | NA | NA | NA | NA | NA | NA | NA | NA | NA | NA |
| A630077J23 | 0 | 0 | 0 | 1 | 2 | 1 | 0 | 0 | 0 | 1 | 2 | 1 | 0.67 | 0 | 1 | 7.1 | 2.828 | 0.25693906 | NA | 0 | 4.9086 | 3.1164 | 3.1164 | TRUE | 0.1607 |
| A630089N07 | 267 | 125 | 144 | 279 | 192 | 132 | 284 | 125 | 145 | 219 | 207 | 144 | 187.24 | 185 | 190 | 1.034 | 0.048 | 0.88015942 | 0.94069759 | 0.1165 | 0.0245 | 0.0664 | 0.0664 | TRUE | 2.3722 |
| A630095E13 | 0 | 0 | 0 | 0 | 0 | 0 | 0 | 0 | 0 | 0 | 0 | 0 | 0 | 0 | 0 | NA | NA | NA | NA | NA | NA | NA | NA | NA | NA |
| A630095N17 | 3 | 2 | 2 | 2 | 0 | 0 | 3 | 2 | 2 | 2 | 0 | 0 | 1.46 | 2 | 1 | 0.246 | -2.023 | 0.25463936 | NA | 0 | 2.2622 | 1.233 | 1.233 | TRUE | 0.9709 |
| A730008H23 | 0 | 0 | 0 | 0 | 0 | 0 | 0 | 0 | 0 | 0 | 0 | 0 | 0 | 0 | 0 | NA | NA | NA | NA | NA | NA | NA | NA | NA | NA |
| A730017C20 | 11 | 9 | 3 | 45 | 30 | 18 | 12 | 9 | 3 | 35 | 32 | 20 | 18.51 | 8 | 29 | 3.719 | 1.895 | 0.000854 | 0.00787808 | 0.0961 | 0.1849 | 0.151 | 0.151 | TRUE | 0.8644 |
| A730017L22 | 29 | 8 | 22 | 25 | 14 | 11 | 31 | 8 | 22 | 20 | 15 | 12 | 17.94 | 20 | 16 | 0.775 | -0.367 | 0.52409184 | 0.71387655 | 0.1752 | 0.1905 | 0.1831 | 0.1831 | TRUE | 0.2923 |
| A730018C14 | 0 | 0 | 1 | 0 | 0 | 0 | 0 | 0 | 1 | 0 | 0 | 0 | 0.17 | 0 | 0 | 0.49 | -1.029 | 0.80087 | NA | 0 | 19.6928 | 10 | 10 | TRUE | 0.4098 |
| A730020E08 | 0 | 2 | 0 | 1 | 1 | 0 | 0 | 2 | 0 | 1 | 1 | 0 | 0.64 | 1 | 1 | 0.935 | -0.096 | 0.97106576 | NA | 0 | 5.1285 | 3.5446 | 3.5446 | TRUE | 0.9738 |
| A730020M07 | 2 | 0 | 1 | 1 | 1 | 0 | 2 | 0 | 1 | 1 | 1 | 0 | 0.83 | 1 | 1 | 0.608 | -0.719 | 0.74058884 | NA | 0 | 3.9678 | 2.1265 | 2.1265 | TRUE | 0.1445 |
| A730036I17 | 19 | 21 | 21 | 43 | 30 | 20 | 20 | 21 | 21 | 34 | 32 | 22 | 25.04 | 21 | 29 | 1.421 | 0.507 | 0.19495837 | 0.40087767 | 0 | 0.1385 | 0.069 | 0.069 | TRUE | 0.2994 |
| A730043L09 | 0 | 1 | 0 | 0 | 0 | 0 | 0 | 1 | 0 | 0 | 0 | 0 | 0.17 | 0 | 0 | 0.49 | -1.029 | 0.80087 | NA | 0 | 19.7991 | 10 | 10 | TRUE | 0.405 |
| A730046J19 | 19 | 0 | 3 | 9 | 7 | 1 | 20 | 0 | 3 | 7 | 8 | 1 | 6.48 | 8 | 5 | 0.684 | -0.549 | 0.6722841 | 0.8172703 | 1.7627 | 0.5151 | 1.0542 | 1.0542 | TRUE | 2.2994 |
| A730056A06 | 0 | 1 | 1 | 0 | 1 | 0 | 0 | 1 | 1 | 0 | 1 | 0 | 0.51 | 1 | 0 | 0.52 | -0.943 | 0.73680291 | NA | 0 | 6.4273 | 3.9121 | 3.9121 | TRUE | 0.5204 |
| A730082K24 | 6 | 7 | 2 | 3 | 5 | 4 | 6 | 7 | 2 | 2 | 5 | 4 | 4.58 | 5 | 4 | 0.776 | -0.366 | 0.6933452 | NA | 0 | 0.726 | 0.3965 | 0.3965 | TRUE | 0.5927 |
| A730085A09 | 61 | 71 | 77 | 225 | 174 | 156 | 65 | 71 | 77 | 177 | 187 | 171 | 124.65 | 71 | 178 | 2.509 | 1.327 | 1.37E-12 | 1.37E-10 | 0 | 0.0333 | 0.0153 | 0.0153 | TRUE | 0.0545 |
| A730085K08 | 0 | 0 | 0 | 0 | 0 | 0 | 0 | 0 | 0 | 0 | 0 | 0 | 0 | 0 | 0 | NA | NA | NA | NA | NA | NA | NA | NA | NA | NA |
| A730090H04 | 6 | 5 | 4 | 4 | 4 | 3 | 6 | 5 | 4 | 3 | 4 | 3 | 4.35 | 5 | 3 | 0.695 | -0.525 | 0.55004483 | NA | 0 | 0.7638 | 0.319 | 0.319 | TRUE | 0.0883 |
| A730090N16 | 4 | 0 | 2 | 5 | 5 | 4 | 4 | 0 | 2 | 4 | 5 | 4 | 3.32 | 2 | 4 | 2.201 | 1.138 | 0.32531367 | NA | 0 | 0.9979 | 0.6098 | 0.6098 | TRUE | 0.422 |
| A730098P11 | 222 | 280 | 264 | 330 | 249 | 245 | 236 | 280 | 265 | 259 | 268 | 268 | 262.67 | 260 | 265 | 1.018 | 0.026 | 0.84859986 | 0.92431747 | 0 | 0.0194 | 0.0093 | 0.0093 | TRUE | 0.0737 |
| A830009L08 | 0 | 0 | 0 | 0 | 0 | 0 | 0 | 0 | 0 | 0 | 0 | 0 | 0 | 0 | 0 | NA | NA | NA | NA | NA | NA | NA | NA | NA | NA |
| A830010M20 | 347 | 465 | 497 | 616 | 475 | 450 | 369 | 464 | 499 | 484 | 511 | 493 | 469.99 | 444 | 496 | 1.116 | 0.158 | 0.25464953 | 0.47273216 | 0.0099 | 0.0139 | 0.0118 | 0.0118 | TRUE | 0.2534 |
| A830018L16 | 81 | 134 | 132 | 156 | 109 | 127 | 86 | 134 | 133 | 123 | 117 | 139 | 121.89 | 118 | 126 | 1.073 | 0.102 | 0.65362023 | 0.80540538 | 0.0235 | 0.0339 | 0.0289 | 0.0289 | TRUE | 0.5413 |
| A830019L24 | 0 | 0 | 0 | 0 | 0 | 0 | 0 | 0 | 0 | 0 | 0 | 0 | 0 | 0 | 0 | NA | NA | NA | NA | NA | NA | NA | NA | NA | NA |
| A830052D11 | 2 | 1 | 0 | 1 | 0 | 0 | 2 | 1 | 0 | 1 | 0 | 0 | 0.65 | 1 | 0 | 0.337 | -1.57 | 0.54866551 | NA | 0 | 5.0634 | 3.4431 | 3.4431 | TRUE | 0.1066 |
| A830080D01 | 1714 | 2267 | 2322 | 2774 | 2063 | 2048 | 1821 | 2264 | 2332 | 2181 | 2220 | 2242 | 2176.4 | 2139 | 2214 | 1.035 | 0.05 | 0.65497632 | 0.80598319 | 0.0084 | 0.0084 | 0.0084 | 0.0084 | TRUE | 0.2044 |
| A830082K12 | 2 | 1 | 1 | 4 | 1 | 1 | 2 | 1 | 1 | 3 | 1 | 1 | 1.57 | 1 | 2 | 1.334 | 0.415 | 0.77612482 | NA | 0 | 2.1003 | 0.8974 | 0.8974 | TRUE | 0.4861 |
| A830082N09 | 32 | 25 | 24 | 51 | 32 | 22 | 34 | 25 | 24 | 40 | 34 | 24 | 30.28 | 28 | 33 | 1.198 | 0.26 | 0.49103263 | 0.68736724 | 2.00E-04 | 0.1157 | 0.0695 | 0.0695 | TRUE | 0.356 |
| A930001A20 | 0 | 0 | 0 | 0 | 0 | 0 | 0 | 0 | 0 | 0 | 0 | 0 | 0 | 0 | 0 | NA | NA | NA | NA | NA | NA | NA | NA | NA | NA |
| A930001C03 | 25 | 40 | 40 | 35 | 27 | 31 | 27 | 40 | 40 | 28 | 29 | 34 | 32.86 | 36 | 30 | 0.844 | -0.245 | 0.48399142 | 0.68159234 | 0 | 0.1072 | 0.0574 | 0.0574 | TRUE | 0.3386 |
| A930003A15 | 0 | 0 | 0 | 0 | 0 | 0 | 0 | 0 | 0 | 0 | 0 | 0 | 0 | 0 | 0 | NA | NA | NA | NA | NA | NA | NA | NA | NA | NA |
| A930003O13 | 0 | 0 | 0 | 0 | 0 | 0 | 0 | 0 | 0 | 0 | 0 | 0 | 0 | 0 | 0 | NA | NA | NA | NA | NA | NA | NA | NA | NA | NA |
| A930004D18 | 125 | 135 | 147 | 330 | 271 | 183 | 133 | 135 | 148 | 259 | 292 | 200 | 194.42 | 139 | 250 | 1.811 | 0.856 | 4.89E-06 | 0.00010833 | 0.0158 | 0.0238 | 0.0197 | 0.0197 | TRUE | 0.2645 |
| A930005H10 | 162 | 125 | 173 | 256 | 157 | 148 | 172 | 125 | 174 | 201 | 169 | 162 | 167.13 | 157 | 177 | 1.135 | 0.182 | 0.35547128 | 0.57387109 | 0.0174 | 0.0266 | 0.022 | 0.022 | TRUE | 0.3394 |
| A930006I01 | 0 | 0 | 0 | 0 | 0 | 0 | 0 | 0 | 0 | 0 | 0 | 0 | 0 | 0 | 0 | NA | NA | NA | NA | NA | NA | NA | NA | NA | NA |
| A930006K02 | 8 | 3 | 3 | 9 | 1 | 9 | 8 | 3 | 3 | 7 | 1 | 10 | 5.42 | 5 | 6 | 1.251 | 0.323 | 0.74207613 | 0.86204793 | 0.3862 | 0.6149 | 0.5065 | 0.5065 | TRUE | 0.4557 |
| A930007I19 | 4 | 6 | 6 | 4 | 4 | 8 | 4 | 6 | 6 | 3 | 4 | 9 | 5.41 | 5 | 5 | 0.977 | -0.033 | 0.96841012 | 0.98419772 | 0 | 0.6157 | 0.3108 | 0.3108 | TRUE | 0.5902 |

| | | | | | | | | | | | | | | | | | | | | | | | | | |
|---|---|---|---|---|---|---|---|---|---|---|---|---|---|---|---|---|---|---|---|---|---|---|---|---|---|
| A930009A15 | 0 | 1 | 0 | 0 | 0 | 0 | 0 | 1 | 0 | 0 | 0 | 0 | 0.17 | 0 | 0 | 0.49 | -1.029 | 0.80087 | NA | 0 | 19.7991 | 10 | 10 | TRUE | 0.405 |
| A930011G23 | 0 | 1 | 0 | 0 | 0 | 0 | 0 | 1 | 0 | 0 | 0 | 0 | 0.17 | 0 | 0 | 0.49 | -1.029 | 0.80087 | NA | 0 | 19.7991 | 10 | 10 | TRUE | 0.405 |
| A930011O12 | 89 | 96 | 96 | 99 | 94 | 74 | 95 | 96 | 96 | 78 | 101 | 81 | 91.13 | 96 | 87 | 0.903 | -0.147 | 0.49093609 | 0.68729309 | 0 | 0.0431 | 0.0217 | 0.0217 | TRUE | 0.1959 |
| A930012L18I | 4 | 2 | 1 | 2 | 4 | 1 | 4 | 2 | 1 | 2 | 4 | 1 | 2.37 | 2 | 2 | 0.956 | -0.065 | 0.95898257 | NA | 0 | 1.3969 | 0.7284 | 0.7284 | TRUE | 0.5234 |
| A930013F10 | 164 | 142 | 164 | 172 | 168 | 147 | 174 | 142 | 165 | 135 | 181 | 161 | 159.6 | 160 | 159 | 0.989 | -0.016 | 0.93068229 | 0.96589597 | 0.0104 | 0.0275 | 0.0188 | 0.0188 | TRUE | 0.1975 |
| A930015D03 | 98 | 70 | 83 | 127 | 99 | 57 | 104 | 70 | 83 | 100 | 107 | 62 | 87.68 | 86 | 90 | 1.048 | 0.068 | 0.80936834 | 0.90182956 | 0.0461 | 0.0445 | 0.0452 | 0.0452 | TRUE | 0.6409 |
| A930016O22 | 24 | 33 | 34 | 19 | 20 | 33 | 25 | 33 | 34 | 15 | 22 | 36 | 27.53 | 31 | 24 | 0.774 | -0.37 | 0.39551074 | 0.60894137 | 0.0705 | 0.1266 | 0.0998 | 0.0998 | TRUE | 0.7281 |
| A930017M01 | 6 | 1 | 3 | 7 | 1 | 7 | 6 | 1 | 3 | 6 | 1 | 8 | 4.1 | 3 | 5 | 1.383 | 0.468 | 0.66856986 | NA | 0.3877 | 0.8096 | 0.6111 | 0.6111 | TRUE | 0.4207 |
| A930018P22 | 0 | 0 | 1 | 1 | 1 | 1 | 0 | 0 | 1 | 1 | 1 | 1 | 0.66 | 0 | 1 | 2.676 | 1.42 | 0.53608725 | NA | 0 | 4.9975 | 2.297 | 2.297 | TRUE | 0.4098 |
| A930019D19 | 0 | 0 | 0 | 0 | 0 | 0 | 0 | 0 | 0 | 0 | 0 | 0 | 0 | 0 | 0 | NA | NA | NA | NA | NA | NA | NA | NA | NA | NA |
| A930024E05 | 70 | 77 | 83 | 66 | 32 | 50 | 74 | 77 | 83 | 52 | 34 | 55 | 62.61 | 78 | 47 | 0.603 | -0.729 | 0.00684978 | 0.03951526 | 0.0028 | 0.0595 | 0.0355 | 0.0355 | TRUE | 0.406 |
| A930041C12 | 0 | 0 | 0 | 0 | 0 | 0 | 0 | 0 | 0 | 0 | 0 | 0 | 0 | 0 | 0 | NA | NA | NA | NA | NA | NA | NA | NA | NA | NA |
| AA387883 | 0 | 0 | 0 | 0 | 0 | 0 | 0 | 0 | 0 | 0 | 0 | 0 | 0 | 0 | 0 | NA | NA | NA | NA | NA | NA | NA | NA | NA | NA |
| AA388235 | 43 | 18 | 20 | 45 | 27 | 16 | 46 | 18 | 20 | 35 | 29 | 18 | 27.61 | 28 | 27 | 0.987 | -0.018 | 0.97071759 | 0.98472179 | 0.1586 | 0.1262 | 0.142 | 0.142 | TRUE | 1.3896 |
| AA414768 | 100 | 61 | 59 | 330 | 160 | 101 | 106 | 61 | 59 | 259 | 172 | 111 | 128.09 | 75 | 181 | 2.405 | 1.266 | 0.00031197 | 0.0035171 | 0.1339 | 0.0326 | 0.0793 | 0.0793 | TRUE | 0.153 |
| AA415398 | 21 | 20 | 21 | 29 | 19 | 10 | 22 | 20 | 21 | 23 | 20 | 11 | 19.59 | 21 | 18 | 0.864 | -0.21 | 0.64412826 | 0.79987331 | 0 | 0.1751 | 0.0986 | 0.0986 | TRUE | 0.5538 |
| AA465934 | 397 | 341 | 272 | 600 | 358 | 308 | 422 | 341 | 273 | 472 | 385 | 337 | 371.56 | 345 | 398 | 1.156 | 0.209 | 0.28852287 | 0.50952666 | 0.0348 | 0.0158 | 0.0252 | 0.0252 | TRUE | 0.2721 |
| AA467197 | 52 | 49 | 36 | 87 | 86 | 67 | 55 | 49 | 36 | 68 | 93 | 73 | 62.43 | 47 | 78 | 1.667 | 0.737 | 0.00856905 | 0.0468461 | 0.0168 | 0.0597 | 0.0394 | 0.0394 | TRUE | 0.316 |
| AA474331 | 66 | 81 | 70 | 84 | 68 | 67 | 70 | 81 | 70 | 66 | 73 | 73 | 72.3 | 74 | 71 | 0.958 | -0.063 | 0.78323151 | 0.88813914 | 0 | 0.0525 | 0.0235 | 0.0235 | TRUE | 0.066 |
| AA536875 | 0 | 0 | 0 | 0 | 0 | 0 | 0 | 0 | 0 | 0 | 0 | 0 | 0 | 0 | 0 | NA | NA | NA | NA | NA | NA | NA | NA | NA | NA |
| AA543186 | 6 | 6 | 0 | 6 | 11 | 6 | 6 | 6 | 0 | 5 | 12 | 7 | 5.91 | 4 | 8 | 1.862 | 0.897 | 0.37076729 | 0.58760362 | 0.4804 | 0.5639 | 0.5346 | 0.5346 | TRUE | 1.3331 |
| AA545190 | 0 | 0 | 0 | 0 | 0 | 0 | 0 | 0 | 0 | 0 | 0 | 0 | 0 | 0 | 0 | NA | NA | NA | NA | NA | NA | NA | NA | NA | NA |
| AA619741 | 3 | 1 | 0 | 4 | 2 | 0 | 3 | 1 | 0 | 3 | 2 | 0 | 1.58 | 1 | 2 | 1.303 | 0.381 | 0.8294413 | NA | 0.8421 | 2.0916 | 1.6187 | 1.6187 | TRUE | 0.6532 |
| AA792892 | 0 | 0 | 0 | 0 | 0 | 0 | 0 | 0 | 0 | 0 | 0 | 0 | 0 | 0 | 0 | NA | NA | NA | NA | NA | NA | NA | NA | NA | NA |
| AA986860 | 57 | 51 | 48 | 62 | 40 | 32 | 61 | 51 | 48 | 49 | 43 | 35 | 47.75 | 53 | 42 | 0.8 | -0.322 | 0.26702726 | 0.48677231 | 0 | 0.0759 | 0.0397 | 0.0397 | TRUE | 0.163 |
| AA987161 | 1671 | 1817 | 1871 | 2096 | 1658 | 1835 | 1775 | 1815 | 1879 | 1648 | 1784 | 2009 | 1818.09 | 1823 | 1814 | 0.994 | -0.009 | 0.93095176 | 0.96610976 | 0.0049 | 0.0087 | 0.0064 | 0.0064 | TRUE | 0.1069 |
| Aaas | 5699 | 5976 | 5878 | 7334 | 5391 | 5583 | 6053 | 5968 | 5902 | 5765 | 5800 | 6111 | 5933.51 | 5974 | 5892 | 0.986 | -0.02 | 0.74172899 | 0.86177162 | 4.00E-04 | 0.0075 | 0.0025 | 0.0025 | TRUE | 0.0128 |
| Aacs | 3427 | 2535 | 2575 | 3177 | 2492 | 2330 | 3640 | 2532 | 2586 | 2498 | 2681 | 2551 | 2747.78 | 2919 | 2577 | 0.883 | -0.18 | 0.2191186 | 0.43131916 | 0.023 | 0.0081 | 0.0151 | 0.0151 | TRUE | 0.5663 |
| Aadac | 2 | 0 | 0 | 1 | 1 | 1 | 2 | 0 | 0 | 1 | 1 | 1 | 0.85 | 1 | 1 | 1.405 | 0.49 | 0.82086568 | NA | 0 | 3.8972 | 2.1473 | 2.1473 | TRUE | 0.9744 |
| Aadacl2 | 1 | 0 | 1 | 6 | 2 | 0 | 1 | 0 | 1 | 5 | 2 | 0 | 1.49 | 1 | 2 | 3.422 | 1.775 | 0.34604143 | NA | 0 | 2.2192 | 1.6012 | 1.6012 | TRUE | 0.1068 |
| Aadacl3 | 1 | 0 | 0 | 2 | 1 | 1 | 1 | 0 | 0 | 2 | 1 | 1 | 0.8 | 0 | 1 | 3.436 | 1.781 | 0.40868918 | NA | 0 | 4.1207 | 1.9586 | 1.9586 | TRUE | 0.4614 |
| Aadat | 18 | 11 | 12 | 27 | 12 | 10 | 19 | 11 | 12 | 21 | 13 | 11 | 14.54 | 14 | 15 | 1.087 | 0.12 | 0.82499629 | 0.91128174 | 0.0498 | 0.2335 | 0.143 | 0.143 | TRUE | 0.7214 |
| Aaed1 | 480 | 635 | 609 | 728 | 568 | 571 | 510 | 634 | 612 | 572 | 611 | 625 | 594 | 585 | 603 | 1.029 | 0.041 | 0.73373122 | 0.85735379 | 0.0059 | 0.0124 | 0.0088 | 0.0088 | TRUE | 0.1475 |
| Aagab | 3237 | 3563 | 3458 | 4236 | 3232 | 3160 | 3438 | 3558 | 3472 | 3330 | 3477 | 3459 | 3455.9 | 3489 | 3422 | 0.98 | -0.029 | 0.64898782 | 0.8032811 | 0 | 0.0079 | 0.0026 | 0.0026 | TRUE | 0.0069 |
| Aak1 | 2184 | 1794 | 1633 | 3444 | 2141 | 1964 | 2320 | 1792 | 1640 | 2707 | 2304 | 2150 | 2152.01 | 1917 | 2387 | 1.246 | 0.317 | 0.03549768 | 0.12941966 | 0.0239 | 0.0084 | 0.0159 | 0.0159 | TRUE | 0.4077 |
| Aamdc | 276 | 383 | 354 | 360 | 283 | 257 | 293 | 383 | 355 | 283 | 304 | 281 | 316.66 | 344 | 289 | 0.842 | -0.249 | 0.08576411 | 0.23605624 | 0.0071 | 0.0173 | 0.0119 | 0.0119 | TRUE | 0.1865 |
| Aamp | 6046 | 5665 | 5739 | 8580 | 6274 | 5995 | 6422 | 5658 | 5763 | 6745 | 6750 | 6562 | 6316.67 | 5948 | 6686 | 1.124 | 0.169 | 0.0278951 | 0.10967118 | 0.0023 | 0.0074 | 0.0041 | 0.0041 | TRUE | 0.0593 |
| Aanat | 0 | 3 | 1 | 4 | 3 | 1 | 0 | 3 | 1 | 3 | 3 | 1 | 1.91 | 1 | 2 | 1.866 | 0.9 | 0.54192208 | NA | 0 | 1.7307 | 0.9931 | 0.9931 | TRUE | 0.7355 |
| Aar2 | 1999 | 1649 | 1621 | 2720 | 1776 | 1505 | 2123 | 1647 | 1628 | 2138 | 1911 | 1647 | 1849.07 | 1799 | 1899 | 1.056 | 0.079 | 0.58390665 | 0.75932524 | 0.0194 | 0.0087 | 0.0143 | 0.0143 | TRUE | 0.2992 |
| Aard | 1390 | 834 | 702 | 3109 | 1754 | 1135 | 1476 | 833 | 705 | 2444 | 1887 | 1242 | 1431.32 | 1005 | 1858 | 1.849 | 0.887 | 0.04164394 | 0.14505482 | 0.1359 | 0.0092 | 0.0583 | 0.1359 | TRUE | 0.2673 |
| Aars | 12395 | 11419 | 11396 | 21778 | 15294 | 13210 | 13166 | 11404 | 11443 | 17120 | 16455 | 14460 | 14008.12 | 12004 | 16012 | 1.334 | 0.416 | 3.31E-05 | 0.00054853 | 0.0071 | 0.0071 | 0.0072 | 0.0072 | TRUE | 0.0878 |
| Aars2 | 3089 | 2829 | 2787 | 3607 | 2728 | 2432 | 3281 | 2825 | 2798 | 2836 | 2935 | 2662 | 2889.63 | 2968 | 2811 | 0.947 | -0.078 | 0.40936228 | 0.62130006 | 0.0049 | 0.008 | 0.0061 | 0.0061 | TRUE | 0.103 |
| Aarsd1 | 3615 | 4779 | 4732 | 5520 | 4072 | 4272 | 3840 | 4773 | 4751 | 4339 | 4381 | 4676 | 4460.17 | 4455 | 4465 | 1.002 | 0.003 | 0.9750478 | 0.98697924 | 0.0081 | 0.0076 | 0.0079 | 0.0079 | TRUE | 0.1773 |
| Aasdh | 1051 | 1622 | 1580 | 1534 | 1250 | 1337 | 1116 | 1620 | 1586 | 1206 | 1345 | 1464 | 1389.52 | 1441 | 1338 | 0.928 | -0.108 | 0.49351446 | 0.68986174 | 0.0247 | 0.0093 | 0.0171 | 0.0171 | TRUE | 0.4662 |
| Aasdhppt | 2926 | 3259 | 3183 | 3846 | 2536 | 2848 | 3108 | 3255 | 3196 | 3023 | 2729 | 3118 | 3071.4 | 3186 | 2957 | 0.928 | -0.108 | 0.18146918 | 0.38330413 | 0.0021 | 0.008 | 0.0044 | 0.0044 | TRUE | 0.0547 |
| Aass | 615 | 944 | 907 | 1027 | 788 | 1149 | 653 | 943 | 911 | 807 | 848 | 1258 | 903.28 | 836 | 971 | 1.161 | 0.215 | 0.28747689 | 0.5087006 | 0.0484 | 0.0106 | 0.0285 | 0.0285 | TRUE | 0.7937 |
| Aatf | 2751 | 2719 | 2795 | 4442 | 3185 | 2744 | 2922 | 2715 | 2806 | 3492 | 3427 | 3004 | 3061.09 | 2814 | 3308 | 1.176 | 0.233 | 0.0083644 | 0.04604652 | 0.0034 | 0.008 | 0.0053 | 0.0053 | TRUE | 0.078 |
| Aatk | 117 | 147 | 126 | 145 | 116 | 101 | 124 | 147 | 127 | 114 | 125 | 111 | 124.49 | 133 | 117 | 0.878 | -0.188 | 0.30353569 | 0.52431834 | 0 | 0.0334 | 0.0161 | 0.0161 | TRUE | 0.092 |
| AB099516 | 0 | 0 | 0 | 0 | 0 | 1 | 0 | 0 | 0 | 0 | 0 | 1 | 0.18 | 0 | 0 | 1.859 | 0.894 | 0.82650734 | NA | 0 | 18.0643 | 10 | 10 | TRUE | 0.5383 |
| AB124611 | 2 | 0 | 2 | 2 | 1 | 2 | 2 | 0 | 2 | 2 | 1 | 2 | 1.5 | 1 | 2 | 1.174 | 0.232 | 0.88198289 | NA | 0 | 2.2105 | 1.0816 | 1.0816 | TRUE | 0.4969 |
| Abat | 357 | 275 | 266 | 505 | 302 | 254 | 379 | 275 | 267 | 397 | 325 | 278 | 320.15 | 307 | 333 | 1.089 | 0.123 | 0.5331578 | 0.72111868 | 0.0324 | 0.0172 | 0.0248 | 0.0248 | TRUE | 0.4689 |
| Abca1 | 1424 | 1515 | 1710 | 1180 | 794 | 1006 | 1513 | 1513 | 1717 | 928 | 854 | 1101 | 1270.96 | 1581 | 961 | 0.608 | -0.719 | 3.24E-09 | 1.73E-07 | 0.0102 | 0.0095 | 0.0098 | 0.0098 | TRUE | 0.1911 |
| Abca12 | 8 | 1 | 2 | 12 | 4 | 9 | 8 | 1 | 2 | 9 | 4 | 10 | 5.85 | 4 | 8 | 2.077 | 1.054 | 0.27076961 | 0.49074002 | 0.3312 | 0.5702 | 0.4642 | 0.4642 | TRUE | 1.7413 |
| Abca13 | 92 | 90 | 52 | 141 | 95 | 87 | 98 | 90 | 52 | 111 | 102 | 95 | 91.35 | 80 | 103 | 1.289 | 0.366 | 0.17707403 | 0.37761108 | 0.0404 | 0.043 | 0.0418 | 0.0418 | TRUE | 0.8619 |
| Abca14 | 1 | 2 | 0 | 5 | 2 | 0 | 1 | 2 | 0 | 4 | 2 | 0 | 1.52 | 1 | 2 | 2.033 | 1.024 | 0.56825104 | NA | 0 | 2.1691 | 1.5832 | 1.5832 | TRUE | 0.1342 |
| Abca15 | 0 | 0 | 0 | 1 | 0 | 0 | 0 | 0 | 0 | 1 | 0 | 0 | 0.13 | 0 | 0 | 1.859 | 0.894 | 0.82650734 | NA | 0 | 25.1515 | 10 | 10 | TRUE | 0.2563 |
| Abca16 | 12 | 8 | 23 | 45 | 48 | 32 | 13 | 8 | 23 | 35 | 52 | 35 | 27.65 | 15 | 41 | 2.777 | 1.473 | 0.00159035 | 0.01283673 | 0.0871 | 0.1261 | 0.1099 | 0.1099 | TRUE | 1.1751 |
| Abca17 | 1 | 4 | 1 | 5 | 6 | 5 | 1 | 4 | 1 | 4 | 6 | 5 | 3.65 | 2 | 5 | 2.583 | 1.369 | 0.19724476 | NA | 0 | 0.9086 | 0.4675 | 0.4675 | TRUE | 0.6742 |
| Abca2 | 1985 | 2474 | 2534 | 2808 | 2109 | 2153 | 2108 | 2471 | 2544 | 2207 | 2269 | 2357 | 2326.17 | 2374 | 2278 | 0.959 | -0.061 | 0.53407984 | 0.72173954 | 0.0045 | 0.0083 | 0.0064 | 0.0064 | TRUE | 0.1161 |
| Abca3 | 1801 | 1920 | 2085 | 3045 | 2161 | 1971 | 1913 | 1918 | 2094 | 2394 | 2325 | 2158 | 2133.41 | 1975 | 2292 | 1.161 | 0.216 | 0.01010643 | 0.0531506 | 0.0021 | 0.0084 | 0.0046 | 0.0046 | TRUE | 0.0336 |
| Abca4 | 145 | 128 | 151 | 230 | 169 | 168 | 154 | 128 | 152 | 181 | 182 | 184 | 163.34 | 145 | 182 | 1.261 | 0.335 | 0.03933156 | 0.13899559 | 0 | 0.0271 | 0.0128 | 0.0128 | TRUE | 0.1071 |
| Abca5 | 1744 | 1602 | 1510 | 1953 | 1480 | 1350 | 1852 | 1600 | 1516 | 1535 | 1592 | 1478 | 1595.67 | 1656 | 1535 | 0.927 | -0.109 | 0.28655948 | 0.50793617 | 0.0053 | 0.009 | 0.0069 | 0.0069 | TRUE | 0.1292 |
| Abca6 | 34 | 19 | 4 | 22 | 18 | 5 | 36 | 19 | 4 | 17 | 19 | 5 | 16.87 | 20 | 14 | 0.717 | -0.479 | 0.53563871 | 0.72271961 | 0.5561 | 0.2021 | 0.3708 | 0.3708 | TRUE | 0.1626 |
| Abca7 | 647 | 994 | 989 | 710 | 682 | 767 | 687 | 993 | 993 | 558 | 734 | 840 | 800.76 | 891 | 711 | 0.796 | -0.329 | 0.08776229 | 0.24001852 | 0.041 | 0.011 | 0.0254 | 0.0254 | TRUE | 0.4752 |
| Abca8a | 1 | 8 | 1 | 12 | 18 | 9 | 1 | 8 | 1 | 9 | 19 | 10 | 8.12 | 3 | 13 | 3.803 | 1.927 | 0.03394319 | 0.12537516 | 0.394 | 0.4127 | 0.4056 | 0.4056 | TRUE | 2.1291 |
| Abca8b | 60 | 32 | 27 | 39 | 24 | 19 | 64 | 32 | 27 | 31 | 26 | 21 | 33.35 | 41 | 26 | 0.634 | -0.657 | 0.13031606 | 0.31086811 | 0.1037 | 0.1057 | 0.1046 | 0.1046 | TRUE | 1.7561 |
| Abca9 | 4 | 1 | 0 | 7 | 1 | 3 | 4 | 1 | 0 | 6 | 1 | 3 | 2.52 | 2 | 3 | 1.928 | 0.947 | 0.50333643 | NA | 0.5948 | 1.3151 | 1.002 | 1.002 | TRUE | 0.4315 |
| Abcb10 | 2659 | 2761 | 2821 | 3363 | 2438 | 2401 | 2824 | 2757 | 2833 | 2644 | 2623 | 2628 | 2718.25 | 2805 | 2632 | 0.938 | -0.092 | 0.13734866 | 0.32135635 | 0 | 0.0081 | 0.0024 | 0.0024 | TRUE | 0.0027 |
| Abcb11 | 1 | 0 | 0 | 11 | 10 | 5 | 1 | 0 | 0 | 9 | 11 | 5 | 4.32 | 0 | 8 | 22.696 | 4.504 | 0.00190229 | NA | 0 | 0.7689 | 0.4575 | 0.4575 | TRUE | 0.43 |

| | | | | | | | | | | | | | | | | | | | | | | | | | |
|---|---|---|---|---|---|---|---|---|---|---|---|---|---|---|---|---|---|---|---|---|---|---|---|---|---|
| Abcb1a | 799 | 611 | 498 | 1385 | 1099 | 948 | 849 | 610 | 500 | 1089 | 1182 | 1038 | 877.98 | 653 | 1103 | 1.69 | 0.757 | 3.56E-05 | 0.00057814 | 0.0376 | 0.0107 | 0.0229 | 0.0229 | TRUE | 0.8077 |
| Abcb1b | 1197 | 1075 | 976 | 2046 | 1216 | 954 | 1271 | 1074 | 980 | 1608 | 1308 | 1044 | 1214.35 | 1108 | 1320 | 1.192 | 0.254 | 0.13731665 | 0.32135635 | 0.0306 | 0.0096 | 0.0202 | 0.0202 | TRUE | 0.1865 |
| Abcb4 | 25 | 40 | 33 | 52 | 32 | 54 | 27 | 40 | 33 | 41 | 34 | 59 | 39.01 | 33 | 45 | 1.344 | 0.426 | 0.2280141 | 0.44159023 | 0.0339 | 0.0914 | 0.0637 | 0.0637 | TRUE | 0.5719 |
| Abcb5 | 46 | 15 | 14 | 133 | 85 | 28 | 49 | 15 | 14 | 105 | 91 | 31 | 50.76 | 26 | 76 | 2.925 | 1.548 | 0.00734777 | 0.04158202 | 0.3891 | 0.0718 | 0.2142 | 0.2142 | TRUE | 1.8533 |
| Abcb6 | 1659 | 1901 | 1885 | 1551 | 1175 | 1245 | 1762 | 1899 | 1893 | 1219 | 1264 | 1363 | 1566.63 | 1851 | 1282 | 0.692 | -0.531 | 4.35E-10 | 2.88E-08 | 0.0018 | 0.009 | 0.0046 | 0.0046 | TRUE | 0.0361 |
| Abcb7 | 4120 | 4371 | 4483 | 5139 | 3621 | 3907 | 4376 | 4365 | 4501 | 4040 | 3896 | 4277 | 4242.6 | 4414 | 4071 | 0.922 | -0.117 | 0.08960577 | 0.24345601 | 9.00E-04 | 0.0077 | 0.0032 | 0.0032 | TRUE | 0.0235 |
| Abcb8 | 991 | 1380 | 1463 | 1564 | 1130 | 1215 | 1053 | 1378 | 1469 | 1229 | 1216 | 1330 | 1279.19 | 1300 | 1258 | 0.968 | -0.047 | 0.72865634 | 0.85400667 | 0.0154 | 0.0095 | 0.0126 | 0.0126 | TRUE | 0.3317 |
| Abcb9 | 36 | 27 | 30 | 35 | 30 | 18 | 38 | 27 | 30 | 28 | 32 | 20 | 29.14 | 32 | 27 | 0.837 | -0.257 | 0.49406116 | 0.69039846 | 0 | 0.12 | 0.0675 | 0.0675 | TRUE | 0.283 |
| Abcc1 | 5295 | 7722 | 8232 | 6027 | 4824 | 6024 | 5624 | 7712 | 8266 | 4738 | 5190 | 6594 | 6354.09 | 7201 | 5507 | 0.765 | -0.387 | 0.02238018 | 0.09422857 | 0.0339 | 0.0074 | 0.0205 | 0.0205 | TRUE | 0.4476 |
| Abcc10 | 893 | 1156 | 1202 | 1274 | 960 | 971 | 949 | 1155 | 1207 | 1002 | 1033 | 1063 | 1067.88 | 1104 | 1033 | 0.935 | -0.097 | 0.40019085 | 0.61321384 | 0.0075 | 0.01 | 0.0086 | 0.0086 | TRUE | 0.1794 |
| Abcc12 | 23 | 14 | 9 | 4 | 2 | 5 | 24 | 14 | 9 | 3 | 2 | 5 | 9.7 | 16 | 3 | 0.226 | -2.149 | 0.0043888 | 0.02832387 | 0 | 0.3464 | 0.2407 | 0.2407 | TRUE | 0.2242 |
| Abcc2 | 255 | 323 | 369 | 283 | 291 | 251 | 271 | 323 | 371 | 222 | 313 | 275 | 295.71 | 322 | 270 | 0.838 | -0.255 | 0.1645338 | 0.36082133 | 0.0235 | 0.018 | 0.0208 | 0.0208 | TRUE | 0.2276 |
| Abcc3 | 81 | 84 | 81 | 108 | 85 | 71 | 86 | 84 | 81 | 85 | 91 | 78 | 84.22 | 84 | 85 | 1.012 | 0.017 | 0.93655126 | 0.96825513 | 0 | 0.046 | 0.02 | 0.02 | TRUE | 0.0444 |
| Abcc4 | 7072 | 7908 | 8789 | 6823 | 5149 | 5793 | 7512 | 7898 | 8825 | 5364 | 5540 | 6341 | 6913.28 | 8078 | 5748 | 0.711 | -0.491 | 1.38E-06 | 3.65E-05 | 0.0074 | 0.0074 | 0.0073 | 0.0073 | TRUE | 0.0991 |
| Abcc5 | 5022 | 4180 | 4326 | 4964 | 3553 | 3509 | 5334 | 4175 | 4344 | 3902 | 3823 | 3841 | 4236.48 | 4618 | 3855 | 0.835 | -0.26 | 0.01658468 | 0.07624971 | 0.0089 | 0.0077 | 0.0083 | 0.0083 | TRUE | 0.2244 |
| Abcc6 | 26 | 25 | 26 | 42 | 34 | 39 | 28 | 25 | 26 | 33 | 37 | 43 | 31.83 | 26 | 38 | 1.421 | 0.507 | 0.13399164 | 0.31649098 | 0 | 0.1104 | 0.05 | 0.05 | TRUE | 0.106 |
| Abcc8 | 442 | 567 | 559 | 493 | 459 | 491 | 469 | 566 | 561 | 388 | 494 | 537 | 502.66 | 532 | 473 | 0.886 | -0.174 | 0.26474466 | 0.48451529 | 0.0175 | 0.0135 | 0.0156 | 0.0156 | TRUE | 0.3018 |
| Abcc9 | 2 | 0 | 0 | 7 | 1 | 0 | 2 | 0 | 0 | 6 | 1 | 0 | 1.45 | 1 | 2 | 3.215 | 1.685 | 0.44317818 | NA | 3.0865 | 2.2781 | 2.5283 | 2.5283 | TRUE | 1.0731 |
| Abcd1 | 475 | 630 | 595 | 780 | 549 | 598 | 505 | 629 | 597 | 613 | 591 | 655 | 598.27 | 577 | 620 | 1.073 | 0.101 | 0.4012852 | 0.6141287 | 0.0059 | 0.0124 | 0.0089 | 0.0089 | TRUE | 0.1405 |
| Abcd2 | 39 | 18 | 9 | 34 | 13 | 13 | 41 | 18 | 9 | 27 | 14 | 14 | 20.56 | 23 | 18 | 0.812 | -0.3 | 0.62830889 | 0.79087692 | 0.2907 | 0.1671 | 0.228 | 0.228 | TRUE | 0.6498 |
| Abcd3 | 2981 | 2774 | 2456 | 3935 | 2761 | 2404 | 3166 | 2770 | 2466 | 3093 | 2971 | 2632 | 2849.74 | 2801 | 2899 | 1.035 | 0.05 | 0.67207616 | 0.81720647 | 0.0109 | 0.0081 | 0.0097 | 0.0097 | TRUE | 0.1578 |
| Abcd4 | 1128 | 1385 | 1399 | 1653 | 1296 | 1369 | 1198 | 1383 | 1405 | 1299 | 1394 | 1499 | 1363.09 | 1329 | 1397 | 1.051 | 0.072 | 0.49318099 | 0.68957895 | 0.0056 | 0.0093 | 0.0071 | 0.0071 | TRUE | 0.0884 |
| Abce1 | 26905 | 28547 | 28602 | 38769 | 26324 | 26580 | 28578 | 28510 | 28719 | 30477 | 28323 | 29096 | 28950.6 | 28602 | 29299 | 1.024 | 0.035 | 0.55407504 | 0.73739528 | 6.00E-04 | 0.007 | 0.0025 | 0.0025 | TRUE | 0.0152 |
| Abcf1 | 15471 | 14013 | 14361 | 16698 | 12748 | 11843 | 16433 | 13995 | 14420 | 13127 | 13716 | 12964 | 14109.09 | 14949 | 13269 | 0.888 | -0.172 | 0.04438759 | 0.15134402 | 0.0041 | 0.0071 | 0.0052 | 0.0052 | TRUE | 0.0921 |
| Abcf2 | 11074 | 9421 | 9298 | 12876 | 9098 | 8042 | 11763 | 9409 | 9336 | 10122 | 9789 | 8803 | 9870.3 | 10169 | 9571 | 0.941 | -0.087 | 0.45003765 | 0.65553889 | 0.0108 | 0.0072 | 0.0095 | 0.0095 | TRUE | 0.2295 |
| Abcf3 | 2213 | 2010 | 1990 | 2909 | 2124 | 1931 | 2351 | 2007 | 1998 | 2287 | 2285 | 2114 | 2173.68 | 2119 | 2229 | 1.052 | 0.073 | 0.44793383 | 0.6536651 | 0.0046 | 0.0084 | 0.0063 | 0.0063 | TRUE | 0.1106 |
| Abcg1 | 1963 | 3336 | 3636 | 1014 | 970 | 1387 | 2085 | 3332 | 3651 | 797 | 1044 | 1518 | 2071.13 | 3023 | 1120 | 0.37 | -1.433 | 5.37E-05 | 0.00082297 | 0.0901 | 0.0085 | 0.0439 | 0.0901 | TRUE | 0.9707 |
| Abcg2 | 1978 | 2072 | 2152 | 3150 | 2050 | 2047 | 2101 | 2069 | 2161 | 2476 | 2206 | 2241 | 2208.98 | 2110 | 2308 | 1.094 | 0.13 | 0.10758738 | 0.27453369 | 0.0019 | 0.0084 | 0.0042 | 0.0042 | TRUE | 0.0506 |
| Abcg3 | 43 | 34 | 38 | 42 | 35 | 31 | 46 | 34 | 38 | 33 | 38 | 34 | 37.07 | 39 | 35 | 0.887 | -0.172 | 0.58561161 | 0.76074968 | 0 | 0.0958 | 0.0449 | 0.0449 | TRUE | 0.1473 |
| Abcg4 | 28 | 44 | 38 | 34 | 26 | 38 | 30 | 44 | 38 | 27 | 28 | 42 | 34.69 | 37 | 32 | 0.855 | -0.227 | 0.52576257 | 0.71510238 | 0.0198 | 0.1019 | 0.0629 | 0.0629 | TRUE | 0.4324 |
| Abcg5 | 10 | 17 | 5 | 16 | 13 | 11 | 11 | 17 | 5 | 13 | 14 | 12 | 11.87 | 11 | 13 | 1.182 | 0.242 | 0.69152124 | 0.83090245 | 0.0584 | 0.2844 | 0.182 | 0.182 | TRUE | 0.2592 |
| Abcg8 | 20 | 18 | 20 | 24 | 11 | 14 | 21 | 18 | 20 | 19 | 12 | 15 | 17.55 | 20 | 15 | 0.784 | -0.351 | 0.44039527 | 0.64746865 | 0 | 0.1946 | 0.0917 | 0.0917 | TRUE | 0.233 |
| Abhd1 | 326 | 328 | 311 | 425 | 265 | 280 | 346 | 328 | 312 | 334 | 285 | 307 | 318.64 | 329 | 309 | 0.941 | -0.088 | 0.49624402 | 0.69196835 | 0 | 0.0172 | 0.0088 | 0.0088 | TRUE | 0.0635 |
| Abhd10 | 1257 | 1399 | 1527 | 1642 | 1084 | 1316 | 1335 | 1397 | 1533 | 1291 | 1166 | 1441 | 1360.55 | 1422 | 1299 | 0.914 | -0.13 | 0.23956263 | 0.45564001 | 0.007 | 0.0093 | 0.0081 | 0.0081 | TRUE | 0.1069 |
| Abhd11 | 1443 | 1597 | 1564 | 1910 | 1442 | 1464 | 1533 | 1595 | 1570 | 1501 | 1552 | 1603 | 1558.94 | 1566 | 1552 | 0.99 | -0.014 | 0.84962207 | 0.92515564 | 0 | 0.009 | 0.0033 | 0.0033 | TRUE | 0.0101 |
| Abhd12 | 1232 | 1655 | 1751 | 1895 | 1495 | 1535 | 1309 | 1653 | 1758 | 1490 | 1609 | 1680 | 1583.03 | 1573 | 1593 | 1.012 | 0.017 | 0.89340433 | 0.94836271 | 0.0129 | 0.009 | 0.0111 | 0.0111 | TRUE | 0.2602 |
| Abhd12b | 1 | 0 | 0 | 1 | 1 | 3 | 1 | 0 | 0 | 1 | 1 | 3 | 1.03 | 0 | 2 | 4.619 | 2.208 | 0.28545434 | NA | 0 | 3.1909 | 1.7882 | 1.7882 | TRUE | 0.4614 |
| Abhd13 | 2924 | 3022 | 3156 | 3896 | 2954 | 2906 | 3106 | 3018 | 3169 | 3063 | 3178 | 3181 | 3119.16 | 3098 | 3141 | 1.014 | 0.02 | 0.76211928 | 0.87457944 | 2.00E-04 | 0.008 | 0.0027 | 0.0027 | TRUE | 0.0062 |
| Abhd14a | 1684 | 1668 | 1565 | 1784 | 1321 | 1320 | 1789 | 1666 | 1571 | 1402 | 1421 | 1445 | 1549.11 | 1675 | 1423 | 0.849 | -0.236 | 0.00489759 | 0.03089793 | 0.0016 | 0.009 | 0.0044 | 0.0044 | TRUE | 0.0421 |
| Abhd14b | 637 | 689 | 710 | 432 | 339 | 468 | 677 | 688 | 713 | 340 | 365 | 512 | 549.05 | 693 | 406 | 0.584 | -0.775 | 2.53E-06 | 6.18E-05 | 0.0223 | 0.0129 | 0.0176 | 0.0176 | TRUE | 0.6045 |
| Abhd15 | 38 | 60 | 53 | 60 | 30 | 34 | 40 | 60 | 53 | 47 | 32 | 37 | 45.03 | 51 | 39 | 0.765 | -0.387 | 0.22152298 | 0.43369669 | 0.0172 | 0.0801 | 0.0498 | 0.0498 | TRUE | 0.2988 |
| Abhd16a | 649 | 699 | 689 | 704 | 572 | 585 | 689 | 698 | 692 | 553 | 615 | 640 | 648.09 | 693 | 603 | 0.868 | -0.203 | 0.04476225 | 0.15235007 | 0.0014 | 0.012 | 0.0059 | 0.0059 | TRUE | 0.0645 |
| Abhd16b | 8 | 10 | 9 | 13 | 9 | 3 | 8 | 10 | 9 | 10 | 10 | 3 | 8.45 | 9 | 8 | 0.853 | -0.229 | 0.73750072 | 0.85945481 | 0 | 0.3967 | 0.2183 | 0.2183 | TRUE | 0.6468 |
| Abhd2 | 1283 | 1341 | 1412 | 1647 | 1221 | 1324 | 1363 | 1339 | 1418 | 1295 | 1314 | 1449 | 1362.94 | 1373 | 1353 | 0.984 | -0.023 | 0.78977844 | 0.89184403 | 0.0016 | 0.0093 | 0.0046 | 0.0046 | TRUE | 0.0465 |
| Abhd3 | 187 | 211 | 195 | 343 | 238 | 244 | 199 | 211 | 196 | 270 | 256 | 267 | 232.99 | 202 | 264 | 1.311 | 0.39 | 0.00359156 | 0.02430119 | 0 | 0.021 | 0.0086 | 0.0086 | TRUE | 0.0168 |
| Abhd4 | 1873 | 2410 | 2327 | 2440 | 1729 | 2126 | 1989 | 2407 | 2337 | 1918 | 1860 | 2327 | 2139.76 | 2244 | 2035 | 0.906 | -0.142 | 0.25018806 | 0.46733938 | 0.0121 | 0.0084 | 0.0105 | 0.0105 | TRUE | 0.1895 |
| Abhd5 | 447 | 440 | 414 | 842 | 510 | 531 | 475 | 439 | 416 | 662 | 549 | 581 | 520.31 | 443 | 597 | 1.35 | 0.433 | 0.00040353 | 0.00437186 | 0.0054 | 0.0132 | 0.0088 | 0.0088 | TRUE | 0.1093 |
| Abhd6 | 340 | 351 | 342 | 431 | 337 | 324 | 361 | 351 | 343 | 339 | 363 | 355 | 351.86 | 352 | 352 | 1 | 0 | 0.99927497 | 0.99950602 | 0 | 0.0163 | 0.0065 | 0.0065 | TRUE | 0.0133 |
| Abhd8 | 830 | 1020 | 1019 | 1055 | 738 | 910 | 882 | 1019 | 1023 | 829 | 794 | 996 | 923.84 | 975 | 873 | 0.895 | -0.159 | 0.19896082 | 0.4062556 | 0.0094 | 0.0105 | 0.01 | 0.01 | TRUE | 0.1779 |
| Abi1 | 2978 | 3300 | 3476 | 4257 | 3096 | 3109 | 3163 | 3296 | 3490 | 3347 | 3331 | 3403 | 3338.35 | 3316 | 3360 | 1.013 | 0.019 | 0.78995273 | 0.89193627 | 0.001 | 0.0079 | 0.0033 | 0.0033 | TRUE | 0.0256 |
| Abi2 | 3964 | 4375 | 4916 | 5051 | 3830 | 3802 | 4210 | 4369 | 4936 | 3971 | 4121 | 4162 | 4294.91 | 4505 | 4085 | 0.906 | -0.142 | 0.09987677 | 0.26096381 | 0.0035 | 0.0077 | 0.0051 | 0.0051 | TRUE | 0.0853 |
| Abi3 | 106 | 132 | 120 | 83 | 73 | 75 | 113 | 132 | 120 | 65 | 79 | 82 | 98.47 | 122 | 75 | 0.615 | -0.701 | 0.00080622 | 0.00753744 | 0 | 0.0404 | 0.0208 | 0.0208 | TRUE | 0.1445 |
| Abi3bp | 29 | 22 | 19 | 44 | 38 | 25 | 31 | 22 | 19 | 35 | 41 | 27 | 29.12 | 24 | 34 | 1.435 | 0.521 | 0.17288567 | 0.37169476 | 0 | 0.1201 | 0.0697 | 0.0697 | TRUE | 0.2877 |
| Abl1 | 5147 | 4696 | 4881 | 6205 | 4819 | 4534 | 5467 | 4690 | 4901 | 4878 | 5185 | 4963 | 5014 | 5019 | 5009 | 0.998 | -0.003 | 0.9707364 | 0.98472179 | 0.0034 | 0.0076 | 0.005 | 0.005 | TRUE | 0.0741 |
| Abl2 | 7966 | 7831 | 8703 | 8812 | 6426 | 6658 | 8461 | 7821 | 8739 | 6927 | 6914 | 7288 | 7691.74 | 8340 | 7043 | 0.844 | -0.244 | 0.00089376 | 0.00815291 | 0.0019 | 0.0073 | 0.0038 | 0.0038 | TRUE | 0.0363 |
| Ablim1 | 663 | 1115 | 1059 | 666 | 579 | 863 | 704 | 1114 | 1063 | 524 | 623 | 945 | 828.72 | 960 | 697 | 0.725 | -0.464 | 0.05057911 | 0.16543627 | 0.075 | 0.0109 | 0.0393 | 0.0393 | TRUE | 1.1375 |
| Ablim2 | 259 | 336 | 328 | 264 | 237 | 299 | 275 | 336 | 329 | 208 | 255 | 327 | 288.31 | 313 | 263 | 0.837 | -0.256 | 0.18415741 | 0.38701276 | 0.0282 | 0.0183 | 0.0233 | 0.0233 | TRUE | 0.3964 |
| Ablim3 | 287 | 271 | 285 | 193 | 168 | 164 | 305 | 271 | 286 | 152 | 181 | 180 | 228.94 | 287 | 171 | 0.592 | -0.756 | 3.91E-07 | 1.20E-05 | 0 | 0.0213 | 0.0114 | 0.0114 | TRUE | 0.1118 |
| Abo | 16 | 30 | 21 | 25 | 18 | 12 | 17 | 30 | 21 | 20 | 19 | 13 | 20.03 | 23 | 17 | 0.769 | -0.378 | 0.40458601 | 0.61732751 | 0 | 0.1714 | 0.098 | 0.098 | TRUE | 0.4604 |
| Abp1 | 17 | 15 | 13 | 34 | 19 | 21 | 18 | 15 | 13 | 27 | 20 | 23 | 19.37 | 15 | 23 | 1.532 | 0.615 | 0.15670917 | 0.34969116 | 0 | 0.1769 | 0.082 | 0.082 | TRUE | 0.1107 |
| Abr | 4636 | 4496 | 4533 | 5570 | 4106 | 3855 | 4924 | 4490 | 4552 | 4379 | 4418 | 4220 | 4497.08 | 4655 | 4339 | 0.932 | -0.101 | 0.15404826 | 0.34595603 | 0.0012 | 0.0076 | 0.0034 | 0.0034 | TRUE | 0.031 |
| Abra | 3 | 2 | 0 | 6 | 1 | 1 | 3 | 2 | 0 | 5 | 1 | 1 | 2.01 | 2 | 2 | 1.376 | 0.46 | 0.75824336 | NA | 0.489 | 1.6444 | 1.1081 | 1.1081 | TRUE | 0.758 |
| Abracl | 1122 | 1133 | 1122 | 1346 | 831 | 905 | 1192 | 1132 | 1127 | 1058 | 894 | 991 | 1065.47 | 1150 | 981 | 0.854 | -0.228 | 0.0196564 | 0.08580701 | 0.0031 | 0.01 | 0.0059 | 0.0059 | TRUE | 0.0712 |
| Abt1 | 1284 | 1193 | 1163 | 1897 | 1360 | 1132 | 1364 | 1191 | 1168 | 1491 | 1463 | 1239 | 1319.46 | 1241 | 1398 | 1.127 | 0.173 | 0.12844791 | 0.30816556 | 0.0075 | 0.0094 | 0.0085 | 0.0085 | TRUE | 0.1176 |
| Abtb1 | 896 | 1050 | 986 | 1455 | 1093 | 1076 | 952 | 1049 | 990 | 1144 | 1176 | 1178 | 1081.34 | 997 | 1166 | 1.169 | 0.225 | 0.00686411 | 0.03956199 | 0 | 0.0099 | 0.0041 | 0.0041 | TRUE | 0.0248 |
| Abtb2 | 546 | 591 | 541 | 624 | 458 | 457 | 580 | 590 | 543 | 491 | 493 | 500 | 532.83 | 571 | 495 | 0.866 | -0.208 | 0.03557147 | 0.12950116 | 0 | 0.0131 | 0.0052 | 0.0052 | TRUE | 0.0216 |
| Acaa1a | 1436 | 1551 | 1506 | 2059 | 1660 | 1397 | 1525 | 1549 | 1512 | 1619 | 1786 | 1529 | 1586.73 | 1529 | 1645 | 1.076 | 0.105 | 0.23507589 | 0.45011738 | 0.0023 | 0.009 | 0.005 | 0.005 | TRUE | 0.0671 |
| Acaa1b | 188 | 231 | 193 | 243 | 191 | 188 | 200 | 231 | 194 | 191 | 206 | 206 | 204.42 | 208 | 201 | 0.963 | -0.054 | 0.72004237 | 0.8481969 | 0 | 0.023 | 0.0115 | 0.0115 | TRUE | 0.0999 |
| Acaa2 | 1465 | 1677 | 1499 | 2153 | 1549 | 1627 | 1556 | 1675 | 1505 | 1693 | 1667 | 1781 | 1646.04 | 1579 | 1714 | 1.085 | 0.118 | 0.15170726 | 0.3428955 | 0.0015 | 0.0089 | 0.0043 | 0.0043 | TRUE | 0.0344 |

| | | | | | | | | | | | | | | | | | | | | | | | | | |
|---|---|---|---|---|---|---|---|---|---|---|---|---|---|---|---|---|---|---|---|---|---|---|---|---|---|
| Acaca | 10861 | 8620 | 9565 | 10467 | 7409 | 7077 | 11536 | 8609 | 9604 | 8228 | 7972 | 7747 | 8949.38 | 9916 | 7982 | 0.805 | -0.313 | 0.00693982 | 0.03986242 | 0.0116 | 0.0073 | 0.0096 | 0.0096 | TRUE | 0.2494 |
| Acacb | 1726 | 1872 | 2191 | 2111 | 1541 | 1601 | 1833 | 1870 | 2200 | 1660 | 1658 | 1753 | 1828.83 | 1968 | 1690 | 0.859 | -0.22 | 0.02699713 | 0.10720009 | 0.005 | 0.0087 | 0.0066 | 0.0066 | TRUE | 0.1292 |
| Acad10 | 714 | 820 | 864 | 1120 | 739 | 846 | 758 | 819 | 868 | 880 | 795 | 926 | 841.09 | 815 | 867 | 1.064 | 0.09 | 0.39697976 | 0.61042847 | 0.0039 | 0.0108 | 0.0069 | 0.0069 | TRUE | 0.0614 |
| Acad11 | 2200 | 2314 | 2290 | 2527 | 1726 | 1938 | 2337 | 2311 | 2299 | 1987 | 1857 | 2121 | 2152.04 | 2316 | 1988 | 0.859 | -0.22 | 0.00606387 | 0.03624074 | 0.0015 | 0.0084 | 0.0042 | 0.0042 | TRUE | 0.0405 |
| Acad12 | 337 | 364 | 391 | 471 | 352 | 247 | 358 | 364 | 393 | 370 | 379 | 270 | 355.58 | 372 | 340 | 0.916 | -0.126 | 0.42809105 | 0.63805066 | 0.0147 | 0.0162 | 0.0154 | 0.0154 | TRUE | 0.3538 |
| Acad8 | 1252 | 1287 | 1257 | 1672 | 1225 | 1214 | 1330 | 1285 | 1262 | 1314 | 1318 | 1329 | 1306.45 | 1292 | 1320 | 1.022 | 0.031 | 0.67621707 | 0.81979608 | 0 | 0.0094 | 0.0032 | 0.0032 | TRUE | 0.0076 |
| Acad9 | 3483 | 3043 | 2984 | 4843 | 3398 | 3002 | 3700 | 3039 | 2996 | 3807 | 3656 | 3286 | 3414.05 | 3245 | 3583 | 1.105 | 0.143 | 0.20308724 | 0.41080946 | 0.0097 | 0.0079 | 0.0089 | 0.0089 | TRUE | 0.1823 |
| Acadl | 1125 | 1505 | 1553 | 1540 | 1148 | 1326 | 1195 | 1503 | 1559 | 1211 | 1235 | 1452 | 1359.12 | 1419 | 1299 | 0.915 | -0.128 | 0.33444174 | 0.5548783 | 0.0137 | 0.0093 | 0.012 | 0.012 | TRUE | 0.229 |
| Acadm | 5360 | 4999 | 4965 | 6986 | 5140 | 4983 | 5693 | 4993 | 4985 | 5492 | 5530 | 5455 | 5358.01 | 5224 | 5492 | 1.051 | 0.072 | 0.36770802 | 0.58492091 | 0.0028 | 0.0075 | 0.0045 | 0.0045 | TRUE | 0.0752 |
| Acads | 5205 | 5510 | 5482 | 5778 | 4223 | 4546 | 5529 | 5503 | 5505 | 4542 | 4544 | 4976 | 5099.71 | 5512 | 4687 | 0.85 | -0.234 | 0.00073182 | 0.00699527 | 0.0011 | 0.0075 | 0.0033 | 0.0033 | TRUE | 0.0352 |
| Acadsb | 1693 | 1750 | 1681 | 2237 | 1602 | 1755 | 1798 | 1748 | 1688 | 1759 | 1724 | 1921 | 1772.87 | 1745 | 1801 | 1.032 | 0.046 | 0.57726719 | 0.75475903 | 0.0016 | 0.0088 | 0.0043 | 0.0043 | TRUE | 0.0404 |
| Acadvl | 3532 | 3135 | 3028 | 4313 | 2861 | 2761 | 3752 | 3131 | 3040 | 3391 | 3078 | 3022 | 3235.69 | 3308 | 3164 | 0.957 | -0.064 | 0.55553661 | 0.73847101 | 0.0075 | 0.0079 | 0.0081 | 0.0081 | TRUE | 0.1673 |
| Acan | 1687 | 1372 | 1382 | 2172 | 1471 | 1207 | 1792 | 1370 | 1388 | 1707 | 1583 | 1321 | 1526.87 | 1517 | 1537 | 1.014 | 0.02 | 0.88841071 | 0.94541621 | 0.0195 | 0.0091 | 0.0145 | 0.0145 | TRUE | 0.3032 |
| Acap1 | 794 | 985 | 958 | 711 | 587 | 571 | 843 | 984 | 962 | 559 | 632 | 625 | 767.43 | 930 | 605 | 0.65 | -0.622 | 1.56E-08 | 7.14E-07 | 0.0045 | 0.0112 | 0.0073 | 0.0073 | TRUE | 0.0781 |
| Acap2 | 3797 | 4515 | 4834 | 5231 | 3855 | 4280 | 4033 | 4509 | 4854 | 4112 | 4148 | 4685 | 4390.2 | 4465 | 4315 | 0.966 | -0.05 | 0.62263329 | 0.7869989 | 0.0058 | 0.0077 | 0.0071 | 0.0071 | TRUE | 0.0872 |
| Acap3 | 547 | 345 | 383 | 579 | 413 | 299 | 581 | 345 | 385 | 455 | 444 | 327 | 422.83 | 437 | 409 | 0.938 | -0.093 | 0.67336396 | 0.81805326 | 0.0525 | 0.0147 | 0.0326 | 0.0326 | TRUE | 0.9632 |
| Acat1 | 1972 | 2369 | 2376 | 2148 | 1610 | 1757 | 2095 | 2366 | 2386 | 1689 | 1732 | 1923 | 2031.75 | 2282 | 1781 | 0.78 | -0.358 | 0.00018356 | 0.00226357 | 0.0042 | 0.0085 | 0.0061 | 0.0061 | TRUE | 0.0624 |
| Acat2 | 6316 | 5365 | 5045 | 6444 | 4643 | 4412 | 6709 | 5358 | 5066 | 5066 | 4996 | 4830 | 5337.25 | 5711 | 4964 | 0.869 | -0.202 | 0.08417373 | 0.2326245 | 0.0118 | 0.0075 | 0.0097 | 0.0097 | TRUE | 0.2847 |
| Acat3 | 14 | 27 | 20 | 17 | 13 | 17 | 15 | 27 | 20 | 13 | 14 | 19 | 17.98 | 21 | 15 | 0.737 | -0.441 | 0.35031517 | 0.56925314 | 0 | 0.1901 | 0.1036 | 0.1036 | TRUE | 0.2144 |
| Acbd3 | 2671 | 3208 | 3113 | 3401 | 2627 | 2723 | 2837 | 3204 | 3126 | 2674 | 2826 | 2981 | 2941.26 | 3056 | 2827 | 0.925 | -0.113 | 0.18990233 | 0.39508927 | 0.003 | 0.008 | 0.005 | 0.005 | TRUE | 0.0474 |
| Acbd4 | 855 | 1122 | 1020 | 1279 | 939 | 1058 | 908 | 1121 | 1024 | 1005 | 1010 | 1158 | 1037.8 | 1018 | 1058 | 1.039 | 0.055 | 0.63617862 | 0.79536224 | 0.0077 | 0.0101 | 0.0088 | 0.0088 | TRUE | 0.1053 |
| Acbd5 | 1399 | 1625 | 1664 | 1919 | 1418 | 1553 | 1486 | 1623 | 1671 | 1509 | 1526 | 1700 | 1585.66 | 1593 | 1578 | 0.99 | -0.015 | 0.87453516 | 0.93811777 | 0.0034 | 0.009 | 0.0056 | 0.0056 | TRUE | 0.0545 |
| Acbd6 | 2141 | 2569 | 2633 | 3104 | 2325 | 2335 | 2274 | 2566 | 2644 | 2440 | 2502 | 2556 | 2496.89 | 2495 | 2499 | 1.002 | 0.002 | 0.97908571 | 0.98867206 | 0.003 | 0.0082 | 0.005 | 0.005 | TRUE | 0.0722 |
| Acbd7 | 106 | 121 | 141 | 153 | 91 | 127 | 113 | 121 | 142 | 120 | 98 | 139 | 122.04 | 125 | 119 | 0.952 | -0.071 | 0.73586232 | 0.8583065 | 0.0127 | 0.0339 | 0.0236 | 0.0236 | TRUE | 0.2298 |
| Accs | 45 | 78 | 72 | 115 | 102 | 78 | 48 | 78 | 72 | 90 | 110 | 85 | 80.59 | 66 | 95 | 1.438 | 0.524 | 0.04531808 | 0.15357903 | 0.0235 | 0.0478 | 0.0366 | 0.0366 | TRUE | 0.5011 |
| Accsl | 256 | 93 | 73 | 222 | 143 | 94 | 272 | 93 | 73 | 175 | 154 | 103 | 144.9 | 146 | 144 | 0.986 | -0.02 | 0.96422948 | 0.98203715 | 0.2713 | 0.0296 | 0.1338 | 0.1338 | TRUE | 5.8922 |
| Acd | 1834 | 1912 | 1943 | 2657 | 1923 | 1879 | 1948 | 1910 | 1951 | 2089 | 2069 | 2057 | 2003.86 | 1936 | 2072 | 1.07 | 0.098 | 0.13496391 | 0.31789886 | 0 | 0.0085 | 0.0026 | 0.0026 | TRUE | 0.0018 |
| Ace | 500 | 597 | 610 | 859 | 796 | 811 | 531 | 596 | 613 | 675 | 856 | 888 | 693.22 | 580 | 806 | 1.388 | 0.473 | 0.0005724 | 0.00578738 | 0.0125 | 0.0117 | 0.0121 | 0.0121 | TRUE | 0.2475 |
| Ace2 | 0 | 0 | 0 | 0 | 0 | 0 | 0 | 0 | 0 | 0 | 0 | 0 | 0 | 0 | 0 | NA | NA | NA | NA | NA | NA | NA | NA | NA | NA |
| Ace3 | 0 | 0 | 3 | 4 | 1 | 2 | 0 | 0 | 3 | 3 | 1 | 2 | 1.57 | 1 | 2 | 2.137 | 1.095 | 0.53448268 | NA | 0.4192 | 2.1048 | 1.5087 | 1.5087 | TRUE | 1.3408 |
| Acer1 | 39 | 39 | 30 | 22 | 13 | 19 | 41 | 39 | 30 | 17 | 14 | 21 | 27.1 | 37 | 17 | 0.471 | -1.085 | 0.005037 | 0.03151273 | 0 | 0.1285 | 0.0659 | 0.0659 | TRUE | 0.1857 |
| Acer2 | 610 | 566 | 534 | 714 | 557 | 492 | 648 | 565 | 536 | 561 | 599 | 539 | 574.76 | 583 | 566 | 0.972 | -0.042 | 0.72134351 | 0.84915882 | 0.0044 | 0.0126 | 0.0081 | 0.0081 | TRUE | 0.11 |
| Acer3 | 712 | 896 | 915 | 1091 | 784 | 819 | 756 | 895 | 919 | 858 | 844 | 897 | 861.26 | 857 | 866 | 1.01 | 0.015 | 0.89068953 | 0.94694677 | 0.0046 | 0.0107 | 0.0073 | 0.0073 | TRUE | 0.124 |
| Ache | 4 | 4 | 0 | 2 | 1 | 1 | 4 | 4 | 0 | 2 | 1 | 1 | 2 | 3 | 1 | 0.461 | -1.119 | 0.45406787 | NA | 0 | 1.656 | 1.0357 | 1.0357 | TRUE | 0.9312 |
| Acin1 | 13297 | 13326 | 14251 | 17640 | 13485 | 12964 | 14124 | 13309 | 14310 | 13867 | 14509 | 14191 | 14051.57 | 13914 | 14189 | 1.02 | 0.028 | 0.65462604 | 0.80586653 | 8.00E-04 | 0.0071 | 0.0028 | 0.0028 | TRUE | 0.0177 |
| Acly | 40485 | 36079 | 36753 | 41911 | 30725 | 32008 | 43002 | 36032 | 36904 | 32947 | 33058 | 35038 | 36163.61 | 38646 | 33681 | 0.872 | -0.198 | 0.02896971 | 0.11226339 | 0.0054 | 0.007 | 0.0059 | 0.0059 | TRUE | 0.119 |
| Acmsd | 0 | 0 | 1 | 1 | 0 | 0 | 0 | 0 | 1 | 1 | 0 | 0 | 0.3 | 0 | 0 | 0.954 | -0.067 | 0.98640617 | NA | 0 | 11.0483 | 9.2783 | 9.2783 | TRUE | 0.4098 |
| Acn9 | 337 | 343 | 344 | 423 | 300 | 300 | 358 | 343 | 345 | 333 | 323 | 328 | 338.27 | 349 | 328 | 0.941 | -0.088 | 0.44037929 | 0.64746865 | 0 | 0.0166 | 0.0063 | 0.0063 | TRUE | 0.0061 |
| Acnat1 | 10 | 4 | 1 | 0 | 3 | 1 | 11 | 4 | 1 | 0 | 3 | 1 | 3.32 | 5 | 1 | 0.266 | -1.912 | 0.17445007 | NA | 0.9488 | 0.9981 | 0.9758 | 0.9758 | TRUE | 0.2912 |
| Acnat2 | 0 | 0 | 0 | 0 | 0 | 0 | 0 | 0 | 0 | 0 | 0 | 0 | 0 | 0 | 0 | NA | NA | NA | NA | NA | NA | NA | NA | NA | NA |
| Aco1 | 4878 | 4987 | 5059 | 4794 | 3530 | 3859 | 5181 | 4981 | 5080 | 3769 | 3798 | 4224 | 4505.44 | 5081 | 3930 | 0.773 | -0.371 | 1.14E-06 | 3.09E-05 | 0.002 | 0.0076 | 0.004 | 0.004 | TRUE | 0.0518 |
| Aco2 | 17672 | 21330 | 22006 | 22995 | 17017 | 18160 | 18771 | 21302 | 22096 | 18077 | 18309 | 19879 | 19739.11 | 20723 | 18755 | 0.905 | -0.144 | 0.10761907 | 0.27453369 | 0.0048 | 0.0071 | 0.0057 | 0.0057 | TRUE | 0.0831 |
| Acot1 | 2034 | 2563 | 2553 | 1533 | 1405 | 1787 | 2160 | 2560 | 2563 | 1205 | 1512 | 1956 | 1992.77 | 2428 | 1558 | 0.641 | -0.641 | 0.00017542 | 0.00218544 | 0.0331 | 0.0086 | 0.0205 | 0.0205 | TRUE | 0.5064 |
| Acot10 | 10 | 8 | 6 | 4 | 5 | 8 | 11 | 8 | 6 | 3 | 5 | 9 | 6.99 | 8 | 6 | 0.689 | -0.538 | 0.47340992 | 0.67430879 | 0 | 0.4785 | 0.2572 | 0.2572 | TRUE | 0.4251 |
| Acot11 | 337 | 359 | 327 | 522 | 315 | 298 | 358 | 359 | 328 | 410 | 339 | 326 | 353.39 | 348 | 358 | 1.033 | 0.046 | 0.73950961 | 0.86066702 | 0.0068 | 0.0162 | 0.0111 | 0.0111 | TRUE | 0.1924 |
| Acot12 | 1 | 0 | 0 | 5 | 8 | 1 | 1 | 0 | 0 | 4 | 9 | 1 | 2.45 | 0 | 5 | 12.401 | 3.632 | 0.03520768 | NA | 0 | 1.352 | 1.0358 | 1.0358 | TRUE | 0.2597 |
| Acot13 | 998 | 1282 | 1325 | 1121 | 799 | 999 | 1060 | 1280 | 1330 | 881 | 860 | 1094 | 1084.22 | 1223 | 945 | 0.772 | -0.374 | 0.00650905 | 0.03815231 | 0.0149 | 0.0099 | 0.0127 | 0.0127 | TRUE | 0.2239 |
| Acot2 | 274 | 340 | 276 | 357 | 262 | 235 | 291 | 340 | 277 | 281 | 282 | 257 | 287.92 | 303 | 273 | 0.904 | -0.146 | 0.29519783 | 0.51607626 | 0.0037 | 0.0183 | 0.0106 | 0.0106 | TRUE | 0.1303 |
| Acot3 | 16 | 28 | 27 | 45 | 25 | 23 | 17 | 28 | 27 | 35 | 27 | 25 | 26.59 | 24 | 29 | 1.219 | 0.286 | 0.47030965 | 0.67209973 | 0 | 0.1308 | 0.0752 | 0.0752 | TRUE | 0.3746 |
| Acot4 | 11 | 4 | 2 | 10 | 10 | 9 | 12 | 4 | 2 | 8 | 11 | 10 | 7.69 | 6 | 10 | 1.614 | 0.69 | 0.38433506 | 0.5985123 | 0.1634 | 0.4351 | 0.3148 | 0.3148 | TRUE | 1.6315 |
| Acot5 | 3 | 0 | 5 | 7 | 6 | 3 | 3 | 0 | 5 | 6 | 6 | 3 | 3.91 | 3 | 5 | 1.866 | 0.9 | 0.41509623 | NA | 0.074 | 0.8499 | 0.5949 | 0.5949 | TRUE | 0.6824 |
| Acot6 | 22 | 14 | 20 | 13 | 16 | 10 | 23 | 14 | 20 | 10 | 17 | 11 | 15.97 | 19 | 13 | 0.664 | -0.591 | 0.2413939 | 0.45809946 | 0 | 0.2132 | 0.118 | 0.118 | TRUE | 0.3531 |
| Acot7 | 2767 | 3217 | 3139 | 3442 | 2566 | 2509 | 2939 | 3213 | 3152 | 2706 | 2761 | 2746 | 2919.49 | 3101 | 2738 | 0.883 | -0.18 | 0.01060331 | 0.05502976 | 8.00E-04 | 0.008 | 0.0032 | 0.0032 | TRUE | 0.0253 |
| Acot8 | 429 | 566 | 530 | 685 | 465 | 441 | 456 | 565 | 532 | 538 | 500 | 483 | 512.44 | 518 | 507 | 0.98 | -0.029 | 0.81730421 | 0.90708637 | 0.0056 | 0.0133 | 0.0091 | 0.0091 | TRUE | 0.127 |
| Acot9 | 3251 | 3746 | 3613 | 4052 | 2953 | 3320 | 3453 | 3741 | 3628 | 3185 | 3177 | 3634 | 3469.83 | 3607 | 3332 | 0.923 | -0.115 | 0.18554753 | 0.3890589 | 0.0035 | 0.0079 | 0.0051 | 0.0051 | TRUE | 0.0759 |
| Acox1 | 1208 | 1236 | 1183 | 1782 | 1260 | 1303 | 1283 | 1234 | 1188 | 1401 | 1356 | 1426 | 1314.71 | 1235 | 1394 | 1.129 | 0.175 | 0.02584869 | 0.10405451 | 0 | 0.0094 | 0.0037 | 0.0037 | TRUE | 0.0138 |
| Acox2 | 0 | 1 | 0 | 2 | 0 | 1 | 0 | 1 | 0 | 2 | 0 | 1 | 0.61 | 0 | 1 | 2.474 | 1.307 | 0.62064351 | NA | 0 | 5.3994 | 3.481 | 3.481 | TRUE | 0.405 |
| Acox3 | 211 | 245 | 254 | 319 | 235 | 265 | 224 | 245 | 255 | 251 | 253 | 290 | 252.92 | 241 | 265 | 1.094 | 0.13 | 0.35557303 | 0.57395331 | 0 | 0.0199 | 0.0103 | 0.0103 | TRUE | 0.0771 |
| Acoxl | 17 | 19 | 35 | 25 | 20 | 16 | 18 | 19 | 35 | 20 | 22 | 18 | 21.81 | 24 | 20 | 0.812 | -0.3 | 0.50572508 | 0.70029387 | 0.0408 | 0.158 | 0.1001 | 0.1001 | TRUE | 0.9765 |
| Acp1 | 1654 | 1918 | 1904 | 2686 | 1884 | 1856 | 1757 | 1916 | 1912 | 2112 | 2027 | 2032 | 1959.08 | 1862 | 2057 | 1.105 | 0.144 | 0.05855002 | 0.18288804 | 9.00E-04 | 0.0086 | 0.0037 | 0.0037 | TRUE | 0.029 |
| Acp2 | 453 | 485 | 534 | 838 | 598 | 467 | 481 | 484 | 536 | 659 | 643 | 511 | 552.52 | 500 | 604 | 1.209 | 0.274 | 0.04109351 | 0.14360584 | 0.0096 | 0.0129 | 0.0111 | 0.0111 | TRUE | 0.2104 |
| Acp5 | 515 | 728 | 794 | 858 | 631 | 612 | 547 | 727 | 797 | 674 | 679 | 670 | 682.45 | 690 | 674 | 0.976 | -0.034 | 0.81723265 | 0.90708637 | 0.017 | 0.0117 | 0.0145 | 0.0145 | TRUE | 0.3878 |
| Acp6 | 7776 | 8947 | 8345 | 11431 | 8739 | 8924 | 8259 | 8935 | 8379 | 8986 | 9403 | 9769 | 8955.28 | 8524 | 9386 | 1.101 | 0.139 | 0.04990277 | 0.1639384 | 0.0016 | 0.0073 | 0.0035 | 0.0035 | TRUE | 0.0217 |
| Acpl2 | 751 | 797 | 726 | 910 | 641 | 750 | 798 | 796 | 729 | 715 | 690 | 821 | 758.11 | 774 | 742 | 0.958 | -0.062 | 0.56784459 | 0.74733351 | 0.0041 | 0.0112 | 0.0072 | 0.0072 | TRUE | 0.1003 |
| Acpp | 192 | 266 | 277 | 384 | 319 | 363 | 204 | 266 | 278 | 302 | 343 | 397 | 298.36 | 249 | 347 | 1.391 | 0.476 | 0.00650564 | 0.03815231 | 0.0192 | 0.0179 | 0.0186 | 0.0186 | TRUE | 0.2776 |
| Acpt | 1 | 2 | 0 | 0 | 0 | 0 | 1 | 2 | 0 | 0 | 0 | 0 | 0.51 | 1 | 0 | 0.176 | -2.508 | 0.44268408 | NA | 0 | 6.4675 | 6.1932 | 6.1932 | TRUE | 0.064 |
| Acr | 1 | 1 | 0 | 2 | 3 | 2 | 1 | 1 | 0 | 2 | 3 | 2 | 1.51 | 1 | 2 | 3.377 | 1.756 | 0.28504553 | NA | 0 | 2.191 | 0.9842 | 0.9842 | TRUE | 0.2535 |
| Acrbp | 685 | 752 | 789 | 532 | 433 | 559 | 728 | 751 | 792 | 418 | 466 | 612 | 627.81 | 757 | 499 | 0.657 | -0.605 | 8.18E-05 | 0.0011734 | 0.0189 | 0.0122 | 0.0153 | 0.0153 | TRUE | 0.4564 |
| Acrv1 | 0 | 0 | 0 | 0 | 0 | 0 | 0 | 0 | 0 | 0 | 0 | 0 | 0 | 0 | 0 | NA | NA | NA | NA | NA | NA | NA | NA | NA | NA |

| | | | | | | | | | | | | | | | | | | | | | | | | | |
|---|---|---|---|---|---|---|---|---|---|---|---|---|---|---|---|---|---|---|---|---|---|---|---|---|---|
| Acsbg1 | 133 | 67 | 64 | 111 | 71 | 59 | 141 | 67 | 64 | 87 | 76 | 65 | 83.45 | 91 | 76 | 0.841 | -0.25 | 0.46916182 | 0.67141562 | 0.1034 | 0.0464 | 0.0739 | 0.0739 | TRUE | 2.233 |
| Acsbg2 | 16 | 17 | 11 | 30 | 21 | 24 | 17 | 17 | 11 | 24 | 23 | 26 | 19.58 | 15 | 24 | 1.609 | 0.687 | 0.11560061 | 0.28713974 | 0 | 0.1752 | 0.0829 | 0.0829 | TRUE | 0.2466 |
| Acsf2 | 2834 | 3122 | 3192 | 2601 | 2084 | 2307 | 3010 | 3118 | 3205 | 2045 | 2242 | 2525 | 2690.94 | 3111 | 2271 | 0.729 | -0.455 | 4.19E-06 | 9.51E-05 | 0.0056 | 0.0081 | 0.0067 | 0.0067 | TRUE | 0.1163 |
| Acsf3 | 1089 | 1053 | 1112 | 1587 | 1133 | 1089 | 1157 | 1052 | 1117 | 1248 | 1219 | 1192 | 1163.94 | 1109 | 1220 | 1.101 | 0.139 | 0.09290719 | 0.24896038 | 5.00E-04 | 0.0097 | 0.0041 | 0.0041 | TRUE | 0.024 |
| Acsl1 | 5921 | 6975 | 7228 | 5776 | 4618 | 5251 | 6289 | 6966 | 7258 | 4541 | 4969 | 5748 | 5961.7 | 6838 | 5086 | 0.744 | -0.427 | 0.00011574 | 0.00154825 | 0.0096 | 0.0075 | 0.0087 | 0.0087 | TRUE | 0.158 |
| Acsl3 | 5829 | 6108 | 5776 | 7573 | 5629 | 6121 | 6191 | 6100 | 5800 | 5953 | 6056 | 6700 | 6133.57 | 6030 | 6236 | 1.034 | 0.048 | 0.53589943 | 0.72271961 | 0.0024 | 0.0074 | 0.0042 | 0.0042 | TRUE | 0.0514 |
| Acsl4 | 4519 | 4247 | 4293 | 5090 | 3474 | 3797 | 4800 | 4242 | 4311 | 4001 | 3738 | 4156 | 4207.95 | 4451 | 3965 | 0.891 | -0.167 | 0.05180343 | 0.16792362 | 0.0034 | 0.0077 | 0.005 | 0.005 | TRUE | 0.0573 |
| Acsl5 | 484 | 652 | 794 | 679 | 504 | 594 | 514 | 651 | 797 | 534 | 542 | 650 | 614.8 | 654 | 575 | 0.879 | -0.186 | 0.28490093 | 0.50627409 | 0.0273 | 0.0123 | 0.0202 | 0.0202 | TRUE | 0.177 |
| Acsl6 | 664 | 851 | 784 | 910 | 650 | 628 | 705 | 850 | 787 | 715 | 699 | 687 | 740.76 | 781 | 700 | 0.897 | -0.156 | 0.13966525 | 0.32422732 | 0.0033 | 0.0114 | 0.0067 | 0.0067 | TRUE | 0.0842 |
| Acsm1 | 0 | 0 | 0 | 0 | 0 | 0 | 0 | 0 | 0 | 0 | 0 | 0 | 0 | 0 | 0 | NA | NA | NA | NA | NA | NA | NA | NA | NA | NA |
| Acsm2 | 0 | 0 | 0 | 0 | 0 | 0 | 0 | 0 | 0 | 0 | 0 | 0 | 0 | 0 | 0 | NA | NA | NA | NA | NA | NA | NA | NA | NA | NA |
| Acsm3 | 2 | 5 | 3 | 4 | 3 | 3 | 2 | 5 | 3 | 3 | 3 | 3 | 3.3 | 3 | 3 | 0.949 | -0.075 | 0.94027256 | NA | 0 | 1.0059 | 0.4253 | 0.4253 | TRUE | 0.2564 |
| Acsm4 | 0 | 0 | 0 | 3 | 4 | 1 | 0 | 0 | 0 | 2 | 4 | 1 | 1.29 | 0 | 2 | 13.751 | 3.781 | 0.06446831 | NA | 0 | 2.5552 | 1.8246 | 1.8246 | TRUE | 0.1011 |
| Acsm5 | 0 | 0 | 0 | 0 | 0 | 0 | 0 | 0 | 0 | 0 | 0 | 0 | 0 | 0 | 0 | NA | NA | NA | NA | NA | NA | NA | NA | NA | NA |
| Acss1 | 1180 | 2471 | 2579 | 699 | 616 | 986 | 1253 | 2468 | 2590 | 549 | 663 | 1079 | 1433.73 | 2104 | 764 | 0.363 | -1.462 | 0.00052082 | 0.00534696 | 0.127 | 0.0092 | 0.0593 | 0.127 | TRUE | 1.5434 |
| Acss2 | 2445 | 2695 | 2552 | 2438 | 1848 | 1909 | 2597 | 2692 | 2562 | 1917 | 1988 | 2090 | 2307.6 | 2617 | 1998 | 0.763 | -0.39 | 1.05E-07 | 3.73E-06 | 8.00E-04 | 0.0083 | 0.0034 | 0.0034 | TRUE | 0.0193 |
| Acss3 | 71 | 65 | 63 | 72 | 44 | 23 | 75 | 65 | 63 | 57 | 47 | 25 | 55.45 | 68 | 43 | 0.64 | -0.643 | 0.054025 | 0.17281409 | 0.0553 | 0.0663 | 0.0615 | 0.0615 | TRUE | 0.9561 |
| Acta1 | 11 | 9 | 11 | 13 | 10 | 11 | 12 | 9 | 11 | 10 | 11 | 12 | 10.79 | 11 | 11 | 1.039 | 0.055 | 0.92185743 | 0.96126008 | 0 | 0.3122 | 0.1323 | 0.1323 | TRUE | 0.0631 |
| Acta2 | 34 | 30 | 19 | 62 | 31 | 27 | 36 | 30 | 19 | 49 | 33 | 30 | 32.8 | 28 | 37 | 1.324 | 0.404 | 0.30686578 | 0.52738472 | 0.0543 | 0.1073 | 0.0818 | 0.0818 | TRUE | 0.5969 |
| Actb | 51206 | 39568 | 38371 | 80203 | 54840 | 42302 | 54390 | 39517 | 38529 | 63049 | 59004 | 46306 | 50132.54 | 44145 | 56120 | 1.271 | 0.346 | 0.0327798 | 0.12222295 | 0.0323 | 0.007 | 0.0189 | 0.0189 | TRUE | 0.5046 |
| Actbl2 | 0 | 0 | 0 | 0 | 0 | 0 | 0 | 0 | 0 | 0 | 0 | 0 | 0 | 0 | 0 | NA | NA | NA | NA | NA | NA | NA | NA | NA | NA |
| Actc1 | 5 | 2 | 1 | 10 | 3 | 2 | 5 | 2 | 1 | 8 | 3 | 2 | 3.6 | 3 | 4 | 1.644 | 0.717 | 0.5169318 | NA | 0.2636 | 0.9224 | 0.5884 | 0.5884 | TRUE | 1.0679 |
| Actg1 | 43362 | 42323 | 44161 | 67970 | 45735 | 44380 | 46058 | 42268 | 44342 | 53433 | 49208 | 48581 | 47315.07 | 44223 | 50407 | 1.14 | 0.189 | 0.00966682 | 0.05131778 | 0.0023 | 0.007 | 0.0038 | 0.0038 | TRUE | 0.0338 |
| Actg2 | 1 | 2 | 0 | 3 | 1 | 0 | 1 | 2 | 0 | 2 | 1 | 0 | 1.08 | 1 | 1 | 1.161 | 0.215 | 0.91303847 | NA | 0 | 3.0508 | 1.8901 | 1.8901 | TRUE | 0.2381 |
| Actl10 | 5 | 8 | 9 | 7 | 3 | 7 | 5 | 8 | 9 | 6 | 3 | 8 | 6.46 | 7 | 6 | 0.732 | -0.45 | 0.55051042 | 0.73491924 | 0 | 0.5173 | 0.2518 | 0.2518 | TRUE | 0.2137 |
| Actl11 | 0 | 0 | 0 | 0 | 0 | 0 | 0 | 0 | 0 | 0 | 0 | 0 | 0 | 0 | 0 | NA | NA | NA | NA | NA | NA | NA | NA | NA | NA |
| Actl6a | 7044 | 7187 | 7035 | 9147 | 6640 | 7084 | 7482 | 7178 | 7064 | 7191 | 7144 | 7755 | 7302.17 | 7241 | 7363 | 1.017 | 0.024 | 0.72790894 | 0.85376514 | 0.0012 | 0.0074 | 0.0033 | 0.0033 | TRUE | 0.0263 |
| Actl6b | 0 | 0 | 1 | 0 | 0 | 0 | 0 | 0 | 1 | 0 | 0 | 0 | 0.17 | 0 | 0 | 0.49 | -1.029 | 0.80087 | NA | 0 | 19.6928 | 10 | 10 | TRUE | 0.4098 |
| Actl7a | 0 | 0 | 0 | 0 | 1 | 0 | 0 | 0 | 0 | 0 | 1 | 0 | 0.18 | 0 | 0 | 1.859 | 0.894 | 0.82650734 | NA | 0 | 18.3785 | 10 | 10 | TRUE | 0.5204 |
| Actl7b | 43 | 39 | 37 | 67 | 42 | 34 | 46 | 39 | 37 | 53 | 45 | 37 | 42.81 | 41 | 45 | 1.117 | 0.16 | 0.60008112 | 0.77212039 | 0 | 0.0839 | 0.0434 | 0.0434 | TRUE | 0.1988 |
| Actl9 | 0 | 0 | 0 | 0 | 0 | 0 | 0 | 0 | 0 | 0 | 0 | 0 | 0 | 0 | 0 | NA | NA | NA | NA | NA | NA | NA | NA | NA | NA |
| Actn1 | 11297 | 11594 | 11811 | 13359 | 10435 | 10749 | 11999 | 11579 | 11859 | 10502 | 11227 | 11766 | 11488.94 | 11812 | 11165 | 0.945 | -0.082 | 0.24760752 | 0.46433167 | 0.0016 | 0.0072 | 0.0035 | 0.0035 | TRUE | 0.0332 |
| Actn2 | 343 | 307 | 250 | 523 | 423 | 294 | 364 | 307 | 251 | 411 | 455 | 322 | 351.67 | 307 | 396 | 1.29 | 0.367 | 0.05134713 | 0.16701811 | 0.0292 | 0.0163 | 0.0227 | 0.0227 | TRUE | 0.2542 |
| Actn3 | 2044 | 1464 | 1619 | 611 | 456 | 410 | 2171 | 1462 | 1626 | 480 | 491 | 449 | 1113.1 | 1753 | 473 | 0.27 | -1.889 | 1.89E-33 | 1.30E-30 | 0.0215 | 0.0099 | 0.0164 | 0.0164 | TRUE | 0.4873 |
| Actn4 | 23418 | 20450 | 20273 | 33580 | 23110 | 20685 | 24874 | 20424 | 20356 | 26398 | 24865 | 22643 | 23259.96 | 21885 | 24635 | 1.126 | 0.171 | 0.11706095 | 0.28934106 | 0.0099 | 0.007 | 0.0085 | 0.0085 | TRUE | 0.1747 |
| Actr10 | 2242 | 2058 | 2025 | 3559 | 2460 | 2276 | 2381 | 2055 | 2033 | 2798 | 2647 | 2491 | 2401.02 | 2156 | 2645 | 1.227 | 0.295 | 0.00245538 | 0.01808793 | 0.0051 | 0.0083 | 0.0064 | 0.0064 | TRUE | 0.1003 |
| Actr1a | 5259 | 5322 | 5360 | 6598 | 4410 | 4508 | 5586 | 5315 | 5382 | 5187 | 4745 | 4935 | 5191.59 | 5428 | 4956 | 0.913 | -0.131 | 0.05730542 | 0.18046194 | 0.001 | 0.0075 | 0.0032 | 0.0032 | TRUE | 0.0206 |
| Actr1b | 3098 | 3269 | 3192 | 4741 | 3465 | 3323 | 3291 | 3265 | 3205 | 3727 | 3728 | 3638 | 3475.53 | 3254 | 3698 | 1.137 | 0.185 | 0.00213274 | 0.0162031 | 0 | 0.0079 | 0.0023 | 0.0023 | TRUE | 0.0024 |
| Actr2 | 19693 | 20963 | 20352 | 23691 | 16974 | 17659 | 20917 | 20936 | 20436 | 18624 | 18263 | 19331 | 19751.08 | 20763 | 18739 | 0.903 | -0.148 | 0.00927339 | 0.04966378 | 4.00E-04 | 0.0071 | 0.0023 | 0.0023 | TRUE | 0.0093 |
| Actr3 | 12601 | 14544 | 14562 | 16933 | 12282 | 13686 | 13384 | 14525 | 14622 | 13311 | 13215 | 14981 | 14006.51 | 14177 | 13836 | 0.976 | -0.035 | 0.67146915 | 0.81667744 | 0.0031 | 0.0071 | 0.0049 | 0.0049 | TRUE | 0.064 |
| Actr3b | 848 | 967 | 918 | 1176 | 866 | 793 | 901 | 966 | 922 | 924 | 932 | 868 | 918.76 | 930 | 908 | 0.977 | -0.033 | 0.70288816 | 0.83762068 | 0 | 0.0105 | 0.0043 | 0.0043 | TRUE | 0.0172 |
| Actr5 | 1157 | 1154 | 1076 | 1362 | 1044 | 974 | 1229 | 1153 | 1080 | 1071 | 1123 | 1066 | 1120.34 | 1154 | 1087 | 0.942 | -0.087 | 0.33162726 | 0.55217325 | 0.0016 | 0.0098 | 0.0048 | 0.0048 | TRUE | 0.0385 |
| Actr6 | 1508 | 1562 | 1420 | 2054 | 1477 | 1447 | 1602 | 1560 | 1426 | 1615 | 1589 | 1584 | 1562.57 | 1529 | 1596 | 1.044 | 0.062 | 0.4460148 | 0.65236459 | 0.0012 | 0.009 | 0.0041 | 0.0041 | TRUE | 0.0422 |
| Actr8 | 2487 | 2717 | 2631 | 2814 | 2154 | 2324 | 2642 | 2713 | 2642 | 2212 | 2318 | 2544 | 2511.77 | 2666 | 2358 | 0.884 | -0.178 | 0.03016214 | 0.11571813 | 0.0021 | 0.0082 | 0.0044 | 0.0044 | TRUE | 0.0575 |
| Actrt1 | 0 | 0 | 0 | 0 | 0 | 0 | 0 | 0 | 0 | 0 | 0 | 0 | 0 | 0 | 0 | NA | NA | NA | NA | NA | NA | NA | NA | NA | NA |
| Actrt2 | 0 | 0 | 1 | 0 | 0 | 0 | 0 | 0 | 1 | 0 | 0 | 0 | 0.17 | 0 | 0 | 0.49 | -1.029 | 0.80087 | NA | 0 | 19.6928 | 10 | 10 | TRUE | 0.4098 |
| Actrt3 | 4 | 9 | 5 | 8 | 3 | 2 | 4 | 9 | 5 | 6 | 3 | 2 | 4.99 | 6 | 4 | 0.657 | -0.606 | 0.49245226 | 0.6887432 | 0 | 0.6666 | 0.3565 | 0.3565 | TRUE | 0.6114 |
| Acvr1 | 303 | 456 | 471 | 578 | 444 | 437 | 322 | 455 | 473 | 454 | 478 | 478 | 443.44 | 417 | 470 | 1.127 | 0.173 | 0.28678878 | 0.50800007 | 0.0189 | 0.0143 | 0.0167 | 0.0167 | TRUE | 0.4541 |
| Acvr1b | 3109 | 3986 | 4077 | 3813 | 2957 | 3549 | 3302 | 3981 | 4094 | 2997 | 3182 | 3885 | 3573.48 | 3792 | 3355 | 0.884 | -0.177 | 0.17512329 | 0.37485771 | 0.0157 | 0.0078 | 0.0121 | 0.0121 | TRUE | 0.2321 |
| Acvr1c | 2 | 4 | 3 | 7 | 9 | 8 | 2 | 4 | 3 | 6 | 10 | 9 | 5.51 | 3 | 8 | 2.588 | 1.372 | 0.10374065 | 0.26796554 | 0 | 0.6045 | 0.2831 | 0.2831 | TRUE | 0.2815 |
| Acvr2a | 864 | 968 | 930 | 1271 | 911 | 897 | 918 | 967 | 934 | 999 | 980 | 982 | 963.26 | 940 | 987 | 1.051 | 0.072 | 0.37291614 | 0.58928297 | 0 | 0.0103 | 0.0036 | 0.0036 | TRUE | 0.0077 |
| Acvr2b | 946 | 1244 | 1379 | 1360 | 1022 | 1049 | 1005 | 1242 | 1385 | 1069 | 1100 | 1148 | 1158.15 | 1211 | 1106 | 0.913 | -0.132 | 0.31320122 | 0.53374497 | 0.0119 | 0.0097 | 0.0114 | 0.0114 | TRUE | 0.2643 |
| Acvrl1 | 164 | 185 | 188 | 131 | 131 | 151 | 174 | 185 | 189 | 103 | 141 | 165 | 159.49 | 183 | 136 | 0.743 | -0.429 | 0.03883137 | 0.13776689 | 0.0218 | 0.0276 | 0.0247 | 0.0247 | TRUE | 0.5065 |
| Acy1 | 2386 | 2787 | 2879 | 2363 | 1958 | 2092 | 2534 | 2783 | 2891 | 1858 | 2107 | 2290 | 2410.48 | 2736 | 2085 | 0.761 | -0.393 | 0.00021755 | 0.00259938 | 0.007 | 0.0083 | 0.0077 | 0.0077 | TRUE | 0.1117 |
| Acy3 | 209 | 267 | 245 | 350 | 244 | 269 | 222 | 267 | 246 | 275 | 263 | 294 | 261.13 | 245 | 277 | 1.132 | 0.179 | 0.20152889 | 0.40908409 | 0.0014 | 0.0195 | 0.0102 | 0.0102 | TRUE | 0.073 |
| Acyp1 | 352 | 601 | 609 | 578 | 439 | 545 | 374 | 600 | 612 | 454 | 472 | 597 | 518.15 | 529 | 508 | 0.96 | -0.059 | 0.77213312 | 0.88112898 | 0.045 | 0.0133 | 0.0284 | 0.0284 | TRUE | 0.7618 |
| Acyp2 | 31 | 23 | 26 | 83 | 53 | 33 | 33 | 23 | 26 | 65 | 57 | 36 | 40.07 | 27 | 53 | 1.945 | 0.96 | 0.00770832 | 0.04320978 | 0.0412 | 0.0891 | 0.0656 | 0.0656 | TRUE | 0.4468 |
| Ada | 1131 | 966 | 868 | 1299 | 968 | 846 | 1201 | 965 | 872 | 1021 | 1042 | 926 | 1004.4 | 1013 | 996 | 0.984 | -0.023 | 0.86826641 | 0.93521122 | 0.0145 | 0.0102 | 0.0125 | 0.0125 | TRUE | 0.3167 |
| Adad1 | 37 | 53 | 48 | 68 | 70 | 64 | 39 | 53 | 48 | 53 | 75 | 70 | 56.54 | 47 | 66 | 1.406 | 0.492 | 0.08083367 | 0.22612783 | 0 | 0.0652 | 0.0389 | 0.0389 | TRUE | 0.2892 |
| Adad2 | 97 | 98 | 112 | 168 | 99 | 92 | 103 | 98 | 112 | 132 | 107 | 101 | 108.78 | 104 | 113 | 1.088 | 0.122 | 0.55259292 | 0.73641702 | 0 | 0.0372 | 0.0212 | 0.0212 | TRUE | 0.2359 |
| Adal | 854 | 806 | 828 | 1056 | 831 | 775 | 907 | 805 | 831 | 830 | 894 | 848 | 852.68 | 848 | 857 | 1.011 | 0.016 | 0.86557345 | 0.93364615 | 0.0015 | 0.0108 | 0.0053 | 0.0053 | TRUE | 0.0441 |
| Adam10 | 3589 | 3544 | 3593 | 4653 | 3254 | 3272 | 3812 | 3539 | 3608 | 3658 | 3501 | 3582 | 3616.66 | 3653 | 3580 | 0.98 | -0.029 | 0.67120927 | 0.81655574 | 6.00E-04 | 0.0078 | 0.003 | 0.003 | TRUE | 0.0176 |
| Adam11 | 274 | 267 | 254 | 235 | 196 | 193 | 291 | 267 | 255 | 185 | 211 | 211 | 236.6 | 271 | 202 | 0.745 | -0.425 | 0.00309512 | 0.02175412 | 0 | 0.0208 | 0.0106 | 0.0106 | TRUE | 0.069 |
| Adam12 | 6 | 5 | 7 | 4 | 5 | 2 | 6 | 5 | 7 | 3 | 5 | 2 | 4.85 | 6 | 3 | 0.58 | -0.786 | 0.36009434 | 0.57852786 | 0 | 0.686 | 0.3114 | 0.3114 | TRUE | 0.2632 |
| Adam15 | 1603 | 1682 | 1575 | 1964 | 1494 | 1405 | 1703 | 1680 | 1581 | 1544 | 1607 | 1538 | 1608.89 | 1655 | 1563 | 0.945 | -0.082 | 0.2778728 | 0.4989878 | 4.00E-04 | 0.009 | 0.0035 | 0.0035 | TRUE | 0.0181 |
| Adam17 | 3547 | 3460 | 3315 | 4007 | 2834 | 2898 | 3768 | 3456 | 3329 | 3150 | 3049 | 3172 | 3320.53 | 3518 | 3124 | 0.888 | -0.171 | 0.02806111 | 0.11003973 | 0.0018 | 0.0079 | 0.0041 | 0.0041 | TRUE | 0.047 |
| Adam18 | 0 | 4 | 0 | 5 | 3 | 1 | 0 | 4 | 0 | 4 | 3 | 1 | 2.04 | 1 | 3 | 2.069 | 1.049 | 0.52660606 | NA | 1.0558 | 1.6208 | 1.4206 | 1.4206 | TRUE | 1.898 |
| Adam19 | 9035 | 13034 | 13578 | 7656 | 6319 | 9163 | 9597 | 13017 | 13634 | 6019 | 6799 | 10030 | 9849.24 | 12083 | 7616 | 0.63 | -0.666 | 0.00075359 | 0.00713849 | 0.055 | 0.0072 | 0.0281 | 0.0281 | TRUE | 0.9387 |
| Adam1a | 11 | 8 | 6 | 12 | 7 | 6 | 12 | 8 | 6 | 9 | 8 | 7 | 8.21 | 9 | 8 | 0.926 | -0.111 | 0.86690447 | 0.93412739 | 0 | 0.4084 | 0.1945 | 0.1945 | TRUE | 0.2992 |

| | | | | | | | | | | | | | | | | | | | | | | | | | |
|---|---|---|---|---|---|---|---|---|---|---|---|---|---|---|---|---|---|---|---|---|---|---|---|---|---|
| Adam1b | 1 | 0 | 0 | 1 | 1 | 0 | 1 | 0 | 0 | 1 | 1 | 0 | 0.49 | 0 | 1 | 1.712 | 0.776 | 0.78488768 | NA | 0 | 6.7665 | 4.0387 | 4.0387 | TRUE | 0.4614 |
| Adam2 | 14 | 14 | 17 | 9 | 3 | 7 | 15 | 14 | 17 | 7 | 3 | 8 | 10.65 | 15 | 6 | 0.395 | -1.342 | 0.02870092 | 0.11155023 | 0 | 0.3163 | 0.157 | 0.157 | TRUE | 0.3372 |
| Adam20 | 0 | 0 | 2 | 0 | 1 | 0 | 0 | 0 | 2 | 0 | 1 | 0 | 0.51 | 1 | 0 | 0.52 | -0.944 | 0.77815412 | NA | 0 | 6.416 | 6.3253 | 6.3253 | TRUE | 0.9739 |
| Adam21 | 1 | 2 | 0 | 0 | 0 | 0 | 1 | 2 | 0 | 0 | 0 | 0 | 0.51 | 1 | 0 | 0.176 | -2.508 | 0.44268408 | NA | 0 | 6.4675 | 6.1932 | 6.1932 | TRUE | 0.064 |
| Adam22 | 597 | 710 | 784 | 750 | 570 | 602 | 634 | 709 | 787 | 590 | 613 | 659 | 665.38 | 710 | 621 | 0.873 | -0.196 | 0.09725975 | 0.25625993 | 0.006 | 0.0119 | 0.0086 | 0.0086 | TRUE | 0.1066 |
| Adam23 | 4339 | 5239 | 5568 | 5022 | 3937 | 4438 | 4609 | 5232 | 5591 | 3948 | 4236 | 4858 | 4745.64 | 5144 | 4347 | 0.845 | -0.243 | 0.03185041 | 0.12006324 | 0.0101 | 0.0076 | 0.009 | 0.009 | TRUE | 0.1285 |
| Adam24 | 1 | 0 | 0 | 0 | 0 | 0 | 1 | 0 | 0 | 0 | 0 | 0 | 0.18 | 0 | 0 | 0.49 | -1.029 | 0.80087 | NA | 0 | 18.6165 | 10 | 10 | TRUE | 0.4614 |
| Adam25 | 2 | 1 | 0 | 2 | 2 | 1 | 2 | 1 | 0 | 2 | 2 | 1 | 1.32 | 1 | 2 | 1.557 | 0.639 | 0.69951162 | NA | 0 | 2.4958 | 1.173 | 1.173 | TRUE | 0.2827 |
| Adam26a | 0 | 0 | 0 | 0 | 0 | 0 | 0 | 0 | 0 | 0 | 0 | 0 | 0 | 0 | 0 | NA | NA | NA | NA | NA | NA | NA | NA | NA | NA |
| Adam26b | 0 | 0 | 0 | 0 | 0 | 0 | 0 | 0 | 0 | 0 | 0 | 0 | 0 | 0 | 0 | NA | NA | NA | NA | NA | NA | NA | NA | NA | NA |
| Adam28 | 0 | 0 | 0 | 1 | 0 | 0 | 0 | 0 | 0 | 1 | 0 | 0 | 0.13 | 0 | 0 | 1.859 | 0.894 | 0.82650734 | NA | 0 | 25.1515 | 10 | 10 | TRUE | 0.2563 |
| Adam29 | 0 | 0 | 0 | 0 | 0 | 0 | 0 | 0 | 0 | 0 | 0 | 0 | 0 | 0 | 0 | NA | NA | NA | NA | NA | NA | NA | NA | NA | NA |
| Adam3 | 4 | 2 | 3 | 8 | 5 | 1 | 4 | 2 | 3 | 6 | 5 | 1 | 3.67 | 3 | 4 | 1.405 | 0.491 | 0.63829365 | NA | 0 | 0.9045 | 0.5072 | 0.5072 | TRUE | 0.6649 |
| Adam30 | 2 | 3 | 6 | 10 | 5 | 3 | 2 | 3 | 6 | 8 | 5 | 3 | 4.61 | 4 | 5 | 1.499 | 0.584 | 0.52307355 | 0.71298212 | 0 | 0.7213 | 0.3789 | 0.3789 | TRUE | 0.3051 |
| Adam32 | 20 | 12 | 14 | 24 | 23 | 17 | 21 | 12 | 14 | 19 | 25 | 19 | 18.25 | 16 | 21 | 1.316 | 0.396 | 0.39408431 | 0.6078132 | 0 | 0.1874 | 0.0995 | 0.0995 | TRUE | 0.4175 |
| Adam33 | 48 | 25 | 19 | 31 | 20 | 22 | 51 | 25 | 19 | 24 | 22 | 24 | 27.5 | 32 | 23 | 0.739 | -0.436 | 0.34500715 | 0.56434898 | 0.1066 | 0.1267 | 0.1167 | 0.1167 | TRUE | 1.5284 |
| Adam34 | 0 | 0 | 0 | 0 | 0 | 0 | 0 | 0 | 0 | 0 | 0 | 0 | 0 | 0 | 0 | NA | NA | NA | NA | NA | NA | NA | NA | NA | NA |
| Adam39 | 0 | 0 | 0 | 0 | 0 | 0 | 0 | 0 | 0 | 0 | 0 | 0 | 0 | 0 | 0 | NA | NA | NA | NA | NA | NA | NA | NA | NA | NA |
| Adam4 | 7 | 5 | 2 | 7 | 1 | 3 | 7 | 5 | 2 | 6 | 1 | 3 | 4.05 | 5 | 3 | 0.703 | -0.509 | 0.61923074 | NA | 0.1456 | 0.8204 | 0.5058 | 0.5058 | TRUE | 0.2402 |
| Adam5 | 1 | 4 | 0 | 2 | 1 | 0 | 1 | 4 | 0 | 2 | 1 | 0 | 1.28 | 2 | 1 | 0.539 | -0.891 | 0.64308139 | NA | 0 | 2.5723 | 1.8277 | 1.8277 | TRUE | 0.5843 |
| Adam6a | 0 | 0 | 0 | 0 | 0 | 0 | 0 | 0 | 0 | 0 | 0 | 0 | 0 | 0 | 0 | NA | NA | NA | NA | NA | NA | NA | NA | NA | NA |
| Adam6b | 0 | 0 | 0 | 0 | 0 | 0 | 0 | 0 | 0 | 0 | 0 | 0 | 0 | 0 | 0 | NA | NA | NA | NA | NA | NA | NA | NA | NA | NA |
| Adam7 | 1 | 1 | 0 | 1 | 0 | 0 | 1 | 1 | 0 | 1 | 0 | 0 | 0.47 | 1 | 0 | 0.51 | -0.973 | 0.73452857 | NA | 0 | 6.9499 | 4.1835 | 4.1835 | TRUE | 0.2562 |
| Adam8 | 66 | 55 | 57 | 78 | 54 | 62 | 70 | 55 | 57 | 61 | 58 | 68 | 61.59 | 61 | 62 | 1.028 | 0.04 | 0.87489615 | 0.93825131 | 0 | 0.0604 | 0.0294 | 0.0294 | TRUE | 0.1525 |
| Adam9 | 2755 | 2835 | 2873 | 4162 | 2958 | 2976 | 2926 | 2831 | 2885 | 3272 | 3183 | 3258 | 3059.1 | 2881 | 3238 | 1.124 | 0.169 | 0.00647273 | 0.03809248 | 0 | 0.008 | 0.0024 | 0.0024 | TRUE | 0.0027 |
| Adamdec1 | 0 | 1 | 0 | 0 | 0 | 0 | 0 | 1 | 0 | 0 | 0 | 0 | 0.17 | 0 | 0 | 0.49 | -1.029 | 0.80087 | NA | 0 | 19.7991 | 10 | 10 | TRUE | 0.405 |
| Adamts1 | 82 | 52 | 62 | 115 | 101 | 78 | 87 | 52 | 62 | 90 | 109 | 85 | 80.96 | 67 | 95 | 1.414 | 0.5 | 0.05924559 | 0.1843701 | 0.027 | 0.0476 | 0.0379 | 0.0379 | TRUE | 0.5921 |
| Adamts10 | 721 | 1036 | 923 | 929 | 832 | 806 | 766 | 1035 | 927 | 730 | 895 | 882 | 872.51 | 909 | 836 | 0.918 | -0.123 | 0.39332459 | 0.60676012 | 0.0165 | 0.0107 | 0.0138 | 0.0138 | TRUE | 0.2255 |
| Adamts12 | 5 | 4 | 4 | 2 | 3 | 1 | 5 | 4 | 4 | 2 | 3 | 1 | 3.2 | 4 | 2 | 0.439 | -1.188 | 0.26552289 | NA | 0 | 1.0355 | 0.4555 | 0.4555 | TRUE | 0.2472 |
| Adamts13 | 1 | 0 | 0 | 0 | 1 | 1 | 1 | 0 | 0 | 0 | 1 | 1 | 0.54 | 0 | 1 | 1.923 | 0.944 | 0.73523723 | NA | 0 | 6.1214 | 3.9144 | 3.9144 | TRUE | 0.4614 |
| Adamts14 | 1008 | 1389 | 1459 | 1042 | 968 | 1056 | 1071 | 1387 | 1465 | 819 | 1042 | 1156 | 1156.58 | 1308 | 1006 | 0.768 | -0.381 | 0.02120651 | 0.09065994 | 0.0259 | 0.0098 | 0.0188 | 0.0188 | TRUE | 0.3197 |
| Adamts15 | 291 | 376 | 387 | 264 | 222 | 204 | 309 | 376 | 389 | 208 | 239 | 223 | 290.48 | 358 | 223 | 0.623 | -0.684 | 4.08E-06 | 9.30E-05 | 0.007 | 0.0182 | 0.0122 | 0.0122 | TRUE | 0.16 |
| Adamts16 | 1 | 1 | 4 | 1 | 4 | 1 | 1 | 1 | 4 | 1 | 4 | 1 | 2.04 | 2 | 2 | 0.994 | -0.009 | 0.99481086 | NA | 0.3171 | 1.619 | 0.9277 | 0.9277 | TRUE | 0.7847 |
| Adamts17 | 90 | 103 | 74 | 106 | 66 | 70 | 96 | 103 | 74 | 83 | 71 | 77 | 83.96 | 91 | 77 | 0.849 | -0.236 | 0.30765331 | 0.5282036 | 0 | 0.0461 | 0.0265 | 0.0265 | TRUE | 0.2474 |
| Adamts18 | 2 | 0 | 1 | 2 | 5 | 0 | 2 | 0 | 1 | 2 | 5 | 0 | 1.68 | 1 | 2 | 2.228 | 1.156 | 0.52044707 | NA | 1.1042 | 1.9678 | 1.6241 | 1.6241 | TRUE | 0.3871 |
| Adamts19 | 77 | 80 | 70 | 104 | 61 | 51 | 82 | 80 | 70 | 82 | 66 | 56 | 72.53 | 77 | 68 | 0.882 | -0.18 | 0.46901032 | 0.67137691 | 0.0093 | 0.0523 | 0.0309 | 0.0309 | TRUE | 0.3274 |
| Adamts2 | 65 | 88 | 110 | 140 | 107 | 80 | 69 | 88 | 110 | 110 | 115 | 88 | 96.69 | 89 | 104 | 1.17 | 0.227 | 0.35846144 | 0.57702121 | 0.0254 | 0.041 | 0.0335 | 0.0335 | TRUE | 0.2731 |
| Adamts20 | 128 | 120 | 110 | 167 | 152 | 127 | 136 | 120 | 110 | 131 | 164 | 139 | 133.35 | 122 | 145 | 1.182 | 0.242 | 0.20120688 | 0.40877592 | 0.0049 | 0.0316 | 0.0182 | 0.0182 | TRUE | 0.1341 |
| Adamts3 | 90 | 106 | 101 | 110 | 77 | 84 | 96 | 106 | 101 | 86 | 83 | 92 | 94.02 | 101 | 87 | 0.862 | -0.214 | 0.28212848 | 0.50353405 | 0 | 0.0419 | 0.0179 | 0.0179 | TRUE | 0.0206 |
| Adamts4 | 49 | 59 | 60 | 88 | 79 | 70 | 52 | 59 | 60 | 69 | 85 | 77 | 67 | 57 | 77 | 1.343 | 0.425 | 0.0777493 | 0.22059811 | 0 | 0.0561 | 0.0265 | 0.0265 | TRUE | 0.0817 |
| Adamts5 | 6 | 11 | 10 | 25 | 30 | 9 | 6 | 11 | 10 | 20 | 32 | 10 | 14.86 | 9 | 21 | 2.249 | 1.169 | 0.05480145 | 0.17448762 | 0.1454 | 0.2285 | 0.1876 | 0.1876 | TRUE | 0.1334 |
| Adamts6 | 5 | 1 | 3 | 3 | 1 | 2 | 5 | 1 | 3 | 2 | 1 | 2 | 2.49 | 3 | 2 | 0.613 | -0.706 | 0.56500434 | NA | 0 | 1.3295 | 0.6621 | 0.6621 | TRUE | 0.1475 |
| Adamts7 | 491 | 475 | 488 | 557 | 448 | 424 | 522 | 474 | 490 | 438 | 482 | 464 | 478.32 | 495 | 461 | 0.93 | -0.104 | 0.33334101 | 0.55395173 | 0 | 0.0138 | 0.0062 | 0.0062 | TRUE | 0.0247 |
| Adamts8 | 61 | 83 | 84 | 210 | 144 | 153 | 65 | 83 | 84 | 165 | 155 | 167 | 119.92 | 77 | 162 | 2.099 | 1.07 | 3.90E-08 | 1.60E-06 | 0 | 0.0344 | 0.0177 | 0.0177 | TRUE | 0.1802 |
| Adamts9 | 597 | 566 | 648 | 806 | 617 | 586 | 634 | 565 | 651 | 634 | 664 | 641 | 631.5 | 617 | 646 | 1.048 | 0.067 | 0.51133211 | 0.7048357 | 0.0014 | 0.0121 | 0.006 | 0.006 | TRUE | 0.063 |
| Adamtsl1 | 209 | 203 | 280 | 265 | 173 | 215 | 222 | 203 | 281 | 208 | 186 | 235 | 222.62 | 235 | 210 | 0.892 | -0.165 | 0.36288324 | 0.58058374 | 0.0169 | 0.0217 | 0.0192 | 0.0192 | TRUE | 0.3232 |
| Adamtsl2 | 1 | 0 | 0 | 0 | 0 | 0 | 1 | 0 | 0 | 0 | 0 | 0 | 0.18 | 0 | 0 | 0.49 | -1.029 | 0.80087 | NA | 0 | 18.6165 | 10 | 10 | TRUE | 0.4614 |
| Adamtsl3 | 13 | 22 | 16 | 31 | 12 | 19 | 14 | 22 | 16 | 24 | 13 | 21 | 18.32 | 17 | 19 | 1.128 | 0.173 | 0.71675175 | 0.84630504 | 0 | 0.1867 | 0.1099 | 0.1099 | TRUE | 0.4251 |
| Adamtsl4 | 316 | 412 | 405 | 351 | 297 | 340 | 336 | 411 | 407 | 276 | 320 | 372 | 353.57 | 385 | 323 | 0.836 | -0.258 | 0.1051392 | 0.27035167 | 0.0147 | 0.0162 | 0.0154 | 0.0154 | TRUE | 0.2047 |
| Adamtsl5 | 114 | 122 | 163 | 132 | 103 | 122 | 121 | 122 | 164 | 104 | 111 | 134 | 125.79 | 136 | 116 | 0.853 | -0.229 | 0.28346041 | 0.50505395 | 0.0166 | 0.0331 | 0.0248 | 0.0248 | TRUE | 0.3419 |
| Adap1 | 930 | 1207 | 1138 | 1381 | 979 | 954 | 988 | 1205 | 1143 | 1086 | 1053 | 1044 | 1086.54 | 1112 | 1061 | 0.954 | -0.068 | 0.5127195 | 0.70600768 | 0.0045 | 0.0099 | 0.0068 | 0.0068 | TRUE | 0.1136 |
| Adap2 | 248 | 371 | 321 | 388 | 283 | 264 | 263 | 371 | 322 | 305 | 304 | 289 | 309.13 | 319 | 299 | 0.939 | -0.09 | 0.56451454 | 0.74512706 | 0.0116 | 0.0176 | 0.0145 | 0.0145 | TRUE | 0.2255 |
| Adar | 3830 | 3458 | 3317 | 4540 | 3309 | 3099 | 4068 | 3454 | 3331 | 3569 | 3560 | 3392 | 3562.32 | 3618 | 3507 | 0.97 | -0.044 | 0.65132201 | 0.80449923 | 0.0052 | 0.0078 | 0.0067 | 0.0067 | TRUE | 0.1443 |
| Adarb1 | 1116 | 1176 | 1141 | 1188 | 952 | 936 | 1185 | 1174 | 1146 | 934 | 1024 | 1025 | 1081.39 | 1168 | 994 | 0.85 | -0.235 | 0.00577007 | 0.03488136 | 7.00E-04 | 0.0099 | 0.0043 | 0.0043 | TRUE | 0.0353 |
| Adarb2 | 3 | 0 | 1 | 5 | 1 | 2 | 3 | 0 | 1 | 4 | 1 | 2 | 1.9 | 1 | 2 | 1.767 | 0.821 | 0.58491712 | NA | 0.1799 | 1.7429 | 1.0584 | 1.0584 | TRUE | 0.6558 |
| Adat1 | 1342 | 1079 | 1073 | 1665 | 1200 | 917 | 1425 | 1078 | 1077 | 1309 | 1291 | 1004 | 1197.38 | 1193 | 1201 | 1.007 | 0.01 | 0.94703144 | 0.97351104 | 0.0232 | 0.0097 | 0.0166 | 0.0166 | TRUE | 0.3459 |
| Adat2 | 462 | 605 | 633 | 678 | 537 | 538 | 491 | 604 | 636 | 533 | 578 | 589 | 571.71 | 577 | 567 | 0.981 | -0.028 | 0.83101429 | 0.91427424 | 0.0087 | 0.0127 | 0.0105 | 0.0105 | TRUE | 0.1984 |
| Adat3 | 341 | 255 | 271 | 342 | 275 | 280 | 362 | 255 | 272 | 269 | 296 | 307 | 293.37 | 296 | 291 | 0.98 | -0.03 | 0.86082651 | 0.93125634 | 0.0169 | 0.0181 | 0.0175 | 0.0175 | TRUE | 0.423 |
| Adc | 868 | 1126 | 1162 | 992 | 813 | 922 | 922 | 1125 | 1167 | 780 | 875 | 1009 | 979.52 | 1071 | 888 | 0.828 | -0.272 | 0.04983258 | 0.16387811 | 0.0143 | 0.0103 | 0.0129 | 0.0129 | TRUE | 0.1768 |
| Adck1 | 853 | 926 | 954 | 1218 | 838 | 905 | 906 | 925 | 958 | 957 | 902 | 991 | 939.76 | 930 | 950 | 1.022 | 0.031 | 0.71932787 | 0.8479218 | 0 | 0.0104 | 0.0044 | 0.0044 | TRUE | 0.0228 |
| Adck2 | 198 | 249 | 249 | 315 | 235 | 250 | 210 | 249 | 250 | 248 | 253 | 274 | 247.19 | 236 | 258 | 1.09 | 0.124 | 0.38457936 | 0.59865631 | 0 | 0.0202 | 0.0106 | 0.0106 | TRUE | 0.1009 |
| Adck3 | 1703 | 2344 | 2542 | 1310 | 1211 | 1440 | 1809 | 2341 | 2552 | 1030 | 1303 | 1576 | 1768.56 | 2234 | 1303 | 0.583 | -0.779 | 1.29E-05 | 0.00024673 | 0.0358 | 0.0088 | 0.0224 | 0.0224 | TRUE | 0.3412 |
| Adck4 | 336 | 331 | 314 | 612 | 440 | 424 | 357 | 331 | 315 | 481 | 473 | 464 | 403.57 | 334 | 473 | 1.416 | 0.502 | 7.77E-06 | 0.00015985 | 0 | 0.0151 | 0.0065 | 0.0065 | TRUE | 0.0394 |
| Adck5 | 569 | 614 | 576 | 878 | 604 | 505 | 604 | 613 | 578 | 690 | 650 | 553 | 614.81 | 598 | 631 | 1.056 | 0.078 | 0.50765347 | 0.70185051 | 0.0054 | 0.0123 | 0.0084 | 0.0084 | TRUE | 0.1361 |
| Adcy1 | 1689 | 1610 | 1540 | 1464 | 1108 | 1171 | 1794 | 1608 | 1546 | 1151 | 1192 | 1282 | 1428.85 | 1649 | 1208 | 0.732 | -0.45 | 3.24E-06 | 7.64E-05 | 0.0035 | 0.0092 | 0.006 | 0.006 | TRUE | 0.0707 |
| Adcy10 | 10 | 7 | 6 | 14 | 13 | 5 | 11 | 7 | 6 | 11 | 14 | 5 | 9.02 | 8 | 10 | 1.297 | 0.375 | 0.57587174 | 0.75380936 | 0 | 0.3723 | 0.2107 | 0.2107 | TRUE | 0.3408 |
| Adcy2 | 71 | 110 | 81 | 186 | 104 | 91 | 75 | 110 | 81 | 146 | 112 | 100 | 104.06 | 89 | 119 | 1.347 | 0.429 | 0.08489159 | 0.23407519 | 0.0313 | 0.0386 | 0.0349 | 0.0349 | TRUE | 0.4243 |
| Adcy3 | 1174 | 1540 | 1562 | 1774 | 1310 | 1392 | 1247 | 1538 | 1568 | 1395 | 1409 | 1524 | 1446.87 | 1451 | 1443 | 0.994 | -0.009 | 0.93570905 | 0.96811881 | 0.0083 | 0.0092 | 0.0087 | 0.0087 | TRUE | 0.1816 |
| Adcy4 | 10 | 23 | 15 | 6 | 19 | 11 | 11 | 23 | 15 | 5 | 20 | 12 | 14.31 | 16 | 12 | 0.751 | -0.413 | 0.52552903 | 0.71494576 | 0.2283 | 0.2371 | 0.233 | 0.233 | TRUE | 0.2966 |

| | | | | | | | | | | | | | | | | | | | | | | | | | |
|---|---|---|---|---|---|---|---|---|---|---|---|---|---|---|---|---|---|---|---|---|---|---|---|---|---|
| Adcy5 | 179 | 168 | 135 | 149 | 123 | 134 | 190 | 168 | 136 | 117 | 132 | 147 | 148.27 | 165 | 132 | 0.801 | -0.32 | 0.1069908 | 0.27340553 | 0.0144 | 0.0291 | 0.0216 | 0.0216 | TRUE | 0.252 |
| Adcy6 | 370 | 580 | 622 | 467 | 359 | 414 | 393 | 579 | 625 | 367 | 386 | 453 | 467.23 | 532 | 402 | 0.755 | -0.406 | 0.03140899 | 0.11909665 | 0.0332 | 0.014 | 0.0235 | 0.0235 | TRUE | 0.609 |
| Adcy7 | 501 | 342 | 414 | 775 | 494 | 388 | 532 | 342 | 416 | 609 | 532 | 425 | 475.82 | 430 | 522 | 1.216 | 0.282 | 0.15190675 | 0.34295352 | 0.0376 | 0.0138 | 0.0258 | 0.0258 | TRUE | 0.387 |
| Adcy8 | 1 | 0 | 1 | 3 | 1 | 1 | 1 | 0 | 1 | 2 | 1 | 1 | 1.1 | 1 | 1 | 2.248 | 1.169 | 0.51947915 | NA | 0 | 3.004 | 1.3093 | 1.3093 | TRUE | 0.2561 |
| Adcy9 | 312 | 311 | 311 | 453 | 351 | 294 | 331 | 311 | 312 | 356 | 378 | 322 | 334.98 | 318 | 352 | 1.107 | 0.147 | 0.23936519 | 0.45534528 | 0 | 0.0167 | 0.0082 | 0.0082 | TRUE | 0.0604 |
| Adcyap1 | 20 | 31 | 26 | 27 | 26 | 20 | 21 | 31 | 26 | 21 | 28 | 22 | 24.9 | 26 | 24 | 0.903 | -0.147 | 0.70573412 | 0.83911335 | 0 | 0.1392 | 0.0691 | 0.0691 | TRUE | 0.1657 |
| Adcyap1r1 | 131 | 126 | 125 | 203 | 127 | 105 | 139 | 126 | 126 | 160 | 137 | 115 | 133.61 | 130 | 137 | 1.058 | 0.082 | 0.67520847 | 0.8190765 | 0.0087 | 0.0316 | 0.0199 | 0.0199 | TRUE | 0.2031 |
| Add1 | 3792 | 3736 | 3663 | 7421 | 5043 | 4298 | 4028 | 3731 | 3678 | 5834 | 5426 | 4705 | 4566.93 | 3812 | 5322 | 1.396 | 0.482 | 2.14E-06 | 5.32E-05 | 0.0067 | 0.0076 | 0.0072 | 0.0072 | TRUE | 0.1251 |
| Add2 | 46 | 58 | 49 | 80 | 59 | 56 | 49 | 58 | 49 | 63 | 63 | 61 | 57.28 | 52 | 62 | 1.203 | 0.266 | 0.29074304 | 0.51144315 | 0 | 0.0644 | 0.0282 | 0.0282 | TRUE | 0.083 |
| Add3 | 3143 | 3144 | 3265 | 4235 | 3196 | 3140 | 3338 | 3140 | 3278 | 3329 | 3439 | 3437 | 3326.99 | 3252 | 3402 | 1.046 | 0.065 | 0.32257506 | 0.54376341 | 4.00E-04 | 0.0079 | 0.0028 | 0.0028 | TRUE | 0.0111 |
| Adgb | 5 | 6 | 4 | 2 | 5 | 4 | 5 | 6 | 4 | 2 | 5 | 4 | 4.44 | 5 | 4 | 0.721 | -0.472 | 0.60522043 | NA | 0 | 0.7486 | 0.3691 | 0.3691 | TRUE | 0.5785 |
| Adh1 | 9 | 9 | 12 | 16 | 15 | 13 | 10 | 9 | 12 | 13 | 16 | 14 | 12.26 | 10 | 14 | 1.397 | 0.482 | 0.36646555 | 0.5839457 | 0 | 0.2757 | 0.121 | 0.121 | TRUE | 0.09 |
| Adh4 | 33 | 39 | 26 | 110 | 83 | 61 | 35 | 39 | 26 | 86 | 89 | 67 | 57.11 | 33 | 81 | 2.428 | 1.28 | 1.07E-05 | 0.00021085 | 0 | 0.0646 | 0.0396 | 0.0396 | TRUE | 0.2567 |
| Adh5 | 14686 | 16873 | 17372 | 17431 | 12778 | 13864 | 15599 | 16851 | 17443 | 13703 | 13748 | 15176 | 15420.2 | 16631 | 14209 | 0.854 | -0.227 | 0.00491549 | 0.03097762 | 0.0032 | 0.0071 | 0.0046 | 0.0046 | TRUE | 0.0433 |
| Adh6-ps1 | 0 | 0 | 0 | 0 | 0 | 0 | 0 | 0 | 0 | 0 | 0 | 0 | 0 | 0 | 0 | NA | NA | NA | NA | NA | NA | NA | NA | NA | NA |
| Adh6a | 0 | 0 | 0 | 0 | 0 | 0 | 0 | 0 | 0 | 0 | 0 | 0 | 0 | 0 | 0 | NA | NA | NA | NA | NA | NA | NA | NA | NA | NA |
| Adh7 | 42 | 15 | 14 | 54 | 46 | 21 | 45 | 15 | 14 | 42 | 49 | 23 | 31.43 | 25 | 38 | 1.568 | 0.649 | 0.22815889 | 0.44159023 | 0.2471 | 0.1117 | 0.1753 | 0.1753 | TRUE | 2.1933 |
| Adhfe1 | 148 | 206 | 189 | 90 | 100 | 132 | 157 | 206 | 190 | 71 | 108 | 144 | 145.93 | 184 | 108 | 0.58 | -0.785 | 0.00342782 | 0.02345689 | 0.0615 | 0.0295 | 0.0445 | 0.0445 | TRUE | 0.353 |
| Adi1 | 2219 | 2322 | 2253 | 2925 | 2051 | 2308 | 2357 | 2319 | 2262 | 2299 | 2207 | 2526 | 2328.47 | 2313 | 2344 | 1.013 | 0.019 | 0.81493494 | 0.9055343 | 0.0022 | 0.0083 | 0.0044 | 0.0044 | TRUE | 0.0553 |
| Adig | 1 | 0 | 0 | 0 | 0 | 0 | 1 | 0 | 0 | 0 | 0 | 0 | 0.18 | 0 | 0 | 0.49 | -1.029 | 0.80087 | NA | 0 | 18.6165 | 10 | 10 | TRUE | 0.4614 |
| Adipoq | 0 | 1 | 0 | 5 | 4 | 1 | 0 | 1 | 0 | 4 | 4 | 1 | 1.72 | 0 | 3 | 8.554 | 3.097 | 0.08391132 | NA | 0 | 1.9208 | 1.1565 | 1.1565 | TRUE | 0.405 |
| Adipor1 | 5361 | 6715 | 6936 | 6998 | 5054 | 5854 | 5694 | 6706 | 6964 | 5501 | 5438 | 6408 | 6118.72 | 6455 | 5782 | 0.896 | -0.159 | 0.15220072 | 0.34327323 | 0.0096 | 0.0074 | 0.0087 | 0.0087 | TRUE | 0.1296 |
| Adipor2 | 15757 | 15629 | 15821 | 14787 | 10780 | 11588 | 16737 | 15609 | 15886 | 11624 | 11599 | 12685 | 14023.23 | 16077 | 11969 | 0.744 | -0.426 | 2.77E-09 | 1.50E-07 | 0.0019 | 0.0071 | 0.0036 | 0.0036 | TRUE | 0.0334 |
| Adk | 3700 | 3027 | 2927 | 3350 | 2312 | 2206 | 3930 | 3023 | 2939 | 2634 | 2488 | 2415 | 2904.68 | 3297 | 2512 | 0.762 | -0.392 | 0.00195669 | 0.01513531 | 0.0143 | 0.008 | 0.0112 | 0.0112 | TRUE | 0.3421 |
| Adm | 93 | 107 | 136 | 133 | 111 | 82 | 99 | 107 | 137 | 105 | 119 | 90 | 109.32 | 114 | 105 | 0.916 | -0.126 | 0.57003851 | 0.74897095 | 0.015 | 0.037 | 0.0262 | 0.0262 | TRUE | 0.2991 |
| Adm2 | 16 | 25 | 40 | 29 | 25 | 34 | 17 | 25 | 40 | 23 | 27 | 37 | 28.17 | 27 | 29 | 1.05 | 0.071 | 0.8725961 | 0.93737812 | 0.085 | 0.1238 | 0.1051 | 0.1051 | TRUE | 0.4962 |
| Adnp | 7457 | 7102 | 7231 | 10241 | 7244 | 7078 | 7921 | 7093 | 7261 | 8051 | 7794 | 7748 | 7644.49 | 7425 | 7864 | 1.059 | 0.083 | 0.24992705 | 0.46722738 | 0.0018 | 0.0073 | 0.0036 | 0.0036 | TRUE | 0.0416 |
| Adnp2 | 3550 | 3740 | 3553 | 4302 | 3264 | 3376 | 3771 | 3735 | 3568 | 3382 | 3512 | 3696 | 3610.47 | 3691 | 3530 | 0.956 | -0.065 | 0.36285036 | 0.58058374 | 0.001 | 0.0078 | 0.0034 | 0.0034 | TRUE | 0.0205 |
| Ado | 2593 | 3061 | 3232 | 3258 | 2554 | 2533 | 2754 | 3057 | 3245 | 2561 | 2748 | 2773 | 2856.41 | 3019 | 2694 | 0.892 | -0.165 | 0.06911417 | 0.20401809 | 0.0038 | 0.0081 | 0.0056 | 0.0056 | TRUE | 0.0712 |
| Adora1 | 37 | 40 | 38 | 22 | 23 | 18 | 39 | 40 | 38 | 17 | 25 | 20 | 29.86 | 39 | 21 | 0.522 | -0.938 | 0.0083796 | 0.04608178 | 0 | 0.1172 | 0.0543 | 0.0543 | TRUE | 0.1652 |
| Adora2a | 39 | 48 | 46 | 71 | 61 | 42 | 41 | 48 | 46 | 56 | 66 | 46 | 50.5 | 45 | 56 | 1.235 | 0.304 | 0.28079173 | 0.50219126 | 0 | 0.0721 | 0.0373 | 0.0373 | TRUE | 0.142 |
| Adora2b | 382 | 340 | 306 | 404 | 279 | 246 | 406 | 340 | 307 | 318 | 300 | 269 | 323.27 | 351 | 296 | 0.845 | -0.244 | 0.10990233 | 0.27827573 | 0.0106 | 0.0171 | 0.0136 | 0.0136 | TRUE | 0.212 |
| Adora3 | 0 | 0 | 0 | 0 | 0 | 0 | 0 | 0 | 0 | 0 | 0 | 0 | 0 | 0 | 0 | NA | NA | NA | NA | NA | NA | NA | NA | NA | NA |
| Adpgk | 4222 | 3980 | 3950 | 5246 | 3788 | 3509 | 4485 | 3975 | 3966 | 4124 | 4076 | 3841 | 4077.73 | 4142 | 4014 | 0.969 | -0.045 | 0.58682429 | 0.76150238 | 0.0027 | 0.0077 | 0.0047 | 0.0047 | TRUE | 0.0636 |
| Adprh | 6620 | 7654 | 7254 | 9752 | 7120 | 7395 | 7032 | 7644 | 7284 | 7666 | 7661 | 8095 | 7563.57 | 7320 | 7807 | 1.066 | 0.093 | 0.17173453 | 0.37062439 | 0.0012 | 0.0073 | 0.0032 | 0.0032 | TRUE | 0.0183 |
| Adprhl1 | 51 | 54 | 45 | 37 | 26 | 12 | 54 | 54 | 45 | 29 | 28 | 13 | 37.25 | 51 | 23 | 0.463 | -1.112 | 0.00279693 | 0.0200423 | 0.0266 | 0.0954 | 0.069 | 0.069 | TRUE | 0.787 |
| Adprhl2 | 1347 | 1853 | 1759 | 2158 | 1527 | 1657 | 1431 | 1851 | 1766 | 1696 | 1643 | 1814 | 1700.14 | 1683 | 1718 | 1.021 | 0.029 | 0.80165504 | 0.89760779 | 0.0097 | 0.0088 | 0.0093 | 0.0093 | TRUE | 0.2061 |
| Adprm | 1276 | 1338 | 1294 | 1503 | 1036 | 1008 | 1355 | 1336 | 1299 | 1182 | 1115 | 1103 | 1231.76 | 1330 | 1133 | 0.853 | -0.23 | 0.00355409 | 0.02409934 | 0 | 0.0096 | 0.0037 | 0.0037 | TRUE | 0.0174 |
| Adra1a | 1 | 0 | 2 | 1 | 0 | 0 | 1 | 0 | 2 | 1 | 0 | 0 | 0.64 | 1 | 0 | 0.341 | -1.551 | 0.55438633 | NA | 0 | 5.1324 | 3.4582 | 3.4582 | TRUE | 0.1148 |
| Adra1b | 0 | 1 | 1 | 0 | 1 | 0 | 0 | 1 | 1 | 0 | 1 | 0 | 0.51 | 1 | 0 | 0.52 | -0.943 | 0.73680291 | NA | 0 | 6.4273 | 3.9121 | 3.9121 | TRUE | 0.5204 |
| Adra1d | 4 | 2 | 2 | 3 | 2 | 2 | 4 | 2 | 2 | 2 | 2 | 2 | 2.49 | 3 | 2 | 0.817 | -0.291 | 0.79998379 | NA | 0 | 1.3287 | 0.5464 | 0.5464 | TRUE | 0.2577 |
| Adra2a | 28 | 37 | 35 | 43 | 36 | 36 | 30 | 37 | 35 | 34 | 39 | 39 | 35.63 | 34 | 37 | 1.094 | 0.13 | 0.68331496 | 0.82451561 | 0 | 0.0994 | 0.0447 | 0.0447 | TRUE | 0.0793 |
| Adra2b | 28 | 34 | 29 | 73 | 55 | 45 | 30 | 34 | 29 | 57 | 59 | 49 | 43.11 | 31 | 55 | 1.788 | 0.839 | 0.00463754 | 0.02958193 | 0 | 0.0833 | 0.0379 | 0.0379 | TRUE | 0.0707 |
| Adra2c | 41 | 70 | 54 | 122 | 87 | 73 | 44 | 70 | 54 | 96 | 94 | 80 | 72.85 | 56 | 90 | 1.607 | 0.684 | 0.00855252 | 0.0468044 | 0.0126 | 0.0521 | 0.0343 | 0.0343 | TRUE | 0.4075 |
| Adrb1 | 100 | 154 | 150 | 102 | 99 | 101 | 106 | 154 | 151 | 80 | 107 | 111 | 117.98 | 137 | 99 | 0.72 | -0.474 | 0.04332076 | 0.14882487 | 0.0272 | 0.0348 | 0.031 | 0.031 | TRUE | 0.3903 |
| Adrb2 | 196 | 210 | 200 | 269 | 218 | 198 | 208 | 210 | 201 | 211 | 235 | 217 | 213.58 | 206 | 221 | 1.07 | 0.097 | 0.48690265 | 0.68383157 | 0 | 0.0223 | 0.0094 | 0.0094 | TRUE | 0.0313 |
| Adrb3 | 1229 | 976 | 927 | 1335 | 994 | 857 | 1305 | 975 | 931 | 1049 | 1069 | 938 | 1044.67 | 1070 | 1019 | 0.953 | -0.07 | 0.63307331 | 0.7935807 | 0.0188 | 0.0101 | 0.0146 | 0.0146 | TRUE | 0.4405 |
| Adrbk1 | 3236 | 3670 | 3622 | 4433 | 3189 | 3238 | 3437 | 3665 | 3637 | 3485 | 3431 | 3545 | 3533.31 | 3580 | 3487 | 0.974 | -0.038 | 0.561325 | 0.74256648 | 4.00E-04 | 0.0078 | 0.0028 | 0.0028 | TRUE | 0.0147 |
| Adrbk2 | 657 | 834 | 822 | 735 | 602 | 594 | 698 | 833 | 825 | 578 | 648 | 650 | 705.31 | 785 | 625 | 0.795 | -0.331 | 0.00435979 | 0.02819293 | 0.0058 | 0.0116 | 0.0083 | 0.0083 | TRUE | 0.112 |
| Adrm1 | 5322 | 4649 | 4725 | 6312 | 4523 | 4129 | 5653 | 4643 | 4744 | 4962 | 4866 | 4520 | 4898.1 | 5013 | 4783 | 0.954 | -0.068 | 0.50397708 | 0.69903496 | 0.0065 | 0.0076 | 0.0072 | 0.0072 | TRUE | 0.1516 |
| Adsl | 5730 | 7277 | 7542 | 8061 | 5800 | 6085 | 6086 | 7268 | 7573 | 6337 | 6240 | 6661 | 6694.2 | 6976 | 6413 | 0.919 | -0.121 | 0.2282146 | 0.44159023 | 0.0067 | 0.0074 | 0.0072 | 0.0072 | TRUE | 0.1517 |
| Adss | 8093 | 8277 | 8105 | 10308 | 7303 | 7447 | 8596 | 8266 | 8138 | 8103 | 7858 | 8152 | 8185.6 | 8333 | 8038 | 0.965 | -0.052 | 0.3845017 | 0.5985945 | 4.00E-04 | 0.0073 | 0.0025 | 0.0025 | TRUE | 0.0093 |
| Adssl1 | 209 | 476 | 461 | 212 | 209 | 261 | 222 | 475 | 463 | 167 | 225 | 286 | 306.25 | 387 | 226 | 0.582 | -0.78 | 0.00878903 | 0.04776723 | 0.114 | 0.0177 | 0.0604 | 0.0604 | TRUE | 0.9277 |
| Adtrp | 23 | 24 | 21 | 58 | 35 | 40 | 24 | 24 | 21 | 46 | 38 | 44 | 32.75 | 23 | 43 | 1.834 | 0.875 | 0.00939881 | 0.05019913 | 0 | 0.1075 | 0.0482 | 0.0482 | TRUE | 0.0687 |
| Aebp1 | 133 | 133 | 97 | 212 | 168 | 118 | 141 | 133 | 97 | 167 | 181 | 129 | 141.35 | 124 | 159 | 1.285 | 0.362 | 0.10110342 | 0.26330963 | 0.0254 | 0.0302 | 0.0279 | 0.0279 | TRUE | 0.3564 |
| Aebp2 | 6833 | 6604 | 6484 | 10778 | 7915 | 7448 | 7258 | 6595 | 6511 | 8473 | 8516 | 8153 | 7584.3 | 6788 | 8381 | 1.235 | 0.304 | 3.32E-05 | 0.00054919 | 0.0019 | 0.0073 | 0.0037 | 0.0037 | TRUE | 0.0447 |
| Aen | 4193 | 4259 | 4395 | 4837 | 3564 | 3734 | 4454 | 4254 | 4413 | 3802 | 3835 | 4087 | 4140.8 | 4374 | 3908 | 0.893 | -0.163 | 0.01625383 | 0.0751555 | 7.00E-04 | 0.0077 | 0.0031 | 0.0031 | TRUE | 0.0195 |
| Aes | 10997 | 11945 | 11503 | 16672 | 11884 | 11863 | 11681 | 11930 | 11550 | 13106 | 12786 | 12986 | 12339.86 | 11720 | 12959 | 1.106 | 0.145 | 0.00695398 | 0.039883 | 1.00E-04 | 0.0072 | 0.002 | 0.002 | TRUE | 0.003 |
| AF067061 | 172 | 99 | 63 | 370 | 272 | 110 | 183 | 99 | 63 | 291 | 293 | 120 | 174.79 | 115 | 235 | 2.045 | 1.032 | 0.01195554 | 0.05999451 | 0.2363 | 0.0258 | 0.115 | 0.115 | TRUE | 1.4752 |
| AF067063 | 4 | 4 | 4 | 29 | 25 | 17 | 4 | 4 | 4 | 23 | 27 | 19 | 13.43 | 4 | 23 | 5.576 | 2.479 | 5.06E-05 | 0.00078456 | 0 | 0.2523 | 0.1232 | 0.1232 | TRUE | 0.0722 |
| AF251705 | 0 | 0 | 0 | 0 | 1 | 0 | 0 | 0 | 0 | 0 | 1 | 0 | 0.18 | 0 | 0 | 1.859 | 0.894 | 0.82650734 | NA | 0 | 18.3785 | 10 | 10 | TRUE | 0.5204 |
| AF357355 | 2 | 0 | 0 | 1 | 1 | 0 | 2 | 0 | 0 | 1 | 1 | 0 | 0.66 | 1 | 1 | 0.896 | -0.159 | 0.95209679 | NA | 0 | 4.9654 | 3.5226 | 3.5226 | TRUE | 0.9743 |
| AF357359 | 79 | 72 | 73 | 70 | 47 | 50 | 84 | 72 | 73 | 55 | 51 | 55 | 64.91 | 76 | 54 | 0.701 | -0.512 | 0.03254245 | 0.121608 | 0 | 0.0577 | 0.0256 | 0.0256 | TRUE | 0.067 |
| AF357399 | 7 | 11 | 6 | 5 | 3 | 5 | 7 | 11 | 6 | 4 | 3 | 5 | 6.18 | 8 | 4 | 0.514 | -0.96 | 0.21497283 | 0.42675287 | 0 | 0.54 | 0.2527 | 0.2527 | TRUE | 0.2824 |
| AF357425 | 5 | 7 | 1 | 2 | 3 | 2 | 5 | 7 | 1 | 2 | 3 | 2 | 3.38 | 4 | 2 | 0.518 | -0.948 | 0.39620103 | NA | 0 | 0.9809 | 0.5726 | 0.5726 | TRUE | 0.5217 |
| AF357426 | 3 | 5 | 6 | 4 | 1 | 1 | 3 | 5 | 6 | 3 | 1 | 1 | 3.25 | 5 | 2 | 0.39 | -1.36 | 0.21346228 | NA | 0 | 1.0196 | 0.49 | 0.49 | TRUE | 0.4941 |
| AF366264 | 0 | 0 | 0 | 0 | 0 | 0 | 0 | 0 | 0 | 0 | 0 | 0 | 0 | 0 | 0 | NA | NA | NA | NA | NA | NA | NA | NA | NA | NA |
| AF529169 | 73 | 79 | 87 | 103 | 73 | 69 | 78 | 79 | 87 | 81 | 79 | 76 | 79.81 | 81 | 79 | 0.965 | -0.051 | 0.81131692 | 0.90291353 | 0 | 0.0482 | 0.0206 | 0.0206 | TRUE | 0.0404 |
| Afap1 | 775 | 900 | 849 | 1369 | 959 | 1013 | 823 | 899 | 852 | 1076 | 1032 | 1109 | 965.24 | 858 | 1072 | 1.249 | 0.321 | 0.00022837 | 0.00270487 | 5.00E-04 | 0.0103 | 0.0044 | 0.0044 | TRUE | 0.0205 |

| | | | | | | | | | | | | | | | | | | | | | | | | | |
|---|---|---|---|---|---|---|---|---|---|---|---|---|---|---|---|---|---|---|---|---|---|---|---|---|---|
| Afap1l1 | 1279 | 1206 | 1161 | 786 | 632 | 668 | 1359 | 1204 | 1166 | 618 | 680 | 731 | 959.64 | 1243 | 676 | 0.543 | -0.88 | 2.25E-15 | 3.22E-13 | 0.0058 | 0.0103 | 0.0077 | 0.0077 | TRUE | 0.0791 |
| Afap1l2 | 231 | 205 | 193 | 271 | 202 | 179 | 245 | 205 | 194 | 213 | 217 | 196 | 211.7 | 215 | 209 | 0.974 | -0.038 | 0.80982786 | 0.90213132 | 0.0043 | 0.0225 | 0.0132 | 0.0132 | TRUE | 0.1695 |
| Aff1 | 3552 | 4502 | 5046 | 4134 | 3179 | 3490 | 3773 | 4496 | 5067 | 3250 | 3420 | 3820 | 3971.06 | 4445 | 3497 | 0.786 | -0.347 | 0.00589692 | 0.03543097 | 0.0138 | 0.0077 | 0.0112 | 0.0112 | TRUE | 0.2036 |
| Aff2 | 6 | 5 | 4 | 14 | 4 | 6 | 6 | 5 | 4 | 11 | 4 | 7 | 6.21 | 5 | 7 | 1.449 | 0.535 | 0.49449659 | 0.69052608 | 0 | 0.5374 | 0.2765 | 0.2765 | TRUE | 0.465 |
| Aff3 | 655 | 755 | 831 | 724 | 520 | 531 | 696 | 754 | 834 | 569 | 559 | 581 | 665.68 | 761 | 570 | 0.748 | -0.418 | 0.00010047 | 0.00138509 | 0.0031 | 0.0119 | 0.0068 | 0.0068 | TRUE | 0.0834 |
| Aff4 | 16182 | 15023 | 17071 | 22205 | 15020 | 14934 | 17188 | 15004 | 17141 | 17456 | 16161 | 16348 | 16549.48 | 16444 | 16655 | 1.013 | 0.018 | 0.82573377 | 0.9116315 | 0.003 | 0.0071 | 0.005 | 0.005 | TRUE | 0.0719 |
| Afg3l1 | 10333 | 10672 | 10678 | 13484 | 9802 | 9148 | 10975 | 10658 | 10722 | 10600 | 10546 | 10014 | 10585.98 | 10785 | 10387 | 0.963 | -0.054 | 0.3615677 | 0.57995094 | 5.00E-04 | 0.0072 | 0.0024 | 0.0024 | TRUE | 0.012 |
| Afg3l2 | 9873 | 9877 | 9581 | 10650 | 8059 | 8493 | 10487 | 9864 | 9620 | 8372 | 8671 | 9297 | 9385.26 | 9990 | 8780 | 0.879 | -0.186 | 0.01354186 | 0.06561143 | 0.0023 | 0.0073 | 0.004 | 0.004 | TRUE | 0.0324 |
| Afm | 0 | 0 | 0 | 0 | 0 | 0 | 0 | 0 | 0 | 0 | 0 | 0 | 0 | 0 | 0 | NA | NA | NA | NA | NA | NA | NA | NA | NA | NA |
| Afmid | 167 | 219 | 207 | 284 | 205 | 196 | 177 | 219 | 208 | 223 | 221 | 215 | 210.39 | 201 | 220 | 1.09 | 0.124 | 0.40783378 | 0.620644 | 0 | 0.0226 | 0.0115 | 0.0115 | TRUE | 0.1155 |
| Afp | 1 | 1 | 0 | 11 | 4 | 1 | 1 | 1 | 0 | 9 | 4 | 1 | 2.68 | 1 | 5 | 6.976 | 2.802 | 0.06847348 | NA | 0 | 1.2341 | 0.8544 | 0.8544 | TRUE | 0.0997 |
| Aftph | 2289 | 2607 | 2694 | 2927 | 2134 | 2244 | 2431 | 2604 | 2705 | 2301 | 2296 | 2456 | 2465.58 | 2580 | 2351 | 0.911 | -0.134 | 0.08939345 | 0.24300475 | 0.0016 | 0.0082 | 0.0041 | 0.0041 | TRUE | 0.0307 |
| Aga | 937 | 1383 | 1206 | 1616 | 1218 | 1356 | 995 | 1381 | 1211 | 1270 | 1310 | 1484 | 1275.44 | 1196 | 1355 | 1.133 | 0.18 | 0.19234965 | 0.39802397 | 0.0154 | 0.0095 | 0.0129 | 0.0129 | TRUE | 0.2572 |
| Agap1 | 2799 | 3129 | 3312 | 3355 | 2549 | 2558 | 2973 | 3125 | 3326 | 2637 | 2743 | 2800 | 2933.95 | 3141 | 2727 | 0.868 | -0.205 | 0.00795976 | 0.04415807 | 0.0015 | 0.008 | 0.0039 | 0.0039 | TRUE | 0.032 |
| Agap2 | 89 | 77 | 57 | 76 | 48 | 39 | 95 | 77 | 57 | 60 | 52 | 43 | 63.79 | 76 | 52 | 0.678 | -0.561 | 0.05427536 | 0.17320019 | 0.0309 | 0.0585 | 0.0449 | 0.0449 | TRUE | 0.152 |
| Agap3 | 2048 | 2400 | 2506 | 2843 | 2149 | 2145 | 2175 | 2397 | 2516 | 2235 | 2312 | 2348 | 2330.62 | 2363 | 2298 | 0.972 | -0.04 | 0.6331819 | 0.79365373 | 0.0026 | 0.0083 | 0.0047 | 0.0047 | TRUE | 0.0582 |
| Agbl1 | 0 | 0 | 0 | 0 | 0 | 0 | 0 | 0 | 0 | 0 | 0 | 0 | 0 | 0 | 0 | NA | NA | NA | NA | NA | NA | NA | NA | NA | NA |
| Agbl2 | 43 | 86 | 64 | 40 | 37 | 59 | 46 | 86 | 64 | 31 | 40 | 65 | 55.28 | 65 | 45 | 0.688 | -0.539 | 0.14751536 | 0.3367022 | 0.0971 | 0.0665 | 0.0812 | 0.0812 | TRUE | 0.2874 |
| Agbl3 | 44 | 38 | 41 | 73 | 56 | 45 | 47 | 38 | 41 | 57 | 60 | 49 | 48.79 | 42 | 55 | 1.329 | 0.41 | 0.1401627 | 0.32504722 | 0 | 0.0744 | 0.0348 | 0.0348 | TRUE | 0.078 |
| Agbl4 | 1 | 0 | 0 | 0 | 0 | 0 | 1 | 0 | 0 | 0 | 0 | 0 | 0.18 | 0 | 0 | 0.49 | -1.029 | 0.80087 | NA | 0 | 18.6165 | 10 | 10 | TRUE | 0.4614 |
| Agbl5 | 1122 | 1283 | 1327 | 1359 | 1053 | 1082 | 1192 | 1281 | 1332 | 1068 | 1133 | 1184 | 1198.55 | 1268 | 1128 | 0.889 | -0.17 | 0.06074783 | 0.18759699 | 0.0021 | 0.0097 | 0.0051 | 0.0051 | TRUE | 0.0334 |
| Ager | 8 | 12 | 9 | 10 | 10 | 10 | 8 | 12 | 9 | 8 | 11 | 11 | 9.85 | 10 | 10 | 0.992 | -0.012 | 0.98430605 | 0.99178911 | 0 | 0.3414 | 0.1527 | 0.1527 | TRUE | 0.1406 |
| Agfg1 | 8279 | 8866 | 8960 | 10103 | 7538 | 7814 | 8794 | 8855 | 8997 | 7942 | 8110 | 8554 | 8541.89 | 8882 | 8202 | 0.923 | -0.115 | 0.06376064 | 0.19406269 | 6.00E-04 | 0.0073 | 0.0027 | 0.0027 | TRUE | 0.0171 |
| Agfg2 | 1145 | 1436 | 1514 | 1416 | 1060 | 1200 | 1216 | 1434 | 1520 | 1113 | 1140 | 1314 | 1289.63 | 1390 | 1189 | 0.855 | -0.226 | 0.05985328 | 0.18557224 | 0.0094 | 0.0095 | 0.0097 | 0.0097 | TRUE | 0.1436 |
| Aggf1 | 5267 | 4717 | 4728 | 7192 | 5061 | 4488 | 5594 | 4711 | 4747 | 5654 | 5445 | 4913 | 5177.46 | 5017 | 5337 | 1.064 | 0.089 | 0.38178913 | 0.59595983 | 0.0069 | 0.0075 | 0.0073 | 0.0073 | TRUE | 0.1232 |
| Agk | 1498 | 1418 | 1286 | 2025 | 1424 | 1432 | 1591 | 1416 | 1291 | 1592 | 1532 | 1568 | 1498.36 | 1433 | 1564 | 1.092 | 0.127 | 0.20834455 | 0.41780168 | 0.0047 | 0.0091 | 0.0066 | 0.0066 | TRUE | 0.1119 |
| Agl | 1402 | 1709 | 1923 | 2272 | 1600 | 1683 | 1489 | 1707 | 1931 | 1786 | 1722 | 1842 | 1746.12 | 1709 | 1783 | 1.043 | 0.061 | 0.58607296 | 0.76109862 | 0.0083 | 0.0088 | 0.0085 | 0.0085 | TRUE | 0.1558 |
| Agmat | 61 | 68 | 57 | 62 | 41 | 44 | 65 | 68 | 57 | 49 | 44 | 48 | 55.16 | 63 | 47 | 0.744 | -0.427 | 0.09764804 | 0.25676975 | 0 | 0.0666 | 0.0295 | 0.0295 | TRUE | 0.0625 |
| Agmo | 0 | 1 | 0 | 0 | 0 | 0 | 0 | 1 | 0 | 0 | 0 | 0 | 0.17 | 0 | 0 | 0.49 | -1.029 | 0.80087 | NA | 0 | 19.7991 | 10 | 10 | TRUE | 0.405 |
| Agpat1 | 3533 | 3876 | 3942 | 4845 | 3743 | 3624 | 3753 | 3871 | 3958 | 3809 | 4027 | 3967 | 3897.48 | 3861 | 3934 | 1.019 | 0.027 | 0.68089097 | 0.82311098 | 5.00E-04 | 0.0077 | 0.0028 | 0.0028 | TRUE | 0.0097 |
| Agpat2 | 1758 | 1910 | 1816 | 2323 | 1770 | 1669 | 1867 | 1908 | 1823 | 1826 | 1904 | 1827 | 1859.31 | 1866 | 1852 | 0.993 | -0.011 | 0.87713118 | 0.93932608 | 0 | 0.0087 | 0.003 | 0.003 | TRUE | 0.0072 |
| Agpat3 | 1633 | 2632 | 2806 | 1791 | 1269 | 1634 | 1735 | 2629 | 2818 | 1408 | 1365 | 1789 | 1957.11 | 2394 | 1521 | 0.635 | -0.655 | 0.00047181 | 0.00496092 | 0.0401 | 0.0086 | 0.0247 | 0.0247 | TRUE | 0.7023 |
| Agpat4 | 761 | 992 | 994 | 912 | 746 | 950 | 808 | 991 | 998 | 717 | 803 | 1040 | 892.77 | 932 | 853 | 0.914 | -0.13 | 0.41983502 | 0.63033203 | 0.0242 | 0.0106 | 0.0175 | 0.0175 | TRUE | 0.4343 |
| Agpat5 | 7011 | 7590 | 7236 | 8246 | 6174 | 6419 | 7447 | 7580 | 7266 | 6482 | 6643 | 7027 | 7074.11 | 7431 | 6717 | 0.904 | -0.146 | 0.02579502 | 0.10394459 | 9.00E-04 | 0.0074 | 0.0029 | 0.0029 | TRUE | 0.0198 |
| Agpat6 | 3536 | 3487 | 3326 | 4804 | 3478 | 3163 | 3756 | 3482 | 3340 | 3777 | 3742 | 3462 | 3593.18 | 3526 | 3660 | 1.038 | 0.054 | 0.50492599 | 0.69967429 | 0.0024 | 0.0078 | 0.0045 | 0.0045 | TRUE | 0.0395 |
| Agpat9 | 20 | 23 | 25 | 256 | 175 | 145 | 21 | 23 | 25 | 201 | 188 | 159 | 102.93 | 23 | 183 | 7.918 | 2.985 | 4.65E-31 | 2.82E-28 | 0 | 0.0389 | 0.0231 | 0.0231 | TRUE | 0.1388 |
| Agphd1 | 576 | 459 | 502 | 544 | 403 | 392 | 612 | 458 | 504 | 428 | 434 | 429 | 477.44 | 525 | 430 | 0.82 | -0.286 | 0.03576994 | 0.12981211 | 0.0092 | 0.0138 | 0.0113 | 0.0113 | TRUE | 0.2444 |
| Agps | 10814 | 12384 | 12747 | 14159 | 10342 | 10669 | 11486 | 12368 | 12799 | 11131 | 11127 | 11679 | 11765.11 | 12218 | 11312 | 0.926 | -0.111 | 0.11918313 | 0.29247802 | 0.0018 | 0.0072 | 0.0036 | 0.0036 | TRUE | 0.0335 |
| Agr2 | 0 | 1 | 0 | 0 | 0 | 0 | 0 | 1 | 0 | 0 | 0 | 0 | 0.17 | 0 | 0 | 0.49 | -1.029 | 0.80087 | NA | 0 | 19.7991 | 10 | 10 | TRUE | 0.405 |
| Agr3 | 0 | 0 | 0 | 0 | 0 | 0 | 0 | 0 | 0 | 0 | 0 | 0 | 0 | 0 | 0 | NA | NA | NA | NA | NA | NA | NA | NA | NA | NA |
| Agrn | 10600 | 12317 | 12555 | 12261 | 9609 | 9103 | 11259 | 12301 | 12607 | 9639 | 10339 | 9965 | 11018.12 | 12056 | 9981 | 0.828 | -0.273 | 0.00027308 | 0.00314868 | 0.0022 | 0.0072 | 0.004 | 0.004 | TRUE | 0.0408 |
| Agrp | 4 | 4 | 8 | 11 | 7 | 4 | 4 | 4 | 8 | 9 | 8 | 4 | 6.14 | 5 | 7 | 1.274 | 0.349 | 0.6542335 | 0.80568968 | 0 | 0.5435 | 0.2728 | 0.2728 | TRUE | 0.3871 |
| Agt | 7 | 3 | 6 | 4 | 4 | 4 | 7 | 3 | 6 | 3 | 4 | 4 | 4.71 | 5 | 4 | 0.714 | -0.486 | 0.57644605 | 0.75418558 | 0 | 0.7058 | 0.3276 | 0.3276 | TRUE | 0.3503 |
| Agtpbp1 | 1132 | 1234 | 1275 | 1317 | 1052 | 1035 | 1202 | 1232 | 1280 | 1035 | 1132 | 1133 | 1169.2 | 1238 | 1100 | 0.887 | -0.172 | 0.0430494 | 0.14822702 | 0.0011 | 0.0097 | 0.0044 | 0.0044 | TRUE | 0.0331 |
| Agtr1a | 79 | 68 | 55 | 117 | 75 | 64 | 84 | 68 | 55 | 92 | 81 | 70 | 74.96 | 69 | 81 | 1.178 | 0.237 | 0.35818161 | 0.57662962 | 0.0167 | 0.0509 | 0.0344 | 0.0344 | TRUE | 0.2859 |
| Agtr1b | 6 | 1 | 3 | 3 | 2 | 3 | 6 | 1 | 3 | 2 | 2 | 3 | 3.03 | 3 | 2 | 0.752 | -0.412 | 0.71483066 | NA | 0 | 1.0943 | 0.5783 | 0.5783 | TRUE | 0.3012 |
| Agtr2 | 0 | 0 | 0 | 0 | 0 | 0 | 0 | 0 | 0 | 0 | 0 | 0 | 0 | 0 | 0 | NA | NA | NA | NA | NA | NA | NA | NA | NA | NA |
| Agtrap | 2933 | 3812 | 3725 | 4431 | 3326 | 3123 | 3115 | 3807 | 3740 | 3483 | 3579 | 3419 | 3523.87 | 3554 | 3494 | 0.983 | -0.025 | 0.79969492 | 0.89674997 | 0.006 | 0.0078 | 0.0067 | 0.0067 | TRUE | 0.1416 |
| Agxt | 2 | 0 | 1 | 7 | 1 | 0 | 2 | 0 | 1 | 6 | 1 | 0 | 1.62 | 1 | 2 | 2.185 | 1.127 | 0.54432462 | NA | 1.4163 | 2.0432 | 1.7811 | 1.7811 | TRUE | 1.3233 |
| Agxt2 | 0 | 0 | 0 | 0 | 0 | 0 | 0 | 0 | 0 | 0 | 0 | 0 | 0 | 0 | 0 | NA | NA | NA | NA | NA | NA | NA | NA | NA | NA |
| Agxt2l1 | 0 | 0 | 0 | 0 | 0 | 1 | 0 | 0 | 0 | 0 | 0 | 1 | 0.18 | 0 | 0 | 1.859 | 0.894 | 0.82650734 | NA | 0 | 18.0643 | 10 | 10 | TRUE | 0.5383 |
| Agxt2l2 | 442 | 464 | 443 | 468 | 349 | 352 | 469 | 463 | 445 | 368 | 376 | 385 | 417.74 | 459 | 376 | 0.819 | -0.288 | 0.00653592 | 0.03822575 | 0 | 0.0148 | 0.0057 | 0.0057 | TRUE | 0.0088 |
| Ahctf1 | 15598 | 13321 | 14168 | 19839 | 13688 | 12755 | 16568 | 13304 | 14226 | 15596 | 14727 | 13962 | 14730.58 | 14699 | 14762 | 1.004 | 0.006 | 0.95159009 | 0.97536556 | 0.0064 | 0.0071 | 0.0075 | 0.0075 | TRUE | 0.1512 |
| Ahcy | 5330 | 5474 | 5299 | 6064 | 4231 | 4437 | 5661 | 5467 | 5321 | 4767 | 4552 | 4857 | 5104.24 | 5483 | 4725 | 0.862 | -0.214 | 0.00120209 | 0.01025873 | 7.00E-04 | 0.0075 | 0.003 | 0.003 | TRUE | 0.0124 |
| Ahcyl1 | 7925 | 9205 | 9223 | 10682 | 7950 | 8796 | 8418 | 9193 | 9261 | 8397 | 8554 | 9629 | 8908.57 | 8957 | 8860 | 0.989 | -0.016 | 0.85331272 | 0.92681753 | 0.0033 | 0.0073 | 0.0053 | 0.0053 | TRUE | 0.0702 |
| Ahcyl2 | 2349 | 2513 | 2580 | 2642 | 1875 | 2026 | 2495 | 2510 | 2591 | 2077 | 2017 | 2218 | 2317.92 | 2532 | 2104 | 0.831 | -0.267 | 0.00029766 | 0.00339318 | 7.00E-04 | 0.0083 | 0.0035 | 0.0035 | TRUE | 0.0265 |
| Ahdc1 | 4687 | 4681 | 4557 | 5462 | 4177 | 3655 | 4978 | 4675 | 4576 | 4294 | 4494 | 4001 | 4503.01 | 4743 | 4263 | 0.899 | -0.154 | 0.05167764 | 0.16770316 | 0.0023 | 0.0076 | 0.0043 | 0.0043 | TRUE | 0.0349 |
| Ahi1 | 731 | 851 | 888 | 914 | 705 | 724 | 776 | 850 | 892 | 719 | 759 | 793 | 797.93 | 839 | 757 | 0.9 | -0.152 | 0.13128682 | 0.31203371 | 0.0024 | 0.011 | 0.006 | 0.006 | TRUE | 0.0506 |
| Ahnak | 2254 | 2107 | 2773 | 3433 | 2345 | 2180 | 2394 | 2104 | 2784 | 2699 | 2523 | 2386 | 2481.83 | 2427 | 2536 | 1.045 | 0.063 | 0.59657797 | 0.76948215 | 0.0113 | 0.0082 | 0.0099 | 0.0099 | TRUE | 0.2007 |
| Ahr | 470 | 551 | 516 | 598 | 460 | 410 | 499 | 550 | 518 | 470 | 495 | 449 | 496.91 | 522 | 471 | 0.902 | -0.149 | 0.16017234 | 0.35474763 | 0 | 0.0135 | 0.0061 | 0.0061 | TRUE | 0.0253 |
| Ahrr | 103 | 135 | 135 | 182 | 141 | 132 | 109 | 135 | 136 | 143 | 152 | 144 | 136.51 | 127 | 146 | 1.155 | 0.208 | 0.24404183 | 0.4605753 | 0 | 0.031 | 0.0155 | 0.0155 | TRUE | 0.1407 |
| Ahsa1 | 7295 | 7215 | 7006 | 11198 | 8132 | 7542 | 7749 | 7206 | 7035 | 8803 | 8750 | 8256 | 7966.24 | 7330 | 8603 | 1.174 | 0.231 | 0.00131949 | 0.01106764 | 0.0017 | 0.0073 | 0.0036 | 0.0036 | TRUE | 0.0305 |
| Ahsa2 | 1319 | 1277 | 1212 | 1938 | 1406 | 1286 | 1401 | 1275 | 1217 | 1524 | 1513 | 1408 | 1389.56 | 1298 | 1482 | 1.142 | 0.192 | 0.03658941 | 0.1319662 | 0.0025 | 0.0093 | 0.0053 | 0.0053 | TRUE | 0.0578 |
| Ahsg | 58 | 53 | 38 | 120 | 101 | 70 | 62 | 53 | 38 | 94 | 109 | 77 | 72.05 | 51 | 93 | 1.835 | 0.875 | 0.00153925 | 0.01250108 | 0.0247 | 0.0526 | 0.0398 | 0.0398 | TRUE | 0.2723 |
| AI115009 | 0 | 0 | 0 | 0 | 0 | 0 | 0 | 0 | 0 | 0 | 0 | 0 | 0 | 0 | 0 | NA | NA | NA | NA | NA | NA | NA | NA | NA | NA |
| AI118078 | 80 | 56 | 69 | 102 | 77 | 49 | 85 | 56 | 69 | 80 | 83 | 54 | 71.14 | 70 | 72 | 1.035 | 0.05 | 0.86042102 | 0.93102087 | 0.0332 | 0.0532 | 0.0436 | 0.0436 | TRUE | 0.3282 |
| AI182371 | 4 | 9 | 6 | 15 | 10 | 8 | 4 | 9 | 6 | 12 | 11 | 9 | 8.43 | 6 | 11 | 1.628 | 0.703 | 0.288153 | 0.50913413 | 0 | 0.3978 | 0.1897 | 0.1897 | TRUE | 0.3068 |
| AI197445 | 3 | 7 | 6 | 11 | 6 | 3 | 3 | 7 | 6 | 9 | 6 | 3 | 5.76 | 5 | 6 | 1.149 | 0.2 | 0.80612437 | 0.90019698 | 0 | 0.5784 | 0.3048 | 0.3048 | TRUE | 0.265 |
| AI314180 | 2381 | 2915 | 3373 | 5305 | 3533 | 3490 | 2529 | 2911 | 3387 | 4170 | 3801 | 3820 | 3436.52 | 2942 | 3930 | 1.336 | 0.418 | 0.00045645 | 0.00484467 | 0.0116 | 0.0079 | 0.0099 | 0.0099 | TRUE | 0.2121 |

| | | | | | | | | | | | | | | | | | | | | | | | | | |
|---|---|---|---|---|---|---|---|---|---|---|---|---|---|---|---|---|---|---|---|---|---|---|---|---|---|
| AI314831 | 4 | 3 | 1 | 3 | 1 | 0 | 4 | 3 | 1 | 2 | 1 | 0 | 1.95 | 3 | 1 | 0.439 | -1.186 | 0.42052199 | NA | 0 | 1.6988 | 0.9715 | 0.9715 | TRUE | 0.4234 |
| AI316807 | 319 | 447 | 487 | 654 | 491 | 435 | 339 | 446 | 489 | 514 | 528 | 476 | 465.47 | 425 | 506 | 1.191 | 0.253 | 0.10294114 | 0.26699353 | 0.0162 | 0.014 | 0.0151 | 0.0151 | TRUE | 0.3586 |
| AI317395 | 39 | 34 | 23 | 67 | 63 | 49 | 41 | 34 | 23 | 53 | 68 | 54 | 45.43 | 33 | 58 | 1.767 | 0.822 | 0.01171978 | 0.05913128 | 0.0188 | 0.0794 | 0.0525 | 0.0525 | TRUE | 0.4684 |
| AI413582 | 341 | 577 | 585 | 492 | 371 | 443 | 362 | 576 | 587 | 387 | 399 | 485 | 466.12 | 508 | 424 | 0.832 | -0.265 | 0.18496641 | 0.38824731 | 0.0403 | 0.014 | 0.0267 | 0.0267 | TRUE | 0.7357 |
| AI414108 | 644 | 789 | 790 | 1011 | 799 | 892 | 684 | 788 | 793 | 795 | 860 | 976 | 816.02 | 755 | 877 | 1.16 | 0.214 | 0.07433007 | 0.21420314 | 0.0077 | 0.0109 | 0.0091 | 0.0091 | TRUE | 0.1166 |
| AI427809 | 198 | 92 | 90 | 163 | 109 | 88 | 210 | 92 | 90 | 128 | 117 | 96 | 122.38 | 131 | 114 | 0.873 | -0.196 | 0.56571351 | 0.74589814 | 0.1242 | 0.0338 | 0.0761 | 0.0761 | TRUE | 2.8688 |
| AI429214 | 30 | 27 | 30 | 47 | 40 | 33 | 32 | 27 | 30 | 37 | 43 | 36 | 34.18 | 30 | 39 | 1.304 | 0.383 | 0.23703098 | 0.45237811 | 0 | 0.1033 | 0.0458 | 0.0458 | TRUE | 0.0667 |
| AI450353 | 78 | 92 | 82 | 115 | 95 | 109 | 83 | 92 | 82 | 90 | 102 | 119 | 94.83 | 86 | 104 | 1.208 | 0.273 | 0.20247252 | 0.4100233 | 0 | 0.0416 | 0.0223 | 0.0223 | TRUE | 0.1609 |
| AI462493 | 457 | 459 | 446 | 710 | 439 | 455 | 485 | 458 | 448 | 558 | 472 | 498 | 486.7 | 464 | 509 | 1.101 | 0.138 | 0.23105696 | 0.44492876 | 0.0029 | 0.0137 | 0.0076 | 0.0076 | TRUE | 0.0854 |
| AI463170 | 3 | 1 | 1 | 2 | 2 | 2 | 3 | 1 | 1 | 2 | 2 | 2 | 1.85 | 2 | 2 | 1.138 | 0.187 | 0.88963194 | NA | 0 | 1.7872 | 0.7557 | 0.7557 | TRUE | 0.4027 |
| AI464131 | 937 | 1200 | 1243 | 1326 | 1059 | 1055 | 995 | 1198 | 1248 | 1042 | 1139 | 1155 | 1129.75 | 1147 | 1112 | 0.969 | -0.046 | 0.68922311 | 0.82917062 | 0.0077 | 0.0098 | 0.0086 | 0.0086 | TRUE | 0.1602 |
| AI467606 | 218 | 294 | 263 | 298 | 205 | 187 | 232 | 294 | 264 | 234 | 221 | 205 | 241.46 | 263 | 220 | 0.837 | -0.257 | 0.09192745 | 0.24742196 | 0.0054 | 0.0205 | 0.0126 | 0.0126 | TRUE | 0.1211 |
| AI480653 | 1760 | 2017 | 2056 | 2209 | 1755 | 1824 | 1869 | 2014 | 2064 | 1737 | 1888 | 1997 | 1928.29 | 1982 | 1874 | 0.944 | -0.083 | 0.35967418 | 0.57800887 | 0.003 | 0.0086 | 0.0053 | 0.0053 | TRUE | 0.0508 |
| AI504432 | 31 | 15 | 25 | 84 | 49 | 24 | 33 | 15 | 25 | 66 | 53 | 26 | 36.34 | 24 | 48 | 2.001 | 1.001 | 0.02887328 | 0.11199935 | 0.1451 | 0.0976 | 0.1201 | 0.1201 | TRUE | 0.2853 |
| AI506816 | 36 | 35 | 25 | 37 | 32 | 22 | 38 | 35 | 25 | 29 | 34 | 24 | 30.98 | 33 | 29 | 0.892 | -0.164 | 0.65007779 | 0.80390676 | 0 | 0.1132 | 0.0623 | 0.0623 | TRUE | 0.2916 |
| AI507597 | 0 | 0 | 0 | 1 | 0 | 0 | 0 | 0 | 0 | 1 | 0 | 0 | 0.13 | 0 | 0 | 1.859 | 0.894 | 0.82650734 | NA | 0 | 25.1515 | 10 | 10 | TRUE | 0.2563 |
| AI593442 | 21 | 13 | 5 | 46 | 36 | 33 | 22 | 13 | 5 | 36 | 39 | 36 | 25.22 | 13 | 37 | 2.766 | 1.468 | 0.00338333 | 0.02323115 | 0.1142 | 0.1375 | 0.1293 | 0.1293 | TRUE | 0.6681 |
| AI597468 | 1733 | 1969 | 1907 | 2447 | 1842 | 1846 | 1841 | 1966 | 1915 | 1924 | 1982 | 2021 | 1941.38 | 1907 | 1976 | 1.035 | 0.05 | 0.48816854 | 0.68476003 | 4.00E-04 | 0.0086 | 0.0032 | 0.0032 | TRUE | 0.0112 |
| AI597479 | 2416 | 2553 | 2606 | 3501 | 2587 | 2552 | 2566 | 2550 | 2617 | 2752 | 2783 | 2794 | 2676.98 | 2578 | 2776 | 1.077 | 0.107 | 0.08417156 | 0.2326245 | 0 | 0.0081 | 0.0024 | 0.0024 | TRUE | 0.0022 |
| AI606473 | 0 | 0 | 0 | 0 | 0 | 0 | 0 | 0 | 0 | 0 | 0 | 0 | 0 | 0 | 0 | NA | NA | NA | NA | NA | NA | NA | NA | NA | NA |
| AI607873 | 0 | 1 | 0 | 0 | 2 | 0 | 0 | 1 | 0 | 0 | 2 | 0 | 0.53 | 0 | 1 | 1.927 | 0.947 | 0.77747358 | NA | 0 | 6.2809 | 6.39 | 6.39 | TRUE | 0.9744 |
| AI646519 | 1 | 1 | 0 | 0 | 1 | 0 | 1 | 1 | 0 | 0 | 1 | 0 | 0.52 | 1 | 0 | 0.51 | -0.972 | 0.72797553 | NA | 0 | 6.3084 | 3.8879 | 3.8879 | TRUE | 0.5204 |
| AI661453 | 247 | 228 | 255 | 398 | 327 | 325 | 262 | 228 | 256 | 313 | 352 | 356 | 294.43 | 249 | 340 | 1.366 | 0.449 | 0.00084291 | 0.00780086 | 0 | 0.0181 | 0.0096 | 0.0096 | TRUE | 0.0611 |
| AI662270 | 41 | 30 | 22 | 1099 | 863 | 720 | 44 | 30 | 22 | 864 | 929 | 788 | 446.04 | 32 | 860 | 27.128 | 4.762 | 1.92E-119 | 3.02E-115 | 0.0111 | 0.0143 | 0.0136 | 0.0136 | TRUE | 0.668 |
| AI747448 | 24 | 17 | 13 | 12 | 10 | 15 | 25 | 17 | 13 | 9 | 11 | 16 | 15.36 | 18 | 12 | 0.654 | -0.612 | 0.24809428 | 0.46483048 | 0.0409 | 0.2214 | 0.1339 | 0.1339 | TRUE | 0.4218 |
| AI836003 | 13 | 11 | 13 | 69 | 49 | 54 | 14 | 11 | 13 | 54 | 53 | 59 | 33.99 | 13 | 55 | 4.391 | 2.134 | 6.43E-09 | 3.19E-07 | 0 | 0.1038 | 0.048 | 0.048 | TRUE | 0.0531 |
| AI837181 | 906 | 994 | 1039 | 1270 | 903 | 945 | 962 | 993 | 1043 | 998 | 972 | 1034 | 1000.45 | 999 | 1001 | 1.002 | 0.003 | 0.97562876 | 0.98697924 | 0 | 0.0102 | 0.0042 | 0.0042 | TRUE | 0.0176 |
| AI846148 | 1802 | 1678 | 1709 | 2778 | 1967 | 1778 | 1914 | 1676 | 1716 | 2184 | 2116 | 1946 | 1925.4 | 1769 | 2082 | 1.178 | 0.236 | 0.01079387 | 0.05571471 | 0.0035 | 0.0086 | 0.0057 | 0.0057 | TRUE | 0.0622 |
| AI847159 | 0 | 0 | 1 | 2 | 2 | 2 | 0 | 0 | 1 | 2 | 2 | 2 | 1.15 | 0 | 2 | 5.357 | 2.422 | 0.20735133 | NA | 0 | 2.8644 | 1.4081 | 1.4081 | TRUE | 0.4098 |
| AI848285 | 307 | 383 | 403 | 382 | 282 | 290 | 326 | 383 | 405 | 300 | 303 | 317 | 339.07 | 371 | 307 | 0.827 | -0.275 | 0.0383221 | 0.13634473 | 0.0038 | 0.0166 | 0.0097 | 0.0097 | TRUE | 0.1274 |
| AI854517 | 50 | 35 | 28 | 108 | 69 | 65 | 53 | 35 | 28 | 85 | 74 | 71 | 57.75 | 39 | 77 | 1.992 | 0.994 | 0.00114402 | 0.00987062 | 0.0263 | 0.064 | 0.0477 | 0.0477 | TRUE | 0.7592 |
| AI854703 | 81 | 141 | 100 | 149 | 118 | 131 | 86 | 141 | 100 | 117 | 127 | 143 | 119.13 | 109 | 129 | 1.18 | 0.239 | 0.30879179 | 0.529319 | 0.0281 | 0.0346 | 0.0314 | 0.0314 | TRUE | 0.6462 |
| AI987944 | 692 | 791 | 830 | 846 | 635 | 718 | 735 | 790 | 833 | 665 | 683 | 786 | 748.78 | 786 | 711 | 0.904 | -0.146 | 0.18839148 | 0.39350861 | 0.0047 | 0.0113 | 0.0075 | 0.0075 | TRUE | 0.098 |
| Aicda | 20 | 12 | 10 | 45 | 27 | 16 | 21 | 12 | 10 | 35 | 29 | 18 | 20.87 | 14 | 27 | 1.913 | 0.936 | 0.06153378 | 0.18933101 | 0.0853 | 0.1648 | 0.1273 | 0.1273 | TRUE | 0.39 |
| Aida | 899 | 1051 | 972 | 1772 | 1263 | 1087 | 955 | 1050 | 976 | 1393 | 1359 | 1190 | 1153.72 | 994 | 1314 | 1.323 | 0.404 | 5.44E-05 | 0.00083133 | 0.004 | 0.0098 | 0.0063 | 0.0063 | TRUE | 0.081 |
| Aif1 | 0 | 0 | 0 | 0 | 0 | 0 | 0 | 0 | 0 | 0 | 0 | 0 | 0 | 0 | 0 | NA | NA | NA | NA | NA | NA | NA | NA | NA | NA |
| Aif1l | 1179 | 1430 | 1405 | 1598 | 1087 | 1148 | 1252 | 1428 | 1411 | 1256 | 1170 | 1257 | 1295.61 | 1364 | 1228 | 0.9 | -0.152 | 0.09958127 | 0.26055007 | 0.0026 | 0.0094 | 0.0053 | 0.0053 | TRUE | 0.0611 |
| Aifm1 | 4017 | 3831 | 3836 | 5052 | 3433 | 3564 | 4267 | 3826 | 3852 | 3971 | 3694 | 3901 | 3918.52 | 3982 | 3855 | 0.969 | -0.046 | 0.56109041 | 0.74256648 | 0.0022 | 0.0077 | 0.0043 | 0.0043 | TRUE | 0.0477 |
| Aifm2 | 1135 | 1277 | 1283 | 1453 | 1052 | 1141 | 1206 | 1275 | 1288 | 1142 | 1132 | 1249 | 1215.39 | 1256 | 1174 | 0.934 | -0.098 | 0.25294388 | 0.47086616 | 0.0012 | 0.0096 | 0.0045 | 0.0045 | TRUE | 0.0363 |
| Aifm3 | 578 | 837 | 772 | 655 | 525 | 564 | 614 | 836 | 775 | 515 | 565 | 617 | 653.7 | 742 | 566 | 0.762 | -0.393 | 0.00694029 | 0.03986242 | 0.0152 | 0.0119 | 0.0137 | 0.0137 | TRUE | 0.2674 |
| Aig1 | 96 | 124 | 158 | 182 | 154 | 142 | 102 | 124 | 159 | 143 | 166 | 155 | 141.44 | 128 | 155 | 1.204 | 0.268 | 0.20110843 | 0.40877592 | 0.0188 | 0.0302 | 0.0246 | 0.0246 | TRUE | 0.4282 |
| Aim1 | 265 | 177 | 201 | 334 | 228 | 198 | 281 | 177 | 202 | 263 | 245 | 217 | 230.78 | 220 | 242 | 1.1 | 0.138 | 0.49629381 | 0.69196835 | 0.0292 | 0.0212 | 0.0252 | 0.0252 | TRUE | 0.6497 |
| Aim1l | 434 | 509 | 496 | 437 | 344 | 343 | 461 | 508 | 498 | 344 | 370 | 375 | 426.08 | 489 | 363 | 0.741 | -0.433 | 0.00011182 | 0.00150346 | 0 | 0.0146 | 0.0066 | 0.0066 | TRUE | 0.0291 |
| Aim2 | 224 | 292 | 283 | 949 | 711 | 760 | 238 | 292 | 284 | 746 | 765 | 832 | 526.11 | 271 | 781 | 2.875 | 1.524 | 3.76E-35 | 2.82E-32 | 0.0037 | 0.0132 | 0.0084 | 0.0084 | TRUE | 0.127 |
| Aimp1 | 4523 | 5531 | 5610 | 6205 | 4515 | 4725 | 4804 | 5524 | 5633 | 4878 | 4858 | 5172 | 5144.85 | 5320 | 4969 | 0.934 | -0.099 | 0.26348393 | 0.48271211 | 0.0039 | 0.0075 | 0.0054 | 0.0054 | TRUE | 0.0877 |
| Aimp2 | 2606 | 2544 | 2415 | 3728 | 2583 | 2382 | 2768 | 2541 | 2425 | 2931 | 2779 | 2607 | 2675.16 | 2578 | 2772 | 1.076 | 0.106 | 0.23531227 | 0.45046063 | 0.0034 | 0.0081 | 0.0053 | 0.0053 | TRUE | 0.0504 |
| Aip | 1168 | 1629 | 1584 | 1735 | 1197 | 1312 | 1241 | 1627 | 1591 | 1364 | 1288 | 1436 | 1424.34 | 1486 | 1363 | 0.917 | -0.125 | 0.31605201 | 0.53684153 | 0.0107 | 0.0092 | 0.0106 | 0.0106 | TRUE | 0.2504 |
| Aipl1 | 45 | 75 | 66 | 48 | 43 | 51 | 48 | 75 | 66 | 38 | 46 | 56 | 54.8 | 63 | 47 | 0.734 | -0.445 | 0.14179696 | 0.32738322 | 0.027 | 0.067 | 0.0476 | 0.0476 | TRUE | 0.3791 |
| Aire | 621 | 846 | 807 | 851 | 841 | 964 | 660 | 845 | 810 | 669 | 905 | 1055 | 823.99 | 772 | 876 | 1.134 | 0.182 | 0.3114342 | 0.53199265 | 0.0334 | 0.0109 | 0.022 | 0.022 | TRUE | 0.3142 |
| Airn | 3 | 1 | 2 | 3 | 0 | 3 | 3 | 1 | 2 | 2 | 0 | 3 | 1.97 | 2 | 2 | 0.921 | -0.118 | 0.93404774 | NA | 0 | 1.677 | 0.962 | 0.962 | TRUE | 0.5766 |
| Ajap1 | 23 | 12 | 14 | 20 | 15 | 11 | 24 | 12 | 14 | 16 | 16 | 12 | 15.73 | 17 | 15 | 0.875 | -0.193 | 0.70670791 | 0.8398044 | 0 | 0.2164 | 0.125 | 0.125 | TRUE | 0.7367 |
| Ajuba | 3718 | 3844 | 3989 | 5827 | 4304 | 4009 | 3949 | 3839 | 4005 | 4581 | 4631 | 4388 | 4232.27 | 3931 | 4533 | 1.153 | 0.206 | 0.00116948 | 0.01001906 | 4.00E-04 | 0.0077 | 0.0027 | 0.0027 | TRUE | 0.0094 |
| AK010878 | 305 | 272 | 355 | 386 | 266 | 242 | 324 | 272 | 356 | 303 | 286 | 265 | 301.1 | 317 | 285 | 0.899 | -0.154 | 0.30631878 | 0.52685494 | 0.0085 | 0.0178 | 0.0129 | 0.0129 | TRUE | 0.1816 |
| Ak1 | 2298 | 2737 | 2520 | 2424 | 1883 | 2080 | 2441 | 2733 | 2530 | 1906 | 2026 | 2277 | 2318.86 | 2568 | 2070 | 0.805 | -0.312 | 0.00151376 | 0.01235125 | 0.0052 | 0.0083 | 0.0066 | 0.0066 | TRUE | 0.0927 |
| AK129341 | 669 | 546 | 615 | 741 | 581 | 545 | 711 | 545 | 618 | 583 | 625 | 597 | 612.94 | 625 | 602 | 0.963 | -0.054 | 0.66606522 | 0.81310515 | 0.0078 | 0.0123 | 0.0098 | 0.0098 | TRUE | 0.1702 |
| Ak2 | 11155 | 11638 | 11675 | 14575 | 10477 | 10257 | 11849 | 11623 | 11723 | 11458 | 11273 | 11228 | 11525.45 | 11732 | 11320 | 0.965 | -0.051 | 0.32475155 | 0.54594394 | 0 | 0.0072 | 0.0019 | 0.0019 | TRUE | 0.0014 |
| Ak3 | 768 | 647 | 704 | 908 | 616 | 526 | 816 | 646 | 707 | 714 | 663 | 576 | 686.86 | 723 | 651 | 0.902 | -0.15 | 0.26509668 | 0.48482952 | 0.0106 | 0.0117 | 0.0115 | 0.0115 | TRUE | 0.1485 |
| Ak4 | 8007 | 9884 | 9083 | 18891 | 14328 | 14733 | 8505 | 9871 | 9120 | 14851 | 15416 | 16128 | 12315.11 | 9165 | 15465 | 1.687 | 0.755 | 7.94E-20 | 1.90E-17 | 0.0035 | 0.0072 | 0.0049 | 0.0049 | TRUE | 0.0554 |
| Ak5 | 5 | 21 | 13 | 30 | 21 | 16 | 5 | 21 | 13 | 24 | 23 | 18 | 17.17 | 13 | 22 | 1.617 | 0.693 | 0.21247128 | 0.4234926 | 0.1097 | 0.1988 | 0.1611 | 0.1611 | TRUE | 0.3096 |
| Ak7 | 250 | 160 | 158 | 629 | 400 | 328 | 266 | 160 | 159 | 494 | 430 | 359 | 311.31 | 195 | 428 | 2.203 | 1.139 | 5.99E-07 | 1.75E-05 | 0.0526 | 0.0175 | 0.0338 | 0.0338 | TRUE | 0.5585 |
| Ak8 | 25 | 51 | 60 | 38 | 30 | 48 | 27 | 51 | 60 | 30 | 32 | 53 | 42.07 | 46 | 38 | 0.827 | -0.274 | 0.50080678 | 0.69665943 | 0.1061 | 0.0852 | 0.0952 | 0.0952 | TRUE | 0.6643 |
| Akap1 | 14220 | 14483 | 14066 | 18861 | 13987 | 13654 | 15104 | 14464 | 14124 | 14827 | 15049 | 14946 | 14752.47 | 14564 | 14941 | 1.026 | 0.037 | 0.52929046 | 0.71762315 | 5.00E-04 | 0.0071 | 0.0024 | 0.0024 | TRUE | 0.0129 |
| Akap10 | 3322 | 3398 | 3395 | 4226 | 3005 | 3058 | 3529 | 3394 | 3409 | 3322 | 3233 | 3347 | 3372.32 | 3444 | 3301 | 0.959 | -0.061 | 0.32982724 | 0.55084296 | 0 | 0.0079 | 0.0025 | 0.0025 | TRUE | 0.0056 |
| Akap11 | 8857 | 6750 | 7339 | 11186 | 7642 | 6857 | 9408 | 6741 | 7369 | 8794 | 8222 | 7506 | 8006.68 | 7839 | 8174 | 1.043 | 0.06 | 0.65421362 | 0.80568968 | 0.0151 | 0.0073 | 0.013 | 0.013 | TRUE | 0.3738 |
| Akap12 | 32369 | 29609 | 31980 | 35783 | 27082 | 26890 | 34382 | 29571 | 32111 | 28130 | 29139 | 29435 | 30461.23 | 32021 | 28901 | 0.903 | -0.148 | 0.06127193 | 0.18874628 | 0.0031 | 0.007 | 0.0045 | 0.0045 | TRUE | 0.0549 |
| Akap13 | 3562 | 2808 | 3196 | 3945 | 2707 | 2512 | 3783 | 2804 | 3209 | 3101 | 2913 | 2750 | 3093.43 | 3265 | 2921 | 0.895 | -0.16 | 0.19257462 | 0.39828047 | 0.0127 | 0.008 | 0.0106 | 0.0106 | TRUE | 0.2336 |
| Akap14 | 0 | 2 | 2 | 3 | 2 | 1 | 0 | 2 | 2 | 2 | 2 | 1 | 1.6 | 1 | 2 | 1.401 | 0.487 | 0.7476056 | NA | 0 | 2.0637 | 1.0108 | 1.0108 | TRUE | 0.4374 |
| Akap17b | 877 | 818 | 782 | 1310 | 878 | 753 | 932 | 817 | 785 | 1030 | 945 | 824 | 888.74 | 845 | 933 | 1.106 | 0.145 | 0.23815391 | 0.45396957 | 0.0086 | 0.0106 | 0.0098 | 0.0098 | TRUE | 0.1228 |
| Akap2 | 3698 | 4678 | 5116 | 3527 | 2710 | 3663 | 3928 | 4672 | 5137 | 2773 | 2916 | 4010 | 3905.84 | 4579 | 3233 | 0.706 | -0.503 | 0.00177058 | 0.01395578 | 0.0304 | 0.0077 | 0.0184 | 0.0184 | TRUE | 0.5365 |

| | | | | | | | | | | | | | | | | | | | | | | | | | |
|---|---|---|---|---|---|---|---|---|---|---|---|---|---|---|---|---|---|---|---|---|---|---|---|---|---|
| Akap3 | 1 | 3 | 4 | 7 | 4 | 2 | 1 | 3 | 4 | 6 | 4 | 2 | 3.35 | 3 | 4 | 1.498 | 0.583 | 0.58030881 | NA | 0 | 0.9918 | 0.4923 | 0.4923 | TRUE | 0.309 |
| Akap4 | 0 | 0 | 0 | 2 | 1 | 1 | 0 | 0 | 0 | 2 | 1 | 1 | 0.62 | 0 | 1 | 6.676 | 2.739 | 0.27800695 | NA | 0 | 5.2881 | 3.2034 | 3.2034 | TRUE | 0.07 |
| Akap5 | 288 | 346 | 326 | 319 | 287 | 299 | 306 | 346 | 327 | 251 | 309 | 327 | 310.94 | 326 | 296 | 0.903 | -0.147 | 0.3214053 | 0.54250904 | 0.0085 | 0.0175 | 0.0127 | 0.0127 | TRUE | 0.2123 |
| Akap6 | 13 | 9 | 10 | 22 | 14 | 10 | 14 | 9 | 10 | 17 | 15 | 11 | 12.69 | 11 | 14 | 1.331 | 0.413 | 0.44465353 | 0.6512281 | 0 | 0.2665 | 0.13 | 0.13 | TRUE | 0.1846 |
| Akap7 | 671 | 1087 | 1227 | 1202 | 911 | 1027 | 713 | 1086 | 1232 | 945 | 980 | 1124 | 1013.28 | 1010 | 1016 | 1.006 | 0.008 | 0.96535782 | 0.98257199 | 0.0417 | 0.0102 | 0.0252 | 0.0252 | TRUE | 0.7602 |
| Akap8 | 7528 | 6133 | 5988 | 8701 | 6416 | 5841 | 7996 | 6125 | 6013 | 6840 | 6903 | 6394 | 6711.82 | 6711 | 6712 | 1 | 0 | 0.99765975 | 0.99854611 | 0.0116 | 0.0074 | 0.0109 | 0.0109 | TRUE | 0.3421 |
| Akap8l | 2772 | 3374 | 3387 | 3237 | 2696 | 2739 | 2944 | 3370 | 3401 | 2545 | 2901 | 2998 | 3026.43 | 3238 | 2815 | 0.869 | -0.203 | 0.04643112 | 0.15587882 | 0.0062 | 0.008 | 0.0072 | 0.0072 | TRUE | 0.0865 |
| Akap9 | 1916 | 2067 | 2607 | 2421 | 1783 | 1712 | 2035 | 2064 | 2618 | 1903 | 1918 | 1874 | 2068.8 | 2239 | 1898 | 0.848 | -0.238 | 0.04089422 | 0.14311971 | 0.0101 | 0.0085 | 0.0093 | 0.0093 | TRUE | 0.2653 |
| Akip1 | 272 | 419 | 491 | 718 | 470 | 511 | 289 | 418 | 493 | 564 | 506 | 559 | 471.65 | 400 | 543 | 1.357 | 0.441 | 0.02034196 | 0.08797186 | 0.0344 | 0.0139 | 0.0238 | 0.0238 | TRUE | 0.5597 |
| Akirin1 | 3395 | 3236 | 3104 | 4431 | 2998 | 2880 | 3606 | 3232 | 3117 | 3483 | 3226 | 3153 | 3302.71 | 3318 | 3287 | 0.991 | -0.013 | 0.88502869 | 0.94366363 | 0.004 | 0.0079 | 0.0055 | 0.0055 | TRUE | 0.0699 |
| Akirin2 | 4715 | 5368 | 5191 | 6338 | 4643 | 4529 | 5008 | 5361 | 5212 | 4982 | 4996 | 4958 | 5086.21 | 5194 | 4979 | 0.959 | -0.061 | 0.32324313 | 0.54444583 | 4.00E-04 | 0.0076 | 0.0026 | 0.0026 | TRUE | 0.0119 |
| Akna | 838 | 930 | 956 | 1240 | 970 | 1036 | 890 | 929 | 960 | 975 | 1044 | 1134 | 988.56 | 926 | 1051 | 1.133 | 0.18 | 0.06293959 | 0.19237538 | 0.0027 | 0.0102 | 0.0058 | 0.0058 | TRUE | 0.0572 |
| Aknad1 | 1 | 0 | 1 | 1 | 4 | 2 | 1 | 0 | 1 | 1 | 4 | 2 | 1.56 | 1 | 2 | 3.482 | 1.8 | 0.28863368 | NA | 0 | 2.122 | 1.1217 | 1.1217 | TRUE | 0.1582 |
| Akp3 | 3 | 4 | 1 | 3 | 2 | 4 | 3 | 4 | 1 | 2 | 2 | 4 | 2.85 | 3 | 3 | 1.077 | 0.107 | 0.92378349 | NA | 0 | 1.1646 | 0.5535 | 0.5535 | TRUE | 0.3714 |
| Akr1a1 | 12800 | 12797 | 12211 | 14920 | 10158 | 9744 | 13596 | 12780 | 12261 | 11729 | 10929 | 10666 | 11993.69 | 12879 | 11108 | 0.863 | -0.213 | 0.00519633 | 0.03212721 | 0.0025 | 0.0072 | 0.0041 | 0.0041 | TRUE | 0.0293 |
| Akr1b10 | 325 | 286 | 320 | 459 | 366 | 280 | 345 | 286 | 321 | 361 | 394 | 307 | 335.55 | 317 | 354 | 1.115 | 0.158 | 0.2867136 | 0.50796477 | 0.0092 | 0.0167 | 0.0128 | 0.0128 | TRUE | 0.1499 |
| Akr1b3 | 0 | 1 | 0 | 0 | 0 | 0 | 0 | 1 | 0 | 0 | 0 | 0 | 0.17 | 0 | 0 | 0.49 | -1.029 | 0.80087 | NA | 0 | 19.7991 | 10 | 10 | TRUE | 0.405 |
| Akr1b7 | 25 | 15 | 31 | 2 | 12 | 16 | 27 | 15 | 31 | 2 | 13 | 18 | 17.44 | 24 | 11 | 0.431 | -1.214 | 0.09184534 | 0.24740985 | 0.4551 | 0.1958 | 0.3055 | 0.3055 | TRUE | 1.8634 |
| Akr1b8 | 2437 | 1411 | 1373 | 1518 | 941 | 929 | 2589 | 1409 | 1379 | 1193 | 1012 | 1017 | 1433.18 | 1792 | 1074 | 0.6 | -0.738 | 0.00114966 | 0.00990303 | 0.0703 | 0.0092 | 0.0364 | 0.0364 | TRUE | 1.8238 |
| Akr1c12 | 0 | 0 | 0 | 12 | 8 | 3 | 0 | 0 | 0 | 9 | 9 | 3 | 3.55 | 0 | 7 | 38.062 | 5.25 | 0.0008171 | NA | 0.1456 | 0.9338 | 0.7048 | 0.7048 | TRUE | 0.5473 |
| Akr1c13 | 14 | 5 | 7 | 68 | 43 | 37 | 15 | 5 | 7 | 53 | 46 | 41 | 27.85 | 9 | 47 | 5.268 | 2.397 | 3.38E-07 | 1.05E-05 | 0.0142 | 0.1252 | 0.0913 | 0.0913 | TRUE | 0.9464 |
| Akr1c14 | 4 | 4 | 3 | 14 | 12 | 7 | 4 | 4 | 3 | 11 | 13 | 8 | 7.14 | 4 | 11 | 2.815 | 1.493 | 0.04488564 | 0.15263826 | 0 | 0.4684 | 0.2166 | 0.2166 | TRUE | 0.1821 |
| Akr1c18 | 5 | 1 | 2 | 6 | 9 | 14 | 5 | 1 | 2 | 5 | 10 | 15 | 6.34 | 3 | 10 | 3.565 | 1.834 | 0.04940562 | 0.16298379 | 0.238 | 0.5265 | 0.3917 | 0.3917 | TRUE | 0.2121 |
| Akr1c19 | 0 | 0 | 0 | 2 | 0 | 0 | 0 | 0 | 0 | 2 | 0 | 0 | 0.26 | 0 | 1 | 2.886 | 1.529 | 0.70676491 | NA | 0 | 12.5792 | 10 | 10 | TRUE | 0.9669 |
| Akr1c20 | 0 | 0 | 0 | 0 | 0 | 0 | 0 | 0 | 0 | 0 | 0 | 0 | 0 | 0 | 0 | NA | NA | NA | NA | NA | NA | NA | NA | NA | NA |
| Akr1c21 | 1 | 1 | 0 | 0 | 1 | 1 | 1 | 1 | 0 | 0 | 1 | 1 | 0.71 | 1 | 1 | 1.016 | 0.022 | 0.99229748 | NA | 0 | 4.6782 | 2.3784 | 2.3784 | TRUE | 0.3742 |
| Akr1c6 | 1 | 1 | 0 | 0 | 0 | 0 | 1 | 1 | 0 | 0 | 0 | 0 | 0.34 | 1 | 0 | 0.261 | -1.936 | 0.60454921 | NA | 0 | 9.5983 | 8.3269 | 8.3269 | TRUE | 0.2507 |
| Akr1cl | 34 | 18 | 14 | 93 | 42 | 20 | 36 | 18 | 14 | 73 | 45 | 22 | 34.72 | 23 | 47 | 2.075 | 1.053 | 0.04913736 | 0.16239647 | 0.2526 | 0.1018 | 0.1736 | 0.1736 | TRUE | 0.1388 |
| Akr1d1 | 0 | 0 | 0 | 0 | 0 | 0 | 0 | 0 | 0 | 0 | 0 | 0 | 0 | 0 | 0 | NA | NA | NA | NA | NA | NA | NA | NA | NA | NA |
| Akr1e1 | 381 | 501 | 526 | 721 | 473 | 584 | 405 | 500 | 528 | 567 | 509 | 639 | 524.7 | 478 | 572 | 1.196 | 0.258 | 0.07807718 | 0.22104348 | 0.0139 | 0.0132 | 0.0135 | 0.0135 | TRUE | 0.2063 |
| Akr7a5 | 1189 | 1559 | 1576 | 1533 | 1239 | 1323 | 1263 | 1557 | 1582 | 1205 | 1333 | 1448 | 1398.14 | 1467 | 1329 | 0.905 | -0.144 | 0.24338439 | 0.46008258 | 0.0104 | 0.0093 | 0.0103 | 0.0103 | TRUE | 0.1784 |
| Akt1 | 2278 | 2775 | 2828 | 3264 | 2517 | 2546 | 2420 | 2771 | 2840 | 2566 | 2708 | 2787 | 2681.95 | 2677 | 2687 | 1.003 | 0.005 | 0.95770859 | 0.97811176 | 0.0041 | 0.0081 | 0.0057 | 0.0057 | TRUE | 0.0855 |
| Akt1s1 | 1839 | 1869 | 1787 | 2684 | 1878 | 1525 | 1953 | 1867 | 1794 | 2110 | 2021 | 1669 | 1902.36 | 1871 | 1933 | 1.034 | 0.048 | 0.66439808 | 0.81221452 | 0.0079 | 0.0086 | 0.0082 | 0.0082 | TRUE | 0.1713 |
| Akt2 | 2815 | 2868 | 2881 | 3971 | 2953 | 2944 | 2990 | 2864 | 2893 | 3122 | 3177 | 3223 | 3044.79 | 2916 | 3174 | 1.088 | 0.122 | 0.05352182 | 0.17185385 | 0 | 0.008 | 0.0026 | 0.0026 | TRUE | 0.006 |
| Akt3 | 1530 | 2050 | 2104 | 2915 | 2085 | 2313 | 1625 | 2047 | 2113 | 2292 | 2243 | 2532 | 2141.99 | 1928 | 2356 | 1.221 | 0.288 | 0.01730924 | 0.07824632 | 0.0109 | 0.0084 | 0.0101 | 0.0101 | TRUE | 0.2282 |
| Aktip | 4082 | 4756 | 4689 | 4662 | 3576 | 3906 | 4336 | 4750 | 4708 | 3665 | 3848 | 4276 | 4263.68 | 4598 | 3930 | 0.854 | -0.227 | 0.01079863 | 0.05571471 | 0.0041 | 0.0077 | 0.0055 | 0.0055 | TRUE | 0.0722 |
| Alad | 1643 | 1644 | 1577 | 2917 | 2037 | 1841 | 1745 | 1642 | 1583 | 2293 | 2192 | 2015 | 1911.76 | 1657 | 2167 | 1.309 | 0.388 | 1.07E-05 | 0.00021082 | 0.0025 | 0.0086 | 0.0051 | 0.0051 | TRUE | 0.045 |
| Alas1 | 2345 | 2150 | 2142 | 2841 | 2071 | 1856 | 2491 | 2147 | 2151 | 2233 | 2228 | 2032 | 2213.69 | 2263 | 2164 | 0.957 | -0.064 | 0.50595033 | 0.70029387 | 0.0044 | 0.0084 | 0.0061 | 0.0061 | TRUE | 0.0937 |
| Alas2 | 3 | 5 | 3 | 6 | 2 | 0 | 3 | 5 | 3 | 5 | 2 | 0 | 3.01 | 4 | 2 | 0.639 | -0.646 | 0.59210212 | NA | 0.1179 | 1.1013 | 0.7068 | 0.7068 | TRUE | 0.4014 |
| Alb | 1 | 0 | 0 | 0 | 0 | 0 | 1 | 0 | 0 | 0 | 0 | 0 | 0.18 | 0 | 0 | 0.49 | -1.029 | 0.80087 | NA | 0 | 18.6165 | 10 | 10 | TRUE | 0.4614 |
| Alcam | 58 | 41 | 33 | 74 | 62 | 61 | 62 | 41 | 33 | 58 | 67 | 67 | 54.56 | 45 | 64 | 1.412 | 0.498 | 0.10598304 | 0.27162244 | 0.0276 | 0.0673 | 0.0494 | 0.0494 | TRUE | 0.7674 |
| Aldh16a1 | 1673 | 1765 | 1618 | 2607 | 1949 | 1766 | 1777 | 1763 | 1625 | 2049 | 2097 | 1933 | 1874 | 1722 | 2026 | 1.177 | 0.236 | 0.00349957 | 0.02385271 | 0.0015 | 0.0087 | 0.0042 | 0.0042 | TRUE | 0.0293 |
| Aldh18a1 | 2588 | 2020 | 1919 | 3107 | 2307 | 2139 | 2749 | 2017 | 1927 | 2442 | 2482 | 2341 | 2326.55 | 2231 | 2422 | 1.086 | 0.119 | 0.400906 | 0.61389115 | 0.0191 | 0.0083 | 0.014 | 0.014 | TRUE | 0.4986 |
| Aldh1a1 | 3 | 3 | 5 | 15 | 7 | 10 | 3 | 3 | 5 | 12 | 8 | 11 | 6.91 | 4 | 10 | 2.712 | 1.439 | 0.05661355 | 0.17876766 | 0 | 0.4835 | 0.226 | 0.226 | TRUE | 0.1604 |
| Aldh1a2 | 264 | 203 | 151 | 319 | 217 | 189 | 280 | 203 | 152 | 251 | 233 | 207 | 220.99 | 212 | 230 | 1.091 | 0.125 | 0.58565956 | 0.76074968 | 0.0462 | 0.0218 | 0.0336 | 0.0336 | TRUE | 0.3605 |
| Aldh1a3 | 33 | 24 | 24 | 31 | 39 | 35 | 35 | 24 | 24 | 24 | 42 | 38 | 31.29 | 28 | 35 | 1.25 | 0.322 | 0.40465891 | 0.61732751 | 0.0372 | 0.1122 | 0.0752 | 0.0752 | TRUE | 0.5842 |
| Aldh1a7 | 8 | 11 | 5 | 28 | 20 | 15 | 8 | 11 | 5 | 22 | 22 | 16 | 14.08 | 8 | 20 | 2.454 | 1.295 | 0.0173812 | 0.07846133 | 0 | 0.241 | 0.1268 | 0.1268 | TRUE | 0.3474 |
| Aldh1b1 | 615 | 647 | 656 | 883 | 648 | 685 | 653 | 646 | 659 | 694 | 697 | 750 | 683.22 | 653 | 714 | 1.093 | 0.128 | 0.16313972 | 0.35866179 | 0 | 0.0117 | 0.0046 | 0.0046 | TRUE | 0.0226 |
| Aldh1l1 | 905 | 1424 | 1294 | 802 | 668 | 940 | 961 | 1422 | 1299 | 630 | 719 | 1029 | 1010.15 | 1227 | 793 | 0.645 | -0.632 | 0.00209078 | 0.01595234 | 0.0522 | 0.0102 | 0.0294 | 0.0294 | TRUE | 0.8039 |
| Aldh1l2 | 98 | 61 | 43 | 128 | 108 | 87 | 104 | 61 | 43 | 101 | 116 | 95 | 86.71 | 69 | 104 | 1.501 | 0.586 | 0.07079811 | 0.20727967 | 0.0858 | 0.0449 | 0.0637 | 0.0637 | TRUE | 1.0463 |
| Aldh2 | 8553 | 11157 | 11882 | 7866 | 6064 | 7109 | 9085 | 11143 | 11931 | 6184 | 6524 | 7782 | 8774.7 | 10720 | 6830 | 0.637 | -0.65 | 8.32E-07 | 2.35E-05 | 0.0168 | 0.0073 | 0.0124 | 0.0124 | TRUE | 0.2174 |
| Aldh3a1 | 64 | 54 | 57 | 60 | 41 | 52 | 68 | 54 | 57 | 47 | 44 | 57 | 54.56 | 60 | 49 | 0.826 | -0.275 | 0.30824047 | 0.52871873 | 0 | 0.0673 | 0.0341 | 0.0341 | TRUE | 0.1324 |
| Aldh3a2 | 1912 | 2198 | 2156 | 3200 | 2288 | 2262 | 2031 | 2195 | 2165 | 2516 | 2462 | 2476 | 2307.39 | 2130 | 2485 | 1.166 | 0.222 | 0.00164119 | 0.01314617 | 4.00E-04 | 0.0083 | 0.0031 | 0.0031 | TRUE | 0.0201 |
| Aldh3b1 | 74 | 97 | 124 | 80 | 56 | 71 | 79 | 97 | 125 | 63 | 60 | 78 | 83.47 | 100 | 67 | 0.668 | -0.583 | 0.02435828 | 0.09979885 | 0.0253 | 0.0464 | 0.0359 | 0.0359 | TRUE | 0.2349 |
| Aldh3b2 | 12 | 8 | 10 | 25 | 16 | 21 | 13 | 8 | 10 | 20 | 17 | 23 | 15.11 | 10 | 20 | 1.948 | 0.962 | 0.05476704 | 0.1744317 | 0 | 0.225 | 0.1063 | 0.1063 | TRUE | 0.1512 |
| Aldh4a1 | 4277 | 4636 | 4657 | 4887 | 3683 | 3909 | 4543 | 4630 | 4676 | 3842 | 3963 | 4279 | 4322.09 | 4616 | 4028 | 0.872 | -0.197 | 0.00626756 | 0.03723458 | 0.0013 | 0.0077 | 0.0035 | 0.0035 | TRUE | 0.0361 |
| Aldh5a1 | 1449 | 1810 | 1864 | 2304 | 1708 | 1705 | 1539 | 1808 | 1872 | 1811 | 1838 | 1866 | 1788.95 | 1740 | 1838 | 1.057 | 0.079 | 0.41940564 | 0.62992722 | 0.0043 | 0.0087 | 0.0064 | 0.0064 | TRUE | 0.1221 |
| Aldh6a1 | 3325 | 4288 | 4354 | 3389 | 2569 | 3153 | 3532 | 4282 | 4372 | 2664 | 2764 | 3451 | 3510.96 | 4062 | 2960 | 0.728 | -0.457 | 0.00057326 | 0.00578738 | 0.0165 | 0.0078 | 0.0124 | 0.0124 | TRUE | 0.2557 |
| Aldh7a1 | 4502 | 4996 | 5190 | 5716 | 4138 | 4392 | 4782 | 4990 | 5211 | 4493 | 4452 | 4808 | 4789.37 | 4994 | 4584 | 0.918 | -0.124 | 0.08951627 | 0.24329667 | 0.0015 | 0.0076 | 0.0036 | 0.0036 | TRUE | 0.0219 |
| Aldh8a1 | 0 | 1 | 0 | 1 | 2 | 3 | 0 | 1 | 0 | 1 | 2 | 3 | 1.2 | 0 | 2 | 5.579 | 2.48 | 0.20613863 | NA | 0 | 2.7444 | 1.5328 | 1.5328 | TRUE | 0.3029 |
| Aldh9a1 | 4932 | 5745 | 5747 | 6406 | 4833 | 5307 | 5239 | 5738 | 5771 | 5036 | 5200 | 5809 | 5465.35 | 5583 | 5348 | 0.958 | -0.062 | 0.48021733 | 0.67946422 | 0.0038 | 0.0075 | 0.0054 | 0.0054 | TRUE | 0.0691 |
| Aldoa | 69970 | 125444 | 133991 | 104176 | 83317 | 109329 | 74321 | 125282 | 134541 | 81895 | 89644 | 119678 | 104226.74 | 111381 | 97072 | 0.872 | -0.198 | 0.35673135 | 0.57524954 | 0.059 | 0.0069 | 0.0334 | 0.0334 | TRUE | 1.0377 |
| Aldoart1 | 13 | 25 | 18 | 21 | 16 | 22 | 14 | 25 | 18 | 17 | 17 | 24 | 19.11 | 19 | 19 | 1.009 | 0.013 | 0.97687256 | 0.98739003 | 0 | 0.1793 | 0.1007 | 0.1007 | TRUE | 0.3707 |
| Aldoart2 | 8 | 6 | 5 | 15 | 2 | 6 | 8 | 6 | 5 | 12 | 2 | 7 | 6.67 | 6 | 7 | 1.076 | 0.106 | 0.89792049 | 0.95044288 | 0.1706 | 0.5008 | 0.3405 | 0.3405 | TRUE | 0.2341 |
| Aldob | 1 | 0 | 0 | 0 | 0 | 0 | 1 | 0 | 0 | 0 | 0 | 0 | 0.18 | 0 | 0 | 0.49 | -1.029 | 0.80087 | NA | 0 | 18.6165 | 10 | 10 | TRUE | 0.4614 |
| Aldoc | 2785 | 4822 | 5181 | 2983 | 2611 | 4103 | 2958 | 4816 | 5202 | 2345 | 2809 | 4491 | 3770.31 | 4325 | 3215 | 0.743 | -0.428 | 0.25052407 | 0.46788225 | 0.0997 | 0.0078 | 0.0458 | 0.0997 | TRUE | 1.4644 |
| Alg1 | 943 | 1029 | 926 | 1176 | 881 | 951 | 1002 | 1028 | 930 | 924 | 948 | 1041 | 978.75 | 987 | 971 | 0.984 | -0.024 | 0.80479907 | 0.89931073 | 0.0023 | 0.0103 | 0.0055 | 0.0055 | TRUE | 0.0467 |
| Alg10b | 2665 | 2763 | 2947 | 4013 | 2799 | 2893 | 2831 | 2759 | 2959 | 3155 | 3012 | 3167 | 2980.39 | 2850 | 3111 | 1.092 | 0.127 | 0.06708528 | 0.20001306 | 7.00E-04 | 0.008 | 0.0031 | 0.0031 | TRUE | 0.0137 |
| Alg11 | 3160 | 3156 | 2937 | 4130 | 2994 | 2845 | 3356 | 3152 | 2949 | 3247 | 3221 | 3114 | 3173.3 | 3152 | 3194 | 1.013 | 0.019 | 0.80640629 | 0.90024179 | 0.002 | 0.0079 | 0.0041 | 0.0041 | TRUE | 0.0388 |

| | | | | | | | | | | | | | | | | | | | | | | | | | |
|---|---|---|---|---|---|---|---|---|---|---|---|---|---|---|---|---|---|---|---|---|---|---|---|---|---|
| Alg12 | 439 | 503 | 541 | 662 | 436 | 501 | 466 | 502 | 543 | 520 | 469 | 548 | 508.3 | 504 | 512 | 1.017 | 0.025 | 0.83572231 | 0.91655742 | 0.0038 | 0.0134 | 0.0081 | 0.0081 | TRUE | 0.0623 |
| Alg13 | 12067 | 14904 | 15301 | 13373 | 10811 | 12187 | 12817 | 14885 | 15364 | 10513 | 11632 | 13341 | 13091.87 | 14355 | 11829 | 0.824 | -0.279 | 0.0162802 | 0.07525537 | 0.0116 | 0.0072 | 0.0097 | 0.0097 | TRUE | 0.1528 |
| Alg14 | 803 | 587 | 550 | 835 | 532 | 474 | 853 | 586 | 552 | 656 | 572 | 519 | 623.18 | 664 | 582 | 0.879 | -0.187 | 0.31273508 | 0.5332037 | 0.0354 | 0.0122 | 0.023 | 0.023 | TRUE | 0.7303 |
| Alg2 | 1347 | 1238 | 1176 | 1817 | 1372 | 1164 | 1431 | 1236 | 1181 | 1428 | 1476 | 1274 | 1337.79 | 1283 | 1393 | 1.086 | 0.12 | 0.28129453 | 0.5026144 | 0.007 | 0.0094 | 0.0081 | 0.0081 | TRUE | 0.122 |
| Alg3 | 1007 | 1384 | 1559 | 1520 | 1129 | 1205 | 1070 | 1382 | 1565 | 1195 | 1215 | 1319 | 1290.99 | 1339 | 1243 | 0.928 | -0.108 | 0.45608819 | 0.66096526 | 0.0189 | 0.0095 | 0.0144 | 0.0144 | TRUE | 0.3686 |
| Alg5 | 504 | 756 | 809 | 1009 | 747 | 741 | 535 | 755 | 812 | 793 | 804 | 811 | 751.79 | 701 | 803 | 1.145 | 0.195 | 0.21764521 | 0.42962457 | 0.0219 | 0.0113 | 0.0167 | 0.0167 | TRUE | 0.502 |
| Alg6 | 1385 | 1233 | 1267 | 1932 | 1338 | 1360 | 1471 | 1231 | 1272 | 1519 | 1440 | 1489 | 1403.64 | 1325 | 1483 | 1.119 | 0.163 | 0.09695674 | 0.25584771 | 0.004 | 0.0093 | 0.0062 | 0.0062 | TRUE | 0.1115 |
| Alg8 | 366 | 570 | 659 | 721 | 509 | 579 | 389 | 569 | 662 | 567 | 548 | 634 | 561.33 | 540 | 583 | 1.079 | 0.109 | 0.56532888 | 0.74551564 | 0.0364 | 0.0128 | 0.0243 | 0.0243 | TRUE | 0.5275 |
| Alg9 | 1390 | 1579 | 1639 | 2019 | 1538 | 1559 | 1476 | 1577 | 1646 | 1587 | 1655 | 1707 | 1607.94 | 1566 | 1650 | 1.052 | 0.074 | 0.3729015 | 0.58928297 | 0.0015 | 0.009 | 0.0043 | 0.0043 | TRUE | 0.0302 |
| Alk | 80 | 96 | 69 | 99 | 75 | 62 | 85 | 96 | 69 | 78 | 81 | 68 | 79.42 | 83 | 76 | 0.906 | -0.142 | 0.54536957 | 0.73043214 | 0 | 0.0484 | 0.0272 | 0.0272 | TRUE | 0.208 |
| Alkbh1 | 1014 | 1171 | 1225 | 1181 | 859 | 1002 | 1077 | 1169 | 1230 | 928 | 924 | 1097 | 1071.01 | 1159 | 983 | 0.848 | -0.238 | 0.02965834 | 0.11414623 | 0.0058 | 0.01 | 0.0077 | 0.0077 | TRUE | 0.1205 |
| Alkbh2 | 719 | 799 | 761 | 875 | 671 | 719 | 764 | 798 | 764 | 688 | 722 | 787 | 753.78 | 775 | 732 | 0.943 | -0.084 | 0.38597737 | 0.59988521 | 0.0014 | 0.0113 | 0.0055 | 0.0055 | TRUE | 0.0502 |
| Alkbh3 | 312 | 367 | 412 | 504 | 402 | 320 | 331 | 367 | 414 | 396 | 433 | 350 | 381.77 | 371 | 393 | 1.061 | 0.085 | 0.55156858 | 0.73561134 | 0.0088 | 0.0155 | 0.012 | 0.012 | TRUE | 0.119 |
| Alkbh4 | 512 | 502 | 481 | 606 | 465 | 433 | 544 | 501 | 483 | 476 | 500 | 474 | 496.47 | 509 | 483 | 0.949 | -0.075 | 0.47884482 | 0.67817611 | 0 | 0.0135 | 0.0061 | 0.0061 | TRUE | 0.0404 |
| Alkbh5 | 9752 | 13789 | 14761 | 16020 | 12987 | 14507 | 10358 | 13771 | 14822 | 12594 | 13973 | 15880 | 13566.37 | 12984 | 14149 | 1.09 | 0.124 | 0.40098628 | 0.61389591 | 0.0199 | 0.0071 | 0.0156 | 0.0156 | TRUE | 0.3824 |
| Alkbh6 | 221 | 253 | 264 | 383 | 269 | 268 | 235 | 253 | 265 | 301 | 289 | 293 | 272.73 | 251 | 294 | 1.175 | 0.233 | 0.0692936 | 0.20428374 | 0 | 0.019 | 0.0081 | 0.0081 | TRUE | 0.0344 |
| Alkbh7 | 1063 | 1196 | 1136 | 1323 | 1011 | 947 | 1129 | 1194 | 1141 | 1040 | 1088 | 1037 | 1104.78 | 1155 | 1055 | 0.913 | -0.131 | 0.10143007 | 0.2638989 | 0 | 0.0099 | 0.0037 | 0.0037 | TRUE | 0.0109 |
| Alkbh8 | 792 | 766 | 818 | 1058 | 811 | 806 | 841 | 765 | 821 | 832 | 873 | 882 | 835.7 | 809 | 862 | 1.065 | 0.091 | 0.31551991 | 0.53622946 | 0 | 0.0108 | 0.0047 | 0.0047 | TRUE | 0.0272 |
| Allc | 1 | 0 | 0 | 2 | 1 | 0 | 1 | 0 | 0 | 2 | 1 | 0 | 0.62 | 0 | 1 | 2.462 | 1.3 | 0.62119797 | NA | 0 | 5.3343 | 3.4416 | 3.4416 | TRUE | 0.4614 |
| Alms1 | 3381 | 2873 | 3678 | 4519 | 3248 | 3052 | 3591 | 2869 | 3693 | 3552 | 3495 | 3341 | 3423.61 | 3384 | 3463 | 1.023 | 0.033 | 0.76786099 | 0.87886092 | 0.0095 | 0.0079 | 0.0088 | 0.0088 | TRUE | 0.2159 |
| Alms1-ps2 | 0 | 1 | 0 | 3 | 0 | 1 | 0 | 1 | 0 | 2 | 0 | 1 | 0.74 | 0 | 1 | 3.216 | 1.685 | 0.50251474 | NA | 0 | 4.4471 | 3.135 | 3.135 | TRUE | 0.1927 |
| Alox12 | 70 | 116 | 92 | 136 | 95 | 71 | 74 | 116 | 92 | 107 | 102 | 78 | 94.9 | 94 | 96 | 1.017 | 0.024 | 0.92237136 | 0.96163677 | 0.0274 | 0.0416 | 0.0347 | 0.0347 | TRUE | 0.269 |
| Alox12b | 92 | 141 | 145 | 105 | 80 | 106 | 98 | 141 | 146 | 83 | 86 | 116 | 111.46 | 128 | 95 | 0.738 | -0.439 | 0.07314736 | 0.21183992 | 0.0314 | 0.0365 | 0.034 | 0.034 | TRUE | 0.4315 |
| Alox12e | 11 | 9 | 6 | 19 | 13 | 12 | 12 | 9 | 6 | 15 | 14 | 13 | 11.46 | 9 | 14 | 1.583 | 0.663 | 0.24160897 | 0.45834215 | 0 | 0.2944 | 0.1387 | 0.1387 | TRUE | 0.2566 |
| Alox15 | 302 | 149 | 123 | 200 | 133 | 90 | 321 | 149 | 124 | 157 | 143 | 99 | 165.32 | 198 | 133 | 0.674 | -0.569 | 0.10808753 | 0.27513825 | 0.1539 | 0.0268 | 0.0841 | 0.0841 | TRUE | 3.0823 |
| Alox5 | 156 | 197 | 162 | 193 | 105 | 122 | 166 | 197 | 163 | 152 | 113 | 134 | 153.89 | 175 | 133 | 0.761 | -0.394 | 0.03684709 | 0.13268317 | 0.0099 | 0.0283 | 0.019 | 0.019 | TRUE | 0.1756 |
| Alox5ap | 93 | 65 | 61 | 72 | 34 | 36 | 99 | 65 | 61 | 57 | 37 | 39 | 59.59 | 75 | 44 | 0.595 | -0.749 | 0.01900289 | 0.08369548 | 0.049 | 0.0622 | 0.0557 | 0.0557 | TRUE | 0.6935 |
| Alox8 | 44 | 29 | 31 | 54 | 39 | 29 | 47 | 29 | 31 | 42 | 42 | 32 | 37.16 | 36 | 39 | 1.093 | 0.128 | 0.71040782 | 0.84213468 | 0.0185 | 0.0955 | 0.0592 | 0.0592 | TRUE | 0.5194 |
| Aloxe3 | 88 | 141 | 124 | 97 | 67 | 113 | 93 | 141 | 125 | 76 | 72 | 124 | 105.14 | 120 | 91 | 0.755 | -0.405 | 0.14716873 | 0.3363323 | 0.0557 | 0.0382 | 0.0466 | 0.0466 | TRUE | 0.9379 |
| Alpi | 0 | 0 | 0 | 0 | 0 | 0 | 0 | 0 | 0 | 0 | 0 | 0 | 0 | 0 | 0 | NA | NA | NA | NA | NA | NA | NA | NA | NA | NA |
| Alpk1 | 58 | 47 | 40 | 57 | 44 | 29 | 62 | 47 | 40 | 45 | 47 | 32 | 45.43 | 50 | 41 | 0.837 | -0.257 | 0.42315821 | 0.63351222 | 0.0216 | 0.0794 | 0.052 | 0.052 | TRUE | 0.3566 |
| Alpk2 | 1 | 0 | 1 | 4 | 3 | 1 | 1 | 0 | 1 | 3 | 3 | 1 | 1.59 | 1 | 2 | 3.67 | 1.876 | 0.25325644 | NA | 0 | 2.0804 | 1.0013 | 1.0013 | TRUE | 0.2567 |
| Alpk3 | 365 | 516 | 512 | 764 | 658 | 670 | 388 | 515 | 514 | 601 | 708 | 733 | 576.52 | 472 | 681 | 1.439 | 0.525 | 0.00048794 | 0.00508071 | 0.0163 | 0.0126 | 0.0145 | 0.0145 | TRUE | 0.2835 |
| Alpl | 5187 | 5401 | 5412 | 6768 | 4767 | 5010 | 5510 | 5394 | 5434 | 5320 | 5129 | 5484 | 5378.58 | 5446 | 5311 | 0.975 | -0.036 | 0.55924083 | 0.74125763 | 3.00E-04 | 0.0075 | 0.0026 | 0.0026 | TRUE | 0.0108 |
| Alppl2 | 63 | 38 | 33 | 154 | 111 | 71 | 67 | 38 | 33 | 121 | 119 | 78 | 76.04 | 46 | 106 | 2.316 | 1.211 | 0.000273 | 0.00314868 | 0.0823 | 0.0502 | 0.0641 | 0.0641 | TRUE | 1.2208 |
| Als2 | 7596 | 5327 | 5174 | 8676 | 5889 | 5372 | 8068 | 5320 | 5195 | 6820 | 6336 | 5880 | 6270.13 | 6194 | 6345 | 1.025 | 0.035 | 0.83588639 | 0.91664769 | 0.0291 | 0.0074 | 0.0203 | 0.0203 | TRUE | 0.8539 |
| Als2cl | 1240 | 1337 | 1323 | 1564 | 1228 | 1122 | 1317 | 1335 | 1328 | 1229 | 1321 | 1228 | 1293.29 | 1327 | 1259 | 0.949 | -0.076 | 0.33061664 | 0.55147568 | 0 | 0.0095 | 0.0036 | 0.0036 | TRUE | 0.0216 |
| Als2cr11 | 12 | 5 | 15 | 35 | 21 | 26 | 13 | 5 | 15 | 28 | 23 | 28 | 18.56 | 11 | 26 | 2.402 | 1.264 | 0.01301589 | 0.0638925 | 0 | 0.1844 | 0.1213 | 0.1213 | TRUE | 0.8556 |
| Als2cr12 | 3 | 7 | 3 | 8 | 6 | 4 | 3 | 7 | 3 | 6 | 6 | 4 | 5.05 | 4 | 5 | 1.3 | 0.379 | 0.65447888 | 0.80574825 | 0 | 0.659 | 0.3149 | 0.3149 | TRUE | 0.4922 |
| Alx1 | 9 | 8 | 21 | 14 | 16 | 18 | 10 | 8 | 21 | 11 | 17 | 20 | 14.43 | 13 | 16 | 1.228 | 0.296 | 0.61139806 | 0.77987759 | 0.111 | 0.2353 | 0.1749 | 0.1749 | TRUE | 1.2974 |
| Alx3 | 8 | 10 | 3 | 11 | 4 | 1 | 8 | 10 | 3 | 9 | 4 | 1 | 5.92 | 7 | 5 | 0.672 | -0.573 | 0.53898911 | 0.7252619 | 0.3298 | 0.563 | 0.4539 | 0.4539 | TRUE | 0.4562 |
| Alx4 | 2 | 10 | 7 | 10 | 4 | 5 | 2 | 10 | 7 | 8 | 4 | 5 | 6.13 | 6 | 6 | 0.928 | -0.108 | 0.89692283 | 0.9501066 | 0 | 0.5444 | 0.3365 | 0.3365 | TRUE | 0.3419 |
| Alyref | 5587 | 5953 | 5663 | 8053 | 6215 | 5171 | 5934 | 5945 | 5686 | 6331 | 6687 | 5660 | 6040.67 | 5855 | 6226 | 1.063 | 0.089 | 0.299539 | 0.51972817 | 0.0036 | 0.0074 | 0.0051 | 0.0051 | TRUE | 0.0766 |
| Alyref2 | 46 | 78 | 102 | 143 | 101 | 90 | 49 | 78 | 102 | 112 | 109 | 99 | 91.46 | 76 | 107 | 1.394 | 0.479 | 0.09162722 | 0.24703325 | 0.0515 | 0.0429 | 0.0468 | 0.0468 | TRUE | 0.3218 |
| Amacr | 735 | 786 | 840 | 939 | 663 | 633 | 781 | 785 | 843 | 738 | 713 | 693 | 758.93 | 803 | 715 | 0.891 | -0.167 | 0.06766725 | 0.20121567 | 0 | 0.0112 | 0.0047 | 0.0047 | TRUE | 0.0231 |
| Ambn | 0 | 0 | 0 | 0 | 1 | 0 | 0 | 0 | 0 | 0 | 1 | 0 | 0.18 | 0 | 0 | 1.859 | 0.894 | 0.82650734 | NA | 0 | 18.3785 | 10 | 10 | TRUE | 0.5204 |
| Ambp | 0 | 0 | 1 | 0 | 0 | 1 | 0 | 0 | 1 | 0 | 0 | 1 | 0.35 | 0 | 0 | 0.954 | -0.067 | 0.98566485 | NA | 0 | 9.4252 | 8.1416 | 8.1416 | TRUE | 0.5383 |
| Ambra1 | 2946 | 2827 | 2941 | 4241 | 3001 | 2691 | 3129 | 2823 | 2953 | 3334 | 3229 | 2946 | 3069.03 | 2968 | 3170 | 1.068 | 0.095 | 0.2635286 | 0.48273788 | 0.0029 | 0.008 | 0.0049 | 0.0049 | TRUE | 0.0461 |
| Amd1 | 0 | 0 | 0 | 0 | 0 | 0 | 0 | 0 | 0 | 0 | 0 | 0 | 0 | 0 | 0 | NA | NA | NA | NA | NA | NA | NA | NA | NA | NA |
| Amd2 | 0 | 0 | 0 | 0 | 0 | 0 | 0 | 0 | 0 | 0 | 0 | 0 | 0 | 0 | 0 | NA | NA | NA | NA | NA | NA | NA | NA | NA | NA |
| Amdhd1 | 58 | 55 | 52 | 43 | 40 | 48 | 62 | 55 | 52 | 34 | 43 | 53 | 49.69 | 56 | 43 | 0.761 | -0.394 | 0.17892312 | 0.38011522 | 0 | 0.0732 | 0.0416 | 0.0416 | TRUE | 0.2782 |
| Amdhd2 | 495 | 777 | 751 | 1071 | 805 | 770 | 526 | 776 | 754 | 842 | 866 | 843 | 767.8 | 685 | 850 | 1.24 | 0.311 | 0.04509535 | 0.15308737 | 0.0206 | 0.0112 | 0.016 | 0.016 | TRUE | 0.487 |
| Amelx | 0 | 0 | 0 | 0 | 0 | 0 | 0 | 0 | 0 | 0 | 0 | 0 | 0 | 0 | 0 | NA | NA | NA | NA | NA | NA | NA | NA | NA | NA |
| Amer1 | 1029 | 824 | 786 | 1444 | 1160 | 968 | 1093 | 823 | 789 | 1135 | 1248 | 1060 | 1024.67 | 902 | 1148 | 1.273 | 0.348 | 0.0170289 | 0.07746751 | 0.019 | 0.0101 | 0.0144 | 0.0144 | TRUE | 0.4088 |
| Amer2 | 8 | 12 | 9 | 24 | 19 | 13 | 8 | 12 | 9 | 19 | 20 | 14 | 13.84 | 10 | 18 | 1.815 | 0.86 | 0.09727544 | 0.25625993 | 0 | 0.2449 | 0.1148 | 0.1148 | TRUE | 0.1438 |
| Amer3 | 2 | 1 | 2 | 5 | 1 | 3 | 2 | 1 | 2 | 4 | 1 | 3 | 2.24 | 2 | 3 | 1.643 | 0.716 | 0.56927432 | NA | 0 | 1.4796 | 0.6683 | 0.6683 | TRUE | 0.3144 |
| Amfr | 3570 | 5226 | 5443 | 5558 | 3957 | 4170 | 3792 | 5219 | 5465 | 4369 | 4257 | 4565 | 4611.34 | 4825 | 4397 | 0.911 | -0.134 | 0.33463622 | 0.55489149 | 0.0178 | 0.0076 | 0.0137 | 0.0137 | TRUE | 0.4274 |
| Amh | 5 | 16 | 17 | 14 | 7 | 16 | 5 | 16 | 17 | 11 | 8 | 18 | 12.4 | 13 | 12 | 0.934 | -0.098 | 0.87852225 | 0.93996524 | 0.1557 | 0.2725 | 0.218 | 0.218 | TRUE | 0.7768 |
| Amhr2 | 303 | 378 | 442 | 387 | 310 | 328 | 322 | 378 | 444 | 304 | 334 | 359 | 356.66 | 381 | 332 | 0.87 | -0.2 | 0.20007922 | 0.40767872 | 0.0136 | 0.0161 | 0.0148 | 0.0148 | TRUE | 0.2051 |
| Amica1 | 5 | 5 | 1 | 8 | 7 | 5 | 5 | 5 | 1 | 6 | 8 | 5 | 5.1 | 4 | 6 | 1.711 | 0.775 | 0.3815786 | 0.59581767 | 0 | 0.6528 | 0.3527 | 0.3527 | TRUE | 0.6653 |
| Amigo1 | 362 | 498 | 508 | 486 | 395 | 445 | 385 | 497 | 510 | 382 | 425 | 487 | 447.69 | 464 | 431 | 0.928 | -0.108 | 0.4933062 | 0.68969289 | 0.0169 | 0.0143 | 0.0156 | 0.0156 | TRUE | 0.2587 |
| Amigo2 | 240 | 282 | 223 | 342 | 237 | 227 | 255 | 282 | 224 | 269 | 255 | 248 | 255.47 | 254 | 257 | 1.017 | 0.024 | 0.86855442 | 0.93527821 | 0.0035 | 0.0198 | 0.0113 | 0.0113 | TRUE | 0.1172 |
| Amigo3 | 11 | 6 | 4 | 18 | 12 | 7 | 12 | 6 | 4 | 14 | 13 | 8 | 9.4 | 7 | 12 | 1.619 | 0.695 | 0.30636104 | 0.52687016 | 0.0537 | 0.3573 | 0.22 | 0.22 | TRUE | 0.7533 |
| Ammecr1 | 844 | 844 | 951 | 1375 | 960 | 916 | 896 | 843 | 955 | 1081 | 1033 | 1003 | 968.47 | 898 | 1039 | 1.157 | 0.211 | 0.02245241 | 0.09436936 | 0.0016 | 0.0103 | 0.0051 | 0.0051 | TRUE | 0.0367 |
| Ammecr1l | 1487 | 1343 | 1313 | 2270 | 1589 | 1377 | 1579 | 1341 | 1318 | 1784 | 1710 | 1507 | 1540.1 | 1413 | 1667 | 1.181 | 0.24 | 0.03228347 | 0.12108797 | 0.008 | 0.009 | 0.0084 | 0.0084 | TRUE | 0.1273 |
| Amn | 310 | 371 | 378 | 320 | 260 | 267 | 329 | 371 | 380 | 252 | 280 | 292 | 317.15 | 360 | 275 | 0.761 | -0.394 | 0.00300454 | 0.02125004 | 0.0028 | 0.0173 | 0.0095 | 0.0095 | TRUE | 0.065 |
| Amn1 | 132 | 193 | 188 | 171 | 112 | 165 | 140 | 193 | 189 | 134 | 121 | 181 | 159.55 | 174 | 145 | 0.833 | -0.264 | 0.2315746 | 0.44541398 | 0.0302 | 0.0276 | 0.0287 | 0.0287 | TRUE | 0.4578 |
| Amot | 570 | 455 | 437 | 966 | 635 | 537 | 605 | 454 | 439 | 759 | 683 | 588 | 588.18 | 499 | 677 | 1.357 | 0.44 | 0.00704282 | 0.04028992 | 0.0224 | 0.0125 | 0.0175 | 0.0175 | TRUE | 0.3987 |
| Amotl1 | 820 | 937 | 1003 | 1939 | 1412 | 1287 | 871 | 936 | 1007 | 1524 | 1519 | 1409 | 1211.04 | 938 | 1484 | 1.582 | 0.662 | 1.50E-12 | 1.49E-10 | 0.0023 | 0.0096 | 0.0054 | 0.0054 | TRUE | 0.0496 |

| | | | | | | | | | | | | | | | | | | | | | | | | | |
|---|---|---|---|---|---|---|---|---|---|---|---|---|---|---|---|---|---|---|---|---|---|---|---|---|---|
| Amotl2 | 1362 | 1204 | 1047 | 2378 | 1520 | 1269 | 1447 | 1202 | 1051 | 1869 | 1635 | 1389 | 1432.4 | 1233 | 1631 | 1.323 | 0.404 | 0.00845764 | 0.04641862 | 0.0229 | 0.0092 | 0.0163 | 0.0163 | TRUE | 0.2049 |
| Ampd1 | 9 | 12 | 15 | 20 | 14 | 17 | 10 | 12 | 15 | 16 | 15 | 19 | 14.33 | 12 | 17 | 1.344 | 0.427 | 0.39316515 | 0.60663283 | 0 | 0.2367 | 0.1087 | 0.1087 | TRUE | 0.168 |
| Ampd2 | 3549 | 2385 | 2195 | 3743 | 2835 | 2290 | 3770 | 2382 | 2204 | 2942 | 3050 | 2507 | 2809.18 | 2785 | 2833 | 1.017 | 0.025 | 0.89775845 | 0.95042599 | 0.0483 | 0.0081 | 0.0266 | 0.0266 | TRUE | 1.1598 |
| Ampd3 | 923 | 1028 | 1054 | 847 | 735 | 867 | 980 | 1027 | 1058 | 666 | 791 | 949 | 911.85 | 1022 | 802 | 0.784 | -0.352 | 0.0118748 | 0.05975174 | 0.0147 | 0.0105 | 0.013 | 0.013 | TRUE | 0.3055 |
| Amph | 1334 | 1550 | 1590 | 1752 | 1265 | 1207 | 1417 | 1548 | 1597 | 1377 | 1361 | 1321 | 1436.84 | 1521 | 1353 | 0.89 | -0.168 | 0.04539109 | 0.15376038 | 0.0014 | 0.0092 | 0.0044 | 0.0044 | TRUE | 0.0425 |
| Amt | 1613 | 2012 | 1850 | 2550 | 1874 | 1851 | 1713 | 2009 | 1858 | 2005 | 2016 | 2026 | 1937.9 | 1860 | 2016 | 1.083 | 0.116 | 0.18253837 | 0.38458391 | 0.0024 | 0.0086 | 0.0049 | 0.0049 | TRUE | 0.0596 |
| Amtn | 0 | 0 | 0 | 0 | 0 | 0 | 0 | 0 | 0 | 0 | 0 | 0 | 0 | 0 | 0 | NA | NA | NA | NA | NA | NA | NA | NA | NA | NA |
| Amy1 | 4 | 5 | 3 | 6 | 3 | 1 | 4 | 5 | 3 | 5 | 3 | 1 | 3.55 | 4 | 3 | 0.757 | -0.403 | 0.69061119 | NA | 0 | 0.9352 | 0.4541 | 0.4541 | TRUE | 0.3725 |
| Amy2a2 | 0 | 0 | 0 | 0 | 0 | 0 | 0 | 0 | 0 | 0 | 0 | 0 | 0 | 0 | 0 | NA | NA | NA | NA | NA | NA | NA | NA | NA | NA |
| Amy2a3 | 0 | 0 | 0 | 0 | 0 | 0 | 0 | 0 | 0 | 0 | 0 | 0 | 0 | 0 | 0 | NA | NA | NA | NA | NA | NA | NA | NA | NA | NA |
| Amy2a4 | 0 | 0 | 0 | 0 | 0 | 0 | 0 | 0 | 0 | 0 | 0 | 0 | 0 | 0 | 0 | NA | NA | NA | NA | NA | NA | NA | NA | NA | NA |
| Amy2a5 | 0 | 0 | 0 | 0 | 0 | 0 | 0 | 0 | 0 | 0 | 0 | 0 | 0 | 0 | 0 | NA | NA | NA | NA | NA | NA | NA | NA | NA | NA |
| Amy2b | 0 | 0 | 0 | 0 | 0 | 0 | 0 | 0 | 0 | 0 | 0 | 0 | 0 | 0 | 0 | NA | NA | NA | NA | NA | NA | NA | NA | NA | NA |
| Amz1 | 130 | 94 | 105 | 105 | 92 | 66 | 138 | 94 | 105 | 83 | 99 | 72 | 98.53 | 112 | 85 | 0.753 | -0.41 | 0.08833269 | 0.24101084 | 0.0221 | 0.0403 | 0.0314 | 0.0314 | TRUE | 0.3909 |
| Amz2 | 2726 | 3329 | 3355 | 3633 | 2713 | 3157 | 2895 | 3325 | 3369 | 2856 | 2919 | 3456 | 3136.63 | 3196 | 3077 | 0.962 | -0.055 | 0.61330493 | 0.78121832 | 0.0086 | 0.008 | 0.0084 | 0.0084 | TRUE | 0.1405 |
| Anapc1 | 11787 | 11051 | 11993 | 15244 | 10727 | 10988 | 12520 | 11037 | 12042 | 11984 | 11542 | 12028 | 11858.7 | 11866 | 11851 | 0.999 | -0.002 | 0.98093576 | 0.98961586 | 0.0018 | 0.0072 | 0.0039 | 0.0039 | TRUE | 0.0457 |
| Anapc10 | 901 | 876 | 830 | 1861 | 1299 | 1094 | 957 | 875 | 833 | 1463 | 1398 | 1198 | 1120.58 | 888 | 1353 | 1.524 | 0.608 | 7.41E-08 | 2.78E-06 | 0.0071 | 0.0098 | 0.0083 | 0.0083 | TRUE | 0.1201 |
| Anapc11 | 3075 | 3700 | 3615 | 4624 | 3204 | 3092 | 3266 | 3695 | 3630 | 3635 | 3447 | 3385 | 3509.71 | 3530 | 3489 | 0.988 | -0.017 | 0.83684988 | 0.91714685 | 0.0026 | 0.0078 | 0.0045 | 0.0045 | TRUE | 0.052 |
| Anapc13 | 907 | 1118 | 1085 | 1261 | 919 | 929 | 963 | 1117 | 1089 | 991 | 989 | 1017 | 1027.74 | 1056 | 999 | 0.945 | -0.081 | 0.38715169 | 0.60099965 | 0.0022 | 0.0101 | 0.0054 | 0.0054 | TRUE | 0.0705 |
| Anapc15 | 1794 | 2088 | 2098 | 2985 | 2158 | 2036 | 1906 | 2085 | 2107 | 2347 | 2322 | 2229 | 2165.77 | 2033 | 2299 | 1.131 | 0.178 | 0.02249913 | 0.09450494 | 0.0011 | 0.0084 | 0.0039 | 0.0039 | TRUE | 0.0359 |
| Anapc16 | 927 | 931 | 930 | 1088 | 794 | 746 | 985 | 930 | 934 | 855 | 854 | 817 | 895.74 | 950 | 842 | 0.887 | -0.172 | 0.04141349 | 0.14437965 | 0 | 0.0106 | 0.004 | 0.004 | TRUE | 0.0125 |
| Anapc2 | 4721 | 4848 | 4673 | 6735 | 4987 | 4742 | 5015 | 4842 | 4692 | 5295 | 5366 | 5191 | 5066.59 | 4850 | 5284 | 1.09 | 0.124 | 0.04901724 | 0.16204149 | 4.00E-04 | 0.0076 | 0.0027 | 0.0027 | TRUE | 0.0108 |
| Anapc4 | 2961 | 3154 | 3239 | 4314 | 3072 | 3057 | 3145 | 3150 | 3252 | 3391 | 3305 | 3346 | 3265.05 | 3182 | 3347 | 1.052 | 0.073 | 0.2338571 | 0.44850881 | 0 | 0.0079 | 0.0024 | 0.0024 | TRUE | 0.0045 |
| Anapc5 | 17113 | 19005 | 19268 | 22986 | 16651 | 17260 | 18177 | 18980 | 19347 | 18070 | 17915 | 18894 | 18563.94 | 18835 | 18293 | 0.971 | -0.042 | 0.49452952 | 0.69052608 | 8.00E-04 | 0.0071 | 0.0027 | 0.0027 | TRUE | 0.0114 |
| Anapc7 | 4134 | 4124 | 4091 | 4676 | 3517 | 3623 | 4391 | 4119 | 4108 | 3676 | 3784 | 3966 | 4007.24 | 4206 | 3809 | 0.905 | -0.143 | 0.04336069 | 0.14892963 | 0.0011 | 0.0077 | 0.0034 | 0.0034 | TRUE | 0.018 |
| Ang | 8 | 11 | 16 | 11 | 6 | 8 | 8 | 11 | 16 | 9 | 6 | 9 | 9.9 | 12 | 8 | 0.672 | -0.573 | 0.35026835 | 0.5692357 | 0 | 0.3396 | 0.1666 | 0.1666 | TRUE | 0.3803 |
| Ang2 | 0 | 0 | 0 | 0 | 0 | 0 | 0 | 0 | 0 | 0 | 0 | 0 | 0 | 0 | 0 | NA | NA | NA | NA | NA | NA | NA | NA | NA | NA |
| Ang3 | 0 | 0 | 0 | 0 | 0 | 0 | 0 | 0 | 0 | 0 | 0 | 0 | 0 | 0 | 0 | NA | NA | NA | NA | NA | NA | NA | NA | NA | NA |
| Ang4 | 0 | 0 | 0 | 0 | 0 | 0 | 0 | 0 | 0 | 0 | 0 | 0 | 0 | 0 | 0 | NA | NA | NA | NA | NA | NA | NA | NA | NA | NA |
| Ang5 | 0 | 0 | 0 | 0 | 0 | 0 | 0 | 0 | 0 | 0 | 0 | 0 | 0 | 0 | 0 | NA | NA | NA | NA | NA | NA | NA | NA | NA | NA |
| Ang6 | 0 | 0 | 0 | 0 | 0 | 0 | 0 | 0 | 0 | 0 | 0 | 0 | 0 | 0 | 0 | NA | NA | NA | NA | NA | NA | NA | NA | NA | NA |
| Angel1 | 1922 | 2141 | 2142 | 2596 | 1935 | 1885 | 2042 | 2138 | 2151 | 2041 | 2082 | 2063 | 2086.11 | 2110 | 2062 | 0.977 | -0.034 | 0.62033005 | 0.78509433 | 0 | 0.0085 | 0.0028 | 0.0028 | TRUE | 0.0097 |
| Angel2 | 1284 | 1329 | 1444 | 2371 | 1647 | 1501 | 1364 | 1327 | 1450 | 1864 | 1772 | 1643 | 1570.02 | 1380 | 1760 | 1.276 | 0.351 | 7.04E-05 | 0.00103709 | 0.0025 | 0.009 | 0.005 | 0.005 | TRUE | 0.0403 |
| Angpt1 | 0 | 1 | 0 | 0 | 0 | 0 | 0 | 1 | 0 | 0 | 0 | 0 | 0.17 | 0 | 0 | 0.49 | -1.029 | 0.80087 | NA | 0 | 19.7991 | 10 | 10 | TRUE | 0.405 |
| Angpt2 | 10 | 7 | 9 | 12 | 8 | 6 | 11 | 7 | 9 | 9 | 9 | 7 | 8.54 | 9 | 8 | 0.931 | -0.103 | 0.87129503 | 0.93670819 | 0 | 0.3925 | 0.1741 | 0.1741 | TRUE | 0.1121 |
| Angpt4 | 1 | 2 | 0 | 2 | 0 | 0 | 1 | 2 | 0 | 2 | 0 | 0 | 0.77 | 1 | 1 | 0.553 | -0.856 | 0.7316688 | NA | 0 | 4.2745 | 3.1222 | 3.1222 | TRUE | 0.4509 |
| Angptl1 | 0 | 0 | 0 | 1 | 0 | 0 | 0 | 0 | 0 | 1 | 0 | 0 | 0.13 | 0 | 0 | 1.859 | 0.894 | 0.82650734 | NA | 0 | 25.1515 | 10 | 10 | TRUE | 0.2563 |
| Angptl2 | 33 | 24 | 30 | 39 | 24 | 25 | 35 | 24 | 30 | 31 | 26 | 27 | 28.83 | 30 | 28 | 0.945 | -0.081 | 0.82199314 | 0.90911907 | 0 | 0.1212 | 0.0587 | 0.0587 | TRUE | 0.1913 |
| Angptl3 | 0 | 0 | 0 | 0 | 0 | 0 | 0 | 0 | 0 | 0 | 0 | 0 | 0 | 0 | 0 | NA | NA | NA | NA | NA | NA | NA | NA | NA | NA |
| Angptl4 | 974 | 752 | 710 | 1212 | 851 | 720 | 1035 | 751 | 713 | 953 | 916 | 788 | 859.18 | 833 | 886 | 1.064 | 0.09 | 0.57994237 | 0.75669188 | 0.0245 | 0.0107 | 0.0177 | 0.0177 | TRUE | 0.532 |
| Angptl6 | 218 | 317 | 321 | 317 | 231 | 275 | 232 | 317 | 322 | 249 | 249 | 301 | 278.21 | 290 | 266 | 0.916 | -0.127 | 0.47224434 | 0.67343922 | 0.0188 | 0.0187 | 0.0188 | 0.0188 | TRUE | 0.3477 |
| Angptl7 | 0 | 0 | 0 | 0 | 0 | 0 | 0 | 0 | 0 | 0 | 0 | 0 | 0 | 0 | 0 | NA | NA | NA | NA | NA | NA | NA | NA | NA | NA |
| Ank | 716 | 883 | 829 | 1054 | 734 | 808 | 761 | 882 | 832 | 829 | 790 | 884 | 829.6 | 825 | 834 | 1.011 | 0.016 | 0.87880755 | 0.94007522 | 0.003 | 0.0109 | 0.0064 | 0.0064 | TRUE | 0.055 |
| Ank1 | 31 | 30 | 26 | 61 | 32 | 32 | 33 | 30 | 26 | 48 | 34 | 35 | 34.4 | 30 | 39 | 1.33 | 0.412 | 0.22596301 | 0.43879447 | 0 | 0.1027 | 0.0537 | 0.0537 | TRUE | 0.3336 |
| Ank2 | 850 | 564 | 651 | 613 | 435 | 433 | 903 | 563 | 654 | 482 | 468 | 474 | 590.62 | 707 | 475 | 0.672 | -0.573 | 0.0011343 | 0.00982978 | 0.0284 | 0.0125 | 0.0206 | 0.0206 | TRUE | 0.6066 |
| Ank3 | 1159 | 1325 | 1402 | 872 | 642 | 831 | 1231 | 1323 | 1408 | 685 | 691 | 910 | 1041.34 | 1321 | 762 | 0.576 | -0.795 | 4.43E-09 | 2.27E-07 | 0.0148 | 0.0101 | 0.0122 | 0.0122 | TRUE | 0.3372 |
| Ankar | 0 | 0 | 0 | 2 | 0 | 1 | 0 | 0 | 0 | 2 | 0 | 1 | 0.44 | 0 | 1 | 4.782 | 2.258 | 0.50653358 | NA | 0 | 7.4188 | 6.7775 | 6.7775 | TRUE | 0.2804 |
| Ankdd1b | 2 | 1 | 2 | 5 | 4 | 5 | 2 | 1 | 2 | 4 | 4 | 5 | 3.14 | 2 | 4 | 2.666 | 1.415 | 0.19366685 | NA | 0 | 1.0562 | 0.4466 | 0.4466 | TRUE | 0.1056 |
| Ankef1 | 17 | 29 | 24 | 41 | 32 | 28 | 18 | 29 | 24 | 32 | 34 | 31 | 28.07 | 24 | 32 | 1.365 | 0.449 | 0.22313659 | 0.4350736 | 0 | 0.1243 | 0.0615 | 0.0615 | TRUE | 0.2465 |
| Ankfn1 | 0 | 0 | 0 | 0 | 0 | 0 | 0 | 0 | 0 | 0 | 0 | 0 | 0 | 0 | 0 | NA | NA | NA | NA | NA | NA | NA | NA | NA | NA |
| Ankfy1 | 13056 | 10798 | 11240 | 16716 | 12052 | 10668 | 13868 | 10784 | 11286 | 13141 | 12967 | 11678 | 12287.3 | 11979 | 12595 | 1.051 | 0.072 | 0.52738293 | 0.71613289 | 0.0096 | 0.0072 | 0.0094 | 0.0094 | TRUE | 0.2324 |
| Ankhd1 | 6524 | 5344 | 5879 | 7177 | 5203 | 5079 | 6930 | 5337 | 5903 | 5642 | 5598 | 5560 | 5828.29 | 6057 | 5600 | 0.925 | -0.113 | 0.29199508 | 0.51281892 | 0.0087 | 0.0075 | 0.0081 | 0.0081 | TRUE | 0.1938 |
| Ankib1 | 1981 | 1956 | 2085 | 2857 | 1989 | 1968 | 2104 | 1953 | 2094 | 2246 | 2140 | 2154 | 2115.25 | 2050 | 2180 | 1.064 | 0.089 | 0.22642736 | 0.43926351 | 7.00E-04 | 0.0085 | 0.0034 | 0.0034 | TRUE | 0.0208 |
| Ankk1 | 10 | 10 | 7 | 12 | 8 | 2 | 11 | 10 | 7 | 9 | 9 | 2 | 7.98 | 9 | 7 | 0.744 | -0.426 | 0.56126255 | 0.74256648 | 0.0022 | 0.4198 | 0.261 | 0.261 | TRUE | 0.8011 |
| Ankle1 | 442 | 511 | 525 | 543 | 507 | 410 | 469 | 510 | 527 | 427 | 546 | 449 | 488.03 | 502 | 474 | 0.941 | -0.087 | 0.51315675 | 0.70632889 | 0.0083 | 0.0137 | 0.0107 | 0.0107 | TRUE | 0.2022 |
| Ankle2 | 3302 | 3447 | 3827 | 4506 | 3329 | 3466 | 3507 | 3443 | 3843 | 3542 | 3582 | 3794 | 3618.45 | 3598 | 3639 | 1.011 | 0.016 | 0.83457668 | 0.91571799 | 0.0022 | 0.0078 | 0.0042 | 0.0042 | TRUE | 0.0433 |
| Ankmy1 | 49 | 64 | 74 | 84 | 52 | 58 | 52 | 64 | 74 | 66 | 56 | 63 | 62.62 | 63 | 62 | 0.976 | -0.035 | 0.89184799 | 0.94753967 | 0 | 0.0595 | 0.0325 | 0.0325 | TRUE | 0.2088 |
| Ankmy2 | 1391 | 2050 | 2080 | 1967 | 1635 | 1804 | 1477 | 2047 | 2089 | 1546 | 1759 | 1975 | 1815.6 | 1871 | 1760 | 0.94 | -0.089 | 0.56694642 | 0.74677459 | 0.0249 | 0.0087 | 0.0169 | 0.0169 | TRUE | 0.4085 |
| Ankra2 | 805 | 1194 | 1206 | 1314 | 960 | 1064 | 855 | 1192 | 1211 | 1033 | 1033 | 1165 | 1081.51 | 1086 | 1077 | 0.991 | -0.013 | 0.92975627 | 0.96550671 | 0.0198 | 0.0099 | 0.0151 | 0.0151 | TRUE | 0.4131 |
| Ankrd1 | 3 | 4 | 2 | 11 | 3 | 0 | 3 | 4 | 2 | 9 | 3 | 0 | 3.51 | 3 | 4 | 1.329 | 0.411 | 0.73933418 | NA | 0.6422 | 0.9453 | 0.8117 | 0.8117 | TRUE | 0.3187 |
| Ankrd10 | 8305 | 6901 | 6505 | 6532 | 5534 | 5657 | 8821 | 6892 | 6532 | 5135 | 5954 | 6192 | 6587.81 | 7415 | 5760 | 0.777 | -0.364 | 0.00670799 | 0.03896808 | 0.0182 | 0.0074 | 0.0129 | 0.0129 | TRUE | 0.3359 |
| Ankrd11 | 14246 | 13197 | 14989 | 18918 | 14101 | 12788 | 15132 | 13180 | 15051 | 14872 | 15172 | 13998 | 14567.4 | 14454 | 14681 | 1.016 | 0.022 | 0.78907025 | 0.891612 | 0.003 | 0.0071 | 0.005 | 0.005 | TRUE | 0.0727 |
| Ankrd12 | 830 | 870 | 863 | 888 | 684 | 723 | 882 | 869 | 867 | 698 | 736 | 791 | 807.08 | 873 | 742 | 0.849 | -0.236 | 0.01101163 | 0.05654356 | 8.00E-04 | 0.011 | 0.0049 | 0.0049 | TRUE | 0.0402 |
| Ankrd13a | 4157 | 4033 | 4211 | 5749 | 4275 | 4281 | 4415 | 4028 | 4228 | 4519 | 4600 | 4686 | 4412.8 | 4224 | 4602 | 1.089 | 0.124 | 0.0712926 | 0.20807308 | 9.00E-04 | 0.0076 | 0.0032 | 0.0032 | TRUE | 0.0201 |
| Ankrd13b | 1522 | 1274 | 1325 | 1267 | 1015 | 946 | 1617 | 1272 | 1330 | 996 | 1092 | 1036 | 1223.84 | 1406 | 1041 | 0.74 | -0.434 | 0.00021341 | 0.00256152 | 0.0084 | 0.0096 | 0.0091 | 0.0091 | TRUE | 0.2049 |
| Ankrd13c | 2039 | 2109 | 2038 | 3144 | 2160 | 2023 | 2166 | 2106 | 2046 | 2472 | 2324 | 2214 | 2221.42 | 2106 | 2337 | 1.11 | 0.151 | 0.0546249 | 0.17410453 | 0.0015 | 0.0084 | 0.004 | 0.004 | TRUE | 0.0315 |
| Ankrd13d | 322 | 547 | 487 | 472 | 393 | 385 | 342 | 546 | 489 | 371 | 423 | 421 | 432.11 | 459 | 405 | 0.881 | -0.182 | 0.31759693 | 0.53856183 | 0.0288 | 0.0145 | 0.0217 | 0.0217 | TRUE | 0.5738 |
| Ankrd16 | 305 | 277 | 253 | 389 | 321 | 305 | 324 | 277 | 254 | 306 | 345 | 334 | 306.62 | 285 | 328 | 1.152 | 0.204 | 0.15622424 | 0.34905281 | 0.0061 | 0.0176 | 0.0116 | 0.0116 | TRUE | 0.16 |
| Ankrd17 | 10761 | 8901 | 9818 | 14889 | 10698 | 9893 | 11430 | 8890 | 9858 | 11705 | 11510 | 10829 | 10703.72 | 10059 | 11348 | 1.128 | 0.174 | 0.10245932 | 0.26598803 | 0.0088 | 0.0072 | 0.0081 | 0.0081 | TRUE | 0.1736 |

| | | | | | | | | | | | | | | | | | | | | | | | | | |
|---|---|---|---|---|---|---|---|---|---|---|---|---|---|---|---|---|---|---|---|---|---|---|---|---|---|
| Ankrd2 | 1 | 2 | 0 | 2 | 1 | 1 | 1 | 2 | 0 | 2 | 1 | 1 | 1.13 | 1 | 1 | 1.235 | 0.305 | 0.86149571 | NA | 0 | 2.9127 | 1.3119 | 1.3119 | TRUE | 0.2553 |
| Ankrd22 | 13 | 3 | 6 | 37 | 27 | 2 | 14 | 3 | 6 | 29 | 29 | 2 | 13.86 | 8 | 20 | 2.664 | 1.413 | 0.11606533 | 0.28769172 | 0.7703 | 0.2446 | 0.4913 | 0.4913 | TRUE | 3.1018 |
| Ankrd23 | 98 | 113 | 95 | 99 | 100 | 89 | 104 | 113 | 95 | 78 | 108 | 97 | 99.2 | 104 | 94 | 0.901 | -0.151 | 0.48775831 | 0.68458483 | 0.0079 | 0.0401 | 0.024 | 0.024 | TRUE | 0.2542 |
| Ankrd24 | 344 | 523 | 528 | 462 | 440 | 441 | 365 | 522 | 530 | 363 | 473 | 483 | 456.2 | 472 | 440 | 0.929 | -0.107 | 0.56411896 | 0.74494404 | 0.031 | 0.0141 | 0.0225 | 0.0225 | TRUE | 0.455 |
| Ankrd26 | 2511 | 2446 | 2605 | 3559 | 2611 | 2514 | 2667 | 2443 | 2616 | 2798 | 2809 | 2752 | 2680.79 | 2575 | 2786 | 1.082 | 0.114 | 0.10759822 | 0.27453369 | 7.00E-04 | 0.0081 | 0.0032 | 0.0032 | TRUE | 0.0246 |
| Ankrd27 | 4946 | 5357 | 5373 | 5643 | 4345 | 4247 | 5254 | 5350 | 5395 | 4436 | 4675 | 4649 | 4959.79 | 5333 | 4587 | 0.86 | -0.218 | 0.0003672 | 0.00405 | 3.00E-04 | 0.0076 | 0.0025 | 0.0025 | TRUE | 0.0102 |
| Ankrd28 | 3935 | 4175 | 4130 | 5652 | 4063 | 3897 | 4180 | 4170 | 4147 | 4443 | 4372 | 4266 | 4262.8 | 4166 | 4360 | 1.047 | 0.066 | 0.26069463 | 0.47999951 | 0 | 0.0077 | 0.0023 | 0.0023 | TRUE | 0.0043 |
| Ankrd29 | 39 | 29 | 21 | 38 | 17 | 24 | 41 | 29 | 21 | 30 | 18 | 26 | 27.65 | 30 | 25 | 0.82 | -0.286 | 0.4950587 | 0.69104043 | 0.0512 | 0.126 | 0.09 | 0.09 | TRUE | 0.2914 |
| Ankrd32 | 709 | 822 | 773 | 1241 | 896 | 806 | 753 | 821 | 776 | 976 | 964 | 882 | 862.02 | 783 | 941 | 1.201 | 0.265 | 0.00467653 | 0.02979091 | 0.0013 | 0.0107 | 0.0051 | 0.0051 | TRUE | 0.0344 |
| Ankrd33 | 0 | 0 | 0 | 0 | 0 | 0 | 0 | 0 | 0 | 0 | 0 | 0 | 0 | 0 | 0 | NA | NA | NA | NA | NA | NA | NA | NA | NA | NA |
| Ankrd33b | 840 | 952 | 987 | 873 | 617 | 583 | 892 | 951 | 991 | 686 | 664 | 638 | 803.73 | 945 | 663 | 0.702 | -0.51 | 4.75E-08 | 1.90E-06 | 9.00E-04 | 0.011 | 0.005 | 0.005 | TRUE | 0.0279 |
| Ankrd34a | 48 | 49 | 33 | 30 | 14 | 24 | 51 | 49 | 33 | 24 | 15 | 26 | 33 | 44 | 22 | 0.49 | -1.03 | 0.00677105 | 0.03913268 | 0.0266 | 0.1067 | 0.0702 | 0.0702 | TRUE | 0.3846 |
| Ankrd34b | 69 | 96 | 122 | 116 | 100 | 106 | 73 | 96 | 123 | 91 | 108 | 116 | 101.08 | 97 | 105 | 1.075 | 0.104 | 0.67404534 | 0.81833909 | 0.0291 | 0.0395 | 0.0344 | 0.0344 | TRUE | 0.2473 |
| Ankrd34c | 0 | 0 | 0 | 0 | 0 | 1 | 0 | 0 | 0 | 0 | 0 | 1 | 0.18 | 0 | 0 | 1.859 | 0.894 | 0.82650734 | NA | 0 | 18.0643 | 10 | 10 | TRUE | 0.5383 |
| Ankrd35 | 318 | 310 | 306 | 393 | 293 | 335 | 338 | 310 | 307 | 309 | 315 | 367 | 324.26 | 318 | 330 | 1.037 | 0.052 | 0.69452171 | 0.83268692 | 0.0028 | 0.0171 | 0.0094 | 0.0094 | TRUE | 0.1031 |
| Ankrd36 | 15 | 10 | 13 | 35 | 14 | 22 | 16 | 10 | 13 | 28 | 15 | 24 | 17.61 | 13 | 22 | 1.725 | 0.787 | 0.10804351 | 0.27511586 | 0 | 0.194 | 0.1114 | 0.1114 | TRUE | 0.4301 |
| Ankrd37 | 837 | 1141 | 1253 | 1098 | 848 | 1081 | 889 | 1140 | 1258 | 863 | 912 | 1183 | 1040.93 | 1096 | 986 | 0.9 | -0.153 | 0.3642107 | 0.58164551 | 0.0289 | 0.0101 | 0.0195 | 0.0195 | TRUE | 0.363 |
| Ankrd39 | 365 | 444 | 481 | 516 | 355 | 375 | 388 | 443 | 483 | 406 | 382 | 410 | 418.7 | 438 | 399 | 0.912 | -0.134 | 0.29070577 | 0.51144315 | 0.0044 | 0.0148 | 0.0091 | 0.0091 | TRUE | 0.1157 |
| Ankrd40 | 3926 | 4035 | 4013 | 4978 | 3566 | 3437 | 4170 | 4030 | 4029 | 3913 | 3837 | 3762 | 3956.97 | 4076 | 3837 | 0.942 | -0.087 | 0.1567665 | 0.34972176 | 1.00E-04 | 0.0077 | 0.0025 | 0.0025 | TRUE | 0.0049 |
| Ankrd42 | 44 | 51 | 64 | 106 | 64 | 51 | 47 | 51 | 64 | 83 | 69 | 56 | 61.66 | 54 | 69 | 1.291 | 0.368 | 0.18879575 | 0.39393017 | 0.0193 | 0.0603 | 0.0401 | 0.0401 | TRUE | 0.1843 |
| Ankrd44 | 422 | 495 | 464 | 838 | 523 | 458 | 448 | 494 | 466 | 659 | 563 | 501 | 521.89 | 469 | 574 | 1.226 | 0.293 | 0.02871195 | 0.11156564 | 0.0092 | 0.0132 | 0.011 | 0.011 | TRUE | 0.2007 |
| Ankrd45 | 116 | 73 | 92 | 198 | 139 | 113 | 123 | 73 | 92 | 156 | 150 | 124 | 119.57 | 96 | 143 | 1.491 | 0.577 | 0.01594359 | 0.07419956 | 0.0303 | 0.0345 | 0.0325 | 0.0325 | TRUE | 0.5188 |
| Ankrd46 | 3237 | 3570 | 3462 | 4590 | 3358 | 3529 | 3438 | 3565 | 3476 | 3608 | 3613 | 3863 | 3594.04 | 3493 | 3695 | 1.057 | 0.081 | 0.23117208 | 0.44494515 | 5.00E-04 | 0.0078 | 0.003 | 0.003 | TRUE | 0.0191 |
| Ankrd49 | 819 | 712 | 734 | 1311 | 899 | 724 | 870 | 711 | 737 | 1031 | 967 | 793 | 851.4 | 773 | 930 | 1.205 | 0.269 | 0.0500776 | 0.16437542 | 0.0131 | 0.0108 | 0.0124 | 0.0124 | TRUE | 0.1979 |
| Ankrd50 | 927 | 1008 | 1068 | 1222 | 833 | 917 | 985 | 1007 | 1072 | 961 | 896 | 1004 | 987.4 | 1021 | 954 | 0.934 | -0.099 | 0.27974518 | 0.50092427 | 0.0015 | 0.0102 | 0.005 | 0.005 | TRUE | 0.032 |
| Ankrd52 | 1614 | 1644 | 1730 | 2017 | 1619 | 1655 | 1714 | 1642 | 1737 | 1586 | 1742 | 1812 | 1705.42 | 1698 | 1713 | 1.008 | 0.012 | 0.89113032 | 0.94728769 | 0.0023 | 0.0088 | 0.0047 | 0.0047 | TRUE | 0.0525 |
| Ankrd53 | 0 | 0 | 0 | 0 | 0 | 0 | 0 | 0 | 0 | 0 | 0 | 0 | 0 | 0 | 0 | NA | NA | NA | NA | NA | NA | NA | NA | NA | NA |
| Ankrd54 | 736 | 712 | 813 | 984 | 692 | 689 | 782 | 711 | 816 | 774 | 745 | 754 | 763.58 | 770 | 758 | 0.985 | -0.022 | 0.81627138 | 0.90643681 | 0.0014 | 0.0112 | 0.0054 | 0.0054 | TRUE | 0.0531 |
| Ankrd55 | 9 | 9 | 14 | 10 | 10 | 8 | 10 | 9 | 14 | 8 | 11 | 9 | 10 | 11 | 9 | 0.834 | -0.262 | 0.6606423 | 0.80885977 | 0 | 0.3364 | 0.1549 | 0.1549 | TRUE | 0.2455 |
| Ankrd6 | 2102 | 1558 | 1504 | 1814 | 1314 | 1282 | 2233 | 1556 | 1510 | 1426 | 1414 | 1403 | 1590.34 | 1766 | 1414 | 0.801 | -0.32 | 0.0384382 | 0.13666468 | 0.0253 | 0.009 | 0.0166 | 0.0166 | TRUE | 0.6434 |
| Ankrd60 | 0 | 0 | 0 | 0 | 0 | 0 | 0 | 0 | 0 | 0 | 0 | 0 | 0 | 0 | 0 | NA | NA | NA | NA | NA | NA | NA | NA | NA | NA |
| Ankrd61 | 2 | 0 | 0 | 3 | 0 | 0 | 2 | 0 | 0 | 2 | 0 | 0 | 0.75 | 1 | 1 | 1.163 | 0.218 | 0.94242245 | NA | 8.7098 | 4.4164 | 5.2331 | 5.2331 | TRUE | 1.4402 |
| Ankrd63 | 13 | 15 | 12 | 11 | 13 | 8 | 14 | 15 | 12 | 9 | 14 | 9 | 12.04 | 14 | 11 | 0.763 | -0.39 | 0.47671829 | 0.67645532 | 0 | 0.2806 | 0.1317 | 0.1317 | TRUE | 0.2886 |
| Ankrd66 | 1 | 0 | 6 | 2 | 2 | 2 | 1 | 0 | 6 | 2 | 2 | 2 | 2.17 | 2 | 2 | 0.827 | -0.274 | 0.85245013 | NA | 0.6313 | 1.5274 | 1.1012 | 1.1012 | TRUE | 1.9456 |
| Ankrd7 | 0 | 0 | 0 | 0 | 0 | 0 | 0 | 0 | 0 | 0 | 0 | 0 | 0 | 0 | 0 | NA | NA | NA | NA | NA | NA | NA | NA | NA | NA |
| Ankrd9 | 547 | 627 | 591 | 692 | 502 | 414 | 581 | 626 | 593 | 544 | 540 | 453 | 556.32 | 600 | 512 | 0.855 | -0.226 | 0.05068607 | 0.16564872 | 0.0038 | 0.0128 | 0.0079 | 0.0079 | TRUE | 0.1162 |
| Anks1 | 2401 | 2407 | 2356 | 3125 | 2293 | 2356 | 2550 | 2404 | 2366 | 2457 | 2467 | 2579 | 2470.44 | 2440 | 2501 | 1.025 | 0.035 | 0.62134826 | 0.78582572 | 7.00E-04 | 0.0082 | 0.0033 | 0.0033 | TRUE | 0.0189 |
| Anks1b | 24 | 28 | 35 | 49 | 22 | 37 | 25 | 28 | 35 | 39 | 24 | 41 | 31.88 | 29 | 35 | 1.162 | 0.217 | 0.56014326 | 0.74190289 | 0.0217 | 0.1102 | 0.0682 | 0.0682 | TRUE | 0.4801 |
| Anks3 | 676 | 730 | 707 | 808 | 661 | 622 | 718 | 729 | 710 | 635 | 711 | 681 | 697.38 | 719 | 676 | 0.939 | -0.091 | 0.33396223 | 0.55468116 | 0 | 0.0116 | 0.005 | 0.005 | TRUE | 0.0345 |
| Anks4b | 0 | 0 | 0 | 1 | 0 | 0 | 0 | 0 | 0 | 1 | 0 | 0 | 0.13 | 0 | 0 | 1.859 | 0.894 | 0.82650734 | NA | 0 | 25.1515 | 10 | 10 | TRUE | 0.2563 |
| Anks6 | 51 | 62 | 59 | 92 | 92 | 71 | 54 | 62 | 59 | 72 | 99 | 78 | 70.73 | 58 | 83 | 1.414 | 0.5 | 0.04324108 | 0.14864827 | 0 | 0.0535 | 0.0295 | 0.0295 | TRUE | 0.2594 |
| Ankzf1 | 1294 | 1711 | 1700 | 1801 | 1674 | 1628 | 1374 | 1709 | 1707 | 1416 | 1801 | 1782 | 1631.54 | 1597 | 1666 | 1.043 | 0.06 | 0.65694938 | 0.80677456 | 0.016 | 0.0089 | 0.0127 | 0.0127 | TRUE | 0.2116 |
| Anln | 14120 | 17251 | 17729 | 19947 | 14661 | 16993 | 14998 | 17229 | 17802 | 15681 | 15774 | 18602 | 16680.84 | 16676 | 16686 | 1 | 0.001 | 0.99458346 | 0.99723905 | 0.0071 | 0.0071 | 0.0081 | 0.0081 | TRUE | 0.1233 |
| Ano1 | 75 | 70 | 49 | 63 | 45 | 55 | 80 | 70 | 49 | 50 | 48 | 60 | 59.49 | 66 | 53 | 0.795 | -0.332 | 0.24698382 | 0.46379674 | 0.0207 | 0.0623 | 0.0421 | 0.0421 | TRUE | 0.452 |
| Ano10 | 376 | 603 | 655 | 499 | 342 | 416 | 399 | 602 | 658 | 392 | 368 | 455 | 479.15 | 553 | 405 | 0.732 | -0.45 | 0.02113292 | 0.09040097 | 0.0373 | 0.0138 | 0.0253 | 0.0253 | TRUE | 0.6886 |
| Ano2 | 11 | 9 | 8 | 14 | 14 | 3 | 12 | 9 | 8 | 11 | 15 | 3 | 9.68 | 10 | 10 | 1.029 | 0.041 | 0.95281767 | 0.97608731 | 0.1149 | 0.3474 | 0.2434 | 0.2434 | TRUE | 0.4245 |
| Ano3 | 343 | 438 | 382 | 454 | 329 | 408 | 364 | 437 | 384 | 357 | 354 | 447 | 390.47 | 395 | 386 | 0.975 | -0.037 | 0.80225236 | 0.89777365 | 0.0107 | 0.0153 | 0.0129 | 0.0129 | TRUE | 0.2137 |
| Ano4 | 58 | 48 | 55 | 25 | 16 | 14 | 62 | 48 | 55 | 20 | 17 | 15 | 36.16 | 55 | 17 | 0.319 | -1.647 | 1.71E-06 | 4.38E-05 | 0 | 0.098 | 0.0483 | 0.0483 | TRUE | 0.1045 |
| Ano5 | 4 | 2 | 2 | 14 | 8 | 7 | 4 | 2 | 2 | 11 | 9 | 8 | 5.92 | 3 | 9 | 3.346 | 1.743 | 0.03806279 | 0.13563631 | 0 | 0.5632 | 0.2684 | 0.2684 | TRUE | 0.2564 |
| Ano6 | 2451 | 2636 | 3000 | 3090 | 2424 | 2364 | 2603 | 2633 | 3012 | 2429 | 2608 | 2588 | 2645.54 | 2749 | 2542 | 0.924 | -0.114 | 0.2049588 | 0.41337964 | 0.0036 | 0.0081 | 0.0054 | 0.0054 | TRUE | 0.085 |
| Ano7 | 28 | 39 | 30 | 39 | 34 | 30 | 30 | 39 | 30 | 31 | 37 | 33 | 33.15 | 33 | 34 | 1.009 | 0.013 | 0.96942134 | 0.98448447 | 0 | 0.1063 | 0.0502 | 0.0502 | TRUE | 0.1788 |
| Ano8 | 644 | 886 | 828 | 773 | 683 | 654 | 684 | 885 | 831 | 608 | 735 | 716 | 743.12 | 800 | 686 | 0.856 | -0.224 | 0.10101633 | 0.26316973 | 0.0127 | 0.0113 | 0.0121 | 0.0121 | TRUE | 0.1901 |
| Ano9 | 855 | 1213 | 1196 | 1149 | 1066 | 1184 | 908 | 1211 | 1201 | 903 | 1147 | 1296 | 1111.13 | 1107 | 1115 | 1.007 | 0.01 | 0.95380755 | 0.97651612 | 0.0284 | 0.0099 | 0.0191 | 0.0191 | TRUE | 0.3368 |
| Anp32a | 14707 | 18227 | 18790 | 16040 | 13140 | 14058 | 15621 | 18203 | 18867 | 12609 | 14138 | 15389 | 15804.66 | 17564 | 14045 | 0.8 | -0.323 | 0.00335544 | 0.02312011 | 0.0096 | 0.0071 | 0.0087 | 0.0087 | TRUE | 0.1145 |
| Anp32b | 13377 | 13466 | 13885 | 22662 | 16568 | 14758 | 14209 | 13449 | 13942 | 17815 | 17826 | 16155 | 15565.92 | 13867 | 17265 | 1.245 | 0.316 | 9.75E-06 | 0.00019504 | 0.0018 | 0.0071 | 0.0036 | 0.0036 | TRUE | 0.0386 |
| Anp32e | 10608 | 10987 | 10588 | 15690 | 11097 | 10327 | 11268 | 10973 | 10631 | 12334 | 11940 | 11305 | 11408.39 | 10957 | 11860 | 1.082 | 0.114 | 0.0884695 | 0.24128324 | 0.0012 | 0.0072 | 0.0032 | 0.0032 | TRUE | 0.0205 |
| Anpep | 2530 | 2891 | 2966 | 3116 | 2140 | 2121 | 2687 | 2887 | 2978 | 2450 | 2303 | 2322 | 2604.43 | 2851 | 2358 | 0.827 | -0.273 | 0.00040604 | 0.00438329 | 0.0016 | 0.0082 | 0.0039 | 0.0039 | TRUE | 0.0305 |
| Antxr1 | 164 | 137 | 129 | 299 | 199 | 152 | 174 | 137 | 130 | 235 | 214 | 166 | 176.02 | 147 | 205 | 1.402 | 0.488 | 0.01664733 | 0.07641362 | 0.0224 | 0.0256 | 0.024 | 0.024 | TRUE | 0.2914 |
| Antxr2 | 20 | 12 | 13 | 36 | 17 | 23 | 21 | 12 | 13 | 28 | 18 | 25 | 19.67 | 15 | 24 | 1.563 | 0.645 | 0.16438737 | 0.36058077 | 0 | 0.1743 | 0.1014 | 0.1014 | TRUE | 0.4841 |
| Antxrl | 0 | 0 | 0 | 0 | 0 | 0 | 0 | 0 | 0 | 0 | 0 | 0 | 0 | 0 | 0 | NA | NA | NA | NA | NA | NA | NA | NA | NA | NA |
| Anxa1 | 81 | 87 | 87 | 275 | 199 | 158 | 86 | 87 | 87 | 216 | 214 | 173 | 143.92 | 87 | 201 | 2.322 | 1.215 | 5.00E-11 | 3.87E-09 | 0 | 0.0298 | 0.0163 | 0.0163 | TRUE | 0.157 |
| Anxa10 | 6 | 8 | 4 | 5 | 1 | 1 | 6 | 8 | 4 | 4 | 1 | 1 | 4.08 | 6 | 2 | 0.349 | -1.519 | 0.13757847 | NA | 0 | 0.8144 | 0.4432 | 0.4432 | TRUE | 0.7395 |
| Anxa11 | 4101 | 3712 | 3983 | 3554 | 2510 | 2796 | 4356 | 3707 | 3999 | 2794 | 2701 | 3061 | 3436.28 | 4021 | 2852 | 0.709 | -0.496 | 1.67E-07 | 5.59E-06 | 0.005 | 0.0079 | 0.0062 | 0.0062 | TRUE | 0.0646 |
| Anxa13 | 0 | 1 | 1 | 1 | 1 | 0 | 0 | 1 | 1 | 1 | 1 | 0 | 0.64 | 1 | 1 | 0.935 | -0.097 | 0.96674856 | NA | 0 | 5.1213 | 2.3899 | 2.3899 | TRUE | 0.2246 |
| Anxa2 | 2470 | 5726 | 6708 | 4399 | 3428 | 4910 | 2624 | 5719 | 6736 | 3458 | 3688 | 5375 | 4599.83 | 5026 | 4174 | 0.83 | -0.268 | 0.54112101 | 0.7269093 | 0.1385 | 0.0076 | 0.0593 | 0.1385 | TRUE | 0.8593 |
| Anxa3 | 100 | 134 | 134 | 407 | 252 | 200 | 106 | 134 | 135 | 320 | 271 | 219 | 197.44 | 125 | 270 | 2.166 | 1.115 | 2.62E-08 | 1.13E-06 | 0.0224 | 0.0236 | 0.023 | 0.023 | TRUE | 0.1076 |
| Anxa4 | 1157 | 1561 | 1548 | 1757 | 1283 | 1376 | 1229 | 1559 | 1554 | 1381 | 1380 | 1506 | 1435.03 | 1447 | 1422 | 0.982 | -0.026 | 0.82893302 | 0.91328449 | 0.0095 | 0.0092 | 0.0094 | 0.0094 | TRUE | 0.2091 |
| Anxa5 | 931 | 1166 | 1262 | 836 | 688 | 870 | 989 | 1164 | 1267 | 657 | 740 | 952 | 961.73 | 1140 | 783 | 0.686 | -0.544 | 0.00074046 | 0.00705114 | 0.0249 | 0.0103 | 0.0176 | 0.0176 | TRUE | 0.4213 |
| Anxa6 | 5956 | 6009 | 5899 | 6939 | 5032 | 4851 | 6326 | 6001 | 5923 | 5455 | 5414 | 5310 | 5738.33 | 6083 | 5393 | 0.887 | -0.174 | 0.00550332 | 0.03354297 | 5.00E-04 | 0.0075 | 0.0026 | 0.0026 | TRUE | 0.0148 |

| | | | | | | | | | | | | | | | | | | | | | | | | | |
|---|---|---|---|---|---|---|---|---|---|---|---|---|---|---|---|---|---|---|---|---|---|---|---|---|---|
| Anxa7 | 5616 | 6904 | 7120 | 6115 | 4525 | 5015 | 5965 | 6895 | 7149 | 4807 | 4869 | 5490 | 5862.49 | 6670 | 5055 | 0.758 | -0.4 | 7.79E-05 | 0.00113151 | 0.0069 | 0.0075 | 0.0072 | 0.0072 | TRUE | 0.1041 |
| Anxa8 | 331 | 469 | 423 | 316 | 272 | 351 | 352 | 468 | 425 | 248 | 293 | 384 | 361.67 | 415 | 308 | 0.741 | -0.432 | 0.02481275 | 0.1012303 | 0.0311 | 0.016 | 0.0239 | 0.0239 | TRUE | 0.5191 |
| Anxa9 | 10 | 13 | 14 | 28 | 18 | 10 | 11 | 13 | 14 | 22 | 19 | 11 | 15 | 13 | 17 | 1.4 | 0.486 | 0.34770192 | 0.56662263 | 0 | 0.2266 | 0.1242 | 0.1242 | TRUE | 0.4883 |
| Aoah | 110 | 21 | 27 | 113 | 79 | 54 | 117 | 21 | 27 | 89 | 85 | 59 | 66.31 | 55 | 78 | 1.416 | 0.502 | 0.37599192 | 0.59147378 | 0.4181 | 0.0566 | 0.2161 | 0.2161 | TRUE | 8.0043 |
| Aoc2 | 248 | 217 | 189 | 228 | 192 | 187 | 263 | 217 | 190 | 179 | 207 | 205 | 210.07 | 223 | 197 | 0.88 | -0.184 | 0.2934588 | 0.5143804 | 0.0124 | 0.0226 | 0.0173 | 0.0173 | TRUE | 0.2685 |
| Aoc3 | 345 | 449 | 480 | 388 | 371 | 518 | 366 | 448 | 482 | 305 | 399 | 567 | 428.01 | 432 | 424 | 0.978 | -0.032 | 0.88726352 | 0.94451405 | 0.0555 | 0.0146 | 0.034 | 0.034 | TRUE | 0.4336 |
| Aox1 | 82 | 121 | 137 | 93 | 62 | 71 | 87 | 121 | 138 | 73 | 67 | 78 | 93.84 | 115 | 73 | 0.629 | -0.669 | 0.00595169 | 0.03570564 | 0.0209 | 0.042 | 0.0314 | 0.0314 | TRUE | 0.348 |
| Aox3 | 15 | 16 | 9 | 48 | 32 | 36 | 16 | 16 | 9 | 38 | 34 | 39 | 25.42 | 14 | 37 | 2.729 | 1.449 | 0.00041561 | 0.00446827 | 0 | 0.1365 | 0.0707 | 0.0707 | TRUE | 0.3812 |
| Aox3l1 | 4 | 5 | 7 | 2 | 3 | 1 | 4 | 5 | 7 | 2 | 3 | 1 | 3.69 | 5 | 2 | 0.358 | -1.481 | 0.14935914 | NA | 0 | 0.8987 | 0.4162 | 0.4162 | TRUE | 0.2465 |
| Aox4 | 97 | 81 | 65 | 112 | 81 | 76 | 103 | 81 | 65 | 88 | 87 | 83 | 84.6 | 83 | 86 | 1.039 | 0.055 | 0.81930362 | 0.90766966 | 0.0134 | 0.0458 | 0.0302 | 0.0302 | TRUE | 0.3041 |
| Ap1ar | 2116 | 2197 | 2231 | 3186 | 2318 | 2175 | 2248 | 2194 | 2240 | 2505 | 2494 | 2381 | 2343.56 | 2227 | 2460 | 1.105 | 0.144 | 0.0315435 | 0.11939144 | 0 | 0.0083 | 0.0028 | 0.0028 | TRUE | 0.0094 |
| Ap1b1 | 6828 | 6437 | 6772 | 7577 | 5455 | 5129 | 7253 | 6429 | 6800 | 5956 | 5869 | 5614 | 6320.2 | 6827 | 5813 | 0.852 | -0.232 | 0.00217673 | 0.01646586 | 0.0021 | 0.0074 | 0.004 | 0.004 | TRUE | 0.0362 |
| Ap1g1 | 10134 | 10849 | 11233 | 14978 | 10494 | 10327 | 10764 | 10835 | 11279 | 11775 | 11291 | 11305 | 11208.03 | 10959 | 11457 | 1.045 | 0.064 | 0.27869876 | 0.49990184 | 5.00E-04 | 0.0072 | 0.0024 | 0.0024 | TRUE | 0.008 |
| Ap1g2 | 576 | 664 | 643 | 795 | 577 | 597 | 612 | 663 | 646 | 625 | 621 | 654 | 636.65 | 640 | 633 | 0.988 | -0.017 | 0.85779411 | 0.92990093 | 0 | 0.0121 | 0.0049 | 0.0049 | TRUE | 0.0175 |
| Ap1m1 | 2619 | 2320 | 2349 | 3456 | 2490 | 2260 | 2782 | 2317 | 2359 | 2717 | 2679 | 2474 | 2554.56 | 2486 | 2623 | 1.056 | 0.078 | 0.43747338 | 0.64548863 | 0.0056 | 0.0082 | 0.0069 | 0.0069 | TRUE | 0.1313 |
| Ap1m2 | 263 | 426 | 446 | 309 | 269 | 343 | 279 | 425 | 448 | 243 | 289 | 375 | 343.41 | 384 | 302 | 0.786 | -0.348 | 0.12253789 | 0.29792934 | 0.0523 | 0.0165 | 0.0336 | 0.0336 | TRUE | 0.6507 |
| Ap1s1 | 1705 | 1861 | 1859 | 2774 | 1874 | 1911 | 1811 | 1859 | 1867 | 2181 | 2016 | 2092 | 1970.86 | 1846 | 2096 | 1.136 | 0.185 | 0.01079656 | 0.05571471 | 5.00E-04 | 0.0086 | 0.0033 | 0.0033 | TRUE | 0.0154 |
| Ap1s2 | 893 | 663 | 635 | 1304 | 804 | 764 | 949 | 662 | 638 | 1025 | 865 | 836 | 829.13 | 750 | 909 | 1.214 | 0.28 | 0.10403627 | 0.26828975 | 0.031 | 0.0109 | 0.0201 | 0.0201 | TRUE | 0.6378 |
| Ap1s3 | 986 | 807 | 844 | 1075 | 792 | 811 | 1047 | 806 | 847 | 845 | 852 | 888 | 880.95 | 900 | 862 | 0.957 | -0.063 | 0.60856051 | 0.77825479 | 0.0091 | 0.0106 | 0.0098 | 0.0098 | TRUE | 0.2422 |
| Ap2a1 | 12253 | 12218 | 12585 | 14493 | 10402 | 10286 | 13015 | 12202 | 12637 | 11393 | 11192 | 11260 | 11949.76 | 12618 | 11282 | 0.894 | -0.161 | 0.00578596 | 0.03491054 | 4.00E-04 | 0.0072 | 0.0024 | 0.0024 | TRUE | 0.0102 |
| Ap2a2 | 10742 | 11264 | 11726 | 15361 | 10959 | 11530 | 11410 | 11249 | 11774 | 12076 | 11791 | 12621 | 11820.29 | 11478 | 12163 | 1.06 | 0.084 | 0.17757545 | 0.37816905 | 7.00E-04 | 0.0072 | 0.0027 | 0.0027 | TRUE | 0.0132 |
| Ap2b1 | 14366 | 15911 | 17259 | 20438 | 14405 | 15342 | 15259 | 15890 | 17330 | 16067 | 15499 | 16794 | 16139.9 | 16160 | 16120 | 0.998 | -0.004 | 0.96332228 | 0.98149348 | 0.0023 | 0.0071 | 0.0044 | 0.0044 | TRUE | 0.0491 |
| Ap2m1 | 4915 | 4693 | 4767 | 7192 | 5020 | 4651 | 5221 | 4687 | 4787 | 5654 | 5401 | 5091 | 5140.06 | 4898 | 5382 | 1.099 | 0.136 | 0.09213929 | 0.24762974 | 0.0028 | 0.0075 | 0.0045 | 0.0045 | TRUE | 0.0404 |
| Ap2s1 | 1944 | 2054 | 2190 | 2938 | 2097 | 2099 | 2065 | 2051 | 2199 | 2310 | 2256 | 2298 | 2196.46 | 2105 | 2288 | 1.087 | 0.12 | 0.08712673 | 0.2388187 | 3.00E-04 | 0.0084 | 0.0031 | 0.0031 | TRUE | 0.0185 |
| Ap3b1 | 5253 | 6063 | 6474 | 8217 | 6118 | 6157 | 5580 | 6055 | 6501 | 6460 | 6583 | 6740 | 6319.55 | 6045 | 6594 | 1.091 | 0.125 | 0.12216734 | 0.2973034 | 0.0029 | 0.0074 | 0.0046 | 0.0046 | TRUE | 0.0553 |
| Ap3b2 | 229 | 234 | 262 | 176 | 137 | 153 | 243 | 234 | 263 | 138 | 147 | 167 | 198.88 | 247 | 151 | 0.61 | -0.712 | 4.76E-06 | 0.00010598 | 0 | 0.0235 | 0.0121 | 0.0121 | TRUE | 0.0921 |
| Ap3d1 | 6905 | 5746 | 5916 | 10098 | 7640 | 6049 | 7334 | 5739 | 5940 | 7938 | 8220 | 6622 | 6965.53 | 6338 | 7593 | 1.198 | 0.261 | 0.04170918 | 0.14515382 | 0.0155 | 0.0074 | 0.0117 | 0.0117 | TRUE | 0.2307 |
| Ap3m1 | 3642 | 3920 | 4032 | 4842 | 3546 | 3600 | 3868 | 3915 | 4049 | 3806 | 3815 | 3941 | 3899.06 | 3944 | 3854 | 0.977 | -0.033 | 0.59053035 | 0.7646207 | 2.00E-04 | 0.0077 | 0.0025 | 0.0025 | TRUE | 0.0066 |
| Ap3m2 | 436 | 707 | 750 | 930 | 842 | 1112 | 463 | 706 | 753 | 731 | 906 | 1217 | 796.09 | 641 | 951 | 1.483 | 0.569 | 0.01121602 | 0.05732309 | 0.061 | 0.011 | 0.035 | 0.035 | TRUE | 0.2978 |
| Ap3s1 | 1422 | 1574 | 1558 | 1908 | 1346 | 1422 | 1510 | 1572 | 1564 | 1500 | 1448 | 1557 | 1525.25 | 1549 | 1502 | 0.969 | -0.045 | 0.54934056 | 0.7339342 | 0 | 0.0091 | 0.0034 | 0.0034 | TRUE | 0.012 |
| Ap3s2 | 4560 | 4349 | 4136 | 5579 | 3863 | 3736 | 4844 | 4343 | 4153 | 4386 | 4156 | 4090 | 4328.61 | 4447 | 4211 | 0.947 | -0.078 | 0.36357411 | 0.58098184 | 0.0036 | 0.0077 | 0.0051 | 0.0051 | TRUE | 0.0741 |
| Ap4b1 | 647 | 820 | 800 | 1188 | 805 | 797 | 687 | 819 | 803 | 934 | 866 | 872 | 830.32 | 770 | 891 | 1.158 | 0.211 | 0.04737424 | 0.15796756 | 0.004 | 0.0109 | 0.007 | 0.007 | TRUE | 0.1036 |
| Ap4e1 | 2193 | 2070 | 2210 | 2898 | 2085 | 2081 | 2329 | 2067 | 2219 | 2278 | 2243 | 2278 | 2235.87 | 2205 | 2266 | 1.028 | 0.04 | 0.607761 | 0.77767374 | 0.0013 | 0.0084 | 0.0039 | 0.0039 | TRUE | 0.0363 |
| Ap4m1 | 517 | 498 | 511 | 756 | 523 | 488 | 549 | 497 | 513 | 594 | 563 | 534 | 541.8 | 520 | 564 | 1.086 | 0.119 | 0.25820246 | 0.47713757 | 0 | 0.013 | 0.0061 | 0.0061 | TRUE | 0.028 |
| Ap4s1 | 186 | 182 | 172 | 343 | 192 | 170 | 198 | 182 | 173 | 270 | 207 | 186 | 202.39 | 184 | 221 | 1.205 | 0.269 | 0.14996564 | 0.34065871 | 0.0175 | 0.0232 | 0.0202 | 0.0202 | TRUE | 0.4338 |
| Ap5b1 | 167 | 155 | 147 | 330 | 184 | 149 | 177 | 155 | 148 | 259 | 198 | 163 | 183.38 | 160 | 207 | 1.299 | 0.377 | 0.07297032 | 0.21158403 | 0.0278 | 0.0249 | 0.0264 | 0.0264 | TRUE | 0.3438 |
| Ap5m1 | 1212 | 1183 | 1156 | 1533 | 1092 | 1120 | 1287 | 1181 | 1161 | 1205 | 1175 | 1226 | 1205.94 | 1210 | 1202 | 0.994 | -0.009 | 0.91438341 | 0.95780402 | 9.00E-04 | 0.0096 | 0.0043 | 0.0043 | TRUE | 0.0374 |
| Ap5s1 | 271 | 310 | 275 | 400 | 277 | 235 | 288 | 310 | 276 | 314 | 298 | 257 | 290.55 | 291 | 290 | 0.998 | -0.003 | 0.98135257 | 0.98988825 | 0.0036 | 0.0182 | 0.0105 | 0.0105 | TRUE | 0.1066 |
| Ap5z1 | 397 | 361 | 375 | 691 | 587 | 430 | 422 | 361 | 377 | 543 | 632 | 471 | 467.37 | 387 | 549 | 1.42 | 0.506 | 0.00047579 | 0.00499686 | 0.012 | 0.014 | 0.0129 | 0.0129 | TRUE | 0.1719 |
| Apaf1 | 3624 | 3819 | 4153 | 5247 | 3655 | 4059 | 3849 | 3814 | 4170 | 4125 | 3933 | 4443 | 4055.67 | 3944 | 4167 | 1.056 | 0.079 | 0.33679958 | 0.55677852 | 0.0028 | 0.0077 | 0.0046 | 0.0046 | TRUE | 0.0406 |
| Apba1 | 36 | 22 | 21 | 47 | 18 | 20 | 38 | 22 | 21 | 37 | 19 | 22 | 26.58 | 27 | 26 | 0.975 | -0.037 | 0.93442769 | 0.96749848 | 0.0863 | 0.1308 | 0.1088 | 0.1088 | TRUE | 0.9639 |
| Apba2 | 249 | 313 | 351 | 362 | 327 | 325 | 264 | 313 | 352 | 285 | 352 | 356 | 320.28 | 310 | 331 | 1.064 | 0.09 | 0.58013684 | 0.75669188 | 0.0149 | 0.0172 | 0.016 | 0.016 | TRUE | 0.1836 |
| Apba3 | 1444 | 1329 | 1388 | 2166 | 1592 | 1463 | 1534 | 1327 | 1394 | 1703 | 1713 | 1601 | 1545.31 | 1418 | 1672 | 1.18 | 0.238 | 0.00800117 | 0.04430987 | 0.0025 | 0.009 | 0.0052 | 0.0052 | TRUE | 0.0608 |
| Apbb1 | 593 | 696 | 759 | 1235 | 932 | 944 | 630 | 695 | 762 | 971 | 1003 | 1033 | 849.01 | 696 | 1002 | 1.44 | 0.526 | 4.53E-07 | 1.37E-05 | 0.0032 | 0.0108 | 0.0066 | 0.0066 | TRUE | 0.0824 |
| Apbb1ip | 21 | 23 | 18 | 40 | 31 | 21 | 22 | 23 | 18 | 31 | 33 | 23 | 25.19 | 21 | 29 | 1.39 | 0.475 | 0.22090573 | 0.43302575 | 0 | 0.1377 | 0.0678 | 0.0678 | TRUE | 0.2113 |
| Apbb2 | 1485 | 1527 | 1589 | 1702 | 1224 | 1223 | 1577 | 1525 | 1596 | 1338 | 1317 | 1339 | 1448.6 | 1566 | 1331 | 0.85 | -0.234 | 0.00116552 | 0.01000526 | 0 | 0.0092 | 0.0031 | 0.0031 | TRUE | 0.0063 |
| Apbb3 | 255 | 290 | 280 | 469 | 364 | 311 | 271 | 290 | 281 | 369 | 392 | 340 | 323.73 | 281 | 367 | 1.308 | 0.387 | 0.00178024 | 0.01401798 | 0 | 0.0171 | 0.0079 | 0.0079 | TRUE | 0.0432 |
| Apc | 8344 | 7278 | 8120 | 9660 | 6867 | 6691 | 8863 | 7269 | 8153 | 7594 | 7388 | 7324 | 7765.25 | 8095 | 7435 | 0.919 | -0.122 | 0.18011864 | 0.38150157 | 0.0049 | 0.0073 | 0.0059 | 0.0059 | TRUE | 0.0974 |
| Apc2 | 146 | 190 | 183 | 155 | 142 | 150 | 155 | 190 | 184 | 122 | 153 | 164 | 161.24 | 176 | 146 | 0.826 | -0.276 | 0.14309611 | 0.3294281 | 0.0115 | 0.0273 | 0.0193 | 0.0193 | TRUE | 0.2448 |
| Apcdd1 | 345 | 380 | 379 | 529 | 407 | 411 | 366 | 380 | 381 | 416 | 438 | 450 | 405.03 | 376 | 435 | 1.156 | 0.209 | 0.05474985 | 0.17441216 | 0 | 0.015 | 0.006 | 0.006 | TRUE | 0.0176 |
| Apcs | 0 | 0 | 0 | 0 | 0 | 0 | 0 | 0 | 0 | 0 | 0 | 0 | 0 | 0 | 0 | NA | NA | NA | NA | NA | NA | NA | NA | NA | NA |
| Apeh | 1865 | 1853 | 1831 | 2385 | 1893 | 1765 | 1981 | 1851 | 1839 | 1875 | 2037 | 1932 | 1918.97 | 1890 | 1948 | 1.03 | 0.043 | 0.58267436 | 0.75868406 | 0.0012 | 0.0086 | 0.0039 | 0.0039 | TRUE | 0.0213 |
| Apex1 | 6593 | 5408 | 5303 | 9544 | 6388 | 5302 | 7003 | 5401 | 5325 | 7503 | 6873 | 5804 | 6318.07 | 5910 | 6727 | 1.138 | 0.187 | 0.18386231 | 0.38659864 | 0.0208 | 0.0074 | 0.0141 | 0.0141 | TRUE | 0.3193 |
| Apex2 | 463 | 360 | 388 | 603 | 420 | 351 | 492 | 360 | 390 | 474 | 452 | 384 | 425.18 | 414 | 437 | 1.058 | 0.081 | 0.61132496 | 0.77987759 | 0.0169 | 0.0147 | 0.0158 | 0.0158 | TRUE | 0.3131 |
| Aph1a | 1335 | 1563 | 1549 | 1753 | 1244 | 1376 | 1418 | 1561 | 1555 | 1378 | 1338 | 1506 | 1459.52 | 1511 | 1407 | 0.931 | -0.103 | 0.25251136 | 0.47053511 | 0.0023 | 0.0092 | 0.0052 | 0.0052 | TRUE | 0.0444 |
| Aph1b | 123 | 142 | 153 | 216 | 152 | 134 | 131 | 142 | 154 | 170 | 164 | 147 | 151.02 | 142 | 160 | 1.128 | 0.174 | 0.30437957 | 0.52512118 | 0 | 0.0287 | 0.0141 | 0.0141 | TRUE | 0.0542 |
| Aph1c | 114 | 160 | 155 | 163 | 125 | 120 | 121 | 160 | 156 | 128 | 134 | 131 | 138.42 | 146 | 131 | 0.901 | -0.15 | 0.41701142 | 0.62807995 | 0.004 | 0.0307 | 0.0175 | 0.0175 | TRUE | 0.2204 |
| Api5 | 10319 | 10398 | 10095 | 15126 | 10950 | 10423 | 10961 | 10385 | 10136 | 11891 | 11782 | 11410 | 11093.94 | 10494 | 11694 | 1.114 | 0.156 | 0.014299 | 0.06836127 | 9.00E-04 | 0.0072 | 0.0028 | 0.0028 | TRUE | 0.0185 |
| Apip | 706 | 920 | 942 | 1126 | 799 | 729 | 750 | 919 | 946 | 885 | 860 | 798 | 859.57 | 872 | 848 | 0.973 | -0.04 | 0.73975127 | 0.86066702 | 0.0079 | 0.0107 | 0.0092 | 0.0092 | TRUE | 0.1763 |
| Apitd1 | 711 | 746 | 757 | 1083 | 776 | 776 | 755 | 745 | 760 | 851 | 835 | 849 | 799.35 | 753 | 845 | 1.122 | 0.166 | 0.04587528 | 0.15496785 | 0 | 0.011 | 0.0037 | 0.0037 | TRUE | 0.0013 |
| Aplf | 697 | 568 | 628 | 868 | 606 | 570 | 740 | 567 | 631 | 682 | 652 | 624 | 649.42 | 646 | 653 | 1.011 | 0.016 | 0.89703508 | 0.9501066 | 0.0084 | 0.012 | 0.01 | 0.01 | TRUE | 0.1909 |
| Apln | 113 | 194 | 155 | 150 | 131 | 144 | 120 | 194 | 156 | 118 | 141 | 158 | 147.65 | 157 | 139 | 0.884 | -0.178 | 0.43483634 | 0.64330164 | 0.0322 | 0.0292 | 0.0307 | 0.0307 | TRUE | 0.1798 |
| Aplnr | 0 | 0 | 0 | 0 | 0 | 0 | 0 | 0 | 0 | 0 | 0 | 0 | 0 | 0 | 0 | NA | NA | NA | NA | NA | NA | NA | NA | NA | NA |
| Aplp1 | 3280 | 4378 | 4577 | 3018 | 2132 | 2674 | 3484 | 4372 | 4596 | 2373 | 2294 | 2927 | 3340.93 | 4151 | 2531 | 0.61 | -0.714 | 3.20E-07 | 1.00E-05 | 0.0189 | 0.0079 | 0.0137 | 0.0137 | TRUE | 0.2402 |
| Aplp2 | 17356 | 18949 | 18996 | 21023 | 15631 | 15769 | 18435 | 18925 | 19074 | 16527 | 16818 | 17262 | 17839.99 | 18811 | 16869 | 0.897 | -0.157 | 0.00463539 | 0.02958193 | 3.00E-04 | 0.0071 | 0.0022 | 0.0022 | TRUE | 0.0051 |
| Apmap | 690 | 618 | 604 | 1059 | 678 | 572 | 733 | 617 | 606 | 833 | 729 | 626 | 690.79 | 652 | 729 | 1.12 | 0.164 | 0.24876469 | 0.46564404 | 0.0136 | 0.0117 | 0.0131 | 0.0131 | TRUE | 0.1692 |
| Apoa1 | 5 | 5 | 9 | 9 | 4 | 4 | 5 | 5 | 9 | 7 | 4 | 4 | 5.85 | 6 | 5 | 0.826 | -0.276 | 0.72414475 | 0.85080534 | 0 | 0.5701 | 0.2697 | 0.2697 | TRUE | 0.311 |
| Apoa1bp | 1667 | 2499 | 2372 | 2653 | 2006 | 2010 | 1771 | 2496 | 2382 | 2086 | 2158 | 2200 | 2182.06 | 2216 | 2148 | 0.969 | -0.045 | 0.7373305 | 0.85938343 | 0.0165 | 0.0084 | 0.0127 | 0.0127 | TRUE | 0.3736 |
| Apoa2 | 7 | 8 | 9 | 6 | 7 | 8 | 7 | 8 | 9 | 5 | 8 | 9 | 7.58 | 8 | 7 | 0.846 | -0.242 | 0.72186719 | 0.84955819 | 0 | 0.4416 | 0.1994 | 0.1994 | TRUE | 0.2904 |

| | | | | | | | | | | | | | | | | | | | | | | | | | |
|---|---|---|---|---|---|---|---|---|---|---|---|---|---|---|---|---|---|---|---|---|---|---|---|---|---|
| Apoa4 | 0 | 0 | 0 | 1 | 0 | 0 | 0 | 0 | 0 | 1 | 0 | 0 | 0.13 | 0 | 0 | 1.859 | 0.894 | 0.82650734 | NA | 0 | 25.1515 | 10 | 10 | TRUE | 0.2563 |
| Apoa5 | 0 | 1 | 0 | 4 | 0 | 3 | 0 | 1 | 0 | 3 | 0 | 3 | 1.24 | 0 | 2 | 5.9 | 2.561 | 0.24029802 | NA | 0 | 2.6683 | 2.2 | 2.2 | TRUE | 0.6744 |
| Apob | 52 | 32 | 27 | 142 | 75 | 45 | 55 | 32 | 27 | 112 | 81 | 49 | 59.31 | 38 | 81 | 2.127 | 1.089 | 0.00632566 | 0.03746464 | 0.1317 | 0.0624 | 0.0952 | 0.0952 | TRUE | 0.1333 |
| Apobec1 | 370 | 492 | 548 | 383 | 288 | 352 | 393 | 491 | 550 | 301 | 310 | 385 | 405.15 | 478 | 332 | 0.693 | -0.529 | 0.00180457 | 0.01417411 | 0.0211 | 0.015 | 0.0181 | 0.0181 | TRUE | 0.2764 |
| Apobec2 | 128 | 148 | 130 | 137 | 100 | 101 | 136 | 148 | 131 | 108 | 108 | 111 | 123.36 | 138 | 109 | 0.786 | -0.347 | 0.04974664 | 0.16373204 | 0 | 0.0336 | 0.0144 | 0.0144 | TRUE | 0.0397 |
| Apobec3 | 4849 | 7112 | 7852 | 7321 | 5817 | 6799 | 5150 | 7103 | 7884 | 5755 | 6259 | 7443 | 6599.01 | 6712 | 6486 | 0.966 | -0.05 | 0.76208659 | 0.87457944 | 0.032 | 0.0074 | 0.0194 | 0.0194 | TRUE | 0.5055 |
| Apobec4 | 0 | 0 | 0 | 0 | 0 | 0 | 0 | 0 | 0 | 0 | 0 | 0 | 0 | 0 | 0 | NA | NA | NA | NA | NA | NA | NA | NA | NA | NA |
| Apobr | 50 | 59 | 71 | 102 | 66 | 51 | 53 | 59 | 71 | 80 | 71 | 56 | 65.06 | 61 | 69 | 1.134 | 0.181 | 0.4962458 | 0.69196835 | 0.013 | 0.0575 | 0.0358 | 0.0358 | TRUE | 0.2397 |
| Apoc1 | 215 | 302 | 305 | 377 | 283 | 347 | 228 | 302 | 306 | 296 | 304 | 380 | 302.82 | 279 | 327 | 1.17 | 0.227 | 0.19067087 | 0.39574431 | 0.0189 | 0.0178 | 0.0183 | 0.0183 | TRUE | 0.2774 |
| Apoc2 | 4 | 6 | 2 | 6 | 4 | 8 | 4 | 6 | 2 | 5 | 4 | 9 | 5 | 4 | 6 | 1.44 | 0.526 | 0.5485056 | 0.73326245 | 0 | 0.6652 | 0.346 | 0.346 | TRUE | 0.3561 |
| Apoc3 | 0 | 0 | 0 | 1 | 1 | 0 | 0 | 0 | 0 | 1 | 1 | 0 | 0.31 | 0 | 1 | 3.32 | 1.731 | 0.65383052 | NA | 0 | 10.6224 | 8.9541 | 8.9541 | TRUE | 0.202 |
| Apoc4 | 1 | 2 | 4 | 1 | 4 | 2 | 1 | 2 | 4 | 1 | 4 | 2 | 2.39 | 2 | 2 | 1 | -0.001 | 0.99963555 | NA | 0 | 1.3839 | 0.7374 | 0.7374 | TRUE | 0.3174 |
| Apod | 216 | 206 | 223 | 211 | 150 | 162 | 229 | 206 | 224 | 166 | 161 | 177 | 193.95 | 220 | 168 | 0.765 | -0.386 | 0.00911377 | 0.04907559 | 0 | 0.0239 | 0.0105 | 0.0105 | TRUE | 0.0342 |
| Apoe | 5562 | 8322 | 9033 | 4131 | 3542 | 4833 | 5908 | 8311 | 9070 | 3247 | 3811 | 5290 | 5939.68 | 7763 | 4116 | 0.53 | -0.916 | 5.25E-06 | 0.00011557 | 0.0552 | 0.0075 | 0.0289 | 0.0289 | TRUE | 0.7577 |
| Apof | 68 | 39 | 35 | 47 | 43 | 25 | 72 | 39 | 35 | 37 | 46 | 27 | 42.82 | 49 | 37 | 0.757 | -0.401 | 0.30580169 | 0.52665845 | 0.09 | 0.0838 | 0.0866 | 0.0866 | TRUE | 0.9084 |
| Apoh | 0 | 0 | 0 | 2 | 0 | 0 | 0 | 0 | 0 | 2 | 0 | 0 | 0.26 | 0 | 1 | 2.886 | 1.529 | 0.70676491 | NA | 0 | 12.5792 | 10 | 10 | TRUE | 0.9669 |
| Apol10a | 0 | 0 | 0 | 0 | 1 | 0 | 0 | 0 | 0 | 0 | 1 | 0 | 0.18 | 0 | 0 | 1.859 | 0.894 | 0.82650734 | NA | 0 | 18.3785 | 10 | 10 | TRUE | 0.5204 |
| Apol10b | 2 | 3 | 4 | 11 | 3 | 6 | 2 | 3 | 4 | 9 | 3 | 7 | 4.6 | 3 | 6 | 2.038 | 1.027 | 0.26261207 | NA | 0 | 0.7236 | 0.3602 | 0.3602 | TRUE | 0.2199 |
| Apol11a | 0 | 0 | 0 | 0 | 0 | 0 | 0 | 0 | 0 | 0 | 0 | 0 | 0 | 0 | 0 | NA | NA | NA | NA | NA | NA | NA | NA | NA | NA |
| Apol11b | 0 | 0 | 0 | 0 | 0 | 0 | 0 | 0 | 0 | 0 | 0 | 0 | 0 | 0 | 0 | NA | NA | NA | NA | NA | NA | NA | NA | NA | NA |
| Apol6 | 6 | 4 | 2 | 12 | 7 | 2 | 6 | 4 | 2 | 9 | 8 | 2 | 5.26 | 4 | 6 | 1.576 | 0.656 | 0.47622192 | 0.67617682 | 0.1799 | 0.6338 | 0.411 | 0.411 | TRUE | 0.5716 |
| Apol7a | 16 | 11 | 10 | 31 | 23 | 21 | 17 | 11 | 10 | 24 | 25 | 23 | 18.35 | 13 | 24 | 1.903 | 0.929 | 0.04261432 | 0.14739322 | 0 | 0.1864 | 0.0904 | 0.0904 | TRUE | 0.3469 |
| Apol7b | 30 | 8 | 4 | 49 | 40 | 14 | 32 | 8 | 4 | 39 | 43 | 15 | 23.46 | 15 | 32 | 2.222 | 1.152 | 0.11215299 | 0.28161615 | 0.5464 | 0.1473 | 0.3284 | 0.3284 | TRUE | 4.6209 |
| Apol7c | 0 | 0 | 0 | 1 | 0 | 0 | 0 | 0 | 0 | 1 | 0 | 0 | 0.13 | 0 | 0 | 1.859 | 0.894 | 0.82650734 | NA | 0 | 25.1515 | 10 | 10 | TRUE | 0.2563 |
| Apol7d | 8 | 23 | 24 | 23 | 24 | 16 | 8 | 23 | 24 | 18 | 26 | 18 | 19.5 | 18 | 21 | 1.099 | 0.136 | 0.79128846 | 0.89271828 | 0.0945 | 0.1759 | 0.139 | 0.139 | TRUE | 1.111 |
| Apol7e | 10 | 5 | 5 | 14 | 11 | 2 | 11 | 5 | 5 | 11 | 12 | 2 | 7.61 | 7 | 8 | 1.228 | 0.296 | 0.71622396 | 0.84578349 | 0.2402 | 0.4398 | 0.3466 | 0.3466 | TRUE | 1.1408 |
| Apol8 | 32 | 32 | 23 | 29 | 18 | 30 | 34 | 32 | 23 | 23 | 19 | 33 | 27.34 | 30 | 25 | 0.84 | -0.252 | 0.52418657 | 0.71388227 | 0.0176 | 0.1274 | 0.0757 | 0.0757 | TRUE | 0.4187 |
| Apol9a | 1 | 0 | 0 | 0 | 0 | 0 | 1 | 0 | 0 | 0 | 0 | 0 | 0.18 | 0 | 0 | 0.49 | -1.029 | 0.80087 | NA | 0 | 18.6165 | 10 | 10 | TRUE | 0.4614 |
| Apol9b | 0 | 0 | 0 | 0 | 0 | 0 | 0 | 0 | 0 | 0 | 0 | 0 | 0 | 0 | 0 | NA | NA | NA | NA | NA | NA | NA | NA | NA | NA |
| Apold1 | 32 | 33 | 41 | 84 | 67 | 58 | 34 | 33 | 41 | 66 | 72 | 63 | 51.62 | 36 | 67 | 1.863 | 0.898 | 0.00105169 | 0.00926659 | 0 | 0.0707 | 0.0327 | 0.0327 | TRUE | 0.113 |
| Apom | 15 | 7 | 12 | 26 | 24 | 20 | 16 | 7 | 12 | 20 | 26 | 22 | 17.19 | 12 | 23 | 1.95 | 0.963 | 0.05114662 | 0.1666015 | 0 | 0.1986 | 0.1103 | 0.1103 | TRUE | 0.46 |
| Apon | 0 | 0 | 0 | 3 | 0 | 1 | 0 | 0 | 0 | 2 | 0 | 1 | 0.58 | 0 | 1 | 6.21 | 2.635 | 0.40034089 | NA | 0 | 5.7314 | 5.6518 | 5.6518 | TRUE | 0.0653 |
| Apoo | 144 | 164 | 150 | 239 | 142 | 150 | 153 | 164 | 151 | 188 | 153 | 164 | 162.04 | 156 | 168 | 1.084 | 0.116 | 0.48680424 | 0.68377024 | 0 | 0.0272 | 0.014 | 0.014 | TRUE | 0.1201 |
| Apoo-ps | 0 | 0 | 0 | 0 | 0 | 0 | 0 | 0 | 0 | 0 | 0 | 0 | 0 | 0 | 0 | NA | NA | NA | NA | NA | NA | NA | NA | NA | NA |
| Apool | 744 | 1004 | 982 | 1214 | 904 | 875 | 790 | 1003 | 986 | 954 | 973 | 958 | 943.97 | 926 | 962 | 1.038 | 0.053 | 0.64691865 | 0.80169726 | 0.0074 | 0.0104 | 0.0087 | 0.0087 | TRUE | 0.1957 |
| Apopt1 | 382 | 716 | 749 | 787 | 622 | 588 | 406 | 715 | 752 | 619 | 669 | 644 | 634.08 | 624 | 644 | 1.031 | 0.044 | 0.83680762 | 0.91714685 | 0.0525 | 0.0121 | 0.0309 | 0.0309 | TRUE | 1.0991 |
| App | 936 | 1453 | 1504 | 1562 | 1137 | 1316 | 994 | 1451 | 1510 | 1228 | 1223 | 1441 | 1307.89 | 1318 | 1297 | 0.983 | -0.024 | 0.88471389 | 0.94351934 | 0.0287 | 0.0094 | 0.019 | 0.019 | TRUE | 0.5549 |
| Appbp2 | 5516 | 5532 | 5478 | 7455 | 5543 | 5119 | 5859 | 5525 | 5500 | 5861 | 5964 | 5604 | 5718.72 | 5628 | 5810 | 1.032 | 0.046 | 0.4913811 | 0.68773282 | 9.00E-04 | 0.0075 | 0.003 | 0.003 | TRUE | 0.0157 |
| Appl1 | 3806 | 3875 | 4232 | 4768 | 3461 | 3914 | 4043 | 3870 | 4249 | 3748 | 3724 | 4284 | 3986.43 | 4054 | 3919 | 0.966 | -0.049 | 0.57921722 | 0.75611769 | 0.0035 | 0.0077 | 0.0054 | 0.0054 | TRUE | 0.0807 |
| Appl2 | 1531 | 1989 | 2015 | 1908 | 1497 | 1719 | 1626 | 1986 | 2023 | 1500 | 1611 | 1882 | 1771.37 | 1878 | 1664 | 0.885 | -0.176 | 0.16971021 | 0.3683321 | 0.0129 | 0.0088 | 0.0113 | 0.0113 | TRUE | 0.1666 |
| Aprt | 6077 | 5874 | 5787 | 7824 | 5587 | 5645 | 6455 | 5866 | 5811 | 6151 | 6011 | 6179 | 6078.87 | 6044 | 6114 | 1.012 | 0.017 | 0.8171648 | 0.90708637 | 0.0013 | 0.0074 | 0.0036 | 0.0036 | TRUE | 0.0431 |
| Aptx | 622 | 770 | 841 | 976 | 655 | 668 | 661 | 769 | 844 | 767 | 705 | 731 | 746.23 | 758 | 734 | 0.969 | -0.045 | 0.70469686 | 0.83847762 | 0.0071 | 0.0113 | 0.009 | 0.009 | TRUE | 0.1486 |
| Aqp1 | 7 | 3 | 2 | 7 | 2 | 1 | 7 | 3 | 2 | 6 | 2 | 1 | 3.53 | 4 | 3 | 0.73 | -0.454 | 0.68133073 | NA | 0.2594 | 0.9398 | 0.5936 | 0.5936 | TRUE | 0.9855 |
| Aqp11 | 288 | 324 | 306 | 378 | 274 | 280 | 306 | 324 | 307 | 297 | 295 | 307 | 305.87 | 312 | 300 | 0.959 | -0.061 | 0.60896008 | 0.77857639 | 0 | 0.0177 | 0.0069 | 0.0069 | TRUE | 0.0115 |
| Aqp12 | 1 | 0 | 0 | 0 | 0 | 0 | 1 | 0 | 0 | 0 | 0 | 0 | 0.18 | 0 | 0 | 0.49 | -1.029 | 0.80087 | NA | 0 | 18.6165 | 10 | 10 | TRUE | 0.4614 |
| Aqp2 | 3 | 6 | 7 | 5 | 4 | 3 | 3 | 6 | 7 | 4 | 4 | 3 | 4.62 | 5 | 4 | 0.71 | -0.495 | 0.56821539 | 0.74742773 | 0 | 0.7198 | 0.3207 | 0.3207 | TRUE | 0.2618 |
| Aqp3 | 3205 | 2880 | 2426 | 7408 | 4546 | 3410 | 3404 | 2876 | 2436 | 5824 | 4891 | 3733 | 3860.68 | 2905 | 4816 | 1.658 | 0.729 | 3.55E-05 | 0.00057741 | 0.0372 | 0.0078 | 0.0221 | 0.0221 | TRUE | 0.1594 |
| Aqp4 | 3 | 0 | 0 | 3 | 2 | 0 | 3 | 0 | 0 | 2 | 2 | 0 | 1.28 | 1 | 1 | 1.449 | 0.535 | 0.80451291 | NA | 2.5227 | 2.5751 | 2.5614 | 2.5614 | TRUE | 1.4429 |
| Aqp5 | 228 | 216 | 220 | 200 | 131 | 111 | 242 | 216 | 221 | 157 | 141 | 122 | 183.08 | 226 | 140 | 0.621 | -0.687 | 3.78E-05 | 0.00061105 | 0.0033 | 0.0249 | 0.0143 | 0.0143 | TRUE | 0.1335 |
| Aqp6 | 60 | 52 | 40 | 57 | 54 | 57 | 64 | 52 | 40 | 45 | 58 | 62 | 53.52 | 52 | 55 | 1.056 | 0.079 | 0.7922971 | 0.89305706 | 0.0224 | 0.0685 | 0.0461 | 0.0461 | TRUE | 0.1945 |
| Aqp7 | 0 | 0 | 0 | 5 | 5 | 2 | 0 | 0 | 0 | 4 | 5 | 2 | 1.92 | 0 | 4 | 20.424 | 4.352 | 0.01483812 | NA | 0 | 1.7258 | 1.1719 | 1.1719 | TRUE | 0.0714 |
| Aqp8 | 1 | 0 | 0 | 10 | 8 | 3 | 1 | 0 | 0 | 8 | 9 | 3 | 3.47 | 0 | 7 | 18.075 | 4.176 | 0.0062104 | NA | 0 | 0.9565 | 0.6041 | 0.6041 | TRUE | 0.4614 |
| Aqp9 | 8 | 8 | 3 | 32 | 21 | 17 | 8 | 8 | 3 | 25 | 23 | 19 | 14.31 | 6 | 22 | 3.425 | 1.776 | 0.0019322 | 0.01496811 | 0 | 0.2371 | 0.1357 | 0.1357 | TRUE | 0.5612 |
| Aqr | 8591 | 7575 | 7872 | 10460 | 7468 | 7510 | 9125 | 7565 | 7904 | 8223 | 8035 | 8221 | 8178.92 | 8198 | 8160 | 0.995 | -0.007 | 0.94079172 | 0.97003791 | 0.004 | 0.0073 | 0.0058 | 0.0058 | TRUE | 0.1194 |
| Ar | 7 | 9 | 16 | 20 | 12 | 14 | 7 | 9 | 16 | 16 | 13 | 15 | 12.74 | 11 | 15 | 1.353 | 0.436 | 0.43216327 | 0.64176836 | 0 | 0.2655 | 0.142 | 0.142 | TRUE | 0.6614 |
| Araf | 2644 | 2989 | 3101 | 3395 | 2659 | 2698 | 2808 | 2985 | 3114 | 2669 | 2861 | 2953 | 2898.41 | 2969 | 2828 | 0.952 | -0.071 | 0.38174079 | 0.59594336 | 0.0021 | 0.008 | 0.0044 | 0.0044 | TRUE | 0.0298 |
| Arap1 | 3237 | 2988 | 3132 | 2915 | 2219 | 2386 | 3438 | 2984 | 3145 | 2292 | 2388 | 2612 | 2809.7 | 3189 | 2431 | 0.762 | -0.392 | 2.51E-05 | 0.00043466 | 0.0044 | 0.0081 | 0.0059 | 0.0059 | TRUE | 0.0567 |
| Arap2 | 396 | 612 | 598 | 770 | 537 | 527 | 421 | 611 | 600 | 605 | 578 | 577 | 565.38 | 544 | 587 | 1.078 | 0.108 | 0.49112022 | 0.68742883 | 0.0193 | 0.0127 | 0.0161 | 0.0161 | TRUE | 0.458 |
| Arap3 | 754 | 688 | 689 | 932 | 625 | 616 | 801 | 687 | 692 | 733 | 672 | 674 | 709.88 | 727 | 693 | 0.955 | -0.066 | 0.54123958 | 0.72699971 | 0.0037 | 0.0115 | 0.007 | 0.007 | TRUE | 0.0939 |
| Arc | 655 | 774 | 733 | 705 | 629 | 666 | 696 | 773 | 736 | 554 | 677 | 729 | 694.13 | 735 | 653 | 0.887 | -0.173 | 0.18284958 | 0.38498245 | 0.0099 | 0.0116 | 0.0107 | 0.0107 | TRUE | 0.2156 |
| Arcn1 | 14206 | 14494 | 14787 | 17551 | 12310 | 12986 | 15089 | 14475 | 14848 | 13797 | 13245 | 14215 | 14278.25 | 14804 | 13752 | 0.929 | -0.106 | 0.08321735 | 0.23091204 | 7.00E-04 | 0.0071 | 0.0026 | 0.0026 | TRUE | 0.0127 |
| Areg | 14 | 14 | 5 | 23 | 17 | 12 | 15 | 14 | 5 | 18 | 18 | 13 | 13.9 | 11 | 16 | 1.469 | 0.555 | 0.33095881 | 0.55190129 | 0.0411 | 0.244 | 0.1597 | 0.1597 | TRUE | 0.904 |
| Arf1 | 18225 | 20054 | 19492 | 25051 | 17709 | 18326 | 19358 | 20028 | 19572 | 19693 | 19054 | 20061 | 19627.65 | 19653 | 19603 | 0.997 | -0.004 | 0.94800149 | 0.97406219 | 3.00E-04 | 0.0071 | 0.0023 | 0.0023 | TRUE | 0.0073 |
| Arf2 | 1830 | 1952 | 1786 | 2627 | 1735 | 1575 | 1944 | 1949 | 1793 | 2065 | 1867 | 1724 | 1890.43 | 1895 | 1885 | 0.995 | -0.007 | 0.94563207 | 0.97276996 | 0.0047 | 0.0086 | 0.0063 | 0.0063 | TRUE | 0.0856 |
| Arf3 | 5491 | 4924 | 4987 | 9526 | 6529 | 5693 | 5832 | 4918 | 5007 | 7489 | 7025 | 6232 | 6083.8 | 5252 | 6915 | 1.317 | 0.397 | 0.0002103 | 0.00253352 | 0.0086 | 0.0074 | 0.0081 | 0.0081 | TRUE | 0.1136 |
| Arf4 | 6471 | 7331 | 7012 | 8914 | 6029 | 6464 | 6873 | 7322 | 7041 | 7007 | 6487 | 7076 | 6967.64 | 7079 | 6857 | 0.969 | -0.046 | 0.51386653 | 0.70666169 | 0.0014 | 0.0074 | 0.0034 | 0.0034 | TRUE | 0.027 |
| Arf5 | 2633 | 3582 | 3474 | 4193 | 3175 | 3539 | 2797 | 3577 | 3488 | 3296 | 3416 | 3874 | 3408.11 | 3287 | 3529 | 1.073 | 0.102 | 0.40025279 | 0.61321853 | 0.0107 | 0.0079 | 0.0103 | 0.0103 | TRUE | 0.207 |
| Arf6 | 5605 | 6236 | 6100 | 8037 | 5889 | 5777 | 5954 | 6228 | 6125 | 6318 | 6336 | 6324 | 6214.1 | 6102 | 6326 | 1.037 | 0.052 | 0.362542 | 0.58035501 | 1.00E-04 | 0.0074 | 0.0022 | 0.0022 | TRUE | 0.0055 |
| Arfgap1 | 1184 | 1214 | 1130 | 2164 | 1435 | 1234 | 1258 | 1212 | 1135 | 1701 | 1544 | 1351 | 1366.77 | 1202 | 1532 | 1.276 | 0.352 | 0.0016052 | 0.01293018 | 0.0068 | 0.0093 | 0.0082 | 0.0082 | TRUE | 0.1284 |

| | | | | | | | | | | | | | | | | | | | | | | | | | |
|---|---|---|---|---|---|---|---|---|---|---|---|---|---|---|---|---|---|---|---|---|---|---|---|---|---|
| Arfgap2 | 1390 | 1524 | 1457 | 2484 | 1773 | 1500 | 1476 | 1522 | 1463 | 1953 | 1908 | 1642 | 1660.63 | 1487 | 1834 | 1.234 | 0.303 | 0.00148499 | 0.01215425 | 0.004 | 0.0089 | 0.006 | 0.006 | TRUE | 0.1005 |
| Arfgap3 | 808 | 910 | 885 | 874 | 732 | 824 | 858 | 909 | 889 | 687 | 788 | 902 | 838.73 | 885 | 792 | 0.893 | -0.163 | 0.18260886 | 0.38465155 | 0.0085 | 0.0108 | 0.0095 | 0.0095 | TRUE | 0.1741 |
| Arfgef1 | 4287 | 5629 | 5944 | 7510 | 5572 | 5912 | 4554 | 5622 | 5968 | 5904 | 5995 | 6472 | 5752.37 | 5381 | 6124 | 1.138 | 0.186 | 0.10543424 | 0.2706117 | 0.0104 | 0.0075 | 0.0094 | 0.0094 | TRUE | 0.2205 |
| Arfgef2 | 6053 | 5677 | 6411 | 7527 | 5533 | 5365 | 6429 | 5670 | 6437 | 5917 | 5953 | 5873 | 6046.58 | 6179 | 5914 | 0.957 | -0.063 | 0.41929285 | 0.62987779 | 0.0023 | 0.0074 | 0.0042 | 0.0042 | TRUE | 0.0635 |
| Arfip1 | 2247 | 2149 | 2089 | 2435 | 1831 | 1707 | 2387 | 2146 | 2098 | 1914 | 1970 | 1869 | 2063.89 | 2210 | 1918 | 0.868 | -0.205 | 0.01488834 | 0.07047385 | 0.0021 | 0.0085 | 0.0046 | 0.0046 | TRUE | 0.0589 |
| Arfip2 | 1096 | 1093 | 1018 | 1427 | 968 | 941 | 1164 | 1092 | 1022 | 1122 | 1042 | 1030 | 1078.55 | 1093 | 1065 | 0.975 | -0.036 | 0.69823875 | 0.83485873 | 0.0023 | 0.01 | 0.0054 | 0.0054 | TRUE | 0.039 |
| Arfrp1 | 871 | 1017 | 999 | 1560 | 1071 | 1062 | 925 | 1016 | 1003 | 1226 | 1152 | 1163 | 1080.86 | 981 | 1180 | 1.203 | 0.267 | 0.00197908 | 0.01525612 | 9.00E-04 | 0.01 | 0.0044 | 0.0044 | TRUE | 0.0297 |
| Arg1 | 20 | 7 | 6 | 14 | 8 | 6 | 21 | 7 | 6 | 11 | 9 | 7 | 10.07 | 11 | 9 | 0.774 | -0.37 | 0.6040022 | 0.77481274 | 0.2005 | 0.3339 | 0.266 | 0.266 | TRUE | 2.077 |
| Arg2 | 73 | 23 | 20 | 114 | 76 | 41 | 78 | 23 | 20 | 90 | 82 | 45 | 56.14 | 40 | 72 | 1.801 | 0.849 | 0.1039658 | 0.26817827 | 0.3131 | 0.0656 | 0.1768 | 0.1768 | TRUE | 4.8656 |
| Arglu1 | 3280 | 2444 | 2308 | 4110 | 3471 | 2881 | 3484 | 2441 | 2317 | 3231 | 3735 | 3154 | 3060.25 | 2747 | 3373 | 1.228 | 0.296 | 0.06653769 | 0.19895694 | 0.0311 | 0.008 | 0.0184 | 0.0184 | TRUE | 0.6669 |
| Arhgap1 | 1690 | 1780 | 1786 | 2583 | 1850 | 1779 | 1795 | 1778 | 1793 | 2031 | 1990 | 1947 | 1889.09 | 1789 | 1989 | 1.113 | 0.154 | 0.02236634 | 0.09421777 | 0 | 0.0086 | 0.0027 | 0.0027 | TRUE | 0.0041 |
| Arhgap10 | 1299 | 1526 | 1570 | 2359 | 1821 | 1847 | 1380 | 1524 | 1576 | 1854 | 1959 | 2022 | 1719.3 | 1493 | 1945 | 1.302 | 0.38 | 1.74E-05 | 0.00031563 | 0.0024 | 0.0088 | 0.0051 | 0.0051 | TRUE | 0.0531 |
| Arhgap11a | 4961 | 5332 | 5600 | 6991 | 5132 | 5179 | 5269 | 5325 | 5623 | 5496 | 5522 | 5669 | 5484.05 | 5406 | 5562 | 1.029 | 0.041 | 0.51651163 | 0.70838447 | 5.00E-04 | 0.0075 | 0.0027 | 0.0027 | TRUE | 0.0151 |
| Arhgap12 | 4925 | 5302 | 5674 | 5507 | 4266 | 4749 | 5231 | 5295 | 5697 | 4329 | 4590 | 5199 | 5056.89 | 5408 | 4706 | 0.87 | -0.201 | 0.03250737 | 0.12155199 | 0.0053 | 0.0076 | 0.0062 | 0.0062 | TRUE | 0.102 |
| Arhgap15 | 0 | 0 | 0 | 0 | 0 | 0 | 0 | 0 | 0 | 0 | 0 | 0 | 0 | 0 | 0 | NA | NA | NA | NA | NA | NA | NA | NA | NA | NA |
| Arhgap17 | 1720 | 1952 | 1876 | 2578 | 1887 | 1863 | 1827 | 1949 | 1884 | 2027 | 2030 | 2039 | 1959.4 | 1887 | 2032 | 1.077 | 0.107 | 0.12277126 | 0.29835875 | 0 | 0.0086 | 0.0029 | 0.0029 | TRUE | 0.0102 |
| Arhgap18 | 284 | 385 | 416 | 514 | 357 | 408 | 302 | 385 | 418 | 404 | 384 | 447 | 389.78 | 368 | 412 | 1.118 | 0.16 | 0.29750345 | 0.5184073 | 0.0137 | 0.0154 | 0.0145 | 0.0145 | TRUE | 0.2817 |
| Arhgap19 | 6850 | 8656 | 9216 | 9397 | 7035 | 7998 | 7276 | 8645 | 9254 | 7387 | 7569 | 8755 | 8147.68 | 8392 | 7904 | 0.942 | -0.087 | 0.45937915 | 0.66378277 | 0.0105 | 0.0073 | 0.0097 | 0.0097 | TRUE | 0.1651 |
| Arhgap20 | 31 | 31 | 39 | 89 | 68 | 79 | 33 | 31 | 39 | 70 | 73 | 86 | 55.44 | 34 | 76 | 2.222 | 1.152 | 2.85E-05 | 0.00048165 | 0 | 0.0663 | 0.0333 | 0.0333 | TRUE | 0.1144 |
| Arhgap21 | 2780 | 2721 | 2956 | 4574 | 3177 | 2742 | 2953 | 2717 | 2968 | 3596 | 3418 | 3002 | 3109 | 2879 | 3339 | 1.16 | 0.214 | 0.02556419 | 0.10326339 | 0.0049 | 0.008 | 0.0063 | 0.0063 | TRUE | 0.0944 |
| Arhgap22 | 433 | 510 | 532 | 370 | 346 | 376 | 460 | 509 | 534 | 291 | 372 | 412 | 429.7 | 501 | 358 | 0.713 | -0.489 | 0.0017626 | 0.01391376 | 0.0158 | 0.0146 | 0.0152 | 0.0152 | TRUE | 0.326 |
| Arhgap23 | 5449 | 4836 | 5150 | 4648 | 3825 | 3741 | 5788 | 4830 | 5171 | 3654 | 4115 | 4095 | 4608.86 | 5263 | 3955 | 0.751 | -0.413 | 2.90E-05 | 0.00048801 | 0.0061 | 0.0076 | 0.0068 | 0.0068 | TRUE | 0.0927 |
| Arhgap24 | 31 | 32 | 38 | 35 | 30 | 26 | 33 | 32 | 38 | 28 | 32 | 28 | 31.88 | 34 | 29 | 0.854 | -0.228 | 0.49481568 | 0.69076234 | 0 | 0.1102 | 0.0486 | 0.0486 | TRUE | 0.0671 |
| Arhgap25 | 36 | 42 | 33 | 50 | 25 | 22 | 38 | 42 | 33 | 39 | 27 | 24 | 33.93 | 38 | 30 | 0.806 | -0.312 | 0.37828264 | 0.5934918 | 0 | 0.104 | 0.0606 | 0.0606 | TRUE | 0.5595 |
| Arhgap26 | 522 | 606 | 538 | 617 | 422 | 429 | 554 | 605 | 540 | 485 | 454 | 470 | 518.1 | 566 | 470 | 0.829 | -0.27 | 0.01040249 | 0.05432589 | 0 | 0.0133 | 0.0061 | 0.0061 | TRUE | 0.0419 |
| Arhgap27 | 812 | 795 | 866 | 1174 | 846 | 810 | 862 | 794 | 870 | 923 | 910 | 887 | 874.31 | 842 | 907 | 1.077 | 0.107 | 0.22311964 | 0.4350736 | 0 | 0.0107 | 0.0045 | 0.0045 | TRUE | 0.0298 |
| Arhgap28 | 1962 | 1901 | 1908 | 1932 | 1385 | 1472 | 2084 | 1899 | 1916 | 1519 | 1490 | 1611 | 1753.11 | 1966 | 1540 | 0.783 | -0.352 | 1.71E-05 | 0.00031227 | 0.0016 | 0.0088 | 0.0043 | 0.0043 | TRUE | 0.0331 |
| Arhgap29 | 2121 | 2062 | 2184 | 3196 | 2509 | 2509 | 2253 | 2059 | 2193 | 2512 | 2700 | 2746 | 2410.61 | 2168 | 2653 | 1.223 | 0.29 | 0.00025176 | 0.00294128 | 0.0018 | 0.0083 | 0.0041 | 0.0041 | TRUE | 0.0265 |
| Arhgap30 | 717 | 666 | 668 | 991 | 760 | 747 | 762 | 665 | 671 | 779 | 818 | 818 | 751.99 | 699 | 805 | 1.151 | 0.203 | 0.04204642 | 0.14610182 | 0.0018 | 0.0113 | 0.0058 | 0.0058 | TRUE | 0.0715 |
| Arhgap31 | 32 | 31 | 24 | 90 | 56 | 50 | 34 | 31 | 24 | 71 | 60 | 55 | 45.8 | 30 | 62 | 2.097 | 1.068 | 0.00046732 | 0.00493162 | 0 | 0.0788 | 0.0421 | 0.0421 | TRUE | 0.1806 |
| Arhgap32 | 3749 | 3791 | 4473 | 4604 | 3552 | 3666 | 3982 | 3786 | 4491 | 3619 | 3822 | 4013 | 3952.27 | 4086 | 3818 | 0.934 | -0.098 | 0.29326763 | 0.51433527 | 0.0049 | 0.0077 | 0.0061 | 0.0061 | TRUE | 0.0915 |
| Arhgap33 | 389 | 633 | 720 | 416 | 386 | 398 | 413 | 632 | 723 | 327 | 415 | 436 | 491.06 | 589 | 393 | 0.665 | -0.588 | 0.00587317 | 0.03535557 | 0.0496 | 0.0136 | 0.0308 | 0.0308 | TRUE | 0.4813 |
| Arhgap36 | 27 | 28 | 15 | 28 | 24 | 11 | 29 | 28 | 15 | 22 | 26 | 12 | 21.93 | 24 | 20 | 0.839 | -0.253 | 0.59649203 | 0.76943423 | 0.0753 | 0.1571 | 0.1193 | 0.1193 | TRUE | 0.6289 |
| Arhgap39 | 1529 | 1709 | 1876 | 1666 | 1380 | 1439 | 1624 | 1707 | 1884 | 1310 | 1485 | 1575 | 1597.37 | 1738 | 1457 | 0.837 | -0.256 | 0.01665189 | 0.07641362 | 0.0063 | 0.009 | 0.0076 | 0.0076 | TRUE | 0.0958 |
| Arhgap4 | 392 | 379 | 387 | 328 | 279 | 298 | 416 | 379 | 389 | 258 | 300 | 326 | 344.62 | 395 | 295 | 0.745 | -0.425 | 0.00184793 | 0.01445319 | 0.0052 | 0.0165 | 0.0105 | 0.0105 | TRUE | 0.1454 |
| Arhgap40 | 19 | 24 | 20 | 29 | 19 | 13 | 20 | 24 | 20 | 23 | 20 | 14 | 20.28 | 21 | 19 | 0.901 | -0.15 | 0.72643398 | 0.85254683 | 0 | 0.1693 | 0.0827 | 0.0827 | TRUE | 0.2459 |
| Arhgap42 | 471 | 414 | 421 | 1176 | 800 | 720 | 500 | 413 | 423 | 924 | 861 | 788 | 651.64 | 445 | 858 | 1.928 | 0.947 | 1.19E-14 | 1.57E-12 | 0.0063 | 0.012 | 0.0091 | 0.0091 | TRUE | 0.1329 |
| Arhgap44 | 2212 | 2111 | 2005 | 2720 | 2075 | 2120 | 2350 | 2108 | 2013 | 2138 | 2233 | 2321 | 2193.76 | 2157 | 2231 | 1.034 | 0.048 | 0.59473503 | 0.7677918 | 0.0035 | 0.0084 | 0.0054 | 0.0054 | TRUE | 0.0736 |
| Arhgap5 | 5198 | 5411 | 5757 | 6826 | 5028 | 4886 | 5521 | 5404 | 5781 | 5366 | 5410 | 5348 | 5471.7 | 5569 | 5375 | 0.965 | -0.051 | 0.40772177 | 0.62062543 | 3.00E-04 | 0.0075 | 0.0026 | 0.0026 | TRUE | 0.0136 |
| Arhgap6 | 27 | 20 | 20 | 59 | 32 | 23 | 29 | 20 | 20 | 46 | 34 | 25 | 29.12 | 23 | 35 | 1.558 | 0.64 | 0.11223303 | 0.2816899 | 0.0407 | 0.12 | 0.0808 | 0.0808 | TRUE | 0.1959 |
| Arhgap8 | 651 | 770 | 845 | 626 | 521 | 637 | 691 | 769 | 848 | 492 | 561 | 697 | 676.49 | 769 | 583 | 0.757 | -0.402 | 0.00905398 | 0.04887042 | 0.019 | 0.0118 | 0.0156 | 0.0156 | TRUE | 0.3412 |
| Arhgap9 | 124 | 155 | 163 | 251 | 165 | 193 | 132 | 155 | 164 | 197 | 178 | 211 | 172.72 | 150 | 195 | 1.301 | 0.38 | 0.02467416 | 0.10081886 | 0.0032 | 0.026 | 0.0147 | 0.0147 | TRUE | 0.1174 |
| Arhgdia | 15619 | 15402 | 16041 | 23829 | 16686 | 15255 | 16590 | 15382 | 16107 | 18732 | 17953 | 16699 | 16910.62 | 16026 | 17795 | 1.11 | 0.151 | 0.04283125 | 0.14775471 | 0.0023 | 0.0071 | 0.004 | 0.004 | TRUE | 0.0355 |
| Arhgdib | 156 | 101 | 97 | 325 | 172 | 131 | 166 | 101 | 97 | 255 | 185 | 143 | 157.99 | 121 | 194 | 1.611 | 0.688 | 0.016555 | 0.07619066 | 0.0826 | 0.0278 | 0.0526 | 0.0526 | TRUE | 0.3465 |
| Arhgdig | 485 | 556 | 546 | 660 | 467 | 425 | 515 | 555 | 548 | 519 | 502 | 465 | 517.54 | 539 | 495 | 0.919 | -0.121 | 0.24794214 | 0.46465582 | 0 | 0.0133 | 0.006 | 0.006 | TRUE | 0.0326 |
| Arhgef1 | 3836 | 4050 | 4078 | 4397 | 3271 | 3494 | 4075 | 4045 | 4095 | 3457 | 3519 | 3825 | 3835.79 | 4072 | 3600 | 0.884 | -0.178 | 0.01272426 | 0.06283249 | 0.0011 | 0.0078 | 0.0034 | 0.0034 | TRUE | 0.036 |
| Arhgef10 | 2368 | 2580 | 2666 | 2723 | 2015 | 2186 | 2515 | 2577 | 2677 | 2141 | 2168 | 2393 | 2411.73 | 2590 | 2234 | 0.862 | -0.214 | 0.00780248 | 0.04356131 | 0.0018 | 0.0083 | 0.0042 | 0.0042 | TRUE | 0.0466 |
| Arhgef10l | 1617 | 2040 | 2044 | 2317 | 1934 | 2089 | 1718 | 2037 | 2052 | 1821 | 2081 | 2287 | 1999.39 | 1936 | 2063 | 1.065 | 0.091 | 0.44911152 | 0.65471734 | 0.0109 | 0.0086 | 0.0099 | 0.0099 | TRUE | 0.1286 |
| Arhgef11 | 1913 | 1693 | 1844 | 2552 | 1846 | 1868 | 2032 | 1691 | 1852 | 2006 | 1986 | 2045 | 1935.25 | 1858 | 2012 | 1.083 | 0.115 | 0.2132996 | 0.42455341 | 0.0036 | 0.0086 | 0.0057 | 0.0057 | TRUE | 0.0807 |
| Arhgef12 | 3715 | 3542 | 3742 | 4564 | 3355 | 3183 | 3946 | 3537 | 3757 | 3588 | 3610 | 3484 | 3653.78 | 3747 | 3561 | 0.95 | -0.073 | 0.3158009 | 0.53660977 | 0.0013 | 0.0078 | 0.0036 | 0.0036 | TRUE | 0.0291 |
| Arhgef15 | 256 | 326 | 345 | 440 | 301 | 351 | 272 | 326 | 346 | 346 | 324 | 384 | 332.98 | 315 | 351 | 1.116 | 0.158 | 0.2774775 | 0.49850497 | 0.008 | 0.0168 | 0.0122 | 0.0122 | TRUE | 0.1579 |
| Arhgef16 | 514 | 615 | 686 | 654 | 503 | 589 | 546 | 614 | 689 | 514 | 541 | 645 | 591.51 | 616 | 567 | 0.918 | -0.123 | 0.37645283 | 0.59184749 | 0.0116 | 0.0125 | 0.0123 | 0.0123 | TRUE | 0.1685 |
| Arhgef17 | 1485 | 1686 | 1735 | 2249 | 1764 | 1776 | 1577 | 1684 | 1742 | 1768 | 1898 | 1944 | 1768.89 | 1668 | 1870 | 1.12 | 0.164 | 0.05026455 | 0.16478332 | 0.0019 | 0.0088 | 0.0045 | 0.0045 | TRUE | 0.0282 |
| Arhgef18 | 4925 | 5873 | 6325 | 6096 | 4959 | 5417 | 5231 | 5865 | 6351 | 4792 | 5336 | 5930 | 5584.19 | 5816 | 5353 | 0.92 | -0.12 | 0.28750679 | 0.5087006 | 0.01 | 0.0075 | 0.009 | 0.009 | TRUE | 0.1085 |
| Arhgef19 | 299 | 393 | 411 | 338 | 299 | 289 | 318 | 392 | 413 | 266 | 322 | 316 | 337.76 | 374 | 301 | 0.803 | -0.317 | 0.04041519 | 0.14199787 | 0.0121 | 0.0167 | 0.0142 | 0.0142 | TRUE | 0.1992 |
| Arhgef2 | 3423 | 4170 | 4730 | 4129 | 3291 | 3534 | 3636 | 4165 | 4749 | 3246 | 3541 | 3869 | 3867.53 | 4183 | 3552 | 0.849 | -0.237 | 0.05133744 | 0.16701811 | 0.0123 | 0.0078 | 0.0104 | 0.0104 | TRUE | 0.1706 |
| Arhgef25 | 2289 | 2236 | 2155 | 1822 | 1519 | 1588 | 2431 | 2233 | 2164 | 1432 | 1634 | 1738 | 1938.88 | 2276 | 1601 | 0.703 | -0.508 | 8.53E-07 | 2.39E-05 | 0.0058 | 0.0086 | 0.0071 | 0.0071 | TRUE | 0.1054 |
| Arhgef26 | 523 | 207 | 140 | 1017 | 634 | 346 | 556 | 207 | 141 | 799 | 682 | 379 | 460.53 | 301 | 620 | 2.061 | 1.044 | 0.11123326 | 0.28048257 | 0.3069 | 0.0141 | 0.1271 | 0.3069 | TRUE | 2.0056 |
| Arhgef28 | 1084 | 994 | 1070 | 524 | 382 | 495 | 1151 | 993 | 1074 | 412 | 411 | 542 | 763.88 | 1073 | 455 | 0.423 | -1.24 | 3.51E-19 | 8.03E-17 | 0.0141 | 0.0112 | 0.0123 | 0.0123 | TRUE | 0.3184 |
| Arhgef3 | 909 | 943 | 1089 | 675 | 618 | 629 | 966 | 942 | 1093 | 531 | 665 | 689 | 814.15 | 1000 | 628 | 0.627 | -0.674 | 3.62E-07 | 1.12E-05 | 0.0116 | 0.0109 | 0.0114 | 0.0114 | TRUE | 0.2249 |
| Arhgef33 | 31 | 39 | 33 | 84 | 60 | 60 | 33 | 39 | 33 | 66 | 65 | 66 | 50.21 | 35 | 66 | 1.868 | 0.902 | 0.00094345 | 0.00850777 | 0 | 0.0725 | 0.0315 | 0.0315 | TRUE | 0.0702 |
| Arhgef37 | 26 | 26 | 25 | 49 | 34 | 32 | 28 | 26 | 25 | 39 | 37 | 35 | 31.47 | 26 | 37 | 1.403 | 0.488 | 0.14216487 | 0.32790646 | 0 | 0.1116 | 0.047 | 0.047 | TRUE | 0.0159 |
| Arhgef38 | 5 | 8 | 9 | 12 | 9 | 17 | 5 | 8 | 9 | 9 | 10 | 19 | 10.01 | 7 | 13 | 1.671 | 0.741 | 0.2467793 | 0.46379674 | 0 | 0.336 | 0.187 | 0.187 | TRUE | 0.6648 |
| Arhgef39 | 914 | 1070 | 1046 | 1168 | 881 | 914 | 971 | 1069 | 1050 | 918 | 948 | 1001 | 992.73 | 1030 | 956 | 0.927 | -0.109 | 0.22320813 | 0.43511464 | 0.0012 | 0.0102 | 0.0048 | 0.0048 | TRUE | 0.0299 |
| Arhgef4 | 436 | 595 | 588 | 596 | 474 | 546 | 463 | 594 | 590 | 469 | 510 | 598 | 537.33 | 549 | 526 | 0.955 | -0.066 | 0.65754932 | 0.80725118 | 0.0155 | 0.013 | 0.0144 | 0.0144 | TRUE | 0.2188 |
| Arhgef40 | 445 | 514 | 590 | 689 | 493 | 440 | 473 | 513 | 592 | 542 | 530 | 482 | 522.03 | 526 | 518 | 0.985 | -0.022 | 0.86152548 | 0.93157685 | 0.0065 | 0.0132 | 0.0096 | 0.0096 | TRUE | 0.142 |
| Arhgef5 | 299 | 357 | 318 | 367 | 310 | 333 | 318 | 357 | 319 | 289 | 334 | 365 | 330 | 331 | 329 | 0.99 | -0.014 | 0.91937102 | 0.96048697 | 0.0064 | 0.0169 | 0.0112 | 0.0112 | TRUE | 0.1396 |
| Arhgef6 | 54 | 51 | 41 | 108 | 63 | 51 | 57 | 51 | 41 | 85 | 68 | 56 | 59.66 | 50 | 70 | 1.405 | 0.49 | 0.08717073 | 0.23889779 | 0.0213 | 0.0621 | 0.0419 | 0.0419 | TRUE | 0.2501 |
| Arhgef7 | 4158 | 3980 | 4004 | 5363 | 3993 | 3810 | 4417 | 3975 | 4020 | 4216 | 4296 | 4171 | 4182.44 | 4137 | 4228 | 1.022 | 0.031 | 0.67004079 | 0.81584564 | 0.0013 | 0.0077 | 0.0036 | 0.0036 | TRUE | 0.0423 |

| | | | | | | | | | | | | | | | | | | | | | | | | | |
|---|---|---|---|---|---|---|---|---|---|---|---|---|---|---|---|---|---|---|---|---|---|---|---|---|---|
| Arhgef9 | 421 | 351 | 349 | 233 | 162 | 150 | 447 | 351 | 350 | 183 | 174 | 164 | 278.3 | 383 | 174 | 0.455 | -1.135 | 6.81E-13 | 7.07E-11 | 0.0097 | 0.0187 | 0.0138 | 0.0138 | TRUE | 0.2473 |
| Arid1a | 14245 | 11227 | 12303 | 18615 | 12677 | 10579 | 15131 | 11213 | 12354 | 14634 | 13640 | 11580 | 13091.74 | 12899 | 13285 | 1.03 | 0.043 | 0.75398507 | 0.87020836 | 0.0153 | 0.0072 | 0.0133 | 0.0133 | TRUE | 0.28 |
| Arid1b | 851 | 1070 | 1145 | 1688 | 1253 | 1113 | 904 | 1069 | 1150 | 1327 | 1348 | 1218 | 1169.28 | 1041 | 1298 | 1.247 | 0.318 | 0.00563839 | 0.03422969 | 0.0074 | 0.0097 | 0.0087 | 0.0087 | TRUE | 0.1573 |
| Arid2 | 4115 | 4377 | 4609 | 5941 | 4167 | 4227 | 4371 | 4371 | 4628 | 4670 | 4483 | 4627 | 4525.17 | 4457 | 4593 | 1.031 | 0.044 | 0.49645243 | 0.69197217 | 4.00E-04 | 0.0076 | 0.0028 | 0.0028 | TRUE | 0.0138 |
| Arid3a | 985 | 1348 | 1394 | 1073 | 958 | 1022 | 1046 | 1346 | 1400 | 844 | 1031 | 1119 | 1130.87 | 1264 | 998 | 0.788 | -0.343 | 0.02431162 | 0.09969912 | 0.0199 | 0.0098 | 0.0158 | 0.0158 | TRUE | 0.2721 |
| Arid3b | 3688 | 3893 | 4001 | 4381 | 3009 | 3102 | 3917 | 3888 | 4017 | 3444 | 3237 | 3396 | 3649.97 | 3941 | 3359 | 0.853 | -0.23 | 0.0003722 | 0.0040937 | 4.00E-04 | 0.0078 | 0.0027 | 0.0027 | TRUE | 0.0121 |
| Arid3c | 58 | 41 | 50 | 44 | 23 | 30 | 62 | 41 | 50 | 35 | 25 | 33 | 40.82 | 51 | 31 | 0.607 | -0.72 | 0.02810966 | 0.11006223 | 0 | 0.0876 | 0.0517 | 0.0517 | TRUE | 0.1617 |
| Arid4a | 1489 | 1540 | 1566 | 2262 | 1703 | 1559 | 1582 | 1538 | 1572 | 1778 | 1832 | 1707 | 1668.19 | 1564 | 1772 | 1.133 | 0.181 | 0.01338863 | 0.06505405 | 0 | 0.0089 | 0.0032 | 0.0032 | TRUE | 0.0124 |
| Arid4b | 2130 | 2148 | 2280 | 2774 | 2118 | 2119 | 2262 | 2145 | 2289 | 2181 | 2279 | 2320 | 2246.02 | 2232 | 2260 | 1.012 | 0.017 | 0.81402043 | 0.90482811 | 7.00E-04 | 0.0084 | 0.0033 | 0.0033 | TRUE | 0.0142 |
| Arid5a | 186 | 206 | 197 | 318 | 215 | 170 | 198 | 206 | 198 | 250 | 231 | 186 | 211.42 | 201 | 222 | 1.114 | 0.155 | 0.34321412 | 0.5633815 | 0.0073 | 0.0225 | 0.0146 | 0.0146 | TRUE | 0.2186 |
| Arid5b | 4842 | 3866 | 4173 | 5641 | 4250 | 4001 | 5143 | 3861 | 4190 | 4435 | 4573 | 4380 | 4430.2 | 4398 | 4463 | 1.015 | 0.021 | 0.85682137 | 0.92916575 | 0.0111 | 0.0076 | 0.0096 | 0.0096 | TRUE | 0.2672 |
| Arih1 | 8741 | 9723 | 10023 | 10952 | 8223 | 8868 | 9284 | 9710 | 10064 | 8610 | 8847 | 9707 | 9370.59 | 9686 | 9055 | 0.935 | -0.097 | 0.21238968 | 0.42338348 | 0.0027 | 0.0073 | 0.0043 | 0.0043 | TRUE | 0.0485 |
| Arih2 | 7025 | 7370 | 6693 | 12398 | 8759 | 8279 | 7462 | 7360 | 6720 | 9746 | 9424 | 9063 | 8295.98 | 7181 | 9411 | 1.311 | 0.39 | 1.68E-07 | 5.61E-06 | 0.002 | 0.0073 | 0.0039 | 0.0039 | TRUE | 0.0384 |
| Arl1 | 1931 | 2151 | 2035 | 2699 | 1903 | 1925 | 2051 | 2148 | 2043 | 2122 | 2048 | 2107 | 2086.52 | 2081 | 2092 | 1.006 | 0.008 | 0.90751149 | 0.95354464 | 0 | 0.0085 | 0.0029 | 0.0029 | TRUE | 0.0097 |
| Arl10 | 347 | 446 | 418 | 406 | 286 | 348 | 369 | 445 | 420 | 319 | 308 | 381 | 373.59 | 411 | 336 | 0.816 | -0.294 | 0.03803456 | 0.13558524 | 0.0082 | 0.0157 | 0.0118 | 0.0118 | TRUE | 0.1516 |
| Arl11 | 3 | 2 | 4 | 2 | 0 | 1 | 3 | 2 | 4 | 2 | 0 | 1 | 1.98 | 3 | 1 | 0.303 | -1.722 | 0.23147412 | NA | 0 | 1.6726 | 0.8029 | 0.8029 | TRUE | 0.2794 |
| Arl13a | 0 | 0 | 0 | 0 | 0 | 0 | 0 | 0 | 0 | 0 | 0 | 0 | 0 | 0 | 0 | NA | NA | NA | NA | NA | NA | NA | NA | NA | NA |
| Arl13b | 1028 | 1213 | 1347 | 1573 | 1107 | 1194 | 1092 | 1211 | 1353 | 1237 | 1191 | 1307 | 1231.76 | 1219 | 1245 | 1.021 | 0.03 | 0.7772713 | 0.88436899 | 0.0059 | 0.0096 | 0.0075 | 0.0075 | TRUE | 0.1108 |
| Arl14 | 0 | 1 | 0 | 1 | 0 | 0 | 0 | 1 | 0 | 1 | 0 | 0 | 0.3 | 0 | 0 | 0.954 | -0.067 | 0.98642001 | NA | 0 | 11.0817 | 9.3014 | 9.3014 | TRUE | 0.405 |
| Arl14ep | 1355 | 1363 | 1383 | 2134 | 1484 | 1401 | 1439 | 1361 | 1389 | 1678 | 1597 | 1534 | 1499.51 | 1396 | 1603 | 1.149 | 0.2 | 0.01166871 | 0.05900575 | 9.00E-04 | 0.0091 | 0.0039 | 0.0039 | TRUE | 0.0208 |
| Arl14epl | 1280 | 1324 | 1240 | 4071 | 2783 | 2494 | 1360 | 1322 | 1245 | 3200 | 2994 | 2730 | 2141.95 | 1309 | 2975 | 2.274 | 1.185 | 2.03E-37 | 2.00E-34 | 0.0039 | 0.0084 | 0.0056 | 0.0056 | TRUE | 0.0628 |
| Arl15 | 1665 | 1766 | 1844 | 2533 | 1730 | 1754 | 1769 | 1764 | 1852 | 1991 | 1861 | 1920 | 1859.41 | 1795 | 1924 | 1.073 | 0.101 | 0.16713853 | 0.3646215 | 5.00E-04 | 0.0087 | 0.0033 | 0.0033 | TRUE | 0.0116 |
| Arl16 | 342 | 284 | 276 | 470 | 347 | 246 | 363 | 284 | 277 | 369 | 373 | 269 | 322.69 | 308 | 337 | 1.097 | 0.134 | 0.46281821 | 0.66632303 | 0.0246 | 0.0171 | 0.0209 | 0.0209 | TRUE | 0.3472 |
| Arl2 | 1130 | 1426 | 1534 | 1528 | 1068 | 1099 | 1200 | 1424 | 1540 | 1201 | 1149 | 1203 | 1286.34 | 1388 | 1184 | 0.853 | -0.229 | 0.04222379 | 0.14652443 | 0.0069 | 0.0095 | 0.0084 | 0.0084 | TRUE | 0.168 |
| Arl2bp | 1808 | 1743 | 1679 | 3098 | 1986 | 1963 | 1920 | 1741 | 1686 | 2435 | 2137 | 2149 | 2011.35 | 1782 | 2240 | 1.258 | 0.331 | 0.00060117 | 0.00601624 | 0.0045 | 0.0085 | 0.0062 | 0.0062 | TRUE | 0.0713 |
| Arl3 | 1234 | 1379 | 1347 | 1547 | 1075 | 1163 | 1311 | 1377 | 1353 | 1216 | 1157 | 1273 | 1281.05 | 1347 | 1215 | 0.902 | -0.148 | 0.06762386 | 0.20113702 | 5.00E-04 | 0.0095 | 0.004 | 0.004 | TRUE | 0.0207 |
| Arl4a | 824 | 797 | 778 | 1108 | 783 | 732 | 875 | 796 | 781 | 871 | 842 | 801 | 827.86 | 817 | 838 | 1.026 | 0.038 | 0.69346414 | 0.83197184 | 0.0015 | 0.0109 | 0.0054 | 0.0054 | TRUE | 0.045 |
| Arl4c | 288 | 361 | 327 | 326 | 259 | 294 | 306 | 361 | 328 | 256 | 279 | 322 | 308.59 | 332 | 286 | 0.859 | -0.22 | 0.13019826 | 0.3107577 | 0.0069 | 0.0176 | 0.0119 | 0.0119 | TRUE | 0.1368 |
| Arl4d | 532 | 460 | 455 | 1178 | 862 | 637 | 565 | 459 | 457 | 926 | 927 | 697 | 672.03 | 494 | 850 | 1.724 | 0.786 | 3.47E-07 | 1.08E-05 | 0.0189 | 0.0118 | 0.0155 | 0.0155 | TRUE | 0.2913 |
| Arl5a | 3251 | 3267 | 3258 | 4296 | 3083 | 3047 | 3453 | 3263 | 3271 | 3377 | 3317 | 3335 | 3336.17 | 3329 | 3343 | 1.004 | 0.006 | 0.9209965 | 0.96116102 | 2.00E-04 | 0.0079 | 0.0027 | 0.0027 | TRUE | 0.0129 |
| Arl5b | 915 | 1012 | 1160 | 1486 | 1095 | 1134 | 972 | 1011 | 1165 | 1168 | 1178 | 1241 | 1122.5 | 1049 | 1196 | 1.139 | 0.188 | 0.06344387 | 0.19335977 | 0.0039 | 0.0098 | 0.0065 | 0.0065 | TRUE | 0.1114 |
| Arl5c | 628 | 595 | 580 | 664 | 485 | 418 | 667 | 594 | 582 | 522 | 522 | 458 | 557.51 | 614 | 501 | 0.815 | -0.294 | 0.01035282 | 0.05416054 | 0.0036 | 0.0128 | 0.0077 | 0.0077 | TRUE | 0.0651 |
| Arl6 | 804 | 816 | 893 | 1254 | 864 | 862 | 854 | 815 | 897 | 986 | 930 | 944 | 904.1 | 855 | 953 | 1.115 | 0.157 | 0.07543522 | 0.21637678 | 0 | 0.0105 | 0.0045 | 0.0045 | TRUE | 0.0216 |
| Arl6ip1 | 16425 | 21303 | 20229 | 23877 | 17041 | 17511 | 17446 | 21275 | 20312 | 18770 | 18335 | 19169 | 19217.94 | 19678 | 18758 | 0.953 | -0.069 | 0.45459232 | 0.65959798 | 0.0051 | 0.0071 | 0.0061 | 0.0061 | TRUE | 0.1204 |
| Arl6ip4 | 2128 | 2188 | 2141 | 2833 | 1979 | 1852 | 2260 | 2185 | 2150 | 2227 | 2129 | 2027 | 2163.16 | 2198 | 2128 | 0.969 | -0.046 | 0.53929274 | 0.72536087 | 0.001 | 0.0084 | 0.0036 | 0.0036 | TRUE | 0.0206 |
| Arl6ip5 | 4208 | 4361 | 4367 | 4650 | 3346 | 3751 | 4470 | 4355 | 4385 | 3655 | 3600 | 4106 | 4095.26 | 4403 | 3787 | 0.86 | -0.218 | 0.00620459 | 0.0369436 | 0.0024 | 0.0077 | 0.0043 | 0.0043 | TRUE | 0.0655 |
| Arl6ip6 | 1974 | 2212 | 2191 | 3137 | 2219 | 2385 | 2097 | 2209 | 2200 | 2466 | 2388 | 2611 | 2328.37 | 2169 | 2488 | 1.147 | 0.198 | 0.00788356 | 0.04384325 | 0.001 | 0.0083 | 0.0036 | 0.0036 | TRUE | 0.0221 |
| Arl8a | 1081 | 1774 | 1750 | 2018 | 1530 | 1741 | 1148 | 1772 | 1757 | 1586 | 1646 | 1906 | 1635.91 | 1559 | 1713 | 1.098 | 0.135 | 0.43088163 | 0.64057692 | 0.0329 | 0.0089 | 0.0206 | 0.0206 | TRUE | 0.6391 |
| Arl8b | 6460 | 7976 | 8298 | 9223 | 6781 | 7222 | 6862 | 7966 | 8332 | 7250 | 7296 | 7906 | 7601.9 | 7720 | 7484 | 0.969 | -0.045 | 0.6420512 | 0.79842557 | 0.0051 | 0.0073 | 0.0066 | 0.0066 | TRUE | 0.1153 |
| Arl9 | 2 | 1 | 0 | 0 | 0 | 0 | 2 | 1 | 0 | 0 | 0 | 0 | 0.52 | 1 | 0 | 0.173 | -2.534 | 0.43568356 | NA | 0 | 6.3362 | 6.1226 | 6.1226 | TRUE | 0.0614 |
| Armc1 | 5115 | 5055 | 4899 | 6647 | 4859 | 4692 | 5433 | 5048 | 4919 | 5225 | 5228 | 5136 | 5165.01 | 5133 | 5196 | 1.012 | 0.018 | 0.79794238 | 0.89608114 | 0.001 | 0.0075 | 0.0033 | 0.0033 | TRUE | 0.0317 |
| Armc10 | 3607 | 3775 | 3853 | 4767 | 3539 | 3712 | 3831 | 3770 | 3869 | 3747 | 3808 | 4063 | 3848.13 | 3823 | 3873 | 1.013 | 0.018 | 0.7879415 | 0.89079015 | 8.00E-04 | 0.0078 | 0.003 | 0.003 | TRUE | 0.0224 |
| Armc12 | 10 | 9 | 0 | 6 | 6 | 8 | 11 | 9 | 0 | 5 | 6 | 9 | 6.59 | 7 | 7 | 1.012 | 0.018 | 0.9861248 | 0.99254373 | 0.7118 | 0.5068 | 0.5818 | 0.5818 | TRUE | 1.9473 |
| Armc2 | 192 | 184 | 158 | 251 | 177 | 151 | 204 | 184 | 159 | 197 | 190 | 165 | 183.23 | 182 | 184 | 1.015 | 0.021 | 0.90076629 | 0.95187048 | 0.0064 | 0.0249 | 0.0155 | 0.0155 | TRUE | 0.1369 |
| Armc3 | 9 | 4 | 9 | 19 | 5 | 12 | 10 | 4 | 9 | 15 | 5 | 13 | 9.34 | 8 | 11 | 1.497 | 0.582 | 0.4050599 | 0.61779371 | 0.1059 | 0.3596 | 0.2408 | 0.2408 | TRUE | 0.7154 |
| Armc4 | 5 | 9 | 11 | 13 | 4 | 13 | 5 | 9 | 11 | 10 | 4 | 14 | 9.02 | 8 | 9 | 1.133 | 0.18 | 0.79654578 | 0.89536272 | 0.0876 | 0.3723 | 0.2412 | 0.2412 | TRUE | 0.266 |
| Armc5 | 609 | 533 | 487 | 785 | 561 | 461 | 647 | 532 | 489 | 617 | 604 | 505 | 565.59 | 556 | 575 | 1.036 | 0.05 | 0.73019478 | 0.85505616 | 0.0145 | 0.0127 | 0.0136 | 0.0136 | TRUE | 0.2376 |
| Armc6 | 1506 | 1529 | 1469 | 2302 | 1629 | 1404 | 1600 | 1527 | 1475 | 1810 | 1753 | 1537 | 1616.82 | 1534 | 1700 | 1.109 | 0.149 | 0.11747311 | 0.29004162 | 0.0039 | 0.0089 | 0.0059 | 0.0059 | TRUE | 0.084 |
| Armc7 | 103 | 92 | 109 | 153 | 135 | 121 | 109 | 92 | 109 | 120 | 145 | 132 | 118.12 | 103 | 132 | 1.278 | 0.354 | 0.06658467 | 0.19904728 | 0 | 0.0348 | 0.0182 | 0.0182 | TRUE | 0.0972 |
| Armc8 | 3483 | 3392 | 3287 | 4616 | 3133 | 2969 | 3700 | 3388 | 3300 | 3629 | 3371 | 3250 | 3439.56 | 3463 | 3417 | 0.987 | -0.019 | 0.82551638 | 0.91151248 | 0.0031 | 0.0079 | 0.0049 | 0.0049 | TRUE | 0.0435 |
| Armc9 | 798 | 895 | 882 | 1178 | 866 | 960 | 848 | 894 | 886 | 926 | 932 | 1051 | 922.63 | 876 | 970 | 1.106 | 0.146 | 0.12483717 | 0.30189081 | 0.0019 | 0.0105 | 0.0054 | 0.0054 | TRUE | 0.0632 |
| Armcx1 | 1363 | 1814 | 2019 | 1781 | 1308 | 1530 | 1448 | 1812 | 2027 | 1400 | 1407 | 1675 | 1628.16 | 1762 | 1494 | 0.847 | -0.239 | 0.09560261 | 0.25355123 | 0.0184 | 0.0089 | 0.0142 | 0.0142 | TRUE | 0.2936 |
| Armcx2 | 3697 | 3694 | 3747 | 3113 | 2226 | 2426 | 3927 | 3689 | 3762 | 2447 | 2395 | 2656 | 3146.06 | 3793 | 2499 | 0.659 | -0.602 | 2.71E-15 | 3.85E-13 | 0.0015 | 0.008 | 0.0038 | 0.0038 | TRUE | 0.0358 |
| Armcx3 | 219 | 227 | 254 | 218 | 164 | 175 | 233 | 227 | 255 | 171 | 176 | 192 | 208.96 | 238 | 180 | 0.754 | -0.408 | 0.00517628 | 0.03204091 | 0 | 0.0227 | 0.0105 | 0.0105 | TRUE | 0.0431 |
| Armcx4 | 2634 | 1831 | 2122 | 2669 | 1757 | 1419 | 2798 | 1829 | 2131 | 2098 | 1890 | 1553 | 2049.84 | 2253 | 1847 | 0.821 | -0.285 | 0.09857603 | 0.25873541 | 0.0346 | 0.0085 | 0.021 | 0.021 | TRUE | 0.4962 |
| Armcx5 | 637 | 657 | 650 | 907 | 628 | 650 | 677 | 656 | 653 | 713 | 676 | 712 | 680.94 | 662 | 700 | 1.058 | 0.082 | 0.36355161 | 0.58098184 | 0 | 0.0117 | 0.0044 | 0.0044 | TRUE | 0.0107 |
| Armcx6 | 203 | 227 | 261 | 204 | 157 | 147 | 216 | 227 | 262 | 160 | 169 | 161 | 199.1 | 235 | 163 | 0.695 | -0.524 | 0.00068262 | 0.00664176 | 0 | 0.0234 | 0.012 | 0.012 | TRUE | 0.1144 |
| Arnt | 5800 | 6171 | 6478 | 6409 | 4706 | 5404 | 6161 | 6163 | 6505 | 5038 | 5063 | 5916 | 5807.57 | 6276 | 5339 | 0.851 | -0.234 | 0.00939789 | 0.05019913 | 0.0046 | 0.0075 | 0.0057 | 0.0057 | TRUE | 0.1084 |
| Arnt2 | 988 | 939 | 960 | 994 | 679 | 671 | 1049 | 938 | 964 | 781 | 731 | 735 | 866.27 | 984 | 749 | 0.762 | -0.392 | 2.93E-05 | 0.00049221 | 0.0014 | 0.0107 | 0.0052 | 0.0052 | TRUE | 0.0405 |
| Arntl | 1273 | 1051 | 972 | 1292 | 889 | 814 | 1352 | 1050 | 976 | 1016 | 957 | 891 | 1040.17 | 1126 | 955 | 0.848 | -0.237 | 0.09202531 | 0.24755639 | 0.0164 | 0.0101 | 0.0133 | 0.0133 | TRUE | 0.369 |
| Arntl2 | 76 | 72 | 73 | 202 | 177 | 147 | 81 | 72 | 73 | 159 | 190 | 161 | 122.68 | 75 | 170 | 2.256 | 1.174 | 1.27E-09 | 7.50E-08 | 0 | 0.0338 | 0.0173 | 0.0173 | TRUE | 0.1143 |
| Arpc1a | 3727 | 4397 | 4436 | 5539 | 3919 | 4070 | 3959 | 4391 | 4454 | 4354 | 4217 | 4455 | 4305.08 | 4268 | 4342 | 1.017 | 0.025 | 0.75050056 | 0.86797274 | 0.0022 | 0.0077 | 0.0041 | 0.0041 | TRUE | 0.0488 |
| Arpc1b | 4033 | 5241 | 4982 | 6845 | 5091 | 5053 | 4284 | 5234 | 5002 | 5381 | 5478 | 5531 | 5151.73 | 4840 | 5463 | 1.129 | 0.175 | 0.06335459 | 0.19312496 | 0.005 | 0.0075 | 0.0062 | 0.0062 | TRUE | 0.123 |
| Arpc2 | 4944 | 5359 | 5235 | 7610 | 5559 | 5262 | 5251 | 5352 | 5256 | 5982 | 5981 | 5760 | 5597.27 | 5286 | 5908 | 1.118 | 0.16 | 0.00570824 | 0.03458377 | 1.00E-04 | 0.0075 | 0.0022 | 0.0022 | TRUE | 0.0058 |
| Arpc3 | 3557 | 4014 | 4079 | 5259 | 3766 | 3928 | 3778 | 4009 | 4096 | 4134 | 4052 | 4300 | 4061.46 | 3961 | 4162 | 1.051 | 0.071 | 0.30708834 | 0.5274878 | 9.00E-04 | 0.0077 | 0.0033 | 0.0033 | TRUE | 0.0198 |
| Arpc4 | 5090 | 4950 | 4665 | 7835 | 5506 | 4783 | 5406 | 4944 | 4684 | 6159 | 5924 | 5236 | 5392.23 | 5011 | 5773 | 1.152 | 0.204 | 0.03515246 | 0.12848773 | 0.0058 | 0.0075 | 0.0066 | 0.0066 | TRUE | 0.0807 |
| Arpc5 | 2652 | 2744 | 2621 | 4618 | 3136 | 2792 | 2817 | 2740 | 2632 | 3630 | 3374 | 3056 | 3041.64 | 2730 | 3353 | 1.229 | 0.298 | 0.00097968 | 0.00876929 | 0.0038 | 0.008 | 0.0055 | 0.0055 | TRUE | 0.0729 |
| Arpc5l | 1693 | 1694 | 1653 | 2321 | 1779 | 1574 | 1798 | 1692 | 1660 | 1825 | 1914 | 1723 | 1768.59 | 1717 | 1821 | 1.061 | 0.085 | 0.30129399 | 0.521912 | 0.0014 | 0.0088 | 0.0043 | 0.0043 | TRUE | 0.0259 |
| Arpp19 | 3265 | 3613 | 3590 | 5309 | 3581 | 3483 | 3468 | 3608 | 3605 | 4174 | 3853 | 3813 | 3753.37 | 3560 | 3947 | 1.109 | 0.149 | 0.03796536 | 0.13544213 | 0.0011 | 0.0078 | 0.0034 | 0.0034 | TRUE | 0.0313 |

| | | | | | | | | | | | | | | | | | | | | | | | | | |
|---|---|---|---|---|---|---|---|---|---|---|---|---|---|---|---|---|---|---|---|---|---|---|---|---|---|
| Arpp21 | 8 | 5 | 4 | 12 | 11 | 10 | 8 | 5 | 4 | 9 | 12 | 11 | 8.29 | 6 | 11 | 1.84 | 0.88 | 0.19050102 | 0.39565211 | 0 | 0.4044 | 0.192 | 0.192 | TRUE | 0.3464 |
| Arr3 | 0 | 1 | 0 | 0 | 0 | 0 | 0 | 1 | 0 | 0 | 0 | 0 | 0.17 | 0 | 0 | 0.49 | -1.029 | 0.80087 | NA | 0 | 19.7991 | 10 | 10 | TRUE | 0.405 |
| Arrb1 | 2551 | 2957 | 3118 | 2247 | 1842 | 2098 | 2710 | 2953 | 3131 | 1766 | 1982 | 2297 | 2473.08 | 2931 | 2015 | 0.687 | -0.542 | 4.72E-06 | 0.0001051 | 0.0107 | 0.0082 | 0.0097 | 0.0097 | TRUE | 0.1806 |
| Arrb2 | 3201 | 2861 | 2913 | 3751 | 2762 | 2707 | 3400 | 2857 | 2925 | 2949 | 2972 | 2963 | 3011 | 3061 | 2961 | 0.968 | -0.048 | 0.60237338 | 0.77379815 | 0.0039 | 0.008 | 0.0057 | 0.0057 | TRUE | 0.114 |
| Arrdc1 | 451 | 602 | 607 | 724 | 543 | 613 | 479 | 601 | 609 | 569 | 584 | 671 | 585.69 | 563 | 608 | 1.078 | 0.109 | 0.42456296 | 0.63471156 | 0.0109 | 0.0125 | 0.0117 | 0.0117 | TRUE | 0.1989 |
| Arrdc2 | 1102 | 1392 | 1617 | 1226 | 1096 | 1204 | 1171 | 1390 | 1624 | 964 | 1179 | 1318 | 1274.22 | 1395 | 1154 | 0.826 | -0.275 | 0.08211627 | 0.22884146 | 0.024 | 0.0095 | 0.0173 | 0.0173 | TRUE | 0.1697 |
| Arrdc3 | 1675 | 3020 | 3291 | 4351 | 3090 | 2795 | 1779 | 3016 | 3305 | 3420 | 3325 | 3060 | 2984.07 | 2700 | 3268 | 1.21 | 0.276 | 0.16671377 | 0.36418584 | 0.0526 | 0.008 | 0.0283 | 0.0283 | TRUE | 1.079 |
| Arrdc4 | 2491 | 4253 | 4153 | 3731 | 2796 | 3222 | 2646 | 4248 | 4170 | 2933 | 3008 | 3527 | 3421.96 | 3688 | 3156 | 0.856 | -0.225 | 0.20445727 | 0.41284088 | 0.0348 | 0.0079 | 0.0223 | 0.0223 | TRUE | 0.7425 |
| Arrdc5 | 1 | 0 | 1 | 5 | 2 | 0 | 1 | 0 | 1 | 4 | 2 | 0 | 1.36 | 1 | 2 | 3.027 | 1.598 | 0.40199684 | NA | 0 | 2.4326 | 1.6383 | 1.6383 | TRUE | 0.1083 |
| Arsa | 842 | 1013 | 1111 | 807 | 583 | 744 | 894 | 1012 | 1116 | 634 | 627 | 814 | 849.62 | 1007 | 692 | 0.686 | -0.543 | 0.0001397 | 0.00180153 | 0.016 | 0.0108 | 0.0134 | 0.0134 | TRUE | 0.2794 |
| Arsb | 89 | 65 | 87 | 192 | 116 | 92 | 95 | 65 | 87 | 151 | 125 | 101 | 103.88 | 82 | 126 | 1.533 | 0.616 | 0.01283538 | 0.06326237 | 0.0294 | 0.0386 | 0.0341 | 0.0341 | TRUE | 0.1494 |
| Arsg | 50 | 120 | 123 | 126 | 109 | 106 | 53 | 120 | 124 | 99 | 117 | 116 | 104.8 | 99 | 111 | 1.118 | 0.161 | 0.61003046 | 0.77891178 | 0.0904 | 0.0383 | 0.0623 | 0.0623 | TRUE | 1.5685 |
| Arsi | 13 | 21 | 12 | 9 | 6 | 9 | 14 | 21 | 12 | 7 | 6 | 10 | 11.7 | 16 | 8 | 0.496 | -1.011 | 0.0824956 | 0.22939361 | 0 | 0.2884 | 0.1485 | 0.1485 | TRUE | 0.4289 |
| Arsj | 1 | 1 | 0 | 3 | 2 | 5 | 1 | 1 | 0 | 2 | 2 | 5 | 2.01 | 1 | 3 | 4.831 | 2.272 | 0.14232082 | NA | 0 | 1.6481 | 0.8322 | 0.8322 | TRUE | 0.4128 |
| Arsk | 80 | 96 | 100 | 107 | 82 | 78 | 85 | 96 | 100 | 84 | 88 | 85 | 89.83 | 94 | 86 | 0.915 | -0.128 | 0.5322975 | 0.72038752 | 0 | 0.0436 | 0.0192 | 0.0192 | TRUE | 0.0631 |
| Art1 | 24 | 33 | 49 | 60 | 47 | 61 | 25 | 33 | 49 | 47 | 51 | 67 | 45.36 | 36 | 55 | 1.52 | 0.605 | 0.08070387 | 0.22592501 | 0.0445 | 0.0795 | 0.0631 | 0.0631 | TRUE | 0.7591 |
| Art2a-ps | 0 | 0 | 0 | 0 | 0 | 0 | 0 | 0 | 0 | 0 | 0 | 0 | 0 | 0 | 0 | NA | NA | NA | NA | NA | NA | NA | NA | NA | NA |
| Art2b | 0 | 0 | 0 | 0 | 0 | 0 | 0 | 0 | 0 | 0 | 0 | 0 | 0 | 0 | 0 | NA | NA | NA | NA | NA | NA | NA | NA | NA | NA |
| Art3 | 8 | 19 | 13 | 32 | 18 | 28 | 8 | 19 | 13 | 25 | 19 | 31 | 19.28 | 13 | 25 | 1.849 | 0.887 | 0.06681019 | 0.1994568 | 0.0253 | 0.1777 | 0.1113 | 0.1113 | TRUE | 0.3264 |
| Art4 | 2 | 4 | 8 | 12 | 8 | 7 | 2 | 4 | 8 | 9 | 9 | 8 | 6.64 | 5 | 9 | 1.814 | 0.859 | 0.26891817 | 0.48891959 | 0 | 0.5029 | 0.2706 | 0.2706 | TRUE | 0.7305 |
| Art5 | 75 | 111 | 111 | 91 | 67 | 71 | 80 | 111 | 111 | 72 | 72 | 78 | 87.22 | 101 | 74 | 0.731 | -0.451 | 0.05071514 | 0.16568278 | 0 | 0.0447 | 0.0268 | 0.0268 | TRUE | 0.3186 |
| Artn | 59 | 88 | 102 | 51 | 42 | 66 | 63 | 88 | 102 | 40 | 45 | 72 | 68.42 | 84 | 52 | 0.618 | -0.694 | 0.03088166 | 0.11770554 | 0.0633 | 0.0551 | 0.059 | 0.059 | TRUE | 0.5337 |
| Arv1 | 652 | 785 | 726 | 693 | 594 | 590 | 693 | 784 | 729 | 545 | 639 | 646 | 672.54 | 735 | 610 | 0.828 | -0.272 | 0.01743278 | 0.07864697 | 0.005 | 0.0118 | 0.008 | 0.008 | TRUE | 0.1076 |
| Arvcf | 3866 | 3990 | 3734 | 4614 | 3386 | 3428 | 4106 | 3985 | 3749 | 3627 | 3643 | 3752 | 3810.56 | 3947 | 3674 | 0.931 | -0.103 | 0.13710028 | 0.32113122 | 8.00E-04 | 0.0078 | 0.0032 | 0.0032 | TRUE | 0.0233 |
| Arx | 3 | 4 | 5 | 9 | 12 | 5 | 3 | 4 | 5 | 7 | 13 | 5 | 6.28 | 4 | 8 | 2.073 | 1.052 | 0.18592468 | 0.38963646 | 0 | 0.5318 | 0.2725 | 0.2725 | TRUE | 0.5902 |
| Arxes1 | 189 | 143 | 150 | 397 | 271 | 188 | 201 | 143 | 151 | 312 | 292 | 206 | 217.27 | 165 | 270 | 1.642 | 0.716 | 0.0009633 | 0.00864718 | 0.0361 | 0.0221 | 0.029 | 0.029 | TRUE | 0.4732 |
| Arxes2 | 242 | 279 | 245 | 580 | 381 | 320 | 257 | 279 | 246 | 456 | 410 | 350 | 332.98 | 261 | 405 | 1.559 | 0.641 | 1.25E-05 | 0.00024051 | 0.0086 | 0.0168 | 0.0123 | 0.0123 | TRUE | 0.1607 |
| As3mt | 646 | 669 | 671 | 1023 | 706 | 654 | 686 | 668 | 674 | 804 | 760 | 716 | 717.96 | 676 | 760 | 1.126 | 0.171 | 0.07113297 | 0.20768405 | 0 | 0.0115 | 0.005 | 0.005 | TRUE | 0.0322 |
| Asah1 | 8046 | 10175 | 9973 | 8635 | 6308 | 7497 | 8546 | 10162 | 10014 | 6788 | 6787 | 8207 | 8417.32 | 9574 | 7261 | 0.758 | -0.399 | 0.00041614 | 0.00447098 | 0.0105 | 0.0073 | 0.0091 | 0.0091 | TRUE | 0.1583 |
| Asah2 | 52 | 76 | 65 | 100 | 52 | 79 | 55 | 76 | 65 | 79 | 56 | 86 | 69.57 | 65 | 74 | 1.126 | 0.171 | 0.52760499 | 0.71625197 | 0.0213 | 0.0543 | 0.0385 | 0.0385 | TRUE | 0.3824 |
| Asap1 | 2574 | 2771 | 3094 | 3830 | 2803 | 2818 | 2734 | 2767 | 3107 | 3011 | 3016 | 3085 | 2953.27 | 2869 | 3037 | 1.058 | 0.082 | 0.31051969 | 0.5310688 | 0.0022 | 0.008 | 0.0043 | 0.0043 | TRUE | 0.0636 |
| Asap2 | 2004 | 2329 | 2562 | 2239 | 1765 | 1953 | 2129 | 2326 | 2573 | 1760 | 1899 | 2138 | 2137.36 | 2343 | 1932 | 0.824 | -0.278 | 0.01346093 | 0.06530475 | 0.0087 | 0.0084 | 0.0087 | 0.0087 | TRUE | 0.1044 |
| Asap3 | 997 | 858 | 877 | 1003 | 782 | 770 | 1059 | 857 | 881 | 788 | 841 | 843 | 878.21 | 932 | 824 | 0.884 | -0.178 | 0.11929097 | 0.29269713 | 0.006 | 0.0107 | 0.0082 | 0.0082 | TRUE | 0.1681 |
| Asb1 | 477 | 658 | 576 | 691 | 439 | 470 | 507 | 657 | 578 | 543 | 472 | 514 | 545.37 | 581 | 510 | 0.879 | -0.186 | 0.16027971 | 0.35474763 | 0.0091 | 0.0129 | 0.0109 | 0.0109 | TRUE | 0.1557 |
| Asb10 | 32 | 34 | 28 | 38 | 19 | 38 | 34 | 34 | 28 | 30 | 20 | 42 | 31.33 | 32 | 31 | 0.956 | -0.065 | 0.86426894 | 0.93287105 | 0.0276 | 0.1121 | 0.0732 | 0.0732 | TRUE | 0.2398 |
| Asb11 | 1 | 2 | 1 | 2 | 3 | 2 | 1 | 2 | 1 | 2 | 3 | 2 | 1.84 | 1 | 2 | 1.695 | 0.761 | 0.57269886 | NA | 0 | 1.7952 | 0.7233 | 0.7233 | TRUE | 0.1152 |
| Asb12 | 4 | 0 | 1 | 3 | 1 | 3 | 4 | 0 | 1 | 2 | 1 | 3 | 2 | 2 | 2 | 1.293 | 0.371 | 0.80308668 | NA | 0.3411 | 1.6581 | 1.0798 | 1.0798 | TRUE | 1.1766 |
| Asb13 | 223 | 259 | 219 | 220 | 187 | 193 | 237 | 259 | 220 | 173 | 201 | 211 | 216.81 | 239 | 195 | 0.815 | -0.294 | 0.05912111 | 0.18413436 | 0.0044 | 0.0221 | 0.0129 | 0.0129 | TRUE | 0.1173 |
| Asb14 | 9 | 4 | 10 | 12 | 10 | 7 | 10 | 4 | 10 | 9 | 11 | 8 | 8.58 | 8 | 9 | 1.184 | 0.244 | 0.71567776 | 0.84567137 | 0 | 0.3911 | 0.2045 | 0.2045 | TRUE | 0.5503 |
| Asb15 | 0 | 0 | 0 | 1 | 0 | 0 | 0 | 0 | 0 | 1 | 0 | 0 | 0.13 | 0 | 0 | 1.859 | 0.894 | 0.82650734 | NA | 0 | 25.1515 | 10 | 10 | TRUE | 0.2563 |
| Asb16 | 58 | 69 | 60 | 102 | 79 | 90 | 62 | 69 | 60 | 80 | 85 | 99 | 75.74 | 64 | 88 | 1.378 | 0.462 | 0.04361832 | 0.14952757 | 0 | 0.0504 | 0.0242 | 0.0242 | TRUE | 0.1026 |
| Asb17 | 1 | 0 | 2 | 5 | 2 | 0 | 1 | 0 | 2 | 4 | 2 | 0 | 1.53 | 1 | 2 | 2.029 | 1.021 | 0.56945318 | NA | 0 | 2.1665 | 1.5843 | 1.5843 | TRUE | 0.1345 |
| Asb18 | 1 | 3 | 2 | 0 | 1 | 0 | 1 | 3 | 2 | 0 | 1 | 0 | 1.19 | 2 | 0 | 0.172 | -2.537 | 0.19183372 | NA | 0 | 2.7744 | 1.472 | 1.472 | TRUE | 0.4602 |
| Asb2 | 6 | 10 | 11 | 17 | 8 | 16 | 6 | 10 | 11 | 13 | 9 | 18 | 11.15 | 9 | 13 | 1.437 | 0.523 | 0.38001507 | 0.59466194 | 0 | 0.3024 | 0.1631 | 0.1631 | TRUE | 0.1538 |
| Asb3 | 817 | 907 | 928 | 1314 | 981 | 933 | 868 | 906 | 932 | 1033 | 1055 | 1021 | 969.2 | 902 | 1036 | 1.149 | 0.201 | 0.0147872 | 0.0701003 | 0 | 0.0103 | 0.0038 | 0.0038 | TRUE | 0.0128 |
| Asb4 | 12 | 6 | 6 | 11 | 20 | 9 | 13 | 6 | 6 | 9 | 22 | 10 | 10.8 | 8 | 14 | 1.609 | 0.686 | 0.30907379 | 0.52957217 | 0.1455 | 0.312 | 0.2285 | 0.2285 | TRUE | 1.1585 |
| Asb5 | 1 | 0 | 1 | 2 | 0 | 1 | 1 | 0 | 1 | 2 | 0 | 1 | 0.79 | 1 | 1 | 1.33 | 0.412 | 0.85027746 | NA | 0 | 4.1831 | 2.149 | 2.149 | TRUE | 0.2882 |
| Asb6 | 1216 | 940 | 779 | 1650 | 1131 | 879 | 1292 | 939 | 782 | 1297 | 1217 | 962 | 1081.46 | 1004 | 1159 | 1.155 | 0.207 | 0.28117846 | 0.50246394 | 0.0433 | 0.0099 | 0.0258 | 0.0258 | TRUE | 0.7152 |
| Asb7 | 773 | 809 | 812 | 1291 | 939 | 821 | 821 | 808 | 815 | 1015 | 1010 | 899 | 894.71 | 815 | 975 | 1.197 | 0.26 | 0.00509807 | 0.03183168 | 0.0014 | 0.0106 | 0.0051 | 0.0051 | TRUE | 0.0543 |
| Asb8 | 598 | 644 | 630 | 1022 | 660 | 724 | 635 | 643 | 633 | 803 | 710 | 793 | 702.83 | 637 | 769 | 1.208 | 0.272 | 0.00517043 | 0.03201727 | 0.0011 | 0.0116 | 0.0054 | 0.0054 | TRUE | 0.0517 |
| Asb9 | 4 | 3 | 0 | 6 | 4 | 5 | 4 | 3 | 0 | 5 | 4 | 5 | 3.62 | 2 | 5 | 2.011 | 1.008 | 0.36088168 | NA | 0 | 0.9162 | 0.5625 | 0.5625 | TRUE | 0.8303 |
| Ascc1 | 1850 | 2488 | 2572 | 2179 | 1737 | 2036 | 1965 | 2485 | 2583 | 1713 | 1869 | 2229 | 2140.49 | 2344 | 1937 | 0.826 | -0.276 | 0.05170652 | 0.16770316 | 0.0186 | 0.0084 | 0.014 | 0.014 | TRUE | 0.2421 |
| Ascc2 | 1698 | 1789 | 1803 | 2683 | 1779 | 1821 | 1804 | 1787 | 1810 | 2109 | 1914 | 1993 | 1902.88 | 1800 | 2005 | 1.115 | 0.157 | 0.03705214 | 0.13320865 | 8.00E-04 | 0.0086 | 0.0035 | 0.0035 | TRUE | 0.0254 |
| Ascc3 | 4381 | 3296 | 3627 | 5994 | 4137 | 3692 | 4653 | 3292 | 3642 | 4712 | 4451 | 4041 | 4131.95 | 3862 | 4401 | 1.14 | 0.189 | 0.17156316 | 0.37051816 | 0.0193 | 0.0077 | 0.0135 | 0.0135 | TRUE | 0.3903 |
| Ascl1 | 6 | 12 | 8 | 16 | 15 | 16 | 6 | 12 | 8 | 13 | 16 | 18 | 12.1 | 9 | 16 | 1.736 | 0.796 | 0.15737322 | 0.35057775 | 0 | 0.2791 | 0.1386 | 0.1386 | TRUE | 0.3215 |
| Ascl2 | 147 | 259 | 236 | 222 | 199 | 200 | 156 | 259 | 237 | 175 | 214 | 219 | 209.89 | 217 | 203 | 0.93 | -0.105 | 0.62887291 | 0.79123697 | 0.0366 | 0.0226 | 0.0295 | 0.0295 | TRUE | 0.6566 |
| Ascl3 | 0 | 1 | 0 | 0 | 0 | 0 | 0 | 1 | 0 | 0 | 0 | 0 | 0.17 | 0 | 0 | 0.49 | -1.029 | 0.80087 | NA | 0 | 19.7991 | 10 | 10 | TRUE | 0.405 |
| Ascl4 | 4 | 2 | 3 | 9 | 6 | 6 | 4 | 2 | 3 | 7 | 6 | 7 | 4.89 | 3 | 7 | 2.184 | 1.127 | 0.19021437 | 0.39517265 | 0 | 0.6802 | 0.2944 | 0.2944 | TRUE | 0.1364 |
| Ascl5 | 3 | 0 | 1 | 0 | 1 | 0 | 3 | 0 | 1 | 0 | 1 | 0 | 0.88 | 1 | 0 | 0.251 | -1.993 | 0.41455294 | NA | 0 | 3.7601 | 2.9308 | 2.9308 | TRUE | 0.3522 |
| Asf1a | 1219 | 1305 | 1303 | 2276 | 1588 | 1287 | 1295 | 1303 | 1308 | 1789 | 1709 | 1409 | 1468.85 | 1302 | 1636 | 1.257 | 0.33 | 0.00278808 | 0.01999712 | 0.0073 | 0.0091 | 0.0081 | 0.0081 | TRUE | 0.176 |
| Asf1b | 2725 | 2694 | 2662 | 3584 | 2554 | 2496 | 2894 | 2691 | 2673 | 2817 | 2748 | 2732 | 2759.26 | 2753 | 2766 | 1.005 | 0.007 | 0.91699658 | 0.95930534 | 7.00E-04 | 0.0081 | 0.0032 | 0.0032 | TRUE | 0.0245 |
| Asgr1 | 14 | 20 | 11 | 18 | 20 | 16 | 15 | 20 | 11 | 14 | 22 | 18 | 16.51 | 15 | 18 | 1.151 | 0.202 | 0.67913728 | 0.82180091 | 0 | 0.2064 | 0.1112 | 0.1112 | TRUE | 0.3112 |
| Asgr2 | 71 | 66 | 40 | 72 | 56 | 62 | 75 | 66 | 40 | 57 | 60 | 68 | 61.04 | 60 | 62 | 1.017 | 0.024 | 0.93663889 | 0.96825513 | 0.0363 | 0.0609 | 0.0492 | 0.0492 | TRUE | 0.7514 |
| Ash1l | 1310 | 1471 | 1678 | 1810 | 1375 | 1406 | 1391 | 1469 | 1685 | 1423 | 1479 | 1539 | 1497.8 | 1515 | 1480 | 0.977 | -0.034 | 0.73559183 | 0.85807931 | 0.005 | 0.0091 | 0.0067 | 0.0067 | TRUE | 0.1158 |
| Ash2l | 26082 | 32867 | 32478 | 33783 | 24428 | 25609 | 27704 | 32825 | 32611 | 26558 | 26283 | 28033 | 29002.21 | 31047 | 26958 | 0.868 | -0.204 | 0.02388611 | 0.09844049 | 0.0049 | 0.007 | 0.0058 | 0.0058 | TRUE | 0.1086 |
| Asic1 | 1572 | 1462 | 1494 | 1598 | 1176 | 1130 | 1670 | 1460 | 1500 | 1256 | 1265 | 1237 | 1398.08 | 1543 | 1253 | 0.812 | -0.3 | 0.00057741 | 0.00581537 | 0.002 | 0.0093 | 0.0048 | 0.0048 | TRUE | 0.0616 |
| Asic2 | 55 | 59 | 73 | 82 | 70 | 53 | 58 | 59 | 73 | 64 | 75 | 58 | 64.74 | 63 | 66 | 1.036 | 0.052 | 0.83812226 | 0.91784921 | 0 | 0.0578 | 0.0307 | 0.0307 | TRUE | 0.159 |
| Asic3 | 6 | 9 | 10 | 4 | 9 | 10 | 6 | 9 | 10 | 3 | 10 | 11 | 8.2 | 8 | 8 | 0.915 | -0.129 | 0.85909141 | 0.93059974 | 0 | 0.4089 | 0.2549 | 0.2549 | TRUE | 1.0772 |
| Asic4 | 3 | 1 | 5 | 1 | 1 | 0 | 3 | 1 | 5 | 1 | 1 | 0 | 1.84 | 3 | 1 | 0.207 | -2.275 | 0.16050203 | NA | 0 | 1.7929 | 0.9606 | 0.9606 | TRUE | 0.1013 |

| | | | | | | | | | | | | | | | | | | | | | | | | | |
|---|---|---|---|---|---|---|---|---|---|---|---|---|---|---|---|---|---|---|---|---|---|---|---|---|---|
| Asic5 | 0 | 0 | 0 | 0 | 0 | 0 | 0 | 0 | 0 | 0 | 0 | 0 | 0 | 0 | 0 | NA | NA | NA | NA | NA | NA | NA | NA | NA | NA |
| Asl | 2255 | 2495 | 2524 | 2347 | 1729 | 1912 | 2395 | 2492 | 2534 | 1845 | 1860 | 2093 | 2203.28 | 2474 | 1933 | 0.781 | -0.357 | 2.30E-05 | 0.00040077 | 0.0023 | 0.0084 | 0.0047 | 0.0047 | TRUE | 0.0629 |
| Asmt | 0 | 0 | 0 | 0 | 0 | 0 | 0 | 0 | 0 | 0 | 0 | 0 | 0 | 0 | 0 | NA | NA | NA | NA | NA | NA | NA | NA | NA | NA |
| Asna1 | 9835 | 9904 | 9700 | 12276 | 8992 | 8632 | 10447 | 9891 | 9740 | 9650 | 9675 | 9449 | 9808.65 | 10026 | 9591 | 0.957 | -0.064 | 0.2976525 | 0.5184073 | 6.00E-04 | 0.0072 | 0.0026 | 0.0026 | TRUE | 0.0164 |
| Asns | 10190 | 10141 | 9164 | 11318 | 8338 | 8310 | 10824 | 10128 | 9202 | 8897 | 8971 | 9097 | 9519.7 | 10051 | 8988 | 0.894 | -0.161 | 0.04846689 | 0.16053487 | 0.0032 | 0.0072 | 0.0047 | 0.0047 | TRUE | 0.0668 |
| Asnsd1 | 6852 | 7725 | 7642 | 10110 | 7351 | 7345 | 7278 | 7715 | 7673 | 7948 | 7909 | 8040 | 7760.6 | 7555 | 7966 | 1.054 | 0.076 | 0.2008357 | 0.40850494 | 4.00E-04 | 0.0073 | 0.0024 | 0.0024 | TRUE | 0.0126 |
| Aspa | 28 | 11 | 6 | 153 | 123 | 77 | 30 | 11 | 6 | 120 | 132 | 84 | 63.94 | 16 | 112 | 7.256 | 2.859 | 8.84E-10 | 5.48E-08 | 0.2439 | 0.0584 | 0.1195 | 0.1195 | TRUE | 2.4604 |
| Aspdh | 0 | 0 | 0 | 0 | 0 | 1 | 0 | 0 | 0 | 0 | 0 | 1 | 0.18 | 0 | 0 | 1.859 | 0.894 | 0.82650734 | NA | 0 | 18.0643 | 10 | 10 | TRUE | 0.5383 |
| Aspg | 486 | 859 | 912 | 553 | 564 | 601 | 516 | 858 | 916 | 435 | 607 | 658 | 664.88 | 763 | 567 | 0.741 | -0.432 | 0.06110673 | 0.18838455 | 0.0666 | 0.0119 | 0.0368 | 0.0368 | TRUE | 0.9475 |
| Asph | 3108 | 3327 | 3512 | 4585 | 3161 | 3228 | 3301 | 3323 | 3526 | 3604 | 3401 | 3534 | 3448.22 | 3383 | 3513 | 1.038 | 0.054 | 0.42965501 | 0.63935454 | 7.00E-04 | 0.0079 | 0.0031 | 0.0031 | TRUE | 0.0166 |
| Asphd1 | 8 | 7 | 10 | 9 | 3 | 7 | 8 | 7 | 10 | 7 | 3 | 8 | 7.25 | 8 | 6 | 0.709 | -0.496 | 0.48556648 | 0.68272174 | 0 | 0.4614 | 0.2243 | 0.2243 | TRUE | 0.3418 |
| Asphd2 | 246 | 253 | 241 | 293 | 212 | 208 | 261 | 253 | 242 | 230 | 228 | 228 | 240.35 | 252 | 229 | 0.908 | -0.139 | 0.28810951 | 0.50913413 | 0 | 0.0206 | 0.0082 | 0.0082 | TRUE | 0.0134 |
| Aspm | 4521 | 4475 | 4931 | 6243 | 4363 | 4353 | 4802 | 4469 | 4951 | 4908 | 4694 | 4765 | 4764.95 | 4741 | 4789 | 1.01 | 0.015 | 0.83601314 | 0.91668515 | 0.0014 | 0.0076 | 0.0035 | 0.0035 | TRUE | 0.0307 |
| Aspn | 0 | 0 | 0 | 0 | 0 | 0 | 0 | 0 | 0 | 0 | 0 | 0 | 0 | 0 | 0 | NA | NA | NA | NA | NA | NA | NA | NA | NA | NA |
| Asprv1 | 38 | 26 | 15 | 40 | 22 | 21 | 40 | 26 | 15 | 31 | 24 | 23 | 26.58 | 27 | 26 | 0.967 | -0.048 | 0.9155579 | 0.95839777 | 0.0907 | 0.1308 | 0.1115 | 0.1115 | TRUE | 0.2572 |
| Aspscr1 | 5090 | 4789 | 4845 | 5919 | 4393 | 4022 | 5406 | 4783 | 4865 | 4653 | 4727 | 4403 | 4806.09 | 5018 | 4594 | 0.916 | -0.127 | 0.11419752 | 0.28498787 | 0.0027 | 0.0076 | 0.0045 | 0.0045 | TRUE | 0.0558 |
| Asrgl1 | 2134 | 2324 | 2385 | 2384 | 1795 | 1803 | 2267 | 2321 | 2395 | 1874 | 1931 | 1974 | 2126.93 | 2328 | 1926 | 0.827 | -0.274 | 9.21E-05 | 0.00129382 | 3.00E-04 | 0.0085 | 0.0031 | 0.0031 | TRUE | 0.0078 |
| Ass1 | 11 | 7 | 8 | 10 | 8 | 5 | 12 | 7 | 8 | 8 | 9 | 5 | 8.11 | 9 | 7 | 0.827 | -0.275 | 0.67806632 | 0.82107193 | 0 | 0.4132 | 0.1917 | 0.1917 | TRUE | 0.2271 |
| Aste1 | 393 | 370 | 394 | 516 | 394 | 400 | 417 | 370 | 396 | 406 | 424 | 438 | 408.33 | 394 | 423 | 1.071 | 0.099 | 0.38334329 | 0.597676 | 0 | 0.015 | 0.0068 | 0.0068 | TRUE | 0.0346 |
| Astl | 2 | 3 | 1 | 2 | 0 | 1 | 2 | 3 | 1 | 2 | 0 | 1 | 1.47 | 2 | 1 | 0.452 | -1.144 | 0.47547291 | NA | 0 | 2.2553 | 1.075 | 1.075 | TRUE | 0.2821 |
| Astn1 | 8 | 8 | 11 | 12 | 15 | 9 | 8 | 8 | 11 | 9 | 16 | 10 | 10.49 | 9 | 12 | 1.277 | 0.353 | 0.55237221 | 0.73624731 | 0 | 0.3209 | 0.1571 | 0.1571 | TRUE | 0.3765 |
| Astn2 | 2 | 3 | 3 | 13 | 5 | 4 | 2 | 3 | 3 | 10 | 5 | 4 | 4.69 | 3 | 6 | 2.492 | 1.317 | 0.15215567 | 0.34326982 | 0 | 0.7101 | 0.3494 | 0.3494 | TRUE | 0.7048 |
| Asun | 1801 | 1560 | 1449 | 2006 | 1386 | 1309 | 1913 | 1558 | 1455 | 1577 | 1491 | 1433 | 1571.17 | 1642 | 1500 | 0.914 | -0.129 | 0.28772422 | 0.50897112 | 0.0109 | 0.009 | 0.01 | 0.01 | TRUE | 0.2508 |
| Asxl1 | 11604 | 10114 | 11438 | 13263 | 9442 | 9789 | 12325 | 10101 | 11485 | 10426 | 10159 | 10716 | 10868.72 | 11304 | 10434 | 0.923 | -0.116 | 0.21036925 | 0.42084523 | 0.0052 | 0.0072 | 0.006 | 0.006 | TRUE | 0.1059 |
| Asxl2 | 2393 | 1739 | 1818 | 2740 | 1967 | 1726 | 2542 | 1737 | 1825 | 2154 | 2116 | 1889 | 2043.96 | 2035 | 2053 | 1.009 | 0.014 | 0.92947325 | 0.9654152 | 0.024 | 0.0085 | 0.0164 | 0.0164 | TRUE | 0.5744 |
| Asxl3 | 286 | 175 | 199 | 323 | 220 | 153 | 304 | 175 | 200 | 254 | 237 | 167 | 222.75 | 226 | 219 | 0.972 | -0.04 | 0.87188219 | 0.93724959 | 0.0612 | 0.0217 | 0.0405 | 0.0405 | TRUE | 0.9869 |
| Asz1 | 121 | 210 | 208 | 686 | 492 | 422 | 129 | 210 | 209 | 539 | 529 | 462 | 346.28 | 183 | 510 | 2.796 | 1.483 | 2.90E-14 | 3.60E-12 | 0.0326 | 0.0164 | 0.0237 | 0.0237 | TRUE | 0.7065 |
| Atad1 | 3642 | 4594 | 4898 | 5006 | 3709 | 4088 | 3868 | 4588 | 4918 | 3935 | 3991 | 4475 | 4295.93 | 4458 | 4134 | 0.927 | -0.109 | 0.33010803 | 0.55093867 | 0.0095 | 0.0077 | 0.0089 | 0.0089 | TRUE | 0.1629 |
| Atad2 | 8937 | 9790 | 10439 | 11571 | 8772 | 8740 | 9493 | 9777 | 10482 | 9096 | 9438 | 9567 | 9642.26 | 9917 | 9367 | 0.944 | -0.082 | 0.2360014 | 0.45112853 | 0.0015 | 0.0072 | 0.0034 | 0.0034 | TRUE | 0.0303 |
| Atad2b | 4962 | 4754 | 5264 | 5893 | 4038 | 3986 | 5271 | 4748 | 5286 | 4633 | 4345 | 4363 | 4774.09 | 5102 | 4447 | 0.872 | -0.198 | 0.0111216 | 0.05695128 | 0.0021 | 0.0076 | 0.0042 | 0.0042 | TRUE | 0.0448 |
| Atad3a | 6762 | 6893 | 6847 | 8574 | 6492 | 5969 | 7182 | 6884 | 6875 | 6740 | 6985 | 6534 | 6866.81 | 6980 | 6753 | 0.967 | -0.048 | 0.45131348 | 0.65683457 | 7.00E-04 | 0.0074 | 0.0028 | 0.0028 | TRUE | 0.0109 |
| Atad5 | 3706 | 3425 | 3730 | 5020 | 3697 | 3712 | 3936 | 3421 | 3745 | 3946 | 3978 | 4063 | 3848.29 | 3701 | 3996 | 1.08 | 0.111 | 0.16346077 | 0.35916735 | 0.0022 | 0.0078 | 0.0043 | 0.0043 | TRUE | 0.0534 |
| Atat1 | 431 | 427 | 401 | 471 | 354 | 393 | 458 | 426 | 403 | 370 | 381 | 430 | 411.37 | 429 | 394 | 0.917 | -0.125 | 0.305008 | 0.52574713 | 0.0028 | 0.0149 | 0.0083 | 0.0083 | TRUE | 0.0737 |
| Atcay | 53 | 66 | 76 | 40 | 28 | 15 | 56 | 66 | 76 | 31 | 30 | 16 | 46.09 | 66 | 26 | 0.396 | -1.335 | 7.90E-05 | 0.00114274 | 0.0241 | 0.0784 | 0.0561 | 0.0561 | TRUE | 0.4401 |
| Ate1 | 3231 | 3214 | 3368 | 4276 | 3023 | 3044 | 3432 | 3210 | 3382 | 3361 | 3253 | 3332 | 3328.29 | 3341 | 3315 | 0.992 | -0.011 | 0.8688959 | 0.93544206 | 5.00E-04 | 0.0079 | 0.0029 | 0.0029 | TRUE | 0.0144 |
| Atf1 | 3406 | 3618 | 3585 | 4765 | 3492 | 3427 | 3618 | 3613 | 3600 | 3746 | 3757 | 3751 | 3680.88 | 3610 | 3751 | 1.039 | 0.055 | 0.33570489 | 0.55587029 | 0 | 0.0078 | 0.0021 | 0.0021 | TRUE | 1.00E-04 |
| Atf2 | 5158 | 4405 | 4556 | 6782 | 4893 | 4541 | 5479 | 4399 | 4575 | 5331 | 5265 | 4971 | 5003.27 | 4818 | 5189 | 1.077 | 0.107 | 0.29865337 | 0.51910525 | 0.0074 | 0.0076 | 0.0075 | 0.0075 | TRUE | 0.1754 |
| Atf3 | 50 | 48 | 41 | 95 | 43 | 45 | 53 | 48 | 41 | 75 | 46 | 49 | 52.07 | 47 | 57 | 1.208 | 0.272 | 0.3792406 | 0.59433453 | 0.0298 | 0.0702 | 0.0498 | 0.0498 | TRUE | 0.7234 |
| Atf4 | 1315 | 939 | 904 | 2309 | 1594 | 1377 | 1397 | 938 | 908 | 1815 | 1715 | 1507 | 1379.97 | 1081 | 1679 | 1.554 | 0.636 | 0.00024069 | 0.00282666 | 0.0354 | 0.0093 | 0.0209 | 0.0209 | TRUE | 0.7813 |
| Atf5 | 349 | 309 | 285 | 612 | 458 | 346 | 371 | 309 | 286 | 481 | 493 | 379 | 386.35 | 322 | 451 | 1.403 | 0.489 | 0.002351 | 0.01749883 | 0.0164 | 0.0154 | 0.0159 | 0.0159 | TRUE | 0.2223 |
| Atf6 | 2488 | 2492 | 2423 | 2861 | 1964 | 2070 | 2643 | 2489 | 2433 | 2249 | 2113 | 2266 | 2365.43 | 2522 | 2209 | 0.876 | -0.19 | 0.01206848 | 0.06036778 | 0.0012 | 0.0083 | 0.0037 | 0.0037 | TRUE | 0.0214 |
| Atf6b | 2350 | 2390 | 2412 | 2841 | 2157 | 2102 | 2496 | 2387 | 2422 | 2233 | 2321 | 2301 | 2360.01 | 2435 | 2285 | 0.938 | -0.092 | 0.16714516 | 0.3646215 | 0 | 0.0083 | 0.0028 | 0.0028 | TRUE | 0.0058 |
| Atf7 | 286 | 258 | 332 | 396 | 268 | 311 | 304 | 258 | 333 | 311 | 288 | 340 | 305.82 | 298 | 313 | 1.05 | 0.071 | 0.63390266 | 0.79407494 | 0.0083 | 0.0177 | 0.0128 | 0.0128 | TRUE | 0.1616 |
| Atf7ip | 8583 | 6797 | 7592 | 10062 | 7429 | 7691 | 9117 | 6788 | 7623 | 7910 | 7993 | 8419 | 7975.03 | 7843 | 8107 | 1.034 | 0.048 | 0.68107063 | 0.82319303 | 0.0094 | 0.0073 | 0.0097 | 0.0097 | TRUE | 0.2464 |
| Atf7ip2 | 749 | 800 | 721 | 983 | 671 | 636 | 796 | 799 | 724 | 773 | 722 | 696 | 751.57 | 773 | 730 | 0.946 | -0.08 | 0.41813953 | 0.62897231 | 0.0017 | 0.0113 | 0.0057 | 0.0057 | TRUE | 0.0364 |
| Atg10 | 276 | 351 | 327 | 403 | 270 | 312 | 293 | 351 | 328 | 317 | 291 | 342 | 320.15 | 324 | 317 | 0.976 | -0.036 | 0.79299837 | 0.89324171 | 0.0038 | 0.0172 | 0.0102 | 0.0102 | TRUE | 0.0778 |
| Atg12 | 2923 | 3690 | 3654 | 3795 | 2791 | 3115 | 3105 | 3685 | 3669 | 2983 | 3003 | 3410 | 3309.19 | 3486 | 3132 | 0.898 | -0.155 | 0.13639485 | 0.31988395 | 0.007 | 0.0079 | 0.0075 | 0.0075 | TRUE | 0.1113 |
| Atg13 | 2362 | 2449 | 2255 | 3441 | 2403 | 2328 | 2509 | 2446 | 2264 | 2705 | 2585 | 2548 | 2509.64 | 2406 | 2613 | 1.086 | 0.119 | 0.12151218 | 0.29644084 | 0.0013 | 0.0082 | 0.0039 | 0.0039 | TRUE | 0.0324 |
| Atg14 | 2299 | 2457 | 2246 | 3231 | 2402 | 2394 | 2442 | 2454 | 2255 | 2540 | 2584 | 2621 | 2482.66 | 2384 | 2582 | 1.083 | 0.115 | 0.11217593 | 0.28161615 | 8.00E-04 | 0.0082 | 0.0034 | 0.0034 | TRUE | 0.027 |
| Atg16l1 | 1992 | 1793 | 1702 | 2644 | 1890 | 1640 | 2116 | 1791 | 1709 | 2079 | 2034 | 1795 | 1920.47 | 1872 | 1969 | 1.052 | 0.074 | 0.51631498 | 0.70829955 | 0.0089 | 0.0086 | 0.0088 | 0.0088 | TRUE | 0.1568 |
| Atg16l2 | 204 | 247 | 225 | 376 | 254 | 216 | 217 | 247 | 226 | 296 | 273 | 236 | 249.1 | 230 | 268 | 1.171 | 0.228 | 0.12830367 | 0.30801003 | 0.0049 | 0.0201 | 0.0121 | 0.0121 | TRUE | 0.119 |
| Atg2a | 3871 | 2962 | 2918 | 4675 | 3421 | 3015 | 4112 | 2958 | 2930 | 3675 | 3681 | 3300 | 3442.69 | 3333 | 3552 | 1.066 | 0.092 | 0.52119146 | 0.71158515 | 0.0207 | 0.0079 | 0.0145 | 0.0145 | TRUE | 0.5069 |
| Atg2b | 7286 | 6500 | 7185 | 7371 | 5411 | 5499 | 7739 | 6492 | 7214 | 5795 | 5822 | 6020 | 6513.51 | 7148 | 5879 | 0.822 | -0.282 | 0.00112743 | 0.00977564 | 0.0038 | 0.0074 | 0.0053 | 0.0053 | TRUE | 0.0789 |
| Atg3 | 951 | 1155 | 1115 | 1611 | 1148 | 1091 | 1010 | 1154 | 1120 | 1266 | 1235 | 1194 | 1163.18 | 1095 | 1232 | 1.126 | 0.171 | 0.05790618 | 0.18185909 | 0.0018 | 0.0097 | 0.005 | 0.005 | TRUE | 0.0539 |
| Atg4a | 125 | 123 | 147 | 215 | 145 | 135 | 133 | 123 | 148 | 169 | 156 | 148 | 146 | 135 | 158 | 1.175 | 0.233 | 0.17772309 | 0.37833021 | 0 | 0.0295 | 0.0146 | 0.0146 | TRUE | 0.077 |
| Atg4b | 1678 | 2211 | 2045 | 2543 | 1982 | 2016 | 1782 | 2208 | 2053 | 1999 | 2133 | 2207 | 2063.72 | 2014 | 2113 | 1.048 | 0.068 | 0.51544881 | 0.70778849 | 0.0066 | 0.0085 | 0.0074 | 0.0074 | TRUE | 0.1227 |
| Atg4c | 816 | 940 | 857 | 845 | 630 | 658 | 867 | 939 | 861 | 664 | 678 | 720 | 788.07 | 889 | 687 | 0.773 | -0.372 | 7.23E-05 | 0.00106402 | 9.00E-04 | 0.0111 | 0.005 | 0.005 | TRUE | 0.0291 |
| Atg4d | 1954 | 1956 | 1827 | 2237 | 1666 | 1640 | 2075 | 1953 | 1835 | 1759 | 1793 | 1795 | 1868.3 | 1954 | 1782 | 0.912 | -0.133 | 0.09640238 | 0.25485389 | 0.0015 | 0.0087 | 0.0041 | 0.0041 | TRUE | 0.0353 |
| Atg5 | 2286 | 2225 | 2126 | 3020 | 2219 | 2192 | 2428 | 2222 | 2135 | 2374 | 2388 | 2399 | 2324.35 | 2262 | 2387 | 1.056 | 0.078 | 0.32710037 | 0.54824943 | 0.0016 | 0.0083 | 0.0041 | 0.0041 | TRUE | 0.05 |
| Atg7 | 1299 | 1322 | 1235 | 1473 | 1141 | 1060 | 1380 | 1320 | 1240 | 1158 | 1228 | 1160 | 1247.68 | 1313 | 1182 | 0.9 | -0.152 | 0.07312994 | 0.21182835 | 0.0012 | 0.0095 | 0.0044 | 0.0044 | TRUE | 0.0287 |
| Atg9a | 1443 | 1325 | 1398 | 1629 | 1219 | 1292 | 1533 | 1323 | 1404 | 1281 | 1312 | 1414 | 1377.7 | 1420 | 1336 | 0.94 | -0.089 | 0.34783416 | 0.56662263 | 0.0031 | 0.0093 | 0.0057 | 0.0057 | TRUE | 0.0579 |
| Atg9b | 328 | 349 | 336 | 324 | 296 | 313 | 348 | 349 | 337 | 255 | 318 | 343 | 325.02 | 345 | 305 | 0.882 | -0.181 | 0.2229232 | 0.43503398 | 0.0089 | 0.017 | 0.0127 | 0.0127 | TRUE | 0.2534 |
| Athl1 | 776 | 741 | 751 | 1016 | 766 | 703 | 824 | 740 | 754 | 799 | 824 | 770 | 785.13 | 773 | 798 | 1.032 | 0.046 | 0.62636573 | 0.78958445 | 8.00E-04 | 0.0111 | 0.0051 | 0.0051 | TRUE | 0.0399 |
| Atic | 11021 | 9952 | 9649 | 14347 | 10256 | 9806 | 11706 | 9939 | 9689 | 11278 | 11035 | 10734 | 10730.26 | 10445 | 11016 | 1.055 | 0.077 | 0.41068532 | 0.62196522 | 0.0052 | 0.0072 | 0.0062 | 0.0062 | TRUE | 0.1364 |
| Atl1 | 251 | 295 | 305 | 294 | 230 | 264 | 267 | 295 | 306 | 231 | 247 | 289 | 272.51 | 289 | 256 | 0.882 | -0.181 | 0.21872695 | 0.4308432 | 0.0054 | 0.019 | 0.0119 | 0.0119 | TRUE | 0.1404 |
| Atl2 | 4379 | 4939 | 5003 | 5539 | 4149 | 4112 | 4651 | 4933 | 5024 | 4354 | 4464 | 4501 | 4654.51 | 4869 | 4440 | 0.912 | -0.133 | 0.04298798 | 0.14813336 | 6.00E-04 | 0.0076 | 0.0029 | 0.0029 | TRUE | 0.0186 |
| Atl3 | 2122 | 3263 | 3520 | 3252 | 2450 | 2780 | 2254 | 3259 | 3534 | 2556 | 2636 | 3043 | 2880.47 | 3016 | 2745 | 0.91 | -0.136 | 0.41165552 | 0.62279432 | 0.0275 | 0.008 | 0.0194 | 0.0194 | TRUE | 0.5925 |
| Atm | 4339 | 3248 | 3650 | 5516 | 3696 | 3139 | 4609 | 3244 | 3665 | 4336 | 3977 | 3436 | 3877.77 | 3839 | 3916 | 1.02 | 0.029 | 0.8433232 | 0.92112915 | 0.0226 | 0.0078 | 0.0153 | 0.0153 | TRUE | 0.3739 |
| Atmin | 7534 | 7213 | 7534 | 9313 | 6924 | 7091 | 8002 | 7204 | 7565 | 7321 | 7450 | 7762 | 7550.7 | 7590 | 7511 | 0.989 | -0.015 | 0.83176554 | 0.91463474 | 0.0014 | 0.0073 | 0.0036 | 0.0036 | TRUE | 0.0275 |

| | | | | | | | | | | | | | | | | | | | | | | | | | |
|---|---|---|---|---|---|---|---|---|---|---|---|---|---|---|---|---|---|---|---|---|---|---|---|---|---|
| Atn1 | 3006 | 2765 | 2731 | 4305 | 3217 | 2567 | 3193 | 2761 | 2742 | 3384 | 3461 | 2810 | 3058.68 | 2899 | 3218 | 1.111 | 0.151 | 0.18338023 | 0.38589366 | 0.0095 | 0.008 | 0.009 | 0.009 | TRUE | 0.1491 |
| Atoh1 | 6 | 11 | 9 | 16 | 17 | 5 | 6 | 11 | 9 | 13 | 18 | 5 | 10.46 | 9 | 12 | 1.377 | 0.461 | 0.48394889 | 0.68159234 | 0.0949 | 0.322 | 0.2147 | 0.2147 | TRUE | 0.176 |
| Atoh7 | 1 | 0 | 0 | 1 | 6 | 0 | 1 | 0 | 0 | 1 | 6 | 0 | 1.38 | 0 | 2 | 6.541 | 2.71 | 0.23373219 | NA | 2.8044 | 2.3873 | 2.5259 | 2.5259 | TRUE | 2.1498 |
| Atoh8 | 7 | 12 | 11 | 13 | 11 | 9 | 7 | 12 | 11 | 10 | 12 | 10 | 10.4 | 10 | 11 | 1.044 | 0.062 | 0.91524597 | 0.95830774 | 0 | 0.3238 | 0.1448 | 0.1448 | TRUE | 0.1826 |
| Atox1 | 4877 | 4859 | 4823 | 5144 | 3674 | 3818 | 5180 | 4853 | 4843 | 4044 | 3953 | 4179 | 4508.66 | 4959 | 4059 | 0.819 | -0.289 | 1.98E-05 | 0.00035389 | 8.00E-04 | 0.0076 | 0.0031 | 0.0031 | TRUE | 0.0186 |
| Atp10a | 670 | 924 | 984 | 764 | 583 | 756 | 712 | 923 | 988 | 601 | 627 | 828 | 779.66 | 874 | 685 | 0.783 | -0.353 | 0.03912458 | 0.13851242 | 0.0281 | 0.0111 | 0.0198 | 0.0198 | TRUE | 0.3878 |
| Atp10b | 1 | 0 | 0 | 3 | 0 | 0 | 1 | 0 | 0 | 2 | 0 | 0 | 0.57 | 0 | 1 | 2.223 | 1.153 | 0.72182708 | NA | 0 | 5.7857 | 5.9519 | 5.9519 | TRUE | 1.4404 |
| Atp10d | 5310 | 4992 | 5410 | 4067 | 3015 | 3281 | 5640 | 4986 | 5432 | 3197 | 3244 | 3592 | 4348.43 | 5353 | 3344 | 0.625 | -0.679 | 6.81E-15 | 9.34E-13 | 0.0037 | 0.0077 | 0.0052 | 0.0052 | TRUE | 0.0506 |
| Atp11a | 2685 | 2654 | 2604 | 4199 | 3030 | 2908 | 2852 | 2651 | 2615 | 3301 | 3260 | 3183 | 2976.92 | 2706 | 3248 | 1.201 | 0.264 | 0.00020842 | 0.00252229 | 8.00E-04 | 0.008 | 0.0033 | 0.0033 | TRUE | 0.027 |
| Atp11b | 8838 | 8733 | 9215 | 10555 | 7789 | 7570 | 9388 | 8722 | 9253 | 8298 | 8380 | 8287 | 8721.11 | 9121 | 8322 | 0.912 | -0.132 | 0.03211132 | 0.1207006 | 6.00E-04 | 0.0073 | 0.0026 | 0.0026 | TRUE | 0.0179 |
| Atp11c | 1243 | 1534 | 1617 | 1930 | 1333 | 1450 | 1320 | 1532 | 1624 | 1517 | 1434 | 1587 | 1502.44 | 1492 | 1513 | 1.014 | 0.02 | 0.85130714 | 0.92587032 | 0.0061 | 0.0091 | 0.0074 | 0.0074 | TRUE | 0.1215 |
| Atp12a | 131 | 366 | 421 | 207 | 159 | 212 | 139 | 366 | 423 | 163 | 171 | 232 | 248.88 | 309 | 189 | 0.609 | -0.715 | 0.04051338 | 0.14224779 | 0.1668 | 0.0201 | 0.0834 | 0.0834 | TRUE | 1.0555 |
| Atp13a1 | 4197 | 4089 | 4095 | 5645 | 4265 | 3705 | 4458 | 4084 | 4112 | 4438 | 4589 | 4056 | 4289.29 | 4218 | 4361 | 1.034 | 0.048 | 0.55886399 | 0.74101748 | 0.0027 | 0.0077 | 0.0047 | 0.0047 | TRUE | 0.0452 |
| Atp13a2 | 2545 | 2365 | 2188 | 2622 | 2012 | 1848 | 2703 | 2362 | 2197 | 2061 | 2165 | 2023 | 2251.85 | 2421 | 2083 | 0.861 | -0.217 | 0.03108122 | 0.11818058 | 0.0056 | 0.0084 | 0.0068 | 0.0068 | TRUE | 0.126 |
| Atp13a3 | 10702 | 10796 | 11047 | 14294 | 10034 | 9894 | 11367 | 10782 | 11092 | 11237 | 10796 | 10831 | 11017.53 | 11080 | 10955 | 0.989 | -0.016 | 0.7820393 | 0.88736142 | 4.00E-04 | 0.0072 | 0.0024 | 0.0024 | TRUE | 0.0068 |
| Atp13a4 | 8 | 7 | 8 | 32 | 23 | 19 | 8 | 7 | 8 | 25 | 25 | 21 | 15.7 | 8 | 24 | 3.016 | 1.592 | 0.00156569 | 0.01268283 | 0 | 0.2167 | 0.0969 | 0.0969 | TRUE | 0.0579 |
| Atp13a5 | 0 | 0 | 0 | 0 | 0 | 0 | 0 | 0 | 0 | 0 | 0 | 0 | 0 | 0 | 0 | NA | NA | NA | NA | NA | NA | NA | NA | NA | NA |
| Atp1a1 | 36305 | 39931 | 39266 | 39704 | 29276 | 31523 | 38562 | 39879 | 39427 | 31212 | 31499 | 34507 | 35847.87 | 39289 | 32406 | 0.825 | -0.278 | 5.05E-05 | 0.00078298 | 0.0017 | 0.007 | 0.0034 | 0.0034 | TRUE | 0.0393 |
| Atp1a2 | 143 | 365 | 359 | 98 | 82 | 177 | 152 | 365 | 360 | 77 | 88 | 194 | 205.99 | 292 | 120 | 0.408 | -1.293 | 0.00114551 | 0.00987744 | 0.2225 | 0.0229 | 0.108 | 0.108 | TRUE | 2.9184 |
| Atp1a3 | 6256 | 7912 | 8644 | 4655 | 3719 | 4826 | 6645 | 7902 | 8679 | 3659 | 4001 | 5283 | 6028.31 | 7742 | 4314 | 0.557 | -0.844 | 6.47E-08 | 2.48E-06 | 0.0282 | 0.0074 | 0.0174 | 0.0174 | TRUE | 0.4689 |
| Atp1a4 | 33 | 20 | 20 | 16 | 11 | 14 | 35 | 20 | 20 | 13 | 12 | 15 | 19.14 | 25 | 13 | 0.529 | -0.919 | 0.05257887 | 0.16979402 | 0 | 0.179 | 0.1045 | 0.1045 | TRUE | 0.7224 |
| Atp1b1 | 6980 | 8973 | 9300 | 5871 | 4733 | 5193 | 7414 | 8961 | 9338 | 4615 | 5092 | 5685 | 6850.98 | 8571 | 5131 | 0.598 | -0.741 | 7.70E-10 | 4.90E-08 | 0.0123 | 0.0074 | 0.0103 | 0.0103 | TRUE | 0.1703 |
| Atp1b2 | 4882 | 5636 | 5364 | 4332 | 3220 | 3818 | 5186 | 5629 | 5386 | 3405 | 3465 | 4179 | 4541.62 | 5400 | 3683 | 0.682 | -0.553 | 6.91E-08 | 2.61E-06 | 0.0073 | 0.0076 | 0.0073 | 0.0073 | TRUE | 0.1685 |
| Atp1b3 | 5631 | 5013 | 4818 | 7830 | 5246 | 5030 | 5981 | 5007 | 4838 | 6155 | 5644 | 5506 | 5521.88 | 5275 | 5768 | 1.094 | 0.129 | 0.22102043 | 0.43311388 | 0.0083 | 0.0075 | 0.0079 | 0.0079 | TRUE | 0.1669 |
| Atp1b4 | 2 | 0 | 0 | 0 | 0 | 1 | 2 | 0 | 0 | 0 | 0 | 1 | 0.54 | 1 | 0 | 0.497 | -1.009 | 0.76081378 | NA | 0 | 6.1475 | 6.1929 | 6.1929 | TRUE | 0.9742 |
| Atp2a1 | 3 | 9 | 13 | 8 | 5 | 7 | 3 | 9 | 13 | 6 | 5 | 8 | 7.43 | 8 | 6 | 0.763 | -0.391 | 0.60563365 | 0.77621309 | 0 | 0.4505 | 0.277 | 0.277 | TRUE | 0.2355 |
| Atp2a2 | 29783 | 30404 | 30210 | 35590 | 25045 | 23785 | 31635 | 30365 | 30334 | 27978 | 26947 | 26036 | 28882.49 | 30778 | 26987 | 0.877 | -0.19 | 0.00204967 | 0.01570053 | 9.00E-04 | 0.007 | 0.0027 | 0.0027 | TRUE | 0.0127 |
| Atp2a3 | 2692 | 3554 | 3456 | 2722 | 2165 | 2151 | 2859 | 3549 | 3470 | 2140 | 2329 | 2355 | 2783.8 | 3293 | 2275 | 0.69 | -0.534 | 7.15E-07 | 2.05E-05 | 0.0073 | 0.0081 | 0.008 | 0.008 | TRUE | 0.161 |
| Atp2b1 | 4695 | 4837 | 5263 | 6088 | 4353 | 4323 | 4987 | 4831 | 5285 | 4786 | 4684 | 4732 | 4883.99 | 5034 | 4734 | 0.94 | -0.089 | 0.18645494 | 0.390281 | 8.00E-04 | 0.0076 | 0.003 | 0.003 | TRUE | 0.0231 |
| Atp2b2 | 179 | 220 | 226 | 177 | 119 | 132 | 190 | 220 | 227 | 139 | 128 | 144 | 174.74 | 212 | 137 | 0.646 | -0.629 | 0.00010709 | 0.00145354 | 0 | 0.0258 | 0.0131 | 0.0131 | TRUE | 0.0895 |
| Atp2b3 | 23 | 20 | 16 | 35 | 19 | 20 | 24 | 20 | 16 | 28 | 20 | 22 | 21.72 | 20 | 23 | 1.165 | 0.22 | 0.59840299 | 0.77076456 | 0 | 0.1586 | 0.0792 | 0.0792 | TRUE | 0.1793 |
| Atp2b4 | 1384 | 1435 | 1353 | 2137 | 1784 | 1851 | 1470 | 1433 | 1359 | 1680 | 1919 | 2026 | 1647.9 | 1421 | 1875 | 1.319 | 0.4 | 7.77E-05 | 0.00112888 | 0.005 | 0.0089 | 0.0067 | 0.0067 | TRUE | 0.1022 |
| Atp2c1 | 4530 | 4312 | 4081 | 5486 | 4223 | 3979 | 4812 | 4306 | 4098 | 4313 | 4544 | 4356 | 4404.64 | 4405 | 4404 | 1 | 0 | 0.99601753 | 0.99810588 | 0.0036 | 0.0077 | 0.0051 | 0.0051 | TRUE | 0.0792 |
| Atp2c2 | 69 | 99 | 75 | 83 | 70 | 88 | 73 | 99 | 75 | 65 | 75 | 96 | 80.73 | 82 | 79 | 0.952 | -0.071 | 0.78174451 | 0.88721842 | 0.0222 | 0.0477 | 0.0351 | 0.0351 | TRUE | 0.3346 |
| Atp4a | 59 | 108 | 104 | 62 | 61 | 70 | 63 | 108 | 104 | 49 | 66 | 77 | 77.66 | 92 | 64 | 0.69 | -0.536 | 0.07536426 | 0.21631395 | 0.0555 | 0.0493 | 0.052 | 0.052 | TRUE | 0.7205 |
| Atp4b | 0 | 0 | 0 | 0 | 0 | 0 | 0 | 0 | 0 | 0 | 0 | 0 | 0 | 0 | 0 | NA | NA | NA | NA | NA | NA | NA | NA | NA | NA |
| Atp5a1 | 56171 | 64134 | 64788 | 72181 | 52773 | 56516 | 59664 | 64051 | 65054 | 56743 | 56781 | 61866 | 60692.99 | 62923 | 58463 | 0.929 | -0.106 | 0.1469056 | 0.33594245 | 0.0023 | 0.007 | 0.0038 | 0.0038 | TRUE | 0.0317 |
| Atp5b | 103317 | 120127 | 119942 | 128724 | 92155 | 98253 | 109741 | 119972 | 120434 | 101193 | 99153 | 107553 | 109674.41 | 116716 | 102633 | 0.879 | -0.186 | 0.01079787 | 0.05571471 | 0.0022 | 0.0069 | 0.0038 | 0.0038 | TRUE | 0.0335 |
| Atp5c1 | 10973 | 13289 | 13093 | 16036 | 11579 | 12718 | 11655 | 13272 | 13147 | 12606 | 12458 | 13922 | 12843.38 | 12691 | 12995 | 1.024 | 0.034 | 0.69659047 | 0.83378245 | 0.0036 | 0.0072 | 0.0054 | 0.0054 | TRUE | 0.0623 |
| Atp5d | 7705 | 8214 | 8426 | 10241 | 7369 | 7463 | 8184 | 8203 | 8461 | 8051 | 7929 | 8169 | 8166.13 | 8283 | 8050 | 0.972 | -0.041 | 0.46268728 | 0.66629195 | 1.00E-04 | 0.0073 | 0.0021 | 0.0021 | TRUE | 0.0043 |
| Atp5e | 2996 | 4130 | 3960 | 4444 | 3088 | 3338 | 3182 | 4125 | 3976 | 3494 | 3322 | 3654 | 3625.53 | 3761 | 3490 | 0.928 | -0.108 | 0.34770646 | 0.56662263 | 0.0098 | 0.0078 | 0.0093 | 0.0093 | TRUE | 0.2202 |
| Atp5f1 | 11959 | 13283 | 13098 | 17192 | 11686 | 11958 | 12703 | 13266 | 13152 | 13515 | 12573 | 13090 | 13049.76 | 13040 | 13059 | 1.002 | 0.002 | 0.971133 | 0.98480349 | 7.00E-04 | 0.0072 | 0.0027 | 0.0027 | TRUE | 0.013 |
| Atp5g1 | 9641 | 9121 | 8917 | 13036 | 8527 | 7796 | 10240 | 9109 | 8954 | 10248 | 9175 | 8534 | 9376.61 | 9434 | 9319 | 0.988 | -0.018 | 0.86006462 | 0.9309173 | 0.0056 | 0.0073 | 0.007 | 0.007 | TRUE | 0.0932 |
| Atp5g2 | 2897 | 3806 | 3970 | 4069 | 3060 | 3265 | 3077 | 3801 | 3986 | 3199 | 3292 | 3574 | 3488.28 | 3621 | 3355 | 0.926 | -0.111 | 0.33991943 | 0.56009289 | 0.0101 | 0.0078 | 0.0094 | 0.0094 | TRUE | 0.21 |
| Atp5g3 | 13274 | 12837 | 11868 | 18992 | 13146 | 11825 | 14099 | 12820 | 11917 | 14930 | 14144 | 12944 | 13475.85 | 12945 | 14006 | 1.082 | 0.114 | 0.23341288 | 0.44812824 | 0.006 | 0.0071 | 0.0065 | 0.0065 | TRUE | 0.0743 |
| Atp5h | 6552 | 7513 | 7423 | 9035 | 6417 | 6636 | 6959 | 7503 | 7453 | 7103 | 6904 | 7264 | 7197.87 | 7305 | 7090 | 0.971 | -0.043 | 0.51497707 | 0.70743379 | 0.001 | 0.0074 | 0.003 | 0.003 | TRUE | 0.0209 |
| Atp5j | 4182 | 4360 | 4283 | 6062 | 4075 | 4189 | 4442 | 4354 | 4301 | 4765 | 4384 | 4586 | 4472.07 | 4366 | 4578 | 1.049 | 0.069 | 0.29998627 | 0.52016089 | 7.00E-04 | 0.0076 | 0.003 | 0.003 | TRUE | 0.0167 |
| Atp5j2 | 5570 | 6096 | 5778 | 7598 | 5579 | 5464 | 5916 | 6088 | 5802 | 5973 | 6003 | 5981 | 5960.5 | 5935 | 5986 | 1.008 | 0.012 | 0.83364128 | 0.91548463 | 1.00E-04 | 0.0075 | 0.0022 | 0.0022 | TRUE | 0.0062 |
| Atp5k | 1685 | 1637 | 1648 | 2738 | 1788 | 1763 | 1790 | 1635 | 1655 | 2152 | 1924 | 1930 | 1847.58 | 1693 | 2002 | 1.183 | 0.243 | 0.00588936 | 0.03541251 | 0.0025 | 0.0087 | 0.0051 | 0.0051 | TRUE | 0.0533 |
| Atp5l | 774 | 662 | 678 | 1044 | 636 | 643 | 822 | 661 | 681 | 821 | 684 | 704 | 728.82 | 721 | 736 | 1.022 | 0.032 | 0.80823653 | 0.90132974 | 0.0107 | 0.0114 | 0.011 | 0.011 | TRUE | 0.1758 |
| Atp5o | 9593 | 10129 | 10026 | 12972 | 9059 | 8889 | 10189 | 10116 | 10067 | 10198 | 9747 | 9730 | 10007.91 | 10124 | 9892 | 0.977 | -0.033 | 0.55635208 | 0.73899149 | 2.00E-04 | 0.0072 | 0.0022 | 0.0022 | TRUE | 0.009 |
| Atp5s | 549 | 573 | 600 | 523 | 423 | 433 | 583 | 572 | 602 | 411 | 455 | 474 | 516.35 | 586 | 447 | 0.761 | -0.394 | 0.00025294 | 0.00294853 | 0 | 0.0133 | 0.0064 | 0.0064 | TRUE | 0.0603 |
| Atp5sl | 1537 | 1391 | 1409 | 1859 | 1416 | 1539 | 1633 | 1389 | 1415 | 1461 | 1524 | 1685 | 1517.69 | 1479 | 1557 | 1.052 | 0.073 | 0.48452693 | 0.68207415 | 0.0055 | 0.0091 | 0.0073 | 0.0073 | TRUE | 0.0992 |
| Atp6ap1 | 1865 | 2474 | 2430 | 2319 | 1782 | 2023 | 1981 | 2471 | 2440 | 1823 | 1917 | 2214 | 2141.1 | 2297 | 1985 | 0.864 | -0.212 | 0.08619332 | 0.23687768 | 0.0117 | 0.0084 | 0.0105 | 0.0105 | TRUE | 0.1754 |
| Atp6ap1l | 13 | 14 | 11 | 27 | 11 | 16 | 14 | 14 | 11 | 21 | 12 | 18 | 14.9 | 13 | 17 | 1.314 | 0.394 | 0.43480172 | 0.64330164 | 0 | 0.228 | 0.1154 | 0.1154 | TRUE | 0.265 |
| Atp6ap2 | 3231 | 3992 | 4034 | 3772 | 2704 | 2980 | 3432 | 3987 | 4051 | 2965 | 2909 | 3262 | 3434.33 | 3823 | 3045 | 0.796 | -0.328 | 0.00076671 | 0.0072281 | 0.0055 | 0.0079 | 0.0066 | 0.0066 | TRUE | 0.0973 |
| Atp6v0a1 | 2783 | 2425 | 2390 | 3403 | 2255 | 2078 | 2956 | 2422 | 2400 | 2675 | 2426 | 2275 | 2525.64 | 2593 | 2459 | 0.949 | -0.076 | 0.51234973 | 0.70574497 | 0.0092 | 0.0082 | 0.0092 | 0.0092 | TRUE | 0.1821 |
| Atp6v0a2 | 809 | 670 | 682 | 916 | 614 | 544 | 859 | 669 | 685 | 720 | 661 | 595 | 698.24 | 738 | 659 | 0.894 | -0.161 | 0.2424938 | 0.45909216 | 0.0126 | 0.0116 | 0.0123 | 0.0123 | TRUE | 0.2449 |
| Atp6v0a4 | 951 | 2087 | 2276 | 1096 | 1080 | 1650 | 1010 | 2084 | 2285 | 862 | 1162 | 1806 | 1534.93 | 1793 | 1277 | 0.712 | -0.491 | 0.272765 | 0.49295859 | 0.1435 | 0.009 | 0.0667 | 0.1435 | TRUE | 0.9156 |
| Atp6v0b | 3073 | 4098 | 3807 | 3970 | 2798 | 3010 | 3264 | 4093 | 3823 | 3121 | 3010 | 3295 | 3434.28 | 3727 | 3142 | 0.843 | -0.246 | 0.01841487 | 0.08197227 | 0.007 | 0.0079 | 0.0076 | 0.0076 | TRUE | 0.1431 |
| Atp6v0c | 0 | 1 | 0 | 1 | 1 | 0 | 0 | 1 | 0 | 1 | 1 | 0 | 0.48 | 0 | 1 | 1.712 | 0.776 | 0.78614772 | NA | 0 | 6.9165 | 4.1067 | 4.1067 | TRUE | 0.405 |
| Atp6v0c-ps2 | 0 | 0 | 0 | 0 | 0 | 0 | 0 | 0 | 0 | 0 | 0 | 0 | 0 | 0 | 0 | NA | NA | NA | NA | NA | NA | NA | NA | NA | NA |
| Atp6v0d1 | 6162 | 7286 | 7777 | 8080 | 5729 | 5971 | 6545 | 7277 | 7809 | 6352 | 6164 | 6536 | 6780.46 | 7210 | 6351 | 0.881 | -0.183 | 0.03689616 | 0.13276893 | 0.0041 | 0.0074 | 0.0054 | 0.0054 | TRUE | 0.0794 |
| Atp6v0d2 | 0 | 1 | 0 | 0 | 0 | 0 | 0 | 1 | 0 | 0 | 0 | 0 | 0.17 | 0 | 0 | 0.49 | -1.029 | 0.80087 | NA | 0 | 19.7991 | 10 | 10 | TRUE | 0.405 |
| Atp6v0e | 1823 | 1748 | 1641 | 2221 | 1447 | 1464 | 1936 | 1746 | 1648 | 1746 | 1557 | 1603 | 1705.88 | 1777 | 1635 | 0.921 | -0.119 | 0.2252238 | 0.43798949 | 0.0044 | 0.0088 | 0.0063 | 0.0063 | TRUE | 0.0744 |
| Atp6v0e2 | 335 | 483 | 498 | 510 | 345 | 420 | 356 | 482 | 500 | 401 | 371 | 460 | 428.35 | 446 | 411 | 0.92 | -0.121 | 0.46555594 | 0.6680084 | 0.02 | 0.0146 | 0.0174 | 0.0174 | TRUE | 0.3602 |
| Atp6v1a | 10504 | 11256 | 11602 | 14109 | 9955 | 9796 | 11157 | 11241 | 11650 | 11091 | 10711 | 10723 | 11095.64 | 11349 | 10842 | 0.955 | -0.066 | 0.25172688 | 0.46946156 | 3.00E-04 | 0.0072 | 0.0023 | 0.0023 | TRUE | 0.0065 |
| Atp6v1b1 | 0 | 0 | 0 | 0 | 0 | 1 | 0 | 0 | 0 | 0 | 0 | 1 | 0.18 | 0 | 0 | 1.859 | 0.894 | 0.82650734 | NA | 0 | 18.0643 | 10 | 10 | TRUE | 0.5383 |
| Atp6v1b2 | 12179 | 12518 | 12941 | 16913 | 12172 | 12221 | 12936 | 12502 | 12994 | 13296 | 13096 | 13378 | 13033.67 | 12811 | 13257 | 1.035 | 0.049 | 0.36574071 | 0.58314421 | 2.00E-04 | 0.0072 | 0.0021 | 0.0021 | TRUE | 0.0054 |

| | | | | | | | | | | | | | | | | | | | | | | | | | |
|---|---|---|---|---|---|---|---|---|---|---|---|---|---|---|---|---|---|---|---|---|---|---|---|---|---|
| Atp6v1c1 | 4070 | 4825 | 4712 | 5649 | 4127 | 4181 | 4323 | 4819 | 4731 | 4441 | 4440 | 4577 | 4555.19 | 4624 | 4486 | 0.97 | -0.044 | 0.54737505 | 0.73246078 | 0.0015 | 0.0076 | 0.0036 | 0.0036 | TRUE | 0.0395 |
| Atp6v1c2 | 124 | 122 | 100 | 146 | 134 | 112 | 132 | 122 | 100 | 115 | 144 | 123 | 122.59 | 118 | 127 | 1.076 | 0.105 | 0.60094323 | 0.77268877 | 0.0083 | 0.0338 | 0.0211 | 0.0211 | TRUE | 0.172 |
| Atp6v1d | 2405 | 2860 | 3092 | 3187 | 2444 | 2546 | 2555 | 2856 | 3105 | 2505 | 2630 | 2787 | 2739.58 | 2839 | 2641 | 0.93 | -0.105 | 0.29145278 | 0.5121693 | 0.0056 | 0.0081 | 0.0067 | 0.0067 | TRUE | 0.0928 |
| Atp6v1e1 | 4862 | 4960 | 4808 | 6715 | 4545 | 4577 | 5164 | 4954 | 4828 | 5279 | 4890 | 5010 | 5020.81 | 4982 | 5060 | 1.016 | 0.023 | 0.7416689 | 0.86176528 | 0.001 | 0.0076 | 0.0032 | 0.0032 | TRUE | 0.0177 |
| Atp6v1e2 | 5 | 1 | 1 | 3 | 2 | 2 | 5 | 1 | 1 | 2 | 2 | 2 | 2.34 | 2 | 2 | 0.926 | -0.111 | 0.93035895 | NA | 0 | 1.4175 | 0.7352 | 0.7352 | TRUE | 1.0891 |
| Atp6v1f | 3368 | 3961 | 3909 | 4338 | 3005 | 3210 | 3577 | 3956 | 3925 | 3410 | 3233 | 3514 | 3602.6 | 3819 | 3386 | 0.886 | -0.174 | 0.02661445 | 0.10621536 | 0.002 | 0.0078 | 0.0042 | 0.0042 | TRUE | 0.0373 |
| Atp6v1g1 | 2337 | 2901 | 2954 | 3382 | 2393 | 2783 | 2482 | 2897 | 2966 | 2659 | 2575 | 3046 | 2770.92 | 2782 | 2760 | 0.992 | -0.012 | 0.91346574 | 0.95735142 | 0.0082 | 0.0081 | 0.0082 | 0.0082 | TRUE | 0.1075 |
| Atp6v1g2 | 12 | 25 | 20 | 22 | 21 | 24 | 13 | 25 | 20 | 17 | 23 | 26 | 20.66 | 19 | 22 | 1.134 | 0.181 | 0.68960597 | 0.82944142 | 0 | 0.1664 | 0.0995 | 0.0995 | TRUE | 0.3494 |
| Atp6v1g3 | 1 | 0 | 0 | 0 | 0 | 0 | 1 | 0 | 0 | 0 | 0 | 0 | 0.18 | 0 | 0 | 0.49 | -1.029 | 0.80087 | NA | 0 | 18.6165 | 10 | 10 | TRUE | 0.4614 |
| Atp6v1h | 2578 | 3182 | 3294 | 4043 | 3014 | 3070 | 2738 | 3178 | 3308 | 3178 | 3243 | 3361 | 3167.58 | 3075 | 3261 | 1.06 | 0.084 | 0.37131524 | 0.58799036 | 0.0046 | 0.0079 | 0.0061 | 0.0061 | TRUE | 0.1109 |
| Atp7a | 1835 | 2053 | 2387 | 2731 | 1788 | 2105 | 1949 | 2050 | 2397 | 2147 | 1924 | 2304 | 2128.53 | 2132 | 2125 | 0.997 | -0.005 | 0.96598989 | 0.98275125 | 0.0095 | 0.0085 | 0.009 | 0.009 | TRUE | 0.1429 |
| Atp7b | 991 | 963 | 949 | 1022 | 797 | 902 | 1053 | 962 | 953 | 803 | 858 | 987 | 935.93 | 989 | 883 | 0.892 | -0.166 | 0.13851988 | 0.3230815 | 0.0057 | 0.0104 | 0.0079 | 0.0079 | TRUE | 0.127 |
| Atp8a1 | 1497 | 1684 | 1889 | 1492 | 1116 | 1326 | 1590 | 1682 | 1897 | 1173 | 1201 | 1452 | 1498.97 | 1723 | 1275 | 0.74 | -0.435 | 0.00030005 | 0.00341688 | 0.0104 | 0.0091 | 0.0098 | 0.0098 | TRUE | 0.1748 |
| Atp8a2 | 82 | 71 | 59 | 87 | 62 | 86 | 87 | 71 | 59 | 68 | 67 | 94 | 74.42 | 72 | 76 | 1.053 | 0.075 | 0.77963502 | 0.88603691 | 0.0236 | 0.0512 | 0.0378 | 0.0378 | TRUE | 0.3618 |
| Atp8b1 | 9 | 13 | 9 | 23 | 18 | 19 | 10 | 13 | 9 | 18 | 19 | 21 | 14.97 | 11 | 19 | 1.838 | 0.878 | 0.07493099 | 0.21543609 | 0 | 0.227 | 0.1015 | 0.1015 | TRUE | 0.1532 |
| Atp8b2 | 2443 | 2043 | 2049 | 3112 | 2298 | 2030 | 2595 | 2040 | 2057 | 2446 | 2473 | 2222 | 2305.62 | 2231 | 2380 | 1.067 | 0.094 | 0.42773445 | 0.6378693 | 0.0097 | 0.0083 | 0.0097 | 0.0097 | TRUE | 0.2463 |
| Atp8b3 | 227 | 331 | 352 | 225 | 177 | 218 | 241 | 331 | 353 | 177 | 190 | 239 | 255.18 | 308 | 202 | 0.653 | -0.614 | 0.0018922 | 0.01473765 | 0.0284 | 0.0198 | 0.0241 | 0.0241 | TRUE | 0.4083 |
| Atp8b4 | 7 | 4 | 2 | 13 | 13 | 10 | 7 | 4 | 2 | 10 | 14 | 11 | 8.1 | 4 | 12 | 2.624 | 1.392 | 0.05698781 | 0.17976235 | 0 | 0.4137 | 0.2285 | 0.2285 | TRUE | 0.5814 |
| Atp8b5 | 114 | 113 | 111 | 63 | 72 | 66 | 121 | 113 | 111 | 50 | 77 | 72 | 90.77 | 115 | 66 | 0.572 | -0.806 | 0.00073848 | 0.0070376 | 0.013 | 0.0432 | 0.0293 | 0.0293 | TRUE | 0.5093 |
| Atp9a | 2103 | 2461 | 2337 | 2465 | 1869 | 2047 | 2234 | 2458 | 2347 | 1938 | 2011 | 2241 | 2204.61 | 2346 | 2063 | 0.879 | -0.186 | 0.0392707 | 0.13887388 | 0.0034 | 0.0084 | 0.0054 | 0.0054 | TRUE | 0.0682 |
| Atp9b | 1495 | 1457 | 1502 | 2108 | 1539 | 1419 | 1588 | 1455 | 1508 | 1657 | 1656 | 1553 | 1569.6 | 1517 | 1622 | 1.07 | 0.097 | 0.22349149 | 0.4354964 | 0.001 | 0.009 | 0.004 | 0.004 | TRUE | 0.02 |
| Atpaf1 | 1094 | 1203 | 1173 | 1558 | 1064 | 1056 | 1162 | 1201 | 1178 | 1225 | 1145 | 1156 | 1177.8 | 1180 | 1175 | 0.996 | -0.005 | 0.94597095 | 0.97280132 | 0 | 0.0097 | 0.0037 | 0.0037 | TRUE | 0.017 |
| Atpaf2 | 1790 | 1703 | 1700 | 2482 | 1749 | 1566 | 1901 | 1701 | 1707 | 1951 | 1882 | 1714 | 1809.38 | 1770 | 1849 | 1.046 | 0.064 | 0.48929157 | 0.68553847 | 0.0034 | 0.0087 | 0.0057 | 0.0057 | TRUE | 0.0509 |
| Atpbd4 | 1714 | 1773 | 1830 | 2476 | 1818 | 1814 | 1821 | 1771 | 1838 | 1946 | 1956 | 1986 | 1886.17 | 1810 | 1963 | 1.085 | 0.117 | 0.08245451 | 0.22936023 | 0 | 0.0086 | 0.0027 | 0.0027 | TRUE | 0.0043 |
| Atpif1 | 6772 | 7528 | 7647 | 7389 | 5410 | 5630 | 7193 | 7518 | 7678 | 5809 | 5821 | 6163 | 6697.02 | 7463 | 5931 | 0.795 | -0.332 | 4.92E-07 | 1.48E-05 | 9.00E-04 | 0.0074 | 0.003 | 0.003 | TRUE | 0.0142 |
| Atr | 4985 | 6315 | 6916 | 7133 | 5566 | 5639 | 5295 | 6307 | 6944 | 5607 | 5989 | 6173 | 6052.51 | 6182 | 5923 | 0.958 | -0.062 | 0.58398524 | 0.75932524 | 0.0088 | 0.0074 | 0.0091 | 0.0091 | TRUE | 0.1921 |
| Atraid | 485 | 666 | 699 | 768 | 470 | 532 | 515 | 665 | 702 | 604 | 506 | 582 | 595.66 | 627 | 564 | 0.899 | -0.153 | 0.30124246 | 0.52188007 | 0.0157 | 0.0124 | 0.0142 | 0.0142 | TRUE | 0.2864 |
| Atrip | 1039 | 1214 | 1122 | 1284 | 1016 | 973 | 1104 | 1212 | 1127 | 1009 | 1093 | 1065 | 1101.71 | 1148 | 1056 | 0.919 | -0.121 | 0.16479287 | 0.36133924 | 0.0012 | 0.0099 | 0.0046 | 0.0046 | TRUE | 0.0294 |
| Atrn | 1617 | 1976 | 2265 | 2426 | 1792 | 1857 | 1718 | 1973 | 2274 | 1907 | 1928 | 2033 | 1972.21 | 1988 | 1956 | 0.983 | -0.024 | 0.83620699 | 0.91680998 | 0.0099 | 0.0086 | 0.0093 | 0.0093 | TRUE | 0.1913 |
| Atrnl1 | 601 | 750 | 806 | 856 | 665 | 702 | 638 | 749 | 809 | 673 | 715 | 768 | 725.6 | 732 | 719 | 0.981 | -0.028 | 0.81933128 | 0.90766966 | 0.008 | 0.0114 | 0.0095 | 0.0095 | TRUE | 0.1478 |
| Atrx | 8377 | 6247 | 6755 | 12109 | 8558 | 7616 | 8898 | 6239 | 6783 | 9519 | 9208 | 8337 | 8163.91 | 7307 | 9021 | 1.235 | 0.304 | 0.02848282 | 0.11100349 | 0.0206 | 0.0073 | 0.0138 | 0.0138 | TRUE | 0.443 |
| Atxn1 | 286 | 286 | 252 | 367 | 242 | 246 | 304 | 286 | 253 | 289 | 260 | 269 | 276.77 | 281 | 273 | 0.973 | -0.039 | 0.77451089 | 0.8825015 | 0.0021 | 0.0188 | 0.01 | 0.01 | TRUE | 0.0844 |
| Atxn10 | 11713 | 14865 | 15649 | 17201 | 12799 | 14259 | 12441 | 14846 | 15713 | 13522 | 13771 | 15609 | 14317.01 | 14333 | 14301 | 0.998 | -0.003 | 0.97565388 | 0.98697924 | 0.0081 | 0.0071 | 0.0089 | 0.0089 | TRUE | 0.163 |
| Atxn1l | 2529 | 2224 | 2406 | 3308 | 2309 | 2266 | 2686 | 2221 | 2416 | 2600 | 2484 | 2480 | 2481.43 | 2441 | 2521 | 1.033 | 0.047 | 0.61414535 | 0.781712 | 0.0045 | 0.0082 | 0.006 | 0.006 | TRUE | 0.0933 |
| Atxn2 | 5025 | 4879 | 4885 | 6251 | 4774 | 4594 | 5337 | 4873 | 4905 | 4914 | 5137 | 5029 | 5032.44 | 5038 | 5027 | 0.998 | -0.003 | 0.9605567 | 0.9800052 | 0.0013 | 0.0076 | 0.0034 | 0.0034 | TRUE | 0.0328 |
| Atxn2l | 12657 | 13912 | 14765 | 15102 | 11787 | 12173 | 13444 | 13894 | 14826 | 11872 | 12682 | 13325 | 13340.5 | 14055 | 12626 | 0.898 | -0.155 | 0.04832501 | 0.16025697 | 0.0028 | 0.0072 | 0.0044 | 0.0044 | TRUE | 0.0335 |
| Atxn3 | 4905 | 4956 | 4943 | 5074 | 3869 | 3944 | 5210 | 4950 | 4963 | 3989 | 4163 | 4317 | 4598.63 | 5041 | 4156 | 0.824 | -0.279 | 4.41E-05 | 0.00069699 | 9.00E-04 | 0.0076 | 0.0031 | 0.0031 | TRUE | 0.0154 |
| Atxn7 | 348 | 407 | 396 | 600 | 438 | 453 | 370 | 406 | 398 | 472 | 471 | 496 | 435.43 | 391 | 480 | 1.225 | 0.293 | 0.00667864 | 0.03884053 | 0 | 0.0145 | 0.0061 | 0.0061 | TRUE | 0.0265 |
| Atxn7l1 | 490 | 517 | 554 | 484 | 322 | 368 | 520 | 516 | 556 | 380 | 346 | 403 | 453.81 | 531 | 376 | 0.709 | -0.496 | 1.23E-05 | 0.00023712 | 7.00E-04 | 0.0142 | 0.007 | 0.007 | TRUE | 0.0534 |
| Atxn7l2 | 489 | 420 | 493 | 509 | 427 | 428 | 519 | 419 | 495 | 400 | 459 | 469 | 460.33 | 478 | 443 | 0.925 | -0.113 | 0.39801568 | 0.61130619 | 0.0077 | 0.0141 | 0.0106 | 0.0106 | TRUE | 0.1339 |
| Atxn7l3 | 6982 | 6290 | 5874 | 8711 | 6220 | 5334 | 7416 | 6282 | 5898 | 6848 | 6692 | 5839 | 6495.88 | 6532 | 6460 | 0.989 | -0.016 | 0.88931664 | 0.94606369 | 0.0106 | 0.0074 | 0.0092 | 0.0092 | TRUE | 0.1709 |
| Atxn7l3b | 2639 | 2777 | 2798 | 5041 | 3576 | 3233 | 2803 | 2773 | 2809 | 3963 | 3848 | 3539 | 3289.23 | 2795 | 3783 | 1.354 | 0.437 | 4.53E-09 | 2.32E-07 | 0.0015 | 0.0079 | 0.0037 | 0.0037 | TRUE | 0.0386 |
| AU015228 | 29 | 10 | 9 | 62 | 41 | 9 | 31 | 10 | 9 | 49 | 44 | 10 | 25.42 | 17 | 34 | 2.076 | 1.054 | 0.12361631 | 0.29971966 | 0.4677 | 0.1365 | 0.2923 | 0.2923 | TRUE | 2.6271 |
| AU015836 | 567 | 293 | 243 | 786 | 501 | 536 | 602 | 293 | 244 | 618 | 539 | 587 | 480.42 | 380 | 581 | 1.532 | 0.616 | 0.03274495 | 0.12215075 | 0.1189 | 0.0138 | 0.0577 | 0.0577 | TRUE | 3.0111 |
| AU016765 | 0 | 0 | 0 | 0 | 0 | 0 | 0 | 0 | 0 | 0 | 0 | 0 | 0 | 0 | 0 | NA | NA | NA | NA | NA | NA | NA | NA | NA | NA |
| AU018091 | 3602 | 4697 | 4744 | 12792 | 8622 | 9727 | 3826 | 4691 | 4763 | 10056 | 9277 | 10648 | 7210.15 | 4427 | 9994 | 2.257 | 1.175 | 1.07E-26 | 4.81E-24 | 0.0091 | 0.0074 | 0.0085 | 0.0085 | TRUE | 0.1715 |
| AU018829 | 0 | 0 | 0 | 0 | 0 | 0 | 0 | 0 | 0 | 0 | 0 | 0 | 0 | 0 | 0 | NA | NA | NA | NA | NA | NA | NA | NA | NA | NA |
| AU019823 | 1388 | 1517 | 1452 | 2322 | 1706 | 1427 | 1474 | 1515 | 1458 | 1825 | 1836 | 1562 | 1611.72 | 1482 | 1741 | 1.175 | 0.233 | 0.01367138 | 0.06606159 | 0.0037 | 0.0089 | 0.0058 | 0.0058 | TRUE | 0.0964 |
| AU019990 | 4 | 1 | 0 | 22 | 19 | 1 | 4 | 1 | 0 | 17 | 20 | 1 | 7.35 | 2 | 13 | 7.514 | 2.91 | 0.01985863 | 0.08641479 | 1.3368 | 0.4553 | 0.7915 | 0.7915 | TRUE | 0.9554 |
| AU021063 | 1 | 0 | 1 | 1 | 0 | 2 | 1 | 0 | 1 | 1 | 0 | 2 | 0.84 | 1 | 1 | 1.434 | 0.52 | 0.81041906 | NA | 0 | 3.9275 | 2.1451 | 2.1451 | TRUE | 0.3074 |
| AU021092 | 0 | 0 | 0 | 7 | 2 | 1 | 0 | 0 | 0 | 6 | 2 | 1 | 1.46 | 0 | 3 | 15.752 | 3.977 | 0.04868368 | NA | 0.6399 | 2.2661 | 1.7679 | 1.7679 | TRUE | 0.4534 |
| AU022252 | 1013 | 1098 | 1069 | 1138 | 870 | 884 | 1076 | 1097 | 1073 | 895 | 936 | 968 | 1007.38 | 1082 | 933 | 0.861 | -0.215 | 0.00865229 | 0.04715409 | 0 | 0.0102 | 0.0039 | 0.0039 | TRUE | 0.0161 |
| AU022751 | 165 | 235 | 187 | 872 | 596 | 511 | 175 | 235 | 188 | 685 | 641 | 559 | 413.98 | 199 | 628 | 3.156 | 1.658 | 7.74E-27 | 3.59E-24 | 0.0126 | 0.0149 | 0.0139 | 0.0139 | TRUE | 0.2646 |
| AU022754 | 0 | 0 | 1 | 0 | 1 | 1 | 0 | 0 | 1 | 0 | 1 | 1 | 0.53 | 0 | 1 | 1.924 | 0.944 | 0.73632075 | NA | 0 | 6.2332 | 3.9677 | 3.9677 | TRUE | 0.4098 |
| AU022793 | 0 | 0 | 0 | 0 | 0 | 0 | 0 | 0 | 0 | 0 | 0 | 0 | 0 | 0 | 0 | NA | NA | NA | NA | NA | NA | NA | NA | NA | NA |
| AU023762 | 0 | 0 | 0 | 1 | 2 | 3 | 0 | 0 | 0 | 1 | 2 | 3 | 1.04 | 0 | 2 | 10.912 | 3.448 | 0.11578645 | NA | 0 | 3.1838 | 2.2241 | 2.2241 | TRUE | 0.0828 |
| AU023871 | 32 | 69 | 61 | 43 | 50 | 72 | 34 | 69 | 61 | 34 | 54 | 79 | 55.09 | 55 | 56 | 1.006 | 0.009 | 0.98272143 | 0.99095214 | 0.1307 | 0.0667 | 0.0974 | 0.0974 | TRUE | 0.2477 |
| AU040320 | 593 | 748 | 794 | 888 | 578 | 619 | 630 | 747 | 797 | 698 | 622 | 678 | 695.29 | 725 | 666 | 0.919 | -0.122 | 0.3204021 | 0.54151113 | 0.0076 | 0.0116 | 0.0094 | 0.0094 | TRUE | 0.154 |
| AU040972 | 0 | 5 | 3 | 18 | 9 | 17 | 0 | 5 | 3 | 14 | 10 | 19 | 8.41 | 3 | 14 | 5.241 | 2.39 | 0.00523196 | 0.03229683 | 0.0655 | 0.3987 | 0.3039 | 0.3039 | TRUE | 0.4231 |
| AU041133 | 63 | 56 | 53 | 88 | 79 | 57 | 67 | 56 | 53 | 69 | 85 | 62 | 65.44 | 59 | 72 | 1.23 | 0.299 | 0.24386637 | 0.46043022 | 0 | 0.0572 | 0.0318 | 0.0318 | TRUE | 0.2057 |
| Auh | 515 | 778 | 811 | 710 | 598 | 712 | 547 | 777 | 814 | 558 | 643 | 779 | 686.55 | 713 | 660 | 0.925 | -0.112 | 0.54339516 | 0.72865992 | 0.0349 | 0.0117 | 0.0231 | 0.0231 | TRUE | 0.4874 |
| Aup1 | 1352 | 1606 | 1664 | 2122 | 1526 | 1623 | 1436 | 1604 | 1671 | 1668 | 1642 | 1777 | 1632.91 | 1570 | 1696 | 1.079 | 0.11 | 0.23029793 | 0.44404144 | 0.0029 | 0.0089 | 0.0055 | 0.0055 | TRUE | 0.067 |
| Aurka | 7093 | 7568 | 7280 | 8674 | 5989 | 6276 | 7534 | 7558 | 7310 | 6819 | 6444 | 6870 | 7089.14 | 7467 | 6711 | 0.899 | -0.154 | 0.01355558 | 0.06564282 | 6.00E-04 | 0.0074 | 0.0027 | 0.0027 | TRUE | 0.0147 |
| Aurkaip1 | 2578 | 2714 | 2756 | 3595 | 2473 | 2512 | 2738 | 2710 | 2767 | 2826 | 2661 | 2750 | 2742.13 | 2738 | 2746 | 1.003 | 0.004 | 0.9501487 | 0.97481103 | 0 | 0.0081 | 0.0027 | 0.0027 | TRUE | 0.0088 |
| Aurkb | 5106 | 6595 | 6616 | 6674 | 4925 | 5622 | 5423 | 6586 | 6643 | 5247 | 5299 | 6154 | 5892.14 | 6217 | 5567 | 0.895 | -0.16 | 0.15775136 | 0.35117213 | 0.0101 | 0.0075 | 0.0091 | 0.0091 | TRUE | 0.1522 |
| Aurkc | 496 | 553 | 480 | 621 | 463 | 485 | 527 | 552 | 482 | 488 | 498 | 531 | 513.06 | 520 | 506 | 0.971 | -0.042 | 0.69822609 | 0.83485873 | 0.0014 | 0.0133 | 0.0066 | 0.0066 | TRUE | 0.049 |
| Auts2 | 611 | 658 | 742 | 852 | 671 | 700 | 649 | 657 | 745 | 670 | 722 | 766 | 701.53 | 684 | 719 | 1.051 | 0.071 | 0.51403584 | 0.70677127 | 0.0039 | 0.0116 | 0.0072 | 0.0072 | TRUE | 0.0729 |
| AV039307 | 2 | 1 | 3 | 2 | 3 | 1 | 2 | 1 | 3 | 2 | 3 | 1 | 2.01 | 2 | 2 | 0.954 | -0.067 | 0.95876719 | NA | 0 | 1.65 | 0.7255 | 0.7255 | TRUE | 0.251 |
| AV051173 | 1 | 0 | 0 | 2 | 2 | 1 | 1 | 0 | 0 | 2 | 2 | 1 | 0.98 | 0 | 2 | 4.39 | 2.134 | 0.29289438 | NA | 0 | 3.368 | 1.662 | 1.662 | TRUE | 0.4614 |

| | | | | | | | | | | | | | | | | | | | | | | | | | |
|---|---|---|---|---|---|---|---|---|---|---|---|---|---|---|---|---|---|---|---|---|---|---|---|---|---|
| AV320801 | 0 | 0 | 0 | 0 | 0 | 0 | 0 | 0 | 0 | 0 | 0 | 0 | 0 | 0 | 0 | NA | NA | NA | NA | NA | NA | NA | NA | NA | NA |
| Aven | 1467 | 1229 | 1098 | 1645 | 1142 | 965 | 1558 | 1227 | 1103 | 1293 | 1229 | 1056 | 1244.4 | 1296 | 1193 | 0.921 | -0.119 | 0.42538991 | 0.63552616 | 0.0195 | 0.0096 | 0.0152 | 0.0152 | TRUE | 0.3753 |
| Avil | 33 | 40 | 26 | 52 | 37 | 39 | 35 | 40 | 26 | 41 | 40 | 43 | 37.41 | 34 | 41 | 1.22 | 0.287 | 0.37326931 | 0.58968554 | 0 | 0.095 | 0.0479 | 0.0479 | TRUE | 0.2762 |
| Avl9 | 2007 | 2258 | 2335 | 2829 | 1962 | 2104 | 2132 | 2255 | 2345 | 2224 | 2111 | 2303 | 2228.26 | 2244 | 2213 | 0.986 | -0.02 | 0.79818945 | 0.89608114 | 0.0016 | 0.0084 | 0.0041 | 0.0041 | TRUE | 0.023 |
| Avp | 1 | 0 | 0 | 0 | 1 | 0 | 1 | 0 | 0 | 0 | 1 | 0 | 0.36 | 0 | 0 | 0.954 | -0.067 | 0.98557668 | NA | 0 | 9.2518 | 8.0179 | 8.0179 | TRUE | 0.5204 |
| Avpi1 | 892 | 979 | 909 | 1542 | 1101 | 1074 | 947 | 978 | 913 | 1212 | 1185 | 1176 | 1068.4 | 946 | 1191 | 1.259 | 0.333 | 3.36E-05 | 0.00055305 | 0 | 0.01 | 0.0037 | 0.0037 | TRUE | 0.0114 |
| Avpr1a | 3 | 1 | 1 | 8 | 2 | 1 | 3 | 1 | 1 | 6 | 2 | 1 | 2.45 | 2 | 3 | 1.903 | 0.929 | 0.47789509 | NA | 0 | 1.3493 | 0.7867 | 0.7867 | TRUE | 1.2795 |
| Avpr1b | 4 | 6 | 3 | 5 | 2 | 4 | 4 | 6 | 3 | 4 | 2 | 4 | 3.95 | 4 | 3 | 0.793 | -0.335 | 0.72032437 | NA | 0 | 0.8404 | 0.3758 | 0.3758 | TRUE | 0.1815 |
| Avpr2 | 1 | 1 | 2 | 1 | 0 | 1 | 1 | 1 | 2 | 1 | 0 | 1 | 0.99 | 1 | 1 | 0.467 | -1.098 | 0.55705614 | NA | 0 | 3.3291 | 1.3772 | 1.3772 | TRUE | 0.1944 |
| AW011738 | 1 | 1 | 2 | 1 | 3 | 3 | 1 | 1 | 2 | 1 | 3 | 3 | 1.89 | 1 | 2 | 1.747 | 0.805 | 0.56085494 | NA | 0 | 1.7459 | 0.7981 | 0.7981 | TRUE | 0.521 |
| AW046200 | 1 | 2 | 4 | 4 | 1 | 0 | 1 | 2 | 4 | 3 | 1 | 0 | 1.88 | 2 | 1 | 0.625 | -0.678 | 0.64973768 | NA | 0 | 1.7567 | 1.0593 | 1.0593 | TRUE | 0.8387 |
| AW112010 | 0 | 0 | 0 | 0 | 0 | 0 | 0 | 0 | 0 | 0 | 0 | 0 | 0 | 0 | 0 | NA | NA | NA | NA | NA | NA | NA | NA | NA | NA |
| AW146154 | 247 | 198 | 172 | 279 | 197 | 155 | 262 | 198 | 173 | 219 | 212 | 170 | 205.63 | 211 | 200 | 0.952 | -0.071 | 0.72828785 | 0.85395791 | 0.0275 | 0.0229 | 0.0252 | 0.0252 | TRUE | 0.4923 |
| AW209491 | 686 | 717 | 690 | 1091 | 748 | 714 | 729 | 716 | 693 | 858 | 805 | 782 | 763.6 | 713 | 815 | 1.145 | 0.195 | 0.03317093 | 0.12331393 | 0 | 0.0112 | 0.0047 | 0.0047 | TRUE | 0.0265 |
| AW495222 | 1 | 1 | 1 | 4 | 3 | 1 | 1 | 1 | 1 | 3 | 3 | 1 | 1.76 | 1 | 2 | 2.462 | 1.3 | 0.36410688 | NA | 0 | 1.8837 | 0.7833 | 0.7833 | TRUE | 0.233 |
| AW549542 | 13 | 20 | 11 | 33 | 33 | 33 | 14 | 20 | 11 | 26 | 36 | 36 | 23.73 | 15 | 33 | 2.162 | 1.113 | 0.00963416 | 0.05119607 | 0 | 0.1457 | 0.0838 | 0.0838 | TRUE | 0.4034 |
| AW549877 | 3677 | 5169 | 5434 | 5202 | 4202 | 4828 | 3906 | 5162 | 5456 | 4089 | 4521 | 5285 | 4736.63 | 4841 | 4632 | 0.957 | -0.064 | 0.66607396 | 0.81310515 | 0.0236 | 0.0076 | 0.0157 | 0.0157 | TRUE | 0.3481 |
| AW551984 | 246 | 247 | 225 | 157 | 130 | 141 | 261 | 247 | 226 | 123 | 140 | 154 | 191.92 | 245 | 139 | 0.567 | -0.819 | 5.01E-07 | 1.50E-05 | 0.0032 | 0.0241 | 0.0135 | 0.0135 | TRUE | 0.114 |
| AW554918 | 356 | 425 | 473 | 585 | 400 | 456 | 378 | 424 | 475 | 460 | 430 | 499 | 444.49 | 426 | 463 | 1.087 | 0.12 | 0.36082818 | 0.57917586 | 0.0067 | 0.0143 | 0.0102 | 0.0102 | TRUE | 0.1175 |
| Awat1 | 0 | 0 | 0 | 0 | 0 | 0 | 0 | 0 | 0 | 0 | 0 | 0 | 0 | 0 | 0 | NA | NA | NA | NA | NA | NA | NA | NA | NA | NA |
| Awat2 | 35 | 35 | 32 | 44 | 24 | 21 | 37 | 35 | 32 | 35 | 26 | 23 | 31.28 | 35 | 28 | 0.808 | -0.307 | 0.38158473 | 0.59581767 | 0 | 0.1122 | 0.0567 | 0.0567 | TRUE | 0.3481 |
| Axin1 | 4105 | 4036 | 4049 | 5002 | 3861 | 3854 | 4360 | 4031 | 4066 | 3932 | 4154 | 4219 | 4126.97 | 4152 | 4102 | 0.988 | -0.018 | 0.80350655 | 0.89855367 | 0.0014 | 0.0077 | 0.0035 | 0.0035 | TRUE | 0.0233 |
| Axin2 | 811 | 769 | 872 | 867 | 638 | 605 | 861 | 768 | 876 | 682 | 686 | 662 | 755.88 | 835 | 677 | 0.811 | -0.303 | 0.0019295 | 0.01496171 | 0.0015 | 0.0113 | 0.0055 | 0.0055 | TRUE | 0.0589 |
| Axl | 279 | 350 | 315 | 253 | 198 | 260 | 296 | 350 | 316 | 199 | 213 | 285 | 276.45 | 321 | 232 | 0.722 | -0.471 | 0.00706369 | 0.04038003 | 0.0175 | 0.0188 | 0.0183 | 0.0183 | TRUE | 0.4231 |
| AY074887 | 1 | 2 | 2 | 3 | 4 | 4 | 1 | 2 | 2 | 2 | 4 | 4 | 2.68 | 2 | 3 | 2.142 | 1.099 | 0.3408132 | NA | 0 | 1.234 | 0.5237 | 0.5237 | TRUE | 0.2075 |
| AY358078 | 1 | 4 | 3 | 3 | 7 | 2 | 1 | 4 | 3 | 2 | 8 | 2 | 3.36 | 3 | 4 | 1.472 | 0.558 | 0.61383291 | NA | 0 | 0.9879 | 0.5682 | 0.5682 | TRUE | 0.9091 |
| AY512915 | 1 | 1 | 2 | 0 | 0 | 0 | 1 | 1 | 2 | 0 | 0 | 0 | 0.68 | 1 | 0 | 0.132 | -2.92 | 0.23843536 | NA | 0 | 4.8647 | 3.0465 | 3.0465 | TRUE | 0.1113 |
| AY512931 | 34 | 20 | 13 | 17 | 20 | 13 | 36 | 20 | 13 | 13 | 22 | 14 | 19.71 | 23 | 16 | 0.708 | -0.497 | 0.34396267 | 0.56375136 | 0.119 | 0.1741 | 0.1467 | 0.1467 | TRUE | 0.4859 |
| AY761184 | 2 | 0 | 0 | 1 | 0 | 3 | 2 | 0 | 0 | 1 | 0 | 3 | 1.03 | 1 | 1 | 1.922 | 0.942 | 0.68925662 | NA | 2.2856 | 3.1979 | 2.9062 | 2.9062 | TRUE | 0.8765 |
| AY761185 | 0 | 0 | 0 | 0 | 0 | 0 | 0 | 0 | 0 | 0 | 0 | 0 | 0 | 0 | 0 | NA | NA | NA | NA | NA | NA | NA | NA | NA | NA |
| Aym1 | 3 | 9 | 6 | 2 | 3 | 10 | 3 | 9 | 6 | 2 | 3 | 11 | 5.66 | 6 | 5 | 0.844 | -0.245 | 0.79893397 | 0.89640568 | 0.3829 | 0.5892 | 0.4846 | 0.4846 | TRUE | 0.8199 |
| Azgp1 | 0 | 0 | 0 | 0 | 0 | 0 | 0 | 0 | 0 | 0 | 0 | 0 | 0 | 0 | 0 | NA | NA | NA | NA | NA | NA | NA | NA | NA | NA |
| Azi1 | 3268 | 3479 | 3624 | 3577 | 2901 | 2960 | 3471 | 3475 | 3639 | 2812 | 3121 | 3240 | 3293 | 3528 | 3058 | 0.866 | -0.207 | 0.01233797 | 0.06139101 | 0.0025 | 0.0079 | 0.0046 | 0.0046 | TRUE | 0.0611 |
| Azi2 | 2227 | 2098 | 1961 | 3052 | 2148 | 2078 | 2365 | 2095 | 1969 | 2399 | 2311 | 2275 | 2235.81 | 2143 | 2328 | 1.087 | 0.12 | 0.20083474 | 0.40850494 | 0.0042 | 0.0084 | 0.0059 | 0.0059 | TRUE | 0.0993 |
| Azin1 | 7293 | 6984 | 6597 | 11537 | 7979 | 6591 | 7746 | 6975 | 6624 | 9069 | 8585 | 7215 | 7702.47 | 7115 | 8290 | 1.165 | 0.221 | 0.04836323 | 0.1603163 | 0.0098 | 0.0073 | 0.0089 | 0.0089 | TRUE | 0.1569 |
| B020004C17 | 0 | 0 | 0 | 1 | 0 | 0 | 0 | 0 | 0 | 1 | 0 | 0 | 0.13 | 0 | 0 | 1.859 | 0.894 | 0.82650734 | NA | 0 | 25.1515 | 10 | 10 | TRUE | 0.2563 |
| B020004J07 | 44 | 7 | 8 | 88 | 67 | 26 | 47 | 7 | 8 | 69 | 72 | 28 | 38.58 | 21 | 56 | 2.762 | 1.465 | 0.0332419 | 0.12344978 | 0.5853 | 0.0923 | 0.3078 | 0.3078 | TRUE | 6.6024 |
| B020014A21 | 0 | 0 | 0 | 0 | 0 | 0 | 0 | 0 | 0 | 0 | 0 | 0 | 0 | 0 | 0 | NA | NA | NA | NA | NA | NA | NA | NA | NA | NA |
| B020018J22 | 0 | 0 | 0 | 1 | 1 | 0 | 0 | 0 | 0 | 1 | 1 | 0 | 0.31 | 0 | 1 | 3.32 | 1.731 | 0.65383052 | NA | 0 | 10.6224 | 8.9541 | 8.9541 | TRUE | 0.202 |
| B020031M1 | 0 | 0 | 0 | 0 | 0 | 0 | 0 | 0 | 0 | 0 | 0 | 0 | 0 | 0 | 0 | NA | NA | NA | NA | NA | NA | NA | NA | NA | NA |
| B130006D01 | 6 | 15 | 20 | 4 | 4 | 9 | 6 | 15 | 20 | 3 | 4 | 10 | 9.79 | 14 | 6 | 0.408 | -1.292 | 0.08378157 | 0.23193226 | 0.2081 | 0.3434 | 0.2788 | 0.2788 | TRUE | 0.2931 |
| B130024G19 | 0 | 0 | 0 | 0 | 1 | 0 | 0 | 0 | 0 | 0 | 1 | 0 | 0.18 | 0 | 0 | 1.859 | 0.894 | 0.82650734 | NA | 0 | 18.3785 | 10 | 10 | TRUE | 0.5204 |
| B130034C11 | 4 | 6 | 5 | 8 | 8 | 1 | 4 | 6 | 5 | 6 | 9 | 1 | 5.21 | 5 | 5 | 1.054 | 0.076 | 0.93320178 | 0.96698367 | 0 | 0.6394 | 0.4024 | 0.4024 | TRUE | 0.5318 |
| B230118H07 | 506 | 565 | 670 | 833 | 559 | 512 | 537 | 564 | 673 | 655 | 601 | 560 | 598.54 | 591 | 605 | 1.025 | 0.035 | 0.78383889 | 0.88838079 | 0.0085 | 0.0124 | 0.0103 | 0.0103 | TRUE | 0.1699 |
| B230119M0 | 0 | 1 | 3 | 3 | 1 | 0 | 0 | 1 | 3 | 2 | 1 | 0 | 1.24 | 1 | 1 | 0.882 | -0.181 | 0.92424706 | NA | 0 | 2.6618 | 1.8311 | 1.8311 | TRUE | 0.3955 |
| B230120H23 | 1021 | 1331 | 1431 | 1572 | 1122 | 1224 | 1084 | 1329 | 1437 | 1236 | 1207 | 1340 | 1272.25 | 1283 | 1261 | 0.982 | -0.026 | 0.83275887 | 0.91523598 | 0.0109 | 0.0095 | 0.0103 | 0.0103 | TRUE | 0.2202 |
| B230206H07 | 4 | 1 | 5 | 2 | 0 | 0 | 4 | 1 | 5 | 2 | 0 | 0 | 1.97 | 3 | 1 | 0.173 | -2.532 | 0.13982677 | NA | 0 | 1.6764 | 1.1476 | 1.1476 | TRUE | 0.9708 |
| B230208H11 | 11 | 6 | 18 | 22 | 15 | 18 | 12 | 6 | 18 | 17 | 16 | 20 | 14.81 | 12 | 18 | 1.485 | 0.571 | 0.29848772 | 0.51905693 | 0.0364 | 0.2293 | 0.1467 | 0.1467 | TRUE | 0.3193 |
| B230209E15 | 0 | 0 | 0 | 0 | 0 | 0 | 0 | 0 | 0 | 0 | 0 | 0 | 0 | 0 | 0 | NA | NA | NA | NA | NA | NA | NA | NA | NA | NA |
| B230209K01 | 7 | 9 | 9 | 8 | 3 | 5 | 7 | 9 | 9 | 6 | 3 | 5 | 6.74 | 8 | 5 | 0.595 | -0.748 | 0.30143894 | 0.52210575 | 0 | 0.4956 | 0.221 | 0.221 | TRUE | 0.1772 |
| B230214G05 | 0 | 0 | 0 | 0 | 0 | 0 | 0 | 0 | 0 | 0 | 0 | 0 | 0 | 0 | 0 | NA | NA | NA | NA | NA | NA | NA | NA | NA | NA |
| B230216G23 | 4 | 1 | 1 | 7 | 5 | 0 | 4 | 1 | 1 | 6 | 5 | 0 | 2.86 | 2 | 4 | 1.774 | 0.827 | 0.54918354 | NA | 0.7703 | 1.1606 | 0.9942 | 0.9942 | TRUE | 1.0572 |
| B230216N24 | 1 | 0 | 3 | 2 | 5 | 2 | 1 | 0 | 3 | 2 | 5 | 2 | 2.2 | 1 | 3 | 2.214 | 1.147 | 0.42343921 | NA | 0.2746 | 1.5026 | 0.9403 | 0.9403 | TRUE | 0.7226 |
| B230217C12 | 400 | 353 | 361 | 463 | 327 | 315 | 425 | 353 | 362 | 364 | 352 | 345 | 366.75 | 380 | 354 | 0.932 | -0.102 | 0.42020194 | 0.63064279 | 0.0029 | 0.0159 | 0.0088 | 0.0088 | TRUE | 0.1214 |
| B230217O12 | 7 | 10 | 11 | 15 | 18 | 12 | 7 | 10 | 11 | 12 | 19 | 13 | 12.13 | 9 | 15 | 1.543 | 0.626 | 0.26245082 | 0.48186844 | 0 | 0.2786 | 0.1377 | 0.1377 | TRUE | 0.3186 |
| B230219D22 | 3866 | 5452 | 5941 | 5474 | 4378 | 4821 | 4106 | 5445 | 5965 | 4303 | 4710 | 5277 | 4967.96 | 5172 | 4763 | 0.921 | -0.119 | 0.42092965 | 0.63116133 | 0.0206 | 0.0076 | 0.0155 | 0.0155 | TRUE | 0.3959 |
| B230312C02 | 0 | 1 | 1 | 1 | 1 | 1 | 0 | 1 | 1 | 1 | 1 | 1 | 0.83 | 1 | 1 | 1.45 | 0.536 | 0.78654731 | NA | 0 | 3.9925 | 1.5656 | 1.5656 | TRUE | 0.2233 |
| B230319C09 | 0 | 0 | 0 | 2 | 1 | 1 | 0 | 0 | 0 | 2 | 1 | 1 | 0.62 | 0 | 1 | 6.676 | 2.739 | 0.27800695 | NA | 0 | 5.2881 | 3.2034 | 3.2034 | TRUE | 0.07 |
| B230323A14 | 0 | 0 | 0 | 0 | 0 | 0 | 0 | 0 | 0 | 0 | 0 | 0 | 0 | 0 | 0 | NA | NA | NA | NA | NA | NA | NA | NA | NA | NA |
| B230378P21 | 0 | 0 | 0 | 0 | 0 | 0 | 0 | 0 | 0 | 0 | 0 | 0 | 0 | 0 | 0 | NA | NA | NA | NA | NA | NA | NA | NA | NA | NA |
| B2m | 453 | 524 | 542 | 680 | 455 | 501 | 481 | 523 | 544 | 535 | 490 | 548 | 520.21 | 516 | 524 | 1.015 | 0.022 | 0.84023082 | 0.91912023 | 0.0016 | 0.0132 | 0.0068 | 0.0068 | TRUE | 0.0408 |
| B330016D10 | 34 | 48 | 53 | 32 | 52 | 49 | 36 | 48 | 53 | 25 | 56 | 54 | 45.34 | 46 | 45 | 0.971 | -0.042 | 0.91271928 | 0.95682342 | 0.0874 | 0.0796 | 0.083 | 0.083 | TRUE | 1.3388 |
| B3galnt1 | 733 | 701 | 738 | 922 | 629 | 703 | 779 | 700 | 741 | 725 | 677 | 770 | 731.8 | 740 | 724 | 0.978 | -0.032 | 0.7533601 | 0.86987269 | 0.0019 | 0.0114 | 0.006 | 0.006 | TRUE | 0.0367 |
| B3galnt2 | 2535 | 2527 | 2617 | 2774 | 2135 | 2204 | 2693 | 2524 | 2628 | 2181 | 2297 | 2413 | 2455.76 | 2615 | 2297 | 0.878 | -0.188 | 0.01443966 | 0.06884592 | 0.0014 | 0.0082 | 0.0038 | 0.0038 | TRUE | 0.0243 |
| B3galt1 | 63 | 50 | 63 | 57 | 33 | 42 | 67 | 50 | 63 | 45 | 36 | 46 | 51.07 | 60 | 42 | 0.703 | -0.508 | 0.07410153 | 0.21385716 | 0 | 0.0714 | 0.0381 | 0.0381 | TRUE | 0.1886 |
| B3galt2 | 0 | 0 | 0 | 0 | 0 | 0 | 0 | 0 | 0 | 0 | 0 | 0 | 0 | 0 | 0 | NA | NA | NA | NA | NA | NA | NA | NA | NA | NA |
| B3galt4 | 65 | 111 | 95 | 138 | 103 | 115 | 69 | 111 | 95 | 108 | 111 | 126 | 103.41 | 92 | 115 | 1.25 | 0.322 | 0.16580055 | 0.36290577 | 0.0185 | 0.0388 | 0.0292 | 0.0292 | TRUE | 0.44 |
| B3galt5 | 95 | 26 | 23 | 16 | 13 | 12 | 101 | 26 | 23 | 13 | 14 | 13 | 31.61 | 50 | 13 | 0.265 | -1.917 | 0.0018172 | 0.01425917 | 0.3388 | 0.1111 | 0.2249 | 0.2249 | TRUE | 6.3487 |
| B3galt6 | 248 | 211 | 185 | 422 | 258 | 224 | 263 | 211 | 186 | 332 | 278 | 245 | 252.41 | 220 | 285 | 1.299 | 0.377 | 0.04576181 | 0.15478347 | 0.0233 | 0.02 | 0.0217 | 0.0217 | TRUE | 0.3245 |

| | | | | | | | | | | | | | | | | | | | | | | | | | |
|---|---|---|---|---|---|---|---|---|---|---|---|---|---|---|---|---|---|---|---|---|---|---|---|---|---|
| B3galtl | 1168 | 1106 | 1067 | 1268 | 910 | 931 | 1241 | 1105 | 1071 | 997 | 979 | 1019 | 1068.6 | 1139 | 998 | 0.877 | -0.19 | 0.04320308 | 0.14861733 | 0.0023 | 0.01 | 0.0054 | 0.0054 | TRUE | 0.0728 |
| B3gat1 | 27 | 30 | 27 | 50 | 33 | 31 | 29 | 30 | 27 | 39 | 36 | 34 | 32.42 | 29 | 36 | 1.272 | 0.347 | 0.29035252 | 0.51138358 | 0 | 0.1085 | 0.0465 | 0.0465 | TRUE | 0.0463 |
| B3gat2 | 4 | 4 | 7 | 6 | 4 | 8 | 4 | 4 | 7 | 5 | 4 | 9 | 5.51 | 5 | 6 | 1.153 | 0.205 | 0.80088497 | 0.89725514 | 0 | 0.605 | 0.2932 | 0.2932 | TRUE | 0.3669 |
| B3gat3 | 706 | 814 | 865 | 1040 | 728 | 725 | 750 | 813 | 869 | 818 | 783 | 794 | 804.31 | 811 | 798 | 0.985 | -0.022 | 0.82205431 | 0.90911907 | 0.0017 | 0.011 | 0.0055 | 0.0055 | TRUE | 0.0503 |
| B3gnt1 | 284 | 315 | 322 | 548 | 407 | 330 | 302 | 315 | 323 | 431 | 438 | 361 | 361.59 | 313 | 410 | 1.31 | 0.39 | 0.00263224 | 0.01912281 | 0.0036 | 0.016 | 0.0093 | 0.0093 | TRUE | 0.1207 |
| B3gnt2 | 1913 | 2387 | 2322 | 2314 | 1795 | 1849 | 2032 | 2384 | 2332 | 1819 | 1931 | 2024 | 2086.97 | 2249 | 1925 | 0.855 | -0.225 | 0.01895716 | 0.08357464 | 0.0043 | 0.0085 | 0.0062 | 0.0062 | TRUE | 0.0861 |
| B3gnt3 | 43 | 45 | 45 | 101 | 82 | 83 | 46 | 45 | 45 | 79 | 88 | 91 | 65.71 | 45 | 86 | 1.899 | 0.925 | 0.00012445 | 0.00164804 | 0 | 0.057 | 0.0249 | 0.0249 | TRUE | 0.0503 |
| B3gnt4 | 20 | 15 | 17 | 22 | 18 | 12 | 21 | 15 | 17 | 17 | 19 | 13 | 17.18 | 18 | 16 | 0.938 | -0.093 | 0.84056202 | 0.91930825 | 0 | 0.1986 | 0.0948 | 0.0948 | TRUE | 0.1435 |
| B3gnt5 | 1020 | 1813 | 1906 | 2068 | 1503 | 1551 | 1083 | 1811 | 1914 | 1626 | 1617 | 1698 | 1624.76 | 1603 | 1647 | 1.027 | 0.039 | 0.83861028 | 0.91812865 | 0.0451 | 0.0089 | 0.0258 | 0.0258 | TRUE | 0.9667 |
| B3gnt6 | 1 | 1 | 3 | 4 | 0 | 4 | 1 | 1 | 3 | 3 | 0 | 4 | 2.1 | 2 | 2 | 1.488 | 0.573 | 0.69340587 | NA | 0 | 1.5761 | 1.0295 | 1.0295 | TRUE | 0.5431 |
| B3gnt7 | 5289 | 6207 | 6613 | 5645 | 4085 | 4127 | 5618 | 6199 | 6640 | 4438 | 4395 | 4518 | 5301.25 | 6152 | 4450 | 0.723 | -0.467 | 3.17E-08 | 1.34E-06 | 0.0033 | 0.0075 | 0.005 | 0.005 | TRUE | 0.0703 |
| B3gnt8 | 18 | 21 | 14 | 24 | 12 | 9 | 19 | 21 | 14 | 19 | 13 | 10 | 15.96 | 18 | 14 | 0.78 | -0.358 | 0.47475186 | 0.67524193 | 0 | 0.2133 | 0.1182 | 0.1182 | TRUE | 0.5295 |
| B3gnt9 | 71 | 104 | 120 | 76 | 65 | 87 | 75 | 104 | 120 | 60 | 70 | 95 | 87.45 | 100 | 75 | 0.746 | -0.423 | 0.1294871 | 0.30966954 | 0.0443 | 0.0446 | 0.0444 | 0.0444 | TRUE | 0.3542 |
| B3gntl1 | 1029 | 1101 | 1168 | 1312 | 1076 | 995 | 1093 | 1100 | 1173 | 1031 | 1158 | 1089 | 1107.27 | 1122 | 1093 | 0.973 | -0.039 | 0.66097619 | 0.80907385 | 0.0016 | 0.0099 | 0.0048 | 0.0048 | TRUE | 0.0324 |
| B430010I23F | 0 | 1 | 0 | 2 | 0 | 0 | 0 | 1 | 0 | 2 | 0 | 0 | 0.43 | 0 | 1 | 1.493 | 0.578 | 0.86962266 | NA | 0 | 7.6954 | 7.0545 | 7.0545 | TRUE | 0.9718 |
| B430212C06 | 5 | 2 | 3 | 2 | 6 | 1 | 5 | 2 | 3 | 2 | 6 | 1 | 3.24 | 3 | 3 | 0.873 | -0.196 | 0.86381622 | NA | 0.2354 | 1.0235 | 0.6257 | 0.6257 | TRUE | 1.1754 |
| B430306N03 | 3 | 4 | 3 | 33 | 19 | 14 | 3 | 4 | 3 | 26 | 20 | 15 | 11.98 | 3 | 20 | 6.088 | 2.606 | 0.00010132 | 0.0013944 | 0 | 0.2818 | 0.1501 | 0.1501 | TRUE | 0.081 |
| B430319G15 | 1 | 3 | 2 | 10 | 5 | 2 | 1 | 3 | 2 | 8 | 5 | 2 | 3.58 | 2 | 5 | 2.571 | 1.362 | 0.20722988 | NA | 0 | 0.9264 | 0.4956 | 0.4956 | TRUE | 0.1308 |
| B4galnt1 | 248 | 355 | 312 | 508 | 394 | 389 | 263 | 355 | 313 | 399 | 424 | 426 | 363.39 | 310 | 416 | 1.339 | 0.422 | 0.00306249 | 0.02159216 | 0.0078 | 0.016 | 0.0118 | 0.0118 | TRUE | 0.196 |
| B4galnt2 | 203 | 409 | 424 | 259 | 173 | 322 | 216 | 408 | 426 | 204 | 186 | 352 | 298.68 | 350 | 247 | 0.706 | -0.502 | 0.0996057 | 0.26057075 | 0.1186 | 0.0179 | 0.0636 | 0.0636 | TRUE | 1.5145 |
| B4galnt3 | 1700 | 1582 | 1585 | 1750 | 1421 | 1467 | 1806 | 1580 | 1592 | 1376 | 1529 | 1606 | 1581.27 | 1659 | 1504 | 0.906 | -0.143 | 0.16063285 | 0.35521608 | 0.005 | 0.009 | 0.0068 | 0.0068 | TRUE | 0.0719 |
| B4galnt4 | 762 | 1127 | 1122 | 1074 | 832 | 832 | 809 | 1126 | 1127 | 844 | 895 | 911 | 951.96 | 1021 | 883 | 0.865 | -0.209 | 0.14690421 | 0.33594245 | 0.0171 | 0.0104 | 0.0139 | 0.0139 | TRUE | 0.3898 |
| B4galt1 | 1483 | 1635 | 1583 | 2314 | 1688 | 1681 | 1575 | 1633 | 1589 | 1819 | 1816 | 1840 | 1712.16 | 1599 | 1825 | 1.141 | 0.19 | 0.00540883 | 0.0331145 | 0 | 0.0088 | 0.0028 | 0.0028 | TRUE | 0.0041 |
| B4galt2 | 520 | 731 | 646 | 763 | 622 | 679 | 552 | 730 | 649 | 600 | 669 | 743 | 657.23 | 644 | 671 | 1.041 | 0.057 | 0.68565369 | 0.82638927 | 0.0139 | 0.0119 | 0.013 | 0.013 | TRUE | 0.1805 |
| B4galt3 | 1213 | 1192 | 1155 | 1611 | 1137 | 1128 | 1288 | 1190 | 1160 | 1266 | 1223 | 1235 | 1227.2 | 1213 | 1241 | 1.024 | 0.034 | 0.67940619 | 0.82193713 | 8.00E-04 | 0.0096 | 0.0042 | 0.0042 | TRUE | 0.0354 |
| B4galt4 | 196 | 387 | 348 | 526 | 343 | 430 | 208 | 387 | 349 | 413 | 369 | 471 | 366.23 | 315 | 418 | 1.326 | 0.408 | 0.07008793 | 0.20596559 | 0.0541 | 0.0159 | 0.0337 | 0.0337 | TRUE | 0.9859 |
| B4galt5 | 3906 | 3842 | 3658 | 5138 | 3717 | 3588 | 4149 | 3837 | 3673 | 4039 | 3999 | 3928 | 3937.48 | 3886 | 3989 | 1.027 | 0.038 | 0.61577889 | 0.78272604 | 0.0015 | 0.0077 | 0.0038 | 0.0038 | TRUE | 0.0424 |
| B4galt6 | 8556 | 8348 | 7554 | 8018 | 5968 | 6313 | 9088 | 8337 | 7585 | 6303 | 6421 | 6911 | 7440.85 | 8337 | 6545 | 0.785 | -0.349 | 0.00015397 | 0.00194735 | 0.0051 | 0.0073 | 0.006 | 0.006 | TRUE | 0.076 |
| B4galt7 | 434 | 502 | 522 | 586 | 432 | 433 | 461 | 501 | 524 | 461 | 465 | 474 | 480.99 | 495 | 467 | 0.941 | -0.088 | 0.40944667 | 0.62130006 | 0 | 0.0138 | 0.0061 | 0.0061 | TRUE | 0.0427 |
| B630005N14 | 1419 | 1738 | 1725 | 2293 | 1683 | 1983 | 1507 | 1736 | 1732 | 1803 | 1811 | 2171 | 1793.19 | 1658 | 1928 | 1.162 | 0.217 | 0.05374403 | 0.17235682 | 0.0082 | 0.0087 | 0.0085 | 0.0085 | TRUE | 0.1457 |
| B630019K06 | 39 | 34 | 42 | 85 | 56 | 49 | 41 | 34 | 42 | 67 | 60 | 54 | 49.71 | 39 | 60 | 1.544 | 0.626 | 0.02543982 | 0.10298688 | 0 | 0.0732 | 0.0355 | 0.0355 | TRUE | 0.1031 |
| B830017H08 | 1 | 4 | 1 | 4 | 6 | 3 | 1 | 4 | 1 | 3 | 6 | 3 | 3.16 | 2 | 4 | 2.099 | 1.07 | 0.34416731 | NA | 0 | 1.0503 | 0.5552 | 0.5552 | TRUE | 0.6739 |
| B930003M22 | 23 | 41 | 52 | 33 | 30 | 46 | 24 | 41 | 52 | 26 | 32 | 50 | 37.69 | 39 | 36 | 0.916 | -0.127 | 0.75860893 | 0.87264094 | 0.0997 | 0.0943 | 0.0967 | 0.0967 | TRUE | 0.3079 |
| B930018H19 | 1 | 0 | 0 | 1 | 0 | 0 | 1 | 0 | 0 | 1 | 0 | 0 | 0.31 | 0 | 0 | 0.954 | -0.067 | 0.98625928 | NA | 0 | 10.7014 | 9.0384 | 9.0384 | TRUE | 0.4614 |
| B930025P03 | 0 | 0 | 0 | 0 | 0 | 0 | 0 | 0 | 0 | 0 | 0 | 0 | 0 | 0 | 0 | NA | NA | NA | NA | NA | NA | NA | NA | NA | NA |
| B930041F14 | 179 | 315 | 340 | 304 | 257 | 295 | 190 | 315 | 341 | 239 | 277 | 323 | 280.76 | 282 | 280 | 0.989 | -0.016 | 0.94482251 | 0.97244458 | 0.0535 | 0.0186 | 0.0352 | 0.0352 | TRUE | 0.9055 |
| B930059L03 | 10 | 9 | 15 | 18 | 13 | 9 | 11 | 9 | 15 | 14 | 14 | 10 | 12.11 | 12 | 13 | 1.1 | 0.138 | 0.80182498 | 0.89762453 | 0 | 0.2789 | 0.1357 | 0.1357 | TRUE | 0.2744 |
| B930092H01 | 13 | 6 | 11 | 4 | 4 | 3 | 14 | 6 | 11 | 3 | 4 | 3 | 6.93 | 10 | 3 | 0.346 | -1.529 | 0.04899249 | 0.16199362 | 0 | 0.4823 | 0.2481 | 0.2481 | TRUE | 0.237 |
| B9d1 | 987 | 1175 | 1072 | 1654 | 1222 | 1133 | 1048 | 1173 | 1076 | 1300 | 1315 | 1240 | 1192.26 | 1099 | 1285 | 1.169 | 0.225 | 0.00947555 | 0.05050637 | 0.0013 | 0.0097 | 0.0046 | 0.0046 | TRUE | 0.0417 |
| B9d2 | 148 | 152 | 146 | 195 | 155 | 157 | 157 | 152 | 147 | 153 | 167 | 172 | 157.92 | 152 | 164 | 1.077 | 0.108 | 0.49987726 | 0.69567318 | 0 | 0.0278 | 0.012 | 0.012 | TRUE | 0.0381 |
| Baalc | 22 | 15 | 16 | 11 | 9 | 17 | 23 | 15 | 16 | 9 | 10 | 19 | 15.23 | 18 | 13 | 0.672 | -0.574 | 0.28546238 | 0.50675641 | 0.046 | 0.2233 | 0.1399 | 0.1399 | TRUE | 0.7451 |
| Baat | 12 | 5 | 2 | 8 | 4 | 3 | 13 | 5 | 2 | 6 | 4 | 3 | 5.6 | 7 | 4 | 0.715 | -0.484 | 0.59931386 | 0.77143379 | 0.2689 | 0.5948 | 0.4292 | 0.4292 | TRUE | 0.5837 |
| Baat1 | 4 | 1 | 7 | 13 | 11 | 6 | 4 | 1 | 7 | 10 | 12 | 7 | 6.82 | 4 | 10 | 2.339 | 1.226 | 0.13951068 | 0.32412923 | 0.0424 | 0.4902 | 0.3198 | 0.3198 | TRUE | 0.4999 |
| Babam1 | 2309 | 2534 | 2525 | 2900 | 2233 | 2345 | 2453 | 2531 | 2535 | 2280 | 2403 | 2567 | 2461.33 | 2506 | 2417 | 0.964 | -0.054 | 0.49087756 | 0.68729309 | 0.0016 | 0.0082 | 0.0039 | 0.0039 | TRUE | 0.0359 |
| Bace1 | 269 | 222 | 305 | 371 | 235 | 222 | 286 | 222 | 306 | 292 | 253 | 243 | 266.87 | 271 | 263 | 0.97 | -0.044 | 0.79645356 | 0.89536272 | 0.015 | 0.0192 | 0.0171 | 0.0171 | TRUE | 0.2878 |
| Bace2 | 7 | 5 | 5 | 13 | 8 | 10 | 7 | 5 | 5 | 10 | 9 | 11 | 7.87 | 6 | 10 | 1.712 | 0.776 | 0.24634194 | 0.46336376 | 0 | 0.4255 | 0.1856 | 0.1856 | TRUE | 0.1284 |
| Bach1 | 2360 | 2952 | 3124 | 2522 | 1908 | 2264 | 2507 | 2948 | 3137 | 1983 | 2053 | 2478 | 2517.59 | 2864 | 2171 | 0.758 | -0.4 | 0.00152513 | 0.01240557 | 0.0134 | 0.0082 | 0.0111 | 0.0111 | TRUE | 0.1846 |
| Bach2 | 225 | 222 | 238 | 193 | 154 | 158 | 239 | 222 | 239 | 152 | 166 | 173 | 198.34 | 233 | 164 | 0.699 | -0.517 | 0.00047743 | 0.00500001 | 0 | 0.0235 | 0.0106 | 0.0106 | TRUE | 0.0466 |
| Bad | 193 | 213 | 185 | 392 | 244 | 259 | 205 | 213 | 186 | 308 | 263 | 284 | 242.95 | 201 | 285 | 1.418 | 0.504 | 0.00046867 | 0.0049378 | 0 | 0.0205 | 0.0107 | 0.0107 | TRUE | 0.0622 |
| Bag1 | 4436 | 4898 | 4658 | 6110 | 4354 | 4388 | 4712 | 4892 | 4677 | 4803 | 4685 | 4803 | 4761.97 | 4760 | 4764 | 1.001 | 0.001 | 0.98503179 | 0.99199132 | 2.00E-04 | 0.0076 | 0.0024 | 0.0024 | TRUE | 0.0071 |
| Bag2 | 420 | 571 | 511 | 620 | 474 | 506 | 446 | 570 | 513 | 487 | 510 | 554 | 513.46 | 510 | 517 | 1.013 | 0.018 | 0.88720425 | 0.94451405 | 0.0076 | 0.0133 | 0.0102 | 0.0102 | TRUE | 0.1382 |
| Bag3 | 3500 | 2956 | 2979 | 3153 | 2326 | 2306 | 3718 | 2952 | 2991 | 2479 | 2503 | 2524 | 2861.1 | 3220 | 2502 | 0.777 | -0.364 | 0.00083701 | 0.00776546 | 0.0085 | 0.0081 | 0.0082 | 0.0082 | TRUE | 0.2214 |
| Bag4 | 8242 | 8836 | 8962 | 9100 | 6916 | 7375 | 8754 | 8825 | 8999 | 7154 | 7441 | 8073 | 8207.64 | 8859 | 7556 | 0.853 | -0.23 | 0.00159707 | 0.01287122 | 0.0019 | 0.0073 | 0.0037 | 0.0037 | TRUE | 0.0437 |
| Bag5 | 408 | 484 | 480 | 721 | 564 | 454 | 433 | 483 | 482 | 567 | 607 | 497 | 511.55 | 466 | 557 | 1.195 | 0.256 | 0.03518077 | 0.12856141 | 0.0049 | 0.0133 | 0.0087 | 0.0087 | TRUE | 0.1002 |
| Bag6 | 14453 | 14000 | 13651 | 19206 | 14136 | 13271 | 15352 | 13982 | 13707 | 15098 | 15209 | 14527 | 14645.92 | 14347 | 14945 | 1.042 | 0.059 | 0.41839282 | 0.62905885 | 0.0021 | 0.0071 | 0.0038 | 0.0038 | TRUE | 0.0459 |
| Bahcc1 | 2853 | 4297 | 4924 | 3155 | 2600 | 3445 | 3030 | 4291 | 4944 | 2480 | 2797 | 3771 | 3552.47 | 4088 | 3016 | 0.738 | -0.439 | 0.02795643 | 0.10975819 | 0.0527 | 0.0078 | 0.0285 | 0.0285 | TRUE | 0.3647 |
| Bahd1 | 979 | 1346 | 1417 | 1574 | 1166 | 1218 | 1040 | 1344 | 1423 | 1237 | 1255 | 1333 | 1272.02 | 1269 | 1275 | 1.004 | 0.006 | 0.96179306 | 0.98075905 | 0.0134 | 0.0095 | 0.0116 | 0.0116 | TRUE | 0.2984 |
| Bai1 | 285 | 403 | 397 | 429 | 343 | 356 | 303 | 402 | 399 | 337 | 369 | 390 | 366.64 | 368 | 365 | 0.991 | -0.013 | 0.9319651 | 0.96628017 | 0.0123 | 0.0159 | 0.014 | 0.014 | TRUE | 0.2727 |
| Bai2 | 1338 | 1521 | 1504 | 1466 | 1233 | 1215 | 1421 | 1519 | 1510 | 1152 | 1327 | 1330 | 1376.58 | 1483 | 1270 | 0.855 | -0.226 | 0.01648779 | 0.07601969 | 0.0033 | 0.0093 | 0.0057 | 0.0057 | TRUE | 0.0808 |
| Bai3 | 21 | 26 | 32 | 64 | 35 | 51 | 22 | 26 | 32 | 50 | 38 | 56 | 37.37 | 27 | 48 | 1.788 | 0.838 | 0.0139404 | 0.06707384 | 0 | 0.0951 | 0.0546 | 0.0546 | TRUE | 0.2641 |
| Baiap2 | 1077 | 1193 | 1082 | 1379 | 1123 | 1144 | 1144 | 1191 | 1086 | 1084 | 1208 | 1252 | 1161.08 | 1140 | 1181 | 1.035 | 0.049 | 0.60739969 | 0.77733755 | 0.0031 | 0.0097 | 0.0058 | 0.0058 | TRUE | 0.0645 |
| Baiap2l1 | 655 | 809 | 863 | 689 | 532 | 706 | 696 | 808 | 867 | 542 | 572 | 773 | 709.51 | 790 | 629 | 0.795 | -0.331 | 0.04136678 | 0.14428059 | 0.0238 | 0.0115 | 0.0176 | 0.0176 | TRUE | 0.4678 |
| Baiap2l2 | 70 | 74 | 77 | 75 | 54 | 44 | 74 | 74 | 77 | 59 | 58 | 48 | 65.13 | 75 | 55 | 0.735 | -0.445 | 0.06362442 | 0.19378103 | 0 | 0.0575 | 0.0258 | 0.0258 | TRUE | 0.0939 |
| Baiap3 | 130 | 156 | 127 | 215 | 143 | 158 | 138 | 156 | 128 | 169 | 154 | 173 | 152.87 | 141 | 165 | 1.177 | 0.235 | 0.16728384 | 0.3648231 | 0 | 0.0285 | 0.0143 | 0.0143 | TRUE | 0.0959 |
| Bak1 | 2512 | 2280 | 2291 | 3614 | 2383 | 2161 | 2668 | 2277 | 2300 | 2841 | 2564 | 2366 | 2502.7 | 2415 | 2590 | 1.073 | 0.102 | 0.34605203 | 0.56535453 | 0.0078 | 0.0082 | 0.008 | 0.008 | TRUE | 0.1016 |
| Bambi | 638 | 693 | 606 | 709 | 452 | 506 | 678 | 692 | 608 | 557 | 486 | 554 | 595.97 | 659 | 532 | 0.808 | -0.307 | 0.00630423 | 0.03736575 | 0.0035 | 0.0124 | 0.0074 | 0.0074 | TRUE | 0.0659 |
| Bambi-ps1 | 48 | 37 | 33 | 78 | 33 | 34 | 51 | 37 | 33 | 61 | 36 | 37 | 42.52 | 40 | 45 | 1.119 | 0.162 | 0.65208268 | 0.80502827 | 0.0545 | 0.0844 | 0.0695 | 0.0695 | TRUE | 0.933 |
| Banf1 | 7843 | 9561 | 9599 | 9864 | 7022 | 7904 | 8331 | 9549 | 9638 | 7754 | 7555 | 8652 | 8579.91 | 9173 | 7987 | 0.871 | -0.2 | 0.03515211 | 0.12848773 | 0.0057 | 0.0073 | 0.0064 | 0.0064 | TRUE | 0.0787 |
| Banf2 | 2 | 0 | 0 | 2 | 0 | 0 | 2 | 0 | 0 | 2 | 0 | 0 | 0.62 | 1 | 1 | 0.783 | -0.353 | 0.91186553 | NA | 0 | 5.3542 | 5.6997 | 5.6997 | TRUE | 0.9742 |

| | | | | | | | | | | | | | | | | | | | | | | | | | |
|---|---|---|---|---|---|---|---|---|---|---|---|---|---|---|---|---|---|---|---|---|---|---|---|---|---|
| Bank1 | 1 | 1 | 0 | 4 | 4 | 1 | 1 | 1 | 0 | 3 | 4 | 1 | 1.77 | 1 | 3 | 4.185 | 2.065 | 0.2002502 | NA | 0 | 1.871 | 0.9546 | 0.9546 | TRUE | 0.2148 |
| Banp | 2315 | 3035 | 2979 | 2953 | 2435 | 2637 | 2459 | 3031 | 2991 | 2321 | 2620 | 2887 | 2718.2 | 2827 | 2609 | 0.923 | -0.116 | 0.34077402 | 0.56103213 | 0.0114 | 0.0081 | 0.0104 | 0.0104 | TRUE | 0.1573 |
| Bap1 | 5399 | 5010 | 5041 | 7356 | 5272 | 4995 | 5735 | 5004 | 5062 | 5783 | 5672 | 5468 | 5453.8 | 5267 | 5641 | 1.071 | 0.099 | 0.22805563 | 0.44159023 | 0.0031 | 0.0075 | 0.0047 | 0.0047 | TRUE | 0.0736 |
| Bard1 | 3214 | 3365 | 3486 | 4245 | 3266 | 3112 | 3414 | 3361 | 3500 | 3337 | 3514 | 3407 | 3422.08 | 3425 | 3419 | 0.998 | -0.003 | 0.96582082 | 0.98274821 | 3.00E-04 | 0.0079 | 0.0027 | 0.0027 | TRUE | 0.0071 |
| Barhl1 | 0 | 0 | 1 | 6 | 3 | 1 | 0 | 0 | 1 | 5 | 3 | 1 | 1.67 | 0 | 3 | 8.341 | 3.06 | 0.09050099 | NA | 0 | 1.975 | 1.1947 | 1.1947 | TRUE | 0.3034 |
| Barhl2 | 8 | 15 | 9 | 27 | 17 | 16 | 8 | 15 | 9 | 21 | 18 | 18 | 14.92 | 11 | 19 | 1.757 | 0.813 | 0.10957459 | 0.27766878 | 0 | 0.2276 | 0.1134 | 0.1134 | TRUE | 0.4171 |
| Barx1 | 51 | 101 | 100 | 149 | 114 | 160 | 54 | 101 | 100 | 117 | 123 | 175 | 111.73 | 85 | 138 | 1.619 | 0.695 | 0.01723664 | 0.07798519 | 0.0696 | 0.0364 | 0.0518 | 0.0518 | TRUE | 0.9367 |
| Barx2 | 92 | 152 | 187 | 109 | 92 | 90 | 98 | 152 | 188 | 86 | 99 | 99 | 120.08 | 146 | 95 | 0.646 | -0.631 | 0.01694589 | 0.07731284 | 0.049 | 0.0343 | 0.0416 | 0.0416 | TRUE | 0.2222 |
| Basp1 | 5372 | 5136 | 4938 | 7063 | 5132 | 4134 | 5706 | 5129 | 4958 | 5552 | 5522 | 4525 | 5232.18 | 5264 | 5200 | 0.988 | -0.018 | 0.87246566 | 0.93737812 | 0.0091 | 0.0075 | 0.0084 | 0.0084 | TRUE | 0.1565 |
| Batf | 23 | 22 | 22 | 28 | 20 | 15 | 24 | 22 | 22 | 22 | 22 | 16 | 21.41 | 23 | 20 | 0.88 | -0.185 | 0.64835767 | 0.80259769 | 0 | 0.1608 | 0.0715 | 0.0715 | TRUE | 0.1198 |
| Batf2 | 7 | 9 | 9 | 5 | 4 | 6 | 7 | 9 | 9 | 4 | 4 | 7 | 6.71 | 8 | 5 | 0.574 | -0.8 | 0.26914125 | 0.48909964 | 0 | 0.4978 | 0.2174 | 0.2174 | TRUE | 0.1519 |
| Batf3 | 65 | 52 | 40 | 59 | 51 | 32 | 69 | 52 | 40 | 46 | 55 | 35 | 49.57 | 54 | 45 | 0.848 | -0.238 | 0.46688043 | 0.66929996 | 0.0393 | 0.0734 | 0.057 | 0.057 | TRUE | 0.2645 |
| Bax | 6032 | 7045 | 7133 | 8207 | 5972 | 6015 | 6407 | 7036 | 7162 | 6452 | 6426 | 6584 | 6677.8 | 6868 | 6487 | 0.944 | -0.082 | 0.25295771 | 0.47086616 | 0.0016 | 0.0074 | 0.0036 | 0.0036 | TRUE | 0.0421 |
| Baz1a | 2905 | 2341 | 2330 | 4071 | 2820 | 2482 | 3086 | 2338 | 2340 | 3200 | 3034 | 2717 | 2785.76 | 2588 | 2984 | 1.153 | 0.206 | 0.12158561 | 0.29652827 | 0.0167 | 0.0081 | 0.0124 | 0.0124 | TRUE | 0.3432 |
| Baz1b | 7234 | 6137 | 6712 | 9934 | 7167 | 6228 | 7684 | 6129 | 6740 | 7809 | 7711 | 6818 | 7148.42 | 6851 | 7446 | 1.087 | 0.12 | 0.2672865 | 0.48702174 | 0.009 | 0.0074 | 0.0083 | 0.0083 | TRUE | 0.138 |
| Baz2a | 29361 | 28915 | 31491 | 35225 | 26648 | 27188 | 31187 | 28878 | 31620 | 27691 | 28672 | 29762 | 29634.8 | 30562 | 28708 | 0.939 | -0.09 | 0.19592592 | 0.40225769 | 0.0018 | 0.007 | 0.0035 | 0.0035 | TRUE | 0.0284 |
| Baz2b | 1745 | 1618 | 1772 | 2287 | 1614 | 1653 | 1853 | 1616 | 1779 | 1798 | 1737 | 1809 | 1765.43 | 1749 | 1781 | 1.018 | 0.026 | 0.75705488 | 0.87167978 | 0.0021 | 0.0088 | 0.0046 | 0.0046 | TRUE | 0.054 |
| BB014433 | 0 | 0 | 0 | 0 | 1 | 0 | 0 | 0 | 0 | 0 | 1 | 0 | 0.18 | 0 | 0 | 1.859 | 0.894 | 0.82650734 | NA | 0 | 18.3785 | 10 | 10 | TRUE | 0.5204 |
| BB019430 | 2 | 0 | 0 | 0 | 0 | 0 | 2 | 0 | 0 | 0 | 0 | 0 | 0.35 | 1 | 0 | 0.254 | -1.975 | 0.62446185 | NA | 10 | 9.3117 | 10 | 10 | TRUE | 0.9741 |
| BB031773 | 0 | 0 | 0 | 0 | 0 | 0 | 0 | 0 | 0 | 0 | 0 | 0 | 0 | 0 | 0 | NA | NA | NA | NA | NA | NA | NA | NA | NA | NA |
| BB123696 | 2 | 1 | 0 | 1 | 0 | 1 | 2 | 1 | 0 | 1 | 0 | 1 | 0.83 | 1 | 1 | 0.612 | -0.709 | 0.7436921 | NA | 0 | 3.9572 | 2.1269 | 2.1269 | TRUE | 0.1474 |
| BB283400 | 18 | 18 | 9 | 12 | 13 | 22 | 19 | 18 | 9 | 9 | 14 | 24 | 15.61 | 15 | 16 | 1.019 | 0.027 | 0.962629 | 0.98130897 | 0.1225 | 0.218 | 0.1712 | 0.1712 | TRUE | 0.571 |
| BB287469 | 82 | 32 | 20 | 92 | 85 | 39 | 87 | 32 | 20 | 72 | 91 | 43 | 57.6 | 46 | 69 | 1.488 | 0.573 | 0.27051928 | 0.49069926 | 0.3132 | 0.0641 | 0.1767 | 0.1767 | TRUE | 1.2525 |
| BB557941 | 4 | 7 | 3 | 9 | 4 | 4 | 4 | 4 | 7 | 3 | 7 | 4 | 5 | 5 | 5 | 1.119 | 0.162 | 0.84790441 | 0.92391533 | 0 | 0.6656 | 0.3178 | 0.3178 | TRUE | 0.3372 |
| Bbc3 | 1552 | 2408 | 2427 | 1617 | 1394 | 1845 | 1648 | 2405 | 2437 | 1271 | 1500 | 2020 | 1880.17 | 2163 | 1597 | 0.738 | -0.439 | 0.02697588 | 0.10720009 | 0.0493 | 0.0087 | 0.0278 | 0.0278 | TRUE | 0.6458 |
| Bbip1 | 401 | 442 | 398 | 587 | 399 | 381 | 426 | 441 | 400 | 461 | 429 | 417 | 429.14 | 422 | 436 | 1.034 | 0.048 | 0.66867865 | 0.81492803 | 0 | 0.0146 | 0.0067 | 0.0067 | TRUE | 0.0322 |
| Bbox1 | 0 | 0 | 0 | 0 | 0 | 0 | 0 | 0 | 0 | 0 | 0 | 0 | 0 | 0 | 0 | NA | NA | NA | NA | NA | NA | NA | NA | NA | NA |
| Bbs1 | 214 | 166 | 193 | 382 | 287 | 218 | 227 | 166 | 194 | 300 | 309 | 239 | 239.1 | 196 | 283 | 1.447 | 0.533 | 0.00300626 | 0.0212527 | 0.0172 | 0.0207 | 0.0189 | 0.0189 | TRUE | 0.215 |
| Bbs10 | 346 | 270 | 322 | 558 | 375 | 346 | 368 | 270 | 323 | 439 | 403 | 379 | 363.56 | 320 | 407 | 1.273 | 0.349 | 0.02072408 | 0.08911126 | 0.0114 | 0.016 | 0.0136 | 0.0136 | TRUE | 0.2169 |
| Bbs12 | 148 | 137 | 118 | 231 | 164 | 120 | 157 | 137 | 118 | 182 | 176 | 131 | 150.32 | 137 | 163 | 1.19 | 0.251 | 0.22042999 | 0.43243666 | 0.0183 | 0.0288 | 0.0236 | 0.0236 | TRUE | 0.2972 |
| Bbs2 | 1252 | 1283 | 1237 | 1458 | 1093 | 1130 | 1330 | 1281 | 1242 | 1146 | 1176 | 1237 | 1235.4 | 1284 | 1186 | 0.923 | -0.115 | 0.15626819 | 0.34905281 | 5.00E-04 | 0.0096 | 0.004 | 0.004 | TRUE | 0.0165 |
| Bbs4 | 736 | 783 | 781 | 867 | 641 | 733 | 782 | 782 | 784 | 682 | 690 | 802 | 753.6 | 783 | 725 | 0.925 | -0.113 | 0.27500152 | 0.49548018 | 0.0028 | 0.0113 | 0.0064 | 0.0064 | TRUE | 0.1026 |
| Bbs5 | 325 | 374 | 363 | 545 | 445 | 408 | 345 | 374 | 364 | 428 | 479 | 447 | 406.18 | 361 | 451 | 1.248 | 0.32 | 0.00485748 | 0.0306817 | 0 | 0.015 | 0.0068 | 0.0068 | TRUE | 0.0327 |
| Bbs7 | 280 | 350 | 300 | 459 | 348 | 380 | 297 | 350 | 301 | 361 | 374 | 416 | 349.9 | 316 | 384 | 1.212 | 0.277 | 0.0356868 | 0.1296297 | 0.004 | 0.0163 | 0.0097 | 0.0097 | TRUE | 0.098 |
| Bbs9 | 1022 | 1200 | 1234 | 1587 | 1136 | 1197 | 1086 | 1198 | 1239 | 1248 | 1222 | 1310 | 1217.2 | 1174 | 1260 | 1.073 | 0.101 | 0.26338407 | 0.48264194 | 0.002 | 0.0096 | 0.0051 | 0.0051 | TRUE | 0.0521 |
| Bbx | 1388 | 1515 | 1643 | 2319 | 1672 | 1655 | 1474 | 1513 | 1650 | 1823 | 1799 | 1812 | 1678.46 | 1546 | 1811 | 1.172 | 0.229 | 0.004141 | 0.0271454 | 9.00E-04 | 0.0089 | 0.004 | 0.004 | TRUE | 0.0418 |
| BC002163 | 745 | 771 | 752 | 934 | 664 | 626 | 791 | 770 | 755 | 734 | 714 | 685 | 741.72 | 772 | 711 | 0.922 | -0.117 | 0.18870815 | 0.39389954 | 0 | 0.0113 | 0.0044 | 0.0044 | TRUE | 0.012 |
| BC003331 | 1145 | 1441 | 1488 | 1590 | 1179 | 1256 | 1216 | 1439 | 1494 | 1250 | 1269 | 1375 | 1340.47 | 1383 | 1298 | 0.938 | -0.093 | 0.39084009 | 0.60464245 | 0.0059 | 0.0094 | 0.0077 | 0.0077 | TRUE | 0.1335 |
| BC003965 | 880 | 870 | 780 | 1319 | 959 | 782 | 935 | 869 | 783 | 1037 | 1032 | 856 | 918.59 | 862 | 975 | 1.132 | 0.178 | 0.13997826 | 0.32471496 | 0.0086 | 0.0105 | 0.0094 | 0.0094 | TRUE | 0.134 |
| BC004004 | 2002 | 2381 | 2459 | 2671 | 1924 | 2100 | 2126 | 2378 | 2469 | 2100 | 2070 | 2299 | 2240.35 | 2324 | 2156 | 0.927 | -0.109 | 0.24223169 | 0.45880608 | 0.0039 | 0.0084 | 0.0058 | 0.0058 | TRUE | 0.0671 |
| BC005537 | 5375 | 4695 | 4685 | 8285 | 6051 | 4719 | 5709 | 4689 | 4704 | 6513 | 6511 | 5166 | 5548.6 | 5034 | 6063 | 1.205 | 0.269 | 0.03499313 | 0.12811271 | 0.0147 | 0.0075 | 0.0115 | 0.0115 | TRUE | 0.2041 |
| BC005561 | 621 | 451 | 488 | 680 | 471 | 441 | 660 | 450 | 490 | 535 | 507 | 483 | 520.68 | 533 | 508 | 0.954 | -0.069 | 0.66956194 | 0.8154066 | 0.02 | 0.0132 | 0.0167 | 0.0167 | TRUE | 0.499 |
| BC005624 | 1358 | 1355 | 1404 | 1941 | 1475 | 1469 | 1442 | 1353 | 1410 | 1526 | 1587 | 1608 | 1487.73 | 1402 | 1574 | 1.122 | 0.166 | 0.02814556 | 0.11009715 | 0 | 0.0091 | 0.0035 | 0.0035 | TRUE | 0.0111 |
| BC005764 | 335 | 406 | 364 | 186 | 179 | 233 | 356 | 405 | 365 | 146 | 193 | 255 | 286.78 | 375 | 198 | 0.525 | -0.931 | 5.62E-06 | 0.00012183 | 0.0347 | 0.0184 | 0.0264 | 0.0264 | TRUE | 0.5651 |
| BC006965 | 0 | 0 | 0 | 0 | 0 | 0 | 0 | 0 | 0 | 0 | 0 | 0 | 0 | 0 | 0 | NA | NA | NA | NA | NA | NA | NA | NA | NA | NA |
| BC016423 | 3457 | 2791 | 2862 | 6002 | 3960 | 3308 | 3672 | 2787 | 2874 | 4718 | 4261 | 3621 | 3655.54 | 3111 | 4200 | 1.35 | 0.433 | 0.00222805 | 0.0167337 | 0.0205 | 0.0078 | 0.0142 | 0.0142 | TRUE | 0.302 |
| BC016579 | 1 | 1 | 2 | 4 | 1 | 0 | 1 | 1 | 2 | 3 | 1 | 0 | 1.38 | 1 | 1 | 1.083 | 0.115 | 0.94463486 | NA | 0 | 2.3914 | 1.2681 | 1.2681 | TRUE | 0.535 |
| BC017158 | 200 | 238 | 239 | 389 | 306 | 226 | 212 | 238 | 240 | 306 | 329 | 247 | 262.09 | 230 | 294 | 1.279 | 0.356 | 0.02440577 | 0.0999318 | 0.0091 | 0.0195 | 0.0141 | 0.0141 | TRUE | 0.2096 |
| BC017612 | 215 | 308 | 287 | 292 | 237 | 251 | 228 | 308 | 288 | 230 | 255 | 275 | 263.91 | 275 | 253 | 0.919 | -0.122 | 0.45721147 | 0.66181528 | 0.0121 | 0.0194 | 0.0156 | 0.0156 | TRUE | 0.2412 |
| BC017643 | 845 | 1046 | 1065 | 1387 | 973 | 948 | 898 | 1045 | 1069 | 1090 | 1047 | 1038 | 1031.09 | 1004 | 1058 | 1.054 | 0.076 | 0.44892748 | 0.65455233 | 0.0037 | 0.0101 | 0.0064 | 0.0064 | TRUE | 0.1018 |
| BC018242 | 239 | 402 | 404 | 348 | 302 | 309 | 254 | 401 | 406 | 274 | 325 | 338 | 332.96 | 354 | 312 | 0.881 | -0.182 | 0.36707655 | 0.58453273 | 0.0365 | 0.0168 | 0.0265 | 0.0265 | TRUE | 0.691 |
| BC018473 | 0 | 0 | 0 | 0 | 0 | 0 | 0 | 0 | 0 | 0 | 0 | 0 | 0 | 0 | 0 | NA | NA | NA | NA | NA | NA | NA | NA | NA | NA |
| BC018507 | 5402 | 5469 | 6121 | 6897 | 5065 | 5302 | 5738 | 5462 | 6146 | 5422 | 5450 | 5804 | 5670.22 | 5782 | 5559 | 0.961 | -0.057 | 0.45984561 | 0.66404367 | 0.0022 | 0.0075 | 0.0041 | 0.0041 | TRUE | 0.0371 |
| BC020402 | 18 | 30 | 17 | 25 | 23 | 28 | 19 | 30 | 17 | 20 | 25 | 31 | 23.53 | 22 | 25 | 1.125 | 0.17 | 0.69215065 | 0.83110773 | 0.0303 | 0.1469 | 0.0906 | 0.0906 | TRUE | 0.4945 |
| BC021614 | 261 | 395 | 338 | 299 | 231 | 287 | 277 | 394 | 339 | 235 | 249 | 314 | 301.48 | 337 | 266 | 0.787 | -0.345 | 0.06049109 | 0.18692505 | 0.0241 | 0.0178 | 0.021 | 0.021 | TRUE | 0.1774 |
| BC021767 | 75 | 66 | 64 | 73 | 50 | 48 | 80 | 66 | 64 | 57 | 54 | 53 | 62.26 | 70 | 55 | 0.783 | -0.354 | 0.15342295 | 0.34502219 | 0 | 0.0598 | 0.028 | 0.028 | TRUE | 0.1295 |
| BC021785 | 1 | 2 | 1 | 3 | 1 | 4 | 1 | 2 | 1 | 2 | 1 | 4 | 1.98 | 1 | 2 | 1.909 | 0.933 | 0.49031454 | NA | 0 | 1.6712 | 0.7534 | 0.7534 | TRUE | 0.2132 |
| BC021891 | 34 | 72 | 60 | 45 | 31 | 41 | 36 | 72 | 60 | 35 | 33 | 45 | 46.98 | 56 | 38 | 0.673 | -0.572 | 0.09927141 | 0.25995495 | 0.0519 | 0.077 | 0.0648 | 0.0648 | TRUE | 0.3813 |
| BC022687 | 172 | 119 | 137 | 215 | 139 | 151 | 183 | 119 | 138 | 169 | 150 | 165 | 153.83 | 147 | 161 | 1.104 | 0.143 | 0.48508703 | 0.68237017 | 0.0191 | 0.0283 | 0.0237 | 0.0237 | TRUE | 0.4855 |
| BC023829 | 876 | 904 | 888 | 1354 | 948 | 830 | 930 | 903 | 892 | 1064 | 1020 | 909 | 952.98 | 908 | 998 | 1.1 | 0.137 | 0.15849697 | 0.35238429 | 0.0026 | 0.0104 | 0.0057 | 0.0057 | TRUE | 0.0719 |
| BC024139 | 4 | 4 | 5 | 2 | 1 | 3 | 4 | 4 | 5 | 2 | 1 | 3 | 3.2 | 4 | 2 | 0.441 | -1.18 | 0.26862509 | NA | 0 | 1.0366 | 0.4553 | 0.4553 | TRUE | 0.2575 |
| BC024386 | 0 | 0 | 0 | 2 | 0 | 1 | 0 | 0 | 0 | 2 | 0 | 1 | 0.44 | 0 | 1 | 4.782 | 2.258 | 0.50653358 | NA | 0 | 7.4188 | 6.7775 | 6.7775 | TRUE | 0.2804 |
| BC024582 | 0 | 1 | 0 | 2 | 0 | 0 | 0 | 1 | 0 | 2 | 0 | 0 | 0.43 | 0 | 1 | 1.493 | 0.578 | 0.86962266 | NA | 0 | 7.6954 | 7.0545 | 7.0545 | TRUE | 0.9718 |
| BC024978 | 160 | 220 | 192 | 272 | 211 | 212 | 170 | 220 | 193 | 214 | 227 | 232 | 209.23 | 194 | 224 | 1.153 | 0.205 | 0.19178827 | 0.39722713 | 0.0036 | 0.0226 | 0.013 | 0.013 | TRUE | 0.1441 |
| BC025920 | 34 | 40 | 37 | 43 | 30 | 29 | 36 | 40 | 37 | 34 | 32 | 32 | 35.17 | 38 | 33 | 0.865 | -0.209 | 0.50455482 | 0.69949949 | 0 | 0.1006 | 0.0422 | 0.0422 | TRUE | 0.0194 |
| BC026585 | 407 | 636 | 617 | 753 | 525 | 593 | 432 | 635 | 620 | 592 | 565 | 649 | 582.16 | 562 | 602 | 1.07 | 0.097 | 0.55150963 | 0.73559491 | 0.0224 | 0.0126 | 0.0176 | 0.0176 | TRUE | 0.4765 |
| BC026590 | 684 | 546 | 523 | 866 | 556 | 471 | 727 | 545 | 525 | 681 | 598 | 516 | 598.59 | 599 | 598 | 1 | 0 | 0.99846972 | 0.99891307 | 0.0241 | 0.0124 | 0.0184 | 0.0184 | TRUE | 0.4059 |
| BC027072 | 12 | 6 | 5 | 21 | 7 | 9 | 13 | 6 | 5 | 17 | 8 | 10 | 9.61 | 8 | 12 | 1.451 | 0.537 | 0.43355472 | 0.64236967 | 0.1082 | 0.3498 | 0.2314 | 0.2314 | TRUE | 0.7927 |
| BC027231 | 2403 | 2220 | 2429 | 2809 | 1945 | 2086 | 2552 | 2217 | 2439 | 2208 | 2093 | 2283 | 2298.81 | 2403 | 2195 | 0.914 | -0.13 | 0.13583141 | 0.31903692 | 0.0028 | 0.0083 | 0.0051 | 0.0051 | TRUE | 0.0555 |

| | | | | | | | | | | | | | | | | | | | | | | | | | |
|---|---|---|---|---|---|---|---|---|---|---|---|---|---|---|---|---|---|---|---|---|---|---|---|---|---|
| BC028528 | 60 | 86 | 81 | 68 | 45 | 52 | 64 | 86 | 81 | 53 | 48 | 57 | 64.96 | 77 | 53 | 0.687 | -0.542 | 0.03238536 | 0.12138353 | 0 | 0.0576 | 0.0303 | 0.0303 | TRUE | 0.2038 |
| BC029214 | 41 | 63 | 63 | 71 | 50 | 35 | 44 | 63 | 63 | 56 | 54 | 38 | 52.94 | 57 | 49 | 0.874 | -0.194 | 0.51903593 | 0.71037146 | 0.0205 | 0.0691 | 0.0462 | 0.0462 | TRUE | 0.3338 |
| BC029722 | 363 | 396 | 414 | 485 | 359 | 385 | 386 | 395 | 416 | 381 | 386 | 421 | 397.62 | 399 | 396 | 0.992 | -0.011 | 0.92036233 | 0.96087606 | 0 | 0.0152 | 0.0067 | 0.0067 | TRUE | 0.0344 |
| BC030307 | 61 | 53 | 48 | 79 | 53 | 42 | 65 | 53 | 48 | 62 | 57 | 46 | 55.17 | 55 | 55 | 1.001 | 0.001 | 0.99708771 | 0.99828918 | 0 | 0.0666 | 0.0371 | 0.0371 | TRUE | 0.1842 |
| BC030336 | 1157 | 1335 | 1334 | 1481 | 1102 | 1086 | 1229 | 1333 | 1339 | 1164 | 1186 | 1189 | 1240.07 | 1300 | 1180 | 0.907 | -0.141 | 0.07920096 | 0.22317416 | 0 | 0.0096 | 0.0039 | 0.0039 | TRUE | 0.0277 |
| BC030476 | 20 | 19 | 23 | 23 | 21 | 12 | 21 | 19 | 23 | 18 | 23 | 13 | 19.52 | 21 | 18 | 0.851 | -0.233 | 0.59463755 | 0.7677918 | 0 | 0.1757 | 0.0869 | 0.0869 | TRUE | 0.2079 |
| BC030500 | 5 | 2 | 0 | 0 | 1 | 0 | 5 | 2 | 0 | 0 | 1 | 0 | 1.4 | 2 | 0 | 0.144 | -2.797 | 0.19876762 | NA | 1.8301 | 2.3645 | 2.2043 | 2.2043 | TRUE | 0.2486 |
| BC030867 | 1973 | 2177 | 2165 | 3254 | 2374 | 2451 | 2096 | 2174 | 2174 | 2558 | 2554 | 2683 | 2373.18 | 2148 | 2598 | 1.209 | 0.274 | 5.37E-05 | 0.00082297 | 0 | 0.0083 | 0.0029 | 0.0029 | TRUE | 0.0097 |
| BC030870 | 1 | 0 | 1 | 1 | 8 | 0 | 1 | 0 | 1 | 1 | 9 | 0 | 1.91 | 1 | 3 | 4.524 | 2.178 | 0.26319287 | NA | 1.9492 | 1.7318 | 1.8261 | 1.8261 | TRUE | 2.9754 |
| BC031181 | 2026 | 2062 | 1960 | 2559 | 1701 | 1633 | 2152 | 2059 | 1968 | 2012 | 1830 | 1788 | 1968.13 | 2060 | 1877 | 0.912 | -0.133 | 0.12026842 | 0.2942289 | 0.0022 | 0.0086 | 0.0048 | 0.0048 | TRUE | 0.0491 |
| BC031361 | 5 | 19 | 14 | 22 | 12 | 12 | 5 | 19 | 14 | 17 | 13 | 13 | 13.61 | 13 | 14 | 1.133 | 0.18 | 0.76117707 | 0.87423727 | 0.0917 | 0.2489 | 0.1781 | 0.1781 | TRUE | 0.4561 |
| BC033916 | 209 | 181 | 167 | 395 | 274 | 258 | 222 | 181 | 168 | 311 | 295 | 282 | 243.03 | 190 | 296 | 1.559 | 0.641 | 4.18E-05 | 0.00066529 | 0.0058 | 0.0205 | 0.0133 | 0.0133 | TRUE | 0.2285 |
| BC035044 | 0 | 0 | 0 | 0 | 0 | 1 | 0 | 0 | 0 | 0 | 0 | 1 | 0.18 | 0 | 0 | 1.859 | 0.894 | 0.82650734 | NA | 0 | 18.0643 | 10 | 10 | TRUE | 0.5383 |
| BC037032 | 0 | 3 | 1 | 0 | 1 | 2 | 0 | 3 | 1 | 0 | 1 | 2 | 1.21 | 1 | 1 | 0.783 | -0.353 | 0.85707656 | NA | 0 | 2.7275 | 1.9263 | 1.9263 | TRUE | 0.3779 |
| BC037034 | 1063 | 1152 | 1045 | 1185 | 1006 | 934 | 1129 | 1151 | 1049 | 932 | 1082 | 1022 | 1060.88 | 1110 | 1012 | 0.911 | -0.134 | 0.16947479 | 0.36794046 | 0.0032 | 0.01 | 0.0059 | 0.0059 | TRUE | 0.0601 |
| BC037704 | 78 | 86 | 99 | 98 | 86 | 73 | 83 | 86 | 99 | 77 | 93 | 80 | 86.27 | 89 | 83 | 0.928 | -0.108 | 0.617412 | 0.78353895 | 0 | 0.0451 | 0.0223 | 0.0223 | TRUE | 0.0923 |
| BC039771 | 6 | 4 | 8 | 8 | 5 | 4 | 6 | 4 | 8 | 6 | 5 | 4 | 5.74 | 6 | 5 | 0.879 | -0.185 | 0.81163625 | 0.90320517 | 0 | 0.5807 | 0.2623 | 0.2623 | TRUE | 0.2282 |
| BC039966 | 1 | 5 | 3 | 15 | 6 | 10 | 1 | 5 | 3 | 12 | 6 | 11 | 6.38 | 3 | 10 | 3.214 | 1.684 | 0.04643157 | 0.15587882 | 0 | 0.5235 | 0.2976 | 0.2976 | TRUE | 0.4279 |
| BC048403 | 1479 | 1483 | 1358 | 1590 | 1293 | 1276 | 1571 | 1481 | 1364 | 1250 | 1391 | 1397 | 1408.92 | 1472 | 1346 | 0.914 | -0.13 | 0.17778254 | 0.37837007 | 0.0038 | 0.0092 | 0.006 | 0.006 | TRUE | 0.0499 |
| BC048502 | 1 | 0 | 2 | 1 | 2 | 3 | 1 | 0 | 2 | 1 | 2 | 3 | 1.55 | 1 | 2 | 1.988 | 0.992 | 0.53786608 | NA | 0 | 2.1341 | 1.1265 | 1.1265 | TRUE | 0.2694 |
| BC048507 | 0 | 0 | 0 | 1 | 1 | 0 | 0 | 0 | 0 | 1 | 1 | 0 | 0.31 | 0 | 1 | 3.32 | 1.731 | 0.65383052 | NA | 0 | 10.6224 | 8.9541 | 8.9541 | TRUE | 0.202 |
| BC048546 | 13 | 11 | 10 | 21 | 18 | 13 | 14 | 11 | 10 | 17 | 19 | 14 | 14.16 | 12 | 17 | 1.441 | 0.527 | 0.29406066 | 0.51492239 | 0 | 0.2396 | 0.1082 | 0.1082 | TRUE | 0.1004 |
| BC048562 | 0 | 0 | 0 | 0 | 0 | 0 | 0 | 0 | 0 | 0 | 0 | 0 | 0 | 0 | 0 | NA | NA | NA | NA | NA | NA | NA | NA | NA | NA |
| BC048602 | 0 | 0 | 0 | 0 | 0 | 0 | 0 | 0 | 0 | 0 | 0 | 0 | 0 | 0 | 0 | NA | NA | NA | NA | NA | NA | NA | NA | NA | NA |
| BC048609 | 0 | 0 | 0 | 1 | 0 | 2 | 0 | 0 | 0 | 1 | 0 | 2 | 0.5 | 0 | 1 | 5.26 | 2.395 | 0.46836297 | NA | 0 | 6.6502 | 6.3639 | 6.3639 | TRUE | 0.1275 |
| BC048644 | 3 | 4 | 3 | 3 | 3 | 3 | 3 | 4 | 3 | 2 | 3 | 3 | 3.18 | 3 | 3 | 0.861 | -0.216 | 0.83062756 | NA | 0 | 1.0438 | 0.4147 | 0.4147 | TRUE | 0.0556 |
| BC048671 | 0 | 0 | 0 | 0 | 0 | 0 | 0 | 0 | 0 | 0 | 0 | 0 | 0 | 0 | 0 | NA | NA | NA | NA | NA | NA | NA | NA | NA | NA |
| BC048679 | 3 | 4 | 2 | 3 | 1 | 5 | 3 | 4 | 2 | 2 | 1 | 5 | 3.02 | 3 | 3 | 0.96 | -0.059 | 0.95767291 | NA | 0 | 1.0992 | 0.5573 | 0.5573 | TRUE | 0.6194 |
| BC048943 | 0 | 0 | 0 | 0 | 0 | 0 | 0 | 0 | 0 | 0 | 0 | 0 | 0 | 0 | 0 | NA | NA | NA | NA | NA | NA | NA | NA | NA | NA |
| BC049352 | 1 | 1 | 3 | 1 | 0 | 0 | 1 | 1 | 3 | 1 | 0 | 0 | 0.98 | 2 | 0 | 0.206 | -2.278 | 0.27214788 | NA | 0 | 3.3804 | 1.8001 | 1.8001 | TRUE | 0.3622 |
| BC049635 | 13 | 11 | 4 | 6 | 8 | 5 | 14 | 11 | 4 | 5 | 9 | 5 | 7.93 | 10 | 6 | 0.648 | -0.626 | 0.40094425 | 0.61389115 | 0 | 0.4221 | 0.2683 | 0.2683 | TRUE | 0.5392 |
| BC049715 | 0 | 0 | 0 | 3 | 0 | 0 | 0 | 0 | 0 | 2 | 0 | 0 | 0.39 | 0 | 1 | 4.278 | 2.097 | 0.60136958 | NA | 10 | 8.3884 | 10 | 10 | TRUE | 1.4411 |
| BC049730 | 0 | 0 | 1 | 1 | 0 | 0 | 0 | 0 | 1 | 1 | 0 | 0 | 0.3 | 0 | 0 | 0.954 | -0.067 | 0.98640617 | NA | 0 | 11.0483 | 9.2783 | 9.2783 | TRUE | 0.4098 |
| BC049762 | 5 | 13 | 6 | 21 | 10 | 8 | 5 | 13 | 6 | 17 | 11 | 9 | 10.06 | 8 | 12 | 1.493 | 0.579 | 0.37975703 | 0.59453654 | 0.0747 | 0.3345 | 0.2097 | 0.2097 | TRUE | 0.8349 |
| BC051019 | 80 | 83 | 60 | 91 | 81 | 72 | 85 | 83 | 60 | 72 | 87 | 79 | 77.6 | 76 | 79 | 1.039 | 0.055 | 0.82136934 | 0.90875327 | 0.0086 | 0.0494 | 0.0298 | 0.0298 | TRUE | 0.3057 |
| BC051070 | 0 | 0 | 0 | 0 | 0 | 0 | 0 | 0 | 0 | 0 | 0 | 0 | 0 | 0 | 0 | NA | NA | NA | NA | NA | NA | NA | NA | NA | NA |
| BC051142 | 8 | 7 | 3 | 4 | 4 | 5 | 8 | 7 | 3 | 3 | 4 | 5 | 5.24 | 6 | 4 | 0.691 | -0.532 | 0.53075818 | 0.71904843 | 0 | 0.636 | 0.3217 | 0.3217 | TRUE | 0.4863 |
| BC051226 | 157 | 180 | 205 | 186 | 157 | 163 | 167 | 180 | 206 | 146 | 169 | 178 | 174.32 | 184 | 164 | 0.89 | -0.168 | 0.32818207 | 0.54918701 | 0.0057 | 0.0258 | 0.0155 | 0.0155 | TRUE | 0.1148 |
| BC051537 | 5 | 9 | 10 | 4 | 5 | 4 | 5 | 9 | 10 | 3 | 5 | 4 | 6.21 | 8 | 4 | 0.522 | -0.938 | 0.22541318 | 0.43815832 | 0 | 0.5377 | 0.2539 | 0.2539 | TRUE | 0.2576 |
| BC051628 | 3 | 5 | 0 | 12 | 4 | 5 | 3 | 5 | 0 | 9 | 4 | 5 | 4.57 | 3 | 6 | 2.375 | 1.248 | 0.23875936 | NA | 0.2139 | 0.7286 | 0.5454 | 0.5454 | TRUE | 0.8106 |
| BC051665 | 0 | 0 | 0 | 0 | 0 | 0 | 0 | 0 | 0 | 0 | 0 | 0 | 0 | 0 | 0 | NA | NA | NA | NA | NA | NA | NA | NA | NA | NA |
| BC052040 | 2985 | 3680 | 3686 | 3628 | 2758 | 3362 | 3171 | 3675 | 3701 | 2852 | 2967 | 3680 | 3341.12 | 3516 | 3166 | 0.9 | -0.151 | 0.22039337 | 0.43243666 | 0.0132 | 0.0079 | 0.0107 | 0.0107 | TRUE | 0.2441 |
| BC052688 | 30 | 31 | 19 | 60 | 56 | 34 | 32 | 31 | 19 | 47 | 60 | 37 | 37.76 | 27 | 48 | 1.767 | 0.822 | 0.02373919 | 0.09803994 | 0.0338 | 0.0942 | 0.0663 | 0.0663 | TRUE | 0.2255 |
| BC053393 | 1 | 0 | 0 | 3 | 3 | 0 | 1 | 0 | 0 | 2 | 3 | 0 | 1.11 | 0 | 2 | 5.12 | 2.356 | 0.29230162 | NA | 0 | 2.9801 | 2.3453 | 2.3453 | TRUE | 0.417 |
| BC053749 | 602 | 515 | 578 | 607 | 514 | 532 | 639 | 514 | 580 | 477 | 553 | 582 | 557.78 | 578 | 537 | 0.929 | -0.107 | 0.42259412 | 0.63284794 | 0.0095 | 0.0128 | 0.011 | 0.011 | TRUE | 0.1177 |
| BC055111 | 85 | 89 | 82 | 127 | 80 | 81 | 90 | 89 | 82 | 100 | 86 | 89 | 89.35 | 87 | 92 | 1.054 | 0.076 | 0.71474903 | 0.8452151 | 0 | 0.0438 | 0.0196 | 0.0196 | TRUE | 0.068 |
| BC055324 | 5315 | 5288 | 5229 | 6449 | 4884 | 4812 | 5645 | 5281 | 5250 | 5070 | 5255 | 5267 | 5294.86 | 5392 | 5197 | 0.964 | -0.053 | 0.42098765 | 0.63116133 | 8.00E-04 | 0.0075 | 0.003 | 0.003 | TRUE | 0.0205 |
| BC055402 | 0 | 0 | 0 | 1 | 2 | 0 | 0 | 0 | 0 | 1 | 2 | 0 | 0.49 | 0 | 1 | 5.204 | 2.38 | 0.47267317 | NA | 0 | 6.7348 | 6.4099 | 6.4099 | TRUE | 0.1222 |
| BC056474 | 1134 | 1232 | 1289 | 1623 | 1136 | 1191 | 1205 | 1230 | 1294 | 1276 | 1222 | 1304 | 1255.18 | 1243 | 1267 | 1.019 | 0.028 | 0.72884224 | 0.85409762 | 3.00E-04 | 0.0095 | 0.0038 | 0.0038 | TRUE | 0.0156 |
| BC061194 | 0 | 0 | 0 | 1 | 0 | 0 | 0 | 0 | 0 | 1 | 0 | 0 | 0.13 | 0 | 0 | 1.859 | 0.894 | 0.82650734 | NA | 0 | 25.1515 | 10 | 10 | TRUE | 0.2563 |
| BC061212 | 0 | 0 | 0 | 0 | 0 | 0 | 0 | 0 | 0 | 0 | 0 | 0 | 0 | 0 | 0 | NA | NA | NA | NA | NA | NA | NA | NA | NA | NA |
| BC061237 | 0 | 0 | 0 | 0 | 0 | 0 | 0 | 0 | 0 | 0 | 0 | 0 | 0 | 0 | 0 | NA | NA | NA | NA | NA | NA | NA | NA | NA | NA |
| BC064078 | 3 | 1 | 0 | 6 | 4 | 3 | 3 | 1 | 0 | 5 | 4 | 3 | 2.75 | 1 | 4 | 2.991 | 1.58 | 0.21561846 | NA | 0 | 1.2056 | 0.6843 | 0.6843 | TRUE | 0.7657 |
| BC065397 | 119 | 159 | 183 | 131 | 136 | 137 | 126 | 159 | 184 | 103 | 146 | 150 | 144.7 | 156 | 133 | 0.847 | -0.24 | 0.2947151 | 0.51566968 | 0.0317 | 0.0297 | 0.0307 | 0.0307 | TRUE | 0.3375 |
| BC068157 | 351 | 414 | 441 | 442 | 403 | 489 | 373 | 413 | 443 | 347 | 434 | 535 | 424.24 | 410 | 439 | 1.068 | 0.095 | 0.58679094 | 0.76150238 | 0.0248 | 0.0147 | 0.0199 | 0.0199 | TRUE | 0.2157 |
| BC068281 | 300 | 265 | 265 | 317 | 251 | 227 | 319 | 265 | 266 | 249 | 270 | 248 | 269.52 | 283 | 256 | 0.904 | -0.146 | 0.30290833 | 0.52367315 | 0.0032 | 0.0191 | 0.0107 | 0.0107 | TRUE | 0.1337 |
| BC080695 | 55 | 25 | 15 | 50 | 53 | 42 | 58 | 25 | 15 | 39 | 57 | 46 | 40.13 | 33 | 47 | 1.446 | 0.532 | 0.27251308 | 0.49272859 | 0.2074 | 0.089 | 0.1437 | 0.1437 | TRUE | 1.728 |
| BC089491 | 1 | 0 | 5 | 2 | 2 | 1 | 1 | 0 | 5 | 2 | 2 | 1 | 1.82 | 2 | 2 | 0.79 | -0.339 | 0.82757656 | NA | 0.513 | 1.8201 | 1.1924 | 1.1924 | TRUE | 1.2785 |
| BC089597 | 2 | 1 | 0 | 1 | 1 | 0 | 2 | 1 | 0 | 1 | 1 | 0 | 0.83 | 1 | 1 | 0.608 | -0.717 | 0.74122071 | NA | 0 | 3.972 | 2.1266 | 2.1266 | TRUE | 0.1465 |
| BC090627 | 0 | 0 | 0 | 1 | 0 | 0 | 0 | 0 | 0 | 1 | 0 | 0 | 0.13 | 0 | 0 | 1.859 | 0.894 | 0.82650734 | NA | 0 | 25.1515 | 10 | 10 | TRUE | 0.2563 |
| BC094916 | 0 | 0 | 0 | 0 | 0 | 0 | 0 | 0 | 0 | 0 | 0 | 0 | 0 | 0 | 0 | NA | NA | NA | NA | NA | NA | NA | NA | NA | NA |
| BC096441 | 10 | 7 | 20 | 16 | 7 | 22 | 11 | 7 | 20 | 13 | 8 | 24 | 13.65 | 13 | 15 | 1.167 | 0.223 | 0.73384944 | 0.85735379 | 0.2213 | 0.2483 | 0.2351 | 0.2351 | TRUE | 0.5239 |
| Bc1 | 0 | 0 | 0 | 0 | 0 | 0 | 0 | 0 | 0 | 0 | 0 | 0 | 0 | 0 | 0 | NA | NA | NA | NA | NA | NA | NA | NA | NA | NA |
| BC100451 | 6 | 5 | 4 | 11 | 12 | 5 | 6 | 5 | 4 | 9 | 13 | 5 | 7.07 | 5 | 9 | 1.76 | 0.816 | 0.27033827 | 0.49054018 | 0 | 0.4729 | 0.2404 | 0.2404 | TRUE | 0.1706 |
| BC100530 | 0 | 0 | 0 | 0 | 0 | 0 | 0 | 0 | 0 | 0 | 0 | 0 | 0 | 0 | 0 | NA | NA | NA | NA | NA | NA | NA | NA | NA | NA |
| BC107364 | 5 | 11 | 12 | 4 | 15 | 11 | 5 | 11 | 12 | 3 | 16 | 12 | 9.95 | 9 | 10 | 1.084 | 0.116 | 0.87712176 | 0.93932608 | 0.2697 | 0.3382 | 0.3044 | 0.3044 | TRUE | 0.6178 |
| BC117090 | 0 | 0 | 0 | 0 | 0 | 0 | 0 | 0 | 0 | 0 | 0 | 0 | 0 | 0 | 0 | NA | NA | NA | NA | NA | NA | NA | NA | NA | NA |
| BC125332 | 30 | 25 | 15 | 39 | 25 | 19 | 32 | 25 | 15 | 31 | 27 | 21 | 25.04 | 24 | 26 | 1.098 | 0.135 | 0.75369464 | 0.8700338 | 0.0406 | 0.1385 | 0.0929 | 0.0929 | TRUE | 0.3314 |
| BC147527 | 65 | 40 | 26 | 96 | 93 | 39 | 69 | 40 | 26 | 75 | 100 | 43 | 58.89 | 45 | 73 | 1.619 | 0.695 | 0.10720567 | 0.27377717 | 0.1779 | 0.0628 | 0.1161 | 0.1161 | TRUE | 0.2931 |

| | | | | | | | | | | | | | | | | | | | | | | | | | |
|---|---|---|---|---|---|---|---|---|---|---|---|---|---|---|---|---|---|---|---|---|---|---|---|---|---|
| Bcam | 2154 | 2075 | 2204 | 2747 | 1747 | 1661 | 2288 | 2072 | 2213 | 2159 | 1880 | 1818 | 2071.78 | 2191 | 1952 | 0.892 | -0.165 | 0.09192827 | 0.24742196 | 0.0049 | 0.0085 | 0.0064 | 0.0064 | TRUE | 0.1057 |
| Bcan | 383 | 495 | 415 | 355 | 337 | 278 | 407 | 494 | 417 | 279 | 363 | 304 | 377.31 | 439 | 315 | 0.716 | -0.482 | 0.00143614 | 0.01184044 | 0.012 | 0.0156 | 0.0137 | 0.0137 | TRUE | 0.1935 |
| Bcap29 | 732 | 1346 | 1370 | 1557 | 1197 | 1276 | 778 | 1344 | 1376 | 1224 | 1288 | 1397 | 1234.35 | 1166 | 1303 | 1.117 | 0.16 | 0.42320595 | 0.63352356 | 0.0491 | 0.0096 | 0.0279 | 0.0279 | TRUE | 1.0152 |
| Bcap31 | 4301 | 5810 | 5648 | 6050 | 4609 | 4885 | 4568 | 5802 | 5671 | 4756 | 4959 | 5347 | 5184.09 | 5347 | 5021 | 0.939 | -0.091 | 0.41817167 | 0.62897231 | 0.0092 | 0.0075 | 0.0089 | 0.0089 | TRUE | 0.1978 |
| Bcar1 | 3863 | 3847 | 3974 | 5169 | 3790 | 3469 | 4103 | 3842 | 3990 | 4063 | 4078 | 3797 | 3979.03 | 3978 | 3979 | 1 | 0.001 | 0.99223016 | 0.99608263 | 0.0011 | 0.0077 | 0.0033 | 0.0033 | TRUE | 0.0193 |
| Bcar3 | 1155 | 1494 | 1618 | 1667 | 1177 | 1426 | 1227 | 1492 | 1625 | 1310 | 1266 | 1561 | 1413.56 | 1448 | 1379 | 0.952 | -0.071 | 0.60446735 | 0.77522029 | 0.0157 | 0.0092 | 0.0127 | 0.0127 | TRUE | 0.214 |
| Bcas1 | 870 | 966 | 915 | 812 | 670 | 742 | 924 | 965 | 919 | 638 | 721 | 812 | 829.84 | 936 | 724 | 0.772 | -0.373 | 0.00115652 | 0.00995671 | 0.0063 | 0.0109 | 0.0083 | 0.0083 | TRUE | 0.1349 |
| Bcas2 | 6697 | 7447 | 7357 | 9040 | 6442 | 6730 | 7113 | 7437 | 7387 | 7107 | 6931 | 7367 | 7223.79 | 7312 | 7135 | 0.976 | -0.036 | 0.56658208 | 0.74648171 | 5.00E-04 | 0.0074 | 0.0026 | 0.0026 | TRUE | 0.0098 |
| Bcas3 | 2692 | 3030 | 3205 | 3954 | 2899 | 3153 | 2859 | 3026 | 3218 | 3108 | 3119 | 3451 | 3130.43 | 3034 | 3226 | 1.063 | 0.088 | 0.30526609 | 0.526077 | 0.0031 | 0.008 | 0.005 | 0.005 | TRUE | 0.045 |
| Bcat1 | 5476 | 6933 | 6736 | 8529 | 6102 | 6059 | 5816 | 6924 | 6764 | 6705 | 6565 | 6633 | 6567.82 | 6501 | 6634 | 1.02 | 0.029 | 0.74132633 | 0.86165152 | 0.0043 | 0.0074 | 0.0055 | 0.0055 | TRUE | 0.1035 |
| Bcat2 | 8292 | 9294 | 9226 | 6656 | 4962 | 5872 | 8808 | 9282 | 9264 | 5232 | 5339 | 6428 | 7392.08 | 9118 | 5666 | 0.621 | -0.687 | 5.49E-12 | 5.07E-10 | 0.007 | 0.0073 | 0.007 | 0.007 | TRUE | 0.1682 |
| Bccip | 4420 | 4855 | 5015 | 5573 | 4148 | 4062 | 4695 | 4849 | 5036 | 4381 | 4463 | 4446 | 4644.95 | 4860 | 4430 | 0.911 | -0.134 | 0.03426974 | 0.12613824 | 5.00E-04 | 0.0076 | 0.0027 | 0.0027 | TRUE | 0.0122 |
| Bcdin3d | 251 | 205 | 226 | 326 | 226 | 209 | 267 | 205 | 227 | 256 | 243 | 229 | 237.75 | 233 | 243 | 1.045 | 0.064 | 0.68200367 | 0.82362649 | 0.0061 | 0.0208 | 0.0132 | 0.0132 | TRUE | 0.1765 |
| Bche | 155 | 105 | 107 | 466 | 282 | 160 | 165 | 105 | 107 | 366 | 303 | 175 | 203.64 | 126 | 281 | 2.247 | 1.168 | 7.02E-05 | 0.00103709 | 0.0955 | 0.0231 | 0.0564 | 0.0564 | TRUE | 0.2749 |
| Bckdha | 3543 | 3838 | 3725 | 3913 | 2946 | 3178 | 3763 | 3833 | 3740 | 3076 | 3170 | 3479 | 3510.21 | 3779 | 3242 | 0.857 | -0.222 | 0.00383877 | 0.02552492 | 0.0018 | 0.0078 | 0.004 | 0.004 | TRUE | 0.0496 |
| Bckdhb | 2193 | 2430 | 2364 | 2547 | 1861 | 2076 | 2329 | 2427 | 2374 | 2002 | 2002 | 2273 | 2234.5 | 2377 | 2092 | 0.88 | -0.184 | 0.02830471 | 0.11058257 | 0.0023 | 0.0084 | 0.0046 | 0.0046 | TRUE | 0.068 |
| Bckdk | 3385 | 3440 | 3437 | 4561 | 3310 | 3156 | 3595 | 3436 | 3451 | 3585 | 3561 | 3455 | 3513.96 | 3494 | 3534 | 1.012 | 0.017 | 0.79275502 | 0.89315895 | 2.00E-04 | 0.0078 | 0.0026 | 0.0026 | TRUE | 0.0078 |
| Bcl10 | 1588 | 1534 | 1474 | 2177 | 1460 | 1524 | 1687 | 1532 | 1480 | 1711 | 1571 | 1668 | 1608.22 | 1566 | 1650 | 1.054 | 0.076 | 0.39260355 | 0.6062924 | 0.0024 | 0.009 | 0.0051 | 0.0051 | TRUE | 0.0543 |
| Bcl11a | 184 | 210 | 244 | 52 | 44 | 60 | 195 | 210 | 245 | 41 | 47 | 66 | 134.01 | 217 | 51 | 0.234 | -2.093 | 7.52E-20 | 1.82E-17 | 0.0184 | 0.0315 | 0.026 | 0.026 | TRUE | 0.4732 |
| Bcl11b | 262 | 409 | 407 | 289 | 226 | 250 | 278 | 408 | 409 | 227 | 243 | 274 | 306.57 | 365 | 248 | 0.678 | -0.561 | 0.0024372 | 0.01800449 | 0.0248 | 0.0176 | 0.0213 | 0.0213 | TRUE | 0.4917 |
| Bcl2 | 12 | 8 | 17 | 40 | 22 | 14 | 13 | 8 | 17 | 31 | 24 | 15 | 18.04 | 13 | 23 | 1.88 | 0.911 | 0.08005479 | 0.2245863 | 0.0727 | 0.1895 | 0.1338 | 0.1338 | TRUE | 0.1267 |
| Bcl2a1a | 0 | 0 | 0 | 0 | 0 | 0 | 0 | 0 | 0 | 0 | 0 | 0 | 0 | 0 | 0 | NA | NA | NA | NA | NA | NA | NA | NA | NA | NA |
| Bcl2a1b | 0 | 0 | 0 | 0 | 0 | 0 | 0 | 0 | 0 | 0 | 0 | 0 | 0 | 0 | 0 | NA | NA | NA | NA | NA | NA | NA | NA | NA | NA |
| Bcl2a1c | 0 | 0 | 0 | 0 | 0 | 0 | 0 | 0 | 0 | 0 | 0 | 0 | 0 | 0 | 0 | NA | NA | NA | NA | NA | NA | NA | NA | NA | NA |
| Bcl2a1d | 0 | 0 | 0 | 0 | 0 | 0 | 0 | 0 | 0 | 0 | 0 | 0 | 0 | 0 | 0 | NA | NA | NA | NA | NA | NA | NA | NA | NA | NA |
| Bcl2l1 | 916 | 879 | 860 | 1542 | 1020 | 955 | 973 | 878 | 864 | 1212 | 1097 | 1045 | 1011.57 | 905 | 1118 | 1.238 | 0.308 | 0.0027592 | 0.01986225 | 0.0041 | 0.0102 | 0.0066 | 0.0066 | TRUE | 0.0663 |
| Bcl2l10 | 26 | 13 | 9 | 14 | 11 | 4 | 28 | 13 | 9 | 11 | 12 | 4 | 12.81 | 17 | 9 | 0.554 | -0.852 | 0.20002585 | 0.40765056 | 0.2017 | 0.2641 | 0.2338 | 0.2338 | TRUE | 0.9304 |
| Bcl2l11 | 1179 | 1364 | 1468 | 1621 | 1233 | 1280 | 1252 | 1362 | 1474 | 1274 | 1327 | 1401 | 1348.44 | 1363 | 1334 | 0.978 | -0.032 | 0.74162734 | 0.86176528 | 0.0037 | 0.0093 | 0.006 | 0.006 | TRUE | 0.0612 |
| Bcl2l12 | 318 | 373 | 382 | 448 | 354 | 361 | 338 | 373 | 384 | 352 | 381 | 395 | 370.35 | 365 | 376 | 1.029 | 0.042 | 0.73104306 | 0.85559596 | 0 | 0.0158 | 0.0079 | 0.0079 | TRUE | 0.0468 |
| Bcl2l13 | 5771 | 6060 | 6209 | 7249 | 5029 | 5546 | 6130 | 6052 | 6234 | 5699 | 5411 | 6071 | 5932.83 | 6139 | 5727 | 0.933 | -0.1 | 0.16230913 | 0.35768332 | 0.0015 | 0.0075 | 0.0035 | 0.0035 | TRUE | 0.0334 |
| Bcl2l14 | 174 | 235 | 201 | 527 | 368 | 382 | 185 | 235 | 202 | 414 | 396 | 418 | 308.29 | 207 | 409 | 1.976 | 0.983 | 9.67E-13 | 9.90E-11 | 0.0015 | 0.0176 | 0.01 | 0.01 | TRUE | 0.1487 |
| Bcl2l15 | 3 | 1 | 4 | 1 | 1 | 1 | 3 | 1 | 4 | 1 | 1 | 1 | 1.86 | 3 | 1 | 0.357 | -1.488 | 0.29888497 | NA | 0 | 1.7784 | 0.7909 | 0.7909 | TRUE | 0.377 |
| Bcl2l2 | 783 | 1099 | 1172 | 929 | 683 | 907 | 832 | 1098 | 1177 | 730 | 735 | 993 | 927.35 | 1036 | 819 | 0.791 | -0.339 | 0.05040606 | 0.16500246 | 0.0304 | 0.0105 | 0.0205 | 0.0205 | TRUE | 0.4049 |
| Bcl3 | 1661 | 1820 | 1935 | 3036 | 2415 | 2480 | 1764 | 1818 | 1943 | 2387 | 2598 | 2715 | 2204.11 | 1842 | 2567 | 1.393 | 0.478 | 4.32E-08 | 1.75E-06 | 0.0026 | 0.0084 | 0.005 | 0.005 | TRUE | 0.0464 |
| Bcl6 | 189 | 215 | 170 | 253 | 177 | 211 | 201 | 215 | 171 | 199 | 190 | 231 | 201.08 | 196 | 207 | 1.057 | 0.08 | 0.62889593 | 0.79123697 | 0.0067 | 0.0233 | 0.0149 | 0.0149 | TRUE | 0.1337 |
| Bcl6b | 400 | 394 | 387 | 450 | 338 | 281 | 425 | 393 | 389 | 354 | 364 | 308 | 372 | 402 | 342 | 0.85 | -0.234 | 0.05947966 | 0.18472911 | 0.0018 | 0.0158 | 0.0084 | 0.0084 | TRUE | 0.083 |
| Bcl7a | 874 | 1088 | 1120 | 1034 | 868 | 916 | 928 | 1087 | 1125 | 813 | 934 | 1003 | 981.5 | 1047 | 917 | 0.875 | -0.193 | 0.11773114 | 0.29026879 | 0.0098 | 0.0103 | 0.01 | 0.01 | TRUE | 0.1199 |
| Bcl7b | 849 | 865 | 907 | 1063 | 765 | 780 | 902 | 864 | 911 | 836 | 823 | 854 | 864.83 | 892 | 838 | 0.939 | -0.091 | 0.27583727 | 0.49651968 | 0 | 0.0107 | 0.0039 | 0.0039 | TRUE | 0.0092 |
| Bcl7c | 886 | 1162 | 1128 | 1271 | 1035 | 965 | 941 | 1160 | 1133 | 999 | 1114 | 1056 | 1067.22 | 1078 | 1056 | 0.979 | -0.031 | 0.78676294 | 0.89028751 | 0.0069 | 0.01 | 0.0082 | 0.0082 | TRUE | 0.147 |
| Bcl9 | 2292 | 1957 | 1934 | 2584 | 1990 | 1785 | 2435 | 1954 | 1942 | 2031 | 2141 | 1954 | 2076.22 | 2110 | 2042 | 0.968 | -0.047 | 0.67755219 | 0.82070141 | 0.009 | 0.0085 | 0.0088 | 0.0088 | TRUE | 0.2181 |
| Bcl9l | 312 | 397 | 425 | 553 | 444 | 428 | 331 | 396 | 427 | 435 | 478 | 469 | 422.6 | 385 | 461 | 1.194 | 0.256 | 0.05480672 | 0.17448762 | 0.0066 | 0.0147 | 0.0104 | 0.0104 | TRUE | 0.1677 |
| Bclaf1 | 11555 | 7037 | 6533 | 16624 | 11233 | 7988 | 12273 | 7028 | 6560 | 13068 | 12086 | 8744 | 9959.97 | 8620 | 11299 | 1.311 | 0.39 | 0.24991102 | 0.46722738 | 0.0829 | 0.0072 | 0.0389 | 0.0829 | TRUE | 1.6783 |
| Bcmo1 | 9 | 4 | 4 | 13 | 6 | 6 | 10 | 4 | 4 | 10 | 6 | 7 | 6.8 | 6 | 8 | 1.344 | 0.426 | 0.57446883 | 0.75272138 | 0 | 0.4912 | 0.2651 | 0.2651 | TRUE | 0.6704 |
| Bco2 | 8 | 9 | 7 | 15 | 9 | 6 | 8 | 9 | 7 | 12 | 10 | 7 | 8.76 | 8 | 10 | 1.156 | 0.21 | 0.74124246 | 0.86165065 | 0 | 0.383 | 0.176 | 0.176 | TRUE | 0.219 |
| Bcor | 1995 | 2305 | 2680 | 2231 | 1847 | 1781 | 2119 | 2302 | 2691 | 1754 | 1987 | 1950 | 2133.79 | 2371 | 1897 | 0.8 | -0.322 | 0.00470531 | 0.02994842 | 0.0091 | 0.0084 | 0.0089 | 0.0089 | TRUE | 0.1694 |
| Bcorl1 | 1307 | 1227 | 1160 | 2098 | 1521 | 1195 | 1388 | 1225 | 1165 | 1649 | 1637 | 1308 | 1395.39 | 1259 | 1531 | 1.217 | 0.283 | 0.02432052 | 0.09970969 | 0.0111 | 0.0093 | 0.0106 | 0.0106 | TRUE | 0.1943 |
| Bcr | 2011 | 1904 | 2093 | 2517 | 1871 | 1856 | 2136 | 1902 | 2102 | 1979 | 2013 | 2032 | 2027.1 | 2047 | 2008 | 0.981 | -0.028 | 0.73004849 | 0.85500295 | 0.0015 | 0.0085 | 0.0041 | 0.0041 | TRUE | 0.0465 |
| Bcs1l | 931 | 1090 | 1072 | 1506 | 1017 | 1054 | 989 | 1089 | 1076 | 1184 | 1094 | 1154 | 1097.63 | 1051 | 1144 | 1.089 | 0.122 | 0.16343272 | 0.35915575 | 0.0012 | 0.0099 | 0.0046 | 0.0046 | TRUE | 0.032 |
| Bdh1 | 1204 | 1398 | 1468 | 1474 | 1061 | 1270 | 1279 | 1396 | 1474 | 1159 | 1142 | 1390 | 1306.6 | 1383 | 1230 | 0.889 | -0.17 | 0.13441773 | 0.31703851 | 0.0077 | 0.0094 | 0.0085 | 0.0085 | TRUE | 0.1538 |
| Bdh2 | 179 | 171 | 182 | 252 | 157 | 182 | 190 | 171 | 183 | 198 | 169 | 199 | 184.98 | 181 | 189 | 1.043 | 0.061 | 0.69574573 | 0.83345494 | 0 | 0.0247 | 0.0123 | 0.0123 | TRUE | 0.0876 |
| Bdkrb1 | 3 | 8 | 4 | 6 | 2 | 4 | 3 | 8 | 4 | 5 | 2 | 4 | 4.41 | 5 | 4 | 0.746 | -0.423 | 0.64524822 | NA | 0 | 0.7545 | 0.3786 | 0.3786 | TRUE | 0.5289 |
| Bdkrb2 | 39 | 44 | 47 | 38 | 31 | 37 | 41 | 44 | 47 | 30 | 33 | 41 | 39.38 | 44 | 35 | 0.777 | -0.363 | 0.23900377 | 0.4549321 | 0 | 0.0906 | 0.0428 | 0.0428 | TRUE | 0.1516 |
| Bdnf | 48 | 38 | 24 | 81 | 60 | 33 | 51 | 38 | 24 | 64 | 65 | 36 | 46.23 | 38 | 55 | 1.461 | 0.547 | 0.15435853 | 0.34635691 | 0.0902 | 0.0782 | 0.0838 | 0.0838 | TRUE | 0.2921 |
| Bdp1 | 3245 | 2874 | 3172 | 3988 | 2972 | 2892 | 3447 | 2870 | 3185 | 3135 | 3198 | 3166 | 3166.76 | 3167 | 3166 | 1 | 0 | 0.99579186 | 0.99794314 | 0.0039 | 0.0079 | 0.0055 | 0.0055 | TRUE | 0.0819 |
| Bean1 | 6 | 5 | 1 | 8 | 6 | 4 | 6 | 5 | 1 | 6 | 6 | 4 | 4.92 | 4 | 5 | 1.395 | 0.48 | 0.59803692 | 0.7706077 | 0 | 0.6771 | 0.3871 | 0.3871 | TRUE | 0.7627 |
| Becn1 | 3325 | 3283 | 3217 | 4704 | 3471 | 3086 | 3532 | 3279 | 3230 | 3698 | 3735 | 3378 | 3475.22 | 3347 | 3604 | 1.077 | 0.107 | 0.18192486 | 0.38376383 | 0.0022 | 0.0079 | 0.0043 | 0.0043 | TRUE | 0.0361 |
| Begain | 103 | 183 | 163 | 118 | 115 | 123 | 109 | 183 | 164 | 93 | 124 | 135 | 134.5 | 152 | 117 | 0.767 | -0.383 | 0.12834113 | 0.30804996 | 0.0454 | 0.0314 | 0.0382 | 0.0382 | TRUE | 0.621 |
| Bend3 | 1777 | 1214 | 1193 | 2398 | 1581 | 1204 | 1887 | 1212 | 1198 | 1885 | 1701 | 1318 | 1533.66 | 1432 | 1635 | 1.142 | 0.191 | 0.33876636 | 0.55879656 | 0.0502 | 0.0091 | 0.0281 | 0.0281 | TRUE | 0.9274 |
| Bend4 | 11036 | 12045 | 12464 | 17263 | 12531 | 12148 | 11722 | 12029 | 12515 | 13571 | 13483 | 13298 | 12769.68 | 12089 | 13451 | 1.113 | 0.154 | 0.00876382 | 0.04766308 | 5.00E-04 | 0.0072 | 0.0024 | 0.0024 | TRUE | 0.0116 |
| Bend5 | 58 | 104 | 91 | 147 | 120 | 114 | 62 | 104 | 91 | 116 | 129 | 125 | 104.38 | 86 | 123 | 1.434 | 0.52 | 0.02836381 | 0.1107586 | 0.0214 | 0.0385 | 0.0307 | 0.0307 | TRUE | 0.5551 |
| Bend6 | 109 | 132 | 136 | 154 | 108 | 126 | 116 | 132 | 137 | 121 | 116 | 138 | 126.56 | 128 | 125 | 0.975 | -0.037 | 0.84139368 | 0.91989887 | 0 | 0.0329 | 0.0164 | 0.0164 | TRUE | 0.0776 |
| Bend7 | 309 | 360 | 371 | 385 | 257 | 283 | 328 | 360 | 373 | 303 | 277 | 310 | 324.87 | 354 | 297 | 0.839 | -0.254 | 0.04338258 | 0.14897236 | 0 | 0.017 | 0.0083 | 0.0083 | TRUE | 0.0439 |
| Best1 | 45 | 51 | 46 | 57 | 36 | 39 | 48 | 51 | 46 | 45 | 39 | 43 | 45.19 | 48 | 42 | 0.873 | -0.195 | 0.48489366 | 0.68236871 | 0 | 0.0798 | 0.0343 | 0.0343 | TRUE | 0.0355 |
| Best2 | 88 | 111 | 116 | 58 | 45 | 63 | 93 | 111 | 116 | 46 | 48 | 69 | 80.63 | 107 | 54 | 0.504 | -0.987 | 9.78E-05 | 0.00135608 | 0.0142 | 0.0478 | 0.0324 | 0.0324 | TRUE | 0.4419 |
| Best3 | 15 | 9 | 5 | 22 | 22 | 10 | 16 | 9 | 5 | 17 | 24 | 11 | 13.64 | 10 | 17 | 1.742 | 0.8 | 0.19218696 | 0.3978439 | 0.1245 | 0.2484 | 0.1917 | 0.1917 | TRUE | 0.2789 |
| Bet1 | 882 | 922 | 943 | 1127 | 802 | 827 | 937 | 921 | 947 | 886 | 863 | 905 | 909.78 | 935 | 885 | 0.946 | -0.079 | 0.32916575 | 0.55024931 | 0 | 0.0105 | 0.0037 | 0.0037 | TRUE | 0.0053 |
| Bet1l | 405 | 519 | 391 | 583 | 414 | 409 | 430 | 518 | 393 | 458 | 445 | 448 | 448.76 | 447 | 450 | 1.008 | 0.011 | 0.93349736 | 0.96713012 | 0.0081 | 0.0142 | 0.0109 | 0.0109 | TRUE | 0.227 |
| Bet3l | 0 | 0 | 0 | 1 | 1 | 0 | 0 | 0 | 0 | 1 | 1 | 0 | 0.31 | 0 | 1 | 3.32 | 1.731 | 0.65383052 | NA | 0 | 10.6224 | 8.9541 | 8.9541 | TRUE | 0.202 |
| Beta-s | 0 | 0 | 0 | 0 | 0 | 0 | 0 | 0 | 0 | 0 | 0 | 0 | 0 | 0 | 0 | NA | NA | NA | NA | NA | NA | NA | NA | NA | NA |

| | | | | | | | | | | | | | | | | | | | | | | | | | |
|---|---|---|---|---|---|---|---|---|---|---|---|---|---|---|---|---|---|---|---|---|---|---|---|---|---|
| Bex1 | 8223 | 8642 | 8310 | 12754 | 9113 | 8861 | 8734 | 8631 | 8344 | 10026 | 9805 | 9700 | 9206.7 | 8570 | 9844 | 1.149 | 0.2 | 0.00050168 | 0.00517876 | 3.00E-04 | 0.0073 | 0.0023 | 0.0023 | TRUE | 0.0065 |
| Bex2 | 1188 | 1274 | 1266 | 2101 | 1544 | 1265 | 1262 | 1272 | 1271 | 1652 | 1661 | 1385 | 1417.18 | 1268 | 1566 | 1.235 | 0.305 | 0.00230328 | 0.01722488 | 0.0046 | 0.0092 | 0.0065 | 0.0065 | TRUE | 0.1219 |
| Bex4 | 3095 | 3460 | 3428 | 4205 | 3148 | 3120 | 3287 | 3456 | 3442 | 3306 | 3387 | 3415 | 3382.18 | 3395 | 3369 | 0.992 | -0.011 | 0.85920577 | 0.93059974 | 2.00E-04 | 0.0079 | 0.0026 | 0.0026 | TRUE | 0.0093 |
| Bex6 | 4 | 1 | 1 | 14 | 21 | 10 | 4 | 1 | 1 | 11 | 23 | 11 | 8.47 | 2 | 15 | 7.186 | 2.845 | 0.00136442 | 0.01136804 | 0.1348 | 0.396 | 0.2875 | 0.2875 | TRUE | 0.8967 |
| Bfar | 1726 | 1578 | 1796 | 2012 | 1447 | 1443 | 1833 | 1576 | 1803 | 1582 | 1557 | 1580 | 1655.13 | 1737 | 1573 | 0.905 | -0.144 | 0.10943828 | 0.27750169 | 0.0028 | 0.0089 | 0.0052 | 0.0052 | TRUE | 0.0801 |
| Bfsp1 | 30 | 22 | 19 | 52 | 41 | 35 | 32 | 22 | 19 | 41 | 44 | 38 | 32.7 | 24 | 41 | 1.695 | 0.762 | 0.03192041 | 0.12026965 | 0 | 0.1076 | 0.0578 | 0.0578 | TRUE | 0.4278 |
| Bfsp2 | 5 | 3 | 2 | 8 | 6 | 3 | 5 | 3 | 2 | 6 | 6 | 3 | 4.39 | 3 | 5 | 1.572 | 0.652 | 0.47739887 | NA | 0 | 0.7572 | 0.3674 | 0.3674 | TRUE | 0.3146 |
| Bglap | 0 | 0 | 0 | 0 | 0 | 0 | 0 | 0 | 0 | 0 | 0 | 0 | 0 | 0 | 0 | NA | NA | NA | NA | NA | NA | NA | NA | NA | NA |
| Bglap2 | 1 | 1 | 0 | 0 | 0 | 0 | 1 | 1 | 0 | 0 | 0 | 0 | 0.34 | 1 | 0 | 0.261 | -1.936 | 0.60454921 | NA | 0 | 9.5983 | 8.3269 | 8.3269 | TRUE | 0.2507 |
| Bglap3 | 22 | 10 | 14 | 10 | 21 | 18 | 23 | 10 | 14 | 8 | 23 | 20 | 16.26 | 16 | 17 | 1.047 | 0.066 | 0.91124025 | 0.95603547 | 0.1696 | 0.2095 | 0.1898 | 0.1898 | TRUE | 0.9959 |
| Bgn | 70 | 73 | 59 | 91 | 54 | 61 | 74 | 73 | 59 | 72 | 58 | 67 | 67.15 | 69 | 66 | 0.955 | -0.067 | 0.78470068 | 0.88897422 | 0 | 0.056 | 0.0284 | 0.0284 | TRUE | 0.1341 |
| Bhlha15 | 643 | 715 | 671 | 410 | 353 | 428 | 683 | 714 | 674 | 322 | 380 | 469 | 540.24 | 690 | 390 | 0.564 | -0.827 | 3.37E-08 | 1.41E-06 | 0.0153 | 0.013 | 0.0142 | 0.0142 | TRUE | 0.3525 |
| Bhlha9 | 1 | 3 | 4 | 5 | 5 | 1 | 1 | 3 | 4 | 4 | 5 | 1 | 3.08 | 3 | 3 | 1.29 | 0.367 | 0.7447374 | NA | 0 | 1.0765 | 0.5889 | 0.5889 | TRUE | 0.3721 |
| Bhlhb9 | 2546 | 3043 | 3058 | 3659 | 2526 | 2409 | 2704 | 3039 | 3071 | 2876 | 2718 | 2637 | 2840.87 | 2938 | 2744 | 0.934 | -0.098 | 0.25167192 | 0.46941457 | 0.0028 | 0.0081 | 0.0049 | 0.0049 | TRUE | 0.0586 |
| Bhlhe22 | 9 | 9 | 16 | 38 | 17 | 12 | 10 | 9 | 16 | 30 | 18 | 13 | 15.99 | 12 | 20 | 1.79 | 0.84 | 0.12814072 | 0.30785004 | 0.09 | 0.213 | 0.1516 | 0.1516 | TRUE | 0.4371 |
| Bhlhe23 | 88 | 37 | 24 | 56 | 27 | 15 | 93 | 37 | 24 | 44 | 29 | 16 | 40.67 | 51 | 30 | 0.584 | -0.776 | 0.16400942 | 0.35997169 | 0.3192 | 0.0879 | 0.1975 | 0.1975 | TRUE | 1.2446 |
| Bhlhe40 | 2517 | 7345 | 9877 | 5256 | 5383 | 7796 | 2673 | 7336 | 9918 | 4132 | 5792 | 8534 | 6397.36 | 6642 | 6153 | 0.926 | -0.111 | 0.8524663 | 0.92628008 | 0.2544 | 0.0074 | 0.0975 | 0.2544 | TRUE | 0.4028 |
| Bhlhe41 | 50 | 98 | 112 | 58 | 65 | 62 | 53 | 98 | 112 | 46 | 70 | 68 | 74.47 | 88 | 61 | 0.692 | -0.531 | 0.11415865 | 0.28497856 | 0.0855 | 0.0511 | 0.0677 | 0.0677 | TRUE | 0.7554 |
| Bhmt | 47 | 41 | 21 | 91 | 59 | 37 | 50 | 41 | 21 | 72 | 63 | 41 | 47.91 | 37 | 59 | 1.576 | 0.656 | 0.09374161 | 0.25046464 | 0.1034 | 0.0757 | 0.0885 | 0.0885 | TRUE | 0.9175 |
| Bhmt2 | 77 | 108 | 76 | 87 | 60 | 67 | 82 | 108 | 76 | 68 | 65 | 73 | 78.71 | 89 | 69 | 0.775 | -0.368 | 0.1263652 | 0.30479156 | 0.0093 | 0.0488 | 0.0288 | 0.0288 | TRUE | 0.3458 |
| Bicc1 | 171 | 250 | 229 | 363 | 278 | 277 | 182 | 250 | 230 | 285 | 299 | 303 | 258.16 | 221 | 296 | 1.34 | 0.423 | 0.00737005 | 0.04167815 | 0.0081 | 0.0197 | 0.0139 | 0.0139 | TRUE | 0.2561 |
| Bicd1 | 156 | 81 | 76 | 206 | 133 | 82 | 166 | 81 | 76 | 162 | 143 | 90 | 119.62 | 108 | 132 | 1.227 | 0.295 | 0.39777659 | 0.61117705 | 0.1314 | 0.0344 | 0.0791 | 0.0791 | TRUE | 2.0892 |
| Bicd2 | 1553 | 1487 | 1462 | 2179 | 1602 | 1500 | 1650 | 1485 | 1468 | 1713 | 1724 | 1642 | 1613.54 | 1534 | 1693 | 1.104 | 0.142 | 0.09056233 | 0.24533651 | 0.0015 | 0.0089 | 0.0045 | 0.0045 | TRUE | 0.0518 |
| Bid | 4799 | 4724 | 4369 | 4075 | 3116 | 3195 | 5097 | 4718 | 4387 | 3203 | 3353 | 3497 | 4042.62 | 4734 | 3351 | 0.708 | -0.499 | 7.61E-09 | 3.73E-07 | 0.0034 | 0.0077 | 0.0051 | 0.0051 | TRUE | 0.0548 |
| Bik | 135 | 179 | 173 | 178 | 142 | 141 | 143 | 179 | 174 | 140 | 153 | 154 | 157.16 | 165 | 149 | 0.899 | -0.153 | 0.37166891 | 0.58818853 | 0 | 0.0279 | 0.0149 | 0.0149 | TRUE | 0.1399 |
| Bin1 | 283 | 370 | 355 | 437 | 328 | 297 | 301 | 370 | 356 | 344 | 353 | 325 | 341.36 | 342 | 341 | 0.995 | -0.008 | 0.95468573 | 0.97698418 | 0.0035 | 0.0166 | 0.0097 | 0.0097 | TRUE | 0.1272 |
| Bin2 | 2214 | 2375 | 2397 | 2428 | 1813 | 1876 | 2352 | 2372 | 2407 | 1909 | 1951 | 2054 | 2173.9 | 2377 | 1971 | 0.829 | -0.271 | 0.00011086 | 0.00149574 | 3.00E-04 | 0.0084 | 0.0031 | 0.0031 | TRUE | 0.0161 |
| Bin3 | 411 | 384 | 354 | 581 | 455 | 409 | 437 | 384 | 355 | 457 | 490 | 448 | 428.25 | 392 | 465 | 1.186 | 0.246 | 0.04767813 | 0.15867893 | 0.0038 | 0.0146 | 0.0088 | 0.0088 | TRUE | 0.1134 |
| Birc2 | 2236 | 2905 | 2789 | 2728 | 2025 | 2286 | 2375 | 2901 | 2800 | 2145 | 2179 | 2502 | 2483.74 | 2692 | 2275 | 0.845 | -0.243 | 0.02923788 | 0.11295336 | 0.0085 | 0.0082 | 0.0086 | 0.0086 | TRUE | 0.1286 |
| Birc3 | 257 | 257 | 276 | 303 | 190 | 186 | 273 | 257 | 277 | 238 | 204 | 204 | 242.17 | 269 | 215 | 0.804 | -0.315 | 0.02608401 | 0.10460134 | 0 | 0.0205 | 0.0103 | 0.0103 | TRUE | 0.1016 |
| Birc5 | 7704 | 7976 | 7839 | 10658 | 7583 | 7289 | 8183 | 7966 | 7871 | 8378 | 8159 | 7979 | 8089.36 | 8007 | 8172 | 1.021 | 0.03 | 0.61356198 | 0.78135644 | 3.00E-04 | 0.0073 | 0.0024 | 0.0024 | TRUE | 0.006 |
| Birc6 | 7255 | 5121 | 6662 | 10221 | 6714 | 5704 | 7706 | 5114 | 6689 | 8035 | 7224 | 6244 | 6835.43 | 6503 | 7168 | 1.102 | 0.141 | 0.37512531 | 0.59102546 | 0.0242 | 0.0074 | 0.0179 | 0.0179 | TRUE | 0.3789 |
| Birc7 | 22 | 18 | 24 | 29 | 21 | 24 | 23 | 18 | 24 | 23 | 23 | 26 | 22.85 | 22 | 24 | 1.094 | 0.129 | 0.74345506 | 0.86314121 | 0 | 0.1511 | 0.0682 | 0.0682 | TRUE | 0.1369 |
| Bivm | 1135 | 1469 | 1573 | 1663 | 1189 | 1374 | 1206 | 1467 | 1579 | 1307 | 1279 | 1504 | 1390.47 | 1417 | 1363 | 0.962 | -0.057 | 0.65904308 | 0.80794771 | 0.0126 | 0.0093 | 0.0111 | 0.0111 | TRUE | 0.2049 |
| Blcap | 1115 | 1178 | 1086 | 1369 | 1021 | 1004 | 1184 | 1176 | 1090 | 1076 | 1099 | 1099 | 1120.84 | 1150 | 1091 | 0.948 | -0.076 | 0.34904672 | 0.56795263 | 0 | 0.0098 | 0.0039 | 0.0039 | TRUE | 0.025 |
| Blk | 0 | 0 | 0 | 0 | 1 | 0 | 0 | 0 | 0 | 0 | 1 | 0 | 0.18 | 0 | 0 | 1.859 | 0.894 | 0.82650734 | NA | 0 | 18.3785 | 10 | 10 | TRUE | 0.5204 |
| Blm | 10237 | 9858 | 10309 | 11850 | 9080 | 9185 | 10874 | 9845 | 10351 | 9316 | 9770 | 10054 | 10034.93 | 10357 | 9713 | 0.938 | -0.093 | 0.19904368 | 0.40627888 | 0.0018 | 0.0072 | 0.0037 | 0.0037 | TRUE | 0.0233 |
| Blmh | 9622 | 9572 | 9625 | 13713 | 9480 | 8690 | 10220 | 9560 | 9665 | 10780 | 10200 | 9513 | 9989.5 | 9815 | 10164 | 1.036 | 0.051 | 0.50878631 | 0.70257778 | 0.0024 | 0.0072 | 0.0041 | 0.0041 | TRUE | 0.0384 |
| Blnk | 100 | 114 | 118 | 123 | 84 | 107 | 106 | 114 | 118 | 97 | 90 | 117 | 107.13 | 113 | 101 | 0.897 | -0.158 | 0.4315177 | 0.64133973 | 0 | 0.0377 | 0.0196 | 0.0196 | TRUE | 0.1692 |
| Bloc1s1 | 556 | 577 | 541 | 959 | 649 | 645 | 591 | 576 | 543 | 754 | 698 | 706 | 644.71 | 570 | 719 | 1.264 | 0.338 | 0.00051 | 0.00525737 | 0 | 0.012 | 0.0052 | 0.0052 | TRUE | 0.0218 |
| Bloc1s2a | 1599 | 1803 | 1716 | 2090 | 1570 | 1497 | 1698 | 1801 | 1723 | 1643 | 1689 | 1639 | 1698.84 | 1741 | 1657 | 0.952 | -0.071 | 0.31955995 | 0.5406844 | 0 | 0.0088 | 0.0031 | 0.0031 | TRUE | 0.011 |
| Bloc1s3 | 431 | 590 | 622 | 580 | 448 | 499 | 458 | 589 | 625 | 456 | 482 | 546 | 525.97 | 557 | 495 | 0.887 | -0.174 | 0.2486821 | 0.46559996 | 0.0155 | 0.0132 | 0.0144 | 0.0144 | TRUE | 0.2833 |
| Bloc1s4 | 394 | 510 | 486 | 697 | 539 | 470 | 418 | 509 | 488 | 548 | 580 | 514 | 509.7 | 472 | 547 | 1.16 | 0.214 | 0.07846523 | 0.22169438 | 0.0046 | 0.0134 | 0.0086 | 0.0086 | TRUE | 0.1128 |
| Bloc1s5 | 488 | 555 | 518 | 686 | 513 | 520 | 518 | 554 | 520 | 539 | 552 | 569 | 542.2 | 531 | 553 | 1.042 | 0.059 | 0.55092229 | 0.73511866 | 0 | 0.013 | 0.0052 | 0.0052 | TRUE | 0.0174 |
| Bloc1s6 | 1177 | 1296 | 1395 | 2279 | 1511 | 1546 | 1250 | 1294 | 1401 | 1792 | 1626 | 1692 | 1509.15 | 1315 | 1703 | 1.296 | 0.374 | 2.09E-05 | 0.00036931 | 0.0022 | 0.0091 | 0.0049 | 0.0049 | TRUE | 0.039 |
| Blvra | 388 | 488 | 467 | 645 | 406 | 419 | 412 | 487 | 469 | 507 | 437 | 459 | 461.83 | 456 | 468 | 1.026 | 0.037 | 0.762604 | 0.87481201 | 0.0046 | 0.014 | 0.0089 | 0.0089 | TRUE | 0.0818 |
| Blvrb | 1028 | 1392 | 1516 | 1153 | 896 | 1003 | 1092 | 1390 | 1522 | 906 | 964 | 1098 | 1162.12 | 1335 | 989 | 0.741 | -0.433 | 0.00279633 | 0.0200423 | 0.0172 | 0.0097 | 0.0142 | 0.0142 | TRUE | 0.3036 |
| Blzf1 | 1284 | 1301 | 1316 | 1777 | 1331 | 1267 | 1364 | 1299 | 1321 | 1397 | 1432 | 1387 | 1366.75 | 1328 | 1405 | 1.058 | 0.081 | 0.26993191 | 0.49014116 | 0 | 0.0093 | 0.0032 | 0.0032 | TRUE | 0.0066 |
| Bmf | 2136 | 2698 | 2719 | 1586 | 1205 | 1657 | 2269 | 2695 | 2730 | 1247 | 1297 | 1814 | 2008.44 | 2565 | 1453 | 0.566 | -0.821 | 1.95E-07 | 6.40E-06 | 0.0276 | 0.0085 | 0.0174 | 0.0174 | TRUE | 0.5685 |
| Bmi1 | 358 | 610 | 490 | 1211 | 915 | 816 | 380 | 609 | 492 | 952 | 984 | 893 | 718.53 | 494 | 943 | 1.909 | 0.933 | 2.32E-08 | 1.01E-06 | 0.0258 | 0.0115 | 0.0185 | 0.0185 | TRUE | 0.4094 |
| Bmp1 | 191 | 222 | 207 | 400 | 255 | 239 | 203 | 222 | 208 | 314 | 274 | 262 | 247.15 | 211 | 283 | 1.348 | 0.431 | 0.00279894 | 0.02004762 | 0 | 0.0202 | 0.0108 | 0.0108 | TRUE | 0.1092 |
| Bmp10 | 0 | 1 | 1 | 0 | 0 | 0 | 0 | 1 | 1 | 0 | 0 | 0 | 0.33 | 1 | 0 | 0.268 | -1.901 | 0.61401143 | NA | 0 | 9.8764 | 8.4772 | 8.4772 | TRUE | 0.2278 |
| Bmp15 | 2 | 0 | 0 | 4 | 2 | 3 | 2 | 0 | 0 | 3 | 2 | 3 | 1.78 | 1 | 3 | 4.141 | 2.05 | 0.21974603 | NA | 0 | 1.8534 | 1.1021 | 1.1021 | TRUE | 0.9746 |
| Bmp2 | 11 | 18 | 16 | 27 | 10 | 22 | 12 | 18 | 16 | 21 | 11 | 24 | 16.97 | 15 | 19 | 1.228 | 0.297 | 0.5587943 | 0.74101748 | 0.0409 | 0.2011 | 0.1263 | 0.1263 | TRUE | 0.6617 |
| Bmp2k | 466 | 565 | 519 | 625 | 528 | 475 | 495 | 564 | 521 | 491 | 568 | 520 | 526.63 | 527 | 526 | 0.998 | -0.003 | 0.9777008 | 0.98797393 | 0.0031 | 0.0132 | 0.0075 | 0.0075 | TRUE | 0.0557 |
| Bmp3 | 0 | 0 | 1 | 1 | 1 | 1 | 0 | 0 | 1 | 1 | 1 | 1 | 0.66 | 0 | 1 | 2.676 | 1.42 | 0.53608725 | NA | 0 | 4.9975 | 2.297 | 2.297 | TRUE | 0.4098 |
| Bmp4 | 2717 | 2205 | 2031 | 1983 | 1502 | 1495 | 2886 | 2202 | 2039 | 1559 | 1616 | 1637 | 1989.81 | 2376 | 1604 | 0.675 | -0.567 | 3.56E-05 | 0.00057836 | 0.0174 | 0.0086 | 0.013 | 0.013 | TRUE | 0.427 |
| Bmp5 | 0 | 0 | 0 | 0 | 0 | 0 | 0 | 0 | 0 | 0 | 0 | 0 | 0 | 0 | 0 | NA | NA | NA | NA | NA | NA | NA | NA | NA | NA |
| Bmp6 | 80 | 94 | 78 | 120 | 82 | 77 | 85 | 94 | 78 | 94 | 88 | 84 | 87.34 | 86 | 89 | 1.04 | 0.056 | 0.78868295 | 0.8913728 | 0 | 0.0446 | 0.0205 | 0.0205 | TRUE | 0.0666 |
| Bmp7 | 511 | 1011 | 1116 | 719 | 619 | 830 | 543 | 1010 | 1121 | 565 | 666 | 909 | 802.14 | 891 | 713 | 0.8 | -0.322 | 0.21774234 | 0.42970873 | 0.0966 | 0.011 | 0.048 | 0.048 | TRUE | 1.3888 |
| Bmp8a | 55 | 66 | 50 | 98 | 91 | 39 | 58 | 66 | 50 | 77 | 98 | 43 | 65.36 | 58 | 73 | 1.249 | 0.32 | 0.341007 | 0.56123995 | 0.0758 | 0.0573 | 0.0661 | 0.0661 | TRUE | 0.2955 |
| Bmp8b | 308 | 312 | 314 | 235 | 185 | 209 | 327 | 312 | 315 | 185 | 199 | 229 | 261.1 | 318 | 204 | 0.64 | -0.644 | 4.92E-06 | 0.00010866 | 0.0011 | 0.0195 | 0.0103 | 0.0103 | TRUE | 0.1185 |
| Bmper | 13 | 10 | 8 | 39 | 20 | 14 | 14 | 10 | 8 | 31 | 22 | 15 | 16.56 | 11 | 23 | 2.147 | 1.102 | 0.03570994 | 0.12965404 | 0 | 0.2059 | 0.1286 | 0.1286 | TRUE | 0.1957 |
| Bmpr1a | 3609 | 4054 | 4159 | 5940 | 4215 | 4364 | 3833 | 4049 | 4176 | 4670 | 4535 | 4777 | 4339.99 | 4019 | 4661 | 1.159 | 0.213 | 0.00198321 | 0.01527738 | 9.00E-04 | 0.0077 | 0.0032 | 0.0032 | TRUE | 0.0199 |
| Bmpr1b | 126 | 132 | 159 | 115 | 86 | 91 | 134 | 132 | 160 | 90 | 93 | 100 | 117.98 | 142 | 94 | 0.663 | -0.593 | 0.00185925 | 0.01451196 | 0 | 0.0348 | 0.0174 | 0.0174 | TRUE | 0.1273 |
| Bmpr2 | 474 | 443 | 553 | 956 | 715 | 628 | 503 | 442 | 555 | 752 | 769 | 687 | 618.24 | 500 | 736 | 1.472 | 0.557 | 4.21E-06 | 9.54E-05 | 0.0061 | 0.0122 | 0.0089 | 0.0089 | TRUE | 0.1207 |
| Bms1 | 19739 | 17530 | 17738 | 22809 | 16605 | 15396 | 20966 | 17507 | 17811 | 17931 | 17866 | 16853 | 18155.74 | 18761 | 17550 | 0.935 | -0.096 | 0.30021905 | 0.52045009 | 0.0057 | 0.0071 | 0.0062 | 0.0062 | TRUE | 0.1293 |
| Bmx | 2 | 2 | 1 | 0 | 0 | 0 | 2 | 2 | 1 | 0 | 0 | 0 | 0.85 | 2 | 0 | 0.105 | -3.251 | 0.15501195 | NA | 0 | 3.8632 | 2.4682 | 2.4682 | TRUE | 0.1019 |
| Bmyc | 434 | 586 | 656 | 790 | 633 | 638 | 461 | 585 | 659 | 621 | 681 | 698 | 617.57 | 568 | 667 | 1.172 | 0.229 | 0.12209902 | 0.29718298 | 0.016 | 0.0122 | 0.0142 | 0.0142 | TRUE | 0.3177 |

| | | | | | | | | | | | | | | | | | | | | | | | | | |
|---|---|---|---|---|---|---|---|---|---|---|---|---|---|---|---|---|---|---|---|---|---|---|---|---|---|
| Bnc1 | 38 | 31 | 34 | 35 | 29 | 24 | 40 | 31 | 34 | 28 | 31 | 26 | 31.74 | 35 | 28 | 0.806 | -0.311 | 0.35789318 | 0.57640061 | 0 | 0.1107 | 0.0508 | 0.0508 | TRUE | 0.1158 |
| Bnc2 | 1384 | 1145 | 1306 | 1034 | 839 | 858 | 1470 | 1144 | 1311 | 813 | 903 | 939 | 1096.62 | 1308 | 885 | 0.676 | -0.565 | 4.00E-06 | 9.15E-05 | 0.0091 | 0.0099 | 0.0099 | 0.0099 | TRUE | 0.1398 |
| Bnip1 | 653 | 931 | 933 | 945 | 714 | 763 | 694 | 930 | 937 | 743 | 768 | 835 | 817.76 | 854 | 782 | 0.916 | -0.127 | 0.3643867 | 0.58180878 | 0.0146 | 0.0109 | 0.0129 | 0.0129 | TRUE | 0.3176 |
| Bnip2 | 1033 | 1137 | 1090 | 1880 | 1376 | 1189 | 1097 | 1136 | 1094 | 1478 | 1480 | 1302 | 1264.53 | 1109 | 1420 | 1.281 | 0.357 | 7.35E-05 | 0.00107706 | 0.0022 | 0.0095 | 0.005 | 0.005 | TRUE | 0.063 |
| Bnip3 | 5960 | 11481 | 12543 | 9654 | 7982 | 11115 | 6331 | 11466 | 12594 | 7589 | 8588 | 12167 | 9789.29 | 10130 | 9448 | 0.933 | -0.101 | 0.77932222 | 0.88599943 | 0.0929 | 0.0072 | 0.0428 | 0.0929 | TRUE | 1.3155 |
| Bnip3l | 6517 | 8595 | 8720 | 8817 | 6897 | 7855 | 6922 | 8584 | 8756 | 6931 | 7421 | 8599 | 7868.74 | 8087 | 7650 | 0.946 | -0.08 | 0.51910582 | 0.71037146 | 0.0122 | 0.0073 | 0.0111 | 0.0111 | TRUE | 0.1938 |
| Bnipl | 35 | 40 | 63 | 34 | 20 | 28 | 37 | 40 | 63 | 27 | 22 | 31 | 36.55 | 47 | 27 | 0.562 | -0.832 | 0.02326636 | 0.09677138 | 0.0374 | 0.097 | 0.0675 | 0.0675 | TRUE | 0.7568 |
| Boc | 566 | 480 | 545 | 382 | 334 | 341 | 601 | 479 | 547 | 300 | 359 | 373 | 443.46 | 542 | 344 | 0.633 | -0.659 | 4.34E-06 | 9.77E-05 | 0.0109 | 0.0143 | 0.0125 | 0.0125 | TRUE | 0.1512 |
| Bod1 | 3416 | 3576 | 3414 | 4450 | 3257 | 3086 | 3628 | 3571 | 3428 | 3498 | 3504 | 3378 | 3501.41 | 3542 | 3460 | 0.977 | -0.034 | 0.60219926 | 0.77369557 | 3.00E-04 | 0.0078 | 0.0027 | 0.0027 | TRUE | 0.0097 |
| Bod1l | 5099 | 4606 | 5211 | 6943 | 4905 | 4428 | 5416 | 4600 | 5232 | 5458 | 5277 | 4847 | 5138.53 | 5083 | 5194 | 1.022 | 0.032 | 0.73716699 | 0.85934705 | 0.0053 | 0.0075 | 0.0062 | 0.0062 | TRUE | 0.0842 |
| Bok | 364 | 460 | 420 | 809 | 540 | 445 | 387 | 459 | 422 | 636 | 581 | 487 | 495.31 | 423 | 568 | 1.346 | 0.429 | 0.00227336 | 0.01703343 | 0.0109 | 0.0136 | 0.0121 | 0.0121 | TRUE | 0.1799 |
| Bola1 | 630 | 722 | 726 | 975 | 695 | 606 | 669 | 721 | 729 | 766 | 748 | 663 | 716.14 | 706 | 726 | 1.028 | 0.04 | 0.69658911 | 0.83378245 | 0.0026 | 0.0115 | 0.0064 | 0.0064 | TRUE | 0.0659 |
| Bola2 | 1467 | 1368 | 1311 | 2296 | 1576 | 1294 | 1558 | 1366 | 1316 | 1805 | 1696 | 1416 | 1526.32 | 1413 | 1639 | 1.16 | 0.215 | 0.08011986 | 0.22472888 | 0.0106 | 0.0091 | 0.0102 | 0.0102 | TRUE | 0.1692 |
| Bola3 | 304 | 418 | 448 | 706 | 467 | 437 | 323 | 417 | 450 | 555 | 502 | 478 | 454.34 | 397 | 512 | 1.291 | 0.369 | 0.01538251 | 0.07209895 | 0.0147 | 0.0142 | 0.0144 | 0.0144 | TRUE | 0.3021 |
| Boll | 33 | 35 | 44 | 42 | 53 | 45 | 35 | 35 | 44 | 33 | 57 | 49 | 42.25 | 38 | 46 | 1.21 | 0.275 | 0.41326698 | 0.62457329 | 0.0293 | 0.0849 | 0.0571 | 0.0571 | TRUE | 0.5401 |
| Bop1 | 7599 | 5917 | 5813 | 9037 | 6572 | 5381 | 8071 | 5909 | 5837 | 7104 | 7071 | 5890 | 6647.22 | 6606 | 6688 | 1.013 | 0.018 | 0.90139046 | 0.95205059 | 0.0188 | 0.0074 | 0.0152 | 0.0152 | TRUE | 0.4595 |
| Bora | 2052 | 2054 | 1939 | 2704 | 1933 | 1656 | 2180 | 2051 | 1947 | 2126 | 2080 | 1813 | 2032.69 | 2059 | 2006 | 0.975 | -0.037 | 0.702337 | 0.83715683 | 0.0047 | 0.0085 | 0.0063 | 0.0063 | TRUE | 0.0853 |
| Bpgm | 1262 | 1136 | 1062 | 1417 | 946 | 805 | 1340 | 1135 | 1066 | 1114 | 1018 | 881 | 1092.39 | 1180 | 1004 | 0.852 | -0.232 | 0.07921183 | 0.22317416 | 0.0127 | 0.0099 | 0.0116 | 0.0116 | TRUE | 0.1681 |
| Bphl | 411 | 498 | 470 | 664 | 464 | 407 | 437 | 497 | 472 | 522 | 499 | 446 | 478.76 | 469 | 489 | 1.045 | 0.063 | 0.59527116 | 0.76817352 | 0.0033 | 0.0138 | 0.008 | 0.008 | TRUE | 0.0689 |
| Bpi | 8 | 17 | 11 | 10 | 6 | 12 | 8 | 17 | 11 | 8 | 6 | 13 | 10.66 | 12 | 9 | 0.745 | -0.424 | 0.49444446 | 0.69052608 | 0 | 0.3159 | 0.1817 | 0.1817 | TRUE | 0.4792 |
| Bpifa1 | 0 | 0 | 0 | 0 | 0 | 0 | 0 | 0 | 0 | 0 | 0 | 0 | 0 | 0 | 0 | NA | NA | NA | NA | NA | NA | NA | NA | NA | NA |
| Bpifa2 | 0 | 0 | 0 | 1 | 0 | 1 | 0 | 0 | 0 | 1 | 0 | 1 | 0.31 | 0 | 1 | 3.348 | 1.743 | 0.65064889 | NA | 0 | 10.5167 | 8.8968 | 8.8968 | TRUE | 0.2082 |
| Bpifa3 | 0 | 0 | 0 | 0 | 0 | 0 | 0 | 0 | 0 | 0 | 0 | 0 | 0 | 0 | 0 | NA | NA | NA | NA | NA | NA | NA | NA | NA | NA |
| Bpifa5 | 0 | 0 | 0 | 0 | 0 | 0 | 0 | 0 | 0 | 0 | 0 | 0 | 0 | 0 | 0 | NA | NA | NA | NA | NA | NA | NA | NA | NA | NA |
| Bpifa6 | 1 | 0 | 0 | 0 | 0 | 0 | 1 | 0 | 0 | 0 | 0 | 0 | 0.18 | 0 | 0 | 0.49 | -1.029 | 0.80087 | NA | 0 | 18.6165 | 10 | 10 | TRUE | 0.4614 |
| Bpifb1 | 3 | 4 | 3 | 8 | 6 | 1 | 3 | 4 | 3 | 6 | 6 | 1 | 4.01 | 3 | 4 | 1.374 | 0.459 | 0.6455553 | NA | 0 | 0.8294 | 0.4623 | 0.4623 | TRUE | 0.7465 |
| Bpifb2 | 0 | 3 | 0 | 1 | 2 | 1 | 0 | 3 | 0 | 1 | 2 | 1 | 1.17 | 1 | 1 | 1.32 | 0.401 | 0.8384527 | NA | 0.6358 | 2.8192 | 1.9105 | 1.9105 | TRUE | 1.4419 |
| Bpifb3 | 0 | 0 | 0 | 0 | 1 | 0 | 0 | 0 | 0 | 0 | 1 | 0 | 0.18 | 0 | 0 | 1.859 | 0.894 | 0.82650734 | NA | 0 | 18.3785 | 10 | 10 | TRUE | 0.5204 |
| Bpifb4 | 87 | 80 | 60 | 110 | 89 | 65 | 92 | 80 | 60 | 86 | 96 | 71 | 80.99 | 77 | 84 | 1.092 | 0.126 | 0.61921507 | 0.78469067 | 0.0197 | 0.0476 | 0.0343 | 0.0343 | TRUE | 0.3524 |
| Bpifb5 | 15 | 22 | 27 | 36 | 20 | 9 | 16 | 22 | 27 | 28 | 22 | 10 | 20.78 | 22 | 20 | 0.928 | -0.108 | 0.8308262 | 0.91425507 | 0.1061 | 0.1654 | 0.1376 | 0.1376 | TRUE | 0.4121 |
| Bpifb6 | 0 | 0 | 0 | 0 | 0 | 0 | 0 | 0 | 0 | 0 | 0 | 0 | 0 | 0 | 0 | NA | NA | NA | NA | NA | NA | NA | NA | NA | NA |
| Bpifb9a | 0 | 0 | 0 | 0 | 0 | 0 | 0 | 0 | 0 | 0 | 0 | 0 | 0 | 0 | 0 | NA | NA | NA | NA | NA | NA | NA | NA | NA | NA |
| Bpifb9b | 0 | 0 | 0 | 1 | 0 | 0 | 0 | 0 | 0 | 1 | 0 | 0 | 0.13 | 0 | 0 | 1.859 | 0.894 | 0.82650734 | NA | 0 | 25.1515 | 10 | 10 | TRUE | 0.2563 |
| Bpifc | 1 | 0 | 0 | 2 | 2 | 0 | 1 | 0 | 0 | 2 | 2 | 0 | 0.8 | 0 | 1 | 3.417 | 1.773 | 0.46651319 | NA | 0 | 4.1368 | 2.8785 | 2.8785 | TRUE | 0.4614 |
| Bpnt1 | 2305 | 1967 | 1821 | 2494 | 1826 | 1673 | 2448 | 1964 | 1828 | 1961 | 1965 | 1831 | 1999.64 | 2080 | 1919 | 0.923 | -0.116 | 0.34666504 | 0.56576997 | 0.0123 | 0.0086 | 0.0105 | 0.0105 | TRUE | 0.289 |
| Bptf | 15486 | 12705 | 14342 | 17150 | 12484 | 11633 | 16449 | 12689 | 14401 | 13482 | 13432 | 12734 | 13864.42 | 14513 | 13216 | 0.911 | -0.135 | 0.20554896 | 0.41419679 | 0.0088 | 0.0071 | 0.0081 | 0.0081 | TRUE | 0.1665 |
| Braf | 2961 | 3203 | 3955 | 4038 | 2919 | 3076 | 3145 | 3199 | 3971 | 3174 | 3141 | 3367 | 3332.9 | 3438 | 3227 | 0.939 | -0.092 | 0.40899078 | 0.62126578 | 0.0091 | 0.0079 | 0.0086 | 0.0086 | TRUE | 0.2236 |
| Brap | 2520 | 2358 | 2368 | 3171 | 2210 | 2147 | 2677 | 2355 | 2378 | 2493 | 2378 | 2350 | 2438.37 | 2470 | 2407 | 0.975 | -0.037 | 0.66558805 | 0.81276361 | 0.0023 | 0.0083 | 0.0047 | 0.0047 | TRUE | 0.0649 |
| Brat1 | 9 | 0 | 1 | 8 | 5 | 6 | 10 | 0 | 1 | 6 | 5 | 7 | 4.8 | 4 | 6 | 1.743 | 0.802 | 0.50020934 | 0.69607391 | 0.9458 | 0.6932 | 0.7922 | 0.7922 | TRUE | 3.2267 |
| Brca1 | 7012 | 7517 | 8118 | 10911 | 7917 | 8194 | 7448 | 7507 | 8151 | 8577 | 8518 | 8970 | 8195.3 | 7702 | 8688 | 1.128 | 0.174 | 0.01308551 | 0.06413445 | 0.0015 | 0.0073 | 0.0034 | 0.0034 | TRUE | 0.0318 |
| Brca2 | 9119 | 6929 | 7625 | 10554 | 7728 | 6795 | 9686 | 6920 | 7656 | 8297 | 8315 | 7438 | 8052.02 | 8087 | 8017 | 0.991 | -0.013 | 0.92405947 | 0.96274037 | 0.0141 | 0.0073 | 0.0124 | 0.0124 | TRUE | 0.365 |
| Brcc3 | 881 | 896 | 1002 | 1484 | 1050 | 1033 | 936 | 895 | 1006 | 1167 | 1130 | 1131 | 1043.98 | 946 | 1143 | 1.209 | 0.273 | 0.00173561 | 0.01372132 | 8.00E-04 | 0.0101 | 0.0045 | 0.0045 | TRUE | 0.0377 |
| Brd1 | 5561 | 7036 | 7225 | 8744 | 6780 | 7115 | 5907 | 7027 | 7255 | 6874 | 7295 | 7788 | 7024.26 | 6730 | 7319 | 1.087 | 0.121 | 0.24349904 | 0.46008258 | 0.0075 | 0.0074 | 0.0076 | 0.0076 | TRUE | 0.1394 |
| Brd2 | 14926 | 14248 | 13575 | 19719 | 14187 | 12825 | 15854 | 14230 | 13631 | 15502 | 15264 | 14039 | 14753.21 | 14572 | 14935 | 1.025 | 0.036 | 0.68285644 | 0.82421457 | 0.0037 | 0.0071 | 0.0054 | 0.0054 | TRUE | 0.0725 |
| Brd3 | 5877 | 6284 | 6593 | 8305 | 6361 | 6048 | 6242 | 6276 | 6620 | 6529 | 6844 | 6620 | 6521.94 | 6379 | 6664 | 1.045 | 0.063 | 0.31990404 | 0.54101722 | 6.00E-04 | 0.0074 | 0.0027 | 0.0027 | TRUE | 0.0133 |
| Brd4 | 11426 | 10274 | 10469 | 13687 | 10195 | 9374 | 12136 | 10261 | 10512 | 10760 | 10969 | 10261 | 10816.55 | 10970 | 10663 | 0.972 | -0.041 | 0.64731547 | 0.80199996 | 0.0041 | 0.0072 | 0.0056 | 0.0056 | TRUE | 0.1057 |
| Brd7 | 6515 | 6813 | 6858 | 9234 | 6863 | 6661 | 6920 | 6804 | 6886 | 7259 | 7384 | 7292 | 7090.86 | 6870 | 7312 | 1.064 | 0.09 | 0.09563634 | 0.25355123 | 0 | 0.0074 | 0.002 | 0.002 | TRUE | 9.00E-04 |
| Brd8 | 6021 | 5930 | 5920 | 7838 | 5842 | 5534 | 6395 | 5922 | 5944 | 6162 | 6286 | 6058 | 6127.85 | 6087 | 6169 | 1.013 | 0.019 | 0.77238723 | 0.88115034 | 8.00E-04 | 0.0074 | 0.003 | 0.003 | TRUE | 0.0239 |
| Brd9 | 855 | 849 | 771 | 1552 | 1076 | 877 | 908 | 848 | 774 | 1220 | 1158 | 960 | 978 | 843 | 1113 | 1.32 | 0.401 | 0.00124224 | 0.01051667 | 0.01 | 0.0103 | 0.0101 | 0.0101 | TRUE | 0.1708 |
| Brdt | 148 | 159 | 163 | 275 | 180 | 170 | 157 | 159 | 164 | 216 | 194 | 186 | 179.27 | 160 | 199 | 1.246 | 0.317 | 0.04022955 | 0.14150323 | 0 | 0.0253 | 0.0116 | 0.0116 | TRUE | 0.0702 |
| Bre | 1739 | 2123 | 2242 | 2759 | 1931 | 2046 | 1847 | 2120 | 2251 | 2169 | 2078 | 2240 | 2117.47 | 2073 | 2162 | 1.043 | 0.061 | 0.53947826 | 0.72541204 | 0.0052 | 0.0085 | 0.0066 | 0.0066 | TRUE | 0.1094 |
| Brf1 | 902 | 947 | 961 | 1258 | 997 | 881 | 958 | 946 | 965 | 989 | 1073 | 964 | 982.48 | 956 | 1009 | 1.054 | 0.077 | 0.37716295 | 0.59226764 | 6.00E-04 | 0.0103 | 0.0044 | 0.0044 | TRUE | 0.0359 |
| Brf2 | 968 | 966 | 918 | 1345 | 910 | 799 | 1028 | 965 | 922 | 1057 | 979 | 875 | 970.96 | 972 | 970 | 1 | 0 | 0.99937928 | 0.99950602 | 0.005 | 0.0103 | 0.0073 | 0.0073 | TRUE | 0.0882 |
| Bri3 | 1370 | 1851 | 1979 | 1848 | 1518 | 1583 | 1455 | 1849 | 1987 | 1453 | 1633 | 1733 | 1684.96 | 1764 | 1606 | 0.91 | -0.136 | 0.31956599 | 0.5406844 | 0.0153 | 0.0089 | 0.0128 | 0.0128 | TRUE | 0.2819 |
| Bri3bp | 3529 | 3835 | 3783 | 4282 | 3280 | 3493 | 3748 | 3830 | 3799 | 3366 | 3529 | 3824 | 3682.65 | 3792 | 3573 | 0.942 | -0.087 | 0.2579503 | 0.47672745 | 0.0018 | 0.0078 | 0.0039 | 0.0039 | TRUE | 0.0457 |
| Bricd5 | 4 | 1 | 2 | 4 | 3 | 3 | 4 | 1 | 2 | 3 | 3 | 3 | 2.82 | 2 | 3 | 1.339 | 0.421 | 0.70611486 | NA | 0 | 1.1757 | 0.5331 | 0.5331 | TRUE | 0.4438 |
| Brip1 | 2715 | 2077 | 2084 | 3518 | 2404 | 2294 | 2884 | 2074 | 2093 | 2766 | 2587 | 2511 | 2485.66 | 2350 | 2621 | 1.116 | 0.158 | 0.26118314 | 0.48061843 | 0.0181 | 0.0082 | 0.0138 | 0.0138 | TRUE | 0.4774 |
| Brix1 | 6133 | 5963 | 6029 | 8062 | 5641 | 5507 | 6514 | 5955 | 6054 | 6338 | 6069 | 6028 | 6159.79 | 6174 | 6145 | 0.995 | -0.007 | 0.92478965 | 0.96292 | 0.0014 | 0.0074 | 0.0033 | 0.0033 | TRUE | 0.0282 |
| Brk1 | 5236 | 5513 | 5260 | 6971 | 4644 | 4822 | 5562 | 5506 | 5282 | 5480 | 4997 | 5278 | 5350.7 | 5450 | 5252 | 0.964 | -0.053 | 0.44668219 | 0.65274451 | 0.0012 | 0.0075 | 0.0033 | 0.0033 | TRUE | 0.022 |
| Brms1 | 718 | 606 | 576 | 1316 | 897 | 680 | 763 | 605 | 578 | 1035 | 965 | 744 | 781.71 | 649 | 915 | 1.411 | 0.497 | 0.00231117 | 0.01726613 | 0.0235 | 0.0111 | 0.0179 | 0.0179 | TRUE | 0.3138 |
| Brms1l | 1701 | 1566 | 1611 | 2272 | 1570 | 1537 | 1807 | 1564 | 1618 | 1786 | 1689 | 1682 | 1691.02 | 1663 | 1719 | 1.035 | 0.049 | 0.58351328 | 0.75896185 | 0.0028 | 0.0089 | 0.0052 | 0.0052 | TRUE | 0.0689 |
| Brox | 1449 | 1475 | 1380 | 2277 | 1620 | 1495 | 1539 | 1473 | 1386 | 1790 | 1743 | 1637 | 1594.56 | 1466 | 1723 | 1.176 | 0.234 | 0.00561212 | 0.03408328 | 0.0018 | 0.009 | 0.0045 | 0.0045 | TRUE | 0.0277 |
| Brpf1 | 3778 | 3283 | 3262 | 5268 | 3703 | 3312 | 4013 | 3279 | 3275 | 4141 | 3984 | 3626 | 3719.68 | 3522 | 3917 | 1.112 | 0.154 | 0.16330101 | 0.35895107 | 0.0091 | 0.0078 | 0.0085 | 0.0085 | TRUE | 0.1803 |
| Brpf3 | 2365 | 2394 | 2616 | 4219 | 3142 | 3276 | 2512 | 2391 | 2627 | 3317 | 3381 | 3586 | 2968.84 | 2510 | 3428 | 1.365 | 0.449 | 3.95E-09 | 2.04E-07 | 0.0014 | 0.008 | 0.0039 | 0.0039 | TRUE | 0.0209 |
| Brs3 | 0 | 0 | 0 | 0 | 0 | 0 | 0 | 0 | 0 | 0 | 0 | 0 | 0 | 0 | 0 | NA | NA | NA | NA | NA | NA | NA | NA | NA | NA |
| Brsk1 | 883 | 978 | 946 | 1115 | 784 | 730 | 938 | 977 | 950 | 877 | 844 | 799 | 897.28 | 955 | 840 | 0.88 | -0.184 | 0.03405605 | 0.12552745 | 0 | 0.0106 | 0.0043 | 0.0043 | TRUE | 0.0211 |
| Brsk2 | 253 | 272 | 310 | 477 | 394 | 355 | 269 | 272 | 311 | 375 | 424 | 389 | 339.86 | 284 | 396 | 1.392 | 0.478 | 0.00021546 | 0.00258221 | 0.0023 | 0.0166 | 0.009 | 0.009 | TRUE | 0.0805 |
| Brwd1 | 4810 | 5396 | 5564 | 6427 | 4854 | 4883 | 5109 | 5389 | 5587 | 5052 | 5223 | 5345 | 5284.19 | 5362 | 5207 | 0.971 | -0.043 | 0.53903923 | 0.7252619 | 0.0011 | 0.0075 | 0.0033 | 0.0033 | TRUE | 0.0207 |
| Brwd3 | 1353 | 1415 | 1563 | 2421 | 1618 | 1555 | 1437 | 1413 | 1569 | 1903 | 1741 | 1702 | 1627.66 | 1473 | 1782 | 1.21 | 0.275 | 0.00212851 | 0.01617872 | 0.0028 | 0.0089 | 0.0052 | 0.0052 | TRUE | 0.0436 |

| | | | | | | | | | | | | | | | | | | | | | | | | |
|---|---|---|---|---|---|---|---|---|---|---|---|---|---|---|---|---|---|---|---|---|---|---|---|---|
| Bscl2 | 519 | 593 | 624 | 1095 | 742 | 679 | 551 | 592 | 627 | 861 | 798 | 743 | 695.41 | 590 | 801 | 1.359 | 0.442 | 4.22E-05 | 0.00067047 | 0.0034 | 0.0116 | 0.0069 | 0.0069 | TRUE | 0.0528 |
| Bsdc1 | 2288 | 2286 | 2306 | 2723 | 1961 | 2013 | 2430 | 2283 | 2315 | 2141 | 2110 | 2204 | 2247.14 | 2343 | 2152 | 0.918 | -0.123 | 0.07781946 | 0.22065963 | 3.00E-04 | 0.0084 | 0.0031 | 0.0031 | TRUE | 0.0128 |
| Bsg | 17015 | 23145 | 23297 | 23220 | 17605 | 20909 | 18073 | 23115 | 23393 | 18254 | 18942 | 22888 | 20777.42 | 21527 | 20028 | 0.93 | -0.104 | 0.43277327 | 0.64199587 | 0.0156 | 0.0071 | 0.0127 | 0.0127 | TRUE | 0.241 |
| Bsn | 75 | 64 | 66 | 130 | 96 | 76 | 80 | 64 | 66 | 102 | 103 | 83 | 83.09 | 70 | 96 | 1.38 | 0.465 | 0.04129285 | 0.14408646 | 0 | 0.0466 | 0.025 | 0.025 | TRUE | 0.1294 |
| Bsnd | 0 | 1 | 2 | 0 | 1 | 1 | 0 | 1 | 2 | 0 | 1 | 1 | 0.86 | 1 | 1 | 0.687 | -0.541 | 0.80218377 | NA | 0 | 3.8247 | 2.1472 | 2.1472 | TRUE | 0.1636 |
| Bsph1 | 0 | 0 | 0 | 0 | 0 | 0 | 0 | 0 | 0 | 0 | 0 | 0 | 0 | 0 | 0 | NA | NA | NA | NA | NA | NA | NA | NA | NA | NA |
| Bsph2 | 0 | 0 | 0 | 0 | 0 | 0 | 0 | 0 | 0 | 0 | 0 | 0 | 0 | 0 | 0 | NA | NA | NA | NA | NA | NA | NA | NA | NA | NA |
| Bspry | 443 | 437 | 464 | 544 | 448 | 442 | 471 | 436 | 466 | 428 | 482 | 484 | 461.07 | 458 | 465 | 1.013 | 0.019 | 0.86615841 | 0.93370586 | 0 | 0.014 | 0.0069 | 0.0069 | TRUE | 0.0596 |
| Bst1 | 126 | 90 | 100 | 137 | 113 | 101 | 134 | 90 | 100 | 108 | 122 | 111 | 110.66 | 108 | 114 | 1.049 | 0.068 | 0.75491402 | 0.87058798 | 0.014 | 0.0367 | 0.0255 | 0.0255 | TRUE | 0.4253 |
| Bst2 | 1490 | 2411 | 2376 | 2178 | 1650 | 2261 | 1583 | 2408 | 2386 | 1712 | 1775 | 2475 | 2056.46 | 2126 | 1987 | 0.935 | -0.097 | 0.61560128 | 0.78256334 | 0.0475 | 0.0085 | 0.0266 | 0.0266 | TRUE | 0.6033 |
| Bsx | 1 | 0 | 0 | 1 | 0 | 0 | 1 | 0 | 0 | 1 | 0 | 0 | 0.31 | 0 | 0 | 0.954 | -0.067 | 0.98625928 | NA | 0 | 10.7014 | 9.0384 | 9.0384 | TRUE | 0.4614 |
| Btaf1 | 7396 | 6163 | 6497 | 8629 | 6035 | 5339 | 7856 | 6155 | 6524 | 6783 | 6493 | 5844 | 6609.28 | 6845 | 6373 | 0.931 | -0.103 | 0.37289327 | 0.58928297 | 0.0108 | 0.0074 | 0.0094 | 0.0094 | TRUE | 0.2036 |
| Btbd1 | 2321 | 3490 | 3493 | 3507 | 2678 | 2921 | 2465 | 3485 | 3507 | 2757 | 2881 | 3197 | 3048.99 | 3152 | 2945 | 0.934 | -0.099 | 0.50127488 | 0.69669611 | 0.0219 | 0.008 | 0.0151 | 0.0151 | TRUE | 0.4415 |
| Btbd10 | 2528 | 2102 | 2016 | 2931 | 2040 | 1951 | 2685 | 2099 | 2024 | 2304 | 2195 | 2136 | 2240.58 | 2269 | 2212 | 0.975 | -0.037 | 0.76545801 | 0.87700158 | 0.0124 | 0.0084 | 0.0106 | 0.0106 | TRUE | 0.3103 |
| Btbd11 | 742 | 740 | 686 | 1157 | 812 | 706 | 788 | 739 | 689 | 910 | 874 | 773 | 795.34 | 739 | 852 | 1.155 | 0.208 | 0.05733807 | 0.18050619 | 0.0046 | 0.011 | 0.0073 | 0.0073 | TRUE | 0.0776 |
| Btbd16 | 2 | 2 | 1 | 3 | 1 | 1 | 2 | 2 | 1 | 2 | 1 | 1 | 1.61 | 2 | 1 | 0.908 | -0.139 | 0.92153658 | NA | 0 | 2.0542 | 0.8218 | 0.8218 | TRUE | 0.2247 |
| Btbd17 | 15 | 29 | 38 | 27 | 28 | 22 | 16 | 29 | 38 | 21 | 30 | 24 | 26.41 | 28 | 25 | 0.902 | -0.148 | 0.73493049 | 0.85772708 | 0.0656 | 0.1316 | 0.1002 | 0.1002 | TRUE | 0.2688 |
| Btbd19 | 425 | 591 | 597 | 859 | 796 | 698 | 451 | 590 | 599 | 675 | 856 | 764 | 656.15 | 547 | 765 | 1.397 | 0.482 | 0.00137648 | 0.01143536 | 0.0175 | 0.0119 | 0.0148 | 0.0148 | TRUE | 0.2717 |
| Btbd2 | 2267 | 4016 | 4238 | 3430 | 2992 | 3427 | 2408 | 4011 | 4255 | 2696 | 3219 | 3751 | 3390.19 | 3558 | 3222 | 0.905 | -0.143 | 0.48773332 | 0.68458483 | 0.0582 | 0.0079 | 0.0304 | 0.0304 | TRUE | 0.8072 |
| Btbd3 | 1262 | 1441 | 1490 | 1813 | 1374 | 1447 | 1340 | 1439 | 1496 | 1425 | 1478 | 1584 | 1460.55 | 1425 | 1496 | 1.049 | 0.069 | 0.43766452 | 0.64554801 | 0.0023 | 0.0092 | 0.005 | 0.005 | TRUE | 0.0324 |
| Btbd6 | 579 | 569 | 564 | 841 | 603 | 499 | 615 | 568 | 566 | 661 | 649 | 546 | 600.96 | 583 | 619 | 1.062 | 0.087 | 0.4539924 | 0.65915199 | 0.0048 | 0.0124 | 0.0081 | 0.0081 | TRUE | 0.121 |
| Btbd7 | 1731 | 1767 | 1864 | 2221 | 1604 | 1784 | 1839 | 1765 | 1872 | 1746 | 1726 | 1953 | 1816.61 | 1825 | 1808 | 0.99 | -0.014 | 0.87065959 | 0.93644593 | 0.0022 | 0.0087 | 0.0047 | 0.0047 | TRUE | 0.0584 |
| Btbd8 | 99 | 155 | 140 | 149 | 117 | 114 | 105 | 155 | 141 | 117 | 126 | 125 | 128.06 | 134 | 123 | 0.916 | -0.126 | 0.53775148 | 0.72419528 | 0.0121 | 0.0326 | 0.0224 | 0.0224 | TRUE | 0.3485 |
| Btbd9 | 2486 | 2366 | 2335 | 2796 | 2022 | 1975 | 2641 | 2363 | 2345 | 2198 | 2176 | 2162 | 2313.93 | 2450 | 2179 | 0.89 | -0.169 | 0.03556347 | 0.12950116 | 0.0017 | 0.0083 | 0.0042 | 0.0042 | TRUE | 0.0563 |
| Btc | 17 | 14 | 12 | 22 | 15 | 18 | 18 | 14 | 12 | 17 | 16 | 20 | 16.2 | 15 | 18 | 1.206 | 0.27 | 0.56426803 | 0.74500513 | 0 | 0.2102 | 0.0961 | 0.0961 | TRUE | 0.1722 |
| Btd | 421 | 392 | 376 | 533 | 375 | 311 | 447 | 391 | 378 | 419 | 403 | 340 | 396.52 | 405 | 387 | 0.958 | -0.062 | 0.65234679 | 0.80502827 | 0.0073 | 0.0152 | 0.011 | 0.011 | TRUE | 0.1275 |
| Btf3 | 14430 | 17230 | 17580 | 19926 | 14228 | 14896 | 15327 | 17208 | 17652 | 15664 | 15308 | 16306 | 16244.31 | 16729 | 15759 | 0.942 | -0.086 | 0.28499915 | 0.50627409 | 0.0031 | 0.0071 | 0.0046 | 0.0046 | TRUE | 0.0657 |
| Btf3l4 | 1134 | 1203 | 1196 | 1669 | 1225 | 1277 | 1205 | 1201 | 1201 | 1312 | 1318 | 1398 | 1272.47 | 1202 | 1343 | 1.116 | 0.159 | 0.03796506 | 0.13544213 | 0 | 0.0095 | 0.0034 | 0.0034 | TRUE | 0.0153 |
| Btg1 | 1498 | 2032 | 2067 | 1879 | 1519 | 1851 | 1591 | 2029 | 2075 | 1477 | 1634 | 2026 | 1805.61 | 1898 | 1712 | 0.901 | -0.15 | 0.32405118 | 0.54535337 | 0.0212 | 0.0087 | 0.0161 | 0.0161 | TRUE | 0.3092 |
| Btg2 | 1740 | 2092 | 1783 | 2407 | 1795 | 1826 | 1848 | 2089 | 1790 | 1892 | 1931 | 1999 | 1925.03 | 1909 | 1941 | 1.016 | 0.023 | 0.79675655 | 0.89536272 | 0.0032 | 0.0086 | 0.0053 | 0.0053 | TRUE | 0.0825 |
| Btg3 | 0 | 1 | 0 | 1 | 0 | 1 | 0 | 1 | 0 | 1 | 0 | 1 | 0.48 | 0 | 1 | 1.723 | 0.785 | 0.78325239 | NA | 0 | 6.8716 | 4.0968 | 4.0968 | TRUE | 0.405 |
| Btg4 | 0 | 0 | 0 | 0 | 0 | 0 | 0 | 0 | 0 | 0 | 0 | 0 | 0 | 0 | 0 | NA | NA | NA | NA | NA | NA | NA | NA | NA | NA |
| Btk | 5 | 8 | 4 | 22 | 9 | 12 | 5 | 8 | 4 | 17 | 10 | 13 | 9.57 | 6 | 13 | 2.334 | 1.222 | 0.06220092 | 0.19086242 | 0 | 0.3511 | 0.185 | 0.185 | TRUE | 0.1681 |
| Btla | 5 | 0 | 0 | 1 | 1 | 1 | 5 | 0 | 0 | 1 | 1 | 1 | 1.38 | 2 | 1 | 0.558 | -0.842 | 0.67211302 | NA | 1.6425 | 2.3978 | 2.0642 | 2.0642 | TRUE | 2.345 |
| Btn1a1 | 13 | 11 | 21 | 14 | 8 | 4 | 14 | 11 | 21 | 11 | 9 | 4 | 11.65 | 15 | 8 | 0.531 | -0.912 | 0.13829397 | 0.32280192 | 0.0506 | 0.2898 | 0.1794 | 0.1794 | TRUE | 0.5127 |
| Btn2a2 | 107 | 80 | 89 | 110 | 81 | 75 | 114 | 80 | 89 | 86 | 87 | 82 | 89.77 | 94 | 85 | 0.905 | -0.143 | 0.52425069 | 0.71390795 | 0.0061 | 0.0436 | 0.0253 | 0.0253 | TRUE | 0.305 |
| Btnl1 | 9 | 21 | 12 | 14 | 14 | 13 | 10 | 21 | 12 | 11 | 15 | 14 | 13.81 | 14 | 13 | 0.938 | -0.092 | 0.86572937 | 0.93364615 | 0 | 0.2454 | 0.1416 | 0.1416 | TRUE | 0.7766 |
| Btnl10 | 0 | 0 | 0 | 0 | 0 | 0 | 0 | 0 | 0 | 0 | 0 | 0 | 0 | 0 | 0 | NA | NA | NA | NA | NA | NA | NA | NA | NA | NA |
| Btnl2 | 0 | 0 | 0 | 0 | 0 | 0 | 0 | 0 | 0 | 0 | 0 | 0 | 0 | 0 | 0 | NA | NA | NA | NA | NA | NA | NA | NA | NA | NA |
| Btnl4 | 4 | 6 | 7 | 4 | 4 | 3 | 4 | 6 | 7 | 3 | 4 | 3 | 4.67 | 6 | 3 | 0.617 | -0.697 | 0.41558128 | 0.62710945 | 0 | 0.7128 | 0.303 | 0.303 | TRUE | 0.1126 |
| Btnl5 | 1 | 1 | 0 | 2 | 0 | 2 | 1 | 1 | 0 | 2 | 0 | 2 | 0.97 | 1 | 1 | 1.844 | 0.883 | 0.66663732 | NA | 0 | 3.4018 | 1.8936 | 1.8936 | TRUE | 0.3972 |
| Btnl6 | 1 | 3 | 2 | 1 | 1 | 0 | 1 | 3 | 2 | 1 | 1 | 0 | 1.32 | 2 | 1 | 0.312 | -1.68 | 0.32925837 | NA | 0 | 2.5 | 1.1176 | 1.1176 | TRUE | 0.1621 |
| Btnl9 | 24 | 15 | 10 | 18 | 10 | 10 | 25 | 15 | 10 | 14 | 11 | 11 | 14.4 | 17 | 12 | 0.717 | -0.481 | 0.38694091 | 0.6009682 | 0.0652 | 0.2358 | 0.1514 | 0.1514 | TRUE | 0.4208 |
| Btrc | 2118 | 2468 | 2418 | 3101 | 2208 | 2293 | 2250 | 2465 | 2428 | 2438 | 2376 | 2510 | 2410.99 | 2381 | 2441 | 1.025 | 0.036 | 0.63210942 | 0.79293961 | 0.0011 | 0.0083 | 0.0036 | 0.0036 | TRUE | 0.028 |
| Bub1 | 10987 | 13400 | 13789 | 14124 | 10330 | 11451 | 11670 | 13383 | 13846 | 11103 | 11114 | 12535 | 12275.17 | 12966 | 11584 | 0.893 | -0.163 | 0.09368282 | 0.25039238 | 0.0063 | 0.0072 | 0.0067 | 0.0067 | TRUE | 0.0934 |
| Bub1b | 15039 | 16525 | 16622 | 18220 | 13506 | 14056 | 15974 | 16504 | 16690 | 14323 | 14532 | 15387 | 15568.21 | 16389 | 14747 | 0.9 | -0.152 | 0.01494295 | 0.07064591 | 9.00E-04 | 0.0071 | 0.0028 | 0.0028 | TRUE | 0.0176 |
| Bub3 | 6097 | 6661 | 6454 | 7989 | 5783 | 5926 | 6476 | 6652 | 6480 | 6280 | 6222 | 6487 | 6433.07 | 6536 | 6330 | 0.968 | -0.046 | 0.42391181 | 0.63416469 | 1.00E-04 | 0.0074 | 0.0023 | 0.0023 | TRUE | 0.0057 |
| Bud13 | 1532 | 1465 | 1597 | 1901 | 1398 | 1344 | 1627 | 1463 | 1604 | 1494 | 1504 | 1471 | 1527.29 | 1565 | 1490 | 0.952 | -0.07 | 0.38324528 | 0.59760644 | 0.001 | 0.0091 | 0.004 | 0.004 | TRUE | 0.039 |
| Bud31 | 1043 | 828 | 784 | 1717 | 1148 | 844 | 1108 | 827 | 787 | 1350 | 1235 | 924 | 1038.47 | 907 | 1170 | 1.29 | 0.367 | 0.04103687 | 0.14344714 | 0.0348 | 0.0101 | 0.0223 | 0.0223 | TRUE | 0.4442 |
| Bves | 77 | 103 | 111 | 153 | 98 | 95 | 82 | 103 | 111 | 120 | 105 | 104 | 104.3 | 99 | 110 | 1.115 | 0.158 | 0.45439006 | 0.65948652 | 0.0049 | 0.0385 | 0.0224 | 0.0224 | TRUE | 0.2138 |
| Bysl | 5052 | 4468 | 4256 | 6758 | 4628 | 3699 | 5366 | 4462 | 4273 | 5313 | 4979 | 4049 | 4740.5 | 4700 | 4780 | 1.017 | 0.025 | 0.85335047 | 0.92681753 | 0.017 | 0.0076 | 0.0126 | 0.0126 | TRUE | 0.2178 |
| Bzrap1 | 119 | 110 | 100 | 73 | 76 | 74 | 126 | 110 | 100 | 57 | 82 | 81 | 92.81 | 112 | 73 | 0.65 | -0.622 | 0.00837772 | 0.04608178 | 0.0146 | 0.0424 | 0.0289 | 0.0289 | TRUE | 0.3799 |
| Bzw1 | 16062 | 13489 | 13663 | 20484 | 14129 | 12613 | 17061 | 13472 | 13719 | 16103 | 15202 | 13807 | 14893.85 | 14751 | 15037 | 1.02 | 0.028 | 0.8105992 | 0.90275177 | 0.0097 | 0.0071 | 0.0097 | 0.0097 | TRUE | 0.2295 |
| Bzw2 | 11727 | 13439 | 13973 | 12381 | 9421 | 10577 | 12456 | 13422 | 14030 | 9733 | 10136 | 11578 | 11892.62 | 13303 | 10482 | 0.788 | -0.344 | 0.00028559 | 0.00327348 | 0.0059 | 0.0072 | 0.0064 | 0.0064 | TRUE | 0.1021 |
| C030006K11I | 233 | 213 | 219 | 279 | 159 | 143 | 247 | 213 | 220 | 219 | 171 | 157 | 204.51 | 227 | 182 | 0.808 | -0.307 | 0.08831544 | 0.24101084 | 0.0141 | 0.023 | 0.0184 | 0.0184 | TRUE | 0.3616 |
| C030007H22 | 0 | 0 | 0 | 0 | 0 | 0 | 0 | 0 | 0 | 0 | 0 | 0 | 0 | 0 | 0 | NA | NA | NA | NA | NA | NA | NA | NA | NA | NA |
| C030013G03 | 0 | 0 | 0 | 0 | 0 | 0 | 0 | 0 | 0 | 0 | 0 | 0 | 0 | 0 | 0 | NA | NA | NA | NA | NA | NA | NA | NA | NA | NA |
| C030016D13 | 5 | 11 | 13 | 4 | 6 | 8 | 5 | 11 | 13 | 3 | 6 | 9 | 7.95 | 10 | 6 | 0.611 | -0.71 | 0.33732219 | 0.5573856 | 0 | 0.4212 | 0.2611 | 0.2611 | TRUE | 0.5652 |
| C030018K13I | 0 | 1 | 4 | 1 | 1 | 2 | 0 | 1 | 4 | 1 | 1 | 2 | 1.51 | 2 | 1 | 0.79 | -0.341 | 0.83722826 | NA | 0 | 2.1871 | 1.308 | 1.308 | TRUE | 0.8847 |
| C030023E24I | 0 | 0 | 0 | 0 | 0 | 0 | 0 | 0 | 0 | 0 | 0 | 0 | 0 | 0 | 0 | NA | NA | NA | NA | NA | NA | NA | NA | NA | NA |
| C030030A07 | 0 | 1 | 0 | 3 | 0 | 0 | 0 | 1 | 0 | 2 | 0 | 0 | 0.56 | 0 | 1 | 2.222 | 1.152 | 0.72351576 | NA | 0 | 5.895 | 6.0404 | 6.0404 | TRUE | 1.4404 |
| C030034I22R | 28 | 54 | 60 | 40 | 39 | 39 | 30 | 54 | 60 | 31 | 42 | 43 | 43.34 | 48 | 39 | 0.801 | -0.32 | 0.37364793 | 0.58999877 | 0.0558 | 0.0829 | 0.0699 | 0.0699 | TRUE | 0.861 |
| C030034L19I | 0 | 1 | 1 | 1 | 0 | 0 | 0 | 1 | 1 | 1 | 0 | 0 | 0.46 | 1 | 0 | 0.52 | -0.943 | 0.74346893 | NA | 0 | 7.0944 | 4.2167 | 4.2167 | TRUE | 0.2562 |
| C030037D09 | 37 | 47 | 55 | 47 | 42 | 48 | 39 | 47 | 55 | 37 | 45 | 53 | 46.02 | 47 | 45 | 0.945 | -0.081 | 0.78920183 | 0.891612 | 0 | 0.0785 | 0.0451 | 0.0451 | TRUE | 0.2203 |
| C030039L03I | 244 | 238 | 212 | 853 | 653 | 461 | 259 | 238 | 213 | 671 | 703 | 505 | 431.25 | 237 | 626 | 2.649 | 1.405 | 1.40E-17 | 2.66E-15 | 0.0185 | 0.0145 | 0.0166 | 0.0166 | TRUE | 0.3328 |
| C030046E11I | 2193 | 2388 | 2370 | 3240 | 2304 | 2266 | 2329 | 2385 | 2380 | 2547 | 2479 | 2480 | 2433.42 | 2365 | 2502 | 1.058 | 0.082 | 0.20035576 | 0.40805928 | 0 | 0.0083 | 0.0025 | 0.0025 | TRUE | 0.0031 |
| C130021I20R | 5 | 9 | 5 | 37 | 26 | 16 | 5 | 9 | 5 | 29 | 28 | 18 | 15.65 | 6 | 25 | 3.869 | 1.952 | 0.00048184 | 0.00503618 | 0 | 0.2174 | 0.1266 | 0.1266 | TRUE | 0.3779 |
| C130026I21R | 9 | 6 | 7 | 20 | 17 | 7 | 10 | 6 | 7 | 16 | 18 | 8 | 10.71 | 8 | 14 | 1.858 | 0.894 | 0.15510741 | 0.34739479 | 0 | 0.3145 | 0.1818 | 0.1818 | TRUE | 0.604 |
| C130026L21I | 8 | 9 | 9 | 2 | 2 | 3 | 8 | 9 | 9 | 2 | 2 | 3 | 5.59 | 9 | 2 | 0.258 | -1.953 | 0.02370357 | 0.09797529 | 0 | 0.5964 | 0.2669 | 0.2669 | TRUE | 0.1231 |

| | | | | | | | | | | | | | | | | | | | | | | | | | |
|---|---|---|---|---|---|---|---|---|---|---|---|---|---|---|---|---|---|---|---|---|---|---|---|---|---|
| C130030K03I | 0 | 0 | 0 | 1 | 0 | 0 | 0 | 0 | 0 | 1 | 0 | 0 | 0.13 | 0 | 0 | 1.859 | 0.894 | 0.82650734 | NA | 0 | 25.1515 | 10 | 10 | TRUE | 0.2563 |
| C130036L24I | 16 | 22 | 19 | 27 | 20 | 12 | 17 | 22 | 19 | 21 | 22 | 13 | 18.99 | 19 | 19 | 0.967 | -0.048 | 0.9148037 | 0.95805338 | 0 | 0.1804 | 0.0911 | 0.0911 | TRUE | 0.313 |
| C130050O18 | 0 | 1 | 1 | 0 | 0 | 1 | 0 | 1 | 1 | 0 | 0 | 1 | 0.52 | 1 | 0 | 0.52 | -0.943 | 0.73639299 | NA | 0 | 6.3885 | 3.8941 | 3.8941 | TRUE | 0.5383 |
| C130060C02I | 6 | 3 | 3 | 4 | 2 | 2 | 6 | 3 | 3 | 3 | 2 | 2 | 3.31 | 4 | 2 | 0.615 | -0.702 | 0.49510149 | NA | 0 | 1.0018 | 0.4458 | 0.4458 | TRUE | 0.3698 |
| C130060K24I | 2 | 1 | 0 | 1 | 2 | 3 | 2 | 1 | 0 | 1 | 2 | 3 | 1.56 | 1 | 2 | 1.971 | 0.979 | 0.54311098 | NA | 0 | 2.1221 | 1.1304 | 1.1304 | TRUE | 0.2769 |
| C130071C03I | 0 | 0 | 0 | 1 | 0 | 0 | 0 | 0 | 0 | 1 | 0 | 0 | 0.13 | 0 | 0 | 1.859 | 0.894 | 0.82650734 | NA | 0 | 25.1515 | 10 | 10 | TRUE | 0.2563 |
| C130074G19 | 354 | 448 | 540 | 598 | 418 | 458 | 376 | 447 | 542 | 470 | 450 | 501 | 464.47 | 455 | 474 | 1.04 | 0.057 | 0.71270496 | 0.84358593 | 0.0158 | 0.014 | 0.0149 | 0.0149 | TRUE | 0.2835 |
| C130079G13 | 0 | 0 | 0 | 0 | 0 | 0 | 0 | 0 | 0 | 0 | 0 | 0 | 0 | 0 | 0 | NA | NA | NA | NA | NA | NA | NA | NA | NA | NA |
| C130080G10 | 0 | 0 | 0 | 0 | 0 | 0 | 0 | 0 | 0 | 0 | 0 | 0 | 0 | 0 | 0 | NA | NA | NA | NA | NA | NA | NA | NA | NA | NA |
| C130083M11 | 1 | 2 | 0 | 3 | 2 | 1 | 1 | 2 | 0 | 2 | 2 | 1 | 1.44 | 1 | 2 | 1.848 | 0.886 | 0.58065249 | NA | 0 | 2.2882 | 1.0946 | 1.0946 | TRUE | 0.3322 |
| C1d | 1332 | 1317 | 1360 | 2157 | 1519 | 1364 | 1415 | 1315 | 1366 | 1696 | 1634 | 1493 | 1486.47 | 1365 | 1608 | 1.179 | 0.237 | 0.0067442 | 0.0390347 | 0.0022 | 0.0091 | 0.0048 | 0.0048 | TRUE | 0.0465 |
| C1galt1 | 945 | 1020 | 852 | 1157 | 785 | 805 | 1004 | 1019 | 855 | 910 | 845 | 881 | 918.88 | 959 | 879 | 0.916 | -0.126 | 0.22957283 | 0.44313092 | 0.0042 | 0.0105 | 0.0069 | 0.0069 | TRUE | 0.1073 |
| C1galt1c1 | 311 | 259 | 219 | 401 | 261 | 277 | 330 | 259 | 220 | 315 | 281 | 303 | 284.7 | 270 | 300 | 1.113 | 0.155 | 0.37737755 | 0.59235754 | 0.0188 | 0.0185 | 0.0187 | 0.0187 | TRUE | 0.4302 |
| C1qa | 0 | 0 | 0 | 0 | 0 | 0 | 0 | 0 | 0 | 0 | 0 | 0 | 0 | 0 | 0 | NA | NA | NA | NA | NA | NA | NA | NA | NA | NA |
| C1qb | 0 | 0 | 0 | 0 | 0 | 0 | 0 | 0 | 0 | 0 | 0 | 0 | 0 | 0 | 0 | NA | NA | NA | NA | NA | NA | NA | NA | NA | NA |
| C1qbp | 20749 | 23679 | 24236 | 28550 | 20071 | 20241 | 22039 | 23648 | 24336 | 22444 | 21595 | 22157 | 22703.16 | 23341 | 22065 | 0.945 | -0.081 | 0.22647929 | 0.43931021 | 0.0014 | 0.007 | 0.0032 | 0.0032 | TRUE | 0.0291 |
| C1qc | 0 | 0 | 0 | 0 | 0 | 0 | 0 | 0 | 0 | 0 | 0 | 0 | 0 | 0 | 0 | NA | NA | NA | NA | NA | NA | NA | NA | NA | NA |
| C1ql1 | 352 | 465 | 467 | 305 | 251 | 291 | 374 | 464 | 469 | 240 | 270 | 319 | 355.93 | 436 | 276 | 0.632 | -0.662 | 3.72E-05 | 0.0006022 | 0.0152 | 0.0162 | 0.0156 | 0.0156 | TRUE | 0.2007 |
| C1ql2 | 1 | 1 | 1 | 11 | 4 | 1 | 1 | 1 | 1 | 9 | 4 | 1 | 2.85 | 1 | 5 | 4.678 | 2.226 | 0.09880667 | NA | 0 | 1.1621 | 0.7077 | 0.7077 | TRUE | 0.1127 |
| C1ql3 | 1 | 8 | 3 | 6 | 9 | 7 | 1 | 8 | 3 | 5 | 10 | 8 | 5.69 | 4 | 8 | 1.801 | 0.849 | 0.35178677 | 0.57063668 | 0.1815 | 0.5861 | 0.4058 | 0.4058 | TRUE | 1.1037 |
| C1ql4 | 71 | 67 | 61 | 95 | 70 | 76 | 75 | 67 | 61 | 75 | 75 | 83 | 72.79 | 68 | 78 | 1.145 | 0.195 | 0.39543099 | 0.60887802 | 0 | 0.0522 | 0.0241 | 0.0241 | TRUE | 0.0823 |
| C1qtnf1 | 16 | 15 | 8 | 7 | 7 | 10 | 17 | 15 | 8 | 6 | 8 | 11 | 10.66 | 13 | 8 | 0.592 | -0.756 | 0.22827956 | 0.44159023 | 0 | 0.3158 | 0.1837 | 0.1837 | TRUE | 0.5167 |
| C1qtnf2 | 0 | 0 | 0 | 3 | 0 | 2 | 0 | 0 | 0 | 2 | 0 | 2 | 0.76 | 0 | 1 | 8.123 | 3.022 | 0.28653847 | NA | 0 | 4.3534 | 4.4726 | 4.4726 | TRUE | 0.4916 |
| C1qtnf3 | 0 | 0 | 0 | 0 | 0 | 0 | 0 | 0 | 0 | 0 | 0 | 0 | 0 | 0 | 0 | NA | NA | NA | NA | NA | NA | NA | NA | NA | NA |
| C1qtnf4 | 61 | 84 | 93 | 90 | 79 | 81 | 65 | 84 | 93 | 71 | 85 | 89 | 81.08 | 81 | 82 | 1.005 | 0.007 | 0.97841776 | 0.98828002 | 0.0115 | 0.0475 | 0.03 | 0.03 | TRUE | 0.2699 |
| C1qtnf5 | 0 | 0 | 0 | 0 | 0 | 0 | 0 | 0 | 0 | 0 | 0 | 0 | 0 | 0 | 0 | NA | NA | NA | NA | NA | NA | NA | NA | NA | NA |
| C1qtnf6 | 21 | 19 | 13 | 16 | 20 | 22 | 22 | 19 | 13 | 13 | 22 | 24 | 18.75 | 18 | 20 | 1.06 | 0.084 | 0.86197839 | 0.93187491 | 0.0409 | 0.1826 | 0.1144 | 0.1144 | TRUE | 0.6225 |
| C1qtnf7 | 1 | 1 | 1 | 0 | 0 | 0 | 1 | 1 | 1 | 0 | 0 | 0 | 0.51 | 1 | 0 | 0.176 | -2.51 | 0.35480135 | NA | 0 | 6.4561 | 3.8046 | 3.8046 | TRUE | 0.0847 |
| C1qtnf9 | 5 | 7 | 6 | 4 | 5 | 3 | 5 | 7 | 6 | 3 | 5 | 3 | 5.02 | 6 | 4 | 0.638 | -0.649 | 0.43270027 | 0.64199587 | 0 | 0.6628 | 0.2847 | 0.2847 | TRUE | 0.156 |
| C1ra | 12 | 10 | 7 | 15 | 16 | 10 | 13 | 10 | 7 | 12 | 17 | 11 | 11.62 | 10 | 13 | 1.341 | 0.423 | 0.45957059 | 0.66382887 | 0 | 0.2904 | 0.1472 | 0.1472 | TRUE | 0.2565 |
| C1rb | 6 | 1 | 3 | 6 | 16 | 5 | 6 | 1 | 3 | 5 | 17 | 5 | 6.3 | 3 | 9 | 2.634 | 1.397 | 0.14627424 | 0.33503301 | 0.3953 | 0.53 | 0.4637 | 0.4637 | TRUE | 1.7429 |
| C1rl | 148 | 186 | 178 | 269 | 197 | 191 | 157 | 186 | 179 | 211 | 212 | 209 | 192.37 | 174 | 211 | 1.212 | 0.277 | 0.06436476 | 0.19526081 | 0 | 0.024 | 0.0109 | 0.0109 | TRUE | 0.0739 |
| C1s | 216 | 213 | 120 | 463 | 364 | 284 | 229 | 213 | 120 | 364 | 392 | 311 | 271.52 | 187 | 356 | 1.897 | 0.924 | 9.15E-05 | 0.00129038 | 0.0564 | 0.019 | 0.0361 | 0.0361 | TRUE | 1.0597 |
| C2 | 50 | 105 | 114 | 42 | 29 | 63 | 53 | 105 | 114 | 33 | 31 | 69 | 67.6 | 91 | 44 | 0.486 | -1.041 | 0.01081405 | 0.05575653 | 0.1595 | 0.0556 | 0.1035 | 0.1035 | TRUE | 1.7534 |
| C230004F18I | 0 | 0 | 0 | 0 | 0 | 0 | 0 | 0 | 0 | 0 | 0 | 0 | 0 | 0 | 0 | NA | NA | NA | NA | NA | NA | NA | NA | NA | NA |
| C230024C17I | 0 | 0 | 0 | 0 | 0 | 1 | 0 | 0 | 0 | 0 | 0 | 1 | 0.18 | 0 | 0 | 1.859 | 0.894 | 0.82650734 | NA | 0 | 18.0643 | 10 | 10 | TRUE | 0.5383 |
| C230035I16R | 7 | 16 | 4 | 9 | 12 | 10 | 7 | 16 | 4 | 7 | 13 | 11 | 9.73 | 9 | 10 | 1.116 | 0.158 | 0.82229101 | 0.90923689 | 0.1592 | 0.3456 | 0.255 | 0.255 | TRUE | 0.817 |
| C230052I12R | 614 | 525 | 453 | 712 | 494 | 362 | 652 | 524 | 455 | 560 | 532 | 396 | 519.81 | 544 | 496 | 0.913 | -0.131 | 0.47259108 | 0.67369003 | 0.031 | 0.0132 | 0.0221 | 0.0221 | TRUE | 0.355 |
| C230079O03 | 0 | 0 | 0 | 0 | 0 | 0 | 0 | 0 | 0 | 0 | 0 | 0 | 0 | 0 | 0 | NA | NA | NA | NA | NA | NA | NA | NA | NA | NA |
| C230081A13 | 2739 | 2151 | 2512 | 3136 | 2152 | 2078 | 2909 | 2148 | 2522 | 2465 | 2315 | 2275 | 2439.21 | 2526 | 2352 | 0.931 | -0.103 | 0.39772665 | 0.61115985 | 0.0115 | 0.0083 | 0.0103 | 0.0103 | TRUE | 0.179 |
| C230091D08 | 0 | 1 | 0 | 1 | 1 | 1 | 0 | 1 | 0 | 1 | 1 | 1 | 0.66 | 0 | 1 | 2.676 | 1.42 | 0.53620179 | NA | 0 | 5.0043 | 2.2992 | 2.2992 | TRUE | 0.405 |
| C2cd2 | 546 | 710 | 662 | 772 | 579 | 533 | 580 | 709 | 665 | 607 | 623 | 583 | 627.84 | 651 | 604 | 0.928 | -0.108 | 0.33933563 | 0.55942318 | 0.0041 | 0.0122 | 0.0076 | 0.0076 | TRUE | 0.1072 |
| C2cd2l | 355 | 354 | 347 | 549 | 381 | 393 | 377 | 354 | 348 | 432 | 410 | 430 | 391.79 | 360 | 424 | 1.179 | 0.238 | 0.03124007 | 0.11858444 | 0 | 0.0153 | 0.0062 | 0.0062 | TRUE | 0.0202 |
| C2cd3 | 2168 | 1846 | 2275 | 2821 | 2052 | 2016 | 2303 | 1844 | 2284 | 2218 | 2208 | 2207 | 2177.18 | 2144 | 2211 | 1.031 | 0.045 | 0.67518356 | 0.8190765 | 0.0072 | 0.0084 | 0.0077 | 0.0077 | TRUE | 0.1817 |
| C2cd4a | 3 | 3 | 1 | 2 | 6 | 2 | 3 | 3 | 1 | 2 | 6 | 2 | 2.9 | 2 | 3 | 1.401 | 0.487 | 0.67779004 | NA | 0 | 1.1427 | 0.6285 | 0.6285 | TRUE | 0.8404 |
| C2cd4b | 45 | 57 | 42 | 78 | 70 | 42 | 48 | 57 | 42 | 61 | 75 | 46 | 54.92 | 49 | 61 | 1.244 | 0.314 | 0.2913776 | 0.51210246 | 0.0223 | 0.0669 | 0.0455 | 0.0455 | TRUE | 0.1605 |
| C2cd4c | 89 | 121 | 104 | 97 | 84 | 91 | 95 | 121 | 104 | 76 | 90 | 100 | 97.68 | 107 | 89 | 0.828 | -0.272 | 0.21219968 | 0.42316517 | 0.0071 | 0.0406 | 0.024 | 0.024 | TRUE | 0.1634 |
| C2cd4d | 0 | 0 | 0 | 0 | 0 | 0 | 0 | 0 | 0 | 0 | 0 | 0 | 0 | 0 | 0 | NA | NA | NA | NA | NA | NA | NA | NA | NA | NA |
| C2cd5 | 3234 | 3627 | 3682 | 3816 | 2891 | 2904 | 3435 | 3622 | 3697 | 3000 | 3111 | 3179 | 3340.63 | 3585 | 3097 | 0.863 | -0.212 | 0.00232492 | 0.01735382 | 9.00E-04 | 0.0079 | 0.0032 | 0.0032 | TRUE | 0.0162 |
| C3 | 207 | 232 | 248 | 160 | 136 | 216 | 220 | 232 | 249 | 126 | 146 | 236 | 201.52 | 234 | 169 | 0.723 | -0.468 | 0.0523131 | 0.16921292 | 0.0516 | 0.0233 | 0.0368 | 0.0368 | TRUE | 1.2531 |
| C330005M16 | 34 | 33 | 35 | 21 | 27 | 29 | 36 | 33 | 35 | 17 | 29 | 32 | 30.25 | 35 | 26 | 0.732 | -0.45 | 0.2395779 | 0.45564001 | 2.00E-04 | 0.1158 | 0.0716 | 0.0716 | TRUE | 0.7526 |
| C330006A16 | 1295 | 1452 | 1407 | 1716 | 1291 | 1322 | 1376 | 1450 | 1413 | 1349 | 1389 | 1447 | 1403.93 | 1413 | 1395 | 0.987 | -0.019 | 0.80314596 | 0.89838427 | 0 | 0.0092 | 0.0036 | 0.0036 | TRUE | 0.0128 |
| C330007P06I | 800 | 810 | 859 | 1346 | 977 | 867 | 850 | 809 | 863 | 1058 | 1051 | 949 | 929.93 | 841 | 1019 | 1.214 | 0.28 | 0.00236922 | 0.01760119 | 0.0014 | 0.0104 | 0.005 | 0.005 | TRUE | 0.0428 |
| C330008G21 | 4 | 4 | 3 | 24 | 19 | 12 | 4 | 4 | 3 | 19 | 20 | 13 | 10.62 | 4 | 17 | 4.678 | 2.226 | 0.00085321 | 0.00787808 | 0 | 0.3172 | 0.1568 | 0.1568 | TRUE | 0.2163 |
| C330011F03I | 1 | 0 | 1 | 0 | 0 | 0 | 1 | 0 | 1 | 0 | 0 | 0 | 0.34 | 1 | 0 | 0.261 | -1.939 | 0.60367425 | NA | 0 | 9.5732 | 8.3134 | 8.3134 | TRUE | 0.2529 |
| C330013E15I | 32 | 37 | 39 | 36 | 31 | 21 | 34 | 37 | 39 | 28 | 33 | 23 | 32.46 | 37 | 28 | 0.769 | -0.379 | 0.26296313 | 0.48235255 | 0 | 0.1084 | 0.0515 | 0.0515 | TRUE | 0.1496 |
| C330013F16I | 101 | 118 | 86 | 201 | 155 | 141 | 107 | 118 | 86 | 158 | 167 | 154 | 131.77 | 104 | 160 | 1.538 | 0.621 | 0.00107163 | 0.00938986 | 8.00E-04 | 0.0319 | 0.018 | 0.018 | TRUE | 0.2161 |
| C330018D20 | 248 | 313 | 276 | 678 | 484 | 459 | 263 | 313 | 277 | 533 | 521 | 502 | 401.56 | 284 | 519 | 1.824 | 0.867 | 4.43E-13 | 4.72E-11 | 3.00E-04 | 0.0151 | 0.0076 | 0.0076 | TRUE | 0.0851 |
| C330021F23I | 329 | 514 | 589 | 449 | 314 | 402 | 349 | 513 | 591 | 353 | 338 | 440 | 430.85 | 484 | 377 | 0.777 | -0.364 | 0.07720227 | 0.21947265 | 0.0429 | 0.0145 | 0.0283 | 0.0283 | TRUE | 0.4352 |
| C330022C24I | 1 | 2 | 3 | 0 | 0 | 0 | 1 | 2 | 3 | 0 | 0 | 0 | 1.01 | 2 | 0 | 0.088 | -3.499 | 0.10850143 | NA | 0 | 3.2623 | 2.174 | 2.174 | TRUE | 0.073 |
| C330024C12I | 3 | 0 | 1 | 1 | 4 | 1 | 3 | 0 | 1 | 1 | 4 | 1 | 1.73 | 1 | 2 | 1.465 | 0.551 | 0.73317067 | NA | 0.5866 | 1.9121 | 1.2736 | 1.2736 | TRUE | 0.7051 |
| C330024D21 | 510 | 539 | 537 | 888 | 676 | 569 | 542 | 538 | 539 | 698 | 727 | 623 | 611.25 | 540 | 683 | 1.266 | 0.34 | 0.00113603 | 0.00983397 | 0.0017 | 0.0123 | 0.0062 | 0.0062 | TRUE | 0.0672 |
| C330027C09I | 6803 | 7181 | 7185 | 8037 | 5723 | 5882 | 7226 | 7172 | 7214 | 6318 | 6158 | 6439 | 6754.44 | 7204 | 6305 | 0.875 | -0.192 | 0.00067166 | 0.0065594 | 0 | 0.0074 | 0.0022 | 0.0022 | TRUE | 0.005 |
| C330046G13 | 454 | 576 | 562 | 402 | 316 | 439 | 482 | 575 | 564 | 316 | 340 | 481 | 459.73 | 540 | 379 | 0.699 | -0.516 | 0.00390909 | 0.02589866 | 0.0286 | 0.0141 | 0.0208 | 0.0208 | TRUE | 0.6262 |
| C3ar1 | 0 | 0 | 0 | 0 | 0 | 0 | 0 | 0 | 0 | 0 | 0 | 0 | 0 | 0 | 0 | NA | NA | NA | NA | NA | NA | NA | NA | NA | NA |
| C430002E04I | 157 | 151 | 152 | 241 | 187 | 186 | 167 | 151 | 153 | 189 | 201 | 204 | 177.41 | 157 | 198 | 1.263 | 0.336 | 0.02697066 | 0.10720009 | 0 | 0.0255 | 0.0109 | 0.0109 | TRUE | 0.0324 |
| C430049B03 | 24 | 29 | 26 | 39 | 44 | 31 | 25 | 29 | 26 | 31 | 47 | 34 | 32.08 | 27 | 37 | 1.381 | 0.466 | 0.18937576 | 0.39456203 | 0 | 0.1096 | 0.0586 | 0.0586 | TRUE | 0.3894 |
| C4a | 45 | 61 | 46 | 56 | 70 | 41 | 48 | 61 | 46 | 44 | 75 | 45 | 53.19 | 52 | 55 | 1.055 | 0.077 | 0.81130142 | 0.90291353 | 0.0429 | 0.0688 | 0.0561 | 0.0561 | TRUE | 0.8752 |
| C4b | 20 | 20 | 13 | 30 | 16 | 18 | 21 | 20 | 13 | 24 | 17 | 20 | 19.13 | 18 | 20 | 1.123 | 0.167 | 0.70832153 | 0.84123849 | 0 | 0.1791 | 0.0916 | 0.0916 | TRUE | 0.3063 |

| | | | | | | | | | | | | | | | | | | | | | | | | | |
|---|---|---|---|---|---|---|---|---|---|---|---|---|---|---|---|---|---|---|---|---|---|---|---|---|---|
| C4bp | 0 | 0 | 0 | 0 | 0 | 0 | 0 | 0 | 0 | 0 | 0 | 0 | 0 | 0 | 0 | NA | NA | NA | NA | NA | NA | NA | NA | NA | NA |
| C4bp-ps1 | 3 | 0 | 1 | 2 | 1 | 1 | 3 | 0 | 1 | 2 | 1 | 1 | 1.32 | 1 | 1 | 0.911 | -0.135 | 0.93641308 | NA | 0 | 2.4985 | 1.2973 | 1.2973 | TRUE | 0.5207 |
| C530005A16 | 5 | 12 | 14 | 13 | 17 | 12 | 5 | 12 | 14 | 10 | 18 | 13 | 12.17 | 10 | 14 | 1.314 | 0.394 | 0.51131121 | 0.7048357 | 0.0528 | 0.2777 | 0.1745 | 0.1745 | TRUE | 0.6285 |
| C530008M17 | 297 | 404 | 406 | 369 | 337 | 288 | 315 | 403 | 408 | 290 | 363 | 315 | 349.09 | 375 | 323 | 0.857 | -0.222 | 0.15799752 | 0.35157152 | 0.0137 | 0.0163 | 0.015 | 0.015 | TRUE | 0.2225 |
| C530044C16I | 1 | 0 | 0 | 1 | 0 | 0 | 1 | 0 | 0 | 1 | 0 | 0 | 0.31 | 0 | 0 | 0.954 | -0.067 | 0.98625928 | NA | 0 | 10.7014 | 9.0384 | 9.0384 | TRUE | 0.4614 |
| C5ar1 | 0 | 0 | 1 | 0 | 0 | 0 | 0 | 0 | 1 | 0 | 0 | 0 | 0.17 | 0 | 0 | 0.49 | -1.029 | 0.80087 | NA | 0 | 19.6928 | 10 | 10 | TRUE | 0.4098 |
| C5ar2 | 1 | 0 | 0 | 2 | 2 | 0 | 1 | 0 | 0 | 2 | 2 | 0 | 0.8 | 0 | 1 | 3.417 | 1.773 | 0.46651319 | NA | 0 | 4.1368 | 2.8785 | 2.8785 | TRUE | 0.4614 |
| C6 | 0 | 2 | 0 | 2 | 0 | 2 | 0 | 2 | 0 | 2 | 0 | 2 | 0.96 | 1 | 1 | 1.877 | 0.908 | 0.6959042 | NA | 0 | 3.4392 | 2.7428 | 2.7428 | TRUE | 0.9738 |
| C630020P19I | 16 | 16 | 18 | 21 | 13 | 17 | 17 | 16 | 18 | 17 | 14 | 19 | 16.69 | 17 | 17 | 0.962 | -0.056 | 0.90188926 | 0.95205059 | 0 | 0.2043 | 0.0883 | 0.0883 | TRUE | 0.0694 |
| C630028M04 | 4 | 0 | 0 | 3 | 1 | 0 | 4 | 0 | 0 | 2 | 1 | 0 | 1.28 | 1 | 1 | 0.836 | -0.258 | 0.90789528 | NA | 3.3118 | 2.5797 | 2.8138 | 2.8138 | TRUE | 1.0684 |
| C630031E19I | 0 | 0 | 0 | 0 | 0 | 0 | 0 | 0 | 0 | 0 | 0 | 0 | 0 | 0 | 0 | NA | NA | NA | NA | NA | NA | NA | NA | NA | NA |
| C630043F03I | 74 | 70 | 75 | 79 | 62 | 40 | 79 | 70 | 75 | 62 | 67 | 44 | 66.07 | 75 | 58 | 0.774 | -0.37 | 0.1529614 | 0.34444707 | 0.0032 | 0.0568 | 0.0329 | 0.0329 | TRUE | 0.3428 |
| C7 | 22 | 4 | 12 | 51 | 26 | 33 | 23 | 4 | 12 | 40 | 28 | 36 | 23.93 | 13 | 35 | 2.662 | 1.413 | 0.01187469 | 0.05975174 | 0.2203 | 0.1446 | 0.174 | 0.174 | TRUE | 0.839 |
| C730002L08I | 0 | 0 | 0 | 0 | 0 | 0 | 0 | 0 | 0 | 0 | 0 | 0 | 0 | 0 | 0 | NA | NA | NA | NA | NA | NA | NA | NA | NA | NA |
| C730027H18 | 1 | 0 | 0 | 2 | 0 | 0 | 1 | 0 | 0 | 2 | 0 | 0 | 0.44 | 0 | 1 | 1.494 | 0.579 | 0.86844164 | NA | 0 | 7.5101 | 6.9157 | 6.9157 | TRUE | 0.9721 |
| C730036E19I | 6 | 1 | 1 | 4 | 1 | 2 | 6 | 1 | 1 | 3 | 1 | 2 | 2.46 | 3 | 2 | 0.784 | -0.35 | 0.78905748 | NA | 0.3625 | 1.3438 | 0.8266 | 0.8266 | TRUE | 1.462 |
| C77080 | 4500 | 5917 | 5905 | 5082 | 4138 | 4868 | 4780 | 5909 | 5929 | 3995 | 4452 | 5329 | 5065.75 | 5539 | 4592 | 0.829 | -0.271 | 0.04505338 | 0.1529778 | 0.0176 | 0.0076 | 0.013 | 0.013 | TRUE | 0.2398 |
| C77370 | 1101 | 953 | 1160 | 1958 | 1322 | 1187 | 1169 | 952 | 1165 | 1539 | 1422 | 1299 | 1257.83 | 1095 | 1420 | 1.298 | 0.376 | 0.00166047 | 0.01326692 | 0.0088 | 0.0095 | 0.0095 | 0.0095 | TRUE | 0.1578 |
| C78339 | 987 | 1225 | 1216 | 1385 | 1012 | 976 | 1048 | 1223 | 1221 | 1089 | 1089 | 1068 | 1123.13 | 1164 | 1082 | 0.929 | -0.106 | 0.27039867 | 0.49059332 | 0.003 | 0.0098 | 0.0058 | 0.0058 | TRUE | 0.0903 |
| C86187 | 1 | 0 | 0 | 1 | 0 | 0 | 1 | 0 | 0 | 1 | 0 | 0 | 0.31 | 0 | 0 | 0.954 | -0.067 | 0.98625928 | NA | 0 | 10.7014 | 9.0384 | 9.0384 | TRUE | 0.4614 |
| C86695 | 7 | 4 | 3 | 21 | 16 | 4 | 7 | 4 | 3 | 17 | 17 | 4 | 8.76 | 5 | 13 | 2.668 | 1.416 | 0.07358162 | 0.212824 | 0.2232 | 0.3831 | 0.3063 | 0.3063 | TRUE | 1.2338 |
| C87198 | 0 | 0 | 0 | 0 | 0 | 0 | 0 | 0 | 0 | 0 | 0 | 0 | 0 | 0 | 0 | NA | NA | NA | NA | NA | NA | NA | NA | NA | NA |
| C87414 | 0 | 0 | 0 | 1 | 0 | 0 | 0 | 0 | 0 | 1 | 0 | 0 | 0.13 | 0 | 0 | 1.859 | 0.894 | 0.82650734 | NA | 0 | 25.1515 | 10 | 10 | TRUE | 0.2563 |
| C87436 | 412 | 394 | 385 | 422 | 380 | 330 | 438 | 393 | 387 | 332 | 409 | 361 | 386.59 | 406 | 367 | 0.903 | -0.147 | 0.26615086 | 0.48579232 | 0.0053 | 0.0154 | 0.01 | 0.01 | TRUE | 0.1119 |
| C87499 | 0 | 1 | 0 | 0 | 2 | 0 | 0 | 1 | 0 | 0 | 2 | 0 | 0.53 | 0 | 1 | 1.927 | 0.947 | 0.77747358 | NA | 0 | 6.2809 | 6.39 | 6.39 | TRUE | 0.9744 |
| C87977 | 0 | 0 | 0 | 0 | 0 | 0 | 0 | 0 | 0 | 0 | 0 | 0 | 0 | 0 | 0 | NA | NA | NA | NA | NA | NA | NA | NA | NA | NA |
| C8a | 11 | 4 | 6 | 6 | 3 | 4 | 12 | 4 | 6 | 5 | 3 | 4 | 5.67 | 7 | 4 | 0.574 | -0.802 | 0.33616499 | 0.55628939 | 0 | 0.5878 | 0.3118 | 0.3118 | TRUE | 0.6077 |
| C8b | 46 | 54 | 58 | 99 | 88 | 84 | 49 | 54 | 58 | 78 | 95 | 92 | 70.91 | 54 | 88 | 1.635 | 0.709 | 0.00296977 | 0.02107029 | 0 | 0.0534 | 0.026 | 0.026 | TRUE | 0.1126 |
| C8g | 60 | 155 | 149 | 121 | 74 | 98 | 64 | 155 | 150 | 95 | 80 | 107 | 108.36 | 123 | 94 | 0.765 | -0.386 | 0.24826942 | 0.46499291 | 0.1098 | 0.0373 | 0.0711 | 0.0711 | TRUE | 1.77 |
| C9 | 0 | 0 | 0 | 1 | 1 | 1 | 0 | 0 | 0 | 1 | 1 | 1 | 0.49 | 0 | 1 | 5.224 | 2.385 | 0.38465 | NA | 0 | 6.6923 | 3.9249 | 3.9249 | TRUE | 0.07 |
| C920006O11 | 71 | 162 | 177 | 153 | 111 | 103 | 75 | 162 | 178 | 120 | 119 | 113 | 127.9 | 138 | 117 | 0.849 | -0.236 | 0.43734264 | 0.64548863 | 0.0887 | 0.0327 | 0.0589 | 0.0589 | TRUE | 1.6162 |
| C920009B18 | 5 | 6 | 4 | 3 | 8 | 5 | 5 | 6 | 4 | 2 | 9 | 5 | 5.29 | 5 | 5 | 1.051 | 0.072 | 0.93285276 | 0.96671764 | 0 | 0.6293 | 0.3386 | 0.3386 | TRUE | 0.2055 |
| C920021L13I | 3 | 5 | 2 | 3 | 2 | 4 | 3 | 5 | 2 | 2 | 2 | 4 | 3.18 | 3 | 3 | 0.863 | -0.212 | 0.83931926 | NA | 0 | 1.043 | 0.473 | 0.473 | TRUE | 0.2517 |
| C920025E04I | 9 | 3 | 4 | 9 | 5 | 5 | 10 | 3 | 4 | 7 | 5 | 5 | 5.75 | 6 | 6 | 1.095 | 0.131 | 0.87330361 | 0.93762803 | 0 | 0.5798 | 0.3111 | 0.3111 | TRUE | 0.8545 |
| Caap1 | 859 | 875 | 856 | 1543 | 1073 | 872 | 912 | 874 | 860 | 1213 | 1154 | 955 | 994.63 | 882 | 1107 | 1.257 | 0.33 | 0.00437833 | 0.02828966 | 0.0074 | 0.0102 | 0.0086 | 0.0086 | TRUE | 0.1727 |
| Cab39 | 5337 | 5103 | 5107 | 7643 | 5549 | 5096 | 5669 | 5096 | 5128 | 6008 | 5970 | 5578 | 5575.05 | 5298 | 5852 | 1.105 | 0.144 | 0.06584653 | 0.1978909 | 0.0024 | 0.0075 | 0.0042 | 0.0042 | TRUE | 0.0457 |
| Cab39l | 3174 | 3454 | 3453 | 3843 | 2767 | 2749 | 3371 | 3450 | 3467 | 3021 | 2977 | 3009 | 3215.91 | 3429 | 3002 | 0.876 | -0.192 | 0.00147763 | 0.01211262 | 0 | 0.0079 | 0.0023 | 0.0023 | TRUE | 0.0026 |
| Cabin1 | 5356 | 4340 | 4455 | 5892 | 4490 | 4001 | 5689 | 4334 | 4473 | 4632 | 4831 | 4380 | 4723.2 | 4832 | 4614 | 0.955 | -0.066 | 0.58009941 | 0.75669188 | 0.0113 | 0.0076 | 0.0102 | 0.0102 | TRUE | 0.2929 |
| Cables1 | 106 | 99 | 102 | 165 | 118 | 111 | 113 | 99 | 102 | 130 | 127 | 122 | 115.34 | 105 | 126 | 1.207 | 0.271 | 0.1377062 | 0.32185866 | 0 | 0.0355 | 0.0153 | 0.0153 | TRUE | 0.0427 |
| Cables2 | 1041 | 1106 | 1104 | 1426 | 1121 | 994 | 1106 | 1105 | 1109 | 1121 | 1206 | 1088 | 1122.34 | 1107 | 1138 | 1.029 | 0.041 | 0.62224036 | 0.78656528 | 5.00E-04 | 0.0098 | 0.0041 | 0.0041 | TRUE | 0.0316 |
| Cabp1 | 418 | 370 | 344 | 348 | 271 | 240 | 444 | 370 | 345 | 274 | 292 | 263 | 331.13 | 386 | 276 | 0.715 | -0.485 | 0.00074079 | 0.00705114 | 0.0074 | 0.0169 | 0.0118 | 0.0118 | TRUE | 0.1943 |
| Cabp2 | 0 | 6 | 9 | 5 | 4 | 2 | 0 | 6 | 9 | 4 | 4 | 2 | 4.24 | 5 | 3 | 0.694 | -0.528 | 0.64632557 | NA | 0.6038 | 0.7835 | 0.7112 | 0.7112 | TRUE | 0.359 |
| Cabp4 | 25 | 18 | 26 | 28 | 20 | 26 | 27 | 18 | 26 | 22 | 22 | 28 | 23.77 | 24 | 24 | 1.017 | 0.024 | 0.9523839 | 0.9757697 | 0 | 0.1455 | 0.0738 | 0.0738 | TRUE | 0.2569 |
| Cabp5 | 0 | 0 | 0 | 1 | 1 | 1 | 0 | 0 | 0 | 1 | 1 | 1 | 0.49 | 0 | 1 | 5.224 | 2.385 | 0.38465 | NA | 0 | 6.6923 | 3.9249 | 3.9249 | TRUE | 0.07 |
| Cabp7 | 29 | 66 | 46 | 64 | 41 | 48 | 31 | 66 | 46 | 50 | 44 | 53 | 48.31 | 48 | 49 | 1.027 | 0.039 | 0.91011381 | 0.95510779 | 0.0504 | 0.0751 | 0.0634 | 0.0634 | TRUE | 0.266 |
| Cabs1 | 3 | 1 | 0 | 2 | 0 | 0 | 3 | 1 | 0 | 2 | 0 | 0 | 0.96 | 1 | 1 | 0.408 | -1.294 | 0.58647705 | NA | 0 | 3.4401 | 2.8489 | 2.8489 | TRUE | 0.5433 |
| Cabyr | 29 | 18 | 16 | 31 | 27 | 13 | 31 | 18 | 16 | 24 | 29 | 14 | 22.08 | 22 | 22 | 1.049 | 0.069 | 0.88538049 | 0.94387096 | 0.075 | 0.1561 | 0.1177 | 0.1177 | TRUE | 0.7038 |
| Cacfd1 | 258 | 368 | 357 | 431 | 333 | 345 | 274 | 368 | 358 | 339 | 358 | 378 | 345.8 | 333 | 358 | 1.073 | 0.102 | 0.50069871 | 0.69657052 | 0.0108 | 0.0164 | 0.0135 | 0.0135 | TRUE | 0.2725 |
| Cachd1 | 610 | 431 | 462 | 459 | 327 | 235 | 648 | 430 | 464 | 361 | 352 | 257 | 418.7 | 514 | 323 | 0.63 | -0.667 | 0.00087896 | 0.00805521 | 0.0379 | 0.0148 | 0.0264 | 0.0264 | TRUE | 0.603 |
| Cacna1a | 2554 | 2619 | 2558 | 2389 | 1906 | 2028 | 2713 | 2616 | 2569 | 1878 | 2051 | 2220 | 2340.94 | 2633 | 2050 | 0.778 | -0.362 | 4.99E-05 | 0.00077404 | 0.0032 | 0.0083 | 0.0053 | 0.0053 | TRUE | 0.0662 |
| Cacna1b | 704 | 689 | 721 | 546 | 436 | 455 | 748 | 688 | 724 | 429 | 469 | 498 | 592.71 | 720 | 465 | 0.645 | -0.632 | 3.31E-09 | 1.75E-07 | 0.0018 | 0.0125 | 0.0065 | 0.0065 | TRUE | 0.0575 |
| Cacna1c | 77 | 98 | 92 | 140 | 81 | 91 | 82 | 98 | 92 | 110 | 87 | 100 | 94.81 | 91 | 99 | 1.094 | 0.13 | 0.53883067 | 0.72512264 | 0 | 0.0417 | 0.0217 | 0.0217 | TRUE | 0.1048 |
| Cacna1d | 124 | 110 | 100 | 112 | 65 | 61 | 132 | 110 | 100 | 88 | 70 | 67 | 94.46 | 114 | 75 | 0.662 | -0.596 | 0.00911033 | 0.04907381 | 0.0112 | 0.0418 | 0.0266 | 0.0266 | TRUE | 0.241 |
| Cacna1e | 8 | 13 | 6 | 33 | 21 | 16 | 8 | 13 | 6 | 26 | 23 | 18 | 15.59 | 9 | 22 | 2.411 | 1.27 | 0.01601121 | 0.07444833 | 0 | 0.2182 | 0.1225 | 0.1225 | TRUE | 0.4426 |
| Cacna1f | 2 | 6 | 7 | 3 | 3 | 6 | 2 | 6 | 7 | 2 | 3 | 7 | 4.55 | 5 | 4 | 0.786 | -0.348 | 0.71123696 | NA | 0 | 0.731 | 0.4133 | 0.4133 | TRUE | 0.4937 |
| Cacna1g | 241 | 249 | 235 | 143 | 123 | 94 | 256 | 249 | 236 | 112 | 132 | 103 | 181.38 | 247 | 116 | 0.469 | -1.092 | 3.12E-11 | 2.53E-09 | 0 | 0.0251 | 0.0132 | 0.0132 | TRUE | 0.1443 |
| Cacna1h | 538 | 811 | 858 | 490 | 429 | 471 | 571 | 810 | 862 | 385 | 462 | 516 | 600.88 | 748 | 454 | 0.606 | -0.722 | 7.82E-05 | 0.0011353 | 0.0324 | 0.0124 | 0.0223 | 0.0223 | TRUE | 0.5014 |
| Cacna1i | 85 | 76 | 75 | 118 | 74 | 68 | 90 | 76 | 75 | 93 | 80 | 74 | 81.39 | 80 | 82 | 1.027 | 0.039 | 0.86275795 | 0.93221844 | 0 | 0.0474 | 0.0245 | 0.0245 | TRUE | 0.1303 |
| Cacna1s | 43 | 40 | 51 | 25 | 23 | 24 | 46 | 40 | 51 | 20 | 25 | 26 | 34.58 | 46 | 24 | 0.512 | -0.965 | 0.00430219 | 0.02791204 | 0 | 0.1022 | 0.0502 | 0.0502 | TRUE | 0.1558 |
| Cacna2d1 | 200 | 204 | 178 | 431 | 320 | 305 | 212 | 204 | 179 | 339 | 344 | 334 | 268.65 | 198 | 339 | 1.71 | 0.774 | 8.27E-09 | 3.99E-07 | 0 | 0.0192 | 0.009 | 0.009 | TRUE | 0.0814 |
| Cacna2d2 | 247 | 330 | 340 | 275 | 258 | 282 | 262 | 330 | 341 | 216 | 278 | 309 | 289.3 | 311 | 268 | 0.857 | -0.223 | 0.22207283 | 0.43428028 | 0.0227 | 0.0183 | 0.0205 | 0.0205 | TRUE | 0.3331 |
| Cacna2d3 | 17 | 20 | 19 | 29 | 20 | 18 | 18 | 20 | 19 | 23 | 22 | 20 | 20.19 | 19 | 22 | 1.124 | 0.168 | 0.68154721 | 0.82351664 | 0 | 0.1701 | 0.0716 | 0.0716 | TRUE | 0.0245 |
| Cacna2d4 | 6 | 7 | 4 | 6 | 3 | 2 | 6 | 7 | 4 | 5 | 3 | 2 | 4.59 | 6 | 3 | 0.595 | -0.749 | 0.3960442 | NA | 0 | 0.7253 | 0.3316 | 0.3316 | TRUE | 0.2314 |
| Cacnb1 | 653 | 823 | 839 | 633 | 540 | 631 | 694 | 822 | 842 | 498 | 581 | 691 | 687.89 | 786 | 590 | 0.749 | -0.417 | 0.00529898 | 0.03266055 | 0.0165 | 0.0117 | 0.0146 | 0.0146 | TRUE | 0.2625 |
| Cacnb2 | 21 | 45 | 48 | 78 | 50 | 57 | 22 | 45 | 48 | 61 | 54 | 62 | 48.83 | 38 | 59 | 1.535 | 0.618 | 0.06958517 | 0.20502472 | 0.0457 | 0.0744 | 0.0621 | 0.0621 | TRUE | 0.9647 |
| Cacnb3 | 1543 | 1939 | 1921 | 1716 | 1442 | 1576 | 1639 | 1936 | 1929 | 1349 | 1552 | 1725 | 1688.33 | 1835 | 1542 | 0.84 | -0.252 | 0.03987255 | 0.14046681 | 0.0108 | 0.0089 | 0.0103 | 0.0103 | TRUE | 0.1468 |
| Cacnb4 | 20 | 26 | 18 | 17 | 16 | 20 | 21 | 26 | 18 | 13 | 17 | 22 | 19.63 | 22 | 17 | 0.796 | -0.33 | 0.45863747 | 0.66320988 | 0 | 0.1748 | 0.0911 | 0.0911 | TRUE | 0.2688 |
| Cacng1 | 1 | 2 | 6 | 12 | 5 | 2 | 1 | 2 | 6 | 9 | 5 | 2 | 4.35 | 3 | 5 | 1.895 | 0.922 | 0.3816919 | NA | 0.3205 | 0.7646 | 0.5476 | 0.5476 | TRUE | 0.3049 |
| Cacng2 | 2 | 2 | 1 | 5 | 0 | 1 | 2 | 2 | 1 | 4 | 0 | 1 | 1.69 | 2 | 2 | 1.027 | 0.038 | 0.98054995 | NA | 0 | 1.9541 | 1.1536 | 1.1536 | TRUE | 0.9346 |

| | | | | | | | | | | | | | | | | | | | | | | | | | |
|---|---|---|---|---|---|---|---|---|---|---|---|---|---|---|---|---|---|---|---|---|---|---|---|---|---|
| Cacng3 | 0 | 0 | 2 | 0 | 1 | 0 | 0 | 0 | 2 | 0 | 1 | 0 | 0.51 | 1 | 0 | 0.52 | -0.944 | 0.77815412 | NA | 0 | 6.416 | 6.3253 | 6.3253 | TRUE | 0.9739 |
| Cacng4 | 82 | 81 | 59 | 104 | 95 | 95 | 87 | 81 | 59 | 82 | 102 | 104 | 85.87 | 76 | 96 | 1.263 | 0.337 | 0.16708224 | 0.3645851 | 0.016 | 0.0453 | 0.0311 | 0.0311 | TRUE | 0.3357 |
| Cacng5 | 468 | 412 | 345 | 275 | 195 | 206 | 497 | 411 | 346 | 216 | 210 | 225 | 317.75 | 418 | 217 | 0.519 | -0.945 | 1.03E-08 | 4.83E-07 | 0.0152 | 0.0173 | 0.0162 | 0.0162 | TRUE | 0.1559 |
| Cacng6 | 110 | 184 | 156 | 146 | 106 | 117 | 117 | 184 | 157 | 115 | 114 | 128 | 135.69 | 153 | 119 | 0.779 | -0.36 | 0.09472829 | 0.25228627 | 0.0209 | 0.0312 | 0.026 | 0.026 | TRUE | 0.2495 |
| Cacng7 | 414 | 402 | 437 | 689 | 478 | 481 | 440 | 401 | 439 | 542 | 514 | 527 | 477.08 | 427 | 528 | 1.237 | 0.307 | 0.00342811 | 0.02345689 | 0 | 0.0138 | 0.0058 | 0.0058 | TRUE | 0.0311 |
| Cacng8 | 59 | 76 | 84 | 92 | 61 | 60 | 63 | 76 | 84 | 72 | 66 | 66 | 71.09 | 74 | 68 | 0.915 | -0.129 | 0.59061377 | 0.76466594 | 0 | 0.0532 | 0.0271 | 0.0271 | TRUE | 0.1668 |
| Cactin | 2651 | 2191 | 2096 | 3718 | 2647 | 2070 | 2816 | 2188 | 2105 | 2923 | 2848 | 2266 | 2524.23 | 2370 | 2679 | 1.131 | 0.178 | 0.22748473 | 0.44077263 | 0.0217 | 0.0082 | 0.0152 | 0.0152 | TRUE | 0.3286 |
| Cacul1 | 3992 | 4291 | 4466 | 5601 | 4031 | 3885 | 4240 | 4285 | 4484 | 4403 | 4337 | 4253 | 4333.82 | 4336 | 4331 | 0.999 | -0.002 | 0.97790576 | 0.98799114 | 4.00E-04 | 0.0077 | 0.0026 | 0.0026 | TRUE | 0.0108 |
| Cacybp | 10057 | 9834 | 9618 | 15218 | 10590 | 10527 | 10682 | 9821 | 9657 | 11963 | 11394 | 11523 | 10840.32 | 10053 | 11627 | 1.157 | 0.21 | 0.00295347 | 0.02098296 | 0.0017 | 0.0072 | 0.0035 | 0.0035 | TRUE | 0.0365 |
| Cad | 28419 | 28427 | 29840 | 32873 | 23549 | 23957 | 30186 | 28390 | 29963 | 25842 | 25337 | 26225 | 27657.16 | 29513 | 25801 | 0.874 | -0.194 | 0.00099674 | 0.00888669 | 6.00E-04 | 0.007 | 0.0025 | 0.0025 | TRUE | 0.0136 |
| Cadm1 | 2290 | 2694 | 2725 | 2682 | 2239 | 2478 | 2432 | 2691 | 2736 | 2108 | 2409 | 2713 | 2514.84 | 2620 | 2410 | 0.919 | -0.121 | 0.28695665 | 0.50821107 | 0.0092 | 0.0082 | 0.009 | 0.009 | TRUE | 0.1471 |
| Cadm2 | 6 | 16 | 12 | 8 | 6 | 6 | 6 | 16 | 12 | 6 | 6 | 7 | 8.95 | 11 | 6 | 0.559 | -0.839 | 0.20794275 | 0.4172612 | 0 | 0.3749 | 0.1992 | 0.1992 | TRUE | 0.1946 |
| Cadm3 | 34 | 23 | 23 | 28 | 14 | 14 | 36 | 23 | 23 | 22 | 15 | 15 | 22.43 | 27 | 17 | 0.646 | -0.631 | 0.1463146 | 0.33505366 | 0 | 0.1538 | 0.0898 | 0.0898 | TRUE | 0.4783 |
| Cadm4 | 690 | 901 | 938 | 606 | 536 | 602 | 733 | 900 | 942 | 476 | 577 | 659 | 714.44 | 858 | 571 | 0.664 | -0.591 | 0.00014746 | 0.00188015 | 0.0189 | 0.0115 | 0.016 | 0.016 | TRUE | 0.2531 |
| Cadps | 39 | 37 | 35 | 35 | 28 | 28 | 41 | 37 | 35 | 28 | 30 | 31 | 33.64 | 38 | 30 | 0.776 | -0.366 | 0.25827299 | 0.47721173 | 0 | 0.1048 | 0.0454 | 0.0454 | TRUE | 0.0487 |
| Cadps2 | 68 | 65 | 64 | 69 | 54 | 43 | 72 | 65 | 64 | 54 | 58 | 47 | 60.14 | 67 | 53 | 0.793 | -0.335 | 0.17948745 | 0.3805964 | 0 | 0.0617 | 0.0281 | 0.0281 | TRUE | 0.0749 |
| Cage1 | 58 | 72 | 74 | 43 | 35 | 32 | 62 | 72 | 74 | 34 | 38 | 35 | 52.38 | 69 | 36 | 0.511 | -0.969 | 0.0003524 | 0.00390866 | 0 | 0.0698 | 0.0318 | 0.0318 | TRUE | 0.0819 |
| Calb1 | 11 | 9 | 4 | 19 | 19 | 17 | 12 | 9 | 4 | 15 | 20 | 19 | 13.11 | 8 | 18 | 2.184 | 1.127 | 0.05088964 | 0.16607313 | 0 | 0.2581 | 0.1506 | 0.1506 | TRUE | 0.6103 |
| Calb2 | 198 | 246 | 247 | 324 | 270 | 217 | 210 | 246 | 248 | 255 | 291 | 238 | 247.79 | 235 | 261 | 1.111 | 0.152 | 0.31542366 | 0.53617717 | 0.005 | 0.0202 | 0.0124 | 0.0124 | TRUE | 0.1054 |
| Calca | 292 | 500 | 525 | 263 | 229 | 304 | 310 | 499 | 527 | 207 | 246 | 333 | 353.76 | 445 | 262 | 0.587 | -0.77 | 0.00146911 | 0.01206724 | 0.0655 | 0.0162 | 0.0392 | 0.0392 | TRUE | 0.8147 |
| Calcb | 1 | 0 | 1 | 0 | 0 | 0 | 1 | 0 | 1 | 0 | 0 | 0 | 0.34 | 1 | 0 | 0.261 | -1.939 | 0.60367425 | NA | 0 | 9.5732 | 8.3134 | 8.3134 | TRUE | 0.2529 |
| Calcoco1 | 1931 | 1884 | 1854 | 2298 | 1817 | 1833 | 2051 | 1882 | 1862 | 1807 | 1955 | 2007 | 1927.04 | 1932 | 1923 | 0.995 | -0.007 | 0.93242804 | 0.9664359 | 0.0024 | 0.0086 | 0.0048 | 0.0048 | TRUE | 0.0354 |
| Calcoco2 | 973 | 553 | 472 | 4991 | 2524 | 1501 | 1033 | 552 | 474 | 3924 | 2716 | 1643 | 1723.67 | 686 | 2761 | 4.022 | 2.008 | 6.56E-05 | 0.00097912 | 0.1814 | 0.0088 | 0.0728 | 0.1814 | TRUE | 0.121 |
| Calcr | 33 | 17 | 12 | 47 | 32 | 28 | 35 | 17 | 12 | 37 | 34 | 31 | 27.68 | 21 | 34 | 1.601 | 0.679 | 0.13884815 | 0.3234564 | 0.0978 | 0.1259 | 0.1133 | 0.1133 | TRUE | 1.706 |
| Calcrl | 8 | 8 | 9 | 18 | 12 | 13 | 8 | 8 | 9 | 14 | 13 | 14 | 11.14 | 8 | 14 | 1.619 | 0.695 | 0.20982748 | 0.42013444 | 0 | 0.3027 | 0.1261 | 0.1261 | TRUE | 0.0117 |
| Cald1 | 2338 | 2312 | 2028 | 3066 | 2029 | 2126 | 2483 | 2309 | 2036 | 2410 | 2183 | 2327 | 2291.55 | 2276 | 2307 | 1.014 | 0.02 | 0.8436924 | 0.92132635 | 0.0058 | 0.0083 | 0.0068 | 0.0068 | TRUE | 0.103 |
| Calhm1 | 0 | 1 | 2 | 1 | 6 | 0 | 0 | 1 | 2 | 1 | 6 | 0 | 1.71 | 1 | 2 | 2.365 | 1.242 | 0.50826734 | NA | 1.6817 | 1.9356 | 1.8285 | 1.8285 | TRUE | 1.8709 |
| Calhm2 | 86 | 85 | 75 | 124 | 94 | 108 | 91 | 85 | 75 | 97 | 101 | 118 | 94.07 | 84 | 105 | 1.257 | 0.33 | 0.11677787 | 0.28895749 | 0 | 0.0417 | 0.0212 | 0.0212 | TRUE | 0.1038 |
| Calm1 | 7443 | 6096 | 5761 | 13992 | 9089 | 7090 | 7906 | 6088 | 5785 | 10999 | 9779 | 7761 | 8053.05 | 6593 | 9513 | 1.443 | 0.529 | 0.00091115 | 0.00828284 | 0.0297 | 0.0073 | 0.0182 | 0.0182 | TRUE | 0.3172 |
| Calm2 | 16167 | 17753 | 16612 | 21312 | 15393 | 15457 | 17172 | 17730 | 16680 | 16754 | 16562 | 16920 | 16969.73 | 17194 | 16745 | 0.974 | -0.038 | 0.50535071 | 0.69995534 | 4.00E-04 | 0.0071 | 0.0023 | 0.0023 | TRUE | 0.0091 |
| Calm3 | 7716 | 8758 | 8553 | 11395 | 8239 | 8057 | 8196 | 8747 | 8588 | 8958 | 8865 | 8820 | 8695.46 | 8510 | 8881 | 1.044 | 0.061 | 0.30276992 | 0.52360605 | 4.00E-04 | 0.0073 | 0.0025 | 0.0025 | TRUE | 0.0127 |
| Calm4 | 0 | 0 | 0 | 0 | 0 | 0 | 0 | 0 | 0 | 0 | 0 | 0 | 0 | 0 | 0 | NA | NA | NA | NA | NA | NA | NA | NA | NA | NA |
| Calm5 | 0 | 0 | 0 | 0 | 0 | 0 | 0 | 0 | 0 | 0 | 0 | 0 | 0 | 0 | 0 | NA | NA | NA | NA | NA | NA | NA | NA | NA | NA |
| Calml3 | 0 | 0 | 0 | 0 | 0 | 1 | 0 | 0 | 0 | 0 | 0 | 1 | 0.18 | 0 | 0 | 1.859 | 0.894 | 0.82650734 | NA | 0 | 18.0643 | 10 | 10 | TRUE | 0.5383 |
| Calml4 | 179 | 179 | 190 | 189 | 110 | 152 | 190 | 179 | 191 | 149 | 118 | 166 | 165.5 | 187 | 144 | 0.775 | -0.368 | 0.04026248 | 0.14155593 | 0.0063 | 0.0268 | 0.0171 | 0.0171 | TRUE | 0.2454 |
| Caln1 | 0 | 6 | 1 | 6 | 5 | 0 | 0 | 6 | 1 | 5 | 5 | 0 | 2.85 | 2 | 3 | 1.447 | 0.533 | 0.73776565 | NA | 1.9889 | 1.1633 | 1.4638 | 1.4638 | TRUE | 1.9698 |
| Calr | 31863 | 35055 | 33122 | 41574 | 29677 | 28175 | 33844 | 35010 | 33258 | 32682 | 31931 | 30842 | 32927.78 | 34037 | 31818 | 0.935 | -0.097 | 0.1021477 | 0.26541573 | 7.00E-04 | 0.007 | 0.0025 | 0.0025 | TRUE | 0.0088 |
| Calr3 | 70 | 89 | 106 | 120 | 91 | 75 | 74 | 89 | 106 | 94 | 98 | 82 | 90.67 | 90 | 91 | 1.017 | 0.025 | 0.91262882 | 0.95679219 | 0.008 | 0.0432 | 0.0263 | 0.0263 | TRUE | 0.2412 |
| Calr4 | 73 | 70 | 84 | 74 | 41 | 42 | 78 | 70 | 84 | 58 | 44 | 46 | 63.34 | 77 | 49 | 0.644 | -0.634 | 0.0132917 | 0.06476267 | 0 | 0.0589 | 0.0309 | 0.0309 | TRUE | 0.225 |
| Calu | 13463 | 15184 | 14406 | 16866 | 11957 | 12604 | 14300 | 15164 | 14465 | 13259 | 12865 | 13797 | 13975.07 | 14643 | 13307 | 0.909 | -0.138 | 0.03101587 | 0.11804593 | 0.001 | 0.0071 | 0.0029 | 0.0029 | TRUE | 0.0126 |
| Caly | 9 | 26 | 28 | 34 | 22 | 33 | 10 | 26 | 28 | 27 | 24 | 36 | 25.03 | 21 | 29 | 1.353 | 0.436 | 0.36310046 | 0.58070747 | 0.1071 | 0.1385 | 0.1246 | 0.1246 | TRUE | 1.2373 |
| Camk1 | 450 | 425 | 393 | 796 | 507 | 419 | 478 | 424 | 395 | 626 | 546 | 459 | 487.83 | 432 | 544 | 1.259 | 0.333 | 0.02740862 | 0.10831591 | 0.0149 | 0.0137 | 0.0143 | 0.0143 | TRUE | 0.164 |
| Camk1d | 764 | 970 | 947 | 996 | 769 | 877 | 812 | 969 | 951 | 783 | 827 | 960 | 883.59 | 911 | 857 | 0.94 | -0.089 | 0.46843564 | 0.67079779 | 0.0091 | 0.0106 | 0.0098 | 0.0098 | TRUE | 0.1311 |
| Camk1g | 1 | 0 | 0 | 0 | 1 | 1 | 1 | 0 | 0 | 0 | 1 | 1 | 0.54 | 0 | 1 | 1.923 | 0.944 | 0.73523723 | NA | 0 | 6.1214 | 3.9144 | 3.9144 | TRUE | 0.4614 |
| Camk2a | 421 | 350 | 341 | 304 | 183 | 208 | 447 | 350 | 342 | 239 | 197 | 228 | 300.45 | 380 | 221 | 0.584 | -0.776 | 1.87E-06 | 4.75E-05 | 0.0135 | 0.0179 | 0.0156 | 0.0156 | TRUE | 0.2753 |
| Camk2b | 2002 | 2201 | 2333 | 2218 | 1685 | 1820 | 2126 | 2198 | 2343 | 1744 | 1813 | 1992 | 2036.01 | 2222 | 1850 | 0.832 | -0.266 | 0.00271614 | 0.01961492 | 0.0029 | 0.0085 | 0.0052 | 0.0052 | TRUE | 0.0547 |
| Camk2d | 199 | 205 | 202 | 416 | 262 | 273 | 211 | 205 | 203 | 327 | 282 | 299 | 254.45 | 206 | 303 | 1.47 | 0.556 | 4.50E-05 | 0.0007102 | 0 | 0.0199 | 0.0093 | 0.0093 | TRUE | 0.0603 |
| Camk2g | 2669 | 2503 | 2514 | 2993 | 2131 | 2153 | 2835 | 2500 | 2524 | 2353 | 2293 | 2357 | 2476.92 | 2620 | 2334 | 0.891 | -0.166 | 0.04219294 | 0.14648186 | 0.0021 | 0.0082 | 0.0044 | 0.0044 | TRUE | 0.0625 |
| Camk2n1 | 430 | 1076 | 1097 | 411 | 378 | 583 | 457 | 1075 | 1102 | 323 | 407 | 638 | 666.81 | 878 | 456 | 0.519 | -0.945 | 0.04915074 | 0.16239647 | 0.1647 | 0.0118 | 0.0741 | 0.1647 | TRUE | 2.092 |
| Camk2n2 | 70 | 76 | 104 | 117 | 86 | 97 | 74 | 76 | 104 | 92 | 93 | 106 | 90.89 | 85 | 97 | 1.139 | 0.188 | 0.41554625 | 0.62710945 | 0.0102 | 0.0431 | 0.0272 | 0.0272 | TRUE | 0.3844 |
| Camk4 | 11 | 14 | 15 | 39 | 24 | 23 | 12 | 14 | 15 | 31 | 26 | 25 | 20.4 | 14 | 27 | 2.011 | 1.008 | 0.01827229 | 0.08151387 | 0 | 0.1684 | 0.076 | 0.076 | TRUE | 0.0921 |
| Camkk1 | 827 | 875 | 847 | 823 | 709 | 705 | 878 | 874 | 850 | 647 | 763 | 772 | 797.39 | 867 | 727 | 0.837 | -0.257 | 0.01503161 | 0.0709179 | 0.0037 | 0.011 | 0.0068 | 0.0068 | TRUE | 0.1153 |
| Camkk2 | 933 | 1128 | 1202 | 1088 | 833 | 912 | 991 | 1127 | 1207 | 855 | 896 | 998 | 1012.4 | 1108 | 916 | 0.826 | -0.275 | 0.01618621 | 0.07497251 | 0.0066 | 0.0102 | 0.0084 | 0.0084 | TRUE | 0.1019 |
| Camkmt | 217 | 172 | 195 | 308 | 223 | 175 | 230 | 172 | 196 | 242 | 240 | 192 | 211.95 | 199 | 225 | 1.129 | 0.175 | 0.32742432 | 0.54873407 | 0.0142 | 0.0224 | 0.0183 | 0.0183 | TRUE | 0.2004 |
| Camkv | 1068 | 1196 | 1185 | 838 | 691 | 727 | 1134 | 1194 | 1190 | 659 | 743 | 796 | 952.8 | 1173 | 733 | 0.623 | -0.682 | 3.22E-11 | 2.58E-09 | 0.0036 | 0.0104 | 0.0065 | 0.0065 | TRUE | 0.0965 |
| Caml | 504 | 521 | 538 | 772 | 593 | 494 | 535 | 520 | 540 | 607 | 638 | 541 | 563.59 | 532 | 595 | 1.12 | 0.163 | 0.13166818 | 0.31261035 | 0.0019 | 0.0127 | 0.0066 | 0.0066 | TRUE | 0.0725 |
| Camp | 0 | 0 | 0 | 2 | 0 | 0 | 0 | 0 | 0 | 2 | 0 | 0 | 0.26 | 0 | 1 | 2.886 | 1.529 | 0.70676491 | NA | 0 | 12.5792 | 10 | 10 | TRUE | 0.9669 |
| Camsap1 | 3947 | 4459 | 4625 | 5469 | 3989 | 4072 | 4192 | 4453 | 4644 | 4299 | 4292 | 4457 | 4389.72 | 4430 | 4349 | 0.982 | -0.027 | 0.70871658 | 0.84126423 | 0.0011 | 0.0077 | 0.0034 | 0.0034 | TRUE | 0.0267 |
| Camsap2 | 2367 | 2143 | 2315 | 3449 | 2408 | 2272 | 2514 | 2140 | 2325 | 2711 | 2591 | 2487 | 2461.36 | 2326 | 2596 | 1.117 | 0.159 | 0.07841406 | 0.2216789 | 0.0036 | 0.0082 | 0.0055 | 0.0055 | TRUE | 0.0603 |
| Camsap3 | 3037 | 2877 | 3157 | 4210 | 3239 | 3134 | 3226 | 2873 | 3170 | 3310 | 3485 | 3431 | 3249.05 | 3090 | 3409 | 1.103 | 0.141 | 0.06952255 | 0.20487849 | 0.0018 | 0.0079 | 0.0041 | 0.0041 | TRUE | 0.0457 |
| Camta1 | 2305 | 2772 | 2855 | 2511 | 1955 | 2318 | 2448 | 2768 | 2867 | 1974 | 2103 | 2537 | 2449.71 | 2694 | 2205 | 0.818 | -0.29 | 0.01629129 | 0.07528456 | 0.0116 | 0.0082 | 0.0101 | 0.0101 | TRUE | 0.2101 |
| Camta2 | 872 | 793 | 835 | 1084 | 818 | 800 | 926 | 792 | 838 | 852 | 880 | 876 | 860.77 | 852 | 869 | 1.02 | 0.029 | 0.76836674 | 0.87924836 | 0.0021 | 0.0107 | 0.0057 | 0.0057 | TRUE | 0.0682 |
| Cand1 | 17109 | 17005 | 18232 | 24463 | 17518 | 17571 | 18173 | 16983 | 18307 | 19231 | 18848 | 19234 | 18462.68 | 17821 | 19104 | 1.072 | 0.1 | 0.10397604 | 0.26817827 | 8.00E-04 | 0.0071 | 0.0027 | 0.0027 | TRUE | 0.0207 |
| Cand2 | 1186 | 901 | 886 | 1798 | 1324 | 1116 | 1260 | 900 | 890 | 1413 | 1425 | 1222 | 1184.81 | 1017 | 1353 | 1.332 | 0.414 | 0.00737991 | 0.04171898 | 0.0238 | 0.0097 | 0.0163 | 0.0163 | TRUE | 0.5227 |
| Cant1 | 2296 | 2377 | 2353 | 2825 | 2041 | 2058 | 2439 | 2374 | 2363 | 2221 | 2196 | 2253 | 2307.49 | 2392 | 2223 | 0.93 | -0.105 | 0.10317091 | 0.26735607 | 0 | 0.0083 | 0.0026 | 0.0026 | TRUE | 0.0036 |
| Canx | 54391 | 65678 | 63307 | 67262 | 49614 | 52957 | 57773 | 65593 | 63567 | 52876 | 53382 | 57970 | 58526.75 | 62311 | 54743 | 0.879 | -0.187 | 0.02081194 | 0.08934293 | 0.0034 | 0.007 | 0.0047 | 0.0047 | TRUE | 0.0497 |
| Cap1 | 10059 | 10021 | 9687 | 14767 | 10490 | 10172 | 10684 | 10008 | 9727 | 11609 | 11287 | 11135 | 10741.56 | 10140 | 11344 | 1.119 | 0.162 | 0.01593866 | 0.0741985 | 0.0013 | 0.0072 | 0.0032 | 0.0032 | TRUE | 0.0269 |
| Cap2 | 117 | 190 | 176 | 164 | 89 | 121 | 124 | 190 | 177 | 129 | 96 | 132 | 141.31 | 164 | 119 | 0.728 | -0.457 | 0.0471929 | 0.15772971 | 0.0321 | 0.0302 | 0.031 | 0.031 | TRUE | 0.462 |
| Capg | 1164 | 1203 | 1296 | 1062 | 814 | 984 | 1236 | 1201 | 1301 | 835 | 876 | 1077 | 1087.83 | 1246 | 929 | 0.745 | -0.425 | 0.00039435 | 0.0042924 | 0.0092 | 0.0099 | 0.0094 | 0.0094 | TRUE | 0.2291 |

| | | | | | | | | | | | | | | | | | | | | | | | | |
|---|---|---|---|---|---|---|---|---|---|---|---|---|---|---|---|---|---|---|---|---|---|---|---|---|
| Capn1 | 1213 | 1233 | 1210 | 1932 | 1386 | 1449 | 1288 | 1231 | 1215 | 1519 | 1491 | 1586 | 1388.5 | 1245 | 1532 | 1.231 | 0.299 | 0.00010707 | 0.00145354 | 0 | 0.0093 | 0.0036 | 0.0036 | TRUE | 0.0112 |
| Capn10 | 1351 | 1197 | 1118 | 1871 | 1405 | 1244 | 1435 | 1195 | 1123 | 1471 | 1512 | 1362 | 1349.55 | 1251 | 1448 | 1.158 | 0.212 | 0.06972648 | 0.20532601 | 0.0089 | 0.0093 | 0.0091 | 0.0091 | TRUE | 0.198 |
| Capn11 | 85 | 130 | 120 | 109 | 84 | 82 | 90 | 130 | 120 | 86 | 90 | 90 | 101.07 | 113 | 89 | 0.779 | -0.361 | 0.09861112 | 0.25874141 | 0.0091 | 0.0395 | 0.0244 | 0.0244 | TRUE | 0.3147 |
| Capn12 | 146 | 182 | 156 | 110 | 67 | 70 | 155 | 182 | 157 | 86 | 72 | 77 | 121.45 | 165 | 78 | 0.479 | -1.063 | 3.69E-08 | 1.53E-06 | 0 | 0.034 | 0.0176 | 0.0176 | TRUE | 0.0902 |
| Capn13 | 0 | 2 | 1 | 1 | 0 | 0 | 0 | 2 | 1 | 1 | 0 | 0 | 0.63 | 1 | 0 | 0.347 | -1.525 | 0.56199923 | NA | 0 | 5.2256 | 3.4788 | 3.4788 | TRUE | 0.1027 |
| Capn2 | 692 | 1083 | 1161 | 1012 | 756 | 958 | 735 | 1082 | 1166 | 796 | 813 | 1049 | 940.01 | 994 | 886 | 0.89 | -0.167 | 0.37426718 | 0.5904186 | 0.0397 | 0.0104 | 0.0245 | 0.0245 | TRUE | 0.619 |
| Capn3 | 117 | 72 | 64 | 120 | 86 | 70 | 124 | 72 | 64 | 94 | 93 | 77 | 87.32 | 87 | 88 | 1.015 | 0.021 | 0.94306949 | 0.97165482 | 0.0593 | 0.0446 | 0.0517 | 0.0517 | TRUE | 1.3304 |
| Capn5 | 1867 | 2970 | 2926 | 2126 | 1737 | 2215 | 1983 | 2966 | 2938 | 1671 | 1869 | 2425 | 2308.69 | 2629 | 1988 | 0.756 | -0.404 | 0.03013227 | 0.1156598 | 0.0413 | 0.0083 | 0.0245 | 0.0245 | TRUE | 0.5599 |
| Capn6 | 53 | 35 | 34 | 226 | 168 | 165 | 56 | 35 | 34 | 178 | 181 | 181 | 110.74 | 42 | 180 | 4.312 | 2.108 | 8.27E-19 | 1.76E-16 | 0 | 0.0367 | 0.0259 | 0.0259 | TRUE | 0.6802 |
| Capn7 | 2503 | 2374 | 2330 | 3368 | 2380 | 2272 | 2659 | 2371 | 2340 | 2648 | 2561 | 2487 | 2510.76 | 2457 | 2565 | 1.045 | 0.063 | 0.45227317 | 0.65768509 | 0.0025 | 0.0082 | 0.0047 | 0.0047 | TRUE | 0.0627 |
| Capn8 | 3 | 0 | 0 | 4 | 1 | 1 | 3 | 0 | 0 | 3 | 1 | 1 | 1.42 | 1 | 2 | 1.724 | 0.786 | 0.67381234 | NA | 0.8744 | 2.332 | 1.7526 | 1.7526 | TRUE | 1.4431 |
| Capn9 | 12 | 19 | 14 | 20 | 9 | 15 | 13 | 19 | 14 | 16 | 10 | 16 | 14.6 | 15 | 14 | 0.917 | -0.126 | 0.80526032 | 0.8995696 | 0 | 0.2325 | 0.119 | 0.119 | TRUE | 0.2843 |
| Capns1 | 8458 | 9339 | 9695 | 12511 | 9059 | 9153 | 8984 | 9327 | 9735 | 9835 | 9747 | 10019 | 9607.86 | 9349 | 9867 | 1.055 | 0.078 | 0.2153461 | 0.42717126 | 8.00E-04 | 0.0072 | 0.0027 | 0.0027 | TRUE | 0.016 |
| Capns2 | 2 | 2 | 2 | 6 | 3 | 2 | 2 | 2 | 2 | 5 | 3 | 2 | 2.71 | 2 | 3 | 1.679 | 0.748 | 0.5029787 | NA | 0 | 1.2223 | 0.5058 | 0.5058 | TRUE | 0.228 |
| Caprin1 | 31786 | 34142 | 36699 | 43107 | 31900 | 32132 | 33762 | 34098 | 36850 | 33887 | 34322 | 35174 | 34682.21 | 34903 | 34461 | 0.987 | -0.018 | 0.77806108 | 0.88507604 | 0.0011 | 0.007 | 0.0031 | 0.0031 | TRUE | 0.0291 |
| Caprin2 | 1348 | 1291 | 1389 | 1714 | 1303 | 1306 | 1432 | 1289 | 1395 | 1347 | 1402 | 1430 | 1382.47 | 1372 | 1393 | 1.015 | 0.021 | 0.79657304 | 0.89536272 | 0.0012 | 0.0093 | 0.0043 | 0.0043 | TRUE | 0.0335 |
| Caps2 | 44 | 54 | 56 | 80 | 93 | 65 | 47 | 54 | 56 | 63 | 100 | 71 | 65.17 | 52 | 78 | 1.484 | 0.57 | 0.04181378 | 0.14545367 | 0.024 | 0.0575 | 0.0404 | 0.0404 | TRUE | 0.5525 |
| Capsl | 258 | 251 | 283 | 442 | 322 | 295 | 274 | 251 | 284 | 347 | 346 | 323 | 304.29 | 270 | 339 | 1.258 | 0.331 | 0.00838249 | 0.04608178 | 0 | 0.0177 | 0.008 | 0.008 | TRUE | 0.0428 |
| Capza1 | 2020 | 2107 | 2163 | 2955 | 1993 | 2045 | 2146 | 2104 | 2172 | 2323 | 2144 | 2239 | 2187.94 | 2141 | 2235 | 1.045 | 0.063 | 0.37625822 | 0.59171848 | 5.00E-04 | 0.0084 | 0.0032 | 0.0032 | TRUE | 0.0153 |
| Capza2 | 5755 | 6079 | 5798 | 8676 | 5953 | 5870 | 6113 | 6071 | 5822 | 6820 | 6405 | 6426 | 6276.15 | 6002 | 6550 | 1.092 | 0.126 | 0.05147621 | 0.16728458 | 7.00E-04 | 0.0074 | 0.0029 | 0.0029 | TRUE | 0.016 |
| Capza3 | 0 | 0 | 0 | 0 | 0 | 0 | 0 | 0 | 0 | 0 | 0 | 0 | 0 | 0 | 0 | NA | NA | NA | NA | NA | NA | NA | NA | NA | NA |
| Capzb | 9299 | 10180 | 9984 | 13196 | 9244 | 9389 | 9877 | 10167 | 10025 | 10374 | 9946 | 10278 | 10111.07 | 10023 | 10199 | 1.018 | 0.025 | 0.65389944 | 0.80556928 | 2.00E-04 | 0.0072 | 0.0022 | 0.0022 | TRUE | 0.0057 |
| Car1 | 0 | 2 | 0 | 0 | 0 | 1 | 0 | 2 | 0 | 0 | 0 | 1 | 0.52 | 1 | 0 | 0.522 | -0.937 | 0.77935356 | NA | 0 | 6.3996 | 6.3045 | 6.3045 | TRUE | 0.9739 |
| Car10 | 2 | 7 | 3 | 8 | 6 | 4 | 2 | 7 | 3 | 6 | 6 | 4 | 4.88 | 4 | 5 | 1.411 | 0.497 | 0.57277407 | 0.7512479 | 0 | 0.6827 | 0.3487 | 0.3487 | TRUE | 0.6964 |
| Car11 | 21 | 30 | 38 | 22 | 20 | 25 | 22 | 30 | 38 | 17 | 22 | 27 | 26.1 | 30 | 22 | 0.725 | -0.464 | 0.25626242 | 0.4749437 | 0.0268 | 0.1331 | 0.0814 | 0.0814 | TRUE | 0.155 |
| Car12 | 547 | 623 | 632 | 797 | 589 | 707 | 581 | 622 | 635 | 627 | 634 | 774 | 645.33 | 613 | 678 | 1.105 | 0.145 | 0.23467325 | 0.44950957 | 0.0068 | 0.012 | 0.0091 | 0.0091 | TRUE | 0.1779 |
| Car13 | 503 | 719 | 668 | 667 | 552 | 661 | 534 | 718 | 671 | 524 | 594 | 724 | 627.49 | 641 | 614 | 0.956 | -0.065 | 0.69423334 | 0.83246774 | 0.0233 | 0.0122 | 0.0178 | 0.0178 | TRUE | 0.2857 |
| Car14 | 61 | 81 | 118 | 62 | 32 | 38 | 65 | 81 | 118 | 49 | 34 | 42 | 64.82 | 88 | 42 | 0.474 | -1.076 | 0.00066564 | 0.00651272 | 0.0513 | 0.0577 | 0.0545 | 0.0545 | TRUE | 0.3672 |
| Car15 | 4 | 1 | 0 | 5 | 3 | 5 | 4 | 1 | 0 | 4 | 3 | 5 | 2.98 | 2 | 4 | 2.432 | 1.282 | 0.30826153 | NA | 0 | 1.1125 | 0.7292 | 0.7292 | TRUE | 1.1774 |
| Car2 | 1072 | 949 | 869 | 2678 | 1772 | 1412 | 1139 | 948 | 873 | 2105 | 1907 | 1546 | 1419.41 | 987 | 1853 | 1.879 | 0.91 | 9.66E-10 | 5.93E-08 | 0.0202 | 0.0092 | 0.0152 | 0.0152 | TRUE | 0.2528 |
| Car3 | 15 | 20 | 21 | 21 | 14 | 9 | 16 | 20 | 21 | 17 | 15 | 10 | 16.4 | 19 | 14 | 0.732 | -0.449 | 0.34664391 | 0.56576997 | 0 | 0.2078 | 0.1022 | 0.1022 | TRUE | 0.2468 |
| Car4 | 249 | 176 | 144 | 709 | 349 | 304 | 264 | 176 | 145 | 557 | 376 | 333 | 308.41 | 195 | 422 | 2.169 | 1.117 | 2.29E-05 | 0.00040039 | 0.0808 | 0.0176 | 0.0464 | 0.0464 | TRUE | 1.0468 |
| Car5a | 30 | 39 | 23 | 30 | 25 | 37 | 32 | 39 | 23 | 24 | 27 | 41 | 30.82 | 31 | 31 | 0.962 | -0.056 | 0.88706073 | 0.94442566 | 0.0432 | 0.1138 | 0.0796 | 0.0796 | TRUE | 0.3741 |
| Car5b | 61 | 46 | 52 | 53 | 27 | 43 | 65 | 46 | 52 | 42 | 29 | 47 | 46.79 | 54 | 39 | 0.725 | -0.464 | 0.14123963 | 0.32677593 | 0.0179 | 0.0773 | 0.0499 | 0.0499 | TRUE | 0.3565 |
| Car6 | 143 | 96 | 80 | 137 | 69 | 64 | 152 | 96 | 80 | 108 | 74 | 70 | 96.68 | 109 | 84 | 0.773 | -0.372 | 0.22464589 | 0.43704002 | 0.0736 | 0.041 | 0.0572 | 0.0572 | TRUE | 1.0837 |
| Car7 | 72 | 53 | 56 | 100 | 70 | 56 | 76 | 53 | 56 | 79 | 75 | 61 | 66.81 | 62 | 72 | 1.165 | 0.22 | 0.40560674 | 0.61838855 | 0.0123 | 0.0562 | 0.0353 | 0.0353 | TRUE | 0.3614 |
| Car8 | 217 | 254 | 218 | 222 | 173 | 205 | 230 | 254 | 219 | 175 | 186 | 224 | 214.69 | 234 | 195 | 0.83 | -0.27 | 0.09630408 | 0.25472211 | 0.0067 | 0.0222 | 0.0142 | 0.0142 | TRUE | 0.184 |
| Car9 | 115 | 178 | 197 | 106 | 71 | 108 | 122 | 178 | 198 | 83 | 76 | 118 | 129.28 | 166 | 92 | 0.557 | -0.845 | 0.00114308 | 0.00987062 | 0.0488 | 0.0324 | 0.0403 | 0.0403 | TRUE | 0.5435 |
| Card10 | 796 | 863 | 880 | 1088 | 873 | 757 | 845 | 862 | 884 | 855 | 939 | 829 | 869.04 | 864 | 874 | 1.012 | 0.017 | 0.85251596 | 0.92628008 | 0.0011 | 0.0107 | 0.0051 | 0.0051 | TRUE | 0.0489 |
| Card11 | 46 | 36 | 45 | 48 | 29 | 32 | 49 | 36 | 45 | 38 | 31 | 35 | 38.99 | 43 | 35 | 0.803 | -0.316 | 0.31236328 | 0.5327199 | 0 | 0.0914 | 0.0447 | 0.0447 | TRUE | 0.1735 |
| Card14 | 16 | 18 | 14 | 17 | 11 | 13 | 17 | 18 | 14 | 13 | 12 | 14 | 14.74 | 16 | 13 | 0.805 | -0.313 | 0.5162703 | 0.70829955 | 0 | 0.2304 | 0.0995 | 0.0995 | TRUE | 0.073 |
| Card6 | 82 | 73 | 78 | 164 | 103 | 109 | 87 | 73 | 78 | 129 | 111 | 119 | 99.56 | 79 | 120 | 1.511 | 0.596 | 0.00327921 | 0.0226841 | 0 | 0.04 | 0.0191 | 0.0191 | TRUE | 0.0644 |
| Card9 | 12 | 24 | 20 | 30 | 33 | 32 | 13 | 24 | 20 | 24 | 36 | 35 | 25.15 | 19 | 32 | 1.642 | 0.715 | 0.09150937 | 0.24684211 | 0.0286 | 0.1379 | 0.0868 | 0.0868 | TRUE | 0.409 |
| Carf | 201 | 298 | 285 | 311 | 227 | 232 | 213 | 298 | 286 | 244 | 244 | 254 | 256.66 | 266 | 247 | 0.931 | -0.104 | 0.52865814 | 0.71718529 | 0.0118 | 0.0197 | 0.0157 | 0.0157 | TRUE | 0.3266 |
| Carhsp1 | 4160 | 4437 | 4298 | 4669 | 3454 | 3496 | 4419 | 4431 | 4316 | 3670 | 3716 | 3827 | 4063.2 | 4389 | 3738 | 0.852 | -0.232 | 0.00012813 | 0.0016868 | 0 | 0.0077 | 0.0024 | 0.0024 | TRUE | 0.0053 |
| Carkd | 1091 | 1277 | 1337 | 1668 | 1231 | 1175 | 1159 | 1275 | 1342 | 1311 | 1324 | 1286 | 1283.1 | 1259 | 1307 | 1.038 | 0.054 | 0.54322413 | 0.72861181 | 0.002 | 0.0095 | 0.005 | 0.005 | TRUE | 0.0576 |
| Carm1 | 9208 | 8790 | 9197 | 9927 | 7128 | 7405 | 9781 | 8779 | 9235 | 7804 | 7669 | 8106 | 8562.17 | 9265 | 7860 | 0.848 | -0.237 | 0.00092666 | 0.00840436 | 0.0017 | 0.0073 | 0.0036 | 0.0036 | TRUE | 0.029 |
| Carns1 | 49 | 37 | 25 | 51 | 35 | 27 | 52 | 37 | 25 | 40 | 38 | 30 | 36.9 | 38 | 36 | 0.946 | -0.081 | 0.82897418 | 0.91328449 | 0.0488 | 0.0962 | 0.0735 | 0.0735 | TRUE | 0.2392 |
| Cars | 2938 | 2888 | 3001 | 4604 | 3400 | 3147 | 3121 | 2884 | 3013 | 3619 | 3658 | 3445 | 3290.11 | 3006 | 3574 | 1.189 | 0.25 | 0.00040775 | 0.00439275 | 8.00E-04 | 0.0079 | 0.0033 | 0.0033 | TRUE | 0.0153 |
| Cars2 | 2440 | 2339 | 2431 | 2822 | 1982 | 2105 | 2592 | 2336 | 2441 | 2218 | 2133 | 2304 | 2337.31 | 2456 | 2218 | 0.903 | -0.147 | 0.06312543 | 0.19253807 | 0.0015 | 0.0083 | 0.0041 | 0.0041 | TRUE | 0.0281 |
| Cartpt | 0 | 0 | 1 | 0 | 0 | 1 | 0 | 0 | 1 | 0 | 0 | 1 | 0.35 | 0 | 0 | 0.954 | -0.067 | 0.98566485 | NA | 0 | 9.4252 | 8.1416 | 8.1416 | TRUE | 0.5383 |
| Casc1 | 2 | 1 | 1 | 2 | 2 | 2 | 2 | 1 | 1 | 2 | 2 | 2 | 1.67 | 1 | 2 | 1.428 | 0.514 | 0.71101204 | NA | 0 | 1.9756 | 0.7648 | 0.7648 | TRUE | 0.1348 |
| Casc3 | 6716 | 7040 | 6990 | 8703 | 6943 | 6435 | 7134 | 7031 | 7019 | 6842 | 7470 | 7044 | 7089.86 | 7061 | 7119 | 1.008 | 0.011 | 0.85999602 | 0.9309173 | 7.00E-04 | 0.0074 | 0.0029 | 0.0029 | TRUE | 0.0228 |
| Casc4 | 2417 | 2777 | 3012 | 3631 | 2617 | 2734 | 2567 | 2773 | 3024 | 2854 | 2816 | 2993 | 2838 | 2788 | 2888 | 1.035 | 0.05 | 0.56854323 | 0.74762914 | 0.003 | 0.0081 | 0.0052 | 0.0052 | TRUE | 0.0665 |
| Casc5 | 3527 | 3711 | 4091 | 4587 | 3251 | 3569 | 3746 | 3706 | 4108 | 3606 | 3498 | 3907 | 3761.83 | 3853 | 3670 | 0.952 | -0.07 | 0.39800402 | 0.61130619 | 0.0029 | 0.0078 | 0.0047 | 0.0047 | TRUE | 0.0406 |
| Casd1 | 495 | 495 | 434 | 770 | 534 | 498 | 526 | 494 | 436 | 605 | 575 | 545 | 530.15 | 485 | 575 | 1.186 | 0.247 | 0.03521799 | 0.12864281 | 0.0038 | 0.0131 | 0.008 | 0.008 | TRUE | 0.093 |
| Cask | 1076 | 1144 | 1106 | 1615 | 1182 | 1135 | 1143 | 1143 | 1111 | 1270 | 1272 | 1242 | 1196.62 | 1132 | 1261 | 1.114 | 0.156 | 0.03656885 | 0.13192221 | 0 | 0.0097 | 0.0032 | 0.0032 | TRUE | 0.0033 |
| Caskin1 | 216 | 239 | 230 | 339 | 300 | 261 | 229 | 239 | 231 | 266 | 323 | 286 | 262.34 | 233 | 292 | 1.249 | 0.321 | 0.02124569 | 0.09076032 | 0 | 0.0195 | 0.0101 | 0.0101 | TRUE | 0.0974 |
| Caskin2 | 1275 | 1133 | 1095 | 1647 | 1250 | 1118 | 1354 | 1132 | 1099 | 1295 | 1345 | 1224 | 1241.47 | 1195 | 1288 | 1.078 | 0.108 | 0.32564755 | 0.54686044 | 0.0066 | 0.0096 | 0.0079 | 0.0079 | TRUE | 0.1621 |
| Casp1 | 0 | 0 | 0 | 0 | 0 | 0 | 0 | 0 | 0 | 0 | 0 | 0 | 0 | 0 | 0 | NA | NA | NA | NA | NA | NA | NA | NA | NA | NA |
| Casp12 | 1 | 0 | 0 | 0 | 0 | 0 | 1 | 0 | 0 | 0 | 0 | 0 | 0.18 | 0 | 0 | 0.49 | -1.029 | 0.80087 | NA | 0 | 18.6165 | 10 | 10 | TRUE | 0.4614 |
| Casp14 | 5 | 19 | 19 | 34 | 19 | 13 | 5 | 19 | 19 | 27 | 20 | 14 | 17.46 | 14 | 20 | 1.421 | 0.507 | 0.38014228 | 0.59468399 | 0.1625 | 0.1956 | 0.181 | 0.181 | TRUE | 0.5375 |
| Casp2 | 676 | 684 | 768 | 1075 | 818 | 785 | 718 | 683 | 771 | 845 | 880 | 859 | 792.8 | 724 | 861 | 1.189 | 0.25 | 0.00724263 | 0.04119391 | 6.00E-04 | 0.0111 | 0.005 | 0.005 | TRUE | 0.0385 |
| Casp3 | 2840 | 3447 | 3668 | 2966 | 2144 | 2620 | 3017 | 3443 | 3683 | 2332 | 2307 | 2868 | 2941.44 | 3381 | 2502 | 0.74 | -0.435 | 0.00037495 | 0.0041153 | 0.0125 | 0.008 | 0.0104 | 0.0104 | TRUE | 0.1975 |
| Casp4 | 0 | 0 | 0 | 0 | 0 | 0 | 0 | 0 | 0 | 0 | 0 | 0 | 0 | 0 | 0 | NA | NA | NA | NA | NA | NA | NA | NA | NA | NA |
| Casp6 | 704 | 849 | 919 | 998 | 784 | 938 | 748 | 848 | 923 | 785 | 844 | 1027 | 862.22 | 840 | 885 | 1.053 | 0.074 | 0.59032393 | 0.76449681 | 0.0142 | 0.0107 | 0.0126 | 0.0126 | TRUE | 0.2327 |
| Casp7 | 160 | 219 | 243 | 379 | 279 | 261 | 170 | 219 | 244 | 298 | 300 | 286 | 252.75 | 211 | 295 | 1.396 | 0.481 | 0.00355036 | 0.02408495 | 0.011 | 0.0199 | 0.0156 | 0.0156 | TRUE | 0.3103 |
| Casp8 | 229 | 283 | 216 | 438 | 304 | 316 | 243 | 283 | 217 | 344 | 327 | 346 | 293.35 | 248 | 339 | 1.37 | 0.454 | 0.00157746 | 0.01275225 | 0.0048 | 0.0181 | 0.0114 | 0.0114 | TRUE | 0.1723 |
| Casp8ap2 | 4616 | 4128 | 4396 | 5706 | 4121 | 4000 | 4903 | 4123 | 4414 | 4486 | 4434 | 4379 | 4456.32 | 4480 | 4433 | 0.99 | -0.015 | 0.86148124 | 0.93157685 | 0.0037 | 0.0076 | 0.0052 | 0.0052 | TRUE | 0.083 |

| | | | | | | | | | | | | | | | | | | | | | | | | | |
|---|---|---|---|---|---|---|---|---|---|---|---|---|---|---|---|---|---|---|---|---|---|---|---|---|---|
| Casp9 | 2052 | 2193 | 2200 | 2749 | 2071 | 2047 | 2180 | 2190 | 2209 | 2161 | 2228 | 2241 | 2201.48 | 2193 | 2210 | 1.007 | 0.011 | 0.86965752 | 0.93600645 | 0 | 0.0084 | 0.0026 | 0.0026 | TRUE | 0.0047 |
| Casq1 | 84 | 53 | 67 | 73 | 47 | 49 | 89 | 53 | 67 | 57 | 51 | 54 | 61.84 | 70 | 54 | 0.774 | -0.369 | 0.19300948 | 0.39856646 | 0.0226 | 0.0602 | 0.0415 | 0.0415 | TRUE | 0.2646 |
| Casq2 | 20 | 9 | 6 | 14 | 6 | 5 | 21 | 9 | 6 | 11 | 6 | 5 | 9.87 | 12 | 7 | 0.644 | -0.634 | 0.37845541 | 0.59351683 | 0.1983 | 0.3408 | 0.2683 | 0.2683 | TRUE | 1.2563 |
| Casr | 12 | 7 | 14 | 19 | 11 | 9 | 13 | 7 | 14 | 15 | 12 | 10 | 11.74 | 11 | 12 | 1.094 | 0.13 | 0.82103666 | 0.90872914 | 0 | 0.2876 | 0.1513 | 0.1513 | TRUE | 0.4247 |
| Cass4 | 430 | 356 | 383 | 185 | 148 | 110 | 457 | 356 | 385 | 145 | 159 | 120 | 270.32 | 399 | 141 | 0.356 | -1.491 | 1.71E-18 | 3.50E-16 | 0.013 | 0.0191 | 0.0161 | 0.0161 | TRUE | 0.1836 |
| Cast | 1265 | 1424 | 1487 | 2139 | 1472 | 1505 | 1344 | 1422 | 1493 | 1682 | 1584 | 1647 | 1528.61 | 1420 | 1638 | 1.154 | 0.206 | 0.01128265 | 0.05757037 | 0.0011 | 0.0091 | 0.0041 | 0.0041 | TRUE | 0.0262 |
| Casz1 | 1485 | 1441 | 1480 | 1415 | 1074 | 1134 | 1577 | 1439 | 1486 | 1112 | 1156 | 1241 | 1335.3 | 1501 | 1170 | 0.779 | -0.361 | 3.99E-05 | 0.00063937 | 0.0019 | 0.0094 | 0.0048 | 0.0048 | TRUE | 0.0341 |
| Cat | 5432 | 5726 | 5791 | 5949 | 4433 | 4563 | 5770 | 5719 | 5815 | 4677 | 4770 | 4995 | 5290.72 | 5768 | 4814 | 0.834 | -0.261 | 2.41E-05 | 0.00041863 | 3.00E-04 | 0.0075 | 0.0026 | 0.0026 | TRUE | 0.0132 |
| Catsper1 | 0 | 0 | 0 | 0 | 0 | 0 | 0 | 0 | 0 | 0 | 0 | 0 | 0 | 0 | 0 | NA | NA | NA | NA | NA | NA | NA | NA | NA | NA |
| Catsper2 | 44 | 79 | 80 | 120 | 83 | 57 | 47 | 79 | 80 | 94 | 89 | 62 | 75.33 | 69 | 82 | 1.197 | 0.259 | 0.38538248 | 0.59920343 | 0.0509 | 0.0506 | 0.0507 | 0.0507 | TRUE | 0.6812 |
| Catsper3 | 3 | 3 | 1 | 7 | 2 | 3 | 3 | 3 | 1 | 6 | 2 | 3 | 3.02 | 2 | 4 | 1.553 | 0.635 | 0.56450932 | NA | 0 | 1.0975 | 0.5292 | 0.5292 | TRUE | 0.3994 |
| Catsper4 | 2 | 3 | 1 | 1 | 1 | 2 | 2 | 3 | 1 | 1 | 1 | 2 | 1.7 | 2 | 1 | 0.647 | -0.629 | 0.65717739 | NA | 0 | 1.9494 | 0.8264 | 0.8264 | TRUE | 0.1844 |
| Catsperb | 0 | 0 | 0 | 1 | 0 | 0 | 0 | 0 | 0 | 1 | 0 | 0 | 0.13 | 0 | 0 | 1.859 | 0.894 | 0.82650734 | NA | 0 | 25.1515 | 10 | 10 | TRUE | 0.2563 |
| Catsperd | 83 | 171 | 210 | 95 | 72 | 108 | 88 | 171 | 211 | 75 | 77 | 118 | 123.36 | 157 | 90 | 0.574 | -0.802 | 0.01713515 | 0.07771795 | 0.1178 | 0.0336 | 0.0728 | 0.0728 | TRUE | 0.3331 |
| Catsperg1 | 113 | 170 | 172 | 161 | 166 | 190 | 120 | 170 | 173 | 127 | 179 | 208 | 162.61 | 154 | 171 | 1.105 | 0.143 | 0.55294928 | 0.73676743 | 0.045 | 0.0272 | 0.0359 | 0.0359 | TRUE | 0.4038 |
| Catsperg2 | 43 | 42 | 43 | 46 | 48 | 49 | 46 | 42 | 43 | 36 | 52 | 54 | 45.37 | 44 | 47 | 1.073 | 0.102 | 0.73560554 | 0.85807931 | 0 | 0.0795 | 0.0433 | 0.0433 | TRUE | 0.3891 |
| Cav1 | 5 | 5 | 5 | 7 | 1 | 4 | 5 | 5 | 5 | 6 | 1 | 4 | 4.38 | 5 | 4 | 0.73 | -0.454 | 0.62391862 | NA | 0 | 0.759 | 0.3873 | 0.3873 | TRUE | 0.5336 |
| Cav2 | 4 | 13 | 10 | 26 | 18 | 23 | 4 | 13 | 10 | 20 | 19 | 25 | 15.38 | 9 | 21 | 2.367 | 1.243 | 0.02240315 | 0.0942498 | 0 | 0.2212 | 0.135 | 0.135 | TRUE | 0.6625 |
| Cav3 | 2 | 1 | 0 | 0 | 1 | 0 | 2 | 1 | 0 | 0 | 1 | 0 | 0.7 | 1 | 0 | 0.337 | -1.569 | 0.54130184 | NA | 0 | 4.7144 | 3.2639 | 3.2639 | TRUE | 0.2797 |
| Cbfa2t2 | 5940 | 6715 | 6904 | 7408 | 5336 | 5911 | 6309 | 6706 | 6932 | 5824 | 5741 | 6471 | 6330.55 | 6649 | 6012 | 0.904 | -0.146 | 0.07544103 | 0.21637678 | 0.0031 | 0.0074 | 0.0047 | 0.0047 | TRUE | 0.0541 |
| Cbfa2t3 | 84 | 136 | 157 | 125 | 97 | 87 | 89 | 136 | 158 | 98 | 104 | 95 | 113.43 | 128 | 99 | 0.777 | -0.363 | 0.14099448 | 0.32640025 | 0.0338 | 0.0359 | 0.035 | 0.035 | TRUE | 0.4473 |
| Cbfb | 2779 | 3335 | 3224 | 5336 | 4031 | 3547 | 2952 | 3331 | 3237 | 4195 | 4337 | 3883 | 3655.72 | 3173 | 4138 | 1.304 | 0.383 | 7.02E-06 | 0.00014677 | 0.0031 | 0.0078 | 0.005 | 0.005 | TRUE | 0.0452 |
| Cbl | 1534 | 1459 | 1695 | 2252 | 1547 | 1479 | 1629 | 1457 | 1702 | 1770 | 1664 | 1619 | 1640.38 | 1596 | 1684 | 1.056 | 0.079 | 0.40149332 | 0.61413564 | 0.0033 | 0.0089 | 0.0057 | 0.0057 | TRUE | 0.0703 |
| Cblb | 1153 | 1322 | 1375 | 1697 | 1242 | 1347 | 1225 | 1320 | 1381 | 1334 | 1336 | 1475 | 1345.08 | 1309 | 1382 | 1.055 | 0.078 | 0.39869807 | 0.61201859 | 0.0027 | 0.0094 | 0.0053 | 0.0053 | TRUE | 0.041 |
| Cblc | 293 | 375 | 418 | 260 | 224 | 303 | 311 | 375 | 420 | 204 | 241 | 332 | 313.76 | 369 | 259 | 0.701 | -0.513 | 0.01314822 | 0.06424219 | 0.038 | 0.0174 | 0.0275 | 0.0275 | TRUE | 0.6655 |
| Cbll1 | 2073 | 2072 | 1965 | 2592 | 1976 | 1872 | 2202 | 2069 | 1973 | 2038 | 2126 | 2049 | 2076.19 | 2081 | 2071 | 0.995 | -0.007 | 0.92490743 | 0.96294409 | 0.0013 | 0.0085 | 0.0039 | 0.0039 | TRUE | 0.0309 |
| Cbln1 | 892 | 1390 | 1398 | 1610 | 1390 | 1461 | 947 | 1388 | 1404 | 1266 | 1496 | 1599 | 1349.99 | 1246 | 1454 | 1.165 | 0.221 | 0.1877392 | 0.39246444 | 0.0274 | 0.0093 | 0.0195 | 0.0195 | TRUE | 0.5272 |
| Cbln2 | 8 | 27 | 18 | 32 | 20 | 20 | 8 | 27 | 18 | 25 | 22 | 22 | 20.35 | 18 | 23 | 1.28 | 0.356 | 0.47564117 | 0.67577395 | 0.0804 | 0.1688 | 0.1296 | 0.1296 | TRUE | 0.2529 |
| Cbln3 | 32 | 40 | 40 | 47 | 44 | 56 | 34 | 40 | 40 | 37 | 47 | 61 | 43.28 | 38 | 48 | 1.266 | 0.34 | 0.2911077 | 0.51183021 | 0.0176 | 0.083 | 0.0511 | 0.0511 | TRUE | 0.2226 |
| Cbln4 | 2 | 1 | 2 | 0 | 1 | 2 | 2 | 1 | 2 | 0 | 1 | 2 | 1.4 | 2 | 1 | 0.607 | -0.721 | 0.66232166 | NA | 0 | 2.3611 | 1.1939 | 1.1939 | TRUE | 0.3613 |
| Cbr1 | 1536 | 1358 | 1325 | 1962 | 1370 | 1394 | 1632 | 1356 | 1330 | 1542 | 1474 | 1526 | 1476.76 | 1439 | 1514 | 1.052 | 0.074 | 0.48511999 | 0.68237017 | 0.0054 | 0.0091 | 0.0073 | 0.0073 | TRUE | 0.1635 |
| Cbr2 | 1 | 1 | 1 | 0 | 2 | 2 | 1 | 1 | 1 | 0 | 2 | 2 | 1.23 | 1 | 1 | 1.366 | 0.45 | 0.79674244 | NA | 0 | 2.6758 | 1.3453 | 1.3453 | TRUE | 0.7169 |
| Cbr3 | 133 | 188 | 164 | 216 | 150 | 165 | 141 | 188 | 165 | 170 | 161 | 181 | 167.59 | 165 | 171 | 1.036 | 0.05 | 0.77156134 | 0.88085888 | 0.005 | 0.0266 | 0.0158 | 0.0158 | TRUE | 0.1615 |
| Cbr4 | 829 | 1174 | 1144 | 1422 | 953 | 1070 | 881 | 1172 | 1149 | 1118 | 1025 | 1171 | 1086.04 | 1067 | 1105 | 1.035 | 0.05 | 0.70901168 | 0.84150527 | 0.0135 | 0.0099 | 0.0118 | 0.0118 | TRUE | 0.279 |
| Cbs | 1971 | 2288 | 2374 | 1394 | 1058 | 1390 | 2094 | 2285 | 2384 | 1096 | 1138 | 1522 | 1753.02 | 2254 | 1252 | 0.555 | -0.85 | 1.26E-09 | 7.49E-08 | 0.0188 | 0.0088 | 0.0135 | 0.0135 | TRUE | 0.424 |
| Cbwd1 | 2139 | 2139 | 2077 | 2684 | 1895 | 1788 | 2272 | 2136 | 2086 | 2110 | 2039 | 1957 | 2099.98 | 2165 | 2035 | 0.941 | -0.088 | 0.25411726 | 0.47207743 | 0.0012 | 0.0085 | 0.0038 | 0.0038 | TRUE | 0.0227 |
| Cbx1 | 8962 | 7487 | 7427 | 8843 | 6604 | 5867 | 9519 | 7477 | 7457 | 6952 | 7105 | 6422 | 7488.93 | 8151 | 6826 | 0.838 | -0.256 | 0.02749587 | 0.10857909 | 0.0117 | 0.0073 | 0.0096 | 0.0096 | TRUE | 0.263 |
| Cbx2 | 1917 | 1950 | 1980 | 3137 | 2353 | 2148 | 2036 | 1947 | 1988 | 2466 | 2532 | 2351 | 2220.15 | 1990 | 2450 | 1.231 | 0.3 | 2.64E-05 | 0.00045224 | 5.00E-04 | 0.0084 | 0.0032 | 0.0032 | TRUE | 0.0147 |
| Cbx3 | 2557 | 3028 | 2974 | 3675 | 2915 | 2948 | 2716 | 3024 | 2986 | 2889 | 3136 | 3227 | 2996.45 | 2909 | 3084 | 1.06 | 0.084 | 0.32468531 | 0.54594394 | 0.0028 | 0.008 | 0.0049 | 0.0049 | TRUE | 0.0407 |
| Cbx4 | 95 | 222 | 215 | 391 | 275 | 275 | 101 | 222 | 216 | 307 | 296 | 301 | 240.47 | 180 | 301 | 1.678 | 0.747 | 0.00514767 | 0.0319768 | 0.0799 | 0.0206 | 0.0469 | 0.0469 | TRUE | 1.5562 |
| Cbx5 | 11891 | 13132 | 15080 | 19692 | 13957 | 13954 | 12630 | 13115 | 15142 | 15480 | 15017 | 15275 | 14443.22 | 13629 | 15257 | 1.119 | 0.163 | 0.06670212 | 0.19928507 | 0.0048 | 0.0071 | 0.0056 | 0.0056 | TRUE | 0.1153 |
| Cbx6 | 2041 | 2118 | 2099 | 2599 | 2017 | 1900 | 2168 | 2115 | 2108 | 2043 | 2170 | 2080 | 2113.99 | 2130 | 2098 | 0.984 | -0.023 | 0.74274671 | 0.86266578 | 0 | 0.0085 | 0.003 | 0.003 | TRUE | 0.011 |
| Cbx7 | 8008 | 8979 | 8754 | 9037 | 6968 | 7194 | 8506 | 8967 | 8790 | 7104 | 7497 | 7875 | 8123.26 | 8754 | 7492 | 0.856 | -0.225 | 0.00137238 | 0.01141626 | 0.0015 | 0.0073 | 0.0034 | 0.0034 | TRUE | 0.0253 |
| Cbx8 | 12 | 25 | 37 | 76 | 71 | 72 | 13 | 25 | 37 | 60 | 76 | 79 | 48.3 | 25 | 72 | 2.855 | 1.513 | 8.19E-05 | 0.0011743 | 0.0853 | 0.0751 | 0.0791 | 0.0791 | TRUE | 0.5958 |
| Cby1 | 834 | 839 | 772 | 1000 | 737 | 674 | 886 | 838 | 775 | 786 | 793 | 738 | 802.64 | 833 | 772 | 0.928 | -0.108 | 0.26579565 | 0.48548107 | 0.0017 | 0.011 | 0.0056 | 0.0056 | TRUE | 0.0439 |
| Cc2d1a | 1430 | 1630 | 1620 | 2391 | 1689 | 1680 | 1519 | 1628 | 1627 | 1880 | 1817 | 1839 | 1718.23 | 1591 | 1845 | 1.16 | 0.214 | 0.00379637 | 0.02534985 | 3.00E-04 | 0.0088 | 0.0033 | 0.0033 | TRUE | 0.0189 |
| Cc2d1b | 3014 | 2923 | 3009 | 4144 | 3165 | 2903 | 3201 | 2919 | 3021 | 3258 | 3405 | 3178 | 3163.8 | 3047 | 3280 | 1.076 | 0.106 | 0.15134787 | 0.34239864 | 0.0012 | 0.0079 | 0.0036 | 0.0036 | TRUE | 0.0237 |
| Cc2d2a | 568 | 772 | 876 | 1691 | 1152 | 1364 | 603 | 771 | 880 | 1329 | 1239 | 1493 | 1052.64 | 751 | 1354 | 1.801 | 0.849 | 2.01E-08 | 9.01E-07 | 0.0201 | 0.01 | 0.0155 | 0.0155 | TRUE | 0.35 |
| Ccar1 | 4267 | 3671 | 3965 | 6030 | 4366 | 3552 | 4532 | 3666 | 3981 | 4740 | 4698 | 3888 | 4250.99 | 4060 | 4442 | 1.094 | 0.13 | 0.27155095 | 0.49163634 | 0.0113 | 0.0077 | 0.0099 | 0.0099 | TRUE | 0.1445 |
| Ccbe1 | 139 | 157 | 135 | 109 | 92 | 90 | 148 | 157 | 136 | 86 | 99 | 99 | 120.53 | 147 | 95 | 0.641 | -0.641 | 0.00058716 | 0.00589103 | 0 | 0.0342 | 0.0164 | 0.0164 | TRUE | 0.072 |
| Ccbl1 | 319 | 434 | 437 | 464 | 388 | 325 | 339 | 433 | 439 | 365 | 417 | 356 | 391.51 | 404 | 379 | 0.939 | -0.091 | 0.53639203 | 0.72319842 | 0.0112 | 0.0153 | 0.0132 | 0.0132 | TRUE | 0.2254 |
| Ccbl2 | 1423 | 1185 | 1160 | 1054 | 691 | 839 | 1511 | 1183 | 1165 | 829 | 743 | 918 | 1058.36 | 1286 | 830 | 0.645 | -0.632 | 5.54E-06 | 0.00012063 | 0.0156 | 0.01 | 0.013 | 0.013 | TRUE | 0.2802 |
| Ccbp2 | 20 | 15 | 11 | 17 | 10 | 11 | 21 | 15 | 11 | 13 | 11 | 12 | 13.91 | 16 | 12 | 0.77 | -0.377 | 0.47234014 | 0.67351493 | 0 | 0.2438 | 0.1252 | 0.1252 | TRUE | 0.2833 |
| Ccdc101 | 1149 | 1096 | 1100 | 1508 | 1039 | 1002 | 1220 | 1095 | 1105 | 1185 | 1118 | 1097 | 1136.63 | 1140 | 1133 | 0.995 | -0.007 | 0.93946165 | 0.96945854 | 0.0018 | 0.0098 | 0.005 | 0.005 | TRUE | 0.0455 |
| Ccdc102a | 464 | 608 | 716 | 628 | 449 | 393 | 493 | 607 | 719 | 494 | 483 | 430 | 537.66 | 606 | 469 | 0.774 | -0.37 | 0.01868615 | 0.08287906 | 0.0186 | 0.013 | 0.016 | 0.016 | TRUE | 0.1486 |
| Ccdc103 | 23 | 25 | 23 | 66 | 31 | 27 | 24 | 25 | 23 | 52 | 33 | 30 | 31.21 | 24 | 38 | 1.599 | 0.677 | 0.07486614 | 0.2153938 | 0 | 0.1124 | 0.0704 | 0.0704 | TRUE | 0.8142 |
| Ccdc104 | 1858 | 2034 | 2013 | 2646 | 2014 | 2067 | 1974 | 2031 | 2021 | 2080 | 2167 | 2263 | 2089.31 | 2009 | 2170 | 1.08 | 0.111 | 0.12724361 | 0.30648843 | 6.00E-04 | 0.0085 | 0.0033 | 0.0033 | TRUE | 0.0169 |
| Ccdc105 | 2 | 3 | 0 | 4 | 1 | 0 | 2 | 3 | 0 | 3 | 1 | 0 | 1.56 | 2 | 1 | 0.857 | -0.223 | 0.89971629 | NA | 0.831 | 2.123 | 1.6264 | 1.6264 | TRUE | 0.6463 |
| Ccdc106 | 273 | 333 | 276 | 381 | 284 | 199 | 290 | 333 | 277 | 300 | 306 | 218 | 287.1 | 300 | 275 | 0.916 | -0.126 | 0.46457192 | 0.66775069 | 0.0176 | 0.0184 | 0.018 | 0.018 | TRUE | 0.3527 |
| Ccdc107 | 408 | 565 | 494 | 695 | 528 | 555 | 433 | 564 | 496 | 546 | 568 | 608 | 535.94 | 498 | 574 | 1.151 | 0.204 | 0.11755335 | 0.29014889 | 0.008 | 0.0131 | 0.0103 | 0.0103 | TRUE | 0.1587 |
| Ccdc108 | 5 | 0 | 4 | 12 | 7 | 1 | 5 | 0 | 4 | 9 | 8 | 1 | 4.56 | 3 | 6 | 1.965 | 0.975 | 0.41540455 | NA | 0.877 | 0.7287 | 0.7868 | 0.7868 | TRUE | 0.8301 |
| Ccdc109b | 2 | 2 | 4 | 20 | 12 | 6 | 2 | 2 | 4 | 16 | 13 | 7 | 7.22 | 3 | 12 | 4.354 | 2.122 | 0.01059756 | 0.05501803 | 0 | 0.463 | 0.2678 | 0.2678 | TRUE | 0.2677 |
| Ccdc11 | 38 | 62 | 67 | 45 | 32 | 38 | 40 | 62 | 67 | 35 | 34 | 42 | 46.83 | 56 | 37 | 0.655 | -0.611 | 0.05396345 | 0.17274437 | 0.0217 | 0.0773 | 0.0502 | 0.0502 | TRUE | 0.5144 |
| Ccdc110 | 4 | 1 | 1 | 3 | 1 | 2 | 4 | 1 | 1 | 2 | 1 | 2 | 1.98 | 2 | 2 | 0.916 | -0.127 | 0.9249302 | NA | 0 | 1.6714 | 0.8068 | 0.8068 | TRUE | 0.7313 |
| Ccdc111 | 1234 | 1741 | 1813 | 1406 | 1204 | 1509 | 1311 | 1739 | 1820 | 1105 | 1295 | 1652 | 1487.08 | 1623 | 1351 | 0.832 | -0.266 | 0.1307154 | 0.31130728 | 0.0343 | 0.0091 | 0.0217 | 0.0217 | TRUE | 0.456 |
| Ccdc112 | 2649 | 2350 | 2227 | 2255 | 1684 | 1400 | 2814 | 2347 | 2236 | 1773 | 1812 | 1533 | 2085.65 | 2466 | 1706 | 0.692 | -0.531 | 9.76E-06 | 0.00019504 | 0.0109 | 0.0085 | 0.0099 | 0.0099 | TRUE | 0.1846 |
| Ccdc113 | 150 | 227 | 218 | 220 | 171 | 205 | 159 | 227 | 219 | 173 | 184 | 224 | 197.71 | 202 | 194 | 0.958 | -0.062 | 0.75435392 | 0.87032896 | 0.022 | 0.0236 | 0.0228 | 0.0228 | TRUE | 0.3617 |
| Ccdc114 | 518 | 492 | 515 | 706 | 472 | 392 | 550 | 491 | 517 | 555 | 508 | 429 | 508.44 | 519 | 497 | 0.959 | -0.06 | 0.64771963 | 0.80218563 | 0.008 | 0.0134 | 0.0105 | 0.0105 | TRUE | 0.1653 |
| Ccdc115 | 1235 | 1413 | 1487 | 1911 | 1438 | 1387 | 1312 | 1411 | 1493 | 1502 | 1547 | 1518 | 1463.97 | 1405 | 1522 | 1.083 | 0.115 | 0.17044195 | 0.36901257 | 0.0014 | 0.0092 | 0.0044 | 0.0044 | TRUE | 0.0405 |

| | | | | | | | | | | | | | | | | | | | | | | | | | |
|---|---|---|---|---|---|---|---|---|---|---|---|---|---|---|---|---|---|---|---|---|---|---|---|---|---|
| Ccdc116 | 125 | 55 | 59 | 115 | 76 | 36 | 133 | 55 | 59 | 90 | 82 | 39 | 76.42 | 82 | 70 | 0.861 | -0.217 | 0.61143412 | 0.77987759 | 0.1971 | 0.05 | 0.1179 | 0.1179 | TRUE | 2.6589 |
| Ccdc117 | 3672 | 4610 | 4620 | 6850 | 5030 | 5234 | 3900 | 4604 | 4639 | 5385 | 5412 | 5729 | 4944.94 | 4381 | 5509 | 1.257 | 0.33 | 0.00038644 | 0.00422385 | 0.0048 | 0.0076 | 0.006 | 0.006 | TRUE | 0.112 |
| Ccdc12 | 1182 | 1145 | 1183 | 1394 | 1087 | 1015 | 1255 | 1144 | 1188 | 1096 | 1170 | 1111 | 1160.56 | 1196 | 1126 | 0.941 | -0.088 | 0.29929375 | 0.51961003 | 8.00E-04 | 0.0097 | 0.0043 | 0.0043 | TRUE | 0.0229 |
| Ccdc120 | 916 | 769 | 872 | 960 | 719 | 747 | 973 | 768 | 876 | 755 | 774 | 818 | 827.09 | 872 | 782 | 0.896 | -0.158 | 0.17411393 | 0.37335066 | 0.0067 | 0.0109 | 0.0085 | 0.0085 | TRUE | 0.1302 |
| Ccdc121 | 9 | 5 | 2 | 13 | 8 | 3 | 10 | 5 | 2 | 10 | 9 | 3 | 6.45 | 6 | 7 | 1.356 | 0.439 | 0.60976809 | 0.77891178 | 0.2154 | 0.518 | 0.3755 | 0.3755 | TRUE | 0.4201 |
| Ccdc122 | 273 | 265 | 247 | 310 | 237 | 216 | 290 | 265 | 248 | 244 | 255 | 236 | 256.3 | 268 | 245 | 0.916 | -0.126 | 0.35216989 | 0.57073249 | 0 | 0.0198 | 0.0094 | 0.0094 | TRUE | 0.0593 |
| Ccdc124 | 4043 | 3497 | 3259 | 5223 | 3695 | 3074 | 4294 | 3492 | 3272 | 4106 | 3976 | 3365 | 3750.95 | 3686 | 3816 | 1.035 | 0.05 | 0.70068737 | 0.83629834 | 0.0155 | 0.0078 | 0.0119 | 0.0119 | TRUE | 0.253 |
| Ccdc125 | 301 | 396 | 335 | 355 | 262 | 293 | 320 | 395 | 336 | 279 | 282 | 321 | 322.21 | 350 | 294 | 0.837 | -0.257 | 0.07048461 | 0.206671 | 0.0064 | 0.0171 | 0.0114 | 0.0114 | TRUE | 0.1443 |
| Ccdc126 | 285 | 275 | 275 | 376 | 261 | 255 | 303 | 275 | 276 | 296 | 281 | 279 | 284.84 | 285 | 285 | 1.004 | 0.006 | 0.96394536 | 0.98187459 | 0 | 0.0185 | 0.0079 | 0.0079 | TRUE | 0.0348 |
| Ccdc127 | 2231 | 2158 | 2047 | 3016 | 2167 | 1967 | 2370 | 2155 | 2055 | 2371 | 2332 | 2153 | 2239.34 | 2193 | 2285 | 1.042 | 0.06 | 0.50619579 | 0.70051066 | 0.0032 | 0.0084 | 0.0054 | 0.0054 | TRUE | 0.0596 |
| Ccdc129 | 2 | 1 | 0 | 1 | 0 | 3 | 2 | 1 | 0 | 1 | 0 | 3 | 1.2 | 1 | 1 | 1.293 | 0.371 | 0.85008668 | NA | 0.7906 | 2.7549 | 1.9195 | 1.9195 | TRUE | 0.5327 |
| Ccdc13 | 16 | 26 | 29 | 44 | 35 | 27 | 17 | 26 | 29 | 35 | 38 | 30 | 28.98 | 24 | 34 | 1.41 | 0.496 | 0.18630082 | 0.3900622 | 0 | 0.1206 | 0.0659 | 0.0659 | TRUE | 0.3742 |
| Ccdc130 | 770 | 974 | 953 | 862 | 673 | 678 | 818 | 973 | 957 | 678 | 724 | 742 | 815.24 | 916 | 715 | 0.779 | -0.36 | 0.00090599 | 0.00825973 | 0.0045 | 0.0109 | 0.0072 | 0.0072 | TRUE | 0.1036 |
| Ccdc132 | 4047 | 3229 | 3052 | 4180 | 3105 | 2799 | 4299 | 3225 | 3065 | 3286 | 3341 | 3064 | 3379.79 | 3530 | 3230 | 0.915 | -0.128 | 0.34988293 | 0.56878516 | 0.0186 | 0.0079 | 0.0131 | 0.0131 | TRUE | 0.4418 |
| Ccdc134 | 401 | 445 | 396 | 483 | 368 | 402 | 426 | 444 | 398 | 380 | 396 | 440 | 413.95 | 423 | 405 | 0.957 | -0.063 | 0.59746597 | 0.77004845 | 0.0022 | 0.0149 | 0.0079 | 0.0079 | TRUE | 0.064 |
| Ccdc135 | 93 | 43 | 39 | 73 | 42 | 52 | 99 | 43 | 39 | 57 | 45 | 57 | 56.73 | 60 | 53 | 0.885 | -0.177 | 0.65504169 | 0.80600074 | 0.128 | 0.065 | 0.0955 | 0.0955 | TRUE | 2.6444 |
| Ccdc136 | 2650 | 2739 | 3047 | 1822 | 1566 | 1794 | 2815 | 2735 | 3060 | 1432 | 1685 | 1964 | 2281.8 | 2870 | 1694 | 0.59 | -0.762 | 1.89E-09 | 1.08E-07 | 0.0133 | 0.0083 | 0.0111 | 0.0111 | TRUE | 0.2348 |
| Ccdc137 | 501 | 597 | 631 | 845 | 658 | 578 | 532 | 596 | 634 | 664 | 708 | 633 | 627.82 | 587 | 668 | 1.137 | 0.186 | 0.09617417 | 0.25449008 | 0.0036 | 0.0122 | 0.0074 | 0.0074 | TRUE | 0.0785 |
| Ccdc138 | 628 | 800 | 864 | 719 | 615 | 703 | 667 | 799 | 868 | 565 | 662 | 770 | 721.67 | 778 | 666 | 0.854 | -0.227 | 0.13798896 | 0.32239185 | 0.0183 | 0.0115 | 0.0155 | 0.0155 | TRUE | 0.2203 |
| Ccdc14 | 1394 | 1443 | 1572 | 1857 | 1390 | 1387 | 1481 | 1441 | 1578 | 1460 | 1496 | 1518 | 1495.66 | 1500 | 1491 | 0.994 | -0.009 | 0.90783834 | 0.95354464 | 6.00E-04 | 0.0091 | 0.0038 | 0.0038 | TRUE | 0.0253 |
| Ccdc141 | 632 | 575 | 634 | 404 | 386 | 512 | 671 | 574 | 637 | 318 | 415 | 560 | 529.25 | 627 | 431 | 0.686 | -0.544 | 0.00606015 | 0.03623226 | 0.0403 | 0.0131 | 0.0264 | 0.0264 | TRUE | 0.4815 |
| Ccdc142 | 64 | 103 | 76 | 108 | 114 | 90 | 68 | 103 | 76 | 85 | 123 | 99 | 92.21 | 82 | 102 | 1.233 | 0.302 | 0.23870096 | 0.45452029 | 0.0299 | 0.0426 | 0.0363 | 0.0363 | TRUE | 0.4455 |
| Ccdc144b | 0 | 0 | 0 | 0 | 0 | 0 | 0 | 0 | 0 | 0 | 0 | 0 | 0 | 0 | 0 | NA | NA | NA | NA | NA | NA | NA | NA | NA | NA |
| Ccdc146 | 3 | 3 | 3 | 8 | 5 | 4 | 3 | 3 | 3 | 6 | 5 | 4 | 4.21 | 3 | 5 | 1.758 | 0.814 | 0.3645632 | NA | 0 | 0.79 | 0.3232 | 0.3232 | TRUE | 0.0659 |
| Ccdc147 | 4 | 0 | 2 | 1 | 1 | 2 | 4 | 0 | 2 | 1 | 1 | 2 | 1.72 | 2 | 1 | 0.64 | -0.644 | 0.68229885 | NA | 0 | 1.9245 | 1.1698 | 1.1698 | TRUE | 0.1486 |
| Ccdc148 | 6 | 0 | 4 | 3 | 0 | 1 | 6 | 0 | 4 | 2 | 0 | 1 | 2.31 | 3 | 1 | 0.347 | -1.528 | 0.34987152 | NA | 1.2495 | 1.4349 | 1.368 | 1.368 | TRUE | 0.1905 |
| Ccdc149 | 15 | 6 | 8 | 20 | 16 | 12 | 16 | 6 | 8 | 16 | 17 | 13 | 12.67 | 10 | 15 | 1.547 | 0.629 | 0.27846693 | 0.49962841 | 0.0206 | 0.2669 | 0.1596 | 0.1596 | TRUE | 0.9001 |
| Ccdc15 | 252 | 367 | 370 | 469 | 322 | 330 | 268 | 367 | 372 | 369 | 346 | 361 | 347.02 | 336 | 359 | 1.07 | 0.098 | 0.53417648 | 0.72175374 | 0.0134 | 0.0164 | 0.0149 | 0.0149 | TRUE | 0.3501 |
| Ccdc150 | 35 | 35 | 35 | 43 | 30 | 21 | 37 | 35 | 35 | 34 | 32 | 23 | 32.72 | 36 | 30 | 0.836 | -0.259 | 0.44433202 | 0.65087811 | 0 | 0.1076 | 0.0522 | 0.0522 | TRUE | 0.2375 |
| Ccdc151 | 88 | 114 | 130 | 113 | 78 | 92 | 93 | 114 | 131 | 89 | 84 | 101 | 101.89 | 113 | 91 | 0.808 | -0.308 | 0.15658046 | 0.34960179 | 0.0085 | 0.0392 | 0.0241 | 0.0241 | TRUE | 0.1971 |
| Ccdc152 | 116 | 189 | 170 | 235 | 165 | 152 | 123 | 189 | 171 | 185 | 178 | 166 | 168.55 | 161 | 176 | 1.095 | 0.131 | 0.50668916 | 0.70098718 | 0.0178 | 0.0264 | 0.0222 | 0.0222 | TRUE | 0.4366 |
| Ccdc153 | 0 | 1 | 0 | 3 | 1 | 0 | 0 | 1 | 0 | 2 | 1 | 0 | 0.74 | 0 | 1 | 3.203 | 1.679 | 0.50423271 | NA | 0 | 4.4659 | 3.139 | 3.139 | TRUE | 0.1967 |
| Ccdc154 | 0 | 1 | 2 | 1 | 1 | 0 | 0 | 1 | 2 | 1 | 1 | 0 | 0.81 | 1 | 1 | 0.624 | -0.68 | 0.75467218 | NA | 0 | 4.0666 | 2.13 | 2.13 | TRUE | 0.1306 |
| Ccdc155 | 46 | 123 | 121 | 102 | 88 | 82 | 49 | 123 | 121 | 80 | 95 | 90 | 92.97 | 98 | 88 | 0.9 | -0.151 | 0.65817976 | 0.80743861 | 0.1098 | 0.0423 | 0.0736 | 0.0736 | TRUE | 1.8304 |
| Ccdc157 | 174 | 192 | 189 | 322 | 239 | 172 | 185 | 192 | 190 | 253 | 257 | 188 | 210.82 | 189 | 233 | 1.236 | 0.305 | 0.07427545 | 0.21412399 | 0.0104 | 0.0225 | 0.0163 | 0.0163 | TRUE | 0.2986 |
| Ccdc158 | 212 | 193 | 201 | 241 | 152 | 196 | 225 | 193 | 202 | 189 | 164 | 215 | 197.88 | 207 | 189 | 0.916 | -0.127 | 0.44760341 | 0.65334741 | 0.0063 | 0.0236 | 0.0149 | 0.0149 | TRUE | 0.1432 |
| Ccdc159 | 48 | 58 | 51 | 47 | 60 | 43 | 51 | 58 | 51 | 37 | 65 | 47 | 51.45 | 53 | 50 | 0.921 | -0.119 | 0.69682633 | 0.83386531 | 0.0241 | 0.0709 | 0.048 | 0.048 | TRUE | 0.4274 |
| Ccdc160 | 183 | 182 | 218 | 301 | 234 | 238 | 194 | 182 | 219 | 237 | 252 | 261 | 223.99 | 198 | 250 | 1.257 | 0.33 | 0.02467156 | 0.10081886 | 0 | 0.0216 | 0.011 | 0.011 | TRUE | 0.0902 |
| Ccdc162 | 149 | 253 | 268 | 135 | 98 | 145 | 158 | 253 | 269 | 106 | 105 | 159 | 175.06 | 227 | 123 | 0.543 | -0.881 | 0.00070671 | 0.00681315 | 0.061 | 0.0257 | 0.0426 | 0.0426 | TRUE | 0.7607 |
| Ccdc163 | 409 | 421 | 393 | 477 | 389 | 411 | 434 | 420 | 395 | 375 | 419 | 450 | 415.49 | 416 | 415 | 0.993 | -0.01 | 0.93504103 | 0.96780858 | 0.0033 | 0.0148 | 0.0084 | 0.0084 | TRUE | 0.0853 |
| Ccdc164 | 164 | 207 | 232 | 231 | 193 | 227 | 174 | 207 | 233 | 182 | 208 | 248 | 208.6 | 205 | 213 | 1.035 | 0.05 | 0.78929029 | 0.891612 | 0.0183 | 0.0227 | 0.0204 | 0.0204 | TRUE | 0.2364 |
| Ccdc166 | 135 | 181 | 214 | 263 | 215 | 210 | 143 | 181 | 215 | 207 | 231 | 230 | 201.17 | 180 | 223 | 1.236 | 0.306 | 0.09789432 | 0.25720594 | 0.0159 | 0.0233 | 0.0196 | 0.0196 | TRUE | 0.2538 |
| Ccdc167 | 266 | 276 | 253 | 402 | 305 | 287 | 283 | 276 | 254 | 316 | 328 | 314 | 295.1 | 271 | 319 | 1.18 | 0.239 | 0.05376601 | 0.1723792 | 0 | 0.0181 | 0.0076 | 0.0076 | TRUE | 0.0328 |
| Ccdc169 | 107 | 84 | 59 | 167 | 124 | 89 | 114 | 84 | 59 | 131 | 133 | 97 | 103.15 | 86 | 120 | 1.414 | 0.5 | 0.07251919 | 0.21075596 | 0.0537 | 0.0388 | 0.0458 | 0.0458 | TRUE | 0.336 |
| Ccdc17 | 10 | 9 | 5 | 9 | 11 | 10 | 11 | 9 | 5 | 7 | 12 | 11 | 9.08 | 8 | 10 | 1.202 | 0.266 | 0.68208761 | 0.82364225 | 0 | 0.3697 | 0.1912 | 0.1912 | TRUE | 0.3505 |
| Ccdc170 | 40 | 59 | 80 | 50 | 35 | 62 | 42 | 59 | 80 | 39 | 38 | 68 | 54.43 | 60 | 48 | 0.793 | -0.335 | 0.3552826 | 0.57377812 | 0.0858 | 0.0674 | 0.0763 | 0.0763 | TRUE | 0.3188 |
| Ccdc171 | 273 | 214 | 233 | 272 | 225 | 180 | 290 | 214 | 234 | 214 | 242 | 197 | 231.77 | 246 | 218 | 0.886 | -0.175 | 0.31023608 | 0.53069885 | 0.0134 | 0.0211 | 0.0172 | 0.0172 | TRUE | 0.2704 |
| Ccdc172 | 437 | 561 | 552 | 703 | 529 | 513 | 464 | 560 | 554 | 553 | 569 | 562 | 543.68 | 526 | 561 | 1.065 | 0.092 | 0.42625437 | 0.63633543 | 0.0034 | 0.013 | 0.0077 | 0.0077 | TRUE | 0.1231 |
| Ccdc173 | 70 | 63 | 64 | 84 | 69 | 69 | 74 | 63 | 64 | 66 | 74 | 76 | 69.56 | 67 | 72 | 1.068 | 0.095 | 0.68297852 | 0.82423576 | 0 | 0.0543 | 0.0249 | 0.0249 | TRUE | 0.0756 |
| Ccdc174 | 394 | 400 | 389 | 692 | 483 | 408 | 418 | 399 | 391 | 544 | 520 | 447 | 453.15 | 403 | 504 | 1.252 | 0.324 | 0.00790663 | 0.04392512 | 0.004 | 0.0142 | 0.0085 | 0.0085 | TRUE | 0.1114 |
| Ccdc175 | 12 | 18 | 7 | 14 | 15 | 10 | 13 | 18 | 7 | 11 | 16 | 11 | 12.64 | 13 | 13 | 1.005 | 0.007 | 0.99076342 | 0.99518028 | 0.0409 | 0.2675 | 0.162 | 0.162 | TRUE | 0.2113 |
| Ccdc176 | 37 | 52 | 56 | 79 | 82 | 57 | 39 | 52 | 56 | 62 | 88 | 62 | 60.03 | 49 | 71 | 1.437 | 0.523 | 0.06883723 | 0.20346716 | 0.0213 | 0.0618 | 0.0421 | 0.0421 | TRUE | 0.3981 |
| Ccdc177 | 6 | 2 | 3 | 8 | 14 | 4 | 6 | 2 | 3 | 6 | 15 | 4 | 6.19 | 4 | 8 | 2.261 | 1.177 | 0.18636226 | 0.39013878 | 0.2178 | 0.5395 | 0.3787 | 0.3787 | TRUE | 1.2672 |
| Ccdc178 | 2 | 0 | 0 | 0 | 1 | 0 | 2 | 0 | 0 | 0 | 1 | 0 | 0.53 | 1 | 0 | 0.497 | -1.009 | 0.76131406 | NA | 0 | 6.1834 | 6.228 | 6.228 | TRUE | 0.9742 |
| Ccdc18 | 1013 | 1167 | 1226 | 1420 | 961 | 923 | 1076 | 1165 | 1231 | 1116 | 1034 | 1010 | 1105.53 | 1157 | 1053 | 0.911 | -0.135 | 0.15772842 | 0.35117062 | 0.0028 | 0.0099 | 0.0057 | 0.0057 | TRUE | 0.0452 |
| Ccdc19 | 100 | 75 | 91 | 53 | 47 | 47 | 106 | 75 | 91 | 42 | 51 | 51 | 69.36 | 91 | 48 | 0.525 | -0.93 | 0.00033924 | 0.00377325 | 0 | 0.0544 | 0.0326 | 0.0326 | TRUE | 0.1285 |
| Ccdc22 | 1308 | 1030 | 996 | 1321 | 1021 | 861 | 1389 | 1029 | 1000 | 1038 | 1099 | 943 | 1082.93 | 1139 | 1027 | 0.901 | -0.15 | 0.30894991 | 0.52947495 | 0.0197 | 0.0099 | 0.0147 | 0.0147 | TRUE | 0.4402 |
| Ccdc23 | 643 | 557 | 489 | 860 | 581 | 547 | 683 | 556 | 491 | 676 | 625 | 599 | 605.04 | 577 | 633 | 1.099 | 0.137 | 0.34235679 | 0.56253908 | 0.0141 | 0.0123 | 0.0133 | 0.0133 | TRUE | 0.3028 |
| Ccdc24 | 56 | 119 | 123 | 93 | 74 | 94 | 59 | 119 | 124 | 73 | 80 | 103 | 92.91 | 101 | 85 | 0.844 | -0.245 | 0.434363 | 0.6429029 | 0.0793 | 0.0424 | 0.0599 | 0.0599 | TRUE | 1.2298 |
| Ccdc25 | 2015 | 1992 | 1950 | 2690 | 1879 | 1911 | 2140 | 1989 | 1958 | 2115 | 2022 | 2092 | 2052.66 | 2029 | 2076 | 1.023 | 0.033 | 0.65764459 | 0.80725118 | 9.00E-04 | 0.0085 | 0.0036 | 0.0036 | TRUE | 0.0276 |
| Ccdc27 | 0 | 0 | 0 | 0 | 1 | 0 | 0 | 0 | 0 | 0 | 1 | 0 | 0.18 | 0 | 0 | 1.859 | 0.894 | 0.82650734 | NA | 0 | 18.3785 | 10 | 10 | TRUE | 0.5204 |
| Ccdc28a | 283 | 382 | 363 | 424 | 302 | 311 | 301 | 382 | 364 | 333 | 325 | 340 | 340.88 | 349 | 333 | 0.954 | -0.068 | 0.61448049 | 0.78183174 | 0.0049 | 0.0166 | 0.0104 | 0.0104 | TRUE | 0.1652 |
| Ccdc28b | 101 | 84 | 100 | 109 | 82 | 90 | 107 | 84 | 100 | 86 | 88 | 99 | 94 | 97 | 91 | 0.933 | -0.1 | 0.63609362 | 0.79536224 | 0 | 0.0419 | 0.0215 | 0.0215 | TRUE | 0.1405 |
| Ccdc3 | 334 | 297 | 274 | 216 | 166 | 165 | 355 | 297 | 275 | 170 | 179 | 181 | 242.59 | 309 | 177 | 0.571 | -0.809 | 1.96E-07 | 6.41E-06 | 0.0066 | 0.0205 | 0.013 | 0.013 | TRUE | 0.1896 |
| Ccdc30 | 139 | 82 | 80 | 160 | 126 | 99 | 148 | 82 | 80 | 126 | 136 | 108 | 113.26 | 103 | 123 | 1.195 | 0.258 | 0.35431995 | 0.57275128 | 0.0585 | 0.036 | 0.0468 | 0.0468 | TRUE | 1.3693 |
| Ccdc32 | 526 | 492 | 472 | 620 | 442 | 422 | 559 | 491 | 474 | 487 | 476 | 462 | 491.49 | 508 | 475 | 0.936 | -0.095 | 0.3992791 | 0.61241174 | 0.0021 | 0.0136 | 0.0072 | 0.0072 | TRUE | 0.088 |
| Ccdc33 | 37 | 48 | 55 | 36 | 23 | 20 | 39 | 48 | 55 | 28 | 25 | 22 | 36.23 | 47 | 25 | 0.529 | -0.919 | 0.0060138 | 0.03600972 | 0 | 0.0978 | 0.0505 | 0.0505 | TRUE | 0.1422 |
| Ccdc34 | 1757 | 2091 | 2112 | 2489 | 1932 | 1911 | 1866 | 2088 | 2121 | 1957 | 2079 | 2092 | 2033.75 | 2025 | 2043 | 1.008 | 0.012 | 0.89320265 | 0.94834594 | 0.0026 | 0.0085 | 0.0048 | 0.0048 | TRUE | 0.0567 |
| Ccdc36 | 163 | 126 | 111 | 209 | 139 | 120 | 173 | 126 | 111 | 164 | 150 | 131 | 142.61 | 137 | 148 | 1.088 | 0.122 | 0.57774257 | 0.7550676 | 0.0253 | 0.03 | 0.0277 | 0.0277 | TRUE | 0.5499 |

| | | | | | | | | | | | | | | | | | | | | | | | | | |
|---|---|---|---|---|---|---|---|---|---|---|---|---|---|---|---|---|---|---|---|---|---|---|---|---|---|
| Ccdc37 | 22 | 20 | 15 | 21 | 32 | 23 | 23 | 20 | 15 | 17 | 34 | 25 | 22.42 | 19 | 25 | 1.292 | 0.37 | 0.41724132 | 0.62823199 | 0.0538 | 0.1538 | 0.1041 | 0.1041 | TRUE | 0.1694 |
| Ccdc38 | 106 | 104 | 102 | 173 | 120 | 120 | 113 | 104 | 102 | 136 | 129 | 131 | 119.22 | 106 | 132 | 1.245 | 0.316 | 0.07626948 | 0.21787537 | 0 | 0.0345 | 0.0144 | 0.0144 | TRUE | 0.0259 |
| Ccdc39 | 106 | 132 | 132 | 131 | 140 | 129 | 113 | 132 | 133 | 103 | 151 | 141 | 128.63 | 126 | 132 | 1.041 | 0.059 | 0.78335038 | 0.88821007 | 0.0176 | 0.0325 | 0.0249 | 0.0249 | TRUE | 0.4086 |
| Ccdc40 | 414 | 360 | 389 | 481 | 342 | 311 | 440 | 360 | 391 | 378 | 368 | 340 | 379.4 | 397 | 362 | 0.915 | -0.129 | 0.31728868 | 0.53815218 | 0.004 | 0.0156 | 0.0093 | 0.0093 | TRUE | 0.1031 |
| Ccdc41 | 2082 | 2362 | 2409 | 2856 | 2231 | 2216 | 2211 | 2359 | 2419 | 2245 | 2400 | 2426 | 2343.44 | 2330 | 2357 | 1.011 | 0.016 | 0.83718793 | 0.91727183 | 0.0015 | 0.0083 | 0.004 | 0.004 | TRUE | 0.0238 |
| Ccdc42 | 1 | 2 | 1 | 2 | 3 | 2 | 1 | 2 | 1 | 2 | 3 | 2 | 1.84 | 1 | 2 | 1.695 | 0.761 | 0.57269886 | NA | 0 | 1.7952 | 0.7233 | 0.7233 | TRUE | 0.1152 |
| Ccdc42b | 0 | 0 | 0 | 0 | 1 | 0 | 0 | 0 | 0 | 0 | 1 | 0 | 0.18 | 0 | 0 | 1.859 | 0.894 | 0.82650734 | NA | 0 | 18.3785 | 10 | 10 | TRUE | 0.5204 |
| Ccdc43 | 5612 | 5805 | 5645 | 7879 | 5847 | 5847 | 5961 | 5798 | 5668 | 6194 | 6291 | 6400 | 6051.99 | 5809 | 6295 | 1.084 | 0.116 | 0.05147631 | 0.16728458 | 3.00E-04 | 0.0074 | 0.0024 | 0.0024 | TRUE | 0.0064 |
| Ccdc47 | 6268 | 6326 | 6504 | 8291 | 5850 | 5652 | 6658 | 6318 | 6531 | 6518 | 6294 | 6187 | 6417.54 | 6502 | 6333 | 0.974 | -0.038 | 0.54316593 | 0.72861181 | 5.00E-04 | 0.0074 | 0.0026 | 0.0026 | TRUE | 0.008 |
| Ccdc50 | 3107 | 3653 | 3814 | 4489 | 3314 | 3448 | 3300 | 3648 | 3830 | 3529 | 3566 | 3774 | 3607.84 | 3593 | 3623 | 1.008 | 0.012 | 0.88988543 | 0.94623031 | 0.0032 | 0.0078 | 0.0049 | 0.0049 | TRUE | 0.0615 |
| Ccdc51 | 447 | 543 | 489 | 604 | 438 | 403 | 475 | 542 | 491 | 475 | 471 | 441 | 482.55 | 503 | 462 | 0.92 | -0.12 | 0.27767686 | 0.49880631 | 0.0011 | 0.0137 | 0.0067 | 0.0067 | TRUE | 0.0556 |
| Ccdc53 | 869 | 1078 | 1097 | 1461 | 1024 | 1054 | 923 | 1077 | 1102 | 1149 | 1102 | 1154 | 1084.2 | 1034 | 1135 | 1.098 | 0.134 | 0.18115915 | 0.38280307 | 0.0037 | 0.0099 | 0.0063 | 0.0063 | TRUE | 0.1042 |
| Ccdc54 | 0 | 0 | 0 | 0 | 0 | 0 | 0 | 0 | 0 | 0 | 0 | 0 | 0 | 0 | 0 | NA | NA | NA | NA | NA | NA | NA | NA | NA | NA |
| Ccdc55 | 2315 | 2357 | 2171 | 3398 | 2561 | 2370 | 2459 | 2354 | 2180 | 2671 | 2755 | 2594 | 2502.31 | 2331 | 2673 | 1.147 | 0.198 | 0.01285611 | 0.06328543 | 0.0017 | 0.0082 | 0.0042 | 0.0042 | TRUE | 0.039 |
| Ccdc57 | 213 | 229 | 231 | 207 | 181 | 228 | 226 | 229 | 232 | 163 | 195 | 250 | 215.66 | 229 | 203 | 0.88 | -0.184 | 0.32293607 | 0.54404482 | 0.0186 | 0.0222 | 0.0204 | 0.0204 | TRUE | 0.4461 |
| Ccdc58 | 3534 | 4763 | 4701 | 4857 | 3451 | 4049 | 3754 | 4757 | 4720 | 3818 | 3713 | 4432 | 4199.07 | 4410 | 3988 | 0.904 | -0.146 | 0.2366149 | 0.45191896 | 0.0127 | 0.0077 | 0.0107 | 0.0107 | TRUE | 0.2063 |
| Ccdc59 | 904 | 823 | 807 | 1323 | 964 | 897 | 960 | 822 | 810 | 1040 | 1037 | 982 | 941.93 | 864 | 1020 | 1.181 | 0.24 | 0.02001422 | 0.08691195 | 0.0036 | 0.0104 | 0.0066 | 0.0066 | TRUE | 0.1118 |
| Ccdc6 | 7569 | 8264 | 8980 | 9732 | 7541 | 7623 | 8040 | 8253 | 9017 | 7651 | 8114 | 8345 | 8236.43 | 8437 | 8037 | 0.952 | -0.07 | 0.3692548 | 0.58663742 | 0.0026 | 0.0073 | 0.0043 | 0.0043 | TRUE | 0.0442 |
| Ccdc60 | 61 | 89 | 83 | 50 | 53 | 49 | 65 | 89 | 83 | 39 | 57 | 54 | 64.5 | 79 | 50 | 0.627 | -0.673 | 0.01433938 | 0.0684732 | 0.0167 | 0.058 | 0.0381 | 0.0381 | TRUE | 0.334 |
| Ccdc61 | 1891 | 3116 | 3388 | 2174 | 1781 | 2421 | 2009 | 3112 | 3402 | 1709 | 1916 | 2650 | 2466.32 | 2841 | 2092 | 0.736 | -0.442 | 0.038701 | 0.13741383 | 0.0608 | 0.0082 | 0.0325 | 0.0325 | TRUE | 0.7967 |
| Ccdc62 | 33 | 52 | 59 | 84 | 54 | 35 | 35 | 52 | 59 | 66 | 58 | 38 | 51.45 | 49 | 54 | 1.115 | 0.158 | 0.63857837 | 0.79637583 | 0.0515 | 0.0709 | 0.0616 | 0.0616 | TRUE | 0.5202 |
| Ccdc63 | 101 | 91 | 83 | 86 | 69 | 60 | 107 | 91 | 83 | 68 | 74 | 66 | 81.5 | 94 | 69 | 0.737 | -0.44 | 0.05022108 | 0.16470925 | 0 | 0.0473 | 0.0238 | 0.0238 | TRUE | 0.1484 |
| Ccdc64 | 750 | 791 | 802 | 1104 | 880 | 778 | 797 | 790 | 805 | 868 | 947 | 852 | 843.04 | 797 | 889 | 1.114 | 0.156 | 0.08300473 | 0.23060605 | 0 | 0.0108 | 0.0047 | 0.0047 | TRUE | 0.0379 |
| Ccdc64b | 65 | 46 | 51 | 48 | 30 | 40 | 69 | 46 | 51 | 38 | 32 | 44 | 46.66 | 55 | 38 | 0.685 | -0.545 | 0.07807211 | 0.22104348 | 0.0139 | 0.0775 | 0.0468 | 0.0468 | TRUE | 0.3811 |
| Ccdc65 | 36 | 29 | 29 | 54 | 38 | 42 | 38 | 29 | 29 | 42 | 41 | 46 | 37.61 | 32 | 43 | 1.344 | 0.426 | 0.17621343 | 0.37648852 | 0 | 0.0945 | 0.0443 | 0.0443 | TRUE | 0.1845 |
| Ccdc66 | 1146 | 1004 | 985 | 1704 | 1252 | 1092 | 1217 | 1003 | 989 | 1340 | 1347 | 1195 | 1181.83 | 1070 | 1294 | 1.21 | 0.276 | 0.01739371 | 0.07849317 | 0.0081 | 0.0097 | 0.0088 | 0.0088 | TRUE | 0.1736 |
| Ccdc67 | 3 | 5 | 1 | 10 | 7 | 3 | 3 | 5 | 1 | 8 | 8 | 3 | 4.64 | 3 | 6 | 2.05 | 1.035 | 0.27848297 | 0.49962841 | 0 | 0.7164 | 0.413 | 0.413 | TRUE | 0.408 |
| Ccdc68 | 152 | 231 | 266 | 294 | 238 | 291 | 161 | 231 | 267 | 231 | 256 | 319 | 244.16 | 220 | 269 | 1.219 | 0.286 | 0.18991501 | 0.39508927 | 0.0406 | 0.0204 | 0.0302 | 0.0302 | TRUE | 0.5692 |
| Ccdc69 | 39 | 67 | 55 | 73 | 39 | 67 | 41 | 67 | 55 | 57 | 42 | 73 | 56.04 | 54 | 57 | 1.054 | 0.076 | 0.81290004 | 0.90416498 | 0.0458 | 0.0657 | 0.0562 | 0.0562 | TRUE | 0.1724 |
| Ccdc7 | 0 | 0 | 0 | 0 | 0 | 0 | 0 | 0 | 0 | 0 | 0 | 0 | 0 | 0 | 0 | NA | NA | NA | NA | NA | NA | NA | NA | NA | NA |
| Ccdc70 | 0 | 0 | 0 | 0 | 0 | 0 | 0 | 0 | 0 | 0 | 0 | 0 | 0 | 0 | 0 | NA | NA | NA | NA | NA | NA | NA | NA | NA | NA |
| Ccdc71 | 1648 | 1544 | 1443 | 2473 | 1774 | 1513 | 1750 | 1542 | 1449 | 1944 | 1909 | 1656 | 1708.4 | 1580 | 1836 | 1.163 | 0.217 | 0.0498628 | 0.16390913 | 0.0075 | 0.0088 | 0.0083 | 0.0083 | TRUE | 0.1064 |
| Ccdc71l | 370 | 357 | 323 | 794 | 559 | 410 | 393 | 357 | 324 | 624 | 601 | 449 | 458.05 | 358 | 558 | 1.562 | 0.643 | 6.27E-05 | 0.00094132 | 0.0183 | 0.0141 | 0.0163 | 0.0163 | TRUE | 0.339 |
| Ccdc73 | 307 | 310 | 299 | 510 | 381 | 368 | 326 | 310 | 300 | 401 | 410 | 403 | 358.27 | 312 | 405 | 1.297 | 0.375 | 0.00090697 | 0.00826119 | 0 | 0.0161 | 0.0064 | 0.0064 | TRUE | 0.0175 |
| Ccdc74a | 15 | 26 | 30 | 36 | 38 | 38 | 16 | 26 | 30 | 28 | 41 | 42 | 30.47 | 24 | 37 | 1.525 | 0.609 | 0.12180928 | 0.29675631 | 0.0333 | 0.115 | 0.077 | 0.077 | TRUE | 0.4947 |
| Ccdc77 | 2174 | 2554 | 2543 | 2941 | 2065 | 2087 | 2309 | 2551 | 2553 | 2312 | 2222 | 2285 | 2371.94 | 2471 | 2273 | 0.92 | -0.121 | 0.1183887 | 0.29119031 | 0.0014 | 0.0083 | 0.0039 | 0.0039 | TRUE | 0.0397 |
| Ccdc78 | 3 | 3 | 0 | 0 | 1 | 0 | 3 | 3 | 0 | 0 | 1 | 0 | 1.21 | 2 | 0 | 0.17 | -2.559 | 0.24614593 | NA | 0 | 2.7301 | 2.2597 | 2.2597 | TRUE | 0.7145 |
| Ccdc79 | 70 | 71 | 64 | 115 | 77 | 59 | 74 | 71 | 64 | 90 | 83 | 65 | 74.56 | 70 | 79 | 1.141 | 0.19 | 0.43088261 | 0.64057692 | 0 | 0.0511 | 0.0285 | 0.0285 | TRUE | 0.2329 |
| Ccdc8 | 100 | 169 | 173 | 156 | 159 | 140 | 106 | 169 | 174 | 123 | 171 | 153 | 149.28 | 150 | 149 | 0.992 | -0.011 | 0.96360435 | 0.98165404 | 0.0421 | 0.029 | 0.0354 | 0.0354 | TRUE | 0.6634 |
| Ccdc80 | 2 | 1 | 1 | 1 | 0 | 0 | 2 | 1 | 1 | 1 | 0 | 0 | 0.82 | 1 | 0 | 0.255 | -1.974 | 0.35972496 | NA | 0 | 4.03 | 1.975 | 1.975 | TRUE | 0.2562 |
| Ccdc81 | 4 | 2 | 4 | 5 | 4 | 8 | 4 | 2 | 4 | 4 | 4 | 9 | 4.54 | 3 | 6 | 1.641 | 0.715 | 0.43357882 | NA | 0 | 0.7322 | 0.3632 | 0.3632 | TRUE | 0.4697 |
| Ccdc82 | 364 | 330 | 322 | 288 | 221 | 193 | 387 | 330 | 323 | 226 | 238 | 211 | 285.83 | 347 | 225 | 0.65 | -0.621 | 9.26E-06 | 0.00018624 | 0.0036 | 0.0184 | 0.0105 | 0.0105 | TRUE | 0.1158 |
| Ccdc83 | 0 | 0 | 0 | 1 | 2 | 0 | 0 | 0 | 0 | 1 | 2 | 0 | 0.49 | 0 | 1 | 5.204 | 2.38 | 0.47267317 | NA | 0 | 6.7348 | 6.4099 | 6.4099 | TRUE | 0.1222 |
| Ccdc84 | 149 | 152 | 141 | 198 | 140 | 145 | 158 | 152 | 142 | 156 | 151 | 159 | 152.78 | 151 | 155 | 1.03 | 0.043 | 0.79010904 | 0.8919706 | 0 | 0.0285 | 0.012 | 0.012 | TRUE | 0.0286 |
| Ccdc85a | 184 | 130 | 155 | 260 | 180 | 195 | 195 | 130 | 156 | 204 | 194 | 213 | 182.07 | 160 | 204 | 1.273 | 0.348 | 0.06528097 | 0.19695312 | 0.015 | 0.025 | 0.0201 | 0.0201 | TRUE | 0.3836 |
| Ccdc85b | 483 | 569 | 662 | 726 | 564 | 586 | 513 | 568 | 665 | 571 | 607 | 641 | 594.17 | 582 | 606 | 1.041 | 0.057 | 0.65618331 | 0.80670515 | 0.0085 | 0.0124 | 0.0103 | 0.0103 | TRUE | 0.1815 |
| Ccdc85c | 1181 | 1226 | 1199 | 1323 | 1102 | 1036 | 1254 | 1224 | 1204 | 1040 | 1186 | 1134 | 1173.76 | 1227 | 1120 | 0.911 | -0.134 | 0.12978778 | 0.31009882 | 0.0017 | 0.0097 | 0.0048 | 0.0048 | TRUE | 0.0485 |
| Ccdc86 | 1219 | 1009 | 889 | 1795 | 1258 | 958 | 1295 | 1008 | 893 | 1411 | 1354 | 1049 | 1168.07 | 1065 | 1271 | 1.194 | 0.256 | 0.12863138 | 0.3083712 | 0.0294 | 0.0097 | 0.0196 | 0.0196 | TRUE | 0.425 |
| Ccdc87 | 6 | 3 | 5 | 4 | 5 | 3 | 6 | 3 | 5 | 3 | 5 | 3 | 4.37 | 5 | 4 | 0.816 | -0.294 | 0.7428702 | NA | 0 | 0.7614 | 0.3452 | 0.3452 | TRUE | 0.2142 |
| Ccdc88a | 4994 | 5919 | 6230 | 7625 | 5387 | 5214 | 5305 | 5911 | 6256 | 5994 | 5796 | 5708 | 5828.21 | 5824 | 5833 | 1.002 | 0.002 | 0.97912869 | 0.98867206 | 0.0036 | 0.0075 | 0.005 | 0.005 | TRUE | 0.0741 |
| Ccdc88b | 144 | 173 | 144 | 156 | 117 | 115 | 153 | 173 | 145 | 123 | 126 | 126 | 140.79 | 157 | 125 | 0.795 | -0.33 | 0.05428155 | 0.17320019 | 0 | 0.0303 | 0.014 | 0.014 | TRUE | 0.0851 |
| Ccdc88c | 3702 | 3933 | 4433 | 4057 | 3115 | 3101 | 3932 | 3928 | 4451 | 3189 | 3352 | 3395 | 3707.78 | 4104 | 3312 | 0.807 | -0.31 | 0.0001948 | 0.00238358 | 0.0028 | 0.0078 | 0.0047 | 0.0047 | TRUE | 0.0668 |
| Ccdc89 | 9 | 12 | 15 | 56 | 41 | 24 | 10 | 12 | 15 | 44 | 44 | 26 | 25.17 | 12 | 38 | 3.132 | 1.647 | 0.00020584 | 0.00250114 | 0 | 0.1378 | 0.0873 | 0.0873 | TRUE | 0.5114 |
| Ccdc9 | 1545 | 1602 | 1682 | 2244 | 1653 | 1574 | 1641 | 1600 | 1689 | 1764 | 1779 | 1723 | 1699.25 | 1643 | 1755 | 1.068 | 0.095 | 0.17964654 | 0.38088253 | 0 | 0.0088 | 0.003 | 0.003 | TRUE | 0.0072 |
| Ccdc90a | 523 | 573 | 528 | 691 | 521 | 487 | 556 | 572 | 530 | 543 | 561 | 533 | 549.14 | 553 | 546 | 0.987 | -0.019 | 0.84990771 | 0.92526408 | 0 | 0.0129 | 0.0051 | 0.0051 | TRUE | 0.015 |
| Ccdc90b | 1318 | 1784 | 1642 | 1749 | 1252 | 1338 | 1400 | 1782 | 1649 | 1375 | 1347 | 1465 | 1502.84 | 1610 | 1396 | 0.866 | -0.207 | 0.06411583 | 0.19484295 | 0.0072 | 0.0091 | 0.0083 | 0.0083 | TRUE | 0.1567 |
| Ccdc91 | 607 | 559 | 586 | 749 | 501 | 517 | 645 | 558 | 588 | 589 | 539 | 566 | 580.87 | 597 | 565 | 0.947 | -0.079 | 0.4604489 | 0.66458277 | 0.002 | 0.0126 | 0.0065 | 0.0065 | TRUE | 0.0566 |
| Ccdc92 | 1259 | 1153 | 1120 | 1494 | 1116 | 992 | 1337 | 1152 | 1125 | 1174 | 1201 | 1086 | 1179.08 | 1205 | 1154 | 0.958 | -0.061 | 0.55477372 | 0.73801375 | 0.0046 | 0.0097 | 0.0069 | 0.0069 | TRUE | 0.1111 |
| Ccdc93 | 2060 | 2159 | 2093 | 2301 | 1745 | 1754 | 2188 | 2156 | 2102 | 1809 | 1878 | 1920 | 2008.72 | 2149 | 1869 | 0.869 | -0.202 | 0.00397445 | 0.02622804 | 0 | 0.0085 | 0.003 | 0.003 | TRUE | 0.0098 |
| Ccdc94 | 288 | 276 | 268 | 489 | 345 | 293 | 306 | 276 | 269 | 384 | 371 | 321 | 321.17 | 284 | 359 | 1.268 | 0.343 | 0.01143227 | 0.05814571 | 0.0039 | 0.0172 | 0.01 | 0.01 | TRUE | 0.096 |
| Ccdc96 | 33 | 14 | 21 | 72 | 47 | 34 | 35 | 14 | 21 | 57 | 51 | 37 | 35.75 | 23 | 48 | 2.073 | 1.052 | 0.01066322 | 0.05526794 | 0.0787 | 0.0991 | 0.0902 | 0.0902 | TRUE | 0.9479 |
| Ccdc97 | 1531 | 1442 | 1512 | 2127 | 1515 | 1432 | 1626 | 1440 | 1518 | 1672 | 1630 | 1568 | 1575.7 | 1528 | 1623 | 1.063 | 0.088 | 0.29726585 | 0.51817827 | 0.0017 | 0.009 | 0.0045 | 0.0045 | TRUE | 0.0377 |
| Ccer1 | 3 | 2 | 2 | 7 | 3 | 0 | 3 | 2 | 2 | 6 | 3 | 0 | 2.65 | 2 | 3 | 1.25 | 0.322 | 0.80323594 | NA | 0 | 1.2483 | 0.8256 | 0.8256 | TRUE | 0.2361 |
| Cchcr1 | 953 | 915 | 862 | 1437 | 1121 | 1080 | 1012 | 914 | 866 | 1130 | 1206 | 1182 | 1051.6 | 931 | 1173 | 1.26 | 0.333 | 0.00051392 | 0.00528397 | 0.0023 | 0.01 | 0.0057 | 0.0057 | TRUE | 0.0699 |
| Ccin | 1 | 0 | 0 | 4 | 1 | 1 | 1 | 0 | 0 | 3 | 1 | 1 | 1.06 | 0 | 2 | 4.937 | 2.304 | 0.25605048 | NA | 0 | 3.1065 | 1.6934 | 1.6934 | TRUE | 0.4788 |
| Cck | 2 | 0 | 1 | 2 | 1 | 2 | 2 | 0 | 1 | 2 | 1 | 2 | 1.33 | 1 | 2 | 1.559 | 0.641 | 0.6985967 | NA | 0 | 2.4883 | 1.1741 | 1.1741 | TRUE | 0.2803 |
| Cckar | 1 | 3 | 0 | 1 | 0 | 2 | 1 | 3 | 0 | 1 | 0 | 2 | 1.17 | 1 | 1 | 0.725 | -0.464 | 0.81300805 | NA | 0 | 2.8172 | 1.9034 | 1.9034 | TRUE | 0.3573 |
| Cckbr | 21 | 12 | 22 | 55 | 36 | 34 | 22 | 12 | 22 | 43 | 39 | 37 | 29.26 | 19 | 40 | 2.123 | 1.086 | 0.00494553 | 0.03107894 | 0 | 0.1195 | 0.0681 | 0.0681 | TRUE | 0.535 |
| Ccl1 | 0 | 0 | 0 | 0 | 0 | 0 | 0 | 0 | 0 | 0 | 0 | 0 | 0 | 0 | 0 | NA | NA | NA | NA | NA | NA | NA | NA | NA | NA |

| | | | | | | | | | | | | | | | | | | | | | | | | | |
|---|---|---|---|---|---|---|---|---|---|---|---|---|---|---|---|---|---|---|---|---|---|---|---|---|---|
| Ccl11 | 0 | 0 | 0 | 0 | 0 | 0 | 0 | 0 | 0 | 0 | 0 | 0 | 0 | 0 | 0 | NA | NA | NA | NA | NA | NA | NA | NA | NA | NA |
| Ccl12 | 0 | 0 | 0 | 0 | 0 | 0 | 0 | 0 | 0 | 0 | 0 | 0 | 0 | 0 | 0 | NA | NA | NA | NA | NA | NA | NA | NA | NA | NA |
| Ccl17 | 0 | 1 | 0 | 0 | 0 | 0 | 0 | 1 | 0 | 0 | 0 | 0 | 0.17 | 0 | 0 | 0.49 | -1.029 | 0.80087 | NA | 0 | 19.7991 | 10 | 10 | TRUE | 0.405 |
| Ccl19 | 0 | 0 | 0 | 0 | 0 | 0 | 0 | 0 | 0 | 0 | 0 | 0 | 0 | 0 | 0 | NA | NA | NA | NA | NA | NA | NA | NA | NA | NA |
| Ccl2 | 1 | 0 | 0 | 1 | 0 | 0 | 1 | 0 | 0 | 1 | 0 | 0 | 0.31 | 0 | 0 | 0.954 | -0.067 | 0.98625928 | NA | 0 | 10.7014 | 9.0384 | 9.0384 | TRUE | 0.4614 |
| Ccl20 | 3 | 0 | 1 | 6 | 4 | 0 | 3 | 0 | 1 | 5 | 4 | 0 | 2.2 | 1 | 3 | 2.197 | 1.136 | 0.48889698 | NA | 1.2419 | 1.5031 | 1.4109 | 1.4109 | TRUE | 0.8993 |
| Ccl21a | 0 | 0 | 0 | 0 | 0 | 0 | 0 | 0 | 0 | 0 | 0 | 0 | 0 | 0 | 0 | NA | NA | NA | NA | NA | NA | NA | NA | NA | NA |
| Ccl21b | 0 | 0 | 0 | 0 | 0 | 0 | 0 | 0 | 0 | 0 | 0 | 0 | 0 | 0 | 0 | NA | NA | NA | NA | NA | NA | NA | NA | NA | NA |
| Ccl21c | 0 | 0 | 0 | 0 | 0 | 0 | 0 | 0 | 0 | 0 | 0 | 0 | 0 | 0 | 0 | NA | NA | NA | NA | NA | NA | NA | NA | NA | NA |
| Ccl22 | 0 | 2 | 1 | 0 | 1 | 0 | 0 | 2 | 1 | 0 | 1 | 0 | 0.68 | 1 | 0 | 0.347 | -1.526 | 0.55436871 | NA | 0 | 4.8547 | 3.291 | 3.291 | TRUE | 0.2722 |
| Ccl24 | 0 | 0 | 0 | 0 | 0 | 0 | 0 | 0 | 0 | 0 | 0 | 0 | 0 | 0 | 0 | NA | NA | NA | NA | NA | NA | NA | NA | NA | NA |
| Ccl25 | 88 | 78 | 88 | 84 | 63 | 66 | 93 | 78 | 88 | 66 | 68 | 72 | 77.63 | 86 | 69 | 0.793 | -0.335 | 0.12980423 | 0.31009882 | 0 | 0.0493 | 0.0223 | 0.0223 | TRUE | 0.0735 |
| Ccl26 | 0 | 0 | 0 | 0 | 0 | 0 | 0 | 0 | 0 | 0 | 0 | 0 | 0 | 0 | 0 | NA | NA | NA | NA | NA | NA | NA | NA | NA | NA |
| Ccl27a | 7 | 8 | 5 | 5 | 3 | 5 | 7 | 8 | 5 | 4 | 3 | 5 | 5.51 | 7 | 4 | 0.616 | -0.7 | 0.38012572 | 0.59468399 | 0 | 0.6045 | 0.2669 | 0.2669 | TRUE | 0.1401 |
| Ccl27b | 0 | 0 | 0 | 0 | 0 | 0 | 0 | 0 | 0 | 0 | 0 | 0 | 0 | 0 | 0 | NA | NA | NA | NA | NA | NA | NA | NA | NA | NA |
| Ccl28 | 8 | 10 | 4 | 2 | 1 | 4 | 8 | 10 | 4 | 2 | 1 | 4 | 4.92 | 7 | 2 | 0.306 | -1.708 | 0.0783217 | 0.2214575 | 0 | 0.6763 | 0.3978 | 0.3978 | TRUE | 0.5301 |
| Ccl3 | 47 | 11 | 6 | 68 | 40 | 8 | 50 | 11 | 6 | 53 | 43 | 9 | 28.7 | 22 | 35 | 1.582 | 0.661 | 0.41317442 | 0.62449324 | 0.7949 | 0.1217 | 0.4338 | 0.4338 | TRUE | 2.0401 |
| Ccl4 | 0 | 0 | 0 | 0 | 0 | 0 | 0 | 0 | 0 | 0 | 0 | 0 | 0 | 0 | 0 | NA | NA | NA | NA | NA | NA | NA | NA | NA | NA |
| Ccl5 | 0 | 0 | 0 | 0 | 0 | 0 | 0 | 0 | 0 | 0 | 0 | 0 | 0 | 0 | 0 | NA | NA | NA | NA | NA | NA | NA | NA | NA | NA |
| Ccl6 | 1 | 0 | 0 | 0 | 0 | 0 | 1 | 0 | 0 | 0 | 0 | 0 | 0.18 | 0 | 0 | 0.49 | -1.029 | 0.80087 | NA | 0 | 18.6165 | 10 | 10 | TRUE | 0.4614 |
| Ccl7 | 0 | 0 | 0 | 1 | 0 | 0 | 0 | 0 | 0 | 1 | 0 | 0 | 0.13 | 0 | 0 | 1.859 | 0.894 | 0.82650734 | NA | 0 | 25.1515 | 10 | 10 | TRUE | 0.2563 |
| Ccl8 | 0 | 0 | 0 | 1 | 0 | 0 | 0 | 0 | 0 | 1 | 0 | 0 | 0.13 | 0 | 0 | 1.859 | 0.894 | 0.82650734 | NA | 0 | 25.1515 | 10 | 10 | TRUE | 0.2563 |
| Ccl9 | 0 | 0 | 0 | 0 | 3 | 1 | 0 | 0 | 0 | 0 | 3 | 1 | 0.72 | 0 | 1 | 7.579 | 2.922 | 0.3263762 | NA | 6.973 | 4.5799 | 5.0433 | 5.0433 | TRUE | 0.1444 |
| Ccm2 | 1633 | 1673 | 1683 | 2305 | 1473 | 1375 | 1735 | 1671 | 1690 | 1812 | 1585 | 1505 | 1666.22 | 1699 | 1634 | 0.963 | -0.054 | 0.57456332 | 0.75272138 | 0.0043 | 0.0089 | 0.0061 | 0.0061 | TRUE | 0.1117 |
| Ccm2l | 44 | 25 | 36 | 29 | 18 | 24 | 47 | 25 | 36 | 23 | 19 | 26 | 29.38 | 36 | 23 | 0.635 | -0.654 | 0.09620022 | 0.25449008 | 0.0304 | 0.119 | 0.0758 | 0.0758 | TRUE | 0.1359 |
| Ccna1 | 77 | 184 | 168 | 130 | 84 | 108 | 82 | 184 | 169 | 102 | 90 | 118 | 124.17 | 145 | 103 | 0.715 | -0.484 | 0.11376587 | 0.28445074 | 0.0882 | 0.0334 | 0.0593 | 0.0593 | TRUE | 1.4899 |
| Ccna2 | 13608 | 17924 | 17900 | 19068 | 13936 | 15333 | 14454 | 17901 | 17973 | 14990 | 14994 | 16784 | 16182.81 | 16776 | 15589 | 0.929 | -0.106 | 0.33265798 | 0.55309738 | 0.0093 | 0.0071 | 0.0086 | 0.0086 | TRUE | 0.1793 |
| Ccnb1 | 15174 | 17006 | 16847 | 20226 | 14096 | 14383 | 16117 | 16984 | 16916 | 15900 | 15166 | 15744 | 16138.11 | 16672 | 15603 | 0.936 | -0.096 | 0.1105705 | 0.27920555 | 6.00E-04 | 0.0071 | 0.0025 | 0.0025 | TRUE | 0.0104 |
| Ccnb1ip1 | 1495 | 1336 | 1266 | 2166 | 1519 | 1419 | 1588 | 1334 | 1271 | 1703 | 1634 | 1553 | 1513.97 | 1398 | 1630 | 1.167 | 0.223 | 0.04278378 | 0.14772018 | 0.0072 | 0.0091 | 0.008 | 0.008 | TRUE | 0.1697 |
| Ccnb2 | 8378 | 11292 | 11255 | 11610 | 8537 | 9612 | 8899 | 11277 | 11301 | 9127 | 9185 | 10522 | 10051.93 | 10492 | 9611 | 0.916 | -0.127 | 0.28553727 | 0.50680776 | 0.0119 | 0.0072 | 0.01 | 0.01 | TRUE | 0.2156 |
| Ccnb3 | 2 | 8 | 2 | 8 | 8 | 3 | 2 | 8 | 2 | 6 | 9 | 3 | 5.05 | 4 | 6 | 1.497 | 0.582 | 0.53674701 | 0.72342966 | 0.1822 | 0.6592 | 0.4324 | 0.4324 | TRUE | 0.7495 |
| Ccnc | 1769 | 1958 | 1834 | 2459 | 1794 | 1696 | 1879 | 1955 | 1842 | 1933 | 1930 | 1857 | 1899.31 | 1892 | 1907 | 1.008 | 0.011 | 0.87310891 | 0.93761484 | 0 | 0.0086 | 0.0031 | 0.0031 | TRUE | 0.0105 |
| Ccnd1 | 2585 | 3165 | 3448 | 3911 | 2924 | 2896 | 2746 | 3161 | 3462 | 3075 | 3146 | 3170 | 3126.58 | 3123 | 3130 | 1.002 | 0.003 | 0.97557744 | 0.98697924 | 0.0065 | 0.008 | 0.0071 | 0.0071 | TRUE | 0.1353 |
| Ccnd2 | 158 | 173 | 166 | 131 | 106 | 78 | 168 | 173 | 167 | 103 | 114 | 85 | 134.95 | 169 | 101 | 0.597 | -0.745 | 4.79E-05 | 0.00075027 | 0 | 0.0313 | 0.0163 | 0.0163 | TRUE | 0.1605 |
| Ccnd3 | 5186 | 6204 | 6449 | 12624 | 8758 | 8868 | 5508 | 6196 | 6475 | 9924 | 9423 | 9707 | 7872.4 | 6060 | 9685 | 1.598 | 0.676 | 9.59E-16 | 1.44E-13 | 0.0035 | 0.0073 | 0.005 | 0.005 | TRUE | 0.0772 |
| Ccndbp1 | 1049 | 1147 | 1175 | 1484 | 1089 | 1094 | 1114 | 1146 | 1180 | 1167 | 1172 | 1198 | 1162.57 | 1147 | 1179 | 1.028 | 0.039 | 0.60987345 | 0.77891178 | 0 | 0.0097 | 0.0034 | 0.0034 | TRUE | 0.0077 |
| Ccne1 | 10170 | 8613 | 8586 | 12226 | 8644 | 7764 | 10802 | 8602 | 8621 | 9611 | 9300 | 8499 | 9239.32 | 9342 | 9137 | 0.978 | -0.032 | 0.77875847 | 0.88548616 | 0.0091 | 0.0073 | 0.0092 | 0.0092 | TRUE | 0.2284 |
| Ccne2 | 515 | 518 | 512 | 1156 | 813 | 650 | 547 | 517 | 514 | 909 | 875 | 712 | 678.91 | 526 | 832 | 1.583 | 0.662 | 1.07E-07 | 3.81E-06 | 0.008 | 0.0118 | 0.0096 | 0.0096 | TRUE | 0.1873 |
| Ccnf | 9737 | 13066 | 13243 | 13456 | 9839 | 10466 | 10342 | 13049 | 13297 | 10578 | 10586 | 11457 | 11551.64 | 12229 | 10874 | 0.889 | -0.17 | 0.1317735 | 0.31279177 | 0.0099 | 0.0072 | 0.009 | 0.009 | TRUE | 0.2227 |
| Ccng1 | 17224 | 24481 | 23186 | 22259 | 15917 | 17667 | 18295 | 24449 | 23281 | 17498 | 17126 | 19339 | 19998.14 | 22008 | 17988 | 0.817 | -0.291 | 0.01721748 | 0.07792082 | 0.0131 | 0.0071 | 0.0107 | 0.0107 | TRUE | 0.2665 |
| Ccng2 | 903 | 2084 | 2496 | 1488 | 1422 | 1956 | 959 | 2081 | 2506 | 1170 | 1530 | 2141 | 1731.26 | 1849 | 1614 | 0.873 | -0.197 | 0.67344684 | 0.81805326 | 0.1562 | 0.0088 | 0.0671 | 0.1562 | TRUE | 0.7065 |
| Ccnh | 2540 | 2844 | 2967 | 3558 | 2591 | 2714 | 2698 | 2840 | 2979 | 2797 | 2788 | 2971 | 2845.32 | 2839 | 2852 | 1.004 | 0.006 | 0.93618463 | 0.96816629 | 0.0015 | 0.0081 | 0.0038 | 0.0038 | TRUE | 0.0229 |
| Ccni | 13217 | 16027 | 15816 | 16548 | 12303 | 13392 | 14039 | 16006 | 15881 | 13009 | 13237 | 14660 | 14471.95 | 15309 | 13635 | 0.891 | -0.167 | 0.05999374 | 0.18578558 | 0.0046 | 0.0071 | 0.0056 | 0.0056 | TRUE | 0.0644 |
| Ccnj | 361 | 424 | 397 | 634 | 423 | 398 | 383 | 423 | 399 | 498 | 455 | 436 | 432.45 | 402 | 463 | 1.154 | 0.207 | 0.07388262 | 0.21342564 | 0.0017 | 0.0145 | 0.0073 | 0.0073 | TRUE | 0.0546 |
| Ccnjl | 887 | 606 | 575 | 754 | 554 | 505 | 942 | 605 | 577 | 593 | 596 | 553 | 644.39 | 708 | 581 | 0.82 | -0.286 | 0.13182392 | 0.31279177 | 0.0396 | 0.012 | 0.0244 | 0.0244 | TRUE | 0.9841 |
| Ccnk | 6564 | 6610 | 6915 | 7195 | 5539 | 5561 | 6972 | 6601 | 6943 | 5656 | 5960 | 6087 | 6370.02 | 6839 | 5901 | 0.863 | -0.213 | 0.00139024 | 0.01154043 | 9.00E-04 | 0.0074 | 0.003 | 0.003 | TRUE | 0.0163 |
| Ccnl1 | 1169 | 1200 | 1239 | 1634 | 1245 | 1245 | 1242 | 1198 | 1244 | 1285 | 1340 | 1363 | 1278.52 | 1228 | 1329 | 1.082 | 0.113 | 0.14068899 | 0.32593224 | 0 | 0.0095 | 0.0035 | 0.0035 | TRUE | 0.0107 |
| Ccnl2 | 2717 | 3096 | 2927 | 3971 | 3310 | 2960 | 2886 | 3092 | 2939 | 3122 | 3561 | 3240 | 3140.03 | 2972 | 3308 | 1.112 | 0.154 | 0.06306348 | 0.19252493 | 0.0025 | 0.008 | 0.0046 | 0.0046 | TRUE | 0.0546 |
| Ccno | 115 | 204 | 194 | 317 | 221 | 207 | 122 | 204 | 195 | 249 | 238 | 227 | 205.71 | 174 | 238 | 1.37 | 0.454 | 0.03036632 | 0.11627521 | 0.0309 | 0.0229 | 0.0267 | 0.0267 | TRUE | 0.707 |
| Ccnt1 | 1386 | 1255 | 1293 | 1807 | 1237 | 1151 | 1472 | 1253 | 1298 | 1421 | 1331 | 1260 | 1339.21 | 1341 | 1337 | 0.998 | -0.003 | 0.97445791 | 0.98671997 | 0.0047 | 0.0094 | 0.0066 | 0.0066 | TRUE | 0.0872 |
| Ccnt2 | 4140 | 4428 | 4714 | 4878 | 3864 | 3859 | 4397 | 4422 | 4733 | 3835 | 4157 | 4224 | 4294.91 | 4517 | 4072 | 0.901 | -0.15 | 0.04833582 | 0.16025909 | 0.0018 | 0.0077 | 0.0039 | 0.0039 | TRUE | 0.0321 |
| Ccny | 3326 | 3363 | 3443 | 4547 | 3276 | 3539 | 3533 | 3359 | 3457 | 3574 | 3525 | 3874 | 3553.64 | 3450 | 3658 | 1.06 | 0.084 | 0.24759795 | 0.46433167 | 0.0012 | 0.0078 | 0.0036 | 0.0036 | TRUE | 0.0322 |
| Ccnyl1 | 987 | 941 | 871 | 1529 | 1129 | 868 | 1048 | 940 | 875 | 1202 | 1215 | 950 | 1038.27 | 954 | 1122 | 1.177 | 0.235 | 0.07266291 | 0.21105697 | 0.0116 | 0.0101 | 0.0114 | 0.0114 | TRUE | 0.2132 |
| Ccp110 | 3737 | 3081 | 3038 | 5036 | 3727 | 3320 | 3969 | 3077 | 3050 | 3959 | 4010 | 3634 | 3616.67 | 3365 | 3868 | 1.149 | 0.201 | 0.09920539 | 0.25984818 | 0.013 | 0.0078 | 0.0104 | 0.0104 | TRUE | 0.299 |
| Ccpg1 | 3473 | 4114 | 4380 | 4403 | 3268 | 3635 | 3689 | 4109 | 4398 | 3461 | 3516 | 3979 | 3858.69 | 4065 | 3652 | 0.898 | -0.155 | 0.1241388 | 0.30066297 | 0.0065 | 0.0078 | 0.0071 | 0.0071 | TRUE | 0.0797 |
| Ccr1 | 0 | 0 | 0 | 0 | 0 | 0 | 0 | 0 | 0 | 0 | 0 | 0 | 0 | 0 | 0 | NA | NA | NA | NA | NA | NA | NA | NA | NA | NA |
| Ccr10 | 0 | 0 | 0 | 0 | 0 | 1 | 0 | 0 | 0 | 0 | 0 | 1 | 0.18 | 0 | 0 | 1.859 | 0.894 | 0.82650734 | NA | 0 | 18.0643 | 10 | 10 | TRUE | 0.5383 |
| Ccr1l1 | 1 | 0 | 0 | 1 | 0 | 1 | 1 | 0 | 0 | 1 | 0 | 1 | 0.49 | 0 | 1 | 1.724 | 0.785 | 0.78200058 | NA | 0 | 6.7235 | 4.0295 | 4.0295 | TRUE | 0.4614 |
| Ccr2 | 0 | 0 | 0 | 0 | 0 | 0 | 0 | 0 | 0 | 0 | 0 | 0 | 0 | 0 | 0 | NA | NA | NA | NA | NA | NA | NA | NA | NA | NA |
| Ccr3 | 0 | 0 | 0 | 0 | 0 | 0 | 0 | 0 | 0 | 0 | 0 | 0 | 0 | 0 | 0 | NA | NA | NA | NA | NA | NA | NA | NA | NA | NA |
| Ccr4 | 15 | 10 | 2 | 27 | 23 | 17 | 16 | 10 | 2 | 21 | 25 | 19 | 15.42 | 9 | 22 | 2.322 | 1.216 | 0.05813926 | 0.18220301 | 0.2157 | 0.2206 | 0.2188 | 0.2188 | TRUE | 0.671 |
| Ccr5 | 0 | 0 | 0 | 0 | 0 | 0 | 0 | 0 | 0 | 0 | 0 | 0 | 0 | 0 | 0 | NA | NA | NA | NA | NA | NA | NA | NA | NA | NA |
| Ccr6 | 0 | 0 | 0 | 0 | 0 | 0 | 0 | 0 | 0 | 0 | 0 | 0 | 0 | 0 | 0 | NA | NA | NA | NA | NA | NA | NA | NA | NA | NA |
| Ccr7 | 33 | 45 | 59 | 28 | 27 | 26 | 35 | 45 | 59 | 22 | 29 | 28 | 36.46 | 46 | 26 | 0.567 | -0.82 | 0.02113086 | 0.09040097 | 0.0268 | 0.0973 | 0.0615 | 0.0615 | TRUE | 0.1982 |
| Ccr8 | 8 | 6 | 4 | 7 | 4 | 2 | 8 | 6 | 4 | 6 | 4 | 2 | 5.08 | 6 | 4 | 0.661 | -0.597 | 0.48566217 | 0.68272174 | 0 | 0.6549 | 0.3249 | 0.3249 | TRUE | 0.1906 |
| Ccr9 | 0 | 0 | 0 | 0 | 0 | 0 | 0 | 0 | 0 | 0 | 0 | 0 | 0 | 0 | 0 | NA | NA | NA | NA | NA | NA | NA | NA | NA | NA |
| Ccrl1 | 5 | 8 | 9 | 4 | 1 | 5 | 5 | 8 | 9 | 3 | 1 | 5 | 5.34 | 7 | 3 | 0.432 | -1.211 | 0.16857225 | 0.36640749 | 0 | 0.624 | 0.3379 | 0.3379 | TRUE | 0.3581 |
| Ccrl2 | 5 | 4 | 6 | 3 | 6 | 2 | 5 | 4 | 6 | 2 | 6 | 2 | 4.39 | 5 | 3 | 0.708 | -0.499 | 0.58873636 | NA | 0 | 0.7575 | 0.3783 | 0.3783 | TRUE | 0.632 |
| Ccrn4l | 3675 | 3029 | 2954 | 5118 | 3432 | 2889 | 3903 | 3025 | 2966 | 4023 | 3693 | 3162 | 3462.2 | 3298 | 3626 | 1.1 | 0.137 | 0.32466289 | 0.54594394 | 0.0179 | 0.0079 | 0.0137 | 0.0137 | TRUE | 0.313 |

| | | | | | | | | | | | | | | | | | | | | | | | | |
|---|---|---|---|---|---|---|---|---|---|---|---|---|---|---|---|---|---|---|---|---|---|---|---|---|
| Ccs | 546 | 708 | 745 | 948 | 629 | 684 | 580 | 707 | 748 | 745 | 677 | 749 | 700.97 | 678 | 724 | 1.067 | 0.093 | 0.45994159 | 0.66406067 | 0.0087 | 0.0116 | 0.01 | 0.01 | TRUE | 0.1888 |
| Ccsap | 813 | 813 | 779 | 1002 | 760 | 742 | 864 | 812 | 782 | 788 | 818 | 812 | 812.56 | 819 | 806 | 0.983 | -0.024 | 0.78825867 | 0.8909931 | 0 | 0.011 | 0.0046 | 0.0046 | TRUE | 0.0264 |
| Ccser1 | 14 | 15 | 9 | 44 | 18 | 11 | 15 | 15 | 9 | 35 | 19 | 12 | 17.48 | 13 | 22 | 1.72 | 0.783 | 0.16141495 | 0.35641001 | 0.1325 | 0.1954 | 0.1637 | 0.1637 | TRUE | 0.3714 |
| Ccser2 | 2959 | 3233 | 3437 | 3584 | 2695 | 2905 | 3143 | 3229 | 3451 | 2817 | 2900 | 3180 | 3120 | 3274 | 2966 | 0.905 | -0.143 | 0.0872406 | 0.23904774 | 0.0028 | 0.008 | 0.0047 | 0.0047 | TRUE | 0.0483 |
| Cct2 | 39746 | 44048 | 45026 | 54421 | 39483 | 40418 | 42217 | 43991 | 45211 | 42782 | 42481 | 44244 | 43487.66 | 43806 | 43169 | 0.985 | -0.021 | 0.72426905 | 0.850888 | 7.00E-04 | 0.007 | 0.0026 | 0.0026 | TRUE | 0.0123 |
| Cct3 | 25947 | 25907 | 25696 | 36410 | 25456 | 23372 | 27560 | 25874 | 25802 | 28623 | 27389 | 25584 | 26805.24 | 26412 | 27199 | 1.03 | 0.042 | 0.56327738 | 0.74417917 | 0.0021 | 0.007 | 0.0038 | 0.0038 | TRUE | 0.033 |
| Cct4 | 27522 | 36370 | 36609 | 38301 | 28786 | 30938 | 29233 | 36323 | 36759 | 30109 | 30972 | 33867 | 32877.22 | 34105 | 31649 | 0.928 | -0.108 | 0.3265546 | 0.54768387 | 0.0096 | 0.007 | 0.0087 | 0.0087 | TRUE | 0.1911 |
| Cct5 | 37321 | 41052 | 40697 | 50808 | 36878 | 37171 | 39642 | 40999 | 40864 | 39941 | 39678 | 40690 | 40302.3 | 40502 | 40103 | 0.99 | -0.014 | 0.78541216 | 0.88971634 | 2.00E-04 | 0.007 | 0.002 | 0.002 | TRUE | 0.0042 |
| Cct6a | 20450 | 21661 | 21625 | 28660 | 19915 | 18957 | 21722 | 21633 | 21714 | 22530 | 21427 | 20751 | 21629.55 | 21690 | 21569 | 0.995 | -0.008 | 0.89727183 | 0.95016592 | 7.00E-04 | 0.0071 | 0.0026 | 0.0026 | TRUE | 0.0186 |
| Cct6b | 44 | 70 | 58 | 115 | 71 | 64 | 47 | 70 | 58 | 90 | 76 | 70 | 68.62 | 58 | 79 | 1.358 | 0.441 | 0.09230223 | 0.24796299 | 0.0125 | 0.0549 | 0.0346 | 0.0346 | TRUE | 0.2484 |
| Cct7 | 38880 | 45648 | 45648 | 51518 | 37322 | 38162 | 41297 | 45589 | 45835 | 40499 | 40156 | 41774 | 42525.3 | 44240 | 40810 | 0.922 | -0.116 | 0.09763368 | 0.25676975 | 0.0018 | 0.007 | 0.0035 | 0.0035 | TRUE | 0.0415 |
| Cct8 | 32543 | 32713 | 32519 | 45760 | 32165 | 32011 | 34566 | 32671 | 32653 | 35973 | 34608 | 35041 | 34251.88 | 33297 | 35207 | 1.057 | 0.081 | 0.17259655 | 0.37130521 | 7.00E-04 | 0.007 | 0.0025 | 0.0025 | TRUE | 0.0136 |
| Cct8l1 | 1 | 1 | 1 | 3 | 2 | 7 | 1 | 1 | 1 | 2 | 2 | 8 | 2.54 | 1 | 4 | 3.923 | 1.972 | 0.14154237 | NA | 0 | 1.3041 | 0.6744 | 0.6744 | TRUE | 0.9387 |
| Ccz1 | 3711 | 3804 | 3668 | 5171 | 3626 | 3471 | 3942 | 3799 | 3683 | 4065 | 3901 | 3800 | 3864.97 | 3808 | 3922 | 1.03 | 0.043 | 0.5313972 | 0.71960132 | 8.00E-04 | 0.0078 | 0.0031 | 0.0031 | TRUE | 0.0126 |
| Cd101 | 2 | 1 | 2 | 1 | 4 | 4 | 2 | 1 | 2 | 1 | 4 | 4 | 2.43 | 2 | 3 | 1.804 | 0.852 | 0.50271939 | NA | 0 | 1.3608 | 0.705 | 0.705 | TRUE | 0.8015 |
| Cd109 | 325 | 484 | 509 | 454 | 369 | 473 | 345 | 483 | 511 | 357 | 397 | 518 | 435.23 | 446 | 424 | 0.948 | -0.078 | 0.69395319 | 0.83228303 | 0.0376 | 0.0145 | 0.0258 | 0.0258 | TRUE | 0.454 |
| Cd14 | 1 | 5 | 7 | 13 | 12 | 6 | 1 | 5 | 7 | 10 | 13 | 7 | 7.13 | 4 | 10 | 2.259 | 1.175 | 0.14721321 | 0.3363323 | 0.0487 | 0.4689 | 0.3082 | 0.3082 | TRUE | 0.5306 |
| Cd151 | 2578 | 2813 | 2869 | 2787 | 1976 | 2049 | 2738 | 2809 | 2881 | 2191 | 2126 | 2243 | 2498.06 | 2809 | 2187 | 0.778 | -0.362 | 9.95E-08 | 3.58E-06 | 2.00E-04 | 0.0082 | 0.0029 | 0.0029 | TRUE | 0.0069 |
| Cd160 | 10 | 4 | 6 | 4 | 0 | 0 | 11 | 4 | 6 | 3 | 0 | 0 | 3.96 | 7 | 1 | 0.17 | -2.554 | 0.04878178 | NA | 0.4093 | 0.8379 | 0.7201 | 0.7201 | TRUE | 1.4082 |
| Cd163 | 1 | 1 | 0 | 9 | 7 | 1 | 1 | 1 | 0 | 7 | 8 | 1 | 2.96 | 1 | 5 | 7.725 | 2.95 | 0.04919198 | NA | 0 | 1.1198 | 0.7798 | 0.7798 | TRUE | 0.9108 |
| Cd163l1 | 4 | 2 | 2 | 6 | 0 | 3 | 4 | 2 | 2 | 5 | 0 | 3 | 2.71 | 3 | 3 | 0.996 | -0.006 | 0.99598555 | NA | 0 | 1.2229 | 0.8089 | 0.8089 | TRUE | 0.5935 |
| Cd164 | 14280 | 16228 | 15540 | 17417 | 12943 | 14032 | 15168 | 16207 | 15604 | 13692 | 13926 | 15360 | 14992.79 | 15660 | 14326 | 0.915 | -0.129 | 0.08826326 | 0.24092906 | 0.0024 | 0.0071 | 0.004 | 0.004 | TRUE | 0.0487 |
| Cd164l2 | 203 | 312 | 293 | 193 | 134 | 174 | 216 | 312 | 294 | 152 | 144 | 190 | 217.96 | 274 | 162 | 0.591 | -0.759 | 0.00013523 | 0.0017612 | 0.0252 | 0.022 | 0.0236 | 0.0236 | TRUE | 0.3835 |
| Cd177 | 5 | 3 | 5 | 7 | 3 | 1 | 5 | 3 | 5 | 6 | 3 | 1 | 3.86 | 4 | 3 | 0.76 | -0.397 | 0.68967425 | NA | 0 | 0.8607 | 0.4508 | 0.4508 | TRUE | 0.2963 |
| Cd180 | 3 | 3 | 1 | 6 | 8 | 0 | 3 | 3 | 1 | 5 | 9 | 0 | 3.42 | 2 | 5 | 1.862 | 0.897 | 0.48425497 | NA | 0.7093 | 0.9706 | 0.8614 | 0.8614 | TRUE | 0.1379 |
| Cd19 | 2 | 2 | 4 | 3 | 4 | 2 | 2 | 2 | 4 | 2 | 4 | 2 | 2.83 | 3 | 3 | 1.078 | 0.108 | 0.92137061 | NA | 0 | 1.1704 | 0.5077 | 0.5077 | TRUE | 0.2125 |
| Cd1d1 | 433 | 464 | 495 | 568 | 463 | 457 | 460 | 463 | 497 | 447 | 498 | 500 | 477.55 | 473 | 482 | 1.015 | 0.022 | 0.84176509 | 0.920175 | 0 | 0.0138 | 0.0067 | 0.0067 | TRUE | 0.0504 |
| Cd1d2 | 33 | 36 | 39 | 60 | 30 | 53 | 35 | 36 | 39 | 47 | 32 | 58 | 41.27 | 37 | 46 | 1.248 | 0.32 | 0.33518418 | 0.55542391 | 0.0192 | 0.0867 | 0.0546 | 0.0546 | TRUE | 0.2218 |
| Cd2 | 94 | 107 | 93 | 59 | 46 | 47 | 100 | 107 | 93 | 46 | 49 | 51 | 74.57 | 100 | 49 | 0.49 | -1.031 | 6.72E-06 | 0.00014214 | 0 | 0.0511 | 0.0227 | 0.0227 | TRUE | 0.0354 |
| Cd200 | 39 | 14 | 26 | 102 | 66 | 49 | 41 | 14 | 26 | 80 | 71 | 54 | 47.72 | 27 | 68 | 2.527 | 1.338 | 0.00088152 | 0.0080693 | 0.109 | 0.0759 | 0.0905 | 0.0905 | TRUE | 0.6849 |
| Cd200r1 | 6 | 4 | 4 | 5 | 2 | 2 | 6 | 4 | 4 | 4 | 2 | 2 | 3.78 | 5 | 3 | 0.589 | -0.764 | 0.42675804 | NA | 0 | 0.8794 | 0.3865 | 0.3865 | TRUE | 0.2246 |
| Cd200r2 | 9 | 6 | 9 | 27 | 17 | 7 | 10 | 6 | 9 | 21 | 18 | 8 | 11.96 | 8 | 16 | 1.94 | 0.956 | 0.12552863 | 0.30336443 | 0 | 0.2823 | 0.1873 | 0.1873 | TRUE | 0.8708 |
| Cd200r3 | 3 | 1 | 0 | 1 | 1 | 0 | 3 | 1 | 0 | 1 | 1 | 0 | 1.01 | 1 | 1 | 0.454 | -1.138 | 0.58578015 | NA | 0 | 3.2756 | 1.9983 | 1.9983 | TRUE | 0.3488 |
| Cd200r4 | 0 | 1 | 0 | 0 | 1 | 0 | 0 | 1 | 0 | 0 | 1 | 0 | 0.35 | 0 | 0 | 0.954 | -0.067 | 0.98571955 | NA | 0 | 9.5346 | 8.2194 | 8.2194 | TRUE | 0.5204 |
| Cd207 | 2 | 1 | 2 | 0 | 0 | 2 | 2 | 1 | 2 | 0 | 0 | 2 | 1.22 | 2 | 1 | 0.405 | -1.306 | 0.49676963 | NA | 0 | 2.7071 | 1.6549 | 1.6549 | TRUE | 0.9754 |
| Cd209a | 0 | 1 | 1 | 0 | 1 | 1 | 0 | 1 | 1 | 0 | 1 | 1 | 0.7 | 1 | 1 | 1.033 | 0.047 | 0.9836759 | NA | 0 | 4.7432 | 2.3808 | 2.3808 | TRUE | 0.3742 |
| Cd209b | 0 | 0 | 0 | 0 | 0 | 0 | 0 | 0 | 0 | 0 | 0 | 0 | 0 | 0 | 0 | NA | NA | NA | NA | NA | NA | NA | NA | NA | NA |
| Cd209c | 0 | 0 | 0 | 0 | 1 | 0 | 0 | 0 | 0 | 0 | 1 | 0 | 0.18 | 0 | 0 | 1.859 | 0.894 | 0.82650734 | NA | 0 | 18.3785 | 10 | 10 | TRUE | 0.5204 |
| Cd209d | 0 | 0 | 0 | 0 | 0 | 0 | 0 | 0 | 0 | 0 | 0 | 0 | 0 | 0 | 0 | NA | NA | NA | NA | NA | NA | NA | NA | NA | NA |
| Cd209e | 1 | 0 | 0 | 6 | 5 | 1 | 1 | 0 | 0 | 5 | 5 | 1 | 2.04 | 0 | 4 | 10.258 | 3.359 | 0.05405063 | NA | 0 | 1.6201 | 1.0599 | 1.0599 | TRUE | 0.5425 |
| Cd209f | 4 | 1 | 3 | 3 | 3 | 2 | 4 | 1 | 3 | 2 | 3 | 2 | 2.67 | 3 | 2 | 0.942 | -0.085 | 0.9403064 | NA | 0 | 1.2396 | 0.5639 | 0.5639 | TRUE | 0.3613 |
| Cd209g | 0 | 0 | 0 | 0 | 1 | 1 | 0 | 0 | 0 | 0 | 1 | 1 | 0.36 | 0 | 1 | 3.794 | 1.924 | 0.60247382 | NA | 0 | 9.1135 | 8.1425 | 8.1425 | TRUE | 0.3482 |
| Cd22 | 17 | 14 | 12 | 23 | 10 | 11 | 18 | 14 | 12 | 18 | 11 | 12 | 14.16 | 15 | 14 | 0.94 | -0.089 | 0.86407952 | 0.93284299 | 0 | 0.2395 | 0.1236 | 0.1236 | TRUE | 0.4342 |
| Cd226 | 0 | 0 | 0 | 0 | 0 | 0 | 0 | 0 | 0 | 0 | 0 | 0 | 0 | 0 | 0 | NA | NA | NA | NA | NA | NA | NA | NA | NA | NA |
| Cd244 | 3 | 3 | 3 | 1 | 1 | 1 | 3 | 3 | 3 | 1 | 1 | 1 | 2.03 | 3 | 1 | 0.317 | -1.656 | 0.21880888 | NA | 0 | 1.6336 | 0.6402 | 0.6402 | TRUE | 0.021 |
| Cd247 | 5 | 8 | 14 | 8 | 4 | 4 | 5 | 8 | 14 | 6 | 4 | 4 | 7.05 | 9 | 5 | 0.552 | -0.856 | 0.25655254 | 0.47519694 | 0 | 0.4739 | 0.2578 | 0.2578 | TRUE | 0.4181 |
| Cd248 | 3 | 8 | 8 | 20 | 11 | 4 | 3 | 8 | 8 | 16 | 12 | 4 | 8.52 | 6 | 11 | 1.677 | 0.746 | 0.32313025 | 0.54431383 | 0.1679 | 0.3934 | 0.2861 | 0.2861 | TRUE | 0.3064 |
| Cd24a | 2596 | 2546 | 2464 | 2076 | 1555 | 1490 | 2757 | 2543 | 2474 | 1632 | 1673 | 1631 | 2118.39 | 2591 | 1645 | 0.635 | -0.655 | 3.95E-17 | 6.92E-15 | 0.0014 | 0.0085 | 0.0039 | 0.0039 | TRUE | 0.0379 |
| Cd27 | 0 | 0 | 1 | 0 | 0 | 0 | 0 | 0 | 1 | 0 | 0 | 0 | 0.17 | 0 | 0 | 0.49 | -1.029 | 0.80087 | NA | 0 | 19.6928 | 10 | 10 | TRUE | 0.4098 |
| Cd274 | 53 | 56 | 70 | 59 | 38 | 36 | 56 | 56 | 70 | 46 | 41 | 39 | 51.53 | 61 | 42 | 0.697 | -0.521 | 0.0580619 | 0.18202192 | 0 | 0.0708 | 0.0346 | 0.0346 | TRUE | 0.161 |
| Cd276 | 2937 | 3133 | 3166 | 3392 | 2485 | 2451 | 3120 | 3129 | 3179 | 2667 | 2674 | 2683 | 2908.47 | 3143 | 2675 | 0.851 | -0.233 | 0.00011707 | 0.00155947 | 0 | 0.008 | 0.0023 | 0.0023 | TRUE | 0.0013 |
| Cd28 | 1 | 0 | 0 | 2 | 3 | 1 | 1 | 0 | 0 | 2 | 3 | 1 | 1.16 | 0 | 2 | 5.347 | 2.419 | 0.21719477 | NA | 0 | 2.8482 | 1.518 | 1.518 | TRUE | 0.4614 |
| Cd2ap | 7201 | 7587 | 8065 | 9272 | 7016 | 7162 | 7649 | 7577 | 8098 | 7289 | 7549 | 7840 | 7666.95 | 7775 | 7559 | 0.972 | -0.041 | 0.54494795 | 0.73005344 | 0.0011 | 0.0073 | 0.0031 | 0.0031 | TRUE | 0.0162 |
| Cd2bp2 | 4031 | 3804 | 3597 | 5821 | 4120 | 3894 | 4282 | 3799 | 3612 | 4576 | 4433 | 4263 | 4160.66 | 3898 | 4424 | 1.135 | 0.183 | 0.04115394 | 0.14372894 | 0.0042 | 0.0077 | 0.0056 | 0.0056 | TRUE | 0.0903 |
| Cd300a | 1 | 7 | 8 | 17 | 15 | 6 | 1 | 7 | 8 | 13 | 16 | 7 | 8.69 | 5 | 12 | 2.236 | 1.161 | 0.14250491 | 0.32840262 | 0.2096 | 0.3859 | 0.3161 | 0.3161 | TRUE | 0.8102 |
| Cd300c | 1 | 0 | 0 | 0 | 0 | 0 | 1 | 0 | 0 | 0 | 0 | 0 | 0.18 | 0 | 0 | 0.49 | -1.029 | 0.80087 | NA | 0 | 18.6165 | 10 | 10 | TRUE | 0.4614 |
| Cd300e | 5 | 1 | 1 | 5 | 0 | 1 | 5 | 1 | 1 | 4 | 0 | 1 | 2.06 | 2 | 2 | 0.721 | -0.472 | 0.76088096 | NA | 0.8442 | 1.6089 | 1.2399 | 1.2399 | TRUE | 1.2298 |
| Cd300lb | 4 | 4 | 5 | 6 | 5 | 6 | 4 | 4 | 5 | 5 | 5 | 7 | 4.99 | 4 | 6 | 1.248 | 0.319 | 0.69494704 | 0.83281701 | 0 | 0.6674 | 0.2742 | 0.2742 | TRUE | 0.052 |
| Cd300ld | 1 | 0 | 0 | 1 | 1 | 1 | 1 | 0 | 0 | 1 | 1 | 1 | 0.67 | 0 | 1 | 2.676 | 1.42 | 0.53486561 | NA | 0 | 4.9253 | 2.274 | 2.274 | TRUE | 0.4614 |
| Cd300lf | 38 | 45 | 30 | 24 | 13 | 23 | 40 | 45 | 30 | 19 | 14 | 25 | 28.91 | 38 | 19 | 0.502 | -0.994 | 0.0117023 | 0.05908087 | 0.0176 | 0.1209 | 0.0736 | 0.0736 | TRUE | 0.3259 |
| Cd300lg | 28 | 41 | 41 | 28 | 24 | 22 | 30 | 41 | 41 | 22 | 26 | 24 | 30.63 | 37 | 24 | 0.64 | -0.644 | 0.06829633 | 0.20245789 | 0 | 0.1145 | 0.0559 | 0.0559 | TRUE | 0.2208 |
| Cd300lh | 0 | 0 | 0 | 0 | 0 | 0 | 0 | 0 | 0 | 0 | 0 | 0 | 0 | 0 | 0 | NA | NA | NA | NA | NA | NA | NA | NA | NA | NA |
| Cd302 | 254 | 376 | 313 | 347 | 268 | 314 | 270 | 376 | 314 | 273 | 288 | 344 | 310.74 | 320 | 302 | 0.941 | -0.088 | 0.6063336 | 0.77676352 | 0.0178 | 0.0175 | 0.0177 | 0.0177 | TRUE | 0.2633 |
| Cd320 | 1736 | 1917 | 1900 | 1893 | 1375 | 1441 | 1844 | 1915 | 1908 | 1488 | 1479 | 1577 | 1701.87 | 1889 | 1515 | 0.802 | -0.319 | 1.41E-05 | 0.00026567 | 0 | 0.0088 | 0.0033 | 0.0033 | TRUE | 0.0153 |
| Cd33 | 1 | 2 | 0 | 0 | 1 | 0 | 1 | 2 | 0 | 0 | 1 | 0 | 0.69 | 1 | 0 | 0.342 | -1.547 | 0.54808115 | NA | 0 | 4.7866 | 3.2778 | 3.2778 | TRUE | 0.3016 |
| Cd34 | 645 | 670 | 722 | 754 | 576 | 534 | 685 | 669 | 725 | 593 | 620 | 585 | 646.04 | 693 | 599 | 0.864 | -0.211 | 0.02599322 | 0.10434335 | 0 | 0.012 | 0.0049 | 0.0049 | TRUE | 0.0193 |
| Cd36 | 0 | 1 | 0 | 2 | 0 | 1 | 0 | 1 | 0 | 2 | 0 | 1 | 0.61 | 0 | 1 | 2.474 | 1.307 | 0.62064351 | NA | 0 | 5.3994 | 3.481 | 3.481 | TRUE | 0.405 |
| Cd37 | 434 | 463 | 434 | 391 | 293 | 268 | 461 | 462 | 436 | 307 | 315 | 293 | 379.19 | 453 | 305 | 0.674 | -0.569 | 3.86E-07 | 1.19E-05 | 0 | 0.0156 | 0.0063 | 0.0063 | TRUE | 0.013 |
| Cd38 | 550 | 671 | 637 | 723 | 577 | 686 | 584 | 670 | 640 | 568 | 621 | 751 | 639.01 | 631 | 647 | 1.023 | 0.032 | 0.81117899 | 0.90290246 | 0.0112 | 0.0121 | 0.0116 | 0.0116 | TRUE | 0.2326 |

| | | | | | | | | | | | | | | | | | | | | | | | | | |
|---|---|---|---|---|---|---|---|---|---|---|---|---|---|---|---|---|---|---|---|---|---|---|---|---|---|
| Cd3d | 2 | 1 | 0 | 2 | 3 | 1 | 2 | 1 | 0 | 2 | 3 | 1 | 1.5 | 1 | 2 | 1.895 | 0.922 | 0.56601369 | NA | 0 | 2.1989 | 1.1137 | 1.1137 | TRUE | 0.3207 |
| Cd3e | 26 | 14 | 17 | 52 | 33 | 35 | 28 | 14 | 17 | 41 | 36 | 38 | 28.89 | 20 | 38 | 1.965 | 0.975 | 0.01300728 | 0.0638925 | 0 | 0.1209 | 0.072 | 0.072 | TRUE | 0.6678 |
| Cd3eap | 1249 | 1134 | 1089 | 1833 | 1319 | 1093 | 1327 | 1133 | 1093 | 1441 | 1419 | 1196 | 1268.21 | 1184 | 1352 | 1.143 | 0.192 | 0.1097913 | 0.27808389 | 0.0094 | 0.0095 | 0.0096 | 0.0096 | TRUE | 0.1323 |
| Cd3g | 1 | 0 | 0 | 5 | 5 | 1 | 1 | 0 | 0 | 4 | 5 | 1 | 1.91 | 0 | 3 | 9.513 | 3.25 | 0.06543483 | NA | 0 | 1.7307 | 1.1018 | 1.1018 | TRUE | 0.3823 |
| Cd4 | 7 | 8 | 4 | 5 | 5 | 2 | 7 | 8 | 4 | 4 | 5 | 2 | 5.16 | 6 | 4 | 0.593 | -0.753 | 0.37576042 | 0.59147378 | 0 | 0.6458 | 0.3151 | 0.3151 | TRUE | 0.2804 |
| Cd40 | 103 | 146 | 147 | 112 | 78 | 77 | 109 | 146 | 148 | 88 | 84 | 84 | 109.85 | 134 | 85 | 0.636 | -0.652 | 0.00176157 | 0.01391264 | 0 | 0.0369 | 0.0218 | 0.0218 | TRUE | 0.2653 |
| Cd40lg | 0 | 0 | 0 | 0 | 0 | 0 | 0 | 0 | 0 | 0 | 0 | 0 | 0 | 0 | 0 | NA | NA | NA | NA | NA | NA | NA | NA | NA | NA |
| Cd44 | 78 | 89 | 68 | 68 | 58 | 44 | 83 | 89 | 68 | 53 | 62 | 48 | 67.34 | 80 | 54 | 0.683 | -0.55 | 0.02864961 | 0.1114332 | 0 | 0.0558 | 0.0302 | 0.0302 | TRUE | 0.1546 |
| Cd46 | 2 | 0 | 0 | 1 | 2 | 3 | 2 | 0 | 0 | 1 | 2 | 3 | 1.39 | 1 | 2 | 2.939 | 1.555 | 0.40577738 | NA | 0 | 2.3752 | 1.5473 | 1.5473 | TRUE | 0.7075 |
| Cd47 | 324 | 530 | 569 | 546 | 406 | 447 | 344 | 529 | 571 | 429 | 437 | 489 | 466.69 | 481 | 452 | 0.937 | -0.093 | 0.62581577 | 0.78918736 | 0.0351 | 0.014 | 0.0243 | 0.0243 | TRUE | 0.7208 |
| Cd48 | 0 | 0 | 0 | 0 | 0 | 0 | 0 | 0 | 0 | 0 | 0 | 0 | 0 | 0 | 0 | NA | NA | NA | NA | NA | NA | NA | NA | NA | NA |
| Cd5 | 1 | 2 | 4 | 1 | 2 | 1 | 1 | 2 | 4 | 1 | 2 | 1 | 1.85 | 2 | 1 | 0.556 | -0.846 | 0.54315866 | NA | 0 | 1.7863 | 0.8078 | 0.8078 | TRUE | 0.397 |
| Cd52 | 0 | 0 | 5 | 2 | 5 | 3 | 0 | 0 | 5 | 2 | 5 | 3 | 2.54 | 2 | 3 | 2.008 | 1.006 | 0.52081167 | NA | 1.3505 | 1.3025 | 1.3199 | 1.3199 | TRUE | 1.3227 |
| Cd53 | 1 | 0 | 0 | 0 | 0 | 0 | 1 | 0 | 0 | 0 | 0 | 0 | 0.18 | 0 | 0 | 0.49 | -1.029 | 0.80087 | NA | 0 | 18.6165 | 10 | 10 | TRUE | 0.4614 |
| Cd55 | 428 | 725 | 662 | 759 | 516 | 651 | 455 | 724 | 665 | 597 | 555 | 713 | 617.98 | 615 | 622 | 1.011 | 0.015 | 0.93405883 | 0.96729979 | 0.0354 | 0.0122 | 0.0235 | 0.0235 | TRUE | 0.6062 |
| Cd59a | 114 | 104 | 81 | 138 | 98 | 112 | 121 | 104 | 81 | 108 | 105 | 123 | 107.14 | 102 | 112 | 1.099 | 0.136 | 0.5357851 | 0.72271961 | 0.0121 | 0.0377 | 0.0253 | 0.0253 | TRUE | 0.2919 |
| Cd59b | 18 | 14 | 11 | 14 | 18 | 7 | 19 | 14 | 11 | 11 | 19 | 8 | 13.7 | 15 | 13 | 0.86 | -0.218 | 0.70222449 | 0.83714926 | 0.0613 | 0.2474 | 0.1602 | 0.1602 | TRUE | 0.7485 |
| Cd5l | 0 | 0 | 0 | 0 | 1 | 1 | 0 | 0 | 0 | 0 | 1 | 1 | 0.36 | 0 | 1 | 3.794 | 1.924 | 0.60247382 | NA | 0 | 9.1135 | 8.1425 | 8.1425 | TRUE | 0.3482 |
| Cd6 | 342 | 361 | 360 | 270 | 237 | 253 | 363 | 361 | 361 | 212 | 255 | 277 | 304.91 | 362 | 248 | 0.683 | -0.55 | 0.00011178 | 0.00150346 | 0.0052 | 0.0177 | 0.0112 | 0.0112 | TRUE | 0.1922 |
| Cd63 | 1140 | 1801 | 1676 | 3260 | 2282 | 2173 | 1211 | 1799 | 1683 | 2563 | 2455 | 2379 | 2014.86 | 1564 | 2466 | 1.576 | 0.656 | 7.97E-06 | 0.00016286 | 0.0197 | 0.0085 | 0.015 | 0.015 | TRUE | 0.4693 |
| Cd68 | 135 | 205 | 235 | 226 | 119 | 203 | 143 | 205 | 236 | 178 | 128 | 222 | 185.33 | 195 | 176 | 0.903 | -0.147 | 0.56808965 | 0.74740657 | 0.0618 | 0.0247 | 0.0422 | 0.0422 | TRUE | 0.2295 |
| Cd69 | 0 | 0 | 0 | 1 | 0 | 1 | 0 | 0 | 0 | 1 | 0 | 1 | 0.31 | 0 | 1 | 3.348 | 1.743 | 0.65064889 | NA | 0 | 10.5167 | 8.8968 | 8.8968 | TRUE | 0.2082 |
| Cd7 | 39 | 93 | 64 | 58 | 51 | 68 | 41 | 93 | 64 | 46 | 55 | 74 | 62.25 | 66 | 58 | 0.876 | -0.191 | 0.59196161 | 0.7655312 | 0.0924 | 0.0598 | 0.0755 | 0.0755 | TRUE | 0.2225 |
| Cd70 | 0 | 0 | 0 | 0 | 0 | 0 | 0 | 0 | 0 | 0 | 0 | 0 | 0 | 0 | 0 | NA | NA | NA | NA | NA | NA | NA | NA | NA | NA |
| Cd72 | 1 | 3 | 0 | 2 | 0 | 0 | 1 | 3 | 0 | 2 | 0 | 0 | 0.94 | 1 | 1 | 0.418 | -1.259 | 0.59753736 | NA | 0 | 3.5175 | 2.8598 | 2.8598 | TRUE | 0.473 |
| Cd74 | 506 | 257 | 235 | 274 | 142 | 139 | 537 | 257 | 236 | 215 | 153 | 152 | 258.41 | 343 | 173 | 0.506 | -0.982 | 0.00172295 | 0.0136623 | 0.1238 | 0.0197 | 0.0664 | 0.0664 | TRUE | 2.7785 |
| Cd79a | 40 | 51 | 42 | 64 | 41 | 32 | 42 | 51 | 42 | 50 | 44 | 35 | 44.17 | 45 | 43 | 0.96 | -0.059 | 0.84584935 | 0.92275081 | 0 | 0.0815 | 0.0428 | 0.0428 | TRUE | 0.1984 |
| Cd79b | 189 | 265 | 260 | 226 | 136 | 168 | 201 | 265 | 261 | 178 | 146 | 184 | 205.73 | 242 | 169 | 0.699 | -0.516 | 0.00415077 | 0.02719816 | 0.0138 | 0.0229 | 0.0183 | 0.0183 | TRUE | 0.2449 |
| Cd80 | 808 | 704 | 608 | 507 | 302 | 335 | 858 | 703 | 610 | 399 | 325 | 367 | 543.67 | 724 | 364 | 0.503 | -0.992 | 3.18E-10 | 2.18E-08 | 0.0185 | 0.013 | 0.0159 | 0.0159 | TRUE | 0.2189 |
| Cd81 | 6493 | 8892 | 8710 | 6944 | 5301 | 6795 | 6897 | 8881 | 8746 | 5459 | 5704 | 7438 | 7187.26 | 8175 | 6200 | 0.758 | -0.399 | 0.00715957 | 0.0407509 | 0.0241 | 0.0074 | 0.0157 | 0.0157 | TRUE | 0.3719 |
| Cd82 | 69 | 115 | 95 | 87 | 87 | 87 | 73 | 115 | 95 | 68 | 94 | 95 | 90.13 | 94 | 86 | 0.902 | -0.149 | 0.56478948 | 0.74534144 | 0.0308 | 0.0435 | 0.0372 | 0.0372 | TRUE | 0.1964 |
| Cd83 | 2 | 15 | 6 | 23 | 10 | 13 | 2 | 15 | 6 | 18 | 11 | 14 | 11.03 | 8 | 14 | 1.863 | 0.897 | 0.21745345 | 0.4294633 | 0.2494 | 0.3055 | 0.2815 | 0.2815 | TRUE | 1.1308 |
| Cd84 | 0 | 0 | 0 | 0 | 1 | 0 | 0 | 0 | 0 | 0 | 1 | 0 | 0.18 | 0 | 0 | 1.859 | 0.894 | 0.82650734 | NA | 0 | 18.3785 | 10 | 10 | TRUE | 0.5204 |
| Cd86 | 8 | 6 | 3 | 3 | 4 | 0 | 8 | 6 | 3 | 2 | 4 | 0 | 4.03 | 6 | 2 | 0.383 | -1.383 | 0.21048264 | NA | 0.1465 | 0.8249 | 0.5726 | 0.5726 | TRUE | 0.3289 |
| Cd8a | 49 | 29 | 21 | 53 | 27 | 26 | 52 | 29 | 21 | 42 | 29 | 28 | 33.55 | 34 | 33 | 0.98 | -0.03 | 0.94458879 | 0.97241382 | 0.1005 | 0.1051 | 0.103 | 0.103 | TRUE | 0.9181 |
| Cd8b1 | 12 | 6 | 10 | 37 | 13 | 12 | 13 | 6 | 10 | 29 | 14 | 13 | 14.16 | 10 | 19 | 1.984 | 0.988 | 0.09782181 | 0.25709809 | 0.1164 | 0.2395 | 0.1775 | 0.1775 | TRUE | 1.4337 |
| Cd9 | 14834 | 20553 | 19239 | 16725 | 12303 | 15097 | 15756 | 20526 | 19318 | 13148 | 13237 | 16526 | 16418.66 | 18533 | 14304 | 0.772 | -0.374 | 0.00512808 | 0.03191795 | 0.0179 | 0.0071 | 0.0128 | 0.0128 | TRUE | 0.2258 |
| Cd93 | 0 | 0 | 1 | 3 | 2 | 1 | 0 | 0 | 1 | 2 | 2 | 1 | 1.1 | 0 | 2 | 5.144 | 2.363 | 0.22933208 | NA | 0 | 2.9978 | 1.5242 | 1.5242 | TRUE | 0.4098 |
| Cd96 | 1 | 5 | 2 | 5 | 1 | 4 | 1 | 5 | 2 | 4 | 1 | 4 | 2.91 | 3 | 3 | 1.167 | 0.223 | 0.84855102 | NA | 0 | 1.1397 | 0.6401 | 0.6401 | TRUE | 0.6705 |
| Cd97 | 7313 | 8876 | 9378 | 7738 | 5871 | 6104 | 7768 | 8865 | 9417 | 6083 | 6317 | 6682 | 7521.73 | 8683 | 6361 | 0.732 | -0.449 | 2.38E-06 | 5.86E-05 | 0.0056 | 0.0073 | 0.0064 | 0.0064 | TRUE | 0.1038 |
| Cd99l2 | 506 | 616 | 598 | 891 | 584 | 682 | 537 | 615 | 600 | 700 | 628 | 747 | 638.08 | 584 | 692 | 1.184 | 0.243 | 0.0339311 | 0.12537411 | 0.0046 | 0.0121 | 0.0079 | 0.0079 | TRUE | 0.0739 |
| Cda | 86 | 71 | 64 | 92 | 42 | 69 | 91 | 71 | 64 | 72 | 45 | 76 | 69.93 | 75 | 64 | 0.855 | -0.225 | 0.43590824 | 0.644102 | 0.0368 | 0.054 | 0.0459 | 0.0459 | TRUE | 0.5669 |
| Cdadc1 | 357 | 298 | 277 | 476 | 349 | 295 | 379 | 298 | 278 | 374 | 376 | 323 | 337.93 | 318 | 358 | 1.125 | 0.17 | 0.28537311 | 0.5067445 | 0.0132 | 0.0167 | 0.0152 | 0.0152 | TRUE | 0.3147 |
| Cdan1 | 1525 | 1538 | 1637 | 2124 | 1572 | 1534 | 1620 | 1536 | 1644 | 1670 | 1691 | 1679 | 1639.98 | 1600 | 1680 | 1.05 | 0.071 | 0.33087522 | 0.55182023 | 0 | 0.0089 | 0.0032 | 0.0032 | TRUE | 0.0148 |
| Cdc123 | 2979 | 2930 | 3011 | 4133 | 2715 | 2604 | 3164 | 2926 | 3023 | 3249 | 2921 | 2850 | 3022.42 | 3038 | 3007 | 0.99 | -0.014 | 0.86856581 | 0.93527821 | 0.0029 | 0.008 | 0.0048 | 0.0048 | TRUE | 0.0611 |
| Cdc14a | 134 | 110 | 118 | 311 | 190 | 180 | 142 | 110 | 118 | 244 | 204 | 197 | 169.44 | 123 | 215 | 1.749 | 0.807 | 1.16E-05 | 0.00022571 | 0.0099 | 0.0263 | 0.018 | 0.018 | TRUE | 0.1763 |
| Cdc14b | 2459 | 3432 | 3614 | 2927 | 2256 | 2597 | 2612 | 3428 | 3629 | 2301 | 2427 | 2843 | 2873.24 | 3223 | 2524 | 0.783 | -0.353 | 0.0140627 | 0.0676004 | 0.0197 | 0.008 | 0.0146 | 0.0146 | TRUE | 0.3337 |
| Cdc16 | 4463 | 5602 | 5330 | 6948 | 4691 | 4936 | 4740 | 5595 | 5352 | 5462 | 5047 | 5403 | 5266.6 | 5229 | 5304 | 1.014 | 0.021 | 0.81745296 | 0.90708637 | 0.0043 | 0.0075 | 0.0056 | 0.0056 | TRUE | 0.0813 |
| Cdc20 | 8991 | 10584 | 10565 | 12629 | 9395 | 9243 | 9550 | 10570 | 10608 | 9928 | 10108 | 10118 | 10147.17 | 10243 | 10051 | 0.981 | -0.027 | 0.6997724 | 0.83581217 | 0.0015 | 0.0072 | 0.0035 | 0.0035 | TRUE | 0.0427 |
| Cdc23 | 5871 | 6021 | 6188 | 8395 | 6100 | 5839 | 6236 | 6013 | 6213 | 6599 | 6563 | 6392 | 6336.18 | 6154 | 6518 | 1.059 | 0.083 | 0.15230822 | 0.34346657 | 2.00E-04 | 0.0074 | 0.0023 | 0.0023 | TRUE | 0.0049 |
| Cdc25a | 4726 | 5456 | 5493 | 6689 | 4828 | 4969 | 5020 | 5449 | 5516 | 5258 | 5195 | 5439 | 5312.78 | 5328 | 5297 | 0.994 | -0.008 | 0.90453595 | 0.95255222 | 0.0014 | 0.0075 | 0.0034 | 0.0034 | TRUE | 0.0312 |
| Cdc25b | 2276 | 2693 | 2721 | 2381 | 1755 | 1912 | 2418 | 2690 | 2732 | 1872 | 1888 | 2093 | 2282.04 | 2613 | 1951 | 0.746 | -0.422 | 3.29E-06 | 7.72E-05 | 0.0035 | 0.0083 | 0.0055 | 0.0055 | TRUE | 0.0518 |
| Cdc25c | 869 | 1370 | 1375 | 1457 | 1080 | 1170 | 923 | 1368 | 1381 | 1145 | 1162 | 1281 | 1210.01 | 1224 | 1196 | 0.977 | -0.034 | 0.8330154 | 0.91523598 | 0.0259 | 0.0096 | 0.0178 | 0.0178 | TRUE | 0.5535 |
| Cdc26 | 889 | 835 | 827 | 1267 | 824 | 821 | 944 | 834 | 830 | 996 | 887 | 899 | 898.32 | 869 | 927 | 1.068 | 0.094 | 0.36068919 | 0.57907063 | 0.0037 | 0.0106 | 0.0066 | 0.0066 | TRUE | 0.0668 |
| Cdc27 | 5453 | 4328 | 4076 | 7199 | 4661 | 3989 | 5792 | 4322 | 4093 | 5659 | 5015 | 4367 | 4874.67 | 4736 | 5014 | 1.059 | 0.083 | 0.5908926 | 0.76478301 | 0.0221 | 0.0076 | 0.0168 | 0.0168 | TRUE | 0.4636 |
| Cdc34 | 10380 | 10608 | 9784 | 15068 | 10557 | 9346 | 11025 | 10594 | 9824 | 11845 | 11359 | 10231 | 10813.08 | 10481 | 11145 | 1.063 | 0.089 | 0.31279336 | 0.53322275 | 0.0043 | 0.0072 | 0.0055 | 0.0055 | TRUE | 0.0629 |
| Cdc37 | 11122 | 10729 | 10579 | 14465 | 10846 | 10306 | 11814 | 10715 | 10622 | 11371 | 11670 | 11282 | 11245.59 | 11050 | 11441 | 1.035 | 0.05 | 0.48310562 | 0.68129391 | 0.0018 | 0.0072 | 0.0036 | 0.0036 | TRUE | 0.0446 |
| Cdc37l1 | 2780 | 2802 | 2734 | 3190 | 2375 | 2470 | 2953 | 2798 | 2745 | 2508 | 2555 | 2704 | 2710.56 | 2832 | 2589 | 0.914 | -0.13 | 0.07776601 | 0.22059811 | 0.001 | 0.0081 | 0.0035 | 0.0035 | TRUE | 0.0181 |
| Cdc40 | 1649 | 1905 | 1853 | 2370 | 1733 | 1780 | 1752 | 1903 | 1861 | 1863 | 1865 | 1948 | 1865.15 | 1839 | 1892 | 1.029 | 0.041 | 0.58397401 | 0.75932524 | 7.00E-04 | 0.0087 | 0.0035 | 0.0035 | TRUE | 0.0204 |
| Cdc42 | 13886 | 14429 | 14065 | 19519 | 13140 | 13502 | 14749 | 14410 | 14123 | 15344 | 14138 | 14780 | 14590.78 | 14427 | 14754 | 1.023 | 0.032 | 0.61084571 | 0.77958142 | 0.001 | 0.0071 | 0.0029 | 0.0029 | TRUE | 0.0163 |
| Cdc42bpa | 1393 | 1541 | 1516 | 2614 | 1835 | 1633 | 1480 | 1539 | 1522 | 2055 | 1974 | 1788 | 1726.28 | 1514 | 1939 | 1.282 | 0.358 | 3.14E-05 | 0.00052293 | 0.0023 | 0.0088 | 0.0047 | 0.0047 | TRUE | 0.056 |
| Cdc42bpb | 4780 | 3926 | 4073 | 6446 | 4731 | 4108 | 5077 | 3921 | 4090 | 5067 | 5090 | 4497 | 4623.72 | 4363 | 4885 | 1.12 | 0.163 | 0.17091527 | 0.36957439 | 0.0123 | 0.0076 | 0.01 | 0.01 | TRUE | 0.2498 |
| Cdc42bpg | 5257 | 5867 | 5832 | 4991 | 4041 | 4240 | 5584 | 5859 | 5856 | 3924 | 4348 | 4641 | 5035.33 | 5766 | 4304 | 0.746 | -0.423 | 9.71E-07 | 2.68E-05 | 0.0036 | 0.0076 | 0.0052 | 0.0052 | TRUE | 0.0737 |
| Cdc42ep1 | 135 | 146 | 133 | 144 | 99 | 102 | 143 | 146 | 134 | 113 | 107 | 112 | 125.69 | 141 | 111 | 0.785 | -0.35 | 0.04483666 | 0.15251702 | 0 | 0.0331 | 0.0139 | 0.0139 | TRUE | 0.022 |
| Cdc42ep2 | 75 | 170 | 212 | 198 | 159 | 232 | 80 | 170 | 213 | 156 | 171 | 254 | 173.83 | 154 | 194 | 1.253 | 0.326 | 0.33681359 | 0.55677852 | 0.1416 | 0.0259 | 0.0771 | 0.0771 | TRUE | 0.6886 |
| Cdc42ep3 | 597 | 496 | 475 | 731 | 554 | 503 | 634 | 495 | 477 | 575 | 596 | 551 | 554.63 | 535 | 574 | 1.072 | 0.1 | 0.46605675 | 0.66857491 | 0.011 | 0.0128 | 0.0119 | 0.0119 | TRUE | 0.3017 |
| Cdc42ep4 | 3917 | 4770 | 4972 | 4888 | 3732 | 3838 | 4161 | 4764 | 4992 | 3843 | 4015 | 4201 | 4329.34 | 4639 | 4020 | 0.866 | -0.207 | 0.027913 | 0.10967118 | 0.0049 | 0.0077 | 0.0062 | 0.0062 | TRUE | 0.099 |
| Cdc42ep5 | 310 | 375 | 357 | 431 | 281 | 357 | 329 | 375 | 358 | 339 | 302 | 391 | 349.03 | 354 | 344 | 0.971 | -0.043 | 0.76138239 | 0.87423727 | 0.0071 | 0.0163 | 0.0115 | 0.0115 | TRUE | 0.1549 |
| Cdc42se1 | 1120 | 1456 | 1422 | 1755 | 1182 | 1217 | 1190 | 1454 | 1428 | 1380 | 1272 | 1332 | 1342.53 | 1357 | 1328 | 0.979 | -0.031 | 0.76876092 | 0.87928573 | 0.006 | 0.0094 | 0.0074 | 0.0074 | TRUE | 0.1396 |

| | | | | | | | | | | | | | | | | | | | | | | | | | |
|---|---|---|---|---|---|---|---|---|---|---|---|---|---|---|---|---|---|---|---|---|---|---|---|---|---|
| Cdc42se2 | 6265 | 7185 | 7209 | 9957 | 7122 | 6641 | 6655 | 7176 | 7239 | 7827 | 7663 | 7270 | 7304.79 | 7023 | 7587 | 1.08 | 0.111 | 0.1179798 | 0.2905195 | 0.0015 | 0.0074 | 0.0035 | 0.0035 | TRUE | 0.0257 |
| Cdc45 | 2031 | 2390 | 2557 | 3386 | 2462 | 2547 | 2157 | 2387 | 2567 | 2662 | 2649 | 2788 | 2535.09 | 2370 | 2700 | 1.139 | 0.187 | 0.03800036 | 0.1355057 | 0.0036 | 0.0082 | 0.0055 | 0.0055 | TRUE | 0.0748 |
| Cdc5l | 13196 | 13645 | 12956 | 20288 | 14155 | 12644 | 14016 | 13627 | 13009 | 15949 | 15230 | 13841 | 14278.78 | 13551 | 15007 | 1.108 | 0.147 | 0.06795749 | 0.20177545 | 0.0031 | 0.0071 | 0.0046 | 0.0046 | TRUE | 0.0564 |
| Cdc6 | 3914 | 3373 | 3456 | 5942 | 4110 | 3908 | 4157 | 3369 | 3470 | 4671 | 4422 | 4278 | 4061.23 | 3665 | 4457 | 1.216 | 0.282 | 0.00633373 | 0.03748426 | 0.0075 | 0.0077 | 0.0075 | 0.0075 | TRUE | 0.1675 |
| Cdc7 | 2613 | 2807 | 2725 | 3150 | 2456 | 2632 | 2775 | 2803 | 2736 | 2476 | 2643 | 2881 | 2719.16 | 2771 | 2667 | 0.962 | -0.057 | 0.49806627 | 0.69358124 | 0.0024 | 0.0081 | 0.0046 | 0.0046 | TRUE | 0.06 |
| Cdc73 | 3837 | 3027 | 3077 | 5248 | 3698 | 3077 | 4076 | 3023 | 3090 | 4126 | 3979 | 3368 | 3610.16 | 3396 | 3824 | 1.126 | 0.172 | 0.22202599 | 0.43425024 | 0.0196 | 0.0078 | 0.014 | 0.014 | TRUE | 0.3721 |
| Cdca2 | 4732 | 6288 | 6223 | 6195 | 4753 | 5000 | 5026 | 6280 | 6249 | 4870 | 5114 | 5473 | 5501.98 | 5852 | 5152 | 0.88 | -0.184 | 0.0955318 | 0.25348715 | 0.0089 | 0.0075 | 0.0086 | 0.0086 | TRUE | 0.1855 |
| Cdca3 | 5730 | 6409 | 6590 | 8165 | 5999 | 6336 | 6086 | 6401 | 6617 | 6419 | 6455 | 6936 | 6485.51 | 6368 | 6603 | 1.037 | 0.052 | 0.46966357 | 0.67170238 | 0.0016 | 0.0074 | 0.0036 | 0.0036 | TRUE | 0.0236 |
| Cdca4 | 2761 | 2956 | 2819 | 3490 | 2506 | 2588 | 2933 | 2952 | 2831 | 2744 | 2696 | 2833 | 2831.38 | 2905 | 2758 | 0.949 | -0.075 | 0.25184412 | 0.46962468 | 2.00E-04 | 0.0081 | 0.0028 | 0.0028 | TRUE | 0.0068 |
| Cdca5 | 2478 | 2436 | 2517 | 3649 | 2567 | 2415 | 2632 | 2433 | 2527 | 2869 | 2762 | 2644 | 2644.39 | 2531 | 2758 | 1.09 | 0.125 | 0.09524163 | 0.25297254 | 0.0012 | 0.0081 | 0.0036 | 0.0036 | TRUE | 0.0159 |
| Cdca7 | 6037 | 7251 | 7673 | 8151 | 6434 | 6977 | 6412 | 7242 | 7705 | 6408 | 6923 | 7637 | 7054.37 | 7120 | 6989 | 0.982 | -0.027 | 0.79714444 | 0.89554542 | 0.008 | 0.0074 | 0.0078 | 0.0078 | TRUE | 0.0921 |
| Cdca7l | 1434 | 1581 | 1702 | 2487 | 1898 | 1824 | 1523 | 1579 | 1709 | 1955 | 2042 | 1997 | 1800.83 | 1604 | 1998 | 1.245 | 0.317 | 8.13E-05 | 0.00117147 | 0.0012 | 0.0087 | 0.0041 | 0.0041 | TRUE | 0.0398 |
| Cdca8 | 6142 | 7068 | 6887 | 9229 | 6674 | 6571 | 6524 | 7059 | 6915 | 7255 | 7181 | 7193 | 7021.16 | 6833 | 7210 | 1.055 | 0.077 | 0.21965715 | 0.43181262 | 6.00E-04 | 0.0074 | 0.0027 | 0.0027 | TRUE | 0.019 |
| Cdcp1 | 164 | 242 | 260 | 229 | 219 | 299 | 174 | 242 | 261 | 180 | 236 | 327 | 236.65 | 226 | 248 | 1.094 | 0.13 | 0.60261709 | 0.77387557 | 0.0622 | 0.0208 | 0.0406 | 0.0406 | TRUE | 0.3442 |
| Cdcp2 | 12 | 5 | 6 | 8 | 3 | 5 | 13 | 5 | 6 | 6 | 3 | 5 | 6.46 | 8 | 5 | 0.64 | -0.644 | 0.41448862 | 0.62611959 | 0 | 0.517 | 0.2878 | 0.2878 | TRUE | 0.7931 |
| Cdh1 | 21363 | 23005 | 23767 | 21555 | 16523 | 18049 | 22691 | 22975 | 23865 | 16945 | 17778 | 19757 | 20668.54 | 23177 | 18160 | 0.783 | -0.352 | 1.49E-05 | 0.00027759 | 0.0034 | 0.0071 | 0.0047 | 0.0047 | TRUE | 0.0724 |
| Cdh10 | 29 | 21 | 19 | 10 | 5 | 9 | 31 | 21 | 19 | 8 | 5 | 10 | 15.66 | 24 | 8 | 0.327 | -1.615 | 0.00248518 | 0.01828184 | 0 | 0.2173 | 0.1204 | 0.1204 | TRUE | 0.4051 |
| Cdh11 | 7 | 8 | 13 | 13 | 6 | 7 | 7 | 8 | 13 | 10 | 6 | 8 | 8.8 | 9 | 8 | 0.862 | -0.214 | 0.74075358 | 0.86146332 | 0 | 0.3812 | 0.1868 | 0.1868 | TRUE | 0.3631 |
| Cdh12 | 1 | 0 | 0 | 4 | 2 | 1 | 1 | 0 | 0 | 3 | 2 | 1 | 1.24 | 0 | 2 | 5.894 | 2.559 | 0.18208358 | NA | 0 | 2.659 | 1.4239 | 1.4239 | TRUE | 0.3994 |
| Cdh13 | 2 | 4 | 2 | 1 | 1 | 2 | 2 | 4 | 2 | 1 | 1 | 2 | 2.03 | 3 | 1 | 0.486 | -1.042 | 0.42985549 | NA | 0 | 1.6299 | 0.6984 | 0.6984 | TRUE | 0.208 |
| Cdh15 | 133 | 117 | 118 | 161 | 127 | 95 | 141 | 117 | 118 | 127 | 137 | 104 | 123.97 | 125 | 123 | 0.977 | -0.034 | 0.8650064 | 0.93316559 | 0.0061 | 0.0335 | 0.0202 | 0.0202 | TRUE | 0.1647 |
| Cdh16 | 0 | 1 | 0 | 1 | 0 | 0 | 0 | 1 | 0 | 1 | 0 | 0 | 0.3 | 0 | 0 | 0.954 | -0.067 | 0.98642001 | NA | 0 | 11.0817 | 9.3014 | 9.3014 | TRUE | 0.405 |
| Cdh17 | 1 | 0 | 0 | 5 | 7 | 4 | 1 | 0 | 0 | 4 | 8 | 4 | 2.82 | 0 | 5 | 14.354 | 3.843 | 0.01402844 | NA | 0 | 1.1763 | 0.6707 | 0.6707 | TRUE | 0.4614 |
| Cdh18 | 2 | 1 | 5 | 13 | 9 | 9 | 2 | 1 | 5 | 10 | 10 | 10 | 6.32 | 3 | 10 | 3.651 | 1.868 | 0.02793758 | 0.10971901 | 0 | 0.5285 | 0.2834 | 0.2834 | TRUE | 0.6692 |
| Cdh19 | 1 | 0 | 0 | 3 | 1 | 0 | 1 | 0 | 0 | 2 | 1 | 0 | 0.75 | 0 | 1 | 3.204 | 1.68 | 0.5025945 | NA | 0 | 4.4029 | 3.1058 | 3.1058 | TRUE | 0.2352 |
| Cdh2 | 120 | 137 | 177 | 293 | 184 | 197 | 127 | 137 | 178 | 230 | 198 | 216 | 180.99 | 147 | 215 | 1.458 | 0.544 | 0.00297721 | 0.02109461 | 0.0115 | 0.0251 | 0.0184 | 0.0184 | TRUE | 0.3414 |
| Cdh20 | 2 | 2 | 0 | 5 | 4 | 5 | 2 | 2 | 0 | 4 | 4 | 5 | 2.97 | 1 | 4 | 3.329 | 1.735 | 0.1515111 | NA | 0 | 1.1156 | 0.5737 | 0.5737 | TRUE | 0.4934 |
| Cdh22 | 148 | 208 | 214 | 47 | 46 | 70 | 157 | 208 | 215 | 37 | 49 | 77 | 123.81 | 193 | 54 | 0.278 | -1.845 | 7.35E-11 | 5.54E-09 | 0.0628 | 0.0335 | 0.0461 | 0.0461 | TRUE | 1.0465 |
| Cdh23 | 117 | 161 | 162 | 103 | 86 | 121 | 124 | 161 | 163 | 81 | 93 | 132 | 125.61 | 149 | 102 | 0.68 | -0.557 | 0.02124035 | 0.09076032 | 0.0349 | 0.0331 | 0.0339 | 0.0339 | TRUE | 0.6545 |
| Cdh24 | 215 | 232 | 178 | 279 | 232 | 176 | 228 | 232 | 179 | 219 | 250 | 193 | 216.73 | 213 | 221 | 1.036 | 0.051 | 0.77196063 | 0.88105963 | 0.0135 | 0.0221 | 0.0177 | 0.0177 | TRUE | 0.2174 |
| Cdh26 | 2 | 2 | 3 | 2 | 0 | 1 | 2 | 2 | 3 | 2 | 0 | 1 | 1.63 | 2 | 1 | 0.389 | -1.361 | 0.3697981 | NA | 0 | 2.0237 | 0.9185 | 0.9185 | TRUE | 0.2806 |
| Cdh3 | 3477 | 3714 | 4168 | 3203 | 2977 | 3487 | 3693 | 3709 | 4185 | 2518 | 3203 | 3817 | 3520.93 | 3862 | 3179 | 0.823 | -0.281 | 0.06013354 | 0.18607537 | 0.0231 | 0.0078 | 0.0159 | 0.0159 | TRUE | 0.263 |
| Cdh4 | 105 | 54 | 53 | 145 | 104 | 105 | 112 | 54 | 53 | 114 | 112 | 115 | 93.25 | 73 | 114 | 1.562 | 0.643 | 0.03902096 | 0.13823868 | 0.0778 | 0.0422 | 0.0586 | 0.0586 | TRUE | 1.9226 |
| Cdh5 | 10 | 12 | 17 | 39 | 21 | 17 | 11 | 12 | 17 | 31 | 23 | 19 | 18.59 | 13 | 24 | 1.824 | 0.867 | 0.06768492 | 0.20123026 | 0 | 0.1841 | 0.1034 | 0.1034 | TRUE | 0.4213 |
| Cdh6 | 13 | 8 | 4 | 27 | 22 | 19 | 14 | 8 | 4 | 21 | 24 | 21 | 15.25 | 9 | 22 | 2.557 | 1.354 | 0.01512011 | 0.07116516 | 0 | 0.2229 | 0.1419 | 0.1419 | TRUE | 0.6905 |
| Cdh7 | 0 | 1 | 1 | 1 | 4 | 1 | 0 | 1 | 1 | 1 | 4 | 1 | 1.36 | 1 | 2 | 3.006 | 1.588 | 0.37276278 | NA | 0 | 2.4212 | 1.2881 | 1.2881 | TRUE | 0.7858 |
| Cdh8 | 1 | 3 | 0 | 12 | 3 | 7 | 1 | 3 | 0 | 9 | 3 | 8 | 4.06 | 1 | 7 | 5.03 | 2.33 | 0.04913773 | NA | 0.147 | 0.8176 | 0.5632 | 0.5632 | TRUE | 0.639 |
| Cdh9 | 66 | 54 | 31 | 23 | 27 | 18 | 70 | 54 | 31 | 18 | 29 | 20 | 37 | 52 | 22 | 0.429 | -1.222 | 0.00335986 | 0.0231247 | 0.0899 | 0.0959 | 0.0931 | 0.0931 | TRUE | 0.1559 |
| Cdhr1 | 318 | 582 | 657 | 352 | 304 | 447 | 338 | 581 | 660 | 277 | 327 | 489 | 445.31 | 526 | 364 | 0.691 | -0.533 | 0.05111707 | 0.16653965 | 0.0939 | 0.0143 | 0.0514 | 0.0514 | TRUE | 0.7609 |
| Cdhr2 | 5 | 5 | 5 | 12 | 4 | 7 | 5 | 5 | 5 | 9 | 4 | 8 | 6.12 | 5 | 7 | 1.413 | 0.499 | 0.51366642 | 0.70657132 | 0 | 0.5451 | 0.2521 | 0.2521 | TRUE | 0.3016 |
| Cdhr3 | 12 | 4 | 14 | 8 | 1 | 5 | 13 | 4 | 14 | 6 | 1 | 5 | 7.27 | 10 | 4 | 0.425 | -1.236 | 0.15281353 | 0.34416322 | 0.2725 | 0.4599 | 0.3764 | 0.3764 | TRUE | 1.0265 |
| Cdhr5 | 3 | 2 | 1 | 6 | 4 | 2 | 3 | 2 | 1 | 5 | 4 | 2 | 2.9 | 2 | 4 | 1.838 | 0.878 | 0.43249829 | NA | 0 | 1.1431 | 0.524 | 0.524 | TRUE | 0.1977 |
| Cdip1 | 1050 | 1219 | 1234 | 1747 | 1252 | 1291 | 1115 | 1217 | 1239 | 1373 | 1347 | 1413 | 1284.24 | 1190 | 1378 | 1.157 | 0.21 | 0.01180726 | 0.05951553 | 9.00E-04 | 0.0095 | 0.0042 | 0.0042 | TRUE | 0.0364 |
| Cdipt | 1502 | 1512 | 1474 | 2295 | 1497 | 1423 | 1595 | 1510 | 1480 | 1804 | 1611 | 1558 | 1593 | 1528 | 1658 | 1.085 | 0.118 | 0.19859152 | 0.4057235 | 0.0033 | 0.009 | 0.0055 | 0.0055 | TRUE | 0.0738 |
| Cdk1 | 6747 | 8278 | 8012 | 10575 | 7520 | 7884 | 7167 | 8267 | 8045 | 8313 | 8091 | 8630 | 8085.55 | 7826 | 8345 | 1.066 | 0.092 | 0.25834632 | 0.47723567 | 0.0031 | 0.0073 | 0.0047 | 0.0047 | TRUE | 0.0663 |
| Cdk10 | 1384 | 1643 | 1506 | 2378 | 1827 | 1619 | 1470 | 1641 | 1512 | 1869 | 1966 | 1772 | 1705.08 | 1541 | 1869 | 1.213 | 0.278 | 0.001345 | 0.01121802 | 0.0021 | 0.0088 | 0.0048 | 0.0048 | TRUE | 0.0388 |
| Cdk11b | 6085 | 6135 | 6196 | 8140 | 6188 | 5605 | 6463 | 6127 | 6221 | 6399 | 6658 | 6136 | 6334.06 | 6270 | 6398 | 1.02 | 0.029 | 0.66531778 | 0.8127356 | 9.00E-04 | 0.0074 | 0.0031 | 0.0031 | TRUE | 0.0155 |
| Cdk12 | 6171 | 5445 | 5769 | 8717 | 5831 | 4864 | 6555 | 5438 | 5793 | 6853 | 6274 | 5324 | 6039.36 | 5929 | 6150 | 1.038 | 0.053 | 0.65672149 | 0.80670515 | 0.0103 | 0.0074 | 0.0102 | 0.0102 | TRUE | 0.1681 |
| Cdk13 | 4358 | 3772 | 3780 | 5573 | 3985 | 3648 | 4629 | 3767 | 3796 | 4381 | 4288 | 3993 | 4142.27 | 4064 | 4221 | 1.039 | 0.055 | 0.60313725 | 0.77412127 | 0.0067 | 0.0077 | 0.0078 | 0.0078 | TRUE | 0.1799 |
| Cdk14 | 77 | 79 | 112 | 186 | 127 | 103 | 82 | 79 | 112 | 146 | 137 | 113 | 111.46 | 91 | 132 | 1.451 | 0.537 | 0.01766473 | 0.07933035 | 0.0183 | 0.0365 | 0.0276 | 0.0276 | TRUE | 0.4084 |
| Cdk15 | 3 | 1 | 4 | 3 | 5 | 1 | 3 | 1 | 4 | 2 | 5 | 1 | 2.84 | 3 | 3 | 1.072 | 0.101 | 0.93162272 | NA | 0 | 1.1673 | 0.6437 | 0.6437 | TRUE | 0.5871 |
| Cdk16 | 5942 | 6933 | 6768 | 8116 | 5997 | 5886 | 6311 | 6924 | 6796 | 6380 | 6452 | 6443 | 6551.17 | 6677 | 6425 | 0.962 | -0.056 | 0.40550101 | 0.61834473 | 0.001 | 0.0074 | 0.0031 | 0.0031 | TRUE | 0.0279 |
| Cdk17 | 2162 | 2238 | 2189 | 2937 | 2082 | 1976 | 2296 | 2235 | 2198 | 2309 | 2240 | 2163 | 2240.25 | 2243 | 2237 | 0.998 | -0.003 | 0.9649643 | 0.98240491 | 4.00E-04 | 0.0084 | 0.0031 | 0.0031 | TRUE | 0.0102 |
| Cdk18 | 1866 | 1568 | 1438 | 1661 | 1232 | 1187 | 1982 | 1566 | 1444 | 1306 | 1326 | 1299 | 1487.09 | 1664 | 1310 | 0.788 | -0.345 | 0.00776441 | 0.04341025 | 0.0134 | 0.0091 | 0.0114 | 0.0114 | TRUE | 0.3366 |
| Cdk19 | 855 | 1071 | 1066 | 1059 | 896 | 898 | 908 | 1070 | 1070 | 833 | 964 | 983 | 971.28 | 1016 | 927 | 0.911 | -0.135 | 0.24500871 | 0.4616907 | 0.0075 | 0.0103 | 0.0087 | 0.0087 | TRUE | 0.1025 |
| Cdk2 | 4088 | 4122 | 4089 | 4903 | 3733 | 3652 | 4342 | 4117 | 4106 | 3854 | 4016 | 3998 | 4072.19 | 4188 | 3956 | 0.944 | -0.082 | 0.20338682 | 0.41115315 | 5.00E-04 | 0.0077 | 0.0028 | 0.0028 | TRUE | 0.0126 |
| Cdk20 | 239 | 240 | 261 | 342 | 247 | 265 | 254 | 240 | 262 | 269 | 266 | 290 | 263.39 | 252 | 275 | 1.091 | 0.125 | 0.33554033 | 0.55585035 | 0 | 0.0194 | 0.0084 | 0.0084 | TRUE | 0.025 |
| Cdk2ap1 | 1361 | 1288 | 1202 | 1495 | 1006 | 885 | 1446 | 1286 | 1207 | 1175 | 1082 | 969 | 1194.22 | 1313 | 1075 | 0.82 | -0.286 | 0.0130993 | 0.06414534 | 0.0076 | 0.0097 | 0.0087 | 0.0087 | TRUE | 0.0935 |
| Cdk2ap2 | 938 | 970 | 988 | 1160 | 839 | 962 | 996 | 969 | 992 | 912 | 903 | 1053 | 970.8 | 986 | 956 | 0.969 | -0.045 | 0.64294861 | 0.79907039 | 0.0028 | 0.0103 | 0.0059 | 0.0059 | TRUE | 0.0926 |
| Cdk3-ps | 40 | 51 | 43 | 66 | 45 | 31 | 42 | 51 | 43 | 52 | 48 | 34 | 45.14 | 45 | 45 | 0.988 | -0.018 | 0.95410918 | 0.97664915 | 0 | 0.0799 | 0.045 | 0.045 | TRUE | 0.3273 |
| Cdk4 | 8405 | 9174 | 9518 | 14948 | 10610 | 9915 | 8928 | 9162 | 9557 | 11751 | 11416 | 10854 | 10277.83 | 9216 | 11340 | 1.231 | 0.299 | 8.91E-06 | 0.00018048 | 0.0012 | 0.0072 | 0.0032 | 0.0032 | TRUE | 0.0172 |
| Cdk5 | 622 | 723 | 669 | 937 | 658 | 694 | 661 | 722 | 672 | 737 | 708 | 760 | 709.79 | 685 | 735 | 1.073 | 0.101 | 0.28179412 | 0.50333599 | 0 | 0.0115 | 0.005 | 0.005 | TRUE | 0.0268 |
| Cdk5r1 | 113 | 95 | 97 | 146 | 146 | 111 | 120 | 95 | 97 | 115 | 157 | 122 | 117.61 | 104 | 131 | 1.257 | 0.33 | 0.12612447 | 0.30435055 | 0.015 | 0.0349 | 0.0249 | 0.0249 | TRUE | 0.3018 |
| Cdk5r2 | 119 | 156 | 145 | 160 | 129 | 114 | 126 | 156 | 146 | 126 | 139 | 125 | 136.19 | 143 | 130 | 0.909 | -0.138 | 0.43880738 | 0.6460857 | 0 | 0.0311 | 0.0156 | 0.0156 | TRUE | 0.1006 |
| Cdk5rap1 | 1227 | 1321 | 1409 | 1650 | 1242 | 1037 | 1303 | 1319 | 1415 | 1297 | 1336 | 1135 | 1300.99 | 1346 | 1256 | 0.934 | -0.099 | 0.31170241 | 0.53217414 | 0.0035 | 0.0094 | 0.0061 | 0.0061 | TRUE | 0.0837 |
| Cdk5rap2 | 2727 | 2830 | 3202 | 4279 | 3194 | 3071 | 2897 | 2826 | 3215 | 3364 | 3437 | 3362 | 3183.35 | 2979 | 3388 | 1.137 | 0.185 | 0.01938433 | 0.0848542 | 0.002 | 0.0079 | 0.0042 | 0.0042 | TRUE | 0.0583 |
| Cdk5rap3 | 6586 | 8683 | 8458 | 7787 | 6320 | 6395 | 6995 | 8672 | 8493 | 6122 | 6800 | 7000 | 7346.97 | 8053 | 6641 | 0.824 | -0.279 | 0.01046166 | 0.05456395 | 0.0087 | 0.0074 | 0.0084 | 0.0084 | TRUE | 0.1611 |
| Cdk6 | 665 | 917 | 934 | 630 | 555 | 613 | 706 | 916 | 938 | 495 | 597 | 671 | 720.57 | 853 | 588 | 0.688 | -0.54 | 0.00070671 | 0.00681315 | 0.022 | 0.0115 | 0.0169 | 0.0169 | TRUE | 0.2689 |
| Cdk7 | 2668 | 2746 | 2668 | 3795 | 2744 | 2601 | 2834 | 2742 | 2679 | 2983 | 2952 | 2847 | 2839.7 | 2752 | 2927 | 1.064 | 0.09 | 0.17935679 | 0.38052398 | 3.00E-04 | 0.0081 | 0.0029 | 0.0029 | TRUE | 0.0082 |

| | | | | | | | | | | | | | | | | | | | | | | | | | |
|---|---|---|---|---|---|---|---|---|---|---|---|---|---|---|---|---|---|---|---|---|---|---|---|---|---|
| Cdk8 | 3856 | 4338 | 4296 | 4879 | 3548 | 3516 | 4096 | 4332 | 4314 | 3835 | 3817 | 3849 | 4040.59 | 4247 | 3834 | 0.903 | -0.148 | 0.01758784 | 0.07914276 | 2.00E-04 | 0.0077 | 0.0025 | 0.0025 | TRUE | 0.0118 |
| Cdk9 | 3922 | 3956 | 3680 | 5489 | 4040 | 3642 | 4166 | 3951 | 3695 | 4315 | 4347 | 3987 | 4076.74 | 3937 | 4216 | 1.071 | 0.099 | 0.22233679 | 0.43444058 | 0.0025 | 0.0077 | 0.0045 | 0.0045 | TRUE | 0.0352 |
| Cdkal1 | 1147 | 1193 | 1187 | 1465 | 1015 | 1075 | 1218 | 1191 | 1192 | 1152 | 1092 | 1177 | 1170.36 | 1200 | 1140 | 0.95 | -0.074 | 0.34695646 | 0.5660113 | 0 | 0.0097 | 0.0036 | 0.0036 | TRUE | 0.0158 |
| Cdkl1 | 9 | 4 | 4 | 18 | 16 | 13 | 10 | 4 | 4 | 14 | 17 | 14 | 10.53 | 6 | 15 | 2.607 | 1.382 | 0.03019917 | 0.11580388 | 0 | 0.3198 | 0.173 | 0.173 | TRUE | 0.6686 |
| Cdkl2 | 1296 | 1189 | 1226 | 1388 | 1075 | 1067 | 1377 | 1187 | 1231 | 1091 | 1157 | 1168 | 1201.81 | 1265 | 1139 | 0.9 | -0.152 | 0.10637509 | 0.27236168 | 0.0029 | 0.0096 | 0.0056 | 0.0056 | TRUE | 0.0711 |
| Cdkl3 | 192 | 159 | 173 | 220 | 136 | 113 | 204 | 159 | 174 | 173 | 146 | 124 | 163.24 | 179 | 148 | 0.83 | -0.269 | 0.16940323 | 0.36787035 | 0.016 | 0.0271 | 0.0215 | 0.0215 | TRUE | 0.2522 |
| Cdkl4 | 254 | 255 | 248 | 206 | 140 | 151 | 270 | 255 | 249 | 162 | 151 | 165 | 208.56 | 258 | 159 | 0.618 | -0.694 | 1.24E-06 | 3.31E-05 | 0 | 0.0227 | 0.0097 | 0.0097 | TRUE | 0.0224 |
| Cdkl5 | 272 | 192 | 240 | 412 | 268 | 233 | 289 | 192 | 241 | 324 | 288 | 255 | 264.82 | 241 | 289 | 1.205 | 0.269 | 0.15141494 | 0.34243138 | 0.0236 | 0.0193 | 0.0215 | 0.0215 | TRUE | 0.2301 |
| Cdkn1a | 3342 | 5901 | 5239 | 3578 | 2735 | 3779 | 3550 | 5893 | 5261 | 2813 | 2943 | 4137 | 4099.31 | 4901 | 3298 | 0.673 | -0.572 | 0.00474383 | 0.03014373 | 0.0547 | 0.0077 | 0.0293 | 0.0293 | TRUE | 0.5865 |
| Cdkn1b | 1798 | 3146 | 3503 | 2606 | 2171 | 2488 | 1910 | 3142 | 3517 | 2049 | 2336 | 2724 | 2612.85 | 2856 | 2370 | 0.829 | -0.27 | 0.19395112 | 0.39960772 | 0.053 | 0.0082 | 0.0308 | 0.0308 | TRUE | 0.9774 |
| Cdkn1c | 263 | 334 | 333 | 354 | 288 | 339 | 279 | 334 | 334 | 278 | 310 | 371 | 317.76 | 316 | 320 | 1.01 | 0.014 | 0.92927623 | 0.96532601 | 0.0128 | 0.0173 | 0.0149 | 0.0149 | TRUE | 0.2217 |
| Cdkn2a | 426 | 553 | 516 | 919 | 621 | 532 | 452 | 552 | 518 | 722 | 668 | 582 | 582.64 | 507 | 657 | 1.297 | 0.375 | 0.00443013 | 0.02855181 | 0.0092 | 0.0126 | 0.0108 | 0.0108 | TRUE | 0.1171 |
| Cdkn2aip | 1302 | 1272 | 1261 | 1837 | 1337 | 1292 | 1383 | 1270 | 1266 | 1444 | 1439 | 1414 | 1369.4 | 1306 | 1432 | 1.097 | 0.133 | 0.09414104 | 0.25112088 | 5.00E-04 | 0.0093 | 0.0038 | 0.0038 | TRUE | 0.0313 |
| Cdkn2aipnl | 3895 | 4278 | 4321 | 6664 | 4639 | 4600 | 4137 | 4272 | 4339 | 5239 | 4991 | 5035 | 4668.97 | 4249 | 5088 | 1.198 | 0.26 | 3.44E-05 | 0.00056367 | 4.00E-04 | 0.0076 | 0.0026 | 0.0026 | TRUE | 0.0083 |
| Cdkn2b | 190 | 151 | 127 | 243 | 138 | 123 | 202 | 151 | 128 | 191 | 148 | 135 | 159.05 | 160 | 158 | 0.992 | -0.012 | 0.9590995 | 0.97898705 | 0.0378 | 0.0276 | 0.0327 | 0.0327 | TRUE | 0.5448 |
| Cdkn2c | 9 | 17 | 17 | 54 | 32 | 31 | 10 | 17 | 17 | 42 | 34 | 34 | 25.74 | 15 | 37 | 2.543 | 1.347 | 0.0010774 | 0.00942472 | 0 | 0.1349 | 0.074 | 0.074 | TRUE | 0.3796 |
| Cdkn2d | 347 | 520 | 561 | 606 | 464 | 526 | 369 | 519 | 563 | 476 | 499 | 576 | 500.44 | 484 | 517 | 1.068 | 0.095 | 0.58889365 | 0.76362992 | 0.0267 | 0.0135 | 0.0201 | 0.0201 | TRUE | 0.5013 |
| Cdkn3 | 342 | 485 | 451 | 636 | 449 | 414 | 363 | 484 | 453 | 500 | 483 | 453 | 456.12 | 433 | 479 | 1.105 | 0.143 | 0.3023791 | 0.52310225 | 0.0096 | 0.0141 | 0.0117 | 0.0117 | TRUE | 0.2301 |
| Cdnf | 55 | 76 | 85 | 69 | 41 | 42 | 58 | 76 | 85 | 54 | 44 | 46 | 60.67 | 73 | 48 | 0.659 | -0.601 | 0.02728044 | 0.10791751 | 0 | 0.0612 | 0.0363 | 0.0363 | TRUE | 0.2775 |
| Cdo1 | 183 | 218 | 179 | 316 | 217 | 223 | 194 | 218 | 180 | 248 | 233 | 244 | 219.64 | 197 | 242 | 1.227 | 0.296 | 0.04327156 | 0.14872066 | 0 | 0.0219 | 0.0108 | 0.0108 | TRUE | 0.0904 |
| Cdon | 1669 | 1567 | 1867 | 2583 | 1866 | 1783 | 1773 | 1565 | 1875 | 2031 | 2008 | 1952 | 1867.07 | 1738 | 1997 | 1.149 | 0.201 | 0.03142031 | 0.11909665 | 0.0035 | 0.0087 | 0.0057 | 0.0057 | TRUE | 0.0914 |
| Cdpf1 | 606 | 771 | 769 | 734 | 593 | 548 | 644 | 770 | 772 | 577 | 638 | 600 | 666.79 | 729 | 605 | 0.829 | -0.27 | 0.01927876 | 0.08453285 | 0.0052 | 0.0118 | 0.0081 | 0.0081 | TRUE | 0.1221 |
| Cdr1 | 0 | 0 | 0 | 0 | 0 | 0 | 0 | 0 | 0 | 0 | 0 | 0 | 0 | 0 | 0 | NA | NA | NA | NA | NA | NA | NA | NA | NA | NA |
| Cdr2 | 2012 | 1951 | 1808 | 2542 | 1683 | 1649 | 2137 | 1948 | 1815 | 1998 | 1811 | 1805 | 1919.2 | 1967 | 1871 | 0.952 | -0.071 | 0.46207076 | 0.66585474 | 0.0043 | 0.0086 | 0.0062 | 0.0062 | TRUE | 0.069 |
| Cdr2l | 1330 | 1759 | 1782 | 1983 | 1641 | 1679 | 1413 | 1757 | 1789 | 1559 | 1766 | 1838 | 1686.86 | 1653 | 1721 | 1.04 | 0.057 | 0.64167409 | 0.79835032 | 0.0114 | 0.0089 | 0.0103 | 0.0103 | TRUE | 0.1944 |
| Cdrt4 | 7 | 4 | 5 | 14 | 13 | 12 | 7 | 4 | 5 | 11 | 14 | 13 | 9.1 | 5 | 13 | 2.316 | 1.212 | 0.06148722 | 0.18922467 | 0 | 0.3691 | 0.1705 | 0.1705 | TRUE | 0.2001 |
| Cds1 | 646 | 998 | 973 | 737 | 554 | 622 | 686 | 997 | 977 | 579 | 596 | 681 | 752.7 | 887 | 619 | 0.697 | -0.52 | 0.00132236 | 0.011082 | 0.0235 | 0.0113 | 0.0175 | 0.0175 | TRUE | 0.4639 |
| Cds2 | 5135 | 5586 | 5986 | 6668 | 5053 | 5020 | 5454 | 5579 | 6011 | 5242 | 5437 | 5495 | 5536.23 | 5681 | 5391 | 0.949 | -0.076 | 0.28437279 | 0.5058795 | 0.0014 | 0.0075 | 0.0034 | 0.0034 | TRUE | 0.0314 |
| Cdsn | 98 | 81 | 75 | 95 | 68 | 83 | 104 | 81 | 75 | 75 | 73 | 91 | 83.17 | 87 | 80 | 0.916 | -0.126 | 0.5950075 | 0.76795894 | 0.01 | 0.0465 | 0.0287 | 0.0287 | TRUE | 0.2835 |
| Cdt1 | 9929 | 10829 | 11665 | 12673 | 10073 | 10444 | 10546 | 10815 | 11713 | 9963 | 10838 | 11433 | 10884.55 | 11025 | 10745 | 0.974 | -0.037 | 0.65859819 | 0.8075619 | 0.0032 | 0.0072 | 0.0051 | 0.0051 | TRUE | 0.0497 |
| Cdv3 | 16392 | 16644 | 16832 | 22343 | 15666 | 15177 | 17411 | 16623 | 16901 | 17564 | 16856 | 16614 | 16994.73 | 16978 | 17011 | 1.002 | 0.003 | 0.96076123 | 0.98008034 | 5.00E-04 | 0.0071 | 0.0025 | 0.0025 | TRUE | 0.0099 |
| Cdx1 | 332 | 288 | 259 | 120 | 89 | 77 | 353 | 288 | 260 | 94 | 96 | 84 | 195.79 | 300 | 91 | 0.305 | -1.711 | 2.45E-20 | 6.13E-18 | 0.0123 | 0.0237 | 0.0177 | 0.0177 | TRUE | 0.2624 |
| Cdx2 | 2 | 8 | 5 | 28 | 12 | 8 | 2 | 8 | 5 | 22 | 13 | 9 | 9.8 | 5 | 15 | 2.903 | 1.538 | 0.03597006 | 0.13026862 | 0.1502 | 0.343 | 0.2534 | 0.2534 | TRUE | 0.3151 |
| Cdx4 | 0 | 0 | 2 | 8 | 2 | 3 | 0 | 0 | 2 | 6 | 2 | 3 | 2.29 | 1 | 4 | 5.852 | 2.549 | 0.11127297 | NA | 0 | 1.4463 | 0.9675 | 0.9675 | TRUE | 0.9598 |
| Cdyl | 2473 | 2642 | 2752 | 3141 | 2427 | 2357 | 2627 | 2639 | 2763 | 2469 | 2611 | 2580 | 2614.88 | 2676 | 2553 | 0.954 | -0.068 | 0.32088266 | 0.54202034 | 5.00E-04 | 0.0082 | 0.003 | 0.003 | TRUE | 0.0104 |
| Cdyl2 | 11879 | 14307 | 16012 | 17839 | 13415 | 14833 | 12618 | 14289 | 16078 | 14024 | 14434 | 16237 | 14613.04 | 14328 | 14898 | 1.04 | 0.056 | 0.61471232 | 0.78200055 | 0.0086 | 0.0071 | 0.0089 | 0.0089 | TRUE | 0.1396 |
| Ceacam-ps1 | 0 | 0 | 0 | 0 | 1 | 0 | 0 | 0 | 0 | 0 | 1 | 0 | 0.18 | 0 | 0 | 1.859 | 0.894 | 0.82650734 | NA | 0 | 18.3785 | 10 | 10 | TRUE | 0.5204 |
| Ceacam1 | 576 | 536 | 572 | 933 | 631 | 638 | 612 | 535 | 574 | 733 | 679 | 698 | 638.71 | 574 | 703 | 1.228 | 0.296 | 0.00376701 | 0.02525689 | 0.0013 | 0.0121 | 0.0059 | 0.0059 | TRUE | 0.0405 |
| Ceacam10 | 41 | 38 | 48 | 45 | 25 | 37 | 44 | 38 | 48 | 35 | 27 | 41 | 38.75 | 43 | 34 | 0.793 | -0.334 | 0.29862181 | 0.51910525 | 0 | 0.0919 | 0.0485 | 0.0485 | TRUE | 0.227 |
| Ceacam11 | 0 | 0 | 0 | 0 | 0 | 0 | 0 | 0 | 0 | 0 | 0 | 0 | 0 | 0 | 0 | NA | NA | NA | NA | NA | NA | NA | NA | NA | NA |
| Ceacam12 | 0 | 0 | 0 | 0 | 0 | 0 | 0 | 0 | 0 | 0 | 0 | 0 | 0 | 0 | 0 | NA | NA | NA | NA | NA | NA | NA | NA | NA | NA |
| Ceacam13 | 0 | 0 | 0 | 1 | 0 | 0 | 0 | 0 | 0 | 1 | 0 | 0 | 0.13 | 0 | 0 | 1.859 | 0.894 | 0.82650734 | NA | 0 | 25.1515 | 10 | 10 | TRUE | 0.2563 |
| Ceacam14 | 0 | 0 | 0 | 0 | 0 | 0 | 0 | 0 | 0 | 0 | 0 | 0 | 0 | 0 | 0 | NA | NA | NA | NA | NA | NA | NA | NA | NA | NA |
| Ceacam15 | 0 | 0 | 0 | 0 | 0 | 0 | 0 | 0 | 0 | 0 | 0 | 0 | 0 | 0 | 0 | NA | NA | NA | NA | NA | NA | NA | NA | NA | NA |
| Ceacam16 | 38 | 55 | 48 | 43 | 40 | 45 | 40 | 55 | 48 | 34 | 43 | 49 | 44.93 | 48 | 42 | 0.872 | -0.198 | 0.51897914 | 0.71037146 | 0 | 0.0802 | 0.0457 | 0.0457 | TRUE | 0.2652 |
| Ceacam18 | 14 | 15 | 16 | 25 | 11 | 10 | 15 | 15 | 16 | 20 | 12 | 11 | 14.73 | 15 | 14 | 0.938 | -0.092 | 0.85546022 | 0.92813638 | 0 | 0.2306 | 0.1179 | 0.1179 | TRUE | 0.6272 |
| Ceacam19 | 16 | 11 | 13 | 40 | 30 | 19 | 17 | 11 | 13 | 31 | 32 | 21 | 20.93 | 14 | 28 | 2.072 | 1.051 | 0.01890148 | 0.0834988 | 0 | 0.1643 | 0.0897 | 0.0897 | TRUE | 0.3128 |
| Ceacam2 | 28 | 5 | 13 | 41 | 30 | 26 | 30 | 5 | 13 | 32 | 32 | 28 | 23.46 | 16 | 31 | 1.955 | 0.967 | 0.09288385 | 0.24896038 | 0.2498 | 0.1473 | 0.1908 | 0.1908 | TRUE | 1.0624 |
| Ceacam20 | 238 | 172 | 141 | 415 | 277 | 230 | 253 | 172 | 142 | 326 | 298 | 252 | 240.37 | 189 | 292 | 1.551 | 0.633 | 0.00549562 | 0.03351776 | 0.0482 | 0.0206 | 0.033 | 0.033 | TRUE | 0.9438 |
| Ceacam3 | 0 | 0 | 0 | 0 | 0 | 0 | 0 | 0 | 0 | 0 | 0 | 0 | 0 | 0 | 0 | NA | NA | NA | NA | NA | NA | NA | NA | NA | NA |
| Ceacam5 | 7 | 2 | 1 | 0 | 1 | 2 | 7 | 2 | 1 | 0 | 1 | 2 | 2.28 | 3 | 1 | 0.298 | -1.745 | 0.25573841 | NA | 0.6694 | 1.4495 | 1.0797 | 1.0797 | TRUE | 1.3999 |
| Ceacam9 | 12 | 10 | 4 | 21 | 11 | 17 | 13 | 10 | 4 | 17 | 12 | 19 | 12.28 | 9 | 16 | 1.763 | 0.818 | 0.17565973 | 0.37576369 | 0.0268 | 0.2751 | 0.1741 | 0.1741 | TRUE | 0.7466 |
| Cebpa | 303 | 369 | 324 | 527 | 435 | 307 | 322 | 369 | 325 | 414 | 468 | 336 | 372.35 | 339 | 406 | 1.2 | 0.263 | 0.09213931 | 0.24762974 | 0.0138 | 0.0158 | 0.0148 | 0.0148 | TRUE | 0.2544 |
| Cebpb | 243 | 229 | 210 | 347 | 304 | 229 | 258 | 229 | 211 | 273 | 327 | 251 | 258.04 | 233 | 284 | 1.219 | 0.286 | 0.077471 | 0.21991948 | 0.0105 | 0.0197 | 0.0149 | 0.0149 | TRUE | 0.1969 |
| Cebpd | 39 | 50 | 68 | 93 | 71 | 53 | 41 | 50 | 68 | 73 | 76 | 58 | 61.19 | 53 | 69 | 1.3 | 0.379 | 0.1877469 | 0.39246444 | 0.0229 | 0.0607 | 0.043 | 0.043 | TRUE | 0.5104 |
| Cebpe | 16 | 17 | 13 | 17 | 14 | 12 | 17 | 17 | 13 | 13 | 15 | 13 | 14.76 | 16 | 14 | 0.883 | -0.179 | 0.71006301 | 0.8420764 | 0 | 0.23 | 0.0998 | 0.0998 | TRUE | 0.1016 |
| Cebpg | 1056 | 1491 | 1537 | 1586 | 1259 | 1291 | 1122 | 1489 | 1543 | 1247 | 1355 | 1413 | 1361.44 | 1385 | 1338 | 0.966 | -0.05 | 0.71483679 | 0.84522122 | 0.0159 | 0.0093 | 0.0128 | 0.0128 | TRUE | 0.331 |
| Cebpz | 11407 | 10644 | 10887 | 13874 | 10311 | 10242 | 12116 | 10630 | 10932 | 10907 | 11094 | 11211 | 11148.39 | 11226 | 11071 | 0.986 | -0.02 | 0.79075401 | 0.89242185 | 0.002 | 0.0072 | 0.004 | 0.004 | TRUE | 0.0588 |
| Cecr2 | 8301 | 7521 | 8331 | 9982 | 7229 | 7266 | 8817 | 7511 | 8365 | 7847 | 7778 | 7954 | 8045.4 | 8231 | 7860 | 0.955 | -0.067 | 0.4162639 | 0.62771911 | 0.0031 | 0.0073 | 0.0047 | 0.0047 | TRUE | 0.0715 |
| Cecr5 | 2525 | 2775 | 2658 | 3183 | 2375 | 2166 | 2682 | 2771 | 2669 | 2502 | 2555 | 2371 | 2591.82 | 2707 | 2476 | 0.915 | -0.129 | 0.06495754 | 0.19626919 | 5.00E-04 | 0.0082 | 0.0031 | 0.0031 | TRUE | 0.0164 |
| Cecr6 | 1 | 7 | 4 | 5 | 3 | 0 | 1 | 7 | 4 | 4 | 3 | 0 | 3.2 | 4 | 2 | 0.605 | -0.725 | 0.56556371 | NA | 0.51 | 1.0349 | 0.8208 | 0.8208 | TRUE | 0.186 |
| Cel | 1 | 1 | 0 | 0 | 0 | 1 | 1 | 1 | 0 | 0 | 0 | 1 | 0.53 | 1 | 0 | 0.51 | -0.972 | 0.72757263 | NA | 0 | 6.271 | 3.8704 | 3.8704 | TRUE | 0.5383 |
| Cela1 | 166 | 212 | 204 | 166 | 113 | 135 | 176 | 212 | 205 | 130 | 122 | 148 | 165.46 | 198 | 133 | 0.673 | -0.57 | 0.00086432 | 0.00794413 | 0.0026 | 0.0268 | 0.0149 | 0.0149 | TRUE | 0.095 |
| Cela2a | 134 | 112 | 88 | 193 | 192 | 158 | 142 | 112 | 88 | 152 | 207 | 173 | 145.63 | 114 | 177 | 1.549 | 0.631 | 0.00577698 | 0.03490977 | 0.0316 | 0.0295 | 0.0305 | 0.0305 | TRUE | 0.4074 |
| Cela3b | 0 | 0 | 0 | 0 | 0 | 0 | 0 | 0 | 0 | 0 | 0 | 0 | 0 | 0 | 0 | NA | NA | NA | NA | NA | NA | NA | NA | NA | NA |
| Celf1 | 17150 | 17276 | 16964 | 23301 | 17015 | 16701 | 18216 | 17254 | 17034 | 18317 | 18307 | 18282 | 17901.68 | 17501 | 18302 | 1.046 | 0.065 | 0.2706058 | 0.49074002 | 6.00E-04 | 0.0071 | 0.0024 | 0.0024 | TRUE | 0.0156 |
| Celf2 | 389 | 636 | 660 | 774 | 593 | 640 | 413 | 635 | 663 | 608 | 638 | 701 | 609.69 | 570 | 649 | 1.137 | 0.185 | 0.30840791 | 0.52894841 | 0.0325 | 0.0123 | 0.0222 | 0.0222 | TRUE | 0.6777 |
| Celf3 | 14 | 8 | 12 | 6 | 10 | 8 | 15 | 8 | 12 | 5 | 11 | 9 | 9.86 | 12 | 8 | 0.684 | -0.549 | 0.39297663 | 0.60646061 | 0 | 0.3411 | 0.1918 | 0.1918 | TRUE | 0.4424 |
| Celf4 | 722 | 671 | 642 | 602 | 497 | 489 | 767 | 670 | 645 | 473 | 535 | 535 | 604.16 | 694 | 514 | 0.74 | -0.434 | 0.00023801 | 0.00280352 | 0.0052 | 0.0124 | 0.0083 | 0.0083 | TRUE | 0.0994 |

| | | | | | | | | | | | | | | | | | | | | | | | | | |
|---|---|---|---|---|---|---|---|---|---|---|---|---|---|---|---|---|---|---|---|---|---|---|---|---|---|
| Celf5 | 739 | 991 | 1138 | 876 | 744 | 742 | 785 | 990 | 1143 | 689 | 800 | 812 | 869.79 | 973 | 767 | 0.788 | -0.344 | 0.02523673 | 0.10245398 | 0.0192 | 0.0107 | 0.0158 | 0.0158 | TRUE | 0.2205 |
| Celf6 | 36 | 23 | 18 | 37 | 25 | 27 | 38 | 23 | 18 | 29 | 27 | 30 | 27.47 | 26 | 29 | 1.082 | 0.114 | 0.77981005 | 0.8861249 | 0.0358 | 0.1268 | 0.084 | 0.084 | TRUE | 0.9259 |
| Celsr1 | 2103 | 2221 | 2712 | 2344 | 1812 | 1746 | 2234 | 2218 | 2723 | 1843 | 1950 | 1911 | 2146.43 | 2392 | 1901 | 0.795 | -0.331 | 0.00170123 | 0.01353095 | 0.0069 | 0.0084 | 0.0076 | 0.0076 | TRUE | 0.1783 |
| Celsr2 | 714 | 467 | 533 | 980 | 696 | 523 | 758 | 466 | 535 | 770 | 749 | 573 | 641.96 | 586 | 697 | 1.19 | 0.251 | 0.2072059 | 0.41652453 | 0.0414 | 0.012 | 0.0269 | 0.0269 | TRUE | 0.7673 |
| Celsr3 | 589 | 538 | 564 | 672 | 534 | 473 | 626 | 537 | 566 | 528 | 575 | 518 | 558.31 | 576 | 540 | 0.937 | -0.094 | 0.39961554 | 0.61268945 | 0.0027 | 0.0128 | 0.0071 | 0.0071 | TRUE | 0.0649 |
| Cend1 | 24 | 10 | 11 | 13 | 10 | 14 | 25 | 10 | 11 | 10 | 11 | 15 | 13.8 | 15 | 12 | 0.779 | -0.361 | 0.53923267 | 0.72535418 | 0.105 | 0.2455 | 0.1752 | 0.1752 | TRUE | 1.424 |
| Cenpa | 5586 | 7611 | 7089 | 7362 | 5546 | 6241 | 5933 | 7601 | 7118 | 5787 | 5967 | 6832 | 6539.82 | 6884 | 6195 | 0.9 | -0.152 | 0.19660743 | 0.40313213 | 0.0116 | 0.0074 | 0.0099 | 0.0099 | TRUE | 0.1781 |
| Cenpb | 2646 | 1965 | 1689 | 4434 | 2897 | 2135 | 2811 | 1962 | 1696 | 3486 | 3117 | 2337 | 2568.11 | 2156 | 2980 | 1.382 | 0.467 | 0.0208487 | 0.08945202 | 0.0552 | 0.0082 | 0.0291 | 0.0291 | TRUE | 0.8187 |
| Cenpc1 | 2170 | 2131 | 2114 | 3179 | 2281 | 2307 | 2305 | 2128 | 2123 | 2499 | 2454 | 2525 | 2339.09 | 2185 | 2493 | 1.141 | 0.19 | 0.00879847 | 0.04780211 | 6.00E-04 | 0.0083 | 0.0034 | 0.0034 | TRUE | 0.0276 |
| Cenpe | 7747 | 6869 | 7474 | 10380 | 7570 | 6718 | 8229 | 6860 | 7505 | 8160 | 8145 | 7354 | 7708.7 | 7531 | 7886 | 1.047 | 0.067 | 0.48287993 | 0.68129391 | 0.0051 | 0.0073 | 0.0064 | 0.0064 | TRUE | 0.0801 |
| Cenpf | 11552 | 12432 | 14708 | 14781 | 10908 | 11291 | 12270 | 12416 | 14768 | 11620 | 11736 | 12360 | 12528.4 | 13151 | 11905 | 0.905 | -0.144 | 0.13064022 | 0.31124736 | 0.0061 | 0.0072 | 0.0064 | 0.0064 | TRUE | 0.1414 |
| Cenph | 1633 | 2001 | 1820 | 2719 | 1984 | 1929 | 1735 | 1998 | 1827 | 2137 | 2135 | 2112 | 1990.69 | 1853 | 2128 | 1.148 | 0.199 | 0.01666773 | 0.07644185 | 0.0019 | 0.0086 | 0.0045 | 0.0045 | TRUE | 0.0566 |
| Cenpi | 1653 | 1580 | 1506 | 2290 | 1632 | 1702 | 1756 | 1578 | 1512 | 1800 | 1756 | 1863 | 1710.86 | 1615 | 1806 | 1.118 | 0.162 | 0.06968538 | 0.20524328 | 0.0026 | 0.0088 | 0.0051 | 0.0051 | TRUE | 0.0695 |
| Cenpj | 1408 | 1056 | 1029 | 1641 | 1116 | 951 | 1496 | 1055 | 1033 | 1290 | 1201 | 1041 | 1185.87 | 1195 | 1177 | 0.986 | -0.02 | 0.90291759 | 0.95205059 | 0.0272 | 0.0097 | 0.0185 | 0.0185 | TRUE | 0.5819 |
| Cenpk | 451 | 709 | 714 | 937 | 667 | 754 | 479 | 708 | 717 | 737 | 718 | 825 | 697.28 | 635 | 760 | 1.197 | 0.259 | 0.12137947 | 0.29630042 | 0.0257 | 0.0116 | 0.0187 | 0.0187 | TRUE | 0.539 |
| Cenpl | 645 | 684 | 642 | 929 | 604 | 624 | 685 | 683 | 645 | 730 | 650 | 683 | 679.35 | 671 | 688 | 1.026 | 0.038 | 0.70102863 | 0.83650923 | 0.0011 | 0.0118 | 0.0055 | 0.0055 | TRUE | 0.0363 |
| Cenpm | 743 | 1034 | 1033 | 2038 | 1281 | 1281 | 789 | 1033 | 1037 | 1602 | 1378 | 1402 | 1206.96 | 953 | 1461 | 1.533 | 0.617 | 4.00E-06 | 9.15E-05 | 0.0133 | 0.0096 | 0.012 | 0.012 | TRUE | 0.2685 |
| Cenpn | 2920 | 2898 | 2860 | 3778 | 2882 | 2626 | 3102 | 2894 | 2872 | 2970 | 3101 | 2875 | 2968.82 | 2956 | 2982 | 1.009 | 0.013 | 0.86351591 | 0.9324082 | 0.0013 | 0.008 | 0.0036 | 0.0036 | TRUE | 0.0225 |
| Cenpo | 1020 | 912 | 919 | 1392 | 1009 | 949 | 1083 | 911 | 923 | 1094 | 1086 | 1039 | 1022.62 | 972 | 1073 | 1.104 | 0.143 | 0.16357785 | 0.35937453 | 0.0038 | 0.0101 | 0.0066 | 0.0066 | TRUE | 0.1185 |
| Cenpp | 715 | 742 | 730 | 1139 | 886 | 748 | 759 | 741 | 733 | 895 | 953 | 819 | 816.83 | 744 | 889 | 1.195 | 0.257 | 0.00845848 | 0.04641862 | 0.0018 | 0.0109 | 0.0056 | 0.0056 | TRUE | 0.0551 |
| Cenpq | 568 | 518 | 528 | 739 | 565 | 563 | 603 | 517 | 530 | 581 | 608 | 616 | 575.99 | 550 | 602 | 1.093 | 0.129 | 0.23467314 | 0.44950957 | 0.0021 | 0.0126 | 0.0067 | 0.0067 | TRUE | 0.0825 |
| Cenpt | 3372 | 3919 | 3828 | 4897 | 3690 | 3263 | 3582 | 3914 | 3844 | 3850 | 3970 | 3572 | 3788.51 | 3780 | 3797 | 1.005 | 0.007 | 0.93236523 | 0.96643431 | 0.0023 | 0.0078 | 0.0042 | 0.0042 | TRUE | 0.0325 |
| Cenpv | 2703 | 3356 | 3323 | 4152 | 2970 | 2782 | 2871 | 3352 | 3337 | 3264 | 3196 | 3045 | 3177.37 | 3187 | 3168 | 0.994 | -0.008 | 0.92810644 | 0.96472155 | 0.0041 | 0.0079 | 0.0056 | 0.0056 | TRUE | 0.0909 |
| Cenpw | 882 | 1090 | 1054 | 1340 | 982 | 1029 | 937 | 1089 | 1058 | 1053 | 1057 | 1126 | 1053.36 | 1028 | 1079 | 1.049 | 0.069 | 0.47606521 | 0.67601516 | 0.0028 | 0.01 | 0.0058 | 0.0058 | TRUE | 0.0713 |
| Cep104 | 894 | 1028 | 1016 | 1500 | 1007 | 1021 | 950 | 1027 | 1020 | 1179 | 1083 | 1118 | 1062.79 | 999 | 1127 | 1.129 | 0.175 | 0.04411787 | 0.15064452 | 0.001 | 0.01 | 0.0045 | 0.0045 | TRUE | 0.022 |
| Cep110 | 3758 | 3726 | 4121 | 4648 | 3544 | 3426 | 3992 | 3721 | 4138 | 3654 | 3813 | 3750 | 3844.68 | 3950 | 3739 | 0.946 | -0.079 | 0.27633016 | 0.49717993 | 0.0013 | 0.0078 | 0.0036 | 0.0036 | TRUE | 0.0314 |
| Cep112 | 131 | 242 | 247 | 297 | 236 | 254 | 139 | 242 | 248 | 233 | 254 | 278 | 232.38 | 210 | 255 | 1.215 | 0.281 | 0.21820676 | 0.43024841 | 0.0466 | 0.0211 | 0.0332 | 0.0332 | TRUE | 0.9342 |
| Cep120 | 2980 | 3243 | 3403 | 4303 | 3139 | 3306 | 3165 | 3239 | 3417 | 3383 | 3377 | 3619 | 3366.68 | 3274 | 3460 | 1.057 | 0.079 | 0.27490771 | 0.49548018 | 0.0011 | 0.0079 | 0.0035 | 0.0035 | TRUE | 0.0196 |
| Cep128 | 1142 | 1042 | 1135 | 1762 | 1276 | 1147 | 1213 | 1041 | 1140 | 1385 | 1373 | 1256 | 1234.49 | 1131 | 1338 | 1.184 | 0.243 | 0.0123421 | 0.06139101 | 0.0035 | 0.0096 | 0.006 | 0.006 | TRUE | 0.0587 |
| Cep135 | 1104 | 853 | 964 | 1710 | 1221 | 1079 | 1173 | 852 | 968 | 1344 | 1314 | 1181 | 1138.6 | 998 | 1280 | 1.284 | 0.36 | 0.0074653 | 0.04211843 | 0.0142 | 0.0098 | 0.0122 | 0.0122 | TRUE | 0.2808 |
| Cep152 | 672 | 669 | 735 | 968 | 805 | 782 | 714 | 668 | 738 | 761 | 866 | 856 | 767.18 | 707 | 828 | 1.17 | 0.226 | 0.02699393 | 0.10720009 | 0.0027 | 0.0112 | 0.0062 | 0.0062 | TRUE | 0.0617 |
| Cep164 | 2678 | 2621 | 2644 | 4322 | 3305 | 3113 | 2845 | 2618 | 2655 | 3398 | 3556 | 3408 | 3079.71 | 2706 | 3454 | 1.276 | 0.352 | 8.11E-07 | 2.30E-05 | 8.00E-04 | 0.008 | 0.0033 | 0.0033 | TRUE | 0.0244 |
| Cep170 | 2349 | 2830 | 2827 | 3171 | 2449 | 2479 | 2495 | 2826 | 2839 | 2493 | 2635 | 2714 | 2666.9 | 2720 | 2614 | 0.961 | -0.058 | 0.5035304 | 0.69859971 | 0.0029 | 0.0081 | 0.0051 | 0.0051 | TRUE | 0.0633 |
| Cep170b | 4919 | 5962 | 6216 | 5811 | 4440 | 4946 | 5225 | 5954 | 6242 | 4568 | 4777 | 5414 | 5363.36 | 5807 | 4920 | 0.847 | -0.239 | 0.0221367 | 0.09357813 | 0.0078 | 0.0075 | 0.0077 | 0.0077 | TRUE | 0.094 |
| Cep19 | 899 | 816 | 849 | 1159 | 808 | 799 | 955 | 815 | 852 | 911 | 869 | 875 | 879.57 | 874 | 885 | 1.013 | 0.019 | 0.84697233 | 0.92352627 | 0.0025 | 0.0106 | 0.0059 | 0.0059 | TRUE | 0.0773 |
| Cep192 | 9727 | 9798 | 10886 | 11469 | 8299 | 8766 | 10332 | 9785 | 10931 | 9016 | 8929 | 9596 | 9764.81 | 10349 | 9180 | 0.887 | -0.173 | 0.02043371 | 0.08822352 | 0.0022 | 0.0072 | 0.0039 | 0.0039 | TRUE | 0.0295 |
| Cep250 | 1557 | 1628 | 1788 | 2473 | 2137 | 1980 | 1654 | 1626 | 1795 | 1944 | 2299 | 2167 | 1914.3 | 1692 | 2137 | 1.262 | 0.336 | 0.00050205 | 0.00517876 | 0.0042 | 0.0086 | 0.0062 | 0.0062 | TRUE | 0.0768 |
| Cep290 | 942 | 751 | 774 | 1636 | 1104 | 918 | 1001 | 750 | 777 | 1286 | 1188 | 1005 | 1001.1 | 843 | 1160 | 1.377 | 0.462 | 0.00189423 | 0.01474623 | 0.0193 | 0.0102 | 0.0149 | 0.0149 | TRUE | 0.3187 |
| Cep350 | 3659 | 3200 | 3997 | 4696 | 3185 | 3134 | 3887 | 3196 | 4013 | 3692 | 3427 | 3431 | 3607.49 | 3699 | 3517 | 0.951 | -0.073 | 0.49711353 | 0.69256025 | 0.0071 | 0.0078 | 0.008 | 0.008 | TRUE | 0.1722 |
| Cep44 | 1017 | 1289 | 1177 | 1651 | 1111 | 1197 | 1080 | 1287 | 1182 | 1298 | 1195 | 1310 | 1225.49 | 1183 | 1268 | 1.072 | 0.1 | 0.31955642 | 0.5406844 | 0.0038 | 0.0096 | 0.0064 | 0.0064 | TRUE | 0.0712 |
| Cep55 | 3566 | 4138 | 3971 | 4895 | 3731 | 3847 | 3788 | 4133 | 3987 | 3848 | 4014 | 4211 | 3996.87 | 3969 | 4024 | 1.014 | 0.019 | 0.79507241 | 0.8946193 | 0.0017 | 0.0077 | 0.0038 | 0.0038 | TRUE | 0.02 |
| Cep57 | 3056 | 3789 | 4001 | 4317 | 3239 | 3559 | 3246 | 3784 | 4017 | 3394 | 3485 | 3896 | 3637.01 | 3682 | 3592 | 0.975 | -0.036 | 0.73537033 | 0.85804993 | 0.0084 | 0.0078 | 0.0081 | 0.0081 | TRUE | 0.1306 |
| Cep57l1 | 1644 | 1572 | 1597 | 2402 | 1884 | 1676 | 1746 | 1570 | 1604 | 1888 | 2027 | 1835 | 1778.29 | 1640 | 1917 | 1.169 | 0.225 | 0.00895742 | 0.04849861 | 0.0021 | 0.0088 | 0.0048 | 0.0048 | TRUE | 0.0386 |
| Cep63 | 925 | 992 | 990 | 1282 | 949 | 923 | 983 | 991 | 994 | 1008 | 1021 | 1010 | 1001.09 | 989 | 1013 | 1.024 | 0.034 | 0.65684171 | 0.80670515 | 0 | 0.0102 | 0.0033 | 0.0033 | TRUE | 6.00E-04 |
| Cep68 | 2815 | 3412 | 3473 | 3764 | 2903 | 2984 | 2990 | 3408 | 3487 | 2959 | 3123 | 3266 | 3205.63 | 3295 | 3116 | 0.945 | -0.081 | 0.37556412 | 0.59147378 | 0.004 | 0.0079 | 0.0057 | 0.0057 | TRUE | 0.0795 |
| Cep70 | 891 | 868 | 910 | 1117 | 814 | 885 | 946 | 867 | 914 | 878 | 876 | 969 | 908.28 | 909 | 908 | 0.998 | -0.003 | 0.9744537 | 0.98671997 | 0.0015 | 0.0105 | 0.0052 | 0.0052 | TRUE | 0.0406 |
| Cep72 | 1479 | 1793 | 1798 | 2086 | 1634 | 1614 | 1571 | 1791 | 1805 | 1640 | 1758 | 1767 | 1721.96 | 1722 | 1722 | 0.999 | -0.002 | 0.98636303 | 0.99265683 | 0.0033 | 0.0088 | 0.0054 | 0.0054 | TRUE | 0.071 |
| Cep76 | 1686 | 1536 | 1568 | 2154 | 1435 | 1312 | 1791 | 1534 | 1574 | 1693 | 1544 | 1436 | 1595.46 | 1633 | 1558 | 0.955 | -0.067 | 0.5269315 | 0.71593432 | 0.0063 | 0.009 | 0.0074 | 0.0074 | TRUE | 0.0858 |
| Cep78 | 2789 | 2459 | 2553 | 3027 | 2228 | 2168 | 2962 | 2456 | 2563 | 2380 | 2397 | 2373 | 2521.95 | 2660 | 2383 | 0.896 | -0.159 | 0.09191449 | 0.24742196 | 0.0044 | 0.0082 | 0.006 | 0.006 | TRUE | 0.1192 |
| Cep85 | 6105 | 6229 | 6142 | 7294 | 5336 | 5318 | 6485 | 6221 | 6167 | 5734 | 5741 | 5821 | 6028.22 | 6291 | 5765 | 0.916 | -0.126 | 0.03256625 | 0.121608 | 2.00E-04 | 0.0074 | 0.0023 | 0.0023 | TRUE | 0.0088 |
| Cep85l | 270 | 252 | 288 | 366 | 258 | 243 | 287 | 252 | 289 | 288 | 278 | 266 | 276.49 | 276 | 277 | 1.006 | 0.008 | 0.94902245 | 0.97441456 | 0 | 0.0188 | 0.009 | 0.009 | TRUE | 0.0667 |
| Cep89 | 780 | 922 | 931 | 1217 | 1013 | 919 | 828 | 921 | 935 | 957 | 1090 | 1006 | 956.13 | 895 | 1018 | 1.136 | 0.184 | 0.06633438 | 0.19871336 | 0.0033 | 0.0103 | 0.0062 | 0.0062 | TRUE | 0.0495 |
| Cep95 | 1252 | 1491 | 1541 | 1608 | 1124 | 1135 | 1330 | 1489 | 1547 | 1264 | 1209 | 1242 | 1347.02 | 1455 | 1238 | 0.851 | -0.233 | 0.01080275 | 0.05571645 | 0.0026 | 0.0093 | 0.0053 | 0.0053 | TRUE | 0.0682 |
| Cep97 | 929 | 985 | 1106 | 1132 | 819 | 844 | 987 | 984 | 1111 | 890 | 881 | 924 | 962.67 | 1027 | 898 | 0.874 | -0.194 | 0.03757163 | 0.13461602 | 0.0018 | 0.0103 | 0.0052 | 0.0052 | TRUE | 0.0607 |
| Cept1 | 1102 | 1475 | 1445 | 1781 | 1273 | 1420 | 1171 | 1473 | 1451 | 1400 | 1370 | 1554 | 1403.12 | 1365 | 1441 | 1.056 | 0.078 | 0.50974901 | 0.70326815 | 0.0095 | 0.0093 | 0.0094 | 0.0094 | TRUE | 0.1858 |
| Cer1 | 0 | 1 | 0 | 0 | 0 | 0 | 0 | 1 | 0 | 0 | 0 | 0 | 0.17 | 0 | 0 | 0.49 | -1.029 | 0.80087 | NA | 0 | 19.7991 | 10 | 10 | TRUE | 0.405 |
| Cercam | 1235 | 1559 | 1547 | 1397 | 1071 | 1222 | 1312 | 1557 | 1553 | 1098 | 1152 | 1338 | 1335.06 | 1474 | 1196 | 0.811 | -0.303 | 0.01048886 | 0.05463353 | 0.0089 | 0.0094 | 0.0093 | 0.0093 | TRUE | 0.1279 |
| Cerk | 352 | 375 | 371 | 440 | 297 | 318 | 374 | 375 | 373 | 346 | 320 | 348 | 355.75 | 374 | 338 | 0.905 | -0.144 | 0.20391536 | 0.41206317 | 0 | 0.0162 | 0.0065 | 0.0065 | TRUE | 0.0246 |
| Cerkl | 22 | 20 | 27 | 32 | 32 | 32 | 23 | 20 | 27 | 25 | 34 | 35 | 27.51 | 23 | 31 | 1.334 | 0.415 | 0.26897167 | 0.48896048 | 0 | 0.1267 | 0.0644 | 0.0644 | TRUE | 0.2556 |
| Cers1 | 0 | 0 | 0 | 0 | 0 | 0 | 0 | 0 | 0 | 0 | 0 | 0 | 0 | 0 | 0 | NA | NA | NA | NA | NA | NA | NA | NA | NA | NA |
| Cers2 | 1569 | 2111 | 2205 | 2824 | 2129 | 2209 | 1667 | 2108 | 2214 | 2220 | 2291 | 2418 | 2152.94 | 1996 | 2310 | 1.157 | 0.21 | 0.08367528 | 0.23178124 | 0.0108 | 0.0084 | 0.0101 | 0.0101 | TRUE | 0.2519 |
| Cers3 | 25 | 39 | 23 | 58 | 28 | 44 | 27 | 39 | 23 | 46 | 30 | 48 | 35.41 | 30 | 41 | 1.401 | 0.486 | 0.19098034 | 0.39611884 | 0.0372 | 0.0999 | 0.0704 | 0.0704 | TRUE | 0.5208 |
| Cers4 | 3945 | 4321 | 4042 | 3808 | 2951 | 3431 | 4190 | 4315 | 4059 | 2994 | 3175 | 3756 | 3748.12 | 4188 | 3308 | 0.79 | -0.341 | 0.00097673 | 0.00874783 | 0.0073 | 0.0078 | 0.0074 | 0.0074 | TRUE | 0.1699 |
| Cers5 | 1763 | 1919 | 2010 | 2720 | 1897 | 1987 | 1873 | 1917 | 2018 | 2138 | 2041 | 2175 | 2026.96 | 1936 | 2118 | 1.094 | 0.13 | 0.08142556 | 0.22746086 | 7.00E-04 | 0.0085 | 0.0035 | 0.0035 | TRUE | 0.0168 |
| Cers6 | 1014 | 1212 | 1371 | 1351 | 942 | 1016 | 1077 | 1210 | 1377 | 1062 | 1014 | 1112 | 1141.98 | 1221 | 1063 | 0.87 | -0.201 | 0.07987626 | 0.22436154 | 0.0073 | 0.0098 | 0.0086 | 0.0086 | TRUE | 0.1484 |
| Ces1a | 1 | 0 | 1 | 2 | 0 | 0 | 1 | 0 | 1 | 2 | 0 | 0 | 0.61 | 1 | 1 | 0.816 | -0.293 | 0.91321207 | NA | 0 | 5.4395 | 3.5982 | 3.5982 | TRUE | 0.9722 |
| Ces1b | 3 | 0 | 1 | 0 | 0 | 0 | 3 | 0 | 1 | 0 | 0 | 0 | 0.7 | 1 | 0 | 0.129 | -2.957 | 0.32154538 | NA | 6.363 | 4.7238 | 5.0467 | 5.0467 | TRUE | 0.1738 |
| Ces1c | 11 | 2 | 3 | 7 | 1 | 0 | 12 | 2 | 3 | 6 | 1 | 0 | 3.88 | 6 | 2 | 0.415 | -1.268 | 0.33460914 | NA | 1.0431 | 0.8563 | 0.9401 | 0.9401 | TRUE | 2.1636 |
| Ces1d | 0 | 0 | 1 | 0 | 0 | 0 | 0 | 0 | 1 | 0 | 0 | 0 | 0.17 | 0 | 0 | 0.49 | -1.029 | 0.80087 | NA | 0 | 19.6928 | 10 | 10 | TRUE | 0.4098 |

| | | | | | | | | | | | | | | | | | | | | | | | | | |
|---|---|---|---|---|---|---|---|---|---|---|---|---|---|---|---|---|---|---|---|---|---|---|---|---|---|
| Ces1e | 0 | 0 | 0 | 0 | 0 | 0 | 0 | 0 | 0 | 0 | 0 | 0 | 0 | 0 | 0 | NA | NA | NA | NA | NA | NA | NA | NA | NA | NA |
| Ces1f | 0 | 0 | 0 | 0 | 0 | 0 | 0 | 0 | 0 | 0 | 0 | 0 | 0 | 0 | 0 | NA | NA | NA | NA | NA | NA | NA | NA | NA | NA |
| Ces1g | 0 | 0 | 0 | 0 | 0 | 0 | 0 | 0 | 0 | 0 | 0 | 0 | 0 | 0 | 0 | NA | NA | NA | NA | NA | NA | NA | NA | NA | NA |
| Ces2a | 0 | 0 | 0 | 0 | 0 | 0 | 0 | 0 | 0 | 0 | 0 | 0 | 0 | 0 | 0 | NA | NA | NA | NA | NA | NA | NA | NA | NA | NA |
| Ces2b | 0 | 0 | 0 | 0 | 0 | 1 | 0 | 0 | 0 | 0 | 0 | 1 | 0.18 | 0 | 0 | 1.859 | 0.894 | 0.82650734 | NA | 0 | 18.0643 | 10 | 10 | TRUE | 0.5383 |
| Ces2c | 0 | 0 | 0 | 0 | 0 | 0 | 0 | 0 | 0 | 0 | 0 | 0 | 0 | 0 | 0 | NA | NA | NA | NA | NA | NA | NA | NA | NA | NA |
| Ces2d-ps | 0 | 0 | 0 | 0 | 0 | 0 | 0 | 0 | 0 | 0 | 0 | 0 | 0 | 0 | 0 | NA | NA | NA | NA | NA | NA | NA | NA | NA | NA |
| Ces2e | 14 | 2 | 6 | 31 | 20 | 15 | 15 | 2 | 6 | 24 | 22 | 16 | 14.2 | 8 | 21 | 2.749 | 1.459 | 0.03005822 | 0.11543176 | 0.2288 | 0.2389 | 0.2351 | 0.2351 | TRUE | 1.4434 |
| Ces2f | 1 | 1 | 0 | 3 | 0 | 1 | 1 | 1 | 0 | 2 | 0 | 1 | 0.92 | 1 | 1 | 1.734 | 0.794 | 0.70563564 | NA | 0 | 3.5918 | 2.0282 | 2.0282 | TRUE | 0.167 |
| Ces2g | 0 | 2 | 0 | 9 | 5 | 3 | 0 | 2 | 0 | 7 | 5 | 3 | 2.96 | 1 | 5 | 7.822 | 2.968 | 0.04402616 | NA | 0 | 1.1214 | 0.7167 | 0.7167 | TRUE | 0.8162 |
| Ces2h | 0 | 0 | 0 | 0 | 0 | 0 | 0 | 0 | 0 | 0 | 0 | 0 | 0 | 0 | 0 | NA | NA | NA | NA | NA | NA | NA | NA | NA | NA |
| Ces3a | 0 | 0 | 0 | 0 | 0 | 0 | 0 | 0 | 0 | 0 | 0 | 0 | 0 | 0 | 0 | NA | NA | NA | NA | NA | NA | NA | NA | NA | NA |
| Ces3b | 0 | 0 | 0 | 0 | 0 | 0 | 0 | 0 | 0 | 0 | 0 | 0 | 0 | 0 | 0 | NA | NA | NA | NA | NA | NA | NA | NA | NA | NA |
| Ces4a | 0 | 1 | 0 | 1 | 2 | 1 | 0 | 1 | 0 | 1 | 2 | 1 | 0.84 | 0 | 1 | 3.633 | 1.861 | 0.38689776 | NA | 0 | 3.9356 | 1.9655 | 1.9655 | TRUE | 0.405 |
| Ces5a | 2 | 1 | 0 | 2 | 3 | 0 | 2 | 1 | 0 | 2 | 3 | 0 | 1.32 | 1 | 2 | 1.55 | 0.632 | 0.73394872 | NA | 0 | 2.5017 | 1.6934 | 1.6934 | TRUE | 0.1309 |
| Cetn1 | 0 | 0 | 0 | 0 | 0 | 0 | 0 | 0 | 0 | 0 | 0 | 0 | 0 | 0 | 0 | NA | NA | NA | NA | NA | NA | NA | NA | NA | NA |
| Cetn2 | 1168 | 1200 | 1094 | 1469 | 1110 | 1039 | 1241 | 1198 | 1098 | 1155 | 1194 | 1137 | 1170.67 | 1179 | 1162 | 0.986 | -0.021 | 0.81085997 | 0.90283686 | 0.0014 | 0.0097 | 0.0046 | 0.0046 | TRUE | 0.043 |
| Cetn3 | 1926 | 2493 | 2492 | 3484 | 2358 | 2675 | 2046 | 2490 | 2502 | 2739 | 2537 | 2928 | 2540.31 | 2346 | 2735 | 1.166 | 0.221 | 0.04520728 | 0.15333532 | 0.0081 | 0.0082 | 0.0084 | 0.0084 | TRUE | 0.1514 |
| Cetn4 | 15 | 17 | 11 | 44 | 23 | 12 | 16 | 17 | 11 | 35 | 25 | 13 | 19.4 | 15 | 24 | 1.667 | 0.737 | 0.15188772 | 0.34295352 | 0.0941 | 0.1767 | 0.1358 | 0.1358 | TRUE | 0.1466 |
| Cfb | 3 | 6 | 1 | 0 | 1 | 1 | 3 | 6 | 1 | 0 | 1 | 1 | 2.06 | 3 | 1 | 0.199 | -2.327 | 0.14564898 | NA | 0 | 1.607 | 0.9744 | 0.9744 | TRUE | 0.163 |
| Cfc1 | 3 | 5 | 4 | 3 | 4 | 3 | 3 | 5 | 4 | 2 | 4 | 3 | 3.69 | 4 | 3 | 0.803 | -0.316 | 0.73993457 | NA | 0 | 0.8996 | 0.3783 | 0.3783 | TRUE | 0.1256 |
| Cfd | 0 | 1 | 0 | 0 | 0 | 0 | 0 | 1 | 0 | 0 | 0 | 0 | 0.17 | 0 | 0 | 0.49 | -1.029 | 0.80087 | NA | 0 | 19.7991 | 10 | 10 | TRUE | 0.405 |
| Cfdp1 | 10646 | 11149 | 11002 | 15312 | 10854 | 10506 | 11308 | 11135 | 11047 | 12037 | 11678 | 11500 | 11450.92 | 11163 | 11738 | 1.052 | 0.073 | 0.19276812 | 0.39842218 | 2.00E-04 | 0.0072 | 0.0022 | 0.0022 | TRUE | 0.0061 |
| Cfh | 3 | 3 | 2 | 6 | 5 | 0 | 3 | 3 | 2 | 5 | 5 | 0 | 3.05 | 3 | 3 | 1.25 | 0.322 | 0.79167403 | NA | 0 | 1.0878 | 0.7398 | 0.7398 | TRUE | 0.9744 |
| Cfhr1 | 0 | 0 | 0 | 0 | 0 | 0 | 0 | 0 | 0 | 0 | 0 | 0 | 0 | 0 | 0 | NA | NA | NA | NA | NA | NA | NA | NA | NA | NA |
| Cfhr2 | 0 | 0 | 0 | 0 | 2 | 0 | 0 | 0 | 0 | 0 | 2 | 0 | 0.36 | 0 | 1 | 3.775 | 1.916 | 0.63423067 | NA | 10 | 9.1927 | 10 | 10 | TRUE | 0.9743 |
| Cfi | 2 | 5 | 2 | 2 | 9 | 2 | 2 | 5 | 2 | 2 | 10 | 2 | 3.76 | 3 | 5 | 1.446 | 0.532 | 0.63564557 | NA | 0.4117 | 0.8827 | 0.6292 | 0.6292 | TRUE | 1.7982 |
| Cfl1 | 26318 | 20841 | 20575 | 35591 | 24604 | 21533 | 27954 | 20814 | 20659 | 27979 | 26472 | 23571 | 24575.07 | 23142 | 26007 | 1.124 | 0.168 | 0.21546969 | 0.42730894 | 0.0189 | 0.007 | 0.0133 | 0.0133 | TRUE | 0.4048 |
| Cfl2 | 1834 | 2394 | 2408 | 3166 | 2254 | 2310 | 1948 | 2391 | 2418 | 2489 | 2425 | 2529 | 2366.59 | 2252 | 2481 | 1.101 | 0.139 | 0.18385593 | 0.38659864 | 0.0064 | 0.0083 | 0.0075 | 0.0075 | TRUE | 0.1686 |
| Cflar | 399 | 261 | 277 | 305 | 218 | 202 | 424 | 261 | 278 | 240 | 235 | 221 | 276.34 | 321 | 232 | 0.723 | -0.467 | 0.0226379 | 0.0949083 | 0.0343 | 0.0188 | 0.0266 | 0.0266 | TRUE | 0.8886 |
| Cfp | 64 | 62 | 57 | 72 | 58 | 56 | 68 | 62 | 57 | 57 | 62 | 61 | 61.24 | 62 | 60 | 0.962 | -0.056 | 0.81889582 | 0.90748106 | 0 | 0.0607 | 0.0268 | 0.0268 | TRUE | 0.0523 |
| Cftr | 6 | 11 | 11 | 38 | 31 | 10 | 6 | 11 | 11 | 30 | 33 | 11 | 17.1 | 9 | 25 | 2.617 | 1.388 | 0.01818849 | 0.08124294 | 0.1492 | 0.1996 | 0.1752 | 0.1752 | TRUE | 1.3381 |
| Cga | 7 | 4 | 4 | 5 | 3 | 3 | 7 | 4 | 4 | 4 | 3 | 3 | 4.31 | 5 | 3 | 0.682 | -0.551 | 0.53752797 | NA | 0 | 0.7704 | 0.3372 | 0.3372 | TRUE | 0.297 |
| Cggbp1 | 6733 | 6597 | 6492 | 8259 | 5861 | 5905 | 7152 | 6588 | 6519 | 6493 | 6306 | 6464 | 6586.9 | 6753 | 6421 | 0.951 | -0.073 | 0.29052692 | 0.5114334 | 0.0012 | 0.0074 | 0.0032 | 0.0032 | TRUE | 0.0325 |
| Cgn | 3171 | 2619 | 2543 | 2786 | 2075 | 2071 | 3368 | 2616 | 2553 | 2190 | 2233 | 2267 | 2537.83 | 2846 | 2230 | 0.784 | -0.352 | 0.00379189 | 0.02533246 | 0.0122 | 0.0082 | 0.0102 | 0.0102 | TRUE | 0.3127 |
| Cgnl1 | 2042 | 1205 | 1314 | 1632 | 1164 | 964 | 2169 | 1203 | 1319 | 1283 | 1252 | 1055 | 1380.4 | 1564 | 1197 | 0.766 | -0.385 | 0.06616143 | 0.19842139 | 0.061 | 0.0093 | 0.031 | 0.031 | TRUE | 1.379 |
| Cgref1 | 307 | 512 | 448 | 562 | 373 | 405 | 326 | 511 | 450 | 442 | 401 | 443 | 428.95 | 429 | 429 | 0.999 | -0.001 | 0.99433662 | 0.99705494 | 0.0247 | 0.0146 | 0.0197 | 0.0197 | TRUE | 0.5061 |
| Cgrrf1 | 251 | 359 | 306 | 260 | 194 | 258 | 267 | 359 | 307 | 204 | 209 | 282 | 271.32 | 311 | 232 | 0.744 | -0.427 | 0.02243384 | 0.09432859 | 0.0232 | 0.019 | 0.0214 | 0.0214 | TRUE | 0.376 |
| Ch25h | 3 | 3 | 0 | 3 | 1 | 3 | 3 | 3 | 0 | 2 | 1 | 3 | 2.15 | 2 | 2 | 1.091 | 0.126 | 0.92768193 | NA | 0 | 1.5391 | 0.9196 | 0.9196 | TRUE | 0.7176 |
| Chac1 | 42 | 46 | 52 | 119 | 84 | 48 | 45 | 46 | 52 | 94 | 90 | 53 | 63.21 | 48 | 79 | 1.663 | 0.734 | 0.01544263 | 0.07233773 | 0.0392 | 0.059 | 0.0492 | 0.0492 | TRUE | 0.7523 |
| Chac2 | 435 | 348 | 338 | 372 | 258 | 247 | 462 | 348 | 339 | 292 | 278 | 270 | 331.57 | 383 | 280 | 0.733 | -0.449 | 0.00475781 | 0.03017855 | 0.0137 | 0.0168 | 0.0152 | 0.0152 | TRUE | 0.3717 |
| Chad | 2 | 2 | 3 | 7 | 5 | 5 | 2 | 2 | 3 | 6 | 5 | 5 | 3.91 | 2 | 5 | 2.291 | 1.196 | 0.20866963 | NA | 0 | 0.8484 | 0.3481 | 0.3481 | TRUE | 0.0583 |
| Chadl | 29 | 59 | 50 | 81 | 44 | 66 | 31 | 59 | 50 | 64 | 47 | 72 | 53.87 | 47 | 61 | 1.308 | 0.388 | 0.24184094 | 0.45850647 | 0.0508 | 0.0681 | 0.0601 | 0.0601 | TRUE | 0.6799 |
| Chaf1a | 5874 | 5057 | 4998 | 8054 | 5871 | 5086 | 6239 | 5050 | 5019 | 6331 | 6317 | 5567 | 5753.98 | 5436 | 6072 | 1.117 | 0.16 | 0.15826736 | 0.35207233 | 0.0105 | 0.0075 | 0.0091 | 0.0091 | TRUE | 0.2035 |
| Chaf1b | 3492 | 3838 | 3950 | 5473 | 4173 | 3884 | 3709 | 3833 | 3966 | 4302 | 4490 | 4252 | 4092.06 | 3836 | 4348 | 1.133 | 0.181 | 0.00683807 | 0.03946215 | 6.00E-04 | 0.0077 | 0.003 | 0.003 | TRUE | 0.0107 |
| Chat | 3 | 4 | 6 | 4 | 3 | 2 | 3 | 4 | 6 | 3 | 3 | 2 | 3.63 | 4 | 3 | 0.651 | -0.62 | 0.52183357 | NA | 0 | 0.915 | 0.3893 | 0.3893 | TRUE | 0.1901 |
| Chchd1 | 1539 | 1919 | 1884 | 2444 | 1601 | 1612 | 1635 | 1917 | 1892 | 1921 | 1723 | 1765 | 1808.57 | 1815 | 1803 | 0.994 | -0.009 | 0.93011389 | 0.96557628 | 0.0049 | 0.0087 | 0.0065 | 0.0065 | TRUE | 0.0903 |
| Chchd10 | 17344 | 22847 | 24198 | 17284 | 13510 | 14584 | 18422 | 22818 | 24297 | 13587 | 14536 | 15964 | 18270.83 | 21846 | 14696 | 0.673 | -0.572 | 2.51E-06 | 6.14E-05 | 0.013 | 0.0071 | 0.0106 | 0.0106 | TRUE | 0.2299 |
| Chchd2 | 8477 | 8245 | 7702 | 10747 | 7607 | 7292 | 9004 | 8234 | 7734 | 8448 | 8185 | 7982 | 8264.57 | 8324 | 8205 | 0.986 | -0.021 | 0.80100492 | 0.89732577 | 0.0027 | 0.0073 | 0.0047 | 0.0047 | TRUE | 0.0623 |
| Chchd3 | 5283 | 5750 | 5724 | 6852 | 4888 | 4635 | 5611 | 5743 | 5748 | 5387 | 5259 | 5074 | 5470.16 | 5701 | 5240 | 0.919 | -0.121 | 0.0468349 | 0.15688648 | 4.00E-04 | 0.0075 | 0.0025 | 0.0025 | TRUE | 0.0093 |
| Chchd4 | 2784 | 3140 | 2986 | 3361 | 2465 | 2596 | 2957 | 3136 | 2998 | 2642 | 2652 | 2842 | 2871.23 | 3030 | 2712 | 0.895 | -0.161 | 0.0254848 | 0.10303752 | 8.00E-04 | 0.0081 | 0.0034 | 0.0034 | TRUE | 0.021 |
| Chchd5 | 829 | 1214 | 1218 | 1425 | 992 | 1174 | 881 | 1212 | 1223 | 1120 | 1067 | 1285 | 1131.44 | 1105 | 1157 | 1.047 | 0.066 | 0.65942114 | 0.80811113 | 0.0204 | 0.0098 | 0.0153 | 0.0153 | TRUE | 0.3776 |
| Chchd6 | 778 | 968 | 993 | 1189 | 918 | 888 | 826 | 967 | 997 | 935 | 988 | 972 | 947.44 | 930 | 965 | 1.037 | 0.052 | 0.61937263 | 0.78476423 | 0.0042 | 0.0104 | 0.0068 | 0.0068 | TRUE | 0.1126 |
| Chchd7 | 706 | 794 | 714 | 889 | 637 | 632 | 750 | 793 | 717 | 699 | 685 | 692 | 722.64 | 753 | 692 | 0.919 | -0.122 | 0.18427763 | 0.38709544 | 0 | 0.0115 | 0.0047 | 0.0047 | TRUE | 0.0254 |
| Chd1 | 7368 | 6533 | 7068 | 10321 | 7237 | 6977 | 7826 | 6525 | 7097 | 8114 | 7787 | 7637 | 7497.54 | 7149 | 7846 | 1.098 | 0.134 | 0.1310522 | 0.31166855 | 0.0044 | 0.0073 | 0.0056 | 0.0056 | TRUE | 0.0837 |
| Chd1l | 3364 | 3720 | 3830 | 4252 | 3337 | 3181 | 3573 | 3715 | 3846 | 3343 | 3590 | 3482 | 3591.53 | 3711 | 3472 | 0.935 | -0.097 | 0.1698743 | 0.36848541 | 9.00E-04 | 0.0078 | 0.0033 | 0.0033 | TRUE | 0.0131 |
| Chd2 | 5894 | 5880 | 7263 | 5883 | 4182 | 4973 | 6260 | 5872 | 7293 | 4625 | 4500 | 5444 | 5665.63 | 6475 | 4856 | 0.75 | -0.415 | 0.00038897 | 0.00423965 | 0.0117 | 0.0075 | 0.0097 | 0.0097 | TRUE | 0.1489 |
| Chd3 | 2036 | 1927 | 2097 | 2859 | 2128 | 2135 | 2163 | 1925 | 2106 | 2248 | 2290 | 2337 | 2177.82 | 2065 | 2292 | 1.11 | 0.151 | 0.05684703 | 0.17939001 | 0.0013 | 0.0084 | 0.004 | 0.004 | TRUE | 0.0425 |
| Chd4 | 43818 | 34493 | 37322 | 50094 | 35623 | 32851 | 46542 | 34448 | 37475 | 39380 | 38328 | 35961 | 38689.15 | 39488 | 37890 | 0.96 | -0.06 | 0.62085858 | 0.78551111 | 0.0119 | 0.007 | 0.0104 | 0.0104 | TRUE | 0.2989 |
| Chd5 | 628 | 727 | 643 | 525 | 418 | 424 | 667 | 726 | 646 | 413 | 450 | 464 | 560.89 | 680 | 442 | 0.649 | -0.623 | 1.09E-08 | 5.11E-07 | 0.0021 | 0.0128 | 0.0067 | 0.0067 | TRUE | 0.0425 |
| Chd6 | 3773 | 3749 | 4077 | 4173 | 3099 | 3254 | 4008 | 3744 | 4094 | 3280 | 3334 | 3562 | 3670.39 | 3949 | 3392 | 0.859 | -0.219 | 0.00379216 | 0.02533246 | 0.0016 | 0.0078 | 0.0039 | 0.0039 | TRUE | 0.025 |
| Chd7 | 4528 | 3961 | 4558 | 4568 | 3252 | 3003 | 4810 | 3956 | 4577 | 3591 | 3499 | 3287 | 3953.22 | 4448 | 3459 | 0.778 | -0.362 | 0.0002087 | 0.00252229 | 0.0056 | 0.0077 | 0.0066 | 0.0066 | TRUE | 0.1139 |
| Chd8 | 7645 | 6358 | 6887 | 9746 | 6665 | 5853 | 8120 | 6350 | 6915 | 7662 | 7171 | 6407 | 7104.18 | 7128 | 7080 | 0.993 | -0.01 | 0.93469161 | 0.9675739 | 0.0118 | 0.0074 | 0.0098 | 0.0098 | TRUE | 0.1807 |
| Chd9 | 10493 | 9849 | 11903 | 9642 | 6937 | 7164 | 11145 | 9836 | 11952 | 7580 | 7464 | 7842 | 9303.21 | 10978 | 7629 | 0.695 | -0.525 | 9.08E-09 | 4.32E-07 | 0.0049 | 0.0073 | 0.0059 | 0.0059 | TRUE | 0.1012 |
| Chdh | 9 | 9 | 6 | 26 | 29 | 12 | 10 | 9 | 6 | 20 | 31 | 13 | 14.89 | 8 | 21 | 2.638 | 1.399 | 0.01436156 | 0.06853571 | 0 | 0.2281 | 0.1503 | 0.1503 | TRUE | 0.1508 |
| Chek1 | 3276 | 3811 | 3820 | 4410 | 3305 | 3608 | 3480 | 3806 | 3836 | 3467 | 3556 | 3950 | 3682.29 | 3707 | 3658 | 0.986 | -0.02 | 0.81824855 | 0.90726388 | 0.0036 | 0.0078 | 0.0052 | 0.0052 | TRUE | 0.0591 |
| Chek2 | 3550 | 3749 | 3698 | 3673 | 2663 | 2822 | 3771 | 3744 | 3713 | 2887 | 2865 | 3089 | 3344.97 | 3743 | 2947 | 0.787 | -0.345 | 2.78E-07 | 8.87E-06 | 5.00E-04 | 0.0079 | 0.0029 | 0.0029 | TRUE | 0.0213 |
| Cherp | 10096 | 8761 | 8932 | 13752 | 10044 | 9060 | 10724 | 8750 | 8969 | 10811 | 10807 | 9918 | 9996.19 | 9481 | 10512 | 1.109 | 0.149 | 0.14181348 | 0.32738322 | 0.0076 | 0.0072 | 0.0073 | 0.0073 | TRUE | 0.1606 |
| Chfr | 2900 | 3138 | 3196 | 4628 | 3221 | 3204 | 3080 | 3134 | 3209 | 3638 | 3466 | 3507 | 3339.07 | 3141 | 3537 | 1.126 | 0.172 | 0.00745252 | 0.0420743 | 3.00E-04 | 0.0079 | 0.0027 | 0.0027 | TRUE | 0.0078 |

| | | | | | | | | | | | | | | | | | | | | | | | | | |
|---|---|---|---|---|---|---|---|---|---|---|---|---|---|---|---|---|---|---|---|---|---|---|---|---|---|
| Chga | 1106 | 1166 | 1166 | 874 | 668 | 704 | 1175 | 1164 | 1171 | 687 | 719 | 771 | 947.75 | 1170 | 726 | 0.619 | -0.691 | 3.28E-15 | 4.57E-13 | 0 | 0.0104 | 0.0044 | 0.0044 | TRUE | 0.0347 |
| Chgb | 14 | 6 | 17 | 15 | 13 | 11 | 15 | 6 | 17 | 12 | 14 | 12 | 12.63 | 13 | 13 | 0.996 | -0.005 | 0.99287941 | 0.99616826 | 0 | 0.2678 | 0.1644 | 0.1644 | TRUE | 0.8822 |
| Chi3l1 | 65 | 95 | 80 | 41 | 36 | 29 | 69 | 95 | 80 | 32 | 39 | 32 | 57.83 | 81 | 34 | 0.419 | -1.255 | 6.46E-06 | 0.00013794 | 0 | 0.0639 | 0.0352 | 0.0352 | TRUE | 0.1411 |
| Chi3l3 | 2 | 1 | 1 | 2 | 2 | 3 | 2 | 1 | 1 | 2 | 2 | 3 | 1.86 | 1 | 2 | 1.685 | 0.752 | 0.57714393 | NA | 0 | 1.782 | 0.7273 | 0.7273 | TRUE | 0.1348 |
| Chi3l4 | 1 | 2 | 0 | 1 | 0 | 1 | 1 | 2 | 0 | 1 | 0 | 1 | 0.82 | 1 | 1 | 0.62 | -0.689 | 0.75103894 | NA | 0 | 4.008 | 2.1286 | 2.1286 | TRUE | 0.1579 |
| Chia | 0 | 0 | 0 | 0 | 0 | 0 | 0 | 0 | 0 | 0 | 0 | 0 | 0 | 0 | 0 | NA | NA | NA | NA | NA | NA | NA | NA | NA | NA |
| Chic1 | 967 | 1040 | 1089 | 1389 | 983 | 1100 | 1027 | 1039 | 1093 | 1092 | 1058 | 1204 | 1085.49 | 1053 | 1118 | 1.061 | 0.085 | 0.35037936 | 0.5692556 | 0.0019 | 0.0099 | 0.0051 | 0.0051 | TRUE | 0.0533 |
| Chic2 | 1058 | 1116 | 1098 | 1562 | 1106 | 968 | 1124 | 1115 | 1103 | 1228 | 1190 | 1060 | 1136.4 | 1114 | 1159 | 1.042 | 0.059 | 0.51913147 | 0.71037146 | 0.0022 | 0.0098 | 0.0052 | 0.0052 | TRUE | 0.0667 |
| Chid1 | 679 | 906 | 926 | 927 | 677 | 610 | 721 | 905 | 930 | 729 | 728 | 668 | 780.12 | 852 | 708 | 0.831 | -0.266 | 0.0348571 | 0.12780763 | 0.0094 | 0.0111 | 0.0102 | 0.0102 | TRUE | 0.213 |
| Chit1 | 27 | 19 | 19 | 24 | 24 | 23 | 29 | 19 | 19 | 19 | 26 | 25 | 22.77 | 22 | 23 | 1.042 | 0.059 | 0.88668186 | 0.94438861 | 0 | 0.1516 | 0.0806 | 0.0806 | TRUE | 0.3536 |
| Chka | 2723 | 1980 | 2166 | 2383 | 1911 | 1960 | 2892 | 1977 | 2175 | 1873 | 2056 | 2146 | 2186.6 | 2348 | 2025 | 0.862 | -0.214 | 0.14905288 | 0.33932765 | 0.0227 | 0.0084 | 0.0154 | 0.0154 | TRUE | 0.4973 |
| Chkb | 11 | 15 | 17 | 17 | 15 | 12 | 12 | 15 | 17 | 13 | 16 | 13 | 14.4 | 15 | 14 | 0.971 | -0.042 | 0.93215526 | 0.96628017 | 0 | 0.2358 | 0.106 | 0.106 | TRUE | 0.1286 |
| Chl1 | 8 | 5 | 3 | 18 | 2 | 5 | 8 | 5 | 3 | 14 | 2 | 5 | 6.38 | 5 | 7 | 1.353 | 0.436 | 0.6329918 | 0.79357325 | 0.3653 | 0.5233 | 0.4425 | 0.4425 | TRUE | 0.6754 |
| Chm | 1752 | 1520 | 1559 | 2118 | 1552 | 1435 | 1861 | 1518 | 1565 | 1665 | 1670 | 1571 | 1641.68 | 1648 | 1635 | 0.993 | -0.011 | 0.91763673 | 0.95968215 | 0.0061 | 0.0089 | 0.0073 | 0.0073 | TRUE | 0.1534 |
| Chml | 1161 | 1170 | 1200 | 1427 | 1116 | 1152 | 1233 | 1168 | 1205 | 1122 | 1201 | 1261 | 1198.37 | 1202 | 1195 | 0.993 | -0.011 | 0.90281543 | 0.95205059 | 0.0014 | 0.0097 | 0.0045 | 0.0045 | TRUE | 0.0353 |
| Chmp1a | 5533 | 5312 | 5023 | 8245 | 5903 | 5427 | 5877 | 5305 | 5044 | 6482 | 6351 | 5941 | 5833.22 | 5409 | 6258 | 1.157 | 0.211 | 0.01505216 | 0.0709936 | 0.0039 | 0.0075 | 0.0052 | 0.0052 | TRUE | 0.0699 |
| Chmp1b | 863 | 811 | 800 | 1506 | 1048 | 945 | 917 | 810 | 803 | 1184 | 1128 | 1034 | 979.3 | 843 | 1115 | 1.324 | 0.405 | 8.87E-05 | 0.00125888 | 0.004 | 0.0103 | 0.0066 | 0.0066 | TRUE | 0.0684 |
| Chmp2a | 3510 | 4716 | 4504 | 4472 | 3378 | 3886 | 3728 | 4710 | 4522 | 3516 | 3635 | 4254 | 4060.76 | 4320 | 3802 | 0.88 | -0.185 | 0.12810321 | 0.30785004 | 0.0123 | 0.0077 | 0.0104 | 0.0104 | TRUE | 0.1748 |
| Chmp2b | 1371 | 1503 | 1504 | 2043 | 1395 | 1456 | 1456 | 1501 | 1510 | 1606 | 1501 | 1594 | 1528.05 | 1489 | 1567 | 1.053 | 0.074 | 0.32555521 | 0.54676357 | 0 | 0.0091 | 0.0034 | 0.0034 | TRUE | 0.0162 |
| Chmp3 | 4763 | 4628 | 4478 | 6005 | 4122 | 4104 | 5059 | 4622 | 4496 | 4721 | 4435 | 4492 | 4637.62 | 4726 | 4549 | 0.963 | -0.055 | 0.48293481 | 0.68129391 | 0.0022 | 0.0076 | 0.0041 | 0.0041 | TRUE | 0.0463 |
| Chmp4b | 3933 | 4040 | 4023 | 6015 | 4458 | 4164 | 4178 | 4035 | 4040 | 4729 | 4797 | 4558 | 4389.18 | 4084 | 4695 | 1.15 | 0.201 | 0.00121827 | 0.0103694 | 3.00E-04 | 0.0077 | 0.0026 | 0.0026 | TRUE | 0.0078 |
| Chmp4c | 861 | 1384 | 1446 | 1371 | 1103 | 1457 | 915 | 1382 | 1452 | 1078 | 1187 | 1595 | 1268.02 | 1250 | 1287 | 1.029 | 0.041 | 0.83830744 | 0.91786582 | 0.0505 | 0.0095 | 0.0285 | 0.0285 | TRUE | 0.659 |
| Chmp5 | 2499 | 2841 | 2751 | 3568 | 2531 | 2616 | 2654 | 2837 | 2762 | 2805 | 2723 | 2864 | 2774.29 | 2751 | 2797 | 1.017 | 0.024 | 0.72881645 | 0.85409762 | 5.00E-04 | 0.0081 | 0.003 | 0.003 | TRUE | 0.0115 |
| Chmp6 | 1817 | 1969 | 1976 | 2883 | 2035 | 1914 | 1930 | 1966 | 1984 | 2266 | 2190 | 2095 | 2071.94 | 1960 | 2184 | 1.115 | 0.156 | 0.02879532 | 0.11183447 | 5.00E-04 | 0.0085 | 0.0032 | 0.0032 | TRUE | 0.0152 |
| Chmp7 | 2671 | 2988 | 3013 | 4234 | 3039 | 2923 | 2837 | 2984 | 3025 | 3328 | 3270 | 3200 | 3107.41 | 2949 | 3266 | 1.108 | 0.148 | 0.02699293 | 0.10720009 | 4.00E-04 | 0.008 | 0.0029 | 0.0029 | TRUE | 0.0133 |
| Chn1 | 984 | 1311 | 1304 | 1605 | 1272 | 1331 | 1045 | 1309 | 1309 | 1262 | 1369 | 1457 | 1291.86 | 1221 | 1363 | 1.115 | 0.157 | 0.19413323 | 0.39972184 | 0.0092 | 0.0095 | 0.0097 | 0.0097 | TRUE | 0.1901 |
| Chn2 | 174 | 170 | 181 | 210 | 112 | 119 | 185 | 170 | 182 | 165 | 121 | 130 | 158.7 | 179 | 139 | 0.78 | -0.359 | 0.05096613 | 0.16620787 | 0.0086 | 0.0277 | 0.018 | 0.018 | TRUE | 0.3139 |
| Chodl | 0 | 0 | 0 | 0 | 1 | 0 | 0 | 0 | 0 | 0 | 1 | 0 | 0.18 | 0 | 0 | 1.859 | 0.894 | 0.82650734 | NA | 0 | 18.3785 | 10 | 10 | TRUE | 0.5204 |
| Chordc1 | 2949 | 2505 | 2429 | 4552 | 3253 | 2791 | 3132 | 2502 | 2439 | 3578 | 3500 | 3055 | 3034.46 | 2691 | 3378 | 1.256 | 0.328 | 0.00788654 | 0.04384439 | 0.0131 | 0.008 | 0.0107 | 0.0107 | TRUE | 0.2494 |
| Chp1 | 2907 | 3217 | 3200 | 4202 | 2766 | 3014 | 3088 | 3213 | 3213 | 3303 | 2976 | 3299 | 3182.06 | 3171 | 3193 | 1.007 | 0.01 | 0.89498099 | 0.94926968 | 0.0017 | 0.0079 | 0.0039 | 0.0039 | TRUE | 0.0426 |
| Chp2 | 4 | 2 | 3 | 6 | 6 | 3 | 4 | 2 | 3 | 5 | 6 | 3 | 3.95 | 3 | 5 | 1.567 | 0.648 | 0.49323205 | NA | 0 | 0.8404 | 0.3769 | 0.3769 | TRUE | 0.1537 |
| Chpf | 1841 | 1813 | 1781 | 2179 | 1603 | 1510 | 1955 | 1811 | 1788 | 1713 | 1725 | 1653 | 1774.18 | 1851 | 1697 | 0.917 | -0.125 | 0.10415522 | 0.26850868 | 9.00E-04 | 0.0088 | 0.0037 | 0.0037 | TRUE | 0.029 |
| Chpf2 | 212 | 229 | 224 | 272 | 214 | 169 | 225 | 229 | 225 | 214 | 230 | 185 | 217.98 | 226 | 210 | 0.927 | -0.109 | 0.46321399 | 0.6665886 | 0 | 0.022 | 0.0112 | 0.0112 | TRUE | 0.1088 |
| Chpt1 | 679 | 691 | 636 | 1040 | 766 | 661 | 721 | 690 | 639 | 818 | 824 | 724 | 735.87 | 683 | 789 | 1.155 | 0.208 | 0.04846842 | 0.16053487 | 0.003 | 0.0114 | 0.0066 | 0.0066 | TRUE | 0.0601 |
| Chrac1 | 997 | 920 | 857 | 1173 | 869 | 891 | 1059 | 919 | 861 | 922 | 935 | 975 | 945.13 | 946 | 944 | 0.998 | -0.003 | 0.97767452 | 0.98797393 | 0.005 | 0.0104 | 0.0073 | 0.0073 | TRUE | 0.1292 |
| Chrd | 188 | 172 | 180 | 162 | 136 | 134 | 200 | 172 | 181 | 127 | 146 | 147 | 162.09 | 184 | 140 | 0.759 | -0.398 | 0.01670574 | 0.07648271 | 0 | 0.0272 | 0.0136 | 0.0136 | TRUE | 0.0736 |
| Chrdl1 | 50 | 42 | 38 | 62 | 54 | 30 | 53 | 42 | 38 | 49 | 58 | 33 | 45.48 | 44 | 47 | 1.052 | 0.073 | 0.82538071 | 0.91142649 | 0.0295 | 0.0793 | 0.056 | 0.056 | TRUE | 0.3511 |
| Chrdl2 | 15 | 9 | 9 | 17 | 15 | 11 | 16 | 9 | 9 | 13 | 16 | 12 | 12.58 | 11 | 14 | 1.226 | 0.294 | 0.59142898 | 0.76507383 | 0 | 0.2687 | 0.1358 | 0.1358 | TRUE | 0.46 |
| Chrm1 | 0 | 0 | 0 | 0 | 0 | 0 | 0 | 0 | 0 | 0 | 0 | 0 | 0 | 0 | 0 | NA | NA | NA | NA | NA | NA | NA | NA | NA | NA |
| Chrm2 | 0 | 0 | 0 | 0 | 0 | 0 | 0 | 0 | 0 | 0 | 0 | 0 | 0 | 0 | 0 | NA | NA | NA | NA | NA | NA | NA | NA | NA | NA |
| Chrm3 | 44 | 26 | 17 | 103 | 82 | 40 | 47 | 26 | 17 | 81 | 88 | 44 | 50.46 | 30 | 71 | 2.383 | 1.253 | 0.00378788 | 0.0253253 | 0.1579 | 0.0722 | 0.1109 | 0.1109 | TRUE | 1.4881 |
| Chrm4 | 340 | 364 | 416 | 508 | 392 | 382 | 361 | 364 | 418 | 399 | 422 | 418 | 396.94 | 381 | 413 | 1.084 | 0.116 | 0.32816707 | 0.54918701 | 0.0014 | 0.0152 | 0.0076 | 0.0076 | TRUE | 0.083 |
| Chrm5 | 0 | 0 | 0 | 0 | 1 | 0 | 0 | 0 | 0 | 0 | 1 | 0 | 0.18 | 0 | 0 | 1.859 | 0.894 | 0.82650734 | NA | 0 | 18.3785 | 10 | 10 | TRUE | 0.5204 |
| Chrna1 | 1 | 3 | 0 | 2 | 3 | 2 | 1 | 3 | 0 | 2 | 3 | 2 | 1.84 | 1 | 2 | 1.701 | 0.766 | 0.60306066 | NA | 0 | 1.7961 | 0.975 | 0.975 | TRUE | 0.6407 |
| Chrna10 | 5 | 7 | 8 | 13 | 10 | 7 | 5 | 7 | 8 | 10 | 11 | 8 | 8.16 | 7 | 10 | 1.41 | 0.496 | 0.44730277 | 0.65333562 | 0 | 0.4105 | 0.1806 | 0.1806 | TRUE | 0.0893 |
| Chrna2 | 5 | 0 | 1 | 1 | 0 | 3 | 5 | 0 | 1 | 1 | 0 | 3 | 1.73 | 2 | 1 | 0.639 | -0.646 | 0.72646303 | NA | 1.7521 | 1.9103 | 1.8443 | 1.8443 | TRUE | 1.4171 |
| Chrna3 | 3 | 0 | 1 | 2 | 0 | 0 | 3 | 0 | 1 | 2 | 0 | 0 | 0.96 | 1 | 1 | 0.407 | -1.296 | 0.5860077 | NA | 0 | 3.4369 | 2.8485 | 2.8485 | TRUE | 0.5389 |
| Chrna4 | 186 | 243 | 239 | 193 | 152 | 165 | 198 | 243 | 240 | 152 | 164 | 181 | 196.02 | 227 | 166 | 0.727 | -0.461 | 0.00516099 | 0.03198394 | 0.0055 | 0.0237 | 0.0143 | 0.0143 | TRUE | 0.1375 |
| Chrna5 | 108 | 107 | 113 | 139 | 110 | 93 | 115 | 107 | 113 | 109 | 118 | 102 | 110.74 | 112 | 110 | 0.983 | -0.024 | 0.89723922 | 0.95016592 | 0 | 0.0367 | 0.0161 | 0.0161 | TRUE | 0.0426 |
| Chrna6 | 0 | 0 | 0 | 1 | 0 | 0 | 0 | 0 | 0 | 1 | 0 | 0 | 0.13 | 0 | 0 | 1.859 | 0.894 | 0.82650734 | NA | 0 | 25.1515 | 10 | 10 | TRUE | 0.2563 |
| Chrna7 | 215 | 221 | 251 | 328 | 268 | 259 | 228 | 221 | 252 | 258 | 288 | 284 | 255.14 | 234 | 277 | 1.181 | 0.24 | 0.0805131 | 0.22559116 | 0 | 0.0198 | 0.0096 | 0.0096 | TRUE | 0.0525 |
| Chrna9 | 151 | 107 | 122 | 214 | 144 | 128 | 160 | 107 | 123 | 168 | 155 | 140 | 142.17 | 130 | 154 | 1.192 | 0.254 | 0.22197973 | 0.43421675 | 0.0178 | 0.0301 | 0.024 | 0.024 | TRUE | 0.425 |
| Chrnb1 | 101 | 194 | 209 | 137 | 135 | 121 | 107 | 194 | 210 | 108 | 145 | 132 | 149.38 | 170 | 128 | 0.752 | -0.411 | 0.13097652 | 0.31162492 | 0.0657 | 0.029 | 0.0466 | 0.0466 | TRUE | 1.1038 |
| Chrnb2 | 47 | 50 | 41 | 40 | 34 | 26 | 50 | 50 | 41 | 31 | 37 | 28 | 39.59 | 47 | 32 | 0.684 | -0.548 | 0.07563866 | 0.21682534 | 0 | 0.0901 | 0.0425 | 0.0425 | TRUE | 0.0949 |
| Chrnb3 | 6 | 2 | 2 | 5 | 3 | 4 | 6 | 2 | 2 | 4 | 3 | 4 | 3.65 | 3 | 4 | 1.12 | 0.163 | 0.87108532 | NA | 0 | 0.9088 | 0.4508 | 0.4508 | TRUE | 0.76 |
| Chrnb4 | 332 | 339 | 371 | 332 | 227 | 265 | 353 | 339 | 373 | 261 | 244 | 290 | 309.84 | 355 | 265 | 0.747 | -0.42 | 0.00123289 | 0.01046004 | 0 | 0.0175 | 0.0089 | 0.0089 | TRUE | 0.0716 |
| Chrnd | 8 | 24 | 16 | 16 | 10 | 8 | 8 | 24 | 16 | 13 | 11 | 9 | 13.44 | 16 | 11 | 0.664 | -0.591 | 0.30961542 | 0.52992476 | 0.0775 | 0.2521 | 0.1668 | 0.1668 | TRUE | 0.1366 |
| Chrne | 7 | 8 | 9 | 10 | 8 | 11 | 7 | 8 | 9 | 8 | 9 | 12 | 8.83 | 8 | 10 | 1.156 | 0.209 | 0.73780856 | 0.85962301 | 0 | 0.3801 | 0.1671 | 0.1671 | TRUE | 0.1682 |
| Chrng | 7 | 15 | 5 | 15 | 7 | 9 | 7 | 15 | 5 | 12 | 8 | 10 | 9.44 | 9 | 10 | 1.069 | 0.096 | 0.88613486 | 0.94407721 | 0.071 | 0.3561 | 0.219 | 0.219 | TRUE | 1.0316 |
| Chst1 | 51 | 40 | 36 | 49 | 38 | 60 | 54 | 40 | 36 | 39 | 41 | 66 | 45.89 | 43 | 49 | 1.109 | 0.149 | 0.66384662 | 0.81170547 | 0.0468 | 0.0787 | 0.063 | 0.063 | TRUE | 0.7558 |
| Chst10 | 360 | 460 | 438 | 480 | 366 | 359 | 382 | 459 | 440 | 377 | 394 | 393 | 407.62 | 427 | 388 | 0.907 | -0.14 | 0.2456226 | 0.46267284 | 0.0024 | 0.015 | 0.0081 | 0.0081 | TRUE | 0.0962 |
| Chst11 | 1000 | 1120 | 1006 | 1058 | 709 | 862 | 1062 | 1119 | 1010 | 832 | 763 | 944 | 954.84 | 1064 | 846 | 0.795 | -0.331 | 0.00274477 | 0.0197854 | 0.0055 | 0.0104 | 0.0077 | 0.0077 | TRUE | 0.1179 |
| Chst12 | 796 | 809 | 803 | 790 | 552 | 646 | 845 | 808 | 806 | 621 | 594 | 707 | 730.31 | 820 | 641 | 0.781 | -0.357 | 0.00065534 | 0.00643468 | 0.0025 | 0.0114 | 0.0065 | 0.0065 | TRUE | 0.094 |
| Chst13 | 43 | 56 | 35 | 58 | 54 | 65 | 46 | 56 | 35 | 46 | 58 | 71 | 51.93 | 46 | 58 | 1.271 | 0.346 | 0.27209497 | 0.49206309 | 0.0327 | 0.0703 | 0.0519 | 0.0519 | TRUE | 0.1676 |
| Chst14 | 120 | 137 | 165 | 236 | 174 | 206 | 127 | 137 | 166 | 186 | 187 | 225 | 171.37 | 143 | 199 | 1.388 | 0.473 | 0.0089985 | 0.04865243 | 0.0092 | 0.0261 | 0.0176 | 0.0176 | TRUE | 0.1944 |
| Chst15 | 86 | 86 | 67 | 120 | 87 | 71 | 91 | 86 | 67 | 94 | 94 | 78 | 85.03 | 81 | 89 | 1.09 | 0.124 | 0.59046307 | 0.76459635 | 0 | 0.0456 | 0.0264 | 0.0264 | TRUE | 0.2194 |
| Chst2 | 83 | 154 | 160 | 294 | 192 | 173 | 88 | 154 | 161 | 231 | 207 | 189 | 171.62 | 134 | 209 | 1.558 | 0.639 | 0.0078055 | 0.04356275 | 0.0464 | 0.0261 | 0.0355 | 0.0355 | TRUE | 0.9128 |
| Chst3 | 171 | 130 | 127 | 147 | 118 | 117 | 182 | 130 | 128 | 116 | 127 | 128 | 134.93 | 147 | 124 | 0.844 | -0.245 | 0.23582664 | 0.45107164 | 0.0157 | 0.0313 | 0.0234 | 0.0234 | TRUE | 0.4586 |
| Chst4 | 0 | 1 | 0 | 1 | 0 | 0 | 0 | 1 | 0 | 1 | 0 | 0 | 0.3 | 0 | 0 | 0.954 | -0.067 | 0.98642001 | NA | 0 | 11.0817 | 9.3014 | 9.3014 | TRUE | 0.405 |

| | | | | | | | | | | | | | | | | | | | | | | | | | |
|---|---|---|---|---|---|---|---|---|---|---|---|---|---|---|---|---|---|---|---|---|---|---|---|---|---|
| Chst5 | 67 | 77 | 67 | 106 | 101 | 66 | 71 | 77 | 67 | 83 | 109 | 72 | 79.93 | 72 | 88 | 1.226 | 0.293 | 0.23047517 | 0.44432886 | 0.0123 | 0.0481 | 0.0305 | 0.0305 | TRUE | 0.3773 |
| Chst7 | 112 | 117 | 116 | 144 | 89 | 113 | 119 | 117 | 116 | 113 | 96 | 124 | 114.16 | 117 | 111 | 0.945 | -0.082 | 0.66901234 | 0.81504759 | 0 | 0.0358 | 0.0177 | 0.0177 | TRUE | 0.1325 |
| Chst8 | 51 | 34 | 50 | 87 | 52 | 56 | 54 | 34 | 50 | 68 | 56 | 61 | 54 | 46 | 62 | 1.348 | 0.431 | 0.1376986 | 0.32185866 | 0.0102 | 0.0679 | 0.0416 | 0.0416 | TRUE | 0.4259 |
| Chst9 | 1 | 0 | 2 | 1 | 1 | 6 | 1 | 0 | 2 | 1 | 1 | 7 | 1.92 | 1 | 3 | 2.699 | 1.433 | 0.38094717 | NA | 0.7788 | 1.7257 | 1.2498 | 1.2498 | TRUE | 1.5552 |
| Chsy1 | 3002 | 3510 | 3326 | 3428 | 2588 | 2670 | 3189 | 3505 | 3340 | 2695 | 2785 | 2923 | 3072.64 | 3345 | 2801 | 0.837 | -0.257 | 0.00076305 | 0.00720645 | 0.0015 | 0.008 | 0.0039 | 0.0039 | TRUE | 0.0215 |
| Chsy3 | 36 | 43 | 38 | 66 | 44 | 28 | 38 | 43 | 38 | 52 | 47 | 31 | 41.54 | 40 | 43 | 1.095 | 0.131 | 0.68668259 | 0.82731325 | 0 | 0.0862 | 0.0521 | 0.0521 | TRUE | 0.4731 |
| Chtf18 | 3075 | 2982 | 2945 | 4766 | 3522 | 3133 | 3266 | 2978 | 2957 | 3747 | 3789 | 3430 | 3361.18 | 3067 | 3655 | 1.192 | 0.253 | 0.00213537 | 0.01620743 | 0.0026 | 0.0079 | 0.0046 | 0.0046 | TRUE | 0.039 |
| Chtf8 | 6971 | 7821 | 8115 | 10410 | 7600 | 7546 | 7404 | 7811 | 8148 | 8184 | 8177 | 8260 | 7997.43 | 7788 | 8207 | 1.054 | 0.076 | 0.2508787 | 0.46837817 | 0.001 | 0.0073 | 0.003 | 0.003 | TRUE | 0.0226 |
| Chtop | 7223 | 7729 | 7541 | 10420 | 7632 | 7152 | 7672 | 7719 | 7572 | 8191 | 8212 | 7829 | 7865.84 | 7654 | 8077 | 1.055 | 0.078 | 0.17907555 | 0.38024408 | 2.00E-04 | 0.0073 | 0.0023 | 0.0023 | TRUE | 0.0088 |
| Chuk | 3855 | 3078 | 2868 | 5048 | 3493 | 3031 | 4095 | 3074 | 2880 | 3968 | 3758 | 3318 | 3515.5 | 3350 | 3681 | 1.099 | 0.137 | 0.34780948 | 0.56662263 | 0.0197 | 0.0078 | 0.015 | 0.015 | TRUE | 0.4601 |
| Churc1 | 1102 | 1442 | 1331 | 1749 | 1214 | 1203 | 1171 | 1440 | 1336 | 1375 | 1306 | 1317 | 1324.19 | 1316 | 1333 | 1.013 | 0.019 | 0.8562241 | 0.928793 | 0.005 | 0.0094 | 0.0068 | 0.0068 | TRUE | 0.1114 |
| Ciao1 | 1968 | 2003 | 1875 | 2895 | 2009 | 1954 | 2090 | 2000 | 1883 | 2276 | 2162 | 2139 | 2091.64 | 1991 | 2192 | 1.101 | 0.139 | 0.0767839 | 0.21883549 | 0.0014 | 0.0085 | 0.004 | 0.004 | TRUE | 0.0275 |
| Ciapin1 | 11205 | 9823 | 9033 | 12304 | 8626 | 7543 | 11902 | 9810 | 9070 | 9672 | 9281 | 8257 | 9665.43 | 10261 | 9070 | 0.884 | -0.178 | 0.1422033 | 0.32793991 | 0.0133 | 0.0072 | 0.0105 | 0.0105 | TRUE | 0.2391 |
| Cib1 | 1262 | 1273 | 1299 | 1945 | 1384 | 1236 | 1340 | 1271 | 1304 | 1529 | 1489 | 1353 | 1381.21 | 1305 | 1457 | 1.117 | 0.16 | 0.06363124 | 0.19378103 | 0.0017 | 0.0093 | 0.0046 | 0.0046 | TRUE | 0.0464 |
| Cib2 | 157 | 198 | 182 | 182 | 130 | 147 | 167 | 198 | 183 | 143 | 140 | 161 | 165.19 | 183 | 148 | 0.81 | -0.305 | 0.06458115 | 0.19578237 | 0 | 0.0268 | 0.0135 | 0.0135 | TRUE | 0.0596 |
| Cib3 | 0 | 1 | 0 | 2 | 1 | 0 | 0 | 1 | 0 | 2 | 1 | 0 | 0.61 | 0 | 1 | 2.461 | 1.299 | 0.62286289 | NA | 0 | 5.4271 | 3.4871 | 3.4871 | TRUE | 0.405 |
| Cib4 | 2 | 2 | 0 | 0 | 0 | 0 | 2 | 2 | 0 | 0 | 0 | 0 | 0.69 | 1 | 0 | 0.131 | -2.936 | 0.31808931 | NA | 0 | 4.8026 | 4.861 | 4.861 | TRUE | 0.4881 |
| Cic | 7607 | 7288 | 7531 | 9547 | 7287 | 7046 | 8080 | 7279 | 7562 | 7505 | 7840 | 7713 | 7663.15 | 7640 | 7686 | 1.006 | 0.009 | 0.90242405 | 0.95205059 | 0.0015 | 0.0073 | 0.0034 | 0.0034 | TRUE | 0.0309 |
| Cidea | 115 | 75 | 83 | 206 | 89 | 77 | 122 | 75 | 83 | 162 | 96 | 84 | 103.73 | 93 | 114 | 1.227 | 0.295 | 0.34974246 | 0.56865085 | 0.0893 | 0.0387 | 0.0622 | 0.0622 | TRUE | 1.4354 |
| Cideb | 18 | 27 | 24 | 22 | 16 | 29 | 19 | 27 | 24 | 17 | 17 | 32 | 22.74 | 23 | 22 | 0.936 | -0.096 | 0.82759429 | 0.91271118 | 0.0357 | 0.1518 | 0.0962 | 0.0962 | TRUE | 0.7808 |
| Cidec | 17 | 10 | 7 | 40 | 25 | 18 | 18 | 10 | 7 | 31 | 27 | 20 | 18.85 | 12 | 26 | 2.246 | 1.168 | 0.02255045 | 0.09466745 | 0.0537 | 0.1816 | 0.1262 | 0.1262 | TRUE | 0.7033 |
| Ciita | 1 | 6 | 4 | 5 | 2 | 6 | 1 | 6 | 4 | 4 | 2 | 7 | 3.95 | 4 | 4 | 1.133 | 0.18 | 0.85980362 | NA | 0 | 0.8402 | 0.4964 | 0.4964 | TRUE | 0.342 |
| Cilp | 6 | 10 | 9 | 23 | 17 | 8 | 6 | 10 | 9 | 18 | 18 | 9 | 11.75 | 8 | 15 | 1.785 | 0.836 | 0.15990525 | 0.35445491 | 0 | 0.2872 | 0.1623 | 0.1623 | TRUE | 0.5563 |
| Cilp2 | 89 | 149 | 131 | 238 | 147 | 129 | 95 | 149 | 132 | 187 | 158 | 141 | 143.56 | 125 | 162 | 1.3 | 0.378 | 0.09098613 | 0.24600797 | 0.028 | 0.0299 | 0.029 | 0.029 | TRUE | 0.4536 |
| Cinp | 2769 | 2794 | 2746 | 3676 | 2608 | 2436 | 2941 | 2790 | 2757 | 2890 | 2806 | 2667 | 2808.54 | 2829 | 2788 | 0.986 | -0.021 | 0.77297381 | 0.88138685 | 0.0011 | 0.0081 | 0.0035 | 0.0035 | TRUE | 0.0174 |
| Cir1 | 545 | 533 | 599 | 720 | 594 | 576 | 579 | 532 | 601 | 566 | 639 | 631 | 591.38 | 571 | 612 | 1.07 | 0.098 | 0.36769656 | 0.58492091 | 0.0026 | 0.0125 | 0.0068 | 0.0068 | TRUE | 0.0532 |
| Cirbp | 2031 | 2248 | 2143 | 2876 | 2304 | 1953 | 2157 | 2245 | 2152 | 2261 | 2479 | 2138 | 2238.65 | 2185 | 2293 | 1.049 | 0.069 | 0.41557287 | 0.62710945 | 0.0025 | 0.0084 | 0.0048 | 0.0048 | TRUE | 0.0604 |
| Cirh1a | 12956 | 11429 | 11135 | 16758 | 11802 | 10287 | 13762 | 11414 | 11181 | 13174 | 12698 | 11261 | 12248.22 | 12119 | 12378 | 1.021 | 0.031 | 0.78184564 | 0.88723778 | 0.0081 | 0.0072 | 0.0087 | 0.0087 | TRUE | 0.1718 |
| Cisd1 | 2157 | 2621 | 2763 | 3058 | 2103 | 2292 | 2291 | 2618 | 2774 | 2404 | 2263 | 2509 | 2476.45 | 2561 | 2392 | 0.934 | -0.099 | 0.3201523 | 0.54120494 | 0.0054 | 0.0082 | 0.0067 | 0.0067 | TRUE | 0.1028 |
| Cisd2 | 820 | 792 | 810 | 1137 | 735 | 689 | 871 | 791 | 813 | 894 | 791 | 754 | 819.02 | 825 | 813 | 0.987 | -0.019 | 0.85883574 | 0.9304418 | 0.0041 | 0.0109 | 0.0069 | 0.0069 | TRUE | 0.093 |
| Cisd3 | 2054 | 1510 | 1449 | 1831 | 1263 | 1283 | 2182 | 1508 | 1455 | 1439 | 1359 | 1404 | 1557.91 | 1715 | 1401 | 0.817 | -0.291 | 0.06641859 | 0.19885665 | 0.0273 | 0.009 | 0.0175 | 0.0175 | TRUE | 0.6834 |
| Cish | 208 | 206 | 212 | 211 | 131 | 182 | 221 | 206 | 213 | 166 | 141 | 199 | 190.93 | 213 | 169 | 0.791 | -0.339 | 0.05098042 | 0.16620787 | 0.0077 | 0.0242 | 0.0164 | 0.0164 | TRUE | 0.2515 |
| Cit | 2675 | 2737 | 2921 | 2974 | 2238 | 2228 | 2841 | 2733 | 2933 | 2338 | 2408 | 2439 | 2615.43 | 2836 | 2395 | 0.844 | -0.244 | 0.00040761 | 0.00439275 | 5.00E-04 | 0.0082 | 0.0031 | 0.0031 | TRUE | 0.0122 |
| Cited1 | 104 | 152 | 154 | 130 | 98 | 95 | 110 | 152 | 155 | 102 | 105 | 104 | 121.42 | 139 | 104 | 0.746 | -0.422 | 0.03953276 | 0.13942429 | 0.0099 | 0.034 | 0.0219 | 0.0219 | TRUE | 0.3272 |
| Cited2 | 967 | 733 | 668 | 1426 | 1108 | 843 | 1027 | 732 | 671 | 1121 | 1192 | 923 | 944.31 | 810 | 1079 | 1.332 | 0.414 | 0.01959799 | 0.08559943 | 0.0344 | 0.0104 | 0.0216 | 0.0216 | TRUE | 0.6514 |
| Cited4 | 91 | 148 | 149 | 126 | 106 | 129 | 97 | 148 | 150 | 99 | 114 | 141 | 124.73 | 132 | 118 | 0.895 | -0.16 | 0.51240023 | 0.7057529 | 0.0366 | 0.0333 | 0.0348 | 0.0348 | TRUE | 0.5394 |
| Ciz1 | 695 | 780 | 930 | 1002 | 842 | 884 | 738 | 779 | 934 | 788 | 906 | 968 | 852.06 | 817 | 887 | 1.084 | 0.117 | 0.37671993 | 0.59206639 | 0.0113 | 0.0108 | 0.0114 | 0.0114 | TRUE | 0.186 |
| CK137956 | 1 | 0 | 1 | 1 | 0 | 0 | 1 | 0 | 1 | 1 | 0 | 0 | 0.48 | 1 | 0 | 0.509 | -0.975 | 0.7336918 | NA | 0 | 6.9367 | 4.1806 | 4.1806 | TRUE | 0.2562 |
| Ckap2 | 7408 | 7768 | 7495 | 9572 | 6968 | 7126 | 7869 | 7758 | 7526 | 7525 | 7497 | 7801 | 7662.46 | 7718 | 7608 | 0.986 | -0.021 | 0.72496433 | 0.85132436 | 3.00E-04 | 0.0073 | 0.0024 | 0.0024 | TRUE | 0.006 |
| Ckap2l | 3335 | 4010 | 3937 | 4398 | 3161 | 3369 | 3542 | 4005 | 3953 | 3457 | 3401 | 3688 | 3674.44 | 3833 | 3515 | 0.917 | -0.125 | 0.13108573 | 0.31169668 | 0.0027 | 0.0078 | 0.0047 | 0.0047 | TRUE | 0.0536 |
| Ckap4 | 3940 | 5114 | 5140 | 5553 | 4202 | 4023 | 4185 | 5107 | 5161 | 4365 | 4521 | 4404 | 4623.95 | 4818 | 4430 | 0.919 | -0.121 | 0.22895672 | 0.44224019 | 0.0063 | 0.0076 | 0.0071 | 0.0071 | TRUE | 0.1607 |
| Ckap5 | 14101 | 15701 | 17403 | 17984 | 13377 | 14019 | 14978 | 15681 | 17474 | 14138 | 14393 | 15346 | 15334.9 | 16044 | 14626 | 0.912 | -0.134 | 0.1169495 | 0.28911091 | 0.0041 | 0.0071 | 0.0052 | 0.0052 | TRUE | 0.0744 |
| Ckb | 4893 | 5077 | 4835 | 8363 | 6445 | 5864 | 5197 | 5070 | 4855 | 6574 | 6934 | 6419 | 5841.73 | 5041 | 6642 | 1.318 | 0.398 | 7.03E-09 | 3.45E-07 | 0.0011 | 0.0075 | 0.0032 | 0.0032 | TRUE | 0.0179 |
| Cklf | 294 | 328 | 271 | 471 | 293 | 246 | 312 | 328 | 272 | 370 | 315 | 269 | 311.13 | 304 | 318 | 1.05 | 0.07 | 0.66496475 | 0.81250574 | 0.0143 | 0.0175 | 0.0158 | 0.0158 | TRUE | 0.2348 |
| Ckm | 11 | 19 | 16 | 8 | 4 | 6 | 12 | 19 | 16 | 6 | 4 | 7 | 10.65 | 16 | 6 | 0.369 | -1.44 | 0.01894646 | 0.08357464 | 0 | 0.3163 | 0.1539 | 0.1539 | TRUE | 0.1823 |
| Ckmt1 | 493 | 653 | 647 | 505 | 417 | 535 | 524 | 652 | 650 | 397 | 449 | 586 | 542.79 | 609 | 477 | 0.783 | -0.354 | 0.03939019 | 0.13910805 | 0.0254 | 0.013 | 0.0194 | 0.0194 | TRUE | 0.4576 |
| Ckmt2 | 1 | 0 | 0 | 0 | 0 | 2 | 1 | 0 | 0 | 0 | 0 | 2 | 0.54 | 0 | 1 | 1.954 | 0.967 | 0.77097429 | NA | 0 | 6.0862 | 6.2584 | 6.2584 | TRUE | 0.9745 |
| Cks1b | 3326 | 3445 | 3336 | 4511 | 3067 | 3108 | 3533 | 3441 | 3350 | 3546 | 3300 | 3402 | 3428.56 | 3441 | 3416 | 0.993 | -0.01 | 0.88407225 | 0.94309013 | 7.00E-04 | 0.0079 | 0.0031 | 0.0031 | TRUE | 0.0138 |
| Cks2 | 3150 | 3464 | 3373 | 4247 | 3016 | 3331 | 3346 | 3460 | 3387 | 3339 | 3245 | 3646 | 3403.7 | 3398 | 3410 | 1.004 | 0.005 | 0.94691026 | 0.97351104 | 0.0017 | 0.0079 | 0.0039 | 0.0039 | TRUE | 0.0442 |
| Clasp1 | 2735 | 2257 | 2324 | 4078 | 2757 | 2400 | 2905 | 2254 | 2334 | 3206 | 2966 | 2627 | 2715.34 | 2498 | 2933 | 1.175 | 0.232 | 0.0692288 | 0.20425473 | 0.0145 | 0.0081 | 0.0114 | 0.0114 | TRUE | 0.2467 |
| Clasp2 | 5759 | 5926 | 6785 | 7481 | 5579 | 5739 | 6117 | 5918 | 6813 | 5881 | 6003 | 6282 | 6169.03 | 6283 | 6055 | 0.964 | -0.053 | 0.51609709 | 0.70824704 | 0.003 | 0.0074 | 0.0047 | 0.0047 | TRUE | 0.0665 |
| Clasrp | 1943 | 2033 | 1875 | 2155 | 1904 | 1753 | 2064 | 2030 | 1883 | 1694 | 2049 | 1919 | 1939.75 | 1992 | 1887 | 0.947 | -0.079 | 0.42750502 | 0.63765927 | 0.0048 | 0.0086 | 0.0067 | 0.0067 | TRUE | 0.0988 |
| Clca1 | 2 | 2 | 2 | 6 | 8 | 3 | 2 | 2 | 2 | 5 | 9 | 3 | 3.79 | 2 | 6 | 2.7 | 1.433 | 0.16082446 | NA | 0 | 0.8762 | 0.4139 | 0.4139 | TRUE | 0.4734 |
| Clca2 | 1 | 0 | 1 | 3 | 0 | 3 | 1 | 0 | 1 | 2 | 0 | 3 | 1.28 | 1 | 2 | 2.76 | 1.465 | 0.44476877 | NA | 0 | 2.5711 | 1.643 | 1.643 | TRUE | 0.4216 |
| Clca3 | 106 | 81 | 61 | 53 | 50 | 54 | 113 | 81 | 61 | 42 | 54 | 59 | 68.22 | 85 | 52 | 0.604 | -0.727 | 0.01818509 | 0.08124294 | 0.0501 | 0.0552 | 0.0527 | 0.0527 | TRUE | 0.2373 |
| Clca4 | 216 | 98 | 107 | 282 | 262 | 228 | 229 | 98 | 107 | 222 | 282 | 250 | 197.98 | 145 | 251 | 1.733 | 0.794 | 0.01007023 | 0.05301324 | 0.1123 | 0.0235 | 0.063 | 0.063 | TRUE | 2.6901 |
| Clca5 | 2 | 1 | 0 | 1 | 1 | 1 | 2 | 1 | 0 | 1 | 1 | 1 | 1.01 | 1 | 1 | 0.946 | -0.08 | 0.96534307 | NA | 0 | 3.2581 | 1.4592 | 1.4592 | TRUE | 0.2006 |
| Clca6 | 0 | 0 | 0 | 0 | 0 | 1 | 0 | 0 | 0 | 0 | 0 | 1 | 0.18 | 0 | 0 | 1.859 | 0.894 | 0.82650734 | NA | 0 | 18.0643 | 10 | 10 | TRUE | 0.5383 |
| Clcc1 | 935 | 1263 | 1280 | 1292 | 939 | 1164 | 993 | 1261 | 1285 | 1016 | 1010 | 1274 | 1139.99 | 1180 | 1100 | 0.932 | -0.102 | 0.47861756 | 0.6781132 | 0.018 | 0.0098 | 0.0141 | 0.0141 | TRUE | 0.2292 |
| Clcf1 | 0 | 15 | 15 | 5 | 7 | 9 | 0 | 15 | 15 | 4 | 8 | 10 | 8.56 | 10 | 7 | 0.701 | -0.512 | 0.6164524 | 0.78309404 | 1.0227 | 0.3918 | 0.631 | 0.631 | TRUE | 2.5443 |
| Clcn1 | 58 | 66 | 68 | 71 | 77 | 66 | 62 | 66 | 68 | 56 | 83 | 72 | 67.78 | 65 | 70 | 1.07 | 0.097 | 0.70338543 | 0.83766892 | 0 | 0.0555 | 0.0324 | 0.0324 | TRUE | 0.3351 |
| Clcn2 | 1192 | 1240 | 1160 | 1171 | 1048 | 1014 | 1266 | 1238 | 1165 | 921 | 1128 | 1110 | 1137.9 | 1223 | 1053 | 0.86 | -0.218 | 0.04674747 | 0.15663929 | 0.0063 | 0.0098 | 0.0078 | 0.0078 | TRUE | 0.1486 |
| Clcn3 | 8521 | 9239 | 9584 | 12179 | 9085 | 9234 | 9051 | 9227 | 9623 | 9574 | 9775 | 10108 | 9559.73 | 9300 | 9819 | 1.056 | 0.078 | 0.21107337 | 0.42177236 | 7.00E-04 | 0.0072 | 0.0027 | 0.0027 | TRUE | 0.0113 |
| Clcn4-2 | 739 | 572 | 500 | 1025 | 686 | 573 | 785 | 571 | 502 | 806 | 738 | 627 | 671.56 | 619 | 724 | 1.17 | 0.226 | 0.2136851 | 0.42515975 | 0.0322 | 0.0118 | 0.0223 | 0.0223 | TRUE | 0.6406 |
| Clcn5 | 618 | 567 | 516 | 1257 | 863 | 738 | 656 | 566 | 518 | 988 | 929 | 808 | 744.23 | 580 | 908 | 1.567 | 0.648 | 8.82E-07 | 2.47E-05 | 0.0103 | 0.0113 | 0.0111 | 0.0111 | TRUE | 0.1537 |
| Clcn6 | 2094 | 2120 | 2145 | 2145 | 1847 | 1782 | 2224 | 2117 | 2154 | 1686 | 1987 | 1951 | 2019.91 | 2165 | 1875 | 0.865 | -0.209 | 0.02389918 | 0.0984429 | 0.0034 | 0.0085 | 0.0057 | 0.0057 | TRUE | 0.0956 |
| Clcn7 | 1207 | 1430 | 1336 | 1692 | 1350 | 1244 | 1282 | 1428 | 1341 | 1330 | 1453 | 1362 | 1366.01 | 1350 | 1382 | 1.022 | 0.032 | 0.71444238 | 0.84513502 | 0.0018 | 0.0093 | 0.0047 | 0.0047 | TRUE | 0.0304 |
| Clcnka | 217 | 218 | 200 | 257 | 207 | 172 | 230 | 218 | 201 | 202 | 223 | 188 | 210.34 | 216 | 204 | 0.945 | -0.082 | 0.58412799 | 0.75938564 | 0 | 0.0226 | 0.0115 | 0.0115 | TRUE | 0.0636 |
| Clcnkb | 81 | 115 | 113 | 161 | 143 | 143 | 86 | 115 | 113 | 127 | 154 | 157 | 125.22 | 105 | 146 | 1.384 | 0.469 | 0.02240132 | 0.0942498 | 0.0107 | 0.0332 | 0.0221 | 0.0221 | TRUE | 0.2361 |

| | | | | | | | | | | | | | | | | | | | | | | | | | |
|---|---|---|---|---|---|---|---|---|---|---|---|---|---|---|---|---|---|---|---|---|---|---|---|---|---|
| Cldn1 | 20 | 13 | 7 | 37 | 18 | 17 | 21 | 13 | 7 | 29 | 19 | 19 | 18.05 | 14 | 22 | 1.643 | 0.716 | 0.17977268 | 0.38109875 | 0.0952 | 0.1894 | 0.1464 | 0.1464 | TRUE | 0.4529 |
| Cldn10 | 37 | 31 | 27 | 27 | 20 | 25 | 39 | 31 | 27 | 21 | 22 | 27 | 27.91 | 32 | 23 | 0.718 | -0.478 | 0.1996028 | 0.40715667 | 0 | 0.1249 | 0.0633 | 0.0633 | TRUE | 0.2266 |
| Cldn11 | 37 | 21 | 28 | 29 | 28 | 32 | 39 | 21 | 28 | 23 | 30 | 35 | 29.39 | 29 | 29 | 0.99 | -0.014 | 0.9712957 | 0.98480349 | 0.0387 | 0.119 | 0.0802 | 0.0802 | TRUE | 0.3386 |
| Cldn12 | 1727 | 1953 | 1913 | 2675 | 1848 | 1791 | 1834 | 1950 | 1921 | 2103 | 1988 | 1961 | 1959.58 | 1902 | 2017 | 1.061 | 0.086 | 0.25092973 | 0.468418 | 7.00E-04 | 0.0086 | 0.0035 | 0.0035 | TRUE | 0.0171 |
| Cldn13 | 0 | 1 | 1 | 0 | 0 | 0 | 0 | 1 | 1 | 0 | 0 | 0 | 0.33 | 1 | 0 | 0.268 | -1.901 | 0.61401143 | NA | 0 | 9.8764 | 8.4772 | 8.4772 | TRUE | 0.2278 |
| Cldn14 | 4 | 5 | 5 | 4 | 3 | 5 | 4 | 5 | 5 | 3 | 3 | 5 | 4.35 | 5 | 4 | 0.821 | -0.284 | 0.74614488 | NA | 0 | 0.764 | 0.322 | 0.322 | TRUE | 0.1698 |
| Cldn15 | 17 | 19 | 9 | 9 | 7 | 5 | 18 | 19 | 9 | 7 | 8 | 5 | 11.02 | 15 | 7 | 0.438 | -1.192 | 0.05333225 | 0.17148954 | 0 | 0.3057 | 0.1684 | 0.1684 | TRUE | 0.6067 |
| Cldn16 | 0 | 1 | 0 | 3 | 3 | 3 | 0 | 1 | 0 | 2 | 3 | 3 | 1.64 | 0 | 3 | 8.039 | 3.007 | 0.08484496 | NA | 0 | 2.0098 | 1.0278 | 1.0278 | TRUE | 0.405 |
| Cldn17 | 0 | 0 | 0 | 0 | 0 | 0 | 0 | 0 | 0 | 0 | 0 | 0 | 0 | 0 | 0 | NA | NA | NA | NA | NA | NA | NA | NA | NA | NA |
| Cldn18 | 34 | 22 | 19 | 25 | 20 | 15 | 36 | 22 | 19 | 20 | 22 | 16 | 22.46 | 26 | 19 | 0.749 | -0.417 | 0.3418138 | 0.56204134 | 0.0298 | 0.1536 | 0.0933 | 0.0933 | TRUE | 0.7467 |
| Cldn19 | 29 | 44 | 30 | 31 | 21 | 24 | 31 | 44 | 30 | 24 | 23 | 26 | 29.68 | 35 | 24 | 0.698 | -0.52 | 0.15253613 | 0.34375257 | 0 | 0.1179 | 0.0604 | 0.0604 | TRUE | 0.3637 |
| Cldn2 | 42 | 7 | 7 | 23 | 12 | 9 | 45 | 7 | 7 | 18 | 13 | 10 | 16.58 | 20 | 14 | 0.703 | -0.509 | 0.5137991 | 0.70663057 | 0.5443 | 0.2056 | 0.376 | 0.376 | TRUE | 6.5362 |
| Cldn20 | 0 | 1 | 0 | 0 | 0 | 1 | 0 | 1 | 0 | 0 | 0 | 1 | 0.35 | 0 | 0 | 0.954 | -0.067 | 0.98567705 | NA | 0 | 9.4494 | 8.1588 | 8.1588 | TRUE | 0.5383 |
| Cldn22 | 0 | 0 | 0 | 0 | 0 | 0 | 0 | 0 | 0 | 0 | 0 | 0 | 0 | 0 | 0 | NA | NA | NA | NA | NA | NA | NA | NA | NA | NA |
| Cldn23 | 7 | 26 | 9 | 20 | 17 | 19 | 7 | 26 | 9 | 16 | 18 | 21 | 16.21 | 14 | 18 | 1.284 | 0.36 | 0.54327129 | 0.72861181 | 0.1682 | 0.2102 | 0.1902 | 0.1902 | TRUE | 2.3709 |
| Cldn24 | 0 | 0 | 0 | 0 | 0 | 0 | 0 | 0 | 0 | 0 | 0 | 0 | 0 | 0 | 0 | NA | NA | NA | NA | NA | NA | NA | NA | NA | NA |
| Cldn25 | 1323 | 1632 | 1716 | 2114 | 1456 | 1616 | 1405 | 1630 | 1723 | 1662 | 1567 | 1769 | 1625.93 | 1586 | 1666 | 1.05 | 0.07 | 0.50905623 | 0.70268115 | 0.0066 | 0.0089 | 0.0076 | 0.0076 | TRUE | 0.1194 |
| Cldn26 | 13 | 5 | 2 | 10 | 7 | 5 | 14 | 5 | 2 | 8 | 8 | 5 | 6.95 | 7 | 7 | 1.012 | 0.017 | 0.9834979 | 0.99129879 | 0.2552 | 0.4812 | 0.3715 | 0.3715 | TRUE | 0.9508 |
| Cldn3 | 72 | 119 | 132 | 106 | 80 | 115 | 76 | 119 | 133 | 83 | 86 | 126 | 103.86 | 109 | 98 | 0.897 | -0.157 | 0.57672551 | 0.75440349 | 0.0562 | 0.0386 | 0.0471 | 0.0471 | TRUE | 0.6742 |
| Cldn4 | 524 | 1576 | 1711 | 869 | 729 | 1140 | 557 | 1574 | 1718 | 683 | 784 | 1248 | 1094 | 1283 | 905 | 0.705 | -0.503 | 0.35112154 | 0.56999652 | 0.2091 | 0.0099 | 0.0871 | 0.2091 | TRUE | 2.9432 |
| Cldn5 | 95 | 142 | 124 | 76 | 65 | 73 | 101 | 142 | 125 | 60 | 70 | 80 | 96.14 | 123 | 70 | 0.567 | -0.818 | 0.00048582 | 0.00506439 | 0.0154 | 0.0412 | 0.0284 | 0.0284 | TRUE | 0.1779 |
| Cldn6 | 1310 | 1305 | 1412 | 900 | 798 | 1137 | 1391 | 1303 | 1418 | 708 | 859 | 1245 | 1153.88 | 1371 | 937 | 0.683 | -0.551 | 0.00355883 | 0.02412116 | 0.0433 | 0.0098 | 0.0248 | 0.0248 | TRUE | 0.9821 |
| Cldn7 | 401 | 1331 | 1545 | 632 | 603 | 1002 | 426 | 1329 | 1551 | 497 | 649 | 1097 | 924.84 | 1102 | 748 | 0.678 | -0.561 | 0.36446585 | 0.58187624 | 0.2742 | 0.0105 | 0.1103 | 0.2742 | TRUE | 1.2012 |
| Cldn8 | 0 | 0 | 0 | 0 | 0 | 1 | 0 | 0 | 0 | 0 | 0 | 1 | 0.18 | 0 | 0 | 1.859 | 0.894 | 0.82650734 | NA | 0 | 18.0643 | 10 | 10 | TRUE | 0.5383 |
| Cldn9 | 59 | 65 | 62 | 54 | 33 | 47 | 63 | 65 | 62 | 42 | 36 | 51 | 53.21 | 63 | 43 | 0.681 | -0.555 | 0.04184128 | 0.14551725 | 0 | 0.0688 | 0.0341 | 0.0341 | TRUE | 0.1969 |
| Cldnd2 | 1 | 2 | 2 | 2 | 4 | 3 | 1 | 2 | 2 | 2 | 4 | 3 | 2.37 | 2 | 3 | 1.769 | 0.823 | 0.49965522 | NA | 0 | 1.3962 | 0.6077 | 0.6077 | TRUE | 0.3254 |
| Clec10a | 7 | 10 | 2 | 71 | 44 | 19 | 7 | 10 | 2 | 56 | 47 | 21 | 23.9 | 6 | 41 | 6.412 | 2.681 | 1.43E-05 | 0.00026785 | 0.2452 | 0.1448 | 0.1843 | 0.1843 | TRUE | 0.49 |
| Clec11a | 37 | 56 | 62 | 61 | 43 | 55 | 39 | 56 | 62 | 48 | 46 | 60 | 51.98 | 52 | 51 | 0.977 | -0.034 | 0.90962577 | 0.95489676 | 0.0163 | 0.0703 | 0.0448 | 0.0448 | TRUE | 0.3875 |
| Clec12a | 0 | 3 | 6 | 0 | 3 | 0 | 0 | 3 | 6 | 0 | 3 | 0 | 2.04 | 3 | 1 | 0.347 | -1.529 | 0.43830888 | NA | 3.6216 | 1.6207 | 2.1658 | 2.1658 | TRUE | 0.3131 |
| Clec12b | 0 | 0 | 0 | 0 | 0 | 0 | 0 | 0 | 0 | 0 | 0 | 0 | 0 | 0 | 0 | NA | NA | NA | NA | NA | NA | NA | NA | NA | NA |
| Clec14a | 2 | 1 | 0 | 1 | 2 | 1 | 2 | 1 | 0 | 1 | 2 | 1 | 1.19 | 1 | 1 | 1.283 | 0.359 | 0.83685447 | NA | 0 | 2.7693 | 1.331 | 1.331 | TRUE | 0.2503 |
| Clec16a | 1297 | 933 | 939 | 1825 | 1203 | 928 | 1378 | 932 | 943 | 1435 | 1294 | 1016 | 1166.19 | 1084 | 1248 | 1.152 | 0.204 | 0.27190119 | 0.49187272 | 0.0375 | 0.0097 | 0.0241 | 0.0241 | TRUE | 0.6702 |
| Clec18a | 1 | 0 | 0 | 1 | 0 | 0 | 1 | 0 | 0 | 1 | 0 | 0 | 0.31 | 0 | 0 | 0.954 | -0.067 | 0.98625928 | NA | 0 | 10.7014 | 9.0384 | 9.0384 | TRUE | 0.4614 |
| Clec1a | 1 | 0 | 1 | 1 | 3 | 0 | 1 | 0 | 1 | 1 | 3 | 0 | 1.01 | 1 | 1 | 1.934 | 0.951 | 0.64880762 | NA | 0 | 3.2579 | 2.0305 | 2.0305 | TRUE | 0.6977 |
| Clec1b | 0 | 0 | 0 | 1 | 1 | 0 | 0 | 0 | 0 | 1 | 1 | 0 | 0.31 | 0 | 1 | 3.32 | 1.731 | 0.65383052 | NA | 0 | 10.6224 | 8.9541 | 8.9541 | TRUE | 0.202 |
| Clec2d | 6 | 6 | 5 | 8 | 6 | 2 | 6 | 6 | 5 | 6 | 6 | 2 | 5.39 | 6 | 5 | 0.87 | -0.201 | 0.80630756 | 0.90019698 | 0 | 0.6185 | 0.3002 | 0.3002 | TRUE | 0.4255 |
| Clec2e | 0 | 1 | 0 | 0 | 0 | 0 | 0 | 1 | 0 | 0 | 0 | 0 | 0.17 | 0 | 0 | 0.49 | -1.029 | 0.80087 | NA | 0 | 19.7991 | 10 | 10 | TRUE | 0.405 |
| Clec2g | 3 | 3 | 0 | 3 | 7 | 2 | 3 | 3 | 0 | 2 | 8 | 2 | 3.04 | 2 | 4 | 1.943 | 0.958 | 0.44815799 | NA | 0.3686 | 1.0893 | 0.7779 | 0.7779 | TRUE | 0.9125 |
| Clec2h | 0 | 0 | 0 | 0 | 0 | 0 | 0 | 0 | 0 | 0 | 0 | 0 | 0 | 0 | 0 | NA | NA | NA | NA | NA | NA | NA | NA | NA | NA |
| Clec2i | 28 | 8 | 10 | 54 | 27 | 28 | 30 | 8 | 10 | 42 | 29 | 31 | 24.99 | 16 | 34 | 2.157 | 1.109 | 0.04121826 | 0.14388985 | 0.2004 | 0.1387 | 0.166 | 0.166 | TRUE | 2.6344 |
| Clec2l | 112 | 172 | 152 | 196 | 135 | 152 | 119 | 172 | 153 | 154 | 145 | 166 | 151.51 | 148 | 155 | 1.049 | 0.069 | 0.72062172 | 0.8486892 | 0.0117 | 0.0286 | 0.0203 | 0.0203 | TRUE | 0.2988 |
| Clec3a | 5 | 0 | 3 | 1 | 2 | 0 | 5 | 0 | 3 | 1 | 2 | 0 | 1.88 | 3 | 1 | 0.352 | -1.505 | 0.38553361 | NA | 1.0131 | 1.7622 | 1.4821 | 1.4821 | TRUE | 0.1397 |
| Clec3b | 1 | 0 | 1 | 2 | 2 | 2 | 1 | 0 | 1 | 2 | 2 | 2 | 1.33 | 1 | 2 | 2.862 | 1.517 | 0.3691033 | NA | 0 | 2.484 | 1.0599 | 1.0599 | TRUE | 0.2566 |
| Clec4a1 | 3 | 5 | 0 | 14 | 12 | 9 | 3 | 5 | 0 | 11 | 13 | 10 | 6.99 | 3 | 11 | 4.131 | 2.046 | 0.0204285 | 0.08822352 | 0 | 0.4781 | 0.3304 | 0.3304 | TRUE | 0.9345 |
| Clec4a2 | 0 | 0 | 0 | 3 | 1 | 2 | 0 | 0 | 0 | 2 | 1 | 2 | 0.94 | 0 | 2 | 10.025 | 3.326 | 0.13309345 | NA | 0 | 3.5218 | 2.2722 | 2.2722 | TRUE | 0.1088 |
| Clec4a3 | 0 | 0 | 0 | 4 | 2 | 2 | 0 | 0 | 0 | 3 | 2 | 2 | 1.25 | 0 | 2 | 13.35 | 3.739 | 0.0610996 | NA | 0 | 2.6475 | 1.6767 | 1.6767 | TRUE | 0.0739 |
| Clec4a4 | 1 | 0 | 0 | 0 | 1 | 0 | 1 | 0 | 0 | 0 | 1 | 0 | 0.36 | 0 | 0 | 0.954 | -0.067 | 0.98557668 | NA | 0 | 9.2518 | 8.0179 | 8.0179 | TRUE | 0.5204 |
| Clec4b1 | 0 | 0 | 0 | 0 | 0 | 0 | 0 | 0 | 0 | 0 | 0 | 0 | 0 | 0 | 0 | NA | NA | NA | NA | NA | NA | NA | NA | NA | NA |
| Clec4b2 | 0 | 0 | 0 | 0 | 0 | 0 | 0 | 0 | 0 | 0 | 0 | 0 | 0 | 0 | 0 | NA | NA | NA | NA | NA | NA | NA | NA | NA | NA |
| Clec4d | 0 | 0 | 0 | 0 | 0 | 0 | 0 | 0 | 0 | 0 | 0 | 0 | 0 | 0 | 0 | NA | NA | NA | NA | NA | NA | NA | NA | NA | NA |
| Clec4e | 0 | 5 | 2 | 5 | 7 | 7 | 0 | 5 | 2 | 4 | 8 | 8 | 4.35 | 2 | 7 | 2.683 | 1.424 | 0.18749271 | NA | 0.1503 | 0.7635 | 0.5464 | 0.5464 | TRUE | 0.7637 |
| Clec4f | 3 | 2 | 2 | 8 | 2 | 5 | 3 | 2 | 2 | 6 | 2 | 5 | 3.52 | 2 | 4 | 1.962 | 0.973 | 0.34550638 | NA | 0 | 0.9434 | 0.4482 | 0.4482 | TRUE | 0.3834 |
| Clec4g | 1 | 1 | 0 | 1 | 2 | 2 | 1 | 1 | 0 | 1 | 2 | 2 | 1.2 | 1 | 2 | 2.455 | 1.296 | 0.46556115 | NA | 0 | 2.7568 | 1.2338 | 1.2338 | TRUE | 0.2531 |
| Clec4n | 0 | 0 | 0 | 0 | 0 | 0 | 0 | 0 | 0 | 0 | 0 | 0 | 0 | 0 | 0 | NA | NA | NA | NA | NA | NA | NA | NA | NA | NA |
| Clec5a | 0 | 0 | 0 | 0 | 0 | 0 | 0 | 0 | 0 | 0 | 0 | 0 | 0 | 0 | 0 | NA | NA | NA | NA | NA | NA | NA | NA | NA | NA |
| Clec7a | 0 | 0 | 0 | 0 | 0 | 0 | 0 | 0 | 0 | 0 | 0 | 0 | 0 | 0 | 0 | NA | NA | NA | NA | NA | NA | NA | NA | NA | NA |
| Clec9a | 0 | 0 | 0 | 0 | 0 | 0 | 0 | 0 | 0 | 0 | 0 | 0 | 0 | 0 | 0 | NA | NA | NA | NA | NA | NA | NA | NA | NA | NA |
| Clgn | 3194 | 3357 | 3320 | 2790 | 1910 | 1975 | 3393 | 3353 | 3334 | 2193 | 2055 | 2162 | 2748.19 | 3360 | 2137 | 0.636 | -0.652 | 1.04E-22 | 3.41E-20 | 1.00E-04 | 0.0081 | 0.0028 | 0.0028 | TRUE | 0.0134 |
| Clhc1 | 1171 | 1182 | 1203 | 1476 | 1047 | 1062 | 1244 | 1180 | 1208 | 1160 | 1127 | 1163 | 1180.26 | 1211 | 1150 | 0.95 | -0.074 | 0.33451887 | 0.5548783 | 0 | 0.0097 | 0.0034 | 0.0034 | TRUE | 0.0068 |
| Clic1 | 1255 | 2187 | 2350 | 2040 | 1641 | 2017 | 1333 | 2184 | 2360 | 1604 | 1766 | 2208 | 1909.01 | 1959 | 1859 | 0.949 | -0.076 | 0.71474212 | 0.8452151 | 0.0573 | 0.0086 | 0.0306 | 0.0306 | TRUE | 0.9435 |
| Clic3 | 10 | 12 | 8 | 16 | 14 | 12 | 11 | 12 | 8 | 13 | 15 | 13 | 11.9 | 10 | 14 | 1.328 | 0.409 | 0.4502687 | 0.65567703 | 0 | 0.2837 | 0.1251 | 0.1251 | TRUE | 0.1252 |
| Clic4 | 9120 | 9371 | 9408 | 12827 | 9238 | 9271 | 9687 | 9359 | 9447 | 10084 | 9940 | 10149 | 9777.37 | 9498 | 10058 | 1.059 | 0.083 | 0.12942771 | 0.30962138 | 1.00E-04 | 0.0072 | 0.002 | 0.002 | TRUE | 0.0037 |
| Clic5 | 7 | 16 | 8 | 11 | 11 | 7 | 7 | 16 | 8 | 9 | 12 | 8 | 9.93 | 10 | 10 | 0.891 | -0.167 | 0.7921913 | 0.89305706 | 0 | 0.3386 | 0.1879 | 0.1879 | TRUE | 0.7654 |
| Clic6 | 133 | 130 | 123 | 143 | 105 | 97 | 141 | 130 | 124 | 112 | 113 | 106 | 121.03 | 132 | 110 | 0.841 | -0.249 | 0.16424144 | 0.36033051 | 0 | 0.0341 | 0.0149 | 0.0149 | TRUE | 0.0422 |
| Clint1 | 8861 | 8315 | 8162 | 10694 | 7668 | 7134 | 9412 | 8304 | 8196 | 8407 | 8250 | 7809 | 8396.35 | 8637 | 8155 | 0.944 | -0.083 | 0.32370086 | 0.54504216 | 0.0036 | 0.0073 | 0.0049 | 0.0049 | TRUE | 0.0751 |
| Clip1 | 939 | 1132 | 1268 | 1190 | 912 | 912 | 997 | 1131 | 1273 | 935 | 981 | 998 | 1052.7 | 1134 | 971 | 0.857 | -0.223 | 0.04956354 | 0.16329979 | 0.0065 | 0.01 | 0.0084 | 0.0084 | TRUE | 0.1388 |
| Clip2 | 657 | 755 | 777 | 729 | 585 | 636 | 698 | 754 | 780 | 573 | 629 | 696 | 688.46 | 744 | 633 | 0.849 | -0.236 | 0.03793258 | 0.13541706 | 0.0049 | 0.0117 | 0.0079 | 0.0079 | TRUE | 0.0898 |
| Clip3 | 1008 | 740 | 774 | 851 | 585 | 610 | 1071 | 739 | 777 | 669 | 629 | 668 | 758.84 | 862 | 655 | 0.76 | -0.395 | 0.01083218 | 0.05581353 | 0.0207 | 0.0112 | 0.016 | 0.016 | TRUE | 0.53 |
| Clip4 | 1179 | 1557 | 1600 | 1533 | 1160 | 1222 | 1252 | 1555 | 1607 | 1205 | 1248 | 1338 | 1367.46 | 1471 | 1264 | 0.858 | -0.22 | 0.06408547 | 0.19484295 | 0.0088 | 0.0093 | 0.0095 | 0.0095 | TRUE | 0.2034 |

| | | | | | | | | | | | | | | | | | | | | | | | | | |
|---|---|---|---|---|---|---|---|---|---|---|---|---|---|---|---|---|---|---|---|---|---|---|---|---|---|
| Clk1 | 5307 | 5882 | 6365 | 4311 | 3755 | 4688 | 5637 | 5874 | 6391 | 3389 | 4040 | 5132 | 5077.23 | 5967 | 4187 | 0.701 | -0.512 | 0.00056286 | 0.0057126 | 0.0239 | 0.0076 | 0.0157 | 0.0157 | TRUE | 0.4742 |
| Clk2 | 2271 | 2585 | 2530 | 3115 | 2582 | 2427 | 2412 | 2582 | 2540 | 2449 | 2778 | 2657 | 2569.64 | 2511 | 2628 | 1.046 | 0.065 | 0.43159408 | 0.6413928 | 0.002 | 0.0082 | 0.0045 | 0.0045 | TRUE | 0.044 |
| Clk3 | 1906 | 2569 | 2598 | 2387 | 1719 | 2056 | 2025 | 2566 | 2609 | 1876 | 1850 | 2251 | 2195.91 | 2400 | 1992 | 0.83 | -0.269 | 0.04170804 | 0.14515382 | 0.0148 | 0.0084 | 0.0121 | 0.0121 | TRUE | 0.2262 |
| Clk4 | 881 | 1091 | 1223 | 1402 | 1081 | 1247 | 936 | 1090 | 1228 | 1102 | 1163 | 1365 | 1147.28 | 1085 | 1210 | 1.115 | 0.157 | 0.24596219 | 0.46298074 | 0.0142 | 0.0098 | 0.0123 | 0.0123 | TRUE | 0.1715 |
| Clmn | 1597 | 2281 | 2477 | 1829 | 1462 | 1649 | 1696 | 2278 | 2487 | 1438 | 1573 | 1805 | 1879.57 | 2154 | 1605 | 0.745 | -0.425 | 0.00662024 | 0.03861402 | 0.024 | 0.0087 | 0.0171 | 0.0171 | TRUE | 0.4174 |
| Clmp | 117 | 73 | 67 | 116 | 79 | 92 | 124 | 73 | 67 | 91 | 85 | 101 | 90.23 | 88 | 92 | 1.048 | 0.068 | 0.80899334 | 0.90174222 | 0.0498 | 0.0434 | 0.0462 | 0.0462 | TRUE | 1.2036 |
| Cln3 | 936 | 894 | 894 | 1009 | 762 | 823 | 994 | 893 | 898 | 793 | 820 | 901 | 883.11 | 928 | 838 | 0.902 | -0.149 | 0.13863503 | 0.3231198 | 0.0029 | 0.0106 | 0.0061 | 0.0061 | TRUE | 0.0507 |
| Cln5 | 1675 | 1802 | 1824 | 2121 | 1512 | 1553 | 1779 | 1800 | 1831 | 1667 | 1627 | 1700 | 1734.08 | 1803 | 1665 | 0.923 | -0.116 | 0.09604616 | 0.25427176 | 0 | 0.0088 | 0.0029 | 0.0029 | TRUE | 0.0046 |
| Cln6 | 1001 | 1839 | 1921 | 1992 | 1564 | 1864 | 1063 | 1837 | 1929 | 1566 | 1683 | 2040 | 1686.32 | 1610 | 1763 | 1.095 | 0.131 | 0.5342471 | 0.72175874 | 0.0587 | 0.0089 | 0.0314 | 0.0314 | TRUE | 1.0615 |
| Cln8 | 2594 | 2741 | 2696 | 3421 | 2539 | 2444 | 2755 | 2737 | 2707 | 2689 | 2732 | 2675 | 2716.05 | 2733 | 2699 | 0.987 | -0.018 | 0.76594476 | 0.87749552 | 0 | 0.0081 | 0.0024 | 0.0024 | TRUE | 0.0014 |
| Clnk | 6 | 4 | 5 | 10 | 8 | 4 | 6 | 4 | 5 | 8 | 9 | 4 | 6.04 | 5 | 7 | 1.365 | 0.449 | 0.5588203 | 0.74101748 | 0 | 0.5524 | 0.2556 | 0.2556 | TRUE | 0.2451 |
| Clns1a | 5543 | 5410 | 5202 | 8362 | 5805 | 5470 | 5888 | 5403 | 5223 | 6574 | 6246 | 5988 | 5886.86 | 5505 | 6269 | 1.139 | 0.188 | 0.01953357 | 0.08541264 | 0.0028 | 0.0075 | 0.0045 | 0.0045 | TRUE | 0.0451 |
| Clock | 1065 | 995 | 1023 | 1488 | 1104 | 1064 | 1131 | 994 | 1027 | 1170 | 1188 | 1165 | 1112.41 | 1051 | 1174 | 1.118 | 0.16 | 0.06915299 | 0.20409448 | 0.0013 | 0.0099 | 0.0047 | 0.0047 | TRUE | 0.0534 |
| Clp1 | 2436 | 1764 | 1546 | 3073 | 2129 | 1521 | 2587 | 1762 | 1552 | 2416 | 2291 | 1665 | 2045.49 | 1967 | 2124 | 1.08 | 0.111 | 0.58662082 | 0.76149659 | 0.0547 | 0.0085 | 0.0295 | 0.0295 | TRUE | 0.9193 |
| Clpb | 5569 | 6096 | 6191 | 6848 | 5079 | 5144 | 5915 | 6088 | 6216 | 5383 | 5465 | 5631 | 5783.13 | 6073 | 5493 | 0.904 | -0.145 | 0.01747782 | 0.07880507 | 3.00E-04 | 0.0075 | 0.0025 | 0.0025 | TRUE | 0.0063 |
| Clpp | 2874 | 2998 | 3008 | 3876 | 2827 | 2541 | 3053 | 2994 | 3020 | 3047 | 3042 | 2782 | 2989.56 | 3022 | 2957 | 0.979 | -0.031 | 0.66506589 | 0.81256637 | 9.00E-04 | 0.008 | 0.0034 | 0.0034 | TRUE | 0.0323 |
| Clps | 5 | 9 | 0 | 4 | 7 | 5 | 5 | 9 | 0 | 3 | 8 | 5 | 5.07 | 5 | 5 | 1.119 | 0.162 | 0.88106016 | 0.94127757 | 0.6365 | 0.6561 | 0.6486 | 0.6486 | TRUE | 0.3486 |
| Clpsl2 | 2 | 1 | 1 | 0 | 4 | 2 | 2 | 1 | 1 | 0 | 4 | 2 | 1.77 | 1 | 2 | 1.535 | 0.618 | 0.6960565 | NA | 0 | 1.8681 | 1.1981 | 1.1981 | TRUE | 0.1549 |
| Clptm1 | 5480 | 5782 | 5950 | 7849 | 5642 | 5541 | 5821 | 5775 | 5974 | 6170 | 6070 | 6065 | 5979.32 | 5857 | 6102 | 1.042 | 0.059 | 0.29396198 | 0.51480884 | 0 | 0.0075 | 0.0021 | 0.0021 | TRUE | 0.0038 |
| Clptm1l | 2549 | 2547 | 2693 | 3634 | 2637 | 2673 | 2707 | 2544 | 2704 | 2857 | 2837 | 2926 | 2762.55 | 2652 | 2873 | 1.084 | 0.116 | 0.08693467 | 0.23841657 | 4.00E-04 | 0.0081 | 0.0029 | 0.0029 | TRUE | 0.0155 |
| Clpx | 2382 | 2693 | 2750 | 3355 | 2462 | 2568 | 2530 | 2690 | 2761 | 2637 | 2649 | 2811 | 2679.73 | 2660 | 2699 | 1.014 | 0.02 | 0.78434784 | 0.88870216 | 0.0012 | 0.0081 | 0.0037 | 0.0037 | TRUE | 0.0221 |
| Clrn1 | 0 | 0 | 0 | 2 | 1 | 1 | 0 | 0 | 0 | 2 | 1 | 1 | 0.62 | 0 | 1 | 6.676 | 2.739 | 0.27800695 | NA | 0 | 5.2881 | 3.2034 | 3.2034 | TRUE | 0.07 |
| Clrn2 | 0 | 0 | 0 | 0 | 0 | 0 | 0 | 0 | 0 | 0 | 0 | 0 | 0 | 0 | 0 | NA | NA | NA | NA | NA | NA | NA | NA | NA | NA |
| Clrn3 | 0 | 0 | 0 | 0 | 0 | 0 | 0 | 0 | 0 | 0 | 0 | 0 | 0 | 0 | 0 | NA | NA | NA | NA | NA | NA | NA | NA | NA | NA |
| Clspn | 3078 | 2701 | 2791 | 4375 | 3038 | 2736 | 3269 | 2698 | 2802 | 3439 | 3269 | 2995 | 3078.72 | 2923 | 3234 | 1.107 | 0.146 | 0.16262996 | 0.35794023 | 0.0073 | 0.008 | 0.0076 | 0.0076 | TRUE | 0.1302 |
| Clstn1 | 5959 | 8334 | 8869 | 8722 | 6614 | 7253 | 6330 | 8323 | 8905 | 6857 | 7116 | 7940 | 7578.42 | 7853 | 7304 | 0.93 | -0.105 | 0.44192417 | 0.6489784 | 0.0164 | 0.0073 | 0.0132 | 0.0132 | TRUE | 0.3514 |
| Clstn2 | 26 | 21 | 17 | 33 | 26 | 22 | 28 | 21 | 17 | 26 | 28 | 24 | 23.94 | 22 | 26 | 1.19 | 0.251 | 0.52665425 | 0.71578825 | 0 | 0.1445 | 0.0714 | 0.0714 | TRUE | 0.2865 |
| Clstn3 | 1642 | 1471 | 1383 | 1950 | 1389 | 1357 | 1744 | 1469 | 1389 | 1533 | 1494 | 1485 | 1519.12 | 1534 | 1504 | 0.981 | -0.028 | 0.79686828 | 0.89536272 | 0.0066 | 0.0091 | 0.0077 | 0.0077 | TRUE | 0.1725 |
| Clta | 8170 | 9123 | 9361 | 10661 | 8187 | 8013 | 8678 | 9111 | 9399 | 8381 | 8809 | 8771 | 8858.28 | 9063 | 8654 | 0.955 | -0.067 | 0.30953281 | 0.52992476 | 0.001 | 0.0073 | 0.003 | 0.003 | TRUE | 0.0168 |
| Cltb | 1526 | 1598 | 1567 | 1382 | 997 | 1070 | 1621 | 1596 | 1573 | 1086 | 1073 | 1171 | 1353.44 | 1597 | 1110 | 0.695 | -0.525 | 3.20E-11 | 2.57E-09 | 1.00E-04 | 0.0093 | 0.0037 | 0.0037 | TRUE | 0.0271 |
| Cltc | 37880 | 34598 | 40073 | 50456 | 34169 | 32936 | 40235 | 34553 | 40238 | 39665 | 36764 | 36054 | 37918.01 | 38342 | 37494 | 0.978 | -0.032 | 0.71980702 | 0.8481969 | 0.0041 | 0.007 | 0.0058 | 0.0058 | TRUE | 0.0915 |
| Clu | 1050 | 1094 | 1039 | 1837 | 1367 | 1288 | 1115 | 1093 | 1043 | 1444 | 1471 | 1410 | 1262.66 | 1084 | 1442 | 1.33 | 0.412 | 1.14E-07 | 3.98E-06 | 0 | 0.0095 | 0.0035 | 0.0035 | TRUE | 0.0128 |
| Cluap1 | 388 | 478 | 540 | 689 | 546 | 592 | 412 | 477 | 542 | 542 | 587 | 648 | 534.81 | 477 | 592 | 1.239 | 0.31 | 0.02641025 | 0.1056142 | 0.0113 | 0.0131 | 0.0121 | 0.0121 | TRUE | 0.1649 |
| Cluh | 16055 | 13628 | 13471 | 17523 | 13242 | 11883 | 17053 | 13610 | 13526 | 13775 | 14248 | 13008 | 14203.43 | 14730 | 13677 | 0.929 | -0.107 | 0.33227802 | 0.55269869 | 0.0103 | 0.0071 | 0.0087 | 0.0087 | TRUE | 0.2327 |
| Clvs1 | 11 | 3 | 11 | 51 | 27 | 25 | 12 | 3 | 11 | 40 | 29 | 27 | 20.37 | 9 | 32 | 3.784 | 1.92 | 0.0003565 | 0.00394582 | 0.0652 | 0.1686 | 0.1334 | 0.1334 | TRUE | 1.029 |
| Clvs2 | 69 | 49 | 43 | 67 | 41 | 30 | 73 | 49 | 43 | 53 | 44 | 33 | 49.17 | 55 | 43 | 0.79 | -0.34 | 0.3115964 | 0.53210246 | 0.0474 | 0.0739 | 0.061 | 0.061 | TRUE | 0.6653 |
| Clybl | 1646 | 2283 | 2248 | 1777 | 1311 | 1688 | 1748 | 2280 | 2257 | 1397 | 1411 | 1848 | 1823.48 | 2095 | 1552 | 0.74 | -0.434 | 0.00430628 | 0.02791559 | 0.0231 | 0.0087 | 0.0161 | 0.0161 | TRUE | 0.3342 |
| Cma1 | 0 | 0 | 0 | 0 | 0 | 1 | 0 | 0 | 0 | 0 | 0 | 1 | 0.18 | 0 | 0 | 1.859 | 0.894 | 0.82650734 | NA | 0 | 18.0643 | 10 | 10 | TRUE | 0.5383 |
| Cma2 | 0 | 0 | 0 | 0 | 0 | 0 | 0 | 0 | 0 | 0 | 0 | 0 | 0 | 0 | 0 | NA | NA | NA | NA | NA | NA | NA | NA | NA | NA |
| Cmah | 7 | 4 | 8 | 19 | 10 | 13 | 7 | 4 | 8 | 15 | 11 | 14 | 9.9 | 6 | 13 | 2.062 | 1.044 | 0.09461299 | 0.25210798 | 0 | 0.3397 | 0.1658 | 0.1658 | TRUE | 0.2898 |
| Cmas | 947 | 969 | 897 | 1156 | 803 | 684 | 1006 | 968 | 901 | 909 | 864 | 749 | 899.3 | 958 | 841 | 0.878 | -0.187 | 0.08843064 | 0.24121897 | 0.0053 | 0.0106 | 0.0076 | 0.0076 | TRUE | 0.107 |
| Cmbl | 57 | 67 | 54 | 47 | 21 | 31 | 61 | 67 | 54 | 37 | 23 | 34 | 45.86 | 61 | 31 | 0.518 | -0.948 | 0.0023309 | 0.01738202 | 0 | 0.0787 | 0.0459 | 0.0459 | TRUE | 0.3665 |
| Cmc1 | 870 | 950 | 941 | 1069 | 730 | 786 | 924 | 949 | 945 | 840 | 785 | 860 | 883.99 | 939 | 828 | 0.883 | -0.18 | 0.03686293 | 0.13270991 | 0 | 0.0106 | 0.0042 | 0.0042 | TRUE | 0.024 |
| Cmc2 | 844 | 721 | 719 | 1204 | 816 | 704 | 896 | 720 | 722 | 946 | 878 | 771 | 822.27 | 779 | 865 | 1.111 | 0.152 | 0.25447207 | 0.47255892 | 0.0119 | 0.0109 | 0.0116 | 0.0116 | TRUE | 0.2035 |
| Cmip | 8608 | 8930 | 9261 | 11125 | 8367 | 8293 | 9143 | 8918 | 9299 | 8746 | 9002 | 9078 | 9031.11 | 9120 | 8942 | 0.98 | -0.029 | 0.61824508 | 0.78428066 | 2.00E-04 | 0.0073 | 0.0023 | 0.0023 | TRUE | 0.0046 |
| Cmklr1 | 1279 | 1134 | 1165 | 1261 | 983 | 1031 | 1359 | 1133 | 1170 | 991 | 1058 | 1129 | 1139.73 | 1221 | 1059 | 0.868 | -0.205 | 0.05933571 | 0.18449436 | 0.0057 | 0.0098 | 0.0076 | 0.0076 | TRUE | 0.1174 |
| Cml1 | 41 | 46 | 64 | 48 | 35 | 45 | 44 | 46 | 64 | 38 | 38 | 49 | 46.4 | 51 | 42 | 0.807 | -0.309 | 0.31915434 | 0.54044491 | 0.0154 | 0.0779 | 0.0475 | 0.0475 | TRUE | 0.4067 |
| Cml2 | 117 | 49 | 35 | 138 | 141 | 69 | 124 | 49 | 35 | 108 | 152 | 76 | 90.68 | 69 | 112 | 1.614 | 0.69 | 0.12781814 | 0.30739486 | 0.2535 | 0.0432 | 0.1363 | 0.1363 | TRUE | 0.853 |
| Cml3 | 0 | 0 | 0 | 0 | 0 | 0 | 0 | 0 | 0 | 0 | 0 | 0 | 0 | 0 | 0 | NA | NA | NA | NA | NA | NA | NA | NA | NA | NA |
| Cml5 | 0 | 0 | 0 | 0 | 1 | 0 | 0 | 0 | 0 | 0 | 1 | 0 | 0.18 | 0 | 0 | 1.859 | 0.894 | 0.82650734 | NA | 0 | 18.3785 | 10 | 10 | TRUE | 0.5204 |
| Cmpk1 | 2993 | 3000 | 2984 | 4521 | 3088 | 3024 | 3179 | 2996 | 2996 | 3554 | 3322 | 3310 | 3226.38 | 3057 | 3395 | 1.111 | 0.152 | 0.03432154 | 0.12615226 | 9.00E-04 | 0.0079 | 0.0034 | 0.0034 | TRUE | 0.0206 |
| Cmpk2 | 77 | 78 | 73 | 156 | 93 | 77 | 82 | 78 | 73 | 123 | 100 | 84 | 90 | 78 | 102 | 1.325 | 0.406 | 0.07869766 | 0.22216207 | 0.0116 | 0.0435 | 0.0271 | 0.0271 | TRUE | 0.2598 |
| Cmss1 | 778 | 661 | 664 | 1135 | 807 | 676 | 826 | 660 | 667 | 892 | 868 | 740 | 775.63 | 718 | 833 | 1.162 | 0.217 | 0.10338542 | 0.26760005 | 0.0115 | 0.0112 | 0.0115 | 0.0115 | TRUE | 0.2063 |
| Cmtm1 | 0 | 0 | 0 | 0 | 0 | 0 | 0 | 0 | 0 | 0 | 0 | 0 | 0 | 0 | 0 | NA | NA | NA | NA | NA | NA | NA | NA | NA | NA |
| Cmtm2a | 0 | 0 | 1 | 0 | 0 | 1 | 0 | 0 | 1 | 0 | 0 | 1 | 0.35 | 0 | 0 | 0.954 | -0.067 | 0.98566485 | NA | 0 | 9.4252 | 8.1416 | 8.1416 | TRUE | 0.5383 |
| Cmtm2b | 4 | 4 | 3 | 1 | 2 | 1 | 4 | 4 | 3 | 1 | 2 | 1 | 2.55 | 4 | 1 | 0.35 | -1.515 | 0.21176627 | NA | 0 | 1.2998 | 0.5541 | 0.5541 | TRUE | 0.1596 |
| Cmtm3 | 198 | 290 | 236 | 334 | 269 | 250 | 210 | 290 | 237 | 263 | 289 | 274 | 260.43 | 246 | 275 | 1.119 | 0.162 | 0.31328374 | 0.53377024 | 0.0101 | 0.0196 | 0.0147 | 0.0147 | TRUE | 0.2732 |
| Cmtm4 | 8863 | 11278 | 12546 | 13007 | 9661 | 10690 | 9414 | 11263 | 12598 | 10225 | 10395 | 11702 | 10932.77 | 11092 | 10774 | 0.971 | -0.042 | 0.72907967 | 0.85431237 | 0.0109 | 0.0072 | 0.0105 | 0.0105 | TRUE | 0.2139 |
| Cmtm5 | 0 | 2 | 1 | 0 | 1 | 0 | 0 | 2 | 1 | 0 | 1 | 0 | 0.68 | 1 | 0 | 0.347 | -1.526 | 0.55436871 | NA | 0 | 4.8547 | 3.291 | 3.291 | TRUE | 0.2722 |
| Cmtm6 | 6690 | 7788 | 7987 | 7681 | 5770 | 6545 | 7106 | 7778 | 8020 | 6038 | 6208 | 7165 | 7052.43 | 7635 | 6470 | 0.847 | -0.239 | 0.01342953 | 0.06517246 | 0.0061 | 0.0074 | 0.0066 | 0.0066 | TRUE | 0.1073 |
| Cmtm7 | 899 | 877 | 845 | 1443 | 989 | 903 | 955 | 876 | 848 | 1134 | 1064 | 988 | 977.7 | 893 | 1062 | 1.191 | 0.252 | 0.01162012 | 0.05883547 | 0.0033 | 0.0103 | 0.0062 | 0.0062 | TRUE | 0.044 |
| Cmtm8 | 82 | 149 | 110 | 103 | 66 | 116 | 87 | 149 | 110 | 81 | 71 | 127 | 104.22 | 115 | 93 | 0.803 | -0.317 | 0.28980277 | 0.51098594 | 0.0729 | 0.0385 | 0.0548 | 0.0548 | TRUE | 0.4703 |
| Cmya5 | 20 | 22 | 7 | 51 | 32 | 20 | 21 | 22 | 7 | 40 | 34 | 22 | 24.44 | 17 | 32 | 1.931 | 0.949 | 0.06228806 | 0.19101833 | 0.1402 | 0.1417 | 0.1411 | 0.1411 | TRUE | 0.9903 |
| Cnbd2 | 40 | 63 | 46 | 43 | 44 | 32 | 42 | 63 | 46 | 34 | 47 | 35 | 44.63 | 50 | 39 | 0.762 | -0.391 | 0.22005368 | 0.43222937 | 0.0187 | 0.0807 | 0.0506 | 0.0506 | TRUE | 0.3782 |
| Cnbp | 20172 | 23879 | 22715 | 27872 | 19464 | 20289 | 21426 | 23848 | 22808 | 21911 | 20942 | 22210 | 22190.84 | 22694 | 21688 | 0.956 | -0.065 | 0.35413158 | 0.57256416 | 0.0018 | 0.0071 | 0.0036 | 0.0036 | TRUE | 0.0292 |
| Cndp1 | 3 | 4 | 2 | 3 | 4 | 2 | 3 | 4 | 2 | 2 | 4 | 2 | 3.01 | 3 | 3 | 0.956 | -0.065 | 0.95067781 | NA | 0 | 1.1026 | 0.4736 | 0.4736 | TRUE | 0.1877 |
| Cndp2 | 3066 | 3392 | 3285 | 5327 | 3755 | 3687 | 3257 | 3388 | 3298 | 4188 | 4040 | 4036 | 3701.09 | 3314 | 4088 | 1.234 | 0.303 | 1.14E-06 | 3.08E-05 | 2.00E-04 | 0.0078 | 0.0025 | 0.0025 | TRUE | 0.0056 |
| Cnep1r1 | 2433 | 2403 | 2339 | 3261 | 2209 | 2296 | 2584 | 2400 | 2349 | 2564 | 2377 | 2513 | 2464.4 | 2444 | 2485 | 1.017 | 0.024 | 0.75736821 | 0.8719099 | 0.0016 | 0.0082 | 0.004 | 0.004 | TRUE | 0.0303 |
| Cnfn | 0 | 1 | 0 | 1 | 1 | 0 | 0 | 1 | 0 | 1 | 1 | 0 | 0.48 | 0 | 1 | 1.712 | 0.776 | 0.78614772 | NA | 0 | 6.9165 | 4.1067 | 4.1067 | TRUE | 0.405 |

| | | | | | | | | | | | | | | | | | | | | | | | | | |
|---|---|---|---|---|---|---|---|---|---|---|---|---|---|---|---|---|---|---|---|---|---|---|---|---|---|
| Cnga1 | 0 | 0 | 0 | 0 | 0 | 0 | 0 | 0 | 0 | 0 | 0 | 0 | 0 | 0 | 0 | NA | NA | NA | NA | NA | NA | NA | NA | NA | NA |
| Cnga2 | 0 | 0 | 0 | 0 | 0 | 0 | 0 | 0 | 0 | 0 | 0 | 0 | 0 | 0 | 0 | NA | NA | NA | NA | NA | NA | NA | NA | NA | NA |
| Cnga3 | 440 | 434 | 346 | 594 | 436 | 362 | 467 | 433 | 347 | 467 | 469 | 396 | 430.09 | 416 | 444 | 1.069 | 0.096 | 0.52473178 | 0.71437805 | 0.0135 | 0.0146 | 0.014 | 0.014 | TRUE | 0.2411 |
| Cnga4 | 2 | 7 | 8 | 12 | 2 | 7 | 2 | 7 | 8 | 9 | 2 | 8 | 6.07 | 6 | 6 | 1.133 | 0.18 | 0.8390947 | 0.91834027 | 0.2238 | 0.55 | 0.399 | 0.399 | TRUE | 0.7742 |
| Cngb1 | 332 | 352 | 355 | 330 | 244 | 249 | 353 | 352 | 356 | 259 | 263 | 273 | 309.19 | 354 | 265 | 0.748 | -0.418 | 0.00038828 | 0.00423523 | 0 | 0.0176 | 0.0067 | 0.0067 | TRUE | 0.0071 |
| Cngb3 | 0 | 0 | 0 | 0 | 0 | 0 | 0 | 0 | 0 | 0 | 0 | 0 | 0 | 0 | 0 | NA | NA | NA | NA | NA | NA | NA | NA | NA | NA |
| Cnih | 3024 | 4115 | 4227 | 4114 | 2978 | 3420 | 3212 | 4110 | 4244 | 3234 | 3204 | 3744 | 3624.67 | 3855 | 3394 | 0.88 | -0.184 | 0.15105001 | 0.34214573 | 0.0142 | 0.0078 | 0.0116 | 0.0116 | TRUE | 0.259 |
| Cnih2 | 92 | 110 | 101 | 205 | 166 | 130 | 98 | 110 | 101 | 161 | 179 | 142 | 131.84 | 103 | 161 | 1.56 | 0.641 | 0.00051376 | 0.00528397 | 0 | 0.0319 | 0.0166 | 0.0166 | TRUE | 0.099 |
| Cnih3 | 2 | 3 | 3 | 6 | 3 | 6 | 2 | 3 | 3 | 5 | 3 | 7 | 3.77 | 3 | 5 | 1.778 | 0.83 | 0.38913573 | NA | 0 | 0.8798 | 0.3803 | 0.3803 | TRUE | 0.1607 |
| Cnih4 | 2169 | 2948 | 2925 | 2930 | 1977 | 2406 | 2304 | 2944 | 2937 | 2303 | 2127 | 2634 | 2541.55 | 2728 | 2355 | 0.863 | -0.213 | 0.10287657 | 0.26686995 | 0.0144 | 0.0082 | 0.0119 | 0.0119 | TRUE | 0.2244 |
| Cnksr1 | 128 | 185 | 148 | 159 | 121 | 146 | 136 | 185 | 149 | 125 | 130 | 160 | 147.39 | 157 | 138 | 0.881 | -0.182 | 0.36245845 | 0.58031617 | 0.0149 | 0.0293 | 0.022 | 0.022 | TRUE | 0.2654 |
| Cnksr2 | 244 | 282 | 314 | 431 | 315 | 263 | 259 | 282 | 315 | 339 | 339 | 288 | 303.62 | 285 | 322 | 1.129 | 0.175 | 0.2206842 | 0.43275286 | 0.0056 | 0.0178 | 0.0114 | 0.0114 | TRUE | 0.0947 |
| Cnksr3 | 1155 | 1134 | 1126 | 1565 | 1064 | 941 | 1227 | 1133 | 1131 | 1230 | 1145 | 1030 | 1149.19 | 1164 | 1135 | 0.977 | -0.034 | 0.73826594 | 0.85986006 | 0.0042 | 0.0098 | 0.0065 | 0.0065 | TRUE | 0.0778 |
| Cnn1 | 190 | 192 | 179 | 66 | 53 | 52 | 202 | 192 | 180 | 52 | 57 | 57 | 123.19 | 191 | 55 | 0.288 | -1.794 | 1.84E-20 | 4.91E-18 | 0 | 0.0336 | 0.0155 | 0.0155 | TRUE | 0.0293 |
| Cnn2 | 1343 | 1693 | 1853 | 2129 | 1533 | 1537 | 1427 | 1691 | 1861 | 1674 | 1649 | 1682 | 1663.91 | 1660 | 1668 | 1.005 | 0.008 | 0.9451864 | 0.97262869 | 0.0083 | 0.0089 | 0.0086 | 0.0086 | TRUE | 0.1811 |
| Cnn3 | 13105 | 14964 | 14356 | 17730 | 12870 | 13418 | 13920 | 14945 | 14415 | 13938 | 13847 | 14688 | 14292.14 | 14427 | 14158 | 0.981 | -0.027 | 0.67327224 | 0.81803018 | 0.001 | 0.0071 | 0.0029 | 0.0029 | TRUE | 0.0131 |
| Cnnm1 | 167 | 135 | 140 | 229 | 198 | 134 | 177 | 135 | 141 | 180 | 213 | 147 | 165.42 | 151 | 180 | 1.193 | 0.255 | 0.21493034 | 0.42675287 | 0.0218 | 0.0268 | 0.0243 | 0.0243 | TRUE | 0.1513 |
| Cnnm2 | 1791 | 2095 | 2156 | 2309 | 1659 | 1838 | 1902 | 2092 | 2165 | 1815 | 1785 | 2012 | 1961.94 | 2053 | 1871 | 0.911 | -0.135 | 0.1456896 | 0.33413063 | 0.0036 | 0.0086 | 0.0057 | 0.0057 | TRUE | 0.0521 |
| Cnnm3 | 1731 | 1726 | 1735 | 2113 | 1728 | 1462 | 1839 | 1724 | 1742 | 1661 | 1859 | 1600 | 1737.53 | 1768 | 1707 | 0.965 | -0.051 | 0.57122824 | 0.74993329 | 0.0027 | 0.0088 | 0.0053 | 0.0053 | TRUE | 0.0726 |
| Cnnm4 | 815 | 798 | 799 | 1327 | 987 | 898 | 866 | 797 | 802 | 1043 | 1062 | 983 | 925.51 | 822 | 1029 | 1.253 | 0.326 | 0.00025795 | 0.00300253 | 6.00E-04 | 0.0105 | 0.0046 | 0.0046 | TRUE | 0.0259 |
| Cnot1 | 50635 | 49515 | 55398 | 59189 | 42857 | 46396 | 53783 | 49451 | 55625 | 46530 | 46111 | 50788 | 50381.47 | 52953 | 47810 | 0.903 | -0.147 | 0.06424333 | 0.19508007 | 0.0032 | 0.007 | 0.0046 | 0.0046 | TRUE | 0.041 |
| Cnot10 | 4217 | 4326 | 4410 | 5431 | 4205 | 3991 | 4479 | 4320 | 4428 | 4269 | 4524 | 4369 | 4398.37 | 4409 | 4387 | 0.995 | -0.007 | 0.90632236 | 0.95350897 | 4.00E-04 | 0.0077 | 0.0026 | 0.0026 | TRUE | 0.0091 |
| Cnot11 | 2476 | 2139 | 2131 | 3531 | 2367 | 2184 | 2630 | 2136 | 2140 | 2776 | 2547 | 2391 | 2436.54 | 2302 | 2571 | 1.117 | 0.16 | 0.16229722 | 0.35768332 | 0.0098 | 0.0083 | 0.0091 | 0.0091 | TRUE | 0.1878 |
| Cnot2 | 5008 | 4802 | 4893 | 5846 | 4361 | 4230 | 5319 | 4796 | 4913 | 4596 | 4692 | 4630 | 4824.42 | 5009 | 4639 | 0.926 | -0.111 | 0.11796695 | 0.2905195 | 0.0012 | 0.0076 | 0.0034 | 0.0034 | TRUE | 0.0356 |
| Cnot3 | 6837 | 6191 | 6166 | 8247 | 6163 | 5910 | 7262 | 6183 | 6191 | 6483 | 6631 | 6469 | 6536.67 | 6545 | 6528 | 0.997 | -0.004 | 0.96544999 | 0.98258226 | 0.0035 | 0.0074 | 0.0055 | 0.0055 | TRUE | 0.1118 |
| Cnot4 | 2342 | 2114 | 2133 | 2783 | 2064 | 1893 | 2488 | 2111 | 2142 | 2188 | 2221 | 2072 | 2203.56 | 2247 | 2160 | 0.962 | -0.056 | 0.55063708 | 0.73491924 | 0.0041 | 0.0084 | 0.006 | 0.006 | TRUE | 0.1061 |
| Cnot6 | 4161 | 3698 | 3670 | 5826 | 4023 | 3600 | 4420 | 3693 | 3685 | 4580 | 4329 | 3941 | 4107.87 | 3933 | 4283 | 1.089 | 0.124 | 0.246808 | 0.46379674 | 0.0082 | 0.0077 | 0.008 | 0.008 | TRUE | 0.1427 |
| Cnot6l | 661 | 1011 | 974 | 1329 | 1079 | 1137 | 702 | 1010 | 978 | 1045 | 1161 | 1245 | 1023.35 | 897 | 1150 | 1.282 | 0.358 | 0.02008186 | 0.08708579 | 0.0219 | 0.0101 | 0.0161 | 0.0161 | TRUE | 0.4271 |
| Cnot7 | 5398 | 5992 | 5956 | 7959 | 5758 | 5879 | 5734 | 5984 | 5980 | 6257 | 6195 | 6435 | 6097.64 | 5899 | 6296 | 1.067 | 0.094 | 0.11782334 | 0.29040627 | 2.00E-04 | 0.0074 | 0.0024 | 0.0024 | TRUE | 0.0074 |
| Cnot8 | 4195 | 5008 | 5017 | 5180 | 3792 | 4043 | 4456 | 5002 | 5038 | 4072 | 4080 | 4426 | 4512.12 | 4832 | 4193 | 0.868 | -0.205 | 0.0147043 | 0.06982452 | 0.0031 | 0.0076 | 0.0049 | 0.0049 | TRUE | 0.0563 |
| Cnp | 3086 | 2592 | 2343 | 3192 | 2170 | 2113 | 3278 | 2589 | 2353 | 2509 | 2335 | 2313 | 2562.71 | 2740 | 2386 | 0.871 | -0.199 | 0.1300236 | 0.31057582 | 0.0158 | 0.0082 | 0.0121 | 0.0121 | TRUE | 0.3579 |
| Cnppd1 | 728 | 1011 | 1115 | 1337 | 976 | 965 | 773 | 1010 | 1120 | 1051 | 1050 | 1056 | 1010.01 | 968 | 1052 | 1.088 | 0.121 | 0.39271387 | 0.6062924 | 0.0164 | 0.0102 | 0.0135 | 0.0135 | TRUE | 0.3665 |
| Cnpy1 | 669 | 685 | 653 | 449 | 467 | 497 | 711 | 684 | 656 | 353 | 502 | 544 | 574.98 | 684 | 466 | 0.681 | -0.555 | 0.00090837 | 0.00826706 | 0.0239 | 0.0126 | 0.0183 | 0.0183 | TRUE | 0.5467 |
| Cnpy2 | 1409 | 1608 | 1591 | 2215 | 1610 | 1406 | 1497 | 1606 | 1598 | 1741 | 1732 | 1539 | 1618.78 | 1567 | 1671 | 1.067 | 0.093 | 0.29012888 | 0.5113596 | 0.0025 | 0.0089 | 0.005 | 0.005 | TRUE | 0.0567 |
| Cnpy3 | 1965 | 2288 | 2205 | 2284 | 1753 | 1877 | 2087 | 2285 | 2214 | 1796 | 1886 | 2055 | 2053.76 | 2195 | 1912 | 0.87 | -0.2 | 0.02218805 | 0.09369474 | 0.0027 | 0.0085 | 0.005 | 0.005 | TRUE | 0.0512 |
| Cnpy4 | 615 | 561 | 508 | 929 | 694 | 641 | 653 | 560 | 510 | 730 | 747 | 702 | 650.38 | 574 | 726 | 1.265 | 0.339 | 0.00494787 | 0.03107894 | 0.0061 | 0.012 | 0.0089 | 0.0089 | TRUE | 0.1672 |
| Cnr1 | 132 | 153 | 146 | 143 | 90 | 81 | 140 | 153 | 147 | 112 | 97 | 89 | 122.92 | 147 | 99 | 0.681 | -0.554 | 0.00312927 | 0.02194525 | 0 | 0.0337 | 0.017 | 0.017 | TRUE | 0.1453 |
| Cnr2 | 3 | 0 | 0 | 3 | 3 | 0 | 3 | 0 | 0 | 2 | 3 | 0 | 1.46 | 1 | 2 | 1.784 | 0.835 | 0.68953422 | NA | 2.7471 | 2.2601 | 2.399 | 2.399 | TRUE | 1.4429 |
| Cnrip1 | 3 | 13 | 3 | 6 | 9 | 2 | 3 | 13 | 3 | 5 | 10 | 2 | 5.96 | 6 | 6 | 0.86 | -0.218 | 0.81969821 | 0.90781402 | 0.4226 | 0.5595 | 0.4896 | 0.4896 | TRUE | 1.123 |
| Cnst | 1092 | 1140 | 1094 | 1400 | 998 | 1005 | 1160 | 1139 | 1098 | 1101 | 1074 | 1100 | 1111.9 | 1132 | 1092 | 0.964 | -0.053 | 0.49924646 | 0.6949847 | 0 | 0.0099 | 0.0035 | 0.0035 | TRUE | 0.0082 |
| Cntd1 | 219 | 544 | 547 | 397 | 317 | 455 | 233 | 543 | 549 | 312 | 341 | 498 | 412.73 | 442 | 384 | 0.868 | -0.205 | 0.50464726 | 0.69953388 | 0.1328 | 0.0149 | 0.0653 | 0.0653 | TRUE | 1.9778 |
| Cntf | 4 | 0 | 4 | 8 | 5 | 6 | 4 | 0 | 4 | 6 | 5 | 7 | 4.42 | 3 | 6 | 2.22 | 1.15 | 0.26107591 | NA | 0 | 0.7528 | 0.4881 | 0.4881 | TRUE | 0.9454 |
| Cntfr | 362 | 435 | 471 | 513 | 382 | 376 | 385 | 434 | 473 | 403 | 411 | 412 | 419.63 | 431 | 409 | 0.948 | -0.077 | 0.52917785 | 0.71760295 | 0.003 | 0.0148 | 0.0083 | 0.0083 | TRUE | 0.1003 |
| Cntln | 94 | 134 | 143 | 327 | 213 | 185 | 100 | 134 | 144 | 257 | 229 | 203 | 177.67 | 126 | 230 | 1.828 | 0.87 | 8.46E-06 | 0.00017193 | 0.0169 | 0.0254 | 0.0213 | 0.0213 | TRUE | 0.3222 |
| Cntn1 | 13 | 12 | 11 | 21 | 11 | 9 | 14 | 12 | 11 | 17 | 12 | 10 | 12.51 | 12 | 13 | 1.05 | 0.071 | 0.89547559 | 0.94945374 | 0 | 0.2703 | 0.1282 | 0.1282 | TRUE | 0.3437 |
| Cntn2 | 954 | 1167 | 1339 | 906 | 715 | 819 | 1013 | 1165 | 1344 | 712 | 769 | 897 | 983.56 | 1174 | 793 | 0.674 | -0.569 | 5.41E-05 | 0.00082712 | 0.0155 | 0.0103 | 0.0132 | 0.0132 | TRUE | 0.1615 |
| Cntn3 | 42 | 36 | 36 | 60 | 28 | 29 | 45 | 36 | 36 | 47 | 30 | 32 | 37.63 | 39 | 36 | 0.944 | -0.082 | 0.80795284 | 0.90112858 | 0.0176 | 0.0945 | 0.0564 | 0.0564 | TRUE | 0.5665 |
| Cntn4 | 5 | 3 | 5 | 5 | 4 | 4 | 5 | 3 | 5 | 4 | 4 | 4 | 4.32 | 4 | 4 | 0.945 | -0.081 | 0.9261365 | NA | 0 | 0.7689 | 0.3208 | 0.3208 | TRUE | 0.1516 |
| Cntn5 | 4 | 1 | 0 | 3 | 1 | 3 | 4 | 1 | 0 | 2 | 1 | 3 | 1.99 | 2 | 2 | 1.294 | 0.372 | 0.80255964 | NA | 0.3391 | 1.6589 | 1.0795 | 1.0795 | TRUE | 1.1786 |
| Cntn6 | 8 | 2 | 2 | 12 | 12 | 12 | 8 | 2 | 2 | 9 | 13 | 13 | 8 | 4 | 12 | 2.851 | 1.512 | 0.05578867 | 0.17697081 | 0.0588 | 0.4188 | 0.2848 | 0.2848 | TRUE | 1.3622 |
| Cntnap1 | 40 | 56 | 47 | 44 | 42 | 45 | 42 | 56 | 47 | 35 | 45 | 49 | 45.77 | 48 | 43 | 0.879 | -0.186 | 0.53778799 | 0.72419528 | 0 | 0.0789 | 0.0437 | 0.0437 | TRUE | 0.2677 |
| Cntnap2 | 808 | 593 | 509 | 898 | 603 | 483 | 858 | 592 | 511 | 706 | 649 | 529 | 640.84 | 654 | 628 | 0.961 | -0.057 | 0.77771806 | 0.88481347 | 0.0454 | 0.012 | 0.028 | 0.028 | TRUE | 0.8789 |
| Cntnap3 | 29 | 18 | 13 | 71 | 43 | 20 | 31 | 18 | 13 | 56 | 46 | 22 | 30.97 | 21 | 41 | 2.021 | 1.015 | 0.03850383 | 0.13680613 | 0.163 | 0.1133 | 0.1368 | 0.1368 | TRUE | 0.4319 |
| Cntnap4 | 4 | 6 | 0 | 8 | 3 | 4 | 4 | 6 | 0 | 6 | 3 | 4 | 4.02 | 3 | 4 | 1.371 | 0.455 | 0.68154993 | NA | 0.2944 | 0.8259 | 0.6309 | 0.6309 | TRUE | 0.8836 |
| Cntnap5a | 0 | 0 | 0 | 1 | 0 | 0 | 0 | 0 | 0 | 1 | 0 | 0 | 0.13 | 0 | 0 | 1.859 | 0.894 | 0.82650734 | NA | 0 | 25.1515 | 10 | 10 | TRUE | 0.2563 |
| Cntnap5b | 0 | 0 | 0 | 0 | 0 | 0 | 0 | 0 | 0 | 0 | 0 | 0 | 0 | 0 | 0 | NA | NA | NA | NA | NA | NA | NA | NA | NA | NA |
| Cntnap5c | 0 | 0 | 0 | 0 | 0 | 0 | 0 | 0 | 0 | 0 | 0 | 0 | 0 | 0 | 0 | NA | NA | NA | NA | NA | NA | NA | NA | NA | NA |
| Cntrob | 2528 | 2799 | 2958 | 3272 | 2448 | 2418 | 2685 | 2795 | 2970 | 2572 | 2634 | 2647 | 2717.28 | 2817 | 2618 | 0.929 | -0.106 | 0.14615764 | 0.33488812 | 0.0011 | 0.0081 | 0.0035 | 0.0035 | TRUE | 0.0275 |
| Coa3 | 1251 | 1382 | 1353 | 2089 | 1354 | 1353 | 1329 | 1380 | 1359 | 1642 | 1457 | 1481 | 1441.28 | 1356 | 1527 | 1.127 | 0.173 | 0.04373035 | 0.14971024 | 0.0018 | 0.0092 | 0.0046 | 0.0046 | TRUE | 0.0542 |
| Coa4 | 328 | 443 | 467 | 509 | 343 | 322 | 348 | 442 | 469 | 400 | 369 | 352 | 396.9 | 420 | 374 | 0.891 | -0.166 | 0.26090684 | 0.48027809 | 0.0116 | 0.0152 | 0.0133 | 0.0133 | TRUE | 0.2539 |
| Coa5 | 5734 | 6085 | 6231 | 7726 | 5477 | 5580 | 6091 | 6077 | 6257 | 6074 | 5893 | 6108 | 6083.15 | 6142 | 6025 | 0.981 | -0.028 | 0.6334776 | 0.79377204 | 1.00E-04 | 0.0074 | 0.0022 | 0.0022 | TRUE | 0.0044 |
| Coa6 | 860 | 752 | 822 | 1151 | 796 | 692 | 913 | 751 | 825 | 905 | 856 | 758 | 834.78 | 830 | 840 | 1.013 | 0.018 | 0.87759241 | 0.93954572 | 0.0077 | 0.0108 | 0.0091 | 0.0091 | TRUE | 0.0916 |
| Coasy | 1758 | 2367 | 2375 | 2418 | 1820 | 1996 | 1867 | 2364 | 2385 | 1901 | 1958 | 2185 | 2110 | 2205 | 2015 | 0.913 | -0.131 | 0.28083301 | 0.50219126 | 0.0109 | 0.0085 | 0.0102 | 0.0102 | TRUE | 0.2172 |
| Cobl | 6825 | 7324 | 8555 | 7403 | 5295 | 5228 | 7249 | 7315 | 8590 | 5820 | 5697 | 5723 | 6732.27 | 7718 | 5747 | 0.745 | -0.426 | 2.15E-06 | 5.32E-05 | 0.0046 | 0.0074 | 0.0057 | 0.0057 | TRUE | 0.1194 |
| Cobll1 | 200 | 164 | 203 | 459 | 319 | 275 | 212 | 164 | 204 | 361 | 343 | 301 | 264.19 | 193 | 335 | 1.736 | 0.796 | 5.38E-07 | 1.60E-05 | 0.0088 | 0.0194 | 0.0141 | 0.0141 | TRUE | 0.1952 |
| Coch | 1730 | 1388 | 1315 | 1277 | 848 | 790 | 1838 | 1386 | 1320 | 1004 | 912 | 865 | 1220.87 | 1515 | 927 | 0.612 | -0.707 | 1.13E-06 | 3.06E-05 | 0.0189 | 0.0096 | 0.0143 | 0.0143 | TRUE | 0.4179 |
| Cog1 | 2546 | 2953 | 2964 | 3602 | 2630 | 2946 | 2704 | 2949 | 2976 | 2832 | 2830 | 3225 | 2919.31 | 2876 | 2962 | 1.029 | 0.042 | 0.64252042 | 0.7988831 | 0.0034 | 0.008 | 0.0055 | 0.0055 | TRUE | 0.0727 |
| Cog2 | 2724 | 2457 | 2425 | 3279 | 2413 | 2285 | 2893 | 2454 | 2435 | 2578 | 2596 | 2501 | 2576.23 | 2594 | 2558 | 0.986 | -0.02 | 0.83480593 | 0.91587083 | 0.0046 | 0.0082 | 0.006 | 0.006 | TRUE | 0.1233 |
| Cog3 | 2581 | 2380 | 2475 | 3159 | 2298 | 2292 | 2741 | 2377 | 2485 | 2483 | 2473 | 2509 | 2511.4 | 2534 | 2488 | 0.982 | -0.026 | 0.74832841 | 0.86655986 | 0.0023 | 0.0082 | 0.0045 | 0.0045 | TRUE | 0.0617 |

| | | | | | | | | | | | | | | | | | | | | | | | | | |
|---|---|---|---|---|---|---|---|---|---|---|---|---|---|---|---|---|---|---|---|---|---|---|---|---|---|
| Cog4 | 7284 | 7925 | 8257 | 9855 | 7423 | 7750 | 7737 | 7915 | 8291 | 7747 | 7987 | 8484 | 8026.68 | 7981 | 8073 | 1.011 | 0.016 | 0.81776199 | 0.90711028 | 0.0013 | 0.0073 | 0.0035 | 0.0035 | TRUE | 0.0242 |
| Cog5 | 1628 | 1988 | 2070 | 2212 | 1619 | 1787 | 1729 | 1985 | 2078 | 1739 | 1742 | 1956 | 1871.69 | 1931 | 1812 | 0.938 | -0.092 | 0.37516778 | 0.59102546 | 0.0059 | 0.0087 | 0.0073 | 0.0073 | TRUE | 0.1007 |
| Cog6 | 3016 | 3455 | 3450 | 3975 | 2759 | 2947 | 3204 | 3451 | 3464 | 3125 | 2969 | 3226 | 3239.59 | 3373 | 3107 | 0.921 | -0.119 | 0.11302109 | 0.28312717 | 0.0014 | 0.0079 | 0.0037 | 0.0037 | TRUE | 0.0233 |
| Cog7 | 1645 | 1659 | 1691 | 2544 | 1879 | 1858 | 1747 | 1657 | 1698 | 2000 | 2022 | 2034 | 1859.59 | 1701 | 2019 | 1.187 | 0.247 | 0.00033023 | 0.00368603 | 0 | 0.0087 | 0.0029 | 0.0029 | TRUE | 0.0069 |
| Cog8 | 2079 | 1979 | 1894 | 3154 | 2086 | 1701 | 2208 | 1976 | 1902 | 2479 | 2244 | 1862 | 2112.06 | 2029 | 2195 | 1.083 | 0.115 | 0.36177417 | 0.58006064 | 0.0124 | 0.0085 | 0.0109 | 0.0109 | TRUE | 0.2128 |
| Coil | 4192 | 4610 | 4567 | 5473 | 3915 | 4102 | 4453 | 4604 | 4586 | 4302 | 4212 | 4490 | 4441.25 | 4548 | 4335 | 0.953 | -0.069 | 0.27698934 | 0.49802507 | 4.00E-04 | 0.0076 | 0.0027 | 0.0027 | TRUE | 0.0118 |
| Col10a1 | 0 | 0 | 0 | 1 | 0 | 0 | 0 | 0 | 0 | 1 | 0 | 0 | 0.13 | 0 | 0 | 1.859 | 0.894 | 0.82650734 | NA | 0 | 25.1515 | 10 | 10 | TRUE | 0.2563 |
| Col11a1 | 351 | 234 | 197 | 551 | 343 | 294 | 373 | 234 | 198 | 433 | 369 | 322 | 321.39 | 268 | 375 | 1.4 | 0.485 | 0.04272578 | 0.1475652 | 0.0621 | 0.0172 | 0.0381 | 0.0381 | TRUE | 1.2715 |
| Col11a2 | 23 | 26 | 17 | 18 | 25 | 19 | 24 | 26 | 17 | 14 | 27 | 21 | 21.55 | 22 | 21 | 0.907 | -0.14 | 0.75416855 | 0.87026239 | 0.0331 | 0.1598 | 0.0977 | 0.0977 | TRUE | 0.2595 |
| Col12a1 | 49 | 49 | 35 | 85 | 52 | 48 | 52 | 49 | 35 | 67 | 56 | 53 | 51.91 | 45 | 59 | 1.295 | 0.373 | 0.1983359 | 0.4053588 | 0 | 0.0704 | 0.0409 | 0.0409 | TRUE | 0.3086 |
| Col13a1 | 136 | 170 | 181 | 214 | 133 | 152 | 144 | 170 | 182 | 168 | 143 | 166 | 162.28 | 165 | 159 | 0.964 | -0.053 | 0.76136941 | 0.87423727 | 0.0041 | 0.0272 | 0.0157 | 0.0157 | TRUE | 0.1271 |
| Col14a1 | 20 | 9 | 15 | 15 | 12 | 13 | 21 | 9 | 15 | 12 | 13 | 14 | 14.04 | 15 | 13 | 0.859 | -0.219 | 0.68384831 | 0.82484365 | 0 | 0.2416 | 0.1374 | 0.1374 | TRUE | 0.164 |
| Col15a1 | 24 | 17 | 17 | 46 | 37 | 35 | 25 | 17 | 17 | 36 | 40 | 38 | 28.97 | 20 | 38 | 1.921 | 0.942 | 0.01068851 | 0.05538083 | 0 | 0.1206 | 0.0595 | 0.0595 | TRUE | 0.3197 |
| Col16a1 | 47 | 28 | 28 | 50 | 43 | 37 | 50 | 28 | 28 | 39 | 46 | 41 | 38.68 | 35 | 42 | 1.19 | 0.251 | 0.4771086 | 0.67670475 | 0.0323 | 0.0921 | 0.0639 | 0.0639 | TRUE | 0.9048 |
| Col17a1 | 137 | 156 | 123 | 176 | 149 | 163 | 146 | 156 | 124 | 138 | 160 | 178 | 150.32 | 142 | 159 | 1.119 | 0.163 | 0.3865329 | 0.60061236 | 0.009 | 0.0288 | 0.0187 | 0.0187 | TRUE | 0.1491 |
| Col18a1 | 14071 | 14763 | 15055 | 11973 | 9100 | 9379 | 14946 | 14744 | 15117 | 9412 | 9791 | 10267 | 12379.45 | 14936 | 9823 | 0.658 | -0.605 | 1.64E-21 | 4.97E-19 | 9.00E-04 | 0.0072 | 0.0028 | 0.0028 | TRUE | 0.019 |
| Col19a1 | 2 | 1 | 0 | 4 | 1 | 1 | 2 | 1 | 0 | 3 | 1 | 1 | 1.41 | 1 | 2 | 1.76 | 0.815 | 0.62311235 | NA | 0 | 2.3494 | 1.2193 | 1.2193 | TRUE | 0.3039 |
| Col1a1 | 276 | 188 | 174 | 182 | 122 | 98 | 293 | 188 | 175 | 143 | 131 | 107 | 172.87 | 219 | 127 | 0.584 | -0.776 | 0.00139193 | 0.0115423 | 0.0469 | 0.026 | 0.0363 | 0.0363 | TRUE | 0.9714 |
| Col1a2 | 91 | 60 | 59 | 81 | 64 | 42 | 97 | 60 | 59 | 64 | 69 | 46 | 65.72 | 72 | 60 | 0.83 | -0.269 | 0.3795225 | 0.59448576 | 0.0469 | 0.057 | 0.0521 | 0.0521 | TRUE | 0.8035 |
| Col20a1 | 35 | 71 | 65 | 59 | 59 | 55 | 37 | 71 | 65 | 46 | 63 | 60 | 57.24 | 58 | 56 | 0.975 | -0.036 | 0.9104826 | 0.95543125 | 0.05 | 0.0645 | 0.0575 | 0.0575 | TRUE | 0.8072 |
| Col22a1 | 308 | 363 | 365 | 585 | 450 | 403 | 327 | 363 | 366 | 460 | 484 | 441 | 406.9 | 352 | 462 | 1.311 | 0.391 | 0.00067473 | 0.00657712 | 0 | 0.015 | 0.007 | 0.007 | TRUE | 0.0431 |
| Col23a1 | 59 | 69 | 64 | 118 | 80 | 77 | 63 | 69 | 64 | 93 | 86 | 84 | 76.49 | 65 | 88 | 1.346 | 0.429 | 0.05102804 | 0.1663183 | 0 | 0.05 | 0.0214 | 0.0214 | TRUE | 0.0271 |
| Col24a1 | 14 | 5 | 6 | 23 | 12 | 15 | 15 | 5 | 6 | 18 | 13 | 16 | 12.22 | 9 | 16 | 1.848 | 0.886 | 0.15000474 | 0.34065871 | 0.0587 | 0.2766 | 0.1828 | 0.1828 | TRUE | 1.1925 |
| Col25a1 | 12 | 19 | 22 | 31 | 24 | 24 | 13 | 19 | 22 | 24 | 26 | 26 | 21.71 | 18 | 25 | 1.415 | 0.501 | 0.23054194 | 0.44434895 | 0 | 0.1586 | 0.0783 | 0.0783 | TRUE | 0.3102 |
| Col26a1 | 92 | 115 | 108 | 62 | 48 | 72 | 98 | 115 | 108 | 49 | 52 | 79 | 83.37 | 107 | 60 | 0.554 | -0.851 | 0.00093715 | 0.00847033 | 0.0206 | 0.0464 | 0.0347 | 0.0347 | TRUE | 0.6423 |
| Col27a1 | 122 | 161 | 139 | 254 | 181 | 170 | 130 | 161 | 140 | 200 | 195 | 186 | 168.41 | 144 | 194 | 1.351 | 0.434 | 0.00811697 | 0.04491959 | 0 | 0.0265 | 0.0132 | 0.0132 | TRUE | 0.119 |
| Col28a1 | 12 | 7 | 6 | 8 | 9 | 6 | 13 | 7 | 6 | 6 | 10 | 7 | 8.05 | 9 | 8 | 0.873 | -0.197 | 0.77448389 | 0.8825015 | 0 | 0.4161 | 0.2143 | 0.2143 | TRUE | 0.5319 |
| Col2a1 | 208 | 157 | 150 | 297 | 246 | 207 | 221 | 157 | 151 | 233 | 265 | 227 | 208.85 | 176 | 242 | 1.373 | 0.457 | 0.01696219 | 0.07731702 | 0.0206 | 0.0227 | 0.0215 | 0.0215 | TRUE | 0.5241 |
| Col3a1 | 0 | 0 | 0 | 1 | 2 | 0 | 0 | 0 | 0 | 1 | 2 | 0 | 0.49 | 0 | 1 | 5.204 | 2.38 | 0.47267317 | NA | 0 | 6.7348 | 6.4099 | 6.4099 | TRUE | 0.1222 |
| Col4a1 | 5623 | 6977 | 6788 | 8951 | 5991 | 5910 | 5973 | 6968 | 6816 | 7037 | 6446 | 6469 | 6618.07 | 6586 | 6651 | 1.01 | 0.014 | 0.87250526 | 0.93737812 | 0.0045 | 0.0074 | 0.0056 | 0.0056 | TRUE | 0.0808 |
| Col4a2 | 1658 | 2159 | 2011 | 3201 | 2066 | 2034 | 1761 | 2156 | 2019 | 2516 | 2223 | 2227 | 2150.39 | 1979 | 2322 | 1.174 | 0.231 | 0.03094491 | 0.11783272 | 0.0071 | 0.0084 | 0.0078 | 0.0078 | TRUE | 0.1117 |
| Col4a3 | 1 | 1 | 0 | 2 | 0 | 0 | 1 | 1 | 0 | 2 | 0 | 0 | 0.61 | 1 | 1 | 0.818 | -0.291 | 0.91402744 | NA | 0 | 5.4476 | 3.5995 | 3.5995 | TRUE | 0.9722 |
| Col4a3bp | 2652 | 2356 | 2583 | 4031 | 2850 | 2639 | 2817 | 2353 | 2594 | 3169 | 3066 | 2889 | 2814.59 | 2588 | 3041 | 1.176 | 0.233 | 0.01335532 | 0.06494505 | 0.0046 | 0.0081 | 0.0061 | 0.0061 | TRUE | 0.0765 |
| Col4a4 | 2 | 4 | 3 | 6 | 6 | 3 | 2 | 4 | 3 | 5 | 6 | 3 | 3.93 | 3 | 5 | 1.578 | 0.658 | 0.4851118 | NA | 0 | 0.8449 | 0.3722 | 0.3722 | TRUE | 0.1536 |
| Col4a5 | 61 | 63 | 45 | 109 | 67 | 61 | 65 | 63 | 45 | 86 | 72 | 67 | 66.24 | 58 | 75 | 1.305 | 0.384 | 0.14533335 | 0.33350757 | 0.0109 | 0.0566 | 0.0348 | 0.0348 | TRUE | 0.3064 |
| Col4a6 | 29 | 25 | 12 | 34 | 28 | 15 | 31 | 25 | 12 | 27 | 30 | 16 | 23.52 | 23 | 24 | 1.085 | 0.118 | 0.80669756 | 0.90035554 | 0.1006 | 0.147 | 0.1254 | 0.1254 | TRUE | 0.6442 |
| Col5a1 | 1104 | 1142 | 1187 | 1908 | 1306 | 1265 | 1173 | 1141 | 1192 | 1500 | 1405 | 1385 | 1299.15 | 1169 | 1430 | 1.225 | 0.293 | 0.00025008 | 0.00292927 | 6.00E-04 | 0.0094 | 0.0038 | 0.0038 | TRUE | 0.0228 |
| Col5a2 | 63 | 38 | 32 | 172 | 103 | 110 | 67 | 38 | 32 | 135 | 111 | 120 | 83.91 | 46 | 122 | 2.686 | 1.426 | 1.93E-06 | 4.88E-05 | 0.0539 | 0.0462 | 0.0494 | 0.0494 | TRUE | 1.2736 |
| Col5a3 | 1008 | 912 | 855 | 618 | 460 | 541 | 1071 | 911 | 859 | 486 | 495 | 592 | 735.5 | 947 | 524 | 0.553 | -0.854 | 1.46E-10 | 1.05E-08 | 0.0109 | 0.0114 | 0.0113 | 0.0113 | TRUE | 0.156 |
| Col6a1 | 46 | 58 | 44 | 33 | 36 | 23 | 49 | 58 | 44 | 26 | 39 | 25 | 40.14 | 50 | 30 | 0.592 | -0.756 | 0.02206288 | 0.09336617 | 0.0093 | 0.089 | 0.0521 | 0.0521 | TRUE | 0.4104 |
| Col6a2 | 174 | 180 | 178 | 208 | 135 | 142 | 185 | 180 | 179 | 164 | 145 | 155 | 167.92 | 181 | 155 | 0.856 | -0.224 | 0.14728932 | 0.3363323 | 0 | 0.0265 | 0.0112 | 0.0112 | TRUE | 0.0298 |
| Col6a3 | 47 | 38 | 40 | 34 | 47 | 50 | 50 | 38 | 40 | 27 | 51 | 55 | 43.34 | 43 | 44 | 1.022 | 0.031 | 0.93133098 | 0.96621594 | 0.06 | 0.0829 | 0.0716 | 0.0716 | TRUE | 1.0664 |
| Col6a4 | 235 | 230 | 224 | 300 | 260 | 243 | 250 | 230 | 225 | 236 | 280 | 266 | 247.64 | 235 | 261 | 1.107 | 0.147 | 0.30010056 | 0.52030187 | 0 | 0.0202 | 0.0105 | 0.0105 | TRUE | 0.0831 |
| Col6a5 | 376 | 213 | 222 | 210 | 138 | 111 | 399 | 213 | 223 | 165 | 148 | 122 | 211.68 | 278 | 145 | 0.522 | -0.937 | 0.00046695 | 0.00493162 | 0.0747 | 0.0225 | 0.0465 | 0.0465 | TRUE | 1.6158 |
| Col6a6 | 238 | 187 | 157 | 187 | 182 | 143 | 253 | 187 | 158 | 147 | 196 | 157 | 182.76 | 199 | 167 | 0.835 | -0.26 | 0.24252827 | 0.45909216 | 0.0353 | 0.0249 | 0.0301 | 0.0301 | TRUE | 0.5268 |
| Col7a1 | 226 | 143 | 137 | 188 | 159 | 120 | 240 | 143 | 138 | 148 | 171 | 131 | 161.78 | 174 | 150 | 0.866 | -0.208 | 0.41154938 | 0.62279432 | 0.0549 | 0.0273 | 0.0399 | 0.0399 | TRUE | 1.1935 |
| Col8a1 | 1 | 0 | 0 | 0 | 1 | 0 | 1 | 0 | 0 | 0 | 1 | 0 | 0.36 | 0 | 0 | 0.954 | -0.067 | 0.98557668 | NA | 0 | 9.2518 | 8.0179 | 8.0179 | TRUE | 0.5204 |
| Col8a2 | 214 | 150 | 179 | 111 | 91 | 82 | 227 | 150 | 180 | 87 | 98 | 90 | 138.63 | 186 | 92 | 0.493 | -1.019 | 1.96E-06 | 4.96E-05 | 0.0197 | 0.0307 | 0.025 | 0.025 | TRUE | 0.2011 |
| Col9a1 | 26 | 10 | 10 | 43 | 28 | 18 | 28 | 10 | 10 | 34 | 30 | 20 | 21.88 | 16 | 28 | 1.77 | 0.824 | 0.121906 | 0.29685061 | 0.1517 | 0.1575 | 0.1547 | 0.1547 | TRUE | 1.9046 |
| Col9a2 | 634 | 662 | 542 | 893 | 639 | 508 | 673 | 661 | 544 | 702 | 688 | 556 | 637.4 | 626 | 649 | 1.037 | 0.052 | 0.71099188 | 0.8426333 | 0.0132 | 0.0121 | 0.0126 | 0.0126 | TRUE | 0.1806 |
| Col9a3 | 517 | 506 | 493 | 399 | 317 | 253 | 549 | 505 | 495 | 314 | 341 | 277 | 413.53 | 516 | 311 | 0.602 | -0.733 | 5.88E-09 | 2.93E-07 | 0.003 | 0.0149 | 0.0089 | 0.0089 | TRUE | 0.0958 |
| Colec10 | 0 | 0 | 0 | 0 | 0 | 0 | 0 | 0 | 0 | 0 | 0 | 0 | 0 | 0 | 0 | NA | NA | NA | NA | NA | NA | NA | NA | NA | NA |
| Colec11 | 0 | 1 | 0 | 1 | 0 | 1 | 0 | 1 | 0 | 1 | 0 | 1 | 0.48 | 0 | 1 | 1.723 | 0.785 | 0.78325239 | NA | 0 | 6.8716 | 4.0968 | 4.0968 | TRUE | 0.405 |
| Colec12 | 352 | 284 | 296 | 566 | 387 | 334 | 374 | 284 | 297 | 445 | 416 | 366 | 363.61 | 318 | 409 | 1.288 | 0.365 | 0.01833073 | 0.08171632 | 0.0123 | 0.016 | 0.0144 | 0.0144 | TRUE | 0.2629 |
| Colq | 20 | 18 | 15 | 16 | 13 | 12 | 21 | 18 | 15 | 13 | 14 | 13 | 15.66 | 18 | 13 | 0.731 | -0.453 | 0.33881382 | 0.55879656 | 0 | 0.2172 | 0.0962 | 0.0962 | TRUE | 0.1136 |
| Commd1 | 1918 | 2531 | 2348 | 2644 | 1982 | 2110 | 2037 | 2528 | 2358 | 2079 | 2133 | 2310 | 2240.56 | 2308 | 2174 | 0.942 | -0.087 | 0.41089702 | 0.62206449 | 0.0067 | 0.0084 | 0.0076 | 0.0076 | TRUE | 0.1269 |
| Commd10 | 2669 | 3137 | 3112 | 3747 | 2667 | 2933 | 2835 | 3133 | 3125 | 2946 | 2870 | 3211 | 3019.74 | 3031 | 3009 | 0.992 | -0.011 | 0.89678774 | 0.95009983 | 0.003 | 0.008 | 0.0049 | 0.0049 | TRUE | 0.0415 |
| Commd2 | 1377 | 1416 | 1300 | 1975 | 1381 | 1307 | 1463 | 1414 | 1305 | 1553 | 1486 | 1431 | 1441.88 | 1394 | 1490 | 1.069 | 0.097 | 0.2615657 | 0.48115398 | 0.0018 | 0.0092 | 0.0047 | 0.0047 | TRUE | 0.0373 |
| Commd3 | 455 | 876 | 722 | 1268 | 874 | 941 | 483 | 875 | 725 | 997 | 940 | 1030 | 841.73 | 694 | 989 | 1.424 | 0.51 | 0.00858057 | 0.04689284 | 0.0428 | 0.0108 | 0.0259 | 0.0259 | TRUE | 0.5313 |
| Commd4 | 1245 | 1434 | 1436 | 1718 | 1340 | 1364 | 1322 | 1432 | 1442 | 1351 | 1442 | 1493 | 1413.65 | 1399 | 1429 | 1.02 | 0.029 | 0.73600617 | 0.8584108 | 0.0018 | 0.0092 | 0.0046 | 0.0046 | TRUE | 0.0283 |
| Commd5 | 602 | 561 | 532 | 1103 | 688 | 601 | 639 | 560 | 534 | 867 | 740 | 658 | 666.52 | 578 | 755 | 1.309 | 0.388 | 0.00513799 | 0.03195814 | 0.0119 | 0.0118 | 0.0123 | 0.0123 | TRUE | 0.2043 |
| Commd6 | 743 | 675 | 687 | 1048 | 717 | 655 | 789 | 674 | 690 | 824 | 771 | 717 | 744.24 | 718 | 771 | 1.075 | 0.105 | 0.34629173 | 0.56542111 | 0.0047 | 0.0113 | 0.0076 | 0.0076 | TRUE | 0.0892 |
| Commd7 | 495 | 419 | 456 | 677 | 509 | 484 | 526 | 418 | 458 | 532 | 548 | 530 | 501.96 | 467 | 537 | 1.148 | 0.2 | 0.09903145 | 0.25958518 | 0.0044 | 0.0135 | 0.0086 | 0.0086 | TRUE | 0.1376 |
| Commd8 | 2296 | 2368 | 2230 | 2902 | 2232 | 2169 | 2439 | 2365 | 2239 | 2281 | 2401 | 2374 | 2350 | 2348 | 2352 | 1.002 | 0.003 | 0.97216319 | 0.98528196 | 9.00E-04 | 0.0083 | 0.0035 | 0.0035 | TRUE | 0.0198 |
| Commd9 | 901 | 693 | 706 | 1106 | 783 | 726 | 957 | 692 | 709 | 869 | 842 | 795 | 810.78 | 786 | 835 | 1.064 | 0.089 | 0.53775013 | 0.72419528 | 0.0165 | 0.011 | 0.0139 | 0.0139 | TRUE | 0.4281 |
| Comp | 35 | 121 | 119 | 68 | 62 | 74 | 37 | 121 | 119 | 53 | 67 | 81 | 79.78 | 92 | 67 | 0.722 | -0.469 | 0.26254939 | 0.48195169 | 0.1911 | 0.0482 | 0.1135 | 0.1135 | TRUE | 2.4157 |
| Comt | 992 | 1147 | 1128 | 1580 | 1060 | 1135 | 1054 | 1146 | 1133 | 1242 | 1140 | 1242 | 1159.47 | 1111 | 1208 | 1.088 | 0.122 | 0.16196604 | 0.3571768 | 0.0013 | 0.0097 | 0.0046 | 0.0046 | TRUE | 0.0285 |
| Comtd1 | 668 | 759 | 834 | 720 | 605 | 616 | 710 | 758 | 837 | 566 | 651 | 674 | 699.37 | 768 | 630 | 0.819 | -0.288 | 0.01506655 | 0.07104027 | 0.0064 | 0.0116 | 0.0087 | 0.0087 | TRUE | 0.0982 |
| Copa | 14240 | 14809 | 14963 | 17368 | 12328 | 12858 | 15125 | 14790 | 15024 | 13653 | 13264 | 14075 | 14322.06 | 14980 | 13664 | 0.912 | -0.133 | 0.02076281 | 0.08921464 | 4.00E-04 | 0.0071 | 0.0023 | 0.0023 | TRUE | 0.0084 |

| | | | | | | | | | | | | | | | | | | | | | | | | | |
|---|---|---|---|---|---|---|---|---|---|---|---|---|---|---|---|---|---|---|---|---|---|---|---|---|---|
| Copb1 | 7425 | 8662 | 8782 | 9975 | 6909 | 7798 | 7887 | 8651 | 8818 | 7842 | 7434 | 8536 | 8194.48 | 8452 | 7937 | 0.939 | -0.091 | 0.29452125 | 0.51538767 | 0.004 | 0.0073 | 0.0053 | 0.0053 | TRUE | 0.053 |
| Copb2 | 12920 | 13071 | 12971 | 16305 | 11556 | 11429 | 13723 | 13054 | 13024 | 12818 | 12434 | 12511 | 12927.3 | 13267 | 12588 | 0.949 | -0.076 | 0.19437307 | 0.40016343 | 5.00E-04 | 0.0072 | 0.0024 | 0.0024 | TRUE | 0.011 |
| Cope | 4192 | 5219 | 5285 | 5137 | 3948 | 4332 | 4453 | 5212 | 5307 | 4038 | 4248 | 4742 | 4666.63 | 4991 | 4343 | 0.87 | -0.201 | 0.05495269 | 0.17484643 | 0.0075 | 0.0076 | 0.0077 | 0.0077 | TRUE | 0.1082 |
| Copg1 | 7065 | 6890 | 7297 | 10499 | 7429 | 7114 | 7504 | 6881 | 7327 | 8253 | 7993 | 7787 | 7624.4 | 7237 | 8011 | 1.107 | 0.147 | 0.03153241 | 0.11937809 | 0.0012 | 0.0073 | 0.0032 | 0.0032 | TRUE | 0.0227 |
| Copg2 | 3775 | 5264 | 5402 | 5309 | 4039 | 4426 | 4010 | 5257 | 5424 | 4174 | 4346 | 4845 | 4675.88 | 4897 | 4455 | 0.91 | -0.137 | 0.29573859 | 0.5165093 | 0.0152 | 0.0076 | 0.0121 | 0.0121 | TRUE | 0.3059 |
| Coprs | 2311 | 2357 | 2292 | 2985 | 2287 | 2100 | 2455 | 2354 | 2301 | 2347 | 2461 | 2299 | 2369.35 | 2370 | 2369 | 0.999 | -0.001 | 0.99087804 | 0.99518186 | 7.00E-04 | 0.0083 | 0.0033 | 0.0033 | TRUE | 0.0138 |
| Cops2 | 11373 | 10871 | 10957 | 14192 | 9811 | 9688 | 12080 | 10857 | 11002 | 11157 | 10556 | 10605 | 11042.8 | 11313 | 10773 | 0.952 | -0.07 | 0.33827731 | 0.5583911 | 0.0021 | 0.0072 | 0.0038 | 0.0038 | TRUE | 0.043 |
| Cops3 | 6247 | 5913 | 6035 | 10109 | 6849 | 6048 | 6635 | 5905 | 6060 | 7947 | 7369 | 6620 | 6756.18 | 6200 | 7312 | 1.18 | 0.238 | 0.01319204 | 0.06441016 | 0.0058 | 0.0074 | 0.0065 | 0.0065 | TRUE | 0.0835 |
| Cops4 | 3561 | 3918 | 3985 | 5471 | 3948 | 3817 | 3782 | 3913 | 4001 | 4301 | 4248 | 4178 | 4070.62 | 3899 | 4242 | 1.088 | 0.122 | 0.05037261 | 0.16493186 | 2.00E-04 | 0.0077 | 0.0025 | 0.0025 | TRUE | 0.0083 |
| Cops5 | 4132 | 4542 | 4695 | 6319 | 4365 | 4358 | 4389 | 4536 | 4714 | 4968 | 4696 | 4771 | 4678.97 | 4546 | 4812 | 1.058 | 0.082 | 0.22054289 | 0.43252954 | 7.00E-04 | 0.0076 | 0.003 | 0.003 | TRUE | 0.0127 |
| Cops6 | 5091 | 5471 | 5520 | 7331 | 5304 | 5182 | 5408 | 5464 | 5543 | 5763 | 5707 | 5673 | 5592.75 | 5472 | 5714 | 1.044 | 0.063 | 0.25995825 | 0.47920307 | 0 | 0.0075 | 0.0021 | 0.0021 | TRUE | 0.0016 |
| Cops7a | 4866 | 5282 | 5419 | 8985 | 6780 | 6532 | 5169 | 5275 | 5441 | 7063 | 7295 | 7150 | 6232.24 | 5295 | 7169 | 1.354 | 0.437 | 2.19E-13 | 2.50E-11 | 2.00E-04 | 0.0074 | 0.0024 | 0.0024 | TRUE | 0.0071 |
| Cops7b | 1899 | 1818 | 1770 | 2658 | 1893 | 1920 | 2017 | 1816 | 1777 | 2090 | 2037 | 2102 | 1973 | 1870 | 2076 | 1.11 | 0.151 | 0.06673304 | 0.1993297 | 0.0017 | 0.0086 | 0.0044 | 0.0044 | TRUE | 0.0569 |
| Cops8 | 2893 | 2815 | 2668 | 4191 | 2932 | 2738 | 3073 | 2811 | 2679 | 3295 | 3155 | 2997 | 3001.61 | 2854 | 3149 | 1.104 | 0.142 | 0.09879375 | 0.25917749 | 0.0031 | 0.008 | 0.005 | 0.005 | TRUE | 0.0543 |
| Copz1 | 6797 | 6783 | 6724 | 8837 | 6106 | 6011 | 7220 | 6774 | 6752 | 6947 | 6570 | 6580 | 6807.01 | 6915 | 6699 | 0.969 | -0.046 | 0.49627961 | 0.69196835 | 0.001 | 0.0074 | 0.0031 | 0.0031 | TRUE | 0.0181 |
| Copz2 | 285 | 539 | 551 | 257 | 172 | 342 | 303 | 538 | 553 | 202 | 185 | 374 | 359.29 | 465 | 254 | 0.545 | -0.875 | 0.00378672 | 0.0253253 | 0.1263 | 0.0161 | 0.0628 | 0.0628 | TRUE | 1.8998 |
| Coq10a | 594 | 868 | 827 | 559 | 437 | 588 | 631 | 867 | 830 | 439 | 470 | 644 | 646.92 | 776 | 518 | 0.666 | -0.586 | 0.00139851 | 0.0115847 | 0.0336 | 0.012 | 0.0226 | 0.0226 | TRUE | 0.5242 |
| Coq10b | 905 | 905 | 814 | 1150 | 782 | 820 | 961 | 904 | 817 | 904 | 841 | 898 | 887.58 | 894 | 881 | 0.986 | -0.02 | 0.83831202 | 0.91786582 | 0.0029 | 0.0106 | 0.0061 | 0.0061 | TRUE | 0.0673 |
| Coq2 | 1547 | 2004 | 2062 | 2163 | 1617 | 1681 | 1643 | 2001 | 2070 | 1700 | 1740 | 1840 | 1832.56 | 1905 | 1760 | 0.924 | -0.115 | 0.29807081 | 0.51872997 | 0.0073 | 0.0087 | 0.0082 | 0.0082 | TRUE | 0.1742 |
| Coq3 | 1731 | 2454 | 2394 | 2061 | 1586 | 1874 | 1839 | 2451 | 2404 | 1620 | 1706 | 2051 | 2011.89 | 2231 | 1792 | 0.803 | -0.316 | 0.02754725 | 0.10870032 | 0.019 | 0.0085 | 0.0143 | 0.0143 | TRUE | 0.2863 |
| Coq4 | 650 | 600 | 559 | 875 | 649 | 560 | 690 | 599 | 561 | 688 | 698 | 613 | 641.68 | 617 | 666 | 1.081 | 0.112 | 0.35185521 | 0.57063668 | 0.0064 | 0.012 | 0.0089 | 0.0089 | TRUE | 0.1266 |
| Coq5 | 2876 | 2994 | 2978 | 3527 | 2551 | 2578 | 3055 | 2990 | 2990 | 2773 | 2745 | 2822 | 2895.76 | 3012 | 2780 | 0.923 | -0.116 | 0.06078629 | 0.18761671 | 0 | 0.008 | 0.0024 | 0.0024 | TRUE | 0.0021 |
| Coq6 | 1533 | 2014 | 1957 | 2185 | 1603 | 1658 | 1628 | 2011 | 1965 | 1718 | 1725 | 1815 | 1810.35 | 1868 | 1753 | 0.938 | -0.093 | 0.37683557 | 0.59206639 | 0.0059 | 0.0087 | 0.0074 | 0.0074 | TRUE | 0.152 |
| Coq7 | 1289 | 1679 | 1537 | 1915 | 1391 | 1410 | 1369 | 1677 | 1543 | 1505 | 1497 | 1543 | 1522.47 | 1530 | 1515 | 0.99 | -0.014 | 0.88586407 | 0.94396169 | 0.0046 | 0.0091 | 0.0064 | 0.0064 | TRUE | 0.1012 |
| Coq9 | 2432 | 2994 | 3194 | 3240 | 2323 | 2392 | 2583 | 2990 | 3207 | 2547 | 2499 | 2618 | 2740.89 | 2927 | 2555 | 0.873 | -0.196 | 0.0492338 | 0.16260219 | 0.0056 | 0.0081 | 0.0068 | 0.0068 | TRUE | 0.1278 |
| Corin | 13 | 4 | 11 | 24 | 16 | 11 | 14 | 4 | 11 | 19 | 17 | 12 | 12.83 | 10 | 16 | 1.682 | 0.75 | 0.21443355 | 0.42616506 | 0.0625 | 0.2637 | 0.1795 | 0.1795 | TRUE | 0.899 |
| Coro1a | 391 | 365 | 372 | 382 | 248 | 263 | 415 | 365 | 374 | 300 | 267 | 288 | 334.73 | 385 | 285 | 0.743 | -0.429 | 0.00068038 | 0.00662405 | 0 | 0.0167 | 0.0084 | 0.0084 | TRUE | 0.056 |
| Coro1b | 3533 | 4054 | 4228 | 5402 | 3804 | 4053 | 3753 | 4049 | 4245 | 4247 | 4093 | 4437 | 4137.16 | 4016 | 4259 | 1.06 | 0.085 | 0.28883659 | 0.50977692 | 0.0024 | 0.0077 | 0.0043 | 0.0043 | TRUE | 0.0398 |
| Coro1c | 13327 | 11485 | 11072 | 15518 | 11391 | 9577 | 14156 | 11470 | 11117 | 12199 | 12256 | 10484 | 11946.97 | 12248 | 11646 | 0.951 | -0.073 | 0.54282021 | 0.7283165 | 0.0113 | 0.0072 | 0.0102 | 0.0102 | TRUE | 0.2269 |
| Coro2a | 446 | 706 | 682 | 637 | 439 | 550 | 474 | 705 | 685 | 501 | 472 | 602 | 573.13 | 621 | 525 | 0.845 | -0.244 | 0.17058875 | 0.36917202 | 0.0295 | 0.0127 | 0.021 | 0.021 | TRUE | 0.5049 |
| Coro2b | 2558 | 3573 | 3908 | 2289 | 1888 | 2363 | 2717 | 3568 | 3924 | 1799 | 2031 | 2587 | 2771.16 | 3403 | 2139 | 0.628 | -0.67 | 9.16E-05 | 0.00129038 | 0.0339 | 0.0081 | 0.0208 | 0.0208 | TRUE | 0.4049 |
| Coro6 | 113 | 116 | 106 | 158 | 120 | 101 | 120 | 116 | 106 | 124 | 129 | 111 | 117.7 | 114 | 121 | 1.064 | 0.09 | 0.62552292 | 0.78901854 | 0 | 0.0349 | 0.016 | 0.016 | TRUE | 0.0566 |
| Coro7 | 853 | 811 | 766 | 1068 | 796 | 754 | 906 | 810 | 769 | 840 | 856 | 825 | 834.42 | 828 | 840 | 1.015 | 0.021 | 0.83026238 | 0.91393762 | 0.0024 | 0.0109 | 0.0059 | 0.0059 | TRUE | 0.0793 |
| Cort | 23 | 28 | 35 | 20 | 20 | 32 | 24 | 28 | 35 | 16 | 22 | 35 | 26.63 | 29 | 24 | 0.816 | -0.294 | 0.49659767 | 0.69198951 | 0.0606 | 0.1306 | 0.0968 | 0.0968 | TRUE | 0.7392 |
| Cotl1 | 7847 | 9061 | 9231 | 7365 | 5253 | 6441 | 8335 | 9049 | 9269 | 5790 | 5652 | 7051 | 7524.25 | 8884 | 6164 | 0.694 | -0.528 | 9.46E-07 | 2.62E-05 | 0.0091 | 0.0073 | 0.0082 | 0.0082 | TRUE | 0.1927 |
| Cox10 | 3072 | 2799 | 2797 | 3529 | 2523 | 2327 | 3263 | 2795 | 2808 | 2774 | 2715 | 2547 | 2817.16 | 2955 | 2679 | 0.907 | -0.141 | 0.1302299 | 0.31078621 | 0.0045 | 0.0081 | 0.0059 | 0.0059 | TRUE | 0.1003 |
| Cox11 | 3001 | 2631 | 2438 | 3264 | 2538 | 2198 | 3188 | 2628 | 2448 | 2566 | 2731 | 2406 | 2660.98 | 2755 | 2568 | 0.932 | -0.101 | 0.39414177 | 0.60784239 | 0.011 | 0.0081 | 0.0098 | 0.0098 | TRUE | 0.2294 |
| Cox14 | 1042 | 1259 | 1270 | 1344 | 995 | 1059 | 1107 | 1257 | 1275 | 1057 | 1071 | 1159 | 1154.29 | 1213 | 1096 | 0.902 | -0.148 | 0.12813272 | 0.30785004 | 0.0034 | 0.0098 | 0.006 | 0.006 | TRUE | 0.0701 |
| Cox15 | 2148 | 2378 | 2474 | 3305 | 2359 | 2317 | 2282 | 2375 | 2484 | 2598 | 2538 | 2536 | 2468.87 | 2380 | 2557 | 1.075 | 0.104 | 0.14152904 | 0.32701377 | 6.00E-04 | 0.0082 | 0.0032 | 0.0032 | TRUE | 0.0177 |
| Cox16 | 523 | 616 | 623 | 805 | 529 | 579 | 556 | 615 | 626 | 633 | 569 | 634 | 605.35 | 599 | 612 | 1.022 | 0.032 | 0.76538197 | 0.87700158 | 0.0021 | 0.0123 | 0.0066 | 0.0066 | TRUE | 0.0464 |
| Cox17 | 3073 | 3042 | 2916 | 3618 | 2604 | 2503 | 3264 | 3038 | 2928 | 2844 | 2802 | 2740 | 2936 | 3077 | 2795 | 0.909 | -0.138 | 0.0648813 | 0.1962003 | 0.0012 | 0.008 | 0.0037 | 0.0037 | TRUE | 0.0344 |
| Cox18 | 1632 | 2295 | 2511 | 2305 | 1735 | 1968 | 1733 | 2292 | 2521 | 1812 | 1867 | 2154 | 2063.31 | 2182 | 1944 | 0.891 | -0.167 | 0.26164442 | 0.48118655 | 0.021 | 0.0085 | 0.0155 | 0.0155 | TRUE | 0.3912 |
| Cox19 | 527 | 481 | 479 | 729 | 491 | 423 | 560 | 480 | 481 | 573 | 528 | 463 | 514.25 | 507 | 521 | 1.03 | 0.043 | 0.7392857 | 0.86064468 | 0.0078 | 0.0133 | 0.0103 | 0.0103 | TRUE | 0.1108 |
| Cox20 | 139 | 149 | 126 | 257 | 158 | 148 | 148 | 149 | 127 | 202 | 170 | 162 | 159.5 | 141 | 178 | 1.267 | 0.342 | 0.05399788 | 0.17278444 | 0.0059 | 0.0276 | 0.0163 | 0.0163 | TRUE | 0.1606 |
| Cox4i1 | 19994 | 24481 | 23886 | 27673 | 20038 | 20779 | 21237 | 24449 | 23984 | 21754 | 21560 | 22746 | 22621.75 | 23223 | 22020 | 0.948 | -0.077 | 0.33835555 | 0.5583911 | 0.0031 | 0.007 | 0.0046 | 0.0046 | TRUE | 0.0685 |
| Cox4i2 | 146 | 235 | 256 | 209 | 156 | 233 | 155 | 235 | 257 | 164 | 168 | 255 | 205.67 | 216 | 196 | 0.906 | -0.143 | 0.56941391 | 0.74821263 | 0.0603 | 0.0229 | 0.0408 | 0.0408 | TRUE | 0.7494 |
| Cox5a | 9902 | 11521 | 11285 | 13288 | 9599 | 10007 | 10518 | 11506 | 11331 | 10446 | 10328 | 10954 | 10847.21 | 11118 | 10576 | 0.951 | -0.072 | 0.29698321 | 0.51785729 | 0.0015 | 0.0072 | 0.0034 | 0.0034 | TRUE | 0.0273 |
| Cox5b | 4169 | 5918 | 6285 | 6509 | 4673 | 4990 | 4428 | 5910 | 6311 | 5117 | 5028 | 5462 | 5376.07 | 5550 | 5202 | 0.937 | -0.093 | 0.48920981 | 0.68548482 | 0.0154 | 0.0075 | 0.013 | 0.013 | TRUE | 0.3808 |
| Cox6a1 | 6046 | 7146 | 7141 | 7464 | 5336 | 5876 | 6422 | 7137 | 7170 | 5868 | 5741 | 6432 | 6461.67 | 6910 | 6014 | 0.87 | -0.201 | 0.01772349 | 0.0795489 | 0.0035 | 0.0074 | 0.005 | 0.005 | TRUE | 0.0465 |
| Cox6a2 | 0 | 0 | 0 | 2 | 1 | 0 | 0 | 0 | 0 | 2 | 1 | 0 | 0.44 | 0 | 1 | 4.754 | 2.249 | 0.50892201 | NA | 0 | 7.4712 | 6.8061 | 6.8061 | TRUE | 0.2361 |
| Cox6b1 | 3439 | 4150 | 4220 | 4463 | 3116 | 3499 | 3653 | 4145 | 4237 | 3508 | 3353 | 3830 | 3787.68 | 4012 | 3564 | 0.888 | -0.171 | 0.07100997 | 0.20748529 | 0.005 | 0.0078 | 0.0062 | 0.0062 | TRUE | 0.0744 |
| Cox6b2 | 938 | 1922 | 1859 | 1244 | 1200 | 1453 | 996 | 1920 | 1867 | 978 | 1291 | 1591 | 1440.35 | 1594 | 1287 | 0.807 | -0.31 | 0.21161634 | 0.42221542 | 0.0898 | 0.0092 | 0.0437 | 0.0437 | TRUE | 0.598 |
| Cox6c | 5603 | 6588 | 6852 | 7223 | 5286 | 5926 | 5951 | 6579 | 6880 | 5678 | 5687 | 6487 | 6210.58 | 6470 | 5951 | 0.92 | -0.121 | 0.20032987 | 0.40805924 | 0.0055 | 0.0074 | 0.0063 | 0.0063 | TRUE | 0.0755 |
| Cox7a1 | 482 | 700 | 609 | 576 | 445 | 513 | 512 | 699 | 612 | 453 | 479 | 562 | 552.62 | 608 | 498 | 0.818 | -0.29 | 0.05659642 | 0.17876766 | 0.0156 | 0.0129 | 0.0148 | 0.0148 | TRUE | 0.1804 |
| Cox7a2 | 5186 | 5908 | 5919 | 6287 | 4382 | 4769 | 5508 | 5900 | 5943 | 4942 | 4715 | 5220 | 5371.61 | 5784 | 4959 | 0.857 | -0.222 | 0.00309125 | 0.02173658 | 0.0019 | 0.0075 | 0.0039 | 0.0039 | TRUE | 0.0256 |
| Cox7a2l | 5015 | 6235 | 6084 | 5775 | 4198 | 4679 | 5327 | 6227 | 6109 | 4540 | 4517 | 5122 | 5306.88 | 5888 | 4726 | 0.803 | -0.317 | 0.00104953 | 0.00925791 | 0.0058 | 0.0075 | 0.0066 | 0.0066 | TRUE | 0.0846 |
| Cox7b | 6385 | 7544 | 7728 | 7440 | 5501 | 5920 | 6782 | 7534 | 7760 | 5849 | 5919 | 6480 | 6720.64 | 7359 | 6083 | 0.826 | -0.275 | 0.0014335 | 0.01183108 | 0.0038 | 0.0074 | 0.0052 | 0.0052 | TRUE | 0.0573 |
| Cox7b2 | 32 | 32 | 30 | 264 | 167 | 154 | 34 | 32 | 30 | 208 | 180 | 169 | 108.64 | 32 | 186 | 5.804 | 2.537 | 1.44E-27 | 7.60E-25 | 0 | 0.0372 | 0.0206 | 0.0206 | TRUE | 0.127 |
| Cox7c | 5347 | 7155 | 6979 | 8346 | 5817 | 6339 | 5679 | 7146 | 7008 | 6561 | 6259 | 6939 | 6598.6 | 6611 | 6586 | 0.996 | -0.005 | 0.95964125 | 0.97926125 | 0.009 | 0.0074 | 0.0083 | 0.0083 | TRUE | 0.1852 |
| Cox8a | 8447 | 10631 | 11236 | 10312 | 7524 | 8480 | 8972 | 10617 | 11282 | 8106 | 8095 | 9283 | 9392.69 | 10290 | 8495 | 0.825 | -0.277 | 0.01263485 | 0.06252481 | 0.0097 | 0.0073 | 0.0088 | 0.0088 | TRUE | 0.1534 |
| Cox8b | 0 | 0 | 0 | 0 | 0 | 0 | 0 | 0 | 0 | 0 | 0 | 0 | 0 | 0 | 0 | NA | NA | NA | NA | NA | NA | NA | NA | NA | NA |
| Cox8c | 65 | 54 | 43 | 97 | 85 | 70 | 69 | 54 | 43 | 76 | 91 | 77 | 68.41 | 55 | 81 | 1.471 | 0.556 | 0.03650909 | 0.13177462 | 0.0131 | 0.0551 | 0.0358 | 0.0358 | TRUE | 0.3811 |
| Cp | 0 | 0 | 0 | 1 | 1 | 0 | 0 | 0 | 0 | 1 | 1 | 0 | 0.31 | 0 | 1 | 3.32 | 1.731 | 0.65383052 | NA | 0 | 10.6224 | 8.9541 | 8.9541 | TRUE | 0.202 |
| Cpa1 | 5 | 2 | 2 | 7 | 3 | 5 | 5 | 2 | 2 | 6 | 3 | 5 | 3.92 | 3 | 5 | 1.542 | 0.625 | 0.51949132 | NA | 0 | 0.8473 | 0.4102 | 0.4102 | TRUE | 0.4897 |
| Cpa2 | 57 | 62 | 70 | 78 | 58 | 56 | 61 | 62 | 70 | 61 | 62 | 61 | 62.96 | 64 | 61 | 0.959 | -0.06 | 0.80248269 | 0.89796077 | 0 | 0.0592 | 0.0255 | 0.0255 | TRUE | 0.0598 |
| Cpa3 | 0 | 0 | 0 | 0 | 0 | 0 | 0 | 0 | 0 | 0 | 0 | 0 | 0 | 0 | 0 | NA | NA | NA | NA | NA | NA | NA | NA | NA | NA |
| Cpa4 | 2 | 0 | 0 | 1 | 0 | 0 | 2 | 0 | 0 | 1 | 0 | 0 | 0.49 | 1 | 0 | 0.497 | -1.01 | 0.76786142 | NA | 0 | 6.7985 | 6.7083 | 6.7083 | TRUE | 0.9742 |
| Cpa5 | 0 | 0 | 1 | 1 | 0 | 0 | 0 | 0 | 1 | 1 | 0 | 0 | 0.3 | 0 | 0 | 0.954 | -0.067 | 0.98640617 | NA | 0 | 11.0483 | 9.2783 | 9.2783 | TRUE | 0.4098 |
| Cpa6 | 0 | 0 | 0 | 0 | 0 | 0 | 0 | 0 | 0 | 0 | 0 | 0 | 0 | 0 | 0 | NA | NA | NA | NA | NA | NA | NA | NA | NA | NA |

| | | | | | | | | | | | | | | | | | | | | | | | | | |
|---|---|---|---|---|---|---|---|---|---|---|---|---|---|---|---|---|---|---|---|---|---|---|---|---|---|
| Cpb1 | 20 | 18 | 9 | 25 | 16 | 12 | 21 | 18 | 9 | 20 | 17 | 13 | 16.38 | 16 | 17 | 1.044 | 0.063 | 0.9029752 | 0.95205059 | 0.0409 | 0.2081 | 0.13 | 0.13 | TRUE | 0.7066 |
| Cpb2 | 29 | 12 | 7 | 41 | 34 | 10 | 31 | 12 | 7 | 32 | 37 | 11 | 21.6 | 17 | 27 | 1.611 | 0.688 | 0.29510208 | 0.51600332 | 0.3757 | 0.1595 | 0.2619 | 0.2619 | TRUE | 2.1851 |
| Cpd | 2148 | 2837 | 3155 | 3818 | 2719 | 2880 | 2282 | 2833 | 3168 | 3001 | 2925 | 3153 | 2893.72 | 2761 | 3026 | 1.096 | 0.132 | 0.29654978 | 0.51727308 | 0.0124 | 0.008 | 0.0112 | 0.0112 | TRUE | 0.2795 |
| Cpe | 1029 | 1254 | 1275 | 1075 | 793 | 873 | 1093 | 1252 | 1280 | 845 | 853 | 956 | 1046.59 | 1208 | 885 | 0.731 | -0.451 | 2.08E-05 | 0.0003682 | 0.005 | 0.0101 | 0.0071 | 0.0071 | TRUE | 0.0835 |
| Cpeb1 | 293 | 467 | 513 | 492 | 347 | 391 | 311 | 466 | 515 | 387 | 373 | 428 | 413.48 | 431 | 396 | 0.918 | -0.123 | 0.52144469 | 0.71165526 | 0.0334 | 0.0149 | 0.024 | 0.024 | TRUE | 0.6782 |
| Cpeb2 | 619 | 883 | 871 | 1164 | 802 | 800 | 657 | 882 | 875 | 915 | 863 | 876 | 844.6 | 805 | 885 | 1.099 | 0.137 | 0.3069653 | 0.52744926 | 0.0124 | 0.0108 | 0.0117 | 0.0117 | TRUE | 0.3021 |
| Cpeb3 | 553 | 577 | 574 | 632 | 462 | 502 | 587 | 576 | 576 | 497 | 497 | 550 | 547.24 | 580 | 515 | 0.886 | -0.174 | 0.08401051 | 0.23229554 | 0 | 0.0129 | 0.0055 | 0.0055 | TRUE | 0.0402 |
| Cpeb4 | 501 | 477 | 538 | 820 | 560 | 506 | 532 | 476 | 540 | 645 | 603 | 554 | 558.3 | 516 | 601 | 1.165 | 0.22 | 0.05381658 | 0.1723792 | 0.0035 | 0.0128 | 0.0076 | 0.0076 | TRUE | 0.0536 |
| Cped1 | 6 | 3 | 2 | 8 | 6 | 1 | 6 | 3 | 2 | 6 | 6 | 1 | 4.2 | 4 | 4 | 1.237 | 0.306 | 0.76462518 | NA | 0.1972 | 0.7908 | 0.5146 | 0.5146 | TRUE | 0.7494 |
| Cphx1 | 0 | 0 | 0 | 0 | 0 | 0 | 0 | 0 | 0 | 0 | 0 | 0 | 0 | 0 | 0 | NA | NA | NA | NA | NA | NA | NA | NA | NA | NA |
| Cphx2 | 0 | 0 | 0 | 0 | 0 | 0 | 0 | 0 | 0 | 0 | 0 | 0 | 0 | 0 | 0 | NA | NA | NA | NA | NA | NA | NA | NA | NA | NA |
| Cplx1 | 168 | 164 | 126 | 135 | 80 | 73 | 178 | 164 | 127 | 106 | 86 | 80 | 123.48 | 156 | 91 | 0.584 | -0.776 | 0.00043287 | 0.00463487 | 0.0193 | 0.0336 | 0.0264 | 0.0264 | TRUE | 0.2932 |
| Cplx2 | 10 | 9 | 14 | 10 | 12 | 11 | 11 | 9 | 14 | 8 | 13 | 12 | 11.08 | 11 | 11 | 0.964 | -0.054 | 0.92589149 | 0.96345983 | 0 | 0.3042 | 0.1487 | 0.1487 | TRUE | 0.291 |
| Cplx3 | 0 | 0 | 0 | 1 | 0 | 0 | 0 | 0 | 0 | 1 | 0 | 0 | 0.13 | 0 | 0 | 1.859 | 0.894 | 0.82650734 | NA | 0 | 25.1515 | 10 | 10 | TRUE | 0.2563 |
| Cplx4 | 0 | 1 | 1 | 0 | 4 | 0 | 0 | 1 | 1 | 0 | 4 | 0 | 1.05 | 1 | 1 | 2.098 | 1.069 | 0.65857658 | NA | 3.0516 | 3.1412 | 3.1111 | 3.1111 | TRUE | 1.9004 |
| Cpm | 252 | 233 | 204 | 377 | 275 | 250 | 268 | 233 | 205 | 296 | 296 | 274 | 261.85 | 235 | 289 | 1.23 | 0.298 | 0.04611738 | 0.15538742 | 0.0052 | 0.0195 | 0.0122 | 0.0122 | TRUE | 0.1605 |
| Cpn1 | 716 | 917 | 924 | 1100 | 831 | 880 | 761 | 916 | 928 | 865 | 894 | 963 | 887.71 | 868 | 907 | 1.044 | 0.063 | 0.58174494 | 0.75797449 | 0.0063 | 0.0106 | 0.0082 | 0.0082 | TRUE | 0.1388 |
| Cpn2 | 5 | 8 | 5 | 16 | 6 | 4 | 5 | 8 | 5 | 13 | 6 | 4 | 6.96 | 6 | 8 | 1.302 | 0.381 | 0.61710132 | 0.7832683 | 0 | 0.4805 | 0.2733 | 0.2733 | TRUE | 1.0342 |
| Cpne1 | 4604 | 4341 | 3998 | 4291 | 2949 | 3226 | 4890 | 4335 | 4014 | 3373 | 3173 | 3531 | 3886.27 | 4413 | 3359 | 0.761 | -0.394 | 7.03E-05 | 0.00103709 | 0.0061 | 0.0078 | 0.0068 | 0.0068 | TRUE | 0.1087 |
| Cpne2 | 239 | 214 | 178 | 353 | 212 | 190 | 254 | 214 | 179 | 278 | 228 | 208 | 226.65 | 216 | 238 | 1.108 | 0.148 | 0.4380792 | 0.64579729 | 0.0219 | 0.0214 | 0.0217 | 0.0217 | TRUE | 0.2412 |
| Cpne3 | 3872 | 4607 | 4719 | 4799 | 3626 | 4239 | 4113 | 4601 | 4738 | 3773 | 3901 | 4640 | 4294.4 | 4484 | 4105 | 0.915 | -0.128 | 0.23989524 | 0.45609524 | 0.0088 | 0.0077 | 0.0083 | 0.0083 | TRUE | 0.1582 |
| Cpne4 | 5 | 2 | 2 | 4 | 4 | 8 | 5 | 2 | 2 | 3 | 4 | 9 | 4.25 | 3 | 5 | 1.726 | 0.787 | 0.42221417 | NA | 0 | 0.7814 | 0.4364 | 0.4364 | TRUE | 0.5916 |
| Cpne5 | 1016 | 1138 | 1126 | 1052 | 812 | 868 | 1079 | 1137 | 1131 | 827 | 874 | 950 | 999.52 | 1116 | 884 | 0.791 | -0.338 | 0.00026828 | 0.00310638 | 0.0018 | 0.0102 | 0.0052 | 0.0052 | TRUE | 0.0513 |
| Cpne6 | 9 | 10 | 4 | 3 | 5 | 5 | 10 | 10 | 4 | 2 | 5 | 5 | 6.13 | 8 | 4 | 0.549 | -0.864 | 0.29173594 | 0.51252275 | 0 | 0.5444 | 0.3055 | 0.3055 | TRUE | 0.5373 |
| Cpne7 | 650 | 508 | 409 | 265 | 263 | 286 | 690 | 507 | 411 | 208 | 283 | 313 | 402.13 | 536 | 268 | 0.499 | -1.003 | 7.32E-06 | 0.00015197 | 0.0511 | 0.0151 | 0.0332 | 0.0332 | TRUE | 0.2641 |
| Cpne8 | 282 | 363 | 397 | 284 | 236 | 233 | 300 | 363 | 399 | 223 | 254 | 255 | 298.82 | 354 | 244 | 0.689 | -0.538 | 0.00052851 | 0.00541274 | 0.0104 | 0.0179 | 0.0139 | 0.0139 | TRUE | 0.202 |
| Cpne9 | 184 | 213 | 235 | 290 | 199 | 241 | 195 | 213 | 236 | 228 | 214 | 264 | 225 | 215 | 235 | 1.094 | 0.13 | 0.40722975 | 0.62008376 | 0.0055 | 0.0215 | 0.0133 | 0.0133 | TRUE | 0.1188 |
| Cpox | 3187 | 3551 | 3667 | 4258 | 3191 | 3221 | 3385 | 3546 | 3682 | 3347 | 3433 | 3526 | 3486.69 | 3538 | 3435 | 0.971 | -0.043 | 0.54048083 | 0.72639311 | 8.00E-04 | 0.0078 | 0.0032 | 0.0032 | TRUE | 0.0173 |
| Cpped1 | 887 | 931 | 874 | 1526 | 1080 | 953 | 942 | 930 | 878 | 1200 | 1162 | 1043 | 1025.73 | 917 | 1135 | 1.239 | 0.31 | 0.00114369 | 0.00987062 | 0.0025 | 0.0101 | 0.0055 | 0.0055 | TRUE | 0.0591 |
| Cpq | 13 | 36 | 42 | 52 | 39 | 44 | 14 | 36 | 42 | 41 | 42 | 48 | 37.16 | 31 | 44 | 1.419 | 0.505 | 0.23053769 | 0.44434895 | 0.1062 | 0.0956 | 0.1001 | 0.1001 | TRUE | 1.4948 |
| Cps1 | 41 | 35 | 45 | 147 | 89 | 73 | 44 | 35 | 45 | 116 | 96 | 80 | 69.15 | 41 | 97 | 2.366 | 1.243 | 4.16E-06 | 9.46E-05 | 0 | 0.0545 | 0.0351 | 0.0351 | TRUE | 0.1398 |
| Cpsf1 | 7136 | 8126 | 8504 | 9191 | 7251 | 7637 | 7580 | 8116 | 8539 | 7225 | 7802 | 8360 | 7936.81 | 8078 | 7796 | 0.965 | -0.052 | 0.55715622 | 0.73979383 | 0.0038 | 0.0073 | 0.0054 | 0.0054 | TRUE | 0.0503 |
| Cpsf2 | 5008 | 4821 | 4784 | 6735 | 4857 | 4697 | 5319 | 4815 | 4804 | 5295 | 5226 | 5142 | 5099.96 | 4979 | 5221 | 1.049 | 0.069 | 0.34614509 | 0.56538943 | 0.0016 | 0.0075 | 0.0036 | 0.0036 | TRUE | 0.0434 |
| Cpsf3 | 5410 | 5759 | 5743 | 7122 | 5346 | 5443 | 5746 | 5752 | 5767 | 5599 | 5752 | 5958 | 5762.24 | 5755 | 5770 | 1.002 | 0.003 | 0.95483141 | 0.97700798 | 3.00E-04 | 0.0075 | 0.0024 | 0.0024 | TRUE | 0.01 |
| Cpsf3l | 4419 | 4608 | 4414 | 5734 | 4167 | 4062 | 4694 | 4602 | 4432 | 4508 | 4483 | 4446 | 4527.58 | 4576 | 4479 | 0.979 | -0.031 | 0.61469162 | 0.78200055 | 2.00E-04 | 0.0076 | 0.0025 | 0.0025 | TRUE | 0.0092 |
| Cpsf4 | 1873 | 1600 | 1625 | 2760 | 2001 | 1904 | 1989 | 1598 | 1632 | 2170 | 2153 | 2084 | 1937.66 | 1740 | 2136 | 1.228 | 0.296 | 0.00556509 | 0.03386288 | 0.0072 | 0.0086 | 0.0077 | 0.0077 | TRUE | 0.1899 |
| Cpsf4l | 383 | 671 | 678 | 738 | 601 | 677 | 407 | 670 | 681 | 580 | 647 | 741 | 620.94 | 586 | 656 | 1.118 | 0.162 | 0.42442612 | 0.63456714 | 0.0452 | 0.0122 | 0.0278 | 0.0278 | TRUE | 0.8354 |
| Cpsf6 | 8974 | 9190 | 9141 | 11842 | 8806 | 8143 | 9532 | 9178 | 9179 | 9309 | 9475 | 8914 | 9264.4 | 9296 | 9233 | 0.993 | -0.01 | 0.87217376 | 0.93737135 | 5.00E-04 | 0.0073 | 0.0026 | 0.0026 | TRUE | 0.0111 |
| Cpsf7 | 3697 | 3425 | 3494 | 4814 | 3594 | 3389 | 3927 | 3421 | 3508 | 3784 | 3867 | 3710 | 3702.82 | 3619 | 3787 | 1.047 | 0.066 | 0.41991507 | 0.63035126 | 0.0026 | 0.0078 | 0.0045 | 0.0045 | TRUE | 0.0674 |
| Cpt1a | 4227 | 7434 | 8043 | 6236 | 5127 | 6039 | 4490 | 7424 | 8076 | 4902 | 5516 | 6611 | 6169.91 | 6663 | 5676 | 0.852 | -0.231 | 0.25678999 | 0.47536288 | 0.0525 | 0.0074 | 0.0299 | 0.0299 | TRUE | 0.9933 |
| Cpt1b | 7 | 10 | 10 | 1 | 2 | 6 | 7 | 10 | 10 | 1 | 2 | 7 | 6.16 | 9 | 3 | 0.332 | -1.59 | 0.07687611 | 0.21894005 | 0.0075 | 0.5416 | 0.366 | 0.366 | TRUE | 1.1283 |
| Cpt1c | 1226 | 1740 | 1567 | 1402 | 1157 | 1395 | 1302 | 1738 | 1573 | 1102 | 1245 | 1527 | 1414.58 | 1538 | 1291 | 0.839 | -0.253 | 0.10122476 | 0.26349512 | 0.0232 | 0.0092 | 0.0165 | 0.0165 | TRUE | 0.3055 |
| Cpt2 | 1482 | 1848 | 1918 | 1894 | 1414 | 1577 | 1574 | 1846 | 1926 | 1489 | 1521 | 1726 | 1680.37 | 1782 | 1579 | 0.886 | -0.175 | 0.11774799 | 0.29026879 | 0.0078 | 0.0089 | 0.0085 | 0.0085 | TRUE | 0.1252 |
| Cpvl | 17 | 8 | 8 | 10 | 4 | 7 | 18 | 8 | 8 | 8 | 4 | 8 | 8.98 | 11 | 7 | 0.588 | -0.766 | 0.268021 | 0.48779451 | 0 | 0.3736 | 0.2263 | 0.2263 | TRUE | 0.9695 |
| Cpxcr1 | 0 | 0 | 0 | 0 | 0 | 0 | 0 | 0 | 0 | 0 | 0 | 0 | 0 | 0 | 0 | NA | NA | NA | NA | NA | NA | NA | NA | NA | NA |
| Cpxm1 | 1062 | 1131 | 1018 | 1516 | 1114 | 1021 | 1128 | 1130 | 1022 | 1192 | 1199 | 1118 | 1131.29 | 1093 | 1170 | 1.07 | 0.098 | 0.26885413 | 0.48886282 | 0.0014 | 0.0098 | 0.0047 | 0.0047 | TRUE | 0.0388 |
| Cpxm2 | 103 | 143 | 127 | 177 | 138 | 127 | 109 | 143 | 128 | 139 | 148 | 139 | 134.4 | 127 | 142 | 1.122 | 0.166 | 0.36407138 | 0.58154079 | 0 | 0.0314 | 0.0166 | 0.0166 | TRUE | 0.1407 |
| Cpz | 30 | 35 | 36 | 69 | 47 | 56 | 32 | 35 | 36 | 54 | 51 | 61 | 44.85 | 34 | 55 | 1.611 | 0.688 | 0.01690415 | 0.07714476 | 0 | 0.0804 | 0.036 | 0.036 | TRUE | 0.068 |
| Cr1l | 467 | 554 | 580 | 1010 | 700 | 655 | 496 | 553 | 582 | 794 | 753 | 717 | 649.31 | 544 | 755 | 1.388 | 0.473 | 1.19E-05 | 0.00022959 | 0.0028 | 0.012 | 0.0068 | 0.0068 | TRUE | 0.0686 |
| Cr2 | 76 | 57 | 45 | 100 | 61 | 56 | 81 | 57 | 45 | 79 | 66 | 61 | 64.73 | 61 | 69 | 1.13 | 0.177 | 0.54768717 | 0.73261374 | 0.0346 | 0.0578 | 0.0466 | 0.0466 | TRUE | 0.5671 |
| Crabp1 | 1880 | 2286 | 2288 | 2201 | 1643 | 1880 | 1997 | 2283 | 2297 | 1730 | 1768 | 2058 | 2022.22 | 2192 | 1852 | 0.844 | -0.244 | 0.02196303 | 0.09310133 | 0.0069 | 0.0085 | 0.0077 | 0.0077 | TRUE | 0.1137 |
| Crabp2 | 454 | 581 | 569 | 192 | 181 | 223 | 482 | 580 | 571 | 151 | 195 | 244 | 370.6 | 544 | 197 | 0.359 | -1.477 | 2.92E-15 | 4.11E-13 | 0.0279 | 0.0158 | 0.0218 | 0.0218 | TRUE | 0.4828 |
| Cradd | 423 | 335 | 348 | 516 | 376 | 374 | 449 | 335 | 349 | 406 | 405 | 409 | 392.15 | 378 | 407 | 1.077 | 0.107 | 0.46161603 | 0.66559865 | 0.0102 | 0.0153 | 0.0126 | 0.0126 | TRUE | 0.3123 |
| Cramp1l | 1925 | 2087 | 2190 | 2435 | 1935 | 1931 | 2045 | 2084 | 2199 | 1914 | 2082 | 2114 | 2072.99 | 2109 | 2037 | 0.965 | -0.052 | 0.52055141 | 0.71109881 | 0.0017 | 0.0085 | 0.0042 | 0.0042 | TRUE | 0.0343 |
| Crat | 613 | 578 | 611 | 798 | 583 | 528 | 651 | 577 | 614 | 627 | 627 | 578 | 612.41 | 614 | 611 | 0.996 | -0.006 | 0.95355659 | 0.97650213 | 0.0012 | 0.0123 | 0.006 | 0.006 | TRUE | 0.0327 |
| Crb1 | 3 | 3 | 5 | 4 | 2 | 6 | 3 | 3 | 5 | 3 | 2 | 7 | 3.84 | 4 | 4 | 1.048 | 0.068 | 0.9442913 | NA | 0 | 0.8638 | 0.4176 | 0.4176 | TRUE | 0.4928 |
| Crb2 | 73 | 82 | 94 | 108 | 58 | 55 | 78 | 82 | 94 | 85 | 62 | 60 | 76.89 | 85 | 69 | 0.823 | -0.28 | 0.25699271 | 0.47568231 | 0.012 | 0.0498 | 0.0311 | 0.0311 | TRUE | 0.3969 |
| Crb3 | 248 | 431 | 501 | 224 | 196 | 338 | 263 | 430 | 503 | 176 | 211 | 370 | 325.65 | 399 | 252 | 0.631 | -0.664 | 0.02967261 | 0.11417333 | 0.1265 | 0.017 | 0.064 | 0.064 | TRUE | 0.7933 |
| Crbn | 4226 | 4346 | 4295 | 5133 | 3697 | 3915 | 4489 | 4340 | 4313 | 4035 | 3978 | 4286 | 4240.05 | 4381 | 4100 | 0.936 | -0.096 | 0.15208513 | 0.3431743 | 7.00E-04 | 0.0077 | 0.003 | 0.003 | TRUE | 0.019 |
| Crcp | 1759 | 1637 | 1591 | 2338 | 1585 | 1320 | 1868 | 1635 | 1598 | 1838 | 1705 | 1445 | 1681.51 | 1700 | 1663 | 0.979 | -0.031 | 0.79519864 | 0.8946193 | 0.0105 | 0.0089 | 0.0097 | 0.0097 | TRUE | 0.1576 |
| Crct1 | 4 | 0 | 2 | 7 | 5 | 3 | 4 | 0 | 2 | 6 | 5 | 3 | 3.4 | 2 | 5 | 2.296 | 1.199 | 0.30035535 | NA | 0 | 0.9747 | 0.616 | 0.616 | TRUE | 0.4344 |
| Creb1 | 1690 | 1928 | 1997 | 2540 | 1722 | 1793 | 1795 | 1926 | 2005 | 1997 | 1853 | 1963 | 1923 | 1909 | 1938 | 1.015 | 0.022 | 0.78941403 | 0.89162403 | 0.0018 | 0.0086 | 0.0043 | 0.0043 | TRUE | 0.0326 |
| Creb3 | 198 | 189 | 204 | 357 | 242 | 181 | 210 | 189 | 205 | 281 | 260 | 198 | 223.84 | 201 | 246 | 1.228 | 0.296 | 0.09083306 | 0.24573226 | 0.0137 | 0.0216 | 0.0175 | 0.0175 | TRUE | 0.3176 |
| Creb3l1 | 710 | 589 | 627 | 613 | 459 | 418 | 754 | 588 | 630 | 482 | 494 | 458 | 567.55 | 657 | 478 | 0.727 | -0.459 | 0.00031715 | 0.00356785 | 0.0077 | 0.0127 | 0.0099 | 0.0099 | TRUE | 0.1947 |
| Creb3l2 | 640 | 818 | 820 | 667 | 558 | 585 | 680 | 817 | 823 | 524 | 600 | 640 | 680.87 | 773 | 588 | 0.759 | -0.397 | 0.00207284 | 0.01583951 | 0.0094 | 0.0117 | 0.0105 | 0.0105 | TRUE | 0.132 |
| Creb3l3 | 79 | 131 | 120 | 87 | 64 | 49 | 84 | 131 | 120 | 68 | 69 | 54 | 87.69 | 112 | 64 | 0.57 | -0.81 | 0.00159638 | 0.01287122 | 0.0257 | 0.0445 | 0.0352 | 0.0352 | TRUE | 0.4653 |
| Creb3l4 | 124 | 116 | 114 | 133 | 94 | 119 | 132 | 116 | 114 | 105 | 101 | 130 | 116.33 | 121 | 112 | 0.926 | -0.11 | 0.5770167 | 0.75455662 | 0.0032 | 0.0352 | 0.0197 | 0.0197 | TRUE | 0.1934 |
| Creb5 | 2 | 3 | 7 | 17 | 10 | 9 | 2 | 3 | 7 | 13 | 11 | 10 | 7.69 | 4 | 11 | 2.799 | 1.485 | 0.04811665 | 0.15980118 | 0 | 0.4355 | 0.2381 | 0.2381 | TRUE | 0.7052 |
| Crebbp | 1753 | 1445 | 1429 | 2472 | 1721 | 1304 | 1862 | 1443 | 1435 | 1943 | 1852 | 1427 | 1660.4 | 1580 | 1741 | 1.102 | 0.141 | 0.36182752 | 0.58006064 | 0.0219 | 0.0089 | 0.0165 | 0.0165 | TRUE | 0.298 |

| | | | | | | | | | | | | | | | | | | | | | | | | | |
|---|---|---|---|---|---|---|---|---|---|---|---|---|---|---|---|---|---|---|---|---|---|---|---|---|---|
| Crebl2 | 857 | 963 | 972 | 1011 | 787 | 773 | 910 | 962 | 976 | 795 | 847 | 846 | 889.29 | 949 | 829 | 0.873 | -0.197 | 0.02428325 | 0.09963462 | 0 | 0.0106 | 0.0044 | 0.0044 | TRUE | 0.0166 |
| Crebrf | 1979 | 2656 | 3132 | 2019 | 1734 | 2208 | 2102 | 2653 | 3145 | 1587 | 1866 | 2417 | 2294.89 | 2633 | 1957 | 0.743 | -0.429 | 0.02037806 | 0.08807968 | 0.0423 | 0.0083 | 0.0242 | 0.0242 | TRUE | 0.2208 |
| Crebzf | 2054 | 2016 | 2045 | 2372 | 2143 | 2045 | 2182 | 2013 | 2053 | 1865 | 2306 | 2239 | 2109.58 | 2083 | 2137 | 1.025 | 0.036 | 0.73631624 | 0.85870895 | 0.007 | 0.0085 | 0.0076 | 0.0076 | TRUE | 0.1522 |
| Creg1 | 855 | 990 | 977 | 858 | 699 | 748 | 908 | 989 | 981 | 674 | 752 | 819 | 853.88 | 959 | 748 | 0.779 | -0.361 | 0.0008392 | 0.00778126 | 0.0046 | 0.0108 | 0.0072 | 0.0072 | TRUE | 0.0923 |
| Creg2 | 120 | 214 | 167 | 127 | 148 | 159 | 127 | 214 | 168 | 100 | 159 | 174 | 157 | 170 | 144 | 0.847 | -0.239 | 0.37947354 | 0.59448576 | 0.0677 | 0.0279 | 0.047 | 0.047 | TRUE | 0.2775 |
| Creld1 | 1319 | 1716 | 1669 | 1471 | 1223 | 1320 | 1401 | 1714 | 1676 | 1156 | 1316 | 1445 | 1451.31 | 1597 | 1306 | 0.817 | -0.292 | 0.01891622 | 0.0835003 | 0.0108 | 0.0092 | 0.0104 | 0.0104 | TRUE | 0.1384 |
| Creld2 | 3467 | 3488 | 3174 | 4550 | 3130 | 2948 | 3683 | 3483 | 3187 | 3577 | 3368 | 3227 | 3420.78 | 3451 | 3391 | 0.983 | -0.025 | 0.77616483 | 0.88373755 | 0.0037 | 0.0079 | 0.0053 | 0.0053 | TRUE | 0.0545 |
| Crem | 391 | 431 | 489 | 642 | 412 | 397 | 415 | 430 | 491 | 505 | 443 | 435 | 453.22 | 445 | 461 | 1.036 | 0.051 | 0.68685765 | 0.82743019 | 0.0053 | 0.0142 | 0.0093 | 0.0093 | TRUE | 0.0924 |
| Crh | 0 | 0 | 1 | 0 | 0 | 0 | 0 | 0 | 1 | 0 | 0 | 0 | 0.17 | 0 | 0 | 0.49 | -1.029 | 0.80087 | NA | 0 | 19.6928 | 10 | 10 | TRUE | 0.4098 |
| Crhbp | 24 | 9 | 10 | 23 | 23 | 17 | 25 | 9 | 10 | 18 | 25 | 19 | 17.66 | 15 | 21 | 1.382 | 0.467 | 0.39288009 | 0.60637095 | 0.1151 | 0.1935 | 0.1568 | 0.1568 | TRUE | 1.7208 |
| Crhr1 | 12 | 10 | 10 | 23 | 16 | 10 | 13 | 10 | 10 | 18 | 17 | 11 | 13.17 | 11 | 15 | 1.422 | 0.508 | 0.33798157 | 0.55812432 | 0 | 0.2571 | 0.1243 | 0.1243 | TRUE | 0.2722 |
| Crhr2 | 152 | 151 | 90 | 103 | 77 | 67 | 161 | 151 | 90 | 81 | 83 | 73 | 106.63 | 134 | 79 | 0.59 | -0.762 | 0.00376366 | 0.02524902 | 0.0418 | 0.0378 | 0.0398 | 0.0398 | TRUE | 0.8452 |
| Crim1 | 606 | 889 | 841 | 1020 | 724 | 707 | 644 | 888 | 844 | 802 | 779 | 774 | 788.46 | 792 | 785 | 0.991 | -0.013 | 0.92323448 | 0.9623458 | 0.0132 | 0.0111 | 0.0122 | 0.0122 | TRUE | 0.3167 |
| Crip1 | 981 | 1276 | 1362 | 1141 | 953 | 1217 | 1042 | 1274 | 1368 | 897 | 1025 | 1332 | 1156.41 | 1228 | 1085 | 0.883 | -0.18 | 0.28323469 | 0.50491557 | 0.0271 | 0.0098 | 0.0194 | 0.0194 | TRUE | 0.4749 |
| Crip2 | 1574 | 2030 | 2147 | 2240 | 1746 | 2031 | 1672 | 2027 | 2156 | 1761 | 1879 | 2223 | 1952.97 | 1952 | 1954 | 1.001 | 0.001 | 0.99326446 | 0.99627909 | 0.0154 | 0.0086 | 0.0122 | 0.0122 | TRUE | 0.1898 |
| Crip3 | 6 | 7 | 5 | 4 | 4 | 7 | 6 | 7 | 5 | 3 | 4 | 8 | 5.58 | 6 | 5 | 0.808 | -0.308 | 0.70208469 | 0.83705948 | 0 | 0.597 | 0.284 | 0.284 | TRUE | 0.392 |
| Cript | 1515 | 1654 | 1603 | 2136 | 1476 | 1485 | 1609 | 1652 | 1610 | 1679 | 1588 | 1626 | 1627.24 | 1624 | 1631 | 1.005 | 0.007 | 0.91998087 | 0.9607958 | 0 | 0.0089 | 0.0031 | 0.0031 | TRUE | 0.0084 |
| Crisp1 | 4 | 1 | 3 | 6 | 6 | 3 | 4 | 1 | 3 | 5 | 6 | 3 | 3.79 | 3 | 5 | 1.759 | 0.815 | 0.41612314 | NA | 0 | 0.8771 | 0.4359 | 0.4359 | TRUE | 0.3819 |
| Crisp2 | 0 | 0 | 0 | 1 | 3 | 1 | 0 | 0 | 0 | 1 | 3 | 1 | 0.85 | 0 | 2 | 8.982 | 3.167 | 0.17867542 | NA | 0 | 3.8762 | 2.7057 | 2.7057 | TRUE | 0.4462 |
| Crisp3 | 0 | 0 | 0 | 0 | 0 | 0 | 0 | 0 | 0 | 0 | 0 | 0 | 0 | 0 | 0 | NA | NA | NA | NA | NA | NA | NA | NA | NA | NA |
| Crisp4 | 0 | 0 | 0 | 0 | 0 | 0 | 0 | 0 | 0 | 0 | 0 | 0 | 0 | 0 | 0 | NA | NA | NA | NA | NA | NA | NA | NA | NA | NA |
| Crispld1 | 65 | 69 | 67 | 99 | 63 | 63 | 69 | 69 | 67 | 78 | 68 | 69 | 69.97 | 68 | 72 | 1.049 | 0.069 | 0.76160582 | 0.87430286 | 0 | 0.054 | 0.0233 | 0.0233 | TRUE | 0.0612 |
| Crispld2 | 563 | 571 | 515 | 549 | 383 | 488 | 598 | 570 | 517 | 432 | 412 | 534 | 510.54 | 562 | 459 | 0.817 | -0.292 | 0.03369756 | 0.12474457 | 0.0099 | 0.0134 | 0.0116 | 0.0116 | TRUE | 0.2308 |
| Crk | 9707 | 10935 | 11108 | 13285 | 9505 | 9544 | 10311 | 10921 | 11154 | 10444 | 10227 | 10447 | 10583.81 | 10795 | 10373 | 0.961 | -0.058 | 0.35685884 | 0.57538108 | 7.00E-04 | 0.0072 | 0.0027 | 0.0027 | TRUE | 0.0188 |
| Crkl | 3054 | 3169 | 3215 | 4404 | 3098 | 3161 | 3244 | 3165 | 3228 | 3462 | 3333 | 3460 | 3315.42 | 3212 | 3418 | 1.064 | 0.09 | 0.14637306 | 0.33511336 | 0 | 0.0079 | 0.0025 | 0.0025 | TRUE | 0.0057 |
| Crlf1 | 103 | 207 | 211 | 178 | 142 | 193 | 109 | 207 | 212 | 140 | 153 | 211 | 172 | 176 | 168 | 0.952 | -0.071 | 0.8012558 | 0.89741637 | 0.0803 | 0.0261 | 0.0514 | 0.0514 | TRUE | 1.16 |
| Crlf2 | 234 | 327 | 330 | 492 | 425 | 394 | 249 | 327 | 331 | 387 | 457 | 431 | 363.64 | 302 | 425 | 1.404 | 0.49 | 0.00146838 | 0.01206724 | 0.0126 | 0.016 | 0.0142 | 0.0142 | TRUE | 0.2689 |
| Crlf3 | 5035 | 5136 | 5108 | 6445 | 4670 | 4877 | 5348 | 5129 | 5129 | 5067 | 5025 | 5339 | 5172.71 | 5202 | 5144 | 0.989 | -0.017 | 0.79703825 | 0.89548991 | 5.00E-04 | 0.0075 | 0.0028 | 0.0028 | TRUE | 0.0133 |
| Crls1 | 1441 | 1742 | 1722 | 1850 | 1450 | 1573 | 1531 | 1740 | 1729 | 1454 | 1560 | 1722 | 1622.62 | 1667 | 1579 | 0.947 | -0.079 | 0.44000829 | 0.64727303 | 0.0051 | 0.0089 | 0.007 | 0.007 | TRUE | 0.0756 |
| Crmp1 | 2096 | 2142 | 2430 | 2078 | 1540 | 1601 | 2226 | 2139 | 2440 | 1634 | 1657 | 1753 | 1974.76 | 2268 | 1681 | 0.741 | -0.433 | 4.83E-07 | 1.46E-05 | 0.0023 | 0.0086 | 0.0048 | 0.0048 | TRUE | 0.0531 |
| Crnkl1 | 2763 | 2496 | 2485 | 3497 | 2475 | 2288 | 2935 | 2493 | 2495 | 2749 | 2663 | 2505 | 2639.89 | 2641 | 2639 | 1 | -0.001 | 0.99513069 | 0.9976609 | 0.0052 | 0.0082 | 0.0064 | 0.0064 | TRUE | 0.1147 |
| Crnn | 0 | 0 | 0 | 1 | 0 | 0 | 0 | 0 | 0 | 1 | 0 | 0 | 0.13 | 0 | 0 | 1.859 | 0.894 | 0.82650734 | NA | 0 | 25.1515 | 10 | 10 | TRUE | 0.2563 |
| Crocc | 1257 | 1312 | 1334 | 1804 | 1617 | 1441 | 1335 | 1310 | 1339 | 1418 | 1740 | 1577 | 1453.38 | 1328 | 1578 | 1.187 | 0.248 | 0.01356119 | 0.06564981 | 0.0048 | 0.0092 | 0.0066 | 0.0066 | TRUE | 0.0974 |
| Crot | 1820 | 2107 | 2010 | 2568 | 1970 | 1981 | 1933 | 2104 | 2018 | 2019 | 2120 | 2169 | 2060.43 | 2018 | 2103 | 1.041 | 0.058 | 0.45004915 | 0.65553889 | 0.0011 | 0.0085 | 0.0037 | 0.0037 | TRUE | 0.0167 |
| Crp | 0 | 0 | 0 | 0 | 0 | 0 | 0 | 0 | 0 | 0 | 0 | 0 | 0 | 0 | 0 | NA | NA | NA | NA | NA | NA | NA | NA | NA | NA |
| Crtac1 | 506 | 789 | 832 | 743 | 562 | 748 | 537 | 788 | 835 | 584 | 605 | 819 | 694.74 | 720 | 669 | 0.928 | -0.108 | 0.58617537 | 0.76113826 | 0.0431 | 0.0116 | 0.0267 | 0.0267 | TRUE | 0.5811 |
| Crtam | 14 | 8 | 7 | 51 | 40 | 23 | 15 | 8 | 7 | 40 | 43 | 25 | 23.03 | 10 | 36 | 3.65 | 1.868 | 0.00013102 | 0.00171482 | 0.0502 | 0.1499 | 0.1069 | 0.1069 | TRUE | 0.6122 |
| Crtap | 1151 | 1162 | 1159 | 2022 | 1389 | 1270 | 1223 | 1160 | 1164 | 1590 | 1494 | 1390 | 1336.84 | 1182 | 1491 | 1.263 | 0.336 | 0.00014156 | 0.00181661 | 0.0021 | 0.0094 | 0.0049 | 0.0049 | TRUE | 0.0422 |
| Crtc1 | 807 | 793 | 791 | 1045 | 755 | 673 | 857 | 792 | 794 | 821 | 812 | 737 | 802.32 | 814 | 790 | 0.971 | -0.042 | 0.66093681 | 0.80907385 | 0.0015 | 0.011 | 0.0054 | 0.0054 | TRUE | 0.0407 |
| Crtc2 | 759 | 637 | 605 | 1087 | 860 | 654 | 806 | 636 | 607 | 855 | 925 | 716 | 757.6 | 683 | 832 | 1.218 | 0.285 | 0.05951332 | 0.18477246 | 0.0186 | 0.0113 | 0.0151 | 0.0151 | TRUE | 0.2911 |
| Crtc3 | 136 | 168 | 170 | 334 | 236 | 206 | 144 | 168 | 171 | 263 | 254 | 225 | 204.15 | 161 | 247 | 1.538 | 0.621 | 6.68E-05 | 0.00099647 | 0 | 0.023 | 0.0123 | 0.0123 | TRUE | 0.0835 |
| Crx | 5 | 1 | 1 | 4 | 9 | 6 | 5 | 1 | 1 | 3 | 10 | 7 | 4.45 | 2 | 7 | 2.648 | 1.405 | 0.18551261 | NA | 0.2641 | 0.747 | 0.5232 | 0.5232 | TRUE | 0.43 |
| Crxos1 | 447 | 509 | 360 | 597 | 500 | 516 | 475 | 508 | 361 | 469 | 538 | 565 | 486.12 | 448 | 524 | 1.168 | 0.224 | 0.15577629 | 0.34834873 | 0.0179 | 0.0137 | 0.0159 | 0.0159 | TRUE | 0.3335 |
| Cry1 | 2523 | 2783 | 2672 | 3094 | 2338 | 2276 | 2680 | 2779 | 2683 | 2432 | 2516 | 2491 | 2596.92 | 2714 | 2480 | 0.913 | -0.131 | 0.04349608 | 0.14928338 | 0 | 0.0082 | 0.0026 | 0.0026 | TRUE | 0.0054 |
| Cry2 | 1885 | 1901 | 1991 | 1720 | 1414 | 1410 | 2002 | 1899 | 1999 | 1352 | 1521 | 1543 | 1719.48 | 1967 | 1472 | 0.748 | -0.419 | 1.59E-06 | 4.11E-05 | 0.0025 | 0.0088 | 0.0049 | 0.0049 | TRUE | 0.0633 |
| Cryaa | 19 | 4 | 5 | 31 | 18 | 4 | 20 | 4 | 5 | 24 | 19 | 4 | 12.89 | 10 | 16 | 1.667 | 0.737 | 0.37656947 | 0.59197185 | 0.581 | 0.2626 | 0.4178 | 0.4178 | TRUE | 1.274 |
| Cryab | 28 | 32 | 35 | 47 | 40 | 46 | 30 | 32 | 35 | 37 | 43 | 50 | 37.86 | 32 | 43 | 1.338 | 0.42 | 0.18614033 | 0.38985171 | 0 | 0.0939 | 0.0456 | 0.0456 | TRUE | 0.1556 |
| Cryba1 | 2 | 4 | 2 | 4 | 1 | 2 | 2 | 4 | 2 | 3 | 1 | 2 | 2.42 | 3 | 2 | 0.804 | -0.316 | 0.78991348 | NA | 0 | 1.3666 | 0.5989 | 0.5989 | TRUE | 0.2188 |
| Cryba2 | 0 | 0 | 0 | 0 | 0 | 0 | 0 | 0 | 0 | 0 | 0 | 0 | 0 | 0 | 0 | NA | NA | NA | NA | NA | NA | NA | NA | NA | NA |
| Cryba4 | 3 | 3 | 4 | 3 | 1 | 6 | 3 | 3 | 4 | 2 | 1 | 7 | 3.37 | 3 | 3 | 0.968 | -0.047 | 0.96483028 | NA | 0 | 0.9854 | 0.5297 | 0.5297 | TRUE | 0.9184 |
| Crybb1 | 3 | 0 | 0 | 1 | 1 | 0 | 3 | 0 | 0 | 1 | 1 | 0 | 0.84 | 1 | 1 | 0.596 | -0.747 | 0.76594026 | NA | 2.0314 | 3.9222 | 3.2678 | 3.2678 | TRUE | 1.4428 |
| Crybb2 | 0 | 0 | 0 | 0 | 0 | 0 | 0 | 0 | 0 | 0 | 0 | 0 | 0 | 0 | 0 | NA | NA | NA | NA | NA | NA | NA | NA | NA | NA |
| Crybb3 | 66 | 72 | 63 | 63 | 51 | 68 | 70 | 72 | 63 | 50 | 55 | 74 | 64.02 | 68 | 60 | 0.866 | -0.207 | 0.43284314 | 0.64199926 | 0.0092 | 0.0584 | 0.0347 | 0.0347 | TRUE | 0.3912 |
| Crybg3 | 390 | 366 | 393 | 515 | 319 | 285 | 414 | 366 | 395 | 405 | 343 | 312 | 372.41 | 392 | 353 | 0.905 | -0.143 | 0.311335 | 0.53199265 | 0.0083 | 0.0157 | 0.0117 | 0.0117 | TRUE | 0.1949 |
| Cryga | 52 | 47 | 37 | 63 | 43 | 30 | 55 | 47 | 37 | 50 | 46 | 33 | 44.66 | 46 | 43 | 0.929 | -0.106 | 0.73973873 | 0.86066702 | 0.0181 | 0.0807 | 0.0511 | 0.0511 | TRUE | 0.2701 |
| Crygb | 3 | 1 | 0 | 10 | 4 | 3 | 3 | 1 | 0 | 8 | 4 | 3 | 3.27 | 1 | 5 | 3.786 | 1.921 | 0.12311821 | NA | 0.112 | 1.0137 | 0.6536 | 0.6536 | TRUE | 0.7656 |
| Crygc | 18 | 14 | 8 | 37 | 23 | 9 | 19 | 14 | 8 | 29 | 25 | 10 | 17.47 | 14 | 21 | 1.564 | 0.645 | 0.2651008 | 0.48482952 | 0.1662 | 0.1955 | 0.1815 | 0.1815 | TRUE | 0.6127 |
| Crygd | 74 | 61 | 44 | 91 | 71 | 61 | 79 | 61 | 44 | 72 | 76 | 67 | 66.4 | 61 | 72 | 1.171 | 0.227 | 0.41672203 | 0.62804967 | 0.0239 | 0.0565 | 0.0413 | 0.0413 | TRUE | 0.2265 |
| Cryge | 69 | 32 | 44 | 96 | 51 | 41 | 73 | 32 | 44 | 75 | 55 | 45 | 54.11 | 50 | 58 | 1.181 | 0.24 | 0.52527479 | 0.7148075 | 0.1016 | 0.0678 | 0.084 | 0.084 | TRUE | 0.776 |
| Crygf | 0 | 2 | 0 | 1 | 2 | 0 | 0 | 2 | 0 | 1 | 2 | 0 | 0.82 | 1 | 1 | 1.455 | 0.541 | 0.82645895 | NA | 0 | 4.012 | 3.1289 | 3.1289 | TRUE | 0.6969 |
| Crygn | 14 | 16 | 14 | 15 | 25 | 20 | 15 | 16 | 14 | 12 | 27 | 22 | 17.58 | 15 | 20 | 1.332 | 0.414 | 0.406668 | 0.61934798 | 0 | 0.1943 | 0.1206 | 0.1206 | TRUE | 0.4779 |
| Crygs | 20 | 9 | 6 | 9 | 8 | 10 | 21 | 9 | 6 | 7 | 9 | 11 | 10.48 | 12 | 9 | 0.732 | -0.45 | 0.5094684 | 0.7030655 | 0.1552 | 0.3212 | 0.2372 | 0.2372 | TRUE | 1.3769 |
| Cryl1 | 535 | 521 | 508 | 827 | 562 | 544 | 568 | 520 | 510 | 650 | 605 | 595 | 574.83 | 533 | 617 | 1.159 | 0.213 | 0.04060516 | 0.14244318 | 0.0011 | 0.0126 | 0.006 | 0.006 | TRUE | 0.039 |
| Crym | 255 | 372 | 303 | 302 | 247 | 254 | 271 | 372 | 304 | 237 | 266 | 278 | 287.97 | 316 | 260 | 0.823 | -0.28 | 0.08494653 | 0.23418574 | 0.0131 | 0.0183 | 0.0156 | 0.0156 | TRUE | 0.2737 |
| Cryz | 1302 | 1892 | 1880 | 2461 | 1665 | 1859 | 1383 | 1890 | 1888 | 1935 | 1791 | 2035 | 1820.21 | 1720 | 1920 | 1.116 | 0.159 | 0.24758344 | 0.46433167 | 0.0152 | 0.0087 | 0.013 | 0.013 | TRUE | 0.3539 |
| Cryzl1 | 508 | 605 | 624 | 889 | 624 | 652 | 540 | 604 | 627 | 699 | 671 | 714 | 642.39 | 590 | 695 | 1.177 | 0.235 | 0.02322171 | 0.09663663 | 0.0015 | 0.012 | 0.0061 | 0.0061 | TRUE | 0.0652 |
| Cs | 17077 | 18723 | 19425 | 20965 | 15133 | 16117 | 18139 | 18699 | 19505 | 16481 | 16282 | 17643 | 17791.36 | 18781 | 16802 | 0.895 | -0.161 | 0.01869904 | 0.0829129 | 0.0015 | 0.0071 | 0.0033 | 0.0033 | TRUE | 0.0234 |
| Csad | 1633 | 1772 | 1829 | 1589 | 1233 | 1137 | 1735 | 1770 | 1837 | 1249 | 1327 | 1245 | 1526.86 | 1781 | 1274 | 0.715 | -0.484 | 2.60E-10 | 1.82E-08 | 3.00E-04 | 0.0091 | 0.0036 | 0.0036 | TRUE | 0.0157 |
| Csda | 31260 | 33543 | 32537 | 38925 | 29221 | 27246 | 33204 | 33500 | 32671 | 30600 | 31440 | 29825 | 31873.12 | 33125 | 30622 | 0.924 | -0.113 | 0.04000981 | 0.14082452 | 4.00E-04 | 0.007 | 0.0022 | 0.0022 | TRUE | 0.0067 |

| | | | | | | | | | | | | | | | | | | | | | | | | | |
|---|---|---|---|---|---|---|---|---|---|---|---|---|---|---|---|---|---|---|---|---|---|---|---|---|---|
| Csdc2 | 91 | 58 | 43 | 86 | 51 | 47 | 97 | 58 | 43 | 68 | 55 | 51 | 61.95 | 66 | 58 | 0.884 | -0.178 | 0.6053269 | 0.77594408 | 0.079 | 0.0601 | 0.0693 | 0.0693 | TRUE | 0.7023 |
| Csde1 | 24400 | 26684 | 26748 | 32625 | 23392 | 23542 | 25917 | 26650 | 26858 | 25647 | 25168 | 25770 | 26001.75 | 26475 | 25528 | 0.964 | -0.053 | 0.32063203 | 0.54182417 | 2.00E-04 | 0.007 | 0.002 | 0.002 | TRUE | 0.0042 |
| Cse1l | 17594 | 16542 | 16578 | 22260 | 15386 | 15497 | 18688 | 16521 | 16646 | 17499 | 16554 | 16964 | 17145.33 | 17285 | 17006 | 0.984 | -0.023 | 0.76226869 | 0.87464971 | 0.0023 | 0.0071 | 0.0043 | 0.0043 | TRUE | 0.0616 |
| Csf1 | 13 | 15 | 12 | 43 | 21 | 22 | 14 | 15 | 12 | 34 | 23 | 24 | 20.22 | 14 | 27 | 1.987 | 0.99 | 0.02541179 | 0.1028998 | 0 | 0.1698 | 0.0859 | 0.0859 | TRUE | 0.384 |
| Csf1r | 0 | 1 | 1 | 1 | 1 | 0 | 0 | 1 | 1 | 1 | 1 | 0 | 0.64 | 1 | 1 | 0.935 | -0.097 | 0.96674856 | NA | 0 | 5.1213 | 2.3899 | 2.3899 | TRUE | 0.2246 |
| Csf2 | 0 | 0 | 0 | 0 | 0 | 0 | 0 | 0 | 0 | 0 | 0 | 0 | 0 | 0 | 0 | NA | NA | NA | NA | NA | NA | NA | NA | NA | NA |
| Csf2ra | 268 | 294 | 287 | 318 | 241 | 249 | 285 | 294 | 288 | 250 | 259 | 273 | 274.72 | 289 | 261 | 0.901 | -0.15 | 0.22890132 | 0.44224019 | 0 | 0.0189 | 0.0076 | 0.0076 | TRUE | 0.0177 |
| Csf2rb | 0 | 1 | 0 | 0 | 3 | 0 | 0 | 1 | 0 | 0 | 3 | 0 | 0.7 | 0 | 1 | 2.901 | 1.536 | 0.62035471 | NA | 8.6404 | 4.6837 | 5.4705 | 5.4705 | TRUE | 1.4431 |
| Csf2rb2 | 0 | 0 | 0 | 0 | 1 | 0 | 0 | 0 | 0 | 0 | 1 | 0 | 0.18 | 0 | 0 | 1.859 | 0.894 | 0.82650734 | NA | 0 | 18.3785 | 10 | 10 | TRUE | 0.5204 |
| Csf3 | 1 | 0 | 0 | 0 | 0 | 0 | 1 | 0 | 0 | 0 | 0 | 0 | 0.18 | 0 | 0 | 0.49 | -1.029 | 0.80087 | NA | 0 | 18.6165 | 10 | 10 | TRUE | 0.4614 |
| Csf3r | 11 | 14 | 7 | 18 | 12 | 2 | 12 | 14 | 7 | 14 | 13 | 2 | 10.32 | 11 | 10 | 0.906 | -0.142 | 0.84736512 | 0.92373809 | 0.2598 | 0.326 | 0.2967 | 0.2967 | TRUE | 1.422 |
| Csgalnact1 | 25 | 12 | 13 | 28 | 21 | 24 | 27 | 12 | 13 | 22 | 23 | 26 | 20.41 | 17 | 24 | 1.376 | 0.461 | 0.32974972 | 0.55084296 | 0.0409 | 0.1683 | 0.1104 | 0.1104 | TRUE | 1.0816 |
| Csgalnact2 | 1096 | 1107 | 979 | 1614 | 1116 | 1013 | 1164 | 1106 | 983 | 1269 | 1201 | 1109 | 1138.53 | 1084 | 1193 | 1.101 | 0.139 | 0.1857845 | 0.38944619 | 0.0051 | 0.0098 | 0.0071 | 0.0071 | TRUE | 0.0799 |
| Csk | 2087 | 2119 | 1986 | 3260 | 2438 | 2479 | 2217 | 2116 | 1994 | 2563 | 2623 | 2714 | 2371.12 | 2109 | 2633 | 1.248 | 0.32 | 3.16E-05 | 0.00052575 | 0.0012 | 0.0083 | 0.0038 | 0.0038 | TRUE | 0.0275 |
| Csl | 3 | 2 | 2 | 3 | 2 | 4 | 3 | 2 | 2 | 2 | 2 | 4 | 2.68 | 2 | 3 | 1.228 | 0.296 | 0.79065877 | NA | 0 | 1.2361 | 0.5178 | 0.5178 | TRUE | 0.2006 |
| Csmd1 | 2 | 0 | 1 | 14 | 7 | 3 | 2 | 0 | 1 | 11 | 8 | 3 | 4.16 | 1 | 7 | 7.15 | 2.838 | 0.02147237 | NA | 0 | 0.7992 | 0.5345 | 0.5345 | TRUE | 0.1825 |
| Csmd3 | 0 | 0 | 0 | 2 | 0 | 0 | 0 | 0 | 0 | 2 | 0 | 0 | 0.26 | 0 | 1 | 2.886 | 1.529 | 0.70676491 | NA | 0 | 12.5792 | 10 | 10 | TRUE | 0.9669 |
| Csn1s1 | 0 | 0 | 0 | 2 | 0 | 0 | 0 | 0 | 0 | 2 | 0 | 0 | 0.26 | 0 | 1 | 2.886 | 1.529 | 0.70676491 | NA | 0 | 12.5792 | 10 | 10 | TRUE | 0.9669 |
| Csn1s2a | 0 | 0 | 0 | 0 | 0 | 0 | 0 | 0 | 0 | 0 | 0 | 0 | 0 | 0 | 0 | NA | NA | NA | NA | NA | NA | NA | NA | NA | NA |
| Csn1s2b | 0 | 0 | 0 | 0 | 0 | 0 | 0 | 0 | 0 | 0 | 0 | 0 | 0 | 0 | 0 | NA | NA | NA | NA | NA | NA | NA | NA | NA | NA |
| Csn2 | 0 | 1 | 0 | 1 | 0 | 0 | 0 | 1 | 0 | 1 | 0 | 0 | 0.3 | 0 | 0 | 0.954 | -0.067 | 0.98642001 | NA | 0 | 11.0817 | 9.3014 | 9.3014 | TRUE | 0.405 |
| Csn3 | 0 | 0 | 0 | 4 | 0 | 1 | 0 | 0 | 0 | 3 | 0 | 1 | 0.71 | 0 | 1 | 7.635 | 2.933 | 0.32080697 | NA | 6.5513 | 4.6698 | 4.9457 | 4.9457 | TRUE | 0.1391 |
| Csnk1a1 | 13406 | 19217 | 19339 | 18778 | 13974 | 15556 | 14240 | 19192 | 19418 | 14762 | 15035 | 17028 | 16612.6 | 17617 | 15608 | 0.886 | -0.175 | 0.19021147 | 0.39517265 | 0.0169 | 0.0071 | 0.0128 | 0.0128 | TRUE | 0.3439 |
| Csnk1d | 7945 | 8561 | 8581 | 11345 | 8153 | 8170 | 8439 | 8550 | 8616 | 8919 | 8772 | 8943 | 8706.53 | 8535 | 8878 | 1.04 | 0.057 | 0.28710898 | 0.50828183 | 0 | 0.0073 | 0.0019 | 0.0019 | TRUE | 0.0013 |
| Csnk1e | 6715 | 10391 | 10865 | 7522 | 5880 | 7326 | 7133 | 10378 | 10910 | 5913 | 6327 | 8019 | 8113.15 | 9474 | 6753 | 0.713 | -0.488 | 0.00530472 | 0.03266927 | 0.0368 | 0.0073 | 0.022 | 0.022 | TRUE | 0.5706 |
| Csnk1g1 | 2185 | 2066 | 2103 | 2780 | 2074 | 1983 | 2321 | 2063 | 2112 | 2185 | 2231 | 2171 | 2180.57 | 2165 | 2196 | 1.014 | 0.02 | 0.79675517 | 0.89536272 | 0.0016 | 0.0084 | 0.0041 | 0.0041 | TRUE | 0.0476 |
| Csnk1g2 | 6915 | 6579 | 6748 | 7716 | 6101 | 5956 | 7345 | 6571 | 6776 | 6066 | 6564 | 6520 | 6640.16 | 6897 | 6383 | 0.925 | -0.112 | 0.14972499 | 0.34036305 | 0.0024 | 0.0074 | 0.0042 | 0.0042 | TRUE | 0.0393 |
| Csnk1g3 | 1637 | 1724 | 1730 | 2306 | 1538 | 1516 | 1739 | 1722 | 1737 | 1813 | 1655 | 1660 | 1720.79 | 1733 | 1709 | 0.987 | -0.019 | 0.81097528 | 0.90283686 | 9.00E-04 | 0.0088 | 0.0037 | 0.0037 | TRUE | 0.035 |
| Csnk2a1 | 12373 | 13094 | 12981 | 14822 | 10476 | 10955 | 13142 | 13077 | 13034 | 11652 | 11272 | 11992 | 12361.52 | 13084 | 11639 | 0.889 | -0.169 | 0.00326699 | 0.02260943 | 4.00E-04 | 0.0072 | 0.0023 | 0.0023 | TRUE | 0.0092 |
| Csnk2a2 | 7075 | 8291 | 8062 | 8983 | 6759 | 6364 | 7515 | 8280 | 8095 | 7062 | 7272 | 6966 | 7531.78 | 7963 | 7100 | 0.892 | -0.166 | 0.01658373 | 0.07624971 | 0.0013 | 0.0073 | 0.0033 | 0.0033 | TRUE | 0.0296 |
| Csnk2b | 4355 | 3783 | 3782 | 5211 | 3524 | 3415 | 4626 | 3778 | 3798 | 4096 | 3792 | 3738 | 3971.29 | 4067 | 3875 | 0.953 | -0.069 | 0.51110149 | 0.70464097 | 0.0073 | 0.0077 | 0.0078 | 0.0078 | TRUE | 0.1755 |
| Csnka2ip | 0 | 0 | 0 | 0 | 0 | 0 | 0 | 0 | 0 | 0 | 0 | 0 | 0 | 0 | 0 | NA | NA | NA | NA | NA | NA | NA | NA | NA | NA |
| Cspg4 | 26 | 70 | 51 | 56 | 50 | 58 | 28 | 70 | 51 | 44 | 54 | 63 | 51.67 | 50 | 54 | 1.079 | 0.109 | 0.77044944 | 0.88022663 | 0.0942 | 0.0707 | 0.0818 | 0.0818 | TRUE | 0.2703 |
| Cspg5 | 143 | 257 | 245 | 175 | 135 | 165 | 152 | 257 | 246 | 138 | 145 | 181 | 186.33 | 218 | 155 | 0.706 | -0.502 | 0.03305686 | 0.12295269 | 0.0447 | 0.0246 | 0.0344 | 0.0344 | TRUE | 0.767 |
| Cspp1 | 1868 | 1667 | 1692 | 2533 | 1845 | 1676 | 1984 | 1665 | 1699 | 1991 | 1985 | 1835 | 1859.82 | 1783 | 1937 | 1.087 | 0.12 | 0.22623243 | 0.43904737 | 0.0051 | 0.0087 | 0.0066 | 0.0066 | TRUE | 0.1177 |
| Csprs | 1 | 1 | 1 | 0 | 0 | 0 | 1 | 1 | 1 | 0 | 0 | 0 | 0.51 | 1 | 0 | 0.176 | -2.51 | 0.35480135 | NA | 0 | 6.4561 | 3.8046 | 3.8046 | TRUE | 0.0847 |
| Csrnp1 | 704 | 641 | 639 | 1286 | 779 | 710 | 748 | 640 | 642 | 1011 | 838 | 777 | 775.98 | 677 | 875 | 1.296 | 0.374 | 0.00549194 | 0.03350826 | 0.0118 | 0.0111 | 0.0118 | 0.0118 | TRUE | 0.2229 |
| Csrnp2 | 2319 | 2017 | 1764 | 2685 | 1903 | 1472 | 2463 | 2014 | 1771 | 2111 | 2048 | 1611 | 2003.07 | 2083 | 1923 | 0.924 | -0.115 | 0.45384591 | 0.65906064 | 0.0239 | 0.0085 | 0.0163 | 0.0163 | TRUE | 0.3082 |
| Csrnp3 | 995 | 947 | 983 | 902 | 694 | 741 | 1057 | 946 | 987 | 709 | 747 | 811 | 876.1 | 997 | 756 | 0.757 | -0.401 | 5.98E-05 | 0.00090389 | 0.0027 | 0.0107 | 0.006 | 0.006 | TRUE | 0.0485 |
| Csrp1 | 1650 | 1826 | 1904 | 2225 | 1569 | 1519 | 1753 | 1824 | 1912 | 1749 | 1688 | 1663 | 1764.68 | 1830 | 1700 | 0.93 | -0.105 | 0.16717542 | 0.36463707 | 7.00E-04 | 0.0088 | 0.0036 | 0.0036 | TRUE | 0.0188 |
| Csrp2 | 6469 | 7485 | 7363 | 7767 | 5934 | 6243 | 6871 | 7475 | 7393 | 6106 | 6385 | 6834 | 6844.03 | 7246 | 6442 | 0.889 | -0.17 | 0.02864673 | 0.1114332 | 0.0024 | 0.0074 | 0.0042 | 0.0042 | TRUE | 0.0346 |
| Csrp2bp | 3242 | 3150 | 3183 | 3617 | 2734 | 2606 | 3444 | 3146 | 3196 | 2843 | 2942 | 2853 | 3070.55 | 3262 | 2879 | 0.883 | -0.18 | 0.0121269 | 0.06060374 | 9.00E-04 | 0.008 | 0.0034 | 0.0034 | TRUE | 0.0288 |
| Csrp3 | 0 | 1 | 0 | 1 | 0 | 0 | 0 | 1 | 0 | 1 | 0 | 0 | 0.3 | 0 | 0 | 0.954 | -0.067 | 0.98642001 | NA | 0 | 11.0817 | 9.3014 | 9.3014 | TRUE | 0.405 |
| Cst10 | 0 | 0 | 0 | 0 | 0 | 0 | 0 | 0 | 0 | 0 | 0 | 0 | 0 | 0 | 0 | NA | NA | NA | NA | NA | NA | NA | NA | NA | NA |
| Cst11 | 0 | 0 | 0 | 0 | 0 | 0 | 0 | 0 | 0 | 0 | 0 | 0 | 0 | 0 | 0 | NA | NA | NA | NA | NA | NA | NA | NA | NA | NA |
| Cst12 | 0 | 0 | 0 | 0 | 0 | 0 | 0 | 0 | 0 | 0 | 0 | 0 | 0 | 0 | 0 | NA | NA | NA | NA | NA | NA | NA | NA | NA | NA |
| Cst13 | 2 | 0 | 0 | 3 | 2 | 2 | 2 | 0 | 0 | 2 | 2 | 2 | 1.47 | 1 | 2 | 3.226 | 1.69 | 0.33871732 | NA | 0 | 2.247 | 1.2922 | 1.2922 | TRUE | 0.9745 |
| Cst3 | 1865 | 2730 | 2631 | 2428 | 1753 | 2075 | 1981 | 2726 | 2642 | 1909 | 1886 | 2271 | 2235.91 | 2450 | 2022 | 0.825 | -0.277 | 0.05316833 | 0.1712068 | 0.0189 | 0.0084 | 0.0144 | 0.0144 | TRUE | 0.3392 |
| Cst6 | 2 | 1 | 1 | 6 | 3 | 3 | 2 | 1 | 1 | 5 | 3 | 3 | 2.56 | 1 | 4 | 2.761 | 1.465 | 0.2249212 | NA | 0 | 1.2942 | 0.5479 | 0.5479 | TRUE | 0.1345 |
| Cst7 | 0 | 0 | 0 | 0 | 1 | 1 | 0 | 0 | 0 | 0 | 1 | 1 | 0.36 | 0 | 1 | 3.794 | 1.924 | 0.60247382 | NA | 0 | 9.1135 | 8.1425 | 8.1425 | TRUE | 0.3482 |
| Cst8 | 0 | 0 | 0 | 0 | 0 | 0 | 0 | 0 | 0 | 0 | 0 | 0 | 0 | 0 | 0 | NA | NA | NA | NA | NA | NA | NA | NA | NA | NA |
| Cst9 | 0 | 0 | 0 | 0 | 0 | 0 | 0 | 0 | 0 | 0 | 0 | 0 | 0 | 0 | 0 | NA | NA | NA | NA | NA | NA | NA | NA | NA | NA |
| Csta | 0 | 0 | 0 | 0 | 0 | 0 | 0 | 0 | 0 | 0 | 0 | 0 | 0 | 0 | 0 | NA | NA | NA | NA | NA | NA | NA | NA | NA | NA |
| Cstad | 0 | 0 | 0 | 1 | 0 | 0 | 0 | 0 | 0 | 1 | 0 | 0 | 0.13 | 0 | 0 | 1.859 | 0.894 | 0.82650734 | NA | 0 | 25.1515 | 10 | 10 | TRUE | 0.2563 |
| Cstb | 1560 | 2401 | 2639 | 2008 | 1530 | 1841 | 1657 | 2398 | 2650 | 1579 | 1646 | 2015 | 1990.79 | 2235 | 1747 | 0.781 | -0.356 | 0.0451238 | 0.15315098 | 0.035 | 0.0086 | 0.0223 | 0.0223 | TRUE | 0.6189 |
| Cstf1 | 3314 | 3350 | 3185 | 4065 | 2967 | 2941 | 3520 | 3346 | 3198 | 3196 | 3192 | 3219 | 3278.51 | 3355 | 3202 | 0.955 | -0.067 | 0.33766072 | 0.55771127 | 7.00E-04 | 0.0079 | 0.0032 | 0.0032 | TRUE | 0.0225 |
| Cstf2 | 4979 | 5729 | 5808 | 7940 | 5773 | 5843 | 5289 | 5722 | 5832 | 6242 | 6211 | 6396 | 5948.55 | 5614 | 6283 | 1.119 | 0.162 | 0.01902747 | 0.08376584 | 0.0012 | 0.0075 | 0.0033 | 0.0033 | TRUE | 0.0313 |
| Cstf2t | 1875 | 1532 | 1604 | 2990 | 2243 | 1835 | 1992 | 1530 | 1611 | 2351 | 2413 | 2009 | 1984.12 | 1711 | 2258 | 1.32 | 0.401 | 0.0019888 | 0.01530149 | 0.0143 | 0.0086 | 0.0116 | 0.0116 | TRUE | 0.2484 |
| Cstf3 | 5584 | 3661 | 3386 | 6064 | 3900 | 3026 | 5931 | 3656 | 3400 | 4767 | 4196 | 3312 | 4210.5 | 4329 | 4092 | 0.945 | -0.081 | 0.70280866 | 0.83759243 | 0.0566 | 0.0077 | 0.0323 | 0.0323 | TRUE | 1.2759 |
| Cstl1 | 0 | 0 | 0 | 0 | 0 | 0 | 0 | 0 | 0 | 0 | 0 | 0 | 0 | 0 | 0 | NA | NA | NA | NA | NA | NA | NA | NA | NA | NA |
| Ctag2 | 3 | 0 | 1 | 17 | 3 | 5 | 3 | 0 | 1 | 13 | 3 | 5 | 4.38 | 1 | 7 | 5.416 | 2.437 | 0.04550936 | NA | 0.4586 | 0.7598 | 0.6339 | 0.6339 | TRUE | 0.6087 |
| Ctage5 | 2498 | 2114 | 2138 | 2980 | 2138 | 1995 | 2653 | 2111 | 2147 | 2343 | 2300 | 2184 | 2289.7 | 2304 | 2276 | 0.988 | -0.017 | 0.87597355 | 0.93888485 | 0.0085 | 0.0083 | 0.0084 | 0.0084 | TRUE | 0.213 |
| Ctbp1 | 5338 | 5204 | 5128 | 6636 | 4817 | 4847 | 5670 | 5197 | 5149 | 5217 | 5183 | 5306 | 5286.92 | 5339 | 5235 | 0.981 | -0.028 | 0.68718744 | 0.8275422 | 0.0012 | 0.0075 | 0.0034 | 0.0034 | TRUE | 0.0358 |
| Ctbp2 | 13251 | 11966 | 12366 | 14329 | 10024 | 9579 | 14075 | 11951 | 12417 | 11264 | 10785 | 10486 | 11829.58 | 12814 | 10845 | 0.846 | -0.241 | 0.00568086 | 0.0344477 | 0.0043 | 0.0072 | 0.0054 | 0.0054 | TRUE | 0.0905 |
| Ctbs | 424 | 581 | 642 | 741 | 513 | 636 | 450 | 580 | 645 | 583 | 552 | 696 | 584.32 | 558 | 610 | 1.092 | 0.127 | 0.43548859 | 0.64367054 | 0.022 | 0.0125 | 0.0173 | 0.0173 | TRUE | 0.3336 |
| Ctc1 | 10400 | 13233 | 13738 | 13877 | 10634 | 11845 | 11047 | 13216 | 13794 | 10909 | 11442 | 12966 | 12228.95 | 12686 | 11772 | 0.928 | -0.108 | 0.34077297 | 0.56103213 | 0.0103 | 0.0072 | 0.0092 | 0.0092 | TRUE | 0.1561 |
| Ctcf | 8569 | 8509 | 8552 | 12495 | 8913 | 8457 | 9102 | 8498 | 8587 | 9823 | 9590 | 9258 | 9142.81 | 8729 | 9557 | 1.095 | 0.131 | 0.04465372 | 0.15211197 | 0.001 | 0.0073 | 0.003 | 0.003 | TRUE | 0.017 |
| Ctcfl | 61 | 54 | 48 | 150 | 134 | 92 | 65 | 54 | 48 | 118 | 144 | 101 | 88.29 | 56 | 121 | 2.175 | 1.121 | 5.42E-06 | 0.00011876 | 0.0162 | 0.0442 | 0.0306 | 0.0306 | TRUE | 0.1779 |
| Ctdnep1 | 1927 | 2055 | 2300 | 2989 | 2053 | 2070 | 2047 | 2052 | 2309 | 2350 | 2209 | 2266 | 2205.53 | 2136 | 2275 | 1.065 | 0.091 | 0.28000638 | 0.50109681 | 0.0023 | 0.0084 | 0.0047 | 0.0047 | TRUE | 0.061 |

| | | | | | | | | | | | | | | | | | | | | | | | | |
|---|---|---|---|---|---|---|---|---|---|---|---|---|---|---|---|---|---|---|---|---|---|---|---|---|
| Ctdp1 | 2962 | 2092 | 2068 | 3564 | 2457 | 1993 | 3146 | 2089 | 2076 | 2802 | 2644 | 2182 | 2489.82 | 2437 | 2543 | 1.043 | 0.061 | 0.7286454 | 0.85400667 | 0.0371 | 0.0082 | 0.0221 | 0.0221 | TRUE | 0.784 |
| Ctdsp1 | 468 | 404 | 436 | 966 | 714 | 504 | 497 | 403 | 438 | 759 | 768 | 552 | 569.61 | 446 | 693 | 1.555 | 0.637 | 7.66E-05 | 0.00111578 | 0.0208 | 0.0127 | 0.0169 | 0.0169 | TRUE | 0.3713 |
| Ctdsp2 | 5326 | 6200 | 6790 | 7687 | 5758 | 5985 | 5657 | 6192 | 6818 | 6043 | 6195 | 6552 | 6242.79 | 6222 | 6263 | 1.006 | 0.009 | 0.92017638 | 0.96082634 | 0.005 | 0.0074 | 0.006 | 0.006 | TRUE | 0.0855 |
| Ctdspl | 495 | 498 | 415 | 713 | 588 | 489 | 526 | 497 | 417 | 561 | 633 | 535 | 528.05 | 480 | 576 | 1.2 | 0.264 | 0.04729934 | 0.15785128 | 0.0088 | 0.0131 | 0.0108 | 0.0108 | TRUE | 0.155 |
| Ctdspl2 | 2443 | 2759 | 2771 | 3689 | 2809 | 2905 | 2595 | 2755 | 2782 | 2900 | 3022 | 3180 | 2872.5 | 2711 | 3034 | 1.119 | 0.162 | 0.03183102 | 0.12006324 | 0.0012 | 0.0081 | 0.0037 | 0.0037 | TRUE | 0.0214 |
| Ctf1 | 89 | 108 | 123 | 114 | 74 | 98 | 95 | 108 | 124 | 90 | 80 | 107 | 100.4 | 109 | 92 | 0.847 | -0.24 | 0.27601817 | 0.49673192 | 0.009 | 0.0397 | 0.0249 | 0.0249 | TRUE | 0.1842 |
| Ctf2 | 6 | 6 | 4 | 29 | 11 | 6 | 6 | 6 | 4 | 23 | 12 | 7 | 9.6 | 5 | 14 | 2.556 | 1.354 | 0.06275301 | 0.19199621 | 0.1632 | 0.3502 | 0.2528 | 0.2528 | TRUE | 0.2921 |
| Ctgf | 434 | 349 | 280 | 1634 | 967 | 837 | 461 | 349 | 281 | 1285 | 1040 | 916 | 721.98 | 364 | 1080 | 2.975 | 1.573 | 1.32E-15 | 1.97E-13 | 0.0435 | 0.0115 | 0.0261 | 0.0261 | TRUE | 0.4336 |
| Cth | 964 | 896 | 840 | 964 | 638 | 636 | 1024 | 895 | 843 | 758 | 686 | 696 | 817.12 | 921 | 713 | 0.776 | -0.366 | 0.00105019 | 0.0092586 | 0.0054 | 0.0109 | 0.0078 | 0.0078 | TRUE | 0.1139 |
| Cthrc1 | 202 | 193 | 172 | 126 | 73 | 110 | 215 | 193 | 173 | 99 | 79 | 120 | 146.34 | 194 | 99 | 0.514 | -0.961 | 3.72E-06 | 8.58E-05 | 0.0166 | 0.0294 | 0.0235 | 0.0235 | TRUE | 0.3109 |
| Ctif | 520 | 465 | 484 | 631 | 471 | 464 | 552 | 464 | 486 | 496 | 507 | 508 | 502.24 | 501 | 504 | 1.005 | 0.008 | 0.94439524 | 0.97238554 | 0.0021 | 0.0135 | 0.0071 | 0.0071 | TRUE | 0.0931 |
| Ctla2a | 0 | 0 | 0 | 0 | 0 | 0 | 0 | 0 | 0 | 0 | 0 | 0 | 0 | 0 | 0 | NA | NA | NA | NA | NA | NA | NA | NA | NA | NA |
| Ctla2b | 0 | 0 | 0 | 0 | 0 | 1 | 0 | 0 | 0 | 0 | 0 | 1 | 0.18 | 0 | 0 | 1.859 | 0.894 | 0.82650734 | NA | 0 | 18.0643 | 10 | 10 | TRUE | 0.5383 |
| Ctla4 | 1 | 0 | 0 | 0 | 0 | 0 | 1 | 0 | 0 | 0 | 0 | 0 | 0.18 | 0 | 0 | 0.49 | -1.029 | 0.80087 | NA | 0 | 18.6165 | 10 | 10 | TRUE | 0.4614 |
| Ctnna1 | 21592 | 22583 | 23533 | 32693 | 22845 | 23381 | 22935 | 22554 | 23630 | 25701 | 24580 | 25594 | 24165.44 | 23040 | 25292 | 1.098 | 0.135 | 0.01930166 | 0.0845862 | 5.00E-04 | 0.007 | 0.0023 | 0.0023 | TRUE | 0.0074 |
| Ctnna2 | 13 | 12 | 7 | 17 | 14 | 10 | 14 | 12 | 7 | 13 | 15 | 11 | 12.03 | 11 | 13 | 1.203 | 0.267 | 0.63262333 | 0.79326881 | 0 | 0.2807 | 0.1404 | 0.1404 | TRUE | 0.3658 |
| Ctnna3 | 1 | 0 | 0 | 1 | 0 | 0 | 1 | 0 | 0 | 1 | 0 | 0 | 0.31 | 0 | 0 | 0.954 | -0.067 | 0.98625928 | NA | 0 | 10.7014 | 9.0384 | 9.0384 | TRUE | 0.4614 |
| Ctnnal1 | 5336 | 4761 | 4675 | 7639 | 5730 | 5888 | 5668 | 4755 | 4694 | 6005 | 6165 | 6445 | 5622.08 | 5039 | 6205 | 1.231 | 0.3 | 0.00201716 | 0.01548915 | 0.0062 | 0.0075 | 0.0066 | 0.0066 | TRUE | 0.1451 |
| Ctnnb1 | 47243 | 44520 | 43431 | 59889 | 43788 | 42299 | 50180 | 44463 | 43609 | 47080 | 47113 | 46303 | 46458.07 | 46084 | 46832 | 1.016 | 0.023 | 0.76544625 | 0.87700158 | 0.0025 | 0.007 | 0.0044 | 0.0044 | TRUE | 0.074 |
| Ctnnbip1 | 2139 | 1973 | 1830 | 2266 | 1663 | 1649 | 2272 | 1970 | 1838 | 1781 | 1789 | 1805 | 1909.28 | 2027 | 1792 | 0.884 | -0.177 | 0.07720196 | 0.21947265 | 0.0053 | 0.0086 | 0.0067 | 0.0067 | TRUE | 0.1353 |
| Ctnnbl1 | 3706 | 3501 | 3618 | 4245 | 3042 | 2857 | 3936 | 3496 | 3633 | 3337 | 3273 | 3127 | 3467.21 | 3688 | 3246 | 0.88 | -0.184 | 0.01906254 | 0.08386453 | 0.002 | 0.0079 | 0.0042 | 0.0042 | TRUE | 0.0419 |
| Ctnnd1 | 9669 | 9578 | 9984 | 14116 | 9932 | 8904 | 10270 | 9566 | 10025 | 11097 | 10686 | 9747 | 10231.79 | 9954 | 10510 | 1.056 | 0.079 | 0.31529222 | 0.53609195 | 0.0027 | 0.0072 | 0.0043 | 0.0043 | TRUE | 0.0492 |
| Ctnnd2 | 291 | 303 | 325 | 324 | 290 | 288 | 309 | 303 | 326 | 255 | 312 | 315 | 303.34 | 313 | 294 | 0.937 | -0.094 | 0.50436302 | 0.69937761 | 0.005 | 0.0178 | 0.0109 | 0.0109 | TRUE | 0.1657 |
| Ctns | 1265 | 1385 | 1405 | 1410 | 1109 | 1180 | 1344 | 1383 | 1411 | 1108 | 1193 | 1292 | 1288.5 | 1379 | 1198 | 0.867 | -0.205 | 0.02454896 | 0.10041135 | 0.0025 | 0.0095 | 0.0052 | 0.0052 | TRUE | 0.0563 |
| Ctps | 9709 | 10120 | 10182 | 11652 | 8410 | 8737 | 10313 | 10107 | 10224 | 9160 | 9049 | 9564 | 9735.99 | 10215 | 9258 | 0.906 | -0.142 | 0.01425318 | 0.06820421 | 3.00E-04 | 0.0072 | 0.0023 | 0.0023 | TRUE | 0.0102 |
| Ctps2 | 1183 | 1442 | 1421 | 1814 | 1256 | 1317 | 1257 | 1440 | 1427 | 1426 | 1351 | 1442 | 1390.43 | 1375 | 1406 | 1.023 | 0.033 | 0.71791501 | 0.84708278 | 0.0026 | 0.0093 | 0.0053 | 0.0053 | TRUE | 0.0674 |
| Ctr9 | 5963 | 4763 | 4683 | 8425 | 5875 | 5026 | 6334 | 4757 | 4702 | 6623 | 6321 | 5502 | 5706.47 | 5264 | 6149 | 1.168 | 0.224 | 0.10335612 | 0.26758579 | 0.0197 | 0.0075 | 0.0135 | 0.0135 | TRUE | 0.3848 |
| Ctrb1 | 3 | 6 | 6 | 10 | 0 | 5 | 3 | 6 | 6 | 8 | 0 | 5 | 4.76 | 5 | 4 | 0.893 | -0.163 | 0.87841815 | 0.93991934 | 0.4682 | 0.6995 | 0.613 | 0.613 | TRUE | 0.6591 |
| Ctrc | 1 | 0 | 2 | 1 | 1 | 1 | 1 | 0 | 2 | 1 | 1 | 1 | 1 | 1 | 1 | 0.956 | -0.066 | 0.97163585 | NA | 0 | 3.2865 | 1.4574 | 1.4574 | TRUE | 0.2133 |
| Ctrl | 18 | 12 | 14 | 9 | 16 | 6 | 19 | 12 | 14 | 7 | 17 | 7 | 12.67 | 15 | 10 | 0.677 | -0.562 | 0.35195063 | 0.57067599 | 0.0832 | 0.2669 | 0.181 | 0.181 | TRUE | 1.1495 |
| Cts3 | 1 | 2 | 1 | 2 | 0 | 1 | 1 | 2 | 1 | 2 | 0 | 1 | 1.12 | 1 | 1 | 0.678 | -0.56 | 0.7486666 | NA | 0 | 2.9437 | 1.2969 | 1.2969 | TRUE | 0.2838 |
| Cts6 | 0 | 0 | 0 | 0 | 0 | 0 | 0 | 0 | 0 | 0 | 0 | 0 | 0 | 0 | 0 | NA | NA | NA | NA | NA | NA | NA | NA | NA | NA |
| Cts7 | 0 | 0 | 0 | 0 | 1 | 0 | 0 | 0 | 0 | 0 | 1 | 0 | 0.18 | 0 | 0 | 1.859 | 0.894 | 0.82650734 | NA | 0 | 18.3785 | 10 | 10 | TRUE | 0.5204 |
| Cts8 | 0 | 0 | 0 | 0 | 0 | 0 | 0 | 0 | 0 | 0 | 0 | 0 | 0 | 0 | 0 | NA | NA | NA | NA | NA | NA | NA | NA | NA | NA |
| Cts8-ps | 0 | 0 | 0 | 0 | 0 | 0 | 0 | 0 | 0 | 0 | 0 | 0 | 0 | 0 | 0 | NA | NA | NA | NA | NA | NA | NA | NA | NA | NA |
| Ctsa | 1389 | 2318 | 2452 | 2205 | 1753 | 2039 | 1475 | 2315 | 2462 | 1733 | 1886 | 2232 | 2017.33 | 2084 | 1950 | 0.936 | -0.096 | 0.61108769 | 0.77977953 | 0.0442 | 0.0085 | 0.0252 | 0.0252 | TRUE | 0.7891 |
| Ctsb | 5663 | 6825 | 6709 | 7089 | 5168 | 5741 | 6015 | 6816 | 6737 | 5573 | 5560 | 6284 | 6164.26 | 6523 | 5806 | 0.89 | -0.168 | 0.06283547 | 0.19213555 | 0.0046 | 0.0074 | 0.0057 | 0.0057 | TRUE | 0.0632 |
| Ctsc | 650 | 868 | 785 | 1086 | 777 | 846 | 690 | 867 | 788 | 854 | 836 | 926 | 826.89 | 782 | 872 | 1.115 | 0.156 | 0.17689484 | 0.37738204 | 0.0061 | 0.0109 | 0.0085 | 0.0085 | TRUE | 0.1233 |
| Ctsd | 14418 | 16350 | 15716 | 15707 | 11273 | 12627 | 15314 | 16329 | 15781 | 12348 | 12129 | 13822 | 14287.13 | 15808 | 12766 | 0.808 | -0.308 | 9.66E-05 | 0.00134008 | 0.003 | 0.0071 | 0.0044 | 0.0044 | TRUE | 0.0639 |
| Ctse | 0 | 0 | 0 | 0 | 0 | 0 | 0 | 0 | 0 | 0 | 0 | 0 | 0 | 0 | 0 | NA | NA | NA | NA | NA | NA | NA | NA | NA | NA |
| Ctsf | 301 | 318 | 240 | 289 | 182 | 220 | 320 | 318 | 241 | 227 | 196 | 241 | 257.02 | 293 | 221 | 0.756 | -0.403 | 0.01780455 | 0.07982188 | 0.0142 | 0.0197 | 0.0169 | 0.0169 | TRUE | 0.271 |
| Ctsg | 1 | 3 | 1 | 1 | 3 | 0 | 1 | 3 | 1 | 1 | 3 | 0 | 1.51 | 2 | 1 | 0.78 | -0.358 | 0.8270085 | NA | 0.2223 | 2.1847 | 1.2646 | 1.2646 | TRUE | 0.8526 |
| Ctsh | 529 | 790 | 735 | 574 | 434 | 504 | 562 | 789 | 738 | 451 | 467 | 552 | 593.13 | 696 | 490 | 0.703 | -0.509 | 0.00128746 | 0.0108182 | 0.0198 | 0.0125 | 0.0163 | 0.0163 | TRUE | 0.3352 |
| Ctsj | 0 | 0 | 0 | 1 | 0 | 0 | 0 | 0 | 0 | 1 | 0 | 0 | 0.13 | 0 | 0 | 1.859 | 0.894 | 0.82650734 | NA | 0 | 25.1515 | 10 | 10 | TRUE | 0.2563 |
| Ctsk | 15 | 5 | 6 | 12 | 7 | 4 | 16 | 5 | 6 | 9 | 8 | 4 | 8.05 | 9 | 7 | 0.804 | -0.314 | 0.67892632 | 0.82160868 | 0.1631 | 0.4162 | 0.2894 | 0.2894 | TRUE | 1.41 |
| Ctsl | 6765 | 6799 | 6561 | 8228 | 5642 | 5605 | 7186 | 6790 | 6588 | 6468 | 6070 | 6136 | 6539.67 | 6855 | 6225 | 0.908 | -0.139 | 0.04715461 | 0.15766855 | 0.0013 | 0.0074 | 0.0034 | 0.0034 | TRUE | 0.0217 |
| Ctsll3 | 0 | 0 | 0 | 1 | 0 | 0 | 0 | 0 | 0 | 1 | 0 | 0 | 0.13 | 0 | 0 | 1.859 | 0.894 | 0.82650734 | NA | 0 | 25.1515 | 10 | 10 | TRUE | 0.2563 |
| Ctsm | 5 | 1 | 1 | 2 | 2 | 1 | 5 | 1 | 1 | 2 | 2 | 1 | 2.02 | 2 | 2 | 0.664 | -0.591 | 0.66790873 | NA | 0 | 1.6361 | 0.8524 | 0.8524 | TRUE | 1.0896 |
| Ctso | 48 | 65 | 65 | 96 | 73 | 71 | 51 | 65 | 65 | 75 | 79 | 78 | 68.82 | 60 | 77 | 1.277 | 0.352 | 0.13806393 | 0.32245868 | 0 | 0.0548 | 0.0259 | 0.0259 | TRUE | 0.1552 |
| Ctsq | 0 | 0 | 0 | 0 | 0 | 0 | 0 | 0 | 0 | 0 | 0 | 0 | 0 | 0 | 0 | NA | NA | NA | NA | NA | NA | NA | NA | NA | NA |
| Ctsr | 0 | 0 | 0 | 0 | 0 | 0 | 0 | 0 | 0 | 0 | 0 | 0 | 0 | 0 | 0 | NA | NA | NA | NA | NA | NA | NA | NA | NA | NA |
| Ctss | 5 | 5 | 4 | 13 | 6 | 4 | 5 | 5 | 4 | 10 | 6 | 4 | 5.9 | 5 | 7 | 1.496 | 0.581 | 0.46155755 | 0.66559865 | 0 | 0.5657 | 0.2726 | 0.2726 | TRUE | 0.4168 |
| Ctsw | 12 | 22 | 24 | 19 | 10 | 18 | 13 | 22 | 24 | 15 | 11 | 20 | 17.37 | 20 | 15 | 0.769 | -0.378 | 0.4474231 | 0.65334295 | 0.0331 | 0.1966 | 0.1201 | 0.1201 | TRUE | 0.3946 |
| Ctsz | 4454 | 4589 | 4415 | 4254 | 3269 | 3506 | 4731 | 4583 | 4433 | 3344 | 3517 | 3838 | 4074.4 | 4582 | 3566 | 0.778 | -0.362 | 8.49E-06 | 0.00017228 | 0.0026 | 0.0077 | 0.0045 | 0.0045 | TRUE | 0.0538 |
| Cttn | 7119 | 6963 | 6760 | 10460 | 7439 | 7073 | 7562 | 6954 | 6788 | 8223 | 8004 | 7743 | 7545.44 | 7101 | 7990 | 1.125 | 0.17 | 0.02040553 | 0.08815008 | 0.0019 | 0.0073 | 0.0038 | 0.0038 | TRUE | 0.0392 |
| Cttnbp2 | 149 | 108 | 91 | 143 | 112 | 108 | 158 | 108 | 91 | 112 | 121 | 118 | 118.11 | 119 | 117 | 0.983 | -0.025 | 0.91717753 | 0.95939276 | 0.0326 | 0.0348 | 0.0338 | 0.0338 | TRUE | 0.8205 |
| Cttnbp2nl | 1682 | 1426 | 1469 | 2054 | 1470 | 1398 | 1787 | 1424 | 1475 | 1615 | 1582 | 1530 | 1568.74 | 1562 | 1576 | 1.009 | 0.013 | 0.90471926 | 0.95255222 | 0.0073 | 0.009 | 0.008 | 0.008 | TRUE | 0.1902 |
| Ctu1 | 674 | 745 | 718 | 803 | 611 | 602 | 716 | 744 | 721 | 631 | 657 | 659 | 688.09 | 727 | 649 | 0.892 | -0.164 | 0.06538346 | 0.19707572 | 0 | 0.0117 | 0.0043 | 0.0043 | TRUE | 0.0074 |
| Ctu2 | 3278 | 3175 | 3187 | 4143 | 3076 | 2604 | 3482 | 3171 | 3200 | 3257 | 3310 | 2850 | 3211.63 | 3284 | 3139 | 0.956 | -0.065 | 0.47797701 | 0.67743419 | 0.0039 | 0.0079 | 0.0057 | 0.0057 | TRUE | 0.0779 |
| Ctxn1 | 518 | 530 | 570 | 493 | 389 | 468 | 550 | 529 | 572 | 388 | 419 | 512 | 495.04 | 550 | 440 | 0.796 | -0.328 | 0.0151552 | 0.0712676 | 0.0086 | 0.0136 | 0.0111 | 0.0111 | TRUE | 0.2405 |
| Ctxn2 | 0 | 0 | 0 | 0 | 0 | 0 | 0 | 0 | 0 | 0 | 0 | 0 | 0 | 0 | 0 | NA | NA | NA | NA | NA | NA | NA | NA | NA | NA |
| Ctxn3 | 1 | 3 | 4 | 0 | 0 | 3 | 1 | 3 | 4 | 0 | 0 | 3 | 1.89 | 3 | 1 | 0.387 | -1.368 | 0.4224925 | NA | 0.866 | 1.7471 | 1.4399 | 1.4399 | TRUE | 1.4132 |
| Cubn | 39 | 13 | 21 | 240 | 107 | 38 | 41 | 13 | 21 | 189 | 115 | 42 | 70.15 | 25 | 115 | 4.598 | 2.201 | 6.76E-05 | 0.00100495 | 0.3788 | 0.0539 | 0.1953 | 0.1953 | TRUE | 0.0688 |
| Cuedc1 | 644 | 827 | 810 | 1409 | 1043 | 1082 | 684 | 826 | 813 | 1108 | 1122 | 1184 | 956.26 | 774 | 1138 | 1.469 | 0.554 | 1.75E-07 | 5.79E-06 | 0.0042 | 0.0103 | 0.007 | 0.007 | TRUE | 0.1226 |
| Cuedc2 | 2324 | 2721 | 2673 | 2720 | 2029 | 1960 | 2468 | 2717 | 2684 | 2138 | 2183 | 2146 | 2389.47 | 2623 | 2156 | 0.822 | -0.284 | 0.00013361 | 0.00174294 | 0.001 | 0.0083 | 0.0035 | 0.0035 | TRUE | 0.0322 |
| Cul1 | 4440 | 4194 | 4429 | 7529 | 4999 | 4540 | 4716 | 4189 | 4447 | 5919 | 5379 | 4970 | 4936.48 | 4451 | 5423 | 1.219 | 0.285 | 0.00259543 | 0.01891642 | 0.0053 | 0.0076 | 0.0063 | 0.0063 | TRUE | 0.0787 |
| Cul2 | 5788 | 5596 | 5403 | 7880 | 5567 | 5690 | 6148 | 5589 | 5425 | 6195 | 5990 | 6229 | 5929.14 | 5721 | 6138 | 1.073 | 0.102 | 0.18234756 | 0.38423323 | 0.0022 | 0.0075 | 0.004 | 0.004 | TRUE | 0.052 |
| Cul3 | 6200 | 6044 | 6097 | 8969 | 6248 | 6060 | 6585 | 6036 | 6122 | 7051 | 6722 | 6634 | 6525.09 | 6248 | 6802 | 1.089 | 0.123 | 0.08061867 | 0.22580672 | 0.0014 | 0.0074 | 0.0034 | 0.0034 | TRUE | 0.0272 |
| Cul4a | 4327 | 5656 | 5838 | 6531 | 4741 | 5256 | 4596 | 5649 | 5862 | 5134 | 5101 | 5754 | 5349.23 | 5369 | 5330 | 0.993 | -0.011 | 0.92446762 | 0.96286755 | 0.0105 | 0.0075 | 0.0092 | 0.0092 | TRUE | 0.1931 |

| | | | | | | | | | | | | | | | | | | | | | | | | | |
|---|---|---|---|---|---|---|---|---|---|---|---|---|---|---|---|---|---|---|---|---|---|---|---|---|---|
| Cul4b | 15620 | 16807 | 17051 | 23846 | 16653 | 17426 | 16591 | 16785 | 17121 | 18746 | 17918 | 19076 | 17706.08 | 16832 | 18580 | 1.104 | 0.142 | 0.01527499 | 0.07168019 | 6.00E-04 | 0.0071 | 0.0024 | 0.0024 | TRUE | 0.0118 |
| Cul5 | 5753 | 4965 | 4694 | 7037 | 4899 | 4487 | 6111 | 4959 | 4713 | 5532 | 5271 | 4912 | 5249.54 | 5261 | 5238 | 0.996 | -0.006 | 0.95918633 | 0.97898705 | 0.0112 | 0.0075 | 0.0096 | 0.0096 | TRUE | 0.2433 |
| Cul7 | 2848 | 3596 | 3892 | 4532 | 3373 | 3503 | 3025 | 3591 | 3908 | 3563 | 3629 | 3835 | 3591.81 | 3508 | 3676 | 1.047 | 0.067 | 0.54110811 | 0.7269093 | 0.0088 | 0.0078 | 0.0084 | 0.0084 | TRUE | 0.1761 |
| Cul9 | 256 | 438 | 427 | 302 | 250 | 304 | 272 | 437 | 429 | 237 | 269 | 333 | 329.55 | 379 | 280 | 0.736 | -0.443 | 0.04079096 | 0.14287252 | 0.0453 | 0.0169 | 0.0306 | 0.0306 | TRUE | 0.7 |
| Cuta | 1311 | 1348 | 1365 | 1508 | 1193 | 1183 | 1393 | 1346 | 1371 | 1185 | 1284 | 1295 | 1312.24 | 1370 | 1255 | 0.915 | -0.128 | 0.11174227 | 0.28113131 | 7.00E-04 | 0.0094 | 0.0039 | 0.0039 | TRUE | 0.0291 |
| Cutc | 319 | 342 | 384 | 484 | 344 | 295 | 339 | 342 | 386 | 380 | 370 | 323 | 356.58 | 356 | 358 | 1.008 | 0.012 | 0.92533674 | 0.96313674 | 0.0036 | 0.0161 | 0.0094 | 0.0094 | TRUE | 0.0811 |
| Cux1 | 1116 | 1592 | 1758 | 1885 | 1428 | 1422 | 1185 | 1590 | 1765 | 1482 | 1536 | 1557 | 1519.24 | 1513 | 1525 | 1.007 | 0.01 | 0.94320309 | 0.9717062 | 0.0201 | 0.0091 | 0.0148 | 0.0148 | TRUE | 0.4321 |
| Cux2 | 347 | 317 | 315 | 394 | 310 | 325 | 369 | 317 | 316 | 310 | 334 | 356 | 333.42 | 334 | 333 | 0.996 | -0.005 | 0.96795569 | 0.98398951 | 0.0034 | 0.0168 | 0.0096 | 0.0096 | TRUE | 0.0933 |
| Cuzd1 | 1 | 0 | 2 | 15 | 3 | 0 | 1 | 0 | 2 | 12 | 3 | 0 | 3.02 | 1 | 5 | 4.983 | 2.317 | 0.16013976 | NA | 1.8016 | 1.0996 | 1.3649 | 1.3649 | TRUE | 0.3156 |
| Cwc15 | 2701 | 2710 | 2687 | 4111 | 2955 | 2712 | 2869 | 2707 | 2698 | 3232 | 3179 | 2969 | 2942.22 | 2758 | 3127 | 1.134 | 0.182 | 0.0141003 | 0.06773983 | 0.0011 | 0.008 | 0.0036 | 0.0036 | TRUE | 0.0235 |
| Cwc22 | 3559 | 3313 | 3482 | 5344 | 3815 | 3423 | 3780 | 3309 | 3496 | 4201 | 4105 | 3747 | 3773.01 | 3528 | 4018 | 1.139 | 0.188 | 0.03256903 | 0.121608 | 0.0036 | 0.0078 | 0.0053 | 0.0053 | TRUE | 0.0473 |
| Cwc25 | 1659 | 1508 | 1287 | 2410 | 1752 | 1539 | 1762 | 1506 | 1292 | 1895 | 1885 | 1685 | 1670.79 | 1520 | 1822 | 1.199 | 0.261 | 0.04225271 | 0.14656032 | 0.0132 | 0.0089 | 0.0113 | 0.0113 | TRUE | 0.233 |
| Cwc27 | 2035 | 2318 | 2348 | 2849 | 2199 | 2303 | 2162 | 2315 | 2358 | 2240 | 2366 | 2521 | 2326.8 | 2278 | 2376 | 1.042 | 0.06 | 0.47487849 | 0.67524193 | 0.0021 | 0.0083 | 0.0046 | 0.0046 | TRUE | 0.0347 |
| Cwf19l1 | 1285 | 1228 | 1272 | 1586 | 1172 | 1138 | 1365 | 1226 | 1277 | 1247 | 1261 | 1246 | 1270.34 | 1289 | 1251 | 0.97 | -0.043 | 0.59598166 | 0.76883877 | 7.00E-04 | 0.0095 | 0.004 | 0.004 | TRUE | 0.0312 |
| Cwf19l2 | 2417 | 2218 | 2304 | 3164 | 2272 | 2460 | 2567 | 2215 | 2313 | 2487 | 2445 | 2693 | 2453.43 | 2365 | 2542 | 1.074 | 0.104 | 0.2533472 | 0.47129891 | 0.0037 | 0.0082 | 0.0055 | 0.0055 | TRUE | 0.0674 |
| Cwh43 | 45 | 54 | 58 | 29 | 21 | 40 | 48 | 54 | 58 | 23 | 23 | 44 | 41.52 | 53 | 30 | 0.552 | -0.856 | 0.01704557 | 0.07749861 | 0.0421 | 0.0862 | 0.0667 | 0.0667 | TRUE | 1.0644 |
| Cx3cl1 | 410 | 252 | 232 | 514 | 338 | 225 | 435 | 252 | 233 | 404 | 364 | 246 | 322.36 | 307 | 338 | 1.104 | 0.143 | 0.59748859 | 0.77004845 | 0.0884 | 0.0171 | 0.0495 | 0.0495 | TRUE | 1.5189 |
| Cx3cr1 | 0 | 0 | 0 | 0 | 0 | 0 | 0 | 0 | 0 | 0 | 0 | 0 | 0 | 0 | 0 | NA | NA | NA | NA | NA | NA | NA | NA | NA | NA |
| Cxadr | 944 | 989 | 947 | 1443 | 1043 | 1111 | 1003 | 988 | 951 | 1134 | 1122 | 1216 | 1069.01 | 981 | 1157 | 1.18 | 0.239 | 0.00430031 | 0.02791204 | 0 | 0.01 | 0.0041 | 0.0041 | TRUE | 0.0229 |
| Cxcl1 | 4 | 2 | 1 | 4 | 0 | 1 | 4 | 2 | 1 | 3 | 0 | 1 | 1.91 | 2 | 1 | 0.616 | -0.7 | 0.63977861 | NA | 0.2746 | 1.7273 | 1.0693 | 1.0693 | TRUE | 0.8211 |
| Cxcl10 | 33 | 26 | 14 | 57 | 35 | 24 | 35 | 26 | 14 | 45 | 38 | 26 | 30.64 | 25 | 36 | 1.459 | 0.545 | 0.21137736 | 0.42189873 | 0.0901 | 0.1144 | 0.1032 | 0.1032 | TRUE | 0.3868 |
| Cxcl11 | 76 | 80 | 82 | 51 | 38 | 40 | 81 | 80 | 82 | 40 | 41 | 44 | 61.29 | 81 | 42 | 0.513 | -0.964 | 8.76E-05 | 0.00124626 | 0 | 0.0607 | 0.0255 | 0.0255 | TRUE | 0.0152 |
| Cxcl12 | 105 | 179 | 202 | 128 | 76 | 114 | 112 | 179 | 203 | 101 | 82 | 125 | 133.39 | 165 | 103 | 0.622 | -0.685 | 0.01177224 | 0.059358 | 0.0602 | 0.0316 | 0.0453 | 0.0453 | TRUE | 0.5895 |
| Cxcl13 | 0 | 0 | 0 | 0 | 0 | 0 | 0 | 0 | 0 | 0 | 0 | 0 | 0 | 0 | 0 | NA | NA | NA | NA | NA | NA | NA | NA | NA | NA |
| Cxcl14 | 153 | 201 | 144 | 333 | 238 | 256 | 163 | 201 | 145 | 262 | 256 | 280 | 217.65 | 170 | 266 | 1.57 | 0.651 | 0.00010313 | 0.00141301 | 0.0082 | 0.022 | 0.0154 | 0.0154 | TRUE | 0.2846 |
| Cxcl16 | 227 | 273 | 246 | 166 | 139 | 161 | 241 | 273 | 247 | 130 | 150 | 176 | 202.84 | 254 | 152 | 0.597 | -0.744 | 1.02E-05 | 0.00020153 | 0.0072 | 0.0231 | 0.0153 | 0.0153 | TRUE | 0.1997 |
| Cxcl17 | 0 | 0 | 0 | 0 | 0 | 0 | 0 | 0 | 0 | 0 | 0 | 0 | 0 | 0 | 0 | NA | NA | NA | NA | NA | NA | NA | NA | NA | NA |
| Cxcl2 | 0 | 0 | 0 | 0 | 0 | 0 | 0 | 0 | 0 | 0 | 0 | 0 | 0 | 0 | 0 | NA | NA | NA | NA | NA | NA | NA | NA | NA | NA |
| Cxcl3 | 2 | 0 | 0 | 1 | 2 | 3 | 2 | 0 | 0 | 1 | 2 | 3 | 1.39 | 1 | 2 | 2.939 | 1.555 | 0.40577738 | NA | 0 | 2.3752 | 1.5473 | 1.5473 | TRUE | 0.7075 |
| Cxcl5 | 7 | 7 | 3 | 6 | 6 | 5 | 7 | 7 | 3 | 5 | 6 | 5 | 5.68 | 6 | 5 | 0.951 | -0.073 | 0.9268687 | 0.96390434 | 0 | 0.5868 | 0.2796 | 0.2796 | TRUE | 0.4122 |
| Cxcl9 | 0 | 1 | 0 | 2 | 3 | 1 | 0 | 1 | 0 | 2 | 3 | 1 | 1.15 | 0 | 2 | 5.347 | 2.419 | 0.21801469 | NA | 0 | 2.8744 | 1.528 | 1.528 | TRUE | 0.405 |
| Cxcr1 | 7 | 1 | 0 | 3 | 2 | 4 | 7 | 1 | 0 | 2 | 2 | 4 | 2.89 | 3 | 3 | 1.057 | 0.08 | 0.95391604 | NA | 0.87 | 1.148 | 1.0217 | 1.0217 | TRUE | 2.4249 |
| Cxcr2 | 0 | 2 | 0 | 1 | 1 | 1 | 0 | 2 | 0 | 1 | 1 | 1 | 0.83 | 1 | 1 | 1.457 | 0.543 | 0.80279454 | NA | 0 | 3.9968 | 2.1438 | 2.1438 | TRUE | 0.9738 |
| Cxcr3 | 0 | 0 | 0 | 2 | 0 | 0 | 0 | 0 | 0 | 2 | 0 | 0 | 0.26 | 0 | 1 | 2.886 | 1.529 | 0.70676491 | NA | 0 | 12.5792 | 10 | 10 | TRUE | 0.9669 |
| Cxcr4 | 66 | 93 | 117 | 81 | 65 | 59 | 70 | 93 | 117 | 64 | 70 | 65 | 79.78 | 93 | 66 | 0.705 | -0.504 | 0.05319052 | 0.17120833 | 0.024 | 0.0482 | 0.0361 | 0.0361 | TRUE | 0.1497 |
| Cxcr5 | 0 | 1 | 1 | 3 | 2 | 2 | 0 | 1 | 1 | 2 | 2 | 2 | 1.45 | 1 | 2 | 3.312 | 1.728 | 0.29265222 | NA | 0 | 2.2783 | 0.9676 | 0.9676 | TRUE | 0.2217 |
| Cxcr6 | 0 | 0 | 0 | 0 | 0 | 0 | 0 | 0 | 0 | 0 | 0 | 0 | 0 | 0 | 0 | NA | NA | NA | NA | NA | NA | NA | NA | NA | NA |
| Cxcr7 | 111 | 140 | 154 | 198 | 158 | 179 | 118 | 140 | 155 | 156 | 170 | 196 | 155.66 | 138 | 174 | 1.261 | 0.334 | 0.07395944 | 0.21352521 | 0.0093 | 0.0281 | 0.0187 | 0.0187 | TRUE | 0.1569 |
| Cxx1a | 78 | 75 | 77 | 141 | 90 | 100 | 83 | 75 | 77 | 111 | 97 | 109 | 92.04 | 78 | 106 | 1.352 | 0.435 | 0.0334135 | 0.12394114 | 0 | 0.0427 | 0.019 | 0.019 | TRUE | 0.0512 |
| Cxx1b | 81 | 84 | 89 | 127 | 89 | 110 | 86 | 84 | 89 | 100 | 96 | 120 | 95.88 | 86 | 105 | 1.216 | 0.283 | 0.1716756 | 0.37055804 | 0 | 0.0413 | 0.0202 | 0.0202 | TRUE | 0.1462 |
| Cxx1c | 17 | 20 | 14 | 23 | 18 | 12 | 18 | 20 | 14 | 18 | 19 | 13 | 17.11 | 17 | 17 | 0.974 | -0.037 | 0.93569067 | 0.96811881 | 0 | 0.1994 | 0.0957 | 0.0957 | TRUE | 0.164 |
| Cxxc1 | 1835 | 1655 | 1672 | 2488 | 1884 | 1725 | 1949 | 1653 | 1679 | 1956 | 2027 | 1888 | 1858.67 | 1760 | 1957 | 1.112 | 0.153 | 0.10804385 | 0.27511586 | 0.0042 | 0.0087 | 0.006 | 0.006 | TRUE | 0.1059 |
| Cxxc4 | 9 | 19 | 9 | 14 | 3 | 6 | 10 | 19 | 9 | 11 | 3 | 7 | 9.73 | 13 | 7 | 0.565 | -0.824 | 0.24198294 | 0.4585617 | 0.141 | 0.3455 | 0.2476 | 0.2476 | TRUE | 0.5369 |
| Cxxc5 | 1511 | 1977 | 2137 | 2005 | 1665 | 1792 | 1605 | 1974 | 2146 | 1576 | 1791 | 1962 | 1842.4 | 1908 | 1776 | 0.93 | -0.104 | 0.44279513 | 0.64973793 | 0.0163 | 0.0087 | 0.0128 | 0.0128 | TRUE | 0.2333 |
| Cyb5 | 2530 | 2470 | 2428 | 2906 | 2024 | 2054 | 2687 | 2467 | 2438 | 2284 | 2178 | 2248 | 2383.78 | 2531 | 2237 | 0.884 | -0.178 | 0.02007619 | 0.08708517 | 0.0013 | 0.0083 | 0.0038 | 0.0038 | TRUE | 0.0354 |
| Cyb561 | 1522 | 2155 | 2249 | 2489 | 1834 | 2107 | 1617 | 2152 | 2258 | 1957 | 1973 | 2306 | 2043.91 | 2009 | 2079 | 1.034 | 0.049 | 0.7328332 | 0.8565538 | 0.0195 | 0.0085 | 0.0142 | 0.0142 | TRUE | 0.3524 |
| Cyb561d1 | 201 | 262 | 237 | 282 | 207 | 215 | 213 | 262 | 238 | 222 | 223 | 235 | 232.15 | 238 | 227 | 0.952 | -0.071 | 0.62444867 | 0.78834114 | 0 | 0.0211 | 0.0109 | 0.0109 | TRUE | 0.0865 |
| Cyb561d2 | 507 | 518 | 463 | 665 | 442 | 412 | 539 | 517 | 465 | 523 | 476 | 451 | 495.02 | 507 | 483 | 0.955 | -0.067 | 0.57449873 | 0.75272138 | 0.0038 | 0.0136 | 0.0081 | 0.0081 | TRUE | 0.063 |
| Cyb5b | 10423 | 8007 | 7659 | 14225 | 9632 | 8226 | 11071 | 7997 | 7690 | 11183 | 10363 | 9005 | 9551.47 | 8919 | 10184 | 1.142 | 0.191 | 0.21149674 | 0.42203316 | 0.0258 | 0.0072 | 0.0168 | 0.0168 | TRUE | 0.5438 |
| Cyb5d1 | 320 | 186 | 197 | 437 | 267 | 201 | 340 | 186 | 198 | 344 | 287 | 220 | 262.38 | 241 | 284 | 1.179 | 0.237 | 0.36833729 | 0.58564866 | 0.0772 | 0.0195 | 0.0462 | 0.0462 | TRUE | 0.6648 |
| Cyb5d2 | 181 | 226 | 180 | 252 | 195 | 182 | 192 | 226 | 181 | 198 | 210 | 199 | 200.97 | 200 | 202 | 1.013 | 0.019 | 0.90296393 | 0.95205059 | 0 | 0.0233 | 0.0125 | 0.0125 | TRUE | 0.1442 |
| Cyb5r1 | 1606 | 2099 | 2041 | 2846 | 2252 | 1941 | 1706 | 2096 | 2049 | 2237 | 2423 | 2125 | 2106.1 | 1950 | 2262 | 1.159 | 0.213 | 0.05085545 | 0.16600416 | 0.0075 | 0.0085 | 0.0081 | 0.0081 | TRUE | 0.1451 |
| Cyb5r2 | 11 | 7 | 10 | 27 | 14 | 18 | 12 | 7 | 10 | 21 | 15 | 20 | 14.12 | 10 | 19 | 1.962 | 0.973 | 0.06261777 | 0.19169399 | 0 | 0.2403 | 0.1171 | 0.1171 | TRUE | 0.1904 |
| Cyb5r3 | 8330 | 9786 | 10348 | 9134 | 7073 | 7879 | 8848 | 9773 | 10390 | 7180 | 7610 | 8625 | 8737.86 | 9670 | 7805 | 0.807 | -0.309 | 0.00252322 | 0.01851845 | 0.0076 | 0.0073 | 0.0074 | 0.0074 | TRUE | 0.103 |
| Cyb5r4 | 2137 | 2234 | 2212 | 3078 | 2257 | 2188 | 2270 | 2231 | 2221 | 2420 | 2428 | 2395 | 2327.54 | 2241 | 2414 | 1.078 | 0.108 | 0.08849094 | 0.24129999 | 0 | 0.0083 | 0.0025 | 0.0025 | TRUE | 0.0016 |
| Cyb5rl | 80 | 83 | 60 | 169 | 121 | 82 | 85 | 83 | 60 | 133 | 130 | 90 | 96.82 | 76 | 118 | 1.552 | 0.634 | 0.01229119 | 0.06128887 | 0.0294 | 0.0409 | 0.0353 | 0.0353 | TRUE | 0.4157 |
| Cyba | 604 | 878 | 903 | 758 | 592 | 660 | 642 | 877 | 907 | 596 | 637 | 722 | 730.07 | 809 | 652 | 0.805 | -0.312 | 0.04621986 | 0.15563032 | 0.0209 | 0.0114 | 0.0163 | 0.0163 | TRUE | 0.3852 |
| Cybasc3 | 635 | 889 | 951 | 1193 | 844 | 915 | 674 | 888 | 955 | 938 | 908 | 1002 | 894.13 | 839 | 949 | 1.131 | 0.177 | 0.21256162 | 0.42356562 | 0.016 | 0.0106 | 0.0134 | 0.0134 | TRUE | 0.3483 |
| Cybb | 422 | 403 | 397 | 614 | 579 | 566 | 448 | 402 | 399 | 483 | 623 | 620 | 495.76 | 416 | 575 | 1.378 | 0.463 | 0.00100106 | 0.00891512 | 0.0111 | 0.0135 | 0.0122 | 0.0122 | TRUE | 0.2409 |
| Cybrd1 | 108 | 128 | 94 | 146 | 80 | 74 | 115 | 128 | 94 | 115 | 86 | 81 | 103.13 | 112 | 94 | 0.842 | -0.249 | 0.28798102 | 0.50903254 | 0.0208 | 0.0388 | 0.0298 | 0.0298 | TRUE | 0.3968 |
| Cyc1 | 11798 | 13444 | 13609 | 16322 | 11668 | 11561 | 12532 | 13427 | 13665 | 12831 | 12554 | 12655 | 12943.92 | 13208 | 12680 | 0.96 | -0.059 | 0.35935825 | 0.57787504 | 0.001 | 0.0072 | 0.0029 | 0.0029 | TRUE | 0.0245 |
| Cycs | 2960 | 2571 | 2531 | 4283 | 2729 | 2413 | 3144 | 2568 | 2541 | 3367 | 2936 | 2641 | 2866.29 | 2751 | 2981 | 1.084 | 0.117 | 0.3624408 | 0.58031617 | 0.0137 | 0.0081 | 0.0115 | 0.0115 | TRUE | 0.1893 |
| Cyct | 382 | 552 | 516 | 826 | 646 | 612 | 406 | 551 | 518 | 649 | 695 | 670 | 581.58 | 492 | 671 | 1.364 | 0.448 | 0.00101289 | 0.00900526 | 0.0108 | 0.0126 | 0.0116 | 0.0116 | TRUE | 0.27 |
| Cyfip1 | 4748 | 4950 | 4982 | 7140 | 4851 | 4911 | 5043 | 4944 | 5002 | 5613 | 5219 | 5376 | 5199.57 | 4996 | 5403 | 1.082 | 0.113 | 0.07356815 | 0.212824 | 6.00E-04 | 0.0075 | 0.0027 | 0.0027 | TRUE | 0.0143 |
| Cyfip2 | 6248 | 7483 | 8018 | 6105 | 4767 | 5510 | 6636 | 7473 | 8051 | 4799 | 5129 | 6032 | 6353.43 | 7387 | 5320 | 0.72 | -0.474 | 5.12E-05 | 0.00079172 | 0.0116 | 0.0074 | 0.0097 | 0.0097 | TRUE | 0.1667 |
| Cygb | 25 | 14 | 15 | 25 | 27 | 18 | 27 | 14 | 15 | 20 | 29 | 20 | 20.67 | 19 | 23 | 1.229 | 0.298 | 0.52107327 | 0.71154697 | 0.0392 | 0.1663 | 0.1061 | 0.1061 | TRUE | 0.7153 |
| Cyhr1 | 2443 | 2889 | 3101 | 2964 | 2420 | 2697 | 2595 | 2885 | 3114 | 2330 | 2604 | 2952 | 2746.67 | 2865 | 2629 | 0.917 | -0.125 | 0.28912613 | 0.50996392 | 0.0106 | 0.0081 | 0.0096 | 0.0096 | TRUE | 0.1406 |
| Cylc1 | 0 | 0 | 0 | 0 | 0 | 0 | 0 | 0 | 0 | 0 | 0 | 0 | 0 | 0 | 0 | NA | NA | NA | NA | NA | NA | NA | NA | NA | NA |
| Cylc2 | 0 | 0 | 0 | 0 | 0 | 0 | 0 | 0 | 0 | 0 | 0 | 0 | 0 | 0 | 0 | NA | NA | NA | NA | NA | NA | NA | NA | NA | NA |

| | | | | | | | | | | | | | | | | | | | | | | | | | |
|---|---|---|---|---|---|---|---|---|---|---|---|---|---|---|---|---|---|---|---|---|---|---|---|---|---|
| Cyld | 1692 | 1940 | 1966 | 2196 | 1732 | 1629 | 1797 | 1937 | 1974 | 1726 | 1864 | 1783 | 1846.97 | 1903 | 1791 | 0.941 | -0.088 | 0.27108046 | 0.49103952 | 0.0014 | 0.0087 | 0.0041 | 0.0041 | TRUE | 0.0284 |
| Cym | 0 | 0 | 0 | 0 | 0 | 0 | 0 | 0 | 0 | 0 | 0 | 0 | 0 | 0 | 0 | NA | NA | NA | NA | NA | NA | NA | NA | NA | NA |
| Cyp11a1 | 399 | 371 | 318 | 393 | 254 | 200 | 424 | 371 | 319 | 309 | 273 | 219 | 319.13 | 371 | 267 | 0.722 | -0.471 | 0.007769 | 0.04342052 | 0.0214 | 0.0172 | 0.0193 | 0.0193 | TRUE | 0.225 |
| Cyp11b1 | 0 | 0 | 0 | 0 | 0 | 0 | 0 | 0 | 0 | 0 | 0 | 0 | 0 | 0 | 0 | NA | NA | NA | NA | NA | NA | NA | NA | NA | NA |
| Cyp11b2 | 1 | 0 | 0 | 0 | 0 | 0 | 1 | 0 | 0 | 0 | 0 | 0 | 0.18 | 0 | 0 | 0.49 | -1.029 | 0.80087 | NA | 0 | 18.6165 | 10 | 10 | TRUE | 0.4614 |
| Cyp17a1 | 1 | 0 | 0 | 0 | 1 | 1 | 1 | 0 | 0 | 0 | 1 | 1 | 0.54 | 0 | 1 | 1.923 | 0.944 | 0.73523723 | NA | 0 | 6.1214 | 3.9144 | 3.9144 | TRUE | 0.4614 |
| Cyp19a1 | 2 | 4 | 2 | 3 | 1 | 3 | 2 | 4 | 2 | 2 | 1 | 3 | 2.47 | 3 | 2 | 0.826 | -0.276 | 0.81427809 | NA | 0 | 1.3384 | 0.5872 | 0.5872 | TRUE | 0.2082 |
| Cyp1a1 | 8 | 9 | 6 | 11 | 7 | 4 | 8 | 9 | 6 | 9 | 8 | 4 | 7.34 | 8 | 7 | 0.885 | -0.176 | 0.80017975 | 0.89686328 | 0 | 0.4555 | 0.2141 | 0.2141 | TRUE | 0.2345 |
| Cyp1a2 | 3 | 0 | 2 | 6 | 6 | 2 | 3 | 0 | 2 | 5 | 6 | 2 | 3.09 | 2 | 4 | 2.599 | 1.378 | 0.26016271 | NA | 0 | 1.0721 | 0.6724 | 0.6724 | TRUE | 0.424 |
| Cyp1b1 | 28 | 14 | 21 | 85 | 37 | 76 | 30 | 14 | 21 | 67 | 40 | 83 | 42.44 | 22 | 63 | 2.938 | 1.555 | 0.00017952 | 0.00222422 | 0.1023 | 0.0845 | 0.0927 | 0.0927 | TRUE | 0.1693 |
| Cyp20a1 | 634 | 770 | 783 | 844 | 620 | 684 | 673 | 769 | 786 | 663 | 667 | 749 | 717.99 | 743 | 693 | 0.932 | -0.102 | 0.35883133 | 0.57738091 | 0.0043 | 0.0115 | 0.0074 | 0.0074 | TRUE | 0.0786 |
| Cyp21a1 | 0 | 2 | 0 | 2 | 0 | 1 | 0 | 2 | 0 | 2 | 0 | 1 | 0.78 | 1 | 1 | 1.351 | 0.434 | 0.86130335 | NA | 0 | 4.2448 | 3.1538 | 3.1538 | TRUE | 0.9738 |
| Cyp24a1 | 25 | 30 | 30 | 57 | 33 | 33 | 27 | 30 | 30 | 45 | 36 | 36 | 33.85 | 29 | 39 | 1.351 | 0.434 | 0.18839611 | 0.39350861 | 0 | 0.1042 | 0.0481 | 0.0481 | TRUE | 0.1578 |
| Cyp26a1 | 17 | 22 | 26 | 40 | 31 | 14 | 18 | 22 | 26 | 31 | 33 | 15 | 24.38 | 22 | 26 | 1.217 | 0.283 | 0.52319247 | 0.71308258 | 0.0528 | 0.1421 | 0.1004 | 0.1004 | TRUE | 0.8054 |
| Cyp26b1 | 17 | 17 | 13 | 19 | 10 | 15 | 18 | 17 | 13 | 15 | 11 | 16 | 15.03 | 16 | 14 | 0.878 | -0.187 | 0.70305532 | 0.83762068 | 0 | 0.226 | 0.1067 | 0.1067 | TRUE | 0.165 |
| Cyp26c1 | 11 | 10 | 15 | 25 | 18 | 14 | 12 | 10 | 15 | 20 | 19 | 15 | 15.18 | 12 | 18 | 1.484 | 0.569 | 0.24438528 | 0.4610511 | 0 | 0.2239 | 0.1039 | 0.1039 | TRUE | 0.1648 |
| Cyp27a1 | 79 | 135 | 112 | 114 | 96 | 86 | 84 | 135 | 112 | 90 | 103 | 94 | 103.04 | 110 | 96 | 0.864 | -0.21 | 0.37018222 | 0.58726979 | 0.0209 | 0.0389 | 0.03 | 0.03 | TRUE | 0.2223 |
| Cyp27b1 | 21 | 13 | 17 | 34 | 26 | 14 | 22 | 13 | 17 | 27 | 28 | 15 | 20.4 | 17 | 23 | 1.347 | 0.429 | 0.35506561 | 0.57354514 | 0.0331 | 0.1684 | 0.1051 | 0.1051 | TRUE | 0.4871 |
| Cyp2a12 | 0 | 0 | 0 | 0 | 0 | 0 | 0 | 0 | 0 | 0 | 0 | 0 | 0 | 0 | 0 | NA | NA | NA | NA | NA | NA | NA | NA | NA | NA |
| Cyp2a22 | 0 | 0 | 0 | 0 | 0 | 0 | 0 | 0 | 0 | 0 | 0 | 0 | 0 | 0 | 0 | NA | NA | NA | NA | NA | NA | NA | NA | NA | NA |
| Cyp2a4 | 2 | 2 | 0 | 4 | 0 | 0 | 2 | 2 | 0 | 3 | 0 | 0 | 1.21 | 1 | 1 | 0.805 | -0.314 | 0.8862134 | NA | 2.4646 | 2.7272 | 2.6479 | 2.6479 | TRUE | 1.8937 |
| Cyp2a5 | 0 | 0 | 0 | 0 | 0 | 0 | 0 | 0 | 0 | 0 | 0 | 0 | 0 | 0 | 0 | NA | NA | NA | NA | NA | NA | NA | NA | NA | NA |
| Cyp2ab1 | 4 | 3 | 1 | 17 | 9 | 9 | 4 | 3 | 1 | 13 | 10 | 10 | 6.86 | 3 | 11 | 4.036 | 2.013 | 0.0143042 | 0.06836539 | 0 | 0.4873 | 0.2554 | 0.2554 | TRUE | 0.3759 |
| Cyp2b10 | 3 | 0 | 0 | 1 | 0 | 0 | 3 | 0 | 0 | 1 | 0 | 0 | 0.66 | 1 | 0 | 0.331 | -1.597 | 0.61164788 | NA | 8.2589 | 4.9826 | 5.6428 | 5.6428 | TRUE | 1.4426 |
| Cyp2b13 | 0 | 0 | 0 | 0 | 0 | 0 | 0 | 0 | 0 | 0 | 0 | 0 | 0 | 0 | 0 | NA | NA | NA | NA | NA | NA | NA | NA | NA | NA |
| Cyp2b19 | 2 | 0 | 0 | 2 | 3 | 0 | 2 | 0 | 0 | 2 | 3 | 0 | 1.15 | 1 | 2 | 2.294 | 1.198 | 0.59102508 | NA | 1.8421 | 2.8615 | 2.5448 | 2.5448 | TRUE | 0.3426 |
| Cyp2b23 | 155 | 159 | 105 | 549 | 411 | 360 | 165 | 159 | 105 | 432 | 442 | 394 | 282.79 | 143 | 423 | 2.959 | 1.565 | 1.44E-17 | 2.71E-15 | 0.0208 | 0.0186 | 0.0195 | 0.0195 | TRUE | 0.5522 |
| Cyp2b9 | 0 | 0 | 0 | 0 | 0 | 0 | 0 | 0 | 0 | 0 | 0 | 0 | 0 | 0 | 0 | NA | NA | NA | NA | NA | NA | NA | NA | NA | NA |
| Cyp2c29 | 1 | 5 | 3 | 8 | 0 | 4 | 1 | 5 | 3 | 6 | 0 | 4 | 3.29 | 3 | 3 | 1.193 | 0.255 | 0.84046909 | NA | 0.6361 | 1.0085 | 0.8545 | 0.8545 | TRUE | 0.5044 |
| Cyp2c37 | 0 | 0 | 0 | 0 | 0 | 0 | 0 | 0 | 0 | 0 | 0 | 0 | 0 | 0 | 0 | NA | NA | NA | NA | NA | NA | NA | NA | NA | NA |
| Cyp2c38 | 0 | 0 | 0 | 0 | 0 | 0 | 0 | 0 | 0 | 0 | 0 | 0 | 0 | 0 | 0 | NA | NA | NA | NA | NA | NA | NA | NA | NA | NA |
| Cyp2c39 | 0 | 0 | 0 | 0 | 3 | 0 | 0 | 0 | 0 | 0 | 3 | 0 | 0.54 | 0 | 1 | 5.683 | 2.507 | 0.52963929 | NA | 10 | 6.1308 | 10 | 10 | TRUE | 1.4428 |
| Cyp2c40 | 1 | 1 | 1 | 1 | 1 | 0 | 1 | 1 | 1 | 1 | 1 | 0 | 0.82 | 1 | 1 | 0.618 | -0.695 | 0.72575676 | NA | 0 | 4.0188 | 1.5537 | 1.5537 | TRUE | 0.1881 |
| Cyp2c44 | 51 | 54 | 36 | 61 | 40 | 60 | 54 | 54 | 36 | 48 | 43 | 66 | 50.15 | 48 | 52 | 1.084 | 0.116 | 0.71194603 | 0.84325407 | 0.0282 | 0.0726 | 0.0514 | 0.0514 | TRUE | 0.3938 |
| Cyp2c50 | 0 | 0 | 0 | 0 | 0 | 0 | 0 | 0 | 0 | 0 | 0 | 0 | 0 | 0 | 0 | NA | NA | NA | NA | NA | NA | NA | NA | NA | NA |
| Cyp2c53-ps | 1 | 0 | 0 | 1 | 2 | 3 | 1 | 0 | 0 | 1 | 2 | 3 | 1.21 | 0 | 2 | 5.578 | 2.48 | 0.20539363 | NA | 0 | 2.7206 | 1.5235 | 1.5235 | TRUE | 0.3524 |
| Cyp2c54 | 0 | 0 | 0 | 0 | 0 | 0 | 0 | 0 | 0 | 0 | 0 | 0 | 0 | 0 | 0 | NA | NA | NA | NA | NA | NA | NA | NA | NA | NA |
| Cyp2c55 | 36 | 32 | 27 | 249 | 196 | 181 | 38 | 32 | 27 | 196 | 211 | 198 | 117.01 | 32 | 202 | 6.225 | 2.638 | 7.72E-32 | 4.87E-29 | 0 | 0.0351 | 0.0182 | 0.0182 | TRUE | 0.1612 |
| Cyp2c65 | 0 | 0 | 0 | 1 | 1 | 0 | 0 | 0 | 0 | 1 | 1 | 0 | 0.31 | 0 | 1 | 3.32 | 1.731 | 0.65383052 | NA | 0 | 10.6224 | 8.9541 | 8.9541 | TRUE | 0.202 |
| Cyp2c66 | 0 | 0 | 0 | 1 | 0 | 0 | 0 | 0 | 0 | 1 | 0 | 0 | 0.13 | 0 | 0 | 1.859 | 0.894 | 0.82650734 | NA | 0 | 25.1515 | 10 | 10 | TRUE | 0.2563 |
| Cyp2c67 | 0 | 0 | 0 | 0 | 0 | 0 | 0 | 0 | 0 | 0 | 0 | 0 | 0 | 0 | 0 | NA | NA | NA | NA | NA | NA | NA | NA | NA | NA |
| Cyp2c68 | 0 | 0 | 0 | 0 | 0 | 0 | 0 | 0 | 0 | 0 | 0 | 0 | 0 | 0 | 0 | NA | NA | NA | NA | NA | NA | NA | NA | NA | NA |
| Cyp2c69 | 17 | 8 | 7 | 30 | 10 | 16 | 18 | 8 | 7 | 24 | 11 | 18 | 14.16 | 11 | 18 | 1.588 | 0.667 | 0.2690219 | 0.48899543 | 0.1312 | 0.2396 | 0.1877 | 0.1877 | TRUE | 0.3785 |
| Cyp2c70 | 13 | 3 | 5 | 1 | 2 | 2 | 14 | 3 | 5 | 1 | 2 | 2 | 4.49 | 7 | 2 | 0.229 | -2.13 | 0.05325276 | NA | 0 | 0.7403 | 0.5104 | 0.5104 | TRUE | 0.8935 |
| Cyp2d10 | 0 | 0 | 0 | 0 | 0 | 0 | 0 | 0 | 0 | 0 | 0 | 0 | 0 | 0 | 0 | NA | NA | NA | NA | NA | NA | NA | NA | NA | NA |
| Cyp2d11 | 0 | 0 | 0 | 0 | 0 | 0 | 0 | 0 | 0 | 0 | 0 | 0 | 0 | 0 | 0 | NA | NA | NA | NA | NA | NA | NA | NA | NA | NA |
| Cyp2d12 | 0 | 1 | 1 | 2 | 2 | 0 | 0 | 1 | 1 | 2 | 2 | 0 | 0.95 | 1 | 1 | 1.861 | 0.896 | 0.66253952 | NA | 0 | 3.4584 | 1.8903 | 1.8903 | TRUE | 0.384 |
| Cyp2d13 | 0 | 0 | 0 | 0 | 0 | 0 | 0 | 0 | 0 | 0 | 0 | 0 | 0 | 0 | 0 | NA | NA | NA | NA | NA | NA | NA | NA | NA | NA |
| Cyp2d22 | 124 | 228 | 193 | 138 | 140 | 153 | 132 | 228 | 194 | 108 | 151 | 167 | 163.3 | 185 | 142 | 0.768 | -0.381 | 0.13656774 | 0.32019421 | 0.0558 | 0.0271 | 0.041 | 0.041 | TRUE | 0.3474 |
| Cyp2d26 | 2 | 3 | 0 | 0 | 2 | 2 | 2 | 3 | 0 | 0 | 2 | 2 | 1.58 | 2 | 1 | 0.821 | -0.284 | 0.87204289 | NA | 0 | 2.096 | 1.5901 | 1.5901 | TRUE | 0.7083 |
| Cyp2d34 | 0 | 0 | 0 | 0 | 0 | 0 | 0 | 0 | 0 | 0 | 0 | 0 | 0 | 0 | 0 | NA | NA | NA | NA | NA | NA | NA | NA | NA | NA |
| Cyp2d37-ps | 0 | 0 | 0 | 0 | 0 | 0 | 0 | 0 | 0 | 0 | 0 | 0 | 0 | 0 | 0 | NA | NA | NA | NA | NA | NA | NA | NA | NA | NA |
| Cyp2d40 | 0 | 0 | 0 | 0 | 0 | 0 | 0 | 0 | 0 | 0 | 0 | 0 | 0 | 0 | 0 | NA | NA | NA | NA | NA | NA | NA | NA | NA | NA |
| Cyp2d9 | 0 | 0 | 0 | 0 | 1 | 0 | 0 | 0 | 0 | 0 | 1 | 0 | 0.18 | 0 | 0 | 1.859 | 0.894 | 0.82650734 | NA | 0 | 18.3785 | 10 | 10 | TRUE | 0.5204 |
| Cyp2e1 | 1 | 0 | 0 | 0 | 0 | 0 | 1 | 0 | 0 | 0 | 0 | 0 | 0.18 | 0 | 0 | 0.49 | -1.029 | 0.80087 | NA | 0 | 18.6165 | 10 | 10 | TRUE | 0.4614 |
| Cyp2f2 | 3 | 1 | 2 | 3 | 1 | 1 | 3 | 1 | 2 | 2 | 1 | 1 | 1.79 | 2 | 1 | 0.754 | -0.408 | 0.76615741 | NA | 0 | 1.8504 | 0.7899 | 0.7899 | TRUE | 0.2056 |
| Cyp2g1 | 0 | 0 | 0 | 0 | 0 | 0 | 0 | 0 | 0 | 0 | 0 | 0 | 0 | 0 | 0 | NA | NA | NA | NA | NA | NA | NA | NA | NA | NA |
| Cyp2j11 | 0 | 0 | 0 | 0 | 0 | 0 | 0 | 0 | 0 | 0 | 0 | 0 | 0 | 0 | 0 | NA | NA | NA | NA | NA | NA | NA | NA | NA | NA |
| Cyp2j12 | 0 | 0 | 0 | 1 | 0 | 0 | 0 | 0 | 0 | 1 | 0 | 0 | 0.13 | 0 | 0 | 1.859 | 0.894 | 0.82650734 | NA | 0 | 25.1515 | 10 | 10 | TRUE | 0.2563 |
| Cyp2j13 | 86 | 56 | 49 | 60 | 45 | 52 | 91 | 56 | 49 | 47 | 48 | 57 | 58.16 | 65 | 51 | 0.776 | -0.366 | 0.24832928 | 0.4650498 | 0.0463 | 0.0635 | 0.055 | 0.055 | TRUE | 1.0334 |
| Cyp2j5 | 0 | 0 | 0 | 0 | 0 | 0 | 0 | 0 | 0 | 0 | 0 | 0 | 0 | 0 | 0 | NA | NA | NA | NA | NA | NA | NA | NA | NA | NA |
| Cyp2j6 | 62 | 80 | 55 | 103 | 64 | 53 | 66 | 80 | 55 | 81 | 69 | 58 | 68.14 | 67 | 69 | 1.039 | 0.055 | 0.83511014 | 0.91601343 | 0.017 | 0.0553 | 0.0365 | 0.0365 | TRUE | 0.2562 |
| Cyp2j8 | 0 | 0 | 0 | 0 | 0 | 0 | 0 | 0 | 0 | 0 | 0 | 0 | 0 | 0 | 0 | NA | NA | NA | NA | NA | NA | NA | NA | NA | NA |
| Cyp2j9 | 97 | 106 | 107 | 135 | 94 | 91 | 103 | 106 | 107 | 106 | 101 | 100 | 103.87 | 105 | 102 | 0.971 | -0.042 | 0.82367489 | 0.91017184 | 0 | 0.0386 | 0.0158 | 0.0158 | TRUE | 0.0119 |
| Cyp2r1 | 13 | 7 | 7 | 33 | 14 | 9 | 14 | 7 | 7 | 26 | 15 | 10 | 13.11 | 9 | 17 | 1.857 | 0.893 | 0.1511027 | 0.34216681 | 0.1318 | 0.2581 | 0.1951 | 0.1951 | TRUE | 0.3688 |
| Cyp2s1 | 1586 | 3109 | 3161 | 2101 | 1897 | 2584 | 1685 | 3105 | 3174 | 1652 | 2041 | 2829 | 2414.14 | 2655 | 2174 | 0.819 | -0.289 | 0.24698087 | 0.46379674 | 0.0847 | 0.0083 | 0.0444 | 0.0444 | TRUE | 0.943 |
| Cyp2t4 | 5 | 9 | 3 | 6 | 4 | 7 | 5 | 9 | 3 | 5 | 4 | 8 | 5.67 | 6 | 6 | 0.957 | -0.064 | 0.93801519 | 0.96893419 | 0 | 0.5884 | 0.3112 | 0.3112 | TRUE | 0.355 |
| Cyp2u1 | 57 | 91 | 84 | 98 | 42 | 37 | 61 | 91 | 84 | 77 | 45 | 41 | 66.42 | 79 | 54 | 0.696 | -0.522 | 0.11141453 | 0.28068465 | 0.068 | 0.0565 | 0.062 | 0.062 | TRUE | 1.2502 |
| Cyp2w1 | 0 | 1 | 1 | 0 | 0 | 1 | 0 | 1 | 1 | 0 | 0 | 1 | 0.52 | 1 | 0 | 0.52 | -0.943 | 0.73639299 | NA | 0 | 6.3885 | 3.8941 | 3.8941 | TRUE | 0.5383 |

| | | | | | | | | | | | | | | | | | | | | | | | | | |
|---|---|---|---|---|---|---|---|---|---|---|---|---|---|---|---|---|---|---|---|---|---|---|---|---|---|
| Cyp39a1 | 171 | 179 | 198 | 187 | 135 | 150 | 182 | 179 | 199 | 147 | 145 | 164 | 169.28 | 187 | 152 | 0.815 | -0.295 | 0.06213222 | 0.19072584 | 0 | 0.0264 | 0.0121 | 0.0121 | TRUE | 0.0477 |
| Cyp3a11 | 0 | 0 | 0 | 0 | 0 | 0 | 0 | 0 | 0 | 0 | 0 | 0 | 0 | 0 | 0 | NA | NA | NA | NA | NA | NA | NA | NA | NA | NA |
| Cyp3a13 | 3 | 9 | 5 | 5 | 6 | 2 | 3 | 9 | 5 | 4 | 6 | 2 | 4.96 | 6 | 4 | 0.728 | -0.458 | 0.60892823 | 0.77857639 | 0 | 0.6709 | 0.3714 | 0.3714 | TRUE | 0.4814 |
| Cyp3a16 | 0 | 0 | 0 | 0 | 0 | 0 | 0 | 0 | 0 | 0 | 0 | 0 | 0 | 0 | 0 | NA | NA | NA | NA | NA | NA | NA | NA | NA | NA |
| Cyp3a25 | 0 | 0 | 0 | 0 | 0 | 0 | 0 | 0 | 0 | 0 | 0 | 0 | 0 | 0 | 0 | NA | NA | NA | NA | NA | NA | NA | NA | NA | NA |
| Cyp3a41a | 0 | 0 | 0 | 0 | 0 | 0 | 0 | 0 | 0 | 0 | 0 | 0 | 0 | 0 | 0 | NA | NA | NA | NA | NA | NA | NA | NA | NA | NA |
| Cyp3a41b | 0 | 0 | 0 | 0 | 0 | 0 | 0 | 0 | 0 | 0 | 0 | 0 | 0 | 0 | 0 | NA | NA | NA | NA | NA | NA | NA | NA | NA | NA |
| Cyp3a44 | 0 | 0 | 0 | 0 | 0 | 0 | 0 | 0 | 0 | 0 | 0 | 0 | 0 | 0 | 0 | NA | NA | NA | NA | NA | NA | NA | NA | NA | NA |
| Cyp3a57 | 0 | 0 | 0 | 0 | 0 | 0 | 0 | 0 | 0 | 0 | 0 | 0 | 0 | 0 | 0 | NA | NA | NA | NA | NA | NA | NA | NA | NA | NA |
| Cyp3a59 | 0 | 0 | 0 | 0 | 0 | 0 | 0 | 0 | 0 | 0 | 0 | 0 | 0 | 0 | 0 | NA | NA | NA | NA | NA | NA | NA | NA | NA | NA |
| Cyp46a1 | 155 | 218 | 191 | 129 | 113 | 140 | 165 | 218 | 192 | 101 | 122 | 153 | 158.4 | 192 | 125 | 0.652 | -0.617 | 0.00365209 | 0.02466839 | 0.0241 | 0.0277 | 0.0259 | 0.0259 | TRUE | 0.3232 |
| Cyp4a10 | 0 | 0 | 0 | 0 | 1 | 0 | 0 | 0 | 0 | 0 | 1 | 0 | 0.18 | 0 | 0 | 1.859 | 0.894 | 0.82650734 | NA | 0 | 18.3785 | 10 | 10 | TRUE | 0.5204 |
| Cyp4a12a | 0 | 0 | 0 | 0 | 0 | 0 | 0 | 0 | 0 | 0 | 0 | 0 | 0 | 0 | 0 | NA | NA | NA | NA | NA | NA | NA | NA | NA | NA |
| Cyp4a12b | 2 | 1 | 4 | 12 | 7 | 3 | 2 | 1 | 4 | 9 | 8 | 3 | 4.56 | 2 | 7 | 2.866 | 1.519 | 0.12531424 | NA | 0 | 0.7286 | 0.4215 | 0.4215 | TRUE | 0.3232 |
| Cyp4a14 | 1 | 0 | 0 | 1 | 1 | 0 | 1 | 0 | 0 | 1 | 1 | 0 | 0.49 | 0 | 1 | 1.712 | 0.776 | 0.78488768 | NA | 0 | 6.7665 | 4.0387 | 4.0387 | TRUE | 0.4614 |
| Cyp4a29-ps | 0 | 0 | 0 | 0 | 0 | 0 | 0 | 0 | 0 | 0 | 0 | 0 | 0 | 0 | 0 | NA | NA | NA | NA | NA | NA | NA | NA | NA | NA |
| Cyp4a30b-ps | 0 | 0 | 0 | 0 | 0 | 1 | 0 | 0 | 0 | 0 | 0 | 1 | 0.18 | 0 | 0 | 1.859 | 0.894 | 0.82650734 | NA | 0 | 18.0643 | 10 | 10 | TRUE | 0.5383 |
| Cyp4a31 | 0 | 0 | 0 | 2 | 0 | 0 | 0 | 0 | 0 | 2 | 0 | 0 | 0.26 | 0 | 1 | 2.886 | 1.529 | 0.70676491 | NA | 0 | 12.5792 | 10 | 10 | TRUE | 0.9669 |
| Cyp4a32 | 0 | 0 | 0 | 0 | 0 | 0 | 0 | 0 | 0 | 0 | 0 | 0 | 0 | 0 | 0 | NA | NA | NA | NA | NA | NA | NA | NA | NA | NA |
| Cyp4b1 | 0 | 0 | 0 | 0 | 2 | 0 | 0 | 0 | 0 | 0 | 2 | 0 | 0.36 | 0 | 1 | 3.775 | 1.916 | 0.63423067 | NA | 10 | 9.1927 | 10 | 10 | TRUE | 0.9743 |
| Cyp4f13 | 307 | 516 | 522 | 384 | 274 | 361 | 326 | 515 | 524 | 302 | 295 | 395 | 392.9 | 455 | 331 | 0.725 | -0.463 | 0.02809798 | 0.11006223 | 0.0447 | 0.0153 | 0.0295 | 0.0295 | TRUE | 0.71 |
| Cyp4f14 | 3 | 4 | 4 | 9 | 2 | 5 | 3 | 4 | 4 | 7 | 2 | 5 | 4.32 | 4 | 5 | 1.331 | 0.413 | 0.653261 | NA | 0 | 0.7702 | 0.374 | 0.374 | TRUE | 0.4255 |
| Cyp4f15 | 6 | 1 | 2 | 4 | 2 | 0 | 6 | 1 | 2 | 3 | 2 | 0 | 2.45 | 3 | 2 | 0.585 | -0.773 | 0.57963051 | NA | 0.5131 | 1.3537 | 0.9723 | 0.9723 | TRUE | 1.0604 |
| Cyp4f16 | 629 | 872 | 819 | 882 | 664 | 778 | 668 | 871 | 822 | 693 | 714 | 852 | 770.13 | 787 | 753 | 0.956 | -0.065 | 0.64189021 | 0.79842557 | 0.0144 | 0.0112 | 0.0129 | 0.0129 | TRUE | 0.2065 |
| Cyp4f17 | 153 | 179 | 202 | 222 | 185 | 210 | 163 | 179 | 203 | 175 | 199 | 230 | 191.26 | 182 | 201 | 1.105 | 0.144 | 0.41538306 | 0.62699029 | 0.0109 | 0.0241 | 0.0173 | 0.0173 | TRUE | 0.1682 |
| Cyp4f18 | 4 | 2 | 5 | 2 | 2 | 4 | 4 | 2 | 5 | 2 | 2 | 4 | 3.23 | 4 | 3 | 0.706 | -0.503 | 0.63621114 | NA | 0 | 1.0274 | 0.4945 | 0.4945 | TRUE | 0.3206 |
| Cyp4f39 | 92 | 144 | 129 | 163 | 108 | 112 | 98 | 144 | 130 | 128 | 116 | 123 | 123 | 124 | 122 | 0.989 | -0.016 | 0.93864274 | 0.96918319 | 0.0114 | 0.0337 | 0.0228 | 0.0228 | TRUE | 0.336 |
| Cyp4f40 | 3 | 2 | 2 | 1 | 3 | 4 | 3 | 2 | 2 | 1 | 3 | 4 | 2.6 | 2 | 3 | 1.137 | 0.185 | 0.87627082 | NA | 0 | 1.2752 | 0.6285 | 0.6285 | TRUE | 0.6706 |
| Cyp4f41-ps | 8 | 12 | 16 | 11 | 8 | 11 | 8 | 12 | 16 | 9 | 9 | 12 | 10.97 | 12 | 10 | 0.796 | -0.33 | 0.57214473 | 0.7508626 | 0 | 0.3071 | 0.1531 | 0.1531 | TRUE | 0.1924 |
| Cyp4v3 | 103 | 114 | 80 | 159 | 94 | 120 | 109 | 114 | 80 | 125 | 101 | 131 | 110.18 | 101 | 119 | 1.179 | 0.238 | 0.28725368 | 0.50848092 | 0.0162 | 0.0368 | 0.0268 | 0.0268 | TRUE | 0.3215 |
| Cyp4x1 | 1 | 0 | 0 | 2 | 2 | 0 | 1 | 0 | 0 | 2 | 2 | 0 | 0.8 | 0 | 1 | 3.417 | 1.773 | 0.46651319 | NA | 0 | 4.1368 | 2.8785 | 2.8785 | TRUE | 0.4614 |
| Cyp51 | 5703 | 6991 | 7283 | 7908 | 5913 | 7014 | 6058 | 6982 | 7313 | 6217 | 6362 | 7678 | 6768.18 | 6784 | 6752 | 0.995 | -0.007 | 0.95177438 | 0.97539867 | 0.0114 | 0.0074 | 0.0096 | 0.0096 | TRUE | 0.1753 |
| Cyp7a1 | 0 | 0 | 0 | 1 | 0 | 0 | 0 | 0 | 0 | 1 | 0 | 0 | 0.13 | 0 | 0 | 1.859 | 0.894 | 0.82650734 | NA | 0 | 25.1515 | 10 | 10 | TRUE | 0.2563 |
| Cyp7b1 | 60 | 45 | 37 | 105 | 90 | 39 | 64 | 45 | 37 | 83 | 97 | 43 | 61.32 | 49 | 74 | 1.528 | 0.611 | 0.09802397 | 0.25741492 | 0.104 | 0.0606 | 0.0813 | 0.0813 | TRUE | 0.5989 |
| Cyp8b1 | 0 | 0 | 0 | 0 | 0 | 0 | 0 | 0 | 0 | 0 | 0 | 0 | 0 | 0 | 0 | NA | NA | NA | NA | NA | NA | NA | NA | NA | NA |
| Cypt1 | 0 | 0 | 0 | 0 | 0 | 0 | 0 | 0 | 0 | 0 | 0 | 0 | 0 | 0 | 0 | NA | NA | NA | NA | NA | NA | NA | NA | NA | NA |
| Cypt10 | 0 | 0 | 0 | 0 | 0 | 0 | 0 | 0 | 0 | 0 | 0 | 0 | 0 | 0 | 0 | NA | NA | NA | NA | NA | NA | NA | NA | NA | NA |
| Cypt12 | 0 | 0 | 0 | 0 | 0 | 0 | 0 | 0 | 0 | 0 | 0 | 0 | 0 | 0 | 0 | NA | NA | NA | NA | NA | NA | NA | NA | NA | NA |
| Cypt14 | 0 | 0 | 0 | 1 | 0 | 0 | 0 | 0 | 0 | 1 | 0 | 0 | 0.13 | 0 | 0 | 1.859 | 0.894 | 0.82650734 | NA | 0 | 25.1515 | 10 | 10 | TRUE | 0.2563 |
| Cypt15 | 0 | 0 | 0 | 0 | 0 | 0 | 0 | 0 | 0 | 0 | 0 | 0 | 0 | 0 | 0 | NA | NA | NA | NA | NA | NA | NA | NA | NA | NA |
| Cypt2 | 1 | 1 | 3 | 5 | 3 | 1 | 1 | 1 | 3 | 4 | 3 | 1 | 2.22 | 2 | 3 | 1.647 | 0.72 | 0.57601288 | NA | 0 | 1.4902 | 0.7189 | 0.7189 | TRUE | 0.3631 |
| Cypt3 | 30 | 27 | 32 | 48 | 36 | 36 | 32 | 27 | 32 | 38 | 39 | 39 | 34.47 | 30 | 39 | 1.273 | 0.349 | 0.27497724 | 0.49548018 | 0 | 0.1025 | 0.044 | 0.044 | TRUE | 0.0637 |
| Cypt4 | 0 | 0 | 0 | 0 | 0 | 0 | 0 | 0 | 0 | 0 | 0 | 0 | 0 | 0 | 0 | NA | NA | NA | NA | NA | NA | NA | NA | NA | NA |
| Cypt7 | 0 | 0 | 0 | 0 | 0 | 0 | 0 | 0 | 0 | 0 | 0 | 0 | 0 | 0 | 0 | NA | NA | NA | NA | NA | NA | NA | NA | NA | NA |
| Cypt8 | 0 | 0 | 0 | 0 | 0 | 0 | 0 | 0 | 0 | 0 | 0 | 0 | 0 | 0 | 0 | NA | NA | NA | NA | NA | NA | NA | NA | NA | NA |
| Cypt9 | 0 | 0 | 0 | 0 | 0 | 0 | 0 | 0 | 0 | 0 | 0 | 0 | 0 | 0 | 0 | NA | NA | NA | NA | NA | NA | NA | NA | NA | NA |
| Cyr61 | 410 | 401 | 259 | 1252 | 804 | 636 | 435 | 400 | 260 | 984 | 865 | 696 | 606.92 | 365 | 848 | 2.325 | 1.217 | 3.92E-09 | 2.03E-07 | 0.0479 | 0.0123 | 0.0288 | 0.0288 | TRUE | 0.3911 |
| Cys1 | 2 | 2 | 4 | 1 | 3 | 2 | 2 | 2 | 4 | 1 | 3 | 2 | 2.39 | 3 | 2 | 0.74 | -0.434 | 0.72160176 | NA | 0 | 1.3852 | 0.6399 | 0.6399 | TRUE | 0.3869 |
| Cysltr1 | 3 | 3 | 0 | 2 | 3 | 1 | 3 | 3 | 0 | 2 | 3 | 1 | 2.01 | 2 | 2 | 0.951 | -0.073 | 0.95938242 | NA | 0 | 1.6436 | 0.9787 | 0.9787 | TRUE | 0.7174 |
| Cysltr2 | 33 | 24 | 16 | 98 | 77 | 46 | 35 | 24 | 16 | 77 | 83 | 50 | 47.55 | 25 | 70 | 2.813 | 1.492 | 6.76E-05 | 0.00100495 | 0.0702 | 0.0762 | 0.0736 | 0.0736 | TRUE | 0.7218 |
| Cystm1 | 534 | 641 | 663 | 725 | 478 | 485 | 567 | 640 | 666 | 570 | 514 | 531 | 581.37 | 624 | 538 | 0.863 | -0.213 | 0.0572121 | 0.18025354 | 0.0032 | 0.0126 | 0.0073 | 0.0073 | TRUE | 0.0748 |
| Cyth1 | 410 | 537 | 486 | 533 | 385 | 421 | 435 | 536 | 488 | 419 | 414 | 461 | 458.98 | 486 | 431 | 0.886 | -0.175 | 0.16064902 | 0.35521608 | 0.0049 | 0.0141 | 0.0091 | 0.0091 | TRUE | 0.0973 |
| Cyth2 | 1660 | 1824 | 1850 | 2424 | 1764 | 1717 | 1763 | 1822 | 1858 | 1906 | 1898 | 1880 | 1854.25 | 1814 | 1895 | 1.044 | 0.062 | 0.36340677 | 0.58083215 | 0 | 0.0087 | 0.0029 | 0.0029 | TRUE | 0.0072 |
| Cyth3 | 1333 | 1295 | 1287 | 1539 | 1102 | 1260 | 1416 | 1293 | 1292 | 1210 | 1186 | 1379 | 1296.05 | 1334 | 1258 | 0.943 | -0.085 | 0.38960642 | 0.60362205 | 0.0038 | 0.0094 | 0.0062 | 0.0062 | TRUE | 0.0837 |
| Cyth4 | 171 | 129 | 121 | 200 | 122 | 108 | 182 | 129 | 121 | 157 | 131 | 118 | 139.78 | 144 | 135 | 0.946 | -0.081 | 0.71984873 | 0.8481969 | 0.0281 | 0.0305 | 0.0293 | 0.0293 | TRUE | 0.5378 |
| Cytip | 0 | 0 | 0 | 0 | 1 | 0 | 0 | 0 | 0 | 0 | 1 | 0 | 0.18 | 0 | 0 | 1.859 | 0.894 | 0.82650734 | NA | 0 | 18.3785 | 10 | 10 | TRUE | 0.5204 |
| Cytl1 | 1 | 2 | 1 | 5 | 1 | 1 | 1 | 2 | 1 | 4 | 1 | 1 | 1.69 | 1 | 2 | 1.549 | 0.631 | 0.66190448 | NA | 0 | 1.9515 | 0.8913 | 0.8913 | TRUE | 0.7909 |
| Cyyr1 | 19 | 23 | 22 | 62 | 37 | 39 | 20 | 23 | 22 | 49 | 40 | 43 | 32.75 | 22 | 44 | 2.018 | 1.013 | 0.00286179 | 0.02040515 | 0 | 0.1075 | 0.0485 | 0.0485 | TRUE | 0.0894 |
| D030018L15I | 1 | 1 | 1 | 2 | 0 | 0 | 1 | 1 | 1 | 2 | 0 | 0 | 0.77 | 1 | 1 | 0.562 | -0.832 | 0.70497892 | NA | 0 | 4.2695 | 2.1321 | 2.1321 | TRUE | 0.9716 |
| D030024E09 | 0 | 0 | 0 | 0 | 0 | 0 | 0 | 0 | 0 | 0 | 0 | 0 | 0 | 0 | 0 | NA | NA | NA | NA | NA | NA | NA | NA | NA | NA |
| D030025E07 | 0 | 0 | 0 | 0 | 0 | 0 | 0 | 0 | 0 | 0 | 0 | 0 | 0 | 0 | 0 | NA | NA | NA | NA | NA | NA | NA | NA | NA | NA |
| D030025P21 | 15 | 25 | 22 | 32 | 36 | 18 | 16 | 25 | 22 | 25 | 39 | 20 | 24.43 | 21 | 28 | 1.322 | 0.403 | 0.35544065 | 0.57387109 | 0.044 | 0.1418 | 0.095 | 0.095 | TRUE | 0.5538 |
| D030028A08 | 74 | 68 | 77 | 130 | 78 | 72 | 79 | 68 | 77 | 102 | 84 | 79 | 81.46 | 75 | 88 | 1.19 | 0.251 | 0.27046217 | 0.49065209 | 0 | 0.0473 | 0.0251 | 0.0251 | TRUE | 0.1997 |
| D030040B21 | 0 | 2 | 1 | 2 | 2 | 1 | 0 | 2 | 1 | 2 | 2 | 1 | 1.3 | 1 | 2 | 1.59 | 0.669 | 0.68609916 | NA | 0 | 2.5345 | 1.1653 | 1.1653 | TRUE | 0.2601 |
| D030045P18 | 0 | 0 | 0 | 2 | 1 | 2 | 0 | 0 | 0 | 2 | 1 | 2 | 0.81 | 0 | 2 | 8.576 | 3.1 | 0.18121756 | NA | 0 | 4.093 | 2.5705 | 2.5705 | TRUE | 0.07 |
| D030047H15 | 7 | 1 | 5 | 3 | 5 | 5 | 7 | 1 | 5 | 2 | 5 | 5 | 4.44 | 4 | 4 | 0.971 | -0.043 | 0.96585279 | NA | 0.1799 | 0.7482 | 0.4838 | 0.4838 | TRUE | 0.3816 |
| D030056L22I | 1977 | 2468 | 2538 | 3244 | 2369 | 2426 | 2100 | 2465 | 2548 | 2550 | 2549 | 2656 | 2477.98 | 2371 | 2585 | 1.09 | 0.124 | 0.19865993 | 0.40581069 | 0.0047 | 0.0082 | 0.0063 | 0.0063 | TRUE | 0.1209 |
| D10Bwg1379 | 234 | 249 | 258 | 584 | 391 | 338 | 249 | 249 | 259 | 459 | 421 | 370 | 334.34 | 252 | 417 | 1.656 | 0.728 | 6.65E-08 | 2.53E-06 | 0.0044 | 0.0168 | 0.0099 | 0.0099 | TRUE | 0.1089 |
| D10Jhu81e | 2427 | 2577 | 2651 | 2901 | 2164 | 2079 | 2578 | 2574 | 2662 | 2281 | 2328 | 2276 | 2449.69 | 2605 | 2295 | 0.881 | -0.183 | 0.00451396 | 0.02894075 | 0 | 0.0082 | 0.0026 | 0.0026 | TRUE | 0.0045 |

| | | | | | | | | | | | | | | | | | | | | | | | | | |
|---|---|---|---|---|---|---|---|---|---|---|---|---|---|---|---|---|---|---|---|---|---|---|---|---|---|
| D10Wsu102e | 3484 | 3463 | 3552 | 4554 | 3314 | 3014 | 3701 | 3459 | 3567 | 3580 | 3566 | 3299 | 3528.45 | 3576 | 3482 | 0.974 | -0.038 | 0.6035228 | 0.77438672 | 0.0012 | 0.0078 | 0.0035 | 0.0035 | TRUE | 0.0253 |
| D10Wsu52e | 8483 | 8518 | 8371 | 12462 | 8800 | 8408 | 9010 | 8507 | 8405 | 9797 | 9468 | 9204 | 9065.27 | 8641 | 9490 | 1.098 | 0.135 | 0.03937059 | 0.13910238 | 0.001 | 0.0073 | 0.003 | 0.003 | TRUE | 0.0171 |
| D11Wsu47e | 2590 | 2662 | 2593 | 3297 | 2517 | 2173 | 2751 | 2659 | 2604 | 2592 | 2708 | 2379 | 2615.32 | 2671 | 2560 | 0.958 | -0.061 | 0.44840354 | 0.65422947 | 0.0019 | 0.0082 | 0.0043 | 0.0043 | TRUE | 0.0456 |
| D130009I18F | 0 | 0 | 0 | 0 | 0 | 0 | 0 | 0 | 0 | 0 | 0 | 0 | 0 | 0 | 0 | NA | NA | NA | NA | NA | NA | NA | NA | NA | NA |
| D130017N08 | 61 | 84 | 78 | 67 | 46 | 46 | 65 | 84 | 78 | 53 | 49 | 50 | 63.25 | 76 | 51 | 0.672 | -0.573 | 0.02141534 | 0.09133658 | 0 | 0.059 | 0.0284 | 0.0284 | TRUE | 0.1413 |
| D130020L05I | 24 | 59 | 60 | 81 | 58 | 52 | 25 | 59 | 60 | 64 | 62 | 57 | 54.61 | 48 | 61 | 1.263 | 0.337 | 0.33906915 | 0.55910074 | 0.076 | 0.0672 | 0.0712 | 0.0712 | TRUE | 1.3263 |
| D130040H23 | 101 | 93 | 73 | 169 | 125 | 91 | 107 | 93 | 73 | 133 | 134 | 100 | 106.74 | 91 | 122 | 1.346 | 0.429 | 0.06632543 | 0.19871336 | 0.021 | 0.0378 | 0.0297 | 0.0297 | TRUE | 0.2825 |
| D130043K22I | 26 | 33 | 27 | 52 | 41 | 30 | 28 | 33 | 27 | 41 | 44 | 33 | 34.25 | 29 | 39 | 1.345 | 0.428 | 0.19707137 | 0.40371603 | 0 | 0.1031 | 0.0495 | 0.0495 | TRUE | 0.14 |
| D130058E03 | 0 | 0 | 0 | 0 | 0 | 0 | 0 | 0 | 0 | 0 | 0 | 0 | 0 | 0 | 0 | NA | NA | NA | NA | NA | NA | NA | NA | NA | NA |
| D14Abb1e | 6102 | 4264 | 4432 | 7200 | 5148 | 4442 | 6481 | 4258 | 4450 | 5660 | 5539 | 4862 | 5208.59 | 5063 | 5354 | 1.057 | 0.081 | 0.61946985 | 0.78482436 | 0.0258 | 0.0075 | 0.0188 | 0.0188 | TRUE | 0.7313 |
| D15Ertd621e | 9438 | 8873 | 9230 | 12406 | 8982 | 8511 | 10025 | 8862 | 9268 | 9753 | 9664 | 9317 | 9481.27 | 9385 | 9578 | 1.021 | 0.03 | 0.69127103 | 0.83074715 | 0.0018 | 0.0073 | 0.0039 | 0.0039 | TRUE | 0.0434 |
| D16Ertd472e | 1329 | 1100 | 1047 | 1488 | 1071 | 942 | 1412 | 1099 | 1051 | 1170 | 1152 | 1031 | 1152.46 | 1187 | 1118 | 0.942 | -0.086 | 0.52349016 | 0.71330339 | 0.0135 | 0.0098 | 0.0123 | 0.0123 | TRUE | 0.3271 |
| D16Ertd519e | 0 | 0 | 0 | 0 | 0 | 0 | 0 | 0 | 0 | 0 | 0 | 0 | 0 | 0 | 0 | NA | NA | NA | NA | NA | NA | NA | NA | NA | NA |
| D17Ertd648e | 1 | 1 | 0 | 3 | 0 | 3 | 1 | 1 | 0 | 2 | 0 | 3 | 1.28 | 1 | 2 | 2.764 | 1.467 | 0.44417354 | NA | 0 | 2.5729 | 1.6424 | 1.6424 | TRUE | 0.4211 |
| D17H6S53E | 74 | 92 | 86 | 136 | 105 | 95 | 79 | 92 | 86 | 107 | 113 | 104 | 96.79 | 86 | 108 | 1.26 | 0.333 | 0.09381158 | 0.25057385 | 0 | 0.0409 | 0.018 | 0.018 | TRUE | 0.0472 |
| D17Wsu104e | 1734 | 2185 | 2145 | 2263 | 1624 | 1706 | 1842 | 2182 | 2154 | 1779 | 1747 | 1867 | 1928.6 | 2059 | 1798 | 0.873 | -0.196 | 0.0407143 | 0.14276255 | 0.004 | 0.0086 | 0.0061 | 0.0061 | TRUE | 0.1029 |
| D17Wsu92e | 8313 | 8409 | 8463 | 11257 | 8069 | 8442 | 8830 | 8398 | 8498 | 8849 | 8682 | 9241 | 8749.66 | 8575 | 8924 | 1.041 | 0.057 | 0.35795736 | 0.57644512 | 7.00E-04 | 0.0073 | 0.0027 | 0.0027 | TRUE | 0.0117 |
| D19Bwg1357 | 7733 | 7465 | 7520 | 10675 | 7416 | 6854 | 8214 | 7455 | 7551 | 8392 | 7979 | 7503 | 7848.97 | 7740 | 7958 | 1.028 | 0.04 | 0.61207477 | 0.78028154 | 0.0025 | 0.0073 | 0.0044 | 0.0044 | TRUE | 0.035 |
| D19Ertd737e | 749 | 898 | 925 | 1414 | 980 | 943 | 796 | 897 | 929 | 1112 | 1054 | 1032 | 969.91 | 874 | 1066 | 1.221 | 0.288 | 0.00341468 | 0.02340565 | 0.0028 | 0.0103 | 0.0059 | 0.0059 | TRUE | 0.0723 |
| D1Ertd622e | 672 | 809 | 845 | 888 | 659 | 655 | 714 | 808 | 848 | 698 | 709 | 717 | 749.05 | 790 | 708 | 0.896 | -0.159 | 0.12296942 | 0.29870225 | 0.0027 | 0.0113 | 0.0063 | 0.0063 | TRUE | 0.0839 |
| D1Pas1 | 938 | 665 | 593 | 1765 | 1238 | 778 | 996 | 664 | 595 | 1388 | 1332 | 852 | 971.18 | 752 | 1191 | 1.584 | 0.664 | 0.00393149 | 0.02599892 | 0.0697 | 0.0103 | 0.0371 | 0.0371 | TRUE | 0.9553 |
| D230025D16 | 3413 | 3154 | 3010 | 5108 | 3603 | 3551 | 3625 | 3150 | 3022 | 4016 | 3877 | 3887 | 3596.12 | 3266 | 3927 | 1.203 | 0.266 | 0.00369208 | 0.02484276 | 0.0045 | 0.0078 | 0.0058 | 0.0058 | TRUE | 0.1124 |
| D230030E09I | 0 | 0 | 0 | 0 | 0 | 1 | 0 | 0 | 0 | 0 | 0 | 1 | 0.18 | 0 | 0 | 1.859 | 0.894 | 0.82650734 | NA | 0 | 18.0643 | 10 | 10 | TRUE | 0.5383 |
| D2hgdh | 297 | 380 | 363 | 447 | 362 | 329 | 315 | 380 | 364 | 351 | 389 | 360 | 360.08 | 353 | 367 | 1.038 | 0.053 | 0.67770383 | 0.82075902 | 0.0031 | 0.0161 | 0.0091 | 0.0091 | TRUE | 0.0983 |
| D2Wsu81e | 2123 | 2037 | 2113 | 2602 | 2088 | 2090 | 2255 | 2034 | 2122 | 2045 | 2247 | 2288 | 2165.15 | 2137 | 2193 | 1.026 | 0.037 | 0.66844191 | 0.81486055 | 0.0027 | 0.0084 | 0.0049 | 0.0049 | TRUE | 0.043 |
| D330022K07I | 0 | 0 | 0 | 1 | 1 | 0 | 0 | 0 | 0 | 1 | 1 | 0 | 0.31 | 0 | 1 | 3.32 | 1.731 | 0.65383052 | NA | 0 | 10.6224 | 8.9541 | 8.9541 | TRUE | 0.202 |
| D330023K18I | 15 | 17 | 8 | 31 | 18 | 22 | 16 | 17 | 8 | 24 | 19 | 24 | 18.13 | 14 | 22 | 1.662 | 0.733 | 0.12586643 | 0.30396038 | 0 | 0.1886 | 0.1061 | 0.1061 | TRUE | 0.5646 |
| D330041H03 | 23 | 30 | 25 | 59 | 36 | 39 | 24 | 30 | 25 | 46 | 39 | 43 | 34.55 | 26 | 43 | 1.611 | 0.688 | 0.03540645 | 0.12914673 | 0 | 0.1023 | 0.0464 | 0.0464 | TRUE | 0.0831 |
| D330045A20 | 147 | 98 | 94 | 143 | 87 | 93 | 156 | 98 | 94 | 112 | 94 | 102 | 109.37 | 116 | 103 | 0.886 | -0.174 | 0.49216194 | 0.6883982 | 0.037 | 0.037 | 0.037 | 0.037 | TRUE | 0.9007 |
| D330050G23 | 4 | 8 | 13 | 17 | 5 | 13 | 4 | 8 | 13 | 13 | 5 | 14 | 9.71 | 8 | 11 | 1.307 | 0.386 | 0.58240938 | 0.7585269 | 0.1406 | 0.3462 | 0.2502 | 0.2502 | TRUE | 0.3519 |
| D330050I16F | 53 | 51 | 32 | 74 | 45 | 51 | 56 | 51 | 32 | 58 | 48 | 56 | 50.3 | 46 | 54 | 1.169 | 0.225 | 0.46500463 | 0.66788976 | 0.0208 | 0.0724 | 0.0486 | 0.0486 | TRUE | 0.5813 |
| D3Bwg0562e | 6 | 5 | 7 | 24 | 19 | 10 | 6 | 5 | 7 | 19 | 20 | 11 | 11.44 | 6 | 17 | 2.745 | 1.457 | 0.01670148 | 0.07648271 | 0 | 0.2948 | 0.1545 | 0.1545 | TRUE | 0.4089 |
| D3Ertd254e | 1644 | 1254 | 1248 | 1921 | 1479 | 1227 | 1746 | 1252 | 1253 | 1510 | 1591 | 1343 | 1449.39 | 1417 | 1481 | 1.046 | 0.064 | 0.66913024 | 0.81506968 | 0.0219 | 0.0092 | 0.0157 | 0.0157 | TRUE | 0.4952 |
| D3Ertd751e | 278 | 329 | 316 | 481 | 339 | 294 | 295 | 329 | 317 | 378 | 365 | 322 | 334.31 | 314 | 355 | 1.133 | 0.18 | 0.16081779 | 0.35543978 | 0.0021 | 0.0168 | 0.0089 | 0.0089 | TRUE | 0.074 |
| D430019H16 | 454 | 505 | 497 | 426 | 368 | 368 | 482 | 504 | 499 | 335 | 396 | 403 | 436.55 | 495 | 378 | 0.761 | -0.394 | 0.00109826 | 0.00957007 | 0.003 | 0.0144 | 0.0082 | 0.0082 | TRUE | 0.1218 |
| D430020J02F | 181 | 172 | 156 | 431 | 335 | 272 | 192 | 172 | 157 | 339 | 360 | 298 | 252.95 | 174 | 332 | 1.917 | 0.939 | 8.85E-10 | 5.48E-08 | 0.0053 | 0.0199 | 0.0125 | 0.0125 | TRUE | 0.0932 |
| D430036J16F | 2 | 2 | 2 | 0 | 0 | 3 | 2 | 2 | 2 | 0 | 0 | 3 | 1.57 | 2 | 1 | 0.512 | -0.965 | 0.58205359 | NA | 0.3393 | 2.1066 | 1.4988 | 1.4988 | TRUE | 1.4447 |
| D430041D05 | 42 | 57 | 56 | 93 | 79 | 51 | 45 | 57 | 56 | 73 | 85 | 56 | 61.95 | 53 | 71 | 1.356 | 0.439 | 0.11059405 | 0.27922033 | 0.014 | 0.0601 | 0.0381 | 0.0381 | TRUE | 0.2256 |
| D430042O09 | 628 | 667 | 729 | 801 | 571 | 569 | 667 | 666 | 732 | 630 | 614 | 623 | 655.35 | 688 | 622 | 0.904 | -0.145 | 0.12779669 | 0.30739486 | 0 | 0.0119 | 0.005 | 0.005 | TRUE | 0.0364 |
| D4Ertd617e | 0 | 0 | 0 | 0 | 0 | 0 | 0 | 0 | 0 | 0 | 0 | 0 | 0 | 0 | 0 | NA | NA | NA | NA | NA | NA | NA | NA | NA | NA |
| D4Wsu53e | 2998 | 3805 | 4285 | 3445 | 3762 | 4090 | 3184 | 3800 | 4303 | 2708 | 4048 | 4477 | 3753.35 | 3762 | 3744 | 0.995 | -0.007 | 0.96830866 | 0.98419772 | 0.0442 | 0.0078 | 0.0248 | 0.0248 | TRUE | 0.7161 |
| D530049I02F | 0 | 0 | 0 | 0 | 0 | 0 | 0 | 0 | 0 | 0 | 0 | 0 | 0 | 0 | 0 | NA | NA | NA | NA | NA | NA | NA | NA | NA | NA |
| D5Ertd577e | 0 | 0 | 0 | 0 | 0 | 0 | 0 | 0 | 0 | 0 | 0 | 0 | 0 | 0 | 0 | NA | NA | NA | NA | NA | NA | NA | NA | NA | NA |
| D5Ertd579e | 1833 | 2001 | 2103 | 2794 | 2076 | 2030 | 1947 | 1998 | 2112 | 2196 | 2234 | 2222 | 2118.21 | 2019 | 2217 | 1.098 | 0.135 | 0.05831541 | 0.18243185 | 4.00E-04 | 0.0085 | 0.0032 | 0.0032 | TRUE | 0.0195 |
| D5Ertd605e | 92 | 107 | 102 | 232 | 166 | 124 | 98 | 107 | 102 | 182 | 179 | 136 | 133.95 | 102 | 166 | 1.622 | 0.698 | 0.00036762 | 0.00405173 | 0.0085 | 0.0315 | 0.0197 | 0.0197 | TRUE | 0.2539 |
| D630003M21 | 52 | 47 | 43 | 87 | 84 | 70 | 55 | 47 | 43 | 68 | 90 | 77 | 63.46 | 48 | 78 | 1.615 | 0.692 | 0.00783886 | 0.04365617 | 0 | 0.0588 | 0.032 | 0.032 | TRUE | 0.1609 |
| D630010B17 | 0 | 0 | 0 | 0 | 0 | 0 | 0 | 0 | 0 | 0 | 0 | 0 | 0 | 0 | 0 | NA | NA | NA | NA | NA | NA | NA | NA | NA | NA |
| D630013N20 | 0 | 0 | 0 | 0 | 0 | 0 | 0 | 0 | 0 | 0 | 0 | 0 | 0 | 0 | 0 | NA | NA | NA | NA | NA | NA | NA | NA | NA | NA |
| D630023F18I | 238 | 251 | 231 | 450 | 265 | 339 | 253 | 251 | 232 | 354 | 285 | 371 | 290.9 | 245 | 337 | 1.375 | 0.459 | 0.00214952 | 0.01629913 | 0.0076 | 0.0182 | 0.0126 | 0.0126 | TRUE | 0.1982 |
| D630029K05I | 1 | 3 | 1 | 1 | 4 | 4 | 1 | 3 | 1 | 1 | 4 | 4 | 2.42 | 2 | 3 | 1.82 | 0.864 | 0.50609899 | NA | 0 | 1.3672 | 0.7564 | 0.7564 | TRUE | 0.7979 |
| D630032N06 | 6 | 20 | 34 | 35 | 30 | 19 | 6 | 20 | 34 | 28 | 32 | 21 | 23.51 | 20 | 27 | 1.33 | 0.411 | 0.47071918 | 0.67223677 | 0.2438 | 0.147 | 0.1904 | 0.1904 | TRUE | 0.2597 |
| D630033O11 | 5 | 11 | 8 | 10 | 8 | 9 | 5 | 11 | 8 | 8 | 9 | 10 | 8.44 | 8 | 9 | 1.074 | 0.103 | 0.8741164 | 0.93791982 | 0 | 0.3972 | 0.187 | 0.187 | TRUE | 0.217 |
| D630037F22I | 976 | 1042 | 1195 | 1381 | 1058 | 1122 | 1037 | 1041 | 1200 | 1086 | 1138 | 1228 | 1121.57 | 1093 | 1151 | 1.053 | 0.074 | 0.47374997 | 0.67458712 | 0.0043 | 0.0098 | 0.0068 | 0.0068 | TRUE | 0.0889 |
| D630039A03 | 86 | 83 | 104 | 126 | 78 | 81 | 91 | 83 | 104 | 99 | 84 | 89 | 91.72 | 93 | 91 | 0.978 | -0.032 | 0.87971894 | 0.94060925 | 0 | 0.0428 | 0.0219 | 0.0219 | TRUE | 0.1151 |
| D630041G03 | 1 | 5 | 4 | 1 | 4 | 5 | 1 | 5 | 4 | 1 | 4 | 5 | 3.44 | 3 | 3 | 1.017 | 0.025 | 0.98247828 | NA | 0 | 0.9648 | 0.6359 | 0.6359 | TRUE | 0.9421 |
| D630045J12F | 1505 | 1883 | 2031 | 2258 | 1761 | 1864 | 1599 | 1881 | 2039 | 1775 | 1895 | 2040 | 1871.45 | 1840 | 1903 | 1.034 | 0.049 | 0.67459314 | 0.81862606 | 0.0094 | 0.0087 | 0.0091 | 0.0091 | TRUE | 0.1581 |
| D630045M09 | 170 | 102 | 72 | 546 | 363 | 282 | 181 | 102 | 72 | 429 | 391 | 309 | 247.2 | 118 | 376 | 3.187 | 1.672 | 1.83E-08 | 8.24E-07 | 0.1095 | 0.0202 | 0.058 | 0.058 | TRUE | 1.895 |
| D6Ertd474e | 1 | 0 | 0 | 0 | 1 | 2 | 1 | 0 | 0 | 0 | 1 | 2 | 0.72 | 0 | 1 | 2.905 | 1.539 | 0.54944501 | NA | 0 | 4.5747 | 3.3003 | 3.3003 | TRUE | 0.2242 |
| D6Ertd527e | 8 | 4 | 9 | 6 | 4 | 4 | 8 | 4 | 9 | 5 | 4 | 4 | 5.82 | 7 | 4 | 0.625 | -0.678 | 0.39097559 | 0.60467415 | 0 | 0.5728 | 0.2702 | 0.2702 | TRUE | 0.4158 |
| D6Wsu163e | 927 | 1007 | 1014 | 1304 | 879 | 906 | 985 | 1006 | 1018 | 1025 | 946 | 992 | 995.19 | 1003 | 988 | 0.985 | -0.021 | 0.79856555 | 0.89619481 | 0 | 0.0102 | 0.004 | 0.004 | TRUE | 0.0163 |
| D730001G18 | 2 | 1 | 3 | 4 | 1 | 1 | 2 | 1 | 3 | 3 | 1 | 1 | 1.91 | 2 | 2 | 0.896 | -0.159 | 0.9058504 | NA | 0 | 1.7332 | 0.7846 | 0.7846 | TRUE | 0.4878 |
| D730005E14I | 26 | 40 | 49 | 54 | 52 | 65 | 28 | 40 | 49 | 42 | 56 | 71 | 47.72 | 39 | 56 | 1.441 | 0.528 | 0.12643761 | 0.30491957 | 0.0513 | 0.0759 | 0.064 | 0.064 | TRUE | 0.1838 |
| D730039F16I | 21 | 31 | 27 | 34 | 15 | 28 | 22 | 31 | 27 | 27 | 16 | 31 | 25.65 | 27 | 25 | 0.916 | -0.127 | 0.75669529 | 0.87155626 | 0.0142 | 0.1353 | 0.0813 | 0.0813 | TRUE | 0.4944 |
| D730045A05 | 2 | 4 | 3 | 7 | 3 | 2 | 2 | 4 | 3 | 6 | 3 | 2 | 3.34 | 3 | 4 | 1.217 | 0.283 | 0.78115122 | NA | 0 | 0.9927 | 0.4498 | 0.4498 | TRUE | 0.4036 |
| D730048I06F | 0 | 1 | 0 | 0 | 1 | 0 | 0 | 1 | 0 | 0 | 1 | 0 | 0.35 | 0 | 0 | 0.954 | -0.067 | 0.98571955 | NA | 0 | 9.5346 | 8.2194 | 8.2194 | TRUE | 0.5204 |
| D730050B12 | 0 | 1 | 1 | 0 | 0 | 0 | 0 | 1 | 1 | 0 | 0 | 0 | 0.33 | 1 | 0 | 0.268 | -1.901 | 0.61401143 | NA | 0 | 9.8764 | 8.4772 | 8.4772 | TRUE | 0.2278 |
| D7Ertd143e | 781 | 564 | 381 | 1142 | 1053 | 529 | 830 | 563 | 383 | 898 | 1133 | 579 | 730.86 | 592 | 870 | 1.47 | 0.556 | 0.17753139 | 0.37816905 | 0.1211 | 0.0114 | 0.0586 | 0.1211 | TRUE | 0.2858 |
| D7Ertd443e | 2 | 2 | 0 | 0 | 2 | 1 | 2 | 2 | 0 | 0 | 2 | 1 | 1.23 | 1 | 1 | 0.762 | -0.392 | 0.83789866 | NA | 0 | 2.6896 | 1.8021 | 1.8021 | TRUE | 0.2983 |
| D7Ertd715e | 649 | 596 | 574 | 641 | 585 | 526 | 689 | 595 | 576 | 504 | 629 | 576 | 595.01 | 620 | 570 | 0.917 | -0.125 | 0.34146697 | 0.5617039 | 0.0093 | 0.0124 | 0.0107 | 0.0107 | TRUE | 0.1248 |

| | | | | | | | | | | | | | | | | | | | | | | | | | |
|---|---|---|---|---|---|---|---|---|---|---|---|---|---|---|---|---|---|---|---|---|---|---|---|---|---|
| D830005E20 | 0 | 0 | 0 | 0 | 0 | 0 | 0 | 0 | 0 | 0 | 0 | 0 | 0 | 0 | 0 | NA | NA | NA | NA | NA | NA | NA | NA | NA | NA |
| D830013O20 | 0 | 0 | 0 | 0 | 0 | 0 | 0 | 0 | 0 | 0 | 0 | 0 | 0 | 0 | 0 | NA | NA | NA | NA | NA | NA | NA | NA | NA | NA |
| D830015G02 | 0 | 0 | 0 | 1 | 0 | 0 | 0 | 0 | 0 | 1 | 0 | 0 | 0.13 | 0 | 0 | 1.859 | 0.894 | 0.82650734 | NA | 0 | 25.1515 | 10 | 10 | TRUE | 0.2563 |
| D830030K20 | 0 | 0 | 0 | 0 | 0 | 1 | 0 | 0 | 0 | 0 | 0 | 1 | 0.18 | 0 | 0 | 1.859 | 0.894 | 0.82650734 | NA | 0 | 18.0643 | 10 | 10 | TRUE | 0.5383 |
| D830031N03 | 1352 | 1319 | 1581 | 1401 | 999 | 983 | 1436 | 1317 | 1587 | 1101 | 1075 | 1076 | 1265.52 | 1447 | 1084 | 0.749 | -0.416 | 2.19E-05 | 0.0003854 | 0.0039 | 0.0095 | 0.0061 | 0.0061 | TRUE | 0.0872 |
| D830046C22 | 0 | 0 | 0 | 0 | 0 | 0 | 0 | 0 | 0 | 0 | 0 | 0 | 0 | 0 | 0 | NA | NA | NA | NA | NA | NA | NA | NA | NA | NA |
| D8Ertd738e | 3163 | 2484 | 2293 | 4303 | 2894 | 2532 | 3360 | 2481 | 2302 | 3383 | 3114 | 2772 | 2901.83 | 2714 | 3090 | 1.139 | 0.187 | 0.22255709 | 0.4346499 | 0.0243 | 0.008 | 0.0166 | 0.0166 | TRUE | 0.5245 |
| D8Ertd82e | 806 | 809 | 775 | 1085 | 949 | 956 | 856 | 808 | 778 | 853 | 1021 | 1046 | 893.79 | 814 | 973 | 1.194 | 0.256 | 0.02434462 | 0.09978259 | 0.0064 | 0.0106 | 0.0082 | 0.0082 | TRUE | 0.1445 |
| D930007P13 | 0 | 0 | 0 | 0 | 1 | 0 | 0 | 0 | 0 | 0 | 1 | 0 | 0.18 | 0 | 0 | 1.859 | 0.894 | 0.82650734 | NA | 0 | 18.3785 | 10 | 10 | TRUE | 0.5204 |
| D930015E06 | 2116 | 2473 | 2381 | 2237 | 1835 | 1912 | 2248 | 2470 | 2391 | 1759 | 1974 | 2093 | 2155.67 | 2370 | 1942 | 0.819 | -0.288 | 0.00254714 | 0.01864798 | 0.0042 | 0.0084 | 0.0061 | 0.0061 | TRUE | 0.0842 |
| D930015M05 | 0 | 0 | 0 | 0 | 0 | 0 | 0 | 0 | 0 | 0 | 0 | 0 | 0 | 0 | 0 | NA | NA | NA | NA | NA | NA | NA | NA | NA | NA |
| D930016D06 | 542 | 394 | 436 | 606 | 448 | 347 | 576 | 393 | 438 | 476 | 482 | 380 | 457.54 | 469 | 446 | 0.952 | -0.071 | 0.68857564 | 0.82864452 | 0.026 | 0.0141 | 0.0201 | 0.0201 | TRUE | 0.458 |
| D930020B18 | 11 | 7 | 12 | 15 | 9 | 6 | 12 | 7 | 12 | 12 | 10 | 7 | 9.79 | 10 | 10 | 0.924 | -0.115 | 0.85269771 | 0.92636518 | 0 | 0.3433 | 0.1732 | 0.1732 | TRUE | 0.2776 |
| D930028M14 | 49 | 70 | 81 | 95 | 53 | 82 | 52 | 70 | 81 | 75 | 57 | 90 | 70.79 | 68 | 74 | 1.088 | 0.122 | 0.66804849 | 0.8144439 | 0.0338 | 0.0534 | 0.0441 | 0.0441 | TRUE | 0.2117 |
| D930032P07 | 0 | 0 | 0 | 0 | 0 | 0 | 0 | 0 | 0 | 0 | 0 | 0 | 0 | 0 | 0 | NA | NA | NA | NA | NA | NA | NA | NA | NA | NA |
| D930048N14 | 421 | 443 | 472 | 430 | 508 | 467 | 447 | 442 | 474 | 338 | 547 | 511 | 459.89 | 454 | 465 | 1.021 | 0.03 | 0.87005226 | 0.93623963 | 0.0308 | 0.0141 | 0.0225 | 0.0225 | TRUE | 0.6879 |
| Daam1 | 1825 | 1649 | 1635 | 2410 | 1634 | 1596 | 1938 | 1647 | 1642 | 1895 | 1758 | 1747 | 1771.13 | 1742 | 1800 | 1.034 | 0.048 | 0.63104398 | 0.79265149 | 0.005 | 0.0088 | 0.0066 | 0.0066 | TRUE | 0.1167 |
| Daam2 | 21 | 15 | 16 | 43 | 29 | 18 | 22 | 15 | 16 | 34 | 31 | 20 | 23.01 | 18 | 28 | 1.601 | 0.679 | 0.1164242 | 0.28830938 | 0 | 0.1501 | 0.0886 | 0.0886 | TRUE | 0.4206 |
| Dab1 | 656 | 784 | 789 | 868 | 599 | 550 | 697 | 783 | 792 | 682 | 644 | 602 | 700.15 | 757 | 643 | 0.85 | -0.235 | 0.0269971 | 0.10720009 | 0.003 | 0.0116 | 0.0067 | 0.0067 | TRUE | 0.0575 |
| Dab2 | 437 | 430 | 360 | 961 | 608 | 424 | 464 | 429 | 361 | 755 | 654 | 464 | 521.48 | 418 | 624 | 1.495 | 0.58 | 0.002508 | 0.01842391 | 0.0363 | 0.0132 | 0.0246 | 0.0246 | TRUE | 0.3204 |
| Dab2ip | 4486 | 5225 | 5358 | 4843 | 3991 | 3995 | 4765 | 5218 | 5380 | 3807 | 4294 | 4373 | 4639.6 | 5121 | 4158 | 0.812 | -0.301 | 0.00092785 | 0.0084055 | 0.0043 | 0.0076 | 0.0057 | 0.0057 | TRUE | 0.067 |
| Dach1 | 6 | 12 | 14 | 31 | 22 | 30 | 6 | 12 | 14 | 24 | 24 | 33 | 18.88 | 11 | 27 | 2.479 | 1.31 | 0.00705336 | 0.04033561 | 0 | 0.1814 | 0.105 | 0.105 | TRUE | 0.4454 |
| Dach2 | 47 | 78 | 98 | 94 | 70 | 57 | 50 | 78 | 98 | 74 | 75 | 62 | 72.97 | 75 | 70 | 0.935 | -0.097 | 0.74366262 | 0.86319155 | 0.0461 | 0.0521 | 0.0492 | 0.0492 | TRUE | 0.2623 |
| Dact1 | 422 | 480 | 552 | 497 | 421 | 439 | 448 | 479 | 554 | 391 | 453 | 481 | 467.69 | 494 | 442 | 0.892 | -0.165 | 0.23057777 | 0.44436371 | 0.0096 | 0.0139 | 0.0116 | 0.0116 | TRUE | 0.133 |
| Dact2 | 962 | 1184 | 1221 | 1253 | 982 | 1029 | 1022 | 1182 | 1226 | 985 | 1057 | 1126 | 1099.71 | 1143 | 1056 | 0.923 | -0.116 | 0.28495591 | 0.50627409 | 0.0059 | 0.0099 | 0.0076 | 0.0076 | TRUE | 0.1032 |
| Dact3 | 1497 | 1948 | 2069 | 1218 | 1047 | 1236 | 1590 | 1945 | 2077 | 957 | 1127 | 1353 | 1508.34 | 1871 | 1146 | 0.612 | -0.709 | 4.34E-06 | 9.77E-05 | 0.023 | 0.0091 | 0.0165 | 0.0165 | TRUE | 0.2999 |
| Dad1 | 3500 | 2961 | 2718 | 3843 | 2728 | 2452 | 3718 | 2957 | 2729 | 3021 | 2935 | 2684 | 3007.38 | 3135 | 2880 | 0.919 | -0.122 | 0.34341485 | 0.56349762 | 0.015 | 0.008 | 0.0116 | 0.0116 | TRUE | 0.3212 |
| Daf2 | 1 | 8 | 3 | 9 | 8 | 8 | 1 | 8 | 3 | 7 | 9 | 9 | 6.08 | 4 | 8 | 2.006 | 1.004 | 0.24578511 | 0.46286838 | 0.0409 | 0.5484 | 0.3528 | 0.3528 | TRUE | 1.2413 |
| Dag1 | 14032 | 13798 | 13793 | 19491 | 14081 | 13835 | 14904 | 13780 | 13850 | 15322 | 15150 | 15145 | 14691.91 | 14178 | 15206 | 1.073 | 0.101 | 0.10803339 | 0.27511586 | 9.00E-04 | 0.0071 | 0.0028 | 0.0028 | TRUE | 0.0245 |
| Dagla | 197 | 201 | 192 | 221 | 160 | 141 | 209 | 201 | 193 | 174 | 172 | 154 | 183.83 | 201 | 167 | 0.831 | -0.266 | 0.0776955 | 0.22051708 | 0 | 0.0248 | 0.0109 | 0.0109 | TRUE | 0.0428 |
| Daglb | 391 | 473 | 451 | 718 | 507 | 439 | 415 | 472 | 453 | 564 | 546 | 481 | 488.51 | 447 | 530 | 1.188 | 0.248 | 0.03674768 | 0.1324159 | 0.0036 | 0.0136 | 0.0081 | 0.0081 | TRUE | 0.0768 |
| Dak | 740 | 1087 | 1115 | 1137 | 813 | 880 | 786 | 1086 | 1120 | 894 | 875 | 963 | 953.84 | 997 | 911 | 0.913 | -0.131 | 0.37245539 | 0.58896796 | 0.0185 | 0.0104 | 0.0146 | 0.0146 | TRUE | 0.4079 |
| Dalrd3 | 1949 | 2127 | 2106 | 2253 | 1989 | 1893 | 2070 | 2124 | 2115 | 1771 | 2140 | 2072 | 2048.74 | 2103 | 1994 | 0.948 | -0.078 | 0.42062066 | 0.63103102 | 0.0047 | 0.0085 | 0.0062 | 0.0062 | TRUE | 0.1184 |
| Dancr | 241 | 234 | 213 | 262 | 185 | 191 | 256 | 234 | 214 | 206 | 199 | 209 | 219.61 | 235 | 205 | 0.873 | -0.195 | 0.17735693 | 0.3779353 | 0 | 0.0219 | 0.0105 | 0.0105 | TRUE | 0.0698 |
| Dand5 | 89 | 91 | 100 | 101 | 77 | 56 | 95 | 91 | 100 | 79 | 83 | 61 | 84.9 | 95 | 74 | 0.784 | -0.351 | 0.11564238 | 0.28713974 | 0 | 0.0457 | 0.0239 | 0.0239 | TRUE | 0.204 |
| Dao | 0 | 1 | 1 | 1 | 0 | 0 | 0 | 1 | 1 | 1 | 0 | 0 | 0.46 | 1 | 0 | 0.52 | -0.943 | 0.74346893 | NA | 0 | 7.0944 | 4.2167 | 4.2167 | TRUE | 0.2562 |
| Dap | 644 | 697 | 700 | 697 | 522 | 582 | 684 | 696 | 703 | 548 | 562 | 637 | 638.28 | 694 | 582 | 0.837 | -0.256 | 0.01332507 | 0.06484511 | 0.0016 | 0.0121 | 0.0061 | 0.0061 | TRUE | 0.0782 |
| Dap3 | 5157 | 7130 | 7174 | 8565 | 6198 | 6687 | 5478 | 7121 | 7203 | 6733 | 6669 | 7320 | 6753.94 | 6601 | 6907 | 1.046 | 0.065 | 0.58764609 | 0.76226286 | 0.0127 | 0.0074 | 0.0103 | 0.0103 | TRUE | 0.2701 |
| Dapk1 | 3539 | 3985 | 4022 | 3648 | 2713 | 3070 | 3759 | 3980 | 4039 | 2868 | 2919 | 3361 | 3487.47 | 3926 | 3049 | 0.776 | -0.365 | 5.49E-05 | 0.00083761 | 0.0042 | 0.0078 | 0.0056 | 0.0056 | TRUE | 0.0965 |
| Dapk2 | 241 | 199 | 158 | 272 | 172 | 175 | 256 | 199 | 159 | 214 | 185 | 192 | 200.64 | 205 | 197 | 0.965 | -0.052 | 0.79949394 | 0.89671586 | 0.0263 | 0.0233 | 0.0248 | 0.0248 | TRUE | 0.29 |
| Dapk3 | 1406 | 1381 | 1396 | 2168 | 1481 | 1211 | 1493 | 1379 | 1402 | 1704 | 1593 | 1326 | 1482.96 | 1425 | 1541 | 1.083 | 0.114 | 0.31774326 | 0.53869147 | 0.0079 | 0.0091 | 0.0088 | 0.0088 | TRUE | 0.1791 |
| Dapl1 | 4 | 5 | 3 | 3 | 4 | 1 | 4 | 5 | 3 | 2 | 4 | 1 | 3.34 | 4 | 2 | 0.631 | -0.663 | 0.52334949 | NA | 0 | 0.9947 | 0.4644 | 0.4644 | TRUE | 0.3359 |
| Dapp1 | 469 | 431 | 485 | 427 | 339 | 315 | 498 | 430 | 487 | 336 | 365 | 345 | 410.14 | 472 | 349 | 0.738 | -0.439 | 0.00021875 | 0.0026118 | 0.0017 | 0.0149 | 0.0077 | 0.0077 | TRUE | 0.0689 |
| Darc | 1 | 1 | 0 | 2 | 0 | 0 | 1 | 1 | 0 | 2 | 0 | 0 | 0.61 | 1 | 1 | 0.818 | -0.291 | 0.91402744 | NA | 0 | 5.4476 | 3.5995 | 3.5995 | TRUE | 0.9722 |
| Dars | 9493 | 11521 | 11717 | 13063 | 9558 | 10004 | 10083 | 11506 | 11765 | 10269 | 10284 | 10951 | 10809.73 | 11118 | 10501 | 0.944 | -0.082 | 0.33558249 | 0.55585035 | 0.0039 | 0.0072 | 0.0052 | 0.0052 | TRUE | 0.081 |
| Dars2 | 4033 | 4135 | 3960 | 4905 | 3352 | 3009 | 4284 | 4130 | 3976 | 3856 | 3607 | 3294 | 3857.66 | 4130 | 3586 | 0.869 | -0.203 | 0.01862523 | 0.08270188 | 0.0034 | 0.0078 | 0.0051 | 0.0051 | TRUE | 0.0614 |
| Daxx | 2783 | 2339 | 2364 | 3611 | 2492 | 2111 | 2956 | 2336 | 2374 | 2839 | 2681 | 2311 | 2582.74 | 2555 | 2610 | 1.022 | 0.031 | 0.80592631 | 0.90019698 | 0.014 | 0.0082 | 0.0113 | 0.0113 | TRUE | 0.228 |
| Dazap1 | 7764 | 8560 | 8243 | 9703 | 7448 | 7814 | 8247 | 8549 | 8277 | 7628 | 8014 | 8554 | 8211.25 | 8358 | 8065 | 0.965 | -0.052 | 0.47065611 | 0.67223251 | 0.0017 | 0.0073 | 0.0036 | 0.0036 | TRUE | 0.0343 |
| Dazap2 | 2540 | 3679 | 3546 | 3666 | 2814 | 3208 | 2698 | 3674 | 3561 | 2882 | 3028 | 3512 | 3225.67 | 3311 | 3141 | 0.948 | -0.077 | 0.58193463 | 0.75815901 | 0.0189 | 0.0079 | 0.0137 | 0.0137 | TRUE | 0.3185 |
| Dazl | 2018 | 2411 | 1768 | 8472 | 6561 | 5270 | 2143 | 2408 | 1775 | 6660 | 7059 | 5769 | 4302.45 | 2109 | 6496 | 3.08 | 1.623 | 1.47E-34 | 1.06E-31 | 0.016 | 0.0077 | 0.0123 | 0.0123 | TRUE | 0.2322 |
| Dbc1 | 242 | 186 | 172 | 250 | 194 | 145 | 257 | 186 | 173 | 197 | 209 | 159 | 196.58 | 205 | 188 | 0.918 | -0.124 | 0.54986822 | 0.73422496 | 0.0278 | 0.0237 | 0.0257 | 0.0257 | TRUE | 0.528 |
| Dbf4 | 6894 | 7778 | 8031 | 8474 | 6140 | 6366 | 7323 | 7768 | 8064 | 6662 | 6606 | 6969 | 7231.84 | 7718 | 6746 | 0.874 | -0.194 | 0.00536408 | 0.03293197 | 0.0014 | 0.0074 | 0.0034 | 0.0034 | TRUE | 0.0245 |
| Dbh | 0 | 0 | 0 | 0 | 0 | 0 | 0 | 0 | 0 | 0 | 0 | 0 | 0 | 0 | 0 | NA | NA | NA | NA | NA | NA | NA | NA | NA | NA |
| Dbi | 4171 | 3913 | 3799 | 5010 | 3438 | 3276 | 4430 | 3908 | 3815 | 3938 | 3699 | 3586 | 3896.09 | 4051 | 3741 | 0.924 | -0.114 | 0.20261988 | 0.41018108 | 0.0041 | 0.0077 | 0.0055 | 0.0055 | TRUE | 0.0816 |
| Dbil5 | 15 | 17 | 20 | 27 | 20 | 24 | 16 | 17 | 20 | 21 | 22 | 26 | 20.33 | 18 | 23 | 1.297 | 0.375 | 0.36835117 | 0.58564866 | 0 | 0.1689 | 0.0753 | 0.0753 | TRUE | 0.0834 |
| Dbn1 | 1578 | 1859 | 1896 | 1772 | 1356 | 1239 | 1676 | 1857 | 1904 | 1393 | 1459 | 1356 | 1607.46 | 1812 | 1403 | 0.774 | -0.37 | 2.30E-05 | 0.00040077 | 0.0023 | 0.009 | 0.0049 | 0.0049 | TRUE | 0.0518 |
| Dbndd1 | 56 | 34 | 42 | 88 | 70 | 44 | 59 | 34 | 42 | 69 | 75 | 48 | 54.71 | 45 | 64 | 1.426 | 0.512 | 0.11389837 | 0.28464666 | 0.045 | 0.0671 | 0.0567 | 0.0567 | TRUE | 0.5845 |
| Dbndd2 | 223 | 315 | 348 | 234 | 158 | 195 | 237 | 315 | 349 | 184 | 170 | 213 | 244.72 | 300 | 189 | 0.629 | -0.669 | 0.00039553 | 0.0042993 | 0.0224 | 0.0204 | 0.0214 | 0.0214 | TRUE | 0.3818 |
| Dbnl | 2364 | 2069 | 1907 | 4024 | 2819 | 2234 | 2511 | 2066 | 1915 | 3163 | 3033 | 2445 | 2522.34 | 2164 | 2880 | 1.332 | 0.413 | 0.00314935 | 0.02204781 | 0.0185 | 0.0082 | 0.0137 | 0.0137 | TRUE | 0.2378 |
| Dbp | 103 | 172 | 130 | 131 | 88 | 105 | 109 | 172 | 131 | 103 | 95 | 115 | 120.72 | 137 | 104 | 0.758 | -0.399 | 0.07902147 | 0.22294674 | 0.0235 | 0.0342 | 0.0288 | 0.0288 | TRUE | 0.3865 |
| Dbpht2 | 0 | 1 | 0 | 1 | 0 | 0 | 0 | 1 | 0 | 1 | 0 | 0 | 0.3 | 0 | 0 | 0.954 | -0.067 | 0.98642001 | NA | 0 | 11.0817 | 9.3014 | 9.3014 | TRUE | 0.405 |
| Dbr1 | 2028 | 1539 | 1397 | 2569 | 1760 | 1473 | 2154 | 1537 | 1403 | 2020 | 1894 | 1612 | 1769.91 | 1698 | 1842 | 1.085 | 0.118 | 0.48681563 | 0.68377024 | 0.0321 | 0.0088 | 0.0202 | 0.0202 | TRUE | 0.6656 |
| Dbt | 3128 | 3655 | 3771 | 3840 | 2753 | 3088 | 3322 | 3650 | 3786 | 3019 | 2962 | 3380 | 3353.39 | 3586 | 3120 | 0.87 | -0.201 | 0.02905381 | 0.1124507 | 0.0044 | 0.0079 | 0.0058 | 0.0058 | TRUE | 0.0641 |
| Dbx1 | 1 | 0 | 1 | 2 | 1 | 0 | 1 | 0 | 1 | 2 | 1 | 0 | 0.79 | 1 | 1 | 1.325 | 0.406 | 0.85251519 | NA | 0 | 4.1997 | 2.1499 | 2.1499 | TRUE | 0.2789 |
| Dbx2 | 235 | 370 | 399 | 216 | 172 | 222 | 250 | 370 | 401 | 170 | 185 | 243 | 269.61 | 340 | 199 | 0.585 | -0.774 | 0.00047038 | 0.0049525 | 0.0443 | 0.0191 | 0.0313 | 0.0313 | TRUE | 0.6111 |
| Dcaf10 | 674 | 762 | 749 | 1118 | 833 | 759 | 716 | 761 | 752 | 879 | 896 | 831 | 805.83 | 743 | 869 | 1.169 | 0.226 | 0.01131428 | 0.05767576 | 0 | 0.011 | 0.0045 | 0.0045 | TRUE | 0.0167 |
| Dcaf11 | 1453 | 1491 | 1468 | 1832 | 1429 | 1294 | 1543 | 1489 | 1474 | 1440 | 1538 | 1416 | 1483.44 | 1502 | 1465 | 0.975 | -0.037 | 0.63841901 | 0.79637583 | 5.00E-04 | 0.0091 | 0.0037 | 0.0037 | TRUE | 0.0223 |
| Dcaf12 | 4699 | 4967 | 4967 | 5881 | 4087 | 4390 | 4991 | 4961 | 4987 | 4623 | 4397 | 4806 | 4794.21 | 4980 | 4609 | 0.925 | -0.112 | 0.09057962 | 0.24534121 | 7.00E-04 | 0.0076 | 0.0029 | 0.0029 | TRUE | 0.0194 |
| Dcaf12l1 | 479 | 555 | 486 | 1775 | 1067 | 974 | 509 | 554 | 488 | 1395 | 1148 | 1066 | 860.11 | 517 | 1203 | 2.33 | 1.22 | 1.98E-20 | 5.13E-18 | 0.0114 | 0.0107 | 0.0111 | 0.0111 | TRUE | 0.238 |

| | | | | | | | | | | | | | | | | | | | | | | | | | |
|---|---|---|---|---|---|---|---|---|---|---|---|---|---|---|---|---|---|---|---|---|---|---|---|---|---|
| Dcaf12l2 | 6 | 4 | 1 | 13 | 15 | 7 | 6 | 4 | 1 | 10 | 16 | 8 | 7.57 | 4 | 11 | 3.004 | 1.587 | 0.05128807 | 0.16695882 | 0.0873 | 0.4424 | 0.299 | 0.299 | TRUE | 0.5582 |
| Dcaf13 | 3680 | 3779 | 3824 | 4741 | 3500 | 3498 | 3909 | 3774 | 3840 | 3727 | 3766 | 3829 | 3807.42 | 3841 | 3774 | 0.982 | -0.025 | 0.67096407 | 0.81648371 | 0 | 0.0078 | 0.0023 | 0.0023 | TRUE | 0.0029 |
| Dcaf15 | 3973 | 4748 | 4617 | 5303 | 4093 | 4152 | 4220 | 4742 | 4636 | 4169 | 4404 | 4545 | 4452.58 | 4533 | 4373 | 0.964 | -0.052 | 0.51477933 | 0.70742351 | 0.0024 | 0.0076 | 0.0044 | 0.0044 | TRUE | 0.0442 |
| Dcaf17 | 2850 | 3154 | 3264 | 3220 | 2368 | 2714 | 3027 | 3150 | 3277 | 2531 | 2548 | 2971 | 2917.43 | 3151 | 2683 | 0.851 | -0.233 | 0.01271258 | 0.06279449 | 0.0046 | 0.008 | 0.0059 | 0.0059 | TRUE | 0.106 |
| Dcaf4 | 3670 | 4867 | 5008 | 4203 | 3177 | 3823 | 3898 | 4861 | 5029 | 3304 | 3418 | 4185 | 4115.78 | 4596 | 3636 | 0.791 | -0.338 | 0.01184689 | 0.05967717 | 0.0171 | 0.0077 | 0.0128 | 0.0128 | TRUE | 0.2146 |
| Dcaf5 | 1549 | 1683 | 1842 | 2334 | 1715 | 1608 | 1645 | 1681 | 1850 | 1835 | 1845 | 1760 | 1769.33 | 1725 | 1813 | 1.051 | 0.072 | 0.38296378 | 0.59743865 | 0.0015 | 0.0088 | 0.0043 | 0.0043 | TRUE | 0.048 |
| Dcaf6 | 925 | 923 | 950 | 1379 | 1053 | 1009 | 983 | 922 | 954 | 1084 | 1133 | 1105 | 1029.96 | 953 | 1107 | 1.162 | 0.216 | 0.00760079 | 0.04273786 | 0 | 0.0101 | 0.0038 | 0.0038 | TRUE | 0.0096 |
| Dcaf7 | 10359 | 10215 | 10834 | 11843 | 8591 | 8798 | 11003 | 10202 | 10878 | 9310 | 9243 | 9631 | 10044.6 | 10694 | 9395 | 0.878 | -0.187 | 0.00366441 | 0.02471785 | 9.00E-04 | 0.0072 | 0.0029 | 0.0029 | TRUE | 0.0199 |
| Dcaf8 | 3134 | 3834 | 3944 | 4709 | 3447 | 3680 | 3329 | 3829 | 3960 | 3702 | 3709 | 4028 | 3759.51 | 3706 | 3813 | 1.029 | 0.041 | 0.66541053 | 0.8127356 | 0.005 | 0.0078 | 0.0061 | 0.0061 | TRUE | 0.0962 |
| Dcakd | 10326 | 8639 | 8708 | 12821 | 9160 | 8411 | 10968 | 8628 | 8744 | 10079 | 9856 | 9207 | 9580.21 | 9447 | 9714 | 1.028 | 0.04 | 0.7189192 | 0.84776027 | 0.0086 | 0.0072 | 0.009 | 0.009 | TRUE | 0.2424 |
| Dcbld1 | 1487 | 1170 | 1126 | 2134 | 1491 | 1303 | 1579 | 1168 | 1131 | 1678 | 1604 | 1426 | 1431.12 | 1293 | 1569 | 1.215 | 0.28 | 0.05618173 | 0.17785995 | 0.0209 | 0.0092 | 0.0148 | 0.0148 | TRUE | 0.4507 |
| Dcbld2 | 3675 | 3331 | 3532 | 4859 | 3453 | 3319 | 3903 | 3327 | 3547 | 3820 | 3715 | 3633 | 3657.47 | 3592 | 3723 | 1.037 | 0.052 | 0.54345508 | 0.7286671 | 0.0029 | 0.0078 | 0.005 | 0.005 | TRUE | 0.0697 |
| Dcc | 42 | 10 | 15 | 116 | 92 | 29 | 45 | 10 | 15 | 91 | 99 | 32 | 48.6 | 23 | 74 | 3.2 | 1.678 | 0.00428006 | 0.02786014 | 0.4054 | 0.0747 | 0.22 | 0.22 | TRUE | 3.7042 |
| Dcdc2a | 31 | 47 | 47 | 60 | 46 | 50 | 33 | 47 | 47 | 47 | 49 | 55 | 46.41 | 42 | 50 | 1.187 | 0.247 | 0.40137686 | 0.6141287 | 0 | 0.0779 | 0.0407 | 0.0407 | TRUE | 0.2805 |
| Dcdc2b | 3 | 6 | 5 | 4 | 5 | 3 | 3 | 6 | 5 | 3 | 5 | 3 | 4.33 | 5 | 4 | 0.822 | -0.282 | 0.75167818 | NA | 0 | 0.7669 | 0.3394 | 0.3394 | TRUE | 0.1565 |
| Dcdc2c | 1 | 0 | 1 | 1 | 2 | 1 | 1 | 0 | 1 | 1 | 2 | 1 | 1.02 | 1 | 1 | 1.936 | 0.953 | 0.61010723 | NA | 0 | 3.2479 | 1.4013 | 1.4013 | TRUE | 0.256 |
| Dchs1 | 60 | 43 | 47 | 135 | 84 | 68 | 64 | 43 | 47 | 106 | 90 | 74 | 70.8 | 51 | 90 | 1.771 | 0.825 | 0.00252672 | 0.01853551 | 0.0221 | 0.0534 | 0.0383 | 0.0383 | TRUE | 0.2031 |
| Dck | 2430 | 2328 | 2145 | 3556 | 2558 | 2385 | 2581 | 2325 | 2154 | 2795 | 2752 | 2611 | 2536.39 | 2353 | 2719 | 1.156 | 0.209 | 0.02473939 | 0.10100689 | 0.0042 | 0.0082 | 0.0059 | 0.0059 | TRUE | 0.0866 |
| Dclk1 | 81 | 107 | 125 | 41 | 34 | 58 | 86 | 107 | 126 | 32 | 37 | 63 | 75.12 | 106 | 44 | 0.412 | -1.28 | 5.24E-05 | 0.00080592 | 0.0628 | 0.0508 | 0.0562 | 0.0562 | TRUE | 0.5223 |
| Dclk2 | 1563 | 2065 | 2017 | 1797 | 1702 | 1831 | 1660 | 2062 | 2025 | 1413 | 1831 | 2004 | 1832.67 | 1916 | 1749 | 0.912 | -0.132 | 0.37909142 | 0.59421883 | 0.0225 | 0.0087 | 0.0158 | 0.0158 | TRUE | 0.3464 |
| Dclk3 | 100 | 118 | 135 | 130 | 102 | 93 | 106 | 118 | 136 | 102 | 110 | 102 | 112.23 | 120 | 105 | 0.871 | -0.199 | 0.30614393 | 0.52666802 | 0 | 0.0363 | 0.0183 | 0.0183 | TRUE | 0.1326 |
| Dclre1a | 559 | 585 | 594 | 923 | 653 | 583 | 594 | 584 | 596 | 726 | 703 | 638 | 640.13 | 591 | 689 | 1.166 | 0.221 | 0.0259351 | 0.10426694 | 0 | 0.012 | 0.0056 | 0.0056 | TRUE | 0.0479 |
| Dclre1b | 472 | 438 | 428 | 763 | 485 | 426 | 501 | 437 | 430 | 600 | 522 | 466 | 492.75 | 456 | 529 | 1.163 | 0.218 | 0.11255013 | 0.28230608 | 0.0098 | 0.0136 | 0.0115 | 0.0115 | TRUE | 0.1641 |
| Dclre1c | 898 | 937 | 933 | 1125 | 897 | 868 | 954 | 936 | 937 | 884 | 965 | 950 | 937.69 | 942 | 933 | 0.99 | -0.015 | 0.85833117 | 0.93028174 | 0 | 0.0104 | 0.0042 | 0.0042 | TRUE | 0.0263 |
| Dcn | 0 | 0 | 0 | 0 | 0 | 0 | 0 | 0 | 0 | 0 | 0 | 0 | 0 | 0 | 0 | NA | NA | NA | NA | NA | NA | NA | NA | NA | NA |
| Dcp1a | 2992 | 3033 | 2989 | 4214 | 2962 | 2991 | 3178 | 3029 | 3001 | 3313 | 3187 | 3274 | 3163.69 | 3069 | 3258 | 1.062 | 0.086 | 0.18915136 | 0.39431214 | 3.00E-04 | 0.0079 | 0.0028 | 0.0028 | TRUE | 0.0116 |
| Dcp1b | 340 | 460 | 501 | 420 | 320 | 359 | 361 | 459 | 503 | 330 | 344 | 393 | 398.51 | 441 | 356 | 0.805 | -0.313 | 0.04862523 | 0.1609142 | 0.016 | 0.0152 | 0.0156 | 0.0156 | TRUE | 0.2894 |
| Dcp2 | 11279 | 8071 | 7440 | 13447 | 9462 | 8194 | 11980 | 8061 | 7471 | 10571 | 10181 | 8970 | 9538.76 | 9171 | 9907 | 1.08 | 0.112 | 0.516723 | 0.70855113 | 0.0312 | 0.0072 | 0.0212 | 0.0212 | TRUE | 0.8775 |
| Dcpp1 | 0 | 0 | 0 | 0 | 0 | 0 | 0 | 0 | 0 | 0 | 0 | 0 | 0 | 0 | 0 | NA | NA | NA | NA | NA | NA | NA | NA | NA | NA |
| Dcpp2 | 0 | 0 | 0 | 0 | 0 | 0 | 0 | 0 | 0 | 0 | 0 | 0 | 0 | 0 | 0 | NA | NA | NA | NA | NA | NA | NA | NA | NA | NA |
| Dcpp3 | 0 | 0 | 0 | 0 | 0 | 0 | 0 | 0 | 0 | 0 | 0 | 0 | 0 | 0 | 0 | NA | NA | NA | NA | NA | NA | NA | NA | NA | NA |
| Dcps | 2554 | 2954 | 2883 | 3158 | 2279 | 2534 | 2713 | 2950 | 2895 | 2483 | 2452 | 2774 | 2711.05 | 2853 | 2570 | 0.9 | -0.151 | 0.07540143 | 0.21635982 | 0.0028 | 0.0081 | 0.0048 | 0.0048 | TRUE | 0.0582 |
| Dcst1 | 0 | 4 | 2 | 8 | 1 | 1 | 0 | 4 | 2 | 6 | 1 | 1 | 2.41 | 2 | 3 | 1.442 | 0.528 | 0.71797949 | NA | 0.8267 | 1.3736 | 1.1171 | 1.1171 | TRUE | 0.452 |
| Dcstamp | 62 | 46 | 32 | 93 | 63 | 53 | 66 | 46 | 32 | 73 | 68 | 58 | 57.14 | 48 | 66 | 1.388 | 0.473 | 0.13960207 | 0.32417231 | 0.0457 | 0.0646 | 0.0558 | 0.0558 | TRUE | 0.4175 |
| Dct | 6 | 4 | 3 | 7 | 8 | 3 | 6 | 4 | 3 | 6 | 9 | 3 | 5.13 | 4 | 6 | 1.303 | 0.382 | 0.65550394 | 0.80619668 | 0 | 0.6492 | 0.3294 | 0.3294 | TRUE | 0.2225 |
| Dctd | 4799 | 5025 | 4706 | 5965 | 4654 | 4818 | 5097 | 5019 | 4725 | 4689 | 5007 | 5274 | 4968.65 | 4947 | 4990 | 1.008 | 0.012 | 0.87602267 | 0.93888485 | 0.0023 | 0.0076 | 0.0042 | 0.0042 | TRUE | 0.0343 |
| Dctn1 | 5115 | 4765 | 4917 | 7365 | 5412 | 4728 | 5433 | 4759 | 4937 | 5790 | 5823 | 5176 | 5319.56 | 5043 | 5596 | 1.11 | 0.15 | 0.09197012 | 0.24745015 | 0.0042 | 0.0075 | 0.0056 | 0.0056 | TRUE | 0.0557 |
| Dctn2 | 5406 | 5355 | 5341 | 6514 | 4704 | 4533 | 5742 | 5348 | 5363 | 5121 | 5061 | 4962 | 5266.21 | 5484 | 5048 | 0.921 | -0.119 | 0.06731552 | 0.20047213 | 7.00E-04 | 0.0075 | 0.0029 | 0.0029 | TRUE | 0.0206 |
| Dctn3 | 1594 | 1768 | 1756 | 2258 | 1652 | 1709 | 1693 | 1766 | 1763 | 1775 | 1777 | 1871 | 1774.22 | 1741 | 1808 | 1.038 | 0.054 | 0.45381751 | 0.65906064 | 0 | 0.0088 | 0.0032 | 0.0032 | TRUE | 0.011 |
| Dctn4 | 2082 | 2557 | 2536 | 2967 | 2277 | 2397 | 2211 | 2554 | 2546 | 2332 | 2450 | 2624 | 2452.97 | 2437 | 2469 | 1.012 | 0.018 | 0.85062319 | 0.92562639 | 0.0047 | 0.0082 | 0.0061 | 0.0061 | TRUE | 0.0793 |
| Dctn5 | 1364 | 1299 | 1286 | 1971 | 1342 | 1257 | 1449 | 1297 | 1291 | 1549 | 1444 | 1376 | 1401.13 | 1346 | 1456 | 1.083 | 0.115 | 0.21944543 | 0.43161158 | 0.0029 | 0.0093 | 0.0056 | 0.0056 | TRUE | 0.0536 |
| Dctn6 | 1720 | 1958 | 2054 | 2084 | 1535 | 1657 | 1827 | 1955 | 2062 | 1638 | 1652 | 1814 | 1824.76 | 1948 | 1701 | 0.873 | -0.196 | 0.02706092 | 0.10723237 | 0.0027 | 0.0087 | 0.0051 | 0.0051 | TRUE | 0.0401 |
| Dctpp1 | 6459 | 7150 | 7048 | 7570 | 5424 | 5730 | 6861 | 7141 | 7077 | 5951 | 5836 | 6272 | 6522.92 | 7026 | 6020 | 0.857 | -0.223 | 0.00049394 | 0.00512241 | 7.00E-04 | 0.0074 | 0.0028 | 0.0028 | TRUE | 0.0163 |
| Dcun1d1 | 1279 | 1415 | 1448 | 1622 | 1150 | 1197 | 1359 | 1413 | 1454 | 1275 | 1237 | 1310 | 1341.39 | 1409 | 1274 | 0.905 | -0.145 | 0.06269589 | 0.19185866 | 0 | 0.0094 | 0.0036 | 0.0036 | TRUE | 0.0115 |
| Dcun1d2 | 1482 | 1249 | 1186 | 1704 | 1178 | 1005 | 1574 | 1247 | 1191 | 1340 | 1267 | 1100 | 1286.59 | 1337 | 1236 | 0.925 | -0.113 | 0.40967044 | 0.62152002 | 0.0151 | 0.0095 | 0.0128 | 0.0128 | TRUE | 0.2873 |
| Dcun1d3 | 1207 | 1198 | 1196 | 1704 | 1137 | 1189 | 1282 | 1196 | 1201 | 1340 | 1223 | 1302 | 1257.31 | 1226 | 1288 | 1.051 | 0.072 | 0.39245697 | 0.60619247 | 0.0011 | 0.0095 | 0.0043 | 0.0043 | TRUE | 0.0231 |
| Dcun1d4 | 3261 | 3364 | 3399 | 3814 | 3113 | 3047 | 3464 | 3360 | 3413 | 2998 | 3349 | 3335 | 3319.91 | 3412 | 3227 | 0.945 | -0.081 | 0.29115779 | 0.51183021 | 0.0016 | 0.0079 | 0.0039 | 0.0039 | TRUE | 0.0478 |
| Dcun1d5 | 2568 | 2907 | 2733 | 4265 | 2952 | 2623 | 2728 | 2903 | 2744 | 3353 | 3176 | 2871 | 2962.57 | 2792 | 3133 | 1.123 | 0.167 | 0.05464428 | 0.17411103 | 0.0031 | 0.008 | 0.0051 | 0.0051 | TRUE | 0.0648 |
| Dcx | 1 | 0 | 0 | 0 | 0 | 1 | 1 | 0 | 0 | 0 | 0 | 1 | 0.36 | 0 | 0 | 0.954 | -0.067 | 0.98553519 | NA | 0 | 9.1716 | 7.9605 | 7.9605 | TRUE | 0.5383 |
| Dcxr | 473 | 901 | 796 | 598 | 462 | 643 | 502 | 900 | 799 | 470 | 497 | 704 | 645.43 | 734 | 557 | 0.758 | -0.399 | 0.08428196 | 0.23276049 | 0.0636 | 0.012 | 0.037 | 0.037 | TRUE | 0.7532 |
| Dda1 | 3313 | 3020 | 3038 | 4267 | 3016 | 2545 | 3519 | 3016 | 3050 | 3354 | 3245 | 2786 | 3161.81 | 3195 | 3128 | 0.979 | -0.03 | 0.78187408 | 0.88723778 | 0.0082 | 0.0079 | 0.0081 | 0.0081 | TRUE | 0.111 |
| Ddah1 | 6164 | 5782 | 5583 | 7712 | 5938 | 5947 | 6547 | 5775 | 5606 | 6063 | 6389 | 6510 | 6148.19 | 5976 | 6321 | 1.058 | 0.081 | 0.35150149 | 0.57030012 | 0.004 | 0.0074 | 0.0053 | 0.0053 | TRUE | 0.0852 |
| Ddah2 | 2447 | 3805 | 3723 | 3694 | 3133 | 4000 | 2599 | 3800 | 3738 | 2904 | 3371 | 4379 | 3465.17 | 3379 | 3551 | 1.051 | 0.071 | 0.700301 | 0.83619323 | 0.0432 | 0.0079 | 0.0244 | 0.0244 | TRUE | 0.505 |
| Ddb1 | 57804 | 61519 | 61182 | 79130 | 56262 | 59584 | 61398 | 61440 | 61433 | 62206 | 60534 | 65224 | 62039.19 | 61424 | 62655 | 1.02 | 0.029 | 0.62449438 | 0.78834114 | 7.00E-04 | 0.007 | 0.0024 | 0.0024 | TRUE | 0.0157 |
| Ddb2 | 458 | 791 | 799 | 684 | 597 | 713 | 486 | 790 | 802 | 538 | 642 | 780 | 673.21 | 693 | 653 | 0.942 | -0.086 | 0.68547826 | 0.82631951 | 0.0531 | 0.0118 | 0.0311 | 0.0311 | TRUE | 0.6733 |
| Ddc | 750 | 543 | 507 | 1116 | 815 | 734 | 797 | 542 | 509 | 877 | 877 | 803 | 734.28 | 616 | 852 | 1.385 | 0.47 | 0.00694648 | 0.03986953 | 0.0318 | 0.0114 | 0.0204 | 0.0204 | TRUE | 0.7711 |
| Ddhd1 | 892 | 1015 | 980 | 1498 | 1166 | 1111 | 947 | 1014 | 984 | 1178 | 1255 | 1216 | 1098.92 | 982 | 1216 | 1.238 | 0.308 | 0.00016899 | 0.00211033 | 0 | 0.0099 | 0.0039 | 0.0039 | TRUE | 0.011 |
| Ddhd2 | 3871 | 3813 | 3786 | 4599 | 3264 | 3190 | 4112 | 3808 | 3802 | 3615 | 3512 | 3492 | 3723.42 | 3907 | 3540 | 0.906 | -0.142 | 0.04013755 | 0.14124261 | 8.00E-04 | 0.0078 | 0.0032 | 0.0032 | TRUE | 0.0255 |
| Ddi1 | 0 | 0 | 0 | 0 | 0 | 0 | 0 | 0 | 0 | 0 | 0 | 0 | 0 | 0 | 0 | NA | NA | NA | NA | NA | NA | NA | NA | NA | NA |
| Ddi2 | 2198 | 1777 | 1791 | 2738 | 1865 | 1526 | 2335 | 1775 | 1798 | 2152 | 2007 | 1670 | 1956.2 | 1969 | 1943 | 0.987 | -0.018 | 0.89820662 | 0.95051289 | 0.0201 | 0.0086 | 0.0146 | 0.0146 | TRUE | 0.3182 |
| Ddit3 | 128 | 223 | 277 | 209 | 165 | 198 | 136 | 223 | 278 | 164 | 178 | 217 | 199.23 | 212 | 186 | 0.875 | -0.192 | 0.46275438 | 0.66629195 | 0.0674 | 0.0234 | 0.0442 | 0.0442 | TRUE | 0.3054 |
| Ddit4 | 2262 | 3796 | 4157 | 4329 | 3529 | 4108 | 2403 | 3791 | 4174 | 3403 | 3797 | 4497 | 3677.46 | 3456 | 3899 | 1.128 | 0.174 | 0.37281305 | 0.58928297 | 0.05 | 0.0078 | 0.0271 | 0.0271 | TRUE | 0.8637 |
| Ddit4l | 718 | 820 | 689 | 1136 | 731 | 612 | 763 | 819 | 692 | 893 | 787 | 670 | 770.48 | 758 | 783 | 1.035 | 0.05 | 0.71512584 | 0.84537286 | 0.0126 | 0.0112 | 0.012 | 0.012 | TRUE | 0.1885 |
| Ddn | 7 | 3 | 7 | 6 | 11 | 6 | 7 | 3 | 7 | 5 | 12 | 7 | 6.76 | 6 | 8 | 1.311 | 0.391 | 0.61704519 | 0.7832683 | 0 | 0.494 | 0.2887 | 0.2887 | TRUE | 0.5988 |
| Ddo | 9 | 7 | 11 | 5 | 11 | 9 | 10 | 7 | 11 | 4 | 12 | 10 | 8.87 | 9 | 9 | 0.911 | -0.135 | 0.84442635 | 0.92161735 | 0 | 0.3784 | 0.2254 | 0.2254 | TRUE | 0.91 |
| Ddost | 8846 | 9666 | 9858 | 10521 | 7728 | 7896 | 9396 | 9654 | 9898 | 8271 | 8315 | 8643 | 9029.51 | 9649 | 8410 | 0.871 | -0.199 | 0.00094008 | 0.00848708 | 5.00E-04 | 0.0073 | 0.0025 | 0.0025 | TRUE | 0.0072 |
| Ddr1 | 1370 | 1389 | 1348 | 1411 | 999 | 1065 | 1455 | 1387 | 1354 | 1109 | 1075 | 1166 | 1257.64 | 1399 | 1117 | 0.798 | -0.325 | 7.44E-05 | 0.00108802 | 6.00E-04 | 0.0095 | 0.004 | 0.004 | TRUE | 0.0172 |
| Ddr2 | 358 | 211 | 172 | 1168 | 964 | 552 | 380 | 211 | 173 | 918 | 1037 | 604 | 553.89 | 255 | 853 | 3.354 | 1.746 | 1.35E-09 | 7.91E-08 | 0.1248 | 0.0129 | 0.0572 | 0.0572 | TRUE | 0.9456 |
| Ddrgk1 | 1114 | 1517 | 1551 | 1653 | 1272 | 1392 | 1183 | 1515 | 1557 | 1299 | 1369 | 1524 | 1407.92 | 1418 | 1397 | 0.984 | -0.023 | 0.86209915 | 0.93192349 | 0.0136 | 0.0092 | 0.0116 | 0.0116 | TRUE | 0.2525 |

| | | | | | | | | | | | | | | | | | | | | | | | | | |
|---|---|---|---|---|---|---|---|---|---|---|---|---|---|---|---|---|---|---|---|---|---|---|---|---|---|
| Ddt | 2528 | 2859 | 2779 | 2501 | 1865 | 2118 | 2685 | 2855 | 2790 | 1966 | 2007 | 2318 | 2437.02 | 2777 | 2097 | 0.755 | -0.406 | 1.09E-05 | 0.00021363 | 0.0041 | 0.0083 | 0.0057 | 0.0057 | TRUE | 0.1025 |
| Ddx1 | 14473 | 15925 | 15771 | 18945 | 13517 | 13373 | 15373 | 15904 | 15836 | 14893 | 14543 | 14639 | 15198.08 | 15704 | 14692 | 0.936 | -0.096 | 0.0735791 | 0.212824 | 1.00E-04 | 0.0071 | 0.002 | 0.002 | TRUE | 0.0042 |
| Ddx10 | 7145 | 6723 | 6901 | 9516 | 6863 | 6306 | 7589 | 6714 | 6929 | 7481 | 7384 | 6903 | 7166.79 | 7077 | 7256 | 1.025 | 0.036 | 0.65035685 | 0.80412061 | 0.0024 | 0.0074 | 0.0044 | 0.0044 | TRUE | 0.0487 |
| Ddx11 | 2407 | 2383 | 2381 | 3166 | 2599 | 2241 | 2557 | 2380 | 2391 | 2489 | 2796 | 2453 | 2510.95 | 2443 | 2579 | 1.056 | 0.079 | 0.36332597 | 0.58083215 | 0.0029 | 0.0082 | 0.005 | 0.005 | TRUE | 0.0651 |
| Ddx17 | 15259 | 18056 | 18708 | 22877 | 17471 | 18535 | 16208 | 18033 | 18785 | 17984 | 18798 | 20289 | 18349.43 | 17675 | 19024 | 1.076 | 0.106 | 0.23109665 | 0.44492876 | 0.0046 | 0.0071 | 0.0056 | 0.0056 | TRUE | 0.0645 |
| Ddx18 | 7105 | 6005 | 5905 | 9192 | 6221 | 5987 | 7547 | 5997 | 5929 | 7226 | 6693 | 6554 | 6657.74 | 6491 | 6824 | 1.052 | 0.072 | 0.52522799 | 0.7148075 | 0.0093 | 0.0074 | 0.0092 | 0.0092 | TRUE | 0.2468 |
| Ddx19a | 4199 | 3696 | 3704 | 4961 | 3553 | 3436 | 4460 | 3691 | 3719 | 3900 | 3823 | 3761 | 3892.42 | 3957 | 3828 | 0.968 | -0.048 | 0.62496987 | 0.78868818 | 0.0052 | 0.0077 | 0.0066 | 0.0066 | TRUE | 0.1504 |
| Ddx19b | 4250 | 4950 | 5100 | 6716 | 4798 | 5525 | 4514 | 4944 | 5121 | 5280 | 5162 | 6048 | 5178.12 | 4860 | 5497 | 1.131 | 0.177 | 0.0630723 | 0.19252493 | 0.0056 | 0.0075 | 0.0064 | 0.0064 | TRUE | 0.0935 |
| Ddx20 | 2880 | 2529 | 2630 | 4043 | 2826 | 2535 | 3059 | 2526 | 2641 | 3178 | 3041 | 2775 | 2869.91 | 2742 | 2998 | 1.094 | 0.129 | 0.21494897 | 0.42675287 | 0.0071 | 0.0081 | 0.0075 | 0.0075 | TRUE | 0.1241 |
| Ddx21 | 30670 | 26750 | 26282 | 32670 | 23610 | 22152 | 32577 | 26715 | 26390 | 25683 | 25403 | 24249 | 26836.11 | 28561 | 25112 | 0.879 | -0.186 | 0.06644962 | 0.19890746 | 0.0078 | 0.007 | 0.0073 | 0.0073 | TRUE | 0.1852 |
| Ddx23 | 7546 | 6365 | 6068 | 8514 | 6452 | 5901 | 8015 | 6357 | 6093 | 6693 | 6942 | 6460 | 6759.91 | 6822 | 6698 | 0.982 | -0.026 | 0.82263028 | 0.90941476 | 0.01 | 0.0074 | 0.0097 | 0.0097 | TRUE | 0.2857 |
| Ddx24 | 7610 | 7141 | 6993 | 9646 | 7069 | 6341 | 8083 | 7132 | 7022 | 7583 | 7606 | 6941 | 7394.44 | 7412 | 7377 | 0.995 | -0.007 | 0.93841739 | 0.9691313 | 0.0034 | 0.0073 | 0.0054 | 0.0054 | TRUE | 0.0765 |
| Ddx25 | 526 | 620 | 589 | 931 | 669 | 608 | 559 | 619 | 591 | 732 | 720 | 666 | 647.76 | 590 | 706 | 1.197 | 0.26 | 0.00951082 | 0.05064301 | 0.001 | 0.012 | 0.0057 | 0.0057 | TRUE | 0.0287 |
| Ddx26b | 447 | 601 | 573 | 619 | 595 | 666 | 475 | 600 | 575 | 487 | 640 | 729 | 584.37 | 550 | 619 | 1.122 | 0.167 | 0.33442953 | 0.5548783 | 0.0267 | 0.0125 | 0.0197 | 0.0197 | TRUE | 0.3721 |
| Ddx27 | 6290 | 6676 | 7056 | 8638 | 6586 | 6401 | 6681 | 6667 | 7085 | 6791 | 7086 | 7007 | 6886.17 | 6811 | 6961 | 1.022 | 0.031 | 0.62112563 | 0.78565991 | 6.00E-04 | 0.0074 | 0.0027 | 0.0027 | TRUE | 0.0151 |
| Ddx28 | 893 | 846 | 898 | 1363 | 951 | 837 | 949 | 845 | 902 | 1071 | 1023 | 916 | 951.01 | 899 | 1003 | 1.118 | 0.161 | 0.11639603 | 0.28828489 | 0.0039 | 0.0104 | 0.0066 | 0.0066 | TRUE | 0.0686 |
| Ddx31 | 1335 | 1258 | 1123 | 1764 | 1238 | 1179 | 1418 | 1256 | 1128 | 1387 | 1332 | 1291 | 1301.89 | 1267 | 1337 | 1.055 | 0.077 | 0.47471873 | 0.67524193 | 0.0064 | 0.0094 | 0.0077 | 0.0077 | TRUE | 0.1293 |
| Ddx39 | 11632 | 10292 | 9923 | 15154 | 10984 | 9463 | 12355 | 10279 | 9964 | 11913 | 11818 | 10359 | 11114.57 | 10866 | 11363 | 1.046 | 0.065 | 0.55774475 | 0.7401735 | 0.0084 | 0.0072 | 0.0087 | 0.0087 | TRUE | 0.1756 |
| Ddx39b | 7327 | 7533 | 7686 | 13896 | 9893 | 7964 | 7783 | 7523 | 7718 | 10924 | 10644 | 8718 | 8884.91 | 7675 | 10095 | 1.316 | 0.396 | 0.00010625 | 0.00144581 | 0.0071 | 0.0073 | 0.0074 | 0.0074 | TRUE | 0.1738 |
| Ddx3x | 39267 | 33083 | 32665 | 49395 | 33487 | 30352 | 41708 | 33040 | 32799 | 38830 | 36030 | 33225 | 35938.89 | 35849 | 36028 | 1.005 | 0.007 | 0.9512254 | 0.97528069 | 0.0104 | 0.007 | 0.0101 | 0.0101 | TRUE | 0.2502 |
| Ddx3y | 3209 | 2753 | 2479 | 6416 | 4598 | 3956 | 3409 | 2749 | 2489 | 5044 | 4947 | 4330 | 3828.09 | 2882 | 4774 | 1.657 | 0.728 | 3.06E-08 | 1.31E-06 | 0.0164 | 0.0078 | 0.0122 | 0.0122 | TRUE | 0.3092 |
| Ddx4 | 544 | 927 | 918 | 1152 | 819 | 873 | 578 | 926 | 922 | 906 | 881 | 956 | 861.31 | 809 | 914 | 1.13 | 0.177 | 0.32280405 | 0.54399673 | 0.0336 | 0.0107 | 0.0219 | 0.0219 | TRUE | 0.7367 |
| Ddx41 | 4741 | 5219 | 5287 | 5893 | 4587 | 4600 | 5036 | 5212 | 5309 | 4633 | 4935 | 5035 | 5026.69 | 5186 | 4868 | 0.938 | -0.092 | 0.18193014 | 0.38376383 | 0.001 | 0.0076 | 0.0032 | 0.0032 | TRUE | 0.0221 |
| Ddx42 | 7248 | 7130 | 7143 | 11777 | 8388 | 7511 | 7699 | 7121 | 7172 | 9258 | 9025 | 8222 | 8082.81 | 7331 | 8835 | 1.205 | 0.27 | 0.00063041 | 0.00623767 | 0.0027 | 0.0073 | 0.0044 | 0.0044 | TRUE | 0.0449 |
| Ddx43 | 252 | 278 | 286 | 475 | 411 | 255 | 268 | 278 | 287 | 373 | 442 | 279 | 321.21 | 278 | 365 | 1.315 | 0.396 | 0.02944167 | 0.11342307 | 0.0239 | 0.0172 | 0.0206 | 0.0206 | TRUE | 0.2166 |
| Ddx46 | 13800 | 11966 | 12153 | 15559 | 12113 | 11587 | 14658 | 11951 | 12203 | 12231 | 13033 | 12684 | 12793.23 | 12937 | 12649 | 0.978 | -0.032 | 0.74277208 | 0.86266578 | 0.0059 | 0.0072 | 0.007 | 0.007 | TRUE | 0.1655 |
| Ddx47 | 6738 | 6198 | 6075 | 7826 | 5653 | 5635 | 7157 | 6190 | 6100 | 6152 | 6082 | 6168 | 6308.29 | 6482 | 6134 | 0.946 | -0.08 | 0.35523677 | 0.57376286 | 0.0039 | 0.0074 | 0.0052 | 0.0052 | TRUE | 0.101 |
| Ddx49 | 5424 | 5723 | 5816 | 7131 | 5468 | 4874 | 5761 | 5716 | 5840 | 5606 | 5883 | 5335 | 5690.19 | 5772 | 5608 | 0.972 | -0.042 | 0.53865551 | 0.72507252 | 0.001 | 0.0075 | 0.0031 | 0.0031 | TRUE | 0.0222 |
| Ddx5 | 64845 | 65460 | 65053 | 79025 | 56409 | 54393 | 68877 | 65375 | 65320 | 62123 | 60693 | 59542 | 63655 | 66524 | 60786 | 0.914 | -0.13 | 0.02528968 | 0.10256332 | 7.00E-04 | 0.007 | 0.0024 | 0.0024 | TRUE | 0.0117 |
| Ddx50 | 3620 | 4742 | 4683 | 4750 | 3471 | 3994 | 3845 | 4736 | 4702 | 3734 | 3735 | 4372 | 4187.32 | 4428 | 3947 | 0.891 | -0.166 | 0.15103347 | 0.34214573 | 0.0105 | 0.0077 | 0.0094 | 0.0094 | TRUE | 0.1612 |
| Ddx51 | 1622 | 1305 | 1238 | 2173 | 1637 | 1290 | 1723 | 1303 | 1243 | 1708 | 1761 | 1412 | 1525.15 | 1423 | 1627 | 1.144 | 0.194 | 0.19893713 | 0.4062556 | 0.0226 | 0.0091 | 0.0158 | 0.0158 | TRUE | 0.4078 |
| Ddx52 | 3874 | 3423 | 3293 | 5341 | 3675 | 3228 | 4115 | 3419 | 3307 | 4199 | 3954 | 3534 | 3754.38 | 3614 | 3896 | 1.078 | 0.109 | 0.34586019 | 0.56517324 | 0.0101 | 0.0078 | 0.0093 | 0.0093 | TRUE | 0.1791 |
| Ddx54 | 12940 | 12520 | 12560 | 14947 | 11357 | 11395 | 13745 | 12504 | 12612 | 11750 | 12219 | 12474 | 12550.53 | 12954 | 12148 | 0.938 | -0.093 | 0.1900632 | 0.39515909 | 0.0017 | 0.0072 | 0.0035 | 0.0035 | TRUE | 0.0349 |
| Ddx55 | 959 | 786 | 833 | 1120 | 802 | 656 | 1019 | 785 | 836 | 880 | 863 | 718 | 850.25 | 880 | 820 | 0.933 | -0.1 | 0.47189401 | 0.67314138 | 0.0143 | 0.0108 | 0.0127 | 0.0127 | TRUE | 0.2252 |
| Ddx56 | 4494 | 4387 | 4486 | 6870 | 4846 | 4154 | 4773 | 4381 | 4504 | 5401 | 5214 | 4547 | 4803.51 | 4553 | 5054 | 1.11 | 0.151 | 0.10092291 | 0.26301324 | 0.0046 | 0.0076 | 0.0059 | 0.0059 | TRUE | 0.0935 |
| Ddx58 | 1565 | 1043 | 1241 | 963 | 787 | 673 | 1662 | 1042 | 1246 | 757 | 847 | 737 | 1048.43 | 1317 | 780 | 0.593 | -0.755 | 1.00E-05 | 0.00019863 | 0.0301 | 0.01 | 0.02 | 0.02 | TRUE | 0.3334 |
| Ddx59 | 1267 | 1341 | 1233 | 1502 | 1023 | 1067 | 1346 | 1339 | 1238 | 1181 | 1101 | 1168 | 1228.76 | 1308 | 1150 | 0.88 | -0.185 | 0.02796148 | 0.10975819 | 0.001 | 0.0096 | 0.0043 | 0.0043 | TRUE | 0.0262 |
| Ddx6 | 10042 | 10019 | 10187 | 13911 | 9624 | 9386 | 10666 | 10006 | 10229 | 10936 | 10355 | 10274 | 10411.05 | 10300 | 10522 | 1.022 | 0.031 | 0.63328252 | 0.79371543 | 9.00E-04 | 0.0072 | 0.0029 | 0.0029 | TRUE | 0.0146 |
| Ddx60 | 488 | 686 | 779 | 302 | 221 | 327 | 518 | 685 | 782 | 237 | 238 | 358 | 469.8 | 662 | 278 | 0.419 | -1.256 | 2.69E-09 | 1.46E-07 | 0.0478 | 0.0139 | 0.0296 | 0.0296 | TRUE | 0.349 |
| Deaf1 | 865 | 1045 | 1018 | 1589 | 1421 | 1258 | 919 | 1044 | 1022 | 1249 | 1529 | 1377 | 1189.96 | 995 | 1385 | 1.391 | 0.476 | 1.71E-05 | 0.00031227 | 0.0067 | 0.0097 | 0.008 | 0.008 | TRUE | 0.0994 |
| Dear1 | 0 | 0 | 0 | 0 | 0 | 0 | 0 | 0 | 0 | 0 | 0 | 0 | 0 | 0 | 0 | NA | NA | NA | NA | NA | NA | NA | NA | NA | NA |
| Deb1 | 265 | 218 | 215 | 385 | 279 | 215 | 281 | 218 | 216 | 303 | 300 | 235 | 258.88 | 238 | 279 | 1.175 | 0.232 | 0.18717254 | 0.39157517 | 0.0173 | 0.0196 | 0.0185 | 0.0185 | TRUE | 0.2743 |
| Decr1 | 1127 | 1507 | 1576 | 1682 | 1153 | 1194 | 1197 | 1505 | 1582 | 1322 | 1241 | 1307 | 1359.07 | 1428 | 1290 | 0.903 | -0.147 | 0.22883216 | 0.44224019 | 0.0096 | 0.0093 | 0.01 | 0.01 | TRUE | 0.2403 |
| Decr2 | 753 | 644 | 622 | 1041 | 776 | 728 | 800 | 643 | 625 | 818 | 835 | 797 | 752.96 | 689 | 817 | 1.186 | 0.246 | 0.04495679 | 0.15281433 | 0.008 | 0.0113 | 0.0095 | 0.0095 | TRUE | 0.2316 |
| Dedd | 1152 | 1396 | 1463 | 1674 | 1199 | 1217 | 1224 | 1394 | 1469 | 1316 | 1290 | 1332 | 1337.51 | 1362 | 1313 | 0.963 | -0.054 | 0.57871916 | 0.75588355 | 0.0037 | 0.0094 | 0.006 | 0.006 | TRUE | 0.0948 |
| Dedd2 | 719 | 721 | 793 | 1028 | 871 | 814 | 764 | 720 | 796 | 808 | 937 | 891 | 819.39 | 760 | 879 | 1.155 | 0.208 | 0.04227414 | 0.1466024 | 0.0031 | 0.0109 | 0.0063 | 0.0063 | TRUE | 0.0613 |
| Def6 | 1106 | 1133 | 1123 | 1468 | 1091 | 1091 | 1175 | 1132 | 1128 | 1154 | 1174 | 1194 | 1159.34 | 1145 | 1174 | 1.025 | 0.036 | 0.63605928 | 0.79536224 | 0 | 0.0097 | 0.0034 | 0.0034 | TRUE | 0.0063 |
| Def8 | 3374 | 3416 | 3605 | 5396 | 3869 | 3560 | 3584 | 3412 | 3620 | 4242 | 4163 | 3897 | 3819.48 | 3539 | 4101 | 1.159 | 0.213 | 0.00288575 | 0.02056668 | 0.0011 | 0.0078 | 0.0034 | 0.0034 | TRUE | 0.0228 |
| Defa-ps1 | 0 | 0 | 0 | 0 | 0 | 0 | 0 | 0 | 0 | 0 | 0 | 0 | 0 | 0 | 0 | NA | NA | NA | NA | NA | NA | NA | NA | NA | NA |
| Defa-ps12 | 0 | 0 | 0 | 0 | 0 | 0 | 0 | 0 | 0 | 0 | 0 | 0 | 0 | 0 | 0 | NA | NA | NA | NA | NA | NA | NA | NA | NA | NA |
| Defa-ps13 | 0 | 0 | 0 | 0 | 0 | 0 | 0 | 0 | 0 | 0 | 0 | 0 | 0 | 0 | 0 | NA | NA | NA | NA | NA | NA | NA | NA | NA | NA |
| Defa-rs1 | 0 | 0 | 0 | 2 | 1 | 0 | 0 | 0 | 0 | 2 | 1 | 0 | 0.44 | 0 | 1 | 4.754 | 2.249 | 0.50892201 | NA | 0 | 7.4712 | 6.8061 | 6.8061 | TRUE | 0.2361 |
| Defa-rs7 | 0 | 0 | 0 | 0 | 0 | 0 | 0 | 0 | 0 | 0 | 0 | 0 | 0 | 0 | 0 | NA | NA | NA | NA | NA | NA | NA | NA | NA | NA |
| Defa17 | 0 | 0 | 0 | 0 | 0 | 0 | 0 | 0 | 0 | 0 | 0 | 0 | 0 | 0 | 0 | NA | NA | NA | NA | NA | NA | NA | NA | NA | NA |
| Defa20 | 0 | 0 | 0 | 0 | 0 | 0 | 0 | 0 | 0 | 0 | 0 | 0 | 0 | 0 | 0 | NA | NA | NA | NA | NA | NA | NA | NA | NA | NA |
| Defa21 | 0 | 0 | 0 | 0 | 0 | 0 | 0 | 0 | 0 | 0 | 0 | 0 | 0 | 0 | 0 | NA | NA | NA | NA | NA | NA | NA | NA | NA | NA |
| Defa22 | 0 | 0 | 0 | 0 | 0 | 0 | 0 | 0 | 0 | 0 | 0 | 0 | 0 | 0 | 0 | NA | NA | NA | NA | NA | NA | NA | NA | NA | NA |
| Defa23 | 0 | 0 | 0 | 0 | 0 | 0 | 0 | 0 | 0 | 0 | 0 | 0 | 0 | 0 | 0 | NA | NA | NA | NA | NA | NA | NA | NA | NA | NA |
| Defa24 | 0 | 0 | 0 | 0 | 0 | 0 | 0 | 0 | 0 | 0 | 0 | 0 | 0 | 0 | 0 | NA | NA | NA | NA | NA | NA | NA | NA | NA | NA |
| Defa25 | 0 | 0 | 0 | 0 | 0 | 0 | 0 | 0 | 0 | 0 | 0 | 0 | 0 | 0 | 0 | NA | NA | NA | NA | NA | NA | NA | NA | NA | NA |
| Defa26 | 0 | 0 | 0 | 0 | 0 | 0 | 0 | 0 | 0 | 0 | 0 | 0 | 0 | 0 | 0 | NA | NA | NA | NA | NA | NA | NA | NA | NA | NA |
| Defa3 | 0 | 0 | 0 | 0 | 0 | 0 | 0 | 0 | 0 | 0 | 0 | 0 | 0 | 0 | 0 | NA | NA | NA | NA | NA | NA | NA | NA | NA | NA |
| Defa4 | 0 | 0 | 0 | 0 | 0 | 0 | 0 | 0 | 0 | 0 | 0 | 0 | 0 | 0 | 0 | NA | NA | NA | NA | NA | NA | NA | NA | NA | NA |
| Defa5 | 0 | 0 | 0 | 0 | 0 | 0 | 0 | 0 | 0 | 0 | 0 | 0 | 0 | 0 | 0 | NA | NA | NA | NA | NA | NA | NA | NA | NA | NA |
| Defa6 | 0 | 0 | 0 | 0 | 0 | 0 | 0 | 0 | 0 | 0 | 0 | 0 | 0 | 0 | 0 | NA | NA | NA | NA | NA | NA | NA | NA | NA | NA |
| Defb1 | 0 | 0 | 2 | 0 | 0 | 1 | 0 | 0 | 2 | 0 | 0 | 1 | 0.52 | 1 | 0 | 0.52 | -0.944 | 0.77774021 | NA | 0 | 6.3774 | 6.2947 | 6.2947 | TRUE | 0.9739 |
| Defb10 | 0 | 0 | 0 | 0 | 0 | 0 | 0 | 0 | 0 | 0 | 0 | 0 | 0 | 0 | 0 | NA | NA | NA | NA | NA | NA | NA | NA | NA | NA |
| Defb11 | 0 | 0 | 0 | 0 | 0 | 0 | 0 | 0 | 0 | 0 | 0 | 0 | 0 | 0 | 0 | NA | NA | NA | NA | NA | NA | NA | NA | NA | NA |
| Defb12 | 0 | 0 | 0 | 0 | 0 | 0 | 0 | 0 | 0 | 0 | 0 | 0 | 0 | 0 | 0 | NA | NA | NA | NA | NA | NA | NA | NA | NA | NA |

| | | | | | | | | | | | | | | | | | | | | | | | | | |
|---|---|---|---|---|---|---|---|---|---|---|---|---|---|---|---|---|---|---|---|---|---|---|---|---|---|
| Defb13 | 0 | 0 | 0 | 0 | 1 | 0 | 0 | 0 | 0 | 0 | 1 | 0 | 0.18 | 0 | 0 | 1.859 | 0.894 | 0.82650734 | NA | 0 | 18.3785 | 10 | 10 | TRUE | 0.5204 |
| Defb14 | 0 | 0 | 0 | 0 | 0 | 0 | 0 | 0 | 0 | 0 | 0 | 0 | 0 | 0 | 0 | NA | NA | NA | NA | NA | NA | NA | NA | NA | NA |
| Defb15 | 0 | 0 | 0 | 0 | 0 | 0 | 0 | 0 | 0 | 0 | 0 | 0 | 0 | 0 | 0 | NA | NA | NA | NA | NA | NA | NA | NA | NA | NA |
| Defb18 | 0 | 0 | 0 | 0 | 0 | 0 | 0 | 0 | 0 | 0 | 0 | 0 | 0 | 0 | 0 | NA | NA | NA | NA | NA | NA | NA | NA | NA | NA |
| Defb19 | 0 | 0 | 0 | 0 | 0 | 0 | 0 | 0 | 0 | 0 | 0 | 0 | 0 | 0 | 0 | NA | NA | NA | NA | NA | NA | NA | NA | NA | NA |
| Defb2 | 0 | 0 | 0 | 0 | 0 | 0 | 0 | 0 | 0 | 0 | 0 | 0 | 0 | 0 | 0 | NA | NA | NA | NA | NA | NA | NA | NA | NA | NA |
| Defb20 | 7 | 10 | 6 | 6 | 4 | 1 | 7 | 10 | 6 | 5 | 4 | 1 | 5.59 | 8 | 3 | 0.44 | -1.183 | 0.16506121 | 0.36167622 | 0 | 0.5959 | 0.3163 | 0.3163 | TRUE | 0.4369 |
| Defb21 | 0 | 0 | 0 | 0 | 0 | 0 | 0 | 0 | 0 | 0 | 0 | 0 | 0 | 0 | 0 | NA | NA | NA | NA | NA | NA | NA | NA | NA | NA |
| Defb22 | 0 | 0 | 0 | 0 | 0 | 0 | 0 | 0 | 0 | 0 | 0 | 0 | 0 | 0 | 0 | NA | NA | NA | NA | NA | NA | NA | NA | NA | NA |
| Defb23 | 0 | 0 | 0 | 4 | 1 | 0 | 0 | 0 | 0 | 3 | 1 | 0 | 0.7 | 0 | 1 | 7.607 | 2.927 | 0.32216877 | NA | 6.5513 | 4.6905 | 4.9568 | 4.9568 | TRUE | 0.1456 |
| Defb25 | 3 | 1 | 2 | 14 | 11 | 1 | 3 | 1 | 2 | 11 | 12 | 1 | 5.02 | 2 | 8 | 3.914 | 1.969 | 0.07616709 | 0.21774648 | 0 | 0.663 | 0.5784 | 0.5784 | TRUE | 1.5271 |
| Defb26 | 0 | 0 | 0 | 0 | 0 | 0 | 0 | 0 | 0 | 0 | 0 | 0 | 0 | 0 | 0 | NA | NA | NA | NA | NA | NA | NA | NA | NA | NA |
| Defb28 | 0 | 0 | 0 | 0 | 0 | 1 | 0 | 0 | 0 | 0 | 0 | 1 | 0.18 | 0 | 0 | 1.859 | 0.894 | 0.82650734 | NA | 0 | 18.0643 | 10 | 10 | TRUE | 0.5383 |
| Defb29 | 2 | 2 | 4 | 3 | 4 | 3 | 2 | 2 | 4 | 2 | 4 | 3 | 3.01 | 3 | 3 | 1.206 | 0.27 | 0.79848667 | NA | 0 | 1.0999 | 0.4717 | 0.4717 | TRUE | 0.2126 |
| Defb3 | 0 | 0 | 0 | 0 | 0 | 0 | 0 | 0 | 0 | 0 | 0 | 0 | 0 | 0 | 0 | NA | NA | NA | NA | NA | NA | NA | NA | NA | NA |
| Defb30 | 35 | 19 | 27 | 40 | 22 | 22 | 37 | 19 | 27 | 31 | 24 | 24 | 27.08 | 28 | 26 | 0.959 | -0.06 | 0.88233109 | 0.94169383 | 0.0321 | 0.1286 | 0.0823 | 0.0823 | TRUE | 0.2311 |
| Defb33 | 0 | 0 | 0 | 0 | 0 | 0 | 0 | 0 | 0 | 0 | 0 | 0 | 0 | 0 | 0 | NA | NA | NA | NA | NA | NA | NA | NA | NA | NA |
| Defb34 | 0 | 0 | 0 | 0 | 0 | 0 | 0 | 0 | 0 | 0 | 0 | 0 | 0 | 0 | 0 | NA | NA | NA | NA | NA | NA | NA | NA | NA | NA |
| Defb35 | 0 | 0 | 0 | 0 | 0 | 0 | 0 | 0 | 0 | 0 | 0 | 0 | 0 | 0 | 0 | NA | NA | NA | NA | NA | NA | NA | NA | NA | NA |
| Defb36 | 4 | 6 | 3 | 6 | 10 | 6 | 4 | 6 | 3 | 5 | 11 | 7 | 5.88 | 4 | 8 | 1.643 | 0.717 | 0.37598009 | 0.59147378 | 0 | 0.5669 | 0.2893 | 0.2893 | TRUE | 0.4341 |
| Defb37 | 0 | 0 | 0 | 0 | 0 | 0 | 0 | 0 | 0 | 0 | 0 | 0 | 0 | 0 | 0 | NA | NA | NA | NA | NA | NA | NA | NA | NA | NA |
| Defb38 | 0 | 0 | 0 | 0 | 0 | 0 | 0 | 0 | 0 | 0 | 0 | 0 | 0 | 0 | 0 | NA | NA | NA | NA | NA | NA | NA | NA | NA | NA |
| Defb39 | 0 | 0 | 0 | 0 | 0 | 0 | 0 | 0 | 0 | 0 | 0 | 0 | 0 | 0 | 0 | NA | NA | NA | NA | NA | NA | NA | NA | NA | NA |
| Defb4 | 0 | 0 | 0 | 0 | 0 | 0 | 0 | 0 | 0 | 0 | 0 | 0 | 0 | 0 | 0 | NA | NA | NA | NA | NA | NA | NA | NA | NA | NA |
| Defb40 | 0 | 0 | 0 | 0 | 0 | 0 | 0 | 0 | 0 | 0 | 0 | 0 | 0 | 0 | 0 | NA | NA | NA | NA | NA | NA | NA | NA | NA | NA |
| Defb41 | 0 | 1 | 0 | 0 | 0 | 0 | 0 | 1 | 0 | 0 | 0 | 0 | 0.17 | 0 | 0 | 0.49 | -1.029 | 0.80087 | NA | 0 | 19.7991 | 10 | 10 | TRUE | 0.405 |
| Defb42 | 100 | 149 | 176 | 118 | 113 | 125 | 106 | 149 | 177 | 93 | 122 | 137 | 130.49 | 144 | 117 | 0.81 | -0.305 | 0.22563779 | 0.43842997 | 0.0438 | 0.0322 | 0.0379 | 0.0379 | TRUE | 0.2742 |
| Defb43 | 0 | 0 | 0 | 0 | 0 | 0 | 0 | 0 | 0 | 0 | 0 | 0 | 0 | 0 | 0 | NA | NA | NA | NA | NA | NA | NA | NA | NA | NA |
| Defb44-ps | 0 | 0 | 0 | 0 | 0 | 0 | 0 | 0 | 0 | 0 | 0 | 0 | 0 | 0 | 0 | NA | NA | NA | NA | NA | NA | NA | NA | NA | NA |
| Defb45 | 0 | 0 | 0 | 0 | 0 | 0 | 0 | 0 | 0 | 0 | 0 | 0 | 0 | 0 | 0 | NA | NA | NA | NA | NA | NA | NA | NA | NA | NA |
| Defb46 | 0 | 0 | 0 | 0 | 0 | 0 | 0 | 0 | 0 | 0 | 0 | 0 | 0 | 0 | 0 | NA | NA | NA | NA | NA | NA | NA | NA | NA | NA |
| Defb47 | 0 | 1 | 0 | 1 | 1 | 0 | 0 | 1 | 0 | 1 | 1 | 0 | 0.48 | 0 | 1 | 1.712 | 0.776 | 0.78614772 | NA | 0 | 6.9165 | 4.1067 | 4.1067 | TRUE | 0.405 |
| Defb48 | 0 | 0 | 0 | 0 | 0 | 0 | 0 | 0 | 0 | 0 | 0 | 0 | 0 | 0 | 0 | NA | NA | NA | NA | NA | NA | NA | NA | NA | NA |
| Defb5 | 0 | 0 | 0 | 0 | 0 | 0 | 0 | 0 | 0 | 0 | 0 | 0 | 0 | 0 | 0 | NA | NA | NA | NA | NA | NA | NA | NA | NA | NA |
| Defb50 | 0 | 0 | 0 | 0 | 0 | 0 | 0 | 0 | 0 | 0 | 0 | 0 | 0 | 0 | 0 | NA | NA | NA | NA | NA | NA | NA | NA | NA | NA |
| Defb6 | 0 | 0 | 0 | 0 | 0 | 0 | 0 | 0 | 0 | 0 | 0 | 0 | 0 | 0 | 0 | NA | NA | NA | NA | NA | NA | NA | NA | NA | NA |
| Defb7 | 0 | 0 | 0 | 0 | 1 | 0 | 0 | 0 | 0 | 0 | 1 | 0 | 0.18 | 0 | 0 | 1.859 | 0.894 | 0.82650734 | NA | 0 | 18.3785 | 10 | 10 | TRUE | 0.5204 |
| Defb8 | 0 | 0 | 0 | 0 | 0 | 0 | 0 | 0 | 0 | 0 | 0 | 0 | 0 | 0 | 0 | NA | NA | NA | NA | NA | NA | NA | NA | NA | NA |
| Defb9 | 0 | 0 | 0 | 0 | 0 | 0 | 0 | 0 | 0 | 0 | 0 | 0 | 0 | 0 | 0 | NA | NA | NA | NA | NA | NA | NA | NA | NA | NA |
| Degs1 | 5957 | 5555 | 5424 | 5933 | 4198 | 4285 | 6327 | 5548 | 5446 | 4664 | 4517 | 4691 | 5198.82 | 5774 | 4624 | 0.801 | -0.32 | 0.00014782 | 0.00188324 | 0.0034 | 0.0075 | 0.0049 | 0.0049 | TRUE | 0.0857 |
| Degs2 | 42 | 38 | 29 | 32 | 18 | 23 | 45 | 38 | 29 | 25 | 19 | 25 | 30.23 | 37 | 23 | 0.627 | -0.674 | 0.06600427 | 0.19817615 | 0 | 0.1159 | 0.0621 | 0.0621 | TRUE | 0.1918 |
| Dek | 7350 | 6935 | 6942 | 14250 | 10436 | 8415 | 7807 | 6926 | 6971 | 11202 | 11228 | 9212 | 8890.97 | 7235 | 10547 | 1.458 | 0.544 | 2.83E-07 | 8.99E-06 | 0.0082 | 0.0073 | 0.008 | 0.008 | TRUE | 0.1498 |
| Dennd1a | 907 | 764 | 818 | 1238 | 854 | 813 | 963 | 763 | 821 | 973 | 919 | 890 | 888.3 | 849 | 927 | 1.093 | 0.128 | 0.26936658 | 0.48939635 | 0.0067 | 0.0106 | 0.0086 | 0.0086 | TRUE | 0.1635 |
| Dennd1b | 1197 | 1470 | 1677 | 2032 | 1511 | 1459 | 1271 | 1468 | 1684 | 1597 | 1626 | 1597 | 1540.61 | 1474 | 1607 | 1.089 | 0.124 | 0.28747447 | 0.5087006 | 0.0088 | 0.009 | 0.0091 | 0.0091 | TRUE | 0.1858 |
| Dennd1c | 96 | 83 | 79 | 93 | 65 | 66 | 102 | 83 | 79 | 73 | 70 | 72 | 79.91 | 88 | 72 | 0.816 | -0.293 | 0.19233758 | 0.39802397 | 0 | 0.0481 | 0.0237 | 0.0237 | TRUE | 0.1776 |
| Dennd2a | 1894 | 2025 | 2063 | 2875 | 2103 | 1980 | 2012 | 2022 | 2071 | 2260 | 2263 | 2167 | 2132.64 | 2035 | 2230 | 1.096 | 0.132 | 0.04978067 | 0.16379495 | 0 | 0.0084 | 0.0028 | 0.0028 | TRUE | 0.0072 |
| Dennd2c | 3491 | 4738 | 5049 | 4489 | 3421 | 3874 | 3708 | 4732 | 5070 | 3529 | 3681 | 4241 | 4160.01 | 4503 | 3817 | 0.847 | -0.239 | 0.07665692 | 0.21859209 | 0.0169 | 0.0077 | 0.0129 | 0.0129 | TRUE | 0.2904 |
| Dennd2d | 54 | 82 | 94 | 101 | 85 | 97 | 57 | 82 | 94 | 79 | 91 | 106 | 85.11 | 78 | 92 | 1.18 | 0.239 | 0.35959081 | 0.57800887 | 0.0281 | 0.0456 | 0.0372 | 0.0372 | TRUE | 0.4792 |
| Dennd3 | 830 | 833 | 799 | 909 | 753 | 802 | 882 | 832 | 802 | 715 | 810 | 878 | 819.75 | 839 | 801 | 0.954 | -0.068 | 0.53948579 | 0.72541204 | 0.0054 | 0.0109 | 0.0077 | 0.0077 | TRUE | 0.1095 |
| Dennd4a | 1949 | 2109 | 2385 | 2464 | 1858 | 2152 | 2070 | 2106 | 2395 | 1937 | 1999 | 2356 | 2143.84 | 2190 | 2097 | 0.957 | -0.063 | 0.56724114 | 0.7471004 | 0.0083 | 0.0084 | 0.0084 | 0.0084 | TRUE | 0.14 |
| Dennd4b | 1442 | 1993 | 2090 | 1955 | 1604 | 1786 | 1532 | 1990 | 2099 | 1537 | 1726 | 1955 | 1806.4 | 1874 | 1739 | 0.928 | -0.108 | 0.45799539 | 0.66258515 | 0.0203 | 0.0087 | 0.0147 | 0.0147 | TRUE | 0.3073 |
| Dennd4c | 2539 | 2285 | 2407 | 3075 | 2210 | 2021 | 2697 | 2282 | 2417 | 2417 | 2378 | 2212 | 2400.54 | 2465 | 2336 | 0.948 | -0.077 | 0.40792333 | 0.62066049 | 0.0041 | 0.0083 | 0.0059 | 0.0059 | TRUE | 0.0817 |
| Dennd5a | 4617 | 5069 | 4894 | 6481 | 4535 | 4678 | 4904 | 5062 | 4914 | 5095 | 4879 | 5121 | 4995.94 | 4960 | 5032 | 1.014 | 0.021 | 0.73433135 | 0.85753055 | 3.00E-04 | 0.0076 | 0.0025 | 0.0025 | TRUE | 0.0085 |
| Dennd5b | 526 | 435 | 460 | 668 | 422 | 353 | 559 | 434 | 462 | 525 | 454 | 386 | 470.1 | 485 | 455 | 0.941 | -0.088 | 0.58003185 | 0.75669188 | 0.0183 | 0.0139 | 0.0162 | 0.0162 | TRUE | 0.2158 |
| Dennd6a | 3792 | 3958 | 3875 | 4537 | 3219 | 3391 | 4028 | 3953 | 3891 | 3567 | 3463 | 3712 | 3768.94 | 3957 | 3581 | 0.905 | -0.144 | 0.02682819 | 0.10687906 | 4.00E-04 | 0.0078 | 0.0028 | 0.0028 | TRUE | 0.0123 |
| Dennd6b | 357 | 425 | 339 | 470 | 486 | 385 | 379 | 424 | 340 | 369 | 523 | 421 | 409.64 | 381 | 438 | 1.146 | 0.197 | 0.23400843 | 0.44867261 | 0.0194 | 0.0149 | 0.0173 | 0.0173 | TRUE | 0.3325 |
| Denr | 6822 | 5847 | 5430 | 7711 | 5578 | 5071 | 7246 | 5839 | 5452 | 6062 | 6002 | 5551 | 6025.38 | 6179 | 5872 | 0.95 | -0.074 | 0.53715688 | 0.72385837 | 0.0113 | 0.0074 | 0.0101 | 0.0101 | TRUE | 0.2781 |
| Depdc1a | 838 | 1158 | 1113 | 1379 | 1078 | 1158 | 890 | 1157 | 1118 | 1084 | 1160 | 1268 | 1112.62 | 1055 | 1171 | 1.109 | 0.149 | 0.24512277 | 0.46184163 | 0.0111 | 0.0099 | 0.011 | 0.011 | TRUE | 0.222 |
| Depdc1b | 1637 | 1554 | 1460 | 1852 | 1295 | 1258 | 1739 | 1552 | 1466 | 1456 | 1393 | 1377 | 1497.18 | 1586 | 1409 | 0.889 | -0.17 | 0.07305958 | 0.21170231 | 0.0033 | 0.0091 | 0.0058 | 0.0058 | TRUE | 0.0858 |
| Depdc5 | 4337 | 3816 | 4040 | 5738 | 4064 | 4042 | 4607 | 3811 | 4057 | 4511 | 4373 | 4425 | 4297.05 | 4158 | 4436 | 1.067 | 0.094 | 0.30563218 | 0.52647781 | 0.0046 | 0.0077 | 0.0058 | 0.0058 | TRUE | 0.1083 |
| Depdc7 | 1046 | 864 | 824 | 1626 | 1034 | 920 | 1111 | 863 | 827 | 1278 | 1113 | 1007 | 1033.19 | 934 | 1133 | 1.214 | 0.28 | 0.05868975 | 0.18315291 | 0.0195 | 0.0101 | 0.0148 | 0.0148 | TRUE | 0.3278 |
| Deptor | 1530 | 1696 | 1756 | 1635 | 1240 | 1250 | 1625 | 1694 | 1763 | 1285 | 1334 | 1368 | 1511.66 | 1694 | 1329 | 0.784 | -0.351 | 7.96E-06 | 0.00016285 | 7.00E-04 | 0.0091 | 0.0038 | 0.0038 | TRUE | 0.0154 |
| Dera | 4077 | 4917 | 4999 | 5420 | 4094 | 4199 | 4330 | 4911 | 5020 | 4261 | 4405 | 4596 | 4587.13 | 4754 | 4421 | 0.93 | -0.105 | 0.22217395 | 0.43437804 | 0.0034 | 0.0076 | 0.0051 | 0.0051 | TRUE | 0.0737 |
| Derl1 | 2374 | 2516 | 2429 | 3299 | 2377 | 2369 | 2522 | 2513 | 2439 | 2593 | 2558 | 2593 | 2536.25 | 2491 | 2581 | 1.036 | 0.051 | 0.41621909 | 0.62771911 | 0 | 0.0082 | 0.0025 | 0.0025 | TRUE | 0.0041 |
| Derl2 | 2226 | 2482 | 2383 | 3006 | 2241 | 2206 | 2364 | 2479 | 2393 | 2363 | 2411 | 2415 | 2404.18 | 2412 | 2396 | 0.993 | -0.01 | 0.8819737 | 0.94167948 | 0 | 0.0083 | 0.0027 | 0.0027 | TRUE | 0.0071 |
| Derl3 | 396 | 451 | 425 | 374 | 259 | 260 | 421 | 450 | 427 | 294 | 279 | 285 | 359.18 | 433 | 286 | 0.661 | -0.597 | 1.65E-07 | 5.52E-06 | 0 | 0.0161 | 0.0065 | 0.0065 | TRUE | 0.0151 |
| Des | 1109 | 1522 | 1546 | 1147 | 901 | 1142 | 1178 | 1520 | 1552 | 902 | 969 | 1250 | 1228.59 | 1417 | 1040 | 0.734 | -0.447 | 0.0052174 | 0.03223222 | 0.0252 | 0.0096 | 0.0176 | 0.0176 | TRUE | 0.3701 |
| Desi1 | 2959 | 2789 | 2817 | 3593 | 2560 | 2550 | 3143 | 2785 | 2829 | 2825 | 2754 | 2791 | 2854.54 | 2919 | 2790 | 0.956 | -0.065 | 0.4100515 | 0.62158382 | 0.0018 | 0.0081 | 0.0041 | 0.0041 | TRUE | 0.0546 |
| Desi2 | 3504 | 3475 | 3437 | 4280 | 2909 | 3079 | 3722 | 3471 | 3451 | 3365 | 3130 | 3370 | 3418.07 | 3548 | 3288 | 0.927 | -0.109 | 0.14103028 | 0.32643523 | 0.0013 | 0.0079 | 0.0037 | 0.0037 | TRUE | 0.0223 |

| | | | | | | | | | | | | | | | | | | | | | | | | | |
|---|---|---|---|---|---|---|---|---|---|---|---|---|---|---|---|---|---|---|---|---|---|---|---|---|---|
| Det1 | 672 | 772 | 749 | 907 | 700 | 706 | 714 | 771 | 752 | 713 | 753 | 773 | 745.98 | 746 | 746 | 1 | 0 | 0.99693003 | 0.99828918 | 0 | 0.0113 | 0.0048 | 0.0048 | TRUE | 0.019 |
| Dexi | 166 | 148 | 189 | 229 | 186 | 180 | 176 | 148 | 190 | 180 | 200 | 197 | 181.85 | 171 | 192 | 1.121 | 0.165 | 0.31915444 | 0.54044491 | 0.0038 | 0.025 | 0.0143 | 0.0143 | TRUE | 0.1555 |
| Dffa | 1357 | 1166 | 1217 | 1407 | 949 | 894 | 1441 | 1164 | 1222 | 1106 | 1021 | 979 | 1155.6 | 1276 | 1035 | 0.812 | -0.3 | 0.00843313 | 0.04631176 | 0.0072 | 0.0098 | 0.0085 | 0.0085 | TRUE | 0.1539 |
| Dffb | 251 | 272 | 237 | 383 | 230 | 230 | 267 | 272 | 238 | 301 | 247 | 252 | 262.76 | 259 | 267 | 1.034 | 0.049 | 0.74144976 | 0.86165152 | 0.0055 | 0.0194 | 0.0119 | 0.0119 | TRUE | 0.1506 |
| Dfna5 | 61 | 82 | 74 | 118 | 70 | 76 | 65 | 82 | 74 | 93 | 75 | 83 | 78.71 | 74 | 84 | 1.14 | 0.189 | 0.40939166 | 0.62130006 | 0 | 0.0488 | 0.0251 | 0.0251 | TRUE | 0.0984 |
| Dfnb59 | 2 | 2 | 1 | 3 | 2 | 4 | 2 | 2 | 1 | 2 | 2 | 4 | 2.34 | 2 | 3 | 1.723 | 0.785 | 0.51821345 | NA | 0 | 1.4173 | 0.5978 | 0.5978 | TRUE | 0.2014 |
| Dgat1 | 727 | 752 | 726 | 1070 | 749 | 623 | 772 | 751 | 729 | 841 | 806 | 682 | 763.54 | 751 | 776 | 1.035 | 0.05 | 0.65362925 | 0.80540538 | 0.0051 | 0.0112 | 0.0078 | 0.0078 | TRUE | 0.132 |
| Dgat2 | 489 | 428 | 417 | 774 | 545 | 491 | 519 | 427 | 419 | 608 | 586 | 537 | 516.31 | 455 | 577 | 1.27 | 0.345 | 0.00707258 | 0.0404016 | 0.0065 | 0.0133 | 0.0099 | 0.0099 | TRUE | 0.1752 |
| Dgat2l6 | 0 | 0 | 0 | 0 | 0 | 0 | 0 | 0 | 0 | 0 | 0 | 0 | 0 | 0 | 0 | NA | NA | NA | NA | NA | NA | NA | NA | NA | NA |
| Dgcr14 | 1365 | 1341 | 1367 | 1866 | 1396 | 1330 | 1450 | 1339 | 1373 | 1467 | 1502 | 1456 | 1431.09 | 1387 | 1475 | 1.063 | 0.089 | 0.24764881 | 0.46433167 | 0 | 0.0092 | 0.0035 | 0.0035 | TRUE | 0.0186 |
| Dgcr2 | 5403 | 5607 | 6012 | 6670 | 4885 | 5098 | 5739 | 5600 | 6037 | 5243 | 5256 | 5581 | 5575.89 | 5792 | 5360 | 0.925 | -0.112 | 0.10351484 | 0.26764507 | 0.0011 | 0.0075 | 0.0032 | 0.0032 | TRUE | 0.0167 |
| Dgcr6 | 969 | 1086 | 1191 | 1343 | 976 | 1023 | 1029 | 1085 | 1196 | 1056 | 1050 | 1120 | 1089.24 | 1103 | 1075 | 0.974 | -0.038 | 0.69063513 | 0.83034167 | 0.0026 | 0.0099 | 0.0056 | 0.0056 | TRUE | 0.0647 |
| Dgcr8 | 2319 | 1807 | 1458 | 3900 | 2794 | 1901 | 2463 | 1805 | 1464 | 3066 | 3006 | 2081 | 2314.14 | 1911 | 2718 | 1.423 | 0.509 | 0.01320973 | 0.06446285 | 0.0553 | 0.0083 | 0.0299 | 0.0299 | TRUE | 0.7071 |
| Dgka | 1476 | 1364 | 1301 | 1331 | 1110 | 1035 | 1568 | 1362 | 1306 | 1046 | 1194 | 1133 | 1268.32 | 1412 | 1124 | 0.796 | -0.329 | 0.00220157 | 0.01662193 | 0.0057 | 0.0095 | 0.0075 | 0.0075 | TRUE | 0.1115 |
| Dgkb | 26 | 7 | 25 | 30 | 19 | 7 | 28 | 7 | 25 | 24 | 20 | 8 | 18.57 | 20 | 17 | 0.874 | -0.195 | 0.76101017 | 0.87423727 | 0.3068 | 0.1843 | 0.2422 | 0.2422 | TRUE | 1.7597 |
| Dgkd | 4881 | 6679 | 7017 | 7503 | 5389 | 6204 | 5184 | 6670 | 7046 | 5898 | 5798 | 6791 | 6231.4 | 6300 | 6162 | 0.978 | -0.032 | 0.80753545 | 0.90093012 | 0.0166 | 0.0074 | 0.0123 | 0.0123 | TRUE | 0.2926 |
| Dgke | 1935 | 1634 | 1496 | 2080 | 1580 | 1427 | 2055 | 1632 | 1502 | 1635 | 1700 | 1562 | 1681.09 | 1730 | 1632 | 0.944 | -0.083 | 0.52013618 | 0.71106768 | 0.0129 | 0.0089 | 0.0115 | 0.0115 | TRUE | 0.3265 |
| Dgkg | 29 | 27 | 33 | 28 | 33 | 28 | 31 | 27 | 33 | 22 | 36 | 31 | 29.85 | 30 | 30 | 0.961 | -0.057 | 0.87647801 | 0.93924522 | 0 | 0.1173 | 0.063 | 0.063 | TRUE | 0.3861 |
| Dgkh | 1085 | 1197 | 1399 | 936 | 695 | 836 | 1152 | 1195 | 1405 | 736 | 748 | 915 | 1025.23 | 1251 | 800 | 0.639 | -0.647 | 6.53E-07 | 1.89E-05 | 0.0121 | 0.0101 | 0.0112 | 0.0112 | TRUE | 0.1878 |
| Dgki | 47 | 33 | 32 | 30 | 37 | 20 | 50 | 33 | 32 | 24 | 40 | 22 | 33.38 | 38 | 29 | 0.739 | -0.436 | 0.26817397 | 0.4879603 | 0.0546 | 0.1056 | 0.0812 | 0.0812 | TRUE | 0.7518 |
| Dgkk | 114 | 73 | 51 | 168 | 116 | 74 | 121 | 73 | 51 | 132 | 125 | 81 | 97.18 | 82 | 113 | 1.383 | 0.467 | 0.17227266 | 0.37111375 | 0.1121 | 0.0408 | 0.0739 | 0.0739 | TRUE | 0.83 |
| Dgkq | 884 | 859 | 795 | 1155 | 829 | 750 | 939 | 858 | 798 | 908 | 892 | 821 | 869.34 | 865 | 874 | 1.011 | 0.016 | 0.8805278 | 0.94090865 | 0.0035 | 0.0107 | 0.0065 | 0.0065 | TRUE | 0.066 |
| Dgkz | 3511 | 3959 | 4060 | 5175 | 3874 | 3605 | 3729 | 3954 | 4077 | 4068 | 4168 | 3946 | 3990.41 | 3920 | 4061 | 1.036 | 0.051 | 0.46994711 | 0.6718642 | 0.001 | 0.0077 | 0.0033 | 0.0033 | TRUE | 0.022 |
| Dguok | 412 | 589 | 555 | 662 | 497 | 539 | 438 | 588 | 557 | 520 | 535 | 590 | 538.05 | 528 | 548 | 1.038 | 0.054 | 0.70345707 | 0.83766892 | 0.0121 | 0.013 | 0.0125 | 0.0125 | TRUE | 0.2586 |
| Dhcr24 | 15472 | 12378 | 12344 | 18190 | 12885 | 11868 | 16434 | 12362 | 12395 | 14300 | 13863 | 12991 | 13724.19 | 13730 | 13718 | 0.999 | -0.001 | 0.99242006 | 0.99616826 | 0.0125 | 0.0071 | 0.0113 | 0.0113 | TRUE | 0.3628 |
| Dhcr7 | 10157 | 8882 | 8413 | 9540 | 6842 | 7229 | 10789 | 8871 | 8448 | 7500 | 7362 | 7913 | 8480.18 | 9369 | 7592 | 0.81 | -0.303 | 0.00510738 | 0.03185068 | 0.0094 | 0.0073 | 0.0083 | 0.0083 | TRUE | 0.2146 |
| Dhdds | 900 | 971 | 935 | 1277 | 969 | 921 | 956 | 970 | 939 | 1004 | 1043 | 1008 | 986.53 | 955 | 1018 | 1.066 | 0.092 | 0.24588271 | 0.46297247 | 0 | 0.0102 | 0.0035 | 0.0035 | TRUE | 0.0051 |
| Dhdh | 665 | 871 | 897 | 1040 | 829 | 891 | 706 | 870 | 901 | 818 | 892 | 975 | 860.29 | 826 | 895 | 1.083 | 0.115 | 0.37594098 | 0.59147378 | 0.011 | 0.0107 | 0.0109 | 0.0109 | TRUE | 0.1887 |
| Dhfr | 2495 | 2759 | 2796 | 4533 | 3149 | 3042 | 2650 | 2755 | 2807 | 3563 | 3388 | 3330 | 3082.44 | 2737 | 3427 | 1.252 | 0.324 | 3.00E-06 | 7.18E-05 | 8.00E-04 | 0.008 | 0.0031 | 0.0031 | TRUE | 0.015 |
| Dhh | 54 | 80 | 72 | 62 | 43 | 58 | 57 | 80 | 72 | 49 | 46 | 63 | 61.34 | 70 | 53 | 0.753 | -0.409 | 0.134005 | 0.31649098 | 0.011 | 0.0606 | 0.037 | 0.037 | TRUE | 0.2411 |
| Dhodh | 3986 | 4302 | 4538 | 4303 | 3431 | 3573 | 4234 | 4296 | 4557 | 3383 | 3692 | 3911 | 4012.06 | 4362 | 3662 | 0.839 | -0.253 | 0.00262717 | 0.01909482 | 0.003 | 0.0077 | 0.0049 | 0.0049 | TRUE | 0.0548 |
| Dhps | 1107 | 1076 | 1177 | 1871 | 1234 | 1236 | 1176 | 1075 | 1182 | 1471 | 1328 | 1353 | 1263.97 | 1144 | 1384 | 1.211 | 0.276 | 0.00221507 | 0.01666687 | 0.0022 | 0.0095 | 0.0051 | 0.0051 | TRUE | 0.0374 |
| Dhrs1 | 1187 | 1211 | 1192 | 1449 | 1040 | 1102 | 1261 | 1209 | 1197 | 1139 | 1119 | 1206 | 1188.59 | 1222 | 1155 | 0.945 | -0.082 | 0.30750076 | 0.52802417 | 0 | 0.0097 | 0.0038 | 0.0038 | TRUE | 0.0177 |
| Dhrs11 | 229 | 243 | 205 | 312 | 232 | 231 | 243 | 243 | 206 | 245 | 250 | 253 | 239.92 | 231 | 249 | 1.081 | 0.112 | 0.42518315 | 0.63533763 | 0 | 0.0206 | 0.01 | 0.01 | TRUE | 0.0979 |
| Dhrs13 | 626 | 833 | 777 | 1132 | 803 | 711 | 665 | 832 | 780 | 890 | 864 | 778 | 801.53 | 759 | 844 | 1.112 | 0.154 | 0.19979618 | 0.40732488 | 0.0076 | 0.011 | 0.0091 | 0.0091 | TRUE | 0.1384 |
| Dhrs2 | 0 | 0 | 2 | 2 | 0 | 1 | 0 | 0 | 2 | 2 | 0 | 1 | 0.78 | 1 | 1 | 1.346 | 0.429 | 0.8629169 | NA | 0 | 4.235 | 3.1534 | 3.1534 | TRUE | 0.9739 |
| Dhrs3 | 32 | 46 | 60 | 54 | 32 | 35 | 34 | 46 | 60 | 42 | 34 | 38 | 42.56 | 47 | 38 | 0.823 | -0.28 | 0.39619969 | 0.60964502 | 0.0241 | 0.0843 | 0.0551 | 0.0551 | TRUE | 0.1971 |
| Dhrs4 | 451 | 366 | 360 | 648 | 389 | 355 | 479 | 366 | 361 | 509 | 419 | 389 | 420.43 | 402 | 439 | 1.094 | 0.13 | 0.43806784 | 0.64579729 | 0.0207 | 0.0147 | 0.0178 | 0.0178 | TRUE | 0.3215 |
| Dhrs7 | 1137 | 3039 | 3179 | 1776 | 1404 | 2197 | 1208 | 3035 | 3192 | 1396 | 1511 | 2405 | 2124.43 | 2478 | 1771 | 0.714 | -0.485 | 0.31701247 | 0.53795592 | 0.169 | 0.0085 | 0.071 | 0.169 | TRUE | 2.4377 |
| Dhrs7b | 2367 | 2884 | 2912 | 3026 | 2170 | 2233 | 2514 | 2880 | 2924 | 2379 | 2335 | 2444 | 2579.39 | 2773 | 2386 | 0.86 | -0.217 | 0.01328947 | 0.06476267 | 0.003 | 0.0082 | 0.0051 | 0.0051 | TRUE | 0.0806 |
| Dhrs7c | 1 | 0 | 1 | 0 | 0 | 0 | 1 | 0 | 1 | 0 | 0 | 0 | 0.34 | 1 | 0 | 0.261 | -1.939 | 0.60367425 | NA | 0 | 9.5732 | 8.3134 | 8.3134 | TRUE | 0.2529 |
| Dhrs9 | 0 | 0 | 0 | 0 | 3 | 0 | 0 | 0 | 0 | 0 | 3 | 0 | 0.54 | 0 | 1 | 5.683 | 2.507 | 0.52963929 | NA | 10 | 6.1308 | 10 | 10 | TRUE | 1.4428 |
| Dhrsx | 231 | 255 | 253 | 419 | 321 | 308 | 245 | 255 | 254 | 329 | 345 | 337 | 294.33 | 251 | 337 | 1.341 | 0.423 | 0.00046118 | 0.00487838 | 0 | 0.0181 | 0.007 | 0.007 | TRUE | 0.0052 |
| Dhtkd1 | 2190 | 2687 | 2797 | 2509 | 1977 | 2004 | 2326 | 2684 | 2808 | 1972 | 2127 | 2194 | 2351.9 | 2606 | 2098 | 0.805 | -0.314 | 0.00179178 | 0.01410179 | 0.0054 | 0.0083 | 0.0068 | 0.0068 | TRUE | 0.1068 |
| Dhx15 | 17801 | 17657 | 17467 | 22466 | 15834 | 16128 | 18908 | 17634 | 17539 | 17661 | 17036 | 17655 | 17738.8 | 18027 | 17451 | 0.968 | -0.047 | 0.4621406 | 0.66589453 | 0.001 | 0.0071 | 0.0029 | 0.0029 | TRUE | 0.0223 |
| Dhx16 | 6457 | 6653 | 6413 | 9533 | 7052 | 6775 | 6858 | 6644 | 6439 | 7494 | 7588 | 7416 | 7073.36 | 6647 | 7499 | 1.128 | 0.174 | 0.00375247 | 0.02520611 | 3.00E-04 | 0.0074 | 0.0025 | 0.0025 | TRUE | 0.0094 |
| Dhx29 | 3990 | 3917 | 4051 | 5511 | 3930 | 3710 | 4238 | 3912 | 4068 | 4332 | 4228 | 4061 | 4139.93 | 4073 | 4207 | 1.033 | 0.047 | 0.49765671 | 0.69319452 | 0.001 | 0.0077 | 0.0033 | 0.0033 | TRUE | 0.0153 |
| Dhx30 | 8495 | 9116 | 9237 | 10124 | 7486 | 7327 | 9023 | 9104 | 9275 | 7959 | 8054 | 8021 | 8572.68 | 9134 | 8011 | 0.877 | -0.189 | 0.00041274 | 0.00444046 | 0 | 0.0073 | 0.002 | 0.002 | TRUE | 0.0022 |
| Dhx32 | 3556 | 3825 | 3776 | 4486 | 3029 | 3120 | 3777 | 3820 | 3792 | 3527 | 3259 | 3415 | 3598.26 | 3796 | 3400 | 0.896 | -0.158 | 0.0164228 | 0.07580343 | 5.00E-04 | 0.0078 | 0.0029 | 0.0029 | TRUE | 0.016 |
| Dhx33 | 3870 | 3175 | 3032 | 5011 | 3461 | 3058 | 4111 | 3171 | 3044 | 3939 | 3724 | 3347 | 3556.09 | 3442 | 3670 | 1.067 | 0.093 | 0.48536249 | 0.68258022 | 0.0145 | 0.0078 | 0.0125 | 0.0125 | TRUE | 0.3508 |
| Dhx34 | 567 | 866 | 862 | 912 | 714 | 736 | 602 | 865 | 866 | 717 | 768 | 806 | 770.58 | 778 | 764 | 0.981 | -0.027 | 0.86121422 | 0.93144485 | 0.021 | 0.0112 | 0.0162 | 0.0162 | TRUE | 0.4591 |
| Dhx35 | 2419 | 2271 | 2240 | 3022 | 2132 | 2154 | 2569 | 2268 | 2249 | 2376 | 2294 | 2358 | 2352.35 | 2362 | 2343 | 0.992 | -0.012 | 0.88931906 | 0.94606369 | 0.0025 | 0.0083 | 0.0047 | 0.0047 | TRUE | 0.0711 |
| Dhx36 | 7369 | 8428 | 8587 | 11027 | 8193 | 8168 | 7827 | 8417 | 8622 | 8669 | 8815 | 8941 | 8548.57 | 8289 | 8808 | 1.063 | 0.088 | 0.19499959 | 0.40087767 | 0.0012 | 0.0073 | 0.0032 | 0.0032 | TRUE | 0.0289 |
| Dhx37 | 2811 | 2146 | 2156 | 3321 | 2364 | 1939 | 2986 | 2143 | 2165 | 2611 | 2544 | 2123 | 2428.44 | 2431 | 2426 | 0.998 | -0.003 | 0.98463973 | 0.9919379 | 0.024 | 0.0083 | 0.0163 | 0.0163 | TRUE | 0.482 |
| Dhx38 | 5164 | 4304 | 4418 | 7675 | 5469 | 4606 | 5485 | 4298 | 4436 | 6033 | 5884 | 5042 | 5196.57 | 4740 | 5653 | 1.193 | 0.254 | 0.03869692 | 0.13741383 | 0.0136 | 0.0075 | 0.0107 | 0.0107 | TRUE | 0.2303 |
| Dhx40 | 5351 | 5712 | 5831 | 7145 | 5374 | 5549 | 5684 | 5705 | 5855 | 5617 | 5782 | 6074 | 5786.08 | 5748 | 5824 | 1.013 | 0.019 | 0.77129296 | 0.88080416 | 6.00E-04 | 0.0075 | 0.0028 | 0.0028 | TRUE | 0.0171 |
| Dhx57 | 1399 | 1331 | 1449 | 1838 | 1362 | 1268 | 1486 | 1329 | 1455 | 1445 | 1465 | 1388 | 1428.09 | 1423 | 1433 | 1.007 | 0.01 | 0.90635344 | 0.95350897 | 0.0014 | 0.0092 | 0.0044 | 0.0044 | TRUE | 0.0404 |
| Dhx58 | 171 | 172 | 181 | 147 | 116 | 132 | 182 | 172 | 182 | 116 | 125 | 144 | 153.34 | 179 | 128 | 0.717 | -0.481 | 0.00474747 | 0.03014373 | 0 | 0.0284 | 0.0142 | 0.0142 | TRUE | 0.1216 |
| Dhx8 | 8021 | 6421 | 6667 | 10975 | 7601 | 6503 | 8520 | 6413 | 6694 | 8628 | 8178 | 7119 | 7591.88 | 7209 | 7975 | 1.106 | 0.146 | 0.2668072 | 0.48648172 | 0.0162 | 0.0073 | 0.0123 | 0.0123 | TRUE | 0.3087 |
| Dhx9 | 22888 | 15121 | 14834 | 32232 | 22223 | 16543 | 24311 | 15101 | 14895 | 25338 | 23911 | 18109 | 20277.54 | 18102 | 22453 | 1.24 | 0.311 | 0.11981687 | 0.29353087 | 0.0562 | 0.0071 | 0.0287 | 0.0287 | TRUE | 1.1011 |
| Diablo | 2272 | 2342 | 2315 | 2861 | 2153 | 2045 | 2413 | 2339 | 2325 | 2249 | 2316 | 2239 | 2313.49 | 2359 | 2268 | 0.961 | -0.057 | 0.38987565 | 0.6037429 | 0 | 0.0083 | 0.0027 | 0.0027 | TRUE | 0.0049 |
| Diap1 | 9132 | 10106 | 10539 | 9681 | 7157 | 7948 | 9700 | 10093 | 10582 | 7610 | 7700 | 8700 | 9064.38 | 10125 | 8003 | 0.79 | -0.339 | 5.13E-05 | 0.0007922 | 0.0036 | 0.0073 | 0.0049 | 0.0049 | TRUE | 0.0706 |
| Diap2 | 1541 | 1748 | 1925 | 1944 | 1472 | 1554 | 1637 | 1746 | 1933 | 1528 | 1584 | 1701 | 1688.09 | 1772 | 1604 | 0.905 | -0.144 | 0.13852248 | 0.3230815 | 0.0042 | 0.0089 | 0.0062 | 0.0062 | TRUE | 0.0764 |
| Diap3 | 2124 | 2406 | 2616 | 3042 | 2151 | 2250 | 2256 | 2403 | 2627 | 2391 | 2314 | 2463 | 2409.07 | 2429 | 2389 | 0.984 | -0.024 | 0.78587838 | 0.89005269 | 0.003 | 0.0083 | 0.005 | 0.005 | TRUE | 0.0618 |
| Dicer1 | 11200 | 9800 | 10943 | 13420 | 9349 | 9320 | 11896 | 9787 | 10988 | 10550 | 10059 | 10202 | 10580.43 | 10890 | 10270 | 0.943 | -0.084 | 0.35189021 | 0.57063668 | 0.0049 | 0.0072 | 0.0058 | 0.0058 | TRUE | 0.096 |
| Dido1 | 7634 | 5731 | 5507 | 8578 | 6253 | 5208 | 8109 | 5724 | 5530 | 6743 | 6728 | 5701 | 6422.34 | 6454 | 6391 | 0.99 | -0.014 | 0.92814385 | 0.96472155 | 0.0233 | 0.0074 | 0.0174 | 0.0174 | TRUE | 0.6136 |
| Diexf | 2530 | 2047 | 2057 | 2797 | 1979 | 1721 | 2687 | 2044 | 2065 | 2199 | 2129 | 1884 | 2168.18 | 2265 | 2071 | 0.914 | -0.129 | 0.32587073 | 0.54706048 | 0.0155 | 0.0084 | 0.012 | 0.012 | TRUE | 0.3205 |
| Dimt1 | 1469 | 1609 | 1464 | 2098 | 1427 | 1402 | 1560 | 1607 | 1470 | 1649 | 1535 | 1535 | 1559.44 | 1546 | 1573 | 1.018 | 0.026 | 0.74779512 | 0.86607612 | 0.0013 | 0.009 | 0.0042 | 0.0042 | TRUE | 0.0223 |
| Dio1 | 0 | 1 | 1 | 0 | 1 | 0 | 0 | 1 | 1 | 0 | 1 | 0 | 0.51 | 1 | 0 | 0.52 | -0.943 | 0.73680291 | NA | 0 | 6.4273 | 3.9121 | 3.9121 | TRUE | 0.5204 |

| | | | | | | | | | | | | | | | | | | | | | | | | | |
|---|---|---|---|---|---|---|---|---|---|---|---|---|---|---|---|---|---|---|---|---|---|---|---|---|---|
| Dio2 | 0 | 0 | 0 | 0 | 1 | 0 | 0 | 0 | 0 | 0 | 1 | 0 | 0.18 | 0 | 0 | 1.859 | 0.894 | 0.82650734 | NA | 0 | 18.3785 | 10 | 10 | TRUE | 0.5204 |
| Dio3 | 29 | 32 | 23 | 54 | 33 | 22 | 31 | 32 | 23 | 42 | 36 | 24 | 31.32 | 29 | 34 | 1.198 | 0.261 | 0.48830328 | 0.6847621 | 0.0246 | 0.1121 | 0.07 | 0.07 | TRUE | 0.3096 |
| Dio3os | 0 | 5 | 3 | 5 | 4 | 3 | 0 | 5 | 3 | 4 | 4 | 3 | 3.25 | 3 | 4 | 1.429 | 0.515 | 0.6556166 | NA | 0 | 1.0193 | 0.6448 | 0.6448 | TRUE | 0.4315 |
| Dip2a | 2440 | 2658 | 2846 | 2945 | 2219 | 2264 | 2592 | 2655 | 2858 | 2315 | 2388 | 2478 | 2547.48 | 2702 | 2394 | 0.886 | -0.175 | 0.02306472 | 0.09626652 | 0.0013 | 0.0082 | 0.0039 | 0.0039 | TRUE | 0.0311 |
| Dip2b | 5131 | 4857 | 5629 | 5602 | 4022 | 3998 | 5450 | 4851 | 5652 | 4404 | 4327 | 4376 | 4843.43 | 5318 | 4369 | 0.822 | -0.283 | 0.00057353 | 0.00578738 | 0.0028 | 0.0076 | 0.0047 | 0.0047 | TRUE | 0.072 |
| Dip2c | 1949 | 2309 | 2444 | 2631 | 1834 | 1812 | 2070 | 2306 | 2454 | 2068 | 1973 | 1984 | 2142.55 | 2277 | 2008 | 0.882 | -0.181 | 0.04605556 | 0.15528248 | 0.0032 | 0.0084 | 0.0054 | 0.0054 | TRUE | 0.076 |
| Diras1 | 77 | 57 | 75 | 89 | 87 | 58 | 82 | 57 | 75 | 70 | 94 | 63 | 73.51 | 71 | 76 | 1.06 | 0.084 | 0.75367017 | 0.8700338 | 0.0236 | 0.0517 | 0.0381 | 0.0381 | TRUE | 0.3758 |
| Diras2 | 31 | 29 | 13 | 64 | 49 | 32 | 33 | 29 | 13 | 50 | 53 | 35 | 35.5 | 25 | 46 | 1.849 | 0.887 | 0.03174855 | 0.11986925 | 0.0813 | 0.0997 | 0.0918 | 0.0918 | TRUE | 1.0722 |
| Dirc2 | 1794 | 1968 | 1855 | 2104 | 1548 | 1558 | 1906 | 1965 | 1863 | 1654 | 1666 | 1705 | 1793.11 | 1911 | 1675 | 0.876 | -0.191 | 0.00662255 | 0.03861402 | 0 | 0.0087 | 0.003 | 0.003 | TRUE | 0.0075 |
| Dis3 | 8470 | 7106 | 7292 | 9226 | 6687 | 6184 | 8997 | 7097 | 7322 | 7253 | 7195 | 6769 | 7438.72 | 7805 | 7072 | 0.906 | -0.142 | 0.18956836 | 0.3947026 | 0.0093 | 0.0073 | 0.0083 | 0.0083 | TRUE | 0.2176 |
| Dis3l | 1493 | 1907 | 2008 | 3334 | 2727 | 2530 | 1586 | 1905 | 2016 | 2621 | 2934 | 2769 | 2305.18 | 1836 | 2775 | 1.511 | 0.596 | 9.47E-08 | 3.42E-06 | 0.0083 | 0.0083 | 0.0085 | 0.0085 | TRUE | 0.1706 |
| Dis3l2 | 1132 | 1529 | 1603 | 1393 | 1056 | 1174 | 1202 | 1527 | 1610 | 1095 | 1136 | 1285 | 1309.23 | 1446 | 1172 | 0.81 | -0.304 | 0.02307642 | 0.09626652 | 0.0136 | 0.0094 | 0.0121 | 0.0121 | TRUE | 0.2612 |
| Disc1 | 33 | 33 | 34 | 61 | 47 | 28 | 35 | 33 | 34 | 48 | 51 | 31 | 38.55 | 34 | 43 | 1.27 | 0.345 | 0.29953482 | 0.51972817 | 0 | 0.0924 | 0.0532 | 0.0532 | TRUE | 0.454 |
| Disp1 | 916 | 793 | 767 | 1003 | 719 | 739 | 973 | 792 | 770 | 788 | 774 | 809 | 817.69 | 845 | 790 | 0.936 | -0.096 | 0.41842863 | 0.62905885 | 0.0071 | 0.0109 | 0.0089 | 0.0089 | TRUE | 0.2075 |
| Disp2 | 39 | 19 | 14 | 51 | 21 | 14 | 41 | 19 | 14 | 40 | 23 | 15 | 25.41 | 25 | 26 | 1.06 | 0.085 | 0.87892145 | 0.94007522 | 0.2312 | 0.1365 | 0.1829 | 0.1829 | TRUE | 0.982 |
| Dixdc1 | 680 | 880 | 915 | 762 | 563 | 690 | 722 | 879 | 919 | 599 | 606 | 755 | 746.67 | 840 | 653 | 0.777 | -0.364 | 0.01093178 | 0.05627155 | 0.0147 | 0.0113 | 0.0134 | 0.0134 | TRUE | 0.2171 |
| Dkc1 | 11431 | 11723 | 11235 | 14091 | 10092 | 11027 | 12142 | 11708 | 11281 | 11077 | 10858 | 12071 | 11522.86 | 11710 | 11335 | 0.968 | -0.047 | 0.52597243 | 0.71529414 | 0.002 | 0.0072 | 0.0039 | 0.0039 | TRUE | 0.0392 |
| Dkk1 | 6 | 0 | 5 | 23 | 7 | 10 | 6 | 0 | 5 | 18 | 8 | 11 | 7.99 | 4 | 12 | 3.253 | 1.702 | 0.05943508 | 0.18463857 | 0.3962 | 0.4191 | 0.4122 | 0.4122 | TRUE | 0.7457 |
| Dkk2 | 0 | 0 | 0 | 0 | 0 | 0 | 0 | 0 | 0 | 0 | 0 | 0 | 0 | 0 | 0 | NA | NA | NA | NA | NA | NA | NA | NA | NA | NA |
| Dkk3 | 811 | 1238 | 1270 | 786 | 624 | 767 | 861 | 1236 | 1275 | 618 | 671 | 840 | 916.99 | 1124 | 710 | 0.631 | -0.665 | 0.00021059 | 0.00253352 | 0.0322 | 0.0105 | 0.0221 | 0.0221 | TRUE | 0.4999 |
| Dkk4 | 0 | 1 | 1 | 0 | 1 | 1 | 0 | 1 | 1 | 0 | 1 | 1 | 0.7 | 1 | 1 | 1.033 | 0.047 | 0.9836759 | NA | 0 | 4.7432 | 2.3808 | 2.3808 | TRUE | 0.3742 |
| Dkkl1 | 108 | 141 | 155 | 123 | 72 | 107 | 115 | 141 | 156 | 97 | 77 | 117 | 117.08 | 137 | 97 | 0.708 | -0.499 | 0.02881716 | 0.11186421 | 0.0219 | 0.035 | 0.0288 | 0.0288 | TRUE | 0.2618 |
| Dlat | 5276 | 5968 | 5790 | 7280 | 5086 | 5154 | 5604 | 5960 | 5814 | 5723 | 5472 | 5642 | 5702.53 | 5793 | 5612 | 0.969 | -0.046 | 0.46928089 | 0.67144012 | 5.00E-04 | 0.0075 | 0.0027 | 0.0027 | TRUE | 0.0099 |
| Dlc1 | 14391 | 16497 | 17521 | 13652 | 10952 | 12050 | 15286 | 16476 | 17593 | 10732 | 11784 | 13191 | 14176.81 | 16452 | 11902 | 0.723 | -0.467 | 5.04E-06 | 0.00011116 | 0.0077 | 0.0071 | 0.0075 | 0.0075 | TRUE | 0.1096 |
| Dld | 6670 | 6767 | 6815 | 8919 | 6280 | 6296 | 7085 | 6758 | 6843 | 7011 | 6757 | 6892 | 6891.04 | 6895 | 6887 | 0.999 | -0.002 | 0.97842171 | 0.98828002 | 3.00E-04 | 0.0074 | 0.0024 | 0.0024 | TRUE | 0.007 |
| Dlec1 | 43 | 56 | 52 | 65 | 47 | 46 | 46 | 56 | 52 | 51 | 51 | 50 | 50.97 | 51 | 51 | 0.988 | -0.018 | 0.94699537 | 0.97351104 | 0 | 0.0715 | 0.0313 | 0.0313 | TRUE | 0.0724 |
| Dleu2 | 9 | 12 | 5 | 16 | 10 | 12 | 10 | 12 | 5 | 13 | 11 | 13 | 10.51 | 9 | 12 | 1.375 | 0.459 | 0.44620996 | 0.65236459 | 0 | 0.3205 | 0.1641 | 0.1641 | TRUE | 0.4643 |
| Dleu7 | 27 | 40 | 28 | 56 | 46 | 45 | 29 | 40 | 28 | 44 | 49 | 49 | 39.92 | 32 | 47 | 1.471 | 0.556 | 0.07693166 | 0.2190587 | 0 | 0.0894 | 0.0452 | 0.0452 | TRUE | 0.3038 |
| Dlg1 | 5945 | 6134 | 6579 | 8013 | 5744 | 5799 | 6315 | 6126 | 6606 | 6299 | 6180 | 6348 | 6312.34 | 6349 | 6276 | 0.988 | -0.017 | 0.79178239 | 0.89301263 | 5.00E-04 | 0.0074 | 0.0027 | 0.0027 | TRUE | 0.0153 |
| Dlg2 | 0 | 2 | 2 | 1 | 2 | 1 | 0 | 2 | 2 | 1 | 2 | 1 | 1.34 | 1 | 1 | 0.983 | -0.024 | 0.988347 | NA | 0 | 2.466 | 1.2122 | 1.2122 | TRUE | 0.4385 |
| Dlg3 | 1716 | 1812 | 1949 | 1915 | 1460 | 1453 | 1823 | 1810 | 1957 | 1505 | 1571 | 1591 | 1709.36 | 1863 | 1556 | 0.835 | -0.261 | 0.00072701 | 0.00698182 | 8.00E-04 | 0.0088 | 0.0037 | 0.0037 | TRUE | 0.0235 |
| Dlg4 | 403 | 412 | 461 | 250 | 183 | 211 | 428 | 411 | 463 | 197 | 197 | 231 | 321.14 | 434 | 208 | 0.478 | -1.064 | 1.93E-15 | 2.82E-13 | 0.0021 | 0.0172 | 0.0094 | 0.0094 | TRUE | 0.0964 |
| Dlg5 | 3184 | 3649 | 3966 | 4328 | 3161 | 3393 | 3382 | 3644 | 3982 | 3402 | 3401 | 3714 | 3587.68 | 3669 | 3506 | 0.955 | -0.066 | 0.46771745 | 0.67001268 | 0.0042 | 0.0078 | 0.0057 | 0.0057 | TRUE | 0.0676 |
| Dlgap1 | 158 | 74 | 66 | 86 | 70 | 55 | 168 | 74 | 66 | 68 | 75 | 60 | 85.19 | 103 | 68 | 0.66 | -0.599 | 0.10659782 | 0.27266637 | 0.1317 | 0.0456 | 0.0869 | 0.0869 | TRUE | 2.9948 |
| Dlgap2 | 36 | 54 | 56 | 73 | 42 | 35 | 38 | 54 | 56 | 57 | 45 | 38 | 48.21 | 49 | 47 | 0.955 | -0.067 | 0.83047122 | 0.91410371 | 0.0217 | 0.0752 | 0.0493 | 0.0493 | TRUE | 0.3413 |
| Dlgap3 | 2574 | 2650 | 2653 | 2219 | 1737 | 1607 | 2734 | 2647 | 2664 | 1744 | 1869 | 1759 | 2236.16 | 2682 | 1791 | 0.668 | -0.583 | 1.31E-16 | 2.18E-14 | 3.00E-04 | 0.0084 | 0.0031 | 0.0031 | TRUE | 0.0174 |
| Dlgap4 | 3183 | 3365 | 3467 | 3593 | 2638 | 2694 | 3381 | 3361 | 3481 | 2825 | 2838 | 2949 | 3139.11 | 3408 | 2871 | 0.842 | -0.248 | 0.0001033 | 0.00141301 | 0 | 0.008 | 0.0026 | 0.0026 | TRUE | 0.0069 |
| Dlgap5 | 7002 | 8487 | 8997 | 9654 | 7116 | 7859 | 7437 | 8476 | 9034 | 7589 | 7656 | 8603 | 8132.64 | 8316 | 7949 | 0.956 | -0.065 | 0.51921163 | 0.71041952 | 0.0064 | 0.0073 | 0.0072 | 0.0072 | TRUE | 0.1042 |
| Dlk1 | 422 | 391 | 386 | 443 | 301 | 242 | 448 | 390 | 388 | 348 | 324 | 265 | 360.56 | 409 | 312 | 0.766 | -0.384 | 0.00918271 | 0.04934573 | 0.0099 | 0.016 | 0.0129 | 0.0129 | TRUE | 0.1954 |
| Dlk2 | 4 | 3 | 7 | 9 | 6 | 7 | 4 | 3 | 7 | 7 | 6 | 8 | 5.91 | 5 | 7 | 1.484 | 0.569 | 0.46407016 | 0.66733357 | 0 | 0.5642 | 0.2591 | 0.2591 | TRUE | 0.3444 |
| Dll1 | 258 | 653 | 647 | 114 | 100 | 165 | 274 | 652 | 650 | 90 | 108 | 181 | 325.61 | 525 | 126 | 0.239 | -2.065 | 2.04E-09 | 1.16E-07 | 0.1688 | 0.017 | 0.0806 | 0.0806 | TRUE | 2.0387 |
| Dll3 | 224 | 239 | 224 | 216 | 137 | 160 | 238 | 239 | 225 | 170 | 147 | 175 | 198.98 | 234 | 164 | 0.703 | -0.509 | 0.00068319 | 0.00664316 | 0 | 0.0235 | 0.011 | 0.011 | TRUE | 0.0796 |
| Dll4 | 6 | 9 | 15 | 10 | 10 | 9 | 6 | 9 | 15 | 8 | 11 | 10 | 9.82 | 10 | 10 | 0.928 | -0.108 | 0.86313626 | 0.9324082 | 0 | 0.3425 | 0.1829 | 0.1829 | TRUE | 0.6213 |
| Dlst | 9987 | 10996 | 11033 | 13097 | 9326 | 9490 | 10608 | 10982 | 11078 | 10296 | 10034 | 10388 | 10564.4 | 10889 | 10239 | 0.94 | -0.089 | 0.11974699 | 0.29345372 | 3.00E-04 | 0.0072 | 0.0023 | 0.0023 | TRUE | 0.0062 |
| Dlx1 | 1 | 2 | 2 | 2 | 9 | 4 | 1 | 2 | 2 | 2 | 10 | 4 | 3.45 | 2 | 5 | 3.022 | 1.595 | 0.17399014 | NA | 0 | 0.9617 | 0.5956 | 0.5956 | TRUE | 0.2234 |
| Dlx1as | 0 | 0 | 0 | 0 | 0 | 0 | 0 | 0 | 0 | 0 | 0 | 0 | 0 | 0 | 0 | NA | NA | NA | NA | NA | NA | NA | NA | NA | NA |
| Dlx2 | 12 | 12 | 12 | 23 | 26 | 16 | 13 | 12 | 12 | 18 | 28 | 18 | 16.72 | 12 | 21 | 1.721 | 0.784 | 0.10320035 | 0.26735607 | 0 | 0.2039 | 0.1018 | 0.1018 | TRUE | 0.4122 |
| Dlx3 | 88 | 143 | 122 | 185 | 125 | 143 | 93 | 143 | 123 | 145 | 134 | 157 | 132.54 | 120 | 145 | 1.215 | 0.281 | 0.17948225 | 0.3805964 | 0.0154 | 0.0318 | 0.0239 | 0.0239 | TRUE | 0.3588 |
| Dlx4 | 34 | 47 | 44 | 20 | 17 | 19 | 36 | 47 | 44 | 16 | 18 | 21 | 30.34 | 42 | 18 | 0.427 | -1.227 | 0.00072775 | 0.00698182 | 0 | 0.1155 | 0.0561 | 0.0561 | TRUE | 0.1254 |
| Dlx5 | 1 | 0 | 0 | 1 | 1 | 0 | 1 | 0 | 0 | 1 | 1 | 0 | 0.49 | 0 | 1 | 1.712 | 0.776 | 0.78488768 | NA | 0 | 6.7665 | 4.0387 | 4.0387 | TRUE | 0.4614 |
| Dlx6 | 0 | 0 | 1 | 9 | 4 | 1 | 0 | 0 | 1 | 7 | 4 | 1 | 2.25 | 0 | 4 | 11.519 | 3.526 | 0.04186847 | NA | 0 | 1.4735 | 1.0474 | 1.0474 | TRUE | 0.2111 |
| Dlx6as1 | 0 | 2 | 0 | 0 | 0 | 0 | 0 | 2 | 0 | 0 | 0 | 0 | 0.33 | 1 | 0 | 0.268 | -1.897 | 0.63866967 | NA | 0 | 9.903 | 10 | 10 | TRUE | 0.9739 |
| Dlx6as2 | 0 | 0 | 0 | 0 | 0 | 0 | 0 | 0 | 0 | 0 | 0 | 0 | 0 | 0 | 0 | NA | NA | NA | NA | NA | NA | NA | NA | NA | NA |
| Dmap1 | 1642 | 1752 | 1702 | 2388 | 1746 | 1727 | 1744 | 1750 | 1709 | 1877 | 1879 | 1890 | 1808.19 | 1734 | 1882 | 1.085 | 0.118 | 0.07704425 | 0.2193397 | 0 | 0.0087 | 0.0027 | 0.0027 | TRUE | 0.002 |
| Dmbt1 | 0 | 0 | 0 | 2 | 5 | 2 | 0 | 0 | 0 | 2 | 5 | 2 | 1.52 | 0 | 3 | 16.091 | 4.008 | 0.03962786 | NA | 0 | 2.1693 | 1.5722 | 1.5722 | TRUE | 0.5576 |
| Dmbx1 | 39 | 75 | 60 | 118 | 90 | 111 | 41 | 75 | 60 | 93 | 97 | 122 | 81.28 | 59 | 104 | 1.755 | 0.811 | 0.00329242 | 0.02274555 | 0.0345 | 0.0474 | 0.0415 | 0.0415 | TRUE | 0.4499 |
| Dmc1 | 406 | 502 | 519 | 526 | 409 | 462 | 431 | 501 | 521 | 413 | 440 | 506 | 468.84 | 484 | 453 | 0.933 | -0.1 | 0.45650748 | 0.66122104 | 0.0081 | 0.0139 | 0.0108 | 0.0108 | TRUE | 0.1184 |
| Dmd | 2004 | 2453 | 2722 | 2653 | 1919 | 2095 | 2129 | 2450 | 2733 | 2086 | 2065 | 2293 | 2292.54 | 2437 | 2148 | 0.881 | -0.183 | 0.10505631 | 0.2702574 | 0.0088 | 0.0083 | 0.0087 | 0.0087 | TRUE | 0.1484 |
| Dmgdh | 78 | 81 | 60 | 86 | 63 | 81 | 83 | 81 | 60 | 68 | 68 | 89 | 74.67 | 75 | 75 | 0.998 | -0.003 | 0.98990905 | 0.99457546 | 0.0135 | 0.051 | 0.0329 | 0.0329 | TRUE | 0.263 |
| Dmkn | 13 | 8 | 4 | 10 | 4 | 9 | 14 | 8 | 4 | 8 | 4 | 10 | 7.97 | 9 | 7 | 0.857 | -0.222 | 0.76603089 | 0.87753045 | 0.1172 | 0.4202 | 0.2753 | 0.2753 | TRUE | 0.27 |
| Dmp1 | 0 | 0 | 0 | 3 | 0 | 0 | 0 | 0 | 0 | 2 | 0 | 0 | 0.39 | 0 | 1 | 4.278 | 2.097 | 0.60136958 | NA | 10 | 8.3884 | 10 | 10 | TRUE | 1.4411 |
| Dmpk | 100 | 82 | 63 | 124 | 101 | 67 | 106 | 82 | 63 | 97 | 109 | 73 | 88.48 | 84 | 93 | 1.114 | 0.156 | 0.5659648 | 0.74604236 | 0.0399 | 0.0441 | 0.0421 | 0.0421 | TRUE | 0.2904 |
| Dmrt1 | 1070 | 2059 | 2212 | 3091 | 2396 | 2501 | 1137 | 2056 | 2221 | 2430 | 2578 | 2738 | 2193.26 | 1805 | 2582 | 1.43 | 0.516 | 0.01542678 | 0.07228497 | 0.0621 | 0.0084 | 0.0323 | 0.0323 | TRUE | 1.2652 |
| Dmrt2 | 24 | 58 | 69 | 79 | 43 | 46 | 25 | 58 | 69 | 62 | 46 | 50 | 51.9 | 51 | 53 | 1.041 | 0.058 | 0.8813797 | 0.94135674 | 0.1089 | 0.0704 | 0.0883 | 0.0883 | TRUE | 0.9311 |
| Dmrt3 | 55 | 104 | 112 | 204 | 167 | 170 | 58 | 104 | 112 | 160 | 180 | 186 | 133.48 | 91 | 175 | 1.91 | 0.933 | 0.00022288 | 0.00264906 | 0.0455 | 0.0316 | 0.0377 | 0.0377 | TRUE | 0.9416 |
| Dmrta1 | 3 | 10 | 4 | 13 | 6 | 1 | 3 | 10 | 4 | 10 | 6 | 1 | 5.83 | 6 | 6 | 1.051 | 0.072 | 0.94035508 | 0.96983421 | 0.4099 | 0.5723 | 0.4943 | 0.4943 | TRUE | 0.276 |
| Dmrta2 | 63 | 99 | 105 | 175 | 113 | 80 | 67 | 99 | 105 | 138 | 122 | 88 | 102.99 | 90 | 116 | 1.281 | 0.358 | 0.17848879 | 0.37940773 | 0.0435 | 0.0389 | 0.0411 | 0.0411 | TRUE | 0.484 |
| Dmrtb1 | 137 | 186 | 164 | 365 | 304 | 218 | 146 | 186 | 165 | 287 | 327 | 239 | 224.77 | 166 | 284 | 1.719 | 0.781 | 1.30E-05 | 0.00024846 | 0.0152 | 0.0216 | 0.0183 | 0.0183 | TRUE | 0.1448 |
| Dmrtc1a | 2 | 0 | 1 | 2 | 0 | 0 | 2 | 0 | 1 | 2 | 0 | 0 | 0.78 | 1 | 1 | 0.543 | -0.88 | 0.72386184 | NA | 0 | 4.212 | 3.1127 | 3.1127 | TRUE | 0.402 |

| | | | | | | | | | | | | | | | | | | | | | | | | | |
|---|---|---|---|---|---|---|---|---|---|---|---|---|---|---|---|---|---|---|---|---|---|---|---|---|---|
| Dmrtc1b | 126 | 83 | 79 | 169 | 171 | 149 | 134 | 83 | 79 | 133 | 184 | 163 | 129.33 | 99 | 160 | 1.62 | 0.696 | 0.00677816 | 0.03915941 | 0.0484 | 0.0324 | 0.0393 | 0.0393 | TRUE | 0.8416 |
| Dmrtc1c1 | 0 | 0 | 0 | 0 | 0 | 0 | 0 | 0 | 0 | 0 | 0 | 0 | 0 | 0 | 0 | NA | NA | NA | NA | NA | NA | NA | NA | NA | NA |
| Dmrtc1c2 | 0 | 0 | 0 | 0 | 0 | 0 | 0 | 0 | 0 | 0 | 0 | 0 | 0 | 0 | 0 | NA | NA | NA | NA | NA | NA | NA | NA | NA | NA |
| Dmrtc2 | 84 | 183 | 178 | 95 | 88 | 135 | 89 | 183 | 179 | 75 | 95 | 148 | 127.98 | 150 | 106 | 0.701 | -0.513 | 0.1310203 | 0.31166855 | 0.1248 | 0.0326 | 0.0751 | 0.0751 | TRUE | 1.2544 |
| Dmtf1 | 4605 | 4218 | 4203 | 5672 | 4222 | 4072 | 4891 | 4213 | 4220 | 4459 | 4543 | 4457 | 4463.85 | 4441 | 4486 | 1.01 | 0.015 | 0.86473722 | 0.93305133 | 0.0036 | 0.0076 | 0.0051 | 0.0051 | TRUE | 0.0955 |
| Dmwd | 2820 | 3002 | 2885 | 3777 | 2960 | 2803 | 2995 | 2998 | 2897 | 2969 | 3185 | 3068 | 3018.77 | 2963 | 3074 | 1.037 | 0.052 | 0.43669784 | 0.64496661 | 5.00E-04 | 0.008 | 0.0029 | 0.0029 | TRUE | 0.0121 |
| Dmxl1 | 4848 | 4354 | 4795 | 5887 | 4171 | 4080 | 5149 | 4348 | 4815 | 4628 | 4488 | 4466 | 4649.06 | 4771 | 4527 | 0.949 | -0.075 | 0.37914242 | 0.59423972 | 0.0034 | 0.0076 | 0.0051 | 0.0051 | TRUE | 0.0732 |
| Dmxl2 | 3403 | 3159 | 3589 | 4195 | 2991 | 3030 | 3615 | 3155 | 3604 | 3298 | 3218 | 3317 | 3367.66 | 3458 | 3278 | 0.948 | -0.077 | 0.35309873 | 0.57159736 | 0.0026 | 0.0079 | 0.0047 | 0.0047 | TRUE | 0.0715 |
| Dna2 | 3858 | 3649 | 3734 | 5237 | 3692 | 3611 | 4098 | 3644 | 3749 | 4117 | 3972 | 3953 | 3922.26 | 3830 | 4014 | 1.048 | 0.068 | 0.37244896 | 0.58896796 | 0.0018 | 0.0077 | 0.0039 | 0.0039 | TRUE | 0.0453 |
| Dnaaf1 | 32 | 34 | 33 | 44 | 42 | 35 | 34 | 34 | 33 | 35 | 45 | 38 | 36.53 | 34 | 39 | 1.163 | 0.218 | 0.48918759 | 0.68548482 | 0 | 0.0971 | 0.044 | 0.044 | TRUE | 0.1215 |
| Dnaaf2 | 1170 | 1280 | 1271 | 1566 | 1175 | 1190 | 1243 | 1278 | 1276 | 1231 | 1264 | 1303 | 1265.87 | 1266 | 1266 | 1 | -0.001 | 0.99301023 | 0.99616826 | 0 | 0.0095 | 0.0033 | 0.0033 | TRUE | 0.0077 |
| Dnaaf3 | 557 | 555 | 540 | 462 | 378 | 461 | 592 | 554 | 542 | 363 | 407 | 505 | 493.78 | 563 | 425 | 0.753 | -0.409 | 0.00493638 | 0.03107714 | 0.0125 | 0.0136 | 0.0132 | 0.0132 | TRUE | 0.3094 |
| Dnahc1 | 249 | 368 | 333 | 252 | 216 | 248 | 264 | 368 | 334 | 198 | 232 | 271 | 278.06 | 322 | 234 | 0.724 | -0.466 | 0.0114239 | 0.05814061 | 0.0227 | 0.0188 | 0.0208 | 0.0208 | TRUE | 0.2757 |
| Dnahc10 | 201 | 230 | 251 | 269 | 191 | 194 | 213 | 230 | 252 | 211 | 206 | 212 | 220.76 | 232 | 210 | 0.905 | -0.144 | 0.31274838 | 0.5332037 | 0 | 0.0218 | 0.0101 | 0.0101 | TRUE | 0.0648 |
| Dnahc11 | 64 | 45 | 37 | 149 | 91 | 63 | 68 | 45 | 37 | 117 | 98 | 69 | 72.35 | 50 | 95 | 1.903 | 0.928 | 0.00388327 | 0.02575565 | 0.0665 | 0.0524 | 0.0591 | 0.0591 | TRUE | 0.3168 |
| Dnahc17 | 183 | 207 | 204 | 238 | 197 | 208 | 194 | 207 | 205 | 187 | 212 | 228 | 205.45 | 202 | 209 | 1.031 | 0.044 | 0.77257714 | 0.88119062 | 0 | 0.0229 | 0.0116 | 0.0116 | TRUE | 0.0996 |
| Dnahc2 | 347 | 383 | 424 | 557 | 470 | 458 | 369 | 383 | 426 | 438 | 506 | 501 | 436.96 | 393 | 482 | 1.225 | 0.293 | 0.01759629 | 0.07915819 | 0.004 | 0.0144 | 0.0087 | 0.0087 | TRUE | 0.0776 |
| Dnahc5 | 22 | 17 | 21 | 45 | 37 | 27 | 23 | 17 | 21 | 35 | 40 | 30 | 27.7 | 20 | 35 | 1.708 | 0.773 | 0.03836558 | 0.13646862 | 0 | 0.1259 | 0.0613 | 0.0613 | TRUE | 0.1242 |
| Dnahc6 | 71 | 101 | 89 | 158 | 99 | 96 | 75 | 101 | 89 | 124 | 107 | 105 | 100.24 | 88 | 112 | 1.267 | 0.341 | 0.10894116 | 0.2765969 | 0 | 0.0398 | 0.0225 | 0.0225 | TRUE | 0.1564 |
| Dnahc7a | 2 | 2 | 1 | 6 | 9 | 1 | 2 | 2 | 1 | 5 | 10 | 1 | 3.44 | 2 | 5 | 3.026 | 1.598 | 0.18000751 | NA | 0 | 0.9655 | 0.6287 | 0.6287 | TRUE | 0.1263 |
| Dnahc7b | 8 | 3 | 4 | 25 | 24 | 16 | 8 | 3 | 4 | 20 | 26 | 18 | 13.08 | 5 | 21 | 4.086 | 2.031 | 0.0010438 | 0.00921772 | 5.00E-04 | 0.2587 | 0.1536 | 0.1536 | TRUE | 0.6243 |
| Dnahc8 | 6001 | 9120 | 11591 | 2769 | 2336 | 3094 | 6374 | 9108 | 11639 | 2177 | 2513 | 3387 | 5866.33 | 9040 | 2692 | 0.298 | -1.748 | 1.44E-15 | 2.12E-13 | 0.069 | 0.0075 | 0.0343 | 0.0343 | TRUE | 0.0859 |
| Dnahc9 | 17 | 11 | 6 | 13 | 7 | 7 | 18 | 11 | 6 | 10 | 8 | 8 | 10.08 | 12 | 9 | 0.733 | -0.449 | 0.4908653 | 0.68729309 | 0 | 0.3337 | 0.2048 | 0.2048 | TRUE | 0.2276 |
| Dnaic1 | 153 | 193 | 168 | 182 | 143 | 137 | 163 | 193 | 169 | 143 | 154 | 150 | 161.81 | 175 | 149 | 0.851 | -0.232 | 0.15457778 | 0.34660912 | 0 | 0.0273 | 0.013 | 0.013 | TRUE | 0.0885 |
| Dnaic2 | 279 | 428 | 438 | 340 | 250 | 309 | 296 | 427 | 440 | 267 | 269 | 338 | 339.68 | 388 | 291 | 0.75 | -0.414 | 0.02872725 | 0.11159757 | 0.0291 | 0.0166 | 0.0229 | 0.0229 | TRUE | 0.486 |
| Dnaja1 | 6013 | 5212 | 5019 | 10878 | 7543 | 6228 | 6387 | 5205 | 5040 | 8551 | 8116 | 6818 | 6686.08 | 5544 | 7828 | 1.412 | 0.498 | 9.34E-05 | 0.00130539 | 0.0152 | 0.0074 | 0.0116 | 0.0116 | TRUE | 0.2156 |
| Dnaja2 | 14930 | 15253 | 14992 | 20189 | 14456 | 14144 | 15858 | 15233 | 15054 | 15871 | 15554 | 15483 | 15508.79 | 15382 | 15636 | 1.017 | 0.024 | 0.67534644 | 0.81916205 | 4.00E-04 | 0.0071 | 0.0022 | 0.0022 | TRUE | 0.009 |
| Dnaja3 | 5249 | 5465 | 5402 | 6510 | 4812 | 4789 | 5575 | 5458 | 5424 | 5118 | 5177 | 5242 | 5332.48 | 5486 | 5179 | 0.944 | -0.083 | 0.14364595 | 0.33045273 | 0 | 0.0075 | 0.0021 | 0.0021 | TRUE | 0.0025 |
| Dnaja4 | 368 | 389 | 377 | 526 | 402 | 375 | 391 | 388 | 379 | 413 | 433 | 410 | 402.41 | 386 | 419 | 1.085 | 0.118 | 0.27178233 | 0.49174612 | 0 | 0.0151 | 0.0057 | 0.0057 | TRUE | 0.0091 |
| Dnajb1 | 2319 | 2913 | 2943 | 4535 | 3457 | 3443 | 2463 | 2909 | 2955 | 3565 | 3720 | 3769 | 3230.17 | 2776 | 3685 | 1.327 | 0.408 | 1.66E-05 | 0.00030445 | 0.0047 | 0.0079 | 0.0062 | 0.0062 | TRUE | 0.1175 |
| Dnajb11 | 3304 | 3137 | 2994 | 4339 | 3060 | 2652 | 3509 | 3133 | 3006 | 3411 | 3292 | 2903 | 3209.18 | 3216 | 3202 | 0.996 | -0.006 | 0.95422498 | 0.97670429 | 0.0065 | 0.0079 | 0.0071 | 0.0071 | TRUE | 0.0808 |
| Dnajb12 | 832 | 814 | 805 | 1440 | 1134 | 1010 | 884 | 813 | 808 | 1132 | 1220 | 1106 | 993.79 | 835 | 1153 | 1.38 | 0.465 | 4.01E-07 | 1.23E-05 | 0.0015 | 0.0102 | 0.005 | 0.005 | TRUE | 0.0309 |
| Dnajb13 | 75 | 83 | 71 | 105 | 61 | 60 | 80 | 83 | 71 | 83 | 66 | 66 | 74.62 | 78 | 72 | 0.92 | -0.121 | 0.60654048 | 0.77676352 | 0 | 0.0511 | 0.0261 | 0.0261 | TRUE | 0.1946 |
| Dnajb14 | 143 | 204 | 207 | 299 | 232 | 244 | 152 | 204 | 208 | 235 | 250 | 267 | 219.21 | 188 | 251 | 1.331 | 0.413 | 0.0155749 | 0.07282754 | 0.0105 | 0.0219 | 0.0163 | 0.0163 | TRUE | 0.2971 |
| Dnajb2 | 447 | 495 | 516 | 772 | 557 | 547 | 475 | 494 | 518 | 607 | 599 | 599 | 548.71 | 496 | 602 | 1.214 | 0.279 | 0.00428284 | 0.02786626 | 0 | 0.0129 | 0.005 | 0.005 | TRUE | 0.0181 |
| Dnajb3 | 19 | 23 | 21 | 73 | 42 | 31 | 20 | 23 | 21 | 57 | 45 | 34 | 33.46 | 21 | 45 | 2.139 | 1.097 | 0.00282804 | 0.02021934 | 0 | 0.1054 | 0.0627 | 0.0627 | TRUE | 0.1216 |
| Dnajb4 | 588 | 402 | 362 | 459 | 346 | 322 | 625 | 401 | 363 | 361 | 372 | 352 | 412.52 | 463 | 362 | 0.782 | -0.355 | 0.08286842 | 0.23026793 | 0.0431 | 0.0149 | 0.0278 | 0.0278 | TRUE | 1.0743 |
| Dnajb5 | 761 | 935 | 937 | 904 | 650 | 721 | 808 | 934 | 941 | 711 | 699 | 789 | 813.7 | 894 | 733 | 0.819 | -0.288 | 0.00798938 | 0.04426009 | 0.0045 | 0.011 | 0.0073 | 0.0073 | TRUE | 0.0837 |
| Dnajb6 | 3342 | 3272 | 3136 | 5107 | 3609 | 3312 | 3550 | 3268 | 3149 | 4015 | 3883 | 3626 | 3581.62 | 3322 | 3841 | 1.157 | 0.21 | 0.01197357 | 0.06006589 | 0.0028 | 0.0078 | 0.0047 | 0.0047 | TRUE | 0.0436 |
| Dnajb7 | 0 | 0 | 0 | 0 | 0 | 0 | 0 | 0 | 0 | 0 | 0 | 0 | 0 | 0 | 0 | NA | NA | NA | NA | NA | NA | NA | NA | NA | NA |
| Dnajb8 | 0 | 0 | 0 | 0 | 0 | 0 | 0 | 0 | 0 | 0 | 0 | 0 | 0 | 0 | 0 | NA | NA | NA | NA | NA | NA | NA | NA | NA | NA |
| Dnajb9 | 556 | 657 | 598 | 920 | 700 | 672 | 591 | 656 | 600 | 723 | 753 | 736 | 676.53 | 616 | 737 | 1.197 | 0.259 | 0.00672329 | 0.03901628 | 0 | 0.0118 | 0.0051 | 0.0051 | TRUE | 0.039 |
| Dnajc1 | 918 | 1081 | 1101 | 1249 | 907 | 962 | 975 | 1080 | 1106 | 982 | 976 | 1053 | 1028.5 | 1054 | 1004 | 0.952 | -0.071 | 0.44891767 | 0.65455233 | 0.0021 | 0.0101 | 0.0053 | 0.0053 | TRUE | 0.0502 |
| Dnajc10 | 5468 | 5135 | 4960 | 7911 | 5450 | 5174 | 5808 | 5128 | 4980 | 6219 | 5864 | 5664 | 5610.56 | 5305 | 5916 | 1.115 | 0.157 | 0.07685841 | 0.21892957 | 0.0043 | 0.0075 | 0.0055 | 0.0055 | TRUE | 0.0835 |
| Dnajc11 | 3805 | 3444 | 3431 | 6386 | 4401 | 3496 | 4042 | 3440 | 3445 | 5020 | 4735 | 3827 | 4084.75 | 3642 | 4527 | 1.243 | 0.314 | 0.01243294 | 0.06171896 | 0.0136 | 0.0077 | 0.0111 | 0.0111 | TRUE | 0.2227 |
| Dnajc12 | 249 | 196 | 166 | 265 | 168 | 165 | 264 | 196 | 167 | 208 | 181 | 181 | 199.43 | 209 | 190 | 0.911 | -0.134 | 0.51026483 | 0.7037951 | 0.0265 | 0.0234 | 0.025 | 0.025 | TRUE | 0.584 |
| Dnajc13 | 10715 | 8979 | 9494 | 14070 | 10195 | 9497 | 11381 | 8967 | 9533 | 11061 | 10969 | 10396 | 10384.58 | 9960 | 10809 | 1.085 | 0.118 | 0.25984908 | 0.47911383 | 0.0083 | 0.0072 | 0.0078 | 0.0078 | TRUE | 0.1901 |
| Dnajc14 | 1719 | 1719 | 1647 | 2749 | 1936 | 1834 | 1826 | 1717 | 1654 | 2161 | 2083 | 2008 | 1908.02 | 1732 | 2084 | 1.204 | 0.268 | 0.00078291 | 0.00736471 | 0.0014 | 0.0086 | 0.004 | 0.004 | TRUE | 0.0269 |
| Dnajc15 | 1081 | 1275 | 1183 | 1585 | 1132 | 1023 | 1148 | 1273 | 1188 | 1246 | 1218 | 1120 | 1198.87 | 1203 | 1195 | 0.993 | -0.01 | 0.9162517 | 0.95891617 | 0.0021 | 0.0097 | 0.0051 | 0.0051 | TRUE | 0.0355 |
| Dnajc16 | 1028 | 1094 | 1162 | 1456 | 1086 | 1063 | 1092 | 1093 | 1167 | 1145 | 1168 | 1164 | 1137.99 | 1117 | 1159 | 1.037 | 0.053 | 0.50775918 | 0.70193512 | 0 | 0.0098 | 0.0037 | 0.0037 | TRUE | 0.0182 |
| Dnajc17 | 614 | 659 | 598 | 802 | 618 | 569 | 652 | 658 | 600 | 630 | 665 | 623 | 638.17 | 637 | 639 | 1.004 | 0.005 | 0.95607688 | 0.97741618 | 0 | 0.0121 | 0.0053 | 0.0053 | TRUE | 0.0297 |
| Dnajc18 | 856 | 1014 | 1006 | 1124 | 840 | 901 | 909 | 1013 | 1010 | 884 | 904 | 986 | 950.95 | 977 | 925 | 0.945 | -0.082 | 0.40027167 | 0.61321853 | 0.0025 | 0.0104 | 0.0057 | 0.0057 | TRUE | 0.0441 |
| Dnajc19 | 1558 | 1901 | 1926 | 2034 | 1370 | 1439 | 1655 | 1899 | 1934 | 1599 | 1474 | 1575 | 1689.26 | 1829 | 1549 | 0.847 | -0.239 | 0.01158025 | 0.05865246 | 0.0039 | 0.0089 | 0.0059 | 0.0059 | TRUE | 0.0835 |
| Dnajc2 | 7299 | 7888 | 7997 | 9310 | 7059 | 6703 | 7753 | 7878 | 8030 | 7319 | 7595 | 7337 | 7651.97 | 7887 | 7417 | 0.94 | -0.089 | 0.12267328 | 0.29821254 | 2.00E-04 | 0.0073 | 0.0022 | 0.0022 | TRUE | 0.0054 |
| Dnajc21 | 5988 | 6456 | 6426 | 7047 | 5436 | 5660 | 6360 | 6448 | 6452 | 5540 | 5849 | 6196 | 6140.79 | 6420 | 5862 | 0.913 | -0.132 | 0.06137482 | 0.1889894 | 0.0014 | 0.0074 | 0.0034 | 0.0034 | TRUE | 0.0303 |
| Dnajc22 | 356 | 443 | 479 | 366 | 278 | 299 | 378 | 442 | 481 | 288 | 299 | 327 | 369.28 | 434 | 305 | 0.701 | -0.512 | 0.00022844 | 0.00270487 | 0.0071 | 0.0158 | 0.0111 | 0.0111 | TRUE | 0.1445 |
| Dnajc24 | 552 | 620 | 630 | 830 | 577 | 554 | 586 | 619 | 633 | 652 | 621 | 606 | 619.64 | 613 | 626 | 1.023 | 0.033 | 0.7284453 | 0.85395791 | 0 | 0.0122 | 0.0051 | 0.0051 | TRUE | 0.0165 |
| Dnajc25 | 203 | 184 | 171 | 335 | 231 | 209 | 216 | 184 | 172 | 263 | 249 | 229 | 218.63 | 191 | 247 | 1.3 | 0.379 | 0.01546873 | 0.07242591 | 0.0042 | 0.022 | 0.013 | 0.013 | TRUE | 0.1434 |
| Dnajc27 | 1410 | 1322 | 1334 | 1641 | 1202 | 1142 | 1498 | 1320 | 1339 | 1290 | 1293 | 1250 | 1331.81 | 1386 | 1278 | 0.922 | -0.117 | 0.18275287 | 0.38488159 | 0.0018 | 0.0094 | 0.0048 | 0.0048 | TRUE | 0.0596 |
| Dnajc28 | 260 | 205 | 195 | 232 | 172 | 179 | 276 | 205 | 196 | 182 | 185 | 196 | 206.68 | 226 | 188 | 0.833 | -0.264 | 0.14280738 | 0.32890741 | 0.0145 | 0.0228 | 0.0185 | 0.0185 | TRUE | 0.4194 |
| Dnajc3 | 2490 | 2800 | 2901 | 2947 | 2065 | 2229 | 2645 | 2796 | 2913 | 2317 | 2222 | 2440 | 2555.44 | 2785 | 2326 | 0.835 | -0.26 | 0.00106193 | 0.00932968 | 0.0017 | 0.0082 | 0.0041 | 0.0041 | TRUE | 0.0234 |
| Dnajc30 | 246 | 243 | 261 | 477 | 346 | 295 | 261 | 243 | 262 | 375 | 372 | 323 | 306.04 | 255 | 357 | 1.399 | 0.485 | 0.00021369 | 0.00256295 | 0 | 0.0177 | 0.009 | 0.009 | TRUE | 0.0761 |
| Dnajc4 | 341 | 482 | 549 | 530 | 420 | 475 | 362 | 481 | 551 | 417 | 452 | 520 | 463.89 | 465 | 463 | 0.994 | -0.009 | 0.96081678 | 0.98008034 | 0.0262 | 0.014 | 0.0201 | 0.0201 | TRUE | 0.3926 |
| Dnajc5 | 4236 | 4741 | 4806 | 6217 | 4680 | 4486 | 4499 | 4735 | 4826 | 4887 | 5035 | 4911 | 4815.56 | 4687 | 4944 | 1.055 | 0.077 | 0.2290449 | 0.44227427 | 5.00E-04 | 0.0076 | 0.0027 | 0.0027 | TRUE | 0.0148 |
| Dnajc5b | 0 | 0 | 1 | 2 | 2 | 1 | 0 | 0 | 1 | 2 | 2 | 1 | 0.97 | 0 | 2 | 4.39 | 2.134 | 0.29386148 | NA | 0 | 3.4016 | 1.6739 | 1.6739 | TRUE | 0.4098 |
| Dnajc5g | 82 | 105 | 101 | 147 | 115 | 112 | 87 | 105 | 101 | 116 | 124 | 123 | 109.21 | 98 | 121 | 1.231 | 0.3 | 0.11686038 | 0.28898118 | 0 | 0.0371 | 0.017 | 0.017 | TRUE | 0.0867 |
| Dnajc6 | 3189 | 4377 | 4530 | 3931 | 3029 | 3932 | 3387 | 4371 | 4549 | 3090 | 3259 | 4304 | 3826.78 | 4102 | 3551 | 0.865 | -0.209 | 0.18904103 | 0.39422915 | 0.0284 | 0.0078 | 0.0179 | 0.0179 | TRUE | 0.418 |
| Dnajc7 | 13461 | 14978 | 14868 | 18414 | 13055 | 12915 | 14298 | 14959 | 14929 | 14476 | 14046 | 14138 | 14474.2 | 14729 | 14220 | 0.965 | -0.051 | 0.37126647 | 0.58797215 | 3.00E-04 | 0.0071 | 0.0022 | 0.0022 | TRUE | 0.008 |
| Dnajc8 | 10113 | 10742 | 10501 | 12702 | 9226 | 9246 | 10742 | 10728 | 10544 | 9985 | 9927 | 10121 | 10341.19 | 10671 | 10011 | 0.938 | -0.092 | 0.07996447 | 0.22447967 | 0 | 0.0072 | 0.0019 | 0.0019 | TRUE | 0.0013 |

| | | | | | | | | | | | | | | | | | | | | | | | | | |
|---|---|---|---|---|---|---|---|---|---|---|---|---|---|---|---|---|---|---|---|---|---|---|---|---|---|
| Dnajc9 | 1867 | 2097 | 2177 | 2950 | 2015 | 2071 | 1983 | 2094 | 2186 | 2319 | 2168 | 2267 | 2169.57 | 2088 | 2251 | 1.079 | 0.109 | 0.15859379 | 0.35254987 | 0.0013 | 0.0084 | 0.0039 | 0.0039 | TRUE | 0.0232 |
| Dnalc1 | 435 | 463 | 453 | 708 | 536 | 495 | 462 | 462 | 455 | 557 | 577 | 542 | 509.07 | 460 | 559 | 1.214 | 0.28 | 0.00447007 | 0.0287294 | 0 | 0.0134 | 0.005 | 0.005 | TRUE | 0.0092 |
| Dnalc4 | 558 | 582 | 531 | 789 | 565 | 562 | 593 | 581 | 533 | 620 | 608 | 615 | 591.75 | 569 | 614 | 1.08 | 0.111 | 0.25653484 | 0.47519694 | 0 | 0.0125 | 0.0052 | 0.0052 | TRUE | 0.0358 |
| Dnali1 | 5 | 2 | 4 | 7 | 3 | 6 | 5 | 2 | 4 | 6 | 3 | 7 | 4.44 | 4 | 5 | 1.359 | 0.442 | 0.62505688 | NA | 0 | 0.7494 | 0.3591 | 0.3591 | TRUE | 0.2751 |
| Dnase1 | 0 | 1 | 0 | 0 | 0 | 0 | 0 | 1 | 0 | 0 | 0 | 0 | 0.17 | 0 | 0 | 0.49 | -1.029 | 0.80087 | NA | 0 | 19.7991 | 10 | 10 | TRUE | 0.405 |
| Dnase1l1 | 49 | 67 | 49 | 52 | 46 | 45 | 52 | 67 | 49 | 41 | 49 | 49 | 51.3 | 56 | 46 | 0.826 | -0.276 | 0.32893688 | 0.55009994 | 0 | 0.0711 | 0.0378 | 0.0378 | TRUE | 0.2454 |
| Dnase1l2 | 11 | 7 | 9 | 7 | 10 | 11 | 12 | 7 | 9 | 6 | 11 | 12 | 9.34 | 9 | 10 | 1.007 | 0.011 | 0.98696077 | 0.99290468 | 0 | 0.3598 | 0.1931 | 0.1931 | TRUE | 0.5789 |
| Dnase1l3 | 3 | 2 | 1 | 2 | 4 | 3 | 3 | 2 | 1 | 2 | 4 | 3 | 2.56 | 2 | 3 | 1.461 | 0.547 | 0.64425221 | NA | 0 | 1.2948 | 0.599 | 0.599 | TRUE | 0.3257 |
| Dnase2a | 231 | 302 | 288 | 539 | 324 | 302 | 245 | 302 | 289 | 424 | 349 | 331 | 323.18 | 279 | 368 | 1.322 | 0.402 | 0.00976398 | 0.05171406 | 0.0118 | 0.0171 | 0.0143 | 0.0143 | TRUE | 0.2127 |
| Dnase2b | 2 | 0 | 0 | 0 | 0 | 0 | 2 | 0 | 0 | 0 | 0 | 0 | 0.35 | 1 | 0 | 0.254 | -1.975 | 0.62446185 | NA | 10 | 9.3117 | 10 | 10 | TRUE | 0.9741 |
| Dnd1 | 734 | 887 | 873 | 1032 | 835 | 854 | 780 | 886 | 877 | 811 | 898 | 935 | 864.43 | 848 | 881 | 1.039 | 0.055 | 0.60172926 | 0.77328069 | 0.004 | 0.0107 | 0.0068 | 0.0068 | TRUE | 0.06 |
| Dner | 137 | 244 | 245 | 118 | 122 | 136 | 146 | 244 | 246 | 93 | 131 | 149 | 168.02 | 212 | 124 | 0.584 | -0.775 | 0.00353198 | 0.02398037 | 0.0641 | 0.0265 | 0.0445 | 0.0445 | TRUE | 0.8107 |
| Dnlz | 216 | 335 | 322 | 544 | 332 | 304 | 229 | 335 | 323 | 428 | 357 | 333 | 334.16 | 296 | 373 | 1.261 | 0.334 | 0.06815374 | 0.20224285 | 0.0256 | 0.0168 | 0.0212 | 0.0212 | TRUE | 0.4299 |
| Dnm1 | 496 | 681 | 683 | 706 | 542 | 631 | 527 | 680 | 686 | 555 | 583 | 691 | 620.28 | 631 | 610 | 0.965 | -0.051 | 0.72751404 | 0.85342889 | 0.0158 | 0.0122 | 0.0141 | 0.0141 | TRUE | 0.2437 |
| Dnm1l | 7089 | 8192 | 8604 | 10234 | 7239 | 8030 | 7530 | 8181 | 8639 | 8045 | 7789 | 8790 | 8162.42 | 8117 | 8208 | 1.011 | 0.016 | 0.85484919 | 0.92766489 | 0.0034 | 0.0073 | 0.0054 | 0.0054 | TRUE | 0.0488 |
| Dnm2 | 5596 | 5543 | 5522 | 7473 | 5574 | 5148 | 5944 | 5536 | 5545 | 5875 | 5997 | 5635 | 5755.29 | 5675 | 5836 | 1.028 | 0.04 | 0.55359346 | 0.73700313 | 0.0011 | 0.0075 | 0.0032 | 0.0032 | TRUE | 0.0209 |
| Dnm3 | 12 | 4 | 5 | 19 | 13 | 11 | 13 | 4 | 5 | 15 | 14 | 12 | 10.45 | 7 | 14 | 1.899 | 0.925 | 0.15668228 | 0.34969116 | 0.0268 | 0.322 | 0.2011 | 0.2011 | TRUE | 1.1432 |
| Dnm3os | 0 | 0 | 0 | 0 | 0 | 0 | 0 | 0 | 0 | 0 | 0 | 0 | 0 | 0 | 0 | NA | NA | NA | NA | NA | NA | NA | NA | NA | NA |
| Dnmbp | 1677 | 1567 | 1721 | 2077 | 1609 | 1648 | 1781 | 1565 | 1728 | 1633 | 1731 | 1804 | 1707.04 | 1691 | 1723 | 1.018 | 0.026 | 0.77414951 | 0.88240848 | 0.0029 | 0.0088 | 0.0052 | 0.0052 | TRUE | 0.0517 |
| Dnmt1 | 16036 | 17809 | 19082 | 22370 | 17014 | 16833 | 17033 | 17786 | 19160 | 17586 | 18306 | 18426 | 18049.56 | 17993 | 18106 | 1.006 | 0.009 | 0.90162225 | 0.95205059 | 0.0016 | 0.0071 | 0.0037 | 0.0037 | TRUE | 0.0394 |
| Dnmt3a | 11589 | 9006 | 9379 | 10505 | 7339 | 6156 | 12310 | 8994 | 9418 | 8258 | 7896 | 6739 | 8935.78 | 10241 | 7631 | 0.745 | -0.424 | 0.00240133 | 0.01779104 | 0.0208 | 0.0073 | 0.014 | 0.014 | TRUE | 0.3817 |
| Dnmt3b | 5154 | 3409 | 3598 | 6706 | 4921 | 3624 | 5474 | 3405 | 3613 | 5272 | 5295 | 3967 | 4504.22 | 4164 | 4845 | 1.164 | 0.219 | 0.25081711 | 0.46831864 | 0.0436 | 0.0076 | 0.0259 | 0.0259 | TRUE | 0.9225 |
| Dnmt3l | 19951 | 16994 | 16575 | 18463 | 12574 | 12687 | 21191 | 16972 | 16643 | 14514 | 13529 | 13888 | 16122.92 | 18269 | 13977 | 0.765 | -0.386 | 0.00043762 | 0.00467622 | 0.0102 | 0.0071 | 0.0086 | 0.0086 | TRUE | 0.2395 |
| Dnpep | 3548 | 4151 | 3987 | 4213 | 3035 | 3536 | 3769 | 4146 | 4003 | 3312 | 3265 | 3871 | 3727.62 | 3973 | 3483 | 0.876 | -0.19 | 0.04688294 | 0.15699314 | 0.0054 | 0.0078 | 0.0063 | 0.0063 | TRUE | 0.1149 |
| Dnph1 | 2531 | 3174 | 3117 | 3167 | 2352 | 2600 | 2688 | 3170 | 3130 | 2490 | 2531 | 2846 | 2809.07 | 2996 | 2622 | 0.875 | -0.193 | 0.05825326 | 0.18235677 | 0.0062 | 0.0081 | 0.0071 | 0.0071 | TRUE | 0.0978 |
| Dntt | 0 | 0 | 0 | 2 | 1 | 0 | 0 | 0 | 0 | 2 | 1 | 0 | 0.44 | 0 | 1 | 4.754 | 2.249 | 0.50892201 | NA | 0 | 7.4712 | 6.8061 | 6.8061 | TRUE | 0.2361 |
| Dnttip1 | 803 | 996 | 929 | 1343 | 923 | 858 | 853 | 995 | 933 | 1056 | 993 | 939 | 961.42 | 927 | 996 | 1.075 | 0.105 | 0.30308887 | 0.52387042 | 0.0036 | 0.0103 | 0.0064 | 0.0064 | TRUE | 0.0576 |
| Dnttip2 | 6906 | 6547 | 6464 | 8572 | 6059 | 5741 | 7335 | 6539 | 6491 | 6739 | 6519 | 6284 | 6651.11 | 6788 | 6514 | 0.96 | -0.059 | 0.45940511 | 0.66378277 | 0.0028 | 0.0074 | 0.0045 | 0.0045 | TRUE | 0.0606 |
| Doc2a | 273 | 320 | 347 | 359 | 287 | 282 | 290 | 320 | 348 | 282 | 309 | 309 | 309.62 | 319 | 300 | 0.937 | -0.093 | 0.48106833 | 0.68018016 | 0.0024 | 0.0175 | 0.0094 | 0.0094 | TRUE | 0.0723 |
| Doc2b | 32 | 42 | 29 | 60 | 41 | 17 | 34 | 42 | 29 | 47 | 44 | 19 | 35.82 | 35 | 37 | 1.053 | 0.074 | 0.85871484 | 0.9304418 | 0.0971 | 0.0989 | 0.0981 | 0.0981 | TRUE | 1.2585 |
| Doc2g | 69 | 90 | 91 | 55 | 59 | 73 | 73 | 90 | 91 | 43 | 63 | 80 | 73.53 | 85 | 62 | 0.727 | -0.461 | 0.10778704 | 0.27486132 | 0.0382 | 0.0517 | 0.0452 | 0.0452 | TRUE | 0.3698 |
| Dock1 | 2440 | 3346 | 3702 | 4006 | 3021 | 3417 | 2592 | 3342 | 3717 | 3149 | 3250 | 3740 | 3298.44 | 3217 | 3380 | 1.05 | 0.071 | 0.61932146 | 0.78476244 | 0.0205 | 0.0079 | 0.0144 | 0.0144 | TRUE | 0.3508 |
| Dock10 | 13 | 15 | 22 | 35 | 27 | 13 | 14 | 15 | 22 | 28 | 29 | 14 | 20.28 | 17 | 24 | 1.397 | 0.482 | 0.31213001 | 0.53255705 | 0.0506 | 0.1694 | 0.1134 | 0.1134 | TRUE | 0.6555 |
| Dock11 | 914 | 944 | 987 | 1682 | 1206 | 1166 | 971 | 943 | 991 | 1322 | 1298 | 1276 | 1133.48 | 968 | 1299 | 1.342 | 0.424 | 4.71E-08 | 1.89E-06 | 0 | 0.0098 | 0.0034 | 0.0034 | TRUE | 0.0064 |
| Dock2 | 57 | 36 | 47 | 70 | 43 | 27 | 61 | 36 | 47 | 55 | 46 | 30 | 45.76 | 48 | 44 | 0.918 | -0.123 | 0.72824991 | 0.85395791 | 0.0577 | 0.0789 | 0.0688 | 0.0688 | TRUE | 0.3007 |
| Dock3 | 1116 | 877 | 893 | 1628 | 1071 | 911 | 1185 | 876 | 897 | 1280 | 1152 | 997 | 1064.55 | 986 | 1143 | 1.16 | 0.215 | 0.15924946 | 0.35340966 | 0.0216 | 0.01 | 0.0158 | 0.0158 | TRUE | 0.3727 |
| Dock4 | 1053 | 1029 | 1125 | 1118 | 850 | 880 | 1118 | 1028 | 1130 | 879 | 915 | 963 | 1005.42 | 1092 | 919 | 0.841 | -0.25 | 0.00581745 | 0.03504689 | 0.0015 | 0.0102 | 0.0049 | 0.0049 | TRUE | 0.0319 |
| Dock5 | 1609 | 1849 | 1915 | 2409 | 1719 | 1870 | 1709 | 1847 | 1923 | 1894 | 1850 | 2047 | 1878.14 | 1826 | 1930 | 1.057 | 0.079 | 0.36217969 | 0.58014481 | 0.0024 | 0.0087 | 0.0049 | 0.0049 | TRUE | 0.0378 |
| Dock6 | 7277 | 9122 | 9809 | 9718 | 8594 | 8686 | 7729 | 9110 | 9849 | 7640 | 9247 | 9508 | 8847.22 | 8896 | 8798 | 0.989 | -0.016 | 0.89636916 | 0.94994561 | 0.0143 | 0.0073 | 0.0111 | 0.0111 | TRUE | 0.1626 |
| Dock7 | 1275 | 1097 | 1261 | 1945 | 1418 | 1151 | 1354 | 1096 | 1266 | 1529 | 1526 | 1260 | 1338.45 | 1239 | 1438 | 1.162 | 0.216 | 0.08017468 | 0.22480268 | 0.0103 | 0.0094 | 0.0103 | 0.0103 | TRUE | 0.1402 |
| Dock8 | 109 | 145 | 126 | 143 | 108 | 111 | 116 | 145 | 127 | 112 | 116 | 122 | 122.87 | 129 | 117 | 0.903 | -0.148 | 0.42566382 | 0.63581483 | 0 | 0.0337 | 0.0167 | 0.0167 | TRUE | 0.1176 |
| Dock9 | 2445 | 2327 | 2480 | 3269 | 2299 | 2397 | 2597 | 2324 | 2490 | 2570 | 2474 | 2624 | 2513.08 | 2470 | 2556 | 1.035 | 0.049 | 0.52687174 | 0.71593432 | 0.0014 | 0.0082 | 0.004 | 0.004 | TRUE | 0.0326 |
| Dohh | 4039 | 4111 | 4102 | 5305 | 3761 | 3660 | 4290 | 4106 | 4119 | 4170 | 4047 | 4006 | 4123.02 | 4172 | 4074 | 0.977 | -0.034 | 0.59033773 | 0.76449681 | 3.00E-04 | 0.0077 | 0.0026 | 0.0026 | TRUE | 0.0075 |
| Dok1 | 355 | 368 | 385 | 454 | 360 | 327 | 377 | 368 | 387 | 357 | 387 | 358 | 372.23 | 377 | 367 | 0.974 | -0.039 | 0.73224981 | 0.85630933 | 0 | 0.0158 | 0.0065 | 0.0065 | TRUE | 0.0251 |
| Dok2 | 43 | 50 | 59 | 94 | 52 | 40 | 46 | 50 | 59 | 74 | 56 | 44 | 54.75 | 52 | 58 | 1.129 | 0.175 | 0.56171898 | 0.74274244 | 0.0276 | 0.0671 | 0.0475 | 0.0475 | TRUE | 0.3107 |
| Dok3 | 32 | 29 | 20 | 35 | 33 | 21 | 34 | 29 | 20 | 28 | 36 | 23 | 28.17 | 28 | 29 | 1.036 | 0.051 | 0.89541568 | 0.94945374 | 0.0176 | 0.1238 | 0.0748 | 0.0748 | TRUE | 0.3715 |
| Dok4 | 554 | 466 | 492 | 478 | 372 | 391 | 588 | 465 | 494 | 376 | 400 | 428 | 458.65 | 516 | 401 | 0.777 | -0.364 | 0.00644409 | 0.03796645 | 0.0078 | 0.0141 | 0.0106 | 0.0106 | TRUE | 0.1751 |
| Dok5 | 31 | 18 | 13 | 13 | 13 | 3 | 33 | 18 | 13 | 10 | 14 | 3 | 15.24 | 21 | 9 | 0.433 | -1.208 | 0.05974283 | 0.1853391 | 0.2156 | 0.2231 | 0.2196 | 0.2196 | TRUE | 0.5619 |
| Dok6 | 4 | 0 | 0 | 1 | 0 | 0 | 4 | 0 | 0 | 1 | 0 | 0 | 0.84 | 1 | 0 | 0.248 | -2.013 | 0.4979877 | NA | 9.4889 | 3.9329 | 4.9952 | 4.9952 | TRUE | 1.8992 |
| Dok7 | 50 | 56 | 65 | 35 | 31 | 33 | 53 | 56 | 65 | 28 | 33 | 36 | 45.22 | 58 | 32 | 0.552 | -0.857 | 0.00384134 | 0.02553125 | 0 | 0.0798 | 0.0393 | 0.0393 | TRUE | 0.1428 |
| Dolk | 219 | 215 | 243 | 447 | 296 | 220 | 233 | 215 | 244 | 351 | 318 | 241 | 267.01 | 231 | 303 | 1.321 | 0.401 | 0.02207668 | 0.09337449 | 0.0175 | 0.0192 | 0.0183 | 0.0183 | TRUE | 0.3622 |
| Dolpp1 | 1408 | 1531 | 1515 | 1582 | 1171 | 1208 | 1496 | 1529 | 1521 | 1244 | 1260 | 1322 | 1395.28 | 1515 | 1275 | 0.841 | -0.25 | 0.00081366 | 0.00758041 | 0 | 0.0093 | 0.0033 | 0.0033 | TRUE | 0.0123 |
| Dom3z | 717 | 574 | 500 | 894 | 695 | 535 | 762 | 573 | 502 | 703 | 748 | 586 | 645.52 | 612 | 679 | 1.109 | 0.15 | 0.39271298 | 0.6062924 | 0.0291 | 0.012 | 0.0205 | 0.0205 | TRUE | 0.5329 |
| Donson | 2824 | 2716 | 2553 | 3689 | 2722 | 2584 | 3000 | 2712 | 2563 | 2900 | 2929 | 2829 | 2822.14 | 2758 | 2886 | 1.046 | 0.065 | 0.44349205 | 0.65031207 | 0.0029 | 0.0081 | 0.0049 | 0.0049 | TRUE | 0.0707 |
| Dopey1 | 1199 | 1012 | 1111 | 1749 | 1210 | 1031 | 1274 | 1011 | 1116 | 1375 | 1302 | 1129 | 1200.87 | 1134 | 1269 | 1.12 | 0.164 | 0.18920177 | 0.39431214 | 0.0105 | 0.0096 | 0.0104 | 0.0104 | TRUE | 0.14 |
| Dopey2 | 3210 | 3444 | 3740 | 3438 | 2576 | 2908 | 3410 | 3440 | 3755 | 2703 | 2772 | 3183 | 3210.34 | 3535 | 2886 | 0.816 | -0.293 | 0.002019 | 0.01549571 | 0.0051 | 0.0079 | 0.0062 | 0.0062 | TRUE | 0.0982 |
| Dos | 605 | 844 | 675 | 1056 | 880 | 541 | 643 | 843 | 678 | 830 | 947 | 592 | 755.41 | 721 | 790 | 1.095 | 0.131 | 0.48093265 | 0.68004928 | 0.037 | 0.0113 | 0.0237 | 0.0237 | TRUE | 0.477 |
| Dot1l | 6007 | 5753 | 6279 | 7406 | 5584 | 5242 | 6380 | 5746 | 6305 | 5822 | 6008 | 5738 | 5999.85 | 6144 | 5856 | 0.953 | -0.069 | 0.34266042 | 0.56278664 | 0.0016 | 0.0075 | 0.0037 | 0.0037 | TRUE | 0.0392 |
| Doxl2 | 0 | 0 | 0 | 0 | 0 | 0 | 0 | 0 | 0 | 0 | 0 | 0 | 0 | 0 | 0 | NA | NA | NA | NA | NA | NA | NA | NA | NA | NA |
| Dpagt1 | 547 | 671 | 692 | 843 | 594 | 579 | 581 | 670 | 695 | 663 | 639 | 634 | 646.93 | 649 | 645 | 0.995 | -0.008 | 0.94469546 | 0.97241382 | 0.003 | 0.012 | 0.0069 | 0.0069 | TRUE | 0.0972 |
| Dpcd | 747 | 769 | 766 | 994 | 801 | 725 | 793 | 768 | 769 | 781 | 862 | 794 | 794.58 | 777 | 812 | 1.045 | 0.063 | 0.48569031 | 0.68272174 | 0 | 0.011 | 0.0047 | 0.0047 | TRUE | 0.0333 |
| Dpcr1 | 60 | 51 | 49 | 66 | 48 | 39 | 64 | 51 | 49 | 52 | 52 | 43 | 51.68 | 55 | 49 | 0.896 | -0.159 | 0.56480748 | 0.74534144 | 0 | 0.0706 | 0.0356 | 0.0356 | TRUE | 0.1733 |
| Dpep1 | 2 | 0 | 2 | 2 | 0 | 0 | 2 | 0 | 2 | 2 | 0 | 0 | 0.95 | 1 | 1 | 0.413 | -1.276 | 0.58424672 | NA | 0 | 3.4718 | 2.6841 | 2.6841 | TRUE | 0.9719 |
| Dpep2 | 5 | 3 | 5 | 4 | 4 | 6 | 5 | 3 | 5 | 3 | 4 | 7 | 4.56 | 4 | 5 | 1.039 | 0.055 | 0.95016145 | NA | 0 | 0.7298 | 0.3331 | 0.3331 | TRUE | 0.2238 |
| Dpep3 | 225 | 272 | 252 | 201 | 162 | 197 | 239 | 272 | 253 | 158 | 174 | 216 | 218.61 | 255 | 183 | 0.715 | -0.485 | 0.00393965 | 0.02604197 | 0.0095 | 0.022 | 0.0157 | 0.0157 | TRUE | 0.2635 |
| Dpf1 | 709 | 751 | 706 | 892 | 713 | 637 | 753 | 750 | 709 | 701 | 767 | 697 | 729.61 | 737 | 722 | 0.979 | -0.031 | 0.74146173 | 0.86165152 | 0 | 0.0114 | 0.0051 | 0.0051 | TRUE | 0.0348 |
| Dpf2 | 3918 | 4354 | 4523 | 5613 | 4130 | 4187 | 4162 | 4348 | 4542 | 4412 | 4444 | 4583 | 4415.17 | 4351 | 4480 | 1.03 | 0.042 | 0.53661532 | 0.72331397 | 8.00E-04 | 0.0076 | 0.0031 | 0.0031 | TRUE | 0.018 |
| Dpf3 | 55 | 47 | 50 | 107 | 96 | 67 | 58 | 47 | 50 | 84 | 103 | 73 | 69.39 | 52 | 87 | 1.677 | 0.746 | 0.00362017 | 0.02447376 | 0 | 0.0544 | 0.0318 | 0.0318 | TRUE | 0.2435 |
| Dph1 | 1370 | 1382 | 1284 | 1839 | 1383 | 1225 | 1455 | 1380 | 1289 | 1446 | 1488 | 1341 | 1399.89 | 1375 | 1425 | 1.037 | 0.052 | 0.56413972 | 0.74494404 | 0.0025 | 0.0093 | 0.0052 | 0.0052 | TRUE | 0.0357 |

| | | | | | | | | | | | | | | | | | | | | | | | | | |
|---|---|---|---|---|---|---|---|---|---|---|---|---|---|---|---|---|---|---|---|---|---|---|---|---|---|
| Dph2 | 920 | 816 | 771 | 1103 | 838 | 757 | 977 | 815 | 774 | 867 | 902 | 829 | 860.62 | 855 | 866 | 1.012 | 0.018 | 0.87887191 | 0.94007522 | 0.0073 | 0.0107 | 0.0088 | 0.0088 | TRUE | 0.1835 |
| Dph3 | 1699 | 1334 | 1202 | 2366 | 1517 | 1321 | 1805 | 1332 | 1207 | 1860 | 1632 | 1446 | 1547.01 | 1448 | 1646 | 1.137 | 0.186 | 0.26298475 | 0.48235255 | 0.0281 | 0.009 | 0.0192 | 0.0192 | TRUE | 0.5581 |
| Dph5 | 829 | 953 | 968 | 1201 | 899 | 957 | 881 | 952 | 972 | 944 | 967 | 1048 | 960.55 | 935 | 986 | 1.054 | 0.076 | 0.41483966 | 0.62638488 | 0.0018 | 0.0103 | 0.0052 | 0.0052 | TRUE | 0.0348 |
| Dpm1 | 1375 | 1841 | 1772 | 1836 | 1349 | 1543 | 1460 | 1839 | 1779 | 1443 | 1451 | 1689 | 1610.37 | 1693 | 1528 | 0.902 | -0.149 | 0.22198134 | 0.43421675 | 0.0105 | 0.0089 | 0.01 | 0.01 | TRUE | 0.1734 |
| Dpm2 | 717 | 530 | 504 | 1042 | 692 | 534 | 762 | 529 | 506 | 819 | 745 | 585 | 657.53 | 599 | 716 | 1.197 | 0.259 | 0.1761349 | 0.37637393 | 0.0395 | 0.0119 | 0.0249 | 0.0249 | TRUE | 0.6602 |
| Dpm3 | 548 | 741 | 719 | 613 | 439 | 536 | 582 | 740 | 722 | 482 | 472 | 587 | 597.51 | 681 | 514 | 0.753 | -0.409 | 0.00443158 | 0.02855181 | 0.0131 | 0.0124 | 0.0132 | 0.0132 | TRUE | 0.1907 |
| Dpp10 | 5 | 0 | 0 | 4 | 2 | 2 | 5 | 0 | 0 | 3 | 2 | 2 | 2.13 | 2 | 2 | 1.434 | 0.52 | 0.75029054 | NA | 1.2367 | 1.5516 | 1.4367 | 1.4367 | TRUE | 2.3454 |
| Dpp3 | 7293 | 7116 | 6853 | 8690 | 6332 | 6283 | 7746 | 7107 | 6881 | 6831 | 6813 | 6878 | 7042.73 | 7245 | 6841 | 0.944 | -0.083 | 0.25291361 | 0.47086616 | 0.0017 | 0.0074 | 0.0036 | 0.0036 | TRUE | 0.0447 |
| Dpp4 | 1111 | 1015 | 982 | 2524 | 1787 | 1503 | 1180 | 1014 | 986 | 1984 | 1923 | 1645 | 1455.32 | 1060 | 1851 | 1.747 | 0.805 | 4.74E-12 | 4.42E-10 | 0.0085 | 0.0092 | 0.009 | 0.009 | TRUE | 0.1171 |
| Dpp6 | 55 | 76 | 67 | 66 | 70 | 67 | 58 | 76 | 67 | 52 | 75 | 73 | 67.02 | 67 | 67 | 0.987 | -0.018 | 0.94468756 | 0.97241382 | 0.016 | 0.0561 | 0.0362 | 0.0362 | TRUE | 0.3898 |
| Dpp7 | 503 | 574 | 579 | 762 | 591 | 632 | 534 | 573 | 581 | 599 | 636 | 692 | 602.61 | 563 | 642 | 1.139 | 0.188 | 0.07802223 | 0.22101584 | 0.0022 | 0.0124 | 0.0065 | 0.0065 | TRUE | 0.0533 |
| Dpp8 | 1379 | 1714 | 1706 | 2082 | 1538 | 1458 | 1465 | 1712 | 1713 | 1637 | 1655 | 1596 | 1629.51 | 1630 | 1629 | 0.999 | -0.001 | 0.99311166 | 0.99620664 | 0.0035 | 0.0089 | 0.0056 | 0.0056 | TRUE | 0.0942 |
| Dpp9 | 2256 | 2108 | 2267 | 3029 | 2116 | 1982 | 2396 | 2105 | 2276 | 2381 | 2277 | 2170 | 2267.55 | 2259 | 2276 | 1.008 | 0.011 | 0.89572414 | 0.94959182 | 0.0028 | 0.0084 | 0.0049 | 0.0049 | TRUE | 0.0431 |
| Dppa1 | 0 | 0 | 0 | 14 | 2 | 4 | 0 | 0 | 0 | 11 | 2 | 4 | 2.92 | 0 | 6 | 31.51 | 4.978 | 0.00350081 | NA | 0.7224 | 1.1341 | 1.0114 | 1.0114 | TRUE | 0.301 |
| Dppa2 | 3071 | 2412 | 2400 | 7012 | 4520 | 2992 | 3262 | 2409 | 2410 | 5512 | 4863 | 3275 | 3621.9 | 2694 | 4550 | 1.69 | 0.757 | 0.0001064 | 0.00144661 | 0.0482 | 0.0078 | 0.0272 | 0.0272 | TRUE | 0.4493 |
| Dppa3 | 250 | 403 | 320 | 1243 | 989 | 1163 | 266 | 402 | 321 | 977 | 1064 | 1273 | 717.28 | 330 | 1105 | 3.346 | 1.743 | 4.66E-24 | 1.75E-21 | 0.0283 | 0.0115 | 0.0194 | 0.0194 | TRUE | 0.4236 |
| Dppa4 | 5307 | 5234 | 5328 | 7858 | 5654 | 4988 | 5637 | 5227 | 5350 | 6177 | 6083 | 5460 | 5655.82 | 5405 | 5907 | 1.093 | 0.129 | 0.10997306 | 0.27832078 | 0.0027 | 0.0075 | 0.0045 | 0.0045 | TRUE | 0.0532 |
| Dppa5a | 42503 | 55997 | 53778 | 60945 | 41970 | 49764 | 45146 | 55925 | 53999 | 47910 | 45157 | 54475 | 50435.17 | 51690 | 49181 | 0.951 | -0.072 | 0.52699722 | 0.71593432 | 0.0095 | 0.007 | 0.0093 | 0.0093 | TRUE | 0.1502 |
| Dpt | 0 | 1 | 0 | 1 | 0 | 0 | 0 | 1 | 0 | 1 | 0 | 0 | 0.3 | 0 | 0 | 0.954 | -0.067 | 0.98642001 | NA | 0 | 11.0817 | 9.3014 | 9.3014 | TRUE | 0.405 |
| Dpy19l1 | 4841 | 5280 | 5353 | 6812 | 5013 | 5194 | 5142 | 5273 | 5375 | 5355 | 5394 | 5686 | 5370.76 | 5263 | 5478 | 1.041 | 0.057 | 0.36658671 | 0.58407976 | 5.00E-04 | 0.0075 | 0.0027 | 0.0027 | TRUE | 0.0133 |
| Dpy19l2 | 0 | 1 | 3 | 2 | 3 | 3 | 0 | 1 | 3 | 2 | 3 | 3 | 2.02 | 1 | 3 | 1.974 | 0.981 | 0.49227556 | NA | 0 | 1.6412 | 0.9075 | 0.9075 | TRUE | 0.7394 |
| Dpy19l3 | 1429 | 1382 | 1498 | 1524 | 1129 | 1055 | 1518 | 1380 | 1504 | 1198 | 1215 | 1155 | 1328.31 | 1467 | 1189 | 0.811 | -0.303 | 0.00024614 | 0.00288633 | 0.001 | 0.0094 | 0.0042 | 0.0042 | TRUE | 0.0326 |
| Dpy19l4 | 1140 | 1352 | 1385 | 1383 | 957 | 1100 | 1211 | 1350 | 1391 | 1087 | 1030 | 1204 | 1212.14 | 1317 | 1107 | 0.84 | -0.252 | 0.01486356 | 0.07037769 | 0.0045 | 0.0096 | 0.0069 | 0.0069 | TRUE | 0.0691 |
| Dpy30 | 3946 | 4373 | 4222 | 5520 | 3872 | 3968 | 4191 | 4367 | 4239 | 4339 | 4166 | 4344 | 4274.51 | 4266 | 4283 | 1.004 | 0.006 | 0.92502125 | 0.96299902 | 3.00E-04 | 0.0077 | 0.0025 | 0.0025 | TRUE | 0.0069 |
| Dpyd | 32 | 29 | 20 | 40 | 31 | 36 | 34 | 29 | 20 | 31 | 33 | 39 | 31.21 | 28 | 34 | 1.253 | 0.325 | 0.37018997 | 0.58726979 | 0 | 0.1125 | 0.0623 | 0.0623 | TRUE | 0.3716 |
| Dpys | 153 | 182 | 183 | 112 | 100 | 115 | 163 | 182 | 184 | 88 | 108 | 126 | 141.59 | 176 | 107 | 0.605 | -0.725 | 0.00020732 | 0.00251338 | 0.0091 | 0.0302 | 0.02 | 0.02 | TRUE | 0.2716 |
| Dpysl2 | 1021 | 1227 | 1023 | 686 | 533 | 563 | 1084 | 1225 | 1027 | 539 | 573 | 616 | 844.36 | 1112 | 576 | 0.517 | -0.951 | 1.86E-17 | 3.41E-15 | 0.0054 | 0.0108 | 0.0077 | 0.0077 | TRUE | 0.095 |
| Dpysl3 | 1229 | 1204 | 1291 | 776 | 682 | 649 | 1305 | 1202 | 1296 | 610 | 734 | 710 | 976.4 | 1268 | 685 | 0.539 | -0.892 | 4.26E-17 | 7.30E-15 | 0.0045 | 0.0103 | 0.007 | 0.007 | TRUE | 0.1128 |
| Dpysl4 | 248 | 180 | 167 | 249 | 188 | 168 | 263 | 180 | 168 | 196 | 202 | 184 | 198.8 | 204 | 194 | 0.954 | -0.068 | 0.73826243 | 0.85986006 | 0.0268 | 0.0235 | 0.0251 | 0.0251 | TRUE | 0.7117 |
| Dpysl5 | 1622 | 1515 | 1544 | 1675 | 1380 | 1393 | 1723 | 1513 | 1550 | 1317 | 1485 | 1525 | 1518.77 | 1595 | 1442 | 0.903 | -0.147 | 0.14222096 | 0.32793991 | 0.0044 | 0.0091 | 0.0065 | 0.0065 | TRUE | 0.0714 |
| DQ267100 | 2 | 1 | 2 | 1 | 1 | 1 | 2 | 1 | 2 | 1 | 1 | 1 | 1.35 | 2 | 1 | 0.571 | -0.809 | 0.60062408 | NA | 0 | 2.4509 | 0.9209 | 0.9209 | TRUE | 0.1051 |
| DQ267101 | 0 | 2 | 2 | 0 | 2 | 0 | 0 | 2 | 2 | 0 | 2 | 0 | 1.03 | 1 | 1 | 0.517 | -0.951 | 0.6798388 | NA | 0 | 3.2171 | 2.7386 | 2.7386 | TRUE | 0.9748 |
| DQ267102 | 6 | 3 | 5 | 5 | 3 | 9 | 6 | 3 | 5 | 4 | 3 | 10 | 5.23 | 5 | 6 | 1.171 | 0.228 | 0.79455188 | 0.89422522 | 0 | 0.6364 | 0.3588 | 0.3588 | TRUE | 0.8495 |
| Dqx1 | 112 | 96 | 86 | 101 | 73 | 59 | 119 | 96 | 86 | 79 | 79 | 65 | 87.29 | 100 | 74 | 0.741 | -0.432 | 0.0631207 | 0.19253807 | 0.0092 | 0.0446 | 0.0272 | 0.0272 | TRUE | 0.2512 |
| Dr1 | 3739 | 3617 | 3424 | 5491 | 3907 | 3526 | 3971 | 3612 | 3438 | 4317 | 4204 | 3860 | 3900.32 | 3674 | 4127 | 1.124 | 0.168 | 0.06038176 | 0.18666034 | 0.004 | 0.0077 | 0.0055 | 0.0055 | TRUE | 0.0609 |
| Dram1 | 86 | 58 | 31 | 124 | 72 | 68 | 91 | 58 | 31 | 97 | 77 | 74 | 71.63 | 60 | 83 | 1.388 | 0.473 | 0.204842 | 0.41330025 | 0.1246 | 0.0529 | 0.0857 | 0.0857 | TRUE | 0.3481 |
| Dram2 | 957 | 1324 | 1398 | 1341 | 932 | 1129 | 1017 | 1322 | 1404 | 1054 | 1003 | 1236 | 1172.56 | 1248 | 1098 | 0.879 | -0.185 | 0.20728356 | 0.41658937 | 0.0183 | 0.0097 | 0.0147 | 0.0147 | TRUE | 0.314 |
| Drap1 | 1779 | 2042 | 2219 | 2711 | 2018 | 2061 | 1890 | 2039 | 2228 | 2131 | 2171 | 2256 | 2119.27 | 2052 | 2186 | 1.065 | 0.091 | 0.3114326 | 0.53199265 | 0.0031 | 0.0085 | 0.0053 | 0.0053 | TRUE | 0.0679 |
| Draxin | 129 | 102 | 100 | 161 | 116 | 89 | 137 | 102 | 100 | 127 | 125 | 97 | 114.68 | 113 | 116 | 1.032 | 0.045 | 0.83931406 | 0.91841152 | 0.017 | 0.0356 | 0.0265 | 0.0265 | TRUE | 0.3369 |
| Drd1a | 7 | 7 | 13 | 1 | 2 | 2 | 7 | 7 | 13 | 1 | 2 | 2 | 5.43 | 9 | 2 | 0.179 | -2.479 | 0.01058796 | 0.05499872 | 0 | 0.6131 | 0.3267 | 0.3267 | TRUE | 0.4546 |
| Drd2 | 23 | 30 | 26 | 31 | 30 | 23 | 24 | 30 | 26 | 24 | 32 | 25 | 27.05 | 27 | 27 | 1.012 | 0.017 | 0.96338479 | 0.98149376 | 0 | 0.1287 | 0.06 | 0.06 | TRUE | 0.155 |
| Drd3 | 5 | 2 | 0 | 16 | 7 | 2 | 5 | 2 | 0 | 13 | 8 | 2 | 4.93 | 2 | 8 | 3.113 | 1.638 | 0.15751037 | 0.35083371 | 0.7169 | 0.6745 | 0.6916 | 0.6916 | TRUE | 0.2206 |
| Drd4 | 23 | 39 | 42 | 54 | 34 | 35 | 24 | 39 | 42 | 42 | 37 | 38 | 37.15 | 35 | 39 | 1.112 | 0.154 | 0.65081732 | 0.80429196 | 0 | 0.0956 | 0.0562 | 0.0562 | TRUE | 0.4908 |
| Drd5 | 15 | 12 | 12 | 27 | 22 | 23 | 16 | 12 | 12 | 21 | 24 | 25 | 18.34 | 13 | 23 | 1.751 | 0.808 | 0.06961118 | 0.20506305 | 0 | 0.1865 | 0.0835 | 0.0835 | TRUE | 0.1183 |
| Drg1 | 6326 | 6840 | 6751 | 9623 | 6941 | 6465 | 6719 | 6831 | 6779 | 7565 | 7468 | 7077 | 7073.19 | 6776 | 7370 | 1.088 | 0.121 | 0.04726122 | 0.15785128 | 4.00E-04 | 0.0074 | 0.0026 | 0.0026 | TRUE | 0.0147 |
| Drg2 | 3402 | 2911 | 3105 | 4931 | 3369 | 3132 | 3614 | 2907 | 3118 | 3876 | 3625 | 3428 | 3428.03 | 3213 | 3643 | 1.134 | 0.182 | 0.0866541 | 0.23793671 | 0.0078 | 0.0079 | 0.0078 | 0.0078 | TRUE | 0.1444 |
| Drosha | 11652 | 11484 | 11962 | 16347 | 11853 | 11400 | 12376 | 11469 | 12011 | 12851 | 12753 | 12479 | 12323.28 | 11952 | 12694 | 1.062 | 0.087 | 0.15838281 | 0.35221777 | 7.00E-04 | 0.0072 | 0.0027 | 0.0027 | TRUE | 0.0153 |
| Drp2 | 55 | 45 | 22 | 98 | 87 | 56 | 58 | 45 | 22 | 77 | 94 | 61 | 59.57 | 42 | 77 | 1.852 | 0.889 | 0.01654678 | 0.07619066 | 0.1038 | 0.0622 | 0.0806 | 0.0806 | TRUE | 0.5598 |
| Dsc1 | 1 | 0 | 0 | 0 | 0 | 0 | 1 | 0 | 0 | 0 | 0 | 0 | 0.18 | 0 | 0 | 0.49 | -1.029 | 0.80087 | NA | 0 | 18.6165 | 10 | 10 | TRUE | 0.4614 |
| Dsc2 | 401 | 504 | 518 | 570 | 407 | 382 | 426 | 503 | 520 | 448 | 438 | 418 | 458.93 | 483 | 435 | 0.9 | -0.152 | 0.21319619 | 0.42440114 | 0.004 | 0.0141 | 0.0086 | 0.0086 | TRUE | 0.1235 |
| Dsc3 | 19 | 47 | 38 | 62 | 43 | 49 | 20 | 47 | 38 | 49 | 46 | 54 | 42.32 | 35 | 50 | 1.408 | 0.493 | 0.17205633 | 0.37089495 | 0.0502 | 0.0847 | 0.0696 | 0.0696 | TRUE | 0.7053 |
| Dscam | 25 | 22 | 27 | 28 | 24 | 19 | 27 | 22 | 27 | 22 | 26 | 21 | 24.04 | 25 | 23 | 0.907 | -0.141 | 0.71199686 | 0.84325407 | 0 | 0.1439 | 0.0642 | 0.0642 | TRUE | 0.0787 |
| Dscaml1 | 76 | 150 | 151 | 78 | 75 | 96 | 81 | 150 | 152 | 61 | 81 | 105 | 104.88 | 128 | 82 | 0.643 | -0.637 | 0.0414443 | 0.14445515 | 0.0844 | 0.0383 | 0.0603 | 0.0603 | TRUE | 0.7304 |
| Dscc1 | 366 | 339 | 363 | 721 | 471 | 407 | 389 | 339 | 364 | 567 | 507 | 446 | 435.15 | 364 | 507 | 1.394 | 0.48 | 0.00038785 | 0.00423523 | 0.0078 | 0.0145 | 0.0108 | 0.0108 | TRUE | 0.1319 |
| Dscr3 | 1771 | 2229 | 2132 | 2339 | 1719 | 1771 | 1881 | 2226 | 2141 | 1839 | 1850 | 1939 | 1979.15 | 2083 | 1876 | 0.9 | -0.152 | 0.10062898 | 0.26242067 | 0.0034 | 0.0086 | 0.0056 | 0.0056 | TRUE | 0.0864 |
| Dse | 2731 | 2779 | 2739 | 4334 | 2881 | 2799 | 2901 | 2775 | 2750 | 3407 | 3100 | 3064 | 2999.54 | 2809 | 3190 | 1.136 | 0.185 | 0.01763208 | 0.07928957 | 0.0017 | 0.008 | 0.004 | 0.004 | TRUE | 0.0436 |
| Dsel | 353 | 296 | 316 | 643 | 450 | 410 | 375 | 296 | 317 | 505 | 484 | 449 | 404.39 | 329 | 479 | 1.458 | 0.544 | 4.88E-05 | 0.00076109 | 0.0062 | 0.015 | 0.0104 | 0.0104 | TRUE | 0.1655 |
| Dsg1a | 0 | 0 | 0 | 0 | 0 | 1 | 0 | 0 | 0 | 0 | 0 | 1 | 0.18 | 0 | 0 | 1.859 | 0.894 | 0.82650734 | NA | 0 | 18.0643 | 10 | 10 | TRUE | 0.5383 |
| Dsg1b | 3 | 0 | 0 | 4 | 3 | 2 | 3 | 0 | 0 | 3 | 3 | 2 | 1.96 | 1 | 3 | 2.742 | 1.455 | 0.36423281 | NA | 0 | 1.6894 | 1.1935 | 1.1935 | TRUE | 1.4433 |
| Dsg1c | 0 | 0 | 0 | 0 | 0 | 0 | 0 | 0 | 0 | 0 | 0 | 0 | 0 | 0 | 0 | NA | NA | NA | NA | NA | NA | NA | NA | NA | NA |
| Dsg2 | 5546 | 6112 | 6685 | 7918 | 5883 | 6004 | 5891 | 6104 | 6712 | 6225 | 6330 | 6572 | 6305.66 | 6236 | 6376 | 1.022 | 0.032 | 0.68434031 | 0.82524776 | 0.0021 | 0.0074 | 0.0042 | 0.0042 | TRUE | 0.0546 |
| Dsg3 | 2 | 2 | 0 | 3 | 3 | 0 | 2 | 2 | 0 | 2 | 3 | 0 | 1.62 | 1 | 2 | 1.371 | 0.456 | 0.78923111 | NA | 0 | 2.043 | 1.4608 | 1.4608 | TRUE | 0.5671 |
| Dsg4 | 0 | 0 | 0 | 1 | 1 | 0 | 0 | 0 | 0 | 1 | 1 | 0 | 0.31 | 0 | 1 | 3.32 | 1.731 | 0.65383052 | NA | 0 | 10.6224 | 8.9541 | 8.9541 | TRUE | 0.202 |
| Dsn1 | 1271 | 1420 | 1428 | 1714 | 1223 | 1216 | 1350 | 1418 | 1434 | 1347 | 1316 | 1331 | 1366.07 | 1401 | 1331 | 0.951 | -0.073 | 0.32936861 | 0.55035509 | 0 | 0.0093 | 0.0033 | 0.0033 | TRUE | 0.0119 |
| Dsp | 2790 | 3263 | 3728 | 4352 | 3179 | 3283 | 2963 | 3259 | 3743 | 3421 | 3420 | 3594 | 3400.16 | 3322 | 3478 | 1.047 | 0.066 | 0.52070242 | 0.71116371 | 0.007 | 0.0079 | 0.0074 | 0.0074 | TRUE | 0.1498 |
| Dspp | 0 | 0 | 0 | 0 | 0 | 0 | 0 | 0 | 0 | 0 | 0 | 0 | 0 | 0 | 0 | NA | NA | NA | NA | NA | NA | NA | NA | NA | NA |
| Dst | 9120 | 8822 | 11710 | 11200 | 7844 | 8028 | 9687 | 8811 | 11758 | 8805 | 8440 | 8788 | 9381.31 | 10085 | 8678 | 0.86 | -0.217 | 0.05936982 | 0.18450897 | 0.0114 | 0.0073 | 0.0094 | 0.0094 | TRUE | 0.2574 |
| Dstn | 8327 | 8346 | 8235 | 10191 | 7489 | 7919 | 8845 | 8335 | 8269 | 8011 | 8058 | 8669 | 8364.41 | 8483 | 8246 | 0.972 | -0.041 | 0.55792938 | 0.74033839 | 0.0015 | 0.0073 | 0.0034 | 0.0034 | TRUE | 0.0245 |
| Dstyk | 761 | 936 | 954 | 1439 | 1004 | 1054 | 808 | 935 | 958 | 1131 | 1080 | 1154 | 1011.04 | 900 | 1122 | 1.246 | 0.317 | 0.00160922 | 0.01295588 | 0.0034 | 0.0102 | 0.0063 | 0.0063 | TRUE | 0.0945 |

| | | | | | | | | | | | | | | | | | | | | | | | | | |
|---|---|---|---|---|---|---|---|---|---|---|---|---|---|---|---|---|---|---|---|---|---|---|---|---|---|
| Dtd1 | 853 | 1070 | 1098 | 1135 | 860 | 921 | 906 | 1069 | 1103 | 892 | 925 | 1008 | 983.82 | 1026 | 942 | 0.917 | -0.124 | 0.26782586 | 0.48755188 | 0.0064 | 0.0103 | 0.008 | 0.008 | TRUE | 0.1238 |
| Dtd2 | 820 | 926 | 887 | 1419 | 1010 | 976 | 871 | 925 | 891 | 1116 | 1087 | 1068 | 992.84 | 896 | 1090 | 1.218 | 0.284 | 0.00048829 | 0.00508071 | 0 | 0.0102 | 0.0038 | 0.0038 | TRUE | 0.0098 |
| Dtl | 4547 | 5130 | 4894 | 5836 | 4353 | 4528 | 4830 | 5123 | 4914 | 4588 | 4684 | 4957 | 4849.19 | 4956 | 4743 | 0.957 | -0.064 | 0.3521866 | 0.57073249 | 0.001 | 0.0076 | 0.0032 | 0.0032 | TRUE | 0.0189 |
| Dtna | 400 | 403 | 453 | 382 | 261 | 340 | 425 | 402 | 455 | 300 | 281 | 372 | 372.59 | 427 | 318 | 0.743 | -0.43 | 0.00307682 | 0.02165876 | 0.0092 | 0.0157 | 0.0124 | 0.0124 | TRUE | 0.2463 |
| Dtnb | 1761 | 2185 | 2342 | 2732 | 2058 | 2134 | 1870 | 2182 | 2352 | 2148 | 2214 | 2336 | 2183.71 | 2135 | 2233 | 1.045 | 0.064 | 0.54530981 | 0.73041412 | 0.0071 | 0.0084 | 0.0077 | 0.0077 | TRUE | 0.1415 |
| Dtnbp1 | 833 | 934 | 940 | 1323 | 946 | 897 | 885 | 933 | 944 | 1040 | 1018 | 982 | 966.87 | 921 | 1013 | 1.101 | 0.139 | 0.09671385 | 0.2552922 | 0 | 0.0103 | 0.004 | 0.004 | TRUE | 0.0136 |
| Dtwd1 | 518 | 578 | 660 | 774 | 553 | 607 | 550 | 577 | 663 | 608 | 595 | 664 | 609.68 | 597 | 622 | 1.043 | 0.06 | 0.60311996 | 0.77412127 | 0.0047 | 0.0123 | 0.0081 | 0.0081 | TRUE | 0.1101 |
| Dtwd2 | 771 | 751 | 852 | 903 | 632 | 710 | 819 | 750 | 855 | 710 | 680 | 777 | 765.26 | 808 | 722 | 0.894 | -0.162 | 0.12096141 | 0.29573762 | 0.0032 | 0.0112 | 0.0066 | 0.0066 | TRUE | 0.0506 |
| Dtx1 | 814 | 1389 | 1487 | 872 | 832 | 1106 | 865 | 1387 | 1493 | 685 | 895 | 1211 | 1089.38 | 1248 | 930 | 0.745 | -0.425 | 0.07569261 | 0.21690316 | 0.0753 | 0.0099 | 0.0404 | 0.0404 | TRUE | 0.4687 |
| Dtx2 | 2130 | 1735 | 1793 | 2153 | 1561 | 1494 | 2262 | 1733 | 1800 | 1693 | 1680 | 1635 | 1800.51 | 1932 | 1669 | 0.864 | -0.21 | 0.07628124 | 0.21787537 | 0.0105 | 0.0087 | 0.0096 | 0.0096 | TRUE | 0.2703 |
| Dtx3 | 1023 | 1372 | 1460 | 935 | 899 | 852 | 1087 | 1370 | 1466 | 735 | 967 | 933 | 1092.96 | 1308 | 878 | 0.671 | -0.576 | 0.00017618 | 0.00219316 | 0.0201 | 0.0099 | 0.016 | 0.016 | TRUE | 0.2619 |
| Dtx3l | 999 | 661 | 705 | 1090 | 795 | 627 | 1061 | 660 | 708 | 857 | 855 | 686 | 804.62 | 810 | 799 | 0.988 | -0.017 | 0.92794556 | 0.96471962 | 0.0404 | 0.011 | 0.0251 | 0.0251 | TRUE | 0.8739 |
| Dtx4 | 79 | 123 | 132 | 173 | 128 | 130 | 84 | 123 | 133 | 136 | 138 | 142 | 125.89 | 113 | 139 | 1.224 | 0.292 | 0.1773938 | 0.3779353 | 0.0173 | 0.0331 | 0.0256 | 0.0256 | TRUE | 0.4992 |
| Dtymk | 3816 | 3704 | 3596 | 5315 | 3649 | 3279 | 4053 | 3699 | 3611 | 4178 | 3926 | 3589 | 3842.83 | 3788 | 3898 | 1.029 | 0.042 | 0.64707593 | 0.80180791 | 0.0045 | 0.0078 | 0.0058 | 0.0058 | TRUE | 0.0582 |
| Duox1 | 32 | 39 | 31 | 47 | 44 | 24 | 34 | 39 | 31 | 37 | 47 | 26 | 35.77 | 35 | 37 | 1.063 | 0.088 | 0.80215721 | 0.89777365 | 0.0165 | 0.099 | 0.0607 | 0.0607 | TRUE | 0.1663 |
| Duox2 | 11 | 14 | 11 | 14 | 5 | 13 | 12 | 14 | 11 | 11 | 5 | 14 | 11.22 | 12 | 10 | 0.835 | -0.259 | 0.66212368 | 0.81016405 | 0 | 0.3005 | 0.1645 | 0.1645 | TRUE | 0.4249 |
| Duoxa1 | 19 | 20 | 20 | 13 | 13 | 15 | 20 | 20 | 20 | 10 | 14 | 16 | 16.81 | 20 | 13 | 0.667 | -0.585 | 0.20814705 | 0.41756491 | 0 | 0.2029 | 0.0938 | 0.0938 | TRUE | 0.2554 |
| Duoxa2 | 14 | 9 | 4 | 15 | 19 | 9 | 15 | 9 | 4 | 12 | 20 | 10 | 11.66 | 9 | 14 | 1.51 | 0.595 | 0.36746691 | 0.58475554 | 0.1481 | 0.2894 | 0.2237 | 0.2237 | TRUE | 0.336 |
| Dupd1 | 0 | 3 | 3 | 0 | 2 | 0 | 0 | 3 | 3 | 0 | 2 | 0 | 1.36 | 2 | 1 | 0.344 | -1.54 | 0.47142161 | NA | 2.2758 | 2.4292 | 2.335 | 2.335 | TRUE | 0.9749 |
| Dus1l | 5745 | 5068 | 4777 | 5814 | 4373 | 3746 | 6102 | 5061 | 4797 | 4571 | 4705 | 4101 | 4889.41 | 5320 | 4459 | 0.838 | -0.255 | 0.02598206 | 0.10433639 | 0.0106 | 0.0076 | 0.0092 | 0.0092 | TRUE | 0.2015 |
| Dus2l | 853 | 972 | 954 | 1287 | 893 | 760 | 906 | 971 | 958 | 1012 | 961 | 832 | 939.86 | 945 | 935 | 0.99 | -0.014 | 0.89551571 | 0.94945374 | 0.0046 | 0.0104 | 0.0071 | 0.0071 | TRUE | 0.1092 |
| Dus3l | 2024 | 1671 | 1552 | 2677 | 2027 | 1618 | 2150 | 1669 | 1558 | 2104 | 2181 | 1771 | 1905.6 | 1792 | 2019 | 1.127 | 0.172 | 0.23443424 | 0.44932517 | 0.0198 | 0.0086 | 0.0146 | 0.0146 | TRUE | 0.3671 |
| Dus4l | 499 | 608 | 587 | 700 | 505 | 436 | 530 | 607 | 589 | 550 | 543 | 477 | 549.59 | 575 | 523 | 0.911 | -0.135 | 0.24023211 | 0.4564989 | 0.0036 | 0.0129 | 0.0077 | 0.0077 | TRUE | 0.0684 |
| Dusp1 | 1018 | 1681 | 1814 | 1557 | 1239 | 1373 | 1081 | 1679 | 1821 | 1224 | 1333 | 1503 | 1440.27 | 1527 | 1353 | 0.886 | -0.175 | 0.34913441 | 0.56797812 | 0.0414 | 0.0092 | 0.0245 | 0.0245 | TRUE | 0.7845 |
| Dusp10 | 163 | 100 | 87 | 158 | 109 | 86 | 173 | 100 | 87 | 124 | 117 | 94 | 116 | 120 | 112 | 0.934 | -0.098 | 0.73449328 | 0.85755524 | 0.0698 | 0.0353 | 0.052 | 0.052 | TRUE | 1.4903 |
| Dusp11 | 3766 | 3251 | 3249 | 4074 | 2857 | 2752 | 4000 | 3247 | 3262 | 3203 | 3074 | 3012 | 3299.74 | 3503 | 3096 | 0.884 | -0.178 | 0.0904684 | 0.24512413 | 0.0076 | 0.0079 | 0.0076 | 0.0076 | TRUE | 0.1871 |
| Dusp12 | 1274 | 1197 | 1121 | 1661 | 1162 | 1154 | 1353 | 1195 | 1126 | 1306 | 1250 | 1263 | 1248.92 | 1225 | 1273 | 1.04 | 0.057 | 0.56530564 | 0.74551564 | 0.0038 | 0.0095 | 0.0062 | 0.0062 | TRUE | 0.1005 |
| Dusp13 | 5 | 7 | 5 | 3 | 9 | 6 | 5 | 7 | 5 | 2 | 10 | 7 | 5.99 | 6 | 6 | 1.051 | 0.072 | 0.93101079 | 0.96610976 | 0 | 0.557 | 0.3236 | 0.3236 | TRUE | 0.3132 |
| Dusp14 | 378 | 295 | 266 | 536 | 373 | 291 | 402 | 295 | 267 | 421 | 401 | 319 | 350.74 | 321 | 380 | 1.187 | 0.247 | 0.19607599 | 0.40230395 | 0.0304 | 0.0163 | 0.0235 | 0.0235 | TRUE | 0.5414 |
| Dusp15 | 26 | 19 | 33 | 17 | 19 | 13 | 28 | 19 | 33 | 13 | 20 | 14 | 21.29 | 27 | 16 | 0.599 | -0.74 | 0.09897394 | 0.25956384 | 0 | 0.1616 | 0.0949 | 0.0949 | TRUE | 0.2429 |
| Dusp16 | 3520 | 3359 | 3510 | 4677 | 3132 | 3092 | 3739 | 3355 | 3524 | 3677 | 3370 | 3385 | 3508.19 | 3539 | 3477 | 0.983 | -0.025 | 0.75607419 | 0.87112296 | 0.0024 | 0.0078 | 0.0044 | 0.0044 | TRUE | 0.0311 |
| Dusp18 | 338 | 299 | 280 | 314 | 216 | 245 | 359 | 299 | 281 | 247 | 232 | 268 | 281.04 | 313 | 249 | 0.796 | -0.328 | 0.0290733 | 0.11249856 | 0.0074 | 0.0186 | 0.0127 | 0.0127 | TRUE | 0.1861 |
| Dusp19 | 460 | 644 | 643 | 633 | 489 | 507 | 489 | 643 | 646 | 498 | 526 | 555 | 559.36 | 593 | 526 | 0.887 | -0.173 | 0.21762958 | 0.42962457 | 0.012 | 0.0128 | 0.0123 | 0.0123 | TRUE | 0.2742 |
| Dusp2 | 38 | 52 | 35 | 38 | 19 | 22 | 40 | 52 | 35 | 30 | 20 | 24 | 33.64 | 42 | 25 | 0.588 | -0.765 | 0.03184896 | 0.12006324 | 0 | 0.1048 | 0.06 | 0.06 | TRUE | 0.295 |
| Dusp21 | 0 | 0 | 0 | 0 | 0 | 0 | 0 | 0 | 0 | 0 | 0 | 0 | 0 | 0 | 0 | NA | NA | NA | NA | NA | NA | NA | NA | NA | NA |
| Dusp22 | 483 | 703 | 725 | 861 | 666 | 660 | 513 | 702 | 728 | 677 | 717 | 722 | 676.5 | 648 | 705 | 1.088 | 0.122 | 0.40962381 | 0.62150907 | 0.0165 | 0.0118 | 0.0142 | 0.0142 | TRUE | 0.3875 |
| Dusp23 | 8 | 20 | 10 | 13 | 14 | 13 | 8 | 20 | 10 | 10 | 15 | 14 | 13 | 13 | 13 | 1.016 | 0.023 | 0.96839935 | 0.98419772 | 0.0556 | 0.2602 | 0.1608 | 0.1608 | TRUE | 0.9866 |
| Dusp26 | 436 | 407 | 328 | 285 | 234 | 269 | 463 | 406 | 329 | 224 | 252 | 294 | 328.2 | 399 | 257 | 0.641 | -0.641 | 0.00026271 | 0.0030511 | 0.021 | 0.0169 | 0.019 | 0.019 | TRUE | 0.1928 |
| Dusp27 | 424 | 290 | 251 | 652 | 508 | 324 | 450 | 290 | 252 | 513 | 547 | 355 | 400.97 | 331 | 472 | 1.427 | 0.513 | 0.0350297 | 0.12812811 | 0.0705 | 0.0151 | 0.04 | 0.04 | TRUE | 1.1348 |
| Dusp28 | 202 | 183 | 128 | 236 | 175 | 173 | 215 | 183 | 129 | 186 | 188 | 189 | 181.51 | 176 | 188 | 1.072 | 0.1 | 0.63140821 | 0.79271062 | 0.0264 | 0.0251 | 0.0256 | 0.0256 | TRUE | 0.4256 |
| Dusp3 | 2729 | 3845 | 4062 | 2836 | 2298 | 2668 | 2899 | 3840 | 4079 | 2229 | 2473 | 2921 | 3073.31 | 3606 | 2541 | 0.704 | -0.506 | 0.00095547 | 0.00859163 | 0.024 | 0.008 | 0.0165 | 0.0165 | TRUE | 0.3576 |
| Dusp4 | 857 | 1290 | 1352 | 1072 | 839 | 811 | 910 | 1288 | 1358 | 843 | 903 | 888 | 1031.56 | 1185 | 878 | 0.74 | -0.434 | 0.00511518 | 0.03186279 | 0.0222 | 0.0101 | 0.0163 | 0.0163 | TRUE | 0.4927 |
| Dusp5 | 213 | 200 | 172 | 292 | 239 | 188 | 226 | 200 | 173 | 230 | 257 | 206 | 215.2 | 200 | 231 | 1.157 | 0.211 | 0.21335158 | 0.42460329 | 0.01 | 0.0222 | 0.016 | 0.016 | TRUE | 0.1509 |
| Dusp6 | 1611 | 1853 | 1838 | 2603 | 1832 | 1601 | 1711 | 1851 | 1846 | 2046 | 1971 | 1753 | 1862.88 | 1803 | 1923 | 1.068 | 0.094 | 0.30808813 | 0.52856814 | 0.0033 | 0.0087 | 0.0056 | 0.0056 | TRUE | 0.0724 |
| Dusp7 | 507 | 720 | 776 | 827 | 699 | 722 | 539 | 719 | 779 | 650 | 752 | 790 | 704.89 | 679 | 731 | 1.075 | 0.105 | 0.50961068 | 0.70313881 | 0.0216 | 0.0116 | 0.0167 | 0.0167 | TRUE | 0.3843 |
| Dusp8 | 76 | 50 | 44 | 159 | 84 | 62 | 81 | 50 | 44 | 125 | 90 | 68 | 76.35 | 58 | 94 | 1.63 | 0.705 | 0.03496443 | 0.12807094 | 0.0825 | 0.0501 | 0.0666 | 0.0666 | TRUE | 0.3032 |
| Dusp9 | 11340 | 9739 | 9086 | 14043 | 8705 | 7441 | 12045 | 9726 | 9123 | 11040 | 9366 | 8145 | 9907.62 | 10298 | 9517 | 0.924 | -0.114 | 0.43072985 | 0.64055214 | 0.0201 | 0.0072 | 0.0149 | 0.0149 | TRUE | 0.269 |
| Dut | 6612 | 6347 | 6326 | 9478 | 6799 | 6569 | 7023 | 6339 | 6352 | 7451 | 7315 | 7191 | 6945.14 | 6571 | 7319 | 1.114 | 0.156 | 0.03130432 | 0.11879974 | 0.0017 | 0.0074 | 0.0036 | 0.0036 | TRUE | 0.0441 |
| Dux | 27 | 10 | 10 | 39 | 30 | 14 | 29 | 10 | 10 | 31 | 32 | 15 | 21.16 | 16 | 26 | 1.619 | 0.695 | 0.22237905 | 0.43445588 | 0.2064 | 0.1626 | 0.1833 | 0.1833 | TRUE | 2.0794 |
| Duxbl1 | 0 | 0 | 0 | 0 | 0 | 0 | 0 | 0 | 0 | 0 | 0 | 0 | 0 | 0 | 0 | NA | NA | NA | NA | NA | NA | NA | NA | NA | NA |
| Duxbl2 | 0 | 0 | 0 | 0 | 0 | 0 | 0 | 0 | 0 | 0 | 0 | 0 | 0 | 0 | 0 | NA | NA | NA | NA | NA | NA | NA | NA | NA | NA |
| Duxbl3 | 0 | 0 | 0 | 0 | 0 | 0 | 0 | 0 | 0 | 0 | 0 | 0 | 0 | 0 | 0 | NA | NA | NA | NA | NA | NA | NA | NA | NA | NA |
| Dvl1 | 1955 | 2359 | 2125 | 2454 | 1944 | 1795 | 2077 | 2356 | 2134 | 1929 | 2092 | 1965 | 2091.99 | 2189 | 1995 | 0.911 | -0.134 | 0.11939122 | 0.29280644 | 0.0025 | 0.0085 | 0.0049 | 0.0049 | TRUE | 0.0542 |
| Dvl2 | 5369 | 5879 | 6039 | 7416 | 5947 | 5799 | 5703 | 5871 | 6064 | 5830 | 6399 | 6348 | 6035.74 | 5879 | 6192 | 1.053 | 0.074 | 0.30202235 | 0.52259973 | 0.0015 | 0.0074 | 0.0036 | 0.0036 | TRUE | 0.0323 |
| Dvl3 | 760 | 838 | 828 | 1305 | 981 | 862 | 807 | 837 | 831 | 1026 | 1055 | 944 | 916.76 | 825 | 1008 | 1.222 | 0.29 | 0.0012205 | 0.01038168 | 8.00E-04 | 0.0105 | 0.0047 | 0.0047 | TRUE | 0.0366 |
| DXBay18 | 0 | 0 | 0 | 0 | 0 | 0 | 0 | 0 | 0 | 0 | 0 | 0 | 0 | 0 | 0 | NA | NA | NA | NA | NA | NA | NA | NA | NA | NA |
| Dydc1 | 0 | 0 | 0 | 1 | 0 | 0 | 0 | 0 | 0 | 1 | 0 | 0 | 0.13 | 0 | 0 | 1.859 | 0.894 | 0.82650734 | NA | 0 | 25.1515 | 10 | 10 | TRUE | 0.2563 |
| Dydc2 | 6 | 1 | 1 | 3 | 2 | 1 | 6 | 1 | 1 | 2 | 2 | 1 | 2.33 | 3 | 2 | 0.682 | -0.552 | 0.67847928 | NA | 0.3393 | 1.4208 | 0.8358 | 0.8358 | TRUE | 1.4621 |
| Dym | 2574 | 2867 | 2858 | 3026 | 2298 | 2510 | 2734 | 2863 | 2870 | 2379 | 2473 | 2748 | 2677.66 | 2822 | 2533 | 0.897 | -0.157 | 0.06381276 | 0.19414637 | 0.0027 | 0.0081 | 0.0048 | 0.0048 | TRUE | 0.0663 |
| Dynap | 0 | 0 | 0 | 0 | 0 | 0 | 0 | 0 | 0 | 0 | 0 | 0 | 0 | 0 | 0 | NA | NA | NA | NA | NA | NA | NA | NA | NA | NA |
| Dync1h1 | 33625 | 30985 | 39959 | 39462 | 28028 | 27642 | 35716 | 30945 | 40123 | 31022 | 30156 | 30259 | 33036.77 | 35595 | 30479 | 0.856 | -0.224 | 0.03250256 | 0.12155199 | 0.0084 | 0.007 | 0.0079 | 0.0079 | TRUE | 0.1599 |
| Dync1i1 | 101 | 65 | 56 | 80 | 65 | 51 | 107 | 65 | 56 | 63 | 70 | 56 | 69.51 | 76 | 63 | 0.827 | -0.273 | 0.37366838 | 0.58999877 | 0.0529 | 0.0543 | 0.0536 | 0.0536 | TRUE | 1.1563 |
| Dync1i2 | 5138 | 5412 | 5602 | 6930 | 5096 | 5345 | 5457 | 5405 | 5625 | 5448 | 5483 | 5851 | 5544.87 | 5496 | 5594 | 1.018 | 0.025 | 0.69987881 | 0.8358692 | 7.00E-04 | 0.0075 | 0.0029 | 0.0029 | TRUE | 0.0196 |
| Dync1li1 | 3255 | 3488 | 3420 | 4043 | 3084 | 3056 | 3457 | 3483 | 3434 | 3178 | 3318 | 3345 | 3369.45 | 3458 | 3280 | 0.948 | -0.077 | 0.22190189 | 0.43416903 | 0 | 0.0079 | 0.0025 | 0.0025 | TRUE | 0.0093 |
| Dync1li2 | 2865 | 3406 | 3462 | 4588 | 3485 | 3424 | 3043 | 3402 | 3476 | 3607 | 3750 | 3748 | 3504.24 | 3307 | 3702 | 1.119 | 0.162 | 0.04416089 | 0.15072615 | 0.0023 | 0.0078 | 0.0044 | 0.0044 | TRUE | 0.059 |
| Dync2h1 | 569 | 505 | 613 | 893 | 606 | 473 | 604 | 504 | 616 | 702 | 652 | 518 | 599.34 | 575 | 624 | 1.087 | 0.121 | 0.42134955 | 0.63146381 | 0.0164 | 0.0124 | 0.0145 | 0.0145 | TRUE | 0.2577 |
| Dync2li1 | 585 | 824 | 776 | 810 | 566 | 673 | 621 | 823 | 779 | 637 | 609 | 737 | 700.99 | 741 | 661 | 0.891 | -0.167 | 0.23896136 | 0.45490629 | 0.0142 | 0.0116 | 0.013 | 0.013 | TRUE | 0.2354 |
| Dynll1 | 6795 | 7220 | 6973 | 10893 | 7683 | 7305 | 7217 | 7211 | 7002 | 8563 | 8266 | 7996 | 7709.32 | 7143 | 8275 | 1.159 | 0.212 | 0.00058094 | 0.00584347 | 5.00E-04 | 0.0073 | 0.0026 | 0.0026 | TRUE | 0.0114 |
| Dynll2 | 6354 | 7490 | 7067 | 10993 | 7869 | 7086 | 6749 | 7480 | 7096 | 8642 | 8467 | 7757 | 7698.43 | 7108 | 8289 | 1.166 | 0.222 | 0.00527697 | 0.03254925 | 0.0027 | 0.0073 | 0.0044 | 0.0044 | TRUE | 0.0384 |

| | | | | | | | | | | | | | | | | | | | | | | | | | |
|---|---|---|---|---|---|---|---|---|---|---|---|---|---|---|---|---|---|---|---|---|---|---|---|---|---|
| Dynlrb1 | 2288 | 2275 | 2101 | 2943 | 1908 | 1920 | 2430 | 2272 | 2110 | 2314 | 2053 | 2102 | 2213.36 | 2271 | 2156 | 0.95 | -0.074 | 0.42766449 | 0.63783681 | 0.0039 | 0.0084 | 0.0058 | 0.0058 | TRUE | 0.0503 |
| Dynlrb2 | 2 | 2 | 0 | 3 | 2 | 0 | 2 | 2 | 0 | 2 | 2 | 0 | 1.44 | 1 | 1 | 1.118 | 0.161 | 0.92689866 | NA | 0 | 2.2968 | 1.534 | 1.534 | TRUE | 0.4905 |
| Dynlt1a | 544 | 684 | 628 | 617 | 440 | 513 | 578 | 683 | 631 | 485 | 473 | 562 | 568.59 | 631 | 507 | 0.803 | -0.317 | 0.00935852 | 0.05003628 | 0.0059 | 0.0127 | 0.009 | 0.009 | TRUE | 0.1021 |
| Dynlt1b | 559 | 615 | 556 | 599 | 483 | 468 | 594 | 614 | 558 | 471 | 520 | 512 | 544.85 | 589 | 501 | 0.85 | -0.235 | 0.02487666 | 0.10140264 | 0 | 0.0129 | 0.0061 | 0.0061 | TRUE | 0.0343 |
| Dynlt1c | 0 | 0 | 0 | 0 | 0 | 0 | 0 | 0 | 0 | 0 | 0 | 0 | 0 | 0 | 0 | NA | NA | NA | NA | NA | NA | NA | NA | NA | NA |
| Dynlt1f | 156 | 181 | 143 | 151 | 131 | 135 | 166 | 181 | 144 | 119 | 141 | 148 | 149.58 | 164 | 136 | 0.828 | -0.272 | 0.14081752 | 0.32618209 | 0.0071 | 0.0289 | 0.0178 | 0.0178 | TRUE | 0.1387 |
| Dynlt3 | 887 | 1563 | 1495 | 1637 | 1206 | 1391 | 942 | 1561 | 1501 | 1287 | 1298 | 1523 | 1351.9 | 1335 | 1369 | 1.025 | 0.036 | 0.84609885 | 0.92282842 | 0.0404 | 0.0093 | 0.0241 | 0.0241 | TRUE | 0.794 |
| Dyrk1a | 2613 | 2845 | 2844 | 4105 | 3056 | 2822 | 2775 | 2841 | 2856 | 3227 | 3288 | 3089 | 3012.78 | 2824 | 3201 | 1.134 | 0.181 | 0.00589477 | 0.03543097 | 3.00E-04 | 0.008 | 0.0028 | 0.0028 | TRUE | 0.0112 |
| Dyrk1b | 425 | 377 | 371 | 723 | 577 | 514 | 451 | 377 | 373 | 568 | 621 | 563 | 492.05 | 400 | 584 | 1.46 | 0.546 | 9.87E-06 | 0.00019646 | 0.0045 | 0.0136 | 0.0089 | 0.0089 | TRUE | 0.1432 |
| Dyrk2 | 507 | 625 | 620 | 809 | 629 | 574 | 539 | 624 | 623 | 636 | 677 | 628 | 621.06 | 595 | 647 | 1.087 | 0.12 | 0.26550164 | 0.48511258 | 0.0024 | 0.0122 | 0.0067 | 0.0067 | TRUE | 0.0804 |
| Dyrk3 | 1920 | 1806 | 1699 | 2458 | 1733 | 1583 | 2039 | 1804 | 1706 | 1932 | 1865 | 1733 | 1846.46 | 1850 | 1843 | 0.997 | -0.004 | 0.96598954 | 0.98275125 | 0.0052 | 0.0087 | 0.0066 | 0.0066 | TRUE | 0.0969 |
| Dyrk4 | 15 | 19 | 8 | 42 | 31 | 14 | 16 | 19 | 8 | 33 | 33 | 15 | 20.77 | 14 | 27 | 1.913 | 0.936 | 0.07305701 | 0.21170231 | 0.1164 | 0.1655 | 0.1427 | 0.1427 | TRUE | 0.8669 |
| Dysf | 602 | 553 | 550 | 606 | 462 | 464 | 639 | 552 | 552 | 476 | 497 | 508 | 537.56 | 581 | 494 | 0.849 | -0.236 | 0.03396929 | 0.12544221 | 0.0025 | 0.013 | 0.0071 | 0.0071 | TRUE | 0.0889 |
| Dytn | 0 | 0 | 0 | 0 | 0 | 0 | 0 | 0 | 0 | 0 | 0 | 0 | 0 | 0 | 0 | NA | NA | NA | NA | NA | NA | NA | NA | NA | NA |
| Dyx1c1 | 177 | 277 | 294 | 235 | 202 | 219 | 188 | 277 | 295 | 185 | 217 | 240 | 233.61 | 253 | 214 | 0.842 | -0.248 | 0.23190363 | 0.44599245 | 0.0323 | 0.021 | 0.0266 | 0.0266 | TRUE | 0.56 |
| Dzank1 | 127 | 124 | 136 | 171 | 177 | 136 | 135 | 124 | 137 | 134 | 190 | 149 | 144.84 | 132 | 158 | 1.195 | 0.256 | 0.19102749 | 0.39611884 | 0.0124 | 0.0296 | 0.0207 | 0.0207 | TRUE | 0.3382 |
| Dzip1 | 637 | 752 | 741 | 875 | 716 | 751 | 677 | 751 | 744 | 688 | 770 | 822 | 742 | 724 | 760 | 1.048 | 0.068 | 0.53748077 | 0.72416259 | 0.0045 | 0.0113 | 0.0074 | 0.0074 | TRUE | 0.0853 |
| Dzip1l | 573 | 562 | 522 | 903 | 650 | 521 | 609 | 561 | 524 | 710 | 699 | 570 | 612.26 | 565 | 660 | 1.17 | 0.227 | 0.07676223 | 0.21881328 | 0.0084 | 0.0123 | 0.0102 | 0.0102 | TRUE | 0.1633 |
| Dzip3 | 3460 | 4070 | 4165 | 5605 | 3848 | 3200 | 3675 | 4065 | 4182 | 4406 | 4140 | 3503 | 3995.22 | 3974 | 4016 | 1.011 | 0.016 | 0.8861991 | 0.94408188 | 0.0089 | 0.0077 | 0.0084 | 0.0084 | TRUE | 0.1519 |
| E030002O03 | 0 | 0 | 0 | 0 | 0 | 0 | 0 | 0 | 0 | 0 | 0 | 0 | 0 | 0 | 0 | NA | NA | NA | NA | NA | NA | NA | NA | NA | NA |
| E030003E18I | 0 | 0 | 0 | 0 | 0 | 0 | 0 | 0 | 0 | 0 | 0 | 0 | 0 | 0 | 0 | NA | NA | NA | NA | NA | NA | NA | NA | NA | NA |
| E030011O05 | 14 | 15 | 14 | 23 | 24 | 20 | 15 | 15 | 14 | 18 | 26 | 22 | 18.28 | 15 | 22 | 1.489 | 0.575 | 0.19605867 | 0.40230395 | 0 | 0.1871 | 0.0851 | 0.0851 | TRUE | 0.1655 |
| E030013I19R | 0 | 0 | 0 | 0 | 0 | 0 | 0 | 0 | 0 | 0 | 0 | 0 | 0 | 0 | 0 | NA | NA | NA | NA | NA | NA | NA | NA | NA | NA |
| E030018B13I | 0 | 0 | 0 | 0 | 0 | 0 | 0 | 0 | 0 | 0 | 0 | 0 | 0 | 0 | 0 | NA | NA | NA | NA | NA | NA | NA | NA | NA | NA |
| E030019B06I | 0 | 0 | 0 | 0 | 1 | 0 | 0 | 0 | 0 | 0 | 1 | 0 | 0.18 | 0 | 0 | 1.859 | 0.894 | 0.82650734 | NA | 0 | 18.3785 | 10 | 10 | TRUE | 0.5204 |
| E030019B13I | 0 | 0 | 1 | 1 | 0 | 0 | 0 | 0 | 1 | 1 | 0 | 0 | 0.3 | 0 | 0 | 0.954 | -0.067 | 0.98640617 | NA | 0 | 11.0483 | 9.2783 | 9.2783 | TRUE | 0.4098 |
| E030024N20 | 2 | 1 | 5 | 1 | 2 | 1 | 2 | 1 | 5 | 1 | 2 | 1 | 2.03 | 3 | 1 | 0.485 | -1.045 | 0.44783797 | NA | 0 | 1.6303 | 0.807 | 0.807 | TRUE | 0.5401 |
| E030025P04I | 0 | 0 | 0 | 0 | 0 | 0 | 0 | 0 | 0 | 0 | 0 | 0 | 0 | 0 | 0 | NA | NA | NA | NA | NA | NA | NA | NA | NA | NA |
| E030030I06R | 29 | 47 | 50 | 37 | 43 | 45 | 31 | 47 | 50 | 29 | 46 | 49 | 42.09 | 43 | 41 | 0.965 | -0.052 | 0.88332084 | 0.94254355 | 0.0478 | 0.0852 | 0.067 | 0.067 | TRUE | 0.6113 |
| E030044B06I | 8 | 3 | 4 | 0 | 1 | 1 | 8 | 3 | 4 | 0 | 1 | 1 | 2.95 | 5 | 1 | 0.13 | -2.946 | 0.03993902 | NA | 0 | 1.1249 | 0.6572 | 0.6572 | TRUE | 0.6293 |
| E130003G02 | 1 | 1 | 0 | 2 | 1 | 1 | 1 | 1 | 0 | 2 | 1 | 1 | 0.97 | 1 | 1 | 1.843 | 0.882 | 0.63765974 | NA | 0 | 3.4128 | 1.3924 | 1.3924 | TRUE | 0.2529 |
| E130006D01I | 10 | 7 | 9 | 8 | 0 | 3 | 11 | 7 | 9 | 6 | 0 | 3 | 6.04 | 9 | 3 | 0.374 | -1.419 | 0.13105413 | 0.31166855 | 0.1407 | 0.5526 | 0.4314 | 0.4314 | TRUE | 0.6165 |
| E130008D07I | 35 | 37 | 40 | 37 | 44 | 45 | 37 | 37 | 40 | 29 | 47 | 49 | 40 | 38 | 42 | 1.089 | 0.123 | 0.71350394 | 0.8442782 | 0 | 0.0893 | 0.0557 | 0.0557 | TRUE | 0.6439 |
| E130012A19I | 19245 | 7819 | 7406 | 25387 | 15281 | 8948 | 20442 | 7809 | 7436 | 19957 | 16441 | 9795 | 13646.76 | 11896 | 15398 | 1.294 | 0.372 | 0.47377641 | 0.67458712 | 0.1946 | 0.0071 | 0.0876 | 0.1946 | TRUE | 1.4292 |
| E130018N17 | 6 | 10 | 6 | 17 | 17 | 26 | 6 | 10 | 6 | 13 | 18 | 28 | 13.75 | 7 | 20 | 2.658 | 1.41 | 0.01618542 | 0.07497251 | 0 | 0.2465 | 0.1549 | 0.1549 | TRUE | 0.2722 |
| E130102H24I | 29 | 56 | 38 | 41 | 53 | 50 | 31 | 56 | 38 | 32 | 57 | 55 | 44.81 | 42 | 48 | 1.143 | 0.193 | 0.60204475 | 0.77356006 | 0.0716 | 0.0804 | 0.0761 | 0.0761 | TRUE | 0.7588 |
| E130112N10 | 45 | 47 | 40 | 48 | 44 | 36 | 48 | 47 | 40 | 38 | 47 | 39 | 43.23 | 45 | 41 | 0.92 | -0.12 | 0.68181794 | 0.82362649 | 0 | 0.0831 | 0.0389 | 0.0389 | TRUE | 0.1107 |
| E130114P18I | 0 | 0 | 0 | 0 | 0 | 0 | 0 | 0 | 0 | 0 | 0 | 0 | 0 | 0 | 0 | NA | NA | NA | NA | NA | NA | NA | NA | NA | NA |
| E130201H02I | 2 | 3 | 3 | 8 | 2 | 2 | 2 | 3 | 3 | 6 | 2 | 2 | 3.13 | 3 | 3 | 1.341 | 0.424 | 0.69376717 | NA | 0 | 1.0604 | 0.5121 | 0.5121 | TRUE | 0.8931 |
| E130215H24I | 0 | 0 | 0 | 0 | 0 | 0 | 0 | 0 | 0 | 0 | 0 | 0 | 0 | 0 | 0 | NA | NA | NA | NA | NA | NA | NA | NA | NA | NA |
| E130218I03R | 2 | 0 | 0 | 3 | 1 | 2 | 2 | 0 | 0 | 2 | 1 | 2 | 1.29 | 1 | 2 | 2.716 | 1.442 | 0.44085401 | NA | 0 | 2.5581 | 1.5265 | 1.5265 | TRUE | 0.9745 |
| E130304I02R | 0 | 2 | 1 | 0 | 0 | 1 | 0 | 2 | 1 | 0 | 0 | 1 | 0.68 | 1 | 0 | 0.347 | -1.526 | 0.55389466 | NA | 0 | 4.8325 | 3.2797 | 3.2797 | TRUE | 0.2854 |
| E130307A14I | 42 | 43 | 47 | 51 | 37 | 40 | 45 | 43 | 47 | 40 | 40 | 44 | 43.07 | 45 | 41 | 0.917 | -0.125 | 0.65947811 | 0.80811815 | 0 | 0.0834 | 0.0352 | 0.0352 | TRUE | 0.0206 |
| E130308A19I | 1264 | 1118 | 1227 | 1467 | 1140 | 1117 | 1343 | 1117 | 1232 | 1153 | 1227 | 1223 | 1215.62 | 1231 | 1201 | 0.976 | -0.035 | 0.72169753 | 0.84944877 | 0.004 | 0.0096 | 0.0063 | 0.0063 | TRUE | 0.0787 |
| E130309D02I | 1629 | 1491 | 1463 | 2049 | 1420 | 1336 | 1730 | 1489 | 1469 | 1611 | 1528 | 1462 | 1548.24 | 1563 | 1534 | 0.982 | -0.026 | 0.79247571 | 0.89309946 | 0.0047 | 0.009 | 0.0065 | 0.0065 | TRUE | 0.1055 |
| E130309D14I | 2059 | 1823 | 1678 | 1067 | 842 | 847 | 2187 | 1821 | 1685 | 839 | 906 | 927 | 1394.08 | 1898 | 891 | 0.469 | -1.092 | 1.70E-19 | 3.93E-17 | 0.0095 | 0.0093 | 0.0097 | 0.0097 | TRUE | 0.2148 |
| E130309F12I | 22 | 22 | 18 | 30 | 16 | 23 | 23 | 22 | 18 | 24 | 17 | 25 | 21.56 | 21 | 22 | 1.044 | 0.062 | 0.88123309 | 0.94135674 | 0 | 0.1597 | 0.0765 | 0.0765 | TRUE | 0.1887 |
| E130310I04R | 2 | 0 | 0 | 0 | 1 | 0 | 2 | 0 | 0 | 0 | 1 | 0 | 0.53 | 1 | 0 | 0.497 | -1.009 | 0.76131406 | NA | 0 | 6.1834 | 6.228 | 6.228 | TRUE | 0.9742 |
| E130311K13I | 120 | 220 | 199 | 260 | 242 | 260 | 127 | 220 | 200 | 204 | 260 | 285 | 216.06 | 182 | 250 | 1.366 | 0.45 | 0.05318206 | 0.17120833 | 0.0476 | 0.0222 | 0.0343 | 0.0343 | TRUE | 0.7359 |
| E130317F20I | 28 | 46 | 41 | 111 | 94 | 62 | 30 | 46 | 41 | 87 | 101 | 68 | 62.19 | 39 | 85 | 2.191 | 1.131 | 0.00011178 | 0.00150346 | 0.0234 | 0.0599 | 0.043 | 0.043 | TRUE | 0.1826 |
| E230008N13 | 4 | 5 | 2 | 6 | 5 | 7 | 4 | 5 | 2 | 5 | 5 | 8 | 4.83 | 4 | 6 | 1.568 | 0.649 | 0.45377649 | 0.65906064 | 0 | 0.6883 | 0.3208 | 0.3208 | TRUE | 0.2663 |
| E230016K23I | 27 | 13 | 9 | 28 | 25 | 19 | 29 | 13 | 9 | 22 | 27 | 21 | 20.07 | 17 | 23 | 1.379 | 0.464 | 0.37585748 | 0.59147378 | 0.117 | 0.1711 | 0.146 | 0.146 | TRUE | 1.7602 |
| E230016M11 | 3 | 8 | 9 | 12 | 4 | 10 | 3 | 8 | 9 | 9 | 4 | 11 | 7.48 | 7 | 8 | 1.222 | 0.289 | 0.7011299 | 0.83650923 | 0 | 0.4472 | 0.2729 | 0.2729 | TRUE | 0.5173 |
| E230019M04 | 3 | 0 | 0 | 0 | 0 | 4 | 3 | 0 | 0 | 0 | 0 | 4 | 1.26 | 1 | 1 | 1.363 | 0.447 | 0.86919973 | NA | 10 | 2.6198 | 4.487 | 4.487 | TRUE | 1.9001 |
| E230025N22 | 17 | 23 | 23 | 30 | 19 | 27 | 18 | 23 | 23 | 24 | 20 | 30 | 22.95 | 21 | 25 | 1.144 | 0.194 | 0.62873206 | 0.79115702 | 0 | 0.1504 | 0.0724 | 0.0724 | TRUE | 0.1752 |
| E230029C05I | 0 | 0 | 0 | 0 | 0 | 0 | 0 | 0 | 0 | 0 | 0 | 0 | 0 | 0 | 0 | NA | NA | NA | NA | NA | NA | NA | NA | NA | NA |
| E2f1 | 2244 | 3013 | 3126 | 3057 | 2265 | 2428 | 2384 | 3009 | 3139 | 2403 | 2437 | 2658 | 2671.58 | 2844 | 2499 | 0.879 | -0.187 | 0.12236784 | 0.29760764 | 0.0111 | 0.0081 | 0.0101 | 0.0101 | TRUE | 0.243 |
| E2f2 | 567 | 618 | 652 | 471 | 347 | 358 | 602 | 617 | 655 | 370 | 373 | 392 | 501.61 | 625 | 378 | 0.605 | -0.724 | 2.61E-12 | 2.55E-10 | 0 | 0.0135 | 0.0056 | 0.0056 | TRUE | 0.0208 |
| E2f3 | 2159 | 1392 | 1299 | 2671 | 1816 | 1510 | 2293 | 1390 | 1304 | 2100 | 1954 | 1653 | 1782.39 | 1662 | 1902 | 1.144 | 0.195 | 0.34628005 | 0.56542111 | 0.056 | 0.0088 | 0.0302 | 0.0302 | TRUE | 1.327 |
| E2f4 | 8319 | 7815 | 7853 | 10174 | 7510 | 7150 | 8836 | 7805 | 7885 | 7998 | 8080 | 7827 | 8071.92 | 8175 | 7968 | 0.975 | -0.037 | 0.62974944 | 0.79193176 | 0.0022 | 0.0073 | 0.0041 | 0.0041 | TRUE | 0.061 |
| E2f5 | 1090 | 1608 | 1596 | 1768 | 1357 | 1592 | 1158 | 1606 | 1603 | 1390 | 1460 | 1743 | 1493.14 | 1456 | 1531 | 1.051 | 0.072 | 0.63889061 | 0.7965678 | 0.0233 | 0.0091 | 0.0164 | 0.0164 | TRUE | 0.3843 |
| E2f6 | 1410 | 1281 | 1410 | 1937 | 1281 | 1171 | 1498 | 1279 | 1416 | 1523 | 1378 | 1282 | 1395.94 | 1398 | 1394 | 0.999 | -0.002 | 0.98470528 | 0.9919379 | 0.0062 | 0.0093 | 0.0075 | 0.0075 | TRUE | 0.0797 |
| E2f7 | 1579 | 1322 | 1307 | 1726 | 1242 | 1219 | 1677 | 1320 | 1312 | 1357 | 1336 | 1334 | 1389.56 | 1436 | 1342 | 0.935 | -0.097 | 0.40996179 | 0.62158382 | 0.0095 | 0.0093 | 0.0093 | 0.0093 | TRUE | 0.2575 |
| E2f8 | 2591 | 2174 | 2227 | 3954 | 2815 | 2858 | 2752 | 2171 | 2236 | 3108 | 3029 | 3129 | 2737.51 | 2386 | 3089 | 1.294 | 0.372 | 0.00061874 | 0.00614921 | 0.0085 | 0.0081 | 0.0082 | 0.0082 | TRUE | 0.2173 |
| E330009J07F | 253 | 263 | 256 | 324 | 245 | 262 | 269 | 263 | 257 | 255 | 264 | 287 | 265.59 | 263 | 269 | 1.02 | 0.028 | 0.82956304 | 0.91355036 | 0 | 0.0193 | 0.0083 | 0.0083 | TRUE | 0.0394 |
| E330011O21 | 2 | 0 | 0 | 1 | 1 | 0 | 2 | 0 | 0 | 1 | 1 | 0 | 0.66 | 1 | 1 | 0.896 | -0.159 | 0.95209679 | NA | 0 | 4.9654 | 3.5226 | 3.5226 | TRUE | 0.9743 |
| E330012B07I | 0 | 0 | 0 | 0 | 0 | 0 | 0 | 0 | 0 | 0 | 0 | 0 | 0 | 0 | 0 | NA | NA | NA | NA | NA | NA | NA | NA | NA | NA |
| E330013P04I | 1 | 0 | 0 | 0 | 1 | 1 | 1 | 0 | 0 | 0 | 1 | 1 | 0.54 | 0 | 1 | 1.923 | 0.944 | 0.73523723 | NA | 0 | 6.1214 | 3.9144 | 3.9144 | TRUE | 0.4614 |
| E330014E10I | 0 | 0 | 0 | 0 | 0 | 0 | 0 | 0 | 0 | 0 | 0 | 0 | 0 | 0 | 0 | NA | NA | NA | NA | NA | NA | NA | NA | NA | NA |

| | | | | | | | | | | | | | | | | | | | | | | | | | |
|---|---|---|---|---|---|---|---|---|---|---|---|---|---|---|---|---|---|---|---|---|---|---|---|---|---|
| E330017A01 | 0 | 0 | 1 | 1 | 0 | 0 | 0 | 0 | 1 | 1 | 0 | 0 | 0.3 | 0 | 0 | 0.954 | -0.067 | 0.98640617 | NA | 0 | 11.0483 | 9.2783 | 9.2783 | TRUE | 0.4098 |
| E330017L17 | 0 | 0 | 0 | 1 | 0 | 0 | 0 | 0 | 0 | 1 | 0 | 0 | 0.13 | 0 | 0 | 1.859 | 0.894 | 0.82650734 | NA | 0 | 25.1515 | 10 | 10 | TRUE | 0.2563 |
| E330020D12 | 229 | 177 | 116 | 112 | 83 | 73 | 243 | 177 | 116 | 88 | 89 | 80 | 132.29 | 179 | 86 | 0.48 | -1.058 | 0.00014607 | 0.00186547 | 0.0633 | 0.0318 | 0.0474 | 0.0474 | TRUE | 0.1149 |
| E330021D16 | 10 | 1 | 1 | 7 | 1 | 3 | 11 | 1 | 1 | 6 | 1 | 3 | 3.75 | 4 | 3 | 0.8 | -0.322 | 0.79820132 | NA | 0.8689 | 0.8859 | 0.8757 | 0.8757 | TRUE | 1.2584 |
| E330023G01 | 0 | 0 | 6 | 5 | 3 | 3 | 0 | 0 | 6 | 4 | 3 | 3 | 2.74 | 2 | 3 | 1.729 | 0.79 | 0.59920939 | NA | 1.2878 | 1.2073 | 1.2345 | 1.2345 | TRUE | 2.7768 |
| E330033B04 | 10 | 10 | 18 | 20 | 21 | 14 | 11 | 10 | 18 | 16 | 23 | 15 | 15.39 | 13 | 18 | 1.38 | 0.465 | 0.36497142 | 0.58244757 | 0 | 0.221 | 0.1228 | 0.1228 | TRUE | 0.5165 |
| E330034G19 | 0 | 1 | 0 | 1 | 0 | 0 | 0 | 1 | 0 | 1 | 0 | 0 | 0.3 | 0 | 0 | 0.954 | -0.067 | 0.98642001 | NA | 0 | 11.0817 | 9.3014 | 9.3014 | TRUE | 0.405 |
| E430016F16 | 0 | 1 | 0 | 0 | 0 | 0 | 0 | 1 | 0 | 0 | 0 | 0 | 0.17 | 0 | 0 | 0.49 | -1.029 | 0.80087 | NA | 0 | 19.7991 | 10 | 10 | TRUE | 0.405 |
| E430018J23 | 57 | 68 | 74 | 104 | 77 | 50 | 61 | 68 | 74 | 82 | 83 | 55 | 70.35 | 68 | 73 | 1.085 | 0.118 | 0.65364568 | 0.80540538 | 0.0155 | 0.0537 | 0.0355 | 0.0355 | TRUE | 0.4144 |
| E430025E21 | 6999 | 5935 | 6124 | 9789 | 6911 | 6463 | 7434 | 5927 | 6149 | 7695 | 7436 | 7075 | 6952.76 | 6503 | 7402 | 1.138 | 0.187 | 0.07662877 | 0.21855135 | 0.0085 | 0.0074 | 0.0079 | 0.0079 | TRUE | 0.1911 |
| E4f1 | 863 | 1026 | 1041 | 1052 | 850 | 897 | 917 | 1025 | 1045 | 827 | 915 | 982 | 951.68 | 996 | 908 | 0.911 | -0.135 | 0.21132293 | 0.42189873 | 0.0052 | 0.0104 | 0.0074 | 0.0074 | TRUE | 0.0748 |
| E530001F21 | 0 | 1 | 2 | 1 | 1 | 0 | 0 | 1 | 2 | 1 | 1 | 0 | 0.81 | 1 | 1 | 0.624 | -0.68 | 0.75467218 | NA | 0 | 4.0666 | 2.13 | 2.13 | TRUE | 0.1306 |
| E530011L22 | 7 | 13 | 13 | 6 | 15 | 6 | 7 | 13 | 13 | 5 | 16 | 7 | 10.15 | 11 | 9 | 0.806 | -0.311 | 0.65188579 | 0.80494346 | 0.1512 | 0.3315 | 0.2435 | 0.2435 | TRUE | 1.3895 |
| Eaf1 | 4595 | 4179 | 4055 | 5296 | 3816 | 3388 | 4881 | 4174 | 4072 | 4163 | 4106 | 3709 | 4183.96 | 4376 | 3993 | 0.913 | -0.132 | 0.19064043 | 0.39573321 | 0.0066 | 0.0077 | 0.0071 | 0.0071 | TRUE | 0.1242 |
| Eaf2 | 67 | 93 | 80 | 86 | 60 | 69 | 71 | 93 | 80 | 68 | 65 | 76 | 75.34 | 81 | 70 | 0.848 | -0.237 | 0.30811916 | 0.52856814 | 0 | 0.0506 | 0.0258 | 0.0258 | TRUE | 0.1412 |
| Eapp | 969 | 1281 | 1313 | 1648 | 1222 | 1125 | 1029 | 1279 | 1318 | 1296 | 1315 | 1231 | 1244.8 | 1209 | 1281 | 1.059 | 0.083 | 0.47686349 | 0.67653962 | 0.0085 | 0.0095 | 0.009 | 0.009 | TRUE | 0.202 |
| Ear1 | 0 | 0 | 0 | 0 | 0 | 0 | 0 | 0 | 0 | 0 | 0 | 0 | 0 | 0 | 0 | NA | NA | NA | NA | NA | NA | NA | NA | NA | NA |
| Ear10 | 0 | 0 | 0 | 0 | 0 | 0 | 0 | 0 | 0 | 0 | 0 | 0 | 0 | 0 | 0 | NA | NA | NA | NA | NA | NA | NA | NA | NA | NA |
| Ear11 | 0 | 0 | 0 | 0 | 0 | 0 | 0 | 0 | 0 | 0 | 0 | 0 | 0 | 0 | 0 | NA | NA | NA | NA | NA | NA | NA | NA | NA | NA |
| Ear12 | 0 | 0 | 0 | 0 | 0 | 0 | 0 | 0 | 0 | 0 | 0 | 0 | 0 | 0 | 0 | NA | NA | NA | NA | NA | NA | NA | NA | NA | NA |
| Ear14 | 0 | 0 | 0 | 0 | 0 | 0 | 0 | 0 | 0 | 0 | 0 | 0 | 0 | 0 | 0 | NA | NA | NA | NA | NA | NA | NA | NA | NA | NA |
| Ear2 | 0 | 0 | 0 | 0 | 0 | 0 | 0 | 0 | 0 | 0 | 0 | 0 | 0 | 0 | 0 | NA | NA | NA | NA | NA | NA | NA | NA | NA | NA |
| Ear3 | 0 | 0 | 0 | 0 | 0 | 0 | 0 | 0 | 0 | 0 | 0 | 0 | 0 | 0 | 0 | NA | NA | NA | NA | NA | NA | NA | NA | NA | NA |
| Ear4 | 0 | 0 | 0 | 0 | 0 | 0 | 0 | 0 | 0 | 0 | 0 | 0 | 0 | 0 | 0 | NA | NA | NA | NA | NA | NA | NA | NA | NA | NA |
| Ear5 | 0 | 0 | 0 | 0 | 0 | 0 | 0 | 0 | 0 | 0 | 0 | 0 | 0 | 0 | 0 | NA | NA | NA | NA | NA | NA | NA | NA | NA | NA |
| Ear6 | 0 | 0 | 0 | 0 | 0 | 0 | 0 | 0 | 0 | 0 | 0 | 0 | 0 | 0 | 0 | NA | NA | NA | NA | NA | NA | NA | NA | NA | NA |
| Ear7 | 0 | 0 | 0 | 0 | 0 | 0 | 0 | 0 | 0 | 0 | 0 | 0 | 0 | 0 | 0 | NA | NA | NA | NA | NA | NA | NA | NA | NA | NA |
| Ears2 | 1115 | 1191 | 1241 | 1564 | 1099 | 1102 | 1184 | 1189 | 1246 | 1229 | 1182 | 1206 | 1206.36 | 1206 | 1206 | 1 | 0 | 0.9979094 | 0.99860587 | 0 | 0.0096 | 0.0034 | 0.0034 | TRUE | 0.0099 |
| Ebag9 | 737 | 747 | 733 | 962 | 741 | 713 | 783 | 746 | 736 | 756 | 797 | 780 | 766.48 | 755 | 778 | 1.03 | 0.043 | 0.62842585 | 0.79094062 | 0 | 0.0112 | 0.0043 | 0.0043 | TRUE | 0.0122 |
| Ebf1 | 119 | 110 | 132 | 59 | 44 | 47 | 126 | 110 | 133 | 46 | 47 | 51 | 85.66 | 123 | 48 | 0.393 | -1.348 | 1.71E-09 | 9.89E-08 | 0 | 0.0454 | 0.0218 | 0.0218 | TRUE | 0.0889 |
| Ebf2 | 6 | 3 | 5 | 6 | 5 | 6 | 6 | 3 | 5 | 5 | 5 | 7 | 5.18 | 5 | 6 | 1.154 | 0.207 | 0.80025366 | 0.89686328 | 0 | 0.6434 | 0.286 | 0.286 | TRUE | 0.2143 |
| Ebf3 | 17 | 17 | 24 | 84 | 51 | 47 | 18 | 17 | 24 | 66 | 55 | 51 | 38.58 | 20 | 57 | 2.925 | 1.548 | 5.30E-06 | 0.0001164 | 0 | 0.0923 | 0.049 | 0.049 | TRUE | 0.2064 |
| Ebf4 | 43 | 62 | 60 | 40 | 35 | 43 | 46 | 62 | 60 | 31 | 38 | 47 | 47.34 | 56 | 39 | 0.686 | -0.543 | 0.07650379 | 0.21834836 | 0.0115 | 0.0765 | 0.0459 | 0.0459 | TRUE | 0.2543 |
| Ebi3 | 26 | 35 | 33 | 11 | 14 | 10 | 28 | 35 | 33 | 9 | 15 | 11 | 21.73 | 32 | 12 | 0.357 | -1.486 | 0.00076527 | 0.00721877 | 0 | 0.1585 | 0.0818 | 0.0818 | TRUE | 0.2605 |
| Ebna1bp2 | 7024 | 6290 | 6295 | 9218 | 6397 | 5980 | 7461 | 6282 | 6321 | 7246 | 6883 | 6546 | 6789.79 | 6688 | 6892 | 1.031 | 0.044 | 0.65097412 | 0.80432149 | 0.0051 | 0.0074 | 0.0065 | 0.0065 | TRUE | 0.1246 |
| Ebp | 673 | 771 | 763 | 939 | 655 | 647 | 715 | 770 | 766 | 738 | 705 | 708 | 733.69 | 750 | 717 | 0.956 | -0.065 | 0.47572395 | 0.67577395 | 0 | 0.0114 | 0.0046 | 0.0046 | TRUE | 0.0201 |
| Ebpl | 1215 | 1320 | 1184 | 1608 | 1056 | 1145 | 1291 | 1318 | 1189 | 1264 | 1136 | 1253 | 1241.89 | 1266 | 1218 | 0.963 | -0.055 | 0.54744152 | 0.73246078 | 0.0024 | 0.0096 | 0.0052 | 0.0052 | TRUE | 0.0408 |
| Ecd | 3446 | 3103 | 3135 | 4551 | 3237 | 3069 | 3660 | 3099 | 3148 | 3578 | 3483 | 3360 | 3387.85 | 3302 | 3474 | 1.052 | 0.073 | 0.42618609 | 0.63629373 | 0.0043 | 0.0079 | 0.0058 | 0.0058 | TRUE | 0.109 |
| Ece1 | 5100 | 4909 | 5198 | 4093 | 3059 | 3511 | 5417 | 4903 | 5219 | 3218 | 3291 | 3843 | 4315.22 | 5180 | 3451 | 0.666 | -0.586 | 1.34E-09 | 7.88E-08 | 0.0058 | 0.0077 | 0.0065 | 0.0065 | TRUE | 0.12 |
| Ece2 | 817 | 874 | 847 | 998 | 755 | 711 | 868 | 873 | 850 | 785 | 812 | 778 | 827.72 | 864 | 792 | 0.916 | -0.126 | 0.13202713 | 0.31299142 | 0 | 0.0109 | 0.0038 | 0.0038 | TRUE | 0.006 |
| Ecel1 | 100 | 150 | 169 | 234 | 143 | 134 | 106 | 150 | 170 | 184 | 154 | 147 | 151.7 | 142 | 162 | 1.14 | 0.189 | 0.39021591 | 0.60395507 | 0.0277 | 0.0286 | 0.0282 | 0.0282 | TRUE | 0.4945 |
| Ech1 | 1963 | 2602 | 2854 | 2469 | 1936 | 2384 | 2085 | 2599 | 2866 | 1941 | 2083 | 2610 | 2363.84 | 2517 | 2211 | 0.878 | -0.187 | 0.22260114 | 0.4346744 | 0.0245 | 0.0083 | 0.0165 | 0.0165 | TRUE | 0.3003 |
| Echdc1 | 1132 | 1260 | 1313 | 1189 | 867 | 936 | 1202 | 1258 | 1318 | 935 | 933 | 1025 | 1111.88 | 1259 | 964 | 0.765 | -0.387 | 1.50E-05 | 0.00027878 | 0.0015 | 0.0099 | 0.0048 | 0.0048 | TRUE | 0.0353 |
| Echdc2 | 1343 | 1383 | 1346 | 1770 | 1286 | 1258 | 1427 | 1381 | 1352 | 1391 | 1384 | 1377 | 1385.24 | 1387 | 1384 | 0.999 | -0.002 | 0.97652755 | 0.98729439 | 0 | 0.0093 | 0.0031 | 0.0031 | TRUE | 0.0076 |
| Echdc3 | 43 | 42 | 42 | 78 | 53 | 59 | 46 | 42 | 42 | 61 | 57 | 65 | 52.12 | 43 | 61 | 1.41 | 0.496 | 0.05934186 | 0.18449436 | 0 | 0.0701 | 0.03 | 0.03 | TRUE | 0.0256 |
| Echs1 | 2685 | 3405 | 3562 | 3173 | 2501 | 2792 | 2852 | 3401 | 3577 | 2494 | 2691 | 3056 | 3011.79 | 3277 | 2747 | 0.838 | -0.255 | 0.03458287 | 0.12693531 | 0.0115 | 0.008 | 0.0101 | 0.0101 | TRUE | 0.1559 |
| Eci1 | 680 | 890 | 894 | 941 | 657 | 670 | 722 | 889 | 898 | 740 | 707 | 733 | 781.48 | 836 | 727 | 0.869 | -0.203 | 0.08081623 | 0.22612512 | 0.0063 | 0.0111 | 0.0084 | 0.0084 | TRUE | 0.1679 |
| Eci2 | 964 | 1183 | 1083 | 1634 | 1128 | 1234 | 1024 | 1181 | 1087 | 1285 | 1214 | 1351 | 1190.31 | 1097 | 1283 | 1.169 | 0.225 | 0.01886992 | 0.08338931 | 0.0028 | 0.0097 | 0.0058 | 0.0058 | TRUE | 0.0535 |
| Eci3 | 18 | 25 | 19 | 27 | 13 | 23 | 19 | 25 | 19 | 21 | 14 | 25 | 20.59 | 21 | 20 | 0.957 | -0.064 | 0.8842546 | 0.94317602 | 0 | 0.1669 | 0.0899 | 0.0899 | TRUE | 0.3517 |
| Ecm1 | 58 | 41 | 50 | 49 | 32 | 28 | 62 | 41 | 50 | 39 | 34 | 31 | 42.73 | 51 | 35 | 0.682 | -0.552 | 0.07869159 | 0.22216207 | 0 | 0.084 | 0.0468 | 0.0468 | TRUE | 0.1755 |
| Ecm2 | 5 | 1 | 1 | 17 | 16 | 6 | 5 | 1 | 1 | 13 | 17 | 7 | 7.41 | 2 | 12 | 5.154 | 2.366 | 0.00988099 | 0.05221089 | 0.1984 | 0.4515 | 0.3543 | 0.3543 | TRUE | 0.6098 |
| Ecscr | 246 | 285 | 331 | 129 | 96 | 148 | 261 | 285 | 332 | 101 | 103 | 162 | 207.5 | 293 | 122 | 0.416 | -1.266 | 6.53E-09 | 3.23E-07 | 0.0352 | 0.0228 | 0.0286 | 0.0286 | TRUE | 0.7977 |
| Ecsit | 2349 | 2524 | 2628 | 2505 | 1833 | 1994 | 2495 | 2521 | 2639 | 1969 | 1972 | 2183 | 2296.46 | 2552 | 2041 | 0.8 | -0.322 | 5.13E-05 | 0.0007922 | 0.0016 | 0.0083 | 0.0041 | 0.0041 | TRUE | 0.0439 |
| Ect2 | 12671 | 13788 | 14051 | 15847 | 11331 | 12453 | 13459 | 13770 | 14109 | 12458 | 12191 | 13632 | 13269.77 | 13779 | 12760 | 0.926 | -0.111 | 0.1245694 | 0.30133568 | 0.002 | 0.0072 | 0.0037 | 0.0037 | TRUE | 0.0435 |
| Ect2l | 3 | 3 | 1 | 7 | 2 | 2 | 3 | 3 | 1 | 6 | 2 | 2 | 2.84 | 2 | 3 | 1.406 | 0.491 | 0.66679262 | NA | 0 | 1.1675 | 0.5794 | 0.5794 | TRUE | 0.6406 |
| Eda | 312 | 263 | 283 | 436 | 297 | 266 | 331 | 263 | 284 | 343 | 320 | 291 | 305.28 | 293 | 318 | 1.088 | 0.122 | 0.40347264 | 0.6162915 | 0.007 | 0.0177 | 0.0121 | 0.0121 | TRUE | 0.1488 |
| Eda2r | 2969 | 3933 | 4051 | 3704 | 2655 | 3272 | 3154 | 3928 | 4068 | 2912 | 2857 | 3582 | 3416.55 | 3717 | 3117 | 0.838 | -0.254 | 0.05801868 | 0.1819496 | 0.0167 | 0.0079 | 0.0127 | 0.0127 | TRUE | 0.2133 |
| Edar | 14 | 12 | 6 | 9 | 9 | 12 | 15 | 12 | 6 | 7 | 10 | 13 | 10.46 | 11 | 10 | 0.901 | -0.15 | 0.8122449 | 0.90375496 | 0 | 0.3218 | 0.1906 | 0.1906 | TRUE | 0.505 |
| Edaradd | 50 | 57 | 59 | 73 | 70 | 78 | 53 | 57 | 59 | 57 | 75 | 85 | 64.56 | 56 | 72 | 1.28 | 0.356 | 0.1766948 | 0.37715933 | 0 | 0.0579 | 0.0341 | 0.0341 | TRUE | 0.3503 |
| Edc3 | 3349 | 3234 | 3317 | 4217 | 3033 | 2960 | 3557 | 3230 | 3331 | 3315 | 3263 | 3240 | 3322.71 | 3373 | 3273 | 0.971 | -0.043 | 0.54324762 | 0.72861181 | 9.00E-04 | 0.0079 | 0.0033 | 0.0033 | TRUE | 0.0278 |
| Edc4 | 10280 | 9207 | 9191 | 11661 | 9091 | 8529 | 10919 | 9195 | 9229 | 9167 | 9781 | 9336 | 9604.61 | 9781 | 9428 | 0.964 | -0.053 | 0.56759569 | 0.74731774 | 0.005 | 0.0072 | 0.0061 | 0.0061 | TRUE | 0.1265 |
| Eddm3b | 0 | 0 | 0 | 0 | 1 | 0 | 0 | 0 | 0 | 0 | 1 | 0 | 0.18 | 0 | 0 | 1.859 | 0.894 | 0.82650734 | NA | 0 | 18.3785 | 10 | 10 | TRUE | 0.5204 |
| Edem1 | 5479 | 5981 | 6080 | 6951 | 5117 | 5274 | 5820 | 5973 | 6105 | 5464 | 5506 | 5773 | 5773.51 | 5966 | 5581 | 0.935 | -0.096 | 0.1242497 | 0.30079299 | 5.00E-04 | 0.0075 | 0.0027 | 0.0027 | TRUE | 0.011 |
| Edem2 | 2764 | 3750 | 3716 | 3439 | 2709 | 2857 | 2936 | 3745 | 3731 | 2703 | 2915 | 3127 | 3192.98 | 3471 | 2915 | 0.84 | -0.252 | 0.03557574 | 0.12950116 | 0.0111 | 0.0079 | 0.0101 | 0.0101 | TRUE | 0.2207 |
| Edem3 | 5744 | 4618 | 4765 | 5770 | 4171 | 4057 | 6101 | 4612 | 4785 | 4536 | 4488 | 4441 | 4827.07 | 5166 | 4488 | 0.869 | -0.203 | 0.08649672 | 0.23754591 | 0.0122 | 0.0076 | 0.0099 | 0.0099 | TRUE | 0.3055 |
| Edf1 | 2386 | 2592 | 2683 | 4047 | 2828 | 2637 | 2534 | 2589 | 2694 | 3181 | 3043 | 2887 | 2821.3 | 2606 | 3037 | 1.166 | 0.221 | 0.00307743 | 0.02165876 | 0.0014 | 0.0081 | 0.0037 | 0.0037 | TRUE | 0.0227 |
| Edil3 | 10 | 8 | 3 | 12 | 23 | 11 | 11 | 8 | 3 | 9 | 25 | 12 | 11.31 | 7 | 15 | 2.128 | 1.089 | 0.11886298 | 0.29199879 | 0.1942 | 0.2983 | 0.2484 | 0.2484 | TRUE | 1.2509 |
| Edn1 | 18 | 8 | 6 | 45 | 28 | 21 | 19 | 8 | 6 | 35 | 30 | 23 | 20.27 | 11 | 29 | 2.697 | 1.431 | 0.0073352 | 0.0415407 | 0.1032 | 0.1694 | 0.1423 | 0.1423 | TRUE | 1.0833 |
| Edn2 | 11 | 18 | 15 | 20 | 8 | 14 | 12 | 18 | 15 | 16 | 9 | 15 | 14.06 | 15 | 13 | 0.891 | -0.166 | 0.75179418 | 0.86898482 | 0 | 0.2412 | 0.1276 | 0.1276 | TRUE | 0.3605 |

| | | | | | | | | | | | | | | | | | | | | | | | | | |
|---|---|---|---|---|---|---|---|---|---|---|---|---|---|---|---|---|---|---|---|---|---|---|---|---|---|
| Edn3 | 7 | 7 | 9 | 9 | 14 | 14 | 7 | 7 | 9 | 7 | 15 | 15 | 10.15 | 8 | 12 | 1.574 | 0.654 | 0.2984777 | 0.51905693 | 0 | 0.3313 | 0.1805 | 0.1805 | TRUE | 0.7425 |
| Ednra | 43 | 30 | 26 | 73 | 53 | 55 | 46 | 30 | 26 | 57 | 57 | 60 | 46.06 | 34 | 58 | 1.72 | 0.782 | 0.01307098 | 0.06408314 | 0.0049 | 0.0784 | 0.0481 | 0.0481 | TRUE | 0.625 |
| Ednrb | 7 | 12 | 6 | 8 | 4 | 10 | 7 | 12 | 6 | 6 | 4 | 11 | 7.83 | 8 | 7 | 0.841 | -0.25 | 0.72695929 | 0.852905 | 0 | 0.4276 | 0.2412 | 0.2412 | TRUE | 0.5078 |
| Eea1 | 1383 | 1509 | 1551 | 2107 | 1537 | 1589 | 1469 | 1507 | 1557 | 1656 | 1654 | 1739 | 1597.15 | 1511 | 1683 | 1.113 | 0.155 | 0.03695367 | 0.13293794 | 0 | 0.009 | 0.0033 | 0.0033 | TRUE | 0.0101 |
| Eed | 5607 | 4971 | 4857 | 6743 | 4904 | 4704 | 5956 | 4965 | 4877 | 5301 | 5276 | 5149 | 5253.94 | 5266 | 5242 | 0.996 | -0.006 | 0.94852553 | 0.9742329 | 0.0061 | 0.0075 | 0.0067 | 0.0067 | TRUE | 0.16 |
| Eef1a1 | 475883 | 584626 | 590739 | 643842 | 450583 | 481952 | 505472 | 583871 | 593164 | 506138 | 484800 | 527572 | 533502.91 | 560836 | 506170 | 0.903 | -0.148 | 0.09139642 | 0.24662179 | 0.0046 | 0.0069 | 0.0055 | 0.0055 | TRUE | 0.0914 |
| Eef1a2 | 210 | 205 | 177 | 214 | 124 | 147 | 223 | 205 | 178 | 168 | 133 | 161 | 178.01 | 202 | 154 | 0.766 | -0.384 | 0.02886826 | 0.11199935 | 0.0079 | 0.0254 | 0.0166 | 0.0166 | TRUE | 0.1418 |
| Eef1b2 | 34012 | 40112 | 39520 | 48161 | 32851 | 34221 | 36127 | 40060 | 39682 | 37860 | 35346 | 37460 | 37755.93 | 38623 | 36889 | 0.955 | -0.066 | 0.36561295 | 0.58306561 | 0.0022 | 0.007 | 0.0038 | 0.0038 | TRUE | 0.0391 |
| Eef1d | 12046 | 13355 | 13660 | 17947 | 13151 | 12685 | 12795 | 13338 | 13716 | 14109 | 14150 | 13886 | 13665.46 | 13283 | 14048 | 1.058 | 0.081 | 0.1735428 | 0.37270111 | 5.00E-04 | 0.0071 | 0.0025 | 0.0025 | TRUE | 0.0126 |
| Eef1e1 | 3490 | 4074 | 3911 | 4545 | 3196 | 3369 | 3707 | 4069 | 3927 | 3573 | 3439 | 3688 | 3733.72 | 3901 | 3567 | 0.914 | -0.129 | 0.07735338 | 0.21970423 | 0.0013 | 0.0078 | 0.0036 | 0.0036 | TRUE | 0.023 |
| Eef1g | 42160 | 53158 | 54052 | 59818 | 44135 | 44846 | 44781 | 53089 | 54274 | 47024 | 47487 | 49091 | 49291.07 | 50715 | 47867 | 0.944 | -0.083 | 0.36800992 | 0.58529007 | 0.0054 | 0.007 | 0.0062 | 0.0062 | TRUE | 0.1282 |
| Eef2 | 229254 | 288502 | 301645 | 293149 | 224883 | 245332 | 243508 | 288130 | 302883 | 230451 | 241960 | 268555 | 262581.13 | 278174 | 246989 | 0.888 | -0.172 | 0.11177397 | 0.28113131 | 0.0093 | 0.0069 | 0.0084 | 0.0084 | TRUE | 0.1456 |
| Eef2k | 983 | 1070 | 1129 | 1064 | 779 | 801 | 1044 | 1069 | 1134 | 836 | 838 | 877 | 966.3 | 1082 | 850 | 0.786 | -0.348 | 4.52E-05 | 0.00071212 | 0 | 0.0103 | 0.0042 | 0.0042 | TRUE | 0.0208 |
| Eefsec | 2123 | 2151 | 2148 | 2914 | 2125 | 2078 | 2255 | 2148 | 2157 | 2291 | 2286 | 2275 | 2235.31 | 2187 | 2284 | 1.045 | 0.063 | 0.34236239 | 0.56253908 | 0 | 0.0084 | 0.0027 | 0.0027 | TRUE | 0.009 |
| Eepd1 | 204 | 183 | 167 | 329 | 198 | 249 | 217 | 183 | 168 | 259 | 213 | 273 | 218.56 | 189 | 248 | 1.314 | 0.394 | 0.02235317 | 0.09421777 | 0.0117 | 0.022 | 0.0168 | 0.0168 | TRUE | 0.1742 |
| Efcab1 | 42 | 79 | 87 | 90 | 81 | 72 | 45 | 79 | 87 | 71 | 87 | 79 | 74.6 | 70 | 79 | 1.118 | 0.161 | 0.58461267 | 0.75995311 | 0.0474 | 0.0511 | 0.0494 | 0.0494 | TRUE | 0.8973 |
| Efcab10 | 42 | 92 | 98 | 56 | 50 | 69 | 45 | 92 | 98 | 44 | 54 | 76 | 68.04 | 78 | 58 | 0.734 | -0.447 | 0.2178519 | 0.42981738 | 0.1063 | 0.0553 | 0.0796 | 0.0796 | TRUE | 1.2859 |
| Efcab11 | 141 | 194 | 211 | 220 | 160 | 193 | 150 | 194 | 212 | 173 | 172 | 211 | 185.29 | 185 | 185 | 0.999 | -0.001 | 0.99657558 | 0.99828918 | 0.0167 | 0.0247 | 0.0207 | 0.0207 | TRUE | 0.2962 |
| Efcab12 | 293 | 212 | 211 | 302 | 249 | 185 | 311 | 212 | 212 | 237 | 268 | 203 | 240.44 | 245 | 236 | 0.964 | -0.053 | 0.79616754 | 0.89534078 | 0.0309 | 0.0206 | 0.0257 | 0.0257 | TRUE | 0.6168 |
| Efcab14 | 2187 | 2381 | 2454 | 2852 | 2030 | 2133 | 2323 | 2378 | 2464 | 2242 | 2184 | 2335 | 2321.01 | 2388 | 2254 | 0.943 | -0.084 | 0.23837268 | 0.45416935 | 5.00E-04 | 0.0083 | 0.0032 | 0.0032 | TRUE | 0.0118 |
| Efcab2 | 149 | 194 | 187 | 281 | 223 | 186 | 158 | 194 | 188 | 221 | 240 | 204 | 200.7 | 180 | 222 | 1.23 | 0.299 | 0.06082988 | 0.18771451 | 0.0031 | 0.0233 | 0.0133 | 0.0133 | TRUE | 0.1169 |
| Efcab3 | 11 | 12 | 12 | 17 | 20 | 8 | 12 | 12 | 12 | 13 | 22 | 9 | 13.23 | 12 | 15 | 1.219 | 0.286 | 0.60666181 | 0.77677139 | 0 | 0.256 | 0.146 | 0.146 | TRUE | 0.3333 |
| Efcab4a | 270 | 321 | 281 | 262 | 241 | 228 | 287 | 321 | 282 | 206 | 259 | 250 | 267.4 | 297 | 238 | 0.801 | -0.321 | 0.03256328 | 0.121608 | 0.0064 | 0.0192 | 0.0124 | 0.0124 | TRUE | 0.1688 |
| Efcab4b | 25 | 26 | 42 | 36 | 32 | 16 | 27 | 26 | 42 | 28 | 34 | 18 | 29.16 | 32 | 27 | 0.849 | -0.237 | 0.56840607 | 0.7475111 | 0.0554 | 0.1199 | 0.0895 | 0.0895 | TRUE | 0.4848 |
| Efcab5 | 37 | 17 | 24 | 24 | 15 | 25 | 39 | 17 | 24 | 19 | 16 | 27 | 23.79 | 27 | 21 | 0.775 | -0.367 | 0.4338553 | 0.64252445 | 0.085 | 0.1454 | 0.1159 | 0.1159 | TRUE | 0.4508 |
| Efcab6 | 55 | 59 | 53 | 35 | 40 | 32 | 58 | 59 | 53 | 28 | 43 | 35 | 46.02 | 57 | 35 | 0.614 | -0.705 | 0.01931222 | 0.08460898 | 0 | 0.0785 | 0.0424 | 0.0424 | TRUE | 0.316 |
| Efcab7 | 178 | 183 | 165 | 312 | 162 | 121 | 189 | 183 | 166 | 245 | 174 | 132 | 181.59 | 179 | 184 | 1.032 | 0.045 | 0.84794834 | 0.92391533 | 0.0448 | 0.025 | 0.0348 | 0.0348 | TRUE | 0.3948 |
| Efcab8 | 14 | 10 | 8 | 8 | 10 | 6 | 15 | 10 | 8 | 6 | 11 | 7 | 9.42 | 11 | 8 | 0.714 | -0.486 | 0.44615545 | 0.65236459 | 0 | 0.3567 | 0.1842 | 0.1842 | TRUE | 0.3442 |
| Efcab9 | 5 | 6 | 6 | 3 | 1 | 1 | 5 | 6 | 6 | 2 | 1 | 1 | 3.64 | 6 | 1 | 0.271 | -1.884 | 0.07375815 | NA | 0 | 0.9113 | 0.4012 | 0.4012 | TRUE | 0.2301 |
| Efcc1 | 0 | 1 | 3 | 3 | 3 | 1 | 0 | 1 | 3 | 2 | 3 | 1 | 1.78 | 1 | 2 | 1.657 | 0.729 | 0.6284253 | NA | 0 | 1.8557 | 1.0362 | 1.0362 | TRUE | 0.7043 |
| Efemp1 | 32 | 34 | 42 | 61 | 43 | 30 | 34 | 34 | 42 | 48 | 46 | 33 | 39.53 | 37 | 42 | 1.159 | 0.213 | 0.5084021 | 0.70245448 | 0 | 0.0902 | 0.0491 | 0.0491 | TRUE | 0.2763 |
| Efemp2 | 126 | 182 | 205 | 243 | 178 | 190 | 134 | 182 | 206 | 191 | 192 | 208 | 185.33 | 174 | 197 | 1.131 | 0.177 | 0.35800503 | 0.57646303 | 0.0179 | 0.0247 | 0.0214 | 0.0214 | TRUE | 0.4257 |
| Efha1 | 503 | 724 | 785 | 923 | 634 | 665 | 534 | 723 | 788 | 726 | 682 | 728 | 696.87 | 682 | 712 | 1.044 | 0.062 | 0.68582406 | 0.8264683 | 0.019 | 0.0116 | 0.0154 | 0.0154 | TRUE | 0.4209 |
| Efha2 | 839 | 846 | 866 | 940 | 599 | 744 | 891 | 845 | 870 | 739 | 644 | 814 | 800.58 | 869 | 732 | 0.844 | -0.245 | 0.03002675 | 0.11536709 | 0.0052 | 0.011 | 0.008 | 0.008 | TRUE | 0.1274 |
| Efhb | 87 | 102 | 109 | 138 | 118 | 124 | 92 | 102 | 109 | 108 | 127 | 136 | 112.48 | 101 | 124 | 1.217 | 0.283 | 0.15275443 | 0.34407924 | 0 | 0.0362 | 0.0192 | 0.0192 | TRUE | 0.1311 |
| Efhc1 | 135 | 201 | 203 | 235 | 157 | 181 | 143 | 201 | 204 | 185 | 169 | 198 | 183.29 | 183 | 184 | 1.006 | 0.009 | 0.96322078 | 0.98149348 | 0.0156 | 0.0249 | 0.0203 | 0.0203 | TRUE | 0.3743 |
| Efhc2 | 443 | 605 | 514 | 2025 | 1520 | 1304 | 471 | 604 | 516 | 1592 | 1635 | 1427 | 1040.94 | 530 | 1551 | 2.925 | 1.549 | 2.62E-37 | 2.43E-34 | 0.0088 | 0.0101 | 0.0093 | 0.0093 | TRUE | 0.1742 |
| Efhd1 | 216 | 172 | 128 | 118 | 91 | 99 | 229 | 172 | 129 | 93 | 98 | 108 | 138.13 | 177 | 100 | 0.564 | -0.826 | 0.00073252 | 0.00699776 | 0.0398 | 0.0308 | 0.0353 | 0.0353 | TRUE | 0.1683 |
| Efhd2 | 1441 | 1900 | 1889 | 1742 | 1310 | 1497 | 1531 | 1898 | 1897 | 1369 | 1409 | 1639 | 1623.75 | 1775 | 1472 | 0.829 | -0.27 | 0.02855173 | 0.11121704 | 0.011 | 0.0089 | 0.0104 | 0.0104 | TRUE | 0.1747 |
| Efna1 | 97 | 133 | 122 | 136 | 106 | 136 | 103 | 133 | 123 | 107 | 114 | 149 | 121.37 | 120 | 123 | 1.028 | 0.039 | 0.85480879 | 0.92766489 | 0.0159 | 0.034 | 0.025 | 0.025 | TRUE | 0.3259 |
| Efna2 | 372 | 406 | 404 | 519 | 363 | 367 | 395 | 405 | 406 | 408 | 391 | 402 | 401.09 | 402 | 400 | 0.996 | -0.006 | 0.95174718 | 0.97539867 | 0 | 0.0151 | 0.0057 | 0.0057 | TRUE | 0.0049 |
| Efna3 | 227 | 337 | 281 | 269 | 223 | 245 | 241 | 337 | 282 | 211 | 240 | 268 | 263.24 | 287 | 240 | 0.835 | -0.261 | 0.13689816 | 0.32082583 | 0.0174 | 0.0194 | 0.0184 | 0.0184 | TRUE | 0.2312 |
| Efna4 | 407 | 316 | 340 | 424 | 312 | 302 | 432 | 316 | 341 | 333 | 336 | 331 | 348.15 | 363 | 333 | 0.918 | -0.123 | 0.41470191 | 0.62626183 | 0.011 | 0.0164 | 0.0135 | 0.0135 | TRUE | 0.3157 |
| Efna5 | 37 | 63 | 57 | 74 | 55 | 60 | 39 | 63 | 57 | 58 | 59 | 66 | 57.08 | 53 | 61 | 1.144 | 0.194 | 0.48820211 | 0.68476003 | 0.0093 | 0.0646 | 0.039 | 0.039 | TRUE | 0.4179 |
| Efnb1 | 140 | 140 | 120 | 171 | 123 | 131 | 149 | 140 | 120 | 134 | 132 | 143 | 136.53 | 136 | 136 | 1.003 | 0.004 | 0.98232318 | 0.99061388 | 0 | 0.031 | 0.0152 | 0.0152 | TRUE | 0.1073 |
| Efnb2 | 463 | 478 | 467 | 629 | 408 | 312 | 492 | 477 | 469 | 494 | 439 | 342 | 452.18 | 479 | 425 | 0.889 | -0.17 | 0.26642129 | 0.48611714 | 0.0151 | 0.0142 | 0.0147 | 0.0147 | TRUE | 0.336 |
| Efnb3 | 527 | 842 | 909 | 1008 | 735 | 846 | 560 | 841 | 913 | 792 | 791 | 926 | 803.79 | 771 | 836 | 1.084 | 0.116 | 0.52227132 | 0.71226036 | 0.0349 | 0.011 | 0.0226 | 0.0226 | TRUE | 0.679 |
| Efr3a | 6559 | 6562 | 6694 | 9223 | 6878 | 6958 | 6967 | 6554 | 6721 | 7250 | 7400 | 7617 | 7084.86 | 6747 | 7422 | 1.1 | 0.137 | 0.02768039 | 0.10914377 | 6.00E-04 | 0.0074 | 0.0027 | 0.0027 | TRUE | 0.0099 |
| Efr3b | 796 | 814 | 843 | 1237 | 815 | 808 | 845 | 813 | 846 | 972 | 877 | 884 | 873.12 | 835 | 911 | 1.093 | 0.128 | 0.16024745 | 0.35474763 | 0.0011 | 0.0107 | 0.0048 | 0.0048 | TRUE | 0.0428 |
| Efs | 528 | 756 | 778 | 807 | 581 | 653 | 561 | 755 | 781 | 634 | 625 | 715 | 678.56 | 699 | 658 | 0.941 | -0.088 | 0.55447763 | 0.73774432 | 0.017 | 0.0118 | 0.0145 | 0.0145 | TRUE | 0.3514 |
| Eftud1 | 2475 | 2097 | 2213 | 2785 | 1966 | 1774 | 2629 | 2094 | 2222 | 2189 | 2115 | 1942 | 2198.64 | 2315 | 2082 | 0.9 | -0.152 | 0.1707633 | 0.36937928 | 0.0084 | 0.0084 | 0.0085 | 0.0085 | TRUE | 0.17 |
| Eftud2 | 27684 | 26068 | 26272 | 34017 | 24952 | 23625 | 29405 | 26034 | 26380 | 26741 | 26847 | 25861 | 26878.18 | 27273 | 26483 | 0.971 | -0.042 | 0.57060488 | 0.74940297 | 0.0023 | 0.007 | 0.004 | 0.004 | TRUE | 0.0572 |
| Egf | 46 | 67 | 68 | 89 | 62 | 65 | 49 | 67 | 68 | 70 | 67 | 71 | 65.31 | 61 | 69 | 1.128 | 0.174 | 0.48955706 | 0.68569518 | 0 | 0.0573 | 0.0301 | 0.0301 | TRUE | 0.2668 |
| Egfbp2 | 0 | 0 | 0 | 0 | 0 | 0 | 0 | 0 | 0 | 0 | 0 | 0 | 0 | 0 | 0 | NA | NA | NA | NA | NA | NA | NA | NA | NA | NA |
| Egfem1 | 0 | 0 | 0 | 0 | 1 | 0 | 0 | 0 | 0 | 0 | 1 | 0 | 0.18 | 0 | 0 | 1.859 | 0.894 | 0.82650734 | NA | 0 | 18.3785 | 10 | 10 | TRUE | 0.5204 |
| Egfl6 | 87 | 100 | 85 | 128 | 72 | 85 | 92 | 100 | 85 | 101 | 77 | 93 | 91.46 | 92 | 90 | 0.98 | -0.029 | 0.89328575 | 0.94834594 | 0 | 0.0429 | 0.0225 | 0.0225 | TRUE | 0.1448 |
| Egfl7 | 1740 | 2263 | 2147 | 2204 | 1610 | 1643 | 1848 | 2260 | 2156 | 1733 | 1732 | 1799 | 1921.25 | 2088 | 1755 | 0.84 | -0.251 | 0.01126665 | 0.05750733 | 0.0047 | 0.0086 | 0.0066 | 0.0066 | TRUE | 0.1218 |
| Egfl8 | 3 | 0 | 6 | 1 | 6 | 4 | 3 | 0 | 6 | 1 | 6 | 4 | 3.47 | 3 | 4 | 1.24 | 0.311 | 0.81202894 | NA | 0.9069 | 0.9558 | 0.9341 | 0.9341 | TRUE | 0.2709 |
| Egflam | 23 | 21 | 22 | 50 | 19 | 21 | 24 | 21 | 22 | 39 | 20 | 23 | 25.04 | 22 | 27 | 1.242 | 0.313 | 0.4609458 | 0.66511381 | 0.0419 | 0.1385 | 0.0892 | 0.0892 | TRUE | 1.0282 |
| Egfr | 152 | 141 | 107 | 208 | 149 | 172 | 161 | 141 | 107 | 164 | 160 | 188 | 153.64 | 136 | 171 | 1.25 | 0.322 | 0.10995455 | 0.27831859 | 0.0165 | 0.0283 | 0.0226 | 0.0226 | TRUE | 0.3611 |
| Egln1 | 2107 | 3952 | 3831 | 5009 | 4089 | 4706 | 2238 | 3947 | 3847 | 3938 | 4400 | 5151 | 3920.05 | 3344 | 4496 | 1.344 | 0.427 | 0.03522168 | 0.12864281 | 0.05 | 0.0077 | 0.0294 | 0.0294 | TRUE | 1.0167 |
| Egln2 | 1923 | 1720 | 1722 | 2419 | 1817 | 1586 | 2043 | 1718 | 1729 | 1902 | 1955 | 1736 | 1847.03 | 1830 | 1864 | 1.019 | 0.027 | 0.79232484 | 0.89305706 | 0.0062 | 0.0087 | 0.0072 | 0.0072 | TRUE | 0.1246 |
| Egln3 | 2213 | 3493 | 4364 | 2435 | 1857 | 2487 | 2351 | 3488 | 4382 | 1914 | 1998 | 2722 | 2809.27 | 3407 | 2211 | 0.649 | -0.624 | 0.00400035 | 0.02634386 | 0.0643 | 0.0081 | 0.0335 | 0.0335 | TRUE | 0.1747 |
| Egr1 | 673 | 1209 | 669 | 766 | 613 | 325 | 715 | 1207 | 672 | 602 | 660 | 356 | 701.92 | 865 | 539 | 0.624 | -0.681 | 0.01123169 | 0.05736599 | 0.1018 | 0.0116 | 0.0505 | 0.0505 | TRUE | 1.4324 |
| Egr2 | 7 | 16 | 10 | 12 | 9 | 8 | 7 | 16 | 10 | 9 | 10 | 9 | 10.22 | 11 | 9 | 0.832 | -0.266 | 0.66067543 | 0.80885977 | 0 | 0.3292 | 0.1661 | 0.1661 | TRUE | 0.543 |
| Egr3 | 0 | 0 | 0 | 1 | 2 | 1 | 0 | 0 | 0 | 1 | 2 | 1 | 0.67 | 0 | 1 | 7.1 | 2.828 | 0.25693906 | NA | 0 | 4.9086 | 3.1164 | 3.1164 | TRUE | 0.1607 |
| Egr4 | 19 | 38 | 41 | 33 | 22 | 21 | 20 | 38 | 41 | 26 | 24 | 23 | 28.65 | 33 | 24 | 0.731 | -0.451 | 0.26264067 | 0.48200706 | 0.0372 | 0.1219 | 0.0814 | 0.0814 | TRUE | 0.7812 |
| Ehbp1 | 1553 | 1555 | 1442 | 1863 | 1379 | 1304 | 1650 | 1553 | 1448 | 1465 | 1484 | 1427 | 1504.36 | 1550 | 1459 | 0.941 | -0.088 | 0.29993464 | 0.52012855 | 0.0016 | 0.0091 | 0.0045 | 0.0045 | TRUE | 0.0401 |
| Ehbp1l1 | 844 | 891 | 793 | 701 | 619 | 656 | 896 | 890 | 796 | 551 | 666 | 718 | 752.96 | 861 | 645 | 0.748 | -0.419 | 0.00109998 | 0.00957448 | 0.0099 | 0.0113 | 0.0105 | 0.0105 | TRUE | 0.1988 |

| | | | | | | | | | | | | | | | | | | | | | | | | | |
|---|---|---|---|---|---|---|---|---|---|---|---|---|---|---|---|---|---|---|---|---|---|---|---|---|---|
| Ehd1 | 8117 | 6744 | 6413 | 9474 | 6545 | 5999 | 8622 | 6735 | 6439 | 7448 | 7042 | 6567 | 7142.15 | 7265 | 7019 | 0.966 | -0.05 | 0.6924484 | 0.83139114 | 0.0127 | 0.0074 | 0.0112 | 0.0112 | TRUE | 0.3254 |
| Ehd2 | 89 | 41 | 37 | 142 | 88 | 52 | 95 | 41 | 37 | 112 | 95 | 57 | 72.64 | 58 | 88 | 1.532 | 0.615 | 0.13662443 | 0.3202795 | 0.1755 | 0.0523 | 0.1086 | 0.1086 | TRUE | 1.336 |
| Ehd3 | 336 | 355 | 338 | 584 | 403 | 349 | 357 | 355 | 339 | 459 | 434 | 382 | 387.59 | 350 | 425 | 1.215 | 0.282 | 0.02222141 | 0.0937905 | 0.0026 | 0.0154 | 0.0083 | 0.0083 | TRUE | 0.0883 |
| Ehd4 | 3151 | 2995 | 3092 | 4385 | 2951 | 2784 | 3347 | 2991 | 3105 | 3447 | 3175 | 3048 | 3185.42 | 3148 | 3223 | 1.025 | 0.035 | 0.68553458 | 0.82631951 | 0.0033 | 0.0079 | 0.0051 | 0.0051 | TRUE | 0.0454 |
| Ehf | 20 | 10 | 6 | 13 | 7 | 6 | 21 | 10 | 6 | 10 | 8 | 7 | 10.26 | 12 | 8 | 0.661 | -0.597 | 0.38374718 | 0.59804511 | 0.1489 | 0.3279 | 0.2373 | 0.2373 | TRUE | 0.6452 |
| Ehhadh | 508 | 408 | 337 | 548 | 386 | 358 | 540 | 407 | 338 | 431 | 415 | 392 | 420.57 | 428 | 413 | 0.964 | -0.053 | 0.76764392 | 0.87867624 | 0.0264 | 0.0147 | 0.0206 | 0.0206 | TRUE | 0.4557 |
| Ehmt1 | 4329 | 4597 | 4990 | 5966 | 4384 | 4444 | 4598 | 4591 | 5010 | 4690 | 4717 | 4865 | 4745.21 | 4733 | 4757 | 1.005 | 0.007 | 0.91932914 | 0.96048697 | 0.0012 | 0.0076 | 0.0033 | 0.0033 | TRUE | 0.032 |
| Ehmt2 | 6578 | 7001 | 7550 | 7288 | 5505 | 5767 | 6987 | 6992 | 7581 | 5729 | 5923 | 6313 | 6587.52 | 7187 | 5988 | 0.833 | -0.263 | 0.00050199 | 0.00517876 | 0.0021 | 0.0074 | 0.004 | 0.004 | TRUE | 0.0281 |
| Ei24 | 7294 | 8369 | 7956 | 8830 | 6411 | 7035 | 7748 | 8358 | 7989 | 6941 | 6898 | 7701 | 7605.76 | 8032 | 7180 | 0.894 | -0.162 | 0.03688715 | 0.1327668 | 0.0025 | 0.0073 | 0.0042 | 0.0042 | TRUE | 0.049 |
| Eid1 | 582 | 666 | 724 | 1575 | 1188 | 911 | 618 | 665 | 727 | 1238 | 1278 | 997 | 920.65 | 670 | 1171 | 1.748 | 0.806 | 4.46E-10 | 2.93E-08 | 0.0112 | 0.0105 | 0.0109 | 0.0109 | TRUE | 0.2 |
| Eid2 | 720 | 859 | 783 | 1458 | 1071 | 890 | 765 | 858 | 786 | 1146 | 1152 | 974 | 946.94 | 803 | 1091 | 1.359 | 0.443 | 4.63E-05 | 0.00072733 | 0.0052 | 0.0104 | 0.0074 | 0.0074 | TRUE | 0.1032 |
| Eid2b | 230 | 274 | 254 | 352 | 260 | 246 | 244 | 274 | 255 | 277 | 280 | 269 | 266.46 | 258 | 275 | 1.068 | 0.095 | 0.45949659 | 0.66378277 | 0 | 0.0193 | 0.0081 | 0.0081 | TRUE | 0.033 |
| Eid3 | 23 | 15 | 20 | 17 | 10 | 12 | 24 | 15 | 20 | 13 | 11 | 13 | 16.13 | 20 | 12 | 0.629 | -0.668 | 0.16626999 | 0.36351681 | 0 | 0.2112 | 0.1029 | 0.1029 | TRUE | 0.1649 |
| Eif1 | 19577 | 20289 | 20230 | 23869 | 16435 | 16892 | 20794 | 20263 | 20313 | 18764 | 17683 | 18491 | 19384.67 | 20457 | 18313 | 0.895 | -0.16 | 0.00571231 | 0.03458377 | 5.00E-04 | 0.0071 | 0.0024 | 0.0024 | TRUE | 0.0111 |
| Eif1a | 9135 | 8879 | 8393 | 13108 | 9521 | 9052 | 9703 | 8868 | 8427 | 10304 | 10244 | 9909 | 9575.89 | 8999 | 10152 | 1.128 | 0.174 | 0.0254719 | 0.10303739 | 0.0027 | 0.0072 | 0.0043 | 0.0043 | TRUE | 0.0571 |
| Eif1ad | 1889 | 1961 | 1839 | 3239 | 2126 | 1856 | 2006 | 1958 | 1847 | 2546 | 2287 | 2032 | 2112.81 | 1937 | 2288 | 1.182 | 0.241 | 0.02203058 | 0.09330457 | 0.0065 | 0.0085 | 0.0075 | 0.0075 | TRUE | 0.1191 |
| Eif1ax | 6432 | 7307 | 7343 | 8802 | 6239 | 6210 | 6832 | 7298 | 7373 | 6919 | 6713 | 6798 | 6988.78 | 7168 | 6810 | 0.95 | -0.074 | 0.25165812 | 0.46941457 | 7.00E-04 | 0.0074 | 0.0028 | 0.0028 | TRUE | 0.0204 |
| Eif1b | 1162 | 1471 | 1545 | 1695 | 1327 | 1346 | 1234 | 1469 | 1551 | 1332 | 1428 | 1473 | 1414.72 | 1418 | 1411 | 0.994 | -0.008 | 0.942197 | 0.97094617 | 0.0075 | 0.0092 | 0.0083 | 0.0083 | TRUE | 0.1543 |
| Eif2a | 7846 | 8170 | 8382 | 9987 | 7080 | 7500 | 8334 | 8159 | 8416 | 7851 | 7618 | 8210 | 8098.05 | 8303 | 7893 | 0.951 | -0.073 | 0.24187986 | 0.45852514 | 7.00E-04 | 0.0073 | 0.0027 | 0.0027 | TRUE | 0.015 |
| Eif2ak1 | 2357 | 1795 | 1818 | 2934 | 1972 | 1777 | 2504 | 1793 | 1825 | 2306 | 2122 | 1945 | 2082.52 | 2041 | 2124 | 1.042 | 0.059 | 0.68920557 | 0.82917062 | 0.0212 | 0.0085 | 0.0151 | 0.0151 | TRUE | 0.476 |
| Eif2ak2 | 1551 | 1460 | 1445 | 2078 | 1437 | 1419 | 1647 | 1458 | 1451 | 1634 | 1546 | 1553 | 1548.25 | 1519 | 1578 | 1.039 | 0.056 | 0.5286302 | 0.71718529 | 0.0022 | 0.009 | 0.005 | 0.005 | TRUE | 0.0658 |
| Eif2ak3 | 2484 | 2611 | 2598 | 2698 | 2118 | 2017 | 2638 | 2608 | 2609 | 2121 | 2279 | 2208 | 2410.41 | 2618 | 2203 | 0.841 | -0.25 | 0.00024461 | 0.00287051 | 3.00E-04 | 0.0083 | 0.0029 | 0.0029 | TRUE | 0.0131 |
| Eif2ak4 | 1735 | 1773 | 1916 | 2436 | 1870 | 1836 | 1843 | 1771 | 1924 | 1915 | 2012 | 2010 | 1912.37 | 1846 | 1979 | 1.072 | 0.1 | 0.18380748 | 0.38659864 | 8.00E-04 | 0.0086 | 0.0035 | 0.0035 | TRUE | 0.0166 |
| Eif2b1 | 4359 | 4034 | 3936 | 5132 | 3725 | 3437 | 4630 | 4029 | 3952 | 4034 | 4008 | 3762 | 4069.26 | 4204 | 3935 | 0.936 | -0.095 | 0.29114574 | 0.51183021 | 0.0043 | 0.0077 | 0.0056 | 0.0056 | TRUE | 0.0957 |
| Eif2b2 | 2229 | 1825 | 1934 | 3194 | 2241 | 1952 | 2368 | 1823 | 1942 | 2511 | 2411 | 2137 | 2198.5 | 2044 | 2353 | 1.152 | 0.204 | 0.10029396 | 0.26180673 | 0.0125 | 0.0084 | 0.0106 | 0.0106 | TRUE | 0.2315 |
| Eif2b3 | 3574 | 4182 | 4102 | 5027 | 3409 | 3220 | 3796 | 4177 | 4119 | 3952 | 3668 | 3525 | 3872.69 | 4031 | 3715 | 0.922 | -0.117 | 0.15242111 | 0.3436229 | 0.0026 | 0.0078 | 0.0046 | 0.0046 | TRUE | 0.0383 |
| Eif2b4 | 2220 | 2017 | 1956 | 3175 | 2258 | 1958 | 2358 | 2014 | 1964 | 2496 | 2429 | 2143 | 2234.2 | 2112 | 2356 | 1.116 | 0.158 | 0.14387299 | 0.33083026 | 0.0076 | 0.0084 | 0.008 | 0.008 | TRUE | 0.1252 |
| Eif2b5 | 7121 | 7669 | 7832 | 10660 | 7928 | 7682 | 7564 | 7659 | 7864 | 8380 | 8530 | 8409 | 8067.71 | 7696 | 8440 | 1.097 | 0.133 | 0.01669769 | 0.07648271 | 1.00E-04 | 0.0073 | 0.0021 | 0.0021 | TRUE | 0.0045 |
| Eif2c1 | 4060 | 4362 | 4744 | 6048 | 4538 | 4661 | 4312 | 4356 | 4763 | 4754 | 4883 | 5102 | 4695.26 | 4477 | 4913 | 1.097 | 0.134 | 0.07659189 | 0.21848569 | 0.0019 | 0.0076 | 0.0039 | 0.0039 | TRUE | 0.0381 |
| Eif2c2 | 5309 | 7099 | 8571 | 7880 | 5937 | 6442 | 5639 | 7090 | 8606 | 6195 | 6388 | 7052 | 6828.23 | 7112 | 6545 | 0.92 | -0.12 | 0.42119149 | 0.6313469 | 0.0225 | 0.0074 | 0.0159 | 0.0159 | TRUE | 0.1798 |
| Eif2c3 | 635 | 553 | 669 | 792 | 495 | 497 | 674 | 552 | 672 | 623 | 533 | 544 | 599.63 | 633 | 567 | 0.897 | -0.158 | 0.22030057 | 0.43242065 | 0.0085 | 0.0124 | 0.0102 | 0.0102 | TRUE | 0.1466 |
| Eif2c4 | 143 | 184 | 187 | 197 | 167 | 170 | 152 | 184 | 188 | 155 | 180 | 186 | 174.01 | 175 | 174 | 0.991 | -0.013 | 0.93943164 | 0.96945854 | 0.0057 | 0.0258 | 0.0156 | 0.0156 | TRUE | 0.1347 |
| Eif2d | 5183 | 5622 | 5630 | 8263 | 5692 | 5495 | 5505 | 5615 | 5653 | 6496 | 6124 | 6015 | 5901.37 | 5591 | 6212 | 1.111 | 0.152 | 0.01840092 | 0.08197227 | 7.00E-04 | 0.0075 | 0.0028 | 0.0028 | TRUE | 0.0197 |
| Eif2s1 | 13118 | 14185 | 13568 | 17699 | 12393 | 12467 | 13934 | 14167 | 13624 | 13914 | 13334 | 13647 | 13769.8 | 13908 | 13632 | 0.98 | -0.029 | 0.6085023 | 0.77824912 | 3.00E-04 | 0.0071 | 0.0022 | 0.0022 | TRUE | 0.0045 |
| Eif2s2 | 19789 | 19833 | 17192 | 47256 | 32648 | 31374 | 21019 | 19807 | 17263 | 37149 | 35127 | 34344 | 27451.56 | 19363 | 35540 | 1.836 | 0.876 | 7.94E-21 | 2.28E-18 | 0.0057 | 0.007 | 0.0063 | 0.0063 | TRUE | 0.1102 |
| Eif2s3x | 4366 | 4623 | 4764 | 6105 | 4332 | 3948 | 4637 | 4617 | 4784 | 4799 | 4661 | 4322 | 4636.67 | 4679 | 4594 | 0.982 | -0.026 | 0.71525628 | 0.84540034 | 0.0014 | 0.0076 | 0.0035 | 0.0035 | TRUE | 0.0326 |
| Eif2s3y | 3095 | 3317 | 3136 | 4330 | 3380 | 3247 | 3287 | 3313 | 3149 | 3404 | 3637 | 3554 | 3390.66 | 3250 | 3532 | 1.086 | 0.12 | 0.07725841 | 0.21955308 | 7.00E-04 | 0.0079 | 0.003 | 0.003 | TRUE | 0.0124 |
| Eif3a | 40625 | 44047 | 48077 | 51977 | 38815 | 40436 | 43151 | 43990 | 48274 | 40860 | 41763 | 44264 | 43716.97 | 45138 | 42296 | 0.937 | -0.094 | 0.21618698 | 0.4282468 | 0.0027 | 0.007 | 0.0041 | 0.0041 | TRUE | 0.0452 |
| Eif3b | 20711 | 19387 | 19608 | 28104 | 20507 | 18852 | 21999 | 19362 | 19689 | 22093 | 22064 | 20636 | 20973.85 | 20350 | 21598 | 1.061 | 0.086 | 0.28066196 | 0.50209438 | 0.0032 | 0.0071 | 0.0045 | 0.0045 | TRUE | 0.0615 |
| Eif3c | 21387 | 23153 | 23986 | 26841 | 19551 | 20070 | 22717 | 23123 | 24084 | 21100 | 21036 | 21970 | 22338.35 | 23308 | 21369 | 0.917 | -0.125 | 0.03538168 | 0.12908624 | 7.00E-04 | 0.0071 | 0.0025 | 0.0025 | TRUE | 0.0104 |
| Eif3d | 16311 | 19197 | 19422 | 25070 | 18226 | 17997 | 17325 | 19172 | 19502 | 19708 | 19610 | 19701 | 19169.63 | 18666 | 19673 | 1.054 | 0.076 | 0.29046899 | 0.51141738 | 0.0019 | 0.0071 | 0.0036 | 0.0036 | TRUE | 0.0483 |
| Eif3e | 12478 | 13242 | 12697 | 17596 | 12201 | 11882 | 13254 | 13225 | 12749 | 13833 | 13128 | 13007 | 13199.12 | 13076 | 13323 | 1.019 | 0.027 | 0.65667998 | 0.80670515 | 7.00E-04 | 0.0072 | 0.0026 | 0.0026 | TRUE | 0.0138 |
| Eif3f | 13371 | 16076 | 16561 | 18158 | 13666 | 14504 | 14202 | 16055 | 16629 | 14274 | 14704 | 15877 | 15290.28 | 15629 | 14952 | 0.957 | -0.064 | 0.47425643 | 0.67490502 | 0.0045 | 0.0071 | 0.0057 | 0.0057 | TRUE | 0.0779 |
| Eif3g | 8059 | 8984 | 9026 | 11215 | 8340 | 7846 | 8560 | 8972 | 9063 | 8816 | 8973 | 8589 | 8828.98 | 8865 | 8793 | 0.992 | -0.012 | 0.84573771 | 0.92273463 | 5.00E-04 | 0.0073 | 0.0026 | 0.0026 | TRUE | 0.0111 |
| Eif3h | 10554 | 12802 | 12689 | 13746 | 9892 | 10704 | 11210 | 12785 | 12741 | 10806 | 10643 | 11717 | 11650.53 | 12245 | 11055 | 0.903 | -0.148 | 0.08464471 | 0.23359865 | 0.0039 | 0.0072 | 0.0052 | 0.0052 | TRUE | 0.0668 |
| Eif3i | 8921 | 10242 | 10545 | 11911 | 8519 | 8948 | 9476 | 10229 | 10588 | 9363 | 9166 | 9795 | 9769.53 | 10098 | 9441 | 0.935 | -0.097 | 0.18810824 | 0.39306349 | 0.002 | 0.0072 | 0.0038 | 0.0038 | TRUE | 0.0354 |
| Eif3j1 | 0 | 0 | 0 | 0 | 0 | 0 | 0 | 0 | 0 | 0 | 0 | 0 | 0 | 0 | 0 | NA | NA | NA | NA | NA | NA | NA | NA | NA | NA |
| Eif3j2 | 3 | 3 | 3 | 4 | 7 | 2 | 3 | 3 | 3 | 3 | 8 | 2 | 3.68 | 3 | 4 | 1.388 | 0.473 | 0.63701768 | NA | 0 | 0.9029 | 0.4423 | 0.4423 | TRUE | 0.7121 |
| Eif3k | 6788 | 8030 | 8246 | 10089 | 7234 | 7592 | 7210 | 8020 | 8280 | 7931 | 7783 | 8311 | 7922.45 | 7837 | 8008 | 1.022 | 0.031 | 0.69898774 | 0.8353743 | 0.0025 | 0.0073 | 0.0046 | 0.0046 | TRUE | 0.0596 |
| Eif3l | 15442 | 15634 | 15903 | 18593 | 13299 | 13380 | 16402 | 15614 | 15968 | 14616 | 14309 | 14647 | 15259.34 | 15995 | 14524 | 0.908 | -0.139 | 0.01251594 | 0.06201743 | 3.00E-04 | 0.0071 | 0.0022 | 0.0022 | TRUE | 0.0061 |
| Eif3m | 8585 | 9892 | 10006 | 12115 | 8428 | 9226 | 9119 | 9879 | 10047 | 9524 | 9068 | 10099 | 9622.71 | 9682 | 9564 | 0.988 | -0.018 | 0.81974703 | 0.90781402 | 0.0022 | 0.0072 | 0.0043 | 0.0043 | TRUE | 0.0316 |
| Eif4a1 | 91749 | 92965 | 90673 | 138527 | 92700 | 86310 | 97454 | 92845 | 91045 | 108899 | 99740 | 94480 | 97410.39 | 93781 | 101040 | 1.077 | 0.108 | 0.17415298 | 0.37335066 | 0.0032 | 0.0069 | 0.0045 | 0.0045 | TRUE | 0.0567 |
| Eif4a2 | 35879 | 58024 | 56562 | 48510 | 38066 | 45529 | 38110 | 57949 | 56794 | 38135 | 40957 | 49839 | 46963.87 | 50951 | 42977 | 0.843 | -0.246 | 0.14851254 | 0.33829287 | 0.034 | 0.007 | 0.0208 | 0.0208 | TRUE | 0.5952 |
| Eif4a3 | 11196 | 10910 | 10846 | 16361 | 11568 | 10424 | 11892 | 10896 | 10891 | 12862 | 12446 | 11411 | 11732.91 | 11226 | 12240 | 1.09 | 0.125 | 0.11974817 | 0.29345372 | 0.0031 | 0.0072 | 0.0046 | 0.0046 | TRUE | 0.0428 |
| Eif4b | 48316 | 59167 | 62807 | 57130 | 43386 | 48818 | 51320 | 59091 | 63065 | 44911 | 46681 | 53439 | 53084.4 | 57825 | 48344 | 0.836 | -0.258 | 0.0175172 | 0.07896006 | 0.0096 | 0.007 | 0.0085 | 0.0085 | TRUE | 0.1186 |
| Eif4e | 8767 | 9482 | 9475 | 11506 | 8226 | 8496 | 9312 | 9470 | 9514 | 9045 | 8851 | 9300 | 9248.63 | 9432 | 9065 | 0.961 | -0.057 | 0.31350459 | 0.5339318 | 2.00E-04 | 0.0073 | 0.0022 | 0.0022 | TRUE | 0.0062 |
| Eif4e1b | 3 | 1 | 2 | 7 | 2 | 1 | 3 | 1 | 2 | 6 | 2 | 1 | 2.49 | 2 | 3 | 1.461 | 0.547 | 0.66032701 | NA | 0 | 1.3297 | 0.7066 | 0.7066 | TRUE | 0.982 |
| Eif4e2 | 4584 | 4406 | 4110 | 6500 | 4466 | 4068 | 4869 | 4400 | 4127 | 5110 | 4805 | 4453 | 4627.37 | 4465 | 4789 | 1.073 | 0.101 | 0.29068748 | 0.51144315 | 0.0056 | 0.0076 | 0.0064 | 0.0064 | TRUE | 0.076 |
| Eif4e3 | 508 | 713 | 737 | 562 | 460 | 584 | 540 | 712 | 740 | 442 | 495 | 639 | 594.62 | 664 | 525 | 0.79 | -0.34 | 0.05793795 | 0.18189239 | 0.0297 | 0.0124 | 0.0215 | 0.0215 | TRUE | 0.4181 |
| Eif4ebp1 | 10819 | 10350 | 10175 | 13999 | 10305 | 9983 | 11492 | 10337 | 10217 | 11005 | 11088 | 10928 | 10844.25 | 10682 | 11007 | 1.03 | 0.043 | 0.55464567 | 0.73790565 | 0.0019 | 0.0072 | 0.0038 | 0.0038 | TRUE | 0.0537 |
| Eif4ebp2 | 11069 | 9730 | 10082 | 13420 | 9767 | 8637 | 11757 | 9717 | 10123 | 10550 | 10509 | 9455 | 10351.84 | 10532 | 10171 | 0.966 | -0.05 | 0.61315033 | 0.78108449 | 0.0061 | 0.0072 | 0.007 | 0.007 | TRUE | 0.1263 |
| Eif4ebp3 | 309 | 292 | 279 | 429 | 288 | 272 | 328 | 292 | 280 | 337 | 310 | 298 | 307.47 | 300 | 315 | 1.052 | 0.074 | 0.57937632 | 0.75620021 | 0.0024 | 0.0176 | 0.0094 | 0.0094 | TRUE | 0.0755 |
| Eif4enif1 | 5261 | 5584 | 5722 | 6605 | 4722 | 4975 | 5588 | 5577 | 5745 | 5192 | 5081 | 5446 | 5438.21 | 5637 | 5240 | 0.929 | -0.106 | 0.09577588 | 0.25370795 | 5.00E-04 | 0.0075 | 0.0027 | 0.0027 | TRUE | 0.0143 |
| Eif4g1 | 41641 | 35499 | 35963 | 58422 | 41444 | 35693 | 44230 | 35453 | 36111 | 45927 | 44591 | 39072 | 40897.25 | 38598 | 43197 | 1.119 | 0.162 | 0.15546572 | 0.34780218 | 0.0115 | 0.007 | 0.0094 | 0.0094 | TRUE | 0.1995 |
| Eif4g2 | 81035 | 72367 | 71561 | 106706 | 71782 | 64684 | 86073 | 72274 | 71855 | 83884 | 77233 | 70807 | 77020.93 | 76734 | 77308 | 1.007 | 0.011 | 0.91903452 | 0.96035815 | 0.0073 | 0.0069 | 0.0081 | 0.0081 | TRUE | 0.1388 |
| Eif4g3 | 1109 | 1588 | 1679 | 1566 | 1290 | 1337 | 1178 | 1586 | 1686 | 1231 | 1388 | 1464 | 1422.06 | 1483 | 1361 | 0.917 | -0.125 | 0.3994534 | 0.61255995 | 0.0208 | 0.0092 | 0.0152 | 0.0152 | TRUE | 0.3894 |
| Eif4h | 17237 | 17569 | 17573 | 24357 | 17340 | 16065 | 18309 | 17546 | 17645 | 19148 | 18657 | 17586 | 18148.37 | 17833 | 18464 | 1.035 | 0.05 | 0.4383901 | 0.64591156 | 0.0011 | 0.0071 | 0.003 | 0.003 | TRUE | 0.0211 |
| Eif5 | 24720 | 21119 | 20180 | 31297 | 21855 | 19928 | 26257 | 21092 | 20263 | 24603 | 23515 | 21814 | 22923.97 | 22537 | 23311 | 1.034 | 0.049 | 0.67423116 | 0.81838909 | 0.0098 | 0.007 | 0.0096 | 0.0096 | TRUE | 0.2551 |
| Eif5a | 63713 | 58156 | 56997 | 81094 | 56410 | 53660 | 67674 | 58081 | 57231 | 63750 | 60694 | 58739 | 61028.19 | 60995 | 61061 | 1.001 | 0.002 | 0.98611274 | 0.99254373 | 0.0043 | 0.007 | 0.0059 | 0.0059 | TRUE | 0.1123 |

| | | | | | | | | | | | | | | | | | | | | | | | | | |
|---|---|---|---|---|---|---|---|---|---|---|---|---|---|---|---|---|---|---|---|---|---|---|---|---|---|
| Eif5a2 | 1264 | 1534 | 1496 | 2045 | 1601 | 1587 | 1343 | 1532 | 1502 | 1608 | 1723 | 1737 | 1574.03 | 1459 | 1689 | 1.157 | 0.21 | 0.01908514 | 0.08390936 | 0.0027 | 0.009 | 0.0052 | 0.0052 | TRUE | 0.0582 |
| Eif5b | 12917 | 12324 | 12122 | 16855 | 12143 | 11333 | 13720 | 12308 | 12172 | 13250 | 13065 | 12406 | 12820.16 | 12733 | 12907 | 1.014 | 0.02 | 0.80030775 | 0.89686328 | 0.0022 | 0.0072 | 0.0043 | 0.0043 | TRUE | 0.0562 |
| Eif6 | 4883 | 4687 | 4236 | 7429 | 5213 | 4236 | 5187 | 4681 | 4253 | 5840 | 5609 | 4637 | 5034.48 | 4707 | 5362 | 1.139 | 0.188 | 0.11484776 | 0.28611259 | 0.0117 | 0.0076 | 0.0101 | 0.0101 | TRUE | 0.1702 |
| Elac1 | 480 | 602 | 600 | 689 | 469 | 533 | 510 | 601 | 602 | 542 | 505 | 583 | 557.21 | 571 | 543 | 0.951 | -0.073 | 0.54244885 | 0.72818991 | 0.0051 | 0.0128 | 0.0086 | 0.0086 | TRUE | 0.1025 |
| Elac2 | 5624 | 4961 | 5140 | 7446 | 5258 | 4640 | 5974 | 4955 | 5161 | 5853 | 5657 | 5079 | 5446.56 | 5363 | 5530 | 1.031 | 0.044 | 0.66555903 | 0.81276361 | 0.0062 | 0.0075 | 0.0075 | 0.0075 | TRUE | 0.1208 |
| Elane | 1 | 0 | 0 | 0 | 0 | 0 | 1 | 0 | 0 | 0 | 0 | 0 | 0.18 | 0 | 0 | 0.49 | -1.029 | 0.80087 | NA | 0 | 18.6165 | 10 | 10 | TRUE | 0.4614 |
| Elavl1 | 5881 | 6189 | 6392 | 8135 | 5950 | 5994 | 6247 | 6181 | 6418 | 6395 | 6402 | 6561 | 6367.37 | 6282 | 6453 | 1.027 | 0.039 | 0.5009002 | 0.69666649 | 1.00E-04 | 0.0074 | 0.0022 | 0.0022 | TRUE | 0.0044 |
| Elavl2 | 1919 | 2439 | 2546 | 2547 | 1923 | 2150 | 2038 | 2436 | 2556 | 2002 | 2069 | 2354 | 2242.57 | 2343 | 2142 | 0.913 | -0.131 | 0.26341503 | 0.48264194 | 0.0098 | 0.0084 | 0.0094 | 0.0094 | TRUE | 0.1569 |
| Elavl3 | 1329 | 1863 | 2204 | 1354 | 1156 | 1444 | 1412 | 1861 | 2213 | 1064 | 1244 | 1581 | 1562.36 | 1829 | 1296 | 0.709 | -0.497 | 0.00911939 | 0.04908907 | 0.0436 | 0.009 | 0.0255 | 0.0255 | TRUE | 0.1876 |
| Elavl4 | 101 | 125 | 135 | 232 | 189 | 182 | 107 | 125 | 136 | 182 | 203 | 199 | 158.77 | 123 | 195 | 1.587 | 0.666 | 0.0001024 | 0.00140568 | 0 | 0.0277 | 0.0145 | 0.0145 | TRUE | 0.1178 |
| Elf1 | 643 | 462 | 464 | 840 | 579 | 540 | 683 | 461 | 466 | 660 | 623 | 591 | 580.79 | 537 | 625 | 1.165 | 0.22 | 0.19402683 | 0.39965928 | 0.0256 | 0.0126 | 0.019 | 0.019 | TRUE | 0.6611 |
| Elf2 | 1830 | 2277 | 2441 | 2919 | 2068 | 2127 | 1944 | 2274 | 2451 | 2295 | 2225 | 2328 | 2252.82 | 2223 | 2283 | 1.027 | 0.038 | 0.71457365 | 0.8452151 | 0.0067 | 0.0084 | 0.0074 | 0.0074 | TRUE | 0.1457 |
| Elf3 | 1455 | 1850 | 1806 | 1627 | 1391 | 1384 | 1545 | 1848 | 1813 | 1279 | 1497 | 1515 | 1582.86 | 1735 | 1430 | 0.823 | -0.28 | 0.01374659 | 0.06630601 | 0.0078 | 0.009 | 0.0087 | 0.0087 | TRUE | 0.1104 |
| Elf4 | 726 | 1137 | 1245 | 1421 | 1159 | 1192 | 771 | 1136 | 1250 | 1117 | 1247 | 1305 | 1137.62 | 1052 | 1223 | 1.162 | 0.216 | 0.21578385 | 0.4277706 | 0.0329 | 0.0098 | 0.0211 | 0.0211 | TRUE | 0.6512 |
| Elf5 | 4 | 5 | 2 | 15 | 18 | 12 | 4 | 5 | 2 | 12 | 19 | 13 | 9.26 | 4 | 15 | 3.922 | 1.971 | 0.005568 | 0.03386755 | 0 | 0.3628 | 0.1948 | 0.1948 | TRUE | 0.3196 |
| Elfn1 | 9 | 8 | 8 | 17 | 8 | 8 | 10 | 8 | 8 | 13 | 9 | 9 | 9.39 | 9 | 10 | 1.217 | 0.283 | 0.64395931 | 0.79977047 | 0 | 0.3579 | 0.1628 | 0.1628 | TRUE | 0.3164 |
| Elfn2 | 42 | 50 | 60 | 62 | 43 | 54 | 45 | 50 | 60 | 49 | 46 | 59 | 51.48 | 52 | 51 | 0.993 | -0.011 | 0.96996273 | 0.98456155 | 0 | 0.0709 | 0.0373 | 0.0373 | TRUE | 0.1772 |
| Elk1 | 1470 | 1525 | 1446 | 1682 | 1199 | 1232 | 1561 | 1523 | 1452 | 1322 | 1290 | 1349 | 1416.22 | 1512 | 1320 | 0.873 | -0.196 | 0.01070768 | 0.0554255 | 0 | 0.0092 | 0.0035 | 0.0035 | TRUE | 0.0146 |
| Elk3 | 530 | 652 | 641 | 784 | 644 | 696 | 563 | 651 | 644 | 616 | 693 | 762 | 654.81 | 619 | 690 | 1.113 | 0.154 | 0.21079372 | 0.4213737 | 0.0074 | 0.0119 | 0.0094 | 0.0094 | TRUE | 0.1079 |
| Elk4 | 374 | 294 | 320 | 367 | 283 | 238 | 397 | 294 | 321 | 289 | 304 | 261 | 310.95 | 337 | 285 | 0.844 | -0.245 | 0.12137893 | 0.29630042 | 0.0122 | 0.0175 | 0.0147 | 0.0147 | TRUE | 0.2721 |
| Ell | 2444 | 2315 | 2243 | 3172 | 2336 | 2204 | 2596 | 2312 | 2252 | 2494 | 2513 | 2413 | 2429.96 | 2387 | 2473 | 1.036 | 0.052 | 0.54261171 | 0.72828457 | 0.0024 | 0.0083 | 0.0048 | 0.0048 | TRUE | 0.071 |
| Ell2 | 4979 | 5178 | 4952 | 4572 | 3531 | 3672 | 5289 | 5171 | 4972 | 3594 | 3799 | 4020 | 4474.18 | 5144 | 3804 | 0.739 | -0.436 | 6.12E-09 | 3.04E-07 | 0.0017 | 0.0076 | 0.0038 | 0.0038 | TRUE | 0.0298 |
| Ell3 | 713 | 895 | 932 | 775 | 615 | 683 | 757 | 894 | 936 | 609 | 662 | 748 | 767.6 | 862 | 673 | 0.779 | -0.36 | 0.00510476 | 0.03184823 | 0.0101 | 0.0112 | 0.0106 | 0.0106 | TRUE | 0.1342 |
| Elmo1 | 791 | 1026 | 1180 | 629 | 475 | 628 | 840 | 1025 | 1185 | 494 | 511 | 687 | 790.45 | 1017 | 564 | 0.554 | -0.851 | 1.01E-06 | 2.78E-05 | 0.0303 | 0.0111 | 0.0204 | 0.0204 | TRUE | 0.2102 |
| Elmo2 | 2492 | 3212 | 3186 | 3073 | 2355 | 2764 | 2647 | 3208 | 3199 | 2416 | 2534 | 3026 | 2838.18 | 3018 | 2659 | 0.881 | -0.184 | 0.13645052 | 0.31996694 | 0.0125 | 0.0081 | 0.0106 | 0.0106 | TRUE | 0.1767 |
| Elmo3 | 321 | 443 | 463 | 466 | 349 | 467 | 341 | 442 | 465 | 366 | 376 | 511 | 416.89 | 416 | 418 | 1.002 | 0.003 | 0.98673551 | 0.9928109 | 0.0285 | 0.0148 | 0.0217 | 0.0217 | TRUE | 0.4379 |
| Elmod1 | 2 | 1 | 2 | 6 | 5 | 2 | 2 | 1 | 2 | 5 | 5 | 2 | 2.9 | 2 | 4 | 2.415 | 1.272 | 0.26414171 | NA | 0 | 1.1418 | 0.5169 | 0.5169 | TRUE | 0.2504 |
| Elmod2 | 2038 | 3020 | 3305 | 2354 | 1916 | 2262 | 2165 | 3016 | 3319 | 1851 | 2061 | 2476 | 2481.25 | 2833 | 2129 | 0.751 | -0.413 | 0.01647367 | 0.07601603 | 0.0329 | 0.0082 | 0.0209 | 0.0209 | TRUE | 0.5165 |
| Elmod3 | 400 | 409 | 396 | 551 | 377 | 365 | 425 | 408 | 398 | 433 | 406 | 400 | 411.55 | 410 | 413 | 1.008 | 0.011 | 0.92033389 | 0.96087606 | 0 | 0.0149 | 0.0062 | 0.0062 | TRUE | 0.023 |
| Elmsan1 | 2732 | 4377 | 4573 | 3049 | 2591 | 2855 | 2902 | 4371 | 4592 | 2397 | 2788 | 3125 | 3362.48 | 3955 | 2770 | 0.7 | -0.514 | 0.0037225 | 0.02503681 | 0.0352 | 0.0079 | 0.0223 | 0.0223 | TRUE | 0.6599 |
| Eln | 103 | 192 | 171 | 27 | 39 | 46 | 109 | 192 | 172 | 21 | 42 | 50 | 97.73 | 158 | 38 | 0.237 | -2.077 | 2.41E-09 | 1.34E-07 | 0.1111 | 0.0406 | 0.071 | 0.071 | TRUE | 0.81 |
| Elof1 | 1092 | 1027 | 979 | 1326 | 940 | 964 | 1160 | 1026 | 983 | 1042 | 1011 | 1055 | 1046.27 | 1056 | 1036 | 0.982 | -0.027 | 0.78234026 | 0.88741185 | 0.003 | 0.0101 | 0.0059 | 0.0059 | TRUE | 0.0877 |
| Elovl1 | 1064 | 1138 | 1175 | 1224 | 837 | 942 | 1130 | 1137 | 1180 | 962 | 901 | 1031 | 1056.74 | 1149 | 965 | 0.84 | -0.252 | 0.00503911 | 0.03151339 | 0.0011 | 0.01 | 0.0049 | 0.0049 | TRUE | 0.0418 |
| Elovl2 | 863 | 1078 | 1109 | 1200 | 863 | 939 | 917 | 1077 | 1114 | 943 | 929 | 1028 | 1001.1 | 1036 | 967 | 0.933 | -0.1 | 0.35960416 | 0.57800887 | 0.0057 | 0.0102 | 0.0076 | 0.0076 | TRUE | 0.12 |
| Elovl3 | 6 | 1 | 2 | 2 | 1 | 3 | 6 | 1 | 2 | 2 | 1 | 3 | 2.55 | 3 | 2 | 0.63 | -0.666 | 0.5989695 | NA | 0 | 1.2978 | 0.7432 | 0.7432 | TRUE | 1.0594 |
| Elovl4 | 1420 | 1649 | 1618 | 1597 | 1177 | 1124 | 1508 | 1647 | 1625 | 1255 | 1266 | 1230 | 1422 | 1593 | 1250 | 0.785 | -0.349 | 9.09E-06 | 0.00018342 | 6.00E-04 | 0.0092 | 0.0038 | 0.0038 | TRUE | 0.0261 |
| Elovl5 | 7195 | 8116 | 8141 | 8964 | 6461 | 6899 | 7642 | 8106 | 8174 | 7047 | 6952 | 7552 | 7578.8 | 7974 | 7184 | 0.901 | -0.151 | 0.03179393 | 0.11996504 | 0.0015 | 0.0073 | 0.0034 | 0.0034 | TRUE | 0.0245 |
| Elovl6 | 17691 | 17374 | 17354 | 19517 | 13067 | 13463 | 18791 | 17352 | 17425 | 15343 | 14059 | 14737 | 16284.53 | 17856 | 14713 | 0.824 | -0.279 | 9.27E-05 | 0.00129815 | 0.0019 | 0.0071 | 0.0036 | 0.0036 | TRUE | 0.0257 |
| Elovl7 | 389 | 584 | 575 | 518 | 456 | 515 | 413 | 583 | 577 | 407 | 491 | 564 | 505.9 | 524 | 487 | 0.927 | -0.109 | 0.5438784 | 0.72899211 | 0.0294 | 0.0134 | 0.0214 | 0.0214 | TRUE | 0.3948 |
| Elp2 | 8601 | 10321 | 10419 | 10789 | 8178 | 8600 | 9136 | 10308 | 10462 | 8481 | 8799 | 9414 | 9433.3 | 9969 | 8898 | 0.893 | -0.164 | 0.05646207 | 0.17849665 | 0.0039 | 0.0073 | 0.0052 | 0.0052 | TRUE | 0.0652 |
| Elp3 | 6977 | 7112 | 7402 | 8960 | 6396 | 6704 | 7411 | 7103 | 7432 | 7044 | 6882 | 7339 | 7201.66 | 7315 | 7088 | 0.969 | -0.046 | 0.46982603 | 0.67180893 | 6.00E-04 | 0.0074 | 0.0027 | 0.0027 | TRUE | 0.0116 |
| Elp4 | 935 | 972 | 1045 | 1418 | 1015 | 918 | 993 | 971 | 1049 | 1115 | 1092 | 1005 | 1037.48 | 1004 | 1071 | 1.067 | 0.093 | 0.29840164 | 0.51905693 | 0.0013 | 0.0101 | 0.0048 | 0.0048 | TRUE | 0.0339 |
| Elp5 | 2781 | 2747 | 2556 | 4107 | 2851 | 2538 | 2954 | 2743 | 2566 | 3229 | 3068 | 2778 | 2889.7 | 2754 | 3025 | 1.099 | 0.136 | 0.15542666 | 0.34776412 | 0.0048 | 0.008 | 0.0062 | 0.0062 | TRUE | 0.0615 |
| Elp6 | 406 | 345 | 394 | 666 | 512 | 415 | 431 | 345 | 396 | 524 | 551 | 454 | 450.02 | 391 | 510 | 1.306 | 0.385 | 0.00493782 | 0.03107714 | 0.0087 | 0.0142 | 0.0113 | 0.0113 | TRUE | 0.1221 |
| Eltd1 | 0 | 0 | 0 | 1 | 0 | 1 | 0 | 0 | 0 | 1 | 0 | 1 | 0.31 | 0 | 1 | 3.348 | 1.743 | 0.65064889 | NA | 0 | 10.5167 | 8.8968 | 8.8968 | TRUE | 0.2082 |
| Emb | 2968 | 3783 | 4273 | 3600 | 2535 | 3086 | 3153 | 3778 | 4291 | 2830 | 2728 | 3378 | 3359.48 | 3741 | 2979 | 0.796 | -0.329 | 0.01657269 | 0.07624971 | 0.0181 | 0.0079 | 0.0133 | 0.0133 | TRUE | 0.195 |
| Emc1 | 2993 | 3185 | 3210 | 3778 | 2887 | 2587 | 3179 | 3181 | 3223 | 2970 | 3106 | 2832 | 3081.87 | 3194 | 2969 | 0.93 | -0.105 | 0.12939631 | 0.30962138 | 6.00E-04 | 0.008 | 0.0032 | 0.0032 | TRUE | 0.0195 |
| Emc10 | 2988 | 3705 | 3632 | 3966 | 2927 | 3037 | 3174 | 3700 | 3647 | 3118 | 3149 | 3324 | 3352.07 | 3507 | 3197 | 0.911 | -0.134 | 0.13148346 | 0.31231281 | 0.0036 | 0.0079 | 0.0054 | 0.0054 | TRUE | 0.0838 |
| Emc2 | 2568 | 2848 | 2933 | 3717 | 2621 | 2812 | 2728 | 2844 | 2945 | 2922 | 2820 | 3078 | 2889.54 | 2839 | 2940 | 1.035 | 0.05 | 0.50103777 | 0.69669611 | 0.0012 | 0.008 | 0.0037 | 0.0037 | TRUE | 0.0202 |
| Emc3 | 1856 | 2294 | 2264 | 2940 | 2190 | 2217 | 1971 | 2291 | 2273 | 2311 | 2356 | 2427 | 2271.68 | 2178 | 2365 | 1.085 | 0.118 | 0.18510736 | 0.38842056 | 0.003 | 0.0084 | 0.0053 | 0.0053 | TRUE | 0.0835 |
| Emc4 | 1233 | 1409 | 1371 | 1996 | 1353 | 1249 | 1310 | 1407 | 1377 | 1569 | 1456 | 1367 | 1414.26 | 1365 | 1464 | 1.074 | 0.103 | 0.2512369 | 0.46875424 | 0.0024 | 0.0092 | 0.0051 | 0.0051 | TRUE | 0.0487 |
| Emc6 | 4648 | 4251 | 4201 | 5269 | 3899 | 3784 | 4937 | 4246 | 4218 | 4142 | 4195 | 4142 | 4313.35 | 4467 | 4160 | 0.931 | -0.103 | 0.24021846 | 0.4564989 | 0.0038 | 0.0077 | 0.0053 | 0.0053 | TRUE | 0.1031 |
| Emc7 | 1012 | 1242 | 1239 | 1512 | 1027 | 1074 | 1075 | 1240 | 1244 | 1189 | 1105 | 1176 | 1171.45 | 1186 | 1157 | 0.975 | -0.037 | 0.70343094 | 0.83766892 | 0.0033 | 0.0097 | 0.0059 | 0.0059 | TRUE | 0.0806 |
| Emc8 | 4372 | 3458 | 3315 | 6121 | 4440 | 3655 | 4644 | 3454 | 3329 | 4812 | 4777 | 4001 | 4169.33 | 3809 | 4530 | 1.19 | 0.25 | 0.0858842 | 0.23631641 | 0.0229 | 0.0077 | 0.0151 | 0.0151 | TRUE | 0.4474 |
| Emc9 | 105 | 124 | 111 | 187 | 152 | 133 | 112 | 124 | 111 | 147 | 164 | 146 | 133.83 | 116 | 152 | 1.314 | 0.393 | 0.02397843 | 0.09871776 | 0 | 0.0315 | 0.0142 | 0.0142 | TRUE | 0.0441 |
| Emcn | 0 | 0 | 0 | 1 | 0 | 0 | 0 | 0 | 0 | 1 | 0 | 0 | 0.13 | 0 | 0 | 1.859 | 0.894 | 0.82650734 | NA | 0 | 25.1515 | 10 | 10 | TRUE | 0.2563 |
| Emd | 2099 | 2596 | 2777 | 3635 | 2631 | 2534 | 2230 | 2593 | 2788 | 2858 | 2831 | 2774 | 2678.79 | 2537 | 2821 | 1.112 | 0.153 | 0.12851969 | 0.308244 | 0.0058 | 0.0081 | 0.0069 | 0.0069 | TRUE | 0.1358 |
| Eme1 | 2285 | 1895 | 1929 | 2830 | 1993 | 1770 | 2427 | 1893 | 1937 | 2225 | 2144 | 1938 | 2093.86 | 2086 | 2102 | 1.008 | 0.012 | 0.92094537 | 0.96116102 | 0.0117 | 0.0085 | 0.0102 | 0.0102 | TRUE | 0.2479 |
| Eme2 | 44 | 39 | 33 | 110 | 77 | 48 | 47 | 39 | 33 | 86 | 83 | 53 | 56.78 | 40 | 74 | 1.877 | 0.908 | 0.00345925 | 0.02362743 | 0.0344 | 0.0649 | 0.05 | 0.05 | TRUE | 0.5582 |
| Emg1 | 6790 | 7212 | 7076 | 9784 | 6817 | 6432 | 7212 | 7203 | 7105 | 7691 | 7335 | 7041 | 7264.47 | 7173 | 7356 | 1.026 | 0.037 | 0.57286701 | 0.7512479 | 8.00E-04 | 0.0074 | 0.0029 | 0.0029 | TRUE | 0.0196 |
| Emid1 | 239 | 377 | 385 | 297 | 199 | 217 | 254 | 377 | 387 | 233 | 214 | 238 | 283.68 | 339 | 228 | 0.674 | -0.57 | 0.00236469 | 0.01758409 | 0.0248 | 0.0185 | 0.0217 | 0.0217 | TRUE | 0.5449 |
| Emilin1 | 168 | 157 | 138 | 277 | 178 | 141 | 178 | 157 | 139 | 218 | 192 | 154 | 172.9 | 158 | 188 | 1.194 | 0.256 | 0.18920583 | 0.39431214 | 0.0173 | 0.026 | 0.0216 | 0.0216 | TRUE | 0.2506 |
| Emilin2 | 234 | 237 | 212 | 310 | 231 | 205 | 249 | 237 | 213 | 244 | 249 | 224 | 235.79 | 233 | 239 | 1.028 | 0.039 | 0.78054349 | 0.88642943 | 0 | 0.0209 | 0.0101 | 0.0101 | TRUE | 0.0616 |
| Emilin3 | 300 | 251 | 268 | 428 | 323 | 279 | 319 | 251 | 269 | 336 | 348 | 305 | 304.64 | 280 | 330 | 1.181 | 0.241 | 0.09536071 | 0.25320356 | 0.0061 | 0.0177 | 0.0117 | 0.0117 | TRUE | 0.1669 |
| Eml1 | 576 | 730 | 810 | 681 | 517 | 585 | 612 | 729 | 813 | 535 | 556 | 640 | 647.7 | 718 | 577 | 0.803 | -0.316 | 0.02399487 | 0.09875967 | 0.0131 | 0.012 | 0.0126 | 0.0126 | TRUE | 0.197 |
| Eml2 | 1854 | 3103 | 3213 | 2439 | 1916 | 2476 | 1969 | 3099 | 3226 | 1917 | 2061 | 2710 | 2497.28 | 2765 | 2229 | 0.806 | -0.311 | 0.11759469 | 0.29016904 | 0.0487 | 0.0082 | 0.028 | 0.028 | TRUE | 0.7683 |
| Eml3 | 950 | 961 | 867 | 1212 | 930 | 788 | 1009 | 960 | 871 | 953 | 1001 | 863 | 942.56 | 947 | 939 | 0.992 | -0.011 | 0.91460737 | 0.95791138 | 0.0044 | 0.0104 | 0.007 | 0.007 | TRUE | 0.0588 |
| Eml4 | 9893 | 8711 | 8804 | 10339 | 7820 | 7407 | 10508 | 8700 | 8840 | 8128 | 8414 | 8108 | 8782.95 | 9349 | 8217 | 0.879 | -0.186 | 0.04855687 | 0.16075543 | 0.0058 | 0.0073 | 0.0063 | 0.0063 | TRUE | 0.1435 |
| Eml5 | 2737 | 2998 | 3232 | 3404 | 2842 | 2856 | 2907 | 2994 | 3245 | 2676 | 3058 | 3126 | 3001.12 | 3049 | 2953 | 0.968 | -0.047 | 0.62041435 | 0.78513802 | 0.0048 | 0.008 | 0.0061 | 0.0061 | TRUE | 0.0831 |

| | | | | | | | | | | | | | | | | | | | | | | | | | |
|---|---|---|---|---|---|---|---|---|---|---|---|---|---|---|---|---|---|---|---|---|---|---|---|---|---|
| Eml6 | 94 | 50 | 57 | 144 | 92 | 61 | 100 | 50 | 57 | 113 | 99 | 67 | 81 | 69 | 93 | 1.354 | 0.437 | 0.19124472 | 0.3962054 | 0.0901 | 0.0476 | 0.068 | 0.068 | TRUE | 1.1322 |
| Emp1 | 777 | 1048 | 1037 | 2043 | 1456 | 1646 | 825 | 1047 | 1041 | 1606 | 1567 | 1802 | 1314.61 | 971 | 1658 | 1.707 | 0.771 | 3.41E-10 | 2.33E-08 | 0.0101 | 0.0094 | 0.01 | 0.01 | TRUE | 0.2047 |
| Emp2 | 275 | 319 | 323 | 354 | 225 | 313 | 292 | 319 | 324 | 278 | 242 | 343 | 299.67 | 312 | 288 | 0.922 | -0.117 | 0.46686485 | 0.66929996 | 0.0129 | 0.0179 | 0.0153 | 0.0153 | TRUE | 0.3029 |
| Emp3 | 140 | 154 | 168 | 149 | 105 | 152 | 149 | 154 | 169 | 117 | 113 | 166 | 144.61 | 157 | 132 | 0.839 | -0.254 | 0.21901573 | 0.43123579 | 0.017 | 0.0297 | 0.0238 | 0.0238 | TRUE | 0.5082 |
| Emr1 | 1 | 0 | 1 | 2 | 0 | 1 | 1 | 0 | 1 | 2 | 0 | 1 | 0.79 | 1 | 1 | 1.33 | 0.412 | 0.85027746 | NA | 0 | 4.1831 | 2.149 | 2.149 | TRUE | 0.2882 |
| Emr4 | 8 | 2 | 2 | 3 | 3 | 1 | 8 | 2 | 2 | 2 | 3 | 1 | 3.2 | 4 | 2 | 0.539 | -0.89 | 0.4420707 | NA | 0.2746 | 1.0373 | 0.6252 | 0.6252 | TRUE | 1.3663 |
| Emx1 | 8 | 26 | 37 | 21 | 15 | 16 | 8 | 26 | 37 | 17 | 16 | 18 | 20.3 | 24 | 17 | 0.698 | -0.518 | 0.35768376 | 0.57623987 | 0.1868 | 0.1692 | 0.1776 | 0.1776 | TRUE | 0.2498 |
| Emx2 | 0 | 1 | 0 | 0 | 1 | 1 | 0 | 1 | 0 | 0 | 1 | 1 | 0.53 | 0 | 1 | 1.924 | 0.944 | 0.7364225 | NA | 0 | 6.2438 | 3.9727 | 3.9727 | TRUE | 0.405 |
| Emx2os | 0 | 0 | 0 | 1 | 0 | 0 | 0 | 0 | 0 | 1 | 0 | 0 | 0.13 | 0 | 0 | 1.859 | 0.894 | 0.82650734 | NA | 0 | 25.1515 | 10 | 10 | TRUE | 0.2563 |
| En1 | 8 | 11 | 12 | 30 | 18 | 13 | 8 | 11 | 12 | 24 | 19 | 14 | 14.79 | 10 | 19 | 1.825 | 0.868 | 0.0906781 | 0.24556577 | 0 | 0.2297 | 0.1155 | 0.1155 | TRUE | 0.1586 |
| En2 | 485 | 536 | 537 | 777 | 611 | 568 | 515 | 535 | 539 | 611 | 657 | 622 | 579.94 | 530 | 630 | 1.188 | 0.248 | 0.01016514 | 0.05344155 | 0 | 0.0126 | 0.005 | 0.005 | TRUE | 0.0168 |
| Enah | 16019 | 14205 | 15322 | 16915 | 12220 | 12083 | 17015 | 14187 | 15385 | 13297 | 13148 | 13227 | 14376.42 | 15529 | 13224 | 0.852 | -0.232 | 0.00688741 | 0.0396598 | 0.0041 | 0.0071 | 0.0052 | 0.0052 | TRUE | 0.0857 |
| Enam | 0 | 0 | 0 | 0 | 0 | 0 | 0 | 0 | 0 | 0 | 0 | 0 | 0 | 0 | 0 | NA | NA | NA | NA | NA | NA | NA | NA | NA | NA |
| Enc1 | 4596 | 4756 | 4954 | 5037 | 3813 | 3494 | 4882 | 4750 | 4974 | 3960 | 4103 | 3825 | 4415.49 | 4869 | 3963 | 0.814 | -0.297 | 5.63E-06 | 0.00012199 | 5.00E-04 | 0.0076 | 0.0029 | 0.0029 | TRUE | 0.0115 |
| Endod1 | 349 | 322 | 294 | 466 | 327 | 265 | 371 | 322 | 295 | 366 | 352 | 290 | 332.62 | 329 | 336 | 1.023 | 0.033 | 0.82827218 | 0.9129575 | 0.0111 | 0.0168 | 0.0138 | 0.0138 | TRUE | 0.1595 |
| Endog | 505 | 434 | 430 | 937 | 681 | 533 | 536 | 433 | 432 | 737 | 733 | 583 | 575.73 | 467 | 684 | 1.466 | 0.552 | 0.00016319 | 0.00204603 | 0.0137 | 0.0126 | 0.0137 | 0.0137 | TRUE | 0.1937 |
| Endou | 0 | 2 | 1 | 7 | 3 | 2 | 0 | 2 | 1 | 6 | 3 | 2 | 2.32 | 1 | 4 | 3.647 | 1.867 | 0.1803142 | NA | 0 | 1.4268 | 0.7615 | 0.7615 | TRUE | 0.3976 |
| Endov | 350 | 362 | 350 | 435 | 350 | 266 | 372 | 362 | 351 | 342 | 377 | 291 | 349.08 | 362 | 337 | 0.931 | -0.102 | 0.45154953 | 0.65705685 | 0.0051 | 0.0163 | 0.0105 | 0.0105 | TRUE | 0.1512 |
| Eng | 2182 | 2266 | 2229 | 2403 | 1831 | 1788 | 2318 | 2263 | 2238 | 1889 | 1970 | 1957 | 2105.87 | 2273 | 1939 | 0.853 | -0.23 | 0.00066168 | 0.00648196 | 0 | 0.0085 | 0.0028 | 0.0028 | TRUE | 0.0063 |
| Engase | 1262 | 1214 | 1227 | 1381 | 1087 | 1110 | 1340 | 1212 | 1232 | 1086 | 1170 | 1215 | 1209.2 | 1261 | 1157 | 0.916 | -0.126 | 0.16741229 | 0.36505276 | 0.0023 | 0.0096 | 0.0052 | 0.0052 | TRUE | 0.0359 |
| Enho | 148 | 163 | 169 | 175 | 144 | 125 | 157 | 163 | 170 | 138 | 155 | 137 | 153.17 | 163 | 143 | 0.875 | -0.192 | 0.23835277 | 0.45416935 | 0 | 0.0284 | 0.0126 | 0.0126 | TRUE | 0.052 |
| Enkd1 | 524 | 776 | 795 | 1036 | 856 | 826 | 557 | 775 | 798 | 814 | 921 | 904 | 794.91 | 710 | 880 | 1.238 | 0.308 | 0.04296341 | 0.1481134 | 0.0196 | 0.011 | 0.0154 | 0.0154 | TRUE | 0.42 |
| Enkur | 18 | 16 | 27 | 18 | 17 | 14 | 19 | 16 | 27 | 14 | 18 | 15 | 18.33 | 21 | 16 | 0.764 | -0.388 | 0.3982102 | 0.6115454 | 0 | 0.1866 | 0.0966 | 0.0966 | TRUE | 0.4042 |
| Eno1 | 866 | 1096 | 1218 | 843 | 629 | 871 | 920 | 1095 | 1223 | 663 | 677 | 953 | 921.72 | 1079 | 764 | 0.708 | -0.499 | 0.00376188 | 0.02524784 | 0.0313 | 0.0105 | 0.0203 | 0.0203 | TRUE | 0.5518 |
| Eno2 | 480 | 739 | 775 | 298 | 257 | 340 | 510 | 738 | 778 | 234 | 277 | 372 | 484.84 | 675 | 294 | 0.435 | -1.202 | 2.26E-08 | 9.87E-07 | 0.0481 | 0.0137 | 0.0309 | 0.0309 | TRUE | 0.5993 |
| Eno3 | 480 | 515 | 475 | 858 | 603 | 579 | 510 | 514 | 477 | 674 | 649 | 634 | 576.37 | 500 | 652 | 1.305 | 0.384 | 9.18E-05 | 0.00129127 | 0 | 0.0126 | 0.0052 | 0.0052 | TRUE | 0.0197 |
| Eno4 | 75 | 45 | 52 | 99 | 67 | 74 | 80 | 45 | 52 | 78 | 72 | 81 | 67.96 | 59 | 77 | 1.309 | 0.388 | 0.16950643 | 0.36794046 | 0.0275 | 0.0554 | 0.0425 | 0.0425 | TRUE | 0.7928 |
| Enoph1 | 3146 | 3260 | 3177 | 3346 | 2568 | 2618 | 3342 | 3256 | 3190 | 2630 | 2763 | 2866 | 3007.77 | 3263 | 2753 | 0.843 | -0.246 | 0.00051176 | 0.00527203 | 9.00E-04 | 0.008 | 0.0033 | 0.0033 | TRUE | 0.0189 |
| Enox1 | 922 | 738 | 886 | 1264 | 914 | 741 | 979 | 737 | 890 | 994 | 983 | 811 | 899.04 | 869 | 929 | 1.071 | 0.099 | 0.48274083 | 0.68126238 | 0.0154 | 0.0106 | 0.0131 | 0.0131 | TRUE | 0.2095 |
| Enox2 | 716 | 833 | 890 | 814 | 594 | 550 | 761 | 832 | 894 | 640 | 639 | 602 | 727.86 | 829 | 627 | 0.757 | -0.402 | 0.0001018 | 0.00139866 | 0.0026 | 0.0114 | 0.0063 | 0.0063 | TRUE | 0.061 |
| Enpep | 12 | 7 | 0 | 22 | 13 | 10 | 13 | 7 | 0 | 17 | 14 | 11 | 10.33 | 7 | 14 | 2.157 | 1.109 | 0.20495998 | 0.41337964 | 0.6968 | 0.3259 | 0.4386 | 0.4386 | TRUE | 0.5484 |
| Enpp1 | 178 | 153 | 144 | 236 | 158 | 175 | 189 | 153 | 145 | 186 | 170 | 192 | 172.26 | 162 | 183 | 1.126 | 0.171 | 0.32286991 | 0.54401187 | 0.0055 | 0.026 | 0.0158 | 0.0158 | TRUE | 0.2198 |
| Enpp2 | 13 | 6 | 4 | 5 | 8 | 11 | 14 | 6 | 4 | 4 | 9 | 12 | 8.07 | 8 | 8 | 1.02 | 0.029 | 0.97041075 | 0.9846576 | 0.2215 | 0.4153 | 0.318 | 0.318 | TRUE | 0.5375 |
| Enpp3 | 893 | 1271 | 1424 | 1355 | 1040 | 1586 | 949 | 1269 | 1430 | 1065 | 1119 | 1736 | 1261.34 | 1216 | 1307 | 1.074 | 0.103 | 0.62622836 | 0.78958445 | 0.0577 | 0.0095 | 0.0315 | 0.0315 | TRUE | 0.99 |
| Enpp4 | 941 | 894 | 909 | 1419 | 935 | 864 | 1000 | 893 | 913 | 1116 | 1006 | 946 | 978.73 | 935 | 1023 | 1.095 | 0.131 | 0.21034924 | 0.42084523 | 0.0044 | 0.0103 | 0.0068 | 0.0068 | TRUE | 0.0779 |
| Enpp5 | 593 | 913 | 971 | 650 | 463 | 681 | 630 | 912 | 975 | 511 | 498 | 745 | 711.88 | 839 | 585 | 0.697 | -0.522 | 0.01232602 | 0.06138491 | 0.0501 | 0.0115 | 0.0299 | 0.0299 | TRUE | 0.6719 |
| Enpp6 | 1 | 0 | 0 | 3 | 3 | 0 | 1 | 0 | 0 | 2 | 3 | 0 | 1.11 | 0 | 2 | 5.12 | 2.356 | 0.29230162 | NA | 0 | 2.9801 | 2.3453 | 2.3453 | TRUE | 0.417 |
| Enpp7 | 0 | 0 | 0 | 0 | 0 | 0 | 0 | 0 | 0 | 0 | 0 | 0 | 0 | 0 | 0 | NA | NA | NA | NA | NA | NA | NA | NA | NA | NA |
| Ensa | 3817 | 4465 | 4500 | 5381 | 3850 | 4234 | 4054 | 4459 | 4518 | 4230 | 4142 | 4635 | 4339.88 | 4344 | 4336 | 0.998 | -0.003 | 0.97151505 | 0.98490859 | 0.0033 | 0.0077 | 0.0049 | 0.0049 | TRUE | 0.044 |
| Enthd1 | 22 | 36 | 27 | 28 | 37 | 50 | 23 | 36 | 27 | 22 | 40 | 55 | 33.83 | 29 | 39 | 1.335 | 0.417 | 0.33164704 | 0.55217325 | 0.0999 | 0.1043 | 0.1022 | 0.1022 | TRUE | 0.1925 |
| Enthd2 | 1565 | 1482 | 1381 | 1965 | 1599 | 1392 | 1662 | 1480 | 1387 | 1545 | 1720 | 1524 | 1553 | 1510 | 1596 | 1.057 | 0.08 | 0.43915734 | 0.64641993 | 0.0055 | 0.009 | 0.0071 | 0.0071 | TRUE | 0.0939 |
| Entpd1 | 38 | 22 | 23 | 26 | 14 | 16 | 40 | 22 | 23 | 20 | 15 | 18 | 23.07 | 28 | 18 | 0.626 | -0.677 | 0.12605395 | 0.30427348 | 0.041 | 0.1497 | 0.0955 | 0.0955 | TRUE | 0.8391 |
| Entpd2 | 221 | 347 | 334 | 202 | 171 | 218 | 235 | 347 | 335 | 159 | 184 | 239 | 249.68 | 306 | 194 | 0.632 | -0.662 | 0.00221704 | 0.01666687 | 0.0393 | 0.0201 | 0.0295 | 0.0295 | TRUE | 0.461 |
| Entpd3 | 41 | 38 | 42 | 42 | 40 | 58 | 44 | 38 | 42 | 33 | 43 | 63 | 43.87 | 41 | 46 | 1.119 | 0.163 | 0.63253075 | 0.79321578 | 0.0391 | 0.082 | 0.0605 | 0.0605 | TRUE | 0.466 |
| Entpd4 | 0 | 0 | 0 | 0 | 0 | 0 | 0 | 0 | 0 | 0 | 0 | 0 | 0 | 0 | 0 | NA | NA | NA | NA | NA | NA | NA | NA | NA | NA |
| Entpd5 | 1227 | 1290 | 1219 | 1401 | 1002 | 1040 | 1303 | 1288 | 1224 | 1101 | 1078 | 1138 | 1188.92 | 1272 | 1106 | 0.87 | -0.202 | 0.01108821 | 0.05680195 | 0 | 0.0097 | 0.0037 | 0.0037 | TRUE | 0.0131 |
| Entpd6 | 2337 | 2400 | 2180 | 2942 | 2022 | 1991 | 2482 | 2397 | 2189 | 2313 | 2176 | 2179 | 2289.32 | 2356 | 2223 | 0.944 | -0.083 | 0.31421428 | 0.5346626 | 0.002 | 0.0083 | 0.0045 | 0.0045 | TRUE | 0.0467 |
| Entpd7 | 437 | 371 | 360 | 463 | 363 | 318 | 464 | 371 | 361 | 364 | 391 | 348 | 383.14 | 399 | 368 | 0.922 | -0.117 | 0.41088244 | 0.62206449 | 0.0088 | 0.0155 | 0.0119 | 0.0119 | TRUE | 0.2354 |
| Entpd8 | 0 | 0 | 0 | 0 | 0 | 0 | 0 | 0 | 0 | 0 | 0 | 0 | 0 | 0 | 0 | NA | NA | NA | NA | NA | NA | NA | NA | NA | NA |
| Eny2 | 1718 | 1404 | 1428 | 2179 | 1522 | 1345 | 1825 | 1402 | 1434 | 1713 | 1638 | 1472 | 1580.62 | 1554 | 1608 | 1.035 | 0.05 | 0.69708495 | 0.83403726 | 0.0132 | 0.009 | 0.0113 | 0.0113 | TRUE | 0.2804 |
| Eogt | 609 | 825 | 885 | 1272 | 929 | 975 | 647 | 824 | 889 | 1000 | 1000 | 1067 | 904.37 | 787 | 1022 | 1.299 | 0.378 | 0.00448244 | 0.02877093 | 0.0117 | 0.0105 | 0.0116 | 0.0116 | TRUE | 0.2844 |
| Eomes | 704 | 738 | 706 | 1038 | 736 | 669 | 748 | 737 | 709 | 816 | 792 | 732 | 755.65 | 731 | 780 | 1.068 | 0.095 | 0.31170585 | 0.53217414 | 0 | 0.0113 | 0.005 | 0.005 | TRUE | 0.0334 |
| Ep300 | 8136 | 7276 | 7791 | 10770 | 7446 | 7027 | 8642 | 7267 | 7823 | 8467 | 8011 | 7692 | 7983.6 | 7911 | 8057 | 1.019 | 0.027 | 0.76859396 | 0.87928573 | 0.0039 | 0.0073 | 0.0058 | 0.0058 | TRUE | 0.0798 |
| Ep400 | 12117 | 10302 | 12355 | 14061 | 10369 | 10098 | 12870 | 10289 | 12406 | 11054 | 11156 | 11054 | 11471.46 | 11855 | 11088 | 0.935 | -0.096 | 0.3325878 | 0.55303898 | 0.0066 | 0.0072 | 0.0071 | 0.0071 | TRUE | 0.1633 |
| Epas1 | 4123 | 4466 | 4859 | 4024 | 2924 | 3129 | 4379 | 4460 | 4879 | 3163 | 3146 | 3425 | 3908.85 | 4573 | 3245 | 0.709 | -0.495 | 8.74E-10 | 5.45E-08 | 0.0025 | 0.0077 | 0.0044 | 0.0044 | TRUE | 0.0418 |
| Epb4.1 | 3530 | 3510 | 3735 | 4401 | 3295 | 3327 | 3749 | 3505 | 3750 | 3460 | 3545 | 3642 | 3608.69 | 3668 | 3549 | 0.967 | -0.048 | 0.48335697 | 0.68146464 | 8.00E-04 | 0.0078 | 0.0031 | 0.0031 | TRUE | 0.0184 |
| Epb4.1l1 | 2249 | 2442 | 2657 | 3100 | 2157 | 2293 | 2389 | 2439 | 2668 | 2437 | 2321 | 2510 | 2460.57 | 2499 | 2423 | 0.97 | -0.045 | 0.58322168 | 0.75875085 | 0.0019 | 0.0082 | 0.0043 | 0.0043 | TRUE | 0.0427 |
| Epb4.1l2 | 7543 | 7742 | 8008 | 9763 | 7215 | 6909 | 8012 | 7732 | 8041 | 7675 | 7763 | 7563 | 7797.61 | 7928 | 7667 | 0.967 | -0.048 | 0.39446782 | 0.60810737 | 2.00E-04 | 0.0073 | 0.0022 | 0.0022 | TRUE | 0.0057 |
| Epb4.1l3 | 590 | 677 | 697 | 1079 | 878 | 810 | 627 | 676 | 700 | 848 | 945 | 887 | 780.37 | 668 | 893 | 1.337 | 0.419 | 2.10E-05 | 0.00036993 | 0.0018 | 0.0111 | 0.0057 | 0.0057 | TRUE | 0.0335 |
| Epb4.1l4a | 2303 | 2560 | 2678 | 3168 | 2276 | 2340 | 2446 | 2557 | 2689 | 2490 | 2449 | 2561 | 2532.11 | 2564 | 2500 | 0.975 | -0.037 | 0.61852235 | 0.78448356 | 0.001 | 0.0082 | 0.0035 | 0.0035 | TRUE | 0.0221 |
| Epb4.1l4b | 830 | 824 | 877 | 1035 | 698 | 730 | 882 | 823 | 881 | 814 | 751 | 799 | 824.81 | 862 | 788 | 0.915 | -0.128 | 0.15656828 | 0.34960179 | 0 | 0.0109 | 0.0047 | 0.0047 | TRUE | 0.0196 |
| Epb4.1l5 | 2115 | 2175 | 2284 | 2367 | 1828 | 1992 | 2247 | 2172 | 2293 | 1861 | 1967 | 2181 | 2120.03 | 2237 | 2003 | 0.895 | -0.161 | 0.06980152 | 0.20540329 | 0.003 | 0.0085 | 0.0052 | 0.0052 | TRUE | 0.0727 |
| Epb4.2 | 1 | 2 | 1 | 1 | 0 | 0 | 1 | 2 | 1 | 1 | 0 | 0 | 0.81 | 1 | 0 | 0.257 | -1.958 | 0.36424053 | NA | 0 | 4.0827 | 1.9787 | 1.9787 | TRUE | 0.2562 |
| Epb4.9 | 1389 | 1814 | 2005 | 2058 | 1618 | 2035 | 1475 | 1812 | 2013 | 1618 | 1741 | 2228 | 1814.43 | 1767 | 1862 | 1.053 | 0.075 | 0.63397106 | 0.79407494 | 0.0259 | 0.0087 | 0.0174 | 0.0174 | TRUE | 0.3555 |
| Epc1 | 644 | 636 | 686 | 1179 | 767 | 684 | 684 | 635 | 689 | 927 | 825 | 749 | 751.48 | 669 | 834 | 1.247 | 0.319 | 0.00534059 | 0.03284711 | 0.0057 | 0.0113 | 0.0081 | 0.0081 | TRUE | 0.1171 |
| Epc2 | 1468 | 1529 | 1606 | 2199 | 1528 | 1544 | 1559 | 1527 | 1613 | 1729 | 1644 | 1690 | 1626.96 | 1566 | 1688 | 1.078 | 0.108 | 0.1388384 | 0.3234564 | 0 | 0.0089 | 0.0032 | 0.0032 | TRUE | 0.0081 |
| Epcam | 4425 | 6294 | 6643 | 5400 | 4419 | 5368 | 4700 | 6286 | 6670 | 4245 | 4755 | 5876 | 5422 | 5885 | 4959 | 0.842 | -0.248 | 0.12621267 | 0.30451678 | 0.0296 | 0.0075 | 0.0187 | 0.0187 | TRUE | 0.3784 |
| Epdr1 | 773 | 1006 | 1001 | 857 | 594 | 649 | 821 | 1005 | 1005 | 674 | 639 | 710 | 809.02 | 944 | 674 | 0.714 | -0.485 | 3.61E-05 | 0.00058477 | 0.0069 | 0.011 | 0.0087 | 0.0087 | TRUE | 0.1531 |
| Epg5 | 1410 | 1140 | 1329 | 1909 | 1284 | 1170 | 1498 | 1139 | 1334 | 1501 | 1382 | 1281 | 1355.6 | 1324 | 1388 | 1.049 | 0.069 | 0.5816088 | 0.75788087 | 0.0117 | 0.0093 | 0.0106 | 0.0106 | TRUE | 0.18 |

| | | | | | | | | | | | | | | | | | | | | | | | | | |
|---|---|---|---|---|---|---|---|---|---|---|---|---|---|---|---|---|---|---|---|---|---|---|---|---|---|
| Epgn | 6 | 0 | 4 | 4 | 3 | 3 | 6 | 0 | 4 | 3 | 3 | 3 | 3.34 | 3 | 3 | 0.934 | -0.099 | 0.93330341 | NA | 0 | 0.993 | 0.7114 | 0.7114 | TRUE | 0.3779 |
| Epha1 | 510 | 566 | 562 | 522 | 393 | 434 | 542 | 565 | 564 | 410 | 423 | 475 | 496.59 | 557 | 436 | 0.781 | -0.356 | 0.00116597 | 0.01000526 | 9.00E-04 | 0.0135 | 0.0066 | 0.0066 | TRUE | 0.0693 |
| Epha10 | 75 | 132 | 106 | 73 | 58 | 72 | 80 | 132 | 106 | 57 | 62 | 79 | 86.09 | 106 | 66 | 0.622 | -0.686 | 0.01076571 | 0.05569518 | 0.0345 | 0.0452 | 0.0399 | 0.0399 | TRUE | 0.1302 |
| Epha2 | 9419 | 10215 | 10659 | 10229 | 7093 | 5759 | 10005 | 10202 | 10703 | 8041 | 7632 | 6304 | 8814.37 | 10303 | 7326 | 0.711 | -0.492 | 3.40E-06 | 7.94E-05 | 0.0081 | 0.0073 | 0.008 | 0.008 | TRUE | 0.1813 |
| Epha3 | 2 | 2 | 0 | 2 | 5 | 2 | 2 | 2 | 0 | 2 | 5 | 2 | 2.21 | 1 | 3 | 2.199 | 1.137 | 0.4157955 | NA | 0 | 1.4973 | 0.87 | 0.87 | TRUE | 0.5566 |
| Epha4 | 5649 | 5732 | 5926 | 4430 | 3312 | 3617 | 6000 | 5725 | 5950 | 3483 | 3564 | 3959 | 4780.09 | 5892 | 3669 | 0.622 | -0.684 | 3.93E-18 | 7.84E-16 | 0.0024 | 0.0076 | 0.0043 | 0.0043 | TRUE | 0.0583 |
| Epha5 | 1 | 0 | 0 | 4 | 2 | 2 | 1 | 0 | 0 | 3 | 2 | 2 | 1.42 | 0 | 2 | 6.866 | 2.779 | 0.12585242 | NA | 0 | 2.3194 | 1.181 | 1.181 | TRUE | 0.4614 |
| Epha6 | 1 | 0 | 1 | 3 | 1 | 0 | 1 | 0 | 1 | 2 | 1 | 0 | 0.92 | 1 | 1 | 1.726 | 0.787 | 0.70823394 | NA | 0 | 3.6005 | 2.0291 | 2.0291 | TRUE | 0.1755 |
| Epha7 | 109 | 188 | 224 | 266 | 188 | 211 | 116 | 188 | 225 | 209 | 202 | 231 | 195.14 | 176 | 214 | 1.214 | 0.28 | 0.24296529 | 0.45958847 | 0.0501 | 0.0238 | 0.0362 | 0.0362 | TRUE | 0.5839 |
| Epha8 | 500 | 463 | 504 | 253 | 219 | 226 | 531 | 462 | 506 | 199 | 236 | 247 | 363.58 | 500 | 227 | 0.453 | -1.142 | 1.02E-16 | 1.71E-14 | 0.0053 | 0.016 | 0.0105 | 0.0105 | TRUE | 0.1441 |
| Ephb1 | 14 | 21 | 10 | 44 | 53 | 29 | 15 | 21 | 10 | 35 | 57 | 32 | 28.21 | 15 | 41 | 2.68 | 1.422 | 0.00157292 | 0.01272272 | 0.0743 | 0.1237 | 0.1005 | 0.1005 | TRUE | 0.8333 |
| Ephb2 | 1076 | 1198 | 1315 | 1494 | 1113 | 1132 | 1143 | 1196 | 1320 | 1174 | 1198 | 1239 | 1211.82 | 1220 | 1204 | 0.986 | -0.02 | 0.82637696 | 0.91211327 | 0.0023 | 0.0096 | 0.0052 | 0.0052 | TRUE | 0.0624 |
| Ephb3 | 1959 | 2356 | 2458 | 1997 | 1668 | 1782 | 2081 | 2353 | 2468 | 1570 | 1795 | 1951 | 2036.18 | 2301 | 1772 | 0.77 | -0.378 | 0.00095938 | 0.00862184 | 0.0088 | 0.0085 | 0.0089 | 0.0089 | TRUE | 0.1219 |
| Ephb4 | 2224 | 3561 | 3552 | 3185 | 2652 | 3285 | 2362 | 3556 | 3567 | 2504 | 2853 | 3596 | 3073.07 | 3162 | 2984 | 0.944 | -0.084 | 0.65087593 | 0.80429196 | 0.0429 | 0.008 | 0.0243 | 0.0243 | TRUE | 0.594 |
| Ephb6 | 52 | 105 | 98 | 174 | 106 | 125 | 55 | 105 | 98 | 137 | 114 | 137 | 107.69 | 86 | 129 | 1.499 | 0.584 | 0.02881423 | 0.11186421 | 0.0466 | 0.0375 | 0.0416 | 0.0416 | TRUE | 0.9131 |
| Ephx1 | 962 | 1603 | 1491 | 1219 | 952 | 1390 | 1022 | 1601 | 1497 | 958 | 1024 | 1522 | 1270.67 | 1373 | 1168 | 0.85 | -0.234 | 0.27079067 | 0.49074002 | 0.0541 | 0.0095 | 0.0319 | 0.0319 | TRUE | 0.838 |
| Ephx2 | 984 | 1394 | 1536 | 691 | 583 | 810 | 1045 | 1392 | 1542 | 543 | 627 | 887 | 1006.14 | 1326 | 686 | 0.516 | -0.954 | 2.89E-06 | 6.92E-05 | 0.0509 | 0.0102 | 0.0288 | 0.0288 | TRUE | 0.6991 |
| Ephx3 | 8 | 3 | 4 | 7 | 2 | 3 | 8 | 3 | 4 | 6 | 2 | 3 | 4.41 | 5 | 4 | 0.723 | -0.468 | 0.618775 | NA | 0 | 0.7543 | 0.407 | 0.407 | TRUE | 0.6279 |
| Ephx4 | 33 | 40 | 48 | 94 | 60 | 54 | 35 | 40 | 48 | 74 | 65 | 59 | 53.46 | 41 | 66 | 1.608 | 0.685 | 0.01378681 | 0.06645647 | 0 | 0.0685 | 0.036 | 0.036 | TRUE | 0.1753 |
| Epm2a | 37 | 34 | 43 | 134 | 120 | 89 | 39 | 34 | 43 | 105 | 129 | 97 | 74.72 | 39 | 110 | 2.848 | 1.51 | 3.29E-09 | 1.75E-07 | 0 | 0.051 | 0.0294 | 0.0294 | TRUE | 0.2041 |
| Epm2aip1 | 1125 | 1275 | 1372 | 1752 | 1418 | 1323 | 1195 | 1273 | 1378 | 1377 | 1526 | 1448 | 1366.19 | 1282 | 1450 | 1.131 | 0.177 | 0.05852754 | 0.18288171 | 0.0031 | 0.0093 | 0.0056 | 0.0056 | TRUE | 0.0512 |
| Epn1 | 4663 | 4270 | 4313 | 5145 | 3866 | 3772 | 4953 | 4264 | 4331 | 4045 | 4160 | 4129 | 4313.56 | 4516 | 4111 | 0.91 | -0.136 | 0.10921806 | 0.2771661 | 0.0033 | 0.0077 | 0.0049 | 0.0049 | TRUE | 0.0871 |
| Epn2 | 5050 | 5624 | 5726 | 6486 | 4768 | 4999 | 5364 | 5617 | 5750 | 5099 | 5130 | 5472 | 5405.22 | 5577 | 5234 | 0.938 | -0.092 | 0.1837475 | 0.38656336 | 0.0011 | 0.0075 | 0.0033 | 0.0033 | TRUE | 0.0193 |
| Epn3 | 8 | 10 | 13 | 14 | 17 | 5 | 8 | 10 | 13 | 11 | 18 | 5 | 11.05 | 10 | 11 | 1.101 | 0.138 | 0.82805251 | 0.91291407 | 0.084 | 0.305 | 0.2012 | 0.2012 | TRUE | 0.2313 |
| Epo | 14 | 22 | 18 | 24 | 9 | 10 | 15 | 22 | 18 | 19 | 10 | 11 | 15.74 | 18 | 13 | 0.73 | -0.453 | 0.37592884 | 0.59147378 | 0 | 0.2163 | 0.1244 | 0.1244 | TRUE | 0.7432 |
| Epor | 173 | 203 | 182 | 167 | 156 | 121 | 184 | 203 | 183 | 131 | 168 | 132 | 166.8 | 190 | 144 | 0.756 | -0.403 | 0.02049561 | 0.08835387 | 0.0046 | 0.0267 | 0.0157 | 0.0157 | TRUE | 0.2145 |
| Eppin | 7 | 4 | 5 | 0 | 2 | 3 | 7 | 4 | 5 | 0 | 2 | 3 | 3.65 | 5 | 2 | 0.313 | -1.674 | 0.14782427 | NA | 0 | 0.9101 | 0.5872 | 0.5872 | TRUE | 0.9267 |
| Eppk1 | 137 | 147 | 155 | 113 | 97 | 97 | 146 | 147 | 156 | 89 | 104 | 106 | 124.56 | 150 | 100 | 0.665 | -0.588 | 0.00125408 | 0.01061121 | 0 | 0.0334 | 0.0156 | 0.0156 | TRUE | 0.103 |
| Eprs | 21640 | 21015 | 21678 | 26271 | 18713 | 18778 | 22986 | 20988 | 21767 | 20652 | 20134 | 20555 | 21180.35 | 21914 | 20447 | 0.933 | -0.1 | 0.11800366 | 0.2905329 | 0.0011 | 0.0071 | 0.0029 | 0.0029 | TRUE | 0.0224 |
| Eps15 | 2979 | 3513 | 3538 | 5127 | 3640 | 3618 | 3164 | 3508 | 3553 | 4030 | 3916 | 3960 | 3688.76 | 3408 | 3969 | 1.164 | 0.22 | 0.00382949 | 0.02548466 | 0.0016 | 0.0078 | 0.0039 | 0.0039 | TRUE | 0.0476 |
| Eps15l1 | 5015 | 5399 | 5593 | 7104 | 5458 | 5170 | 5327 | 5392 | 5616 | 5585 | 5872 | 5659 | 5575.21 | 5445 | 5705 | 1.048 | 0.067 | 0.28482246 | 0.50626428 | 5.00E-04 | 0.0075 | 0.0027 | 0.0027 | TRUE | 0.0092 |
| Eps8 | 5474 | 4380 | 4381 | 5988 | 4268 | 3586 | 5814 | 4374 | 4399 | 4707 | 4592 | 3925 | 4635.42 | 4862 | 4408 | 0.907 | -0.141 | 0.29930619 | 0.51961003 | 0.0186 | 0.0076 | 0.0131 | 0.0131 | TRUE | 0.3571 |
| Eps8l1 | 6 | 2 | 2 | 4 | 4 | 3 | 6 | 2 | 2 | 3 | 4 | 3 | 3.52 | 3 | 3 | 1.036 | 0.051 | 0.95999509 | NA | 0 | 0.9432 | 0.4668 | 0.4668 | TRUE | 0.7601 |
| Eps8l2 | 602 | 903 | 951 | 726 | 569 | 707 | 639 | 902 | 955 | 571 | 612 | 774 | 742.17 | 832 | 652 | 0.783 | -0.353 | 0.05211297 | 0.16873862 | 0.0321 | 0.0113 | 0.0224 | 0.0224 | TRUE | 0.4857 |
| Eps8l3 | 0 | 0 | 0 | 1 | 0 | 0 | 0 | 0 | 0 | 1 | 0 | 0 | 0.13 | 0 | 0 | 1.859 | 0.894 | 0.82650734 | NA | 0 | 25.1515 | 10 | 10 | TRUE | 0.2563 |
| Epsti1 | 0 | 0 | 1 | 0 | 0 | 0 | 0 | 0 | 1 | 0 | 0 | 0 | 0.17 | 0 | 0 | 0.49 | -1.029 | 0.80087 | NA | 0 | 19.6928 | 10 | 10 | TRUE | 0.4098 |
| Ept1 | 3608 | 2659 | 2677 | 4164 | 2802 | 2702 | 3832 | 2656 | 2688 | 3273 | 3015 | 2958 | 3070.31 | 3059 | 3082 | 1.008 | 0.011 | 0.93989611 | 0.96977897 | 0.0235 | 0.008 | 0.0159 | 0.0159 | TRUE | 0.5944 |
| Epx | 80 | 55 | 52 | 51 | 33 | 37 | 85 | 55 | 52 | 40 | 36 | 41 | 51.37 | 64 | 39 | 0.606 | -0.723 | 0.01997361 | 0.08675953 | 0.0273 | 0.071 | 0.0489 | 0.0489 | TRUE | 0.7032 |
| Epyc | 0 | 0 | 0 | 0 | 0 | 0 | 0 | 0 | 0 | 0 | 0 | 0 | 0 | 0 | 0 | NA | NA | NA | NA | NA | NA | NA | NA | NA | NA |
| Eqtn | 1 | 0 | 0 | 0 | 0 | 0 | 1 | 0 | 0 | 0 | 0 | 0 | 0.18 | 0 | 0 | 0.49 | -1.029 | 0.80087 | NA | 0 | 18.6165 | 10 | 10 | TRUE | 0.4614 |
| Eral1 | 1986 | 2239 | 2291 | 2662 | 2011 | 2115 | 2109 | 2236 | 2300 | 2093 | 2164 | 2315 | 2202.93 | 2215 | 2191 | 0.988 | -0.017 | 0.83289493 | 0.91523598 | 0.0019 | 0.0084 | 0.0042 | 0.0042 | TRUE | 0.0299 |
| Erap1 | 138 | 116 | 146 | 174 | 119 | 103 | 147 | 116 | 147 | 137 | 128 | 113 | 131.1 | 137 | 126 | 0.926 | -0.111 | 0.56486598 | 0.74534144 | 0.0061 | 0.032 | 0.0191 | 0.0191 | TRUE | 0.1803 |
| Eras | 1248 | 979 | 956 | 1747 | 1251 | 1269 | 1326 | 978 | 960 | 1373 | 1346 | 1389 | 1228.62 | 1088 | 1369 | 1.259 | 0.333 | 0.01616475 | 0.07496338 | 0.0169 | 0.0096 | 0.013 | 0.013 | TRUE | 0.4366 |
| Erbb2 | 755 | 764 | 706 | 1454 | 995 | 893 | 802 | 763 | 709 | 1143 | 1071 | 978 | 910.83 | 758 | 1064 | 1.405 | 0.491 | 2.58E-06 | 6.27E-05 | 0.004 | 0.0105 | 0.0067 | 0.0067 | TRUE | 0.0597 |
| Erbb2ip | 1175 | 1595 | 1722 | 2310 | 1678 | 1515 | 1248 | 1593 | 1729 | 1816 | 1805 | 1658 | 1641.64 | 1523 | 1760 | 1.155 | 0.208 | 0.11373901 | 0.28442867 | 0.0136 | 0.0089 | 0.0119 | 0.0119 | TRUE | 0.3002 |
| Erbb3 | 873 | 1128 | 1137 | 1769 | 1226 | 1242 | 927 | 1127 | 1142 | 1391 | 1319 | 1360 | 1210.8 | 1065 | 1357 | 1.273 | 0.349 | 0.0011642 | 0.01000428 | 0.0054 | 0.0096 | 0.0075 | 0.0075 | TRUE | 0.1525 |
| Erbb4 | 1 | 1 | 1 | 1 | 0 | 0 | 1 | 1 | 1 | 1 | 0 | 0 | 0.64 | 1 | 0 | 0.342 | -1.549 | 0.50187412 | NA | 0 | 5.1396 | 2.332 | 2.332 | TRUE | 0.2563 |
| Erc1 | 7909 | 6006 | 6026 | 10575 | 7323 | 6340 | 8401 | 5998 | 6051 | 8313 | 7879 | 6940 | 7263.7 | 6817 | 7711 | 1.131 | 0.178 | 0.22441771 | 0.43681551 | 0.0224 | 0.0074 | 0.0153 | 0.0153 | TRUE | 0.5042 |
| Erc2 | 445 | 450 | 479 | 633 | 436 | 419 | 473 | 449 | 481 | 498 | 469 | 459 | 471.41 | 468 | 475 | 1.017 | 0.025 | 0.81333993 | 0.90450545 | 0 | 0.0139 | 0.0058 | 0.0058 | TRUE | 0.0212 |
| Ercc1 | 553 | 528 | 524 | 876 | 645 | 514 | 587 | 527 | 526 | 689 | 694 | 563 | 597.69 | 547 | 649 | 1.187 | 0.247 | 0.04756155 | 0.15835777 | 0.0072 | 0.0124 | 0.0095 | 0.0095 | TRUE | 0.1546 |
| Ercc2 | 2168 | 1924 | 1886 | 2710 | 2045 | 1752 | 2303 | 1922 | 1894 | 2130 | 2200 | 1918 | 2061.1 | 2040 | 2083 | 1.022 | 0.031 | 0.77921008 | 0.8859358 | 0.0081 | 0.0085 | 0.0082 | 0.0082 | TRUE | 0.1541 |
| Ercc3 | 4013 | 3584 | 3534 | 5416 | 3862 | 3464 | 4263 | 3579 | 3549 | 4258 | 4155 | 3792 | 3932.53 | 3797 | 4068 | 1.072 | 0.1 | 0.3292485 | 0.5502542 | 0.0069 | 0.0077 | 0.0073 | 0.0073 | TRUE | 0.1399 |
| Ercc4 | 1222 | 915 | 900 | 1829 | 1254 | 997 | 1298 | 914 | 904 | 1438 | 1349 | 1091 | 1165.65 | 1039 | 1293 | 1.246 | 0.317 | 0.06370703 | 0.19393693 | 0.0318 | 0.0097 | 0.0202 | 0.0202 | TRUE | 0.5699 |
| Ercc5 | 2059 | 2510 | 2438 | 2801 | 1959 | 2143 | 2187 | 2507 | 2448 | 2202 | 2108 | 2346 | 2299.56 | 2381 | 2219 | 0.932 | -0.102 | 0.25838013 | 0.47724223 | 0.0034 | 0.0083 | 0.0054 | 0.0054 | TRUE | 0.0611 |
| Ercc6 | 2246 | 2310 | 2341 | 2627 | 1979 | 1901 | 2386 | 2307 | 2351 | 2065 | 2129 | 2081 | 2219.77 | 2348 | 2092 | 0.891 | -0.167 | 0.01095941 | 0.05634023 | 0 | 0.0084 | 0.0026 | 0.0026 | TRUE | 0.003 |
| Ercc6l | 3110 | 2302 | 2285 | 4260 | 2719 | 2233 | 3303 | 2299 | 2294 | 3349 | 2925 | 2444 | 2769.25 | 2632 | 2906 | 1.104 | 0.143 | 0.40352742 | 0.6162915 | 0.0309 | 0.0081 | 0.0209 | 0.0209 | TRUE | 0.5104 |
| Ercc6l2 | 4079 | 3325 | 3634 | 4860 | 3368 | 3360 | 4333 | 3321 | 3649 | 3821 | 3624 | 3678 | 3737.43 | 3768 | 3708 | 0.984 | -0.023 | 0.83688617 | 0.91714685 | 0.0093 | 0.0078 | 0.0086 | 0.0086 | TRUE | 0.2093 |
| Ercc8 | 1135 | 1098 | 1095 | 1348 | 1030 | 948 | 1206 | 1097 | 1099 | 1060 | 1108 | 1038 | 1101.22 | 1134 | 1069 | 0.942 | -0.085 | 0.32594899 | 0.5471171 | 0.0011 | 0.0099 | 0.0045 | 0.0045 | TRUE | 0.0364 |
| Erdr1 | 1249 | 1432 | 1427 | 1319 | 1214 | 1268 | 1327 | 1430 | 1433 | 1037 | 1306 | 1388 | 1320.13 | 1397 | 1244 | 0.889 | -0.169 | 0.18013202 | 0.38150157 | 0.0107 | 0.0094 | 0.0107 | 0.0107 | TRUE | 0.2593 |
| Ereg | 0 | 1 | 1 | 0 | 2 | 0 | 0 | 1 | 1 | 0 | 2 | 0 | 0.69 | 1 | 1 | 1.039 | 0.055 | 0.98329309 | NA | 0 | 4.7646 | 3.5085 | 3.5085 | TRUE | 0.9747 |
| Erf | 8065 | 5441 | 5276 | 13433 | 8521 | 5801 | 8566 | 5434 | 5298 | 10560 | 9168 | 6350 | 7562.71 | 6433 | 8693 | 1.351 | 0.435 | 0.04742214 | 0.15806925 | 0.0716 | 0.0073 | 0.0345 | 0.0345 | TRUE | 0.5967 |
| Erg | 22 | 16 | 10 | 27 | 22 | 10 | 23 | 16 | 10 | 21 | 24 | 11 | 17.54 | 16 | 19 | 1.138 | 0.187 | 0.72596304 | 0.85230677 | 0.0941 | 0.1947 | 0.1476 | 0.1476 | TRUE | 0.2711 |
| Ergic1 | 3381 | 4413 | 4523 | 3507 | 2658 | 3227 | 3591 | 4407 | 4542 | 2757 | 2860 | 3532 | 3614.89 | 4180 | 3050 | 0.729 | -0.455 | 0.00064686 | 0.00636848 | 0.0167 | 0.0078 | 0.0126 | 0.0126 | TRUE | 0.2324 |
| Ergic2 | 1723 | 1712 | 1703 | 2477 | 1770 | 1683 | 1830 | 1710 | 1710 | 1947 | 1904 | 1842 | 1823.98 | 1750 | 1898 | 1.085 | 0.118 | 0.11575387 | 0.2873261 | 6.00E-04 | 0.0087 | 0.0035 | 0.0035 | TRUE | 0.0193 |
| Ergic3 | 3163 | 3891 | 4003 | 4208 | 3193 | 3253 | 3360 | 3886 | 4019 | 3308 | 3435 | 3561 | 3594.91 | 3755 | 3435 | 0.914 | -0.129 | 0.16766975 | 0.36546259 | 0.0046 | 0.0078 | 0.006 | 0.006 | TRUE | 0.103 |
| Erh | 4048 | 3574 | 3421 | 6014 | 3896 | 3813 | 4300 | 3569 | 3435 | 4728 | 4192 | 4174 | 4066.27 | 3768 | 4365 | 1.159 | 0.212 | 0.05780545 | 0.18161504 | 0.0099 | 0.0077 | 0.0088 | 0.0088 | TRUE | 0.1851 |
| Eri1 | 7627 | 7219 | 7116 | 9792 | 7116 | 6877 | 8101 | 7210 | 7145 | 7698 | 7656 | 7528 | 7556.36 | 7485 | 7627 | 1.019 | 0.027 | 0.72312241 | 0.85005881 | 0.002 | 0.0073 | 0.0041 | 0.0041 | TRUE | 0.0632 |
| Eri2 | 1289 | 1398 | 1369 | 1798 | 1322 | 1266 | 1369 | 1396 | 1375 | 1413 | 1422 | 1386 | 1393.61 | 1380 | 1407 | 1.02 | 0.028 | 0.69176489 | 0.83095304 | 0 | 0.0093 | 0.003 | 0.003 | TRUE | 0.0021 |
| Eri3 | 2372 | 2388 | 2223 | 2470 | 1833 | 1961 | 2519 | 2385 | 2232 | 1942 | 1972 | 2147 | 2199.51 | 2379 | 2020 | 0.849 | -0.236 | 0.00639825 | 0.03775281 | 0.0026 | 0.0084 | 0.0049 | 0.0049 | TRUE | 0.036 |

| | | | | | | | | | | | | | | | | | | | | | | | | | |
|---|---|---|---|---|---|---|---|---|---|---|---|---|---|---|---|---|---|---|---|---|---|---|---|---|---|
| Erich1 | 1241 | 1476 | 1423 | 1783 | 1366 | 1354 | 1318 | 1474 | 1429 | 1402 | 1470 | 1482 | 1429.11 | 1407 | 1451 | 1.031 | 0.044 | 0.60253896 | 0.77387557 | 0.0014 | 0.0092 | 0.0044 | 0.0044 | TRUE | 0.0365 |
| Erich2 | 4 | 3 | 3 | 7 | 6 | 10 | 4 | 3 | 3 | 6 | 6 | 11 | 5.53 | 3 | 8 | 2.217 | 1.149 | 0.16810742 | 0.36589195 | 0 | 0.603 | 0.286 | 0.286 | TRUE | 0.3789 |
| Erlec1 | 1331 | 1787 | 1840 | 1981 | 1389 | 1560 | 1414 | 1785 | 1848 | 1557 | 1494 | 1708 | 1634.24 | 1682 | 1586 | 0.943 | -0.085 | 0.49620121 | 0.69196835 | 0.0119 | 0.0089 | 0.0106 | 0.0106 | TRUE | 0.2341 |
| Erlin1 | 2365 | 2698 | 2685 | 3242 | 2308 | 2385 | 2512 | 2695 | 2696 | 2549 | 2483 | 2611 | 2590.87 | 2634 | 2548 | 0.967 | -0.048 | 0.49645287 | 0.69197217 | 7.00E-04 | 0.0082 | 0.0033 | 0.0033 | TRUE | 0.0199 |
| Erlin2 | 4460 | 4225 | 4131 | 6085 | 4424 | 4233 | 4737 | 4220 | 4148 | 4784 | 4760 | 4634 | 4547 | 4368 | 4726 | 1.082 | 0.114 | 0.15420213 | 0.34613091 | 0.0025 | 0.0076 | 0.0044 | 0.0044 | TRUE | 0.0664 |
| Ermap | 0 | 1 | 3 | 1 | 0 | 1 | 0 | 1 | 3 | 1 | 0 | 1 | 0.98 | 1 | 1 | 0.471 | -1.086 | 0.60350353 | NA | 0 | 3.3618 | 2.0009 | 2.0009 | TRUE | 0.3174 |
| Ermn | 0 | 0 | 0 | 0 | 0 | 0 | 0 | 0 | 0 | 0 | 0 | 0 | 0 | 0 | 0 | NA | NA | NA | NA | NA | NA | NA | NA | NA | NA |
| Ermp1 | 4901 | 5235 | 5568 | 6835 | 4764 | 5122 | 5206 | 5228 | 5591 | 5373 | 5126 | 5607 | 5355.1 | 5342 | 5369 | 1.005 | 0.007 | 0.92063749 | 0.96109972 | 0.0016 | 0.0075 | 0.0036 | 0.0036 | TRUE | 0.0203 |
| Ern1 | 848 | 956 | 1046 | 1182 | 847 | 830 | 901 | 955 | 1050 | 929 | 911 | 909 | 942.48 | 969 | 916 | 0.946 | -0.08 | 0.40012928 | 0.61317907 | 0.0021 | 0.0104 | 0.0054 | 0.0054 | TRUE | 0.065 |
| Ern2 | 8 | 4 | 8 | 11 | 6 | 6 | 8 | 4 | 8 | 9 | 6 | 7 | 7.03 | 7 | 7 | 1.066 | 0.093 | 0.89694811 | 0.9501066 | 0 | 0.4754 | 0.226 | 0.226 | TRUE | 0.3527 |
| Ero1l | 3339 | 5241 | 5867 | 6326 | 4798 | 6331 | 3547 | 5234 | 5891 | 4973 | 5162 | 6930 | 5289.59 | 4891 | 5688 | 1.163 | 0.218 | 0.26066691 | 0.47999951 | 0.0442 | 0.0075 | 0.0268 | 0.0268 | TRUE | 0.7039 |
| Ero1lb | 527 | 534 | 511 | 586 | 452 | 472 | 560 | 533 | 513 | 461 | 486 | 517 | 511.64 | 535 | 488 | 0.91 | -0.136 | 0.20193272 | 0.40947324 | 0 | 0.0133 | 0.0062 | 0.0062 | TRUE | 0.0308 |
| Erp27 | 0 | 0 | 0 | 1 | 0 | 0 | 0 | 0 | 0 | 1 | 0 | 0 | 0.13 | 0 | 0 | 1.859 | 0.894 | 0.82650734 | NA | 0 | 25.1515 | 10 | 10 | TRUE | 0.2563 |
| Erp29 | 4464 | 5250 | 5400 | 5948 | 4347 | 4550 | 4742 | 5243 | 5422 | 4676 | 4677 | 4981 | 4956.77 | 5136 | 4778 | 0.93 | -0.104 | 0.19924594 | 0.40658649 | 0.0027 | 0.0076 | 0.0046 | 0.0046 | TRUE | 0.0548 |
| Erp44 | 2325 | 2838 | 2843 | 3180 | 2263 | 2481 | 2470 | 2834 | 2855 | 2500 | 2435 | 2716 | 2634.85 | 2720 | 2550 | 0.937 | -0.093 | 0.32020504 | 0.54123609 | 0.0043 | 0.0082 | 0.0059 | 0.0059 | TRUE | 0.0782 |
| Errfi1 | 992 | 1232 | 1187 | 1617 | 1168 | 1106 | 1054 | 1230 | 1192 | 1271 | 1257 | 1211 | 1202.42 | 1159 | 1246 | 1.076 | 0.105 | 0.26195792 | 0.48137098 | 0.0027 | 0.0096 | 0.0055 | 0.0055 | TRUE | 0.0748 |
| Erv3 | 0 | 0 | 0 | 0 | 0 | 0 | 0 | 0 | 0 | 0 | 0 | 0 | 0 | 0 | 0 | NA | NA | NA | NA | NA | NA | NA | NA | NA | NA |
| Esam | 176 | 123 | 126 | 245 | 164 | 206 | 187 | 123 | 127 | 193 | 176 | 225 | 171.81 | 146 | 198 | 1.364 | 0.447 | 0.03795202 | 0.13544213 | 0.0296 | 0.0261 | 0.0275 | 0.0275 | TRUE | 0.642 |
| Esco1 | 2243 | 2348 | 2367 | 2967 | 2133 | 2061 | 2382 | 2345 | 2377 | 2332 | 2295 | 2256 | 2331.27 | 2368 | 2294 | 0.969 | -0.045 | 0.48142067 | 0.68055631 | 0 | 0.0083 | 0.0025 | 0.0025 | TRUE | 0.0026 |
| Esco2 | 7305 | 8131 | 7736 | 8195 | 6284 | 7076 | 7759 | 8121 | 7768 | 6442 | 6761 | 7746 | 7432.79 | 7883 | 6983 | 0.886 | -0.175 | 0.05346838 | 0.17175218 | 0.0049 | 0.0073 | 0.0058 | 0.0058 | TRUE | 0.1113 |
| Esd | 2319 | 3306 | 3183 | 3835 | 2602 | 3096 | 2463 | 3302 | 3196 | 3015 | 2800 | 3389 | 3027.4 | 2987 | 3068 | 1.027 | 0.038 | 0.77508548 | 0.88302862 | 0.0167 | 0.008 | 0.0126 | 0.0126 | TRUE | 0.2854 |
| Esf1 | 5725 | 4835 | 4517 | 7519 | 5405 | 4507 | 6081 | 4829 | 4536 | 5911 | 5815 | 4934 | 5350.86 | 5149 | 5553 | 1.079 | 0.109 | 0.40868021 | 0.62120237 | 0.015 | 0.0075 | 0.0125 | 0.0125 | TRUE | 0.3058 |
| Esm1 | 0 | 0 | 0 | 0 | 0 | 0 | 0 | 0 | 0 | 0 | 0 | 0 | 0 | 0 | 0 | NA | NA | NA | NA | NA | NA | NA | NA | NA | NA |
| Espl1 | 15480 | 16282 | 17361 | 16605 | 12473 | 12522 | 16442 | 16261 | 17432 | 13054 | 13420 | 13707 | 15052.8 | 16712 | 13394 | 0.801 | -0.319 | 3.93E-07 | 1.20E-05 | 9.00E-04 | 0.0071 | 0.0028 | 0.0028 | TRUE | 0.0174 |
| Espn | 362 | 567 | 581 | 511 | 450 | 583 | 385 | 566 | 583 | 402 | 484 | 638 | 509.7 | 511 | 508 | 0.992 | -0.012 | 0.95579624 | 0.97741618 | 0.0505 | 0.0134 | 0.031 | 0.031 | TRUE | 0.4592 |
| Espnl | 7 | 2 | 5 | 7 | 3 | 8 | 7 | 2 | 5 | 6 | 3 | 9 | 5.32 | 5 | 6 | 1.212 | 0.277 | 0.7528864 | 0.86954631 | 0 | 0.6258 | 0.3672 | 0.3672 | TRUE | 0.2951 |
| Esr1 | 9 | 6 | 1 | 3 | 10 | 3 | 10 | 6 | 1 | 2 | 11 | 3 | 5.49 | 6 | 5 | 0.981 | -0.028 | 0.97842242 | 0.98828002 | 0.5507 | 0.6067 | 0.5787 | 0.5787 | TRUE | 0.4035 |
| Esr2 | 53 | 33 | 41 | 46 | 25 | 25 | 56 | 33 | 41 | 36 | 27 | 27 | 36.81 | 43 | 30 | 0.699 | -0.516 | 0.14632574 | 0.33505366 | 0.0284 | 0.0964 | 0.063 | 0.063 | TRUE | 0.3277 |
| Esrp1 | 499 | 828 | 977 | 961 | 842 | 1061 | 530 | 827 | 981 | 755 | 906 | 1161 | 860.13 | 779 | 941 | 1.206 | 0.271 | 0.23860455 | 0.45439156 | 0.0689 | 0.0107 | 0.0369 | 0.0369 | TRUE | 0.5833 |
| Esrp2 | 200 | 163 | 183 | 354 | 302 | 201 | 212 | 163 | 184 | 278 | 325 | 220 | 230.37 | 186 | 274 | 1.474 | 0.56 | 0.00367466 | 0.02476783 | 0.0233 | 0.0212 | 0.0222 | 0.0222 | TRUE | 0.172 |
| Esrra | 1299 | 1315 | 1227 | 1510 | 1136 | 1037 | 1380 | 1313 | 1232 | 1187 | 1222 | 1135 | 1244.93 | 1308 | 1181 | 0.903 | -0.147 | 0.09025335 | 0.24458348 | 0.0015 | 0.0095 | 0.0046 | 0.0046 | TRUE | 0.0313 |
| Esrrb | 28373 | 37865 | 38643 | 35661 | 27283 | 29821 | 30137 | 37816 | 38802 | 28034 | 29355 | 32644 | 32797.9 | 35585 | 30011 | 0.843 | -0.246 | 0.03717634 | 0.13359428 | 0.012 | 0.007 | 0.01 | 0.01 | TRUE | 0.2195 |
| Esrrg | 50 | 41 | 22 | 24 | 33 | 16 | 53 | 41 | 22 | 19 | 36 | 18 | 31.34 | 39 | 24 | 0.617 | -0.698 | 0.13506761 | 0.31796579 | 0.1354 | 0.112 | 0.1233 | 0.1233 | TRUE | 0.2669 |
| Esx1 | 595 | 735 | 559 | 641 | 407 | 370 | 632 | 734 | 561 | 504 | 438 | 405 | 545.7 | 642 | 449 | 0.7 | -0.515 | 0.0003943 | 0.0042924 | 0.0136 | 0.0129 | 0.0133 | 0.0133 | TRUE | 0.1842 |
| Esyt1 | 2933 | 3073 | 3168 | 3876 | 2756 | 2827 | 3115 | 3069 | 3181 | 3047 | 2965 | 3095 | 3078.72 | 3122 | 3036 | 0.972 | -0.04 | 0.52389084 | 0.71372603 | 0 | 0.008 | 0.0026 | 0.0026 | TRUE | 0.0049 |
| Esyt2 | 2265 | 2805 | 3208 | 3777 | 2715 | 2887 | 2406 | 2801 | 3221 | 2969 | 2921 | 3160 | 2913.17 | 2809 | 3017 | 1.074 | 0.103 | 0.3855224 | 0.59935527 | 0.0098 | 0.008 | 0.0097 | 0.0097 | TRUE | 0.1995 |
| Esyt3 | 1476 | 1618 | 1607 | 1513 | 1138 | 1205 | 1568 | 1616 | 1614 | 1189 | 1224 | 1319 | 1421.69 | 1599 | 1244 | 0.777 | -0.363 | 6.65E-06 | 0.00014114 | 8.00E-04 | 0.0092 | 0.004 | 0.004 | TRUE | 0.0328 |
| Etaa1 | 3419 | 2822 | 2656 | 3714 | 2726 | 2706 | 3632 | 2818 | 2667 | 2920 | 2933 | 2962 | 2988.61 | 3039 | 2938 | 0.967 | -0.048 | 0.69947793 | 0.83558024 | 0.0121 | 0.008 | 0.011 | 0.011 | TRUE | 0.3531 |
| Etd | 0 | 0 | 0 | 0 | 0 | 0 | 0 | 0 | 0 | 0 | 0 | 0 | 0 | 0 | 0 | NA | NA | NA | NA | NA | NA | NA | NA | NA | NA |
| Etf1 | 26548 | 22277 | 21480 | 32341 | 22472 | 19362 | 28199 | 22248 | 21568 | 25424 | 24178 | 21195 | 23802.05 | 24005 | 23599 | 0.983 | -0.025 | 0.84605869 | 0.92282842 | 0.0128 | 0.007 | 0.0115 | 0.0115 | TRUE | 0.2858 |
| Etfa | 7566 | 8470 | 8283 | 9986 | 6969 | 7011 | 8036 | 8459 | 8317 | 7850 | 7498 | 7675 | 7972.59 | 8271 | 7674 | 0.928 | -0.108 | 0.07281931 | 0.21131669 | 4.00E-04 | 0.0073 | 0.0025 | 0.0025 | TRUE | 0.0075 |
| Etfb | 1888 | 2355 | 2332 | 3556 | 2488 | 2533 | 2005 | 2352 | 2342 | 2795 | 2677 | 2773 | 2490.68 | 2233 | 2748 | 1.231 | 0.3 | 0.0009601 | 0.00862336 | 0.0036 | 0.0082 | 0.0055 | 0.0055 | TRUE | 0.0959 |
| Etfdh | 1393 | 1840 | 1976 | 2069 | 1409 | 1402 | 1480 | 1838 | 1984 | 1626 | 1516 | 1535 | 1663.09 | 1767 | 1559 | 0.882 | -0.181 | 0.14149396 | 0.32701377 | 0.0106 | 0.0089 | 0.0103 | 0.0103 | TRUE | 0.2439 |
| Ethe1 | 435 | 450 | 435 | 483 | 289 | 332 | 462 | 449 | 437 | 380 | 311 | 363 | 400.39 | 449 | 351 | 0.783 | -0.352 | 0.00446852 | 0.0287294 | 0.0027 | 0.0151 | 0.0085 | 0.0085 | TRUE | 0.1133 |
| Etl4 | 8531 | 7199 | 7725 | 9575 | 6711 | 6307 | 9061 | 7190 | 7757 | 7527 | 7221 | 6904 | 7609.93 | 8003 | 7217 | 0.902 | -0.149 | 0.15135149 | 0.34239864 | 0.0079 | 0.0073 | 0.0076 | 0.0076 | TRUE | 0.1635 |
| Etnk1 | 8659 | 8525 | 9292 | 10930 | 7917 | 7651 | 9197 | 8514 | 9330 | 8592 | 8518 | 8375 | 8754.54 | 9014 | 8495 | 0.943 | -0.085 | 0.20069555 | 0.40847133 | 0.0011 | 0.0073 | 0.0031 | 0.0031 | TRUE | 0.0288 |
| Etnk2 | 517 | 624 | 731 | 661 | 467 | 545 | 549 | 623 | 734 | 520 | 502 | 597 | 587.5 | 635 | 540 | 0.848 | -0.237 | 0.09487775 | 0.25238857 | 0.0125 | 0.0125 | 0.0128 | 0.0128 | TRUE | 0.2169 |
| Etohd2 | 13 | 23 | 18 | 24 | 20 | 22 | 14 | 23 | 18 | 19 | 22 | 24 | 19.89 | 18 | 22 | 1.168 | 0.224 | 0.60786383 | 0.77774221 | 0 | 0.1726 | 0.086 | 0.086 | TRUE | 0.2506 |
| Etohi1 | 358 | 235 | 255 | 483 | 384 | 290 | 380 | 235 | 256 | 380 | 413 | 317 | 330.22 | 290 | 370 | 1.276 | 0.352 | 0.08890945 | 0.24218996 | 0.0407 | 0.0169 | 0.0277 | 0.0277 | TRUE | 0.8215 |
| Ets1 | 186 | 176 | 143 | 348 | 271 | 243 | 198 | 176 | 144 | 274 | 292 | 266 | 224.68 | 173 | 277 | 1.609 | 0.686 | 2.64E-05 | 0.00045224 | 0.0065 | 0.0216 | 0.0145 | 0.0145 | TRUE | 0.2278 |
| Ets2 | 681 | 738 | 716 | 1268 | 944 | 888 | 723 | 737 | 719 | 997 | 1016 | 972 | 860.65 | 726 | 995 | 1.37 | 0.454 | 4.43E-08 | 1.79E-06 | 0 | 0.0107 | 0.0038 | 0.0038 | TRUE | 0.0046 |
| Etv1 | 328 | 258 | 288 | 448 | 323 | 284 | 348 | 258 | 289 | 352 | 348 | 311 | 317.64 | 298 | 337 | 1.131 | 0.177 | 0.24720533 | 0.46398502 | 0.0104 | 0.0173 | 0.0137 | 0.0137 | TRUE | 0.2399 |
| Etv2 | 19 | 21 | 18 | 47 | 33 | 16 | 20 | 21 | 18 | 37 | 36 | 18 | 24.87 | 20 | 30 | 1.529 | 0.613 | 0.15870278 | 0.35263911 | 0.0458 | 0.1394 | 0.0942 | 0.0942 | TRUE | 0.8177 |
| Etv3 | 984 | 1137 | 1030 | 1635 | 1088 | 1035 | 1045 | 1136 | 1034 | 1285 | 1171 | 1133 | 1133.97 | 1072 | 1196 | 1.117 | 0.16 | 0.0902531 | 0.24458348 | 0.0027 | 0.0098 | 0.0055 | 0.0055 | TRUE | 0.0523 |
| Etv4 | 10741 | 10523 | 10717 | 10885 | 7885 | 7067 | 11409 | 10509 | 10761 | 8557 | 8484 | 7736 | 9575.98 | 10893 | 8259 | 0.758 | -0.399 | 1.30E-07 | 4.47E-06 | 0.0023 | 0.0072 | 0.004 | 0.004 | TRUE | 0.0373 |
| Etv5 | 14612 | 15197 | 15914 | 19360 | 14294 | 14042 | 15521 | 15177 | 15979 | 15219 | 15379 | 15371 | 15441.2 | 15559 | 15323 | 0.985 | -0.022 | 0.68939541 | 0.82925141 | 3.00E-04 | 0.0071 | 0.0021 | 0.0021 | TRUE | 0.0068 |
| Etv6 | 1894 | 1614 | 1665 | 2115 | 1609 | 1502 | 2012 | 1612 | 1672 | 1663 | 1731 | 1644 | 1722.25 | 1765 | 1679 | 0.952 | -0.072 | 0.50475544 | 0.69962236 | 0.0067 | 0.0088 | 0.0078 | 0.0078 | TRUE | 0.1798 |
| EU599041 | 979 | 771 | 742 | 1081 | 899 | 749 | 1040 | 770 | 745 | 850 | 967 | 820 | 865.31 | 852 | 879 | 1.032 | 0.046 | 0.76323213 | 0.87540522 | 0.0199 | 0.0107 | 0.0155 | 0.0155 | TRUE | 0.4431 |
| Eva1a | 50 | 71 | 51 | 129 | 109 | 76 | 53 | 71 | 51 | 101 | 117 | 83 | 79.52 | 58 | 100 | 1.722 | 0.784 | 0.0021199 | 0.01613567 | 0.0169 | 0.0483 | 0.0333 | 0.0333 | TRUE | 0.2017 |
| Eva1b | 510 | 681 | 749 | 730 | 556 | 593 | 542 | 680 | 752 | 574 | 598 | 649 | 632.52 | 658 | 607 | 0.922 | -0.118 | 0.41109587 | 0.62224822 | 0.0141 | 0.0121 | 0.0132 | 0.0132 | TRUE | 0.2799 |
| Eva1c | 27 | 32 | 26 | 35 | 24 | 34 | 29 | 32 | 26 | 28 | 26 | 37 | 29.55 | 29 | 30 | 1.04 | 0.056 | 0.87484481 | 0.93825131 | 0 | 0.1184 | 0.0587 | 0.0587 | TRUE | 0.2553 |
| Evc | 533 | 541 | 569 | 711 | 601 | 533 | 566 | 540 | 571 | 559 | 647 | 583 | 577.8 | 559 | 596 | 1.065 | 0.091 | 0.39160398 | 0.60534847 | 0.0018 | 0.0126 | 0.0064 | 0.0064 | TRUE | 0.0634 |
| Evc2 | 1464 | 1226 | 1356 | 1715 | 1226 | 1211 | 1555 | 1224 | 1362 | 1348 | 1319 | 1326 | 1355.66 | 1380 | 1331 | 0.965 | -0.052 | 0.63063273 | 0.79261236 | 0.0065 | 0.0093 | 0.0077 | 0.0077 | TRUE | 0.1469 |
| Evi2a | 0 | 0 | 1 | 0 | 0 | 1 | 0 | 0 | 1 | 0 | 0 | 1 | 0.35 | 0 | 0 | 0.954 | -0.067 | 0.98566485 | NA | 0 | 9.4252 | 8.1416 | 8.1416 | TRUE | 0.5383 |
| Evi2b | 0 | 0 | 0 | 0 | 0 | 0 | 0 | 0 | 0 | 0 | 0 | 0 | 0 | 0 | 0 | NA | NA | NA | NA | NA | NA | NA | NA | NA | NA |
| Evi5 | 4269 | 4694 | 5000 | 5474 | 3722 | 3844 | 4534 | 4688 | 5021 | 4303 | 4005 | 4208 | 4459.77 | 4748 | 4172 | 0.879 | -0.186 | 0.01283987 | 0.06326472 | 0.0017 | 0.0076 | 0.0038 | 0.0038 | TRUE | 0.0308 |
| Evi5l | 133 | 167 | 177 | 214 | 156 | 199 | 141 | 167 | 178 | 168 | 168 | 218 | 173.28 | 162 | 185 | 1.137 | 0.185 | 0.31933668 | 0.54057938 | 0.0125 | 0.0259 | 0.0191 | 0.0191 | TRUE | 0.2585 |
| Evl | 1889 | 2188 | 2336 | 2454 | 1772 | 1761 | 2006 | 2185 | 2346 | 1929 | 1907 | 1928 | 2050.1 | 2179 | 1921 | 0.882 | -0.182 | 0.03430098 | 0.12614807 | 0.0023 | 0.0085 | 0.0048 | 0.0048 | TRUE | 0.0579 |
| Evpl | 796 | 738 | 747 | 760 | 526 | 538 | 845 | 737 | 750 | 597 | 566 | 589 | 680.82 | 777 | 584 | 0.752 | -0.412 | 5.79E-05 | 0.00087833 | 0.002 | 0.0117 | 0.0061 | 0.0061 | TRUE | 0.0688 |

| | | | | | | | | | | | | | | | | | | | | | | | | | |
|---|---|---|---|---|---|---|---|---|---|---|---|---|---|---|---|---|---|---|---|---|---|---|---|---|---|
| Evx1 | 5 | 4 | 6 | 22 | 14 | 7 | 5 | 4 | 6 | 17 | 15 | 8 | 9.23 | 5 | 13 | 2.635 | 1.398 | 0.04097642 | 0.14332177 | 0 | 0.364 | 0.2006 | 0.2006 | TRUE | 0.5391 |
| Evx2 | 0 | 0 | 0 | 1 | 0 | 0 | 0 | 0 | 0 | 1 | 0 | 0 | 0.13 | 0 | 0 | 1.859 | 0.894 | 0.82650734 | NA | 0 | 25.1515 | 10 | 10 | TRUE | 0.2563 |
| Ewsr1 | 16513 | 18244 | 18306 | 23830 | 17995 | 17826 | 17540 | 18220 | 18381 | 18733 | 19362 | 19513 | 18624.91 | 18047 | 19203 | 1.064 | 0.089 | 0.11912685 | 0.29243092 | 5.00E-04 | 0.0071 | 0.0023 | 0.0023 | TRUE | 0.0074 |
| Exd1 | 126 | 135 | 144 | 208 | 163 | 131 | 134 | 135 | 145 | 164 | 175 | 143 | 149.26 | 138 | 161 | 1.168 | 0.224 | 0.18898638 | 0.39421944 | 0 | 0.029 | 0.0141 | 0.0141 | TRUE | 0.0883 |
| Exd2 | 1740 | 2248 | 2188 | 2421 | 1754 | 1864 | 1848 | 2245 | 2197 | 1903 | 1887 | 2040 | 2020.18 | 2097 | 1943 | 0.927 | -0.11 | 0.28202319 | 0.5034031 | 0.0055 | 0.0085 | 0.007 | 0.007 | TRUE | 0.1297 |
| Exo1 | 3577 | 3100 | 3116 | 4506 | 3155 | 2963 | 3799 | 3096 | 3129 | 3542 | 3395 | 3243 | 3367.42 | 3341 | 3393 | 1.016 | 0.023 | 0.82863243 | 0.91322447 | 0.0074 | 0.0079 | 0.0076 | 0.0076 | TRUE | 0.1745 |
| Exo5 | 928 | 770 | 752 | 1195 | 766 | 706 | 986 | 769 | 755 | 939 | 824 | 773 | 841.03 | 837 | 845 | 1.012 | 0.017 | 0.90401353 | 0.95248677 | 0.0152 | 0.0108 | 0.0131 | 0.0131 | TRUE | 0.2878 |
| Exoc1 | 2687 | 2366 | 2400 | 3232 | 2477 | 2245 | 2854 | 2363 | 2410 | 2541 | 2665 | 2458 | 2548.37 | 2542 | 2555 | 1.005 | 0.007 | 0.94381617 | 0.97197967 | 0.0059 | 0.0082 | 0.0068 | 0.0068 | TRUE | 0.1393 |
| Exoc2 | 2234 | 2848 | 3158 | 3333 | 2506 | 2646 | 2373 | 2844 | 3171 | 2620 | 2696 | 2896 | 2766.85 | 2796 | 2737 | 0.979 | -0.031 | 0.79684199 | 0.89536272 | 0.0115 | 0.0081 | 0.01 | 0.01 | TRUE | 0.2124 |
| Exoc3 | 2690 | 2752 | 2727 | 4008 | 2797 | 2730 | 2857 | 2748 | 2738 | 3151 | 3009 | 2988 | 2915.41 | 2781 | 3049 | 1.097 | 0.133 | 0.04556578 | 0.15421962 | 4.00E-04 | 0.008 | 0.0029 | 0.0029 | TRUE | 0.0105 |
| Exoc3l | 276 | 274 | 286 | 235 | 155 | 163 | 293 | 274 | 287 | 185 | 167 | 178 | 230.65 | 285 | 177 | 0.622 | -0.686 | 6.00E-07 | 1.75E-05 | 0 | 0.0212 | 0.009 | 0.009 | TRUE | 0.025 |
| Exoc3l4 | 17 | 20 | 30 | 36 | 19 | 26 | 18 | 20 | 30 | 28 | 20 | 28 | 24.23 | 23 | 25 | 1.135 | 0.182 | 0.65845674 | 0.80755675 | 0 | 0.1429 | 0.0813 | 0.0813 | TRUE | 0.4742 |
| Exoc4 | 5255 | 7522 | 7958 | 7228 | 5295 | 6151 | 5582 | 7512 | 7991 | 5682 | 5697 | 6733 | 6532.85 | 7028 | 6037 | 0.859 | -0.219 | 0.13052396 | 0.31106435 | 0.0212 | 0.0074 | 0.015 | 0.015 | TRUE | 0.3954 |
| Exoc5 | 2910 | 2903 | 2831 | 3827 | 2677 | 2594 | 3091 | 2899 | 2843 | 3008 | 2880 | 2840 | 2926.85 | 2944 | 2909 | 0.989 | -0.017 | 0.81930656 | 0.90766966 | 0.0011 | 0.008 | 0.0035 | 0.0035 | TRUE | 0.023 |
| Exoc6 | 1597 | 1672 | 1735 | 2034 | 1438 | 1502 | 1696 | 1670 | 1742 | 1599 | 1547 | 1644 | 1649.77 | 1703 | 1597 | 0.938 | -0.093 | 0.20212316 | 0.40964874 | 0 | 0.0089 | 0.0032 | 0.0032 | TRUE | 0.0086 |
| Exoc6b | 1740 | 1866 | 1891 | 2497 | 1775 | 1897 | 1848 | 1864 | 1899 | 1963 | 1910 | 2077 | 1926.64 | 1870 | 1983 | 1.06 | 0.084 | 0.24917701 | 0.46630515 | 4.00E-04 | 0.0086 | 0.0033 | 0.0033 | TRUE | 0.0201 |
| Exoc7 | 928 | 1191 | 1227 | 1569 | 1226 | 1150 | 986 | 1189 | 1232 | 1233 | 1319 | 1259 | 1203.1 | 1136 | 1270 | 1.118 | 0.161 | 0.14407668 | 0.33120207 | 0.0061 | 0.0096 | 0.0079 | 0.0079 | TRUE | 0.1591 |
| Exoc8 | 1404 | 1292 | 1188 | 2352 | 1683 | 1297 | 1491 | 1290 | 1193 | 1849 | 1811 | 1420 | 1509.01 | 1325 | 1693 | 1.279 | 0.355 | 0.00963948 | 0.0512071 | 0.0154 | 0.0091 | 0.0129 | 0.0129 | TRUE | 0.2393 |
| Exog | 1048 | 962 | 863 | 1139 | 862 | 791 | 1113 | 961 | 867 | 895 | 927 | 866 | 938.2 | 980 | 896 | 0.915 | -0.129 | 0.27069064 | 0.49074002 | 0.007 | 0.0104 | 0.0088 | 0.0088 | TRUE | 0.1674 |
| Exosc1 | 1509 | 1423 | 1390 | 1883 | 1332 | 1232 | 1603 | 1421 | 1396 | 1480 | 1433 | 1349 | 1446.95 | 1473 | 1421 | 0.965 | -0.051 | 0.58288013 | 0.75875085 | 0.003 | 0.0092 | 0.0056 | 0.0056 | TRUE | 0.071 |
| Exosc10 | 3335 | 3769 | 4028 | 5461 | 4003 | 3647 | 3542 | 3764 | 4045 | 4293 | 4307 | 3992 | 3990.54 | 3784 | 4197 | 1.109 | 0.15 | 0.068717 | 0.20330228 | 0.0027 | 0.0077 | 0.0046 | 0.0046 | TRUE | 0.0443 |
| Exosc2 | 3868 | 4010 | 3947 | 5193 | 3681 | 3499 | 4108 | 4005 | 3963 | 4082 | 3961 | 3830 | 3991.6 | 4025 | 3958 | 0.983 | -0.024 | 0.70793004 | 0.84083686 | 4.00E-04 | 0.0077 | 0.0027 | 0.0027 | TRUE | 0.0096 |
| Exosc3 | 1386 | 1485 | 1475 | 2226 | 1601 | 1490 | 1472 | 1483 | 1481 | 1750 | 1723 | 1631 | 1589.97 | 1479 | 1701 | 1.151 | 0.203 | 0.00574858 | 0.0347648 | 0 | 0.009 | 0.0033 | 0.0033 | TRUE | 0.0155 |
| Exosc4 | 989 | 896 | 836 | 1522 | 1100 | 950 | 1050 | 895 | 839 | 1196 | 1184 | 1040 | 1034.12 | 928 | 1140 | 1.229 | 0.297 | 0.01268452 | 0.0627148 | 0.0084 | 0.0101 | 0.0093 | 0.0093 | TRUE | 0.1575 |
| Exosc5 | 3184 | 3851 | 3879 | 4444 | 3322 | 2984 | 3382 | 3846 | 3895 | 3494 | 3574 | 3266 | 3576.2 | 3708 | 3445 | 0.929 | -0.106 | 0.22893872 | 0.44224019 | 0.0036 | 0.0078 | 0.0053 | 0.0053 | TRUE | 0.0717 |
| Exosc6 | 1489 | 1664 | 1595 | 2238 | 1625 | 1446 | 1582 | 1662 | 1602 | 1759 | 1748 | 1583 | 1655.93 | 1615 | 1697 | 1.051 | 0.072 | 0.37956489 | 0.59448576 | 0.0014 | 0.0089 | 0.0042 | 0.0042 | TRUE | 0.0411 |
| Exosc7 | 2138 | 2178 | 2174 | 3151 | 2364 | 2182 | 2271 | 2175 | 2183 | 2477 | 2544 | 2389 | 2339.7 | 2210 | 2470 | 1.118 | 0.161 | 0.02051151 | 0.08835612 | 3.00E-04 | 0.0083 | 0.003 | 0.003 | TRUE | 0.0098 |
| Exosc8 | 2485 | 3183 | 3216 | 3582 | 2578 | 2872 | 2640 | 3179 | 3229 | 2816 | 2774 | 3144 | 2963.52 | 3016 | 2911 | 0.965 | -0.051 | 0.63482196 | 0.79482512 | 0.0081 | 0.008 | 0.0081 | 0.0081 | TRUE | 0.1445 |
| Exosc9 | 2383 | 2377 | 2486 | 3404 | 2375 | 2218 | 2531 | 2374 | 2496 | 2676 | 2555 | 2428 | 2510.09 | 2467 | 2553 | 1.035 | 0.05 | 0.51009102 | 0.7036169 | 0.0014 | 0.0082 | 0.0038 | 0.0038 | TRUE | 0.0222 |
| Exph5 | 16 | 22 | 15 | 16 | 6 | 12 | 17 | 22 | 15 | 13 | 6 | 13 | 14.37 | 18 | 11 | 0.599 | -0.739 | 0.16234146 | 0.3577046 | 0 | 0.2362 | 0.1279 | 0.1279 | TRUE | 0.4003 |
| Ext1 | 1646 | 1888 | 1889 | 1932 | 1337 | 1425 | 1748 | 1886 | 1897 | 1519 | 1439 | 1560 | 1674.64 | 1844 | 1506 | 0.817 | -0.292 | 0.00028527 | 0.00327226 | 0.0012 | 0.0089 | 0.0041 | 0.0041 | TRUE | 0.0245 |
| Ext2 | 3137 | 3413 | 3406 | 3526 | 2534 | 2615 | 3332 | 3409 | 3420 | 2772 | 2726 | 2863 | 3086.91 | 3387 | 2787 | 0.823 | -0.281 | 8.96E-06 | 0.0001811 | 0 | 0.008 | 0.0026 | 0.0026 | TRUE | 0.0067 |
| Extl1 | 974 | 958 | 895 | 796 | 610 | 611 | 1035 | 957 | 899 | 626 | 656 | 669 | 806.82 | 964 | 650 | 0.675 | -0.568 | 8.28E-09 | 3.99E-07 | 0.0021 | 0.011 | 0.0057 | 0.0057 | TRUE | 0.0496 |
| Extl2 | 463 | 424 | 393 | 451 | 327 | 335 | 492 | 423 | 395 | 355 | 352 | 367 | 397.16 | 437 | 358 | 0.819 | -0.287 | 0.02490077 | 0.10143269 | 0.0044 | 0.0152 | 0.0093 | 0.0093 | TRUE | 0.1401 |
| Extl3 | 2397 | 2886 | 2731 | 3343 | 2497 | 2576 | 2546 | 2882 | 2742 | 2628 | 2687 | 2820 | 2717.5 | 2723 | 2712 | 0.995 | -0.007 | 0.9321424 | 0.96628017 | 0.0022 | 0.0081 | 0.0044 | 0.0044 | TRUE | 0.0393 |
| Eya1 | 138 | 74 | 91 | 158 | 118 | 111 | 147 | 74 | 91 | 124 | 127 | 122 | 114.09 | 104 | 124 | 1.197 | 0.259 | 0.33661642 | 0.55668597 | 0.0521 | 0.0358 | 0.0436 | 0.0436 | TRUE | 1.2501 |
| Eya2 | 83 | 104 | 93 | 77 | 77 | 101 | 88 | 104 | 93 | 61 | 83 | 111 | 89.89 | 95 | 85 | 0.883 | -0.179 | 0.5037881 | 0.69883429 | 0.0372 | 0.0436 | 0.0404 | 0.0404 | TRUE | 0.2774 |
| Eya3 | 2349 | 2543 | 2654 | 3126 | 2395 | 2085 | 2495 | 2540 | 2665 | 2457 | 2577 | 2282 | 2502.72 | 2567 | 2439 | 0.95 | -0.074 | 0.36092284 | 0.57926885 | 0.0018 | 0.0082 | 0.0043 | 0.0043 | TRUE | 0.0376 |
| Eya4 | 1 | 2 | 5 | 13 | 10 | 14 | 1 | 2 | 5 | 10 | 11 | 15 | 7.4 | 3 | 12 | 4.446 | 2.153 | 0.00903258 | 0.04878832 | 0 | 0.4523 | 0.2608 | 0.2608 | TRUE | 0.6768 |
| Ezh1 | 5346 | 7408 | 7842 | 4554 | 3585 | 4830 | 5678 | 7398 | 7874 | 3580 | 3857 | 5287 | 5612.58 | 6983 | 4241 | 0.607 | -0.72 | 2.91E-05 | 0.00048979 | 0.0367 | 0.0075 | 0.0212 | 0.0212 | TRUE | 0.5659 |
| Ezh2 | 6317 | 6855 | 7222 | 10960 | 8066 | 8253 | 6710 | 6846 | 7252 | 8616 | 8679 | 9034 | 7856.03 | 6936 | 8776 | 1.265 | 0.339 | 2.38E-07 | 7.65E-06 | 0.001 | 0.0073 | 0.003 | 0.003 | TRUE | 0.0194 |
| Ezr | 13533 | 14347 | 14850 | 16531 | 12808 | 12744 | 14374 | 14328 | 14911 | 12995 | 13781 | 13950 | 14056.7 | 14538 | 13575 | 0.934 | -0.099 | 0.11400418 | 0.28478082 | 8.00E-04 | 0.0071 | 0.0028 | 0.0028 | TRUE | 0.0172 |
| F10 | 0 | 1 | 0 | 0 | 0 | 0 | 0 | 1 | 0 | 0 | 0 | 0 | 0.17 | 0 | 0 | 0.49 | -1.029 | 0.80087 | NA | 0 | 19.7991 | 10 | 10 | TRUE | 0.405 |
| F11 | 0 | 0 | 1 | 1 | 0 | 0 | 0 | 0 | 1 | 1 | 0 | 0 | 0.3 | 0 | 0 | 0.954 | -0.067 | 0.98640617 | NA | 0 | 11.0483 | 9.2783 | 9.2783 | TRUE | 0.4098 |
| F11r | 2216 | 2359 | 2149 | 3098 | 2170 | 2180 | 2354 | 2356 | 2158 | 2435 | 2335 | 2386 | 2337.35 | 2289 | 2385 | 1.042 | 0.06 | 0.42329834 | 0.6335416 | 0.001 | 0.0083 | 0.0036 | 0.0036 | TRUE | 0.0306 |
| F12 | 0 | 3 | 1 | 1 | 2 | 3 | 0 | 3 | 1 | 1 | 2 | 3 | 1.7 | 1 | 2 | 1.512 | 0.596 | 0.70221358 | NA | 0 | 1.9406 | 1.1313 | 1.1313 | TRUE | 0.5451 |
| F13a1 | 0 | 0 | 0 | 0 | 0 | 0 | 0 | 0 | 0 | 0 | 0 | 0 | 0 | 0 | 0 | NA | NA | NA | NA | NA | NA | NA | NA | NA | NA |
| F13b | 1 | 1 | 0 | 0 | 0 | 0 | 1 | 1 | 0 | 0 | 0 | 0 | 0.34 | 1 | 0 | 0.261 | -1.936 | 0.60454921 | NA | 0 | 9.5983 | 8.3269 | 8.3269 | TRUE | 0.2507 |
| F2 | 32 | 52 | 53 | 75 | 64 | 81 | 34 | 52 | 53 | 59 | 69 | 89 | 59.27 | 46 | 72 | 1.546 | 0.629 | 0.037159 | 0.13356238 | 0.0319 | 0.0625 | 0.0478 | 0.0478 | TRUE | 0.4187 |
| F2r | 742 | 748 | 719 | 1542 | 1045 | 909 | 788 | 747 | 722 | 1212 | 1124 | 995 | 931.45 | 752 | 1110 | 1.478 | 0.563 | 1.96E-07 | 6.41E-06 | 0.0051 | 0.0104 | 0.0073 | 0.0073 | TRUE | 0.0983 |
| F2rl1 | 3956 | 4128 | 4287 | 5312 | 3901 | 4021 | 4202 | 4123 | 4305 | 4176 | 4197 | 4402 | 4234 | 4210 | 4258 | 1.011 | 0.016 | 0.79796834 | 0.89608114 | 4.00E-04 | 0.0077 | 0.0027 | 0.0027 | TRUE | 0.0105 |
| F2rl2 | 4 | 0 | 1 | 5 | 1 | 1 | 4 | 0 | 1 | 4 | 1 | 1 | 1.89 | 2 | 2 | 1.204 | 0.268 | 0.86422573 | NA | 0.6703 | 1.7478 | 1.2357 | 1.2357 | TRUE | 1.1663 |
| F2rl3 | 2 | 1 | 2 | 4 | 1 | 3 | 2 | 1 | 2 | 3 | 1 | 3 | 2.11 | 2 | 2 | 1.479 | 0.565 | 0.65802498 | NA | 0 | 1.5713 | 0.6802 | 0.6802 | TRUE | 0.2466 |
| F3 | 325 | 431 | 400 | 326 | 301 | 352 | 345 | 430 | 402 | 256 | 324 | 385 | 357.12 | 392 | 322 | 0.817 | -0.291 | 0.10321363 | 0.26735607 | 0.024 | 0.0161 | 0.0201 | 0.0201 | TRUE | 0.2673 |
| F420014N23 | 0 | 1 | 0 | 0 | 5 | 0 | 0 | 1 | 0 | 0 | 5 | 0 | 1.06 | 0 | 2 | 4.851 | 2.278 | 0.42092096 | NA | 10 | 3.1059 | 4.498 | 4.498 | TRUE | 2.3453 |
| F5 | 68 | 87 | 98 | 224 | 163 | 124 | 72 | 87 | 98 | 176 | 175 | 136 | 124.12 | 86 | 162 | 1.894 | 0.921 | 1.48E-05 | 0.00027657 | 0.0132 | 0.0334 | 0.0236 | 0.0236 | TRUE | 0.2107 |
| F630028O10 | 0 | 0 | 0 | 0 | 0 | 0 | 0 | 0 | 0 | 0 | 0 | 0 | 0 | 0 | 0 | NA | NA | NA | NA | NA | NA | NA | NA | NA | NA |
| F630042J09F | 31 | 45 | 62 | 45 | 49 | 62 | 33 | 45 | 62 | 35 | 53 | 68 | 49.35 | 47 | 52 | 1.104 | 0.143 | 0.6973832 | 0.83408871 | 0.0814 | 0.0737 | 0.0771 | 0.0771 | TRUE | 0.2156 |
| F630111L10F | 0 | 0 | 0 | 0 | 0 | 0 | 0 | 0 | 0 | 0 | 0 | 0 | 0 | 0 | 0 | NA | NA | NA | NA | NA | NA | NA | NA | NA | NA |
| F630206G17 | 0 | 0 | 0 | 0 | 0 | 0 | 0 | 0 | 0 | 0 | 0 | 0 | 0 | 0 | 0 | NA | NA | NA | NA | NA | NA | NA | NA | NA | NA |
| F7 | 1 | 6 | 0 | 5 | 2 | 4 | 1 | 6 | 0 | 4 | 2 | 4 | 2.92 | 2 | 3 | 1.481 | 0.566 | 0.66767741 | NA | 0.5691 | 1.1354 | 0.8969 | 0.8969 | TRUE | 1.943 |
| F730035M05 | 2 | 0 | 1 | 0 | 0 | 1 | 2 | 0 | 1 | 0 | 0 | 1 | 0.7 | 1 | 0 | 0.337 | -1.571 | 0.54026722 | NA | 0 | 4.6875 | 3.2519 | 3.2519 | TRUE | 0.2914 |
| F730043M19 | 59 | 78 | 56 | 69 | 76 | 66 | 63 | 78 | 56 | 54 | 82 | 72 | 67.51 | 66 | 69 | 1.052 | 0.073 | 0.79110866 | 0.89260914 | 0.0226 | 0.0557 | 0.0392 | 0.0392 | TRUE | 0.373 |
| F8 | 37 | 27 | 20 | 75 | 71 | 31 | 39 | 27 | 20 | 59 | 76 | 34 | 42.61 | 29 | 56 | 1.966 | 0.975 | 0.01781075 | 0.079827 | 0.1084 | 0.0842 | 0.0957 | 0.0957 | TRUE | 0.1711 |
| F830002L21F | 2 | 2 | 0 | 1 | 0 | 0 | 2 | 2 | 0 | 1 | 0 | 0 | 0.82 | 1 | 0 | 0.255 | -1.973 | 0.41705779 | NA | 0 | 4.0344 | 2.8891 | 2.8891 | TRUE | 0.489 |
| F830016B08 | 0 | 0 | 0 | 0 | 0 | 0 | 0 | 0 | 0 | 0 | 0 | 0 | 0 | 0 | 0 | NA | NA | NA | NA | NA | NA | NA | NA | NA | NA |
| F830045P16I | 0 | 2 | 1 | 0 | 1 | 0 | 0 | 2 | 1 | 0 | 1 | 0 | 0.68 | 1 | 0 | 0.347 | -1.526 | 0.55436871 | NA | 0 | 4.8547 | 3.291 | 3.291 | TRUE | 0.2722 |
| F8a | 256 | 261 | 245 | 356 | 240 | 228 | 272 | 261 | 246 | 280 | 258 | 250 | 261.04 | 260 | 263 | 1.014 | 0.02 | 0.87985976 | 0.94062783 | 0 | 0.0195 | 0.0089 | 0.0089 | TRUE | 0.04 |
| F9 | 0 | 0 | 0 | 0 | 0 | 0 | 0 | 0 | 0 | 0 | 0 | 0 | 0 | 0 | 0 | NA | NA | NA | NA | NA | NA | NA | NA | NA | NA |

| | | | | | | | | | | | | | | | | | | | | | | | | | |
|---|---|---|---|---|---|---|---|---|---|---|---|---|---|---|---|---|---|---|---|---|---|---|---|---|---|
| F930015N05 | 0 | 0 | 3 | 1 | 2 | 0 | 0 | 0 | 3 | 1 | 2 | 0 | 0.99 | 1 | 1 | 0.968 | -0.047 | 0.98406172 | NA | 2.3792 | 3.3289 | 3.0202 | 3.0202 | TRUE | 1.1768 |
| Fa2h | 94 | 133 | 131 | 98 | 67 | 89 | 100 | 133 | 132 | 77 | 72 | 97 | 101.79 | 122 | 82 | 0.675 | -0.567 | 0.01270919 | 0.06279449 | 0.0144 | 0.0393 | 0.0272 | 0.0272 | TRUE | 0.24 |
| Faah | 317 | 454 | 430 | 537 | 425 | 414 | 337 | 453 | 432 | 422 | 457 | 453 | 425.75 | 407 | 444 | 1.089 | 0.123 | 0.38993672 | 0.60377793 | 0.0104 | 0.0146 | 0.0124 | 0.0124 | TRUE | 0.2623 |
| Fabp1 | 0 | 0 | 0 | 0 | 0 | 0 | 0 | 0 | 0 | 0 | 0 | 0 | 0 | 0 | 0 | NA | NA | NA | NA | NA | NA | NA | NA | NA | NA |
| Fabp12 | 0 | 3 | 1 | 2 | 4 | 4 | 0 | 3 | 1 | 2 | 4 | 4 | 2.38 | 1 | 3 | 2.501 | 1.323 | 0.33598224 | NA | 0 | 1.3936 | 0.8373 | 0.8373 | TRUE | 0.7351 |
| Fabp2 | 0 | 0 | 0 | 0 | 0 | 0 | 0 | 0 | 0 | 0 | 0 | 0 | 0 | 0 | 0 | NA | NA | NA | NA | NA | NA | NA | NA | NA | NA |
| Fabp3 | 4236 | 9466 | 9311 | 5032 | 4463 | 7246 | 4499 | 9454 | 9349 | 3956 | 4802 | 7932 | 6665.33 | 7767 | 5563 | 0.716 | -0.482 | 0.2696985 | 0.48977372 | 0.137 | 0.0074 | 0.0608 | 0.137 | TRUE | 1.6912 |
| Fabp4 | 1 | 2 | 1 | 5 | 3 | 3 | 1 | 2 | 1 | 4 | 3 | 3 | 2.42 | 1 | 3 | 2.574 | 1.364 | 0.26375818 | NA | 0 | 1.3696 | 0.5578 | 0.5578 | TRUE | 0.1098 |
| Fabp5 | 930 | 1102 | 1015 | 1897 | 1521 | 1524 | 988 | 1101 | 1019 | 1491 | 1637 | 1668 | 1317.27 | 1036 | 1599 | 1.542 | 0.625 | 1.14E-11 | 9.89E-10 | 0.0027 | 0.0094 | 0.0053 | 0.0053 | TRUE | 0.0427 |
| Fabp6 | 1 | 1 | 4 | 15 | 3 | 6 | 1 | 1 | 4 | 12 | 3 | 7 | 4.61 | 2 | 7 | 3.587 | 1.843 | 0.08192658 | 0.22845159 | 0.2382 | 0.7214 | 0.4905 | 0.4905 | TRUE | 0.269 |
| Fabp7 | 3 | 0 | 0 | 4 | 0 | 1 | 3 | 0 | 0 | 3 | 0 | 1 | 1.24 | 1 | 1 | 1.377 | 0.461 | 0.83657965 | NA | 3.0229 | 2.6688 | 2.7812 | 2.7812 | TRUE | 0.8427 |
| Fabp9 | 2 | 3 | 1 | 16 | 10 | 6 | 2 | 3 | 1 | 13 | 11 | 7 | 6 | 2 | 10 | 4.917 | 2.298 | 0.01116874 | 0.05715554 | 0 | 0.5555 | 0.2922 | 0.2922 | TRUE | 0.2842 |
| Fadd | 304 | 249 | 225 | 423 | 272 | 191 | 323 | 249 | 226 | 333 | 293 | 209 | 271.96 | 266 | 278 | 1.049 | 0.069 | 0.74888837 | 0.86684463 | 0.0409 | 0.019 | 0.0297 | 0.0297 | TRUE | 0.5182 |
| Fads1 | 5911 | 7706 | 8027 | 7914 | 5894 | 6849 | 6279 | 7696 | 8060 | 6221 | 6342 | 7497 | 7015.8 | 7345 | 6687 | 0.91 | -0.136 | 0.27254402 | 0.49272859 | 0.0135 | 0.0074 | 0.0109 | 0.0109 | TRUE | 0.1968 |
| Fads2 | 3287 | 3821 | 4098 | 4621 | 3196 | 3279 | 3491 | 3816 | 4115 | 3633 | 3439 | 3589 | 3680.5 | 3807 | 3554 | 0.933 | -0.099 | 0.25011221 | 0.46733938 | 0.0034 | 0.0078 | 0.0051 | 0.0051 | TRUE | 0.064 |
| Fads3 | 156 | 167 | 155 | 201 | 119 | 128 | 166 | 167 | 156 | 158 | 128 | 140 | 152.38 | 163 | 142 | 0.876 | -0.19 | 0.26209607 | 0.48147442 | 0 | 0.0285 | 0.0142 | 0.0142 | TRUE | 0.1104 |
| Fads6 | 216 | 331 | 388 | 288 | 237 | 267 | 229 | 331 | 390 | 226 | 255 | 292 | 287.21 | 317 | 258 | 0.813 | -0.299 | 0.15891202 | 0.35285941 | 0.0397 | 0.0184 | 0.0288 | 0.0288 | TRUE | 0.3284 |
| Faf1 | 5773 | 7624 | 8139 | 8696 | 6076 | 6667 | 6132 | 7614 | 8172 | 6836 | 6537 | 7298 | 7098.35 | 7306 | 6890 | 0.943 | -0.085 | 0.47772906 | 0.67729589 | 0.0109 | 0.0074 | 0.0101 | 0.0101 | TRUE | 0.2411 |
| Faf2 | 2814 | 2667 | 2833 | 4091 | 3016 | 2740 | 2989 | 2664 | 2845 | 3216 | 3245 | 2999 | 2992.93 | 2833 | 3153 | 1.114 | 0.155 | 0.05338981 | 0.17158711 | 0.0021 | 0.008 | 0.0043 | 0.0043 | TRUE | 0.033 |
| Fah | 1054 | 1526 | 1585 | 971 | 788 | 1051 | 1120 | 1524 | 1592 | 763 | 848 | 1150 | 1166.12 | 1412 | 920 | 0.652 | -0.618 | 0.00089152 | 0.00813721 | 0.0395 | 0.0097 | 0.024 | 0.024 | TRUE | 0.5671 |
| Fahd1 | 271 | 361 | 364 | 349 | 295 | 312 | 288 | 361 | 365 | 274 | 317 | 342 | 324.53 | 338 | 311 | 0.918 | -0.123 | 0.42665762 | 0.63685362 | 0.0118 | 0.0171 | 0.0143 | 0.0143 | TRUE | 0.1896 |
| Fahd2a | 295 | 341 | 295 | 388 | 327 | 307 | 313 | 341 | 296 | 305 | 352 | 336 | 323.84 | 317 | 331 | 1.043 | 0.061 | 0.64136631 | 0.79835032 | 0.0024 | 0.0171 | 0.0091 | 0.0091 | TRUE | 0.0575 |
| Faim | 1082 | 966 | 951 | 1568 | 1059 | 1024 | 1149 | 965 | 955 | 1233 | 1139 | 1121 | 1093.65 | 1023 | 1164 | 1.139 | 0.188 | 0.08337282 | 0.23130274 | 0.0057 | 0.0099 | 0.0076 | 0.0076 | TRUE | 0.1387 |
| Faim2 | 123 | 106 | 76 | 127 | 74 | 75 | 131 | 106 | 76 | 100 | 80 | 82 | 95.73 | 104 | 87 | 0.84 | -0.252 | 0.32908246 | 0.55024931 | 0.0336 | 0.0413 | 0.0375 | 0.0375 | TRUE | 0.1987 |
| Faim3 | 0 | 0 | 0 | 3 | 2 | 2 | 0 | 0 | 0 | 2 | 2 | 2 | 1.12 | 0 | 2 | 11.901 | 3.573 | 0.08259001 | NA | 0 | 2.9574 | 1.8339 | 1.8339 | TRUE | 0.07 |
| Fam101a | 26 | 23 | 14 | 35 | 23 | 19 | 28 | 23 | 14 | 28 | 25 | 21 | 22.95 | 22 | 25 | 1.137 | 0.186 | 0.66441743 | 0.81221452 | 0 | 0.1504 | 0.088 | 0.088 | TRUE | 0.5252 |
| Fam101b | 478 | 460 | 371 | 748 | 534 | 472 | 508 | 459 | 373 | 588 | 575 | 517 | 503.15 | 447 | 560 | 1.255 | 0.327 | 0.02205548 | 0.09335992 | 0.0121 | 0.0135 | 0.0127 | 0.0127 | TRUE | 0.2442 |
| Fam102a | 743 | 1196 | 1154 | 919 | 752 | 865 | 789 | 1194 | 1159 | 722 | 809 | 947 | 936.8 | 1047 | 826 | 0.788 | -0.344 | 0.05307863 | 0.17105764 | 0.0333 | 0.0104 | 0.0217 | 0.0217 | TRUE | 0.5547 |
| Fam102b | 734 | 749 | 775 | 1022 | 791 | 731 | 780 | 748 | 778 | 803 | 851 | 800 | 793.42 | 769 | 818 | 1.064 | 0.09 | 0.30370961 | 0.52448352 | 0 | 0.0111 | 0.0042 | 0.0042 | TRUE | 0.0143 |
| Fam103a1 | 1509 | 1478 | 1414 | 1756 | 1236 | 1368 | 1603 | 1476 | 1420 | 1380 | 1330 | 1497 | 1451.08 | 1500 | 1402 | 0.935 | -0.097 | 0.29628842 | 0.51704591 | 0.0028 | 0.0092 | 0.0055 | 0.0055 | TRUE | 0.0435 |
| Fam104a | 3202 | 3263 | 3304 | 3853 | 2816 | 2765 | 3401 | 3259 | 3318 | 3029 | 3030 | 3027 | 3177.16 | 3326 | 3029 | 0.911 | -0.135 | 0.02785063 | 0.10959798 | 0 | 0.0079 | 0.0024 | 0.0024 | TRUE | 0.0047 |
| Fam105a | 95 | 99 | 104 | 124 | 105 | 102 | 101 | 99 | 104 | 97 | 113 | 112 | 104.39 | 101 | 107 | 1.055 | 0.078 | 0.68694879 | 0.82743019 | 0 | 0.0385 | 0.0171 | 0.0171 | TRUE | 0.0726 |
| Fam105b | 979 | 845 | 836 | 1457 | 1002 | 837 | 1040 | 844 | 839 | 1145 | 1078 | 916 | 977.15 | 908 | 1046 | 1.154 | 0.207 | 0.12101023 | 0.29581113 | 0.0127 | 0.0103 | 0.0118 | 0.0118 | TRUE | 0.1924 |
| Fam107a | 3 | 5 | 5 | 3 | 10 | 3 | 3 | 5 | 5 | 2 | 11 | 3 | 4.93 | 4 | 5 | 1.22 | 0.286 | 0.76049895 | 0.87392272 | 0.2049 | 0.6746 | 0.4295 | 0.4295 | TRUE | 1.478 |
| Fam107b | 566 | 684 | 630 | 1104 | 725 | 726 | 601 | 683 | 633 | 868 | 780 | 795 | 726.59 | 639 | 814 | 1.275 | 0.351 | 0.00063593 | 0.00627653 | 0.0024 | 0.0114 | 0.0062 | 0.0062 | TRUE | 0.0431 |
| Fam108a | 2778 | 2574 | 2614 | 3321 | 2452 | 2285 | 2951 | 2571 | 2625 | 2611 | 2638 | 2501 | 2649.39 | 2716 | 2583 | 0.952 | -0.072 | 0.39860282 | 0.6119696 | 0.0027 | 0.0081 | 0.0048 | 0.0048 | TRUE | 0.0696 |
| Fam108b | 1729 | 1753 | 1609 | 2227 | 1551 | 1566 | 1837 | 1751 | 1616 | 1751 | 1669 | 1714 | 1722.76 | 1735 | 1711 | 0.987 | -0.019 | 0.82208066 | 0.90911907 | 0.0018 | 0.0088 | 0.0044 | 0.0044 | TRUE | 0.0433 |
| Fam108c | 3329 | 3570 | 3514 | 4111 | 3143 | 3445 | 3536 | 3565 | 3528 | 3232 | 3382 | 3771 | 3502.39 | 3543 | 3462 | 0.977 | -0.034 | 0.6811706 | 0.82325078 | 0.0029 | 0.0078 | 0.0047 | 0.0047 | TRUE | 0.0742 |
| Fam109a | 488 | 582 | 501 | 696 | 521 | 466 | 518 | 581 | 503 | 547 | 561 | 510 | 536.74 | 534 | 539 | 1.01 | 0.014 | 0.90008638 | 0.95154919 | 0.0022 | 0.013 | 0.007 | 0.007 | TRUE | 0.0698 |
| Fam109b | 56 | 49 | 59 | 75 | 53 | 44 | 59 | 49 | 59 | 59 | 57 | 48 | 55.3 | 56 | 55 | 0.983 | -0.025 | 0.92341151 | 0.96235363 | 0 | 0.0665 | 0.032 | 0.032 | TRUE | 0.1011 |
| Fam110a | 413 | 512 | 510 | 658 | 450 | 568 | 439 | 511 | 512 | 517 | 484 | 622 | 514.22 | 487 | 541 | 1.109 | 0.149 | 0.27806293 | 0.49924225 | 0.0103 | 0.0133 | 0.0117 | 0.0117 | TRUE | 0.1947 |
| Fam110b | 552 | 649 | 683 | 710 | 526 | 669 | 586 | 648 | 686 | 558 | 566 | 732 | 629.45 | 640 | 619 | 0.965 | -0.051 | 0.72122825 | 0.8490865 | 0.0135 | 0.0121 | 0.013 | 0.013 | TRUE | 0.2994 |
| Fam110c | 28 | 55 | 44 | 42 | 41 | 54 | 30 | 55 | 44 | 33 | 44 | 59 | 44.18 | 43 | 45 | 1.049 | 0.069 | 0.85114406 | 0.92587032 | 0.0648 | 0.0815 | 0.0734 | 0.0734 | TRUE | 0.268 |
| Fam111a | 1485 | 1347 | 1439 | 2577 | 1731 | 1560 | 1577 | 1345 | 1445 | 2026 | 1862 | 1708 | 1660.58 | 1456 | 1865 | 1.282 | 0.359 | 0.00064285 | 0.00633292 | 0.0059 | 0.0089 | 0.0073 | 0.0073 | TRUE | 0.0695 |
| Fam114a1 | 414 | 734 | 728 | 379 | 324 | 404 | 440 | 733 | 731 | 298 | 349 | 442 | 498.76 | 635 | 363 | 0.571 | -0.809 | 0.0003189 | 0.0035799 | 0.0578 | 0.0135 | 0.0342 | 0.0342 | TRUE | 0.8464 |
| Fam114a2 | 6075 | 7121 | 6976 | 7743 | 5974 | 5894 | 6453 | 7112 | 7005 | 6087 | 6428 | 6452 | 6589.28 | 6857 | 6322 | 0.922 | -0.117 | 0.10531989 | 0.27039976 | 0.0016 | 0.0074 | 0.0036 | 0.0036 | TRUE | 0.0323 |
| Fam115a | 464 | 437 | 494 | 534 | 355 | 341 | 493 | 436 | 496 | 420 | 382 | 373 | 433.39 | 475 | 392 | 0.826 | -0.276 | 0.0200885 | 0.08709066 | 0.0025 | 0.0145 | 0.0078 | 0.0078 | TRUE | 0.0593 |
| Fam115c | 497 | 410 | 386 | 624 | 477 | 397 | 528 | 409 | 388 | 491 | 513 | 435 | 460.55 | 442 | 480 | 1.087 | 0.12 | 0.43285718 | 0.64199926 | 0.0151 | 0.0141 | 0.0146 | 0.0146 | TRUE | 0.3358 |
| Fam115e | 0 | 0 | 0 | 0 | 0 | 0 | 0 | 0 | 0 | 0 | 0 | 0 | 0 | 0 | 0 | NA | NA | NA | NA | NA | NA | NA | NA | NA | NA |
| Fam117a | 3066 | 2829 | 2767 | 3236 | 2348 | 2344 | 3257 | 2825 | 2778 | 2544 | 2526 | 2566 | 2749.4 | 2953 | 2545 | 0.862 | -0.214 | 0.01554155 | 0.07269316 | 0.0035 | 0.0081 | 0.0053 | 0.0053 | TRUE | 0.0977 |
| Fam117b | 3757 | 4149 | 4230 | 5286 | 4022 | 3957 | 3991 | 4144 | 4247 | 4155 | 4327 | 4332 | 4199.34 | 4127 | 4271 | 1.035 | 0.049 | 0.4476941 | 0.65337572 | 5.00E-04 | 0.0077 | 0.0028 | 0.0028 | TRUE | 0.0102 |
| Fam118a | 1391 | 1460 | 1387 | 1993 | 1400 | 1350 | 1477 | 1458 | 1393 | 1567 | 1506 | 1478 | 1479.86 | 1443 | 1517 | 1.052 | 0.073 | 0.33566599 | 0.55587029 | 0 | 0.0091 | 0.0035 | 0.0035 | TRUE | 0.0111 |
| Fam118b | 1464 | 1106 | 1000 | 1748 | 1222 | 1009 | 1555 | 1105 | 1004 | 1374 | 1315 | 1105 | 1242.86 | 1221 | 1265 | 1.036 | 0.051 | 0.76894547 | 0.87928573 | 0.0327 | 0.0096 | 0.0209 | 0.0209 | TRUE | 0.6847 |
| Fam120a | 7912 | 9421 | 9464 | 11034 | 8695 | 8638 | 8404 | 9409 | 9503 | 8674 | 9355 | 9456 | 9133.44 | 9105 | 9162 | 1.006 | 0.009 | 0.91543544 | 0.95835171 | 0.0033 | 0.0073 | 0.0047 | 0.0047 | TRUE | 0.0554 |
| Fam120aos | 113 | 172 | 195 | 174 | 160 | 158 | 120 | 172 | 196 | 137 | 172 | 173 | 161.58 | 163 | 161 | 0.985 | -0.022 | 0.92157499 | 0.96126008 | 0.0326 | 0.0273 | 0.0299 | 0.0299 | TRUE | 0.5464 |
| Fam120b | 732 | 730 | 652 | 938 | 622 | 557 | 778 | 729 | 655 | 737 | 669 | 610 | 696.26 | 721 | 672 | 0.934 | -0.098 | 0.416001 | 0.62756268 | 0.0069 | 0.0116 | 0.009 | 0.009 | TRUE | 0.0881 |
| Fam120c | 100 | 95 | 119 | 193 | 123 | 108 | 106 | 95 | 119 | 152 | 132 | 118 | 120.48 | 107 | 134 | 1.259 | 0.333 | 0.09680614 | 0.25549305 | 0.0071 | 0.0342 | 0.0205 | 0.0205 | TRUE | 0.1478 |
| Fam122a | 186 | 203 | 160 | 346 | 223 | 209 | 198 | 203 | 161 | 272 | 240 | 229 | 216.95 | 187 | 247 | 1.324 | 0.405 | 0.01295836 | 0.06372909 | 0.0071 | 0.0221 | 0.0144 | 0.0144 | TRUE | 0.1649 |
| Fam122b | 1927 | 2512 | 2715 | 3155 | 2320 | 2521 | 2047 | 2509 | 2726 | 2480 | 2496 | 2760 | 2502.96 | 2427 | 2579 | 1.062 | 0.087 | 0.47524925 | 0.67546567 | 0.012 | 0.0082 | 0.0103 | 0.0103 | TRUE | 0.2274 |
| Fam122c | 0 | 0 | 0 | 0 | 0 | 0 | 0 | 0 | 0 | 0 | 0 | 0 | 0 | 0 | 0 | NA | NA | NA | NA | NA | NA | NA | NA | NA | NA |
| Fam124a | 226 | 140 | 117 | 485 | 341 | 168 | 240 | 140 | 117 | 381 | 367 | 184 | 238.24 | 166 | 311 | 1.877 | 0.909 | 0.00551788 | 0.03361449 | 0.1392 | 0.0207 | 0.0726 | 0.0726 | TRUE | 1.6187 |
| Fam124b | 8 | 3 | 2 | 10 | 9 | 7 | 8 | 3 | 2 | 8 | 10 | 8 | 6.45 | 4 | 9 | 1.878 | 0.909 | 0.26393348 | 0.48331114 | 0 | 0.5176 | 0.3044 | 0.3044 | TRUE | 1.0622 |
| Fam126a | 1323 | 1554 | 1666 | 1798 | 1310 | 1213 | 1405 | 1552 | 1673 | 1413 | 1409 | 1328 | 1463.47 | 1543 | 1383 | 0.897 | -0.158 | 0.09958018 | 0.26055007 | 0.0034 | 0.0092 | 0.0059 | 0.0059 | TRUE | 0.0735 |
| Fam126b | 1542 | 1566 | 1558 | 1842 | 1418 | 1436 | 1638 | 1564 | 1564 | 1448 | 1526 | 1572 | 1551.98 | 1589 | 1515 | 0.953 | -0.069 | 0.37082323 | 0.58760362 | 6.00E-04 | 0.009 | 0.0037 | 0.0037 | TRUE | 0.0188 |
| Fam129a | 1214 | 1152 | 1124 | 1850 | 1425 | 1315 | 1289 | 1151 | 1129 | 1454 | 1533 | 1439 | 1332.6 | 1190 | 1475 | 1.241 | 0.311 | 0.00057335 | 0.00578738 | 0.0021 | 0.0094 | 0.0051 | 0.0051 | TRUE | 0.0644 |
| Fam129b | 9301 | 10073 | 10417 | 7576 | 6242 | 6998 | 9879 | 10060 | 10460 | 5956 | 6716 | 7660 | 8455.19 | 10133 | 6777 | 0.669 | -0.581 | 3.16E-08 | 1.34E-06 | 0.0082 | 0.0073 | 0.0078 | 0.0078 | TRUE | 0.1585 |
| Fam129c | 140 | 157 | 161 | 150 | 142 | 120 | 149 | 157 | 162 | 118 | 153 | 131 | 144.87 | 156 | 134 | 0.857 | -0.222 | 0.21314177 | 0.42434637 | 0 | 0.0296 | 0.016 | 0.016 | TRUE | 0.1531 |
| Fam131a | 253 | 222 | 230 | 222 | 130 | 171 | 269 | 222 | 231 | 175 | 140 | 187 | 203.83 | 241 | 167 | 0.696 | -0.522 | 0.00260017 | 0.01893359 | 0.01 | 0.0231 | 0.0166 | 0.0166 | TRUE | 0.2072 |
| Fam131b | 86 | 76 | 64 | 110 | 74 | 63 | 91 | 76 | 64 | 86 | 80 | 69 | 77.76 | 77 | 78 | 1.02 | 0.028 | 0.90728165 | 0.95354464 | 0.0087 | 0.0493 | 0.0296 | 0.0296 | TRUE | 0.2329 |

| | | | | | | | | | | | | | | | | | | | | | | | | | |
|---|---|---|---|---|---|---|---|---|---|---|---|---|---|---|---|---|---|---|---|---|---|---|---|---|---|
| Fam131c | 1146 | 1072 | 1011 | 913 | 685 | 713 | 1217 | 1071 | 1015 | 718 | 737 | 780 | 923.04 | 1101 | 745 | 0.676 | -0.564 | 1.00E-07 | 3.59E-06 | 0.0045 | 0.0105 | 0.007 | 0.007 | TRUE | 0.1016 |
| Fam132a | 520 | 847 | 825 | 522 | 459 | 615 | 552 | 846 | 828 | 410 | 494 | 673 | 634.01 | 742 | 526 | 0.707 | -0.499 | 0.02236057 | 0.09421777 | 0.0548 | 0.0121 | 0.0328 | 0.0328 | TRUE | 0.7 |
| Fam132b | 144 | 197 | 151 | 268 | 240 | 226 | 153 | 197 | 152 | 211 | 258 | 247 | 202.94 | 167 | 239 | 1.425 | 0.511 | 0.00379904 | 0.02535696 | 0.0117 | 0.0231 | 0.0173 | 0.0173 | TRUE | 0.2579 |
| Fam133b | 1055 | 1082 | 971 | 1569 | 1224 | 975 | 1121 | 1081 | 975 | 1233 | 1317 | 1067 | 1132.31 | 1059 | 1206 | 1.139 | 0.188 | 0.09712738 | 0.25620253 | 0.0067 | 0.0098 | 0.0084 | 0.0084 | TRUE | 0.1192 |
| Fam134a | 1151 | 1457 | 1522 | 1967 | 1444 | 1412 | 1223 | 1455 | 1528 | 1546 | 1554 | 1546 | 1475.26 | 1402 | 1549 | 1.104 | 0.143 | 0.17311308 | 0.37203155 | 0.0055 | 0.0091 | 0.0073 | 0.0073 | TRUE | 0.1501 |
| Fam134b | 1229 | 1885 | 1952 | 1184 | 955 | 1315 | 1305 | 1883 | 1960 | 931 | 1028 | 1439 | 1424.29 | 1716 | 1133 | 0.66 | -0.6 | 0.00258314 | 0.01884424 | 0.0491 | 0.0092 | 0.0279 | 0.0279 | TRUE | 0.6712 |
| Fam134c | 3447 | 3775 | 3799 | 5033 | 3587 | 3524 | 3661 | 3770 | 3815 | 3957 | 3859 | 3858 | 3819.93 | 3749 | 3891 | 1.038 | 0.054 | 0.37521798 | 0.59102546 | 0 | 0.0078 | 0.0024 | 0.0024 | TRUE | 0.005 |
| Fam135a | 1755 | 1662 | 1749 | 2529 | 1844 | 1739 | 1864 | 1660 | 1756 | 1988 | 1984 | 1904 | 1859.32 | 1760 | 1959 | 1.113 | 0.155 | 0.05336619 | 0.17156371 | 0.0014 | 0.0087 | 0.0041 | 0.0041 | TRUE | 0.0321 |
| Fam135b | 0 | 0 | 1 | 0 | 1 | 0 | 0 | 0 | 1 | 0 | 1 | 0 | 0.35 | 0 | 0 | 0.954 | -0.067 | 0.98570726 | NA | 0 | 9.5099 | 8.2019 | 8.2019 | TRUE | 0.5204 |
| Fam136a | 2275 | 2168 | 1979 | 4148 | 2899 | 2287 | 2416 | 2165 | 1987 | 3261 | 3119 | 2503 | 2575.37 | 2189 | 2961 | 1.353 | 0.436 | 0.00066011 | 0.0064706 | 0.0135 | 0.0082 | 0.0114 | 0.0114 | TRUE | 0.2213 |
| Fam13a | 76 | 74 | 60 | 85 | 68 | 62 | 81 | 74 | 60 | 67 | 73 | 68 | 70.45 | 72 | 69 | 0.967 | -0.049 | 0.83895347 | 0.91831315 | 0 | 0.0537 | 0.0269 | 0.0269 | TRUE | 0.1757 |
| Fam13b | 778 | 829 | 768 | 1075 | 800 | 715 | 826 | 828 | 771 | 845 | 861 | 783 | 818.99 | 808 | 830 | 1.027 | 0.038 | 0.68295076 | 0.82423576 | 7.00E-04 | 0.0109 | 0.0049 | 0.0049 | TRUE | 0.0281 |
| Fam13c | 517 | 568 | 576 | 529 | 385 | 452 | 549 | 567 | 578 | 416 | 414 | 495 | 503.28 | 565 | 442 | 0.781 | -0.357 | 0.00220339 | 0.01662318 | 0.0031 | 0.0134 | 0.0078 | 0.0078 | TRUE | 0.125 |
| Fam149a | 111 | 115 | 113 | 247 | 183 | 172 | 118 | 115 | 113 | 194 | 197 | 188 | 154.26 | 115 | 193 | 1.674 | 0.743 | 3.20E-06 | 7.58E-05 | 0 | 0.0283 | 0.0114 | 0.0114 | TRUE | 0.0049 |
| Fam149b | 638 | 866 | 912 | 1116 | 805 | 858 | 678 | 865 | 916 | 877 | 866 | 939 | 856.83 | 820 | 894 | 1.091 | 0.125 | 0.34244449 | 0.56254916 | 0.0119 | 0.0107 | 0.0114 | 0.0114 | TRUE | 0.2705 |
| Fam150a | 18 | 12 | 10 | 23 | 17 | 14 | 19 | 12 | 10 | 18 | 18 | 15 | 15.47 | 14 | 17 | 1.263 | 0.336 | 0.49961643 | 0.69537154 | 0 | 0.2198 | 0.1133 | 0.1133 | TRUE | 0.4899 |
| Fam150b | 1 | 0 | 1 | 2 | 2 | 1 | 1 | 0 | 1 | 2 | 2 | 1 | 1.15 | 1 | 2 | 2.347 | 1.231 | 0.48826222 | NA | 0 | 2.8779 | 1.2226 | 1.2226 | TRUE | 0.2563 |
| Fam151a | 14 | 16 | 9 | 27 | 23 | 15 | 15 | 16 | 9 | 21 | 25 | 16 | 17.05 | 13 | 21 | 1.567 | 0.648 | 0.17999912 | 0.38137375 | 0 | 0.2002 | 0.1064 | 0.1064 | TRUE | 0.3365 |
| Fam151b | 19 | 12 | 23 | 44 | 17 | 22 | 20 | 12 | 23 | 35 | 18 | 24 | 22.04 | 18 | 26 | 1.407 | 0.492 | 0.29201286 | 0.51281892 | 0.0615 | 0.1564 | 0.1105 | 0.1105 | TRUE | 0.3741 |
| Fam154a | 47 | 33 | 29 | 37 | 26 | 41 | 50 | 33 | 29 | 29 | 28 | 45 | 35.66 | 37 | 34 | 0.908 | -0.14 | 0.71139468 | 0.84292829 | 0.0488 | 0.0993 | 0.0748 | 0.0748 | TRUE | 0.618 |
| Fam154b | 3 | 6 | 11 | 12 | 4 | 6 | 3 | 6 | 11 | 9 | 4 | 7 | 6.75 | 7 | 7 | 1.014 | 0.02 | 0.98018953 | 0.98922122 | 0.099 | 0.4946 | 0.3038 | 0.3038 | TRUE | 0.4315 |
| Fam155a | 0 | 0 | 0 | 3 | 0 | 2 | 0 | 0 | 0 | 2 | 0 | 2 | 0.76 | 0 | 1 | 8.123 | 3.022 | 0.28653847 | NA | 0 | 4.3534 | 4.4726 | 4.4726 | TRUE | 0.4916 |
| Fam159a | 102 | 138 | 162 | 105 | 66 | 78 | 108 | 138 | 163 | 83 | 71 | 85 | 107.96 | 136 | 80 | 0.585 | -0.775 | 0.00066705 | 0.00652245 | 0.0177 | 0.0374 | 0.0275 | 0.0275 | TRUE | 0.1463 |
| Fam159b | 26 | 62 | 29 | 61 | 45 | 57 | 28 | 62 | 29 | 48 | 48 | 62 | 46.24 | 40 | 53 | 1.333 | 0.415 | 0.28104691 | 0.50229789 | 0.092 | 0.0782 | 0.0847 | 0.0847 | TRUE | 1.8425 |
| Fam160a1 | 208 | 157 | 167 | 283 | 195 | 213 | 221 | 157 | 168 | 222 | 210 | 233 | 201.81 | 182 | 222 | 1.221 | 0.288 | 0.10475233 | 0.26969536 | 0.0117 | 0.0232 | 0.0178 | 0.0178 | TRUE | 0.3761 |
| Fam160a2 | 2674 | 3057 | 3087 | 3956 | 2962 | 2971 | 2840 | 3053 | 3100 | 3110 | 3187 | 3252 | 3090.34 | 2998 | 3183 | 1.062 | 0.086 | 0.2281477 | 0.44159023 | 8.00E-04 | 0.008 | 0.0034 | 0.0034 | TRUE | 0.0255 |
| Fam160b1 | 1615 | 1925 | 2016 | 2230 | 1691 | 1835 | 1715 | 1923 | 2024 | 1753 | 1819 | 2009 | 1873.89 | 1887 | 1860 | 0.985 | -0.022 | 0.83010745 | 0.9138554 | 0.0055 | 0.0087 | 0.0068 | 0.0068 | TRUE | 0.0765 |
| Fam160b2 | 1279 | 1176 | 1161 | 1888 | 1390 | 1225 | 1359 | 1174 | 1166 | 1484 | 1496 | 1341 | 1336.58 | 1233 | 1440 | 1.169 | 0.225 | 0.02671732 | 0.1065181 | 0.0046 | 0.0094 | 0.0067 | 0.0067 | TRUE | 0.0949 |
| Fam161a | 64 | 92 | 88 | 192 | 128 | 141 | 68 | 92 | 88 | 151 | 138 | 154 | 115.2 | 83 | 148 | 1.783 | 0.834 | 3.43E-05 | 0.00056194 | 0 | 0.0355 | 0.0198 | 0.0198 | TRUE | 0.222 |
| Fam161b | 147 | 169 | 181 | 269 | 173 | 183 | 156 | 169 | 182 | 211 | 186 | 200 | 184.1 | 169 | 199 | 1.182 | 0.241 | 0.12127062 | 0.29612636 | 0 | 0.0248 | 0.0119 | 0.0119 | TRUE | 0.0472 |
| Fam162a | 4143 | 5886 | 5920 | 4802 | 3418 | 4235 | 4401 | 5878 | 5944 | 3775 | 3678 | 4636 | 4718.62 | 5408 | 4030 | 0.745 | -0.425 | 0.00340354 | 0.02334954 | 0.0213 | 0.0076 | 0.0149 | 0.0149 | TRUE | 0.3234 |
| Fam162b | 3 | 4 | 3 | 7 | 2 | 2 | 3 | 4 | 3 | 6 | 2 | 2 | 3.34 | 3 | 3 | 0.992 | -0.012 | 0.99094171 | NA | 0 | 0.9934 | 0.4561 | 0.4561 | TRUE | 0.6448 |
| Fam163a | 8 | 6 | 2 | 15 | 8 | 6 | 8 | 6 | 2 | 12 | 9 | 7 | 7.24 | 5 | 9 | 1.656 | 0.728 | 0.34419408 | 0.5640134 | 1.00E-04 | 0.4617 | 0.2797 | 0.2797 | TRUE | 0.5272 |
| Fam163b | 97 | 164 | 158 | 150 | 143 | 163 | 103 | 164 | 159 | 118 | 154 | 178 | 145.95 | 142 | 150 | 1.054 | 0.075 | 0.76061388 | 0.87399112 | 0.0454 | 0.0295 | 0.0372 | 0.0372 | TRUE | 0.4761 |
| Fam166a | 3 | 5 | 5 | 4 | 3 | 2 | 3 | 5 | 5 | 3 | 3 | 2 | 3.63 | 4 | 3 | 0.651 | -0.62 | 0.51909296 | NA | 0 | 0.9152 | 0.3796 | 0.3796 | TRUE | 0.0996 |
| Fam166b | 64 | 69 | 74 | 38 | 23 | 25 | 68 | 69 | 74 | 30 | 25 | 27 | 48.86 | 70 | 27 | 0.39 | -1.358 | 1.76E-06 | 4.49E-05 | 0 | 0.0743 | 0.0332 | 0.0332 | TRUE | 0.0523 |
| Fam167a | 204 | 215 | 217 | 176 | 170 | 200 | 217 | 215 | 218 | 138 | 183 | 219 | 198.25 | 217 | 180 | 0.828 | -0.272 | 0.16170113 | 0.35683099 | 0.0209 | 0.0235 | 0.0222 | 0.0222 | TRUE | 0.3168 |
| Fam167b | 0 | 0 | 1 | 1 | 0 | 0 | 0 | 0 | 1 | 1 | 0 | 0 | 0.3 | 0 | 0 | 0.954 | -0.067 | 0.98640617 | NA | 0 | 11.0483 | 9.2783 | 9.2783 | TRUE | 0.4098 |
| Fam168a | 2581 | 2470 | 2381 | 3566 | 2487 | 2478 | 2741 | 2467 | 2391 | 2803 | 2676 | 2713 | 2631.8 | 2533 | 2731 | 1.078 | 0.109 | 0.1896053 | 0.39472739 | 0.0024 | 0.0082 | 0.0046 | 0.0046 | TRUE | 0.0626 |
| Fam168b | 3414 | 3816 | 3736 | 5183 | 3944 | 3830 | 3626 | 3811 | 3751 | 4074 | 4244 | 4193 | 3949.87 | 3729 | 4170 | 1.118 | 0.161 | 0.01057675 | 0.05499872 | 3.00E-04 | 0.0077 | 0.0026 | 0.0026 | TRUE | 0.0071 |
| Fam169a | 9507 | 8304 | 8893 | 14279 | 9697 | 9017 | 10098 | 8293 | 8930 | 11225 | 10433 | 9871 | 9808.31 | 9107 | 10510 | 1.154 | 0.207 | 0.03808863 | 0.13569774 | 0.007 | 0.0072 | 0.0071 | 0.0071 | TRUE | 0.1107 |
| Fam169b | 227 | 240 | 282 | 256 | 176 | 244 | 241 | 240 | 283 | 201 | 189 | 267 | 236.95 | 255 | 219 | 0.859 | -0.219 | 0.22160716 | 0.43380763 | 0.0166 | 0.0208 | 0.0189 | 0.0189 | TRUE | 0.3881 |
| Fam170a | 0 | 0 | 0 | 1 | 0 | 0 | 0 | 0 | 0 | 1 | 0 | 0 | 0.13 | 0 | 0 | 1.859 | 0.894 | 0.82650734 | NA | 0 | 25.1515 | 10 | 10 | TRUE | 0.2563 |
| Fam170b | 3 | 4 | 5 | 5 | 7 | 1 | 3 | 4 | 5 | 4 | 8 | 1 | 4.13 | 4 | 4 | 1.027 | 0.038 | 0.96895221 | NA | 0 | 0.8053 | 0.4595 | 0.4595 | TRUE | 0.2301 |
| Fam171a1 | 500 | 651 | 713 | 1147 | 891 | 845 | 531 | 650 | 716 | 902 | 959 | 925 | 780.42 | 632 | 929 | 1.467 | 0.553 | 1.54E-05 | 0.00028511 | 0.0095 | 0.0111 | 0.0105 | 0.0105 | TRUE | 0.2294 |
| Fam171a2 | 411 | 356 | 344 | 612 | 473 | 409 | 437 | 356 | 345 | 481 | 509 | 448 | 429.21 | 379 | 479 | 1.265 | 0.339 | 0.01232116 | 0.06138491 | 0.0072 | 0.0146 | 0.0108 | 0.0108 | TRUE | 0.1993 |
| Fam171b | 391 | 332 | 355 | 506 | 373 | 320 | 415 | 332 | 356 | 398 | 401 | 350 | 375.46 | 368 | 383 | 1.043 | 0.061 | 0.65641734 | 0.80670515 | 0.0066 | 0.0157 | 0.0109 | 0.0109 | TRUE | 0.145 |
| Fam172a | 869 | 955 | 942 | 1488 | 1028 | 1065 | 923 | 954 | 946 | 1170 | 1106 | 1166 | 1044.05 | 941 | 1147 | 1.22 | 0.286 | 0.0003568 | 0.00394635 | 0 | 0.0101 | 0.0037 | 0.0037 | TRUE | 0.0116 |
| Fam173a | 245 | 376 | 354 | 356 | 288 | 299 | 260 | 376 | 355 | 280 | 310 | 327 | 318.04 | 330 | 306 | 0.923 | -0.115 | 0.50088515 | 0.69666649 | 0.0185 | 0.0173 | 0.0179 | 0.0179 | TRUE | 0.3886 |
| Fam173b | 126 | 107 | 99 | 174 | 119 | 93 | 134 | 107 | 99 | 137 | 128 | 102 | 117.79 | 113 | 122 | 1.082 | 0.114 | 0.59684516 | 0.76965176 | 0.0151 | 0.0349 | 0.0251 | 0.0251 | TRUE | 0.2454 |
| Fam174a | 232 | 403 | 433 | 438 | 360 | 382 | 246 | 402 | 435 | 344 | 387 | 418 | 372.25 | 361 | 383 | 1.06 | 0.083 | 0.69461042 | 0.83270212 | 0.0453 | 0.0158 | 0.0299 | 0.0299 | TRUE | 0.8781 |
| Fam174b | 205 | 230 | 207 | 389 | 285 | 213 | 218 | 230 | 208 | 306 | 307 | 233 | 250.15 | 219 | 282 | 1.293 | 0.37 | 0.02119185 | 0.09062847 | 0.0095 | 0.0201 | 0.0145 | 0.0145 | TRUE | 0.2487 |
| Fam175a | 1219 | 1242 | 1158 | 1631 | 1185 | 1118 | 1295 | 1240 | 1163 | 1282 | 1275 | 1224 | 1246.49 | 1233 | 1260 | 1.023 | 0.033 | 0.69611531 | 0.83364698 | 9.00E-04 | 0.0095 | 0.0042 | 0.0042 | TRUE | 0.0296 |
| Fam175b | 1740 | 1993 | 2088 | 1985 | 1520 | 1732 | 1848 | 1990 | 2097 | 1560 | 1635 | 1896 | 1837.84 | 1978 | 1697 | 0.857 | -0.222 | 0.03566242 | 0.12960084 | 0.0065 | 0.0087 | 0.0075 | 0.0075 | TRUE | 0.1259 |
| Fam177a | 0 | 0 | 0 | 0 | 0 | 0 | 0 | 0 | 0 | 0 | 0 | 0 | 0 | 0 | 0 | NA | NA | NA | NA | NA | NA | NA | NA | NA | NA |
| Fam178a | 5779 | 5691 | 6007 | 7526 | 5625 | 5447 | 6138 | 5684 | 6032 | 5916 | 6052 | 5963 | 5964.12 | 5951 | 5977 | 1.004 | 0.006 | 0.922576 | 0.96178655 | 7.00E-04 | 0.0075 | 0.0028 | 0.0028 | TRUE | 0.0189 |
| Fam178b | 311 | 435 | 460 | 652 | 549 | 540 | 330 | 434 | 462 | 513 | 591 | 591 | 486.84 | 409 | 565 | 1.379 | 0.464 | 0.00247472 | 0.01822188 | 0.0159 | 0.0137 | 0.0148 | 0.0148 | TRUE | 0.3227 |
| Fam179a | 12 | 4 | 7 | 11 | 9 | 13 | 13 | 4 | 7 | 9 | 10 | 14 | 9.39 | 8 | 11 | 1.367 | 0.451 | 0.50522121 | 0.69983743 | 0.0601 | 0.3578 | 0.2202 | 0.2202 | TRUE | 0.5731 |
| Fam179b | 3867 | 3270 | 3293 | 4604 | 3203 | 2843 | 4107 | 3266 | 3307 | 3619 | 3446 | 3112 | 3476.23 | 3560 | 3392 | 0.953 | -0.069 | 0.55719065 | 0.73979383 | 0.0101 | 0.0079 | 0.0097 | 0.0097 | TRUE | 0.2199 |
| Fam180a | 0 | 1 | 0 | 3 | 1 | 0 | 0 | 1 | 0 | 2 | 1 | 0 | 0.74 | 0 | 1 | 3.203 | 1.679 | 0.50423271 | NA | 0 | 4.4659 | 3.139 | 3.139 | TRUE | 0.1967 |
| Fam181a | 5 | 3 | 5 | 3 | 7 | 4 | 5 | 3 | 5 | 2 | 8 | 4 | 4.6 | 4 | 5 | 1.052 | 0.074 | 0.93499035 | NA | 0 | 0.7232 | 0.3703 | 0.3703 | TRUE | 0.3235 |
| Fam181b | 46 | 86 | 64 | 79 | 87 | 73 | 49 | 86 | 64 | 62 | 94 | 80 | 72.44 | 66 | 79 | 1.177 | 0.235 | 0.42606011 | 0.63621327 | 0.0457 | 0.0524 | 0.0491 | 0.0491 | TRUE | 0.3482 |
| Fam183b | 30 | 52 | 75 | 71 | 68 | 71 | 32 | 52 | 75 | 56 | 73 | 78 | 60.97 | 53 | 69 | 1.293 | 0.37 | 0.28673385 | 0.50796477 | 0.0811 | 0.0609 | 0.0704 | 0.0704 | TRUE | 0.2948 |
| Fam184a | 473 | 429 | 410 | 631 | 452 | 420 | 502 | 428 | 412 | 496 | 486 | 460 | 464.11 | 447 | 481 | 1.075 | 0.105 | 0.39003835 | 0.60381673 | 0.0041 | 0.014 | 0.0086 | 0.0086 | TRUE | 0.1322 |
| Fam184b | 65 | 80 | 63 | 131 | 88 | 80 | 69 | 80 | 63 | 103 | 95 | 88 | 82.91 | 71 | 95 | 1.348 | 0.43 | 0.05287237 | 0.1705323 | 0 | 0.0466 | 0.0233 | 0.0233 | TRUE | 0.1178 |
| Fam185a | 1193 | 1388 | 1430 | 1545 | 1125 | 1225 | 1267 | 1386 | 1436 | 1215 | 1210 | 1341 | 1309.2 | 1363 | 1255 | 0.92 | -0.12 | 0.20060883 | 0.40841648 | 0.0029 | 0.0094 | 0.0055 | 0.0055 | TRUE | 0.0453 |
| Fam186b | 15 | 9 | 9 | 14 | 6 | 8 | 16 | 9 | 9 | 11 | 6 | 9 | 10.03 | 11 | 9 | 0.782 | -0.354 | 0.56670677 | 0.74658363 | 0 | 0.3354 | 0.1752 | 0.1752 | TRUE | 0.4608 |
| Fam187a | 0 | 0 | 0 | 0 | 0 | 0 | 0 | 0 | 0 | 0 | 0 | 0 | 0 | 0 | 0 | NA | NA | NA | NA | NA | NA | NA | NA | NA | NA |
| Fam187b | 16 | 12 | 9 | 14 | 9 | 6 | 17 | 12 | 9 | 11 | 10 | 7 | 10.88 | 13 | 9 | 0.725 | -0.463 | 0.43527573 | 0.64352914 | 0 | 0.3097 | 0.1606 | 0.1606 | TRUE | 0.3048 |

| | | | | | | | | | | | | | | | | | | | | | | | | | |
|---|---|---|---|---|---|---|---|---|---|---|---|---|---|---|---|---|---|---|---|---|---|---|---|---|---|
| Fam188a | 2714 | 2805 | 2931 | 2957 | 2212 | 2299 | 2883 | 2801 | 2943 | 2325 | 2380 | 2517 | 2641.39 | 2876 | 2407 | 0.837 | -0.257 | 0.0003014 | 0.00342732 | 7.00E-04 | 0.0082 | 0.0033 | 0.0033 | TRUE | 0.0191 |
| Fam188b | 130 | 151 | 152 | 159 | 134 | 155 | 138 | 151 | 153 | 125 | 144 | 170 | 146.73 | 147 | 146 | 0.989 | -0.015 | 0.93376803 | 0.96713012 | 0.0072 | 0.0294 | 0.018 | 0.018 | TRUE | 0.2017 |
| Fam189a1 | 15 | 16 | 16 | 27 | 17 | 13 | 16 | 16 | 16 | 21 | 18 | 14 | 16.95 | 16 | 18 | 1.129 | 0.175 | 0.70201548 | 0.83705948 | 0 | 0.2012 | 0.0911 | 0.0911 | TRUE | 0.1623 |
| Fam189a2 | 388 | 578 | 482 | 413 | 320 | 381 | 412 | 577 | 484 | 325 | 344 | 417 | 426.57 | 491 | 362 | 0.736 | -0.443 | 0.00802176 | 0.04440829 | 0.0197 | 0.0146 | 0.0177 | 0.0177 | TRUE | 0.2003 |
| Fam189b | 900 | 839 | 915 | 1022 | 773 | 771 | 956 | 838 | 919 | 803 | 832 | 844 | 865.29 | 904 | 826 | 0.914 | -0.13 | 0.16615548 | 0.36339067 | 0.0015 | 0.0107 | 0.0052 | 0.0052 | TRUE | 0.0492 |
| Fam192a | 2929 | 2980 | 2898 | 4452 | 3111 | 2949 | 3111 | 2976 | 2910 | 3500 | 3347 | 3228 | 3178.73 | 2999 | 3358 | 1.12 | 0.164 | 0.02280501 | 0.09545021 | 9.00E-04 | 0.0079 | 0.0034 | 0.0034 | TRUE | 0.0168 |
| Fam193a | 2522 | 2565 | 2690 | 3417 | 2480 | 2405 | 2679 | 2562 | 2701 | 2686 | 2668 | 2633 | 2654.78 | 2647 | 2662 | 1.006 | 0.008 | 0.89667636 | 0.95009983 | 0 | 0.0081 | 0.0027 | 0.0027 | TRUE | 0.0097 |
| Fam193b | 1991 | 2394 | 2144 | 2299 | 2172 | 1953 | 2115 | 2391 | 2153 | 1807 | 2337 | 2138 | 2156.77 | 2220 | 2094 | 0.943 | -0.085 | 0.46810323 | 0.67044353 | 0.0102 | 0.0084 | 0.0094 | 0.0094 | TRUE | 0.1761 |
| Fam194a | 2 | 1 | 2 | 4 | 5 | 3 | 2 | 1 | 2 | 3 | 5 | 3 | 2.82 | 2 | 4 | 2.293 | 1.197 | 0.29123481 | NA | 0 | 1.1738 | 0.5027 | 0.5027 | TRUE | 0.1575 |
| Fam195a | 282 | 419 | 430 | 324 | 242 | 295 | 300 | 418 | 432 | 255 | 260 | 323 | 331.29 | 383 | 279 | 0.727 | -0.459 | 0.01225523 | 0.06114822 | 0.0253 | 0.0168 | 0.0211 | 0.0211 | TRUE | 0.4161 |
| Fam195b | 1553 | 2226 | 2321 | 1956 | 1454 | 1678 | 1650 | 2223 | 2331 | 1538 | 1564 | 1837 | 1857.02 | 2068 | 1646 | 0.796 | -0.329 | 0.0254699 | 0.10303739 | 0.0199 | 0.0087 | 0.0151 | 0.0151 | TRUE | 0.378 |
| Fam196a | 37 | 25 | 25 | 56 | 31 | 28 | 39 | 25 | 25 | 44 | 33 | 31 | 32.9 | 30 | 36 | 1.219 | 0.286 | 0.43759419 | 0.64550463 | 0.0247 | 0.107 | 0.0673 | 0.0673 | TRUE | 0.5001 |
| Fam196b | 48 | 28 | 38 | 20 | 19 | 18 | 51 | 28 | 38 | 16 | 20 | 20 | 28.83 | 39 | 19 | 0.475 | -1.075 | 0.00735269 | 0.04159492 | 0 | 0.1212 | 0.0764 | 0.0764 | TRUE | 0.141 |
| Fam198a | 8 | 5 | 9 | 6 | 2 | 3 | 8 | 5 | 9 | 5 | 2 | 3 | 5.45 | 7 | 3 | 0.46 | -1.12 | 0.18037404 | 0.38170661 | 0 | 0.6118 | 0.2941 | 0.2941 | TRUE | 0.2456 |
| Fam198b | 402 | 342 | 278 | 471 | 332 | 366 | 427 | 342 | 279 | 370 | 357 | 401 | 362.64 | 349 | 376 | 1.077 | 0.107 | 0.53333827 | 0.72124217 | 0.0211 | 0.016 | 0.0186 | 0.0186 | TRUE | 0.3048 |
| Fam199x | 1130 | 1272 | 1362 | 1380 | 969 | 1061 | 1200 | 1270 | 1368 | 1085 | 1043 | 1161 | 1187.85 | 1279 | 1096 | 0.857 | -0.223 | 0.01764057 | 0.07928957 | 0.0027 | 0.0097 | 0.0055 | 0.0055 | TRUE | 0.0437 |
| Fam19a1 | 3 | 4 | 5 | 6 | 6 | 1 | 3 | 4 | 5 | 5 | 6 | 1 | 4.08 | 4 | 4 | 1.011 | 0.016 | 0.98715305 | NA | 0 | 0.8147 | 0.4396 | 0.4396 | TRUE | 0.4763 |
| Fam19a2 | 2 | 3 | 1 | 1 | 2 | 1 | 2 | 3 | 1 | 1 | 2 | 1 | 1.69 | 2 | 1 | 0.645 | -0.632 | 0.65547609 | NA | 0 | 1.953 | 0.8254 | 0.8254 | TRUE | 0.1844 |
| Fam19a3 | 22 | 30 | 28 | 20 | 22 | 8 | 23 | 30 | 28 | 16 | 24 | 9 | 21.6 | 27 | 16 | 0.591 | -0.759 | 0.10393016 | 0.26817827 | 0.0332 | 0.1594 | 0.1078 | 0.1078 | TRUE | 0.338 |
| Fam19a4 | 167 | 181 | 211 | 225 | 157 | 122 | 177 | 181 | 212 | 177 | 169 | 134 | 174.89 | 190 | 160 | 0.843 | -0.246 | 0.17163437 | 0.37055804 | 0.0095 | 0.0257 | 0.0176 | 0.0176 | TRUE | 0.2099 |
| Fam19a5 | 270 | 325 | 324 | 532 | 397 | 348 | 287 | 325 | 325 | 418 | 427 | 381 | 360.5 | 312 | 409 | 1.309 | 0.389 | 0.00162954 | 0.01307428 | 0.0011 | 0.016 | 0.0081 | 0.0081 | TRUE | 0.0567 |
| Fam203a | 934 | 981 | 934 | 1153 | 867 | 649 | 992 | 980 | 938 | 906 | 933 | 710 | 909.89 | 970 | 850 | 0.877 | -0.189 | 0.12964553 | 0.30981353 | 0.0098 | 0.0105 | 0.0101 | 0.0101 | TRUE | 0.2409 |
| Fam207a | 1701 | 1511 | 1471 | 2375 | 1722 | 1505 | 1807 | 1509 | 1477 | 1867 | 1853 | 1647 | 1693.35 | 1598 | 1789 | 1.12 | 0.164 | 0.14134499 | 0.32680725 | 0.0079 | 0.0088 | 0.0084 | 0.0084 | TRUE | 0.1576 |
| Fam209 | 1 | 0 | 1 | 0 | 0 | 0 | 1 | 0 | 1 | 0 | 0 | 0 | 0.34 | 1 | 0 | 0.261 | -1.939 | 0.60367425 | NA | 0 | 9.5732 | 8.3134 | 8.3134 | TRUE | 0.2529 |
| Fam20a | 14 | 18 | 37 | 22 | 25 | 23 | 15 | 18 | 37 | 17 | 27 | 25 | 23.23 | 23 | 23 | 0.983 | -0.025 | 0.95945097 | 0.97919379 | 0.1069 | 0.1487 | 0.1282 | 0.1282 | TRUE | 1.5852 |
| Fam20b | 2026 | 2302 | 2248 | 2995 | 2091 | 2236 | 2152 | 2299 | 2257 | 2354 | 2250 | 2448 | 2293.35 | 2236 | 2351 | 1.051 | 0.072 | 0.33565024 | 0.55587029 | 0.001 | 0.0083 | 0.0036 | 0.0036 | TRUE | 0.0167 |
| Fam20c | 104 | 162 | 159 | 127 | 113 | 117 | 110 | 162 | 160 | 100 | 122 | 128 | 130.23 | 144 | 117 | 0.806 | -0.312 | 0.15963306 | 0.35411148 | 0.0231 | 0.0322 | 0.0277 | 0.0277 | TRUE | 0.4245 |
| Fam21 | 6667 | 6807 | 7229 | 10660 | 7691 | 7152 | 7082 | 6798 | 7259 | 8380 | 8275 | 7829 | 7603.75 | 7046 | 8161 | 1.158 | 0.212 | 0.00137582 | 0.01143536 | 0.001 | 0.0073 | 0.003 | 0.003 | TRUE | 0.0154 |
| Fam210a | 2327 | 2686 | 2807 | 3626 | 2565 | 2811 | 2472 | 2683 | 2819 | 2850 | 2760 | 3077 | 2776.68 | 2658 | 2896 | 1.089 | 0.124 | 0.15816029 | 0.35188378 | 0.0032 | 0.0081 | 0.0052 | 0.0052 | TRUE | 0.0453 |
| Fam210b | 724 | 814 | 766 | 903 | 686 | 646 | 769 | 813 | 769 | 710 | 738 | 707 | 751.04 | 784 | 718 | 0.916 | -0.126 | 0.15323988 | 0.34475084 | 0 | 0.0113 | 0.0043 | 0.0043 | TRUE | 0.0127 |
| Fam211a | 476 | 561 | 538 | 801 | 601 | 624 | 506 | 560 | 540 | 630 | 647 | 683 | 594.24 | 535 | 653 | 1.219 | 0.285 | 0.00462438 | 0.02952862 | 0 | 0.0124 | 0.0056 | 0.0056 | TRUE | 0.0273 |
| Fam211b | 37 | 62 | 48 | 50 | 45 | 41 | 39 | 62 | 48 | 39 | 48 | 45 | 47 | 50 | 44 | 0.883 | -0.18 | 0.55485434 | 0.73805875 | 0 | 0.077 | 0.0453 | 0.0453 | TRUE | 0.3088 |
| Fam212a | 67 | 74 | 70 | 74 | 57 | 56 | 71 | 74 | 70 | 58 | 61 | 61 | 66.03 | 72 | 60 | 0.838 | -0.255 | 0.27176133 | 0.49174612 | 0 | 0.0568 | 0.0235 | 0.0235 | TRUE | 0.0093 |
| Fam212b | 24 | 26 | 27 | 83 | 34 | 20 | 25 | 26 | 27 | 65 | 37 | 22 | 33.72 | 26 | 41 | 1.591 | 0.67 | 0.13756289 | 0.3217144 | 0.126 | 0.1046 | 0.1153 | 0.1153 | TRUE | 0.342 |
| Fam213a | 1365 | 1554 | 1571 | 1626 | 1215 | 1450 | 1450 | 1552 | 1577 | 1278 | 1307 | 1587 | 1458.68 | 1526 | 1391 | 0.911 | -0.135 | 0.22328418 | 0.43519995 | 0.0075 | 0.0092 | 0.0082 | 0.0082 | TRUE | 0.182 |
| Fam213b | 419 | 508 | 463 | 580 | 454 | 456 | 445 | 507 | 465 | 456 | 488 | 499 | 476.81 | 472 | 481 | 1.017 | 0.024 | 0.82747507 | 0.91265292 | 0.0013 | 0.0138 | 0.0068 | 0.0068 | TRUE | 0.0487 |
| Fam214a | 388 | 452 | 436 | 587 | 434 | 409 | 412 | 451 | 438 | 461 | 467 | 448 | 446.24 | 434 | 459 | 1.057 | 0.08 | 0.44618761 | 0.65236459 | 0 | 0.0143 | 0.0058 | 0.0058 | TRUE | 0.0218 |
| Fam214b | 398 | 384 | 405 | 505 | 377 | 373 | 423 | 384 | 407 | 397 | 406 | 408 | 403.97 | 405 | 404 | 0.998 | -0.003 | 0.9804649 | 0.98937252 | 0 | 0.0151 | 0.0061 | 0.0061 | TRUE | 0.0234 |
| Fam216a | 606 | 1031 | 1005 | 1224 | 848 | 946 | 644 | 1030 | 1009 | 962 | 912 | 1036 | 932.1 | 894 | 970 | 1.085 | 0.117 | 0.51071114 | 0.70422593 | 0.0337 | 0.0104 | 0.0217 | 0.0217 | TRUE | 0.7127 |
| Fam216b | 0 | 0 | 0 | 0 | 0 | 0 | 0 | 0 | 0 | 0 | 0 | 0 | 0 | 0 | 0 | NA | NA | NA | NA | NA | NA | NA | NA | NA | NA |
| Fam217a | 14 | 16 | 11 | 15 | 12 | 10 | 15 | 16 | 11 | 12 | 13 | 11 | 12.92 | 14 | 12 | 0.851 | -0.233 | 0.65359438 | 0.80540538 | 0 | 0.2618 | 0.1158 | 0.1158 | TRUE | 0.148 |
| Fam217b | 346 | 451 | 421 | 359 | 277 | 332 | 368 | 450 | 423 | 282 | 298 | 363 | 364.06 | 414 | 314 | 0.759 | -0.398 | 0.00818491 | 0.04521626 | 0.0114 | 0.016 | 0.0136 | 0.0136 | TRUE | 0.2058 |
| Fam219a | 150 | 116 | 128 | 230 | 161 | 132 | 159 | 116 | 129 | 181 | 173 | 144 | 150.37 | 135 | 166 | 1.239 | 0.309 | 0.11491997 | 0.28620404 | 0.0129 | 0.0288 | 0.0209 | 0.0209 | TRUE | 0.2628 |
| Fam219b | 618 | 946 | 981 | 1017 | 790 | 861 | 656 | 945 | 985 | 799 | 850 | 943 | 863.03 | 862 | 864 | 1.001 | 0.002 | 0.98971323 | 0.99456878 | 0.0258 | 0.0107 | 0.0183 | 0.0183 | TRUE | 0.516 |
| Fam220a | 1288 | 1292 | 1336 | 1465 | 1083 | 1053 | 1368 | 1290 | 1341 | 1152 | 1165 | 1153 | 1244.91 | 1333 | 1157 | 0.867 | -0.205 | 0.00673174 | 0.03902025 | 0 | 0.0095 | 0.0033 | 0.0033 | TRUE | 0.0096 |
| Fam221a | 333 | 338 | 325 | 364 | 242 | 283 | 354 | 338 | 326 | 286 | 260 | 310 | 312.32 | 339 | 285 | 0.842 | -0.249 | 0.0531464 | 0.17117112 | 0 | 0.0175 | 0.0087 | 0.0087 | TRUE | 0.0625 |
| Fam221b | 1 | 8 | 6 | 6 | 4 | 6 | 1 | 8 | 6 | 5 | 4 | 7 | 5.11 | 5 | 5 | 1.025 | 0.036 | 0.9689796 | 0.98446557 | 0 | 0.6515 | 0.4062 | 0.4062 | TRUE | 0.7206 |
| Fam222a | 155 | 135 | 114 | 187 | 146 | 109 | 165 | 135 | 114 | 147 | 157 | 119 | 139.56 | 138 | 141 | 1.025 | 0.036 | 0.86574921 | 0.93364615 | 0.0186 | 0.0305 | 0.0246 | 0.0246 | TRUE | 0.2912 |
| Fam222b | 2506 | 2078 | 1912 | 3299 | 2240 | 1816 | 2662 | 2075 | 1920 | 2593 | 2410 | 1988 | 2274.73 | 2219 | 2330 | 1.051 | 0.071 | 0.63768692 | 0.79603827 | 0.0236 | 0.0084 | 0.0161 | 0.0161 | TRUE | 0.3686 |
| Fam227a | 52 | 51 | 32 | 44 | 42 | 33 | 55 | 51 | 32 | 35 | 45 | 36 | 42.37 | 46 | 39 | 0.836 | -0.258 | 0.44040348 | 0.64746865 | 0.0267 | 0.0847 | 0.0568 | 0.0568 | TRUE | 0.5598 |
| Fam227b | 0 | 0 | 0 | 1 | 0 | 0 | 0 | 0 | 0 | 1 | 0 | 0 | 0.13 | 0 | 0 | 1.859 | 0.894 | 0.82650734 | NA | 0 | 25.1515 | 10 | 10 | TRUE | 0.2563 |
| Fam228a | 16 | 11 | 11 | 11 | 10 | 10 | 17 | 11 | 11 | 9 | 11 | 11 | 11.56 | 13 | 10 | 0.774 | -0.369 | 0.50889225 | 0.70257778 | 0 | 0.2918 | 0.1366 | 0.1366 | TRUE | 0.2861 |
| Fam228b | 119 | 187 | 178 | 231 | 200 | 164 | 126 | 187 | 179 | 182 | 215 | 180 | 178.03 | 164 | 192 | 1.169 | 0.226 | 0.25525321 | 0.47357413 | 0.0199 | 0.0254 | 0.0227 | 0.0227 | TRUE | 0.4191 |
| Fam229a | 10 | 5 | 8 | 8 | 8 | 4 | 11 | 5 | 8 | 6 | 9 | 4 | 7.15 | 8 | 6 | 0.817 | -0.291 | 0.68695031 | 0.82743019 | 0 | 0.4674 | 0.236 | 0.236 | TRUE | 0.1994 |
| Fam229b | 83 | 134 | 127 | 160 | 103 | 85 | 88 | 134 | 128 | 126 | 111 | 93 | 113.19 | 117 | 110 | 0.945 | -0.081 | 0.73160967 | 0.85592014 | 0.0265 | 0.036 | 0.0314 | 0.0314 | TRUE | 0.4471 |
| Fam24a | 0 | 0 | 0 | 0 | 0 | 0 | 0 | 0 | 0 | 0 | 0 | 0 | 0 | 0 | 0 | NA | NA | NA | NA | NA | NA | NA | NA | NA | NA |
| Fam25c | 710 | 596 | 505 | 1052 | 743 | 709 | 754 | 595 | 507 | 827 | 799 | 776 | 709.83 | 619 | 801 | 1.295 | 0.373 | 0.01413726 | 0.0678554 | 0.0188 | 0.0115 | 0.0152 | 0.0152 | TRUE | 0.4285 |
| Fam26d | 4 | 1 | 4 | 2 | 2 | 2 | 4 | 1 | 4 | 2 | 2 | 2 | 2.53 | 3 | 2 | 0.634 | -0.658 | 0.58120433 | NA | 0 | 1.3093 | 0.6081 | 0.6081 | TRUE | 0.4773 |
| Fam26e | 53 | 88 | 69 | 184 | 116 | 94 | 56 | 88 | 69 | 145 | 125 | 103 | 97.64 | 71 | 124 | 1.748 | 0.806 | 0.00128533 | 0.01080612 | 0.0269 | 0.0406 | 0.034 | 0.034 | TRUE | 0.2356 |
| Fam26f | 52 | 46 | 47 | 149 | 101 | 65 | 55 | 46 | 47 | 117 | 109 | 71 | 74.22 | 49 | 99 | 2.011 | 1.008 | 0.00029095 | 0.00332771 | 0.0298 | 0.0513 | 0.0405 | 0.0405 | TRUE | 0.5672 |
| Fam32a | 2887 | 2507 | 2328 | 4356 | 3199 | 2704 | 3067 | 2504 | 2338 | 3424 | 3442 | 2960 | 2955.68 | 2636 | 3275 | 1.243 | 0.314 | 0.01202641 | 0.0602013 | 0.0135 | 0.008 | 0.0109 | 0.0109 | TRUE | 0.2474 |
| Fam35a | 1447 | 1668 | 1679 | 1813 | 1200 | 1319 | 1537 | 1666 | 1686 | 1425 | 1291 | 1444 | 1508.15 | 1630 | 1387 | 0.851 | -0.233 | 0.00862753 | 0.04706791 | 0.0023 | 0.0091 | 0.005 | 0.005 | TRUE | 0.0431 |
| Fam3a | 280 | 438 | 452 | 565 | 390 | 453 | 297 | 437 | 454 | 444 | 420 | 496 | 424.73 | 396 | 453 | 1.143 | 0.193 | 0.27965 | 0.5008107 | 0.0265 | 0.0147 | 0.0205 | 0.0205 | TRUE | 0.5435 |
| Fam3b | 74 | 90 | 55 | 160 | 89 | 105 | 79 | 90 | 55 | 126 | 96 | 115 | 93.36 | 75 | 112 | 1.508 | 0.593 | 0.01847052 | 0.08210741 | 0.0257 | 0.0422 | 0.0344 | 0.0344 | TRUE | 0.4761 |
| Fam3c | 1837 | 2141 | 1962 | 2820 | 2146 | 2253 | 1951 | 2138 | 1970 | 2217 | 2309 | 2466 | 2175.27 | 2020 | 2331 | 1.153 | 0.206 | 0.01376678 | 0.0663802 | 0.0021 | 0.0084 | 0.0046 | 0.0046 | TRUE | 0.0319 |
| Fam43a | 841 | 821 | 718 | 1244 | 862 | 773 | 893 | 820 | 721 | 978 | 927 | 846 | 864.29 | 811 | 917 | 1.132 | 0.178 | 0.12819996 | 0.30794544 | 0.0072 | 0.0107 | 0.0087 | 0.0087 | TRUE | 0.113 |
| Fam43b | 107 | 107 | 140 | 108 | 83 | 99 | 114 | 107 | 141 | 85 | 89 | 108 | 107.28 | 121 | 94 | 0.78 | -0.359 | 0.09572696 | 0.25362095 | 0.0103 | 0.0376 | 0.024 | 0.024 | TRUE | 0.22 |
| Fam45a | 1605 | 2073 | 2197 | 2520 | 1684 | 1810 | 1705 | 2070 | 2206 | 1981 | 1812 | 1981 | 1959.23 | 1994 | 1925 | 0.965 | -0.051 | 0.66060546 | 0.80885977 | 0.0096 | 0.0086 | 0.0091 | 0.0091 | TRUE | 0.1938 |
| Fam46a | 111 | 133 | 139 | 217 | 154 | 159 | 118 | 133 | 140 | 171 | 166 | 174 | 150.11 | 130 | 170 | 1.307 | 0.386 | 0.01980444 | 0.08626224 | 0 | 0.0289 | 0.013 | 0.013 | TRUE | 0.0673 |

| | | | | | | | | | | | | | | | | | | | | | | | | | |
|---|---|---|---|---|---|---|---|---|---|---|---|---|---|---|---|---|---|---|---|---|---|---|---|---|---|
| Fam46b | 306 | 244 | 268 | 293 | 220 | 206 | 325 | 244 | 269 | 230 | 237 | 225 | 255.06 | 279 | 231 | 0.827 | -0.274 | 0.07783231 | 0.22065963 | 0.0077 | 0.0198 | 0.0134 | 0.0134 | TRUE | 0.2282 |
| Fam46c | 212 | 270 | 234 | 250 | 207 | 225 | 225 | 270 | 235 | 197 | 223 | 246 | 232.56 | 243 | 222 | 0.909 | -0.138 | 0.38259813 | 0.59692727 | 0.0069 | 0.0211 | 0.0136 | 0.0136 | TRUE | 0.1184 |
| Fam46d | 5 | 5 | 1 | 9 | 5 | 4 | 5 | 5 | 1 | 7 | 5 | 4 | 4.69 | 4 | 5 | 1.506 | 0.59 | 0.5215373 | 0.71165526 | 0 | 0.7093 | 0.3892 | 0.3892 | TRUE | 0.6649 |
| Fam47c | 0 | 0 | 0 | 0 | 0 | 0 | 0 | 0 | 0 | 0 | 0 | 0 | 0 | 0 | 0 | NA | NA | NA | NA | NA | NA | NA | NA | NA | NA |
| Fam47e | 1 | 0 | 1 | 0 | 1 | 3 | 1 | 0 | 1 | 0 | 1 | 3 | 1.07 | 1 | 1 | 2.062 | 1.044 | 0.61619838 | NA | 0 | 3.0829 | 2.0349 | 2.0349 | TRUE | 0.3266 |
| Fam49a | 593 | 488 | 525 | 737 | 538 | 465 | 630 | 487 | 527 | 579 | 579 | 509 | 551.94 | 548 | 556 | 1.015 | 0.021 | 0.87241327 | 0.93737812 | 0.0095 | 0.0129 | 0.0111 | 0.0111 | TRUE | 0.1979 |
| Fam49b | 6326 | 8098 | 7912 | 8612 | 6285 | 6667 | 6719 | 8088 | 7944 | 6770 | 6762 | 7298 | 7263.63 | 7584 | 6943 | 0.915 | -0.127 | 0.18429493 | 0.38709544 | 0.0057 | 0.0074 | 0.0065 | 0.0065 | TRUE | 0.1213 |
| Fam50a | 2126 | 2195 | 2261 | 3164 | 2328 | 2215 | 2258 | 2192 | 2270 | 2487 | 2505 | 2425 | 2356.23 | 2240 | 2472 | 1.104 | 0.142 | 0.02932291 | 0.11318673 | 0 | 0.0083 | 0.0026 | 0.0026 | TRUE | 0.0043 |
| Fam50b | 2 | 6 | 1 | 2 | 0 | 2 | 2 | 6 | 1 | 2 | 0 | 2 | 2.15 | 3 | 1 | 0.416 | -1.267 | 0.38414992 | NA | 0 | 1.5413 | 0.9734 | 0.9734 | TRUE | 0.8501 |
| Fam53a | 1462 | 1393 | 1410 | 2054 | 1361 | 1338 | 1553 | 1391 | 1416 | 1615 | 1464 | 1465 | 1483.93 | 1453 | 1515 | 1.043 | 0.061 | 0.50121662 | 0.69669611 | 0.0028 | 0.0091 | 0.0052 | 0.0052 | TRUE | 0.0431 |
| Fam53b | 1194 | 1358 | 1287 | 1059 | 862 | 917 | 1268 | 1356 | 1292 | 833 | 927 | 1004 | 1113.42 | 1305 | 921 | 0.705 | -0.505 | 5.63E-07 | 1.66E-05 | 0.004 | 0.0099 | 0.0064 | 0.0064 | TRUE | 0.0879 |
| Fam53c | 2582 | 1914 | 1731 | 3411 | 2407 | 1948 | 2743 | 1912 | 1738 | 2681 | 2590 | 2132 | 2299.3 | 2131 | 2468 | 1.159 | 0.212 | 0.22743273 | 0.44072728 | 0.0344 | 0.0083 | 0.0219 | 0.0219 | TRUE | 0.763 |
| Fam57a | 207 | 323 | 344 | 371 | 282 | 297 | 220 | 323 | 345 | 292 | 303 | 325 | 301.34 | 296 | 307 | 1.035 | 0.05 | 0.78997323 | 0.89193627 | 0.0255 | 0.0178 | 0.0217 | 0.0217 | TRUE | 0.5647 |
| Fam57b | 235 | 197 | 195 | 122 | 88 | 110 | 250 | 197 | 196 | 96 | 95 | 120 | 158.86 | 214 | 104 | 0.483 | -1.049 | 6.79E-08 | 2.58E-06 | 0.0126 | 0.0276 | 0.0201 | 0.0201 | TRUE | 0.2287 |
| Fam58b | 846 | 1139 | 1159 | 1402 | 1065 | 1025 | 899 | 1138 | 1164 | 1102 | 1146 | 1122 | 1094.99 | 1067 | 1123 | 1.053 | 0.074 | 0.53618451 | 0.72304225 | 0.009 | 0.0099 | 0.0094 | 0.0094 | TRUE | 0.226 |
| Fam5b | 67 | 42 | 36 | 97 | 64 | 52 | 71 | 42 | 36 | 76 | 69 | 57 | 58.55 | 50 | 67 | 1.36 | 0.444 | 0.17414694 | 0.37335066 | 0.0554 | 0.0632 | 0.0594 | 0.0594 | TRUE | 1.1246 |
| Fam5c | 2 | 0 | 0 | 1 | 0 | 0 | 2 | 0 | 0 | 1 | 0 | 0 | 0.49 | 1 | 0 | 0.497 | -1.01 | 0.76786142 | NA | 0 | 6.7985 | 6.7083 | 6.7083 | TRUE | 0.9742 |
| Fam60a | 927 | 1067 | 1091 | 1016 | 734 | 836 | 985 | 1066 | 1095 | 799 | 790 | 915 | 941.55 | 1049 | 835 | 0.795 | -0.331 | 0.00128364 | 0.0107976 | 0.0034 | 0.0104 | 0.0065 | 0.0065 | TRUE | 0.0832 |
| Fam63a | 1359 | 1458 | 1585 | 2121 | 1563 | 1645 | 1443 | 1456 | 1592 | 1667 | 1682 | 1801 | 1606.82 | 1497 | 1717 | 1.146 | 0.197 | 0.01915859 | 0.08408354 | 0.0017 | 0.009 | 0.0045 | 0.0045 | TRUE | 0.0368 |
| Fam63b | 1108 | 1551 | 1747 | 961 | 832 | 897 | 1177 | 1549 | 1754 | 755 | 895 | 982 | 1185.44 | 1493 | 877 | 0.587 | -0.768 | 3.55E-06 | 8.24E-05 | 0.0263 | 0.0097 | 0.0189 | 0.0189 | TRUE | 0.2757 |
| Fam64a | 2767 | 2983 | 3046 | 3142 | 2352 | 2427 | 2939 | 2979 | 3059 | 2470 | 2531 | 2657 | 2772.34 | 2992 | 2553 | 0.853 | -0.23 | 0.00082977 | 0.00771188 | 5.00E-04 | 0.0081 | 0.003 | 0.003 | TRUE | 0.0154 |
| Fam65a | 2369 | 2563 | 2555 | 3012 | 2266 | 2189 | 2516 | 2560 | 2565 | 2368 | 2438 | 2396 | 2473.93 | 2547 | 2401 | 0.942 | -0.086 | 0.17537107 | 0.37524793 | 0 | 0.0082 | 0.0025 | 0.0025 | TRUE | 0.0022 |
| Fam65b | 92 | 99 | 103 | 74 | 66 | 85 | 98 | 99 | 103 | 58 | 71 | 93 | 87.04 | 100 | 74 | 0.735 | -0.443 | 0.06701736 | 0.19992393 | 0.0149 | 0.0448 | 0.0305 | 0.0305 | TRUE | 0.4478 |
| Fam65c | 7 | 10 | 7 | 19 | 11 | 10 | 7 | 10 | 7 | 15 | 12 | 11 | 10.36 | 8 | 13 | 1.552 | 0.634 | 0.27918676 | 0.50026513 | 0 | 0.3249 | 0.1461 | 0.1461 | TRUE | 0.1373 |
| Fam69a | 436 | 407 | 368 | 658 | 452 | 405 | 463 | 406 | 370 | 517 | 486 | 443 | 447.67 | 413 | 482 | 1.17 | 0.226 | 0.08745341 | 0.23950603 | 0.007 | 0.0143 | 0.0104 | 0.0104 | TRUE | 0.1286 |
| Fam69b | 510 | 673 | 616 | 645 | 559 | 537 | 542 | 672 | 619 | 507 | 601 | 588 | 588.12 | 611 | 565 | 0.924 | -0.114 | 0.37688403 | 0.59206639 | 0.0085 | 0.0125 | 0.0103 | 0.0103 | TRUE | 0.1142 |
| Fam69c | 8 | 4 | 8 | 7 | 0 | 4 | 8 | 4 | 8 | 6 | 0 | 4 | 5.07 | 7 | 3 | 0.494 | -1.018 | 0.31223402 | 0.53255705 | 0.2255 | 0.657 | 0.5122 | 0.5122 | TRUE | 0.9739 |
| Fam71a | 0 | 0 | 0 | 0 | 0 | 0 | 0 | 0 | 0 | 0 | 0 | 0 | 0 | 0 | 0 | NA | NA | NA | NA | NA | NA | NA | NA | NA | NA |
| Fam71b | 0 | 1 | 0 | 2 | 2 | 0 | 0 | 1 | 0 | 2 | 2 | 0 | 0.79 | 0 | 1 | 3.417 | 1.773 | 0.46807648 | NA | 0 | 4.1923 | 2.9081 | 2.9081 | TRUE | 0.405 |
| Fam71d | 0 | 1 | 0 | 2 | 1 | 1 | 0 | 1 | 0 | 2 | 1 | 1 | 0.79 | 0 | 1 | 3.436 | 1.781 | 0.41000155 | NA | 0 | 4.1757 | 1.9771 | 1.9771 | TRUE | 0.405 |
| Fam71e1 | 14 | 18 | 25 | 12 | 12 | 15 | 15 | 18 | 25 | 9 | 13 | 16 | 16.12 | 19 | 13 | 0.66 | -0.599 | 0.23386464 | 0.44850881 | 0 | 0.2113 | 0.1176 | 0.1176 | TRUE | 0.3658 |
| Fam71e2 | 7 | 4 | 4 | 3 | 3 | 2 | 7 | 4 | 4 | 2 | 3 | 2 | 3.87 | 5 | 2 | 0.503 | -0.992 | 0.30281825 | NA | 0 | 0.8581 | 0.3797 | 0.3797 | TRUE | 0.2972 |
| Fam71f1 | 1 | 0 | 2 | 0 | 0 | 2 | 1 | 0 | 2 | 0 | 0 | 2 | 0.88 | 1 | 1 | 0.688 | -0.539 | 0.82569208 | NA | 0 | 3.765 | 3.1169 | 3.1169 | TRUE | 0.5573 |
| Fam71f2 | 9 | 15 | 12 | 15 | 4 | 8 | 10 | 15 | 12 | 12 | 4 | 9 | 10.24 | 12 | 8 | 0.69 | -0.536 | 0.39521914 | 0.60867066 | 0 | 0.3286 | 0.1869 | 0.1869 | TRUE | 0.4034 |
| Fam72a | 485 | 473 | 488 | 582 | 368 | 387 | 515 | 472 | 490 | 458 | 396 | 424 | 459.11 | 492 | 426 | 0.866 | -0.207 | 0.06720269 | 0.20026653 | 0.0015 | 0.0141 | 0.0071 | 0.0071 | TRUE | 0.0525 |
| Fam73a | 3179 | 3882 | 3935 | 4137 | 2961 | 3004 | 3377 | 3877 | 3951 | 3252 | 3186 | 3288 | 3488.53 | 3735 | 3242 | 0.868 | -0.204 | 0.01827498 | 0.08151387 | 0.0032 | 0.0078 | 0.0051 | 0.0051 | TRUE | 0.0855 |
| Fam73b | 684 | 698 | 609 | 885 | 686 | 585 | 727 | 697 | 612 | 696 | 738 | 640 | 684.89 | 679 | 691 | 1.02 | 0.028 | 0.80615033 | 0.90019698 | 0.0049 | 0.0117 | 0.0079 | 0.0079 | TRUE | 0.088 |
| Fam76a | 1631 | 1883 | 1869 | 2419 | 1692 | 1777 | 1732 | 1881 | 1877 | 1902 | 1820 | 1945 | 1859.5 | 1830 | 1889 | 1.032 | 0.046 | 0.55628563 | 0.73896547 | 0.001 | 0.0087 | 0.0038 | 0.0038 | TRUE | 0.0261 |
| Fam76b | 2516 | 2672 | 2733 | 3090 | 2365 | 2430 | 2672 | 2669 | 2744 | 2429 | 2545 | 2660 | 2619.82 | 2695 | 2545 | 0.944 | -0.084 | 0.24159408 | 0.45834215 | 8.00E-04 | 0.0082 | 0.0033 | 0.0033 | TRUE | 0.0196 |
| Fam78a | 538 | 663 | 667 | 637 | 524 | 534 | 571 | 662 | 670 | 501 | 564 | 585 | 592.07 | 634 | 550 | 0.865 | -0.21 | 0.07882322 | 0.2224767 | 0.0055 | 0.0125 | 0.0086 | 0.0086 | TRUE | 0.0881 |
| Fam78b | 13 | 6 | 7 | 29 | 14 | 12 | 14 | 6 | 7 | 23 | 15 | 13 | 12.97 | 9 | 17 | 1.927 | 0.946 | 0.10873406 | 0.27620449 | 0.0569 | 0.2609 | 0.1649 | 0.1649 | TRUE | 0.6917 |
| Fam81a | 2124 | 2569 | 2495 | 2088 | 1616 | 1802 | 2256 | 2566 | 2505 | 1641 | 1739 | 1973 | 2113.28 | 2442 | 1784 | 0.73 | -0.454 | 1.05E-05 | 0.00020784 | 0.0061 | 0.0085 | 0.0072 | 0.0072 | TRUE | 0.1026 |
| Fam83a | 34 | 23 | 13 | 35 | 23 | 19 | 36 | 23 | 13 | 28 | 25 | 21 | 24.2 | 24 | 25 | 1.019 | 0.027 | 0.9534367 | 0.97650213 | 0.0849 | 0.143 | 0.1154 | 0.1154 | TRUE | 0.2717 |
| Fam83b | 62 | 77 | 55 | 119 | 104 | 96 | 66 | 77 | 55 | 94 | 112 | 105 | 84.75 | 66 | 104 | 1.564 | 0.645 | 0.00514224 | 0.03195814 | 0 | 0.0458 | 0.0259 | 0.0259 | TRUE | 0.1886 |
| Fam83c | 2 | 1 | 2 | 1 | 4 | 0 | 2 | 1 | 2 | 1 | 4 | 0 | 1.7 | 2 | 2 | 0.978 | -0.032 | 0.98355619 | NA | 0.3696 | 1.9408 | 1.1991 | 1.1991 | TRUE | 1.2803 |
| Fam83d | 2201 | 2716 | 2533 | 3049 | 2229 | 2213 | 2338 | 2712 | 2543 | 2397 | 2398 | 2422 | 2468.56 | 2531 | 2406 | 0.95 | -0.074 | 0.37484593 | 0.59085248 | 0.0022 | 0.0082 | 0.0045 | 0.0045 | TRUE | 0.054 |
| Fam83e | 5 | 6 | 2 | 7 | 5 | 4 | 5 | 6 | 2 | 6 | 5 | 4 | 4.76 | 4 | 5 | 1.152 | 0.204 | 0.81406993 | 0.90482811 | 0 | 0.6987 | 0.3307 | 0.3307 | TRUE | 0.4265 |
| Fam83f | 59 | 82 | 83 | 83 | 63 | 59 | 63 | 82 | 83 | 65 | 68 | 65 | 70.92 | 76 | 66 | 0.866 | -0.208 | 0.38468995 | 0.59871028 | 0 | 0.0534 | 0.027 | 0.027 | TRUE | 0.2097 |
| Fam83g | 893 | 867 | 746 | 897 | 682 | 650 | 949 | 866 | 749 | 705 | 734 | 712 | 785.66 | 855 | 717 | 0.839 | -0.253 | 0.02753196 | 0.10866719 | 0.0061 | 0.0111 | 0.0082 | 0.0082 | TRUE | 0.1387 |
| Fam83h | 220 | 246 | 217 | 325 | 242 | 189 | 234 | 246 | 218 | 255 | 260 | 207 | 236.67 | 233 | 241 | 1.038 | 0.054 | 0.72370023 | 0.85044852 | 0.0051 | 0.0208 | 0.0127 | 0.0127 | TRUE | 0.1623 |
| Fam84a | 57 | 73 | 47 | 127 | 79 | 71 | 61 | 73 | 47 | 100 | 85 | 78 | 73.87 | 60 | 88 | 1.459 | 0.545 | 0.03592204 | 0.13015446 | 0.0159 | 0.0515 | 0.0345 | 0.0345 | TRUE | 0.2944 |
| Fam84b | 26 | 55 | 48 | 91 | 58 | 56 | 28 | 55 | 48 | 72 | 62 | 61 | 54.33 | 44 | 65 | 1.493 | 0.579 | 0.06690037 | 0.19961269 | 0.0336 | 0.0675 | 0.0526 | 0.0526 | TRUE | 0.7696 |
| Fam86 | 710 | 865 | 774 | 1027 | 791 | 764 | 754 | 864 | 777 | 807 | 851 | 836 | 814.99 | 798 | 831 | 1.041 | 0.058 | 0.55107925 | 0.73520741 | 0.0017 | 0.0109 | 0.0055 | 0.0055 | TRUE | 0.0613 |
| Fam89a | 475 | 437 | 484 | 427 | 315 | 315 | 505 | 436 | 486 | 336 | 339 | 345 | 407.73 | 476 | 340 | 0.714 | -0.485 | 2.70E-05 | 0.00045953 | 0 | 0.015 | 0.0071 | 0.0071 | TRUE | 0.0608 |
| Fam89b | 463 | 438 | 401 | 515 | 357 | 380 | 492 | 437 | 403 | 405 | 384 | 416 | 422.8 | 444 | 402 | 0.905 | -0.144 | 0.24387169 | 0.46043022 | 0.0035 | 0.0147 | 0.0086 | 0.0086 | TRUE | 0.1019 |
| Fam92a | 429 | 689 | 684 | 1124 | 817 | 911 | 456 | 688 | 687 | 884 | 879 | 997 | 765.08 | 610 | 920 | 1.507 | 0.591 | 0.00043869 | 0.00468451 | 0.027 | 0.0112 | 0.019 | 0.019 | TRUE | 0.5738 |
| Fam92b | 4 | 4 | 4 | 7 | 4 | 7 | 4 | 4 | 4 | 6 | 4 | 8 | 4.95 | 4 | 6 | 1.421 | 0.507 | 0.54032599 | 0.72632929 | 0 | 0.6718 | 0.2856 | 0.2856 | TRUE | 0.1539 |
| Fam96a | 1324 | 1472 | 1413 | 2178 | 1523 | 1429 | 1406 | 1470 | 1419 | 1712 | 1639 | 1564 | 1535.05 | 1432 | 1638 | 1.145 | 0.195 | 0.01250465 | 0.06198093 | 8.00E-04 | 0.009 | 0.0038 | 0.0038 | TRUE | 0.0193 |
| Fam96b | 1861 | 2125 | 2210 | 3246 | 2198 | 2137 | 1977 | 2122 | 2219 | 2552 | 2365 | 2339 | 2262.33 | 2106 | 2419 | 1.149 | 0.2 | 0.01684451 | 0.07693937 | 0.0022 | 0.0084 | 0.0046 | 0.0046 | TRUE | 0.0348 |
| Fam98a | 5034 | 4514 | 4234 | 5068 | 3711 | 3816 | 5347 | 4508 | 4251 | 3984 | 3993 | 4177 | 4376.77 | 4702 | 4051 | 0.862 | -0.215 | 0.03779547 | 0.13505193 | 0.0074 | 0.0077 | 0.0075 | 0.0075 | TRUE | 0.1751 |
| Fam98b | 1608 | 1561 | 1454 | 2494 | 1705 | 1541 | 1708 | 1559 | 1460 | 1961 | 1834 | 1687 | 1701.48 | 1576 | 1827 | 1.161 | 0.215 | 0.03374219 | 0.1248667 | 0.0051 | 0.0088 | 0.0068 | 0.0068 | TRUE | 0.0648 |
| Fam98c | 644 | 740 | 742 | 957 | 743 | 593 | 684 | 739 | 745 | 752 | 799 | 649 | 728.17 | 723 | 733 | 1.015 | 0.022 | 0.84587441 | 0.92275081 | 0.0052 | 0.0114 | 0.0079 | 0.0079 | TRUE | 0.1169 |
| Fan1 | 535 | 519 | 509 | 695 | 518 | 451 | 568 | 518 | 511 | 546 | 557 | 494 | 532.51 | 532 | 532 | 1.001 | 0.001 | 0.99199943 | 0.99591438 | 0.0017 | 0.0131 | 0.0067 | 0.0067 | TRUE | 0.0458 |
| Fanca | 5417 | 5892 | 6137 | 7037 | 5485 | 5292 | 5754 | 5884 | 6162 | 5532 | 5902 | 5793 | 5837.8 | 5933 | 5742 | 0.968 | -0.048 | 0.47677193 | 0.67647057 | 9.00E-04 | 0.0075 | 0.0031 | 0.0031 | TRUE | 0.0139 |
| Fancb | 837 | 783 | 757 | 1061 | 719 | 655 | 889 | 782 | 760 | 834 | 774 | 717 | 792.64 | 810 | 775 | 0.958 | -0.063 | 0.57475868 | 0.75283624 | 0.0052 | 0.0111 | 0.0077 | 0.0077 | TRUE | 0.0851 |
| Fancc | 808 | 971 | 932 | 995 | 793 | 791 | 858 | 970 | 936 | 782 | 853 | 866 | 877.52 | 921 | 834 | 0.904 | -0.146 | 0.13636212 | 0.31988395 | 0.0024 | 0.0107 | 0.0058 | 0.0058 | TRUE | 0.0424 |
| Fancd2 | 5775 | 7311 | 7595 | 8427 | 6419 | 6773 | 6134 | 7302 | 7626 | 6625 | 6906 | 7414 | 7001.17 | 7021 | 6982 | 0.994 | -0.008 | 0.93730288 | 0.96864531 | 0.008 | 0.0074 | 0.0077 | 0.0077 | TRUE | 0.1488 |
| Fance | 2109 | 2329 | 2346 | 2439 | 1863 | 1864 | 2240 | 2326 | 2356 | 1917 | 2004 | 2040 | 2147.34 | 2307 | 1987 | 0.861 | -0.216 | 0.00230762 | 0.01724919 | 4.00E-04 | 0.0084 | 0.0032 | 0.0032 | TRUE | 0.0119 |
| Fancf | 292 | 317 | 344 | 504 | 407 | 324 | 310 | 317 | 345 | 396 | 438 | 355 | 360.16 | 324 | 396 | 1.223 | 0.29 | 0.02811854 | 0.11006223 | 0.0043 | 0.0161 | 0.0098 | 0.0098 | TRUE | 0.0931 |

| | | | | | | | | | | | | | | | | | | | | | | | | |
|---|---|---|---|---|---|---|---|---|---|---|---|---|---|---|---|---|---|---|---|---|---|---|---|---|
| Fancg | 803 | 822 | 823 | 1103 | 800 | 804 | 853 | 821 | 826 | 867 | 861 | 880 | 851.37 | 833 | 869 | 1.043 | 0.061 | 0.46001525 | 0.66410622 | 0 | 0.0108 | 0.0037 | 0.0037 | TRUE | 0.0049 |
| Fanci | 3480 | 4090 | 4243 | 5223 | 4110 | 4312 | 3696 | 4085 | 4260 | 4106 | 4422 | 4720 | 4214.95 | 4014 | 4416 | 1.1 | 0.137 | 0.13808689 | 0.32245868 | 0.0046 | 0.0077 | 0.0059 | 0.0059 | TRUE | 0.0582 |
| Fancl | 1966 | 2004 | 1958 | 3061 | 2179 | 2028 | 2088 | 2001 | 1966 | 2406 | 2344 | 2220 | 2171.07 | 2018 | 2323 | 1.152 | 0.204 | 0.0061695 | 0.03677448 | 9.00E-04 | 0.0084 | 0.0035 | 0.0035 | TRUE | 0.0183 |
| Fancm | 4454 | 3361 | 3562 | 6074 | 4243 | 3637 | 4731 | 3357 | 3577 | 4775 | 4565 | 3981 | 4164.27 | 3888 | 4440 | 1.142 | 0.192 | 0.18162108 | 0.38350003 | 0.0215 | 0.0077 | 0.0146 | 0.0146 | TRUE | 0.4373 |
| Fank1 | 203 | 285 | 223 | 332 | 269 | 308 | 216 | 285 | 224 | 261 | 289 | 337 | 268.62 | 242 | 296 | 1.223 | 0.29 | 0.09274071 | 0.24873893 | 0.0163 | 0.0192 | 0.0177 | 0.0177 | TRUE | 0.2734 |
| Fap | 0 | 0 | 0 | 0 | 1 | 1 | 0 | 0 | 0 | 0 | 1 | 1 | 0.36 | 0 | 1 | 3.794 | 1.924 | 0.60247382 | NA | 0 | 9.1135 | 8.1425 | 8.1425 | TRUE | 0.3482 |
| Far1 | 5538 | 5714 | 5504 | 6480 | 4797 | 4938 | 5882 | 5707 | 5527 | 5094 | 5161 | 5405 | 5462.72 | 5705 | 5220 | 0.915 | -0.128 | 0.04961788 | 0.16342811 | 7.00E-04 | 0.0075 | 0.0029 | 0.0029 | TRUE | 0.0117 |
| Far2 | 10 | 5 | 3 | 18 | 11 | 8 | 11 | 5 | 3 | 14 | 12 | 9 | 8.9 | 6 | 12 | 1.888 | 0.917 | 0.19309259 | 0.39859714 | 0.0438 | 0.3773 | 0.2327 | 0.2327 | TRUE | 0.8883 |
| Farp1 | 1676 | 2023 | 2138 | 2295 | 1796 | 1703 | 1780 | 2020 | 2147 | 1804 | 1932 | 1864 | 1924.69 | 1982 | 1867 | 0.941 | -0.087 | 0.36595624 | 0.5833699 | 0.0042 | 0.0086 | 0.0062 | 0.0062 | TRUE | 0.096 |
| Farp2 | 146 | 144 | 146 | 156 | 118 | 106 | 155 | 144 | 147 | 123 | 127 | 116 | 135.19 | 149 | 122 | 0.821 | -0.284 | 0.09263975 | 0.24858829 | 0 | 0.0313 | 0.0131 | 0.0131 | TRUE | 0.0163 |
| Fars2 | 929 | 1008 | 1030 | 1257 | 902 | 896 | 987 | 1007 | 1034 | 988 | 970 | 981 | 994.53 | 1009 | 980 | 0.971 | -0.043 | 0.59107159 | 0.76481919 | 0 | 0.0102 | 0.0035 | 0.0035 | TRUE | 0.0056 |
| Farsa | 6705 | 6482 | 6420 | 7858 | 5900 | 5340 | 7122 | 6474 | 6446 | 6177 | 6348 | 5845 | 6402.12 | 6681 | 6123 | 0.917 | -0.125 | 0.10219962 | 0.26550691 | 0.0023 | 0.0074 | 0.0041 | 0.0041 | TRUE | 0.0407 |
| Farsb | 7352 | 7420 | 7375 | 9283 | 6570 | 6654 | 7809 | 7410 | 7405 | 7298 | 7069 | 7284 | 7379.19 | 7541 | 7217 | 0.957 | -0.063 | 0.29575168 | 0.5165093 | 4.00E-04 | 0.0073 | 0.0025 | 0.0025 | TRUE | 0.0117 |
| Fas | 42 | 55 | 35 | 48 | 31 | 45 | 45 | 55 | 35 | 38 | 33 | 49 | 42.51 | 45 | 40 | 0.891 | -0.166 | 0.61266735 | 0.78070375 | 0.0203 | 0.0844 | 0.0538 | 0.0538 | TRUE | 0.1897 |
| Fasl | 2 | 0 | 0 | 0 | 0 | 0 | 2 | 0 | 0 | 0 | 0 | 0 | 0.35 | 1 | 0 | 0.254 | -1.975 | 0.62446185 | NA | 10 | 9.3117 | 10 | 10 | TRUE | 0.9741 |
| Fasn | 59140 | 45649 | 49557 | 62124 | 43193 | 41106 | 62817 | 45590 | 49760 | 48837 | 46473 | 44997 | 49745.78 | 52722 | 46769 | 0.887 | -0.173 | 0.16591689 | 0.36299641 | 0.0154 | 0.007 | 0.0112 | 0.0112 | TRUE | 0.3435 |
| Fastk | 1527 | 2473 | 2483 | 2818 | 2329 | 2321 | 1622 | 2470 | 2493 | 2215 | 2506 | 2541 | 2307.8 | 2195 | 2421 | 1.102 | 0.141 | 0.39468913 | 0.60827017 | 0.0306 | 0.0083 | 0.0193 | 0.0193 | TRUE | 0.6306 |
| Fastkd1 | 3064 | 3228 | 3257 | 3464 | 2540 | 2617 | 3255 | 3224 | 3270 | 2723 | 2733 | 2865 | 3011.57 | 3250 | 2774 | 0.853 | -0.229 | 0.0003352 | 0.00373354 | 0 | 0.008 | 0.0026 | 0.0026 | TRUE | 0.0099 |
| Fastkd2 | 1678 | 1896 | 1867 | 2800 | 1989 | 1964 | 1782 | 1894 | 1875 | 2201 | 2140 | 2150 | 2006.94 | 1850 | 2164 | 1.17 | 0.226 | 0.00121295 | 0.01032972 | 0 | 0.0085 | 0.003 | 0.003 | TRUE | 0.0124 |
| Fastkd3 | 1100 | 854 | 754 | 1231 | 899 | 774 | 1168 | 853 | 757 | 968 | 967 | 847 | 926.77 | 926 | 927 | 1.002 | 0.003 | 0.98670039 | 0.9928109 | 0.0274 | 0.0105 | 0.0189 | 0.0189 | TRUE | 0.6226 |
| Fastkd5 | 1101 | 804 | 749 | 1259 | 880 | 787 | 1169 | 803 | 752 | 990 | 947 | 861 | 920.42 | 908 | 933 | 1.028 | 0.039 | 0.82003498 | 0.90793905 | 0.0306 | 0.0105 | 0.0204 | 0.0204 | TRUE | 0.7529 |
| Fat1 | 10297 | 8534 | 10262 | 13496 | 9133 | 8226 | 10937 | 8523 | 10304 | 10609 | 9827 | 9005 | 9867.51 | 9921 | 9814 | 0.989 | -0.016 | 0.89313615 | 0.94834594 | 0.0115 | 0.0072 | 0.0096 | 0.0096 | TRUE | 0.1858 |
| Fat2 | 52 | 36 | 42 | 75 | 54 | 39 | 55 | 36 | 42 | 59 | 58 | 43 | 48.85 | 44 | 53 | 1.204 | 0.268 | 0.38321033 | 0.59760644 | 0.0169 | 0.0743 | 0.0472 | 0.0472 | TRUE | 0.3417 |
| Fat3 | 31 | 30 | 28 | 76 | 56 | 32 | 33 | 30 | 28 | 60 | 60 | 35 | 41.01 | 30 | 52 | 1.712 | 0.776 | 0.02365316 | 0.09783886 | 0.0284 | 0.0872 | 0.0584 | 0.0584 | TRUE | 0.6136 |
| Fat4 | 0 | 2 | 3 | 2 | 1 | 1 | 0 | 2 | 3 | 2 | 1 | 1 | 1.46 | 2 | 1 | 0.75 | -0.414 | 0.79550055 | NA | 0 | 2.2653 | 1.1497 | 1.1497 | TRUE | 0.3908 |
| Fate1 | 0 | 0 | 0 | 0 | 0 | 0 | 0 | 0 | 0 | 0 | 0 | 0 | 0 | 0 | 0 | NA | NA | NA | NA | NA | NA | NA | NA | NA | NA |
| Fau | 10934 | 15008 | 15704 | 16681 | 12639 | 13675 | 11614 | 14989 | 15768 | 13113 | 13599 | 14969 | 14008.74 | 14124 | 13894 | 0.984 | -0.024 | 0.85137568 | 0.92587032 | 0.0118 | 0.0071 | 0.0115 | 0.0115 | TRUE | 0.2954 |
| Faxc | 720 | 880 | 987 | 987 | 790 | 827 | 765 | 879 | 991 | 776 | 850 | 905 | 860.98 | 878 | 844 | 0.96 | -0.059 | 0.63955432 | 0.79689135 | 0.0102 | 0.0107 | 0.0104 | 0.0104 | TRUE | 0.1512 |
| Fbf1 | 1738 | 2039 | 2027 | 2549 | 2013 | 1814 | 1846 | 2036 | 2035 | 2004 | 2166 | 1986 | 2012.19 | 1972 | 2052 | 1.04 | 0.056 | 0.50197011 | 0.69723229 | 0.002 | 0.0085 | 0.0046 | 0.0046 | TRUE | 0.0379 |
| Fbl | 8083 | 7751 | 7830 | 10951 | 7863 | 7509 | 8586 | 7741 | 7862 | 8609 | 8460 | 8220 | 8246.24 | 8063 | 8430 | 1.046 | 0.064 | 0.36850951 | 0.58584135 | 0.0017 | 0.0073 | 0.0036 | 0.0036 | TRUE | 0.0392 |
| Fblim1 | 7809 | 8319 | 8403 | 9026 | 7066 | 7298 | 8295 | 8308 | 8438 | 7096 | 7603 | 7989 | 7954.54 | 8347 | 7563 | 0.906 | -0.143 | 0.04563185 | 0.15441011 | 0.0016 | 0.0073 | 0.0035 | 0.0035 | TRUE | 0.0359 |
| Fbll1 | 9 | 13 | 6 | 16 | 13 | 3 | 10 | 13 | 6 | 13 | 14 | 3 | 9.74 | 10 | 10 | 1.053 | 0.075 | 0.91642382 | 0.95892234 | 0.1712 | 0.3453 | 0.2656 | 0.2656 | TRUE | 0.617 |
| Fbln1 | 2216 | 3224 | 3399 | 2194 | 1785 | 2359 | 2354 | 3220 | 3413 | 1725 | 1921 | 2582 | 2535.7 | 2996 | 2076 | 0.693 | -0.53 | 0.00369169 | 0.02484276 | 0.0405 | 0.0082 | 0.0236 | 0.0236 | TRUE | 0.5503 |
| Fbln2 | 306 | 333 | 328 | 396 | 262 | 317 | 325 | 333 | 329 | 311 | 282 | 347 | 321.19 | 329 | 313 | 0.952 | -0.071 | 0.58760145 | 0.76226286 | 0.0016 | 0.0172 | 0.0091 | 0.0091 | TRUE | 0.0942 |
| Fbln5 | 8 | 7 | 8 | 23 | 21 | 7 | 8 | 7 | 8 | 18 | 23 | 8 | 11.98 | 8 | 16 | 2.065 | 1.046 | 0.09258951 | 0.2485231 | 0 | 0.282 | 0.1832 | 0.1832 | TRUE | 0.3583 |
| Fbln7 | 20 | 19 | 27 | 25 | 17 | 21 | 21 | 19 | 27 | 20 | 18 | 23 | 21.38 | 22 | 20 | 0.903 | -0.147 | 0.72203206 | 0.8495889 | 0 | 0.161 | 0.0758 | 0.0758 | TRUE | 0.1933 |
| Fbn1 | 678 | 613 | 694 | 1088 | 637 | 533 | 720 | 612 | 697 | 855 | 685 | 583 | 692.22 | 676 | 708 | 1.049 | 0.069 | 0.66335108 | 0.81128824 | 0.0208 | 0.0117 | 0.0164 | 0.0164 | TRUE | 0.3741 |
| Fbn2 | 427 | 413 | 454 | 551 | 375 | 304 | 454 | 412 | 456 | 433 | 403 | 333 | 415.21 | 441 | 390 | 0.887 | -0.173 | 0.20680003 | 0.41592069 | 0.008 | 0.0148 | 0.0112 | 0.0112 | TRUE | 0.1846 |
| Fbp1 | 5 | 15 | 13 | 23 | 11 | 6 | 5 | 15 | 13 | 18 | 12 | 7 | 11.64 | 11 | 12 | 1.106 | 0.145 | 0.82515645 | 0.91134777 | 0.153 | 0.29 | 0.2247 | 0.2247 | TRUE | 0.2352 |
| Fbp2 | 45 | 39 | 41 | 95 | 55 | 46 | 48 | 39 | 41 | 75 | 59 | 50 | 52.02 | 43 | 61 | 1.451 | 0.537 | 0.06471548 | 0.19598551 | 0 | 0.0702 | 0.0409 | 0.0409 | TRUE | 0.3471 |
| Fbrs | 1066 | 1054 | 1045 | 1338 | 1061 | 988 | 1132 | 1053 | 1049 | 1052 | 1142 | 1082 | 1084.86 | 1078 | 1092 | 1.012 | 0.017 | 0.83964236 | 0.91868465 | 9.00E-04 | 0.0099 | 0.0044 | 0.0044 | TRUE | 0.023 |
| Fbrsl1 | 1835 | 1820 | 1640 | 2238 | 1854 | 1637 | 1949 | 1818 | 1647 | 1759 | 1995 | 1792 | 1826.59 | 1805 | 1849 | 1.024 | 0.035 | 0.73026458 | 0.85505616 | 0.0053 | 0.0087 | 0.0067 | 0.0067 | TRUE | 0.0707 |
| Fbxl12 | 438 | 794 | 748 | 622 | 517 | 586 | 465 | 793 | 751 | 489 | 556 | 641 | 616 | 670 | 562 | 0.839 | -0.254 | 0.21743937 | 0.4294633 | 0.0474 | 0.0123 | 0.0289 | 0.0289 | TRUE | 0.8387 |
| Fbxl13 | 7 | 17 | 11 | 9 | 5 | 7 | 7 | 17 | 11 | 7 | 5 | 8 | 9.26 | 12 | 7 | 0.567 | -0.818 | 0.21122324 | 0.42189873 | 0 | 0.3626 | 0.1929 | 0.1929 | TRUE | 0.2491 |
| Fbxl14 | 2183 | 2779 | 3071 | 3560 | 2786 | 2975 | 2319 | 2775 | 3084 | 2799 | 2998 | 3257 | 2871.75 | 2726 | 3018 | 1.107 | 0.146 | 0.23567269 | 0.45087696 | 0.0123 | 0.0081 | 0.0106 | 0.0106 | TRUE | 0.2068 |
| Fbxl15 | 158 | 137 | 120 | 237 | 170 | 136 | 168 | 137 | 120 | 186 | 183 | 149 | 157.21 | 142 | 173 | 1.222 | 0.289 | 0.13969026 | 0.32423765 | 0.0144 | 0.0279 | 0.0212 | 0.0212 | TRUE | 0.2657 |
| Fbxl16 | 244 | 237 | 222 | 429 | 315 | 302 | 259 | 237 | 223 | 337 | 339 | 331 | 287.59 | 240 | 336 | 1.402 | 0.487 | 0.00014274 | 0.00183038 | 0 | 0.0184 | 0.0082 | 0.0082 | TRUE | 0.0556 |
| Fbxl17 | 371 | 402 | 470 | 702 | 602 | 510 | 394 | 401 | 472 | 552 | 648 | 558 | 504.22 | 422 | 586 | 1.385 | 0.47 | 0.00026235 | 0.00304916 | 0.007 | 0.0134 | 0.0099 | 0.0099 | TRUE | 0.1214 |
| Fbxl18 | 1037 | 880 | 855 | 1222 | 938 | 769 | 1101 | 879 | 859 | 961 | 1009 | 842 | 941.75 | 946 | 937 | 0.991 | -0.013 | 0.92153131 | 0.96126008 | 0.0129 | 0.0104 | 0.0117 | 0.0117 | TRUE | 0.2446 |
| Fbxl19 | 4113 | 4128 | 4012 | 5563 | 4074 | 3602 | 4369 | 4123 | 4028 | 4373 | 4383 | 3943 | 4203.23 | 4173 | 4233 | 1.015 | 0.021 | 0.79245914 | 0.89309946 | 0.0024 | 0.0077 | 0.0043 | 0.0043 | TRUE | 0.0435 |
| Fbxl2 | 463 | 508 | 527 | 503 | 360 | 443 | 492 | 507 | 529 | 395 | 387 | 485 | 466 | 509 | 422 | 0.828 | -0.272 | 0.03376431 | 0.12491925 | 0.0056 | 0.014 | 0.0097 | 0.0097 | TRUE | 0.1877 |
| Fbxl20 | 3330 | 4310 | 4434 | 4130 | 3199 | 3581 | 3537 | 4304 | 4452 | 3247 | 3442 | 3920 | 3817.04 | 4098 | 3536 | 0.863 | -0.213 | 0.07524605 | 0.21609262 | 0.0116 | 0.0078 | 0.0101 | 0.0101 | TRUE | 0.1743 |
| Fbxl21 | 57 | 45 | 52 | 89 | 80 | 78 | 61 | 45 | 52 | 70 | 86 | 85 | 66.52 | 53 | 80 | 1.527 | 0.61 | 0.01609136 | 0.07471091 | 0 | 0.0564 | 0.0305 | 0.0305 | TRUE | 0.1416 |
| Fbxl22 | 1 | 1 | 1 | 5 | 6 | 1 | 1 | 1 | 1 | 4 | 6 | 1 | 2.42 | 1 | 4 | 3.755 | 1.909 | 0.16097148 | NA | 0 | 1.3658 | 0.7082 | 0.7082 | TRUE | 0.0864 |
| Fbxl3 | 1656 | 2563 | 2474 | 2083 | 1780 | 1984 | 1759 | 2560 | 2484 | 1637 | 1915 | 2172 | 2087.88 | 2268 | 1908 | 0.841 | -0.25 | 0.1301006 | 0.31070082 | 0.0281 | 0.0085 | 0.0191 | 0.0191 | TRUE | 0.4657 |
| Fbxl4 | 483 | 507 | 536 | 691 | 466 | 463 | 513 | 506 | 538 | 543 | 501 | 507 | 518.17 | 519 | 517 | 0.997 | -0.004 | 0.96975341 | 0.98448447 | 0 | 0.0133 | 0.0055 | 0.0055 | TRUE | 0.0241 |
| Fbxl5 | 4625 | 5322 | 5173 | 6007 | 4378 | 4257 | 4913 | 5315 | 5194 | 4722 | 4710 | 4660 | 4919.1 | 5141 | 4697 | 0.914 | -0.13 | 0.04404106 | 0.15049954 | 5.00E-04 | 0.0076 | 0.0028 | 0.0028 | TRUE | 0.0183 |
| Fbxl6 | 1497 | 1949 | 1907 | 2036 | 1610 | 1536 | 1590 | 1946 | 1915 | 1601 | 1732 | 1681 | 1744.27 | 1817 | 1671 | 0.919 | -0.121 | 0.24822744 | 0.4649695 | 0.0058 | 0.0088 | 0.0074 | 0.0074 | TRUE | 0.1438 |
| Fbxl7 | 20 | 10 | 14 | 39 | 26 | 13 | 21 | 10 | 14 | 31 | 28 | 14 | 19.69 | 15 | 24 | 1.624 | 0.7 | 0.17185874 | 0.37064898 | 0.0913 | 0.1742 | 0.135 | 0.135 | TRUE | 0.7336 |
| Fbxl8 | 138 | 134 | 138 | 145 | 98 | 115 | 147 | 134 | 139 | 114 | 105 | 126 | 127.38 | 140 | 115 | 0.824 | -0.279 | 0.11750606 | 0.29007757 | 0 | 0.0328 | 0.015 | 0.015 | TRUE | 0.0619 |
| Fbxo10 | 546 | 555 | 547 | 837 | 678 | 642 | 580 | 554 | 549 | 658 | 729 | 703 | 628.95 | 561 | 697 | 1.24 | 0.311 | 0.00152267 | 0.01239195 | 0 | 0.0121 | 0.0053 | 0.0053 | TRUE | 0.0295 |
| Fbxo11 | 2279 | 2427 | 2349 | 3180 | 2356 | 2252 | 2421 | 2424 | 2359 | 2500 | 2535 | 2465 | 2450.53 | 2401 | 2500 | 1.041 | 0.058 | 0.36141677 | 0.57982558 | 0 | 0.0082 | 0.0025 | 0.0025 | TRUE | 0.0029 |
| Fbxo15 | 10316 | 9665 | 8500 | 23227 | 16618 | 14233 | 10957 | 9653 | 8535 | 18259 | 17880 | 15580 | 13477.38 | 9715 | 17240 | 1.775 | 0.828 | 6.75E-13 | 7.05E-11 | 0.0112 | 0.0071 | 0.0095 | 0.0095 | TRUE | 0.1529 |
| Fbxo16 | 220 | 236 | 241 | 226 | 181 | 152 | 234 | 236 | 242 | 178 | 195 | 166 | 208.36 | 237 | 180 | 0.757 | -0.401 | 0.00539941 | 0.03309738 | 0 | 0.0227 | 0.0101 | 0.0101 | TRUE | 0.0545 |
| Fbxo17 | 336 | 288 | 283 | 414 | 296 | 268 | 357 | 288 | 284 | 325 | 318 | 293 | 311 | 310 | 312 | 1.011 | 0.016 | 0.91332925 | 0.95732555 | 0.0066 | 0.0175 | 0.0117 | 0.0117 | TRUE | 0.2003 |
| Fbxo18 | 3960 | 5008 | 5179 | 5198 | 3841 | 3964 | 4206 | 5002 | 5200 | 4086 | 4133 | 4339 | 4494.36 | 4803 | 4186 | 0.871 | -0.198 | 0.04712862 | 0.15766097 | 0.0062 | 0.0076 | 0.007 | 0.007 | TRUE | 0.1436 |
| Fbxo2 | 2476 | 2199 | 2088 | 1631 | 1344 | 1281 | 2630 | 2196 | 2097 | 1282 | 1446 | 1402 | 1842.19 | 2308 | 1377 | 0.596 | -0.746 | 5.80E-11 | 4.44E-09 | 0.0086 | 0.0087 | 0.0088 | 0.0088 | TRUE | 0.1806 |
| Fbxo21 | 4763 | 4804 | 4779 | 5464 | 4126 | 4255 | 5059 | 4798 | 4799 | 4295 | 4439 | 4658 | 4674.67 | 4885 | 4464 | 0.914 | -0.13 | 0.05888221 | 0.18363368 | 0.001 | 0.0076 | 0.0032 | 0.0032 | TRUE | 0.0175 |
| Fbxo22 | 5793 | 5962 | 6128 | 6378 | 4696 | 5329 | 6153 | 5954 | 6153 | 5014 | 5053 | 5833 | 5693.43 | 6087 | 5300 | 0.871 | -0.2 | 0.0192979 | 0.0845862 | 0.0037 | 0.0075 | 0.0051 | 0.0051 | TRUE | 0.0942 |

| | | | | | | | | | | | | | | | | | | | | | | | | | |
|---|---|---|---|---|---|---|---|---|---|---|---|---|---|---|---|---|---|---|---|---|---|---|---|---|---|
| Fbxo24 | 38 | 41 | 43 | 58 | 51 | 43 | 40 | 41 | 43 | 46 | 55 | 47 | 45.34 | 41 | 49 | 1.182 | 0.241 | 0.39266369 | 0.6062924 | 0 | 0.0796 | 0.035 | 0.035 | TRUE | 0.0795 |
| Fbxo25 | 1554 | 1581 | 1609 | 1866 | 1384 | 1369 | 1651 | 1579 | 1616 | 1467 | 1489 | 1499 | 1549.96 | 1615 | 1485 | 0.919 | -0.121 | 0.08641316 | 0.23735777 | 0 | 0.009 | 0.003 | 0.003 | TRUE | 0.0046 |
| Fbxo27 | 160 | 245 | 278 | 321 | 278 | 251 | 170 | 245 | 279 | 252 | 299 | 275 | 253.33 | 231 | 275 | 1.189 | 0.249 | 0.21208692 | 0.42299379 | 0.0299 | 0.0199 | 0.0248 | 0.0248 | TRUE | 0.5866 |
| Fbxo28 | 3307 | 3616 | 3559 | 3958 | 2837 | 2990 | 3513 | 3611 | 3574 | 3111 | 3052 | 3273 | 3355.75 | 3566 | 3145 | 0.882 | -0.181 | 0.00598947 | 0.03587769 | 3.00E-04 | 0.0079 | 0.0028 | 0.0028 | TRUE | 0.015 |
| Fbxo3 | 1152 | 1631 | 1712 | 1965 | 1443 | 1636 | 1224 | 1629 | 1719 | 1545 | 1553 | 1791 | 1576.62 | 1524 | 1630 | 1.069 | 0.096 | 0.50238894 | 0.69762972 | 0.0189 | 0.009 | 0.0141 | 0.0141 | TRUE | 0.3569 |
| Fbxo30 | 2411 | 2422 | 2401 | 3145 | 2308 | 2321 | 2561 | 2419 | 2411 | 2472 | 2483 | 2541 | 2481.16 | 2464 | 2499 | 1.014 | 0.02 | 0.7637941 | 0.87579502 | 3.00E-04 | 0.0082 | 0.0029 | 0.0029 | TRUE | 0.0144 |
| Fbxo31 | 2918 | 3218 | 3157 | 3574 | 2836 | 2797 | 3099 | 3214 | 3170 | 2810 | 3051 | 3062 | 3067.66 | 3161 | 2974 | 0.94 | -0.089 | 0.21670795 | 0.42872021 | 0.0011 | 0.008 | 0.0034 | 0.0034 | TRUE | 0.0291 |
| Fbxo32 | 190 | 77 | 94 | 131 | 73 | 83 | 202 | 77 | 94 | 103 | 79 | 91 | 107.58 | 124 | 91 | 0.732 | -0.45 | 0.20897765 | 0.41875181 | 0.1341 | 0.0375 | 0.083 | 0.083 | TRUE | 2.7328 |
| Fbxo33 | 1259 | 1129 | 1065 | 1752 | 1305 | 1108 | 1337 | 1128 | 1069 | 1377 | 1404 | 1213 | 1254.74 | 1178 | 1331 | 1.131 | 0.177 | 0.13447211 | 0.31711933 | 0.009 | 0.0095 | 0.0093 | 0.0093 | TRUE | 0.167 |
| Fbxo34 | 1594 | 1556 | 1589 | 2211 | 1622 | 1564 | 1693 | 1554 | 1596 | 1738 | 1745 | 1712 | 1672.99 | 1614 | 1732 | 1.073 | 0.102 | 0.17408852 | 0.37335066 | 3.00E-04 | 0.0089 | 0.0034 | 0.0034 | TRUE | 0.0219 |
| Fbxo36 | 67 | 103 | 117 | 193 | 136 | 113 | 71 | 103 | 117 | 152 | 146 | 124 | 118.88 | 97 | 141 | 1.447 | 0.533 | 0.0220246 | 0.09330433 | 0.0255 | 0.0346 | 0.0303 | 0.0303 | TRUE | 0.5206 |
| Fbxo38 | 4588 | 4796 | 5075 | 6044 | 4322 | 4596 | 4873 | 4790 | 5096 | 4751 | 4650 | 5031 | 4865.25 | 4920 | 4811 | 0.978 | -0.032 | 0.63922841 | 0.79669776 | 0.001 | 0.0076 | 0.0032 | 0.0032 | TRUE | 0.0193 |
| Fbxo39 | 24 | 33 | 21 | 31 | 21 | 16 | 25 | 33 | 21 | 24 | 23 | 18 | 24 | 26 | 22 | 0.815 | -0.295 | 0.46508139 | 0.66788976 | 0 | 0.1442 | 0.0759 | 0.0759 | TRUE | 0.2886 |
| Fbxo4 | 45 | 53 | 63 | 71 | 44 | 53 | 48 | 53 | 63 | 56 | 47 | 58 | 54.19 | 55 | 54 | 0.983 | -0.024 | 0.92801966 | 0.96471962 | 0 | 0.0677 | 0.0341 | 0.0341 | TRUE | 0.1589 |
| Fbxo40 | 4 | 2 | 1 | 4 | 4 | 2 | 4 | 2 | 1 | 3 | 4 | 2 | 2.81 | 2 | 3 | 1.338 | 0.42 | 0.7108148 | NA | 0 | 1.1774 | 0.5581 | 0.5581 | TRUE | 0.4446 |
| Fbxo41 | 181 | 285 | 257 | 243 | 185 | 205 | 192 | 285 | 258 | 191 | 199 | 224 | 224.9 | 245 | 205 | 0.834 | -0.261 | 0.15910498 | 0.35313872 | 0.0192 | 0.0216 | 0.0203 | 0.0203 | TRUE | 0.3887 |
| Fbxo42 | 4372 | 4348 | 4292 | 5488 | 4147 | 3859 | 4644 | 4342 | 4310 | 4314 | 4462 | 4224 | 4382.71 | 4432 | 4333 | 0.978 | -0.032 | 0.63747204 | 0.79603827 | 9.00E-04 | 0.0077 | 0.0032 | 0.0032 | TRUE | 0.0212 |
| Fbxo43 | 64 | 111 | 99 | 59 | 52 | 62 | 68 | 111 | 99 | 46 | 56 | 68 | 74.74 | 93 | 57 | 0.608 | -0.719 | 0.01135778 | 0.05786997 | 0.0366 | 0.051 | 0.0439 | 0.0439 | TRUE | 0.5148 |
| Fbxo44 | 255 | 313 | 316 | 257 | 212 | 237 | 271 | 313 | 317 | 202 | 228 | 259 | 265.05 | 300 | 230 | 0.763 | -0.391 | 0.01119991 | 0.0572779 | 0.0077 | 0.0193 | 0.0133 | 0.0133 | TRUE | 0.1386 |
| Fbxo45 | 1712 | 1813 | 1730 | 2215 | 1584 | 1588 | 1818 | 1811 | 1737 | 1741 | 1704 | 1738 | 1758.35 | 1789 | 1728 | 0.966 | -0.05 | 0.47565855 | 0.67577395 | 0 | 0.0088 | 0.0029 | 0.0029 | TRUE | 0.0077 |
| Fbxo46 | 456 | 456 | 414 | 748 | 531 | 489 | 484 | 455 | 416 | 588 | 571 | 535 | 508.35 | 452 | 565 | 1.252 | 0.324 | 0.00378733 | 0.0253253 | 0.0019 | 0.0134 | 0.007 | 0.007 | TRUE | 0.0568 |
| Fbxo47 | 482 | 560 | 554 | 367 | 281 | 314 | 512 | 559 | 556 | 289 | 302 | 344 | 427.01 | 542 | 312 | 0.573 | -0.804 | 3.40E-11 | 2.71E-09 | 0.0023 | 0.0146 | 0.0081 | 0.0081 | TRUE | 0.0906 |
| Fbxo48 | 84 | 88 | 93 | 130 | 108 | 99 | 89 | 88 | 93 | 102 | 116 | 108 | 99.54 | 90 | 109 | 1.205 | 0.269 | 0.16697123 | 0.36449415 | 0 | 0.04 | 0.0172 | 0.0172 | TRUE | 0.0336 |
| Fbxo5 | 3169 | 3926 | 4007 | 4360 | 3165 | 3447 | 3366 | 3921 | 4023 | 3427 | 3405 | 3773 | 3652.76 | 3770 | 3535 | 0.938 | -0.093 | 0.34371032 | 0.56363061 | 0.0057 | 0.0078 | 0.0067 | 0.0067 | TRUE | 0.1068 |
| Fbxo6 | 1659 | 2203 | 2224 | 2422 | 1754 | 1974 | 1762 | 2200 | 2233 | 1904 | 1887 | 2161 | 2024.58 | 2065 | 1984 | 0.96 | -0.058 | 0.62628718 | 0.78958445 | 0.0109 | 0.0085 | 0.0098 | 0.0098 | TRUE | 0.1988 |
| Fbxo7 | 1318 | 1529 | 1442 | 1535 | 1179 | 1289 | 1400 | 1527 | 1448 | 1207 | 1269 | 1411 | 1376.86 | 1458 | 1296 | 0.887 | -0.172 | 0.07010038 | 0.20596559 | 0.0031 | 0.0093 | 0.0058 | 0.0058 | TRUE | 0.0727 |
| Fbxo8 | 1120 | 1049 | 1096 | 1492 | 1113 | 980 | 1190 | 1048 | 1100 | 1173 | 1198 | 1073 | 1130.16 | 1113 | 1148 | 1.032 | 0.046 | 0.63163205 | 0.79278217 | 0.0028 | 0.0098 | 0.0056 | 0.0056 | TRUE | 0.0437 |
| Fbxo9 | 769 | 725 | 684 | 1392 | 939 | 798 | 817 | 724 | 687 | 1094 | 1010 | 874 | 867.64 | 743 | 993 | 1.339 | 0.421 | 0.00070944 | 0.00682693 | 0.0093 | 0.0107 | 0.0099 | 0.0099 | TRUE | 0.1307 |
| Fbxw10 | 86 | 71 | 90 | 111 | 78 | 62 | 91 | 71 | 90 | 87 | 84 | 68 | 81.95 | 84 | 80 | 0.95 | -0.074 | 0.75131604 | 0.86862312 | 0 | 0.0471 | 0.0272 | 0.0272 | TRUE | 0.1817 |
| Fbxw11 | 6547 | 6568 | 6611 | 8995 | 6429 | 6258 | 6954 | 6560 | 6638 | 7071 | 6917 | 6850 | 6831.75 | 6717 | 6946 | 1.034 | 0.049 | 0.42411472 | 0.63431045 | 4.00E-04 | 0.0074 | 0.0025 | 0.0025 | TRUE | 0.0116 |
| Fbxw13 | 0 | 0 | 0 | 1 | 2 | 0 | 0 | 0 | 0 | 1 | 2 | 0 | 0.49 | 0 | 1 | 5.204 | 2.38 | 0.47267317 | NA | 0 | 6.7348 | 6.4099 | 6.4099 | TRUE | 0.1222 |
| Fbxw14 | 0 | 0 | 0 | 0 | 0 | 0 | 0 | 0 | 0 | 0 | 0 | 0 | 0 | 0 | 0 | NA | NA | NA | NA | NA | NA | NA | NA | NA | NA |
| Fbxw15 | 0 | 0 | 0 | 1 | 0 | 0 | 0 | 0 | 0 | 1 | 0 | 0 | 0.13 | 0 | 0 | 1.859 | 0.894 | 0.82650734 | NA | 0 | 25.1515 | 10 | 10 | TRUE | 0.2563 |
| Fbxw16 | 0 | 0 | 1 | 0 | 0 | 0 | 0 | 0 | 1 | 0 | 0 | 0 | 0.17 | 0 | 0 | 0.49 | -1.029 | 0.80087 | NA | 0 | 19.6928 | 10 | 10 | TRUE | 0.4098 |
| Fbxw17 | 283 | 270 | 268 | 366 | 277 | 261 | 301 | 270 | 269 | 288 | 298 | 286 | 285.13 | 280 | 291 | 1.038 | 0.054 | 0.66692238 | 0.81379599 | 0 | 0.0185 | 0.008 | 0.008 | TRUE | 0.0468 |
| Fbxw18 | 0 | 0 | 0 | 0 | 0 | 0 | 0 | 0 | 0 | 0 | 0 | 0 | 0 | 0 | 0 | NA | NA | NA | NA | NA | NA | NA | NA | NA | NA |
| Fbxw19 | 0 | 0 | 0 | 0 | 0 | 1 | 0 | 0 | 0 | 0 | 0 | 1 | 0.18 | 0 | 0 | 1.859 | 0.894 | 0.82650734 | NA | 0 | 18.0643 | 10 | 10 | TRUE | 0.5383 |
| Fbxw2 | 2512 | 2978 | 2998 | 3905 | 2774 | 2776 | 2668 | 2974 | 3010 | 3070 | 2985 | 3039 | 2957.65 | 2884 | 3031 | 1.051 | 0.072 | 0.36119984 | 0.57953971 | 0.0017 | 0.008 | 0.0041 | 0.0041 | TRUE | 0.0519 |
| Fbxw20 | 0 | 0 | 0 | 1 | 0 | 1 | 0 | 0 | 0 | 1 | 0 | 1 | 0.31 | 0 | 1 | 3.348 | 1.743 | 0.65064889 | NA | 0 | 10.5167 | 8.8968 | 8.8968 | TRUE | 0.2082 |
| Fbxw21 | 0 | 0 | 0 | 1 | 0 | 0 | 0 | 0 | 0 | 1 | 0 | 0 | 0.13 | 0 | 0 | 1.859 | 0.894 | 0.82650734 | NA | 0 | 25.1515 | 10 | 10 | TRUE | 0.2563 |
| Fbxw22 | 0 | 0 | 0 | 0 | 0 | 0 | 0 | 0 | 0 | 0 | 0 | 0 | 0 | 0 | 0 | NA | NA | NA | NA | NA | NA | NA | NA | NA | NA |
| Fbxw24 | 0 | 0 | 0 | 0 | 0 | 0 | 0 | 0 | 0 | 0 | 0 | 0 | 0 | 0 | 0 | NA | NA | NA | NA | NA | NA | NA | NA | NA | NA |
| Fbxw26 | 0 | 0 | 0 | 0 | 0 | 0 | 0 | 0 | 0 | 0 | 0 | 0 | 0 | 0 | 0 | NA | NA | NA | NA | NA | NA | NA | NA | NA | NA |
| Fbxw28 | 0 | 0 | 0 | 0 | 0 | 0 | 0 | 0 | 0 | 0 | 0 | 0 | 0 | 0 | 0 | NA | NA | NA | NA | NA | NA | NA | NA | NA | NA |
| Fbxw4 | 115 | 265 | 324 | 236 | 169 | 196 | 122 | 265 | 325 | 186 | 182 | 215 | 215.67 | 237 | 194 | 0.816 | -0.293 | 0.34354858 | 0.56358897 | 0.1156 | 0.0222 | 0.0642 | 0.0642 | TRUE | 0.5459 |
| Fbxw5 | 880 | 1019 | 1070 | 1241 | 966 | 932 | 935 | 1018 | 1074 | 976 | 1039 | 1020 | 1010.32 | 1009 | 1012 | 1.002 | 0.003 | 0.97570002 | 0.98697924 | 0.002 | 0.0102 | 0.0053 | 0.0053 | TRUE | 0.0491 |
| Fbxw7 | 1514 | 1223 | 1230 | 2211 | 1513 | 1275 | 1608 | 1221 | 1235 | 1738 | 1628 | 1396 | 1471.05 | 1355 | 1587 | 1.172 | 0.229 | 0.10516152 | 0.27035167 | 0.0182 | 0.0091 | 0.0137 | 0.0137 | TRUE | 0.3208 |
| Fbxw8 | 5488 | 5877 | 6055 | 7416 | 5137 | 4931 | 5829 | 5869 | 6080 | 5830 | 5527 | 5398 | 5755.54 | 5926 | 5585 | 0.943 | -0.085 | 0.19550213 | 0.40149221 | 8.00E-04 | 0.0075 | 0.0029 | 0.0029 | TRUE | 0.0182 |
| Fbxw9 | 1353 | 1595 | 1543 | 2368 | 1870 | 1625 | 1437 | 1593 | 1549 | 1862 | 2012 | 1779 | 1705.29 | 1526 | 1884 | 1.234 | 0.303 | 0.00063949 | 0.00630772 | 0.0025 | 0.0088 | 0.0051 | 0.0051 | TRUE | 0.042 |
| Fcamr | 0 | 1 | 1 | 2 | 0 | 0 | 0 | 1 | 1 | 2 | 0 | 0 | 0.6 | 1 | 1 | 0.834 | -0.262 | 0.92267544 | NA | 0 | 5.536 | 3.6136 | 3.6136 | TRUE | 0.9722 |
| Fcer1a | 0 | 0 | 0 | 0 | 0 | 0 | 0 | 0 | 0 | 0 | 0 | 0 | 0 | 0 | 0 | NA | NA | NA | NA | NA | NA | NA | NA | NA | NA |
| Fcer1g | 22 | 26 | 22 | 15 | 10 | 21 | 23 | 26 | 22 | 12 | 11 | 23 | 19.49 | 24 | 15 | 0.631 | -0.664 | 0.15976369 | 0.35430158 | 0.0182 | 0.1759 | 0.1067 | 0.1067 | TRUE | 0.8537 |
| Fcer2a | 0 | 0 | 0 | 0 | 0 | 0 | 0 | 0 | 0 | 0 | 0 | 0 | 0 | 0 | 0 | NA | NA | NA | NA | NA | NA | NA | NA | NA | NA |
| Fcf1 | 2434 | 2509 | 2450 | 3556 | 2504 | 2419 | 2585 | 2506 | 2460 | 2795 | 2694 | 2648 | 2614.79 | 2517 | 2712 | 1.078 | 0.109 | 0.10906461 | 0.27686578 | 4.00E-04 | 0.0082 | 0.0029 | 0.0029 | TRUE | 0.0089 |
| Fcgbp | 204 | 294 | 275 | 339 | 236 | 234 | 217 | 294 | 276 | 266 | 254 | 256 | 260.5 | 262 | 259 | 0.987 | -0.018 | 0.90674049 | 0.95352883 | 0.0086 | 0.0195 | 0.0139 | 0.0139 | TRUE | 0.2535 |
| Fcgr1 | 1 | 0 | 0 | 0 | 0 | 0 | 1 | 0 | 0 | 0 | 0 | 0 | 0.18 | 0 | 0 | 0.49 | -1.029 | 0.80087 | NA | 0 | 18.6165 | 10 | 10 | TRUE | 0.4614 |
| Fcgr2b | 79 | 43 | 35 | 25 | 30 | 27 | 84 | 43 | 35 | 20 | 32 | 30 | 40.58 | 54 | 27 | 0.5 | -1 | 0.01977129 | 0.08617669 | 0.122 | 0.0881 | 0.1049 | 0.1049 | TRUE | 1.6289 |
| Fcgr3 | 6 | 6 | 3 | 10 | 4 | 11 | 6 | 6 | 3 | 8 | 4 | 12 | 6.6 | 5 | 8 | 1.574 | 0.655 | 0.40414947 | 0.61693403 | 0 | 0.5063 | 0.2833 | 0.2833 | TRUE | 0.1485 |
| Fcgr4 | 0 | 0 | 0 | 0 | 0 | 0 | 0 | 0 | 0 | 0 | 0 | 0 | 0 | 0 | 0 | NA | NA | NA | NA | NA | NA | NA | NA | NA | NA |
| Fcgrt | 263 | 300 | 320 | 359 | 250 | 268 | 279 | 300 | 321 | 282 | 269 | 293 | 290.81 | 300 | 281 | 0.938 | -0.093 | 0.47165941 | 0.67309195 | 0 | 0.0182 | 0.0085 | 0.0085 | TRUE | 0.0432 |
| Fcho1 | 1702 | 1742 | 1805 | 1955 | 1687 | 1532 | 1808 | 1740 | 1812 | 1537 | 1815 | 1677 | 1731.5 | 1787 | 1676 | 0.938 | -0.093 | 0.30555924 | 0.52646716 | 0.0032 | 0.0088 | 0.0054 | 0.0054 | TRUE | 0.0656 |
| Fcho2 | 3018 | 3494 | 3431 | 3403 | 2647 | 2724 | 3206 | 3489 | 3445 | 2675 | 2848 | 2982 | 3107.54 | 3380 | 2835 | 0.838 | -0.254 | 0.00144706 | 0.01191806 | 0.002 | 0.008 | 0.0043 | 0.0043 | TRUE | 0.0301 |
| Fchsd1 | 837 | 768 | 743 | 941 | 760 | 720 | 889 | 767 | 746 | 740 | 818 | 788 | 791.28 | 801 | 782 | 0.976 | -0.035 | 0.74851597 | 0.8665787 | 0.0045 | 0.0111 | 0.0073 | 0.0073 | TRUE | 0.1099 |
| Fchsd2 | 1075 | 1250 | 1339 | 1548 | 1168 | 1233 | 1142 | 1248 | 1344 | 1217 | 1257 | 1350 | 1259.67 | 1245 | 1275 | 1.023 | 0.033 | 0.73940529 | 0.86065684 | 0.0039 | 0.0095 | 0.0062 | 0.0062 | TRUE | 0.0626 |
| Fcna | 0 | 3 | 1 | 1 | 1 | 2 | 0 | 3 | 1 | 1 | 1 | 2 | 1.34 | 1 | 1 | 0.988 | -0.017 | 0.99188734 | NA | 0 | 2.4619 | 1.3077 | 1.3077 | TRUE | 0.5056 |
| Fcnb | 0 | 0 | 0 | 1 | 0 | 1 | 0 | 0 | 0 | 1 | 0 | 1 | 0.31 | 0 | 1 | 3.348 | 1.743 | 0.65064889 | NA | 0 | 10.5167 | 8.8968 | 8.8968 | TRUE | 0.2082 |
| Fcrl1 | 1 | 1 | 1 | 1 | 0 | 0 | 1 | 1 | 1 | 1 | 0 | 0 | 0.64 | 1 | 0 | 0.342 | -1.549 | 0.50187412 | NA | 0 | 5.1396 | 2.332 | 2.332 | TRUE | 0.2563 |
| Fcrl5 | 2 | 0 | 1 | 0 | 1 | 1 | 2 | 0 | 1 | 0 | 1 | 1 | 0.88 | 1 | 1 | 0.669 | -0.58 | 0.78787572 | NA | 0 | 3.7371 | 2.1463 | 2.1463 | TRUE | 0.1679 |
| Fcrl6 | 0 | 0 | 0 | 0 | 0 | 0 | 0 | 0 | 0 | 0 | 0 | 0 | 0 | 0 | 0 | NA | NA | NA | NA | NA | NA | NA | NA | NA | NA |

| | | | | | | | | | | | | | | | | | | | | | | | | | |
|---|---|---|---|---|---|---|---|---|---|---|---|---|---|---|---|---|---|---|---|---|---|---|---|---|---|
| Fcrla | 1 | 0 | 1 | 0 | 0 | 1 | 1 | 0 | 1 | 0 | 0 | 1 | 0.53 | 1 | 0 | 0.509 | -0.975 | 0.72674596 | NA | 0 | 6.2603 | 3.8683 | 3.8683 | TRUE | 0.5383 |
| Fcrlb | 5 | 2 | 3 | 1 | 3 | 0 | 5 | 2 | 3 | 1 | 3 | 0 | 2.39 | 3 | 1 | 0.383 | -1.384 | 0.31486224 | NA | 0.0945 | 1.3859 | 0.8457 | 0.8457 | TRUE | 0.8434 |
| Fcrls | 0 | 0 | 1 | 2 | 0 | 0 | 0 | 0 | 1 | 2 | 0 | 0 | 0.43 | 0 | 1 | 1.493 | 0.579 | 0.86952142 | NA | 0 | 7.6793 | 7.0425 | 7.0425 | TRUE | 0.9718 |
| Fdft1 | 8992 | 9639 | 9625 | 9400 | 7193 | 8185 | 9551 | 9627 | 9665 | 7390 | 7739 | 8960 | 8821.78 | 9614 | 8030 | 0.835 | -0.26 | 0.00437689 | 0.02828966 | 0.0051 | 0.0073 | 0.0059 | 0.0059 | TRUE | 0.1253 |
| Fdps | 11619 | 10817 | 11335 | 10996 | 7966 | 8607 | 12341 | 10803 | 11382 | 8644 | 8571 | 9422 | 10193.81 | 11509 | 8879 | 0.771 | -0.374 | 6.78E-06 | 0.00014306 | 0.0035 | 0.0072 | 0.0049 | 0.0049 | TRUE | 0.0489 |
| Fdx1 | 1020 | 1041 | 1041 | 1483 | 1008 | 911 | 1083 | 1040 | 1045 | 1166 | 1085 | 997 | 1069.32 | 1056 | 1083 | 1.026 | 0.037 | 0.69132364 | 0.83074715 | 0.0025 | 0.01 | 0.0055 | 0.0055 | TRUE | 0.0565 |
| Fdx1l | 110 | 170 | 175 | 164 | 113 | 122 | 117 | 170 | 176 | 129 | 122 | 134 | 141.07 | 154 | 128 | 0.83 | -0.269 | 0.19753993 | 0.40430829 | 0.0182 | 0.0303 | 0.0243 | 0.0243 | TRUE | 0.4629 |
| Fdxacb1 | 431 | 470 | 485 | 691 | 473 | 459 | 458 | 469 | 487 | 543 | 509 | 502 | 494.79 | 471 | 518 | 1.101 | 0.138 | 0.17933558 | 0.38052398 | 0 | 0.0136 | 0.0056 | 0.0056 | TRUE | 0.0221 |
| Fdxr | 1800 | 2045 | 2028 | 2027 | 1527 | 1650 | 1912 | 2042 | 2036 | 1593 | 1643 | 1806 | 1838.87 | 1997 | 1681 | 0.841 | -0.25 | 0.00346052 | 0.02362743 | 0.0021 | 0.0087 | 0.0047 | 0.0047 | TRUE | 0.0509 |
| Fech | 1932 | 2044 | 2127 | 2746 | 2054 | 1971 | 2052 | 2041 | 2136 | 2159 | 2210 | 2158 | 2125.91 | 2076 | 2176 | 1.048 | 0.067 | 0.31874696 | 0.53988745 | 0 | 0.0085 | 0.0028 | 0.0028 | TRUE | 0.0076 |
| Fem1a | 1137 | 1082 | 1010 | 1974 | 1403 | 1083 | 1208 | 1081 | 1014 | 1552 | 1510 | 1186 | 1258.22 | 1101 | 1416 | 1.287 | 0.364 | 0.00596009 | 0.03574242 | 0.0128 | 0.0095 | 0.0118 | 0.0118 | TRUE | 0.2415 |
| Fem1b | 14592 | 15716 | 16707 | 18422 | 13577 | 13969 | 15499 | 15696 | 16776 | 14482 | 14608 | 15291 | 15391.97 | 15990 | 14794 | 0.925 | -0.112 | 0.08968355 | 0.24349956 | 0.0013 | 0.0071 | 0.0031 | 0.0031 | TRUE | 0.0226 |
| Fem1c | 2418 | 2603 | 2619 | 2851 | 2185 | 2203 | 2568 | 2600 | 2630 | 2241 | 2351 | 2412 | 2466.9 | 2599 | 2335 | 0.898 | -0.156 | 0.0235006 | 0.09746293 | 4.00E-04 | 0.0082 | 0.003 | 0.003 | TRUE | 0.0152 |
| Fen1 | 4070 | 3330 | 3152 | 6019 | 4068 | 3329 | 4323 | 3326 | 3165 | 4732 | 4377 | 3644 | 3927.73 | 3605 | 4251 | 1.18 | 0.238 | 0.10646872 | 0.27255715 | 0.0233 | 0.0077 | 0.0154 | 0.0154 | TRUE | 0.3695 |
| Fendrr | 2 | 1 | 1 | 0 | 0 | 1 | 2 | 1 | 1 | 0 | 0 | 1 | 0.87 | 1 | 0 | 0.255 | -1.974 | 0.35376185 | NA | 0 | 3.7923 | 1.8925 | 1.8925 | TRUE | 0.5383 |
| Fer1l4 | 30 | 22 | 13 | 25 | 23 | 15 | 32 | 22 | 13 | 20 | 25 | 16 | 21.28 | 22 | 20 | 0.911 | -0.135 | 0.77682046 | 0.88398357 | 0.0653 | 0.1617 | 0.1158 | 0.1158 | TRUE | 0.1911 |
| Ferd3l | 0 | 0 | 0 | 0 | 0 | 0 | 0 | 0 | 0 | 0 | 0 | 0 | 0 | 0 | 0 | NA | NA | NA | NA | NA | NA | NA | NA | NA | NA |
| Fermt1 | 118 | 98 | 112 | 152 | 96 | 103 | 125 | 98 | 112 | 119 | 103 | 113 | 111.87 | 112 | 112 | 1.003 | 0.004 | 0.98507279 | 0.99199132 | 0 | 0.0364 | 0.0191 | 0.0191 | TRUE | 0.1206 |
| Fermt2 | 11660 | 11646 | 11898 | 14800 | 10887 | 10679 | 12385 | 11631 | 11947 | 11635 | 11714 | 11690 | 11833.5 | 11988 | 11680 | 0.974 | -0.038 | 0.51564138 | 0.70792965 | 4.00E-04 | 0.0072 | 0.0023 | 0.0023 | TRUE | 0.0103 |
| Fermt3 | 28 | 40 | 47 | 44 | 41 | 59 | 30 | 40 | 47 | 35 | 44 | 65 | 43.36 | 39 | 48 | 1.216 | 0.282 | 0.43172534 | 0.64146699 | 0.0554 | 0.0829 | 0.0694 | 0.0694 | TRUE | 0.4728 |
| Fert2 | 1238 | 1471 | 1516 | 1903 | 1321 | 1397 | 1315 | 1469 | 1522 | 1496 | 1421 | 1529 | 1458.81 | 1435 | 1482 | 1.032 | 0.046 | 0.61395106 | 0.78159949 | 0.0028 | 0.0092 | 0.0053 | 0.0053 | TRUE | 0.0645 |
| Fes | 122 | 166 | 156 | 136 | 114 | 120 | 130 | 166 | 157 | 107 | 123 | 131 | 135.49 | 151 | 120 | 0.795 | -0.331 | 0.08306242 | 0.23068521 | 0.0066 | 0.0312 | 0.0188 | 0.0188 | TRUE | 0.1543 |
| Fetub | 335 | 388 | 354 | 505 | 374 | 350 | 356 | 387 | 355 | 397 | 402 | 383 | 380.22 | 366 | 394 | 1.076 | 0.106 | 0.34571369 | 0.56516549 | 0 | 0.0156 | 0.0065 | 0.0065 | TRUE | 0.0297 |
| Fev | 0 | 0 | 0 | 4 | 0 | 0 | 0 | 0 | 0 | 3 | 0 | 0 | 0.52 | 0 | 1 | 5.676 | 2.505 | 0.52992595 | NA | 10 | 6.293 | 10 | 10 | TRUE | 1.8971 |
| Fez1 | 299 | 338 | 306 | 414 | 265 | 237 | 318 | 338 | 307 | 325 | 285 | 259 | 305.4 | 321 | 290 | 0.907 | -0.141 | 0.31065127 | 0.53123624 | 0.0047 | 0.0177 | 0.0107 | 0.0107 | TRUE | 0.1368 |
| Fez2 | 1568 | 1684 | 1625 | 2157 | 1631 | 1563 | 1665 | 1682 | 1632 | 1696 | 1755 | 1711 | 1690.08 | 1660 | 1721 | 1.036 | 0.052 | 0.45645348 | 0.66122104 | 0 | 0.0089 | 0.0029 | 0.0029 | TRUE | 0.0037 |
| Fezf1 | 1 | 2 | 2 | 2 | 0 | 0 | 1 | 2 | 2 | 2 | 0 | 0 | 1.11 | 2 | 1 | 0.344 | -1.541 | 0.42887349 | NA | 0 | 2.9838 | 1.5946 | 1.5946 | TRUE | 0.9712 |
| Fezf2 | 0 | 0 | 0 | 0 | 0 | 0 | 0 | 0 | 0 | 0 | 0 | 0 | 0 | 0 | 0 | NA | NA | NA | NA | NA | NA | NA | NA | NA | NA |
| Ffar1 | 0 | 1 | 0 | 0 | 1 | 1 | 0 | 1 | 0 | 0 | 1 | 1 | 0.53 | 0 | 1 | 1.924 | 0.944 | 0.7364225 | NA | 0 | 6.2438 | 3.9727 | 3.9727 | TRUE | 0.405 |
| Ffar2 | 1 | 2 | 1 | 6 | 4 | 1 | 1 | 2 | 1 | 5 | 4 | 1 | 2.36 | 1 | 3 | 2.516 | 1.331 | 0.30280998 | NA | 0 | 1.401 | 0.6829 | 0.6829 | TRUE | 0.4491 |
| Ffar3 | 4 | 8 | 6 | 16 | 8 | 10 | 4 | 8 | 6 | 13 | 9 | 11 | 8.4 | 6 | 11 | 1.764 | 0.819 | 0.21592578 | 0.42789061 | 0 | 0.3991 | 0.1868 | 0.1868 | TRUE | 0.1792 |
| Ffar4 | 10 | 20 | 13 | 14 | 16 | 15 | 11 | 20 | 13 | 11 | 17 | 16 | 14.71 | 15 | 15 | 1.011 | 0.016 | 0.97553092 | 0.98697924 | 0 | 0.2308 | 0.1286 | 0.1286 | TRUE | 0.4808 |
| Fga | 0 | 0 | 0 | 1 | 0 | 0 | 0 | 0 | 0 | 1 | 0 | 0 | 0.13 | 0 | 0 | 1.859 | 0.894 | 0.82650734 | NA | 0 | 25.1515 | 10 | 10 | TRUE | 0.2563 |
| Fgb | 0 | 0 | 0 | 0 | 0 | 0 | 0 | 0 | 0 | 0 | 0 | 0 | 0 | 0 | 0 | NA | NA | NA | NA | NA | NA | NA | NA | NA | NA |
| Fgd1 | 4438 | 5991 | 5952 | 7720 | 5620 | 5607 | 4714 | 5983 | 5976 | 6069 | 6047 | 6138 | 5821.17 | 5558 | 6085 | 1.095 | 0.13 | 0.22951278 | 0.44306922 | 0.0084 | 0.0075 | 0.0083 | 0.0083 | TRUE | 0.2149 |
| Fgd2 | 72 | 56 | 58 | 58 | 45 | 45 | 76 | 56 | 58 | 46 | 48 | 49 | 55.65 | 63 | 48 | 0.751 | -0.413 | 0.12608042 | 0.3042908 | 0 | 0.0661 | 0.0342 | 0.0342 | TRUE | 0.2709 |
| Fgd3 | 6 | 2 | 0 | 8 | 2 | 5 | 6 | 2 | 0 | 6 | 2 | 5 | 3.71 | 3 | 4 | 1.689 | 0.756 | 0.53034117 | NA | 0.5511 | 0.8939 | 0.7559 | 0.7559 | TRUE | 0.8802 |
| Fgd4 | 86 | 91 | 72 | 208 | 153 | 108 | 91 | 91 | 72 | 164 | 165 | 118 | 116.81 | 85 | 149 | 1.758 | 0.814 | 0.00025287 | 0.00294853 | 0.0175 | 0.0351 | 0.0264 | 0.0264 | TRUE | 0.325 |
| Fgd5 | 2978 | 3430 | 3347 | 2817 | 2274 | 2291 | 3163 | 3426 | 3361 | 2215 | 2447 | 2508 | 2853.09 | 3317 | 2390 | 0.72 | -0.474 | 1.31E-08 | 6.10E-07 | 0.0024 | 0.0081 | 0.0046 | 0.0046 | TRUE | 0.0509 |
| Fgd6 | 1800 | 1615 | 1629 | 1991 | 1344 | 1348 | 1912 | 1613 | 1636 | 1565 | 1446 | 1476 | 1607.89 | 1720 | 1496 | 0.87 | -0.201 | 0.04315389 | 0.1485104 | 0.0046 | 0.009 | 0.0065 | 0.0065 | TRUE | 0.1144 |
| Fgf1 | 113 | 63 | 61 | 236 | 168 | 84 | 120 | 63 | 61 | 186 | 181 | 92 | 117.07 | 81 | 153 | 1.882 | 0.912 | 0.00979658 | 0.05184987 | 0.1345 | 0.035 | 0.0804 | 0.0804 | TRUE | 1.5869 |
| Fgf10 | 116 | 207 | 221 | 353 | 316 | 434 | 123 | 207 | 222 | 278 | 340 | 475 | 274.07 | 184 | 364 | 1.976 | 0.982 | 0.00020655 | 0.00250763 | 0.0791 | 0.0189 | 0.0464 | 0.0464 | TRUE | 0.3757 |
| Fgf11 | 34 | 71 | 110 | 39 | 30 | 47 | 36 | 71 | 110 | 31 | 32 | 51 | 55.31 | 72 | 38 | 0.523 | -0.934 | 0.03104064 | 0.1180925 | 0.1687 | 0.0665 | 0.1153 | 0.1153 | TRUE | 0.1278 |
| Fgf12 | 1 | 6 | 5 | 4 | 4 | 2 | 1 | 6 | 5 | 3 | 4 | 2 | 3.62 | 4 | 3 | 0.794 | -0.333 | 0.74853994 | NA | 0 | 0.9173 | 0.4973 | 0.4973 | TRUE | 0.6562 |
| Fgf13 | 245 | 331 | 391 | 603 | 434 | 427 | 260 | 331 | 393 | 474 | 467 | 467 | 398.64 | 328 | 469 | 1.431 | 0.517 | 0.00133254 | 0.01113768 | 0.0166 | 0.0152 | 0.0161 | 0.0161 | TRUE | 0.3153 |
| Fgf14 | 8 | 8 | 4 | 15 | 8 | 4 | 8 | 8 | 4 | 12 | 9 | 4 | 7.55 | 7 | 8 | 1.228 | 0.296 | 0.68698949 | 0.82743019 | 0 | 0.4434 | 0.2564 | 0.2564 | TRUE | 0.2202 |
| Fgf15 | 197 | 397 | 417 | 243 | 184 | 257 | 209 | 396 | 419 | 191 | 198 | 281 | 282.46 | 341 | 223 | 0.653 | -0.614 | 0.0239582 | 0.09866026 | 0.0865 | 0.0186 | 0.0497 | 0.0497 | TRUE | 1.3003 |
| Fgf16 | 0 | 0 | 0 | 0 | 0 | 1 | 0 | 0 | 0 | 0 | 0 | 1 | 0.18 | 0 | 0 | 1.859 | 0.894 | 0.82650734 | NA | 0 | 18.0643 | 10 | 10 | TRUE | 0.5383 |
| Fgf17 | 114 | 118 | 157 | 134 | 73 | 104 | 121 | 118 | 158 | 105 | 79 | 114 | 115.72 | 132 | 99 | 0.752 | -0.412 | 0.07070301 | 0.20711884 | 0.0216 | 0.0354 | 0.0287 | 0.0287 | TRUE | 0.3088 |
| Fgf18 | 53 | 50 | 56 | 72 | 55 | 51 | 56 | 50 | 56 | 57 | 59 | 56 | 55.68 | 54 | 57 | 1.056 | 0.079 | 0.75461731 | 0.8704618 | 0 | 0.0661 | 0.0281 | 0.0281 | TRUE | 0.0393 |
| Fgf2 | 2 | 4 | 4 | 5 | 2 | 3 | 2 | 4 | 4 | 4 | 2 | 3 | 3.25 | 3 | 3 | 0.931 | -0.102 | 0.91951203 | NA | 0 | 1.0205 | 0.4344 | 0.4344 | TRUE | 0.143 |
| Fgf20 | 2 | 0 | 1 | 3 | 0 | 0 | 2 | 0 | 1 | 2 | 0 | 0 | 0.91 | 1 | 1 | 0.802 | -0.318 | 0.89399339 | NA | 1.8421 | 3.6095 | 3.0187 | 3.0187 | TRUE | 0.6132 |
| Fgf21 | 0 | 2 | 0 | 2 | 1 | 1 | 0 | 2 | 0 | 2 | 1 | 1 | 0.96 | 1 | 1 | 1.87 | 0.903 | 0.66006362 | NA | 0 | 3.4504 | 1.8897 | 1.8897 | TRUE | 0.9738 |
| Fgf22 | 10 | 16 | 10 | 18 | 16 | 14 | 11 | 16 | 10 | 14 | 17 | 15 | 13.89 | 12 | 15 | 1.268 | 0.343 | 0.50164415 | 0.69705269 | 0 | 0.2441 | 0.1146 | 0.1146 | TRUE | 0.2953 |
| Fgf23 | 8 | 15 | 5 | 7 | 4 | 4 | 8 | 15 | 5 | 6 | 4 | 4 | 7.11 | 9 | 5 | 0.501 | -0.997 | 0.19271163 | 0.39842218 | 0 | 0.47 | 0.2659 | 0.2659 | TRUE | 0.2786 |
| Fgf3 | 78 | 143 | 136 | 98 | 94 | 99 | 83 | 143 | 137 | 77 | 101 | 108 | 108.13 | 121 | 95 | 0.787 | -0.345 | 0.19960275 | 0.40715667 | 0.0486 | 0.0374 | 0.0428 | 0.0428 | TRUE | 0.7505 |
| Fgf4 | 9905 | 15676 | 14715 | 12069 | 8773 | 11072 | 10521 | 15656 | 14775 | 9488 | 9439 | 12120 | 11999.83 | 13651 | 10349 | 0.758 | -0.4 | 0.01490075 | 0.07051142 | 0.0311 | 0.0072 | 0.0193 | 0.0193 | TRUE | 0.4918 |
| Fgf5 | 8 | 5 | 7 | 8 | 6 | 6 | 8 | 5 | 7 | 6 | 6 | 7 | 6.64 | 7 | 6 | 0.942 | -0.086 | 0.90403054 | 0.95248677 | 0 | 0.5032 | 0.2158 | 0.2158 | TRUE | 0.1477 |
| Fgf6 | 1 | 1 | 1 | 0 | 0 | 0 | 1 | 1 | 1 | 0 | 0 | 0 | 0.51 | 1 | 0 | 0.176 | -2.51 | 0.35480135 | NA | 0 | 6.4561 | 3.8046 | 3.8046 | TRUE | 0.0847 |
| Fgf7 | 3 | 0 | 0 | 2 | 0 | 4 | 3 | 0 | 0 | 2 | 0 | 4 | 1.52 | 1 | 2 | 1.88 | 0.91 | 0.66568111 | NA | 3.1725 | 2.1702 | 2.4649 | 2.4649 | TRUE | 0.5998 |
| Fgf8 | 49 | 84 | 77 | 38 | 28 | 33 | 52 | 84 | 77 | 30 | 30 | 36 | 51.56 | 71 | 32 | 0.449 | -1.156 | 0.00019985 | 0.00244155 | 0.0217 | 0.0708 | 0.0471 | 0.0471 | TRUE | 0.4841 |
| Fgf9 | 2 | 5 | 3 | 2 | 8 | 3 | 2 | 5 | 3 | 2 | 9 | 3 | 3.93 | 3 | 5 | 1.3 | 0.378 | 0.71725533 | NA | 0.2334 | 0.8447 | 0.5244 | 0.5244 | TRUE | 0.8583 |
| Fgfbp1 | 466 | 231 | 206 | 908 | 479 | 478 | 495 | 231 | 207 | 714 | 515 | 523 | 447.49 | 311 | 584 | 1.881 | 0.912 | 0.00253332 | 0.0185753 | 0.1299 | 0.0143 | 0.0632 | 0.0632 | TRUE | 3.0282 |
| Fgfbp3 | 87 | 116 | 140 | 56 | 44 | 59 | 92 | 116 | 141 | 44 | 47 | 65 | 84.13 | 116 | 52 | 0.444 | -1.171 | 1.39E-05 | 0.0002627 | 0.0303 | 0.0461 | 0.0385 | 0.0385 | TRUE | 0.1291 |
| Fgfr1 | 8811 | 8355 | 7990 | 8988 | 6823 | 6588 | 9359 | 8344 | 8023 | 7066 | 7341 | 7212 | 7890.71 | 8575 | 7206 | 0.84 | -0.251 | 0.00230262 | 0.01722488 | 0.0033 | 0.0073 | 0.0048 | 0.0048 | TRUE | 0.078 |
| Fgfr1op | 2336 | 2540 | 2583 | 3032 | 2202 | 2469 | 2481 | 2537 | 2594 | 2384 | 2369 | 2703 | 2511.17 | 2537 | 2485 | 0.979 | -0.03 | 0.71752307 | 0.84681037 | 0.0026 | 0.0082 | 0.0047 | 0.0047 | TRUE | 0.0705 |
| Fgfr1op2 | 967 | 1130 | 1055 | 1240 | 996 | 937 | 1027 | 1129 | 1059 | 975 | 1072 | 1026 | 1047.85 | 1072 | 1024 | 0.955 | -0.067 | 0.45352061 | 0.65889159 | 0.0014 | 0.01 | 0.0048 | 0.0048 | TRUE | 0.0258 |
| Fgfr2 | 397 | 250 | 222 | 338 | 181 | 156 | 422 | 250 | 223 | 266 | 195 | 171 | 254.25 | 298 | 211 | 0.708 | -0.498 | 0.06579983 | 0.19786368 | 0.0821 | 0.0199 | 0.0488 | 0.0488 | TRUE | 1.4774 |
| Fgfr3 | 252 | 406 | 418 | 672 | 559 | 586 | 268 | 405 | 420 | 528 | 601 | 641 | 477.34 | 364 | 590 | 1.618 | 0.695 | 0.00015559 | 0.00196315 | 0.0309 | 0.0138 | 0.0221 | 0.0221 | TRUE | 0.6112 |

| | | | | | | | | | | | | | | | | | | | | | | | | |
|---|---|---|---|---|---|---|---|---|---|---|---|---|---|---|---|---|---|---|---|---|---|---|---|---|
| Fgfr4 | 320 | 445 | 401 | 387 | 257 | 355 | 340 | 444 | 403 | 304 | 277 | 389 | 359.39 | 396 | 323 | 0.816 | -0.294 | 0.08931257 | 0.24293248 | 0.0209 | 0.0161 | 0.0188 | 0.0188 | TRUE | 0.3481 |
| Fgfrl1 | 893 | 1242 | 1191 | 968 | 772 | 915 | 949 | 1240 | 1196 | 761 | 831 | 1002 | 996.34 | 1128 | 865 | 0.765 | -0.386 | 0.00935378 | 0.05003628 | 0.0184 | 0.0102 | 0.0149 | 0.0149 | TRUE | 0.2319 |
| Fgg | 15 | 19 | 20 | 17 | 16 | 9 | 16 | 19 | 20 | 13 | 17 | 10 | 15.9 | 18 | 13 | 0.735 | -0.445 | 0.35663389 | 0.57524954 | 0 | 0.2141 | 0.1036 | 0.1036 | TRUE | 0.2215 |
| Fggy | 542 | 821 | 929 | 724 | 527 | 573 | 576 | 820 | 933 | 569 | 567 | 627 | 681.98 | 776 | 588 | 0.757 | -0.402 | 0.02261447 | 0.09488573 | 0.0301 | 0.0117 | 0.0209 | 0.0209 | TRUE | 0.447 |
| Fgl1 | 12 | 7 | 7 | 26 | 24 | 26 | 13 | 7 | 7 | 20 | 26 | 28 | 16.91 | 9 | 25 | 2.79 | 1.48 | 0.00363591 | 0.02456967 | 0 | 0.2017 | 0.1094 | 0.1094 | TRUE | 0.4268 |
| Fgl2 | 164 | 153 | 160 | 170 | 109 | 140 | 174 | 153 | 161 | 134 | 117 | 153 | 148.64 | 163 | 135 | 0.829 | -0.271 | 0.12953844 | 0.3096984 | 0.0026 | 0.0291 | 0.0163 | 0.0163 | TRUE | 0.1385 |
| Fgr | 106 | 127 | 155 | 120 | 94 | 77 | 113 | 127 | 156 | 94 | 101 | 84 | 112.47 | 132 | 93 | 0.708 | -0.498 | 0.01821391 | 0.08131559 | 0.0098 | 0.0362 | 0.0229 | 0.0229 | TRUE | 0.2604 |
| Fh1 | 9210 | 11187 | 11049 | 10969 | 7999 | 8526 | 9783 | 11173 | 11094 | 8623 | 8606 | 9333 | 9768.67 | 10683 | 8854 | 0.829 | -0.271 | 0.00130427 | 0.01095363 | 0.0036 | 0.0072 | 0.005 | 0.005 | TRUE | 0.0664 |
| Fhad1 | 123 | 181 | 185 | 168 | 165 | 186 | 131 | 181 | 186 | 132 | 178 | 204 | 168.4 | 166 | 171 | 1.028 | 0.039 | 0.86229012 | 0.93194197 | 0.036 | 0.0265 | 0.0312 | 0.0312 | TRUE | 0.4256 |
| Fhdc1 | 278 | 305 | 318 | 301 | 223 | 263 | 295 | 305 | 319 | 237 | 240 | 288 | 280.61 | 306 | 255 | 0.83 | -0.269 | 0.05339523 | 0.17158711 | 0.0028 | 0.0186 | 0.0104 | 0.0104 | TRUE | 0.1392 |
| Fhit | 42 | 57 | 46 | 29 | 19 | 19 | 45 | 57 | 46 | 23 | 20 | 21 | 35.29 | 49 | 21 | 0.435 | -1.202 | 0.00032322 | 0.00362315 | 0 | 0.1002 | 0.0474 | 0.0474 | TRUE | 0.1522 |
| Fhl1 | 1960 | 2206 | 2166 | 2225 | 1795 | 1823 | 2082 | 2203 | 2175 | 1749 | 1931 | 1996 | 2022.65 | 2153 | 1892 | 0.878 | -0.188 | 0.02595289 | 0.10428766 | 0.0023 | 0.0085 | 0.0046 | 0.0046 | TRUE | 0.0541 |
| Fhl2 | 40 | 39 | 25 | 64 | 54 | 62 | 42 | 39 | 25 | 50 | 58 | 68 | 47.14 | 35 | 59 | 1.65 | 0.723 | 0.02344529 | 0.09731031 | 0.0193 | 0.0768 | 0.0505 | 0.0505 | TRUE | 0.4747 |
| Fhl3 | 271 | 296 | 331 | 352 | 297 | 247 | 288 | 296 | 332 | 277 | 320 | 270 | 297.08 | 305 | 289 | 0.945 | -0.081 | 0.55613856 | 0.73889464 | 0.0036 | 0.018 | 0.0104 | 0.0104 | TRUE | 0.0941 |
| Fhl4 | 89 | 179 | 147 | 290 | 178 | 134 | 95 | 179 | 148 | 228 | 192 | 147 | 164.51 | 141 | 189 | 1.347 | 0.43 | 0.10872779 | 0.27620449 | 0.0664 | 0.0269 | 0.0455 | 0.0455 | TRUE | 0.3897 |
| Fhl5 | 0 | 0 | 0 | 2 | 0 | 0 | 0 | 0 | 0 | 2 | 0 | 0 | 0.26 | 0 | 1 | 2.886 | 1.529 | 0.70676491 | NA | 0 | 12.5792 | 10 | 10 | TRUE | 0.9669 |
| Fhod1 | 1156 | 988 | 939 | 1497 | 1069 | 896 | 1228 | 987 | 943 | 1177 | 1150 | 981 | 1077.55 | 1053 | 1103 | 1.048 | 0.068 | 0.61239093 | 0.78054559 | 0.014 | 0.01 | 0.0121 | 0.0121 | TRUE | 0.2533 |
| Fhod3 | 1863 | 1976 | 2058 | 1574 | 1251 | 1271 | 1979 | 1973 | 2066 | 1237 | 1346 | 1391 | 1665.57 | 2006 | 1325 | 0.66 | -0.6 | 3.30E-13 | 3.59E-11 | 0.0015 | 0.0089 | 0.0043 | 0.0043 | TRUE | 0.0416 |
| Fibcd1 | 7 | 1 | 2 | 4 | 5 | 4 | 7 | 1 | 2 | 3 | 5 | 4 | 3.89 | 3 | 4 | 1.235 | 0.305 | 0.77208734 | NA | 0.1954 | 0.8536 | 0.5363 | 0.5363 | TRUE | 1.3955 |
| Fibin | 0 | 0 | 0 | 0 | 0 | 1 | 0 | 0 | 0 | 0 | 0 | 1 | 0.18 | 0 | 0 | 1.859 | 0.894 | 0.82650734 | NA | 0 | 18.0643 | 10 | 10 | TRUE | 0.5383 |
| Fibp | 745 | 874 | 828 | 1052 | 771 | 760 | 791 | 873 | 831 | 827 | 830 | 832 | 830.68 | 832 | 830 | 0.997 | -0.005 | 0.95890434 | 0.97898705 | 0 | 0.0109 | 0.0044 | 0.0044 | TRUE | 0.0221 |
| Ficd | 335 | 478 | 390 | 537 | 336 | 304 | 356 | 477 | 392 | 422 | 362 | 333 | 390.21 | 408 | 372 | 0.913 | -0.131 | 0.41174782 | 0.62279981 | 0.0164 | 0.0153 | 0.0159 | 0.0159 | TRUE | 0.2534 |
| Fig4 | 4450 | 4359 | 4342 | 5673 | 4246 | 4242 | 4727 | 4353 | 4360 | 4460 | 4568 | 4644 | 4518.59 | 4480 | 4557 | 1.017 | 0.025 | 0.72297435 | 0.85005881 | 9.00E-04 | 0.0076 | 0.0032 | 0.0032 | TRUE | 0.0282 |
| Figf | 209 | 281 | 287 | 208 | 150 | 159 | 222 | 281 | 288 | 164 | 161 | 174 | 214.96 | 264 | 166 | 0.63 | -0.666 | 4.45E-05 | 0.00070286 | 0.0072 | 0.0222 | 0.0143 | 0.0143 | TRUE | 0.2101 |
| Figla | 3 | 7 | 11 | 7 | 3 | 4 | 3 | 7 | 11 | 6 | 3 | 4 | 5.72 | 7 | 4 | 0.622 | -0.684 | 0.41448474 | 0.62611959 | 0 | 0.5827 | 0.3238 | 0.3238 | TRUE | 0.1351 |
| Fign | 103 | 95 | 105 | 155 | 104 | 93 | 109 | 95 | 105 | 122 | 112 | 102 | 107.54 | 103 | 112 | 1.087 | 0.121 | 0.53522781 | 0.72254971 | 0 | 0.0375 | 0.018 | 0.018 | TRUE | 0.0676 |
| Fignl1 | 4168 | 4568 | 4686 | 7157 | 4974 | 4882 | 4427 | 4562 | 4705 | 5626 | 5352 | 5344 | 5002.77 | 4565 | 5441 | 1.192 | 0.253 | 9.88E-05 | 0.00136634 | 6.00E-04 | 0.0076 | 0.0029 | 0.0029 | TRUE | 0.011 |
| Fignl2 | 488 | 786 | 746 | 750 | 566 | 749 | 518 | 785 | 749 | 590 | 609 | 820 | 678.48 | 684 | 673 | 0.982 | -0.026 | 0.8945939 | 0.9490474 | 0.0397 | 0.0118 | 0.0252 | 0.0252 | TRUE | 0.5285 |
| Filip1 | 17 | 33 | 32 | 66 | 50 | 32 | 18 | 33 | 32 | 52 | 54 | 35 | 37.31 | 28 | 47 | 1.693 | 0.76 | 0.04077903 | 0.14287252 | 0.0404 | 0.0952 | 0.0706 | 0.0706 | TRUE | 0.5716 |
| Filip1l | 24 | 16 | 9 | 28 | 13 | 12 | 25 | 16 | 9 | 22 | 14 | 13 | 16.61 | 17 | 16 | 0.985 | -0.022 | 0.96850011 | 0.98422576 | 0.1071 | 0.2053 | 0.1577 | 0.1577 | TRUE | 0.2741 |
| Fip1l1 | 3314 | 3821 | 3828 | 4684 | 3589 | 3578 | 3520 | 3816 | 3844 | 3682 | 3862 | 3917 | 3773.38 | 3727 | 3820 | 1.025 | 0.035 | 0.6296515 | 0.79187174 | 0.0012 | 0.0078 | 0.0036 | 0.0036 | TRUE | 0.0285 |
| Fis1 | 1660 | 1649 | 1591 | 2379 | 1626 | 1657 | 1763 | 1647 | 1598 | 1870 | 1749 | 1814 | 1740.19 | 1669 | 1811 | 1.086 | 0.119 | 0.13833102 | 0.3228406 | 0.0012 | 0.0088 | 0.004 | 0.004 | TRUE | 0.0291 |
| Fitm1 | 32 | 54 | 44 | 43 | 19 | 48 | 34 | 54 | 44 | 34 | 20 | 53 | 39.82 | 44 | 36 | 0.806 | -0.311 | 0.44382946 | 0.65050444 | 0.0977 | 0.0896 | 0.0934 | 0.0934 | TRUE | 0.3467 |
| Fitm2 | 532 | 556 | 487 | 668 | 470 | 459 | 565 | 555 | 489 | 525 | 506 | 502 | 523.77 | 536 | 511 | 0.954 | -0.069 | 0.52737227 | 0.71613289 | 0.0014 | 0.0132 | 0.0066 | 0.0066 | TRUE | 0.0703 |
| Fiz1 | 2858 | 2743 | 2898 | 3686 | 2730 | 2349 | 3036 | 2739 | 2910 | 2898 | 2937 | 2571 | 2848.56 | 2895 | 2802 | 0.968 | -0.047 | 0.59833717 | 0.77074272 | 0.0036 | 0.0081 | 0.0053 | 0.0053 | TRUE | 0.0624 |
| Fjx1 | 251 | 279 | 293 | 364 | 287 | 183 | 267 | 279 | 294 | 286 | 309 | 200 | 272.45 | 280 | 265 | 0.949 | -0.076 | 0.67728732 | 0.82056964 | 0.0219 | 0.019 | 0.0204 | 0.0204 | TRUE | 0.4945 |
| Fkbp10 | 890 | 999 | 1157 | 920 | 607 | 767 | 945 | 998 | 1162 | 723 | 653 | 840 | 886.79 | 1035 | 739 | 0.713 | -0.487 | 0.00025064 | 0.00293261 | 0.0123 | 0.0106 | 0.0116 | 0.0116 | TRUE | 0.1659 |
| Fkbp11 | 1196 | 1447 | 1295 | 1150 | 912 | 952 | 1270 | 1445 | 1300 | 904 | 981 | 1042 | 1157.2 | 1338 | 976 | 0.728 | -0.458 | 5.45E-06 | 0.00011922 | 0.0041 | 0.0098 | 0.0064 | 0.0064 | TRUE | 0.0584 |
| Fkbp14 | 239 | 270 | 264 | 249 | 189 | 198 | 254 | 270 | 265 | 196 | 203 | 217 | 234.07 | 263 | 205 | 0.779 | -0.36 | 0.00783917 | 0.04365617 | 0 | 0.021 | 0.0089 | 0.0089 | TRUE | 0.0254 |
| Fkbp15 | 1495 | 1568 | 1681 | 2178 | 1581 | 1566 | 1588 | 1566 | 1688 | 1712 | 1701 | 1714 | 1661.55 | 1614 | 1709 | 1.059 | 0.083 | 0.26079234 | 0.48012336 | 0 | 0.0089 | 0.0033 | 0.0033 | TRUE | 0.0195 |
| Fkbp1a | 10421 | 10416 | 10032 | 12196 | 8477 | 8515 | 11069 | 10403 | 10073 | 9588 | 9121 | 9321 | 9928.99 | 10515 | 9343 | 0.889 | -0.17 | 0.01235039 | 0.06140932 | 0.0013 | 0.0072 | 0.0032 | 0.0032 | TRUE | 0.0259 |
| Fkbp1b | 262 | 419 | 432 | 463 | 341 | 420 | 278 | 418 | 434 | 364 | 367 | 460 | 386.86 | 377 | 397 | 1.052 | 0.073 | 0.70868687 | 0.84126423 | 0.0345 | 0.0154 | 0.0248 | 0.0248 | TRUE | 0.596 |
| Fkbp2 | 713 | 826 | 768 | 916 | 723 | 716 | 757 | 825 | 771 | 720 | 778 | 784 | 772.53 | 784 | 761 | 0.968 | -0.046 | 0.62215184 | 0.78656528 | 0.001 | 0.0112 | 0.0051 | 0.0051 | TRUE | 0.0271 |
| Fkbp3 | 9179 | 10803 | 10811 | 12287 | 8997 | 9869 | 9750 | 10789 | 10855 | 9659 | 9680 | 10803 | 10256.1 | 10465 | 10047 | 0.96 | -0.059 | 0.48572284 | 0.68272174 | 0.0035 | 0.0072 | 0.005 | 0.005 | TRUE | 0.0528 |
| Fkbp4 | 42325 | 44285 | 43512 | 54481 | 41130 | 40629 | 44957 | 44228 | 43691 | 42829 | 44253 | 44475 | 44072 | 44292 | 43852 | 0.99 | -0.014 | 0.78654105 | 0.89023346 | 3.00E-04 | 0.007 | 0.002 | 0.002 | TRUE | 0.0051 |
| Fkbp5 | 11430 | 11333 | 11367 | 13569 | 9798 | 9705 | 12141 | 11318 | 11414 | 10667 | 10542 | 10624 | 11117.55 | 11624 | 10611 | 0.913 | -0.132 | 0.03079103 | 0.11742794 | 6.00E-04 | 0.0072 | 0.0026 | 0.0026 | TRUE | 0.0184 |
| Fkbp6 | 348 | 545 | 522 | 727 | 562 | 515 | 370 | 544 | 524 | 572 | 605 | 564 | 529.67 | 479 | 580 | 1.209 | 0.274 | 0.08633874 | 0.23719466 | 0.0197 | 0.0131 | 0.0165 | 0.0165 | TRUE | 0.4628 |
| Fkbp7 | 25 | 54 | 59 | 54 | 23 | 33 | 27 | 54 | 59 | 42 | 25 | 36 | 40.51 | 47 | 34 | 0.742 | -0.43 | 0.2844439 | 0.50589188 | 0.0948 | 0.0882 | 0.0911 | 0.0911 | TRUE | 1.0919 |
| Fkbp8 | 8147 | 8540 | 8438 | 11578 | 8485 | 8003 | 8654 | 8529 | 8473 | 9102 | 9129 | 8761 | 8774.46 | 8552 | 8997 | 1.052 | 0.073 | 0.19279928 | 0.39842218 | 1.00E-04 | 0.0073 | 0.0022 | 0.0022 | TRUE | 0.0064 |
| Fkbp9 | 6281 | 7035 | 7112 | 10088 | 6636 | 6533 | 6672 | 7026 | 7141 | 7930 | 7140 | 7151 | 7176.73 | 6946 | 7407 | 1.067 | 0.093 | 0.22389941 | 0.43607575 | 0.0023 | 0.0074 | 0.0041 | 0.0041 | TRUE | 0.0469 |
| Fkbpl | 182 | 172 | 171 | 268 | 195 | 153 | 193 | 172 | 172 | 211 | 210 | 167 | 187.46 | 179 | 196 | 1.098 | 0.135 | 0.41743011 | 0.6283363 | 0.0051 | 0.0245 | 0.0147 | 0.0147 | TRUE | 0.1685 |
| Fkrp | 837 | 719 | 757 | 1080 | 747 | 699 | 889 | 718 | 760 | 849 | 804 | 765 | 797.52 | 789 | 806 | 1.022 | 0.032 | 0.78103182 | 0.88660096 | 0.0063 | 0.011 | 0.0083 | 0.0083 | TRUE | 0.1449 |
| Fktn | 1030 | 1066 | 1153 | 1267 | 900 | 946 | 1094 | 1065 | 1158 | 996 | 968 | 1036 | 1052.72 | 1106 | 1000 | 0.904 | -0.145 | 0.0874689 | 0.23950685 | 5.00E-04 | 0.01 | 0.0042 | 0.0042 | TRUE | 0.0206 |
| Flad1 | 1173 | 1332 | 1298 | 1732 | 1364 | 1335 | 1246 | 1330 | 1303 | 1362 | 1468 | 1461 | 1361.68 | 1293 | 1430 | 1.105 | 0.144 | 0.07427462 | 0.21412399 | 8.00E-04 | 0.0093 | 0.004 | 0.004 | TRUE | 0.022 |
| Flcn | 2629 | 2730 | 2700 | 2938 | 2244 | 2125 | 2792 | 2726 | 2711 | 2310 | 2414 | 2326 | 2546.7 | 2743 | 2350 | 0.856 | -0.223 | 0.00061714 | 0.00613718 | 0 | 0.0082 | 0.0027 | 0.0027 | TRUE | 0.0069 |
| Flg2 | 0 | 0 | 0 | 0 | 0 | 0 | 0 | 0 | 0 | 0 | 0 | 0 | 0 | 0 | 0 | NA | NA | NA | NA | NA | NA | NA | NA | NA | NA |
| Fli1 | 0 | 3 | 1 | 5 | 5 | 4 | 0 | 3 | 1 | 4 | 5 | 4 | 2.95 | 1 | 4 | 3.371 | 1.753 | 0.15729435 | NA | 0 | 1.1243 | 0.6212 | 0.6212 | TRUE | 0.7345 |
| Flii | 9070 | 9171 | 9361 | 10325 | 7877 | 8073 | 9634 | 9159 | 9399 | 8117 | 8475 | 8837 | 8936.93 | 9397 | 8476 | 0.902 | -0.149 | 0.02417467 | 0.09929244 | 0.0011 | 0.0073 | 0.003 | 0.003 | TRUE | 0.017 |
| Flna | 18587 | 16544 | 18166 | 23848 | 16203 | 15111 | 19743 | 16523 | 18241 | 18747 | 17433 | 16541 | 17871.35 | 18169 | 17574 | 0.967 | -0.048 | 0.61170098 | 0.78005723 | 0.0051 | 0.0071 | 0.0064 | 0.0064 | TRUE | 0.0768 |
| Flnb | 19387 | 20505 | 24125 | 26235 | 18847 | 21842 | 20592 | 20479 | 24224 | 20624 | 20278 | 23910 | 21684.44 | 21765 | 21604 | 0.993 | -0.011 | 0.91897183 | 0.96035815 | 0.0072 | 0.0071 | 0.008 | 0.008 | TRUE | 0.1195 |
| Flnc | 3148 | 3040 | 3353 | 2682 | 1988 | 1945 | 3344 | 3036 | 3367 | 2108 | 2139 | 2129 | 2687.17 | 3249 | 2125 | 0.654 | -0.612 | 6.01E-16 | 9.29E-14 | 0.0014 | 0.0081 | 0.0037 | 0.0037 | TRUE | 0.04 |
| Flot1 | 1923 | 2446 | 2424 | 1849 | 1572 | 1926 | 2043 | 2443 | 2434 | 1454 | 1691 | 2108 | 2028.76 | 2307 | 1751 | 0.759 | -0.398 | 0.00746659 | 0.04211843 | 0.0221 | 0.0085 | 0.0155 | 0.0155 | TRUE | 0.384 |
| Flot2 | 4479 | 5607 | 5607 | 5711 | 4182 | 4912 | 4757 | 5600 | 5630 | 4490 | 4500 | 5377 | 5058.89 | 5329 | 4789 | 0.898 | -0.155 | 0.16616621 | 0.36339067 | 0.0097 | 0.0076 | 0.0088 | 0.0088 | TRUE | 0.1404 |
| Flrt1 | 95 | 112 | 86 | 177 | 117 | 99 | 101 | 112 | 86 | 139 | 126 | 108 | 112.09 | 100 | 124 | 1.253 | 0.325 | 0.11761087 | 0.29016904 | 0.0075 | 0.0363 | 0.022 | 0.022 | TRUE | 0.1358 |
| Flrt2 | 76 | 55 | 55 | 154 | 97 | 76 | 81 | 55 | 55 | 121 | 104 | 83 | 83.25 | 64 | 103 | 1.625 | 0.701 | 0.00846991 | 0.04644901 | 0.0295 | 0.0465 | 0.0383 | 0.0383 | TRUE | 0.3095 |
| Flrt3 | 358 | 242 | 188 | 317 | 182 | 165 | 380 | 242 | 189 | 249 | 196 | 181 | 239.39 | 270 | 209 | 0.774 | -0.37 | 0.15996767 | 0.35445491 | 0.0755 | 0.0207 | 0.0458 | 0.0458 | TRUE | 0.6239 |
| Flt1 | 94 | 117 | 114 | 98 | 91 | 75 | 100 | 117 | 114 | 77 | 98 | 82 | 98.04 | 110 | 86 | 0.773 | -0.371 | 0.07597251 | 0.21734486 | 0 | 0.0405 | 0.0212 | 0.0212 | TRUE | 0.1435 |
| Flt3 | 15 | 22 | 12 | 29 | 14 | 9 | 16 | 22 | 12 | 23 | 15 | 10 | 16.28 | 17 | 16 | 0.968 | -0.047 | 0.92960694 | 0.9654152 | 0.0718 | 0.2093 | 0.1425 | 0.1425 | TRUE | 0.3703 |
| Flt3l | 15 | 14 | 16 | 17 | 12 | 11 | 16 | 14 | 16 | 13 | 13 | 12 | 14.05 | 15 | 13 | 0.835 | -0.259 | 0.59590314 | 0.76880035 | 0 | 0.2414 | 0.1009 | 0.1009 | TRUE | 0.0277 |

| | | | | | | | | | | | | | | | | | | | | | | | | | |
|---|---|---|---|---|---|---|---|---|---|---|---|---|---|---|---|---|---|---|---|---|---|---|---|---|---|
| Flt4 | 828 | 848 | 844 | 1275 | 990 | 761 | 879 | 847 | 847 | 1002 | 1065 | 833 | 912.4 | 858 | 967 | 1.128 | 0.173 | 0.1353187 | 0.31824306 | 0.0071 | 0.0105 | 0.0086 | 0.0086 | TRUE | 0.1715 |
| Flywch1 | 743 | 691 | 698 | 1052 | 716 | 609 | 789 | 690 | 701 | 827 | 770 | 667 | 740.7 | 727 | 755 | 1.04 | 0.057 | 0.63846471 | 0.79637583 | 0.0073 | 0.0114 | 0.0091 | 0.0091 | TRUE | 0.122 |
| Flywch2 | 74 | 91 | 114 | 77 | 50 | 64 | 79 | 91 | 114 | 61 | 54 | 70 | 78.06 | 95 | 62 | 0.648 | -0.625 | 0.01300661 | 0.0638925 | 0.015 | 0.0491 | 0.0323 | 0.0323 | TRUE | 0.3247 |
| Fmn1 | 69 | 78 | 96 | 131 | 121 | 88 | 73 | 78 | 96 | 103 | 130 | 96 | 96.18 | 82 | 110 | 1.328 | 0.409 | 0.07415454 | 0.213971 | 0.0128 | 0.0412 | 0.0272 | 0.0272 | TRUE | 0.251 |
| Fmn2 | 474 | 413 | 424 | 473 | 328 | 377 | 503 | 412 | 426 | 372 | 353 | 413 | 413.18 | 447 | 379 | 0.848 | -0.238 | 0.07393102 | 0.21348225 | 0.0065 | 0.0149 | 0.0104 | 0.0104 | TRUE | 0.1391 |
| Fmnl1 | 939 | 1327 | 1448 | 901 | 746 | 839 | 997 | 1325 | 1454 | 708 | 803 | 918 | 1034.33 | 1259 | 810 | 0.643 | -0.638 | 9.17E-05 | 0.00129108 | 0.0244 | 0.0101 | 0.0182 | 0.0182 | TRUE | 0.3953 |
| Fmnl2 | 1738 | 1774 | 1811 | 2386 | 1666 | 1583 | 1846 | 1772 | 1818 | 1876 | 1793 | 1733 | 1806.21 | 1812 | 1801 | 0.994 | -0.008 | 0.90967054 | 0.95489676 | 5.00E-04 | 0.0087 | 0.0034 | 0.0034 | TRUE | 0.0167 |
| Fmnl3 | 1199 | 909 | 911 | 1693 | 1194 | 986 | 1274 | 908 | 915 | 1331 | 1285 | 1079 | 1131.84 | 1032 | 1232 | 1.194 | 0.256 | 0.10591466 | 0.27152998 | 0.0251 | 0.0098 | 0.0172 | 0.0172 | TRUE | 0.4997 |
| Fmo1 | 6 | 3 | 1 | 19 | 14 | 7 | 6 | 3 | 1 | 15 | 15 | 8 | 8.01 | 3 | 13 | 3.677 | 1.879 | 0.02094383 | 0.08978691 | 0.0838 | 0.4184 | 0.2893 | 0.2893 | TRUE | 0.7589 |
| Fmo2 | 0 | 0 | 0 | 0 | 0 | 0 | 0 | 0 | 0 | 0 | 0 | 0 | 0 | 0 | 0 | NA | NA | NA | NA | NA | NA | NA | NA | NA | NA |
| Fmo3 | 0 | 0 | 0 | 0 | 0 | 0 | 0 | 0 | 0 | 0 | 0 | 0 | 0 | 0 | 0 | NA | NA | NA | NA | NA | NA | NA | NA | NA | NA |
| Fmo4 | 20 | 6 | 6 | 19 | 8 | 4 | 21 | 6 | 6 | 15 | 9 | 4 | 10.2 | 11 | 9 | 0.855 | -0.225 | 0.77263254 | 0.88119062 | 0.3495 | 0.33 | 0.3398 | 0.3398 | TRUE | 0.9012 |
| Fmo5 | 165 | 138 | 136 | 175 | 143 | 137 | 175 | 138 | 137 | 138 | 154 | 150 | 148.51 | 150 | 147 | 0.981 | -0.028 | 0.87755524 | 0.93954572 | 0.0054 | 0.0291 | 0.0171 | 0.0171 | TRUE | 0.2262 |
| Fmo6 | 0 | 0 | 0 | 0 | 0 | 0 | 0 | 0 | 0 | 0 | 0 | 0 | 0 | 0 | 0 | NA | NA | NA | NA | NA | NA | NA | NA | NA | NA |
| Fmo9 | 0 | 0 | 0 | 0 | 0 | 0 | 0 | 0 | 0 | 0 | 0 | 0 | 0 | 0 | 0 | NA | NA | NA | NA | NA | NA | NA | NA | NA | NA |
| Fmod | 0 | 0 | 1 | 1 | 0 | 0 | 0 | 0 | 1 | 1 | 0 | 0 | 0.3 | 0 | 0 | 0.954 | -0.067 | 0.98640617 | NA | 0 | 11.0483 | 9.2783 | 9.2783 | TRUE | 0.4098 |
| Fmr1 | 4149 | 4772 | 5058 | 6635 | 4856 | 5184 | 4407 | 4766 | 5079 | 5216 | 5225 | 5675 | 5061.16 | 4751 | 5372 | 1.131 | 0.177 | 0.03679881 | 0.13253958 | 0.0034 | 0.0076 | 0.005 | 0.005 | TRUE | 0.0487 |
| Fmr1nb | 1352 | 1625 | 1577 | 2191 | 1496 | 1640 | 1436 | 1623 | 1583 | 1722 | 1610 | 1795 | 1628.28 | 1547 | 1709 | 1.104 | 0.143 | 0.11286365 | 0.28291251 | 0.0026 | 0.0089 | 0.0053 | 0.0053 | TRUE | 0.0476 |
| Fn1 | 57945 | 68826 | 80898 | 47322 | 36646 | 45048 | 61548 | 68737 | 81230 | 37201 | 39429 | 49312 | 56242.82 | 70505 | 41981 | 0.595 | -0.748 | 1.01E-07 | 3.61E-06 | 0.0217 | 0.007 | 0.0142 | 0.0142 | TRUE | 0.2857 |
| Fn3k | 50 | 103 | 107 | 33 | 20 | 46 | 53 | 103 | 107 | 26 | 22 | 50 | 60.21 | 88 | 33 | 0.369 | -1.438 | 0.00043399 | 0.0046409 | 0.1452 | 0.0616 | 0.0995 | 0.0995 | TRUE | 1.4748 |
| Fn3krp | 763 | 970 | 923 | 1010 | 759 | 823 | 810 | 969 | 927 | 794 | 817 | 901 | 869.58 | 902 | 837 | 0.927 | -0.109 | 0.32252592 | 0.54376341 | 0.0053 | 0.0107 | 0.0076 | 0.0076 | TRUE | 0.0933 |
| Fnbp1 | 2046 | 2105 | 2272 | 2591 | 2034 | 2043 | 2173 | 2102 | 2281 | 2037 | 2188 | 2236 | 2169.75 | 2185 | 2154 | 0.985 | -0.022 | 0.78236507 | 0.88741185 | 0.0016 | 0.0084 | 0.0041 | 0.0041 | TRUE | 0.028 |
| Fnbp1l | 6326 | 6191 | 6486 | 8412 | 6151 | 6131 | 6719 | 6183 | 6513 | 6613 | 6618 | 6711 | 6559.54 | 6472 | 6647 | 1.027 | 0.039 | 0.545806 | 0.73089253 | 7.00E-04 | 0.0074 | 0.0028 | 0.0028 | TRUE | 0.0186 |
| Fnbp4 | 4549 | 4922 | 5058 | 5876 | 4477 | 4655 | 4832 | 4916 | 5079 | 4619 | 4817 | 5096 | 4893.02 | 4942 | 4844 | 0.98 | -0.029 | 0.67813709 | 0.82109459 | 0.0011 | 0.0076 | 0.0034 | 0.0034 | TRUE | 0.0251 |
| Fnd3c2 | 0 | 0 | 0 | 0 | 0 | 0 | 0 | 0 | 0 | 0 | 0 | 0 | 0 | 0 | 0 | NA | NA | NA | NA | NA | NA | NA | NA | NA | NA |
| Fndc1 | 114 | 123 | 125 | 204 | 186 | 182 | 121 | 123 | 126 | 160 | 200 | 199 | 154.86 | 123 | 186 | 1.509 | 0.594 | 0.00073362 | 0.00700402 | 0 | 0.0282 | 0.0155 | 0.0155 | TRUE | 0.1774 |
| Fndc3a | 3818 | 4080 | 4431 | 5349 | 3773 | 3882 | 4055 | 4075 | 4449 | 4205 | 4060 | 4249 | 4182.21 | 4193 | 4171 | 0.995 | -0.008 | 0.9173546 | 0.95951434 | 0.0014 | 0.0077 | 0.0035 | 0.0035 | TRUE | 0.0348 |
| Fndc3b | 1931 | 2085 | 2174 | 2212 | 1598 | 1608 | 2051 | 2082 | 2183 | 1739 | 1719 | 1760 | 1922.46 | 2105 | 1739 | 0.826 | -0.276 | 8.69E-05 | 0.00123868 | 0 | 0.0086 | 0.003 | 0.003 | TRUE | 0.0126 |
| Fndc3c1 | 157 | 97 | 88 | 136 | 81 | 77 | 167 | 97 | 88 | 107 | 87 | 84 | 105.06 | 117 | 93 | 0.794 | -0.333 | 0.2465731 | 0.46362157 | 0.0615 | 0.0383 | 0.0498 | 0.0498 | TRUE | 1.3517 |
| Fndc4 | 217 | 241 | 239 | 359 | 277 | 258 | 230 | 241 | 240 | 282 | 298 | 282 | 262.31 | 237 | 287 | 1.212 | 0.278 | 0.02860289 | 0.11136132 | 0 | 0.0195 | 0.0077 | 0.0077 | TRUE | 0.0111 |
| Fndc5 | 290 | 363 | 314 | 270 | 162 | 168 | 308 | 363 | 315 | 212 | 174 | 184 | 259.38 | 329 | 190 | 0.581 | -0.785 | 1.90E-07 | 6.25E-06 | 0.0055 | 0.0196 | 0.0122 | 0.0122 | TRUE | 0.1212 |
| Fndc7 | 8 | 6 | 2 | 7 | 11 | 8 | 8 | 6 | 2 | 6 | 12 | 9 | 7.1 | 5 | 9 | 1.572 | 0.652 | 0.40848336 | 0.62109317 | 0.0935 | 0.471 | 0.2985 | 0.2985 | TRUE | 0.3578 |
| Fndc8 | 1 | 1 | 0 | 1 | 0 | 1 | 1 | 1 | 0 | 1 | 0 | 1 | 0.66 | 1 | 1 | 0.924 | -0.114 | 0.96080322 | NA | 0 | 5.0217 | 2.3843 | 2.3843 | TRUE | 0.2521 |
| Fndc9 | 1 | 0 | 0 | 1 | 1 | 2 | 1 | 0 | 0 | 1 | 1 | 2 | 0.85 | 0 | 1 | 3.644 | 1.866 | 0.38437313 | NA | 0 | 3.8725 | 1.9476 | 1.9476 | TRUE | 0.4614 |
| Fnip1 | 3903 | 4954 | 5253 | 4283 | 3356 | 3939 | 4146 | 4948 | 5275 | 3367 | 3611 | 4312 | 4276.25 | 4790 | 3763 | 0.786 | -0.348 | 0.00732822 | 0.04152402 | 0.0155 | 0.0077 | 0.0119 | 0.0119 | TRUE | 0.1976 |
| Fnip2 | 3886 | 3557 | 3902 | 4186 | 2989 | 3071 | 4128 | 3552 | 3918 | 3291 | 3216 | 3362 | 3577.74 | 3866 | 3290 | 0.851 | -0.233 | 0.0050005 | 0.03134655 | 0.0027 | 0.0078 | 0.0047 | 0.0047 | TRUE | 0.0613 |
| Fnta | 8279 | 8860 | 8706 | 9824 | 7198 | 7676 | 8794 | 8849 | 8742 | 7723 | 7745 | 8403 | 8375.69 | 8795 | 7957 | 0.905 | -0.145 | 0.02786507 | 0.10959798 | 0.001 | 0.0073 | 0.003 | 0.003 | TRUE | 0.0292 |
| Fntb | 2454 | 2469 | 2554 | 3234 | 2378 | 2311 | 2607 | 2466 | 2564 | 2542 | 2559 | 2530 | 2544.59 | 2546 | 2544 | 0.999 | -0.001 | 0.98676792 | 0.9928109 | 0 | 0.0082 | 0.0027 | 0.0027 | TRUE | 0.0091 |
| Focad | 3207 | 4351 | 4760 | 4556 | 3569 | 3774 | 3406 | 4345 | 4780 | 3582 | 3840 | 4131 | 4014.03 | 4177 | 3851 | 0.922 | -0.118 | 0.38009543 | 0.59468399 | 0.0158 | 0.0077 | 0.0127 | 0.0127 | TRUE | 0.3169 |
| Folh1 | 10 | 7 | 5 | 20 | 14 | 16 | 11 | 7 | 5 | 16 | 15 | 18 | 11.82 | 8 | 16 | 2.14 | 1.098 | 0.05604407 | 0.17757093 | 0 | 0.2856 | 0.1395 | 0.1395 | TRUE | 0.3398 |
| Folr1 | 402 | 322 | 291 | 1371 | 809 | 715 | 427 | 322 | 292 | 1078 | 870 | 783 | 628.61 | 347 | 910 | 2.628 | 1.394 | 9.42E-15 | 1.28E-12 | 0.0317 | 0.0121 | 0.0213 | 0.0213 | TRUE | 0.4621 |
| Folr2 | 3 | 1 | 4 | 15 | 7 | 1 | 3 | 1 | 4 | 12 | 8 | 1 | 4.77 | 3 | 7 | 2.532 | 1.34 | 0.21637685 | 0.42835392 | 0.4721 | 0.6976 | 0.5888 | 0.5888 | TRUE | 0.1613 |
| Folr4 | 2 | 1 | 0 | 2 | 0 | 1 | 2 | 1 | 0 | 2 | 0 | 1 | 0.96 | 1 | 1 | 0.883 | -0.179 | 0.92987033 | NA | 0 | 3.4209 | 1.9665 | 1.9665 | TRUE | 0.1865 |
| Fopnl | 1248 | 1449 | 1299 | 1696 | 1206 | 1190 | 1326 | 1447 | 1304 | 1333 | 1298 | 1303 | 1335.09 | 1359 | 1311 | 0.965 | -0.051 | 0.53206185 | 0.7201304 | 0.001 | 0.0094 | 0.0041 | 0.0041 | TRUE | 0.0388 |
| Fos | 92 | 147 | 148 | 131 | 92 | 106 | 98 | 147 | 149 | 103 | 99 | 116 | 118.52 | 131 | 106 | 0.807 | -0.309 | 0.1717614 | 0.37062439 | 0.0217 | 0.0347 | 0.0283 | 0.0283 | TRUE | 0.4988 |
| Fosb | 4 | 6 | 3 | 8 | 11 | 3 | 4 | 6 | 3 | 6 | 12 | 3 | 5.78 | 4 | 7 | 1.61 | 0.687 | 0.41915606 | 0.62979226 | 0 | 0.5772 | 0.3373 | 0.3373 | TRUE | 0.294 |
| Fosl1 | 11 | 10 | 10 | 18 | 8 | 4 | 12 | 10 | 10 | 14 | 9 | 4 | 9.81 | 11 | 9 | 0.874 | -0.194 | 0.76529975 | 0.87700158 | 0 | 0.3428 | 0.2033 | 0.2033 | TRUE | 0.3385 |
| Fosl2 | 25 | 61 | 64 | 113 | 121 | 111 | 27 | 61 | 64 | 89 | 130 | 122 | 82.04 | 51 | 114 | 2.234 | 1.16 | 0.00067285 | 0.00656698 | 0.0988 | 0.0471 | 0.0694 | 0.0694 | TRUE | 1.3698 |
| Foxa1 | 130 | 152 | 138 | 266 | 187 | 160 | 138 | 152 | 139 | 209 | 201 | 175 | 168.98 | 143 | 195 | 1.369 | 0.453 | 0.00550401 | 0.03354297 | 0 | 0.0264 | 0.0131 | 0.0131 | TRUE | 0.0843 |
| Foxa2 | 3 | 3 | 4 | 18 | 12 | 1 | 3 | 3 | 4 | 14 | 13 | 1 | 6.39 | 3 | 9 | 2.787 | 1.479 | 0.1303527 | 0.31086811 | 0.4479 | 0.5223 | 0.4866 | 0.4866 | TRUE | 1.8135 |
| Foxa3 | 123 | 134 | 124 | 125 | 100 | 81 | 131 | 134 | 125 | 98 | 108 | 89 | 113.92 | 130 | 98 | 0.757 | -0.401 | 0.03226265 | 0.12103864 | 0 | 0.0358 | 0.0164 | 0.0164 | TRUE | 0.0633 |
| Foxb1 | 0 | 0 | 0 | 0 | 0 | 0 | 0 | 0 | 0 | 0 | 0 | 0 | 0 | 0 | 0 | NA | NA | NA | NA | NA | NA | NA | NA | NA | NA |
| Foxb2 | 1 | 2 | 0 | 0 | 0 | 0 | 1 | 2 | 0 | 0 | 0 | 0 | 0.51 | 1 | 0 | 0.176 | -2.508 | 0.44268408 | NA | 0 | 6.4675 | 6.1932 | 6.1932 | TRUE | 0.064 |
| Foxc1 | 8 | 29 | 21 | 46 | 27 | 30 | 8 | 29 | 21 | 36 | 29 | 33 | 26.1 | 19 | 33 | 1.673 | 0.742 | 0.11517738 | 0.28657165 | 0.0987 | 0.1331 | 0.1188 | 0.1188 | TRUE | 0.5816 |
| Foxc2 | 29 | 54 | 65 | 102 | 64 | 95 | 31 | 54 | 65 | 80 | 69 | 104 | 67.17 | 50 | 84 | 1.682 | 0.75 | 0.02204459 | 0.09333886 | 0.0678 | 0.0559 | 0.0614 | 0.0614 | TRUE | 0.8463 |
| Foxd1 | 84 | 62 | 70 | 40 | 40 | 39 | 89 | 62 | 70 | 31 | 43 | 43 | 56.43 | 74 | 39 | 0.526 | -0.928 | 0.00144827 | 0.01192174 | 0.0169 | 0.0653 | 0.0416 | 0.0416 | TRUE | 0.2983 |
| Foxd2 | 5 | 15 | 12 | 45 | 27 | 21 | 5 | 15 | 12 | 35 | 29 | 23 | 19.96 | 11 | 29 | 2.704 | 1.435 | 0.00428726 | 0.02786626 | 0.0381 | 0.172 | 0.1181 | 0.1181 | TRUE | 0.3981 |
| Foxd3 | 1218 | 697 | 655 | 928 | 690 | 501 | 1294 | 696 | 658 | 730 | 742 | 548 | 777.98 | 883 | 673 | 0.764 | -0.389 | 0.11998806 | 0.29385896 | 0.0855 | 0.0111 | 0.0439 | 0.0439 | TRUE | 1.975 |
| Foxd4 | 10 | 3 | 9 | 13 | 14 | 5 | 11 | 3 | 9 | 10 | 15 | 5 | 8.9 | 8 | 10 | 1.362 | 0.446 | 0.54383634 | 0.72899211 | 0.1517 | 0.377 | 0.2737 | 0.2737 | TRUE | 0.2299 |
| Foxe1 | 6 | 11 | 8 | 29 | 17 | 16 | 6 | 11 | 8 | 23 | 18 | 18 | 14 | 8 | 20 | 2.314 | 1.21 | 0.02237071 | 0.09421777 | 0 | 0.2422 | 0.1173 | 0.1173 | TRUE | 0.2111 |
| Foxe3 | 0 | 0 | 0 | 3 | 5 | 2 | 0 | 0 | 0 | 2 | 5 | 2 | 1.65 | 0 | 3 | 17.532 | 4.132 | 0.02746524 | NA | 0 | 1.998 | 1.3819 | 1.3819 | TRUE | 0.3969 |
| Foxf1 | 3 | 7 | 8 | 15 | 5 | 18 | 3 | 7 | 8 | 12 | 5 | 20 | 9.18 | 6 | 12 | 2.014 | 1.01 | 0.17406045 | 0.37335066 | 0.1756 | 0.3657 | 0.2744 | 0.2744 | TRUE | 0.1341 |
| Foxf2 | 0 | 0 | 0 | 5 | 4 | 0 | 0 | 0 | 0 | 4 | 4 | 0 | 1.37 | 0 | 3 | 14.689 | 3.877 | 0.1038995 | NA | 6.3016 | 2.4074 | 2.9173 | 2.9173 | TRUE | 0.838 |
| Foxg1 | 43 | 50 | 47 | 148 | 96 | 86 | 46 | 50 | 47 | 116 | 103 | 94 | 76.1 | 48 | 104 | 2.204 | 1.14 | 1.17E-06 | 3.16E-05 | 0 | 0.0502 | 0.0242 | 0.0242 | TRUE | 0.1047 |
| Foxh1 | 1214 | 1635 | 1755 | 1787 | 1308 | 1392 | 1289 | 1633 | 1762 | 1405 | 1407 | 1524 | 1503.41 | 1561 | 1445 | 0.925 | -0.112 | 0.38525788 | 0.59920343 | 0.0123 | 0.0091 | 0.0113 | 0.0113 | TRUE | 0.2791 |
| Foxi1 | 1 | 0 | 0 | 0 | 1 | 0 | 1 | 0 | 0 | 0 | 1 | 0 | 0.36 | 0 | 0 | 0.954 | -0.067 | 0.98557668 | NA | 0 | 9.2518 | 8.0179 | 8.0179 | TRUE | 0.5204 |
| Foxi2 | 1 | 2 | 0 | 4 | 2 | 2 | 1 | 2 | 0 | 3 | 2 | 2 | 1.76 | 1 | 2 | 2.466 | 1.302 | 0.38149857 | NA | 0 | 1.8814 | 0.8999 | 0.8999 | TRUE | 0.3728 |
| Foxi3 | 36 | 29 | 23 | 27 | 24 | 11 | 38 | 29 | 23 | 21 | 26 | 12 | 24.9 | 30 | 20 | 0.658 | -0.605 | 0.16857146 | 0.36640749 | 0.0465 | 0.1392 | 0.0973 | 0.0973 | TRUE | 0.4123 |
| Foxj1 | 96 | 155 | 157 | 134 | 104 | 152 | 102 | 155 | 158 | 105 | 112 | 166 | 133.01 | 138 | 128 | 0.922 | -0.117 | 0.65444972 | 0.80574825 | 0.0522 | 0.0317 | 0.0416 | 0.0416 | TRUE | 0.6938 |

| | | | | | | | | | | | | | | | | | | | | | | | | | |
|---|---|---|---|---|---|---|---|---|---|---|---|---|---|---|---|---|---|---|---|---|---|---|---|---|---|
| Foxj2 | 1132 | 1293 | 1391 | 1299 | 983 | 1102 | 1202 | 1291 | 1397 | 1021 | 1058 | 1206 | 1195.93 | 1297 | 1095 | 0.844 | -0.245 | 0.02225808 | 0.09391504 | 0.0056 | 0.0097 | 0.0075 | 0.0075 | TRUE | 0.0938 |
| Foxj3 | 2845 | 3156 | 3285 | 3834 | 2870 | 2887 | 3022 | 3152 | 3298 | 3014 | 3088 | 3160 | 3122.42 | 3157 | 3087 | 0.978 | -0.033 | 0.64303207 | 0.79907134 | 9.00E-04 | 0.008 | 0.0033 | 0.0033 | TRUE | 0.0186 |
| Foxk1 | 1701 | 1932 | 1945 | 1930 | 1497 | 1570 | 1807 | 1930 | 1953 | 1517 | 1611 | 1719 | 1755.96 | 1897 | 1616 | 0.851 | -0.232 | 0.00669987 | 0.03893525 | 0.0022 | 0.0088 | 0.0047 | 0.0047 | TRUE | 0.0375 |
| Foxk2 | 8041 | 8975 | 9184 | 9063 | 6709 | 6817 | 8541 | 8963 | 9222 | 7125 | 7218 | 7462 | 8088.58 | 8909 | 7268 | 0.816 | -0.294 | 5.48E-06 | 0.0001197 | 8.00E-04 | 0.0073 | 0.0029 | 0.0029 | TRUE | 0.0159 |
| Foxl1 | 1 | 0 | 0 | 1 | 0 | 0 | 1 | 0 | 0 | 1 | 0 | 0 | 0.31 | 0 | 0 | 0.954 | -0.067 | 0.98625928 | NA | 0 | 10.7014 | 9.0384 | 9.0384 | TRUE | 0.4614 |
| Foxl2 | 5 | 9 | 19 | 30 | 29 | 19 | 5 | 9 | 19 | 24 | 31 | 21 | 18.16 | 11 | 25 | 2.254 | 1.173 | 0.03399065 | 0.12549168 | 0.1089 | 0.1883 | 0.1552 | 0.1552 | TRUE | 1.4682 |
| Foxl2os | 8 | 20 | 26 | 49 | 44 | 34 | 8 | 20 | 26 | 39 | 47 | 37 | 29.61 | 18 | 41 | 2.244 | 1.166 | 0.00788961 | 0.04384602 | 0.068 | 0.1182 | 0.0989 | 0.0989 | TRUE | 0.9309 |
| Foxm1 | 4574 | 4818 | 5014 | 5842 | 4297 | 4004 | 4858 | 4812 | 5035 | 4593 | 4623 | 4383 | 4717.27 | 4902 | 4533 | 0.925 | -0.113 | 0.07574207 | 0.21700345 | 4.00E-04 | 0.0076 | 0.0027 | 0.0027 | TRUE | 0.0101 |
| Foxn1 | 6 | 3 | 3 | 0 | 1 | 2 | 6 | 3 | 3 | 0 | 1 | 2 | 2.61 | 4 | 1 | 0.248 | -2.012 | 0.13987447 | NA | 0 | 1.2702 | 0.7377 | 0.7377 | TRUE | 0.5885 |
| Foxn2 | 2882 | 3270 | 3278 | 3827 | 2855 | 2807 | 3061 | 3266 | 3291 | 3008 | 3072 | 3073 | 3128.57 | 3206 | 3051 | 0.951 | -0.072 | 0.28700456 | 0.50821107 | 5.00E-04 | 0.008 | 0.003 | 0.003 | TRUE | 0.0189 |
| Foxn3 | 545 | 770 | 822 | 660 | 527 | 625 | 579 | 769 | 825 | 519 | 567 | 684 | 657.21 | 724 | 590 | 0.813 | -0.298 | 0.07584599 | 0.21720036 | 0.024 | 0.0119 | 0.0188 | 0.0188 | TRUE | 0.3638 |
| Foxn4 | 233 | 239 | 213 | 281 | 211 | 214 | 247 | 239 | 214 | 221 | 227 | 234 | 230.37 | 233 | 227 | 0.974 | -0.038 | 0.78415755 | 0.88855038 | 0 | 0.0212 | 0.0097 | 0.0097 | TRUE | 0.0591 |
| Foxo1 | 2310 | 2273 | 2446 | 2582 | 2062 | 2012 | 2454 | 2270 | 2456 | 2030 | 2219 | 2202 | 2271.76 | 2393 | 2150 | 0.898 | -0.155 | 0.05277561 | 0.17032473 | 0.0018 | 0.0084 | 0.0042 | 0.0042 | TRUE | 0.0298 |
| Foxo3 | 2099 | 2441 | 2565 | 2987 | 2371 | 2423 | 2230 | 2438 | 2576 | 2348 | 2551 | 2652 | 2465.74 | 2415 | 2517 | 1.042 | 0.059 | 0.520298 | 0.71109881 | 0.0037 | 0.0082 | 0.0057 | 0.0057 | TRUE | 0.0542 |
| Foxo4 | 2150 | 2320 | 2340 | 2695 | 1975 | 2093 | 2284 | 2317 | 2350 | 2119 | 2125 | 2291 | 2247.5 | 2317 | 2178 | 0.94 | -0.09 | 0.21467995 | 0.42654727 | 6.00E-04 | 0.0084 | 0.0033 | 0.0033 | TRUE | 0.0246 |
| Foxo6 | 465 | 376 | 427 | 387 | 319 | 332 | 494 | 376 | 429 | 304 | 343 | 363 | 384.84 | 433 | 337 | 0.778 | -0.363 | 0.01418894 | 0.06793806 | 0.0111 | 0.0155 | 0.0131 | 0.0131 | TRUE | 0.1757 |
| Foxp1 | 3318 | 3091 | 3309 | 2655 | 1943 | 2026 | 3524 | 3087 | 3323 | 2087 | 2091 | 2218 | 2721.56 | 3311 | 2132 | 0.644 | -0.635 | 1.56E-14 | 2.03E-12 | 0.0022 | 0.0081 | 0.0045 | 0.0045 | TRUE | 0.0427 |
| Foxp2 | 2 | 1 | 0 | 11 | 5 | 3 | 2 | 1 | 0 | 9 | 5 | 3 | 3.41 | 1 | 6 | 5.679 | 2.506 | 0.04832853 | NA | 0 | 0.9742 | 0.58 | 0.58 | TRUE | 0.2683 |
| Foxp3 | 0 | 0 | 1 | 5 | 3 | 0 | 0 | 0 | 1 | 4 | 3 | 0 | 1.36 | 0 | 2 | 6.597 | 2.722 | 0.2026911 | NA | 0 | 2.4285 | 2.0864 | 2.0864 | TRUE | 0.7324 |
| Foxp4 | 2167 | 2116 | 2226 | 2159 | 1719 | 1561 | 2302 | 2113 | 2235 | 1697 | 1850 | 1709 | 1984.28 | 2217 | 1752 | 0.79 | -0.34 | 2.29E-05 | 0.00039928 | 0.0016 | 0.0086 | 0.0041 | 0.0041 | TRUE | 0.0285 |
| Foxq1 | 79 | 141 | 141 | 208 | 192 | 172 | 84 | 141 | 142 | 164 | 207 | 188 | 154.11 | 122 | 186 | 1.52 | 0.604 | 0.01018588 | 0.05349496 | 0.0384 | 0.0283 | 0.033 | 0.033 | TRUE | 0.7456 |
| Foxr1 | 105 | 119 | 116 | 146 | 129 | 122 | 112 | 119 | 116 | 115 | 139 | 134 | 122.33 | 116 | 129 | 1.112 | 0.153 | 0.40576163 | 0.61850512 | 0 | 0.0338 | 0.0161 | 0.0161 | TRUE | 0.107 |
| Foxr2 | 179 | 212 | 236 | 186 | 126 | 128 | 190 | 212 | 237 | 146 | 136 | 140 | 176.79 | 213 | 141 | 0.661 | -0.597 | 0.00026678 | 0.00309157 | 0 | 0.0255 | 0.0135 | 0.0135 | TRUE | 0.1066 |
| Foxred1 | 2037 | 2114 | 2215 | 2620 | 1864 | 1808 | 2164 | 2111 | 2224 | 2060 | 2006 | 1979 | 2090.56 | 2166 | 2015 | 0.93 | -0.104 | 0.12961287 | 0.30978243 | 0 | 0.0085 | 0.0029 | 0.0029 | TRUE | 0.0066 |
| Foxred2 | 769 | 947 | 972 | 1075 | 821 | 818 | 817 | 946 | 976 | 845 | 883 | 895 | 893.74 | 913 | 874 | 0.957 | -0.063 | 0.54357753 | 0.72871269 | 0.0037 | 0.0106 | 0.0066 | 0.0066 | TRUE | 0.1002 |
| Foxs1 | 1 | 1 | 1 | 2 | 1 | 2 | 1 | 1 | 1 | 2 | 1 | 2 | 1.32 | 1 | 2 | 1.576 | 0.656 | 0.6715149 | NA | 0 | 2.5082 | 0.9225 | 0.9225 | TRUE | 0.0617 |
| Fpgs | 4505 | 4254 | 4394 | 4815 | 3862 | 3966 | 4785 | 4249 | 4412 | 3785 | 4155 | 4341 | 4287.92 | 4482 | 4094 | 0.913 | -0.131 | 0.13978778 | 0.32441625 | 0.0039 | 0.0077 | 0.0054 | 0.0054 | TRUE | 0.0536 |
| Fpgt | 878 | 869 | 837 | 1352 | 913 | 872 | 933 | 868 | 840 | 1063 | 982 | 955 | 940.1 | 880 | 1000 | 1.137 | 0.186 | 0.05017497 | 0.1646265 | 0.002 | 0.0104 | 0.0054 | 0.0054 | TRUE | 0.0375 |
| Fpr-rs3 | 0 | 0 | 0 | 0 | 0 | 0 | 0 | 0 | 0 | 0 | 0 | 0 | 0 | 0 | 0 | NA | NA | NA | NA | NA | NA | NA | NA | NA | NA |
| Fpr-rs4 | 0 | 0 | 0 | 0 | 0 | 0 | 0 | 0 | 0 | 0 | 0 | 0 | 0 | 0 | 0 | NA | NA | NA | NA | NA | NA | NA | NA | NA | NA |
| Fpr-rs6 | 0 | 0 | 0 | 0 | 0 | 0 | 0 | 0 | 0 | 0 | 0 | 0 | 0 | 0 | 0 | NA | NA | NA | NA | NA | NA | NA | NA | NA | NA |
| Fpr1 | 0 | 0 | 0 | 0 | 0 | 0 | 0 | 0 | 0 | 0 | 0 | 0 | 0 | 0 | 0 | NA | NA | NA | NA | NA | NA | NA | NA | NA | NA |
| Fpr2 | 0 | 0 | 0 | 0 | 0 | 0 | 0 | 0 | 0 | 0 | 0 | 0 | 0 | 0 | 0 | NA | NA | NA | NA | NA | NA | NA | NA | NA | NA |
| Fpr3 | 0 | 0 | 0 | 0 | 0 | 0 | 0 | 0 | 0 | 0 | 0 | 0 | 0 | 0 | 0 | NA | NA | NA | NA | NA | NA | NA | NA | NA | NA |
| Fra10ac1 | 1207 | 1055 | 1088 | 1418 | 1064 | 1029 | 1282 | 1054 | 1092 | 1115 | 1145 | 1126 | 1135.68 | 1143 | 1129 | 0.988 | -0.018 | 0.86444861 | 0.93294221 | 0.0047 | 0.0098 | 0.0068 | 0.0068 | TRUE | 0.1357 |
| Fras1 | 52 | 37 | 41 | 64 | 62 | 42 | 55 | 37 | 41 | 50 | 67 | 46 | 49.39 | 44 | 54 | 1.222 | 0.289 | 0.34827502 | 0.56702733 | 0.02 | 0.0736 | 0.048 | 0.048 | TRUE | 0.3418 |
| Frat1 | 147 | 162 | 158 | 214 | 129 | 124 | 156 | 162 | 159 | 168 | 139 | 136 | 153.22 | 159 | 148 | 0.933 | -0.1 | 0.56131062 | 0.74256648 | 0 | 0.0284 | 0.0147 | 0.0147 | TRUE | 0.1708 |
| Frat2 | 417 | 565 | 618 | 493 | 433 | 450 | 443 | 564 | 621 | 388 | 466 | 493 | 495.63 | 543 | 449 | 0.825 | -0.277 | 0.08909965 | 0.24249864 | 0.0204 | 0.0136 | 0.0171 | 0.0171 | TRUE | 0.3001 |
| Frem1 | 41 | 41 | 20 | 12 | 16 | 14 | 44 | 41 | 20 | 9 | 17 | 15 | 24.43 | 35 | 14 | 0.397 | -1.334 | 0.00571077 | 0.03458377 | 0.0926 | 0.1418 | 0.1179 | 0.1179 | TRUE | 0.9834 |
| Frem2 | 1254 | 1583 | 1932 | 770 | 603 | 686 | 1332 | 1581 | 1940 | 605 | 649 | 751 | 1142.98 | 1618 | 668 | 0.413 | -1.277 | 3.57E-16 | 5.63E-14 | 0.0225 | 0.0098 | 0.0166 | 0.0166 | TRUE | 0.1542 |
| Frem3 | 1 | 0 | 0 | 5 | 7 | 2 | 1 | 0 | 0 | 4 | 8 | 2 | 2.45 | 0 | 5 | 12.41 | 3.633 | 0.02744661 | NA | 0 | 1.3503 | 0.8487 | 0.8487 | TRUE | 0.3854 |
| Frg1 | 2857 | 3191 | 3073 | 4391 | 3361 | 3268 | 3035 | 3187 | 3086 | 3452 | 3616 | 3577 | 3325.43 | 3103 | 3548 | 1.143 | 0.193 | 0.00285159 | 0.02035077 | 3.00E-04 | 0.0079 | 0.0027 | 0.0027 | TRUE | 0.007 |
| Frk | 38 | 33 | 36 | 102 | 50 | 37 | 40 | 33 | 36 | 80 | 54 | 41 | 47.33 | 36 | 58 | 1.608 | 0.685 | 0.0476723 | 0.15867893 | 0.0519 | 0.0765 | 0.064 | 0.064 | TRUE | 0.3596 |
| Frmd3 | 5 | 3 | 4 | 16 | 7 | 5 | 5 | 3 | 4 | 13 | 8 | 5 | 6.32 | 4 | 9 | 2.113 | 1.079 | 0.17449383 | 0.37392889 | 0 | 0.5284 | 0.2738 | 0.2738 | TRUE | 0.5843 |
| Frmd4a | 795 | 840 | 886 | 1197 | 873 | 829 | 844 | 839 | 890 | 941 | 939 | 907 | 893.46 | 858 | 929 | 1.084 | 0.116 | 0.16642455 | 0.36370349 | 0 | 0.0106 | 0.0039 | 0.0039 | TRUE | 0.0127 |
| Frmd4b | 551 | 561 | 504 | 645 | 478 | 517 | 585 | 560 | 506 | 507 | 514 | 566 | 539.81 | 550 | 529 | 0.96 | -0.058 | 0.60300177 | 0.77412127 | 0.0027 | 0.013 | 0.0072 | 0.0072 | TRUE | 0.0586 |
| Frmd5 | 455 | 446 | 451 | 614 | 397 | 413 | 483 | 445 | 453 | 483 | 427 | 452 | 457.25 | 460 | 454 | 0.988 | -0.018 | 0.87058705 | 0.93643176 | 0 | 0.0141 | 0.0066 | 0.0066 | TRUE | 0.0377 |
| Frmd6 | 1320 | 1204 | 1191 | 1923 | 1392 | 1404 | 1402 | 1202 | 1196 | 1512 | 1498 | 1537 | 1391.12 | 1267 | 1516 | 1.197 | 0.259 | 0.00632074 | 0.03744955 | 0.0033 | 0.0093 | 0.0057 | 0.0057 | TRUE | 0.1042 |
| Frmd7 | 0 | 1 | 0 | 0 | 0 | 0 | 0 | 1 | 0 | 0 | 0 | 0 | 0.17 | 0 | 0 | 0.49 | -1.029 | 0.80087 | NA | 0 | 19.7991 | 10 | 10 | TRUE | 0.405 |
| Frmd8 | 2429 | 2115 | 2032 | 3220 | 2151 | 1930 | 2580 | 2112 | 2040 | 2531 | 2314 | 2113 | 2281.83 | 2244 | 2319 | 1.034 | 0.048 | 0.69176441 | 0.83095304 | 0.0118 | 0.0083 | 0.0103 | 0.0103 | TRUE | 0.2072 |
| Frmpd1 | 198 | 189 | 176 | 215 | 176 | 157 | 210 | 189 | 177 | 169 | 189 | 172 | 184.34 | 192 | 177 | 0.92 | -0.12 | 0.44487468 | 0.65143101 | 0 | 0.0248 | 0.0123 | 0.0123 | TRUE | 0.0747 |
| Frmpd3 | 1 | 2 | 1 | 2 | 1 | 0 | 1 | 2 | 1 | 2 | 1 | 0 | 1.12 | 1 | 1 | 0.676 | -0.565 | 0.74666147 | NA | 0 | 2.9519 | 1.2963 | 1.2963 | TRUE | 0.2738 |
| Frmpd4 | 0 | 0 | 1 | 2 | 1 | 0 | 0 | 0 | 1 | 2 | 1 | 0 | 0.61 | 0 | 1 | 2.461 | 1.299 | 0.62272026 | NA | 0 | 5.419 | 3.4832 | 3.4832 | TRUE | 0.4098 |
| Frrs1 | 2471 | 2576 | 2749 | 3194 | 2196 | 2225 | 2625 | 2573 | 2760 | 2511 | 2363 | 2436 | 2544.47 | 2653 | 2437 | 0.919 | -0.122 | 0.08727525 | 0.2390596 | 8.00E-04 | 0.0082 | 0.0033 | 0.0033 | TRUE | 0.0154 |
| Frrs1l | 130 | 114 | 86 | 171 | 111 | 103 | 138 | 114 | 86 | 134 | 119 | 113 | 117.48 | 113 | 122 | 1.087 | 0.121 | 0.59412849 | 0.76757819 | 0.0218 | 0.0349 | 0.0285 | 0.0285 | TRUE | 0.2363 |
| Frs2 | 2876 | 2691 | 2782 | 3187 | 2351 | 2425 | 3055 | 2688 | 2793 | 2505 | 2530 | 2655 | 2704.2 | 2845 | 2563 | 0.901 | -0.151 | 0.06450514 | 0.19561145 | 0.0021 | 0.0081 | 0.0044 | 0.0044 | TRUE | 0.0503 |
| Frs3 | 282 | 508 | 511 | 416 | 366 | 480 | 300 | 507 | 513 | 327 | 394 | 525 | 427.71 | 440 | 415 | 0.943 | -0.085 | 0.72436098 | 0.85090198 | 0.0693 | 0.0146 | 0.0398 | 0.0398 | TRUE | 0.7654 |
| Fry | 1776 | 1909 | 2242 | 2057 | 1599 | 1650 | 1886 | 1907 | 2251 | 1617 | 1720 | 1806 | 1864.64 | 2015 | 1714 | 0.851 | -0.233 | 0.02365328 | 0.09783886 | 0.0059 | 0.0087 | 0.0071 | 0.0071 | TRUE | 0.1277 |
| Fryl | 2399 | 2549 | 2830 | 3169 | 2253 | 2368 | 2548 | 2546 | 2842 | 2491 | 2424 | 2592 | 2573.82 | 2645 | 2502 | 0.946 | -0.08 | 0.32491716 | 0.54594394 | 0.0021 | 0.0082 | 0.0044 | 0.0044 | TRUE | 0.0513 |
| Frzb | 1 | 8 | 5 | 19 | 8 | 12 | 1 | 8 | 5 | 15 | 9 | 13 | 8.46 | 5 | 12 | 2.601 | 1.379 | 0.07287108 | 0.211428 | 0.0721 | 0.3964 | 0.2781 | 0.2781 | TRUE | 0.608 |
| Fsbp | 32 | 25 | 19 | 51 | 33 | 39 | 34 | 25 | 19 | 40 | 36 | 43 | 32.72 | 26 | 40 | 1.52 | 0.604 | 0.09291027 | 0.24896038 | 0 | 0.1076 | 0.061 | 0.061 | TRUE | 0.4309 |
| Fscb | 0 | 0 | 0 | 2 | 0 | 0 | 0 | 0 | 0 | 2 | 0 | 0 | 0.26 | 0 | 1 | 2.886 | 1.529 | 0.70676491 | NA | 0 | 12.5792 | 10 | 10 | TRUE | 0.9669 |
| Fscn1 | 18761 | 21015 | 21874 | 24741 | 19052 | 18451 | 19927 | 20988 | 21964 | 19449 | 20499 | 20198 | 20504.15 | 20960 | 20049 | 0.956 | -0.064 | 0.34300908 | 0.56312473 | 0.0015 | 0.0071 | 0.0033 | 0.0033 | TRUE | 0.0227 |
| Fscn2 | 5 | 4 | 11 | 11 | 7 | 8 | 5 | 4 | 11 | 9 | 8 | 9 | 7.55 | 7 | 9 | 1.225 | 0.293 | 0.68192042 | 0.82362649 | 0 | 0.4434 | 0.2346 | 0.2346 | TRUE | 0.7931 |
| Fscn3 | 0 | 0 | 0 | 1 | 0 | 0 | 0 | 0 | 0 | 1 | 0 | 0 | 0.13 | 0 | 0 | 1.859 | 0.894 | 0.82650734 | NA | 0 | 25.1515 | 10 | 10 | TRUE | 0.2563 |
| Fsd1 | 1257 | 1316 | 1245 | 978 | 833 | 908 | 1335 | 1314 | 1250 | 769 | 896 | 994 | 1093.1 | 1300 | 886 | 0.681 | -0.555 | 1.64E-06 | 4.22E-05 | 0.0073 | 0.0099 | 0.0087 | 0.0087 | TRUE | 0.1654 |
| Fsd1l | 1453 | 1365 | 1374 | 1702 | 1314 | 1158 | 1543 | 1363 | 1380 | 1338 | 1414 | 1268 | 1384.27 | 1429 | 1340 | 0.938 | -0.092 | 0.32206697 | 0.54330584 | 0.0028 | 0.0093 | 0.0056 | 0.0056 | TRUE | 0.059 |
| Fsd2 | 21 | 23 | 23 | 25 | 16 | 24 | 22 | 23 | 23 | 20 | 17 | 26 | 21.92 | 23 | 21 | 0.92 | -0.12 | 0.76858331 | 0.87928573 | 0 | 0.1572 | 0.0735 | 0.0735 | TRUE | 0.2368 |
| Fshb | 0 | 0 | 0 | 0 | 0 | 0 | 0 | 0 | 0 | 0 | 0 | 0 | 0 | 0 | 0 | NA | NA | NA | NA | NA | NA | NA | NA | NA | NA |

| | | | | | | | | | | | | | | | | | | | | | | | | | |
|---|---|---|---|---|---|---|---|---|---|---|---|---|---|---|---|---|---|---|---|---|---|---|---|---|---|
| Fshr | 2 | 0 | 1 | 1 | 1 | 1 | 2 | 0 | 1 | 1 | 1 | 1 | 1.01 | 1 | 1 | 0.945 | -0.082 | 0.96468043 | NA | 0 | 3.2552 | 1.4594 | 1.4594 | TRUE | 0.198 |
| Fsip1 | 63 | 78 | 80 | 129 | 122 | 98 | 67 | 78 | 80 | 101 | 131 | 107 | 94.18 | 75 | 113 | 1.505 | 0.589 | 0.00715243 | 0.04072497 | 0 | 0.0419 | 0.0235 | 0.0235 | TRUE | 0.1879 |
| Fst | 31 | 28 | 25 | 38 | 32 | 32 | 33 | 28 | 25 | 30 | 34 | 35 | 30.89 | 29 | 33 | 1.152 | 0.204 | 0.55096612 | 0.73511866 | 0 | 0.1136 | 0.0519 | 0.0519 | TRUE | 0.106 |
| Fstl1 | 3207 | 3463 | 3387 | 2262 | 1683 | 1893 | 3406 | 3459 | 3401 | 1778 | 1811 | 2072 | 2654.51 | 3422 | 1887 | 0.551 | -0.86 | 2.43E-23 | 8.33E-21 | 0.0031 | 0.0081 | 0.005 | 0.005 | TRUE | 0.0882 |
| Fstl3 | 349 | 560 | 609 | 458 | 342 | 439 | 371 | 559 | 612 | 360 | 368 | 481 | 458.34 | 514 | 403 | 0.783 | -0.353 | 0.08825653 | 0.24092906 | 0.0444 | 0.0141 | 0.0287 | 0.0287 | TRUE | 0.6894 |
| Fstl4 | 42 | 57 | 61 | 88 | 56 | 65 | 45 | 57 | 61 | 69 | 60 | 71 | 60.56 | 54 | 67 | 1.232 | 0.301 | 0.24590448 | 0.46297247 | 0 | 0.0613 | 0.0317 | 0.0317 | TRUE | 0.1957 |
| Fstl5 | 291 | 200 | 121 | 88 | 72 | 78 | 309 | 200 | 121 | 69 | 77 | 85 | 143.73 | 210 | 77 | 0.368 | -1.444 | 7.70E-06 | 0.00015872 | 0.1065 | 0.0298 | 0.0662 | 0.0662 | TRUE | 0.1379 |
| Ftcd | 6 | 17 | 6 | 8 | 7 | 12 | 6 | 17 | 6 | 6 | 8 | 13 | 9.39 | 10 | 9 | 0.906 | -0.142 | 0.84235285 | 0.92050088 | 0.1608 | 0.3578 | 0.2586 | 0.2586 | TRUE | 1.4278 |
| Fth1 | 33697 | 30553 | 30583 | 30630 | 22000 | 20307 | 35792 | 30514 | 30709 | 24079 | 23671 | 22229 | 27832.17 | 32338 | 23326 | 0.721 | -0.471 | 1.51E-07 | 5.10E-06 | 0.0051 | 0.007 | 0.0058 | 0.0058 | TRUE | 0.1068 |
| Fthl17 | 5 | 6 | 9 | 39 | 32 | 19 | 5 | 6 | 9 | 31 | 34 | 21 | 17.7 | 7 | 29 | 4.222 | 2.078 | 8.98E-05 | 0.00126984 | 0 | 0.193 | 0.1106 | 0.1106 | TRUE | 0.344 |
| Ftl1 | 104841 | 116481 | 118413 | 111474 | 79120 | 79943 | 111360 | 116331 | 118899 | 87632 | 85128 | 87510 | 101143.34 | 115530 | 86757 | 0.751 | -0.413 | 1.03E-12 | 1.05E-10 | 7.00E-04 | 0.0069 | 0.0024 | 0.0024 | TRUE | 0.0122 |
| Ftmt | 0 | 0 | 0 | 0 | 0 | 0 | 0 | 0 | 0 | 0 | 0 | 0 | 0 | 0 | 0 | NA | NA | NA | NA | NA | NA | NA | NA | NA | NA |
| Fto | 5550 | 5993 | 6259 | 6974 | 4802 | 5173 | 5895 | 5985 | 6285 | 5482 | 5167 | 5663 | 5746.13 | 6055 | 5437 | 0.898 | -0.155 | 0.02840635 | 0.11086984 | 0.0014 | 0.0075 | 0.0034 | 0.0034 | TRUE | 0.0229 |
| Ftsj1 | 905 | 912 | 910 | 1236 | 834 | 787 | 961 | 911 | 914 | 972 | 897 | 861 | 919.38 | 929 | 910 | 0.981 | -0.027 | 0.76952714 | 0.87955519 | 0.0014 | 0.0105 | 0.005 | 0.005 | TRUE | 0.0433 |
| Ftsj2 | 471 | 488 | 395 | 769 | 503 | 412 | 500 | 487 | 397 | 605 | 541 | 451 | 496.83 | 461 | 532 | 1.156 | 0.209 | 0.17680008 | 0.37733299 | 0.0166 | 0.0135 | 0.0151 | 0.0151 | TRUE | 0.2063 |
| Ftsj3 | 15974 | 13485 | 13288 | 21409 | 15285 | 12864 | 16967 | 13468 | 13343 | 16830 | 16446 | 14082 | 15189.14 | 14593 | 15786 | 1.082 | 0.113 | 0.35673215 | 0.57524954 | 0.013 | 0.0071 | 0.0109 | 0.0109 | TRUE | 0.2478 |
| Ftsjd1 | 677 | 962 | 886 | 895 | 679 | 680 | 719 | 961 | 890 | 704 | 731 | 744 | 791.33 | 857 | 726 | 0.847 | -0.239 | 0.06221774 | 0.19087691 | 0.0102 | 0.0111 | 0.0106 | 0.0106 | TRUE | 0.233 |
| Ftsjd2 | 4433 | 4598 | 4492 | 5780 | 4343 | 4262 | 4709 | 4592 | 4510 | 4544 | 4673 | 4665 | 4615.52 | 4604 | 4627 | 1.005 | 0.007 | 0.90406147 | 0.95248677 | 2.00E-04 | 0.0076 | 0.0024 | 0.0024 | TRUE | 0.0048 |
| Ftx | 89 | 108 | 78 | 129 | 126 | 95 | 95 | 108 | 78 | 101 | 136 | 104 | 103.61 | 94 | 114 | 1.212 | 0.277 | 0.22457813 | 0.43701051 | 0.0165 | 0.0387 | 0.0277 | 0.0277 | TRUE | 0.2774 |
| Fubp1 | 9889 | 13260 | 13522 | 15785 | 12598 | 14279 | 10504 | 13243 | 13578 | 12409 | 13555 | 15631 | 13153.08 | 12442 | 13865 | 1.114 | 0.156 | 0.22961185 | 0.44315199 | 0.016 | 0.0072 | 0.0121 | 0.0121 | TRUE | 0.2268 |
| Fubp3 | 7204 | 5616 | 5679 | 9978 | 6905 | 5865 | 7652 | 5609 | 5702 | 7844 | 7429 | 6420 | 6776.07 | 6321 | 7231 | 1.144 | 0.194 | 0.16999687 | 0.36864988 | 0.0215 | 0.0074 | 0.0143 | 0.0143 | TRUE | 0.4137 |
| Fuca1 | 706 | 824 | 836 | 958 | 722 | 766 | 750 | 823 | 839 | 753 | 777 | 839 | 796.78 | 804 | 790 | 0.981 | -0.028 | 0.77654724 | 0.88386398 | 0.0021 | 0.011 | 0.0058 | 0.0058 | TRUE | 0.0409 |
| Fuca2 | 515 | 793 | 814 | 865 | 615 | 747 | 547 | 792 | 817 | 680 | 662 | 818 | 719.29 | 719 | 720 | 1.001 | 0.001 | 0.99528138 | 0.99768514 | 0.0287 | 0.0115 | 0.0201 | 0.0201 | TRUE | 0.5146 |
| Fuk | 3571 | 3939 | 3731 | 3790 | 2994 | 3110 | 3793 | 3934 | 3746 | 2979 | 3221 | 3404 | 3513.07 | 3824 | 3201 | 0.837 | -0.257 | 0.00122103 | 0.01038168 | 0.0021 | 0.0078 | 0.0043 | 0.0043 | TRUE | 0.0456 |
| Fundc1 | 875 | 854 | 742 | 1169 | 845 | 775 | 929 | 853 | 745 | 919 | 909 | 848 | 867.31 | 842 | 892 | 1.06 | 0.084 | 0.45620538 | 0.6610263 | 0.0058 | 0.0107 | 0.0079 | 0.0079 | TRUE | 0.1218 |
| Fundc2 | 1244 | 1922 | 1886 | 2143 | 1496 | 1761 | 1321 | 1920 | 1894 | 1685 | 1610 | 1928 | 1726.09 | 1712 | 1741 | 1.017 | 0.024 | 0.87821474 | 0.93976544 | 0.025 | 0.0088 | 0.017 | 0.017 | TRUE | 0.479 |
| Fuom | 71 | 86 | 60 | 140 | 95 | 101 | 75 | 86 | 60 | 110 | 102 | 111 | 90.73 | 74 | 108 | 1.458 | 0.544 | 0.01418815 | 0.06793806 | 0 | 0.0432 | 0.024 | 0.024 | TRUE | 0.2403 |
| Furin | 1387 | 1690 | 1758 | 1851 | 1390 | 1519 | 1473 | 1688 | 1765 | 1455 | 1496 | 1663 | 1589.95 | 1642 | 1538 | 0.936 | -0.096 | 0.36454483 | 0.58188453 | 0.0059 | 0.009 | 0.0074 | 0.0074 | TRUE | 0.0972 |
| Fus | 17465 | 14231 | 12779 | 22140 | 15645 | 13017 | 18551 | 14213 | 12831 | 17405 | 16833 | 14249 | 15680.33 | 15198 | 16162 | 1.063 | 0.089 | 0.54731074 | 0.73246078 | 0.0202 | 0.0071 | 0.0156 | 0.0156 | TRUE | 0.4553 |
| Fut1 | 1 | 2 | 1 | 1 | 1 | 0 | 1 | 2 | 1 | 1 | 1 | 0 | 0.99 | 1 | 1 | 0.466 | -1.103 | 0.55558731 | NA | 0 | 3.3426 | 1.3764 | 1.3764 | TRUE | 0.1869 |
| Fut10 | 839 | 750 | 730 | 503 | 400 | 413 | 891 | 749 | 733 | 395 | 430 | 452 | 608.51 | 791 | 426 | 0.538 | -0.895 | 5.10E-13 | 5.36E-11 | 0.0069 | 0.0123 | 0.0093 | 0.0093 | TRUE | 0.1446 |
| Fut11 | 275 | 263 | 245 | 399 | 312 | 251 | 292 | 263 | 246 | 314 | 336 | 275 | 287.48 | 267 | 308 | 1.155 | 0.208 | 0.14817623 | 0.33786837 | 0.005 | 0.0184 | 0.0114 | 0.0114 | TRUE | 0.0965 |
| Fut2 | 11 | 4 | 8 | 16 | 7 | 7 | 12 | 4 | 8 | 13 | 8 | 8 | 8.58 | 8 | 10 | 1.189 | 0.25 | 0.71826938 | 0.84737412 | 0 | 0.3908 | 0.2292 | 0.2292 | TRUE | 0.2438 |
| Fut4 | 170 | 185 | 151 | 226 | 165 | 130 | 181 | 185 | 152 | 178 | 178 | 142 | 169.07 | 173 | 166 | 0.965 | -0.052 | 0.7693689 | 0.87946546 | 0.0071 | 0.0264 | 0.0168 | 0.0168 | TRUE | 0.1566 |
| Fut4-ps1 | 0 | 0 | 0 | 0 | 1 | 0 | 0 | 0 | 0 | 0 | 1 | 0 | 0.18 | 0 | 0 | 1.859 | 0.894 | 0.82650734 | NA | 0 | 18.3785 | 10 | 10 | TRUE | 0.5204 |
| Fut7 | 3 | 7 | 5 | 4 | 2 | 10 | 3 | 7 | 5 | 3 | 2 | 11 | 5.24 | 5 | 5 | 1.051 | 0.072 | 0.93906668 | 0.96943053 | 0.2488 | 0.6356 | 0.44 | 0.44 | TRUE | 1.4667 |
| Fut8 | 1063 | 1106 | 1066 | 1667 | 1104 | 1099 | 1129 | 1105 | 1070 | 1310 | 1188 | 1203 | 1167.56 | 1101 | 1234 | 1.121 | 0.165 | 0.0528117 | 0.17037147 | 0.0012 | 0.0097 | 0.0044 | 0.0044 | TRUE | 0.0367 |
| Fut9 | 1626 | 2245 | 2586 | 1612 | 1412 | 1956 | 1727 | 2242 | 2597 | 1267 | 1519 | 2141 | 1915.57 | 2189 | 1642 | 0.75 | -0.415 | 0.04391517 | 0.15014741 | 0.0567 | 0.0086 | 0.03 | 0.03 | TRUE | 0.4934 |
| Fuz | 94 | 183 | 259 | 210 | 188 | 195 | 100 | 183 | 260 | 165 | 202 | 213 | 187.25 | 181 | 193 | 1.068 | 0.095 | 0.75678282 | 0.87155626 | 0.1095 | 0.0245 | 0.063 | 0.063 | TRUE | 0.2152 |
| Fv1 | 8 | 5 | 13 | 20 | 16 | 11 | 8 | 5 | 13 | 16 | 17 | 12 | 11.92 | 9 | 15 | 1.698 | 0.763 | 0.19527671 | 0.40118591 | 0 | 0.2833 | 0.1601 | 0.1601 | TRUE | 0.4174 |
| Fxn | 744 | 620 | 580 | 853 | 585 | 532 | 790 | 619 | 582 | 671 | 629 | 582 | 645.7 | 664 | 627 | 0.946 | -0.08 | 0.57676334 | 0.75440349 | 0.0143 | 0.012 | 0.0132 | 0.0132 | TRUE | 0.3251 |
| Fxr1 | 21063 | 26322 | 26657 | 25985 | 20343 | 22335 | 22373 | 26288 | 26766 | 20427 | 21888 | 24449 | 23698.57 | 25142 | 22255 | 0.885 | -0.176 | 0.09736522 | 0.2563981 | 0.0087 | 0.007 | 0.0081 | 0.0081 | TRUE | 0.1136 |
| Fxr2 | 15299 | 18339 | 18114 | 18922 | 13777 | 14785 | 16250 | 18315 | 18188 | 14875 | 14823 | 16185 | 16439.44 | 17584 | 15294 | 0.87 | -0.201 | 0.01357765 | 0.06568916 | 0.0033 | 0.0071 | 0.0047 | 0.0047 | TRUE | 0.0539 |
| Fxyd1 | 18 | 26 | 34 | 11 | 9 | 16 | 19 | 26 | 34 | 9 | 10 | 18 | 19.18 | 26 | 12 | 0.446 | -1.164 | 0.0201824 | 0.08737766 | 0.0527 | 0.1787 | 0.1204 | 0.1204 | TRUE | 0.2667 |
| Fxyd2 | 26 | 26 | 18 | 23 | 14 | 18 | 28 | 26 | 18 | 18 | 15 | 20 | 20.75 | 24 | 18 | 0.739 | -0.437 | 0.30599807 | 0.52665845 | 0 | 0.1657 | 0.0821 | 0.0821 | TRUE | 0.272 |
| Fxyd3 | 15 | 15 | 23 | 9 | 9 | 18 | 16 | 15 | 23 | 7 | 10 | 20 | 15.08 | 18 | 12 | 0.664 | -0.59 | 0.29774356 | 0.51843801 | 0.092 | 0.2254 | 0.1622 | 0.1622 | TRUE | 1.1048 |
| Fxyd4 | 5 | 8 | 12 | 14 | 8 | 18 | 5 | 8 | 12 | 11 | 9 | 20 | 10.78 | 8 | 13 | 1.539 | 0.622 | 0.33334744 | 0.55395173 | 0.0785 | 0.3126 | 0.1999 | 0.1999 | TRUE | 0.7434 |
| Fxyd5 | 89 | 124 | 109 | 188 | 147 | 173 | 95 | 124 | 109 | 148 | 158 | 189 | 137.19 | 109 | 165 | 1.506 | 0.59 | 0.00278344 | 0.01998971 | 0.01 | 0.0309 | 0.0204 | 0.0204 | TRUE | 0.1717 |
| Fxyd6 | 1351 | 2476 | 2599 | 1435 | 1358 | 1991 | 1435 | 2473 | 2610 | 1128 | 1461 | 2179 | 1881.02 | 2173 | 1589 | 0.732 | -0.451 | 0.22727953 | 0.44059153 | 0.1 | 0.0087 | 0.0482 | 0.1 | TRUE | 0.8057 |
| Fxyd7 | 9 | 16 | 6 | 5 | 4 | 1 | 10 | 16 | 6 | 4 | 4 | 1 | 6.82 | 11 | 3 | 0.3 | -1.737 | 0.03995661 | 0.14068851 | 0 | 0.4903 | 0.3125 | 0.3125 | TRUE | 0.2045 |
| Fyb | 3 | 4 | 3 | 7 | 4 | 5 | 3 | 4 | 3 | 6 | 4 | 5 | 4.25 | 3 | 5 | 1.502 | 0.587 | 0.50700528 | NA | 0 | 0.7829 | 0.3185 | 0.3185 | TRUE | 0.0351 |
| Fyco1 | 1608 | 1416 | 1469 | 1723 | 1282 | 1287 | 1708 | 1414 | 1475 | 1354 | 1379 | 1409 | 1456.64 | 1532 | 1381 | 0.901 | -0.15 | 0.13016604 | 0.31072784 | 0.0043 | 0.0092 | 0.0064 | 0.0064 | TRUE | 0.1206 |
| Fyn | 1426 | 1508 | 1536 | 1853 | 1367 | 1378 | 1515 | 1506 | 1542 | 1457 | 1471 | 1508 | 1499.83 | 1521 | 1479 | 0.972 | -0.041 | 0.56090346 | 0.74256648 | 0 | 0.0091 | 0.0029 | 0.0029 | TRUE | 0.0037 |
| Fyttd1 | 9212 | 10016 | 9467 | 11975 | 8996 | 9271 | 9785 | 10003 | 9506 | 9414 | 9679 | 10149 | 9755.87 | 9765 | 9747 | 0.998 | -0.003 | 0.96586228 | 0.98274821 | 7.00E-04 | 0.0072 | 0.0029 | 0.0029 | TRUE | 0.0158 |
| Fzd1 | 47 | 93 | 84 | 188 | 143 | 129 | 50 | 93 | 84 | 148 | 154 | 141 | 111.67 | 76 | 148 | 1.946 | 0.961 | 0.00011691 | 0.00155861 | 0.0325 | 0.0364 | 0.0347 | 0.0347 | TRUE | 0.7941 |
| Fzd10 | 661 | 749 | 752 | 885 | 655 | 602 | 702 | 748 | 755 | 696 | 705 | 659 | 710.78 | 735 | 687 | 0.934 | -0.098 | 0.28706421 | 0.50825962 | 0 | 0.0115 | 0.0048 | 0.0048 | TRUE | 0.018 |
| Fzd2 | 304 | 317 | 357 | 501 | 393 | 300 | 323 | 317 | 358 | 394 | 423 | 328 | 357.17 | 333 | 382 | 1.148 | 0.199 | 0.15891144 | 0.35285941 | 0.0076 | 0.0161 | 0.0116 | 0.0116 | TRUE | 0.1652 |
| Fzd3 | 1195 | 1216 | 1537 | 1672 | 1219 | 1246 | 1269 | 1214 | 1543 | 1314 | 1312 | 1364 | 1336.16 | 1342 | 1330 | 0.991 | -0.014 | 0.90417807 | 0.95248677 | 0.0078 | 0.0094 | 0.0085 | 0.0085 | TRUE | 0.2066 |
| Fzd4 | 225 | 272 | 279 | 314 | 239 | 220 | 239 | 272 | 280 | 247 | 257 | 241 | 255.93 | 264 | 248 | 0.941 | -0.087 | 0.52129035 | 0.71159696 | 0 | 0.0198 | 0.0095 | 0.0095 | TRUE | 0.0733 |
| Fzd5 | 10662 | 9906 | 9853 | 10620 | 8335 | 8276 | 11325 | 9893 | 9893 | 8349 | 8968 | 9059 | 9581.26 | 10370 | 8792 | 0.848 | -0.238 | 0.00542348 | 0.03318044 | 0.004 | 0.0072 | 0.0052 | 0.0052 | TRUE | 0.0791 |
| Fzd6 | 774 | 557 | 492 | 915 | 584 | 505 | 822 | 556 | 494 | 719 | 628 | 553 | 628.81 | 624 | 633 | 1.016 | 0.023 | 0.90804067 | 0.95356664 | 0.0433 | 0.0121 | 0.0271 | 0.0271 | TRUE | 0.9034 |
| Fzd7 | 3393 | 3395 | 4118 | 3102 | 2543 | 2865 | 3604 | 3391 | 4135 | 2439 | 2736 | 3136 | 3240.06 | 3710 | 2770 | 0.746 | -0.422 | 0.00061233 | 0.00609706 | 0.0128 | 0.0079 | 0.0106 | 0.0106 | TRUE | 0.1619 |
| Fzd8 | 126 | 131 | 120 | 216 | 145 | 136 | 134 | 131 | 120 | 170 | 156 | 149 | 143.31 | 128 | 158 | 1.236 | 0.305 | 0.07097826 | 0.20748529 | 0 | 0.0299 | 0.0136 | 0.0136 | TRUE | 0.0473 |
| Fzd9 | 477 | 575 | 595 | 547 | 433 | 418 | 507 | 574 | 597 | 430 | 466 | 458 | 505.3 | 559 | 451 | 0.805 | -0.313 | 0.00623696 | 0.0371065 | 0.0027 | 0.0134 | 0.0074 | 0.0074 | TRUE | 0.079 |
| Fzr1 | 3956 | 4594 | 4478 | 5991 | 4682 | 4528 | 4202 | 4588 | 4496 | 4710 | 5038 | 4957 | 4665.04 | 4429 | 4902 | 1.106 | 0.146 | 0.04265107 | 0.14744498 | 0.0013 | 0.0076 | 0.0035 | 0.0035 | TRUE | 0.0244 |
| G0s2 | 36 | 78 | 93 | 57 | 27 | 39 | 38 | 78 | 93 | 45 | 29 | 43 | 54.35 | 70 | 39 | 0.557 | -0.843 | 0.02844851 | 0.11092646 | 0.1077 | 0.0675 | 0.0868 | 0.0868 | TRUE | 0.4689 |
| G2e3 | 7625 | 10863 | 12266 | 7798 | 6608 | 8767 | 8099 | 10849 | 12316 | 6130 | 7110 | 9597 | 9016.88 | 10421 | 7612 | 0.73 | -0.453 | 0.01733856 | 0.07832875 | 0.0485 | 0.0073 | 0.026 | 0.026 | TRUE | 0.4714 |
| G3bp1 | 34959 | 35706 | 34476 | 44403 | 33335 | 31941 | 37133 | 35660 | 34618 | 34906 | 35866 | 34964 | 35524.52 | 35804 | 35245 | 0.984 | -0.023 | 0.70128303 | 0.83650923 | 6.00E-04 | 0.007 | 0.0025 | 0.0025 | TRUE | 0.0129 |
| G3bp2 | 21302 | 16440 | 15829 | 34321 | 23406 | 19308 | 22626 | 16419 | 15894 | 26980 | 25183 | 21136 | 21373.13 | 18313 | 24433 | 1.334 | 0.416 | 0.00696902 | 0.0399402 | 0.0283 | 0.0071 | 0.0171 | 0.0171 | TRUE | 0.5193 |

| | | | | | | | | | | | | | | | | | | | | | | | | | |
|---|---|---|---|---|---|---|---|---|---|---|---|---|---|---|---|---|---|---|---|---|---|---|---|---|---|
| G530011O06 | 0 | 2 | 0 | 0 | 0 | 0 | 0 | 2 | 0 | 0 | 0 | 0 | 0.33 | 1 | 0 | 0.268 | -1.897 | 0.63866967 | NA | 0 | 9.903 | 10 | 10 | TRUE | 0.9739 |
| G630025P09 | 3 | 5 | 4 | 24 | 7 | 4 | 3 | 5 | 4 | 19 | 8 | 4 | 7.16 | 4 | 10 | 2.566 | 1.36 | 0.10785913 | 0.27490073 | 0.2352 | 0.4669 | 0.3431 | 0.3431 | TRUE | 0.8023 |
| G630055G22 | 0 | 0 | 0 | 0 | 0 | 0 | 0 | 0 | 0 | 0 | 0 | 0 | 0 | 0 | 0 | NA | NA | NA | NA | NA | NA | NA | NA | NA | NA |
| G630071F17 | 0 | 0 | 0 | 0 | 1 | 0 | 0 | 0 | 0 | 0 | 1 | 0 | 0.18 | 0 | 0 | 1.859 | 0.894 | 0.82650734 | NA | 0 | 18.3785 | 10 | 10 | TRUE | 0.5204 |
| G630090E17 | 4 | 2 | 0 | 6 | 4 | 5 | 4 | 2 | 0 | 5 | 4 | 5 | 3.46 | 2 | 5 | 2.34 | 1.226 | 0.28133897 | NA | 0 | 0.96 | 0.5847 | 0.5847 | TRUE | 0.4475 |
| G630093K05 | 7 | 0 | 3 | 5 | 2 | 3 | 7 | 0 | 3 | 4 | 2 | 3 | 3.3 | 3 | 3 | 0.909 | -0.138 | 0.91103592 | NA | 0.5137 | 1.0045 | 0.7963 | 0.7963 | TRUE | 0.2845 |
| G6b | 4 | 2 | 3 | 2 | 1 | 1 | 4 | 2 | 3 | 2 | 1 | 1 | 2.17 | 3 | 1 | 0.412 | -1.281 | 0.31669158 | NA | 0 | 1.5273 | 0.6289 | 0.6289 | TRUE | 0.1371 |
| G6pc | 2 | 1 | 1 | 1 | 0 | 0 | 2 | 1 | 1 | 1 | 0 | 0 | 0.82 | 1 | 0 | 0.255 | -1.974 | 0.35972496 | NA | 0 | 4.03 | 1.975 | 1.975 | TRUE | 0.2562 |
| G6pc2 | 6 | 4 | 3 | 3 | 4 | 1 | 6 | 4 | 3 | 2 | 4 | 1 | 3.52 | 4 | 2 | 0.58 | -0.786 | 0.44622645 | NA | 0 | 0.9421 | 0.464 | 0.464 | TRUE | 0.3358 |
| G6pc3 | 837 | 1307 | 1393 | 1262 | 995 | 1269 | 889 | 1305 | 1399 | 992 | 1071 | 1389 | 1174.14 | 1198 | 1151 | 0.96 | -0.059 | 0.75833243 | 0.87245014 | 0.043 | 0.0097 | 0.0255 | 0.0255 | TRUE | 0.6081 |
| G6pd2 | 0 | 1 | 0 | 1 | 0 | 0 | 0 | 1 | 0 | 1 | 0 | 0 | 0.3 | 0 | 0 | 0.954 | -0.067 | 0.98642001 | NA | 0 | 11.0817 | 9.3014 | 9.3014 | TRUE | 0.405 |
| G6pdx | 2198 | 2054 | 2059 | 2973 | 2081 | 2192 | 2335 | 2051 | 2067 | 2337 | 2239 | 2399 | 2238.19 | 2151 | 2325 | 1.081 | 0.113 | 0.19168906 | 0.39707379 | 0.0027 | 0.0084 | 0.0049 | 0.0049 | TRUE | 0.0672 |
| G730013B05 | 197 | 168 | 194 | 267 | 204 | 229 | 209 | 168 | 195 | 210 | 219 | 251 | 208.65 | 191 | 227 | 1.188 | 0.248 | 0.12418302 | 0.30072124 | 0.0056 | 0.0227 | 0.0139 | 0.0139 | TRUE | 0.1193 |
| Gaa | 6286 | 7243 | 7327 | 6768 | 5162 | 5551 | 6677 | 7234 | 7357 | 5320 | 5554 | 6076 | 6369.75 | 7089 | 5650 | 0.797 | -0.328 | 9.61E-05 | 0.00133652 | 0.0034 | 0.0074 | 0.0049 | 0.0049 | TRUE | 0.053 |
| Gab1 | 19873 | 20249 | 20179 | 28806 | 20368 | 19851 | 21109 | 20223 | 20262 | 22645 | 21915 | 21730 | 21313.85 | 20531 | 22097 | 1.076 | 0.106 | 0.06303879 | 0.19249716 | 5.00E-04 | 0.0071 | 0.0023 | 0.0023 | TRUE | 0.0074 |
| Gab2 | 111 | 184 | 157 | 186 | 141 | 135 | 118 | 184 | 158 | 146 | 152 | 148 | 150.84 | 153 | 149 | 0.969 | -0.045 | 0.82338809 | 0.9101083 | 0.0175 | 0.0287 | 0.0232 | 0.0232 | TRUE | 0.3191 |
| Gab3 | 37 | 40 | 50 | 54 | 31 | 34 | 39 | 40 | 50 | 42 | 33 | 37 | 40.41 | 43 | 37 | 0.877 | -0.19 | 0.53786565 | 0.72419528 | 0 | 0.0884 | 0.0435 | 0.0435 | TRUE | 0.1591 |
| Gabarap | 2308 | 2966 | 3133 | 3486 | 2475 | 2835 | 2452 | 2962 | 3146 | 2740 | 2663 | 3103 | 2844.38 | 2853 | 2835 | 0.994 | -0.009 | 0.93786775 | 0.96893419 | 0.0112 | 0.0081 | 0.0098 | 0.0098 | TRUE | 0.1838 |
| Gabarapl1 | 2271 | 3052 | 3181 | 3845 | 2919 | 3183 | 2412 | 3048 | 3194 | 3023 | 3141 | 3484 | 3050.32 | 2885 | 3216 | 1.114 | 0.156 | 0.21002782 | 0.42042884 | 0.0127 | 0.008 | 0.0109 | 0.0109 | TRUE | 0.2489 |
| Gabarapl2 | 2560 | 2329 | 2103 | 6680 | 4135 | 3264 | 2719 | 2326 | 2112 | 5251 | 4449 | 3573 | 3405.01 | 2386 | 4424 | 1.855 | 0.892 | 9.61E-09 | 4.55E-07 | 0.0259 | 0.0079 | 0.0171 | 0.0171 | TRUE | 0.1226 |
| Gabbr1 | 1909 | 2363 | 2452 | 2926 | 2676 | 2762 | 2028 | 2360 | 2462 | 2300 | 2879 | 3023 | 2508.76 | 2283 | 2734 | 1.197 | 0.259 | 0.04729396 | 0.15785128 | 0.0141 | 0.0082 | 0.0119 | 0.0119 | TRUE | 0.2362 |
| Gabbr2 | 8 | 3 | 3 | 8 | 8 | 9 | 8 | 3 | 3 | 6 | 9 | 10 | 6.54 | 5 | 8 | 1.704 | 0.769 | 0.33029216 | 0.55109763 | 0 | 0.5105 | 0.2836 | 0.2836 | TRUE | 0.8205 |
| Gabpa | 9438 | 9002 | 8987 | 12395 | 9059 | 9177 | 10025 | 8990 | 9024 | 9744 | 9747 | 10046 | 9595.95 | 9346 | 9846 | 1.053 | 0.075 | 0.30404753 | 0.52500959 | 0.002 | 0.0072 | 0.0037 | 0.0037 | TRUE | 0.0492 |
| Gabpb1 | 2779 | 2953 | 2966 | 3788 | 2787 | 2936 | 2952 | 2949 | 2978 | 2978 | 2999 | 3214 | 3011.59 | 2960 | 3064 | 1.035 | 0.049 | 0.47051733 | 0.6721364 | 6.00E-04 | 0.008 | 0.003 | 0.003 | TRUE | 0.0222 |
| Gabpb2 | 3048 | 3495 | 3565 | 5147 | 3999 | 3754 | 3238 | 3490 | 3580 | 4046 | 4303 | 4109 | 3794.3 | 3436 | 4153 | 1.208 | 0.273 | 0.00023792 | 0.00280352 | 0.0015 | 0.0078 | 0.0037 | 0.0037 | TRUE | 0.0309 |
| Gabra1 | 64 | 27 | 18 | 153 | 115 | 38 | 68 | 27 | 18 | 120 | 124 | 42 | 66.44 | 38 | 95 | 2.536 | 1.343 | 0.01187373 | 0.05975174 | 0.3521 | 0.0565 | 0.1865 | 0.1865 | TRUE | 3.5519 |
| Gabra2 | 2 | 3 | 1 | 3 | 2 | 2 | 2 | 3 | 1 | 2 | 2 | 2 | 2.14 | 2 | 2 | 1.096 | 0.133 | 0.91421649 | NA | 0 | 1.5483 | 0.6285 | 0.6285 | TRUE | 0.1848 |
| Gabra3 | 12 | 4 | 6 | 11 | 3 | 2 | 13 | 4 | 6 | 9 | 3 | 2 | 6.14 | 8 | 5 | 0.639 | -0.647 | 0.46737186 | 0.66963926 | 0.263 | 0.5436 | 0.4027 | 0.4027 | TRUE | 1.2097 |
| Gabra4 | 173 | 158 | 163 | 181 | 135 | 140 | 184 | 158 | 164 | 142 | 145 | 153 | 157.67 | 169 | 147 | 0.872 | -0.198 | 0.22436477 | 0.43681551 | 0 | 0.0278 | 0.0127 | 0.0127 | TRUE | 0.0665 |
| Gabra5 | 12 | 16 | 20 | 44 | 26 | 14 | 13 | 16 | 20 | 35 | 28 | 15 | 21.12 | 16 | 26 | 1.608 | 0.685 | 0.15256581 | 0.34375257 | 0.065 | 0.1629 | 0.1152 | 0.1152 | TRUE | 0.3137 |
| Gabra6 | 0 | 0 | 0 | 0 | 0 | 0 | 0 | 0 | 0 | 0 | 0 | 0 | 0 | 0 | 0 | NA | NA | NA | NA | NA | NA | NA | NA | NA | NA |
| Gabrb1 | 2 | 1 | 1 | 3 | 1 | 1 | 2 | 1 | 1 | 2 | 1 | 1 | 1.44 | 1 | 1 | 1.13 | 0.176 | 0.906047 | NA | 0 | 2.2905 | 0.9103 | 0.9103 | TRUE | 0.2239 |
| Gabrb2 | 8 | 9 | 6 | 6 | 13 | 15 | 8 | 9 | 6 | 5 | 14 | 16 | 9.77 | 8 | 12 | 1.47 | 0.556 | 0.42243547 | 0.63267043 | 0 | 0.344 | 0.2381 | 0.2381 | TRUE | 1.3464 |
| Gabrb3 | 487 | 677 | 720 | 816 | 674 | 689 | 517 | 676 | 723 | 641 | 725 | 754 | 672.87 | 639 | 707 | 1.105 | 0.145 | 0.33149389 | 0.55215482 | 0.0169 | 0.0118 | 0.0145 | 0.0145 | TRUE | 0.3241 |
| Gabrd | 6 | 6 | 3 | 2 | 5 | 4 | 6 | 6 | 3 | 2 | 5 | 4 | 4.45 | 5 | 4 | 0.72 | -0.474 | 0.61055934 | NA | 0 | 0.747 | 0.3932 | 0.3932 | TRUE | 0.5759 |
| Gabre | 46 | 51 | 36 | 32 | 37 | 32 | 49 | 51 | 36 | 25 | 40 | 35 | 39.32 | 45 | 33 | 0.729 | -0.455 | 0.17768993 | 0.37833021 | 0.0176 | 0.0907 | 0.0561 | 0.0561 | TRUE | 0.3866 |
| Gabrg1 | 7 | 7 | 7 | 5 | 5 | 7 | 7 | 7 | 7 | 4 | 5 | 8 | 6.4 | 7 | 6 | 0.78 | -0.359 | 0.62553182 | 0.78901854 | 0 | 0.5213 | 0.2305 | 0.2305 | TRUE | 0.2163 |
| Gabrg2 | 5 | 12 | 6 | 18 | 12 | 8 | 5 | 12 | 6 | 14 | 13 | 9 | 9.86 | 8 | 12 | 1.541 | 0.624 | 0.33007125 | 0.55093867 | 0 | 0.3411 | 0.1895 | 0.1895 | TRUE | 0.6561 |
| Gabrg3 | 17 | 2 | 11 | 23 | 11 | 5 | 18 | 2 | 11 | 18 | 12 | 5 | 11.08 | 10 | 12 | 1.153 | 0.205 | 0.79976148 | 0.89676085 | 0.4701 | 0.3042 | 0.3821 | 0.3821 | TRUE | 0.2992 |
| Gabrp | 36 | 112 | 97 | 31 | 34 | 44 | 38 | 112 | 97 | 24 | 37 | 48 | 59.43 | 82 | 36 | 0.438 | -1.192 | 0.00650069 | 0.03815231 | 0.182 | 0.0623 | 0.1186 | 0.1186 | TRUE | 0.9193 |
| Gabrq | 3 | 4 | 2 | 10 | 5 | 5 | 3 | 4 | 2 | 8 | 5 | 5 | 4.65 | 3 | 6 | 2.055 | 1.039 | 0.23907521 | 0.45501317 | 0 | 0.7153 | 0.317 | 0.317 | TRUE | 0.1617 |
| Gabrr1 | 70 | 47 | 56 | 79 | 57 | 26 | 74 | 47 | 56 | 62 | 61 | 28 | 54.9 | 59 | 50 | 0.861 | -0.216 | 0.55321205 | 0.73680632 | 0.0898 | 0.0669 | 0.0776 | 0.0776 | TRUE | 1.1282 |
| Gabrr2 | 8 | 2 | 0 | 1 | 0 | 2 | 8 | 2 | 0 | 1 | 0 | 2 | 2.25 | 3 | 1 | 0.282 | -1.828 | 0.29551932 | NA | 1.6818 | 1.4743 | 1.5564 | 1.5564 | TRUE | 0.5207 |
| Gabrr3 | 3 | 0 | 0 | 3 | 3 | 1 | 3 | 0 | 0 | 2 | 3 | 1 | 1.64 | 1 | 2 | 2.134 | 1.094 | 0.53078608 | NA | 0.4769 | 2.0101 | 1.4861 | 1.4861 | TRUE | 1.4432 |
| Gad1 | 264 | 710 | 814 | 277 | 289 | 517 | 280 | 709 | 817 | 218 | 311 | 566 | 483.58 | 602 | 365 | 0.606 | -0.724 | 0.22001603 | 0.43222937 | 0.2488 | 0.0137 | 0.1068 | 0.2488 | TRUE | 1.1038 |
| Gad2 | 2 | 4 | 3 | 4 | 3 | 6 | 2 | 4 | 3 | 3 | 3 | 7 | 3.68 | 3 | 4 | 1.398 | 0.483 | 0.62227181 | NA | 0 | 0.9025 | 0.4104 | 0.4104 | TRUE | 0.3391 |
| Gadd45a | 1075 | 994 | 948 | 1281 | 878 | 850 | 1142 | 993 | 952 | 1007 | 945 | 930 | 994.77 | 1029 | 961 | 0.935 | -0.097 | 0.35214526 | 0.57073249 | 0.0045 | 0.0102 | 0.0069 | 0.0069 | TRUE | 0.1098 |
| Gadd45b | 596 | 850 | 1060 | 951 | 796 | 979 | 633 | 849 | 1064 | 748 | 856 | 1072 | 870.34 | 849 | 892 | 1.05 | 0.07 | 0.72834375 | 0.85395791 | 0.0483 | 0.0107 | 0.0284 | 0.0284 | TRUE | 0.1934 |
| Gadd45g | 19 | 65 | 86 | 70 | 64 | 115 | 20 | 65 | 86 | 55 | 69 | 126 | 70.2 | 57 | 83 | 1.451 | 0.537 | 0.27833744 | 0.49954274 | 0.2993 | 0.0538 | 0.1623 | 0.1623 | TRUE | 0.8014 |
| Gadd45gip1 | 3238 | 3411 | 3295 | 4202 | 3152 | 3154 | 3439 | 3407 | 3309 | 3303 | 3391 | 3453 | 3383.61 | 3385 | 3382 | 0.999 | -0.001 | 0.98297831 | 0.99106917 | 2.00E-04 | 0.0079 | 0.0026 | 0.0026 | TRUE | 0.0052 |
| Gadl1 | 16 | 15 | 13 | 35 | 21 | 18 | 17 | 15 | 13 | 28 | 23 | 20 | 19.14 | 15 | 24 | 1.562 | 0.644 | 0.14255668 | 0.32842594 | 0 | 0.179 | 0.0839 | 0.0839 | TRUE | 0.1788 |
| Gak | 4101 | 4512 | 4695 | 6189 | 4679 | 4560 | 4356 | 4506 | 4714 | 4865 | 5034 | 4992 | 4744.62 | 4525 | 4964 | 1.097 | 0.133 | 0.04281969 | 0.14775471 | 6.00E-04 | 0.0076 | 0.0029 | 0.0029 | TRUE | 0.0162 |
| Gal | 5 | 8 | 6 | 13 | 9 | 12 | 5 | 8 | 6 | 10 | 10 | 13 | 8.73 | 6 | 11 | 1.702 | 0.767 | 0.23010823 | 0.44377649 | 0 | 0.3844 | 0.1706 | 0.1706 | TRUE | 0.1115 |
| Gal3st1 | 134 | 146 | 163 | 190 | 138 | 143 | 142 | 146 | 164 | 149 | 148 | 157 | 151.03 | 151 | 151 | 1.005 | 0.007 | 0.96642126 | 0.98304089 | 0 | 0.0287 | 0.0127 | 0.0127 | TRUE | 0.0607 |
| Gal3st2 | 0 | 1 | 0 | 1 | 0 | 1 | 0 | 1 | 0 | 1 | 0 | 1 | 0.48 | 0 | 1 | 1.723 | 0.785 | 0.78325239 | NA | 0 | 6.8716 | 4.0968 | 4.0968 | TRUE | 0.405 |
| Gal3st3 | 64 | 65 | 72 | 138 | 97 | 99 | 68 | 65 | 72 | 108 | 104 | 108 | 87.74 | 68 | 107 | 1.566 | 0.647 | 0.00171641 | 0.01362413 | 0 | 0.0445 | 0.0187 | 0.0187 | TRUE | 0.0226 |
| Gal3st4 | 5 | 5 | 4 | 4 | 4 | 1 | 5 | 5 | 4 | 3 | 4 | 1 | 3.81 | 5 | 3 | 0.601 | -0.735 | 0.44955178 | NA | 0 | 0.8714 | 0.4033 | 0.4033 | TRUE | 0.3053 |
| Galc | 616 | 823 | 771 | 870 | 594 | 644 | 654 | 822 | 774 | 684 | 639 | 705 | 713.07 | 750 | 676 | 0.901 | -0.15 | 0.20626084 | 0.41518859 | 0.0066 | 0.0115 | 0.0088 | 0.0088 | TRUE | 0.1469 |
| Gale | 656 | 558 | 530 | 742 | 591 | 429 | 697 | 557 | 532 | 583 | 636 | 470 | 579.17 | 595 | 563 | 0.946 | -0.08 | 0.61705398 | 0.7832683 | 0.0205 | 0.0126 | 0.0166 | 0.0166 | TRUE | 0.2593 |
| Galk1 | 7842 | 10229 | 10627 | 10541 | 8082 | 8936 | 8330 | 10216 | 10671 | 8287 | 8696 | 9782 | 9330.02 | 9739 | 8922 | 0.916 | -0.127 | 0.28320857 | 0.50491557 | 0.0117 | 0.0073 | 0.0099 | 0.0099 | TRUE | 0.1956 |
| Galk2 | 1216 | 1161 | 1173 | 1389 | 1025 | 963 | 1292 | 1160 | 1178 | 1092 | 1103 | 1054 | 1146.31 | 1210 | 1083 | 0.896 | -0.159 | 0.06542609 | 0.1971533 | 0.0011 | 0.0098 | 0.0045 | 0.0045 | TRUE | 0.0419 |
| Galm | 679 | 696 | 746 | 660 | 517 | 521 | 721 | 695 | 749 | 519 | 556 | 570 | 635.13 | 722 | 548 | 0.759 | -0.398 | 4.79E-05 | 0.00075052 | 0 | 0.0121 | 0.0053 | 0.0053 | TRUE | 0.0279 |
| Galns | 515 | 609 | 606 | 664 | 492 | 484 | 547 | 608 | 608 | 522 | 529 | 530 | 557.48 | 588 | 527 | 0.896 | -0.158 | 0.11720178 | 0.28955302 | 0 | 0.0128 | 0.0056 | 0.0056 | TRUE | 0.0429 |
| Galnt1 | 3979 | 4633 | 4554 | 5917 | 4308 | 4196 | 4226 | 4627 | 4573 | 4651 | 4635 | 4593 | 4550.99 | 4475 | 4626 | 1.034 | 0.048 | 0.48350903 | 0.68146464 | 9.00E-04 | 0.0076 | 0.0031 | 0.0031 | TRUE | 0.0288 |
| Galnt10 | 1532 | 1038 | 978 | 1511 | 1077 | 957 | 1627 | 1037 | 982 | 1188 | 1159 | 1048 | 1173.36 | 1215 | 1132 | 0.931 | -0.103 | 0.58581417 | 0.76088785 | 0.0395 | 0.0097 | 0.0247 | 0.0247 | TRUE | 1.0532 |
| Galnt11 | 2541 | 2685 | 2577 | 3407 | 2533 | 2692 | 2699 | 2682 | 2588 | 2678 | 2725 | 2947 | 2719.77 | 2656 | 2783 | 1.048 | 0.067 | 0.36561748 | 0.58306561 | 0.001 | 0.0081 | 0.0036 | 0.0036 | TRUE | 0.0317 |
| Galnt12 | 463 | 655 | 676 | 653 | 516 | 632 | 492 | 654 | 679 | 513 | 555 | 692 | 597.51 | 608 | 587 | 0.963 | -0.054 | 0.75018504 | 0.86769716 | 0.0253 | 0.0124 | 0.0189 | 0.0189 | TRUE | 0.328 |
| Galnt13 | 16 | 18 | 15 | 19 | 10 | 10 | 17 | 18 | 15 | 15 | 11 | 11 | 14.45 | 17 | 12 | 0.74 | -0.435 | 0.3783864 | 0.5934918 | 0 | 0.235 | 0.1053 | 0.1053 | TRUE | 0.1921 |
| Galnt14 | 71 | 79 | 79 | 80 | 53 | 65 | 75 | 79 | 79 | 63 | 57 | 71 | 70.78 | 78 | 64 | 0.817 | -0.292 | 0.20609117 | 0.41502674 | 0 | 0.0534 | 0.0241 | 0.0241 | TRUE | 0.0835 |

| | | | | | | | | | | | | | | | | | | | | | | | | | |
|---|---|---|---|---|---|---|---|---|---|---|---|---|---|---|---|---|---|---|---|---|---|---|---|---|---|
| Galnt15 | 0 | 1 | 1 | 1 | 1 | 0 | 0 | 1 | 1 | 1 | 1 | 0 | 0.64 | 1 | 1 | 0.935 | -0.097 | 0.96674856 | NA | 0 | 5.1213 | 2.3899 | 2.3899 | TRUE | 0.2246 |
| Galnt16 | 69 | 126 | 136 | 272 | 169 | 176 | 73 | 126 | 137 | 214 | 182 | 193 | 154 | 112 | 196 | 1.752 | 0.809 | 0.00083239 | 0.00772715 | 0.0435 | 0.0283 | 0.0352 | 0.0352 | TRUE | 0.8922 |
| Galnt18 | 30 | 45 | 41 | 47 | 45 | 44 | 32 | 45 | 41 | 37 | 48 | 48 | 41.92 | 39 | 44 | 1.124 | 0.168 | 0.59020474 | 0.76449681 | 0 | 0.0855 | 0.0464 | 0.0464 | TRUE | 0.2024 |
| Galnt2 | 5834 | 6783 | 6467 | 7787 | 5468 | 5526 | 6197 | 6774 | 6494 | 6122 | 5883 | 6049 | 6253.06 | 6488 | 6018 | 0.928 | -0.108 | 0.1051077 | 0.2703455 | 0.001 | 0.0074 | 0.0031 | 0.0031 | TRUE | 0.0188 |
| Galnt3 | 238 | 312 | 314 | 300 | 205 | 232 | 253 | 312 | 315 | 236 | 221 | 254 | 265.01 | 293 | 237 | 0.807 | -0.31 | 0.03920046 | 0.13865656 | 0.0061 | 0.0193 | 0.0124 | 0.0124 | TRUE | 0.1618 |
| Galnt4 | 189 | 155 | 146 | 303 | 219 | 142 | 201 | 155 | 147 | 238 | 236 | 155 | 188.57 | 168 | 210 | 1.256 | 0.329 | 0.1392875 | 0.32392251 | 0.0366 | 0.0244 | 0.0304 | 0.0304 | TRUE | 0.546 |
| Galnt5 | 1 | 0 | 0 | 0 | 0 | 0 | 1 | 0 | 0 | 0 | 0 | 0 | 0.18 | 0 | 0 | 0.49 | -1.029 | 0.80087 | NA | 0 | 18.6165 | 10 | 10 | TRUE | 0.4614 |
| Galnt6 | 813 | 772 | 693 | 1164 | 830 | 818 | 864 | 771 | 696 | 915 | 893 | 895 | 838.98 | 777 | 901 | 1.161 | 0.215 | 0.04684114 | 0.15688648 | 0.0041 | 0.0108 | 0.0072 | 0.0072 | TRUE | 0.1124 |
| Galnt7 | 1157 | 1539 | 1618 | 2578 | 1716 | 1770 | 1229 | 1537 | 1625 | 2027 | 1846 | 1938 | 1700.18 | 1464 | 1937 | 1.323 | 0.404 | 0.00084043 | 0.00778346 | 0.0103 | 0.0088 | 0.01 | 0.01 | TRUE | 0.2359 |
| Galnt9 | 14 | 25 | 31 | 36 | 31 | 33 | 15 | 25 | 31 | 28 | 33 | 36 | 28.12 | 24 | 32 | 1.369 | 0.453 | 0.25772745 | 0.47651994 | 0.0222 | 0.124 | 0.0789 | 0.0789 | TRUE | 0.4736 |
| Galntl5 | 1 | 2 | 3 | 2 | 4 | 5 | 1 | 2 | 3 | 2 | 4 | 5 | 2.9 | 2 | 4 | 1.825 | 0.868 | 0.45031944 | NA | 0 | 1.1415 | 0.5687 | 0.5687 | TRUE | 0.5636 |
| Galntl6 | 5 | 1 | 1 | 11 | 7 | 2 | 5 | 1 | 1 | 9 | 8 | 2 | 4.28 | 2 | 6 | 2.561 | 1.357 | 0.2153316 | NA | 0.3333 | 0.7766 | 0.5735 | 0.5735 | TRUE | 1.0881 |
| Galp | 1 | 2 | 1 | 2 | 1 | 0 | 1 | 2 | 1 | 2 | 1 | 0 | 1.12 | 1 | 1 | 0.676 | -0.565 | 0.74666147 | NA | 0 | 2.9519 | 1.2963 | 1.2963 | TRUE | 0.2738 |
| Galr1 | 2 | 1 | 0 | 3 | 0 | 1 | 2 | 1 | 0 | 2 | 0 | 1 | 1.1 | 1 | 1 | 1.149 | 0.201 | 0.9188536 | NA | 0 | 3.0127 | 1.8934 | 1.8934 | TRUE | 0.2311 |
| Galr2 | 53 | 74 | 75 | 67 | 31 | 44 | 56 | 74 | 75 | 53 | 33 | 48 | 56.62 | 68 | 45 | 0.656 | -0.608 | 0.037674 | 0.13482965 | 0.0181 | 0.0651 | 0.0433 | 0.0433 | TRUE | 0.3681 |
| Galr3 | 3 | 1 | 3 | 2 | 1 | 2 | 3 | 1 | 3 | 2 | 1 | 2 | 2.01 | 2 | 2 | 0.672 | -0.572 | 0.66028922 | NA | 0 | 1.6493 | 0.7058 | 0.7058 | TRUE | 0.283 |
| Galt | 964 | 1131 | 1093 | 1285 | 992 | 1056 | 1024 | 1130 | 1097 | 1010 | 1067 | 1156 | 1080.74 | 1084 | 1078 | 0.994 | -0.009 | 0.92153582 | 0.96126008 | 0.0027 | 0.01 | 0.0056 | 0.0056 | TRUE | 0.0479 |
| Gamt | 53 | 66 | 70 | 64 | 64 | 49 | 56 | 66 | 70 | 50 | 69 | 54 | 60.88 | 64 | 58 | 0.893 | -0.163 | 0.53680903 | 0.72345143 | 0 | 0.061 | 0.0336 | 0.0336 | TRUE | 0.2403 |
| Gan | 434 | 297 | 310 | 633 | 423 | 303 | 461 | 297 | 311 | 498 | 455 | 332 | 392.22 | 356 | 428 | 1.204 | 0.267 | 0.21794433 | 0.42994595 | 0.0483 | 0.0153 | 0.0314 | 0.0314 | TRUE | 0.7516 |
| Ganab | 14955 | 17365 | 17398 | 19416 | 13886 | 13912 | 15885 | 17343 | 17469 | 15263 | 14940 | 15229 | 16021.59 | 16899 | 15144 | 0.896 | -0.158 | 0.01849994 | 0.08221502 | 0.0013 | 0.0071 | 0.0032 | 0.0032 | TRUE | 0.0337 |
| Ganc | 245 | 263 | 266 | 361 | 247 | 201 | 260 | 263 | 267 | 284 | 266 | 220 | 259.93 | 263 | 257 | 0.977 | -0.034 | 0.81870332 | 0.90735638 | 0.0047 | 0.0196 | 0.0118 | 0.0118 | TRUE | 0.1679 |
| Gap43 | 426 | 264 | 278 | 180 | 100 | 107 | 452 | 264 | 279 | 142 | 108 | 117 | 226.92 | 332 | 122 | 0.369 | -1.437 | 3.44E-09 | 1.81E-07 | 0.0518 | 0.0214 | 0.0371 | 0.0371 | TRUE | 1.1472 |
| Gapdh | 123685 | 154396 | 161665 | 173020 | 128554 | 144585 | 131375 | 154197 | 162329 | 136015 | 138316 | 158271 | 146750.45 | 149300 | 144201 | 0.966 | -0.05 | 0.6420216 | 0.79842557 | 0.0079 | 0.0069 | 0.0084 | 0.0084 | TRUE | 0.1351 |
| Gapdhs | 0 | 0 | 0 | 0 | 0 | 0 | 0 | 0 | 0 | 0 | 0 | 0 | 0 | 0 | 0 | NA | NA | NA | NA | NA | NA | NA | NA | NA | NA |
| Gapt | 0 | 0 | 0 | 0 | 0 | 0 | 0 | 0 | 0 | 0 | 0 | 0 | 0 | 0 | 0 | NA | NA | NA | NA | NA | NA | NA | NA | NA | NA |
| Gapvd1 | 6531 | 6640 | 6804 | 8340 | 5820 | 5793 | 6937 | 6631 | 6832 | 6556 | 6262 | 6341 | 6593.33 | 6800 | 6386 | 0.939 | -0.09 | 0.13375596 | 0.31609216 | 3.00E-04 | 0.0074 | 0.0025 | 0.0025 | TRUE | 0.0067 |
| Gar1 | 3395 | 3209 | 3231 | 5040 | 3657 | 3217 | 3606 | 3205 | 3244 | 3962 | 3935 | 3522 | 3578.91 | 3352 | 3806 | 1.136 | 0.184 | 0.03939866 | 0.13910805 | 0.0039 | 0.0078 | 0.0055 | 0.0055 | TRUE | 0.0535 |
| Garem | 1745 | 1425 | 1587 | 1831 | 1336 | 1342 | 1853 | 1423 | 1594 | 1439 | 1437 | 1469 | 1536.01 | 1623 | 1448 | 0.892 | -0.164 | 0.14765316 | 0.33696796 | 0.008 | 0.009 | 0.0086 | 0.0086 | TRUE | 0.1849 |
| Gareml | 403 | 357 | 337 | 515 | 404 | 369 | 428 | 357 | 338 | 405 | 435 | 404 | 394.41 | 374 | 415 | 1.107 | 0.147 | 0.26796067 | 0.48774099 | 0.0058 | 0.0153 | 0.0102 | 0.0102 | TRUE | 0.179 |
| Garnl3 | 1060 | 1113 | 1163 | 1052 | 787 | 804 | 1126 | 1112 | 1168 | 827 | 847 | 880 | 993.19 | 1135 | 851 | 0.749 | -0.416 | 4.35E-07 | 1.33E-05 | 0 | 0.0102 | 0.0039 | 0.0039 | TRUE | 0.0103 |
| Gars | 12870 | 13179 | 13442 | 18367 | 12852 | 12514 | 13670 | 13162 | 13497 | 14439 | 13828 | 13699 | 13715.76 | 13443 | 13989 | 1.041 | 0.058 | 0.325072 | 0.54606833 | 5.00E-04 | 0.0071 | 0.0024 | 0.0024 | TRUE | 0.0098 |
| Gart | 30936 | 32479 | 33549 | 33796 | 25111 | 24951 | 32860 | 32437 | 33687 | 26568 | 27018 | 27313 | 29980.29 | 32995 | 26966 | 0.817 | -0.291 | 4.33E-08 | 1.75E-06 | 2.00E-04 | 0.007 | 0.002 | 0.002 | TRUE | 0.0041 |
| Gas1 | 99 | 206 | 206 | 258 | 215 | 182 | 105 | 206 | 207 | 203 | 231 | 199 | 191.85 | 173 | 211 | 1.222 | 0.289 | 0.26124273 | 0.48067109 | 0.0636 | 0.0241 | 0.0424 | 0.0424 | TRUE | 1.2319 |
| Gas2 | 503 | 580 | 611 | 933 | 672 | 658 | 534 | 579 | 614 | 733 | 723 | 720 | 650.63 | 576 | 725 | 1.26 | 0.334 | 0.00074287 | 0.00706608 | 4.00E-04 | 0.012 | 0.0055 | 0.0055 | TRUE | 0.0459 |
| Gas2l1 | 1000 | 703 | 676 | 1631 | 1177 | 805 | 1062 | 702 | 679 | 1282 | 1266 | 881 | 978.8 | 814 | 1143 | 1.405 | 0.49 | 0.01765341 | 0.07932466 | 0.053 | 0.0103 | 0.0297 | 0.0297 | TRUE | 0.8399 |
| Gas2l2 | 3 | 5 | 2 | 8 | 2 | 1 | 3 | 5 | 2 | 6 | 2 | 1 | 3.29 | 3 | 3 | 0.968 | -0.047 | 0.96624088 | NA | 0.1799 | 1.0091 | 0.5877 | 0.5877 | TRUE | 1.2859 |
| Gas2l3 | 2725 | 3086 | 3176 | 3158 | 2255 | 2683 | 2894 | 3082 | 3189 | 2483 | 2426 | 2937 | 2835.21 | 3055 | 2615 | 0.856 | -0.225 | 0.02643937 | 0.10567707 | 0.0064 | 0.0081 | 0.007 | 0.007 | TRUE | 0.1396 |
| Gas5 | 8633 | 13426 | 13296 | 11498 | 9257 | 10327 | 9170 | 13409 | 13351 | 9039 | 9960 | 11305 | 11038.72 | 11977 | 10101 | 0.843 | -0.246 | 0.11899192 | 0.29214516 | 0.0268 | 0.0072 | 0.0178 | 0.0178 | TRUE | 0.5136 |
| Gas6 | 2236 | 2609 | 2476 | 2162 | 1710 | 1807 | 2375 | 2606 | 2486 | 1700 | 1840 | 1978 | 2164.05 | 2489 | 1839 | 0.738 | -0.438 | 1.21E-06 | 3.25E-05 | 0.0032 | 0.0084 | 0.0054 | 0.0054 | TRUE | 0.0545 |
| Gas7 | 1184 | 1093 | 1112 | 1170 | 854 | 1016 | 1258 | 1092 | 1117 | 920 | 919 | 1112 | 1069.43 | 1156 | 984 | 0.851 | -0.233 | 0.04630008 | 0.15573574 | 0.008 | 0.01 | 0.0089 | 0.0089 | TRUE | 0.1543 |
| Gas8 | 279 | 209 | 170 | 513 | 389 | 222 | 296 | 209 | 171 | 403 | 419 | 243 | 290.1 | 225 | 355 | 1.578 | 0.658 | 0.01166752 | 0.05900575 | 0.0773 | 0.0183 | 0.0454 | 0.0454 | TRUE | 0.8579 |
| Gast | 4 | 6 | 5 | 2 | 2 | 0 | 4 | 6 | 5 | 2 | 2 | 0 | 3.16 | 5 | 1 | 0.249 | -2.008 | 0.09011516 | NA | 0 | 1.048 | 0.527 | 0.527 | TRUE | 0.3605 |
| Gata1 | 169 | 184 | 170 | 182 | 156 | 177 | 180 | 184 | 171 | 143 | 168 | 194 | 173.11 | 178 | 168 | 0.941 | -0.088 | 0.61589137 | 0.78280592 | 0.007 | 0.0259 | 0.0161 | 0.0161 | TRUE | 0.1993 |
| Gata2 | 54 | 91 | 91 | 138 | 123 | 103 | 57 | 91 | 91 | 108 | 132 | 113 | 98.86 | 80 | 118 | 1.471 | 0.557 | 0.02278824 | 0.09541176 | 0.0236 | 0.0402 | 0.0325 | 0.0325 | TRUE | 0.5513 |
| Gata3 | 37 | 17 | 8 | 112 | 67 | 40 | 39 | 17 | 8 | 88 | 72 | 44 | 44.71 | 21 | 68 | 3.19 | 1.674 | 0.0013967 | 0.01157583 | 0.2853 | 0.0806 | 0.1666 | 0.1666 | TRUE | 0.9047 |
| Gata4 | 76 | 114 | 91 | 260 | 184 | 130 | 81 | 114 | 91 | 204 | 198 | 142 | 138.44 | 95 | 181 | 1.907 | 0.932 | 3.79E-05 | 0.00061168 | 0.0268 | 0.0307 | 0.0288 | 0.0288 | TRUE | 0.3735 |
| Gata5 | 8 | 11 | 7 | 9 | 6 | 3 | 8 | 11 | 7 | 7 | 6 | 3 | 7.22 | 9 | 5 | 0.642 | -0.639 | 0.37049817 | 0.58743656 | 0 | 0.4631 | 0.2232 | 0.2232 | TRUE | 0.2587 |
| Gata6 | 63 | 126 | 111 | 358 | 200 | 180 | 67 | 126 | 111 | 281 | 215 | 197 | 166.31 | 101 | 231 | 2.282 | 1.19 | 4.25E-06 | 9.61E-05 | 0.0584 | 0.0267 | 0.0411 | 0.0411 | TRUE | 0.8507 |
| Gatad1 | 3674 | 4722 | 4482 | 5580 | 3993 | 3934 | 3902 | 4716 | 4500 | 4387 | 4296 | 4306 | 4351.32 | 4373 | 4330 | 0.99 | -0.014 | 0.87503534 | 0.93827298 | 0.0046 | 0.0077 | 0.0058 | 0.0058 | TRUE | 0.1077 |
| Gatad2a | 17882 | 16480 | 16052 | 22929 | 16759 | 15420 | 18994 | 16459 | 16118 | 18025 | 18032 | 16880 | 17417.79 | 17190 | 17646 | 1.027 | 0.038 | 0.67015464 | 0.81587642 | 0.004 | 0.0071 | 0.0056 | 0.0056 | TRUE | 0.103 |
| Gatad2b | 1766 | 1597 | 1807 | 2150 | 1556 | 1582 | 1876 | 1595 | 1814 | 1690 | 1674 | 1732 | 1730.21 | 1762 | 1699 | 0.964 | -0.052 | 0.56389046 | 0.74480197 | 0.0032 | 0.0088 | 0.0054 | 0.0054 | TRUE | 0.083 |
| Gatc | 414 | 436 | 434 | 511 | 409 | 384 | 440 | 435 | 436 | 402 | 440 | 420 | 428.85 | 437 | 421 | 0.962 | -0.057 | 0.59478064 | 0.7677918 | 0 | 0.0146 | 0.0058 | 0.0058 | TRUE | 0.0194 |
| Gatm | 330 | 486 | 425 | 557 | 382 | 413 | 351 | 485 | 427 | 438 | 411 | 452 | 427.27 | 421 | 434 | 1.03 | 0.043 | 0.77145756 | 0.88080416 | 0.0117 | 0.0146 | 0.0131 | 0.0131 | TRUE | 0.2447 |
| Gatsl2 | 465 | 387 | 401 | 427 | 303 | 253 | 494 | 387 | 403 | 336 | 326 | 277 | 370.28 | 428 | 313 | 0.733 | -0.448 | 0.00283246 | 0.02024174 | 0.0114 | 0.0158 | 0.0135 | 0.0135 | TRUE | 0.2106 |
| Gatsl3 | 1363 | 1647 | 1820 | 2039 | 1390 | 1657 | 1448 | 1645 | 1827 | 1603 | 1496 | 1814 | 1638.73 | 1640 | 1638 | 0.998 | -0.003 | 0.98225156 | 0.990605 | 0.0109 | 0.0089 | 0.01 | 0.01 | TRUE | 0.1265 |
| Gba | 578 | 582 | 584 | 1070 | 774 | 730 | 614 | 581 | 586 | 841 | 833 | 799 | 709.1 | 594 | 824 | 1.389 | 0.474 | 1.45E-07 | 4.94E-06 | 0 | 0.0115 | 0.0044 | 0.0044 | TRUE | 0.0101 |
| Gba2 | 1132 | 864 | 864 | 1171 | 845 | 638 | 1202 | 863 | 868 | 921 | 909 | 698 | 910.15 | 978 | 843 | 0.863 | -0.213 | 0.21066694 | 0.42117366 | 0.0292 | 0.0105 | 0.0198 | 0.0198 | TRUE | 0.4818 |
| Gbas | 1756 | 2271 | 2410 | 2594 | 1892 | 1950 | 1865 | 2268 | 2420 | 2039 | 2036 | 2135 | 2127.1 | 2184 | 2070 | 0.947 | -0.078 | 0.48825585 | 0.68476003 | 0.0079 | 0.0085 | 0.0087 | 0.0087 | TRUE | 0.1973 |
| Gbe1 | 286 | 408 | 427 | 683 | 424 | 441 | 304 | 407 | 429 | 537 | 456 | 483 | 435.98 | 380 | 492 | 1.295 | 0.373 | 0.01863151 | 0.08270648 | 0.0171 | 0.0145 | 0.0158 | 0.0158 | TRUE | 0.3501 |
| Gbf1 | 3823 | 3936 | 4404 | 5186 | 3809 | 3852 | 4061 | 3931 | 4422 | 4077 | 4098 | 4217 | 4134.23 | 4138 | 4131 | 0.998 | -0.003 | 0.97131646 | 0.98480349 | 0.0018 | 0.0077 | 0.0038 | 0.0038 | TRUE | 0.044 |
| Gbgt1 | 41 | 32 | 13 | 55 | 43 | 33 | 44 | 32 | 13 | 43 | 46 | 36 | 35.7 | 30 | 42 | 1.423 | 0.508 | 0.24955876 | 0.466798 | 0.1268 | 0.0992 | 0.1115 | 0.1115 | TRUE | 0.5302 |
| Gbp1 | 24 | 34 | 51 | 52 | 25 | 35 | 25 | 34 | 51 | 41 | 27 | 38 | 36.12 | 37 | 35 | 0.961 | -0.057 | 0.88134075 | 0.94135674 | 0.0568 | 0.0981 | 0.0782 | 0.0782 | TRUE | 0.5063 |
| Gbp10 | 0 | 0 | 1 | 0 | 0 | 0 | 0 | 0 | 1 | 0 | 0 | 0 | 0.17 | 0 | 0 | 0.49 | -1.029 | 0.80087 | NA | 0 | 19.6928 | 10 | 10 | TRUE | 0.4098 |
| Gbp11 | 2 | 1 | 4 | 2 | 0 | 0 | 2 | 1 | 4 | 2 | 0 | 0 | 1.45 | 2 | 1 | 0.246 | -2.023 | 0.27141076 | NA | 0 | 2.2759 | 1.3932 | 1.3932 | TRUE | 0.7141 |
| Gbp2 | 43 | 60 | 54 | 79 | 41 | 29 | 46 | 60 | 54 | 62 | 44 | 32 | 49.63 | 53 | 46 | 0.871 | -0.199 | 0.5481805 | 0.73305889 | 0.0439 | 0.0733 | 0.0591 | 0.0591 | TRUE | 0.3854 |
| Gbp3 | 9 | 12 | 11 | 22 | 15 | 14 | 10 | 12 | 11 | 17 | 16 | 15 | 13.56 | 11 | 16 | 1.498 | 0.583 | 0.24708903 | 0.46388344 | 0 | 0.2499 | 0.106 | 0.106 | TRUE | 0.0384 |
| Gbp4 | 97 | 66 | 80 | 87 | 63 | 51 | 103 | 66 | 80 | 68 | 68 | 56 | 73.55 | 83 | 64 | 0.773 | -0.371 | 0.15590652 | 0.34854113 | 0.019 | 0.0517 | 0.0356 | 0.0356 | TRUE | 0.2764 |
| Gbp5 | 0 | 0 | 1 | 4 | 1 | 1 | 0 | 0 | 1 | 3 | 1 | 1 | 1.05 | 0 | 2 | 4.937 | 2.303 | 0.25694648 | NA | 0 | 3.1349 | 1.7042 | 1.7042 | TRUE | 0.4788 |
| Gbp6 | 3 | 4 | 5 | 3 | 0 | 1 | 3 | 4 | 5 | 2 | 0 | 1 | 2.61 | 4 | 1 | 0.3 | -1.738 | 0.17550002 | NA | 0 | 1.2696 | 0.6622 | 0.6622 | TRUE | 0.6014 |

| | | | | | | | | | | | | | | | | | | | | | | | | | |
|---|---|---|---|---|---|---|---|---|---|---|---|---|---|---|---|---|---|---|---|---|---|---|---|---|---|
| Gbp7 | 33 | 48 | 49 | 43 | 35 | 43 | 35 | 48 | 49 | 34 | 38 | 47 | 41.79 | 44 | 40 | 0.891 | -0.167 | 0.59820248 | 0.77063216 | 0 | 0.0857 | 0.0484 | 0.0484 | TRUE | 0.241 |
| Gbp8 | 4 | 4 | 0 | 9 | 2 | 3 | 4 | 4 | 0 | 7 | 2 | 3 | 3.46 | 3 | 4 | 1.549 | 0.631 | 0.59785794 | NA | 0.395 | 0.9593 | 0.7298 | 0.7298 | TRUE | 0.933 |
| Gbp9 | 51 | 38 | 39 | 64 | 60 | 65 | 54 | 38 | 39 | 50 | 65 | 71 | 52.88 | 44 | 62 | 1.412 | 0.497 | 0.09410297 | 0.25112088 | 0.0175 | 0.0692 | 0.044 | 0.044 | TRUE | 0.3266 |
| Gbx1 | 23 | 28 | 24 | 26 | 16 | 14 | 24 | 28 | 24 | 20 | 17 | 15 | 21.58 | 25 | 17 | 0.698 | -0.52 | 0.2006026 | 0.40841648 | 0 | 0.1596 | 0.0714 | 0.0714 | TRUE | 0.1186 |
| Gbx2 | 1590 | 1162 | 1144 | 1291 | 896 | 619 | 1689 | 1160 | 1149 | 1015 | 964 | 678 | 1109.1 | 1333 | 886 | 0.665 | -0.589 | 0.00285295 | 0.02035127 | 0.0465 | 0.0099 | 0.0271 | 0.0271 | TRUE | 0.656 |
| Gc | 6 | 0 | 2 | 2 | 4 | 2 | 6 | 0 | 2 | 2 | 4 | 2 | 2.74 | 3 | 3 | 0.958 | -0.062 | 0.96341897 | NA | 0.6028 | 1.2088 | 0.9355 | 0.9355 | TRUE | 0.5551 |
| Gca | 1525 | 1287 | 1266 | 2679 | 1853 | 1512 | 1620 | 1285 | 1271 | 2106 | 1994 | 1655 | 1655.2 | 1392 | 1918 | 1.379 | 0.463 | 0.00075592 | 0.00715627 | 0.0166 | 0.0089 | 0.013 | 0.013 | TRUE | 0.2457 |
| Gcat | 2686 | 4128 | 4464 | 3068 | 2383 | 3008 | 2853 | 4123 | 4482 | 2412 | 2564 | 3293 | 3287.75 | 3819 | 2756 | 0.721 | -0.471 | 0.00947105 | 0.05049944 | 0.0392 | 0.0079 | 0.0234 | 0.0234 | TRUE | 0.5956 |
| Gcc1 | 834 | 805 | 890 | 1092 | 829 | 837 | 886 | 804 | 894 | 858 | 892 | 916 | 875.02 | 861 | 889 | 1.032 | 0.045 | 0.62569903 | 0.78915254 | 0.0011 | 0.0107 | 0.005 | 0.005 | TRUE | 0.0404 |
| Gcc2 | 1371 | 1111 | 1295 | 2225 | 1539 | 1466 | 1456 | 1110 | 1300 | 1749 | 1656 | 1605 | 1479.31 | 1289 | 1670 | 1.296 | 0.375 | 0.00140933 | 0.01166208 | 0.009 | 0.0091 | 0.0092 | 0.0092 | TRUE | 0.1779 |
| Gcdh | 3686 | 4416 | 4240 | 4305 | 3280 | 3436 | 3915 | 4410 | 4257 | 3384 | 3529 | 3761 | 3876.24 | 4194 | 3558 | 0.848 | -0.238 | 0.00438087 | 0.02829446 | 0.0029 | 0.0078 | 0.0048 | 0.0048 | TRUE | 0.0412 |
| Gcfc2 | 1129 | 1052 | 1069 | 1278 | 1087 | 1094 | 1199 | 1051 | 1073 | 1005 | 1170 | 1198 | 1115.83 | 1108 | 1124 | 1.014 | 0.02 | 0.85949088 | 0.9307807 | 0.0062 | 0.0099 | 0.0079 | 0.0079 | TRUE | 0.1061 |
| Gcg | 0 | 0 | 0 | 0 | 0 | 0 | 0 | 0 | 0 | 0 | 0 | 0 | 0 | 0 | 0 | NA | NA | NA | NA | NA | NA | NA | NA | NA | NA |
| Gcgr | 23 | 14 | 11 | 21 | 25 | 13 | 24 | 14 | 11 | 17 | 27 | 14 | 17.85 | 16 | 19 | 1.165 | 0.22 | 0.67312479 | 0.81799428 | 0.0833 | 0.1915 | 0.1394 | 0.1394 | TRUE | 0.8269 |
| Gch1 | 803 | 883 | 820 | 784 | 542 | 632 | 853 | 882 | 823 | 616 | 583 | 692 | 741.58 | 853 | 630 | 0.739 | -0.437 | 2.70E-05 | 0.00045953 | 0.0024 | 0.0113 | 0.0064 | 0.0064 | TRUE | 0.0825 |
| Gchfr | 397 | 414 | 365 | 311 | 207 | 304 | 422 | 413 | 366 | 244 | 223 | 333 | 333.6 | 400 | 267 | 0.665 | -0.589 | 0.00074771 | 0.00709557 | 0.022 | 0.0168 | 0.0189 | 0.0189 | TRUE | 0.5119 |
| Gck | 13 | 24 | 25 | 22 | 30 | 43 | 14 | 24 | 25 | 17 | 32 | 47 | 26.59 | 21 | 32 | 1.52 | 0.604 | 0.21677436 | 0.42872021 | 0.1349 | 0.1308 | 0.1326 | 0.1326 | TRUE | 0.1375 |
| Gckr | 1 | 7 | 7 | 9 | 6 | 4 | 1 | 7 | 7 | 7 | 6 | 4 | 5.5 | 5 | 6 | 1.188 | 0.249 | 0.77820596 | 0.88511424 | 0 | 0.6061 | 0.3806 | 0.3806 | TRUE | 0.91 |
| Gclc | 3945 | 3844 | 3747 | 4410 | 3336 | 2964 | 4190 | 3839 | 3762 | 3467 | 3589 | 3245 | 3682.07 | 3930 | 3434 | 0.874 | -0.195 | 0.01709969 | 0.07763281 | 0.0026 | 0.0078 | 0.0045 | 0.0045 | TRUE | 0.0406 |
| Gclm | 3149 | 3309 | 3470 | 3743 | 2835 | 2643 | 3345 | 3305 | 3484 | 2942 | 3050 | 2893 | 3169.95 | 3378 | 2962 | 0.877 | -0.19 | 0.00430628 | 0.02791559 | 4.00E-04 | 0.0079 | 0.0029 | 0.0029 | TRUE | 0.0092 |
| Gcm1 | 18 | 11 | 8 | 42 | 28 | 19 | 19 | 11 | 8 | 33 | 30 | 21 | 20.35 | 13 | 28 | 2.22 | 1.15 | 0.01911619 | 0.08400686 | 0.0476 | 0.1688 | 0.116 | 0.116 | TRUE | 0.7622 |
| Gcm2 | 36 | 58 | 45 | 131 | 115 | 118 | 38 | 58 | 45 | 103 | 124 | 129 | 82.87 | 47 | 119 | 2.506 | 1.325 | 1.24E-07 | 4.30E-06 | 0.011 | 0.0467 | 0.0303 | 0.0303 | TRUE | 0.323 |
| Gcn1l1 | 11415 | 13689 | 14215 | 14859 | 11450 | 10908 | 12125 | 13671 | 14273 | 11681 | 12319 | 11941 | 12668.41 | 13356 | 11980 | 0.897 | -0.157 | 0.06255151 | 0.19156552 | 0.0037 | 0.0072 | 0.005 | 0.005 | TRUE | 0.0795 |
| Gcnt1 | 2085 | 2037 | 2106 | 2117 | 1525 | 1602 | 2215 | 2034 | 2115 | 1664 | 1641 | 1754 | 1903.72 | 2121 | 1686 | 0.795 | -0.331 | 1.69E-05 | 0.00030877 | 0.001 | 0.0086 | 0.0037 | 0.0037 | TRUE | 0.0179 |
| Gcnt2 | 2692 | 3054 | 3009 | 2859 | 2432 | 2775 | 2859 | 3050 | 3021 | 2248 | 2617 | 3038 | 2805.44 | 2977 | 2634 | 0.884 | -0.177 | 0.13886148 | 0.3234564 | 0.0113 | 0.0081 | 0.01 | 0.01 | TRUE | 0.2181 |
| Gcnt3 | 14 | 9 | 15 | 59 | 36 | 22 | 15 | 9 | 15 | 46 | 39 | 24 | 24.69 | 13 | 36 | 2.827 | 1.5 | 0.00109627 | 0.00956324 | 0.0545 | 0.1404 | 0.0992 | 0.0992 | TRUE | 0.2265 |
| Gcnt4 | 214 | 209 | 221 | 224 | 188 | 212 | 227 | 209 | 222 | 176 | 202 | 232 | 211.4 | 219 | 203 | 0.925 | -0.113 | 0.4820031 | 0.68083046 | 0.0061 | 0.0225 | 0.0139 | 0.0139 | TRUE | 0.1641 |
| Gcnt7 | 0 | 0 | 0 | 0 | 0 | 0 | 0 | 0 | 0 | 0 | 0 | 0 | 0 | 0 | 0 | NA | NA | NA | NA | NA | NA | NA | NA | NA | NA |
| Gcsam | 0 | 0 | 0 | 0 | 0 | 0 | 0 | 0 | 0 | 0 | 0 | 0 | 0 | 0 | 0 | NA | NA | NA | NA | NA | NA | NA | NA | NA | NA |
| Gcsh | 9644 | 8543 | 8125 | 10699 | 7456 | 7060 | 10244 | 8532 | 8158 | 8411 | 8022 | 7728 | 8515.86 | 8978 | 8054 | 0.897 | -0.157 | 0.13542653 | 0.31837044 | 0.0084 | 0.0073 | 0.0078 | 0.0078 | TRUE | 0.1857 |
| Gda | 37 | 28 | 31 | 75 | 39 | 34 | 39 | 28 | 31 | 59 | 42 | 37 | 39.42 | 33 | 46 | 1.417 | 0.503 | 0.14122521 | 0.32677593 | 0.0255 | 0.0905 | 0.058 | 0.058 | TRUE | 0.5379 |
| Gdap1 | 14 | 18 | 19 | 11 | 8 | 8 | 15 | 18 | 19 | 9 | 9 | 9 | 12.99 | 17 | 9 | 0.5 | -0.999 | 0.05681634 | 0.17932906 | 0 | 0.2605 | 0.1125 | 0.1125 | TRUE | 0.0723 |
| Gdap10 | 165 | 213 | 216 | 185 | 196 | 217 | 175 | 213 | 217 | 145 | 211 | 238 | 199.79 | 202 | 198 | 0.977 | -0.033 | 0.87791868 | 0.93970448 | 0.0333 | 0.0234 | 0.0284 | 0.0284 | TRUE | 0.6234 |
| Gdap1l1 | 151 | 162 | 181 | 84 | 63 | 53 | 160 | 162 | 182 | 66 | 68 | 58 | 115.96 | 168 | 64 | 0.381 | -1.391 | 1.06E-12 | 1.07E-10 | 0 | 0.0353 | 0.0168 | 0.0168 | TRUE | 0.055 |
| Gdap2 | 818 | 822 | 904 | 1147 | 830 | 874 | 869 | 821 | 908 | 902 | 893 | 957 | 891.49 | 866 | 917 | 1.059 | 0.083 | 0.36059281 | 0.57903378 | 8.00E-04 | 0.0106 | 0.0048 | 0.0048 | TRUE | 0.0247 |
| Gde1 | 2218 | 2766 | 2647 | 3302 | 2419 | 2673 | 2356 | 2762 | 2658 | 2596 | 2603 | 2926 | 2650.12 | 2592 | 2708 | 1.044 | 0.063 | 0.52040139 | 0.71109881 | 0.0048 | 0.0081 | 0.0065 | 0.0065 | TRUE | 0.0768 |
| Gdf1 | 0 | 0 | 0 | 0 | 0 | 0 | 0 | 0 | 0 | 0 | 0 | 0 | 0 | 0 | 0 | NA | NA | NA | NA | NA | NA | NA | NA | NA | NA |
| Gdf10 | 10 | 7 | 2 | 13 | 6 | 2 | 11 | 7 | 2 | 10 | 6 | 2 | 6.41 | 7 | 6 | 0.981 | -0.027 | 0.97603587 | 0.98709087 | 0.3408 | 0.5205 | 0.434 | 0.434 | TRUE | 0.2587 |
| Gdf11 | 38 | 51 | 41 | 66 | 76 | 42 | 40 | 51 | 41 | 52 | 82 | 46 | 52.02 | 44 | 60 | 1.352 | 0.435 | 0.17810971 | 0.37884644 | 0.0401 | 0.0702 | 0.0553 | 0.0553 | TRUE | 0.8445 |
| Gdf15 | 64 | 123 | 106 | 84 | 58 | 96 | 68 | 123 | 106 | 66 | 62 | 105 | 88.46 | 99 | 78 | 0.782 | -0.354 | 0.25587336 | 0.47450143 | 0.0741 | 0.0441 | 0.0585 | 0.0585 | TRUE | 0.6194 |
| Gdf2 | 0 | 0 | 0 | 1 | 0 | 3 | 0 | 0 | 0 | 1 | 0 | 3 | 0.68 | 0 | 1 | 7.177 | 2.843 | 0.34581154 | NA | 6.5474 | 4.8635 | 5.1913 | 5.1913 | TRUE | 0.362 |
| Gdf3 | 1871 | 2295 | 2532 | 4248 | 2977 | 2962 | 1987 | 2292 | 2542 | 3339 | 3203 | 3242 | 2767.78 | 2274 | 3261 | 1.434 | 0.52 | 7.80E-07 | 2.21E-05 | 0.0071 | 0.0081 | 0.0076 | 0.0076 | TRUE | 0.1468 |
| Gdf5 | 2 | 0 | 2 | 6 | 3 | 3 | 2 | 0 | 2 | 5 | 3 | 3 | 2.56 | 1 | 4 | 2.755 | 1.462 | 0.25055011 | NA | 0 | 1.2937 | 0.6642 | 0.6642 | TRUE | 0.4989 |
| Gdf6 | 1 | 3 | 3 | 4 | 1 | 1 | 1 | 3 | 3 | 3 | 1 | 1 | 2.06 | 2 | 2 | 0.775 | -0.369 | 0.77717249 | NA | 0 | 1.6028 | 0.7404 | 0.7404 | TRUE | 0.4886 |
| Gdf7 | 12 | 11 | 13 | 22 | 22 | 9 | 13 | 11 | 13 | 17 | 24 | 10 | 14.6 | 12 | 17 | 1.384 | 0.469 | 0.38099029 | 0.59530204 | 0 | 0.2325 | 0.1357 | 0.1357 | TRUE | 0.1598 |
| Gdf9 | 23 | 8 | 7 | 29 | 10 | 17 | 24 | 8 | 7 | 23 | 11 | 19 | 15.27 | 13 | 18 | 1.336 | 0.418 | 0.51343096 | 0.70639477 | 0.2345 | 0.2227 | 0.228 | 0.228 | TRUE | 1.6293 |
| Gdi1 | 2193 | 2986 | 2918 | 3525 | 2714 | 2960 | 2329 | 2982 | 2930 | 2771 | 2920 | 3240 | 2862.14 | 2747 | 2977 | 1.083 | 0.115 | 0.3404962 | 0.56075035 | 0.0108 | 0.0081 | 0.0102 | 0.0102 | TRUE | 0.2144 |
| Gdi2 | 21358 | 24183 | 24311 | 30800 | 21972 | 22680 | 22686 | 24152 | 24411 | 24213 | 23641 | 24827 | 23988.08 | 23750 | 24227 | 1.02 | 0.029 | 0.64957918 | 0.80354219 | 9.00E-04 | 0.007 | 0.0028 | 0.0028 | TRUE | 0.0188 |
| Gdnf | 1 | 0 | 7 | 6 | 5 | 4 | 1 | 0 | 7 | 5 | 5 | 4 | 3.76 | 3 | 5 | 1.783 | 0.834 | 0.49215947 | NA | 0.5963 | 0.8829 | 0.7723 | 0.7723 | TRUE | 2.3552 |
| Gdpd1 | 665 | 711 | 710 | 993 | 662 | 731 | 706 | 710 | 713 | 781 | 712 | 800 | 737.07 | 710 | 764 | 1.077 | 0.107 | 0.25240448 | 0.47049751 | 0 | 0.0114 | 0.005 | 0.005 | TRUE | 0.0409 |
| Gdpd2 | 19 | 24 | 28 | 47 | 35 | 29 | 20 | 24 | 28 | 37 | 38 | 32 | 29.77 | 24 | 36 | 1.472 | 0.558 | 0.11432282 | 0.28516518 | 0 | 0.1176 | 0.055 | 0.055 | TRUE | 0.1279 |
| Gdpd3 | 5 | 8 | 19 | 17 | 4 | 12 | 5 | 8 | 19 | 13 | 4 | 13 | 10.53 | 11 | 10 | 0.956 | -0.065 | 0.92954499 | 0.9654152 | 0.2699 | 0.3198 | 0.2955 | 0.2955 | TRUE | 1.6605 |
| Gdpd4 | 2 | 0 | 1 | 3 | 3 | 0 | 2 | 0 | 1 | 2 | 3 | 0 | 1.45 | 1 | 2 | 1.815 | 0.86 | 0.63307917 | NA | 0 | 2.2751 | 1.5839 | 1.5839 | TRUE | 0.4076 |
| Gdpd5 | 187 | 272 | 264 | 273 | 179 | 199 | 199 | 272 | 265 | 215 | 193 | 218 | 226.73 | 245 | 209 | 0.85 | -0.235 | 0.17083193 | 0.36944484 | 0.0123 | 0.0214 | 0.0167 | 0.0167 | TRUE | 0.3015 |
| Gdpgp1 | 531 | 428 | 436 | 812 | 544 | 450 | 564 | 427 | 438 | 638 | 585 | 493 | 524.25 | 476 | 572 | 1.203 | 0.266 | 0.09129114 | 0.24659083 | 0.0182 | 0.0132 | 0.016 | 0.016 | TRUE | 0.2991 |
| Gem | 119 | 87 | 80 | 151 | 98 | 123 | 126 | 87 | 80 | 119 | 105 | 135 | 108.73 | 98 | 120 | 1.223 | 0.291 | 0.22585848 | 0.43864548 | 0.0269 | 0.0372 | 0.0322 | 0.0322 | TRUE | 0.6219 |
| Gemin2 | 1356 | 1733 | 1736 | 1957 | 1428 | 1504 | 1440 | 1731 | 1743 | 1538 | 1536 | 1646 | 1605.91 | 1638 | 1573 | 0.96 | -0.058 | 0.57370321 | 0.75215686 | 0.0058 | 0.009 | 0.0071 | 0.0071 | TRUE | 0.134 |
| Gemin4 | 5914 | 4496 | 4592 | 7647 | 5522 | 4308 | 6282 | 4490 | 4611 | 6011 | 5941 | 4716 | 5341.89 | 5128 | 5556 | 1.084 | 0.116 | 0.45378757 | 0.65906064 | 0.0232 | 0.0075 | 0.0171 | 0.0171 | TRUE | 0.4723 |
| Gemin5 | 22796 | 20315 | 21456 | 23680 | 17197 | 15905 | 24213 | 20289 | 21544 | 18615 | 18503 | 17411 | 20095.84 | 22015 | 18176 | 0.826 | -0.276 | 0.00180044 | 0.01416289 | 0.0047 | 0.0071 | 0.0056 | 0.0056 | TRUE | 0.0933 |
| Gemin6 | 2096 | 1919 | 2009 | 2674 | 1813 | 1749 | 2226 | 1917 | 2017 | 2102 | 1951 | 1915 | 2021.24 | 2053 | 1989 | 0.969 | -0.045 | 0.62399406 | 0.78821354 | 0.0037 | 0.0085 | 0.0056 | 0.0056 | TRUE | 0.0655 |
| Gemin7 | 1135 | 1259 | 1273 | 1441 | 944 | 1029 | 1206 | 1257 | 1278 | 1133 | 1016 | 1126 | 1169.34 | 1247 | 1092 | 0.876 | -0.191 | 0.0286881 | 0.11152789 | 0.0013 | 0.0097 | 0.0047 | 0.0047 | TRUE | 0.0436 |
| Gemin8 | 406 | 455 | 504 | 526 | 408 | 442 | 431 | 454 | 506 | 413 | 439 | 484 | 454.67 | 464 | 445 | 0.958 | -0.061 | 0.61982186 | 0.78489204 | 0.0044 | 0.0141 | 0.0088 | 0.0088 | TRUE | 0.0735 |
| Gen1 | 746 | 524 | 484 | 1203 | 863 | 586 | 792 | 523 | 486 | 946 | 929 | 641 | 719.57 | 600 | 839 | 1.397 | 0.483 | 0.02570647 | 0.1036672 | 0.0574 | 0.0115 | 0.0323 | 0.0323 | TRUE | 0.9146 |
| Get4 | 3869 | 4072 | 4061 | 4791 | 3470 | 3366 | 4110 | 4067 | 4078 | 3766 | 3734 | 3685 | 3906.4 | 4085 | 3728 | 0.913 | -0.132 | 0.02289086 | 0.09571438 | 0 | 0.0077 | 0.0022 | 0.0022 | TRUE | 0.0013 |
| Gfap | 142 | 71 | 95 | 54 | 40 | 30 | 151 | 71 | 95 | 42 | 43 | 33 | 72.58 | 106 | 39 | 0.374 | -1.417 | 2.52E-05 | 0.00043613 | 0.0764 | 0.0523 | 0.0643 | 0.0643 | TRUE | 0.2983 |
| Gfer | 1787 | 2589 | 2670 | 3010 | 2379 | 2628 | 1898 | 2586 | 2681 | 2366 | 2560 | 2877 | 2494.56 | 2388 | 2601 | 1.089 | 0.123 | 0.40333828 | 0.61618088 | 0.0216 | 0.0082 | 0.0151 | 0.0151 | TRUE | 0.39 |
| Gfi1 | 0 | 1 | 2 | 3 | 1 | 2 | 0 | 1 | 2 | 2 | 1 | 2 | 1.44 | 1 | 2 | 1.869 | 0.902 | 0.57403764 | NA | 0 | 2.2972 | 1.0918 | 1.0918 | TRUE | 0.2891 |
| Gfi1b | 7 | 4 | 2 | 5 | 3 | 3 | 7 | 4 | 2 | 4 | 3 | 3 | 3.98 | 4 | 3 | 0.785 | -0.349 | 0.716245 | NA | 0 | 0.8346 | 0.412 | 0.412 | TRUE | 0.3279 |
| Gfm1 | 4471 | 4413 | 4412 | 6836 | 4615 | 4466 | 4749 | 4407 | 4430 | 5374 | 4965 | 4889 | 4802.42 | 4529 | 5076 | 1.121 | 0.165 | 0.02920604 | 0.11290139 | 0.0019 | 0.0076 | 0.0039 | 0.0039 | TRUE | 0.0324 |

| | | | | | | | | | | | | | | | | | | | | | | | | | |
|---|---|---|---|---|---|---|---|---|---|---|---|---|---|---|---|---|---|---|---|---|---|---|---|---|---|
| Gfm2 | 4106 | 3893 | 3889 | 4929 | 3333 | 2856 | 4361 | 3888 | 3905 | 3875 | 3586 | 3126 | 3790.25 | 4051 | 3529 | 0.871 | -0.199 | 0.05940538 | 0.18458266 | 0.0075 | 0.0078 | 0.0077 | 0.0077 | TRUE | 0.1209 |
| Gfod1 | 935 | 1140 | 1257 | 1035 | 898 | 1008 | 993 | 1139 | 1262 | 814 | 966 | 1103 | 1046.18 | 1131 | 961 | 0.849 | -0.237 | 0.10109105 | 0.26330963 | 0.017 | 0.0101 | 0.0141 | 0.0141 | TRUE | 0.2196 |
| Gfod2 | 833 | 620 | 540 | 864 | 589 | 542 | 885 | 619 | 542 | 679 | 634 | 593 | 658.74 | 682 | 635 | 0.932 | -0.101 | 0.58065036 | 0.75711123 | 0.0315 | 0.0119 | 0.0226 | 0.0226 | TRUE | 0.7957 |
| Gfpt1 | 3039 | 3443 | 3660 | 3829 | 2667 | 2896 | 3228 | 3439 | 3675 | 3010 | 2870 | 3170 | 3231.88 | 3447 | 3017 | 0.875 | -0.193 | 0.02270606 | 0.09511819 | 0.0029 | 0.0079 | 0.0048 | 0.0048 | TRUE | 0.0407 |
| Gfpt2 | 6059 | 7587 | 8337 | 5311 | 4012 | 5269 | 6436 | 7577 | 8371 | 4175 | 4317 | 5768 | 6107.28 | 7461 | 4753 | 0.637 | -0.651 | 1.49E-05 | 0.00027752 | 0.0254 | 0.0074 | 0.0161 | 0.0161 | TRUE | 0.4243 |
| Gfra1 | 185 | 139 | 147 | 137 | 90 | 106 | 197 | 139 | 148 | 108 | 97 | 116 | 133.92 | 161 | 107 | 0.664 | -0.59 | 0.0045677 | 0.02922584 | 0.0156 | 0.0315 | 0.0234 | 0.0234 | TRUE | 0.3894 |
| Gfra2 | 5 | 16 | 19 | 7 | 7 | 18 | 5 | 16 | 19 | 6 | 8 | 20 | 12.18 | 13 | 11 | 0.799 | -0.324 | 0.65567491 | 0.80633954 | 0.3193 | 0.2773 | 0.2979 | 0.2979 | TRUE | 1.6136 |
| Gfra3 | 23 | 17 | 17 | 49 | 31 | 25 | 24 | 17 | 17 | 39 | 33 | 27 | 26.29 | 19 | 33 | 1.71 | 0.774 | 0.04820798 | 0.16003096 | 0 | 0.1322 | 0.0696 | 0.0696 | TRUE | 0.2491 |
| Gfra4 | 171 | 139 | 181 | 295 | 189 | 173 | 182 | 139 | 182 | 232 | 203 | 189 | 187.8 | 168 | 208 | 1.248 | 0.319 | 0.07521677 | 0.21604787 | 0.0113 | 0.0244 | 0.0178 | 0.0178 | TRUE | 0.2394 |
| Gfral | 0 | 0 | 0 | 0 | 0 | 0 | 0 | 0 | 0 | 0 | 0 | 0 | 0 | 0 | 0 | NA | NA | NA | NA | NA | NA | NA | NA | NA | NA |
| Gga1 | 3149 | 3361 | 3255 | 3759 | 2819 | 2871 | 3345 | 3357 | 3268 | 2955 | 3033 | 3143 | 3183.45 | 3323 | 3044 | 0.916 | -0.127 | 0.04985796 | 0.16390913 | 3.00E-04 | 0.0079 | 0.0027 | 0.0027 | TRUE | 0.0098 |
| Gga2 | 12911 | 14217 | 14271 | 13987 | 10873 | 11517 | 13714 | 14199 | 14330 | 10995 | 11699 | 12607 | 12923.89 | 14081 | 11767 | 0.836 | -0.259 | 0.00069294 | 0.00671321 | 0.0025 | 0.0072 | 0.0041 | 0.0041 | TRUE | 0.0477 |
| Gga3 | 6928 | 5902 | 5803 | 7647 | 5692 | 5334 | 7359 | 5894 | 5827 | 6011 | 6124 | 5839 | 6175.76 | 6360 | 5991 | 0.942 | -0.086 | 0.42787074 | 0.63790314 | 0.009 | 0.0074 | 0.0083 | 0.0083 | TRUE | 0.2301 |
| Ggact | 550 | 584 | 618 | 593 | 458 | 444 | 584 | 583 | 621 | 466 | 493 | 486 | 538.83 | 596 | 482 | 0.807 | -0.309 | 0.00180424 | 0.01417411 | 0 | 0.013 | 0.0052 | 0.0052 | TRUE | 0.0153 |
| Ggct | 667 | 689 | 677 | 847 | 605 | 549 | 708 | 688 | 680 | 666 | 651 | 601 | 665.69 | 692 | 639 | 0.925 | -0.113 | 0.23586025 | 0.45107164 | 0 | 0.0119 | 0.005 | 0.005 | TRUE | 0.0316 |
| Ggcx | 322 | 316 | 261 | 357 | 352 | 307 | 342 | 316 | 262 | 281 | 379 | 336 | 319.19 | 307 | 332 | 1.08 | 0.111 | 0.50839691 | 0.70245448 | 0.0174 | 0.0172 | 0.0173 | 0.0173 | TRUE | 0.2179 |
| Ggh | 279 | 275 | 232 | 489 | 299 | 303 | 296 | 275 | 233 | 384 | 322 | 332 | 306.96 | 268 | 346 | 1.294 | 0.372 | 0.01332469 | 0.06484511 | 0.0087 | 0.0176 | 0.013 | 0.013 | TRUE | 0.1467 |
| Ggn | 5 | 3 | 3 | 10 | 7 | 0 | 5 | 3 | 3 | 8 | 8 | 0 | 4.45 | 4 | 5 | 1.381 | 0.466 | 0.67711016 | NA | 0.5531 | 0.7469 | 0.671 | 0.671 | TRUE | 1.4239 |
| Ggnbp1 | 197 | 265 | 285 | 308 | 280 | 247 | 209 | 265 | 286 | 242 | 301 | 270 | 262.31 | 253 | 271 | 1.068 | 0.095 | 0.57746088 | 0.75490686 | 0.0148 | 0.0195 | 0.0171 | 0.0171 | TRUE | 0.2549 |
| Ggnbp2 | 5229 | 6481 | 6807 | 6901 | 5007 | 5583 | 5554 | 6473 | 6835 | 5425 | 5387 | 6111 | 5964.24 | 6287 | 5641 | 0.897 | -0.157 | 0.13638085 | 0.31988395 | 0.0079 | 0.0075 | 0.0078 | 0.0078 | TRUE | 0.1268 |
| Ggps1 | 1260 | 1353 | 1282 | 1437 | 1038 | 1041 | 1338 | 1351 | 1287 | 1130 | 1117 | 1140 | 1227.15 | 1325 | 1129 | 0.851 | -0.232 | 0.00209845 | 0.0159888 | 0 | 0.0096 | 0.0033 | 0.0033 | TRUE | 0.0077 |
| Ggt1 | 567 | 701 | 644 | 593 | 467 | 549 | 602 | 700 | 647 | 466 | 502 | 601 | 586.43 | 650 | 523 | 0.804 | -0.315 | 0.01753705 | 0.07902697 | 0.0091 | 0.0125 | 0.011 | 0.011 | TRUE | 0.1956 |
| Ggt5 | 38 | 31 | 33 | 33 | 29 | 37 | 40 | 31 | 33 | 26 | 31 | 41 | 33.68 | 35 | 33 | 0.929 | -0.107 | 0.75946267 | 0.87317723 | 0 | 0.1047 | 0.0578 | 0.0578 | TRUE | 0.3024 |
| Ggt6 | 121 | 124 | 105 | 113 | 72 | 71 | 129 | 124 | 105 | 89 | 77 | 78 | 100.3 | 119 | 81 | 0.685 | -0.546 | 0.00763733 | 0.04291273 | 0 | 0.0397 | 0.0199 | 0.0199 | TRUE | 0.1046 |
| Ggt7 | 188 | 238 | 229 | 226 | 182 | 211 | 200 | 238 | 230 | 178 | 196 | 231 | 211.96 | 223 | 202 | 0.903 | -0.148 | 0.37376927 | 0.590099 | 0.0084 | 0.0224 | 0.0152 | 0.0152 | TRUE | 0.1755 |
| Ggta1 | 1546 | 1993 | 2173 | 1736 | 1387 | 1604 | 1642 | 1990 | 2182 | 1365 | 1492 | 1756 | 1737.89 | 1938 | 1538 | 0.793 | -0.335 | 0.01696894 | 0.07731702 | 0.0175 | 0.0088 | 0.0136 | 0.0136 | TRUE | 0.2154 |
| Gh | 37 | 35 | 52 | 56 | 43 | 53 | 39 | 35 | 52 | 44 | 46 | 58 | 45.8 | 42 | 49 | 1.168 | 0.224 | 0.46694624 | 0.66933346 | 0.0115 | 0.0788 | 0.0463 | 0.0463 | TRUE | 0.3372 |
| Ghdc | 937 | 1151 | 1212 | 1173 | 871 | 966 | 995 | 1150 | 1217 | 922 | 937 | 1057 | 1046.41 | 1121 | 972 | 0.867 | -0.206 | 0.07063153 | 0.20698633 | 0.0066 | 0.0101 | 0.0084 | 0.0084 | TRUE | 0.114 |
| Ghitm | 8729 | 9062 | 8934 | 11931 | 8282 | 8433 | 9272 | 9050 | 8971 | 9379 | 8911 | 9231 | 9135.68 | 9098 | 9174 | 1.008 | 0.012 | 0.83436685 | 0.91571799 | 3.00E-04 | 0.0073 | 0.0023 | 0.0023 | TRUE | 0.0077 |
| Ghr | 127 | 138 | 118 | 127 | 132 | 123 | 135 | 138 | 118 | 100 | 142 | 135 | 127.95 | 130 | 126 | 0.957 | -0.063 | 0.76177925 | 0.87437467 | 0.0139 | 0.0327 | 0.0231 | 0.0231 | TRUE | 0.3606 |
| Ghrh | 0 | 1 | 1 | 1 | 0 | 0 | 0 | 1 | 1 | 1 | 0 | 0 | 0.46 | 1 | 0 | 0.52 | -0.943 | 0.74346893 | NA | 0 | 7.0944 | 4.2167 | 4.2167 | TRUE | 0.2562 |
| Ghrhr | 1 | 0 | 0 | 0 | 1 | 0 | 1 | 0 | 0 | 0 | 1 | 0 | 0.36 | 0 | 0 | 0.954 | -0.067 | 0.98557668 | NA | 0 | 9.2518 | 8.0179 | 8.0179 | TRUE | 0.5204 |
| Ghrl | 0 | 0 | 1 | 1 | 0 | 0 | 0 | 0 | 1 | 1 | 0 | 0 | 0.3 | 0 | 0 | 0.954 | -0.067 | 0.98640617 | NA | 0 | 11.0483 | 9.2783 | 9.2783 | TRUE | 0.4098 |
| Ghsr | 7 | 10 | 4 | 8 | 7 | 3 | 7 | 10 | 4 | 6 | 8 | 3 | 6.42 | 7 | 6 | 0.802 | -0.319 | 0.68194722 | 0.82362649 | 0 | 0.5197 | 0.279 | 0.279 | TRUE | 0.2372 |
| Gid4 | 8097 | 10058 | 10713 | 8936 | 6984 | 8062 | 8600 | 10045 | 10757 | 7025 | 7514 | 8825 | 8794.45 | 9801 | 7788 | 0.795 | -0.332 | 0.00641208 | 0.03780613 | 0.0133 | 0.0073 | 0.0106 | 0.0106 | TRUE | 0.1656 |
| Gid8 | 3123 | 2520 | 2512 | 4215 | 2945 | 2615 | 3317 | 2517 | 2522 | 3314 | 3169 | 2863 | 2950.15 | 2785 | 3115 | 1.119 | 0.162 | 0.21537791 | 0.42718064 | 0.0156 | 0.008 | 0.0119 | 0.0119 | TRUE | 0.3381 |
| Gif | 0 | 0 | 0 | 0 | 0 | 0 | 0 | 0 | 0 | 0 | 0 | 0 | 0 | 0 | 0 | NA | NA | NA | NA | NA | NA | NA | NA | NA | NA |
| Gigyf1 | 3407 | 3910 | 3782 | 3804 | 3362 | 3492 | 3619 | 3905 | 3798 | 2990 | 3617 | 3823 | 3625.26 | 3774 | 3477 | 0.921 | -0.119 | 0.27429487 | 0.49481628 | 0.0081 | 0.0078 | 0.0083 | 0.0083 | TRUE | 0.1838 |
| Gigyf2 | 7977 | 8195 | 8563 | 9285 | 7440 | 7390 | 8473 | 8184 | 8598 | 7299 | 8005 | 8090 | 8108.2 | 8418 | 7798 | 0.926 | -0.111 | 0.12437373 | 0.30099106 | 0.0017 | 0.0073 | 0.0036 | 0.0036 | TRUE | 0.0385 |
| Gimap1 | 0 | 3 | 9 | 7 | 3 | 2 | 0 | 3 | 9 | 6 | 3 | 2 | 3.83 | 4 | 4 | 0.916 | -0.127 | 0.91687624 | NA | 0.7327 | 0.8681 | 0.8102 | 0.8102 | TRUE | 0.4715 |
| Gimap3 | 0 | 0 | 0 | 0 | 0 | 0 | 0 | 0 | 0 | 0 | 0 | 0 | 0 | 0 | 0 | NA | NA | NA | NA | NA | NA | NA | NA | NA | NA |
| Gimap4 | 0 | 2 | 0 | 0 | 0 | 1 | 0 | 2 | 0 | 0 | 0 | 1 | 0.52 | 1 | 0 | 0.522 | -0.937 | 0.77935356 | NA | 0 | 6.3996 | 6.3045 | 6.3045 | TRUE | 0.9739 |
| Gimap5 | 2 | 0 | 2 | 1 | 1 | 1 | 2 | 0 | 2 | 1 | 1 | 1 | 1.18 | 1 | 1 | 0.712 | -0.49 | 0.77859548 | NA | 0 | 2.7952 | 1.3192 | 1.3192 | TRUE | 0.4961 |
| Gimap6 | 0 | 0 | 0 | 2 | 0 | 0 | 0 | 0 | 0 | 2 | 0 | 0 | 0.26 | 0 | 1 | 2.886 | 1.529 | 0.70676491 | NA | 0 | 12.5792 | 10 | 10 | TRUE | 0.9669 |
| Gimap7 | 0 | 0 | 0 | 1 | 0 | 0 | 0 | 0 | 0 | 1 | 0 | 0 | 0.13 | 0 | 0 | 1.859 | 0.894 | 0.82650734 | NA | 0 | 25.1515 | 10 | 10 | TRUE | 0.2563 |
| Gimap8 | 11 | 9 | 4 | 21 | 17 | 12 | 12 | 9 | 4 | 17 | 18 | 13 | 12.1 | 8 | 16 | 1.951 | 0.964 | 0.10421519 | 0.26861939 | 0 | 0.2791 | 0.1607 | 0.1607 | TRUE | 0.6102 |
| Gimap9 | 111 | 129 | 148 | 172 | 111 | 111 | 118 | 129 | 149 | 135 | 119 | 122 | 128.58 | 132 | 125 | 0.953 | -0.069 | 0.71064671 | 0.84228751 | 0 | 0.0325 | 0.0172 | 0.0172 | TRUE | 0.1284 |
| Gin1 | 472 | 357 | 352 | 597 | 424 | 329 | 501 | 357 | 353 | 469 | 456 | 360 | 416.16 | 404 | 428 | 1.063 | 0.088 | 0.62845979 | 0.79094062 | 0.0281 | 0.0148 | 0.0215 | 0.0215 | TRUE | 0.512 |
| Ginm1 | 791 | 1016 | 957 | 1186 | 842 | 819 | 840 | 1015 | 961 | 932 | 906 | 897 | 925.1 | 939 | 912 | 0.971 | -0.042 | 0.68271938 | 0.82414825 | 0.0037 | 0.0105 | 0.0065 | 0.0065 | TRUE | 0.0996 |
| Gins1 | 1875 | 1954 | 1924 | 2543 | 1809 | 1861 | 1992 | 1951 | 1932 | 1999 | 1946 | 2037 | 1976.27 | 1958 | 1994 | 1.018 | 0.026 | 0.69855617 | 0.83505557 | 0 | 0.0086 | 0.0028 | 0.0028 | TRUE | 0.0052 |
| Gins2 | 4099 | 4672 | 4646 | 4709 | 3353 | 3579 | 4354 | 4666 | 4665 | 3702 | 3608 | 3918 | 4152.03 | 4562 | 3743 | 0.82 | -0.286 | 8.10E-05 | 0.00116834 | 0.0013 | 0.0077 | 0.0035 | 0.0035 | TRUE | 0.0202 |
| Gins3 | 1062 | 918 | 846 | 1421 | 984 | 849 | 1128 | 917 | 849 | 1117 | 1059 | 929 | 999.91 | 965 | 1035 | 1.074 | 0.103 | 0.45114699 | 0.65677407 | 0.0143 | 0.0102 | 0.0124 | 0.0124 | TRUE | 0.2606 |
| Gins4 | 3948 | 4238 | 4179 | 6027 | 4245 | 4305 | 4193 | 4233 | 4196 | 4738 | 4567 | 4713 | 4440 | 4207 | 4673 | 1.111 | 0.151 | 0.00958519 | 0.05097018 | 0 | 0.0076 | 0.0022 | 0.0022 | TRUE | 0.0047 |
| Gip | 2 | 2 | 2 | 5 | 5 | 1 | 2 | 2 | 2 | 4 | 5 | 1 | 2.76 | 2 | 3 | 1.708 | 0.772 | 0.50437325 | NA | 0 | 1.2024 | 0.5739 | 0.5739 | TRUE | 0.318 |
| Gipc1 | 5165 | 6087 | 6118 | 6749 | 5080 | 4650 | 5486 | 6079 | 6143 | 5306 | 5466 | 5090 | 5594.98 | 5903 | 5287 | 0.896 | -0.159 | 0.04082865 | 0.1429728 | 0.0022 | 0.0075 | 0.0042 | 0.0042 | TRUE | 0.0464 |
| Gipc2 | 104 | 156 | 165 | 160 | 118 | 125 | 110 | 156 | 166 | 126 | 127 | 137 | 136.92 | 144 | 130 | 0.9 | -0.152 | 0.46353539 | 0.66686857 | 0.016 | 0.031 | 0.0235 | 0.0235 | TRUE | 0.4241 |
| Gipc3 | 13 | 15 | 13 | 18 | 8 | 11 | 14 | 15 | 13 | 14 | 9 | 12 | 12.77 | 14 | 12 | 0.839 | -0.253 | 0.63051304 | 0.79257604 | 0 | 0.2648 | 0.1209 | 0.1209 | TRUE | 0.1839 |
| Gipr | 69 | 87 | 77 | 62 | 72 | 64 | 73 | 87 | 77 | 49 | 77 | 70 | 72.29 | 79 | 65 | 0.82 | -0.287 | 0.27526201 | 0.49588035 | 0.0185 | 0.0525 | 0.0359 | 0.0359 | TRUE | 0.5052 |
| Git1 | 9891 | 10148 | 10010 | 12138 | 9369 | 9432 | 10506 | 10135 | 10051 | 9542 | 10080 | 10325 | 10106.53 | 10231 | 9982 | 0.976 | -0.036 | 0.58125483 | 0.75771147 | 9.00E-04 | 0.0072 | 0.0029 | 0.0029 | TRUE | 0.0183 |
| Git2 | 1614 | 1582 | 1729 | 2121 | 1560 | 1584 | 1714 | 1580 | 1736 | 1667 | 1678 | 1734 | 1685.03 | 1677 | 1693 | 1.01 | 0.014 | 0.85927013 | 0.93060554 | 9.00E-04 | 0.0089 | 0.0038 | 0.0038 | TRUE | 0.0309 |
| Gja1 | 4184 | 3971 | 4319 | 3427 | 2408 | 2563 | 4444 | 3966 | 4337 | 2694 | 2591 | 2806 | 3472.88 | 4249 | 2697 | 0.635 | -0.656 | 2.09E-16 | 3.40E-14 | 0.0021 | 0.0079 | 0.0043 | 0.0043 | TRUE | 0.0414 |
| Gja10 | 0 | 0 | 0 | 2 | 0 | 1 | 0 | 0 | 0 | 2 | 0 | 1 | 0.44 | 0 | 1 | 4.782 | 2.258 | 0.50653358 | NA | 0 | 7.4188 | 6.7775 | 6.7775 | TRUE | 0.2804 |
| Gja3 | 173 | 138 | 103 | 119 | 90 | 90 | 184 | 138 | 103 | 94 | 97 | 99 | 118.98 | 142 | 97 | 0.68 | -0.556 | 0.0233929 | 0.09714756 | 0.0349 | 0.0346 | 0.0347 | 0.0347 | TRUE | 0.1942 |
| Gja4 | 80 | 94 | 70 | 99 | 88 | 91 | 85 | 94 | 70 | 78 | 95 | 100 | 86.88 | 83 | 91 | 1.088 | 0.121 | 0.59920428 | 0.77143379 | 0.0075 | 0.0448 | 0.0267 | 0.0267 | TRUE | 0.1716 |
| Gja5 | 16 | 26 | 8 | 14 | 11 | 13 | 17 | 26 | 8 | 11 | 12 | 14 | 14.68 | 17 | 12 | 0.724 | -0.466 | 0.41901051 | 0.62963355 | 0.1062 | 0.2314 | 0.1698 | 0.1698 | TRUE | 0.1404 |
| Gja6 | 6 | 8 | 7 | 6 | 2 | 7 | 6 | 8 | 7 | 5 | 2 | 8 | 5.99 | 7 | 5 | 0.677 | -0.563 | 0.47880139 | 0.67817611 | 0 | 0.5572 | 0.2822 | 0.2822 | TRUE | 0.2694 |
| Gja8 | 3 | 1 | 2 | 6 | 3 | 2 | 3 | 1 | 2 | 5 | 3 | 2 | 2.72 | 2 | 3 | 1.669 | 0.739 | 0.51821312 | NA | 0 | 1.2176 | 0.551 | 0.551 | TRUE | 0.2273 |
| Gjb1 | 83 | 70 | 80 | 62 | 60 | 47 | 88 | 70 | 80 | 49 | 65 | 51 | 67.19 | 79 | 55 | 0.689 | -0.538 | 0.03276367 | 0.12219167 | 0 | 0.0559 | 0.0304 | 0.0304 | TRUE | 0.1904 |
| Gjb2 | 28 | 32 | 41 | 61 | 51 | 39 | 30 | 32 | 41 | 48 | 55 | 43 | 41.4 | 34 | 49 | 1.413 | 0.499 | 0.10822203 | 0.27543616 | 0 | 0.0865 | 0.0445 | 0.0445 | TRUE | 0.2184 |

| | | | | | | | | | | | | | | | | | | | | | | | | | |
|---|---|---|---|---|---|---|---|---|---|---|---|---|---|---|---|---|---|---|---|---|---|---|---|---|---|
| Gjb3 | 508 | 627 | 618 | 1923 | 1455 | 1668 | 540 | 626 | 621 | 1512 | 1565 | 1826 | 1114.9 | 596 | 1634 | 2.742 | 1.455 | 6.97E-36 | 5.79E-33 | 0.0076 | 0.0099 | 0.0086 | 0.0086 | TRUE | 0.1261 |
| Gjb4 | 78 | 68 | 62 | 114 | 81 | 89 | 83 | 68 | 62 | 90 | 87 | 97 | 81.2 | 71 | 91 | 1.288 | 0.365 | 0.1062561 | 0.27210118 | 0 | 0.0475 | 0.0242 | 0.0242 | TRUE | 0.1874 |
| Gjb5 | 496 | 679 | 673 | 805 | 754 | 924 | 527 | 678 | 676 | 633 | 811 | 1011 | 722.71 | 627 | 818 | 1.304 | 0.383 | 0.03943009 | 0.13918785 | 0.0342 | 0.0115 | 0.0235 | 0.0235 | TRUE | 0.1679 |
| Gjb6 | 2 | 2 | 2 | 0 | 0 | 0 | 2 | 2 | 2 | 0 | 0 | 0 | 1.02 | 2 | 0 | 0.088 | -3.51 | 0.09851441 | NA | 0 | 3.2315 | 2.0038 | 2.0038 | TRUE | 0.0847 |
| Gjc1 | 5309 | 5104 | 4968 | 5552 | 4219 | 4286 | 5639 | 5097 | 4988 | 4365 | 4539 | 4692 | 4886.76 | 5241 | 4532 | 0.864 | -0.21 | 0.00853494 | 0.0467569 | 0.0026 | 0.0076 | 0.0044 | 0.0044 | TRUE | 0.0535 |
| Gjc2 | 1 | 0 | 1 | 3 | 4 | 3 | 1 | 0 | 1 | 2 | 4 | 3 | 2 | 1 | 3 | 4.812 | 2.267 | 0.13623186 | NA | 0 | 1.6524 | 0.7718 | 0.7718 | TRUE | 0.2572 |
| Gjc3 | 0 | 1 | 4 | 1 | 1 | 1 | 0 | 1 | 4 | 1 | 1 | 1 | 1.33 | 2 | 1 | 0.581 | -0.783 | 0.65389631 | NA | 0 | 2.4865 | 1.3954 | 1.3954 | TRUE | 0.7334 |
| Gjd2 | 0 | 0 | 0 | 1 | 1 | 1 | 0 | 0 | 0 | 1 | 1 | 1 | 0.49 | 0 | 1 | 5.224 | 2.385 | 0.38465 | NA | 0 | 6.6923 | 3.9249 | 3.9249 | TRUE | 0.07 |
| Gjd3 | 9 | 21 | 32 | 16 | 13 | 16 | 10 | 21 | 32 | 13 | 14 | 18 | 17.79 | 21 | 15 | 0.698 | -0.518 | 0.34728854 | 0.56625901 | 0.128 | 0.1921 | 0.1608 | 0.1608 | TRUE | 0.1364 |
| Gjd4 | 0 | 0 | 2 | 1 | 2 | 2 | 0 | 0 | 2 | 1 | 2 | 2 | 1.19 | 1 | 2 | 2.495 | 1.319 | 0.49531723 | NA | 0 | 2.7771 | 1.65 | 1.65 | TRUE | 0.9738 |
| Gje1 | 0 | 0 | 0 | 0 | 0 | 0 | 0 | 0 | 0 | 0 | 0 | 0 | 0 | 0 | 0 | NA | NA | NA | NA | NA | NA | NA | NA | NA | NA |
| Gk2 | 0 | 0 | 0 | 0 | 0 | 0 | 0 | 0 | 0 | 0 | 0 | 0 | 0 | 0 | 0 | NA | NA | NA | NA | NA | NA | NA | NA | NA | NA |
| Gk5 | 702 | 732 | 750 | 989 | 717 | 709 | 746 | 731 | 753 | 777 | 771 | 776 | 759.14 | 743 | 775 | 1.043 | 0.06 | 0.47443295 | 0.67497352 | 0 | 0.0112 | 0.0038 | 0.0038 | TRUE | 0.0025 |
| Gkap1 | 940 | 1324 | 1314 | 1615 | 1148 | 1225 | 998 | 1322 | 1319 | 1270 | 1235 | 1341 | 1247.64 | 1213 | 1282 | 1.056 | 0.079 | 0.54046359 | 0.72639311 | 0.0125 | 0.0095 | 0.0111 | 0.0111 | TRUE | 0.2869 |
| Gkn1 | 0 | 0 | 0 | 0 | 1 | 0 | 0 | 0 | 0 | 0 | 1 | 0 | 0.18 | 0 | 0 | 1.859 | 0.894 | 0.82650734 | NA | 0 | 18.3785 | 10 | 10 | TRUE | 0.5204 |
| Gkn2 | 0 | 0 | 0 | 0 | 1 | 0 | 0 | 0 | 0 | 0 | 1 | 0 | 0.18 | 0 | 0 | 1.859 | 0.894 | 0.82650734 | NA | 0 | 18.3785 | 10 | 10 | TRUE | 0.5204 |
| Gkn3 | 0 | 1 | 1 | 2 | 1 | 1 | 0 | 1 | 1 | 2 | 1 | 1 | 0.96 | 1 | 1 | 1.868 | 0.902 | 0.63074385 | NA | 0 | 3.4472 | 1.3898 | 1.3898 | TRUE | 0.2229 |
| Gla | 633 | 816 | 776 | 822 | 656 | 743 | 672 | 815 | 779 | 646 | 706 | 813 | 738.64 | 755 | 722 | 0.954 | -0.068 | 0.60097973 | 0.77268877 | 0.0104 | 0.0114 | 0.0108 | 0.0108 | TRUE | 0.1449 |
| Glb1 | 4827 | 5060 | 5028 | 5076 | 3717 | 3988 | 5127 | 5053 | 5049 | 3990 | 3999 | 4365 | 4597.39 | 5076 | 4118 | 0.811 | -0.302 | 1.35E-05 | 0.00025817 | 0.0011 | 0.0076 | 0.0033 | 0.0033 | TRUE | 0.0333 |
| Glb1l | 20 | 18 | 19 | 51 | 41 | 32 | 21 | 18 | 19 | 40 | 44 | 35 | 29.59 | 19 | 40 | 2.048 | 1.034 | 0.00385269 | 0.02559586 | 0 | 0.1182 | 0.0537 | 0.0537 | TRUE | 0.0733 |
| Glb1l2 | 209 | 325 | 380 | 390 | 323 | 378 | 222 | 325 | 382 | 307 | 348 | 414 | 332.67 | 310 | 356 | 1.149 | 0.2 | 0.35130671 | 0.57017785 | 0.0444 | 0.0168 | 0.0301 | 0.0301 | TRUE | 0.4938 |
| Glb1l3 | 0 | 1 | 3 | 1 | 0 | 1 | 0 | 1 | 3 | 1 | 0 | 1 | 0.98 | 1 | 1 | 0.471 | -1.086 | 0.60350353 | NA | 0 | 3.3618 | 2.0009 | 2.0009 | TRUE | 0.3174 |
| Glcci1 | 1041 | 737 | 622 | 1147 | 785 | 522 | 1106 | 736 | 625 | 902 | 845 | 571 | 797.34 | 822 | 773 | 0.94 | -0.089 | 0.70428261 | 0.83814573 | 0.0709 | 0.011 | 0.0379 | 0.0379 | TRUE | 1.0799 |
| Glce | 2733 | 2344 | 2326 | 3533 | 2379 | 2277 | 2903 | 2341 | 2336 | 2777 | 2560 | 2493 | 2568.17 | 2527 | 2610 | 1.033 | 0.047 | 0.67355099 | 0.818099 | 0.0092 | 0.0082 | 0.0087 | 0.0087 | TRUE | 0.2057 |
| Gldc | 13607 | 14917 | 15547 | 14591 | 11466 | 12883 | 14453 | 14898 | 15611 | 11470 | 12337 | 14102 | 13811.85 | 14987 | 12636 | 0.843 | -0.246 | 0.01035916 | 0.05416765 | 0.0063 | 0.0071 | 0.0066 | 0.0066 | TRUE | 0.1259 |
| Gldn | 5 | 2 | 1 | 5 | 1 | 2 | 5 | 2 | 1 | 4 | 1 | 2 | 2.58 | 3 | 2 | 0.892 | -0.165 | 0.89310833 | NA | 0 | 1.2815 | 0.706 | 0.706 | TRUE | 0.7391 |
| Gle1 | 3714 | 3457 | 3534 | 4989 | 3701 | 3299 | 3945 | 3453 | 3549 | 3922 | 3982 | 3611 | 3743.54 | 3649 | 3838 | 1.052 | 0.073 | 0.39767844 | 0.61114531 | 0.0035 | 0.0078 | 0.0052 | 0.0052 | TRUE | 0.0613 |
| Glg1 | 7705 | 9404 | 10564 | 12335 | 9013 | 9712 | 8184 | 9392 | 10607 | 9697 | 9697 | 10631 | 9701.48 | 9394 | 10008 | 1.065 | 0.091 | 0.40707662 | 0.61991044 | 0.0086 | 0.0072 | 0.0086 | 0.0086 | TRUE | 0.1558 |
| Gli1 | 455 | 585 | 557 | 305 | 279 | 350 | 483 | 584 | 559 | 240 | 300 | 383 | 424.98 | 542 | 308 | 0.566 | -0.822 | 7.23E-06 | 0.00015024 | 0.0282 | 0.0147 | 0.0217 | 0.0217 | TRUE | 0.5182 |
| Gli2 | 1065 | 1445 | 1793 | 1528 | 1138 | 1501 | 1131 | 1443 | 1800 | 1201 | 1224 | 1643 | 1407.23 | 1458 | 1356 | 0.93 | -0.105 | 0.57406514 | 0.75247257 | 0.0415 | 0.0092 | 0.0246 | 0.0246 | TRUE | 0.2059 |
| Gli3 | 475 | 395 | 460 | 1038 | 647 | 595 | 505 | 394 | 462 | 816 | 696 | 651 | 587.39 | 454 | 721 | 1.592 | 0.671 | 2.10E-06 | 5.26E-05 | 0.0127 | 0.0125 | 0.0126 | 0.0126 | TRUE | 0.1605 |
| Glipr1 | 4 | 6 | 3 | 12 | 9 | 2 | 4 | 6 | 3 | 9 | 10 | 2 | 5.76 | 4 | 7 | 1.623 | 0.699 | 0.42340679 | 0.63364379 | 0 | 0.5789 | 0.3653 | 0.3653 | TRUE | 0.8836 |
| Glipr1l1 | 0 | 0 | 0 | 0 | 0 | 0 | 0 | 0 | 0 | 0 | 0 | 0 | 0 | 0 | 0 | NA | NA | NA | NA | NA | NA | NA | NA | NA | NA |
| Glipr1l2 | 0 | 1 | 1 | 3 | 2 | 2 | 0 | 1 | 1 | 2 | 2 | 2 | 1.45 | 1 | 2 | 3.312 | 1.728 | 0.29265222 | NA | 0 | 2.2783 | 0.9676 | 0.9676 | TRUE | 0.2217 |
| Glipr2 | 742 | 743 | 737 | 1243 | 892 | 676 | 788 | 742 | 740 | 977 | 960 | 740 | 824.51 | 757 | 892 | 1.18 | 0.239 | 0.06689619 | 0.19961269 | 0.0111 | 0.0109 | 0.0111 | 0.0111 | TRUE | 0.2621 |
| Glis1 | 114 | 104 | 107 | 129 | 101 | 99 | 121 | 104 | 107 | 101 | 109 | 108 | 108.47 | 111 | 106 | 0.957 | -0.063 | 0.73839119 | 0.85992069 | 0 | 0.0373 | 0.0167 | 0.0167 | TRUE | 0.0644 |
| Glis2 | 1537 | 1052 | 1043 | 1707 | 1172 | 1116 | 1633 | 1051 | 1047 | 1342 | 1261 | 1222 | 1259.17 | 1244 | 1275 | 1.026 | 0.037 | 0.83548832 | 0.91636452 | 0.0341 | 0.0095 | 0.0215 | 0.0215 | TRUE | 0.8977 |
| Glis3 | 157 | 173 | 186 | 216 | 176 | 181 | 167 | 173 | 187 | 170 | 189 | 198 | 180.6 | 176 | 186 | 1.056 | 0.078 | 0.61781819 | 0.78392828 | 0 | 0.0251 | 0.0121 | 0.0121 | TRUE | 0.0669 |
| Glmn | 1298 | 1348 | 1295 | 1946 | 1284 | 1202 | 1379 | 1346 | 1300 | 1530 | 1382 | 1316 | 1375.39 | 1342 | 1409 | 1.051 | 0.072 | 0.43193829 | 0.6416625 | 0.0028 | 0.0093 | 0.0053 | 0.0053 | TRUE | 0.0694 |
| Glo1 | 11242 | 12510 | 12561 | 12900 | 9580 | 10164 | 11941 | 12494 | 12613 | 10141 | 10307 | 11126 | 11436.99 | 12349 | 10525 | 0.852 | -0.231 | 0.00088088 | 0.0080681 | 0.0015 | 0.0072 | 0.0034 | 0.0034 | TRUE | 0.0305 |
| Glod4 | 4720 | 6490 | 6562 | 6271 | 4641 | 5098 | 5013 | 6482 | 6589 | 4930 | 4993 | 5581 | 5597.97 | 6028 | 5168 | 0.857 | -0.222 | 0.07141085 | 0.20823529 | 0.0128 | 0.0075 | 0.0108 | 0.0108 | TRUE | 0.2642 |
| Glod5 | 57 | 45 | 45 | 105 | 97 | 110 | 61 | 45 | 45 | 83 | 104 | 120 | 76.33 | 50 | 102 | 2.034 | 1.024 | 0.0001134 | 0.00152211 | 0.0212 | 0.0501 | 0.0357 | 0.0357 | TRUE | 0.1756 |
| Glp1r | 285 | 363 | 443 | 309 | 180 | 212 | 303 | 363 | 445 | 243 | 194 | 232 | 296.45 | 370 | 223 | 0.603 | -0.73 | 6.17E-05 | 0.00093049 | 0.0225 | 0.018 | 0.0203 | 0.0203 | TRUE | 0.1859 |
| Glp2r | 3 | 2 | 1 | 5 | 2 | 2 | 3 | 2 | 1 | 4 | 2 | 2 | 2.41 | 2 | 3 | 1.365 | 0.449 | 0.70661582 | NA | 0 | 1.3739 | 0.6023 | 0.6023 | TRUE | 0.221 |
| Glra1 | 0 | 0 | 0 | 1 | 0 | 0 | 0 | 0 | 0 | 1 | 0 | 0 | 0.13 | 0 | 0 | 1.859 | 0.894 | 0.82650734 | NA | 0 | 25.1515 | 10 | 10 | TRUE | 0.2563 |
| Glra2 | 2 | 1 | 0 | 4 | 1 | 0 | 2 | 1 | 0 | 3 | 1 | 0 | 1.22 | 1 | 1 | 1.408 | 0.494 | 0.79785456 | NA | 0 | 2.6986 | 1.8503 | 1.8503 | TRUE | 0.4335 |
| Glra3 | 0 | 0 | 0 | 0 | 0 | 0 | 0 | 0 | 0 | 0 | 0 | 0 | 0 | 0 | 0 | NA | NA | NA | NA | NA | NA | NA | NA | NA | NA |
| Glra4 | 0 | 0 | 0 | 0 | 0 | 0 | 0 | 0 | 0 | 0 | 0 | 0 | 0 | 0 | 0 | NA | NA | NA | NA | NA | NA | NA | NA | NA | NA |
| Glrb | 118 | 160 | 180 | 259 | 155 | 176 | 125 | 160 | 181 | 204 | 167 | 193 | 171.48 | 155 | 188 | 1.21 | 0.274 | 0.15298919 | 0.34446045 | 0.0151 | 0.0261 | 0.0207 | 0.0207 | TRUE | 0.2944 |
| Glrp1 | 1 | 0 | 0 | 2 | 0 | 0 | 1 | 0 | 0 | 2 | 0 | 0 | 0.44 | 0 | 1 | 1.494 | 0.579 | 0.86844164 | NA | 0 | 7.5101 | 6.9157 | 6.9157 | TRUE | 0.9721 |
| Glrx | 728 | 696 | 649 | 1122 | 732 | 746 | 773 | 695 | 652 | 882 | 788 | 817 | 767.71 | 707 | 829 | 1.174 | 0.232 | 0.03240591 | 0.12140286 | 0.0041 | 0.0112 | 0.0071 | 0.0071 | TRUE | 0.0797 |
| Glrx2 | 567 | 534 | 585 | 851 | 581 | 564 | 602 | 533 | 587 | 669 | 625 | 617 | 605.74 | 574 | 637 | 1.111 | 0.152 | 0.14253756 | 0.32842594 | 0.0014 | 0.0123 | 0.0061 | 0.0061 | TRUE | 0.046 |
| Glrx3 | 1060 | 1484 | 1347 | 1478 | 1009 | 1185 | 1126 | 1482 | 1353 | 1162 | 1086 | 1297 | 1250.87 | 1320 | 1182 | 0.895 | -0.161 | 0.21552422 | 0.42736335 | 0.0122 | 0.0095 | 0.0113 | 0.0113 | TRUE | 0.1984 |
| Glrx5 | 1793 | 2073 | 1976 | 2364 | 1699 | 1566 | 1904 | 2070 | 1984 | 1858 | 1828 | 1714 | 1893.26 | 1986 | 1800 | 0.907 | -0.141 | 0.07222893 | 0.21010599 | 0.0012 | 0.0086 | 0.0039 | 0.0039 | TRUE | 0.0208 |
| Gls | 2190 | 2813 | 2935 | 3936 | 3140 | 3170 | 2326 | 2809 | 2947 | 3094 | 3378 | 3470 | 3004.21 | 2694 | 3314 | 1.23 | 0.298 | 0.00731938 | 0.0415107 | 0.0086 | 0.008 | 0.0086 | 0.0086 | TRUE | 0.1729 |
| Gls2 | 1611 | 2391 | 2599 | 2247 | 1902 | 1977 | 1711 | 2388 | 2610 | 1766 | 2046 | 2164 | 2114.29 | 2236 | 1992 | 0.891 | -0.167 | 0.3017916 | 0.52237237 | 0.0265 | 0.0085 | 0.0184 | 0.0184 | TRUE | 0.5102 |
| Glt1d1 | 889 | 904 | 1014 | 1076 | 836 | 895 | 944 | 903 | 1018 | 846 | 899 | 980 | 931.72 | 955 | 908 | 0.95 | -0.074 | 0.46836524 | 0.67075788 | 0.0036 | 0.0104 | 0.0064 | 0.0064 | TRUE | 0.056 |
| Glt25d1 | 8453 | 10738 | 10940 | 11204 | 8478 | 9008 | 8979 | 10724 | 10985 | 8808 | 9122 | 9861 | 9746.3 | 10229 | 9264 | 0.905 | -0.143 | 0.16022329 | 0.35474763 | 0.0072 | 0.0072 | 0.0074 | 0.0074 | TRUE | 0.1397 |
| Glt25d2 | 34 | 38 | 29 | 70 | 46 | 46 | 36 | 38 | 29 | 55 | 49 | 50 | 43.01 | 34 | 51 | 1.505 | 0.59 | 0.04578021 | 0.15481248 | 0 | 0.0835 | 0.0385 | 0.0385 | TRUE | 0.1289 |
| Glt28d2 | 21 | 40 | 30 | 49 | 33 | 26 | 22 | 40 | 30 | 39 | 36 | 28 | 32.48 | 31 | 34 | 1.112 | 0.153 | 0.67717137 | 0.82056964 | 0.0189 | 0.1083 | 0.0667 | 0.0667 | TRUE | 0.2763 |
| Glt6d1 | 0 | 0 | 0 | 0 | 0 | 0 | 0 | 0 | 0 | 0 | 0 | 0 | 0 | 0 | 0 | NA | NA | NA | NA | NA | NA | NA | NA | NA | NA |
| Glt8d1 | 438 | 611 | 622 | 644 | 503 | 564 | 465 | 610 | 625 | 506 | 541 | 617 | 560.81 | 567 | 555 | 0.978 | -0.032 | 0.83061655 | 0.91419987 | 0.0161 | 0.0128 | 0.0145 | 0.0145 | TRUE | 0.2855 |
| Glt8d2 | 17 | 24 | 23 | 33 | 15 | 22 | 18 | 24 | 23 | 26 | 16 | 24 | 21.88 | 22 | 22 | 1.021 | 0.03 | 0.9429943 | 0.97164083 | 0 | 0.1575 | 0.0814 | 0.0814 | TRUE | 0.2944 |
| Gltp | 1402 | 1620 | 1649 | 2194 | 1614 | 1707 | 1489 | 1618 | 1656 | 1725 | 1737 | 1869 | 1682.12 | 1588 | 1777 | 1.118 | 0.162 | 0.0556002 | 0.1765505 | 0.0019 | 0.0089 | 0.0045 | 0.0045 | TRUE | 0.0353 |
| Gltpd1 | 233 | 213 | 187 | 403 | 265 | 222 | 247 | 213 | 188 | 317 | 285 | 243 | 248.82 | 216 | 282 | 1.308 | 0.387 | 0.02411777 | 0.09911035 | 0.0143 | 0.0201 | 0.0171 | 0.0171 | TRUE | 0.1761 |
| Gltpd2 | 15 | 27 | 24 | 15 | 17 | 18 | 16 | 27 | 24 | 12 | 18 | 20 | 19.46 | 22 | 17 | 0.733 | -0.447 | 0.33513443 | 0.55539983 | 0 | 0.1762 | 0.1026 | 0.1026 | TRUE | 0.3938 |
| Gltscr1 | 392 | 400 | 368 | 614 | 458 | 468 | 416 | 399 | 370 | 483 | 493 | 512 | 445.52 | 395 | 496 | 1.255 | 0.327 | 0.00276556 | 0.01989892 | 0 | 0.0143 | 0.0063 | 0.0063 | TRUE | 0.0371 |
| Gltscr1l | 2409 | 2467 | 2657 | 3413 | 2530 | 2487 | 2559 | 2464 | 2668 | 2683 | 2722 | 2722 | 2636.35 | 2564 | 2709 | 1.057 | 0.08 | 0.24611213 | 0.46320719 | 4.00E-04 | 0.0082 | 0.003 | 0.003 | TRUE | 0.0155 |
| Gltscr2 | 8953 | 11828 | 12045 | 12618 | 9847 | 9859 | 9510 | 11813 | 12094 | 9919 | 10595 | 10792 | 10787.19 | 11139 | 10435 | 0.937 | -0.094 | 0.38723434 | 0.60106879 | 0.0088 | 0.0072 | 0.0085 | 0.0085 | TRUE | 0.2 |

| | | | | | | | | | | | | | | | | | | | | | | | | | |
|---|---|---|---|---|---|---|---|---|---|---|---|---|---|---|---|---|---|---|---|---|---|---|---|---|---|
| Glud1 | 23369 | 28274 | 27902 | 33771 | 23740 | 25341 | 24822 | 28237 | 28017 | 26548 | 25543 | 27740 | 26817.78 | 27025 | 26610 | 0.985 | -0.022 | 0.78242284 | 0.88741357 | 0.0027 | 0.007 | 0.0047 | 0.0047 | TRUE | 0.0622 |
| Glul | 7911 | 6730 | 6560 | 8903 | 6107 | 5678 | 8403 | 6721 | 6587 | 6999 | 6571 | 6215 | 6916.03 | 7237 | 6595 | 0.911 | -0.134 | 0.24456123 | 0.4611329 | 0.0113 | 0.0074 | 0.0094 | 0.0094 | TRUE | 0.2423 |
| Glyat | 0 | 0 | 0 | 0 | 0 | 0 | 0 | 0 | 0 | 0 | 0 | 0 | 0 | 0 | 0 | NA | NA | NA | NA | NA | NA | NA | NA | NA | NA |
| Glyatl3 | 0 | 0 | 0 | 0 | 0 | 1 | 0 | 0 | 0 | 0 | 0 | 1 | 0.18 | 0 | 0 | 1.859 | 0.894 | 0.82650734 | NA | 0 | 18.0643 | 10 | 10 | TRUE | 0.5383 |
| Glycam1 | 0 | 0 | 0 | 0 | 0 | 0 | 0 | 0 | 0 | 0 | 0 | 0 | 0 | 0 | 0 | NA | NA | NA | NA | NA | NA | NA | NA | NA | NA |
| Glyctk | 434 | 499 | 499 | 523 | 390 | 367 | 461 | 498 | 501 | 411 | 420 | 402 | 448.81 | 487 | 411 | 0.844 | -0.245 | 0.02047935 | 0.08834803 | 0 | 0.0142 | 0.0058 | 0.0058 | TRUE | 0.0247 |
| Glyr1 | 6057 | 5506 | 5612 | 9635 | 6732 | 5694 | 6434 | 5499 | 5635 | 7574 | 7243 | 6233 | 6436.34 | 5856 | 7017 | 1.198 | 0.261 | 0.01457047 | 0.06930203 | 0.0084 | 0.0074 | 0.0081 | 0.0081 | TRUE | 0.1164 |
| Gm10007 | 0 | 0 | 0 | 0 | 0 | 0 | 0 | 0 | 0 | 0 | 0 | 0 | 0 | 0 | 0 | NA | NA | NA | NA | NA | NA | NA | NA | NA | NA |
| Gm10012 | 0 | 1 | 3 | 0 | 2 | 1 | 0 | 1 | 3 | 0 | 2 | 1 | 1.21 | 1 | 1 | 0.778 | -0.362 | 0.85374568 | NA | 0 | 2.7305 | 1.9255 | 1.9255 | TRUE | 0.3946 |
| Gm10024 | 0 | 0 | 0 | 0 | 0 | 0 | 0 | 0 | 0 | 0 | 0 | 0 | 0 | 0 | 0 | NA | NA | NA | NA | NA | NA | NA | NA | NA | NA |
| Gm10033 | 323 | 228 | 304 | 360 | 263 | 219 | 343 | 228 | 305 | 283 | 283 | 240 | 280.29 | 292 | 269 | 0.921 | -0.119 | 0.51600347 | 0.70818019 | 0.0223 | 0.0187 | 0.0205 | 0.0205 | TRUE | 0.42 |
| Gm10046 | 2 | 11 | 12 | 12 | 9 | 7 | 2 | 11 | 12 | 9 | 10 | 8 | 8.66 | 8 | 9 | 1.062 | 0.086 | 0.90708284 | 0.95354464 | 0.1332 | 0.3875 | 0.2791 | 0.2791 | TRUE | 1.2327 |
| Gm10052 | 0 | 0 | 0 | 0 | 0 | 0 | 0 | 0 | 0 | 0 | 0 | 0 | 0 | 0 | 0 | NA | NA | NA | NA | NA | NA | NA | NA | NA | NA |
| Gm10057 | 0 | 0 | 0 | 0 | 0 | 0 | 0 | 0 | 0 | 0 | 0 | 0 | 0 | 0 | 0 | NA | NA | NA | NA | NA | NA | NA | NA | NA | NA |
| Gm10058 | 0 | 0 | 0 | 0 | 0 | 0 | 0 | 0 | 0 | 0 | 0 | 0 | 0 | 0 | 0 | NA | NA | NA | NA | NA | NA | NA | NA | NA | NA |
| Gm10069 | 85 | 123 | 103 | 141 | 106 | 123 | 90 | 123 | 103 | 111 | 114 | 135 | 112.68 | 105 | 120 | 1.132 | 0.179 | 0.38999804 | 0.6038136 | 0.0081 | 0.0361 | 0.0224 | 0.0224 | TRUE | 0.2045 |
| Gm10081 | 0 | 0 | 0 | 0 | 0 | 0 | 0 | 0 | 0 | 0 | 0 | 0 | 0 | 0 | 0 | NA | NA | NA | NA | NA | NA | NA | NA | NA | NA |
| Gm10094 | 0 | 0 | 0 | 0 | 0 | 0 | 0 | 0 | 0 | 0 | 0 | 0 | 0 | 0 | 0 | NA | NA | NA | NA | NA | NA | NA | NA | NA | NA |
| Gm10096 | 0 | 0 | 0 | 0 | 0 | 0 | 0 | 0 | 0 | 0 | 0 | 0 | 0 | 0 | 0 | NA | NA | NA | NA | NA | NA | NA | NA | NA | NA |
| Gm101 | 7 | 7 | 6 | 6 | 8 | 10 | 7 | 7 | 6 | 5 | 9 | 11 | 7.45 | 7 | 8 | 1.169 | 0.225 | 0.74963712 | 0.8673408 | 0 | 0.4489 | 0.221 | 0.221 | TRUE | 0.437 |
| Gm10100 | 0 | 0 | 0 | 0 | 0 | 0 | 0 | 0 | 0 | 0 | 0 | 0 | 0 | 0 | 0 | NA | NA | NA | NA | NA | NA | NA | NA | NA | NA |
| Gm10104 | 0 | 0 | 0 | 0 | 0 | 0 | 0 | 0 | 0 | 0 | 0 | 0 | 0 | 0 | 0 | NA | NA | NA | NA | NA | NA | NA | NA | NA | NA |
| Gm10125 | 1 | 0 | 0 | 2 | 1 | 0 | 1 | 0 | 0 | 2 | 1 | 0 | 0.62 | 0 | 1 | 2.462 | 1.3 | 0.62119797 | NA | 0 | 5.3343 | 3.4416 | 3.4416 | TRUE | 0.4614 |
| Gm10142 | 0 | 0 | 0 | 0 | 0 | 0 | 0 | 0 | 0 | 0 | 0 | 0 | 0 | 0 | 0 | NA | NA | NA | NA | NA | NA | NA | NA | NA | NA |
| Gm10147 | 0 | 0 | 0 | 0 | 0 | 0 | 0 | 0 | 0 | 0 | 0 | 0 | 0 | 0 | 0 | NA | NA | NA | NA | NA | NA | NA | NA | NA | NA |
| Gm10190 | 9 | 1 | 4 | 6 | 5 | 0 | 10 | 1 | 4 | 5 | 5 | 0 | 4.11 | 5 | 3 | 0.704 | -0.507 | 0.68182971 | NA | 0.9178 | 0.8081 | 0.8531 | 0.8531 | TRUE | 0.3267 |
| Gm10220 | 1 | 3 | 0 | 1 | 3 | 0 | 1 | 3 | 0 | 1 | 3 | 0 | 1.35 | 1 | 1 | 0.977 | -0.033 | 0.98609604 | NA | 1.0346 | 2.4556 | 1.869 | 1.869 | TRUE | 0.5472 |
| Gm10228 | 0 | 0 | 0 | 0 | 0 | 0 | 0 | 0 | 0 | 0 | 0 | 0 | 0 | 0 | 0 | NA | NA | NA | NA | NA | NA | NA | NA | NA | NA |
| Gm10229 | 0 | 0 | 0 | 0 | 0 | 0 | 0 | 0 | 0 | 0 | 0 | 0 | 0 | 0 | 0 | NA | NA | NA | NA | NA | NA | NA | NA | NA | NA |
| Gm10230 | 0 | 0 | 0 | 0 | 0 | 0 | 0 | 0 | 0 | 0 | 0 | 0 | 0 | 0 | 0 | NA | NA | NA | NA | NA | NA | NA | NA | NA | NA |
| Gm10248 | 0 | 0 | 0 | 0 | 0 | 0 | 0 | 0 | 0 | 0 | 0 | 0 | 0 | 0 | 0 | NA | NA | NA | NA | NA | NA | NA | NA | NA | NA |
| Gm10256 | 0 | 0 | 0 | 0 | 0 | 0 | 0 | 0 | 0 | 0 | 0 | 0 | 0 | 0 | 0 | NA | NA | NA | NA | NA | NA | NA | NA | NA | NA |
| Gm10267 | 0 | 0 | 0 | 0 | 0 | 0 | 0 | 0 | 0 | 0 | 0 | 0 | 0 | 0 | 0 | NA | NA | NA | NA | NA | NA | NA | NA | NA | NA |
| Gm10272 | 0 | 0 | 0 | 0 | 0 | 0 | 0 | 0 | 0 | 0 | 0 | 0 | 0 | 0 | 0 | NA | NA | NA | NA | NA | NA | NA | NA | NA | NA |
| Gm10280 | 0 | 0 | 0 | 0 | 0 | 0 | 0 | 0 | 0 | 0 | 0 | 0 | 0 | 0 | 0 | NA | NA | NA | NA | NA | NA | NA | NA | NA | NA |
| Gm10318 | 0 | 0 | 0 | 0 | 0 | 0 | 0 | 0 | 0 | 0 | 0 | 0 | 0 | 0 | 0 | NA | NA | NA | NA | NA | NA | NA | NA | NA | NA |
| Gm10319 | 0 | 0 | 0 | 0 | 0 | 0 | 0 | 0 | 0 | 0 | 0 | 0 | 0 | 0 | 0 | NA | NA | NA | NA | NA | NA | NA | NA | NA | NA |
| Gm10324 | 295 | 210 | 178 | 487 | 349 | 257 | 313 | 210 | 179 | 383 | 376 | 281 | 290.25 | 234 | 347 | 1.484 | 0.569 | 0.01313088 | 0.0641972 | 0.0534 | 0.0183 | 0.0344 | 0.0344 | TRUE | 0.9671 |
| Gm10334 | 0 | 0 | 0 | 0 | 0 | 0 | 0 | 0 | 0 | 0 | 0 | 0 | 0 | 0 | 0 | NA | NA | NA | NA | NA | NA | NA | NA | NA | NA |
| Gm10336 | 1371 | 1127 | 1276 | 1703 | 1036 | 1163 | 1456 | 1126 | 1281 | 1339 | 1115 | 1273 | 1264.93 | 1288 | 1242 | 0.965 | -0.051 | 0.68813486 | 0.82833091 | 0.0119 | 0.0095 | 0.0108 | 0.0108 | TRUE | 0.1571 |
| Gm10352 | 0 | 0 | 0 | 0 | 0 | 0 | 0 | 0 | 0 | 0 | 0 | 0 | 0 | 0 | 0 | NA | NA | NA | NA | NA | NA | NA | NA | NA | NA |
| Gm10364 | 4 | 1 | 1 | 2 | 1 | 2 | 4 | 1 | 1 | 2 | 1 | 2 | 1.85 | 2 | 2 | 0.778 | -0.362 | 0.7947159 | NA | 0 | 1.7894 | 0.8433 | 0.8433 | TRUE | 0.7315 |
| Gm10373 | 1 | 0 | 0 | 4 | 1 | 1 | 1 | 0 | 0 | 3 | 1 | 1 | 1.06 | 0 | 2 | 4.937 | 2.304 | 0.25605048 | NA | 0 | 3.1065 | 1.6934 | 1.6934 | TRUE | 0.4788 |
| Gm10375 | 0 | 0 | 0 | 0 | 0 | 0 | 0 | 0 | 0 | 0 | 0 | 0 | 0 | 0 | 0 | NA | NA | NA | NA | NA | NA | NA | NA | NA | NA |
| Gm10377 | 0 | 0 | 0 | 0 | 0 | 0 | 0 | 0 | 0 | 0 | 0 | 0 | 0 | 0 | 0 | NA | NA | NA | NA | NA | NA | NA | NA | NA | NA |
| Gm10389 | 0 | 0 | 0 | 0 | 0 | 0 | 0 | 0 | 0 | 0 | 0 | 0 | 0 | 0 | 0 | NA | NA | NA | NA | NA | NA | NA | NA | NA | NA |
| Gm10390 | 0 | 1 | 1 | 1 | 2 | 0 | 0 | 1 | 1 | 1 | 2 | 0 | 0.82 | 1 | 1 | 1.448 | 0.535 | 0.80575974 | NA | 0 | 4.0076 | 2.145 | 2.145 | TRUE | 0.2939 |
| Gm10400 | 0 | 0 | 0 | 0 | 0 | 0 | 0 | 0 | 0 | 0 | 0 | 0 | 0 | 0 | 0 | NA | NA | NA | NA | NA | NA | NA | NA | NA | NA |
| Gm10406 | 47 | 56 | 76 | 87 | 54 | 54 | 50 | 56 | 76 | 68 | 58 | 59 | 61.29 | 61 | 62 | 1.021 | 0.03 | 0.91177585 | 0.95644617 | 0.0129 | 0.0607 | 0.0375 | 0.0375 | TRUE | 0.4378 |
| Gm10408 | 0 | 0 | 0 | 0 | 0 | 0 | 0 | 0 | 0 | 0 | 0 | 0 | 0 | 0 | 0 | NA | NA | NA | NA | NA | NA | NA | NA | NA | NA |
| Gm10409 | 0 | 0 | 0 | 0 | 0 | 0 | 0 | 0 | 0 | 0 | 0 | 0 | 0 | 0 | 0 | NA | NA | NA | NA | NA | NA | NA | NA | NA | NA |
| Gm10413 | 0 | 0 | 0 | 0 | 0 | 0 | 0 | 0 | 0 | 0 | 0 | 0 | 0 | 0 | 0 | NA | NA | NA | NA | NA | NA | NA | NA | NA | NA |
| Gm10415 | 0 | 0 | 0 | 0 | 0 | 1 | 0 | 0 | 0 | 0 | 0 | 1 | 0.18 | 0 | 0 | 1.859 | 0.894 | 0.82650734 | NA | 0 | 18.0643 | 10 | 10 | TRUE | 0.5383 |
| Gm10416 | 108 | 71 | 78 | 130 | 86 | 56 | 115 | 71 | 78 | 102 | 93 | 61 | 86.66 | 88 | 85 | 0.975 | -0.036 | 0.90057134 | 0.95174291 | 0.0552 | 0.0449 | 0.0499 | 0.0499 | TRUE | 0.6614 |
| Gm10421 | 0 | 0 | 0 | 0 | 0 | 0 | 0 | 0 | 0 | 0 | 0 | 0 | 0 | 0 | 0 | NA | NA | NA | NA | NA | NA | NA | NA | NA | NA |
| Gm10432 | 24 | 19 | 6 | 24 | 15 | 16 | 25 | 19 | 6 | 19 | 16 | 18 | 17.17 | 17 | 18 | 1.045 | 0.063 | 0.91214456 | 0.95666604 | 0.1508 | 0.1988 | 0.1769 | 0.1769 | TRUE | 0.5263 |
| Gm10433 | 9 | 8 | 4 | 26 | 12 | 9 | 10 | 8 | 4 | 20 | 13 | 10 | 10.79 | 7 | 14 | 2.031 | 1.023 | 0.11175111 | 0.28113131 | 0 | 0.3121 | 0.193 | 0.193 | TRUE | 0.5421 |
| Gm10436 | 1 | 0 | 0 | 2 | 1 | 0 | 1 | 0 | 0 | 2 | 1 | 0 | 0.62 | 0 | 1 | 2.462 | 1.3 | 0.62119797 | NA | 0 | 5.3343 | 3.4416 | 3.4416 | TRUE | 0.4614 |
| Gm10439 | 13 | 5 | 7 | 77 | 54 | 51 | 14 | 5 | 7 | 61 | 58 | 56 | 33.38 | 9 | 58 | 6.812 | 2.768 | 1.63E-10 | 1.16E-08 | 0 | 0.1056 | 0.0669 | 0.0669 | TRUE | 0.8239 |
| Gm10440 | 0 | 0 | 0 | 0 | 0 | 0 | 0 | 0 | 0 | 0 | 0 | 0 | 0 | 0 | 0 | NA | NA | NA | NA | NA | NA | NA | NA | NA | NA |
| Gm10445 | 0 | 0 | 0 | 0 | 0 | 0 | 0 | 0 | 0 | 0 | 0 | 0 | 0 | 0 | 0 | NA | NA | NA | NA | NA | NA | NA | NA | NA | NA |
| Gm1045 | 14 | 17 | 18 | 32 | 10 | 13 | 15 | 17 | 18 | 25 | 11 | 14 | 16.68 | 17 | 17 | 1.02 | 0.029 | 0.95536534 | 0.97727168 | 0.0527 | 0.2044 | 0.1291 | 0.1291 | TRUE | 1.1437 |
| Gm10451 | 195 | 191 | 182 | 368 | 266 | 239 | 207 | 191 | 183 | 289 | 286 | 262 | 236.29 | 194 | 279 | 1.444 | 0.53 | 0.00014549 | 0.00186037 | 0 | 0.0208 | 0.0096 | 0.0096 | TRUE | 0.0401 |
| Gm10466 | 0 | 0 | 0 | 0 | 0 | 0 | 0 | 0 | 0 | 0 | 0 | 0 | 0 | 0 | 0 | NA | NA | NA | NA | NA | NA | NA | NA | NA | NA |
| Gm10471 | 1 | 0 | 0 | 2 | 1 | 2 | 1 | 0 | 0 | 2 | 1 | 2 | 0.98 | 0 | 2 | 4.402 | 2.138 | 0.29192655 | NA | 0 | 3.3574 | 1.6617 | 1.6617 | TRUE | 0.4614 |
| Gm10474 | 5 | 9 | 5 | 5 | 10 | 8 | 5 | 9 | 5 | 4 | 11 | 9 | 7.13 | 6 | 8 | 1.192 | 0.253 | 0.73685359 | 0.85914503 | 0 | 0.4691 | 0.2653 | 0.2653 | TRUE | 0.7202 |
| Gm10485 | 0 | 0 | 0 | 0 | 0 | 0 | 0 | 0 | 0 | 0 | 0 | 0 | 0 | 0 | 0 | NA | NA | NA | NA | NA | NA | NA | NA | NA | NA |
| Gm10486 | 0 | 0 | 0 | 0 | 0 | 0 | 0 | 0 | 0 | 0 | 0 | 0 | 0 | 0 | 0 | NA | NA | NA | NA | NA | NA | NA | NA | NA | NA |
| Gm10487 | 0 | 0 | 0 | 0 | 0 | 0 | 0 | 0 | 0 | 0 | 0 | 0 | 0 | 0 | 0 | NA | NA | NA | NA | NA | NA | NA | NA | NA | NA |

| | | | | | | | | | | | | | | | | | | | | | | | | | |
|---|---|---|---|---|---|---|---|---|---|---|---|---|---|---|---|---|---|---|---|---|---|---|---|---|---|
| Gm10488 | 0 | 0 | 0 | 0 | 0 | 0 | 0 | 0 | 0 | 0 | 0 | 0 | 0 | 0 | 0 | NA | NA | NA | NA | NA | NA | NA | NA | NA | NA |
| Gm10494 | 0 | 0 | 0 | 0 | 1 | 0 | 0 | 0 | 0 | 0 | 1 | 0 | 0.18 | 0 | 0 | 1.859 | 0.894 | 0.82650734 | NA | 0 | 18.3785 | 10 | 10 | TRUE | 0.5204 |
| Gm10509 | 125 | 151 | 151 | 197 | 138 | 118 | 133 | 151 | 152 | 155 | 148 | 129 | 144.62 | 145 | 144 | 0.996 | -0.006 | 0.97247016 | 0.98541137 | 0 | 0.0297 | 0.015 | 0.015 | TRUE | 0.083 |
| Gm10510 | 0 | 0 | 0 | 0 | 0 | 0 | 0 | 0 | 0 | 0 | 0 | 0 | 0 | 0 | 0 | NA | NA | NA | NA | NA | NA | NA | NA | NA | NA |
| Gm10512 | 0 | 0 | 0 | 0 | 1 | 0 | 0 | 0 | 0 | 0 | 1 | 0 | 0.18 | 0 | 0 | 1.859 | 0.894 | 0.82650734 | NA | 0 | 18.3785 | 10 | 10 | TRUE | 0.5204 |
| Gm10516 | 0 | 3 | 5 | 4 | 4 | 4 | 0 | 3 | 5 | 3 | 4 | 4 | 3.31 | 3 | 4 | 1.456 | 0.542 | 0.63769264 | NA | 0 | 1.003 | 0.6406 | 0.6406 | TRUE | 0.4251 |
| Gm10532 | 22 | 19 | 22 | 77 | 48 | 37 | 23 | 19 | 22 | 61 | 52 | 41 | 36.19 | 21 | 51 | 2.384 | 1.254 | 0.00030847 | 0.00349005 | 0 | 0.0979 | 0.0536 | 0.0536 | TRUE | 0.1314 |
| Gm10536 | 0 | 0 | 1 | 1 | 0 | 0 | 0 | 0 | 0 | 1 | 1 | 0 | 0.3 | 0 | 0 | 0.954 | -0.067 | 0.98640617 | NA | 0 | 11.0483 | 9.2783 | 9.2783 | TRUE | 0.4098 |
| Gm10538 | 0 | 0 | 0 | 0 | 0 | 0 | 0 | 0 | 0 | 0 | 0 | 0 | 0 | 0 | 0 | NA | NA | NA | NA | NA | NA | NA | NA | NA | NA |
| Gm10548 | 1 | 1 | 1 | 2 | 2 | 0 | 1 | 1 | 1 | 2 | 2 | 0 | 1.13 | 1 | 1 | 1.231 | 0.299 | 0.86401656 | NA | 0 | 2.9184 | 1.3116 | 1.3116 | TRUE | 0.3778 |
| Gm10549 | 0 | 0 | 0 | 1 | 0 | 0 | 0 | 0 | 0 | 1 | 0 | 0 | 0.13 | 0 | 0 | 1.859 | 0.894 | 0.82650734 | NA | 0 | 25.1515 | 10 | 10 | TRUE | 0.2563 |
| Gm10556 | 1 | 0 | 1 | 2 | 1 | 0 | 1 | 0 | 1 | 2 | 1 | 0 | 0.79 | 1 | 1 | 1.325 | 0.406 | 0.85251519 | NA | 0 | 4.1997 | 2.1499 | 2.1499 | TRUE | 0.2789 |
| Gm10560 | 33 | 46 | 41 | 47 | 28 | 34 | 35 | 46 | 41 | 37 | 30 | 37 | 37.74 | 41 | 35 | 0.855 | -0.226 | 0.4730125 | 0.67398619 | 0 | 0.0942 | 0.0451 | 0.0451 | TRUE | 0.1083 |
| Gm10578 | 0 | 0 | 0 | 0 | 0 | 0 | 0 | 0 | 0 | 0 | 0 | 0 | 0 | 0 | 0 | NA | NA | NA | NA | NA | NA | NA | NA | NA | NA |
| Gm10584 | 0 | 2 | 0 | 1 | 0 | 2 | 0 | 2 | 0 | 1 | 0 | 2 | 0.83 | 1 | 1 | 1.469 | 0.555 | 0.82210875 | NA | 0 | 3.9818 | 3.125 | 3.125 | TRUE | 0.7031 |
| Gm10591 | 0 | 0 | 0 | 0 | 0 | 0 | 0 | 0 | 0 | 0 | 0 | 0 | 0 | 0 | 0 | NA | NA | NA | NA | NA | NA | NA | NA | NA | NA |
| Gm10619 | 0 | 0 | 0 | 0 | 0 | 0 | 0 | 0 | 0 | 0 | 0 | 0 | 0 | 0 | 0 | NA | NA | NA | NA | NA | NA | NA | NA | NA | NA |
| Gm10635 | 3 | 2 | 0 | 4 | 2 | 1 | 3 | 2 | 0 | 3 | 2 | 1 | 1.93 | 2 | 2 | 1.26 | 0.334 | 0.81686255 | NA | 0 | 1.7146 | 0.9747 | 0.9747 | TRUE | 0.4515 |
| Gm10636 | 1 | 0 | 0 | 0 | 0 | 0 | 1 | 0 | 0 | 0 | 0 | 0 | 0.18 | 0 | 0 | 0.49 | -1.029 | 0.80087 | NA | 0 | 18.6165 | 10 | 10 | TRUE | 0.4614 |
| Gm10637 | 0 | 0 | 0 | 0 | 0 | 0 | 0 | 0 | 0 | 0 | 0 | 0 | 0 | 0 | 0 | NA | NA | NA | NA | NA | NA | NA | NA | NA | NA |
| Gm10638 | 6 | 7 | 3 | 11 | 7 | 10 | 6 | 7 | 3 | 9 | 8 | 11 | 7.25 | 5 | 9 | 1.655 | 0.727 | 0.31089523 | 0.53148055 | 0 | 0.4613 | 0.2231 | 0.2231 | TRUE | 0.3402 |
| Gm10639 | 0 | 1 | 0 | 2 | 1 | 0 | 0 | 1 | 0 | 2 | 1 | 0 | 0.61 | 0 | 1 | 2.461 | 1.299 | 0.62286289 | NA | 0 | 5.4271 | 3.4871 | 3.4871 | TRUE | 0.405 |
| Gm10640 | 5 | 0 | 3 | 4 | 1 | 6 | 5 | 0 | 3 | 3 | 1 | 7 | 3.19 | 3 | 4 | 1.298 | 0.376 | 0.77111577 | NA | 0.6649 | 1.0412 | 0.8839 | 0.8839 | TRUE | 0.3422 |
| Gm10649 | 0 | 0 | 0 | 0 | 0 | 0 | 0 | 0 | 0 | 0 | 0 | 0 | 0 | 0 | 0 | NA | NA | NA | NA | NA | NA | NA | NA | NA | NA |
| Gm10653 | 3 | 3 | 7 | 4 | 3 | 2 | 3 | 3 | 7 | 3 | 3 | 2 | 3.63 | 4 | 3 | 0.65 | -0.621 | 0.53082755 | NA | 0 | 0.9148 | 0.4206 | 0.4206 | TRUE | 0.4995 |
| Gm10658 | 47 | 81 | 76 | 74 | 43 | 91 | 50 | 81 | 76 | 58 | 46 | 100 | 68.53 | 69 | 68 | 0.981 | -0.027 | 0.93846959 | 0.9691313 | 0.0944 | 0.055 | 0.0738 | 0.0738 | TRUE | 1.2575 |
| Gm10662 | 0 | 0 | 0 | 0 | 0 | 0 | 0 | 0 | 0 | 0 | 0 | 0 | 0 | 0 | 0 | NA | NA | NA | NA | NA | NA | NA | NA | NA | NA |
| Gm10664 | 1314 | 1554 | 1546 | 921 | 731 | 946 | 1396 | 1552 | 1552 | 724 | 787 | 1036 | 1174.35 | 1500 | 849 | 0.565 | -0.823 | 1.37E-08 | 6.34E-07 | 0.0193 | 0.0097 | 0.0142 | 0.0142 | TRUE | 0.4387 |
| Gm10665 | 0 | 0 | 0 | 0 | 0 | 0 | 0 | 0 | 0 | 0 | 0 | 0 | 0 | 0 | 0 | NA | NA | NA | NA | NA | NA | NA | NA | NA | NA |
| Gm10666 | 0 | 0 | 0 | 0 | 0 | 0 | 0 | 0 | 0 | 0 | 0 | 0 | 0 | 0 | 0 | NA | NA | NA | NA | NA | NA | NA | NA | NA | NA |
| Gm10670 | 0 | 0 | 0 | 0 | 0 | 0 | 0 | 0 | 0 | 0 | 0 | 0 | 0 | 0 | 0 | NA | NA | NA | NA | NA | NA | NA | NA | NA | NA |
| Gm10677 | 0 | 0 | 2 | 1 | 0 | 1 | 0 | 0 | 2 | 1 | 0 | 1 | 0.65 | 1 | 1 | 0.938 | -0.093 | 0.9721749 | NA | 0 | 5.0896 | 3.5394 | 3.5394 | TRUE | 0.9739 |
| Gm10681 | 0 | 0 | 0 | 0 | 0 | 0 | 0 | 0 | 0 | 0 | 0 | 0 | 0 | 0 | 0 | NA | NA | NA | NA | NA | NA | NA | NA | NA | NA |
| Gm10684 | 0 | 0 | 0 | 0 | 0 | 0 | 0 | 0 | 0 | 0 | 0 | 0 | 0 | 0 | 0 | NA | NA | NA | NA | NA | NA | NA | NA | NA | NA |
| Gm10696 | 0 | 1 | 0 | 7 | 14 | 1 | 0 | 1 | 0 | 6 | 15 | 1 | 3.78 | 0 | 7 | 19.687 | 4.299 | 0.00998515 | NA | 1.0337 | 0.8792 | 0.9386 | 0.9386 | TRUE | 0.2139 |
| Gm10697 | 0 | 0 | 0 | 0 | 0 | 0 | 0 | 0 | 0 | 0 | 0 | 0 | 0 | 0 | 0 | NA | NA | NA | NA | NA | NA | NA | NA | NA | NA |
| Gm10714 | 3 | 5 | 9 | 3 | 4 | 7 | 3 | 5 | 9 | 2 | 4 | 8 | 5.26 | 6 | 5 | 0.813 | -0.298 | 0.73746965 | 0.85945481 | 0.1179 | 0.6336 | 0.3762 | 0.3762 | TRUE | 0.4681 |
| Gm10731 | 0 | 0 | 0 | 0 | 0 | 0 | 0 | 0 | 0 | 0 | 0 | 0 | 0 | 0 | 0 | NA | NA | NA | NA | NA | NA | NA | NA | NA | NA |
| Gm10745 | 0 | 0 | 0 | 0 | 1 | 0 | 0 | 0 | 0 | 0 | 1 | 0 | 0.18 | 0 | 0 | 1.859 | 0.894 | 0.82650734 | NA | 0 | 18.3785 | 10 | 10 | TRUE | 0.5204 |
| Gm10754 | 0 | 1 | 1 | 1 | 1 | 2 | 0 | 1 | 1 | 1 | 1 | 2 | 1.01 | 1 | 1 | 1.97 | 0.978 | 0.60105973 | NA | 0 | 3.2719 | 1.3982 | 1.3982 | TRUE | 0.2229 |
| Gm10767 | 9 | 9 | 0 | 6 | 12 | 9 | 10 | 9 | 0 | 5 | 13 | 10 | 7.67 | 6 | 9 | 1.473 | 0.559 | 0.56864585 | 0.74770173 | 0.7673 | 0.4364 | 0.5554 | 0.5554 | TRUE | 1.0591 |
| Gm10768 | 81 | 129 | 120 | 95 | 90 | 103 | 86 | 129 | 120 | 75 | 97 | 113 | 103.27 | 112 | 95 | 0.843 | -0.247 | 0.32845574 | 0.54950119 | 0.0337 | 0.0388 | 0.0363 | 0.0363 | TRUE | 0.3982 |
| Gm10778 | 15 | 21 | 16 | 14 | 14 | 10 | 16 | 21 | 16 | 11 | 15 | 11 | 15 | 18 | 12 | 0.695 | -0.525 | 0.28484588 | 0.50626428 | 0 | 0.2266 | 0.1049 | 0.1049 | TRUE | 0.1384 |
| Gm1078 | 0 | 2 | 1 | 1 | 1 | 0 | 0 | 2 | 1 | 1 | 1 | 0 | 0.81 | 1 | 1 | 0.625 | -0.678 | 0.7553135 | NA | 0 | 4.0711 | 2.1302 | 2.1302 | TRUE | 0.1304 |
| Gm10782 | 0 | 0 | 2 | 2 | 0 | 0 | 0 | 0 | 2 | 2 | 0 | 0 | 0.6 | 1 | 1 | 0.818 | -0.289 | 0.92823606 | NA | 0 | 5.5276 | 5.7664 | 5.7664 | TRUE | 0.9739 |
| Gm10785 | 3 | 0 | 2 | 2 | 0 | 1 | 3 | 0 | 2 | 2 | 0 | 1 | 1.31 | 2 | 1 | 0.532 | -0.912 | 0.62342598 | NA | 0 | 2.5212 | 1.6559 | 1.6559 | TRUE | 0.2588 |
| Gm10787 | 0 | 0 | 0 | 1 | 0 | 3 | 0 | 0 | 0 | 1 | 0 | 3 | 0.68 | 0 | 1 | 7.177 | 2.843 | 0.34581154 | NA | 6.5474 | 4.8635 | 5.1913 | 5.1913 | TRUE | 0.362 |
| Gm10789 | 21 | 33 | 46 | 31 | 29 | 26 | 22 | 33 | 46 | 24 | 31 | 28 | 30.91 | 34 | 28 | 0.823 | -0.28 | 0.475703 | 0.67577395 | 0.0417 | 0.1135 | 0.0787 | 0.0787 | TRUE | 0.2088 |
| Gm10790 | 0 | 0 | 0 | 0 | 0 | 0 | 0 | 0 | 0 | 0 | 0 | 0 | 0 | 0 | 0 | NA | NA | NA | NA | NA | NA | NA | NA | NA | NA |
| Gm10791 | 0 | 0 | 0 | 1 | 0 | 0 | 0 | 0 | 0 | 1 | 0 | 0 | 0.13 | 0 | 0 | 1.859 | 0.894 | 0.82650734 | NA | 0 | 25.1515 | 10 | 10 | TRUE | 0.2563 |
| Gm10804 | 0 | 0 | 0 | 1 | 0 | 0 | 0 | 0 | 0 | 1 | 0 | 0 | 0.13 | 0 | 0 | 1.859 | 0.894 | 0.82650734 | NA | 0 | 25.1515 | 10 | 10 | TRUE | 0.2563 |
| Gm10814 | 0 | 1 | 3 | 1 | 1 | 1 | 0 | 1 | 3 | 1 | 1 | 1 | 1.16 | 1 | 1 | 0.726 | -0.462 | 0.79558366 | NA | 0 | 2.8438 | 1.4137 | 1.4137 | TRUE | 0.4177 |
| Gm10818 | 2 | 1 | 2 | 1 | 0 | 1 | 2 | 1 | 2 | 1 | 0 | 1 | 1.17 | 2 | 1 | 0.372 | -1.427 | 0.42109589 | NA | 0 | 2.8259 | 1.2028 | 1.2028 | TRUE | 0.1932 |
| Gm1082 | 28 | 39 | 41 | 33 | 26 | 26 | 30 | 39 | 41 | 26 | 28 | 28 | 32.04 | 37 | 27 | 0.747 | -0.421 | 0.21689513 | 0.42884167 | 0 | 0.1097 | 0.0519 | 0.0519 | TRUE | 0.1887 |
| Gm10823 | 0 | 0 | 0 | 0 | 0 | 0 | 0 | 0 | 0 | 0 | 0 | 0 | 0 | 0 | 0 | NA | NA | NA | NA | NA | NA | NA | NA | NA | NA |
| Gm10825 | 2 | 3 | 7 | 9 | 5 | 5 | 2 | 3 | 7 | 7 | 5 | 5 | 5.01 | 4 | 6 | 1.48 | 0.565 | 0.51583908 | 0.70801619 | 0 | 0.6641 | 0.3382 | 0.3382 | TRUE | 0.7051 |
| Gm10845 | 0 | 0 | 0 | 0 | 0 | 0 | 0 | 0 | 0 | 0 | 0 | 0 | 0 | 0 | 0 | NA | NA | NA | NA | NA | NA | NA | NA | NA | NA |
| Gm10857 | 2 | 0 | 0 | 4 | 1 | 0 | 2 | 0 | 0 | 3 | 1 | 0 | 1.06 | 1 | 1 | 2.065 | 1.046 | 0.65153896 | NA | 2.081 | 3.1223 | 2.7969 | 2.7969 | TRUE | 0.5659 |
| Gm10863 | 12 | 13 | 19 | 30 | 26 | 19 | 13 | 13 | 19 | 24 | 28 | 21 | 19.53 | 15 | 24 | 1.612 | 0.689 | 0.11950432 | 0.29294718 | 0 | 0.1756 | 0.0867 | 0.0867 | TRUE | 0.2704 |
| Gm10865 | 0 | 0 | 0 | 1 | 0 | 0 | 0 | 0 | 0 | 1 | 0 | 0 | 0.13 | 0 | 0 | 1.859 | 0.894 | 0.82650734 | NA | 0 | 25.1515 | 10 | 10 | TRUE | 0.2563 |
| Gm10872 | 0 | 0 | 0 | 1 | 0 | 0 | 0 | 0 | 0 | 1 | 0 | 0 | 0.13 | 0 | 0 | 1.859 | 0.894 | 0.82650734 | NA | 0 | 25.1515 | 10 | 10 | TRUE | 0.2563 |
| Gm10921 | 0 | 0 | 0 | 0 | 0 | 0 | 0 | 0 | 0 | 0 | 0 | 0 | 0 | 0 | 0 | NA | NA | NA | NA | NA | NA | NA | NA | NA | NA |
| Gm10922 | 0 | 0 | 0 | 0 | 0 | 0 | 0 | 0 | 0 | 0 | 0 | 0 | 0 | 0 | 0 | NA | NA | NA | NA | NA | NA | NA | NA | NA | NA |
| Gm10941 | 12 | 22 | 13 | 17 | 3 | 14 | 13 | 22 | 13 | 13 | 3 | 15 | 13.28 | 16 | 10 | 0.672 | -0.573 | 0.37427653 | 0.5904186 | 0.1792 | 0.255 | 0.2224 | 0.2224 | TRUE | 1.2324 |
| Gm1110 | 0 | 0 | 0 | 0 | 0 | 1 | 0 | 0 | 0 | 0 | 0 | 1 | 0.18 | 0 | 0 | 1.859 | 0.894 | 0.82650734 | NA | 0 | 18.0643 | 10 | 10 | TRUE | 0.5383 |
| Gm11110 | 13 | 25 | 36 | 26 | 15 | 20 | 14 | 25 | 36 | 20 | 16 | 22 | 22.23 | 25 | 19 | 0.78 | -0.359 | 0.4500016 | 0.65553889 | 0.0766 | 0.1551 | 0.1173 | 0.1173 | TRUE | 0.1467 |
| Gm11127 | 3 | 2 | 0 | 3 | 0 | 2 | 3 | 2 | 0 | 2 | 0 | 2 | 1.62 | 2 | 1 | 0.895 | -0.159 | 0.92562072 | NA | 0 | 2.0381 | 1.4841 | 1.4841 | TRUE | 0.356 |
| Gm11128 | 0 | 0 | 0 | 0 | 0 | 0 | 0 | 0 | 0 | 0 | 0 | 0 | 0 | 0 | 0 | NA | NA | NA | NA | NA | NA | NA | NA | NA | NA |
| Gm11149 | 0 | 0 | 0 | 0 | 0 | 0 | 0 | 0 | 0 | 0 | 0 | 0 | 0 | 0 | 0 | NA | NA | NA | NA | NA | NA | NA | NA | NA | NA |

| | | | | | | | | | | | | | | | | | | | | | | | | | |
|---|---|---|---|---|---|---|---|---|---|---|---|---|---|---|---|---|---|---|---|---|---|---|---|---|---|
| Gm11166 | 0 | 0 | 0 | 2 | 0 | 0 | 0 | 0 | 0 | 2 | 0 | 0 | 0.26 | 0 | 1 | 2.886 | 1.529 | 0.70676491 | NA | 0 | 12.5792 | 10 | 10 | TRUE | 0.9669 |
| Gm11186 | 0 | 0 | 0 | 0 | 0 | 0 | 0 | 0 | 0 | 0 | 0 | 0 | 0 | 0 | 0 | NA | NA | NA | NA | NA | NA | NA | NA | NA | NA |
| Gm11190 | 3 | 16 | 10 | 17 | 23 | 10 | 3 | 16 | 10 | 13 | 25 | 11 | 13.04 | 10 | 16 | 1.667 | 0.738 | 0.27664395 | 0.49765942 | 0.236 | 0.2595 | 0.249 | 0.249 | TRUE | 0.366 |
| Gm11194 | 0 | 1 | 1 | 2 | 5 | 1 | 0 | 1 | 1 | 2 | 5 | 1 | 1.67 | 1 | 3 | 3.934 | 1.976 | 0.23853617 | NA | 0 | 1.9739 | 1.0846 | 1.0846 | TRUE | 0.844 |
| Gm11201 | 1 | 0 | 0 | 1 | 1 | 1 | 1 | 0 | 0 | 1 | 1 | 1 | 0.67 | 0 | 1 | 2.676 | 1.42 | 0.53486561 | NA | 0 | 4.9253 | 2.274 | 2.274 | TRUE | 0.4614 |
| Gm11202 | 0 | 0 | 0 | 0 | 2 | 1 | 0 | 0 | 0 | 0 | 2 | 1 | 0.54 | 0 | 1 | 5.685 | 2.507 | 0.43809847 | NA | 0 | 6.0955 | 6.0745 | 6.0745 | TRUE | 0.0593 |
| Gm11213 | 0 | 0 | 0 | 0 | 0 | 0 | 0 | 0 | 0 | 0 | 0 | 0 | 0 | 0 | 0 | NA | NA | NA | NA | NA | NA | NA | NA | NA | NA |
| Gm1123 | 1 | 1 | 8 | 3 | 1 | 2 | 1 | 1 | 8 | 2 | 1 | 2 | 2.62 | 3 | 2 | 0.561 | -0.834 | 0.52887081 | NA | 0.5136 | 1.2645 | 0.8558 | 0.8558 | TRUE | 2.1355 |
| Gm11240 | 1 | 1 | 0 | 0 | 0 | 0 | 1 | 1 | 0 | 0 | 0 | 0 | 0.34 | 1 | 0 | 0.261 | -1.936 | 0.60454921 | NA | 0 | 9.5983 | 8.3269 | 8.3269 | TRUE | 0.2507 |
| Gm11346 | 2 | 7 | 9 | 7 | 5 | 6 | 2 | 7 | 9 | 6 | 5 | 7 | 5.93 | 6 | 6 | 0.955 | -0.066 | 0.93515729 | 0.96786539 | 0 | 0.5621 | 0.3113 | 0.3113 | TRUE | 0.6203 |
| Gm11351 | 0 | 0 | 0 | 0 | 0 | 0 | 0 | 0 | 0 | 0 | 0 | 0 | 0 | 0 | 0 | NA | NA | NA | NA | NA | NA | NA | NA | NA | NA |
| Gm1140 | 0 | 0 | 0 | 0 | 0 | 0 | 0 | 0 | 0 | 0 | 0 | 0 | 0 | 0 | 0 | NA | NA | NA | NA | NA | NA | NA | NA | NA | NA |
| Gm1141 | 5 | 6 | 2 | 53 | 37 | 16 | 5 | 6 | 2 | 42 | 40 | 18 | 18.72 | 4 | 33 | 7.483 | 2.904 | 5.59E-06 | 0.00012154 | 0.1404 | 0.1829 | 0.1645 | 0.1645 | TRUE | 1.1032 |
| Gm11413 | 70 | 74 | 66 | 72 | 47 | 65 | 74 | 74 | 66 | 57 | 51 | 71 | 65.48 | 71 | 60 | 0.83 | -0.269 | 0.28360668 | 0.50514336 | 0 | 0.0572 | 0.0301 | 0.0301 | TRUE | 0.2356 |
| Gm11426 | 4 | 7 | 1 | 5 | 2 | 3 | 4 | 7 | 1 | 4 | 2 | 3 | 3.6 | 4 | 3 | 0.772 | -0.373 | 0.72465057 | NA | 0 | 0.9216 | 0.5267 | 0.5267 | TRUE | 0.1873 |
| Gm11435 | 0 | 0 | 1 | 2 | 2 | 0 | 0 | 0 | 1 | 2 | 2 | 0 | 0.79 | 0 | 1 | 3.417 | 1.773 | 0.4679427 | NA | 0 | 4.1875 | 2.9056 | 2.9056 | TRUE | 0.4098 |
| Gm11437 | 0 | 0 | 0 | 0 | 0 | 0 | 0 | 0 | 0 | 0 | 0 | 0 | 0 | 0 | 0 | NA | NA | NA | NA | NA | NA | NA | NA | NA | NA |
| Gm11468 | 0 | 0 | 0 | 0 | 0 | 0 | 0 | 0 | 0 | 0 | 0 | 0 | 0 | 0 | 0 | NA | NA | NA | NA | NA | NA | NA | NA | NA | NA |
| Gm11487 | 6 | 6 | 1 | 11 | 9 | 3 | 6 | 6 | 1 | 9 | 10 | 3 | 5.83 | 4 | 7 | 1.631 | 0.705 | 0.43687257 | 0.64516425 | 0.1976 | 0.5719 | 0.4106 | 0.4106 | TRUE | 0.8554 |
| Gm11517 | 6 | 1 | 4 | 41 | 17 | 5 | 6 | 1 | 4 | 32 | 18 | 5 | 11.23 | 4 | 18 | 4.991 | 2.319 | 0.00773543 | 0.04330961 | 0.4867 | 0.3002 | 0.3851 | 0.3851 | TRUE | 0.0763 |
| Gm11529 | 1 | 2 | 7 | 3 | 1 | 4 | 1 | 2 | 7 | 2 | 1 | 4 | 2.98 | 3 | 2 | 0.768 | -0.381 | 0.75206999 | NA | 0.3235 | 1.1111 | 0.7076 | 0.7076 | TRUE | 1.3088 |
| Gm11541 | 1 | 2 | 0 | 1 | 3 | 1 | 1 | 2 | 0 | 1 | 3 | 1 | 1.36 | 1 | 2 | 1.641 | 0.715 | 0.67317265 | NA | 0 | 2.4269 | 1.2773 | 1.2773 | TRUE | 0.3487 |
| Gm11544 | 4 | 3 | 4 | 31 | 20 | 5 | 4 | 3 | 4 | 24 | 22 | 5 | 10.44 | 4 | 17 | 4.603 | 2.203 | 0.00514196 | 0.03195814 | 0 | 0.3225 | 0.2819 | 0.2819 | TRUE | 1.4046 |
| Gm11545 | 104 | 70 | 51 | 276 | 181 | 98 | 110 | 70 | 51 | 217 | 195 | 107 | 125.1 | 77 | 173 | 2.248 | 1.169 | 0.00068617 | 0.006664 | 0.1276 | 0.0332 | 0.0759 | 0.0759 | TRUE | 0.9892 |
| Gm11548 | 0 | 0 | 0 | 0 | 0 | 0 | 0 | 0 | 0 | 0 | 0 | 0 | 0 | 0 | 0 | NA | NA | NA | NA | NA | NA | NA | NA | NA | NA |
| Gm11549 | 7 | 7 | 2 | 3 | 3 | 0 | 7 | 7 | 2 | 2 | 3 | 0 | 3.67 | 5 | 2 | 0.346 | -1.533 | 0.18859545 | NA | 0.2223 | 0.9045 | 0.6284 | 0.6284 | TRUE | 0.6772 |
| Gm11554 | 0 | 0 | 0 | 0 | 0 | 0 | 0 | 0 | 0 | 0 | 0 | 0 | 0 | 0 | 0 | NA | NA | NA | NA | NA | NA | NA | NA | NA | NA |
| Gm11559 | 0 | 0 | 0 | 0 | 0 | 0 | 0 | 0 | 0 | 0 | 0 | 0 | 0 | 0 | 0 | NA | NA | NA | NA | NA | NA | NA | NA | NA | NA |
| Gm11562 | 0 | 0 | 0 | 0 | 0 | 0 | 0 | 0 | 0 | 0 | 0 | 0 | 0 | 0 | 0 | NA | NA | NA | NA | NA | NA | NA | NA | NA | NA |
| Gm11563 | 0 | 0 | 0 | 0 | 0 | 0 | 0 | 0 | 0 | 0 | 0 | 0 | 0 | 0 | 0 | NA | NA | NA | NA | NA | NA | NA | NA | NA | NA |
| Gm11564 | 8 | 1 | 1 | 6 | 5 | 2 | 8 | 1 | 1 | 5 | 5 | 2 | 3.8 | 3 | 4 | 1.185 | 0.245 | 0.83171542 | NA | 0.5135 | 0.8744 | 0.6947 | 0.6947 | TRUE | 2.2203 |
| Gm11565 | 0 | 0 | 0 | 0 | 0 | 0 | 0 | 0 | 0 | 0 | 0 | 0 | 0 | 0 | 0 | NA | NA | NA | NA | NA | NA | NA | NA | NA | NA |
| Gm11567 | 0 | 0 | 0 | 0 | 0 | 0 | 0 | 0 | 0 | 0 | 0 | 0 | 0 | 0 | 0 | NA | NA | NA | NA | NA | NA | NA | NA | NA | NA |
| Gm11568 | 0 | 0 | 0 | 0 | 0 | 0 | 0 | 0 | 0 | 0 | 0 | 0 | 0 | 0 | 0 | NA | NA | NA | NA | NA | NA | NA | NA | NA | NA |
| Gm11569 | 0 | 0 | 0 | 0 | 0 | 0 | 0 | 0 | 0 | 0 | 0 | 0 | 0 | 0 | 0 | NA | NA | NA | NA | NA | NA | NA | NA | NA | NA |
| Gm11570 | 0 | 0 | 0 | 0 | 0 | 0 | 0 | 0 | 0 | 0 | 0 | 0 | 0 | 0 | 0 | NA | NA | NA | NA | NA | NA | NA | NA | NA | NA |
| Gm11595 | 0 | 0 | 0 | 0 | 0 | 0 | 0 | 0 | 0 | 0 | 0 | 0 | 0 | 0 | 0 | NA | NA | NA | NA | NA | NA | NA | NA | NA | NA |
| Gm11596 | 0 | 0 | 0 | 0 | 0 | 0 | 0 | 0 | 0 | 0 | 0 | 0 | 0 | 0 | 0 | NA | NA | NA | NA | NA | NA | NA | NA | NA | NA |
| Gm11602 | 14 | 14 | 13 | 39 | 13 | 9 | 15 | 14 | 13 | 31 | 14 | 10 | 16.07 | 14 | 18 | 1.323 | 0.404 | 0.47887695 | 0.67817611 | 0.133 | 0.2119 | 0.1715 | 0.1715 | TRUE | 1.5574 |
| Gm11627 | 63 | 71 | 58 | 87 | 68 | 41 | 67 | 71 | 58 | 68 | 73 | 45 | 63.75 | 65 | 62 | 0.954 | -0.068 | 0.80722682 | 0.90068918 | 0.0182 | 0.0586 | 0.0397 | 0.0397 | TRUE | 0.4796 |
| Gm11648 | 6 | 2 | 2 | 8 | 3 | 2 | 6 | 2 | 2 | 6 | 3 | 2 | 3.68 | 3 | 4 | 1.159 | 0.213 | 0.83852363 | NA | 0.1456 | 0.9019 | 0.5127 | 0.5127 | TRUE | 0.7605 |
| Gm11696 | 38 | 46 | 42 | 43 | 33 | 39 | 40 | 46 | 42 | 34 | 36 | 43 | 40.08 | 43 | 38 | 0.868 | -0.205 | 0.49737059 | 0.69285717 | 0 | 0.0891 | 0.0405 | 0.0405 | TRUE | 0.1084 |
| Gm11710 | 0 | 0 | 0 | 0 | 0 | 0 | 0 | 0 | 0 | 0 | 0 | 0 | 0 | 0 | 0 | NA | NA | NA | NA | NA | NA | NA | NA | NA | NA |
| Gm11711 | 0 | 0 | 0 | 0 | 0 | 0 | 0 | 0 | 0 | 0 | 0 | 0 | 0 | 0 | 0 | NA | NA | NA | NA | NA | NA | NA | NA | NA | NA |
| Gm11744 | 33 | 24 | 41 | 49 | 27 | 33 | 35 | 24 | 41 | 39 | 29 | 36 | 33.98 | 33 | 35 | 1.039 | 0.055 | 0.87664021 | 0.93932608 | 1.00E-04 | 0.1039 | 0.0624 | 0.0624 | TRUE | 0.4327 |
| Gm11747 | 0 | 0 | 0 | 0 | 0 | 0 | 0 | 0 | 0 | 0 | 0 | 0 | 0 | 0 | 0 | NA | NA | NA | NA | NA | NA | NA | NA | NA | NA |
| Gm11756 | 2 | 0 | 0 | 4 | 5 | 1 | 2 | 0 | 0 | 3 | 5 | 1 | 1.96 | 1 | 3 | 4.629 | 2.211 | 0.19572379 | NA | 0.3393 | 1.6902 | 1.2143 | 1.2143 | TRUE | 0.4703 |
| Gm11757 | 0 | 0 | 0 | 0 | 0 | 0 | 0 | 0 | 0 | 0 | 0 | 0 | 0 | 0 | 0 | NA | NA | NA | NA | NA | NA | NA | NA | NA | NA |
| Gm11758 | 0 | 0 | 0 | 0 | 0 | 0 | 0 | 0 | 0 | 0 | 0 | 0 | 0 | 0 | 0 | NA | NA | NA | NA | NA | NA | NA | NA | NA | NA |
| Gm11762 | 0 | 2 | 0 | 1 | 0 | 0 | 0 | 2 | 0 | 1 | 0 | 0 | 0.46 | 1 | 0 | 0.522 | -0.937 | 0.78646161 | NA | 0 | 7.1082 | 6.8573 | 6.8573 | TRUE | 0.9739 |
| Gm11837 | 222 | 279 | 284 | 399 | 267 | 291 | 236 | 279 | 285 | 314 | 287 | 319 | 286.52 | 267 | 307 | 1.15 | 0.201 | 0.14497796 | 0.33274045 | 0.0026 | 0.0184 | 0.0102 | 0.0102 | TRUE | 0.112 |
| Gm11937 | 0 | 0 | 0 | 0 | 0 | 0 | 0 | 0 | 0 | 0 | 0 | 0 | 0 | 0 | 0 | NA | NA | NA | NA | NA | NA | NA | NA | NA | NA |
| Gm11938 | 0 | 0 | 0 | 0 | 0 | 0 | 0 | 0 | 0 | 0 | 0 | 0 | 0 | 0 | 0 | NA | NA | NA | NA | NA | NA | NA | NA | NA | NA |
| Gm11944 | 3 | 7 | 17 | 15 | 10 | 11 | 3 | 7 | 17 | 12 | 11 | 12 | 10.31 | 9 | 12 | 1.266 | 0.34 | 0.6293472 | 0.79155215 | 0.1795 | 0.3265 | 0.259 | 0.259 | TRUE | 0.9914 |
| Gm11961 | 23 | 14 | 18 | 22 | 14 | 18 | 24 | 14 | 18 | 17 | 15 | 20 | 18.09 | 19 | 17 | 0.923 | -0.115 | 0.80214765 | 0.89777365 | 0 | 0.189 | 0.0968 | 0.0968 | TRUE | 0.3113 |
| Gm11974 | 918 | 1148 | 1100 | 1683 | 1131 | 1000 | 975 | 1147 | 1105 | 1323 | 1217 | 1095 | 1143.45 | 1076 | 1212 | 1.127 | 0.173 | 0.12525784 | 0.30276883 | 0.0066 | 0.0098 | 0.0083 | 0.0083 | TRUE | 0.085 |
| Gm11978 | 0 | 0 | 0 | 2 | 1 | 0 | 0 | 0 | 0 | 2 | 1 | 0 | 0.44 | 0 | 1 | 4.754 | 2.249 | 0.50892201 | NA | 0 | 7.4712 | 6.8061 | 6.8061 | TRUE | 0.2361 |
| Gm11981 | 1 | 0 | 0 | 0 | 0 | 0 | 1 | 0 | 0 | 0 | 0 | 0 | 0.18 | 0 | 0 | 0.49 | -1.029 | 0.80087 | NA | 0 | 18.6165 | 10 | 10 | TRUE | 0.4614 |
| Gm11985 | 3 | 1 | 2 | 5 | 7 | 2 | 3 | 1 | 2 | 4 | 8 | 2 | 3.31 | 2 | 5 | 2.209 | 1.144 | 0.30147666 | NA | 0 | 1.003 | 0.5261 | 0.5261 | TRUE | 0.309 |
| Gm11992 | 43 | 69 | 76 | 115 | 78 | 91 | 46 | 69 | 76 | 90 | 84 | 100 | 77.47 | 64 | 91 | 1.432 | 0.518 | 0.04634757 | 0.15580006 | 0.0181 | 0.0494 | 0.0354 | 0.0354 | TRUE | 0.5188 |
| Gm12 | 0 | 0 | 0 | 0 | 0 | 0 | 0 | 0 | 0 | 0 | 0 | 0 | 0 | 0 | 0 | NA | NA | NA | NA | NA | NA | NA | NA | NA | NA |
| Gm12060 | 0 | 0 | 0 | 1 | 1 | 0 | 0 | 0 | 0 | 1 | 1 | 0 | 0.31 | 0 | 1 | 3.32 | 1.731 | 0.65383052 | NA | 0 | 10.6224 | 8.9541 | 8.9541 | TRUE | 0.202 |
| Gm12070 | 14 | 14 | 10 | 8 | 7 | 8 | 15 | 14 | 10 | 6 | 8 | 9 | 10.25 | 13 | 8 | 0.576 | -0.796 | 0.18031347 | 0.38162964 | 0 | 0.3285 | 0.1493 | 0.1493 | TRUE | 0.1631 |
| Gm12130 | 0 | 0 | 0 | 0 | 0 | 0 | 0 | 0 | 0 | 0 | 0 | 0 | 0 | 0 | 0 | NA | NA | NA | NA | NA | NA | NA | NA | NA | NA |
| Gm12159 | 0 | 0 | 0 | 0 | 0 | 0 | 0 | 0 | 0 | 0 | 0 | 0 | 0 | 0 | 0 | NA | NA | NA | NA | NA | NA | NA | NA | NA | NA |
| Gm12169 | 0 | 0 | 0 | 1 | 0 | 0 | 0 | 0 | 0 | 1 | 0 | 0 | 0.13 | 0 | 0 | 1.859 | 0.894 | 0.82650734 | NA | 0 | 25.1515 | 10 | 10 | TRUE | 0.2563 |
| Gm12171 | 0 | 0 | 0 | 0 | 0 | 0 | 0 | 0 | 0 | 0 | 0 | 0 | 0 | 0 | 0 | NA | NA | NA | NA | NA | NA | NA | NA | NA | NA |
| Gm12185 | 0 | 1 | 0 | 0 | 0 | 0 | 0 | 1 | 0 | 0 | 0 | 0 | 0.17 | 0 | 0 | 0.49 | -1.029 | 0.80087 | NA | 0 | 19.7991 | 10 | 10 | TRUE | 0.405 |
| Gm12191 | 0 | 0 | 0 | 0 | 0 | 0 | 0 | 0 | 0 | 0 | 0 | 0 | 0 | 0 | 0 | NA | NA | NA | NA | NA | NA | NA | NA | NA | NA |

| | | | | | | | | | | | | | | | | | | | | | | | | | |
|---|---|---|---|---|---|---|---|---|---|---|---|---|---|---|---|---|---|---|---|---|---|---|---|---|---|
| Gm12216 | 0 | 2 | 1 | 1 | 0 | 0 | 0 | 2 | 1 | 1 | 0 | 0 | 0.63 | 1 | 0 | 0.347 | -1.525 | 0.56199923 | NA | 0 | 5.2256 | 3.4788 | 3.4788 | TRUE | 0.1027 |
| Gm12228 | 0 | 0 | 0 | 0 | 1 | 0 | 0 | 0 | 0 | 0 | 1 | 0 | 0.18 | 0 | 0 | 1.859 | 0.894 | 0.82650734 | NA | 0 | 18.3785 | 10 | 10 | TRUE | 0.5204 |
| Gm12238 | 2 | 8 | 4 | 5 | 3 | 7 | 2 | 8 | 4 | 4 | 3 | 8 | 4.83 | 5 | 5 | 1.036 | 0.051 | 0.95539893 | 0.97727168 | 0.0954 | 0.6897 | 0.3984 | 0.3984 | TRUE | 0.5701 |
| Gm12250 | 70 | 65 | 59 | 65 | 43 | 34 | 74 | 65 | 59 | 51 | 46 | 37 | 55.52 | 66 | 45 | 0.682 | -0.551 | 0.04313788 | 0.14848768 | 0 | 0.0662 | 0.035 | 0.035 | TRUE | 0.164 |
| Gm12253 | 0 | 0 | 0 | 0 | 0 | 0 | 0 | 0 | 0 | 0 | 0 | 0 | 0 | 0 | 0 | NA | NA | NA | NA | NA | NA | NA | NA | NA | NA |
| Gm12295 | 0 | 0 | 0 | 0 | 0 | 0 | 0 | 0 | 0 | 0 | 0 | 0 | 0 | 0 | 0 | NA | NA | NA | NA | NA | NA | NA | NA | NA | NA |
| Gm12298 | 0 | 0 | 0 | 1 | 0 | 1 | 0 | 0 | 0 | 1 | 0 | 1 | 0.31 | 0 | 1 | 3.348 | 1.743 | 0.65064889 | NA | 0 | 10.5167 | 8.8968 | 8.8968 | TRUE | 0.2082 |
| Gm12359 | 12 | 4 | 4 | 14 | 13 | 3 | 13 | 4 | 4 | 11 | 14 | 3 | 8.17 | 7 | 9 | 1.375 | 0.46 | 0.57763759 | 0.75499296 | 0.3172 | 0.41 | 0.3649 | 0.3649 | TRUE | 0.875 |
| Gm12360 | 0 | 0 | 0 | 0 | 1 | 0 | 0 | 0 | 0 | 0 | 1 | 0 | 0.18 | 0 | 0 | 1.859 | 0.894 | 0.82650734 | NA | 0 | 18.3785 | 10 | 10 | TRUE | 0.5204 |
| Gm12409 | 0 | 0 | 0 | 2 | 1 | 0 | 0 | 0 | 0 | 2 | 1 | 0 | 0.44 | 0 | 1 | 4.754 | 2.249 | 0.50892201 | NA | 0 | 7.4712 | 6.8061 | 6.8061 | TRUE | 0.2361 |
| Gm12504 | 13 | 11 | 5 | 11 | 11 | 7 | 14 | 11 | 5 | 9 | 12 | 8 | 9.66 | 10 | 10 | 0.944 | -0.083 | 0.89811455 | 0.95048397 | 0 | 0.3479 | 0.1995 | 0.1995 | TRUE | 0.5376 |
| Gm12505 | 0 | 0 | 0 | 1 | 2 | 2 | 0 | 0 | 0 | 1 | 2 | 2 | 0.85 | 0 | 2 | 9.003 | 3.17 | 0.16844964 | NA | 0 | 3.8621 | 2.5291 | 2.5291 | TRUE | 0.2298 |
| Gm12522 | 35 | 48 | 49 | 29 | 25 | 34 | 37 | 48 | 49 | 23 | 27 | 37 | 36.87 | 45 | 29 | 0.641 | -0.642 | 0.06237849 | 0.19118414 | 0.0115 | 0.0963 | 0.0571 | 0.0571 | TRUE | 0.3861 |
| Gm12530 | 0 | 0 | 0 | 0 | 0 | 0 | 0 | 0 | 0 | 0 | 0 | 0 | 0 | 0 | 0 | NA | NA | NA | NA | NA | NA | NA | NA | NA | NA |
| Gm12597 | 0 | 0 | 0 | 0 | 0 | 0 | 0 | 0 | 0 | 0 | 0 | 0 | 0 | 0 | 0 | NA | NA | NA | NA | NA | NA | NA | NA | NA | NA |
| Gm12603 | 0 | 0 | 0 | 0 | 0 | 0 | 0 | 0 | 0 | 0 | 0 | 0 | 0 | 0 | 0 | NA | NA | NA | NA | NA | NA | NA | NA | NA | NA |
| Gm12633 | 1 | 0 | 0 | 0 | 0 | 0 | 1 | 0 | 0 | 0 | 0 | 0 | 0.18 | 0 | 0 | 0.49 | -1.029 | 0.80087 | NA | 0 | 18.6165 | 10 | 10 | TRUE | 0.4614 |
| Gm12657 | 1 | 11 | 8 | 8 | 1 | 8 | 1 | 11 | 8 | 6 | 1 | 9 | 6.03 | 7 | 5 | 0.803 | -0.317 | 0.75689235 | 0.87155626 | 0.6334 | 0.5529 | 0.5899 | 0.5899 | TRUE | 0.5437 |
| Gm12669 | 0 | 0 | 0 | 0 | 0 | 0 | 0 | 0 | 0 | 0 | 0 | 0 | 0 | 0 | 0 | NA | NA | NA | NA | NA | NA | NA | NA | NA | NA |
| Gm12695 | 0 | 0 | 0 | 0 | 0 | 1 | 0 | 0 | 0 | 0 | 0 | 1 | 0.18 | 0 | 0 | 1.859 | 0.894 | 0.82650734 | NA | 0 | 18.0643 | 10 | 10 | TRUE | 0.5383 |
| Gm12709 | 0 | 0 | 0 | 0 | 0 | 0 | 0 | 0 | 0 | 0 | 0 | 0 | 0 | 0 | 0 | NA | NA | NA | NA | NA | NA | NA | NA | NA | NA |
| Gm12718 | 0 | 0 | 1 | 3 | 0 | 1 | 0 | 0 | 1 | 2 | 0 | 1 | 0.74 | 0 | 1 | 3.216 | 1.685 | 0.50237532 | NA | 0 | 4.4418 | 3.1322 | 3.1322 | TRUE | 0.1958 |
| Gm12789 | 1 | 0 | 0 | 1 | 1 | 0 | 1 | 0 | 0 | 1 | 1 | 0 | 0.49 | 0 | 1 | 1.712 | 0.776 | 0.78488768 | NA | 0 | 6.7665 | 4.0387 | 4.0387 | TRUE | 0.4614 |
| Gm12794 | 202 | 71 | 46 | 384 | 272 | 114 | 215 | 71 | 46 | 302 | 293 | 125 | 175.16 | 111 | 240 | 2.17 | 1.118 | 0.13419257 | 0.31679168 | 0.3948 | 0.0257 | 0.1739 | 0.3948 | TRUE | 6.6897 |
| Gm128 | 64 | 97 | 125 | 107 | 101 | 95 | 68 | 97 | 126 | 84 | 109 | 104 | 97.86 | 97 | 99 | 1.018 | 0.026 | 0.92416237 | 0.96274037 | 0.0437 | 0.0406 | 0.042 | 0.042 | TRUE | 0.1883 |
| Gm12830 | 1 | 0 | 0 | 1 | 0 | 0 | 1 | 0 | 0 | 1 | 0 | 0 | 0.31 | 0 | 0 | 0.954 | -0.067 | 0.98625928 | NA | 0 | 10.7014 | 9.0384 | 9.0384 | TRUE | 0.4614 |
| Gm12839 | 0 | 0 | 0 | 0 | 0 | 0 | 0 | 0 | 0 | 0 | 0 | 0 | 0 | 0 | 0 | NA | NA | NA | NA | NA | NA | NA | NA | NA | NA |
| Gm12886 | 0 | 0 | 0 | 0 | 0 | 0 | 0 | 0 | 0 | 0 | 0 | 0 | 0 | 0 | 0 | NA | NA | NA | NA | NA | NA | NA | NA | NA | NA |
| Gm12887 | 0 | 0 | 0 | 0 | 0 | 0 | 0 | 0 | 0 | 0 | 0 | 0 | 0 | 0 | 0 | NA | NA | NA | NA | NA | NA | NA | NA | NA | NA |
| Gm12888 | 0 | 0 | 0 | 0 | 0 | 0 | 0 | 0 | 0 | 0 | 0 | 0 | 0 | 0 | 0 | NA | NA | NA | NA | NA | NA | NA | NA | NA | NA |
| Gm129 | 85 | 132 | 110 | 151 | 151 | 138 | 90 | 132 | 110 | 119 | 162 | 151 | 127.47 | 111 | 144 | 1.294 | 0.372 | 0.09565479 | 0.25355511 | 0.0229 | 0.0327 | 0.0278 | 0.0278 | TRUE | 0.2569 |
| Gm12942 | 123 | 196 | 212 | 119 | 101 | 188 | 131 | 196 | 213 | 94 | 109 | 206 | 157.88 | 180 | 136 | 0.754 | -0.408 | 0.20294555 | 0.41057686 | 0.1143 | 0.0278 | 0.0675 | 0.0675 | TRUE | 2.0495 |
| Gm13003 | 1 | 3 | 1 | 0 | 2 | 0 | 1 | 3 | 1 | 0 | 2 | 0 | 1.2 | 2 | 1 | 0.406 | -1.301 | 0.50823204 | NA | 0 | 2.7468 | 1.7756 | 1.7756 | TRUE | 0.9751 |
| Gm13011 | 0 | 0 | 0 | 0 | 0 | 0 | 0 | 0 | 0 | 0 | 0 | 0 | 0 | 0 | 0 | NA | NA | NA | NA | NA | NA | NA | NA | NA | NA |
| Gm13023 | 1 | 0 | 0 | 0 | 0 | 0 | 1 | 0 | 0 | 0 | 0 | 0 | 0.18 | 0 | 0 | 0.49 | -1.029 | 0.80087 | NA | 0 | 18.6165 | 10 | 10 | TRUE | 0.4614 |
| Gm13031 | 0 | 4 | 3 | 4 | 2 | 4 | 0 | 4 | 3 | 3 | 2 | 4 | 2.78 | 2 | 3 | 1.369 | 0.453 | 0.71311064 | NA | 0 | 1.1918 | 0.723 | 0.723 | TRUE | 0.7409 |
| Gm13032 | 12 | 18 | 15 | 11 | 13 | 19 | 13 | 18 | 15 | 9 | 14 | 21 | 14.87 | 15 | 15 | 0.934 | -0.099 | 0.85407875 | 0.9274031 | 0.0409 | 0.2285 | 0.1395 | 0.1395 | TRUE | 0.2672 |
| Gm13034 | 38 | 26 | 20 | 57 | 43 | 25 | 40 | 26 | 20 | 45 | 46 | 27 | 34.14 | 29 | 39 | 1.378 | 0.463 | 0.25014539 | 0.46733938 | 0.0679 | 0.1034 | 0.0867 | 0.0867 | TRUE | 0.779 |
| Gm13037 | 0 | 0 | 0 | 0 | 0 | 0 | 0 | 0 | 0 | 0 | 0 | 0 | 0 | 0 | 0 | NA | NA | NA | NA | NA | NA | NA | NA | NA | NA |
| Gm13040 | 0 | 0 | 0 | 0 | 0 | 0 | 0 | 0 | 0 | 0 | 0 | 0 | 0 | 0 | 0 | NA | NA | NA | NA | NA | NA | NA | NA | NA | NA |
| Gm13043 | 0 | 0 | 0 | 0 | 0 | 0 | 0 | 0 | 0 | 0 | 0 | 0 | 0 | 0 | 0 | NA | NA | NA | NA | NA | NA | NA | NA | NA | NA |
| Gm13051 | 378 | 279 | 278 | 534 | 397 | 329 | 402 | 279 | 279 | 420 | 427 | 360 | 361.06 | 320 | 402 | 1.26 | 0.333 | 0.06068516 | 0.18748801 | 0.0245 | 0.016 | 0.0199 | 0.0199 | TRUE | 0.5633 |
| Gm13057 | 0 | 0 | 0 | 0 | 0 | 0 | 0 | 0 | 0 | 0 | 0 | 0 | 0 | 0 | 0 | NA | NA | NA | NA | NA | NA | NA | NA | NA | NA |
| Gm13078 | 24 | 21 | 17 | 48 | 43 | 19 | 25 | 21 | 17 | 38 | 46 | 21 | 28.06 | 21 | 35 | 1.656 | 0.727 | 0.09103827 | 0.24607637 | 0.064 | 0.1243 | 0.0953 | 0.0953 | TRUE | 0.3187 |
| Gm13083 | 11 | 6 | 4 | 18 | 10 | 7 | 12 | 6 | 4 | 14 | 11 | 8 | 9.04 | 7 | 11 | 1.522 | 0.606 | 0.37798349 | 0.59318961 | 0.056 | 0.3712 | 0.2253 | 0.2253 | TRUE | 0.5117 |
| Gm13084 | 0 | 0 | 0 | 0 | 0 | 0 | 0 | 0 | 0 | 0 | 0 | 0 | 0 | 0 | 0 | NA | NA | NA | NA | NA | NA | NA | NA | NA | NA |
| Gm13088 | 0 | 0 | 0 | 0 | 0 | 0 | 0 | 0 | 0 | 0 | 0 | 0 | 0 | 0 | 0 | NA | NA | NA | NA | NA | NA | NA | NA | NA | NA |
| Gm13102 | 0 | 0 | 0 | 0 | 0 | 0 | 0 | 0 | 0 | 0 | 0 | 0 | 0 | 0 | 0 | NA | NA | NA | NA | NA | NA | NA | NA | NA | NA |
| Gm13103 | 0 | 0 | 0 | 0 | 0 | 0 | 0 | 0 | 0 | 0 | 0 | 0 | 0 | 0 | 0 | NA | NA | NA | NA | NA | NA | NA | NA | NA | NA |
| Gm13109 | 90 | 42 | 30 | 155 | 99 | 45 | 96 | 42 | 30 | 122 | 107 | 49 | 74.22 | 56 | 93 | 1.662 | 0.733 | 0.11826098 | 0.29103015 | 0.2544 | 0.0513 | 0.1443 | 0.1443 | TRUE | 2.0411 |
| Gm13119 | 213 | 98 | 78 | 336 | 216 | 126 | 226 | 98 | 78 | 264 | 232 | 138 | 172.82 | 134 | 211 | 1.58 | 0.66 | 0.08903378 | 0.24240305 | 0.1985 | 0.026 | 0.1024 | 0.1024 | TRUE | 2.4899 |
| Gm13124 | 1 | 1 | 0 | 4 | 5 | 0 | 1 | 1 | 0 | 3 | 5 | 0 | 1.76 | 1 | 3 | 4.17 | 2.06 | 0.25462646 | NA | 0 | 1.8743 | 1.4374 | 1.4374 | TRUE | 0.1161 |
| Gm13125 | 0 | 0 | 0 | 2 | 0 | 1 | 0 | 0 | 0 | 2 | 0 | 1 | 0.44 | 0 | 1 | 4.782 | 2.258 | 0.50653358 | NA | 0 | 7.4188 | 6.7775 | 6.7775 | TRUE | 0.2804 |
| Gm13128 | 13 | 2 | 8 | 28 | 18 | 13 | 14 | 2 | 8 | 22 | 19 | 14 | 13.24 | 8 | 18 | 2.355 | 1.235 | 0.06318753 | 0.19265295 | 0.1813 | 0.2557 | 0.2277 | 0.2277 | TRUE | 0.4848 |
| Gm13139 | 31 | 18 | 31 | 41 | 35 | 23 | 33 | 18 | 31 | 32 | 38 | 25 | 29.52 | 27 | 32 | 1.162 | 0.216 | 0.58291959 | 0.75875085 | 0.0287 | 0.1185 | 0.0774 | 0.0774 | TRUE | 0.5799 |
| Gm13152 | 713 | 679 | 674 | 897 | 667 | 625 | 757 | 678 | 677 | 705 | 718 | 684 | 703.2 | 704 | 702 | 0.998 | -0.003 | 0.97560411 | 0.98697924 | 9.00E-04 | 0.0116 | 0.0054 | 0.0054 | TRUE | 0.0513 |
| Gm13154 | 2760 | 2417 | 2459 | 3155 | 2287 | 2206 | 2932 | 2414 | 2469 | 2480 | 2461 | 2415 | 2528.38 | 2605 | 2452 | 0.941 | -0.087 | 0.37597593 | 0.59147378 | 0.0055 | 0.0082 | 0.0066 | 0.0066 | TRUE | 0.1457 |
| Gm13157 | 2643 | 2663 | 2838 | 2719 | 2060 | 2037 | 2807 | 2660 | 2850 | 2137 | 2216 | 2230 | 2483.38 | 2772 | 2194 | 0.791 | -0.338 | 1.48E-06 | 3.86E-05 | 6.00E-04 | 0.0082 | 0.0031 | 0.0031 | TRUE | 0.0154 |
| Gm13177 | 0 | 0 | 0 | 0 | 0 | 0 | 0 | 0 | 0 | 0 | 0 | 0 | 0 | 0 | 0 | NA | NA | NA | NA | NA | NA | NA | NA | NA | NA |
| Gm13178 | 4 | 0 | 0 | 0 | 0 | 0 | 4 | 0 | 0 | 0 | 0 | 0 | 0.71 | 1 | 0 | 0.127 | -2.977 | 0.42703158 | NA | 10 | 4.6593 | 8.7652 | 8.7652 | TRUE | 1.899 |
| Gm13212 | 367 | 354 | 359 | 491 | 354 | 297 | 390 | 354 | 360 | 386 | 381 | 325 | 365.97 | 368 | 364 | 0.991 | -0.013 | 0.91907469 | 0.96035815 | 0.0029 | 0.0159 | 0.0089 | 0.0089 | TRUE | 0.0969 |
| Gm1322 | 7 | 3 | 6 | 12 | 21 | 14 | 7 | 3 | 6 | 9 | 23 | 15 | 10.63 | 5 | 16 | 2.861 | 1.516 | 0.02587835 | 0.10412075 | 0 | 0.3167 | 0.2091 | 0.2091 | TRUE | 0.1182 |
| Gm13238 | 2 | 1 | 1 | 4 | 4 | 3 | 2 | 1 | 1 | 3 | 4 | 3 | 2.48 | 1 | 3 | 2.606 | 1.382 | 0.25589788 | NA | 0 | 1.3372 | 0.5554 | 0.5554 | TRUE | 0.1345 |
| Gm13242 | 2453 | 2157 | 2200 | 2514 | 1884 | 1725 | 2606 | 2154 | 2209 | 1976 | 2027 | 1888 | 2143.4 | 2323 | 1964 | 0.846 | -0.242 | 0.01585211 | 0.07388445 | 0.0056 | 0.0084 | 0.0068 | 0.0068 | TRUE | 0.1369 |
| Gm13247 | 2611 | 2327 | 2402 | 4247 | 3149 | 2803 | 2773 | 2324 | 2412 | 3339 | 3388 | 3068 | 2884.05 | 2503 | 3265 | 1.305 | 0.384 | 7.99E-05 | 0.00115354 | 0.0054 | 0.008 | 0.0065 | 0.0065 | TRUE | 0.1079 |
| Gm13251 | 56 | 33 | 31 | 125 | 75 | 42 | 59 | 33 | 31 | 98 | 81 | 46 | 58.08 | 41 | 75 | 1.832 | 0.873 | 0.02440637 | 0.0999318 | 0.1182 | 0.0636 | 0.0895 | 0.0895 | TRUE | 0.3769 |
| Gm13271 | 0 | 0 | 0 | 0 | 0 | 0 | 0 | 0 | 0 | 0 | 0 | 0 | 0 | 0 | 0 | NA | NA | NA | NA | NA | NA | NA | NA | NA | NA |
| Gm13272 | 0 | 0 | 0 | 0 | 0 | 0 | 0 | 0 | 0 | 0 | 0 | 0 | 0 | 0 | 0 | NA | NA | NA | NA | NA | NA | NA | NA | NA | NA |
| Gm13275 | 0 | 0 | 0 | 0 | 0 | 0 | 0 | 0 | 0 | 0 | 0 | 0 | 0 | 0 | 0 | NA | NA | NA | NA | NA | NA | NA | NA | NA | NA |

| | | | | | | | | | | | | | | | | | | | | | | | | | |
|---|---|---|---|---|---|---|---|---|---|---|---|---|---|---|---|---|---|---|---|---|---|---|---|---|---|
| Gm13276 | 0 | 0 | 0 | 0 | 0 | 0 | 0 | 0 | 0 | 0 | 0 | 0 | 0 | 0 | 0 | NA | NA | NA | NA | NA | NA | NA | NA | NA | NA |
| Gm13277 | 0 | 0 | 0 | 0 | 0 | 0 | 0 | 0 | 0 | 0 | 0 | 0 | 0 | 0 | 0 | NA | NA | NA | NA | NA | NA | NA | NA | NA | NA |
| Gm13278 | 0 | 0 | 0 | 0 | 0 | 0 | 0 | 0 | 0 | 0 | 0 | 0 | 0 | 0 | 0 | NA | NA | NA | NA | NA | NA | NA | NA | NA | NA |
| Gm13279 | 0 | 0 | 0 | 0 | 0 | 0 | 0 | 0 | 0 | 0 | 0 | 0 | 0 | 0 | 0 | NA | NA | NA | NA | NA | NA | NA | NA | NA | NA |
| Gm13280 | 0 | 0 | 0 | 0 | 0 | 0 | 0 | 0 | 0 | 0 | 0 | 0 | 0 | 0 | 0 | NA | NA | NA | NA | NA | NA | NA | NA | NA | NA |
| Gm13283 | 0 | 0 | 0 | 0 | 0 | 0 | 0 | 0 | 0 | 0 | 0 | 0 | 0 | 0 | 0 | NA | NA | NA | NA | NA | NA | NA | NA | NA | NA |
| Gm13285 | 0 | 0 | 0 | 0 | 0 | 0 | 0 | 0 | 0 | 0 | 0 | 0 | 0 | 0 | 0 | NA | NA | NA | NA | NA | NA | NA | NA | NA | NA |
| Gm13286 | 0 | 0 | 0 | 0 | 0 | 0 | 0 | 0 | 0 | 0 | 0 | 0 | 0 | 0 | 0 | NA | NA | NA | NA | NA | NA | NA | NA | NA | NA |
| Gm13287 | 0 | 0 | 0 | 0 | 0 | 0 | 0 | 0 | 0 | 0 | 0 | 0 | 0 | 0 | 0 | NA | NA | NA | NA | NA | NA | NA | NA | NA | NA |
| Gm13288 | 0 | 0 | 0 | 0 | 0 | 0 | 0 | 0 | 0 | 0 | 0 | 0 | 0 | 0 | 0 | NA | NA | NA | NA | NA | NA | NA | NA | NA | NA |
| Gm13289 | 0 | 0 | 0 | 0 | 0 | 0 | 0 | 0 | 0 | 0 | 0 | 0 | 0 | 0 | 0 | NA | NA | NA | NA | NA | NA | NA | NA | NA | NA |
| Gm13290 | 0 | 0 | 0 | 0 | 0 | 0 | 0 | 0 | 0 | 0 | 0 | 0 | 0 | 0 | 0 | NA | NA | NA | NA | NA | NA | NA | NA | NA | NA |
| Gm13293 | 0 | 0 | 0 | 0 | 0 | 0 | 0 | 0 | 0 | 0 | 0 | 0 | 0 | 0 | 0 | NA | NA | NA | NA | NA | NA | NA | NA | NA | NA |
| Gm13298 | 0 | 0 | 0 | 0 | 0 | 0 | 0 | 0 | 0 | 0 | 0 | 0 | 0 | 0 | 0 | NA | NA | NA | NA | NA | NA | NA | NA | NA | NA |
| Gm13304 | 0 | 0 | 0 | 0 | 0 | 0 | 0 | 0 | 0 | 0 | 0 | 0 | 0 | 0 | 0 | NA | NA | NA | NA | NA | NA | NA | NA | NA | NA |
| Gm13305 | 0 | 0 | 0 | 0 | 0 | 0 | 0 | 0 | 0 | 0 | 0 | 0 | 0 | 0 | 0 | NA | NA | NA | NA | NA | NA | NA | NA | NA | NA |
| Gm13306 | 0 | 0 | 0 | 0 | 0 | 0 | 0 | 0 | 0 | 0 | 0 | 0 | 0 | 0 | 0 | NA | NA | NA | NA | NA | NA | NA | NA | NA | NA |
| Gm13308 | 0 | 0 | 0 | 0 | 0 | 0 | 0 | 0 | 0 | 0 | 0 | 0 | 0 | 0 | 0 | NA | NA | NA | NA | NA | NA | NA | NA | NA | NA |
| Gm13315 | 1 | 0 | 0 | 1 | 0 | 0 | 1 | 0 | 0 | 1 | 0 | 0 | 0.31 | 0 | 0 | 0.954 | -0.067 | 0.98625928 | NA | 0 | 10.7014 | 9.0384 | 9.0384 | TRUE | 0.4614 |
| Gm1332 | 0 | 0 | 0 | 0 | 1 | 0 | 0 | 0 | 0 | 0 | 1 | 0 | 0.18 | 0 | 0 | 1.859 | 0.894 | 0.82650734 | NA | 0 | 18.3785 | 10 | 10 | TRUE | 0.5204 |
| Gm13363 | 0 | 0 | 0 | 0 | 0 | 0 | 0 | 0 | 0 | 0 | 0 | 0 | 0 | 0 | 0 | NA | NA | NA | NA | NA | NA | NA | NA | NA | NA |
| Gm13375 | 52 | 114 | 110 | 107 | 91 | 104 | 55 | 114 | 110 | 84 | 98 | 114 | 95.9 | 93 | 99 | 1.055 | 0.077 | 0.80084837 | 0.89725514 | 0.0725 | 0.0413 | 0.056 | 0.056 | TRUE | 1.1993 |
| Gm13399 | 0 | 1 | 0 | 0 | 0 | 0 | 0 | 1 | 0 | 0 | 0 | 0 | 0.17 | 0 | 0 | 0.49 | -1.029 | 0.80087 | NA | 0 | 19.7991 | 10 | 10 | TRUE | 0.405 |
| Gm13446 | 2 | 1 | 1 | 3 | 2 | 2 | 2 | 1 | 1 | 2 | 2 | 2 | 1.8 | 1 | 2 | 1.636 | 0.71 | 0.59760572 | NA | 0 | 1.8326 | 0.7125 | 0.7125 | TRUE | 0.1348 |
| Gm13476 | 0 | 1 | 0 | 1 | 0 | 1 | 0 | 1 | 0 | 1 | 0 | 1 | 0.48 | 0 | 1 | 1.723 | 0.785 | 0.78325239 | NA | 0 | 6.8716 | 4.0968 | 4.0968 | TRUE | 0.405 |
| Gm13483 | 0 | 0 | 0 | 0 | 0 | 0 | 0 | 0 | 0 | 0 | 0 | 0 | 0 | 0 | 0 | NA | NA | NA | NA | NA | NA | NA | NA | NA | NA |
| Gm13490 | 0 | 0 | 0 | 0 | 0 | 0 | 0 | 0 | 0 | 0 | 0 | 0 | 0 | 0 | 0 | NA | NA | NA | NA | NA | NA | NA | NA | NA | NA |
| Gm13497 | 0 | 0 | 0 | 0 | 0 | 0 | 0 | 0 | 0 | 0 | 0 | 0 | 0 | 0 | 0 | NA | NA | NA | NA | NA | NA | NA | NA | NA | NA |
| Gm13498 | 24 | 15 | 13 | 74 | 71 | 20 | 25 | 15 | 13 | 58 | 76 | 22 | 35 | 18 | 52 | 2.934 | 1.553 | 0.00274971 | 0.0198029 | 0.2162 | 0.101 | 0.1558 | 0.1558 | TRUE | 0.2266 |
| Gm13539 | 0 | 0 | 0 | 0 | 0 | 0 | 0 | 0 | 0 | 0 | 0 | 0 | 0 | 0 | 0 | NA | NA | NA | NA | NA | NA | NA | NA | NA | NA |
| Gm13544 | 2 | 2 | 4 | 14 | 2 | 7 | 2 | 2 | 4 | 11 | 2 | 8 | 4.83 | 3 | 7 | 2.59 | 1.373 | 0.163865 | 0.35980492 | 0 | 0.6895 | 0.4432 | 0.4432 | TRUE | 0.1868 |
| Gm13546 | 0 | 0 | 0 | 0 | 2 | 0 | 0 | 0 | 0 | 0 | 2 | 0 | 0.36 | 0 | 1 | 3.775 | 1.916 | 0.63423067 | NA | 10 | 9.1927 | 10 | 10 | TRUE | 0.9743 |
| Gm13547 | 0 | 0 | 0 | 0 | 0 | 0 | 0 | 0 | 0 | 0 | 0 | 0 | 0 | 0 | 0 | NA | NA | NA | NA | NA | NA | NA | NA | NA | NA |
| Gm13580 | 11 | 9 | 5 | 20 | 25 | 21 | 12 | 9 | 5 | 16 | 27 | 23 | 15.22 | 9 | 22 | 2.543 | 1.347 | 0.01451588 | 0.06906319 | 0 | 0.2234 | 0.1364 | 0.1364 | TRUE | 0.3464 |
| Gm13582 | 0 | 0 | 0 | 0 | 0 | 0 | 0 | 0 | 0 | 0 | 0 | 0 | 0 | 0 | 0 | NA | NA | NA | NA | NA | NA | NA | NA | NA | NA |
| Gm136 | 0 | 0 | 0 | 0 | 2 | 0 | 0 | 0 | 0 | 0 | 2 | 0 | 0.36 | 0 | 1 | 3.775 | 1.916 | 0.63423067 | NA | 10 | 9.1927 | 10 | 10 | TRUE | 0.9743 |
| Gm13629 | 0 | 0 | 0 | 0 | 0 | 0 | 0 | 0 | 0 | 0 | 0 | 0 | 0 | 0 | 0 | NA | NA | NA | NA | NA | NA | NA | NA | NA | NA |
| Gm13704 | 4 | 2 | 3 | 5 | 5 | 2 | 4 | 2 | 3 | 4 | 5 | 2 | 3.46 | 3 | 4 | 1.249 | 0.32 | 0.74988628 | NA | 0 | 0.9591 | 0.4342 | 0.4342 | TRUE | 0.1799 |
| Gm13710 | 1 | 2 | 1 | 4 | 3 | 1 | 1 | 2 | 1 | 3 | 3 | 1 | 1.92 | 1 | 2 | 1.852 | 0.889 | 0.50929609 | NA | 0 | 1.7212 | 0.7378 | 0.7378 | TRUE | 0.2325 |
| Gm13718 | 121 | 94 | 86 | 164 | 144 | 124 | 129 | 94 | 86 | 129 | 155 | 136 | 121.39 | 103 | 140 | 1.359 | 0.442 | 0.04260026 | 0.14737689 | 0.017 | 0.034 | 0.0258 | 0.0258 | TRUE | 0.4572 |
| Gm13749 | 0 | 0 | 0 | 0 | 0 | 0 | 0 | 0 | 0 | 0 | 0 | 0 | 0 | 0 | 0 | NA | NA | NA | NA | NA | NA | NA | NA | NA | NA |
| Gm13752 | 8 | 4 | 5 | 12 | 8 | 9 | 8 | 4 | 5 | 9 | 9 | 10 | 7.57 | 6 | 9 | 1.6 | 0.678 | 0.32779198 | 0.54900652 | 0 | 0.4422 | 0.2056 | 0.2056 | TRUE | 0.3462 |
| Gm13769 | 0 | 0 | 0 | 0 | 0 | 0 | 0 | 0 | 0 | 0 | 0 | 0 | 0 | 0 | 0 | NA | NA | NA | NA | NA | NA | NA | NA | NA | NA |
| Gm13807 | 0 | 0 | 0 | 0 | 0 | 0 | 0 | 0 | 0 | 0 | 0 | 0 | 0 | 0 | 0 | NA | NA | NA | NA | NA | NA | NA | NA | NA | NA |
| Gm13826 | 270 | 337 | 340 | 403 | 257 | 303 | 287 | 337 | 341 | 317 | 277 | 332 | 314.96 | 322 | 309 | 0.959 | -0.061 | 0.66650842 | 0.81350958 | 0.0054 | 0.0174 | 0.0111 | 0.0111 | TRUE | 0.1004 |
| Gm13845 | 4 | 3 | 0 | 5 | 1 | 3 | 4 | 3 | 0 | 4 | 1 | 3 | 2.59 | 2 | 3 | 1.163 | 0.218 | 0.86891073 | NA | 0 | 1.2793 | 0.8726 | 0.8726 | TRUE | 0.7452 |
| Gm13871 | 0 | 0 | 0 | 0 | 0 | 0 | 0 | 0 | 0 | 0 | 0 | 0 | 0 | 0 | 0 | NA | NA | NA | NA | NA | NA | NA | NA | NA | NA |
| Gm13889 | 2 | 10 | 16 | 18 | 9 | 16 | 2 | 10 | 16 | 14 | 10 | 18 | 11.59 | 9 | 14 | 1.462 | 0.548 | 0.44341778 | 0.65026362 | 0.2606 | 0.2912 | 0.2783 | 0.2783 | TRUE | 0.3746 |
| Gm13939 | 26 | 23 | 19 | 35 | 26 | 27 | 28 | 23 | 19 | 28 | 28 | 30 | 25.78 | 23 | 29 | 1.221 | 0.288 | 0.4416362 | 0.6486159 | 0 | 0.1347 | 0.0618 | 0.0618 | TRUE | 0.1543 |
| Gm13944 | 0 | 0 | 0 | 0 | 0 | 0 | 0 | 0 | 0 | 0 | 0 | 0 | 0 | 0 | 0 | NA | NA | NA | NA | NA | NA | NA | NA | NA | NA |
| Gm14005 | 14 | 14 | 8 | 18 | 9 | 13 | 15 | 14 | 8 | 14 | 10 | 14 | 12.49 | 12 | 13 | 1.037 | 0.053 | 0.92339975 | 0.96235363 | 0 | 0.2706 | 0.1394 | 0.1394 | TRUE | 0.3728 |
| Gm14015 | 0 | 0 | 0 | 0 | 0 | 0 | 0 | 0 | 0 | 0 | 0 | 0 | 0 | 0 | 0 | NA | NA | NA | NA | NA | NA | NA | NA | NA | NA |
| Gm14023 | 21 | 32 | 22 | 36 | 24 | 25 | 22 | 32 | 22 | 28 | 26 | 27 | 26.31 | 25 | 27 | 1.068 | 0.095 | 0.80156748 | 0.89757347 | 0 | 0.1321 | 0.064 | 0.064 | TRUE | 0.3116 |
| Gm14047 | 1 | 1 | 1 | 0 | 2 | 2 | 1 | 1 | 1 | 0 | 2 | 2 | 1.23 | 1 | 1 | 1.366 | 0.45 | 0.79674244 | NA | 0 | 2.6758 | 1.3453 | 1.3453 | TRUE | 0.7169 |
| Gm14057 | 77 | 79 | 64 | 111 | 75 | 66 | 82 | 79 | 64 | 87 | 81 | 72 | 77.53 | 75 | 80 | 1.072 | 0.1 | 0.66449211 | 0.81224285 | 0 | 0.0494 | 0.0254 | 0.0254 | TRUE | 0.1443 |
| Gm14085 | 2 | 1 | 0 | 2 | 0 | 0 | 2 | 1 | 0 | 2 | 0 | 0 | 0.78 | 1 | 1 | 0.544 | -0.878 | 0.72447748 | NA | 0 | 4.2168 | 3.1134 | 3.1134 | TRUE | 0.4055 |
| Gm14092 | 0 | 0 | 0 | 0 | 0 | 0 | 0 | 0 | 0 | 0 | 0 | 0 | 0 | 0 | 0 | NA | NA | NA | NA | NA | NA | NA | NA | NA | NA |
| Gm14124 | 766 | 403 | 366 | 927 | 644 | 429 | 814 | 402 | 368 | 729 | 693 | 470 | 579.14 | 528 | 631 | 1.195 | 0.257 | 0.37631385 | 0.59174697 | 0.1222 | 0.0126 | 0.0592 | 0.0592 | TRUE | 2.6142 |
| Gm14137 | 5 | 6 | 4 | 3 | 3 | 2 | 5 | 6 | 4 | 2 | 3 | 2 | 3.85 | 5 | 2 | 0.505 | -0.984 | 0.29917847 | NA | 0 | 0.8628 | 0.3591 | 0.3591 | TRUE | 0.073 |
| Gm14139 | 487 | 269 | 266 | 637 | 433 | 273 | 517 | 269 | 267 | 501 | 466 | 299 | 386.42 | 351 | 422 | 1.203 | 0.267 | 0.34925394 | 0.5680554 | 0.1076 | 0.0154 | 0.056 | 0.056 | TRUE | 1.9536 |
| Gm14151 | 0 | 1 | 2 | 0 | 1 | 3 | 0 | 1 | 2 | 0 | 1 | 3 | 1.23 | 1 | 1 | 1.399 | 0.484 | 0.80516646 | NA | 0 | 2.6901 | 1.9261 | 1.9261 | TRUE | 0.4068 |
| Gm14164 | 37 | 31 | 21 | 23 | 16 | 18 | 39 | 31 | 21 | 18 | 17 | 20 | 24.39 | 30 | 18 | 0.603 | -0.731 | 0.07966222 | 0.22396301 | 0 | 0.142 | 0.0818 | 0.0818 | TRUE | 0.1542 |
| Gm14169 | 6 | 18 | 18 | 8 | 5 | 10 | 6 | 18 | 18 | 6 | 5 | 11 | 10.84 | 14 | 7 | 0.528 | -0.922 | 0.16823261 | 0.36603183 | 0.1236 | 0.3108 | 0.2213 | 0.2213 | TRUE | 0.97 |
| Gm14204 | 4 | 2 | 3 | 2 | 1 | 1 | 4 | 2 | 3 | 2 | 1 | 1 | 2.17 | 3 | 1 | 0.412 | -1.281 | 0.31669158 | NA | 0 | 1.5273 | 0.6289 | 0.6289 | TRUE | 0.1371 |
| Gm14207 | 0 | 4 | 5 | 1 | 1 | 2 | 0 | 4 | 5 | 1 | 1 | 2 | 2.18 | 3 | 1 | 0.439 | -1.189 | 0.42068711 | NA | 0 | 1.5197 | 1.034 | 1.034 | TRUE | 0.9387 |
| Gm14288 | 0 | 0 | 0 | 0 | 0 | 0 | 0 | 0 | 0 | 0 | 0 | 0 | 0 | 0 | 0 | NA | NA | NA | NA | NA | NA | NA | NA | NA | NA |
| Gm14295 | 202 | 223 | 228 | 238 | 129 | 195 | 215 | 223 | 229 | 187 | 139 | 213 | 200.93 | 222 | 180 | 0.81 | -0.304 | 0.10351073 | 0.26764507 | 0.0167 | 0.0233 | 0.0201 | 0.0201 | TRUE | 0.4088 |
| Gm14305 | 141 | 163 | 178 | 231 | 173 | 172 | 150 | 163 | 179 | 182 | 186 | 188 | 174.55 | 164 | 185 | 1.13 | 0.177 | 0.25775172 | 0.47651994 | 0 | 0.0258 | 0.0118 | 0.0118 | TRUE | 0.0678 |
| Gm14306 | 0 | 0 | 0 | 0 | 0 | 0 | 0 | 0 | 0 | 0 | 0 | 0 | 0 | 0 | 0 | NA | NA | NA | NA | NA | NA | NA | NA | NA | NA |

| | | | | | | | | | | | | | | | | | | | | | | | | | |
|---|---|---|---|---|---|---|---|---|---|---|---|---|---|---|---|---|---|---|---|---|---|---|---|---|---|
| Gm14308 | 0 | 0 | 0 | 0 | 0 | 0 | 0 | 0 | 0 | 0 | 0 | 0 | 0 | 0 | 0 | NA | NA | NA | NA | NA | NA | NA | NA | NA | NA |
| Gm14318 | 0 | 0 | 1 | 0 | 0 | 0 | 0 | 0 | 1 | 0 | 0 | 0 | 0.17 | 0 | 0 | 0.49 | -1.029 | 0.80087 | NA | 0 | 19.6928 | 10 | 10 | TRUE | 0.4098 |
| Gm14322 | 295 | 252 | 250 | 427 | 268 | 233 | 313 | 252 | 251 | 336 | 288 | 255 | 282.52 | 272 | 293 | 1.081 | 0.112 | 0.50142599 | 0.69684472 | 0.0144 | 0.0186 | 0.0164 | 0.0164 | TRUE | 0.1959 |
| Gm14325 | 500 | 560 | 607 | 853 | 649 | 574 | 531 | 559 | 609 | 671 | 698 | 628 | 616.17 | 566 | 666 | 1.175 | 0.232 | 0.02850586 | 0.11106583 | 0.002 | 0.0122 | 0.0065 | 0.0065 | TRUE | 0.0514 |
| Gm14326 | 940 | 900 | 837 | 1282 | 950 | 818 | 998 | 899 | 840 | 1008 | 1022 | 895 | 943.85 | 912 | 975 | 1.069 | 0.097 | 0.37357099 | 0.58997634 | 0.0048 | 0.0104 | 0.0074 | 0.0074 | TRUE | 0.0803 |
| Gm14327 | 4 | 1 | 0 | 1 | 2 | 2 | 4 | 1 | 0 | 1 | 2 | 2 | 1.73 | 2 | 2 | 0.968 | -0.047 | 0.9761174 | NA | 0.4192 | 1.9122 | 1.2074 | 1.2074 | TRUE | 1.0763 |
| Gm14345 | 0 | 0 | 0 | 0 | 0 | 0 | 0 | 0 | 0 | 0 | 0 | 0 | 0 | 0 | 0 | NA | NA | NA | NA | NA | NA | NA | NA | NA | NA |
| Gm14346 | 0 | 0 | 0 | 0 | 0 | 0 | 0 | 0 | 0 | 0 | 0 | 0 | 0 | 0 | 0 | NA | NA | NA | NA | NA | NA | NA | NA | NA | NA |
| Gm14347 | 0 | 0 | 0 | 0 | 0 | 0 | 0 | 0 | 0 | 0 | 0 | 0 | 0 | 0 | 0 | NA | NA | NA | NA | NA | NA | NA | NA | NA | NA |
| Gm14351 | 0 | 0 | 0 | 0 | 0 | 0 | 0 | 0 | 0 | 0 | 0 | 0 | 0 | 0 | 0 | NA | NA | NA | NA | NA | NA | NA | NA | NA | NA |
| Gm14354 | 0 | 0 | 0 | 0 | 0 | 0 | 0 | 0 | 0 | 0 | 0 | 0 | 0 | 0 | 0 | NA | NA | NA | NA | NA | NA | NA | NA | NA | NA |
| Gm14374 | 0 | 0 | 0 | 0 | 0 | 0 | 0 | 0 | 0 | 0 | 0 | 0 | 0 | 0 | 0 | NA | NA | NA | NA | NA | NA | NA | NA | NA | NA |
| Gm14378 | 9 | 9 | 15 | 17 | 11 | 22 | 10 | 9 | 15 | 13 | 12 | 24 | 13.82 | 11 | 16 | 1.456 | 0.542 | 0.33424587 | 0.5548783 | 0.0534 | 0.2454 | 0.1515 | 0.1515 | TRUE | 0.7411 |
| Gm14379 | 0 | 0 | 0 | 5 | 0 | 2 | 0 | 0 | 0 | 4 | 0 | 2 | 1.02 | 0 | 2 | 10.974 | 3.456 | 0.18527553 | NA | 5.9789 | 3.2368 | 3.6669 | 3.6669 | TRUE | 0.0669 |
| Gm14391 | 0 | 0 | 0 | 0 | 0 | 0 | 0 | 0 | 0 | 0 | 0 | 0 | 0 | 0 | 0 | NA | NA | NA | NA | NA | NA | NA | NA | NA | NA |
| Gm14393 | 76 | 57 | 52 | 88 | 76 | 64 | 81 | 57 | 52 | 69 | 82 | 70 | 68.48 | 63 | 74 | 1.164 | 0.219 | 0.40909643 | 0.62130006 | 0.0153 | 0.055 | 0.0361 | 0.0361 | TRUE | 0.4965 |
| Gm14403 | 188 | 140 | 132 | 386 | 268 | 202 | 200 | 140 | 133 | 303 | 288 | 221 | 214.16 | 158 | 271 | 1.726 | 0.788 | 0.00022224 | 0.00264542 | 0.033 | 0.0223 | 0.0277 | 0.0277 | TRUE | 0.5774 |
| Gm14405 | 0 | 0 | 0 | 1 | 0 | 0 | 0 | 0 | 0 | 1 | 0 | 0 | 0.13 | 0 | 0 | 1.859 | 0.894 | 0.82650734 | NA | 0 | 25.1515 | 10 | 10 | TRUE | 0.2563 |
| Gm14420 | 296 | 182 | 228 | 393 | 290 | 251 | 314 | 182 | 229 | 309 | 312 | 275 | 270.14 | 242 | 299 | 1.237 | 0.307 | 0.13855699 | 0.3230815 | 0.0355 | 0.0191 | 0.0271 | 0.0271 | TRUE | 0.5375 |
| Gm14430 | 0 | 0 | 0 | 0 | 0 | 0 | 0 | 0 | 0 | 0 | 0 | 0 | 0 | 0 | 0 | NA | NA | NA | NA | NA | NA | NA | NA | NA | NA |
| Gm14431 | 0 | 0 | 0 | 0 | 0 | 0 | 0 | 0 | 0 | 0 | 0 | 0 | 0 | 0 | 0 | NA | NA | NA | NA | NA | NA | NA | NA | NA | NA |
| Gm14434 | 0 | 0 | 0 | 0 | 0 | 0 | 0 | 0 | 0 | 0 | 0 | 0 | 0 | 0 | 0 | NA | NA | NA | NA | NA | NA | NA | NA | NA | NA |
| Gm14436 | 0 | 0 | 0 | 1 | 1 | 0 | 0 | 0 | 0 | 1 | 1 | 0 | 0.31 | 0 | 1 | 3.32 | 1.731 | 0.65383052 | NA | 0 | 10.6224 | 8.9541 | 8.9541 | TRUE | 0.202 |
| Gm14440 | 1 | 0 | 1 | 4 | 2 | 0 | 1 | 0 | 1 | 3 | 2 | 0 | 1.23 | 1 | 2 | 2.631 | 1.396 | 0.4715886 | NA | 0 | 2.6916 | 1.6909 | 1.6909 | TRUE | 0.3147 |
| Gm14446 | 11 | 9 | 7 | 5 | 5 | 3 | 12 | 9 | 7 | 4 | 5 | 3 | 6.72 | 9 | 4 | 0.454 | -1.139 | 0.12634692 | 0.30479156 | 0 | 0.4974 | 0.2281 | 0.2281 | TRUE | 0.1684 |
| Gm14458 | 53 | 52 | 39 | 203 | 162 | 114 | 56 | 52 | 39 | 160 | 174 | 125 | 101.01 | 49 | 153 | 3.118 | 1.641 | 2.24E-11 | 1.85E-09 | 0.0187 | 0.0395 | 0.0297 | 0.0297 | TRUE | 0.2604 |
| Gm14459 | 0 | 0 | 0 | 0 | 0 | 0 | 0 | 0 | 0 | 0 | 0 | 0 | 0 | 0 | 0 | NA | NA | NA | NA | NA | NA | NA | NA | NA | NA |
| Gm14461 | 0 | 0 | 0 | 0 | 0 | 0 | 0 | 0 | 0 | 0 | 0 | 0 | 0 | 0 | 0 | NA | NA | NA | NA | NA | NA | NA | NA | NA | NA |
| Gm14474 | 0 | 0 | 0 | 0 | 0 | 0 | 0 | 0 | 0 | 0 | 0 | 0 | 0 | 0 | 0 | NA | NA | NA | NA | NA | NA | NA | NA | NA | NA |
| Gm14475 | 0 | 0 | 0 | 0 | 0 | 0 | 0 | 0 | 0 | 0 | 0 | 0 | 0 | 0 | 0 | NA | NA | NA | NA | NA | NA | NA | NA | NA | NA |
| Gm14476 | 0 | 0 | 0 | 0 | 0 | 0 | 0 | 0 | 0 | 0 | 0 | 0 | 0 | 0 | 0 | NA | NA | NA | NA | NA | NA | NA | NA | NA | NA |
| Gm14477 | 0 | 0 | 0 | 0 | 0 | 0 | 0 | 0 | 0 | 0 | 0 | 0 | 0 | 0 | 0 | NA | NA | NA | NA | NA | NA | NA | NA | NA | NA |
| Gm14478 | 0 | 0 | 0 | 0 | 0 | 0 | 0 | 0 | 0 | 0 | 0 | 0 | 0 | 0 | 0 | NA | NA | NA | NA | NA | NA | NA | NA | NA | NA |
| Gm14479 | 0 | 0 | 0 | 0 | 0 | 0 | 0 | 0 | 0 | 0 | 0 | 0 | 0 | 0 | 0 | NA | NA | NA | NA | NA | NA | NA | NA | NA | NA |
| Gm14482 | 0 | 0 | 0 | 0 | 0 | 0 | 0 | 0 | 0 | 0 | 0 | 0 | 0 | 0 | 0 | NA | NA | NA | NA | NA | NA | NA | NA | NA | NA |
| Gm14483 | 0 | 0 | 0 | 0 | 0 | 0 | 0 | 0 | 0 | 0 | 0 | 0 | 0 | 0 | 0 | NA | NA | NA | NA | NA | NA | NA | NA | NA | NA |
| Gm14484 | 0 | 0 | 0 | 0 | 0 | 0 | 0 | 0 | 0 | 0 | 0 | 0 | 0 | 0 | 0 | NA | NA | NA | NA | NA | NA | NA | NA | NA | NA |
| Gm14492 | 0 | 0 | 1 | 0 | 2 | 0 | 0 | 0 | 1 | 0 | 2 | 0 | 0.53 | 0 | 1 | 1.927 | 0.947 | 0.77736506 | NA | 0 | 6.2701 | 6.3814 | 6.3814 | TRUE | 0.9744 |
| Gm14496 | 0 | 0 | 0 | 0 | 0 | 0 | 0 | 0 | 0 | 0 | 0 | 0 | 0 | 0 | 0 | NA | NA | NA | NA | NA | NA | NA | NA | NA | NA |
| Gm14501 | 0 | 2 | 3 | 0 | 0 | 2 | 0 | 2 | 3 | 0 | 0 | 2 | 1.2 | 2 | 1 | 0.418 | -1.259 | 0.56996925 | NA | 1.8421 | 2.7526 | 2.5385 | 2.5385 | TRUE | 0.714 |
| Gm14511 | 0 | 7 | 4 | 8 | 9 | 8 | 0 | 7 | 4 | 6 | 10 | 9 | 5.96 | 4 | 8 | 2.218 | 1.149 | 0.22627917 | 0.43908403 | 0.1878 | 0.56 | 0.4523 | 0.4523 | TRUE | 0.5794 |
| Gm14525 | 0 | 0 | 0 | 0 | 1 | 0 | 0 | 0 | 0 | 0 | 1 | 0 | 0.18 | 0 | 0 | 1.859 | 0.894 | 0.82650734 | NA | 0 | 18.3785 | 10 | 10 | TRUE | 0.5204 |
| Gm14548 | 0 | 0 | 0 | 0 | 0 | 0 | 0 | 0 | 0 | 0 | 0 | 0 | 0 | 0 | 0 | NA | NA | NA | NA | NA | NA | NA | NA | NA | NA |
| Gm14625 | 0 | 0 | 0 | 0 | 1 | 0 | 0 | 0 | 0 | 0 | 1 | 0 | 0.18 | 0 | 0 | 1.859 | 0.894 | 0.82650734 | NA | 0 | 18.3785 | 10 | 10 | TRUE | 0.5204 |
| Gm14632 | 0 | 0 | 0 | 0 | 0 | 0 | 0 | 0 | 0 | 0 | 0 | 0 | 0 | 0 | 0 | NA | NA | NA | NA | NA | NA | NA | NA | NA | NA |
| Gm14634 | 1 | 6 | 2 | 5 | 7 | 3 | 1 | 6 | 2 | 4 | 8 | 3 | 3.97 | 3 | 5 | 1.61 | 0.687 | 0.5067017 | NA | 0.1286 | 0.8371 | 0.5026 | 0.5026 | TRUE | 0.9793 |
| Gm14635 | 0 | 0 | 0 | 1 | 0 | 0 | 0 | 0 | 0 | 1 | 0 | 0 | 0.13 | 0 | 0 | 1.859 | 0.894 | 0.82650734 | NA | 0 | 25.1515 | 10 | 10 | TRUE | 0.2563 |
| Gm14685 | 0 | 0 | 0 | 0 | 0 | 0 | 0 | 0 | 0 | 0 | 0 | 0 | 0 | 0 | 0 | NA | NA | NA | NA | NA | NA | NA | NA | NA | NA |
| Gm14692 | 0 | 0 | 0 | 0 | 0 | 0 | 0 | 0 | 0 | 0 | 0 | 0 | 0 | 0 | 0 | NA | NA | NA | NA | NA | NA | NA | NA | NA | NA |
| Gm14718 | 0 | 0 | 0 | 0 | 0 | 0 | 0 | 0 | 0 | 0 | 0 | 0 | 0 | 0 | 0 | NA | NA | NA | NA | NA | NA | NA | NA | NA | NA |
| Gm14725 | 0 | 0 | 0 | 0 | 0 | 0 | 0 | 0 | 0 | 0 | 0 | 0 | 0 | 0 | 0 | NA | NA | NA | NA | NA | NA | NA | NA | NA | NA |
| Gm14743 | 0 | 0 | 0 | 0 | 0 | 0 | 0 | 0 | 0 | 0 | 0 | 0 | 0 | 0 | 0 | NA | NA | NA | NA | NA | NA | NA | NA | NA | NA |
| Gm14744 | 0 | 0 | 0 | 0 | 0 | 0 | 0 | 0 | 0 | 0 | 0 | 0 | 0 | 0 | 0 | NA | NA | NA | NA | NA | NA | NA | NA | NA | NA |
| Gm14781 | 0 | 0 | 0 | 0 | 0 | 0 | 0 | 0 | 0 | 0 | 0 | 0 | 0 | 0 | 0 | NA | NA | NA | NA | NA | NA | NA | NA | NA | NA |
| Gm14812 | 0 | 0 | 0 | 0 | 0 | 0 | 0 | 0 | 0 | 0 | 0 | 0 | 0 | 0 | 0 | NA | NA | NA | NA | NA | NA | NA | NA | NA | NA |
| Gm14819 | 0 | 0 | 0 | 0 | 0 | 0 | 0 | 0 | 0 | 0 | 0 | 0 | 0 | 0 | 0 | NA | NA | NA | NA | NA | NA | NA | NA | NA | NA |
| Gm14827 | 3 | 4 | 0 | 8 | 7 | 4 | 3 | 4 | 0 | 6 | 8 | 4 | 4.23 | 2 | 6 | 2.543 | 1.347 | 0.20078331 | NA | 0 | 0.7857 | 0.5041 | 0.5041 | TRUE | 0.8272 |
| Gm14850 | 0 | 0 | 0 | 0 | 0 | 0 | 0 | 0 | 0 | 0 | 0 | 0 | 0 | 0 | 0 | NA | NA | NA | NA | NA | NA | NA | NA | NA | NA |
| Gm14851 | 0 | 0 | 0 | 0 | 0 | 0 | 0 | 0 | 0 | 0 | 0 | 0 | 0 | 0 | 0 | NA | NA | NA | NA | NA | NA | NA | NA | NA | NA |
| Gm14858 | 0 | 0 | 0 | 1 | 0 | 0 | 0 | 0 | 0 | 1 | 0 | 0 | 0.13 | 0 | 0 | 1.859 | 0.894 | 0.82650734 | NA | 0 | 25.1515 | 10 | 10 | TRUE | 0.2563 |
| Gm14872 | 4 | 10 | 8 | 9 | 9 | 3 | 4 | 10 | 8 | 7 | 10 | 3 | 7.05 | 7 | 7 | 0.9 | -0.152 | 0.84197538 | 0.92021591 | 0 | 0.4741 | 0.275 | 0.275 | TRUE | 0.2996 |
| Gm14873 | 52 | 62 | 55 | 65 | 62 | 45 | 55 | 62 | 55 | 51 | 67 | 49 | 56.57 | 57 | 56 | 0.966 | -0.049 | 0.85246463 | 0.92628008 | 0 | 0.0651 | 0.0333 | 0.0333 | TRUE | 0.2389 |
| Gm14920 | 0 | 0 | 0 | 0 | 0 | 0 | 0 | 0 | 0 | 0 | 0 | 0 | 0 | 0 | 0 | NA | NA | NA | NA | NA | NA | NA | NA | NA | NA |
| Gm15008 | 0 | 0 | 0 | 0 | 0 | 1 | 0 | 0 | 0 | 0 | 0 | 1 | 0.18 | 0 | 0 | 1.859 | 0.894 | 0.82650734 | NA | 0 | 18.0643 | 10 | 10 | TRUE | 0.5383 |
| Gm15023 | 0 | 0 | 0 | 0 | 0 | 0 | 0 | 0 | 0 | 0 | 0 | 0 | 0 | 0 | 0 | NA | NA | NA | NA | NA | NA | NA | NA | NA | NA |
| Gm15056 | 0 | 0 | 0 | 0 | 0 | 0 | 0 | 0 | 0 | 0 | 0 | 0 | 0 | 0 | 0 | NA | NA | NA | NA | NA | NA | NA | NA | NA | NA |
| Gm15070 | 14 | 4 | 7 | 13 | 13 | 6 | 15 | 4 | 7 | 10 | 14 | 7 | 9.44 | 9 | 10 | 1.195 | 0.256 | 0.71847952 | 0.84755863 | 0.1491 | 0.3557 | 0.2576 | 0.2576 | TRUE | 0.6054 |
| Gm15080 | 0 | 0 | 0 | 0 | 0 | 0 | 0 | 0 | 0 | 0 | 0 | 0 | 0 | 0 | 0 | NA | NA | NA | NA | NA | NA | NA | NA | NA | NA |
| Gm15085 | 0 | 0 | 0 | 0 | 0 | 0 | 0 | 0 | 0 | 0 | 0 | 0 | 0 | 0 | 0 | NA | NA | NA | NA | NA | NA | NA | NA | NA | NA |

| | | | | | | | | | | | | | | | | | | | | | | | | |
|---|---|---|---|---|---|---|---|---|---|---|---|---|---|---|---|---|---|---|---|---|---|---|---|---|
| Gm15091 | 2 | 6 | 3 | 24 | 21 | 19 | 2 | 6 | 3 | 19 | 23 | 21 | 12.23 | 4 | 21 | 5.549 | 2.472 | 0.0001487 | 0.00188841 | 0 | 0.2762 | 0.1456 | 0.1456 | TRUE | 0.4605 |
| Gm15093 | 0 | 0 | 0 | 0 | 0 | 0 | 0 | 0 | 0 | 0 | 0 | 0 | 0 | 0 | 0 | NA | NA | NA | NA | NA | NA | NA | NA | NA | NA |
| Gm15097 | 23 | 27 | 15 | 83 | 67 | 114 | 24 | 27 | 15 | 65 | 72 | 125 | 54.76 | 22 | 87 | 3.933 | 1.975 | 2.62E-07 | 8.41E-06 | 0.0888 | 0.0671 | 0.0774 | 0.0774 | TRUE | 1.3015 |
| Gm15104 | 1 | 0 | 0 | 2 | 0 | 0 | 1 | 0 | 0 | 2 | 0 | 0 | 0.44 | 0 | 1 | 1.494 | 0.579 | 0.86844164 | NA | 0 | 7.5101 | 6.9157 | 6.9157 | TRUE | 0.9721 |
| Gm15107 | 0 | 2 | 2 | 10 | 11 | 8 | 0 | 2 | 2 | 8 | 12 | 9 | 5.41 | 1 | 10 | 6.981 | 2.803 | 0.006562 | 0.03834601 | 0 | 0.6159 | 0.3373 | 0.3373 | TRUE | 0.4306 |
| Gm15114 | 1 | 0 | 0 | 2 | 0 | 1 | 1 | 0 | 0 | 2 | 0 | 1 | 0.62 | 0 | 1 | 2.474 | 1.307 | 0.61898859 | NA | 0 | 5.3076 | 3.4358 | 3.4358 | TRUE | 0.4614 |
| Gm15127 | 2 | 1 | 0 | 9 | 6 | 8 | 2 | 1 | 0 | 7 | 6 | 9 | 4.24 | 1 | 7 | 7.226 | 2.853 | 0.01441723 | NA | 0 | 0.7848 | 0.4167 | 0.4167 | TRUE | 0.3774 |
| Gm15133 | 0 | 0 | 0 | 0 | 0 | 0 | 0 | 0 | 0 | 0 | 0 | 0 | 0 | 0 | 0 | NA | NA | NA | NA | NA | NA | NA | NA | NA | NA |
| Gm15140 | 0 | 0 | 0 | 0 | 0 | 0 | 0 | 0 | 0 | 0 | 0 | 0 | 0 | 0 | 0 | NA | NA | NA | NA | NA | NA | NA | NA | NA | NA |
| Gm15179 | 1 | 2 | 1 | 0 | 0 | 1 | 1 | 2 | 1 | 0 | 0 | 1 | 0.86 | 1 | 0 | 0.257 | -1.958 | 0.35814405 | NA | 0 | 3.8389 | 1.8948 | 1.8948 | TRUE | 0.5383 |
| Gm15217 | 1 | 1 | 1 | 2 | 0 | 1 | 1 | 1 | 1 | 2 | 0 | 1 | 0.96 | 1 | 1 | 0.899 | -0.153 | 0.93410618 | NA | 0 | 3.4554 | 1.4437 | 1.4437 | TRUE | 0.2848 |
| Gm1527 | 0 | 0 | 0 | 1 | 1 | 0 | 0 | 0 | 0 | 1 | 1 | 0 | 0.31 | 0 | 1 | 3.32 | 1.731 | 0.65383052 | NA | 0 | 10.6224 | 8.9541 | 8.9541 | TRUE | 0.202 |
| Gm15284 | 1 | 1 | 1 | 3 | 2 | 3 | 1 | 1 | 1 | 2 | 2 | 3 | 1.81 | 1 | 2 | 2.534 | 1.342 | 0.3349048 | NA | 0 | 1.8272 | 0.7061 | 0.7061 | TRUE | 0.0541 |
| Gm15292 | 0 | 0 | 0 | 0 | 0 | 0 | 0 | 0 | 0 | 0 | 0 | 0 | 0 | 0 | 0 | NA | NA | NA | NA | NA | NA | NA | NA | NA | NA |
| Gm15293 | 0 | 0 | 0 | 0 | 0 | 0 | 0 | 0 | 0 | 0 | 0 | 0 | 0 | 0 | 0 | NA | NA | NA | NA | NA | NA | NA | NA | NA | NA |
| Gm15299 | 0 | 0 | 0 | 0 | 0 | 0 | 0 | 0 | 0 | 0 | 0 | 0 | 0 | 0 | 0 | NA | NA | NA | NA | NA | NA | NA | NA | NA | NA |
| Gm15308 | 0 | 0 | 0 | 0 | 0 | 0 | 0 | 0 | 0 | 0 | 0 | 0 | 0 | 0 | 0 | NA | NA | NA | NA | NA | NA | NA | NA | NA | NA |
| Gm15315 | 0 | 0 | 0 | 0 | 0 | 0 | 0 | 0 | 0 | 0 | 0 | 0 | 0 | 0 | 0 | NA | NA | NA | NA | NA | NA | NA | NA | NA | NA |
| Gm15319 | 0 | 0 | 0 | 0 | 0 | 0 | 0 | 0 | 0 | 0 | 0 | 0 | 0 | 0 | 0 | NA | NA | NA | NA | NA | NA | NA | NA | NA | NA |
| Gm15328 | 0 | 0 | 1 | 1 | 0 | 0 | 0 | 0 | 1 | 1 | 0 | 0 | 0.3 | 0 | 0 | 0.954 | -0.067 | 0.98640617 | NA | 0 | 11.0483 | 9.2783 | 9.2783 | TRUE | 0.4098 |
| Gm15348 | 0 | 0 | 0 | 0 | 0 | 0 | 0 | 0 | 0 | 0 | 0 | 0 | 0 | 0 | 0 | NA | NA | NA | NA | NA | NA | NA | NA | NA | NA |
| Gm15350 | 0 | 0 | 0 | 0 | 0 | 0 | 0 | 0 | 0 | 0 | 0 | 0 | 0 | 0 | 0 | NA | NA | NA | NA | NA | NA | NA | NA | NA | NA |
| Gm15386 | 0 | 0 | 0 | 0 | 0 | 0 | 0 | 0 | 0 | 0 | 0 | 0 | 0 | 0 | 0 | NA | NA | NA | NA | NA | NA | NA | NA | NA | NA |
| Gm15401 | 0 | 5 | 4 | 0 | 0 | 2 | 0 | 5 | 4 | 0 | 0 | 2 | 1.87 | 3 | 1 | 0.231 | -2.115 | 0.28932596 | NA | 2.6535 | 1.7719 | 2 | 2 | TRUE | 0.9752 |
| Gm15408 | 15 | 9 | 11 | 16 | 6 | 14 | 16 | 9 | 11 | 13 | 6 | 15 | 11.72 | 12 | 11 | 0.96 | -0.059 | 0.92111599 | 0.96116102 | 0 | 0.288 | 0.1694 | 0.1694 | TRUE | 0.504 |
| Gm15412 | 0 | 0 | 0 | 0 | 0 | 0 | 0 | 0 | 0 | 0 | 0 | 0 | 0 | 0 | 0 | NA | NA | NA | NA | NA | NA | NA | NA | NA | NA |
| Gm15413 | 0 | 1 | 0 | 0 | 1 | 0 | 0 | 1 | 0 | 0 | 1 | 0 | 0.35 | 0 | 0 | 0.954 | -0.067 | 0.98571955 | NA | 0 | 9.5346 | 8.2194 | 8.2194 | TRUE | 0.5204 |
| Gm15417 | 15 | 27 | 23 | 23 | 4 | 12 | 16 | 27 | 23 | 18 | 4 | 13 | 16.92 | 22 | 12 | 0.547 | -0.871 | 0.13197437 | 0.31297158 | 0.1444 | 0.2016 | 0.1773 | 0.1773 | TRUE | 0.7123 |
| Gm15421 | 55 | 73 | 71 | 73 | 54 | 64 | 58 | 73 | 71 | 57 | 58 | 70 | 64.69 | 67 | 62 | 0.912 | -0.132 | 0.59462239 | 0.7677918 | 0 | 0.0578 | 0.0292 | 0.0292 | TRUE | 0.1209 |
| Gm15441 | 17 | 21 | 16 | 14 | 16 | 19 | 18 | 21 | 16 | 11 | 17 | 21 | 17.35 | 18 | 16 | 0.878 | -0.187 | 0.69384373 | 0.83228303 | 0 | 0.1968 | 0.1054 | 0.1054 | TRUE | 0.497 |
| Gm15446 | 31 | 36 | 38 | 44 | 33 | 19 | 33 | 36 | 38 | 35 | 36 | 21 | 32.99 | 36 | 31 | 0.854 | -0.228 | 0.52024216 | 0.71109881 | 0 | 0.1068 | 0.0606 | 0.0606 | TRUE | 0.4595 |
| Gm15455 | 46 | 50 | 21 | 153 | 108 | 57 | 49 | 50 | 21 | 120 | 116 | 62 | 69.79 | 40 | 99 | 2.501 | 1.322 | 0.00070742 | 0.00681501 | 0.1396 | 0.0541 | 0.0921 | 0.0921 | TRUE | 1.2913 |
| Gm15471 | 0 | 0 | 0 | 0 | 0 | 0 | 0 | 0 | 0 | 0 | 0 | 0 | 0 | 0 | 0 | NA | NA | NA | NA | NA | NA | NA | NA | NA | NA |
| Gm1553 | 0 | 0 | 0 | 1 | 0 | 0 | 0 | 0 | 0 | 1 | 0 | 0 | 0.13 | 0 | 0 | 1.859 | 0.894 | 0.82650734 | NA | 0 | 25.1515 | 10 | 10 | TRUE | 0.2563 |
| Gm15545 | 25 | 25 | 28 | 32 | 27 | 16 | 27 | 25 | 28 | 25 | 29 | 18 | 25.23 | 27 | 24 | 0.903 | -0.147 | 0.70434171 | 0.83815282 | 0 | 0.1375 | 0.0689 | 0.0689 | TRUE | 0.2913 |
| Gm156 | 0 | 0 | 0 | 0 | 0 | 0 | 0 | 0 | 0 | 0 | 0 | 0 | 0 | 0 | 0 | NA | NA | NA | NA | NA | NA | NA | NA | NA | NA |
| Gm15612 | 0 | 1 | 1 | 4 | 3 | 1 | 0 | 1 | 1 | 3 | 3 | 1 | 1.58 | 1 | 2 | 3.716 | 1.894 | 0.24934509 | NA | 0 | 2.0942 | 0.9967 | 0.9967 | TRUE | 0.2349 |
| Gm1564 | 2866 | 3119 | 2857 | 3109 | 1990 | 2138 | 3044 | 3115 | 2869 | 2444 | 2141 | 2340 | 2658.91 | 3009 | 2308 | 0.767 | -0.382 | 6.27E-06 | 0.00013428 | 0.0025 | 0.0081 | 0.0048 | 0.0048 | TRUE | 0.0486 |
| Gm15645 | 0 | 2 | 0 | 0 | 0 | 0 | 0 | 2 | 0 | 0 | 0 | 0 | 0.33 | 1 | 0 | 0.268 | -1.897 | 0.63866967 | NA | 0 | 9.903 | 10 | 10 | TRUE | 0.9739 |
| Gm15663 | 1 | 0 | 2 | 0 | 1 | 0 | 1 | 0 | 2 | 0 | 1 | 0 | 0.69 | 1 | 0 | 0.341 | -1.551 | 0.54690874 | NA | 0 | 4.7742 | 3.2754 | 3.2754 | TRUE | 0.2976 |
| Gm15698 | 14 | 6 | 5 | 34 | 17 | 14 | 15 | 6 | 5 | 27 | 18 | 15 | 14.37 | 9 | 20 | 2.362 | 1.24 | 0.03996224 | 0.14068851 | 0.1071 | 0.2361 | 0.1789 | 0.1789 | TRUE | 1.193 |
| Gm15706 | 43 | 23 | 24 | 55 | 51 | 36 | 46 | 23 | 24 | 43 | 55 | 39 | 38.38 | 31 | 46 | 1.485 | 0.571 | 0.1326612 | 0.31402258 | 0.0579 | 0.0927 | 0.0766 | 0.0766 | TRUE | 1.1409 |
| Gm15708 | 5 | 3 | 4 | 5 | 3 | 3 | 5 | 3 | 4 | 4 | 3 | 3 | 3.79 | 4 | 3 | 0.855 | -0.227 | 0.80761726 | NA | 0 | 0.8752 | 0.3614 | 0.3614 | TRUE | 0.1064 |
| Gm15713 | 0 | 0 | 0 | 0 | 0 | 0 | 0 | 0 | 0 | 0 | 0 | 0 | 0 | 0 | 0 | NA | NA | NA | NA | NA | NA | NA | NA | NA | NA |
| Gm15760 | 9 | 18 | 19 | 10 | 15 | 7 | 10 | 18 | 19 | 8 | 16 | 8 | 13.05 | 16 | 11 | 0.672 | -0.574 | 0.33144893 | 0.55215482 | 0.0732 | 0.2594 | 0.1718 | 0.1718 | TRUE | 0.7216 |
| Gm15772 | 0 | 0 | 0 | 0 | 0 | 0 | 0 | 0 | 0 | 0 | 0 | 0 | 0 | 0 | 0 | NA | NA | NA | NA | NA | NA | NA | NA | NA | NA |
| Gm15787 | 1 | 2 | 5 | 6 | 5 | 3 | 1 | 2 | 5 | 5 | 5 | 3 | 3.58 | 3 | 4 | 1.652 | 0.724 | 0.48341039 | NA | 0 | 0.928 | 0.4712 | 0.4712 | TRUE | 0.6772 |
| Gm15800 | 2172 | 1622 | 1877 | 2663 | 1830 | 1624 | 2307 | 1620 | 1885 | 2093 | 1969 | 1778 | 1941.97 | 1937 | 1947 | 1.005 | 0.008 | 0.95669587 | 0.97766996 | 0.0186 | 0.0086 | 0.0138 | 0.0138 | TRUE | 0.3367 |
| Gm15850 | 0 | 1 | 0 | 0 | 0 | 0 | 0 | 1 | 0 | 0 | 0 | 0 | 0.17 | 0 | 0 | 0.49 | -1.029 | 0.80087 | NA | 0 | 19.7991 | 10 | 10 | TRUE | 0.405 |
| Gm1587 | 0 | 0 | 0 | 0 | 0 | 0 | 0 | 0 | 0 | 0 | 0 | 0 | 0 | 0 | 0 | NA | NA | NA | NA | NA | NA | NA | NA | NA | NA |
| Gm15880 | 0 | 0 | 0 | 0 | 0 | 0 | 0 | 0 | 0 | 0 | 0 | 0 | 0 | 0 | 0 | NA | NA | NA | NA | NA | NA | NA | NA | NA | NA |
| Gm15881 | 0 | 0 | 0 | 0 | 0 | 0 | 0 | 0 | 0 | 0 | 0 | 0 | 0 | 0 | 0 | NA | NA | NA | NA | NA | NA | NA | NA | NA | NA |
| Gm15910 | 38 | 22 | 18 | 66 | 39 | 34 | 40 | 22 | 18 | 52 | 42 | 37 | 35.25 | 27 | 44 | 1.642 | 0.715 | 0.07290975 | 0.2115013 | 0.0654 | 0.1004 | 0.0843 | 0.0843 | TRUE | 1.1959 |
| Gm15915 | 120 | 108 | 108 | 73 | 55 | 73 | 127 | 108 | 108 | 57 | 59 | 80 | 90.04 | 114 | 65 | 0.569 | -0.814 | 0.00042807 | 0.00458658 | 0.0069 | 0.0435 | 0.0265 | 0.0265 | TRUE | 0.3113 |
| Gm15941 | 0 | 0 | 0 | 0 | 0 | 0 | 0 | 0 | 0 | 0 | 0 | 0 | 0 | 0 | 0 | NA | NA | NA | NA | NA | NA | NA | NA | NA | NA |
| Gm15987 | 1 | 1 | 0 | 5 | 5 | 4 | 1 | 1 | 0 | 4 | 5 | 4 | 2.62 | 1 | 4 | 6.662 | 2.736 | 0.05396719 | NA | 0 | 1.2619 | 0.5968 | 0.5968 | TRUE | 0.2541 |
| Gm15997 | 0 | 0 | 0 | 0 | 0 | 0 | 0 | 0 | 0 | 0 | 0 | 0 | 0 | 0 | 0 | NA | NA | NA | NA | NA | NA | NA | NA | NA | NA |
| Gm16023 | 8 | 7 | 10 | 22 | 11 | 9 | 8 | 7 | 10 | 17 | 12 | 10 | 10.75 | 8 | 13 | 1.545 | 0.627 | 0.28881043 | 0.50977692 | 0 | 0.3133 | 0.1548 | 0.1548 | TRUE | 0.4294 |
| Gm16039 | 83 | 75 | 68 | 126 | 63 | 76 | 88 | 75 | 68 | 99 | 68 | 83 | 80.23 | 77 | 83 | 1.087 | 0.121 | 0.62573812 | 0.78915254 | 0.015 | 0.048 | 0.0316 | 0.0316 | TRUE | 0.1933 |
| Gm1604b | 0 | 0 | 0 | 0 | 0 | 0 | 0 | 0 | 0 | 0 | 0 | 0 | 0 | 0 | 0 | NA | NA | NA | NA | NA | NA | NA | NA | NA | NA |
| Gm16062 | 45 | 43 | 45 | 49 | 31 | 35 | 48 | 43 | 45 | 39 | 33 | 38 | 41.02 | 45 | 37 | 0.813 | -0.299 | 0.30792388 | 0.52834806 | 0 | 0.0872 | 0.0376 | 0.0376 | TRUE | 0.0438 |
| Gm16063 | 23 | 16 | 18 | 51 | 35 | 35 | 24 | 16 | 18 | 40 | 38 | 38 | 29.09 | 19 | 39 | 1.992 | 0.994 | 0.00661539 | 0.03861402 | 0 | 0.1201 | 0.0576 | 0.0576 | TRUE | 0.2506 |
| Gm16065 | 12 | 20 | 18 | 23 | 19 | 23 | 13 | 20 | 18 | 18 | 20 | 25 | 19.08 | 17 | 21 | 1.245 | 0.316 | 0.4780524 | 0.67743419 | 0 | 0.1795 | 0.0895 | 0.0895 | TRUE | 0.219 |
| Gm16119 | 2 | 0 | 0 | 2 | 2 | 3 | 2 | 0 | 0 | 2 | 2 | 3 | 1.52 | 1 | 2 | 3.34 | 1.74 | 0.32519469 | NA | 0 | 2.1713 | 1.3054 | 1.3054 | TRUE | 0.9745 |
| Gm16130 | 0 | 0 | 0 | 0 | 0 | 0 | 0 | 0 | 0 | 0 | 0 | 0 | 0 | 0 | 0 | NA | NA | NA | NA | NA | NA | NA | NA | NA | NA |
| Gm16157 | 0 | 0 | 0 | 4 | 1 | 1 | 0 | 0 | 0 | 3 | 1 | 1 | 0.89 | 0 | 2 | 9.564 | 3.258 | 0.15728766 | NA | 0 | 3.7259 | 2.5507 | 2.5507 | TRUE | 0.4752 |
| Gm16197 | 29 | 38 | 42 | 49 | 39 | 21 | 31 | 38 | 42 | 39 | 42 | 23 | 35.73 | 37 | 35 | 0.936 | -0.096 | 0.79017329 | 0.8919706 | 0.0252 | 0.0991 | 0.0653 | 0.0653 | TRUE | 0.5527 |
| Gm16287 | 1 | 0 | 0 | 0 | 0 | 0 | 1 | 0 | 0 | 0 | 0 | 0 | 0.18 | 0 | 0 | 0.49 | -1.029 | 0.80087 | NA | 0 | 18.6165 | 10 | 10 | TRUE | 0.4614 |
| Gm16291 | 0 | 0 | 0 | 0 | 0 | 0 | 0 | 0 | 0 | 0 | 0 | 0 | 0 | 0 | 0 | NA | NA | NA | NA | NA | NA | NA | NA | NA | NA |

| | | | | | | | | | | | | | | | | | | | | | | | | | |
|---|---|---|---|---|---|---|---|---|---|---|---|---|---|---|---|---|---|---|---|---|---|---|---|---|---|
| Gm16294 | 0 | 0 | 0 | 0 | 0 | 0 | 0 | 0 | 0 | 0 | 0 | 0 | 0 | 0 | 0 | NA | NA | NA | NA | NA | NA | NA | NA | NA | NA |
| Gm1631 | 235 | 280 | 267 | 271 | 187 | 177 | 250 | 280 | 268 | 213 | 201 | 194 | 234.22 | 266 | 203 | 0.764 | -0.389 | 0.00481463 | 0.03045982 | 0 | 0.021 | 0.0094 | 0.0094 | TRUE | 0.0312 |
| Gm16325 | 0 | 0 | 0 | 0 | 0 | 0 | 0 | 0 | 0 | 0 | 0 | 0 | 0 | 0 | 0 | NA | NA | NA | NA | NA | NA | NA | NA | NA | NA |
| Gm16336 | 0 | 0 | 0 | 0 | 0 | 0 | 0 | 0 | 0 | 0 | 0 | 0 | 0 | 0 | 0 | NA | NA | NA | NA | NA | NA | NA | NA | NA | NA |
| Gm16367 | 0 | 0 | 0 | 0 | 0 | 0 | 0 | 0 | 0 | 0 | 0 | 0 | 0 | 0 | 0 | NA | NA | NA | NA | NA | NA | NA | NA | NA | NA |
| Gm16381 | 0 | 0 | 0 | 0 | 0 | 0 | 0 | 0 | 0 | 0 | 0 | 0 | 0 | 0 | 0 | NA | NA | NA | NA | NA | NA | NA | NA | NA | NA |
| Gm16386 | 34 | 16 | 25 | 40 | 27 | 22 | 36 | 16 | 25 | 31 | 29 | 24 | 26.96 | 26 | 28 | 1.102 | 0.14 | 0.73828461 | 0.85986006 | 0.0456 | 0.1291 | 0.0901 | 0.0901 | TRUE | 0.2549 |
| Gm16390 | 0 | 0 | 0 | 2 | 0 | 0 | 0 | 0 | 0 | 2 | 0 | 0 | 0.26 | 0 | 1 | 2.886 | 1.529 | 0.70676491 | NA | 0 | 12.5792 | 10 | 10 | TRUE | 0.9669 |
| Gm16404 | 0 | 0 | 0 | 0 | 0 | 0 | 0 | 0 | 0 | 0 | 0 | 0 | 0 | 0 | 0 | NA | NA | NA | NA | NA | NA | NA | NA | NA | NA |
| Gm16405 | 0 | 0 | 0 | 0 | 0 | 0 | 0 | 0 | 0 | 0 | 0 | 0 | 0 | 0 | 0 | NA | NA | NA | NA | NA | NA | NA | NA | NA | NA |
| Gm16430 | 0 | 0 | 0 | 0 | 0 | 0 | 0 | 0 | 0 | 0 | 0 | 0 | 0 | 0 | 0 | NA | NA | NA | NA | NA | NA | NA | NA | NA | NA |
| Gm16432 | 0 | 0 | 0 | 2 | 1 | 1 | 0 | 0 | 0 | 2 | 1 | 1 | 0.62 | 0 | 1 | 6.676 | 2.739 | 0.27800695 | NA | 0 | 5.2881 | 3.2034 | 3.2034 | TRUE | 0.07 |
| Gm16442 | 0 | 0 | 0 | 0 | 0 | 0 | 0 | 0 | 0 | 0 | 0 | 0 | 0 | 0 | 0 | NA | NA | NA | NA | NA | NA | NA | NA | NA | NA |
| Gm16445 | 0 | 0 | 0 | 0 | 0 | 0 | 0 | 0 | 0 | 0 | 0 | 0 | 0 | 0 | 0 | NA | NA | NA | NA | NA | NA | NA | NA | NA | NA |
| Gm16451 | 0 | 0 | 0 | 0 | 0 | 0 | 0 | 0 | 0 | 0 | 0 | 0 | 0 | 0 | 0 | NA | NA | NA | NA | NA | NA | NA | NA | NA | NA |
| Gm1647 | 0 | 0 | 0 | 1 | 0 | 0 | 0 | 0 | 0 | 1 | 0 | 0 | 0.13 | 0 | 0 | 1.859 | 0.894 | 0.82650734 | NA | 0 | 25.1515 | 10 | 10 | TRUE | 0.2563 |
| Gm16497 | 0 | 0 | 0 | 0 | 0 | 0 | 0 | 0 | 0 | 0 | 0 | 0 | 0 | 0 | 0 | NA | NA | NA | NA | NA | NA | NA | NA | NA | NA |
| Gm16501 | 0 | 0 | 0 | 0 | 0 | 0 | 0 | 0 | 0 | 0 | 0 | 0 | 0 | 0 | 0 | NA | NA | NA | NA | NA | NA | NA | NA | NA | NA |
| Gm16515 | 2206 | 2422 | 2417 | 3870 | 2729 | 2313 | 2343 | 2419 | 2427 | 3042 | 2936 | 2532 | 2616.57 | 2396 | 2837 | 1.184 | 0.244 | 0.00920057 | 0.04940801 | 0.0041 | 0.0082 | 0.0059 | 0.0059 | TRUE | 0.1067 |
| Gm16516 | 351 | 312 | 321 | 271 | 196 | 236 | 373 | 312 | 322 | 213 | 211 | 258 | 281.5 | 336 | 227 | 0.677 | -0.563 | 0.0001895 | 0.0023277 | 0.0073 | 0.0186 | 0.0127 | 0.0127 | TRUE | 0.1505 |
| Gm16523 | 1 | 2 | 1 | 3 | 1 | 0 | 1 | 2 | 1 | 2 | 1 | 0 | 1.25 | 1 | 1 | 0.882 | -0.182 | 0.91426124 | NA | 0 | 2.6432 | 1.2732 | 1.2732 | TRUE | 0.3046 |
| Gm1653 | 4 | 8 | 11 | 14 | 9 | 10 | 4 | 8 | 11 | 11 | 10 | 11 | 9.15 | 8 | 11 | 1.356 | 0.439 | 0.49447509 | 0.69052608 | 0 | 0.3668 | 0.1859 | 0.1859 | TRUE | 0.2947 |
| Gm16532 | 31 | 33 | 55 | 39 | 33 | 44 | 33 | 33 | 55 | 31 | 36 | 48 | 39.24 | 40 | 38 | 0.938 | -0.093 | 0.80011688 | 0.89686328 | 0.0501 | 0.0909 | 0.0708 | 0.0708 | TRUE | 0.7852 |
| Gm16548 | 1 | 1 | 0 | 3 | 1 | 2 | 1 | 1 | 0 | 2 | 1 | 2 | 1.28 | 1 | 2 | 2.765 | 1.467 | 0.39470822 | NA | 0 | 2.5792 | 1.1358 | 1.1358 | TRUE | 0.2532 |
| Gm16551 | 4 | 4 | 1 | 2 | 4 | 4 | 4 | 4 | 1 | 2 | 4 | 4 | 3.25 | 3 | 3 | 1.09 | 0.124 | 0.90959496 | NA | 0 | 1.0205 | 0.5429 | 0.5429 | TRUE | 0.47 |
| Gm16576 | 72 | 74 | 73 | 74 | 51 | 53 | 76 | 74 | 73 | 58 | 55 | 58 | 65.79 | 74 | 57 | 0.766 | -0.385 | 0.09726409 | 0.25625993 | 0 | 0.057 | 0.0235 | 0.0235 | TRUE | 0.0087 |
| Gm16596 | 1 | 2 | 2 | 0 | 2 | 2 | 1 | 2 | 2 | 0 | 2 | 2 | 1.57 | 2 | 1 | 0.819 | -0.288 | 0.85415907 | NA | 0 | 2.1077 | 1.1101 | 1.1101 | TRUE | 0.7276 |
| Gm166 | 104 | 125 | 143 | 188 | 180 | 165 | 110 | 125 | 144 | 148 | 194 | 181 | 150.16 | 126 | 174 | 1.372 | 0.457 | 0.01877883 | 0.08314447 | 0.0122 | 0.0288 | 0.0203 | 0.0203 | TRUE | 0.2015 |
| Gm1661 | 4 | 15 | 12 | 9 | 7 | 8 | 4 | 15 | 12 | 7 | 8 | 9 | 9.11 | 10 | 8 | 0.742 | -0.431 | 0.52995306 | 0.7181798 | 0 | 0.3686 | 0.2279 | 0.2279 | TRUE | 0.6362 |
| Gm16617 | 0 | 0 | 0 | 0 | 0 | 0 | 0 | 0 | 0 | 0 | 0 | 0 | 0 | 0 | 0 | NA | NA | NA | NA | NA | NA | NA | NA | NA | NA |
| Gm16675 | 1 | 0 | 0 | 1 | 0 | 1 | 1 | 0 | 0 | 1 | 0 | 1 | 0.49 | 0 | 1 | 1.724 | 0.785 | 0.78200058 | NA | 0 | 6.7235 | 4.0295 | 4.0295 | TRUE | 0.4614 |
| Gm16677 | 0 | 0 | 0 | 0 | 0 | 0 | 0 | 0 | 0 | 0 | 0 | 0 | 0 | 0 | 0 | NA | NA | NA | NA | NA | NA | NA | NA | NA | NA |
| Gm16701 | 0 | 0 | 0 | 0 | 0 | 1 | 0 | 0 | 0 | 0 | 0 | 1 | 0.18 | 0 | 0 | 1.859 | 0.894 | 0.82650734 | NA | 0 | 18.0643 | 10 | 10 | TRUE | 0.5383 |
| Gm16702 | 9 | 4 | 5 | 5 | 4 | 10 | 10 | 4 | 5 | 4 | 4 | 11 | 6.29 | 6 | 6 | 1.022 | 0.031 | 0.9697234 | 0.98448447 | 0.1308 | 0.5304 | 0.3301 | 0.3301 | TRUE | 0.8844 |
| Gm1673 | 91 | 169 | 194 | 165 | 132 | 159 | 97 | 169 | 195 | 130 | 142 | 174 | 151 | 154 | 149 | 0.966 | -0.05 | 0.85457912 | 0.92756327 | 0.0663 | 0.0287 | 0.0465 | 0.0465 | TRUE | 0.8378 |
| Gm16793 | 2 | 1 | 2 | 2 | 1 | 0 | 2 | 1 | 2 | 2 | 1 | 0 | 1.3 | 2 | 1 | 0.539 | -0.892 | 0.58964573 | NA | 0 | 2.5478 | 1.1441 | 1.1441 | TRUE | 0.2725 |
| Gm16796 | 2 | 0 | 1 | 2 | 1 | 1 | 2 | 0 | 1 | 2 | 1 | 1 | 1.15 | 1 | 1 | 1.22 | 0.287 | 0.86966868 | NA | 0 | 2.8836 | 1.3154 | 1.3154 | TRUE | 0.2346 |
| Gm16833 | 0 | 0 | 0 | 0 | 0 | 0 | 0 | 0 | 0 | 0 | 0 | 0 | 0 | 0 | 0 | NA | NA | NA | NA | NA | NA | NA | NA | NA | NA |
| Gm16845 | 23 | 19 | 20 | 53 | 37 | 12 | 24 | 19 | 20 | 42 | 40 | 13 | 26.35 | 21 | 32 | 1.501 | 0.586 | 0.23667702 | 0.45197602 | 0.143 | 0.1319 | 0.1372 | 0.1372 | TRUE | 1.6597 |
| Gm16853 | 1 | 0 | 0 | 2 | 1 | 0 | 1 | 0 | 0 | 2 | 1 | 0 | 0.62 | 0 | 1 | 2.462 | 1.3 | 0.62119797 | NA | 0 | 5.3343 | 3.4416 | 3.4416 | TRUE | 0.4614 |
| Gm16861 | 7 | 3 | 1 | 10 | 6 | 4 | 7 | 3 | 1 | 8 | 6 | 4 | 5.02 | 4 | 6 | 1.659 | 0.73 | 0.43640125 | 0.64470968 | 0.1071 | 0.6629 | 0.4219 | 0.4219 | TRUE | 0.8405 |
| Gm16863 | 1 | 2 | 1 | 0 | 0 | 1 | 1 | 2 | 1 | 0 | 0 | 1 | 0.86 | 1 | 0 | 0.257 | -1.958 | 0.35814405 | NA | 0 | 3.8389 | 1.8948 | 1.8948 | TRUE | 0.5383 |
| Gm16880 | 35 | 35 | 51 | 70 | 48 | 35 | 37 | 35 | 51 | 55 | 52 | 38 | 44.72 | 41 | 48 | 1.18 | 0.239 | 0.45167142 | 0.65711296 | 0.0166 | 0.0806 | 0.0501 | 0.0501 | TRUE | 0.3532 |
| Gm16894 | 0 | 0 | 0 | 0 | 0 | 0 | 0 | 0 | 0 | 0 | 0 | 0 | 0 | 0 | 0 | NA | NA | NA | NA | NA | NA | NA | NA | NA | NA |
| Gm16897 | 34 | 32 | 21 | 19 | 30 | 16 | 36 | 32 | 21 | 15 | 32 | 18 | 25.65 | 30 | 22 | 0.72 | -0.474 | 0.29820194 | 0.51889243 | 0.0808 | 0.1354 | 0.1092 | 0.1092 | TRUE | 1.0182 |
| Gm16907 | 6 | 0 | 2 | 5 | 1 | 0 | 6 | 0 | 2 | 4 | 1 | 0 | 2.23 | 3 | 2 | 0.623 | -0.683 | 0.68619094 | NA | 1.766 | 1.4834 | 1.5933 | 1.5933 | TRUE | 0.4066 |
| Gm16938 | 30 | 25 | 19 | 38 | 33 | 27 | 32 | 25 | 19 | 30 | 36 | 30 | 28.47 | 25 | 32 | 1.251 | 0.323 | 0.38736001 | 0.60114554 | 0 | 0.1226 | 0.0648 | 0.0648 | TRUE | 0.2644 |
| Gm16973 | 166 | 114 | 147 | 167 | 112 | 129 | 176 | 114 | 148 | 131 | 121 | 141 | 138.46 | 146 | 131 | 0.899 | -0.154 | 0.46615577 | 0.66857491 | 0.0195 | 0.0307 | 0.0251 | 0.0251 | TRUE | 0.2079 |
| Gm16982 | 1 | 1 | 0 | 3 | 0 | 0 | 1 | 1 | 0 | 2 | 0 | 0 | 0.74 | 1 | 1 | 1.21 | 0.275 | 0.91432274 | NA | 0 | 4.4798 | 3.3685 | 3.3685 | TRUE | 1.4393 |
| Gm16998 | 2 | 1 | 1 | 1 | 1 | 2 | 2 | 1 | 1 | 1 | 1 | 2 | 1.36 | 1 | 1 | 0.966 | -0.05 | 0.97423616 | NA | 0 | 2.4238 | 0.9611 | 0.9611 | TRUE | 0.1676 |
| Gm17019 | 0 | 0 | 1 | 0 | 0 | 0 | 0 | 0 | 1 | 0 | 0 | 0 | 0.17 | 0 | 0 | 0.49 | -1.029 | 0.80087 | NA | 0 | 19.6928 | 10 | 10 | TRUE | 0.4098 |
| Gm17066 | 2123 | 1899 | 1834 | 2163 | 1721 | 1682 | 2255 | 1897 | 1842 | 1700 | 1852 | 1841 | 1897.73 | 1998 | 1798 | 0.9 | -0.152 | 0.14926028 | 0.33965275 | 0.0066 | 0.0086 | 0.0075 | 0.0075 | TRUE | 0.1532 |
| Gm1715 | 0 | 0 | 0 | 0 | 0 | 0 | 0 | 0 | 0 | 0 | 0 | 0 | 0 | 0 | 0 | NA | NA | NA | NA | NA | NA | NA | NA | NA | NA |
| Gm1720 | 1 | 0 | 0 | 0 | 0 | 0 | 1 | 0 | 0 | 0 | 0 | 0 | 0.18 | 0 | 0 | 0.49 | -1.029 | 0.80087 | NA | 0 | 18.6165 | 10 | 10 | TRUE | 0.4614 |
| Gm17252 | 0 | 0 | 0 | 0 | 0 | 0 | 0 | 0 | 0 | 0 | 0 | 0 | 0 | 0 | 0 | NA | NA | NA | NA | NA | NA | NA | NA | NA | NA |
| Gm17296 | 1762 | 1800 | 1916 | 2438 | 1884 | 1704 | 1872 | 1798 | 1924 | 1917 | 2027 | 1865 | 1900.34 | 1865 | 1936 | 1.039 | 0.055 | 0.47705232 | 0.67668578 | 9.00E-04 | 0.0086 | 0.0037 | 0.0037 | TRUE | 0.0199 |
| Gm17359 | 2 | 0 | 0 | 0 | 1 | 0 | 2 | 0 | 0 | 0 | 1 | 0 | 0.53 | 1 | 0 | 0.497 | -1.009 | 0.76131406 | NA | 0 | 6.1834 | 6.228 | 6.228 | TRUE | 0.9742 |
| Gm17365 | 0 | 0 | 0 | 0 | 0 | 0 | 0 | 0 | 0 | 0 | 0 | 0 | 0 | 0 | 0 | NA | NA | NA | NA | NA | NA | NA | NA | NA | NA |
| Gm17384 | 0 | 0 | 0 | 0 | 0 | 0 | 0 | 0 | 0 | 0 | 0 | 0 | 0 | 0 | 0 | NA | NA | NA | NA | NA | NA | NA | NA | NA | NA |
| Gm17455 | 2 | 0 | 0 | 1 | 4 | 0 | 2 | 0 | 0 | 1 | 4 | 0 | 1.2 | 1 | 2 | 2.409 | 1.268 | 0.57993696 | NA | 2.789 | 2.7469 | 2.7617 | 2.7617 | TRUE | 1.2955 |
| Gm17495 | 2 | 1 | 0 | 2 | 2 | 3 | 2 | 1 | 0 | 2 | 2 | 3 | 1.69 | 1 | 2 | 2.242 | 1.165 | 0.44453988 | NA | 0 | 1.958 | 0.9649 | 0.9649 | TRUE | 0.3522 |
| Gm17644 | 0 | 0 | 0 | 0 | 0 | 0 | 0 | 0 | 0 | 0 | 0 | 0 | 0 | 0 | 0 | NA | NA | NA | NA | NA | NA | NA | NA | NA | NA |
| Gm17660 | 0 | 0 | 0 | 0 | 0 | 0 | 0 | 0 | 0 | 0 | 0 | 0 | 0 | 0 | 0 | NA | NA | NA | NA | NA | NA | NA | NA | NA | NA |
| Gm17677 | 0 | 0 | 0 | 0 | 0 | 0 | 0 | 0 | 0 | 0 | 0 | 0 | 0 | 0 | 0 | NA | NA | NA | NA | NA | NA | NA | NA | NA | NA |
| Gm17689 | 0 | 0 | 0 | 0 | 0 | 0 | 0 | 0 | 0 | 0 | 0 | 0 | 0 | 0 | 0 | NA | NA | NA | NA | NA | NA | NA | NA | NA | NA |
| Gm17727 | 0 | 0 | 0 | 0 | 1 | 0 | 0 | 0 | 0 | 0 | 1 | 0 | 0.18 | 0 | 0 | 1.859 | 0.894 | 0.82650734 | NA | 0 | 18.3785 | 10 | 10 | TRUE | 0.5204 |
| Gm17745 | 14 | 4 | 7 | 20 | 12 | 8 | 15 | 4 | 7 | 16 | 13 | 9 | 10.55 | 9 | 13 | 1.462 | 0.548 | 0.41660172 | 0.62792836 | 0.1235 | 0.3192 | 0.229 | 0.229 | TRUE | 0.9881 |
| Gm17746 | 0 | 0 | 0 | 0 | 0 | 0 | 0 | 0 | 0 | 0 | 0 | 0 | 0 | 0 | 0 | NA | NA | NA | NA | NA | NA | NA | NA | NA | NA |
| Gm17751 | 0 | 0 | 0 | 0 | 0 | 0 | 0 | 0 | 0 | 0 | 0 | 0 | 0 | 0 | 0 | NA | NA | NA | NA | NA | NA | NA | NA | NA | NA |

| | | | | | | | | | | | | | | | | | | | | | | | | | |
|---|---|---|---|---|---|---|---|---|---|---|---|---|---|---|---|---|---|---|---|---|---|---|---|---|---|
| Gm17757 | 0 | 0 | 0 | 0 | 0 | 0 | 0 | 0 | 0 | 0 | 0 | 0 | 0 | 0 | 0 | NA | NA | NA | NA | NA | NA | NA | NA | NA | NA |
| Gm17762 | 63 | 127 | 120 | 124 | 99 | 125 | 67 | 127 | 120 | 97 | 107 | 137 | 109.18 | 105 | 114 | 1.081 | 0.112 | 0.69269898 | 0.83157621 | 0.0617 | 0.0371 | 0.0487 | 0.0487 | TRUE | 0.9671 |
| Gm17769 | 1 | 1 | 0 | 2 | 4 | 2 | 1 | 1 | 0 | 2 | 4 | 2 | 1.69 | 1 | 3 | 3.891 | 1.96 | 0.22456476 | NA | 0 | 1.9589 | 0.9453 | 0.9453 | TRUE | 0.3065 |
| Gm17801 | 1 | 3 | 2 | 2 | 3 | 0 | 1 | 3 | 2 | 2 | 3 | 0 | 1.81 | 2 | 2 | 0.79 | -0.34 | 0.8168866 | NA | 0 | 1.8259 | 0.9998 | 0.9998 | TRUE | 0.2163 |
| Gm17821 | 0 | 0 | 0 | 0 | 0 | 0 | 0 | 0 | 0 | 0 | 0 | 0 | 0 | 0 | 0 | NA | NA | NA | NA | NA | NA | NA | NA | NA | NA |
| Gm17830 | 0 | 0 | 0 | 0 | 0 | 0 | 0 | 0 | 0 | 0 | 0 | 0 | 0 | 0 | 0 | NA | NA | NA | NA | NA | NA | NA | NA | NA | NA |
| Gm1821 | 293 | 357 | 369 | 436 | 335 | 340 | 311 | 357 | 371 | 343 | 360 | 372 | 352.27 | 346 | 358 | 1.034 | 0.048 | 0.70136384 | 0.83650923 | 0.0019 | 0.0163 | 0.0086 | 0.0086 | TRUE | 0.0874 |
| Gm18409 | 0 | 0 | 0 | 0 | 0 | 0 | 0 | 0 | 0 | 0 | 0 | 0 | 0 | 0 | 0 | NA | NA | NA | NA | NA | NA | NA | NA | NA | NA |
| Gm18853 | 0 | 0 | 0 | 0 | 0 | 0 | 0 | 0 | 0 | 0 | 0 | 0 | 0 | 0 | 0 | NA | NA | NA | NA | NA | NA | NA | NA | NA | NA |
| Gm19276 | 0 | 0 | 0 | 0 | 0 | 0 | 0 | 0 | 0 | 0 | 0 | 0 | 0 | 0 | 0 | NA | NA | NA | NA | NA | NA | NA | NA | NA | NA |
| Gm19277 | 0 | 0 | 0 | 0 | 0 | 0 | 0 | 0 | 0 | 0 | 0 | 0 | 0 | 0 | 0 | NA | NA | NA | NA | NA | NA | NA | NA | NA | NA |
| Gm19299 | 0 | 0 | 1 | 0 | 0 | 1 | 0 | 0 | 1 | 0 | 0 | 1 | 0.35 | 0 | 0 | 0.954 | -0.067 | 0.98566485 | NA | 0 | 9.4252 | 8.1416 | 8.1416 | TRUE | 0.5383 |
| Gm19303 | 0 | 0 | 0 | 0 | 0 | 0 | 0 | 0 | 0 | 0 | 0 | 0 | 0 | 0 | 0 | NA | NA | NA | NA | NA | NA | NA | NA | NA | NA |
| Gm19345 | 52 | 41 | 35 | 36 | 38 | 39 | 55 | 41 | 35 | 28 | 41 | 43 | 40.53 | 44 | 37 | 0.846 | -0.241 | 0.48092792 | 0.68004928 | 0.0284 | 0.0882 | 0.059 | 0.059 | TRUE | 0.3962 |
| Gm19395 | 0 | 0 | 0 | 0 | 0 | 0 | 0 | 0 | 0 | 0 | 0 | 0 | 0 | 0 | 0 | NA | NA | NA | NA | NA | NA | NA | NA | NA | NA |
| Gm19402 | 0 | 0 | 0 | 0 | 0 | 0 | 0 | 0 | 0 | 0 | 0 | 0 | 0 | 0 | 0 | NA | NA | NA | NA | NA | NA | NA | NA | NA | NA |
| Gm19424 | 0 | 0 | 0 | 0 | 0 | 0 | 0 | 0 | 0 | 0 | 0 | 0 | 0 | 0 | 0 | NA | NA | NA | NA | NA | NA | NA | NA | NA | NA |
| Gm1943 | 71 | 87 | 94 | 126 | 79 | 79 | 75 | 87 | 94 | 99 | 85 | 86 | 87.87 | 85 | 90 | 1.057 | 0.079 | 0.71288943 | 0.84374097 | 0 | 0.0444 | 0.0222 | 0.0222 | TRUE | 0.0991 |
| Gm19434 | 1 | 0 | 1 | 0 | 0 | 1 | 1 | 0 | 1 | 0 | 0 | 1 | 0.53 | 1 | 0 | 0.509 | -0.975 | 0.72674596 | NA | 0 | 6.2603 | 3.8683 | 3.8683 | TRUE | 0.5383 |
| Gm19461 | 1 | 4 | 2 | 3 | 1 | 2 | 1 | 4 | 2 | 2 | 1 | 2 | 2.11 | 2 | 2 | 0.801 | -0.32 | 0.80202923 | NA | 0 | 1.5647 | 0.7 | 0.7 | TRUE | 0.3916 |
| Gm19466 | 0 | 0 | 0 | 0 | 0 | 0 | 0 | 0 | 0 | 0 | 0 | 0 | 0 | 0 | 0 | NA | NA | NA | NA | NA | NA | NA | NA | NA | NA |
| Gm19510 | 1 | 0 | 0 | 4 | 1 | 0 | 1 | 0 | 0 | 3 | 1 | 0 | 0.88 | 0 | 1 | 3.941 | 1.978 | 0.4137088 | NA | 0 | 3.7488 | 2.8817 | 2.8817 | TRUE | 0.2821 |
| Gm19522 | 0 | 0 | 0 | 0 | 0 | 0 | 0 | 0 | 0 | 0 | 0 | 0 | 0 | 0 | 0 | NA | NA | NA | NA | NA | NA | NA | NA | NA | NA |
| Gm19557 | 0 | 0 | 0 | 0 | 0 | 0 | 0 | 0 | 0 | 0 | 0 | 0 | 0 | 0 | 0 | NA | NA | NA | NA | NA | NA | NA | NA | NA | NA |
| Gm19583 | 14 | 7 | 20 | 20 | 19 | 12 | 15 | 7 | 20 | 16 | 20 | 13 | 15.21 | 14 | 16 | 1.175 | 0.233 | 0.67289164 | 0.81781975 | 0.0692 | 0.2235 | 0.1528 | 0.1528 | TRUE | 0.3872 |
| Gm19589 | 0 | 0 | 0 | 0 | 0 | 0 | 0 | 0 | 0 | 0 | 0 | 0 | 0 | 0 | 0 | NA | NA | NA | NA | NA | NA | NA | NA | NA | NA |
| Gm19619 | 0 | 0 | 0 | 0 | 0 | 0 | 0 | 0 | 0 | 0 | 0 | 0 | 0 | 0 | 0 | NA | NA | NA | NA | NA | NA | NA | NA | NA | NA |
| Gm1965 | 0 | 1 | 0 | 0 | 0 | 0 | 0 | 1 | 0 | 0 | 0 | 0 | 0.17 | 0 | 0 | 0.49 | -1.029 | 0.80087 | NA | 0 | 19.7991 | 10 | 10 | TRUE | 0.405 |
| Gm19668 | 0 | 0 | 0 | 0 | 0 | 0 | 0 | 0 | 0 | 0 | 0 | 0 | 0 | 0 | 0 | NA | NA | NA | NA | NA | NA | NA | NA | NA | NA |
| Gm1968 | 0 | 0 | 0 | 0 | 0 | 2 | 0 | 0 | 0 | 0 | 0 | 2 | 0.36 | 0 | 1 | 3.832 | 1.938 | 0.63022034 | NA | 10 | 9.0356 | 10 | 10 | TRUE | 0.9744 |
| Gm19689 | 0 | 0 | 0 | 0 | 0 | 0 | 0 | 0 | 0 | 0 | 0 | 0 | 0 | 0 | 0 | NA | NA | NA | NA | NA | NA | NA | NA | NA | NA |
| Gm19705 | 47 | 45 | 40 | 107 | 87 | 63 | 50 | 45 | 40 | 84 | 94 | 69 | 63.62 | 45 | 82 | 1.83 | 0.872 | 0.00084013 | 0.00778346 | 0 | 0.0587 | 0.0318 | 0.0318 | TRUE | 0.1737 |
| Gm19710 | 2 | 4 | 2 | 4 | 3 | 4 | 2 | 4 | 2 | 3 | 3 | 4 | 3.15 | 3 | 3 | 1.313 | 0.393 | 0.70362226 | NA | 0 | 1.054 | 0.4425 | 0.4425 | TRUE | 0.2085 |
| Gm19757 | 1 | 1 | 0 | 4 | 2 | 0 | 1 | 1 | 0 | 3 | 2 | 0 | 1.23 | 1 | 2 | 2.635 | 1.398 | 0.4709707 | NA | 0 | 2.6936 | 1.6903 | 1.6903 | TRUE | 0.3143 |
| Gm1976 | 60 | 92 | 87 | 60 | 50 | 56 | 64 | 92 | 87 | 47 | 54 | 61 | 67.54 | 81 | 54 | 0.664 | -0.591 | 0.02564764 | 0.10353584 | 0.0137 | 0.0557 | 0.0351 | 0.0351 | TRUE | 0.3167 |
| Gm19782 | 0 | 0 | 0 | 0 | 0 | 0 | 0 | 0 | 0 | 0 | 0 | 0 | 0 | 0 | 0 | NA | NA | NA | NA | NA | NA | NA | NA | NA | NA |
| Gm19784 | 56 | 18 | 18 | 60 | 42 | 19 | 59 | 18 | 18 | 47 | 45 | 21 | 34.78 | 32 | 38 | 1.191 | 0.252 | 0.65081491 | 0.80429196 | 0.2987 | 0.1016 | 0.1951 | 0.1951 | TRUE | 3.8436 |
| Gm1987 | 0 | 0 | 0 | 0 | 0 | 0 | 0 | 0 | 0 | 0 | 0 | 0 | 0 | 0 | 0 | NA | NA | NA | NA | NA | NA | NA | NA | NA | NA |
| Gm19897 | 0 | 1 | 1 | 5 | 0 | 0 | 0 | 1 | 1 | 4 | 0 | 0 | 0.99 | 1 | 1 | 2.027 | 1.019 | 0.67267051 | NA | 2.5641 | 3.3383 | 3.0722 | 3.0722 | TRUE | 2.338 |
| Gm1993 | 0 | 0 | 0 | 0 | 0 | 0 | 0 | 0 | 0 | 0 | 0 | 0 | 0 | 0 | 0 | NA | NA | NA | NA | NA | NA | NA | NA | NA | NA |
| Gm1995 | 193 | 60 | 55 | 302 | 269 | 90 | 205 | 60 | 55 | 237 | 289 | 99 | 157.58 | 107 | 208 | 1.955 | 0.967 | 0.19408499 | 0.39967468 | 0.3924 | 0.0278 | 0.1771 | 0.3924 | TRUE | 1.4079 |
| Gm19990 | 0 | 0 | 0 | 0 | 0 | 0 | 0 | 0 | 0 | 0 | 0 | 0 | 0 | 0 | 0 | NA | NA | NA | NA | NA | NA | NA | NA | NA | NA |
| Gm2002 | 0 | 0 | 0 | 0 | 0 | 0 | 0 | 0 | 0 | 0 | 0 | 0 | 0 | 0 | 0 | NA | NA | NA | NA | NA | NA | NA | NA | NA | NA |
| Gm20063 | 0 | 0 | 0 | 0 | 0 | 0 | 0 | 0 | 0 | 0 | 0 | 0 | 0 | 0 | 0 | NA | NA | NA | NA | NA | NA | NA | NA | NA | NA |
| Gm20098 | 0 | 0 | 0 | 0 | 1 | 0 | 0 | 0 | 0 | 0 | 1 | 0 | 0.18 | 0 | 0 | 1.859 | 0.894 | 0.82650734 | NA | 0 | 18.3785 | 10 | 10 | TRUE | 0.5204 |
| Gm2011 | 0 | 2 | 3 | 5 | 3 | 1 | 0 | 2 | 3 | 4 | 3 | 1 | 2.21 | 2 | 3 | 1.657 | 0.728 | 0.5965999 | NA | 0 | 1.4973 | 0.8851 | 0.8851 | TRUE | 0.5411 |
| Gm20110 | 0 | 0 | 0 | 0 | 0 | 0 | 0 | 0 | 0 | 0 | 0 | 0 | 0 | 0 | 0 | NA | NA | NA | NA | NA | NA | NA | NA | NA | NA |
| Gm2012 | 0 | 0 | 0 | 2 | 0 | 1 | 0 | 0 | 0 | 2 | 0 | 1 | 0.44 | 0 | 1 | 4.782 | 2.258 | 0.50653358 | NA | 0 | 7.4188 | 6.7775 | 6.7775 | TRUE | 0.2804 |
| Gm20125 | 0 | 0 | 0 | 1 | 0 | 0 | 0 | 0 | 0 | 1 | 0 | 0 | 0.13 | 0 | 0 | 1.859 | 0.894 | 0.82650734 | NA | 0 | 25.1515 | 10 | 10 | TRUE | 0.2563 |
| Gm20139 | 0 | 0 | 0 | 0 | 0 | 0 | 0 | 0 | 0 | 0 | 0 | 0 | 0 | 0 | 0 | NA | NA | NA | NA | NA | NA | NA | NA | NA | NA |
| Gm2016 | 966 | 310 | 184 | 1457 | 1038 | 447 | 1026 | 310 | 185 | 1145 | 1117 | 489 | 711.99 | 507 | 917 | 1.81 | 0.856 | 0.27995803 | 0.50109666 | 0.4508 | 0.0115 | 0.1718 | 0.4508 | TRUE | 6.2077 |
| Gm20172 | 0 | 0 | 0 | 0 | 0 | 0 | 0 | 0 | 0 | 0 | 0 | 0 | 0 | 0 | 0 | NA | NA | NA | NA | NA | NA | NA | NA | NA | NA |
| Gm20187 | 1 | 1 | 1 | 0 | 1 | 2 | 1 | 1 | 1 | 0 | 1 | 2 | 1.06 | 1 | 1 | 1.021 | 0.029 | 0.98726074 | NA | 0 | 3.1295 | 1.4749 | 1.4749 | TRUE | 0.229 |
| Gm20199 | 2 | 12 | 8 | 2 | 5 | 3 | 2 | 12 | 8 | 2 | 5 | 3 | 5.4 | 7 | 3 | 0.451 | -1.15 | 0.23331986 | 0.44800594 | 0.2875 | 0.6174 | 0.4551 | 0.4551 | TRUE | 0.178 |
| Gm2022 | 940 | 334 | 174 | 1328 | 1027 | 446 | 998 | 334 | 175 | 1044 | 1105 | 488 | 690.65 | 502 | 879 | 1.751 | 0.808 | 0.29883967 | 0.51926751 | 0.4341 | 0.0117 | 0.1665 | 0.4341 | TRUE | 3.3708 |
| Gm20257 | 1 | 0 | 0 | 1 | 0 | 0 | 0 | 1 | 0 | 0 | 1 | 0 | 0.31 | 0 | 0 | 0.954 | -0.067 | 0.98625928 | NA | 0 | 10.7014 | 9.0384 | 9.0384 | TRUE | 0.4614 |
| Gm20268 | 0 | 0 | 0 | 0 | 0 | 0 | 0 | 0 | 0 | 0 | 0 | 0 | 0 | 0 | 0 | NA | NA | NA | NA | NA | NA | NA | NA | NA | NA |
| Gm2027 | 121 | 39 | 36 | 263 | 199 | 61 | 129 | 39 | 36 | 207 | 214 | 67 | 115.21 | 68 | 163 | 2.397 | 1.261 | 0.10044388 | 0.26204412 | 0.4144 | 0.0355 | 0.1917 | 0.4144 | TRUE | 5.3788 |
| Gm2030 | 0 | 0 | 1 | 1 | 2 | 0 | 0 | 0 | 1 | 1 | 2 | 0 | 0.66 | 0 | 1 | 2.668 | 1.416 | 0.58743033 | NA | 0 | 5.0212 | 3.4053 | 3.4053 | TRUE | 0.2754 |
| Gm20300 | 312 | 223 | 234 | 353 | 279 | 275 | 331 | 223 | 235 | 278 | 300 | 301 | 277.97 | 263 | 293 | 1.114 | 0.155 | 0.38986724 | 0.6037429 | 0.021 | 0.0188 | 0.0199 | 0.0199 | TRUE | 0.5727 |
| Gm20324 | 6 | 9 | 7 | 16 | 6 | 9 | 6 | 9 | 7 | 13 | 6 | 10 | 8.55 | 7 | 10 | 1.302 | 0.381 | 0.55956212 | 0.74136428 | 0 | 0.3924 | 0.1889 | 0.1889 | TRUE | 0.2162 |
| Gm20337 | 17 | 24 | 18 | 37 | 28 | 21 | 18 | 24 | 18 | 29 | 30 | 23 | 23.72 | 20 | 27 | 1.37 | 0.454 | 0.25120223 | 0.46875424 | 0 | 0.1458 | 0.0695 | 0.0695 | TRUE | 0.1624 |
| Gm20356 | 0 | 0 | 0 | 0 | 0 | 0 | 0 | 0 | 0 | 0 | 0 | 0 | 0 | 0 | 0 | NA | NA | NA | NA | NA | NA | NA | NA | NA | NA |
| Gm20362 | 0 | 1 | 0 | 1 | 0 | 0 | 0 | 1 | 0 | 1 | 0 | 0 | 0.3 | 0 | 0 | 0.954 | -0.067 | 0.98642001 | NA | 0 | 11.0817 | 9.3014 | 9.3014 | TRUE | 0.405 |
| Gm20408 | 0 | 0 | 0 | 0 | 0 | 0 | 0 | 0 | 0 | 0 | 0 | 0 | 0 | 0 | 0 | NA | NA | NA | NA | NA | NA | NA | NA | NA | NA |
| Gm20410 | 0 | 0 | 0 | 0 | 0 | 0 | 0 | 0 | 0 | 0 | 0 | 0 | 0 | 0 | 0 | NA | NA | NA | NA | NA | NA | NA | NA | NA | NA |
| Gm2042 | 0 | 0 | 0 | 0 | 0 | 0 | 0 | 0 | 0 | 0 | 0 | 0 | 0 | 0 | 0 | NA | NA | NA | NA | NA | NA | NA | NA | NA | NA |
| Gm20516 | 0 | 0 | 0 | 0 | 0 | 0 | 0 | 0 | 0 | 0 | 0 | 0 | 0 | 0 | 0 | NA | NA | NA | NA | NA | NA | NA | NA | NA | NA |
| Gm20554 | 6 | 14 | 10 | 16 | 13 | 9 | 6 | 14 | 10 | 13 | 14 | 10 | 11.14 | 10 | 12 | 1.196 | 0.259 | 0.65856354 | 0.8075619 | 0 | 0.3028 | 0.1567 | 0.1567 | TRUE | 0.2346 |

| | | | | | | | | | | | | | | | | | | | | | | | | | |
|---|---|---|---|---|---|---|---|---|---|---|---|---|---|---|---|---|---|---|---|---|---|---|---|---|---|
| Gm20556 | 1 | 2 | 2 | 2 | 3 | 3 | 1 | 2 | 2 | 2 | 3 | 3 | 2.19 | 2 | 3 | 1.563 | 0.645 | 0.60331534 | NA | 0 | 1.5099 | 0.6238 | 0.6238 | TRUE | 0.2155 |
| Gm20579 | 0 | 0 | 0 | 0 | 0 | 0 | 0 | 0 | 0 | 0 | 0 | 0 | 0 | 0 | 0 | NA | NA | NA | NA | NA | NA | NA | NA | NA | NA |
| Gm20580 | 1 | 0 | 0 | 1 | 0 | 0 | 1 | 0 | 0 | 1 | 0 | 0 | 0.31 | 0 | 0 | 0.954 | -0.067 | 0.98625928 | NA | 0 | 10.7014 | 9.0384 | 9.0384 | TRUE | 0.4614 |
| Gm20581 | 0 | 0 | 0 | 0 | 0 | 0 | 0 | 0 | 0 | 0 | 0 | 0 | 0 | 0 | 0 | NA | NA | NA | NA | NA | NA | NA | NA | NA | NA |
| Gm20594 | 0 | 0 | 0 | 0 | 0 | 0 | 0 | 0 | 0 | 0 | 0 | 0 | 0 | 0 | 0 | NA | NA | NA | NA | NA | NA | NA | NA | NA | NA |
| Gm20597 | 1 | 6 | 0 | 2 | 6 | 6 | 1 | 6 | 0 | 2 | 6 | 7 | 3.61 | 2 | 5 | 2.033 | 1.024 | 0.43024744 | NA | 0.8648 | 0.9199 | 0.8959 | 0.8959 | TRUE | 1.943 |
| Gm20604 | 40 | 39 | 29 | 44 | 36 | 36 | 42 | 39 | 29 | 35 | 39 | 39 | 37.21 | 37 | 38 | 1.018 | 0.025 | 0.93740582 | 0.96865027 | 0 | 0.0954 | 0.0477 | 0.0477 | TRUE | 0.2466 |
| Gm20605 | 59 | 86 | 72 | 59 | 61 | 57 | 63 | 86 | 72 | 46 | 66 | 62 | 65.88 | 74 | 58 | 0.784 | -0.352 | 0.18995018 | 0.39508927 | 0.0151 | 0.0569 | 0.0364 | 0.0364 | TRUE | 0.3093 |
| Gm2061 | 7 | 12 | 26 | 25 | 17 | 29 | 7 | 12 | 26 | 20 | 18 | 32 | 19.2 | 15 | 23 | 1.522 | 0.606 | 0.28342027 | 0.50503949 | 0.1706 | 0.1785 | 0.1749 | 0.1749 | TRUE | 1.511 |
| Gm20611 | 0 | 0 | 0 | 0 | 0 | 0 | 0 | 0 | 0 | 0 | 0 | 0 | 0 | 0 | 0 | NA | NA | NA | NA | NA | NA | NA | NA | NA | NA |
| Gm20735 | 1 | 0 | 0 | 0 | 0 | 1 | 1 | 0 | 0 | 0 | 0 | 1 | 0.36 | 0 | 0 | 0.954 | -0.067 | 0.98553519 | NA | 0 | 9.1716 | 7.9605 | 7.9605 | TRUE | 0.5383 |
| Gm20736 | 0 | 0 | 0 | 0 | 0 | 0 | 0 | 0 | 0 | 0 | 0 | 0 | 0 | 0 | 0 | NA | NA | NA | NA | NA | NA | NA | NA | NA | NA |
| Gm20738 | 0 | 0 | 0 | 0 | 0 | 0 | 0 | 0 | 0 | 0 | 0 | 0 | 0 | 0 | 0 | NA | NA | NA | NA | NA | NA | NA | NA | NA | NA |
| Gm20740 | 0 | 0 | 0 | 0 | 0 | 0 | 0 | 0 | 0 | 0 | 0 | 0 | 0 | 0 | 0 | NA | NA | NA | NA | NA | NA | NA | NA | NA | NA |
| Gm20741 | 0 | 0 | 0 | 0 | 0 | 0 | 0 | 0 | 0 | 0 | 0 | 0 | 0 | 0 | 0 | NA | NA | NA | NA | NA | NA | NA | NA | NA | NA |
| Gm20743 | 0 | 0 | 0 | 0 | 0 | 0 | 0 | 0 | 0 | 0 | 0 | 0 | 0 | 0 | 0 | NA | NA | NA | NA | NA | NA | NA | NA | NA | NA |
| Gm20744 | 0 | 1 | 0 | 0 | 0 | 0 | 0 | 1 | 0 | 0 | 0 | 0 | 0.17 | 0 | 0 | 0.49 | -1.029 | 0.80087 | NA | 0 | 19.7991 | 10 | 10 | TRUE | 0.405 |
| Gm20745 | 0 | 0 | 0 | 0 | 0 | 0 | 0 | 0 | 0 | 0 | 0 | 0 | 0 | 0 | 0 | NA | NA | NA | NA | NA | NA | NA | NA | NA | NA |
| Gm20747 | 0 | 0 | 0 | 0 | 0 | 0 | 0 | 0 | 0 | 0 | 0 | 0 | 0 | 0 | 0 | NA | NA | NA | NA | NA | NA | NA | NA | NA | NA |
| Gm20748 | 139 | 231 | 220 | 194 | 146 | 176 | 148 | 231 | 221 | 153 | 157 | 193 | 183.58 | 200 | 168 | 0.836 | -0.258 | 0.23013238 | 0.44377649 | 0.0311 | 0.0248 | 0.0279 | 0.0279 | TRUE | 0.5588 |
| Gm20750 | 3 | 0 | 1 | 2 | 0 | 0 | 3 | 0 | 1 | 2 | 0 | 0 | 0.96 | 1 | 1 | 0.407 | -1.296 | 0.5860077 | NA | 0 | 3.4369 | 2.8485 | 2.8485 | TRUE | 0.5389 |
| Gm20751 | 10 | 11 | 5 | 10 | 10 | 17 | 11 | 11 | 5 | 8 | 11 | 19 | 10.64 | 9 | 13 | 1.384 | 0.469 | 0.46986586 | 0.67180893 | 0.0902 | 0.3164 | 0.2058 | 0.2058 | TRUE | 0.6408 |
| Gm20752 | 0 | 1 | 0 | 0 | 0 | 0 | 0 | 1 | 0 | 0 | 0 | 0 | 0.17 | 0 | 0 | 0.49 | -1.029 | 0.80087 | NA | 0 | 19.7991 | 10 | 10 | TRUE | 0.405 |
| Gm20753 | 0 | 0 | 0 | 0 | 0 | 0 | 0 | 0 | 0 | 0 | 0 | 0 | 0 | 0 | 0 | NA | NA | NA | NA | NA | NA | NA | NA | NA | NA |
| Gm20754 | 0 | 0 | 0 | 0 | 0 | 0 | 0 | 0 | 0 | 0 | 0 | 0 | 0 | 0 | 0 | NA | NA | NA | NA | NA | NA | NA | NA | NA | NA |
| Gm20755 | 0 | 0 | 0 | 1 | 0 | 0 | 0 | 0 | 0 | 1 | 0 | 0 | 0.13 | 0 | 0 | 1.859 | 0.894 | 0.82650734 | NA | 0 | 25.1515 | 10 | 10 | TRUE | 0.2563 |
| Gm20756 | 3 | 3 | 2 | 8 | 7 | 1 | 3 | 3 | 2 | 6 | 8 | 1 | 3.85 | 3 | 5 | 1.839 | 0.879 | 0.40182701 | NA | 0 | 0.8624 | 0.5075 | 0.5075 | TRUE | 0.8329 |
| Gm20757 | 0 | 0 | 0 | 1 | 0 | 0 | 0 | 0 | 0 | 1 | 0 | 0 | 0.13 | 0 | 0 | 1.859 | 0.894 | 0.82650734 | NA | 0 | 25.1515 | 10 | 10 | TRUE | 0.2563 |
| Gm20758 | 0 | 1 | 0 | 1 | 0 | 0 | 0 | 1 | 0 | 1 | 0 | 0 | 0.3 | 0 | 0 | 0.954 | -0.067 | 0.98642001 | NA | 0 | 11.0817 | 9.3014 | 9.3014 | TRUE | 0.405 |
| Gm20759 | 0 | 0 | 0 | 0 | 0 | 0 | 0 | 0 | 0 | 0 | 0 | 0 | 0 | 0 | 0 | NA | NA | NA | NA | NA | NA | NA | NA | NA | NA |
| Gm20765 | 0 | 0 | 0 | 0 | 0 | 0 | 0 | 0 | 0 | 0 | 0 | 0 | 0 | 0 | 0 | NA | NA | NA | NA | NA | NA | NA | NA | NA | NA |
| Gm20767 | 92 | 38 | 30 | 161 | 123 | 61 | 98 | 38 | 30 | 127 | 132 | 67 | 81.91 | 55 | 109 | 1.97 | 0.978 | 0.03014386 | 0.11567613 | 0.2408 | 0.0471 | 0.133 | 0.133 | TRUE | 3.7304 |
| Gm20806 | 0 | 0 | 0 | 0 | 0 | 0 | 0 | 0 | 0 | 0 | 0 | 0 | 0 | 0 | 0 | NA | NA | NA | NA | NA | NA | NA | NA | NA | NA |
| Gm20809 | 0 | 0 | 0 | 0 | 0 | 0 | 0 | 0 | 0 | 0 | 0 | 0 | 0 | 0 | 0 | NA | NA | NA | NA | NA | NA | NA | NA | NA | NA |
| Gm20815 | 0 | 0 | 0 | 0 | 0 | 0 | 0 | 0 | 0 | 0 | 0 | 0 | 0 | 0 | 0 | NA | NA | NA | NA | NA | NA | NA | NA | NA | NA |
| Gm20816 | 0 | 0 | 0 | 0 | 0 | 0 | 0 | 0 | 0 | 0 | 0 | 0 | 0 | 0 | 0 | NA | NA | NA | NA | NA | NA | NA | NA | NA | NA |
| Gm20822 | 0 | 0 | 0 | 0 | 0 | 0 | 0 | 0 | 0 | 0 | 0 | 0 | 0 | 0 | 0 | NA | NA | NA | NA | NA | NA | NA | NA | NA | NA |
| Gm20823 | 0 | 0 | 0 | 0 | 0 | 0 | 0 | 0 | 0 | 0 | 0 | 0 | 0 | 0 | 0 | NA | NA | NA | NA | NA | NA | NA | NA | NA | NA |
| Gm20826 | 0 | 0 | 0 | 0 | 0 | 0 | 0 | 0 | 0 | 0 | 0 | 0 | 0 | 0 | 0 | NA | NA | NA | NA | NA | NA | NA | NA | NA | NA |
| Gm2083 | 0 | 0 | 0 | 0 | 0 | 0 | 0 | 0 | 0 | 0 | 0 | 0 | 0 | 0 | 0 | NA | NA | NA | NA | NA | NA | NA | NA | NA | NA |
| Gm20831 | 0 | 0 | 0 | 0 | 0 | 0 | 0 | 0 | 0 | 0 | 0 | 0 | 0 | 0 | 0 | NA | NA | NA | NA | NA | NA | NA | NA | NA | NA |
| Gm20854 | 0 | 0 | 0 | 0 | 0 | 0 | 0 | 0 | 0 | 0 | 0 | 0 | 0 | 0 | 0 | NA | NA | NA | NA | NA | NA | NA | NA | NA | NA |
| Gm20857 | 0 | 0 | 0 | 0 | 0 | 0 | 0 | 0 | 0 | 0 | 0 | 0 | 0 | 0 | 0 | NA | NA | NA | NA | NA | NA | NA | NA | NA | NA |
| Gm20858 | 0 | 0 | 0 | 0 | 0 | 0 | 0 | 0 | 0 | 0 | 0 | 0 | 0 | 0 | 0 | NA | NA | NA | NA | NA | NA | NA | NA | NA | NA |
| Gm20865 | 0 | 0 | 0 | 0 | 0 | 0 | 0 | 0 | 0 | 0 | 0 | 0 | 0 | 0 | 0 | NA | NA | NA | NA | NA | NA | NA | NA | NA | NA |
| Gm20867 | 0 | 0 | 0 | 0 | 0 | 0 | 0 | 0 | 0 | 0 | 0 | 0 | 0 | 0 | 0 | NA | NA | NA | NA | NA | NA | NA | NA | NA | NA |
| Gm20871 | 1 | 0 | 1 | 0 | 0 | 0 | 1 | 0 | 1 | 0 | 0 | 0 | 0.34 | 1 | 0 | 0.261 | -1.939 | 0.60367425 | NA | 0 | 9.5732 | 8.3134 | 8.3134 | TRUE | 0.2529 |
| Gm20877 | 0 | 0 | 0 | 0 | 0 | 0 | 0 | 0 | 0 | 0 | 0 | 0 | 0 | 0 | 0 | NA | NA | NA | NA | NA | NA | NA | NA | NA | NA |
| Gm20878 | 0 | 0 | 0 | 0 | 0 | 0 | 0 | 0 | 0 | 0 | 0 | 0 | 0 | 0 | 0 | NA | NA | NA | NA | NA | NA | NA | NA | NA | NA |
| Gm20917 | 0 | 0 | 0 | 0 | 0 | 0 | 0 | 0 | 0 | 0 | 0 | 0 | 0 | 0 | 0 | NA | NA | NA | NA | NA | NA | NA | NA | NA | NA |
| Gm20939 | 1003 | 583 | 640 | 761 | 577 | 450 | 1065 | 582 | 643 | 598 | 621 | 493 | 666.98 | 763 | 571 | 0.748 | -0.419 | 0.06006356 | 0.18589531 | 0.0603 | 0.0118 | 0.0343 | 0.0343 | TRUE | 1.4161 |
| Gm21002 | 0 | 0 | 0 | 0 | 0 | 0 | 0 | 0 | 0 | 0 | 0 | 0 | 0 | 0 | 0 | NA | NA | NA | NA | NA | NA | NA | NA | NA | NA |
| Gm21057 | 0 | 0 | 0 | 0 | 0 | 0 | 0 | 0 | 0 | 0 | 0 | 0 | 0 | 0 | 0 | NA | NA | NA | NA | NA | NA | NA | NA | NA | NA |
| Gm2109 | 1 | 1 | 2 | 4 | 3 | 1 | 1 | 1 | 2 | 3 | 3 | 1 | 1.92 | 1 | 2 | 1.851 | 0.888 | 0.5097652 | NA | 0 | 1.7204 | 0.7381 | 0.7381 | TRUE | 0.2325 |
| Gm21119 | 2 | 0 | 0 | 1 | 0 | 3 | 2 | 0 | 0 | 1 | 0 | 3 | 1.03 | 1 | 1 | 1.922 | 0.942 | 0.68925662 | NA | 2.2856 | 3.1979 | 2.9062 | 2.9062 | TRUE | 0.8765 |
| Gm2115 | 0 | 0 | 0 | 0 | 0 | 0 | 0 | 0 | 0 | 0 | 0 | 0 | 0 | 0 | 0 | NA | NA | NA | NA | NA | NA | NA | NA | NA | NA |
| Gm21276 | 0 | 0 | 0 | 0 | 0 | 0 | 0 | 0 | 0 | 0 | 0 | 0 | 0 | 0 | 0 | NA | NA | NA | NA | NA | NA | NA | NA | NA | NA |
| Gm21293 | 0 | 0 | 0 | 0 | 0 | 0 | 0 | 0 | 0 | 0 | 0 | 0 | 0 | 0 | 0 | NA | NA | NA | NA | NA | NA | NA | NA | NA | NA |
| Gm21304 | 0 | 0 | 0 | 0 | 0 | 0 | 0 | 0 | 0 | 0 | 0 | 0 | 0 | 0 | 0 | NA | NA | NA | NA | NA | NA | NA | NA | NA | NA |
| Gm21312 | 0 | 0 | 0 | 0 | 0 | 0 | 0 | 0 | 0 | 0 | 0 | 0 | 0 | 0 | 0 | NA | NA | NA | NA | NA | NA | NA | NA | NA | NA |
| Gm21319 | 9 | 2 | 1 | 13 | 13 | 5 | 10 | 2 | 1 | 10 | 14 | 5 | 7.04 | 4 | 10 | 2.387 | 1.255 | 0.17719974 | 0.37777701 | 0.4188 | 0.4749 | 0.4507 | 0.4507 | TRUE | 0.8876 |
| Gm21498 | 0 | 0 | 0 | 0 | 0 | 0 | 0 | 0 | 0 | 0 | 0 | 0 | 0 | 0 | 0 | NA | NA | NA | NA | NA | NA | NA | NA | NA | NA |
| Gm21541 | 0 | 0 | 0 | 0 | 0 | 0 | 0 | 0 | 0 | 0 | 0 | 0 | 0 | 0 | 0 | NA | NA | NA | NA | NA | NA | NA | NA | NA | NA |
| Gm21586 | 0 | 0 | 0 | 0 | 0 | 0 | 0 | 0 | 0 | 0 | 0 | 0 | 0 | 0 | 0 | NA | NA | NA | NA | NA | NA | NA | NA | NA | NA |
| Gm216 | 34 | 31 | 23 | 38 | 35 | 37 | 36 | 31 | 23 | 30 | 38 | 41 | 33.03 | 30 | 36 | 1.193 | 0.254 | 0.47096555 | 0.67235914 | 0 | 0.1066 | 0.0591 | 0.0591 | TRUE | 0.2748 |
| Gm21637 | 0 | 0 | 0 | 0 | 0 | 0 | 0 | 0 | 0 | 0 | 0 | 0 | 0 | 0 | 0 | NA | NA | NA | NA | NA | NA | NA | NA | NA | NA |
| Gm21677 | 0 | 0 | 0 | 0 | 0 | 0 | 0 | 0 | 0 | 0 | 0 | 0 | 0 | 0 | 0 | NA | NA | NA | NA | NA | NA | NA | NA | NA | NA |
| Gm21693 | 0 | 0 | 0 | 0 | 0 | 0 | 0 | 0 | 0 | 0 | 0 | 0 | 0 | 0 | 0 | NA | NA | NA | NA | NA | NA | NA | NA | NA | NA |
| Gm21704 | 0 | 0 | 0 | 0 | 0 | 0 | 0 | 0 | 0 | 0 | 0 | 0 | 0 | 0 | 0 | NA | NA | NA | NA | NA | NA | NA | NA | NA | NA |

| | | | | | | | | | | | | | | | | | | | | | | | | | |
|---|---|---|---|---|---|---|---|---|---|---|---|---|---|---|---|---|---|---|---|---|---|---|---|---|---|
| Gm21708 | 0 | 0 | 0 | 0 | 0 | 0 | 0 | 0 | 0 | 0 | 0 | 0 | 0 | 0 | 0 | NA | NA | NA | NA | NA | NA | NA | NA | NA | NA |
| Gm2176 | 0 | 0 | 0 | 0 | 0 | 0 | 0 | 0 | 0 | 0 | 0 | 0 | 0 | 0 | 0 | NA | NA | NA | NA | NA | NA | NA | NA | NA | NA |
| Gm21943 | 0 | 0 | 0 | 0 | 0 | 0 | 0 | 0 | 0 | 0 | 0 | 0 | 0 | 0 | 0 | NA | NA | NA | NA | NA | NA | NA | NA | NA | NA |
| Gm21944 | 0 | 0 | 0 | 0 | 0 | 0 | 0 | 0 | 0 | 0 | 0 | 0 | 0 | 0 | 0 | NA | NA | NA | NA | NA | NA | NA | NA | NA | NA |
| Gm21945 | 0 | 0 | 0 | 0 | 0 | 0 | 0 | 0 | 0 | 0 | 0 | 0 | 0 | 0 | 0 | NA | NA | NA | NA | NA | NA | NA | NA | NA | NA |
| Gm21946 | 0 | 0 | 0 | 0 | 0 | 0 | 0 | 0 | 0 | 0 | 0 | 0 | 0 | 0 | 0 | NA | NA | NA | NA | NA | NA | NA | NA | NA | NA |
| Gm21947 | 4 | 0 | 1 | 8 | 8 | 1 | 4 | 0 | 1 | 6 | 9 | 1 | 3.54 | 2 | 5 | 3.092 | 1.628 | 0.22014567 | NA | 0.8074 | 0.9374 | 0.8843 | 0.8843 | TRUE | 0.6375 |
| Gm21949 | 0 | 0 | 0 | 0 | 0 | 0 | 0 | 0 | 0 | 0 | 0 | 0 | 0 | 0 | 0 | NA | NA | NA | NA | NA | NA | NA | NA | NA | NA |
| Gm21950 | 0 | 0 | 0 | 0 | 0 | 0 | 0 | 0 | 0 | 0 | 0 | 0 | 0 | 0 | 0 | NA | NA | NA | NA | NA | NA | NA | NA | NA | NA |
| Gm21951 | 0 | 0 | 0 | 0 | 0 | 0 | 0 | 0 | 0 | 0 | 0 | 0 | 0 | 0 | 0 | NA | NA | NA | NA | NA | NA | NA | NA | NA | NA |
| Gm2381 | 798 | 574 | 578 | 1151 | 930 | 721 | 848 | 573 | 580 | 905 | 1001 | 789 | 782.66 | 667 | 898 | 1.347 | 0.43 | 0.01426082 | 0.06822009 | 0.0325 | 0.0111 | 0.0209 | 0.0209 | TRUE | 0.659 |
| Gm2382 | 131 | 114 | 112 | 135 | 90 | 97 | 139 | 114 | 112 | 106 | 97 | 106 | 112.43 | 122 | 103 | 0.847 | -0.239 | 0.22035029 | 0.43242065 | 0 | 0.0362 | 0.0184 | 0.0184 | TRUE | 0.154 |
| Gm2447 | 0 | 0 | 0 | 0 | 0 | 0 | 0 | 0 | 0 | 0 | 0 | 0 | 0 | 0 | 0 | NA | NA | NA | NA | NA | NA | NA | NA | NA | NA |
| Gm2506 | 0 | 0 | 0 | 0 | 0 | 0 | 0 | 0 | 0 | 0 | 0 | 0 | 0 | 0 | 0 | NA | NA | NA | NA | NA | NA | NA | NA | NA | NA |
| Gm2516 | 0 | 0 | 0 | 0 | 0 | 0 | 0 | 0 | 0 | 0 | 0 | 0 | 0 | 0 | 0 | NA | NA | NA | NA | NA | NA | NA | NA | NA | NA |
| Gm2518 | 3 | 4 | 3 | 9 | 5 | 2 | 3 | 4 | 3 | 7 | 5 | 2 | 4.14 | 3 | 5 | 1.459 | 0.545 | 0.56161102 | NA | 0 | 0.8027 | 0.3877 | 0.3877 | TRUE | 0.3374 |
| Gm266 | 13 | 26 | 24 | 35 | 35 | 42 | 14 | 26 | 24 | 28 | 38 | 46 | 29.17 | 21 | 37 | 1.724 | 0.786 | 0.05609506 | 0.17765687 | 0.045 | 0.1198 | 0.0848 | 0.0848 | TRUE | 0.2506 |
| Gm2663 | 0 | 0 | 0 | 0 | 0 | 0 | 0 | 0 | 0 | 0 | 0 | 0 | 0 | 0 | 0 | NA | NA | NA | NA | NA | NA | NA | NA | NA | NA |
| Gm2694 | 709 | 746 | 632 | 847 | 635 | 677 | 753 | 745 | 635 | 666 | 683 | 741 | 703.81 | 711 | 697 | 0.979 | -0.03 | 0.7895518 | 0.8916894 | 0.0046 | 0.0116 | 0.0076 | 0.0076 | TRUE | 0.1046 |
| Gm2696 | 0 | 0 | 0 | 0 | 0 | 0 | 0 | 0 | 0 | 0 | 0 | 0 | 0 | 0 | 0 | NA | NA | NA | NA | NA | NA | NA | NA | NA | NA |
| Gm2721 | 0 | 0 | 1 | 0 | 0 | 0 | 0 | 0 | 1 | 0 | 0 | 0 | 0.17 | 0 | 0 | 0.49 | -1.029 | 0.80087 | NA | 0 | 19.6928 | 10 | 10 | TRUE | 0.4098 |
| Gm2762 | 48 | 31 | 48 | 50 | 29 | 26 | 51 | 31 | 48 | 39 | 31 | 28 | 38.19 | 43 | 33 | 0.766 | -0.384 | 0.26841922 | 0.48829387 | 0.025 | 0.0932 | 0.0602 | 0.0602 | TRUE | 0.4911 |
| Gm2799 | 0 | 0 | 0 | 0 | 0 | 0 | 0 | 0 | 0 | 0 | 0 | 0 | 0 | 0 | 0 | NA | NA | NA | NA | NA | NA | NA | NA | NA | NA |
| Gm2825 | 0 | 0 | 0 | 0 | 0 | 0 | 0 | 0 | 0 | 0 | 0 | 0 | 0 | 0 | 0 | NA | NA | NA | NA | NA | NA | NA | NA | NA | NA |
| Gm2837 | 0 | 0 | 0 | 0 | 0 | 0 | 0 | 0 | 0 | 0 | 0 | 0 | 0 | 0 | 0 | NA | NA | NA | NA | NA | NA | NA | NA | NA | NA |
| Gm2848 | 17 | 13 | 8 | 9 | 7 | 9 | 18 | 13 | 8 | 7 | 8 | 10 | 10.59 | 13 | 8 | 0.624 | -0.681 | 0.26646567 | 0.48614188 | 0 | 0.318 | 0.1714 | 0.1714 | TRUE | 0.1289 |
| Gm2863 | 0 | 0 | 0 | 0 | 0 | 0 | 0 | 0 | 0 | 0 | 0 | 0 | 0 | 0 | 0 | NA | NA | NA | NA | NA | NA | NA | NA | NA | NA |
| Gm2897 | 25 | 37 | 46 | 45 | 31 | 51 | 27 | 37 | 46 | 35 | 33 | 56 | 39.04 | 37 | 41 | 1.129 | 0.175 | 0.63494571 | 0.79485386 | 0.0517 | 0.0913 | 0.0723 | 0.0723 | TRUE | 0.3782 |
| Gm2913 | 0 | 0 | 0 | 0 | 0 | 0 | 0 | 0 | 0 | 0 | 0 | 0 | 0 | 0 | 0 | NA | NA | NA | NA | NA | NA | NA | NA | NA | NA |
| Gm2933 | 0 | 0 | 0 | 0 | 0 | 0 | 0 | 0 | 0 | 0 | 0 | 0 | 0 | 0 | 0 | NA | NA | NA | NA | NA | NA | NA | NA | NA | NA |
| Gm2a | 2508 | 4024 | 4328 | 3509 | 2721 | 3398 | 2664 | 4019 | 4346 | 2759 | 2928 | 3720 | 3405.71 | 3676 | 3136 | 0.853 | -0.23 | 0.22022234 | 0.43242065 | 0.0424 | 0.0079 | 0.0251 | 0.0251 | TRUE | 0.7053 |
| Gm3002 | 3 | 7 | 5 | 11 | 8 | 2 | 3 | 7 | 5 | 9 | 9 | 2 | 5.77 | 5 | 7 | 1.29 | 0.368 | 0.66729874 | 0.81392029 | 0 | 0.5775 | 0.352 | 0.352 | TRUE | 0.7506 |
| Gm3020 | 0 | 0 | 0 | 0 | 0 | 0 | 0 | 0 | 0 | 0 | 0 | 0 | 0 | 0 | 0 | NA | NA | NA | NA | NA | NA | NA | NA | NA | NA |
| Gm3086 | 1 | 0 | 0 | 1 | 0 | 0 | 1 | 0 | 0 | 1 | 0 | 0 | 0.31 | 0 | 0 | 0.954 | -0.067 | 0.98625928 | NA | 0 | 10.7014 | 9.0384 | 9.0384 | TRUE | 0.4614 |
| Gm3139 | 0 | 0 | 0 | 0 | 0 | 0 | 0 | 0 | 0 | 0 | 0 | 0 | 0 | 0 | 0 | NA | NA | NA | NA | NA | NA | NA | NA | NA | NA |
| Gm3143 | 12 | 7 | 3 | 17 | 12 | 8 | 13 | 7 | 3 | 13 | 13 | 9 | 9.63 | 8 | 12 | 1.554 | 0.636 | 0.35715182 | 0.57567691 | 0.0839 | 0.349 | 0.2342 | 0.2342 | TRUE | 0.4489 |
| Gm3219 | 43 | 41 | 28 | 48 | 41 | 28 | 46 | 41 | 28 | 38 | 44 | 31 | 37.87 | 38 | 38 | 0.982 | -0.026 | 0.93935859 | 0.96945854 | 0.0166 | 0.0939 | 0.058 | 0.058 | TRUE | 0.3987 |
| Gm3230 | 3 | 3 | 2 | 2 | 2 | 2 | 3 | 3 | 2 | 2 | 2 | 2 | 2.35 | 3 | 2 | 0.715 | -0.483 | 0.67988015 | NA | 0 | 1.4084 | 0.5512 | 0.5512 | TRUE | 0.0647 |
| Gm3238 | 0 | 0 | 0 | 0 | 0 | 0 | 0 | 0 | 0 | 0 | 0 | 0 | 0 | 0 | 0 | NA | NA | NA | NA | NA | NA | NA | NA | NA | NA |
| Gm3258 | 13 | 7 | 6 | 10 | 3 | 8 | 14 | 7 | 6 | 8 | 3 | 9 | 7.78 | 9 | 7 | 0.747 | -0.421 | 0.56931673 | 0.74821263 | 0.0772 | 0.4304 | 0.2638 | 0.2638 | TRUE | 0.6941 |
| Gm3259 | 0 | 0 | 0 | 0 | 0 | 0 | 0 | 0 | 0 | 0 | 0 | 0 | 0 | 0 | 0 | NA | NA | NA | NA | NA | NA | NA | NA | NA | NA |
| Gm3264 | 1 | 0 | 0 | 1 | 1 | 0 | 1 | 0 | 0 | 1 | 1 | 0 | 0.49 | 0 | 1 | 1.712 | 0.776 | 0.78488768 | NA | 0 | 6.7665 | 4.0387 | 4.0387 | TRUE | 0.4614 |
| Gm3279 | 0 | 0 | 0 | 0 | 0 | 0 | 0 | 0 | 0 | 0 | 0 | 0 | 0 | 0 | 0 | NA | NA | NA | NA | NA | NA | NA | NA | NA | NA |
| Gm3285 | 0 | 0 | 0 | 0 | 0 | 0 | 0 | 0 | 0 | 0 | 0 | 0 | 0 | 0 | 0 | NA | NA | NA | NA | NA | NA | NA | NA | NA | NA |
| Gm3286 | 0 | 0 | 0 | 0 | 0 | 0 | 0 | 0 | 0 | 0 | 0 | 0 | 0 | 0 | 0 | NA | NA | NA | NA | NA | NA | NA | NA | NA | NA |
| Gm3317 | 0 | 0 | 0 | 0 | 0 | 0 | 0 | 0 | 0 | 0 | 0 | 0 | 0 | 0 | 0 | NA | NA | NA | NA | NA | NA | NA | NA | NA | NA |
| Gm3336 | 95 | 111 | 133 | 109 | 93 | 113 | 101 | 111 | 134 | 86 | 100 | 124 | 109.13 | 115 | 103 | 0.891 | -0.166 | 0.4648311 | 0.66787976 | 0.0186 | 0.0371 | 0.0279 | 0.0279 | TRUE | 0.2931 |
| Gm3376 | 0 | 0 | 0 | 0 | 0 | 0 | 0 | 0 | 0 | 0 | 0 | 0 | 0 | 0 | 0 | NA | NA | NA | NA | NA | NA | NA | NA | NA | NA |
| Gm3383 | 0 | 0 | 0 | 0 | 0 | 0 | 0 | 0 | 0 | 0 | 0 | 0 | 0 | 0 | 0 | NA | NA | NA | NA | NA | NA | NA | NA | NA | NA |
| Gm3402 | 0 | 0 | 0 | 0 | 0 | 0 | 0 | 0 | 0 | 0 | 0 | 0 | 0 | 0 | 0 | NA | NA | NA | NA | NA | NA | NA | NA | NA | NA |
| Gm3404 | 0 | 0 | 0 | 0 | 0 | 0 | 0 | 0 | 0 | 0 | 0 | 0 | 0 | 0 | 0 | NA | NA | NA | NA | NA | NA | NA | NA | NA | NA |
| Gm3409 | 0 | 0 | 0 | 0 | 0 | 0 | 0 | 0 | 0 | 0 | 0 | 0 | 0 | 0 | 0 | NA | NA | NA | NA | NA | NA | NA | NA | NA | NA |
| Gm3414 | 157 | 134 | 160 | 184 | 117 | 109 | 167 | 134 | 161 | 145 | 126 | 119 | 141.85 | 154 | 130 | 0.849 | -0.237 | 0.20042155 | 0.40814056 | 0.0053 | 0.0301 | 0.0176 | 0.0176 | TRUE | 0.1362 |
| Gm3415 | 0 | 0 | 0 | 0 | 0 | 0 | 0 | 0 | 0 | 0 | 0 | 0 | 0 | 0 | 0 | NA | NA | NA | NA | NA | NA | NA | NA | NA | NA |
| Gm3417 | 0 | 0 | 0 | 0 | 0 | 0 | 0 | 0 | 0 | 0 | 0 | 0 | 0 | 0 | 0 | NA | NA | NA | NA | NA | NA | NA | NA | NA | NA |
| Gm3428 | 0 | 0 | 0 | 0 | 0 | 0 | 0 | 0 | 0 | 0 | 0 | 0 | 0 | 0 | 0 | NA | NA | NA | NA | NA | NA | NA | NA | NA | NA |
| Gm3434 | 0 | 0 | 0 | 0 | 0 | 0 | 0 | 0 | 0 | 0 | 0 | 0 | 0 | 0 | 0 | NA | NA | NA | NA | NA | NA | NA | NA | NA | NA |
| Gm3435 | 14 | 11 | 13 | 18 | 13 | 8 | 15 | 11 | 13 | 14 | 14 | 9 | 12.63 | 13 | 12 | 0.956 | -0.066 | 0.90231463 | 0.95205059 | 0 | 0.2677 | 0.1267 | 0.1267 | TRUE | 0.2288 |
| Gm3448 | 0 | 0 | 0 | 0 | 0 | 0 | 0 | 0 | 0 | 0 | 0 | 0 | 0 | 0 | 0 | NA | NA | NA | NA | NA | NA | NA | NA | NA | NA |
| Gm3494 | 0 | 0 | 0 | 0 | 0 | 0 | 0 | 0 | 0 | 0 | 0 | 0 | 0 | 0 | 0 | NA | NA | NA | NA | NA | NA | NA | NA | NA | NA |
| Gm3500 | 18 | 28 | 30 | 52 | 23 | 27 | 19 | 28 | 30 | 41 | 25 | 30 | 28.73 | 26 | 32 | 1.241 | 0.312 | 0.42863416 | 0.63849811 | 0.029 | 0.1216 | 0.0766 | 0.0766 | TRUE | 0.504 |
| Gm3558 | 151 | 192 | 187 | 282 | 189 | 221 | 160 | 192 | 188 | 222 | 203 | 242 | 201.14 | 180 | 222 | 1.234 | 0.304 | 0.05446935 | 0.17369401 | 0.0026 | 0.0233 | 0.0129 | 0.0129 | TRUE | 0.0954 |
| Gm3604 | 372 | 365 | 375 | 654 | 548 | 461 | 395 | 365 | 377 | 514 | 590 | 505 | 457.43 | 379 | 536 | 1.415 | 0.501 | 2.02E-05 | 0.00036103 | 0.0026 | 0.0141 | 0.0077 | 0.0077 | TRUE | 0.0871 |
| Gm362 | 0 | 0 | 0 | 0 | 0 | 0 | 0 | 0 | 0 | 0 | 0 | 0 | 0 | 0 | 0 | NA | NA | NA | NA | NA | NA | NA | NA | NA | NA |
| Gm364 | 73 | 110 | 85 | 854 | 544 | 471 | 78 | 110 | 85 | 671 | 585 | 516 | 340.83 | 91 | 591 | 6.498 | 2.7 | 2.11E-50 | 3.03E-47 | 0.0178 | 0.0166 | 0.0171 | 0.0171 | TRUE | 0.1748 |
| Gm3646 | 0 | 0 | 0 | 0 | 0 | 0 | 0 | 0 | 0 | 0 | 0 | 0 | 0 | 0 | 0 | NA | NA | NA | NA | NA | NA | NA | NA | NA | NA |
| Gm3696 | 16 | 17 | 17 | 22 | 25 | 23 | 17 | 17 | 17 | 17 | 27 | 25 | 20.07 | 17 | 23 | 1.347 | 0.43 | 0.32008556 | 0.54115843 | 0 | 0.1711 | 0.0832 | 0.0832 | TRUE | 0.35 |
| Gm3701 | 0 | 0 | 0 | 0 | 0 | 0 | 0 | 0 | 0 | 0 | 0 | 0 | 0 | 0 | 0 | NA | NA | NA | NA | NA | NA | NA | NA | NA | NA |
| Gm3706 | 0 | 0 | 0 | 0 | 0 | 0 | 0 | 0 | 0 | 0 | 0 | 0 | 0 | 0 | 0 | NA | NA | NA | NA | NA | NA | NA | NA | NA | NA |

| | | | | | | | | | | | | | | | | | | | | | | | | | |
|---|---|---|---|---|---|---|---|---|---|---|---|---|---|---|---|---|---|---|---|---|---|---|---|---|---|
| Gm3716 | 229 | 303 | 275 | 247 | 187 | 239 | 243 | 303 | 276 | 194 | 201 | 262 | 246.5 | 274 | 219 | 0.797 | -0.327 | 0.05936996 | 0.18450897 | 0.0151 | 0.0203 | 0.0176 | 0.0176 | TRUE | 0.311 |
| Gm3750 | 0 | 0 | 0 | 0 | 0 | 0 | 0 | 0 | 0 | 0 | 0 | 0 | 0 | 0 | 0 | NA | NA | NA | NA | NA | NA | NA | NA | NA | NA |
| Gm3763 | 0 | 0 | 0 | 0 | 0 | 0 | 0 | 0 | 0 | 0 | 0 | 0 | 0 | 0 | 0 | NA | NA | NA | NA | NA | NA | NA | NA | NA | NA |
| Gm3776 | 0 | 0 | 0 | 0 | 0 | 0 | 0 | 0 | 0 | 0 | 0 | 0 | 0 | 0 | 0 | NA | NA | NA | NA | NA | NA | NA | NA | NA | NA |
| Gm382 | 0 | 0 | 0 | 0 | 0 | 0 | 0 | 0 | 0 | 0 | 0 | 0 | 0 | 0 | 0 | NA | NA | NA | NA | NA | NA | NA | NA | NA | NA |
| Gm3833 | 1 | 4 | 5 | 2 | 1 | 5 | 1 | 4 | 5 | 2 | 1 | 5 | 3.03 | 3 | 3 | 0.789 | -0.342 | 0.77210399 | NA | 0.2309 | 1.093 | 0.6723 | 0.6723 | TRUE | 0.8562 |
| Gm3893 | 0 | 0 | 0 | 0 | 0 | 0 | 0 | 0 | 0 | 0 | 0 | 0 | 0 | 0 | 0 | NA | NA | NA | NA | NA | NA | NA | NA | NA | NA |
| Gm3925 | 55 | 106 | 100 | 69 | 34 | 45 | 58 | 106 | 100 | 54 | 37 | 49 | 67.46 | 88 | 47 | 0.531 | -0.913 | 0.00387937 | 0.02574953 | 0.056 | 0.0557 | 0.0559 | 0.0559 | TRUE | 0.8155 |
| Gm3985 | 0 | 1 | 0 | 2 | 4 | 3 | 0 | 1 | 0 | 2 | 4 | 3 | 1.69 | 0 | 3 | 8.25 | 3.044 | 0.08485592 | NA | 0 | 1.9527 | 1.0871 | 1.0871 | TRUE | 0.3896 |
| Gm4013 | 10 | 23 | 14 | 19 | 12 | 17 | 11 | 23 | 14 | 15 | 13 | 19 | 15.68 | 16 | 16 | 0.971 | -0.042 | 0.93467086 | 0.9675739 | 0.0268 | 0.2169 | 0.1279 | 0.1279 | TRUE | 0.7265 |
| Gm4027 | 0 | 0 | 0 | 0 | 0 | 0 | 0 | 0 | 0 | 0 | 0 | 0 | 0 | 0 | 0 | NA | NA | NA | NA | NA | NA | NA | NA | NA | NA |
| Gm4064 | 0 | 0 | 0 | 0 | 0 | 0 | 0 | 0 | 0 | 0 | 0 | 0 | 0 | 0 | 0 | NA | NA | NA | NA | NA | NA | NA | NA | NA | NA |
| Gm4070 | 0 | 0 | 0 | 0 | 0 | 0 | 0 | 0 | 0 | 0 | 0 | 0 | 0 | 0 | 0 | NA | NA | NA | NA | NA | NA | NA | NA | NA | NA |
| Gm41 | 20 | 27 | 27 | 63 | 53 | 53 | 21 | 27 | 27 | 50 | 57 | 58 | 39.98 | 25 | 55 | 2.175 | 1.121 | 0.00036181 | 0.00399613 | 0 | 0.0893 | 0.0419 | 0.0419 | TRUE | 0.1055 |
| Gm410 | 0 | 0 | 0 | 0 | 0 | 0 | 0 | 0 | 0 | 0 | 0 | 0 | 0 | 0 | 0 | NA | NA | NA | NA | NA | NA | NA | NA | NA | NA |
| Gm4133 | 0 | 0 | 0 | 0 | 0 | 0 | 0 | 0 | 0 | 0 | 0 | 0 | 0 | 0 | 0 | NA | NA | NA | NA | NA | NA | NA | NA | NA | NA |
| Gm4141 | 0 | 0 | 0 | 0 | 0 | 0 | 0 | 0 | 0 | 0 | 0 | 0 | 0 | 0 | 0 | NA | NA | NA | NA | NA | NA | NA | NA | NA | NA |
| Gm4172 | 0 | 0 | 0 | 0 | 0 | 0 | 0 | 0 | 0 | 0 | 0 | 0 | 0 | 0 | 0 | NA | NA | NA | NA | NA | NA | NA | NA | NA | NA |
| Gm4175 | 0 | 0 | 0 | 0 | 0 | 0 | 0 | 0 | 0 | 0 | 0 | 0 | 0 | 0 | 0 | NA | NA | NA | NA | NA | NA | NA | NA | NA | NA |
| Gm4177 | 0 | 0 | 0 | 0 | 0 | 0 | 0 | 0 | 0 | 0 | 0 | 0 | 0 | 0 | 0 | NA | NA | NA | NA | NA | NA | NA | NA | NA | NA |
| Gm4179 | 0 | 0 | 0 | 0 | 0 | 0 | 0 | 0 | 0 | 0 | 0 | 0 | 0 | 0 | 0 | NA | NA | NA | NA | NA | NA | NA | NA | NA | NA |
| Gm4187 | 0 | 0 | 0 | 0 | 0 | 0 | 0 | 0 | 0 | 0 | 0 | 0 | 0 | 0 | 0 | NA | NA | NA | NA | NA | NA | NA | NA | NA | NA |
| Gm4201 | 0 | 0 | 0 | 0 | 0 | 0 | 0 | 0 | 0 | 0 | 0 | 0 | 0 | 0 | 0 | NA | NA | NA | NA | NA | NA | NA | NA | NA | NA |
| Gm4214 | 0 | 0 | 0 | 0 | 0 | 0 | 0 | 0 | 0 | 0 | 0 | 0 | 0 | 0 | 0 | NA | NA | NA | NA | NA | NA | NA | NA | NA | NA |
| Gm4216 | 0 | 0 | 0 | 0 | 0 | 0 | 0 | 0 | 0 | 0 | 0 | 0 | 0 | 0 | 0 | NA | NA | NA | NA | NA | NA | NA | NA | NA | NA |
| Gm4220 | 0 | 0 | 0 | 0 | 0 | 0 | 0 | 0 | 0 | 0 | 0 | 0 | 0 | 0 | 0 | NA | NA | NA | NA | NA | NA | NA | NA | NA | NA |
| Gm4224 | 0 | 0 | 0 | 0 | 0 | 0 | 0 | 0 | 0 | 0 | 0 | 0 | 0 | 0 | 0 | NA | NA | NA | NA | NA | NA | NA | NA | NA | NA |
| Gm4251 | 0 | 0 | 0 | 0 | 0 | 0 | 0 | 0 | 0 | 0 | 0 | 0 | 0 | 0 | 0 | NA | NA | NA | NA | NA | NA | NA | NA | NA | NA |
| Gm4262 | 34 | 54 | 50 | 37 | 30 | 31 | 36 | 54 | 50 | 29 | 32 | 34 | 39.26 | 47 | 32 | 0.676 | -0.565 | 0.07960152 | 0.22383228 | 0 | 0.0908 | 0.0484 | 0.0484 | TRUE | 0.305 |
| Gm4265 | 0 | 0 | 0 | 0 | 0 | 0 | 0 | 0 | 0 | 0 | 0 | 0 | 0 | 0 | 0 | NA | NA | NA | NA | NA | NA | NA | NA | NA | NA |
| Gm4278 | 0 | 0 | 0 | 0 | 0 | 0 | 0 | 0 | 0 | 0 | 0 | 0 | 0 | 0 | 0 | NA | NA | NA | NA | NA | NA | NA | NA | NA | NA |
| Gm428 | 0 | 0 | 0 | 1 | 2 | 2 | 0 | 0 | 0 | 1 | 2 | 2 | 0.85 | 0 | 2 | 9.003 | 3.17 | 0.16844964 | NA | 0 | 3.8621 | 2.5291 | 2.5291 | TRUE | 0.2298 |
| Gm4285 | 4 | 13 | 11 | 17 | 11 | 14 | 4 | 13 | 11 | 13 | 12 | 15 | 11.47 | 9 | 13 | 1.426 | 0.512 | 0.39652641 | 0.60990975 | 0 | 0.2942 | 0.1729 | 0.1729 | TRUE | 0.726 |
| Gm4297 | 0 | 0 | 0 | 0 | 0 | 1 | 0 | 0 | 0 | 0 | 0 | 1 | 0.18 | 0 | 0 | 1.859 | 0.894 | 0.82650734 | NA | 0 | 18.0643 | 10 | 10 | TRUE | 0.5383 |
| Gm4301 | 0 | 0 | 0 | 0 | 0 | 0 | 0 | 0 | 0 | 0 | 0 | 0 | 0 | 0 | 0 | NA | NA | NA | NA | NA | NA | NA | NA | NA | NA |
| Gm4302 | 0 | 0 | 0 | 0 | 0 | 0 | 0 | 0 | 0 | 0 | 0 | 0 | 0 | 0 | 0 | NA | NA | NA | NA | NA | NA | NA | NA | NA | NA |
| Gm4303 | 0 | 0 | 0 | 0 | 0 | 0 | 0 | 0 | 0 | 0 | 0 | 0 | 0 | 0 | 0 | NA | NA | NA | NA | NA | NA | NA | NA | NA | NA |
| Gm4305 | 0 | 0 | 0 | 0 | 0 | 0 | 0 | 0 | 0 | 0 | 0 | 0 | 0 | 0 | 0 | NA | NA | NA | NA | NA | NA | NA | NA | NA | NA |
| Gm4307 | 0 | 0 | 0 | 0 | 0 | 0 | 0 | 0 | 0 | 0 | 0 | 0 | 0 | 0 | 0 | NA | NA | NA | NA | NA | NA | NA | NA | NA | NA |
| Gm4312 | 0 | 0 | 0 | 0 | 0 | 0 | 0 | 0 | 0 | 0 | 0 | 0 | 0 | 0 | 0 | NA | NA | NA | NA | NA | NA | NA | NA | NA | NA |
| Gm4340 | 0 | 0 | 0 | 0 | 0 | 0 | 0 | 0 | 0 | 0 | 0 | 0 | 0 | 0 | 0 | NA | NA | NA | NA | NA | NA | NA | NA | NA | NA |
| Gm4345 | 0 | 0 | 0 | 0 | 0 | 0 | 0 | 0 | 0 | 0 | 0 | 0 | 0 | 0 | 0 | NA | NA | NA | NA | NA | NA | NA | NA | NA | NA |
| Gm4349 | 127 | 141 | 126 | 134 | 107 | 143 | 135 | 141 | 127 | 105 | 115 | 157 | 129.87 | 134 | 126 | 0.933 | -0.1 | 0.63516466 | 0.79491607 | 0.0161 | 0.0323 | 0.0241 | 0.0241 | TRUE | 0.4588 |
| Gm436 | 0 | 0 | 1 | 0 | 0 | 0 | 0 | 0 | 1 | 0 | 0 | 0 | 0.17 | 0 | 0 | 0.49 | -1.029 | 0.80087 | NA | 0 | 19.6928 | 10 | 10 | TRUE | 0.4098 |
| Gm4371 | 99 | 97 | 79 | 121 | 86 | 52 | 105 | 97 | 79 | 95 | 93 | 57 | 87.65 | 94 | 82 | 0.873 | -0.195 | 0.46382057 | 0.66715714 | 0.0345 | 0.0445 | 0.0397 | 0.0397 | TRUE | 0.6193 |
| Gm438 | 0 | 0 | 0 | 0 | 0 | 2 | 0 | 0 | 0 | 0 | 0 | 2 | 0.36 | 0 | 1 | 3.832 | 1.938 | 0.63022034 | NA | 10 | 9.0356 | 10 | 10 | TRUE | 0.9744 |
| Gm44 | 39 | 35 | 23 | 94 | 56 | 44 | 41 | 35 | 23 | 74 | 60 | 48 | 46.96 | 33 | 61 | 1.845 | 0.883 | 0.00898426 | 0.04861056 | 0.0382 | 0.0771 | 0.0589 | 0.0589 | TRUE | 0.2407 |
| Gm4461 | 0 | 0 | 0 | 3 | 1 | 0 | 0 | 0 | 0 | 2 | 1 | 0 | 0.57 | 0 | 1 | 6.182 | 2.628 | 0.40213576 | NA | 0 | 5.7626 | 5.6688 | 5.6688 | TRUE | 0.0684 |
| Gm4477 | 0 | 0 | 0 | 0 | 0 | 0 | 0 | 0 | 0 | 0 | 0 | 0 | 0 | 0 | 0 | NA | NA | NA | NA | NA | NA | NA | NA | NA | NA |
| Gm4489 | 1 | 2 | 2 | 5 | 3 | 1 | 1 | 2 | 2 | 4 | 3 | 1 | 2.22 | 2 | 3 | 1.65 | 0.722 | 0.56588262 | NA | 0 | 1.4908 | 0.6636 | 0.6636 | TRUE | 0.2973 |
| Gm4498 | 0 | 0 | 0 | 0 | 0 | 0 | 0 | 0 | 0 | 0 | 0 | 0 | 0 | 0 | 0 | NA | NA | NA | NA | NA | NA | NA | NA | NA | NA |
| Gm4532 | 30 | 29 | 33 | 29 | 30 | 34 | 32 | 29 | 33 | 23 | 32 | 37 | 31.04 | 31 | 31 | 0.973 | -0.04 | 0.91236085 | 0.95675253 | 0 | 0.113 | 0.0607 | 0.0607 | TRUE | 0.4175 |
| Gm4541 | 0 | 0 | 0 | 0 | 0 | 0 | 0 | 0 | 0 | 0 | 0 | 0 | 0 | 0 | 0 | NA | NA | NA | NA | NA | NA | NA | NA | NA | NA |
| Gm4559 | 0 | 0 | 0 | 0 | 0 | 0 | 0 | 0 | 0 | 0 | 0 | 0 | 0 | 0 | 0 | NA | NA | NA | NA | NA | NA | NA | NA | NA | NA |
| Gm4566 | 0 | 0 | 0 | 0 | 0 | 0 | 0 | 0 | 0 | 0 | 0 | 0 | 0 | 0 | 0 | NA | NA | NA | NA | NA | NA | NA | NA | NA | NA |
| Gm4567 | 0 | 0 | 0 | 0 | 0 | 0 | 0 | 0 | 0 | 0 | 0 | 0 | 0 | 0 | 0 | NA | NA | NA | NA | NA | NA | NA | NA | NA | NA |
| Gm4598 | 7 | 2 | 3 | 11 | 2 | 3 | 7 | 2 | 3 | 9 | 2 | 3 | 4.42 | 4 | 5 | 1.166 | 0.221 | 0.8264284 | NA | 0.2824 | 0.752 | 0.5095 | 0.5095 | TRUE | 1.3195 |
| Gm4636 | 2363 | 1940 | 2015 | 3206 | 2159 | 1890 | 2510 | 1937 | 2023 | 2520 | 2323 | 2069 | 2230.48 | 2157 | 2304 | 1.069 | 0.096 | 0.45696892 | 0.66158562 | 0.0126 | 0.0084 | 0.0115 | 0.0115 | TRUE | 0.2478 |
| Gm4710 | 0 | 0 | 0 | 0 | 0 | 0 | 0 | 0 | 0 | 0 | 0 | 0 | 0 | 0 | 0 | NA | NA | NA | NA | NA | NA | NA | NA | NA | NA |
| Gm4719 | 0 | 0 | 0 | 0 | 0 | 0 | 0 | 0 | 0 | 0 | 0 | 0 | 0 | 0 | 0 | NA | NA | NA | NA | NA | NA | NA | NA | NA | NA |
| Gm4724 | 0 | 0 | 0 | 0 | 0 | 0 | 0 | 0 | 0 | 0 | 0 | 0 | 0 | 0 | 0 | NA | NA | NA | NA | NA | NA | NA | NA | NA | NA |
| Gm4736 | 0 | 0 | 0 | 0 | 0 | 0 | 0 | 0 | 0 | 0 | 0 | 0 | 0 | 0 | 0 | NA | NA | NA | NA | NA | NA | NA | NA | NA | NA |
| Gm4745 | 1 | 0 | 0 | 1 | 1 | 0 | 1 | 0 | 0 | 1 | 1 | 0 | 0.49 | 0 | 1 | 1.712 | 0.776 | 0.78488768 | NA | 0 | 6.7665 | 4.0387 | 4.0387 | TRUE | 0.4614 |
| Gm4759 | 0 | 0 | 0 | 0 | 0 | 0 | 0 | 0 | 0 | 0 | 0 | 0 | 0 | 0 | 0 | NA | NA | NA | NA | NA | NA | NA | NA | NA | NA |
| Gm4763 | 0 | 0 | 0 | 0 | 0 | 0 | 0 | 0 | 0 | 0 | 0 | 0 | 0 | 0 | 0 | NA | NA | NA | NA | NA | NA | NA | NA | NA | NA |
| Gm4776 | 0 | 0 | 0 | 0 | 0 | 0 | 0 | 0 | 0 | 0 | 0 | 0 | 0 | 0 | 0 | NA | NA | NA | NA | NA | NA | NA | NA | NA | NA |
| Gm4787 | 1 | 4 | 8 | 6 | 2 | 3 | 1 | 4 | 8 | 5 | 2 | 3 | 3.87 | 4 | 3 | 0.783 | -0.353 | 0.7363212 | NA | 3.00E-04 | 0.8574 | 0.5308 | 0.5308 | TRUE | 0.2307 |
| Gm4788 | 0 | 0 | 0 | 0 | 0 | 0 | 0 | 0 | 0 | 0 | 0 | 0 | 0 | 0 | 0 | NA | NA | NA | NA | NA | NA | NA | NA | NA | NA |
| Gm4791 | 0 | 0 | 0 | 0 | 0 | 0 | 0 | 0 | 0 | 0 | 0 | 0 | 0 | 0 | 0 | NA | NA | NA | NA | NA | NA | NA | NA | NA | NA |

| | | | | | | | | | | | | | | | | | | | | | | | | | |
|---|---|---|---|---|---|---|---|---|---|---|---|---|---|---|---|---|---|---|---|---|---|---|---|---|---|
| Gm4792 | 0 | 0 | 0 | 0 | 0 | 0 | 0 | 0 | 0 | 0 | 0 | 0 | 0 | 0 | 0 | NA | NA | NA | NA | NA | NA | NA | NA | NA | NA |
| Gm4794 | 0 | 0 | 0 | 0 | 0 | 0 | 0 | 0 | 0 | 0 | 0 | 0 | 0 | 0 | 0 | NA | NA | NA | NA | NA | NA | NA | NA | NA | NA |
| Gm4814 | 0 | 0 | 0 | 0 | 0 | 0 | 0 | 0 | 0 | 0 | 0 | 0 | 0 | 0 | 0 | NA | NA | NA | NA | NA | NA | NA | NA | NA | NA |
| Gm4827 | 9 | 5 | 4 | 5 | 2 | 3 | 10 | 5 | 4 | 4 | 2 | 3 | 4.66 | 6 | 3 | 0.512 | -0.965 | 0.28572645 | 0.50703825 | 0 | 0.7145 | 0.3513 | 0.3513 | TRUE | 0.5206 |
| Gm4832 | 0 | 0 | 0 | 1 | 0 | 0 | 0 | 0 | 0 | 1 | 0 | 0 | 0.13 | 0 | 0 | 1.859 | 0.894 | 0.82650734 | NA | 0 | 25.1515 | 10 | 10 | TRUE | 0.2563 |
| Gm4836 | 0 | 0 | 0 | 0 | 0 | 0 | 0 | 0 | 0 | 0 | 0 | 0 | 0 | 0 | 0 | NA | NA | NA | NA | NA | NA | NA | NA | NA | NA |
| Gm4841 | 0 | 0 | 0 | 0 | 0 | 0 | 0 | 0 | 0 | 0 | 0 | 0 | 0 | 0 | 0 | NA | NA | NA | NA | NA | NA | NA | NA | NA | NA |
| Gm4846 | 0 | 0 | 0 | 0 | 0 | 0 | 0 | 0 | 0 | 0 | 0 | 0 | 0 | 0 | 0 | NA | NA | NA | NA | NA | NA | NA | NA | NA | NA |
| Gm4847 | 0 | 1 | 0 | 0 | 0 | 0 | 0 | 1 | 0 | 0 | 0 | 0 | 0.17 | 0 | 0 | 0.49 | -1.029 | 0.80087 | NA | 0 | 19.7991 | 10 | 10 | TRUE | 0.405 |
| Gm4850 | 0 | 0 | 0 | 0 | 0 | 0 | 0 | 0 | 0 | 0 | 0 | 0 | 0 | 0 | 0 | NA | NA | NA | NA | NA | NA | NA | NA | NA | NA |
| Gm4858 | 133 | 30 | 23 | 644 | 390 | 118 | 141 | 30 | 23 | 506 | 420 | 129 | 208.23 | 65 | 352 | 5.434 | 2.442 | 0.00959654 | 0.05101337 | 0.6313 | 0.0227 | 0.2541 | 0.6313 | TRUE | 1.4337 |
| Gm4861 | 0 | 0 | 0 | 0 | 0 | 0 | 0 | 0 | 0 | 0 | 0 | 0 | 0 | 0 | 0 | NA | NA | NA | NA | NA | NA | NA | NA | NA | NA |
| Gm4871 | 0 | 0 | 0 | 0 | 0 | 0 | 0 | 0 | 0 | 0 | 0 | 0 | 0 | 0 | 0 | NA | NA | NA | NA | NA | NA | NA | NA | NA | NA |
| Gm4872 | 0 | 0 | 3 | 1 | 0 | 0 | 0 | 0 | 3 | 1 | 0 | 0 | 0.63 | 1 | 0 | 0.347 | -1.528 | 0.63042337 | NA | 7.8967 | 5.2108 | 5.757 | 5.757 | TRUE | 1.4422 |
| Gm4884 | 0 | 0 | 0 | 0 | 0 | 0 | 0 | 0 | 0 | 0 | 0 | 0 | 0 | 0 | 0 | NA | NA | NA | NA | NA | NA | NA | NA | NA | NA |
| Gm4890 | 7 | 8 | 11 | 35 | 23 | 6 | 7 | 8 | 11 | 28 | 25 | 7 | 14.22 | 9 | 20 | 2.238 | 1.162 | 0.07476775 | 0.21514997 | 0.2088 | 0.2386 | 0.224 | 0.224 | TRUE | 1.6828 |
| Gm4894 | 0 | 0 | 0 | 0 | 0 | 0 | 0 | 0 | 0 | 0 | 0 | 0 | 0 | 0 | 0 | NA | NA | NA | NA | NA | NA | NA | NA | NA | NA |
| Gm4906 | 0 | 0 | 0 | 0 | 0 | 0 | 0 | 0 | 0 | 0 | 0 | 0 | 0 | 0 | 0 | NA | NA | NA | NA | NA | NA | NA | NA | NA | NA |
| Gm4907 | 0 | 0 | 0 | 0 | 0 | 0 | 0 | 0 | 0 | 0 | 0 | 0 | 0 | 0 | 0 | NA | NA | NA | NA | NA | NA | NA | NA | NA | NA |
| Gm4922 | 6 | 5 | 1 | 2 | 0 | 3 | 6 | 5 | 1 | 2 | 0 | 3 | 2.87 | 4 | 2 | 0.392 | -1.351 | 0.30785689 | NA | 0.3918 | 1.1543 | 0.8384 | 0.8384 | TRUE | 0.4256 |
| Gm4925 | 0 | 2 | 1 | 5 | 1 | 1 | 0 | 2 | 1 | 4 | 1 | 1 | 1.52 | 1 | 2 | 2.069 | 1.049 | 0.52182209 | NA | 0 | 2.1784 | 1.193 | 1.193 | TRUE | 0.5532 |
| Gm4926 | 0 | 0 | 0 | 1 | 1 | 1 | 0 | 0 | 0 | 1 | 1 | 1 | 0.49 | 0 | 1 | 5.224 | 2.385 | 0.38465 | NA | 0 | 6.6923 | 3.9249 | 3.9249 | TRUE | 0.07 |
| Gm4937 | 0 | 0 | 0 | 0 | 0 | 0 | 0 | 0 | 0 | 0 | 0 | 0 | 0 | 0 | 0 | NA | NA | NA | NA | NA | NA | NA | NA | NA | NA |
| Gm4944 | 156 | 103 | 124 | 289 | 209 | 149 | 166 | 103 | 125 | 227 | 225 | 163 | 168.04 | 131 | 205 | 1.569 | 0.65 | 0.00494518 | 0.03107894 | 0.0384 | 0.0265 | 0.0322 | 0.0322 | TRUE | 0.5429 |
| Gm4951 | 0 | 1 | 0 | 0 | 1 | 0 | 0 | 1 | 0 | 0 | 1 | 0 | 0.35 | 0 | 0 | 0.954 | -0.067 | 0.98571955 | NA | 0 | 9.5346 | 8.2194 | 8.2194 | TRUE | 0.5204 |
| Gm4952 | 0 | 0 | 0 | 1 | 1 | 0 | 0 | 0 | 0 | 1 | 1 | 0 | 0.31 | 0 | 1 | 3.32 | 1.731 | 0.65383052 | NA | 0 | 10.6224 | 8.9541 | 8.9541 | TRUE | 0.202 |
| Gm4956 | 0 | 0 | 0 | 0 | 0 | 0 | 0 | 0 | 0 | 0 | 0 | 0 | 0 | 0 | 0 | NA | NA | NA | NA | NA | NA | NA | NA | NA | NA |
| Gm4961 | 226 | 183 | 139 | 216 | 163 | 152 | 240 | 183 | 140 | 170 | 175 | 166 | 178.99 | 188 | 170 | 0.91 | -0.135 | 0.53167773 | 0.71979581 | 0.0311 | 0.0253 | 0.0282 | 0.0282 | TRUE | 0.247 |
| Gm4971 | 253 | 96 | 50 | 443 | 404 | 139 | 269 | 96 | 50 | 348 | 435 | 152 | 224.98 | 138 | 312 | 2.255 | 1.173 | 0.13747229 | 0.32159789 | 0.4444 | 0.0215 | 0.1858 | 0.4444 | TRUE | 1.097 |
| Gm4975 | 186 | 244 | 236 | 207 | 225 | 282 | 198 | 244 | 237 | 163 | 242 | 309 | 231.95 | 226 | 238 | 1.048 | 0.068 | 0.77521905 | 0.88305321 | 0.0525 | 0.0211 | 0.0364 | 0.0364 | TRUE | 0.2298 |
| Gm4980 | 12 | 14 | 13 | 21 | 24 | 12 | 13 | 14 | 13 | 17 | 26 | 13 | 15.87 | 13 | 19 | 1.39 | 0.475 | 0.34227543 | 0.56253908 | 0 | 0.2144 | 0.1151 | 0.1151 | TRUE | 0.576 |
| Gm4981 | 4 | 1 | 3 | 9 | 2 | 2 | 4 | 1 | 3 | 7 | 2 | 2 | 3.28 | 3 | 4 | 1.425 | 0.511 | 0.64821704 | NA | 0 | 1.0115 | 0.5935 | 0.5935 | TRUE | 1.1565 |
| Gm4984 | 29 | 51 | 41 | 47 | 42 | 46 | 31 | 51 | 41 | 37 | 45 | 50 | 42.57 | 41 | 44 | 1.071 | 0.099 | 0.76247032 | 0.87472228 | 0.0187 | 0.0843 | 0.053 | 0.053 | TRUE | 0.2071 |
| Gm5 | 4 | 7 | 3 | 13 | 5 | 6 | 4 | 7 | 3 | 10 | 5 | 7 | 6.07 | 5 | 7 | 1.573 | 0.654 | 0.40930632 | 0.62130006 | 0 | 0.5497 | 0.2795 | 0.2795 | TRUE | 0.395 |
| Gm5039 | 828 | 262 | 141 | 1007 | 765 | 301 | 879 | 262 | 142 | 792 | 823 | 329 | 537.82 | 428 | 648 | 1.516 | 0.6 | 0.46930991 | 0.67144012 | 0.4939 | 0.013 | 0.1902 | 0.4939 | TRUE | 4.0939 |
| Gm5065 | 0 | 0 | 0 | 1 | 0 | 0 | 0 | 0 | 0 | 1 | 0 | 0 | 0.13 | 0 | 0 | 1.859 | 0.894 | 0.82650734 | NA | 0 | 25.1515 | 10 | 10 | TRUE | 0.2563 |
| Gm5069 | 11 | 9 | 13 | 5 | 7 | 3 | 12 | 9 | 13 | 4 | 8 | 3 | 8.08 | 11 | 5 | 0.433 | -1.207 | 0.08471794 | 0.23375985 | 0 | 0.4147 | 0.2074 | 0.2074 | TRUE | 0.3896 |
| Gm5071 | 0 | 0 | 0 | 1 | 2 | 0 | 0 | 0 | 0 | 1 | 2 | 0 | 0.49 | 0 | 1 | 5.204 | 2.38 | 0.47267317 | NA | 0 | 6.7348 | 6.4099 | 6.4099 | TRUE | 0.1222 |
| Gm5072 | 0 | 0 | 0 | 0 | 0 | 0 | 0 | 0 | 0 | 0 | 0 | 0 | 0 | 0 | 0 | NA | NA | NA | NA | NA | NA | NA | NA | NA | NA |
| Gm5077 | 39 | 38 | 20 | 49 | 46 | 41 | 41 | 38 | 20 | 39 | 49 | 45 | 38.73 | 33 | 44 | 1.334 | 0.415 | 0.25343895 | 0.47131696 | 0.0406 | 0.092 | 0.0688 | 0.0688 | TRUE | 0.834 |
| Gm5082 | 0 | 0 | 0 | 0 | 0 | 0 | 0 | 0 | 0 | 0 | 0 | 0 | 0 | 0 | 0 | NA | NA | NA | NA | NA | NA | NA | NA | NA | NA |
| Gm5083 | 12 | 13 | 20 | 10 | 12 | 10 | 13 | 13 | 20 | 8 | 13 | 11 | 12.92 | 15 | 11 | 0.684 | -0.548 | 0.31609041 | 0.53684153 | 0 | 0.2619 | 0.1349 | 0.1349 | TRUE | 0.3535 |
| Gm5084 | 0 | 0 | 0 | 0 | 0 | 0 | 0 | 0 | 0 | 0 | 0 | 0 | 0 | 0 | 0 | NA | NA | NA | NA | NA | NA | NA | NA | NA | NA |
| Gm5086 | 6 | 7 | 11 | 4 | 6 | 2 | 6 | 7 | 11 | 3 | 6 | 2 | 6.03 | 8 | 4 | 0.478 | -1.064 | 0.19216674 | 0.3978439 | 0 | 0.553 | 0.2916 | 0.2916 | TRUE | 0.467 |
| Gm5087 | 0 | 0 | 0 | 0 | 0 | 0 | 0 | 0 | 0 | 0 | 0 | 0 | 0 | 0 | 0 | NA | NA | NA | NA | NA | NA | NA | NA | NA | NA |
| Gm5088 | 1 | 1 | 2 | 1 | 1 | 0 | 1 | 1 | 2 | 1 | 1 | 0 | 0.99 | 1 | 1 | 0.465 | -1.104 | 0.55512019 | NA | 0 | 3.3396 | 1.3765 | 1.3765 | TRUE | 0.1869 |
| Gm5089 | 1 | 3 | 2 | 3 | 4 | 4 | 1 | 3 | 2 | 2 | 4 | 4 | 2.85 | 2 | 3 | 1.788 | 0.838 | 0.45309688 | NA | 0 | 1.1624 | 0.5131 | 0.5131 | TRUE | 0.2077 |
| Gm5091 | 286 | 653 | 655 | 521 | 428 | 534 | 304 | 652 | 658 | 410 | 461 | 585 | 511.37 | 538 | 485 | 0.901 | -0.151 | 0.57904161 | 0.7560136 | 0.1006 | 0.0133 | 0.0514 | 0.0514 | TRUE | 1.6888 |
| Gm5095 | 0 | 0 | 0 | 1 | 1 | 0 | 0 | 0 | 0 | 1 | 1 | 0 | 0.31 | 0 | 1 | 3.32 | 1.731 | 0.65383052 | NA | 0 | 10.6224 | 8.9541 | 8.9541 | TRUE | 0.202 |
| Gm5105 | 0 | 0 | 0 | 0 | 1 | 1 | 0 | 0 | 0 | 0 | 1 | 1 | 0.36 | 0 | 1 | 3.794 | 1.924 | 0.60247382 | NA | 0 | 9.1135 | 8.1425 | 8.1425 | TRUE | 0.3482 |
| Gm5108 | 0 | 0 | 0 | 0 | 0 | 0 | 0 | 0 | 0 | 0 | 0 | 0 | 0 | 0 | 0 | NA | NA | NA | NA | NA | NA | NA | NA | NA | NA |
| Gm5111 | 1 | 0 | 0 | 0 | 0 | 0 | 1 | 0 | 0 | 0 | 0 | 0 | 0.18 | 0 | 0 | 0.49 | -1.029 | 0.80087 | NA | 0 | 18.6165 | 10 | 10 | TRUE | 0.4614 |
| Gm5113 | 176 | 180 | 187 | 269 | 175 | 167 | 187 | 180 | 188 | 211 | 188 | 183 | 189.51 | 185 | 194 | 1.054 | 0.076 | 0.61556641 | 0.78256334 | 0 | 0.0243 | 0.0111 | 0.0111 | TRUE | 0.0714 |
| Gm5114 | 0 | 0 | 0 | 0 | 0 | 0 | 0 | 0 | 0 | 0 | 0 | 0 | 0 | 0 | 0 | NA | NA | NA | NA | NA | NA | NA | NA | NA | NA |
| Gm5122 | 3 | 2 | 2 | 4 | 5 | 1 | 3 | 2 | 2 | 3 | 5 | 1 | 2.8 | 2 | 3 | 1.342 | 0.424 | 0.70987246 | NA | 0 | 1.1827 | 0.5718 | 0.5718 | TRUE | 0.15 |
| Gm5124 | 35 | 27 | 30 | 44 | 36 | 23 | 37 | 27 | 30 | 35 | 39 | 25 | 32.13 | 31 | 33 | 1.049 | 0.069 | 0.84699492 | 0.92352627 | 0 | 0.1094 | 0.0599 | 0.0599 | TRUE | 0.2618 |
| Gm5126 | 7 | 4 | 8 | 14 | 5 | 8 | 7 | 4 | 8 | 11 | 5 | 9 | 7.43 | 6 | 8 | 1.307 | 0.387 | 0.58898092 | 0.76368029 | 0 | 0.45 | 0.2323 | 0.2323 | TRUE | 0.2721 |
| Gm5127 | 0 | 0 | 0 | 0 | 1 | 0 | 0 | 0 | 0 | 0 | 1 | 0 | 0.18 | 0 | 0 | 1.859 | 0.894 | 0.82650734 | NA | 0 | 18.3785 | 10 | 10 | TRUE | 0.5204 |
| Gm5128 | 0 | 0 | 0 | 0 | 0 | 0 | 0 | 0 | 0 | 0 | 0 | 0 | 0 | 0 | 0 | NA | NA | NA | NA | NA | NA | NA | NA | NA | NA |
| Gm5129 | 0 | 0 | 0 | 1 | 0 | 1 | 0 | 0 | 0 | 1 | 0 | 1 | 0.31 | 0 | 1 | 3.348 | 1.743 | 0.65064889 | NA | 0 | 10.5167 | 8.8968 | 8.8968 | TRUE | 0.2082 |
| Gm5132 | 0 | 0 | 1 | 0 | 0 | 0 | 0 | 0 | 1 | 0 | 0 | 0 | 0.17 | 0 | 0 | 0.49 | -1.029 | 0.80087 | NA | 0 | 19.6928 | 10 | 10 | TRUE | 0.4098 |
| Gm5134 | 145 | 134 | 113 | 341 | 255 | 232 | 154 | 134 | 113 | 268 | 274 | 254 | 199.62 | 134 | 265 | 1.987 | 0.991 | 2.13E-09 | 1.20E-07 | 0.0024 | 0.0234 | 0.0141 | 0.0141 | TRUE | 0.182 |
| Gm5136 | 3 | 3 | 1 | 19 | 14 | 12 | 3 | 3 | 1 | 15 | 15 | 13 | 8.39 | 2 | 14 | 6.032 | 2.593 | 0.00097672 | 0.00874783 | 0 | 0.3997 | 0.1976 | 0.1976 | TRUE | 0.2794 |
| Gm514 | 5 | 12 | 12 | 28 | 29 | 22 | 5 | 12 | 12 | 22 | 31 | 24 | 17.77 | 10 | 26 | 2.613 | 1.386 | 0.00612175 | 0.03651746 | 0 | 0.1923 | 0.1131 | 0.1131 | TRUE | 0.5145 |
| Gm5141 | 203 | 187 | 226 | 298 | 268 | 212 | 216 | 187 | 227 | 234 | 288 | 232 | 230.67 | 210 | 251 | 1.198 | 0.26 | 0.10847 | 0.27588919 | 0.0084 | 0.0212 | 0.0146 | 0.0146 | TRUE | 0.1778 |
| Gm5142 | 0 | 1 | 0 | 2 | 0 | 0 | 0 | 1 | 0 | 2 | 0 | 0 | 0.43 | 0 | 1 | 1.493 | 0.578 | 0.86962266 | NA | 0 | 7.6954 | 7.0545 | 7.0545 | TRUE | 0.9718 |
| Gm5148 | 0 | 0 | 0 | 1 | 0 | 0 | 0 | 0 | 0 | 1 | 0 | 0 | 0.13 | 0 | 0 | 1.859 | 0.894 | 0.82650734 | NA | 0 | 25.1515 | 10 | 10 | TRUE | 0.2563 |
| Gm5150 | 0 | 0 | 0 | 1 | 0 | 0 | 0 | 0 | 0 | 1 | 0 | 0 | 0.13 | 0 | 0 | 1.859 | 0.894 | 0.82650734 | NA | 0 | 25.1515 | 10 | 10 | TRUE | 0.2563 |
| Gm5166 | 1 | 1 | 0 | 5 | 4 | 0 | 1 | 1 | 0 | 4 | 4 | 0 | 1.72 | 1 | 3 | 4.054 | 2.019 | 0.26365108 | NA | 0 | 1.9269 | 1.4284 | 1.4284 | TRUE | 0.8243 |

| | | | | | | | | | | | | | | | | | | | | | | | | | |
|---|---|---|---|---|---|---|---|---|---|---|---|---|---|---|---|---|---|---|---|---|---|---|---|---|---|
| Gm5168 | 0 | 0 | 0 | 0 | 0 | 0 | 0 | 0 | 0 | 0 | 0 | 0 | 0 | 0 | 0 | NA | NA | NA | NA | NA | NA | NA | NA | NA | NA |
| Gm5169 | 0 | 0 | 0 | 0 | 0 | 0 | 0 | 0 | 0 | 0 | 0 | 0 | 0 | 0 | 0 | NA | NA | NA | NA | NA | NA | NA | NA | NA | NA |
| Gm5176 | 14 | 6 | 4 | 11 | 10 | 1 | 15 | 6 | 4 | 9 | 11 | 1 | 7.56 | 8 | 7 | 0.833 | -0.264 | 0.7711947 | 0.88078194 | 0.4787 | 0.4425 | 0.4592 | 0.4592 | TRUE | 1.3972 |
| Gm5177 | 2 | 0 | 1 | 1 | 5 | 0 | 2 | 0 | 1 | 1 | 5 | 0 | 1.55 | 1 | 2 | 1.963 | 0.973 | 0.60813863 | NA | 1.4824 | 2.1337 | 1.8591 | 1.8591 | TRUE | 1.3215 |
| Gm525 | 3 | 6 | 1 | 18 | 6 | 5 | 3 | 6 | 1 | 14 | 6 | 5 | 6.04 | 3 | 8 | 2.595 | 1.376 | 0.13097264 | 0.31162492 | 0.2111 | 0.552 | 0.3923 | 0.3923 | TRUE | 0.7915 |
| Gm527 | 151 | 203 | 167 | 221 | 165 | 191 | 160 | 203 | 168 | 174 | 178 | 209 | 181.86 | 177 | 187 | 1.053 | 0.075 | 0.66700106 | 0.81379599 | 0.0076 | 0.025 | 0.0161 | 0.0161 | TRUE | 0.1753 |
| Gm5294 | 1 | 0 | 1 | 0 | 0 | 0 | 1 | 0 | 1 | 0 | 0 | 0 | 0.34 | 1 | 0 | 0.261 | -1.939 | 0.60367425 | NA | 0 | 9.5732 | 8.3134 | 8.3134 | TRUE | 0.2529 |
| Gm53 | 85 | 107 | 87 | 192 | 176 | 143 | 90 | 107 | 87 | 151 | 189 | 157 | 130.22 | 95 | 166 | 1.742 | 0.801 | 4.34E-05 | 0.00068731 | 0.0067 | 0.0322 | 0.0193 | 0.0193 | TRUE | 0.1638 |
| Gm5334 | 0 | 0 | 0 | 0 | 0 | 0 | 0 | 0 | 0 | 0 | 0 | 0 | 0 | 0 | 0 | NA | NA | NA | NA | NA | NA | NA | NA | NA | NA |
| Gm5346 | 0 | 0 | 0 | 0 | 0 | 0 | 0 | 0 | 0 | 0 | 0 | 0 | 0 | 0 | 0 | NA | NA | NA | NA | NA | NA | NA | NA | NA | NA |
| Gm5347 | 0 | 0 | 0 | 0 | 0 | 0 | 0 | 0 | 0 | 0 | 0 | 0 | 0 | 0 | 0 | NA | NA | NA | NA | NA | NA | NA | NA | NA | NA |
| Gm5382 | 0 | 0 | 0 | 0 | 0 | 0 | 0 | 0 | 0 | 0 | 0 | 0 | 0 | 0 | 0 | NA | NA | NA | NA | NA | NA | NA | NA | NA | NA |
| Gm5409 | 0 | 0 | 0 | 0 | 0 | 0 | 0 | 0 | 0 | 0 | 0 | 0 | 0 | 0 | 0 | NA | NA | NA | NA | NA | NA | NA | NA | NA | NA |
| Gm5414 | 0 | 0 | 0 | 0 | 0 | 0 | 0 | 0 | 0 | 0 | 0 | 0 | 0 | 0 | 0 | NA | NA | NA | NA | NA | NA | NA | NA | NA | NA |
| Gm5415 | 0 | 0 | 0 | 0 | 0 | 0 | 0 | 0 | 0 | 0 | 0 | 0 | 0 | 0 | 0 | NA | NA | NA | NA | NA | NA | NA | NA | NA | NA |
| Gm5416 | 0 | 0 | 0 | 0 | 0 | 0 | 0 | 0 | 0 | 0 | 0 | 0 | 0 | 0 | 0 | NA | NA | NA | NA | NA | NA | NA | NA | NA | NA |
| Gm5420 | 0 | 0 | 0 | 0 | 0 | 0 | 0 | 0 | 0 | 0 | 0 | 0 | 0 | 0 | 0 | NA | NA | NA | NA | NA | NA | NA | NA | NA | NA |
| Gm5424 | 0 | 0 | 0 | 0 | 0 | 0 | 0 | 0 | 0 | 0 | 0 | 0 | 0 | 0 | 0 | NA | NA | NA | NA | NA | NA | NA | NA | NA | NA |
| Gm5431 | 0 | 0 | 0 | 0 | 0 | 0 | 0 | 0 | 0 | 0 | 0 | 0 | 0 | 0 | 0 | NA | NA | NA | NA | NA | NA | NA | NA | NA | NA |
| Gm5434 | 0 | 0 | 0 | 0 | 0 | 0 | 0 | 0 | 0 | 0 | 0 | 0 | 0 | 0 | 0 | NA | NA | NA | NA | NA | NA | NA | NA | NA | NA |
| Gm5441 | 0 | 0 | 0 | 0 | 0 | 0 | 0 | 0 | 0 | 0 | 0 | 0 | 0 | 0 | 0 | NA | NA | NA | NA | NA | NA | NA | NA | NA | NA |
| Gm5458 | 11 | 2 | 8 | 5 | 7 | 10 | 12 | 2 | 8 | 4 | 8 | 11 | 7.35 | 7 | 8 | 1.022 | 0.031 | 0.97048582 | 0.9846576 | 0.2667 | 0.4549 | 0.3646 | 0.3646 | TRUE | 0.3928 |
| Gm5460 | 0 | 0 | 0 | 0 | 0 | 0 | 0 | 0 | 0 | 0 | 0 | 0 | 0 | 0 | 0 | NA | NA | NA | NA | NA | NA | NA | NA | NA | NA |
| Gm5464 | 1 | 1 | 0 | 2 | 1 | 2 | 1 | 1 | 0 | 2 | 1 | 2 | 1.15 | 1 | 2 | 2.356 | 1.236 | 0.48643677 | NA | 0 | 2.8723 | 1.2227 | 1.2227 | TRUE | 0.2531 |
| Gm5468 | 5 | 3 | 1 | 4 | 2 | 1 | 5 | 3 | 1 | 3 | 2 | 1 | 2.62 | 3 | 2 | 0.704 | -0.506 | 0.67405715 | NA | 0 | 1.2658 | 0.6541 | 0.6541 | TRUE | 0.165 |
| Gm5475 | 4 | 1 | 2 | 1 | 1 | 1 | 4 | 1 | 2 | 1 | 1 | 1 | 1.7 | 2 | 1 | 0.405 | -1.305 | 0.37620901 | NA | 0 | 1.9425 | 0.8548 | 0.8548 | TRUE | 0.3747 |
| Gm5476 | 0 | 0 | 0 | 0 | 1 | 0 | 0 | 0 | 0 | 0 | 1 | 0 | 0.18 | 0 | 0 | 1.859 | 0.894 | 0.82650734 | NA | 0 | 18.3785 | 10 | 10 | TRUE | 0.5204 |
| Gm5477 | 0 | 0 | 0 | 0 | 0 | 0 | 0 | 0 | 0 | 0 | 0 | 0 | 0 | 0 | 0 | NA | NA | NA | NA | NA | NA | NA | NA | NA | NA |
| Gm5478 | 0 | 0 | 0 | 0 | 0 | 0 | 0 | 0 | 0 | 0 | 0 | 0 | 0 | 0 | 0 | NA | NA | NA | NA | NA | NA | NA | NA | NA | NA |
| Gm5480 | 15 | 39 | 44 | 43 | 40 | 49 | 16 | 39 | 44 | 34 | 43 | 54 | 38.26 | 33 | 44 | 1.309 | 0.388 | 0.3625931 | 0.58035501 | 0.1173 | 0.093 | 0.1041 | 0.1041 | TRUE | 0.7936 |
| Gm5483 | 0 | 1 | 0 | 0 | 0 | 0 | 0 | 1 | 0 | 0 | 0 | 0 | 0.17 | 0 | 0 | 0.49 | -1.029 | 0.80087 | NA | 0 | 19.7991 | 10 | 10 | TRUE | 0.405 |
| Gm5485 | 0 | 0 | 0 | 0 | 0 | 0 | 0 | 0 | 0 | 0 | 0 | 0 | 0 | 0 | 0 | NA | NA | NA | NA | NA | NA | NA | NA | NA | NA |
| Gm5494 | 0 | 0 | 2 | 5 | 2 | 1 | 0 | 0 | 2 | 4 | 2 | 1 | 1.53 | 1 | 2 | 3.597 | 1.847 | 0.30588298 | NA | 0 | 2.1589 | 1.3866 | 1.3866 | TRUE | 0.823 |
| Gm5506 | 5966 | 7459 | 7989 | 6788 | 4958 | 5956 | 6337 | 7449 | 8022 | 5336 | 5335 | 6520 | 6499.77 | 7269 | 5730 | 0.788 | -0.343 | 0.00552617 | 0.033652 | 0.0138 | 0.0074 | 0.0109 | 0.0109 | TRUE | 0.1769 |
| Gm5512 | 20 | 7 | 10 | 12 | 9 | 11 | 21 | 7 | 10 | 9 | 10 | 12 | 11.57 | 13 | 10 | 0.814 | -0.296 | 0.63641352 | 0.79543785 | 0.0977 | 0.2916 | 0.1952 | 0.1952 | TRUE | 1.19 |
| Gm5523 | 16 | 19 | 9 | 20 | 7 | 22 | 17 | 19 | 9 | 16 | 8 | 24 | 15.39 | 15 | 16 | 1.051 | 0.072 | 0.90232992 | 0.95205059 | 0.1446 | 0.221 | 0.185 | 0.185 | TRUE | 0.1693 |
| Gm5531 | 0 | 0 | 0 | 0 | 2 | 0 | 0 | 0 | 0 | 0 | 2 | 0 | 0.36 | 0 | 1 | 3.775 | 1.916 | 0.63423067 | NA | 10 | 9.1927 | 10 | 10 | TRUE | 0.9743 |
| Gm5535 | 2 | 0 | 2 | 1 | 0 | 1 | 2 | 0 | 2 | 1 | 0 | 1 | 1 | 1 | 1 | 0.461 | -1.116 | 0.58487872 | NA | 0 | 3.294 | 1.8677 | 1.8677 | TRUE | 0.4949 |
| Gm5538 | 0 | 0 | 0 | 0 | 0 | 0 | 0 | 0 | 0 | 0 | 0 | 0 | 0 | 0 | 0 | NA | NA | NA | NA | NA | NA | NA | NA | NA | NA |
| Gm5544 | 0 | 1 | 0 | 0 | 1 | 0 | 0 | 1 | 0 | 0 | 1 | 0 | 0.35 | 0 | 0 | 0.954 | -0.067 | 0.98571955 | NA | 0 | 9.5346 | 8.2194 | 8.2194 | TRUE | 0.5204 |
| Gm5547 | 0 | 0 | 0 | 0 | 0 | 0 | 0 | 0 | 0 | 0 | 0 | 0 | 0 | 0 | 0 | NA | NA | NA | NA | NA | NA | NA | NA | NA | NA |
| Gm5549 | 0 | 0 | 0 | 0 | 0 | 0 | 0 | 0 | 0 | 0 | 0 | 0 | 0 | 0 | 0 | NA | NA | NA | NA | NA | NA | NA | NA | NA | NA |
| Gm5577 | 25 | 30 | 29 | 42 | 33 | 35 | 27 | 30 | 29 | 33 | 36 | 38 | 32.08 | 29 | 36 | 1.243 | 0.314 | 0.34202554 | 0.56227088 | 0 | 0.1096 | 0.0471 | 0.0471 | TRUE | 0.0347 |
| Gm5591 | 0 | 0 | 0 | 0 | 0 | 0 | 0 | 0 | 0 | 0 | 0 | 0 | 0 | 0 | 0 | NA | NA | NA | NA | NA | NA | NA | NA | NA | NA |
| Gm5592 | 0 | 0 | 0 | 0 | 0 | 0 | 0 | 0 | 0 | 0 | 0 | 0 | 0 | 0 | 0 | NA | NA | NA | NA | NA | NA | NA | NA | NA | NA |
| Gm5595 | 1021 | 771 | 730 | 1279 | 973 | 931 | 1084 | 770 | 733 | 1005 | 1047 | 1019 | 943.16 | 862 | 1024 | 1.187 | 0.248 | 0.11270039 | 0.28259308 | 0.0237 | 0.0104 | 0.0165 | 0.0165 | TRUE | 0.6013 |
| Gm5607 | 2 | 3 | 5 | 8 | 1 | 3 | 2 | 3 | 5 | 6 | 1 | 3 | 3.46 | 3 | 3 | 1.075 | 0.104 | 0.92224842 | NA | 0 | 0.9577 | 0.541 | 0.541 | TRUE | 0.3007 |
| Gm561 | 296 | 389 | 388 | 385 | 258 | 265 | 314 | 388 | 390 | 303 | 278 | 290 | 327.14 | 364 | 290 | 0.797 | -0.327 | 0.01840445 | 0.08197227 | 0.0055 | 0.017 | 0.0108 | 0.0108 | TRUE | 0.1616 |
| Gm5615 | 0 | 0 | 0 | 0 | 0 | 0 | 0 | 0 | 0 | 0 | 0 | 0 | 0 | 0 | 0 | NA | NA | NA | NA | NA | NA | NA | NA | NA | NA |
| Gm5617 | 120 | 214 | 236 | 302 | 275 | 203 | 127 | 214 | 237 | 237 | 296 | 222 | 222.28 | 193 | 252 | 1.305 | 0.384 | 0.11294451 | 0.28302522 | 0.0558 | 0.0217 | 0.0377 | 0.0377 | TRUE | 0.9386 |
| Gm5622 | 0 | 1 | 0 | 3 | 1 | 0 | 0 | 1 | 0 | 2 | 1 | 0 | 0.74 | 0 | 1 | 3.203 | 1.679 | 0.50423271 | NA | 0 | 4.4659 | 3.139 | 3.139 | TRUE | 0.1967 |
| Gm5627 | 0 | 0 | 0 | 0 | 0 | 0 | 0 | 0 | 0 | 0 | 0 | 0 | 0 | 0 | 0 | NA | NA | NA | NA | NA | NA | NA | NA | NA | NA |
| Gm5634 | 0 | 0 | 0 | 1 | 0 | 1 | 0 | 0 | 0 | 1 | 0 | 1 | 0.31 | 0 | 1 | 3.348 | 1.743 | 0.65064889 | NA | 0 | 10.5167 | 8.8968 | 8.8968 | TRUE | 0.2082 |
| Gm5635 | 55 | 72 | 75 | 212 | 166 | 198 | 58 | 72 | 75 | 167 | 179 | 217 | 127.94 | 68 | 188 | 2.723 | 1.445 | 5.61E-12 | 5.14E-10 | 0.0106 | 0.0327 | 0.0217 | 0.0217 | TRUE | 0.1996 |
| Gm5640 | 0 | 0 | 0 | 0 | 0 | 1 | 0 | 0 | 0 | 0 | 0 | 1 | 0.18 | 0 | 0 | 1.859 | 0.894 | 0.82650734 | NA | 0 | 18.0643 | 10 | 10 | TRUE | 0.5383 |
| Gm5643 | 11 | 19 | 21 | 27 | 11 | 17 | 12 | 19 | 21 | 21 | 12 | 19 | 17.24 | 17 | 17 | 1.004 | 0.006 | 0.99074753 | 0.99518028 | 0 | 0.198 | 0.1174 | 0.1174 | TRUE | 0.378 |
| Gm5662 | 1476 | 514 | 298 | 2342 | 1528 | 665 | 1568 | 513 | 299 | 1841 | 1644 | 728 | 1098.9 | 793 | 1404 | 1.77 | 0.824 | 0.2878172 | 0.5090214 | 0.432 | 0.0099 | 0.1615 | 0.432 | TRUE | 4.5862 |
| Gm5712 | 1 | 0 | 1 | 0 | 0 | 0 | 1 | 0 | 1 | 0 | 0 | 0 | 0.34 | 1 | 0 | 0.261 | -1.939 | 0.60367425 | NA | 0 | 9.5732 | 8.3134 | 8.3134 | TRUE | 0.2529 |
| Gm572 | 14 | 14 | 11 | 22 | 17 | 9 | 15 | 14 | 11 | 17 | 18 | 10 | 14.22 | 13 | 15 | 1.146 | 0.197 | 0.70391927 | 0.83796624 | 0 | 0.2385 | 0.123 | 0.123 | TRUE | 0.3865 |
| Gm5725 | 0 | 0 | 0 | 0 | 0 | 0 | 0 | 0 | 0 | 0 | 0 | 0 | 0 | 0 | 0 | NA | NA | NA | NA | NA | NA | NA | NA | NA | NA |
| Gm5726 | 0 | 0 | 0 | 0 | 0 | 0 | 0 | 0 | 0 | 0 | 0 | 0 | 0 | 0 | 0 | NA | NA | NA | NA | NA | NA | NA | NA | NA | NA |
| Gm5728 | 0 | 0 | 0 | 0 | 0 | 0 | 0 | 0 | 0 | 0 | 0 | 0 | 0 | 0 | 0 | NA | NA | NA | NA | NA | NA | NA | NA | NA | NA |
| Gm5741 | 5 | 3 | 2 | 7 | 6 | 1 | 5 | 3 | 2 | 6 | 6 | 1 | 3.89 | 3 | 4 | 1.283 | 0.359 | 0.72786431 | NA | 0 | 0.8528 | 0.5066 | 0.5066 | TRUE | 0.6816 |
| Gm5766 | 0 | 0 | 0 | 0 | 0 | 0 | 0 | 0 | 0 | 0 | 0 | 0 | 0 | 0 | 0 | NA | NA | NA | NA | NA | NA | NA | NA | NA | NA |
| Gm5771 | 0 | 0 | 0 | 0 | 0 | 0 | 0 | 0 | 0 | 0 | 0 | 0 | 0 | 0 | 0 | NA | NA | NA | NA | NA | NA | NA | NA | NA | NA |
| Gm5779 | 0 | 0 | 0 | 1 | 0 | 1 | 0 | 0 | 0 | 1 | 0 | 1 | 0.31 | 0 | 1 | 3.348 | 1.743 | 0.65064889 | NA | 0 | 10.5167 | 8.8968 | 8.8968 | TRUE | 0.2082 |
| Gm5795 | 0 | 0 | 1 | 1 | 0 | 2 | 0 | 0 | 1 | 1 | 0 | 2 | 0.66 | 0 | 1 | 2.693 | 1.429 | 0.58329919 | NA | 0 | 4.974 | 3.3955 | 3.3955 | TRUE | 0.2781 |
| Gm5796 | 27 | 41 | 30 | 59 | 52 | 29 | 29 | 41 | 30 | 46 | 56 | 32 | 38.97 | 33 | 45 | 1.345 | 0.427 | 0.22524301 | 0.43798949 | 0.0319 | 0.0914 | 0.0632 | 0.0632 | TRUE | 0.2455 |
| Gm5797 | 2 | 0 | 0 | 1 | 0 | 3 | 2 | 0 | 0 | 1 | 0 | 3 | 1.03 | 1 | 1 | 1.922 | 0.942 | 0.68925662 | NA | 2.2856 | 3.1979 | 2.9062 | 2.9062 | TRUE | 0.8765 |

| | | | | | | | | | | | | | | | | | | | | | | | | | |
|---|---|---|---|---|---|---|---|---|---|---|---|---|---|---|---|---|---|---|---|---|---|---|---|---|---|
| Gm5800 | 0 | 0 | 0 | 0 | 0 | 0 | 0 | 0 | 0 | 0 | 0 | 0 | 0 | 0 | 0 | NA | NA | NA | NA | NA | NA | NA | NA | NA | NA |
| Gm5801 | 6 | 11 | 12 | 12 | 9 | 4 | 6 | 11 | 12 | 9 | 10 | 4 | 8.82 | 10 | 8 | 0.804 | -0.314 | 0.6374972 | 0.79603827 | 0 | 0.3805 | 0.207 | 0.207 | TRUE | 0.383 |
| Gm5803 | 6 | 12 | 8 | 7 | 7 | 6 | 6 | 12 | 8 | 6 | 8 | 7 | 7.67 | 9 | 7 | 0.736 | -0.442 | 0.51791537 | 0.709446 | 0 | 0.4367 | 0.2037 | 0.2037 | TRUE | 0.3213 |
| Gm581 | 5 | 11 | 13 | 4 | 5 | 5 | 5 | 11 | 13 | 3 | 5 | 5 | 7.22 | 10 | 4 | 0.468 | -1.095 | 0.14620904 | 0.33493231 | 0 | 0.4629 | 0.2511 | 0.2511 | TRUE | 0.5196 |
| Gm5820 | 4 | 4 | 4 | 2 | 3 | 2 | 4 | 4 | 4 | 2 | 3 | 2 | 3.21 | 4 | 2 | 0.561 | -0.835 | 0.41718729 | NA | 0 | 1.0338 | 0.4251 | 0.4251 | TRUE | 0.119 |
| Gm5833 | 0 | 0 | 0 | 0 | 0 | 0 | 0 | 0 | 0 | 0 | 0 | 0 | 0 | 0 | 0 | NA | NA | NA | NA | NA | NA | NA | NA | NA | NA |
| Gm5860 | 0 | 0 | 0 | 0 | 0 | 0 | 0 | 0 | 0 | 0 | 0 | 0 | 0 | 0 | 0 | NA | NA | NA | NA | NA | NA | NA | NA | NA | NA |
| Gm5868 | 1 | 1 | 0 | 5 | 3 | 1 | 1 | 1 | 0 | 4 | 3 | 1 | 1.72 | 1 | 3 | 4.078 | 2.028 | 0.21054998 | NA | 0 | 1.9234 | 0.9672 | 0.9672 | TRUE | 0.3003 |
| Gm5878 | 40 | 35 | 25 | 39 | 35 | 22 | 42 | 35 | 25 | 31 | 38 | 24 | 32.49 | 34 | 31 | 0.903 | -0.148 | 0.69092789 | 0.830485 | 0.0242 | 0.1083 | 0.0689 | 0.0689 | TRUE | 0.245 |
| Gm5885 | 0 | 0 | 0 | 0 | 0 | 0 | 0 | 0 | 0 | 0 | 0 | 0 | 0 | 0 | 0 | NA | NA | NA | NA | NA | NA | NA | NA | NA | NA |
| Gm5886 | 0 | 0 | 0 | 0 | 0 | 0 | 0 | 0 | 0 | 0 | 0 | 0 | 0 | 0 | 0 | NA | NA | NA | NA | NA | NA | NA | NA | NA | NA |
| Gm5891 | 0 | 0 | 0 | 0 | 0 | 0 | 0 | 0 | 0 | 0 | 0 | 0 | 0 | 0 | 0 | NA | NA | NA | NA | NA | NA | NA | NA | NA | NA |
| Gm590 | 8 | 12 | 7 | 11 | 9 | 3 | 8 | 12 | 7 | 9 | 10 | 3 | 8.19 | 9 | 7 | 0.792 | -0.336 | 0.6311585 | 0.79265149 | 0 | 0.4093 | 0.2307 | 0.2307 | TRUE | 0.5431 |
| Gm5901 | 0 | 0 | 0 | 1 | 0 | 0 | 0 | 0 | 0 | 1 | 0 | 0 | 0.13 | 0 | 0 | 1.859 | 0.894 | 0.82650734 | NA | 0 | 25.1515 | 10 | 10 | TRUE | 0.2563 |
| Gm5916 | 0 | 0 | 0 | 0 | 0 | 0 | 0 | 0 | 0 | 0 | 0 | 0 | 0 | 0 | 0 | NA | NA | NA | NA | NA | NA | NA | NA | NA | NA |
| Gm5918 | 159 | 162 | 131 | 214 | 138 | 165 | 169 | 162 | 132 | 168 | 148 | 181 | 159.92 | 154 | 166 | 1.077 | 0.107 | 0.55318 | 0.73680632 | 0.0064 | 0.0275 | 0.0171 | 0.0171 | TRUE | 0.1731 |
| Gm5925 | 0 | 0 | 0 | 0 | 0 | 0 | 0 | 0 | 0 | 0 | 0 | 0 | 0 | 0 | 0 | NA | NA | NA | NA | NA | NA | NA | NA | NA | NA |
| Gm5934 | 0 | 0 | 0 | 0 | 0 | 0 | 0 | 0 | 0 | 0 | 0 | 0 | 0 | 0 | 0 | NA | NA | NA | NA | NA | NA | NA | NA | NA | NA |
| Gm5935 | 0 | 0 | 0 | 0 | 0 | 0 | 0 | 0 | 0 | 0 | 0 | 0 | 0 | 0 | 0 | NA | NA | NA | NA | NA | NA | NA | NA | NA | NA |
| Gm5936 | 0 | 0 | 0 | 0 | 0 | 0 | 0 | 0 | 0 | 0 | 0 | 0 | 0 | 0 | 0 | NA | NA | NA | NA | NA | NA | NA | NA | NA | NA |
| Gm5938 | 0 | 0 | 0 | 0 | 0 | 0 | 0 | 0 | 0 | 0 | 0 | 0 | 0 | 0 | 0 | NA | NA | NA | NA | NA | NA | NA | NA | NA | NA |
| Gm5941 | 0 | 0 | 0 | 0 | 0 | 0 | 0 | 0 | 0 | 0 | 0 | 0 | 0 | 0 | 0 | NA | NA | NA | NA | NA | NA | NA | NA | NA | NA |
| Gm595 | 80 | 115 | 105 | 233 | 159 | 149 | 85 | 115 | 105 | 183 | 171 | 163 | 137.1 | 102 | 172 | 1.695 | 0.762 | 5.18E-05 | 0.00079969 | 0.0021 | 0.0309 | 0.0177 | 0.0177 | TRUE | 0.1989 |
| Gm597 | 0 | 0 | 0 | 0 | 0 | 0 | 0 | 0 | 0 | 0 | 0 | 0 | 0 | 0 | 0 | NA | NA | NA | NA | NA | NA | NA | NA | NA | NA |
| Gm6026 | 0 | 0 | 0 | 0 | 0 | 0 | 0 | 0 | 0 | 0 | 0 | 0 | 0 | 0 | 0 | NA | NA | NA | NA | NA | NA | NA | NA | NA | NA |
| Gm6034 | 0 | 0 | 0 | 1 | 0 | 0 | 0 | 0 | 0 | 1 | 0 | 0 | 0.13 | 0 | 0 | 1.859 | 0.894 | 0.82650734 | NA | 0 | 25.1515 | 10 | 10 | TRUE | 0.2563 |
| Gm6040 | 0 | 0 | 0 | 0 | 0 | 0 | 0 | 0 | 0 | 0 | 0 | 0 | 0 | 0 | 0 | NA | NA | NA | NA | NA | NA | NA | NA | NA | NA |
| Gm6042 | 0 | 0 | 0 | 0 | 0 | 0 | 0 | 0 | 0 | 0 | 0 | 0 | 0 | 0 | 0 | NA | NA | NA | NA | NA | NA | NA | NA | NA | NA |
| Gm608 | 1562 | 1254 | 1419 | 2119 | 1492 | 1320 | 1659 | 1252 | 1425 | 1666 | 1605 | 1445 | 1508.73 | 1445 | 1572 | 1.088 | 0.122 | 0.32979793 | 0.55084296 | 0.0111 | 0.0091 | 0.0106 | 0.0106 | TRUE | 0.2008 |
| Gm6084 | 0 | 0 | 0 | 0 | 0 | 0 | 0 | 0 | 0 | 0 | 0 | 0 | 0 | 0 | 0 | NA | NA | NA | NA | NA | NA | NA | NA | NA | NA |
| Gm6086 | 0 | 1 | 0 | 5 | 3 | 1 | 0 | 1 | 0 | 4 | 3 | 1 | 1.54 | 0 | 3 | 7.597 | 2.925 | 0.10986546 | NA | 0 | 2.1434 | 1.2365 | 1.2365 | TRUE | 0.405 |
| Gm609 | 0 | 0 | 0 | 0 | 0 | 0 | 0 | 0 | 0 | 0 | 0 | 0 | 0 | 0 | 0 | NA | NA | NA | NA | NA | NA | NA | NA | NA | NA |
| Gm6116 | 0 | 0 | 0 | 0 | 0 | 0 | 0 | 0 | 0 | 0 | 0 | 0 | 0 | 0 | 0 | NA | NA | NA | NA | NA | NA | NA | NA | NA | NA |
| Gm6121 | 0 | 0 | 0 | 0 | 0 | 0 | 0 | 0 | 0 | 0 | 0 | 0 | 0 | 0 | 0 | NA | NA | NA | NA | NA | NA | NA | NA | NA | NA |
| Gm614 | 0 | 1 | 1 | 0 | 1 | 0 | 0 | 1 | 1 | 0 | 1 | 0 | 0.51 | 1 | 0 | 0.52 | -0.943 | 0.73680291 | NA | 0 | 6.4273 | 3.9121 | 3.9121 | TRUE | 0.5204 |
| Gm6150 | 0 | 0 | 0 | 0 | 0 | 0 | 0 | 0 | 0 | 0 | 0 | 0 | 0 | 0 | 0 | NA | NA | NA | NA | NA | NA | NA | NA | NA | NA |
| Gm6164 | 0 | 0 | 0 | 0 | 0 | 0 | 0 | 0 | 0 | 0 | 0 | 0 | 0 | 0 | 0 | NA | NA | NA | NA | NA | NA | NA | NA | NA | NA |
| Gm6194 | 1 | 2 | 0 | 0 | 0 | 2 | 1 | 2 | 0 | 0 | 0 | 2 | 0.87 | 1 | 1 | 0.69 | -0.535 | 0.82697486 | NA | 0 | 3.7728 | 3.1177 | 3.1177 | TRUE | 0.5667 |
| Gm6213 | 1 | 0 | 0 | 0 | 0 | 0 | 1 | 0 | 0 | 0 | 0 | 0 | 0.18 | 0 | 0 | 0.49 | -1.029 | 0.80087 | NA | 0 | 18.6165 | 10 | 10 | TRUE | 0.4614 |
| Gm6225 | 2 | 1 | 0 | 2 | 2 | 0 | 2 | 1 | 0 | 2 | 2 | 0 | 1.14 | 1 | 1 | 1.213 | 0.279 | 0.8846965 | NA | 0 | 2.8937 | 1.7759 | 1.7759 | TRUE | 0.2368 |
| Gm6249 | 0 | 0 | 0 | 0 | 0 | 0 | 0 | 0 | 0 | 0 | 0 | 0 | 0 | 0 | 0 | NA | NA | NA | NA | NA | NA | NA | NA | NA | NA |
| Gm6251 | 15 | 25 | 28 | 25 | 14 | 28 | 16 | 25 | 28 | 20 | 15 | 31 | 22.4 | 23 | 22 | 0.942 | -0.086 | 0.85135781 | 0.92587032 | 0.0551 | 0.154 | 0.1076 | 0.1076 | TRUE | 0.6502 |
| Gm6260 | 0 | 0 | 0 | 0 | 0 | 0 | 0 | 0 | 0 | 0 | 0 | 0 | 0 | 0 | 0 | NA | NA | NA | NA | NA | NA | NA | NA | NA | NA |
| Gm6268 | 1 | 0 | 0 | 2 | 0 | 1 | 1 | 0 | 0 | 2 | 0 | 1 | 0.62 | 0 | 1 | 2.474 | 1.307 | 0.61898859 | NA | 0 | 5.3076 | 3.4358 | 3.4358 | TRUE | 0.4614 |
| Gm6277 | 2 | 1 | 5 | 3 | 0 | 0 | 2 | 1 | 5 | 2 | 0 | 0 | 1.75 | 3 | 1 | 0.317 | -1.658 | 0.33869122 | NA | 0.6569 | 1.8891 | 1.4088 | 1.4088 | TRUE | 1.1554 |
| Gm6289 | 0 | 0 | 1 | 0 | 0 | 1 | 0 | 0 | 1 | 0 | 0 | 1 | 0.35 | 0 | 0 | 0.954 | -0.067 | 0.98566485 | NA | 0 | 9.4252 | 8.1416 | 8.1416 | TRUE | 0.5383 |
| Gm6297 | 27 | 45 | 41 | 49 | 37 | 54 | 29 | 45 | 41 | 39 | 40 | 59 | 42.04 | 38 | 46 | 1.19 | 0.251 | 0.46163985 | 0.66559865 | 0.0316 | 0.0853 | 0.0595 | 0.0595 | TRUE | 0.4846 |
| Gm6300 | 0 | 0 | 0 | 0 | 0 | 0 | 0 | 0 | 0 | 0 | 0 | 0 | 0 | 0 | 0 | NA | NA | NA | NA | NA | NA | NA | NA | NA | NA |
| Gm6307 | 0 | 0 | 0 | 0 | 0 | 0 | 0 | 0 | 0 | 0 | 0 | 0 | 0 | 0 | 0 | NA | NA | NA | NA | NA | NA | NA | NA | NA | NA |
| Gm6313 | 0 | 0 | 0 | 0 | 0 | 0 | 0 | 0 | 0 | 0 | 0 | 0 | 0 | 0 | 0 | NA | NA | NA | NA | NA | NA | NA | NA | NA | NA |
| Gm6329 | 85 | 111 | 78 | 162 | 140 | 146 | 90 | 111 | 78 | 127 | 151 | 160 | 119.54 | 93 | 146 | 1.562 | 0.643 | 0.00254324 | 0.01863073 | 0.0128 | 0.0345 | 0.0239 | 0.0239 | TRUE | 0.2698 |
| Gm6367 | 0 | 0 | 0 | 0 | 0 | 0 | 0 | 0 | 0 | 0 | 0 | 0 | 0 | 0 | 0 | NA | NA | NA | NA | NA | NA | NA | NA | NA | NA |
| Gm6370 | 0 | 0 | 0 | 0 | 0 | 0 | 0 | 0 | 0 | 0 | 0 | 0 | 0 | 0 | 0 | NA | NA | NA | NA | NA | NA | NA | NA | NA | NA |
| Gm6377 | 0 | 0 | 0 | 1 | 1 | 3 | 0 | 0 | 0 | 1 | 1 | 3 | 0.86 | 0 | 2 | 9.035 | 3.175 | 0.17710044 | NA | 0 | 3.8481 | 2.6998 | 2.6998 | TRUE | 0.4599 |
| Gm6402 | 34 | 27 | 36 | 38 | 23 | 25 | 36 | 27 | 36 | 30 | 25 | 27 | 30.2 | 33 | 27 | 0.83 | -0.269 | 0.44294075 | 0.64974531 | 0 | 0.116 | 0.0553 | 0.0553 | TRUE | 0.184 |
| Gm6406 | 7 | 17 | 11 | 42 | 33 | 17 | 7 | 17 | 11 | 33 | 36 | 19 | 20.43 | 12 | 29 | 2.458 | 1.297 | 0.01038496 | 0.05425358 | 0.0721 | 0.1681 | 0.1248 | 0.1248 | TRUE | 0.602 |
| Gm6408 | 0 | 0 | 0 | 0 | 0 | 0 | 0 | 0 | 0 | 0 | 0 | 0 | 0 | 0 | 0 | NA | NA | NA | NA | NA | NA | NA | NA | NA | NA |
| Gm6416 | 0 | 0 | 0 | 0 | 0 | 0 | 0 | 0 | 0 | 0 | 0 | 0 | 0 | 0 | 0 | NA | NA | NA | NA | NA | NA | NA | NA | NA | NA |
| Gm6432 | 31 | 30 | 21 | 31 | 32 | 12 | 33 | 30 | 21 | 24 | 34 | 13 | 25.99 | 28 | 24 | 0.858 | -0.22 | 0.62731968 | 0.79013624 | 0.0817 | 0.1337 | 0.1098 | 0.1098 | TRUE | 0.231 |
| Gm6455 | 0 | 0 | 0 | 0 | 0 | 0 | 0 | 0 | 0 | 0 | 0 | 0 | 0 | 0 | 0 | NA | NA | NA | NA | NA | NA | NA | NA | NA | NA |
| Gm6460 | 0 | 0 | 0 | 0 | 0 | 0 | 0 | 0 | 0 | 0 | 0 | 0 | 0 | 0 | 0 | NA | NA | NA | NA | NA | NA | NA | NA | NA | NA |
| Gm648 | 1 | 0 | 3 | 4 | 3 | 2 | 1 | 0 | 3 | 3 | 3 | 2 | 2.11 | 1 | 3 | 2.106 | 1.075 | 0.439907 | NA | 0 | 1.5712 | 0.8484 | 0.8484 | TRUE | 0.7193 |
| Gm6484 | 1 | 0 | 0 | 1 | 0 | 0 | 1 | 0 | 0 | 1 | 0 | 0 | 0.31 | 0 | 0 | 0.954 | -0.067 | 0.98625928 | NA | 0 | 10.7014 | 9.0384 | 9.0384 | TRUE | 0.4614 |
| Gm6498 | 0 | 1 | 5 | 5 | 1 | 0 | 0 | 1 | 5 | 4 | 1 | 0 | 1.84 | 2 | 2 | 0.857 | -0.222 | 0.90134398 | NA | 1.7604 | 1.7997 | 1.7839 | 1.7839 | TRUE | 1.2874 |
| Gm6524 | 3 | 2 | 1 | 4 | 2 | 6 | 3 | 2 | 1 | 3 | 2 | 7 | 3.01 | 2 | 4 | 1.912 | 0.935 | 0.40913953 | NA | 0 | 1.1019 | 0.5511 | 0.5511 | TRUE | 0.4969 |
| Gm6525 | 1 | 1 | 2 | 1 | 0 | 0 | 1 | 1 | 2 | 1 | 0 | 0 | 0.81 | 1 | 0 | 0.257 | -1.959 | 0.36384929 | NA | 0 | 4.0781 | 1.9784 | 1.9784 | TRUE | 0.2562 |
| Gm6531 | 0 | 0 | 1 | 0 | 0 | 0 | 0 | 0 | 1 | 0 | 0 | 0 | 0.17 | 0 | 0 | 0.49 | -1.029 | 0.80087 | NA | 0 | 19.6928 | 10 | 10 | TRUE | 0.4098 |
| Gm6537 | 2 | 2 | 1 | 1 | 1 | 1 | 2 | 2 | 1 | 1 | 1 | 1 | 1.35 | 2 | 1 | 0.571 | -0.808 | 0.60104378 | NA | 0 | 2.4525 | 0.9207 | 0.9207 | TRUE | 0.1023 |
| Gm6548 | 538 | 427 | 440 | 671 | 448 | 349 | 571 | 426 | 442 | 527 | 482 | 382 | 471.87 | 480 | 464 | 0.968 | -0.046 | 0.78710693 | 0.89032073 | 0.0243 | 0.0139 | 0.0192 | 0.0192 | TRUE | 0.3223 |

| | | | | | | | | | | | | | | | | | | | | | | | | | |
|---|---|---|---|---|---|---|---|---|---|---|---|---|---|---|---|---|---|---|---|---|---|---|---|---|---|
| Gm6567 | 0 | 0 | 0 | 0 | 0 | 0 | 0 | 0 | 0 | 0 | 0 | 0 | 0 | 0 | 0 | NA | NA | NA | NA | NA | NA | NA | NA | NA | NA |
| Gm6568 | 0 | 0 | 0 | 0 | 0 | 1 | 0 | 0 | 0 | 0 | 0 | 1 | 0.18 | 0 | 0 | 1.859 | 0.894 | 0.82650734 | NA | | 0 | 18.0643 | 10 | 10 | TRUE | 0.5383 |
| Gm6578 | 0 | 1 | 1 | 0 | 0 | 0 | 0 | 1 | 1 | 0 | 0 | 0 | 0.33 | 1 | 0 | 0.268 | -1.901 | 0.61401143 | NA | | 0 | 9.8764 | 8.4772 | 8.4772 | TRUE | 0.2278 |
| Gm6583 | 4 | 3 | 3 | 8 | 4 | 1 | 4 | 3 | 3 | 6 | 4 | 1 | 3.66 | 3 | 4 | 1.167 | 0.223 | 0.82706439 | NA | | 0 | 0.9077 | 0.4808 | 0.4808 | TRUE | 0.303 |
| Gm6588 | 12 | 14 | 9 | 12 | 12 | 7 | 13 | 14 | 9 | 9 | 13 | 8 | 10.96 | 12 | 10 | 0.838 | -0.255 | 0.65810459 | 0.80743861 | | 0 | 0.3074 | 0.1481 | 0.1481 | TRUE | 0.2017 |
| Gm6592 | 0 | 0 | 0 | 1 | 0 | 1 | 0 | 0 | 0 | 1 | 0 | 1 | 0.31 | 0 | 1 | 3.348 | 1.743 | 0.65064889 | NA | | 0 | 10.5167 | 8.8968 | 8.8968 | TRUE | 0.2082 |
| Gm6602 | 0 | 0 | 0 | 0 | 0 | 0 | 0 | 0 | 0 | 0 | 0 | 0 | 0 | 0 | 0 | NA | NA | NA | NA | NA | NA | NA | NA | NA | NA |
| Gm6607 | 1 | 1 | 2 | 4 | 2 | 3 | 1 | 1 | 2 | 3 | 2 | 3 | 2.11 | 1 | 3 | 2.111 | 1.078 | 0.3992365 | NA | | 0 | 1.5696 | 0.6328 | 0.6328 | TRUE | 0.1119 |
| Gm6614 | 0 | 0 | 0 | 0 | 0 | 0 | 0 | 0 | 0 | 0 | 0 | 0 | 0 | 0 | 0 | NA | NA | NA | NA | NA | NA | NA | NA | NA | NA |
| Gm6623 | 3 | 0 | 0 | 2 | 2 | 0 | 3 | 0 | 0 | 2 | 2 | 0 | 1.15 | 1 | 1 | 1.19 | 0.251 | 0.91066667 | NA | 2.3687 | | 2.8672 | 2.7172 | 2.7172 | TRUE | 1.4429 |
| Gm6634 | 0 | 0 | 0 | 0 | 0 | 0 | 0 | 0 | 0 | 0 | 0 | 0 | 0 | 0 | 0 | NA | NA | NA | NA | NA | NA | NA | NA | NA | NA |
| Gm6639 | 9 | 4 | 3 | 13 | 12 | 6 | 10 | 4 | 3 | 10 | 13 | 7 | 7.71 | 6 | 10 | 1.806 | 0.853 | 0.25717759 | 0.47585698 | 0.0554 | | 0.4341 | 0.2657 | 0.2657 | TRUE | 0.6386 |
| Gm6642 | 0 | 1 | 0 | 0 | 1 | 0 | 0 | 1 | 0 | 0 | 1 | 0 | 0.35 | 0 | 0 | 0.954 | -0.067 | 0.98571955 | NA | | 0 | 9.5346 | 8.2194 | 8.2194 | TRUE | 0.5204 |
| Gm6644 | 0 | 0 | 0 | 0 | 0 | 0 | 0 | 0 | 0 | 0 | 0 | 0 | 0 | 0 | 0 | NA | NA | NA | NA | NA | NA | NA | NA | NA | NA |
| Gm6654 | 0 | 1 | 1 | 1 | 0 | 0 | 0 | 1 | 1 | 1 | 0 | 0 | 0.46 | 1 | 0 | 0.52 | -0.943 | 0.74346893 | NA | | 0 | 7.0944 | 4.2167 | 4.2167 | TRUE | 0.2562 |
| Gm6682 | 258 | 240 | 247 | 384 | 245 | 258 | 274 | 240 | 248 | 302 | 264 | 282 | 268.27 | 254 | 283 | 1.116 | 0.158 | 0.24695868 | 0.46379674 | | 0 | 0.0192 | 0.0097 | 0.0097 | TRUE | 0.0528 |
| Gm6696 | 0 | 0 | 0 | 0 | 0 | 0 | 0 | 0 | 0 | 0 | 0 | 0 | 0 | 0 | 0 | NA | NA | NA | NA | NA | NA | NA | NA | NA | NA |
| Gm6710 | 0 | 0 | 0 | 0 | 0 | 0 | 0 | 0 | 0 | 0 | 0 | 0 | 0 | 0 | 0 | NA | NA | NA | NA | NA | NA | NA | NA | NA | NA |
| Gm6756 | 6 | 7 | 7 | 15 | 8 | 13 | 6 | 7 | 7 | 12 | 9 | 14 | 9.17 | 7 | 12 | 1.698 | 0.764 | 0.22209642 | 0.43428028 | | 0 | 0.3661 | 0.1644 | 0.1644 | TRUE | 0.1708 |
| Gm6760 | 0 | 0 | 0 | 0 | 0 | 0 | 0 | 0 | 0 | 0 | 0 | 0 | 0 | 0 | 0 | NA | NA | NA | NA | NA | NA | NA | NA | NA | NA |
| Gm6763 | 0 | 0 | 0 | 0 | 0 | 0 | 0 | 0 | 0 | 0 | 0 | 0 | 0 | 0 | 0 | NA | NA | NA | NA | NA | NA | NA | NA | NA | NA |
| Gm6772 | 14 | 11 | 14 | 21 | 23 | 15 | 15 | 11 | 14 | 17 | 25 | 16 | 16.26 | 13 | 19 | 1.44 | 0.526 | 0.2753281 | 0.49589028 | | 0 | 0.2095 | 0.1036 | 0.1036 | TRUE | 0.3139 |
| Gm6787 | 4 | 3 | 6 | 11 | 4 | 5 | 4 | 3 | 6 | 9 | 4 | 5 | 5.28 | 4 | 6 | 1.408 | 0.494 | 0.55195233 | 0.73599865 | | 0 | 0.6306 | 0.3033 | 0.3033 | TRUE | 0.3917 |
| Gm6792 | 112 | 136 | 104 | 81 | 55 | 71 | 119 | 136 | 104 | 64 | 59 | 78 | 93.3 | 120 | 67 | 0.557 | -0.844 | 0.00018843 | 0.00231643 | 0.0072 | | 0.0422 | 0.0252 | 0.0252 | TRUE | 0.1663 |
| Gm6793 | 0 | 0 | 0 | 0 | 0 | 0 | 0 | 0 | 0 | 0 | 0 | 0 | 0 | 0 | 0 | NA | NA | NA | NA | NA | NA | NA | NA | NA | NA |
| Gm6812 | 7 | 3 | 2 | 7 | 1 | 5 | 7 | 3 | 2 | 6 | 1 | 5 | 4.08 | 4 | 4 | 0.984 | -0.023 | 0.98258699 | NA | 0.219 | | 0.814 | 0.5307 | 0.5307 | TRUE | 0.7908 |
| Gm684 | 34 | 54 | 49 | 63 | 30 | 34 | 36 | 54 | 49 | 50 | 32 | 37 | 43.04 | 46 | 40 | 0.86 | -0.217 | 0.50867236 | 0.70252018 | 0.0233 | | 0.0834 | 0.0543 | 0.0543 | TRUE | 0.4067 |
| Gm6878 | 0 | 0 | 1 | 0 | 1 | 0 | 0 | 0 | 1 | 0 | 1 | 0 | 0.35 | 0 | 0 | 0.954 | -0.067 | 0.98570726 | NA | | 0 | 9.5099 | 8.2019 | 8.2019 | TRUE | 0.5204 |
| Gm6880 | 142 | 194 | 143 | 349 | 225 | 219 | 151 | 194 | 144 | 274 | 242 | 240 | 207.39 | 163 | 252 | 1.551 | 0.633 | 0.00025114 | 0.00293618 | 0.0101 | | 0.0228 | 0.0165 | 0.0165 | TRUE | 0.2974 |
| Gm6890 | 6 | 7 | 6 | 22 | 19 | 14 | 6 | 7 | 6 | 17 | 20 | 15 | 12.08 | 6 | 17 | 2.735 | 1.452 | 0.01040569 | 0.05432589 | | 0 | 0.2797 | 0.1246 | 0.1246 | TRUE | 0.0856 |
| Gm6902 | 0 | 0 | 0 | 1 | 0 | 0 | 0 | 0 | 0 | 1 | 0 | 0 | 0.13 | 0 | 0 | 1.859 | 0.894 | 0.82650734 | NA | | 0 | 25.1515 | 10 | 10 | TRUE | 0.2563 |
| Gm6904 | 0 | 0 | 0 | 0 | 0 | 0 | 0 | 0 | 0 | 0 | 0 | 0 | 0 | 0 | 0 | NA | NA | NA | NA | NA | NA | NA | NA | NA | NA |
| Gm6927 | 0 | 0 | 0 | 0 | 0 | 0 | 0 | 0 | 0 | 0 | 0 | 0 | 0 | 0 | 0 | NA | NA | NA | NA | NA | NA | NA | NA | NA | NA |
| Gm6936 | 0 | 0 | 0 | 0 | 0 | 0 | 0 | 0 | 0 | 0 | 0 | 0 | 0 | 0 | 0 | NA | NA | NA | NA | NA | NA | NA | NA | NA | NA |
| Gm6938 | 2 | 6 | 6 | 3 | 3 | 3 | 2 | 6 | 6 | 2 | 3 | 3 | 3.84 | 5 | 3 | 0.619 | -0.692 | 0.47668676 | NA | | 0 | 0.8659 | 0.4076 | 0.4076 | TRUE | 0.4189 |
| Gm694 | 1 | 2 | 0 | 2 | 2 | 3 | 1 | 2 | 0 | 2 | 2 | 3 | 1.68 | 1 | 2 | 2.264 | 1.179 | 0.43894473 | NA | | 0 | 1.9703 | 0.9603 | 0.9603 | TRUE | 0.3726 |
| Gm6981 | 28 | 13 | 19 | 56 | 41 | 20 | 30 | 13 | 19 | 44 | 44 | 22 | 28.64 | 21 | 37 | 1.793 | 0.842 | 0.07037185 | 0.20653235 | 0.113 | | 0.1219 | 0.1179 | 0.1179 | TRUE | 0.6784 |
| Gm6994 | 0 | 0 | 0 | 0 | 0 | 0 | 0 | 0 | 0 | 0 | 0 | 0 | 0 | 0 | 0 | NA | NA | NA | NA | NA | NA | NA | NA | NA | NA |
| Gm7008 | 1 | 1 | 2 | 4 | 1 | 4 | 1 | 1 | 2 | 3 | 1 | 4 | 2.11 | 1 | 3 | 2.114 | 1.08 | 0.41480717 | NA | | 0 | 1.5672 | 0.7179 | 0.7179 | TRUE | 0.2306 |
| Gm7030 | 1 | 0 | 3 | 3 | 1 | 0 | 1 | 0 | 3 | 2 | 1 | 0 | 1.25 | 1 | 1 | 0.872 | -0.197 | 0.91760008 | NA | | 0 | 2.6394 | 1.8338 | 1.8338 | TRUE | 0.3871 |
| Gm7056 | 0 | 0 | 0 | 0 | 0 | 0 | 0 | 0 | 0 | 0 | 0 | 0 | 0 | 0 | 0 | NA | NA | NA | NA | NA | NA | NA | NA | NA | NA |
| Gm7073 | 0 | 0 | 0 | 0 | 0 | 0 | 0 | 0 | 0 | 0 | 0 | 0 | 0 | 0 | 0 | NA | NA | NA | NA | NA | NA | NA | NA | NA | NA |
| Gm7092 | 0 | 0 | 1 | 0 | 0 | 1 | 0 | 0 | 1 | 0 | 0 | 1 | 0.35 | 0 | 0 | 0.954 | -0.067 | 0.98566485 | NA | | 0 | 9.4252 | 8.1416 | 8.1416 | TRUE | 0.5383 |
| Gm7102 | 5 | 4 | 2 | 17 | 7 | 6 | 5 | 4 | 2 | 13 | 8 | 7 | 6.46 | 4 | 9 | 2.468 | 1.304 | 0.10693381 | 0.27330417 | | 0 | 0.5166 | 0.2825 | 0.2825 | TRUE | 0.6541 |
| Gm7104 | 26 | 10 | 3 | 48 | 37 | 22 | 28 | 10 | 3 | 38 | 40 | 24 | 23.71 | 14 | 34 | 2.517 | 1.332 | 0.04158718 | 0.14488911 | 0.4099 | | 0.1459 | 0.2554 | 0.2554 | TRUE | 1.9016 |
| Gm711 | 1 | 3 | 3 | 8 | 6 | 3 | 1 | 3 | 3 | 6 | 6 | 3 | 3.85 | 2 | 5 | 2.269 | 1.182 | 0.23632983 | NA | | 0 | 0.8626 | 0.4112 | 0.4112 | TRUE | 0.2246 |
| Gm7120 | 120 | 64 | 82 | 95 | 57 | 68 | 127 | 64 | 82 | 75 | 61 | 74 | 80.69 | 91 | 70 | 0.771 | -0.375 | 0.21112943 | 0.42183093 | 0.0571 | | 0.0477 | 0.0523 | 0.0523 | TRUE | 0.5577 |
| Gm7134 | 0 | 0 | 0 | 0 | 0 | 0 | 0 | 0 | 0 | 0 | 0 | 0 | 0 | 0 | 0 | NA | NA | NA | NA | NA | NA | NA | NA | NA | NA |
| Gm715 | 19 | 18 | 12 | 35 | 23 | 13 | 20 | 18 | 12 | 28 | 25 | 14 | 19.45 | 17 | 22 | 1.336 | 0.418 | 0.37959845 | 0.59448576 | 0.0409 | | 0.1763 | 0.1107 | 0.1107 | TRUE | 0.5196 |
| Gm7157 | 0 | 0 | 0 | 1 | 0 | 0 | 0 | 0 | 0 | 1 | 0 | 0 | 0.13 | 0 | 0 | 1.859 | 0.894 | 0.82650734 | NA | | 0 | 25.1515 | 10 | 10 | TRUE | 0.2563 |
| Gm7168 | 0 | 0 | 1 | 0 | 0 | 0 | 0 | 0 | 1 | 0 | 0 | 0 | 0.17 | 0 | 0 | 0.49 | -1.029 | 0.80087 | NA | | 0 | 19.6928 | 10 | 10 | TRUE | 0.4098 |
| Gm7173 | 15 | 15 | 13 | 33 | 17 | 22 | 16 | 15 | 13 | 26 | 18 | 24 | 18.71 | 15 | 23 | 1.562 | 0.644 | 0.14584735 | 0.33434656 | | 0 | 0.1829 | 0.0849 | 0.0849 | TRUE | 0.1623 |
| Gm7244 | 0 | 0 | 1 | 3 | 0 | 0 | 0 | 0 | 1 | 2 | 0 | 0 | 0.56 | 0 | 1 | 2.223 | 1.152 | 0.72337121 | NA | | 0 | 5.8855 | 6.0328 | 6.0328 | TRUE | 1.4404 |
| Gm7257 | 0 | 0 | 0 | 0 | 0 | 0 | 0 | 0 | 0 | 0 | 0 | 0 | 0 | 0 | 0 | NA | NA | NA | NA | NA | NA | NA | NA | NA | NA |
| Gm7271 | 0 | 0 | 0 | 0 | 0 | 0 | 0 | 0 | 0 | 0 | 0 | 0 | 0 | 0 | 0 | NA | NA | NA | NA | NA | NA | NA | NA | NA | NA |
| Gm732 | 0 | 0 | 0 | 0 | 0 | 0 | 0 | 0 | 0 | 0 | 0 | 0 | 0 | 0 | 0 | NA | NA | NA | NA | NA | NA | NA | NA | NA | NA |
| Gm7325 | 1696 | 2734 | 3164 | 2943 | 2394 | 2520 | 1801 | 2730 | 3177 | 2314 | 2576 | 2759 | 2559.47 | 2569 | 2550 | 0.992 | -0.012 | 0.94981587 | 0.97481103 | 0.0441 | | 0.0082 | 0.025 | 0.025 | TRUE | 0.5391 |
| Gm7334 | 0 | 0 | 0 | 0 | 0 | 0 | 0 | 0 | 0 | 0 | 0 | 0 | 0 | 0 | 0 | NA | NA | NA | NA | NA | NA | NA | NA | NA | NA |
| Gm7337 | 0 | 0 | 0 | 0 | 0 | 0 | 0 | 0 | 0 | 0 | 0 | 0 | 0 | 0 | 0 | NA | NA | NA | NA | NA | NA | NA | NA | NA | NA |
| Gm7367 | 0 | 0 | 0 | 0 | 0 | 0 | 0 | 0 | 0 | 0 | 0 | 0 | 0 | 0 | 0 | NA | NA | NA | NA | NA | NA | NA | NA | NA | NA |
| Gm7444 | 1 | 0 | 3 | 2 | 0 | 0 | 1 | 0 | 3 | 2 | 0 | 0 | 0.94 | 1 | 1 | 0.416 | -1.264 | 0.59609826 | NA | | 0 | 3.5075 | 2.8584 | 2.8584 | TRUE | 0.4749 |
| Gm7457 | 0 | 0 | 0 | 0 | 0 | 0 | 0 | 0 | 0 | 0 | 0 | 0 | 0 | 0 | 0 | NA | NA | NA | NA | NA | NA | NA | NA | NA | NA |
| Gm7534 | 7 | 3 | 9 | 3 | 5 | 4 | 7 | 3 | 9 | 2 | 5 | 4 | 5.26 | 6 | 4 | 0.612 | -0.709 | 0.41702894 | 0.62807995 | | 0 | 0.6327 | 0.3476 | 0.3476 | TRUE | 0.5667 |
| Gm7538 | 0 | 0 | 0 | 2 | 1 | 0 | 0 | 0 | 0 | 2 | 1 | 0 | 0.44 | 0 | 1 | 4.754 | 2.249 | 0.50892201 | NA | | 0 | 7.4712 | 6.8061 | 6.8061 | TRUE | 0.2361 |
| Gm7550 | 0 | 0 | 0 | 0 | 0 | 0 | 0 | 0 | 0 | 0 | 0 | 0 | 0 | 0 | 0 | NA | NA | NA | NA | NA | NA | NA | NA | NA | NA |
| Gm757 | 12 | 19 | 16 | 7 | 9 | 3 | 13 | 19 | 16 | 6 | 10 | 3 | 11.04 | 16 | 6 | 0.385 | -1.379 | 0.02810678 | 0.11006223 | | 0 | 0.3052 | 0.1726 | 0.1726 | TRUE | 0.505 |
| Gm7609 | 0 | 0 | 0 | 0 | 0 | 0 | 0 | 0 | 0 | 0 | 0 | 0 | 0 | 0 | 0 | NA | NA | NA | NA | NA | NA | NA | NA | NA | NA |
| Gm7616 | 0 | 0 | 0 | 0 | 0 | 0 | 0 | 0 | 0 | 0 | 0 | 0 | 0 | 0 | 0 | NA | NA | NA | NA | NA | NA | NA | NA | NA | NA |
| Gm765 | 0 | 1 | 0 | 0 | 0 | 0 | 0 | 1 | 0 | 0 | 0 | 0 | 0.17 | 0 | 0 | 0.49 | -1.029 | 0.80087 | NA | | 0 | 19.7991 | 10 | 10 | TRUE | 0.405 |

| | | | | | | | | | | | | | | | | | | | | | | | | | |
|---|---|---|---|---|---|---|---|---|---|---|---|---|---|---|---|---|---|---|---|---|---|---|---|---|---|
| Gm766 | 12 | 15 | 13 | 18 | 10 | 14 | 13 | 15 | 13 | 14 | 11 | 15 | 13.5 | 14 | 13 | 0.988 | -0.017 | 0.97247832 | 0.98541137 | 0 | 0.2509 | 0.1103 | 0.1103 | TRUE | 0.1148 |
| Gm7694 | 117 | 116 | 140 | 142 | 110 | 116 | 124 | 116 | 141 | 112 | 118 | 127 | 122.94 | 127 | 119 | 0.935 | -0.096 | 0.60321775 | 0.77412127 | 0 | 0.0337 | 0.0165 | 0.0165 | TRUE | 0.0918 |
| Gm7714 | 0 | 0 | 0 | 0 | 0 | 0 | 0 | 0 | 0 | 0 | 0 | 0 | 0 | 0 | 0 | NA | NA | NA | NA | NA | NA | NA | NA | NA | NA |
| Gm773 | 0 | 0 | 3 | 44 | 38 | 15 | 0 | 0 | 3 | 35 | 41 | 16 | 15.82 | 1 | 31 | 30.153 | 4.914 | 1.28E-06 | 3.41E-05 | 0.2553 | 0.2152 | 0.2246 | 0.2246 | TRUE | 1.1871 |
| Gm7849 | 0 | 0 | 0 | 0 | 0 | 0 | 0 | 0 | 0 | 0 | 0 | 0 | 0 | 0 | 0 | NA | NA | NA | NA | NA | NA | NA | NA | NA | NA |
| Gm7854 | 1 | 1 | 1 | 2 | 0 | 0 | 1 | 1 | 1 | 2 | 0 | 0 | 0.77 | 1 | 1 | 0.562 | -0.832 | 0.70497892 | NA | 0 | 4.2695 | 2.1321 | 2.1321 | TRUE | 0.9716 |
| Gm7861 | 0 | 0 | 0 | 0 | 0 | 0 | 0 | 0 | 0 | 0 | 0 | 0 | 0 | 0 | 0 | NA | NA | NA | NA | NA | NA | NA | NA | NA | NA |
| Gm7903 | 0 | 0 | 0 | 0 | 0 | 0 | 0 | 0 | 0 | 0 | 0 | 0 | 0 | 0 | 0 | NA | NA | NA | NA | NA | NA | NA | NA | NA | NA |
| Gm7904 | 1 | 3 | 0 | 0 | 2 | 2 | 1 | 3 | 0 | 0 | 2 | 2 | 1.4 | 1 | 1 | 1.036 | 0.051 | 0.97805171 | NA | 0 | 2.3602 | 1.7701 | 1.7701 | TRUE | 0.4654 |
| Gm7977 | 0 | 1 | 1 | 0 | 1 | 0 | 0 | 1 | 1 | 0 | 1 | 0 | 0.51 | 1 | 0 | 0.52 | -0.943 | 0.73680291 | NA | 0 | 6.4273 | 3.9121 | 3.9121 | TRUE | 0.5204 |
| Gm7978 | 0 | 0 | 0 | 0 | 0 | 0 | 0 | 0 | 0 | 0 | 0 | 0 | 0 | 0 | 0 | NA | NA | NA | NA | NA | NA | NA | NA | NA | NA |
| Gm805 | 7 | 10 | 9 | 20 | 20 | 13 | 7 | 10 | 9 | 16 | 22 | 14 | 12.99 | 9 | 17 | 1.938 | 0.954 | 0.07651606 | 0.21834836 | 0 | 0.2605 | 0.1228 | 0.1228 | TRUE | 0.2268 |
| Gm806 | 0 | 2 | 0 | 1 | 3 | 3 | 0 | 2 | 0 | 1 | 3 | 3 | 1.55 | 1 | 2 | 3.555 | 1.83 | 0.31377818 | NA | 0 | 2.1334 | 1.4149 | 1.4149 | TRUE | 0.9737 |
| Gm8096 | 0 | 0 | 0 | 2 | 1 | 0 | 0 | 0 | 0 | 2 | 1 | 0 | 0.44 | 0 | 1 | 4.754 | 2.249 | 0.50892201 | NA | 0 | 7.4712 | 6.8061 | 6.8061 | TRUE | 0.2361 |
| Gm813 | 0 | 0 | 0 | 0 | 0 | 0 | 0 | 0 | 0 | 0 | 0 | 0 | 0 | 0 | 0 | NA | NA | NA | NA | NA | NA | NA | NA | NA | NA |
| Gm815 | 0 | 0 | 0 | 0 | 0 | 0 | 0 | 0 | 0 | 0 | 0 | 0 | 0 | 0 | 0 | NA | NA | NA | NA | NA | NA | NA | NA | NA | NA |
| Gm8179 | 0 | 0 | 0 | 0 | 0 | 0 | 0 | 0 | 0 | 0 | 0 | 0 | 0 | 0 | 0 | NA | NA | NA | NA | NA | NA | NA | NA | NA | NA |
| Gm8221 | 8 | 2 | 0 | 7 | 5 | 4 | 8 | 2 | 0 | 6 | 5 | 4 | 4.29 | 3 | 5 | 1.469 | 0.555 | 0.62877561 | NA | 0.5999 | 0.7744 | 0.7056 | 0.7056 | TRUE | 1.4941 |
| Gm8234 | 1 | 1 | 0 | 2 | 2 | 4 | 1 | 1 | 0 | 2 | 2 | 4 | 1.69 | 1 | 3 | 3.903 | 1.965 | 0.22355965 | NA | 0 | 1.9517 | 0.9466 | 0.9466 | TRUE | 0.3222 |
| Gm826 | 0 | 0 | 1 | 0 | 1 | 0 | 0 | 0 | 1 | 0 | 1 | 0 | 0.35 | 0 | 0 | 0.954 | -0.067 | 0.98570726 | NA | 0 | 9.5099 | 8.2019 | 8.2019 | TRUE | 0.5204 |
| Gm8267 | 1 | 0 | 0 | 1 | 1 | 0 | 1 | 0 | 0 | 1 | 1 | 0 | 0.49 | 0 | 1 | 1.712 | 0.776 | 0.78488768 | NA | 0 | 6.7665 | 4.0387 | 4.0387 | TRUE | 0.4614 |
| Gm829 | 0 | 0 | 0 | 0 | 0 | 0 | 0 | 0 | 0 | 0 | 0 | 0 | 0 | 0 | 0 | NA | NA | NA | NA | NA | NA | NA | NA | NA | NA |
| Gm8298 | 0 | 0 | 0 | 0 | 0 | 0 | 0 | 0 | 0 | 0 | 0 | 0 | 0 | 0 | 0 | NA | NA | NA | NA | NA | NA | NA | NA | NA | NA |
| Gm8300 | 1611 | 551 | 356 | 2212 | 1551 | 620 | 1711 | 550 | 357 | 1739 | 1669 | 679 | 1117.55 | 873 | 1362 | 1.56 | 0.642 | 0.40125574 | 0.6141287 | 0.4207 | 0.0099 | 0.1577 | 0.4207 | TRUE | 5.6623 |
| Gm833 | 0 | 2 | 0 | 1 | 0 | 0 | 0 | 2 | 0 | 1 | 0 | 0 | 0.46 | 1 | 0 | 0.522 | -0.937 | 0.78646161 | NA | 0 | 7.1082 | 6.8573 | 6.8573 | TRUE | 0.9739 |
| Gm8363 | 0 | 0 | 0 | 0 | 0 | 0 | 0 | 0 | 0 | 0 | 0 | 0 | 0 | 0 | 0 | NA | NA | NA | NA | NA | NA | NA | NA | NA | NA |
| Gm8369 | 0 | 0 | 0 | 0 | 0 | 0 | 0 | 0 | 0 | 0 | 0 | 0 | 0 | 0 | 0 | NA | NA | NA | NA | NA | NA | NA | NA | NA | NA |
| Gm839 | 0 | 0 | 0 | 0 | 0 | 0 | 0 | 0 | 0 | 0 | 0 | 0 | 0 | 0 | 0 | NA | NA | NA | NA | NA | NA | NA | NA | NA | NA |
| Gm8439 | 0 | 0 | 0 | 0 | 0 | 0 | 0 | 0 | 0 | 0 | 0 | 0 | 0 | 0 | 0 | NA | NA | NA | NA | NA | NA | NA | NA | NA | NA |
| Gm8453 | 0 | 0 | 0 | 0 | 0 | 0 | 0 | 0 | 0 | 0 | 0 | 0 | 0 | 0 | 0 | NA | NA | NA | NA | NA | NA | NA | NA | NA | NA |
| Gm853 | 29 | 88 | 78 | 34 | 40 | 48 | 31 | 88 | 78 | 27 | 43 | 53 | 53.22 | 66 | 41 | 0.616 | -0.698 | 0.1112145 | 0.28048257 | 0.1744 | 0.0688 | 0.1187 | 0.1187 | TRUE | 1.659 |
| Gm8579 | 2 | 0 | 0 | 1 | 1 | 0 | 2 | 0 | 0 | 1 | 1 | 0 | 0.66 | 1 | 1 | 0.896 | -0.159 | 0.95209679 | NA | 0 | 4.9654 | 3.5226 | 3.5226 | TRUE | 0.9743 |
| Gm8580 | 3 | 1 | 0 | 1 | 0 | 2 | 3 | 1 | 0 | 1 | 0 | 2 | 1.19 | 1 | 1 | 0.709 | -0.496 | 0.80026663 | NA | 0.6399 | 2.7674 | 1.9072 | 1.9072 | TRUE | 0.4567 |
| Gm8615 | 481 | 544 | 516 | 535 | 408 | 449 | 511 | 543 | 518 | 421 | 439 | 492 | 487.23 | 524 | 451 | 0.858 | -0.221 | 0.04814465 | 0.15986051 | 0.0016 | 0.0137 | 0.007 | 0.007 | TRUE | 0.0728 |
| Gm8633 | 0 | 0 | 0 | 0 | 0 | 0 | 0 | 0 | 0 | 0 | 0 | 0 | 0 | 0 | 0 | NA | NA | NA | NA | NA | NA | NA | NA | NA | NA |
| Gm8660 | 0 | 0 | 0 | 0 | 0 | 0 | 0 | 0 | 0 | 0 | 0 | 0 | 0 | 0 | 0 | NA | NA | NA | NA | NA | NA | NA | NA | NA | NA |
| Gm8677 | 0 | 0 | 0 | 0 | 0 | 0 | 0 | 0 | 0 | 0 | 0 | 0 | 0 | 0 | 0 | NA | NA | NA | NA | NA | NA | NA | NA | NA | NA |
| Gm8693 | 0 | 0 | 0 | 0 | 0 | 0 | 0 | 0 | 0 | 0 | 0 | 0 | 0 | 0 | 0 | NA | NA | NA | NA | NA | NA | NA | NA | NA | NA |
| Gm8709 | 0 | 0 | 1 | 0 | 0 | 0 | 0 | 0 | 1 | 0 | 0 | 0 | 0.17 | 0 | 0 | 0.49 | -1.029 | 0.80087 | NA | 0 | 19.6928 | 10 | 10 | TRUE | 0.4098 |
| Gm872 | 1 | 4 | 2 | 1 | 0 | 2 | 1 | 4 | 2 | 1 | 0 | 2 | 1.67 | 2 | 1 | 0.414 | -1.273 | 0.41787787 | NA | 0 | 1.9756 | 1.0635 | 1.0635 | TRUE | 0.4377 |
| Gm8720 | 0 | 0 | 0 | 0 | 0 | 0 | 0 | 0 | 0 | 0 | 0 | 0 | 0 | 0 | 0 | NA | NA | NA | NA | NA | NA | NA | NA | NA | NA |
| Gm8764 | 0 | 0 | 0 | 0 | 0 | 0 | 0 | 0 | 0 | 0 | 0 | 0 | 0 | 0 | 0 | NA | NA | NA | NA | NA | NA | NA | NA | NA | NA |
| Gm8765 | 0 | 0 | 1 | 0 | 0 | 0 | 0 | 0 | 1 | 0 | 0 | 0 | 0.17 | 0 | 0 | 0.49 | -1.029 | 0.80087 | NA | 0 | 19.6928 | 10 | 10 | TRUE | 0.4098 |
| Gm8773 | 4 | 8 | 5 | 11 | 6 | 10 | 4 | 8 | 5 | 9 | 6 | 11 | 7.22 | 6 | 9 | 1.505 | 0.589 | 0.40819314 | 0.62087325 | 0 | 0.4633 | 0.221 | 0.221 | TRUE | 0.255 |
| Gm8787 | 11 | 15 | 7 | 13 | 12 | 17 | 12 | 15 | 7 | 10 | 13 | 19 | 12.57 | 11 | 14 | 1.228 | 0.296 | 0.60659544 | 0.77676352 | 0 | 0.2689 | 0.157 | 0.157 | TRUE | 0.354 |
| Gm8801 | 247 | 162 | 121 | 220 | 137 | 95 | 262 | 162 | 121 | 173 | 147 | 104 | 161.66 | 182 | 141 | 0.78 | -0.358 | 0.24359874 | 0.46017958 | 0.1005 | 0.0273 | 0.0617 | 0.0617 | TRUE | 0.5797 |
| Gm8817 | 5 | 2 | 2 | 13 | 5 | 3 | 5 | 2 | 2 | 10 | 5 | 3 | 4.7 | 3 | 6 | 2.075 | 1.053 | 0.27194586 | 0.49187272 | 0 | 0.7079 | 0.4206 | 0.4206 | TRUE | 0.4707 |
| Gm884 | 21 | 16 | 6 | 31 | 24 | 16 | 22 | 16 | 6 | 24 | 26 | 18 | 18.67 | 15 | 23 | 1.536 | 0.619 | 0.25335433 | 0.47129891 | 0.118 | 0.1834 | 0.1552 | 0.1552 | TRUE | 0.5317 |
| Gm8882 | 0 | 0 | 0 | 0 | 0 | 0 | 0 | 0 | 0 | 0 | 0 | 0 | 0 | 0 | 0 | NA | NA | NA | NA | NA | NA | NA | NA | NA | NA |
| Gm8883 | 0 | 0 | 0 | 0 | 0 | 0 | 0 | 0 | 0 | 0 | 0 | 0 | 0 | 0 | 0 | NA | NA | NA | NA | NA | NA | NA | NA | NA | NA |
| Gm8884 | 0 | 0 | 0 | 0 | 0 | 0 | 0 | 0 | 0 | 0 | 0 | 0 | 0 | 0 | 0 | NA | NA | NA | NA | NA | NA | NA | NA | NA | NA |
| Gm889 | 0 | 0 | 0 | 0 | 0 | 0 | 0 | 0 | 0 | 0 | 0 | 0 | 0 | 0 | 0 | NA | NA | NA | NA | NA | NA | NA | NA | NA | NA |
| Gm8898 | 0 | 0 | 0 | 0 | 0 | 0 | 0 | 0 | 0 | 0 | 0 | 0 | 0 | 0 | 0 | NA | NA | NA | NA | NA | NA | NA | NA | NA | NA |
| Gm8909 | 1 | 7 | 6 | 10 | 6 | 4 | 1 | 7 | 6 | 8 | 6 | 4 | 5.46 | 5 | 6 | 1.332 | 0.413 | 0.64049685 | 0.79767036 | 0 | 0.61 | 0.3778 | 0.3778 | TRUE | 0.8254 |
| Gm8979 | 0 | 0 | 0 | 0 | 0 | 0 | 0 | 0 | 0 | 0 | 0 | 0 | 0 | 0 | 0 | NA | NA | NA | NA | NA | NA | NA | NA | NA | NA |
| Gm8989 | 0 | 0 | 0 | 0 | 0 | 0 | 0 | 0 | 0 | 0 | 0 | 0 | 0 | 0 | 0 | NA | NA | NA | NA | NA | NA | NA | NA | NA | NA |
| Gm8994 | 109 | 40 | 38 | 127 | 120 | 45 | 116 | 40 | 38 | 100 | 129 | 49 | 78.68 | 65 | 93 | 1.438 | 0.524 | 0.28093865 | 0.50220609 | 0.2906 | 0.0488 | 0.157 | 0.157 | TRUE | 0.8203 |
| Gm9 | 163 | 188 | 195 | 548 | 388 | 284 | 173 | 188 | 196 | 431 | 417 | 311 | 285.97 | 186 | 386 | 2.085 | 1.06 | 4.41E-10 | 2.91E-08 | 0.0153 | 0.0184 | 0.0168 | 0.0168 | TRUE | 0.329 |
| Gm904 | 0 | 0 | 0 | 3 | 2 | 3 | 0 | 0 | 0 | 2 | 2 | 3 | 1.3 | 0 | 2 | 13.802 | 3.787 | 0.05455258 | NA | 0 | 2.543 | 1.607 | 1.607 | TRUE | 0.07 |
| Gm9041 | 0 | 0 | 0 | 0 | 0 | 0 | 0 | 0 | 0 | 0 | 0 | 0 | 0 | 0 | 0 | NA | NA | NA | NA | NA | NA | NA | NA | NA | NA |
| Gm9047 | 0 | 0 | 0 | 0 | 0 | 1 | 0 | 0 | 0 | 0 | 0 | 1 | 0.18 | 0 | 0 | 1.859 | 0.894 | 0.82650734 | NA | 0 | 18.0643 | 10 | 10 | TRUE | 0.5383 |
| Gm9054 | 1 | 1 | 1 | 3 | 1 | 1 | 1 | 1 | 1 | 2 | 1 | 1 | 1.27 | 1 | 1 | 1.513 | 0.597 | 0.70531437 | NA | 0 | 2.6099 | 0.9808 | 0.9808 | TRUE | 0.2233 |
| Gm906 | 22 | 11 | 10 | 5 | 5 | 4 | 23 | 11 | 10 | 4 | 5 | 4 | 9.68 | 15 | 4 | 0.307 | -1.704 | 0.01521649 | 0.07149076 | 0 | 0.3472 | 0.2129 | 0.2129 | TRUE | 1.1184 |
| Gm9079 | 30 | 14 | 35 | 31 | 27 | 21 | 32 | 14 | 35 | 24 | 29 | 23 | 26.23 | 27 | 25 | 0.944 | -0.084 | 0.8505787 | 0.92562639 | 0.0732 | 0.1325 | 0.1047 | 0.1047 | TRUE | 1.1421 |
| Gm9112 | 144 | 123 | 122 | 195 | 175 | 200 | 153 | 123 | 123 | 153 | 188 | 219 | 159.8 | 133 | 187 | 1.403 | 0.489 | 0.015813 | 0.07376595 | 0.0189 | 0.0275 | 0.0231 | 0.0231 | TRUE | 0.1731 |
| Gm9125 | 1 | 0 | 0 | 6 | 1 | 1 | 1 | 0 | 0 | 5 | 1 | 1 | 1.32 | 0 | 2 | 6.419 | 2.682 | 0.17226133 | NA | 0 | 2.4934 | 1.574 | 1.574 | TRUE | 1.1097 |
| Gm9159 | 0 | 0 | 0 | 0 | 0 | 0 | 0 | 0 | 0 | 0 | 0 | 0 | 0 | 0 | 0 | NA | NA | NA | NA | NA | NA | NA | NA | NA | NA |
| Gm9199 | 1 | 0 | 0 | 0 | 0 | 0 | 1 | 0 | 0 | 0 | 0 | 0 | 0.18 | 0 | 0 | 0.49 | -1.029 | 0.80087 | NA | 0 | 18.6165 | 10 | 10 | TRUE | 0.4614 |
| Gm9268 | 0 | 0 | 0 | 0 | 0 | 0 | 0 | 0 | 0 | 0 | 0 | 0 | 0 | 0 | 0 | NA | NA | NA | NA | NA | NA | NA | NA | NA | NA |

| | | | | | | | | | | | | | | | | | | | | | | | | | |
|---|---|---|---|---|---|---|---|---|---|---|---|---|---|---|---|---|---|---|---|---|---|---|---|---|---|
| Gm933 | 4 | 5 | 3 | 2 | 4 | 10 | 4 | 5 | 3 | 2 | 4 | 11 | 4.85 | 4 | 6 | 1.343 | 0.426 | 0.66306997 | 0.81107023 | 0.2811 | 0.6867 | 0.4741 | 0.4741 | TRUE | 0.5441 |
| Gm9376 | 2 | 0 | 0 | 0 | 1 | 0 | 2 | 0 | 0 | 0 | 1 | 0 | 0.53 | 1 | 0 | 0.497 | -1.009 | 0.76131406 | NA | 0 | 6.1834 | 6.228 | 6.228 | TRUE | 0.9742 |
| Gm94 | 0 | 0 | 0 | 0 | 0 | 0 | 0 | 0 | 0 | 0 | 0 | 0 | 0 | 0 | 0 | NA | NA | NA | NA | NA | NA | NA | NA | NA | NA |
| Gm9513 | 0 | 0 | 0 | 0 | 0 | 0 | 0 | 0 | 0 | 0 | 0 | 0 | 0 | 0 | 0 | NA | NA | NA | NA | NA | NA | NA | NA | NA | NA |
| Gm9573 | 0 | 0 | 0 | 0 | 0 | 0 | 0 | 0 | 0 | 0 | 0 | 0 | 0 | 0 | 0 | NA | NA | NA | NA | NA | NA | NA | NA | NA | NA |
| Gm960 | 6 | 7 | 10 | 9 | 7 | 3 | 6 | 7 | 10 | 7 | 8 | 3 | 6.88 | 8 | 6 | 0.77 | -0.377 | 0.60650908 | 0.77676352 | 0 | 0.4856 | 0.2386 | 0.2386 | TRUE | 0.3174 |
| Gm9696 | 0 | 0 | 1 | 0 | 0 | 0 | 0 | 0 | 1 | 0 | 0 | 0 | 0.17 | 0 | 0 | 0.49 | -1.029 | 0.80087 | NA | 0 | 19.6928 | 10 | 10 | TRUE | 0.4098 |
| Gm97 | 0 | 0 | 0 | 0 | 0 | 0 | 0 | 0 | 0 | 0 | 0 | 0 | 0 | 0 | 0 | NA | NA | NA | NA | NA | NA | NA | NA | NA | NA |
| Gm9705 | 2 | 5 | 3 | 3 | 5 | 6 | 2 | 5 | 3 | 2 | 5 | 7 | 4.07 | 3 | 5 | 1.382 | 0.467 | 0.62750087 | NA | 0 | 0.8158 | 0.4115 | 0.4115 | TRUE | 0.5114 |
| Gm973 | 62 | 64 | 49 | 175 | 111 | 85 | 66 | 64 | 49 | 138 | 119 | 93 | 88.17 | 60 | 117 | 1.965 | 0.974 | 0.00010776 | 0.00146142 | 0.0213 | 0.0443 | 0.0329 | 0.0329 | TRUE | 0.1796 |
| Gm9731 | 0 | 0 | 0 | 0 | 0 | 0 | 0 | 0 | 0 | 0 | 0 | 0 | 0 | 0 | 0 | NA | NA | NA | NA | NA | NA | NA | NA | NA | NA |
| Gm9733 | 0 | 0 | 0 | 0 | 0 | 0 | 0 | 0 | 0 | 0 | 0 | 0 | 0 | 0 | 0 | NA | NA | NA | NA | NA | NA | NA | NA | NA | NA |
| Gm9758 | 0 | 0 | 0 | 0 | 0 | 0 | 0 | 0 | 0 | 0 | 0 | 0 | 0 | 0 | 0 | NA | NA | NA | NA | NA | NA | NA | NA | NA | NA |
| Gm9767 | 0 | 0 | 0 | 0 | 1 | 0 | 0 | 0 | 0 | 0 | 1 | 0 | 0.18 | 0 | 0 | 1.859 | 0.894 | 0.82650734 | NA | 0 | 18.3785 | 10 | 10 | TRUE | 0.5204 |
| Gm9776 | 21 | 39 | 28 | 69 | 50 | 31 | 22 | 39 | 28 | 54 | 54 | 34 | 38.56 | 30 | 47 | 1.592 | 0.671 | 0.06687783 | 0.19961269 | 0.0428 | 0.0923 | 0.0691 | 0.0691 | TRUE | 0.4825 |
| Gm9833 | 613 | 586 | 546 | 878 | 643 | 588 | 651 | 585 | 548 | 690 | 692 | 644 | 635.05 | 595 | 675 | 1.136 | 0.184 | 0.08956368 | 0.24338359 | 0.0028 | 0.0121 | 0.0069 | 0.0069 | TRUE | 0.0796 |
| Gm9839 | 0 | 0 | 0 | 0 | 0 | 0 | 0 | 0 | 0 | 0 | 0 | 0 | 0 | 0 | 0 | NA | NA | NA | NA | NA | NA | NA | NA | NA | NA |
| Gm9846 | 0 | 0 | 0 | 0 | 0 | 0 | 0 | 0 | 0 | 0 | 0 | 0 | 0 | 0 | 0 | NA | NA | NA | NA | NA | NA | NA | NA | NA | NA |
| Gm9855 | 0 | 0 | 0 | 0 | 0 | 0 | 0 | 0 | 0 | 0 | 0 | 0 | 0 | 0 | 0 | NA | NA | NA | NA | NA | NA | NA | NA | NA | NA |
| Gm9866 | 8 | 13 | 15 | 3 | 7 | 10 | 8 | 13 | 15 | 2 | 8 | 11 | 9.56 | 12 | 7 | 0.555 | -0.85 | 0.23471278 | 0.44950957 | 0.1355 | 0.3514 | 0.2548 | 0.2548 | TRUE | 0.6815 |
| Gm9871 | 0 | 0 | 0 | 0 | 0 | 0 | 0 | 0 | 0 | 0 | 0 | 0 | 0 | 0 | 0 | NA | NA | NA | NA | NA | NA | NA | NA | NA | NA |
| Gm9895 | 57 | 96 | 99 | 55 | 33 | 75 | 61 | 96 | 99 | 43 | 36 | 82 | 69.44 | 85 | 54 | 0.626 | -0.677 | 0.06524469 | 0.19690761 | 0.1137 | 0.0543 | 0.0819 | 0.0819 | TRUE | 1.7193 |
| Gm9897 | 62 | 65 | 51 | 88 | 50 | 55 | 66 | 65 | 51 | 69 | 54 | 60 | 60.86 | 61 | 61 | 1.012 | 0.017 | 0.94887758 | 0.9743734 | 0 | 0.061 | 0.0325 | 0.0325 | TRUE | 0.1619 |
| Gm9899 | 7 | 4 | 7 | 6 | 6 | 8 | 7 | 4 | 7 | 5 | 6 | 9 | 6.4 | 6 | 7 | 1.069 | 0.096 | 0.89826241 | 0.95051289 | 0 | 0.5218 | 0.2488 | 0.2488 | TRUE | 0.2302 |
| Gm9920 | 5 | 11 | 15 | 19 | 10 | 6 | 5 | 11 | 15 | 15 | 11 | 7 | 10.6 | 10 | 11 | 1.039 | 0.055 | 0.93346175 | 0.96713012 | 0.1 | 0.3176 | 0.2146 | 0.2146 | TRUE | 0.2753 |
| Gm9926 | 10 | 8 | 8 | 9 | 10 | 12 | 11 | 8 | 8 | 7 | 11 | 13 | 9.6 | 9 | 10 | 1.15 | 0.201 | 0.74372051 | 0.86319522 | 0 | 0.35 | 0.1681 | 0.1681 | TRUE | 0.3494 |
| Gm9958 | 21 | 32 | 24 | 32 | 43 | 31 | 22 | 32 | 24 | 25 | 46 | 34 | 30.62 | 26 | 35 | 1.333 | 0.414 | 0.28987294 | 0.51105254 | 0.039 | 0.1145 | 0.0769 | 0.0769 | TRUE | 0.2304 |
| Gm996 | 47 | 43 | 55 | 51 | 49 | 43 | 50 | 43 | 55 | 40 | 53 | 47 | 48 | 49 | 47 | 0.94 | -0.089 | 0.75500269 | 0.87058798 | 0 | 0.0755 | 0.0383 | 0.0383 | TRUE | 0.1408 |
| Gm9961 | 0 | 0 | 0 | 0 | 0 | 0 | 0 | 0 | 0 | 0 | 0 | 0 | 0 | 0 | 0 | NA | NA | NA | NA | NA | NA | NA | NA | NA | NA |
| Gm9962 | 2 | 0 | 0 | 3 | 0 | 0 | 2 | 0 | 0 | 2 | 0 | 0 | 0.75 | 1 | 1 | 1.163 | 0.218 | 0.94242245 | NA | 8.7098 | 4.4164 | 5.2331 | 5.2331 | TRUE | 1.4402 |
| Gm9992 | 9 | 4 | 2 | 24 | 21 | 18 | 10 | 4 | 2 | 19 | 23 | 20 | 12.79 | 5 | 21 | 3.958 | 1.985 | 0.0020881 | 0.01594383 | 0 | 0.2645 | 0.1764 | 0.1764 | TRUE | 1.0759 |
| Gm9994 | 4 | 2 | 2 | 1 | 0 | 0 | 4 | 2 | 2 | 1 | 0 | 0 | 1.51 | 3 | 0 | 0.127 | -2.974 | 0.10094653 | NA | 0 | 2.1934 | 1.1837 | 1.1837 | TRUE | 0.2589 |
| Gm9999 | 0 | 0 | 0 | 0 | 0 | 0 | 0 | 0 | 0 | 0 | 0 | 0 | 0 | 0 | 0 | NA | NA | NA | NA | NA | NA | NA | NA | NA | NA |
| Gmcl1 | 2520 | 2944 | 2776 | 3329 | 2500 | 2403 | 2677 | 2940 | 2787 | 2617 | 2690 | 2630 | 2723.6 | 2801 | 2646 | 0.944 | -0.083 | 0.24757772 | 0.46433167 | 9.00E-04 | 0.0081 | 0.0033 | 0.0033 | TRUE | 0.0228 |
| Gmcl1l | 0 | 0 | 0 | 0 | 0 | 0 | 0 | 0 | 0 | 0 | 0 | 0 | 0 | 0 | 0 | NA | NA | NA | NA | NA | NA | NA | NA | NA | NA |
| Gmds | 497 | 606 | 611 | 809 | 548 | 553 | 528 | 605 | 614 | 636 | 590 | 605 | 596.26 | 582 | 610 | 1.049 | 0.069 | 0.52527121 | 0.7148075 | 0.0023 | 0.0124 | 0.0067 | 0.0067 | TRUE | 0.0773 |
| Gmeb1 | 1473 | 1248 | 1223 | 1702 | 1175 | 1099 | 1565 | 1246 | 1228 | 1338 | 1264 | 1203 | 1307.37 | 1346 | 1268 | 0.943 | -0.085 | 0.48226629 | 0.68089732 | 0.0097 | 0.0094 | 0.0098 | 0.0098 | TRUE | 0.241 |
| Gmeb2 | 1312 | 1531 | 1552 | 1924 | 1490 | 1496 | 1394 | 1529 | 1558 | 1512 | 1603 | 1638 | 1539.04 | 1494 | 1584 | 1.06 | 0.084 | 0.32751604 | 0.54874958 | 0.0019 | 0.009 | 0.0047 | 0.0047 | TRUE | 0.0411 |
| Gmfb | 4905 | 5989 | 6148 | 7084 | 5118 | 5186 | 5210 | 5981 | 6173 | 5569 | 5507 | 5677 | 5686.15 | 5788 | 5584 | 0.965 | -0.052 | 0.55060886 | 0.73491924 | 0.0035 | 0.0075 | 0.0053 | 0.0053 | TRUE | 0.093 |
| Gmfg | 354 | 625 | 616 | 320 | 205 | 304 | 376 | 624 | 619 | 252 | 221 | 333 | 403.94 | 540 | 269 | 0.497 | -1.009 | 1.14E-05 | 0.00022161 | 0.0584 | 0.0151 | 0.0353 | 0.0353 | TRUE | 0.8189 |
| Gmip | 167 | 168 | 195 | 216 | 168 | 180 | 177 | 168 | 196 | 170 | 181 | 197 | 181.43 | 180 | 183 | 1.01 | 0.014 | 0.92849594 | 0.96476961 | 0 | 0.0251 | 0.0126 | 0.0126 | TRUE | 0.0612 |
| Gml | 3 | 2 | 4 | 23 | 12 | 15 | 3 | 2 | 4 | 18 | 13 | 16 | 9.44 | 3 | 16 | 5.177 | 2.372 | 0.00096392 | 0.0086479 | 0 | 0.3561 | 0.1747 | 0.1747 | TRUE | 0.1279 |
| Gmnc | 0 | 0 | 0 | 0 | 0 | 0 | 0 | 0 | 0 | 0 | 0 | 0 | 0 | 0 | 0 | NA | NA | NA | NA | NA | NA | NA | NA | NA | NA |
| Gmnn | 3574 | 4171 | 4203 | 5880 | 4311 | 4454 | 3796 | 4166 | 4220 | 4622 | 4638 | 4876 | 4386.41 | 4061 | 4712 | 1.16 | 0.214 | 0.00438901 | 0.02832387 | 0.0017 | 0.0077 | 0.0039 | 0.0039 | TRUE | 0.0394 |
| Gmppa | 2034 | 2153 | 2156 | 2260 | 1682 | 1698 | 2160 | 2150 | 2165 | 1777 | 1810 | 1859 | 1986.77 | 2158 | 1815 | 0.841 | -0.251 | 0.00018075 | 0.00223589 | 0 | 0.0086 | 0.0027 | 0.0027 | TRUE | 0.0053 |
| Gmppb | 637 | 498 | 400 | 1060 | 708 | 456 | 677 | 497 | 402 | 833 | 762 | 499 | 611.64 | 525 | 698 | 1.331 | 0.412 | 0.0764862 | 0.21834214 | 0.0646 | 0.0123 | 0.0373 | 0.0373 | TRUE | 0.6181 |
| Gmpr | 868 | 1102 | 1126 | 1098 | 855 | 881 | 922 | 1101 | 1131 | 863 | 920 | 964 | 983.44 | 1051 | 916 | 0.871 | -0.2 | 0.07599904 | 0.21734486 | 0.0065 | 0.0103 | 0.0081 | 0.0081 | TRUE | 0.1373 |
| Gmpr2 | 1157 | 1169 | 1071 | 1339 | 971 | 942 | 1229 | 1167 | 1075 | 1053 | 1045 | 1031 | 1100.06 | 1157 | 1043 | 0.901 | -0.15 | 0.09134739 | 0.24661325 | 0.0014 | 0.0099 | 0.0048 | 0.0048 | TRUE | 0.046 |
| Gmps | 5843 | 6900 | 7209 | 9745 | 6890 | 6938 | 6206 | 6891 | 7239 | 7661 | 7413 | 7595 | 7167.45 | 6779 | 7556 | 1.115 | 0.157 | 0.05330938 | 0.17145098 | 0.0029 | 0.0074 | 0.0046 | 0.0046 | TRUE | 0.0665 |
| Gna11 | 3893 | 4572 | 4471 | 5279 | 3824 | 3717 | 4135 | 4566 | 4489 | 4150 | 4114 | 4069 | 4253.95 | 4397 | 4111 | 0.935 | -0.097 | 0.16940416 | 0.36787035 | 0.0011 | 0.0077 | 0.0034 | 0.0034 | TRUE | 0.0329 |
| Gna12 | 2137 | 2051 | 2119 | 2413 | 1702 | 1863 | 2270 | 2048 | 2128 | 1897 | 1831 | 2039 | 2035.57 | 2149 | 1922 | 0.895 | -0.161 | 0.05772255 | 0.18146304 | 0.0022 | 0.0085 | 0.0047 | 0.0047 | TRUE | 0.0336 |
| Gna13 | 7374 | 8309 | 8911 | 10646 | 7922 | 8959 | 7832 | 8298 | 8948 | 8369 | 8524 | 9807 | 8629.67 | 8359 | 8900 | 1.065 | 0.09 | 0.34533537 | 0.5647687 | 0.006 | 0.0073 | 0.0065 | 0.0065 | TRUE | 0.097 |
| Gna14 | 45 | 103 | 154 | 40 | 35 | 87 | 48 | 103 | 155 | 31 | 38 | 95 | 78.27 | 102 | 55 | 0.536 | -0.901 | 0.07214136 | 0.2099287 | 0.3131 | 0.049 | 0.1669 | 0.1669 | TRUE | 0.2273 |
| Gna15 | 632 | 1041 | 1115 | 600 | 486 | 675 | 671 | 1040 | 1120 | 472 | 523 | 739 | 760.67 | 944 | 578 | 0.612 | -0.709 | 0.0014962 | 0.01223964 | 0.0601 | 0.0112 | 0.0345 | 0.0345 | TRUE | 0.7578 |
| Gnai1 | 817 | 903 | 914 | 1755 | 1240 | 1138 | 868 | 902 | 918 | 1380 | 1334 | 1246 | 1107.82 | 896 | 1320 | 1.474 | 0.56 | 8.38E-11 | 6.20E-09 | 0.001 | 0.0099 | 0.0044 | 0.0044 | TRUE | 0.0287 |
| Gnai2 | 7967 | 8157 | 8659 | 9038 | 7083 | 7124 | 8462 | 8146 | 8695 | 7105 | 7621 | 7798 | 7971.26 | 8434 | 7508 | 0.89 | -0.168 | 0.01714698 | 0.0777356 | 0.0015 | 0.0073 | 0.0035 | 0.0035 | TRUE | 0.0272 |
| Gnai3 | 7774 | 7733 | 7941 | 10243 | 7292 | 7164 | 8257 | 7723 | 7974 | 8052 | 7846 | 7842 | 7949.02 | 7985 | 7913 | 0.991 | -0.013 | 0.83399132 | 0.91571799 | 4.00E-04 | 0.0073 | 0.0025 | 0.0025 | TRUE | 0.0109 |
| Gnal | 159 | 194 | 165 | 521 | 342 | 383 | 169 | 194 | 166 | 410 | 368 | 419 | 287.52 | 176 | 399 | 2.266 | 1.18 | 2.88E-17 | 5.16E-15 | 0 | 0.0184 | 0.0099 | 0.0099 | TRUE | 0.0832 |
| Gnao1 | 330 | 312 | 327 | 459 | 284 | 319 | 351 | 312 | 328 | 361 | 306 | 349 | 334.34 | 330 | 339 | 1.027 | 0.039 | 0.76454662 | 0.87640303 | 0.0025 | 0.0168 | 0.0091 | 0.0091 | TRUE | 0.0809 |
| Gnaq | 1659 | 2014 | 2111 | 2440 | 1756 | 1751 | 1762 | 2011 | 2120 | 1918 | 1889 | 1917 | 1936.24 | 1964 | 1908 | 0.971 | -0.042 | 0.65242268 | 0.80502827 | 0.0039 | 0.0086 | 0.0058 | 0.0058 | TRUE | 0.0977 |
| Gnas | 7114 | 9296 | 9742 | 9956 | 7132 | 7352 | 7556 | 9284 | 9782 | 7827 | 7674 | 8048 | 8361.74 | 8874 | 7850 | 0.884 | -0.177 | 0.10360342 | 0.26778648 | 0.0088 | 0.0073 | 0.0084 | 0.0084 | TRUE | 0.206 |
| Gnat1 | 28 | 14 | 15 | 23 | 13 | 13 | 30 | 14 | 15 | 18 | 14 | 14 | 17.51 | 20 | 15 | 0.795 | -0.332 | 0.51191405 | 0.70537329 | 0.0566 | 0.195 | 0.1265 | 0.1265 | TRUE | 1.0595 |
| Gnat2 | 274 | 207 | 193 | 187 | 113 | 111 | 291 | 207 | 194 | 147 | 122 | 122 | 180.28 | 231 | 130 | 0.566 | -0.821 | 0.00010033 | 0.00138446 | 0.027 | 0.0252 | 0.0261 | 0.0261 | TRUE | 0.5768 |
| Gnat3 | 0 | 0 | 0 | 0 | 0 | 0 | 0 | 0 | 0 | 0 | 0 | 0 | 0 | 0 | 0 | NA | NA | NA | NA | NA | NA | NA | NA | NA | NA |
| Gnaz | 373 | 425 | 394 | 478 | 413 | 415 | 396 | 424 | 396 | 376 | 444 | 454 | 415.11 | 405 | 425 | 1.045 | 0.064 | 0.61176266 | 0.78007281 | 0.0042 | 0.0148 | 0.0089 | 0.0089 | TRUE | 0.1251 |
| Gnb1 | 14993 | 17642 | 17840 | 23462 | 16771 | 17423 | 15925 | 17619 | 17913 | 18444 | 18045 | 19072 | 17836.41 | 17152 | 18520 | 1.08 | 0.111 | 0.13725232 | 0.32132159 | 0.0023 | 0.0071 | 0.0039 | 0.0039 | TRUE | 0.0479 |
| Gnb1l | 423 | 714 | 609 | 744 | 563 | 550 | 449 | 713 | 612 | 585 | 606 | 602 | 594.43 | 591 | 598 | 1.01 | 0.014 | 0.93198592 | 0.96628017 | 0.0249 | 0.0124 | 0.0187 | 0.0187 | TRUE | 0.3609 |
| Gnb2 | 5770 | 6860 | 6900 | 8292 | 6024 | 6159 | 6129 | 6851 | 6928 | 6519 | 6481 | 6742 | 6608.37 | 6636 | 6581 | 0.992 | -0.012 | 0.87337096 | 0.93762803 | 0.0023 | 0.0074 | 0.0041 | 0.0041 | TRUE | 0.0545 |
| Gnb2l1 | 95376 | 123920 | 125970 | 131223 | 95141 | 103031 | 101306 | 123760 | 126487 | 103157 | 102366 | 112784 | 111643.35 | 117184 | 106102 | 0.905 | -0.143 | 0.17346804 | 0.37263482 | 0.0084 | 0.0069 | 0.008 | 0.008 | TRUE | 0.1721 |

| | | | | | | | | | | | | | | | | | | | | | | | | | |
|---|---|---|---|---|---|---|---|---|---|---|---|---|---|---|---|---|---|---|---|---|---|---|---|---|---|
| Gnb3 | 0 | 4 | 5 | 1 | 2 | 1 | 0 | 4 | 5 | 1 | 2 | 1 | 2.17 | 3 | 1 | 0.437 | -1.193 | 0.41916673 | NA | 0 | 1.5218 | 1.0326 | 1.0326 | TRUE | 0.9387 |
| Gnb4 | 741 | 1039 | 998 | 1256 | 878 | 901 | 787 | 1038 | 1002 | 987 | 945 | 986 | 957.53 | 942 | 973 | 1.032 | 0.046 | 0.71409333 | 0.84484885 | 0.0101 | 0.0103 | 0.0102 | 0.0102 | TRUE | 0.2463 |
| Gnb5 | 282 | 262 | 233 | 305 | 235 | 234 | 300 | 262 | 234 | 240 | 253 | 256 | 257.32 | 265 | 250 | 0.941 | -0.087 | 0.55272767 | 0.73653437 | 0.0044 | 0.0197 | 0.0117 | 0.0117 | TRUE | 0.1429 |
| Gne | 882 | 1171 | 1246 | 982 | 734 | 859 | 937 | 1169 | 1251 | 772 | 790 | 940 | 976.58 | 1119 | 834 | 0.745 | -0.426 | 0.00257834 | 0.01883541 | 0.0152 | 0.0103 | 0.0133 | 0.0133 | TRUE | 0.2423 |
| Gng10 | 315 | 370 | 316 | 568 | 396 | 384 | 335 | 370 | 317 | 447 | 426 | 420 | 385.72 | 341 | 431 | 1.267 | 0.341 | 0.00396319 | 0.02617567 | 0 | 0.0154 | 0.0075 | 0.0075 | TRUE | 0.0642 |
| Gng11 | 6 | 8 | 9 | 33 | 21 | 26 | 6 | 8 | 9 | 26 | 23 | 28 | 16.73 | 8 | 26 | 3.284 | 1.716 | 0.00057249 | 0.00578738 | 0 | 0.2038 | 0.0944 | 0.0944 | TRUE | 0.0694 |
| Gng12 | 1191 | 1816 | 1877 | 1731 | 1329 | 1685 | 1265 | 1814 | 1885 | 1361 | 1430 | 1844 | 1599.77 | 1655 | 1545 | 0.933 | -0.099 | 0.57413485 | 0.75247257 | 0.0358 | 0.009 | 0.0219 | 0.0219 | TRUE | 0.5105 |
| Gng13 | 9 | 8 | 11 | 17 | 11 | 6 | 10 | 8 | 11 | 13 | 12 | 7 | 10.06 | 10 | 11 | 1.123 | 0.167 | 0.78379268 | 0.88838079 | 0 | 0.3344 | 0.167 | 0.167 | TRUE | 0.362 |
| Gng2 | 1807 | 2273 | 2451 | 1655 | 1242 | 1495 | 1919 | 2270 | 2461 | 1301 | 1336 | 1637 | 1820.72 | 2217 | 1425 | 0.642 | -0.639 | 1.66E-06 | 4.27E-05 | 0.0152 | 0.0087 | 0.0122 | 0.0122 | TRUE | 0.2027 |
| Gng3 | 308 | 293 | 321 | 662 | 519 | 531 | 327 | 293 | 322 | 520 | 558 | 581 | 433.7 | 314 | 553 | 1.76 | 0.816 | 1.71E-12 | 1.68E-10 | 0 | 0.0145 | 0.0071 | 0.0071 | TRUE | 0.0406 |
| Gng4 | 5 | 7 | 3 | 3 | 6 | 2 | 5 | 7 | 3 | 2 | 6 | 2 | 4.39 | 5 | 3 | 0.709 | -0.497 | 0.59893571 | NA | 0 | 0.758 | 0.408 | 0.408 | TRUE | 0.5791 |
| Gng5 | 2163 | 2203 | 2199 | 2808 | 1927 | 1983 | 2297 | 2200 | 2208 | 2207 | 2073 | 2171 | 2192.86 | 2235 | 2150 | 0.962 | -0.055 | 0.43257142 | 0.64193794 | 4.00E-04 | 0.0084 | 0.0031 | 0.0031 | TRUE | 0.0118 |
| Gng7 | 205 | 204 | 233 | 242 | 165 | 145 | 218 | 204 | 234 | 190 | 178 | 159 | 196.99 | 219 | 176 | 0.806 | -0.312 | 0.04467301 | 0.15211197 | 0 | 0.0236 | 0.0123 | 0.0123 | TRUE | 0.0728 |
| Gng8 | 35 | 34 | 41 | 26 | 35 | 26 | 37 | 34 | 41 | 20 | 38 | 28 | 33.14 | 37 | 29 | 0.763 | -0.391 | 0.28355564 | 0.50512347 | 0.0176 | 0.1063 | 0.0648 | 0.0648 | TRUE | 0.2996 |
| Gngt1 | 0 | 0 | 0 | 0 | 0 | 0 | 0 | 0 | 0 | 0 | 0 | 0 | 0 | 0 | 0 | NA | NA | NA | NA | NA | NA | NA | NA | NA | NA |
| Gngt2 | 186 | 309 | 329 | 173 | 121 | 184 | 198 | 309 | 330 | 136 | 130 | 201 | 217.35 | 279 | 156 | 0.558 | -0.843 | 0.00078452 | 0.00737389 | 0.0607 | 0.0221 | 0.0404 | 0.0404 | TRUE | 0.7241 |
| Gnl1 | 4479 | 4408 | 4462 | 6274 | 4527 | 4396 | 4757 | 4402 | 4480 | 4932 | 4871 | 4812 | 4709.19 | 4546 | 4872 | 1.072 | 0.1 | 0.12690495 | 0.30585955 | 6.00E-04 | 0.0076 | 0.0029 | 0.0029 | TRUE | 0.02 |
| Gnl2 | 4941 | 5462 | 5675 | 6710 | 4945 | 4857 | 5248 | 5455 | 5698 | 5275 | 5321 | 5317 | 5385.6 | 5467 | 5304 | 0.97 | -0.044 | 0.4957766 | 0.69173643 | 6.00E-04 | 0.0075 | 0.0028 | 0.0028 | TRUE | 0.0167 |
| Gnl3 | 9874 | 10240 | 10533 | 11690 | 8566 | 8812 | 10488 | 10227 | 10576 | 9190 | 9216 | 9646 | 9890.55 | 10430 | 9351 | 0.896 | -0.158 | 0.00700629 | 0.04011004 | 4.00E-04 | 0.0072 | 0.0024 | 0.0024 | TRUE | 0.0093 |
| Gnl3l | 11419 | 11988 | 11765 | 12117 | 8817 | 9170 | 12129 | 11973 | 11813 | 9525 | 9487 | 10038 | 10827.47 | 11972 | 9683 | 0.809 | -0.306 | 2.08E-07 | 6.78E-06 | 5.00E-04 | 0.0072 | 0.0024 | 0.0024 | TRUE | 0.0125 |
| Gnmt | 112 | 195 | 186 | 127 | 101 | 144 | 119 | 195 | 187 | 100 | 109 | 158 | 144.44 | 167 | 122 | 0.729 | -0.456 | 0.08378557 | 0.23193226 | 0.0576 | 0.0297 | 0.0431 | 0.0431 | TRUE | 0.6616 |
| Gnpat | 10281 | 11195 | 11675 | 12949 | 9601 | 10098 | 10920 | 11181 | 11723 | 10179 | 10330 | 11054 | 10897.86 | 11275 | 10521 | 0.933 | -0.1 | 0.14889254 | 0.33883279 | 0.0015 | 0.0072 | 0.0034 | 0.0034 | TRUE | 0.0239 |
| Gnpda1 | 4602 | 5813 | 5895 | 5723 | 4448 | 4966 | 4888 | 5805 | 5919 | 4499 | 4786 | 5436 | 5222.28 | 5537 | 4907 | 0.886 | -0.175 | 0.11934795 | 0.29279139 | 0.0097 | 0.0075 | 0.0089 | 0.0089 | TRUE | 0.1282 |
| Gnpda2 | 1004 | 1180 | 1081 | 1667 | 1114 | 1273 | 1066 | 1178 | 1085 | 1310 | 1199 | 1393 | 1205.48 | 1110 | 1301 | 1.172 | 0.229 | 0.01841933 | 0.08197227 | 0.0034 | 0.0096 | 0.0059 | 0.0059 | TRUE | 0.0556 |
| Gnpnat1 | 2109 | 2319 | 2139 | 3014 | 2069 | 2225 | 2240 | 2316 | 2148 | 2369 | 2226 | 2436 | 2289.17 | 2235 | 2344 | 1.049 | 0.069 | 0.3702931 | 0.58728707 | 0.0013 | 0.0083 | 0.0038 | 0.0038 | TRUE | 0.023 |
| Gnptab | 2516 | 2610 | 2699 | 3210 | 2286 | 2373 | 2672 | 2607 | 2710 | 2523 | 2460 | 2598 | 2594.97 | 2663 | 2527 | 0.949 | -0.076 | 0.25435452 | 0.47240692 | 0 | 0.0082 | 0.0028 | 0.0028 | TRUE | 0.0071 |
| Gnptg | 346 | 429 | 433 | 413 | 328 | 356 | 368 | 428 | 435 | 325 | 353 | 390 | 383 | 410 | 356 | 0.865 | -0.209 | 0.1184612 | 0.29125021 | 0.0058 | 0.0155 | 0.0103 | 0.0103 | TRUE | 0.0947 |
| Gnrh1 | 1 | 1 | 0 | 3 | 0 | 1 | 1 | 1 | 0 | 2 | 0 | 1 | 0.92 | 1 | 1 | 1.734 | 0.794 | 0.70563564 | NA | 0 | 3.5918 | 2.0282 | 2.0282 | TRUE | 0.167 |
| Gnrhr | 0 | 1 | 0 | 0 | 0 | 0 | 0 | 1 | 0 | 0 | 0 | 0 | 0.17 | 0 | 0 | 0.49 | -1.029 | 0.80087 | NA | 0 | 19.7991 | 10 | 10 | TRUE | 0.405 |
| Gns | 3786 | 3739 | 3909 | 4299 | 3313 | 3181 | 4021 | 3734 | 3925 | 3380 | 3565 | 3482 | 3684.47 | 3893 | 3476 | 0.892 | -0.164 | 0.01624677 | 0.07514487 | 8.00E-04 | 0.0078 | 0.0031 | 0.0031 | TRUE | 0.0156 |
| Golga1 | 432 | 440 | 411 | 661 | 508 | 516 | 459 | 439 | 413 | 520 | 547 | 565 | 490.34 | 437 | 544 | 1.243 | 0.314 | 0.00323901 | 0.02247502 | 0 | 0.0136 | 0.0061 | 0.0061 | TRUE | 0.0274 |
| Golga2 | 2234 | 2207 | 2212 | 2467 | 1919 | 1885 | 2373 | 2204 | 2221 | 1939 | 2065 | 2063 | 2144.28 | 2266 | 2022 | 0.892 | -0.165 | 0.02941918 | 0.11340213 | 0.001 | 0.0084 | 0.0036 | 0.0036 | TRUE | 0.0205 |
| Golga3 | 3446 | 3186 | 3415 | 4065 | 2899 | 2742 | 3660 | 3182 | 3429 | 3196 | 3119 | 3002 | 3264.57 | 3424 | 3106 | 0.907 | -0.14 | 0.08808531 | 0.24056822 | 0.0025 | 0.0079 | 0.0046 | 0.0046 | TRUE | 0.0465 |
| Golga4 | 7955 | 7421 | 8107 | 8896 | 6623 | 6464 | 8450 | 7411 | 8140 | 6993 | 7126 | 7076 | 7532.74 | 8000 | 7065 | 0.883 | -0.179 | 0.01709374 | 0.07762908 | 0.0021 | 0.0073 | 0.0039 | 0.0039 | TRUE | 0.0507 |
| Golga5 | 2312 | 2458 | 2447 | 3046 | 2268 | 2245 | 2456 | 2455 | 2457 | 2395 | 2440 | 2458 | 2443.31 | 2456 | 2431 | 0.99 | -0.015 | 0.80907601 | 0.90175584 | 0 | 0.0083 | 0.0024 | 0.0024 | TRUE | 0.0021 |
| Golga7 | 5316 | 5732 | 5561 | 7044 | 5180 | 5170 | 5647 | 5725 | 5584 | 5537 | 5573 | 5659 | 5620.86 | 5652 | 5590 | 0.989 | -0.016 | 0.77626807 | 0.88373755 | 0 | 0.0075 | 0.0021 | 0.0021 | TRUE | 0.0016 |
| Golga7b | 23 | 20 | 17 | 19 | 13 | 14 | 24 | 20 | 17 | 15 | 14 | 15 | 17.62 | 20 | 15 | 0.721 | -0.472 | 0.29348904 | 0.5143804 | 0 | 0.1939 | 0.0873 | 0.0873 | TRUE | 0.1489 |
| Golgb1 | 4379 | 3820 | 4550 | 5793 | 4184 | 4024 | 4651 | 3815 | 4569 | 4554 | 4502 | 4405 | 4415.94 | 4345 | 4487 | 1.033 | 0.047 | 0.63079551 | 0.79261527 | 0.0058 | 0.0076 | 0.0065 | 0.0065 | TRUE | 0.1388 |
| Golim4 | 1335 | 1568 | 1489 | 1790 | 1273 | 1347 | 1418 | 1566 | 1495 | 1407 | 1370 | 1475 | 1455.07 | 1493 | 1417 | 0.949 | -0.076 | 0.35873477 | 0.57728444 | 0.0012 | 0.0092 | 0.0042 | 0.0042 | TRUE | 0.0231 |
| Golm1 | 5577 | 6294 | 6476 | 6396 | 4459 | 4996 | 5924 | 6286 | 6503 | 5028 | 4798 | 5469 | 5667.8 | 6238 | 5098 | 0.817 | -0.291 | 0.00040545 | 0.00438297 | 0.0031 | 0.0075 | 0.0047 | 0.0047 | TRUE | 0.049 |
| Golph3 | 6058 | 5902 | 5734 | 7562 | 5436 | 5513 | 6435 | 5894 | 5758 | 5945 | 5849 | 6035 | 5985.81 | 6029 | 5943 | 0.986 | -0.021 | 0.77543381 | 0.88317029 | 0.0014 | 0.0075 | 0.0036 | 0.0036 | TRUE | 0.0422 |
| Golph3l | 1497 | 1483 | 1473 | 1944 | 1355 | 1333 | 1590 | 1481 | 1479 | 1528 | 1458 | 1459 | 1499.25 | 1517 | 1482 | 0.978 | -0.033 | 0.67424219 | 0.81838909 | 6.00E-04 | 0.0091 | 0.0037 | 0.0037 | TRUE | 0.0214 |
| Golt1a | 2 | 1 | 0 | 2 | 1 | 3 | 2 | 1 | 0 | 2 | 1 | 3 | 1.51 | 1 | 2 | 1.902 | 0.928 | 0.56355174 | NA | 0 | 2.1898 | 1.1152 | 1.1152 | TRUE | 0.3219 |
| Golt1b | 1738 | 1932 | 1857 | 2378 | 1866 | 1758 | 1846 | 1930 | 1865 | 1869 | 2008 | 1924 | 1906.95 | 1880 | 1934 | 1.028 | 0.04 | 0.58255661 | 0.75865601 | 4.00E-04 | 0.0086 | 0.0033 | 0.0033 | TRUE | 0.0135 |
| Gon4l | 2985 | 2636 | 2798 | 4558 | 3140 | 2774 | 3171 | 2633 | 2809 | 3583 | 3378 | 3037 | 3101.81 | 2871 | 3333 | 1.161 | 0.216 | 0.04181085 | 0.14545367 | 0.0075 | 0.008 | 0.0078 | 0.0078 | TRUE | 0.1011 |
| Gopc | 1649 | 1451 | 1429 | 2368 | 1661 | 1483 | 1752 | 1449 | 1435 | 1862 | 1787 | 1623 | 1651.26 | 1545 | 1757 | 1.138 | 0.186 | 0.09744944 | 0.2564613 | 0.0082 | 0.0089 | 0.0085 | 0.0085 | TRUE | 0.164 |
| Gorab | 687 | 758 | 781 | 1087 | 731 | 717 | 730 | 757 | 784 | 855 | 787 | 785 | 782.81 | 757 | 809 | 1.069 | 0.097 | 0.29715045 | 0.51809163 | 0 | 0.0111 | 0.0049 | 0.0049 | TRUE | 0.0306 |
| Gorasp1 | 1162 | 1260 | 1287 | 1335 | 1067 | 1050 | 1234 | 1258 | 1292 | 1049 | 1148 | 1149 | 1188.63 | 1261 | 1115 | 0.883 | -0.179 | 0.03246358 | 0.1215323 | 9.00E-04 | 0.0097 | 0.0042 | 0.0042 | TRUE | 0.0336 |
| Gorasp2 | 4649 | 4839 | 4921 | 6020 | 4317 | 4390 | 4938 | 4833 | 4941 | 4732 | 4645 | 4806 | 4815.81 | 4904 | 4728 | 0.964 | -0.053 | 0.36270396 | 0.58047356 | 0 | 0.0076 | 0.0022 | 0.0022 | TRUE | 0.0028 |
| Gosr1 | 2109 | 2232 | 2142 | 2916 | 2080 | 2077 | 2240 | 2229 | 2151 | 2292 | 2238 | 2274 | 2237.32 | 2207 | 2268 | 1.028 | 0.04 | 0.54600977 | 0.73106369 | 0 | 0.0084 | 0.0027 | 0.0027 | TRUE | 0.006 |
| Gosr2 | 4328 | 4283 | 4145 | 6526 | 4508 | 4341 | 4597 | 4277 | 4162 | 5130 | 4850 | 4752 | 4628.18 | 4345 | 4911 | 1.13 | 0.177 | 0.01890529 | 0.0834988 | 0.0018 | 0.0076 | 0.0039 | 0.0039 | TRUE | 0.0312 |
| Got1 | 10166 | 10276 | 10250 | 10127 | 7278 | 7925 | 10798 | 10263 | 10292 | 7961 | 7831 | 8675 | 9303.3 | 10451 | 8156 | 0.78 | -0.358 | 6.46E-07 | 1.88E-05 | 0.0018 | 0.0073 | 0.0036 | 0.0036 | TRUE | 0.0378 |
| Got1l1 | 1509 | 2441 | 2051 | 1744 | 1294 | 1548 | 1603 | 2438 | 2059 | 1371 | 1392 | 1695 | 1759.65 | 2033 | 1486 | 0.73 | -0.453 | 0.00535438 | 0.0329055 | 0.0277 | 0.0088 | 0.0185 | 0.0185 | TRUE | 0.1781 |
| Got2 | 19127 | 20681 | 20985 | 25879 | 18297 | 19030 | 20316 | 20654 | 21071 | 20344 | 19686 | 20831 | 20483.92 | 20680 | 20287 | 0.981 | -0.028 | 0.63028513 | 0.79241587 | 5.00E-04 | 0.0071 | 0.0023 | 0.0023 | TRUE | 0.0082 |
| Gp1ba | 0 | 1 | 1 | 0 | 1 | 0 | 0 | 1 | 1 | 0 | 1 | 0 | 0.51 | 1 | 0 | 0.52 | -0.943 | 0.73680291 | NA | 0 | 6.4273 | 3.9121 | 3.9121 | TRUE | 0.5204 |
| Gp1bb | 1 | 1 | 1 | 6 | 3 | 3 | 1 | 1 | 1 | 5 | 3 | 3 | 2.38 | 1 | 4 | 3.696 | 1.886 | 0.14398988 | NA | 0 | 1.3898 | 0.5713 | 0.5713 | TRUE | 0.1072 |
| Gp2 | 0 | 0 | 0 | 0 | 0 | 0 | 0 | 0 | 0 | 0 | 0 | 0 | 0 | 0 | 0 | NA | NA | NA | NA | NA | NA | NA | NA | NA | NA |
| Gp49a | 0 | 0 | 0 | 0 | 0 | 0 | 0 | 0 | 0 | 0 | 0 | 0 | 0 | 0 | 0 | NA | NA | NA | NA | NA | NA | NA | NA | NA | NA |
| Gp5 | 8 | 7 | 10 | 8 | 2 | 3 | 8 | 7 | 10 | 6 | 2 | 3 | 6.21 | 8 | 4 | 0.473 | -1.079 | 0.17358775 | 0.37274186 | 0 | 0.5375 | 0.2724 | 0.2724 | TRUE | 0.6142 |
| Gp6 | 2 | 1 | 0 | 1 | 1 | 2 | 2 | 1 | 0 | 1 | 1 | 2 | 1.2 | 1 | 1 | 1.287 | 0.363 | 0.83501143 | NA | 0 | 2.7621 | 1.3317 | 1.3317 | TRUE | 0.2511 |
| Gp9 | 0 | 0 | 0 | 0 | 0 | 0 | 0 | 0 | 0 | 0 | 0 | 0 | 0 | 0 | 0 | NA | NA | NA | NA | NA | NA | NA | NA | NA | NA |
| Gpa33 | 1188 | 1303 | 1244 | 1056 | 974 | 1083 | 1262 | 1301 | 1249 | 830 | 1048 | 1186 | 1145.99 | 1271 | 1021 | 0.803 | -0.317 | 0.02087814 | 0.08955401 | 0.0143 | 0.0098 | 0.0127 | 0.0127 | TRUE | 0.3274 |
| Gpaa1 | 1285 | 1306 | 1334 | 1786 | 1275 | 1179 | 1365 | 1304 | 1339 | 1404 | 1372 | 1291 | 1345.85 | 1336 | 1356 | 1.015 | 0.022 | 0.78459936 | 0.88892329 | 5.00E-04 | 0.0094 | 0.0038 | 0.0038 | TRUE | 0.0208 |
| Gpam | 1259 | 1253 | 1282 | 1511 | 1138 | 1087 | 1337 | 1251 | 1287 | 1188 | 1224 | 1190 | 1246.34 | 1292 | 1201 | 0.929 | -0.106 | 0.17048669 | 0.36901257 | 0 | 0.0095 | 0.0035 | 0.0035 | TRUE | 0.0112 |
| Gpank1 | 885 | 851 | 781 | 1156 | 822 | 679 | 940 | 850 | 784 | 909 | 884 | 743 | 851.76 | 858 | 845 | 0.986 | -0.02 | 0.87202724 | 0.93734162 | 0.0087 | 0.0108 | 0.0096 | 0.0096 | TRUE | 0.1312 |
| Gpat2 | 783 | 1118 | 1073 | 1220 | 1073 | 979 | 832 | 1117 | 1077 | 959 | 1154 | 1072 | 1035.14 | 1009 | 1062 | 1.052 | 0.073 | 0.60168574 | 0.77328069 | 0.0157 | 0.0101 | 0.0131 | 0.0131 | TRUE | 0.28 |
| Gpatch1 | 1480 | 1416 | 1460 | 2148 | 1673 | 1514 | 1572 | 1414 | 1466 | 1689 | 1800 | 1657 | 1599.69 | 1484 | 1715 | 1.156 | 0.209 | 0.01301302 | 0.0638925 | 0.0017 | 0.009 | 0.0045 | 0.0045 | TRUE | 0.0322 |
| Gpatch11 | 865 | 926 | 925 | 1276 | 890 | 888 | 919 | 925 | 929 | 1003 | 958 | 972 | 950.85 | 924 | 978 | 1.058 | 0.082 | 0.30659635 | 0.52707865 | 0 | 0.0104 | 0.0036 | 0.0036 | TRUE | 0.0065 |
| Gpatch2 | 1457 | 1357 | 1291 | 2025 | 1451 | 1272 | 1548 | 1355 | 1296 | 1592 | 1561 | 1392 | 1457.44 | 1400 | 1515 | 1.083 | 0.115 | 0.27712052 | 0.49805749 | 0.0059 | 0.0092 | 0.0074 | 0.0074 | TRUE | 0.1024 |

| | | | | | | | | | | | | | | | | | | | | | | |
|---|---|---|---|---|---|---|---|---|---|---|---|---|---|---|---|---|---|---|---|---|---|---|
| Gpatch2l | 1155 | 1155 | 1115 | 1506 | 1093 | 1082 | 1227 | 1154 | 1120 | 1184 | 1176 | 1184 | 1174.04 | 1167 | 1181 | 1.013 | 0.019 | 0.81838377 | 0.90726388 | 0 | 0.0097 | 0.0038 | 0.0038 | TRUE | 0.0242 |
| Gpatch3 | 423 | 249 | 233 | 694 | 435 | 289 | 449 | 249 | 234 | 546 | 468 | 316 | 376.98 | 311 | 443 | 1.429 | 0.515 | 0.06491568 | 0.19625266 | 0.1055 | 0.0156 | 0.0534 | 0.0534 | TRUE | 0.857 |
| Gpatch4 | 973 | 831 | 780 | 1425 | 1037 | 754 | 1033 | 830 | 783 | 1120 | 1116 | 825 | 951.33 | 882 | 1020 | 1.158 | 0.211 | 0.18830788 | 0.39342852 | 0.0235 | 0.0104 | 0.0175 | 0.0175 | TRUE | 0.3314 |
| Gpatch8 | 5665 | 4765 | 5000 | 6827 | 5040 | 4771 | 6017 | 4759 | 5021 | 5367 | 5423 | 5223 | 5301.47 | 5266 | 5338 | 1.014 | 0.02 | 0.84962095 | 0.92515564 | 0.0077 | 0.0075 | 0.0076 | 0.0076 | TRUE | 0.1899 |
| Gpbar1 | 31 | 16 | 21 | 16 | 8 | 10 | 33 | 16 | 21 | 13 | 9 | 11 | 17.02 | 23 | 11 | 0.464 | -1.109 | 0.03107007 | 0.11816667 | 0 | 0.2005 | 0.1235 | 0.1235 | TRUE | 0.3699 |
| Gpbp1 | 8939 | 8952 | 8606 | 11779 | 8351 | 8549 | 9495 | 8940 | 8641 | 9260 | 8985 | 9358 | 9113.28 | 9025 | 9201 | 1.019 | 0.028 | 0.67885752 | 0.82158846 | 0.0011 | 0.0073 | 0.0032 | 0.0032 | TRUE | 0.0252 |
| Gpbp1l1 | 6106 | 6239 | 5976 | 7508 | 5605 | 5615 | 6486 | 6231 | 6001 | 5902 | 6031 | 6147 | 6132.74 | 6239 | 6027 | 0.966 | -0.05 | 0.43912256 | 0.64641993 | 7.00E-04 | 0.0074 | 0.0029 | 0.0029 | TRUE | 0.0146 |
| Gpc1 | 1062 | 1175 | 1073 | 1675 | 1155 | 1073 | 1128 | 1173 | 1077 | 1317 | 1243 | 1175 | 1185.49 | 1126 | 1245 | 1.106 | 0.145 | 0.10229708 | 0.26571633 | 0.0018 | 0.0097 | 0.0049 | 0.0049 | TRUE | 0.0318 |
| Gpc2 | 719 | 965 | 913 | 721 | 560 | 695 | 764 | 964 | 917 | 567 | 603 | 761 | 762.39 | 882 | 644 | 0.729 | -0.456 | 0.00232808 | 0.01736916 | 0.0179 | 0.0112 | 0.0148 | 0.0148 | TRUE | 0.2979 |
| Gpc3 | 2377 | 3430 | 3789 | 3376 | 2656 | 3601 | 2525 | 3426 | 3805 | 2654 | 2858 | 3942 | 3201.4 | 3252 | 3151 | 0.969 | -0.046 | 0.80652647 | 0.90025021 | 0.0439 | 0.0079 | 0.0247 | 0.0247 | TRUE | 0.5848 |
| Gpc4 | 235 | 214 | 252 | 585 | 351 | 357 | 250 | 214 | 253 | 460 | 378 | 391 | 324.12 | 239 | 410 | 1.719 | 0.781 | 7.29E-08 | 2.74E-06 | 0.0074 | 0.0171 | 0.0118 | 0.0118 | TRUE | 0.1399 |
| Gpc5 | 4 | 2 | 2 | 7 | 3 | 2 | 4 | 2 | 2 | 6 | 3 | 2 | 3.2 | 3 | 4 | 1.355 | 0.438 | 0.67873083 | NA | 0 | 1.0378 | 0.4874 | 0.4874 | TRUE | 0.4026 |
| Gpc6 | 30 | 62 | 74 | 87 | 80 | 73 | 32 | 62 | 74 | 68 | 86 | 80 | 67.08 | 56 | 78 | 1.389 | 0.474 | 0.1509082 | 0.34192273 | 0.0715 | 0.056 | 0.0631 | 0.0631 | TRUE | 0.9363 |
| Gpcpd1 | 726 | 757 | 703 | 1270 | 908 | 773 | 771 | 756 | 706 | 998 | 977 | 846 | 842.42 | 744 | 940 | 1.265 | 0.339 | 0.00132763 | 0.01110842 | 0.0039 | 0.0108 | 0.0068 | 0.0068 | TRUE | 0.0904 |
| Gpd1 | 71 | 34 | 45 | 68 | 53 | 50 | 75 | 34 | 45 | 53 | 57 | 55 | 53.29 | 51 | 55 | 1.071 | 0.098 | 0.77358833 | 0.88192776 | 0.0621 | 0.0687 | 0.0656 | 0.0656 | TRUE | 0.8686 |
| Gpd1l | 1848 | 1840 | 1896 | 2667 | 1794 | 1999 | 1963 | 1838 | 1904 | 2097 | 1930 | 2188 | 1986.56 | 1902 | 2072 | 1.09 | 0.124 | 0.13569837 | 0.31886691 | 0.0018 | 0.0086 | 0.0045 | 0.0045 | TRUE | 0.0425 |
| Gpd2 | 6880 | 7212 | 7817 | 8117 | 5701 | 5785 | 7308 | 7203 | 7849 | 6381 | 6134 | 6333 | 6867.84 | 7453 | 6283 | 0.843 | -0.246 | 0.00025914 | 0.00301413 | 0.0011 | 0.0074 | 0.0031 | 0.0031 | TRUE | 0.0264 |
| Gpha2 | 39 | 30 | 24 | 45 | 26 | 35 | 41 | 30 | 24 | 35 | 28 | 38 | 32.86 | 32 | 34 | 1.068 | 0.095 | 0.79417969 | 0.89406153 | 0.0176 | 0.1072 | 0.0648 | 0.0648 | TRUE | 0.4598 |
| Gphb5 | 0 | 0 | 0 | 0 | 0 | 0 | 0 | 0 | 0 | 0 | 0 | 0 | 0 | 0 | 0 | NA | NA | NA | NA | NA | NA | NA | NA | NA | NA |
| Gphn | 436 | 538 | 545 | 795 | 514 | 511 | 463 | 537 | 547 | 625 | 553 | 559 | 547.5 | 516 | 579 | 1.124 | 0.168 | 0.15555589 | 0.34795454 | 0.0046 | 0.0129 | 0.0083 | 0.0083 | TRUE | 0.0925 |
| Gpi1 | 30105 | 38947 | 39998 | 43970 | 33701 | 38104 | 31977 | 38897 | 40162 | 34566 | 36260 | 41711 | 37262.09 | 37012 | 37512 | 1.013 | 0.019 | 0.86908124 | 0.93551388 | 0.0096 | 0.007 | 0.0099 | 0.0099 | TRUE | 0.1734 |
| Gpihbp1 | 0 | 0 | 1 | 0 | 0 | 0 | 0 | 0 | 1 | 0 | 0 | 0 | 0.17 | 0 | 0 | 0.49 | -1.029 | 0.80087 | NA | 0 | 19.6928 | 10 | 10 | TRUE | 0.4098 |
| Gpkow | 993 | 937 | 874 | 1297 | 1009 | 907 | 1055 | 936 | 878 | 1020 | 1086 | 993 | 994.37 | 956 | 1033 | 1.08 | 0.111 | 0.28452441 | 0.50597801 | 0.004 | 0.0102 | 0.0068 | 0.0068 | TRUE | 0.0967 |
| Gpld1 | 10 | 8 | 6 | 17 | 9 | 9 | 11 | 8 | 6 | 13 | 10 | 10 | 9.59 | 8 | 11 | 1.349 | 0.432 | 0.48112893 | 0.68020487 | 0 | 0.3505 | 0.1648 | 0.1648 | TRUE | 0.1879 |
| Gpm6a | 35 | 24 | 19 | 36 | 13 | 12 | 37 | 24 | 19 | 28 | 14 | 13 | 22.61 | 27 | 18 | 0.703 | -0.509 | 0.30603758 | 0.52665845 | 0.1118 | 0.1526 | 0.1327 | 0.1327 | TRUE | 1.1689 |
| Gpm6b | 67 | 31 | 43 | 30 | 14 | 24 | 71 | 31 | 43 | 24 | 15 | 26 | 35.04 | 48 | 22 | 0.449 | -1.156 | 0.00742263 | 0.04194545 | 0.1019 | 0.1009 | 0.1013 | 0.1013 | TRUE | 0.341 |
| Gpn1 | 2941 | 2936 | 2861 | 3398 | 2540 | 2513 | 3124 | 2932 | 2873 | 2671 | 2733 | 2751 | 2847.3 | 2976 | 2718 | 0.913 | -0.131 | 0.06193578 | 0.19023391 | 7.00E-04 | 0.0081 | 0.0032 | 0.0032 | TRUE | 0.0228 |
| Gpn2 | 1862 | 1633 | 1598 | 2193 | 1555 | 1400 | 1978 | 1631 | 1605 | 1724 | 1673 | 1533 | 1690.47 | 1738 | 1643 | 0.946 | -0.08 | 0.47548126 | 0.67567262 | 0.0078 | 0.0089 | 0.0085 | 0.0085 | TRUE | 0.1758 |
| Gpn3 | 2863 | 2994 | 2864 | 3477 | 2447 | 2533 | 3041 | 2990 | 2876 | 2733 | 2633 | 2773 | 2840.97 | 2969 | 2713 | 0.914 | -0.13 | 0.05395669 | 0.17274437 | 4.00E-04 | 0.0081 | 0.0029 | 0.0029 | TRUE | 0.0092 |
| Gpnmb | 66 | 60 | 55 | 67 | 51 | 53 | 70 | 60 | 55 | 53 | 55 | 58 | 58.47 | 62 | 55 | 0.893 | -0.163 | 0.52003038 | 0.71102318 | 0 | 0.0632 | 0.0294 | 0.0294 | TRUE | 0.1184 |
| Gpr1 | 5 | 8 | 5 | 10 | 10 | 5 | 5 | 8 | 5 | 8 | 11 | 5 | 7.07 | 6 | 8 | 1.313 | 0.393 | 0.58327019 | 0.75875085 | 0 | 0.4729 | 0.2257 | 0.2257 | TRUE | 0.1796 |
| Gpr101 | 3 | 1 | 0 | 1 | 0 | 0 | 3 | 1 | 0 | 1 | 0 | 0 | 0.83 | 1 | 0 | 0.252 | -1.991 | 0.4214787 | NA | 0 | 3.983 | 3.0547 | 3.0547 | TRUE | 0.2456 |
| Gpr107 | 2167 | 2738 | 2723 | 3407 | 2447 | 2525 | 2302 | 2734 | 2734 | 2678 | 2633 | 2764 | 2640.92 | 2590 | 2692 | 1.039 | 0.055 | 0.5593741 | 0.74125763 | 0.0047 | 0.0082 | 0.0061 | 0.0061 | TRUE | 0.1147 |
| Gpr108 | 851 | 1166 | 1047 | 1196 | 959 | 1028 | 904 | 1164 | 1051 | 940 | 1032 | 1125 | 1036.17 | 1040 | 1032 | 0.992 | -0.012 | 0.92568387 | 0.96333323 | 0.0111 | 0.0101 | 0.0106 | 0.0106 | TRUE | 0.1558 |
| Gpr110 | 0 | 0 | 1 | 5 | 4 | 5 | 0 | 0 | 1 | 4 | 4 | 5 | 2.45 | 0 | 4 | 12.444 | 3.637 | 0.0222212 | NA | 0 | 1.3505 | 0.716 | 0.716 | TRUE | 0.4098 |
| Gpr111 | 2 | 1 | 0 | 3 | 2 | 1 | 2 | 1 | 0 | 2 | 2 | 1 | 1.45 | 1 | 2 | 1.828 | 0.871 | 0.58722174 | NA | 0 | 2.2717 | 1.0991 | 1.0991 | TRUE | 0.3111 |
| Gpr113 | 49 | 65 | 46 | 56 | 57 | 50 | 52 | 65 | 46 | 44 | 61 | 55 | 53.87 | 54 | 53 | 0.975 | -0.036 | 0.90016878 | 0.95157252 | 0.0115 | 0.0681 | 0.0408 | 0.0408 | TRUE | 0.2433 |
| Gpr114 | 86 | 119 | 148 | 72 | 52 | 61 | 91 | 119 | 149 | 57 | 56 | 67 | 89.69 | 120 | 60 | 0.498 | -1.005 | 9.08E-05 | 0.00128229 | 0.0266 | 0.0436 | 0.035 | 0.035 | TRUE | 0.1164 |
| Gpr115 | 8 | 2 | 0 | 0 | 1 | 3 | 8 | 2 | 0 | 0 | 1 | 3 | 2.48 | 3 | 1 | 0.404 | -1.309 | 0.44648292 | NA | 2.0992 | 1.3376 | 1.6361 | 1.6361 | TRUE | 0.6246 |
| Gpr116 | 204 | 166 | 196 | 212 | 187 | 134 | 217 | 166 | 197 | 167 | 201 | 147 | 182.3 | 193 | 172 | 0.888 | -0.171 | 0.36615171 | 0.58362252 | 0.0155 | 0.025 | 0.0202 | 0.0202 | TRUE | 0.2342 |
| Gpr119 | 0 | 0 | 0 | 0 | 0 | 0 | 0 | 0 | 0 | 0 | 0 | 0 | 0 | 0 | 0 | NA | NA | NA | NA | NA | NA | NA | NA | NA | NA |
| Gpr12 | 13 | 22 | 19 | 13 | 19 | 17 | 14 | 22 | 19 | 10 | 20 | 19 | 17.35 | 18 | 16 | 0.885 | -0.176 | 0.72212737 | 0.84963766 | 0 | 0.1967 | 0.1194 | 0.1194 | TRUE | 0.6653 |
| Gpr123 | 1 | 0 | 0 | 4 | 2 | 3 | 1 | 0 | 0 | 3 | 2 | 3 | 1.61 | 0 | 3 | 7.834 | 2.97 | 0.09140814 | NA | 0 | 2.0569 | 1.0597 | 1.0597 | TRUE | 0.4614 |
| Gpr124 | 208 | 242 | 256 | 398 | 245 | 181 | 221 | 242 | 257 | 313 | 264 | 198 | 249.05 | 240 | 258 | 1.08 | 0.11 | 0.56522861 | 0.74551564 | 0.025 | 0.0201 | 0.0226 | 0.0226 | TRUE | 0.2507 |
| Gpr125 | 2464 | 3006 | 2974 | 4528 | 3320 | 2979 | 2617 | 3002 | 2986 | 3560 | 3572 | 3261 | 3166.37 | 2868 | 3464 | 1.208 | 0.272 | 0.00238086 | 0.01766269 | 0.0037 | 0.0079 | 0.0055 | 0.0055 | TRUE | 0.0711 |
| Gpr126 | 32 | 53 | 50 | 183 | 111 | 107 | 34 | 53 | 50 | 144 | 119 | 117 | 86.26 | 46 | 127 | 2.776 | 1.473 | 4.75E-09 | 2.42E-07 | 0.012 | 0.0451 | 0.0306 | 0.0306 | TRUE | 0.3821 |
| Gpr128 | 0 | 2 | 0 | 0 | 0 | 0 | 0 | 2 | 0 | 0 | 0 | 0 | 0.33 | 1 | 0 | 0.268 | -1.897 | 0.63866967 | NA | 0 | 9.903 | 10 | 10 | TRUE | 0.9739 |
| Gpr132 | 2 | 4 | 2 | 3 | 3 | 5 | 2 | 4 | 2 | 2 | 3 | 5 | 3.2 | 3 | 3 | 1.338 | 0.42 | 0.68874356 | NA | 0 | 1.0371 | 0.4705 | 0.4705 | TRUE | 0.2593 |
| Gpr133 | 119 | 56 | 47 | 79 | 48 | 51 | 126 | 56 | 47 | 62 | 52 | 56 | 66.52 | 76 | 57 | 0.741 | -0.432 | 0.26927739 | 0.48929066 | 0.136 | 0.0564 | 0.0949 | 0.0949 | TRUE | 2.9533 |
| Gpr135 | 60 | 72 | 71 | 65 | 70 | 52 | 64 | 72 | 71 | 51 | 75 | 57 | 65.04 | 69 | 61 | 0.881 | -0.183 | 0.4816753 | 0.68073323 | 0.0075 | 0.0576 | 0.0333 | 0.0333 | TRUE | 0.3486 |
| Gpr137 | 351 | 447 | 519 | 671 | 494 | 480 | 373 | 446 | 521 | 527 | 532 | 525 | 487.47 | 447 | 528 | 1.181 | 0.241 | 0.08990914 | 0.24390214 | 0.0107 | 0.0137 | 0.0124 | 0.0124 | TRUE | 0.222 |
| Gpr137b | 377 | 657 | 687 | 687 | 551 | 646 | 400 | 656 | 690 | 540 | 593 | 707 | 597.74 | 582 | 613 | 1.053 | 0.074 | 0.7232973 | 0.85006834 | 0.0492 | 0.0124 | 0.0297 | 0.0297 | TRUE | 0.8709 |
| Gpr137b-ps | 29 | 63 | 58 | 68 | 52 | 54 | 31 | 63 | 58 | 53 | 56 | 59 | 53.41 | 51 | 56 | 1.105 | 0.145 | 0.65682116 | 0.80670515 | 0.0444 | 0.0686 | 0.0575 | 0.0575 | TRUE | 0.9319 |
| Gpr137c | 18 | 33 | 37 | 28 | 25 | 37 | 19 | 33 | 37 | 22 | 27 | 41 | 29.77 | 30 | 30 | 0.993 | -0.01 | 0.98037579 | 0.9893459 | 0.0708 | 0.1176 | 0.0951 | 0.0951 | TRUE | 0.6255 |
| Gpr139 | 1 | 1 | 2 | 2 | 0 | 0 | 1 | 1 | 2 | 2 | 0 | 0 | 0.94 | 1 | 1 | 0.425 | -1.233 | 0.55053282 | NA | 0 | 3.5108 | 1.8559 | 1.8559 | TRUE | 0.9714 |
| Gpr141 | 0 | 0 | 0 | 1 | 0 | 0 | 0 | 0 | 0 | 1 | 0 | 0 | 0.13 | 0 | 0 | 1.859 | 0.894 | 0.82650734 | NA | 0 | 25.1515 | 10 | 10 | TRUE | 0.2563 |
| Gpr142 | 2 | 8 | 2 | 9 | 9 | 3 | 2 | 8 | 2 | 7 | 10 | 3 | 5.36 | 4 | 7 | 1.651 | 0.723 | 0.43869748 | 0.64604451 | 0.2156 | 0.6214 | 0.4292 | 0.4292 | TRUE | 0.6604 |
| Gpr143 | 22 | 15 | 9 | 21 | 9 | 7 | 23 | 15 | 9 | 17 | 10 | 8 | 13.54 | 16 | 12 | 0.727 | -0.46 | 0.444852 | 0.65143101 | 0.1185 | 0.2502 | 0.1856 | 0.1856 | TRUE | 0.2216 |
| Gpr146 | 82 | 125 | 120 | 64 | 66 | 97 | 87 | 125 | 120 | 50 | 71 | 106 | 93.32 | 111 | 76 | 0.679 | -0.558 | 0.07107656 | 0.20759622 | 0.0747 | 0.0422 | 0.0577 | 0.0577 | TRUE | 0.5792 |
| Gpr149 | 0 | 0 | 1 | 3 | 1 | 1 | 0 | 0 | 1 | 2 | 1 | 1 | 0.92 | 0 | 1 | 4.189 | 2.067 | 0.32121847 | NA | 0 | 3.5794 | 1.8085 | 1.8085 | TRUE | 0.4098 |
| Gpr15 | 0 | 0 | 0 | 0 | 0 | 0 | 0 | 0 | 0 | 0 | 0 | 0 | 0 | 0 | 0 | NA | NA | NA | NA | NA | NA | NA | NA | NA | NA |
| Gpr150 | 5 | 1 | 1 | 8 | 12 | 4 | 5 | 1 | 1 | 6 | 13 | 4 | 5.15 | 2 | 8 | 3.248 | 1.699 | 0.09647758 | 0.25496722 | 0.27 | 0.6468 | 0.4789 | 0.4789 | TRUE | 0.9913 |
| Gpr151 | 0 | 0 | 0 | 0 | 0 | 0 | 0 | 0 | 0 | 0 | 0 | 0 | 0 | 0 | 0 | NA | NA | NA | NA | NA | NA | NA | NA | NA | NA |
| Gpr152 | 0 | 0 | 0 | 0 | 0 | 0 | 0 | 0 | 0 | 0 | 0 | 0 | 0 | 0 | 0 | NA | NA | NA | NA | NA | NA | NA | NA | NA | NA |
| Gpr153 | 436 | 467 | 410 | 261 | 197 | 216 | 463 | 466 | 412 | 205 | 212 | 236 | 332.46 | 447 | 218 | 0.486 | -1.04 | 2.07E-15 | 2.99E-13 | 0.0021 | 0.0168 | 0.009 | 0.009 | TRUE | 0.0593 |
| Gpr155 | 275 | 206 | 200 | 279 | 199 | 156 | 292 | 206 | 201 | 219 | 214 | 171 | 217.14 | 233 | 201 | 0.867 | -0.206 | 0.30712151 | 0.5274878 | 0.0274 | 0.0221 | 0.0247 | 0.0247 | TRUE | 0.5416 |
| Gpr156 | 42 | 35 | 36 | 62 | 41 | 41 | 45 | 35 | 36 | 49 | 44 | 45 | 42.24 | 39 | 46 | 1.195 | 0.256 | 0.38480192 | 0.59882544 | 0 | 0.0849 | 0.0389 | 0.0389 | TRUE | 0.1343 |
| Gpr157 | 168 | 237 | 204 | 218 | 156 | 150 | 178 | 237 | 205 | 171 | 168 | 164 | 187.23 | 207 | 168 | 0.812 | -0.301 | 0.07161196 | 0.20861909 | 0.0055 | 0.0245 | 0.0147 | 0.0147 | TRUE | 0.1767 |
| Gpr158 | 1 | 1 | 0 | 3 | 1 | 1 | 1 | 1 | 0 | 2 | 1 | 1 | 1.1 | 1 | 1 | 2.251 | 1.171 | 0.51887941 | NA | 0 | 3.0065 | 1.3089 | 1.3089 | TRUE | 0.253 |

| | | | | | | | | | | | | | | | | | | | | | | | | | |
|---|---|---|---|---|---|---|---|---|---|---|---|---|---|---|---|---|---|---|---|---|---|---|---|---|---|
| Gpr160 | 765 | 841 | 726 | 875 | 632 | 673 | 813 | 840 | 729 | 688 | 680 | 737 | 747.67 | 794 | 702 | 0.883 | -0.179 | 0.07847242 | 0.22169438 | 0.0023 | 0.0113 | 0.0061 | 0.0061 | TRUE | 0.0609 |
| Gpr161 | 84 | 88 | 87 | 141 | 118 | 94 | 89 | 88 | 87 | 111 | 127 | 103 | 100.86 | 88 | 114 | 1.288 | 0.365 | 0.06715084 | 0.20017068 | 0 | 0.0396 | 0.0185 | 0.0185 | TRUE | 0.1003 |
| Gpr162 | 197 | 176 | 167 | 150 | 136 | 129 | 209 | 176 | 168 | 118 | 146 | 141 | 159.69 | 184 | 135 | 0.731 | -0.452 | 0.01301315 | 0.0638925 | 0.0078 | 0.0275 | 0.0174 | 0.0174 | TRUE | 0.1498 |
| Gpr165 | 0 | 0 | 0 | 3 | 3 | 1 | 0 | 0 | 0 | 2 | 3 | 1 | 1.11 | 0 | 2 | 11.874 | 3.57 | 0.0910322 | NA | 0 | 2.9656 | 1.9955 | 1.9955 | TRUE | 0.1116 |
| Gpr17 | 0 | 0 | 0 | 0 | 1 | 0 | 0 | 0 | 0 | 0 | 1 | 0 | 0.18 | 0 | 0 | 1.859 | 0.894 | 0.82650734 | NA | 0 | 18.3785 | 10 | 10 | TRUE | 0.5204 |
| Gpr171 | 0 | 0 | 0 | 0 | 0 | 0 | 0 | 0 | 0 | 0 | 0 | 0 | 0 | 0 | 0 | NA | NA | NA | NA | NA | NA | NA | NA | NA | NA |
| Gpr173 | 23 | 18 | 15 | 29 | 18 | 16 | 24 | 18 | 15 | 23 | 19 | 18 | 19.52 | 19 | 20 | 1.046 | 0.065 | 0.88172457 | 0.94154096 | 0 | 0.1756 | 0.088 | 0.088 | TRUE | 0.2941 |
| Gpr174 | 0 | 0 | 0 | 0 | 0 | 0 | 0 | 0 | 0 | 0 | 0 | 0 | 0 | 0 | 0 | NA | NA | NA | NA | NA | NA | NA | NA | NA | NA |
| Gpr176 | 10 | 1 | 3 | 18 | 14 | 5 | 11 | 1 | 3 | 14 | 15 | 5 | 8.22 | 5 | 11 | 2.399 | 1.263 | 0.1568968 | 0.34981288 | 0.4494 | 0.4077 | 0.4255 | 0.4255 | TRUE | 2.0098 |
| Gpr179 | 293 | 304 | 299 | 443 | 339 | 324 | 311 | 304 | 300 | 348 | 365 | 355 | 330.45 | 305 | 356 | 1.166 | 0.222 | 0.05357467 | 0.1719584 | 0 | 0.0169 | 0.0065 | 0.0065 | TRUE | 0.0054 |
| Gpr18 | 0 | 0 | 0 | 0 | 0 | 0 | 0 | 0 | 0 | 0 | 0 | 0 | 0 | 0 | 0 | NA | NA | NA | NA | NA | NA | NA | NA | NA | NA |
| Gpr180 | 833 | 977 | 1023 | 1328 | 884 | 910 | 885 | 976 | 1027 | 1044 | 951 | 996 | 979.83 | 963 | 997 | 1.036 | 0.051 | 0.60016475 | 0.77215094 | 0.0029 | 0.0103 | 0.0059 | 0.0059 | TRUE | 0.0592 |
| Gpr182 | 8 | 10 | 5 | 16 | 8 | 13 | 8 | 10 | 5 | 13 | 9 | 14 | 9.82 | 8 | 12 | 1.51 | 0.595 | 0.34002027 | 0.56014202 | 0 | 0.3424 | 0.1733 | 0.1733 | TRUE | 0.2914 |
| Gpr183 | 0 | 0 | 0 | 0 | 0 | 0 | 0 | 0 | 0 | 0 | 0 | 0 | 0 | 0 | 0 | NA | NA | NA | NA | NA | NA | NA | NA | NA | NA |
| Gpr19 | 1018 | 1413 | 1494 | 1483 | 1085 | 1246 | 1081 | 1411 | 1500 | 1166 | 1167 | 1364 | 1281.63 | 1331 | 1232 | 0.926 | -0.112 | 0.43394917 | 0.6425633 | 0.0179 | 0.0095 | 0.0139 | 0.0139 | TRUE | 0.3224 |
| Gpr20 | 82 | 80 | 87 | 119 | 72 | 77 | 87 | 80 | 87 | 94 | 77 | 84 | 84.94 | 85 | 85 | 1.008 | 0.011 | 0.95915718 | 0.97898705 | 0 | 0.0457 | 0.021 | 0.021 | TRUE | 0.0802 |
| Gpr21 | 9 | 17 | 10 | 12 | 9 | 8 | 10 | 17 | 10 | 9 | 10 | 9 | 10.74 | 12 | 9 | 0.761 | -0.393 | 0.50113889 | 0.69669611 | 0 | 0.3136 | 0.1521 | 0.1521 | TRUE | 0.4761 |
| Gpr22 | 0 | 0 | 0 | 0 | 0 | 0 | 0 | 0 | 0 | 0 | 0 | 0 | 0 | 0 | 0 | NA | NA | NA | NA | NA | NA | NA | NA | NA | NA |
| Gpr25 | 1 | 19 | 13 | 21 | 18 | 21 | 1 | 19 | 13 | 17 | 19 | 23 | 15.33 | 11 | 20 | 1.767 | 0.821 | 0.25904103 | 0.47795918 | 0.4588 | 0.2219 | 0.3106 | 0.3106 | TRUE | 0.8979 |
| Gpr26 | 1 | 2 | 2 | 0 | 1 | 1 | 1 | 2 | 2 | 0 | 1 | 1 | 1.21 | 2 | 1 | 0.403 | -1.313 | 0.45680201 | NA | 0 | 2.7377 | 1.2272 | 1.2272 | TRUE | 0.3951 |
| Gpr27 | 48 | 135 | 148 | 140 | 102 | 118 | 51 | 135 | 149 | 110 | 110 | 129 | 113.9 | 112 | 116 | 1.042 | 0.059 | 0.86768744 | 0.93471527 | 0.1382 | 0.0358 | 0.0819 | 0.0819 | TRUE | 2.2161 |
| Gpr3 | 89 | 103 | 87 | 66 | 50 | 38 | 95 | 103 | 87 | 52 | 54 | 42 | 72.01 | 95 | 49 | 0.519 | -0.947 | 7.93E-05 | 0.00114641 | 0 | 0.0527 | 0.0262 | 0.0262 | TRUE | 0.1313 |
| Gpr30 | 0 | 0 | 0 | 0 | 1 | 0 | 0 | 0 | 0 | 0 | 1 | 0 | 0.18 | 0 | 0 | 1.859 | 0.894 | 0.82650734 | NA | 0 | 18.3785 | 10 | 10 | TRUE | 0.5204 |
| Gpr31b | 0 | 0 | 0 | 0 | 0 | 0 | 0 | 0 | 0 | 0 | 0 | 0 | 0 | 0 | 0 | NA | NA | NA | NA | NA | NA | NA | NA | NA | NA |
| Gpr33 | 0 | 0 | 0 | 0 | 0 | 0 | 0 | 0 | 0 | 0 | 0 | 0 | 0 | 0 | 0 | NA | NA | NA | NA | NA | NA | NA | NA | NA | NA |
| Gpr34 | 0 | 0 | 0 | 0 | 0 | 0 | 0 | 0 | 0 | 0 | 0 | 0 | 0 | 0 | 0 | NA | NA | NA | NA | NA | NA | NA | NA | NA | NA |
| Gpr35 | 4 | 5 | 3 | 5 | 4 | 2 | 4 | 5 | 3 | 4 | 4 | 2 | 3.78 | 4 | 3 | 0.856 | -0.224 | 0.81262446 | NA | 0 | 0.8785 | 0.3768 | 0.3768 | TRUE | 0.1347 |
| Gpr37 | 1 | 0 | 0 | 2 | 2 | 0 | 1 | 0 | 0 | 2 | 2 | 0 | 0.8 | 0 | 1 | 3.417 | 1.773 | 0.46651319 | NA | 0 | 4.1368 | 2.8785 | 2.8785 | TRUE | 0.4614 |
| Gpr37l1 | 34 | 35 | 35 | 33 | 16 | 29 | 36 | 35 | 35 | 26 | 17 | 32 | 30.19 | 35 | 25 | 0.706 | -0.502 | 0.17161489 | 0.37055804 | 0 | 0.116 | 0.0633 | 0.0633 | TRUE | 0.395 |
| Gpr39 | 25 | 10 | 5 | 18 | 19 | 9 | 27 | 10 | 5 | 14 | 20 | 10 | 14.33 | 14 | 15 | 1.073 | 0.102 | 0.88234539 | 0.94169383 | 0.3041 | 0.2367 | 0.2694 | 0.2694 | TRUE | 1.2488 |
| Gpr4 | 391 | 308 | 316 | 343 | 293 | 273 | 415 | 308 | 317 | 270 | 315 | 299 | 320.66 | 347 | 295 | 0.849 | -0.236 | 0.14342164 | 0.33004787 | 0.0143 | 0.0172 | 0.0157 | 0.0157 | TRUE | 0.3389 |
| Gpr44 | 0 | 0 | 1 | 0 | 0 | 1 | 0 | 0 | 1 | 0 | 0 | 1 | 0.35 | 0 | 0 | 0.954 | -0.067 | 0.98566485 | NA | 0 | 9.4252 | 8.1416 | 8.1416 | TRUE | 0.5383 |
| Gpr45 | 4 | 6 | 5 | 3 | 4 | 3 | 4 | 6 | 5 | 2 | 4 | 3 | 4.2 | 5 | 3 | 0.642 | -0.64 | 0.47729022 | NA | 0 | 0.7911 | 0.3351 | 0.3351 | TRUE | 0.1265 |
| Gpr50 | 3 | 14 | 0 | 26 | 18 | 14 | 3 | 14 | 0 | 20 | 19 | 15 | 12.05 | 6 | 18 | 3.206 | 1.681 | 0.06078336 | 0.18761671 | 0.879 | 0.2803 | 0.4632 | 0.4632 | TRUE | 3.35 |
| Gpr52 | 2 | 0 | 2 | 0 | 0 | 0 | 2 | 0 | 2 | 0 | 0 | 0 | 0.69 | 1 | 0 | 0.13 | -2.939 | 0.31726474 | NA | 0 | 4.7901 | 4.8546 | 4.8546 | TRUE | 0.4924 |
| Gpr55 | 4 | 0 | 4 | 3 | 3 | 2 | 4 | 0 | 4 | 2 | 3 | 2 | 2.67 | 3 | 2 | 0.942 | -0.086 | 0.94560151 | NA | 0 | 1.2392 | 0.7806 | 0.7806 | TRUE | 0.9419 |
| Gpr56 | 876 | 1183 | 1263 | 410 | 375 | 623 | 930 | 1181 | 1268 | 322 | 403 | 682 | 797.98 | 1126 | 469 | 0.416 | -1.266 | 3.69E-07 | 1.14E-05 | 0.0907 | 0.011 | 0.0432 | 0.0432 | TRUE | 1.8196 |
| Gpr6 | 0 | 0 | 0 | 0 | 0 | 0 | 0 | 0 | 0 | 0 | 0 | 0 | 0 | 0 | 0 | NA | NA | NA | NA | NA | NA | NA | NA | NA | NA |
| Gpr61 | 15 | 11 | 8 | 8 | 11 | 8 | 16 | 11 | 8 | 6 | 12 | 9 | 10.31 | 12 | 9 | 0.762 | -0.392 | 0.52667529 | 0.71578825 | 0 | 0.3266 | 0.1777 | 0.1777 | TRUE | 0.3103 |
| Gpr62 | 8 | 15 | 16 | 19 | 12 | 16 | 8 | 15 | 16 | 15 | 13 | 18 | 14.15 | 13 | 15 | 1.143 | 0.193 | 0.70941941 | 0.84171221 | 0 | 0.2397 | 0.1225 | 0.1225 | TRUE | 0.3853 |
| Gpr63 | 35 | 21 | 16 | 83 | 41 | 43 | 37 | 21 | 16 | 65 | 44 | 47 | 38.44 | 25 | 52 | 2.124 | 1.087 | 0.0064709 | 0.03809248 | 0.0753 | 0.0926 | 0.0848 | 0.0848 | TRUE | 1.1333 |
| Gpr64 | 16 | 31 | 22 | 53 | 25 | 25 | 17 | 31 | 22 | 42 | 27 | 27 | 27.66 | 23 | 32 | 1.378 | 0.463 | 0.26217457 | 0.48154389 | 0.0429 | 0.126 | 0.0857 | 0.0857 | TRUE | 0.5547 |
| Gpr65 | 0 | 1 | 0 | 0 | 0 | 0 | 0 | 1 | 0 | 0 | 0 | 0 | 0.17 | 0 | 0 | 0.49 | -1.029 | 0.80087 | NA | 0 | 19.7991 | 10 | 10 | TRUE | 0.405 |
| Gpr68 | 46 | 88 | 94 | 34 | 38 | 47 | 49 | 88 | 94 | 27 | 41 | 51 | 58.37 | 77 | 40 | 0.51 | -0.971 | 0.00797134 | 0.04419123 | 0.0928 | 0.0633 | 0.0774 | 0.0774 | TRUE | 0.7845 |
| Gpr75 | 29 | 65 | 73 | 96 | 93 | 103 | 31 | 65 | 73 | 75 | 100 | 113 | 76.22 | 56 | 96 | 1.698 | 0.764 | 0.02540073 | 0.10288145 | 0.0941 | 0.0501 | 0.0699 | 0.0699 | TRUE | 1.2234 |
| Gpr81 | 0 | 7 | 8 | 10 | 12 | 6 | 0 | 7 | 8 | 8 | 13 | 7 | 7.06 | 5 | 9 | 1.805 | 0.852 | 0.36215302 | 0.58014481 | 0.4705 | 0.4735 | 0.4748 | 0.4748 | TRUE | 1.4599 |
| Gpr82 | 0 | 0 | 0 | 0 | 0 | 0 | 0 | 0 | 0 | 0 | 0 | 0 | 0 | 0 | 0 | NA | NA | NA | NA | NA | NA | NA | NA | NA | NA |
| Gpr83 | 25 | 11 | 11 | 18 | 12 | 4 | 27 | 11 | 11 | 14 | 13 | 4 | 13.34 | 16 | 10 | 0.656 | -0.608 | 0.35440818 | 0.57279009 | 0.2099 | 0.2539 | 0.2329 | 0.2329 | TRUE | 1.4463 |
| Gpr84 | 0 | 4 | 0 | 4 | 2 | 0 | 0 | 4 | 0 | 3 | 2 | 0 | 1.55 | 1 | 2 | 1.339 | 0.421 | 0.83935129 | NA | 3.233 | 2.1344 | 2.4569 | 2.4569 | TRUE | 1.5899 |
| Gpr85 | 13 | 12 | 8 | 39 | 23 | 25 | 14 | 12 | 8 | 31 | 25 | 27 | 19.43 | 11 | 28 | 2.46 | 1.299 | 0.00441448 | 0.02846493 | 0 | 0.1764 | 0.0869 | 0.0869 | TRUE | 0.2416 |
| Gpr87 | 0 | 0 | 0 | 0 | 0 | 1 | 0 | 0 | 0 | 0 | 0 | 1 | 0.18 | 0 | 0 | 1.859 | 0.894 | 0.82650734 | NA | 0 | 18.0643 | 10 | 10 | TRUE | 0.5383 |
| Gpr88 | 0 | 0 | 0 | 0 | 0 | 0 | 0 | 0 | 0 | 0 | 0 | 0 | 0 | 0 | 0 | NA | NA | NA | NA | NA | NA | NA | NA | NA | NA |
| Gpr89 | 1128 | 1152 | 1114 | 1347 | 959 | 884 | 1198 | 1151 | 1119 | 1059 | 1032 | 968 | 1087.6 | 1156 | 1020 | 0.883 | -0.18 | 0.03498722 | 0.12811271 | 7.00E-04 | 0.0099 | 0.0043 | 0.0043 | TRUE | 0.0233 |
| Gpr97 | 11 | 8 | 4 | 9 | 0 | 0 | 12 | 8 | 4 | 7 | 0 | 0 | 5.13 | 8 | 2 | 0.317 | -1.656 | 0.20122599 | 0.40877592 | 1.6668 | 0.6494 | 0.9499 | 0.9499 | TRUE | 1.1999 |
| Gpr98 | 423 | 396 | 390 | 598 | 393 | 380 | 449 | 395 | 392 | 470 | 423 | 416 | 424.22 | 412 | 436 | 1.061 | 0.085 | 0.48179106 | 0.68077623 | 0.003 | 0.0147 | 0.0082 | 0.0082 | TRUE | 0.071 |
| Gprasp1 | 6408 | 6140 | 5975 | 8821 | 6227 | 5794 | 6806 | 6132 | 6000 | 6934 | 6700 | 6342 | 6485.79 | 6313 | 6659 | 1.055 | 0.077 | 0.3458694 | 0.56517324 | 0.0031 | 0.0074 | 0.0047 | 0.0047 | TRUE | 0.057 |
| Gprasp2 | 1864 | 2366 | 2409 | 2514 | 2035 | 2059 | 1980 | 2363 | 2419 | 1976 | 2190 | 2254 | 2196.91 | 2254 | 2140 | 0.949 | -0.076 | 0.48550402 | 0.68271838 | 0.0078 | 0.0084 | 0.008 | 0.008 | TRUE | 0.1366 |
| Gprc5a | 2510 | 5255 | 6065 | 2971 | 2578 | 3173 | 2666 | 5248 | 6090 | 2336 | 2774 | 3473 | 3764.48 | 4668 | 2861 | 0.613 | -0.706 | 0.0492384 | 0.16260219 | 0.0927 | 0.0078 | 0.048 | 0.0927 | TRUE | 0.6765 |
| Gprc5b | 829 | 974 | 948 | 960 | 693 | 785 | 881 | 973 | 952 | 755 | 746 | 859 | 860.8 | 935 | 787 | 0.84 | -0.251 | 0.01416764 | 0.0679328 | 0.0032 | 0.0107 | 0.0064 | 0.0064 | TRUE | 0.0761 |
| Gprc5c | 181 | 309 | 300 | 573 | 431 | 453 | 192 | 309 | 301 | 450 | 464 | 496 | 368.69 | 267 | 470 | 1.756 | 0.812 | 1.38E-05 | 0.00026131 | 0.0292 | 0.0158 | 0.0222 | 0.0222 | TRUE | 0.6681 |
| Gprc5d | 4 | 3 | 3 | 4 | 0 | 4 | 4 | 3 | 3 | 3 | 0 | 4 | 2.96 | 3 | 2 | 0.74 | -0.434 | 0.71512324 | NA | 0 | 1.1186 | 0.6748 | 0.6748 | TRUE | 0.7502 |
| Gprc6a | 0 | 0 | 0 | 0 | 0 | 0 | 0 | 0 | 0 | 0 | 0 | 0 | 0 | 0 | 0 | NA | NA | NA | NA | NA | NA | NA | NA | NA | NA |
| Gprin1 | 80 | 52 | 50 | 66 | 57 | 38 | 85 | 52 | 50 | 52 | 61 | 42 | 56.99 | 62 | 52 | 0.829 | -0.27 | 0.39724298 | 0.61077366 | 0.0466 | 0.0647 | 0.0559 | 0.0559 | TRUE | 0.8582 |
| Gprin2 | 87 | 76 | 101 | 105 | 99 | 98 | 92 | 76 | 101 | 83 | 107 | 107 | 94.34 | 90 | 99 | 1.094 | 0.129 | 0.57093952 | 0.74978003 | 0.0119 | 0.0418 | 0.0269 | 0.0269 | TRUE | 0.2251 |
| Gprin3 | 5 | 6 | 2 | 19 | 10 | 4 | 5 | 6 | 2 | 15 | 11 | 4 | 7.23 | 4 | 10 | 2.29 | 1.195 | 0.14367814 | 0.33047859 | 0.1597 | 0.4625 | 0.3181 | 0.3181 | TRUE | 0.1768 |
| Gps1 | 6481 | 6919 | 7084 | 9202 | 6902 | 6597 | 6884 | 6910 | 7113 | 7234 | 7426 | 7221 | 7131.43 | 6969 | 7294 | 1.047 | 0.066 | 0.24688011 | 0.46379674 | 1.00E-04 | 0.0074 | 0.0022 | 0.0022 | TRUE | 0.004 |
| Gps2 | 1150 | 951 | 1039 | 1514 | 1167 | 1010 | 1222 | 950 | 1043 | 1190 | 1256 | 1106 | 1127.66 | 1072 | 1184 | 1.105 | 0.144 | 0.22864669 | 0.44204654 | 0.009 | 0.0098 | 0.0095 | 0.0095 | TRUE | 0.1787 |
| Gpsm1 | 448 | 448 | 435 | 514 | 417 | 384 | 476 | 447 | 437 | 404 | 449 | 420 | 438.86 | 453 | 424 | 0.935 | -0.097 | 0.38041871 | 0.59482144 | 0 | 0.0144 | 0.0065 | 0.0065 | TRUE | 0.0288 |
| Gpsm2 | 1778 | 2015 | 2012 | 2987 | 2038 | 2311 | 1889 | 2012 | 2020 | 2348 | 2193 | 2530 | 2165.31 | 1974 | 2357 | 1.194 | 0.256 | 0.00311702 | 0.02186905 | 0.0026 | 0.0084 | 0.0049 | 0.0049 | TRUE | 0.0491 |

| | | | | | | | | | | | | | | | | | | | | | | | | | |
|---|---|---|---|---|---|---|---|---|---|---|---|---|---|---|---|---|---|---|---|---|---|---|---|---|---|
| Gpsm3 | 184 | 145 | 155 | 192 | 147 | 132 | 195 | 145 | 156 | 151 | 158 | 144 | 158.25 | 165 | 151 | 0.916 | -0.127 | 0.48350515 | 0.68146464 | 0.0074 | 0.0277 | 0.0174 | 0.0174 | TRUE | 0.2664 |
| Gpt | 182 | 193 | 199 | 190 | 157 | 132 | 193 | 193 | 200 | 149 | 169 | 144 | 174.78 | 195 | 154 | 0.789 | -0.342 | 0.02751573 | 0.10863033 | 0 | 0.0258 | 0.0115 | 0.0115 | TRUE | 0.0689 |
| Gpt2 | 1870 | 2174 | 2180 | 2277 | 1705 | 1746 | 1986 | 2171 | 2189 | 1790 | 1834 | 1911 | 1980.36 | 2115 | 1845 | 0.872 | -0.198 | 0.01302869 | 0.06391556 | 0.0015 | 0.0086 | 0.0041 | 0.0041 | TRUE | 0.0344 |
| Gpx1 | 6220 | 6353 | 6078 | 12428 | 8393 | 7564 | 6607 | 6345 | 6103 | 9770 | 9030 | 8280 | 7689.13 | 6352 | 9027 | 1.421 | 0.507 | 5.32E-09 | 2.69E-07 | 0.004 | 0.0073 | 0.0053 | 0.0053 | TRUE | 0.0639 |
| Gpx2 | 507 | 554 | 504 | 156 | 131 | 160 | 539 | 553 | 506 | 123 | 141 | 175 | 339.43 | 533 | 146 | 0.273 | -1.873 | 3.89E-33 | 2.55E-30 | 0.0089 | 0.0166 | 0.0133 | 0.0133 | TRUE | 0.3063 |
| Gpx2-ps1 | 4 | 1 | 5 | 5 | 4 | 3 | 4 | 1 | 5 | 4 | 4 | 3 | 3.63 | 3 | 4 | 1.126 | 0.171 | 0.86545846 | NA | 0 | 0.9142 | 0.4557 | 0.4557 | TRUE | 0.5747 |
| Gpx3 | 1801 | 2117 | 2115 | 2111 | 1443 | 1617 | 1913 | 2114 | 2124 | 1660 | 1553 | 1770 | 1855.51 | 2050 | 1661 | 0.81 | -0.304 | 0.00078697 | 0.00738191 | 0.003 | 0.0087 | 0.0054 | 0.0054 | TRUE | 0.0414 |
| Gpx4 | 7893 | 11941 | 12162 | 13454 | 9388 | 11040 | 8384 | 11926 | 12212 | 10576 | 10101 | 12085 | 10880.61 | 10841 | 10921 | 1.007 | 0.011 | 0.9440522 | 0.97215927 | 0.0253 | 0.0072 | 0.0163 | 0.0163 | TRUE | 0.4802 |
| Gpx5 | 0 | 1 | 0 | 0 | 1 | 0 | 0 | 1 | 0 | 0 | 1 | 0 | 0.35 | 0 | 0 | 0.954 | -0.067 | 0.98571955 | NA | 0 | 9.5346 | 8.2194 | 8.2194 | TRUE | 0.5204 |
| Gpx6 | 16 | 45 | 37 | 30 | 24 | 13 | 17 | 45 | 37 | 24 | 26 | 14 | 27.12 | 33 | 21 | 0.643 | -0.636 | 0.18189803 | 0.38376383 | 0.1219 | 0.1284 | 0.1254 | 0.1254 | TRUE | 0.5343 |
| Gpx7 | 222 | 363 | 356 | 338 | 233 | 243 | 236 | 363 | 357 | 266 | 251 | 266 | 289.7 | 319 | 261 | 0.818 | -0.289 | 0.12235163 | 0.29760764 | 0.0253 | 0.0183 | 0.0218 | 0.0218 | TRUE | 0.5807 |
| Gpx8 | 116 | 156 | 162 | 161 | 111 | 108 | 123 | 156 | 163 | 127 | 119 | 118 | 134.32 | 147 | 121 | 0.825 | -0.277 | 0.13921379 | 0.32391948 | 0 | 0.0314 | 0.0179 | 0.0179 | TRUE | 0.2086 |
| Gramd1a | 1205 | 1657 | 1628 | 1727 | 1451 | 1770 | 1280 | 1655 | 1635 | 1358 | 1561 | 1938 | 1570.97 | 1523 | 1619 | 1.062 | 0.087 | 0.58296955 | 0.75875085 | 0.0256 | 0.009 | 0.0174 | 0.0174 | TRUE | 0.3573 |
| Gramd1b | 804 | 849 | 770 | 907 | 676 | 740 | 854 | 848 | 773 | 713 | 727 | 810 | 787.57 | 825 | 750 | 0.908 | -0.139 | 0.17268154 | 0.37140778 | 0.0026 | 0.0111 | 0.0062 | 0.0062 | TRUE | 0.0569 |
| Gramd1c | 69 | 67 | 60 | 102 | 61 | 56 | 73 | 67 | 60 | 80 | 66 | 61 | 67.93 | 67 | 69 | 1.04 | 0.056 | 0.81995845 | 0.90791804 | 0 | 0.0554 | 0.0292 | 0.0292 | TRUE | 0.201 |
| Gramd2 | 60 | 76 | 72 | 105 | 74 | 82 | 64 | 76 | 72 | 83 | 80 | 90 | 77.31 | 71 | 84 | 1.187 | 0.247 | 0.26485094 | 0.48459733 | 0 | 0.0495 | 0.0224 | 0.0224 | TRUE | 0.0642 |
| Gramd3 | 312 | 397 | 383 | 466 | 304 | 343 | 331 | 396 | 385 | 366 | 327 | 375 | 363.56 | 371 | 356 | 0.961 | -0.058 | 0.66118887 | 0.80914578 | 0.0041 | 0.016 | 0.0097 | 0.0097 | TRUE | 0.0979 |
| Gramd4 | 1934 | 2416 | 2397 | 1821 | 1456 | 1690 | 2054 | 2413 | 2407 | 1432 | 1567 | 1850 | 1953.67 | 2291 | 1616 | 0.705 | -0.505 | 4.57E-05 | 0.00071888 | 0.012 | 0.0086 | 0.0105 | 0.0105 | TRUE | 0.1929 |
| Grap | 2 | 2 | 2 | 3 | 2 | 4 | 2 | 2 | 2 | 2 | 2 | 4 | 2.5 | 2 | 3 | 1.437 | 0.523 | 0.6492352 | NA | 0 | 1.323 | 0.5382 | 0.5382 | TRUE | 0.2008 |
| Grap2 | 9 | 6 | 5 | 8 | 5 | 3 | 10 | 6 | 5 | 6 | 5 | 3 | 5.92 | 7 | 5 | 0.738 | -0.439 | 0.57453449 | 0.75272138 | 0 | 0.5633 | 0.2691 | 0.2691 | TRUE | 0.2955 |
| Grasp | 1141 | 1640 | 1798 | 1369 | 1047 | 1134 | 1212 | 1638 | 1805 | 1076 | 1127 | 1241 | 1349.88 | 1552 | 1148 | 0.739 | -0.435 | 0.00454036 | 0.02908637 | 0.021 | 0.0093 | 0.0162 | 0.0162 | TRUE | 0.4411 |
| Grb10 | 523 | 773 | 795 | 1664 | 1143 | 1030 | 556 | 772 | 798 | 1308 | 1230 | 1127 | 965.2 | 709 | 1222 | 1.724 | 0.786 | 1.94E-07 | 6.37E-06 | 0.0187 | 0.0103 | 0.0153 | 0.0153 | TRUE | 0.42 |
| Grb14 | 345 | 401 | 405 | 530 | 401 | 448 | 366 | 400 | 407 | 417 | 431 | 490 | 418.68 | 391 | 446 | 1.138 | 0.187 | 0.12502125 | 0.30224327 | 0.003 | 0.0148 | 0.0083 | 0.0083 | TRUE | 0.0856 |
| Grb2 | 7597 | 7050 | 6534 | 9970 | 7349 | 6698 | 8069 | 7041 | 6561 | 7838 | 7907 | 7332 | 7457.97 | 7224 | 7692 | 1.065 | 0.091 | 0.35133601 | 0.57017785 | 0.006 | 0.0073 | 0.0067 | 0.0067 | TRUE | 0.1279 |
| Grb7 | 2445 | 3340 | 3560 | 3127 | 2509 | 3199 | 2597 | 3336 | 3575 | 2458 | 2700 | 3502 | 3027.81 | 3169 | 2887 | 0.911 | -0.135 | 0.40867337 | 0.62120237 | 0.0266 | 0.008 | 0.019 | 0.019 | TRUE | 0.4217 |
| Grcc10 | 1433 | 2114 | 2145 | 1922 | 1458 | 1712 | 1522 | 2111 | 2154 | 1511 | 1569 | 1874 | 1790.15 | 1929 | 1651 | 0.856 | -0.225 | 0.14495371 | 0.3327332 | 0.0229 | 0.0087 | 0.0166 | 0.0166 | TRUE | 0.411 |
| Greb1 | 687 | 760 | 763 | 455 | 339 | 355 | 730 | 759 | 766 | 358 | 365 | 389 | 560.98 | 752 | 371 | 0.492 | -1.023 | 1.31E-24 | 5.17E-22 | 0 | 0.0128 | 0.0052 | 0.0052 | TRUE | 0.0207 |
| Greb1l | 1220 | 1029 | 1130 | 1672 | 1137 | 1008 | 1296 | 1028 | 1135 | 1314 | 1223 | 1103 | 1183.22 | 1153 | 1213 | 1.054 | 0.076 | 0.53200253 | 0.7201304 | 0.0098 | 0.0097 | 0.0097 | 0.0097 | TRUE | 0.1409 |
| Grem1 | 2 | 1 | 4 | 2 | 0 | 2 | 2 | 1 | 4 | 2 | 0 | 2 | 1.82 | 2 | 1 | 0.532 | -0.911 | 0.53930838 | NA | 0 | 1.8202 | 0.9874 | 0.9874 | TRUE | 0.3178 |
| Grem2 | 16 | 15 | 6 | 30 | 9 | 11 | 17 | 15 | 6 | 24 | 10 | 12 | 13.88 | 13 | 15 | 1.211 | 0.276 | 0.65825436 | 0.80743861 | 0.1696 | 0.2442 | 0.2078 | 0.2078 | TRUE | 1.3221 |
| Grhl1 | 196 | 188 | 155 | 257 | 199 | 187 | 208 | 188 | 156 | 202 | 214 | 205 | 195.4 | 184 | 207 | 1.126 | 0.171 | 0.29978455 | 0.51992544 | 0.005 | 0.0238 | 0.0144 | 0.0144 | TRUE | 0.1947 |
| Grhl2 | 475 | 336 | 390 | 625 | 473 | 487 | 505 | 336 | 392 | 491 | 509 | 533 | 460.84 | 411 | 511 | 1.245 | 0.316 | 0.05151199 | 0.16729608 | 0.0195 | 0.0141 | 0.0168 | 0.0168 | TRUE | 0.4595 |
| Grhl3 | 2174 | 2925 | 2950 | 2629 | 2085 | 2378 | 2309 | 2921 | 2962 | 2067 | 2243 | 2603 | 2517.61 | 2731 | 2304 | 0.843 | -0.246 | 0.06475265 | 0.19603458 | 0.0154 | 0.0082 | 0.0123 | 0.0123 | TRUE | 0.2211 |
| Grhpr | 2661 | 3150 | 3116 | 3742 | 2800 | 2956 | 2826 | 3146 | 3129 | 2942 | 3013 | 3236 | 3048.55 | 3034 | 3064 | 1.009 | 0.013 | 0.87092868 | 0.93648357 | 0.0027 | 0.008 | 0.0046 | 0.0046 | TRUE | 0.0433 |
| Gria1 | 5 | 3 | 1 | 14 | 11 | 4 | 5 | 3 | 1 | 11 | 12 | 4 | 6.09 | 3 | 9 | 2.959 | 1.565 | 0.08157797 | 0.22771085 | 0.1179 | 0.548 | 0.3626 | 0.3626 | TRUE | 0.5532 |
| Gria2 | 6 | 2 | 3 | 6 | 1 | 2 | 6 | 2 | 3 | 5 | 1 | 2 | 3.23 | 4 | 3 | 0.726 | -0.462 | 0.67711237 | NA | 0 | 1.0277 | 0.5731 | 0.5731 | TRUE | 0.6984 |
| Gria3 | 28 | 34 | 26 | 91 | 57 | 56 | 30 | 34 | 26 | 72 | 61 | 61 | 47.33 | 30 | 65 | 2.169 | 1.117 | 0.00012939 | 0.00170067 | 0 | 0.0765 | 0.0368 | 0.0368 | TRUE | 0.094 |
| Gria4 | 27 | 26 | 30 | 67 | 36 | 34 | 29 | 26 | 30 | 53 | 39 | 37 | 35.57 | 28 | 43 | 1.529 | 0.612 | 0.06796016 | 0.20177545 | 0 | 0.0995 | 0.0517 | 0.0517 | TRUE | 0.3526 |
| Grid1 | 32 | 61 | 60 | 125 | 105 | 101 | 34 | 61 | 60 | 98 | 113 | 111 | 79.49 | 52 | 107 | 2.066 | 1.047 | 0.00011686 | 0.00155861 | 0.0253 | 0.0483 | 0.0388 | 0.0388 | TRUE | 0.7156 |
| Grid2 | 4 | 1 | 0 | 13 | 11 | 13 | 4 | 1 | 0 | 10 | 12 | 14 | 6.92 | 2 | 12 | 7.016 | 2.811 | 0.00365909 | 0.02470515 | 0 | 0.4828 | 0.3362 | 0.3362 | TRUE | 1.1749 |
| Grid2ip | 51 | 95 | 94 | 70 | 63 | 74 | 54 | 95 | 94 | 55 | 68 | 81 | 74.54 | 81 | 68 | 0.832 | -0.265 | 0.37868448 | 0.593758 | 0.052 | 0.0511 | 0.0516 | 0.0516 | TRUE | 0.6905 |
| Grifin | 5 | 4 | 3 | 7 | 7 | 2 | 5 | 4 | 3 | 6 | 8 | 2 | 4.59 | 4 | 5 | 1.244 | 0.315 | 0.72744332 | NA | 0 | 0.7246 | 0.3671 | 0.3671 | TRUE | 0.2851 |
| Grik1 | 0 | 0 | 0 | 0 | 1 | 1 | 0 | 0 | 0 | 0 | 1 | 1 | 0.36 | 0 | 1 | 3.794 | 1.924 | 0.60247382 | NA | 0 | 9.1135 | 8.1425 | 8.1425 | TRUE | 0.3482 |
| Grik2 | 28 | 21 | 14 | 38 | 26 | 14 | 30 | 21 | 14 | 30 | 28 | 15 | 22.99 | 22 | 24 | 1.139 | 0.188 | 0.68995317 | 0.82966922 | 0.079 | 0.1502 | 0.1166 | 0.1166 | TRUE | 0.2785 |
| Grik3 | 1432 | 1685 | 1985 | 2261 | 1720 | 1731 | 1521 | 1683 | 1993 | 1777 | 1851 | 1895 | 1786.65 | 1732 | 1841 | 1.062 | 0.087 | 0.45022709 | 0.65567698 | 0.0093 | 0.0087 | 0.0091 | 0.0091 | TRUE | 0.2095 |
| Grik4 | 83 | 94 | 80 | 80 | 60 | 64 | 88 | 94 | 80 | 63 | 65 | 70 | 76.65 | 87 | 66 | 0.751 | -0.413 | 0.06310713 | 0.19253807 | 0 | 0.0499 | 0.0222 | 0.0222 | TRUE | 0.0489 |
| Grik5 | 682 | 757 | 722 | 705 | 526 | 585 | 724 | 756 | 725 | 554 | 566 | 640 | 660.99 | 735 | 587 | 0.797 | -0.327 | 0.00143938 | 0.01186094 | 0.0016 | 0.0119 | 0.0061 | 0.0061 | TRUE | 0.0732 |
| Grin1 | 592 | 2340 | 2358 | 1590 | 1995 | 2988 | 629 | 2337 | 2368 | 1250 | 2146 | 3271 | 2000.12 | 1778 | 2222 | 1.25 | 0.322 | 0.63180188 | 0.79278407 | 0.3247 | 0.0086 | 0.1229 | 0.3247 | TRUE | 0.382 |
| Grin2a | 5 | 11 | 14 | 7 | 13 | 8 | 5 | 11 | 14 | 6 | 14 | 9 | 9.77 | 10 | 10 | 0.916 | -0.127 | 0.85237299 | 0.92628008 | 0.1115 | 0.3442 | 0.2316 | 0.2316 | TRUE | 0.3757 |
| Grin2b | 6 | 8 | 8 | 12 | 5 | 2 | 6 | 8 | 8 | 9 | 5 | 2 | 6.57 | 7 | 5 | 0.778 | -0.361 | 0.64558525 | 0.80086288 | 0 | 0.5086 | 0.2928 | 0.2928 | TRUE | 0.3639 |
| Grin2c | 72 | 263 | 322 | 58 | 67 | 99 | 76 | 263 | 323 | 46 | 72 | 108 | 148.09 | 221 | 75 | 0.341 | -1.554 | 0.02017943 | 0.08737766 | 0.3136 | 0.0292 | 0.1494 | 0.3136 | TRUE | 0.4703 |
| Grin2d | 63 | 96 | 79 | 98 | 70 | 69 | 67 | 96 | 79 | 77 | 75 | 76 | 78.33 | 81 | 76 | 0.941 | -0.088 | 0.70660373 | 0.8398044 | 0 | 0.049 | 0.0269 | 0.0269 | TRUE | 0.253 |
| Grin3a | 0 | 0 | 0 | 3 | 0 | 3 | 0 | 0 | 0 | 2 | 0 | 3 | 0.94 | 0 | 2 | 10.037 | 3.327 | 0.21038399 | NA | 0.0077 | 3.5102 | 3.8249 | 3.8249 | TRUE | 0.2471 |
| Grin3b | 2 | 1 | 1 | 0 | 3 | 2 | 2 | 1 | 1 | 0 | 3 | 2 | 1.59 | 1 | 2 | 1.275 | 0.35 | 0.82744695 | NA | 0 | 2.078 | 1.2078 | 1.2078 | TRUE | 0.6194 |
| Grina | 2769 | 3583 | 3658 | 2521 | 2078 | 2317 | 2941 | 3578 | 3673 | 1982 | 2236 | 2536 | 2824.42 | 3397 | 2251 | 0.662 | -0.594 | 3.58E-06 | 8.30E-05 | 0.0139 | 0.0081 | 0.0115 | 0.0115 | TRUE | 0.1676 |
| Grip1 | 122 | 124 | 143 | 334 | 227 | 213 | 130 | 124 | 144 | 263 | 244 | 233 | 189.5 | 133 | 247 | 1.867 | 0.901 | 7.66E-09 | 3.74E-07 | 0 | 0.0243 | 0.0117 | 0.0117 | TRUE | 0.0576 |
| Grip2 | 72 | 93 | 84 | 94 | 64 | 76 | 76 | 93 | 84 | 74 | 69 | 83 | 79.94 | 84 | 75 | 0.889 | -0.169 | 0.44729006 | 0.65333562 | 0 | 0.0481 | 0.0233 | 0.0233 | TRUE | 0.0706 |
| Gripap1 | 362 | 282 | 306 | 488 | 333 | 284 | 385 | 282 | 307 | 384 | 358 | 311 | 337.7 | 325 | 351 | 1.084 | 0.116 | 0.47582505 | 0.67583364 | 0.0157 | 0.0167 | 0.0161 | 0.0161 | TRUE | 0.2952 |
| Grk1 | 6 | 7 | 8 | 12 | 11 | 9 | 6 | 7 | 8 | 9 | 12 | 10 | 8.75 | 7 | 10 | 1.448 | 0.534 | 0.39198089 | 0.60563505 | 0 | 0.3833 | 0.162 | 0.162 | TRUE | 0.0507 |
| Grk4 | 7 | 6 | 8 | 18 | 13 | 13 | 7 | 6 | 8 | 14 | 14 | 14 | 10.64 | 7 | 14 | 1.975 | 0.982 | 0.08920532 | 0.24274436 | 0 | 0.3166 | 0.1346 | 0.1346 | TRUE | 0.056 |
| Grk5 | 649 | 490 | 467 | 707 | 480 | 461 | 689 | 489 | 469 | 556 | 516 | 505 | 537.42 | 549 | 526 | 0.958 | -0.062 | 0.70863503 | 0.84126423 | 0.0225 | 0.013 | 0.0179 | 0.0179 | TRUE | 0.5806 |
| Grk6 | 3715 | 4580 | 4652 | 5057 | 3904 | 4229 | 3946 | 4574 | 4671 | 3975 | 4200 | 4629 | 4332.73 | 4397 | 4268 | 0.97 | -0.043 | 0.67031705 | 0.81594208 | 0.0069 | 0.0077 | 0.0072 | 0.0072 | TRUE | 0.0979 |
| Grlf1 | 6012 | 5138 | 5275 | 8275 | 5523 | 5053 | 6386 | 5131 | 5297 | 6505 | 5942 | 5531 | 5798.78 | 5605 | 5993 | 1.07 | 0.097 | 0.39094665 | 0.60467415 | 0.0095 | 0.0075 | 0.009 | 0.009 | TRUE | 0.1812 |
| Grm1 | 0 | 0 | 0 | 0 | 0 | 0 | 0 | 0 | 0 | 0 | 0 | 0 | 0 | 0 | 0 | NA | NA | NA | NA | NA | NA | NA | NA | NA | NA |
| Grm2 | 16 | 5 | 15 | 21 | 9 | 7 | 17 | 5 | 15 | 17 | 10 | 8 | 11.82 | 12 | 12 | 0.928 | -0.108 | 0.87001307 | 0.93623963 | 0.168 | 0.2857 | 0.2291 | 0.2291 | TRUE | 1.1091 |
| Grm3 | 1 | 1 | 1 | 9 | 0 | 6 | 1 | 1 | 1 | 7 | 0 | 7 | 2.78 | 1 | 5 | 4.496 | 2.169 | 0.1425255 | NA | 0 | 1.19 | 0.969 | 0.969 | TRUE | 1.321 |
| Grm4 | 51 | 34 | 29 | 56 | 43 | 49 | 54 | 34 | 29 | 44 | 46 | 54 | 43.53 | 39 | 48 | 1.228 | 0.296 | 0.38112151 | 0.59544809 | 0.032 | 0.0826 | 0.0588 | 0.0588 | TRUE | 0.8249 |
| Grm5 | 1 | 0 | 0 | 3 | 3 | 0 | 1 | 0 | 0 | 2 | 3 | 0 | 1.11 | 0 | 2 | 5.12 | 2.356 | 0.29230162 | NA | 0 | 2.9801 | 2.3453 | 2.3453 | TRUE | 0.417 |
| Grm6 | 0 | 0 | 0 | 3 | 0 | 0 | 0 | 0 | 0 | 2 | 0 | 0 | 0.39 | 0 | 1 | 4.278 | 2.097 | 0.60136958 | NA | 10 | 8.3884 | 10 | 10 | TRUE | 1.4411 |

| | | | | | | | | | | | | | | | | | | | | | | | | | |
|---|---|---|---|---|---|---|---|---|---|---|---|---|---|---|---|---|---|---|---|---|---|---|---|---|---|
| Grm7 | 163 | 116 | 106 | 191 | 127 | 133 | 173 | 116 | 106 | 150 | 137 | 146 | 137.97 | 132 | 144 | 1.096 | 0.132 | 0.55912386 | 0.74117512 | 0.0281 | 0.0308 | 0.0295 | 0.0295 | TRUE | 0.7625 |
| Grm8 | 0 | 1 | 1 | 1 | 1 | 0 | 0 | 1 | 1 | 1 | 1 | 0 | 0.64 | 1 | 1 | 0.935 | -0.097 | 0.96674856 | NA | 0 | 5.1213 | 2.3899 | 2.3899 | TRUE | 0.2246 |
| Grn | 4643 | 5336 | 5337 | 6717 | 4761 | 5008 | 4932 | 5329 | 5359 | 5280 | 5123 | 5482 | 5250.78 | 5207 | 5295 | 1.017 | 0.024 | 0.73431259 | 0.85753055 | 0.0012 | 0.0075 | 0.0035 | 0.0035 | TRUE | 0.0259 |
| Grp | 1 | 8 | 1 | 6 | 4 | 7 | 1 | 8 | 1 | 5 | 4 | 8 | 4.46 | 3 | 6 | 1.643 | 0.717 | 0.49996477 | NA | 0.3729 | 0.7461 | 0.5735 | 0.5735 | TRUE | 2.1281 |
| Grpel1 | 5327 | 5173 | 4959 | 6771 | 4691 | 4218 | 5658 | 5166 | 4979 | 5323 | 5047 | 4617 | 5131.87 | 5268 | 4996 | 0.949 | -0.076 | 0.3999206 | 0.61294737 | 0.0044 | 0.0075 | 0.0057 | 0.0057 | TRUE | 0.0534 |
| Grpel2 | 2716 | 3125 | 3221 | 2621 | 2028 | 2324 | 2885 | 3121 | 3234 | 2060 | 2182 | 2544 | 2671.08 | 3080 | 2262 | 0.734 | -0.446 | 2.35E-05 | 0.00040838 | 0.0073 | 0.0081 | 0.0076 | 0.0076 | TRUE | 0.1434 |
| Grpr | 1 | 0 | 0 | 0 | 1 | 0 | 1 | 0 | 0 | 0 | 1 | 0 | 0.36 | 0 | 0 | 0.954 | -0.067 | 0.98557668 | NA | 0 | 9.2518 | 8.0179 | 8.0179 | TRUE | 0.5204 |
| Grrp1 | 35 | 28 | 47 | 43 | 54 | 46 | 37 | 28 | 47 | 34 | 58 | 50 | 42.43 | 37 | 47 | 1.258 | 0.331 | 0.35266169 | 0.57124167 | 0.0493 | 0.0845 | 0.0672 | 0.0672 | TRUE | 0.3688 |
| Grsf1 | 5253 | 6522 | 6562 | 7512 | 5440 | 5978 | 5580 | 6514 | 6589 | 5905 | 5853 | 6544 | 6164.07 | 6228 | 6101 | 0.98 | -0.03 | 0.75758565 | 0.8719099 | 0.006 | 0.0074 | 0.0066 | 0.0066 | TRUE | 0.1009 |
| Grtp1 | 1703 | 1996 | 1868 | 1829 | 1520 | 1553 | 1809 | 1993 | 1876 | 1438 | 1635 | 1700 | 1741.87 | 1893 | 1591 | 0.84 | -0.252 | 0.00938619 | 0.0501657 | 0.004 | 0.0088 | 0.0062 | 0.0062 | TRUE | 0.0877 |
| Grwd1 | 4867 | 3566 | 3468 | 6352 | 4429 | 3490 | 5170 | 3561 | 3482 | 4993 | 4765 | 3820 | 4298.73 | 4071 | 4526 | 1.112 | 0.153 | 0.36975394 | 0.58696949 | 0.0311 | 0.0077 | 0.0208 | 0.0208 | TRUE | 0.678 |
| Grxcr1 | 0 | 0 | 0 | 0 | 0 | 0 | 0 | 0 | 0 | 0 | 0 | 0 | 0 | 0 | 0 | NA | NA | NA | NA | NA | NA | NA | NA | NA | NA |
| Grxcr2 | 0 | 0 | 1 | 1 | 0 | 0 | 0 | 0 | 1 | 1 | 0 | 0 | 0.3 | 0 | 0 | 0.954 | -0.067 | 0.98640617 | NA | 0 | 11.0483 | 9.2783 | 9.2783 | TRUE | 0.4098 |
| Gsc | 31 | 74 | 59 | 80 | 69 | 72 | 33 | 74 | 59 | 63 | 74 | 79 | 63.67 | 55 | 72 | 1.295 | 0.373 | 0.25340249 | 0.47130473 | 0.0631 | 0.0586 | 0.0606 | 0.0606 | TRUE | 0.5445 |
| Gsc2 | 11 | 8 | 11 | 30 | 15 | 8 | 12 | 8 | 11 | 24 | 16 | 9 | 13.2 | 10 | 16 | 1.601 | 0.679 | 0.24659659 | 0.46362157 | 0.0777 | 0.2565 | 0.1673 | 0.1673 | TRUE | 0.1331 |
| Gsdma | 30 | 17 | 14 | 32 | 21 | 20 | 32 | 17 | 14 | 25 | 23 | 22 | 22.09 | 21 | 23 | 1.114 | 0.155 | 0.73425231 | 0.85753055 | 0.0485 | 0.156 | 0.1051 | 0.1051 | TRUE | 1.1085 |
| Gsdma2 | 1 | 1 | 0 | 0 | 0 | 0 | 1 | 1 | 0 | 0 | 0 | 0 | 0.34 | 1 | 0 | 0.261 | -1.936 | 0.60454921 | NA | 0 | 9.5983 | 8.3269 | 8.3269 | TRUE | 0.2507 |
| Gsdma3 | 0 | 0 | 1 | 0 | 1 | 0 | 0 | 0 | 1 | 0 | 1 | 0 | 0.35 | 0 | 0 | 0.954 | -0.067 | 0.98570726 | NA | 0 | 9.5099 | 8.2019 | 8.2019 | TRUE | 0.5204 |
| Gsdmc | 1 | 0 | 1 | 2 | 3 | 1 | 1 | 0 | 1 | 2 | 3 | 1 | 1.33 | 1 | 2 | 2.857 | 1.514 | 0.37992559 | NA | 0 | 2.4898 | 1.1469 | 1.1469 | TRUE | 0.2564 |
| Gsdmc2 | 1 | 0 | 0 | 1 | 1 | 1 | 1 | 0 | 0 | 1 | 1 | 1 | 0.67 | 0 | 1 | 2.676 | 1.42 | 0.53486561 | NA | 0 | 4.9253 | 2.274 | 2.274 | TRUE | 0.4614 |
| Gsdmc3 | 0 | 0 | 0 | 0 | 0 | 0 | 0 | 0 | 0 | 0 | 0 | 0 | 0 | 0 | 0 | NA | NA | NA | NA | NA | NA | NA | NA | NA | NA |
| Gsdmc4 | 2 | 1 | 1 | 4 | 5 | 5 | 2 | 1 | 1 | 3 | 5 | 5 | 3.02 | 1 | 4 | 3.378 | 1.756 | 0.13009175 | NA | 0 | 1.0975 | 0.4905 | 0.4905 | TRUE | 0.2058 |
| Gsdmcl-ps | 0 | 0 | 0 | 0 | 0 | 0 | 0 | 0 | 0 | 0 | 0 | 0 | 0 | 0 | 0 | NA | NA | NA | NA | NA | NA | NA | NA | NA | NA |
| Gsdmcl1 | 0 | 0 | 0 | 0 | 0 | 0 | 0 | 0 | 0 | 0 | 0 | 0 | 0 | 0 | 0 | NA | NA | NA | NA | NA | NA | NA | NA | NA | NA |
| Gsdmd | 115 | 102 | 107 | 156 | 104 | 127 | 122 | 102 | 107 | 123 | 112 | 139 | 117.5 | 110 | 125 | 1.127 | 0.173 | 0.37286124 | 0.58928297 | 0 | 0.0349 | 0.0185 | 0.0185 | TRUE | 0.0986 |
| Gse1 | 4780 | 5804 | 6173 | 6029 | 4977 | 4527 | 5077 | 5797 | 6198 | 4740 | 5355 | 4956 | 5353.67 | 5691 | 5017 | 0.881 | -0.182 | 0.07017075 | 0.20605716 | 0.0066 | 0.0075 | 0.0071 | 0.0071 | TRUE | 0.1083 |
| Gsg1 | 38 | 42 | 46 | 57 | 49 | 38 | 40 | 42 | 46 | 45 | 53 | 42 | 44.6 | 43 | 47 | 1.081 | 0.113 | 0.69517726 | 0.83296633 | 0 | 0.0808 | 0.037 | 0.037 | TRUE | 0.1053 |
| Gsg1l | 7 | 21 | 10 | 8 | 5 | 8 | 7 | 21 | 10 | 6 | 5 | 9 | 9.81 | 13 | 7 | 0.528 | -0.921 | 0.17660719 | 0.37707441 | 0.1088 | 0.3426 | 0.2233 | 0.2233 | TRUE | 1.2683 |
| Gsg2 | 2112 | 1662 | 1520 | 3043 | 2125 | 1660 | 2243 | 1660 | 1526 | 2392 | 2286 | 1817 | 1987.51 | 1810 | 2165 | 1.197 | 0.259 | 0.11993993 | 0.29378671 | 0.0314 | 0.0086 | 0.0195 | 0.0195 | TRUE | 0.5297 |
| Gsk3a | 4557 | 4528 | 4471 | 6057 | 4327 | 4305 | 4840 | 4522 | 4489 | 4762 | 4656 | 4713 | 4663.58 | 4617 | 4710 | 1.02 | 0.029 | 0.66048339 | 0.80885221 | 6.00E-04 | 0.0076 | 0.0029 | 0.0029 | TRUE | 0.0217 |
| Gsk3b | 2646 | 2627 | 2644 | 4029 | 2784 | 2589 | 2811 | 2624 | 2655 | 3167 | 2995 | 2834 | 2847.62 | 2697 | 2999 | 1.113 | 0.154 | 0.04979691 | 0.16379495 | 0.0017 | 0.0081 | 0.0041 | 0.0041 | TRUE | 0.0298 |
| Gskip | 372 | 384 | 373 | 451 | 291 | 284 | 395 | 384 | 375 | 355 | 313 | 311 | 355.28 | 385 | 326 | 0.851 | -0.233 | 0.0532692 | 0.17139165 | 0 | 0.0162 | 0.0076 | 0.0076 | TRUE | 0.0705 |
| Gsn | 524 | 590 | 571 | 690 | 453 | 539 | 557 | 589 | 573 | 542 | 487 | 590 | 556.5 | 573 | 540 | 0.942 | -0.086 | 0.44337368 | 0.65025941 | 0.0028 | 0.0128 | 0.0073 | 0.0073 | TRUE | 0.0814 |
| Gspt1 | 17943 | 16908 | 16617 | 23059 | 16130 | 14398 | 19059 | 16886 | 16685 | 18127 | 17355 | 15761 | 17312.16 | 17543 | 17081 | 0.974 | -0.038 | 0.67367821 | 0.818099 | 0.0044 | 0.0071 | 0.0059 | 0.0059 | TRUE | 0.0698 |
| Gspt2 | 475 | 424 | 431 | 742 | 589 | 506 | 505 | 423 | 433 | 583 | 634 | 554 | 521.95 | 454 | 590 | 1.302 | 0.381 | 0.00163726 | 0.01312139 | 0.0047 | 0.0132 | 0.0086 | 0.0086 | TRUE | 0.1109 |
| Gsr | 4584 | 5676 | 5599 | 7275 | 5428 | 5545 | 4869 | 5669 | 5622 | 5719 | 5840 | 6070 | 5631.46 | 5387 | 5876 | 1.091 | 0.125 | 0.14725442 | 0.3363323 | 0.0037 | 0.0075 | 0.0052 | 0.0052 | TRUE | 0.086 |
| Gss | 1590 | 1465 | 1358 | 1682 | 1224 | 1189 | 1689 | 1463 | 1364 | 1322 | 1317 | 1302 | 1409.38 | 1505 | 1314 | 0.873 | -0.196 | 0.05769451 | 0.18143868 | 0.0051 | 0.0092 | 0.007 | 0.007 | TRUE | 0.1368 |
| Gsta1 | 0 | 0 | 0 | 0 | 0 | 0 | 0 | 0 | 0 | 0 | 0 | 0 | 0 | 0 | 0 | NA | NA | NA | NA | NA | NA | NA | NA | NA | NA |
| Gsta2 | 2 | 1 | 2 | 0 | 1 | 3 | 2 | 1 | 2 | 0 | 1 | 3 | 1.58 | 2 | 1 | 0.817 | -0.292 | 0.85543989 | NA | 0 | 2.0895 | 1.2038 | 1.2038 | TRUE | 0.5769 |
| Gsta3 | 159 | 46 | 45 | 125 | 85 | 58 | 169 | 46 | 45 | 98 | 91 | 63 | 85.54 | 87 | 84 | 0.976 | -0.035 | 0.94345138 | 0.97171381 | 0.302 | 0.0454 | 0.1602 | 0.1602 | TRUE | 6.4547 |
| Gsta4 | 12017 | 11829 | 11948 | 13142 | 9675 | 9872 | 12764 | 11814 | 11997 | 10331 | 10410 | 10806 | 11353.72 | 12192 | 10516 | 0.863 | -0.213 | 0.00098419 | 0.00880175 | 0.001 | 0.0072 | 0.0029 | 0.0029 | TRUE | 0.0206 |
| Gstcd | 1662 | 1722 | 1711 | 2275 | 1543 | 1477 | 1765 | 1720 | 1718 | 1788 | 1660 | 1617 | 1711.42 | 1734 | 1688 | 0.974 | -0.037 | 0.6318849 | 0.79278407 | 0.001 | 0.0088 | 0.0038 | 0.0038 | TRUE | 0.0333 |
| Gstk1 | 185 | 209 | 197 | 280 | 164 | 218 | 197 | 209 | 198 | 220 | 176 | 239 | 206.37 | 201 | 212 | 1.054 | 0.076 | 0.64607788 | 0.80125503 | 0.007 | 0.0229 | 0.0148 | 0.0148 | TRUE | 0.2224 |
| Gstm1 | 1476 | 1713 | 1754 | 1379 | 1189 | 1431 | 1568 | 1711 | 1761 | 1084 | 1279 | 1566 | 1494.93 | 1680 | 1310 | 0.779 | -0.36 | 0.0110133 | 0.05654356 | 0.0179 | 0.0091 | 0.0138 | 0.0138 | TRUE | 0.3521 |
| Gstm2 | 149 | 179 | 180 | 252 | 212 | 266 | 158 | 179 | 181 | 198 | 228 | 291 | 205.86 | 173 | 239 | 1.381 | 0.465 | 0.01350254 | 0.06546634 | 0.0184 | 0.0229 | 0.0206 | 0.0206 | TRUE | 0.3972 |
| Gstm3 | 200 | 265 | 260 | 109 | 84 | 152 | 212 | 265 | 261 | 86 | 90 | 166 | 180.1 | 246 | 114 | 0.462 | -1.115 | 2.83E-05 | 0.00047853 | 0.0682 | 0.0252 | 0.0447 | 0.0447 | TRUE | 1.5697 |
| Gstm4 | 598 | 839 | 801 | 957 | 702 | 747 | 635 | 838 | 804 | 752 | 755 | 818 | 767.12 | 759 | 775 | 1.02 | 0.029 | 0.82263634 | 0.90941476 | 0.0104 | 0.0112 | 0.0108 | 0.0108 | TRUE | 0.2403 |
| Gstm5 | 1472 | 1534 | 1482 | 1574 | 1140 | 1216 | 1564 | 1532 | 1488 | 1237 | 1227 | 1331 | 1396.44 | 1528 | 1265 | 0.828 | -0.273 | 0.00057232 | 0.00578738 | 5.00E-04 | 0.0093 | 0.0038 | 0.0038 | TRUE | 0.0245 |
| Gstm6 | 112 | 125 | 97 | 143 | 103 | 121 | 119 | 125 | 97 | 112 | 111 | 132 | 116.15 | 114 | 118 | 1.041 | 0.058 | 0.77213199 | 0.88112898 | 0.004 | 0.0353 | 0.0201 | 0.0201 | TRUE | 0.1597 |
| Gstm7 | 123 | 158 | 171 | 172 | 112 | 147 | 131 | 158 | 172 | 135 | 121 | 161 | 146.13 | 154 | 139 | 0.904 | -0.146 | 0.45937862 | 0.66378277 | 0.0125 | 0.0294 | 0.0211 | 0.0211 | TRUE | 0.1887 |
| Gsto1 | 879 | 1024 | 985 | 1165 | 815 | 868 | 934 | 1023 | 989 | 916 | 877 | 950 | 948.04 | 982 | 914 | 0.931 | -0.103 | 0.2446005 | 0.4611329 | 7.00E-04 | 0.0104 | 0.0046 | 0.0046 | TRUE | 0.0218 |
| Gsto2 | 15 | 27 | 25 | 31 | 20 | 13 | 16 | 27 | 25 | 24 | 22 | 14 | 21.35 | 23 | 20 | 0.889 | -0.169 | 0.70482731 | 0.83847762 | 0 | 0.1612 | 0.0967 | 0.0967 | TRUE | 0.3753 |
| Gstp1 | 1848 | 1780 | 1672 | 2294 | 1626 | 1651 | 1963 | 1778 | 1679 | 1803 | 1749 | 1807 | 1796.6 | 1807 | 1786 | 0.989 | -0.016 | 0.86022455 | 0.9309173 | 0.0027 | 0.0087 | 0.005 | 0.005 | TRUE | 0.069 |
| Gstp2 | 238 | 310 | 300 | 397 | 309 | 309 | 253 | 310 | 301 | 312 | 332 | 338 | 307.74 | 288 | 327 | 1.136 | 0.184 | 0.17494181 | 0.37459602 | 0.003 | 0.0176 | 0.01 | 0.01 | TRUE | 0.126 |
| Gstt1 | 184 | 192 | 155 | 123 | 78 | 107 | 195 | 192 | 156 | 97 | 84 | 117 | 140.1 | 181 | 99 | 0.548 | -0.868 | 1.75E-05 | 0.00031761 | 0.0122 | 0.0304 | 0.0217 | 0.0217 | TRUE | 0.2253 |
| Gstt2 | 176 | 314 | 381 | 245 | 188 | 216 | 187 | 314 | 383 | 193 | 202 | 236 | 252.4 | 295 | 210 | 0.714 | -0.486 | 0.05301008 | 0.17090657 | 0.0654 | 0.02 | 0.0415 | 0.0415 | TRUE | 0.3272 |
| Gstt3 | 379 | 363 | 374 | 373 | 309 | 296 | 403 | 363 | 376 | 293 | 332 | 324 | 348.39 | 381 | 316 | 0.831 | -0.267 | 0.02968268 | 0.11418419 | 0 | 0.0164 | 0.008 | 0.008 | TRUE | 0.0511 |
| Gstt4 | 21 | 30 | 14 | 12 | 17 | 5 | 22 | 30 | 14 | 9 | 18 | 5 | 16.59 | 22 | 11 | 0.498 | -1.004 | 0.08607637 | 0.23672128 | 0.1478 | 0.2055 | 0.1793 | 0.1793 | TRUE | 0.3242 |
| Gstz1 | 1125 | 1364 | 1376 | 1568 | 1077 | 1093 | 1195 | 1362 | 1382 | 1233 | 1159 | 1196 | 1254.45 | 1313 | 1196 | 0.911 | -0.134 | 0.15070992 | 0.3416207 | 0.0028 | 0.0095 | 0.0055 | 0.0055 | TRUE | 0.0738 |
| Gsx1 | 5 | 5 | 8 | 24 | 16 | 12 | 5 | 5 | 8 | 19 | 17 | 13 | 11.26 | 6 | 16 | 2.694 | 1.43 | 0.01673484 | 0.07657149 | 0 | 0.2995 | 0.1442 | 0.1442 | TRUE | 0.1829 |
| Gsx2 | 11 | 8 | 5 | 22 | 22 | 15 | 12 | 8 | 5 | 17 | 24 | 16 | 13.68 | 8 | 19 | 2.327 | 1.218 | 0.02924097 | 0.11295336 | 0 | 0.2477 | 0.1371 | 0.1371 | TRUE | 0.3845 |
| Gt(ROSA)26 | 269 | 328 | 305 | 307 | 253 | 271 | 286 | 328 | 306 | 241 | 272 | 297 | 288.29 | 307 | 270 | 0.878 | -0.187 | 0.18532005 | 0.38873093 | 0.0044 | 0.0183 | 0.0109 | 0.0109 | TRUE | 0.1045 |
| Gtdc1 | 711 | 825 | 816 | 1079 | 706 | 803 | 755 | 824 | 819 | 848 | 760 | 879 | 814.22 | 799 | 829 | 1.037 | 0.052 | 0.60519149 | 0.77583356 | 0.0027 | 0.0109 | 0.0062 | 0.0062 | TRUE | 0.0621 |
| Gtdc2 | 497 | 352 | 289 | 467 | 338 | 310 | 528 | 352 | 290 | 367 | 364 | 339 | 373.29 | 390 | 357 | 0.916 | -0.127 | 0.55205839 | 0.73607785 | 0.0441 | 0.0157 | 0.0302 | 0.0302 | TRUE | 0.8502 |
| Gtf2a1 | 2402 | 2840 | 2648 | 3768 | 2675 | 2579 | 2551 | 2836 | 2659 | 2962 | 2878 | 2823 | 2784.99 | 2682 | 2888 | 1.077 | 0.107 | 0.15526288 | 0.34749626 | 0.0012 | 0.0081 | 0.0037 | 0.0037 | TRUE | 0.0307 |
| Gtf2a1l | 197 | 279 | 279 | 276 | 215 | 232 | 209 | 279 | 280 | 217 | 231 | 254 | 245.05 | 256 | 234 | 0.912 | -0.133 | 0.42955869 | 0.63927153 | 0.0123 | 0.0203 | 0.0162 | 0.0162 | TRUE | 0.2808 |
| Gtf2a2 | 1788 | 1821 | 1661 | 2564 | 1750 | 1671 | 1899 | 1819 | 1668 | 2016 | 1883 | 1829 | 1852.22 | 1795 | 1909 | 1.064 | 0.09 | 0.31110874 | 0.53173029 | 0.0026 | 0.0087 | 0.0051 | 0.0051 | TRUE | 0.0466 |
| Gtf2b | 1844 | 1570 | 1414 | 2971 | 2027 | 1793 | 1959 | 1568 | 1420 | 2336 | 2181 | 1963 | 1904.27 | 1649 | 2160 | 1.31 | 0.39 | 0.00443482 | 0.02856101 | 0.0172 | 0.0086 | 0.013 | 0.013 | TRUE | 0.3253 |
| Gtf2e1 | 4634 | 5191 | 4976 | 6899 | 4765 | 4580 | 4922 | 5184 | 4996 | 5423 | 5127 | 5014 | 5111.11 | 5034 | 5188 | 1.031 | 0.044 | 0.51809952 | 0.70963662 | 9.00E-04 | 0.0075 | 0.0031 | 0.0031 | TRUE | 0.0195 |

| | | | | | | | | | | | | | | | | | | | | | | | | |
|---|---|---|---|---|---|---|---|---|---|---|---|---|---|---|---|---|---|---|---|---|---|---|---|---|
| Gtf2e2 | 6186 | 6876 | 6878 | 8788 | 6647 | 6692 | 6571 | 6867 | 6906 | 6908 | 7152 | 7325 | 6954.94 | 6781 | 7128 | 1.051 | 0.072 | 0.25289678 | 0.47086616 | 6.00E-04 | 0.0074 | 0.0027 | 0.0027 | TRUE | 0.009 |
| Gtf2f1 | 3492 | 3335 | 3337 | 5485 | 4069 | 3440 | 3709 | 3331 | 3351 | 4312 | 4378 | 3766 | 3807.67 | 3464 | 4152 | 1.199 | 0.262 | 0.00519262 | 0.03211744 | 0.0048 | 0.0078 | 0.0061 | 0.0061 | TRUE | 0.0802 |
| Gtf2f2 | 2472 | 2748 | 2680 | 3606 | 2647 | 2406 | 2626 | 2744 | 2691 | 2835 | 2848 | 2634 | 2729.61 | 2687 | 2772 | 1.032 | 0.045 | 0.52546436 | 0.71494339 | 8.00E-04 | 0.0081 | 0.0033 | 0.0033 | TRUE | 0.0229 |
| Gtf2h1 | 1779 | 2172 | 2368 | 4318 | 2860 | 2800 | 1890 | 2169 | 2378 | 3394 | 3077 | 3065 | 2662.2 | 2146 | 3179 | 1.482 | 0.567 | 1.34E-07 | 4.59E-06 | 0.0077 | 0.0081 | 0.008 | 0.008 | TRUE | 0.1314 |
| Gtf2h2 | 1743 | 1791 | 1737 | 2199 | 1598 | 1598 | 1851 | 1789 | 1744 | 1729 | 1719 | 1749 | 1763.58 | 1795 | 1732 | 0.965 | -0.051 | 0.4691417 | 0.67141562 | 0 | 0.0088 | 0.003 | 0.003 | TRUE | 0.0091 |
| Gtf2h3 | 2365 | 2266 | 2132 | 3086 | 2219 | 2028 | 2512 | 2263 | 2141 | 2426 | 2388 | 2220 | 2324.89 | 2305 | 2345 | 1.017 | 0.025 | 0.78696431 | 0.89032073 | 0.004 | 0.0083 | 0.0057 | 0.0057 | TRUE | 0.0744 |
| Gtf2h4 | 2284 | 2301 | 2368 | 2976 | 2143 | 2117 | 2426 | 2298 | 2378 | 2339 | 2306 | 2317 | 2344.06 | 2367 | 2321 | 0.981 | -0.028 | 0.66870394 | 0.81492803 | 0 | 0.0083 | 0.0027 | 0.0027 | TRUE | 0.0079 |
| Gtf2h5 | 2234 | 2126 | 1929 | 2718 | 1842 | 1738 | 2373 | 2123 | 1937 | 2137 | 1982 | 1903 | 2075.69 | 2144 | 2007 | 0.936 | -0.095 | 0.36202335 | 0.58006064 | 0.0062 | 0.0085 | 0.0073 | 0.0073 | TRUE | 0.105 |
| Gtf2i | 8947 | 8541 | 9120 | 11370 | 7739 | 7412 | 9503 | 8530 | 9157 | 8938 | 8327 | 8114 | 8761.53 | 9063 | 8460 | 0.934 | -0.099 | 0.20336292 | 0.41115315 | 0.0026 | 0.0073 | 0.0043 | 0.0043 | TRUE | 0.0324 |
| Gtf2ird1 | 2697 | 3035 | 3041 | 3895 | 2870 | 3067 | 2865 | 3031 | 3053 | 3062 | 3088 | 3357 | 3076.08 | 2983 | 3169 | 1.062 | 0.087 | 0.25270574 | 0.47078608 | 0.0014 | 0.008 | 0.0038 | 0.0038 | TRUE | 0.0326 |
| Gtf2ird2 | 81 | 66 | 57 | 161 | 120 | 68 | 86 | 66 | 57 | 127 | 129 | 74 | 89.88 | 70 | 110 | 1.584 | 0.663 | 0.02245551 | 0.09436936 | 0.055 | 0.0436 | 0.0491 | 0.0491 | TRUE | 0.7679 |
| Gtf3a | 3449 | 4150 | 3941 | 4664 | 3443 | 3641 | 3663 | 4145 | 3957 | 3666 | 3704 | 3986 | 3853.64 | 3922 | 3785 | 0.965 | -0.051 | 0.52907814 | 0.71756991 | 0.0025 | 0.0078 | 0.0045 | 0.0045 | TRUE | 0.0403 |
| Gtf3c1 | 7841 | 7691 | 7985 | 11042 | 7789 | 7628 | 8329 | 7681 | 8018 | 8680 | 8380 | 8350 | 8239.71 | 8009 | 8470 | 1.058 | 0.081 | 0.21059099 | 0.42112857 | 9.00E-04 | 0.0073 | 0.0029 | 0.0029 | TRUE | 0.0157 |
| Gtf3c2 | 6828 | 6038 | 5720 | 8685 | 6305 | 5376 | 7253 | 6030 | 5743 | 6827 | 6784 | 5885 | 6420.39 | 6342 | 6499 | 1.025 | 0.035 | 0.75877826 | 0.87277207 | 0.011 | 0.0074 | 0.0094 | 0.0094 | TRUE | 0.1923 |
| Gtf3c3 | 2558 | 2526 | 2612 | 3769 | 2814 | 2586 | 2717 | 2523 | 2623 | 2963 | 3028 | 2831 | 2780.65 | 2621 | 2941 | 1.122 | 0.166 | 0.02049751 | 0.08835387 | 7.00E-04 | 0.0081 | 0.0033 | 0.0033 | TRUE | 0.013 |
| Gtf3c4 | 3761 | 4544 | 4562 | 4982 | 3679 | 3854 | 3995 | 4538 | 4581 | 3916 | 3958 | 4219 | 4201.23 | 4371 | 4031 | 0.922 | -0.117 | 0.17049167 | 0.36901257 | 0.0033 | 0.0077 | 0.005 | 0.005 | TRUE | 0.069 |
| Gtf3c5 | 2436 | 2305 | 2161 | 3142 | 2345 | 2029 | 2587 | 2302 | 2170 | 2470 | 2523 | 2221 | 2378.91 | 2353 | 2405 | 1.022 | 0.032 | 0.75256659 | 0.86948897 | 0.0059 | 0.0083 | 0.0069 | 0.0069 | TRUE | 0.0917 |
| Gtf3c6 | 821 | 724 | 714 | 1470 | 931 | 785 | 872 | 723 | 717 | 1156 | 1002 | 859 | 888.11 | 771 | 1006 | 1.306 | 0.386 | 0.00690681 | 0.03974249 | 0.0156 | 0.0106 | 0.0135 | 0.0135 | TRUE | 0.1621 |
| Gtl3 | 3490 | 4394 | 4417 | 4968 | 3788 | 4458 | 3707 | 4388 | 4435 | 3905 | 4076 | 4880 | 4231.92 | 4177 | 4287 | 1.026 | 0.037 | 0.75469779 | 0.87049097 | 0.0118 | 0.0077 | 0.01 | 0.01 | TRUE | 0.178 |
| Gtpbp1 | 4045 | 4486 | 4590 | 5301 | 4064 | 4213 | 4297 | 4480 | 4609 | 4167 | 4373 | 4612 | 4422.87 | 4462 | 4384 | 0.982 | -0.026 | 0.72623451 | 0.85249856 | 0.0017 | 0.0076 | 0.0037 | 0.0037 | TRUE | 0.0252 |
| Gtpbp10 | 2732 | 2898 | 2705 | 3350 | 2416 | 2476 | 2902 | 2894 | 2716 | 2634 | 2599 | 2710 | 2742.6 | 2837 | 2648 | 0.933 | -0.1 | 0.14858355 | 0.33840575 | 5.00E-04 | 0.0081 | 0.0031 | 0.0031 | TRUE | 0.017 |
| Gtpbp2 | 965 | 1037 | 990 | 988 | 805 | 856 | 1025 | 1036 | 994 | 777 | 866 | 937 | 939.1 | 1018 | 860 | 0.843 | -0.246 | 0.01587116 | 0.07394975 | 0.0036 | 0.0104 | 0.0064 | 0.0064 | TRUE | 0.0888 |
| Gtpbp3 | 2275 | 2708 | 2820 | 3010 | 2297 | 2310 | 2416 | 2705 | 2832 | 2366 | 2471 | 2529 | 2553.14 | 2651 | 2455 | 0.926 | -0.111 | 0.21050425 | 0.42100851 | 0.0032 | 0.0082 | 0.0053 | 0.0053 | TRUE | 0.0724 |
| Gtpbp4 | 12440 | 12843 | 12905 | 15881 | 11068 | 10393 | 13213 | 12826 | 12958 | 12484 | 11908 | 11377 | 12461.26 | 12999 | 11923 | 0.917 | -0.124 | 0.05606663 | 0.17760251 | 0.0011 | 0.0072 | 0.003 | 0.003 | TRUE | 0.0208 |
| Gtpbp5 | 465 | 445 | 454 | 744 | 535 | 431 | 494 | 444 | 456 | 585 | 576 | 472 | 504.42 | 465 | 544 | 1.173 | 0.23 | 0.07095907 | 0.20748529 | 0.0065 | 0.0134 | 0.0097 | 0.0097 | TRUE | 0.1547 |
| Gtpbp6 | 719 | 958 | 866 | 1008 | 857 | 706 | 764 | 957 | 870 | 792 | 922 | 773 | 846.22 | 864 | 829 | 0.96 | -0.059 | 0.63692548 | 0.79575322 | 0.0097 | 0.0108 | 0.0102 | 0.0102 | TRUE | 0.1206 |
| Gtpbp8 | 189 | 217 | 247 | 327 | 209 | 201 | 201 | 217 | 248 | 257 | 225 | 220 | 227.91 | 222 | 234 | 1.057 | 0.08 | 0.60647344 | 0.77676352 | 0.0053 | 0.0214 | 0.013 | 0.013 | TRUE | 0.1174 |
| Gtse1 | 8210 | 8527 | 8544 | 9937 | 7108 | 7225 | 8720 | 8516 | 8579 | 7812 | 7648 | 7909 | 8197.32 | 8605 | 7790 | 0.905 | -0.144 | 0.00924468 | 0.04954371 | 1.00E-04 | 0.0073 | 0.0021 | 0.0021 | TRUE | 0.0031 |
| Gtsf1 | 183 | 295 | 267 | 713 | 551 | 494 | 194 | 295 | 268 | 561 | 593 | 541 | 408.53 | 252 | 565 | 2.236 | 1.161 | 1.55E-12 | 1.53E-10 | 0.018 | 0.015 | 0.0165 | 0.0165 | TRUE | 0.4437 |
| Gtsf1l | 477 | 519 | 520 | 799 | 568 | 636 | 507 | 518 | 522 | 628 | 611 | 696 | 580.43 | 516 | 645 | 1.25 | 0.322 | 0.00176681 | 0.01394002 | 0 | 0.0126 | 0.0059 | 0.0059 | TRUE | 0.0547 |
| Guca1a | 425 | 541 | 514 | 459 | 395 | 368 | 451 | 540 | 516 | 361 | 425 | 403 | 449.42 | 502 | 396 | 0.787 | -0.346 | 0.00674689 | 0.03903594 | 0.0056 | 0.0142 | 0.0095 | 0.0095 | TRUE | 0.0917 |
| Guca1b | 9 | 7 | 5 | 16 | 2 | 3 | 10 | 7 | 5 | 13 | 2 | 3 | 6.6 | 7 | 6 | 0.863 | -0.213 | 0.80934612 | 0.90182956 | 0.313 | 0.5063 | 0.4092 | 0.4092 | TRUE | 2.6779 |
| Guca2a | 1 | 1 | 0 | 1 | 2 | 1 | 1 | 1 | 0 | 1 | 2 | 1 | 1.02 | 1 | 1 | 1.938 | 0.955 | 0.60944978 | NA | 0 | 3.2508 | 1.401 | 1.401 | TRUE | 0.2529 |
| Guca2b | 0 | 0 | 0 | 1 | 0 | 0 | 0 | 0 | 0 | 1 | 0 | 0 | 0.13 | 0 | 0 | 1.859 | 0.894 | 0.82650734 | NA | 0 | 25.1515 | 10 | 10 | TRUE | 0.2563 |
| Gucd1 | 664 | 646 | 548 | 993 | 698 | 590 | 705 | 645 | 550 | 781 | 751 | 646 | 679.7 | 633 | 726 | 1.147 | 0.198 | 0.13908657 | 0.32374165 | 0.0111 | 0.0118 | 0.0114 | 0.0114 | TRUE | 0.1561 |
| Gucy1a2 | 122 | 70 | 77 | 130 | 93 | 66 | 130 | 70 | 77 | 102 | 100 | 72 | 91.89 | 92 | 91 | 0.995 | -0.007 | 0.98151611 | 0.9899899 | 0.0634 | 0.0428 | 0.0528 | 0.0528 | TRUE | 1.1855 |
| Gucy1a3 | 2 | 5 | 3 | 8 | 7 | 10 | 2 | 5 | 3 | 6 | 8 | 11 | 5.82 | 3 | 8 | 2.416 | 1.273 | 0.12306124 | 0.29887926 | 0 | 0.5733 | 0.2804 | 0.2804 | TRUE | 0.2565 |
| Gucy1b2 | 1 | 1 | 0 | 2 | 1 | 0 | 1 | 1 | 0 | 2 | 1 | 0 | 0.78 | 1 | 1 | 1.327 | 0.408 | 0.85171912 | NA | 0 | 4.2045 | 2.1498 | 2.1498 | TRUE | 0.2789 |
| Gucy1b3 | 125 | 102 | 105 | 156 | 109 | 102 | 133 | 102 | 105 | 123 | 117 | 112 | 115.27 | 113 | 117 | 1.036 | 0.052 | 0.79336556 | 0.89352768 | 0 | 0.0355 | 0.0194 | 0.0194 | TRUE | 0.2205 |
| Gucy2c | 21 | 7 | 6 | 17 | 9 | 11 | 22 | 7 | 6 | 13 | 10 | 12 | 11.74 | 12 | 12 | 1.002 | 0.002 | 0.99708415 | 0.99828918 | 0.1957 | 0.2876 | 0.2421 | 0.2421 | TRUE | 2.2942 |
| Gucy2d | 0 | 0 | 0 | 0 | 2 | 0 | 0 | 0 | 0 | 0 | 2 | 0 | 0.36 | 0 | 1 | 3.775 | 1.916 | 0.63423067 | NA | 10 | 9.1927 | 10 | 10 | TRUE | 0.9743 |
| Gucy2e | 59 | 42 | 38 | 37 | 35 | 37 | 63 | 42 | 38 | 29 | 38 | 41 | 41.67 | 48 | 36 | 0.748 | -0.42 | 0.21635301 | 0.42835392 | 0.0303 | 0.086 | 0.0585 | 0.0585 | TRUE | 0.5992 |
| Gucy2f | 0 | 0 | 0 | 0 | 0 | 0 | 0 | 0 | 0 | 0 | 0 | 0 | 0 | 0 | 0 | NA | NA | NA | NA | NA | NA | NA | NA | NA | NA |
| Gucy2g | 32 | 43 | 34 | 34 | 34 | 29 | 34 | 43 | 34 | 27 | 37 | 32 | 34.35 | 37 | 32 | 0.85 | -0.234 | 0.48335778 | 0.68146464 | 0 | 0.1028 | 0.0513 | 0.0513 | TRUE | 0.1548 |
| Guf1 | 2294 | 2175 | 2159 | 2571 | 1897 | 1888 | 2437 | 2172 | 2168 | 2021 | 2041 | 2067 | 2150.93 | 2259 | 2043 | 0.904 | -0.145 | 0.07424628 | 0.21411818 | 0.0017 | 0.0084 | 0.0043 | 0.0043 | TRUE | 0.0572 |
| Guk1 | 3461 | 3544 | 3501 | 5253 | 3572 | 3170 | 3676 | 3539 | 3515 | 4129 | 3843 | 3470 | 3695.63 | 3577 | 3814 | 1.067 | 0.093 | 0.29055741 | 0.5114334 | 0.0036 | 0.0078 | 0.0053 | 0.0053 | TRUE | 0.0756 |
| Gulo | 172 | 291 | 263 | 295 | 253 | 251 | 183 | 291 | 264 | 232 | 272 | 275 | 252.71 | 246 | 260 | 1.054 | 0.076 | 0.7012258 | 0.83650923 | 0.0283 | 0.0199 | 0.0241 | 0.0241 | TRUE | 0.5537 |
| Gulp1 | 192 | 180 | 159 | 282 | 200 | 144 | 204 | 180 | 160 | 222 | 215 | 158 | 189.64 | 181 | 198 | 1.097 | 0.134 | 0.4875051 | 0.68450195 | 0.0189 | 0.0243 | 0.0216 | 0.0216 | TRUE | 0.3369 |
| Gusb | 1420 | 1784 | 1792 | 1851 | 1354 | 1310 | 1508 | 1782 | 1799 | 1455 | 1457 | 1434 | 1572.55 | 1696 | 1449 | 0.854 | -0.228 | 0.0188481 | 0.08333957 | 0.0042 | 0.009 | 0.0062 | 0.0062 | TRUE | 0.1131 |
| Gvin1 | 0 | 0 | 0 | 0 | 0 | 0 | 0 | 0 | 0 | 0 | 0 | 0 | 0 | 0 | 0 | NA | NA | NA | NA | NA | NA | NA | NA | NA | NA |
| Gxylt1 | 2741 | 2563 | 2474 | 3430 | 2497 | 2457 | 2911 | 2560 | 2484 | 2696 | 2687 | 2690 | 2671.31 | 2652 | 2691 | 1.015 | 0.021 | 0.8062146 | 0.90019698 | 0.0032 | 0.0081 | 0.0051 | 0.0051 | TRUE | 0.0888 |
| Gxylt2 | 36 | 31 | 31 | 41 | 34 | 23 | 38 | 31 | 31 | 32 | 37 | 25 | 32.39 | 33 | 31 | 0.939 | -0.09 | 0.79230401 | 0.89305706 | 0 | 0.1086 | 0.0542 | 0.0542 | TRUE | 0.1798 |
| Gyg | 1149 | 1573 | 1575 | 1516 | 1143 | 1255 | 1220 | 1571 | 1581 | 1192 | 1230 | 1374 | 1361.37 | 1457 | 1265 | 0.867 | -0.205 | 0.10953789 | 0.27762038 | 0.0117 | 0.0093 | 0.0111 | 0.0111 | TRUE | 0.2431 |
| Gyk | 1485 | 1821 | 1716 | 1865 | 1410 | 1684 | 1577 | 1819 | 1723 | 1466 | 1517 | 1843 | 1657.6 | 1706 | 1609 | 0.942 | -0.086 | 0.46504664 | 0.66788976 | 0.0097 | 0.0089 | 0.0093 | 0.0093 | TRUE | 0.195 |
| Gykl1 | 0 | 0 | 0 | 0 | 0 | 0 | 0 | 0 | 0 | 0 | 0 | 0 | 0 | 0 | 0 | NA | NA | NA | NA | NA | NA | NA | NA | NA | NA |
| Gyltl1b | 811 | 916 | 1028 | 1023 | 842 | 937 | 861 | 915 | 1032 | 804 | 906 | 1026 | 924.05 | 936 | 912 | 0.973 | -0.04 | 0.75500163 | 0.87058798 | 0.0106 | 0.0105 | 0.0106 | 0.0106 | TRUE | 0.1416 |
| Gypa | 0 | 1 | 1 | 2 | 1 | 0 | 0 | 1 | 1 | 2 | 1 | 0 | 0.78 | 1 | 1 | 1.349 | 0.432 | 0.84339822 | NA | 0 | 4.2569 | 2.1495 | 2.1495 | TRUE | 0.2789 |
| Gypc | 62 | 60 | 39 | 125 | 76 | 52 | 66 | 60 | 39 | 98 | 82 | 57 | 66.98 | 55 | 79 | 1.445 | 0.531 | 0.0925044 | 0.24833692 | 0.0564 | 0.0561 | 0.0563 | 0.0563 | TRUE | 0.2454 |
| Gys1 | 1215 | 1350 | 1387 | 2107 | 1489 | 1500 | 1291 | 1348 | 1393 | 1656 | 1602 | 1642 | 1488.65 | 1344 | 1633 | 1.216 | 0.282 | 0.00019009 | 0.00233317 | 0 | 0.0091 | 0.0034 | 0.0034 | TRUE | 0.0143 |
| Gys2 | 0 | 0 | 0 | 1 | 1 | 1 | 0 | 0 | 0 | 1 | 1 | 1 | 0.49 | 0 | 1 | 5.224 | 2.385 | 0.38465 | NA | 0 | 6.6923 | 3.9249 | 3.9249 | TRUE | 0.07 |
| Gzf1 | 2373 | 3109 | 3223 | 3350 | 2466 | 2847 | 2521 | 3105 | 3236 | 2634 | 2653 | 3116 | 2877.5 | 2954 | 2801 | 0.948 | -0.077 | 0.53412021 | 0.72173954 | 0.0129 | 0.008 | 0.0107 | 0.0107 | TRUE | 0.1997 |
| Gzma | 0 | 0 | 0 | 0 | 0 | 0 | 0 | 0 | 0 | 0 | 0 | 0 | 0 | 0 | 0 | NA | NA | NA | NA | NA | NA | NA | NA | NA | NA |
| Gzmb | 0 | 1 | 1 | 2 | 0 | 0 | 0 | 1 | 1 | 2 | 0 | 0 | 0.6 | 1 | 1 | 0.834 | -0.262 | 0.92267544 | NA | 0 | 5.536 | 3.6136 | 3.6136 | TRUE | 0.9722 |
| Gzmc | 1 | 0 | 0 | 0 | 0 | 1 | 0 | 1 | 0 | 0 | 1 | 0 | 0.36 | 0 | 0 | 0.954 | -0.067 | 0.98557668 | NA | 0 | 9.2518 | 8.0179 | 8.0179 | TRUE | 0.5204 |
| Gzmd | 0 | 0 | 0 | 0 | 0 | 0 | 0 | 0 | 0 | 0 | 0 | 0 | 0 | 0 | 0 | NA | NA | NA | NA | NA | NA | NA | NA | NA | NA |
| Gzme | 0 | 0 | 0 | 0 | 0 | 0 | 0 | 0 | 0 | 0 | 0 | 0 | 0 | 0 | 0 | NA | NA | NA | NA | NA | NA | NA | NA | NA | NA |
| Gzmf | 0 | 0 | 0 | 0 | 0 | 0 | 0 | 0 | 0 | 0 | 0 | 0 | 0 | 0 | 0 | NA | NA | NA | NA | NA | NA | NA | NA | NA | NA |
| Gzmg | 0 | 0 | 0 | 0 | 0 | 0 | 0 | 0 | 0 | 0 | 0 | 0 | 0 | 0 | 0 | NA | NA | NA | NA | NA | NA | NA | NA | NA | NA |
| Gzmk | 0 | 0 | 0 | 0 | 0 | 0 | 0 | 0 | 0 | 0 | 0 | 0 | 0 | 0 | 0 | NA | NA | NA | NA | NA | NA | NA | NA | NA | NA |

| | | | | | | | | | | | | | | | | | | | | | | | | | |
|---|---|---|---|---|---|---|---|---|---|---|---|---|---|---|---|---|---|---|---|---|---|---|---|---|---|
| Gzmm | 27 | 45 | 43 | 29 | 35 | 49 | 29 | 45 | 43 | 23 | 38 | 54 | 38.48 | 39 | 38 | 0.966 | -0.049 | 0.90314979 | 0.95212067 | 0.0911 | 0.0925 | 0.0917 | 0.0917 | TRUE | 0.2141 |
| Gzmn | 2 | 3 | 3 | 1 | 1 | 0 | 2 | 3 | 3 | 1 | 1 | 0 | 1.67 | 3 | 1 | 0.234 | -2.098 | 0.18714036 | NA | 0 | 1.9846 | 0.8686 | 0.8686 | TRUE | 0.1825 |
| H13 | 1050 | 1054 | 1061 | 1780 | 1325 | 1168 | 1115 | 1053 | 1065 | 1399 | 1426 | 1279 | 1222.79 | 1078 | 1368 | 1.27 | 0.345 | 6.55E-05 | 0.00097883 | 0.0013 | 0.0096 | 0.0045 | 0.0045 | TRUE | 0.0384 |
| H19 | 1156 | 1231 | 1145 | 4559 | 3491 | 2728 | 1228 | 1229 | 1150 | 3584 | 3756 | 2986 | 2322.21 | 1202 | 3442 | 2.863 | 1.518 | 3.85E-44 | 4.67E-41 | 0.0077 | 0.0083 | 0.008 | 0.008 | TRUE | 0.1622 |
| H1f0 | 1705 | 1085 | 1168 | 2419 | 1556 | 1036 | 1811 | 1084 | 1173 | 1902 | 1674 | 1134 | 1462.88 | 1356 | 1570 | 1.159 | 0.212 | 0.35345915 | 0.57188733 | 0.0724 | 0.0092 | 0.037 | 0.037 | TRUE | 0.8549 |
| H1fnt | 0 | 0 | 0 | 1 | 0 | 0 | 0 | 0 | 0 | 1 | 0 | 0 | 0.13 | 0 | 0 | 1.859 | 0.894 | 0.82650734 | NA | 0 | 25.1515 | 10 | 10 | TRUE | 0.2563 |
| H1foo | 0 | 2 | 1 | 2 | 1 | 3 | 0 | 2 | 1 | 2 | 1 | 3 | 1.49 | 1 | 2 | 1.942 | 0.958 | 0.55128086 | NA | 0 | 2.2195 | 1.1066 | 1.1066 | TRUE | 0.2986 |
| H1fx | 2751 | 2841 | 2721 | 4589 | 3668 | 2880 | 2922 | 2837 | 2732 | 3608 | 3947 | 3153 | 3199.7 | 2830 | 3569 | 1.261 | 0.335 | 0.00100667 | 0.00895503 | 0.0063 | 0.0079 | 0.0071 | 0.0071 | TRUE | 0.1257 |
| H2-Aa | 10 | 9 | 9 | 8 | 8 | 8 | 11 | 9 | 9 | 6 | 9 | 9 | 8.72 | 10 | 8 | 0.819 | -0.289 | 0.64381171 | 0.79973208 | 0 | 0.3848 | 0.1647 | 0.1647 | TRUE | 0.1214 |
| H2-Ab1 | 208 | 305 | 270 | 245 | 209 | 206 | 221 | 305 | 271 | 193 | 225 | 225 | 239.94 | 266 | 214 | 0.805 | -0.313 | 0.0660616 | 0.19823509 | 0.0131 | 0.0206 | 0.0167 | 0.0167 | TRUE | 0.2386 |
| H2-Bl | 92 | 94 | 80 | 231 | 163 | 153 | 98 | 94 | 80 | 182 | 175 | 167 | 132.73 | 91 | 175 | 1.932 | 0.95 | 1.34E-07 | 4.59E-06 | 0 | 0.0317 | 0.015 | 0.015 | TRUE | 0.0955 |
| H2-D1 | 805 | 737 | 758 | 576 | 435 | 380 | 855 | 736 | 761 | 453 | 468 | 416 | 614.84 | 784 | 446 | 0.569 | -0.814 | 3.12E-13 | 3.45E-11 | 0.0034 | 0.0123 | 0.0072 | 0.0072 | TRUE | 0.0739 |
| H2-DMa | 333 | 351 | 300 | 262 | 174 | 202 | 354 | 351 | 301 | 206 | 187 | 221 | 269.96 | 335 | 205 | 0.611 | -0.711 | 7.68E-07 | 2.19E-05 | 0.0035 | 0.0191 | 0.011 | 0.011 | TRUE | 0.0897 |
| H2-DMb1 | 79 | 112 | 121 | 82 | 70 | 89 | 84 | 112 | 121 | 64 | 75 | 97 | 92.41 | 106 | 79 | 0.743 | -0.428 | 0.09324043 | 0.24963301 | 0.0289 | 0.0426 | 0.0359 | 0.0359 | TRUE | 0.3734 |
| H2-DMb2 | 2 | 0 | 0 | 0 | 1 | 3 | 2 | 0 | 0 | 0 | 1 | 3 | 1.08 | 1 | 1 | 2.037 | 1.026 | 0.66198198 | NA | 2.552 | 3.0553 | 2.8946 | 2.8946 | TRUE | 0.5809 |
| H2-Ea-ps | 0 | 0 | 0 | 0 | 0 | 0 | 0 | 0 | 0 | 0 | 0 | 0 | 0 | 0 | 0 | NA | NA | NA | NA | NA | NA | NA | NA | NA | NA |
| H2-Eb1 | 78 | 59 | 50 | 127 | 85 | 73 | 83 | 59 | 50 | 100 | 91 | 80 | 77.2 | 64 | 90 | 1.419 | 0.505 | 0.05655731 | 0.17876189 | 0.0229 | 0.0496 | 0.0371 | 0.0371 | TRUE | 0.57 |
| H2-Eb2 | 7 | 3 | 4 | 6 | 4 | 3 | 7 | 3 | 4 | 5 | 4 | 3 | 4.46 | 5 | 4 | 0.861 | -0.216 | 0.8082718 | NA | 0 | 0.7458 | 0.346 | 0.346 | TRUE | 0.4215 |
| H2-K1 | 3478 | 3930 | 3693 | 3067 | 2549 | 2375 | 3694 | 3925 | 3708 | 2411 | 2743 | 2600 | 3180.13 | 3776 | 2585 | 0.684 | -0.548 | 1.12E-11 | 9.83E-10 | 0.0022 | 0.0079 | 0.0044 | 0.0044 | TRUE | 0.0427 |
| H2-K2 | 55 | 64 | 73 | 56 | 55 | 59 | 58 | 64 | 73 | 44 | 59 | 65 | 60.57 | 65 | 56 | 0.851 | -0.233 | 0.39083986 | 0.60464245 | 0 | 0.0613 | 0.0365 | 0.0365 | TRUE | 0.3399 |
| H2-Ke2 | 3200 | 3202 | 3155 | 4933 | 3557 | 3165 | 3399 | 3198 | 3168 | 3878 | 3827 | 3465 | 3489.07 | 3255 | 3723 | 1.144 | 0.194 | 0.01493196 | 0.07063792 | 0.0021 | 0.0078 | 0.0043 | 0.0043 | TRUE | 0.0446 |
| H2-Ke6 | 754 | 1017 | 950 | 612 | 606 | 643 | 801 | 1016 | 954 | 481 | 652 | 704 | 767.91 | 924 | 612 | 0.662 | -0.596 | 0.00031618 | 0.00355942 | 0.0254 | 0.0112 | 0.0183 | 0.0183 | TRUE | 0.4254 |
| H2-L | 0 | 0 | 0 | 0 | 0 | 0 | 0 | 0 | 0 | 0 | 0 | 0 | 0 | 0 | 0 | NA | NA | NA | NA | NA | NA | NA | NA | NA | NA |
| H2-M1 | 0 | 0 | 0 | 0 | 0 | 0 | 0 | 0 | 0 | 0 | 0 | 0 | 0 | 0 | 0 | NA | NA | NA | NA | NA | NA | NA | NA | NA | NA |
| H2-M10.1 | 0 | 0 | 0 | 0 | 1 | 0 | 0 | 0 | 0 | 0 | 1 | 0 | 0.18 | 0 | 0 | 1.859 | 0.894 | 0.82650734 | NA | 0 | 18.3785 | 10 | 10 | TRUE | 0.5204 |
| H2-M10.2 | 0 | 0 | 0 | 0 | 0 | 0 | 0 | 0 | 0 | 0 | 0 | 0 | 0 | 0 | 0 | NA | NA | NA | NA | NA | NA | NA | NA | NA | NA |
| H2-M10.3 | 0 | 0 | 0 | 0 | 0 | 0 | 0 | 0 | 0 | 0 | 0 | 0 | 0 | 0 | 0 | NA | NA | NA | NA | NA | NA | NA | NA | NA | NA |
| H2-M10.4 | 0 | 0 | 0 | 0 | 1 | 0 | 0 | 0 | 0 | 0 | 1 | 0 | 0.18 | 0 | 0 | 1.859 | 0.894 | 0.82650734 | NA | 0 | 18.3785 | 10 | 10 | TRUE | 0.5204 |
| H2-M10.5 | 0 | 0 | 0 | 0 | 0 | 0 | 0 | 0 | 0 | 0 | 0 | 0 | 0 | 0 | 0 | NA | NA | NA | NA | NA | NA | NA | NA | NA | NA |
| H2-M10.6 | 0 | 0 | 0 | 0 | 1 | 0 | 0 | 0 | 0 | 0 | 1 | 0 | 0.18 | 0 | 0 | 1.859 | 0.894 | 0.82650734 | NA | 0 | 18.3785 | 10 | 10 | TRUE | 0.5204 |
| H2-M11 | 0 | 0 | 0 | 0 | 0 | 0 | 0 | 0 | 0 | 0 | 0 | 0 | 0 | 0 | 0 | NA | NA | NA | NA | NA | NA | NA | NA | NA | NA |
| H2-M2 | 0 | 0 | 0 | 1 | 1 | 0 | 0 | 0 | 0 | 1 | 1 | 0 | 0.31 | 0 | 1 | 3.32 | 1.731 | 0.65383052 | NA | 0 | 10.6224 | 8.9541 | 8.9541 | TRUE | 0.202 |
| H2-M3 | 154 | 190 | 180 | 202 | 141 | 134 | 164 | 190 | 181 | 159 | 152 | 147 | 165.21 | 178 | 153 | 0.857 | -0.223 | 0.16171874 | 0.35683099 | 0 | 0.0268 | 0.0122 | 0.0122 | TRUE | 0.053 |
| H2-M5 | 179 | 129 | 128 | 394 | 261 | 190 | 190 | 129 | 129 | 310 | 281 | 208 | 207.67 | 149 | 266 | 1.789 | 0.84 | 0.00020899 | 0.00252387 | 0.0398 | 0.0228 | 0.0317 | 0.0317 | TRUE | 0.5971 |
| H2-M9 | 0 | 0 | 0 | 0 | 0 | 0 | 0 | 0 | 0 | 0 | 0 | 0 | 0 | 0 | 0 | NA | NA | NA | NA | NA | NA | NA | NA | NA | NA |
| H2-Oa | 79 | 88 | 79 | 41 | 42 | 42 | 84 | 88 | 79 | 32 | 45 | 46 | 62.42 | 84 | 41 | 0.487 | -1.039 | 8.10E-05 | 0.00116834 | 0 | 0.0597 | 0.032 | 0.032 | TRUE | 0.3295 |
| H2-Ob | 0 | 1 | 0 | 1 | 0 | 1 | 0 | 1 | 0 | 1 | 0 | 1 | 0.48 | 0 | 1 | 1.723 | 0.785 | 0.78325239 | NA | 0 | 6.8716 | 4.0968 | 4.0968 | TRUE | 0.405 |
| H2-Q1 | 4 | 5 | 2 | 5 | 8 | 3 | 4 | 5 | 2 | 4 | 9 | 3 | 4.51 | 4 | 5 | 1.397 | 0.482 | 0.60273287 | NA | 0 | 0.737 | 0.388 | 0.388 | TRUE | 0.5961 |
| H2-Q10 | 462 | 377 | 374 | 336 | 293 | 270 | 491 | 377 | 376 | 264 | 315 | 296 | 352.95 | 415 | 292 | 0.703 | -0.508 | 0.0012408 | 0.01051017 | 0.0138 | 0.0162 | 0.0149 | 0.0149 | TRUE | 0.2988 |
| H2-Q2 | 70 | 154 | 153 | 63 | 52 | 59 | 74 | 154 | 154 | 50 | 56 | 65 | 91.97 | 127 | 57 | 0.444 | -1.172 | 0.00023987 | 0.00282119 | 0.0797 | 0.0427 | 0.0608 | 0.0608 | TRUE | 1.3322 |
| H2-Q4 | 429 | 373 | 317 | 332 | 288 | 249 | 456 | 373 | 318 | 261 | 310 | 273 | 331.65 | 382 | 281 | 0.735 | -0.444 | 0.00820003 | 0.045272 | 0.0176 | 0.0168 | 0.0172 | 0.0172 | TRUE | 0.2544 |
| H2-Q5 | 10 | 11 | 2 | 4 | 3 | 0 | 11 | 11 | 2 | 3 | 3 | 0 | 5 | 8 | 2 | 0.277 | -1.855 | 0.09551536 | 0.25348616 | 0.5058 | 0.6661 | 0.6025 | 0.6025 | TRUE | 1.2475 |
| H2-Q6 | 0 | 0 | 0 | 0 | 0 | 0 | 0 | 0 | 0 | 0 | 0 | 0 | 0 | 0 | 0 | NA | NA | NA | NA | NA | NA | NA | NA | NA | NA |
| H2-Q7 | 0 | 0 | 0 | 0 | 0 | 0 | 0 | 0 | 0 | 0 | 0 | 0 | 0 | 0 | 0 | NA | NA | NA | NA | NA | NA | NA | NA | NA | NA |
| H2-Q8 | 0 | 0 | 0 | 0 | 0 | 0 | 0 | 0 | 0 | 0 | 0 | 0 | 0 | 0 | 0 | NA | NA | NA | NA | NA | NA | NA | NA | NA | NA |
| H2-Q9 | 0 | 0 | 0 | 0 | 0 | 0 | 0 | 0 | 0 | 0 | 0 | 0 | 0 | 0 | 0 | NA | NA | NA | NA | NA | NA | NA | NA | NA | NA |
| H2-T10 | 8 | 11 | 6 | 7 | 3 | 11 | 8 | 11 | 6 | 6 | 3 | 12 | 7.71 | 8 | 7 | 0.807 | -0.309 | 0.68045743 | 0.82283023 | 0.0937 | 0.434 | 0.2747 | 0.2747 | TRUE | 0.8236 |
| H2-T22 | 1 | 0 | 8 | 4 | 1 | 2 | 1 | 0 | 8 | 3 | 1 | 2 | 2.58 | 3 | 2 | 0.712 | -0.489 | 0.74087798 | NA | 1.0744 | 1.2818 | 1.1828 | 1.1828 | TRUE | 2.7551 |
| H2-T23 | 3 | 10 | 7 | 8 | 4 | 6 | 3 | 10 | 7 | 6 | 4 | 7 | 6.23 | 7 | 6 | 0.849 | -0.236 | 0.76385474 | 0.87580088 | 0 | 0.5359 | 0.2831 | 0.2831 | TRUE | 0.2231 |
| H2-T24 | 0 | 0 | 0 | 0 | 1 | 1 | 0 | 0 | 0 | 0 | 1 | 1 | 0.36 | 0 | 1 | 3.794 | 1.924 | 0.60247382 | NA | 0 | 9.1135 | 8.1425 | 8.1425 | TRUE | 0.3482 |
| H2-T3 | 0 | 0 | 0 | 5 | 0 | 0 | 0 | 0 | 0 | 4 | 0 | 0 | 0.66 | 0 | 1 | 7.081 | 2.824 | 0.45988427 | NA | 10 | 5.0358 | 9.153 | 9.153 | TRUE | 2.3415 |
| H2-T9 | 0 | 0 | 0 | 0 | 0 | 0 | 0 | 0 | 0 | 0 | 0 | 0 | 0 | 0 | 0 | NA | NA | NA | NA | NA | NA | NA | NA | NA | NA |
| H2afb1 | 0 | 0 | 0 | 0 | 0 | 0 | 0 | 0 | 0 | 0 | 0 | 0 | 0 | 0 | 0 | NA | NA | NA | NA | NA | NA | NA | NA | NA | NA |
| H2afj | 1893 | 2183 | 2184 | 2480 | 1951 | 1925 | 2011 | 2180 | 2193 | 1950 | 2099 | 2107 | 2089.97 | 2128 | 2052 | 0.964 | -0.053 | 0.50511415 | 0.69975058 | 0.0017 | 0.0085 | 0.0041 | 0.0041 | TRUE | 0.028 |
| H2afv | 2963 | 3566 | 3612 | 3871 | 2840 | 3128 | 3147 | 3561 | 3627 | 3043 | 3056 | 3424 | 3309.71 | 3445 | 3174 | 0.921 | -0.119 | 0.20729098 | 0.41658937 | 0.0047 | 0.0079 | 0.0061 | 0.0061 | TRUE | 0.0694 |
| H2afx | 4857 | 5357 | 5407 | 4737 | 3545 | 4022 | 5159 | 5350 | 5429 | 3724 | 3814 | 4403 | 4646.51 | 5313 | 3980 | 0.749 | -0.417 | 3.03E-06 | 7.23E-05 | 0.0043 | 0.0076 | 0.0055 | 0.0055 | TRUE | 0.1045 |
| H2afy | 5165 | 8073 | 8051 | 9454 | 7662 | 9301 | 5486 | 8063 | 8084 | 7432 | 8244 | 10181 | 7915 | 7211 | 8619 | 1.195 | 0.257 | 0.134335 | 0.31691521 | 0.0347 | 0.0073 | 0.0211 | 0.0211 | TRUE | 0.5342 |
| H2afy2 | 1199 | 1559 | 1520 | 2404 | 1814 | 1680 | 1274 | 1557 | 1526 | 1890 | 1952 | 1839 | 1672.9 | 1452 | 1894 | 1.304 | 0.382 | 0.00019317 | 0.00236574 | 0.0053 | 0.0089 | 0.007 | 0.007 | TRUE | 0.1389 |
| H2afy3 | 0 | 0 | 1 | 0 | 0 | 1 | 0 | 0 | 1 | 0 | 0 | 1 | 0.35 | 0 | 0 | 0.954 | -0.067 | 0.98566485 | NA | 0 | 9.4252 | 8.1416 | 8.1416 | TRUE | 0.5383 |
| H2afz | 16622 | 14673 | 13969 | 23762 | 16157 | 14329 | 17656 | 14654 | 14026 | 18680 | 17384 | 15685 | 16347.5 | 15445 | 17250 | 1.117 | 0.16 | 0.1656536 | 0.36267194 | 0.0116 | 0.0071 | 0.0095 | 0.0095 | TRUE | 0.1916 |
| H2bfm | 3 | 3 | 5 | 49 | 23 | 15 | 3 | 3 | 5 | 39 | 25 | 16 | 15.15 | 4 | 27 | 7.164 | 2.841 | 1.97E-05 | 0.00035252 | 0 | 0.2244 | 0.1644 | 0.1644 | TRUE | 0.1336 |
| H3f3a | 6032 | 6603 | 6938 | 8876 | 6407 | 6592 | 6407 | 6594 | 6966 | 6978 | 6894 | 7216 | 6842.52 | 6656 | 7029 | 1.056 | 0.079 | 0.2373895 | 0.45289793 | 0.001 | 0.0074 | 0.003 | 0.003 | TRUE | 0.0203 |
| H3f3b | 23616 | 27012 | 25483 | 41503 | 28790 | 24976 | 25084 | 26977 | 25588 | 32626 | 30976 | 27340 | 28098.66 | 25883 | 30314 | 1.171 | 0.228 | 0.0099811 | 0.05261429 | 0.0046 | 0.007 | 0.0056 | 0.0056 | TRUE | 0.0901 |
| H60b | 74 | 78 | 69 | 80 | 56 | 67 | 79 | 78 | 69 | 63 | 60 | 73 | 70.38 | 75 | 65 | 0.869 | -0.203 | 0.38467456 | 0.59871028 | 0 | 0.0537 | 0.0249 | 0.0249 | TRUE | 0.09 |
| H60c | 2 | 2 | 3 | 3 | 2 | 2 | 2 | 2 | 3 | 2 | 2 | 2 | 2.31 | 2 | 2 | 0.941 | -0.088 | 0.93998552 | NA | 0 | 1.4358 | 0.5515 | 0.5515 | TRUE | 0.0582 |
| H6pd | 2156 | 2231 | 2328 | 2533 | 1860 | 1728 | 2290 | 2228 | 2338 | 1991 | 2001 | 1892 | 2123.3 | 2285 | 1961 | 0.858 | -0.22 | 0.00173027 | 0.01370663 | 3.00E-04 | 0.0085 | 0.0031 | 0.0031 | TRUE | 0.0115 |
| Haao | 10 | 32 | 28 | 14 | 19 | 21 | 11 | 32 | 28 | 11 | 20 | 23 | 20.86 | 24 | 18 | 0.76 | -0.395 | 0.46916507 | 0.67141562 | 0.1672 | 0.1649 | 0.1659 | 0.1659 | TRUE | 1.3147 |
| Habp2 | 2 | 2 | 0 | 0 | 3 | 1 | 2 | 2 | 0 | 0 | 3 | 1 | 1.41 | 1 | 1 | 1.022 | 0.031 | 0.98664418 | NA | 0 | 2.3478 | 1.7728 | 1.7728 | TRUE | 0.4686 |
| Habp4 | 369 | 328 | 323 | 589 | 471 | 374 | 392 | 328 | 324 | 463 | 507 | 409 | 403.84 | 348 | 460 | 1.322 | 0.403 | 0.00413313 | 0.02711639 | 0.0087 | 0.0151 | 0.0117 | 0.0117 | TRUE | 0.1383 |

| | | | | | | | | | | | | | | | | | | | | | | | | | |
|---|---|---|---|---|---|---|---|---|---|---|---|---|---|---|---|---|---|---|---|---|---|---|---|---|---|
| Hace1 | 1246 | 1304 | 1336 | 1728 | 1189 | 1160 | 1323 | 1302 | 1341 | 1358 | 1279 | 1270 | 1312.46 | 1322 | 1302 | 0.986 | -0.021 | 0.78790251 | 0.89079015 | 0 | 0.0094 | 0.0035 | 0.0035 | TRUE | 0.0176 |
| Hacl1 | 1099 | 1620 | 1638 | 1561 | 1163 | 1305 | 1167 | 1618 | 1645 | 1227 | 1251 | 1429 | 1389.49 | 1477 | 1302 | 0.882 | -0.182 | 0.21960548 | 0.43181262 | 0.019 | 0.0093 | 0.0151 | 0.0151 | TRUE | 0.4033 |
| Hadh | 5820 | 6886 | 6701 | 8916 | 6583 | 7034 | 6182 | 6877 | 6729 | 7009 | 7083 | 7700 | 6929.88 | 6596 | 7264 | 1.101 | 0.139 | 0.08000487 | 0.22448618 | 0.0027 | 0.0074 | 0.0044 | 0.0044 | TRUE | 0.0367 |
| Hadha | 9422 | 9288 | 9521 | 10627 | 7563 | 7284 | 10008 | 9276 | 9560 | 8354 | 8137 | 7973 | 8884.81 | 9615 | 8155 | 0.848 | -0.237 | 0.00021272 | 0.00255644 | 8.00E-04 | 0.0073 | 0.0028 | 0.0028 | TRUE | 0.0156 |
| Hadhb | 2048 | 1957 | 1962 | 2879 | 1991 | 1810 | 2175 | 1954 | 1970 | 2263 | 2142 | 1981 | 2081.11 | 2033 | 2129 | 1.048 | 0.067 | 0.45937414 | 0.66378277 | 0.0033 | 0.0085 | 0.0055 | 0.0055 | TRUE | 0.045 |
| Hagh | 1750 | 2122 | 2071 | 1811 | 1340 | 1502 | 1859 | 2119 | 2080 | 1424 | 1442 | 1644 | 1761.2 | 2019 | 1503 | 0.744 | -0.427 | 1.88E-05 | 0.00033635 | 0.0049 | 0.0088 | 0.0066 | 0.0066 | TRUE | 0.0803 |
| Haghl | 235 | 333 | 259 | 347 | 300 | 299 | 250 | 333 | 260 | 273 | 323 | 327 | 294.19 | 281 | 308 | 1.093 | 0.128 | 0.4339706 | 0.6425633 | 0.0141 | 0.0181 | 0.016 | 0.016 | TRUE | 0.2941 |
| Hal | 2146 | 1952 | 1762 | 1855 | 1416 | 1285 | 2279 | 1949 | 1769 | 1458 | 1524 | 1407 | 1731.09 | 1999 | 1463 | 0.732 | -0.451 | 6.96E-05 | 0.00103042 | 0.0083 | 0.0088 | 0.0087 | 0.0087 | TRUE | 0.1812 |
| Hamp | 0 | 0 | 0 | 1 | 0 | 0 | 0 | 0 | 0 | 1 | 0 | 0 | 0.13 | 0 | 0 | 1.859 | 0.894 | 0.82650734 | NA | 0 | 25.1515 | 10 | 10 | TRUE | 0.2563 |
| Hamp2 | 0 | 0 | 0 | 0 | 0 | 0 | 0 | 0 | 0 | 0 | 0 | 0 | 0 | 0 | 0 | NA | NA | NA | NA | NA | NA | NA | NA | NA | NA |
| Hand1 | 62 | 96 | 91 | 574 | 343 | 403 | 66 | 96 | 91 | 451 | 369 | 441 | 252.42 | 84 | 420 | 4.979 | 2.316 | 4.43E-36 | 3.88E-33 | 0.0134 | 0.02 | 0.0174 | 0.0174 | TRUE | 0.3368 |
| Hand2 | 1 | 1 | 0 | 10 | 8 | 5 | 1 | 1 | 0 | 8 | 9 | 5 | 4 | 1 | 7 | 10.748 | 3.426 | 0.00903247 | NA | 0 | 0.8304 | 0.4255 | 0.4255 | TRUE | 0.2545 |
| Hao1 | 0 | 0 | 0 | 0 | 0 | 0 | 0 | 0 | 0 | 0 | 0 | 0 | 0 | 0 | 0 | NA | NA | NA | NA | NA | NA | NA | NA | NA | NA |
| Hao2 | 0 | 0 | 0 | 0 | 1 | 1 | 0 | 0 | 0 | 0 | 1 | 1 | 0.36 | 0 | 1 | 3.794 | 1.924 | 0.60247382 | NA | 0 | 9.1135 | 8.1425 | 8.1425 | TRUE | 0.3482 |
| Hap1 | 6323 | 4867 | 4643 | 8102 | 6394 | 5039 | 6716 | 4861 | 4662 | 6369 | 6880 | 5516 | 5833.93 | 5413 | 6255 | 1.156 | 0.209 | 0.17598238 | 0.37630071 | 0.0267 | 0.0075 | 0.017 | 0.017 | TRUE | 0.5405 |
| Hapln1 | 1 | 0 | 0 | 1 | 0 | 3 | 1 | 0 | 0 | 1 | 0 | 3 | 0.86 | 0 | 1 | 3.671 | 1.876 | 0.4462535 | NA | 0 | 3.8584 | 3.0074 | 3.0074 | TRUE | 0.5472 |
| Hapln2 | 0 | 0 | 0 | 0 | 0 | 0 | 0 | 0 | 0 | 0 | 0 | 0 | 0 | 0 | 0 | NA | NA | NA | NA | NA | NA | NA | NA | NA | NA |
| Hapln3 | 39 | 52 | 55 | 56 | 33 | 27 | 41 | 52 | 55 | 44 | 36 | 30 | 42.94 | 49 | 37 | 0.739 | -0.435 | 0.16545394 | 0.36228509 | 0 | 0.0836 | 0.0474 | 0.0474 | TRUE | 0.2863 |
| Hapln4 | 570 | 714 | 721 | 621 | 486 | 429 | 605 | 713 | 724 | 488 | 523 | 470 | 587.2 | 681 | 494 | 0.725 | -0.465 | 7.07E-05 | 0.00104051 | 0.0047 | 0.0125 | 0.0081 | 0.0081 | TRUE | 0.1099 |
| Harbi1 | 868 | 999 | 957 | 1060 | 782 | 823 | 922 | 998 | 961 | 833 | 841 | 901 | 909.36 | 960 | 858 | 0.893 | -0.163 | 0.06631675 | 0.19871336 | 6.00E-04 | 0.0105 | 0.0046 | 0.0046 | TRUE | 0.0217 |
| Hars | 4746 | 4202 | 4166 | 6670 | 4471 | 4252 | 5041 | 4197 | 4183 | 5243 | 4811 | 4654 | 4688.2 | 4474 | 4903 | 1.096 | 0.133 | 0.20120994 | 0.40877592 | 0.0077 | 0.0076 | 0.0075 | 0.0075 | TRUE | 0.1498 |
| Hars2 | 1063 | 1095 | 1061 | 1375 | 1054 | 1023 | 1129 | 1094 | 1065 | 1081 | 1134 | 1120 | 1103.8 | 1096 | 1112 | 1.014 | 0.02 | 0.80407667 | 0.89891531 | 0 | 0.0099 | 0.0037 | 0.0037 | TRUE | 0.0083 |
| Has1 | 1 | 5 | 4 | 8 | 10 | 2 | 1 | 5 | 4 | 6 | 11 | 2 | 4.88 | 3 | 6 | 1.901 | 0.927 | 0.34824243 | 0.56702733 | 0.2389 | 0.6813 | 0.4756 | 0.4756 | TRUE | 0.1172 |
| Has2 | 40 | 31 | 36 | 17 | 7 | 10 | 42 | 31 | 36 | 13 | 8 | 11 | 23.57 | 36 | 11 | 0.295 | -1.761 | 5.86E-05 | 0.00088718 | 0 | 0.1467 | 0.0792 | 0.0792 | TRUE | 0.2116 |
| Has2as | 0 | 0 | 0 | 1 | 1 | 0 | 0 | 0 | 0 | 1 | 1 | 0 | 0.31 | 0 | 1 | 3.32 | 1.731 | 0.65383052 | NA | 0 | 10.6224 | 8.9541 | 8.9541 | TRUE | 0.202 |
| Has3 | 55 | 77 | 67 | 88 | 58 | 46 | 58 | 77 | 67 | 69 | 62 | 50 | 64.09 | 67 | 60 | 0.902 | -0.149 | 0.56680116 | 0.7466456 | 0 | 0.0583 | 0.0333 | 0.0333 | TRUE | 0.1815 |
| Hat1 | 14596 | 15568 | 15006 | 22729 | 16314 | 17975 | 15504 | 15548 | 15068 | 17868 | 17553 | 19676 | 16869.35 | 15373 | 18366 | 1.195 | 0.257 | 0.00037171 | 0.0040911 | 0.002 | 0.0071 | 0.0037 | 0.0037 | TRUE | 0.0476 |
| Haus1 | 1623 | 1800 | 1784 | 2491 | 1716 | 1714 | 1724 | 1798 | 1791 | 1958 | 1846 | 1876 | 1832.28 | 1771 | 1893 | 1.07 | 0.097 | 0.17626848 | 0.37655513 | 0 | 0.0087 | 0.0032 | 0.0032 | TRUE | 0.0111 |
| Haus2 | 2533 | 2594 | 2481 | 3676 | 2574 | 2534 | 2690 | 2591 | 2491 | 2890 | 2769 | 2774 | 2700.91 | 2591 | 2811 | 1.085 | 0.118 | 0.0918378 | 0.24740985 | 7.00E-04 | 0.0081 | 0.0032 | 0.0032 | TRUE | 0.0137 |
| Haus3 | 2034 | 1827 | 1863 | 2701 | 1954 | 1734 | 2160 | 1825 | 1871 | 2123 | 2102 | 1898 | 1996.6 | 1952 | 2041 | 1.046 | 0.065 | 0.51711284 | 0.70874821 | 0.005 | 0.0086 | 0.0068 | 0.0068 | TRUE | 0.1053 |
| Haus4 | 1558 | 1341 | 1394 | 1944 | 1326 | 1221 | 1655 | 1339 | 1400 | 1528 | 1427 | 1337 | 1447.56 | 1465 | 1431 | 0.977 | -0.033 | 0.76982863 | 0.87970853 | 0.0079 | 0.0092 | 0.0084 | 0.0084 | TRUE | 0.155 |
| Haus5 | 1551 | 1642 | 1652 | 2485 | 1833 | 1613 | 1647 | 1640 | 1659 | 1954 | 1972 | 1766 | 1772.91 | 1649 | 1897 | 1.151 | 0.203 | 0.01123126 | 0.05736599 | 0.0013 | 0.0088 | 0.0041 | 0.0041 | TRUE | 0.0437 |
| Haus6 | 4100 | 3693 | 3666 | 4667 | 3301 | 3405 | 4355 | 3688 | 3681 | 3669 | 3552 | 3727 | 3778.67 | 3908 | 3649 | 0.934 | -0.099 | 0.2878649 | 0.50903254 | 0.0048 | 0.0078 | 0.0059 | 0.0059 | TRUE | 0.1216 |
| Haus7 | 2082 | 2140 | 2188 | 2026 | 1512 | 1585 | 2211 | 2137 | 2197 | 1593 | 1627 | 1735 | 1916.7 | 2182 | 1652 | 0.757 | -0.403 | 6.05E-08 | 2.34E-06 | 6.00E-04 | 0.0086 | 0.0034 | 0.0034 | TRUE | 0.0234 |
| Haus8 | 1708 | 1814 | 1849 | 2783 | 2057 | 1953 | 1814 | 1812 | 1857 | 2188 | 2213 | 2138 | 2003.55 | 1828 | 2180 | 1.193 | 0.254 | 0.00013843 | 0.00178965 | 0 | 0.0085 | 0.0027 | 0.0027 | TRUE | 0.0033 |
| Havcr1 | 3 | 0 | 2 | 8 | 3 | 5 | 3 | 0 | 2 | 6 | 3 | 5 | 3.36 | 2 | 5 | 2.93 | 1.551 | 0.18280494 | NA | 0 | 0.9862 | 0.5819 | 0.5819 | TRUE | 0.6167 |
| Havcr2 | 22 | 21 | 16 | 4 | 8 | 10 | 23 | 21 | 16 | 3 | 9 | 11 | 13.85 | 20 | 8 | 0.365 | -1.452 | 0.01495156 | 0.07064591 | 0.0375 | 0.2448 | 0.1649 | 0.1649 | TRUE | 1.0093 |
| Hax1 | 2246 | 2331 | 2146 | 3163 | 2233 | 2046 | 2386 | 2328 | 2155 | 2487 | 2403 | 2240 | 2332.87 | 2290 | 2377 | 1.038 | 0.054 | 0.51461166 | 0.70725471 | 0.0021 | 0.0083 | 0.0046 | 0.0046 | TRUE | 0.0321 |
| Hba-a1 | 0 | 0 | 0 | 0 | 0 | 0 | 0 | 0 | 0 | 0 | 0 | 0 | 0 | 0 | 0 | NA | NA | NA | NA | NA | NA | NA | NA | NA | NA |
| Hba-a2 | 0 | 0 | 0 | 0 | 0 | 0 | 0 | 0 | 0 | 0 | 0 | 0 | 0 | 0 | 0 | NA | NA | NA | NA | NA | NA | NA | NA | NA | NA |
| Hba-x | 0 | 0 | 0 | 0 | 0 | 1 | 0 | 0 | 0 | 0 | 0 | 1 | 0.18 | 0 | 0 | 1.859 | 0.894 | 0.82650734 | NA | 0 | 18.0643 | 10 | 10 | TRUE | 0.5383 |
| Hbb-b1 | 0 | 0 | 0 | 0 | 0 | 0 | 0 | 0 | 0 | 0 | 0 | 0 | 0 | 0 | 0 | NA | NA | NA | NA | NA | NA | NA | NA | NA | NA |
| Hbb-b2 | 0 | 0 | 0 | 0 | 0 | 0 | 0 | 0 | 0 | 0 | 0 | 0 | 0 | 0 | 0 | NA | NA | NA | NA | NA | NA | NA | NA | NA | NA |
| Hbb-bh1 | 0 | 0 | 0 | 1 | 0 | 1 | 0 | 0 | 0 | 1 | 0 | 1 | 0.31 | 0 | 1 | 3.348 | 1.743 | 0.65064889 | NA | 0 | 10.5167 | 8.8968 | 8.8968 | TRUE | 0.2082 |
| Hbb-bh2 | 0 | 0 | 0 | 0 | 0 | 0 | 0 | 0 | 0 | 0 | 0 | 0 | 0 | 0 | 0 | NA | NA | NA | NA | NA | NA | NA | NA | NA | NA |
| Hbb-y | 0 | 0 | 3 | 1 | 1 | 0 | 0 | 0 | 3 | 1 | 1 | 0 | 0.81 | 1 | 1 | 0.622 | -0.684 | 0.78589133 | NA | 1.8306 | 4.0621 | 3.2884 | 3.2884 | TRUE | 1.4421 |
| Hbegf | 771 | 794 | 803 | 1035 | 806 | 777 | 819 | 793 | 806 | 814 | 867 | 851 | 824.93 | 806 | 844 | 1.046 | 0.065 | 0.44748725 | 0.65334741 | 0 | 0.0109 | 0.004 | 0.004 | TRUE | 0.0123 |
| Hbp1 | 3706 | 4585 | 4855 | 4624 | 3233 | 3991 | 3936 | 4579 | 4875 | 3635 | 3479 | 4369 | 4145.46 | 4463 | 3828 | 0.857 | -0.222 | 0.07084999 | 0.20734866 | 0.0131 | 0.0077 | 0.0106 | 0.0106 | TRUE | 0.1856 |
| Hbq1a | 12 | 19 | 12 | 27 | 19 | 18 | 13 | 19 | 12 | 21 | 20 | 20 | 17.52 | 15 | 20 | 1.402 | 0.488 | 0.28624881 | 0.5074591 | 0 | 0.1949 | 0.092 | 0.092 | TRUE | 0.3152 |
| Hbq1b | 33 | 35 | 37 | 55 | 41 | 29 | 35 | 35 | 37 | 43 | 44 | 32 | 37.71 | 36 | 40 | 1.115 | 0.157 | 0.61855424 | 0.78448356 | 0 | 0.0943 | 0.0455 | 0.0455 | TRUE | 0.2118 |
| Hbs1l | 6668 | 7227 | 7379 | 8114 | 5660 | 5943 | 7083 | 7218 | 7409 | 6379 | 6090 | 6506 | 6780.59 | 7237 | 6325 | 0.874 | -0.194 | 0.00205457 | 0.01573045 | 6.00E-04 | 0.0074 | 0.0027 | 0.0027 | TRUE | 0.0128 |
| Hc | 5 | 3 | 6 | 18 | 6 | 6 | 5 | 3 | 6 | 14 | 6 | 7 | 6.92 | 5 | 9 | 1.93 | 0.948 | 0.22106053 | 0.43311388 | 0 | 0.4831 | 0.2726 | 0.2726 | TRUE | 0.9686 |
| Hccs | 1761 | 2125 | 2123 | 2523 | 1797 | 1841 | 1870 | 2122 | 2132 | 1983 | 1933 | 2015 | 2009.43 | 2041 | 1977 | 0.968 | -0.046 | 0.58680996 | 0.76150238 | 0.0021 | 0.0085 | 0.0047 | 0.0047 | TRUE | 0.0647 |
| Hcfc1 | 24171 | 20815 | 22622 | 30441 | 21885 | 20977 | 25674 | 20788 | 22715 | 23930 | 23547 | 22963 | 23269.46 | 23059 | 23480 | 1.018 | 0.026 | 0.78036556 | 0.88629118 | 0.0047 | 0.007 | 0.0063 | 0.0063 | TRUE | 0.1204 |
| Hcfc1r1 | 832 | 822 | 928 | 998 | 754 | 774 | 884 | 821 | 932 | 785 | 811 | 847 | 846.59 | 879 | 814 | 0.926 | -0.111 | 0.24501339 | 0.4616907 | 0.0017 | 0.0108 | 0.0054 | 0.0054 | TRUE | 0.0398 |
| Hcfc2 | 413 | 490 | 458 | 645 | 464 | 482 | 439 | 489 | 460 | 507 | 499 | 528 | 486.97 | 463 | 511 | 1.105 | 0.143 | 0.17197576 | 0.37079995 | 0 | 0.0137 | 0.0059 | 0.0059 | TRUE | 0.0297 |
| Hck | 156 | 182 | 156 | 209 | 148 | 158 | 166 | 182 | 157 | 164 | 159 | 173 | 166.77 | 168 | 165 | 0.984 | -0.023 | 0.88646056 | 0.94423286 | 0 | 0.0267 | 0.0121 | 0.0121 | TRUE | 0.0551 |
| Hcls1 | 3 | 4 | 4 | 7 | 5 | 0 | 3 | 4 | 4 | 6 | 5 | 0 | 3.68 | 4 | 4 | 0.988 | -0.018 | 0.9873713 | NA | 0 | 0.9021 | 0.6345 | 0.6345 | TRUE | 1.0414 |
| Hcn1 | 29 | 6 | 11 | 36 | 23 | 9 | 31 | 6 | 11 | 28 | 25 | 10 | 18.46 | 16 | 21 | 1.327 | 0.408 | 0.55060043 | 0.73491924 | 0.3839 | 0.1854 | 0.2807 | 0.2807 | TRUE | 1.9652 |
| Hcn2 | 1723 | 2704 | 2573 | 1937 | 1731 | 1709 | 1830 | 2701 | 2584 | 1523 | 1862 | 1871 | 2061.69 | 2372 | 1752 | 0.738 | -0.437 | 0.00652315 | 0.03818975 | 0.0255 | 0.0085 | 0.0181 | 0.0181 | TRUE | 0.4825 |
| Hcn3 | 269 | 403 | 431 | 324 | 308 | 311 | 286 | 402 | 433 | 255 | 331 | 340 | 341.25 | 374 | 309 | 0.824 | -0.278 | 0.15481829 | 0.34699361 | 0.0328 | 0.0166 | 0.0247 | 0.0247 | TRUE | 0.4824 |
| Hcn4 | 34 | 51 | 41 | 28 | 23 | 33 | 36 | 51 | 41 | 22 | 25 | 36 | 35.18 | 43 | 28 | 0.64 | -0.643 | 0.07158372 | 0.20857536 | 0.0182 | 0.1005 | 0.0619 | 0.0619 | TRUE | 0.4373 |
| Hcrt | 2 | 1 | 0 | 0 | 1 | 3 | 2 | 1 | 0 | 0 | 1 | 3 | 1.25 | 1 | 1 | 1.365 | 0.449 | 0.81908948 | NA | 0 | 2.6485 | 1.934 | 1.934 | TRUE | 0.4165 |
| Hcrtr1 | 9 | 8 | 6 | 10 | 11 | 9 | 10 | 8 | 6 | 8 | 12 | 10 | 8.85 | 8 | 10 | 1.246 | 0.317 | 0.61531863 | 0.78236493 | 0 | 0.379 | 0.1724 | 0.1724 | TRUE | 0.1392 |
| Hcrtr2 | 1 | 0 | 1 | 10 | 0 | 3 | 1 | 0 | 1 | 8 | 0 | 3 | 2.2 | 1 | 4 | 5.523 | 2.466 | 0.16455565 | NA | 0 | 1.5031 | 1.3918 | 1.3918 | TRUE | 0.1086 |
| Hcst | 0 | 3 | 1 | 1 | 1 | 1 | 0 | 3 | 1 | 1 | 1 | 1 | 1.16 | 1 | 1 | 0.727 | -0.46 | 0.7966897 | NA | 0 | 2.8482 | 1.4133 | 1.4133 | TRUE | 0.4088 |
| Hdac1 | 6051 | 5162 | 5255 | 8726 | 6047 | 5449 | 6427 | 5155 | 5277 | 6860 | 6506 | 5965 | 6031.64 | 5620 | 6444 | 1.147 | 0.198 | 0.07590997 | 0.21724631 | 0.01 | 0.0074 | 0.0088 | 0.0088 | TRUE | 0.1925 |
| Hdac10 | 151 | 166 | 158 | 258 | 200 | 169 | 160 | 166 | 159 | 203 | 215 | 185 | 181.3 | 162 | 201 | 1.244 | 0.315 | 0.0390493 | 0.13830801 | 0 | 0.0251 | 0.0112 | 0.0112 | TRUE | 0.0493 |
| Hdac11 | 563 | 814 | 913 | 502 | 485 | 580 | 598 | 813 | 917 | 395 | 522 | 635 | 646.51 | 776 | 517 | 0.665 | -0.588 | 0.00473656 | 0.03011087 | 0.0465 | 0.012 | 0.0296 | 0.0296 | TRUE | 0.3424 |

| | | | | | | | | | | | | | | | | | | | | | | | | | |
|---|---|---|---|---|---|---|---|---|---|---|---|---|---|---|---|---|---|---|---|---|---|---|---|---|---|
| Hdac2 | 6174 | 6409 | 6126 | 8790 | 6444 | 6346 | 6558 | 6401 | 6151 | 6910 | 6933 | 6947 | 6649.97 | 6370 | 6930 | 1.088 | 0.122 | 0.04222268 | 0.14652443 | 3.00E-04 | 0.0074 | 0.0024 | 0.0024 | TRUE | 0.011 |
| Hdac3 | 2355 | 2288 | 2292 | 3481 | 2511 | 2549 | 2501 | 2285 | 2301 | 2736 | 2702 | 2790 | 2552.72 | 2362 | 2743 | 1.161 | 0.215 | 0.00331836 | 0.02290465 | 8.00E-04 | 0.0082 | 0.0035 | 0.0035 | TRUE | 0.0319 |
| Hdac4 | 1476 | 1797 | 1954 | 1628 | 1249 | 1392 | 1568 | 1795 | 1962 | 1280 | 1344 | 1524 | 1578.65 | 1775 | 1383 | 0.778 | -0.361 | 0.00222751 | 0.0167337 | 0.0094 | 0.009 | 0.0094 | 0.0094 | TRUE | 0.1254 |
| Hdac5 | 3864 | 4742 | 4750 | 4956 | 3773 | 4166 | 4104 | 4736 | 4770 | 3896 | 4060 | 4560 | 4354.25 | 4537 | 4172 | 0.919 | -0.121 | 0.22803524 | 0.44159023 | 0.0065 | 0.0077 | 0.0071 | 0.0071 | TRUE | 0.0845 |
| Hdac6 | 5508 | 5614 | 5438 | 7440 | 5882 | 6307 | 5850 | 5607 | 5460 | 5849 | 6329 | 6904 | 5999.83 | 5639 | 6361 | 1.128 | 0.173 | 0.04527868 | 0.15351147 | 0.0038 | 0.0075 | 0.0052 | 0.0052 | TRUE | 0.0682 |
| Hdac7 | 375 | 785 | 873 | 561 | 496 | 592 | 398 | 784 | 877 | 441 | 534 | 648 | 613.6 | 686 | 541 | 0.787 | -0.345 | 0.19517094 | 0.40102085 | 0.0974 | 0.0123 | 0.0494 | 0.0494 | TRUE | 1.4198 |
| Hdac8 | 239 | 222 | 205 | 307 | 215 | 228 | 254 | 222 | 206 | 241 | 231 | 250 | 233.94 | 227 | 241 | 1.061 | 0.085 | 0.56105236 | 0.74256648 | 0.0017 | 0.021 | 0.0111 | 0.0111 | TRUE | 0.115 |
| Hdac9 | 169 | 222 | 222 | 251 | 215 | 207 | 180 | 222 | 223 | 197 | 231 | 227 | 213.23 | 208 | 218 | 1.047 | 0.066 | 0.68428033 | 0.82523852 | 0.0063 | 0.0224 | 0.0141 | 0.0141 | TRUE | 0.1546 |
| Hdc | 51 | 46 | 33 | 77 | 53 | 32 | 54 | 46 | 33 | 61 | 57 | 35 | 47.64 | 44 | 51 | 1.152 | 0.204 | 0.54955397 | 0.73404177 | 0.0482 | 0.0761 | 0.0628 | 0.0628 | TRUE | 0.5733 |
| Hddc2 | 1139 | 1288 | 1286 | 1854 | 1427 | 1371 | 1210 | 1286 | 1291 | 1457 | 1535 | 1501 | 1380.17 | 1262 | 1498 | 1.186 | 0.246 | 0.00151931 | 0.01237097 | 0 | 0.0093 | 0.0036 | 0.0036 | TRUE | 0.0159 |
| Hddc3 | 188 | 290 | 302 | 384 | 273 | 299 | 200 | 290 | 303 | 302 | 294 | 327 | 285.91 | 264 | 308 | 1.163 | 0.218 | 0.22825175 | 0.44159023 | 0.0218 | 0.0184 | 0.0201 | 0.0201 | TRUE | 0.5037 |
| Hdgf | 24249 | 26284 | 26259 | 31759 | 23168 | 21936 | 25757 | 26250 | 26367 | 24966 | 24927 | 24012 | 25379.96 | 26125 | 24635 | 0.943 | -0.085 | 0.1163199 | 0.28814159 | 3.00E-04 | 0.007 | 0.0021 | 0.0021 | TRUE | 0.006 |
| Hdgfl1 | 52 | 41 | 39 | 95 | 98 | 63 | 55 | 41 | 39 | 75 | 105 | 69 | 64.07 | 45 | 83 | 1.84 | 0.879 | 0.00260078 | 0.01893359 | 0.0295 | 0.0583 | 0.0442 | 0.0442 | TRUE | 0.5045 |
| Hdgfrp2 | 5178 | 5042 | 5144 | 6912 | 5603 | 4859 | 5500 | 5035 | 5165 | 5434 | 6028 | 5319 | 5413.61 | 5233 | 5594 | 1.069 | 0.096 | 0.24450527 | 0.46111886 | 0.0031 | 0.0075 | 0.0047 | 0.0047 | TRUE | 0.0561 |
| Hdgfrp3 | 581 | 849 | 894 | 1566 | 1177 | 1027 | 617 | 848 | 898 | 1231 | 1266 | 1124 | 997.39 | 788 | 1207 | 1.533 | 0.616 | 3.88E-05 | 0.00062306 | 0.0184 | 0.0102 | 0.0151 | 0.0151 | TRUE | 0.4231 |
| Hdhd1a | 9 | 1 | 0 | 3 | 0 | 3 | 10 | 1 | 0 | 2 | 0 | 3 | 2.7 | 4 | 2 | 0.54 | -0.889 | 0.59761258 | NA | 2.283 | 1.227 | 1.6407 | 1.6407 | TRUE | 3.2304 |
| Hdhd2 | 1052 | 992 | 989 | 1503 | 1058 | 995 | 1117 | 991 | 993 | 1182 | 1138 | 1089 | 1085.04 | 1034 | 1136 | 1.1 | 0.138 | 0.13991693 | 0.32462044 | 0.0023 | 0.0099 | 0.0054 | 0.0054 | TRUE | 0.0596 |
| Hdhd3 | 55 | 37 | 25 | 86 | 56 | 36 | 58 | 37 | 25 | 68 | 60 | 39 | 47.96 | 40 | 56 | 1.397 | 0.482 | 0.2136043 | 0.42505262 | 0.0991 | 0.0756 | 0.0867 | 0.0867 | TRUE | 0.5924 |
| Hdlbp | 32147 | 27233 | 27208 | 41990 | 28437 | 25172 | 34146 | 27198 | 27320 | 33009 | 30596 | 27555 | 29970.63 | 29555 | 30387 | 1.028 | 0.04 | 0.73561314 | 0.85807931 | 0.0106 | 0.007 | 0.0101 | 0.0101 | TRUE | 0.226 |
| Hdx | 134 | 107 | 102 | 234 | 143 | 114 | 142 | 107 | 102 | 184 | 154 | 125 | 135.7 | 117 | 154 | 1.322 | 0.402 | 0.07620583 | 0.21781776 | 0.028 | 0.0312 | 0.0296 | 0.0296 | TRUE | 0.1796 |
| Heatr1 | 17492 | 13600 | 14411 | 18966 | 13654 | 11998 | 18580 | 13582 | 14470 | 14910 | 14691 | 13134 | 14894.39 | 15544 | 14245 | 0.916 | -0.126 | 0.33117849 | 0.55212268 | 0.0167 | 0.0071 | 0.012 | 0.012 | TRUE | 0.3569 |
| Heatr2 | 2750 | 3014 | 3234 | 4182 | 3319 | 3319 | 2921 | 3010 | 3247 | 3288 | 3571 | 3633 | 3278.36 | 3059 | 3497 | 1.143 | 0.192 | 0.01916563 | 0.08408354 | 0.0024 | 0.0079 | 0.0046 | 0.0046 | TRUE | 0.0351 |
| Heatr3 | 9563 | 6409 | 6398 | 11052 | 7821 | 6017 | 10158 | 6401 | 6424 | 8688 | 8415 | 6587 | 7778.71 | 7661 | 7897 | 1.031 | 0.044 | 0.81677941 | 0.90681718 | 0.0397 | 0.0073 | 0.0257 | 0.0257 | TRUE | 0.9922 |
| Heatr5a | 4515 | 4329 | 4845 | 5232 | 3947 | 3881 | 4796 | 4323 | 4865 | 4113 | 4247 | 4248 | 4432.02 | 4661 | 4203 | 0.902 | -0.15 | 0.04990288 | 0.1639384 | 0.0018 | 0.0076 | 0.004 | 0.004 | TRUE | 0.0491 |
| Heatr5b | 3238 | 3346 | 3839 | 3870 | 2791 | 3023 | 3439 | 3342 | 3855 | 3042 | 3003 | 3309 | 3331.69 | 3545 | 3118 | 0.879 | -0.185 | 0.03846616 | 0.1367031 | 0.0039 | 0.0079 | 0.0055 | 0.0055 | TRUE | 0.071 |
| Heatr6 | 2619 | 2904 | 2948 | 3664 | 2626 | 2787 | 2782 | 2900 | 2960 | 2880 | 2825 | 3051 | 2899.79 | 2881 | 2919 | 1.013 | 0.019 | 0.7944126 | 0.8942097 | 9.00E-04 | 0.008 | 0.0034 | 0.0034 | TRUE | 0.0187 |
| Heatr8 | 37 | 47 | 52 | 53 | 44 | 48 | 39 | 47 | 52 | 42 | 47 | 53 | 46.67 | 46 | 47 | 1.017 | 0.024 | 0.93373866 | 0.96713012 | 0 | 0.0775 | 0.0387 | 0.0387 | TRUE | 0.129 |
| Hebp1 | 572 | 730 | 697 | 778 | 515 | 580 | 608 | 729 | 700 | 612 | 554 | 635 | 639.52 | 679 | 600 | 0.884 | -0.178 | 0.12813702 | 0.30785004 | 0.0052 | 0.0121 | 0.0083 | 0.0083 | TRUE | 0.0988 |
| Hebp2 | 288 | 309 | 313 | 476 | 352 | 349 | 306 | 309 | 314 | 374 | 379 | 382 | 343.96 | 310 | 378 | 1.221 | 0.289 | 0.0101734 | 0.0534493 | 0 | 0.0165 | 0.0061 | 0.0061 | TRUE | 0.002 |
| Heca | 1930 | 2110 | 2090 | 2416 | 1786 | 1933 | 2050 | 2107 | 2099 | 1899 | 1922 | 2116 | 2032.12 | 2085 | 1979 | 0.949 | -0.076 | 0.3324138 | 0.55286623 | 0.0012 | 0.0085 | 0.004 | 0.004 | TRUE | 0.0439 |
| Hectd1 | 5777 | 6294 | 7189 | 8053 | 5412 | 5539 | 6136 | 6286 | 7219 | 6331 | 5823 | 6063 | 6309.58 | 6547 | 6072 | 0.928 | -0.108 | 0.22897118 | 0.44224019 | 0.0046 | 0.0074 | 0.0057 | 0.0057 | TRUE | 0.0983 |
| Hectd2 | 477 | 658 | 615 | 748 | 531 | 552 | 507 | 657 | 618 | 588 | 571 | 604 | 590.82 | 594 | 588 | 0.99 | -0.015 | 0.90462738 | 0.95255222 | 0.0075 | 0.0125 | 0.0098 | 0.0098 | TRUE | 0.1919 |
| Hectd3 | 1996 | 2169 | 2208 | 2567 | 1847 | 1869 | 2120 | 2166 | 2217 | 2018 | 1987 | 2046 | 2092.42 | 2168 | 2017 | 0.93 | -0.104 | 0.12037496 | 0.29444073 | 0 | 0.0085 | 0.0028 | 0.0028 | TRUE | 0.0048 |
| Hecw1 | 18 | 10 | 8 | 20 | 6 | 8 | 19 | 10 | 8 | 16 | 6 | 9 | 11.35 | 12 | 10 | 0.849 | -0.236 | 0.71576781 | 0.8456753 | 0.1298 | 0.2973 | 0.2141 | 0.2141 | TRUE | 0.925 |
| Hecw2 | 92 | 70 | 62 | 317 | 187 | 191 | 98 | 70 | 62 | 249 | 201 | 209 | 148.23 | 77 | 220 | 2.88 | 1.526 | 3.23E-12 | 3.07E-10 | 0.0214 | 0.0291 | 0.0257 | 0.0257 | TRUE | 0.5219 |
| Heg1 | 453 | 501 | 526 | 537 | 379 | 413 | 481 | 500 | 528 | 422 | 408 | 452 | 465.28 | 503 | 427 | 0.849 | -0.237 | 0.02923864 | 0.11295336 | 0 | 0.014 | 0.0063 | 0.0063 | TRUE | 0.0283 |
| Helb | 6056 | 5683 | 6000 | 5676 | 4623 | 4653 | 6433 | 5676 | 6025 | 4462 | 4974 | 5093 | 5443.73 | 6045 | 4843 | 0.801 | -0.32 | 0.00030946 | 0.00349383 | 0.0041 | 0.0075 | 0.0055 | 0.0055 | TRUE | 0.0583 |
| Hells | 15624 | 14240 | 14792 | 21184 | 15321 | 14463 | 16595 | 14222 | 14853 | 16653 | 16484 | 15832 | 15773.25 | 15223 | 16323 | 1.072 | 0.101 | 0.21994587 | 0.43222937 | 0.0035 | 0.0071 | 0.0048 | 0.0048 | TRUE | 0.076 |
| Helq | 1168 | 1076 | 1143 | 1607 | 1120 | 1010 | 1241 | 1075 | 1148 | 1263 | 1205 | 1106 | 1172.81 | 1155 | 1191 | 1.033 | 0.047 | 0.63909667 | 0.79663579 | 0.004 | 0.0097 | 0.0063 | 0.0063 | TRUE | 0.051 |
| Helt | 2 | 3 | 2 | 1 | 2 | 1 | 2 | 3 | 2 | 1 | 2 | 1 | 1.86 | 2 | 1 | 0.553 | -0.855 | 0.52517718 | NA | 0 | 1.7779 | 0.7184 | 0.7184 | TRUE | 0.1598 |
| Helz | 7525 | 6533 | 7687 | 8735 | 6253 | 6147 | 7993 | 6525 | 7719 | 6867 | 6728 | 6729 | 7093.25 | 7412 | 6775 | 0.914 | -0.13 | 0.17217735 | 0.37108242 | 0.0054 | 0.0074 | 0.0064 | 0.0064 | TRUE | 0.134 |
| Helz2 | 689 | 495 | 554 | 567 | 385 | 350 | 732 | 494 | 556 | 446 | 414 | 383 | 504.26 | 594 | 414 | 0.698 | -0.518 | 0.00181116 | 0.01421879 | 0.0213 | 0.0134 | 0.0178 | 0.0178 | TRUE | 0.4806 |
| Hemgn | 0 | 0 | 1 | 0 | 2 | 0 | 0 | 0 | 1 | 0 | 2 | 0 | 0.53 | 0 | 1 | 1.927 | 0.947 | 0.77736506 | NA | 0 | 6.2701 | 6.3814 | 6.3814 | TRUE | 0.9744 |
| Hemk1 | 1073 | 1244 | 1285 | 1568 | 1084 | 1057 | 1140 | 1242 | 1290 | 1233 | 1166 | 1157 | 1204.73 | 1224 | 1185 | 0.969 | -0.046 | 0.6056352 | 0.77621309 | 0.0018 | 0.0096 | 0.0049 | 0.0049 | TRUE | 0.0434 |
| Hemt1 | 2 | 4 | 7 | 41 | 21 | 20 | 2 | 4 | 7 | 32 | 23 | 22 | 14.98 | 4 | 26 | 5.834 | 2.545 | 3.87E-05 | 0.00062287 | 0 | 0.2269 | 0.1407 | 0.1407 | TRUE | 0.5052 |
| Henmt1 | 42 | 34 | 20 | 88 | 48 | 49 | 45 | 34 | 20 | 69 | 52 | 54 | 45.52 | 33 | 58 | 1.779 | 0.831 | 0.0199387 | 0.08663172 | 0.0538 | 0.0793 | 0.0679 | 0.0679 | TRUE | 0.4739 |
| Hepacam | 18 | 11 | 18 | 34 | 18 | 16 | 19 | 11 | 18 | 27 | 19 | 18 | 18.63 | 16 | 21 | 1.334 | 0.415 | 0.37511343 | 0.59102546 | 0 | 0.1837 | 0.1033 | 0.1033 | TRUE | 0.3697 |
| Hepacam2 | 12 | 16 | 21 | 22 | 9 | 12 | 13 | 16 | 21 | 17 | 10 | 13 | 14.99 | 17 | 13 | 0.813 | -0.298 | 0.56120749 | 0.74256648 | 0 | 0.2267 | 0.1232 | 0.1232 | TRUE | 0.3422 |
| Heph | 12 | 10 | 7 | 43 | 18 | 20 | 13 | 10 | 7 | 34 | 19 | 22 | 17.47 | 10 | 25 | 2.551 | 1.351 | 0.00856569 | 0.04684396 | 0 | 0.1955 | 0.1193 | 0.1193 | TRUE | 0.6709 |
| Hephl1 | 5 | 0 | 1 | 16 | 10 | 4 | 5 | 0 | 1 | 13 | 11 | 4 | 5.67 | 2 | 9 | 4.48 | 2.163 | 0.04790427 | 0.15919999 | 0.5125 | 0.5877 | 0.563 | 0.563 | TRUE | 1.5925 |
| Herc1 | 6006 | 4369 | 5533 | 6835 | 4580 | 4089 | 6379 | 4363 | 5556 | 5373 | 4928 | 4476 | 5179.25 | 5433 | 4926 | 0.907 | -0.141 | 0.33013704 | 0.55093867 | 0.021 | 0.0075 | 0.0149 | 0.0149 | TRUE | 0.2817 |
| Herc2 | 7863 | 7188 | 9397 | 9968 | 6897 | 6598 | 8352 | 7179 | 9436 | 7836 | 7421 | 7223 | 7907.59 | 8322 | 7493 | 0.9 | -0.151 | 0.17565976 | 0.37576369 | 0.0099 | 0.0073 | 0.0089 | 0.0089 | TRUE | 0.1765 |
| Herc3 | 1044 | 1002 | 904 | 1481 | 1067 | 1118 | 1109 | 1001 | 908 | 1164 | 1148 | 1224 | 1092.24 | 1006 | 1179 | 1.172 | 0.229 | 0.02668475 | 0.10641514 | 0.0042 | 0.0099 | 0.0068 | 0.0068 | TRUE | 0.0956 |
| Herc4 | 9763 | 9983 | 9959 | 12201 | 8795 | 8916 | 10370 | 9970 | 10000 | 9591 | 9463 | 9760 | 9859.06 | 10113 | 9605 | 0.95 | -0.074 | 0.18742418 | 0.39199763 | 2.00E-04 | 0.0072 | 0.0022 | 0.0022 | TRUE | 0.006 |
| Herc6 | 66 | 41 | 46 | 159 | 125 | 104 | 70 | 41 | 46 | 125 | 134 | 114 | 88.43 | 52 | 124 | 2.381 | 1.251 | 1.25E-06 | 3.35E-05 | 0.0206 | 0.0442 | 0.0343 | 0.0343 | TRUE | 0.7017 |
| Herpud1 | 348 | 426 | 383 | 501 | 338 | 348 | 370 | 425 | 385 | 394 | 364 | 381 | 386.35 | 393 | 380 | 0.966 | -0.05 | 0.67036377 | 0.81594208 | 0 | 0.0154 | 0.0075 | 0.0075 | TRUE | 0.0594 |
| Herpud2 | 1846 | 2035 | 1920 | 2792 | 2080 | 1867 | 1961 | 2032 | 1928 | 2195 | 2238 | 2044 | 2066.26 | 1974 | 2159 | 1.094 | 0.13 | 0.08779122 | 0.24005601 | 0.001 | 0.0085 | 0.0037 | 0.0037 | TRUE | 0.0259 |
| Hes1 | 746 | 1285 | 1296 | 1075 | 898 | 927 | 792 | 1283 | 1301 | 845 | 966 | 1015 | 1033.84 | 1125 | 942 | 0.836 | -0.258 | 0.17141482 | 0.37034626 | 0.0405 | 0.0101 | 0.0247 | 0.0247 | TRUE | 0.8015 |
| Hes2 | 48 | 65 | 68 | 37 | 37 | 35 | 51 | 65 | 68 | 29 | 40 | 38 | 48.56 | 61 | 36 | 0.577 | -0.793 | 0.00770851 | 0.04320978 | 0 | 0.0747 | 0.0418 | 0.0418 | TRUE | 0.2301 |
| Hes3 | 4 | 4 | 0 | 1 | 2 | 1 | 4 | 4 | 0 | 1 | 2 | 1 | 2.05 | 3 | 1 | 0.482 | -1.053 | 0.4821618 | NA | 0 | 1.6171 | 1.0615 | 1.0615 | TRUE | 0.9311 |
| Hes5 | 18 | 41 | 29 | 26 | 19 | 22 | 19 | 41 | 29 | 20 | 20 | 24 | 25.69 | 30 | 21 | 0.725 | -0.463 | 0.2737945 | 0.49433176 | 0.0414 | 0.1351 | 0.0893 | 0.0893 | TRUE | 0.1754 |
| Hes6 | 397 | 497 | 469 | 562 | 413 | 452 | 422 | 496 | 471 | 442 | 444 | 495 | 461.65 | 463 | 460 | 0.993 | -0.01 | 0.93135816 | 0.96621594 | 0.0032 | 0.014 | 0.0081 | 0.0081 | TRUE | 0.0699 |
| Hes7 | 46 | 45 | 36 | 15 | 19 | 15 | 49 | 45 | 36 | 12 | 20 | 16 | 29.77 | 43 | 16 | 0.37 | -1.436 | 0.00022451 | 0.00266645 | 0 | 0.1176 | 0.0669 | 0.0669 | TRUE | 0.3558 |
| Hesx1 | 30 | 22 | 12 | 90 | 59 | 69 | 32 | 22 | 12 | 71 | 63 | 76 | 45.94 | 22 | 70 | 3.195 | 1.676 | 4.59E-06 | 0.00010254 | 0.0415 | 0.0786 | 0.066 | 0.066 | TRUE | 0.647 |
| Hexa | 1625 | 3723 | 4024 | 2549 | 2005 | 2600 | 1726 | 3718 | 4041 | 2004 | 2157 | 2846 | 2748.66 | 3162 | 2336 | 0.739 | -0.437 | 0.2703359 | 0.49054018 | 0.1128 | 0.0081 | 0.051 | 0.1128 | TRUE | 1.9164 |
| Hexb | 4697 | 6265 | 6640 | 4150 | 3270 | 4132 | 4989 | 6257 | 6667 | 3262 | 3518 | 4523 | 4869.51 | 5971 | 3768 | 0.631 | -0.665 | 1.49E-05 | 0.00027752 | 0.026 | 0.0076 | 0.0168 | 0.0168 | TRUE | 0.3736 |
| Hexdc | 351 | 470 | 493 | 692 | 508 | 443 | 373 | 469 | 495 | 544 | 547 | 485 | 485.46 | 446 | 525 | 1.178 | 0.237 | 0.09177199 | 0.24729673 | 0.0107 | 0.0137 | 0.0121 | 0.0121 | TRUE | 0.2351 |
| Hexim1 | 3046 | 2210 | 1964 | 4170 | 3005 | 2319 | 3235 | 2207 | 1972 | 3278 | 3233 | 2539 | 2744.07 | 2471 | 3017 | 1.221 | 0.288 | 0.12383469 | 0.30006463 | 0.0446 | 0.0081 | 0.0248 | 0.0248 | TRUE | 0.8856 |
| Hexim2 | 318 | 367 | 408 | 472 | 363 | 344 | 338 | 367 | 410 | 371 | 391 | 377 | 375.36 | 372 | 380 | 1.021 | 0.03 | 0.81099499 | 0.90283686 | 0.0022 | 0.0157 | 0.0084 | 0.0084 | TRUE | 0.0936 |

| | | | | | | | | | | | | | | | | | | | | | | | | |
|---|---|---|---|---|---|---|---|---|---|---|---|---|---|---|---|---|---|---|---|---|---|---|---|---|
| Hey1 | 223 | 275 | 273 | 271 | 253 | 248 | 237 | 275 | 274 | 213 | 272 | 271 | 257.06 | 262 | 252 | 0.96 | -0.06 | 0.70851801 | 0.84126423 | 0.0097 | 0.0197 | 0.0144 | 0.0144 | TRUE | 0.2208 |
| Hey2 | 62 | 105 | 73 | 69 | 55 | 65 | 66 | 105 | 73 | 54 | 59 | 71 | 71.43 | 81 | 61 | 0.753 | -0.409 | 0.13837711 | 0.3229004 | 0.0285 | 0.053 | 0.0408 | 0.0408 | TRUE | 0.6013 |
| Heyl | 4 | 3 | 2 | 4 | 3 | 3 | 4 | 3 | 2 | 3 | 3 | 3 | 3.15 | 3 | 3 | 1.046 | 0.065 | 0.94952946 | NA | 0 | 1.0522 | 0.4311 | 0.4311 | TRUE | 0.137 |
| Hfe | 25 | 21 | 18 | 43 | 24 | 21 | 27 | 21 | 18 | 34 | 26 | 23 | 24.7 | 22 | 28 | 1.271 | 0.346 | 0.38713471 | 0.60099965 | 0 | 0.1403 | 0.0744 | 0.0744 | TRUE | 0.2984 |
| Hfe2 | 109 | 116 | 105 | 149 | 92 | 79 | 116 | 116 | 105 | 117 | 99 | 86 | 106.61 | 112 | 101 | 0.903 | -0.148 | 0.4731351 | 0.67409998 | 0 | 0.0378 | 0.0211 | 0.0211 | TRUE | 0.215 |
| Hfm1 | 67 | 124 | 114 | 163 | 148 | 115 | 71 | 124 | 114 | 128 | 159 | 126 | 120.46 | 103 | 138 | 1.332 | 0.414 | 0.09715691 | 0.25620253 | 0.0387 | 0.0343 | 0.0363 | 0.0363 | TRUE | 0.7094 |
| Hgd | 0 | 1 | 0 | 3 | 0 | 1 | 0 | 1 | 0 | 2 | 0 | 1 | 0.74 | 0 | 1 | 3.216 | 1.685 | 0.50251474 | NA | 0 | 4.4471 | 3.135 | 3.135 | TRUE | 0.1927 |
| Hgf | 1 | 0 | 1 | 4 | 0 | 2 | 1 | 0 | 1 | 3 | 0 | 2 | 1.23 | 1 | 2 | 2.644 | 1.403 | 0.46925634 | NA | 0 | 2.678 | 1.6912 | 1.6912 | TRUE | 0.3556 |
| Hgfac | 1 | 1 | 0 | 2 | 2 | 6 | 1 | 1 | 0 | 2 | 2 | 7 | 2.06 | 1 | 4 | 4.951 | 2.308 | 0.14558731 | NA | 0 | 1.6071 | 0.9194 | 0.9194 | TRUE | 0.8673 |
| Hgs | 4252 | 4367 | 4639 | 5826 | 4356 | 4303 | 4516 | 4361 | 4658 | 4580 | 4687 | 4710 | 4585.47 | 4512 | 4659 | 1.032 | 0.046 | 0.4650706 | 0.66788976 | 4.00E-04 | 0.0076 | 0.0027 | 0.0027 | TRUE | 0.0104 |
| Hgsnat | 1578 | 2006 | 1912 | 2208 | 1668 | 1844 | 1676 | 2003 | 1920 | 1736 | 1795 | 2019 | 1858.06 | 1866 | 1850 | 0.99 | -0.014 | 0.89640125 | 0.94994561 | 0.0069 | 0.0087 | 0.0076 | 0.0076 | TRUE | 0.0959 |
| Hhat | 51 | 88 | 86 | 93 | 53 | 64 | 54 | 88 | 86 | 73 | 57 | 70 | 71.43 | 76 | 67 | 0.877 | -0.189 | 0.4953934 | 0.69138525 | 0.0284 | 0.053 | 0.0412 | 0.0412 | TRUE | 0.5716 |
| Hhatl | 0 | 0 | 0 | 0 | 0 | 0 | 0 | 0 | 0 | 0 | 0 | 0 | 0 | 0 | 0 | NA | NA | NA | NA | NA | NA | NA | NA | NA | NA |
| Hhex | 305 | 458 | 461 | 467 | 396 | 412 | 324 | 457 | 463 | 367 | 426 | 451 | 414.74 | 415 | 415 | 0.998 | -0.003 | 0.98715468 | 0.99294666 | 0.0225 | 0.0148 | 0.0187 | 0.0187 | TRUE | 0.4198 |
| Hhip | 11 | 11 | 7 | 27 | 20 | 9 | 12 | 11 | 7 | 21 | 22 | 10 | 13.72 | 10 | 18 | 1.785 | 0.836 | 0.14187814 | 0.32748457 | 0 | 0.2471 | 0.1538 | 0.1538 | TRUE | 0.6747 |
| Hhipl1 | 26 | 31 | 18 | 22 | 16 | 8 | 28 | 31 | 18 | 17 | 17 | 9 | 19.99 | 26 | 14 | 0.569 | -0.812 | 0.08932066 | 0.24293248 | 0.0374 | 0.1717 | 0.1111 | 0.1111 | TRUE | 0.4724 |
| Hhipl2 | 5 | 5 | 5 | 12 | 7 | 8 | 5 | 5 | 5 | 9 | 8 | 9 | 6.84 | 5 | 9 | 1.685 | 0.753 | 0.28614286 | 0.50736877 | 0 | 0.4885 | 0.2023 | 0.2023 | TRUE | 0.033 |
| Hhla1 | 0 | 0 | 0 | 1 | 0 | 1 | 0 | 0 | 0 | 1 | 0 | 1 | 0.31 | 0 | 1 | 3.348 | 1.743 | 0.65064889 | NA | 0 | 10.5167 | 8.8968 | 8.8968 | TRUE | 0.2082 |
| Hiat1 | 4090 | 4402 | 4425 | 6476 | 4718 | 4734 | 4344 | 4396 | 4443 | 5091 | 5076 | 5182 | 4755.52 | 4394 | 5116 | 1.164 | 0.219 | 0.00011789 | 0.00156779 | 0 | 0.0076 | 0.0021 | 0.0021 | TRUE | 0.0015 |
| Hiatl1 | 2720 | 2955 | 3091 | 3965 | 3020 | 3123 | 2889 | 2951 | 3104 | 3117 | 3249 | 3419 | 3121.49 | 2981 | 3262 | 1.094 | 0.129 | 0.08407864 | 0.23244317 | 0.0013 | 0.008 | 0.0037 | 0.0037 | TRUE | 0.0215 |
| Hibadh | 4154 | 5512 | 5568 | 4504 | 3511 | 4293 | 4412 | 5505 | 5591 | 3541 | 3778 | 4699 | 4587.62 | 5169 | 4006 | 0.775 | -0.368 | 0.00824829 | 0.04543906 | 0.0194 | 0.0076 | 0.0138 | 0.0138 | TRUE | 0.2788 |
| Hibch | 1608 | 1888 | 1795 | 1876 | 1284 | 1351 | 1708 | 1886 | 1802 | 1475 | 1382 | 1479 | 1621.84 | 1799 | 1445 | 0.804 | -0.316 | 0.00010901 | 0.00147575 | 0.0013 | 0.0089 | 0.0042 | 0.0042 | TRUE | 0.0234 |
| Hic1 | 123 | 226 | 168 | 250 | 186 | 167 | 131 | 226 | 169 | 197 | 200 | 183 | 184.08 | 175 | 193 | 1.103 | 0.141 | 0.51326355 | 0.70632889 | 0.0319 | 0.0248 | 0.0283 | 0.0283 | TRUE | 0.3599 |
| Hic2 | 4396 | 4036 | 3821 | 5019 | 3642 | 3305 | 4669 | 4031 | 3837 | 3946 | 3919 | 3618 | 4003.13 | 4179 | 3828 | 0.916 | -0.127 | 0.2011586 | 0.40877592 | 0.0062 | 0.0077 | 0.0068 | 0.0068 | TRUE | 0.1281 |
| Hid1 | 587 | 583 | 570 | 798 | 670 | 655 | 623 | 582 | 572 | 627 | 721 | 717 | 640.55 | 592 | 688 | 1.16 | 0.214 | 0.04608175 | 0.15533262 | 0.0029 | 0.012 | 0.0067 | 0.0067 | TRUE | 0.0746 |
| Hif1a | 5839 | 5641 | 5368 | 8326 | 5567 | 4800 | 6202 | 5634 | 5390 | 6545 | 5990 | 5254 | 5835.86 | 5742 | 5930 | 1.033 | 0.047 | 0.66116266 | 0.80914578 | 0.0071 | 0.0075 | 0.008 | 0.008 | TRUE | 0.1209 |
| Hif1an | 3849 | 3778 | 3836 | 5241 | 3815 | 3768 | 4088 | 3773 | 3852 | 4120 | 4105 | 4125 | 4010.44 | 3904 | 4117 | 1.054 | 0.076 | 0.24650184 | 0.46355396 | 5.00E-04 | 0.0077 | 0.0029 | 0.0029 | TRUE | 0.0206 |
| Hif3a | 917 | 1381 | 1764 | 937 | 836 | 1128 | 974 | 1379 | 1771 | 737 | 899 | 1235 | 1165.89 | 1375 | 957 | 0.696 | -0.524 | 0.02594113 | 0.10426694 | 0.0759 | 0.0097 | 0.039 | 0.039 | TRUE | 0.1381 |
| Higd1a | 1210 | 1805 | 1867 | 2547 | 1922 | 2060 | 1285 | 1803 | 1875 | 2002 | 2068 | 2255 | 1881.29 | 1654 | 2108 | 1.274 | 0.35 | 0.01846151 | 0.08209049 | 0.02 | 0.0087 | 0.0153 | 0.0153 | TRUE | 0.4582 |
| Higd1b | 9 | 2 | 4 | 2 | 2 | 1 | 10 | 2 | 4 | 2 | 2 | 1 | 3.4 | 5 | 2 | 0.311 | -1.686 | 0.14441079 | NA | 0 | 0.9762 | 0.5597 | 0.5597 | TRUE | 0.4202 |
| Higd1c | 0 | 0 | 0 | 0 | 0 | 0 | 0 | 0 | 0 | 0 | 0 | 0 | 0 | 0 | 0 | NA | NA | NA | NA | NA | NA | NA | NA | NA | NA |
| Higd2a | 2497 | 2759 | 2728 | 2426 | 1747 | 2114 | 2652 | 2755 | 2739 | 1907 | 1880 | 2314 | 2374.63 | 2715 | 2034 | 0.749 | -0.418 | 5.19E-05 | 0.00079989 | 0.0067 | 0.0083 | 0.0073 | 0.0073 | TRUE | 0.1748 |
| Hilpda | 272 | 345 | 355 | 460 | 341 | 317 | 289 | 345 | 356 | 362 | 367 | 347 | 344.24 | 330 | 359 | 1.086 | 0.119 | 0.36336189 | 0.58083215 | 0.0031 | 0.0165 | 0.0095 | 0.0095 | TRUE | 0.1329 |
| Hils1 | 1 | 0 | 0 | 1 | 2 | 0 | 1 | 0 | 0 | 1 | 2 | 0 | 0.67 | 0 | 1 | 2.668 | 1.416 | 0.58602256 | NA | 0 | 4.9484 | 3.3683 | 3.3683 | TRUE | 0.3203 |
| Hinfp | 753 | 715 | 689 | 936 | 735 | 713 | 800 | 714 | 692 | 736 | 791 | 780 | 752.14 | 735 | 769 | 1.045 | 0.064 | 0.52703394 | 0.71593432 | 0.0023 | 0.0113 | 0.0061 | 0.0061 | TRUE | 0.0693 |
| Hint1 | 7395 | 8788 | 8629 | 11300 | 7859 | 7970 | 7855 | 8777 | 8664 | 8883 | 8456 | 8724 | 8559.88 | 8432 | 8688 | 1.03 | 0.043 | 0.55764879 | 0.7401735 | 0.0018 | 0.0073 | 0.0038 | 0.0038 | TRUE | 0.0437 |
| Hint2 | 505 | 741 | 718 | 807 | 557 | 628 | 536 | 740 | 721 | 634 | 599 | 687 | 653.09 | 666 | 640 | 0.961 | -0.057 | 0.69891169 | 0.8353743 | 0.016 | 0.0119 | 0.0141 | 0.0141 | TRUE | 0.339 |
| Hint3 | 243 | 303 | 280 | 345 | 272 | 284 | 258 | 303 | 281 | 271 | 293 | 311 | 286.1 | 281 | 292 | 1.037 | 0.052 | 0.70209611 | 0.83705948 | 0.0021 | 0.0184 | 0.0097 | 0.0097 | TRUE | 0.0545 |
| Hip1 | 1556 | 1393 | 1410 | 2355 | 1675 | 1549 | 1653 | 1391 | 1416 | 1851 | 1802 | 1696 | 1634.81 | 1487 | 1783 | 1.2 | 0.263 | 0.00824515 | 0.04543864 | 0.0048 | 0.0089 | 0.0065 | 0.0065 | TRUE | 0.1147 |
| Hip1r | 2793 | 2794 | 2981 | 2533 | 2045 | 2104 | 2967 | 2790 | 2993 | 1991 | 2200 | 2303 | 2540.83 | 2917 | 2165 | 0.742 | -0.431 | 6.50E-07 | 1.89E-05 | 0.0028 | 0.0082 | 0.005 | 0.005 | TRUE | 0.0609 |
| Hipk1 | 3151 | 3185 | 3592 | 3931 | 2793 | 2810 | 3347 | 3181 | 3607 | 3090 | 3005 | 3076 | 3217.65 | 3378 | 3057 | 0.905 | -0.144 | 0.06188952 | 0.19012885 | 0.0017 | 0.0079 | 0.004 | 0.004 | TRUE | 0.0427 |
| Hipk2 | 639 | 593 | 560 | 1163 | 793 | 722 | 679 | 592 | 562 | 914 | 853 | 790 | 731.85 | 611 | 852 | 1.397 | 0.482 | 3.36E-05 | 0.00055305 | 0.0059 | 0.0114 | 0.0083 | 0.0083 | TRUE | 0.1094 |
| Hipk3 | 675 | 1065 | 1181 | 1578 | 1178 | 1292 | 717 | 1064 | 1186 | 1240 | 1267 | 1414 | 1148.12 | 989 | 1307 | 1.322 | 0.402 | 0.02292496 | 0.09578079 | 0.0343 | 0.0098 | 0.0217 | 0.0217 | TRUE | 0.6884 |
| Hipk4 | 40 | 22 | 38 | 37 | 38 | 19 | 42 | 22 | 38 | 29 | 41 | 21 | 32.23 | 34 | 30 | 0.885 | -0.176 | 0.67263235 | 0.81756762 | 0.0766 | 0.1091 | 0.0937 | 0.0937 | TRUE | 0.3627 |
| Hira | 5966 | 4637 | 4601 | 7493 | 5128 | 4239 | 6337 | 4631 | 4620 | 5890 | 5517 | 4640 | 5272.65 | 5196 | 5349 | 1.03 | 0.042 | 0.7778177 | 0.88486302 | 0.0242 | 0.0075 | 0.016 | 0.016 | TRUE | 0.4496 |
| Hirip3 | 2993 | 2717 | 2897 | 4436 | 3442 | 3104 | 3179 | 2713 | 2909 | 3487 | 3703 | 3398 | 3231.65 | 2934 | 3529 | 1.203 | 0.267 | 0.00265351 | 0.01925966 | 0.0037 | 0.0079 | 0.0054 | 0.0054 | TRUE | 0.0648 |
| Hist1h1a | 17 | 28 | 23 | 51 | 39 | 29 | 18 | 28 | 23 | 40 | 42 | 32 | 30.49 | 23 | 38 | 1.647 | 0.72 | 0.04786905 | 0.15916553 | 0 | 0.115 | 0.0603 | 0.0603 | TRUE | 0.2053 |
| Hist1h1b | 6 | 13 | 14 | 14 | 9 | 5 | 6 | 13 | 14 | 11 | 10 | 5 | 9.93 | 11 | 9 | 0.789 | -0.342 | 0.59438406 | 0.7677918 | 0 | 0.3387 | 0.1955 | 0.1955 | TRUE | 0.4989 |
| Hist1h1c | 134 | 130 | 116 | 211 | 145 | 125 | 142 | 130 | 116 | 166 | 156 | 137 | 141.23 | 129 | 153 | 1.184 | 0.244 | 0.17785608 | 0.37846007 | 0 | 0.0302 | 0.0164 | 0.0164 | TRUE | 0.0861 |
| Hist1h1d | 13 | 13 | 13 | 19 | 11 | 14 | 14 | 13 | 13 | 15 | 12 | 15 | 13.66 | 13 | 14 | 1.059 | 0.083 | 0.86723278 | 0.93428931 | 0 | 0.2481 | 0.1057 | 0.1057 | TRUE | 0.0749 |
| Hist1h1e | 3 | 7 | 10 | 15 | 9 | 7 | 3 | 7 | 10 | 12 | 10 | 8 | 8.23 | 7 | 10 | 1.446 | 0.532 | 0.4463151 | 0.65236459 | 0 | 0.4074 | 0.2257 | 0.2257 | TRUE | 0.322 |
| Hist1h1t | 1 | 1 | 1 | 9 | 4 | 1 | 1 | 1 | 1 | 7 | 4 | 1 | 2.59 | 1 | 4 | 4.148 | 2.052 | 0.12937655 | NA | 0 | 1.279 | 0.704 | 0.704 | TRUE | 0.0883 |
| Hist1h2aa | 0 | 0 | 0 | 2 | 3 | 0 | 0 | 0 | 0 | 2 | 3 | 0 | 0.8 | 0 | 2 | 8.522 | 3.091 | 0.27029335 | NA | 5.3026 | 4.1249 | 4.3597 | 4.3597 | TRUE | 0.0518 |
| Hist1h2ab | 2 | 10 | 12 | 3 | 9 | 3 | 2 | 10 | 12 | 2 | 10 | 3 | 6.58 | 8 | 5 | 0.623 | -0.684 | 0.46184059 | 0.66571988 | 0.4148 | 0.5075 | 0.4619 | 0.4619 | TRUE | 1.1569 |
| Hist1h2ac | 9 | 5 | 8 | 9 | 8 | 7 | 10 | 5 | 8 | 7 | 9 | 8 | 7.66 | 8 | 8 | 1.032 | 0.046 | 0.94591757 | 0.97280132 | 0 | 0.4373 | 0.1979 | 0.1979 | TRUE | 0.2481 |
| Hist1h2ad | 0 | 2 | 1 | 7 | 1 | 2 | 0 | 2 | 1 | 6 | 1 | 2 | 1.96 | 1 | 3 | 2.956 | 1.564 | 0.30523978 | NA | 0 | 1.6864 | 1.0071 | 1.0071 | TRUE | 0.7165 |
| Hist1h2ae | 27 | 25 | 27 | 56 | 38 | 22 | 29 | 25 | 27 | 44 | 41 | 24 | 31.62 | 27 | 36 | 1.359 | 0.443 | 0.23751776 | 0.45308781 | 0 | 0.1111 | 0.0689 | 0.0689 | TRUE | 0.5865 |
| Hist1h2af | 10 | 11 | 27 | 30 | 26 | 24 | 11 | 11 | 27 | 24 | 28 | 26 | 21.09 | 16 | 26 | 1.591 | 0.67 | 0.17716496 | 0.37775391 | 0.0834 | 0.1631 | 0.1273 | 0.1273 | TRUE | 1.6547 |
| Hist1h2ag | 7 | 3 | 3 | 9 | 4 | 4 | 7 | 3 | 3 | 7 | 4 | 4 | 4.87 | 4 | 5 | 1.193 | 0.255 | 0.77121472 | 0.88078194 | 0 | 0.6838 | 0.3465 | 0.3465 | TRUE | 0.5824 |
| Hist1h2ah | 6 | 5 | 8 | 5 | 12 | 3 | 6 | 5 | 8 | 4 | 13 | 3 | 6.59 | 6 | 7 | 1.025 | 0.036 | 0.9653776 | 0.98257199 | 0.1743 | 0.507 | 0.3397 | 0.3397 | TRUE | 1.5704 |
| Hist1h2ai | 4 | 5 | 5 | 7 | 9 | 7 | 4 | 5 | 5 | 6 | 10 | 8 | 6.19 | 5 | 8 | 1.584 | 0.664 | 0.37884935 | 0.59395745 | 0 | 0.5395 | 0.238 | 0.238 | TRUE | 0.1978 |
| Hist1h2ak | 9 | 3 | 10 | 15 | 9 | 9 | 10 | 3 | 10 | 12 | 10 | 10 | 8.99 | 8 | 11 | 1.395 | 0.481 | 0.47803124 | 0.67743419 | 0 | 0.3735 | 0.216 | 0.216 | TRUE | 0.7978 |
| Hist1h2an | 0 | 1 | 2 | 3 | 2 | 0 | 0 | 1 | 2 | 2 | 2 | 0 | 1.25 | 1 | 1 | 1.516 | 0.6 | 0.74724324 | NA | 0 | 2.6364 | 1.6718 | 1.6718 | TRUE | 0.2901 |
| Hist1h2ao | 0 | 0 | 0 | 0 | 0 | 0 | 0 | 0 | 0 | 0 | 0 | 0 | 0 | 0 | 0 | NA | NA | NA | NA | NA | NA | NA | NA | NA | NA |
| Hist1h2ap | 0 | 0 | 0 | 0 | 0 | 0 | 0 | 0 | 0 | 0 | 0 | 0 | 0 | 0 | 0 | NA | NA | NA | NA | NA | NA | NA | NA | NA | NA |
| Hist1h2ba | 0 | 0 | 1 | 1 | 1 | 0 | 0 | 0 | 1 | 1 | 1 | 0 | 0.48 | 0 | 1 | 1.712 | 0.776 | 0.78603958 | NA | 0 | 6.9035 | 4.1008 | 4.1008 | TRUE | 0.4098 |
| Hist1h2bb | 5 | 6 | 6 | 4 | 7 | 1 | 5 | 6 | 6 | 3 | 8 | 1 | 4.85 | 6 | 4 | 0.675 | -0.568 | 0.53786606 | 0.72419528 | 0.0193 | 0.6862 | 0.3982 | 0.3982 | TRUE | 0.6952 |
| Hist1h2bc | 79 | 64 | 53 | 93 | 46 | 39 | 84 | 64 | 53 | 73 | 49 | 43 | 61.06 | 67 | 55 | 0.83 | -0.269 | 0.39564537 | 0.60902975 | 0.0507 | 0.0609 | 0.0559 | 0.0559 | TRUE | 0.6846 |
| Hist1h2be | 4 | 3 | 0 | 6 | 9 | 9 | 4 | 3 | 0 | 5 | 10 | 10 | 5.25 | 2 | 8 | 3.339 | 1.739 | 0.08368044 | 0.23178124 | 0.1188 | 0.6345 | 0.4547 | 0.4547 | TRUE | 0.8312 |

| | | | | | | | | | | | | | | | | | | | | | | | | |
|---|---|---|---|---|---|---|---|---|---|---|---|---|---|---|---|---|---|---|---|---|---|---|---|---|
| Hist1h2bf | 2 | 3 | 2 | 3 | 3 | 5 | 2 | 3 | 2 | 2 | 3 | 5 | 3.03 | 2 | 3 | 1.527 | 0.611 | 0.56758562 | NA | 0 | 1.0937 | 0.474 | 0.474 | TRUE | 0.2593 |
| Hist1h2bg | 13 | 14 | 12 | 24 | 15 | 13 | 14 | 14 | 12 | 19 | 16 | 14 | 14.85 | 13 | 16 | 1.244 | 0.315 | 0.51447641 | 0.70719208 | 0 | 0.2288 | 0.0998 | 0.0998 | TRUE | 0.1019 |
| Hist1h2bh | 2 | 3 | 2 | 9 | 3 | 3 | 2 | 3 | 2 | 7 | 3 | 3 | 3.45 | 2 | 4 | 1.94 | 0.956 | 0.35288434 | NA | 0 | 0.9611 | 0.4427 | 0.4427 | TRUE | 0.5795 |
| Hist1h2bj | 13 | 8 | 10 | 12 | 15 | 11 | 14 | 8 | 10 | 9 | 16 | 12 | 11.58 | 11 | 12 | 1.174 | 0.231 | 0.68650395 | 0.82716121 | 0 | 0.2915 | 0.1492 | 0.1492 | TRUE | 0.2469 |
| Hist1h2bk | 5 | 18 | 8 | 16 | 11 | 6 | 5 | 18 | 8 | 13 | 12 | 7 | 10.38 | 10 | 11 | 0.993 | -0.01 | 0.98803892 | 0.99364598 | 0.149 | 0.3242 | 0.239 | 0.239 | TRUE | 1.4011 |
| Hist1h2bl | 3 | 1 | 5 | 5 | 6 | 4 | 3 | 1 | 5 | 4 | 6 | 4 | 4 | 3 | 5 | 1.595 | 0.674 | 0.49245359 | NA | 0 | 0.8315 | 0.4274 | 0.4274 | TRUE | 0.344 |
| Hist1h2bm | 4 | 2 | 2 | 0 | 1 | 3 | 4 | 2 | 2 | 0 | 1 | 3 | 2.1 | 3 | 1 | 0.505 | -0.985 | 0.49621037 | NA | 0 | 1.5739 | 0.9723 | 0.9723 | TRUE | 0.7524 |
| Hist1h2bn | 0 | 3 | 5 | 2 | 1 | 1 | 0 | 3 | 5 | 2 | 1 | 1 | 1.96 | 3 | 1 | 0.47 | -1.089 | 0.46841852 | NA | 0 | 1.6878 | 1.0498 | 1.0498 | TRUE | 0.1781 |
| Hist1h2bp | 6 | 5 | 8 | 13 | 11 | 13 | 6 | 5 | 8 | 10 | 12 | 14 | 9.28 | 6 | 12 | 1.86 | 0.895 | 0.15362157 | 0.34534199 | 0 | 0.3619 | 0.1641 | 0.1641 | TRUE | 0.1146 |
| Hist1h2bq | 0 | 0 | 0 | 0 | 0 | 0 | 0 | 0 | 0 | 0 | 0 | 0 | 0 | 0 | 0 | NA | NA | NA | NA | NA | NA | NA | NA | NA | NA |
| Hist1h2br | 0 | 0 | 0 | 0 | 0 | 0 | 0 | 0 | 0 | 0 | 0 | 0 | 0 | 0 | 0 | NA | NA | NA | NA | NA | NA | NA | NA | NA | NA |
| Hist1h3a | 8 | 8 | 4 | 14 | 19 | 10 | 8 | 8 | 4 | 11 | 20 | 11 | 10.48 | 7 | 14 | 2.06 | 1.043 | 0.10560167 | 0.27095324 | 0 | 0.3212 | 0.1894 | 0.1894 | TRUE | 0.6321 |
| Hist1h3b | 2 | 2 | 3 | 4 | 4 | 3 | 2 | 2 | 3 | 3 | 4 | 3 | 2.98 | 2 | 3 | 1.496 | 0.581 | 0.58052421 | NA | 0 | 1.1133 | 0.4457 | 0.4457 | TRUE | 0.0583 |
| Hist1h3c | 32 | 22 | 33 | 44 | 42 | 40 | 34 | 22 | 33 | 35 | 45 | 44 | 35.44 | 30 | 41 | 1.382 | 0.466 | 0.1757788 | 0.37591637 | 0 | 0.0999 | 0.0564 | 0.0564 | TRUE | 0.3481 |
| Hist1h3d | 4 | 6 | 3 | 14 | 5 | 6 | 4 | 6 | 3 | 11 | 5 | 7 | 6.03 | 4 | 8 | 1.755 | 0.812 | 0.3078592 | 0.52834806 | 0 | 0.5528 | 0.2782 | 0.2782 | TRUE | 0.5288 |
| Hist1h3e | 13 | 16 | 14 | 12 | 21 | 14 | 14 | 16 | 14 | 9 | 23 | 15 | 15.2 | 15 | 16 | 1.065 | 0.091 | 0.86220058 | 0.93192349 | 0 | 0.2236 | 0.1338 | 0.1338 | TRUE | 0.2309 |
| Hist1h3f | 3 | 4 | 1 | 3 | 10 | 7 | 3 | 4 | 1 | 2 | 11 | 8 | 4.83 | 3 | 7 | 2.501 | 1.323 | 0.18877413 | 0.39393017 | 0 | 0.6893 | 0.4707 | 0.4707 | TRUE | 0.2364 |
| Hist1h3g | 6 | 7 | 14 | 28 | 17 | 9 | 6 | 7 | 14 | 22 | 18 | 10 | 12.93 | 9 | 17 | 1.841 | 0.88 | 0.14803022 | 0.3376819 | 0.0931 | 0.2617 | 0.182 | 0.182 | TRUE | 0.5401 |
| Hist1h3h | 8 | 8 | 7 | 6 | 8 | 4 | 8 | 8 | 7 | 5 | 9 | 4 | 6.87 | 8 | 6 | 0.747 | -0.421 | 0.5582428 | 0.74059171 | 0 | 0.4865 | 0.2236 | 0.2236 | TRUE | 0.3414 |
| Hist1h3i | 5 | 4 | 6 | 6 | 5 | 5 | 5 | 4 | 6 | 5 | 5 | 5 | 5.15 | 5 | 5 | 1.012 | 0.017 | 0.9828315 | 0.99099977 | 0 | 0.6466 | 0.2667 | 0.2667 | TRUE | 0.077 |
| Hist1h4a | 5 | 12 | 7 | 10 | 7 | 9 | 5 | 12 | 7 | 8 | 8 | 10 | 8.26 | 8 | 9 | 1.032 | 0.045 | 0.94591673 | 0.97280132 | 0 | 0.4057 | 0.2024 | 0.2024 | TRUE | 0.5263 |
| Hist1h4b | 8 | 12 | 4 | 21 | 19 | 16 | 8 | 12 | 4 | 17 | 20 | 18 | 13.16 | 8 | 18 | 2.218 | 1.149 | 0.04554136 | 0.15417008 | 0 | 0.2572 | 0.1484 | 0.1484 | TRUE | 0.5171 |
| Hist1h4c | 0 | 0 | 2 | 4 | 2 | 2 | 0 | 0 | 2 | 3 | 2 | 2 | 1.58 | 1 | 2 | 3.717 | 1.894 | 0.27339521 | NA | 0 | 2.0889 | 1.2047 | 1.2047 | TRUE | 0.9738 |
| Hist1h4d | 13 | 19 | 23 | 26 | 20 | 22 | 14 | 19 | 23 | 20 | 22 | 24 | 20.32 | 19 | 22 | 1.176 | 0.234 | 0.58474456 | 0.75998583 | 0 | 0.169 | 0.0828 | 0.0828 | TRUE | 0.2537 |
| Hist1h4f | 2 | 6 | 6 | 2 | 3 | 1 | 2 | 6 | 6 | 2 | 3 | 1 | 3.34 | 5 | 2 | 0.412 | -1.28 | 0.24222868 | NA | 0 | 0.9935 | 0.5054 | 0.5054 | TRUE | 0.4199 |
| Hist1h4h | 3 | 5 | 9 | 13 | 6 | 6 | 3 | 5 | 9 | 10 | 6 | 7 | 6.74 | 6 | 8 | 1.36 | 0.444 | 0.5593435 | 0.74125763 | 0 | 0.4955 | 0.265 | 0.265 | TRUE | 0.5752 |
| Hist1h4i | 86 | 69 | 55 | 214 | 144 | 61 | 91 | 69 | 55 | 168 | 155 | 67 | 100.9 | 72 | 130 | 1.816 | 0.861 | 0.01738168 | 0.07846133 | 0.134 | 0.0396 | 0.0835 | 0.0835 | TRUE | 1.8086 |
| Hist1h4j | 0 | 2 | 2 | 3 | 2 | 3 | 0 | 2 | 2 | 2 | 2 | 3 | 1.97 | 1 | 2 | 1.921 | 0.942 | 0.49685421 | NA | 0 | 1.6821 | 0.8167 | 0.8167 | TRUE | 0.436 |
| Hist1h4k | 6 | 14 | 10 | 11 | 16 | 13 | 6 | 14 | 10 | 9 | 17 | 14 | 11.75 | 10 | 13 | 1.301 | 0.38 | 0.52700816 | 0.71593432 | 0 | 0.2873 | 0.1721 | 0.1721 | TRUE | 0.2934 |
| Hist1h4m | 0 | 0 | 0 | 0 | 0 | 0 | 0 | 0 | 0 | 0 | 0 | 0 | 0 | 0 | 0 | NA | NA | NA | NA | NA | NA | NA | NA | NA | NA |
| Hist1h4n | 2 | 8 | 7 | 10 | 7 | 6 | 2 | 8 | 7 | 8 | 8 | 7 | 6.52 | 6 | 8 | 1.278 | 0.354 | 0.64663331 | 0.80159447 | 0 | 0.5124 | 0.2743 | 0.2743 | TRUE | 0.6415 |
| Hist2h2aa1 | 0 | 0 | 0 | 0 | 0 | 0 | 0 | 0 | 0 | 0 | 0 | 0 | 0 | 0 | 0 | NA | NA | NA | NA | NA | NA | NA | NA | NA | NA |
| Hist2h2aa2 | 0 | 0 | 0 | 0 | 1 | 0 | 0 | 0 | 0 | 0 | 1 | 0 | 0.18 | 0 | 0 | 1.859 | 0.894 | 0.82650734 | NA | 0 | 18.3785 | 10 | 10 | TRUE | 0.5204 |
| Hist2h2ab | 0 | 0 | 1 | 0 | 0 | 0 | 0 | 0 | 1 | 0 | 0 | 0 | 0.17 | 0 | 0 | 0.49 | -1.029 | 0.80087 | NA | 0 | 19.6928 | 10 | 10 | TRUE | 0.4098 |
| Hist2h2ac | 0 | 1 | 0 | 3 | 4 | 0 | 0 | 1 | 0 | 2 | 4 | 0 | 1.28 | 0 | 2 | 6.082 | 2.605 | 0.23147909 | NA | 0 | 2.5871 | 2.1939 | 2.1939 | TRUE | 0.1679 |
| Hist2h2bb | 3 | 11 | 13 | 12 | 8 | 11 | 3 | 11 | 13 | 9 | 9 | 12 | 9.55 | 9 | 10 | 1.098 | 0.135 | 0.84346285 | 0.92120325 | 0.0741 | 0.3518 | 0.2315 | 0.2315 | TRUE | 0.9872 |
| Hist2h2be | 29 | 28 | 42 | 44 | 19 | 32 | 31 | 28 | 42 | 35 | 20 | 35 | 31.83 | 34 | 30 | 0.896 | -0.159 | 0.67722866 | 0.82056964 | 0.0311 | 0.1104 | 0.0733 | 0.0733 | TRUE | 0.4844 |
| Hist2h3b | 3 | 2 | 2 | 5 | 4 | 2 | 3 | 2 | 2 | 4 | 4 | 2 | 2.94 | 2 | 3 | 1.462 | 0.548 | 0.60875593 | NA | 0 | 1.129 | 0.4731 | 0.4731 | TRUE | 0.136 |
| Hist2h3c1 | 30 | 12 | 13 | 19 | 11 | 20 | 32 | 12 | 13 | 15 | 12 | 22 | 17.59 | 19 | 16 | 0.856 | -0.225 | 0.68992919 | 0.82966922 | 0.1484 | 0.1941 | 0.1715 | 0.1715 | TRUE | 1.7973 |
| Hist2h3c2 | 3 | 4 | 2 | 6 | 4 | 4 | 3 | 4 | 2 | 5 | 4 | 4 | 3.76 | 3 | 4 | 1.461 | 0.547 | 0.56185433 | NA | 0 | 0.882 | 0.3658 | 0.3658 | TRUE | 0.1228 |
| Hist2h4 | 3 | 2 | 1 | 3 | 2 | 2 | 3 | 2 | 1 | 2 | 2 | 2 | 2.15 | 2 | 2 | 1.09 | 0.125 | 0.91960055 | NA | 0 | 1.5407 | 0.6314 | 0.6314 | TRUE | 0.1979 |
| Hist3h2a | 32 | 28 | 37 | 44 | 22 | 24 | 34 | 28 | 37 | 35 | 24 | 26 | 30.61 | 33 | 28 | 0.861 | -0.217 | 0.54571659 | 0.73083485 | 0 | 0.1145 | 0.06 | 0.06 | TRUE | 0.3041 |
| Hist3h2ba | 21 | 24 | 34 | 46 | 37 | 26 | 22 | 24 | 34 | 36 | 40 | 28 | 30.81 | 27 | 35 | 1.299 | 0.378 | 0.3019733 | 0.52259973 | 0 | 0.1138 | 0.0634 | 0.0634 | TRUE | 0.3636 |
| Hist3h2bb-ps | 0 | 0 | 0 | 0 | 1 | 0 | 0 | 0 | 0 | 0 | 1 | 0 | 0.18 | 0 | 0 | 1.859 | 0.894 | 0.82650734 | NA | 0 | 18.3785 | 10 | 10 | TRUE | 0.5204 |
| Hist4h4 | 10 | 7 | 3 | 15 | 9 | 12 | 11 | 7 | 3 | 12 | 10 | 13 | 9.21 | 7 | 12 | 1.686 | 0.754 | 0.26329176 | 0.48264194 | 0 | 0.3648 | 0.21 | 0.21 | TRUE | 0.3356 |
| Hivep1 | 881 | 918 | 1065 | 1413 | 1061 | 1063 | 936 | 917 | 1069 | 1111 | 1142 | 1164 | 1056.32 | 974 | 1139 | 1.169 | 0.225 | 0.01970985 | 0.08601559 | 0.0025 | 0.01 | 0.0057 | 0.0057 | TRUE | 0.0879 |
| Hivep2 | 17 | 27 | 38 | 67 | 36 | 38 | 18 | 27 | 38 | 53 | 39 | 42 | 36.03 | 28 | 45 | 1.603 | 0.68 | 0.0704691 | 0.20666393 | 0.0415 | 0.0983 | 0.0726 | 0.0726 | TRUE | 0.3969 |
| Hivep3 | 49 | 21 | 23 | 60 | 43 | 27 | 52 | 21 | 23 | 47 | 46 | 30 | 36.52 | 32 | 41 | 1.287 | 0.364 | 0.41005692 | 0.62158382 | 0.1299 | 0.0971 | 0.1127 | 0.1127 | TRUE | 1.9905 |
| Hjurp | 685 | 841 | 678 | 891 | 736 | 737 | 728 | 840 | 681 | 700 | 792 | 807 | 757.9 | 750 | 766 | 1.021 | 0.031 | 0.80017816 | 0.89686328 | 0.0075 | 0.0113 | 0.0091 | 0.0091 | TRUE | 0.1328 |
| Hk1 | 5882 | 9699 | 9717 | 6472 | 5571 | 7680 | 6248 | 9686 | 9757 | 5088 | 5994 | 8407 | 7529.98 | 8564 | 6496 | 0.758 | -0.399 | 0.05868164 | 0.18315291 | 0.0621 | 0.0073 | 0.0319 | 0.0319 | TRUE | 0.8075 |
| Hk2 | 16995 | 21402 | 22664 | 21413 | 18706 | 21561 | 18052 | 21374 | 22757 | 16833 | 20127 | 23602 | 20457.46 | 20728 | 20187 | 0.974 | -0.038 | 0.78651733 | 0.89023346 | 0.0167 | 0.0071 | 0.0143 | 0.0143 | TRUE | 0.2029 |
| Hk3 | 7 | 8 | 5 | 8 | 6 | 5 | 7 | 8 | 5 | 6 | 6 | 5 | 6.44 | 7 | 6 | 0.893 | -0.163 | 0.82151796 | 0.90875327 | 0 | 0.5182 | 0.2211 | 0.2211 | TRUE | 0.1402 |
| Hkdc1 | 32 | 20 | 21 | 37 | 30 | 16 | 34 | 20 | 21 | 29 | 32 | 18 | 25.65 | 25 | 26 | 1.057 | 0.08 | 0.85280592 | 0.92640367 | 0.0499 | 0.1353 | 0.0951 | 0.0951 | TRUE | 0.58 |
| Hlcs | 690 | 827 | 890 | 964 | 732 | 676 | 733 | 826 | 894 | 758 | 788 | 740 | 789.65 | 818 | 762 | 0.932 | -0.102 | 0.34243499 | 0.56254916 | 0.0041 | 0.0111 | 0.0071 | 0.0071 | TRUE | 0.0966 |
| Hlf | 41 | 42 | 39 | 55 | 35 | 36 | 44 | 42 | 39 | 43 | 38 | 39 | 40.83 | 42 | 40 | 0.969 | -0.046 | 0.87497181 | 0.93826865 | 0 | 0.0876 | 0.0376 | 0.0376 | TRUE | 0.0415 |
| Hltf | 4386 | 4905 | 5299 | 4946 | 3743 | 4380 | 4659 | 4899 | 5321 | 3888 | 4027 | 4795 | 4598.02 | 4960 | 4237 | 0.854 | -0.228 | 0.03393495 | 0.12537411 | 0.0086 | 0.0076 | 0.0081 | 0.0081 | TRUE | 0.1612 |
| Hlx | 5 | 6 | 2 | 12 | 14 | 11 | 5 | 6 | 2 | 9 | 15 | 12 | 8.31 | 4 | 12 | 2.732 | 1.45 | 0.04363466 | 0.14952757 | 0 | 0.4034 | 0.2165 | 0.2165 | TRUE | 0.3712 |
| Hmbox1 | 579 | 670 | 671 | 871 | 673 | 654 | 615 | 669 | 674 | 685 | 724 | 716 | 680.44 | 653 | 708 | 1.084 | 0.117 | 0.22003682 | 0.43222937 | 0 | 0.0117 | 0.005 | 0.005 | TRUE | 0.0297 |
| Hmbs | 3020 | 4069 | 4279 | 4190 | 3055 | 3422 | 3208 | 4064 | 4297 | 3294 | 3287 | 3746 | 3649.14 | 3856 | 3442 | 0.893 | -0.164 | 0.1950951 | 0.40091726 | 0.0136 | 0.0078 | 0.0113 | 0.0113 | TRUE | 0.2629 |
| Hmcn1 | 103 | 80 | 63 | 139 | 60 | 61 | 109 | 80 | 63 | 109 | 65 | 67 | 82.19 | 84 | 80 | 0.96 | -0.059 | 0.85017187 | 0.92539067 | 0.0717 | 0.047 | 0.0591 | 0.0591 | TRUE | 0.6206 |
| Hmg20a | 2664 | 2696 | 2844 | 3138 | 2434 | 2519 | 2830 | 2693 | 2856 | 2467 | 2619 | 2757 | 2703.49 | 2793 | 2614 | 0.936 | -0.096 | 0.21737158 | 0.4294633 | 0.0015 | 0.0081 | 0.004 | 0.004 | TRUE | 0.0302 |
| Hmg20b | 826 | 857 | 797 | 1327 | 935 | 886 | 877 | 856 | 800 | 1043 | 1006 | 970 | 925.43 | 844 | 1006 | 1.193 | 0.254 | 0.00422615 | 0.02755469 | 7.00E-04 | 0.0105 | 0.0046 | 0.0046 | TRUE | 0.025 |
| Hmga1 | 132 | 207 | 195 | 234 | 185 | 178 | 140 | 207 | 196 | 184 | 199 | 195 | 186.77 | 181 | 193 | 1.063 | 0.088 | 0.64032688 | 0.79767036 | 0.0154 | 0.0245 | 0.02 | 0.02 | TRUE | 0.4096 |
| Hmga1-rs1 | 0 | 0 | 0 | 0 | 0 | 0 | 0 | 0 | 0 | 0 | 0 | 0 | 0 | 0 | 0 | NA | NA | NA | NA | NA | NA | NA | NA | NA | NA |
| Hmga2 | 1513 | 1362 | 1355 | 2787 | 1952 | 1599 | 1607 | 1360 | 1361 | 2191 | 2100 | 1750 | 1728.23 | 1443 | 2014 | 1.397 | 0.482 | 7.34E-05 | 0.00107696 | 0.0107 | 0.0088 | 0.0101 | 0.0101 | TRUE | 0.1576 |
| Hmga2-ps1 | 46 | 73 | 70 | 45 | 55 | 46 | 49 | 73 | 70 | 35 | 59 | 50 | 56.16 | 64 | 48 | 0.748 | -0.419 | 0.18019116 | 0.38152437 | 0.0382 | 0.0656 | 0.0523 | 0.0523 | TRUE | 0.5127 |
| Hmgb1 | 5070 | 5092 | 4810 | 7194 | 5005 | 4702 | 5385 | 5085 | 4830 | 5655 | 5385 | 5147 | 5247.99 | 5100 | 5396 | 1.058 | 0.082 | 0.2958812 | 0.51662108 | 0.0023 | 0.0075 | 0.0042 | 0.0042 | TRUE | 0.0291 |
| Hmgb1-rs17 | 1 | 0 | 3 | 2 | 0 | 0 | 1 | 0 | 3 | 2 | 0 | 0 | 0.94 | 1 | 1 | 0.416 | -1.264 | 0.59609826 | NA | 0 | 3.5075 | 2.8584 | 2.8584 | TRUE | 0.4749 |
| Hmgb2 | 11899 | 16951 | 17048 | 25104 | 17825 | 16796 | 12639 | 16929 | 17118 | 19735 | 19179 | 18386 | 17330.87 | 15562 | 19100 | 1.227 | 0.296 | 0.01830166 | 0.08160978 | 0.0138 | 0.0071 | 0.0112 | 0.0112 | TRUE | 0.3301 |

| | | | | | | | | | | | | | | | | | | | | | | | | | |
|---|---|---|---|---|---|---|---|---|---|---|---|---|---|---|---|---|---|---|---|---|---|---|---|---|---|
| Hmgb3 | 823 | 1054 | 1008 | 1884 | 1320 | 1201 | 874 | 1053 | 1012 | 1481 | 1420 | 1315 | 1192.49 | 980 | 1405 | 1.435 | 0.521 | 9.35E-07 | 2.60E-05 | 0.0051 | 0.0097 | 0.0073 | 0.0073 | TRUE | 0.1052 |
| Hmgb4 | 0 | 0 | 1 | 1 | 0 | 0 | 0 | 0 | 1 | 1 | 0 | 0 | 0.3 | 0 | 0 | 0.954 | -0.067 | 0.98640617 | NA | 0 | 11.0483 | 9.2783 | 9.2783 | TRUE | 0.4098 |
| Hmgcl | 1377 | 1536 | 1501 | 1872 | 1333 | 1384 | 1463 | 1534 | 1507 | 1472 | 1434 | 1515 | 1487.44 | 1501 | 1474 | 0.981 | -0.027 | 0.71494136 | 0.8452815 | 0 | 0.0091 | 0.0033 | 0.0033 | TRUE | 0.007 |
| Hmgcll1 | 207 | 211 | 227 | 303 | 211 | 211 | 220 | 211 | 228 | 238 | 227 | 231 | 225.79 | 220 | 232 | 1.058 | 0.082 | 0.54602652 | 0.73106369 | 0 | 0.0215 | 0.0087 | 0.0087 | TRUE | 0.0137 |
| Hmgcr | 15266 | 14330 | 14590 | 16315 | 12222 | 12902 | 16215 | 14311 | 14650 | 12826 | 13150 | 14123 | 14212.59 | 15059 | 13366 | 0.888 | -0.172 | 0.03549469 | 0.12941966 | 0.0034 | 0.0071 | 0.0048 | 0.0048 | TRUE | 0.0552 |
| Hmgcs1 | 19856 | 18129 | 18367 | 22117 | 16201 | 17109 | 21091 | 18106 | 18442 | 17387 | 17431 | 18728 | 18530.83 | 19213 | 17849 | 0.929 | -0.106 | 0.22097832 | 0.43311388 | 0.0045 | 0.0071 | 0.0054 | 0.0054 | TRUE | 0.0894 |
| Hmgcs2 | 98 | 91 | 53 | 130 | 90 | 92 | 104 | 91 | 53 | 102 | 97 | 101 | 91.32 | 83 | 100 | 1.209 | 0.274 | 0.31659017 | 0.53748764 | 0.043 | 0.043 | 0.043 | 0.043 | TRUE | 0.9186 |
| Hmgn1 | 23371 | 24141 | 23973 | 30708 | 21843 | 20235 | 24824 | 24110 | 24071 | 24140 | 23502 | 22150 | 23799.62 | 24335 | 23264 | 0.956 | -0.065 | 0.3071944 | 0.52755554 | 0.001 | 0.007 | 0.0029 | 0.0029 | TRUE | 0.0214 |
| Hmgn2 | 1607 | 1630 | 1559 | 3000 | 2080 | 1686 | 1707 | 1628 | 1565 | 2358 | 2238 | 1846 | 1890.35 | 1633 | 2147 | 1.315 | 0.395 | 0.00045697 | 0.00484685 | 0.0079 | 0.0086 | 0.0086 | 0.0086 | TRUE | 0.1819 |
| Hmgn3 | 352 | 367 | 336 | 614 | 491 | 419 | 374 | 367 | 337 | 483 | 528 | 459 | 424.57 | 359 | 490 | 1.364 | 0.447 | 0.00014079 | 0.00181033 | 0.0016 | 0.0147 | 0.0075 | 0.0075 | TRUE | 0.0528 |
| Hmgn5 | 1391 | 1149 | 1136 | 1818 | 1219 | 1054 | 1477 | 1148 | 1141 | 1429 | 1312 | 1154 | 1276.7 | 1255 | 1298 | 1.035 | 0.05 | 0.71943251 | 0.8479218 | 0.0161 | 0.0095 | 0.013 | 0.013 | TRUE | 0.2873 |
| Hmgxb3 | 4849 | 4752 | 4623 | 5674 | 4071 | 4069 | 5150 | 4746 | 4642 | 4460 | 4380 | 4454 | 4638.85 | 4846 | 4431 | 0.915 | -0.129 | 0.07025889 | 0.20627759 | 0.0013 | 0.0076 | 0.0034 | 0.0034 | TRUE | 0.0367 |
| Hmgxb4 | 3581 | 3977 | 4231 | 4397 | 3226 | 3255 | 3804 | 3972 | 4248 | 3457 | 3471 | 3563 | 3752.43 | 4008 | 3497 | 0.872 | -0.197 | 0.0071423 | 0.04068194 | 0.0013 | 0.0078 | 0.0036 | 0.0036 | TRUE | 0.0335 |
| Hmha1 | 1201 | 1819 | 1904 | 873 | 832 | 1162 | 1276 | 1817 | 1912 | 686 | 895 | 1272 | 1309.6 | 1668 | 951 | 0.57 | -0.812 | 0.00032725 | 0.00365534 | 0.0686 | 0.0094 | 0.036 | 0.036 | TRUE | 0.8162 |
| Hmmr | 13340 | 16629 | 17189 | 16353 | 11889 | 13620 | 14169 | 16608 | 17260 | 12855 | 12792 | 14909 | 14765.51 | 16012 | 13519 | 0.844 | -0.244 | 0.02318447 | 0.0965516 | 0.009 | 0.0071 | 0.0083 | 0.0083 | TRUE | 0.1239 |
| Hmox1 | 3354 | 2915 | 2806 | 4716 | 3352 | 2875 | 3563 | 2911 | 2818 | 3707 | 3607 | 3147 | 3292.06 | 3097 | 3487 | 1.126 | 0.171 | 0.15135705 | 0.34239864 | 0.0118 | 0.0079 | 0.01 | 0.01 | TRUE | 0.2097 |
| Hmox2 | 1645 | 1609 | 1628 | 2313 | 1603 | 1515 | 1747 | 1607 | 1635 | 1818 | 1725 | 1658 | 1698.39 | 1663 | 1734 | 1.043 | 0.061 | 0.45334305 | 0.65875499 | 0.0015 | 0.0088 | 0.0042 | 0.0042 | TRUE | 0.0236 |
| Hmx1 | 36 | 48 | 64 | 120 | 98 | 74 | 38 | 48 | 64 | 94 | 105 | 81 | 71.87 | 50 | 93 | 1.864 | 0.899 | 0.00100632 | 0.00895503 | 0.0208 | 0.0527 | 0.0384 | 0.0384 | TRUE | 0.4943 |
| Hmx2 | 2 | 11 | 5 | 6 | 10 | 12 | 2 | 11 | 5 | 5 | 11 | 13 | 7.79 | 6 | 10 | 1.552 | 0.634 | 0.4380758 | 0.64579729 | 0.2536 | 0.4298 | 0.3453 | 0.3453 | TRUE | 0.7843 |
| Hmx3 | 1 | 3 | 1 | 4 | 5 | 3 | 1 | 3 | 1 | 3 | 5 | 3 | 2.81 | 2 | 4 | 2.308 | 1.207 | 0.2978694 | NA | 0 | 1.1786 | 0.5418 | 0.5418 | TRUE | 0.3596 |
| Hn1 | 7292 | 8372 | 7926 | 9769 | 6917 | 6728 | 7745 | 8361 | 7959 | 7680 | 7442 | 7365 | 7758.64 | 8022 | 7496 | 0.934 | -0.098 | 0.12861104 | 0.3083693 | 8.00E-04 | 0.0073 | 0.0029 | 0.0029 | TRUE | 0.0167 |
| Hn1l | 11223 | 12070 | 11808 | 14067 | 9874 | 10256 | 11921 | 12054 | 11856 | 11058 | 10624 | 11227 | 11456.78 | 11944 | 10970 | 0.918 | -0.123 | 0.03143567 | 0.11909762 | 3.00E-04 | 0.0072 | 0.0023 | 0.0023 | TRUE | 0.0092 |
| Hnf1a | 298 | 316 | 324 | 299 | 252 | 253 | 317 | 316 | 325 | 235 | 271 | 277 | 290.1 | 319 | 261 | 0.815 | -0.294 | 0.02481336 | 0.1012303 | 0 | 0.0183 | 0.0089 | 0.0089 | TRUE | 0.0923 |
| Hnf1b | 94 | 90 | 90 | 105 | 82 | 50 | 100 | 90 | 90 | 83 | 88 | 55 | 84.27 | 93 | 75 | 0.808 | -0.307 | 0.21376655 | 0.42521453 | 0.0157 | 0.046 | 0.0321 | 0.0321 | TRUE | 0.4846 |
| Hnf4a | 217 | 252 | 263 | 252 | 238 | 315 | 230 | 252 | 264 | 198 | 256 | 345 | 257.54 | 249 | 266 | 1.067 | 0.094 | 0.65830923 | 0.80743861 | 0.0373 | 0.0197 | 0.0285 | 0.0285 | TRUE | 0.3149 |
| Hnf4g | 0 | 0 | 0 | 0 | 1 | 0 | 0 | 0 | 0 | 0 | 1 | 0 | 0.18 | 0 | 0 | 1.859 | 0.894 | 0.82650734 | NA | 0 | 18.3785 | 10 | 10 | TRUE | 0.5204 |
| Hnmt | 3 | 2 | 1 | 2 | 8 | 3 | 3 | 2 | 1 | 2 | 9 | 3 | 3.28 | 2 | 5 | 2.149 | 1.103 | 0.34411932 | NA | 0 | 1.0128 | 0.6207 | 0.6207 | TRUE | 0.6504 |
| Hnrnpa0 | 14184 | 14664 | 13691 | 21582 | 16561 | 14137 | 15066 | 14645 | 13747 | 16966 | 17819 | 15475 | 15619.68 | 14486 | 16753 | 1.157 | 0.21 | 0.01108881 | 0.05680195 | 0.0035 | 0.0071 | 0.0049 | 0.0049 | TRUE | 0.0544 |
| Hnrnpa1 | 1643 | 1989 | 1957 | 2252 | 1741 | 1940 | 1745 | 1986 | 1965 | 1770 | 1873 | 2124 | 1910.64 | 1899 | 1922 | 1.012 | 0.017 | 0.87209783 | 0.93735362 | 0.0064 | 0.0086 | 0.0073 | 0.0073 | TRUE | 0.101 |
| Hnrnpa2b1 | 56926 | 53908 | 53495 | 80246 | 58021 | 55252 | 60465 | 53838 | 53715 | 63083 | 62427 | 60482 | 59001.78 | 56006 | 61997 | 1.107 | 0.147 | 0.05054128 | 0.16534682 | 0.0026 | 0.007 | 0.004 | 0.004 | TRUE | 0.0594 |
| Hnrnpa3 | 1932 | 2286 | 2227 | 2642 | 1816 | 2047 | 2052 | 2283 | 2236 | 2077 | 1954 | 2241 | 2140.49 | 2190 | 2091 | 0.954 | -0.068 | 0.45231625 | 0.6576871 | 0.0032 | 0.0084 | 0.0054 | 0.0054 | TRUE | 0.0471 |
| Hnrnpab | 43875 | 41208 | 39517 | 60083 | 42991 | 38621 | 46603 | 41155 | 39679 | 47233 | 46256 | 42277 | 43867 | 42479 | 45255 | 1.065 | 0.091 | 0.31505599 | 0.5358636 | 0.0053 | 0.007 | 0.0059 | 0.0059 | TRUE | 0.0883 |
| Hnrnpc | 22070 | 25647 | 24810 | 33098 | 22633 | 21619 | 23442 | 25614 | 24912 | 26019 | 24352 | 23665 | 24667.36 | 24656 | 24679 | 1.001 | 0.001 | 0.98470234 | 0.9919379 | 0.0017 | 0.007 | 0.0038 | 0.0038 | TRUE | 0.0277 |
| Hnrnpd | 17458 | 12563 | 12107 | 21747 | 16441 | 13121 | 18543 | 12547 | 12157 | 17096 | 17690 | 14363 | 15399.21 | 14416 | 16383 | 1.136 | 0.185 | 0.2716577 | 0.49168984 | 0.0318 | 0.0071 | 0.0203 | 0.0203 | TRUE | 0.7672 |
| Hnrnpf | 17661 | 18965 | 18922 | 25410 | 18768 | 17536 | 18759 | 18941 | 19000 | 19975 | 20193 | 19196 | 19343.96 | 18900 | 19788 | 1.047 | 0.066 | 0.22893943 | 0.44224019 | 3.00E-04 | 0.0071 | 0.0021 | 0.0021 | TRUE | 0.0084 |
| Hnrnph1 | 12783 | 9459 | 9010 | 15090 | 11293 | 9155 | 13578 | 9447 | 9047 | 11863 | 12151 | 10022 | 11017.72 | 10691 | 11345 | 1.061 | 0.086 | 0.5955389 | 0.76839329 | 0.0261 | 0.0072 | 0.0187 | 0.0187 | TRUE | 0.682 |
| Hnrnph2 | 3513 | 3347 | 3278 | 4224 | 2992 | 3001 | 3731 | 3343 | 3291 | 3321 | 3219 | 3285 | 3365.07 | 3455 | 3275 | 0.948 | -0.077 | 0.33125404 | 0.55212268 | 0.0021 | 0.0079 | 0.0042 | 0.0042 | TRUE | 0.0593 |
| Hnrnph3 | 2728 | 3525 | 3656 | 3706 | 2977 | 2905 | 2898 | 3520 | 3671 | 2913 | 3203 | 3180 | 3230.92 | 3363 | 3099 | 0.921 | -0.118 | 0.28414387 | 0.50575748 | 0.0083 | 0.0079 | 0.0085 | 0.0085 | TRUE | 0.1779 |
| Hnrnpk | 33774 | 34757 | 34035 | 45246 | 32727 | 31826 | 35874 | 34712 | 34175 | 35569 | 35212 | 34839 | 35063.41 | 34920 | 35207 | 1.008 | 0.012 | 0.82738969 | 0.91262257 | 3.00E-04 | 0.007 | 0.0021 | 0.0021 | TRUE | 0.007 |
| Hnrnpl | 27166 | 27853 | 27911 | 33792 | 25772 | 26097 | 28855 | 27817 | 28026 | 26565 | 27729 | 28567 | 27926.45 | 28233 | 27620 | 0.978 | -0.032 | 0.60103249 | 0.77268877 | 8.00E-04 | 0.007 | 0.0026 | 0.0026 | TRUE | 0.0137 |
| Hnrnpm | 18037 | 16008 | 16318 | 25887 | 19357 | 16867 | 19158 | 15987 | 16385 | 20350 | 20827 | 18464 | 18528.61 | 17177 | 19880 | 1.157 | 0.211 | 0.03235572 | 0.12130127 | 0.007 | 0.0071 | 0.0069 | 0.0069 | TRUE | 0.1245 |
| Hnrnpr | 3166 | 2901 | 3114 | 5150 | 3800 | 3387 | 3363 | 2897 | 3127 | 4049 | 4089 | 3708 | 3538.6 | 3129 | 3949 | 1.262 | 0.336 | 0.00015515 | 0.00195939 | 0.0037 | 0.0078 | 0.0054 | 0.0054 | TRUE | 0.0518 |
| Hnrnpu | 47318 | 47176 | 48762 | 68929 | 52065 | 47556 | 50260 | 47115 | 48962 | 54187 | 56019 | 52058 | 51433.37 | 48779 | 54088 | 1.109 | 0.149 | 0.01957541 | 0.08552446 | 0.0012 | 0.007 | 0.0029 | 0.0029 | TRUE | 0.0132 |
| Hnrnpul1 | 8145 | 9595 | 10159 | 11866 | 9263 | 9522 | 8651 | 9583 | 10201 | 9328 | 9966 | 10423 | 9692.1 | 9478 | 9906 | 1.045 | 0.063 | 0.482476 | 0.68094957 | 0.0044 | 0.0072 | 0.0058 | 0.0058 | TRUE | 0.0711 |
| Hnrnpul2 | 14041 | 14116 | 14320 | 18976 | 13820 | 13815 | 14914 | 14098 | 14379 | 14917 | 14869 | 15123 | 14716.7 | 14464 | 14970 | 1.035 | 0.05 | 0.38026274 | 0.59475446 | 3.00E-04 | 0.0071 | 0.0022 | 0.0022 | TRUE | 0.0091 |
| Hnrpdl | 7273 | 6531 | 6112 | 9319 | 7124 | 6515 | 7725 | 6523 | 6137 | 7326 | 7665 | 7132 | 7084.57 | 6795 | 7374 | 1.085 | 0.118 | 0.2535542 | 0.4714757 | 0.0079 | 0.0074 | 0.0076 | 0.0076 | TRUE | 0.1749 |
| Hnrpll | 3652 | 4536 | 4728 | 5769 | 4357 | 4364 | 3879 | 4530 | 4747 | 4535 | 4688 | 4777 | 4526.12 | 4385 | 4667 | 1.064 | 0.089 | 0.35087211 | 0.56972959 | 0.0052 | 0.0076 | 0.0064 | 0.0064 | TRUE | 0.1241 |
| Hoga1 | 6 | 13 | 11 | 38 | 21 | 18 | 6 | 13 | 11 | 30 | 23 | 20 | 17.1 | 10 | 24 | 2.382 | 1.252 | 0.01312349 | 0.0641972 | 0 | 0.1996 | 0.1131 | 0.1131 | TRUE | 0.3486 |
| Homer1 | 1452 | 1598 | 1633 | 2445 | 1834 | 1755 | 1542 | 1596 | 1640 | 1922 | 1973 | 1921 | 1765.73 | 1593 | 1939 | 1.217 | 0.283 | 6.64E-05 | 0.00099089 | 0 | 0.0088 | 0.0031 | 0.0031 | TRUE | 0.0092 |
| Homer2 | 815 | 1118 | 1200 | 1328 | 1019 | 974 | 866 | 1117 | 1205 | 1044 | 1096 | 1066 | 1065.62 | 1063 | 1069 | 1.006 | 0.008 | 0.95161839 | 0.97536556 | 0.0137 | 0.01 | 0.012 | 0.012 | TRUE | 0.3125 |
| Homer3 | 507 | 524 | 470 | 894 | 607 | 635 | 539 | 523 | 472 | 703 | 653 | 695 | 597.46 | 511 | 684 | 1.338 | 0.42 | 5.49E-05 | 0.00083794 | 0.0011 | 0.0124 | 0.0061 | 0.0061 | TRUE | 0.053 |
| Homez | 389 | 298 | 318 | 407 | 283 | 200 | 413 | 298 | 319 | 320 | 304 | 219 | 312.25 | 343 | 281 | 0.821 | -0.285 | 0.14641747 | 0.33511773 | 0.0318 | 0.0175 | 0.0246 | 0.0246 | TRUE | 0.4121 |
| Hook1 | 8208 | 7814 | 7948 | 10083 | 7249 | 7193 | 8718 | 7804 | 7981 | 7926 | 7799 | 7874 | 8017.12 | 8168 | 7866 | 0.963 | -0.054 | 0.44617723 | 0.65236459 | 0.0016 | 0.0073 | 0.0035 | 0.0035 | TRUE | 0.0424 |
| Hook2 | 3481 | 4027 | 3962 | 3644 | 3109 | 3584 | 3697 | 4022 | 3978 | 2865 | 3345 | 3923 | 3638.41 | 3899 | 3378 | 0.866 | -0.208 | 0.09213279 | 0.24762974 | 0.0129 | 0.0078 | 0.0107 | 0.0107 | TRUE | 0.2427 |
| Hook3 | 1203 | 1237 | 1283 | 1998 | 1561 | 1419 | 1278 | 1235 | 1288 | 1571 | 1680 | 1553 | 1434.17 | 1267 | 1601 | 1.263 | 0.337 | 1.58E-05 | 0.00029165 | 5.00E-04 | 0.0092 | 0.0037 | 0.0037 | TRUE | 0.0217 |
| Hopx | 158 | 207 | 175 | 159 | 137 | 152 | 168 | 207 | 176 | 125 | 147 | 166 | 164.84 | 184 | 146 | 0.794 | -0.333 | 0.07180465 | 0.20910321 | 0.0105 | 0.0269 | 0.0185 | 0.0185 | TRUE | 0.1856 |
| Hormad1 | 296 | 434 | 396 | 1056 | 805 | 896 | 314 | 433 | 398 | 830 | 866 | 981 | 637.09 | 382 | 892 | 2.334 | 1.223 | 2.83E-17 | 5.13E-15 | 0.0139 | 0.0121 | 0.0132 | 0.0132 | TRUE | 0.2712 |
| Hormad2 | 90 | 141 | 143 | 329 | 225 | 246 | 96 | 141 | 144 | 259 | 242 | 269 | 191.67 | 127 | 257 | 2.024 | 1.017 | 8.75E-08 | 3.20E-06 | 0.0152 | 0.0241 | 0.0201 | 0.0201 | TRUE | 0.4599 |
| Hotair | 0 | 0 | 0 | 0 | 0 | 0 | 0 | 0 | 0 | 0 | 0 | 0 | 0 | 0 | 0 | NA | NA | NA | NA | NA | NA | NA | NA | NA | NA |
| Hoxa1 | 116 | 120 | 104 | 293 | 190 | 150 | 123 | 120 | 104 | 230 | 204 | 164 | 157.74 | 116 | 199 | 1.73 | 0.791 | 4.92E-05 | 0.0007658 | 0.0137 | 0.0278 | 0.0205 | 0.0205 | TRUE | 0.2153 |
| Hoxa10 | 0 | 3 | 2 | 16 | 12 | 6 | 0 | 3 | 2 | 13 | 13 | 7 | 6.18 | 2 | 11 | 6.352 | 2.667 | 0.00660589 | 0.03858817 | 0 | 0.5403 | 0.347 | 0.347 | TRUE | 0.5335 |
| Hoxa11 | 0 | 0 | 1 | 3 | 3 | 1 | 0 | 0 | 1 | 2 | 3 | 1 | 1.28 | 0 | 2 | 6.1 | 2.609 | 0.16990347 | NA | 0 | 2.579 | 1.3839 | 1.3839 | TRUE | 0.393 |
| Hoxa11as | 0 | 3 | 0 | 4 | 2 | 1 | 0 | 3 | 0 | 3 | 2 | 1 | 1.56 | 1 | 2 | 2.139 | 1.097 | 0.53383026 | NA | 0.4192 | 2.1126 | 1.5062 | 1.5062 | TRUE | 1.2673 |
| Hoxa13 | 3 | 10 | 7 | 16 | 13 | 13 | 3 | 10 | 7 | 13 | 14 | 14 | 10.17 | 7 | 14 | 2.005 | 1.004 | 0.11358644 | 0.28422741 | 0 | 0.331 | 0.1783 | 0.1783 | TRUE | 0.4377 |
| Hoxa2 | 0 | 0 | 0 | 1 | 2 | 1 | 0 | 0 | 0 | 1 | 2 | 1 | 0.67 | 0 | 1 | 7.1 | 2.828 | 0.25693906 | NA | 0 | 4.9086 | 3.1164 | 3.1164 | TRUE | 0.1607 |
| Hoxa3 | 5 | 6 | 1 | 27 | 10 | 5 | 5 | 6 | 1 | 21 | 11 | 5 | 8.29 | 4 | 12 | 3.088 | 1.627 | 0.05906111 | 0.1840205 | 0.3335 | 0.4041 | 0.3716 | 0.3716 | TRUE | 0.2407 |
| Hoxa4 | 11 | 19 | 12 | 25 | 15 | 15 | 12 | 19 | 12 | 20 | 16 | 16 | 15.82 | 14 | 17 | 1.226 | 0.294 | 0.54411516 | 0.72918551 | 0 | 0.2151 | 0.1058 | 0.1058 | TRUE | 0.3796 |
| Hoxa5 | 6 | 8 | 5 | 21 | 7 | 7 | 6 | 8 | 5 | 17 | 8 | 8 | 8.51 | 6 | 11 | 1.663 | 0.734 | 0.29190546 | 0.51274451 | 0 | 0.3938 | 0.2246 | 0.2246 | TRUE | 1.0741 |
| Hoxa6 | 0 | 0 | 0 | 1 | 1 | 0 | 0 | 0 | 0 | 1 | 1 | 0 | 0.31 | 0 | 1 | 3.32 | 1.731 | 0.65383052 | NA | 0 | 10.6224 | 8.9541 | 8.9541 | TRUE | 0.202 |

| | | | | | | | | | | | | | | | | | | | | | | | | | |
|---|---|---|---|---|---|---|---|---|---|---|---|---|---|---|---|---|---|---|---|---|---|---|---|---|---|
| Hoxa7 | 3 | 6 | 10 | 24 | 18 | 8 | 3 | 6 | 10 | 19 | 19 | 9 | 11.04 | 6 | 16 | 2.448 | 1.292 | 0.05492207 | 0.17478428 | 0.1047 | 0.3054 | 0.2158 | 0.2158 | TRUE | 0.6338 |
| Hoxa9 | 5 | 6 | 3 | 13 | 9 | 9 | 5 | 6 | 3 | 10 | 10 | 10 | 7.35 | 5 | 10 | 2.082 | 1.058 | 0.13422612 | 0.31680062 | 0 | 0.4554 | 0.2035 | 0.2035 | TRUE | 0.2064 |
| Hoxb1 | 4 | 1 | 0 | 6 | 6 | 2 | 4 | 1 | 0 | 5 | 6 | 2 | 3.1 | 2 | 4 | 2.582 | 1.369 | 0.28451715 | NA | 0.2875 | 1.0691 | 0.7732 | 0.7732 | TRUE | 0.8258 |
| Hoxb13 | 58 | 85 | 61 | 79 | 49 | 41 | 62 | 85 | 61 | 62 | 53 | 45 | 61.24 | 69 | 53 | 0.773 | -0.372 | 0.18017514 | 0.38152437 | 0.0167 | 0.0607 | 0.0391 | 0.0391 | TRUE | 0.3533 |
| Hoxb2 | 3 | 2 | 1 | 12 | 2 | 5 | 3 | 2 | 1 | 9 | 2 | 5 | 3.87 | 2 | 5 | 2.817 | 1.494 | 0.16747692 | NA | 0 | 0.8572 | 0.5108 | 0.5108 | TRUE | 0.1205 |
| Hoxb3 | 1 | 1 | 0 | 2 | 2 | 1 | 1 | 1 | 0 | 2 | 2 | 1 | 1.15 | 1 | 2 | 2.35 | 1.233 | 0.4876953 | NA | 0 | 2.8801 | 1.2223 | 1.2223 | TRUE | 0.2531 |
| Hoxb4 | 19 | 15 | 17 | 53 | 17 | 18 | 20 | 15 | 17 | 42 | 18 | 20 | 21.98 | 17 | 27 | 1.547 | 0.629 | 0.19897126 | 0.4062556 | 0.0962 | 0.1568 | 0.1255 | 0.1255 | TRUE | 1.7892 |
| Hoxb5 | 1 | 0 | 0 | 4 | 0 | 0 | 1 | 0 | 0 | 3 | 0 | 0 | 0.7 | 0 | 1 | 2.951 | 1.561 | 0.61103313 | NA | 7.9145 | 4.7058 | 5.3344 | 5.3344 | TRUE | 1.8958 |
| Hoxb6 | 0 | 1 | 1 | 1 | 1 | 0 | 0 | 1 | 1 | 1 | 1 | 0 | 0.64 | 1 | 1 | 0.935 | -0.097 | 0.96674856 | NA | 0 | 5.1213 | 2.3899 | 2.3899 | TRUE | 0.2246 |
| Hoxb7 | 17 | 38 | 33 | 64 | 21 | 30 | 18 | 38 | 33 | 50 | 23 | 33 | 32.48 | 30 | 35 | 1.194 | 0.256 | 0.56527945 | 0.74551564 | 0.1165 | 0.1083 | 0.112 | 0.112 | TRUE | 0.361 |
| Hoxb8 | 1 | 2 | 0 | 1 | 0 | 2 | 1 | 2 | 0 | 1 | 0 | 2 | 1.01 | 1 | 1 | 0.963 | -0.054 | 0.97862174 | NA | 0 | 3.2822 | 1.9784 | 1.9784 | TRUE | 0.2782 |
| Hoxb9 | 104 | 105 | 80 | 102 | 71 | 77 | 110 | 105 | 80 | 80 | 76 | 84 | 89.42 | 98 | 80 | 0.815 | -0.295 | 0.18605331 | 0.38980244 | 0 | 0.0437 | 0.0246 | 0.0246 | TRUE | 0.2566 |
| Hoxc10 | 11 | 11 | 3 | 7 | 4 | 6 | 12 | 11 | 3 | 6 | 4 | 7 | 7.01 | 9 | 6 | 0.639 | -0.647 | 0.41120183 | 0.62234673 | 0 | 0.4769 | 0.2972 | 0.2972 | TRUE | 1.0079 |
| Hoxc11 | 0 | 0 | 1 | 0 | 0 | 0 | 0 | 0 | 1 | 0 | 0 | 0 | 0.17 | 0 | 0 | 0.49 | -1.029 | 0.80087 | NA | 0 | 19.6928 | 10 | 10 | TRUE | 0.4098 |
| Hoxc12 | 3 | 9 | 7 | 9 | 9 | 12 | 3 | 9 | 7 | 7 | 10 | 13 | 8.18 | 6 | 10 | 1.536 | 0.619 | 0.38022449 | 0.5947536 | 0 | 0.4095 | 0.2293 | 0.2293 | TRUE | 0.3688 |
| Hoxc13 | 76 | 124 | 122 | 193 | 183 | 157 | 81 | 124 | 123 | 152 | 197 | 172 | 141.26 | 109 | 174 | 1.586 | 0.665 | 0.00284965 | 0.02034616 | 0.0262 | 0.0302 | 0.0283 | 0.0283 | TRUE | 0.5017 |
| Hoxc4 | 37 | 43 | 39 | 55 | 39 | 38 | 39 | 43 | 39 | 43 | 42 | 42 | 41.37 | 40 | 42 | 1.045 | 0.064 | 0.82580258 | 0.91163704 | 0 | 0.0865 | 0.0361 | 0.0361 | TRUE | 0.0219 |
| Hoxc5 | 0 | 0 | 1 | 4 | 2 | 3 | 0 | 0 | 1 | 3 | 2 | 3 | 1.6 | 0 | 3 | 7.834 | 2.97 | 0.09176525 | NA | 0 | 2.0693 | 1.0646 | 1.0646 | TRUE | 0.4098 |
| Hoxc6 | 3 | 1 | 2 | 6 | 3 | 4 | 3 | 1 | 2 | 5 | 3 | 4 | 3.09 | 2 | 4 | 2.011 | 1.008 | 0.3503985 | NA | 0 | 1.0744 | 0.4723 | 0.4723 | TRUE | 0.1971 |
| Hoxc8 | 0 | 1 | 0 | 1 | 0 | 0 | 0 | 1 | 0 | 1 | 0 | 0 | 0.3 | 0 | 0 | 0.954 | -0.067 | 0.98642001 | NA | 0 | 11.0817 | 9.3014 | 9.3014 | TRUE | 0.405 |
| Hoxc9 | 3 | 0 | 1 | 5 | 5 | 3 | 3 | 0 | 1 | 4 | 5 | 3 | 2.8 | 1 | 4 | 3.042 | 1.605 | 0.20781553 | NA | 0 | 1.1845 | 0.6829 | 0.6829 | TRUE | 0.7645 |
| Hoxd1 | 55 | 82 | 85 | 96 | 63 | 47 | 58 | 82 | 85 | 75 | 68 | 51 | 70.06 | 75 | 65 | 0.866 | -0.207 | 0.44757511 | 0.65334741 | 0.0239 | 0.0539 | 0.0395 | 0.0395 | TRUE | 0.3415 |
| Hoxd10 | 2 | 2 | 2 | 6 | 4 | 4 | 2 | 2 | 2 | 5 | 4 | 4 | 3.25 | 2 | 4 | 2.191 | 1.132 | 0.27184031 | NA | 0 | 1.0191 | 0.4052 | 0.4052 | TRUE | 0.0058 |
| Hoxd11 | 0 | 7 | 3 | 13 | 3 | 6 | 0 | 7 | 3 | 10 | 3 | 7 | 5 | 3 | 7 | 2.008 | 1.005 | 0.35324103 | 0.57170972 | 0.5475 | 0.6654 | 0.6204 | 0.6204 | TRUE | 0.4428 |
| Hoxd12 | 2 | 0 | 0 | 1 | 0 | 1 | 2 | 0 | 0 | 1 | 0 | 1 | 0.67 | 1 | 1 | 0.902 | -0.15 | 0.95485009 | NA | 0 | 4.9422 | 3.5197 | 3.5197 | TRUE | 0.9743 |
| Hoxd13 | 45 | 109 | 102 | 203 | 139 | 136 | 48 | 109 | 102 | 160 | 150 | 149 | 119.51 | 86 | 153 | 1.766 | 0.82 | 0.00497793 | 0.03122989 | 0.0754 | 0.0345 | 0.0524 | 0.0524 | TRUE | 1.4152 |
| Hoxd3 | 2 | 0 | 1 | 1 | 0 | 1 | 2 | 0 | 1 | 1 | 0 | 1 | 0.83 | 1 | 1 | 0.611 | -0.711 | 0.74305956 | NA | 0 | 3.953 | 2.1268 | 2.1268 | TRUE | 0.1454 |
| Hoxd4 | 9 | 14 | 8 | 9 | 10 | 19 | 10 | 14 | 8 | 7 | 11 | 21 | 11.7 | 11 | 13 | 1.206 | 0.27 | 0.67032642 | 0.81594208 | 0.12 | 0.2885 | 0.2034 | 0.2034 | TRUE | 0.7049 |
| Hoxd8 | 18 | 58 | 59 | 113 | 74 | 66 | 19 | 58 | 59 | 89 | 80 | 72 | 62.83 | 45 | 80 | 1.765 | 0.819 | 0.03677392 | 0.13248017 | 0.1399 | 0.0593 | 0.0936 | 0.0936 | TRUE | 1.9622 |
| Hoxd9 | 7 | 18 | 15 | 40 | 44 | 32 | 7 | 18 | 15 | 31 | 47 | 35 | 25.71 | 13 | 38 | 2.791 | 1.481 | 0.00097541 | 0.008746 | 0.0384 | 0.135 | 0.0946 | 0.0946 | TRUE | 0.6222 |
| Hp | 0 | 0 | 1 | 3 | 1 | 1 | 0 | 0 | 1 | 2 | 1 | 1 | 0.92 | 0 | 1 | 4.189 | 2.067 | 0.32121847 | NA | 0 | 3.5794 | 1.8085 | 1.8085 | TRUE | 0.4098 |
| Hp1bp3 | 1551 | 1869 | 1932 | 2311 | 1849 | 1802 | 1647 | 1867 | 1940 | 1817 | 1989 | 1973 | 1872.11 | 1818 | 1926 | 1.059 | 0.083 | 0.38705701 | 0.60099965 | 0.0039 | 0.0087 | 0.006 | 0.006 | TRUE | 0.081 |
| Hpca | 165 | 172 | 143 | 264 | 206 | 160 | 175 | 172 | 144 | 208 | 222 | 175 | 182.49 | 164 | 202 | 1.233 | 0.303 | 0.07998962 | 0.22448332 | 0.007 | 0.025 | 0.016 | 0.016 | TRUE | 0.135 |
| Hpcal1 | 2052 | 3673 | 3902 | 2164 | 1761 | 2490 | 2180 | 3668 | 3918 | 1701 | 1895 | 2726 | 2681.24 | 3255 | 2107 | 0.647 | -0.628 | 0.00664898 | 0.0387538 | 0.0745 | 0.0081 | 0.0382 | 0.0382 | TRUE | 1.015 |
| Hpcal4 | 236 | 219 | 206 | 230 | 161 | 138 | 251 | 219 | 207 | 181 | 173 | 151 | 196.89 | 226 | 168 | 0.749 | -0.417 | 0.00994905 | 0.05249795 | 0.0042 | 0.0236 | 0.0137 | 0.0137 | TRUE | 0.1044 |
| Hpd | 7 | 9 | 10 | 5 | 6 | 2 | 7 | 9 | 10 | 4 | 6 | 2 | 6.51 | 9 | 4 | 0.474 | -1.077 | 0.16048676 | 0.35495684 | 0 | 0.5132 | 0.2504 | 0.2504 | TRUE | 0.338 |
| Hpdl | 422 | 363 | 363 | 499 | 345 | 329 | 448 | 363 | 364 | 392 | 371 | 360 | 383.15 | 392 | 374 | 0.958 | -0.063 | 0.64040767 | 0.79767036 | 0.0059 | 0.0155 | 0.0103 | 0.0103 | TRUE | 0.1813 |
| Hpgd | 88 | 111 | 87 | 274 | 197 | 143 | 93 | 111 | 87 | 215 | 212 | 157 | 145.93 | 97 | 195 | 2.005 | 1.004 | 1.29E-06 | 3.43E-05 | 0.0171 | 0.0295 | 0.0232 | 0.0232 | TRUE | 0.3078 |
| Hpgds | 0 | 0 | 0 | 0 | 0 | 0 | 0 | 0 | 0 | 0 | 0 | 0 | 0 | 0 | 0 | NA | NA | NA | NA | NA | NA | NA | NA | NA | NA |
| Hpn | 928 | 1466 | 1561 | 865 | 676 | 1061 | 986 | 1464 | 1567 | 680 | 727 | 1161 | 1097.66 | 1339 | 856 | 0.639 | -0.646 | 0.00509349 | 0.03181566 | 0.0734 | 0.0099 | 0.0374 | 0.0374 | TRUE | 1.1522 |
| Hprt | 7486 | 8775 | 8518 | 9447 | 6903 | 7691 | 7951 | 8764 | 8553 | 7426 | 7427 | 8419 | 8090.13 | 8423 | 7757 | 0.921 | -0.119 | 0.1624269 | 0.35779288 | 0.0038 | 0.0073 | 0.0051 | 0.0051 | TRUE | 0.0677 |
| Hps1 | 1432 | 1632 | 1562 | 2547 | 2054 | 1970 | 1521 | 1630 | 1568 | 2002 | 2210 | 2156 | 1848.01 | 1573 | 2123 | 1.349 | 0.431 | 7.75E-08 | 2.91E-06 | 0.0015 | 0.0087 | 0.0041 | 0.0041 | TRUE | 0.0307 |
| Hps3 | 1491 | 1501 | 1535 | 1543 | 1062 | 1182 | 1584 | 1499 | 1541 | 1213 | 1143 | 1294 | 1378.93 | 1541 | 1217 | 0.789 | -0.342 | 5.85E-05 | 0.00088647 | 0.0012 | 0.0093 | 0.0045 | 0.0045 | TRUE | 0.036 |
| Hps4 | 903 | 974 | 928 | 1103 | 885 | 870 | 959 | 973 | 932 | 867 | 952 | 952 | 939.22 | 955 | 924 | 0.967 | -0.049 | 0.57997549 | 0.75669188 | 9.00E-04 | 0.0104 | 0.0046 | 0.0046 | TRUE | 0.0361 |
| Hps5 | 464 | 486 | 530 | 1132 | 713 | 629 | 493 | 485 | 532 | 890 | 767 | 689 | 642.66 | 503 | 782 | 1.555 | 0.637 | 6.06E-07 | 1.77E-05 | 0.0086 | 0.012 | 0.0101 | 0.0101 | TRUE | 0.1779 |
| Hps6 | 127 | 151 | 160 | 280 | 197 | 147 | 135 | 151 | 161 | 220 | 212 | 161 | 173.22 | 149 | 198 | 1.331 | 0.413 | 0.0266443 | 0.10630758 | 0.0125 | 0.0259 | 0.0191 | 0.0191 | TRUE | 0.2779 |
| Hpse | 71 | 135 | 98 | 161 | 130 | 124 | 75 | 135 | 98 | 127 | 140 | 136 | 118.47 | 103 | 134 | 1.3 | 0.379 | 0.11716483 | 0.28950709 | 0.032 | 0.0347 | 0.0335 | 0.0335 | TRUE | 0.4629 |
| Hpse2 | 9 | 6 | 8 | 13 | 14 | 5 | 10 | 6 | 8 | 10 | 15 | 5 | 9.06 | 8 | 10 | 1.307 | 0.387 | 0.56320757 | 0.74414927 | 0 | 0.3707 | 0.2095 | 0.2095 | TRUE | 0.1395 |
| Hpx | 3 | 5 | 4 | 6 | 3 | 2 | 3 | 5 | 4 | 5 | 3 | 2 | 3.72 | 4 | 3 | 0.845 | -0.244 | 0.79797995 | NA | 0 | 0.8921 | 0.3863 | 0.3863 | TRUE | 0.2301 |
| Hr | 717 | 724 | 706 | 781 | 651 | 652 | 762 | 723 | 709 | 614 | 700 | 714 | 703.61 | 731 | 676 | 0.923 | -0.115 | 0.26960623 | 0.48966253 | 0.0028 | 0.0116 | 0.0065 | 0.0065 | TRUE | 0.08 |
| Hras1 | 1523 | 1432 | 1379 | 2425 | 1619 | 1356 | 1618 | 1430 | 1385 | 1906 | 1742 | 1484 | 1594.19 | 1478 | 1711 | 1.159 | 0.213 | 0.07909328 | 0.22307927 | 0.0103 | 0.009 | 0.0099 | 0.0099 | TRUE | 0.1612 |
| Hrasls | 5 | 9 | 13 | 19 | 18 | 12 | 5 | 9 | 13 | 15 | 19 | 13 | 12.47 | 9 | 16 | 1.726 | 0.788 | 0.1703631 | 0.36901257 | 0 | 0.2712 | 0.151 | 0.151 | TRUE | 0.3349 |
| Hrasls5 | 1 | 0 | 0 | 1 | 0 | 1 | 1 | 0 | 0 | 1 | 0 | 1 | 0.49 | 0 | 1 | 1.724 | 0.785 | 0.78200058 | NA | 0 | 6.7235 | 4.0295 | 4.0295 | TRUE | 0.4614 |
| Hrc | 36 | 124 | 110 | 52 | 45 | 69 | 38 | 124 | 110 | 41 | 48 | 76 | 72.89 | 91 | 55 | 0.602 | -0.732 | 0.09412353 | 0.25112088 | 0.2065 | 0.0521 | 0.1232 | 0.1232 | TRUE | 1.8732 |
| Hrct1 | 2 | 2 | 3 | 12 | 5 | 7 | 2 | 2 | 3 | 9 | 5 | 8 | 4.93 | 2 | 7 | 3.172 | 1.665 | 0.06494955 | 0.19626919 | 0 | 0.6745 | 0.3074 | 0.3074 | TRUE | 0.1608 |
| Hrg | 5 | 10 | 5 | 11 | 9 | 8 | 5 | 10 | 5 | 9 | 10 | 9 | 7.9 | 7 | 9 | 1.328 | 0.41 | 0.54481373 | 0.72993562 | 0 | 0.4239 | 0.2005 | 0.2005 | TRUE | 0.454 |
| Hrh1 | 244 | 184 | 188 | 186 | 159 | 169 | 259 | 184 | 189 | 146 | 171 | 185 | 189 | 211 | 167 | 0.794 | -0.334 | 0.09136198 | 0.24661325 | 0.0213 | 0.0243 | 0.0228 | 0.0228 | TRUE | 0.4411 |
| Hrh2 | 33 | 28 | 37 | 38 | 18 | 18 | 35 | 28 | 37 | 30 | 19 | 20 | 28.19 | 33 | 23 | 0.697 | -0.521 | 0.17251355 | 0.37119832 | 0 | 0.1238 | 0.0689 | 0.0689 | TRUE | 0.482 |
| Hrh3 | 37 | 45 | 48 | 19 | 16 | 22 | 39 | 45 | 48 | 15 | 17 | 24 | 31.45 | 44 | 19 | 0.42 | -1.252 | 0.00062762 | 0.00621789 | 0 | 0.1117 | 0.0588 | 0.0588 | TRUE | 0.3027 |
| Hrh4 | 0 | 0 | 0 | 0 | 0 | 0 | 0 | 0 | 0 | 0 | 0 | 0 | 0 | 0 | 0 | NA | NA | NA | NA | NA | NA | NA | NA | NA | NA |
| Hrk | 105 | 180 | 216 | 140 | 107 | 110 | 112 | 180 | 217 | 110 | 115 | 120 | 142.3 | 170 | 115 | 0.679 | -0.559 | 0.02896982 | 0.11226339 | 0.0501 | 0.0301 | 0.0399 | 0.0399 | TRUE | 0.3147 |
| Hrnr | 0 | 0 | 0 | 0 | 0 | 0 | 0 | 0 | 0 | 0 | 0 | 0 | 0 | 0 | 0 | NA | NA | NA | NA | NA | NA | NA | NA | NA | NA |
| Hrsp12 | 1366 | 1446 | 1450 | 1534 | 1061 | 1134 | 1451 | 1444 | 1456 | 1206 | 1142 | 1241 | 1323.31 | 1450 | 1196 | 0.825 | -0.278 | 0.00032474 | 0.00363247 | 0 | 0.0094 | 0.0035 | 0.0035 | TRUE | 0.0186 |
| Hs1bp3 | 1923 | 2366 | 2602 | 2851 | 2148 | 2219 | 2043 | 2363 | 2613 | 2241 | 2311 | 2429 | 2333.26 | 2340 | 2327 | 0.994 | -0.008 | 0.94107026 | 0.97010197 | 0.0079 | 0.0083 | 0.0081 | 0.0081 | TRUE | 0.1489 |
| Hs2st1 | 2196 | 1805 | 1642 | 3308 | 2234 | 2043 | 2333 | 1803 | 1649 | 2600 | 2404 | 2236 | 2170.75 | 1928 | 2413 | 1.252 | 0.325 | 0.02126427 | 0.09081507 | 0.0195 | 0.0084 | 0.0138 | 0.0138 | TRUE | 0.4067 |
| Hs3st1 | 17 | 22 | 12 | 62 | 37 | 33 | 18 | 22 | 12 | 49 | 40 | 36 | 29.46 | 17 | 42 | 2.405 | 1.266 | 0.0012109 | 0.01031786 | 0 | 0.1187 | 0.0692 | 0.0692 | TRUE | 0.364 |
| Hs3st2 | 2 | 1 | 1 | 4 | 2 | 0 | 2 | 1 | 1 | 3 | 2 | 0 | 1.57 | 1 | 2 | 1.324 | 0.405 | 0.79598963 | NA | 0 | 2.1045 | 1.1297 | 1.1297 | TRUE | 0.4456 |
| Hs3st3a1 | 18 | 36 | 19 | 48 | 21 | 45 | 19 | 36 | 19 | 38 | 23 | 49 | 30.62 | 25 | 37 | 1.476 | 0.562 | 0.21014512 | 0.42061026 | 0.1065 | 0.1145 | 0.1108 | 0.1108 | TRUE | 0.2621 |
| Hs3st3b1 | 941 | 1201 | 1306 | 1166 | 1038 | 1156 | 1000 | 1199 | 1311 | 917 | 1117 | 1265 | 1134.86 | 1170 | 1100 | 0.939 | -0.091 | 0.54938036 | 0.7339342 | 0.0215 | 0.0098 | 0.0158 | 0.0158 | TRUE | 0.2583 |

| | | | | | | | | | | | | | | | | | | | | | | | | | |
|---|---|---|---|---|---|---|---|---|---|---|---|---|---|---|---|---|---|---|---|---|---|---|---|---|---|
| Hs3st4 | 0 | 4 | 9 | 11 | 4 | 3 | 0 | 4 | 9 | 9 | 4 | 3 | 4.88 | 4 | 5 | 1.256 | 0.329 | 0.76733405 | 0.8785766 | 0.6857 | 0.6823 | 0.6837 | 0.6837 | TRUE | 0.3111 |
| Hs3st5 | 7 | 3 | 2 | 7 | 6 | 3 | 7 | 3 | 2 | 6 | 6 | 3 | 4.61 | 4 | 5 | 1.237 | 0.307 | 0.73963167 | 0.86066702 | 0 | 0.721 | 0.3941 | 0.3941 | TRUE | 0.7896 |
| Hs3st6 | 97 | 121 | 143 | 117 | 84 | 96 | 103 | 121 | 144 | 92 | 90 | 105 | 109.15 | 123 | 96 | 0.78 | -0.358 | 0.09074354 | 0.24565862 | 0.0087 | 0.0371 | 0.0229 | 0.0229 | TRUE | 0.2082 |
| Hs6st1 | 3614 | 3565 | 3331 | 4230 | 3413 | 3106 | 3839 | 3560 | 3345 | 3325 | 3672 | 3400 | 3523.54 | 3581 | 3466 | 0.968 | -0.047 | 0.58314739 | 0.75875085 | 0.0031 | 0.0078 | 0.0051 | 0.0051 | TRUE | 0.048 |
| Hs6st2 | 78 | 77 | 96 | 200 | 130 | 106 | 83 | 77 | 96 | 157 | 140 | 116 | 111.55 | 85 | 138 | 1.618 | 0.695 | 0.00116884 | 0.01001899 | 0.0107 | 0.0364 | 0.0235 | 0.0235 | TRUE | 0.1894 |
| Hs6st3 | 6 | 1 | 4 | 20 | 12 | 5 | 6 | 1 | 4 | 16 | 13 | 5 | 7.58 | 4 | 11 | 3.039 | 1.604 | 0.05800596 | 0.1819496 | 0.1991 | 0.4414 | 0.3376 | 0.3376 | TRUE | 0.5004 |
| Hsbp1 | 3105 | 3303 | 3308 | 4175 | 2978 | 3125 | 3298 | 3299 | 3322 | 3282 | 3204 | 3421 | 3304.23 | 3306 | 3302 | 0.999 | -0.002 | 0.9778814 | 0.98799114 | 2.00E-04 | 0.0079 | 0.0027 | 0.0027 | TRUE | 0.0118 |
| Hsbp1l1 | 14 | 26 | 24 | 25 | 12 | 11 | 15 | 26 | 24 | 20 | 13 | 12 | 18.26 | 22 | 15 | 0.694 | -0.526 | 0.27407506 | 0.49464606 | 0 | 0.1874 | 0.1109 | 0.1109 | TRUE | 0.4425 |
| Hscb | 446 | 760 | 788 | 669 | 501 | 567 | 474 | 759 | 791 | 526 | 539 | 621 | 618.27 | 675 | 562 | 0.832 | -0.265 | 0.17360903 | 0.37274186 | 0.0401 | 0.0122 | 0.0257 | 0.0257 | TRUE | 0.7978 |
| Hsd11b1 | 1 | 0 | 1 | 1 | 0 | 0 | 1 | 0 | 1 | 1 | 0 | 0 | 0.48 | 1 | 0 | 0.509 | -0.975 | 0.7336918 | NA | 0 | 6.9367 | 4.1806 | 4.1806 | TRUE | 0.2562 |
| Hsd11b2 | 131 | 156 | 128 | 150 | 107 | 93 | 139 | 156 | 129 | 118 | 115 | 102 | 126.39 | 141 | 112 | 0.792 | -0.336 | 0.06929483 | 0.20428374 | 0 | 0.033 | 0.0166 | 0.0166 | TRUE | 0.0866 |
| Hsd17b1 | 12 | 28 | 24 | 23 | 16 | 17 | 13 | 28 | 24 | 18 | 17 | 19 | 19.79 | 22 | 18 | 0.83 | -0.269 | 0.56122388 | 0.74256648 | 0 | 0.1734 | 0.104 | 0.104 | TRUE | 0.6958 |
| Hsd17b10 | 2187 | 2727 | 2761 | 2548 | 1891 | 2180 | 2323 | 2723 | 2772 | 2003 | 2035 | 2386 | 2373.8 | 2606 | 2141 | 0.821 | -0.284 | 0.01142809 | 0.05814319 | 0.0088 | 0.0083 | 0.0087 | 0.0087 | TRUE | 0.1207 |
| Hsd17b11 | 1241 | 1388 | 1408 | 1481 | 1070 | 1169 | 1318 | 1386 | 1414 | 1164 | 1151 | 1280 | 1285.55 | 1373 | 1198 | 0.872 | -0.197 | 0.0226692 | 0.09501424 | 0.0015 | 0.0095 | 0.0046 | 0.0046 | TRUE | 0.0414 |
| Hsd17b12 | 5868 | 6825 | 6907 | 6681 | 4716 | 5335 | 6233 | 6816 | 6935 | 5252 | 5074 | 5840 | 6025.1 | 6661 | 5389 | 0.809 | -0.306 | 0.00049399 | 0.00512241 | 0.0041 | 0.0075 | 0.0054 | 0.0054 | TRUE | 0.0651 |
| Hsd17b13 | 2 | 1 | 1 | 2 | 3 | 2 | 2 | 1 | 1 | 2 | 3 | 2 | 1.85 | 1 | 2 | 1.682 | 0.75 | 0.57825256 | NA | 0 | 1.785 | 0.7265 | 0.7265 | TRUE | 0.1348 |
| Hsd17b14 | 2753 | 3038 | 2954 | 3520 | 2505 | 2743 | 2924 | 3034 | 2966 | 2767 | 2695 | 3003 | 2898.23 | 2975 | 2822 | 0.948 | -0.077 | 0.30777771 | 0.52830165 | 0.0013 | 0.008 | 0.0037 | 0.0037 | TRUE | 0.0377 |
| Hsd17b2 | 2 | 2 | 1 | 2 | 0 | 1 | 2 | 2 | 1 | 2 | 0 | 1 | 1.3 | 2 | 1 | 0.541 | -0.887 | 0.5918554 | NA | 0 | 2.5434 | 1.1447 | 1.1447 | TRUE | 0.2827 |
| Hsd17b3 | 1 | 1 | 1 | 2 | 4 | 2 | 1 | 1 | 1 | 2 | 4 | 2 | 1.86 | 1 | 3 | 2.597 | 1.377 | 0.32907341 | NA | 0 | 1.7828 | 0.7494 | 0.7494 | TRUE | 0.3063 |
| Hsd17b4 | 4636 | 5104 | 5159 | 5684 | 3926 | 4282 | 4924 | 5097 | 5180 | 4468 | 4224 | 4687 | 4763.6 | 5067 | 4460 | 0.88 | -0.184 | 0.01034791 | 0.05416054 | 0.0014 | 0.0076 | 0.0035 | 0.0035 | TRUE | 0.0258 |
| Hsd17b6 | 2 | 1 | 0 | 2 | 2 | 2 | 2 | 1 | 0 | 2 | 2 | 2 | 1.51 | 1 | 2 | 1.898 | 0.925 | 0.55550267 | NA | 0 | 2.1943 | 1.0284 | 1.0284 | TRUE | 0.3212 |
| Hsd17b7 | 1623 | 1605 | 1458 | 1888 | 1388 | 1333 | 1724 | 1603 | 1464 | 1484 | 1493 | 1459 | 1537.93 | 1597 | 1479 | 0.926 | -0.11 | 0.22087863 | 0.43302575 | 0.0028 | 0.009 | 0.0052 | 0.0052 | TRUE | 0.064 |
| Hsd3b1 | 0 | 0 | 0 | 0 | 0 | 0 | 0 | 0 | 0 | 0 | 0 | 0 | 0 | 0 | 0 | NA | NA | NA | NA | NA | NA | NA | NA | NA | NA |
| Hsd3b2 | 0 | 1 | 0 | 1 | 0 | 0 | 0 | 1 | 0 | 1 | 0 | 0 | 0.3 | 0 | 0 | 0.954 | -0.067 | 0.98642001 | NA | 0 | 11.0817 | 9.3014 | 9.3014 | TRUE | 0.405 |
| Hsd3b3 | 0 | 0 | 0 | 0 | 0 | 0 | 0 | 0 | 0 | 0 | 0 | 0 | 0 | 0 | 0 | NA | NA | NA | NA | NA | NA | NA | NA | NA | NA |
| Hsd3b4 | 0 | 0 | 0 | 0 | 0 | 0 | 0 | 0 | 0 | 0 | 0 | 0 | 0 | 0 | 0 | NA | NA | NA | NA | NA | NA | NA | NA | NA | NA |
| Hsd3b5 | 0 | 1 | 0 | 0 | 0 | 0 | 0 | 1 | 0 | 0 | 0 | 0 | 0.17 | 0 | 0 | 0.49 | -1.029 | 0.80087 | NA | 0 | 19.7991 | 10 | 10 | TRUE | 0.405 |
| Hsd3b6 | 0 | 0 | 0 | 0 | 0 | 0 | 0 | 0 | 0 | 0 | 0 | 0 | 0 | 0 | 0 | NA | NA | NA | NA | NA | NA | NA | NA | NA | NA |
| Hsd3b7 | 258 | 440 | 402 | 425 | 322 | 366 | 274 | 439 | 404 | 334 | 346 | 401 | 366.39 | 372 | 360 | 0.967 | -0.049 | 0.79720859 | 0.8955537 | 0.0315 | 0.0159 | 0.0236 | 0.0236 | TRUE | 0.6071 |
| Hsdl1 | 1523 | 2040 | 1972 | 2583 | 1896 | 2030 | 1618 | 2037 | 1980 | 2031 | 2040 | 2222 | 1987.97 | 1878 | 2098 | 1.116 | 0.159 | 0.15488648 | 0.34704782 | 0.0079 | 0.0086 | 0.0085 | 0.0085 | TRUE | 0.1776 |
| Hsdl2 | 1117 | 1086 | 1100 | 1264 | 901 | 880 | 1186 | 1085 | 1105 | 994 | 969 | 963 | 1050.32 | 1125 | 975 | 0.867 | -0.205 | 0.01434373 | 0.0684732 | 0 | 0.01 | 0.0041 | 0.0041 | TRUE | 0.027 |
| Hsf1 | 1710 | 1728 | 1710 | 2321 | 1843 | 1686 | 1816 | 1726 | 1717 | 1825 | 1983 | 1846 | 1818.71 | 1753 | 1885 | 1.075 | 0.104 | 0.17992521 | 0.38129377 | 0.001 | 0.0087 | 0.0038 | 0.0038 | TRUE | 0.0251 |
| Hsf2 | 2711 | 2498 | 2573 | 3243 | 2332 | 2348 | 2880 | 2495 | 2584 | 2549 | 2509 | 2570 | 2597.77 | 2653 | 2543 | 0.959 | -0.061 | 0.4655098 | 0.6680031 | 0.0024 | 0.0082 | 0.0046 | 0.0046 | TRUE | 0.0677 |
| Hsf2bp | 1523 | 2083 | 2237 | 3747 | 2797 | 2971 | 1618 | 2080 | 2246 | 2946 | 3009 | 3252 | 2525.24 | 1981 | 3069 | 1.549 | 0.631 | 1.31E-06 | 3.47E-05 | 0.0143 | 0.0082 | 0.0118 | 0.0118 | TRUE | 0.3111 |
| Hsf3 | 5 | 11 | 2 | 35 | 22 | 25 | 5 | 11 | 2 | 28 | 24 | 27 | 16.14 | 6 | 26 | 4.287 | 2.1 | 0.00045277 | 0.00481533 | 0 | 0.211 | 0.1561 | 0.1561 | TRUE | 1.1983 |
| Hsf4 | 56 | 132 | 157 | 97 | 91 | 110 | 59 | 132 | 158 | 76 | 98 | 120 | 107.26 | 116 | 98 | 0.841 | -0.25 | 0.48084355 | 0.68004523 | 0.1314 | 0.0376 | 0.0809 | 0.0809 | TRUE | 0.7545 |
| Hsf5 | 54 | 134 | 125 | 129 | 81 | 114 | 57 | 134 | 126 | 101 | 87 | 125 | 105.01 | 106 | 104 | 0.988 | -0.018 | 0.9567801 | 0.97766996 | 0.1009 | 0.0383 | 0.0673 | 0.0673 | TRUE | 1.5513 |
| Hsfy2 | 0 | 0 | 0 | 0 | 0 | 0 | 0 | 0 | 0 | 0 | 0 | 0 | 0 | 0 | 0 | NA | NA | NA | NA | NA | NA | NA | NA | NA | NA |
| Hsh2d | 73 | 98 | 94 | 86 | 64 | 93 | 78 | 98 | 94 | 68 | 69 | 102 | 84.68 | 90 | 80 | 0.879 | -0.187 | 0.46201391 | 0.66583365 | 0.0227 | 0.0458 | 0.0345 | 0.0345 | TRUE | 0.542 |
| Hsp90aa1 | 125838 | 128296 | 128839 | 220315 | 153880 | 148792 | 133662 | 128130 | 129368 | 173194 | 165565 | 162876 | 148799.43 | 130387 | 167212 | 1.282 | 0.359 | 9.86E-10 | 6.03E-08 | 7.00E-04 | 0.0069 | 0.0025 | 0.0025 | TRUE | 0.012 |
| Hsp90ab1 | 299851 | 314473 | 316081 | 401502 | 287955 | 275022 | 318495 | 314067 | 317379 | 315629 | 309822 | 301055 | 312741.1 | 316647 | 308835 | 0.975 | -0.036 | 0.49281631 | 0.68913013 | 3.00E-04 | 0.0069 | 0.002 | 0.002 | TRUE | 0.0059 |
| Hsp90b1 | 43859 | 49326 | 47330 | 50164 | 37581 | 39391 | 46586 | 49262 | 47524 | 39435 | 40435 | 43120 | 44393.69 | 47791 | 40997 | 0.858 | -0.221 | 0.00086965 | 0.00798376 | 0.0014 | 0.007 | 0.0032 | 0.0032 | TRUE | 0.0251 |
| Hspa12a | 932 | 1126 | 1198 | 780 | 605 | 711 | 990 | 1125 | 1203 | 613 | 651 | 778 | 893.31 | 1106 | 681 | 0.615 | -0.702 | 6.64E-08 | 2.53E-06 | 0.011 | 0.0106 | 0.011 | 0.011 | TRUE | 0.1837 |
| Hspa12b | 35 | 43 | 37 | 64 | 41 | 36 | 37 | 43 | 37 | 50 | 44 | 39 | 41.85 | 39 | 44 | 1.146 | 0.197 | 0.50818576 | 0.70234015 | 0 | 0.0856 | 0.0399 | 0.0399 | TRUE | 0.1131 |
| Hspa13 | 1084 | 1026 | 946 | 1596 | 1067 | 947 | 1151 | 1025 | 950 | 1255 | 1148 | 1037 | 1094.21 | 1042 | 1147 | 1.101 | 0.139 | 0.23585493 | 0.45107164 | 0.0079 | 0.0099 | 0.009 | 0.009 | TRUE | 0.1004 |
| Hspa14 | 4909 | 5719 | 5829 | 9031 | 6414 | 5951 | 5214 | 5712 | 5853 | 7099 | 6901 | 6514 | 6215.6 | 5593 | 6838 | 1.223 | 0.29 | 0.00023454 | 0.00277092 | 0.0025 | 0.0074 | 0.0043 | 0.0043 | TRUE | 0.0427 |
| Hspa1a | 65 | 32 | 41 | 72 | 76 | 51 | 69 | 32 | 41 | 57 | 82 | 56 | 56.06 | 47 | 65 | 1.366 | 0.45 | 0.20548909 | 0.41412906 | 0.0816 | 0.0657 | 0.0727 | 0.0727 | TRUE | 1.2474 |
| Hspa1b | 138 | 107 | 107 | 341 | 310 | 239 | 147 | 107 | 107 | 268 | 334 | 262 | 204.02 | 120 | 288 | 2.393 | 1.259 | 7.99E-11 | 5.95E-09 | 0.0186 | 0.0231 | 0.0211 | 0.0211 | TRUE | 0.3634 |
| Hspa1l | 3 | 2 | 4 | 7 | 3 | 2 | 3 | 2 | 4 | 6 | 3 | 2 | 3.35 | 3 | 4 | 1.212 | 0.277 | 0.78617288 | NA | 0 | 0.9893 | 0.4525 | 0.4525 | TRUE | 0.4036 |
| Hspa2 | 308 | 375 | 388 | 576 | 436 | 451 | 327 | 375 | 390 | 453 | 469 | 494 | 417.81 | 364 | 472 | 1.295 | 0.373 | 0.00192609 | 0.01494262 | 0.0021 | 0.0148 | 0.008 | 0.008 | TRUE | 0.0875 |
| Hspa4 | 41260 | 31791 | 30054 | 50412 | 35317 | 30785 | 43825 | 31750 | 30177 | 39630 | 37999 | 33699 | 36180.11 | 35251 | 37109 | 1.053 | 0.074 | 0.61689263 | 0.7832683 | 0.0207 | 0.007 | 0.0158 | 0.0158 | TRUE | 0.5542 |
| Hspa4l | 906 | 1264 | 1352 | 2135 | 1451 | 1626 | 962 | 1262 | 1358 | 1678 | 1561 | 1780 | 1433.62 | 1194 | 1673 | 1.401 | 0.486 | 0.00048293 | 0.00503754 | 0.016 | 0.0092 | 0.0133 | 0.0133 | TRUE | 0.3445 |
| Hspa5 | 23503 | 21965 | 19312 | 28473 | 19735 | 17446 | 24964 | 21937 | 19391 | 22383 | 21234 | 19097 | 21501.09 | 22097 | 20905 | 0.946 | -0.08 | 0.48369913 | 0.68148223 | 0.01 | 0.0071 | 0.0094 | 0.0094 | TRUE | 0.1576 |
| Hspa8 | 89077 | 85818 | 85253 | 125417 | 86703 | 78463 | 94616 | 85707 | 85603 | 98593 | 93287 | 85890 | 90615.99 | 88642 | 92590 | 1.045 | 0.063 | 0.45561711 | 0.66053461 | 0.0038 | 0.0069 | 0.0051 | 0.0051 | TRUE | 0.0491 |
| Hspa9 | 47177 | 48952 | 49064 | 57160 | 40908 | 41525 | 50110 | 48889 | 49265 | 44935 | 44014 | 45456 | 47111.57 | 49421 | 44802 | 0.907 | -0.142 | 0.00607891 | 0.03630315 | 2.00E-04 | 0.007 | 0.0019 | 0.0019 | TRUE | 0.0029 |
| Hspb1 | 16511 | 14697 | 14556 | 18080 | 13272 | 12120 | 17538 | 14678 | 14616 | 14213 | 14280 | 13267 | 14765.26 | 15611 | 13920 | 0.892 | -0.165 | 0.08614107 | 0.23681663 | 0.0064 | 0.0071 | 0.0066 | 0.0066 | TRUE | 0.1426 |
| Hspb11 | 209 | 355 | 363 | 317 | 212 | 249 | 222 | 355 | 364 | 249 | 228 | 273 | 281.82 | 314 | 250 | 0.796 | -0.329 | 0.1130558 | 0.28316914 | 0.0365 | 0.0186 | 0.0274 | 0.0274 | TRUE | 0.7344 |
| Hspb2 | 57 | 94 | 97 | 79 | 43 | 56 | 61 | 94 | 97 | 62 | 46 | 61 | 70.25 | 84 | 56 | 0.675 | -0.567 | 0.044995 | 0.15291126 | 0.0316 | 0.0538 | 0.043 | 0.043 | TRUE | 0.5473 |
| Hspb3 | 1 | 3 | 2 | 6 | 2 | 5 | 1 | 3 | 2 | 5 | 2 | 5 | 3.07 | 2 | 4 | 2.033 | 1.024 | 0.3533943 | NA | 0 | 1.0807 | 0.507 | 0.507 | TRUE | 0.2694 |
| Hspb6 | 50 | 68 | 74 | 88 | 66 | 73 | 53 | 68 | 74 | 69 | 71 | 80 | 69.24 | 65 | 73 | 1.123 | 0.168 | 0.49661611 | 0.69198951 | 0 | 0.0545 | 0.0293 | 0.0293 | TRUE | 0.2225 |
| Hspb7 | 8 | 6 | 9 | 13 | 11 | 5 | 8 | 6 | 9 | 10 | 12 | 5 | 8.51 | 8 | 9 | 1.177 | 0.235 | 0.72261445 | 0.84995712 | 0 | 0.3941 | 0.1972 | 0.1972 | TRUE | 0.3602 |
| Hspb8 | 971 | 1658 | 1739 | 1394 | 1076 | 1464 | 1031 | 1656 | 1746 | 1096 | 1158 | 1603 | 1381.59 | 1478 | 1286 | 0.869 | -0.202 | 0.34368282 | 0.56363061 | 0.0596 | 0.0093 | 0.0321 | 0.0321 | TRUE | 0.8395 |
| Hspb9 | 5 | 4 | 5 | 9 | 8 | 10 | 5 | 4 | 5 | 7 | 9 | 11 | 6.83 | 5 | 9 | 1.847 | 0.885 | 0.2212095 | 0.43313671 | 0 | 0.4895 | 0.2147 | 0.2147 | TRUE | 0.1297 |
| Hspbap1 | 4598 | 4693 | 4797 | 5196 | 3918 | 3839 | 4884 | 4687 | 4817 | 4085 | 4216 | 4202 | 4481.69 | 4796 | 4168 | 0.869 | -0.203 | 0.00076471 | 0.00721786 | 2.00E-04 | 0.0076 | 0.0024 | 0.0024 | TRUE | 0.0048 |
| Hspbp1 | 2114 | 2032 | 1879 | 2871 | 2199 | 1961 | 2245 | 2029 | 1887 | 2257 | 2366 | 2147 | 2155.18 | 2054 | 2257 | 1.099 | 0.136 | 0.15307617 | 0.34455792 | 0.0044 | 0.0084 | 0.0061 | 0.0061 | TRUE | 0.0803 |
| Hspd1 | 35300 | 31640 | 30604 | 54063 | 36548 | 32767 | 37495 | 31599 | 30730 | 42500 | 39323 | 35869 | 36252.63 | 33275 | 39231 | 1.179 | 0.238 | 0.02805399 | 0.11003973 | 0.0096 | 0.007 | 0.0084 | 0.0084 | TRUE | 0.1507 |
| Hspe1 | 7401 | 8761 | 8360 | 11974 | 8536 | 8212 | 7861 | 8750 | 8394 | 9413 | 9184 | 8989 | 8765.29 | 8335 | 9195 | 1.103 | 0.142 | 0.04387318 | 0.15007283 | 0.0015 | 0.0073 | 0.0035 | 0.0035 | TRUE | 0.0302 |
| Hspg2 | 15220 | 16717 | 20600 | 19013 | 12944 | 13534 | 16166 | 16695 | 20685 | 14947 | 13927 | 14815 | 16205.82 | 17849 | 14563 | 0.816 | -0.294 | 0.0076476 | 0.04295512 | 0.0102 | 0.0071 | 0.0087 | 0.0087 | TRUE | 0.2363 |
| Hsph1 | 4858 | 3156 | 3177 | 7562 | 5405 | 3850 | 5160 | 3152 | 3190 | 5945 | 5815 | 4214 | 4579.43 | 3834 | 5325 | 1.389 | 0.474 | 0.02097815 | 0.08988517 | 0.0609 | 0.0076 | 0.0302 | 0.0302 | TRUE | 1.1135 |

| | | | | | | | | | | | | | | | | | | | | | | | | | |
|---|---|---|---|---|---|---|---|---|---|---|---|---|---|---|---|---|---|---|---|---|---|---|---|---|---|
| Htatip2 | 527 | 585 | 552 | 836 | 526 | 553 | 560 | 584 | 554 | 657 | 566 | 605 | 587.79 | 566 | 609 | 1.078 | 0.109 | 0.30288983 | 0.52367315 | 0.0019 | 0.0125 | 0.0063 | 0.0063 | TRUE | 0.0578 |
| Htatsf1 | 3819 | 3962 | 3981 | 4930 | 3553 | 3596 | 4056 | 3957 | 3997 | 3876 | 3823 | 3936 | 3940.91 | 4003 | 3878 | 0.969 | -0.046 | 0.43711179 | 0.64533625 | 0 | 0.0077 | 0.0023 | 0.0023 | TRUE | 0.002 |
| Htr1a | 0 | 0 | 0 | 0 | 0 | 0 | 0 | 0 | 0 | 0 | 0 | 0 | 0 | 0 | 0 | NA | NA | NA | NA | NA | NA | NA | NA | NA | NA |
| Htr1b | 6 | 4 | 6 | 10 | 9 | 6 | 6 | 4 | 6 | 8 | 10 | 7 | 6.75 | 5 | 8 | 1.473 | 0.559 | 0.43705122 | 0.64530723 | 0 | 0.4949 | 0.2178 | 0.2178 | TRUE | 0.123 |
| Htr1d | 14 | 10 | 6 | 10 | 16 | 8 | 15 | 10 | 6 | 8 | 17 | 9 | 10.79 | 10 | 11 | 1.089 | 0.123 | 0.84939132 | 0.92505178 | 0.1001 | 0.3123 | 0.2086 | 0.2086 | TRUE | 0.3703 |
| Htr1f | 0 | 0 | 0 | 1 | 0 | 0 | 0 | 0 | 0 | 1 | 0 | 0 | 0.13 | 0 | 0 | 1.859 | 0.894 | 0.82650734 | NA | 0 | 25.1515 | 10 | 10 | TRUE | 0.2563 |
| Htr2a | 0 | 1 | 0 | 0 | 1 | 1 | 0 | 1 | 0 | 0 | 1 | 1 | 0.53 | 0 | 1 | 1.924 | 0.944 | 0.7364225 | NA | 0 | 6.2438 | 3.9727 | 3.9727 | TRUE | 0.405 |
| Htr2b | 0 | 0 | 0 | 0 | 0 | 0 | 0 | 0 | 0 | 0 | 0 | 0 | 0 | 0 | 0 | NA | NA | NA | NA | NA | NA | NA | NA | NA | NA |
| Htr2c | 0 | 1 | 0 | 0 | 0 | 0 | 0 | 1 | 0 | 0 | 0 | 0 | 0.17 | 0 | 0 | 0.49 | -1.029 | 0.80087 | NA | 0 | 19.7991 | 10 | 10 | TRUE | 0.405 |
| Htr3a | 61 | 80 | 68 | 74 | 43 | 63 | 65 | 80 | 68 | 58 | 46 | 69 | 64.4 | 71 | 58 | 0.814 | -0.298 | 0.25705086 | 0.47573411 | 0.0061 | 0.0581 | 0.034 | 0.034 | TRUE | 0.2317 |
| Htr3b | 3 | 2 | 0 | 3 | 0 | 1 | 3 | 2 | 0 | 2 | 0 | 1 | 1.44 | 2 | 1 | 0.693 | -0.529 | 0.76905053 | NA | 0 | 2.2955 | 1.634 | 1.634 | TRUE | 0.3039 |
| Htr4 | 24 | 15 | 24 | 26 | 15 | 17 | 25 | 15 | 24 | 20 | 16 | 19 | 19.96 | 21 | 18 | 0.859 | -0.218 | 0.62012953 | 0.78498274 | 0 | 0.172 | 0.0898 | 0.0898 | TRUE | 0.4045 |
| Htr5a | 0 | 0 | 1 | 1 | 2 | 1 | 0 | 0 | 1 | 1 | 2 | 1 | 0.84 | 0 | 1 | 3.633 | 1.861 | 0.38679592 | NA | 0 | 3.9314 | 1.964 | 1.964 | TRUE | 0.4098 |
| Htr5b | 41 | 28 | 22 | 92 | 64 | 46 | 44 | 28 | 22 | 72 | 69 | 50 | 47.52 | 31 | 64 | 2.058 | 1.041 | 0.00269574 | 0.01950482 | 0.0459 | 0.0762 | 0.0627 | 0.0627 | TRUE | 0.7854 |
| Htr6 | 53 | 76 | 46 | 48 | 37 | 37 | 56 | 76 | 46 | 38 | 40 | 41 | 49.41 | 59 | 40 | 0.66 | -0.598 | 0.04962321 | 0.16342811 | 0.0187 | 0.0736 | 0.0459 | 0.0459 | TRUE | 0.3529 |
| Htr7 | 2 | 1 | 0 | 7 | 5 | 0 | 2 | 1 | 0 | 6 | 5 | 0 | 2.33 | 1 | 4 | 3.549 | 1.827 | 0.26478016 | NA | 0 | 1.4182 | 1.3064 | 1.3064 | TRUE | 1.0651 |
| Htra1 | 1563 | 1304 | 1090 | 2511 | 1753 | 1727 | 1660 | 1302 | 1094 | 1974 | 1886 | 1890 | 1634.59 | 1352 | 1917 | 1.418 | 0.504 | 0.00070493 | 0.00680839 | 0.0218 | 0.0089 | 0.0153 | 0.0153 | TRUE | 0.4758 |
| Htra2 | 1536 | 1306 | 1271 | 2325 | 1725 | 1344 | 1632 | 1304 | 1276 | 1828 | 1856 | 1471 | 1561.16 | 1404 | 1718 | 1.224 | 0.292 | 0.03500971 | 0.12811442 | 0.0167 | 0.009 | 0.0132 | 0.0132 | TRUE | 0.2411 |
| Htra3 | 50 | 73 | 47 | 48 | 29 | 32 | 53 | 73 | 47 | 38 | 31 | 35 | 46.2 | 58 | 35 | 0.602 | -0.732 | 0.01866946 | 0.08285163 | 0 | 0.0782 | 0.047 | 0.047 | TRUE | 0.4551 |
| Htra4 | 13 | 19 | 13 | 18 | 7 | 9 | 14 | 19 | 13 | 14 | 8 | 10 | 12.9 | 15 | 11 | 0.697 | -0.52 | 0.34184546 | 0.56204134 | 0 | 0.2624 | 0.1364 | 0.1364 | TRUE | 0.4174 |
| Htt | 4949 | 5385 | 6603 | 7124 | 5262 | 5426 | 5257 | 5378 | 6630 | 5600 | 5662 | 5940 | 5744.4 | 5755 | 5734 | 0.996 | -0.005 | 0.95978372 | 0.97934326 | 0.0072 | 0.0075 | 0.0082 | 0.0082 | TRUE | 0.2159 |
| Hunk | 2658 | 2875 | 2743 | 4437 | 3083 | 3167 | 2823 | 2871 | 2754 | 3488 | 3317 | 3467 | 3120.12 | 2816 | 3424 | 1.216 | 0.282 | 1.52E-05 | 0.00028153 | 3.00E-04 | 0.008 | 0.0027 | 0.0027 | TRUE | 0.009 |
| Hus1 | 4155 | 4309 | 4365 | 5048 | 3691 | 3704 | 4413 | 4303 | 4383 | 3968 | 3971 | 4055 | 4182.32 | 4366 | 3998 | 0.916 | -0.127 | 0.02895117 | 0.11224628 | 0 | 0.0077 | 0.0022 | 0.0022 | TRUE | 0.002 |
| Hus1b | 147 | 142 | 170 | 173 | 151 | 147 | 156 | 142 | 171 | 136 | 162 | 161 | 154.67 | 156 | 153 | 0.977 | -0.034 | 0.84704834 | 0.92352627 | 0 | 0.0282 | 0.0155 | 0.0155 | TRUE | 0.1111 |
| Huwe1 | 39790 | 31759 | 38945 | 46341 | 31794 | 29072 | 42264 | 31718 | 39105 | 36430 | 34208 | 31824 | 35924.81 | 37696 | 34154 | 0.906 | -0.142 | 0.23454841 | 0.44941796 | 0.0125 | 0.007 | 0.0103 | 0.0103 | TRUE | 0.2356 |
| Hvcn1 | 955 | 884 | 892 | 1020 | 662 | 615 | 1014 | 883 | 896 | 802 | 712 | 673 | 830.04 | 931 | 729 | 0.785 | -0.35 | 0.00197356 | 0.01522845 | 0.0058 | 0.0109 | 0.008 | 0.008 | TRUE | 0.0932 |
| Hyal1 | 188 | 215 | 216 | 210 | 188 | 199 | 200 | 215 | 217 | 165 | 202 | 218 | 202.75 | 211 | 195 | 0.923 | -0.116 | 0.48166136 | 0.68073323 | 0.0067 | 0.0232 | 0.0146 | 0.0146 | TRUE | 0.2135 |
| Hyal2 | 1378 | 1126 | 1080 | 1195 | 939 | 853 | 1464 | 1125 | 1084 | 939 | 1010 | 934 | 1092.69 | 1224 | 961 | 0.785 | -0.349 | 0.0094146 | 0.05026644 | 0.0141 | 0.0099 | 0.0121 | 0.0121 | TRUE | 0.3503 |
| Hyal3 | 7 | 21 | 10 | 8 | 12 | 11 | 7 | 21 | 10 | 6 | 13 | 12 | 11.62 | 13 | 10 | 0.8 | -0.321 | 0.61972343 | 0.78489204 | 0.1392 | 0.2905 | 0.215 | 0.215 | TRUE | 1.2894 |
| Hyal4 | 0 | 0 | 0 | 0 | 0 | 0 | 0 | 0 | 0 | 0 | 0 | 0 | 0 | 0 | 0 | NA | NA | NA | NA | NA | NA | NA | NA | NA | NA |
| Hyal5 | 0 | 0 | 0 | 0 | 0 | 0 | 0 | 0 | 0 | 0 | 0 | 0 | 0 | 0 | 0 | NA | NA | NA | NA | NA | NA | NA | NA | NA | NA |
| Hyal6 | 3 | 0 | 0 | 2 | 1 | 1 | 3 | 0 | 0 | 2 | 1 | 1 | 1.15 | 1 | 1 | 1.2 | 0.263 | 0.89366021 | NA | 0.5179 | 2.8595 | 1.909 | 1.909 | TRUE | 1.443 |
| Hydin | 2017 | 1925 | 1964 | 2396 | 1923 | 1846 | 2142 | 1923 | 1972 | 1884 | 2069 | 2021 | 2001.72 | 2012 | 1991 | 0.989 | -0.016 | 0.85121409 | 0.92587032 | 0.0023 | 0.0085 | 0.0047 | 0.0047 | TRUE | 0.0385 |
| Hyi | 119 | 186 | 181 | 196 | 131 | 148 | 126 | 186 | 182 | 154 | 141 | 162 | 158.49 | 165 | 152 | 0.925 | -0.113 | 0.57303855 | 0.75135563 | 0.0176 | 0.0277 | 0.0227 | 0.0227 | TRUE | 0.431 |
| Hyls1 | 796 | 1024 | 922 | 1278 | 883 | 825 | 845 | 1023 | 926 | 1005 | 950 | 903 | 941.96 | 931 | 953 | 1.023 | 0.033 | 0.75619096 | 0.87119385 | 0.0049 | 0.0104 | 0.0072 | 0.0072 | TRUE | 0.0879 |
| Hyou1 | 16295 | 16327 | 16148 | 15326 | 11463 | 11692 | 17308 | 16306 | 16214 | 12048 | 12333 | 12799 | 14501.45 | 16609 | 12393 | 0.746 | -0.423 | 4.78E-11 | 3.72E-09 | 0.001 | 0.0071 | 0.0029 | 0.0029 | TRUE | 0.0166 |
| Hypk | 3412 | 3541 | 3545 | 5011 | 3531 | 3379 | 3624 | 3536 | 3560 | 3939 | 3799 | 3699 | 3692.9 | 3573 | 3812 | 1.067 | 0.094 | 0.1413374 | 0.32680725 | 4.00E-04 | 0.0078 | 0.0027 | 0.0027 | TRUE | 0.0105 |
| I730028E13R | 0 | 0 | 0 | 0 | 0 | 0 | 0 | 0 | 0 | 0 | 0 | 0 | 0 | 0 | 0 | NA | NA | NA | NA | NA | NA | NA | NA | NA | NA |
| I730030J21R | 1 | 3 | 1 | 0 | 0 | 0 | 1 | 3 | 1 | 0 | 0 | 0 | 0.84 | 2 | 0 | 0.106 | -3.237 | 0.1670717 | NA | 0 | 3.9115 | 2.653 | 2.653 | TRUE | 0.3579 |
| I830012O16F | 0 | 0 | 0 | 0 | 0 | 0 | 0 | 0 | 0 | 0 | 0 | 0 | 0 | 0 | 0 | NA | NA | NA | NA | NA | NA | NA | NA | NA | NA |
| I830077J02R | 2 | 0 | 0 | 1 | 5 | 2 | 2 | 0 | 0 | 1 | 5 | 2 | 1.75 | 1 | 3 | 3.956 | 1.984 | 0.27247598 | NA | 0.7617 | 1.8931 | 1.4414 | 1.4414 | TRUE | 0.6728 |
| Iah1 | 454 | 589 | 632 | 772 | 567 | 507 | 482 | 588 | 635 | 607 | 610 | 555 | 579.5 | 568 | 591 | 1.039 | 0.055 | 0.67359993 | 0.818099 | 0.0092 | 0.0126 | 0.0108 | 0.0108 | TRUE | 0.2043 |
| Iapp | 0 | 0 | 0 | 1 | 0 | 0 | 0 | 0 | 0 | 1 | 0 | 0 | 0.13 | 0 | 0 | 1.859 | 0.894 | 0.82650734 | NA | 0 | 25.1515 | 10 | 10 | TRUE | 0.2563 |
| Iars | 13481 | 12239 | 12895 | 17806 | 12774 | 12099 | 14319 | 12223 | 12948 | 13998 | 13744 | 13244 | 13412.72 | 13163 | 13662 | 1.038 | 0.054 | 0.51583704 | 0.70801619 | 0.0032 | 0.0071 | 0.0049 | 0.0049 | TRUE | 0.0721 |
| Iars2 | 7482 | 8877 | 9155 | 10510 | 7615 | 8366 | 7947 | 8866 | 9193 | 8262 | 8193 | 9158 | 8603.11 | 8669 | 8538 | 0.985 | -0.022 | 0.80528932 | 0.8995696 | 0.0038 | 0.0073 | 0.0057 | 0.0057 | TRUE | 0.0646 |
| Iba57 | 791 | 634 | 529 | 919 | 638 | 562 | 840 | 633 | 531 | 722 | 686 | 615 | 671.44 | 668 | 674 | 1.011 | 0.015 | 0.93006218 | 0.96557628 | 0.0287 | 0.0118 | 0.0202 | 0.0202 | TRUE | 0.5255 |
| Ibsp | 0 | 0 | 0 | 0 | 0 | 0 | 0 | 0 | 0 | 0 | 0 | 0 | 0 | 0 | 0 | NA | NA | NA | NA | NA | NA | NA | NA | NA | NA |
| Ibtk | 8391 | 7590 | 8063 | 9787 | 6867 | 6698 | 8913 | 7580 | 8096 | 7694 | 7388 | 7332 | 7833.88 | 8196 | 7471 | 0.912 | -0.133 | 0.11123298 | 0.28048257 | 0.0035 | 0.0073 | 0.0049 | 0.0049 | TRUE | 0.0713 |
| Ica1 | 307 | 438 | 482 | 515 | 420 | 411 | 326 | 437 | 484 | 405 | 452 | 450 | 425.69 | 416 | 436 | 1.046 | 0.065 | 0.6929399 | 0.83159723 | 0.0194 | 0.0146 | 0.0171 | 0.0171 | TRUE | 0.408 |
| Ica1l | 16 | 14 | 12 | 35 | 21 | 14 | 17 | 14 | 12 | 28 | 23 | 15 | 18.08 | 14 | 22 | 1.536 | 0.619 | 0.19002298 | 0.39514156 | 0 | 0.1892 | 0.103 | 0.103 | TRUE | 0.2213 |
| Icam1 | 4690 | 6240 | 6735 | 4769 | 3788 | 4529 | 4982 | 6232 | 6763 | 3749 | 4076 | 4958 | 5126.43 | 5992 | 4261 | 0.711 | -0.492 | 0.00073553 | 0.00701776 | 0.0221 | 0.0075 | 0.0151 | 0.0151 | TRUE | 0.2653 |
| Icam2 | 1 | 0 | 2 | 8 | 1 | 2 | 1 | 0 | 2 | 6 | 1 | 2 | 2.1 | 1 | 3 | 3.187 | 1.672 | 0.27051046 | NA | 0 | 1.5726 | 1.01 | 1.01 | TRUE | 1.0583 |
| Icam4 | 24 | 38 | 29 | 51 | 40 | 36 | 25 | 38 | 29 | 40 | 43 | 39 | 35.85 | 31 | 41 | 1.321 | 0.402 | 0.21982308 | 0.43208494 | 0 | 0.0988 | 0.0488 | 0.0488 | TRUE | 0.2749 |
| Icam5 | 135 | 160 | 97 | 99 | 82 | 103 | 143 | 160 | 97 | 78 | 88 | 113 | 113.23 | 133 | 93 | 0.693 | -0.528 | 0.04049682 | 0.14222128 | 0.0416 | 0.036 | 0.0388 | 0.0388 | TRUE | 0.5805 |
| Ick | 997 | 1038 | 1044 | 1393 | 1093 | 1030 | 1059 | 1037 | 1048 | 1095 | 1176 | 1127 | 1090.42 | 1048 | 1133 | 1.08 | 0.112 | 0.16212819 | 0.35742865 | 0 | 0.0099 | 0.0037 | 0.0037 | TRUE | 0.0133 |
| Icmt | 3507 | 3327 | 3257 | 3816 | 2830 | 2956 | 3725 | 3323 | 3270 | 3000 | 3045 | 3236 | 3266.45 | 3439 | 3094 | 0.899 | -0.153 | 0.07076085 | 0.20721131 | 0.003 | 0.0079 | 0.0049 | 0.0049 | TRUE | 0.064 |
| Icos | 0 | 0 | 0 | 0 | 0 | 1 | 0 | 0 | 0 | 0 | 0 | 1 | 0.18 | 0 | 0 | 1.859 | 0.894 | 0.82650734 | NA | 0 | 18.0643 | 10 | 10 | TRUE | 0.5383 |
| Icosl | 732 | 735 | 725 | 824 | 585 | 540 | 778 | 734 | 728 | 648 | 629 | 591 | 684.64 | 747 | 623 | 0.835 | -0.26 | 0.00628187 | 0.03728928 | 0 | 0.0117 | 0.005 | 0.005 | TRUE | 0.0228 |
| Ict1 | 1780 | 1988 | 2079 | 2667 | 1883 | 1869 | 1891 | 1985 | 2088 | 2097 | 2026 | 2046 | 2022.02 | 1988 | 2056 | 1.034 | 0.049 | 0.51643861 | 0.70838447 | 9.00E-04 | 0.0085 | 0.0036 | 0.0036 | TRUE | 0.0233 |
| Id1 | 1331 | 1092 | 899 | 1488 | 874 | 880 | 1414 | 1091 | 903 | 1170 | 940 | 963 | 1080.08 | 1136 | 1024 | 0.903 | -0.147 | 0.39483346 | 0.60835724 | 0.0306 | 0.01 | 0.0207 | 0.0207 | TRUE | 0.3636 |
| Id2 | 506 | 463 | 420 | 417 | 338 | 373 | 537 | 462 | 422 | 328 | 364 | 408 | 420.23 | 474 | 367 | 0.772 | -0.373 | 0.01063673 | 0.05514874 | 0.0113 | 0.0147 | 0.0129 | 0.0129 | TRUE | 0.1591 |
| Id3 | 573 | 253 | 183 | 688 | 485 | 303 | 609 | 253 | 184 | 541 | 522 | 332 | 406.57 | 349 | 465 | 1.335 | 0.417 | 0.45474707 | 0.65976182 | 0.2213 | 0.015 | 0.0983 | 0.2213 | TRUE | 3.5349 |
| Id4 | 88 | 99 | 103 | 142 | 143 | 104 | 93 | 99 | 103 | 112 | 154 | 114 | 112.52 | 98 | 127 | 1.279 | 0.355 | 0.09452592 | 0.25193271 | 0.011 | 0.0362 | 0.0233 | 0.0233 | TRUE | 0.3573 |
| Ide | 10304 | 10720 | 11100 | 11627 | 8321 | 8985 | 10945 | 10706 | 11146 | 9140 | 8953 | 9835 | 10120.84 | 10932 | 9309 | 0.852 | -0.232 | 0.00060509 | 0.00604779 | 0.0013 | 0.0072 | 0.0032 | 0.0032 | TRUE | 0.0297 |
| Idh1 | 9667 | 8805 | 8203 | 11663 | 7894 | 8040 | 10268 | 8794 | 8237 | 9169 | 8493 | 8801 | 8960.24 | 9100 | 8821 | 0.969 | -0.045 | 0.6565714 | 0.80670515 | 0.0062 | 0.0073 | 0.0072 | 0.0072 | TRUE | 0.1541 |
| Idh2 | 7984 | 10167 | 10237 | 8691 | 6707 | 7470 | 8480 | 10154 | 10279 | 6832 | 7216 | 8177 | 8523.15 | 9638 | 7408 | 0.769 | -0.38 | 0.0006096 | 0.00608908 | 0.0096 | 0.0073 | 0.0087 | 0.0087 | TRUE | 0.1347 |
| Idh3a | 8831 | 9251 | 9172 | 10452 | 7363 | 7746 | 9380 | 9239 | 9210 | 8217 | 7922 | 8479 | 8741.12 | 9276 | 8206 | 0.885 | -0.177 | 0.0031978 | 0.02229698 | 5.00E-04 | 0.0073 | 0.0025 | 0.0025 | TRUE | 0.0111 |
| Idh3b | 4207 | 5010 | 4949 | 5894 | 4139 | 4473 | 4469 | 5004 | 4969 | 4633 | 4453 | 4896 | 4737.42 | 4814 | 4661 | 0.968 | -0.047 | 0.56778609 | 0.74733351 | 0.0026 | 0.0076 | 0.0046 | 0.0046 | TRUE | 0.0479 |
| Idh3g | 5653 | 6844 | 6854 | 7618 | 5662 | 5766 | 6004 | 6835 | 6882 | 5989 | 6092 | 6312 | 6352.37 | 6574 | 6131 | 0.932 | -0.101 | 0.217597 | 0.42962457 | 0.0029 | 0.0074 | 0.0047 | 0.0047 | TRUE | 0.07 |

| | | | | | | | | | | | | | | | | | | | | | | | | | |
|---|---|---|---|---|---|---|---|---|---|---|---|---|---|---|---|---|---|---|---|---|---|---|---|---|---|
| Idi1 | 2136 | 1733 | 1705 | 2204 | 1525 | 1579 | 2269 | 1731 | 1712 | 1733 | 1641 | 1728 | 1802.24 | 1904 | 1701 | 0.893 | -0.162 | 0.20071347 | 0.40847133 | 0.0134 | 0.0087 | 0.0111 | 0.0111 | TRUE | 0.3392 |
| Idi2 | 0 | 0 | 2 | 0 | 0 | 0 | 0 | 0 | 2 | 0 | 0 | 0 | 0.33 | 1 | 0 | 0.267 | -1.904 | 0.63742563 | NA | 0.0046 | 9.8498 | 10 | 10 | TRUE | 0.974 |
| Idnk | 483 | 496 | 419 | 557 | 398 | 368 | 513 | 495 | 421 | 438 | 428 | 403 | 449.67 | 476 | 423 | 0.889 | -0.17 | 0.16847767 | 0.36640749 | 0.0043 | 0.0142 | 0.0087 | 0.0087 | TRUE | 0.1219 |
| Ido1 | 5 | 8 | 6 | 11 | 0 | 2 | 5 | 8 | 6 | 9 | 0 | 2 | 5.03 | 6 | 4 | 0.584 | -0.777 | 0.47398043 | 0.67469488 | 0.6009 | 0.6623 | 0.6385 | 0.6385 | TRUE | 1.7955 |
| Ido2 | 53 | 48 | 47 | 57 | 41 | 35 | 56 | 48 | 47 | 45 | 44 | 38 | 46.44 | 50 | 42 | 0.843 | -0.246 | 0.38083359 | 0.59526263 | 0 | 0.0778 | 0.0353 | 0.0353 | TRUE | 0.0806 |
| Ids | 975 | 966 | 950 | 1345 | 1013 | 1005 | 1036 | 965 | 954 | 1057 | 1090 | 1100 | 1033.61 | 985 | 1082 | 1.099 | 0.136 | 0.10324897 | 0.2673962 | 0 | 0.0101 | 0.0041 | 0.0041 | TRUE | 0.0242 |
| Idua | 251 | 506 | 533 | 274 | 242 | 336 | 267 | 505 | 535 | 215 | 260 | 368 | 358.45 | 436 | 281 | 0.644 | -0.634 | 0.02373632 | 0.09803994 | 0.1007 | 0.0161 | 0.0538 | 0.0538 | TRUE | 1.328 |
| Ier2 | 5481 | 4895 | 4328 | 6941 | 4830 | 3916 | 5822 | 4889 | 4346 | 5456 | 5197 | 4287 | 4999.36 | 5019 | 4980 | 0.992 | -0.011 | 0.93658098 | 0.96825513 | 0.0186 | 0.0076 | 0.0133 | 0.0133 | TRUE | 0.2386 |
| Ier3 | 612 | 1125 | 1414 | 907 | 934 | 1196 | 650 | 1124 | 1420 | 713 | 1005 | 1309 | 1036.76 | 1065 | 1009 | 0.948 | -0.078 | 0.84768106 | 0.9238415 | 0.1166 | 0.0101 | 0.0546 | 0.1166 | TRUE | 0.341 |
| Ier3ip1 | 1346 | 1485 | 1507 | 1927 | 1340 | 1442 | 1430 | 1483 | 1513 | 1515 | 1442 | 1578 | 1493.51 | 1475 | 1512 | 1.025 | 0.035 | 0.65800789 | 0.80743861 | 7.00E-04 | 0.0091 | 0.0038 | 0.0038 | TRUE | 0.0192 |
| Ier5 | 204 | 286 | 294 | 213 | 175 | 187 | 217 | 286 | 295 | 167 | 188 | 205 | 226.33 | 266 | 187 | 0.701 | -0.513 | 0.00375473 | 0.02521059 | 0.0149 | 0.0215 | 0.0181 | 0.0181 | TRUE | 0.289 |
| Ier5l | 432 | 543 | 530 | 733 | 570 | 483 | 459 | 542 | 532 | 576 | 613 | 529 | 541.93 | 511 | 573 | 1.12 | 0.164 | 0.17047949 | 0.36901257 | 0.0047 | 0.013 | 0.0085 | 0.0085 | TRUE | 0.0924 |
| Iffo1 | 567 | 519 | 509 | 616 | 510 | 478 | 602 | 518 | 511 | 484 | 549 | 523 | 531.32 | 544 | 519 | 0.953 | -0.07 | 0.5573551 | 0.73994989 | 0.0044 | 0.0131 | 0.0082 | 0.0082 | TRUE | 0.1022 |
| Iffo2 | 1110 | 1510 | 1582 | 1614 | 1336 | 1512 | 1179 | 1508 | 1588 | 1269 | 1437 | 1655 | 1439.49 | 1425 | 1454 | 1.019 | 0.028 | 0.8514654 | 0.92588509 | 0.0203 | 0.0092 | 0.015 | 0.015 | TRUE | 0.274 |
| Ifi202b | 0 | 0 | 0 | 0 | 0 | 0 | 0 | 0 | 0 | 0 | 0 | 0 | 0 | 0 | 0 | NA | NA | NA | NA | NA | NA | NA | NA | NA | NA |
| Ifi203 | 0 | 0 | 0 | 0 | 0 | 1 | 0 | 0 | 0 | 0 | 0 | 1 | 0.18 | 0 | 0 | 1.859 | 0.894 | 0.82650734 | NA | 0 | 18.0643 | 10 | 10 | TRUE | 0.5383 |
| Ifi204 | 0 | 0 | 0 | 0 | 0 | 0 | 0 | 0 | 0 | 0 | 0 | 0 | 0 | 0 | 0 | NA | NA | NA | NA | NA | NA | NA | NA | NA | NA |
| Ifi205 | 0 | 0 | 0 | 0 | 0 | 0 | 0 | 0 | 0 | 0 | 0 | 0 | 0 | 0 | 0 | NA | NA | NA | NA | NA | NA | NA | NA | NA | NA |
| Ifi27l1 | 13 | 24 | 28 | 45 | 40 | 35 | 14 | 24 | 28 | 35 | 43 | 38 | 30.44 | 22 | 39 | 1.762 | 0.817 | 0.03330462 | 0.12362448 | 0.0068 | 0.1151 | 0.0705 | 0.0705 | TRUE | 0.5735 |
| Ifi27l2a | 7 | 10 | 6 | 10 | 6 | 12 | 7 | 10 | 6 | 8 | 6 | 13 | 8.48 | 8 | 9 | 1.163 | 0.218 | 0.74344764 | 0.86314121 | 0 | 0.3952 | 0.2012 | 0.2012 | TRUE | 0.4257 |
| Ifi27l2b | 0 | 0 | 0 | 0 | 0 | 0 | 0 | 0 | 0 | 0 | 0 | 0 | 0 | 0 | 0 | NA | NA | NA | NA | NA | NA | NA | NA | NA | NA |
| Ifi30 | 3114 | 4578 | 5085 | 2990 | 2327 | 3144 | 3308 | 4572 | 5106 | 2351 | 2504 | 3442 | 3546.9 | 4329 | 2766 | 0.639 | -0.647 | 0.00058378 | 0.0058683 | 0.0451 | 0.0078 | 0.0252 | 0.0252 | TRUE | 0.4839 |
| Ifi35 | 159 | 178 | 107 | 286 | 179 | 135 | 169 | 178 | 107 | 225 | 193 | 148 | 169.88 | 151 | 189 | 1.248 | 0.32 | 0.19604641 | 0.40230395 | 0.0509 | 0.0263 | 0.0381 | 0.0381 | TRUE | 0.4272 |
| Ifi44 | 0 | 0 | 0 | 0 | 0 | 0 | 0 | 0 | 0 | 0 | 0 | 0 | 0 | 0 | 0 | NA | NA | NA | NA | NA | NA | NA | NA | NA | NA |
| Ifi44l | 0 | 0 | 0 | 0 | 0 | 0 | 0 | 0 | 0 | 0 | 0 | 0 | 0 | 0 | 0 | NA | NA | NA | NA | NA | NA | NA | NA | NA | NA |
| Ifi47 | 1 | 4 | 0 | 3 | 1 | 0 | 1 | 4 | 0 | 2 | 1 | 0 | 1.42 | 2 | 1 | 0.702 | -0.511 | 0.78454392 | NA | 1.0416 | 2.3348 | 1.8009 | 1.8009 | TRUE | 0.6637 |
| Ifih1 | 2 | 1 | 2 | 7 | 8 | 6 | 2 | 1 | 2 | 6 | 9 | 7 | 4.3 | 2 | 7 | 4.02 | 2.007 | 0.04505073 | NA | 0 | 0.7728 | 0.3518 | 0.3518 | TRUE | 0.1181 |
| Ifit1 | 0 | 3 | 6 | 2 | 3 | 4 | 0 | 3 | 6 | 2 | 3 | 4 | 3.03 | 3 | 3 | 0.997 | -0.005 | 0.99706361 | NA | 0.4219 | 1.093 | 0.808 | 0.808 | TRUE | 0.1873 |
| Ifit2 | 107 | 139 | 148 | 116 | 80 | 106 | 114 | 139 | 149 | 91 | 86 | 116 | 115.73 | 134 | 98 | 0.729 | -0.456 | 0.03322841 | 0.12342876 | 0.0124 | 0.0354 | 0.0242 | 0.0242 | TRUE | 0.2462 |
| Ifit3 | 1 | 1 | 1 | 3 | 1 | 1 | 1 | 1 | 1 | 2 | 1 | 1 | 1.27 | 1 | 1 | 1.513 | 0.597 | 0.70531437 | NA | 0 | 2.6099 | 0.9808 | 0.9808 | TRUE | 0.2233 |
| Ifitm1 | 1285 | 1427 | 1415 | 938 | 963 | 1541 | 1365 | 1425 | 1421 | 737 | 1036 | 1687 | 1278.54 | 1404 | 1153 | 0.821 | -0.284 | 0.24403392 | 0.4605753 | 0.0849 | 0.0095 | 0.0421 | 0.0421 | TRUE | 0.7192 |
| Ifitm10 | 3 | 6 | 4 | 1 | 3 | 3 | 3 | 6 | 4 | 1 | 3 | 3 | 3.42 | 4 | 2 | 0.534 | -0.906 | 0.3920668 | NA | 0 | 0.9715 | 0.4844 | 0.4844 | TRUE | 0.5395 |
| Ifitm2 | 6273 | 7169 | 7490 | 8259 | 6786 | 8228 | 6663 | 7160 | 7521 | 6493 | 7301 | 9007 | 7357.38 | 7115 | 7600 | 1.068 | 0.095 | 0.45992603 | 0.66406067 | 0.0147 | 0.0074 | 0.0118 | 0.0118 | TRUE | 0.3199 |
| Ifitm3 | 779 | 751 | 777 | 1324 | 1104 | 1414 | 827 | 750 | 780 | 1041 | 1188 | 1548 | 1022.36 | 786 | 1259 | 1.601 | 0.679 | 1.00E-05 | 0.00019932 | 0.0212 | 0.0101 | 0.016 | 0.016 | TRUE | 0.4693 |
| Ifitm5 | 22 | 24 | 10 | 12 | 23 | 16 | 23 | 24 | 10 | 9 | 25 | 18 | 18.18 | 19 | 17 | 0.891 | -0.166 | 0.76813465 | 0.87911035 | 0.1562 | 0.1881 | 0.1726 | 0.1726 | TRUE | 0.2657 |
| Ifitm6 | 1 | 1 | 0 | 2 | 1 | 6 | 1 | 1 | 0 | 2 | 1 | 7 | 1.88 | 1 | 3 | 4.438 | 2.15 | 0.19592819 | NA | 0 | 1.7597 | 1.0828 | 1.0828 | TRUE | 1.2067 |
| Ifitm7 | 12 | 13 | 4 | 9 | 5 | 10 | 13 | 13 | 4 | 7 | 5 | 11 | 8.86 | 10 | 8 | 0.785 | -0.35 | 0.62461521 | 0.78842986 | 0.1138 | 0.3788 | 0.2537 | 0.2537 | TRUE | 0.9461 |
| Ifltd1 | 0 | 0 | 0 | 0 | 0 | 0 | 0 | 0 | 0 | 0 | 0 | 0 | 0 | 0 | 0 | NA | NA | NA | NA | NA | NA | NA | NA | NA | NA |
| Ifna1 | 0 | 0 | 0 | 0 | 0 | 0 | 0 | 0 | 0 | 0 | 0 | 0 | 0 | 0 | 0 | NA | NA | NA | NA | NA | NA | NA | NA | NA | NA |
| Ifna11 | 0 | 0 | 0 | 0 | 0 | 0 | 0 | 0 | 0 | 0 | 0 | 0 | 0 | 0 | 0 | NA | NA | NA | NA | NA | NA | NA | NA | NA | NA |
| Ifna12 | 0 | 0 | 0 | 0 | 0 | 0 | 0 | 0 | 0 | 0 | 0 | 0 | 0 | 0 | 0 | NA | NA | NA | NA | NA | NA | NA | NA | NA | NA |
| Ifna13 | 0 | 0 | 0 | 0 | 0 | 0 | 0 | 0 | 0 | 0 | 0 | 0 | 0 | 0 | 0 | NA | NA | NA | NA | NA | NA | NA | NA | NA | NA |
| Ifna14 | 0 | 0 | 0 | 0 | 0 | 0 | 0 | 0 | 0 | 0 | 0 | 0 | 0 | 0 | 0 | NA | NA | NA | NA | NA | NA | NA | NA | NA | NA |
| Ifna2 | 0 | 0 | 0 | 0 | 0 | 0 | 0 | 0 | 0 | 0 | 0 | 0 | 0 | 0 | 0 | NA | NA | NA | NA | NA | NA | NA | NA | NA | NA |
| Ifna4 | 0 | 0 | 0 | 0 | 0 | 0 | 0 | 0 | 0 | 0 | 0 | 0 | 0 | 0 | 0 | NA | NA | NA | NA | NA | NA | NA | NA | NA | NA |
| Ifna5 | 0 | 0 | 0 | 0 | 0 | 0 | 0 | 0 | 0 | 0 | 0 | 0 | 0 | 0 | 0 | NA | NA | NA | NA | NA | NA | NA | NA | NA | NA |
| Ifna6 | 0 | 0 | 0 | 0 | 0 | 0 | 0 | 0 | 0 | 0 | 0 | 0 | 0 | 0 | 0 | NA | NA | NA | NA | NA | NA | NA | NA | NA | NA |
| Ifna7 | 0 | 0 | 0 | 0 | 0 | 0 | 0 | 0 | 0 | 0 | 0 | 0 | 0 | 0 | 0 | NA | NA | NA | NA | NA | NA | NA | NA | NA | NA |
| Ifna9 | 0 | 0 | 0 | 0 | 0 | 0 | 0 | 0 | 0 | 0 | 0 | 0 | 0 | 0 | 0 | NA | NA | NA | NA | NA | NA | NA | NA | NA | NA |
| Ifnab | 0 | 0 | 0 | 0 | 0 | 0 | 0 | 0 | 0 | 0 | 0 | 0 | 0 | 0 | 0 | NA | NA | NA | NA | NA | NA | NA | NA | NA | NA |
| Ifnar1 | 2584 | 2350 | 2453 | 2737 | 1924 | 1917 | 2745 | 2347 | 2463 | 2152 | 2070 | 2098 | 2312.48 | 2518 | 2107 | 0.837 | -0.257 | 0.00321216 | 0.02233786 | 0.0028 | 0.0083 | 0.005 | 0.005 | TRUE | 0.0748 |
| Ifnar2 | 430 | 678 | 649 | 523 | 387 | 463 | 457 | 677 | 652 | 411 | 416 | 507 | 519.98 | 595 | 445 | 0.746 | -0.422 | 0.01660491 | 0.07628681 | 0.0275 | 0.0132 | 0.0204 | 0.0204 | TRUE | 0.4837 |
| Ifnb1 | 0 | 0 | 0 | 0 | 0 | 0 | 0 | 0 | 0 | 0 | 0 | 0 | 0 | 0 | 0 | NA | NA | NA | NA | NA | NA | NA | NA | NA | NA |
| Ifne | 0 | 0 | 0 | 0 | 0 | 0 | 0 | 0 | 0 | 0 | 0 | 0 | 0 | 0 | 0 | NA | NA | NA | NA | NA | NA | NA | NA | NA | NA |
| Ifng | 0 | 0 | 0 | 0 | 1 | 0 | 0 | 0 | 0 | 0 | 1 | 0 | 0.18 | 0 | 0 | 1.859 | 0.894 | 0.82650734 | NA | 0 | 18.3785 | 10 | 10 | TRUE | 0.5204 |
| Ifngr1 | 313 | 391 | 354 | 526 | 399 | 426 | 332 | 390 | 355 | 413 | 429 | 466 | 397.92 | 359 | 436 | 1.212 | 0.277 | 0.0238975 | 0.0984429 | 0.0026 | 0.0152 | 0.0083 | 0.0083 | TRUE | 0.0654 |
| Ifngr2 | 488 | 496 | 502 | 308 | 277 | 279 | 518 | 495 | 504 | 242 | 298 | 305 | 393.89 | 506 | 282 | 0.555 | -0.849 | 1.13E-10 | 8.23E-09 | 0.0046 | 0.0153 | 0.0097 | 0.0097 | TRUE | 0.1851 |
| Ifnk | 0 | 0 | 0 | 0 | 0 | 0 | 0 | 0 | 0 | 0 | 0 | 0 | 0 | 0 | 0 | NA | NA | NA | NA | NA | NA | NA | NA | NA | NA |
| Ifnl2 | 0 | 0 | 0 | 1 | 0 | 0 | 0 | 0 | 0 | 1 | 0 | 0 | 0.13 | 0 | 0 | 1.859 | 0.894 | 0.82650734 | NA | 0 | 25.1515 | 10 | 10 | TRUE | 0.2563 |
| Ifnl3 | 0 | 0 | 0 | 0 | 0 | 0 | 0 | 0 | 0 | 0 | 0 | 0 | 0 | 0 | 0 | NA | NA | NA | NA | NA | NA | NA | NA | NA | NA |
| Ifnlr1 | 353 | 461 | 434 | 448 | 297 | 355 | 375 | 460 | 436 | 352 | 320 | 389 | 388.58 | 424 | 354 | 0.834 | -0.262 | 0.05746268 | 0.18082631 | 0.0074 | 0.0154 | 0.0112 | 0.0112 | TRUE | 0.116 |
| Ifnz | 0 | 0 | 0 | 0 | 0 | 0 | 0 | 0 | 0 | 0 | 0 | 0 | 0 | 0 | 0 | NA | NA | NA | NA | NA | NA | NA | NA | NA | NA |
| Ifrd1 | 5457 | 6750 | 6707 | 7980 | 5541 | 5012 | 5796 | 6741 | 6735 | 6273 | 5962 | 5486 | 6165.59 | 6424 | 5907 | 0.92 | -0.121 | 0.20738871 | 0.41661646 | 0.0056 | 0.0074 | 0.0065 | 0.0065 | TRUE | 0.0891 |
| Ifrd2 | 4578 | 4151 | 4184 | 4643 | 3363 | 3242 | 4863 | 4146 | 4201 | 3650 | 3618 | 3549 | 4004.45 | 4403 | 3606 | 0.819 | -0.288 | 0.00105802 | 0.00930642 | 0.0038 | 0.0077 | 0.0053 | 0.0053 | TRUE | 0.1013 |
| Ift122 | 826 | 939 | 1043 | 1256 | 933 | 1047 | 877 | 938 | 1047 | 987 | 1004 | 1146 | 999.96 | 954 | 1046 | 1.095 | 0.131 | 0.24127127 | 0.45797694 | 0.0061 | 0.0102 | 0.008 | 0.008 | TRUE | 0.0871 |
| Ift140 | 1418 | 1779 | 1852 | 2089 | 1544 | 1781 | 1506 | 1777 | 1860 | 1642 | 1661 | 1950 | 1732.59 | 1714 | 1751 | 1.021 | 0.03 | 0.8002439 | 0.89686328 | 0.0101 | 0.0088 | 0.0095 | 0.0095 | TRUE | 0.1356 |
| Ift172 | 979 | 1254 | 1333 | 1534 | 1132 | 1171 | 1040 | 1252 | 1338 | 1206 | 1218 | 1282 | 1222.74 | 1210 | 1235 | 1.02 | 0.029 | 0.80183063 | 0.89762453 | 0.008 | 0.0096 | 0.0087 | 0.0087 | TRUE | 0.1811 |
| Ift20 | 1994 | 1742 | 1504 | 2168 | 1599 | 1436 | 2118 | 1740 | 1510 | 1704 | 1720 | 1572 | 1727.43 | 1789 | 1665 | 0.931 | -0.103 | 0.43913662 | 0.64641993 | 0.0145 | 0.0088 | 0.0122 | 0.0122 | TRUE | 0.3112 |
| Ift27 | 409 | 373 | 386 | 575 | 398 | 366 | 434 | 373 | 388 | 452 | 428 | 401 | 412.57 | 398 | 427 | 1.074 | 0.103 | 0.39427555 | 0.60787039 | 0.0026 | 0.0149 | 0.0082 | 0.0082 | TRUE | 0.0722 |

| | | | | | | | | | | | | | | | | | | | | | | | | | |
|---|---|---|---|---|---|---|---|---|---|---|---|---|---|---|---|---|---|---|---|---|---|---|---|---|---|
| Ift43 | 324 | 466 | 450 | 484 | 361 | 413 | 344 | 465 | 452 | 380 | 388 | 452 | 413.73 | 420 | 407 | 0.967 | -0.049 | 0.75237536 | 0.86946543 | 0.0152 | 0.0149 | 0.015 | 0.015 | TRUE | 0.2887 |
| Ift46 | 3345 | 3080 | 3238 | 4562 | 3189 | 3065 | 3553 | 3076 | 3251 | 3586 | 3431 | 3355 | 3375.48 | 3293 | 3457 | 1.05 | 0.071 | 0.39913829 | 0.61225531 | 0.0028 | 0.0079 | 0.0048 | 0.0048 | TRUE | 0.0576 |
| Ift52 | 2040 | 2628 | 2733 | 2714 | 2006 | 2225 | 2167 | 2625 | 2744 | 2134 | 2158 | 2436 | 2377.19 | 2512 | 2243 | 0.892 | -0.164 | 0.15602936 | 0.3487095 | 0.0095 | 0.0083 | 0.0092 | 0.0092 | TRUE | 0.1747 |
| Ift57 | 565 | 829 | 807 | 1054 | 807 | 874 | 600 | 828 | 810 | 829 | 868 | 957 | 815.33 | 746 | 885 | 1.184 | 0.244 | 0.09225877 | 0.24788847 | 0.0167 | 0.0109 | 0.0139 | 0.0139 | TRUE | 0.3451 |
| Ift74 | 522 | 496 | 555 | 1129 | 733 | 741 | 554 | 495 | 557 | 888 | 789 | 811 | 682.4 | 535 | 829 | 1.55 | 0.632 | 2.47E-09 | 1.36E-07 | 0.0027 | 0.0117 | 0.0065 | 0.0065 | TRUE | 0.0511 |
| Ift80 | 1148 | 1114 | 1178 | 1641 | 1129 | 1214 | 1219 | 1113 | 1183 | 1290 | 1215 | 1329 | 1224.74 | 1172 | 1278 | 1.091 | 0.126 | 0.1434281 | 0.33004787 | 0.0012 | 0.0096 | 0.0045 | 0.0045 | TRUE | 0.0234 |
| Ift81 | 765 | 1029 | 1125 | 1339 | 917 | 930 | 813 | 1028 | 1130 | 1053 | 987 | 1018 | 1004.52 | 990 | 1019 | 1.029 | 0.042 | 0.75455749 | 0.87045646 | 0.0134 | 0.0102 | 0.0119 | 0.0119 | TRUE | 0.292 |
| Ift88 | 616 | 678 | 729 | 876 | 720 | 616 | 654 | 677 | 732 | 689 | 775 | 674 | 700.17 | 688 | 713 | 1.035 | 0.05 | 0.63769386 | 0.79603827 | 0.003 | 0.0116 | 0.0067 | 0.0067 | TRUE | 0.0674 |
| Igbp1 | 2242 | 2859 | 3081 | 3117 | 2237 | 2501 | 2381 | 2855 | 3094 | 2450 | 2407 | 2738 | 2654.22 | 2777 | 2532 | 0.911 | -0.134 | 0.25739268 | 0.47607984 | 0.0105 | 0.0081 | 0.0097 | 0.0097 | TRUE | 0.188 |
| Igbp1b | 26 | 26 | 23 | 80 | 60 | 64 | 28 | 26 | 23 | 63 | 65 | 70 | 45.7 | 26 | 66 | 2.574 | 1.364 | 3.77E-06 | 8.69E-05 | 0 | 0.079 | 0.0353 | 0.0353 | TRUE | 0.0441 |
| Igdcc3 | 555 | 648 | 664 | 855 | 633 | 521 | 590 | 647 | 667 | 672 | 681 | 570 | 637.82 | 635 | 641 | 1.011 | 0.016 | 0.88989565 | 0.94623031 | 0.0051 | 0.0121 | 0.0082 | 0.0082 | TRUE | 0.1075 |
| Igdcc4 | 1023 | 1199 | 1166 | 968 | 699 | 744 | 1087 | 1197 | 1171 | 761 | 752 | 814 | 963.72 | 1152 | 776 | 0.673 | -0.571 | 2.87E-10 | 1.99E-08 | 0.0011 | 0.0103 | 0.0048 | 0.0048 | TRUE | 0.029 |
| Igf1 | 1 | 3 | 1 | 3 | 3 | 6 | 1 | 3 | 1 | 2 | 3 | 7 | 2.87 | 2 | 4 | 2.358 | 1.238 | 0.29565995 | NA | 0 | 1.155 | 0.5855 | 0.5855 | TRUE | 0.4625 |
| Igf1r | 2709 | 2986 | 3301 | 3666 | 2659 | 2738 | 2877 | 2982 | 3315 | 2882 | 2861 | 2997 | 2985.69 | 3058 | 2913 | 0.953 | -0.07 | 0.39958984 | 0.61268945 | 0.0026 | 0.008 | 0.0047 | 0.0047 | TRUE | 0.0655 |
| Igf2 | 31 | 43 | 55 | 59 | 49 | 57 | 33 | 43 | 55 | 46 | 53 | 62 | 48.77 | 44 | 54 | 1.225 | 0.293 | 0.34711068 | 0.56607908 | 0.0209 | 0.0745 | 0.0491 | 0.0491 | TRUE | 0.2938 |
| Igf2as | 1 | 0 | 1 | 4 | 0 | 1 | 1 | 0 | 1 | 3 | 0 | 1 | 1.05 | 1 | 1 | 2.129 | 1.09 | 0.59527377 | NA | 0 | 3.1418 | 1.9522 | 1.9522 | TRUE | 0.3095 |
| Igf2bp1 | 20604 | 17933 | 19091 | 34752 | 23738 | 20038 | 21885 | 17910 | 19169 | 27319 | 25541 | 21935 | 22293.16 | 19655 | 24932 | 1.269 | 0.343 | 0.00284641 | 0.02033221 | 0.0113 | 0.0071 | 0.0095 | 0.0095 | TRUE | 0.1353 |
| Igf2bp2 | 4381 | 6347 | 6642 | 7393 | 5980 | 6880 | 4653 | 6339 | 6669 | 5812 | 6434 | 7531 | 6239.77 | 5887 | 6592 | 1.12 | 0.163 | 0.29014437 | 0.5113596 | 0.0235 | 0.0074 | 0.017 | 0.017 | TRUE | 0.4097 |
| Igf2bp3 | 4141 | 4308 | 4462 | 5287 | 4110 | 3787 | 4398 | 4302 | 4480 | 4156 | 4422 | 4145 | 4317.51 | 4393 | 4241 | 0.965 | -0.051 | 0.43549381 | 0.64367054 | 5.00E-04 | 0.0077 | 0.0029 | 0.0029 | TRUE | 0.0168 |
| Igf2r | 15017 | 15004 | 16762 | 17191 | 12724 | 12928 | 15951 | 14985 | 16831 | 13514 | 13690 | 14152 | 14853.72 | 15922 | 13785 | 0.866 | -0.208 | 0.00366422 | 0.02471785 | 0.0019 | 0.0071 | 0.0036 | 0.0036 | TRUE | 0.0325 |
| Igfals | 64 | 77 | 81 | 82 | 63 | 72 | 68 | 77 | 81 | 64 | 68 | 79 | 72.88 | 75 | 70 | 0.929 | -0.106 | 0.64919934 | 0.8033859 | 0 | 0.0521 | 0.0252 | 0.0252 | TRUE | 0.0955 |
| Igfbp1 | 0 | 0 | 0 | 0 | 1 | 0 | 0 | 0 | 0 | 0 | 1 | 0 | 0.18 | 0 | 0 | 1.859 | 0.894 | 0.82650734 | NA | 0 | 18.3785 | 10 | 10 | TRUE | 0.5204 |
| Igfbp2 | 23467 | 30761 | 30354 | 20904 | 16035 | 19064 | 24926 | 30721 | 30479 | 16433 | 17253 | 20869 | 23446.72 | 28709 | 18185 | 0.633 | -0.659 | 1.47E-07 | 4.98E-06 | 0.0149 | 0.007 | 0.0113 | 0.0113 | TRUE | 0.2038 |
| Igfbp3 | 987 | 758 | 659 | 1235 | 855 | 642 | 1048 | 757 | 662 | 971 | 920 | 703 | 843.44 | 822 | 865 | 1.052 | 0.073 | 0.7032998 | 0.83766892 | 0.0415 | 0.0108 | 0.0255 | 0.0255 | TRUE | 0.685 |
| Igfbp4 | 1477 | 3828 | 4215 | 1462 | 1233 | 1913 | 1569 | 3823 | 4232 | 1149 | 1327 | 2094 | 2365.7 | 3208 | 1523 | 0.475 | -1.075 | 0.02908501 | 0.11251627 | 0.1743 | 0.0083 | 0.0726 | 0.1743 | TRUE | 1.606 |
| Igfbp5 | 76 | 87 | 97 | 76 | 60 | 77 | 81 | 87 | 97 | 60 | 65 | 84 | 78.93 | 88 | 70 | 0.783 | -0.353 | 0.14169142 | 0.32724514 | 0.0074 | 0.0486 | 0.0288 | 0.0288 | TRUE | 0.2975 |
| Igfbp6 | 13 | 21 | 18 | 28 | 8 | 12 | 14 | 21 | 18 | 22 | 9 | 13 | 16.1 | 18 | 15 | 0.841 | -0.251 | 0.63768999 | 0.79603827 | 0.0668 | 0.2115 | 0.1421 | 0.1421 | TRUE | 0.6597 |
| Igfbp7 | 66 | 44 | 40 | 96 | 46 | 40 | 70 | 44 | 40 | 75 | 49 | 44 | 53.83 | 51 | 56 | 1.104 | 0.142 | 0.6838215 | 0.82484365 | 0.0714 | 0.0681 | 0.0696 | 0.0696 | TRUE | 0.8339 |
| Igfbpl1 | 301 | 250 | 240 | 313 | 189 | 201 | 320 | 250 | 241 | 246 | 203 | 220 | 246.64 | 270 | 223 | 0.828 | -0.272 | 0.10859386 | 0.27606395 | 0.0131 | 0.0203 | 0.0166 | 0.0166 | TRUE | 0.2859 |
| Igfl3 | 0 | 0 | 0 | 0 | 0 | 0 | 0 | 0 | 0 | 0 | 0 | 0 | 0 | 0 | 0 | NA | NA | NA | NA | NA | NA | NA | NA | NA | NA |
| Igflr1 | 74 | 74 | 59 | 83 | 65 | 78 | 79 | 74 | 59 | 65 | 70 | 85 | 72.05 | 71 | 73 | 1.038 | 0.054 | 0.82653333 | 0.91211327 | 0 | 0.0526 | 0.0304 | 0.0304 | TRUE | 0.1792 |
| Igfn1 | 7 | 7 | 11 | 4 | 5 | 3 | 7 | 7 | 11 | 3 | 5 | 3 | 6.21 | 8 | 4 | 0.459 | -1.125 | 0.14742693 | 0.33659779 | 0 | 0.5371 | 0.2491 | 0.2491 | TRUE | 0.2158 |
| Ighmbp2 | 739 | 807 | 799 | 1073 | 813 | 742 | 785 | 806 | 802 | 844 | 875 | 812 | 820.61 | 798 | 844 | 1.057 | 0.08 | 0.34962602 | 0.56854332 | 0 | 0.0109 | 0.0041 | 0.0041 | TRUE | 0.0119 |
| Igj | 0 | 0 | 0 | 1 | 1 | 2 | 0 | 0 | 0 | 1 | 1 | 2 | 0.68 | 0 | 1 | 7.126 | 2.833 | 0.25569871 | NA | 0 | 4.8859 | 3.111 | 3.111 | TRUE | 0.1691 |
| Igll1 | 2 | 4 | 2 | 8 | 5 | 8 | 2 | 4 | 2 | 6 | 5 | 9 | 4.76 | 3 | 7 | 2.499 | 1.321 | 0.14106916 | 0.3264773 | 0 | 0.6992 | 0.3209 | 0.3209 | TRUE | 0.2088 |
| Iglon5 | 244 | 245 | 214 | 264 | 199 | 209 | 259 | 245 | 215 | 208 | 214 | 229 | 228.19 | 240 | 217 | 0.904 | -0.145 | 0.32009047 | 0.54115843 | 0 | 0.0213 | 0.011 | 0.011 | TRUE | 0.0904 |
| Igsf1 | 429 | 377 | 387 | 340 | 260 | 279 | 456 | 377 | 389 | 267 | 280 | 305 | 345.54 | 407 | 284 | 0.697 | -0.52 | 0.00012789 | 0.00168512 | 0.0051 | 0.0164 | 0.0103 | 0.0103 | TRUE | 0.1254 |
| Igsf10 | 336 | 317 | 304 | 581 | 362 | 326 | 357 | 317 | 305 | 457 | 389 | 357 | 363.64 | 326 | 401 | 1.233 | 0.302 | 0.03651123 | 0.13177462 | 0.009 | 0.016 | 0.0122 | 0.0122 | TRUE | 0.1774 |
| Igsf11 | 81 | 94 | 100 | 150 | 105 | 133 | 86 | 94 | 100 | 118 | 113 | 146 | 109.47 | 93 | 126 | 1.34 | 0.422 | 0.03843559 | 0.13666468 | 0 | 0.037 | 0.0206 | 0.0206 | TRUE | 0.1906 |
| Igsf21 | 543 | 837 | 902 | 672 | 574 | 632 | 577 | 836 | 906 | 528 | 618 | 692 | 692.68 | 773 | 613 | 0.792 | -0.337 | 0.06811378 | 0.20216232 | 0.035 | 0.0117 | 0.0231 | 0.0231 | TRUE | 0.5816 |
| Igsf23 | 54 | 46 | 38 | 101 | 65 | 60 | 57 | 46 | 38 | 79 | 70 | 66 | 59.41 | 47 | 72 | 1.527 | 0.611 | 0.02463381 | 0.1007062 | 0 | 0.0624 | 0.0356 | 0.0356 | TRUE | 0.2774 |
| Igsf3 | 3381 | 3304 | 3425 | 3684 | 2832 | 2932 | 3591 | 3300 | 3439 | 2896 | 3047 | 3210 | 3247.11 | 3443 | 3051 | 0.886 | -0.175 | 0.0242119 | 0.09939355 | 0.0018 | 0.0079 | 0.004 | 0.004 | TRUE | 0.0251 |
| Igsf5 | 22 | 24 | 11 | 34 | 26 | 14 | 23 | 24 | 11 | 27 | 28 | 15 | 21.4 | 19 | 23 | 1.206 | 0.27 | 0.57721546 | 0.75475396 | 0.0764 | 0.1608 | 0.1219 | 0.1219 | TRUE | 0.771 |
| Igsf6 | 0 | 0 | 0 | 0 | 0 | 0 | 0 | 0 | 0 | 0 | 0 | 0 | 0 | 0 | 0 | NA | NA | NA | NA | NA | NA | NA | NA | NA | NA |
| Igsf8 | 3764 | 4526 | 4754 | 3133 | 2486 | 2836 | 3998 | 4520 | 4774 | 2463 | 2675 | 3104 | 3588.98 | 4431 | 2747 | 0.62 | -0.69 | 2.78E-09 | 1.50E-07 | 0.0106 | 0.0078 | 0.0094 | 0.0094 | TRUE | 0.1566 |
| Igsf9 | 1526 | 1411 | 1446 | 1460 | 1234 | 1242 | 1621 | 1409 | 1452 | 1148 | 1328 | 1360 | 1386.17 | 1494 | 1279 | 0.855 | -0.226 | 0.03425122 | 0.12612895 | 0.0057 | 0.0093 | 0.0075 | 0.0075 | TRUE | 0.0984 |
| Igsf9b | 401 | 458 | 540 | 592 | 459 | 474 | 426 | 457 | 542 | 465 | 494 | 519 | 483.94 | 475 | 493 | 1.036 | 0.05 | 0.69896926 | 0.8353743 | 0.0072 | 0.0137 | 0.0101 | 0.0101 | TRUE | 0.1773 |
| Igtp | 238 | 260 | 277 | 187 | 133 | 125 | 253 | 260 | 278 | 147 | 143 | 137 | 202.92 | 264 | 142 | 0.541 | -0.887 | 1.17E-09 | 7.07E-08 | 0 | 0.0231 | 0.01 | 0.01 | TRUE | 0.0263 |
| Ihh | 8 | 4 | 3 | 9 | 2 | 4 | 8 | 4 | 3 | 7 | 2 | 4 | 4.85 | 5 | 4 | 0.899 | -0.154 | 0.86649375 | 0.93389495 | 0 | 0.6859 | 0.4003 | 0.4003 | TRUE | 0.3601 |
| Iigp1 | 0 | 0 | 0 | 0 | 0 | 0 | 0 | 0 | 0 | 0 | 0 | 0 | 0 | 0 | 0 | NA | NA | NA | NA | NA | NA | NA | NA | NA | NA |
| Ik | 5386 | 5738 | 5712 | 7252 | 5310 | 5385 | 5721 | 5731 | 5735 | 5701 | 5713 | 5895 | 5749.31 | 5729 | 5770 | 1.007 | 0.01 | 0.85838131 | 0.93028174 | 0 | 0.0075 | 0.0021 | 0.0021 | TRUE | 0.0044 |
| Ikbip | 768 | 989 | 952 | 1155 | 802 | 916 | 816 | 988 | 956 | 908 | 863 | 1003 | 922.16 | 920 | 925 | 1.005 | 0.007 | 0.95385997 | 0.97651612 | 0.0068 | 0.0105 | 0.0084 | 0.0084 | TRUE | 0.116 |
| Ikbkap | 7473 | 6301 | 6701 | 10670 | 7324 | 6749 | 7938 | 6293 | 6729 | 8388 | 7880 | 7388 | 7435.83 | 6987 | 7885 | 1.129 | 0.175 | 0.10704758 | 0.27341772 | 0.0092 | 0.0073 | 0.0083 | 0.0083 | TRUE | 0.1731 |
| Ikbkb | 1836 | 1874 | 1872 | 2036 | 1587 | 1465 | 1950 | 1872 | 1880 | 1601 | 1708 | 1604 | 1768.86 | 1901 | 1638 | 0.861 | -0.215 | 0.00342469 | 0.02345689 | 3.00E-04 | 0.0088 | 0.0033 | 0.0033 | TRUE | 0.0168 |
| Ikbke | 40 | 42 | 41 | 151 | 98 | 97 | 42 | 42 | 41 | 119 | 105 | 106 | 75.99 | 42 | 110 | 2.636 | 1.399 | 1.89E-09 | 1.08E-07 | 0 | 0.0503 | 0.0224 | 0.0224 | TRUE | 0.0511 |
| Ikbkg | 1210 | 1404 | 1454 | 1847 | 1363 | 1364 | 1285 | 1402 | 1460 | 1452 | 1467 | 1493 | 1426.5 | 1382 | 1471 | 1.063 | 0.089 | 0.29308553 | 0.51418305 | 0.0014 | 0.0092 | 0.0044 | 0.0044 | TRUE | 0.0452 |
| Ikzf1 | 96 | 131 | 138 | 184 | 132 | 115 | 102 | 131 | 139 | 145 | 142 | 126 | 130.65 | 124 | 138 | 1.111 | 0.152 | 0.43386288 | 0.64252445 | 0.0064 | 0.0321 | 0.0197 | 0.0197 | TRUE | 0.2366 |
| Ikzf2 | 30 | 43 | 45 | 79 | 42 | 62 | 32 | 43 | 45 | 62 | 45 | 68 | 49.19 | 40 | 58 | 1.46 | 0.546 | 0.0751853 | 0.21599682 | 0.0158 | 0.0739 | 0.0467 | 0.0467 | TRUE | 0.315 |
| Ikzf3 | 40 | 30 | 32 | 50 | 33 | 20 | 42 | 30 | 32 | 39 | 36 | 22 | 33.55 | 35 | 32 | 0.933 | -0.1 | 0.78641791 | 0.89023346 | 0.029 | 0.1051 | 0.0694 | 0.0694 | TRUE | 0.5038 |
| Ikzf4 | 133 | 150 | 150 | 132 | 97 | 93 | 141 | 150 | 151 | 104 | 104 | 102 | 125.27 | 147 | 103 | 0.702 | -0.511 | 0.00325955 | 0.02256787 | 0 | 0.0332 | 0.0136 | 0.0136 | TRUE | 0.0128 |
| Ikzf5 | 975 | 1060 | 1029 | 1435 | 1116 | 1055 | 1036 | 1059 | 1033 | 1128 | 1201 | 1155 | 1101.86 | 1043 | 1161 | 1.113 | 0.155 | 0.04968247 | 0.16358911 | 0 | 0.0099 | 0.0036 | 0.0036 | TRUE | 0.0106 |
| Il10 | 0 | 1 | 0 | 0 | 0 | 0 | 0 | 1 | 0 | 0 | 0 | 0 | 0.17 | 0 | 0 | 0.49 | -1.029 | 0.80087 | NA | 0 | 19.7991 | 10 | 10 | TRUE | 0.405 |
| Il10ra | 7 | 5 | 2 | 7 | 11 | 6 | 7 | 5 | 2 | 6 | 12 | 7 | 6.39 | 5 | 8 | 1.648 | 0.721 | 0.37734953 | 0.59235754 | 1.00E-04 | 0.5224 | 0.3122 | 0.3122 | TRUE | 0.3658 |
| Il10rb | 98 | 139 | 151 | 144 | 115 | 129 | 104 | 139 | 152 | 113 | 124 | 141 | 128.78 | 132 | 126 | 0.955 | -0.066 | 0.75401491 | 0.87020836 | 0.0166 | 0.0325 | 0.0246 | 0.0246 | TRUE | 0.3352 |
| Il11 | 12 | 20 | 24 | 19 | 12 | 14 | 13 | 20 | 24 | 15 | 13 | 15 | 16.67 | 19 | 14 | 0.759 | -0.397 | 0.41003087 | 0.62158382 | 0 | 0.2046 | 0.1068 | 0.1068 | TRUE | 0.3258 |
| Il11ra1 | 108 | 131 | 129 | 152 | 117 | 103 | 115 | 131 | 130 | 119 | 126 | 113 | 122.2 | 125 | 119 | 0.954 | -0.067 | 0.70784835 | 0.84126423 | 0 | 0.0339 | 0.0152 | 0.0152 | TRUE | 0.0518 |
| Il11ra2 | 0 | 0 | 0 | 0 | 0 | 0 | 0 | 0 | 0 | 0 | 0 | 0 | 0 | 0 | 0 | NA | NA | NA | NA | NA | NA | NA | NA | NA | NA |
| Il12a | 0 | 0 | 0 | 0 | 0 | 0 | 0 | 0 | 0 | 0 | 0 | 0 | 0 | 0 | 0 | NA | NA | NA | NA | NA | NA | NA | NA | NA | NA |
| Il12b | 0 | 0 | 0 | 1 | 1 | 1 | 0 | 0 | 0 | 1 | 1 | 1 | 0.49 | 0 | 1 | 5.224 | 2.385 | 0.38465 | NA | 0 | 6.6923 | 3.9249 | 3.9249 | TRUE | 0.07 |

| | | | | | | | | | | | | | | | | | | | | | | | | | |
|---|---|---|---|---|---|---|---|---|---|---|---|---|---|---|---|---|---|---|---|---|---|---|---|---|---|
| Il12rb1 | 197 | 272 | 307 | 197 | 168 | 196 | 209 | 272 | 308 | 155 | 181 | 215 | 223.22 | 263 | 184 | 0.695 | -0.525 | 0.00923749 | 0.04952201 | 0.0278 | 0.0217 | 0.0247 | 0.0247 | TRUE | 0.2863 |
| Il12rb2 | 6 | 4 | 8 | 6 | 3 | 5 | 6 | 4 | 8 | 5 | 3 | 5 | 5.3 | 6 | 4 | 0.731 | -0.452 | 0.57934137 | 0.75620021 | 0 | 0.6281 | 0.2873 | 0.2873 | TRUE | 0.228 |
| Il13 | 1 | 1 | 0 | 0 | 0 | 0 | 1 | 1 | 0 | 0 | 0 | 0 | 0.34 | 1 | 0 | 0.261 | -1.936 | 0.60454921 | NA | 0 | 9.5983 | 8.3269 | 8.3269 | TRUE | 0.2507 |
| Il13ra1 | 189 | 264 | 250 | 354 | 249 | 318 | 201 | 264 | 251 | 278 | 268 | 348 | 268.29 | 239 | 298 | 1.248 | 0.319 | 0.06462597 | 0.19582871 | 0.0163 | 0.0192 | 0.0177 | 0.0177 | TRUE | 0.2371 |
| Il13ra2 | 0 | 0 | 0 | 0 | 0 | 0 | 0 | 0 | 0 | 0 | 0 | 0 | 0 | 0 | 0 | NA | NA | NA | NA | NA | NA | NA | NA | NA | NA |
| Il15 | 20 | 22 | 23 | 28 | 14 | 24 | 21 | 22 | 23 | 22 | 15 | 26 | 21.61 | 22 | 21 | 0.956 | -0.065 | 0.87695213 | 0.93932608 | 0 | 0.1594 | 0.08 | 0.08 | TRUE | 0.3173 |
| Il15ra | 7 | 6 | 9 | 1 | 3 | 3 | 7 | 6 | 9 | 1 | 3 | 3 | 4.96 | 7 | 2 | 0.312 | -1.681 | 0.07056998 | 0.20684439 | 0 | 0.6711 | 0.3459 | 0.3459 | TRUE | 0.5531 |
| Il16 | 2 | 8 | 5 | 6 | 6 | 3 | 2 | 8 | 5 | 5 | 6 | 3 | 4.93 | 5 | 5 | 0.951 | -0.072 | 0.93482241 | 0.9676458 | 0 | 0.6749 | 0.363 | 0.363 | TRUE | 0.1818 |
| Il17a | 0 | 0 | 0 | 0 | 0 | 0 | 0 | 0 | 0 | 0 | 0 | 0 | 0 | 0 | 0 | NA | NA | NA | NA | NA | NA | NA | NA | NA | NA |
| Il17b | 65 | 88 | 87 | 87 | 71 | 81 | 69 | 88 | 87 | 68 | 76 | 89 | 79.62 | 81 | 78 | 0.951 | -0.073 | 0.75687173 | 0.87155626 | 0 | 0.0483 | 0.0273 | 0.0273 | TRUE | 0.1613 |
| Il17c | 0 | 1 | 0 | 2 | 1 | 1 | 0 | 1 | 0 | 2 | 1 | 1 | 0.79 | 0 | 1 | 3.436 | 1.781 | 0.41000155 | NA | 0 | 4.1757 | 1.9771 | 1.9771 | TRUE | 0.405 |
| Il17d | 118 | 87 | 91 | 178 | 123 | 101 | 125 | 87 | 91 | 140 | 132 | 111 | 114.4 | 101 | 128 | 1.266 | 0.34 | 0.12772633 | 0.3073226 | 0.0178 | 0.0357 | 0.027 | 0.027 | TRUE | 0.4189 |
| Il17f | 65 | 49 | 34 | 40 | 28 | 31 | 69 | 49 | 34 | 31 | 30 | 34 | 41.27 | 51 | 32 | 0.629 | -0.67 | 0.06000455 | 0.18578558 | 0.0453 | 0.0867 | 0.066 | 0.066 | TRUE | 0.1881 |
| Il17ra | 1611 | 1801 | 1957 | 1632 | 1426 | 1368 | 1711 | 1799 | 1965 | 1283 | 1534 | 1497 | 1631.6 | 1825 | 1438 | 0.787 | -0.345 | 0.00116769 | 0.01001457 | 0.006 | 0.0089 | 0.0075 | 0.0075 | TRUE | 0.1099 |
| Il17rb | 8 | 7 | 4 | 8 | 3 | 1 | 8 | 7 | 4 | 6 | 3 | 1 | 5.02 | 6 | 3 | 0.563 | -0.828 | 0.36873873 | 0.58608759 | 0.0772 | 0.6632 | 0.3992 | 0.3992 | TRUE | 0.5986 |
| Il17rc | 528 | 566 | 545 | 656 | 444 | 501 | 561 | 565 | 547 | 516 | 478 | 548 | 535.86 | 558 | 514 | 0.921 | -0.118 | 0.25723946 | 0.47591564 | 0 | 0.0131 | 0.006 | 0.006 | TRUE | 0.0422 |
| Il17rd | 2974 | 3022 | 3819 | 2991 | 2203 | 2505 | 3159 | 3018 | 3835 | 2351 | 2370 | 2742 | 2912.57 | 3337 | 2488 | 0.745 | -0.424 | 0.00042 | 0.00450936 | 0.012 | 0.008 | 0.0101 | 0.0101 | TRUE | 0.2069 |
| Il17re | 13 | 13 | 9 | 25 | 25 | 17 | 14 | 13 | 9 | 20 | 27 | 19 | 16.83 | 12 | 22 | 1.816 | 0.861 | 0.07350863 | 0.21272994 | 0 | 0.2026 | 0.1013 | 0.1013 | TRUE | 0.2304 |
| Il18 | 1 | 2 | 3 | 2 | 0 | 2 | 1 | 2 | 3 | 2 | 0 | 2 | 1.64 | 2 | 1 | 0.624 | -0.681 | 0.65218974 | NA | 0 | 2.017 | 1.0101 | 1.0101 | TRUE | 0.3809 |
| Il18bp | 54 | 50 | 39 | 50 | 58 | 45 | 57 | 50 | 39 | 39 | 62 | 49 | 49.57 | 49 | 50 | 1.025 | 0.036 | 0.90778036 | 0.95354464 | 0.026 | 0.0734 | 0.0502 | 0.0502 | TRUE | 0.2611 |
| Il18r1 | 5 | 4 | 6 | 3 | 1 | 6 | 5 | 4 | 6 | 2 | 1 | 7 | 4.22 | 5 | 3 | 0.643 | -0.636 | 0.51611364 | NA | 0.0183 | 0.7872 | 0.4437 | 0.4437 | TRUE | 0.9137 |
| Il18rap | 0 | 0 | 0 | 2 | 0 | 0 | 0 | 0 | 0 | 2 | 0 | 0 | 0.26 | 0 | 1 | 2.886 | 1.529 | 0.70676491 | NA | 0 | 12.5792 | 10 | 10 | TRUE | 0.9669 |
| Il19 | 0 | 0 | 1 | 1 | 0 | 2 | 0 | 0 | 1 | 1 | 0 | 2 | 0.66 | 0 | 1 | 2.693 | 1.429 | 0.58329919 | NA | 0 | 4.974 | 3.3955 | 3.3955 | TRUE | 0.2781 |
| Il1a | 0 | 0 | 0 | 0 | 0 | 0 | 0 | 0 | 0 | 0 | 0 | 0 | 0 | 0 | 0 | NA | NA | NA | NA | NA | NA | NA | NA | NA | NA |
| Il1b | 0 | 0 | 0 | 0 | 0 | 0 | 0 | 0 | 0 | 0 | 0 | 0 | 0 | 0 | 0 | NA | NA | NA | NA | NA | NA | NA | NA | NA | NA |
| Il1f10 | 0 | 0 | 0 | 0 | 0 | 0 | 0 | 0 | 0 | 0 | 0 | 0 | 0 | 0 | 0 | NA | NA | NA | NA | NA | NA | NA | NA | NA | NA |
| Il1f5 | 0 | 0 | 0 | 0 | 0 | 0 | 0 | 0 | 0 | 0 | 0 | 0 | 0 | 0 | 0 | NA | NA | NA | NA | NA | NA | NA | NA | NA | NA |
| Il1f6 | 0 | 0 | 0 | 0 | 0 | 0 | 0 | 0 | 0 | 0 | 0 | 0 | 0 | 0 | 0 | NA | NA | NA | NA | NA | NA | NA | NA | NA | NA |
| Il1f8 | 1 | 0 | 0 | 0 | 0 | 0 | 1 | 0 | 0 | 0 | 0 | 0 | 0.18 | 0 | 0 | 0.49 | -1.029 | 0.80087 | NA | 0 | 18.6165 | 10 | 10 | TRUE | 0.4614 |
| Il1f9 | 0 | 0 | 0 | 0 | 0 | 0 | 0 | 0 | 0 | 0 | 0 | 0 | 0 | 0 | 0 | NA | NA | NA | NA | NA | NA | NA | NA | NA | NA |
| Il1r1 | 91 | 92 | 71 | 225 | 127 | 126 | 97 | 92 | 71 | 177 | 137 | 138 | 118.55 | 87 | 151 | 1.745 | 0.803 | 0.00023149 | 0.00273897 | 0.0156 | 0.0347 | 0.0252 | 0.0252 | TRUE | 0.2641 |
| Il1r2 | 76 | 82 | 88 | 117 | 92 | 80 | 81 | 82 | 88 | 92 | 99 | 88 | 88.25 | 84 | 93 | 1.109 | 0.15 | 0.46674794 | 0.66923168 | 0 | 0.0442 | 0.0191 | 0.0191 | TRUE | 0.0299 |
| Il1rap | 298 | 342 | 298 | 573 | 347 | 344 | 317 | 342 | 299 | 450 | 373 | 377 | 359.61 | 319 | 400 | 1.257 | 0.33 | 0.01516581 | 0.07129512 | 0.0059 | 0.0161 | 0.0105 | 0.0105 | TRUE | 0.1462 |
| Il1rapl1 | 0 | 0 | 0 | 0 | 0 | 0 | 0 | 0 | 0 | 0 | 0 | 0 | 0 | 0 | 0 | NA | NA | NA | NA | NA | NA | NA | NA | NA | NA |
| Il1rapl2 | 0 | 0 | 0 | 0 | 0 | 0 | 0 | 0 | 0 | 0 | 0 | 0 | 0 | 0 | 0 | NA | NA | NA | NA | NA | NA | NA | NA | NA | NA |
| Il1rl1 | 2 | 5 | 3 | 3 | 5 | 3 | 2 | 5 | 3 | 2 | 5 | 3 | 3.53 | 3 | 3 | 1.07 | 0.097 | 0.92280605 | NA | 0 | 0.9414 | 0.4375 | 0.4375 | TRUE | 0.2564 |
| Il1rl2 | 39 | 17 | 14 | 56 | 29 | 31 | 41 | 17 | 14 | 44 | 31 | 34 | 30.27 | 24 | 36 | 1.517 | 0.602 | 0.20648897 | 0.41545401 | 0.1437 | 0.1157 | 0.1287 | 0.1287 | TRUE | 2.2725 |
| Il1rn | 4 | 0 | 1 | 0 | 0 | 1 | 4 | 0 | 1 | 0 | 0 | 1 | 1.06 | 2 | 0 | 0.2 | -2.319 | 0.32545953 | NA | 1.897 | 3.121 | 2.7141 | 2.7141 | TRUE | 0.5097 |
| Il2 | 0 | 0 | 0 | 0 | 0 | 0 | 0 | 0 | 0 | 0 | 0 | 0 | 0 | 0 | 0 | NA | NA | NA | NA | NA | NA | NA | NA | NA | NA |
| Il20 | 0 | 0 | 0 | 0 | 0 | 0 | 0 | 0 | 0 | 0 | 0 | 0 | 0 | 0 | 0 | NA | NA | NA | NA | NA | NA | NA | NA | NA | NA |
| Il20ra | 6 | 4 | 4 | 6 | 7 | 1 | 6 | 4 | 4 | 5 | 8 | 1 | 4.62 | 5 | 5 | 0.933 | -0.1 | 0.91545313 | 0.95835171 | 0 | 0.7198 | 0.4236 | 0.4236 | TRUE | 0.2514 |
| Il20rb | 62 | 19 | 8 | 50 | 14 | 10 | 66 | 19 | 8 | 39 | 15 | 11 | 26.36 | 31 | 22 | 0.711 | -0.492 | 0.51718128 | 0.70874821 | 0.6445 | 0.1319 | 0.3777 | 0.3777 | TRUE | 1.3706 |
| Il21 | 0 | 1 | 0 | 1 | 1 | 0 | 0 | 1 | 0 | 1 | 1 | 0 | 0.48 | 0 | 1 | 1.712 | 0.776 | 0.78614772 | NA | 0 | 6.9165 | 4.1067 | 4.1067 | TRUE | 0.405 |
| Il21r | 1 | 5 | 1 | 0 | 0 | 4 | 1 | 5 | 1 | 0 | 0 | 4 | 1.91 | 2 | 1 | 0.599 | -0.739 | 0.68406305 | NA | 1.921 | 1.735 | 1.8115 | 1.8115 | TRUE | 1.9017 |
| Il22 | 0 | 0 | 0 | 0 | 1 | 0 | 0 | 0 | 0 | 0 | 1 | 0 | 0.18 | 0 | 0 | 1.859 | 0.894 | 0.82650734 | NA | 0 | 18.3785 | 10 | 10 | TRUE | 0.5204 |
| Il22ra1 | 68 | 33 | 29 | 73 | 52 | 37 | 72 | 33 | 29 | 57 | 56 | 41 | 48.02 | 45 | 51 | 1.151 | 0.202 | 0.61708294 | 0.7832683 | 0.1207 | 0.0755 | 0.0971 | 0.0971 | TRUE | 2.2054 |
| Il22ra2 | 0 | 0 | 0 | 1 | 0 | 0 | 0 | 0 | 0 | 1 | 0 | 0 | 0.13 | 0 | 0 | 1.859 | 0.894 | 0.82650734 | NA | 0 | 25.1515 | 10 | 10 | TRUE | 0.2563 |
| Il23a | 47 | 42 | 60 | 38 | 30 | 48 | 50 | 42 | 60 | 30 | 32 | 53 | 44.47 | 51 | 38 | 0.749 | -0.418 | 0.22270817 | 0.43477574 | 0.0406 | 0.081 | 0.0616 | 0.0616 | TRUE | 0.7194 |
| Il23r | 1 | 0 | 0 | 0 | 0 | 0 | 1 | 0 | 0 | 0 | 0 | 0 | 0.18 | 0 | 0 | 0.49 | -1.029 | 0.80087 | NA | 0 | 18.6165 | 10 | 10 | TRUE | 0.4614 |
| Il24 | 0 | 0 | 0 | 0 | 2 | 0 | 0 | 0 | 0 | 0 | 2 | 0 | 0.36 | 0 | 1 | 3.775 | 1.916 | 0.63423067 | NA | 10 | 9.1927 | 10 | 10 | TRUE | 0.9743 |
| Il25 | 3 | 4 | 1 | 1 | 2 | 5 | 3 | 4 | 1 | 1 | 2 | 5 | 2.77 | 3 | 3 | 1.001 | 0.002 | 0.99882715 | NA | 0 | 1.1979 | 0.7171 | 0.7171 | TRUE | 0.6537 |
| Il27 | 0 | 0 | 0 | 0 | 0 | 0 | 0 | 0 | 0 | 0 | 0 | 0 | 0 | 0 | 0 | NA | NA | NA | NA | NA | NA | NA | NA | NA | NA |
| Il27ra | 111 | 242 | 256 | 205 | 156 | 190 | 118 | 242 | 257 | 161 | 168 | 208 | 192.27 | 206 | 179 | 0.869 | -0.202 | 0.4771679 | 0.67672014 | 0.0866 | 0.024 | 0.053 | 0.053 | TRUE | 1.5023 |
| Il2ra | 8 | 3 | 5 | 1 | 2 | 1 | 8 | 3 | 5 | 1 | 2 | 1 | 3.42 | 5 | 1 | 0.239 | -2.067 | 0.07414812 | NA | 0 | 0.9689 | 0.5012 | 0.5012 | TRUE | 0.1796 |
| Il2rb | 5 | 6 | 12 | 1 | 2 | 3 | 5 | 6 | 12 | 1 | 2 | 3 | 4.93 | 8 | 2 | 0.256 | -1.966 | 0.04602904 | 0.15528248 | 0 | 0.6753 | 0.3904 | 0.3904 | TRUE | 0.6673 |
| Il2rg | 62 | 65 | 57 | 93 | 57 | 52 | 66 | 65 | 57 | 73 | 61 | 57 | 63.23 | 63 | 64 | 1.024 | 0.034 | 0.89252582 | 0.94794056 | 0 | 0.059 | 0.0291 | 0.0291 | TRUE | 0.1625 |
| Il3 | 0 | 0 | 0 | 0 | 0 | 0 | 0 | 0 | 0 | 0 | 0 | 0 | 0 | 0 | 0 | NA | NA | NA | NA | NA | NA | NA | NA | NA | NA |
| Il31 | 0 | 0 | 1 | 0 | 0 | 0 | 0 | 0 | 1 | 0 | 0 | 0 | 0.17 | 0 | 0 | 0.49 | -1.029 | 0.80087 | NA | 0 | 19.6928 | 10 | 10 | TRUE | 0.4098 |
| Il31ra | 3 | 4 | 5 | 2 | 3 | 1 | 3 | 4 | 5 | 2 | 3 | 1 | 3.02 | 4 | 2 | 0.478 | -1.065 | 0.32965843 | NA | 0 | 1.0992 | 0.4846 | 0.4846 | TRUE | 0.2477 |
| Il33 | 29 | 8 | 8 | 17 | 18 | 6 | 31 | 8 | 8 | 13 | 19 | 7 | 14.35 | 16 | 13 | 0.843 | -0.246 | 0.73144014 | 0.85590756 | 0.3662 | 0.2364 | 0.3011 | 0.3011 | TRUE | 0.9997 |
| Il34 | 35 | 38 | 39 | 73 | 69 | 43 | 37 | 38 | 39 | 57 | 74 | 47 | 48.83 | 38 | 59 | 1.562 | 0.644 | 0.03159635 | 0.11948077 | 0 | 0.0744 | 0.043 | 0.043 | TRUE | 0.2451 |
| Il3ra | 213 | 291 | 255 | 401 | 259 | 283 | 226 | 291 | 256 | 315 | 279 | 310 | 279.43 | 258 | 301 | 1.17 | 0.226 | 0.12563742 | 0.30350029 | 0.0059 | 0.0187 | 0.0121 | 0.0121 | TRUE | 0.1401 |
| Il4 | 0 | 0 | 0 | 0 | 0 | 0 | 0 | 0 | 0 | 0 | 0 | 0 | 0 | 0 | 0 | NA | NA | NA | NA | NA | NA | NA | NA | NA | NA |
| Il4i1 | 0 | 0 | 0 | 1 | 0 | 0 | 0 | 0 | 0 | 1 | 0 | 0 | 0.13 | 0 | 0 | 1.859 | 0.894 | 0.82650734 | NA | 0 | 25.1515 | 10 | 10 | TRUE | 0.2563 |
| Il4ra | 208 | 277 | 268 | 327 | 223 | 241 | 221 | 277 | 269 | 257 | 240 | 264 | 254.58 | 256 | 254 | 0.992 | -0.012 | 0.93692272 | 0.96848507 | 0.0041 | 0.0198 | 0.0117 | 0.0117 | TRUE | 0.1542 |
| Il5 | 0 | 0 | 0 | 0 | 1 | 0 | 0 | 0 | 0 | 0 | 1 | 0 | 0.18 | 0 | 0 | 1.859 | 0.894 | 0.82650734 | NA | 0 | 18.3785 | 10 | 10 | TRUE | 0.5204 |
| Il5ra | 0 | 0 | 0 | 0 | 0 | 0 | 0 | 0 | 0 | 0 | 0 | 0 | 0 | 0 | 0 | NA | NA | NA | NA | NA | NA | NA | NA | NA | NA |
| Il6 | 2 | 7 | 5 | 6 | 3 | 1 | 2 | 7 | 5 | 5 | 3 | 1 | 3.86 | 5 | 3 | 0.653 | -0.615 | 0.54676581 | NA | 0 | 0.8598 | 0.4847 | 0.4847 | TRUE | 0.2343 |
| Il6ra | 72 | 37 | 29 | 139 | 112 | 64 | 76 | 37 | 29 | 109 | 121 | 70 | 73.73 | 47 | 100 | 2.11 | 1.077 | 0.00566873 | 0.03440063 | 0.145 | 0.0516 | 0.0936 | 0.0936 | TRUE | 2.2255 |

| | | | | | | | | | | | | | | | | | | | | | | | | | |
|---|---|---|---|---|---|---|---|---|---|---|---|---|---|---|---|---|---|---|---|---|---|---|---|---|---|
| Il6st | 5944 | 6025 | 6442 | 5378 | 4209 | 4575 | 6314 | 6017 | 6468 | 4228 | 4529 | 5008 | 5427.28 | 6266 | 4588 | 0.732 | -0.45 | 3.18E-07 | 9.99E-06 | 0.0041 | 0.0075 | 0.0054 | 0.0054 | TRUE | 0.078 |
| Il7 | 22 | 9 | 17 | 22 | 18 | 9 | 23 | 9 | 17 | 17 | 19 | 10 | 15.99 | 16 | 15 | 0.947 | -0.078 | 0.88705771 | 0.94442566 | 0.0927 | 0.2129 | 0.1565 | 0.1565 | TRUE | 0.2477 |
| Il7r | 0 | 0 | 0 | 0 | 0 | 0 | 0 | 0 | 0 | 0 | 0 | 0 | 0 | 0 | 0 | NA | NA | NA | NA | NA | NA | NA | NA | NA | NA |
| Il8 | 0 | 0 | 0 | 0 | 0 | 0 | 0 | 0 | 0 | 0 | 0 | 0 | 0 | 0 | 0 | NA | NA | NA | NA | NA | NA | NA | NA | NA | NA |
| Il9 | 0 | 0 | 0 | 0 | 0 | 0 | 0 | 0 | 0 | 0 | 0 | 0 | 0 | 0 | 0 | NA | NA | NA | NA | NA | NA | NA | NA | NA | NA |
| Il9r | 13 | 19 | 16 | 26 | 14 | 17 | 14 | 19 | 16 | 20 | 15 | 19 | 17.16 | 16 | 18 | 1.112 | 0.153 | 0.73719031 | 0.85934705 | 0 | 0.1989 | 0.0914 | 0.0914 | TRUE | 0.1013 |
| Ildr1 | 1172 | 1797 | 1588 | 1197 | 968 | 1078 | 1245 | 1795 | 1595 | 941 | 1042 | 1180 | 1299.43 | 1545 | 1054 | 0.682 | -0.552 | 0.00032428 | 0.00363072 | 0.0215 | 0.0094 | 0.0162 | 0.0162 | TRUE | 0.2918 |
| Ildr2 | 256 | 209 | 191 | 260 | 189 | 142 | 272 | 209 | 192 | 204 | 203 | 155 | 205.94 | 224 | 187 | 0.84 | -0.252 | 0.20199259 | 0.40954199 | 0.0236 | 0.0229 | 0.0233 | 0.0233 | TRUE | 0.3788 |
| Ilf2 | 8711 | 8667 | 8452 | 11171 | 7839 | 7644 | 9253 | 8656 | 8487 | 8782 | 8434 | 8368 | 8663.12 | 8799 | 8528 | 0.969 | -0.045 | 0.50646944 | 0.70082786 | 0.0012 | 0.0073 | 0.0032 | 0.0032 | TRUE | 0.0249 |
| Ilf3 | 14433 | 16479 | 17156 | 17882 | 14054 | 14475 | 15330 | 16458 | 17226 | 14057 | 15121 | 15845 | 15673.06 | 16338 | 15008 | 0.918 | -0.123 | 0.13469597 | 0.31740995 | 0.0034 | 0.0071 | 0.0048 | 0.0048 | TRUE | 0.0377 |
| Ilk | 1295 | 1304 | 1266 | 2248 | 1501 | 1359 | 1376 | 1302 | 1271 | 1767 | 1615 | 1488 | 1469.81 | 1316 | 1623 | 1.234 | 0.304 | 0.00167466 | 0.01336677 | 0.004 | 0.0091 | 0.006 | 0.006 | TRUE | 0.074 |
| Ilkap | 1436 | 1682 | 1862 | 2216 | 1676 | 1743 | 1525 | 1680 | 1870 | 1742 | 1803 | 1908 | 1754.68 | 1692 | 1818 | 1.074 | 0.103 | 0.31213996 | 0.53255705 | 0.0054 | 0.0088 | 0.0069 | 0.0069 | TRUE | 0.1023 |
| Iltifb | 0 | 0 | 0 | 0 | 0 | 0 | 0 | 0 | 0 | 0 | 0 | 0 | 0 | 0 | 0 | NA | NA | NA | NA | NA | NA | NA | NA | NA | NA |
| Ilvbl | 286 | 384 | 393 | 460 | 353 | 395 | 304 | 384 | 395 | 362 | 380 | 432 | 375.95 | 361 | 391 | 1.083 | 0.115 | 0.44112981 | 0.64823436 | 0.0113 | 0.0157 | 0.0134 | 0.0134 | TRUE | 0.2153 |
| Immp1l | 453 | 570 | 539 | 703 | 464 | 502 | 481 | 569 | 541 | 553 | 499 | 550 | 532.17 | 530 | 534 | 1.006 | 0.009 | 0.93612749 | 0.96816629 | 0.0032 | 0.0131 | 0.0076 | 0.0076 | TRUE | 0.0767 |
| Immp2l | 46 | 99 | 104 | 103 | 65 | 85 | 49 | 99 | 104 | 81 | 70 | 93 | 82.69 | 84 | 81 | 0.966 | -0.05 | 0.87417077 | 0.93791982 | 0.0746 | 0.0467 | 0.0599 | 0.0599 | TRUE | 1.2376 |
| Immt | 15237 | 14319 | 14402 | 17585 | 12391 | 12278 | 16184 | 14301 | 14461 | 13824 | 13332 | 13440 | 14257.02 | 14982 | 13532 | 0.903 | -0.147 | 0.05306338 | 0.17104346 | 0.0025 | 0.0071 | 0.0041 | 0.0041 | TRUE | 0.0602 |
| Imp3 | 2164 | 2598 | 2729 | 2796 | 2143 | 2224 | 2299 | 2595 | 2740 | 2198 | 2306 | 2435 | 2428.61 | 2545 | 2313 | 0.909 | -0.138 | 0.1496661 | 0.34028164 | 0.0046 | 0.0083 | 0.0062 | 0.0062 | TRUE | 0.0864 |
| Imp4 | 2818 | 2306 | 2295 | 4153 | 2846 | 2319 | 2993 | 2303 | 2304 | 3265 | 3062 | 2539 | 2744.34 | 2533 | 2955 | 1.167 | 0.223 | 0.11627404 | 0.2881185 | 0.0198 | 0.0081 | 0.0141 | 0.0141 | TRUE | 0.3051 |
| Impa1 | 3331 | 2977 | 2941 | 5127 | 3684 | 3457 | 3538 | 2973 | 2953 | 4030 | 3964 | 3784 | 3540.46 | 3155 | 3926 | 1.245 | 0.316 | 0.00108831 | 0.00950437 | 0.0056 | 0.0078 | 0.0065 | 0.0065 | TRUE | 0.137 |
| Impa2 | 2699 | 3685 | 3738 | 4639 | 3789 | 3965 | 2867 | 3680 | 3753 | 3647 | 4077 | 4340 | 3727.38 | 3433 | 4021 | 1.171 | 0.228 | 0.07346915 | 0.2126547 | 0.0137 | 0.0078 | 0.0114 | 0.0114 | TRUE | 0.2531 |
| Impact | 7727 | 8182 | 8510 | 9768 | 6652 | 6704 | 8207 | 8171 | 8545 | 7679 | 7157 | 7339 | 7849.73 | 8308 | 7392 | 0.89 | -0.168 | 0.00832794 | 0.04586183 | 8.00E-04 | 0.0073 | 0.0028 | 0.0028 | TRUE | 0.0143 |
| Impad1 | 3170 | 3774 | 3834 | 5068 | 3742 | 3545 | 3367 | 3769 | 3850 | 3984 | 4026 | 3881 | 3812.79 | 3662 | 3964 | 1.082 | 0.114 | 0.15421392 | 0.34613091 | 0.0023 | 0.0078 | 0.0044 | 0.0044 | TRUE | 0.0602 |
| Impdh1 | 5334 | 4838 | 4722 | 6294 | 4485 | 4284 | 5666 | 4832 | 4741 | 4948 | 4826 | 4690 | 4950.29 | 5080 | 4821 | 0.949 | -0.075 | 0.41854722 | 0.62917715 | 0.0051 | 0.0076 | 0.006 | 0.006 | TRUE | 0.124 |
| Impdh2 | 20805 | 24426 | 25142 | 28064 | 20870 | 22582 | 22099 | 24394 | 25245 | 22062 | 22455 | 24720 | 23495.73 | 23913 | 23079 | 0.965 | -0.051 | 0.54993593 | 0.73424105 | 0.0037 | 0.007 | 0.0053 | 0.0053 | TRUE | 0.0539 |
| Impg1 | 0 | 0 | 0 | 0 | 1 | 0 | 0 | 0 | 0 | 0 | 1 | 0 | 0.18 | 0 | 0 | 1.859 | 0.894 | 0.82650734 | NA | 0 | 18.3785 | 10 | 10 | TRUE | 0.5204 |
| Impg2 | 2 | 2 | 1 | 1 | 1 | 1 | 2 | 2 | 1 | 1 | 1 | 1 | 1.35 | 2 | 1 | 0.571 | -0.808 | 0.60104378 | NA | 0 | 2.4525 | 0.9207 | 0.9207 | TRUE | 0.1023 |
| Ina | 2211 | 1406 | 1391 | 1806 | 1238 | 908 | 2348 | 1404 | 1397 | 1420 | 1332 | 994 | 1482.51 | 1716 | 1249 | 0.728 | -0.458 | 0.03437823 | 0.12630177 | 0.066 | 0.0091 | 0.0332 | 0.0332 | TRUE | 1.2511 |
| Inadl | 403 | 603 | 629 | 774 | 552 | 667 | 428 | 602 | 632 | 608 | 594 | 730 | 599.06 | 554 | 644 | 1.162 | 0.216 | 0.20269406 | 0.41027859 | 0.0258 | 0.0124 | 0.0191 | 0.0191 | TRUE | 0.4601 |
| Inca1 | 106 | 224 | 182 | 171 | 121 | 161 | 113 | 224 | 183 | 134 | 130 | 176 | 159.98 | 173 | 147 | 0.848 | -0.239 | 0.37390747 | 0.59019901 | 0.0654 | 0.0275 | 0.0456 | 0.0456 | TRUE | 0.3402 |
| Incenp | 9112 | 9407 | 9778 | 12047 | 8986 | 8390 | 9679 | 9395 | 9818 | 9470 | 9668 | 9184 | 9535.75 | 9631 | 9441 | 0.98 | -0.029 | 0.62926249 | 0.79150874 | 4.00E-04 | 0.0072 | 0.0024 | 0.0024 | TRUE | 0.0069 |
| Inf2 | 1307 | 1595 | 1710 | 1541 | 1320 | 1247 | 1388 | 1593 | 1717 | 1211 | 1420 | 1365 | 1449.15 | 1566 | 1332 | 0.85 | -0.235 | 0.0406988 | 0.14273993 | 0.0079 | 0.0092 | 0.0088 | 0.0088 | TRUE | 0.1185 |
| Ing1 | 2874 | 2990 | 3077 | 4011 | 3000 | 2961 | 3053 | 2986 | 3090 | 3153 | 3228 | 3241 | 3125.12 | 3043 | 3207 | 1.054 | 0.076 | 0.22082778 | 0.43298057 | 0 | 0.008 | 0.0024 | 0.0024 | TRUE | 0.0032 |
| Ing2 | 1022 | 1414 | 1378 | 1940 | 1499 | 1494 | 1086 | 1412 | 1384 | 1525 | 1613 | 1635 | 1442.45 | 1294 | 1591 | 1.229 | 0.298 | 0.013228 | 0.06451205 | 0.0094 | 0.0092 | 0.0097 | 0.0097 | TRUE | 0.2372 |
| Ing3 | 1137 | 1136 | 1243 | 1563 | 1168 | 1074 | 1208 | 1135 | 1248 | 1229 | 1257 | 1176 | 1208.57 | 1197 | 1221 | 1.02 | 0.028 | 0.73486774 | 0.85771731 | 9.00E-04 | 0.0096 | 0.0042 | 0.0042 | TRUE | 0.025 |
| Ing4 | 1950 | 1924 | 1915 | 2671 | 1985 | 2157 | 2071 | 1922 | 1923 | 2100 | 2136 | 2361 | 2085.38 | 1972 | 2199 | 1.115 | 0.157 | 0.0658278 | 0.19787232 | 0.0023 | 0.0085 | 0.0047 | 0.0047 | TRUE | 0.0501 |
| Ing5 | 3927 | 4058 | 4088 | 5268 | 3875 | 3691 | 4171 | 4053 | 4105 | 4141 | 4169 | 4040 | 4113.27 | 4110 | 4117 | 1.002 | 0.003 | 0.96320781 | 0.98149348 | 0 | 0.0077 | 0.0023 | 0.0023 | TRUE | 0.0032 |
| Inha | 13 | 38 | 29 | 18 | 17 | 11 | 14 | 38 | 29 | 14 | 18 | 12 | 20.89 | 27 | 15 | 0.548 | -0.867 | 0.08768403 | 0.23988769 | 0.102 | 0.1646 | 0.134 | 0.134 | TRUE | 0.2475 |
| Inhba | 1 | 0 | 0 | 0 | 0 | 0 | 1 | 0 | 0 | 0 | 0 | 0 | 0.18 | 0 | 0 | 0.49 | -1.029 | 0.80087 | NA | 0 | 18.6165 | 10 | 10 | TRUE | 0.4614 |
| Inhbb | 283 | 611 | 668 | 1499 | 1168 | 1125 | 301 | 610 | 671 | 1178 | 1257 | 1231 | 874.69 | 527 | 1222 | 2.317 | 1.212 | 7.03E-07 | 2.02E-05 | 0.0831 | 0.0107 | 0.0417 | 0.0417 | TRUE | 1.6447 |
| Inhbc | 46 | 29 | 22 | 39 | 25 | 7 | 49 | 29 | 22 | 31 | 27 | 8 | 27.52 | 33 | 22 | 0.659 | -0.601 | 0.27500846 | 0.49548018 | 0.2452 | 0.1266 | 0.1804 | 0.1804 | TRUE | 1.0783 |
| Inhbe | 4 | 5 | 3 | 2 | 2 | 4 | 4 | 5 | 3 | 2 | 2 | 4 | 3.39 | 4 | 3 | 0.647 | -0.628 | 0.5394722 | NA | 0 | 0.9779 | 0.4446 | 0.4446 | TRUE | 0.3203 |
| Inip | 1642 | 1642 | 1624 | 2097 | 1453 | 1364 | 1744 | 1640 | 1631 | 1648 | 1563 | 1493 | 1619.93 | 1672 | 1568 | 0.939 | -0.091 | 0.26481944 | 0.48459591 | 0.0013 | 0.0089 | 0.0042 | 0.0042 | TRUE | 0.0248 |
| Inmt | 0 | 0 | 0 | 0 | 0 | 0 | 0 | 0 | 0 | 0 | 0 | 0 | 0 | 0 | 0 | NA | NA | NA | NA | NA | NA | NA | NA | NA | NA |
| Ino80 | 3513 | 3235 | 3374 | 5491 | 3757 | 3493 | 3731 | 3231 | 3388 | 4317 | 4042 | 3824 | 3755.44 | 3450 | 4061 | 1.177 | 0.236 | 0.00920808 | 0.04943151 | 0.0042 | 0.0078 | 0.0056 | 0.0056 | TRUE | 0.0618 |
| Ino80b | 394 | 316 | 308 | 585 | 424 | 315 | 418 | 316 | 309 | 460 | 456 | 345 | 384.04 | 348 | 420 | 1.21 | 0.276 | 0.12115354 | 0.29597972 | 0.0237 | 0.0155 | 0.0201 | 0.0201 | TRUE | 0.3586 |
| Ino80c | 858 | 870 | 823 | 1307 | 935 | 796 | 911 | 869 | 826 | 1027 | 1006 | 871 | 918.57 | 869 | 968 | 1.116 | 0.158 | 0.1307166 | 0.31130728 | 0.004 | 0.0105 | 0.0068 | 0.0068 | TRUE | 0.0901 |
| Ino80d | 1328 | 1118 | 1254 | 2250 | 1545 | 1289 | 1411 | 1117 | 1259 | 1769 | 1662 | 1411 | 1438.06 | 1262 | 1614 | 1.28 | 0.356 | 0.00549159 | 0.03350826 | 0.0122 | 0.0092 | 0.0111 | 0.0111 | TRUE | 0.1453 |
| Ino80e | 1243 | 1039 | 1040 | 1807 | 1239 | 1010 | 1320 | 1038 | 1044 | 1421 | 1333 | 1106 | 1210.24 | 1134 | 1287 | 1.135 | 0.183 | 0.19506063 | 0.40091206 | 0.0169 | 0.0096 | 0.0135 | 0.0135 | TRUE | 0.2465 |
| Inpp1 | 110 | 148 | 128 | 182 | 158 | 145 | 117 | 148 | 129 | 143 | 170 | 159 | 144.16 | 131 | 157 | 1.196 | 0.258 | 0.15476576 | 0.34692519 | 0.004 | 0.0298 | 0.0167 | 0.0167 | TRUE | 0.1288 |
| Inpp4a | 837 | 950 | 895 | 974 | 743 | 745 | 889 | 949 | 899 | 766 | 799 | 816 | 852.85 | 912 | 794 | 0.869 | -0.202 | 0.02027867 | 0.08772219 | 0 | 0.0108 | 0.0043 | 0.0043 | TRUE | 0.0147 |
| Inpp4b | 44 | 13 | 8 | 87 | 53 | 17 | 47 | 13 | 8 | 68 | 57 | 19 | 35.3 | 23 | 48 | 2.138 | 1.096 | 0.10454168 | 0.26925895 | 0.5304 | 0.1002 | 0.2958 | 0.2958 | TRUE | 1.8616 |
| Inpp5a | 2731 | 2649 | 2599 | 3548 | 2477 | 2350 | 2901 | 2646 | 2610 | 2789 | 2665 | 2572 | 2697.13 | 2719 | 2675 | 0.985 | -0.022 | 0.78088254 | 0.88649531 | 0.0021 | 0.0081 | 0.0043 | 0.0043 | TRUE | 0.0415 |
| Inpp5b | 2049 | 1868 | 1941 | 2431 | 1730 | 1632 | 2176 | 1866 | 1949 | 1911 | 1861 | 1786 | 1924.98 | 1997 | 1853 | 0.928 | -0.107 | 0.23349696 | 0.44818232 | 0.003 | 0.0086 | 0.0053 | 0.0053 | TRUE | 0.0744 |
| Inpp5d | 10683 | 12078 | 10977 | 14370 | 10830 | 12210 | 11347 | 12062 | 11022 | 11297 | 11652 | 13366 | 11791.08 | 11477 | 12105 | 1.055 | 0.077 | 0.39880067 | 0.61201859 | 0.0051 | 0.0072 | 0.0059 | 0.0059 | TRUE | 0.1014 |
| Inpp5e | 387 | 425 | 430 | 712 | 483 | 473 | 411 | 424 | 432 | 560 | 520 | 518 | 477.41 | 422 | 533 | 1.262 | 0.335 | 0.0012768 | 0.01075522 | 0 | 0.0138 | 0.0057 | 0.0057 | TRUE | 0.025 |
| Inpp5f | 2861 | 2654 | 2532 | 2628 | 1919 | 1894 | 3039 | 2651 | 2542 | 2066 | 2065 | 2073 | 2405.97 | 2744 | 2068 | 0.754 | -0.408 | 9.77E-06 | 0.00019509 | 0.0039 | 0.0083 | 0.0057 | 0.0057 | TRUE | 0.107 |
| Inpp5j | 10 | 7 | 5 | 6 | 8 | 2 | 11 | 7 | 5 | 5 | 9 | 2 | 6.36 | 8 | 5 | 0.686 | -0.545 | 0.5024943 | 0.69771459 | 0.0625 | 0.5251 | 0.3127 | 0.3127 | TRUE | 0.4162 |
| Inpp5k | 751 | 1106 | 1108 | 1457 | 1159 | 1133 | 798 | 1105 | 1113 | 1145 | 1247 | 1240 | 1107.91 | 1005 | 1211 | 1.204 | 0.268 | 0.05967098 | 0.18518904 | 0.0172 | 0.0099 | 0.0137 | 0.0137 | TRUE | 0.3872 |
| Inppl1 | 3704 | 4090 | 4262 | 5187 | 3923 | 4118 | 3934 | 4085 | 4279 | 4078 | 4221 | 4508 | 4184.14 | 4099 | 4269 | 1.041 | 0.058 | 0.44762118 | 0.65334741 | 0.0018 | 0.0077 | 0.004 | 0.004 | TRUE | 0.0291 |
| Ins1 | 0 | 0 | 0 | 0 | 0 | 0 | 0 | 0 | 0 | 0 | 0 | 0 | 0 | 0 | 0 | NA | NA | NA | NA | NA | NA | NA | NA | NA | NA |
| Ins2 | 0 | 0 | 0 | 0 | 0 | 0 | 0 | 0 | 0 | 0 | 0 | 0 | 0 | 0 | 0 | NA | NA | NA | NA | NA | NA | NA | NA | NA | NA |
| Insc | 115 | 134 | 138 | 90 | 79 | 96 | 122 | 134 | 139 | 71 | 85 | 105 | 109.23 | 132 | 87 | 0.657 | -0.607 | 0.0051026 | 0.03184735 | 0.0103 | 0.0371 | 0.0243 | 0.0243 | TRUE | 0.3102 |
| Insig1 | 8386 | 9761 | 9624 | 8965 | 7143 | 8067 | 8907 | 9748 | 9664 | 7048 | 7685 | 8831 | 8647.16 | 9440 | 7855 | 0.832 | -0.266 | 0.00968478 | 0.0513958 | 0.0076 | 0.0073 | 0.0075 | 0.0075 | TRUE | 0.1442 |
| Insig2 | 1272 | 1298 | 1319 | 1719 | 1232 | 1296 | 1351 | 1296 | 1324 | 1351 | 1326 | 1419 | 1344.57 | 1324 | 1365 | 1.031 | 0.044 | 0.56549099 | 0.74566707 | 0 | 0.0094 | 0.0035 | 0.0035 | TRUE | 0.0137 |
| Insl3 | 10 | 15 | 23 | 12 | 16 | 20 | 11 | 15 | 23 | 9 | 17 | 22 | 16.21 | 16 | 16 | 0.983 | -0.025 | 0.96328906 | 0.98149348 | 0.1015 | 0.2102 | 0.1569 | 0.1569 | TRUE | 0.449 |
| Insl5 | 0 | 0 | 0 | 1 | 2 | 0 | 0 | 0 | 0 | 1 | 2 | 0 | 0.49 | 0 | 1 | 5.204 | 2.38 | 0.47267317 | NA | 0 | 6.7348 | 6.4099 | 6.4099 | TRUE | 0.1222 |
| Insl6 | 112 | 157 | 148 | 142 | 65 | 113 | 119 | 157 | 149 | 112 | 70 | 124 | 121.61 | 142 | 102 | 0.72 | -0.474 | 0.06018136 | 0.18615151 | 0.0411 | 0.034 | 0.0373 | 0.0373 | TRUE | 0.7029 |
| Insm1 | 15 | 29 | 17 | 42 | 23 | 22 | 16 | 29 | 17 | 33 | 25 | 24 | 23.97 | 21 | 27 | 1.327 | 0.408 | 0.34035141 | 0.56057044 | 0.0283 | 0.1444 | 0.0894 | 0.0894 | TRUE | 0.6897 |

| | | | | | | | | | | | | | | | | | | | | | | | | | |
|---|---|---|---|---|---|---|---|---|---|---|---|---|---|---|---|---|---|---|---|---|---|---|---|---|---|
| Insm2 | 2 | 1 | 2 | 2 | 2 | 2 | 2 | 1 | 2 | 2 | 2 | 2 | 1.84 | 2 | 2 | 1.145 | 0.195 | 0.88196197 | NA | 0 | 1.7966 | 0.6956 | 0.6956 | TRUE | 0.1053 |
| Insr | 1142 | 1290 | 1404 | 1824 | 1248 | 1361 | 1213 | 1288 | 1410 | 1434 | 1343 | 1490 | 1362.93 | 1304 | 1422 | 1.091 | 0.125 | 0.18850663 | 0.39358308 | 0.0032 | 0.0093 | 0.0058 | 0.0058 | TRUE | 0.0609 |
| Insrr | 7 | 11 | 4 | 13 | 4 | 7 | 7 | 11 | 4 | 10 | 4 | 8 | 7.44 | 7 | 7 | 1.001 | 0.002 | 0.99828319 | 0.99885317 | 0 | 0.4499 | 0.266 | 0.266 | TRUE | 0.1942 |
| Ints1 | 9039 | 7913 | 8312 | 10278 | 7614 | 7185 | 9601 | 7903 | 8346 | 8080 | 8192 | 7865 | 8331.17 | 8617 | 8046 | 0.934 | -0.099 | 0.28392523 | 0.50542537 | 0.0053 | 0.0073 | 0.006 | 0.006 | TRUE | 0.1219 |
| Ints10 | 4267 | 4391 | 4461 | 6003 | 4313 | 4348 | 4532 | 4385 | 4479 | 4719 | 4641 | 4760 | 4586.02 | 4465 | 4707 | 1.054 | 0.076 | 0.19381703 | 0.39954891 | 0 | 0.0076 | 0.0022 | 0.0022 | TRUE | 0.003 |
| Ints12 | 1151 | 1311 | 1270 | 1817 | 1289 | 1197 | 1223 | 1309 | 1275 | 1428 | 1387 | 1310 | 1322.11 | 1269 | 1375 | 1.084 | 0.117 | 0.15324931 | 0.34475084 | 8.00E-04 | 0.0094 | 0.004 | 0.004 | TRUE | 0.0203 |
| Ints2 | 7048 | 6626 | 6760 | 8920 | 6244 | 6182 | 7486 | 6617 | 6788 | 7012 | 6718 | 6767 | 6898.16 | 6964 | 6832 | 0.981 | -0.027 | 0.7202389 | 0.84830169 | 0.002 | 0.0074 | 0.004 | 0.004 | TRUE | 0.0525 |
| Ints3 | 10210 | 8975 | 8701 | 11850 | 8453 | 8074 | 10845 | 8963 | 8737 | 9316 | 9095 | 8838 | 9298.95 | 9515 | 9083 | 0.955 | -0.067 | 0.50933316 | 0.70294037 | 0.007 | 0.0073 | 0.0073 | 0.0073 | TRUE | 0.1825 |
| Ints4 | 5055 | 5319 | 5782 | 6482 | 4745 | 4808 | 5369 | 5312 | 5806 | 5096 | 5105 | 5263 | 5325.21 | 5496 | 5155 | 0.938 | -0.093 | 0.17908912 | 0.38024408 | 0.0011 | 0.0075 | 0.0032 | 0.0032 | TRUE | 0.0297 |
| Ints5 | 2883 | 2094 | 1945 | 4158 | 2833 | 2015 | 3062 | 2091 | 1953 | 3269 | 3048 | 2206 | 2604.85 | 2369 | 2841 | 1.2 | 0.263 | 0.18284044 | 0.38498245 | 0.0496 | 0.0082 | 0.0276 | 0.0276 | TRUE | 0.794 |
| Ints6 | 1747 | 1483 | 1471 | 2620 | 1694 | 1476 | 1856 | 1481 | 1477 | 2060 | 1823 | 1616 | 1718.62 | 1605 | 1833 | 1.143 | 0.193 | 0.15107696 | 0.34215764 | 0.0156 | 0.0088 | 0.0124 | 0.0124 | TRUE | 0.2254 |
| Ints7 | 3946 | 3627 | 3608 | 4679 | 3442 | 3255 | 4191 | 3622 | 3623 | 3678 | 3703 | 3563 | 3730.21 | 3812 | 3648 | 0.957 | -0.063 | 0.4655099 | 0.6680031 | 0.0035 | 0.0078 | 0.0051 | 0.0051 | TRUE | 0.092 |
| Ints8 | 4633 | 4934 | 4901 | 6145 | 4476 | 4464 | 4921 | 4928 | 4921 | 4831 | 4816 | 4887 | 4883.83 | 4923 | 4845 | 0.984 | -0.023 | 0.67393963 | 0.81827374 | 0 | 0.0076 | 0.002 | 0.002 | TRUE | 7.00E-04 |
| Ints9 | 2955 | 2416 | 2415 | 3627 | 2543 | 2298 | 3139 | 2413 | 2425 | 2851 | 2736 | 2516 | 2679.9 | 2659 | 2701 | 1.016 | 0.023 | 0.85336318 | 0.92681753 | 0.0132 | 0.0081 | 0.0109 | 0.0109 | TRUE | 0.302 |
| Intu | 1045 | 1044 | 1125 | 1468 | 1097 | 1086 | 1110 | 1043 | 1130 | 1154 | 1180 | 1189 | 1134.23 | 1094 | 1174 | 1.073 | 0.102 | 0.20688423 | 0.41598394 | 0 | 0.0098 | 0.0038 | 0.0038 | TRUE | 0.0204 |
| Invs | 145 | 104 | 112 | 265 | 148 | 78 | 154 | 104 | 112 | 208 | 159 | 85 | 137.21 | 123 | 151 | 1.228 | 0.296 | 0.35737331 | 0.57585736 | 0.1093 | 0.0309 | 0.0673 | 0.0673 | TRUE | 0.2684 |
| Ip6k1 | 4132 | 5213 | 4786 | 4843 | 3797 | 4028 | 4389 | 5206 | 4806 | 3807 | 4085 | 4409 | 4450.44 | 4800 | 4100 | 0.854 | -0.228 | 0.02045074 | 0.08827288 | 0.006 | 0.0076 | 0.0067 | 0.0067 | TRUE | 0.0684 |
| Ip6k2 | 413 | 459 | 403 | 591 | 365 | 410 | 439 | 458 | 405 | 465 | 393 | 449 | 434.64 | 434 | 436 | 1.005 | 0.007 | 0.95524553 | 0.97724151 | 0.0036 | 0.0145 | 0.0085 | 0.0085 | TRUE | 0.0835 |
| Ip6k3 | 2 | 0 | 0 | 2 | 2 | 0 | 2 | 0 | 0 | 2 | 2 | 0 | 0.97 | 1 | 1 | 1.79 | 0.84 | 0.71717794 | NA | 0 | 3.3867 | 2.7541 | 2.7541 | TRUE | 0.9744 |
| Ipcef1 | 11 | 5 | 2 | 18 | 5 | 5 | 12 | 5 | 2 | 14 | 5 | 5 | 7.28 | 6 | 8 | 1.367 | 0.451 | 0.60263376 | 0.77387557 | 0.339 | 0.4593 | 0.3996 | 0.3996 | TRUE | 0.7616 |
| Ipmk | 4126 | 4412 | 4558 | 5398 | 4073 | 3979 | 4383 | 4406 | 4577 | 4243 | 4382 | 4356 | 4391.16 | 4455 | 4327 | 0.971 | -0.042 | 0.48756474 | 0.68451764 | 2.00E-04 | 0.0077 | 0.0025 | 0.0025 | TRUE | 0.0069 |
| Ipo11 | 5413 | 6046 | 6263 | 7362 | 5318 | 5387 | 5750 | 6038 | 6289 | 5787 | 5722 | 5897 | 5913.78 | 6026 | 5802 | 0.963 | -0.055 | 0.41089193 | 0.62206449 | 9.00E-04 | 0.0075 | 0.003 | 0.003 | TRUE | 0.0195 |
| Ipo13 | 1739 | 1105 | 1062 | 2423 | 1518 | 1104 | 1847 | 1104 | 1066 | 1905 | 1633 | 1209 | 1460.6 | 1339 | 1582 | 1.182 | 0.241 | 0.29434996 | 0.51515728 | 0.0737 | 0.0092 | 0.0375 | 0.0375 | TRUE | 0.7645 |
| Ipo4 | 7066 | 5631 | 5709 | 9276 | 6694 | 5774 | 7505 | 5624 | 5732 | 7292 | 7202 | 6321 | 6612.74 | 6287 | 6938 | 1.104 | 0.142 | 0.27545761 | 0.49602508 | 0.0156 | 0.0074 | 0.0121 | 0.0121 | TRUE | 0.3503 |
| Ipo5 | 40144 | 40784 | 42627 | 47801 | 35496 | 35959 | 42640 | 40731 | 42802 | 37577 | 38192 | 39363 | 40217.51 | 42058 | 38377 | 0.912 | -0.132 | 0.02272609 | 0.09517683 | 6.00E-04 | 0.007 | 0.0024 | 0.0024 | TRUE | 0.0093 |
| Ipo7 | 18443 | 20008 | 20219 | 22247 | 16348 | 17415 | 19590 | 19982 | 20302 | 17489 | 17589 | 19063 | 19002.61 | 19958 | 18047 | 0.904 | -0.145 | 0.02727161 | 0.10790965 | 0.0013 | 0.0071 | 0.0031 | 0.0031 | TRUE | 0.0296 |
| Ipo8 | 2140 | 2199 | 2262 | 2907 | 2017 | 2067 | 2273 | 2196 | 2271 | 2285 | 2170 | 2263 | 2243.1 | 2247 | 2239 | 0.997 | -0.004 | 0.94866021 | 0.9742329 | 0 | 0.0084 | 0.0029 | 0.0029 | TRUE | 0.0088 |
| Ipo9 | 9032 | 8384 | 8637 | 12546 | 8759 | 8432 | 9594 | 8373 | 8672 | 9863 | 9424 | 9230 | 9192.7 | 8880 | 9506 | 1.071 | 0.098 | 0.21844765 | 0.43051243 | 0.003 | 0.0073 | 0.0045 | 0.0045 | TRUE | 0.0604 |
| Ipp | 634 | 782 | 855 | 948 | 700 | 705 | 673 | 781 | 859 | 745 | 753 | 772 | 763.84 | 771 | 757 | 0.981 | -0.028 | 0.81101906 | 0.90283686 | 0.0062 | 0.0112 | 0.0084 | 0.0084 | TRUE | 0.1442 |
| Ippk | 1303 | 1393 | 1375 | 1757 | 1239 | 1198 | 1384 | 1391 | 1381 | 1381 | 1333 | 1311 | 1363.59 | 1385 | 1342 | 0.969 | -0.045 | 0.5391718 | 0.72533416 | 0 | 0.0093 | 0.0031 | 0.0031 | TRUE | 0.0082 |
| Ipw | 1 | 2 | 0 | 1 | 1 | 0 | 1 | 2 | 0 | 1 | 1 | 0 | 0.82 | 1 | 1 | 0.617 | -0.697 | 0.74855989 | NA | 0 | 4.0232 | 2.1283 | 2.1283 | TRUE | 0.1568 |
| Iqca | 82 | 59 | 54 | 59 | 66 | 75 | 87 | 59 | 54 | 46 | 71 | 82 | 66.62 | 67 | 66 | 0.991 | -0.013 | 0.9674373 | 0.98366749 | 0.0604 | 0.0564 | 0.0582 | 0.0582 | TRUE | 0.6782 |
| Iqcb1 | 1290 | 2140 | 2218 | 2424 | 1864 | 2170 | 1370 | 2137 | 2227 | 1906 | 2006 | 2375 | 2003.51 | 1911 | 2096 | 1.096 | 0.132 | 0.46877273 | 0.67115863 | 0.04 | 0.0085 | 0.0235 | 0.0235 | TRUE | 0.7407 |
| Iqcc | 664 | 786 | 794 | 965 | 711 | 677 | 705 | 785 | 797 | 759 | 765 | 741 | 758.7 | 762 | 755 | 0.99 | -0.015 | 0.87698374 | 0.93932608 | 9.00E-04 | 0.0112 | 0.0052 | 0.0052 | TRUE | 0.0506 |
| Iqcd | 4 | 4 | 3 | 4 | 5 | 4 | 4 | 4 | 3 | 3 | 5 | 4 | 4.03 | 4 | 4 | 1.135 | 0.182 | 0.84104809 | NA | 0 | 0.8251 | 0.3449 | 0.3449 | TRUE | 0.1341 |
| Iqce | 250 | 357 | 332 | 525 | 376 | 362 | 266 | 357 | 333 | 413 | 405 | 396 | 361.5 | 319 | 405 | 1.27 | 0.344 | 0.01578445 | 0.07365454 | 0.0079 | 0.016 | 0.0119 | 0.0119 | TRUE | 0.2368 |
| Iqcf1 | 0 | 0 | 0 | 0 | 0 | 0 | 0 | 0 | 0 | 0 | 0 | 0 | 0 | 0 | 0 | NA | NA | NA | NA | NA | NA | NA | NA | NA | NA |
| Iqcf3 | 0 | 1 | 0 | 0 | 0 | 0 | 0 | 1 | 0 | 0 | 0 | 0 | 0.17 | 0 | 0 | 0.49 | -1.029 | 0.80087 | NA | 0 | 19.7991 | 10 | 10 | TRUE | 0.405 |
| Iqcf4 | 0 | 0 | 1 | 0 | 0 | 0 | 0 | 0 | 1 | 0 | 0 | 0 | 0.17 | 0 | 0 | 0.49 | -1.029 | 0.80087 | NA | 0 | 19.6928 | 10 | 10 | TRUE | 0.4098 |
| Iqcf5 | 2 | 5 | 1 | 9 | 1 | 2 | 2 | 5 | 1 | 7 | 1 | 2 | 3.08 | 3 | 3 | 1.309 | 0.388 | 0.74920462 | NA | 0.4232 | 1.0776 | 0.7351 | 0.7351 | TRUE | 1.585 |
| Iqcf6 | 0 | 0 | 0 | 0 | 0 | 0 | 0 | 0 | 0 | 0 | 0 | 0 | 0 | 0 | 0 | NA | NA | NA | NA | NA | NA | NA | NA | NA | NA |
| Iqcg | 512 | 683 | 653 | 784 | 648 | 719 | 544 | 682 | 656 | 616 | 697 | 787 | 663.7 | 627 | 700 | 1.115 | 0.157 | 0.26208057 | 0.48147442 | 0.0131 | 0.0119 | 0.0125 | 0.0125 | TRUE | 0.1581 |
| Iqch | 1 | 1 | 3 | 14 | 8 | 5 | 1 | 1 | 3 | 11 | 9 | 5 | 5.03 | 2 | 8 | 4.967 | 2.312 | 0.02015722 | 0.08731656 | 0 | 0.6623 | 0.355 | 0.355 | TRUE | 0.2681 |
| Iqcj | 0 | 0 | 0 | 0 | 1 | 0 | 0 | 0 | 0 | 0 | 1 | 0 | 0.18 | 0 | 0 | 1.859 | 0.894 | 0.82650734 | NA | 0 | 18.3785 | 10 | 10 | TRUE | 0.5204 |
| Iqck | 5 | 4 | 3 | 11 | 10 | 2 | 5 | 4 | 3 | 9 | 11 | 2 | 5.65 | 4 | 7 | 1.769 | 0.823 | 0.35113404 | 0.56999652 | 0 | 0.5897 | 0.3688 | 0.3688 | TRUE | 0.6634 |
| Iqgap1 | 12512 | 13081 | 14317 | 17033 | 12215 | 12286 | 13290 | 13064 | 14376 | 13390 | 13143 | 13449 | 13451.9 | 13577 | 13327 | 0.982 | -0.027 | 0.68880739 | 0.82886016 | 0.0012 | 0.0071 | 0.0031 | 0.0031 | TRUE | 0.0324 |
| Iqgap2 | 1648 | 1482 | 1653 | 2541 | 1773 | 1759 | 1750 | 1480 | 1660 | 1998 | 1908 | 1926 | 1786.84 | 1630 | 1944 | 1.193 | 0.254 | 0.00529916 | 0.03266055 | 0.003 | 0.0087 | 0.0054 | 0.0054 | TRUE | 0.0783 |
| Iqgap3 | 2687 | 2845 | 2937 | 3899 | 2874 | 2865 | 2854 | 2841 | 2949 | 3065 | 3092 | 3136 | 2989.66 | 2881 | 3098 | 1.075 | 0.104 | 0.09480411 | 0.25228627 | 0 | 0.008 | 0.0025 | 0.0025 | TRUE | 0.0051 |
| Iqsec1 | 1638 | 1615 | 1688 | 2468 | 1825 | 1608 | 1740 | 1613 | 1695 | 1940 | 1964 | 1760 | 1785.27 | 1683 | 1888 | 1.123 | 0.167 | 0.04652043 | 0.15607749 | 0.0019 | 0.0087 | 0.0045 | 0.0045 | TRUE | 0.0417 |
| Iqsec2 | 323 | 383 | 400 | 427 | 332 | 333 | 343 | 383 | 402 | 336 | 357 | 365 | 364.11 | 376 | 353 | 0.936 | -0.095 | 0.43943574 | 0.64664868 | 0 | 0.016 | 0.0081 | 0.0081 | TRUE | 0.0654 |
| Iqsec3 | 100 | 129 | 124 | 84 | 59 | 73 | 106 | 129 | 125 | 66 | 63 | 80 | 94.83 | 120 | 70 | 0.581 | -0.784 | 0.0002731 | 0.00314868 | 0 | 0.0416 | 0.0222 | 0.0222 | TRUE | 0.134 |
| Iqub | 28 | 22 | 21 | 50 | 41 | 37 | 30 | 22 | 21 | 39 | 44 | 41 | 32.79 | 24 | 41 | 1.703 | 0.768 | 0.02462561 | 0.10069875 | 0 | 0.1074 | 0.0512 | 0.0512 | TRUE | 0.2244 |
| Irak1 | 974 | 1077 | 1026 | 1710 | 1372 | 1160 | 1035 | 1076 | 1030 | 1344 | 1476 | 1270 | 1205.11 | 1047 | 1363 | 1.302 | 0.381 | 3.58E-05 | 0.00058105 | 0.0024 | 0.0096 | 0.0053 | 0.0053 | TRUE | 0.0617 |
| Irak1bp1 | 272 | 375 | 403 | 659 | 533 | 563 | 289 | 375 | 405 | 518 | 573 | 616 | 462.65 | 356 | 569 | 1.596 | 0.675 | 1.22E-05 | 0.00023645 | 0.0157 | 0.014 | 0.0149 | 0.0149 | TRUE | 0.3077 |
| Irak2 | 601 | 728 | 722 | 818 | 641 | 643 | 638 | 727 | 725 | 643 | 690 | 704 | 687.83 | 697 | 679 | 0.973 | -0.04 | 0.70364162 | 0.83769895 | 0.0024 | 0.0117 | 0.0063 | 0.0063 | TRUE | 0.063 |
| Irak3 | 915 | 777 | 869 | 910 | 655 | 559 | 972 | 776 | 873 | 715 | 705 | 612 | 775.41 | 874 | 677 | 0.776 | -0.365 | 0.00321183 | 0.02233786 | 0.0087 | 0.0112 | 0.0098 | 0.0098 | TRUE | 0.1151 |
| Irak4 | 167 | 147 | 162 | 231 | 170 | 160 | 177 | 147 | 163 | 182 | 183 | 175 | 171.08 | 162 | 180 | 1.11 | 0.15 | 0.34475816 | 0.56423428 | 0 | 0.0262 | 0.0123 | 0.0123 | TRUE | 0.0741 |
| Ireb2 | 17857 | 15997 | 16336 | 20964 | 15225 | 15271 | 18967 | 15976 | 16403 | 16480 | 16381 | 16717 | 16820.77 | 17115 | 16526 | 0.966 | -0.051 | 0.56009645 | 0.74190289 | 0.0041 | 0.0071 | 0.0054 | 0.0054 | TRUE | 0.1095 |
| Irf1 | 1301 | 1507 | 1635 | 1752 | 1456 | 1314 | 1382 | 1505 | 1642 | 1377 | 1567 | 1438 | 1485.15 | 1510 | 1461 | 0.967 | -0.048 | 0.63517601 | 0.79491607 | 0.0052 | 0.0091 | 0.0068 | 0.0068 | TRUE | 0.0706 |
| Irf2 | 16 | 55 | 54 | 56 | 44 | 59 | 17 | 55 | 54 | 44 | 47 | 65 | 47.02 | 42 | 52 | 1.231 | 0.3 | 0.49619423 | 0.69196835 | 0.1685 | 0.077 | 0.1182 | 0.1182 | TRUE | 2.0197 |
| Irf2bp1 | 781 | 787 | 734 | 1267 | 993 | 802 | 830 | 786 | 737 | 996 | 1068 | 878 | 882.48 | 784 | 981 | 1.251 | 0.323 | 0.00351395 | 0.02390938 | 0.0055 | 0.0106 | 0.0077 | 0.0077 | TRUE | 0.0982 |
| Irf2bp2 | 1290 | 2086 | 2060 | 2587 | 2272 | 2214 | 1370 | 2083 | 2068 | 2034 | 2445 | 2424 | 2070.63 | 1840 | 2301 | 1.249 | 0.321 | 0.05572416 | 0.17690852 | 0.0288 | 0.0085 | 0.0198 | 0.0198 | TRUE | 0.6029 |
| Irf2bpl | 3173 | 4278 | 4305 | 5468 | 3913 | 3804 | 3370 | 4272 | 4323 | 4299 | 4210 | 4164 | 4106.36 | 3988 | 4224 | 1.059 | 0.083 | 0.45655549 | 0.66122986 | 0.0094 | 0.0077 | 0.0087 | 0.0087 | TRUE | 0.2234 |
| Irf3 | 933 | 993 | 884 | 1342 | 1049 | 845 | 991 | 992 | 888 | 1055 | 1129 | 925 | 996.5 | 957 | 1036 | 1.083 | 0.116 | 0.29564037 | 0.5165093 | 0.006 | 0.0102 | 0.0078 | 0.0078 | TRUE | 0.1032 |
| Irf4 | 8 | 1 | 0 | 7 | 5 | 2 | 8 | 1 | 0 | 6 | 5 | 2 | 3.76 | 3 | 4 | 1.396 | 0.481 | 0.71102476 | NA | 1.0214 | 0.8828 | 0.9423 | 0.9423 | TRUE | 2.8308 |
| Irf5 | 186 | 212 | 163 | 253 | 181 | 179 | 198 | 212 | 164 | 199 | 195 | 196 | 193.76 | 191 | 197 | 1.029 | 0.042 | 0.79405234 | 0.8940332 | 0.0032 | 0.0239 | 0.0134 | 0.0134 | TRUE | 0.1709 |
| Irf6 | 662 | 1027 | 977 | 827 | 658 | 855 | 703 | 1026 | 981 | 650 | 708 | 936 | 833.98 | 903 | 765 | 0.846 | -0.242 | 0.19368921 | 0.39947733 | 0.0364 | 0.0109 | 0.0237 | 0.0237 | TRUE | 0.4529 |
| Irf7 | 9 | 8 | 9 | 10 | 10 | 5 | 10 | 8 | 9 | 8 | 11 | 5 | 8.45 | 9 | 8 | 0.907 | -0.141 | 0.82681082 | 0.91230308 | 0 | 0.3969 | 0.1819 | 0.1819 | TRUE | 0.2242 |
| Irf8 | 560 | 737 | 698 | 474 | 356 | 419 | 595 | 736 | 701 | 373 | 383 | 459 | 541.01 | 677 | 405 | 0.597 | -0.745 | 6.12E-08 | 2.37E-06 | 0.0105 | 0.013 | 0.0117 | 0.0117 | TRUE | 0.1536 |

| | | | | | | | | | | | | | | | | | | | | | | | | | |
|---|---|---|---|---|---|---|---|---|---|---|---|---|---|---|---|---|---|---|---|---|---|---|---|---|---|
| Irf9 | 335 | 390 | 395 | 451 | 364 | 422 | 356 | 389 | 397 | 355 | 392 | 462 | 391.68 | 381 | 403 | 1.055 | 0.078 | 0.57917409 | 0.75611769 | 0.0084 | 0.0153 | 0.0116 | 0.0116 | TRUE | 0.1891 |
| Irg1 | 0 | 0 | 0 | 0 | 1 | 0 | 0 | 0 | 0 | 0 | 1 | 0 | 0.18 | 0 | 0 | 1.859 | 0.894 | 0.82650734 | NA | 0 | 18.3785 | 10 | 10 | TRUE | 0.5204 |
| Irgc1 | 3 | 1 | 1 | 1 | 2 | 1 | 3 | 1 | 1 | 1 | 2 | 1 | 1.54 | 2 | 1 | 0.768 | -0.381 | 0.79807331 | NA | 0 | 2.1503 | 0.9352 | 0.9352 | TRUE | 0.4032 |
| Irgm1 | 321 | 105 | 100 | 144 | 75 | 39 | 341 | 105 | 100 | 113 | 81 | 43 | 130.47 | 182 | 79 | 0.434 | -1.204 | 0.08353636 | 0.23155281 | 0.34 | 0.0322 | 0.1637 | 0.34 | TRUE | 2.2878 |
| Irgm2 | 173 | 205 | 177 | 169 | 116 | 92 | 184 | 205 | 178 | 133 | 125 | 101 | 154.1 | 189 | 120 | 0.635 | -0.654 | 0.00030027 | 0.00341688 | 0.0046 | 0.0283 | 0.0168 | 0.0168 | TRUE | 0.1821 |
| Irgq | 612 | 687 | 703 | 546 | 376 | 431 | 650 | 686 | 706 | 429 | 405 | 472 | 557.94 | 681 | 435 | 0.639 | -0.646 | 2.43E-09 | 1.35E-07 | 0.0013 | 0.0128 | 0.0066 | 0.0066 | TRUE | 0.0596 |
| Irs1 | 46 | 72 | 80 | 104 | 61 | 83 | 49 | 72 | 80 | 82 | 66 | 91 | 73.22 | 67 | 80 | 1.184 | 0.243 | 0.37684151 | 0.59206639 | 0.0284 | 0.0519 | 0.0408 | 0.0408 | TRUE | 0.4869 |
| Irs2 | 251 | 306 | 302 | 642 | 502 | 387 | 267 | 306 | 303 | 505 | 540 | 424 | 390.65 | 292 | 490 | 1.678 | 0.746 | 1.10E-07 | 3.90E-06 | 0.0082 | 0.0153 | 0.0115 | 0.0115 | TRUE | 0.1566 |
| Irs3 | 11 | 14 | 10 | 10 | 6 | 10 | 12 | 14 | 10 | 8 | 6 | 11 | 10.16 | 12 | 8 | 0.705 | -0.505 | 0.39526028 | 0.60867459 | 0 | 0.3311 | 0.1531 | 0.1531 | TRUE | 0.1848 |
| Irs4 | 115 | 109 | 119 | 129 | 126 | 104 | 122 | 109 | 119 | 101 | 136 | 114 | 116.89 | 117 | 117 | 0.997 | -0.004 | 0.98408726 | 0.99163201 | 0 | 0.0351 | 0.0199 | 0.0199 | TRUE | 0.1933 |
| Irx1 | 11 | 11 | 16 | 41 | 25 | 31 | 12 | 11 | 16 | 32 | 27 | 34 | 21.97 | 13 | 31 | 2.403 | 1.265 | 0.00300219 | 0.02124296 | 0 | 0.1569 | 0.0757 | 0.0757 | TRUE | 0.1919 |
| Irx2 | 132 | 249 | 281 | 321 | 279 | 258 | 140 | 249 | 282 | 252 | 300 | 282 | 251 | 224 | 278 | 1.242 | 0.313 | 0.20130587 | 0.40888554 | 0.0614 | 0.02 | 0.0392 | 0.0392 | TRUE | 1.1605 |
| Irx3 | 138 | 240 | 213 | 209 | 193 | 209 | 147 | 240 | 214 | 164 | 208 | 229 | 200.15 | 200 | 200 | 0.998 | -0.003 | 0.98823329 | 0.99377809 | 0.0401 | 0.0234 | 0.0315 | 0.0315 | TRUE | 0.5872 |
| Irx4 | 9 | 11 | 15 | 31 | 20 | 20 | 10 | 11 | 15 | 24 | 22 | 22 | 17.23 | 12 | 23 | 1.905 | 0.93 | 0.04597126 | 0.15519235 | 0 | 0.1981 | 0.092 | 0.092 | TRUE | 0.2181 |
| Irx5 | 37 | 58 | 65 | 76 | 55 | 72 | 39 | 58 | 65 | 60 | 59 | 79 | 60.04 | 54 | 66 | 1.212 | 0.277 | 0.34488751 | 0.56427418 | 0.0265 | 0.0618 | 0.0452 | 0.0452 | TRUE | 0.4669 |
| Irx6 | 3 | 4 | 2 | 14 | 6 | 5 | 3 | 4 | 2 | 11 | 6 | 5 | 5.35 | 3 | 7 | 2.53 | 1.339 | 0.12122383 | 0.29610381 | 0 | 0.6222 | 0.3081 | 0.3081 | TRUE | 0.5289 |
| Isca1 | 1792 | 1642 | 1712 | 2257 | 1712 | 1624 | 1903 | 1640 | 1719 | 1774 | 1842 | 1778 | 1776.06 | 1754 | 1798 | 1.025 | 0.036 | 0.68237368 | 0.82388408 | 0.0025 | 0.0088 | 0.0049 | 0.0049 | TRUE | 0.0667 |
| Isca2 | 555 | 786 | 816 | 867 | 626 | 746 | 590 | 785 | 819 | 682 | 674 | 817 | 727.59 | 731 | 724 | 0.989 | -0.016 | 0.91819559 | 0.95991788 | 0.0198 | 0.0114 | 0.0157 | 0.0157 | TRUE | 0.3387 |
| Iscu | 1357 | 1160 | 1144 | 1622 | 1159 | 984 | 1441 | 1159 | 1149 | 1275 | 1247 | 1077 | 1224.64 | 1250 | 1200 | 0.961 | -0.058 | 0.64926441 | 0.8033859 | 0.0117 | 0.0096 | 0.0108 | 0.0108 | TRUE | 0.2159 |
| Isg15 | 1 | 3 | 3 | 1 | 2 | 0 | 1 | 3 | 3 | 1 | 2 | 0 | 1.67 | 2 | 1 | 0.411 | -1.282 | 0.4088876 | NA | 0 | 1.9819 | 1.02 | 1.02 | TRUE | 0.4365 |
| Isg20 | 9 | 10 | 8 | 9 | 6 | 6 | 10 | 10 | 8 | 7 | 6 | 7 | 7.95 | 9 | 7 | 0.731 | -0.452 | 0.48534384 | 0.68258022 | 0 | 0.4215 | 0.1746 | 0.1746 | TRUE | 0.0404 |
| Isg20l2 | 3840 | 3167 | 3080 | 4714 | 3340 | 3135 | 4079 | 3163 | 3093 | 3706 | 3594 | 3432 | 3510.91 | 3445 | 3577 | 1.039 | 0.055 | 0.65398792 | 0.80558399 | 0.0125 | 0.0078 | 0.0104 | 0.0104 | TRUE | 0.3148 |
| Isl1 | 14 | 14 | 19 | 38 | 26 | 11 | 15 | 14 | 19 | 30 | 28 | 12 | 19.64 | 16 | 23 | 1.469 | 0.555 | 0.26631377 | 0.48597718 | 0.0747 | 0.1747 | 0.1267 | 0.1267 | TRUE | 0.9709 |
| Isl2 | 14 | 40 | 43 | 32 | 36 | 29 | 15 | 40 | 43 | 25 | 39 | 32 | 32.27 | 33 | 32 | 0.969 | -0.045 | 0.92176666 | 0.96126008 | 0.1293 | 0.109 | 0.1184 | 0.1184 | TRUE | 1.2702 |
| Islr | 29 | 45 | 35 | 26 | 20 | 24 | 31 | 45 | 35 | 20 | 22 | 26 | 29.85 | 37 | 23 | 0.612 | -0.709 | 0.05188603 | 0.16814187 | 0 | 0.1173 | 0.0605 | 0.0605 | TRUE | 0.2613 |
| Islr2 | 1532 | 1847 | 1792 | 2054 | 1553 | 1462 | 1627 | 1845 | 1799 | 1615 | 1671 | 1600 | 1692.87 | 1757 | 1629 | 0.927 | -0.11 | 0.19020458 | 0.39517265 | 0.0018 | 0.0088 | 0.0045 | 0.0045 | TRUE | 0.0502 |
| Ism1 | 3 | 4 | 4 | 9 | 5 | 1 | 3 | 4 | 4 | 7 | 5 | 1 | 4.12 | 4 | 4 | 1.233 | 0.302 | 0.75853671 | NA | 0 | 0.8056 | 0.4523 | 0.4523 | TRUE | 0.596 |
| Isoc1 | 3464 | 4218 | 4312 | 4792 | 3679 | 4132 | 3679 | 4213 | 4330 | 3767 | 3958 | 4523 | 4078.37 | 4074 | 4083 | 1.002 | 0.003 | 0.97941711 | 0.98867206 | 0.008 | 0.0077 | 0.0079 | 0.0079 | TRUE | 0.1081 |
| Isoc2a | 2312 | 2394 | 2327 | 2950 | 2053 | 2254 | 2456 | 2391 | 2337 | 2319 | 2209 | 2467 | 2363.09 | 2395 | 2332 | 0.974 | -0.038 | 0.61919126 | 0.78469067 | 0.0012 | 0.0083 | 0.0039 | 0.0039 | TRUE | 0.0307 |
| Isoc2b | 21 | 42 | 34 | 40 | 37 | 28 | 22 | 42 | 34 | 31 | 40 | 31 | 33.38 | 33 | 34 | 1.031 | 0.045 | 0.90355707 | 0.95228213 | 0.0249 | 0.1056 | 0.068 | 0.068 | TRUE | 0.3315 |
| Ispd | 27 | 38 | 29 | 71 | 50 | 48 | 29 | 38 | 29 | 56 | 54 | 53 | 42.98 | 32 | 54 | 1.694 | 0.76 | 0.01077308 | 0.05570918 | 0 | 0.0835 | 0.039 | 0.039 | TRUE | 0.1895 |
| Ist1 | 3031 | 3855 | 4026 | 5302 | 3880 | 3864 | 3219 | 3850 | 4043 | 4168 | 4175 | 4230 | 3947.4 | 3704 | 4191 | 1.131 | 0.178 | 0.07917842 | 0.22317416 | 0.0064 | 0.0077 | 0.0072 | 0.0072 | TRUE | 0.159 |
| Isx | 0 | 0 | 0 | 0 | 0 | 0 | 0 | 0 | 0 | 0 | 0 | 0 | 0 | 0 | 0 | NA | NA | NA | NA | NA | NA | NA | NA | NA | NA |
| Isy1 | 1883 | 1838 | 1945 | 2620 | 1904 | 1686 | 2000 | 1836 | 1953 | 2060 | 2049 | 1846 | 1957.08 | 1930 | 1985 | 1.029 | 0.041 | 0.62715657 | 0.78999388 | 0.0023 | 0.0086 | 0.0047 | 0.0047 | TRUE | 0.0449 |
| Isyna1 | 9116 | 7700 | 7604 | 8673 | 6608 | 6272 | 9683 | 7690 | 7635 | 6818 | 7110 | 6866 | 7633.6 | 8336 | 6931 | 0.831 | -0.266 | 0.01532408 | 0.07184643 | 0.0099 | 0.0073 | 0.0086 | 0.0086 | TRUE | 0.2438 |
| Itch | 7017 | 6770 | 7016 | 7557 | 5148 | 5849 | 7453 | 6761 | 7045 | 5941 | 5539 | 6403 | 6523.61 | 7086 | 5961 | 0.841 | -0.25 | 0.00328415 | 0.02270828 | 0.0036 | 0.0074 | 0.005 | 0.005 | TRUE | 0.051 |
| Itfg1 | 1819 | 2685 | 2921 | 3204 | 2272 | 2598 | 1932 | 2682 | 2933 | 2519 | 2445 | 2844 | 2558.97 | 2516 | 2603 | 1.034 | 0.049 | 0.7542373 | 0.87027805 | 0.0258 | 0.0082 | 0.017 | 0.017 | TRUE | 0.4985 |
| Itfg2 | 1499 | 1612 | 1684 | 2157 | 1623 | 1593 | 1592 | 1610 | 1691 | 1696 | 1746 | 1744 | 1679.79 | 1631 | 1729 | 1.059 | 0.083 | 0.24911701 | 0.46624819 | 0 | 0.0089 | 0.0032 | 0.0032 | TRUE | 0.0125 |
| Itfg3 | 1905 | 2216 | 2178 | 2941 | 2016 | 2175 | 2023 | 2213 | 2187 | 2312 | 2169 | 2381 | 2214.25 | 2141 | 2287 | 1.068 | 0.095 | 0.23718313 | 0.45255895 | 0.0016 | 0.0084 | 0.0042 | 0.0042 | TRUE | 0.0279 |
| Itga1 | 55 | 41 | 27 | 143 | 119 | 114 | 58 | 41 | 27 | 112 | 128 | 125 | 81.95 | 42 | 122 | 2.892 | 1.532 | 1.13E-07 | 3.97E-06 | 0.0389 | 0.0471 | 0.0439 | 0.0439 | TRUE | 0.7761 |
| Itga10 | 72 | 69 | 57 | 78 | 93 | 55 | 76 | 69 | 57 | 61 | 100 | 60 | 70.7 | 67 | 74 | 1.09 | 0.125 | 0.6686744 | 0.81492803 | 0.0405 | 0.0535 | 0.0471 | 0.0471 | TRUE | 0.851 |
| Itga11 | 21 | 15 | 15 | 49 | 29 | 31 | 22 | 15 | 15 | 39 | 31 | 34 | 26 | 17 | 35 | 1.992 | 0.994 | 0.01108884 | 0.05680195 | 0 | 0.1336 | 0.067 | 0.067 | TRUE | 0.2837 |
| Itga2 | 1 | 0 | 0 | 6 | 4 | 2 | 1 | 0 | 0 | 5 | 4 | 2 | 2.05 | 0 | 4 | 10.279 | 3.362 | 0.04510609 | NA | 0 | 1.6176 | 0.8982 | 0.8982 | TRUE | 0.4614 |
| Itga2b | 341 | 444 | 392 | 488 | 323 | 382 | 362 | 443 | 394 | 384 | 348 | 418 | 391.43 | 400 | 383 | 0.958 | -0.062 | 0.64754541 | 0.80215228 | 0.0067 | 0.0153 | 0.0107 | 0.0107 | TRUE | 0.1055 |
| Itga3 | 5117 | 5789 | 5767 | 8749 | 6313 | 6255 | 5435 | 5782 | 5791 | 6878 | 6792 | 6847 | 6254.1 | 5669 | 6839 | 1.206 | 0.271 | 1.14E-05 | 0.0002219 | 4.00E-04 | 0.0074 | 0.0026 | 0.0026 | TRUE | 0.0158 |
| Itga4 | 11 | 20 | 29 | 36 | 28 | 18 | 12 | 20 | 29 | 28 | 30 | 20 | 23.15 | 20 | 26 | 1.286 | 0.363 | 0.43415851 | 0.6427102 | 0.0651 | 0.1492 | 0.1109 | 0.1109 | TRUE | 0.2663 |
| Itga5 | 3084 | 3020 | 3156 | 3786 | 2554 | 2396 | 3276 | 3016 | 3169 | 2976 | 2748 | 2623 | 2967.97 | 3154 | 2782 | 0.883 | -0.18 | 0.02949225 | 0.11359016 | 0.0024 | 0.008 | 0.0046 | 0.0046 | TRUE | 0.0458 |
| Itga6 | 7607 | 7983 | 8369 | 8961 | 6399 | 6820 | 8080 | 7973 | 8403 | 7044 | 6885 | 7466 | 7641.83 | 8152 | 7132 | 0.875 | -0.193 | 0.00380427 | 0.02538111 | 0.0011 | 0.0073 | 0.0031 | 0.0031 | TRUE | 0.0203 |
| Itga7 | 178 | 135 | 94 | 288 | 224 | 170 | 189 | 135 | 94 | 226 | 241 | 186 | 178.63 | 139 | 218 | 1.565 | 0.646 | 0.01138748 | 0.05797398 | 0.0578 | 0.0253 | 0.041 | 0.041 | TRUE | 0.4442 |
| Itga8 | 38 | 45 | 36 | 126 | 76 | 81 | 40 | 45 | 36 | 99 | 82 | 89 | 65.16 | 40 | 90 | 2.225 | 1.154 | 5.31E-06 | 0.0001164 | 0 | 0.0575 | 0.0283 | 0.0283 | TRUE | 0.0854 |
| Itga9 | 1366 | 1225 | 1139 | 1270 | 926 | 959 | 1451 | 1223 | 1144 | 998 | 996 | 1050 | 1143.75 | 1273 | 1015 | 0.797 | -0.327 | 0.00395884 | 0.02615789 | 0.007 | 0.0098 | 0.0084 | 0.0084 | TRUE | 0.1796 |
| Itgad | 1 | 0 | 0 | 1 | 0 | 0 | 1 | 0 | 0 | 1 | 0 | 0 | 0.31 | 0 | 0 | 0.954 | -0.067 | 0.98625928 | NA | 0 | 10.7014 | 9.0384 | 9.0384 | TRUE | 0.4614 |
| Itgae | 222 | 252 | 201 | 277 | 202 | 209 | 236 | 252 | 202 | 218 | 217 | 229 | 225.53 | 230 | 221 | 0.963 | -0.055 | 0.71295331 | 0.84375326 | 0 | 0.0215 | 0.0116 | 0.0116 | TRUE | 0.126 |
| Itgal | 505 | 346 | 271 | 769 | 582 | 513 | 536 | 346 | 272 | 605 | 626 | 562 | 491.06 | 385 | 598 | 1.554 | 0.636 | 0.00455941 | 0.0291965 | 0.0612 | 0.0136 | 0.0341 | 0.0341 | TRUE | 1.2559 |
| Itgam | 141 | 113 | 99 | 229 | 168 | 90 | 150 | 113 | 99 | 180 | 181 | 99 | 136.89 | 121 | 153 | 1.272 | 0.348 | 0.21563737 | 0.42753395 | 0.0696 | 0.031 | 0.0494 | 0.0494 | TRUE | 0.9935 |
| Itgav | 3451 | 3768 | 3885 | 3568 | 2565 | 2712 | 3666 | 3763 | 3901 | 2805 | 2760 | 2969 | 3310.51 | 3777 | 2845 | 0.753 | -0.409 | 5.48E-09 | 2.76E-07 | 8.00E-04 | 0.0079 | 0.0032 | 0.0032 | TRUE | 0.0175 |
| Itgax | 0 | 1 | 0 | 0 | 0 | 0 | 0 | 1 | 0 | 0 | 0 | 0 | 0.17 | 0 | 0 | 0.49 | -1.029 | 0.80087 | NA | 0 | 19.7991 | 10 | 10 | TRUE | 0.405 |
| Itgb1 | 15008 | 18733 | 19419 | 22102 | 15606 | 16944 | 15941 | 18709 | 19499 | 17375 | 16791 | 18548 | 17810.42 | 18050 | 17571 | 0.973 | -0.039 | 0.69294736 | 0.83159723 | 0.0056 | 0.0071 | 0.0069 | 0.0069 | TRUE | 0.1277 |
| Itgb1bp1 | 611 | 683 | 633 | 937 | 620 | 630 | 649 | 682 | 636 | 737 | 667 | 690 | 676.67 | 656 | 698 | 1.066 | 0.092 | 0.3427034 | 0.56279863 | 0 | 0.0118 | 0.0052 | 0.0052 | TRUE | 0.0294 |
| Itgb1bp2 | 16 | 17 | 17 | 26 | 19 | 21 | 17 | 17 | 17 | 20 | 20 | 23 | 19.15 | 17 | 21 | 1.249 | 0.321 | 0.44593229 | 0.65236459 | 0 | 0.1789 | 0.0747 | 0.0747 | TRUE | 0.025 |
| Itgb2 | 13 | 8 | 7 | 16 | 6 | 6 | 14 | 8 | 7 | 13 | 6 | 7 | 9.07 | 10 | 9 | 0.906 | -0.142 | 0.83140028 | 0.91442435 | 0 | 0.3701 | 0.2124 | 0.2124 | TRUE | 0.6689 |
| Itgb2l | 0 | 0 | 0 | 0 | 0 | 0 | 0 | 0 | 0 | 0 | 0 | 0 | 0 | 0 | 0 | NA | NA | NA | NA | NA | NA | NA | NA | NA | NA |
| Itgb3 | 1061 | 1128 | 959 | 1323 | 974 | 956 | 1127 | 1127 | 963 | 1040 | 1048 | 1046 | 1058.49 | 1072 | 1045 | 0.975 | -0.037 | 0.70328853 | 0.83766892 | 0.0031 | 0.01 | 0.0059 | 0.0059 | TRUE | 0.0953 |
| Itgb3bp | 309 | 373 | 372 | 470 | 336 | 311 | 328 | 373 | 374 | 369 | 362 | 340 | 357.61 | 358 | 357 | 0.998 | -0.003 | 0.98013223 | 0.98922122 | 0 | 0.0161 | 0.0078 | 0.0078 | TRUE | 0.06 |
| Itgb4 | 392 | 532 | 526 | 602 | 459 | 508 | 416 | 531 | 528 | 473 | 494 | 556 | 499.84 | 492 | 508 | 1.031 | 0.043 | 0.75539402 | 0.87078457 | 0.0107 | 0.0135 | 0.012 | 0.012 | TRUE | 0.2085 |
| Itgb5 | 1536 | 1649 | 1656 | 1949 | 1460 | 1677 | 1632 | 1647 | 1663 | 1532 | 1571 | 1836 | 1646.66 | 1647 | 1646 | 0.999 | -0.002 | 0.98603367 | 0.99254373 | 0.0043 | 0.0089 | 0.0062 | 0.0062 | TRUE | 0.1213 |
| Itgb6 | 0 | 1 | 0 | 4 | 2 | 0 | 0 | 1 | 0 | 3 | 2 | 0 | 1.05 | 0 | 2 | 4.9 | 2.293 | 0.31302341 | NA | 0 | 3.1469 | 2.4479 | 2.4479 | TRUE | 0.3114 |
| Itgb7 | 1264 | 1428 | 1524 | 1017 | 823 | 848 | 1343 | 1426 | 1530 | 799 | 885 | 928 | 1152.04 | 1433 | 871 | 0.607 | -0.72 | 1.16E-12 | 1.16E-10 | 0.0042 | 0.0098 | 0.0065 | 0.0065 | TRUE | 0.064 |
| Itgb8 | 0 | 0 | 0 | 2 | 1 | 0 | 0 | 0 | 0 | 2 | 1 | 0 | 0.44 | 0 | 1 | 4.754 | 2.249 | 0.50892201 | NA | 0 | 7.4712 | 6.8061 | 6.8061 | TRUE | 0.2361 |

| | | | | | | | | | | | | | | | | | | | | | | | | | |
|---|---|---|---|---|---|---|---|---|---|---|---|---|---|---|---|---|---|---|---|---|---|---|---|---|---|
| Itgbl1 | 2 | 1 | 1 | 4 | 2 | 5 | 2 | 1 | 1 | 3 | 2 | 5 | 2.48 | 1 | 3 | 2.612 | 1.385 | 0.26483075 | NA | 0 | 1.3338 | 0.6017 | 0.6017 | TRUE | 0.2854 |
| Itih1 | 1 | 0 | 0 | 0 | 0 | 1 | 1 | 0 | 0 | 0 | 0 | 1 | 0.36 | 0 | 0 | 0.954 | -0.067 | 0.98553519 | NA | 0 | 9.1716 | 7.9605 | 7.9605 | TRUE | 0.5383 |
| Itih2 | 2 | 0 | 3 | 4 | 1 | 4 | 2 | 0 | 3 | 3 | 1 | 4 | 2.29 | 2 | 3 | 1.681 | 0.75 | 0.58500186 | NA | 0 | 1.446 | 0.8886 | 0.8886 | TRUE | 0.5154 |
| Itih3 | 2 | 4 | 1 | 2 | 1 | 3 | 2 | 4 | 1 | 2 | 1 | 3 | 2.18 | 2 | 2 | 0.823 | -0.281 | 0.82677389 | NA | 0 | 1.521 | 0.7201 | 0.7201 | TRUE | 0.3522 |
| Itih4 | 12 | 12 | 4 | 7 | 1 | 4 | 13 | 12 | 4 | 6 | 1 | 4 | 6.62 | 10 | 4 | 0.39 | -1.359 | 0.12007934 | 0.2939912 | 0.1979 | 0.5047 | 0.3667 | 0.3667 | TRUE | 0.8781 |
| Itih5 | 237 | 208 | 166 | 356 | 226 | 206 | 252 | 208 | 167 | 280 | 243 | 225 | 229.11 | 209 | 249 | 1.199 | 0.261 | 0.16983031 | 0.36844068 | 0.0222 | 0.0213 | 0.0217 | 0.0217 | TRUE | 0.2391 |
| Itk | 0 | 0 | 0 | 2 | 1 | 0 | 0 | 0 | 0 | 2 | 1 | 0 | 0.44 | 0 | 1 | 4.754 | 2.249 | 0.50892201 | NA | 0 | 7.4712 | 6.8061 | 6.8061 | TRUE | 0.2361 |
| Itln1 | 0 | 0 | 0 | 0 | 1 | 0 | 0 | 0 | 0 | 0 | 1 | 0 | 0.18 | 0 | 0 | 1.859 | 0.894 | 0.82650734 | NA | 0 | 18.3785 | 10 | 10 | TRUE | 0.5204 |
| Itm2a | 536 | 816 | 772 | 894 | 612 | 651 | 569 | 815 | 775 | 703 | 658 | 713 | 705.56 | 720 | 691 | 0.96 | -0.059 | 0.6938978 | 0.83228303 | 0.0174 | 0.0116 | 0.0146 | 0.0146 | TRUE | 0.3935 |
| Itm2b | 3565 | 4823 | 4843 | 4949 | 3632 | 3916 | 3787 | 4817 | 4863 | 3891 | 3908 | 4287 | 4258.55 | 4489 | 4029 | 0.897 | -0.156 | 0.17921455 | 0.3804269 | 0.0104 | 0.0077 | 0.0095 | 0.0095 | TRUE | 0.2278 |
| Itm2c | 2055 | 2352 | 2354 | 2319 | 1683 | 1853 | 2183 | 2349 | 2364 | 1823 | 1811 | 2028 | 2092.94 | 2299 | 1887 | 0.821 | -0.285 | 0.00082657 | 0.00769075 | 0.0023 | 0.0085 | 0.0048 | 0.0048 | TRUE | 0.051 |
| Itpa | 3031 | 2370 | 2312 | 4149 | 2678 | 2478 | 3219 | 2367 | 2321 | 3262 | 2881 | 2713 | 2793.91 | 2636 | 2952 | 1.12 | 0.164 | 0.26049589 | 0.47988003 | 0.0206 | 0.0081 | 0.0149 | 0.0149 | TRUE | 0.4544 |
| Itpk1 | 7062 | 6379 | 6523 | 7535 | 5365 | 5203 | 7501 | 6371 | 6550 | 5923 | 5772 | 5696 | 6302.16 | 6807 | 5797 | 0.852 | -0.232 | 0.00745344 | 0.0420743 | 0.0039 | 0.0074 | 0.0052 | 0.0052 | TRUE | 0.0969 |
| Itpka | 581 | 853 | 867 | 1123 | 917 | 915 | 617 | 852 | 871 | 883 | 987 | 1002 | 868.44 | 780 | 957 | 1.226 | 0.294 | 0.04797548 | 0.15936591 | 0.0186 | 0.0107 | 0.0148 | 0.0148 | TRUE | 0.3932 |
| Itpkb | 100 | 124 | 168 | 106 | 103 | 84 | 106 | 124 | 169 | 83 | 111 | 92 | 114.14 | 133 | 95 | 0.715 | -0.484 | 0.04615024 | 0.15546381 | 0.0312 | 0.0358 | 0.0335 | 0.0335 | TRUE | 0.5713 |
| Itpkc | 558 | 609 | 560 | 812 | 642 | 618 | 593 | 608 | 562 | 638 | 691 | 676 | 628.13 | 588 | 668 | 1.136 | 0.184 | 0.05731257 | 0.18046194 | 0 | 0.0121 | 0.0052 | 0.0052 | TRUE | 0.0195 |
| Itpr1 | 11480 | 10633 | 11958 | 10475 | 7846 | 8139 | 12194 | 10619 | 12007 | 8235 | 8442 | 8909 | 10067.67 | 11607 | 8529 | 0.735 | -0.445 | 9.24E-08 | 3.36E-06 | 0.0034 | 0.0072 | 0.0049 | 0.0049 | TRUE | 0.0677 |
| Itpr2 | 199 | 232 | 234 | 175 | 117 | 122 | 211 | 232 | 235 | 138 | 126 | 134 | 179.17 | 226 | 133 | 0.586 | -0.77 | 5.77E-07 | 1.70E-05 | 0 | 0.0253 | 0.0112 | 0.0112 | TRUE | 0.0348 |
| Itpr3 | 5643 | 6389 | 7543 | 7646 | 5903 | 6308 | 5994 | 6381 | 7574 | 6011 | 6351 | 6905 | 6535.94 | 6650 | 6422 | 0.966 | -0.05 | 0.65007753 | 0.80390676 | 0.0085 | 0.0074 | 0.0087 | 0.0087 | TRUE | 0.1805 |
| Itprip | 153 | 163 | 147 | 209 | 166 | 145 | 163 | 163 | 148 | 164 | 179 | 159 | 162.42 | 158 | 167 | 1.06 | 0.085 | 0.59538157 | 0.76825314 | 0 | 0.0272 | 0.0121 | 0.0121 | TRUE | 0.0358 |
| Itpripl1 | 607 | 632 | 612 | 798 | 604 | 621 | 645 | 631 | 615 | 627 | 650 | 680 | 641.24 | 630 | 652 | 1.034 | 0.049 | 0.60464941 | 0.77532769 | 0 | 0.012 | 0.0048 | 0.0048 | TRUE | 0.0158 |
| Itpripl2 | 51 | 125 | 147 | 180 | 129 | 104 | 54 | 125 | 148 | 142 | 139 | 114 | 120.13 | 109 | 132 | 1.206 | 0.271 | 0.41841727 | 0.62905885 | 0.1184 | 0.0343 | 0.0722 | 0.0722 | TRUE | 1.2588 |
| Itsn1 | 3438 | 3245 | 3357 | 4257 | 3161 | 3226 | 3652 | 3241 | 3371 | 3347 | 3401 | 3531 | 3423.71 | 3421 | 3426 | 1.001 | 0.002 | 0.97854989 | 0.98832536 | 0.0019 | 0.0079 | 0.004 | 0.004 | TRUE | 0.0422 |
| Itsn2 | 1604 | 1741 | 1808 | 2634 | 1888 | 1635 | 1704 | 1739 | 1815 | 2071 | 2031 | 1790 | 1858.28 | 1753 | 1964 | 1.121 | 0.165 | 0.06585994 | 0.19789351 | 0.0031 | 0.0087 | 0.0052 | 0.0052 | TRUE | 0.0721 |
| Ivd | 1410 | 1361 | 1368 | 1570 | 1112 | 1107 | 1498 | 1359 | 1374 | 1234 | 1196 | 1212 | 1312.16 | 1410 | 1214 | 0.861 | -0.215 | 0.00855128 | 0.0468044 | 9.00E-04 | 0.0094 | 0.0041 | 0.0041 | TRUE | 0.0352 |
| Ivl | 0 | 0 | 0 | 0 | 0 | 0 | 0 | 0 | 0 | 0 | 0 | 0 | 0 | 0 | 0 | NA | NA | NA | NA | NA | NA | NA | NA | NA | NA |
| Ivns1abp | 7986 | 8144 | 8536 | 9561 | 6884 | 7025 | 8483 | 8133 | 8571 | 7516 | 7407 | 7690 | 7966.65 | 8396 | 7538 | 0.898 | -0.156 | 0.00910777 | 0.04907381 | 4.00E-04 | 0.0073 | 0.0024 | 0.0024 | TRUE | 0.0091 |
| Iws1 | 3989 | 3579 | 3705 | 5778 | 4253 | 3783 | 4237 | 3574 | 3720 | 4542 | 4576 | 4141 | 4131.81 | 3844 | 4420 | 1.15 | 0.202 | 0.03326602 | 0.1235103 | 0.0052 | 0.0077 | 0.0062 | 0.0062 | TRUE | 0.0973 |
| Iyd | 2 | 1 | 2 | 4 | 1 | 3 | 2 | 1 | 2 | 3 | 1 | 3 | 2.11 | 2 | 2 | 1.479 | 0.565 | 0.65802498 | NA | 0 | 1.5713 | 0.6802 | 0.6802 | TRUE | 0.2466 |
| Izumo1 | 59 | 84 | 54 | 25 | 45 | 36 | 63 | 84 | 54 | 20 | 48 | 39 | 51.38 | 67 | 36 | 0.529 | -0.919 | 0.01599111 | 0.0743768 | 0.0975 | 0.071 | 0.0833 | 0.0833 | TRUE | 1.2579 |
| Izumo2 | 5 | 9 | 14 | 13 | 12 | 10 | 5 | 9 | 14 | 10 | 13 | 11 | 10.41 | 9 | 11 | 1.194 | 0.256 | 0.67764811 | 0.82075457 | 0 | 0.3235 | 0.176 | 0.176 | TRUE | 0.3344 |
| Izumo3 | 0 | 0 | 0 | 0 | 0 | 0 | 0 | 0 | 0 | 0 | 0 | 0 | 0 | 0 | 0 | NA | NA | NA | NA | NA | NA | NA | NA | NA | NA |
| Izumo4 | 38 | 76 | 85 | 78 | 90 | 57 | 40 | 76 | 85 | 61 | 97 | 62 | 70.36 | 67 | 73 | 1.09 | 0.124 | 0.71857456 | 0.84760736 | 0.0901 | 0.0537 | 0.0709 | 0.0709 | TRUE | 1.0619 |
| Jag1 | 1179 | 1367 | 1361 | 1114 | 873 | 1087 | 1252 | 1365 | 1367 | 876 | 939 | 1190 | 1164.84 | 1328 | 1002 | 0.753 | -0.409 | 0.00196487 | 0.01517625 | 0.0137 | 0.0097 | 0.0117 | 0.0117 | TRUE | 0.321 |
| Jag2 | 1767 | 2295 | 2401 | 2519 | 1984 | 1935 | 1877 | 2292 | 2411 | 1980 | 2135 | 2118 | 2135.47 | 2193 | 2078 | 0.947 | -0.079 | 0.48569784 | 0.68272174 | 0.0079 | 0.0084 | 0.0087 | 0.0087 | TRUE | 0.1923 |
| Jagn1 | 1881 | 1670 | 1495 | 2395 | 1570 | 1469 | 1998 | 1668 | 1501 | 1883 | 1689 | 1608 | 1724.5 | 1722 | 1727 | 1.003 | 0.005 | 0.97176587 | 0.98494257 | 0.0132 | 0.0088 | 0.0112 | 0.0112 | TRUE | 0.2361 |
| Jak1 | 3534 | 4173 | 4458 | 5998 | 4329 | 4358 | 3754 | 4168 | 4476 | 4715 | 4658 | 4771 | 4423.51 | 4133 | 4715 | 1.141 | 0.19 | 0.02836262 | 0.1107586 | 0.0036 | 0.0076 | 0.0052 | 0.0052 | TRUE | 0.0781 |
| Jak2 | 578 | 732 | 798 | 985 | 671 | 756 | 614 | 731 | 801 | 774 | 722 | 828 | 745.02 | 715 | 775 | 1.082 | 0.114 | 0.37273646 | 0.58928297 | 0.0097 | 0.0113 | 0.0105 | 0.0105 | TRUE | 0.1809 |
| Jak3 | 972 | 1540 | 1590 | 2408 | 2311 | 2272 | 1032 | 1538 | 1597 | 1893 | 2486 | 2487 | 1838.92 | 1389 | 2289 | 1.647 | 0.72 | 6.72E-05 | 0.00100091 | 0.0353 | 0.0087 | 0.0229 | 0.0229 | TRUE | 0.6052 |
| Jakmip1 | 516 | 671 | 649 | 489 | 412 | 505 | 548 | 670 | 652 | 384 | 443 | 553 | 541.73 | 623 | 460 | 0.737 | -0.441 | 0.00629789 | 0.03735628 | 0.02 | 0.013 | 0.0169 | 0.0169 | TRUE | 0.358 |
| Jakmip2 | 33 | 18 | 16 | 24 | 14 | 16 | 35 | 18 | 16 | 19 | 15 | 18 | 20.09 | 23 | 17 | 0.749 | -0.417 | 0.38700214 | 0.60099965 | 0.0618 | 0.1709 | 0.1165 | 0.1165 | TRUE | 1.178 |
| Jakmip3 | 3 | 3 | 7 | 6 | 4 | 3 | 3 | 3 | 7 | 5 | 4 | 3 | 4.25 | 4 | 4 | 0.936 | -0.096 | 0.91625153 | NA | 0 | 0.7816 | 0.3636 | 0.3636 | TRUE | 0.4997 |
| Jam2 | 4372 | 3444 | 3196 | 5659 | 3808 | 3042 | 4644 | 3440 | 3209 | 4449 | 4097 | 3330 | 3861.38 | 3764 | 3959 | 1.052 | 0.073 | 0.65368972 | 0.80540538 | 0.0303 | 0.0078 | 0.0188 | 0.0188 | TRUE | 0.5082 |
| Jam3 | 813 | 600 | 524 | 957 | 662 | 564 | 864 | 599 | 526 | 752 | 712 | 617 | 678.48 | 663 | 694 | 1.048 | 0.067 | 0.72156276 | 0.84935352 | 0.0372 | 0.0118 | 0.0241 | 0.0241 | TRUE | 0.8236 |
| Jarid2 | 22339 | 16381 | 16893 | 26688 | 17518 | 14495 | 23728 | 16360 | 16962 | 20980 | 18848 | 15867 | 18790.92 | 19017 | 18565 | 0.976 | -0.035 | 0.8308406 | 0.91425507 | 0.0266 | 0.0071 | 0.0189 | 0.0189 | TRUE | 0.5746 |
| Jazf1 | 55 | 75 | 121 | 111 | 80 | 89 | 58 | 75 | 121 | 87 | 86 | 97 | 87.6 | 85 | 90 | 1.061 | 0.085 | 0.77433228 | 0.88248925 | 0.06 | 0.0445 | 0.052 | 0.052 | TRUE | 0.9557 |
| Jdp2 | 173 | 334 | 315 | 321 | 282 | 386 | 184 | 334 | 316 | 252 | 303 | 423 | 301.99 | 278 | 326 | 1.171 | 0.228 | 0.38502507 | 0.59905449 | 0.08 | 0.0178 | 0.0463 | 0.0463 | TRUE | 0.7282 |
| Jhdm1d | 1487 | 2116 | 2343 | 2235 | 1587 | 1726 | 1579 | 2113 | 2353 | 1757 | 1708 | 1889 | 1899.87 | 2015 | 1785 | 0.885 | -0.176 | 0.23421781 | 0.44901949 | 0.0199 | 0.0086 | 0.0152 | 0.0152 | TRUE | 0.4318 |
| Jkamp | 2006 | 2113 | 2003 | 2348 | 1745 | 1694 | 2131 | 2110 | 2011 | 1846 | 1878 | 1854 | 1971.65 | 2084 | 1859 | 0.892 | -0.165 | 0.01717189 | 0.07778476 | 0 | 0.0086 | 0.0029 | 0.0029 | TRUE | 0.0113 |
| Jmjd1c | 9509 | 6756 | 8015 | 10399 | 6951 | 6028 | 10100 | 6747 | 8048 | 8175 | 7479 | 6599 | 7857.96 | 8298 | 7418 | 0.894 | -0.162 | 0.28964157 | 0.51075882 | 0.0262 | 0.0073 | 0.0167 | 0.0167 | TRUE | 0.3162 |
| Jmjd4 | 1442 | 1214 | 1117 | 2108 | 1400 | 1117 | 1532 | 1212 | 1122 | 1657 | 1506 | 1223 | 1375.31 | 1289 | 1462 | 1.135 | 0.183 | 0.24204017 | 0.45860854 | 0.0241 | 0.0093 | 0.017 | 0.017 | TRUE | 0.3265 |
| Jmjd6 | 3503 | 3990 | 4265 | 4878 | 3706 | 3849 | 3721 | 3985 | 4283 | 3835 | 3987 | 4213 | 4003.94 | 3996 | 4012 | 1.004 | 0.005 | 0.95060475 | 0.97507515 | 0.0033 | 0.0077 | 0.005 | 0.005 | TRUE | 0.0479 |
| Jmjd7 | 727 | 865 | 857 | 790 | 652 | 675 | 772 | 864 | 861 | 621 | 702 | 739 | 759.67 | 832 | 687 | 0.824 | -0.279 | 0.01163919 | 0.05891313 | 0.0047 | 0.0112 | 0.0075 | 0.0075 | TRUE | 0.0874 |
| Jmjd8 | 462 | 443 | 440 | 764 | 532 | 488 | 491 | 442 | 442 | 601 | 572 | 534 | 513.69 | 458 | 569 | 1.244 | 0.314 | 0.00428807 | 0.02786626 | 0.0017 | 0.0133 | 0.0068 | 0.0068 | TRUE | 0.0439 |
| Jmy | 1869 | 1809 | 2012 | 2759 | 2116 | 2034 | 1985 | 1807 | 2020 | 2169 | 2277 | 2227 | 2080.71 | 1937 | 2224 | 1.148 | 0.199 | 0.01261357 | 0.06244231 | 0.0015 | 0.0085 | 0.0041 | 0.0041 | TRUE | 0.0423 |
| Josd1 | 1639 | 1959 | 1980 | 2195 | 1695 | 1695 | 1741 | 1956 | 1988 | 1726 | 1824 | 1855 | 1848.37 | 1895 | 1802 | 0.95 | -0.074 | 0.40066148 | 0.61357733 | 0.0028 | 0.0087 | 0.005 | 0.005 | TRUE | 0.061 |
| Josd2 | 159 | 134 | 141 | 265 | 184 | 118 | 169 | 134 | 142 | 208 | 198 | 129 | 163.29 | 148 | 178 | 1.209 | 0.274 | 0.2227889 | 0.4348722 | 0.0335 | 0.0271 | 0.0302 | 0.0302 | TRUE | 0.6086 |
| Jph1 | 299 | 284 | 328 | 546 | 381 | 355 | 318 | 284 | 329 | 429 | 410 | 389 | 359.72 | 310 | 409 | 1.321 | 0.402 | 0.0011159 | 0.00969164 | 0 | 0.0161 | 0.0081 | 0.0081 | TRUE | 0.0642 |
| Jph2 | 80 | 85 | 52 | 98 | 52 | 59 | 85 | 85 | 52 | 77 | 56 | 65 | 69.94 | 74 | 66 | 0.894 | -0.162 | 0.57273681 | 0.7512479 | 0.0353 | 0.054 | 0.0449 | 0.0449 | TRUE | 0.6109 |
| Jph3 | 211 | 220 | 258 | 240 | 155 | 186 | 224 | 220 | 259 | 189 | 167 | 204 | 210.32 | 234 | 187 | 0.795 | -0.33 | 0.03591739 | 0.13015446 | 0.0039 | 0.0226 | 0.0131 | 0.0131 | TRUE | 0.095 |
| Jph4 | 126 | 125 | 129 | 131 | 93 | 81 | 134 | 125 | 130 | 103 | 100 | 89 | 113.32 | 130 | 97 | 0.754 | -0.408 | 0.02782417 | 0.1095464 | 0 | 0.036 | 0.0159 | 0.0159 | TRUE | 0.0548 |
| Jpx | 1 | 2 | 2 | 10 | 5 | 3 | 1 | 2 | 2 | 8 | 5 | 3 | 3.6 | 2 | 5 | 3.29 | 1.718 | 0.10890386 | NA | 0 | 0.9223 | 0.4388 | 0.4388 | TRUE | 0.1271 |
| Jrk | 538 | 649 | 484 | 946 | 685 | 502 | 571 | 648 | 486 | 744 | 737 | 550 | 622.63 | 568 | 677 | 1.191 | 0.253 | 0.12209088 | 0.29718298 | 0.0228 | 0.0122 | 0.0176 | 0.0176 | TRUE | 0.3153 |
| Jrkl | 158 | 199 | 160 | 257 | 192 | 153 | 168 | 199 | 161 | 202 | 207 | 167 | 183.89 | 176 | 192 | 1.094 | 0.13 | 0.45131258 | 0.65683457 | 0.0071 | 0.0248 | 0.0159 | 0.0159 | TRUE | 0.1416 |
| Jsrp1 | 37 | 82 | 74 | 66 | 69 | 86 | 39 | 82 | 74 | 52 | 74 | 94 | 69.29 | 65 | 73 | 1.12 | 0.163 | 0.64307714 | 0.79907134 | 0.0975 | 0.0544 | 0.0749 | 0.0749 | TRUE | 0.344 |
| Jtb | 906 | 869 | 848 | 1224 | 817 | 815 | 962 | 868 | 851 | 962 | 879 | 892 | 902.52 | 894 | 911 | 1.02 | 0.029 | 0.76336488 | 0.87543015 | 0.0023 | 0.0106 | 0.0057 | 0.0057 | TRUE | 0.053 |
| Jun | 307 | 346 | 227 | 766 | 505 | 374 | 326 | 346 | 228 | 602 | 543 | 409 | 409.08 | 300 | 518 | 1.731 | 0.791 | 9.93E-05 | 0.0013718 | 0.0402 | 0.015 | 0.0271 | 0.0271 | TRUE | 0.4991 |
| Junb | 556 | 942 | 1134 | 1088 | 1040 | 1029 | 591 | 941 | 1139 | 855 | 1119 | 1126 | 961.78 | 890 | 1033 | 1.161 | 0.215 | 0.3331835 | 0.55385436 | 0.0637 | 0.0103 | 0.0344 | 0.0344 | TRUE | 0.4899 |
| Jund | 2653 | 3285 | 2826 | 6758 | 4504 | 3824 | 2818 | 3281 | 2838 | 5313 | 4846 | 4186 | 3880.15 | 2979 | 4782 | 1.606 | 0.683 | 3.58E-09 | 1.88E-07 | 0.0105 | 0.0078 | 0.0094 | 0.0094 | TRUE | 0.1444 |

| | | | | | | | | | | | | | | | | | | | | | | | |
|---|---|---|---|---|---|---|---|---|---|---|---|---|---|---|---|---|---|---|---|---|---|---|---|
| Jup | 7932 | 7435 | 8095 | 7855 | 6127 | 6611 | 8425 | 7425 | 8128 | 6175 | 6592 | 7237 | 7330.48 | 7993 | 6668 | 0.834 | -0.262 | 0.0045949 | 0.02937605 | 0.0051 | 0.0074 | 0.006 | 0.006 | TRUE | 0.0679 |
| Kalrn | 887 | 950 | 939 | 1083 | 812 | 847 | 942 | 949 | 943 | 851 | 874 | 927 | 914.33 | 945 | 884 | 0.935 | -0.097 | 0.25293391 | 0.47086616 | 0 | 0.0105 | 0.0041 | 0.0041 | TRUE | 0.0214 |
| Kank1 | 1778 | 1774 | 1866 | 2512 | 1796 | 1802 | 1889 | 1772 | 1874 | 1975 | 1932 | 1973 | 1902.27 | 1845 | 1960 | 1.063 | 0.088 | 0.21532299 | 0.42717126 | 0 | 0.0086 | 0.0031 | 0.0031 | TRUE | 0.0145 |
| Kank2 | 6155 | 8490 | 8740 | 5909 | 4836 | 6206 | 6538 | 8479 | 8776 | 4645 | 5203 | 6793 | 6739.08 | 7931 | 5547 | 0.699 | -0.516 | 0.00162973 | 0.01307428 | 0.0317 | 0.0074 | 0.0192 | 0.0192 | TRUE | 0.4706 |
| Kank3 | 2313 | 2515 | 2440 | 2816 | 2284 | 2236 | 2457 | 2512 | 2450 | 2214 | 2457 | 2448 | 2422.9 | 2473 | 2373 | 0.959 | -0.06 | 0.43273411 | 0.64199587 | 0.0015 | 0.0083 | 0.0039 | 0.0039 | TRUE | 0.0427 |
| Kank4 | 611 | 561 | 580 | 1082 | 886 | 724 | 649 | 560 | 582 | 851 | 953 | 793 | 731.34 | 597 | 866 | 1.449 | 0.535 | 4.03E-06 | 9.21E-05 | 0.0059 | 0.0114 | 0.0083 | 0.0083 | TRUE | 0.0918 |
| Kansl1 | 6480 | 6779 | 6848 | 8041 | 5939 | 5991 | 6883 | 6770 | 6876 | 6321 | 6390 | 6558 | 6633.09 | 6843 | 6423 | 0.939 | -0.092 | 0.10347851 | 0.26763907 | 0 | 0.0074 | 0.0021 | 0.0021 | TRUE | 0.0041 |
| Kansl1l | 233 | 265 | 252 | 301 | 245 | 292 | 247 | 265 | 253 | 237 | 264 | 320 | 264.17 | 255 | 274 | 1.068 | 0.095 | 0.54448137 | 0.72961429 | 0.0091 | 0.0194 | 0.0139 | 0.0139 | TRUE | 0.2437 |
| Kansl2 | 2456 | 2103 | 2035 | 2717 | 1928 | 1876 | 2609 | 2100 | 2043 | 2136 | 2074 | 2054 | 2169.37 | 2251 | 2088 | 0.928 | -0.108 | 0.33835219 | 0.5583911 | 0.0091 | 0.0084 | 0.0087 | 0.0087 | TRUE | 0.2339 |
| Kansl3 | 3140 | 3571 | 3686 | 4336 | 3220 | 3403 | 3335 | 3566 | 3701 | 3409 | 3465 | 3725 | 3533.5 | 3534 | 3533 | 0.999 | -0.001 | 0.98938936 | 0.99437047 | 0.0022 | 0.0078 | 0.0042 | 0.0042 | TRUE | 0.0294 |
| Kap | 0 | 0 | 0 | 0 | 0 | 0 | 0 | 0 | 0 | 0 | 0 | 0 | 0 | 0 | 0 | NA | NA | NA | NA | NA | NA | NA | NA | NA | NA |
| Kars | 32688 | 33083 | 33780 | 36242 | 26374 | 26810 | 34720 | 33040 | 33919 | 28491 | 28377 | 29348 | 31315.77 | 33893 | 28739 | 0.848 | -0.238 | 2.06E-05 | 0.00036544 | 4.00E-04 | 0.007 | 0.0022 | 0.0022 | TRUE | 0.0059 |
| Kat2a | 11485 | 11568 | 11955 | 14568 | 10729 | 11018 | 12199 | 11553 | 12004 | 11452 | 11544 | 12061 | 11802.19 | 11919 | 11686 | 0.98 | -0.029 | 0.63952126 | 0.79689135 | 7.00E-04 | 0.0072 | 0.0026 | 0.0026 | TRUE | 0.0096 |
| Kat2b | 2672 | 2751 | 2927 | 4131 | 2926 | 3141 | 2838 | 2747 | 2939 | 3247 | 3148 | 3438 | 3059.76 | 2841 | 3278 | 1.153 | 0.206 | 0.00501216 | 0.03138958 | 0.0011 | 0.008 | 0.0036 | 0.0036 | TRUE | 0.0219 |
| Kat5 | 1878 | 2113 | 2063 | 2394 | 1798 | 1847 | 1995 | 2110 | 2071 | 1882 | 1935 | 2022 | 2002.48 | 2059 | 1946 | 0.945 | -0.082 | 0.26272169 | 0.48209965 | 6.00E-04 | 0.0085 | 0.0034 | 0.0034 | TRUE | 0.0139 |
| Kat6a | 5655 | 5803 | 6462 | 8204 | 6231 | 6263 | 6007 | 5796 | 6489 | 6449 | 6704 | 6856 | 6383.33 | 6097 | 6670 | 1.094 | 0.129 | 0.08251631 | 0.22941076 | 0.0019 | 0.0074 | 0.0038 | 0.0038 | TRUE | 0.0386 |
| Kat6b | 7789 | 7146 | 7498 | 11974 | 8900 | 7910 | 8273 | 7137 | 7529 | 9413 | 9576 | 8659 | 8431.08 | 7646 | 9216 | 1.205 | 0.269 | 0.00190018 | 0.01478521 | 0.0041 | 0.0073 | 0.0053 | 0.0053 | TRUE | 0.0628 |
| Kat7 | 16236 | 16231 | 17249 | 19676 | 14390 | 14777 | 17246 | 16210 | 17320 | 15468 | 15483 | 16176 | 16316.93 | 16925 | 15709 | 0.928 | -0.108 | 0.08695278 | 0.23842477 | 9.00E-04 | 0.0071 | 0.0028 | 0.0028 | TRUE | 0.0167 |
| Kat8 | 1399 | 1532 | 1554 | 1608 | 1156 | 1232 | 1486 | 1530 | 1560 | 1264 | 1244 | 1349 | 1405.48 | 1525 | 1286 | 0.842 | -0.248 | 0.00161285 | 0.01297853 | 4.00E-04 | 0.0092 | 0.0037 | 0.0037 | TRUE | 0.0216 |
| Katna1 | 2681 | 3134 | 3209 | 3163 | 2310 | 2628 | 2848 | 3130 | 3222 | 2487 | 2485 | 2877 | 2841.42 | 3067 | 2616 | 0.853 | -0.23 | 0.01763748 | 0.07928957 | 0.0052 | 0.0081 | 0.0064 | 0.0064 | TRUE | 0.0915 |
| Katnal1 | 422 | 566 | 720 | 501 | 356 | 453 | 448 | 565 | 723 | 394 | 383 | 496 | 501.54 | 579 | 424 | 0.732 | -0.45 | 0.01986138 | 0.08641479 | 0.0361 | 0.0135 | 0.0248 | 0.0248 | TRUE | 0.2097 |
| Katnal2 | 182 | 214 | 239 | 178 | 117 | 154 | 193 | 214 | 240 | 140 | 126 | 169 | 180.24 | 216 | 145 | 0.67 | -0.577 | 0.0013594 | 0.01133214 | 0.0099 | 0.0252 | 0.0176 | 0.0176 | TRUE | 0.2031 |
| Katnb1 | 2350 | 2333 | 2449 | 3220 | 2396 | 2317 | 2496 | 2330 | 2459 | 2531 | 2578 | 2536 | 2488.46 | 2428 | 2548 | 1.049 | 0.07 | 0.30607433 | 0.52666423 | 3.00E-04 | 0.0082 | 0.0029 | 0.0029 | TRUE | 0.0153 |
| Katnbl1 | 2230 | 2096 | 1916 | 2700 | 1914 | 1912 | 2369 | 2093 | 1924 | 2123 | 2059 | 2093 | 2110.11 | 2129 | 2092 | 0.983 | -0.025 | 0.7995774 | 0.8967457 | 0.0051 | 0.0085 | 0.0065 | 0.0065 | TRUE | 0.1175 |
| Kazald1 | 11 | 16 | 11 | 19 | 19 | 14 | 12 | 16 | 11 | 15 | 20 | 15 | 14.9 | 13 | 17 | 1.303 | 0.382 | 0.43981333 | 0.64708356 | 0 | 0.228 | 0.108 | 0.108 | TRUE | 0.1834 |
| Kazn | 3257 | 4730 | 4775 | 3160 | 2646 | 3147 | 3460 | 4724 | 4795 | 2484 | 2847 | 3445 | 3625.66 | 4326 | 2925 | 0.676 | -0.565 | 0.00046308 | 0.00489525 | 0.0284 | 0.0078 | 0.0185 | 0.0185 | TRUE | 0.3736 |
| Kbtbd11 | 488 | 763 | 738 | 569 | 452 | 509 | 518 | 762 | 741 | 447 | 486 | 557 | 585.37 | 674 | 497 | 0.737 | -0.441 | 0.0103182 | 0.05403009 | 0.0264 | 0.0125 | 0.0196 | 0.0196 | TRUE | 0.4784 |
| Kbtbd12 | 1 | 2 | 0 | 4 | 3 | 2 | 1 | 2 | 0 | 3 | 3 | 2 | 1.94 | 1 | 3 | 2.806 | 1.489 | 0.30154579 | NA | 0 | 1.7078 | 0.8251 | 0.8251 | TRUE | 0.373 |
| Kbtbd13 | 29 | 30 | 47 | 109 | 67 | 50 | 31 | 30 | 47 | 86 | 72 | 55 | 53.41 | 36 | 71 | 1.976 | 0.982 | 0.00234054 | 0.01743744 | 0.0381 | 0.0686 | 0.054 | 0.054 | TRUE | 0.2584 |
| Kbtbd2 | 1310 | 1513 | 1467 | 1678 | 1265 | 1193 | 1391 | 1511 | 1473 | 1319 | 1361 | 1306 | 1393.6 | 1458 | 1329 | 0.911 | -0.135 | 0.08425962 | 0.23276049 | 4.00E-04 | 0.0093 | 0.0037 | 0.0037 | TRUE | 0.0193 |
| Kbtbd3 | 97 | 98 | 106 | 119 | 104 | 76 | 103 | 98 | 106 | 94 | 112 | 83 | 99.33 | 102 | 96 | 0.939 | -0.091 | 0.660412 | 0.80885221 | 0 | 0.0401 | 0.0211 | 0.0211 | TRUE | 0.1845 |
| Kbtbd4 | 718 | 687 | 625 | 1000 | 626 | 599 | 763 | 686 | 628 | 786 | 674 | 656 | 698.61 | 692 | 705 | 1.021 | 0.029 | 0.8155129 | 0.90592684 | 0.0083 | 0.0116 | 0.01 | 0.01 | TRUE | 0.123 |
| Kbtbd7 | 705 | 670 | 594 | 1186 | 914 | 733 | 749 | 669 | 596 | 932 | 983 | 802 | 788.76 | 671 | 906 | 1.35 | 0.433 | 0.00080606 | 0.00753744 | 0.0098 | 0.0111 | 0.0107 | 0.0107 | TRUE | 0.1192 |
| Kbtbd8 | 2886 | 2609 | 2668 | 3904 | 2758 | 2516 | 3065 | 2606 | 2679 | 3069 | 2967 | 2754 | 2856.77 | 2783 | 2930 | 1.053 | 0.075 | 0.43349729 | 0.64234492 | 0.0047 | 0.0081 | 0.0062 | 0.0062 | TRUE | 0.0952 |
| Kcmf1 | 4115 | 4772 | 4650 | 6838 | 4599 | 4671 | 4371 | 4766 | 4669 | 5375 | 4948 | 5113 | 4873.78 | 4602 | 5145 | 1.118 | 0.161 | 0.02810164 | 0.11006223 | 0.0016 | 0.0076 | 0.0037 | 0.0037 | TRUE | 0.0234 |
| Kcna1 | 8 | 12 | 6 | 34 | 16 | 8 | 8 | 12 | 6 | 27 | 17 | 9 | 13.2 | 9 | 18 | 2.011 | 1.008 | 0.1083919 | 0.27573501 | 0.1407 | 0.2565 | 0.1988 | 0.1988 | TRUE | 0.129 |
| Kcna10 | 0 | 0 | 0 | 0 | 0 | 0 | 0 | 0 | 0 | 0 | 0 | 0 | 0 | 0 | 0 | NA | NA | NA | NA | NA | NA | NA | NA | NA | NA |
| Kcna2 | 3 | 12 | 7 | 14 | 5 | 9 | 3 | 12 | 7 | 11 | 5 | 10 | 8.07 | 7 | 9 | 1.186 | 0.246 | 0.73978058 | 0.86066702 | 0.1048 | 0.415 | 0.2711 | 0.2711 | TRUE | 0.3028 |
| Kcna3 | 11 | 15 | 9 | 28 | 24 | 11 | 12 | 15 | 9 | 22 | 26 | 12 | 15.93 | 12 | 20 | 1.682 | 0.751 | 0.15391648 | 0.34575855 | 0 | 0.2137 | 0.1323 | 0.1323 | TRUE | 0.5161 |
| Kcna4 | 1 | 0 | 0 | 1 | 2 | 2 | 1 | 0 | 0 | 1 | 2 | 2 | 1.03 | 0 | 2 | 4.603 | 2.203 | 0.27660303 | NA | 0 | 3.2005 | 1.6625 | 1.6625 | TRUE | 0.4614 |
| Kcna5 | 10 | 7 | 7 | 15 | 8 | 5 | 11 | 7 | 7 | 12 | 9 | 5 | 8.42 | 8 | 9 | 1.067 | 0.094 | 0.88817339 | 0.94527988 | 0 | 0.3982 | 0.2025 | 0.2025 | TRUE | 0.2079 |
| Kcna6 | 27 | 41 | 20 | 56 | 32 | 34 | 29 | 41 | 20 | 44 | 34 | 37 | 34.23 | 30 | 38 | 1.294 | 0.372 | 0.32123686 | 0.54237525 | 0.0349 | 0.1031 | 0.0714 | 0.0714 | TRUE | 0.3894 |
| Kcna7 | 25 | 32 | 36 | 49 | 40 | 25 | 27 | 32 | 36 | 39 | 43 | 27 | 33.93 | 32 | 36 | 1.152 | 0.204 | 0.55826115 | 0.74059171 | 0 | 0.104 | 0.0583 | 0.0583 | TRUE | 0.3064 |
| Kcnab1 | 0 | 0 | 0 | 6 | 2 | 0 | 0 | 0 | 0 | 5 | 2 | 0 | 1.14 | 0 | 2 | 12.338 | 3.625 | 0.15273868 | NA | 6.5132 | 2.8847 | 3.4196 | 3.4196 | TRUE | 0.063 |
| Kcnab2 | 233 | 205 | 188 | 236 | 166 | 201 | 247 | 205 | 189 | 186 | 179 | 220 | 204.19 | 214 | 195 | 0.911 | -0.135 | 0.43813137 | 0.64581383 | 0.011 | 0.023 | 0.0169 | 0.0169 | TRUE | 0.2074 |
| Kcnab3 | 550 | 567 | 476 | 465 | 412 | 353 | 584 | 566 | 478 | 366 | 443 | 386 | 470.61 | 543 | 398 | 0.733 | -0.448 | 0.0009339 | 0.0084458 | 0.0086 | 0.0139 | 0.011 | 0.011 | TRUE | 0.1282 |
| Kcnb1 | 174 | 147 | 180 | 250 | 213 | 198 | 185 | 147 | 181 | 197 | 229 | 217 | 192.47 | 171 | 214 | 1.252 | 0.324 | 0.05019766 | 0.16466672 | 0.0052 | 0.024 | 0.0145 | 0.0145 | TRUE | 0.1623 |
| Kcnb2 | 0 | 1 | 0 | 2 | 2 | 2 | 0 | 1 | 0 | 2 | 2 | 2 | 1.15 | 0 | 2 | 5.357 | 2.422 | 0.20741993 | NA | 0 | 2.8666 | 1.409 | 1.409 | TRUE | 0.405 |
| Kcnc1 | 13 | 32 | 18 | 21 | 10 | 16 | 14 | 32 | 18 | 17 | 11 | 18 | 18.1 | 21 | 15 | 0.702 | -0.51 | 0.32171759 | 0.54291994 | 0.0772 | 0.1889 | 0.1341 | 0.1341 | TRUE | 0.9005 |
| Kcnc2 | 5 | 6 | 8 | 13 | 7 | 15 | 5 | 6 | 8 | 10 | 8 | 16 | 8.92 | 6 | 11 | 1.758 | 0.814 | 0.22183536 | 0.43411891 | 0 | 0.3763 | 0.1974 | 0.1974 | TRUE | 0.4125 |
| Kcnc3 | 245 | 231 | 312 | 374 | 326 | 235 | 260 | 231 | 313 | 294 | 351 | 257 | 284.37 | 268 | 301 | 1.121 | 0.165 | 0.35441659 | 0.57279009 | 0.0203 | 0.0185 | 0.0194 | 0.0194 | TRUE | 0.245 |
| Kcnc4 | 17 | 31 | 54 | 43 | 38 | 41 | 18 | 31 | 54 | 34 | 41 | 45 | 37.13 | 34 | 40 | 1.153 | 0.205 | 0.63493481 | 0.79485386 | 0.1196 | 0.0956 | 0.1069 | 0.1069 | TRUE | 0.4985 |
| Kcnd1 | 383 | 425 | 441 | 395 | 346 | 404 | 407 | 424 | 443 | 311 | 372 | 442 | 399.85 | 425 | 375 | 0.88 | -0.184 | 0.23234489 | 0.44654725 | 0.014 | 0.0151 | 0.0145 | 0.0145 | TRUE | 0.246 |
| Kcnd2 | 7 | 5 | 12 | 20 | 22 | 16 | 7 | 5 | 12 | 16 | 24 | 18 | 13.56 | 8 | 19 | 2.313 | 1.21 | 0.03320915 | 0.12341534 | 0 | 0.2498 | 0.1442 | 0.1442 | TRUE | 0.5297 |
| Kcnd3 | 8 | 3 | 2 | 15 | 9 | 10 | 8 | 3 | 2 | 12 | 10 | 11 | 7.65 | 4 | 11 | 2.426 | 1.279 | 0.09510749 | 0.25278654 | 0 | 0.4373 | 0.2634 | 0.2634 | TRUE | 1.0615 |
| Kcne1 | 9 | 10 | 6 | 7 | 9 | 9 | 10 | 10 | 6 | 6 | 10 | 10 | 8.43 | 9 | 9 | 0.967 | -0.049 | 0.9408236 | 0.97003791 | 0 | 0.3975 | 0.1959 | 0.1959 | TRUE | 0.3564 |
| Kcne1l | 4 | 6 | 3 | 15 | 10 | 4 | 4 | 6 | 3 | 12 | 11 | 4 | 6.7 | 4 | 9 | 2.05 | 1.036 | 0.1898161 | 0.39506196 | 0 | 0.4988 | 0.2803 | 0.2803 | TRUE | 0.5849 |
| Kcne2 | 3 | 0 | 0 | 2 | 1 | 0 | 3 | 0 | 0 | 2 | 1 | 0 | 0.97 | 1 | 1 | 0.858 | -0.221 | 0.92570942 | NA | 2.2264 | 3.3947 | 3.0145 | 3.0145 | TRUE | 1.4428 |
| Kcne3 | 3 | 1 | 1 | 4 | 0 | 0 | 3 | 1 | 1 | 3 | 0 | 0 | 1.39 | 2 | 1 | 0.651 | -0.62 | 0.73998811 | NA | 0.9569 | 2.3788 | 1.7849 | 1.7849 | TRUE | 1.8921 |
| Kcne4 | 0 | 0 | 0 | 0 | 0 | 0 | 0 | 0 | 0 | 0 | 0 | 0 | 0 | 0 | 0 | NA | NA | NA | NA | NA | NA | NA | NA | NA | NA |
| Kcnf1 | 40 | 22 | 37 | 103 | 71 | 66 | 42 | 22 | 37 | 81 | 76 | 72 | 55.2 | 34 | 76 | 2.268 | 1.181 | 0.00011389 | 0.00152744 | 0.0118 | 0.0666 | 0.046 | 0.046 | TRUE | 0.6719 |
| Kcng1 | 177 | 201 | 194 | 229 | 183 | 196 | 188 | 201 | 195 | 180 | 197 | 215 | 195.84 | 195 | 197 | 1.01 | 0.015 | 0.92080705 | 0.96114949 | 0 | 0.0237 | 0.0113 | 0.0113 | TRUE | 0.0688 |
| Kcng2 | 35 | 29 | 47 | 34 | 34 | 20 | 37 | 29 | 47 | 27 | 37 | 22 | 33.09 | 38 | 29 | 0.751 | -0.414 | 0.26946342 | 0.48946478 | 0.0304 | 0.1065 | 0.0704 | 0.0704 | TRUE | 0.2219 |
| Kcng3 | 12 | 11 | 12 | 22 | 23 | 12 | 13 | 11 | 12 | 17 | 25 | 13 | 15.16 | 12 | 18 | 1.541 | 0.624 | 0.21854528 | 0.43064669 | 0 | 0.2242 | 0.1157 | 0.1157 | TRUE | 0.2913 |
| Kcng4 | 14 | 11 | 9 | 18 | 12 | 6 | 15 | 11 | 9 | 14 | 13 | 7 | 11.42 | 12 | 11 | 0.975 | -0.036 | 0.95135409 | 0.97533692 | 0 | 0.2954 | 0.1631 | 0.1631 | TRUE | 0.4469 |
| Kcnh1 | 125 | 153 | 132 | 162 | 135 | 112 | 133 | 153 | 133 | 127 | 145 | 123 | 135.55 | 140 | 132 | 0.944 | -0.083 | 0.64165955 | 0.79835032 | 0 | 0.0312 | 0.0156 | 0.0156 | TRUE | 0.0786 |
| Kcnh2 | 1086 | 1419 | 1466 | 1159 | 939 | 1174 | 1154 | 1417 | 1472 | 911 | 1010 | 1285 | 1208.21 | 1348 | 1069 | 0.792 | -0.336 | 0.03080979 | 0.11745999 | 0.0232 | 0.0096 | 0.0166 | 0.0166 | TRUE | 0.3736 |
| Kcnh3 | 403 | 611 | 624 | 291 | 316 | 346 | 428 | 610 | 627 | 229 | 340 | 379 | 435.39 | 555 | 316 | 0.568 | -0.817 | 0.0001932 | 0.00236574 | 0.0519 | 0.0145 | 0.0321 | 0.0321 | TRUE | 0.6862 |
| Kcnh4 | 139 | 119 | 122 | 118 | 104 | 108 | 148 | 119 | 123 | 93 | 112 | 118 | 118.65 | 130 | 108 | 0.827 | -0.274 | 0.17460752 | 0.37396928 | 0.0072 | 0.0347 | 0.0209 | 0.0209 | TRUE | 0.1621 |

| | | | | | | | | | | | | | | | | | | | | | | | | | |
|---|---|---|---|---|---|---|---|---|---|---|---|---|---|---|---|---|---|---|---|---|---|---|---|---|---|
| Kcnh5 | 12 | 8 | 9 | 13 | 9 | 10 | 13 | 8 | 9 | 10 | 10 | 11 | 10.1 | 10 | 10 | 1.038 | 0.054 | 0.92640192 | 0.9637313 | 0 | 0.333 | 0.1478 | 0.1478 | TRUE | 0.2035 |
| Kcnh6 | 230 | 241 | 187 | 262 | 204 | 198 | 244 | 241 | 188 | 206 | 219 | 217 | 219.16 | 224 | 214 | 0.954 | -0.068 | 0.67197767 | 0.81714972 | 0.0064 | 0.0219 | 0.014 | 0.014 | TRUE | 0.2242 |
| Kcnh7 | 2 | 2 | 1 | 0 | 3 | 0 | 2 | 2 | 1 | 0 | 3 | 0 | 1.39 | 2 | 1 | 0.609 | -0.716 | 0.69971547 | NA | 0.6399 | 2.3731 | 1.7131 | 1.7131 | TRUE | 1.4442 |
| Kcnh8 | 20 | 21 | 7 | 3 | 3 | 4 | 21 | 21 | 7 | 2 | 3 | 4 | 9.87 | 16 | 3 | 0.199 | -2.329 | 0.00259229 | 0.01890228 | 0 | 0.3407 | 0.2505 | 0.2505 | TRUE | 1.2176 |
| Kcnip1 | 3 | 3 | 5 | 2 | 6 | 5 | 3 | 3 | 5 | 2 | 6 | 5 | 4.12 | 4 | 4 | 1.176 | 0.234 | 0.80958579 | NA | 0 | 0.807 | 0.433 | 0.433 | TRUE | 0.8151 |
| Kcnip2 | 324 | 594 | 671 | 299 | 270 | 352 | 344 | 593 | 674 | 235 | 291 | 385 | 420.33 | 537 | 304 | 0.564 | -0.825 | 0.0016294 | 0.01307428 | 0.0861 | 0.0147 | 0.0469 | 0.0469 | TRUE | 0.6484 |
| Kcnip3 | 247 | 337 | 339 | 335 | 218 | 230 | 262 | 337 | 340 | 263 | 235 | 252 | 281.5 | 313 | 250 | 0.799 | -0.324 | 0.03486105 | 0.12780763 | 0.0088 | 0.0186 | 0.0135 | 0.0135 | TRUE | 0.225 |
| Kcnip4 | 1 | 1 | 0 | 0 | 0 | 0 | 1 | 1 | 0 | 0 | 0 | 0 | 0.34 | 1 | 0 | 0.261 | -1.936 | 0.60454921 | NA | 0 | 9.5983 | 8.3269 | 8.3269 | TRUE | 0.2507 |
| Kcnj1 | 0 | 0 | 0 | 0 | 0 | 0 | 0 | 0 | 0 | 0 | 0 | 0 | 0 | 0 | 0 | NA | NA | NA | NA | NA | NA | NA | NA | NA | NA |
| Kcnj10 | 323 | 249 | 216 | 233 | 176 | 147 | 343 | 249 | 217 | 183 | 189 | 161 | 223.68 | 270 | 178 | 0.661 | -0.598 | 0.00323247 | 0.02246913 | 0.0284 | 0.0216 | 0.0251 | 0.0251 | TRUE | 0.6326 |
| Kcnj11 | 35 | 24 | 27 | 34 | 32 | 37 | 37 | 24 | 27 | 27 | 34 | 41 | 31.65 | 29 | 34 | 1.146 | 0.197 | 0.59411725 | 0.76757819 | 0.0176 | 0.111 | 0.0661 | 0.0661 | TRUE | 0.3242 |
| Kcnj12 | 551 | 575 | 554 | 679 | 480 | 498 | 585 | 574 | 556 | 534 | 516 | 545 | 551.86 | 572 | 532 | 0.93 | -0.105 | 0.27716527 | 0.49805749 | 0 | 0.0129 | 0.0049 | 0.0049 | TRUE | 0.0071 |
| Kcnj13 | 0 | 0 | 0 | 0 | 0 | 0 | 0 | 0 | 0 | 0 | 0 | 0 | 0 | 0 | 0 | NA | NA | NA | NA | NA | NA | NA | NA | NA | NA |
| Kcnj14 | 6 | 17 | 23 | 16 | 16 | 12 | 6 | 17 | 23 | 13 | 17 | 13 | 14.9 | 15 | 14 | 0.918 | -0.123 | 0.83282137 | 0.91523598 | 0.1174 | 0.2281 | 0.1766 | 0.1766 | TRUE | 0.3696 |
| Kcnj15 | 10 | 2 | 2 | 13 | 6 | 4 | 11 | 2 | 2 | 10 | 6 | 4 | 5.95 | 5 | 7 | 1.466 | 0.552 | 0.55532985 | 0.73844216 | 0.3481 | 0.5609 | 0.457 | 0.457 | TRUE | 1.739 |
| Kcnj16 | 15 | 7 | 4 | 3 | 5 | 2 | 16 | 7 | 4 | 2 | 5 | 2 | 6.14 | 9 | 3 | 0.365 | -1.453 | 0.10860872 | 0.27606395 | 0.2328 | 0.5431 | 0.3846 | 0.3846 | TRUE | 0.5167 |
| Kcnj2 | 41 | 28 | 17 | 36 | 27 | 19 | 44 | 28 | 17 | 28 | 29 | 21 | 27.79 | 30 | 26 | 0.887 | -0.174 | 0.69501939 | 0.83284043 | 0.0841 | 0.1255 | 0.1055 | 0.1055 | TRUE | 0.2681 |
| Kcnj3 | 84 | 86 | 82 | 172 | 120 | 98 | 89 | 86 | 82 | 135 | 129 | 107 | 104.84 | 86 | 124 | 1.448 | 0.534 | 0.00795537 | 0.04414924 | 0 | 0.0383 | 0.0193 | 0.0193 | TRUE | 0.1339 |
| Kcnj4 | 301 | 330 | 344 | 321 | 236 | 240 | 320 | 330 | 345 | 252 | 254 | 263 | 293.95 | 332 | 256 | 0.772 | -0.372 | 0.00227702 | 0.01704465 | 0 | 0.0181 | 0.0073 | 0.0073 | TRUE | 0.0152 |
| Kcnj5 | 2 | 0 | 0 | 1 | 3 | 2 | 2 | 0 | 0 | 1 | 3 | 2 | 1.39 | 1 | 2 | 2.933 | 1.552 | 0.40676253 | NA | 0 | 2.3805 | 1.5468 | 1.5468 | TRUE | 0.7839 |
| Kcnj6 | 1 | 0 | 0 | 0 | 0 | 0 | 1 | 0 | 0 | 0 | 0 | 0 | 0.18 | 0 | 0 | 0.49 | -1.029 | 0.80087 | NA | 0 | 18.6165 | 10 | 10 | TRUE | 0.4614 |
| Kcnj8 | 2 | 0 | 2 | 1 | 1 | 0 | 2 | 0 | 2 | 1 | 1 | 0 | 1 | 1 | 1 | 0.459 | -1.123 | 0.58268644 | NA | 0 | 3.3043 | 1.8666 | 1.8666 | TRUE | 0.4949 |
| Kcnj9 | 15 | 7 | 9 | 14 | 7 | 13 | 16 | 7 | 9 | 11 | 8 | 14 | 10.79 | 11 | 11 | 1.028 | 0.04 | 0.94795748 | 0.97406219 | 0.0409 | 0.3123 | 0.1837 | 0.1837 | TRUE | 0.4824 |
| Kcnk1 | 960 | 1250 | 1435 | 1455 | 1136 | 1281 | 1020 | 1248 | 1441 | 1144 | 1222 | 1402 | 1246.22 | 1236 | 1256 | 1.015 | 0.022 | 0.88095665 | 0.94123075 | 0.0194 | 0.0095 | 0.0147 | 0.0147 | TRUE | 0.2357 |
| Kcnk10 | 0 | 3 | 1 | 0 | 0 | 0 | 0 | 3 | 1 | 0 | 0 | 0 | 0.67 | 1 | 0 | 0.134 | -2.899 | 0.33619836 | NA | 6.5513 | 4.9483 | 5.1656 | 5.1656 | TRUE | 0.1499 |
| Kcnk12 | 26 | 18 | 13 | 34 | 30 | 21 | 28 | 18 | 13 | 27 | 32 | 23 | 23.44 | 20 | 27 | 1.402 | 0.487 | 0.26523883 | 0.48485708 | 0.0278 | 0.1475 | 0.0936 | 0.0936 | TRUE | 0.6216 |
| Kcnk13 | 0 | 1 | 0 | 2 | 1 | 1 | 0 | 1 | 0 | 2 | 1 | 1 | 0.79 | 0 | 1 | 3.436 | 1.781 | 0.41000155 | NA | 0 | 4.1757 | 1.9771 | 1.9771 | TRUE | 0.405 |
| Kcnk15 | 0 | 0 | 1 | 0 | 0 | 0 | 0 | 0 | 1 | 0 | 0 | 0 | 0.17 | 0 | 0 | 0.49 | -1.029 | 0.80087 | NA | 0 | 19.6928 | 10 | 10 | TRUE | 0.4098 |
| Kcnk16 | 0 | 0 | 1 | 1 | 0 | 1 | 0 | 0 | 1 | 1 | 0 | 1 | 0.48 | 0 | 1 | 1.723 | 0.785 | 0.78314495 | NA | 0 | 6.8587 | 4.091 | 4.091 | TRUE | 0.4098 |
| Kcnk18 | 11 | 14 | 12 | 7 | 3 | 7 | 12 | 14 | 12 | 6 | 3 | 8 | 9.02 | 13 | 6 | 0.434 | -1.204 | 0.0654766 | 0.19715481 | 0 | 0.3722 | 0.178 | 0.178 | TRUE | 0.2394 |
| Kcnk2 | 17 | 12 | 10 | 49 | 19 | 14 | 18 | 12 | 10 | 39 | 20 | 15 | 19.06 | 13 | 25 | 1.88 | 0.911 | 0.09072474 | 0.24564988 | 0.1229 | 0.1797 | 0.151 | 0.151 | TRUE | 0.9352 |
| Kcnk3 | 14 | 10 | 7 | 6 | 0 | 8 | 15 | 10 | 7 | 5 | 0 | 9 | 7.56 | 11 | 5 | 0.423 | -1.241 | 0.17528235 | 0.37510898 | 0.4563 | 0.4427 | 0.4471 | 0.4471 | TRUE | 0.5101 |
| Kcnk4 | 9 | 6 | 7 | 11 | 12 | 12 | 10 | 6 | 7 | 9 | 13 | 13 | 9.55 | 8 | 12 | 1.526 | 0.61 | 0.32482414 | 0.54594394 | 0 | 0.352 | 0.166 | 0.166 | TRUE | 0.2398 |
| Kcnk5 | 4099 | 3635 | 3752 | 4038 | 3124 | 2903 | 4354 | 3630 | 3767 | 3174 | 3361 | 3178 | 3577.49 | 3917 | 3238 | 0.827 | -0.275 | 0.00346008 | 0.02362743 | 0.0049 | 0.0078 | 0.0061 | 0.0061 | TRUE | 0.1156 |
| Kcnk6 | 588 | 864 | 995 | 659 | 494 | 555 | 625 | 863 | 999 | 518 | 532 | 608 | 690.61 | 829 | 553 | 0.666 | -0.587 | 0.00088837 | 0.0081131 | 0.0302 | 0.0117 | 0.0209 | 0.0209 | TRUE | 0.2869 |
| Kcnk7 | 10 | 8 | 11 | 5 | 10 | 6 | 11 | 8 | 11 | 4 | 11 | 7 | 8.49 | 10 | 7 | 0.704 | -0.506 | 0.45816672 | 0.66271143 | 0 | 0.3951 | 0.2131 | 0.2131 | TRUE | 0.4429 |
| Kcnk9 | 0 | 0 | 0 | 0 | 0 | 0 | 0 | 0 | 0 | 0 | 0 | 0 | 0 | 0 | 0 | NA | NA | NA | NA | NA | NA | NA | NA | NA | NA |
| Kcnma1 | 78 | 85 | 104 | 135 | 95 | 82 | 83 | 85 | 104 | 106 | 102 | 90 | 95.04 | 91 | 99 | 1.097 | 0.134 | 0.52853447 | 0.71718529 | 0 | 0.0416 | 0.022 | 0.022 | TRUE | 0.1682 |
| Kcnmb1 | 21 | 26 | 25 | 28 | 13 | 11 | 22 | 26 | 25 | 22 | 14 | 12 | 20.24 | 24 | 16 | 0.665 | -0.589 | 0.18465104 | 0.38768853 | 0 | 0.1697 | 0.0912 | 0.0912 | TRUE | 0.6253 |
| Kcnmb2 | 3 | 4 | 4 | 2 | 2 | 3 | 3 | 4 | 4 | 2 | 2 | 3 | 3.03 | 4 | 2 | 0.614 | -0.703 | 0.50424506 | NA | 0 | 1.0926 | 0.4503 | 0.4503 | TRUE | 0.1204 |
| Kcnmb3 | 0 | 0 | 0 | 0 | 1 | 1 | 0 | 0 | 0 | 0 | 1 | 1 | 0.36 | 0 | 1 | 3.794 | 1.924 | 0.60247382 | NA | 0 | 9.1135 | 8.1425 | 8.1425 | TRUE | 0.3482 |
| Kcnmb4 | 31 | 23 | 25 | 31 | 23 | 15 | 33 | 23 | 25 | 24 | 25 | 16 | 24.42 | 27 | 22 | 0.814 | -0.296 | 0.46111593 | 0.66520813 | 0 | 0.1418 | 0.0753 | 0.0753 | TRUE | 0.2434 |
| Kcnn1 | 319 | 455 | 436 | 386 | 291 | 308 | 339 | 454 | 438 | 303 | 313 | 337 | 364.12 | 410 | 318 | 0.774 | -0.37 | 0.01363313 | 0.06589961 | 0.0113 | 0.016 | 0.0135 | 0.0135 | TRUE | 0.2655 |
| Kcnn2 | 279 | 283 | 260 | 404 | 309 | 301 | 296 | 283 | 261 | 318 | 332 | 329 | 303.27 | 280 | 326 | 1.166 | 0.221 | 0.07469509 | 0.2149801 | 0 | 0.0178 | 0.0078 | 0.0078 | TRUE | 0.0395 |
| Kcnn3 | 42 | 57 | 55 | 53 | 41 | 42 | 45 | 57 | 55 | 42 | 44 | 46 | 48.09 | 52 | 44 | 0.838 | -0.255 | 0.35891771 | 0.57740209 | 0 | 0.0754 | 0.0349 | 0.0349 | TRUE | 0.1309 |
| Kcnn4 | 29 | 82 | 52 | 41 | 35 | 39 | 31 | 82 | 52 | 32 | 38 | 43 | 46.25 | 55 | 38 | 0.679 | -0.558 | 0.15996671 | 0.35445491 | 0.1048 | 0.0781 | 0.0912 | 0.0912 | TRUE | 0.207 |
| Kcnq1 | 0 | 8 | 18 | 1 | 2 | 2 | 0 | 8 | 18 | 1 | 2 | 2 | 5.2 | 9 | 2 | 0.192 | -2.382 | 0.07102543 | 0.20748529 | 1.2531 | 0.6406 | 0.9032 | 0.9032 | TRUE | 0.0919 |
| Kcnq1ot1 | 503 | 378 | 588 | 579 | 473 | 459 | 534 | 378 | 590 | 455 | 509 | 502 | 494.79 | 501 | 489 | 0.976 | -0.035 | 0.83903262 | 0.91833605 | 0.026 | 0.0136 | 0.0198 | 0.0198 | TRUE | 0.5425 |
| Kcnq2 | 204 | 269 | 220 | 329 | 215 | 224 | 217 | 269 | 221 | 259 | 231 | 245 | 240.23 | 236 | 245 | 1.042 | 0.059 | 0.69525425 | 0.8329953 | 0.0047 | 0.0206 | 0.0123 | 0.0123 | TRUE | 0.1697 |
| Kcnq3 | 5 | 5 | 0 | 6 | 4 | 4 | 5 | 5 | 0 | 5 | 4 | 4 | 3.95 | 3 | 4 | 1.306 | 0.386 | 0.72281188 | NA | 0 | 0.8408 | 0.5935 | 0.5935 | TRUE | 1.1381 |
| Kcnq4 | 54 | 66 | 74 | 51 | 36 | 39 | 57 | 66 | 74 | 40 | 39 | 43 | 53.18 | 66 | 41 | 0.614 | -0.703 | 0.00915823 | 0.04926454 | 0 | 0.0688 | 0.0327 | 0.0327 | TRUE | 0.1113 |
| Kcnq5 | 39 | 49 | 62 | 76 | 55 | 55 | 41 | 49 | 62 | 60 | 59 | 60 | 55.29 | 51 | 60 | 1.172 | 0.229 | 0.40115087 | 0.61408828 | 0 | 0.0665 | 0.0355 | 0.0355 | TRUE | 0.314 |
| Kcnrg | 18 | 25 | 9 | 19 | 7 | 8 | 19 | 25 | 9 | 15 | 8 | 9 | 14.06 | 18 | 11 | 0.597 | -0.745 | 0.21496899 | 0.42675287 | 0.1261 | 0.2412 | 0.1847 | 0.1847 | TRUE | 0.2676 |
| Kcns1 | 63 | 63 | 62 | 122 | 89 | 83 | 67 | 63 | 62 | 96 | 96 | 91 | 79.1 | 64 | 94 | 1.472 | 0.558 | 0.00951891 | 0.05066899 | 0 | 0.0486 | 0.0202 | 0.0202 | TRUE | 0.0132 |
| Kcns2 | 30 | 29 | 27 | 42 | 31 | 20 | 32 | 29 | 27 | 33 | 33 | 22 | 29.37 | 29 | 29 | 1.009 | 0.013 | 0.97066958 | 0.98472179 | 0 | 0.1191 | 0.0607 | 0.0607 | TRUE | 0.3024 |
| Kcns3 | 160 | 252 | 256 | 449 | 361 | 336 | 170 | 252 | 257 | 353 | 388 | 368 | 297.98 | 226 | 370 | 1.632 | 0.707 | 7.44E-05 | 0.00108802 | 0.0208 | 0.018 | 0.0193 | 0.0193 | TRUE | 0.5144 |
| Kcnt1 | 51 | 63 | 56 | 102 | 86 | 80 | 54 | 63 | 56 | 80 | 93 | 88 | 72.27 | 58 | 87 | 1.497 | 0.582 | 0.01153492 | 0.058498 | 0 | 0.0525 | 0.0238 | 0.0238 | TRUE | 0.0522 |
| Kcnt2 | 35 | 28 | 31 | 19 | 16 | 11 | 37 | 28 | 31 | 15 | 17 | 12 | 23.41 | 32 | 15 | 0.46 | -1.12 | 0.0061031 | 0.03643378 | 0 | 0.1476 | 0.0712 | 0.0712 | TRUE | 0.1276 |
| Kcnu1 | 103 | 161 | 142 | 125 | 121 | 149 | 109 | 161 | 143 | 98 | 130 | 163 | 134.06 | 138 | 130 | 0.944 | -0.084 | 0.73700141 | 0.85925387 | 0.043 | 0.0315 | 0.0371 | 0.0371 | TRUE | 0.209 |
| Kcnv1 | 1 | 3 | 5 | 9 | 10 | 6 | 1 | 3 | 5 | 7 | 11 | 7 | 5.58 | 3 | 8 | 2.663 | 1.413 | 0.10603588 | 0.27171372 | 0 | 0.5973 | 0.3225 | 0.3225 | TRUE | 0.406 |
| Kcnv2 | 24 | 16 | 8 | 29 | 24 | 15 | 25 | 16 | 8 | 23 | 26 | 16 | 19.09 | 16 | 22 | 1.32 | 0.401 | 0.44606053 | 0.65236459 | 0.1049 | 0.1795 | 0.1455 | 0.1455 | TRUE | 0.3026 |
| Kcp | 379 | 341 | 277 | 253 | 241 | 181 | 403 | 341 | 278 | 199 | 259 | 198 | 279.6 | 341 | 219 | 0.642 | -0.639 | 0.00073595 | 0.00701776 | 0.0255 | 0.0187 | 0.0221 | 0.0221 | TRUE | 0.126 |
| Kctd1 | 147 | 147 | 157 | 305 | 213 | 179 | 156 | 147 | 158 | 240 | 229 | 196 | 187.58 | 154 | 222 | 1.447 | 0.533 | 0.00081232 | 0.00758002 | 0 | 0.0245 | 0.0127 | 0.0127 | TRUE | 0.1095 |
| Kctd10 | 1806 | 1958 | 2019 | 2624 | 1854 | 1778 | 1918 | 1955 | 2027 | 2063 | 1995 | 1946 | 1984.15 | 1967 | 2001 | 1.018 | 0.025 | 0.72192889 | 0.84955819 | 4.00E-04 | 0.0086 | 0.0032 | 0.0032 | TRUE | 0.009 |
| Kctd11 | 69 | 76 | 69 | 72 | 56 | 50 | 73 | 76 | 69 | 57 | 60 | 55 | 65.01 | 73 | 57 | 0.785 | -0.349 | 0.13799565 | 0.32239185 | 0 | 0.0576 | 0.0243 | 0.0243 | TRUE | 0.017 |
| Kctd12 | 263 | 301 | 292 | 687 | 562 | 404 | 279 | 301 | 293 | 540 | 605 | 442 | 410.02 | 291 | 529 | 1.818 | 0.862 | 4.97E-09 | 2.53E-07 | 0.0113 | 0.0149 | 0.013 | 0.013 | TRUE | 0.1958 |
| Kctd12b | 9 | 5 | 5 | 8 | 8 | 5 | 10 | 5 | 5 | 6 | 9 | 5 | 6.66 | 7 | 7 | 1.042 | 0.059 | 0.93594755 | 0.96816629 | 0 | 0.5018 | 0.2411 | 0.2411 | TRUE | 0.3956 |
| Kctd13 | 1018 | 1311 | 1292 | 1158 | 908 | 1044 | 1081 | 1309 | 1297 | 910 | 977 | 1143 | 1119.67 | 1229 | 1010 | 0.821 | -0.285 | 0.02536203 | 0.10277746 | 0.0111 | 0.0098 | 0.0108 | 0.0108 | TRUE | 0.1572 |
| Kctd14 | 16 | 5 | 12 | 31 | 16 | 14 | 17 | 5 | 12 | 24 | 17 | 15 | 15.16 | 11 | 19 | 1.69 | 0.757 | 0.18671766 | 0.39077904 | 0.0921 | 0.2242 | 0.166 | 0.166 | TRUE | 0.4957 |

| | | | | | | | | | | | | | | | | | | | | | | | | | |
|---|---|---|---|---|---|---|---|---|---|---|---|---|---|---|---|---|---|---|---|---|---|---|---|---|---|
| Kctd15 | 310 | 431 | 401 | 831 | 646 | 653 | 329 | 430 | 403 | 653 | 695 | 715 | 537.58 | 387 | 688 | 1.773 | 0.826 | 1.94E-10 | 1.38E-08 | 0.0071 | 0.013 | 0.0101 | 0.0101 | TRUE | 0.196 |
| Kctd16 | 1 | 0 | 0 | 0 | 1 | 0 | 1 | 0 | 0 | 0 | 1 | 0 | 0.36 | 0 | 0 | 0.954 | -0.067 | 0.98557668 | NA | 0 | 9.2518 | 8.0179 | 8.0179 | TRUE | 0.5204 |
| Kctd17 | 808 | 728 | 803 | 956 | 721 | 691 | 858 | 727 | 806 | 752 | 776 | 756 | 779.21 | 797 | 761 | 0.955 | -0.067 | 0.50857936 | 0.70252018 | 0.0023 | 0.0111 | 0.006 | 0.006 | TRUE | 0.0705 |
| Kctd18 | 505 | 622 | 637 | 831 | 612 | 629 | 536 | 621 | 640 | 653 | 658 | 689 | 632.92 | 599 | 667 | 1.112 | 0.153 | 0.15884877 | 0.35281832 | 0.0029 | 0.0121 | 0.0069 | 0.0069 | TRUE | 0.0974 |
| Kctd19 | 102 | 136 | 122 | 57 | 41 | 51 | 108 | 136 | 123 | 45 | 44 | 56 | 85.24 | 122 | 48 | 0.393 | -1.347 | 7.95E-09 | 3.87E-07 | 0 | 0.0456 | 0.025 | 0.025 | TRUE | 0.1402 |
| Kctd2 | 1821 | 2270 | 2271 | 2311 | 1780 | 2030 | 1934 | 2267 | 2280 | 1817 | 1915 | 2222 | 2072.61 | 2160 | 1985 | 0.918 | -0.123 | 0.28093417 | 0.50220609 | 0.0091 | 0.0085 | 0.0089 | 0.0089 | TRUE | 0.132 |
| Kctd20 | 3456 | 3537 | 3448 | 4573 | 3147 | 3089 | 3671 | 3532 | 3462 | 3595 | 3386 | 3381 | 3504.63 | 3555 | 3454 | 0.972 | -0.041 | 0.54975708 | 0.73418872 | 8.00E-04 | 0.0078 | 0.0031 | 0.0031 | TRUE | 0.0158 |
| Kctd21 | 220 | 194 | 195 | 234 | 177 | 158 | 234 | 194 | 196 | 184 | 190 | 173 | 195.1 | 208 | 182 | 0.879 | -0.186 | 0.23518021 | 0.45026247 | 0 | 0.0238 | 0.0125 | 0.0125 | TRUE | 0.1283 |
| Kctd3 | 4028 | 4642 | 4896 | 5257 | 4037 | 3851 | 4278 | 4636 | 4916 | 4133 | 4344 | 4216 | 4420.38 | 4610 | 4231 | 0.917 | -0.124 | 0.11885433 | 0.29199879 | 0.0024 | 0.0076 | 0.0043 | 0.0043 | TRUE | 0.0482 |
| Kctd4 | 0 | 0 | 0 | 0 | 0 | 0 | 0 | 0 | 0 | 0 | 0 | 0 | 0 | 0 | 0 | NA | NA | NA | NA | NA | NA | NA | NA | NA | NA |
| Kctd5 | 3633 | 3761 | 3749 | 4792 | 3486 | 3308 | 3859 | 3756 | 3764 | 3767 | 3751 | 3621 | 3753.06 | 3793 | 3713 | 0.979 | -0.031 | 0.61772415 | 0.78387201 | 0 | 0.0078 | 0.0024 | 0.0024 | TRUE | 0.0056 |
| Kctd6 | 689 | 785 | 768 | 839 | 618 | 664 | 732 | 784 | 771 | 660 | 665 | 727 | 723.05 | 762 | 684 | 0.896 | -0.158 | 0.09662529 | 0.25520989 | 6.00E-04 | 0.0115 | 0.0052 | 0.0052 | TRUE | 0.035 |
| Kctd7 | 225 | 238 | 240 | 349 | 218 | 220 | 239 | 238 | 241 | 274 | 235 | 241 | 244.57 | 239 | 250 | 1.048 | 0.067 | 0.62639449 | 0.78958445 | 0 | 0.0204 | 0.0097 | 0.0097 | TRUE | 0.0877 |
| Kctd8 | 9 | 6 | 2 | 8 | 9 | 5 | 10 | 6 | 2 | 6 | 10 | 5 | 6.5 | 6 | 7 | 1.222 | 0.29 | 0.72115429 | 0.84906281 | 0.0772 | 0.5137 | 0.3186 | 0.3186 | TRUE | 0.2577 |
| Kctd9 | 1504 | 1742 | 1775 | 2078 | 1426 | 1528 | 1598 | 1740 | 1782 | 1634 | 1534 | 1673 | 1660.01 | 1707 | 1614 | 0.945 | -0.081 | 0.34136454 | 0.56159399 | 0.002 | 0.0089 | 0.0046 | 0.0046 | TRUE | 0.0375 |
| Kdelc1 | 1446 | 1669 | 1779 | 1621 | 1121 | 1340 | 1536 | 1667 | 1786 | 1274 | 1206 | 1467 | 1489.39 | 1663 | 1316 | 0.791 | -0.339 | 0.00206453 | 0.01579135 | 0.0071 | 0.0091 | 0.008 | 0.008 | TRUE | 0.1196 |
| Kdelc2 | 283 | 447 | 501 | 477 | 380 | 379 | 301 | 446 | 503 | 375 | 409 | 415 | 408.13 | 417 | 400 | 0.958 | -0.062 | 0.74524169 | 0.86438829 | 0.033 | 0.015 | 0.0239 | 0.0239 | TRUE | 0.6809 |
| Kdelr1 | 2443 | 3006 | 2979 | 3379 | 2440 | 2830 | 2595 | 3002 | 2991 | 2656 | 2625 | 3098 | 2827.95 | 2863 | 2793 | 0.975 | -0.036 | 0.73274927 | 0.8565538 | 0.0073 | 0.0081 | 0.0076 | 0.0076 | TRUE | 0.11 |
| Kdelr2 | 4388 | 4510 | 4489 | 6252 | 4362 | 4466 | 4661 | 4504 | 4507 | 4915 | 4693 | 4889 | 4694.88 | 4557 | 4832 | 1.06 | 0.085 | 0.16814512 | 0.36589195 | 3.00E-04 | 0.0076 | 0.0025 | 0.0025 | TRUE | 0.0077 |
| Kdelr3 | 1880 | 3164 | 2980 | 2022 | 1519 | 2108 | 1997 | 3160 | 2992 | 1590 | 1634 | 2308 | 2280.08 | 2716 | 1844 | 0.679 | -0.559 | 0.00481812 | 0.03046967 | 0.0499 | 0.0083 | 0.0279 | 0.0279 | TRUE | 0.6509 |
| Kdm1a | 19459 | 21067 | 20271 | 23239 | 17310 | 17725 | 20669 | 21040 | 20354 | 18269 | 18624 | 19403 | 19726.48 | 20688 | 18765 | 0.907 | -0.141 | 0.01550982 | 0.07256625 | 5.00E-04 | 0.0071 | 0.0024 | 0.0024 | TRUE | 0.0108 |
| Kdm1b | 1736 | 1616 | 1678 | 2226 | 1713 | 1581 | 1844 | 1614 | 1685 | 1750 | 1843 | 1731 | 1744.4 | 1714 | 1775 | 1.035 | 0.05 | 0.56189644 | 0.7429148 | 0.0021 | 0.0088 | 0.0048 | 0.0048 | TRUE | 0.0526 |
| Kdm2a | 3938 | 3950 | 4232 | 5850 | 4380 | 4034 | 4183 | 3945 | 4249 | 4599 | 4713 | 4416 | 4350.73 | 4126 | 4576 | 1.109 | 0.149 | 0.03091851 | 0.11776063 | 9.00E-04 | 0.0077 | 0.0032 | 0.0032 | TRUE | 0.0179 |
| Kdm2b | 4309 | 4356 | 4800 | 5965 | 4351 | 4151 | 4577 | 4350 | 4820 | 4689 | 4681 | 4544 | 4610.26 | 4582 | 4638 | 1.012 | 0.018 | 0.80277587 | 0.89812791 | 0.0012 | 0.0076 | 0.0034 | 0.0034 | TRUE | 0.0251 |
| Kdm3a | 6507 | 4641 | 5034 | 7444 | 5380 | 5007 | 6912 | 4635 | 5055 | 5852 | 5789 | 5481 | 5620.44 | 5534 | 5707 | 1.031 | 0.045 | 0.76145567 | 0.87425776 | 0.023 | 0.0075 | 0.0154 | 0.0154 | TRUE | 0.5782 |
| Kdm3b | 7107 | 7757 | 8238 | 10251 | 7532 | 7486 | 7549 | 7747 | 8272 | 8059 | 8104 | 8195 | 7987.47 | 7856 | 8119 | 1.033 | 0.047 | 0.47038976 | 0.67209973 | 0.001 | 0.0073 | 0.003 | 0.003 | TRUE | 0.0262 |
| Kdm4a | 3420 | 3183 | 3272 | 4828 | 3467 | 3210 | 3633 | 3179 | 3285 | 3795 | 3730 | 3514 | 3522.75 | 3366 | 3680 | 1.094 | 0.129 | 0.12239951 | 0.29763877 | 0.0029 | 0.0078 | 0.0048 | 0.0048 | TRUE | 0.0584 |
| Kdm4b | 1922 | 2201 | 2344 | 2625 | 2201 | 2460 | 2042 | 2198 | 2354 | 2064 | 2368 | 2693 | 2286.31 | 2198 | 2375 | 1.08 | 0.111 | 0.35213252 | 0.57073249 | 0.0106 | 0.0083 | 0.0098 | 0.0098 | TRUE | 0.1662 |
| Kdm4c | 6372 | 7028 | 7822 | 6434 | 4780 | 6064 | 6768 | 7019 | 7854 | 5058 | 5143 | 6638 | 6413.35 | 7214 | 5613 | 0.778 | -0.362 | 0.00417458 | 0.02730876 | 0.0154 | 0.0074 | 0.0114 | 0.0114 | TRUE | 0.3108 |
| Kdm4d | 6 | 5 | 2 | 6 | 11 | 5 | 6 | 5 | 2 | 5 | 12 | 5 | 5.9 | 4 | 7 | 1.636 | 0.71 | 0.40418332 | 0.61693403 | 0 | 0.5653 | 0.3396 | 0.3396 | TRUE | 0.7452 |
| Kdm5a | 7936 | 7263 | 7874 | 10227 | 7248 | 7100 | 8429 | 7254 | 7906 | 8040 | 7798 | 7772 | 7866.59 | 7863 | 7870 | 1.001 | 0.001 | 0.98582319 | 0.99253326 | 0.0029 | 0.0073 | 0.0045 | 0.0045 | TRUE | 0.0562 |
| Kdm5b | 23964 | 22693 | 24228 | 30468 | 22305 | 21403 | 25454 | 22664 | 24327 | 23952 | 23999 | 23429 | 23970.75 | 24148 | 23793 | 0.985 | -0.021 | 0.75910718 | 0.87295942 | 0.0015 | 0.007 | 0.0035 | 0.0035 | TRUE | 0.0354 |
| Kdm5c | 9938 | 9848 | 10617 | 13314 | 9696 | 9836 | 10556 | 9835 | 10661 | 10466 | 10432 | 10767 | 10452.93 | 10351 | 10555 | 1.02 | 0.028 | 0.6623901 | 0.81042717 | 9.00E-04 | 0.0072 | 0.0029 | 0.0029 | TRUE | 0.0232 |
| Kdm5d | 63 | 96 | 71 | 729 | 515 | 430 | 67 | 96 | 71 | 573 | 554 | 471 | 305.33 | 78 | 533 | 6.823 | 2.77 | 9.07E-54 | 1.59E-50 | 0.0131 | 0.0177 | 0.0157 | 0.0157 | TRUE | 0.3737 |
| Kdm6a | 3085 | 3096 | 3070 | 3937 | 2806 | 2710 | 3277 | 3092 | 3083 | 3095 | 3019 | 2967 | 3088.66 | 3151 | 3027 | 0.961 | -0.057 | 0.39475117 | 0.60830634 | 5.00E-04 | 0.008 | 0.0029 | 0.0029 | TRUE | 0.0149 |
| Kdm6b | 3698 | 3423 | 3557 | 4312 | 3592 | 3474 | 3928 | 3419 | 3572 | 3390 | 3865 | 3803 | 3662.58 | 3640 | 3686 | 1.012 | 0.018 | 0.84769428 | 0.9238415 | 0.0048 | 0.0078 | 0.006 | 0.006 | TRUE | 0.0607 |
| Kdm8 | 146 | 197 | 178 | 250 | 177 | 143 | 155 | 197 | 179 | 197 | 190 | 157 | 179.01 | 177 | 181 | 1.026 | 0.037 | 0.83313053 | 0.91523598 | 0.0085 | 0.0253 | 0.0168 | 0.0168 | TRUE | 0.1467 |
| Kdr | 83 | 61 | 44 | 106 | 75 | 58 | 88 | 61 | 44 | 83 | 81 | 63 | 70.13 | 64 | 76 | 1.182 | 0.241 | 0.43165323 | 0.64142027 | 0.0525 | 0.0539 | 0.0532 | 0.0532 | TRUE | 0.4271 |
| Kdsr | 1863 | 2375 | 2474 | 2344 | 1841 | 1988 | 1979 | 2372 | 2484 | 1843 | 1981 | 2176 | 2139.1 | 2278 | 2000 | 0.877 | -0.189 | 0.10588902 | 0.27151385 | 0.0096 | 0.0084 | 0.0094 | 0.0094 | TRUE | 0.1598 |
| Keap1 | 571 | 699 | 603 | 1041 | 669 | 680 | 607 | 698 | 605 | 818 | 720 | 744 | 698.77 | 637 | 761 | 1.196 | 0.258 | 0.01991535 | 0.08657796 | 0.0043 | 0.0116 | 0.0074 | 0.0074 | TRUE | 0.0842 |
| Keg1 | 0 | 0 | 0 | 0 | 0 | 0 | 0 | 0 | 0 | 0 | 0 | 0 | 0 | 0 | 0 | NA | NA | NA | NA | NA | NA | NA | NA | NA | NA |
| Kel | 3 | 0 | 0 | 1 | 0 | 0 | 3 | 0 | 0 | 1 | 0 | 0 | 0.66 | 1 | 0 | 0.331 | -1.597 | 0.61164788 | NA | 8.2589 | 4.9826 | 5.6428 | 5.6428 | TRUE | 1.4426 |
| Kera | 0 | 0 | 0 | 0 | 0 | 0 | 0 | 0 | 0 | 0 | 0 | 0 | 0 | 0 | 0 | NA | NA | NA | NA | NA | NA | NA | NA | NA | NA |
| Khdc1a | 0 | 0 | 0 | 0 | 0 | 1 | 0 | 0 | 0 | 0 | 0 | 1 | 0.18 | 0 | 0 | 1.859 | 0.894 | 0.82650734 | NA | 0 | 18.0643 | 10 | 10 | TRUE | 0.5383 |
| Khdc1b | 0 | 0 | 0 | 0 | 0 | 0 | 0 | 0 | 0 | 0 | 0 | 0 | 0 | 0 | 0 | NA | NA | NA | NA | NA | NA | NA | NA | NA | NA |
| Khdc1c | 0 | 0 | 0 | 0 | 0 | 0 | 0 | 0 | 0 | 0 | 0 | 0 | 0 | 0 | 0 | NA | NA | NA | NA | NA | NA | NA | NA | NA | NA |
| Khdc3 | 2574 | 2957 | 2700 | 6388 | 5386 | 5522 | 2734 | 2953 | 2711 | 5022 | 5795 | 6045 | 4209.96 | 2799 | 5621 | 2.007 | 1.005 | 2.19E-25 | 9.61E-23 | 0.0052 | 0.0077 | 0.0064 | 0.0064 | TRUE | 0.1066 |
| Khdrbs1 | 10038 | 9820 | 10080 | 14651 | 10556 | 9887 | 10662 | 9807 | 10121 | 11517 | 11358 | 10823 | 10714.8 | 10197 | 11233 | 1.102 | 0.14 | 0.03844669 | 0.13666468 | 0.0013 | 0.0072 | 0.0032 | 0.0032 | TRUE | 0.0194 |
| Khdrbs2 | 8 | 9 | 6 | 20 | 13 | 11 | 8 | 9 | 6 | 16 | 14 | 12 | 10.88 | 8 | 14 | 1.785 | 0.836 | 0.14650729 | 0.33522602 | 0 | 0.3098 | 0.1388 | 0.1388 | TRUE | 0.1207 |
| Khdrbs3 | 1857 | 2119 | 2139 | 2764 | 1984 | 1949 | 1972 | 2116 | 2148 | 2173 | 2135 | 2133 | 2112.92 | 2079 | 2147 | 1.033 | 0.047 | 0.52155866 | 0.71165526 | 6.00E-04 | 0.0085 | 0.0033 | 0.0033 | TRUE | 0.0241 |
| Khk | 149 | 155 | 164 | 175 | 124 | 136 | 158 | 155 | 165 | 138 | 133 | 149 | 149.6 | 159 | 140 | 0.878 | -0.187 | 0.2477053 | 0.46437751 | 0 | 0.0289 | 0.0122 | 0.0122 | TRUE | 0.0301 |
| Khnyn | 1200 | 1753 | 1951 | 2073 | 1685 | 2014 | 1275 | 1751 | 1959 | 1630 | 1813 | 2205 | 1771.93 | 1662 | 1883 | 1.132 | 0.179 | 0.3043742 | 0.52512118 | 0.0349 | 0.0088 | 0.0214 | 0.0214 | TRUE | 0.4995 |
| Khsrp | 13872 | 13164 | 12777 | 17029 | 13006 | 12144 | 14735 | 13147 | 12829 | 13387 | 13994 | 13294 | 13564.17 | 13570 | 13558 | 0.999 | -0.001 | 0.98704999 | 0.99290468 | 0.0025 | 0.0071 | 0.0045 | 0.0045 | TRUE | 0.0688 |
| Kidins220 | 3465 | 3234 | 3748 | 4226 | 3186 | 3107 | 3680 | 3230 | 3763 | 3322 | 3428 | 3401 | 3470.81 | 3558 | 3384 | 0.951 | -0.072 | 0.39459223 | 0.60818026 | 0.003 | 0.0079 | 0.0049 | 0.0049 | TRUE | 0.0793 |
| Kif11 | 9423 | 9723 | 10250 | 11985 | 8519 | 9114 | 10009 | 9710 | 10292 | 9422 | 9166 | 9977 | 9762.62 | 10004 | 9522 | 0.952 | -0.071 | 0.2876605 | 0.50891548 | 0.0012 | 0.0072 | 0.0031 | 0.0031 | TRUE | 0.0213 |
| Kif12 | 1 | 2 | 0 | 0 | 0 | 0 | 1 | 2 | 0 | 0 | 0 | 0 | 0.51 | 1 | 0 | 0.176 | -2.508 | 0.44268408 | NA | 0 | 6.4675 | 6.1932 | 6.1932 | TRUE | 0.064 |
| Kif13a | 2803 | 3053 | 3520 | 4382 | 3056 | 3113 | 2977 | 3049 | 3534 | 3445 | 3288 | 3408 | 3283.55 | 3187 | 3380 | 1.061 | 0.085 | 0.35353103 | 0.57194495 | 0.0044 | 0.0079 | 0.0057 | 0.0057 | TRUE | 0.1107 |
| Kif13b | 4588 | 4229 | 4792 | 4897 | 3579 | 3707 | 4873 | 4224 | 4812 | 3850 | 3851 | 4058 | 4277.8 | 4636 | 3920 | 0.845 | -0.242 | 0.0042252 | 0.02755469 | 0.0031 | 0.0077 | 0.0049 | 0.0049 | TRUE | 0.0739 |
| Kif14 | 3142 | 3332 | 3517 | 3885 | 2827 | 2962 | 3337 | 3328 | 3531 | 3054 | 3042 | 3242 | 3255.77 | 3399 | 3113 | 0.916 | -0.127 | 0.06987051 | 0.20548156 | 8.00E-04 | 0.0079 | 0.0032 | 0.0032 | TRUE | 0.016 |
| Kif15 | 3360 | 3502 | 3926 | 5012 | 3762 | 3908 | 3569 | 3497 | 3942 | 3940 | 4048 | 4278 | 3879.03 | 3669 | 4089 | 1.114 | 0.156 | 0.05615985 | 0.17782637 | 0.0026 | 0.0078 | 0.0045 | 0.0045 | TRUE | 0.0515 |
| Kif16b | 1359 | 1504 | 1577 | 2475 | 1680 | 1857 | 1443 | 1502 | 1583 | 1946 | 1808 | 2033 | 1719.17 | 1509 | 1929 | 1.277 | 0.353 | 3.99E-05 | 0.00063937 | 0.0022 | 0.0088 | 0.0047 | 0.0047 | TRUE | 0.0358 |
| Kif17 | 128 | 235 | 239 | 212 | 196 | 170 | 136 | 235 | 240 | 167 | 211 | 186 | 195.71 | 204 | 188 | 0.921 | -0.119 | 0.61802394 | 0.78412283 | 0.0487 | 0.0237 | 0.0358 | 0.0358 | TRUE | 0.9085 |
| Kif18a | 2513 | 3187 | 3352 | 3177 | 2359 | 2730 | 2669 | 3183 | 3366 | 2498 | 2538 | 2988 | 2873.66 | 3073 | 2675 | 0.87 | -0.201 | 0.09610506 | 0.25436633 | 0.0116 | 0.008 | 0.0101 | 0.0101 | TRUE | 0.16 |
| Kif18b | 5439 | 5909 | 5599 | 6989 | 5142 | 5094 | 5777 | 5901 | 5622 | 5494 | 5532 | 5576 | 5650.57 | 5767 | 5534 | 0.96 | -0.059 | 0.30791153 | 0.52834806 | 2.00E-04 | 0.0075 | 0.0023 | 0.0023 | TRUE | 0.0059 |
| Kif19a | 118 | 130 | 124 | 127 | 108 | 106 | 125 | 130 | 125 | 100 | 116 | 116 | 118.63 | 127 | 111 | 0.871 | -0.199 | 0.27667706 | 0.49765942 | 0 | 0.0347 | 0.0156 | 0.0156 | TRUE | 0.083 |
| Kif1a | 1300 | 1689 | 1828 | 768 | 569 | 682 | 1381 | 1687 | 1836 | 604 | 612 | 747 | 1144.28 | 1635 | 654 | 0.4 | -1.322 | 1.03E-20 | 2.86E-18 | 0.016 | 0.0098 | 0.0134 | 0.0134 | TRUE | 0.2216 |
| Kif1b | 5439 | 5255 | 5931 | 7583 | 5074 | 4832 | 5777 | 5248 | 5955 | 5961 | 5459 | 5289 | 5615.1 | 5660 | 5570 | 0.984 | -0.023 | 0.79267064 | 0.89313448 | 0.0039 | 0.0075 | 0.0053 | 0.0053 | TRUE | 0.0495 |
| Kif1c | 4966 | 6131 | 6352 | 5960 | 4339 | 4635 | 5275 | 6123 | 6378 | 4685 | 4668 | 5074 | 5367.24 | 5925 | 4809 | 0.812 | -0.301 | 0.00177579 | 0.01398987 | 0.0056 | 0.0075 | 0.0065 | 0.0065 | TRUE | 0.1124 |
| Kif20a | 10298 | 11174 | 11108 | 13295 | 9338 | 9844 | 10938 | 11160 | 11154 | 10451 | 10047 | 10776 | 10754.31 | 11084 | 10425 | 0.941 | -0.088 | 0.14087733 | 0.32627272 | 5.00E-04 | 0.0072 | 0.0025 | 0.0025 | TRUE | 0.0122 |

| | | | | | | | | | | | | | | | | | | | | | | | | |
|---|---|---|---|---|---|---|---|---|---|---|---|---|---|---|---|---|---|---|---|---|---|---|---|---|
| Kif20b | 10805 | 11515 | 11937 | 11224 | 8768 | 9270 | 11477 | 11500 | 11986 | 8823 | 9434 | 10147 | 10561.28 | 11654 | 9468 | 0.812 | -0.3 | 0.0001099 | 0.0014853 | 0.0026 | 0.0072 | 0.0042 | 0.0042 | TRUE | 0.0481 |
| Kif21a | 888 | 995 | 965 | 1361 | 982 | 928 | 943 | 994 | 969 | 1070 | 1057 | 1016 | 1008.04 | 969 | 1048 | 1.082 | 0.113 | 0.16258241 | 0.35788551 | 0 | 0.0102 | 0.0038 | 0.0038 | TRUE | 0.0082 |
| Kif21b | 2123 | 2665 | 3043 | 2110 | 1714 | 2163 | 2255 | 2662 | 3055 | 1659 | 1844 | 2368 | 2307.11 | 2657 | 1957 | 0.736 | -0.442 | 0.00578435 | 0.03491054 | 0.0283 | 0.0083 | 0.0181 | 0.0181 | TRUE | 0.2771 |
| Kif22 | 5425 | 6053 | 6480 | 7167 | 5228 | 5386 | 5762 | 6045 | 6507 | 5634 | 5625 | 5896 | 5911.51 | 6105 | 5718 | 0.937 | -0.094 | 0.20970719 | 0.42000023 | 0.002 | 0.0075 | 0.0039 | 0.0039 | TRUE | 0.0405 |
| Kif23 | 6873 | 9296 | 9739 | 9364 | 7414 | 8396 | 7300 | 9284 | 9779 | 7361 | 7977 | 9191 | 8482.05 | 8788 | 8176 | 0.93 | -0.104 | 0.4385097 | 0.64591156 | 0.0158 | 0.0073 | 0.0129 | 0.0129 | TRUE | 0.2677 |
| Kif24 | 1514 | 1372 | 1580 | 2059 | 1464 | 1400 | 1608 | 1370 | 1586 | 1619 | 1575 | 1533 | 1548.53 | 1521 | 1576 | 1.036 | 0.051 | 0.5898701 | 0.76439333 | 0.0036 | 0.009 | 0.0057 | 0.0057 | TRUE | 0.0915 |
| Kif26a | 2676 | 2950 | 3217 | 2468 | 2175 | 2346 | 2842 | 2946 | 3230 | 1940 | 2340 | 2568 | 2644.53 | 3006 | 2283 | 0.759 | -0.398 | 0.00099987 | 0.00890956 | 0.0113 | 0.0081 | 0.0102 | 0.0102 | TRUE | 0.2114 |
| Kif26b | 41 | 35 | 41 | 62 | 42 | 40 | 44 | 35 | 41 | 49 | 45 | 44 | 42.9 | 40 | 46 | 1.154 | 0.207 | 0.47585167 | 0.67583364 | 0 | 0.0837 | 0.0374 | 0.0374 | TRUE | 0.0889 |
| Kif27 | 81 | 49 | 66 | 93 | 72 | 47 | 86 | 49 | 66 | 73 | 77 | 51 | 67.21 | 67 | 67 | 1.007 | 0.011 | 0.97161364 | 0.98491487 | 0.0454 | 0.0559 | 0.0509 | 0.0509 | TRUE | 0.2178 |
| Kif2a | 4883 | 4928 | 5026 | 7204 | 5105 | 4922 | 5187 | 4922 | 5047 | 5663 | 5493 | 5388 | 5283.11 | 5052 | 5515 | 1.092 | 0.127 | 0.04231558 | 0.14671385 | 4.00E-04 | 0.0075 | 0.0026 | 0.0026 | TRUE | 0.0069 |
| Kif2b | 0 | 0 | 0 | 2 | 0 | 0 | 0 | 0 | 0 | 2 | 0 | 0 | 0.26 | 0 | 1 | 2.886 | 1.529 | 0.70676491 | NA | 0 | 12.5792 | 10 | 10 | TRUE | 0.9669 |
| Kif2c | 5707 | 6303 | 6183 | 7579 | 5511 | 5599 | 6062 | 6295 | 6208 | 5958 | 5929 | 6129 | 6096.93 | 6188 | 6005 | 0.97 | -0.043 | 0.45526695 | 0.66027313 | 2.00E-04 | 0.0074 | 0.0023 | 0.0023 | TRUE | 0.0039 |
| Kif3a | 3536 | 3002 | 2937 | 4328 | 3082 | 2864 | 3756 | 2998 | 2949 | 3402 | 3316 | 3135 | 3259.42 | 3234 | 3284 | 1.016 | 0.023 | 0.84305318 | 0.92105202 | 0.0099 | 0.0079 | 0.009 | 0.009 | TRUE | 0.2415 |
| Kif3b | 2578 | 2309 | 2362 | 2910 | 2071 | 1973 | 2738 | 2306 | 2372 | 2288 | 2228 | 2160 | 2348.61 | 2472 | 2225 | 0.9 | -0.151 | 0.10610861 | 0.27181176 | 0.0042 | 0.0083 | 0.0059 | 0.0059 | TRUE | 0.1074 |
| Kif3c | 561 | 725 | 724 | 887 | 723 | 806 | 596 | 724 | 727 | 697 | 778 | 882 | 734.07 | 682 | 786 | 1.15 | 0.202 | 0.13423653 | 0.31680062 | 0.0119 | 0.0114 | 0.0117 | 0.0117 | TRUE | 0.144 |
| Kif4 | 2976 | 3518 | 3922 | 4346 | 3096 | 3386 | 3161 | 3513 | 3938 | 3416 | 3331 | 3707 | 3511.12 | 3537 | 3485 | 0.985 | -0.022 | 0.83309001 | 0.91523598 | 0.0073 | 0.0078 | 0.0075 | 0.0075 | TRUE | 0.1193 |
| Kif4-ps | 1 | 2 | 1 | 7 | 2 | 4 | 1 | 2 | 1 | 6 | 2 | 4 | 2.68 | 1 | 4 | 2.986 | 1.578 | 0.19343922 | NA | 0 | 1.2349 | 0.5647 | 0.5647 | TRUE | 0.2556 |
| Kif5a | 563 | 649 | 758 | 507 | 459 | 573 | 598 | 648 | 761 | 399 | 494 | 627 | 587.82 | 669 | 507 | 0.756 | -0.404 | 0.02532367 | 0.10267478 | 0.0309 | 0.0125 | 0.0218 | 0.0218 | TRUE | 0.394 |
| Kif5b | 13600 | 12734 | 12856 | 16762 | 11898 | 11500 | 14446 | 12718 | 12909 | 13177 | 12802 | 12589 | 13106.5 | 13358 | 12856 | 0.963 | -0.055 | 0.4747108 | 0.67524193 | 0.0027 | 0.0072 | 0.0042 | 0.0042 | TRUE | 0.0621 |
| Kif5c | 1445 | 1384 | 1404 | 1842 | 1373 | 1396 | 1535 | 1382 | 1410 | 1448 | 1477 | 1528 | 1463.38 | 1442 | 1484 | 1.029 | 0.041 | 0.61602409 | 0.78291152 | 0.0012 | 0.0092 | 0.0042 | 0.0042 | TRUE | 0.0377 |
| Kif6 | 54 | 62 | 52 | 70 | 61 | 47 | 57 | 62 | 52 | 55 | 66 | 51 | 57.27 | 57 | 57 | 1.002 | 0.003 | 0.98979104 | 0.99457546 | 0 | 0.0644 | 0.031 | 0.031 | TRUE | 0.126 |
| Kif7 | 1031 | 1077 | 1021 | 1273 | 932 | 815 | 1095 | 1076 | 1025 | 1001 | 1003 | 892 | 1015.26 | 1065 | 965 | 0.907 | -0.141 | 0.12452555 | 0.30127581 | 0.0016 | 0.0101 | 0.0051 | 0.0051 | TRUE | 0.0512 |
| Kif9 | 83 | 112 | 91 | 136 | 125 | 81 | 88 | 112 | 91 | 107 | 134 | 89 | 103.58 | 97 | 110 | 1.132 | 0.179 | 0.44425851 | 0.65087811 | 0.02 | 0.0387 | 0.0295 | 0.0295 | TRUE | 0.281 |
| Kifap3 | 2340 | 2530 | 2626 | 3839 | 2811 | 2697 | 2485 | 2527 | 2637 | 3018 | 3024 | 2952 | 2773.95 | 2550 | 2998 | 1.176 | 0.234 | 0.00037716 | 0.0041338 | 1.00E-04 | 0.0081 | 0.0028 | 0.0028 | TRUE | 0.0108 |
| Kifc1 | 4240 | 5228 | 5103 | 6119 | 4415 | 4453 | 4504 | 5221 | 5124 | 4810 | 4750 | 4875 | 4880.65 | 4950 | 4812 | 0.972 | -0.041 | 0.62028475 | 0.78509433 | 0.0026 | 0.0076 | 0.0047 | 0.0047 | TRUE | 0.0756 |
| Kifc2 | 470 | 314 | 326 | 370 | 315 | 303 | 499 | 314 | 327 | 291 | 339 | 332 | 350.27 | 380 | 321 | 0.843 | -0.247 | 0.2151073 | 0.42696606 | 0.036 | 0.0163 | 0.0256 | 0.0256 | TRUE | 0.8593 |
| Kifc3 | 3336 | 2731 | 2902 | 3608 | 2933 | 2879 | 3543 | 2727 | 2914 | 2836 | 3156 | 3152 | 3054.73 | 3061 | 3048 | 0.995 | -0.007 | 0.95474621 | 0.97698418 | 0.0109 | 0.008 | 0.0096 | 0.0096 | TRUE | 0.2299 |
| Kifc5b | 1298 | 1934 | 1774 | 1764 | 1417 | 1418 | 1379 | 1932 | 1781 | 1387 | 1525 | 1552 | 1592.51 | 1697 | 1488 | 0.876 | -0.191 | 0.16265452 | 0.35794434 | 0.015 | 0.009 | 0.0128 | 0.0128 | TRUE | 0.3243 |
| Kin | 644 | 568 | 518 | 858 | 564 | 463 | 684 | 567 | 520 | 674 | 607 | 507 | 593.26 | 590 | 596 | 1.011 | 0.016 | 0.91809814 | 0.95991788 | 0.0185 | 0.0125 | 0.0156 | 0.0156 | TRUE | 0.2245 |
| Kir3dl1 | 0 | 0 | 0 | 0 | 0 | 0 | 0 | 0 | 0 | 0 | 0 | 0 | 0 | 0 | 0 | NA | NA | NA | NA | NA | NA | NA | NA | NA | NA |
| Kir3dl2 | 0 | 0 | 0 | 0 | 0 | 0 | 0 | 0 | 0 | 0 | 0 | 0 | 0 | 0 | 0 | NA | NA | NA | NA | NA | NA | NA | NA | NA | NA |
| Kirrel | 1243 | 977 | 899 | 2476 | 1594 | 1322 | 1320 | 976 | 903 | 1946 | 1715 | 1447 | 1384.56 | 1066 | 1703 | 1.598 | 0.676 | 6.19E-05 | 0.00093196 | 0.0315 | 0.0093 | 0.0198 | 0.0198 | TRUE | 0.3713 |
| Kirrel2 | 437 | 627 | 754 | 300 | 266 | 308 | 464 | 626 | 757 | 236 | 286 | 337 | 451.11 | 616 | 286 | 0.464 | -1.108 | 8.25E-08 | 3.05E-06 | 0.0432 | 0.0142 | 0.0282 | 0.0282 | TRUE | 0.1324 |
| Kirrel3 | 9 | 18 | 12 | 13 | 5 | 8 | 10 | 18 | 12 | 10 | 5 | 9 | 10.66 | 13 | 8 | 0.621 | -0.688 | 0.26052142 | 0.47988003 | 0 | 0.316 | 0.1712 | 0.1712 | TRUE | 0.4263 |
| Kis2 | 1 | 0 | 2 | 2 | 2 | 0 | 1 | 0 | 2 | 2 | 2 | 0 | 1.13 | 1 | 1 | 1.226 | 0.294 | 0.8783029 | NA | 0 | 2.9161 | 1.7728 | 1.7728 | TRUE | 0.2511 |
| Kiss1 | 3 | 0 | 2 | 3 | 8 | 1 | 3 | 0 | 2 | 2 | 9 | 1 | 2.88 | 2 | 4 | 2.32 | 1.214 | 0.37826192 | NA | 0.7019 | 1.1524 | 0.9479 | 0.9479 | TRUE | 1.5299 |
| Kiss1r | 28 | 51 | 42 | 29 | 27 | 36 | 30 | 51 | 42 | 23 | 29 | 39 | 35.68 | 41 | 30 | 0.736 | -0.443 | 0.23854439 | 0.45438673 | 0.0452 | 0.0992 | 0.0732 | 0.0732 | TRUE | 0.2501 |
| Kit | 7212 | 8453 | 8659 | 8755 | 6781 | 7333 | 7660 | 8442 | 8695 | 6882 | 7296 | 8027 | 7833.77 | 8266 | 7402 | 0.895 | -0.16 | 0.08225264 | 0.22891982 | 0.005 | 0.0073 | 0.0059 | 0.0059 | TRUE | 0.0666 |
| Kitl | 485 | 527 | 533 | 549 | 420 | 453 | 515 | 526 | 535 | 432 | 452 | 496 | 492.67 | 525 | 460 | 0.873 | -0.195 | 0.06982291 | 0.20540329 | 0 | 0.0136 | 0.0063 | 0.0063 | TRUE | 0.0539 |
| Kl | 18 | 4 | 4 | 5 | 8 | 10 | 19 | 4 | 4 | 4 | 9 | 11 | 8.44 | 9 | 8 | 0.859 | -0.219 | 0.79812903 | 0.89608114 | 0.4219 | 0.3974 | 0.41 | 0.41 | TRUE | 2.9487 |
| Klb | 21 | 15 | 6 | 43 | 27 | 20 | 22 | 15 | 6 | 34 | 29 | 22 | 21.34 | 14 | 28 | 1.971 | 0.979 | 0.06075611 | 0.18759699 | 0.1193 | 0.1613 | 0.1437 | 0.1437 | TRUE | 0.4868 |
| Klc1 | 1509 | 1722 | 1641 | 2254 | 1854 | 1759 | 1603 | 1720 | 1648 | 1772 | 1995 | 1926 | 1777.09 | 1657 | 1898 | 1.144 | 0.195 | 0.0205037 | 0.08835612 | 0.002 | 0.0088 | 0.0045 | 0.0045 | TRUE | 0.0415 |
| Klc2 | 2564 | 2701 | 2651 | 2592 | 2061 | 2137 | 2723 | 2698 | 2662 | 2038 | 2218 | 2339 | 2446.21 | 2694 | 2198 | 0.815 | -0.295 | 0.00027807 | 0.00319897 | 0.0018 | 0.0082 | 0.0043 | 0.0043 | TRUE | 0.0505 |
| Klc3 | 573 | 822 | 864 | 562 | 516 | 591 | 609 | 821 | 868 | 442 | 555 | 647 | 656.84 | 766 | 548 | 0.714 | -0.485 | 0.00820361 | 0.045272 | 0.0338 | 0.0119 | 0.0227 | 0.0227 | TRUE | 0.3803 |
| Klc4 | 1231 | 1375 | 1318 | 1482 | 1104 | 1155 | 1308 | 1373 | 1323 | 1165 | 1188 | 1264 | 1270.23 | 1335 | 1206 | 0.903 | -0.148 | 0.06591012 | 0.19796884 | 5.00E-04 | 0.0095 | 0.0039 | 0.0039 | TRUE | 0.0214 |
| Klf1 | 13 | 21 | 7 | 32 | 20 | 19 | 14 | 21 | 7 | 25 | 22 | 21 | 18.21 | 14 | 23 | 1.618 | 0.694 | 0.17301599 | 0.37187356 | 0.049 | 0.1878 | 0.1288 | 0.1288 | TRUE | 0.3238 |
| Klf10 | 3694 | 6127 | 6894 | 5483 | 4431 | 5244 | 3924 | 6119 | 6922 | 4310 | 4767 | 5740 | 5297.21 | 5655 | 4939 | 0.873 | -0.195 | 0.32077448 | 0.54196626 | 0.0472 | 0.0075 | 0.0277 | 0.0277 | TRUE | 0.7492 |
| Klf11 | 1759 | 2298 | 2235 | 2857 | 2039 | 2015 | 1868 | 2295 | 2244 | 2246 | 2194 | 2206 | 2175.52 | 2136 | 2215 | 1.037 | 0.052 | 0.60237723 | 0.77379815 | 0.0058 | 0.0084 | 0.0069 | 0.0069 | TRUE | 0.1448 |
| Klf12 | 682 | 650 | 735 | 950 | 680 | 614 | 724 | 649 | 738 | 747 | 732 | 672 | 710.36 | 704 | 717 | 1.019 | 0.028 | 0.7887588 | 0.89139466 | 0.0025 | 0.0115 | 0.0063 | 0.0063 | TRUE | 0.055 |
| Klf13 | 2784 | 2409 | 2225 | 3376 | 2532 | 1905 | 2957 | 2406 | 2234 | 2654 | 2724 | 2085 | 2510.11 | 2532 | 2488 | 0.983 | -0.025 | 0.86184128 | 0.93179056 | 0.0207 | 0.0082 | 0.0147 | 0.0147 | TRUE | 0.2607 |
| Klf14 | 13 | 9 | 9 | 25 | 15 | 9 | 14 | 9 | 9 | 20 | 16 | 10 | 12.91 | 11 | 15 | 1.451 | 0.537 | 0.33707099 | 0.55714563 | 0 | 0.262 | 0.1456 | 0.1456 | TRUE | 0.3106 |
| Klf15 | 896 | 1324 | 1294 | 882 | 808 | 953 | 952 | 1322 | 1299 | 693 | 869 | 1043 | 1029.87 | 1191 | 868 | 0.728 | -0.457 | 0.01155583 | 0.05856637 | 0.034 | 0.0101 | 0.0226 | 0.0226 | TRUE | 0.266 |
| Klf16 | 2176 | 2266 | 2208 | 3103 | 2444 | 2349 | 2311 | 2263 | 2217 | 2439 | 2630 | 2571 | 2405.29 | 2264 | 2547 | 1.125 | 0.169 | 0.0166902 | 0.07648271 | 6.00E-04 | 0.0083 | 0.0032 | 0.0032 | TRUE | 0.0169 |
| Klf17 | 203 | 203 | 155 | 290 | 245 | 204 | 216 | 203 | 156 | 228 | 264 | 223 | 214.81 | 192 | 238 | 1.245 | 0.316 | 0.0721048 | 0.20986102 | 0.0129 | 0.0222 | 0.0176 | 0.0176 | TRUE | 0.2894 |
| Klf2 | 9489 | 12413 | 11447 | 12469 | 9791 | 8795 | 10079 | 12397 | 11494 | 9802 | 10535 | 9628 | 10655.69 | 11323 | 9988 | 0.882 | -0.181 | 0.06348474 | 0.19340955 | 0.0063 | 0.0072 | 0.0068 | 0.0068 | TRUE | 0.1129 |
| Klf3 | 3751 | 5324 | 5218 | 5268 | 3972 | 4473 | 3984 | 5317 | 5239 | 4141 | 4274 | 4896 | 4642.02 | 4847 | 4437 | 0.915 | -0.128 | 0.33326437 | 0.55393041 | 0.0155 | 0.0076 | 0.0123 | 0.0123 | TRUE | 0.2952 |
| Klf4 | 19525 | 18291 | 17144 | 26157 | 20226 | 18311 | 20739 | 18267 | 17214 | 20563 | 21762 | 20044 | 19764.93 | 18740 | 20790 | 1.109 | 0.15 | 0.10353174 | 0.26764507 | 0.0055 | 0.0071 | 0.006 | 0.006 | TRUE | 0.1065 |
| Klf5 | 5393 | 3928 | 3811 | 5279 | 4250 | 3706 | 5728 | 3923 | 3827 | 4150 | 4573 | 4057 | 4376.23 | 4493 | 4260 | 0.948 | -0.077 | 0.62790876 | 0.79062566 | 0.0262 | 0.0077 | 0.0178 | 0.0178 | TRUE | 0.7048 |
| Klf6 | 2273 | 2637 | 2424 | 3326 | 2199 | 2216 | 2414 | 2634 | 2434 | 2615 | 2366 | 2426 | 2481.38 | 2494 | 2469 | 0.99 | -0.014 | 0.86350679 | 0.9324082 | 0.0022 | 0.0082 | 0.0044 | 0.0044 | TRUE | 0.033 |
| Klf7 | 219 | 237 | 244 | 344 | 254 | 249 | 233 | 237 | 245 | 270 | 273 | 273 | 255.1 | 238 | 272 | 1.142 | 0.192 | 0.13029561 | 0.31086811 | 0 | 0.0198 | 0.0077 | 0.0077 | TRUE | 0.0071 |
| Klf8 | 1628 | 1503 | 1575 | 1950 | 1378 | 1410 | 1729 | 1501 | 1581 | 1533 | 1483 | 1543 | 1561.8 | 1604 | 1520 | 0.948 | -0.077 | 0.37234585 | 0.58896764 | 0.0022 | 0.009 | 0.0048 | 0.0048 | TRUE | 0.0561 |
| Klf9 | 7688 | 7584 | 7035 | 10485 | 7604 | 6773 | 8166 | 7574 | 7064 | 8242 | 8181 | 7414 | 7773.69 | 7601 | 7946 | 1.045 | 0.064 | 0.46346569 | 0.66686857 | 0.0039 | 0.0073 | 0.0054 | 0.0054 | TRUE | 0.0515 |
| Klhdc1 | 88 | 137 | 169 | 108 | 80 | 95 | 93 | 137 | 170 | 85 | 86 | 104 | 112.49 | 133 | 92 | 0.686 | -0.545 | 0.03564432 | 0.12960084 | 0.0424 | 0.0362 | 0.0392 | 0.0392 | TRUE | 0.1918 |
| Klhdc10 | 6968 | 6203 | 6208 | 8969 | 6102 | 5423 | 7401 | 6195 | 6233 | 7051 | 6565 | 5936 | 6563.69 | 6610 | 6517 | 0.986 | -0.02 | 0.85218515 | 0.92628008 | 0.0087 | 0.0074 | 0.0082 | 0.0082 | TRUE | 0.1337 |
| Klhdc2 | 2590 | 3739 | 3707 | 2939 | 2163 | 2401 | 2751 | 3734 | 3722 | 2310 | 2327 | 2628 | 2912.23 | 3402 | 2422 | 0.712 | -0.491 | 0.00028785 | 0.00329695 | 0.0159 | 0.008 | 0.0128 | 0.0128 | TRUE | 0.3406 |
| Klhdc3 | 7072 | 6873 | 6821 | 6959 | 5090 | 5140 | 7512 | 6864 | 6849 | 5471 | 5477 | 5627 | 6299.76 | 7075 | 5525 | 0.781 | -0.357 | 3.25E-07 | 1.02E-05 | 0.0013 | 0.0074 | 0.0034 | 0.0034 | TRUE | 0.0355 |
| Klhdc4 | 1542 | 1419 | 1412 | 2686 | 2005 | 1631 | 1638 | 1417 | 1418 | 2112 | 2157 | 1785 | 1754.5 | 1491 | 2018 | 1.354 | 0.437 | 9.27E-05 | 0.00129815 | 0.0078 | 0.0088 | 0.0084 | 0.0084 | TRUE | 0.122 |
| Klhdc5 | 185 | 237 | 231 | 326 | 299 | 218 | 197 | 237 | 232 | 256 | 322 | 239 | 246.96 | 222 | 272 | 1.226 | 0.294 | 0.0837909 | 0.23193226 | 0.0134 | 0.0202 | 0.0167 | 0.0167 | TRUE | 0.2761 |
| Klhdc7a | 58 | 192 | 168 | 109 | 105 | 131 | 62 | 192 | 169 | 86 | 113 | 143 | 127.35 | 141 | 114 | 0.808 | -0.307 | 0.42570407 | 0.63581483 | 0.182 | 0.0328 | 0.0993 | 0.0993 | TRUE | 1.264 |

| | | | | | | | | | | | | | | | | | | | | | | | | |
|---|---|---|---|---|---|---|---|---|---|---|---|---|---|---|---|---|---|---|---|---|---|---|---|---|
| Klhdc7b | 0 | 0 | 0 | 1 | 2 | 2 | 0 | 0 | 0 | 1 | 2 | 2 | 0.85 | 0 | 2 | 9.003 | 3.17 | 0.16844964 | NA | 0 | 3.8621 | 2.5291 | 2.5291 | TRUE | 0.2298 |
| Klhdc8a | 10 | 15 | 4 | 22 | 7 | 18 | 11 | 15 | 4 | 17 | 8 | 20 | 12.36 | 10 | 15 | 1.508 | 0.593 | 0.37482123 | 0.59085248 | 0.1949 | 0.2735 | 0.2372 | 0.2372 | TRUE | 0.4854 |
| Klhdc8b | 257 | 217 | 166 | 277 | 188 | 188 | 273 | 217 | 167 | 218 | 202 | 206 | 213.7 | 219 | 209 | 0.955 | -0.067 | 0.73894519 | 0.86031177 | 0.0259 | 0.0223 | 0.0241 | 0.0241 | TRUE | 0.2043 |
| Klhdc9 | 42 | 75 | 97 | 60 | 49 | 75 | 45 | 75 | 97 | 47 | 53 | 82 | 66.48 | 72 | 61 | 0.835 | -0.261 | 0.46763326 | 0.66995293 | 0.1012 | 0.0565 | 0.0779 | 0.0779 | TRUE | 0.2339 |
| Klhl1 | 0 | 1 | 0 | 1 | 1 | 0 | 0 | 1 | 0 | 1 | 1 | 0 | 0.48 | 0 | 1 | 1.712 | 0.776 | 0.78614772 | NA | 0 | 6.9165 | 4.1067 | 4.1067 | TRUE | 0.405 |
| Klhl10 | 53 | 80 | 73 | 81 | 66 | 78 | 56 | 80 | 73 | 64 | 71 | 85 | 71.59 | 70 | 73 | 1.045 | 0.063 | 0.80484071 | 0.89931073 | 0.0128 | 0.0529 | 0.0335 | 0.0335 | TRUE | 0.2517 |
| Klhl11 | 1850 | 1744 | 1764 | 2317 | 1679 | 1647 | 1965 | 1742 | 1771 | 1821 | 1807 | 1803 | 1818.14 | 1826 | 1810 | 0.992 | -0.012 | 0.88190113 | 0.94166574 | 0.0016 | 0.0087 | 0.0042 | 0.0042 | TRUE | 0.0533 |
| Klhl12 | 3270 | 3174 | 3020 | 3902 | 2846 | 2717 | 3473 | 3170 | 3032 | 3067 | 3062 | 2974 | 3129.9 | 3225 | 3034 | 0.941 | -0.088 | 0.27390612 | 0.4944543 | 0.0021 | 0.008 | 0.0043 | 0.0043 | TRUE | 0.0549 |
| Klhl13 | 1829 | 2097 | 2004 | 6847 | 5160 | 5150 | 1943 | 2094 | 2012 | 5383 | 5552 | 5637 | 3770.19 | 2016 | 5524 | 2.739 | 1.454 | 3.46E-101 | 1.81E-97 | 5.00E-04 | 0.0078 | 0.003 | 0.003 | TRUE | 0.0138 |
| Klhl14 | 3 | 7 | 6 | 11 | 1 | 4 | 3 | 7 | 6 | 9 | 1 | 4 | 5.05 | 5 | 5 | 0.891 | -0.166 | 0.86126533 | 0.93144485 | 0.2237 | 0.6592 | 0.4541 | 0.4541 | TRUE | 0.3196 |
| Klhl15 | 276 | 299 | 323 | 386 | 287 | 337 | 293 | 299 | 324 | 303 | 309 | 369 | 316.21 | 305 | 327 | 1.069 | 0.096 | 0.48380399 | 0.68148223 | 0.0044 | 0.0173 | 0.0104 | 0.0104 | TRUE | 0.139 |
| Klhl17 | 259 | 309 | 293 | 491 | 449 | 376 | 275 | 309 | 294 | 386 | 483 | 412 | 359.76 | 293 | 427 | 1.456 | 0.542 | 8.17E-05 | 0.00117306 | 0.0064 | 0.0161 | 0.0107 | 0.0107 | TRUE | 0.1521 |
| Klhl18 | 444 | 345 | 349 | 616 | 416 | 326 | 472 | 345 | 350 | 484 | 448 | 357 | 409.22 | 389 | 430 | 1.107 | 0.146 | 0.41071315 | 0.62196522 | 0.0256 | 0.015 | 0.0203 | 0.0203 | TRUE | 0.3959 |
| Klhl2 | 2461 | 2832 | 2769 | 3407 | 2482 | 2636 | 2614 | 2828 | 2780 | 2678 | 2670 | 2886 | 2742.84 | 2741 | 2745 | 1.001 | 0.002 | 0.9834865 | 0.99129879 | 0.0014 | 0.0081 | 0.0038 | 0.0038 | TRUE | 0.0242 |
| Klhl20 | 1420 | 1447 | 1481 | 1955 | 1565 | 1368 | 1508 | 1445 | 1487 | 1537 | 1684 | 1497 | 1526.45 | 1480 | 1573 | 1.062 | 0.087 | 0.29489927 | 0.51584451 | 0.0015 | 0.0091 | 0.0043 | 0.0043 | TRUE | 0.0453 |
| Klhl21 | 1677 | 2018 | 2003 | 2108 | 1541 | 1662 | 1781 | 2015 | 2011 | 1657 | 1658 | 1819 | 1823.73 | 1936 | 1711 | 0.884 | -0.178 | 0.050354 | 0.16493186 | 0.0031 | 0.0087 | 0.0054 | 0.0054 | TRUE | 0.0588 |
| Klhl22 | 1658 | 1499 | 1461 | 3439 | 2334 | 2004 | 1761 | 1497 | 1467 | 2703 | 2511 | 2194 | 2022.26 | 1575 | 2469 | 1.569 | 0.65 | 3.20E-08 | 1.35E-06 | 0.0099 | 0.0085 | 0.0094 | 0.0094 | TRUE | 0.1283 |
| Klhl23 | 261 | 280 | 273 | 454 | 305 | 248 | 277 | 280 | 274 | 357 | 328 | 271 | 297.92 | 277 | 319 | 1.154 | 0.207 | 0.1566296 | 0.349662 | 0.0068 | 0.018 | 0.012 | 0.012 | TRUE | 0.1875 |
| Klhl24 | 4337 | 5338 | 5778 | 4392 | 3588 | 4463 | 4607 | 5331 | 5802 | 3453 | 3860 | 4885 | 4656.34 | 5247 | 4066 | 0.775 | -0.368 | 0.0117105 | 0.05910336 | 0.0227 | 0.0076 | 0.0151 | 0.0151 | TRUE | 0.3779 |
| Klhl25 | 706 | 646 | 640 | 870 | 668 | 648 | 750 | 645 | 643 | 684 | 719 | 709 | 691.61 | 679 | 704 | 1.036 | 0.051 | 0.62538568 | 0.78901854 | 0.0027 | 0.0117 | 0.0066 | 0.0066 | TRUE | 0.097 |
| Klhl26 | 1216 | 1729 | 1666 | 2201 | 1628 | 1886 | 1292 | 1727 | 1673 | 1730 | 1752 | 2065 | 1706.27 | 1564 | 1849 | 1.182 | 0.241 | 0.07799929 | 0.22101584 | 0.0157 | 0.0088 | 0.0129 | 0.0129 | TRUE | 0.2786 |
| Klhl28 | 488 | 472 | 472 | 656 | 475 | 493 | 518 | 471 | 474 | 516 | 511 | 540 | 505.02 | 488 | 522 | 1.07 | 0.098 | 0.34554306 | 0.56504978 | 0 | 0.0134 | 0.0058 | 0.0058 | TRUE | 0.0343 |
| Klhl29 | 66 | 43 | 37 | 170 | 109 | 106 | 70 | 43 | 37 | 134 | 117 | 116 | 86.19 | 50 | 122 | 2.453 | 1.295 | 2.85E-06 | 6.84E-05 | 0.035 | 0.0451 | 0.0409 | 0.0409 | TRUE | 0.9723 |
| Klhl3 | 130 | 121 | 102 | 199 | 153 | 130 | 138 | 121 | 102 | 156 | 165 | 142 | 137.45 | 120 | 154 | 1.284 | 0.361 | 0.05589713 | 0.17727921 | 0.0045 | 0.0309 | 0.0182 | 0.0182 | TRUE | 0.174 |
| Klhl30 | 11 | 12 | 7 | 18 | 3 | 11 | 12 | 12 | 7 | 14 | 3 | 12 | 10.02 | 10 | 10 | 0.971 | -0.043 | 0.95113723 | 0.97528069 | 0.1463 | 0.3357 | 0.2497 | 0.2497 | TRUE | 1.0902 |
| Klhl31 | 16 | 4 | 7 | 3 | 2 | 4 | 17 | 4 | 7 | 2 | 2 | 4 | 6.15 | 9 | 3 | 0.315 | -1.667 | 0.06909073 | 0.20400524 | 0.2405 | 0.5425 | 0.3848 | 0.3848 | TRUE | 0.5619 |
| Klhl32 | 32 | 20 | 21 | 43 | 38 | 35 | 34 | 20 | 21 | 34 | 41 | 38 | 31.34 | 25 | 38 | 1.505 | 0.59 | 0.11015134 | 0.27854848 | 0.0061 | 0.112 | 0.0648 | 0.0648 | TRUE | 0.5786 |
| Klhl33 | 3 | 9 | 6 | 8 | 4 | 4 | 3 | 9 | 6 | 6 | 4 | 4 | 5.53 | 6 | 5 | 0.829 | -0.271 | 0.74006781 | 0.86085622 | 0 | 0.6028 | 0.3011 | 0.3011 | TRUE | 0.1661 |
| Klhl34 | 11 | 15 | 8 | 39 | 39 | 18 | 12 | 15 | 8 | 31 | 42 | 20 | 21.17 | 12 | 31 | 2.661 | 1.412 | 0.00447584 | 0.02874304 | 0.0684 | 0.1625 | 0.1178 | 0.1178 | TRUE | 0.0889 |
| Klhl35 | 0 | 2 | 3 | 3 | 2 | 0 | 0 | 2 | 3 | 2 | 2 | 0 | 1.59 | 2 | 1 | 0.913 | -0.132 | 0.9383116 | NA | 0 | 2.0832 | 1.4712 | 1.4712 | TRUE | 0.4378 |
| Klhl36 | 601 | 628 | 593 | 852 | 608 | 525 | 638 | 627 | 595 | 670 | 654 | 575 | 626.61 | 620 | 633 | 1.022 | 0.031 | 0.77242575 | 0.88115034 | 0.0024 | 0.0122 | 0.0066 | 0.0066 | TRUE | 0.0745 |
| Klhl38 | 1 | 0 | 0 | 2 | 1 | 0 | 1 | 0 | 0 | 2 | 1 | 0 | 0.62 | 0 | 1 | 2.462 | 1.3 | 0.62119797 | NA | 0 | 5.3343 | 3.4416 | 3.4416 | TRUE | 0.4614 |
| Klhl4 | 3 | 1 | 0 | 1 | 1 | 1 | 3 | 1 | 0 | 1 | 1 | 1 | 1.19 | 1 | 1 | 0.706 | -0.502 | 0.77820686 | NA | 0 | 2.7746 | 1.4201 | 1.4201 | TRUE | 0.4548 |
| Klhl40 | 1 | 2 | 2 | 7 | 5 | 4 | 1 | 2 | 2 | 6 | 5 | 4 | 3.39 | 2 | 5 | 3.007 | 1.588 | 0.13396901 | NA | 0 | 0.9792 | 0.4127 | 0.4127 | TRUE | 0.0751 |
| Klhl41 | 5 | 6 | 4 | 5 | 10 | 4 | 5 | 6 | 4 | 4 | 11 | 4 | 5.73 | 5 | 6 | 1.23 | 0.298 | 0.71777556 | 0.8469816 | 0 | 0.5817 | 0.3123 | 0.3123 | TRUE | 0.8478 |
| Klhl5 | 1351 | 958 | 902 | 1124 | 714 | 689 | 1435 | 957 | 906 | 884 | 768 | 754 | 950.58 | 1099 | 802 | 0.73 | -0.454 | 0.01253128 | 0.06207393 | 0.0376 | 0.0104 | 0.0227 | 0.0227 | TRUE | 0.8537 |
| Klhl6 | 0 | 0 | 0 | 1 | 0 | 0 | 0 | 0 | 0 | 1 | 0 | 0 | 0.13 | 0 | 0 | 1.859 | 0.894 | 0.82650734 | NA | 0 | 25.1515 | 10 | 10 | TRUE | 0.2563 |
| Klhl7 | 2071 | 2023 | 1996 | 2137 | 1559 | 1691 | 2200 | 2020 | 2004 | 1680 | 1677 | 1851 | 1905.46 | 2075 | 1736 | 0.837 | -0.257 | 0.00265523 | 0.01926324 | 0.0022 | 0.0086 | 0.0048 | 0.0048 | TRUE | 0.04 |
| Klhl8 | 1428 | 1539 | 1498 | 1792 | 1290 | 1377 | 1517 | 1537 | 1504 | 1409 | 1388 | 1507 | 1477 | 1519 | 1435 | 0.944 | -0.083 | 0.27714423 | 0.49805749 | 3.00E-04 | 0.0091 | 0.0035 | 0.0035 | TRUE | 0.0231 |
| Klhl9 | 1693 | 1540 | 1351 | 2884 | 2130 | 1828 | 1798 | 1538 | 1357 | 2267 | 2292 | 2001 | 1875.46 | 1564 | 2187 | 1.398 | 0.484 | 9.58E-05 | 0.0013341 | 0.012 | 0.0087 | 0.0105 | 0.0105 | TRUE | 0.2058 |
| Klk1 | 22 | 72 | 60 | 13 | 12 | 28 | 23 | 72 | 60 | 10 | 13 | 31 | 34.88 | 52 | 18 | 0.342 | -1.55 | 0.0051561 | 0.03198394 | 0.2794 | 0.1013 | 0.1837 | 0.1837 | TRUE | 0.7241 |
| Klk10 | 321 | 295 | 314 | 297 | 244 | 246 | 341 | 295 | 315 | 233 | 263 | 269 | 286.03 | 317 | 255 | 0.803 | -0.316 | 0.02014768 | 0.08729923 | 0.0021 | 0.0184 | 0.0098 | 0.0098 | TRUE | 0.0665 |
| Klk11 | 70 | 77 | 73 | 61 | 59 | 71 | 74 | 77 | 73 | 48 | 63 | 78 | 68.95 | 75 | 63 | 0.836 | -0.259 | 0.32436549 | 0.54575342 | 0.0146 | 0.0547 | 0.0352 | 0.0352 | TRUE | 0.2729 |
| Klk12 | 7 | 5 | 1 | 2 | 3 | 0 | 7 | 5 | 1 | 2 | 3 | 0 | 3.04 | 4 | 2 | 0.359 | -1.478 | 0.26151019 | NA | 0.4709 | 1.091 | 0.8303 | 0.8303 | TRUE | 0.2138 |
| Klk13 | 167 | 256 | 237 | 105 | 115 | 122 | 177 | 256 | 238 | 83 | 124 | 134 | 168.47 | 224 | 114 | 0.503 | -0.991 | 3.07E-05 | 0.00051335 | 0.0423 | 0.0265 | 0.0341 | 0.0341 | TRUE | 0.6192 |
| Klk14 | 41 | 21 | 18 | 12 | 22 | 11 | 44 | 21 | 18 | 9 | 24 | 12 | 21.29 | 28 | 15 | 0.542 | -0.883 | 0.11282295 | 0.28285545 | 0.1831 | 0.1616 | 0.1722 | 0.1722 | TRUE | 1.6087 |
| Klk15 | 17 | 9 | 4 | 12 | 11 | 8 | 18 | 9 | 4 | 9 | 12 | 9 | 10.18 | 10 | 10 | 0.969 | -0.045 | 0.94793314 | 0.97406219 | 0.1526 | 0.3305 | 0.2453 | 0.2453 | TRUE | 0.421 |
| Klk1b1 | 0 | 0 | 0 | 0 | 0 | 0 | 0 | 0 | 0 | 0 | 0 | 0 | 0 | 0 | 0 | NA | NA | NA | NA | NA | NA | NA | NA | NA | NA |
| Klk1b11 | 0 | 0 | 0 | 0 | 0 | 0 | 0 | 0 | 0 | 0 | 0 | 0 | 0 | 0 | 0 | NA | NA | NA | NA | NA | NA | NA | NA | NA | NA |
| Klk1b16 | 0 | 0 | 0 | 0 | 0 | 0 | 0 | 0 | 0 | 0 | 0 | 0 | 0 | 0 | 0 | NA | NA | NA | NA | NA | NA | NA | NA | NA | NA |
| Klk1b21 | 0 | 0 | 0 | 0 | 0 | 0 | 0 | 0 | 0 | 0 | 0 | 0 | 0 | 0 | 0 | NA | NA | NA | NA | NA | NA | NA | NA | NA | NA |
| Klk1b22 | 0 | 1 | 0 | 0 | 0 | 0 | 0 | 1 | 0 | 0 | 0 | 0 | 0.17 | 0 | 0 | 0.49 | -1.029 | 0.80087 | NA | 0 | 19.7991 | 10 | 10 | TRUE | 0.405 |
| Klk1b24 | 0 | 0 | 0 | 0 | 0 | 0 | 0 | 0 | 0 | 0 | 0 | 0 | 0 | 0 | 0 | NA | NA | NA | NA | NA | NA | NA | NA | NA | NA |
| Klk1b26 | 0 | 0 | 0 | 0 | 0 | 0 | 0 | 0 | 0 | 0 | 0 | 0 | 0 | 0 | 0 | NA | NA | NA | NA | NA | NA | NA | NA | NA | NA |
| Klk1b27 | 0 | 1 | 0 | 0 | 0 | 0 | 0 | 1 | 0 | 0 | 0 | 0 | 0.17 | 0 | 0 | 0.49 | -1.029 | 0.80087 | NA | 0 | 19.7991 | 10 | 10 | TRUE | 0.405 |
| Klk1b3 | 0 | 0 | 0 | 0 | 0 | 0 | 0 | 0 | 0 | 0 | 0 | 0 | 0 | 0 | 0 | NA | NA | NA | NA | NA | NA | NA | NA | NA | NA |
| Klk1b4 | 0 | 0 | 0 | 0 | 0 | 0 | 0 | 0 | 0 | 0 | 0 | 0 | 0 | 0 | 0 | NA | NA | NA | NA | NA | NA | NA | NA | NA | NA |
| Klk1b5 | 2 | 5 | 2 | 0 | 1 | 2 | 2 | 5 | 2 | 0 | 1 | 2 | 2.07 | 3 | 1 | 0.337 | -1.569 | 0.28571513 | NA | 0 | 1.6021 | 0.9143 | 0.9143 | TRUE | 0.5576 |
| Klk1b7-ps | 0 | 0 | 0 | 0 | 0 | 0 | 0 | 0 | 0 | 0 | 0 | 0 | 0 | 0 | 0 | NA | NA | NA | NA | NA | NA | NA | NA | NA | NA |
| Klk1b8 | 0 | 0 | 0 | 0 | 0 | 0 | 0 | 0 | 0 | 0 | 0 | 0 | 0 | 0 | 0 | NA | NA | NA | NA | NA | NA | NA | NA | NA | NA |
| Klk1b9 | 2 | 0 | 1 | 2 | 1 | 1 | 2 | 0 | 1 | 2 | 1 | 1 | 1.15 | 1 | 1 | 1.22 | 0.287 | 0.86966868 | NA | 0 | 2.8836 | 1.3154 | 1.3154 | TRUE | 0.2346 |
| Klk4 | 10 | 2 | 5 | 9 | 9 | 4 | 11 | 2 | 5 | 7 | 10 | 4 | 6.46 | 6 | 7 | 1.206 | 0.27 | 0.7489773 | 0.86687239 | 0.1727 | 0.5167 | 0.3557 | 0.3557 | TRUE | 0.5144 |
| Klk5 | 21 | 19 | 17 | 13 | 10 | 4 | 22 | 19 | 17 | 10 | 11 | 4 | 13.95 | 19 | 8 | 0.44 | -1.185 | 0.03048755 | 0.11662616 | 0 | 0.243 | 0.1328 | 0.1328 | TRUE | 0.4768 |
| Klk6 | 2 | 1 | 3 | 3 | 1 | 0 | 2 | 1 | 3 | 2 | 1 | 0 | 1.59 | 2 | 1 | 0.588 | -0.766 | 0.62086489 | NA | 0 | 2.0725 | 1.0738 | 1.0738 | TRUE | 0.4359 |
| Klk7 | 35 | 24 | 25 | 17 | 10 | 17 | 37 | 24 | 25 | 13 | 11 | 19 | 21.5 | 29 | 14 | 0.495 | -1.016 | 0.02384006 | 0.09832776 | 0 | 0.1602 | 0.0935 | 0.0935 | TRUE | 0.4152 |
| Klk8 | 1025 | 1162 | 1199 | 1028 | 766 | 884 | 1089 | 1160 | 1204 | 808 | 824 | 968 | 1008.86 | 1151 | 867 | 0.752 | -0.411 | 0.00012373 | 0.00163995 | 0.0049 | 0.0102 | 0.0073 | 0.0073 | TRUE | 0.1221 |
| Klk9 | 3 | 3 | 0 | 1 | 0 | 2 | 3 | 3 | 0 | 1 | 0 | 2 | 1.53 | 2 | 1 | 0.477 | -1.068 | 0.55237163 | NA | 0 | 2.1653 | 1.5806 | 1.5806 | TRUE | 0.7156 |
| Klkb1 | 0 | 0 | 0 | 0 | 0 | 0 | 0 | 0 | 0 | 0 | 0 | 0 | 0 | 0 | 0 | NA | NA | NA | NA | NA | NA | NA | NA | NA | NA |
| Klra1 | 0 | 0 | 0 | 0 | 0 | 0 | 0 | 0 | 0 | 0 | 0 | 0 | 0 | 0 | 0 | NA | NA | NA | NA | NA | NA | NA | NA | NA | NA |

| | | | | | | | | | | | | | | | | | | | | | | | | | |
|---|---|---|---|---|---|---|---|---|---|---|---|---|---|---|---|---|---|---|---|---|---|---|---|---|---|
| Klra10 | 0 | 0 | 0 | 0 | 0 | 0 | 0 | 0 | 0 | 0 | 0 | 0 | 0 | 0 | 0 | NA | NA | NA | NA | NA | NA | NA | NA | NA | NA |
| Klra12 | 0 | 0 | 0 | 0 | 0 | 0 | 0 | 0 | 0 | 0 | 0 | 0 | 0 | 0 | 0 | NA | NA | NA | NA | NA | NA | NA | NA | NA | NA |
| Klra13-ps | 0 | 0 | 0 | 0 | 0 | 0 | 0 | 0 | 0 | 0 | 0 | 0 | 0 | 0 | 0 | NA | NA | NA | NA | NA | NA | NA | NA | NA | NA |
| Klra14 | 0 | 0 | 0 | 0 | 0 | 0 | 0 | 0 | 0 | 0 | 0 | 0 | 0 | 0 | 0 | NA | NA | NA | NA | NA | NA | NA | NA | NA | NA |
| Klra15 | 0 | 0 | 0 | 0 | 0 | 0 | 0 | 0 | 0 | 0 | 0 | 0 | 0 | 0 | 0 | NA | NA | NA | NA | NA | NA | NA | NA | NA | NA |
| Klra17 | 0 | 0 | 0 | 0 | 0 | 1 | 0 | 0 | 0 | 0 | 0 | 1 | 0.18 | 0 | 0 | 1.859 | 0.894 | 0.82650734 | NA | 0 | 18.0643 | 10 | 10 | TRUE | 0.5383 |
| Klra18 | 0 | 0 | 0 | 0 | 0 | 0 | 0 | 0 | 0 | 0 | 0 | 0 | 0 | 0 | 0 | NA | NA | NA | NA | NA | NA | NA | NA | NA | NA |
| Klra19 | 0 | 0 | 0 | 0 | 0 | 0 | 0 | 0 | 0 | 0 | 0 | 0 | 0 | 0 | 0 | NA | NA | NA | NA | NA | NA | NA | NA | NA | NA |
| Klra2 | 1 | 0 | 0 | 0 | 0 | 0 | 1 | 0 | 0 | 0 | 0 | 0 | 0.18 | 0 | 0 | 0.49 | -1.029 | 0.80087 | NA | 0 | 18.6165 | 10 | 10 | TRUE | 0.4614 |
| Klra21 | 0 | 0 | 0 | 0 | 0 | 0 | 0 | 0 | 0 | 0 | 0 | 0 | 0 | 0 | 0 | NA | NA | NA | NA | NA | NA | NA | NA | NA | NA |
| Klra22 | 0 | 0 | 0 | 0 | 0 | 0 | 0 | 0 | 0 | 0 | 0 | 0 | 0 | 0 | 0 | NA | NA | NA | NA | NA | NA | NA | NA | NA | NA |
| Klra23 | 0 | 0 | 0 | 0 | 0 | 0 | 0 | 0 | 0 | 0 | 0 | 0 | 0 | 0 | 0 | NA | NA | NA | NA | NA | NA | NA | NA | NA | NA |
| Klra3 | 0 | 0 | 0 | 0 | 0 | 0 | 0 | 0 | 0 | 0 | 0 | 0 | 0 | 0 | 0 | NA | NA | NA | NA | NA | NA | NA | NA | NA | NA |
| Klra33 | 0 | 0 | 0 | 0 | 0 | 0 | 0 | 0 | 0 | 0 | 0 | 0 | 0 | 0 | 0 | NA | NA | NA | NA | NA | NA | NA | NA | NA | NA |
| Klra4 | 0 | 0 | 0 | 0 | 0 | 0 | 0 | 0 | 0 | 0 | 0 | 0 | 0 | 0 | 0 | NA | NA | NA | NA | NA | NA | NA | NA | NA | NA |
| Klra5 | 0 | 0 | 0 | 0 | 0 | 0 | 0 | 0 | 0 | 0 | 0 | 0 | 0 | 0 | 0 | NA | NA | NA | NA | NA | NA | NA | NA | NA | NA |
| Klra6 | 0 | 0 | 0 | 0 | 0 | 0 | 0 | 0 | 0 | 0 | 0 | 0 | 0 | 0 | 0 | NA | NA | NA | NA | NA | NA | NA | NA | NA | NA |
| Klra7 | 0 | 0 | 0 | 0 | 0 | 0 | 0 | 0 | 0 | 0 | 0 | 0 | 0 | 0 | 0 | NA | NA | NA | NA | NA | NA | NA | NA | NA | NA |
| Klra8 | 0 | 0 | 0 | 0 | 0 | 0 | 0 | 0 | 0 | 0 | 0 | 0 | 0 | 0 | 0 | NA | NA | NA | NA | NA | NA | NA | NA | NA | NA |
| Klra9 | 0 | 0 | 0 | 0 | 0 | 0 | 0 | 0 | 0 | 0 | 0 | 0 | 0 | 0 | 0 | NA | NA | NA | NA | NA | NA | NA | NA | NA | NA |
| Klrb1 | 0 | 0 | 0 | 0 | 0 | 0 | 0 | 0 | 0 | 0 | 0 | 0 | 0 | 0 | 0 | NA | NA | NA | NA | NA | NA | NA | NA | NA | NA |
| Klrb1-ps1 | 0 | 0 | 0 | 0 | 0 | 0 | 0 | 0 | 0 | 0 | 0 | 0 | 0 | 0 | 0 | NA | NA | NA | NA | NA | NA | NA | NA | NA | NA |
| Klrb1a | 2 | 0 | 0 | 1 | 2 | 2 | 2 | 0 | 0 | 1 | 2 | 2 | 1.21 | 1 | 2 | 2.423 | 1.277 | 0.50849417 | NA | 0 | 2.7327 | 1.6616 | 1.6616 | TRUE | 0.9745 |
| Klrb1b | 0 | 0 | 0 | 0 | 0 | 0 | 0 | 0 | 0 | 0 | 0 | 0 | 0 | 0 | 0 | NA | NA | NA | NA | NA | NA | NA | NA | NA | NA |
| Klrb1c | 0 | 1 | 0 | 0 | 0 | 0 | 0 | 1 | 0 | 0 | 0 | 0 | 0.17 | 0 | 0 | 0.49 | -1.029 | 0.80087 | NA | 0 | 19.7991 | 10 | 10 | TRUE | 0.405 |
| Klrb1f | 6 | 3 | 6 | 7 | 5 | 3 | 6 | 3 | 6 | 6 | 5 | 3 | 4.93 | 5 | 5 | 0.929 | -0.106 | 0.90001649 | 0.95153909 | 0 | 0.6756 | 0.312 | 0.312 | TRUE | 0.2795 |
| Klrc1 | 0 | 0 | 0 | 0 | 0 | 0 | 0 | 0 | 0 | 0 | 0 | 0 | 0 | 0 | 0 | NA | NA | NA | NA | NA | NA | NA | NA | NA | NA |
| Klrc2 | 0 | 0 | 0 | 0 | 0 | 0 | 0 | 0 | 0 | 0 | 0 | 0 | 0 | 0 | 0 | NA | NA | NA | NA | NA | NA | NA | NA | NA | NA |
| Klrc3 | 0 | 0 | 0 | 0 | 0 | 0 | 0 | 0 | 0 | 0 | 0 | 0 | 0 | 0 | 0 | NA | NA | NA | NA | NA | NA | NA | NA | NA | NA |
| Klrd1 | 1 | 0 | 0 | 0 | 0 | 0 | 1 | 0 | 0 | 0 | 0 | 0 | 0.18 | 0 | 0 | 0.49 | -1.029 | 0.80087 | NA | 0 | 18.6165 | 10 | 10 | TRUE | 0.4614 |
| Klre1 | 1 | 0 | 0 | 0 | 0 | 0 | 1 | 0 | 0 | 0 | 0 | 0 | 0.18 | 0 | 0 | 0.49 | -1.029 | 0.80087 | NA | 0 | 18.6165 | 10 | 10 | TRUE | 0.4614 |
| Klrg1 | 0 | 2 | 1 | 3 | 5 | 0 | 0 | 2 | 1 | 2 | 5 | 0 | 1.79 | 1 | 2 | 2.555 | 1.354 | 0.43674306 | NA | 0.9768 | 1.8474 | 1.4866 | 1.4866 | TRUE | 0.1253 |
| Klrg2 | 160 | 159 | 137 | 178 | 112 | 104 | 170 | 159 | 138 | 140 | 121 | 114 | 140.1 | 156 | 125 | 0.806 | -0.311 | 0.09343206 | 0.24989155 | 0.0048 | 0.0304 | 0.0175 | 0.0175 | TRUE | 0.1264 |
| Klri1 | 0 | 0 | 0 | 0 | 0 | 0 | 0 | 0 | 0 | 0 | 0 | 0 | 0 | 0 | 0 | NA | NA | NA | NA | NA | NA | NA | NA | NA | NA |
| Klri2 | 0 | 0 | 0 | 0 | 0 | 0 | 0 | 0 | 0 | 0 | 0 | 0 | 0 | 0 | 0 | NA | NA | NA | NA | NA | NA | NA | NA | NA | NA |
| Klrk1 | 0 | 0 | 0 | 0 | 0 | 0 | 0 | 0 | 0 | 0 | 0 | 0 | 0 | 0 | 0 | NA | NA | NA | NA | NA | NA | NA | NA | NA | NA |
| Kmo | 0 | 0 | 0 | 0 | 0 | 0 | 0 | 0 | 0 | 0 | 0 | 0 | 0 | 0 | 0 | NA | NA | NA | NA | NA | NA | NA | NA | NA | NA |
| Kncn | 0 | 0 | 0 | 0 | 0 | 0 | 0 | 0 | 0 | 0 | 0 | 0 | 0 | 0 | 0 | NA | NA | NA | NA | NA | NA | NA | NA | NA | NA |
| Kndc1 | 334 | 342 | 389 | 474 | 422 | 440 | 355 | 342 | 391 | 373 | 454 | 482 | 399.21 | 363 | 436 | 1.201 | 0.264 | 0.06299686 | 0.1924239 | 0.0093 | 0.0152 | 0.012 | 0.012 | TRUE | 0.1972 |
| Kng1 | 8 | 3 | 10 | 17 | 21 | 12 | 8 | 3 | 10 | 13 | 23 | 13 | 11.77 | 7 | 16 | 2.275 | 1.186 | 0.06387454 | 0.19428502 | 0.0718 | 0.2868 | 0.1937 | 0.1937 | TRUE | 0.7211 |
| Kng2 | 0 | 0 | 1 | 4 | 5 | 3 | 0 | 0 | 1 | 3 | 5 | 3 | 2.14 | 0 | 4 | 10.71 | 3.421 | 0.03818519 | NA | 0 | 1.5497 | 0.8377 | 0.8377 | TRUE | 0.4098 |
| Knstrn | 2516 | 2861 | 2874 | 3119 | 2276 | 2286 | 2672 | 2857 | 2886 | 2452 | 2449 | 2502 | 2636.45 | 2805 | 2468 | 0.88 | -0.185 | 0.00781804 | 0.04358645 | 6.00E-04 | 0.0082 | 0.0031 | 0.0031 | TRUE | 0.0207 |
| Kntc1 | 6240 | 7633 | 8192 | 9440 | 6722 | 6940 | 6628 | 7623 | 8226 | 7421 | 7232 | 7597 | 7454.52 | 7492 | 7417 | 0.99 | -0.015 | 0.87922259 | 0.94020603 | 0.0061 | 0.0073 | 0.0066 | 0.0066 | TRUE | 0.1242 |
| Kpna1 | 7297 | 6850 | 6707 | 8483 | 5741 | 5975 | 7751 | 6841 | 6735 | 6669 | 6177 | 6541 | 6785.43 | 7109 | 6462 | 0.909 | -0.137 | 0.10500884 | 0.27022343 | 0.0036 | 0.0074 | 0.005 | 0.005 | TRUE | 0.0761 |
| Kpna2 | 18097 | 19199 | 18258 | 21228 | 14442 | 15057 | 19222 | 19174 | 18333 | 16688 | 15539 | 16482 | 17573.02 | 18910 | 16236 | 0.859 | -0.22 | 0.00052956 | 0.00542003 | 0.001 | 0.0071 | 0.0028 | 0.0028 | TRUE | 0.0173 |
| Kpna3 | 12515 | 12263 | 12224 | 14789 | 10939 | 10900 | 13293 | 12247 | 12274 | 11626 | 11770 | 11932 | 12190.32 | 12605 | 11776 | 0.934 | -0.098 | 0.13147789 | 0.31231281 | 0.0011 | 0.0072 | 0.003 | 0.003 | TRUE | 0.0279 |
| Kpna4 | 6194 | 5802 | 5807 | 8317 | 5663 | 5196 | 6579 | 5795 | 5831 | 6538 | 6093 | 5688 | 6087.25 | 6068 | 6106 | 1.007 | 0.009 | 0.91806233 | 0.95991788 | 0.0049 | 0.0074 | 0.0059 | 0.0059 | TRUE | 0.0661 |
| Kpna6 | 8318 | 9330 | 9299 | 12453 | 8869 | 8659 | 8835 | 9318 | 9337 | 9790 | 9542 | 9479 | 9383.5 | 9163 | 9604 | 1.048 | 0.068 | 0.25830311 | 0.47721173 | 5.00E-04 | 0.0073 | 0.0025 | 0.0025 | TRUE | 0.012 |
| Kpna7 | 19 | 11 | 18 | 45 | 28 | 34 | 20 | 11 | 18 | 35 | 30 | 37 | 25.33 | 16 | 34 | 2.092 | 1.065 | 0.0088817 | 0.04818309 | 0 | 0.137 | 0.074 | 0.074 | TRUE | 0.4107 |
| Kpnb1 | 90972 | 87918 | 93113 | 107874 | 77220 | 76385 | 96628 | 87804 | 93495 | 84802 | 83084 | 83615 | 88238.26 | 92642 | 83834 | 0.905 | -0.144 | 0.02430134 | 0.0996829 | 0.0012 | 0.0069 | 0.0029 | 0.0029 | TRUE | 0.0256 |
| Kprp | 0 | 0 | 0 | 0 | 0 | 0 | 0 | 0 | 0 | 0 | 0 | 0 | 0 | 0 | 0 | NA | NA | NA | NA | NA | NA | NA | NA | NA | NA |
| Kptn | 444 | 578 | 567 | 729 | 608 | 513 | 472 | 577 | 569 | 573 | 654 | 562 | 567.83 | 539 | 596 | 1.104 | 0.143 | 0.26178156 | 0.48126204 | 0.0077 | 0.0127 | 0.01 | 0.01 | TRUE | 0.1401 |
| Kras | 2104 | 2769 | 2848 | 3258 | 2530 | 2628 | 2235 | 2765 | 2860 | 2561 | 2722 | 2877 | 2670 | 2620 | 2720 | 1.038 | 0.054 | 0.64150732 | 0.79835032 | 0.01 | 0.0081 | 0.0092 | 0.0092 | TRUE | 0.2002 |
| Krba1 | 1159 | 1014 | 1012 | 1798 | 1357 | 1193 | 1231 | 1013 | 1016 | 1413 | 1460 | 1306 | 1239.89 | 1087 | 1393 | 1.283 | 0.359 | 0.00126309 | 0.01066455 | 0.007 | 0.0096 | 0.0081 | 0.0081 | TRUE | 0.1611 |
| Krcc1 | 1170 | 1353 | 1255 | 1947 | 1364 | 1127 | 1243 | 1351 | 1260 | 1531 | 1468 | 1234 | 1347.67 | 1285 | 1411 | 1.099 | 0.136 | 0.21252393 | 0.42354402 | 0.0061 | 0.0093 | 0.0078 | 0.0078 | TRUE | 0.1436 |
| Kremen1 | 1877 | 1988 | 1914 | 2257 | 1593 | 1651 | 1994 | 1985 | 1922 | 1774 | 1714 | 1807 | 1866.09 | 1967 | 1765 | 0.897 | -0.156 | 0.02598487 | 0.10433639 | 0 | 0.0087 | 0.003 | 0.003 | TRUE | 0.0076 |
| Kremen2 | 90 | 75 | 91 | 84 | 60 | 61 | 96 | 75 | 91 | 66 | 65 | 67 | 76.54 | 87 | 66 | 0.754 | -0.407 | 0.07384261 | 0.21338322 | 0 | 0.0499 | 0.0241 | 0.0241 | TRUE | 0.1478 |
| Kri1 | 2895 | 2651 | 2752 | 3739 | 2823 | 2544 | 3075 | 2648 | 2763 | 2939 | 3037 | 2785 | 2874.56 | 2829 | 2920 | 1.033 | 0.046 | 0.60009205 | 0.77212039 | 0.0031 | 0.008 | 0.0053 | 0.0053 | TRUE | 0.0703 |
| Krit1 | 2174 | 2077 | 2095 | 2711 | 1847 | 1874 | 2309 | 2074 | 2104 | 2131 | 1987 | 2051 | 2109.49 | 2162 | 2056 | 0.952 | -0.072 | 0.37840182 | 0.5934918 | 0.0017 | 0.0085 | 0.0043 | 0.0043 | TRUE | 0.0425 |
| Krr1 | 5130 | 4744 | 4573 | 6304 | 4478 | 4313 | 5449 | 4738 | 4592 | 4956 | 4818 | 4721 | 4878.94 | 4926 | 4832 | 0.981 | -0.028 | 0.75654668 | 0.87147635 | 0.0037 | 0.0076 | 0.0056 | 0.0056 | TRUE | 0.1049 |
| Krt1 | 4 | 6 | 6 | 11 | 16 | 12 | 4 | 6 | 6 | 9 | 17 | 13 | 9.21 | 5 | 13 | 2.369 | 1.244 | 0.06165328 | 0.18955078 | 0 | 0.3646 | 0.1874 | 0.1874 | TRUE | 0.1291 |
| Krt10 | 59 | 62 | 50 | 88 | 63 | 55 | 63 | 62 | 50 | 69 | 68 | 60 | 61.99 | 58 | 66 | 1.131 | 0.177 | 0.4766887 | 0.67645532 | 0 | 0.06 | 0.0286 | 0.0286 | TRUE | 0.1262 |
| Krt12 | 43 | 52 | 43 | 96 | 69 | 54 | 46 | 52 | 43 | 75 | 74 | 59 | 58.27 | 47 | 69 | 1.487 | 0.573 | 0.02929152 | 0.11309323 | 0 | 0.0634 | 0.0318 | 0.0318 | TRUE | 0.1474 |
| Krt13 | 3 | 2 | 2 | 3 | 1 | 2 | 3 | 2 | 2 | 2 | 1 | 2 | 2.14 | 2 | 2 | 0.793 | -0.334 | 0.78610422 | NA | 0 | 1.5493 | 0.6233 | 0.6233 | TRUE | 0.1057 |
| Krt14 | 39 | 26 | 15 | 29 | 20 | 11 | 41 | 26 | 15 | 23 | 22 | 12 | 23.14 | 27 | 19 | 0.689 | -0.537 | 0.29031725 | 0.51138358 | 0.1326 | 0.1493 | 0.1412 | 0.1412 | TRUE | 0.214 |
| Krt15 | 5 | 1 | 1 | 5 | 1 | 4 | 5 | 1 | 1 | 4 | 1 | 4 | 2.78 | 2 | 3 | 1.303 | 0.382 | 0.75821416 | NA | 0.2706 | 1.1906 | 0.7432 | 0.7432 | TRUE | 1.0891 |
| Krt16 | 34 | 32 | 23 | 35 | 25 | 25 | 36 | 32 | 23 | 28 | 27 | 27 | 28.82 | 30 | 27 | 0.898 | -0.154 | 0.67051155 | 0.81605896 | 0 | 0.1212 | 0.0603 | 0.0603 | TRUE | 0.2969 |
| Krt17 | 272 | 239 | 217 | 242 | 209 | 217 | 289 | 239 | 218 | 190 | 225 | 238 | 233.02 | 249 | 218 | 0.874 | -0.195 | 0.2558318 | 0.47448015 | 0.0131 | 0.021 | 0.0169 | 0.0169 | TRUE | 0.2225 |
| Krt18 | 367 | 623 | 589 | 749 | 526 | 608 | 390 | 622 | 591 | 589 | 566 | 666 | 570.62 | 534 | 607 | 1.134 | 0.182 | 0.31875678 | 0.53988745 | 0.0322 | 0.0127 | 0.0223 | 0.0223 | TRUE | 0.6517 |

| | | | | | | | | | | | | | | | | | | | | | | | | | |
|---|---|---|---|---|---|---|---|---|---|---|---|---|---|---|---|---|---|---|---|---|---|---|---|---|---|
| Krt19 | 807 | 1512 | 1716 | 1455 | 1254 | 1615 | 857 | 1510 | 1723 | 1144 | 1349 | 1768 | 1391.86 | 1363 | 1420 | 1.041 | 0.058 | 0.81119248 | 0.90290246 | 0.0857 | 0.0093 | 0.0422 | 0.0422 | TRUE | 1.0798 |
| Krt2 | 0 | 0 | 1 | 1 | 0 | 0 | 0 | 0 | 1 | 1 | 0 | 0 | 0.3 | 0 | 0 | 0.954 | -0.067 | 0.98640617 | NA | 0 | 11.0483 | 9.2783 | 9.2783 | TRUE | 0.4098 |
| Krt20 | 10 | 7 | 11 | 41 | 25 | 19 | 11 | 7 | 11 | 32 | 27 | 21 | 18.1 | 10 | 27 | 2.81 | 1.49 | 0.00231208 | 0.01726613 | 0 | 0.1889 | 0.1006 | 0.1006 | TRUE | 0.1209 |
| Krt222 | 79 | 64 | 65 | 66 | 42 | 47 | 84 | 64 | 65 | 52 | 45 | 51 | 60.27 | 71 | 49 | 0.699 | -0.517 | 0.04720756 | 0.15774527 | 0 | 0.0616 | 0.0318 | 0.0318 | TRUE | 0.222 |
| Krt23 | 103 | 123 | 123 | 175 | 121 | 144 | 109 | 123 | 124 | 138 | 130 | 158 | 130.19 | 119 | 142 | 1.194 | 0.256 | 0.15909243 | 0.35313872 | 0 | 0.0322 | 0.016 | 0.016 | TRUE | 0.0935 |
| Krt24 | 5 | 1 | 2 | 3 | 0 | 1 | 5 | 1 | 2 | 2 | 0 | 1 | 1.96 | 3 | 1 | 0.437 | -1.194 | 0.42706905 | NA | 0 | 1.6862 | 1.0393 | 1.0393 | TRUE | 0.6155 |
| Krt25 | 1 | 0 | 1 | 4 | 3 | 0 | 1 | 0 | 1 | 3 | 3 | 0 | 1.41 | 1 | 2 | 3.14 | 1.651 | 0.37876312 | NA | 0 | 2.3493 | 1.5616 | 1.5616 | TRUE | 0.6488 |
| Krt26 | 2 | 0 | 0 | 2 | 3 | 0 | 2 | 0 | 0 | 2 | 3 | 0 | 1.15 | 1 | 2 | 2.294 | 1.198 | 0.59102508 | NA | 1.8421 | 2.8615 | 2.5448 | 2.5448 | TRUE | 0.3426 |
| Krt27 | 23 | 16 | 15 | 23 | 17 | 12 | 24 | 16 | 15 | 18 | 18 | 13 | 17.5 | 18 | 16 | 0.898 | -0.155 | 0.74120052 | 0.86165065 | 0 | 0.1952 | 0.1017 | 0.1017 | TRUE | 0.3928 |
| Krt28 | 9 | 7 | 4 | 8 | 9 | 5 | 10 | 7 | 4 | 6 | 10 | 5 | 7 | 7 | 7 | 1.042 | 0.059 | 0.93618395 | 0.96816629 | 0 | 0.4774 | 0.244 | 0.244 | TRUE | 0.232 |
| Krt31 | 0 | 0 | 0 | 0 | 0 | 0 | 0 | 0 | 0 | 0 | 0 | 0 | 0 | 0 | 0 | NA | NA | NA | NA | NA | NA | NA | NA | NA | NA |
| Krt32 | 0 | 0 | 0 | 0 | 0 | 0 | 0 | 0 | 0 | 0 | 0 | 0 | 0 | 0 | 0 | NA | NA | NA | NA | NA | NA | NA | NA | NA | NA |
| Krt33a | 1 | 1 | 0 | 8 | 1 | 3 | 1 | 1 | 0 | 6 | 1 | 3 | 2.12 | 1 | 3 | 5.29 | 2.403 | 0.12914151 | NA | 0 | 1.5621 | 0.9241 | 0.9241 | TRUE | 0.1191 |
| Krt33b | 0 | 0 | 0 | 0 | 0 | 0 | 0 | 0 | 0 | 0 | 0 | 0 | 0 | 0 | 0 | NA | NA | NA | NA | NA | NA | NA | NA | NA | NA |
| Krt34 | 0 | 0 | 0 | 0 | 0 | 1 | 0 | 0 | 0 | 0 | 0 | 1 | 0.18 | 0 | 0 | 1.859 | 0.894 | 0.82650734 | NA | 0 | 18.0643 | 10 | 10 | TRUE | 0.5383 |
| Krt35 | 0 | 0 | 0 | 0 | 1 | 0 | 0 | 0 | 0 | 0 | 1 | 0 | 0.18 | 0 | 0 | 1.859 | 0.894 | 0.82650734 | NA | 0 | 18.3785 | 10 | 10 | TRUE | 0.5204 |
| Krt36 | 1 | 1 | 2 | 2 | 0 | 0 | 1 | 1 | 2 | 2 | 0 | 0 | 0.94 | 1 | 1 | 0.425 | -1.233 | 0.55053282 | NA | 0 | 3.5108 | 1.8559 | 1.8559 | TRUE | 0.9714 |
| Krt39 | 0 | 0 | 0 | 0 | 0 | 0 | 0 | 0 | 0 | 0 | 0 | 0 | 0 | 0 | 0 | NA | NA | NA | NA | NA | NA | NA | NA | NA | NA |
| Krt4 | 2 | 0 | 2 | 5 | 2 | 2 | 2 | 0 | 2 | 4 | 2 | 2 | 2.07 | 1 | 3 | 2.04 | 1.028 | 0.45645137 | NA | 0 | 1.6004 | 0.8263 | 0.8263 | TRUE | 0.498 |
| Krt40 | 0 | 4 | 1 | 4 | 4 | 2 | 0 | 4 | 1 | 3 | 4 | 2 | 2.44 | 2 | 3 | 1.911 | 0.934 | 0.48663849 | NA | 0.0604 | 1.3574 | 0.8432 | 0.8432 | TRUE | 1.1413 |
| Krt42 | 344 | 340 | 323 | 431 | 362 | 374 | 365 | 340 | 324 | 339 | 389 | 409 | 361.16 | 343 | 379 | 1.103 | 0.141 | 0.27981083 | 0.50098495 | 0.0043 | 0.016 | 0.0096 | 0.0096 | TRUE | 0.1062 |
| Krt5 | 11 | 9 | 10 | 9 | 14 | 8 | 12 | 9 | 10 | 7 | 15 | 9 | 10.27 | 10 | 10 | 0.996 | -0.006 | 0.99174783 | 0.99572517 | 0 | 0.3277 | 0.1698 | 0.1698 | TRUE | 0.5514 |
| Krt6a | 0 | 0 | 0 | 0 | 0 | 0 | 0 | 0 | 0 | 0 | 0 | 0 | 0 | 0 | 0 | NA | NA | NA | NA | NA | NA | NA | NA | NA | NA |
| Krt6b | 1 | 0 | 0 | 2 | 2 | 0 | 1 | 0 | 0 | 2 | 2 | 0 | 0.8 | 0 | 1 | 3.417 | 1.773 | 0.46651319 | NA | 0 | 4.1368 | 2.8785 | 2.8785 | TRUE | 0.4614 |
| Krt7 | 137 | 375 | 452 | 125 | 141 | 219 | 146 | 375 | 454 | 98 | 152 | 240 | 243.93 | 325 | 163 | 0.502 | -0.994 | 0.08900908 | 0.24237762 | 0.2414 | 0.0204 | 0.1124 | 0.2414 | TRUE | 0.5306 |
| Krt71 | 6 | 9 | 6 | 24 | 12 | 13 | 6 | 9 | 6 | 19 | 13 | 14 | 11.23 | 7 | 15 | 2.164 | 1.114 | 0.05661292 | 0.17876766 | 0 | 0.3002 | 0.1429 | 0.1429 | TRUE | 0.2296 |
| Krt72 | 0 | 0 | 0 | 3 | 0 | 0 | 0 | 0 | 0 | 2 | 0 | 0 | 0.39 | 0 | 1 | 4.278 | 2.097 | 0.60136958 | NA | 10 | 8.3884 | 10 | 10 | TRUE | 1.4411 |
| Krt73 | 0 | 2 | 1 | 3 | 2 | 0 | 0 | 2 | 1 | 2 | 2 | 0 | 1.25 | 1 | 1 | 1.517 | 0.602 | 0.74659055 | NA | 0 | 2.6383 | 1.6713 | 1.6713 | TRUE | 0.2931 |
| Krt74 | 6 | 0 | 0 | 2 | 0 | 1 | 6 | 0 | 0 | 2 | 0 | 1 | 1.51 | 2 | 1 | 0.43 | -1.217 | 0.5817883 | NA | 3.9027 | 2.1935 | 2.7392 | 2.7392 | TRUE | 2.779 |
| Krt75 | 8 | 11 | 2 | 8 | 7 | 6 | 8 | 11 | 2 | 6 | 8 | 7 | 6.98 | 7 | 7 | 0.947 | -0.078 | 0.92147485 | 0.96126008 | 0 | 0.4789 | 0.3105 | 0.3105 | TRUE | 0.7655 |
| Krt76 | 0 | 1 | 0 | 0 | 0 | 0 | 0 | 1 | 0 | 0 | 0 | 0 | 0.17 | 0 | 0 | 0.49 | -1.029 | 0.80087 | NA | 0 | 19.7991 | 10 | 10 | TRUE | 0.405 |
| Krt77 | 0 | 0 | 1 | 0 | 0 | 0 | 0 | 0 | 1 | 0 | 0 | 0 | 0.17 | 0 | 0 | 0.49 | -1.029 | 0.80087 | NA | 0 | 19.6928 | 10 | 10 | TRUE | 0.4098 |
| Krt78 | 14 | 14 | 12 | 7 | 5 | 7 | 15 | 14 | 12 | 6 | 5 | 8 | 9.91 | 14 | 6 | 0.451 | -1.15 | 0.05853328 | 0.18288171 | 0 | 0.3394 | 0.1495 | 0.1495 | TRUE | 0.0935 |
| Krt79 | 113 | 90 | 96 | 107 | 78 | 81 | 120 | 90 | 96 | 84 | 84 | 89 | 93.83 | 102 | 86 | 0.838 | -0.254 | 0.23609022 | 0.45112853 | 0 | 0.042 | 0.0225 | 0.0225 | TRUE | 0.227 |
| Krt8 | 507 | 656 | 663 | 682 | 570 | 532 | 539 | 655 | 666 | 536 | 613 | 582 | 598.53 | 620 | 577 | 0.93 | -0.105 | 0.40352792 | 0.6162915 | 0.0074 | 0.0124 | 0.0096 | 0.0096 | TRUE | 0.1536 |
| Krt80 | 5 | 25 | 28 | 7 | 11 | 8 | 5 | 25 | 28 | 6 | 12 | 9 | 14.08 | 19 | 9 | 0.441 | -1.181 | 0.09393457 | 0.2508527 | 0.3116 | 0.2409 | 0.2753 | 0.2753 | TRUE | 2.0765 |
| Krt81 | 1 | 1 | 1 | 0 | 1 | 2 | 1 | 1 | 1 | 0 | 1 | 2 | 1.06 | 1 | 1 | 1.021 | 0.029 | 0.98726074 | NA | 0 | 3.1295 | 1.4749 | 1.4749 | TRUE | 0.229 |
| Krt82 | 0 | 0 | 0 | 0 | 1 | 1 | 0 | 0 | 0 | 0 | 1 | 1 | 0.36 | 0 | 1 | 3.794 | 1.924 | 0.60247382 | NA | 0 | 9.1135 | 8.1425 | 8.1425 | TRUE | 0.3482 |
| Krt83 | 2 | 1 | 2 | 5 | 1 | 3 | 2 | 1 | 2 | 4 | 1 | 3 | 2.24 | 2 | 3 | 1.643 | 0.716 | 0.56927432 | NA | 0 | 1.4796 | 0.6683 | 0.6683 | TRUE | 0.3144 |
| Krt84 | 0 | 1 | 0 | 2 | 0 | 3 | 0 | 1 | 0 | 2 | 0 | 3 | 0.98 | 0 | 2 | 4.416 | 2.143 | 0.35716898 | NA | 0 | 3.383 | 2.5996 | 2.5996 | TRUE | 0.1602 |
| Krt85 | 1 | 1 | 0 | 2 | 0 | 1 | 1 | 1 | 0 | 2 | 0 | 1 | 0.79 | 1 | 1 | 1.332 | 0.414 | 0.8494818 | NA | 0 | 4.1879 | 2.1489 | 2.1489 | TRUE | 0.2882 |
| Krt86 | 0 | 0 | 1 | 3 | 0 | 0 | 0 | 0 | 1 | 2 | 0 | 0 | 0.56 | 0 | 1 | 2.223 | 1.152 | 0.72337121 | NA | 0 | 5.8855 | 6.0328 | 6.0328 | TRUE | 1.4404 |
| Krt9 | 1 | 1 | 0 | 4 | 3 | 0 | 1 | 1 | 0 | 3 | 3 | 0 | 1.41 | 1 | 2 | 3.145 | 1.653 | 0.37822297 | NA | 0 | 2.3508 | 1.5609 | 1.5609 | TRUE | 0.6488 |
| Krtap1-3 | 0 | 0 | 0 | 0 | 0 | 0 | 0 | 0 | 0 | 0 | 0 | 0 | 0 | 0 | 0 | NA | NA | NA | NA | NA | NA | NA | NA | NA | NA |
| Krtap1-4 | 0 | 0 | 0 | 0 | 0 | 0 | 0 | 0 | 0 | 0 | 0 | 0 | 0 | 0 | 0 | NA | NA | NA | NA | NA | NA | NA | NA | NA | NA |
| Krtap1-5 | 0 | 0 | 0 | 0 | 0 | 0 | 0 | 0 | 0 | 0 | 0 | 0 | 0 | 0 | 0 | NA | NA | NA | NA | NA | NA | NA | NA | NA | NA |
| Krtap10-10 | 1 | 3 | 0 | 4 | 2 | 6 | 1 | 3 | 0 | 3 | 2 | 7 | 2.65 | 1 | 4 | 2.897 | 1.535 | 0.25106506 | NA | 0.1456 | 1.2483 | 0.7875 | 0.7875 | TRUE | 0.7208 |
| Krtap10-4 | 0 | 0 | 0 | 0 | 0 | 0 | 0 | 0 | 0 | 0 | 0 | 0 | 0 | 0 | 0 | NA | NA | NA | NA | NA | NA | NA | NA | NA | NA |
| Krtap11-1 | 0 | 0 | 0 | 0 | 0 | 0 | 0 | 0 | 0 | 0 | 0 | 0 | 0 | 0 | 0 | NA | NA | NA | NA | NA | NA | NA | NA | NA | NA |
| Krtap12-1 | 0 | 0 | 0 | 0 | 0 | 0 | 0 | 0 | 0 | 0 | 0 | 0 | 0 | 0 | 0 | NA | NA | NA | NA | NA | NA | NA | NA | NA | NA |
| Krtap13 | 0 | 0 | 0 | 0 | 0 | 0 | 0 | 0 | 0 | 0 | 0 | 0 | 0 | 0 | 0 | NA | NA | NA | NA | NA | NA | NA | NA | NA | NA |
| Krtap13-1 | 0 | 0 | 0 | 0 | 0 | 0 | 0 | 0 | 0 | 0 | 0 | 0 | 0 | 0 | 0 | NA | NA | NA | NA | NA | NA | NA | NA | NA | NA |
| Krtap14 | 0 | 0 | 0 | 0 | 0 | 0 | 0 | 0 | 0 | 0 | 0 | 0 | 0 | 0 | 0 | NA | NA | NA | NA | NA | NA | NA | NA | NA | NA |
| Krtap15 | 0 | 0 | 0 | 0 | 0 | 0 | 0 | 0 | 0 | 0 | 0 | 0 | 0 | 0 | 0 | NA | NA | NA | NA | NA | NA | NA | NA | NA | NA |
| Krtap16-1 | 0 | 0 | 0 | 0 | 0 | 0 | 0 | 0 | 0 | 0 | 0 | 0 | 0 | 0 | 0 | NA | NA | NA | NA | NA | NA | NA | NA | NA | NA |
| Krtap16-3 | 0 | 0 | 0 | 0 | 0 | 0 | 0 | 0 | 0 | 0 | 0 | 0 | 0 | 0 | 0 | NA | NA | NA | NA | NA | NA | NA | NA | NA | NA |
| Krtap17-1 | 0 | 0 | 0 | 0 | 0 | 0 | 0 | 0 | 0 | 0 | 0 | 0 | 0 | 0 | 0 | NA | NA | NA | NA | NA | NA | NA | NA | NA | NA |
| Krtap19-1 | 0 | 0 | 0 | 0 | 0 | 0 | 0 | 0 | 0 | 0 | 0 | 0 | 0 | 0 | 0 | NA | NA | NA | NA | NA | NA | NA | NA | NA | NA |
| Krtap19-3 | 0 | 0 | 0 | 0 | 0 | 0 | 0 | 0 | 0 | 0 | 0 | 0 | 0 | 0 | 0 | NA | NA | NA | NA | NA | NA | NA | NA | NA | NA |
| Krtap19-4 | 0 | 0 | 0 | 0 | 0 | 0 | 0 | 0 | 0 | 0 | 0 | 0 | 0 | 0 | 0 | NA | NA | NA | NA | NA | NA | NA | NA | NA | NA |
| Krtap19-5 | 0 | 0 | 0 | 0 | 0 | 0 | 0 | 0 | 0 | 0 | 0 | 0 | 0 | 0 | 0 | NA | NA | NA | NA | NA | NA | NA | NA | NA | NA |
| Krtap19-9b | 0 | 0 | 0 | 0 | 0 | 0 | 0 | 0 | 0 | 0 | 0 | 0 | 0 | 0 | 0 | NA | NA | NA | NA | NA | NA | NA | NA | NA | NA |
| Krtap2-4 | 0 | 0 | 0 | 0 | 0 | 0 | 0 | 0 | 0 | 0 | 0 | 0 | 0 | 0 | 0 | NA | NA | NA | NA | NA | NA | NA | NA | NA | NA |
| Krtap20-2 | 0 | 0 | 0 | 0 | 0 | 0 | 0 | 0 | 0 | 0 | 0 | 0 | 0 | 0 | 0 | NA | NA | NA | NA | NA | NA | NA | NA | NA | NA |
| Krtap21-1 | 0 | 0 | 0 | 0 | 0 | 0 | 0 | 0 | 0 | 0 | 0 | 0 | 0 | 0 | 0 | NA | NA | NA | NA | NA | NA | NA | NA | NA | NA |
| Krtap22-2 | 0 | 0 | 0 | 0 | 0 | 0 | 0 | 0 | 0 | 0 | 0 | 0 | 0 | 0 | 0 | NA | NA | NA | NA | NA | NA | NA | NA | NA | NA |
| Krtap24-1 | 0 | 0 | 0 | 0 | 0 | 0 | 0 | 0 | 0 | 0 | 0 | 0 | 0 | 0 | 0 | NA | NA | NA | NA | NA | NA | NA | NA | NA | NA |
| Krtap26-1 | 0 | 0 | 0 | 0 | 0 | 0 | 0 | 0 | 0 | 0 | 0 | 0 | 0 | 0 | 0 | NA | NA | NA | NA | NA | NA | NA | NA | NA | NA |
| Krtap27-1 | 0 | 0 | 0 | 0 | 0 | 0 | 0 | 0 | 0 | 0 | 0 | 0 | 0 | 0 | 0 | NA | NA | NA | NA | NA | NA | NA | NA | NA | NA |

| | | | | | | | | | | | | | | | | | | | | | | | | | |
|---|---|---|---|---|---|---|---|---|---|---|---|---|---|---|---|---|---|---|---|---|---|---|---|---|---|
| Krtap3-1 | 0 | 0 | 0 | 0 | 0 | 0 | 0 | 0 | 0 | 0 | 0 | 0 | 0 | 0 | 0 | NA | NA | NA | NA | NA | NA | NA | NA | NA | NA |
| Krtap3-2 | 0 | 0 | 0 | 0 | 0 | 0 | 0 | 0 | 0 | 0 | 0 | 0 | 0 | 0 | 0 | NA | NA | NA | NA | NA | NA | NA | NA | NA | NA |
| Krtap3-3 | 0 | 0 | 0 | 0 | 0 | 0 | 0 | 0 | 0 | 0 | 0 | 0 | 0 | 0 | 0 | NA | NA | NA | NA | NA | NA | NA | NA | NA | NA |
| Krtap31-1 | 0 | 0 | 0 | 0 | 0 | 0 | 0 | 0 | 0 | 0 | 0 | 0 | 0 | 0 | 0 | NA | NA | NA | NA | NA | NA | NA | NA | NA | NA |
| Krtap31-2 | 0 | 0 | 0 | 0 | 0 | 0 | 0 | 0 | 0 | 0 | 0 | 0 | 0 | 0 | 0 | NA | NA | NA | NA | NA | NA | NA | NA | NA | NA |
| Krtap4-1 | 0 | 0 | 0 | 0 | 0 | 0 | 0 | 0 | 0 | 0 | 0 | 0 | 0 | 0 | 0 | NA | NA | NA | NA | NA | NA | NA | NA | NA | NA |
| Krtap4-13 | 0 | 0 | 0 | 5 | 0 | 1 | 0 | 0 | 0 | 4 | 0 | 1 | 0.84 | 0 | 2 | 9.056 | 3.179 | 0.26036982 | NA | 6.8671 | 3.9404 | 4.4601 | 4.4601 | TRUE | 0.2452 |
| Krtap4-16 | 0 | 0 | 0 | 1 | 1 | 0 | 0 | 0 | 0 | 1 | 1 | 0 | 0.31 | 0 | 1 | 3.32 | 1.731 | 0.65383052 | NA | 0 | 10.6224 | 8.9541 | 8.9541 | TRUE | 0.202 |
| Krtap4-2 | 0 | 0 | 0 | 0 | 0 | 0 | 0 | 0 | 0 | 0 | 0 | 0 | 0 | 0 | 0 | NA | NA | NA | NA | NA | NA | NA | NA | NA | NA |
| Krtap4-6 | 0 | 0 | 0 | 0 | 0 | 0 | 0 | 0 | 0 | 0 | 0 | 0 | 0 | 0 | 0 | NA | NA | NA | NA | NA | NA | NA | NA | NA | NA |
| Krtap4-7 | 0 | 0 | 0 | 0 | 0 | 0 | 0 | 0 | 0 | 0 | 0 | 0 | 0 | 0 | 0 | NA | NA | NA | NA | NA | NA | NA | NA | NA | NA |
| Krtap4-8 | 0 | 0 | 0 | 0 | 0 | 0 | 0 | 0 | 0 | 0 | 0 | 0 | 0 | 0 | 0 | NA | NA | NA | NA | NA | NA | NA | NA | NA | NA |
| Krtap4-9 | 0 | 0 | 0 | 0 | 0 | 0 | 0 | 0 | 0 | 0 | 0 | 0 | 0 | 0 | 0 | NA | NA | NA | NA | NA | NA | NA | NA | NA | NA |
| Krtap5-1 | 0 | 0 | 0 | 0 | 0 | 0 | 0 | 0 | 0 | 0 | 0 | 0 | 0 | 0 | 0 | NA | NA | NA | NA | NA | NA | NA | NA | NA | NA |
| Krtap5-2 | 0 | 0 | 0 | 0 | 0 | 0 | 0 | 0 | 0 | 0 | 0 | 0 | 0 | 0 | 0 | NA | NA | NA | NA | NA | NA | NA | NA | NA | NA |
| Krtap5-3 | 0 | 0 | 0 | 0 | 0 | 0 | 0 | 0 | 0 | 0 | 0 | 0 | 0 | 0 | 0 | NA | NA | NA | NA | NA | NA | NA | NA | NA | NA |
| Krtap5-4 | 0 | 0 | 0 | 0 | 0 | 0 | 0 | 0 | 0 | 0 | 0 | 0 | 0 | 0 | 0 | NA | NA | NA | NA | NA | NA | NA | NA | NA | NA |
| Krtap5-5 | 0 | 0 | 0 | 0 | 0 | 0 | 0 | 0 | 0 | 0 | 0 | 0 | 0 | 0 | 0 | NA | NA | NA | NA | NA | NA | NA | NA | NA | NA |
| Krtap6-1 | 0 | 0 | 0 | 0 | 0 | 0 | 0 | 0 | 0 | 0 | 0 | 0 | 0 | 0 | 0 | NA | NA | NA | NA | NA | NA | NA | NA | NA | NA |
| Krtap6-2 | 0 | 0 | 0 | 0 | 0 | 0 | 0 | 0 | 0 | 0 | 0 | 0 | 0 | 0 | 0 | NA | NA | NA | NA | NA | NA | NA | NA | NA | NA |
| Krtap6-5 | 0 | 0 | 0 | 1 | 0 | 0 | 0 | 0 | 0 | 1 | 0 | 0 | 0.13 | 0 | 0 | 1.859 | 0.894 | 0.82650734 | NA | 0 | 25.1515 | 10 | 10 | TRUE | 0.2563 |
| Krtap7-1 | 0 | 0 | 0 | 0 | 0 | 0 | 0 | 0 | 0 | 0 | 0 | 0 | 0 | 0 | 0 | NA | NA | NA | NA | NA | NA | NA | NA | NA | NA |
| Krtap8-1 | 0 | 0 | 0 | 0 | 0 | 0 | 0 | 0 | 0 | 0 | 0 | 0 | 0 | 0 | 0 | NA | NA | NA | NA | NA | NA | NA | NA | NA | NA |
| Krtap9-1 | 0 | 0 | 0 | 0 | 0 | 0 | 0 | 0 | 0 | 0 | 0 | 0 | 0 | 0 | 0 | NA | NA | NA | NA | NA | NA | NA | NA | NA | NA |
| Krtap9-3 | 0 | 0 | 0 | 0 | 0 | 0 | 0 | 0 | 0 | 0 | 0 | 0 | 0 | 0 | 0 | NA | NA | NA | NA | NA | NA | NA | NA | NA | NA |
| Krtap9-5 | 0 | 0 | 0 | 0 | 0 | 0 | 0 | 0 | 0 | 0 | 0 | 0 | 0 | 0 | 0 | NA | NA | NA | NA | NA | NA | NA | NA | NA | NA |
| Krtcap2 | 1570 | 1754 | 1799 | 2421 | 1719 | 1681 | 1668 | 1752 | 1806 | 1903 | 1850 | 1840 | 1803.1 | 1742 | 1864 | 1.07 | 0.098 | 0.18261925 | 0.38465155 | 4.00E-04 | 0.0087 | 0.0034 | 0.0034 | TRUE | 0.0167 |
| Krtcap3 | 262 | 386 | 382 | 226 | 196 | 225 | 278 | 386 | 384 | 178 | 211 | 246 | 280.37 | 349 | 212 | 0.604 | -0.727 | 0.00014067 | 0.00181033 | 0.0263 | 0.0187 | 0.0225 | 0.0225 | TRUE | 0.3567 |
| Krtdap | 3 | 0 | 0 | 7 | 2 | 0 | 3 | 0 | 0 | 6 | 2 | 0 | 1.81 | 1 | 3 | 2.473 | 1.306 | 0.51874749 | NA | 3.4073 | 1.8302 | 2.2776 | 2.2776 | TRUE | 0.4528 |
| Ksr1 | 1148 | 917 | 935 | 1522 | 1065 | 980 | 1219 | 916 | 939 | 1196 | 1146 | 1073 | 1081.52 | 1025 | 1138 | 1.112 | 0.153 | 0.25020376 | 0.46733938 | 0.0135 | 0.0099 | 0.0118 | 0.0118 | TRUE | 0.3291 |
| Ksr2 | 79 | 48 | 43 | 75 | 59 | 52 | 84 | 48 | 43 | 59 | 63 | 57 | 59.06 | 58 | 60 | 1.027 | 0.038 | 0.90580644 | 0.953297 | 0.0506 | 0.0627 | 0.0568 | 0.0568 | TRUE | 1.2301 |
| Kti12 | 1116 | 958 | 957 | 1387 | 1021 | 951 | 1185 | 957 | 961 | 1090 | 1099 | 1041 | 1055.5 | 1034 | 1077 | 1.041 | 0.058 | 0.60687576 | 0.77685614 | 0.0071 | 0.01 | 0.0083 | 0.0083 | TRUE | 0.1942 |
| Ktn1 | 4701 | 5363 | 5643 | 7599 | 5249 | 5155 | 4993 | 5356 | 5666 | 5974 | 5648 | 5643 | 5546.64 | 5338 | 5755 | 1.078 | 0.108 | 0.161929 | 0.35714503 | 0.0023 | 0.0075 | 0.0042 | 0.0042 | TRUE | 0.0389 |
| Kxd1 | 2070 | 3099 | 3404 | 2960 | 2346 | 2485 | 2199 | 3095 | 3418 | 2327 | 2524 | 2720 | 2713.83 | 2904 | 2524 | 0.869 | -0.203 | 0.20510023 | 0.41355675 | 0.0261 | 0.0081 | 0.0181 | 0.0181 | TRUE | 0.5473 |
| Ky | 114 | 86 | 98 | 114 | 90 | 70 | 121 | 86 | 98 | 90 | 97 | 77 | 94.74 | 102 | 88 | 0.863 | -0.212 | 0.35089737 | 0.56972959 | 0.0111 | 0.0417 | 0.0267 | 0.0267 | TRUE | 0.2649 |
| Kynu | 0 | 0 | 2 | 2 | 2 | 1 | 0 | 0 | 2 | 2 | 2 | 1 | 1.14 | 1 | 2 | 2.38 | 1.251 | 0.51829667 | NA | 0 | 2.9023 | 1.6411 | 1.6411 | TRUE | 0.9738 |
| L1cam | 608 | 702 | 740 | 766 | 520 | 545 | 646 | 701 | 743 | 602 | 559 | 597 | 641.36 | 697 | 586 | 0.842 | -0.249 | 0.01602311 | 0.07448174 | 0.0017 | 0.012 | 0.0061 | 0.0061 | TRUE | 0.0477 |
| L1td1 | 20823 | 15025 | 15075 | 31694 | 21781 | 16206 | 22118 | 15006 | 15137 | 24915 | 23435 | 17740 | 19725.1 | 17420 | 22030 | 1.265 | 0.339 | 0.05741352 | 0.18070764 | 0.042 | 0.0071 | 0.0229 | 0.0229 | TRUE | 0.6807 |
| L2hgdh | 2985 | 3271 | 3025 | 3981 | 3154 | 3276 | 3171 | 3267 | 3037 | 3130 | 3394 | 3586 | 3263.99 | 3158 | 3370 | 1.066 | 0.093 | 0.26183804 | 0.48126204 | 0.0025 | 0.0079 | 0.0046 | 0.0046 | TRUE | 0.0479 |
| L3hypdh | 102 | 136 | 162 | 137 | 108 | 109 | 108 | 136 | 163 | 108 | 116 | 119 | 125.01 | 136 | 114 | 0.841 | -0.249 | 0.23605116 | 0.45112853 | 0.0144 | 0.0333 | 0.0238 | 0.0238 | TRUE | 0.1596 |
| L3mbtl1 | 61 | 74 | 58 | 51 | 68 | 56 | 65 | 74 | 58 | 40 | 73 | 61 | 61.92 | 66 | 58 | 0.879 | -0.186 | 0.53570193 | 0.72271961 | 0.0371 | 0.0601 | 0.0488 | 0.0488 | TRUE | 0.5375 |
| L3mbtl2 | 8034 | 8005 | 8134 | 10443 | 7605 | 7131 | 8534 | 7995 | 8167 | 8209 | 8183 | 7806 | 8148.93 | 8232 | 8066 | 0.98 | -0.029 | 0.64668324 | 0.80159447 | 8.00E-04 | 0.0073 | 0.0028 | 0.0028 | TRUE | 0.0125 |
| L3mbtl3 | 264 | 235 | 235 | 407 | 316 | 298 | 280 | 235 | 236 | 320 | 340 | 326 | 289.54 | 250 | 329 | 1.313 | 0.393 | 0.00358774 | 0.02428573 | 0.0011 | 0.0183 | 0.0096 | 0.0096 | TRUE | 0.1209 |
| L3mbtl4 | 9 | 3 | 4 | 5 | 6 | 3 | 10 | 3 | 4 | 4 | 6 | 3 | 5.04 | 6 | 4 | 0.825 | -0.277 | 0.75475506 | 0.87049337 | 0 | 0.6605 | 0.3673 | 0.3673 | TRUE | 0.8552 |
| l7Rn6 | 2180 | 2655 | 2600 | 3087 | 2335 | 2445 | 2316 | 2652 | 2611 | 2427 | 2512 | 2676 | 2532.22 | 2526 | 2538 | 1.005 | 0.007 | 0.94154914 | 0.97046876 | 0.0035 | 0.0082 | 0.0053 | 0.0053 | TRUE | 0.0642 |
| Lacc1 | 482 | 409 | 377 | 774 | 475 | 461 | 512 | 408 | 379 | 608 | 511 | 505 | 487.19 | 433 | 541 | 1.253 | 0.325 | 0.03402524 | 0.12550866 | 0.0157 | 0.0137 | 0.0149 | 0.0149 | TRUE | 0.2925 |
| Lace1 | 545 | 906 | 824 | 722 | 526 | 632 | 579 | 905 | 827 | 568 | 566 | 692 | 689.41 | 770 | 609 | 0.789 | -0.342 | 0.05641492 | 0.17838333 | 0.0318 | 0.0117 | 0.0216 | 0.0216 | TRUE | 0.5583 |
| Lactb | 831 | 725 | 713 | 1127 | 786 | 686 | 883 | 724 | 716 | 886 | 846 | 751 | 800.87 | 774 | 828 | 1.07 | 0.098 | 0.43648311 | 0.64477022 | 0.0094 | 0.011 | 0.0101 | 0.0101 | TRUE | 0.1771 |
| Lactb2 | 4170 | 4662 | 4533 | 4926 | 3836 | 3976 | 4429 | 4656 | 4552 | 3872 | 4127 | 4352 | 4331.49 | 4546 | 4117 | 0.905 | -0.143 | 0.05577452 | 0.17697081 | 0.0017 | 0.0077 | 0.0038 | 0.0038 | TRUE | 0.0334 |
| Lactbl1 | 0 | 0 | 0 | 0 | 1 | 0 | 0 | 0 | 0 | 0 | 1 | 0 | 0.18 | 0 | 0 | 1.859 | 0.894 | 0.82650734 | NA | 0 | 18.3785 | 10 | 10 | TRUE | 0.5204 |
| Lad1 | 405 | 555 | 583 | 298 | 250 | 359 | 430 | 554 | 585 | 234 | 269 | 393 | 411.01 | 523 | 299 | 0.569 | -0.812 | 0.00012986 | 0.00170389 | 0.0474 | 0.0149 | 0.0299 | 0.0299 | TRUE | 0.852 |
| Lag3 | 297 | 422 | 399 | 240 | 168 | 244 | 315 | 421 | 401 | 189 | 181 | 267 | 295.68 | 379 | 212 | 0.558 | -0.841 | 1.86E-05 | 0.00033359 | 0.0308 | 0.018 | 0.0242 | 0.0242 | TRUE | 0.5472 |
| Lage3 | 860 | 1114 | 1057 | 1598 | 1091 | 1143 | 913 | 1113 | 1061 | 1256 | 1174 | 1251 | 1128.11 | 1029 | 1227 | 1.192 | 0.254 | 0.01498417 | 0.07071523 | 0.0044 | 0.0098 | 0.0069 | 0.0069 | TRUE | 0.1148 |
| Lair1 | 10 | 2 | 5 | 3 | 7 | 3 | 11 | 2 | 5 | 2 | 8 | 3 | 5.14 | 6 | 4 | 0.739 | -0.436 | 0.65049192 | 0.80416668 | 0.2936 | 0.6484 | 0.4682 | 0.4682 | TRUE | 0.3511 |
| Lalba | 3 | 2 | 0 | 0 | 1 | 2 | 3 | 2 | 0 | 0 | 1 | 2 | 1.41 | 2 | 1 | 0.608 | -0.717 | 0.6991424 | NA | 0 | 2.3464 | 1.7183 | 1.7183 | TRUE | 0.2906 |
| Lama1 | 6161 | 5301 | 6179 | 8208 | 5787 | 6067 | 6544 | 5294 | 6204 | 6452 | 6226 | 6641 | 6227.14 | 6014 | 6440 | 1.071 | 0.099 | 0.31351648 | 0.5339318 | 0.006 | 0.0074 | 0.0068 | 0.0068 | TRUE | 0.1339 |
| Lama2 | 8 | 15 | 20 | 29 | 20 | 14 | 8 | 15 | 20 | 23 | 22 | 15 | 17.2 | 14 | 20 | 1.372 | 0.456 | 0.36502257 | 0.58247025 | 0.0268 | 0.1984 | 0.1229 | 0.1229 | TRUE | 0.3253 |
| Lama3 | 887 | 777 | 763 | 1063 | 773 | 765 | 942 | 776 | 766 | 836 | 832 | 837 | 831.51 | 828 | 835 | 1.009 | 0.012 | 0.91213312 | 0.95666604 | 0.0056 | 0.0109 | 0.0079 | 0.0079 | TRUE | 0.1715 |
| Lama4 | 57 | 35 | 32 | 173 | 104 | 58 | 61 | 35 | 32 | 136 | 112 | 63 | 73.17 | 43 | 104 | 2.453 | 1.294 | 0.00047618 | 0.00499686 | 0.1166 | 0.0519 | 0.0822 | 0.0822 | TRUE | 0.287 |
| Lama5 | 10054 | 11185 | 12500 | 9269 | 7062 | 6910 | 10679 | 11171 | 12551 | 7287 | 7598 | 7564 | 9474.99 | 11467 | 7483 | 0.653 | -0.616 | 2.60E-13 | 2.93E-11 | 0.0037 | 0.0073 | 0.005 | 0.005 | TRUE | 0.0837 |
| Lamb1 | 4272 | 5080 | 5736 | 6223 | 4265 | 4025 | 4538 | 5073 | 5760 | 4892 | 4589 | 4406 | 4876.25 | 5124 | 4629 | 0.904 | -0.146 | 0.17042141 | 0.36901257 | 0.0082 | 0.0076 | 0.008 | 0.008 | TRUE | 0.1437 |
| Lamb2 | 1498 | 1540 | 1548 | 1886 | 1353 | 1367 | 1591 | 1538 | 1554 | 1483 | 1456 | 1496 | 1519.71 | 1561 | 1478 | 0.947 | -0.079 | 0.26624601 | 0.48590976 | 0 | 0.0091 | 0.0029 | 0.0029 | TRUE | 0.0034 |
| Lamb3 | 179 | 273 | 281 | 220 | 144 | 208 | 190 | 273 | 282 | 173 | 155 | 228 | 216.75 | 248 | 185 | 0.744 | -0.426 | 0.05085798 | 0.16600416 | 0.0373 | 0.0221 | 0.0295 | 0.0295 | TRUE | 0.4622 |
| Lamc1 | 21150 | 19069 | 21002 | 23175 | 16316 | 16795 | 22465 | 19044 | 21088 | 18218 | 17555 | 18385 | 19459.3 | 20866 | 18053 | 0.865 | -0.209 | 0.01193652 | 0.0599563 | 0.0036 | 0.0071 | 0.0049 | 0.0049 | TRUE | 0.0714 |
| Lamc2 | 1751 | 1466 | 1504 | 2179 | 1663 | 1577 | 1860 | 1464 | 1510 | 1713 | 1789 | 1726 | 1677.11 | 1611 | 1743 | 1.082 | 0.113 | 0.31367052 | 0.53402541 | 0.0081 | 0.0089 | 0.0085 | 0.0085 | TRUE | 0.2188 |
| Lamc3 | 16 | 19 | 13 | 12 | 11 | 10 | 17 | 19 | 13 | 9 | 12 | 11 | 13.54 | 16 | 11 | 0.654 | -0.614 | 0.23241861 | 0.44654725 | 0 | 0.2502 | 0.1134 | 0.1134 | TRUE | 0.1512 |
| Lamp1 | 23224 | 27752 | 27204 | 34142 | 24049 | 24146 | 24668 | 27716 | 27316 | 26840 | 25875 | 26432 | 26474.41 | 26567 | 26382 | 0.993 | -0.01 | 0.88969057 | 0.94620362 | 0.0017 | 0.007 | 0.0037 | 0.0037 | TRUE | 0.0478 |
| Lamp2 | 2117 | 4149 | 4663 | 2790 | 2180 | 3166 | 2249 | 4144 | 4682 | 2193 | 2346 | 3466 | 3179.82 | 3692 | 2668 | 0.723 | -0.468 | 0.19384623 | 0.39954891 | 0.0934 | 0.0079 | 0.0462 | 0.0934 | TRUE | 0.9795 |
| Lamp3 | 1 | 1 | 2 | 6 | 2 | 3 | 1 | 1 | 2 | 5 | 2 | 3 | 2.37 | 1 | 3 | 2.522 | 1.335 | 0.28483082 | NA | 0 | 1.3968 | 0.604 | 0.604 | TRUE | 0.2221 |

| | | | | | | | | | | | | | | | | | | | | | | | | | |
|---|---|---|---|---|---|---|---|---|---|---|---|---|---|---|---|---|---|---|---|---|---|---|---|---|---|
| Lamp5 | 0 | 0 | 0 | 1 | 0 | 0 | 0 | 0 | 0 | 1 | 0 | 0 | 0.13 | 0 | 0 | 1.859 | 0.894 | 0.82650734 | NA | 0 | 25.1515 | 10 | 10 | TRUE | 0.2563 |
| Lamtor1 | 2658 | 2817 | 2764 | 4061 | 2891 | 2538 | 2823 | 2813 | 2775 | 3192 | 3111 | 2778 | 2915.53 | 2804 | 3027 | 1.08 | 0.111 | 0.17363294 | 0.37274245 | 0.0022 | 0.008 | 0.0045 | 0.0045 | TRUE | 0.0624 |
| Lamtor2 | 2893 | 3582 | 3439 | 3765 | 2639 | 2848 | 3073 | 3577 | 3453 | 2960 | 2839 | 3118 | 3170.02 | 3368 | 2972 | 0.882 | -0.18 | 0.04353782 | 0.14937535 | 0.0037 | 0.0079 | 0.0054 | 0.0054 | TRUE | 0.0712 |
| Lamtor3 | 1372 | 1193 | 1336 | 1431 | 985 | 1024 | 1457 | 1191 | 1341 | 1125 | 1060 | 1121 | 1215.99 | 1330 | 1102 | 0.829 | -0.271 | 0.00864208 | 0.04711474 | 0.0049 | 0.0096 | 0.0068 | 0.0068 | TRUE | 0.1001 |
| Lamtor4 | 731 | 672 | 644 | 991 | 643 | 567 | 776 | 671 | 647 | 779 | 692 | 621 | 697.63 | 698 | 697 | 1 | 0.001 | 0.99620365 | 0.99816555 | 0.0098 | 0.0116 | 0.0106 | 0.0106 | TRUE | 0.1283 |
| Lamtor5 | 1523 | 1827 | 1749 | 2210 | 1573 | 1629 | 1618 | 1825 | 1756 | 1737 | 1692 | 1783 | 1735.25 | 1733 | 1737 | 1.003 | 0.004 | 0.96369748 | 0.98168551 | 0.0016 | 0.0088 | 0.0043 | 0.0043 | TRUE | 0.0406 |
| Lancl1 | 1951 | 2255 | 2236 | 2768 | 2015 | 2148 | 2072 | 2252 | 2245 | 2176 | 2168 | 2351 | 2210.82 | 2190 | 2232 | 1.019 | 0.027 | 0.73668931 | 0.85901699 | 0.0017 | 0.0084 | 0.0041 | 0.0041 | TRUE | 0.0266 |
| Lancl2 | 1046 | 1161 | 1253 | 1921 | 1402 | 1298 | 1111 | 1160 | 1258 | 1510 | 1508 | 1421 | 1328.03 | 1176 | 1480 | 1.258 | 0.331 | 0.00014871 | 0.00188841 | 0.0017 | 0.0094 | 0.0047 | 0.0047 | TRUE | 0.0445 |
| Lancl3 | 33 | 14 | 12 | 38 | 26 | 9 | 35 | 14 | 12 | 30 | 28 | 10 | 21.46 | 20 | 23 | 1.118 | 0.161 | 0.78933876 | 0.891612 | 0.273 | 0.1604 | 0.2146 | 0.2146 | TRUE | 2.1103 |
| Lao1 | 0 | 0 | 0 | 0 | 0 | 0 | 0 | 0 | 0 | 0 | 0 | 0 | 0 | 0 | 0 | NA | NA | NA | NA | NA | NA | NA | NA | NA | NA |
| Lap3 | 14563 | 16701 | 17124 | 16623 | 12693 | 13788 | 15468 | 16679 | 17194 | 13068 | 13657 | 15093 | 15193.32 | 16447 | 13939 | 0.847 | -0.239 | 0.00543568 | 0.03324219 | 0.0041 | 0.0071 | 0.0053 | 0.0053 | TRUE | 0.0641 |
| Laptm4a | 7492 | 8889 | 8523 | 9960 | 7085 | 7663 | 7958 | 8878 | 8558 | 7830 | 7623 | 8388 | 8205.75 | 8465 | 7947 | 0.939 | -0.091 | 0.24323335 | 0.45993003 | 0.0026 | 0.0073 | 0.0043 | 0.0043 | TRUE | 0.0334 |
| Laptm4b | 6115 | 7944 | 8481 | 6758 | 5192 | 5816 | 6495 | 7934 | 8516 | 5313 | 5586 | 6367 | 6701.7 | 7648 | 5755 | 0.752 | -0.411 | 0.00093032 | 0.00841826 | 0.0135 | 0.0074 | 0.0109 | 0.0109 | TRUE | 0.2122 |
| Laptm5 | 1295 | 1774 | 1824 | 1517 | 1203 | 1262 | 1376 | 1772 | 1831 | 1193 | 1294 | 1381 | 1474.51 | 1660 | 1289 | 0.776 | -0.365 | 0.00540853 | 0.0331145 | 0.0128 | 0.0091 | 0.0117 | 0.0117 | TRUE | 0.2697 |
| Large | 972 | 1011 | 1022 | 1432 | 995 | 1007 | 1032 | 1010 | 1026 | 1126 | 1071 | 1102 | 1061.16 | 1023 | 1100 | 1.076 | 0.105 | 0.17910432 | 0.38024408 | 0 | 0.01 | 0.0035 | 0.0035 | TRUE | 0.0063 |
| Larp1 | 30835 | 30920 | 32926 | 36478 | 26578 | 25314 | 32752 | 30880 | 33061 | 28676 | 28596 | 27710 | 30279.35 | 32231 | 28327 | 0.879 | -0.186 | 0.0021243 | 0.01615455 | 8.00E-04 | 0.007 | 0.0026 | 0.0026 | TRUE | 0.0165 |
| Larp1b | 1158 | 1496 | 1506 | 1413 | 1073 | 1149 | 1230 | 1494 | 1512 | 1111 | 1154 | 1258 | 1293.21 | 1412 | 1174 | 0.831 | -0.267 | 0.01878289 | 0.08314447 | 0.0072 | 0.0095 | 0.0085 | 0.0085 | TRUE | 0.1525 |
| Larp4 | 9970 | 10698 | 10673 | 12678 | 9101 | 9179 | 10590 | 10684 | 10717 | 9966 | 9792 | 10048 | 10299.56 | 10664 | 9935 | 0.932 | -0.102 | 0.05279661 | 0.17035764 | 0 | 0.0072 | 0.0019 | 0.0019 | TRUE | 0.0019 |
| Larp4b | 5238 | 5067 | 5161 | 7209 | 5220 | 4900 | 5564 | 5060 | 5182 | 5667 | 5616 | 5364 | 5408.95 | 5269 | 5549 | 1.053 | 0.075 | 0.29230415 | 0.51317582 | 0.0014 | 0.0075 | 0.0035 | 0.0035 | TRUE | 0.0292 |
| Larp6 | 8 | 9 | 6 | 37 | 11 | 16 | 8 | 9 | 6 | 29 | 12 | 18 | 13.66 | 8 | 20 | 2.516 | 1.331 | 0.02534176 | 0.10272172 | 0 | 0.2481 | 0.1653 | 0.1653 | TRUE | 0.2928 |
| Larp7 | 2895 | 3018 | 2832 | 4953 | 3691 | 3134 | 3075 | 3014 | 2844 | 3894 | 3971 | 3431 | 3371.39 | 2978 | 3765 | 1.265 | 0.339 | 0.00011123 | 0.00149947 | 0.0034 | 0.0079 | 0.0052 | 0.0052 | TRUE | 0.073 |
| Lars | 10762 | 10039 | 10321 | 14367 | 10079 | 9643 | 11431 | 10026 | 10363 | 11294 | 10844 | 10556 | 10752.49 | 10607 | 10898 | 1.028 | 0.039 | 0.61807095 | 0.78412283 | 0.0025 | 0.0072 | 0.0044 | 0.0044 | TRUE | 0.0564 |
| Lars2 | 17751 | 20154 | 21989 | 18302 | 13496 | 16662 | 18855 | 20128 | 22079 | 14388 | 14521 | 18239 | 18034.93 | 20354 | 15716 | 0.772 | -0.373 | 0.00156077 | 0.01265626 | 0.0128 | 0.0071 | 0.01 | 0.01 | TRUE | 0.2411 |
| Las1l | 6580 | 6608 | 6667 | 8218 | 5986 | 5793 | 6989 | 6599 | 6694 | 6460 | 6441 | 6341 | 6587.54 | 6761 | 6414 | 0.949 | -0.076 | 0.20329741 | 0.41107778 | 3.00E-04 | 0.0074 | 0.0024 | 0.0024 | TRUE | 0.0106 |
| Lasp1 | 13383 | 15069 | 16093 | 18844 | 13473 | 13008 | 14215 | 15050 | 16159 | 14814 | 14496 | 14239 | 14828.8 | 15141 | 14516 | 0.959 | -0.061 | 0.40975165 | 0.62158344 | 0.0022 | 0.0071 | 0.0039 | 0.0039 | TRUE | 0.0423 |
| Lat | 240 | 327 | 309 | 363 | 254 | 223 | 255 | 327 | 310 | 285 | 273 | 244 | 282.42 | 297 | 267 | 0.901 | -0.15 | 0.32111481 | 0.54230889 | 0.0079 | 0.0186 | 0.013 | 0.013 | TRUE | 0.1729 |
| Lat2 | 52 | 36 | 27 | 41 | 34 | 28 | 55 | 36 | 27 | 32 | 37 | 31 | 36.29 | 39 | 33 | 0.842 | -0.248 | 0.50305757 | 0.69812793 | 0.0436 | 0.0977 | 0.0712 | 0.0712 | TRUE | 0.4962 |
| Lats1 | 1169 | 1375 | 1315 | 1397 | 1032 | 1123 | 1242 | 1373 | 1320 | 1098 | 1110 | 1229 | 1228.86 | 1312 | 1146 | 0.873 | -0.196 | 0.0324818 | 0.12155199 | 0.0024 | 0.0096 | 0.0052 | 0.0052 | TRUE | 0.0478 |
| Lats2 | 113 | 142 | 128 | 419 | 282 | 202 | 120 | 142 | 129 | 329 | 303 | 221 | 207.38 | 130 | 284 | 2.192 | 1.132 | 8.90E-09 | 4.26E-07 | 0.0218 | 0.0228 | 0.0223 | 0.0223 | TRUE | 0.42 |
| Lax1 | 11 | 6 | 7 | 25 | 17 | 13 | 12 | 6 | 7 | 20 | 18 | 14 | 12.81 | 8 | 17 | 2.129 | 1.09 | 0.0512841 | 0.16695882 | 0 | 0.264 | 0.1352 | 0.1352 | TRUE | 0.3839 |
| Layn | 86 | 106 | 92 | 93 | 90 | 83 | 91 | 106 | 92 | 73 | 97 | 91 | 91.73 | 96 | 87 | 0.895 | -0.159 | 0.4672077 | 0.66963926 | 0 | 0.0428 | 0.0237 | 0.0237 | TRUE | 0.2087 |
| Lbh | 211 | 200 | 195 | 345 | 291 | 292 | 224 | 200 | 196 | 271 | 313 | 320 | 253.94 | 207 | 301 | 1.456 | 0.542 | 0.00018562 | 0.00228537 | 0.0034 | 0.0199 | 0.0111 | 0.0111 | TRUE | 0.0924 |
| Lbp | 38 | 60 | 38 | 35 | 42 | 31 | 40 | 60 | 38 | 28 | 45 | 34 | 40.85 | 46 | 36 | 0.764 | -0.388 | 0.27097432 | 0.49096002 | 0.0403 | 0.0876 | 0.0644 | 0.0644 | TRUE | 0.545 |
| Lbr | 13734 | 15590 | 15455 | 16117 | 11660 | 12371 | 14588 | 15570 | 15518 | 12670 | 12545 | 13542 | 14072.27 | 15225 | 12919 | 0.848 | -0.237 | 0.00049286 | 0.00511741 | 0.0014 | 0.0071 | 0.0033 | 0.0033 | TRUE | 0.0217 |
| Lbx1 | 6 | 6 | 9 | 29 | 14 | 13 | 6 | 6 | 9 | 23 | 15 | 14 | 12.25 | 7 | 17 | 2.454 | 1.295 | 0.02528752 | 0.10256332 | 0 | 0.2759 | 0.1422 | 0.1422 | TRUE | 0.4498 |
| Lbx2 | 2 | 2 | 3 | 8 | 1 | 1 | 2 | 2 | 3 | 6 | 1 | 1 | 2.6 | 2 | 3 | 1.233 | 0.302 | 0.8086335 | NA | 0.2746 | 1.2745 | 0.7378 | 0.7378 | TRUE | 1.8144 |
| Lca5 | 704 | 833 | 853 | 1054 | 723 | 819 | 748 | 832 | 857 | 829 | 778 | 897 | 823.2 | 812 | 835 | 1.027 | 0.038 | 0.71504515 | 0.84534084 | 0.0037 | 0.0109 | 0.0068 | 0.0068 | TRUE | 0.0565 |
| Lca5l | 3 | 11 | 3 | 9 | 7 | 10 | 3 | 11 | 3 | 7 | 8 | 11 | 7.12 | 6 | 9 | 1.474 | 0.56 | 0.48181753 | 0.68077623 | 0.128 | 0.4694 | 0.3097 | 0.3097 | TRUE | 1.4798 |
| Lcat | 31 | 20 | 16 | 13 | 20 | 16 | 33 | 20 | 16 | 10 | 22 | 18 | 19.7 | 23 | 17 | 0.707 | -0.5 | 0.3235538 | 0.54485272 | 0.0895 | 0.1741 | 0.1327 | 0.1327 | TRUE | 0.8073 |
| Lce1a1 | 0 | 0 | 0 | 0 | 0 | 0 | 0 | 0 | 0 | 0 | 0 | 0 | 0 | 0 | 0 | NA | NA | NA | NA | NA | NA | NA | NA | NA | NA |
| Lce1a2 | 0 | 0 | 0 | 0 | 0 | 0 | 0 | 0 | 0 | 0 | 0 | 0 | 0 | 0 | 0 | NA | NA | NA | NA | NA | NA | NA | NA | NA | NA |
| Lce1b | 0 | 0 | 0 | 0 | 0 | 0 | 0 | 0 | 0 | 0 | 0 | 0 | 0 | 0 | 0 | NA | NA | NA | NA | NA | NA | NA | NA | NA | NA |
| Lce1c | 0 | 0 | 0 | 0 | 0 | 0 | 0 | 0 | 0 | 0 | 0 | 0 | 0 | 0 | 0 | NA | NA | NA | NA | NA | NA | NA | NA | NA | NA |
| Lce1d | 0 | 0 | 0 | 0 | 0 | 0 | 0 | 0 | 0 | 0 | 0 | 0 | 0 | 0 | 0 | NA | NA | NA | NA | NA | NA | NA | NA | NA | NA |
| Lce1e | 0 | 0 | 0 | 0 | 0 | 0 | 0 | 0 | 0 | 0 | 0 | 0 | 0 | 0 | 0 | NA | NA | NA | NA | NA | NA | NA | NA | NA | NA |
| Lce1f | 0 | 0 | 0 | 0 | 1 | 0 | 0 | 0 | 0 | 0 | 1 | 0 | 0.18 | 0 | 0 | 1.859 | 0.894 | 0.82650734 | NA | 0 | 18.3785 | 10 | 10 | TRUE | 0.5204 |
| Lce1g | 0 | 0 | 0 | 0 | 0 | 0 | 0 | 0 | 0 | 0 | 0 | 0 | 0 | 0 | 0 | NA | NA | NA | NA | NA | NA | NA | NA | NA | NA |
| Lce1h | 0 | 0 | 0 | 0 | 0 | 0 | 0 | 0 | 0 | 0 | 0 | 0 | 0 | 0 | 0 | NA | NA | NA | NA | NA | NA | NA | NA | NA | NA |
| Lce1i | 0 | 0 | 0 | 0 | 0 | 0 | 0 | 0 | 0 | 0 | 0 | 0 | 0 | 0 | 0 | NA | NA | NA | NA | NA | NA | NA | NA | NA | NA |
| Lce1k | 0 | 0 | 0 | 0 | 0 | 0 | 0 | 0 | 0 | 0 | 0 | 0 | 0 | 0 | 0 | NA | NA | NA | NA | NA | NA | NA | NA | NA | NA |
| Lce1l | 0 | 0 | 0 | 0 | 0 | 0 | 0 | 0 | 0 | 0 | 0 | 0 | 0 | 0 | 0 | NA | NA | NA | NA | NA | NA | NA | NA | NA | NA |
| Lce1m | 0 | 0 | 0 | 0 | 0 | 0 | 0 | 0 | 0 | 0 | 0 | 0 | 0 | 0 | 0 | NA | NA | NA | NA | NA | NA | NA | NA | NA | NA |
| Lce3a | 0 | 0 | 0 | 0 | 0 | 0 | 0 | 0 | 0 | 0 | 0 | 0 | 0 | 0 | 0 | NA | NA | NA | NA | NA | NA | NA | NA | NA | NA |
| Lce3b | 0 | 0 | 0 | 0 | 0 | 0 | 0 | 0 | 0 | 0 | 0 | 0 | 0 | 0 | 0 | NA | NA | NA | NA | NA | NA | NA | NA | NA | NA |
| Lce3c | 0 | 0 | 0 | 0 | 0 | 0 | 0 | 0 | 0 | 0 | 0 | 0 | 0 | 0 | 0 | NA | NA | NA | NA | NA | NA | NA | NA | NA | NA |
| Lce3d | 0 | 0 | 0 | 0 | 0 | 0 | 0 | 0 | 0 | 0 | 0 | 0 | 0 | 0 | 0 | NA | NA | NA | NA | NA | NA | NA | NA | NA | NA |
| Lce3e | 0 | 0 | 0 | 0 | 0 | 0 | 0 | 0 | 0 | 0 | 0 | 0 | 0 | 0 | 0 | NA | NA | NA | NA | NA | NA | NA | NA | NA | NA |
| Lce3f | 0 | 0 | 0 | 0 | 0 | 0 | 0 | 0 | 0 | 0 | 0 | 0 | 0 | 0 | 0 | NA | NA | NA | NA | NA | NA | NA | NA | NA | NA |
| Lce6a | 0 | 0 | 0 | 0 | 0 | 0 | 0 | 0 | 0 | 0 | 0 | 0 | 0 | 0 | 0 | NA | NA | NA | NA | NA | NA | NA | NA | NA | NA |
| Lck | 494 | 534 | 586 | 1062 | 836 | 665 | 525 | 533 | 588 | 835 | 899 | 728 | 684.79 | 549 | 821 | 1.496 | 0.581 | 1.02E-06 | 2.80E-05 | 0.0062 | 0.0117 | 0.0086 | 0.0086 | TRUE | 0.1134 |
| Lclat1 | 941 | 1012 | 1008 | 1313 | 988 | 1062 | 1000 | 1011 | 1012 | 1032 | 1063 | 1163 | 1046.68 | 1008 | 1086 | 1.077 | 0.107 | 0.22321282 | 0.43511464 | 0.0011 | 0.0101 | 0.0046 | 0.0046 | TRUE | 0.0451 |
| Lcmt1 | 2404 | 2659 | 2591 | 2835 | 2230 | 2257 | 2553 | 2656 | 2602 | 2229 | 2399 | 2471 | 2484.89 | 2604 | 2366 | 0.908 | -0.139 | 0.06479052 | 0.19609981 | 0.0012 | 0.0082 | 0.0037 | 0.0037 | TRUE | 0.0321 |
| Lcmt2 | 366 | 353 | 245 | 638 | 447 | 285 | 389 | 353 | 246 | 502 | 481 | 312 | 380.3 | 329 | 432 | 1.313 | 0.393 | 0.08540101 | 0.23523311 | 0.0567 | 0.0156 | 0.0349 | 0.0349 | TRUE | 0.671 |
| Lcn10 | 0 | 0 | 0 | 0 | 0 | 0 | 0 | 0 | 0 | 0 | 0 | 0 | 0 | 0 | 0 | NA | NA | NA | NA | NA | NA | NA | NA | NA | NA |
| Lcn11 | 0 | 0 | 0 | 0 | 0 | 0 | 0 | 0 | 0 | 0 | 0 | 0 | 0 | 0 | 0 | NA | NA | NA | NA | NA | NA | NA | NA | NA | NA |
| Lcn12 | 0 | 0 | 0 | 0 | 0 | 0 | 0 | 0 | 0 | 0 | 0 | 0 | 0 | 0 | 0 | NA | NA | NA | NA | NA | NA | NA | NA | NA | NA |
| Lcn2 | 0 | 0 | 0 | 0 | 0 | 0 | 0 | 0 | 0 | 0 | 0 | 0 | 0 | 0 | 0 | NA | NA | NA | NA | NA | NA | NA | NA | NA | NA |
| Lcn3 | 0 | 0 | 0 | 0 | 0 | 0 | 0 | 0 | 0 | 0 | 0 | 0 | 0 | 0 | 0 | NA | NA | NA | NA | NA | NA | NA | NA | NA | NA |
| Lcn4 | 0 | 0 | 0 | 0 | 0 | 0 | 0 | 0 | 0 | 0 | 0 | 0 | 0 | 0 | 0 | NA | NA | NA | NA | NA | NA | NA | NA | NA | NA |

| | | | | | | | | | | | | | | | | | | | | | | | | | |
|---|---|---|---|---|---|---|---|---|---|---|---|---|---|---|---|---|---|---|---|---|---|---|---|---|---|
| Lcn5 | 0 | 0 | 0 | 0 | 0 | 0 | 0 | 0 | 0 | 0 | 0 | 0 | 0 | 0 | 0 | NA | NA | NA | NA | NA | NA | NA | NA | NA | NA |
| Lcn6 | 0 | 0 | 0 | 0 | 0 | 0 | 0 | 0 | 0 | 0 | 0 | 0 | 0 | 0 | 0 | NA | NA | NA | NA | NA | NA | NA | NA | NA | NA |
| Lcn8 | 0 | 0 | 0 | 0 | 0 | 0 | 0 | 0 | 0 | 0 | 0 | 0 | 0 | 0 | 0 | NA | NA | NA | NA | NA | NA | NA | NA | NA | NA |
| Lcn9 | 0 | 0 | 0 | 2 | 0 | 0 | 0 | 0 | 0 | 2 | 0 | 0 | 0.26 | 0 | 1 | 2.886 | 1.529 | 0.70676491 | NA | 0 | 12.5792 | 10 | 10 | TRUE | 0.9669 |
| Lcor | 203 | 173 | 185 | 248 | 179 | 182 | 216 | 173 | 186 | 195 | 193 | 199 | 193.49 | 192 | 196 | 1.023 | 0.032 | 0.83592085 | 0.91664769 | 0 | 0.0239 | 0.0124 | 0.0124 | TRUE | 0.1307 |
| Lcorl | 709 | 1347 | 1479 | 821 | 723 | 869 | 753 | 1345 | 1485 | 645 | 778 | 951 | 993 | 1194 | 791 | 0.662 | -0.595 | 0.01309737 | 0.06414534 | 0.0786 | 0.0102 | 0.0404 | 0.0404 | TRUE | 1.2509 |
| Lcp1 | 73 | 50 | 38 | 178 | 158 | 125 | 78 | 50 | 38 | 140 | 170 | 137 | 102.07 | 55 | 149 | 2.702 | 1.434 | 2.69E-07 | 8.62E-06 | 0.0492 | 0.0392 | 0.0434 | 0.0434 | TRUE | 1.0269 |
| Lcp2 | 2 | 1 | 0 | 5 | 2 | 1 | 2 | 1 | 0 | 4 | 2 | 1 | 1.72 | 1 | 2 | 2.365 | 1.242 | 0.42319483 | NA | 0 | 1.926 | 1.0362 | 1.0362 | TRUE | 0.3322 |
| Lct | 30 | 27 | 18 | 42 | 41 | 30 | 32 | 27 | 18 | 33 | 44 | 33 | 31.15 | 26 | 37 | 1.428 | 0.514 | 0.16974769 | 0.36836277 | 0.0142 | 0.1127 | 0.0678 | 0.0678 | TRUE | 0.4136 |
| Lctl | 35 | 55 | 47 | 91 | 71 | 65 | 37 | 55 | 47 | 72 | 76 | 71 | 59.73 | 46 | 73 | 1.569 | 0.65 | 0.01359811 | 0.06576799 | 0 | 0.0621 | 0.0323 | 0.0323 | TRUE | 0.2328 |
| Ldb1 | 5098 | 5860 | 6010 | 5905 | 4748 | 4977 | 5415 | 5852 | 6035 | 4642 | 5109 | 5448 | 5416.8 | 5767 | 5066 | 0.878 | -0.187 | 0.03829011 | 0.13632318 | 0.0044 | 0.0075 | 0.0057 | 0.0057 | TRUE | 0.0659 |
| Ldb2 | 2 | 0 | 0 | 0 | 0 | 1 | 2 | 0 | 0 | 0 | 0 | 1 | 0.54 | 1 | 0 | 0.497 | -1.009 | 0.76081378 | NA | 0 | 6.1475 | 6.1929 | 6.1929 | TRUE | 0.9742 |
| Ldb3 | 739 | 987 | 1004 | 583 | 414 | 565 | 785 | 986 | 1008 | 458 | 445 | 618 | 716.84 | 926 | 507 | 0.547 | -0.87 | 1.20E-07 | 4.17E-06 | 0.0246 | 0.0115 | 0.0179 | 0.0179 | TRUE | 0.422 |
| Ldha | 62137 | 83858 | 85033 | 83026 | 64933 | 76682 | 66000 | 83750 | 85382 | 65269 | 69864 | 83941 | 75700.88 | 78377 | 73025 | 0.932 | -0.102 | 0.44763328 | 0.65334741 | 0.0162 | 0.0069 | 0.013 | 0.013 | TRUE | 0.2337 |
| Ldhal6b | 22 | 26 | 25 | 21 | 19 | 25 | 23 | 26 | 25 | 17 | 20 | 27 | 23.13 | 25 | 21 | 0.855 | -0.225 | 0.57868565 | 0.75588355 | 0 | 0.1494 | 0.0751 | 0.0751 | TRUE | 0.3131 |
| Ldhb | 2721 | 4527 | 4825 | 3194 | 2763 | 3516 | 2890 | 4521 | 4845 | 2511 | 2973 | 3849 | 3598.11 | 4085 | 3111 | 0.761 | -0.394 | 0.05826121 | 0.18235677 | 0.0563 | 0.0078 | 0.0308 | 0.0308 | TRUE | 0.7969 |
| Ldhc | 51 | 54 | 51 | 118 | 84 | 94 | 54 | 54 | 51 | 93 | 90 | 103 | 74.22 | 53 | 95 | 1.794 | 0.843 | 0.00020276 | 0.00247138 | 0 | 0.0513 | 0.0224 | 0.0224 | TRUE | 0.0437 |
| Ldhd | 47 | 117 | 110 | 108 | 82 | 126 | 50 | 117 | 110 | 85 | 88 | 138 | 98.05 | 92 | 104 | 1.118 | 0.161 | 0.64503191 | 0.80042827 | 0.1221 | 0.0405 | 0.0781 | 0.0781 | TRUE | 1.5288 |
| Ldlr | 7253 | 7721 | 8119 | 8226 | 6196 | 6304 | 7704 | 7711 | 8152 | 6467 | 6667 | 6901 | 7266.87 | 7856 | 6678 | 0.85 | -0.235 | 0.00031739 | 0.00356794 | 9.00E-04 | 0.0074 | 0.0029 | 0.0029 | TRUE | 0.0133 |
| Ldlrad1 | 0 | 0 | 0 | 0 | 0 | 0 | 0 | 0 | 0 | 0 | 0 | 0 | 0 | 0 | 0 | NA | NA | NA | NA | NA | NA | NA | NA | NA | NA |
| Ldlrad2 | 0 | 1 | 1 | 0 | 0 | 0 | 0 | 1 | 1 | 0 | 0 | 0 | 0.33 | 1 | 0 | 0.268 | -1.901 | 0.61401143 | NA | 0 | 9.8764 | 8.4772 | 8.4772 | TRUE | 0.2278 |
| Ldlrad3 | 4279 | 4621 | 4854 | 4280 | 3125 | 3382 | 4545 | 4615 | 4874 | 3365 | 3362 | 3702 | 4077.18 | 4678 | 3476 | 0.743 | -0.429 | 2.10E-08 | 9.33E-07 | 0.0019 | 0.0077 | 0.004 | 0.004 | TRUE | 0.039 |
| Ldlrad4 | 40 | 46 | 42 | 50 | 28 | 30 | 42 | 46 | 42 | 39 | 30 | 33 | 38.81 | 43 | 34 | 0.788 | -0.344 | 0.26302423 | 0.48235255 | 0 | 0.0918 | 0.042 | 0.042 | TRUE | 0.1497 |
| Ldlrap1 | 1464 | 1473 | 1535 | 1574 | 1211 | 1234 | 1555 | 1471 | 1541 | 1237 | 1303 | 1351 | 1409.76 | 1522 | 1297 | 0.851 | -0.232 | 0.00366568 | 0.02471785 | 7.00E-04 | 0.0092 | 0.0039 | 0.0039 | TRUE | 0.0202 |
| Ldoc1 | 208 | 132 | 108 | 186 | 152 | 126 | 221 | 132 | 108 | 146 | 164 | 138 | 151.48 | 154 | 149 | 0.972 | -0.042 | 0.8782133 | 0.93976544 | 0.0661 | 0.0287 | 0.0466 | 0.0466 | TRUE | 1.4598 |
| Ldoc1l | 320 | 296 | 268 | 515 | 389 | 339 | 340 | 296 | 269 | 405 | 419 | 371 | 349.85 | 302 | 398 | 1.322 | 0.402 | 0.00325507 | 0.02255713 | 0.0052 | 0.0163 | 0.0106 | 0.0106 | TRUE | 0.1382 |
| Leap2 | 0 | 1 | 1 | 0 | 0 | 0 | 0 | 1 | 1 | 0 | 0 | 0 | 0.33 | 1 | 0 | 0.268 | -1.901 | 0.61401143 | NA | 0 | 9.8764 | 8.4772 | 8.4772 | TRUE | 0.2278 |
| Lect1 | 0 | 4 | 2 | 0 | 2 | 0 | 0 | 4 | 2 | 0 | 2 | 0 | 1.36 | 2 | 1 | 0.344 | -1.538 | 0.47633942 | NA | 2.3147 | 2.4308 | 2.3998 | 2.3998 | TRUE | 0.3004 |
| Lect2 | 0 | 0 | 0 | 0 | 0 | 0 | 0 | 0 | 0 | 0 | 0 | 0 | 0 | 0 | 0 | NA | NA | NA | NA | NA | NA | NA | NA | NA | NA |
| Lef1 | 1179 | 1373 | 1869 | 891 | 709 | 896 | 1252 | 1371 | 1877 | 700 | 763 | 981 | 1157.38 | 1500 | 815 | 0.543 | -0.882 | 1.47E-06 | 3.83E-05 | 0.0393 | 0.0097 | 0.0232 | 0.0232 | TRUE | 0.5813 |
| Lefty1 | 659 | 816 | 1019 | 1382 | 829 | 641 | 700 | 815 | 1023 | 1086 | 892 | 702 | 869.69 | 846 | 893 | 1.057 | 0.08 | 0.67614493 | 0.81979608 | 0.0406 | 0.0107 | 0.0251 | 0.0251 | TRUE | 0.181 |
| Lefty2 | 1733 | 2314 | 2183 | 5136 | 3940 | 2876 | 1841 | 2311 | 2192 | 4038 | 4239 | 3148 | 2961.45 | 2115 | 3808 | 1.801 | 0.849 | 1.22E-09 | 7.31E-08 | 0.0177 | 0.008 | 0.0137 | 0.0137 | TRUE | 0.2788 |
| Lekr1 | 30 | 46 | 59 | 65 | 72 | 63 | 32 | 46 | 59 | 51 | 77 | 69 | 55.76 | 46 | 66 | 1.432 | 0.518 | 0.1103459 | 0.27886164 | 0.0477 | 0.066 | 0.0573 | 0.0573 | TRUE | 0.2785 |
| Lelp1 | 0 | 0 | 0 | 0 | 0 | 0 | 0 | 0 | 0 | 0 | 0 | 0 | 0 | 0 | 0 | NA | NA | NA | NA | NA | NA | NA | NA | NA | NA |
| Lemd1 | 5 | 7 | 8 | 9 | 2 | 3 | 5 | 7 | 8 | 7 | 2 | 3 | 5.47 | 7 | 4 | 0.633 | -0.659 | 0.43248324 | 0.64192787 | 0 | 0.6087 | 0.3197 | 0.3197 | TRUE | 0.8373 |
| Lemd2 | 885 | 1520 | 1406 | 1459 | 1255 | 1274 | 940 | 1518 | 1412 | 1147 | 1350 | 1395 | 1293.61 | 1290 | 1297 | 1.005 | 0.007 | 0.96719969 | 0.98366479 | 0.0359 | 0.0094 | 0.0222 | 0.0222 | TRUE | 0.6748 |
| Lemd3 | 1770 | 1768 | 1759 | 2449 | 1744 | 1674 | 1880 | 1766 | 1766 | 1925 | 1876 | 1832 | 1841.02 | 1804 | 1878 | 1.042 | 0.059 | 0.42361515 | 0.63389547 | 4.00E-04 | 0.0087 | 0.0033 | 0.0033 | TRUE | 0.0163 |
| Lenep | 41 | 47 | 38 | 69 | 52 | 46 | 44 | 47 | 38 | 54 | 56 | 50 | 48.2 | 43 | 53 | 1.249 | 0.321 | 0.24370881 | 0.46027726 | 0 | 0.0753 | 0.0334 | 0.0334 | TRUE | 0.0728 |
| Leng1 | 290 | 323 | 337 | 414 | 295 | 289 | 308 | 323 | 338 | 325 | 317 | 316 | 321.37 | 323 | 319 | 0.99 | -0.014 | 0.90440037 | 0.9525344 | 0 | 0.0172 | 0.007 | 0.007 | TRUE | 0.0197 |
| Leng8 | 5760 | 5942 | 5928 | 6172 | 6066 | 5610 | 6118 | 5934 | 5952 | 4852 | 6527 | 6141 | 5920.74 | 6001 | 5840 | 0.973 | -0.04 | 0.73661396 | 0.85899263 | 0.0119 | 0.0075 | 0.0099 | 0.0099 | TRUE | 0.2685 |
| Leng9 | 13 | 18 | 15 | 15 | 18 | 18 | 14 | 18 | 15 | 12 | 19 | 20 | 16.28 | 16 | 17 | 1.072 | 0.101 | 0.8348789 | 0.91588718 | 0 | 0.2092 | 0.1067 | 0.1067 | TRUE | 0.4398 |
| Leo1 | 3891 | 3194 | 3321 | 4828 | 3492 | 3039 | 4133 | 3190 | 3335 | 3795 | 3757 | 3327 | 3589.45 | 3553 | 3626 | 1.021 | 0.03 | 0.80329976 | 0.89842886 | 0.0121 | 0.0078 | 0.0102 | 0.0102 | TRUE | 0.2482 |
| Lep | 0 | 1 | 0 | 0 | 0 | 0 | 0 | 1 | 0 | 0 | 0 | 0 | 0.17 | 0 | 0 | 0.49 | -1.029 | 0.80087 | NA | 0 | 19.7991 | 10 | 10 | TRUE | 0.405 |
| Lepr | 138 | 235 | 239 | 285 | 244 | 263 | 147 | 235 | 240 | 224 | 263 | 288 | 232.62 | 207 | 258 | 1.244 | 0.315 | 0.14606864 | 0.33475656 | 0.0383 | 0.0211 | 0.0294 | 0.0294 | TRUE | 0.7042 |
| Lepre1 | 2820 | 2840 | 2635 | 2814 | 2020 | 2096 | 2995 | 2836 | 2646 | 2212 | 2173 | 2294 | 2526.24 | 2826 | 2226 | 0.788 | -0.344 | 1.70E-05 | 0.00031185 | 0.0017 | 0.0082 | 0.0042 | 0.0042 | TRUE | 0.0377 |
| Leprel1 | 28 | 27 | 19 | 70 | 37 | 21 | 30 | 27 | 19 | 55 | 40 | 23 | 32.27 | 25 | 39 | 1.569 | 0.65 | 0.12895094 | 0.30890254 | 0.0899 | 0.109 | 0.0996 | 0.0996 | TRUE | 0.1402 |
| Leprel2 | 197 | 185 | 168 | 277 | 232 | 155 | 209 | 185 | 169 | 218 | 250 | 170 | 199.96 | 188 | 213 | 1.133 | 0.181 | 0.34352627 | 0.56358897 | 0.019 | 0.0234 | 0.0212 | 0.0212 | TRUE | 0.2815 |
| Leprel4 | 634 | 717 | 733 | 477 | 374 | 398 | 673 | 716 | 736 | 375 | 402 | 436 | 556.43 | 708 | 404 | 0.569 | -0.812 | 1.05E-13 | 1.25E-11 | 0.0017 | 0.0128 | 0.0067 | 0.0067 | TRUE | 0.0525 |
| Leprot | 2726 | 3698 | 3693 | 4447 | 3399 | 3870 | 2895 | 3693 | 3708 | 3496 | 3657 | 4236 | 3614.37 | 3432 | 3796 | 1.106 | 0.145 | 0.25496358 | 0.47316067 | 0.014 | 0.0078 | 0.0114 | 0.0114 | TRUE | 0.2273 |
| Leprotl1 | 1717 | 1981 | 1879 | 2484 | 1788 | 1831 | 1824 | 1978 | 1887 | 1953 | 1924 | 2004 | 1928.29 | 1896 | 1960 | 1.034 | 0.048 | 0.5168578 | 0.70855117 | 5.00E-04 | 0.0086 | 0.0034 | 0.0034 | TRUE | 0.0174 |
| Letm1 | 3433 | 3257 | 3171 | 4575 | 3264 | 3001 | 3646 | 3253 | 3184 | 3597 | 3512 | 3285 | 3412.78 | 3361 | 3465 | 1.031 | 0.044 | 0.61088301 | 0.77958142 | 0.003 | 0.0079 | 0.0051 | 0.0051 | TRUE | 0.0669 |
| Letm2 | 73 | 143 | 196 | 128 | 92 | 92 | 78 | 143 | 197 | 101 | 99 | 101 | 119.58 | 139 | 100 | 0.719 | -0.475 | 0.13191136 | 0.31290508 | 0.095 | 0.0345 | 0.0631 | 0.0631 | TRUE | 0.1757 |
| Letmd1 | 317 | 368 | 383 | 482 | 352 | 349 | 337 | 368 | 385 | 379 | 379 | 382 | 371.41 | 363 | 380 | 1.046 | 0.065 | 0.57261512 | 0.7512479 | 0 | 0.0158 | 0.0069 | 0.0069 | TRUE | 0.045 |
| Lfng | 85 | 130 | 158 | 106 | 88 | 109 | 90 | 130 | 159 | 83 | 95 | 119 | 112.68 | 126 | 99 | 0.782 | -0.355 | 0.17931164 | 0.38052398 | 0.047 | 0.0361 | 0.0415 | 0.0415 | TRUE | 0.227 |
| Lgals1 | 1235 | 1363 | 1393 | 1367 | 975 | 1015 | 1312 | 1361 | 1399 | 1075 | 1049 | 1111 | 1217.75 | 1357 | 1078 | 0.794 | -0.332 | 2.64E-05 | 0.00045224 | 0 | 0.0096 | 0.0037 | 0.0037 | TRUE | 0.0102 |
| Lgals12 | 69 | 71 | 73 | 53 | 40 | 42 | 73 | 71 | 73 | 42 | 43 | 46 | 58.03 | 72 | 44 | 0.6 | -0.738 | 0.00313871 | 0.02200168 | 0 | 0.0637 | 0.0267 | 0.0267 | TRUE | 0.0173 |
| Lgals2 | 3 | 2 | 2 | 2 | 1 | 2 | 3 | 2 | 2 | 2 | 1 | 2 | 2 | 2 | 2 | 0.673 | -0.572 | 0.65339145 | NA | 0 | 1.6501 | 0.6514 | 0.6514 | TRUE | 0.083 |
| Lgals3 | 445 | 419 | 418 | 444 | 312 | 339 | 473 | 418 | 420 | 349 | 336 | 371 | 394.44 | 437 | 352 | 0.806 | -0.312 | 0.00846979 | 0.04644901 | 0 | 0.0153 | 0.0076 | 0.0076 | TRUE | 0.0586 |
| Lgals3bp | 28 | 30 | 27 | 39 | 38 | 26 | 30 | 30 | 27 | 31 | 41 | 28 | 31.14 | 29 | 33 | 1.149 | 0.201 | 0.56395283 | 0.74482198 | 0 | 0.1127 | 0.0548 | 0.0548 | TRUE | 0.2554 |
| Lgals4 | 6 | 8 | 2 | 5 | 10 | 14 | 6 | 8 | 2 | 4 | 11 | 15 | 7.73 | 5 | 10 | 1.809 | 0.855 | 0.30269725 | 0.5235952 | 0.2691 | 0.433 | 0.3527 | 0.3527 | TRUE | 0.2245 |
| Lgals6 | 0 | 0 | 0 | 0 | 0 | 0 | 0 | 0 | 0 | 0 | 0 | 0 | 0 | 0 | 0 | NA | NA | NA | NA | NA | NA | NA | NA | NA | NA |
| Lgals7 | 13 | 11 | 15 | 28 | 27 | 23 | 14 | 11 | 15 | 22 | 29 | 25 | 19.35 | 13 | 25 | 1.906 | 0.93 | 0.03505277 | 0.12818279 | 0 | 0.1772 | 0.0826 | 0.0826 | TRUE | 0.102 |
| Lgals8 | 384 | 470 | 473 | 548 | 393 | 449 | 408 | 469 | 475 | 431 | 423 | 492 | 449.56 | 451 | 449 | 0.994 | -0.009 | 0.93985086 | 0.96977897 | 0.0045 | 0.0142 | 0.0089 | 0.0089 | TRUE | 0.0798 |
| Lgals9 | 140 | 92 | 120 | 163 | 136 | 103 | 149 | 92 | 120 | 128 | 146 | 113 | 124.72 | 120 | 129 | 1.074 | 0.102 | 0.65861456 | 0.8075619 | 0.0278 | 0.0333 | 0.0306 | 0.0306 | TRUE | 0.2011 |
| Lgalsl | 621 | 1188 | 1044 | 1451 | 1012 | 1166 | 660 | 1186 | 1048 | 1141 | 1089 | 1276 | 1066.71 | 965 | 1169 | 1.211 | 0.276 | 0.16214824 | 0.35742865 | 0.0467 | 0.01 | 0.0272 | 0.0272 | TRUE | 0.9111 |
| Lgi1 | 0 | 1 | 0 | 4 | 2 | 0 | 0 | 1 | 0 | 3 | 2 | 0 | 1.05 | 0 | 2 | 4.9 | 2.293 | 0.31302341 | NA | 0 | 3.1469 | 2.4479 | 2.4479 | TRUE | 0.3114 |
| Lgi2 | 342 | 505 | 462 | 574 | 406 | 448 | 363 | 504 | 464 | 451 | 437 | 490 | 451.66 | 444 | 459 | 1.034 | 0.049 | 0.7428415 | 0.86268287 | 0.0133 | 0.0142 | 0.0137 | 0.0137 | TRUE | 0.2896 |
| Lgi3 | 48 | 55 | 42 | 68 | 51 | 52 | 51 | 55 | 42 | 53 | 55 | 57 | 52.22 | 49 | 55 | 1.115 | 0.157 | 0.55773536 | 0.7401735 | 0 | 0.07 | 0.0323 | 0.0323 | TRUE | 0.1333 |
| Lgi4 | 3 | 7 | 5 | 4 | 3 | 2 | 3 | 7 | 5 | 3 | 3 | 2 | 3.96 | 5 | 3 | 0.565 | -0.824 | 0.38313395 | NA | 0 | 0.8388 | 0.3722 | 0.3722 | TRUE | 0.1413 |

| | | | | | | | | | | | | | | | | | | | | | | | | | |
|---|---|---|---|---|---|---|---|---|---|---|---|---|---|---|---|---|---|---|---|---|---|---|---|---|---|
| Lgmn | 3201 | 3447 | 3341 | 3691 | 2586 | 2729 | 3400 | 3443 | 3355 | 2902 | 2782 | 2987 | 3144.76 | 3399 | 2890 | 0.85 | -0.234 | 0.00040257 | 0.00436675 | 2.00E-04 | 0.008 | 0.0028 | 0.0028 | TRUE | 0.0127 |
| Lgr4 | 4902 | 4327 | 4825 | 5793 | 4323 | 4282 | 5207 | 4321 | 4845 | 4554 | 4651 | 4687 | 4710.94 | 4791 | 4631 | 0.967 | -0.049 | 0.58330294 | 0.75875085 | 0.0038 | 0.0076 | 0.0055 | 0.0055 | TRUE | 0.0896 |
| Lgr5 | 4 | 1 | 1 | 10 | 8 | 3 | 4 | 1 | 1 | 8 | 9 | 3 | 4.33 | 2 | 7 | 3.21 | 1.683 | 0.10833968 | NA | 0.0772 | 0.767 | 0.4698 | 0.4698 | TRUE | 0.7296 |
| Lgr6 | 190 | 197 | 167 | 175 | 120 | 141 | 202 | 197 | 168 | 138 | 129 | 154 | 164.55 | 189 | 140 | 0.743 | -0.428 | 0.01198225 | 0.06007058 | 0 | 0.0269 | 0.0147 | 0.0147 | TRUE | 0.1034 |
| Lgsn | 0 | 1 | 0 | 2 | 0 | 0 | 0 | 0 | 1 | 0 | 2 | 0 | 0.43 | 0 | 1 | 1.493 | 0.578 | 0.86962266 | NA | 0 | 7.6954 | 7.0545 | 7.0545 | TRUE | 0.9718 |
| Lhb | 2 | 3 | 2 | 0 | 1 | 0 | 2 | 3 | 2 | 0 | 1 | 0 | 1.37 | 2 | 0 | 0.147 | -2.768 | 0.13272706 | NA | 0 | 2.4161 | 1.2273 | 1.2273 | TRUE | 0.5204 |
| Lhcgr | 0 | 1 | 0 | 0 | 0 | 0 | 0 | 1 | 0 | 0 | 0 | 0 | 0.17 | 0 | 0 | 0.49 | -1.029 | 0.80087 | NA | 0 | 19.7991 | 10 | 10 | TRUE | 0.405 |
| Lhfp | 238 | 247 | 252 | 346 | 233 | 213 | 253 | 247 | 253 | 272 | 251 | 233 | 251.39 | 251 | 252 | 1.007 | 0.01 | 0.94021716 | 0.96979647 | 0 | 0.02 | 0.0092 | 0.0092 | TRUE | 0.0581 |
| Lhfpl1 | 0 | 0 | 0 | 0 | 2 | 2 | 0 | 0 | 0 | 0 | 2 | 2 | 0.72 | 0 | 1 | 7.604 | 2.927 | 0.31659722 | NA | 0 | 4.5602 | 4.8112 | 4.8112 | TRUE | 0.6686 |
| Lhfpl2 | 263 | 261 | 264 | 257 | 183 | 231 | 279 | 261 | 265 | 202 | 197 | 253 | 242.82 | 268 | 217 | 0.808 | -0.308 | 0.04469912 | 0.15216804 | 0.0052 | 0.0205 | 0.0127 | 0.0127 | TRUE | 0.217 |
| Lhfpl3 | 1 | 0 | 2 | 1 | 0 | 1 | 1 | 0 | 2 | 1 | 0 | 1 | 0.83 | 1 | 1 | 0.619 | -0.693 | 0.74976439 | NA | 0 | 3.9992 | 2.1283 | 2.1283 | TRUE | 0.1555 |
| Lhfpl4 | 51 | 86 | 79 | 82 | 49 | 98 | 54 | 86 | 79 | 64 | 53 | 107 | 73.97 | 73 | 75 | 1.019 | 0.028 | 0.93352753 | 0.96713012 | 0.0819 | 0.0514 | 0.066 | 0.066 | TRUE | 1.266 |
| Lhfpl5 | 61 | 108 | 92 | 84 | 60 | 60 | 65 | 108 | 92 | 66 | 65 | 66 | 76.88 | 88 | 66 | 0.74 | -0.435 | 0.09529864 | 0.25308136 | 0.0214 | 0.0498 | 0.0357 | 0.0357 | TRUE | 0.3222 |
| Lhpp | 1111 | 1032 | 1105 | 889 | 626 | 571 | 1180 | 1031 | 1110 | 699 | 674 | 625 | 886.29 | 1107 | 666 | 0.602 | -0.731 | 3.05E-13 | 3.38E-11 | 0.0028 | 0.0106 | 0.0061 | 0.0061 | TRUE | 0.0434 |
| Lhx1 | 8 | 2 | 3 | 4 | 8 | 1 | 8 | 2 | 3 | 3 | 9 | 1 | 4.39 | 4 | 4 | 0.95 | -0.074 | 0.944853 | NA | 0.4506 | 0.757 | 0.6026 | 0.6026 | TRUE | 0.7922 |
| Lhx2 | 15 | 54 | 43 | 126 | 92 | 73 | 16 | 54 | 43 | 99 | 99 | 80 | 65.16 | 38 | 93 | 2.454 | 1.295 | 0.00084966 | 0.00785052 | 0.138 | 0.0575 | 0.0898 | 0.0898 | TRUE | 1.8022 |
| Lhx3 | 6 | 8 | 9 | 3 | 10 | 2 | 6 | 8 | 9 | 2 | 11 | 2 | 6.45 | 8 | 5 | 0.642 | -0.639 | 0.45947883 | 0.66378277 | 0.2135 | 0.5176 | 0.3753 | 0.3753 | TRUE | 1.8119 |
| Lhx4 | 9 | 4 | 8 | 6 | 3 | 9 | 10 | 4 | 8 | 5 | 3 | 10 | 6.56 | 7 | 6 | 0.819 | -0.288 | 0.7182166 | 0.84737412 | 0.0772 | 0.5088 | 0.3052 | 0.3052 | TRUE | 0.6976 |
| Lhx5 | 8 | 14 | 14 | 14 | 13 | 13 | 8 | 14 | 14 | 11 | 14 | 14 | 12.63 | 12 | 13 | 1.064 | 0.09 | 0.86672229 | 0.93405835 | 0 | 0.2678 | 0.1277 | 0.1277 | TRUE | 0.2643 |
| Lhx6 | 94 | 116 | 139 | 179 | 157 | 131 | 100 | 116 | 140 | 141 | 169 | 143 | 134.72 | 119 | 151 | 1.272 | 0.347 | 0.08219187 | 0.22887185 | 0.0107 | 0.0314 | 0.0212 | 0.0212 | TRUE | 0.2464 |
| Lhx8 | 2 | 9 | 10 | 8 | 4 | 6 | 2 | 9 | 10 | 6 | 4 | 7 | 6.39 | 7 | 6 | 0.81 | -0.303 | 0.71111394 | 0.84271464 | 0 | 0.5228 | 0.3257 | 0.3257 | TRUE | 0.9439 |
| Lhx9 | 0 | 1 | 2 | 2 | 1 | 3 | 0 | 1 | 2 | 2 | 1 | 3 | 1.49 | 1 | 2 | 1.94 | 0.956 | 0.55183554 | NA | 0 | 2.2182 | 1.1069 | 1.1069 | TRUE | 0.2991 |
| Lias | 2261 | 2489 | 2469 | 2826 | 2058 | 2098 | 2402 | 2486 | 2479 | 2222 | 2214 | 2297 | 2349.83 | 2456 | 2244 | 0.914 | -0.13 | 0.04830663 | 0.16025697 | 0 | 0.0083 | 0.0027 | 0.0027 | TRUE | 0.005 |
| Lif | 69 | 115 | 110 | 148 | 81 | 80 | 73 | 115 | 110 | 116 | 87 | 88 | 98.28 | 99 | 97 | 0.978 | -0.032 | 0.89990754 | 0.95148767 | 0.0328 | 0.0404 | 0.0367 | 0.0367 | TRUE | 0.509 |
| Lifr | 7327 | 7410 | 7989 | 7023 | 5383 | 5706 | 7783 | 7400 | 8022 | 5521 | 5792 | 6246 | 6793.94 | 7735 | 5853 | 0.757 | -0.403 | 2.99E-07 | 9.47E-06 | 0.0026 | 0.0074 | 0.0043 | 0.0043 | TRUE | 0.0421 |
| Lig1 | 12903 | 14615 | 15047 | 20084 | 15438 | 15197 | 13705 | 14596 | 15109 | 15788 | 16610 | 16636 | 15407.41 | 14470 | 16345 | 1.129 | 0.176 | 0.01082385 | 0.05578884 | 0.0015 | 0.0071 | 0.0034 | 0.0034 | TRUE | 0.0261 |
| Lig3 | 6329 | 6153 | 6582 | 9727 | 6961 | 6462 | 6723 | 6145 | 6609 | 7647 | 7490 | 7074 | 6947.75 | 6492 | 7404 | 1.141 | 0.19 | 0.00904928 | 0.04886175 | 0.0017 | 0.0074 | 0.0037 | 0.0037 | TRUE | 0.0267 |
| Lig4 | 1542 | 1407 | 1329 | 1840 | 1349 | 1223 | 1638 | 1405 | 1334 | 1446 | 1451 | 1339 | 1435.7 | 1459 | 1412 | 0.968 | -0.047 | 0.66024022 | 0.80873787 | 0.0061 | 0.0092 | 0.0074 | 0.0074 | TRUE | 0.1377 |
| Lilra5 | 0 | 1 | 0 | 1 | 3 | 0 | 0 | 1 | 0 | 1 | 3 | 0 | 0.84 | 0 | 1 | 3.632 | 1.861 | 0.45208766 | NA | 0 | 3.9503 | 3.0431 | 3.0431 | TRUE | 0.5202 |
| Lilra6 | 0 | 0 | 0 | 0 | 0 | 0 | 0 | 0 | 0 | 0 | 0 | 0 | 0 | 0 | 0 | NA | NA | NA | NA | NA | NA | NA | NA | NA | NA |
| Lilrb3 | 0 | 0 | 0 | 1 | 0 | 1 | 0 | 0 | 0 | 1 | 0 | 1 | 0.31 | 0 | 1 | 3.348 | 1.743 | 0.65064889 | NA | 0 | 10.5167 | 8.8968 | 8.8968 | TRUE | 0.2082 |
| Lilrb4 | 0 | 0 | 0 | 0 | 0 | 0 | 0 | 0 | 0 | 0 | 0 | 0 | 0 | 0 | 0 | NA | NA | NA | NA | NA | NA | NA | NA | NA | NA |
| Lim2 | 10 | 2 | 5 | 4 | 6 | 4 | 11 | 2 | 5 | 3 | 6 | 4 | 5.27 | 6 | 4 | 0.787 | -0.345 | 0.70572779 | 0.83911335 | 0.1832 | 0.6321 | 0.4077 | 0.4077 | TRUE | 0.3671 |
| Lima1 | 12537 | 17146 | 18081 | 8920 | 7083 | 9713 | 13317 | 17124 | 18155 | 7012 | 7621 | 10632 | 12310.18 | 16199 | 8422 | 0.52 | -0.944 | 5.46E-08 | 2.14E-06 | 0.0387 | 0.0072 | 0.0216 | 0.0216 | TRUE | 0.6437 |
| Limch1 | 162 | 203 | 189 | 319 | 221 | 212 | 172 | 203 | 190 | 251 | 238 | 232 | 214.2 | 188 | 240 | 1.277 | 0.353 | 0.01510038 | 0.07112887 | 0 | 0.0223 | 0.0105 | 0.0105 | TRUE | 0.0595 |
| Limd1 | 1101 | 1254 | 1188 | 1857 | 1359 | 1321 | 1169 | 1252 | 1193 | 1460 | 1462 | 1446 | 1330.46 | 1205 | 1456 | 1.208 | 0.273 | 0.00030621 | 0.00347198 | 0 | 0.0094 | 0.0034 | 0.0034 | TRUE | 0.0143 |
| Limd2 | 1893 | 2447 | 2487 | 2368 | 1559 | 1681 | 2011 | 2444 | 2497 | 1862 | 1677 | 1840 | 2055.13 | 2317 | 1793 | 0.774 | -0.37 | 0.00079395 | 0.00744035 | 0.0075 | 0.0085 | 0.0083 | 0.0083 | TRUE | 0.1618 |
| Lime1 | 421 | 400 | 357 | 545 | 475 | 362 | 447 | 399 | 358 | 428 | 511 | 396 | 423.48 | 401 | 445 | 1.108 | 0.148 | 0.31522768 | 0.53603999 | 0.0122 | 0.0147 | 0.0133 | 0.0133 | TRUE | 0.1894 |
| Limk1 | 2307 | 2145 | 2117 | 3113 | 2250 | 2090 | 2450 | 2142 | 2126 | 2447 | 2421 | 2288 | 2312.38 | 2239 | 2385 | 1.065 | 0.092 | 0.30547062 | 0.52637196 | 0.0033 | 0.0083 | 0.0053 | 0.0053 | TRUE | 0.082 |
| Limk2 | 4314 | 4883 | 5037 | 4697 | 3489 | 3865 | 4582 | 4877 | 5058 | 3692 | 3754 | 4231 | 4365.64 | 4839 | 3892 | 0.804 | -0.314 | 0.00030845 | 0.00349005 | 0.0037 | 0.0077 | 0.0052 | 0.0052 | TRUE | 0.0701 |
| Lims1 | 3692 | 4518 | 4848 | 4717 | 3503 | 3995 | 3922 | 4512 | 4868 | 3708 | 3769 | 4373 | 4191.99 | 4434 | 3950 | 0.891 | -0.167 | 0.13859033 | 0.3230815 | 0.0098 | 0.0077 | 0.0089 | 0.0089 | TRUE | 0.1243 |
| Lims2 | 53 | 109 | 81 | 72 | 56 | 61 | 56 | 109 | 81 | 57 | 60 | 67 | 71.69 | 82 | 61 | 0.742 | -0.43 | 0.14580926 | 0.33433441 | 0.0443 | 0.0529 | 0.0486 | 0.0486 | TRUE | 0.16 |
| Lin28a | 14159 | 15766 | 17159 | 10619 | 7992 | 8824 | 15039 | 15746 | 17229 | 8348 | 8599 | 9659 | 12436.74 | 16005 | 8869 | 0.554 | -0.852 | 2.44E-20 | 6.13E-18 | 0.0054 | 0.0072 | 0.006 | 0.006 | TRUE | 0.0742 |
| Lin28b | 1546 | 1358 | 1492 | 2150 | 1487 | 1312 | 1642 | 1356 | 1498 | 1690 | 1600 | 1436 | 1537.13 | 1499 | 1575 | 1.052 | 0.073 | 0.50662548 | 0.70098228 | 0.0073 | 0.009 | 0.008 | 0.008 | TRUE | 0.084 |
| Lin37 | 415 | 464 | 470 | 621 | 426 | 401 | 441 | 463 | 472 | 488 | 458 | 439 | 460.27 | 459 | 462 | 1.008 | 0.012 | 0.91338035 | 0.95732555 | 0 | 0.0141 | 0.0062 | 0.0062 | TRUE | 0.0307 |
| Lin52 | 653 | 989 | 908 | 716 | 586 | 667 | 694 | 988 | 912 | 563 | 630 | 730 | 752.76 | 865 | 641 | 0.741 | -0.433 | 0.00770938 | 0.04320978 | 0.0222 | 0.0113 | 0.0177 | 0.0177 | TRUE | 0.3539 |
| Lin54 | 2253 | 2429 | 2638 | 2618 | 2000 | 2212 | 2393 | 2426 | 2649 | 2058 | 2152 | 2421 | 2349.85 | 2489 | 2210 | 0.888 | -0.172 | 0.07068914 | 0.20711668 | 0.0046 | 0.0083 | 0.0061 | 0.0061 | TRUE | 0.084 |
| Lin7a | 5 | 4 | 1 | 4 | 7 | 9 | 5 | 4 | 1 | 3 | 8 | 10 | 5.14 | 3 | 7 | 1.97 | 0.978 | 0.30070082 | 0.52111343 | 0.1799 | 0.6479 | 0.4219 | 0.4219 | TRUE | 0.369 |
| Lin7b | 22 | 30 | 33 | 46 | 38 | 35 | 23 | 30 | 33 | 36 | 41 | 38 | 33.64 | 29 | 38 | 1.329 | 0.411 | 0.21721691 | 0.42935653 | 0 | 0.1048 | 0.0494 | 0.0494 | TRUE | 0.1714 |
| Lin7c | 6190 | 6655 | 6553 | 8659 | 6103 | 6156 | 6575 | 6646 | 6580 | 6807 | 6566 | 6739 | 6652.23 | 6600 | 6704 | 1.016 | 0.023 | 0.6852196 | 0.82611973 | 0 | 0.0074 | 0.0021 | 0.0021 | TRUE | 0.0039 |
| Lin9 | 828 | 803 | 905 | 1281 | 934 | 875 | 879 | 802 | 909 | 1007 | 1005 | 958 | 926.66 | 863 | 990 | 1.147 | 0.198 | 0.03160498 | 0.11948077 | 0.0013 | 0.0105 | 0.005 | 0.005 | TRUE | 0.0464 |
| Lingo1 | 161 | 200 | 230 | 240 | 210 | 198 | 171 | 200 | 231 | 189 | 226 | 217 | 205.51 | 201 | 211 | 1.046 | 0.065 | 0.70635165 | 0.8396577 | 0.0107 | 0.0229 | 0.0167 | 0.0167 | TRUE | 0.1908 |
| Lingo2 | 0 | 0 | 0 | 0 | 1 | 0 | 0 | 0 | 0 | 0 | 1 | 0 | 0.18 | 0 | 0 | 1.859 | 0.894 | 0.82650734 | NA | 0 | 18.3785 | 10 | 10 | TRUE | 0.5204 |
| Lingo3 | 55 | 90 | 65 | 78 | 56 | 79 | 58 | 90 | 65 | 61 | 60 | 86 | 70.27 | 71 | 69 | 0.97 | -0.044 | 0.87690791 | 0.93932608 | 0.0328 | 0.0538 | 0.0435 | 0.0435 | TRUE | 0.4796 |
| Lingo4 | 0 | 0 | 0 | 0 | 0 | 0 | 0 | 0 | 0 | 0 | 0 | 0 | 0 | 0 | 0 | NA | NA | NA | NA | NA | NA | NA | NA | NA | NA |
| Lins | 413 | 485 | 441 | 610 | 456 | 432 | 439 | 484 | 443 | 480 | 491 | 473 | 468.15 | 455 | 481 | 1.056 | 0.079 | 0.45457936 | 0.65959798 | 0 | 0.0139 | 0.0059 | 0.0059 | TRUE | 0.0365 |
| Lipa | 1520 | 1635 | 1656 | 1531 | 1064 | 1231 | 1615 | 1633 | 1663 | 1204 | 1145 | 1348 | 1434.35 | 1637 | 1232 | 0.752 | -0.41 | 6.84E-06 | 0.0001441 | 0.0026 | 0.0092 | 0.0053 | 0.0053 | TRUE | 0.0791 |
| Lipc | 7 | 8 | 12 | 23 | 10 | 11 | 7 | 8 | 12 | 18 | 11 | 12 | 11.39 | 9 | 14 | 1.502 | 0.587 | 0.31358567 | 0.53393859 | 0 | 0.2961 | 0.1532 | 0.1532 | TRUE | 0.431 |
| Lipe | 382 | 403 | 432 | 686 | 544 | 484 | 406 | 402 | 434 | 539 | 585 | 530 | 482.73 | 414 | 551 | 1.331 | 0.413 | 0.00011912 | 0.00158273 | 0 | 0.0137 | 0.0062 | 0.0062 | TRUE | 0.0328 |
| Lipf | 0 | 0 | 0 | 0 | 1 | 0 | 0 | 0 | 0 | 0 | 1 | 0 | 0.18 | 0 | 0 | 1.859 | 0.894 | 0.82650734 | NA | 0 | 18.3785 | 10 | 10 | TRUE | 0.5204 |
| Lipg | 104 | 122 | 136 | 93 | 94 | 80 | 110 | 122 | 137 | 73 | 101 | 88 | 105.11 | 123 | 87 | 0.706 | -0.503 | 0.01992601 | 0.08660045 | 0.0089 | 0.0382 | 0.0239 | 0.0239 | TRUE | 0.2177 |
| Liph | 1807 | 2307 | 2237 | 2239 | 1870 | 2054 | 1919 | 2304 | 2246 | 1760 | 2012 | 2248 | 2081.69 | 2156 | 2007 | 0.93 | -0.105 | 0.39193612 | 0.60563505 | 0.0105 | 0.0085 | 0.0103 | 0.0103 | TRUE | 0.1418 |
| Lipi | 0 | 0 | 0 | 0 | 0 | 0 | 0 | 0 | 0 | 0 | 0 | 0 | 0 | 0 | 0 | NA | NA | NA | NA | NA | NA | NA | NA | NA | NA |
| Lipk | 0 | 0 | 0 | 1 | 0 | 0 | 0 | 0 | 0 | 1 | 0 | 0 | 0.13 | 0 | 0 | 1.859 | 0.894 | 0.82650734 | NA | 0 | 25.1515 | 10 | 10 | TRUE | 0.2563 |
| Lipm | 1 | 1 | 2 | 1 | 1 | 0 | 1 | 1 | 2 | 1 | 1 | 0 | 0.99 | 1 | 1 | 0.465 | -1.104 | 0.55512019 | NA | 0 | 3.3396 | 1.3765 | 1.3765 | TRUE | 0.1869 |
| Lipn | 1 | 0 | 0 | 0 | 0 | 0 | 1 | 0 | 0 | 0 | 0 | 0 | 0.18 | 0 | 0 | 0.49 | -1.029 | 0.80087 | NA | 0 | 18.6165 | 10 | 10 | TRUE | 0.4614 |
| Lipo1 | 195 | 213 | 172 | 533 | 349 | 297 | 207 | 213 | 173 | 419 | 376 | 325 | 285.36 | 198 | 373 | 1.894 | 0.921 | 8.49E-09 | 4.08E-07 | 0.011 | 0.0184 | 0.0145 | 0.0145 | TRUE | 0.132 |
| Lipt1 | 511 | 592 | 564 | 534 | 465 | 466 | 543 | 591 | 566 | 420 | 500 | 510 | 521.76 | 567 | 477 | 0.839 | -0.253 | 0.03526556 | 0.12872214 | 0.0048 | 0.0132 | 0.0085 | 0.0085 | TRUE | 0.1343 |
| Lipt2 | 515 | 551 | 471 | 611 | 424 | 382 | 547 | 550 | 473 | 480 | 456 | 418 | 487.49 | 523 | 451 | 0.864 | -0.211 | 0.07931671 | 0.22334997 | 0.0041 | 0.0137 | 0.0083 | 0.0083 | TRUE | 0.0835 |

| | | | | | | | | | | | | | | | | | | | | | | | | | |
|---|---|---|---|---|---|---|---|---|---|---|---|---|---|---|---|---|---|---|---|---|---|---|---|---|---|
| Litaf | 2087 | 2295 | 2392 | 2371 | 1746 | 1946 | 2217 | 2292 | 2402 | 1864 | 1879 | 2130 | 2130.55 | 2304 | 1958 | 0.849 | -0.236 | 0.00783819 | 0.04365617 | 0.003 | 0.0084 | 0.0052 | 0.0052 | TRUE | 0.0713 |
| Lix1 | 23 | 22 | 23 | 30 | 21 | 20 | 24 | 22 | 23 | 24 | 23 | 22 | 22.93 | 23 | 23 | 0.982 | -0.027 | 0.94456057 | 0.97241382 | 0 | 0.1506 | 0.0628 | 0.0628 | TRUE | 0.0128 |
| Lix1l | 1044 | 1223 | 1250 | 979 | 737 | 863 | 1109 | 1221 | 1255 | 770 | 793 | 945 | 1015.46 | 1195 | 836 | 0.698 | -0.518 | 6.29E-06 | 0.00013447 | 0.007 | 0.0101 | 0.0085 | 0.0085 | TRUE | 0.1528 |
| Llgl1 | 7629 | 7692 | 7762 | 10560 | 7798 | 7824 | 8103 | 7682 | 7794 | 8301 | 8390 | 8565 | 8139.25 | 7860 | 8419 | 1.071 | 0.099 | 0.09276454 | 0.24873893 | 3.00E-04 | 0.0073 | 0.0024 | 0.0024 | TRUE | 0.009 |
| Llgl2 | 15604 | 15181 | 15465 | 16334 | 12007 | 11706 | 16574 | 15161 | 15528 | 12841 | 12919 | 12814 | 14306.24 | 15754 | 12858 | 0.816 | -0.293 | 4.38E-06 | 9.84E-05 | 0.001 | 0.0071 | 0.0029 | 0.0029 | TRUE | 0.0253 |
| Llph | 1032 | 908 | 869 | 1355 | 989 | 870 | 1096 | 907 | 873 | 1065 | 1064 | 952 | 992.87 | 959 | 1027 | 1.072 | 0.101 | 0.39882328 | 0.61201859 | 0.008 | 0.0102 | 0.0093 | 0.0093 | TRUE | 0.1874 |
| Lman1 | 8159 | 8637 | 8698 | 9698 | 7210 | 7789 | 8666 | 8626 | 8734 | 7624 | 7758 | 8526 | 8322.25 | 8675 | 7969 | 0.918 | -0.123 | 0.08629031 | 0.23710292 | 0.0017 | 0.0073 | 0.0036 | 0.0036 | TRUE | 0.0455 |
| Lman1l | 0 | 0 | 0 | 0 | 1 | 0 | 0 | 0 | 0 | 0 | 1 | 0 | 0.18 | 0 | 0 | 1.859 | 0.894 | 0.82650734 | NA | 0 | 18.3785 | 10 | 10 | TRUE | 0.5204 |
| Lman2 | 5310 | 5597 | 5504 | 7012 | 5078 | 5123 | 5640 | 5590 | 5527 | 5512 | 5464 | 5608 | 5556.73 | 5586 | 5528 | 0.99 | -0.015 | 0.78958497 | 0.8916894 | 0 | 0.0075 | 0.0021 | 0.0021 | TRUE | 0.0019 |
| Lman2l | 478 | 507 | 538 | 751 | 585 | 559 | 508 | 506 | 540 | 590 | 629 | 612 | 564.33 | 518 | 610 | 1.177 | 0.236 | 0.01623068 | 0.07511451 | 0 | 0.0127 | 0.0051 | 0.0051 | TRUE | 0.0164 |
| Lmbr1 | 601 | 590 | 556 | 998 | 681 | 678 | 638 | 589 | 558 | 785 | 733 | 742 | 674.22 | 595 | 753 | 1.267 | 0.341 | 0.000703 | 0.00679398 | 0.0013 | 0.0118 | 0.0058 | 0.0058 | TRUE | 0.0466 |
| Lmbr1l | 1627 | 1508 | 1448 | 1945 | 1497 | 1396 | 1728 | 1506 | 1454 | 1529 | 1611 | 1528 | 1559.33 | 1563 | 1556 | 0.996 | -0.006 | 0.94861741 | 0.9742329 | 0.004 | 0.009 | 0.006 | 0.006 | TRUE | 0.103 |
| Lmbrd1 | 751 | 735 | 772 | 1054 | 752 | 740 | 798 | 734 | 775 | 829 | 809 | 810 | 792.44 | 769 | 816 | 1.062 | 0.086 | 0.3261499 | 0.54728555 | 0 | 0.0111 | 0.0043 | 0.0043 | TRUE | 0.0188 |
| Lmbrd2 | 1031 | 933 | 1206 | 1277 | 935 | 899 | 1095 | 932 | 1211 | 1004 | 1006 | 984 | 1038.64 | 1079 | 998 | 0.925 | -0.113 | 0.33210711 | 0.55255612 | 0.0078 | 0.0101 | 0.0088 | 0.0088 | TRUE | 0.1718 |
| Lmcd1 | 359 | 473 | 430 | 181 | 122 | 195 | 381 | 472 | 432 | 142 | 131 | 213 | 295.41 | 428 | 162 | 0.378 | -1.405 | 3.45E-12 | 3.26E-10 | 0.0335 | 0.0181 | 0.0251 | 0.0251 | TRUE | 0.7789 |
| Lmf1 | 386 | 562 | 564 | 520 | 407 | 482 | 410 | 561 | 566 | 409 | 438 | 528 | 485.32 | 512 | 458 | 0.892 | -0.164 | 0.32915535 | 0.55024931 | 0.0228 | 0.0137 | 0.0184 | 0.0184 | TRUE | 0.355 |
| Lmf2 | 1664 | 1630 | 1698 | 2244 | 1799 | 1662 | 1767 | 1628 | 1705 | 1764 | 1936 | 1819 | 1769.89 | 1700 | 1840 | 1.082 | 0.113 | 0.16039244 | 0.3548001 | 0.0014 | 0.0088 | 0.0041 | 0.0041 | TRUE | 0.0251 |
| Lmln | 323 | 384 | 397 | 452 | 337 | 371 | 343 | 384 | 399 | 355 | 363 | 406 | 374.88 | 375 | 375 | 0.997 | -0.004 | 0.97477191 | 0.98685208 | 0.0028 | 0.0157 | 0.0088 | 0.0088 | TRUE | 0.063 |
| Lmna | 706 | 904 | 1036 | 1113 | 798 | 936 | 750 | 903 | 1040 | 875 | 859 | 1025 | 908.52 | 898 | 920 | 1.024 | 0.034 | 0.81567486 | 0.90603732 | 0.0169 | 0.0105 | 0.0139 | 0.0139 | TRUE | 0.2165 |
| Lmnb1 | 6587 | 6741 | 6932 | 10454 | 7760 | 7149 | 6997 | 6732 | 6960 | 8218 | 8349 | 7826 | 7513.74 | 6896 | 8131 | 1.179 | 0.238 | 0.00013575 | 0.00176514 | 6.00E-04 | 0.0073 | 0.0027 | 0.0027 | TRUE | 0.0131 |
| Lmnb2 | 4522 | 4068 | 4229 | 5456 | 4277 | 3911 | 4803 | 4063 | 4246 | 4289 | 4602 | 4281 | 4380.72 | 4371 | 4391 | 1.005 | 0.007 | 0.94215487 | 0.97094617 | 0.0044 | 0.0077 | 0.0057 | 0.0057 | TRUE | 0.0911 |
| Lmo1 | 86 | 115 | 101 | 90 | 89 | 80 | 91 | 115 | 101 | 71 | 96 | 88 | 93.62 | 102 | 85 | 0.821 | -0.284 | 0.20422984 | 0.41254009 | 0.0084 | 0.0421 | 0.0254 | 0.0254 | TRUE | 0.2222 |
| Lmo2 | 56 | 113 | 94 | 121 | 92 | 52 | 59 | 113 | 94 | 95 | 99 | 57 | 86.29 | 89 | 84 | 0.943 | -0.085 | 0.78906372 | 0.891612 | 0.0806 | 0.0451 | 0.0619 | 0.0619 | TRUE | 0.4344 |
| Lmo3 | 1 | 1 | 0 | 0 | 1 | 0 | 1 | 1 | 0 | 0 | 1 | 0 | 0.52 | 1 | 0 | 0.51 | -0.972 | 0.72797553 | NA | 0 | 6.3084 | 3.8879 | 3.8879 | TRUE | 0.5204 |
| Lmo4 | 254 | 237 | 216 | 508 | 328 | 242 | 270 | 237 | 217 | 399 | 353 | 265 | 290.09 | 241 | 339 | 1.41 | 0.496 | 0.0077789 | 0.04346046 | 0.0245 | 0.0183 | 0.0214 | 0.0214 | TRUE | 0.3409 |
| Lmo7 | 2051 | 2042 | 2101 | 2275 | 1656 | 1464 | 2179 | 2039 | 2110 | 1788 | 1782 | 1603 | 1916.71 | 2109 | 1724 | 0.818 | -0.29 | 0.00044864 | 0.00477783 | 0.0016 | 0.0086 | 0.0044 | 0.0044 | TRUE | 0.0453 |
| Lmod1 | 137 | 132 | 114 | 99 | 75 | 79 | 146 | 132 | 114 | 78 | 81 | 86 | 106.14 | 131 | 82 | 0.624 | -0.679 | 0.00075788 | 0.00717053 | 0 | 0.0379 | 0.0194 | 0.0194 | TRUE | 0.1202 |
| Lmod2 | 1 | 0 | 1 | 1 | 2 | 2 | 1 | 0 | 1 | 1 | 2 | 2 | 1.2 | 1 | 2 | 2.452 | 1.294 | 0.46611437 | NA | 0 | 2.7547 | 1.2342 | 1.2342 | TRUE | 0.2562 |
| Lmod3 | 4 | 2 | 2 | 11 | 3 | 5 | 4 | 2 | 2 | 9 | 3 | 5 | 4.27 | 3 | 6 | 2.144 | 1.1 | 0.25416226 | NA | 0 | 0.779 | 0.4011 | 0.4011 | TRUE | 0.2044 |
| Lmtk2 | 6024 | 4734 | 4964 | 6946 | 4791 | 4527 | 6399 | 4728 | 4984 | 5460 | 5155 | 4956 | 5280.26 | 5370 | 5190 | 0.967 | -0.049 | 0.6972046 | 0.83403726 | 0.0127 | 0.0075 | 0.0112 | 0.0112 | TRUE | 0.3418 |
| Lmtk3 | 414 | 304 | 313 | 417 | 326 | 326 | 440 | 304 | 314 | 328 | 351 | 357 | 348.84 | 353 | 345 | 0.979 | -0.031 | 0.85732019 | 0.92964277 | 0.0197 | 0.0163 | 0.0181 | 0.0181 | TRUE | 0.5311 |
| Lmx1a | 6 | 19 | 19 | 18 | 24 | 18 | 6 | 19 | 19 | 14 | 26 | 20 | 17.35 | 15 | 20 | 1.33 | 0.411 | 0.45783915 | 0.66241987 | 0.1196 | 0.1968 | 0.1617 | 0.1617 | TRUE | 0.536 |
| Lmx1b | 22 | 44 | 40 | 94 | 77 | 61 | 23 | 44 | 40 | 74 | 83 | 67 | 55.17 | 36 | 75 | 2.074 | 1.053 | 0.00064142 | 0.00632282 | 0.0199 | 0.0666 | 0.0477 | 0.0477 | TRUE | 0.639 |
| Lnp | 2045 | 2430 | 2584 | 3245 | 2363 | 2535 | 2172 | 2427 | 2595 | 2551 | 2542 | 2775 | 2510.33 | 2398 | 2623 | 1.093 | 0.129 | 0.17642163 | 0.3767831 | 0.0046 | 0.0082 | 0.0062 | 0.0062 | TRUE | 0.0819 |
| Lnpep | 1013 | 842 | 950 | 1740 | 1213 | 1169 | 1076 | 841 | 954 | 1368 | 1305 | 1280 | 1137.24 | 957 | 1318 | 1.378 | 0.462 | 3.96E-05 | 0.00063518 | 0.0068 | 0.0098 | 0.0082 | 0.0082 | TRUE | 0.1407 |
| Lnx1 | 62 | 98 | 112 | 63 | 75 | 97 | 66 | 98 | 112 | 50 | 81 | 106 | 85.43 | 92 | 79 | 0.85 | -0.234 | 0.48310874 | 0.68129391 | 0.0939 | 0.0455 | 0.0686 | 0.0686 | TRUE | 0.3003 |
| Lnx2 | 1169 | 1260 | 1258 | 1294 | 1003 | 1017 | 1242 | 1258 | 1263 | 1017 | 1079 | 1113 | 1162.15 | 1254 | 1070 | 0.852 | -0.231 | 0.00427945 | 0.02786014 | 0 | 0.0097 | 0.0038 | 0.0038 | TRUE | 0.0232 |
| LOC1000389 | 1 | 0 | 0 | 2 | 2 | 0 | 1 | 0 | 0 | 2 | 2 | 0 | 0.8 | 0 | 1 | 3.417 | 1.773 | 0.46651319 | NA | 0 | 4.1368 | 2.8785 | 2.8785 | TRUE | 0.4614 |
| LOC1000407 | 0 | 0 | 0 | 0 | 0 | 0 | 0 | 0 | 0 | 0 | 0 | 0 | 0 | 0 | 0 | NA | NA | NA | NA | NA | NA | NA | NA | NA | NA |
| LOC1000433 | 0 | 0 | 0 | 0 | 0 | 0 | 0 | 0 | 0 | 0 | 0 | 0 | 0 | 0 | 0 | NA | NA | NA | NA | NA | NA | NA | NA | NA | NA |
| LOC1000488 | 0 | 0 | 0 | 0 | 0 | 0 | 0 | 0 | 0 | 0 | 0 | 0 | 0 | 0 | 0 | NA | NA | NA | NA | NA | NA | NA | NA | NA | NA |
| LOC1005029 | 0 | 0 | 0 | 0 | 0 | 0 | 0 | 0 | 0 | 0 | 0 | 0 | 0 | 0 | 0 | NA | NA | NA | NA | NA | NA | NA | NA | NA | NA |
| LOC1005032 | 18 | 10 | 13 | 62 | 31 | 27 | 19 | 10 | 13 | 49 | 33 | 30 | 25.63 | 14 | 37 | 2.676 | 1.42 | 0.00139165 | 0.0115423 | 0.0454 | 0.1354 | 0.0927 | 0.0927 | TRUE | 0.6076 |
| LOC1005034 | 0 | 2 | 1 | 7 | 4 | 7 | 0 | 2 | 1 | 6 | 4 | 8 | 3.41 | 1 | 6 | 5.732 | 2.519 | 0.04179691 | NA | 0 | 0.9725 | 0.5158 | 0.5158 | TRUE | 0.3544 |
| LOC1005046 | 5 | 4 | 8 | 7 | 4 | 3 | 5 | 4 | 8 | 6 | 4 | 3 | 5.07 | 6 | 4 | 0.764 | -0.389 | 0.64058899 | 0.79767643 | 0 | 0.6565 | 0.3006 | 0.3006 | TRUE | 0.2695 |
| LOC1005047 | 14 | 14 | 15 | 21 | 2 | 10 | 15 | 14 | 15 | 17 | 2 | 11 | 12.25 | 15 | 10 | 0.687 | -0.542 | 0.42826807 | 0.63819387 | 0.2217 | 0.2758 | 0.2535 | 0.2535 | TRUE | 0.4928 |
| LOC1008616 | 14 | 17 | 22 | 36 | 22 | 18 | 15 | 17 | 22 | 28 | 24 | 20 | 20.94 | 18 | 24 | 1.336 | 0.418 | 0.32809577 | 0.54915913 | 0 | 0.1643 | 0.0828 | 0.0828 | TRUE | 0.2047 |
| LOC1008619 | 0 | 0 | 0 | 0 | 0 | 0 | 0 | 0 | 0 | 0 | 0 | 0 | 0 | 0 | 0 | NA | NA | NA | NA | NA | NA | NA | NA | NA | NA |
| LOC106740 | 184 | 256 | 244 | 257 | 213 | 204 | 195 | 256 | 245 | 202 | 229 | 223 | 225.11 | 232 | 218 | 0.938 | -0.093 | 0.56734847 | 0.74717936 | 0.0077 | 0.0215 | 0.0144 | 0.0144 | TRUE | 0.2071 |
| LOC171588 | 0 | 1 | 0 | 6 | 2 | 1 | 0 | 1 | 0 | 5 | 2 | 1 | 1.49 | 0 | 3 | 7.382 | 2.884 | 0.12222921 | NA | 0 | 2.2125 | 1.3281 | 1.3281 | TRUE | 0.364 |
| LOC666331 | 80 | 78 | 92 | 69 | 62 | 93 | 85 | 78 | 92 | 54 | 67 | 102 | 79.67 | 85 | 74 | 0.867 | -0.206 | 0.46956221 | 0.67161831 | 0.0433 | 0.0483 | 0.0458 | 0.0458 | TRUE | 0.9393 |
| Loh12cr1 | 420 | 464 | 431 | 608 | 426 | 372 | 446 | 463 | 433 | 478 | 458 | 407 | 447.63 | 447 | 448 | 1.003 | 0.004 | 0.97430868 | 0.98671997 | 0.0018 | 0.0143 | 0.0074 | 0.0074 | TRUE | 0.0714 |
| Lonp1 | 14363 | 16025 | 16389 | 16708 | 13404 | 13921 | 15256 | 16004 | 16456 | 13135 | 14422 | 15239 | 15085.29 | 15905 | 14265 | 0.897 | -0.157 | 0.05539474 | 0.17607534 | 0.0034 | 0.0071 | 0.0048 | 0.0048 | TRUE | 0.059 |
| Lonp2 | 2561 | 3303 | 3354 | 4548 | 3249 | 3214 | 2720 | 3299 | 3368 | 3575 | 3496 | 3518 | 3329.33 | 3129 | 3530 | 1.128 | 0.174 | 0.08519172 | 0.23469762 | 0.0061 | 0.0079 | 0.007 | 0.007 | TRUE | 0.1583 |
| Lonrf1 | 1584 | 2108 | 2172 | 2859 | 2139 | 2186 | 1682 | 2105 | 2181 | 2248 | 2301 | 2393 | 2151.76 | 1989 | 2314 | 1.163 | 0.217 | 0.05836423 | 0.18250795 | 0.0089 | 0.0084 | 0.009 | 0.009 | TRUE | 0.2198 |
| Lonrf2 | 130 | 66 | 75 | 121 | 81 | 46 | 138 | 66 | 75 | 95 | 87 | 50 | 85.32 | 93 | 77 | 0.837 | -0.257 | 0.47370234 | 0.67458712 | 0.1211 | 0.0455 | 0.0812 | 0.0812 | TRUE | 1.6982 |
| Lonrf3 | 685 | 881 | 987 | 1365 | 1096 | 1028 | 728 | 880 | 991 | 1073 | 1179 | 1125 | 996.02 | 866 | 1126 | 1.299 | 0.377 | 0.0035131 | 0.02390938 | 0.0112 | 0.0102 | 0.011 | 0.011 | TRUE | 0.2318 |
| Lor | 4 | 7 | 6 | 16 | 20 | 9 | 4 | 7 | 6 | 13 | 22 | 10 | 10.2 | 6 | 15 | 2.531 | 1.34 | 0.04110048 | 0.14360584 | 0 | 0.3298 | 0.1879 | 0.1879 | TRUE | 0.6596 |
| Lox | 6 | 9 | 13 | 17 | 13 | 14 | 6 | 9 | 13 | 13 | 14 | 15 | 11.85 | 9 | 14 | 1.495 | 0.58 | 0.30162046 | 0.5222481 | 0 | 0.2849 | 0.1388 | 0.1388 | TRUE | 0.3663 |
| Loxhd1 | 4 | 8 | 11 | 7 | 5 | 6 | 4 | 8 | 11 | 6 | 5 | 7 | 6.79 | 8 | 6 | 0.745 | -0.424 | 0.56650891 | 0.74644767 | 0 | 0.4922 | 0.2447 | 0.2447 | TRUE | 0.187 |
| Loxl1 | 131 | 210 | 215 | 303 | 227 | 220 | 139 | 210 | 216 | 238 | 244 | 241 | 214.67 | 188 | 241 | 1.279 | 0.355 | 0.06465855 | 0.19588834 | 0.0215 | 0.0222 | 0.0219 | 0.0219 | TRUE | 0.5536 |
| Loxl2 | 1582 | 2061 | 2327 | 2240 | 1672 | 1884 | 1680 | 2058 | 2337 | 1761 | 1799 | 2062 | 1949.58 | 2025 | 1874 | 0.925 | -0.112 | 0.41053927 | 0.62194077 | 0.0155 | 0.0086 | 0.0129 | 0.0129 | TRUE | 0.261 |
| Loxl3 | 79 | 67 | 65 | 55 | 39 | 40 | 84 | 67 | 65 | 43 | 42 | 44 | 57.51 | 72 | 43 | 0.598 | -0.743 | 0.00467679 | 0.02979091 | 0 | 0.0642 | 0.0312 | 0.0312 | TRUE | 0.1844 |
| Loxl4 | 60 | 90 | 50 | 85 | 62 | 61 | 64 | 90 | 50 | 67 | 67 | 67 | 67.35 | 68 | 67 | 0.982 | -0.026 | 0.92728501 | 0.96421012 | 0.0285 | 0.0558 | 0.0425 | 0.0425 | TRUE | 0.4803 |
| Lpar1 | 1046 | 771 | 654 | 1855 | 1234 | 1007 | 1111 | 770 | 657 | 1458 | 1328 | 1102 | 1071 | 846 | 1296 | 1.533 | 0.617 | 0.00164273 | 0.01315186 | 0.0473 | 0.01 | 0.0267 | 0.0267 | TRUE | 0.8918 |
| Lpar2 | 1517 | 1939 | 2091 | 1597 | 1259 | 1570 | 1611 | 1936 | 2100 | 1255 | 1355 | 1719 | 1662.68 | 1882 | 1443 | 0.766 | -0.385 | 0.01035341 | 0.05416054 | 0.0221 | 0.0089 | 0.0156 | 0.0156 | TRUE | 0.3353 |
| Lpar3 | 74 | 69 | 58 | 94 | 80 | 63 | 79 | 69 | 58 | 74 | 86 | 69 | 72.45 | 69 | 76 | 1.113 | 0.154 | 0.52555675 | 0.71494576 | 0 | 0.0524 | 0.0287 | 0.0287 | TRUE | 0.1563 |
| Lpar4 | 273 | 121 | 111 | 237 | 144 | 105 | 290 | 121 | 111 | 186 | 155 | 115 | 163.08 | 174 | 152 | 0.876 | -0.192 | 0.59915229 | 0.77143379 | 0.1704 | 0.0271 | 0.0897 | 0.0897 | TRUE | 2.1992 |
| Lpar5 | 197 | 199 | 157 | 313 | 229 | 211 | 209 | 199 | 158 | 246 | 246 | 231 | 214.84 | 189 | 241 | 1.28 | 0.357 | 0.02684881 | 0.10693422 | 0.0054 | 0.0222 | 0.0139 | 0.0139 | TRUE | 0.2231 |

| | | | | | | | | | | | | | | | | | | | | | | | | |
|---|---|---|---|---|---|---|---|---|---|---|---|---|---|---|---|---|---|---|---|---|---|---|---|---|
| Lpar6 | 71 | 58 | 68 | 193 | 131 | 109 | 75 | 58 | 68 | 152 | 141 | 119 | 102.27 | 67 | 137 | 2.051 | 1.036 | 1.91E-06 | 4.85E-05 | 0 | 0.0391 | 0.023 | 0.023 | TRUE | 0.1321 |
| Lpcat1 | 2853 | 2739 | 2506 | 3201 | 2323 | 2333 | 3030 | 2735 | 2516 | 2516 | 2499 | 2554 | 2641.96 | 2760 | 2523 | 0.914 | -0.13 | 0.15567615 | 0.34817417 | 0.0039 | 0.0082 | 0.0056 | 0.0056 | TRUE | 0.0884 |
| Lpcat2 | 432 | 429 | 349 | 333 | 222 | 318 | 459 | 428 | 350 | 262 | 239 | 348 | 347.75 | 412 | 283 | 0.685 | -0.546 | 0.00274838 | 0.0198024 | 0.0254 | 0.0164 | 0.021 | 0.021 | TRUE | 0.4455 |
| Lpcat2b | 0 | 0 | 0 | 0 | 0 | 0 | 0 | 0 | 0 | 0 | 0 | 0 | 0 | 0 | 0 | NA | NA | NA | NA | NA | NA | NA | NA | NA | NA |
| Lpcat3 | 6097 | 6089 | 5994 | 6556 | 4758 | 4993 | 6476 | 6081 | 6019 | 5154 | 5119 | 5466 | 5719.1 | 6192 | 5246 | 0.847 | -0.239 | 0.00056127 | 0.00570012 | 0.0012 | 0.0075 | 0.0033 | 0.0033 | TRUE | 0.0196 |
| Lpcat4 | 876 | 970 | 921 | 1151 | 884 | 948 | 930 | 969 | 925 | 905 | 951 | 1038 | 952.95 | 941 | 965 | 1.024 | 0.034 | 0.71922027 | 0.8479218 | 0.0018 | 0.0104 | 0.0052 | 0.0052 | TRUE | 0.0521 |
| Lpgat1 | 3139 | 4279 | 4745 | 5151 | 3882 | 3959 | 3334 | 4273 | 4764 | 4049 | 4177 | 4334 | 4155.33 | 4124 | 4187 | 1.015 | 0.022 | 0.8709314 | 0.93648357 | 0.0166 | 0.0077 | 0.0124 | 0.0124 | TRUE | 0.3414 |
| Lphn1 | 14684 | 14307 | 14517 | 12483 | 9818 | 9364 | 15597 | 14289 | 14577 | 9813 | 10564 | 10250 | 12514.87 | 14821 | 10209 | 0.689 | -0.538 | 1.60E-14 | 2.07E-12 | 0.0016 | 0.0072 | 0.0035 | 0.0035 | TRUE | 0.0257 |
| Lphn2 | 1004 | 1107 | 1223 | 1501 | 1059 | 1100 | 1066 | 1106 | 1228 | 1180 | 1139 | 1204 | 1153.92 | 1133 | 1174 | 1.036 | 0.051 | 0.57637709 | 0.75418558 | 0.0022 | 0.0098 | 0.0052 | 0.0052 | TRUE | 0.0641 |
| Lphn3 | 109 | 56 | 50 | 216 | 129 | 78 | 116 | 56 | 50 | 170 | 139 | 85 | 102.65 | 74 | 131 | 1.782 | 0.834 | 0.02565828 | 0.10355231 | 0.151 | 0.039 | 0.09 | 0.09 | TRUE | 0.6265 |
| Lpin1 | 2844 | 3914 | 4067 | 2239 | 1868 | 2462 | 3021 | 3909 | 4084 | 1760 | 2010 | 2695 | 2913.08 | 3671 | 2155 | 0.587 | -0.769 | 1.06E-05 | 0.00020969 | 0.0367 | 0.008 | 0.0216 | 0.0216 | TRUE | 0.581 |
| Lpin2 | 1174 | 1285 | 1283 | 1900 | 1382 | 1359 | 1247 | 1283 | 1288 | 1494 | 1487 | 1488 | 1381.14 | 1273 | 1490 | 1.17 | 0.227 | 0.001573 | 0.01272272 | 0 | 0.0093 | 0.003 | 0.003 | TRUE | 0.0038 |
| Lpin3 | 682 | 888 | 940 | 1161 | 925 | 887 | 724 | 887 | 944 | 913 | 995 | 971 | 905.67 | 852 | 960 | 1.126 | 0.171 | 0.16255271 | 0.35788551 | 0.009 | 0.0105 | 0.0097 | 0.0097 | TRUE | 0.2021 |
| Lpl | 1087 | 1701 | 1752 | 2226 | 1674 | 1646 | 1155 | 1699 | 1759 | 1750 | 1801 | 1802 | 1660.9 | 1538 | 1784 | 1.16 | 0.214 | 0.17100538 | 0.36966788 | 0.0252 | 0.0089 | 0.0171 | 0.0171 | TRUE | 0.5706 |
| Lpo | 8 | 5 | 9 | 19 | 5 | 6 | 8 | 5 | 9 | 15 | 5 | 7 | 8.24 | 7 | 9 | 1.219 | 0.286 | 0.69306904 | 0.83159723 | 0.1108 | 0.4069 | 0.2563 | 0.2563 | TRUE | 1.3499 |
| Lpp | 629 | 864 | 994 | 1221 | 833 | 845 | 668 | 863 | 998 | 960 | 896 | 925 | 885.03 | 843 | 927 | 1.1 | 0.137 | 0.36190743 | 0.58006064 | 0.0193 | 0.0106 | 0.0151 | 0.0151 | TRUE | 0.337 |
| Lpxn | 8 | 9 | 15 | 20 | 14 | 9 | 8 | 9 | 15 | 16 | 15 | 10 | 12.2 | 11 | 14 | 1.254 | 0.327 | 0.56240597 | 0.74333922 | 0 | 0.277 | 0.1463 | 0.1463 | TRUE | 0.4281 |
| Lrat | 17 | 16 | 18 | 29 | 24 | 31 | 18 | 16 | 18 | 23 | 26 | 34 | 22.44 | 17 | 28 | 1.575 | 0.655 | 0.11100752 | 0.28012971 | 0 | 0.1537 | 0.0746 | 0.0746 | TRUE | 0.2522 |
| Lrba | 2884 | 3070 | 3434 | 4397 | 3035 | 2945 | 3063 | 3066 | 3448 | 3457 | 3265 | 3224 | 3253.88 | 3192 | 3315 | 1.039 | 0.055 | 0.50860978 | 0.70252018 | 0.0026 | 0.0079 | 0.0046 | 0.0046 | TRUE | 0.0597 |
| Lrch1 | 1067 | 1185 | 1235 | 1498 | 1140 | 1175 | 1133 | 1183 | 1240 | 1178 | 1227 | 1286 | 1207.88 | 1185 | 1230 | 1.037 | 0.052 | 0.54339957 | 0.72865992 | 0.0012 | 0.0096 | 0.0044 | 0.0044 | TRUE | 0.0193 |
| Lrch2 | 1053 | 1083 | 1016 | 1107 | 778 | 916 | 1118 | 1082 | 1020 | 870 | 837 | 1003 | 988.38 | 1073 | 903 | 0.841 | -0.25 | 0.01729365 | 0.07819824 | 0.0042 | 0.0102 | 0.0069 | 0.0069 | TRUE | 0.1081 |
| Lrch3 | 2769 | 2817 | 2926 | 3074 | 2264 | 2426 | 2941 | 2813 | 2938 | 2417 | 2436 | 2656 | 2700.11 | 2897 | 2503 | 0.863 | -0.212 | 0.00479958 | 0.030389 | 0.0012 | 0.0081 | 0.0037 | 0.0037 | TRUE | 0.0343 |
| Lrch4 | 0 | 0 | 0 | 0 | 0 | 0 | 0 | 0 | 0 | 0 | 0 | 0 | 0 | 0 | 0 | NA | NA | NA | NA | NA | NA | NA | NA | NA | NA |
| Lrcol1 | 67 | 90 | 113 | 52 | 57 | 63 | 71 | 90 | 113 | 41 | 61 | 69 | 74.28 | 91 | 57 | 0.618 | -0.694 | 0.02011637 | 0.08718751 | 0.0485 | 0.0513 | 0.05 | 0.05 | TRUE | 0.2481 |
| Lrdd | 1119 | 1087 | 1009 | 1383 | 1047 | 932 | 1189 | 1086 | 1013 | 1087 | 1127 | 1020 | 1086.87 | 1096 | 1078 | 0.984 | -0.023 | 0.81551796 | 0.90592684 | 0.0035 | 0.0099 | 0.0061 | 0.0061 | TRUE | 0.0653 |
| Lrfn1 | 856 | 857 | 840 | 1024 | 737 | 746 | 909 | 856 | 843 | 805 | 793 | 817 | 837.19 | 869 | 805 | 0.926 | -0.111 | 0.1971442 | 0.40376033 | 0 | 0.0108 | 0.0042 | 0.0042 | TRUE | 0.0188 |
| Lrfn2 | 7 | 3 | 6 | 3 | 1 | 5 | 7 | 3 | 6 | 2 | 1 | 5 | 4.23 | 5 | 3 | 0.536 | -0.898 | 0.36499314 | NA | 0 | 0.7862 | 0.4493 | 0.4493 | TRUE | 0.6146 |
| Lrfn3 | 225 | 288 | 283 | 293 | 191 | 275 | 239 | 288 | 284 | 230 | 206 | 301 | 257.94 | 270 | 246 | 0.907 | -0.14 | 0.44055551 | 0.64757144 | 0.0202 | 0.0197 | 0.0199 | 0.0199 | TRUE | 0.4191 |
| Lrfn4 | 1051 | 1489 | 1531 | 1057 | 922 | 924 | 1116 | 1487 | 1537 | 831 | 992 | 1011 | 1162.52 | 1380 | 945 | 0.684 | -0.548 | 0.00022069 | 0.00262889 | 0.0198 | 0.0097 | 0.015 | 0.015 | TRUE | 0.3355 |
| Lrfn5 | 10 | 1 | 4 | 18 | 5 | 4 | 11 | 1 | 4 | 14 | 5 | 4 | 6.59 | 5 | 8 | 1.562 | 0.643 | 0.50364155 | 0.69869245 | 0.5085 | 0.5068 | 0.5077 | 0.5077 | TRUE | 0.8407 |
| Lrg1 | 0 | 0 | 0 | 0 | 0 | 0 | 0 | 0 | 0 | 0 | 0 | 0 | 0 | 0 | 0 | NA | NA | NA | NA | NA | NA | NA | NA | NA | NA |
| Lrguk | 72 | 64 | 71 | 87 | 80 | 51 | 76 | 64 | 71 | 68 | 86 | 56 | 70.33 | 70 | 70 | 0.994 | -0.009 | 0.97122798 | 0.98480349 | 0.011 | 0.0537 | 0.0335 | 0.0335 | TRUE | 0.279 |
| Lrif1 | 445 | 285 | 314 | 725 | 529 | 298 | 473 | 285 | 315 | 570 | 569 | 326 | 422.98 | 358 | 488 | 1.368 | 0.452 | 0.07745873 | 0.21991948 | 0.0775 | 0.0147 | 0.0448 | 0.0448 | TRUE | 0.9732 |
| Lrig1 | 2163 | 2477 | 2392 | 2546 | 1969 | 1958 | 2297 | 2474 | 2402 | 2001 | 2119 | 2143 | 2239.41 | 2391 | 2088 | 0.873 | -0.197 | 0.00822417 | 0.04536958 | 9.00E-04 | 0.0084 | 0.0035 | 0.0035 | TRUE | 0.0163 |
| Lrig2 | 1747 | 1790 | 1931 | 1888 | 1375 | 1382 | 1856 | 1788 | 1939 | 1484 | 1479 | 1513 | 1676.44 | 1861 | 1492 | 0.802 | -0.319 | 1.82E-05 | 0.00032682 | 0 | 0.0089 | 0.0034 | 0.0034 | TRUE | 0.0164 |
| Lrig3 | 986 | 962 | 1008 | 1036 | 778 | 780 | 1047 | 961 | 1012 | 814 | 837 | 854 | 920.92 | 1007 | 835 | 0.829 | -0.27 | 0.00169862 | 0.01351695 | 0 | 0.0105 | 0.0042 | 0.0042 | TRUE | 0.0191 |
| Lrit1 | 0 | 0 | 0 | 0 | 0 | 0 | 0 | 0 | 0 | 0 | 0 | 0 | 0 | 0 | 0 | NA | NA | NA | NA | NA | NA | NA | NA | NA | NA |
| Lrit2 | 0 | 0 | 0 | 0 | 0 | 0 | 0 | 0 | 0 | 0 | 0 | 0 | 0 | 0 | 0 | NA | NA | NA | NA | NA | NA | NA | NA | NA | NA |
| Lrit3 | 0 | 2 | 0 | 0 | 2 | 0 | 0 | 2 | 0 | 0 | 2 | 0 | 0.69 | 1 | 1 | 1.052 | 0.073 | 0.98136341 | NA | 8.094 | 4.7708 | 5.501 | 5.501 | TRUE | 0.9745 |
| Lrmp | 224 | 201 | 199 | 181 | 207 | 178 | 238 | 201 | 200 | 142 | 223 | 195 | 199.72 | 213 | 187 | 0.873 | -0.195 | 0.33557339 | 0.55585035 | 0.0257 | 0.0234 | 0.0246 | 0.0246 | TRUE | 0.5005 |
| Lrp1 | 4737 | 5120 | 6272 | 4672 | 3531 | 4496 | 5032 | 5113 | 6298 | 3673 | 3799 | 4922 | 4806.03 | 5481 | 4131 | 0.754 | -0.408 | 0.00419537 | 0.0274107 | 0.0219 | 0.0076 | 0.0144 | 0.0144 | TRUE | 0.3405 |
| Lrp10 | 2979 | 3166 | 3232 | 3462 | 2532 | 2716 | 3164 | 3162 | 3245 | 2722 | 2724 | 2973 | 2998.39 | 3190 | 2806 | 0.879 | -0.186 | 0.01016879 | 0.05344289 | 9.00E-04 | 0.008 | 0.0034 | 0.0034 | TRUE | 0.0325 |
| Lrp11 | 3617 | 4661 | 4814 | 4846 | 3525 | 4029 | 3842 | 4655 | 4834 | 3810 | 3793 | 4410 | 4223.87 | 4444 | 4004 | 0.901 | -0.15 | 0.19478659 | 0.40071236 | 0.0106 | 0.0077 | 0.0095 | 0.0095 | TRUE | 0.1707 |
| Lrp12 | 511 | 605 | 611 | 881 | 637 | 587 | 543 | 604 | 614 | 693 | 685 | 643 | 630.17 | 587 | 674 | 1.148 | 0.199 | 0.05146966 | 0.16728458 | 0.0012 | 0.0121 | 0.0059 | 0.0059 | TRUE | 0.0501 |
| Lrp1b | 13 | 5 | 4 | 25 | 12 | 7 | 14 | 5 | 4 | 20 | 13 | 8 | 10.51 | 8 | 14 | 1.791 | 0.841 | 0.24229755 | 0.45887572 | 0.213 | 0.3204 | 0.2689 | 0.2689 | TRUE | 0.4487 |
| Lrp2 | 4272 | 2219 | 2845 | 2862 | 1685 | 1254 | 4538 | 2216 | 2857 | 2250 | 1813 | 1373 | 2507.66 | 3204 | 1812 | 0.566 | -0.822 | 0.02667118 | 0.10638791 | 0.0987 | 0.0082 | 0.0447 | 0.0987 | TRUE | 0.4889 |
| Lrp2bp | 36 | 40 | 28 | 38 | 38 | 33 | 38 | 40 | 28 | 30 | 41 | 36 | 35.53 | 35 | 36 | 1 | 0 | 0.99928161 | 0.99950602 | 0 | 0.0996 | 0.0535 | 0.0535 | TRUE | 0.2361 |
| Lrp3 | 414 | 602 | 595 | 619 | 491 | 496 | 440 | 601 | 597 | 487 | 528 | 543 | 532.71 | 546 | 519 | 0.95 | -0.075 | 0.61607386 | 0.78291168 | 0.0149 | 0.0131 | 0.014 | 0.014 | TRUE | 0.3374 |
| Lrp4 | 1174 | 1421 | 1418 | 1308 | 1046 | 1179 | 1247 | 1419 | 1424 | 1028 | 1125 | 1291 | 1255.71 | 1363 | 1148 | 0.841 | -0.249 | 0.03232991 | 0.12123331 | 0.0082 | 0.0095 | 0.009 | 0.009 | TRUE | 0.1408 |
| Lrp5 | 3757 | 4759 | 5235 | 4212 | 3325 | 3575 | 3991 | 4753 | 5256 | 3311 | 3577 | 3913 | 4133.66 | 4667 | 3600 | 0.771 | -0.375 | 0.00218277 | 0.01650363 | 0.0125 | 0.0077 | 0.0105 | 0.0105 | TRUE | 0.1955 |
| Lrp6 | 7659 | 7677 | 8820 | 9421 | 6775 | 7009 | 8135 | 7667 | 8856 | 7406 | 7289 | 7672 | 7837.75 | 8219 | 7456 | 0.907 | -0.141 | 0.07727416 | 0.21955724 | 0.0028 | 0.0073 | 0.0044 | 0.0044 | TRUE | 0.0561 |
| Lrp8 | 543 | 1033 | 1171 | 1033 | 800 | 953 | 577 | 1032 | 1176 | 812 | 861 | 1043 | 916.71 | 928 | 905 | 0.975 | -0.037 | 0.87565519 | 0.93865053 | 0.0727 | 0.0105 | 0.0382 | 0.0382 | TRUE | 1.0626 |
| Lrpap1 | 1939 | 2098 | 2081 | 3521 | 2357 | 2139 | 2060 | 2095 | 2090 | 2768 | 2536 | 2341 | 2314.96 | 2082 | 2548 | 1.225 | 0.293 | 0.00088733 | 0.0081083 | 0.0029 | 0.0083 | 0.0052 | 0.0052 | TRUE | 0.0699 |
| Lrpprc | 14588 | 12329 | 12430 | 16132 | 12059 | 10607 | 15495 | 12313 | 12481 | 12682 | 12975 | 11611 | 12926.11 | 13430 | 12423 | 0.925 | -0.112 | 0.31214727 | 0.53255705 | 0.0104 | 0.0072 | 0.0088 | 0.0088 | TRUE | 0.2212 |
| Lrr1 | 588 | 538 | 531 | 858 | 537 | 446 | 625 | 537 | 533 | 674 | 578 | 488 | 572.59 | 565 | 580 | 1.029 | 0.041 | 0.78133815 | 0.88682105 | 0.0154 | 0.0127 | 0.0141 | 0.0141 | TRUE | 0.2037 |
| Lrrc1 | 1472 | 1326 | 1343 | 1514 | 1029 | 1043 | 1564 | 1324 | 1349 | 1190 | 1107 | 1142 | 1279.23 | 1412 | 1146 | 0.812 | -0.3 | 0.00256938 | 0.0187873 | 0.0038 | 0.0095 | 0.0063 | 0.0063 | TRUE | 0.1054 |
| Lrrc10 | 4 | 3 | 2 | 6 | 10 | 5 | 4 | 3 | 2 | 5 | 11 | 5 | 5.03 | 3 | 7 | 2.251 | 1.17 | 0.19041156 | 0.39551839 | 0 | 0.6614 | 0.3389 | 0.3389 | TRUE | 0.5588 |
| Lrrc10b | 9 | 9 | 10 | 20 | 7 | 13 | 10 | 9 | 10 | 16 | 8 | 14 | 11.01 | 10 | 13 | 1.322 | 0.403 | 0.49187061 | 0.6881738 | 0 | 0.3061 | 0.1555 | 0.1555 | TRUE | 0.4539 |
| Lrrc14 | 282 | 226 | 226 | 605 | 446 | 293 | 300 | 226 | 227 | 476 | 480 | 321 | 338.06 | 251 | 426 | 1.699 | 0.765 | 0.00013911 | 0.00179543 | 0.0354 | 0.0166 | 0.0259 | 0.0259 | TRUE | 0.5271 |
| Lrrc14b | 3 | 0 | 0 | 0 | 0 | 0 | 3 | 0 | 0 | 0 | 0 | 0 | 0.53 | 1 | 0 | 0.169 | -2.562 | 0.52101829 | NA | 10 | 6.2101 | 10 | 10 | TRUE | 1.4425 |
| Lrrc15 | 147 | 126 | 85 | 249 | 159 | 144 | 156 | 126 | 85 | 196 | 171 | 158 | 148.63 | 122 | 175 | 1.432 | 0.518 | 0.03155238 | 0.11939639 | 0.0412 | 0.0291 | 0.0348 | 0.0348 | TRUE | 0.3997 |
| Lrrc16a | 1332 | 1258 | 1293 | 1923 | 1378 | 1329 | 1415 | 1256 | 1298 | 1512 | 1483 | 1455 | 1403.11 | 1323 | 1483 | 1.121 | 0.165 | 0.04844423 | 0.16053487 | 0.0013 | 0.0093 | 0.0043 | 0.0043 | TRUE | 0.0439 |
| Lrrc16b | 159 | 181 | 159 | 118 | 133 | 151 | 169 | 181 | 160 | 93 | 143 | 165 | 151.74 | 170 | 134 | 0.783 | -0.353 | 0.13218028 | 0.31326032 | 0.0374 | 0.0286 | 0.0329 | 0.0329 | TRUE | 0.8042 |
| Lrrc17 | 2 | 1 | 0 | 8 | 3 | 1 | 2 | 1 | 0 | 6 | 3 | 1 | 2.29 | 1 | 3 | 3.497 | 1.806 | 0.21876488 | NA | 0 | 1.4461 | 0.9229 | 0.9229 | TRUE | 0.1715 |
| Lrrc18 | 4 | 6 | 20 | 2 | 4 | 8 | 4 | 6 | 20 | 2 | 4 | 9 | 7.49 | 10 | 5 | 0.471 | -1.086 | 0.24351822 | 0.46008258 | 0.497 | 0.4466 | 0.4723 | 0.4723 | TRUE | 1.5401 |
| Lrrc19 | 10 | 1 | 1 | 3 | 4 | 2 | 11 | 1 | 1 | 2 | 4 | 2 | 3.58 | 4 | 3 | 0.702 | -0.51 | 0.67850036 | NA | 0.6931 | 0.9273 | 0.8023 | 0.8023 | TRUE | 2.9751 |
| Lrrc2 | 962 | 1427 | 1474 | 1700 | 1230 | 1398 | 1022 | 1425 | 1480 | 1336 | 1323 | 1530 | 1352.86 | 1309 | 1396 | 1.067 | 0.093 | 0.54112618 | 0.7269093 | 0.0223 | 0.0093 | 0.0159 | 0.0159 | TRUE | 0.4413 |
| Lrrc20 | 619 | 452 | 411 | 736 | 479 | 383 | 657 | 451 | 413 | 579 | 515 | 419 | 505.8 | 507 | 504 | 0.996 | -0.006 | 0.97669308 | 0.98733516 | 0.0423 | 0.0134 | 0.0274 | 0.0274 | TRUE | 0.7807 |
| Lrrc23 | 10 | 12 | 6 | 11 | 7 | 9 | 11 | 12 | 6 | 9 | 8 | 10 | 9.11 | 10 | 9 | 0.909 | -0.137 | 0.82792597 | 0.91291407 | 0 | 0.3685 | 0.1768 | 0.1768 | TRUE | 0.3547 |

| | | | | | | | | | | | | | | | | | | | | | | | | | |
|---|---|---|---|---|---|---|---|---|---|---|---|---|---|---|---|---|---|---|---|---|---|---|---|---|---|
| Lrrc24 | 14 | 21 | 24 | 17 | 19 | 22 | 15 | 21 | 24 | 13 | 20 | 24 | 19.64 | 20 | 19 | 0.954 | -0.069 | 0.88210494 | 0.9416921 | 0 | 0.1747 | 0.1027 | 0.1027 | TRUE | 0.477 |
| Lrrc25 | 13 | 12 | 2 | 4 | 2 | 14 | 14 | 12 | 2 | 3 | 2 | 15 | 8.07 | 9 | 7 | 0.733 | -0.447 | 0.64980575 | 0.80369639 | 0.7284 | 0.4151 | 0.5715 | 0.5715 | TRUE | 2.816 |
| Lrrc26 | 1 | 0 | 0 | 1 | 0 | 0 | 1 | 0 | 0 | 1 | 0 | 0 | 0.31 | 0 | 0 | 0.954 | -0.067 | 0.98625928 | NA | 0 | 10.7014 | 9.0384 | 9.0384 | TRUE | 0.4614 |
| Lrrc27 | 319 | 372 | 325 | 299 | 268 | 290 | 339 | 372 | 326 | 235 | 288 | 317 | 312.92 | 346 | 280 | 0.808 | -0.307 | 0.04603621 | 0.15528248 | 0.0106 | 0.0174 | 0.0138 | 0.0138 | TRUE | 0.2391 |
| Lrrc28 | 926 | 1047 | 993 | 1471 | 911 | 1029 | 984 | 1046 | 997 | 1156 | 980 | 1126 | 1048.21 | 1009 | 1087 | 1.079 | 0.11 | 0.27016411 | 0.4903393 | 0.0035 | 0.01 | 0.0062 | 0.0062 | TRUE | 0.0885 |
| Lrrc29 | 116 | 159 | 142 | 184 | 147 | 133 | 123 | 159 | 143 | 145 | 158 | 146 | 145.5 | 142 | 150 | 1.054 | 0.076 | 0.66760711 | 0.81403165 | 0 | 0.0295 | 0.0158 | 0.0158 | TRUE | 0.1305 |
| Lrrc3 | 290 | 359 | 286 | 327 | 232 | 208 | 308 | 359 | 287 | 257 | 250 | 228 | 281.35 | 318 | 245 | 0.771 | -0.375 | 0.00970246 | 0.05145502 | 0.0054 | 0.0186 | 0.0116 | 0.0116 | TRUE | 0.1429 |
| Lrrc30 | 0 | 0 | 0 | 1 | 0 | 0 | 0 | 0 | 0 | 1 | 0 | 0 | 0.13 | 0 | 0 | 1.859 | 0.894 | 0.82650734 | NA | 0 | 25.1515 | 10 | 10 | TRUE | 0.2563 |
| Lrrc32 | 49 | 34 | 34 | 63 | 37 | 45 | 52 | 34 | 34 | 50 | 40 | 49 | 43.12 | 40 | 46 | 1.158 | 0.212 | 0.50732304 | 0.70163969 | 0.0128 | 0.0833 | 0.0503 | 0.0503 | TRUE | 0.501 |
| Lrrc33 | 205 | 146 | 152 | 239 | 173 | 107 | 218 | 146 | 153 | 188 | 186 | 117 | 167.89 | 172 | 164 | 0.954 | -0.068 | 0.78632327 | 0.89023346 | 0.0519 | 0.0265 | 0.0387 | 0.0387 | TRUE | 0.6384 |
| Lrrc34 | 1469 | 2669 | 2902 | 1623 | 1370 | 1991 | 1560 | 2666 | 2914 | 1276 | 1474 | 2179 | 2011.53 | 2380 | 1643 | 0.69 | -0.535 | 0.02987448 | 0.11486598 | 0.0868 | 0.0085 | 0.0432 | 0.0432 | TRUE | 1.099 |
| Lrrc36 | 27 | 28 | 22 | 13 | 13 | 8 | 29 | 28 | 22 | 10 | 14 | 9 | 18.62 | 26 | 11 | 0.418 | -1.26 | 0.00639243 | 0.03773258 | 0 | 0.1839 | 0.0898 | 0.0898 | TRUE | 0.1884 |
| Lrrc38 | 18 | 25 | 16 | 9 | 9 | 25 | 19 | 25 | 16 | 7 | 10 | 27 | 17.38 | 20 | 15 | 0.723 | -0.468 | 0.43745656 | 0.64548863 | 0.2109 | 0.1965 | 0.2029 | 0.2029 | TRUE | 2.4176 |
| Lrrc39 | 1 | 6 | 1 | 5 | 5 | 1 | 1 | 6 | 1 | 4 | 5 | 1 | 3.08 | 3 | 3 | 1.292 | 0.369 | 0.76237325 | NA | 0.3996 | 1.0775 | 0.7439 | 0.7439 | TRUE | 1.1401 |
| Lrrc3b | 74 | 67 | 58 | 121 | 100 | 83 | 79 | 67 | 58 | 95 | 108 | 91 | 82.89 | 68 | 98 | 1.441 | 0.528 | 0.02076509 | 0.08921464 | 0 | 0.0466 | 0.025 | 0.025 | TRUE | 0.1645 |
| Lrrc4 | 43 | 47 | 55 | 74 | 64 | 42 | 46 | 47 | 55 | 58 | 69 | 46 | 53.47 | 49 | 58 | 1.17 | 0.227 | 0.42188396 | 0.63202449 | 0 | 0.0685 | 0.0388 | 0.0388 | TRUE | 0.1768 |
| Lrrc40 | 3643 | 3992 | 4012 | 5147 | 3623 | 3587 | 3870 | 3987 | 4028 | 4046 | 3898 | 3927 | 3959.28 | 3962 | 3957 | 0.999 | -0.001 | 0.98106496 | 0.98966143 | 2.00E-04 | 0.0077 | 0.0025 | 0.0025 | TRUE | 0.005 |
| Lrrc41 | 1194 | 1733 | 1669 | 2126 | 1400 | 1572 | 1268 | 1731 | 1676 | 1671 | 1506 | 1721 | 1595.54 | 1558 | 1633 | 1.048 | 0.067 | 0.61914228 | 0.78469067 | 0.0157 | 0.009 | 0.0126 | 0.0126 | TRUE | 0.3186 |
| Lrrc42 | 2701 | 2788 | 2745 | 3768 | 2672 | 2716 | 2869 | 2784 | 2756 | 2962 | 2875 | 2973 | 2869.95 | 2803 | 2937 | 1.048 | 0.067 | 0.29174672 | 0.51252275 | 0 | 0.0081 | 0.0026 | 0.0026 | TRUE | 0.0051 |
| Lrrc43 | 0 | 3 | 2 | 3 | 1 | 2 | 0 | 3 | 2 | 2 | 1 | 2 | 1.77 | 2 | 2 | 1.125 | 0.17 | 0.90789638 | NA | 0 | 1.8668 | 0.9901 | 0.9901 | TRUE | 0.5185 |
| Lrrc45 | 719 | 704 | 657 | 777 | 723 | 654 | 764 | 703 | 660 | 611 | 778 | 716 | 705.19 | 709 | 702 | 0.988 | -0.017 | 0.89332841 | 0.94834594 | 0.0089 | 0.0116 | 0.0101 | 0.0101 | TRUE | 0.1569 |
| Lrrc46 | 48 | 58 | 46 | 55 | 49 | 50 | 51 | 58 | 46 | 43 | 53 | 55 | 50.96 | 52 | 50 | 0.966 | -0.049 | 0.85801697 | 0.93001469 | 0 | 0.0715 | 0.0354 | 0.0354 | TRUE | 0.1416 |
| Lrrc47 | 4724 | 4527 | 4519 | 6567 | 4934 | 4479 | 5018 | 4521 | 4538 | 5162 | 5309 | 4903 | 4908.42 | 4692 | 5125 | 1.092 | 0.127 | 0.10315322 | 0.26735607 | 0.0023 | 0.0076 | 0.0042 | 0.0042 | TRUE | 0.0448 |
| Lrrc48 | 173 | 337 | 354 | 282 | 220 | 292 | 184 | 337 | 355 | 222 | 237 | 320 | 275.63 | 292 | 260 | 0.887 | -0.173 | 0.50584102 | 0.70029387 | 0.0759 | 0.0189 | 0.0452 | 0.0452 | TRUE | 1.1753 |
| Lrrc49 | 300 | 289 | 244 | 422 | 265 | 257 | 319 | 289 | 245 | 332 | 285 | 281 | 291.75 | 284 | 299 | 1.057 | 0.08 | 0.60588738 | 0.77641012 | 0.0096 | 0.0182 | 0.0137 | 0.0137 | TRUE | 0.1633 |
| Lrrc4b | 17 | 20 | 18 | 9 | 5 | 5 | 18 | 20 | 18 | 7 | 5 | 5 | 12.34 | 19 | 6 | 0.323 | -1.63 | 0.00382528 | 0.02546744 | 0 | 0.2739 | 0.1211 | 0.1211 | TRUE | 0.0819 |
| Lrrc4c | 0 | 0 | 0 | 0 | 0 | 0 | 0 | 0 | 0 | 0 | 0 | 0 | 0 | 0 | 0 | NA | NA | NA | NA | NA | NA | NA | NA | NA | NA |
| Lrrc51 | 97 | 288 | 340 | 116 | 84 | 117 | 103 | 288 | 341 | 91 | 90 | 128 | 173.62 | 244 | 103 | 0.422 | -1.244 | 0.00105429 | 0.00928433 | 0.1881 | 0.0259 | 0.097 | 0.097 | TRUE | 0.6428 |
| Lrrc52 | 2 | 4 | 1 | 1 | 5 | 2 | 2 | 4 | 1 | 1 | 5 | 2 | 2.58 | 2 | 3 | 1.144 | 0.194 | 0.87748995 | NA | 0 | 1.284 | 0.7512 | 0.7512 | TRUE | 0.5508 |
| Lrrc55 | 9 | 11 | 5 | 32 | 14 | 16 | 10 | 11 | 5 | 25 | 15 | 18 | 13.88 | 9 | 19 | 2.279 | 1.188 | 0.03485548 | 0.12780763 | 0 | 0.2442 | 0.1433 | 0.1433 | TRUE | 0.4561 |
| Lrrc56 | 199 | 207 | 200 | 285 | 234 | 218 | 211 | 207 | 201 | 224 | 252 | 239 | 222.23 | 206 | 238 | 1.152 | 0.205 | 0.14051533 | 0.32572131 | 0 | 0.0217 | 0.0094 | 0.0094 | TRUE | 0.0325 |
| Lrrc57 | 937 | 966 | 930 | 1430 | 1006 | 975 | 995 | 965 | 934 | 1124 | 1082 | 1067 | 1027.94 | 965 | 1091 | 1.132 | 0.179 | 0.02866333 | 0.1114591 | 0 | 0.0101 | 0.0038 | 0.0038 | TRUE | 0.0093 |
| Lrrc58 | 3445 | 3651 | 3648 | 4740 | 3584 | 3444 | 3659 | 3646 | 3663 | 3726 | 3856 | 3770 | 3720.14 | 3656 | 3784 | 1.035 | 0.049 | 0.4037537 | 0.61651693 | 0 | 0.0078 | 0.0023 | 0.0023 | TRUE | 0.0034 |
| Lrrc59 | 20263 | 21177 | 20571 | 28828 | 20574 | 18869 | 21523 | 21150 | 20655 | 22662 | 22136 | 20655 | 21463.63 | 21109 | 21818 | 1.034 | 0.048 | 0.46981336 | 0.67180893 | 0.0013 | 0.0071 | 0.0031 | 0.0031 | TRUE | 0.0266 |
| Lrrc6 | 11 | 18 | 20 | 16 | 16 | 22 | 12 | 18 | 20 | 13 | 17 | 24 | 17.27 | 17 | 18 | 1.07 | 0.097 | 0.84441819 | 0.92161735 | 0 | 0.1977 | 0.1193 | 0.1193 | TRUE | 0.2915 |
| Lrrc61 | 473 | 713 | 834 | 563 | 453 | 450 | 502 | 712 | 837 | 443 | 487 | 493 | 579.08 | 684 | 474 | 0.693 | -0.53 | 0.00404733 | 0.02660877 | 0.0329 | 0.0126 | 0.0227 | 0.0227 | TRUE | 0.2761 |
| Lrrc63 | 3 | 3 | 1 | 1 | 1 | 2 | 3 | 3 | 1 | 1 | 1 | 2 | 1.87 | 2 | 1 | 0.552 | -0.857 | 0.53313239 | NA | 0 | 1.7658 | 0.7809 | 0.7809 | TRUE | 0.2775 |
| Lrrc66 | 2 | 0 | 0 | 2 | 2 | 2 | 2 | 0 | 0 | 2 | 2 | 2 | 1.34 | 1 | 2 | 2.825 | 1.498 | 0.41193911 | NA | 0 | 2.4661 | 1.4169 | 1.4169 | TRUE | 0.9745 |
| Lrrc69 | 2 | 1 | 1 | 1 | 0 | 1 | 2 | 1 | 1 | 1 | 0 | 1 | 1 | 1 | 1 | 0.463 | -1.112 | 0.55213889 | NA | 0 | 3.297 | 1.3782 | 1.3782 | TRUE | 0.1944 |
| Lrrc7 | 0 | 1 | 0 | 0 | 0 | 0 | 0 | 1 | 0 | 0 | 0 | 0 | 0.17 | 0 | 0 | 0.49 | -1.029 | 0.80087 | NA | 0 | 19.7991 | 10 | 10 | TRUE | 0.405 |
| Lrrc71 | 12 | 20 | 18 | 11 | 11 | 9 | 13 | 20 | 18 | 9 | 12 | 10 | 13.52 | 17 | 10 | 0.592 | -0.756 | 0.14960704 | 0.34028031 | 0 | 0.2505 | 0.1196 | 0.1196 | TRUE | 0.2196 |
| Lrrc72 | 0 | 0 | 0 | 1 | 1 | 1 | 0 | 0 | 0 | 1 | 1 | 1 | 0.49 | 0 | 1 | 5.224 | 2.385 | 0.38465 | NA | 0 | 6.6923 | 3.9249 | 3.9249 | TRUE | 0.07 |
| Lrrc73 | 21 | 17 | 23 | 41 | 29 | 19 | 22 | 17 | 23 | 32 | 31 | 21 | 24.43 | 21 | 28 | 1.358 | 0.442 | 0.27374167 | 0.49433176 | 0 | 0.1417 | 0.0755 | 0.0755 | TRUE | 0.3066 |
| Lrrc8a | 1100 | 1074 | 1057 | 1223 | 951 | 874 | 1168 | 1073 | 1061 | 961 | 1023 | 957 | 1040.62 | 1101 | 980 | 0.891 | -0.167 | 0.05858169 | 0.18288804 | 0.0011 | 0.0101 | 0.0047 | 0.0047 | TRUE | 0.0343 |
| Lrrc8b | 425 | 630 | 637 | 616 | 468 | 489 | 451 | 629 | 640 | 484 | 504 | 535 | 540.55 | 573 | 508 | 0.885 | -0.177 | 0.25390458 | 0.47188737 | 0.0178 | 0.013 | 0.0155 | 0.0155 | TRUE | 0.4035 |
| Lrrc8c | 282 | 292 | 341 | 444 | 305 | 334 | 300 | 292 | 342 | 349 | 328 | 366 | 329.4 | 311 | 348 | 1.117 | 0.159 | 0.21650382 | 0.4284977 | 0.0019 | 0.0169 | 0.0089 | 0.0089 | TRUE | 0.0878 |
| Lrrc8d | 1253 | 1942 | 2170 | 2606 | 1870 | 1786 | 1331 | 1939 | 2179 | 2049 | 2012 | 1955 | 1910.83 | 1816 | 2005 | 1.104 | 0.143 | 0.39423278 | 0.60787039 | 0.0311 | 0.0086 | 0.0196 | 0.0196 | TRUE | 0.6592 |
| Lrrc8e | 119 | 127 | 121 | 166 | 121 | 144 | 126 | 127 | 121 | 130 | 130 | 158 | 132.17 | 125 | 139 | 1.114 | 0.156 | 0.38574329 | 0.59963958 | 0 | 0.0318 | 0.0156 | 0.0156 | TRUE | 0.1291 |
| Lrrc9 | 169 | 179 | 205 | 266 | 218 | 197 | 180 | 179 | 206 | 209 | 235 | 216 | 203.9 | 188 | 220 | 1.167 | 0.223 | 0.13810603 | 0.32245868 | 0 | 0.0231 | 0.0113 | 0.0113 | TRUE | 0.0745 |
| Lrrcc1 | 468 | 616 | 669 | 1125 | 887 | 783 | 497 | 615 | 672 | 884 | 954 | 857 | 746.65 | 595 | 898 | 1.51 | 0.595 | 7.18E-06 | 0.00014949 | 0.0107 | 0.0113 | 0.0113 | 0.0113 | TRUE | 0.2402 |
| Lrrd1 | 3 | 1 | 1 | 2 | 2 | 2 | 3 | 1 | 1 | 2 | 2 | 2 | 1.85 | 2 | 2 | 1.138 | 0.187 | 0.88963194 | NA | 0 | 1.7872 | 0.7557 | 0.7557 | TRUE | 0.4027 |
| Lrrfip1 | 2348 | 2552 | 2421 | 2523 | 1993 | 1888 | 2494 | 2549 | 2431 | 1983 | 2144 | 2067 | 2278.01 | 2491 | 2065 | 0.828 | -0.272 | 0.00015517 | 0.00195939 | 6.00E-04 | 0.0083 | 0.0033 | 0.0033 | TRUE | 0.0148 |
| Lrrfip2 | 2491 | 2482 | 2415 | 3159 | 2248 | 2076 | 2646 | 2479 | 2425 | 2483 | 2419 | 2273 | 2454.03 | 2517 | 2392 | 0.951 | -0.073 | 0.35297126 | 0.57159736 | 0.0015 | 0.0082 | 0.004 | 0.004 | TRUE | 0.0244 |
| Lrriq1 | 12 | 21 | 19 | 45 | 25 | 10 | 13 | 21 | 19 | 35 | 27 | 11 | 21 | 18 | 24 | 1.399 | 0.484 | 0.36156845 | 0.57995094 | 0.1433 | 0.1638 | 0.1539 | 0.1539 | TRUE | 0.3361 |
| Lrriq3 | 14 | 15 | 11 | 23 | 24 | 18 | 15 | 15 | 11 | 18 | 26 | 20 | 17.42 | 14 | 21 | 1.549 | 0.631 | 0.17315076 | 0.37206182 | 0 | 0.1961 | 0.0941 | 0.0941 | TRUE | 0.1937 |
| Lrriq4 | 281 | 311 | 324 | 247 | 232 | 267 | 298 | 311 | 325 | 194 | 250 | 292 | 278.41 | 311 | 245 | 0.785 | -0.35 | 0.04502908 | 0.15296115 | 0.0179 | 0.0187 | 0.0183 | 0.0183 | TRUE | 0.3587 |
| Lrrk1 | 167 | 183 | 167 | 263 | 224 | 195 | 177 | 183 | 168 | 207 | 241 | 213 | 198.18 | 176 | 220 | 1.251 | 0.323 | 0.03223131 | 0.12094983 | 0 | 0.0235 | 0.0112 | 0.0112 | TRUE | 0.072 |
| Lrrk2 | 106 | 99 | 90 | 125 | 123 | 75 | 113 | 99 | 90 | 98 | 132 | 82 | 102.42 | 101 | 104 | 1.036 | 0.05 | 0.8345051 | 0.91571799 | 0.0248 | 0.0391 | 0.0321 | 0.0321 | TRUE | 0.513 |
| Lrrn1 | 376 | 393 | 395 | 413 | 312 | 337 | 399 | 392 | 397 | 325 | 336 | 369 | 369.63 | 396 | 343 | 0.865 | -0.21 | 0.06980209 | 0.20540329 | 0 | 0.0158 | 0.0069 | 0.0069 | TRUE | 0.0483 |
| Lrrn2 | 88 | 90 | 80 | 70 | 50 | 51 | 93 | 90 | 80 | 55 | 54 | 56 | 71.39 | 88 | 55 | 0.625 | -0.679 | 0.00307574 | 0.02165876 | 0 | 0.0531 | 0.0232 | 0.0232 | TRUE | 0.0542 |
| Lrrn3 | 2 | 1 | 0 | 4 | 0 | 0 | 2 | 1 | 0 | 3 | 0 | 0 | 1.04 | 1 | 1 | 1.061 | 0.085 | 0.97074604 | NA | 2.5033 | 3.1607 | 2.946 | 2.946 | TRUE | 0.8817 |
| Lrrn4 | 272 | 201 | 236 | 707 | 494 | 368 | 289 | 201 | 237 | 556 | 532 | 403 | 369.46 | 242 | 497 | 2.054 | 1.039 | 2.23E-08 | 9.74E-07 | 0.0278 | 0.0158 | 0.0218 | 0.0218 | TRUE | 0.3144 |
| Lrrn4cl | 69 | 86 | 86 | 105 | 79 | 86 | 73 | 86 | 86 | 83 | 85 | 94 | 84.54 | 82 | 87 | 1.063 | 0.088 | 0.68197745 | 0.82362649 | 0 | 0.0459 | 0.0213 | 0.0213 | TRUE | 0.0761 |
| Lrrtm1 | 0 | 0 | 0 | 0 | 1 | 0 | 0 | 0 | 0 | 0 | 1 | 0 | 0.18 | 0 | 0 | 1.859 | 0.894 | 0.82650734 | NA | 0 | 18.3785 | 10 | 10 | TRUE | 0.5204 |
| Lrrtm2 | 0 | 0 | 0 | 0 | 0 | 0 | 0 | 0 | 0 | 0 | 0 | 0 | 0 | 0 | 0 | NA | NA | NA | NA | NA | NA | NA | NA | NA | NA |
| Lrrtm3 | 2 | 0 | 1 | 2 | 1 | 0 | 2 | 0 | 1 | 2 | 1 | 0 | 0.96 | 1 | 1 | 0.878 | -0.187 | 0.92696056 | NA | 0 | 3.4287 | 1.9664 | 1.9664 | TRUE | 0.1831 |
| Lrrtm4 | 0 | 0 | 0 | 0 | 0 | 1 | 0 | 0 | 0 | 0 | 0 | 1 | 0.18 | 0 | 0 | 1.859 | 0.894 | 0.82650734 | NA | 0 | 18.0643 | 10 | 10 | TRUE | 0.5383 |
| Lrsam1 | 767 | 840 | 712 | 958 | 775 | 683 | 815 | 839 | 715 | 753 | 834 | 748 | 783.86 | 790 | 778 | 0.985 | -0.021 | 0.8431492 | 0.92105202 | 0.0041 | 0.0111 | 0.0071 | 0.0071 | TRUE | 0.0814 |
| Lrtm1 | 0 | 0 | 1 | 1 | 0 | 0 | 0 | 0 | 1 | 1 | 0 | 0 | 0.3 | 0 | 0 | 0.954 | -0.067 | 0.98640617 | NA | 0 | 11.0483 | 9.2783 | 9.2783 | TRUE | 0.4098 |

| | | | | | | | | | | | | | | | | | | | | | | | | | |
|---|---|---|---|---|---|---|---|---|---|---|---|---|---|---|---|---|---|---|---|---|---|---|---|---|---|
| Lrtm2 | 64 | 85 | 83 | 37 | 32 | 34 | 68 | 85 | 83 | 29 | 34 | 37 | 56.16 | 79 | 33 | 0.423 | -1.24 | 6.84E-06 | 0.0001441 | 0 | 0.0656 | 0.0337 | 0.0337 | TRUE | 0.1293 |
| Lrwd1 | 2951 | 2980 | 3049 | 3828 | 2833 | 2712 | 3134 | 2976 | 3062 | 3009 | 3048 | 2969 | 3033.05 | 3057 | 3009 | 0.984 | -0.023 | 0.71752203 | 0.84681037 | 0 | 0.008 | 0.0026 | 0.0026 | TRUE | 0.0066 |
| Lsamp | 0 | 0 | 0 | 0 | 0 | 0 | 0 | 0 | 0 | 0 | 0 | 0 | 0 | 0 | 0 | NA | NA | NA | NA | NA | NA | NA | NA | NA | NA |
| Lsg1 | 4593 | 4238 | 3993 | 5614 | 3840 | 3525 | 4879 | 4233 | 4009 | 4413 | 4132 | 3859 | 4254.01 | 4374 | 4135 | 0.946 | -0.081 | 0.43342459 | 0.64234492 | 0.0069 | 0.0077 | 0.0074 | 0.0074 | TRUE | 0.1241 |
| Lsm1 | 1968 | 1892 | 1810 | 2552 | 1708 | 1750 | 2090 | 1890 | 1817 | 2006 | 1838 | 1916 | 1926.15 | 1932 | 1920 | 0.994 | -0.009 | 0.92270045 | 0.96185271 | 0.0031 | 0.0086 | 0.0052 | 0.0052 | TRUE | 0.0615 |
| Lsm10 | 465 | 494 | 515 | 675 | 482 | 476 | 494 | 493 | 517 | 531 | 519 | 521 | 512.45 | 501 | 524 | 1.044 | 0.062 | 0.52394287 | 0.71373526 | 0 | 0.0133 | 0.0049 | 0.0049 | TRUE | 0.0087 |
| Lsm11 | 2002 | 2548 | 2502 | 2301 | 1770 | 1919 | 2126 | 2545 | 2512 | 1809 | 1904 | 2101 | 2166.23 | 2394 | 1938 | 0.809 | -0.306 | 0.00428644 | 0.02786626 | 0.007 | 0.0084 | 0.0078 | 0.0078 | TRUE | 0.1159 |
| Lsm12 | 3012 | 3437 | 3356 | 4317 | 3267 | 3322 | 3199 | 3433 | 3370 | 3394 | 3515 | 3636 | 3424.47 | 3334 | 3515 | 1.054 | 0.076 | 0.28068515 | 0.50209438 | 8.00E-04 | 0.0079 | 0.0033 | 0.0033 | TRUE | 0.0151 |
| Lsm14a | 7834 | 8439 | 8599 | 11595 | 8344 | 8773 | 8321 | 8428 | 8634 | 9115 | 8978 | 9603 | 8846.61 | 8461 | 9232 | 1.091 | 0.126 | 0.04255901 | 0.14726651 | 6.00E-04 | 0.0073 | 0.0027 | 0.0027 | TRUE | 0.015 |
| Lsm14b | 3707 | 4771 | 4957 | 5428 | 4300 | 4706 | 3937 | 4765 | 4977 | 4267 | 4627 | 5151 | 4620.79 | 4560 | 4682 | 1.026 | 0.038 | 0.75114197 | 0.86854921 | 0.0118 | 0.0076 | 0.0099 | 0.0099 | TRUE | 0.1735 |
| Lsm2 | 2120 | 2353 | 2390 | 3064 | 2243 | 2190 | 2252 | 2350 | 2400 | 2409 | 2413 | 2397 | 2370.15 | 2334 | 2406 | 1.031 | 0.044 | 0.51109822 | 0.70464097 | 0 | 0.0083 | 0.0028 | 0.0028 | TRUE | 0.0114 |
| Lsm3 | 2114 | 2193 | 2218 | 3214 | 2124 | 2066 | 2245 | 2190 | 2227 | 2527 | 2285 | 2262 | 2289.36 | 2221 | 2358 | 1.062 | 0.087 | 0.26717691 | 0.48698882 | 0.0017 | 0.0083 | 0.004 | 0.004 | TRUE | 0.0485 |
| Lsm4 | 7049 | 9170 | 8918 | 9010 | 6945 | 7210 | 7487 | 9158 | 8955 | 7083 | 7472 | 7892 | 8007.98 | 8533 | 7482 | 0.877 | -0.19 | 0.06156109 | 0.18937808 | 0.007 | 0.0073 | 0.0073 | 0.0073 | TRUE | 0.1403 |
| Lsm5 | 1500 | 1698 | 1590 | 2164 | 1402 | 1460 | 1593 | 1696 | 1597 | 1701 | 1508 | 1598 | 1615.57 | 1629 | 1602 | 0.985 | -0.022 | 0.79316863 | 0.89336968 | 0.0019 | 0.0089 | 0.0045 | 0.0045 | TRUE | 0.0359 |
| Lsm6 | 3079 | 3022 | 2976 | 4561 | 3019 | 2954 | 3270 | 3018 | 2988 | 3585 | 3248 | 3234 | 3224.02 | 3092 | 3356 | 1.086 | 0.119 | 0.15029843 | 0.34103105 | 0.0026 | 0.0079 | 0.0046 | 0.0046 | TRUE | 0.0442 |
| Lsm7 | 788 | 1022 | 1041 | 1261 | 892 | 912 | 837 | 1021 | 1045 | 991 | 960 | 998 | 975.39 | 968 | 983 | 1.016 | 0.023 | 0.84050557 | 0.91930825 | 0.0063 | 0.0103 | 0.008 | 0.008 | TRUE | 0.1655 |
| Lsmd1 | 1970 | 2141 | 2232 | 3142 | 2217 | 2152 | 2092 | 2138 | 2241 | 2470 | 2385 | 2356 | 2280.49 | 2157 | 2404 | 1.114 | 0.156 | 0.02717344 | 0.10757516 | 5.00E-04 | 0.0083 | 0.0032 | 0.0032 | TRUE | 0.014 |
| Lsp1 | 25 | 33 | 28 | 36 | 11 | 14 | 27 | 33 | 28 | 28 | 12 | 15 | 23.85 | 29 | 18 | 0.644 | -0.634 | 0.16181458 | 0.35694259 | 0.0583 | 0.145 | 0.1047 | 0.1047 | TRUE | 1.3343 |
| Lsr | 2079 | 2202 | 2190 | 2582 | 2012 | 1992 | 2208 | 2199 | 2199 | 2030 | 2165 | 2181 | 2163.59 | 2202 | 2125 | 0.964 | -0.052 | 0.45707055 | 0.66167202 | 4.00E-04 | 0.0084 | 0.0031 | 0.0031 | TRUE | 0.0192 |
| Lss | 8623 | 7022 | 7247 | 7702 | 5649 | 6009 | 9159 | 7013 | 7277 | 6055 | 6078 | 6578 | 7026.55 | 7816 | 6237 | 0.798 | -0.326 | 0.00547922 | 0.03345709 | 0.0121 | 0.0074 | 0.0098 | 0.0098 | TRUE | 0.2757 |
| Lst1 | 7 | 4 | 17 | 8 | 3 | 8 | 7 | 4 | 17 | 6 | 3 | 9 | 7.8 | 9 | 6 | 0.641 | -0.642 | 0.42669679 | 0.63685362 | 0.2431 | 0.4295 | 0.336 | 0.336 | TRUE | 0.6386 |
| Lta | 3 | 8 | 7 | 5 | 3 | 1 | 3 | 8 | 7 | 4 | 3 | 1 | 4.41 | 6 | 3 | 0.462 | -1.114 | 0.24442659 | NA | 0 | 0.754 | 0.4023 | 0.4023 | TRUE | 0.3875 |
| Lta4h | 5214 | 5325 | 5563 | 6821 | 4669 | 4698 | 5538 | 5318 | 5586 | 5362 | 5024 | 5143 | 5328.42 | 5481 | 5176 | 0.945 | -0.082 | 0.20450625 | 0.41288692 | 6.00E-04 | 0.0075 | 0.0028 | 0.0028 | TRUE | 0.0122 |
| Ltb | 95 | 114 | 150 | 58 | 41 | 57 | 101 | 114 | 151 | 46 | 44 | 62 | 86.25 | 122 | 51 | 0.414 | -1.271 | 1.70E-06 | 4.35E-05 | 0.0283 | 0.0451 | 0.0369 | 0.0369 | TRUE | 0.4355 |
| Ltb4r1 | 10 | 13 | 11 | 13 | 7 | 4 | 11 | 13 | 11 | 10 | 8 | 4 | 9.46 | 12 | 7 | 0.65 | -0.622 | 0.32288202 | 0.54401187 | 0 | 0.355 | 0.1762 | 0.1762 | TRUE | 0.3572 |
| Ltb4r2 | 10 | 10 | 6 | 25 | 12 | 11 | 11 | 10 | 6 | 20 | 13 | 12 | 11.87 | 9 | 15 | 1.695 | 0.761 | 0.18752185 | 0.3921499 | 0 | 0.2844 | 0.15 | 0.15 | TRUE | 0.4381 |
| Ltbp1 | 870 | 1226 | 1207 | 1199 | 835 | 1032 | 924 | 1224 | 1212 | 943 | 898 | 1130 | 1055.19 | 1120 | 990 | 0.883 | -0.179 | 0.22245211 | 0.4345448 | 0.0178 | 0.01 | 0.0145 | 0.0145 | TRUE | 0.2798 |
| Ltbp2 | 78 | 101 | 104 | 96 | 93 | 94 | 83 | 101 | 104 | 75 | 100 | 103 | 94.43 | 96 | 93 | 0.96 | -0.059 | 0.79750476 | 0.89574083 | 0.0121 | 0.0418 | 0.027 | 0.027 | TRUE | 0.2889 |
| Ltbp3 | 223 | 342 | 350 | 206 | 191 | 251 | 237 | 342 | 351 | 162 | 206 | 275 | 262.01 | 310 | 214 | 0.688 | -0.539 | 0.02161034 | 0.09194449 | 0.0534 | 0.0195 | 0.0357 | 0.0357 | TRUE | 0.523 |
| Ltbp4 | 3636 | 4137 | 4480 | 5337 | 3964 | 3802 | 3862 | 4132 | 4498 | 4196 | 4265 | 4162 | 4185.76 | 4164 | 4208 | 1.01 | 0.015 | 0.85529582 | 0.92802185 | 0.0027 | 0.0077 | 0.0045 | 0.0045 | TRUE | 0.0601 |
| Ltbr | 221 | 295 | 289 | 306 | 240 | 219 | 235 | 295 | 290 | 241 | 258 | 240 | 259.68 | 273 | 246 | 0.9 | -0.152 | 0.30339718 | 0.5242845 | 0.0047 | 0.0196 | 0.0119 | 0.0119 | TRUE | 0.1674 |
| Ltc4s | 3 | 4 | 5 | 9 | 10 | 5 | 3 | 4 | 5 | 7 | 11 | 5 | 5.92 | 4 | 8 | 1.902 | 0.927 | 0.24149803 | 0.45824194 | 0 | 0.5636 | 0.2634 | 0.2634 | TRUE | 0.2968 |
| Ltf | 7 | 7 | 5 | 13 | 12 | 10 | 7 | 7 | 5 | 10 | 13 | 11 | 8.92 | 6 | 11 | 1.749 | 0.806 | 0.20085949 | 0.40850494 | 0 | 0.3762 | 0.1642 | 0.1642 | TRUE | 0.0985 |
| Ltk | 74 | 159 | 149 | 177 | 168 | 170 | 79 | 159 | 150 | 139 | 181 | 186 | 148.83 | 129 | 169 | 1.303 | 0.382 | 0.16757021 | 0.36534661 | 0.0709 | 0.029 | 0.0485 | 0.0485 | TRUE | 1.1779 |
| Ltn1 | 5438 | 4802 | 5249 | 7136 | 5043 | 4827 | 5776 | 4796 | 5271 | 5610 | 5426 | 5284 | 5360.35 | 5281 | 5440 | 1.03 | 0.043 | 0.63395373 | 0.79407494 | 0.0046 | 0.0075 | 0.0057 | 0.0057 | TRUE | 0.082 |
| Ltv1 | 4999 | 4437 | 4515 | 6438 | 4539 | 4342 | 5310 | 4431 | 4534 | 5061 | 4884 | 4753 | 4828.73 | 4758 | 4899 | 1.03 | 0.042 | 0.65141401 | 0.80454668 | 0.0044 | 0.0076 | 0.0061 | 0.0061 | TRUE | 0.1251 |
| Luc7l | 4056 | 4620 | 4793 | 6002 | 4713 | 4693 | 4308 | 4614 | 4813 | 4718 | 5071 | 5137 | 4776.89 | 4578 | 4975 | 1.086 | 0.12 | 0.12750417 | 0.30697539 | 0.0023 | 0.0076 | 0.0042 | 0.0042 | TRUE | 0.0324 |
| Luc7l2 | 6369 | 5080 | 5310 | 7573 | 5681 | 5401 | 6765 | 5073 | 5332 | 5953 | 6112 | 5912 | 5858.03 | 5723 | 5992 | 1.047 | 0.066 | 0.57634616 | 0.75418558 | 0.0103 | 0.0075 | 0.01 | 0.01 | TRUE | 0.3089 |
| Luc7l3 | 5685 | 5580 | 5271 | 7085 | 6015 | 5443 | 6038 | 5573 | 5293 | 5570 | 6472 | 5958 | 5817.26 | 5635 | 6000 | 1.065 | 0.09 | 0.32405857 | 0.54535337 | 0.0048 | 0.0075 | 0.0059 | 0.0059 | TRUE | 0.0577 |
| Lum | 0 | 0 | 0 | 0 | 1 | 0 | 0 | 0 | 0 | 0 | 1 | 0 | 0.18 | 0 | 0 | 1.859 | 0.894 | 0.82650734 | NA | 0 | 18.3785 | 10 | 10 | TRUE | 0.5204 |
| Lurap1 | 77 | 91 | 98 | 113 | 61 | 77 | 82 | 91 | 98 | 89 | 66 | 84 | 84.97 | 90 | 80 | 0.884 | -0.179 | 0.43330011 | 0.64227707 | 0 | 0.0457 | 0.0257 | 0.0257 | TRUE | 0.2096 |
| Lurap1l | 162 | 163 | 136 | 164 | 129 | 125 | 172 | 163 | 137 | 129 | 139 | 137 | 146 | 157 | 135 | 0.857 | -0.222 | 0.20597272 | 0.41489166 | 0 | 0.0295 | 0.0153 | 0.0153 | TRUE | 0.1391 |
| Luzp1 | 13404 | 12915 | 13617 | 13230 | 9980 | 9732 | 14237 | 12898 | 13673 | 10400 | 10738 | 10653 | 12100.02 | 13603 | 10597 | 0.779 | -0.36 | 7.08E-08 | 2.67E-06 | 0.0012 | 0.0072 | 0.0031 | 0.0031 | TRUE | 0.0251 |
| Luzp2 | 1 | 1 | 0 | 2 | 2 | 0 | 1 | 1 | 0 | 2 | 2 | 0 | 0.96 | 1 | 1 | 1.832 | 0.874 | 0.67015343 | NA | 0 | 3.4238 | 1.894 | 1.894 | TRUE | 0.3841 |
| Luzp4 | 24 | 18 | 21 | 69 | 42 | 54 | 25 | 18 | 21 | 54 | 45 | 59 | 37.18 | 21 | 53 | 2.462 | 1.3 | 0.0001175 | 0.00156388 | 0 | 0.0955 | 0.049 | 0.049 | TRUE | 0.1408 |
| Lxn | 45 | 65 | 59 | 106 | 91 | 69 | 48 | 65 | 59 | 83 | 98 | 76 | 71.45 | 57 | 86 | 1.491 | 0.576 | 0.02079017 | 0.08929808 | 0 | 0.053 | 0.0301 | 0.0301 | TRUE | 0.1739 |
| Ly6a | 196 | 112 | 75 | 624 | 552 | 514 | 208 | 112 | 75 | 491 | 594 | 563 | 340.41 | 132 | 549 | 4.17 | 2.06 | 5.98E-12 | 5.46E-10 | 0.1233 | 0.0166 | 0.0598 | 0.0598 | TRUE | 2.6167 |
| Ly6c1 | 13 | 9 | 5 | 4 | 6 | 6 | 14 | 9 | 5 | 3 | 6 | 7 | 7.33 | 9 | 5 | 0.572 | -0.805 | 0.28953711 | 0.5106317 | 0 | 0.4563 | 0.269 | 0.269 | TRUE | 0.1547 |
| Ly6c2 | 0 | 1 | 0 | 2 | 1 | 0 | 0 | 1 | 0 | 2 | 1 | 0 | 0.61 | 0 | 1 | 2.461 | 1.299 | 0.62286289 | NA | 0 | 5.4271 | 3.4871 | 3.4871 | TRUE | 0.405 |
| Ly6d | 1 | 0 | 0 | 0 | 0 | 1 | 1 | 0 | 0 | 0 | 0 | 1 | 0.36 | 0 | 0 | 0.954 | -0.067 | 0.98553519 | NA | 0 | 9.1716 | 7.9605 | 7.9605 | TRUE | 0.5383 |
| Ly6e | 4590 | 4662 | 4424 | 4947 | 3796 | 4350 | 4875 | 4656 | 4442 | 3889 | 4084 | 4762 | 4451.42 | 4658 | 4245 | 0.911 | -0.134 | 0.17886935 | 0.38005219 | 0.0066 | 0.0076 | 0.007 | 0.007 | TRUE | 0.1378 |
| Ly6f | 1 | 0 | 0 | 3 | 2 | 2 | 1 | 0 | 0 | 2 | 2 | 2 | 1.29 | 0 | 2 | 6.113 | 2.612 | 0.15938094 | NA | 0 | 2.5536 | 1.2636 | 1.2636 | TRUE | 0.4614 |
| Ly6g5b | 8 | 4 | 1 | 6 | 8 | 2 | 8 | 4 | 1 | 5 | 9 | 2 | 4.83 | 4 | 5 | 1.154 | 0.206 | 0.83885524 | 0.91826936 | 0.3622 | 0.6883 | 0.5323 | 0.5323 | TRUE | 0.3645 |
| Ly6g5c | 0 | 1 | 0 | 2 | 0 | 0 | 0 | 1 | 0 | 2 | 0 | 0 | 0.43 | 0 | 1 | 1.493 | 0.578 | 0.86962266 | NA | 0 | 7.6954 | 7.0545 | 7.0545 | TRUE | 0.9718 |
| Ly6g6c | 19 | 38 | 38 | 21 | 30 | 41 | 20 | 38 | 38 | 17 | 32 | 45 | 31.66 | 32 | 31 | 0.962 | -0.056 | 0.90433736 | 0.95253164 | 0.1443 | 0.111 | 0.1271 | 0.1271 | TRUE | 0.2788 |
| Ly6g6d | 71 | 67 | 59 | 82 | 95 | 80 | 75 | 67 | 59 | 64 | 102 | 88 | 75.97 | 67 | 85 | 1.255 | 0.327 | 0.21531656 | 0.42717126 | 0.0239 | 0.0503 | 0.0369 | 0.0369 | TRUE | 0.3177 |
| Ly6g6e | 1278 | 2478 | 2231 | 1650 | 1482 | 2164 | 1357 | 2475 | 2240 | 1297 | 1595 | 2369 | 1888.82 | 2024 | 1754 | 0.866 | -0.207 | 0.56620789 | 0.74617571 | 0.0936 | 0.0086 | 0.0446 | 0.0936 | TRUE | 0.9111 |
| Ly6g6f | 2 | 0 | 0 | 0 | 0 | 4 | 2 | 0 | 0 | 0 | 0 | 4 | 1.08 | 1 | 1 | 2.051 | 1.036 | 0.71218494 | NA | 10 | 3.0465 | 4.6154 | 4.6154 | TRUE | 1.9001 |
| Ly6h | 70 | 76 | 64 | 67 | 30 | 31 | 74 | 76 | 64 | 53 | 32 | 34 | 55.57 | 71 | 40 | 0.561 | -0.834 | 0.00508141 | 0.03176535 | 0.0192 | 0.0662 | 0.0445 | 0.0445 | TRUE | 0.7151 |
| Ly6i | 0 | 0 | 0 | 0 | 0 | 0 | 0 | 0 | 0 | 0 | 0 | 0 | 0 | 0 | 0 | NA | NA | NA | NA | NA | NA | NA | NA | NA | NA |
| Ly6k | 231 | 282 | 222 | 386 | 281 | 290 | 245 | 282 | 223 | 303 | 302 | 317 | 278.86 | 250 | 307 | 1.23 | 0.299 | 0.03349047 | 0.12413905 | 0.003 | 0.0187 | 0.0106 | 0.0106 | TRUE | 0.1381 |
| Ly75 | 2420 | 2734 | 3045 | 2426 | 1870 | 2324 | 2570 | 2730 | 3058 | 1907 | 2012 | 2544 | 2470.26 | 2786 | 2154 | 0.773 | -0.372 | 0.00475143 | 0.03015678 | 0.0159 | 0.0082 | 0.0121 | 0.0121 | TRUE | 0.3021 |
| Ly86 | 0 | 0 | 1 | 3 | 1 | 1 | 0 | 0 | 1 | 2 | 1 | 1 | 0.92 | 0 | 1 | 4.189 | 2.067 | 0.32121847 | NA | 0 | 3.5794 | 1.8085 | 1.8085 | TRUE | 0.4098 |
| Ly9 | 1 | 0 | 0 | 1 | 4 | 1 | 1 | 0 | 0 | 1 | 4 | 1 | 1.21 | 0 | 2 | 5.56 | 2.475 | 0.2201387 | NA | 0 | 2.7346 | 1.6954 | 1.6954 | TRUE | 0.7865 |
| Ly96 | 0 | 0 | 4 | 7 | 4 | 0 | 0 | 0 | 4 | 6 | 4 | 0 | 2.3 | 1 | 3 | 2.453 | 1.295 | 0.49401375 | NA | 3.7884 | 1.4369 | 2.054 | 2.054 | TRUE | 1.7259 |
| Lyar | 9845 | 9495 | 9256 | 11115 | 8512 | 7896 | 10457 | 9483 | 9294 | 8738 | 9158 | 8643 | 9295.57 | 9745 | 8846 | 0.908 | -0.139 | 0.06648295 | 0.1989088 | 0.0024 | 0.0073 | 0.0041 | 0.0041 | TRUE | 0.0499 |
| Lyg1 | 0 | 0 | 0 | 0 | 0 | 0 | 0 | 0 | 0 | 0 | 0 | 0 | 0 | 0 | 0 | NA | NA | NA | NA | NA | NA | NA | NA | NA | NA |
| Lyg2 | 2 | 2 | 4 | 3 | 4 | 1 | 2 | 2 | 4 | 2 | 4 | 1 | 2.65 | 3 | 2 | 0.95 | -0.075 | 0.94857782 | NA | 0 | 1.2505 | 0.5844 | 0.5844 | TRUE | 0.3078 |
| Lyl1 | 5 | 7 | 13 | 3 | 6 | 5 | 5 | 7 | 13 | 2 | 6 | 5 | 6.61 | 8 | 4 | 0.55 | -0.862 | 0.28378191 | 0.50528429 | 0 | 0.5055 | 0.301 | 0.301 | TRUE | 0.7002 |

| | | | | | | | | | | | | | | | | | | | | | | | | | |
|---|---|---|---|---|---|---|---|---|---|---|---|---|---|---|---|---|---|---|---|---|---|---|---|---|---|
| Lyn | 226 | 282 | 226 | 445 | 290 | 264 | 240 | 282 | 227 | 350 | 312 | 289 | 283.24 | 250 | 317 | 1.273 | 0.348 | 0.02109775 | 0.09034854 | 0.0076 | 0.0185 | 0.0128 | 0.0128 | TRUE | 0.142 |
| Lynx1 | 68 | 92 | 106 | 99 | 58 | 90 | 72 | 92 | 106 | 78 | 62 | 99 | 84.88 | 90 | 80 | 0.881 | -0.183 | 0.48902155 | 0.68540227 | 0.0308 | 0.0457 | 0.0385 | 0.0385 | TRUE | 0.2925 |
| Lypd1 | 26 | 20 | 14 | 53 | 41 | 25 | 28 | 20 | 14 | 42 | 44 | 27 | 29.13 | 21 | 38 | 1.845 | 0.884 | 0.03187177 | 0.12011508 | 0.0414 | 0.12 | 0.0843 | 0.0843 | TRUE | 0.4319 |
| Lypd2 | 4 | 6 | 3 | 1 | 3 | 1 | 4 | 6 | 3 | 1 | 3 | 1 | 3.06 | 4 | 2 | 0.374 | -1.417 | 0.21647991 | NA | 0 | 1.0834 | 0.5379 | 0.5379 | TRUE | 0.4426 |
| Lypd3 | 13 | 11 | 14 | 8 | 7 | 7 | 14 | 11 | 14 | 6 | 8 | 8 | 10.06 | 13 | 7 | 0.55 | -0.863 | 0.14447782 | 0.331834 | 0 | 0.3345 | 0.1442 | 0.1442 | TRUE | 0.0744 |
| Lypd4 | 19 | 24 | 25 | 30 | 23 | 23 | 20 | 24 | 25 | 24 | 25 | 25 | 23.79 | 23 | 25 | 1.059 | 0.082 | 0.82890602 | 0.91328449 | 0 | 0.1454 | 0.0624 | 0.0624 | TRUE | 0.0677 |
| Lypd5 | 2 | 4 | 3 | 4 | 1 | 1 | 2 | 4 | 3 | 3 | 1 | 1 | 2.41 | 3 | 2 | 0.603 | -0.73 | 0.54570475 | NA | 0 | 1.3752 | 0.619 | 0.619 | TRUE | 0.491 |
| Lypd6 | 241 | 189 | 110 | 217 | 185 | 130 | 256 | 189 | 110 | 171 | 199 | 142 | 177.86 | 185 | 171 | 0.923 | -0.116 | 0.69200541 | 0.83105988 | 0.0916 | 0.0254 | 0.0561 | 0.0561 | TRUE | 0.2095 |
| Lypd6b | 17 | 19 | 14 | 9 | 11 | 9 | 18 | 19 | 14 | 7 | 12 | 10 | 13.31 | 17 | 10 | 0.556 | -0.847 | 0.11122962 | 0.28048257 | 0 | 0.2544 | 0.1221 | 0.1221 | TRUE | 0.2361 |
| Lypd8 | 0 | 0 | 0 | 0 | 0 | 0 | 0 | 0 | 0 | 0 | 0 | 0 | 0 | 0 | 0 | NA | NA | NA | NA | NA | NA | NA | NA | NA | NA |
| Lypla1 | 5209 | 5830 | 5678 | 7896 | 5547 | 5745 | 5533 | 5822 | 5701 | 6207 | 5968 | 6289 | 5920.15 | 5685 | 6155 | 1.083 | 0.114 | 0.06610222 | 0.19828153 | 4.00E-04 | 0.0075 | 0.0026 | 0.0026 | TRUE | 0.0085 |
| Lypla2 | 1854 | 2205 | 1912 | 2356 | 1721 | 1715 | 1969 | 2202 | 1920 | 1852 | 1852 | 1877 | 1945.4 | 2030 | 1860 | 0.916 | -0.126 | 0.13464686 | 0.31738906 | 0.002 | 0.0086 | 0.0046 | 0.0046 | TRUE | 0.0664 |
| Lyplal1 | 693 | 696 | 654 | 1338 | 847 | 817 | 736 | 695 | 657 | 1052 | 911 | 894 | 824.23 | 696 | 952 | 1.371 | 0.455 | 2.98E-05 | 0.00050078 | 0.0047 | 0.0109 | 0.0073 | 0.0073 | TRUE | 0.1022 |
| Lyrm1 | 80 | 81 | 61 | 96 | 74 | 63 | 85 | 81 | 61 | 75 | 80 | 69 | 75.2 | 76 | 75 | 0.988 | -0.018 | 0.94038008 | 0.96983421 | 0 | 0.0507 | 0.028 | 0.028 | TRUE | 0.258 |
| Lyrm2 | 317 | 405 | 389 | 485 | 331 | 314 | 337 | 404 | 391 | 381 | 356 | 344 | 368.82 | 377 | 360 | 0.956 | -0.065 | 0.61276254 | 0.78070959 | 0.0033 | 0.0158 | 0.0091 | 0.0091 | TRUE | 0.1002 |
| Lyrm4 | 429 | 552 | 536 | 656 | 479 | 466 | 456 | 551 | 538 | 516 | 515 | 510 | 514.39 | 515 | 514 | 0.997 | -0.004 | 0.97047615 | 0.9846576 | 0.0031 | 0.0133 | 0.0077 | 0.0077 | TRUE | 0.1174 |
| Lyrm5 | 612 | 579 | 597 | 719 | 490 | 508 | 650 | 578 | 599 | 565 | 527 | 556 | 579.38 | 609 | 549 | 0.903 | -0.148 | 0.14998575 | 0.34065871 | 0 | 0.0126 | 0.0058 | 0.0058 | TRUE | 0.04 |
| Lyrm7 | 123 | 205 | 259 | 199 | 118 | 125 | 131 | 205 | 260 | 156 | 127 | 137 | 169.28 | 199 | 140 | 0.706 | -0.501 | 0.05168982 | 0.16770316 | 0.0581 | 0.0264 | 0.0418 | 0.0418 | TRUE | 0.2003 |
| Lyrm9 | 1070 | 1347 | 1265 | 1207 | 921 | 975 | 1137 | 1345 | 1270 | 949 | 991 | 1067 | 1126.51 | 1251 | 1002 | 0.801 | -0.321 | 0.00184945 | 0.01445466 | 0.0046 | 0.0098 | 0.0067 | 0.0067 | TRUE | 0.0761 |
| Lysmd1 | 27 | 26 | 18 | 42 | 27 | 27 | 29 | 26 | 18 | 33 | 29 | 30 | 27.39 | 24 | 31 | 1.265 | 0.339 | 0.36378835 | 0.58120643 | 0 | 0.1272 | 0.0636 | 0.0636 | TRUE | 0.2995 |
| Lysmd2 | 124 | 157 | 151 | 245 | 164 | 137 | 132 | 157 | 152 | 193 | 176 | 150 | 159.86 | 147 | 173 | 1.182 | 0.241 | 0.17611755 | 0.37637393 | 0.0062 | 0.0275 | 0.0167 | 0.0167 | TRUE | 0.1411 |
| Lysmd3 | 745 | 644 | 662 | 1014 | 738 | 707 | 791 | 643 | 665 | 797 | 794 | 774 | 744.05 | 700 | 788 | 1.127 | 0.173 | 0.12309241 | 0.29890892 | 0.0046 | 0.0113 | 0.0077 | 0.0077 | TRUE | 0.1539 |
| Lysmd4 | 230 | 359 | 370 | 344 | 274 | 284 | 244 | 359 | 372 | 270 | 295 | 311 | 308.41 | 325 | 292 | 0.898 | -0.156 | 0.39667549 | 0.61002008 | 0.0245 | 0.0176 | 0.0211 | 0.0211 | TRUE | 0.5288 |
| Lyst | 1072 | 737 | 905 | 1161 | 802 | 793 | 1139 | 736 | 909 | 913 | 863 | 868 | 904.51 | 928 | 881 | 0.95 | -0.074 | 0.64169878 | 0.79835032 | 0.023 | 0.0105 | 0.0169 | 0.0169 | TRUE | 0.2599 |
| Lyve1 | 1 | 1 | 0 | 0 | 0 | 0 | 1 | 1 | 0 | 0 | 0 | 0 | 0.34 | 1 | 0 | 0.261 | -1.936 | 0.60454921 | NA | 0 | 9.5983 | 8.3269 | 8.3269 | TRUE | 0.2507 |
| Lyz1 | 0 | 0 | 0 | 0 | 0 | 0 | 0 | 0 | 0 | 0 | 0 | 0 | 0 | 0 | 0 | NA | NA | NA | NA | NA | NA | NA | NA | NA | NA |
| Lyz2 | 0 | 0 | 0 | 0 | 0 | 0 | 0 | 0 | 0 | 0 | 0 | 0 | 0 | 0 | 0 | NA | NA | NA | NA | NA | NA | NA | NA | NA | NA |
| Lyzl1 | 0 | 0 | 0 | 0 | 0 | 0 | 0 | 0 | 0 | 0 | 0 | 0 | 0 | 0 | 0 | NA | NA | NA | NA | NA | NA | NA | NA | NA | NA |
| Lyzl4 | 1 | 3 | 1 | 6 | 0 | 0 | 1 | 3 | 1 | 5 | 0 | 0 | 1.63 | 2 | 2 | 0.977 | -0.033 | 0.98564522 | NA | 1.5281 | 2.0282 | 1.8191 | 1.8191 | TRUE | 2.7673 |
| Lyzl6 | 0 | 0 | 0 | 0 | 0 | 0 | 0 | 0 | 0 | 0 | 0 | 0 | 0 | 0 | 0 | NA | NA | NA | NA | NA | NA | NA | NA | NA | NA |
| Lzic | 1032 | 1101 | 1109 | 1531 | 1031 | 1156 | 1096 | 1100 | 1114 | 1204 | 1109 | 1265 | 1147.93 | 1103 | 1193 | 1.081 | 0.113 | 0.19668167 | 0.40323193 | 0.0013 | 0.0098 | 0.0046 | 0.0046 | TRUE | 0.0437 |
| Lztfl1 | 592 | 759 | 818 | 1046 | 762 | 880 | 629 | 758 | 821 | 822 | 820 | 963 | 802.27 | 736 | 868 | 1.179 | 0.237 | 0.0751837 | 0.21599682 | 0.0115 | 0.011 | 0.0116 | 0.0116 | TRUE | 0.1912 |
| Lztr1 | 1258 | 1445 | 1503 | 2052 | 1518 | 1527 | 1336 | 1443 | 1509 | 1613 | 1633 | 1672 | 1534.41 | 1429 | 1639 | 1.146 | 0.197 | 0.01681522 | 0.07687235 | 0.0013 | 0.0091 | 0.0042 | 0.0042 | TRUE | 0.039 |
| Lzts1 | 10 | 6 | 7 | 18 | 17 | 12 | 11 | 6 | 7 | 14 | 18 | 13 | 11.54 | 8 | 15 | 1.93 | 0.949 | 0.09944874 | 0.26028966 | 0 | 0.2925 | 0.1418 | 0.1418 | TRUE | 0.2558 |
| Lzts2 | 2155 | 2450 | 2421 | 1769 | 1571 | 1688 | 2289 | 2447 | 2431 | 1391 | 1690 | 1848 | 2015.92 | 2389 | 1643 | 0.687 | -0.542 | 4.80E-06 | 0.00010658 | 0.0099 | 0.0085 | 0.0096 | 0.0096 | TRUE | 0.2214 |
| M1ap | 671 | 739 | 677 | 1004 | 749 | 798 | 713 | 738 | 680 | 789 | 806 | 874 | 766.54 | 710 | 823 | 1.158 | 0.211 | 0.02649593 | 0.10579589 | 0.0011 | 0.0112 | 0.0052 | 0.0052 | TRUE | 0.0339 |
| M6pr | 11553 | 12299 | 12059 | 13662 | 9826 | 10014 | 12271 | 12283 | 12109 | 10740 | 10572 | 10962 | 11489.5 | 12221 | 10758 | 0.88 | -0.184 | 0.00061208 | 0.00609706 | 1.00E-04 | 0.0072 | 0.002 | 0.002 | TRUE | 0.0033 |
| Maats1 | 121 | 126 | 128 | 190 | 173 | 149 | 129 | 126 | 129 | 149 | 186 | 163 | 146.92 | 128 | 166 | 1.299 | 0.377 | 0.03002251 | 0.11536709 | 0 | 0.0293 | 0.0148 | 0.0148 | TRUE | 0.1159 |
| Mab21l1 | 1 | 0 | 0 | 0 | 0 | 0 | 1 | 0 | 0 | 0 | 0 | 0 | 0.18 | 0 | 0 | 0.49 | -1.029 | 0.80087 | NA | 0 | 18.6165 | 10 | 10 | TRUE | 0.4614 |
| Mab21l2 | 6 | 2 | 4 | 4 | 1 | 4 | 6 | 2 | 4 | 3 | 1 | 4 | 3.5 | 4 | 3 | 0.699 | -0.517 | 0.62321663 | NA | 0 | 0.9488 | 0.5047 | 0.5047 | TRUE | 0.1766 |
| Mab21l3 | 347 | 484 | 464 | 370 | 316 | 378 | 369 | 483 | 466 | 291 | 340 | 414 | 393.75 | 439 | 348 | 0.791 | -0.339 | 0.0523625 | 0.16923389 | 0.0234 | 0.0153 | 0.0194 | 0.0194 | TRUE | 0.308 |
| Macc1 | 0 | 0 | 2 | 0 | 0 | 0 | 0 | 0 | 2 | 0 | 0 | 0 | 0.33 | 1 | 0 | 0.267 | -1.904 | 0.63742563 | NA | 0.0046 | 9.8498 | 10 | 10 | TRUE | 0.974 |
| Macf1 | 14381 | 14193 | 18109 | 17729 | 11604 | 11859 | 15275 | 14175 | 18183 | 13937 | 12485 | 12982 | 14506.18 | 15878 | 13135 | 0.827 | -0.274 | 0.0128036 | 0.06316496 | 0.01 | 0.0071 | 0.0086 | 0.0086 | TRUE | 0.1974 |
| Macrod1 | 157 | 421 | 408 | 201 | 136 | 183 | 167 | 420 | 410 | 158 | 146 | 200 | 250.26 | 332 | 168 | 0.506 | -0.984 | 0.00195413 | 0.0151229 | 0.1285 | 0.0201 | 0.0682 | 0.0682 | TRUE | 2.1489 |
| Macrod2 | 433 | 410 | 421 | 549 | 359 | 360 | 460 | 409 | 423 | 432 | 386 | 394 | 417.34 | 431 | 404 | 0.94 | -0.089 | 0.44217932 | 0.64905093 | 0.0014 | 0.0148 | 0.0074 | 0.0074 | TRUE | 0.0438 |
| Mad1l1 | 1938 | 1626 | 1595 | 3116 | 2148 | 1789 | 2058 | 1624 | 1602 | 2450 | 2311 | 1958 | 2000.49 | 1761 | 2240 | 1.272 | 0.347 | 0.01009711 | 0.05311929 | 0.0163 | 0.0086 | 0.0126 | 0.0126 | TRUE | 0.2626 |
| Mad2l1 | 4873 | 5242 | 5046 | 6817 | 4863 | 4871 | 5176 | 5235 | 5067 | 5359 | 5232 | 5332 | 5233.55 | 5159 | 5308 | 1.029 | 0.041 | 0.47506541 | 0.67538594 | 0 | 0.0075 | 0.0022 | 0.0022 | TRUE | 0.003 |
| Mad2l1bp | 554 | 447 | 418 | 824 | 515 | 421 | 588 | 446 | 420 | 648 | 554 | 461 | 519.55 | 485 | 554 | 1.145 | 0.195 | 0.2744569 | 0.49488215 | 0.0285 | 0.0132 | 0.0211 | 0.0211 | TRUE | 0.2586 |
| Mad2l2 | 1934 | 2265 | 2238 | 2353 | 1622 | 1841 | 2054 | 2262 | 2247 | 1850 | 1745 | 2015 | 2028.95 | 2188 | 1870 | 0.855 | -0.227 | 0.01260877 | 0.06243815 | 0.0033 | 0.0085 | 0.0055 | 0.0055 | TRUE | 0.0548 |
| Madcam1 | 49 | 48 | 55 | 42 | 38 | 33 | 52 | 48 | 55 | 33 | 41 | 36 | 44.21 | 52 | 37 | 0.706 | -0.502 | 0.08166528 | 0.2278077 | 0 | 0.0814 | 0.0368 | 0.0368 | TRUE | 0.0711 |
| Madd | 927 | 627 | 655 | 995 | 717 | 647 | 985 | 626 | 658 | 782 | 771 | 708 | 755.07 | 756 | 754 | 0.998 | -0.003 | 0.98486631 | 0.99199132 | 0.0319 | 0.0113 | 0.0215 | 0.0215 | TRUE | 0.8254 |
| Maea | 8784 | 8973 | 9017 | 9592 | 7149 | 7327 | 9330 | 8961 | 9054 | 7540 | 7692 | 8021 | 8433.09 | 9115 | 7751 | 0.85 | -0.234 | 0.00013112 | 0.00171482 | 5.00E-04 | 0.0073 | 0.0026 | 0.0026 | TRUE | 0.0113 |
| Mael | 464 | 639 | 581 | 699 | 509 | 519 | 493 | 638 | 583 | 549 | 548 | 568 | 563.28 | 571 | 555 | 0.971 | -0.043 | 0.73164566 | 0.85592014 | 0.0068 | 0.0128 | 0.0095 | 0.0095 | TRUE | 0.1684 |
| Maf | 3 | 10 | 4 | 2 | 4 | 3 | 3 | 10 | 4 | 2 | 4 | 3 | 4.39 | 6 | 3 | 0.52 | -0.944 | 0.33423244 | NA | 0 | 0.7571 | 0.4362 | 0.4362 | TRUE | 0.9679 |
| Maf1 | 2995 | 3394 | 3122 | 3981 | 2914 | 3170 | 3181 | 3390 | 3135 | 3130 | 3135 | 3470 | 3240.09 | 3235 | 3245 | 1.003 | 0.004 | 0.96099248 | 0.98013965 | 0.0023 | 0.0079 | 0.0043 | 0.0043 | TRUE | 0.0444 |
| Mafa | 167 | 145 | 159 | 235 | 167 | 178 | 177 | 145 | 160 | 185 | 180 | 195 | 173.52 | 161 | 187 | 1.161 | 0.215 | 0.17930152 | 0.38052398 | 0 | 0.0259 | 0.0127 | 0.0127 | TRUE | 0.0867 |
| Mafb | 21 | 31 | 18 | 37 | 21 | 29 | 22 | 31 | 18 | 29 | 23 | 32 | 25.79 | 24 | 28 | 1.17 | 0.227 | 0.57124521 | 0.74993329 | 0 | 0.1346 | 0.0764 | 0.0764 | TRUE | 0.4216 |
| Maff | 473 | 892 | 1007 | 611 | 499 | 638 | 502 | 891 | 1011 | 480 | 537 | 698 | 686.67 | 801 | 572 | 0.713 | -0.488 | 0.04728655 | 0.15785128 | 0.0802 | 0.0117 | 0.0422 | 0.0422 | TRUE | 0.8345 |
| Mafg | 1371 | 1318 | 1259 | 1897 | 1405 | 1103 | 1456 | 1316 | 1264 | 1491 | 1512 | 1207 | 1374.51 | 1345 | 1403 | 1.044 | 0.062 | 0.60420619 | 0.77501139 | 0.0096 | 0.0093 | 0.0095 | 0.0095 | TRUE | 0.1778 |
| Mafk | 512 | 623 | 561 | 687 | 492 | 491 | 544 | 622 | 563 | 540 | 529 | 537 | 556.04 | 576 | 535 | 0.929 | -0.106 | 0.30704664 | 0.5274741 | 0 | 0.0128 | 0.006 | 0.006 | TRUE | 0.0568 |
| Mag | 31 | 58 | 43 | 13 | 19 | 24 | 33 | 58 | 43 | 10 | 20 | 26 | 31.83 | 45 | 19 | 0.417 | -1.26 | 0.00468548 | 0.02983424 | 0.0995 | 0.1104 | 0.1055 | 0.1055 | TRUE | 0.3969 |
| Magea1 | 0 | 0 | 0 | 0 | 0 | 0 | 0 | 0 | 0 | 0 | 0 | 0 | 0 | 0 | 0 | NA | NA | NA | NA | NA | NA | NA | NA | NA | NA |
| Magea10 | 2 | 3 | 3 | 26 | 26 | 17 | 2 | 3 | 3 | 20 | 28 | 19 | 12.53 | 3 | 22 | 8.201 | 3.036 | 1.37E-05 | 0.00026067 | 0 | 0.2699 | 0.1415 | 0.1415 | TRUE | 0.2629 |
| Magea2 | 1 | 2 | 0 | 21 | 20 | 12 | 1 | 2 | 0 | 17 | 22 | 13 | 9.04 | 1 | 17 | 16.703 | 4.062 | 6.95E-05 | 0.00102987 | 0 | 0.3714 | 0.2226 | 0.2226 | TRUE | 0.3331 |
| Magea3 | 1 | 3 | 1 | 27 | 26 | 11 | 1 | 3 | 1 | 21 | 28 | 12 | 11.05 | 2 | 20 | 12.038 | 3.59 | 3.36E-05 | 0.00055332 | 0 | 0.305 | 0.2151 | 0.2151 | TRUE | 0.18 |
| Magea4 | 14 | 5 | 12 | 168 | 109 | 111 | 15 | 5 | 12 | 132 | 117 | 122 | 67.13 | 11 | 124 | 11.705 | 3.549 | 1.06E-23 | 3.81E-21 | 0 | 0.056 | 0.0379 | 0.0379 | TRUE | 0.802 |
| Magea5 | 7 | 13 | 16 | 165 | 119 | 89 | 7 | 13 | 16 | 130 | 128 | 97 | 65.28 | 12 | 118 | 9.704 | 3.279 | 6.25E-21 | 1.82E-18 | 0.0172 | 0.0574 | 0.042 | 0.042 | TRUE | 0.4301 |
| Magea6 | 2 | 3 | 5 | 38 | 22 | 14 | 2 | 3 | 5 | 30 | 24 | 15 | 13.17 | 3 | 23 | 6.807 | 2.767 | 4.87E-05 | 0.0007602 | 0 | 0.2571 | 0.1628 | 0.1628 | TRUE | 0.1307 |
| Magea8 | 5 | 6 | 2 | 143 | 108 | 56 | 5 | 6 | 2 | 112 | 116 | 61 | 50.54 | 4 | 96 | 21.873 | 4.451 | 4.09E-17 | 7.09E-15 | 0 | 0.0721 | 0.0815 | 0.0815 | TRUE | 0.9602 |

| | | | | | | | | | | | | | | | | | | | | | | | | | |
|---|---|---|---|---|---|---|---|---|---|---|---|---|---|---|---|---|---|---|---|---|---|---|---|---|---|
| Mageb1 | 0 | 0 | 0 | 0 | 0 | 0 | 0 | 0 | 0 | 0 | 0 | 0 | 0 | 0 | 0 | NA | NA | NA | NA | NA | NA | NA | NA | NA | NA |
| Mageb16 | 669 | 648 | 604 | 977 | 668 | 738 | 711 | 647 | 606 | 768 | 719 | 808 | 709.81 | 655 | 765 | 1.169 | 0.225 | 0.0361013 | 0.13062393 | 0.0033 | 0.0115 | 0.0069 | 0.0069 | TRUE | 0.0651 |
| Mageb16-ps | 0 | 0 | 0 | 0 | 1 | 0 | 0 | 0 | 0 | 0 | 1 | 0 | 0.18 | 0 | 0 | 1.859 | 0.894 | 0.82650734 | NA | 0 | 18.3785 | 10 | 10 | TRUE | 0.5204 |
| Mageb18 | 4 | 1 | 4 | 22 | 11 | 5 | 4 | 1 | 4 | 17 | 12 | 5 | 7.31 | 3 | 11 | 3.792 | 1.923 | 0.02579273 | 0.10394459 | 0 | 0.4575 | 0.328 | 0.328 | TRUE | 0.1319 |
| Mageb2 | 0 | 0 | 0 | 0 | 0 | 0 | 0 | 0 | 0 | 0 | 0 | 0 | 0 | 0 | 0 | NA | NA | NA | NA | NA | NA | NA | NA | NA | NA |
| Mageb3 | 5 | 8 | 3 | 4 | 3 | 7 | 5 | 8 | 3 | 3 | 3 | 8 | 5.06 | 5 | 5 | 0.848 | -0.238 | 0.78785189 | 0.89079015 | 0 | 0.6582 | 0.3624 | 0.3624 | TRUE | 0.3264 |
| Mageb4 | 75 | 99 | 66 | 153 | 109 | 91 | 80 | 99 | 66 | 120 | 117 | 100 | 97 | 82 | 112 | 1.38 | 0.464 | 0.04320381 | 0.14861733 | 0.0125 | 0.0409 | 0.0274 | 0.0274 | TRUE | 0.3251 |
| Mageb5 | 0 | 0 | 0 | 0 | 0 | 0 | 0 | 0 | 0 | 0 | 0 | 0 | 0 | 0 | 0 | NA | NA | NA | NA | NA | NA | NA | NA | NA | NA |
| Maged1 | 6841 | 6468 | 6538 | 9886 | 7127 | 6528 | 7266 | 6460 | 6565 | 7772 | 7668 | 7146 | 7146.1 | 6764 | 7529 | 1.113 | 0.155 | 0.0538451 | 0.17243553 | 0.0029 | 0.0074 | 0.0045 | 0.0045 | TRUE | 0.0515 |
| Maged2 | 871 | 913 | 870 | 1000 | 732 | 730 | 925 | 912 | 874 | 786 | 788 | 799 | 847.23 | 904 | 791 | 0.875 | -0.192 | 0.02189779 | 0.09291686 | 0 | 0.0108 | 0.0039 | 0.0039 | TRUE | 0.01 |
| Magee1 | 157 | 144 | 156 | 465 | 298 | 281 | 167 | 144 | 157 | 366 | 321 | 308 | 243.5 | 156 | 332 | 2.133 | 1.093 | 3.66E-13 | 3.95E-11 | 0 | 0.0204 | 0.0115 | 0.0115 | TRUE | 0.0984 |
| Magee2 | 96 | 56 | 58 | 168 | 108 | 78 | 102 | 56 | 58 | 132 | 116 | 85 | 91.63 | 72 | 111 | 1.551 | 0.633 | 0.0377348 | 0.13498494 | 0.0704 | 0.0429 | 0.0553 | 0.0553 | TRUE | 0.9903 |
| Mageh1 | 170 | 258 | 261 | 314 | 238 | 207 | 181 | 258 | 262 | 247 | 256 | 227 | 238.3 | 234 | 243 | 1.041 | 0.058 | 0.74714343 | 0.8655848 | 0.0179 | 0.0207 | 0.0193 | 0.0193 | TRUE | 0.4279 |
| Magel2 | 130 | 60 | 34 | 272 | 245 | 91 | 138 | 60 | 34 | 214 | 264 | 100 | 134.86 | 77 | 193 | 2.49 | 1.316 | 0.00574746 | 0.0347648 | 0.3266 | 0.0313 | 0.1545 | 0.1545 | TRUE | 0.8111 |
| Magi1 | 2091 | 2024 | 2064 | 2363 | 1770 | 1522 | 2221 | 2021 | 2072 | 1858 | 1904 | 1666 | 1957.16 | 2105 | 1809 | 0.86 | -0.218 | 0.01472715 | 0.06989126 | 0.0029 | 0.0086 | 0.0052 | 0.0052 | TRUE | 0.0571 |
| Magi2 | 80 | 89 | 81 | 85 | 81 | 68 | 85 | 89 | 81 | 67 | 87 | 74 | 80.6 | 85 | 76 | 0.891 | -0.166 | 0.45730263 | 0.66182576 | 0 | 0.0478 | 0.0236 | 0.0236 | TRUE | 0.1448 |
| Magi3 | 1006 | 749 | 726 | 1740 | 1225 | 959 | 1069 | 748 | 729 | 1368 | 1318 | 1050 | 1046.87 | 849 | 1245 | 1.468 | 0.554 | 0.00147731 | 0.01211262 | 0.0332 | 0.0101 | 0.0209 | 0.0209 | TRUE | 0.6103 |
| Magix | 5 | 2 | 2 | 3 | 4 | 5 | 5 | 2 | 2 | 2 | 4 | 5 | 3.58 | 3 | 4 | 1.29 | 0.367 | 0.71902182 | NA | 0 | 0.9283 | 0.4625 | 0.4625 | TRUE | 0.4901 |
| Magoh | 2378 | 2220 | 2065 | 4084 | 2770 | 2317 | 2526 | 2217 | 2073 | 3211 | 2980 | 2536 | 2590.61 | 2272 | 2909 | 1.281 | 0.357 | 0.00320724 | 0.02232327 | 0.0115 | 0.0082 | 0.0102 | 0.0102 | TRUE | 0.1521 |
| Magohb | 836 | 903 | 804 | 1409 | 971 | 949 | 888 | 902 | 807 | 1108 | 1045 | 1039 | 964.72 | 866 | 1064 | 1.23 | 0.298 | 0.00108014 | 0.00944344 | 0.0013 | 0.0103 | 0.0049 | 0.0049 | TRUE | 0.0416 |
| Magt1 | 2251 | 3093 | 2990 | 2978 | 2110 | 2365 | 2391 | 3089 | 3002 | 2341 | 2270 | 2589 | 2613.73 | 2827 | 2400 | 0.849 | -0.237 | 0.04896921 | 0.16195059 | 0.0109 | 0.0082 | 0.01 | 0.01 | TRUE | 0.221 |
| Mak | 178 | 178 | 167 | 203 | 186 | 176 | 189 | 178 | 168 | 160 | 200 | 193 | 181.15 | 178 | 184 | 1.03 | 0.042 | 0.79818868 | 0.89608114 | 0.0044 | 0.0251 | 0.0143 | 0.0143 | TRUE | 0.1602 |
| Mak16 | 8128 | 7394 | 6914 | 10581 | 6953 | 6315 | 8633 | 7384 | 6942 | 8318 | 7481 | 6913 | 7611.99 | 7653 | 7571 | 0.989 | -0.015 | 0.89184324 | 0.94753967 | 0.0087 | 0.0073 | 0.0092 | 0.0092 | TRUE | 0.1531 |
| Mal | 56 | 52 | 32 | 30 | 24 | 41 | 59 | 52 | 32 | 24 | 26 | 45 | 39.64 | 48 | 32 | 0.653 | -0.616 | 0.12081894 | 0.29543509 | 0.0838 | 0.09 | 0.0872 | 0.0872 | TRUE | 0.8101 |
| Mal2 | 80 | 144 | 124 | 134 | 113 | 135 | 85 | 144 | 125 | 105 | 122 | 148 | 121.33 | 118 | 125 | 1.056 | 0.079 | 0.7527265 | 0.86948897 | 0.0403 | 0.0341 | 0.0371 | 0.0371 | TRUE | 0.5495 |
| Malat1 | 16353 | 16335 | 16060 | 13726 | 11817 | 9745 | 17370 | 16314 | 16126 | 10790 | 12714 | 10667 | 13996.96 | 16603 | 11390 | 0.686 | -0.544 | 5.77E-09 | 2.89E-07 | 0.0058 | 0.0071 | 0.0062 | 0.0062 | TRUE | 0.1261 |
| Mall | 97 | 98 | 104 | 127 | 85 | 103 | 103 | 98 | 104 | 100 | 91 | 113 | 101.56 | 102 | 101 | 0.995 | -0.007 | 0.97154501 | 0.98490859 | 0 | 0.0393 | 0.0183 | 0.0183 | TRUE | 0.0876 |
| Malsu1 | 1067 | 1327 | 1260 | 1440 | 1117 | 1102 | 1133 | 1325 | 1265 | 1132 | 1202 | 1206 | 1210.66 | 1241 | 1180 | 0.95 | -0.074 | 0.43496224 | 0.6433672 | 0.003 | 0.0096 | 0.0057 | 0.0057 | TRUE | 0.069 |
| Malt1 | 995 | 1142 | 1228 | 1042 | 776 | 966 | 1057 | 1141 | 1233 | 819 | 835 | 1057 | 1023.66 | 1144 | 904 | 0.79 | -0.341 | 0.00888387 | 0.04818309 | 0.0124 | 0.0101 | 0.0112 | 0.0112 | TRUE | 0.2609 |
| Mamdc2 | 16 | 18 | 11 | 14 | 11 | 6 | 17 | 18 | 11 | 11 | 12 | 7 | 12.57 | 15 | 10 | 0.643 | -0.636 | 0.2525844 | 0.47061562 | 0 | 0.269 | 0.1402 | 0.1402 | TRUE | 0.2808 |
| Mamdc4 | 38 | 38 | 31 | 45 | 40 | 32 | 40 | 38 | 31 | 35 | 43 | 35 | 37.15 | 36 | 38 | 1.035 | 0.05 | 0.875685 | 0.93865053 | 0 | 0.0956 | 0.0455 | 0.0455 | TRUE | 0.1201 |
| Maml1 | 1853 | 2179 | 2157 | 2999 | 2057 | 1962 | 1968 | 2176 | 2166 | 2358 | 2213 | 2148 | 2171.46 | 2103 | 2240 | 1.065 | 0.091 | 0.27577004 | 0.49645532 | 0.002 | 0.0084 | 0.0046 | 0.0046 | TRUE | 0.0381 |
| Maml2 | 49 | 14 | 25 | 25 | 35 | 30 | 52 | 14 | 25 | 20 | 38 | 33 | 30.21 | 30 | 30 | 0.986 | -0.02 | 0.96898576 | 0.98446557 | 0.2215 | 0.1159 | 0.1669 | 0.1669 | TRUE | 0.6845 |
| Maml3 | 40 | 39 | 48 | 92 | 35 | 43 | 42 | 39 | 48 | 72 | 38 | 47 | 47.78 | 43 | 52 | 1.222 | 0.29 | 0.39153121 | 0.60534847 | 0.0471 | 0.0759 | 0.0613 | 0.0613 | TRUE | 0.7691 |
| Mamld1 | 7 | 1 | 1 | 8 | 2 | 3 | 7 | 1 | 1 | 6 | 2 | 3 | 3.53 | 3 | 4 | 1.273 | 0.348 | 0.76685762 | NA | 0.4816 | 0.9409 | 0.7085 | 0.7085 | TRUE | 1.8402 |
| Mamstr | 61 | 102 | 91 | 49 | 60 | 75 | 65 | 102 | 91 | 39 | 65 | 82 | 73.87 | 86 | 62 | 0.711 | -0.491 | 0.13904556 | 0.32369396 | 0.0808 | 0.0515 | 0.0655 | 0.0655 | TRUE | 0.4895 |
| Man1a | 419 | 492 | 543 | 575 | 423 | 510 | 445 | 491 | 545 | 452 | 455 | 558 | 491.18 | 494 | 488 | 0.988 | -0.018 | 0.89676059 | 0.95009983 | 0.0104 | 0.0136 | 0.0119 | 0.0119 | TRUE | 0.1789 |
| Man1a2 | 2687 | 2391 | 2372 | 4178 | 2834 | 2697 | 2854 | 2388 | 2382 | 3284 | 3049 | 2952 | 2818.27 | 2541 | 3095 | 1.219 | 0.285 | 0.00535911 | 0.03291426 | 0.0067 | 0.0081 | 0.0072 | 0.0072 | TRUE | 0.1403 |
| Man1b1 | 3530 | 4260 | 4224 | 5127 | 3879 | 3922 | 3749 | 4254 | 4241 | 4030 | 4174 | 4293 | 4123.76 | 4081 | 4166 | 1.02 | 0.029 | 0.72288572 | 0.85005881 | 0.0028 | 0.0077 | 0.0046 | 0.0046 | TRUE | 0.0616 |
| Man1c1 | 739 | 669 | 669 | 582 | 482 | 597 | 785 | 668 | 672 | 458 | 519 | 654 | 625.74 | 708 | 544 | 0.766 | -0.385 | 0.01303959 | 0.06394915 | 0.019 | 0.0122 | 0.0157 | 0.0157 | TRUE | 0.367 |
| Man2a1 | 3162 | 3254 | 3465 | 5486 | 3495 | 3463 | 3359 | 3250 | 3479 | 4313 | 3760 | 3791 | 3658.58 | 3363 | 3955 | 1.177 | 0.235 | 0.00621091 | 0.03696547 | 0.0033 | 0.0078 | 0.005 | 0.005 | TRUE | 0.0771 |
| Man2a2 | 1824 | 1939 | 1963 | 2020 | 1707 | 1615 | 1937 | 1936 | 1971 | 1588 | 1837 | 1768 | 1839.57 | 1948 | 1731 | 0.888 | -0.172 | 0.04479141 | 0.15241642 | 0.0024 | 0.0087 | 0.0048 | 0.0048 | TRUE | 0.0647 |
| Man2b1 | 3780 | 4346 | 4418 | 4279 | 3227 | 3529 | 4015 | 4340 | 4436 | 3364 | 3472 | 3863 | 3915.08 | 4264 | 3566 | 0.836 | -0.258 | 0.00315571 | 0.02207175 | 0.0037 | 0.0077 | 0.0053 | 0.0053 | TRUE | 0.0642 |
| Man2b2 | 681 | 946 | 988 | 1101 | 866 | 978 | 723 | 945 | 992 | 866 | 932 | 1071 | 921.34 | 887 | 956 | 1.077 | 0.107 | 0.46921687 | 0.67142882 | 0.0186 | 0.0105 | 0.0147 | 0.0147 | TRUE | 0.308 |
| Man2c1 | 2072 | 2409 | 2519 | 2716 | 2154 | 1947 | 2201 | 2406 | 2529 | 2135 | 2318 | 2131 | 2286.67 | 2379 | 2195 | 0.922 | -0.117 | 0.18524929 | 0.38863419 | 0.003 | 0.0083 | 0.0051 | 0.0051 | TRUE | 0.0517 |
| Manba | 5422 | 4917 | 4962 | 9755 | 7127 | 6741 | 5759 | 4911 | 4982 | 7669 | 7668 | 7379 | 6394.68 | 5217 | 7572 | 1.451 | 0.537 | 6.00E-10 | 3.86E-08 | 0.0041 | 0.0074 | 0.0053 | 0.0053 | TRUE | 0.1004 |
| Manbal | 842 | 828 | 764 | 1182 | 770 | 787 | 894 | 827 | 767 | 929 | 828 | 861 | 851.26 | 829 | 873 | 1.054 | 0.076 | 0.46548166 | 0.6680031 | 0.0035 | 0.0108 | 0.0066 | 0.0066 | TRUE | 0.0552 |
| Manea | 858 | 915 | 904 | 1116 | 751 | 788 | 911 | 914 | 908 | 877 | 808 | 863 | 880.13 | 911 | 849 | 0.933 | -0.1 | 0.24065785 | 0.45703258 | 0 | 0.0106 | 0.0041 | 0.0041 | TRUE | 0.0212 |
| Maneal | 258 | 370 | 354 | 421 | 319 | 347 | 274 | 370 | 355 | 331 | 343 | 380 | 342.17 | 333 | 351 | 1.053 | 0.075 | 0.62653648 | 0.78965426 | 0.0119 | 0.0165 | 0.0142 | 0.0142 | TRUE | 0.27 |
| Manf | 5813 | 5859 | 5097 | 6097 | 4440 | 4239 | 6174 | 5851 | 5118 | 4793 | 4777 | 4640 | 5225.7 | 5714 | 4737 | 0.829 | -0.271 | 0.00281813 | 0.02016187 | 0.0044 | 0.0075 | 0.0057 | 0.0057 | TRUE | 0.1018 |
| Mansc1 | 98 | 146 | 129 | 172 | 140 | 160 | 104 | 146 | 130 | 135 | 151 | 175 | 140.07 | 127 | 154 | 1.21 | 0.275 | 0.17730609 | 0.37791649 | 0.0151 | 0.0304 | 0.0228 | 0.0228 | TRUE | 0.2399 |
| Mansc4 | 11 | 13 | 12 | 9 | 10 | 9 | 12 | 13 | 12 | 7 | 11 | 10 | 10.73 | 12 | 9 | 0.746 | -0.423 | 0.45643547 | 0.66122104 | 0 | 0.3138 | 0.1377 | 0.1377 | TRUE | 0.1811 |
| Maoa | 2219 | 2491 | 2437 | 2892 | 2046 | 1962 | 2357 | 2488 | 2447 | 2273 | 2201 | 2148 | 2319.05 | 2431 | 2207 | 0.909 | -0.138 | 0.04669871 | 0.15657569 | 4.00E-04 | 0.0083 | 0.0031 | 0.0031 | TRUE | 0.0085 |
| Maob | 193 | 187 | 216 | 179 | 98 | 119 | 205 | 187 | 217 | 141 | 105 | 130 | 164.18 | 203 | 125 | 0.621 | -0.688 | 0.00012177 | 0.00161521 | 0.0058 | 0.027 | 0.0167 | 0.0167 | TRUE | 0.1928 |
| Map1a | 4783 | 4289 | 4830 | 5104 | 3741 | 3291 | 5080 | 4283 | 4850 | 4012 | 4025 | 3603 | 4308.94 | 4738 | 3880 | 0.819 | -0.288 | 0.00270149 | 0.019527 | 0.0053 | 0.0077 | 0.0064 | 0.0064 | TRUE | 0.0858 |
| Map1b | 8460 | 7103 | 8639 | 7246 | 5601 | 4933 | 8986 | 7094 | 8674 | 5696 | 6026 | 5400 | 6979.47 | 8251 | 5707 | 0.692 | -0.532 | 1.17E-06 | 3.17E-05 | 0.0088 | 0.0074 | 0.0085 | 0.0085 | TRUE | 0.184 |
| Map1lc3a | 1319 | 1397 | 1392 | 2043 | 1334 | 1383 | 1401 | 1395 | 1398 | 1606 | 1435 | 1514 | 1458.2 | 1398 | 1518 | 1.087 | 0.12 | 0.13636003 | 0.31988395 | 0.0011 | 0.0092 | 0.004 | 0.004 | TRUE | 0.0317 |
| Map1lc3b | 4323 | 4825 | 4944 | 5677 | 4208 | 4545 | 4592 | 4819 | 4964 | 4463 | 4528 | 4975 | 4723.41 | 4792 | 4655 | 0.971 | -0.042 | 0.59013083 | 0.76449681 | 0.0021 | 0.0076 | 0.0042 | 0.0042 | TRUE | 0.0439 |
| Map1s | 4668 | 4184 | 3998 | 6452 | 4846 | 4306 | 4958 | 4179 | 4014 | 5072 | 5214 | 4714 | 4691.82 | 4384 | 5000 | 1.141 | 0.19 | 0.06708122 | 0.20001306 | 0.0076 | 0.0076 | 0.0075 | 0.0075 | TRUE | 0.1599 |
| Map2 | 597 | 764 | 748 | 737 | 576 | 666 | 634 | 763 | 751 | 579 | 620 | 729 | 679.39 | 716 | 643 | 0.896 | -0.158 | 0.23274093 | 0.44700316 | 0.01 | 0.0118 | 0.0112 | 0.0112 | TRUE | 0.1613 |
| Map2k1 | 6417 | 7380 | 7411 | 9431 | 7015 | 7387 | 6816 | 7370 | 7441 | 7414 | 7548 | 8086 | 7445.96 | 7209 | 7683 | 1.065 | 0.092 | 0.21919478 | 0.43131916 | 0.002 | 0.0073 | 0.0039 | 0.0039 | TRUE | 0.0278 |
| Map2k2 | 3331 | 3459 | 3465 | 4884 | 3596 | 3135 | 3538 | 3455 | 3479 | 3839 | 3869 | 3432 | 3602.02 | 3491 | 3713 | 1.064 | 0.09 | 0.24814353 | 0.46486754 | 0.0018 | 0.0078 | 0.0041 | 0.0041 | TRUE | 0.0531 |
| Map2k3 | 3142 | 3077 | 2967 | 4099 | 3127 | 2886 | 3337 | 3073 | 2979 | 3222 | 3364 | 3159 | 3189.25 | 3130 | 3248 | 1.038 | 0.054 | 0.48957731 | 0.68569518 | 0.0018 | 0.0079 | 0.0041 | 0.0041 | TRUE | 0.0407 |
| Map2k4 | 4996 | 5436 | 5816 | 7448 | 5249 | 5492 | 5307 | 5429 | 5840 | 5855 | 5648 | 6012 | 5681.66 | 5525 | 5838 | 1.057 | 0.079 | 0.26853775 | 0.48845317 | 0.0015 | 0.0075 | 0.0035 | 0.0035 | TRUE | 0.0303 |
| Map2k5 | 462 | 584 | 595 | 759 | 478 | 551 | 491 | 583 | 597 | 597 | 514 | 603 | 564.26 | 557 | 571 | 1.026 | 0.037 | 0.7728997 | 0.88136607 | 0.0075 | 0.0127 | 0.0099 | 0.0099 | TRUE | 0.1263 |
| Map2k6 | 881 | 1101 | 1115 | 1285 | 952 | 1062 | 936 | 1100 | 1120 | 1010 | 1024 | 1163 | 1058.65 | 1052 | 1066 | 1.013 | 0.018 | 0.87342322 | 0.93762803 | 0.0068 | 0.01 | 0.0082 | 0.0082 | TRUE | 0.1105 |
| Map2k7 | 3216 | 4221 | 4307 | 4483 | 3456 | 3748 | 3416 | 4216 | 4325 | 3524 | 3718 | 4103 | 3883.6 | 3986 | 3782 | 0.949 | -0.076 | 0.51327019 | 0.70632889 | 0.0094 | 0.0078 | 0.0095 | 0.0095 | TRUE | 0.1899 |
| Map3k1 | 2881 | 3586 | 4201 | 3530 | 2988 | 3234 | 3060 | 3581 | 4218 | 2775 | 3215 | 3540 | 3398.3 | 3620 | 3177 | 0.877 | -0.189 | 0.18200821 | 0.38384382 | 0.0197 | 0.0079 | 0.0141 | 0.0141 | TRUE | 0.2135 |
| Map3k10 | 739 | 958 | 952 | 1478 | 1081 | 1064 | 785 | 957 | 956 | 1162 | 1163 | 1165 | 1031.22 | 899 | 1163 | 1.293 | 0.371 | 0.00049836 | 0.00516095 | 0.0046 | 0.0101 | 0.0072 | 0.0072 | TRUE | 0.1461 |

| | | | | | | | | | | | | | | | | | | | | | | | | | |
|---|---|---|---|---|---|---|---|---|---|---|---|---|---|---|---|---|---|---|---|---|---|---|---|---|---|
| Map3k11 | 1013 | 947 | 869 | 1284 | 924 | 826 | 1076 | 946 | 873 | 1009 | 994 | 904 | 967.01 | 965 | 969 | 1.005 | 0.008 | 0.94473097 | 0.97241382 | 0.0063 | 0.0103 | 0.008 | 0.008 | TRUE | 0.1207 |
| Map3k12 | 1762 | 2016 | 2054 | 1732 | 1591 | 1748 | 1872 | 2013 | 2062 | 1362 | 1712 | 1913 | 1822.37 | 1982 | 1662 | 0.838 | -0.255 | 0.05573788 | 0.17691646 | 0.0145 | 0.0087 | 0.0123 | 0.0123 | TRUE | 0.3065 |
| Map3k13 | 387 | 285 | 279 | 238 | 153 | 190 | 411 | 285 | 280 | 187 | 165 | 208 | 255.92 | 325 | 187 | 0.574 | -0.801 | 4.98E-05 | 0.00077404 | 0.0279 | 0.0198 | 0.0239 | 0.0239 | TRUE | 0.6007 |
| Map3k14 | 222 | 245 | 216 | 255 | 201 | 203 | 236 | 245 | 217 | 200 | 216 | 222 | 222.72 | 233 | 213 | 0.914 | -0.129 | 0.36193258 | 0.58006064 | 0 | 0.0217 | 0.0099 | 0.0099 | TRUE | 0.0383 |
| Map3k15 | 604 | 683 | 682 | 1205 | 917 | 780 | 642 | 682 | 685 | 947 | 987 | 854 | 799.37 | 670 | 929 | 1.388 | 0.473 | 2.07E-06 | 5.21E-05 | 0.0022 | 0.011 | 0.0059 | 0.0059 | TRUE | 0.0584 |
| Map3k19 | 1 | 0 | 1 | 2 | 1 | 2 | 1 | 0 | 1 | 2 | 1 | 2 | 1.15 | 1 | 2 | 2.353 | 1.234 | 0.4870029 | NA | 0 | 2.8701 | 1.2231 | 1.2231 | TRUE | 0.2563 |
| Map3k2 | 745 | 938 | 1029 | 1226 | 945 | 1015 | 791 | 937 | 1033 | 964 | 1017 | 1111 | 975.49 | 920 | 1031 | 1.119 | 0.162 | 0.1978389 | 0.40450028 | 0.0098 | 0.0103 | 0.0103 | 0.0103 | TRUE | 0.1785 |
| Map3k3 | 3039 | 3660 | 3778 | 4626 | 3253 | 3278 | 3228 | 3655 | 3794 | 3637 | 3500 | 3588 | 3566.94 | 3559 | 3575 | 1.005 | 0.006 | 0.9400439 | 0.96979647 | 0.0034 | 0.0078 | 0.0051 | 0.0051 | TRUE | 0.0803 |
| Map3k4 | 4835 | 6069 | 6367 | 6868 | 5141 | 5766 | 5136 | 6061 | 6393 | 5399 | 5531 | 6312 | 5805.37 | 5863 | 5747 | 0.98 | -0.029 | 0.7944247 | 0.8942097 | 0.0098 | 0.0075 | 0.0088 | 0.0088 | TRUE | 0.1436 |
| Map3k5 | 770 | 876 | 929 | 1201 | 914 | 846 | 818 | 875 | 933 | 944 | 983 | 926 | 913.2 | 875 | 951 | 1.087 | 0.12 | 0.19810511 | 0.40499208 | 0.0014 | 0.0105 | 0.0051 | 0.0051 | TRUE | 0.0395 |
| Map3k6 | 313 | 296 | 299 | 517 | 377 | 349 | 332 | 296 | 300 | 406 | 406 | 382 | 353.73 | 309 | 398 | 1.288 | 0.365 | 0.00214271 | 0.01625534 | 0 | 0.0162 | 0.0073 | 0.0073 | TRUE | 0.0473 |
| Map3k7 | 4329 | 3431 | 3522 | 6108 | 4174 | 3624 | 4598 | 3427 | 3536 | 4802 | 4491 | 3967 | 4136.8 | 3854 | 4420 | 1.147 | 0.198 | 0.14358999 | 0.33037219 | 0.0183 | 0.0077 | 0.013 | 0.013 | TRUE | 0.3472 |
| Map3k8 | 46 | 74 | 58 | 64 | 57 | 61 | 49 | 74 | 58 | 50 | 61 | 67 | 59.9 | 60 | 59 | 0.98 | -0.03 | 0.91587587 | 0.958667 | 0.016 | 0.0619 | 0.0395 | 0.0395 | TRUE | 0.3357 |
| Map3k9 | 1328 | 1904 | 1866 | 1771 | 1352 | 1472 | 1411 | 1902 | 1874 | 1392 | 1455 | 1611 | 1607.33 | 1729 | 1486 | 0.859 | -0.219 | 0.10649375 | 0.27257698 | 0.0147 | 0.009 | 0.0126 | 0.0126 | TRUE | 0.3119 |
| Map4 | 14274 | 14836 | 15139 | 21715 | 15674 | 15742 | 15162 | 14817 | 15201 | 17071 | 16864 | 17232 | 16057.75 | 15060 | 17056 | 1.133 | 0.18 | 0.00058541 | 0.00588096 | 1.00E-04 | 0.0071 | 0.0019 | 0.0019 | TRUE | 0.0024 |
| Map4k1 | 1033 | 1221 | 1249 | 1490 | 1120 | 1111 | 1097 | 1219 | 1254 | 1171 | 1205 | 1216 | 1193.89 | 1190 | 1197 | 1.006 | 0.008 | 0.92872579 | 0.96488131 | 0.0017 | 0.0097 | 0.0049 | 0.0049 | TRUE | 0.0557 |
| Map4k2 | 665 | 707 | 679 | 776 | 626 | 581 | 706 | 706 | 682 | 610 | 674 | 636 | 668.96 | 698 | 640 | 0.916 | -0.127 | 0.17778979 | 0.37837007 | 0 | 0.0118 | 0.0049 | 0.0049 | TRUE | 0.0249 |
| Map4k3 | 2175 | 2421 | 2656 | 3052 | 2403 | 2437 | 2310 | 2418 | 2667 | 2399 | 2585 | 2668 | 2507.9 | 2465 | 2551 | 1.034 | 0.049 | 0.59084697 | 0.76478301 | 0.0038 | 0.0082 | 0.0055 | 0.0055 | TRUE | 0.0623 |
| Map4k4 | 14803 | 14516 | 14316 | 16224 | 12066 | 13212 | 15723 | 14497 | 14375 | 12754 | 12982 | 14463 | 14132.39 | 14865 | 13400 | 0.901 | -0.15 | 0.0689009 | 0.20357906 | 0.0035 | 0.0071 | 0.0048 | 0.0048 | TRUE | 0.0588 |
| Map4k5 | 1633 | 1647 | 1681 | 2470 | 1759 | 1793 | 1735 | 1645 | 1688 | 1942 | 1893 | 1963 | 1810.72 | 1689 | 1933 | 1.144 | 0.194 | 0.00567425 | 0.03442085 | 0 | 0.0087 | 0.003 | 0.003 | TRUE | 0.0066 |
| Map6 | 467 | 454 | 467 | 682 | 466 | 453 | 496 | 453 | 469 | 536 | 501 | 496 | 491.96 | 473 | 511 | 1.083 | 0.115 | 0.27546815 | 0.49602508 | 0 | 0.0136 | 0.0059 | 0.0059 | TRUE | 0.0226 |
| Map6d1 | 112 | 145 | 134 | 146 | 115 | 141 | 119 | 145 | 135 | 115 | 124 | 154 | 131.86 | 133 | 131 | 0.982 | -0.026 | 0.89549447 | 0.94945374 | 0.0096 | 0.0319 | 0.0207 | 0.0207 | TRUE | 0.2441 |
| Map7 | 2754 | 2815 | 2916 | 4311 | 3247 | 3209 | 2925 | 2811 | 2928 | 3389 | 3494 | 3513 | 3176.65 | 2888 | 3465 | 1.2 | 0.262 | 3.81E-05 | 0.00061422 | 0 | 0.0079 | 0.0026 | 0.0026 | TRUE | 0.0066 |
| Map7d1 | 1814 | 1781 | 1698 | 1875 | 1454 | 1465 | 1927 | 1779 | 1705 | 1474 | 1564 | 1604 | 1675.42 | 1804 | 1547 | 0.858 | -0.222 | 0.01054264 | 0.05487741 | 0.0023 | 0.0089 | 0.0048 | 0.0048 | TRUE | 0.043 |
| Map7d2 | 3670 | 3991 | 4032 | 3721 | 2777 | 2902 | 3898 | 3986 | 4049 | 2925 | 2988 | 3177 | 3503.72 | 3978 | 3030 | 0.761 | -0.393 | 1.09E-08 | 5.09E-07 | 7.00E-04 | 0.0078 | 0.0031 | 0.0031 | TRUE | 0.0217 |
| Map9 | 729 | 883 | 972 | 802 | 687 | 699 | 774 | 882 | 976 | 630 | 739 | 765 | 794.5 | 877 | 711 | 0.81 | -0.304 | 0.0188647 | 0.08338931 | 0.0107 | 0.011 | 0.0108 | 0.0108 | TRUE | 0.125 |
| Mapk1 | 16215 | 16795 | 16599 | 21747 | 15626 | 16123 | 17223 | 16773 | 16667 | 17096 | 16813 | 17649 | 17036.87 | 16888 | 17186 | 1.018 | 0.025 | 0.65479758 | 0.80595188 | 4.00E-04 | 0.0071 | 0.0022 | 0.0022 | TRUE | 0.0068 |
| Mapk10 | 15 | 21 | 11 | 72 | 59 | 47 | 16 | 21 | 11 | 57 | 63 | 51 | 36.58 | 16 | 57 | 3.576 | 1.838 | 5.04E-07 | 1.51E-05 | 0 | 0.097 | 0.0559 | 0.0559 | TRUE | 0.3642 |
| Mapk11 | 486 | 532 | 517 | 614 | 451 | 501 | 516 | 531 | 519 | 483 | 485 | 548 | 513.83 | 522 | 505 | 0.967 | -0.049 | 0.64680022 | 0.80161347 | 0 | 0.0133 | 0.0063 | 0.0063 | TRUE | 0.0628 |
| Mapk12 | 291 | 408 | 436 | 406 | 272 | 314 | 309 | 407 | 438 | 319 | 293 | 344 | 351.65 | 385 | 319 | 0.827 | -0.274 | 0.09479301 | 0.25228627 | 0.0166 | 0.0163 | 0.0164 | 0.0164 | TRUE | 0.3371 |
| Mapk13 | 494 | 687 | 698 | 749 | 639 | 833 | 525 | 686 | 701 | 589 | 688 | 912 | 683.31 | 637 | 730 | 1.143 | 0.193 | 0.30056472 | 0.52099206 | 0.0353 | 0.0117 | 0.0236 | 0.0236 | TRUE | 0.5501 |
| Mapk14 | 6193 | 5419 | 5310 | 9475 | 6355 | 5570 | 6578 | 5412 | 5332 | 7449 | 6838 | 6097 | 6284.2 | 5774 | 6795 | 1.177 | 0.235 | 0.04636018 | 0.15580499 | 0.012 | 0.0074 | 0.0099 | 0.0099 | TRUE | 0.1809 |
| Mapk15 | 68 | 55 | 40 | 57 | 72 | 43 | 72 | 55 | 40 | 45 | 77 | 47 | 56.11 | 56 | 56 | 1.009 | 0.013 | 0.96942634 | 0.98448447 | 0.0716 | 0.0656 | 0.0685 | 0.0685 | TRUE | 0.3614 |
| Mapk1ip1 | 552 | 417 | 412 | 493 | 393 | 362 | 586 | 416 | 414 | 388 | 423 | 396 | 437.19 | 472 | 402 | 0.852 | -0.231 | 0.15872337 | 0.35263911 | 0.0195 | 0.0144 | 0.0171 | 0.0171 | TRUE | 0.5171 |
| Mapk1ip1l | 7362 | 8244 | 8626 | 11571 | 8374 | 7829 | 7820 | 8233 | 8661 | 9096 | 9010 | 8570 | 8565.12 | 8238 | 8892 | 1.079 | 0.11 | 0.12146392 | 0.29636897 | 0.0016 | 0.0073 | 0.0035 | 0.0035 | TRUE | 0.0247 |
| Mapk3 | 4535 | 4341 | 4685 | 4233 | 3172 | 3537 | 4817 | 4335 | 4704 | 3328 | 3413 | 3872 | 4078.16 | 4619 | 3538 | 0.766 | -0.385 | 2.59E-05 | 0.00044528 | 0.0045 | 0.0077 | 0.0058 | 0.0058 | TRUE | 0.0828 |
| Mapk4 | 280 | 400 | 491 | 700 | 552 | 466 | 297 | 399 | 493 | 550 | 594 | 510 | 474.04 | 396 | 551 | 1.39 | 0.475 | 0.01002044 | 0.05276874 | 0.0301 | 0.0139 | 0.0223 | 0.0223 | TRUE | 0.2919 |
| Mapk6 | 11348 | 12137 | 12287 | 12326 | 9141 | 9717 | 12054 | 12121 | 12337 | 9690 | 9835 | 10637 | 11112.34 | 12171 | 10054 | 0.826 | -0.276 | 3.48E-05 | 0.00056705 | 0.0012 | 0.0072 | 0.0031 | 0.0031 | TRUE | 0.0314 |
| Mapk7 | 3271 | 3396 | 3454 | 4838 | 3562 | 3304 | 3474 | 3392 | 3468 | 3803 | 3832 | 3617 | 3597.78 | 3445 | 3751 | 1.089 | 0.123 | 0.0542517 | 0.17317504 | 3.00E-04 | 0.0078 | 0.0027 | 0.0027 | TRUE | 0.0117 |
| Mapk8 | 3024 | 3435 | 3434 | 3985 | 2753 | 3105 | 3212 | 3431 | 3448 | 3133 | 2962 | 3399 | 3264.06 | 3364 | 3165 | 0.941 | -0.088 | 0.29039792 | 0.51139353 | 0.0027 | 0.0079 | 0.0047 | 0.0047 | TRUE | 0.0504 |
| Mapk8ip1 | 2045 | 2139 | 2259 | 2454 | 1861 | 1941 | 2172 | 2136 | 2268 | 1929 | 2002 | 2125 | 2105.48 | 2192 | 2019 | 0.92 | -0.12 | 0.12021404 | 0.29418399 | 0.0012 | 0.0085 | 0.0038 | 0.0038 | TRUE | 0.0254 |
| Mapk8ip2 | 466 | 646 | 621 | 521 | 487 | 494 | 495 | 645 | 624 | 410 | 524 | 541 | 539.67 | 588 | 492 | 0.834 | -0.262 | 0.09975077 | 0.26082048 | 0.0194 | 0.013 | 0.0163 | 0.0163 | TRUE | 0.2567 |
| Mapk8ip3 | 1747 | 2219 | 2293 | 2275 | 1791 | 1989 | 1856 | 2216 | 2302 | 1788 | 1927 | 2177 | 2044.48 | 2125 | 1964 | 0.924 | -0.114 | 0.34086229 | 0.56111888 | 0.0105 | 0.0085 | 0.0099 | 0.0099 | TRUE | 0.1482 |
| Mapk9 | 4816 | 5492 | 5776 | 6560 | 4761 | 5128 | 5115 | 5485 | 5800 | 5157 | 5123 | 5613 | 5382.16 | 5467 | 5298 | 0.969 | -0.046 | 0.58007503 | 0.75669188 | 0.0028 | 0.0075 | 0.0047 | 0.0047 | TRUE | 0.0385 |
| Mapkap1 | 2314 | 2395 | 2395 | 3342 | 2385 | 2193 | 2458 | 2392 | 2405 | 2627 | 2566 | 2401 | 2474.76 | 2418 | 2531 | 1.047 | 0.067 | 0.35601911 | 0.57449697 | 8.00E-04 | 0.0082 | 0.0033 | 0.0033 | TRUE | 0.0245 |
| Mapkapk2 | 1593 | 1629 | 1654 | 2614 | 1890 | 1832 | 1692 | 1627 | 1661 | 2055 | 2034 | 2005 | 1845.6 | 1660 | 2031 | 1.224 | 0.292 | 1.79E-05 | 0.00032377 | 0 | 0.0087 | 0.0028 | 0.0028 | TRUE | 0.0037 |
| Mapkapk3 | 3038 | 3527 | 3615 | 2703 | 2213 | 2648 | 3227 | 3522 | 3630 | 2125 | 2381 | 2899 | 2963.96 | 3460 | 2468 | 0.713 | -0.488 | 0.00011527 | 0.00154332 | 0.0141 | 0.008 | 0.0112 | 0.0112 | TRUE | 0.2818 |
| Mapkapk5 | 1830 | 1954 | 1925 | 2727 | 1906 | 1874 | 1944 | 1951 | 1933 | 2144 | 2051 | 2051 | 2012.34 | 1943 | 2082 | 1.072 | 0.101 | 0.13665904 | 0.32031302 | 0 | 0.0085 | 0.0028 | 0.0028 | TRUE | 0.0084 |
| Mapkbp1 | 1456 | 1314 | 1242 | 1447 | 1116 | 1142 | 1547 | 1312 | 1247 | 1138 | 1201 | 1250 | 1282.38 | 1369 | 1196 | 0.874 | -0.195 | 0.07671603 | 0.21872111 | 0.0066 | 0.0095 | 0.0079 | 0.0079 | TRUE | 0.1549 |
| Mapre1 | 7545 | 6494 | 6370 | 11099 | 7536 | 6520 | 8014 | 6486 | 6396 | 8725 | 8108 | 7137 | 7477.75 | 6965 | 7990 | 1.147 | 0.198 | 0.10271007 | 0.26648186 | 0.0133 | 0.0073 | 0.0105 | 0.0105 | TRUE | 0.2117 |
| Mapre2 | 4725 | 4456 | 4572 | 4085 | 3089 | 2958 | 5019 | 4450 | 4591 | 3211 | 3324 | 3238 | 3972.11 | 4687 | 3258 | 0.695 | -0.525 | 6.15E-12 | 5.58E-10 | 0.0018 | 0.0077 | 0.0039 | 0.0039 | TRUE | 0.0467 |
| Mapre3 | 245 | 277 | 251 | 382 | 247 | 279 | 260 | 277 | 252 | 300 | 266 | 305 | 276.73 | 263 | 290 | 1.105 | 0.145 | 0.27533041 | 0.49589028 | 0 | 0.0188 | 0.0091 | 0.0091 | TRUE | 0.0601 |
| Mapt | 1162 | 1133 | 1027 | 1383 | 1025 | 1046 | 1234 | 1132 | 1031 | 1087 | 1103 | 1145 | 1122.01 | 1132 | 1112 | 0.982 | -0.027 | 0.78618272 | 0.89023346 | 0.0035 | 0.0098 | 0.0061 | 0.0061 | TRUE | 0.0738 |
| Mar-01 | 3 | 1 | 0 | 3 | 7 | 5 | 3 | 1 | 0 | 2 | 8 | 5 | 3.26 | 1 | 5 | 3.673 | 1.877 | 0.13850306 | NA | 0.2192 | 1.0181 | 0.6895 | 0.6895 | TRUE | 0.4859 |
| Mar-02 | 1403 | 1735 | 1613 | 1969 | 1406 | 1602 | 1490 | 1733 | 1620 | 1548 | 1513 | 1754 | 1609.48 | 1614 | 1605 | 0.994 | -0.009 | 0.92844658 | 0.96476961 | 0.0054 | 0.009 | 0.0068 | 0.0068 | TRUE | 0.0785 |
| Mar-01 | 5 | 0 | 1 | 9 | 5 | 6 | 5 | 0 | 1 | 7 | 5 | 7 | 4.22 | 2 | 6 | 3.061 | 1.614 | 0.15323465 | NA | 0 | 0.787 | 0.5974 | 0.5974 | TRUE | 1.5926 |
| Mar-10 | 8 | 4 | 7 | 4 | 2 | 8 | 8 | 4 | 7 | 3 | 2 | 9 | 5.6 | 6 | 5 | 0.711 | -0.492 | 0.57410053 | 0.75247257 | 0.1133 | 0.5956 | 0.3676 | 0.3676 | TRUE | 0.9998 |
| Mar-11 | 68 | 117 | 135 | 83 | 89 | 88 | 72 | 117 | 136 | 65 | 96 | 96 | 96.99 | 108 | 86 | 0.788 | -0.343 | 0.2430792 | 0.45974879 | 0.0634 | 0.0409 | 0.0518 | 0.0518 | TRUE | 0.5457 |
| Mar-02 | 1312 | 1454 | 1310 | 1488 | 1071 | 1152 | 1394 | 1452 | 1315 | 1170 | 1152 | 1261 | 1290.7 | 1387 | 1194 | 0.861 | -0.216 | 0.01269837 | 0.06276363 | 0.0016 | 0.0095 | 0.0046 | 0.0046 | TRUE | 0.028 |
| Mar-03 | 29 | 29 | 33 | 42 | 43 | 39 | 31 | 29 | 33 | 33 | 46 | 43 | 35.81 | 31 | 41 | 1.304 | 0.383 | 0.24177785 | 0.45849708 | 0 | 0.0989 | 0.0487 | 0.0487 | TRUE | 0.2417 |
| Mar-04 | 2 | 0 | 1 | 1 | 0 | 1 | 2 | 0 | 1 | 1 | 0 | 1 | 0.83 | 1 | 1 | 0.611 | -0.711 | 0.74305956 | NA | 0 | 3.953 | 2.1268 | 2.1268 | TRUE | 0.1454 |
| Mar-05 | 4759 | 5154 | 5152 | 5883 | 4333 | 4338 | 5055 | 5147 | 5173 | 4625 | 4662 | 4749 | 4901.8 | 5125 | 4679 | 0.913 | -0.132 | 0.02134117 | 0.09106953 | 0 | 0.0076 | 0.0022 | 0.0022 | TRUE | 0.0021 |
| Mar-06 | 5642 | 6194 | 6741 | 7417 | 5321 | 5495 | 5993 | 6186 | 6769 | 5831 | 5725 | 6015 | 6086.39 | 6316 | 5857 | 0.927 | -0.109 | 0.1511282 | 0.34217543 | 0.0021 | 0.0074 | 0.004 | 0.004 | TRUE | 0.048 |
| Mar-07 | 5440 | 5219 | 5349 | 6624 | 5002 | 4932 | 5778 | 5212 | 5371 | 5207 | 5382 | 5399 | 5391.57 | 5454 | 5329 | 0.977 | -0.033 | 0.63789834 | 0.79611447 | 0.0013 | 0.0075 | 0.0034 | 0.0034 | TRUE | 0.033 |
| Mar-08 | 1465 | 1835 | 1816 | 2143 | 1620 | 1739 | 1556 | 1833 | 1823 | 1685 | 1743 | 1904 | 1757.24 | 1737 | 1777 | 1.022 | 0.032 | 0.75554254 | 0.87082848 | 0.0056 | 0.0088 | 0.0069 | 0.0069 | TRUE | 0.1002 |
| Mar-09 | 849 | 911 | 915 | 1105 | 807 | 799 | 902 | 910 | 919 | 869 | 868 | 875 | 890.32 | 910 | 871 | 0.956 | -0.064 | 0.42208054 | 0.63219889 | 0 | 0.0106 | 0.0035 | 0.0035 | TRUE | 8.00E-04 |
| Marcks | 3112 | 3108 | 3100 | 3864 | 3043 | 2531 | 3305 | 3104 | 3113 | 3038 | 3274 | 2771 | 3100.74 | 3174 | 3028 | 0.954 | -0.068 | 0.44306758 | 0.6498709 | 0.0035 | 0.008 | 0.0054 | 0.0054 | TRUE | 0.0661 |
| Marcksl1 | 5858 | 5424 | 5158 | 14232 | 10389 | 9475 | 6222 | 5417 | 5179 | 11188 | 11178 | 10372 | 8259.38 | 5606 | 10913 | 1.947 | 0.961 | 6.46E-25 | 2.68E-22 | 0.0055 | 0.0073 | 0.0061 | 0.0061 | TRUE | 0.1126 |
| Marcksl1-ps4 | 0 | 0 | 0 | 0 | 0 | 0 | 0 | 0 | 0 | 0 | 0 | 0 | 0 | 0 | 0 | NA | NA | NA | NA | NA | NA | NA | NA | NA | NA |

| | | | | | | | | | | | | | | | | | | | | | | | | | |
|---|---|---|---|---|---|---|---|---|---|---|---|---|---|---|---|---|---|---|---|---|---|---|---|---|---|
| Marco | 0 | 0 | 0 | 0 | 1 | 0 | 0 | 0 | 0 | 0 | 1 | 0 | 0.18 | 0 | 0 | 1.859 | 0.894 | 0.82650734 | NA | 0 | 18.3785 | 10 | 10 | TRUE | 0.5204 |
| Marf1 | 3962 | 4378 | 4549 | 4844 | 4139 | 4332 | 4208 | 4372 | 4568 | 3808 | 4453 | 4742 | 4358.62 | 4383 | 4334 | 0.989 | -0.017 | 0.87040712 | 0.93630208 | 0.0069 | 0.0077 | 0.0072 | 0.0072 | TRUE | 0.1386 |
| Mark1 | 679 | 816 | 939 | 934 | 701 | 694 | 721 | 815 | 943 | 734 | 754 | 760 | 787.86 | 826 | 749 | 0.906 | -0.142 | 0.24248195 | 0.45909216 | 0.0079 | 0.0111 | 0.0093 | 0.0093 | TRUE | 0.1808 |
| Mark2 | 2832 | 2675 | 2718 | 3687 | 2914 | 2657 | 3008 | 2672 | 2729 | 2898 | 3135 | 2909 | 2891.83 | 2803 | 2981 | 1.063 | 0.089 | 0.28666211 | 0.50796477 | 0.0026 | 0.008 | 0.0046 | 0.0046 | TRUE | 0.0496 |
| Mark3 | 4978 | 4884 | 4820 | 6914 | 4938 | 4632 | 5288 | 4878 | 4840 | 5435 | 5313 | 5070 | 5137.28 | 5002 | 5273 | 1.054 | 0.077 | 0.29354865 | 0.5144277 | 0.0016 | 0.0075 | 0.0036 | 0.0036 | TRUE | 0.0304 |
| Mark4 | 2227 | 2890 | 2971 | 2411 | 2001 | 2360 | 2365 | 2886 | 2983 | 1895 | 2153 | 2583 | 2477.77 | 2745 | 2210 | 0.805 | -0.313 | 0.02703268 | 0.10723237 | 0.0191 | 0.0082 | 0.014 | 0.014 | TRUE | 0.2633 |
| Mars | 8789 | 7537 | 7649 | 11085 | 8147 | 7257 | 9335 | 7527 | 7680 | 8714 | 8766 | 7944 | 8327.82 | 8181 | 8475 | 1.036 | 0.051 | 0.63184946 | 0.79278407 | 0.0073 | 0.0073 | 0.008 | 0.008 | TRUE | 0.186 |
| Mars2 | 624 | 485 | 469 | 961 | 698 | 491 | 663 | 484 | 471 | 755 | 751 | 537 | 610.34 | 539 | 681 | 1.264 | 0.339 | 0.06878628 | 0.20341504 | 0.0339 | 0.0123 | 0.0233 | 0.0233 | TRUE | 0.4668 |
| Marveld1 | 234 | 328 | 328 | 804 | 611 | 488 | 249 | 328 | 329 | 632 | 657 | 534 | 454.85 | 302 | 608 | 2.014 | 1.01 | 5.36E-11 | 4.13E-09 | 0.015 | 0.0141 | 0.0146 | 0.0146 | TRUE | 0.2661 |
| Marveld2 | 428 | 510 | 512 | 589 | 412 | 437 | 455 | 509 | 514 | 463 | 443 | 478 | 477.12 | 493 | 461 | 0.937 | -0.095 | 0.38884881 | 0.60282666 | 0 | 0.0138 | 0.0066 | 0.0066 | TRUE | 0.0526 |
| Marveld3 | 159 | 140 | 130 | 183 | 165 | 146 | 169 | 140 | 131 | 144 | 178 | 160 | 153.41 | 147 | 161 | 1.094 | 0.129 | 0.48582902 | 0.6827493 | 0.0082 | 0.0284 | 0.0182 | 0.0182 | TRUE | 0.1852 |
| Mas1 | 1 | 2 | 1 | 3 | 3 | 1 | 1 | 2 | 1 | 2 | 3 | 1 | 1.79 | 1 | 2 | 1.645 | 0.718 | 0.6012616 | NA | 0 | 1.8466 | 0.7673 | 0.7673 | TRUE | 0.1693 |
| Masp1 | 5 | 3 | 3 | 2 | 5 | 1 | 5 | 3 | 3 | 2 | 5 | 1 | 3.23 | 4 | 3 | 0.702 | -0.511 | 0.64195626 | NA | 0 | 1.0276 | 0.5511 | 0.5511 | TRUE | 0.8342 |
| Masp2 | 28 | 44 | 36 | 24 | 27 | 41 | 30 | 44 | 36 | 19 | 29 | 45 | 33.77 | 37 | 31 | 0.835 | -0.26 | 0.52791969 | 0.7165533 | 0.0798 | 0.1045 | 0.0925 | 0.0925 | TRUE | 0.4022 |
| Mast1 | 623 | 551 | 497 | 684 | 517 | 527 | 662 | 550 | 499 | 538 | 556 | 577 | 563.65 | 570 | 557 | 0.976 | -0.034 | 0.79451578 | 0.89422522 | 0.0092 | 0.0127 | 0.0108 | 0.0108 | TRUE | 0.2288 |
| Mast2 | 2738 | 2981 | 3040 | 4646 | 3456 | 3253 | 2908 | 2977 | 3052 | 3652 | 3718 | 3561 | 3311.59 | 2979 | 3644 | 1.223 | 0.29 | 5.74E-06 | 0.00012423 | 2.00E-04 | 0.0079 | 0.0026 | 0.0026 | TRUE | 0.0056 |
| Mast3 | 1703 | 1872 | 1870 | 2306 | 1932 | 1969 | 1809 | 1870 | 1878 | 1813 | 2079 | 2155 | 1933.88 | 1852 | 2016 | 1.087 | 0.121 | 0.19479214 | 0.40071236 | 0.0039 | 0.0086 | 0.0057 | 0.0057 | TRUE | 0.0957 |
| Mast4 | 655 | 514 | 580 | 623 | 433 | 419 | 696 | 513 | 582 | 490 | 466 | 459 | 534.29 | 597 | 472 | 0.79 | -0.34 | 0.0137995 | 0.06649734 | 0.0107 | 0.0131 | 0.0118 | 0.0118 | TRUE | 0.2436 |
| Mastl | 2441 | 2456 | 2548 | 3438 | 2460 | 2475 | 2593 | 2453 | 2558 | 2703 | 2647 | 2709 | 2610.47 | 2535 | 2686 | 1.06 | 0.084 | 0.20208391 | 0.40962182 | 0 | 0.0082 | 0.0027 | 0.0027 | TRUE | 0.0097 |
| Mat1a | 10 | 6 | 7 | 24 | 13 | 8 | 11 | 6 | 7 | 19 | 14 | 9 | 10.88 | 8 | 14 | 1.785 | 0.836 | 0.17756659 | 0.37816905 | 0 | 0.3098 | 0.1769 | 0.1769 | TRUE | 0.1343 |
| Mat2a | 15475 | 11175 | 8814 | 20684 | 13793 | 9357 | 16437 | 11161 | 8850 | 16260 | 14840 | 10243 | 12965.2 | 12149 | 13781 | 1.134 | 0.182 | 0.42177474 | 0.63192089 | 0.0642 | 0.0072 | 0.0369 | 0.0369 | TRUE | 0.7209 |
| Mat2b | 5643 | 6353 | 6181 | 6911 | 5329 | 5626 | 5994 | 6345 | 6206 | 5433 | 5734 | 6159 | 5978.36 | 6182 | 5775 | 0.934 | -0.099 | 0.19703822 | 0.40371603 | 0.0022 | 0.0075 | 0.004 | 0.004 | TRUE | 0.0411 |
| Matk | 41 | 103 | 85 | 53 | 60 | 63 | 44 | 103 | 85 | 42 | 65 | 69 | 67.82 | 77 | 59 | 0.751 | -0.413 | 0.25988587 | 0.47912565 | 0.1106 | 0.0555 | 0.0818 | 0.0818 | TRUE | 0.5243 |
| Matn1 | 29 | 17 | 27 | 15 | 18 | 15 | 31 | 17 | 27 | 12 | 19 | 16 | 20.41 | 25 | 16 | 0.63 | -0.667 | 0.14901431 | 0.33928882 | 0 | 0.1683 | 0.1024 | 0.1024 | TRUE | 0.4826 |
| Matn2 | 138 | 197 | 182 | 167 | 102 | 107 | 147 | 197 | 183 | 131 | 110 | 117 | 147.37 | 176 | 119 | 0.682 | -0.551 | 0.00388071 | 0.02574953 | 0.0097 | 0.0293 | 0.0193 | 0.0193 | TRUE | 0.217 |
| Matn3 | 711 | 1078 | 1111 | 743 | 544 | 821 | 755 | 1077 | 1116 | 584 | 585 | 899 | 835.91 | 983 | 689 | 0.701 | -0.512 | 0.01400677 | 0.06735205 | 0.053 | 0.0108 | 0.0301 | 0.0301 | TRUE | 0.8286 |
| Matn4 | 16 | 3 | 6 | 13 | 9 | 12 | 17 | 3 | 6 | 10 | 10 | 13 | 9.84 | 9 | 11 | 1.275 | 0.35 | 0.63519591 | 0.79491607 | 0.2285 | 0.3416 | 0.2884 | 0.2884 | TRUE | 1.6086 |
| Matr3 | 24576 | 27442 | 26715 | 33333 | 24853 | 25064 | 26104 | 27407 | 26825 | 26204 | 26740 | 27436 | 26785.98 | 26779 | 26793 | 1.001 | 0.001 | 0.98955465 | 0.99447279 | 4.00E-04 | 0.007 | 0.0023 | 0.0023 | TRUE | 0.0059 |
| Mau2 | 3538 | 3981 | 4067 | 5467 | 4139 | 4083 | 3758 | 3976 | 4084 | 4298 | 4453 | 4469 | 4173.01 | 3939 | 4407 | 1.118 | 0.162 | 0.01756389 | 0.07908012 | 8.00E-04 | 0.0077 | 0.0031 | 0.0031 | TRUE | 0.0196 |
| Mavs | 1403 | 1460 | 1607 | 1754 | 1278 | 1433 | 1490 | 1458 | 1614 | 1379 | 1375 | 1569 | 1480.75 | 1521 | 1441 | 0.947 | -0.079 | 0.40778223 | 0.62062543 | 0.0034 | 0.0091 | 0.0058 | 0.0058 | TRUE | 0.0716 |
| Max | 4445 | 4105 | 3947 | 4857 | 3513 | 3339 | 4721 | 4100 | 3963 | 3818 | 3780 | 3655 | 4006.22 | 4261 | 3751 | 0.88 | -0.184 | 0.04272539 | 0.1475652 | 0.0044 | 0.0077 | 0.0057 | 0.0057 | TRUE | 0.1084 |
| Maz | 8718 | 9709 | 10037 | 10207 | 7852 | 7932 | 9260 | 9696 | 10078 | 8024 | 8448 | 8683 | 9031.63 | 9678 | 8385 | 0.866 | -0.207 | 0.0031634 | 0.02211577 | 0.0015 | 0.0073 | 0.0034 | 0.0034 | TRUE | 0.0175 |
| Mb | 7 | 14 | 13 | 18 | 8 | 14 | 7 | 14 | 13 | 14 | 9 | 15 | 12.09 | 11 | 13 | 1.106 | 0.145 | 0.79922324 | 0.89649805 | 0 | 0.2793 | 0.1519 | 0.1519 | TRUE | 0.3464 |
| Mb21d1 | 85 | 124 | 85 | 235 | 154 | 126 | 90 | 124 | 85 | 185 | 166 | 138 | 131.31 | 100 | 163 | 1.634 | 0.708 | 0.00143267 | 0.0118304 | 0.0227 | 0.032 | 0.0275 | 0.0275 | TRUE | 0.4367 |
| Mb21d2 | 501 | 575 | 521 | 1209 | 961 | 939 | 532 | 574 | 523 | 950 | 1034 | 1028 | 773.64 | 543 | 1004 | 1.847 | 0.885 | 4.16E-20 | 1.03E-17 | 0.0012 | 0.0112 | 0.0053 | 0.0053 | TRUE | 0.0295 |
| Mbd1 | 631 | 552 | 535 | 952 | 734 | 597 | 670 | 551 | 537 | 748 | 790 | 654 | 658.39 | 586 | 731 | 1.247 | 0.319 | 0.01616015 | 0.07496338 | 0.01 | 0.0119 | 0.0111 | 0.0111 | TRUE | 0.1833 |
| Mbd2 | 610 | 594 | 524 | 992 | 678 | 594 | 648 | 593 | 526 | 780 | 729 | 650 | 654.48 | 589 | 720 | 1.224 | 0.291 | 0.02018978 | 0.08738564 | 0.008 | 0.0119 | 0.0098 | 0.0098 | TRUE | 0.1027 |
| Mbd3 | 5273 | 5167 | 5175 | 7258 | 5239 | 4711 | 5601 | 5160 | 5196 | 5706 | 5637 | 5157 | 5409.48 | 5319 | 5500 | 1.034 | 0.048 | 0.53252012 | 0.7204764 | 0.0022 | 0.0075 | 0.0042 | 0.0042 | TRUE | 0.0361 |
| Mbd3l1 | 0 | 0 | 0 | 0 | 0 | 0 | 0 | 0 | 0 | 0 | 0 | 0 | 0 | 0 | 0 | NA | NA | NA | NA | NA | NA | NA | NA | NA | NA |
| Mbd3l2 | 0 | 0 | 0 | 1 | 0 | 0 | 0 | 0 | 0 | 1 | 0 | 0 | 0.13 | 0 | 0 | 1.859 | 0.894 | 0.82650734 | NA | 0 | 25.1515 | 10 | 10 | TRUE | 0.2563 |
| Mbd4 | 1542 | 1559 | 1542 | 2027 | 1511 | 1420 | 1638 | 1557 | 1548 | 1593 | 1626 | 1554 | 1586.14 | 1581 | 1591 | 1.007 | 0.009 | 0.89768588 | 0.95041294 | 0 | 0.009 | 0.0033 | 0.0033 | TRUE | 0.0118 |
| Mbd5 | 521 | 437 | 503 | 812 | 568 | 482 | 564 | 436 | 505 | 638 | 611 | 528 | 547.1 | 502 | 592 | 1.182 | 0.241 | 0.08106625 | 0.22666213 | 0.0111 | 0.0129 | 0.0119 | 0.0119 | TRUE | 0.1519 |
| Mbd6 | 2268 | 2148 | 2164 | 2552 | 1956 | 1865 | 2409 | 2145 | 2173 | 2006 | 2105 | 2042 | 2146.56 | 2242 | 2051 | 0.914 | -0.129 | 0.11231121 | 0.28184127 | 0.0017 | 0.0084 | 0.0043 | 0.0043 | TRUE | 0.051 |
| Mbip | 943 | 963 | 872 | 1236 | 858 | 868 | 1002 | 962 | 876 | 972 | 923 | 950 | 947.32 | 947 | 948 | 1.003 | 0.004 | 0.96645721 | 0.98304089 | 0.0016 | 0.0104 | 0.0052 | 0.0052 | TRUE | 0.0511 |
| Mbl1 | 0 | 0 | 0 | 1 | 0 | 0 | 0 | 0 | 0 | 1 | 0 | 0 | 0.13 | 0 | 0 | 1.859 | 0.894 | 0.82650734 | NA | 0 | 25.1515 | 10 | 10 | TRUE | 0.2563 |
| Mbl2 | 7 | 10 | 18 | 29 | 21 | 11 | 7 | 10 | 18 | 23 | 23 | 12 | 15.49 | 12 | 19 | 1.623 | 0.698 | 0.21177436 | 0.42244319 | 0.084 | 0.2196 | 0.1569 | 0.1569 | TRUE | 0.8357 |
| Mblac1 | 34 | 26 | 22 | 68 | 44 | 25 | 36 | 26 | 22 | 53 | 47 | 27 | 35.39 | 28 | 42 | 1.534 | 0.618 | 0.11592544 | 0.28752771 | 0.0619 | 0.1 | 0.0816 | 0.0816 | TRUE | 0.7193 |
| Mblac2 | 61 | 47 | 46 | 99 | 67 | 51 | 65 | 47 | 46 | 78 | 72 | 56 | 60.61 | 53 | 69 | 1.31 | 0.389 | 0.16392582 | 0.35985398 | 0.016 | 0.0613 | 0.0396 | 0.0396 | TRUE | 0.3286 |
| Mbnl1 | 305 | 410 | 381 | 712 | 495 | 384 | 324 | 409 | 383 | 560 | 533 | 420 | 438.11 | 372 | 504 | 1.357 | 0.44 | 0.00490114 | 0.03090794 | 0.0161 | 0.0144 | 0.0153 | 0.0153 | TRUE | 0.2432 |
| Mbnl2 | 1335 | 1341 | 1400 | 1775 | 1285 | 1330 | 1418 | 1339 | 1406 | 1395 | 1383 | 1456 | 1399.48 | 1388 | 1411 | 1.017 | 0.024 | 0.75153338 | 0.86874703 | 0 | 0.0093 | 0.0035 | 0.0035 | TRUE | 0.0112 |
| Mbnl3 | 374 | 398 | 453 | 788 | 564 | 517 | 397 | 397 | 455 | 619 | 607 | 566 | 506.97 | 416 | 597 | 1.435 | 0.521 | 3.64E-06 | 8.41E-05 | 0.0019 | 0.0134 | 0.0071 | 0.0071 | TRUE | 0.0752 |
| Mboat1 | 148 | 180 | 169 | 323 | 240 | 245 | 157 | 180 | 170 | 254 | 258 | 268 | 214.5 | 169 | 260 | 1.538 | 0.621 | 1.28E-05 | 0.00024631 | 0 | 0.0223 | 0.0097 | 0.0097 | TRUE | 0.0382 |
| Mboat2 | 2066 | 2494 | 2724 | 2392 | 1810 | 2091 | 2194 | 2491 | 2735 | 1880 | 1947 | 2289 | 2256.2 | 2473 | 2039 | 0.824 | -0.279 | 0.01985498 | 0.08641479 | 0.0111 | 0.0084 | 0.0099 | 0.0099 | TRUE | 0.1386 |
| Mboat4 | 14 | 33 | 33 | 23 | 18 | 22 | 15 | 33 | 33 | 18 | 19 | 24 | 23.75 | 27 | 20 | 0.755 | -0.405 | 0.37082121 | 0.58760362 | 0.0601 | 0.1456 | 0.105 | 0.105 | TRUE | 0.9394 |
| Mboat7 | 1219 | 1509 | 1558 | 1739 | 1316 | 1396 | 1295 | 1507 | 1564 | 1367 | 1416 | 1528 | 1446.23 | 1455 | 1437 | 0.987 | -0.019 | 0.8543103 | 0.92752251 | 0.0058 | 0.0092 | 0.0072 | 0.0072 | TRUE | 0.1117 |
| Mbp | 219 | 262 | 259 | 257 | 232 | 299 | 233 | 262 | 260 | 202 | 250 | 327 | 255.55 | 252 | 260 | 1.029 | 0.041 | 0.83362801 | 0.91548463 | 0.028 | 0.0198 | 0.0239 | 0.0239 | TRUE | 0.3585 |
| Mbtd1 | 10335 | 9738 | 9817 | 9953 | 7656 | 7765 | 10978 | 9725 | 9857 | 7824 | 8237 | 8500 | 9187 | 10187 | 8187 | 0.804 | -0.315 | 8.26E-05 | 0.00118362 | 0.003 | 0.0073 | 0.0045 | 0.0045 | TRUE | 0.0563 |
| Mbtps1 | 12822 | 14272 | 15007 | 20081 | 14224 | 13377 | 13619 | 14254 | 15069 | 15786 | 15304 | 14643 | 14779.15 | 14314 | 15244 | 1.065 | 0.091 | 0.20479293 | 0.41330025 | 0.0019 | 0.0071 | 0.0036 | 0.0036 | TRUE | 0.026 |
| Mbtps2 | 1648 | 2025 | 2213 | 2817 | 2052 | 2336 | 1750 | 2022 | 2222 | 2215 | 2208 | 2557 | 2162.4 | 1998 | 2327 | 1.164 | 0.219 | 0.06147713 | 0.18922467 | 0.01 | 0.0084 | 0.0094 | 0.0094 | TRUE | 0.142 |
| Mc1r | 5 | 1 | 4 | 10 | 3 | 3 | 5 | 1 | 4 | 8 | 3 | 3 | 4.12 | 3 | 5 | 1.426 | 0.511 | 0.61211547 | NA | 0 | 0.8072 | 0.484 | 0.484 | TRUE | 0.7825 |
| Mc2r | 0 | 0 | 0 | 0 | 0 | 0 | 0 | 0 | 0 | 0 | 0 | 0 | 0 | 0 | 0 | NA | NA | NA | NA | NA | NA | NA | NA | NA | NA |
| Mc3r | 0 | 0 | 0 | 1 | 1 | 0 | 0 | 0 | 0 | 1 | 1 | 0 | 0.31 | 0 | 1 | 3.32 | 1.731 | 0.65383052 | NA | 0 | 10.6224 | 8.9541 | 8.9541 | TRUE | 0.202 |
| Mc4r | 1 | 2 | 0 | 1 | 2 | 2 | 1 | 2 | 0 | 1 | 2 | 2 | 1.36 | 1 | 2 | 1.644 | 0.717 | 0.66514503 | NA | 0 | 2.4213 | 1.185 | 1.185 | TRUE | 0.3149 |
| Mc5r | 34 | 7 | 11 | 35 | 17 | 10 | 36 | 7 | 11 | 28 | 18 | 11 | 18.48 | 18 | 19 | 1.059 | 0.083 | 0.90322276 | 0.95212067 | 0.3845 | 0.1851 | 0.283 | 0.283 | TRUE | 1.1486 |
| Mcam | 3331 | 4611 | 4923 | 3590 | 2975 | 4113 | 3538 | 4605 | 4943 | 2822 | 3201 | 4502 | 3935.3 | 4362 | 3508 | 0.804 | -0.315 | 0.09138977 | 0.24662179 | 0.0453 | 0.0077 | 0.0248 | 0.0248 | TRUE | 0.7462 |
| Mcat | 1515 | 1966 | 1860 | 2229 | 1581 | 1684 | 1609 | 1963 | 1868 | 1752 | 1701 | 1843 | 1789.5 | 1813 | 1765 | 0.973 | -0.039 | 0.70046823 | 0.83619323 | 0.0055 | 0.0087 | 0.0068 | 0.0068 | TRUE | 0.1168 |
| Mcc | 553 | 401 | 363 | 448 | 307 | 256 | 587 | 400 | 364 | 352 | 330 | 280 | 385.85 | 450 | 321 | 0.713 | -0.488 | 0.01496851 | 0.0706643 | 0.0378 | 0.0154 | 0.0263 | 0.0263 | TRUE | 0.8111 |
| Mccc1 | 502 | 493 | 478 | 763 | 610 | 504 | 533 | 492 | 480 | 600 | 656 | 552 | 552.23 | 502 | 603 | 1.201 | 0.264 | 0.0206984 | 0.08904943 | 0.0034 | 0.0129 | 0.0076 | 0.0076 | TRUE | 0.0689 |
| Mccc2 | 1336 | 1416 | 1485 | 1638 | 1142 | 1117 | 1419 | 1414 | 1491 | 1288 | 1229 | 1223 | 1343.91 | 1441 | 1247 | 0.865 | -0.209 | 0.00671994 | 0.03901628 | 0 | 0.0094 | 0.0035 | 0.0035 | TRUE | 0.011 |

| | | | | | | | | | | | | | | | | | | | | | | | | | |
|---|---|---|---|---|---|---|---|---|---|---|---|---|---|---|---|---|---|---|---|---|---|---|---|---|---|
| Mcee | 549 | 656 | 629 | 587 | 446 | 436 | 583 | 655 | 632 | 461 | 480 | 477 | 548.08 | 623 | 473 | 0.758 | -0.4 | 9.64E-05 | 0.00133898 | 0 | 0.0129 | 0.0057 | 0.0057 | TRUE | 0.037 |
| Mcf2 | 298 | 265 | 230 | 185 | 122 | 121 | 317 | 265 | 231 | 145 | 131 | 132 | 203.55 | 271 | 136 | 0.505 | -0.985 | 2.13E-08 | 9.43E-07 | 0.0113 | 0.0231 | 0.0168 | 0.0168 | TRUE | 0.1942 |
| Mcf2l | 1886 | 2458 | 2667 | 1607 | 1291 | 1656 | 2003 | 2455 | 2678 | 1263 | 1389 | 1813 | 1933.52 | 2379 | 1488 | 0.625 | -0.677 | 2.74E-05 | 0.00046564 | 0.0283 | 0.0086 | 0.0183 | 0.0183 | TRUE | 0.4367 |
| Mcfd2 | 1422 | 1372 | 1348 | 1798 | 1296 | 1294 | 1510 | 1370 | 1354 | 1413 | 1394 | 1416 | 1409.75 | 1411 | 1408 | 0.998 | -0.003 | 0.97109202 | 0.98480349 | 0.0011 | 0.0092 | 0.0042 | 0.0042 | TRUE | 0.045 |
| Mchr1 | 7 | 2 | 5 | 3 | 0 | 3 | 7 | 2 | 5 | 2 | 0 | 3 | 3.35 | 5 | 2 | 0.395 | -1.339 | 0.26172405 | NA | 0.1799 | 0.9905 | 0.6629 | 0.6629 | TRUE | 0.2591 |
| Mcin | 4 | 8 | 6 | 6 | 8 | 7 | 4 | 8 | 6 | 5 | 9 | 8 | 6.54 | 6 | 7 | 1.132 | 0.179 | 0.80987049 | 0.90213132 | 0 | 0.5105 | 0.2433 | 0.2433 | TRUE | 0.2862 |
| Mcl1 | 30004 | 37580 | 39749 | 36527 | 28672 | 30107 | 31870 | 37531 | 39912 | 28715 | 30849 | 32957 | 33639.01 | 36438 | 30840 | 0.846 | -0.241 | 0.02379399 | 0.0982148 | 0.0087 | 0.007 | 0.0081 | 0.0081 | TRUE | 0.1472 |
| Mcm10 | 5799 | 5861 | 5927 | 8187 | 5879 | 5693 | 6160 | 5853 | 5951 | 6436 | 6325 | 6232 | 6159.61 | 5988 | 6331 | 1.057 | 0.081 | 0.17648158 | 0.37685723 | 2.00E-04 | 0.0074 | 0.0024 | 0.0024 | TRUE | 0.0076 |
| Mcm2 | 17400 | 17744 | 18591 | 24209 | 17741 | 17309 | 18482 | 17721 | 18667 | 19031 | 19088 | 18947 | 18656.19 | 18290 | 19022 | 1.04 | 0.057 | 0.30583943 | 0.52665845 | 3.00E-04 | 0.0071 | 0.0022 | 0.0022 | TRUE | 0.0091 |
| Mcm3 | 16539 | 18188 | 18840 | 21579 | 16302 | 16771 | 17567 | 18165 | 18917 | 16964 | 17540 | 18359 | 17918.56 | 18216 | 17621 | 0.967 | -0.048 | 0.47438771 | 0.67497005 | 0.0014 | 0.0071 | 0.0032 | 0.0032 | TRUE | 0.0164 |
| Mcm3ap | 8088 | 7723 | 7755 | 12310 | 8580 | 8496 | 8591 | 7713 | 7787 | 9677 | 9232 | 9300 | 8716.61 | 8030 | 9403 | 1.171 | 0.228 | 0.00193223 | 0.01496811 | 0.002 | 0.0073 | 0.0038 | 0.0038 | TRUE | 0.0455 |
| Mcm4 | 11213 | 11774 | 12398 | 18312 | 13055 | 13165 | 11910 | 11759 | 12449 | 14395 | 14046 | 14411 | 13161.81 | 12039 | 14284 | 1.186 | 0.247 | 2.22E-05 | 0.00038966 | 4.00E-04 | 0.0072 | 0.0024 | 0.0024 | TRUE | 0.0108 |
| Mcm5 | 19404 | 23059 | 23338 | 31477 | 23301 | 22581 | 20610 | 23029 | 23434 | 24745 | 25070 | 24718 | 23601.2 | 22358 | 24844 | 1.111 | 0.152 | 0.04100133 | 0.1433547 | 0.0023 | 0.007 | 0.004 | 0.004 | TRUE | 0.0572 |
| Mcm6 | 19792 | 23574 | 24617 | 27580 | 20570 | 22744 | 21023 | 23544 | 24718 | 21681 | 22132 | 24897 | 22999.07 | 23095 | 22903 | 0.992 | -0.012 | 0.89933912 | 0.95120551 | 0.005 | 0.007 | 0.0065 | 0.0065 | TRUE | 0.0754 |
| Mcm7 | 16592 | 19628 | 19849 | 23098 | 17441 | 18007 | 17624 | 19603 | 19930 | 18158 | 18765 | 19711 | 18965.26 | 19052 | 18878 | 0.991 | -0.013 | 0.86580704 | 0.93364615 | 0.0024 | 0.0071 | 0.0044 | 0.0044 | TRUE | 0.0526 |
| Mcm8 | 1230 | 1222 | 1309 | 1802 | 1289 | 1075 | 1306 | 1220 | 1314 | 1417 | 1387 | 1177 | 1303.58 | 1280 | 1327 | 1.037 | 0.052 | 0.61146121 | 0.77987759 | 0.0051 | 0.0094 | 0.0069 | 0.0069 | TRUE | 0.1162 |
| Mcm9 | 888 | 781 | 744 | 1257 | 893 | 713 | 943 | 780 | 747 | 988 | 961 | 780 | 866.62 | 823 | 910 | 1.106 | 0.145 | 0.29641356 | 0.51709265 | 0.0141 | 0.0107 | 0.0128 | 0.0128 | TRUE | 0.1916 |
| Mcmbp | 6426 | 7279 | 7804 | 9584 | 7009 | 7500 | 6826 | 7270 | 7836 | 7534 | 7541 | 8210 | 7536.09 | 7311 | 7762 | 1.062 | 0.086 | 0.30151752 | 0.52212718 | 0.0035 | 0.0073 | 0.0049 | 0.0049 | TRUE | 0.0483 |
| Mcmdc2 | 116 | 205 | 182 | 144 | 122 | 157 | 123 | 205 | 183 | 113 | 131 | 172 | 154.5 | 170 | 139 | 0.812 | -0.3 | 0.22974083 | 0.44334667 | 0.0494 | 0.0282 | 0.0385 | 0.0385 | TRUE | 0.6146 |
| Mcoln1 | 1531 | 2054 | 2173 | 2502 | 1909 | 2049 | 1626 | 2051 | 2182 | 1967 | 2054 | 2243 | 2020.54 | 1953 | 2088 | 1.069 | 0.096 | 0.45136147 | 0.65684381 | 0.0133 | 0.0085 | 0.0111 | 0.0111 | TRUE | 0.2587 |
| Mcoln2 | 4 | 3 | 0 | 5 | 3 | 2 | 4 | 3 | 0 | 4 | 3 | 2 | 2.77 | 2 | 3 | 1.308 | 0.387 | 0.75510832 | NA | 0 | 1.1982 | 0.7413 | 0.7413 | TRUE | 0.7996 |
| Mcoln3 | 91 | 126 | 121 | 72 | 45 | 80 | 97 | 126 | 121 | 57 | 48 | 88 | 89.43 | 115 | 64 | 0.557 | -0.843 | 0.00215541 | 0.01633597 | 0.0405 | 0.0437 | 0.0423 | 0.0423 | TRUE | 0.8428 |
| Mcph1 | 6081 | 5536 | 5385 | 7774 | 5513 | 5215 | 6459 | 5529 | 5407 | 6111 | 5932 | 5709 | 5857.78 | 5798 | 5917 | 1.021 | 0.029 | 0.75120867 | 0.86856265 | 0.0043 | 0.0075 | 0.0061 | 0.0061 | TRUE | 0.1211 |
| Mcpt-ps1 | 0 | 0 | 0 | 0 | 0 | 0 | 0 | 0 | 0 | 0 | 0 | 0 | 0 | 0 | 0 | NA | NA | NA | NA | NA | NA | NA | NA | NA | NA |
| Mcpt1 | 0 | 0 | 0 | 0 | 0 | 0 | 0 | 0 | 0 | 0 | 0 | 0 | 0 | 0 | 0 | NA | NA | NA | NA | NA | NA | NA | NA | NA | NA |
| Mcpt2 | 0 | 0 | 0 | 0 | 0 | 0 | 0 | 0 | 0 | 0 | 0 | 0 | 0 | 0 | 0 | NA | NA | NA | NA | NA | NA | NA | NA | NA | NA |
| Mcpt4 | 0 | 0 | 0 | 0 | 0 | 0 | 0 | 0 | 0 | 0 | 0 | 0 | 0 | 0 | 0 | NA | NA | NA | NA | NA | NA | NA | NA | NA | NA |
| Mcpt8 | 0 | 0 | 0 | 0 | 0 | 0 | 0 | 0 | 0 | 0 | 0 | 0 | 0 | 0 | 0 | NA | NA | NA | NA | NA | NA | NA | NA | NA | NA |
| Mcpt9 | 0 | 0 | 0 | 0 | 0 | 0 | 0 | 0 | 0 | 0 | 0 | 0 | 0 | 0 | 0 | NA | NA | NA | NA | NA | NA | NA | NA | NA | NA |
| Mcrs1 | 4053 | 5076 | 5048 | 5833 | 4535 | 4582 | 4305 | 5069 | 5069 | 4585 | 4879 | 5016 | 4820.62 | 4814 | 4827 | 1.002 | 0.003 | 0.97119199 | 0.98480349 | 0.0052 | 0.0076 | 0.0061 | 0.0061 | TRUE | 0.1042 |
| Mctp1 | 2 | 0 | 0 | 4 | 2 | 1 | 2 | 0 | 0 | 3 | 2 | 1 | 1.42 | 1 | 2 | 3.107 | 1.635 | 0.37150105 | NA | 0 | 2.3282 | 1.4502 | 1.4502 | TRUE | 0.8392 |
| Mctp2 | 558 | 656 | 678 | 465 | 355 | 471 | 593 | 655 | 681 | 366 | 382 | 516 | 531.95 | 643 | 421 | 0.654 | -0.614 | 8.46E-05 | 0.00120841 | 0.0186 | 0.0131 | 0.0156 | 0.0156 | TRUE | 0.4401 |
| Mcts1 | 618 | 666 | 682 | 920 | 613 | 654 | 656 | 665 | 685 | 723 | 660 | 716 | 684.18 | 669 | 700 | 1.047 | 0.066 | 0.48360249 | 0.68148223 | 0 | 0.0117 | 0.0049 | 0.0049 | TRUE | 0.029 |
| Mcts2 | 542 | 480 | 490 | 747 | 593 | 471 | 576 | 479 | 492 | 587 | 638 | 516 | 547.99 | 516 | 580 | 1.126 | 0.171 | 0.19301614 | 0.39856646 | 0.0087 | 0.0129 | 0.0106 | 0.0106 | TRUE | 0.12 |
| Mcu | 1453 | 1655 | 1740 | 1591 | 1160 | 1318 | 1543 | 1653 | 1747 | 1251 | 1248 | 1443 | 1480.82 | 1648 | 1314 | 0.797 | -0.328 | 0.00102945 | 0.00911648 | 0.0045 | 0.0091 | 0.0065 | 0.0065 | TRUE | 0.0875 |
| Mdc1 | 13846 | 18197 | 19873 | 15888 | 13034 | 15459 | 14707 | 18174 | 19955 | 12490 | 14024 | 16922 | 16045.16 | 17612 | 14479 | 0.822 | -0.283 | 0.05431593 | 0.17323981 | 0.0235 | 0.0071 | 0.0155 | 0.0155 | TRUE | 0.2665 |
| Mdfi | 769 | 829 | 772 | 607 | 478 | 498 | 817 | 828 | 775 | 477 | 514 | 545 | 659.42 | 807 | 512 | 0.634 | -0.658 | 7.22E-11 | 5.47E-09 | 0.0011 | 0.0119 | 0.0058 | 0.0058 | TRUE | 0.0445 |
| Mdfic | 329 | 371 | 364 | 434 | 331 | 300 | 349 | 371 | 365 | 341 | 356 | 328 | 351.86 | 362 | 342 | 0.945 | -0.082 | 0.47404243 | 0.67472225 | 0 | 0.0163 | 0.0066 | 0.0066 | TRUE | 0.0142 |
| Mdga1 | 135 | 139 | 127 | 167 | 141 | 136 | 143 | 139 | 128 | 131 | 152 | 149 | 140.27 | 137 | 144 | 1.051 | 0.072 | 0.67608782 | 0.81979608 | 0 | 0.0304 | 0.0143 | 0.0143 | TRUE | 0.0687 |
| Mdga2 | 23 | 8 | 6 | 63 | 43 | 14 | 24 | 8 | 6 | 50 | 46 | 15 | 24.93 | 13 | 37 | 2.913 | 1.543 | 0.01699615 | 0.0773408 | 0.3771 | 0.1391 | 0.2478 | 0.2478 | TRUE | 2.497 |
| Mdh1 | 4779 | 5019 | 5036 | 7212 | 5011 | 5046 | 5076 | 5013 | 5057 | 5670 | 5392 | 5524 | 5288.34 | 5049 | 5529 | 1.095 | 0.131 | 0.02591983 | 0.10426103 | 2.00E-04 | 0.0075 | 0.0023 | 0.0023 | TRUE | 0.0062 |
| Mdh1b | 1 | 4 | 5 | 6 | 6 | 0 | 1 | 4 | 5 | 5 | 6 | 0 | 3.54 | 3 | 4 | 1.114 | 0.155 | 0.89974629 | NA | 0.6186 | 0.9371 | 0.8112 | 0.8112 | TRUE | 0.7082 |
| Mdh2 | 16085 | 13958 | 13516 | 20381 | 13914 | 13037 | 17085 | 13940 | 13571 | 16022 | 14971 | 14271 | 14976.7 | 14865 | 15088 | 1.015 | 0.022 | 0.84439581 | 0.92161735 | 0.008 | 0.0071 | 0.0086 | 0.0086 | TRUE | 0.2086 |
| Mdk | 5113 | 6317 | 6605 | 8078 | 5425 | 5125 | 5431 | 6309 | 6632 | 6350 | 5837 | 5610 | 6028.21 | 6124 | 5932 | 0.969 | -0.046 | 0.65316578 | 0.80540538 | 0.006 | 0.0074 | 0.0073 | 0.0073 | TRUE | 0.1194 |
| Mdm1 | 1954 | 1893 | 1842 | 2299 | 1745 | 1812 | 2075 | 1891 | 1850 | 1807 | 1878 | 1984 | 1913.99 | 1939 | 1890 | 0.974 | -0.037 | 0.66289629 | 0.81098358 | 0.0025 | 0.0086 | 0.0048 | 0.0048 | TRUE | 0.046 |
| Mdm2 | 5880 | 6749 | 6023 | 8388 | 5844 | 6223 | 6246 | 6740 | 6048 | 6594 | 6288 | 6812 | 6454.57 | 6345 | 6565 | 1.035 | 0.049 | 0.51740857 | 0.70899808 | 0.0021 | 0.0074 | 0.004 | 0.004 | TRUE | 0.0364 |
| Mdm4 | 5726 | 6852 | 6627 | 5872 | 4955 | 5677 | 6082 | 6843 | 6654 | 4616 | 5331 | 6214 | 5956.86 | 6526 | 5387 | 0.825 | -0.277 | 0.02180448 | 0.09264555 | 0.0126 | 0.0075 | 0.0104 | 0.0104 | TRUE | 0.2199 |
| Mdn1 | 19148 | 12106 | 15235 | 19479 | 13224 | 10815 | 20339 | 12090 | 15298 | 15313 | 14228 | 11839 | 14851.05 | 15909 | 13793 | 0.867 | -0.206 | 0.25452814 | 0.47256243 | 0.0424 | 0.0071 | 0.0234 | 0.0234 | TRUE | 0.3056 |
| Mdp1 | 1123 | 1348 | 1387 | 1681 | 1254 | 1357 | 1193 | 1346 | 1393 | 1321 | 1349 | 1485 | 1347.99 | 1311 | 1385 | 1.056 | 0.079 | 0.42886052 | 0.63860634 | 0.0041 | 0.0093 | 0.0065 | 0.0065 | TRUE | 0.0738 |
| Me1 | 2623 | 3483 | 3635 | 3715 | 2728 | 3091 | 2786 | 3479 | 3650 | 2920 | 2935 | 3384 | 3192.28 | 3305 | 3080 | 0.932 | -0.102 | 0.41190903 | 0.62293884 | 0.0121 | 0.0079 | 0.0109 | 0.0109 | TRUE | 0.2289 |
| Me2 | 6288 | 6722 | 6958 | 8572 | 6205 | 6224 | 6679 | 6713 | 6987 | 6739 | 6676 | 6813 | 6767.81 | 6793 | 6743 | 0.993 | -0.011 | 0.85242181 | 0.92628008 | 2.00E-04 | 0.0074 | 0.0023 | 0.0023 | TRUE | 0.0076 |
| Me3 | 4 | 7 | 5 | 16 | 5 | 6 | 4 | 7 | 5 | 13 | 5 | 7 | 6.8 | 5 | 8 | 1.532 | 0.615 | 0.41701027 | 0.62807995 | 0 | 0.4915 | 0.2611 | 0.2611 | TRUE | 0.8325 |
| Mea1 | 1475 | 1966 | 1952 | 1771 | 1424 | 1452 | 1567 | 1963 | 1960 | 1392 | 1532 | 1589 | 1667.33 | 1830 | 1504 | 0.822 | -0.283 | 0.01648893 | 0.07601969 | 0.0091 | 0.0089 | 0.0095 | 0.0095 | TRUE | 0.1909 |
| Meaf6 | 621 | 665 | 602 | 859 | 606 | 586 | 660 | 664 | 604 | 675 | 652 | 641 | 649.5 | 643 | 656 | 1.022 | 0.031 | 0.74607183 | 0.86509667 | 0 | 0.012 | 0.0051 | 0.0051 | TRUE | 0.0321 |
| Mecom | 8 | 3 | 11 | 7 | 7 | 7 | 8 | 3 | 11 | 6 | 8 | 8 | 7.21 | 7 | 7 | 0.913 | -0.132 | 0.8608984 | 0.93125634 | 0 | 0.4641 | 0.2694 | 0.2694 | TRUE | 0.551 |
| Mecp2 | 1114 | 1320 | 1246 | 1427 | 1106 | 1070 | 1183 | 1318 | 1251 | 1122 | 1190 | 1171 | 1205.96 | 1251 | 1161 | 0.927 | -0.109 | 0.20237702 | 0.40995781 | 0.0011 | 0.0096 | 0.0044 | 0.0044 | TRUE | 0.0267 |
| Mecr | 1152 | 1149 | 1197 | 1305 | 909 | 938 | 1224 | 1148 | 1202 | 1026 | 978 | 1027 | 1100.63 | 1191 | 1010 | 0.849 | -0.237 | 0.00338316 | 0.02323115 | 0 | 0.0099 | 0.0038 | 0.0038 | TRUE | 0.0123 |
| Med1 | 8832 | 6937 | 6857 | 9952 | 6578 | 5942 | 9381 | 6928 | 6885 | 7823 | 7078 | 6504 | 7433.3 | 7731 | 7135 | 0.923 | -0.116 | 0.40919456 | 0.62130006 | 0.0197 | 0.0073 | 0.014 | 0.014 | TRUE | 0.4252 |
| Med10 | 1244 | 1354 | 1305 | 2181 | 1535 | 1377 | 1321 | 1352 | 1310 | 1715 | 1652 | 1507 | 1476.23 | 1328 | 1625 | 1.224 | 0.292 | 0.00060463 | 0.00604709 | 0.0018 | 0.0091 | 0.0045 | 0.0045 | TRUE | 0.0475 |
| Med11 | 975 | 1014 | 975 | 1365 | 918 | 908 | 1036 | 1013 | 979 | 1073 | 988 | 994 | 1013.67 | 1009 | 1018 | 1.01 | 0.015 | 0.86478217 | 0.93305133 | 7.00E-04 | 0.0102 | 0.0043 | 0.0043 | TRUE | 0.0276 |
| Med12 | 3085 | 2886 | 3382 | 3302 | 2390 | 2434 | 3277 | 2882 | 3396 | 2596 | 2571 | 2664 | 2897.77 | 3185 | 2610 | 0.82 | -0.287 | 0.00112003 | 0.00971681 | 0.0032 | 0.008 | 0.0052 | 0.0052 | TRUE | 0.0842 |
| Med12l | 635 | 866 | 901 | 1207 | 872 | 861 | 674 | 865 | 905 | 949 | 938 | 943 | 878.94 | 815 | 943 | 1.157 | 0.211 | 0.09909919 | 0.25971958 | 0.0106 | 0.0107 | 0.0106 | 0.0106 | TRUE | 0.2676 |
| Med13 | 6430 | 6349 | 6809 | 8425 | 5719 | 5533 | 6830 | 6341 | 6837 | 6623 | 6153 | 6057 | 6473.44 | 6669 | 6278 | 0.942 | -0.087 | 0.23982801 | 0.45606069 | 0.0018 | 0.0074 | 0.0038 | 0.0038 | TRUE | 0.0285 |
| Med13l | 6361 | 6835 | 7141 | 9465 | 6494 | 6403 | 6757 | 6826 | 7170 | 7441 | 6987 | 7009 | 7031.65 | 6918 | 7146 | 1.033 | 0.047 | 0.47669559 | 0.67645532 | 9.00E-04 | 0.0074 | 0.003 | 0.003 | TRUE | 0.0161 |
| Med14 | 2323 | 2087 | 2101 | 2868 | 2019 | 1843 | 2467 | 2084 | 2110 | 2255 | 2172 | 2017 | 2184.29 | 2220 | 2148 | 0.968 | -0.047 | 0.63808603 | 0.79613107 | 0.0056 | 0.0084 | 0.0068 | 0.0068 | TRUE | 0.1144 |
| Med15 | 3154 | 2965 | 3208 | 4074 | 2990 | 2966 | 3350 | 2961 | 3221 | 3203 | 3217 | 3247 | 3199.82 | 3177 | 3222 | 1.014 | 0.02 | 0.79080146 | 0.89242185 | 0.0017 | 0.0079 | 0.0039 | 0.0039 | TRUE | 0.0432 |
| Med16 | 2380 | 2331 | 2400 | 3063 | 2255 | 2137 | 2528 | 2328 | 2410 | 2408 | 2426 | 2339 | 2406.54 | 2422 | 2391 | 0.987 | -0.018 | 0.7983336 | 0.89617918 | 6.00E-04 | 0.0083 | 0.0032 | 0.0032 | TRUE | 0.0177 |
| Med17 | 3686 | 3406 | 3256 | 4526 | 3244 | 2989 | 3915 | 3402 | 3269 | 3558 | 3490 | 3272 | 3484.4 | 3529 | 3440 | 0.975 | -0.036 | 0.70299903 | 0.83762068 | 0.0052 | 0.0078 | 0.0063 | 0.0063 | TRUE | 0.1114 |
| Med18 | 607 | 594 | 571 | 789 | 575 | 522 | 645 | 593 | 573 | 620 | 619 | 571 | 603.61 | 604 | 603 | 1 | 0.001 | 0.99544082 | 0.99778155 | 0.0011 | 0.0124 | 0.006 | 0.006 | TRUE | 0.041 |

| | | | | | | | | | | | | | | | | | | | | | | | | |
|---|---|---|---|---|---|---|---|---|---|---|---|---|---|---|---|---|---|---|---|---|---|---|---|---|
| Med19 | 2693 | 2722 | 2695 | 3873 | 2724 | 2711 | 2860 | 2718 | 2706 | 3045 | 2931 | 2968 | 2871.35 | 2761 | 2981 | 1.08 | 0.111 | 0.09591418 | 0.25398899 | 3.00E-04 | 0.0081 | 0.0028 | 0.0028 | TRUE | 0.0118 |
| Med20 | 1562 | 1344 | 1375 | 1897 | 1331 | 1209 | 1659 | 1342 | 1381 | 1491 | 1432 | 1323 | 1438.14 | 1461 | 1415 | 0.97 | -0.044 | 0.69402653 | 0.83228303 | 0.0079 | 0.0092 | 0.0084 | 0.0084 | TRUE | 0.1695 |
| Med21 | 824 | 884 | 754 | 1285 | 791 | 807 | 875 | 883 | 757 | 1010 | 851 | 883 | 876.64 | 838 | 915 | 1.093 | 0.128 | 0.2677913 | 0.48754524 | 0.0067 | 0.0107 | 0.0084 | 0.0084 | TRUE | 0.1017 |
| Med22 | 1783 | 1581 | 1478 | 2468 | 1639 | 1415 | 1894 | 1579 | 1484 | 1940 | 1763 | 1549 | 1701.57 | 1652 | 1751 | 1.06 | 0.085 | 0.51499278 | 0.70743379 | 0.0139 | 0.0088 | 0.0116 | 0.0116 | TRUE | 0.1969 |
| Med23 | 3134 | 2508 | 2653 | 3736 | 2762 | 2576 | 3329 | 2505 | 2664 | 2937 | 2972 | 2820 | 2871.01 | 2833 | 2910 | 1.027 | 0.039 | 0.74429428 | 0.86347991 | 0.0114 | 0.0081 | 0.0099 | 0.0099 | TRUE | 0.2849 |
| Med24 | 7766 | 6825 | 6859 | 8696 | 6544 | 6330 | 8249 | 6816 | 6887 | 6836 | 7041 | 6929 | 7126.41 | 7317 | 6935 | 0.948 | -0.077 | 0.41840988 | 0.62905885 | 0.0059 | 0.0074 | 0.0064 | 0.0064 | TRUE | 0.1512 |
| Med25 | 5085 | 5465 | 5551 | 6280 | 4785 | 5070 | 5401 | 5458 | 5574 | 4937 | 5148 | 5550 | 5344.67 | 5478 | 5212 | 0.951 | -0.072 | 0.32385707 | 0.54518878 | 0.0016 | 0.0075 | 0.0037 | 0.0037 | TRUE | 0.0392 |
| Med26 | 2128 | 2168 | 2135 | 2882 | 2242 | 2105 | 2260 | 2165 | 2144 | 2266 | 2412 | 2304 | 2258.56 | 2190 | 2327 | 1.063 | 0.088 | 0.21673609 | 0.42872021 | 5.00E-04 | 0.0084 | 0.0032 | 0.0032 | TRUE | 0.0123 |
| Med27 | 598 | 609 | 605 | 939 | 665 | 633 | 635 | 608 | 607 | 738 | 715 | 693 | 666.24 | 617 | 715 | 1.161 | 0.215 | 0.01897019 | 0.08359818 | 0 | 0.0118 | 0.0046 | 0.0046 | TRUE | 0.0095 |
| Med28 | 1711 | 1933 | 1864 | 2990 | 1987 | 1977 | 1817 | 1931 | 1872 | 2351 | 2138 | 2164 | 2045.35 | 1873 | 2218 | 1.184 | 0.244 | 0.00186543 | 0.01455073 | 0.0014 | 0.0085 | 0.0039 | 0.0039 | TRUE | 0.0341 |
| Med29 | 594 | 530 | 506 | 959 | 672 | 530 | 631 | 529 | 508 | 754 | 723 | 580 | 620.9 | 556 | 686 | 1.235 | 0.304 | 0.03588348 | 0.13007452 | 0.0147 | 0.0122 | 0.0135 | 0.0135 | TRUE | 0.2114 |
| Med30 | 1222 | 1561 | 1574 | 1601 | 1160 | 1288 | 1298 | 1559 | 1580 | 1259 | 1248 | 1410 | 1392.34 | 1479 | 1306 | 0.882 | -0.181 | 0.1050525 | 0.2702574 | 0.007 | 0.0093 | 0.0083 | 0.0083 | TRUE | 0.1377 |
| Med31 | 765 | 835 | 782 | 1284 | 851 | 906 | 813 | 834 | 785 | 1009 | 916 | 992 | 891.41 | 811 | 972 | 1.2 | 0.263 | 0.00336197 | 0.0231247 | 7.00E-04 | 0.0106 | 0.0047 | 0.0047 | TRUE | 0.0306 |
| Med4 | 2523 | 2293 | 2153 | 3392 | 2334 | 2202 | 2680 | 2290 | 2162 | 2667 | 2511 | 2410 | 2453.33 | 2377 | 2529 | 1.064 | 0.09 | 0.39254018 | 0.60626162 | 0.0067 | 0.0082 | 0.0076 | 0.0076 | TRUE | 0.15 |
| Med6 | 1093 | 1062 | 1017 | 1756 | 1207 | 1121 | 1161 | 1061 | 1021 | 1380 | 1299 | 1227 | 1191.49 | 1081 | 1302 | 1.206 | 0.27 | 0.00475869 | 0.03017855 | 0.0031 | 0.0097 | 0.0057 | 0.0057 | TRUE | 0.0499 |
| Med7 | 1774 | 1935 | 1907 | 2864 | 1978 | 1816 | 1884 | 1933 | 1915 | 2251 | 2128 | 1988 | 2016.53 | 1911 | 2122 | 1.112 | 0.153 | 0.05630251 | 0.17814528 | 0.0016 | 0.0085 | 0.0041 | 0.0041 | TRUE | 0.0371 |
| Med8 | 1756 | 1678 | 1684 | 2320 | 1583 | 1489 | 1865 | 1676 | 1691 | 1824 | 1703 | 1630 | 1731.48 | 1744 | 1719 | 0.986 | -0.02 | 0.82501851 | 0.91128174 | 0.0028 | 0.0088 | 0.0051 | 0.0051 | TRUE | 0.0444 |
| Med9 | 2547 | 2368 | 2322 | 3425 | 2388 | 2199 | 2705 | 2365 | 2332 | 2692 | 2569 | 2407 | 2511.8 | 2467 | 2556 | 1.037 | 0.052 | 0.58304147 | 0.75875085 | 0.0046 | 0.0082 | 0.006 | 0.006 | TRUE | 0.0862 |
| Medag | 12 | 8 | 3 | 12 | 8 | 6 | 13 | 8 | 3 | 9 | 9 | 7 | 8.06 | 8 | 8 | 1.046 | 0.064 | 0.93012937 | 0.96557628 | 0.0801 | 0.4157 | 0.2626 | 0.2626 | TRUE | 0.2029 |
| Mef2a | 402 | 392 | 469 | 662 | 458 | 422 | 427 | 391 | 471 | 520 | 493 | 462 | 460.76 | 430 | 492 | 1.145 | 0.196 | 0.10673633 | 0.27288789 | 0.0038 | 0.0141 | 0.0084 | 0.0084 | TRUE | 0.0817 |
| Mef2b | 118 | 187 | 179 | 139 | 116 | 119 | 125 | 187 | 180 | 109 | 125 | 130 | 142.7 | 164 | 121 | 0.738 | -0.438 | 0.03909761 | 0.13844802 | 0.0207 | 0.03 | 0.0253 | 0.0253 | TRUE | 0.4432 |
| Mef2c | 8 | 5 | 8 | 8 | 2 | 4 | 8 | 5 | 8 | 6 | 2 | 4 | 5.72 | 7 | 4 | 0.609 | -0.716 | 0.37955817 | 0.59448576 | 0 | 0.5825 | 0.295 | 0.295 | TRUE | 0.3767 |
| Mef2d | 1748 | 2061 | 2077 | 2036 | 1711 | 1804 | 1857 | 2058 | 2086 | 1601 | 1841 | 1975 | 1902.8 | 2000 | 1806 | 0.902 | -0.149 | 0.16490235 | 0.36152904 | 0.0067 | 0.0086 | 0.0078 | 0.0078 | TRUE | 0.1212 |
| Mefv | 0 | 0 | 0 | 0 | 0 | 0 | 0 | 0 | 0 | 0 | 0 | 0 | 0 | 0 | 0 | NA | NA | NA | NA | NA | NA | NA | NA | NA | NA |
| Meg3 | 10588 | 11184 | 10556 | 7978 | 8266 | 6929 | 11246 | 11170 | 10599 | 6272 | 8894 | 7585 | 9294.25 | 11005 | 7584 | 0.689 | -0.538 | 2.66E-05 | 0.00045462 | 0.0153 | 0.0073 | 0.0117 | 0.0117 | TRUE | 0.2781 |
| Megf10 | 6 | 2 | 3 | 6 | 9 | 4 | 6 | 2 | 3 | 5 | 10 | 4 | 5.03 | 4 | 6 | 1.648 | 0.721 | 0.4240544 | 0.63431045 | 0 | 0.6623 | 0.3713 | 0.3713 | TRUE | 0.5361 |
| Megf11 | 332 | 444 | 408 | 221 | 208 | 220 | 353 | 443 | 410 | 174 | 224 | 241 | 307.35 | 402 | 213 | 0.527 | -0.924 | 5.89E-08 | 2.29E-06 | 0.017 | 0.0176 | 0.0173 | 0.0173 | TRUE | 0.3039 |
| Megf6 | 938 | 1181 | 1034 | 964 | 674 | 717 | 996 | 1179 | 1038 | 758 | 725 | 785 | 913.65 | 1071 | 756 | 0.705 | -0.503 | 1.10E-06 | 3.00E-05 | 0.0038 | 0.0105 | 0.0066 | 0.0066 | TRUE | 0.0935 |
| Megf8 | 1814 | 2446 | 2342 | 3474 | 2503 | 2342 | 1927 | 2443 | 2352 | 2731 | 2693 | 2564 | 2451.5 | 2241 | 2663 | 1.188 | 0.249 | 0.02171624 | 0.09232034 | 0.0075 | 0.0082 | 0.0081 | 0.0081 | TRUE | 0.1811 |
| Megf9 | 104 | 92 | 123 | 199 | 125 | 136 | 110 | 92 | 124 | 156 | 134 | 149 | 127.61 | 109 | 146 | 1.352 | 0.435 | 0.02490435 | 0.10143269 | 0.0045 | 0.0327 | 0.0191 | 0.0191 | TRUE | 0.1833 |
| Mei1 | 38 | 78 | 80 | 86 | 73 | 64 | 40 | 78 | 80 | 68 | 79 | 70 | 69.13 | 66 | 72 | 1.085 | 0.118 | 0.69924684 | 0.83549402 | 0.0514 | 0.0546 | 0.0531 | 0.0531 | TRUE | 1.007 |
| Mei4 | 93 | 89 | 78 | 168 | 103 | 100 | 99 | 89 | 78 | 132 | 111 | 109 | 103.06 | 89 | 117 | 1.331 | 0.412 | 0.04754112 | 0.1583566 | 0 | 0.0389 | 0.0212 | 0.0212 | TRUE | 0.1308 |
| Meig1 | 0 | 0 | 0 | 0 | 0 | 1 | 0 | 0 | 0 | 0 | 0 | 1 | 0.18 | 0 | 0 | 1.859 | 0.894 | 0.82650734 | NA | 0 | 18.0643 | 10 | 10 | TRUE | 0.5383 |
| Meiob | 110 | 257 | 305 | 70 | 50 | 100 | 117 | 257 | 306 | 55 | 54 | 109 | 149.68 | 227 | 73 | 0.32 | -1.644 | 2.04E-05 | 0.00036348 | 0.1878 | 0.0289 | 0.0983 | 0.0983 | TRUE | 0.4077 |
| Meis1 | 27 | 24 | 23 | 84 | 48 | 49 | 29 | 24 | 23 | 66 | 52 | 54 | 41.18 | 25 | 57 | 2.275 | 1.186 | 0.00018021 | 0.00223103 | 0 | 0.0869 | 0.0435 | 0.0435 | TRUE | 0.1792 |
| Meis2 | 49 | 27 | 16 | 130 | 55 | 51 | 52 | 27 | 16 | 102 | 59 | 56 | 52.05 | 32 | 72 | 2.3 | 1.202 | 0.0078424 | 0.04365617 | 0.1901 | 0.0702 | 0.1243 | 0.1243 | TRUE | 1.4845 |
| Meis3 | 69 | 61 | 39 | 87 | 63 | 42 | 73 | 61 | 39 | 68 | 68 | 46 | 59.25 | 58 | 61 | 1.055 | 0.077 | 0.81043347 | 0.90263093 | 0.0533 | 0.0625 | 0.0581 | 0.0581 | TRUE | 0.4372 |
| Melk | 5059 | 5267 | 5368 | 6660 | 4782 | 5107 | 5374 | 5260 | 5390 | 5236 | 5145 | 5590 | 5332.49 | 5341 | 5324 | 0.997 | -0.005 | 0.94092256 | 0.97007652 | 8.00E-04 | 0.0075 | 0.003 | 0.003 | TRUE | 0.0232 |
| Memo1 | 1391 | 1777 | 1764 | 2224 | 1594 | 1643 | 1477 | 1775 | 1771 | 1748 | 1715 | 1799 | 1714.22 | 1674 | 1754 | 1.047 | 0.067 | 0.50439182 | 0.69937761 | 0.005 | 0.0088 | 0.0066 | 0.0066 | TRUE | 0.1272 |
| Men1 | 3369 | 3016 | 2906 | 4513 | 3278 | 2915 | 3578 | 3012 | 2918 | 3548 | 3527 | 3191 | 3295.69 | 3169 | 3422 | 1.08 | 0.111 | 0.29259656 | 0.51361523 | 0.0076 | 0.0079 | 0.0077 | 0.0077 | TRUE | 0.1546 |
| Meox1 | 86 | 58 | 57 | 49 | 39 | 29 | 91 | 58 | 57 | 39 | 42 | 32 | 53.12 | 69 | 38 | 0.545 | -0.876 | 0.00541061 | 0.0331145 | 0.033 | 0.0689 | 0.0509 | 0.0509 | TRUE | 0.7189 |
| Meox2 | 1 | 2 | 1 | 5 | 2 | 3 | 1 | 2 | 1 | 4 | 2 | 3 | 2.24 | 1 | 3 | 2.319 | 1.213 | 0.33570669 | NA | 0 | 1.4787 | 0.6138 | 0.6138 | TRUE | 0.1098 |
| Mep1a | 0 | 1 | 0 | 0 | 0 | 0 | 0 | 1 | 0 | 0 | 0 | 0 | 0.17 | 0 | 0 | 0.49 | -1.029 | 0.80087 | NA | 0 | 19.7991 | 10 | 10 | TRUE | 0.405 |
| Mep1b | 487 | 394 | 349 | 1024 | 849 | 886 | 517 | 393 | 350 | 805 | 913 | 970 | 658.26 | 420 | 896 | 2.131 | 1.092 | 1.36E-11 | 1.18E-09 | 0.023 | 0.0119 | 0.017 | 0.017 | TRUE | 0.4664 |
| Mepce | 1743 | 1895 | 1757 | 2467 | 1834 | 1616 | 1851 | 1893 | 1764 | 1939 | 1973 | 1769 | 1864.96 | 1836 | 1894 | 1.032 | 0.045 | 0.58315077 | 0.75875085 | 0.0018 | 0.0087 | 0.0043 | 0.0043 | TRUE | 0.0396 |
| Mepe | 0 | 0 | 1 | 0 | 0 | 0 | 0 | 0 | 1 | 0 | 0 | 0 | 0.17 | 0 | 0 | 0.49 | -1.029 | 0.80087 | NA | 0 | 19.6928 | 10 | 10 | TRUE | 0.4098 |
| Mertk | 377 | 351 | 348 | 486 | 312 | 310 | 400 | 351 | 349 | 382 | 336 | 339 | 359.58 | 367 | 352 | 0.963 | -0.054 | 0.67113315 | 0.81655574 | 0.0032 | 0.0161 | 0.009 | 0.009 | TRUE | 0.0729 |
| Mesdc1 | 541 | 754 | 706 | 1075 | 793 | 709 | 575 | 753 | 709 | 845 | 853 | 776 | 751.83 | 679 | 825 | 1.215 | 0.281 | 0.0270664 | 0.10723237 | 0.0094 | 0.0113 | 0.0103 | 0.0103 | TRUE | 0.2114 |
| Mesdc2 | 5163 | 5618 | 5532 | 7188 | 5298 | 4899 | 5484 | 5611 | 5555 | 5651 | 5700 | 5363 | 5560.53 | 5550 | 5571 | 1.004 | 0.006 | 0.9264579 | 0.9637313 | 4.00E-04 | 0.0075 | 0.0026 | 0.0026 | TRUE | 0.013 |
| Mesp1 | 12 | 18 | 23 | 22 | 14 | 7 | 13 | 18 | 23 | 17 | 15 | 8 | 15.64 | 18 | 13 | 0.751 | -0.413 | 0.43702954 | 0.64530723 | 0.0467 | 0.2176 | 0.1393 | 0.1393 | TRUE | 0.2745 |
| Mesp2 | 120 | 80 | 93 | 197 | 125 | 98 | 127 | 80 | 93 | 155 | 134 | 107 | 116.23 | 100 | 132 | 1.325 | 0.406 | 0.10246914 | 0.26598803 | 0.036 | 0.0352 | 0.0357 | 0.0357 | TRUE | 0.4246 |
| Mest | 10569 | 9351 | 9684 | 11946 | 8513 | 7848 | 11226 | 9339 | 9724 | 9391 | 9159 | 8591 | 9571.7 | 10096 | 9047 | 0.896 | -0.158 | 0.09345196 | 0.24990239 | 0.0058 | 0.0072 | 0.0063 | 0.0063 | TRUE | 0.117 |
| Met | 638 | 766 | 810 | 1078 | 817 | 893 | 678 | 765 | 813 | 847 | 879 | 978 | 826.67 | 752 | 901 | 1.197 | 0.26 | 0.02145326 | 0.09144889 | 0.0057 | 0.0109 | 0.008 | 0.008 | TRUE | 0.0879 |
| Metap1 | 3700 | 4174 | 4335 | 5256 | 3827 | 4022 | 3930 | 4169 | 4353 | 4132 | 4118 | 4403 | 4183.94 | 4151 | 4218 | 1.016 | 0.023 | 0.76126463 | 0.87423727 | 0.0018 | 0.0077 | 0.0038 | 0.0038 | TRUE | 0.0262 |
| Metap1d | 324 | 242 | 268 | 383 | 304 | 256 | 344 | 242 | 269 | 301 | 327 | 280 | 293.89 | 285 | 303 | 1.063 | 0.089 | 0.59872287 | 0.7711136 | 0.0159 | 0.0181 | 0.017 | 0.017 | TRUE | 0.3677 |
| Metap2 | 12649 | 13050 | 13119 | 17184 | 12544 | 11843 | 13435 | 13033 | 13173 | 13509 | 13497 | 12964 | 13268.47 | 13214 | 13323 | 1.008 | 0.012 | 0.83100621 | 0.91427424 | 3.00E-04 | 0.0072 | 0.0022 | 0.0022 | TRUE | 0.0068 |
| Metrn | 495 | 506 | 503 | 668 | 505 | 387 | 526 | 505 | 505 | 525 | 543 | 424 | 504.72 | 512 | 497 | 0.972 | -0.04 | 0.7528491 | 0.86954631 | 0.0067 | 0.0134 | 0.0098 | 0.0098 | TRUE | 0.1905 |
| Metrnl | 92 | 129 | 115 | 109 | 89 | 73 | 98 | 129 | 115 | 86 | 96 | 80 | 100.56 | 114 | 87 | 0.763 | -0.39 | 0.06505143 | 0.19649029 | 0 | 0.0397 | 0.0221 | 0.0221 | TRUE | 0.1533 |
| Mettl1 | 685 | 800 | 717 | 1057 | 795 | 664 | 728 | 799 | 720 | 831 | 855 | 727 | 776.61 | 749 | 804 | 1.075 | 0.104 | 0.33439819 | 0.5548783 | 0.004 | 0.0111 | 0.0071 | 0.0071 | TRUE | 0.0824 |
| Mettl10 | 412 | 402 | 504 | 646 | 474 | 507 | 438 | 401 | 506 | 508 | 510 | 555 | 486.33 | 448 | 524 | 1.168 | 0.225 | 0.07559032 | 0.21672616 | 0.0059 | 0.0137 | 0.0094 | 0.0094 | TRUE | 0.1478 |
| Mettl11b | 1 | 0 | 0 | 0 | 0 | 0 | 1 | 0 | 0 | 0 | 0 | 0 | 0.18 | 0 | 0 | 0.49 | -1.029 | 0.80087 | NA | 0 | 18.6165 | 10 | 10 | TRUE | 0.4614 |
| Mettl13 | 1438 | 1538 | 1489 | 2129 | 1461 | 1349 | 1527 | 1536 | 1495 | 1674 | 1572 | 1477 | 1546.8 | 1519 | 1574 | 1.037 | 0.052 | 0.5292994 | 0.71762315 | 0.0015 | 0.009 | 0.0043 | 0.0043 | TRUE | 0.0379 |
| Mettl14 | 2445 | 2353 | 2403 | 2850 | 2143 | 2177 | 2597 | 2350 | 2413 | 2240 | 2306 | 2383 | 2381.52 | 2453 | 2310 | 0.941 | -0.087 | 0.25748065 | 0.47613844 | 0.0012 | 0.0083 | 0.0039 | 0.0039 | TRUE | 0.0317 |
| Mettl15 | 260 | 328 | 321 | 391 | 286 | 284 | 276 | 328 | 322 | 307 | 308 | 311 | 308.67 | 309 | 309 | 0.999 | -0.001 | 0.99293495 | 0.99616826 | 0 | 0.0176 | 0.0087 | 0.0087 | TRUE | 0.0947 |
| Mettl16 | 6228 | 5797 | 5896 | 7980 | 5803 | 5570 | 6615 | 5790 | 5920 | 6273 | 6244 | 6097 | 6156.52 | 6108 | 6205 | 1.016 | 0.023 | 0.77144048 | 0.88080416 | 0.0021 | 0.0074 | 0.0043 | 0.0043 | TRUE | 0.0642 |
| Mettl17 | 556 | 390 | 383 | 625 | 505 | 415 | 591 | 389 | 385 | 491 | 543 | 454 | 475.6 | 455 | 496 | 1.092 | 0.126 | 0.49857294 | 0.69422552 | 0.0325 | 0.0138 | 0.0231 | 0.0231 | TRUE | 0.7862 |
| Mettl18 | 425 | 386 | 380 | 650 | 416 | 358 | 451 | 386 | 382 | 511 | 448 | 392 | 428.16 | 406 | 450 | 1.111 | 0.152 | 0.29409212 | 0.51492239 | 0.0112 | 0.0146 | 0.0128 | 0.0128 | TRUE | 0.1681 |
| Mettl2 | 2363 | 3011 | 2902 | 3498 | 2571 | 2440 | 2510 | 3007 | 2914 | 2750 | 2766 | 2671 | 2769.67 | 2810 | 2729 | 0.971 | -0.042 | 0.6467576 | 0.80161347 | 0.0044 | 0.0081 | 0.0058 | 0.0058 | TRUE | 0.1058 |
| Mettl20 | 312 | 295 | 307 | 260 | 200 | 240 | 331 | 295 | 308 | 204 | 215 | 263 | 269.43 | 311 | 227 | 0.728 | -0.457 | 0.00240154 | 0.01779104 | 0.0063 | 0.0191 | 0.0125 | 0.0125 | TRUE | 0.1982 |

| | | | | | | | | | | | | | | | | | | | | | | | | | |
|---|---|---|---|---|---|---|---|---|---|---|---|---|---|---|---|---|---|---|---|---|---|---|---|---|---|
| Mettl21a | 428 | 568 | 577 | 719 | 536 | 547 | 455 | 567 | 579 | 565 | 577 | 599 | 556.99 | 534 | 580 | 1.086 | 0.119 | 0.34497615 | 0.56434898 | 0.007 | 0.0128 | 0.0097 | 0.0097 | TRUE | 0.1949 |
| Mettl21c | 0 | 0 | 0 | 0 | 0 | 0 | 0 | 0 | 0 | 0 | 0 | 0 | 0 | 0 | 0 | NA | NA | NA | NA | NA | NA | NA | NA | NA | NA |
| Mettl21d | 881 | 1054 | 1021 | 1252 | 867 | 953 | 936 | 1053 | 1025 | 984 | 933 | 1043 | 995.65 | 1005 | 987 | 0.982 | -0.026 | 0.78369633 | 0.88834728 | 0.0023 | 0.0102 | 0.0056 | 0.0056 | TRUE | 0.0425 |
| Mettl21e | 1 | 2 | 1 | 2 | 0 | 1 | 1 | 2 | 1 | 2 | 0 | 1 | 1.12 | 1 | 1 | 0.678 | -0.56 | 0.7486666 | NA | 0 | 2.9437 | 1.2969 | 1.2969 | TRUE | 0.2838 |
| Mettl22 | 500 | 853 | 898 | 853 | 618 | 753 | 531 | 852 | 902 | 671 | 665 | 824 | 740.74 | 762 | 720 | 0.945 | -0.082 | 0.68128022 | 0.82332018 | 0.0452 | 0.0114 | 0.0274 | 0.0274 | TRUE | 0.8277 |
| Mettl23 | 616 | 710 | 687 | 780 | 588 | 617 | 654 | 709 | 690 | 613 | 633 | 675 | 662.41 | 684 | 640 | 0.935 | -0.098 | 0.31554309 | 0.53622946 | 0 | 0.0119 | 0.0053 | 0.0053 | TRUE | 0.0265 |
| Mettl24 | 9 | 6 | 6 | 8 | 10 | 6 | 10 | 6 | 6 | 6 | 11 | 7 | 7.53 | 7 | 8 | 1.089 | 0.124 | 0.85866685 | 0.9304418 | 0 | 0.4443 | 0.2131 | 0.2131 | TRUE | 0.2788 |
| Mettl25 | 426 | 344 | 332 | 618 | 412 | 409 | 452 | 344 | 333 | 486 | 443 | 448 | 417.71 | 376 | 459 | 1.221 | 0.288 | 0.0560454 | 0.17757093 | 0.0131 | 0.0148 | 0.0139 | 0.0139 | TRUE | 0.3552 |
| Mettl3 | 1350 | 1175 | 1192 | 1859 | 1273 | 1180 | 1434 | 1173 | 1197 | 1461 | 1370 | 1292 | 1321.18 | 1268 | 1374 | 1.085 | 0.117 | 0.29421394 | 0.51502133 | 0.0072 | 0.0094 | 0.0082 | 0.0082 | TRUE | 0.1566 |
| Mettl4 | 1440 | 1313 | 1442 | 1713 | 1256 | 1337 | 1530 | 1311 | 1448 | 1347 | 1351 | 1464 | 1408.39 | 1430 | 1387 | 0.97 | -0.044 | 0.64399474 | 0.79977047 | 0.0034 | 0.0092 | 0.0057 | 0.0057 | TRUE | 0.0632 |
| Mettl5 | 1489 | 1774 | 1690 | 2034 | 1516 | 1581 | 1582 | 1772 | 1697 | 1599 | 1631 | 1731 | 1668.5 | 1684 | 1654 | 0.982 | -0.027 | 0.75228748 | 0.86942757 | 0.0019 | 0.0089 | 0.0045 | 0.0045 | TRUE | 0.0336 |
| Mettl6 | 991 | 971 | 886 | 1624 | 1077 | 983 | 1053 | 970 | 890 | 1277 | 1159 | 1076 | 1070.58 | 971 | 1171 | 1.207 | 0.272 | 0.01408605 | 0.067692 | 0.0058 | 0.01 | 0.0079 | 0.0079 | TRUE | 0.0772 |
| Mettl7a1 | 404 | 686 | 745 | 466 | 354 | 417 | 429 | 685 | 748 | 366 | 381 | 456 | 511 | 621 | 401 | 0.646 | -0.631 | 0.00249729 | 0.01835379 | 0.0467 | 0.0134 | 0.0293 | 0.0293 | TRUE | 0.8545 |
| Mettl7a2 | 14 | 13 | 19 | 22 | 13 | 12 | 15 | 13 | 19 | 17 | 14 | 13 | 15.22 | 16 | 15 | 0.953 | -0.07 | 0.88555514 | 0.94387096 | 0 | 0.2233 | 0.104 | 0.104 | TRUE | 0.1753 |
| Mettl7a3 | 11 | 13 | 5 | 9 | 4 | 8 | 12 | 13 | 5 | 7 | 4 | 9 | 8.3 | 10 | 7 | 0.68 | -0.557 | 0.43036589 | 0.64017078 | 0 | 0.4036 | 0.2349 | 0.2349 | TRUE | 0.6486 |
| Mettl7b | 0 | 0 | 0 | 3 | 3 | 2 | 0 | 0 | 0 | 2 | 3 | 2 | 1.3 | 0 | 2 | 13.776 | 3.784 | 0.05475068 | NA | 0 | 2.5491 | 1.6072 | 1.6072 | TRUE | 0.07 |
| Mettl8 | 1257 | 860 | 772 | 1411 | 901 | 732 | 1335 | 859 | 775 | 1109 | 969 | 801 | 974.86 | 990 | 960 | 0.971 | -0.043 | 0.83819508 | 0.91786523 | 0.0549 | 0.0103 | 0.0309 | 0.0309 | TRUE | 1.0866 |
| Mettl9 | 2079 | 2775 | 2851 | 3294 | 2456 | 2738 | 2208 | 2771 | 2863 | 2589 | 2643 | 2997 | 2678.59 | 2614 | 2743 | 1.049 | 0.069 | 0.57431831 | 0.75265047 | 0.0124 | 0.0081 | 0.0105 | 0.0105 | TRUE | 0.2233 |
| Mex3a | 1906 | 2817 | 2987 | 3457 | 2720 | 2850 | 2025 | 2813 | 2999 | 2718 | 2927 | 3120 | 2766.85 | 2612 | 2922 | 1.118 | 0.161 | 0.28200204 | 0.5034031 | 0.0231 | 0.0081 | 0.0158 | 0.0158 | TRUE | 0.4694 |
| Mex3b | 110 | 110 | 100 | 165 | 141 | 113 | 117 | 110 | 100 | 130 | 152 | 124 | 122.04 | 109 | 135 | 1.238 | 0.308 | 0.10321653 | 0.26735607 | 0 | 0.0339 | 0.0174 | 0.0174 | TRUE | 0.1147 |
| Mex3c | 2809 | 2921 | 3150 | 4143 | 3005 | 2994 | 2984 | 2917 | 3163 | 3257 | 3233 | 3277 | 3138.55 | 3021 | 3256 | 1.078 | 0.108 | 0.11176305 | 0.28113131 | 6.00E-04 | 0.008 | 0.003 | 0.003 | TRUE | 0.0205 |
| Mex3d | 1791 | 2018 | 2027 | 2139 | 1702 | 1886 | 1902 | 2015 | 2035 | 1682 | 1831 | 2065 | 1921.73 | 1984 | 1859 | 0.936 | -0.095 | 0.34382091 | 0.56363615 | 0.0053 | 0.0086 | 0.0068 | 0.0068 | TRUE | 0.1126 |
| Mfap1a | 591 | 484 | 469 | 735 | 541 | 473 | 628 | 483 | 471 | 578 | 582 | 518 | 543.28 | 527 | 559 | 1.061 | 0.086 | 0.54923371 | 0.7338626 | 0.0131 | 0.013 | 0.013 | 0.013 | TRUE | 0.3221 |
| Mfap1b | 218 | 203 | 213 | 313 | 189 | 182 | 232 | 203 | 214 | 246 | 203 | 199 | 216.13 | 216 | 216 | 1.005 | 0.007 | 0.96522573 | 0.98254422 | 0.0054 | 0.0221 | 0.0133 | 0.0133 | TRUE | 0.1712 |
| Mfap2 | 132 | 111 | 102 | 65 | 47 | 56 | 140 | 111 | 102 | 51 | 51 | 61 | 86.08 | 118 | 54 | 0.46 | -1.12 | 2.81E-06 | 6.76E-05 | 0.0115 | 0.0452 | 0.0278 | 0.0278 | TRUE | 0.2738 |
| Mfap3 | 3350 | 2916 | 3057 | 4294 | 3279 | 2886 | 3558 | 2912 | 3070 | 3376 | 3528 | 3159 | 3267.14 | 3180 | 3354 | 1.055 | 0.077 | 0.44886586 | 0.65455233 | 0.0061 | 0.0079 | 0.0072 | 0.0072 | TRUE | 0.1314 |
| Mfap3l | 363 | 282 | 268 | 383 | 281 | 277 | 386 | 282 | 269 | 301 | 302 | 303 | 307.16 | 312 | 302 | 0.969 | -0.045 | 0.78654599 | 0.89023346 | 0.0165 | 0.0176 | 0.017 | 0.017 | TRUE | 0.4764 |
| Mfap4 | 100 | 103 | 80 | 101 | 60 | 53 | 106 | 103 | 80 | 79 | 65 | 58 | 81.9 | 96 | 67 | 0.703 | -0.509 | 0.03632278 | 0.13124466 | 0.0126 | 0.0471 | 0.0302 | 0.0302 | TRUE | 0.2463 |
| Mfap5 | 108 | 160 | 173 | 282 | 188 | 172 | 115 | 160 | 174 | 222 | 202 | 188 | 176.74 | 150 | 204 | 1.367 | 0.451 | 0.02199846 | 0.09321862 | 0.0184 | 0.0255 | 0.0221 | 0.0221 | TRUE | 0.4243 |
| Mff | 3013 | 3341 | 3540 | 3787 | 2684 | 3140 | 3200 | 3337 | 3555 | 2977 | 2888 | 3437 | 3232.27 | 3364 | 3101 | 0.922 | -0.118 | 0.2229988 | 0.43505698 | 0.0055 | 0.0079 | 0.0064 | 0.0064 | TRUE | 0.1088 |
| Mfge8 | 6092 | 8857 | 8061 | 9504 | 7358 | 8897 | 6471 | 8846 | 8094 | 7471 | 7917 | 9739 | 8089.61 | 7804 | 8376 | 1.073 | 0.102 | 0.48162096 | 0.68073323 | 0.0192 | 0.0073 | 0.015 | 0.015 | TRUE | 0.2724 |
| Mfhas1 | 402 | 427 | 398 | 775 | 629 | 561 | 427 | 426 | 400 | 609 | 677 | 614 | 525.53 | 418 | 633 | 1.515 | 0.6 | 1.58E-08 | 7.22E-07 | 0 | 0.0132 | 0.0061 | 0.0061 | TRUE | 0.0416 |
| Mfi2 | 78 | 138 | 135 | 70 | 55 | 87 | 83 | 138 | 136 | 55 | 59 | 95 | 94.28 | 119 | 70 | 0.585 | -0.774 | 0.01151192 | 0.05843773 | 0.0722 | 0.0418 | 0.0563 | 0.0563 | TRUE | 0.8828 |
| Mfn1 | 3319 | 2899 | 2952 | 4480 | 3105 | 2936 | 3525 | 2895 | 2964 | 3522 | 3341 | 3214 | 3243.54 | 3128 | 3359 | 1.074 | 0.103 | 0.31103122 | 0.5316554 | 0.0067 | 0.0079 | 0.0071 | 0.0071 | TRUE | 0.1496 |
| Mfn2 | 1674 | 1470 | 1505 | 2337 | 1672 | 1378 | 1778 | 1468 | 1511 | 1837 | 1799 | 1508 | 1650.32 | 1586 | 1715 | 1.082 | 0.114 | 0.34361264 | 0.56358897 | 0.0101 | 0.0089 | 0.0098 | 0.0098 | TRUE | 0.1353 |
| Mfng | 800 | 819 | 770 | 853 | 623 | 717 | 850 | 818 | 773 | 671 | 670 | 785 | 761.1 | 814 | 709 | 0.87 | -0.201 | 0.06242461 | 0.19128831 | 0.0039 | 0.0112 | 0.007 | 0.007 | TRUE | 0.1029 |
| Mfrp | 0 | 0 | 0 | 0 | 0 | 0 | 0 | 0 | 0 | 0 | 0 | 0 | 0 | 0 | 0 | NA | NA | NA | NA | NA | NA | NA | NA | NA | NA |
| Mfsd1 | 2898 | 3055 | 2967 | 4942 | 3417 | 3253 | 3078 | 3051 | 2979 | 3885 | 3676 | 3561 | 3371.81 | 3036 | 3707 | 1.222 | 0.289 | 3.25E-05 | 0.00053939 | 9.00E-04 | 0.0079 | 0.0032 | 0.0032 | TRUE | 0.0217 |
| Mfsd10 | 1439 | 1343 | 1299 | 2043 | 1448 | 1307 | 1528 | 1341 | 1304 | 1606 | 1558 | 1431 | 1461.47 | 1391 | 1532 | 1.102 | 0.139 | 0.16297718 | 0.35840437 | 0.0045 | 0.0092 | 0.0065 | 0.0065 | TRUE | 0.089 |
| Mfsd11 | 811 | 918 | 856 | 1001 | 751 | 745 | 861 | 917 | 860 | 787 | 808 | 816 | 841.37 | 879 | 804 | 0.913 | -0.131 | 0.12784018 | 0.30739486 | 0 | 0.0108 | 0.0041 | 0.0041 | TRUE | 0.0167 |
| Mfsd12 | 158 | 219 | 215 | 349 | 234 | 171 | 168 | 219 | 216 | 274 | 252 | 187 | 219.29 | 201 | 238 | 1.186 | 0.246 | 0.21274571 | 0.42383894 | 0.0253 | 0.0219 | 0.0236 | 0.0236 | TRUE | 0.3749 |
| Mfsd2a | 70 | 36 | 38 | 118 | 65 | 50 | 74 | 36 | 38 | 93 | 70 | 55 | 60.98 | 49 | 73 | 1.474 | 0.56 | 0.12652874 | 0.30509268 | 0.1002 | 0.0609 | 0.0796 | 0.0796 | TRUE | 0.7149 |
| Mfsd2b | 74 | 68 | 66 | 97 | 59 | 73 | 79 | 68 | 66 | 76 | 63 | 80 | 72.07 | 71 | 73 | 1.034 | 0.049 | 0.83624323 | 0.91680998 | 0 | 0.0526 | 0.0261 | 0.0261 | TRUE | 0.1154 |
| Mfsd3 | 298 | 296 | 302 | 379 | 275 | 264 | 317 | 296 | 303 | 298 | 296 | 289 | 299.7 | 305 | 294 | 0.965 | -0.051 | 0.66892327 | 0.8150064 | 0 | 0.0179 | 0.007 | 0.007 | TRUE | 0.0119 |
| Mfsd4 | 97 | 119 | 117 | 98 | 92 | 82 | 103 | 119 | 117 | 77 | 99 | 90 | 100.86 | 113 | 89 | 0.779 | -0.36 | 0.0815902 | 0.22771085 | 0 | 0.0396 | 0.0207 | 0.0207 | TRUE | 0.1416 |
| Mfsd5 | 2145 | 1866 | 1701 | 2448 | 1817 | 1593 | 2278 | 1864 | 1708 | 1924 | 1955 | 1744 | 1912.19 | 1950 | 1874 | 0.962 | -0.057 | 0.65016235 | 0.8039483 | 0.0124 | 0.0086 | 0.0107 | 0.0107 | TRUE | 0.2619 |
| Mfsd6 | 174 | 254 | 291 | 228 | 155 | 216 | 185 | 254 | 292 | 179 | 167 | 236 | 218.86 | 244 | 194 | 0.796 | -0.33 | 0.13351003 | 0.31560554 | 0.0388 | 0.022 | 0.0302 | 0.0302 | TRUE | 0.3442 |
| Mfsd6l | 12 | 24 | 27 | 20 | 28 | 28 | 13 | 24 | 27 | 16 | 30 | 31 | 23.39 | 21 | 26 | 1.184 | 0.244 | 0.6095675 | 0.77891178 | 0.0926 | 0.1478 | 0.1212 | 0.1212 | TRUE | 0.8302 |
| Mfsd7a | 26 | 27 | 37 | 45 | 44 | 31 | 28 | 27 | 37 | 35 | 47 | 34 | 34.73 | 31 | 39 | 1.268 | 0.342 | 0.31975257 | 0.54087704 | 0 | 0.1018 | 0.0558 | 0.0558 | TRUE | 0.254 |
| Mfsd7b | 334 | 253 | 215 | 490 | 349 | 257 | 355 | 253 | 216 | 385 | 376 | 281 | 310.89 | 275 | 347 | 1.268 | 0.342 | 0.11137073 | 0.28068465 | 0.0431 | 0.0175 | 0.03 | 0.03 | TRUE | 0.7299 |
| Mfsd7c | 311 | 379 | 399 | 333 | 204 | 277 | 330 | 379 | 401 | 262 | 219 | 303 | 315.66 | 370 | 261 | 0.707 | -0.501 | 0.00184654 | 0.01445319 | 0.0135 | 0.0173 | 0.0154 | 0.0154 | TRUE | 0.211 |
| Mfsd8 | 158 | 168 | 172 | 224 | 141 | 171 | 168 | 168 | 173 | 176 | 152 | 187 | 170.55 | 170 | 172 | 1.014 | 0.02 | 0.90286595 | 0.95205059 | 0 | 0.0262 | 0.0128 | 0.0128 | TRUE | 0.1042 |
| Mfsd9 | 342 | 385 | 379 | 466 | 364 | 377 | 363 | 385 | 381 | 366 | 392 | 413 | 383.16 | 376 | 390 | 1.036 | 0.05 | 0.66106326 | 0.80911762 | 0 | 0.0155 | 0.0069 | 0.0069 | TRUE | 0.0354 |
| Mga | 12505 | 12163 | 14933 | 15219 | 11142 | 10559 | 13283 | 12147 | 14994 | 11964 | 11988 | 11558 | 12655.78 | 13475 | 11837 | 0.878 | -0.187 | 0.04691817 | 0.15705929 | 0.0057 | 0.0072 | 0.0063 | 0.0063 | TRUE | 0.1191 |
| Mgam | 0 | 0 | 0 | 3 | 3 | 1 | 0 | 0 | 0 | 2 | 3 | 1 | 1.11 | 0 | 2 | 11.874 | 3.57 | 0.0910322 | NA | 0 | 2.9656 | 1.9955 | 1.9955 | TRUE | 0.1116 |
| Mgarp | 688 | 863 | 929 | 1096 | 884 | 961 | 731 | 862 | 933 | 862 | 951 | 1052 | 898.36 | 842 | 955 | 1.133 | 0.18 | 0.16532058 | 0.36204334 | 0.0107 | 0.0106 | 0.011 | 0.011 | TRUE | 0.1574 |
| Mgat1 | 954 | 1110 | 1086 | 1251 | 934 | 900 | 1013 | 1109 | 1090 | 983 | 1005 | 985 | 1030.98 | 1071 | 991 | 0.925 | -0.112 | 0.18057812 | 0.38193351 | 0 | 0.0101 | 0.0041 | 0.0041 | TRUE | 0.0261 |
| Mgat2 | 1431 | 1425 | 1271 | 2344 | 1653 | 1284 | 1520 | 1423 | 1276 | 1843 | 1779 | 1406 | 1541.01 | 1406 | 1676 | 1.192 | 0.253 | 0.05151896 | 0.16729608 | 0.0127 | 0.009 | 0.0116 | 0.0116 | TRUE | 0.2383 |
| Mgat3 | 660 | 1021 | 1164 | 1187 | 832 | 1001 | 701 | 1020 | 1169 | 933 | 895 | 1096 | 968.93 | 963 | 975 | 1.012 | 0.017 | 0.92868016 | 0.96488131 | 0.0378 | 0.0103 | 0.0235 | 0.0235 | TRUE | 0.5729 |
| Mgat4a | 889 | 936 | 893 | 1257 | 919 | 893 | 944 | 935 | 897 | 988 | 989 | 978 | 955.03 | 925 | 985 | 1.065 | 0.09 | 0.26182572 | 0.48126204 | 0 | 0.0104 | 0.0036 | 0.0036 | TRUE | 0.0087 |
| Mgat4b | 2116 | 2803 | 2674 | 3429 | 2495 | 2742 | 2248 | 2799 | 2685 | 2696 | 2684 | 3002 | 2685.59 | 2577 | 2794 | 1.084 | 0.116 | 0.28735681 | 0.5086064 | 0.0079 | 0.0081 | 0.0082 | 0.0082 | TRUE | 0.1516 |
| Mgat4c | 8 | 2 | 3 | 10 | 14 | 7 | 8 | 2 | 3 | 8 | 15 | 8 | 7.35 | 4 | 10 | 2.27 | 1.183 | 0.14616852 | 0.33488812 | 0.147 | 0.4552 | 0.3147 | 0.3147 | TRUE | 1.0616 |
| Mgat5 | 280 | 303 | 319 | 508 | 361 | 345 | 297 | 303 | 320 | 399 | 388 | 378 | 347.62 | 307 | 388 | 1.267 | 0.342 | 0.0029923 | 0.0211825 | 0 | 0.0164 | 0.0066 | 0.0066 | TRUE | 0.0169 |
| Mgat5b | 1206 | 1189 | 1177 | 1418 | 1107 | 1135 | 1281 | 1187 | 1182 | 1115 | 1191 | 1242 | 1199.75 | 1217 | 1183 | 0.971 | -0.042 | 0.63555904 | 0.79511677 | 0.0018 | 0.0096 | 0.0048 | 0.0048 | TRUE | 0.0315 |
| Mgea5 | 11283 | 11489 | 11985 | 14549 | 11501 | 11367 | 11985 | 11474 | 12034 | 11437 | 12374 | 12443 | 11957.92 | 11831 | 12085 | 1.021 | 0.03 | 0.65229173 | 0.80502827 | 0.0012 | 0.0072 | 0.0032 | 0.0032 | TRUE | 0.027 |
| Mgl2 | 150 | 126 | 120 | 178 | 135 | 124 | 159 | 126 | 120 | 140 | 145 | 136 | 137.76 | 135 | 140 | 1.039 | 0.055 | 0.7687716 | 0.87928573 | 0.0043 | 0.0308 | 0.0177 | 0.0177 | TRUE | 0.2466 |
| Mgll | 183 | 157 | 139 | 305 | 231 | 194 | 194 | 157 | 140 | 240 | 249 | 212 | 198.57 | 164 | 234 | 1.43 | 0.516 | 0.00352628 | 0.02397008 | 0.0109 | 0.0235 | 0.0173 | 0.0173 | TRUE | 0.2833 |
| Mgme1 | 1738 | 2188 | 2253 | 2397 | 1806 | 2040 | 1846 | 2185 | 2262 | 1884 | 1943 | 2233 | 2059.01 | 2098 | 2020 | 0.963 | -0.055 | 0.63127665 | 0.79265149 | 0.0094 | 0.0085 | 0.009 | 0.009 | TRUE | 0.133 |
| Mgmt | 567 | 852 | 855 | 843 | 640 | 786 | 602 | 851 | 859 | 663 | 689 | 860 | 753.89 | 771 | 737 | 0.956 | -0.065 | 0.70269883 | 0.83752481 | 0.0281 | 0.0113 | 0.0197 | 0.0197 | TRUE | 0.431 |

| | | | | | | | | | | | | | | | | | | | | | | | | |
|---|---|---|---|---|---|---|---|---|---|---|---|---|---|---|---|---|---|---|---|---|---|---|---|---|
| Mgp | 1 | 1 | 0 | 1 | 2 | 3 | 1 | 1 | 0 | 1 | 2 | 3 | 1.38 | 1 | 2 | 2.976 | 1.573 | 0.36210123 | NA | 0 | 2.3934 | 1.1595 | 1.1595 | TRUE | 0.1805 |
| Mgrn1 | 2032 | 2367 | 2406 | 2672 | 2005 | 2150 | 2158 | 2364 | 2416 | 2101 | 2157 | 2354 | 2258.24 | 2313 | 2204 | 0.952 | -0.07 | 0.42269781 | 0.63294312 | 0.0029 | 0.0084 | 0.0051 | 0.0051 | TRUE | 0.0425 |
| Mgst1 | 212 | 269 | 254 | 301 | 197 | 222 | 225 | 269 | 255 | 237 | 212 | 243 | 240.08 | 250 | 231 | 0.924 | -0.115 | 0.43082937 | 0.64057692 | 0.0021 | 0.0206 | 0.0111 | 0.0111 | TRUE | 0.0803 |
| Mgst2 | 7 | 6 | 2 | 12 | 4 | 6 | 7 | 6 | 2 | 9 | 4 | 7 | 5.96 | 5 | 7 | 1.336 | 0.418 | 0.61500294 | 0.78215958 | 0 | 0.56 | 0.3259 | 0.3259 | TRUE | 0.436 |
| Mgst3 | 846 | 1112 | 1149 | 926 | 659 | 800 | 899 | 1111 | 1154 | 728 | 709 | 876 | 912.6 | 1055 | 771 | 0.731 | -0.453 | 0.00101449 | 0.0090144 | 0.0137 | 0.0105 | 0.0126 | 0.0126 | TRUE | 0.1988 |
| Mia | 20 | 32 | 30 | 28 | 24 | 19 | 21 | 32 | 30 | 22 | 26 | 21 | 25.33 | 28 | 23 | 0.821 | -0.284 | 0.46290022 | 0.66638027 | 0 | 0.137 | 0.0682 | 0.0682 | TRUE | 0.2608 |
| Mia2 | 0 | 0 | 2 | 4 | 1 | 2 | 0 | 0 | 2 | 3 | 1 | 2 | 1.4 | 1 | 2 | 3.2 | 1.678 | 0.35981347 | NA | 0 | 2.3551 | 1.4375 | 1.4375 | TRUE | 0.8646 |
| Mia3 | 4438 | 3827 | 4083 | 5967 | 4317 | 4013 | 4714 | 3822 | 4100 | 4691 | 4645 | 4393 | 4394.04 | 4212 | 4576 | 1.087 | 0.12 | 0.2226338 | 0.43468437 | 0.0061 | 0.0077 | 0.0067 | 0.0067 | TRUE | 0.1322 |
| Miat | 94 | 49 | 50 | 62 | 74 | 40 | 100 | 49 | 50 | 49 | 80 | 44 | 61.86 | 66 | 58 | 0.864 | -0.21 | 0.58323268 | 0.75875085 | 0.1209 | 0.0602 | 0.0894 | 0.0894 | TRUE | 1.7037 |
| Mib1 | 2373 | 2615 | 2921 | 3156 | 2292 | 2504 | 2521 | 2612 | 2933 | 2481 | 2466 | 2741 | 2625.54 | 2689 | 2563 | 0.953 | -0.069 | 0.45964605 | 0.66387707 | 0.0044 | 0.0082 | 0.006 | 0.006 | TRUE | 0.0769 |
| Mib2 | 1374 | 1672 | 1637 | 2428 | 1961 | 1728 | 1459 | 1670 | 1644 | 1909 | 2110 | 1892 | 1780.53 | 1591 | 1970 | 1.238 | 0.308 | 0.00109605 | 0.00956324 | 0.0036 | 0.0088 | 0.0058 | 0.0058 | TRUE | 0.0628 |
| Mical1 | 1642 | 1670 | 1795 | 1497 | 1213 | 1180 | 1744 | 1668 | 1802 | 1177 | 1305 | 1292 | 1497.99 | 1738 | 1258 | 0.723 | -0.468 | 3.71E-08 | 1.54E-06 | 0.0017 | 0.0091 | 0.0045 | 0.0045 | TRUE | 0.0396 |
| Mical2 | 271 | 210 | 202 | 201 | 179 | 153 | 288 | 210 | 203 | 158 | 193 | 167 | 203.08 | 234 | 173 | 0.739 | -0.436 | 0.02335856 | 0.09707807 | 0.0203 | 0.0231 | 0.0217 | 0.0217 | TRUE | 0.4543 |
| Mical3 | 5252 | 5003 | 5228 | 5754 | 4191 | 4081 | 5579 | 4997 | 5249 | 4523 | 4509 | 4467 | 4887.41 | 5275 | 4500 | 0.853 | -0.229 | 0.00123157 | 0.01045441 | 0.0012 | 0.0076 | 0.0034 | 0.0034 | TRUE | 0.0309 |
| Micalcl | 5 | 4 | 5 | 5 | 6 | 2 | 5 | 4 | 5 | 4 | 6 | 2 | 4.48 | 5 | 4 | 0.877 | -0.189 | 0.83156861 | NA | 0 | 0.7417 | 0.3443 | 0.3443 | TRUE | 0.2982 |
| Micall1 | 1618 | 2007 | 2067 | 1605 | 1229 | 1229 | 1719 | 2004 | 2075 | 1262 | 1322 | 1345 | 1621.31 | 1933 | 1310 | 0.677 | -0.562 | 2.01E-08 | 9.01E-07 | 0.005 | 0.0089 | 0.0066 | 0.0066 | TRUE | 0.1132 |
| Micall2 | 348 | 610 | 610 | 406 | 317 | 454 | 370 | 609 | 613 | 319 | 341 | 497 | 458.09 | 531 | 386 | 0.726 | -0.462 | 0.05035074 | 0.16493186 | 0.0627 | 0.0141 | 0.0379 | 0.0379 | TRUE | 0.8185 |
| Micu1 | 685 | 677 | 671 | 969 | 637 | 617 | 728 | 676 | 674 | 762 | 685 | 675 | 700 | 693 | 707 | 1.023 | 0.033 | 0.7410655 | 0.86159073 | 0.0019 | 0.0116 | 0.0059 | 0.0059 | TRUE | 0.0555 |
| Mid1 | 16 | 11 | 6 | 14 | 12 | 6 | 17 | 11 | 6 | 11 | 13 | 7 | 10.75 | 11 | 10 | 0.902 | -0.149 | 0.8165583 | 0.90663552 | 0.0774 | 0.3134 | 0.2023 | 0.2023 | TRUE | 0.2132 |
| Mid1ip1 | 633 | 751 | 651 | 908 | 729 | 758 | 672 | 750 | 654 | 714 | 784 | 830 | 733.99 | 692 | 776 | 1.12 | 0.163 | 0.13734593 | 0.32135635 | 0.0043 | 0.0114 | 0.0073 | 0.0073 | TRUE | 0.0639 |
| Mid2 | 48 | 101 | 95 | 86 | 58 | 56 | 51 | 101 | 95 | 68 | 62 | 61 | 73.09 | 82 | 64 | 0.774 | -0.37 | 0.22175122 | 0.43403577 | 0.0526 | 0.052 | 0.0522 | 0.0522 | TRUE | 1.0209 |
| Midn | 3635 | 3634 | 3589 | 4948 | 3629 | 3006 | 3861 | 3629 | 3604 | 3890 | 3905 | 3291 | 3696.48 | 3698 | 3695 | 0.999 | -0.001 | 0.99331022 | 0.99627909 | 0.0051 | 0.0078 | 0.0062 | 0.0062 | TRUE | 0.1109 |
| Mien1 | 1937 | 2030 | 1761 | 2218 | 1696 | 1703 | 2057 | 2027 | 1768 | 1744 | 1825 | 1864 | 1880.94 | 1951 | 1811 | 0.928 | -0.108 | 0.23849985 | 0.45435676 | 0.0031 | 0.0087 | 0.0055 | 0.0055 | TRUE | 0.0814 |
| Mier1 | 3088 | 2894 | 3042 | 4215 | 3080 | 3185 | 3280 | 2890 | 3054 | 3314 | 3314 | 3486 | 3223.11 | 3075 | 3371 | 1.096 | 0.133 | 0.09329885 | 0.24974703 | 0.002 | 0.0079 | 0.0042 | 0.0042 | TRUE | 0.0412 |
| Mier2 | 2839 | 2825 | 2726 | 3246 | 2542 | 2455 | 3016 | 2821 | 2737 | 2552 | 2735 | 2687 | 2758.04 | 2858 | 2658 | 0.93 | -0.105 | 0.16923065 | 0.36769608 | 0.0013 | 0.0081 | 0.0038 | 0.0038 | TRUE | 0.0281 |
| Mier3 | 1301 | 1490 | 1421 | 2113 | 1631 | 1529 | 1382 | 1488 | 1427 | 1661 | 1755 | 1674 | 1564.41 | 1432 | 1697 | 1.184 | 0.244 | 0.00146977 | 0.01206724 | 5.00E-04 | 0.009 | 0.0036 | 0.0036 | TRUE | 0.014 |
| Mif | 17977 | 23198 | 24392 | 22599 | 17136 | 20157 | 19095 | 23168 | 24492 | 17766 | 18437 | 22065 | 20837.13 | 22252 | 19423 | 0.873 | -0.196 | 0.11936866 | 0.29279665 | 0.015 | 0.0071 | 0.0114 | 0.0114 | TRUE | 0.1884 |
| Mif4gd | 2474 | 2258 | 2106 | 3429 | 2310 | 2072 | 2628 | 2255 | 2115 | 2696 | 2485 | 2268 | 2407.79 | 2333 | 2483 | 1.065 | 0.091 | 0.42672261 | 0.63685362 | 0.0088 | 0.0083 | 0.009 | 0.009 | TRUE | 0.1483 |
| Miip | 341 | 403 | 413 | 506 | 387 | 429 | 362 | 402 | 415 | 398 | 416 | 470 | 410.53 | 393 | 428 | 1.086 | 0.12 | 0.33967398 | 0.55980544 | 0.0038 | 0.0149 | 0.0089 | 0.0089 | TRUE | 0.0827 |
| Mill1 | 0 | 0 | 0 | 0 | 0 | 0 | 0 | 0 | 0 | 0 | 0 | 0 | 0 | 0 | 0 | NA | NA | NA | NA | NA | NA | NA | NA | NA | NA |
| Mill2 | 0 | 1 | 0 | 2 | 1 | 1 | 0 | 1 | 0 | 2 | 1 | 1 | 0.79 | 0 | 1 | 3.436 | 1.781 | 0.41000155 | NA | 0 | 4.1757 | 1.9771 | 1.9771 | TRUE | 0.405 |
| Milr1 | 0 | 1 | 1 | 5 | 4 | 0 | 0 | 1 | 1 | 4 | 4 | 0 | 1.71 | 1 | 3 | 4.11 | 2.039 | 0.25931071 | NA | 0 | 1.9378 | 1.4212 | 1.4212 | TRUE | 0.8242 |
| Mina | 3265 | 2996 | 2887 | 4252 | 3082 | 2860 | 3468 | 2992 | 2899 | 3343 | 3316 | 3131 | 3191.39 | 3120 | 3263 | 1.046 | 0.065 | 0.4903413 | 0.68670423 | 0.0045 | 0.0079 | 0.0061 | 0.0061 | TRUE | 0.1157 |
| Mink1 | 5687 | 5191 | 4865 | 6864 | 5164 | 4593 | 6041 | 5184 | 4885 | 5396 | 5556 | 5028 | 5348.29 | 5370 | 5327 | 0.992 | -0.012 | 0.90994057 | 0.95498953 | 0.0072 | 0.0075 | 0.0073 | 0.0073 | TRUE | 0.1454 |
| Minos1 | 2570 | 2958 | 2696 | 3684 | 2516 | 2526 | 2730 | 2954 | 2707 | 2896 | 2707 | 2765 | 2793.21 | 2797 | 2789 | 0.998 | -0.004 | 0.9628301 | 0.98130897 | 0.0014 | 0.0081 | 0.0038 | 0.0038 | TRUE | 0.0294 |
| Minpp1 | 3466 | 3643 | 3803 | 4601 | 3226 | 3197 | 3682 | 3638 | 3819 | 3617 | 3471 | 3500 | 3620.99 | 3713 | 3529 | 0.951 | -0.073 | 0.25288845 | 0.47086616 | 3.00E-04 | 0.0078 | 0.0026 | 0.0026 | TRUE | 0.0076 |
| Mios | 1274 | 1449 | 1426 | 1789 | 1239 | 1354 | 1353 | 1447 | 1432 | 1406 | 1333 | 1482 | 1408.97 | 1411 | 1407 | 0.997 | -0.004 | 0.96316229 | 0.98149348 | 0.0012 | 0.0092 | 0.0043 | 0.0043 | TRUE | 0.0253 |
| Miox | 11 | 22 | 9 | 20 | 11 | 11 | 12 | 22 | 9 | 16 | 12 | 12 | 13.72 | 14 | 13 | 0.932 | -0.101 | 0.85679221 | 0.92916575 | 0.0564 | 0.2471 | 0.1538 | 0.1538 | TRUE | 1.0144 |
| Mip | 82 | 70 | 70 | 87 | 59 | 53 | 87 | 70 | 70 | 68 | 63 | 58 | 69.53 | 76 | 63 | 0.839 | -0.253 | 0.29095675 | 0.51170493 | 0 | 0.0543 | 0.0271 | 0.0271 | TRUE | 0.1536 |
| Mipep | 2003 | 2366 | 2481 | 2612 | 1979 | 2224 | 2128 | 2363 | 2491 | 2053 | 2129 | 2435 | 2266.47 | 2327 | 2206 | 0.947 | -0.078 | 0.45559615 | 0.66053461 | 0.0062 | 0.0084 | 0.0075 | 0.0075 | TRUE | 0.0993 |
| Mipol1 | 592 | 740 | 789 | 758 | 524 | 603 | 629 | 739 | 792 | 596 | 564 | 660 | 663.31 | 720 | 607 | 0.842 | -0.248 | 0.04990103 | 0.1639384 | 0.0085 | 0.0119 | 0.01 | 0.01 | TRUE | 0.1443 |
| Mir100 | 0 | 0 | 0 | 0 | 0 | 0 | 0 | 0 | 0 | 0 | 0 | 0 | 0 | 0 | 0 | NA | NA | NA | NA | NA | NA | NA | NA | NA | NA |
| Mir101a | 0 | 0 | 0 | 0 | 0 | 0 | 0 | 0 | 0 | 0 | 0 | 0 | 0 | 0 | 0 | NA | NA | NA | NA | NA | NA | NA | NA | NA | NA |
| Mir101b | 0 | 0 | 0 | 0 | 0 | 0 | 0 | 0 | 0 | 0 | 0 | 0 | 0 | 0 | 0 | NA | NA | NA | NA | NA | NA | NA | NA | NA | NA |
| Mir101c | 0 | 0 | 0 | 0 | 0 | 0 | 0 | 0 | 0 | 0 | 0 | 0 | 0 | 0 | 0 | NA | NA | NA | NA | NA | NA | NA | NA | NA | NA |
| Mir103-1 | 0 | 0 | 0 | 0 | 0 | 0 | 0 | 0 | 0 | 0 | 0 | 0 | 0 | 0 | 0 | NA | NA | NA | NA | NA | NA | NA | NA | NA | NA |
| Mir103-2 | 1 | 0 | 0 | 0 | 0 | 0 | 1 | 0 | 0 | 0 | 0 | 0 | 0.18 | 0 | 0 | 0.49 | -1.029 | 0.80087 | NA | 0 | 18.6165 | 10 | 10 | TRUE | 0.4614 |
| Mir105 | 0 | 0 | 0 | 0 | 0 | 0 | 0 | 0 | 0 | 0 | 0 | 0 | 0 | 0 | 0 | NA | NA | NA | NA | NA | NA | NA | NA | NA | NA |
| Mir106a | 0 | 0 | 0 | 0 | 0 | 0 | 0 | 0 | 0 | 0 | 0 | 0 | 0 | 0 | 0 | NA | NA | NA | NA | NA | NA | NA | NA | NA | NA |
| Mir106b | 2 | 1 | 0 | 0 | 0 | 0 | 2 | 1 | 0 | 0 | 0 | 0 | 0.52 | 1 | 0 | 0.173 | -2.534 | 0.43568356 | NA | 0 | 6.3362 | 6.1226 | 6.1226 | TRUE | 0.0614 |
| Mir107 | 0 | 0 | 0 | 0 | 0 | 0 | 0 | 0 | 0 | 0 | 0 | 0 | 0 | 0 | 0 | NA | NA | NA | NA | NA | NA | NA | NA | NA | NA |
| Mir10a | 0 | 0 | 0 | 0 | 0 | 0 | 0 | 0 | 0 | 0 | 0 | 0 | 0 | 0 | 0 | NA | NA | NA | NA | NA | NA | NA | NA | NA | NA |
| Mir10b | 0 | 0 | 0 | 0 | 0 | 0 | 0 | 0 | 0 | 0 | 0 | 0 | 0 | 0 | 0 | NA | NA | NA | NA | NA | NA | NA | NA | NA | NA |
| Mir1186 | 0 | 0 | 0 | 0 | 0 | 0 | 0 | 0 | 0 | 0 | 0 | 0 | 0 | 0 | 0 | NA | NA | NA | NA | NA | NA | NA | NA | NA | NA |
| Mir1186b | 0 | 0 | 0 | 0 | 0 | 0 | 0 | 0 | 0 | 0 | 0 | 0 | 0 | 0 | 0 | NA | NA | NA | NA | NA | NA | NA | NA | NA | NA |
| Mir1187 | 0 | 0 | 0 | 0 | 0 | 0 | 0 | 0 | 0 | 0 | 0 | 0 | 0 | 0 | 0 | NA | NA | NA | NA | NA | NA | NA | NA | NA | NA |
| Mir1188 | 0 | 0 | 1 | 3 | 0 | 1 | 0 | 0 | 1 | 2 | 0 | 1 | 0.74 | 0 | 1 | 3.216 | 1.685 | 0.50237532 | NA | 0 | 4.4418 | 3.1322 | 3.1322 | TRUE | 0.1958 |
| Mir1190 | 1 | 0 | 0 | 0 | 0 | 0 | 1 | 0 | 0 | 0 | 0 | 0 | 0.18 | 0 | 0 | 0.49 | -1.029 | 0.80087 | NA | 0 | 18.6165 | 10 | 10 | TRUE | 0.4614 |
| Mir1191 | 0 | 0 | 0 | 0 | 0 | 0 | 0 | 0 | 0 | 0 | 0 | 0 | 0 | 0 | 0 | NA | NA | NA | NA | NA | NA | NA | NA | NA | NA |
| Mir1192 | 3 | 0 | 0 | 1 | 1 | 1 | 3 | 0 | 0 | 1 | 1 | 1 | 1.02 | 1 | 1 | 0.933 | -0.1 | 0.9616121 | NA | 0.6399 | 3.2245 | 2.1155 | 2.1155 | TRUE | 1.443 |
| Mir1193 | 0 | 0 | 1 | 0 | 0 | 0 | 0 | 0 | 1 | 0 | 0 | 0 | 0.17 | 0 | 0 | 0.49 | -1.029 | 0.80087 | NA | 0 | 19.6928 | 10 | 10 | TRUE | 0.4098 |
| Mir1195 | 0 | 0 | 0 | 0 | 0 | 0 | 0 | 0 | 0 | 0 | 0 | 0 | 0 | 0 | 0 | NA | NA | NA | NA | NA | NA | NA | NA | NA | NA |
| Mir1196 | 0 | 0 | 2 | 0 | 1 | 0 | 0 | 0 | 2 | 0 | 1 | 0 | 0.51 | 1 | 0 | 0.52 | -0.944 | 0.77815412 | NA | 0 | 6.416 | 6.3253 | 6.3253 | TRUE | 0.9739 |
| Mir1197 | 0 | 0 | 0 | 0 | 0 | 0 | 0 | 0 | 0 | 0 | 0 | 0 | 0 | 0 | 0 | NA | NA | NA | NA | NA | NA | NA | NA | NA | NA |
| Mir1198 | 0 | 0 | 0 | 0 | 0 | 0 | 0 | 0 | 0 | 0 | 0 | 0 | 0 | 0 | 0 | NA | NA | NA | NA | NA | NA | NA | NA | NA | NA |
| Mir1199 | 9 | 32 | 30 | 21 | 36 | 33 | 10 | 32 | 30 | 17 | 39 | 36 | 27.17 | 24 | 31 | 1.263 | 0.337 | 0.53582796 | 0.72271961 | 0.2299 | 0.1282 | 0.1758 | 0.1758 | TRUE | 1.5787 |
| Mir1224 | 0 | 0 | 0 | 0 | 0 | 0 | 0 | 0 | 0 | 0 | 0 | 0 | 0 | 0 | 0 | NA | NA | NA | NA | NA | NA | NA | NA | NA | NA |
| Mir122a | 0 | 0 | 0 | 0 | 0 | 0 | 0 | 0 | 0 | 0 | 0 | 0 | 0 | 0 | 0 | NA | NA | NA | NA | NA | NA | NA | NA | NA | NA |
| Mir1247 | 0 | 0 | 0 | 0 | 0 | 0 | 0 | 0 | 0 | 0 | 0 | 0 | 0 | 0 | 0 | NA | NA | NA | NA | NA | NA | NA | NA | NA | NA |

| | | | | | | | | | | | | | | | | | | | | | | | | | |
|---|---|---|---|---|---|---|---|---|---|---|---|---|---|---|---|---|---|---|---|---|---|---|---|---|---|
| Mir1249 | 7 | 0 | 0 | 0 | 2 | 2 | 7 | 0 | 0 | 0 | 2 | 2 | 1.96 | 2 | 1 | 0.574 | -0.8 | 0.70264313 | NA | 4.7339 | 1.6854 | 2.6127 | 2.6127 | TRUE | 3.2027 |
| Mir124a-1 | 0 | 0 | 0 | 0 | 0 | 0 | 0 | 0 | 0 | 0 | 0 | 0 | 0 | 0 | 0 | NA | NA | NA | NA | NA | NA | NA | NA | NA | NA |
| Mir124a-2 | 0 | 0 | 0 | 0 | 0 | 0 | 0 | 0 | 0 | 0 | 0 | 0 | 0 | 0 | 0 | NA | NA | NA | NA | NA | NA | NA | NA | NA | NA |
| Mir124a-3 | 0 | 0 | 0 | 0 | 0 | 0 | 0 | 0 | 0 | 0 | 0 | 0 | 0 | 0 | 0 | NA | NA | NA | NA | NA | NA | NA | NA | NA | NA |
| Mir1251 | 0 | 0 | 0 | 0 | 0 | 0 | 0 | 0 | 0 | 0 | 0 | 0 | 0 | 0 | 0 | NA | NA | NA | NA | NA | NA | NA | NA | NA | NA |
| Mir125a | 0 | 0 | 0 | 0 | 0 | 0 | 0 | 0 | 0 | 0 | 0 | 0 | 0 | 0 | 0 | NA | NA | NA | NA | NA | NA | NA | NA | NA | NA |
| Mir125b-1 | 0 | 0 | 0 | 0 | 0 | 0 | 0 | 0 | 0 | 0 | 0 | 0 | 0 | 0 | 0 | NA | NA | NA | NA | NA | NA | NA | NA | NA | NA |
| Mir125b-2 | 0 | 0 | 0 | 0 | 0 | 0 | 0 | 0 | 0 | 0 | 0 | 0 | 0 | 0 | 0 | NA | NA | NA | NA | NA | NA | NA | NA | NA | NA |
| Mir126 | 0 | 0 | 0 | 0 | 0 | 0 | 0 | 0 | 0 | 0 | 0 | 0 | 0 | 0 | 0 | NA | NA | NA | NA | NA | NA | NA | NA | NA | NA |
| Mir1264 | 0 | 0 | 0 | 0 | 0 | 0 | 0 | 0 | 0 | 0 | 0 | 0 | 0 | 0 | 0 | NA | NA | NA | NA | NA | NA | NA | NA | NA | NA |
| Mir127 | 1 | 0 | 0 | 0 | 0 | 0 | 1 | 0 | 0 | 0 | 0 | 0 | 0.18 | 0 | 0 | 0.49 | -1.029 | 0.80087 | NA | 0 | 18.6165 | 10 | 10 | TRUE | 0.4614 |
| Mir128-1 | 0 | 0 | 0 | 0 | 0 | 0 | 0 | 0 | 0 | 0 | 0 | 0 | 0 | 0 | 0 | NA | NA | NA | NA | NA | NA | NA | NA | NA | NA |
| Mir128-2 | 0 | 0 | 0 | 0 | 0 | 0 | 0 | 0 | 0 | 0 | 0 | 0 | 0 | 0 | 0 | NA | NA | NA | NA | NA | NA | NA | NA | NA | NA |
| Mir129-1 | 0 | 0 | 0 | 0 | 0 | 0 | 0 | 0 | 0 | 0 | 0 | 0 | 0 | 0 | 0 | NA | NA | NA | NA | NA | NA | NA | NA | NA | NA |
| Mir129-2 | 0 | 0 | 0 | 0 | 0 | 0 | 0 | 0 | 0 | 0 | 0 | 0 | 0 | 0 | 0 | NA | NA | NA | NA | NA | NA | NA | NA | NA | NA |
| Mir1298 | 0 | 0 | 0 | 0 | 0 | 0 | 0 | 0 | 0 | 0 | 0 | 0 | 0 | 0 | 0 | NA | NA | NA | NA | NA | NA | NA | NA | NA | NA |
| Mir1306 | 0 | 0 | 0 | 0 | 0 | 0 | 0 | 0 | 0 | 0 | 0 | 0 | 0 | 0 | 0 | NA | NA | NA | NA | NA | NA | NA | NA | NA | NA |
| Mir130a | 0 | 0 | 0 | 0 | 0 | 0 | 0 | 0 | 0 | 0 | 0 | 0 | 0 | 0 | 0 | NA | NA | NA | NA | NA | NA | NA | NA | NA | NA |
| Mir130b | 0 | 0 | 0 | 0 | 0 | 0 | 0 | 0 | 0 | 0 | 0 | 0 | 0 | 0 | 0 | NA | NA | NA | NA | NA | NA | NA | NA | NA | NA |
| Mir132 | 0 | 0 | 0 | 0 | 0 | 0 | 0 | 0 | 0 | 0 | 0 | 0 | 0 | 0 | 0 | NA | NA | NA | NA | NA | NA | NA | NA | NA | NA |
| Mir133a-1 | 0 | 0 | 0 | 0 | 0 | 0 | 0 | 0 | 0 | 0 | 0 | 0 | 0 | 0 | 0 | NA | NA | NA | NA | NA | NA | NA | NA | NA | NA |
| Mir133a-2 | 0 | 0 | 0 | 0 | 0 | 0 | 0 | 0 | 0 | 0 | 0 | 0 | 0 | 0 | 0 | NA | NA | NA | NA | NA | NA | NA | NA | NA | NA |
| Mir133b | 0 | 0 | 0 | 0 | 0 | 0 | 0 | 0 | 0 | 0 | 0 | 0 | 0 | 0 | 0 | NA | NA | NA | NA | NA | NA | NA | NA | NA | NA |
| Mir134 | 1 | 0 | 0 | 0 | 0 | 0 | 1 | 0 | 0 | 0 | 0 | 0 | 0.18 | 0 | 0 | 0.49 | -1.029 | 0.80087 | NA | 0 | 18.6165 | 10 | 10 | TRUE | 0.4614 |
| Mir135a-1 | 0 | 0 | 0 | 0 | 0 | 0 | 0 | 0 | 0 | 0 | 0 | 0 | 0 | 0 | 0 | NA | NA | NA | NA | NA | NA | NA | NA | NA | NA |
| Mir135a-2 | 0 | 0 | 0 | 0 | 0 | 0 | 0 | 0 | 0 | 0 | 0 | 0 | 0 | 0 | 0 | NA | NA | NA | NA | NA | NA | NA | NA | NA | NA |
| Mir135b | 0 | 0 | 0 | 0 | 0 | 0 | 0 | 0 | 0 | 0 | 0 | 0 | 0 | 0 | 0 | NA | NA | NA | NA | NA | NA | NA | NA | NA | NA |
| Mir136 | 0 | 0 | 0 | 0 | 0 | 0 | 0 | 0 | 0 | 0 | 0 | 0 | 0 | 0 | 0 | NA | NA | NA | NA | NA | NA | NA | NA | NA | NA |
| Mir137 | 0 | 0 | 0 | 0 | 0 | 0 | 0 | 0 | 0 | 0 | 0 | 0 | 0 | 0 | 0 | NA | NA | NA | NA | NA | NA | NA | NA | NA | NA |
| Mir138-1 | 0 | 0 | 0 | 1 | 0 | 0 | 0 | 0 | 0 | 1 | 0 | 0 | 0.13 | 0 | 0 | 1.859 | 0.894 | 0.82650734 | NA | 0 | 25.1515 | 10 | 10 | TRUE | 0.2563 |
| Mir138-2 | 0 | 0 | 0 | 0 | 0 | 0 | 0 | 0 | 0 | 0 | 0 | 0 | 0 | 0 | 0 | NA | NA | NA | NA | NA | NA | NA | NA | NA | NA |
| Mir139 | 0 | 0 | 0 | 0 | 0 | 0 | 0 | 0 | 0 | 0 | 0 | 0 | 0 | 0 | 0 | NA | NA | NA | NA | NA | NA | NA | NA | NA | NA |
| Mir140 | 0 | 0 | 0 | 0 | 0 | 0 | 0 | 0 | 0 | 0 | 0 | 0 | 0 | 0 | 0 | NA | NA | NA | NA | NA | NA | NA | NA | NA | NA |
| Mir141 | 0 | 0 | 0 | 0 | 0 | 0 | 0 | 0 | 0 | 0 | 0 | 0 | 0 | 0 | 0 | NA | NA | NA | NA | NA | NA | NA | NA | NA | NA |
| Mir142 | 0 | 0 | 0 | 0 | 0 | 0 | 0 | 0 | 0 | 0 | 0 | 0 | 0 | 0 | 0 | NA | NA | NA | NA | NA | NA | NA | NA | NA | NA |
| Mir143 | 0 | 0 | 0 | 1 | 0 | 0 | 0 | 0 | 0 | 1 | 0 | 0 | 0.13 | 0 | 0 | 1.859 | 0.894 | 0.82650734 | NA | 0 | 25.1515 | 10 | 10 | TRUE | 0.2563 |
| Mir143hg | 1 | 2 | 2 | 6 | 3 | 3 | 1 | 2 | 2 | 5 | 3 | 3 | 2.72 | 2 | 4 | 2.227 | 1.155 | 0.31043049 | NA | 0 | 1.2199 | 0.5045 | 0.5045 | TRUE | 0.1077 |
| Mir144 | 0 | 0 | 0 | 0 | 0 | 0 | 0 | 0 | 0 | 0 | 0 | 0 | 0 | 0 | 0 | NA | NA | NA | NA | NA | NA | NA | NA | NA | NA |
| Mir145 | 0 | 0 | 0 | 0 | 0 | 0 | 0 | 0 | 0 | 0 | 0 | 0 | 0 | 0 | 0 | NA | NA | NA | NA | NA | NA | NA | NA | NA | NA |
| Mir146 | 0 | 0 | 0 | 0 | 0 | 0 | 0 | 0 | 0 | 0 | 0 | 0 | 0 | 0 | 0 | NA | NA | NA | NA | NA | NA | NA | NA | NA | NA |
| Mir146b | 0 | 0 | 0 | 0 | 0 | 0 | 0 | 0 | 0 | 0 | 0 | 0 | 0 | 0 | 0 | NA | NA | NA | NA | NA | NA | NA | NA | NA | NA |
| Mir147 | 0 | 0 | 0 | 0 | 0 | 0 | 0 | 0 | 0 | 0 | 0 | 0 | 0 | 0 | 0 | NA | NA | NA | NA | NA | NA | NA | NA | NA | NA |
| Mir148a | 0 | 0 | 0 | 0 | 0 | 0 | 0 | 0 | 0 | 0 | 0 | 0 | 0 | 0 | 0 | NA | NA | NA | NA | NA | NA | NA | NA | NA | NA |
| Mir148b | 0 | 0 | 0 | 0 | 0 | 0 | 0 | 0 | 0 | 0 | 0 | 0 | 0 | 0 | 0 | NA | NA | NA | NA | NA | NA | NA | NA | NA | NA |
| Mir149 | 0 | 0 | 0 | 0 | 0 | 0 | 0 | 0 | 0 | 0 | 0 | 0 | 0 | 0 | 0 | NA | NA | NA | NA | NA | NA | NA | NA | NA | NA |
| Mir150 | 0 | 0 | 0 | 0 | 0 | 0 | 0 | 0 | 0 | 0 | 0 | 0 | 0 | 0 | 0 | NA | NA | NA | NA | NA | NA | NA | NA | NA | NA |
| Mir152 | 0 | 0 | 0 | 0 | 0 | 0 | 0 | 0 | 0 | 0 | 0 | 0 | 0 | 0 | 0 | NA | NA | NA | NA | NA | NA | NA | NA | NA | NA |
| Mir153 | 0 | 0 | 0 | 0 | 0 | 0 | 0 | 0 | 0 | 0 | 0 | 0 | 0 | 0 | 0 | NA | NA | NA | NA | NA | NA | NA | NA | NA | NA |
| Mir154 | 0 | 0 | 0 | 0 | 0 | 0 | 0 | 0 | 0 | 0 | 0 | 0 | 0 | 0 | 0 | NA | NA | NA | NA | NA | NA | NA | NA | NA | NA |
| Mir155 | 0 | 0 | 0 | 0 | 0 | 0 | 0 | 0 | 0 | 0 | 0 | 0 | 0 | 0 | 0 | NA | NA | NA | NA | NA | NA | NA | NA | NA | NA |
| Mir15a | 0 | 0 | 1 | 0 | 0 | 0 | 0 | 0 | 1 | 0 | 0 | 0 | 0.17 | 0 | 0 | 0.49 | -1.029 | 0.80087 | NA | 0 | 19.6928 | 10 | 10 | TRUE | 0.4098 |
| Mir15b | 0 | 0 | 0 | 0 | 0 | 0 | 0 | 0 | 0 | 0 | 0 | 0 | 0 | 0 | 0 | NA | NA | NA | NA | NA | NA | NA | NA | NA | NA |
| Mir16-1 | 0 | 0 | 0 | 0 | 0 | 0 | 0 | 0 | 0 | 0 | 0 | 0 | 0 | 0 | 0 | NA | NA | NA | NA | NA | NA | NA | NA | NA | NA |
| Mir16-2 | 0 | 0 | 0 | 0 | 0 | 0 | 0 | 0 | 0 | 0 | 0 | 0 | 0 | 0 | 0 | NA | NA | NA | NA | NA | NA | NA | NA | NA | NA |
| Mir17 | 0 | 0 | 1 | 1 | 0 | 0 | 0 | 0 | 1 | 1 | 0 | 0 | 0.3 | 0 | 0 | 0.954 | -0.067 | 0.98640617 | NA | 0 | 11.0483 | 9.2783 | 9.2783 | TRUE | 0.4098 |
| Mir17hg | 598 | 524 | 362 | 676 | 786 | 506 | 635 | 523 | 363 | 531 | 846 | 554 | 575.5 | 507 | 644 | 1.268 | 0.343 | 0.14848464 | 0.33827816 | 0.0707 | 0.0126 | 0.0388 | 0.0388 | TRUE | 0.4788 |
| Mir18 | 1 | 1 | 0 | 1 | 1 | 0 | 1 | 1 | 0 | 1 | 1 | 0 | 0.65 | 1 | 1 | 0.919 | -0.122 | 0.95812821 | NA | 0 | 5.0456 | 2.3857 | 2.3857 | TRUE | 0.2521 |
| Mir181a-1 | 0 | 0 | 0 | 0 | 0 | 0 | 0 | 0 | 0 | 0 | 0 | 0 | 0 | 0 | 0 | NA | NA | NA | NA | NA | NA | NA | NA | NA | NA |
| Mir181a-2 | 0 | 0 | 0 | 0 | 0 | 0 | 0 | 0 | 0 | 0 | 0 | 0 | 0 | 0 | 0 | NA | NA | NA | NA | NA | NA | NA | NA | NA | NA |
| Mir181b-1 | 0 | 0 | 0 | 0 | 0 | 0 | 0 | 0 | 0 | 0 | 0 | 0 | 0 | 0 | 0 | NA | NA | NA | NA | NA | NA | NA | NA | NA | NA |
| Mir181b-2 | 0 | 0 | 0 | 0 | 0 | 0 | 0 | 0 | 0 | 0 | 0 | 0 | 0 | 0 | 0 | NA | NA | NA | NA | NA | NA | NA | NA | NA | NA |
| Mir181c | 0 | 0 | 0 | 1 | 0 | 0 | 0 | 0 | 0 | 1 | 0 | 0 | 0.13 | 0 | 0 | 1.859 | 0.894 | 0.82650734 | NA | 0 | 25.1515 | 10 | 10 | TRUE | 0.2563 |
| Mir181d | 0 | 0 | 1 | 0 | 0 | 0 | 0 | 0 | 1 | 0 | 0 | 0 | 0.17 | 0 | 0 | 0.49 | -1.029 | 0.80087 | NA | 0 | 19.6928 | 10 | 10 | TRUE | 0.4098 |
| Mir182 | 0 | 0 | 0 | 0 | 0 | 0 | 0 | 0 | 0 | 0 | 0 | 0 | 0 | 0 | 0 | NA | NA | NA | NA | NA | NA | NA | NA | NA | NA |
| Mir183 | 0 | 0 | 0 | 0 | 0 | 0 | 0 | 0 | 0 | 0 | 0 | 0 | 0 | 0 | 0 | NA | NA | NA | NA | NA | NA | NA | NA | NA | NA |
| Mir1839 | 2 | 2 | 2 | 1 | 3 | 3 | 2 | 2 | 2 | 1 | 3 | 3 | 2.24 | 2 | 2 | 1.158 | 0.212 | 0.86494851 | NA | 0 | 1.479 | 0.66 | 0.66 | TRUE | 0.5277 |
| Mir184 | 0 | 0 | 0 | 0 | 0 | 0 | 0 | 0 | 0 | 0 | 0 | 0 | 0 | 0 | 0 | NA | NA | NA | NA | NA | NA | NA | NA | NA | NA |
| Mir1843 | 0 | 0 | 0 | 0 | 0 | 0 | 0 | 0 | 0 | 0 | 0 | 0 | 0 | 0 | 0 | NA | NA | NA | NA | NA | NA | NA | NA | NA | NA |
| Mir1843b | 0 | 0 | 0 | 0 | 0 | 0 | 0 | 0 | 0 | 0 | 0 | 0 | 0 | 0 | 0 | NA | NA | NA | NA | NA | NA | NA | NA | NA | NA |

| | | | | | | | | | | | | | | | | | | | | | | | | | |
|---|---|---|---|---|---|---|---|---|---|---|---|---|---|---|---|---|---|---|---|---|---|---|---|---|---|
| Mir185 | 0 | 0 | 0 | 0 | 0 | 0 | 0 | 0 | 0 | 0 | 0 | 0 | 0 | 0 | 0 | NA | NA | NA | NA | NA | NA | NA | NA | NA | NA |
| Mir186 | 3 | 3 | 3 | 2 | 2 | 1 | 3 | 3 | 3 | 2 | 2 | 1 | 2.34 | 3 | 2 | 0.524 | -0.932 | 0.43494862 | NA | 0 | 1.4174 | 0.557 | 0.557 | TRUE | 0.055 |
| Mir187 | 0 | 0 | 0 | 0 | 0 | 0 | 0 | 0 | 0 | 0 | 0 | 0 | 0 | 0 | 0 | NA | NA | NA | NA | NA | NA | NA | NA | NA | NA |
| Mir188 | 0 | 0 | 0 | 0 | 0 | 0 | 0 | 0 | 0 | 0 | 0 | 0 | 0 | 0 | 0 | NA | NA | NA | NA | NA | NA | NA | NA | NA | NA |
| Mir1892 | 0 | 0 | 0 | 0 | 0 | 0 | 0 | 0 | 0 | 0 | 0 | 0 | 0 | 0 | 0 | NA | NA | NA | NA | NA | NA | NA | NA | NA | NA |
| Mir1893 | 0 | 0 | 0 | 0 | 0 | 0 | 0 | 0 | 0 | 0 | 0 | 0 | 0 | 0 | 0 | NA | NA | NA | NA | NA | NA | NA | NA | NA | NA |
| Mir1894 | 2 | 0 | 0 | 2 | 4 | 1 | 2 | 0 | 0 | 2 | 4 | 1 | 1.52 | 1 | 2 | 3.325 | 1.733 | 0.3440356 | NA | 0.3393 | 2.1803 | 1.4704 | 1.4704 | TRUE | 0.9745 |
| Mir1895 | 0 | 0 | 0 | 0 | 0 | 0 | 0 | 0 | 0 | 0 | 0 | 0 | 0 | 0 | 0 | NA | NA | NA | NA | NA | NA | NA | NA | NA | NA |
| Mir1896 | 1 | 0 | 0 | 0 | 0 | 0 | 1 | 0 | 0 | 0 | 0 | 0 | 0.18 | 0 | 0 | 0.49 | -1.029 | 0.80087 | NA | 0 | 18.6165 | 10 | 10 | TRUE | 0.4614 |
| Mir1897 | 0 | 0 | 0 | 0 | 0 | 0 | 0 | 0 | 0 | 0 | 0 | 0 | 0 | 0 | 0 | NA | NA | NA | NA | NA | NA | NA | NA | NA | NA |
| Mir1898 | 2 | 1 | 4 | 3 | 1 | 0 | 2 | 1 | 4 | 2 | 1 | 0 | 1.76 | 2 | 1 | 0.505 | -0.985 | 0.5168979 | NA | 0 | 1.8763 | 1.0443 | 1.0443 | TRUE | 0.4534 |
| Mir1899 | 0 | 0 | 0 | 0 | 0 | 0 | 0 | 0 | 0 | 0 | 0 | 0 | 0 | 0 | 0 | NA | NA | NA | NA | NA | NA | NA | NA | NA | NA |
| Mir18b | 0 | 0 | 0 | 0 | 0 | 0 | 0 | 0 | 0 | 0 | 0 | 0 | 0 | 0 | 0 | NA | NA | NA | NA | NA | NA | NA | NA | NA | NA |
| Mir190 | 0 | 0 | 0 | 0 | 0 | 0 | 0 | 0 | 0 | 0 | 0 | 0 | 0 | 0 | 0 | NA | NA | NA | NA | NA | NA | NA | NA | NA | NA |
| Mir1900 | 0 | 0 | 0 | 0 | 0 | 0 | 0 | 0 | 0 | 0 | 0 | 0 | 0 | 0 | 0 | NA | NA | NA | NA | NA | NA | NA | NA | NA | NA |
| Mir1901 | 0 | 1 | 1 | 0 | 0 | 0 | 0 | 1 | 1 | 0 | 0 | 0 | 0.33 | 1 | 0 | 0.268 | -1.901 | 0.61401143 | NA | 0 | 9.8764 | 8.4772 | 8.4772 | TRUE | 0.2278 |
| Mir1902 | 0 | 0 | 0 | 0 | 0 | 0 | 0 | 0 | 0 | 0 | 0 | 0 | 0 | 0 | 0 | NA | NA | NA | NA | NA | NA | NA | NA | NA | NA |
| Mir1903 | 0 | 0 | 0 | 0 | 0 | 0 | 0 | 0 | 0 | 0 | 0 | 0 | 0 | 0 | 0 | NA | NA | NA | NA | NA | NA | NA | NA | NA | NA |
| Mir1904 | 0 | 1 | 0 | 0 | 0 | 0 | 0 | 1 | 0 | 0 | 0 | 0 | 0.17 | 0 | 0 | 0.49 | -1.029 | 0.80087 | NA | 0 | 19.7991 | 10 | 10 | TRUE | 0.405 |
| Mir1905 | 0 | 1 | 0 | 0 | 0 | 0 | 0 | 1 | 0 | 0 | 0 | 0 | 0.17 | 0 | 0 | 0.49 | -1.029 | 0.80087 | NA | 0 | 19.7991 | 10 | 10 | TRUE | 0.405 |
| Mir1906-1 | 0 | 0 | 0 | 0 | 0 | 0 | 0 | 0 | 0 | 0 | 0 | 0 | 0 | 0 | 0 | NA | NA | NA | NA | NA | NA | NA | NA | NA | NA |
| Mir1906-2 | 0 | 0 | 0 | 0 | 0 | 0 | 0 | 0 | 0 | 0 | 0 | 0 | 0 | 0 | 0 | NA | NA | NA | NA | NA | NA | NA | NA | NA | NA |
| Mir1907 | 0 | 1 | 0 | 2 | 2 | 0 | 0 | 1 | 0 | 2 | 2 | 0 | 0.79 | 0 | 1 | 3.417 | 1.773 | 0.46807648 | NA | 0 | 4.1923 | 2.9081 | 2.9081 | TRUE | 0.405 |
| Mir190b | 0 | 0 | 0 | 0 | 0 | 0 | 0 | 0 | 0 | 0 | 0 | 0 | 0 | 0 | 0 | NA | NA | NA | NA | NA | NA | NA | NA | NA | NA |
| Mir191 | 0 | 0 | 1 | 1 | 0 | 0 | 0 | 0 | 1 | 1 | 0 | 0 | 0.3 | 0 | 0 | 0.954 | -0.067 | 0.98640617 | NA | 0 | 11.0483 | 9.2783 | 9.2783 | TRUE | 0.4098 |
| Mir1912 | 0 | 0 | 0 | 0 | 0 | 0 | 0 | 0 | 0 | 0 | 0 | 0 | 0 | 0 | 0 | NA | NA | NA | NA | NA | NA | NA | NA | NA | NA |
| Mir192 | 0 | 0 | 0 | 0 | 0 | 0 | 0 | 0 | 0 | 0 | 0 | 0 | 0 | 0 | 0 | NA | NA | NA | NA | NA | NA | NA | NA | NA | NA |
| Mir1928 | 0 | 0 | 0 | 0 | 0 | 0 | 0 | 0 | 0 | 0 | 0 | 0 | 0 | 0 | 0 | NA | NA | NA | NA | NA | NA | NA | NA | NA | NA |
| Mir1929 | 1 | 0 | 0 | 1 | 1 | 0 | 1 | 0 | 0 | 1 | 1 | 0 | 0.49 | 0 | 1 | 1.712 | 0.776 | 0.78488768 | NA | 0 | 6.7665 | 4.0387 | 4.0387 | TRUE | 0.4614 |
| Mir193 | 0 | 0 | 0 | 0 | 0 | 0 | 0 | 0 | 0 | 0 | 0 | 0 | 0 | 0 | 0 | NA | NA | NA | NA | NA | NA | NA | NA | NA | NA |
| Mir1930 | 0 | 0 | 0 | 0 | 0 | 0 | 0 | 0 | 0 | 0 | 0 | 0 | 0 | 0 | 0 | NA | NA | NA | NA | NA | NA | NA | NA | NA | NA |
| Mir1931 | 0 | 0 | 0 | 0 | 0 | 0 | 0 | 0 | 0 | 0 | 0 | 0 | 0 | 0 | 0 | NA | NA | NA | NA | NA | NA | NA | NA | NA | NA |
| Mir1932 | 0 | 0 | 0 | 0 | 0 | 0 | 0 | 0 | 0 | 0 | 0 | 0 | 0 | 0 | 0 | NA | NA | NA | NA | NA | NA | NA | NA | NA | NA |
| Mir1933 | 0 | 0 | 0 | 0 | 0 | 0 | 0 | 0 | 0 | 0 | 0 | 0 | 0 | 0 | 0 | NA | NA | NA | NA | NA | NA | NA | NA | NA | NA |
| Mir1934 | 0 | 0 | 0 | 1 | 0 | 3 | 0 | 0 | 0 | 1 | 0 | 3 | 0.68 | 0 | 1 | 7.177 | 2.843 | 0.34581154 | NA | 6.5474 | 4.8635 | 5.1913 | 5.1913 | TRUE | 0.362 |
| Mir1935 | 0 | 1 | 0 | 0 | 0 | 0 | 0 | 1 | 0 | 0 | 0 | 0 | 0.17 | 0 | 0 | 0.49 | -1.029 | 0.80087 | NA | 0 | 19.7991 | 10 | 10 | TRUE | 0.405 |
| Mir1936 | 0 | 0 | 0 | 0 | 0 | 0 | 0 | 0 | 0 | 0 | 0 | 0 | 0 | 0 | 0 | NA | NA | NA | NA | NA | NA | NA | NA | NA | NA |
| Mir1938 | 0 | 1 | 0 | 2 | 0 | 2 | 0 | 1 | 0 | 2 | 0 | 2 | 0.79 | 0 | 1 | 3.442 | 1.783 | 0.46489926 | NA | 0 | 4.1593 | 2.9019 | 2.9019 | TRUE | 0.405 |
| Mir193b | 0 | 0 | 0 | 0 | 0 | 0 | 0 | 0 | 0 | 0 | 0 | 0 | 0 | 0 | 0 | NA | NA | NA | NA | NA | NA | NA | NA | NA | NA |
| Mir194-1 | 0 | 0 | 0 | 0 | 0 | 0 | 0 | 0 | 0 | 0 | 0 | 0 | 0 | 0 | 0 | NA | NA | NA | NA | NA | NA | NA | NA | NA | NA |
| Mir194-2 | 0 | 0 | 0 | 0 | 0 | 0 | 0 | 0 | 0 | 0 | 0 | 0 | 0 | 0 | 0 | NA | NA | NA | NA | NA | NA | NA | NA | NA | NA |
| Mir1940 | 0 | 0 | 0 | 0 | 0 | 0 | 0 | 0 | 0 | 0 | 0 | 0 | 0 | 0 | 0 | NA | NA | NA | NA | NA | NA | NA | NA | NA | NA |
| Mir1941 | 0 | 0 | 0 | 0 | 0 | 0 | 0 | 0 | 0 | 0 | 0 | 0 | 0 | 0 | 0 | NA | NA | NA | NA | NA | NA | NA | NA | NA | NA |
| Mir1942 | 0 | 0 | 0 | 0 | 0 | 0 | 0 | 0 | 0 | 0 | 0 | 0 | 0 | 0 | 0 | NA | NA | NA | NA | NA | NA | NA | NA | NA | NA |
| Mir1943 | 2 | 1 | 1 | 2 | 2 | 1 | 2 | 1 | 1 | 2 | 2 | 1 | 1.49 | 1 | 2 | 1.173 | 0.231 | 0.87447545 | NA | 0 | 2.2165 | 0.8574 | 0.8574 | TRUE | 0.1349 |
| Mir1945 | 0 | 0 | 0 | 4 | 0 | 1 | 0 | 0 | 0 | 3 | 0 | 1 | 0.71 | 0 | 1 | 7.635 | 2.933 | 0.32080697 | NA | 6.5513 | 4.6698 | 4.9457 | 4.9457 | TRUE | 0.1391 |
| Mir1946a | 0 | 0 | 0 | 0 | 0 | 0 | 0 | 0 | 0 | 0 | 0 | 0 | 0 | 0 | 0 | NA | NA | NA | NA | NA | NA | NA | NA | NA | NA |
| Mir1946b | 0 | 0 | 0 | 1 | 0 | 0 | 0 | 0 | 0 | 1 | 0 | 0 | 0.13 | 0 | 0 | 1.859 | 0.894 | 0.82650734 | NA | 0 | 25.1515 | 10 | 10 | TRUE | 0.2563 |
| Mir1947 | 0 | 0 | 0 | 0 | 0 | 0 | 0 | 0 | 0 | 0 | 0 | 0 | 0 | 0 | 0 | NA | NA | NA | NA | NA | NA | NA | NA | NA | NA |
| Mir1948 | 0 | 0 | 0 | 0 | 0 | 0 | 0 | 0 | 0 | 0 | 0 | 0 | 0 | 0 | 0 | NA | NA | NA | NA | NA | NA | NA | NA | NA | NA |
| Mir1949 | 13 | 8 | 18 | 10 | 16 | 6 | 14 | 8 | 18 | 8 | 17 | 7 | 11.92 | 13 | 11 | 0.788 | -0.344 | 0.58943682 | 0.76400346 | 0.125 | 0.2833 | 0.2069 | 0.2069 | TRUE | 0.5232 |
| Mir195 | 0 | 0 | 0 | 0 | 0 | 0 | 0 | 0 | 0 | 0 | 0 | 0 | 0 | 0 | 0 | NA | NA | NA | NA | NA | NA | NA | NA | NA | NA |
| Mir1950 | 0 | 0 | 0 | 0 | 0 | 0 | 0 | 0 | 0 | 0 | 0 | 0 | 0 | 0 | 0 | NA | NA | NA | NA | NA | NA | NA | NA | NA | NA |
| Mir1951 | 0 | 0 | 0 | 0 | 0 | 0 | 0 | 0 | 0 | 0 | 0 | 0 | 0 | 0 | 0 | NA | NA | NA | NA | NA | NA | NA | NA | NA | NA |
| Mir1952 | 0 | 0 | 0 | 0 | 0 | 0 | 0 | 0 | 0 | 0 | 0 | 0 | 0 | 0 | 0 | NA | NA | NA | NA | NA | NA | NA | NA | NA | NA |
| Mir1953 | 0 | 0 | 0 | 0 | 0 | 0 | 0 | 0 | 0 | 0 | 0 | 0 | 0 | 0 | 0 | NA | NA | NA | NA | NA | NA | NA | NA | NA | NA |
| Mir1954 | 0 | 0 | 0 | 0 | 0 | 0 | 0 | 0 | 0 | 0 | 0 | 0 | 0 | 0 | 0 | NA | NA | NA | NA | NA | NA | NA | NA | NA | NA |
| Mir1955 | 0 | 0 | 1 | 0 | 0 | 0 | 0 | 0 | 1 | 0 | 0 | 0 | 0.17 | 0 | 0 | 0.49 | -1.029 | 0.80087 | NA | 0 | 19.6928 | 10 | 10 | TRUE | 0.4098 |
| Mir1956 | 0 | 0 | 0 | 0 | 0 | 0 | 0 | 0 | 0 | 0 | 0 | 0 | 0 | 0 | 0 | NA | NA | NA | NA | NA | NA | NA | NA | NA | NA |
| Mir1957 | 0 | 0 | 0 | 0 | 0 | 0 | 0 | 0 | 0 | 0 | 0 | 0 | 0 | 0 | 0 | NA | NA | NA | NA | NA | NA | NA | NA | NA | NA |
| Mir1958 | 0 | 0 | 0 | 0 | 0 | 0 | 0 | 0 | 0 | 0 | 0 | 0 | 0 | 0 | 0 | NA | NA | NA | NA | NA | NA | NA | NA | NA | NA |
| Mir1960 | 0 | 0 | 0 | 0 | 0 | 0 | 0 | 0 | 0 | 0 | 0 | 0 | 0 | 0 | 0 | NA | NA | NA | NA | NA | NA | NA | NA | NA | NA |
| Mir1961 | 0 | 0 | 0 | 0 | 0 | 0 | 0 | 0 | 0 | 0 | 0 | 0 | 0 | 0 | 0 | NA | NA | NA | NA | NA | NA | NA | NA | NA | NA |
| Mir1962 | 0 | 0 | 0 | 0 | 0 | 0 | 0 | 0 | 0 | 0 | 0 | 0 | 0 | 0 | 0 | NA | NA | NA | NA | NA | NA | NA | NA | NA | NA |
| Mir1963 | 0 | 0 | 0 | 0 | 1 | 0 | 0 | 0 | 0 | 0 | 1 | 0 | 0.18 | 0 | 0 | 1.859 | 0.894 | 0.82650734 | NA | 0 | 18.3785 | 10 | 10 | TRUE | 0.5204 |
| Mir1964 | 1 | 0 | 0 | 0 | 1 | 0 | 1 | 0 | 0 | 0 | 1 | 0 | 0.36 | 0 | 0 | 0.954 | -0.067 | 0.98557668 | NA | 0 | 9.2518 | 8.0179 | 8.0179 | TRUE | 0.5204 |
| Mir1965 | 0 | 0 | 0 | 0 | 0 | 0 | 0 | 0 | 0 | 0 | 0 | 0 | 0 | 0 | 0 | NA | NA | NA | NA | NA | NA | NA | NA | NA | NA |
| Mir1966 | 0 | 0 | 0 | 0 | 0 | 1 | 0 | 0 | 0 | 0 | 0 | 1 | 0.18 | 0 | 0 | 1.859 | 0.894 | 0.82650734 | NA | 0 | 18.0643 | 10 | 10 | TRUE | 0.5383 |
| Mir1967 | 0 | 1 | 0 | 0 | 0 | 0 | 0 | 1 | 0 | 0 | 0 | 0 | 0.17 | 0 | 0 | 0.49 | -1.029 | 0.80087 | NA | 0 | 19.7991 | 10 | 10 | TRUE | 0.405 |

| | | | | | | | | | | | | | | | | | | | | | | | | | |
|---|---|---|---|---|---|---|---|---|---|---|---|---|---|---|---|---|---|---|---|---|---|---|---|---|---|
| Mir1968 | 4 | 6 | 7 | 5 | 3 | 3 | 4 | 6 | 7 | 4 | 3 | 3 | 4.62 | 6 | 3 | 0.608 | -0.719 | 0.40244191 | 0.61528827 | 0 | 0.7202 | 0.3037 | 0.3037 | TRUE | 0.1126 |
| Mir1969 | 0 | 0 | 0 | 0 | 0 | 0 | 0 | 0 | 0 | 0 | 0 | 0 | 0 | 0 | 0 | NA | NA | NA | NA | NA | NA | NA | NA | NA | NA |
| Mir196a-1 | 0 | 3 | 7 | 7 | 13 | 6 | 0 | 3 | 7 | 6 | 14 | 7 | 6.01 | 3 | 9 | 2.566 | 1.359 | 0.18024619 | 0.3815587 | 0.4946 | 0.5547 | 0.5331 | 0.5331 | TRUE | 0.7055 |
| Mir196a-2 | 0 | 0 | 0 | 0 | 0 | 0 | 0 | 0 | 0 | 0 | 0 | 0 | 0 | 0 | 0 | NA | NA | NA | NA | NA | NA | NA | NA | NA | NA |
| Mir196b | 0 | 0 | 0 | 0 | 0 | 0 | 0 | 0 | 0 | 0 | 0 | 0 | 0 | 0 | 0 | NA | NA | NA | NA | NA | NA | NA | NA | NA | NA |
| Mir1970 | 0 | 0 | 0 | 0 | 0 | 0 | 0 | 0 | 0 | 0 | 0 | 0 | 0 | 0 | 0 | NA | NA | NA | NA | NA | NA | NA | NA | NA | NA |
| Mir1971 | 0 | 0 | 0 | 0 | 0 | 0 | 0 | 0 | 0 | 0 | 0 | 0 | 0 | 0 | 0 | NA | NA | NA | NA | NA | NA | NA | NA | NA | NA |
| Mir1981 | 0 | 0 | 0 | 0 | 0 | 0 | 0 | 0 | 0 | 0 | 0 | 0 | 0 | 0 | 0 | NA | NA | NA | NA | NA | NA | NA | NA | NA | NA |
| Mir1982 | 0 | 0 | 0 | 0 | 0 | 0 | 0 | 0 | 0 | 0 | 0 | 0 | 0 | 0 | 0 | NA | NA | NA | NA | NA | NA | NA | NA | NA | NA |
| Mir1983 | 0 | 1 | 0 | 0 | 1 | 0 | 0 | 1 | 0 | 0 | 1 | 0 | 0.35 | 0 | 0 | 0.954 | -0.067 | 0.98571955 | NA | 0 | 9.5346 | 8.2194 | 8.2194 | TRUE | 0.5204 |
| Mir199a-1 | 0 | 0 | 0 | 0 | 0 | 0 | 0 | 0 | 0 | 0 | 0 | 0 | 0 | 0 | 0 | NA | NA | NA | NA | NA | NA | NA | NA | NA | NA |
| Mir199a-2 | 0 | 0 | 0 | 0 | 0 | 0 | 0 | 0 | 0 | 0 | 0 | 0 | 0 | 0 | 0 | NA | NA | NA | NA | NA | NA | NA | NA | NA | NA |
| Mir199b | 0 | 0 | 0 | 0 | 0 | 0 | 0 | 0 | 0 | 0 | 0 | 0 | 0 | 0 | 0 | NA | NA | NA | NA | NA | NA | NA | NA | NA | NA |
| Mir19a | 0 | 1 | 0 | 0 | 0 | 0 | 0 | 1 | 0 | 0 | 0 | 0 | 0.17 | 0 | 0 | 0.49 | -1.029 | 0.80087 | NA | 0 | 19.7991 | 10 | 10 | TRUE | 0.405 |
| Mir19b-1 | 0 | 0 | 0 | 1 | 1 | 1 | 0 | 0 | 0 | 1 | 1 | 1 | 0.49 | 0 | 1 | 5.224 | 2.385 | 0.38465 | NA | 0 | 6.6923 | 3.9249 | 3.9249 | TRUE | 0.07 |
| Mir19b-2 | 0 | 0 | 0 | 0 | 0 | 0 | 0 | 0 | 0 | 0 | 0 | 0 | 0 | 0 | 0 | NA | NA | NA | NA | NA | NA | NA | NA | NA | NA |
| Mir1a-1 | 0 | 0 | 0 | 0 | 0 | 0 | 0 | 0 | 0 | 0 | 0 | 0 | 0 | 0 | 0 | NA | NA | NA | NA | NA | NA | NA | NA | NA | NA |
| Mir1a-2 | 0 | 0 | 0 | 0 | 0 | 0 | 0 | 0 | 0 | 0 | 0 | 0 | 0 | 0 | 0 | NA | NA | NA | NA | NA | NA | NA | NA | NA | NA |
| Mir1b | 0 | 0 | 0 | 0 | 0 | 0 | 0 | 0 | 0 | 0 | 0 | 0 | 0 | 0 | 0 | NA | NA | NA | NA | NA | NA | NA | NA | NA | NA |
| Mir200a | 0 | 0 | 0 | 0 | 0 | 0 | 0 | 0 | 0 | 0 | 0 | 0 | 0 | 0 | 0 | NA | NA | NA | NA | NA | NA | NA | NA | NA | NA |
| Mir200b | 0 | 0 | 0 | 1 | 0 | 0 | 0 | 0 | 0 | 1 | 0 | 0 | 0.13 | 0 | 0 | 1.859 | 0.894 | 0.82650734 | NA | 0 | 25.1515 | 10 | 10 | TRUE | 0.2563 |
| Mir200c | 0 | 0 | 0 | 0 | 0 | 0 | 0 | 0 | 0 | 0 | 0 | 0 | 0 | 0 | 0 | NA | NA | NA | NA | NA | NA | NA | NA | NA | NA |
| Mir201 | 0 | 0 | 0 | 0 | 0 | 0 | 0 | 0 | 0 | 0 | 0 | 0 | 0 | 0 | 0 | NA | NA | NA | NA | NA | NA | NA | NA | NA | NA |
| Mir202 | 0 | 0 | 0 | 0 | 0 | 0 | 0 | 0 | 0 | 0 | 0 | 0 | 0 | 0 | 0 | NA | NA | NA | NA | NA | NA | NA | NA | NA | NA |
| Mir203 | 0 | 0 | 0 | 0 | 0 | 0 | 0 | 0 | 0 | 0 | 0 | 0 | 0 | 0 | 0 | NA | NA | NA | NA | NA | NA | NA | NA | NA | NA |
| Mir204 | 0 | 0 | 0 | 0 | 0 | 0 | 0 | 0 | 0 | 0 | 0 | 0 | 0 | 0 | 0 | NA | NA | NA | NA | NA | NA | NA | NA | NA | NA |
| Mir205 | 0 | 0 | 0 | 0 | 0 | 0 | 0 | 0 | 0 | 0 | 0 | 0 | 0 | 0 | 0 | NA | NA | NA | NA | NA | NA | NA | NA | NA | NA |
| Mir206 | 0 | 0 | 0 | 0 | 0 | 0 | 0 | 0 | 0 | 0 | 0 | 0 | 0 | 0 | 0 | NA | NA | NA | NA | NA | NA | NA | NA | NA | NA |
| Mir207 | 0 | 0 | 0 | 0 | 0 | 0 | 0 | 0 | 0 | 0 | 0 | 0 | 0 | 0 | 0 | NA | NA | NA | NA | NA | NA | NA | NA | NA | NA |
| Mir208a | 0 | 0 | 0 | 0 | 0 | 0 | 0 | 0 | 0 | 0 | 0 | 0 | 0 | 0 | 0 | NA | NA | NA | NA | NA | NA | NA | NA | NA | NA |
| Mir208b | 0 | 0 | 0 | 0 | 0 | 0 | 0 | 0 | 0 | 0 | 0 | 0 | 0 | 0 | 0 | NA | NA | NA | NA | NA | NA | NA | NA | NA | NA |
| Mir20a | 0 | 0 | 0 | 1 | 4 | 0 | 0 | 0 | 0 | 1 | 4 | 0 | 0.85 | 0 | 2 | 8.983 | 3.167 | 0.26555076 | NA | 7.4659 | 3.8904 | 4.5408 | 4.5408 | TRUE | 0.7289 |
| Mir20b | 0 | 0 | 0 | 0 | 0 | 0 | 0 | 0 | 0 | 0 | 0 | 0 | 0 | 0 | 0 | NA | NA | NA | NA | NA | NA | NA | NA | NA | NA |
| Mir21 | 0 | 0 | 0 | 0 | 0 | 0 | 0 | 0 | 0 | 0 | 0 | 0 | 0 | 0 | 0 | NA | NA | NA | NA | NA | NA | NA | NA | NA | NA |
| Mir210 | 0 | 1 | 0 | 0 | 0 | 0 | 0 | 1 | 0 | 0 | 0 | 0 | 0.17 | 0 | 0 | 0.49 | -1.029 | 0.80087 | NA | 0 | 19.7991 | 10 | 10 | TRUE | 0.405 |
| Mir211 | 0 | 0 | 0 | 0 | 0 | 0 | 0 | 0 | 0 | 0 | 0 | 0 | 0 | 0 | 0 | NA | NA | NA | NA | NA | NA | NA | NA | NA | NA |
| Mir212 | 0 | 0 | 0 | 0 | 0 | 0 | 0 | 0 | 0 | 0 | 0 | 0 | 0 | 0 | 0 | NA | NA | NA | NA | NA | NA | NA | NA | NA | NA |
| Mir2136 | 0 | 0 | 0 | 0 | 0 | 0 | 0 | 0 | 0 | 0 | 0 | 0 | 0 | 0 | 0 | NA | NA | NA | NA | NA | NA | NA | NA | NA | NA |
| Mir2137 | 0 | 0 | 0 | 0 | 0 | 0 | 0 | 0 | 0 | 0 | 0 | 0 | 0 | 0 | 0 | NA | NA | NA | NA | NA | NA | NA | NA | NA | NA |
| Mir214 | 0 | 0 | 0 | 0 | 0 | 0 | 0 | 0 | 0 | 0 | 0 | 0 | 0 | 0 | 0 | NA | NA | NA | NA | NA | NA | NA | NA | NA | NA |
| Mir215 | 0 | 0 | 0 | 0 | 0 | 0 | 0 | 0 | 0 | 0 | 0 | 0 | 0 | 0 | 0 | NA | NA | NA | NA | NA | NA | NA | NA | NA | NA |
| Mir216a | 0 | 0 | 0 | 0 | 0 | 0 | 0 | 0 | 0 | 0 | 0 | 0 | 0 | 0 | 0 | NA | NA | NA | NA | NA | NA | NA | NA | NA | NA |
| Mir216b | 0 | 0 | 0 | 0 | 0 | 0 | 0 | 0 | 0 | 0 | 0 | 0 | 0 | 0 | 0 | NA | NA | NA | NA | NA | NA | NA | NA | NA | NA |
| Mir217 | 0 | 0 | 0 | 0 | 0 | 0 | 0 | 0 | 0 | 0 | 0 | 0 | 0 | 0 | 0 | NA | NA | NA | NA | NA | NA | NA | NA | NA | NA |
| Mir218-1 | 0 | 0 | 0 | 0 | 0 | 0 | 0 | 0 | 0 | 0 | 0 | 0 | 0 | 0 | 0 | NA | NA | NA | NA | NA | NA | NA | NA | NA | NA |
| Mir218-2 | 0 | 0 | 0 | 0 | 0 | 0 | 0 | 0 | 0 | 0 | 0 | 0 | 0 | 0 | 0 | NA | NA | NA | NA | NA | NA | NA | NA | NA | NA |
| Mir219-1 | 0 | 0 | 0 | 0 | 0 | 0 | 0 | 0 | 0 | 0 | 0 | 0 | 0 | 0 | 0 | NA | NA | NA | NA | NA | NA | NA | NA | NA | NA |
| Mir219-2 | 0 | 0 | 0 | 0 | 0 | 0 | 0 | 0 | 0 | 0 | 0 | 0 | 0 | 0 | 0 | NA | NA | NA | NA | NA | NA | NA | NA | NA | NA |
| Mir22 | 0 | 0 | 0 | 0 | 0 | 0 | 0 | 0 | 0 | 0 | 0 | 0 | 0 | 0 | 0 | NA | NA | NA | NA | NA | NA | NA | NA | NA | NA |
| Mir221 | 0 | 0 | 0 | 0 | 0 | 0 | 0 | 0 | 0 | 0 | 0 | 0 | 0 | 0 | 0 | NA | NA | NA | NA | NA | NA | NA | NA | NA | NA |
| Mir222 | 0 | 0 | 0 | 0 | 0 | 0 | 0 | 0 | 0 | 0 | 0 | 0 | 0 | 0 | 0 | NA | NA | NA | NA | NA | NA | NA | NA | NA | NA |
| Mir223 | 0 | 0 | 0 | 0 | 0 | 0 | 0 | 0 | 0 | 0 | 0 | 0 | 0 | 0 | 0 | NA | NA | NA | NA | NA | NA | NA | NA | NA | NA |
| Mir22hg | 110 | 100 | 64 | 156 | 103 | 88 | 117 | 100 | 64 | 123 | 111 | 96 | 101.79 | 94 | 110 | 1.178 | 0.236 | 0.36896511 | 0.58632924 | 0.0402 | 0.0393 | 0.0397 | 0.0397 | TRUE | 0.6433 |
| Mir23a | 0 | 0 | 0 | 0 | 0 | 0 | 0 | 0 | 0 | 0 | 0 | 0 | 0 | 0 | 0 | NA | NA | NA | NA | NA | NA | NA | NA | NA | NA |
| Mir23b | 0 | 0 | 0 | 0 | 0 | 0 | 0 | 0 | 0 | 0 | 0 | 0 | 0 | 0 | 0 | NA | NA | NA | NA | NA | NA | NA | NA | NA | NA |
| Mir24-1 | 0 | 1 | 0 | 1 | 0 | 0 | 0 | 1 | 0 | 1 | 0 | 0 | 0.3 | 0 | 0 | 0.954 | -0.067 | 0.98642001 | NA | 0 | 11.0817 | 9.3014 | 9.3014 | TRUE | 0.405 |
| Mir24-2 | 0 | 0 | 0 | 0 | 0 | 0 | 0 | 0 | 0 | 0 | 0 | 0 | 0 | 0 | 0 | NA | NA | NA | NA | NA | NA | NA | NA | NA | NA |
| Mir25 | 5 | 6 | 2 | 5 | 6 | 6 | 5 | 6 | 2 | 4 | 6 | 7 | 5.04 | 4 | 6 | 1.261 | 0.335 | 0.69700181 | 0.83401203 | 0 | 0.66 | 0.3295 | 0.3295 | TRUE | 0.4266 |
| Mir26a-1 | 0 | 0 | 0 | 0 | 0 | 0 | 0 | 0 | 0 | 0 | 0 | 0 | 0 | 0 | 0 | NA | NA | NA | NA | NA | NA | NA | NA | NA | NA |
| Mir26a-2 | 0 | 0 | 0 | 0 | 0 | 0 | 0 | 0 | 0 | 0 | 0 | 0 | 0 | 0 | 0 | NA | NA | NA | NA | NA | NA | NA | NA | NA | NA |
| Mir26b | 0 | 0 | 0 | 0 | 0 | 0 | 0 | 0 | 0 | 0 | 0 | 0 | 0 | 0 | 0 | NA | NA | NA | NA | NA | NA | NA | NA | NA | NA |
| Mir27a | 0 | 1 | 0 | 0 | 0 | 0 | 0 | 1 | 0 | 0 | 0 | 0 | 0.17 | 0 | 0 | 0.49 | -1.029 | 0.80087 | NA | 0 | 19.7991 | 10 | 10 | TRUE | 0.405 |
| Mir27b | 0 | 0 | 0 | 0 | 0 | 0 | 0 | 0 | 0 | 0 | 0 | 0 | 0 | 0 | 0 | NA | NA | NA | NA | NA | NA | NA | NA | NA | NA |
| Mir28 | 0 | 0 | 0 | 0 | 0 | 0 | 0 | 0 | 0 | 0 | 0 | 0 | 0 | 0 | 0 | NA | NA | NA | NA | NA | NA | NA | NA | NA | NA |
| Mir2861 | 4 | 4 | 4 | 4 | 5 | 2 | 4 | 4 | 4 | 3 | 5 | 2 | 3.83 | 4 | 3 | 0.87 | -0.2 | 0.8314008 | NA | 0 | 0.8673 | 0.3752 | 0.3752 | TRUE | 0.265 |
| Mir28b | 0 | 0 | 0 | 0 | 0 | 0 | 0 | 0 | 0 | 0 | 0 | 0 | 0 | 0 | 0 | NA | NA | NA | NA | NA | NA | NA | NA | NA | NA |
| Mir28c | 0 | 0 | 0 | 0 | 0 | 0 | 0 | 0 | 0 | 0 | 0 | 0 | 0 | 0 | 0 | NA | NA | NA | NA | NA | NA | NA | NA | NA | NA |
| Mir290 | 0 | 0 | 0 | 2 | 1 | 1 | 0 | 0 | 0 | 2 | 1 | 1 | 0.62 | 0 | 1 | 6.676 | 2.739 | 0.27800695 | NA | 0 | 5.2881 | 3.2034 | 3.2034 | TRUE | 0.07 |
| Mir291a | 0 | 0 | 0 | 0 | 0 | 0 | 0 | 0 | 0 | 0 | 0 | 0 | 0 | 0 | 0 | NA | NA | NA | NA | NA | NA | NA | NA | NA | NA |

| | | | | | | | | | | | | | | | | | | | | | | | | | |
|---|---|---|---|---|---|---|---|---|---|---|---|---|---|---|---|---|---|---|---|---|---|---|---|---|---|
| Mir291b | 0 | 0 | 0 | 0 | 0 | 0 | 0 | 0 | 0 | 0 | 0 | 0 | 0 | 0 | 0 | NA | NA | NA | NA | NA | NA | NA | NA | NA | NA |
| Mir292 | 0 | 0 | 0 | 0 | 0 | 0 | 0 | 0 | 0 | 0 | 0 | 0 | 0 | 0 | 0 | NA | NA | NA | NA | NA | NA | NA | NA | NA | NA |
| Mir293 | 0 | 0 | 0 | 0 | 0 | 0 | 0 | 0 | 0 | 0 | 0 | 0 | 0 | 0 | 0 | NA | NA | NA | NA | NA | NA | NA | NA | NA | NA |
| Mir294 | 0 | 0 | 0 | 0 | 0 | 0 | 0 | 0 | 0 | 0 | 0 | 0 | 0 | 0 | 0 | NA | NA | NA | NA | NA | NA | NA | NA | NA | NA |
| Mir295 | 0 | 0 | 0 | 0 | 0 | 0 | 0 | 0 | 0 | 0 | 0 | 0 | 0 | 0 | 0 | NA | NA | NA | NA | NA | NA | NA | NA | NA | NA |
| Mir296 | 0 | 0 | 0 | 0 | 0 | 0 | 0 | 0 | 0 | 0 | 0 | 0 | 0 | 0 | 0 | NA | NA | NA | NA | NA | NA | NA | NA | NA | NA |
| Mir297-1 | 0 | 0 | 0 | 0 | 0 | 0 | 0 | 0 | 0 | 0 | 0 | 0 | 0 | 0 | 0 | NA | NA | NA | NA | NA | NA | NA | NA | NA | NA |
| Mir297-2 | 0 | 0 | 0 | 0 | 0 | 0 | 0 | 0 | 0 | 0 | 0 | 0 | 0 | 0 | 0 | NA | NA | NA | NA | NA | NA | NA | NA | NA | NA |
| Mir297a-3 | 0 | 0 | 0 | 0 | 0 | 0 | 0 | 0 | 0 | 0 | 0 | 0 | 0 | 0 | 0 | NA | NA | NA | NA | NA | NA | NA | NA | NA | NA |
| Mir297a-4 | 0 | 0 | 0 | 0 | 0 | 0 | 0 | 0 | 0 | 0 | 0 | 0 | 0 | 0 | 0 | NA | NA | NA | NA | NA | NA | NA | NA | NA | NA |
| Mir297b | 0 | 0 | 0 | 0 | 0 | 0 | 0 | 0 | 0 | 0 | 0 | 0 | 0 | 0 | 0 | NA | NA | NA | NA | NA | NA | NA | NA | NA | NA |
| Mir297c | 0 | 0 | 0 | 0 | 0 | 0 | 0 | 0 | 0 | 0 | 0 | 0 | 0 | 0 | 0 | NA | NA | NA | NA | NA | NA | NA | NA | NA | NA |
| Mir298 | 0 | 0 | 0 | 0 | 0 | 0 | 0 | 0 | 0 | 0 | 0 | 0 | 0 | 0 | 0 | NA | NA | NA | NA | NA | NA | NA | NA | NA | NA |
| Mir299 | 0 | 0 | 0 | 0 | 0 | 0 | 0 | 0 | 0 | 0 | 0 | 0 | 0 | 0 | 0 | NA | NA | NA | NA | NA | NA | NA | NA | NA | NA |
| Mir29a | 0 | 0 | 0 | 0 | 0 | 0 | 0 | 0 | 0 | 0 | 0 | 0 | 0 | 0 | 0 | NA | NA | NA | NA | NA | NA | NA | NA | NA | NA |
| Mir29b-1 | 0 | 0 | 0 | 0 | 0 | 0 | 0 | 0 | 0 | 0 | 0 | 0 | 0 | 0 | 0 | NA | NA | NA | NA | NA | NA | NA | NA | NA | NA |
| Mir29b-2 | 0 | 0 | 0 | 0 | 0 | 0 | 0 | 0 | 0 | 0 | 0 | 0 | 0 | 0 | 0 | NA | NA | NA | NA | NA | NA | NA | NA | NA | NA |
| Mir29c | 0 | 0 | 0 | 0 | 0 | 0 | 0 | 0 | 0 | 0 | 0 | 0 | 0 | 0 | 0 | NA | NA | NA | NA | NA | NA | NA | NA | NA | NA |
| Mir300 | 0 | 0 | 0 | 0 | 0 | 0 | 0 | 0 | 0 | 0 | 0 | 0 | 0 | 0 | 0 | NA | NA | NA | NA | NA | NA | NA | NA | NA | NA |
| Mir301 | 0 | 0 | 0 | 0 | 0 | 0 | 0 | 0 | 0 | 0 | 0 | 0 | 0 | 0 | 0 | NA | NA | NA | NA | NA | NA | NA | NA | NA | NA |
| Mir301b | 0 | 1 | 0 | 0 | 0 | 0 | 0 | 1 | 0 | 0 | 0 | 0 | 0.17 | 0 | 0 | 0.49 | -1.029 | 0.80087 | NA | 0 | 19.7991 | 10 | 10 | TRUE | 0.405 |
| Mir302a | 0 | 0 | 0 | 0 | 0 | 0 | 0 | 0 | 0 | 0 | 0 | 0 | 0 | 0 | 0 | NA | NA | NA | NA | NA | NA | NA | NA | NA | NA |
| Mir302b | 0 | 0 | 0 | 0 | 0 | 0 | 0 | 0 | 0 | 0 | 0 | 0 | 0 | 0 | 0 | NA | NA | NA | NA | NA | NA | NA | NA | NA | NA |
| Mir302c | 0 | 0 | 0 | 0 | 0 | 0 | 0 | 0 | 0 | 0 | 0 | 0 | 0 | 0 | 0 | NA | NA | NA | NA | NA | NA | NA | NA | NA | NA |
| Mir302d | 0 | 0 | 0 | 0 | 0 | 0 | 0 | 0 | 0 | 0 | 0 | 0 | 0 | 0 | 0 | NA | NA | NA | NA | NA | NA | NA | NA | NA | NA |
| Mir3057 | 2 | 2 | 1 | 1 | 2 | 3 | 2 | 2 | 1 | 1 | 2 | 3 | 1.89 | 2 | 2 | 1.184 | 0.244 | 0.85627192 | NA | 0 | 1.7488 | 0.7695 | 0.7695 | TRUE | 0.2782 |
| Mir3058 | 3 | 5 | 2 | 3 | 1 | 4 | 3 | 5 | 2 | 2 | 1 | 4 | 3 | 3 | 2 | 0.762 | -0.392 | 0.7228735 | NA | 0 | 1.105 | 0.5424 | 0.5424 | TRUE | 0.3518 |
| Mir3059 | 0 | 0 | 0 | 0 | 0 | 0 | 0 | 0 | 0 | 0 | 0 | 0 | 0 | 0 | 0 | NA | NA | NA | NA | NA | NA | NA | NA | NA | NA |
| Mir3060 | 4 | 3 | 0 | 2 | 0 | 0 | 4 | 3 | 0 | 2 | 0 | 0 | 1.47 | 2 | 1 | 0.238 | -2.07 | 0.32363595 | NA | 0 | 2.2488 | 2.1063 | 2.1063 | TRUE | 0.6308 |
| Mir3061 | 14 | 10 | 7 | 7 | 8 | 7 | 15 | 10 | 7 | 6 | 9 | 8 | 8.94 | 11 | 8 | 0.677 | -0.562 | 0.39111071 | 0.60481742 | 0 | 0.3753 | 0.1932 | 0.1932 | TRUE | 0.2799 |
| Mir3062 | 1 | 0 | 2 | 2 | 1 | 1 | 1 | 0 | 2 | 2 | 1 | 1 | 1.14 | 1 | 1 | 1.233 | 0.302 | 0.86280404 | NA | 0 | 2.9081 | 1.3124 | 1.3124 | TRUE | 0.2528 |
| Mir3063 | 0 | 0 | 0 | 0 | 0 | 0 | 0 | 0 | 0 | 0 | 0 | 0 | 0 | 0 | 0 | NA | NA | NA | NA | NA | NA | NA | NA | NA | NA |
| Mir3064 | 82 | 105 | 106 | 105 | 124 | 85 | 87 | 105 | 106 | 83 | 133 | 93 | 101.23 | 99 | 103 | 1.031 | 0.043 | 0.8607493 | 0.93125634 | 0.0292 | 0.0394 | 0.0343 | 0.0343 | TRUE | 0.6447 |
| Mir3065 | 0 | 0 | 0 | 0 | 0 | 0 | 0 | 0 | 0 | 0 | 0 | 0 | 0 | 0 | 0 | NA | NA | NA | NA | NA | NA | NA | NA | NA | NA |
| Mir3066 | 0 | 0 | 0 | 0 | 0 | 0 | 0 | 0 | 0 | 0 | 0 | 0 | 0 | 0 | 0 | NA | NA | NA | NA | NA | NA | NA | NA | NA | NA |
| Mir3067 | 0 | 0 | 0 | 0 | 0 | 0 | 0 | 0 | 0 | 0 | 0 | 0 | 0 | 0 | 0 | NA | NA | NA | NA | NA | NA | NA | NA | NA | NA |
| Mir3068 | 3 | 7 | 2 | 8 | 1 | 1 | 3 | 7 | 2 | 6 | 1 | 1 | 3.44 | 4 | 3 | 0.723 | -0.469 | 0.6872855 | NA | 0.4206 | 0.9643 | 0.6856 | 0.6856 | TRUE | 1.816 |
| Mir3069 | 0 | 0 | 0 | 0 | 0 | 0 | 0 | 0 | 0 | 0 | 0 | 0 | 0 | 0 | 0 | NA | NA | NA | NA | NA | NA | NA | NA | NA | NA |
| Mir3070a | 1 | 0 | 0 | 0 | 0 | 0 | 1 | 0 | 0 | 0 | 0 | 0 | 0.18 | 0 | 0 | 0.49 | -1.029 | 0.80087 | NA | 0 | 18.6165 | 10 | 10 | TRUE | 0.4614 |
| Mir3070b | 1 | 0 | 0 | 1 | 0 | 0 | 1 | 0 | 0 | 1 | 0 | 0 | 0.31 | 0 | 0 | 0.954 | -0.067 | 0.98625928 | NA | 0 | 10.7014 | 9.0384 | 9.0384 | TRUE | 0.4614 |
| Mir3071 | 0 | 0 | 0 | 0 | 0 | 0 | 0 | 0 | 0 | 0 | 0 | 0 | 0 | 0 | 0 | NA | NA | NA | NA | NA | NA | NA | NA | NA | NA |
| Mir3072 | 9 | 9 | 4 | 4 | 6 | 5 | 10 | 9 | 4 | 3 | 6 | 5 | 6.27 | 8 | 5 | 0.658 | -0.603 | 0.44429548 | 0.65087811 | 0 | 0.5321 | 0.2814 | 0.2814 | TRUE | 0.4724 |
| Mir3073 | 0 | 0 | 0 | 0 | 0 | 0 | 0 | 0 | 0 | 0 | 0 | 0 | 0 | 0 | 0 | NA | NA | NA | NA | NA | NA | NA | NA | NA | NA |
| Mir3074-1 | 0 | 0 | 0 | 0 | 0 | 0 | 0 | 0 | 0 | 0 | 0 | 0 | 0 | 0 | 0 | NA | NA | NA | NA | NA | NA | NA | NA | NA | NA |
| Mir3074-2 | 0 | 0 | 0 | 0 | 0 | 0 | 0 | 0 | 0 | 0 | 0 | 0 | 0 | 0 | 0 | NA | NA | NA | NA | NA | NA | NA | NA | NA | NA |
| Mir3075 | 0 | 0 | 0 | 0 | 0 | 0 | 0 | 0 | 0 | 0 | 0 | 0 | 0 | 0 | 0 | NA | NA | NA | NA | NA | NA | NA | NA | NA | NA |
| Mir3076 | 0 | 1 | 0 | 0 | 0 | 1 | 0 | 1 | 0 | 0 | 0 | 1 | 0.35 | 0 | 0 | 0.954 | -0.067 | 0.98567705 | NA | 0 | 9.4494 | 8.1588 | 8.1588 | TRUE | 0.5383 |
| Mir3077 | 0 | 0 | 0 | 0 | 0 | 0 | 0 | 0 | 0 | 0 | 0 | 0 | 0 | 0 | 0 | NA | NA | NA | NA | NA | NA | NA | NA | NA | NA |
| Mir3078 | 0 | 0 | 0 | 0 | 0 | 0 | 0 | 0 | 0 | 0 | 0 | 0 | 0 | 0 | 0 | NA | NA | NA | NA | NA | NA | NA | NA | NA | NA |
| Mir3079 | 0 | 0 | 0 | 0 | 0 | 0 | 0 | 0 | 0 | 0 | 0 | 0 | 0 | 0 | 0 | NA | NA | NA | NA | NA | NA | NA | NA | NA | NA |
| Mir3081 | 0 | 0 | 0 | 0 | 0 | 0 | 0 | 0 | 0 | 0 | 0 | 0 | 0 | 0 | 0 | NA | NA | NA | NA | NA | NA | NA | NA | NA | NA |
| Mir3082 | 21 | 16 | 23 | 20 | 19 | 16 | 22 | 16 | 23 | 16 | 20 | 18 | 19.18 | 20 | 18 | 0.871 | -0.199 | 0.64802457 | 0.80243071 | 0 | 0.1787 | 0.0849 | 0.0849 | TRUE | 0.2046 |
| Mir3083 | 0 | 0 | 0 | 0 | 0 | 0 | 0 | 0 | 0 | 0 | 0 | 0 | 0 | 0 | 0 | NA | NA | NA | NA | NA | NA | NA | NA | NA | NA |
| Mir3085 | 0 | 0 | 0 | 0 | 0 | 0 | 0 | 0 | 0 | 0 | 0 | 0 | 0 | 0 | 0 | NA | NA | NA | NA | NA | NA | NA | NA | NA | NA |
| Mir3086 | 0 | 0 | 0 | 0 | 0 | 0 | 0 | 0 | 0 | 0 | 0 | 0 | 0 | 0 | 0 | NA | NA | NA | NA | NA | NA | NA | NA | NA | NA |
| Mir3087 | 0 | 0 | 0 | 0 | 0 | 0 | 0 | 0 | 0 | 0 | 0 | 0 | 0 | 0 | 0 | NA | NA | NA | NA | NA | NA | NA | NA | NA | NA |
| Mir3088 | 0 | 0 | 0 | 0 | 0 | 0 | 0 | 0 | 0 | 0 | 0 | 0 | 0 | 0 | 0 | NA | NA | NA | NA | NA | NA | NA | NA | NA | NA |
| Mir3089 | 0 | 0 | 0 | 0 | 0 | 0 | 0 | 0 | 0 | 0 | 0 | 0 | 0 | 0 | 0 | NA | NA | NA | NA | NA | NA | NA | NA | NA | NA |
| Mir3091 | 49 | 41 | 34 | 72 | 50 | 28 | 52 | 41 | 34 | 57 | 54 | 31 | 44.7 | 42 | 47 | 1.117 | 0.159 | 0.64957325 | 0.80354219 | 0.0503 | 0.0806 | 0.0662 | 0.0662 | TRUE | 0.6953 |
| Mir3092 | 0 | 0 | 0 | 0 | 0 | 0 | 0 | 0 | 0 | 0 | 0 | 0 | 0 | 0 | 0 | NA | NA | NA | NA | NA | NA | NA | NA | NA | NA |
| Mir3093 | 0 | 0 | 0 | 0 | 0 | 0 | 0 | 0 | 0 | 0 | 0 | 0 | 0 | 0 | 0 | NA | NA | NA | NA | NA | NA | NA | NA | NA | NA |
| Mir3094 | 0 | 1 | 0 | 0 | 0 | 0 | 0 | 1 | 0 | 0 | 0 | 0 | 0.17 | 0 | 0 | 0.49 | -1.029 | 0.80087 | NA | 0 | 19.7991 | 10 | 10 | TRUE | 0.405 |
| Mir3095 | 0 | 0 | 0 | 0 | 0 | 0 | 0 | 0 | 0 | 0 | 0 | 0 | 0 | 0 | 0 | NA | NA | NA | NA | NA | NA | NA | NA | NA | NA |
| Mir3096 | 0 | 0 | 0 | 1 | 0 | 0 | 0 | 0 | 0 | 1 | 0 | 0 | 0.13 | 0 | 0 | 1.859 | 0.894 | 0.82650734 | NA | 0 | 25.1515 | 10 | 10 | TRUE | 0.2563 |
| Mir3096b | 1 | 2 | 0 | 1 | 2 | 0 | 1 | 2 | 0 | 1 | 2 | 0 | 1 | 1 | 1 | 0.955 | -0.066 | 0.9741481 | NA | 0 | 3.3027 | 1.9774 | 1.9774 | TRUE | 0.2713 |
| Mir3097 | 0 | 0 | 0 | 0 | 0 | 0 | 0 | 0 | 0 | 0 | 0 | 0 | 0 | 0 | 0 | NA | NA | NA | NA | NA | NA | NA | NA | NA | NA |
| Mir3098 | 0 | 0 | 0 | 0 | 0 | 0 | 0 | 0 | 0 | 0 | 0 | 0 | 0 | 0 | 0 | NA | NA | NA | NA | NA | NA | NA | NA | NA | NA |
| Mir3099 | 0 | 0 | 0 | 0 | 0 | 0 | 0 | 0 | 0 | 0 | 0 | 0 | 0 | 0 | 0 | NA | NA | NA | NA | NA | NA | NA | NA | NA | NA |
| Mir30a | 0 | 0 | 0 | 0 | 0 | 0 | 0 | 0 | 0 | 0 | 0 | 0 | 0 | 0 | 0 | NA | NA | NA | NA | NA | NA | NA | NA | NA | NA |

| | | | | | | | | | | | | | | | | | | | | | | | | | | |
|---|---|---|---|---|---|---|---|---|---|---|---|---|---|---|---|---|---|---|---|---|---|---|---|---|---|---|
| Mir30b | 0 | 0 | 0 | 0 | 0 | 0 | 0 | 0 | 0 | 0 | 0 | 0 | 0 | 0 | 0 | NA | NA | NA | NA | NA | NA | NA | NA | NA | NA |
| Mir30c-1 | 0 | 0 | 0 | 0 | 0 | 0 | 0 | 0 | 0 | 0 | 0 | 0 | 0 | 0 | 0 | NA | NA | NA | NA | NA | NA | NA | NA | NA | NA |
| Mir30c-2 | 0 | 0 | 0 | 0 | 0 | 0 | 0 | 0 | 0 | 0 | 0 | 0 | 0 | 0 | 0 | NA | NA | NA | NA | NA | NA | NA | NA | NA | NA |
| Mir30d | 0 | 0 | 0 | 0 | 0 | 0 | 0 | 0 | 0 | 0 | 0 | 0 | 0 | 0 | 0 | NA | NA | NA | NA | NA | NA | NA | NA | NA | NA |
| Mir30e | 0 | 0 | 0 | 0 | 0 | 0 | 0 | 0 | 0 | 0 | 0 | 0 | 0 | 0 | 0 | NA | NA | NA | NA | NA | NA | NA | NA | NA | NA |
| Mir31 | 0 | 0 | 0 | 0 | 0 | 0 | 0 | 0 | 0 | 0 | 0 | 0 | 0 | 0 | 0 | NA | NA | NA | NA | NA | NA | NA | NA | NA | NA |
| Mir3100 | 1 | 0 | 0 | 0 | 0 | 0 | 1 | 0 | 0 | 0 | 0 | 0 | 0.18 | 0 | 0 | 0.49 | -1.029 | 0.80087 | NA | | 0 | 18.6165 | 10 | 10 | TRUE | 0.4614 |
| Mir3101 | 4 | 4 | 5 | 4 | 9 | 10 | 4 | 4 | 5 | 3 | 10 | 11 | 6.17 | 4 | 8 | 1.759 | 0.815 | 0.32402701 | 0.54535337 | | 0 | 0.5406 | 0.3138 | 0.3138 | TRUE | 1.0634 |
| Mir3102 | 1 | 0 | 0 | 0 | 1 | 0 | 1 | 0 | 0 | 0 | 1 | 0 | 0.36 | 0 | 0 | 0.954 | -0.067 | 0.98557668 | NA | | 0 | 9.2518 | 8.0179 | 8.0179 | TRUE | 0.5204 |
| Mir3103 | 0 | 0 | 0 | 0 | 0 | 0 | 0 | 0 | 0 | 0 | 0 | 0 | 0 | 0 | 0 | NA | NA | NA | NA | NA | NA | NA | NA | NA | NA |
| Mir3104 | 0 | 0 | 0 | 0 | 0 | 0 | 0 | 0 | 0 | 0 | 0 | 0 | 0 | 0 | 0 | NA | NA | NA | NA | NA | NA | NA | NA | NA | NA |
| Mir3106 | 0 | 0 | 0 | 0 | 0 | 0 | 0 | 0 | 0 | 0 | 0 | 0 | 0 | 0 | 0 | NA | NA | NA | NA | NA | NA | NA | NA | NA | NA |
| Mir3107 | 0 | 0 | 0 | 0 | 0 | 0 | 0 | 0 | 0 | 0 | 0 | 0 | 0 | 0 | 0 | NA | NA | NA | NA | NA | NA | NA | NA | NA | NA |
| Mir3108 | 0 | 0 | 0 | 0 | 0 | 0 | 0 | 0 | 0 | 0 | 0 | 0 | 0 | 0 | 0 | NA | NA | NA | NA | NA | NA | NA | NA | NA | NA |
| Mir3109 | 0 | 0 | 0 | 0 | 0 | 0 | 0 | 0 | 0 | 0 | 0 | 0 | 0 | 0 | 0 | NA | NA | NA | NA | NA | NA | NA | NA | NA | NA |
| Mir3110 | 0 | 0 | 0 | 0 | 0 | 0 | 0 | 0 | 0 | 0 | 0 | 0 | 0 | 0 | 0 | NA | NA | NA | NA | NA | NA | NA | NA | NA | NA |
| Mir3112 | 17 | 18 | 13 | 18 | 16 | 22 | 18 | 18 | 13 | 14 | 17 | 24 | 17.42 | 16 | 18 | 1.12 | 0.164 | 0.72847381 | 0.85395791 | | 0 | 0.196 | 0.1028 | 0.1028 | TRUE | 0.3439 |
| Mir32 | 0 | 0 | 0 | 0 | 0 | 0 | 0 | 0 | 0 | 0 | 0 | 0 | 0 | 0 | 0 | NA | NA | NA | NA | NA | NA | NA | NA | NA | NA |
| Mir320 | 0 | 0 | 1 | 0 | 0 | 1 | 0 | 0 | 1 | 0 | 0 | 1 | 0.35 | 0 | 0 | 0.954 | -0.067 | 0.98566485 | NA | | 0 | 9.4252 | 8.1416 | 8.1416 | TRUE | 0.5383 |
| Mir322 | 0 | 0 | 0 | 0 | 0 | 0 | 0 | 0 | 0 | 0 | 0 | 0 | 0 | 0 | 0 | NA | NA | NA | NA | NA | NA | NA | NA | NA | NA |
| Mir323 | 0 | 0 | 0 | 0 | 0 | 0 | 0 | 0 | 0 | 0 | 0 | 0 | 0 | 0 | 0 | NA | NA | NA | NA | NA | NA | NA | NA | NA | NA |
| Mir324 | 0 | 0 | 0 | 0 | 1 | 0 | 0 | 0 | 0 | 0 | 1 | 0 | 0.18 | 0 | 0 | 1.859 | 0.894 | 0.82650734 | NA | | 0 | 18.3785 | 10 | 10 | TRUE | 0.5204 |
| Mir325 | 0 | 0 | 0 | 0 | 0 | 0 | 0 | 0 | 0 | 0 | 0 | 0 | 0 | 0 | 0 | NA | NA | NA | NA | NA | NA | NA | NA | NA | NA |
| Mir326 | 0 | 0 | 0 | 0 | 0 | 0 | 0 | 0 | 0 | 0 | 0 | 0 | 0 | 0 | 0 | NA | NA | NA | NA | NA | NA | NA | NA | NA | NA |
| Mir328 | 1 | 0 | 0 | 0 | 0 | 0 | 1 | 0 | 0 | 0 | 0 | 0 | 0.18 | 0 | 0 | 0.49 | -1.029 | 0.80087 | NA | | 0 | 18.6165 | 10 | 10 | TRUE | 0.4614 |
| Mir329 | 0 | 0 | 0 | 0 | 0 | 0 | 0 | 0 | 0 | 0 | 0 | 0 | 0 | 0 | 0 | NA | NA | NA | NA | NA | NA | NA | NA | NA | NA |
| Mir33 | 0 | 0 | 0 | 0 | 0 | 0 | 0 | 0 | 0 | 0 | 0 | 0 | 0 | 0 | 0 | NA | NA | NA | NA | NA | NA | NA | NA | NA | NA |
| Mir330 | 1 | 0 | 0 | 0 | 0 | 1 | 1 | 0 | 0 | 0 | 0 | 1 | 0.36 | 0 | 0 | 0.954 | -0.067 | 0.98553519 | NA | | 0 | 9.1716 | 7.9605 | 7.9605 | TRUE | 0.5383 |
| Mir331 | 0 | 1 | 0 | 1 | 0 | 0 | 0 | 1 | 0 | 1 | 0 | 0 | 0.3 | 0 | 0 | 0.954 | -0.067 | 0.98642001 | NA | | 0 | 11.0817 | 9.3014 | 9.3014 | TRUE | 0.405 |
| Mir335 | 1 | 0 | 0 | 0 | 0 | 0 | 1 | 0 | 0 | 0 | 0 | 0 | 0.18 | 0 | 0 | 0.49 | -1.029 | 0.80087 | NA | | 0 | 18.6165 | 10 | 10 | TRUE | 0.4614 |
| Mir337 | 0 | 0 | 0 | 0 | 0 | 0 | 0 | 0 | 0 | 0 | 0 | 0 | 0 | 0 | 0 | NA | NA | NA | NA | NA | NA | NA | NA | NA | NA |
| Mir338 | 0 | 0 | 0 | 0 | 0 | 0 | 0 | 0 | 0 | 0 | 0 | 0 | 0 | 0 | 0 | NA | NA | NA | NA | NA | NA | NA | NA | NA | NA |
| Mir339 | 0 | 0 | 0 | 1 | 0 | 0 | 0 | 0 | 0 | 1 | 0 | 0 | 0.13 | 0 | 0 | 1.859 | 0.894 | 0.82650734 | NA | | 0 | 25.1515 | 10 | 10 | TRUE | 0.2563 |
| Mir340 | 0 | 0 | 0 | 0 | 0 | 0 | 0 | 0 | 0 | 0 | 0 | 0 | 0 | 0 | 0 | NA | NA | NA | NA | NA | NA | NA | NA | NA | NA |
| Mir341 | 0 | 0 | 3 | 0 | 0 | 0 | 0 | 0 | 3 | 0 | 0 | 0 | 0.5 | 1 | 0 | 0.178 | -2.488 | 0.53354604 | NA | | 10 | 6.5689 | 10 | 10 | TRUE | 1.4422 |
| Mir343 | 2 | 1 | 0 | 1 | 2 | 0 | 2 | 1 | 0 | 1 | 2 | 0 | 1.01 | 1 | 1 | 0.942 | -0.086 | 0.96639571 | NA | | 0 | 3.2682 | 1.9792 | 1.9792 | TRUE | 0.2432 |
| Mir344 | 0 | 0 | 0 | 0 | 0 | 0 | 0 | 0 | 0 | 0 | 0 | 0 | 0 | 0 | 0 | NA | NA | NA | NA | NA | NA | NA | NA | NA | NA |
| Mir344-2 | 0 | 0 | 0 | 1 | 0 | 0 | 0 | 0 | 0 | 1 | 0 | 0 | 0.13 | 0 | 0 | 1.859 | 0.894 | 0.82650734 | NA | | 0 | 25.1515 | 10 | 10 | TRUE | 0.2563 |
| Mir344b | 0 | 0 | 0 | 0 | 0 | 0 | 0 | 0 | 0 | 0 | 0 | 0 | 0 | 0 | 0 | NA | NA | NA | NA | NA | NA | NA | NA | NA | NA |
| Mir344c | 0 | 0 | 0 | 0 | 0 | 0 | 0 | 0 | 0 | 0 | 0 | 0 | 0 | 0 | 0 | NA | NA | NA | NA | NA | NA | NA | NA | NA | NA |
| Mir344d-1 | 0 | 0 | 0 | 0 | 0 | 0 | 0 | 0 | 0 | 0 | 0 | 0 | 0 | 0 | 0 | NA | NA | NA | NA | NA | NA | NA | NA | NA | NA |
| Mir344d-2 | 0 | 0 | 0 | 0 | 0 | 0 | 0 | 0 | 0 | 0 | 0 | 0 | 0 | 0 | 0 | NA | NA | NA | NA | NA | NA | NA | NA | NA | NA |
| Mir344d-3 | 0 | 0 | 0 | 0 | 0 | 0 | 0 | 0 | 0 | 0 | 0 | 0 | 0 | 0 | 0 | NA | NA | NA | NA | NA | NA | NA | NA | NA | NA |
| Mir344e | 0 | 0 | 0 | 0 | 0 | 0 | 0 | 0 | 0 | 0 | 0 | 0 | 0 | 0 | 0 | NA | NA | NA | NA | NA | NA | NA | NA | NA | NA |
| Mir344f | 0 | 0 | 0 | 0 | 0 | 0 | 0 | 0 | 0 | 0 | 0 | 0 | 0 | 0 | 0 | NA | NA | NA | NA | NA | NA | NA | NA | NA | NA |
| Mir344g | 0 | 0 | 0 | 0 | 0 | 0 | 0 | 0 | 0 | 0 | 0 | 0 | 0 | 0 | 0 | NA | NA | NA | NA | NA | NA | NA | NA | NA | NA |
| Mir345 | 0 | 0 | 0 | 0 | 0 | 0 | 0 | 0 | 0 | 0 | 0 | 0 | 0 | 0 | 0 | NA | NA | NA | NA | NA | NA | NA | NA | NA | NA |
| Mir346 | 0 | 0 | 0 | 0 | 0 | 0 | 0 | 0 | 0 | 0 | 0 | 0 | 0 | 0 | 0 | NA | NA | NA | NA | NA | NA | NA | NA | NA | NA |
| Mir3470a | 0 | 0 | 0 | 0 | 0 | 0 | 0 | 0 | 0 | 0 | 0 | 0 | 0 | 0 | 0 | NA | NA | NA | NA | NA | NA | NA | NA | NA | NA |
| Mir3470b | 0 | 0 | 0 | 0 | 0 | 0 | 0 | 0 | 0 | 0 | 0 | 0 | 0 | 0 | 0 | NA | NA | NA | NA | NA | NA | NA | NA | NA | NA |
| Mir3471-1 | 0 | 0 | 0 | 0 | 0 | 0 | 0 | 0 | 0 | 0 | 0 | 0 | 0 | 0 | 0 | NA | NA | NA | NA | NA | NA | NA | NA | NA | NA |
| Mir3473 | 0 | 0 | 0 | 0 | 0 | 0 | 0 | 0 | 0 | 0 | 0 | 0 | 0 | 0 | 0 | NA | NA | NA | NA | NA | NA | NA | NA | NA | NA |
| Mir3473c | 0 | 0 | 0 | 0 | 0 | 0 | 0 | 0 | 0 | 0 | 0 | 0 | 0 | 0 | 0 | NA | NA | NA | NA | NA | NA | NA | NA | NA | NA |
| Mir3473d | 0 | 1 | 0 | 0 | 1 | 1 | 0 | 1 | 0 | 0 | 1 | 1 | 0.53 | 0 | 1 | 1.924 | 0.944 | 0.7364225 | NA | | 0 | 6.2438 | 3.9727 | 3.9727 | TRUE | 0.405 |
| Mir3474 | 0 | 0 | 0 | 1 | 0 | 0 | 0 | 0 | 0 | 1 | 0 | 0 | 0.13 | 0 | 0 | 1.859 | 0.894 | 0.82650734 | NA | | 0 | 25.1515 | 10 | 10 | TRUE | 0.2563 |
| Mir3475 | 0 | 0 | 0 | 0 | 0 | 0 | 0 | 0 | 0 | 0 | 0 | 0 | 0 | 0 | 0 | NA | NA | NA | NA | NA | NA | NA | NA | NA | NA |
| Mir34a | 0 | 0 | 0 | 0 | 0 | 0 | 0 | 0 | 0 | 0 | 0 | 0 | 0 | 0 | 0 | NA | NA | NA | NA | NA | NA | NA | NA | NA | NA |
| Mir34b | 0 | 0 | 0 | 0 | 0 | 0 | 0 | 0 | 0 | 0 | 0 | 0 | 0 | 0 | 0 | NA | NA | NA | NA | NA | NA | NA | NA | NA | NA |
| Mir34c | 0 | 0 | 0 | 0 | 0 | 0 | 0 | 0 | 0 | 0 | 0 | 0 | 0 | 0 | 0 | NA | NA | NA | NA | NA | NA | NA | NA | NA | NA |
| Mir350 | 0 | 0 | 0 | 0 | 0 | 0 | 0 | 0 | 0 | 0 | 0 | 0 | 0 | 0 | 0 | NA | NA | NA | NA | NA | NA | NA | NA | NA | NA |
| Mir351 | 0 | 0 | 0 | 0 | 0 | 0 | 0 | 0 | 0 | 0 | 0 | 0 | 0 | 0 | 0 | NA | NA | NA | NA | NA | NA | NA | NA | NA | NA |
| Mir3572 | 0 | 0 | 0 | 0 | 0 | 0 | 0 | 0 | 0 | 0 | 0 | 0 | 0 | 0 | 0 | NA | NA | NA | NA | NA | NA | NA | NA | NA | NA |
| Mir362 | 0 | 0 | 0 | 0 | 0 | 0 | 0 | 0 | 0 | 0 | 0 | 0 | 0 | 0 | 0 | NA | NA | NA | NA | NA | NA | NA | NA | NA | NA |
| Mir363 | 1 | 2 | 1 | 4 | 2 | 1 | 1 | 2 | 1 | 3 | 2 | 1 | 1.74 | 1 | 2 | 1.599 | 0.677 | 0.62576406 | NA | | 0 | 1.8976 | 0.7863 | 0.7863 | TRUE | 0.1953 |
| Mir365-1 | 0 | 0 | 0 | 0 | 0 | 0 | 0 | 0 | 0 | 0 | 0 | 0 | 0 | 0 | 0 | NA | NA | NA | NA | NA | NA | NA | NA | NA | NA |
| Mir365-2 | 0 | 0 | 0 | 0 | 0 | 0 | 0 | 0 | 0 | 0 | 0 | 0 | 0 | 0 | 0 | NA | NA | NA | NA | NA | NA | NA | NA | NA | NA |
| Mir367 | 0 | 0 | 0 | 0 | 0 | 0 | 0 | 0 | 0 | 0 | 0 | 0 | 0 | 0 | 0 | NA | NA | NA | NA | NA | NA | NA | NA | NA | NA |
| Mir369 | 0 | 0 | 0 | 0 | 0 | 0 | 0 | 0 | 0 | 0 | 0 | 0 | 0 | 0 | 0 | NA | NA | NA | NA | NA | NA | NA | NA | NA | NA |
| Mir370 | 0 | 1 | 0 | 0 | 0 | 0 | 0 | 1 | 0 | 0 | 0 | 0 | 0.17 | 0 | 0 | 0.49 | -1.029 | 0.80087 | NA | | 0 | 19.7991 | 10 | 10 | TRUE | 0.405 |

| | | | | | | | | | | | | | | | | | | | | | | | | | |
|---|---|---|---|---|---|---|---|---|---|---|---|---|---|---|---|---|---|---|---|---|---|---|---|---|---|
| Mir374 | 0 | 0 | 0 | 0 | 0 | 0 | 0 | 0 | 0 | 0 | 0 | 0 | 0 | 0 | 0 | NA | NA | NA | NA | NA | NA | NA | NA | NA | NA |
| Mir374c | 0 | 0 | 0 | 0 | 0 | 0 | 0 | 0 | 0 | 0 | 0 | 0 | 0 | 0 | 0 | NA | NA | NA | NA | NA | NA | NA | NA | NA | NA |
| Mir375 | 0 | 0 | 0 | 0 | 0 | 0 | 0 | 0 | 0 | 0 | 0 | 0 | 0 | 0 | 0 | NA | NA | NA | NA | NA | NA | NA | NA | NA | NA |
| Mir376a | 0 | 0 | 0 | 0 | 0 | 0 | 0 | 0 | 0 | 0 | 0 | 0 | 0 | 0 | 0 | NA | NA | NA | NA | NA | NA | NA | NA | NA | NA |
| Mir376b | 0 | 0 | 0 | 0 | 0 | 0 | 0 | 0 | 0 | 0 | 0 | 0 | 0 | 0 | 0 | NA | NA | NA | NA | NA | NA | NA | NA | NA | NA |
| Mir376c | 2 | 0 | 0 | 0 | 0 | 0 | 2 | 0 | 0 | 0 | 0 | 0 | 0.35 | 1 | 0 | 0.254 | -1.975 | 0.62446185 | NA | 10 | 9.3117 | 10 | 10 | TRUE | 0.9741 |
| Mir377 | 1 | 0 | 0 | 1 | 1 | 1 | 1 | 0 | 0 | 1 | 1 | 1 | 0.67 | 0 | 1 | 2.676 | 1.42 | 0.53486561 | NA | 0 | 4.9253 | 2.274 | 2.274 | TRUE | 0.4614 |
| Mir378 | 0 | 0 | 0 | 0 | 0 | 0 | 0 | 0 | 0 | 0 | 0 | 0 | 0 | 0 | 0 | NA | NA | NA | NA | NA | NA | NA | NA | NA | NA |
| Mir378b | 0 | 0 | 0 | 0 | 0 | 0 | 0 | 0 | 0 | 0 | 0 | 0 | 0 | 0 | 0 | NA | NA | NA | NA | NA | NA | NA | NA | NA | NA |
| Mir379 | 0 | 0 | 0 | 0 | 0 | 0 | 0 | 0 | 0 | 0 | 0 | 0 | 0 | 0 | 0 | NA | NA | NA | NA | NA | NA | NA | NA | NA | NA |
| Mir380 | 0 | 0 | 0 | 0 | 0 | 0 | 0 | 0 | 0 | 0 | 0 | 0 | 0 | 0 | 0 | NA | NA | NA | NA | NA | NA | NA | NA | NA | NA |
| Mir381 | 0 | 0 | 0 | 0 | 0 | 0 | 0 | 0 | 0 | 0 | 0 | 0 | 0 | 0 | 0 | NA | NA | NA | NA | NA | NA | NA | NA | NA | NA |
| Mir382 | 0 | 0 | 0 | 0 | 0 | 0 | 0 | 0 | 0 | 0 | 0 | 0 | 0 | 0 | 0 | NA | NA | NA | NA | NA | NA | NA | NA | NA | NA |
| Mir383 | 0 | 0 | 0 | 0 | 0 | 0 | 0 | 0 | 0 | 0 | 0 | 0 | 0 | 0 | 0 | NA | NA | NA | NA | NA | NA | NA | NA | NA | NA |
| Mir384 | 0 | 0 | 0 | 0 | 0 | 0 | 0 | 0 | 0 | 0 | 0 | 0 | 0 | 0 | 0 | NA | NA | NA | NA | NA | NA | NA | NA | NA | NA |
| Mir3960 | 2 | 4 | 0 | 7 | 5 | 3 | 2 | 4 | 0 | 6 | 5 | 3 | 3.38 | 2 | 5 | 2.325 | 1.217 | 0.29175905 | NA | 0 | 0.9813 | 0.6072 | 0.6072 | TRUE | 0.464 |
| Mir3962 | 0 | 0 | 0 | 0 | 0 | 0 | 0 | 0 | 0 | 0 | 0 | 0 | 0 | 0 | 0 | NA | NA | NA | NA | NA | NA | NA | NA | NA | NA |
| Mir3963 | 0 | 0 | 0 | 0 | 0 | 0 | 0 | 0 | 0 | 0 | 0 | 0 | 0 | 0 | 0 | NA | NA | NA | NA | NA | NA | NA | NA | NA | NA |
| Mir3964 | 0 | 0 | 0 | 0 | 0 | 0 | 0 | 0 | 0 | 0 | 0 | 0 | 0 | 0 | 0 | NA | NA | NA | NA | NA | NA | NA | NA | NA | NA |
| Mir3965 | 0 | 0 | 0 | 0 | 0 | 0 | 0 | 0 | 0 | 0 | 0 | 0 | 0 | 0 | 0 | NA | NA | NA | NA | NA | NA | NA | NA | NA | NA |
| Mir3966 | 0 | 0 | 0 | 0 | 0 | 0 | 0 | 0 | 0 | 0 | 0 | 0 | 0 | 0 | 0 | NA | NA | NA | NA | NA | NA | NA | NA | NA | NA |
| Mir3967 | 0 | 0 | 0 | 0 | 0 | 0 | 0 | 0 | 0 | 0 | 0 | 0 | 0 | 0 | 0 | NA | NA | NA | NA | NA | NA | NA | NA | NA | NA |
| Mir3968 | 0 | 0 | 0 | 0 | 0 | 0 | 0 | 0 | 0 | 0 | 0 | 0 | 0 | 0 | 0 | NA | NA | NA | NA | NA | NA | NA | NA | NA | NA |
| Mir3969 | 0 | 0 | 0 | 0 | 0 | 0 | 0 | 0 | 0 | 0 | 0 | 0 | 0 | 0 | 0 | NA | NA | NA | NA | NA | NA | NA | NA | NA | NA |
| Mir3970 | 0 | 0 | 0 | 0 | 0 | 0 | 0 | 0 | 0 | 0 | 0 | 0 | 0 | 0 | 0 | NA | NA | NA | NA | NA | NA | NA | NA | NA | NA |
| Mir3971 | 0 | 0 | 0 | 2 | 0 | 0 | 0 | 0 | 0 | 2 | 0 | 0 | 0.26 | 0 | 1 | 2.886 | 1.529 | 0.70676491 | NA | 0 | 12.5792 | 10 | 10 | TRUE | 0.9669 |
| Mir409 | 1 | 0 | 1 | 2 | 0 | 0 | 1 | 0 | 1 | 2 | 0 | 0 | 0.61 | 1 | 1 | 0.816 | -0.293 | 0.91321207 | NA | 0 | 5.4395 | 3.5982 | 3.5982 | TRUE | 0.9722 |
| Mir410 | 0 | 0 | 0 | 0 | 0 | 0 | 0 | 0 | 0 | 0 | 0 | 0 | 0 | 0 | 0 | NA | NA | NA | NA | NA | NA | NA | NA | NA | NA |
| Mir411 | 0 | 0 | 0 | 0 | 0 | 0 | 0 | 0 | 0 | 0 | 0 | 0 | 0 | 0 | 0 | NA | NA | NA | NA | NA | NA | NA | NA | NA | NA |
| Mir412 | 0 | 0 | 0 | 0 | 0 | 0 | 0 | 0 | 0 | 0 | 0 | 0 | 0 | 0 | 0 | NA | NA | NA | NA | NA | NA | NA | NA | NA | NA |
| Mir421 | 0 | 0 | 0 | 0 | 0 | 0 | 0 | 0 | 0 | 0 | 0 | 0 | 0 | 0 | 0 | NA | NA | NA | NA | NA | NA | NA | NA | NA | NA |
| Mir423 | 0 | 0 | 0 | 1 | 0 | 0 | 0 | 0 | 0 | 1 | 0 | 0 | 0.13 | 0 | 0 | 1.859 | 0.894 | 0.82650734 | NA | 0 | 25.1515 | 10 | 10 | TRUE | 0.2563 |
| Mir425 | 3 | 0 | 1 | 2 | 0 | 1 | 3 | 0 | 1 | 2 | 0 | 1 | 1.14 | 1 | 1 | 0.659 | -0.601 | 0.75967187 | NA | 0 | 2.8894 | 1.8829 | 1.8829 | TRUE | 0.4216 |
| Mir429 | 0 | 0 | 0 | 0 | 0 | 0 | 0 | 0 | 0 | 0 | 0 | 0 | 0 | 0 | 0 | NA | NA | NA | NA | NA | NA | NA | NA | NA | NA |
| Mir431 | 0 | 0 | 0 | 0 | 0 | 0 | 0 | 0 | 0 | 0 | 0 | 0 | 0 | 0 | 0 | NA | NA | NA | NA | NA | NA | NA | NA | NA | NA |
| Mir432 | 0 | 0 | 0 | 0 | 0 | 0 | 0 | 0 | 0 | 0 | 0 | 0 | 0 | 0 | 0 | NA | NA | NA | NA | NA | NA | NA | NA | NA | NA |
| Mir433 | 0 | 0 | 0 | 0 | 0 | 0 | 0 | 0 | 0 | 0 | 0 | 0 | 0 | 0 | 0 | NA | NA | NA | NA | NA | NA | NA | NA | NA | NA |
| Mir434 | 0 | 0 | 0 | 0 | 0 | 0 | 0 | 0 | 0 | 0 | 0 | 0 | 0 | 0 | 0 | NA | NA | NA | NA | NA | NA | NA | NA | NA | NA |
| Mir448 | 0 | 0 | 0 | 0 | 0 | 0 | 0 | 0 | 0 | 0 | 0 | 0 | 0 | 0 | 0 | NA | NA | NA | NA | NA | NA | NA | NA | NA | NA |
| Mir449a | 0 | 0 | 0 | 0 | 0 | 0 | 0 | 0 | 0 | 0 | 0 | 0 | 0 | 0 | 0 | NA | NA | NA | NA | NA | NA | NA | NA | NA | NA |
| Mir449b | 0 | 0 | 0 | 0 | 0 | 0 | 0 | 0 | 0 | 0 | 0 | 0 | 0 | 0 | 0 | NA | NA | NA | NA | NA | NA | NA | NA | NA | NA |
| Mir449c | 0 | 0 | 0 | 0 | 0 | 0 | 0 | 0 | 0 | 0 | 0 | 0 | 0 | 0 | 0 | NA | NA | NA | NA | NA | NA | NA | NA | NA | NA |
| Mir450-1 | 0 | 0 | 0 | 0 | 0 | 0 | 0 | 0 | 0 | 0 | 0 | 0 | 0 | 0 | 0 | NA | NA | NA | NA | NA | NA | NA | NA | NA | NA |
| Mir450-2 | 0 | 0 | 0 | 0 | 0 | 0 | 0 | 0 | 0 | 0 | 0 | 0 | 0 | 0 | 0 | NA | NA | NA | NA | NA | NA | NA | NA | NA | NA |
| Mir450b | 0 | 0 | 0 | 0 | 0 | 0 | 0 | 0 | 0 | 0 | 0 | 0 | 0 | 0 | 0 | NA | NA | NA | NA | NA | NA | NA | NA | NA | NA |
| Mir451 | 0 | 0 | 0 | 0 | 0 | 0 | 0 | 0 | 0 | 0 | 0 | 0 | 0 | 0 | 0 | NA | NA | NA | NA | NA | NA | NA | NA | NA | NA |
| Mir452 | 0 | 0 | 0 | 0 | 0 | 0 | 0 | 0 | 0 | 0 | 0 | 0 | 0 | 0 | 0 | NA | NA | NA | NA | NA | NA | NA | NA | NA | NA |
| Mir453 | 3 | 2 | 2 | 2 | 0 | 0 | 3 | 2 | 2 | 2 | 0 | 0 | 1.46 | 2 | 1 | 0.246 | -2.023 | 0.25463936 | NA | 0 | 2.2622 | 1.233 | 1.233 | TRUE | 0.9709 |
| Mir455 | 0 | 0 | 0 | 0 | 0 | 0 | 0 | 0 | 0 | 0 | 0 | 0 | 0 | 0 | 0 | NA | NA | NA | NA | NA | NA | NA | NA | NA | NA |
| Mir463 | 0 | 0 | 0 | 0 | 0 | 0 | 0 | 0 | 0 | 0 | 0 | 0 | 0 | 0 | 0 | NA | NA | NA | NA | NA | NA | NA | NA | NA | NA |
| Mir465 | 0 | 0 | 0 | 0 | 0 | 0 | 0 | 0 | 0 | 0 | 0 | 0 | 0 | 0 | 0 | NA | NA | NA | NA | NA | NA | NA | NA | NA | NA |
| Mir465b-1 | 0 | 0 | 0 | 0 | 0 | 0 | 0 | 0 | 0 | 0 | 0 | 0 | 0 | 0 | 0 | NA | NA | NA | NA | NA | NA | NA | NA | NA | NA |
| Mir465b-2 | 0 | 0 | 0 | 0 | 0 | 0 | 0 | 0 | 0 | 0 | 0 | 0 | 0 | 0 | 0 | NA | NA | NA | NA | NA | NA | NA | NA | NA | NA |
| Mir465c-1 | 0 | 0 | 0 | 0 | 0 | 0 | 0 | 0 | 0 | 0 | 0 | 0 | 0 | 0 | 0 | NA | NA | NA | NA | NA | NA | NA | NA | NA | NA |
| Mir465c-2 | 0 | 0 | 0 | 0 | 0 | 0 | 0 | 0 | 0 | 0 | 0 | 0 | 0 | 0 | 0 | NA | NA | NA | NA | NA | NA | NA | NA | NA | NA |
| Mir466 | 0 | 0 | 0 | 0 | 0 | 0 | 0 | 0 | 0 | 0 | 0 | 0 | 0 | 0 | 0 | NA | NA | NA | NA | NA | NA | NA | NA | NA | NA |
| Mir4660 | 0 | 0 | 0 | 0 | 0 | 0 | 0 | 0 | 0 | 0 | 0 | 0 | 0 | 0 | 0 | NA | NA | NA | NA | NA | NA | NA | NA | NA | NA |
| Mir466b-2 | 0 | 0 | 0 | 0 | 0 | 0 | 0 | 0 | 0 | 0 | 0 | 0 | 0 | 0 | 0 | NA | NA | NA | NA | NA | NA | NA | NA | NA | NA |
| Mir466b-3 | 0 | 0 | 0 | 0 | 0 | 0 | 0 | 0 | 0 | 0 | 0 | 0 | 0 | 0 | 0 | NA | NA | NA | NA | NA | NA | NA | NA | NA | NA |
| Mir466d | 0 | 0 | 0 | 0 | 0 | 0 | 0 | 0 | 0 | 0 | 0 | 0 | 0 | 0 | 0 | NA | NA | NA | NA | NA | NA | NA | NA | NA | NA |
| Mir466f-1 | 2 | 0 | 0 | 0 | 0 | 0 | 2 | 0 | 0 | 0 | 0 | 0 | 0.35 | 1 | 0 | 0.254 | -1.975 | 0.62446185 | NA | 10 | 9.3117 | 10 | 10 | TRUE | 0.9741 |
| Mir466f-2 | 0 | 1 | 0 | 0 | 0 | 0 | 0 | 1 | 0 | 0 | 0 | 0 | 0.17 | 0 | 0 | 0.49 | -1.029 | 0.80087 | NA | 0 | 19.7991 | 10 | 10 | TRUE | 0.405 |
| Mir466f-3 | 0 | 0 | 0 | 0 | 0 | 0 | 0 | 0 | 0 | 0 | 0 | 0 | 0 | 0 | 0 | NA | NA | NA | NA | NA | NA | NA | NA | NA | NA |
| Mir466g | 0 | 0 | 0 | 0 | 0 | 0 | 0 | 0 | 0 | 0 | 0 | 0 | 0 | 0 | 0 | NA | NA | NA | NA | NA | NA | NA | NA | NA | NA |
| Mir466h | 0 | 0 | 0 | 0 | 0 | 0 | 0 | 0 | 0 | 0 | 0 | 0 | 0 | 0 | 0 | NA | NA | NA | NA | NA | NA | NA | NA | NA | NA |
| Mir466i | 0 | 0 | 0 | 0 | 0 | 0 | 0 | 0 | 0 | 0 | 0 | 0 | 0 | 0 | 0 | NA | NA | NA | NA | NA | NA | NA | NA | NA | NA |
| Mir466n | 0 | 0 | 0 | 0 | 0 | 0 | 0 | 0 | 0 | 0 | 0 | 0 | 0 | 0 | 0 | NA | NA | NA | NA | NA | NA | NA | NA | NA | NA |
| Mir467a-1 | 0 | 0 | 0 | 0 | 0 | 0 | 0 | 0 | 0 | 0 | 0 | 0 | 0 | 0 | 0 | NA | NA | NA | NA | NA | NA | NA | NA | NA | NA |
| Mir467a-10 | 0 | 0 | 0 | 0 | 0 | 0 | 0 | 0 | 0 | 0 | 0 | 0 | 0 | 0 | 0 | NA | NA | NA | NA | NA | NA | NA | NA | NA | NA |

| | | | | | | | | | | | | | | | | | | | | | | | | | |
|---|---|---|---|---|---|---|---|---|---|---|---|---|---|---|---|---|---|---|---|---|---|---|---|---|---|
| Mir467a-2 | 0 | 0 | 0 | 0 | 0 | 0 | 0 | 0 | 0 | 0 | 0 | 0 | 0 | 0 | 0 | NA | NA | NA | NA | NA | NA | NA | NA | NA | NA |
| Mir467a-3 | 0 | 0 | 0 | 0 | 0 | 0 | 0 | 0 | 0 | 0 | 0 | 0 | 0 | 0 | 0 | NA | NA | NA | NA | NA | NA | NA | NA | NA | NA |
| Mir467a-4 | 0 | 0 | 0 | 0 | 0 | 0 | 0 | 0 | 0 | 0 | 0 | 0 | 0 | 0 | 0 | NA | NA | NA | NA | NA | NA | NA | NA | NA | NA |
| Mir467a-5 | 0 | 0 | 0 | 0 | 0 | 0 | 0 | 0 | 0 | 0 | 0 | 0 | 0 | 0 | 0 | NA | NA | NA | NA | NA | NA | NA | NA | NA | NA |
| Mir467a-6 | 0 | 0 | 0 | 0 | 0 | 0 | 0 | 0 | 0 | 0 | 0 | 0 | 0 | 0 | 0 | NA | NA | NA | NA | NA | NA | NA | NA | NA | NA |
| Mir467a-7 | 0 | 0 | 0 | 0 | 0 | 0 | 0 | 0 | 0 | 0 | 0 | 0 | 0 | 0 | 0 | NA | NA | NA | NA | NA | NA | NA | NA | NA | NA |
| Mir467a-8 | 0 | 0 | 0 | 0 | 0 | 0 | 0 | 0 | 0 | 0 | 0 | 0 | 0 | 0 | 0 | NA | NA | NA | NA | NA | NA | NA | NA | NA | NA |
| Mir467a-9 | 0 | 0 | 0 | 0 | 0 | 0 | 0 | 0 | 0 | 0 | 0 | 0 | 0 | 0 | 0 | NA | NA | NA | NA | NA | NA | NA | NA | NA | NA |
| Mir467b | 0 | 0 | 0 | 0 | 0 | 0 | 0 | 0 | 0 | 0 | 0 | 0 | 0 | 0 | 0 | NA | NA | NA | NA | NA | NA | NA | NA | NA | NA |
| Mir467c | 0 | 0 | 0 | 0 | 0 | 0 | 0 | 0 | 0 | 0 | 0 | 0 | 0 | 0 | 0 | NA | NA | NA | NA | NA | NA | NA | NA | NA | NA |
| Mir467d | 0 | 0 | 0 | 0 | 0 | 0 | 0 | 0 | 0 | 0 | 0 | 0 | 0 | 0 | 0 | NA | NA | NA | NA | NA | NA | NA | NA | NA | NA |
| Mir467e | 0 | 0 | 0 | 0 | 0 | 0 | 0 | 0 | 0 | 0 | 0 | 0 | 0 | 0 | 0 | NA | NA | NA | NA | NA | NA | NA | NA | NA | NA |
| Mir467f | 0 | 0 | 0 | 0 | 0 | 0 | 0 | 0 | 0 | 0 | 0 | 0 | 0 | 0 | 0 | NA | NA | NA | NA | NA | NA | NA | NA | NA | NA |
| Mir468 | 0 | 0 | 0 | 0 | 0 | 0 | 0 | 0 | 0 | 0 | 0 | 0 | 0 | 0 | 0 | NA | NA | NA | NA | NA | NA | NA | NA | NA | NA |
| Mir470 | 0 | 0 | 0 | 0 | 0 | 0 | 0 | 0 | 0 | 0 | 0 | 0 | 0 | 0 | 0 | NA | NA | NA | NA | NA | NA | NA | NA | NA | NA |
| Mir471 | 0 | 0 | 0 | 0 | 0 | 0 | 0 | 0 | 0 | 0 | 0 | 0 | 0 | 0 | 0 | NA | NA | NA | NA | NA | NA | NA | NA | NA | NA |
| Mir483 | 0 | 0 | 0 | 0 | 0 | 0 | 0 | 0 | 0 | 0 | 0 | 0 | 0 | 0 | 0 | NA | NA | NA | NA | NA | NA | NA | NA | NA | NA |
| Mir484 | 0 | 0 | 0 | 1 | 0 | 0 | 0 | 0 | 0 | 1 | 0 | 0 | 0.13 | 0 | 0 | 1.859 | 0.894 | 0.82650734 | NA | 0 | 25.1515 | 10 | 10 | TRUE | 0.2563 |
| Mir485 | 1 | 0 | 0 | 1 | 0 | 0 | 1 | 0 | 0 | 1 | 0 | 0 | 0.31 | 0 | 0 | 0.954 | -0.067 | 0.98625928 | NA | 0 | 10.7014 | 9.0384 | 9.0384 | TRUE | 0.4614 |
| Mir486 | 0 | 0 | 0 | 0 | 0 | 0 | 0 | 0 | 0 | 0 | 0 | 0 | 0 | 0 | 0 | NA | NA | NA | NA | NA | NA | NA | NA | NA | NA |
| Mir487b | 0 | 0 | 0 | 0 | 0 | 0 | 0 | 0 | 0 | 0 | 0 | 0 | 0 | 0 | 0 | NA | NA | NA | NA | NA | NA | NA | NA | NA | NA |
| Mir488 | 0 | 0 | 0 | 0 | 0 | 0 | 0 | 0 | 0 | 0 | 0 | 0 | 0 | 0 | 0 | NA | NA | NA | NA | NA | NA | NA | NA | NA | NA |
| Mir489 | 0 | 0 | 0 | 0 | 0 | 0 | 0 | 0 | 0 | 0 | 0 | 0 | 0 | 0 | 0 | NA | NA | NA | NA | NA | NA | NA | NA | NA | NA |
| Mir490 | 0 | 0 | 0 | 0 | 0 | 0 | 0 | 0 | 0 | 0 | 0 | 0 | 0 | 0 | 0 | NA | NA | NA | NA | NA | NA | NA | NA | NA | NA |
| Mir491 | 0 | 0 | 0 | 0 | 0 | 0 | 0 | 0 | 0 | 0 | 0 | 0 | 0 | 0 | 0 | NA | NA | NA | NA | NA | NA | NA | NA | NA | NA |
| Mir493 | 0 | 0 | 0 | 0 | 0 | 0 | 0 | 0 | 0 | 0 | 0 | 0 | 0 | 0 | 0 | NA | NA | NA | NA | NA | NA | NA | NA | NA | NA |
| Mir494 | 0 | 0 | 1 | 0 | 0 | 0 | 0 | 0 | 1 | 0 | 0 | 0 | 0.17 | 0 | 0 | 0.49 | -1.029 | 0.80087 | NA | 0 | 19.6928 | 10 | 10 | TRUE | 0.4098 |
| Mir495 | 0 | 0 | 0 | 0 | 0 | 0 | 0 | 0 | 0 | 0 | 0 | 0 | 0 | 0 | 0 | NA | NA | NA | NA | NA | NA | NA | NA | NA | NA |
| Mir496 | 0 | 0 | 0 | 0 | 1 | 0 | 0 | 0 | 0 | 0 | 1 | 0 | 0.18 | 0 | 0 | 1.859 | 0.894 | 0.82650734 | NA | 0 | 18.3785 | 10 | 10 | TRUE | 0.5204 |
| Mir497 | 0 | 0 | 0 | 0 | 0 | 0 | 0 | 0 | 0 | 0 | 0 | 0 | 0 | 0 | 0 | NA | NA | NA | NA | NA | NA | NA | NA | NA | NA |
| Mir499 | 0 | 0 | 0 | 0 | 0 | 0 | 0 | 0 | 0 | 0 | 0 | 0 | 0 | 0 | 0 | NA | NA | NA | NA | NA | NA | NA | NA | NA | NA |
| Mir500 | 0 | 0 | 0 | 0 | 0 | 0 | 0 | 0 | 0 | 0 | 0 | 0 | 0 | 0 | 0 | NA | NA | NA | NA | NA | NA | NA | NA | NA | NA |
| Mir501 | 0 | 0 | 0 | 0 | 0 | 0 | 0 | 0 | 0 | 0 | 0 | 0 | 0 | 0 | 0 | NA | NA | NA | NA | NA | NA | NA | NA | NA | NA |
| Mir503 | 0 | 0 | 0 | 0 | 0 | 0 | 0 | 0 | 0 | 0 | 0 | 0 | 0 | 0 | 0 | NA | NA | NA | NA | NA | NA | NA | NA | NA | NA |
| Mir504 | 0 | 0 | 0 | 0 | 0 | 0 | 0 | 0 | 0 | 0 | 0 | 0 | 0 | 0 | 0 | NA | NA | NA | NA | NA | NA | NA | NA | NA | NA |
| Mir5046 | 0 | 0 | 0 | 0 | 0 | 0 | 0 | 0 | 0 | 0 | 0 | 0 | 0 | 0 | 0 | NA | NA | NA | NA | NA | NA | NA | NA | NA | NA |
| Mir505 | 0 | 0 | 0 | 0 | 0 | 0 | 0 | 0 | 0 | 0 | 0 | 0 | 0 | 0 | 0 | NA | NA | NA | NA | NA | NA | NA | NA | NA | NA |
| Mir509 | 0 | 0 | 0 | 0 | 0 | 0 | 0 | 0 | 0 | 0 | 0 | 0 | 0 | 0 | 0 | NA | NA | NA | NA | NA | NA | NA | NA | NA | NA |
| Mir5097 | 0 | 0 | 0 | 0 | 0 | 0 | 0 | 0 | 0 | 0 | 0 | 0 | 0 | 0 | 0 | NA | NA | NA | NA | NA | NA | NA | NA | NA | NA |
| Mir5098 | 0 | 0 | 0 | 0 | 0 | 0 | 0 | 0 | 0 | 0 | 0 | 0 | 0 | 0 | 0 | NA | NA | NA | NA | NA | NA | NA | NA | NA | NA |
| Mir5100 | 0 | 0 | 0 | 0 | 0 | 0 | 0 | 0 | 0 | 0 | 0 | 0 | 0 | 0 | 0 | NA | NA | NA | NA | NA | NA | NA | NA | NA | NA |
| Mir5101 | 0 | 0 | 0 | 0 | 0 | 0 | 0 | 0 | 0 | 0 | 0 | 0 | 0 | 0 | 0 | NA | NA | NA | NA | NA | NA | NA | NA | NA | NA |
| Mir5103 | 0 | 0 | 0 | 0 | 0 | 0 | 0 | 0 | 0 | 0 | 0 | 0 | 0 | 0 | 0 | NA | NA | NA | NA | NA | NA | NA | NA | NA | NA |
| Mir5104 | 0 | 0 | 0 | 0 | 0 | 0 | 0 | 0 | 0 | 0 | 0 | 0 | 0 | 0 | 0 | NA | NA | NA | NA | NA | NA | NA | NA | NA | NA |
| Mir5106 | 1 | 0 | 1 | 1 | 0 | 0 | 1 | 0 | 1 | 1 | 0 | 0 | 0.48 | 1 | 0 | 0.509 | -0.975 | 0.7336918 | NA | 0 | 6.9367 | 4.1806 | 4.1806 | TRUE | 0.2562 |
| Mir5107 | 0 | 0 | 0 | 0 | 0 | 0 | 0 | 0 | 0 | 0 | 0 | 0 | 0 | 0 | 0 | NA | NA | NA | NA | NA | NA | NA | NA | NA | NA |
| Mir5108 | 0 | 0 | 0 | 0 | 0 | 0 | 0 | 0 | 0 | 0 | 0 | 0 | 0 | 0 | 0 | NA | NA | NA | NA | NA | NA | NA | NA | NA | NA |
| Mir5109 | 2 | 0 | 0 | 2 | 0 | 0 | 2 | 0 | 0 | 2 | 0 | 0 | 0.62 | 1 | 1 | 0.783 | -0.353 | 0.91186553 | NA | 0 | 5.3542 | 5.6997 | 5.6997 | TRUE | 0.9742 |
| Mir511 | 0 | 0 | 0 | 0 | 0 | 0 | 0 | 0 | 0 | 0 | 0 | 0 | 0 | 0 | 0 | NA | NA | NA | NA | NA | NA | NA | NA | NA | NA |
| Mir5112 | 0 | 0 | 0 | 0 | 0 | 0 | 0 | 0 | 0 | 0 | 0 | 0 | 0 | 0 | 0 | NA | NA | NA | NA | NA | NA | NA | NA | NA | NA |
| Mir5113 | 0 | 0 | 0 | 0 | 0 | 0 | 0 | 0 | 0 | 0 | 0 | 0 | 0 | 0 | 0 | NA | NA | NA | NA | NA | NA | NA | NA | NA | NA |
| Mir5114 | 0 | 0 | 0 | 0 | 0 | 0 | 0 | 0 | 0 | 0 | 0 | 0 | 0 | 0 | 0 | NA | NA | NA | NA | NA | NA | NA | NA | NA | NA |
| Mir5115 | 0 | 0 | 0 | 0 | 0 | 0 | 0 | 0 | 0 | 0 | 0 | 0 | 0 | 0 | 0 | NA | NA | NA | NA | NA | NA | NA | NA | NA | NA |
| Mir5116 | 0 | 0 | 0 | 0 | 0 | 0 | 0 | 0 | 0 | 0 | 0 | 0 | 0 | 0 | 0 | NA | NA | NA | NA | NA | NA | NA | NA | NA | NA |
| Mir5117 | 0 | 0 | 0 | 0 | 0 | 0 | 0 | 0 | 0 | 0 | 0 | 0 | 0 | 0 | 0 | NA | NA | NA | NA | NA | NA | NA | NA | NA | NA |
| Mir5119 | 0 | 1 | 0 | 1 | 0 | 0 | 0 | 1 | 0 | 1 | 0 | 0 | 0.3 | 0 | 0 | 0.954 | -0.067 | 0.98642001 | NA | 0 | 11.0817 | 9.3014 | 9.3014 | TRUE | 0.405 |
| Mir5120 | 0 | 0 | 0 | 0 | 0 | 0 | 0 | 0 | 0 | 0 | 0 | 0 | 0 | 0 | 0 | NA | NA | NA | NA | NA | NA | NA | NA | NA | NA |
| Mir5121 | 6 | 6 | 2 | 5 | 5 | 0 | 6 | 6 | 2 | 4 | 5 | 0 | 3.95 | 5 | 3 | 0.656 | -0.609 | 0.59019822 | NA | 0.3527 | 0.8415 | 0.6593 | 0.6593 | TRUE | 0.8956 |
| Mir5122 | 0 | 0 | 0 | 0 | 0 | 0 | 0 | 0 | 0 | 0 | 0 | 0 | 0 | 0 | 0 | NA | NA | NA | NA | NA | NA | NA | NA | NA | NA |
| Mir5123 | 0 | 0 | 0 | 0 | 0 | 0 | 0 | 0 | 0 | 0 | 0 | 0 | 0 | 0 | 0 | NA | NA | NA | NA | NA | NA | NA | NA | NA | NA |
| Mir5124 | 0 | 0 | 0 | 0 | 0 | 0 | 0 | 0 | 0 | 0 | 0 | 0 | 0 | 0 | 0 | NA | NA | NA | NA | NA | NA | NA | NA | NA | NA |
| Mir5125 | 0 | 0 | 0 | 0 | 0 | 0 | 0 | 0 | 0 | 0 | 0 | 0 | 0 | 0 | 0 | NA | NA | NA | NA | NA | NA | NA | NA | NA | NA |
| Mir5126 | 0 | 0 | 0 | 0 | 0 | 0 | 0 | 0 | 0 | 0 | 0 | 0 | 0 | 0 | 0 | NA | NA | NA | NA | NA | NA | NA | NA | NA | NA |
| Mir5127 | 0 | 0 | 0 | 0 | 0 | 0 | 0 | 0 | 0 | 0 | 0 | 0 | 0 | 0 | 0 | NA | NA | NA | NA | NA | NA | NA | NA | NA | NA |
| Mir5128 | 0 | 0 | 0 | 0 | 0 | 0 | 0 | 0 | 0 | 0 | 0 | 0 | 0 | 0 | 0 | NA | NA | NA | NA | NA | NA | NA | NA | NA | NA |
| Mir5129 | 0 | 0 | 0 | 0 | 0 | 0 | 0 | 0 | 0 | 0 | 0 | 0 | 0 | 0 | 0 | NA | NA | NA | NA | NA | NA | NA | NA | NA | NA |
| Mir5130 | 0 | 0 | 0 | 0 | 0 | 0 | 0 | 0 | 0 | 0 | 0 | 0 | 0 | 0 | 0 | NA | NA | NA | NA | NA | NA | NA | NA | NA | NA |
| Mir5131 | 0 | 0 | 0 | 0 | 0 | 0 | 0 | 0 | 0 | 0 | 0 | 0 | 0 | 0 | 0 | NA | NA | NA | NA | NA | NA | NA | NA | NA | NA |
| Mir5132 | 0 | 0 | 0 | 0 | 0 | 0 | 0 | 0 | 0 | 0 | 0 | 0 | 0 | 0 | 0 | NA | NA | NA | NA | NA | NA | NA | NA | NA | NA |

| | | | | | | | | | | | | | | | | | | | | | | | | | |
|---|---|---|---|---|---|---|---|---|---|---|---|---|---|---|---|---|---|---|---|---|---|---|---|---|---|
| Mir5133 | 17 | 25 | 15 | 32 | 13 | 16 | 18 | 25 | 15 | 25 | 14 | 18 | 19.12 | 19 | 19 | 0.986 | -0.021 | 0.96461223 | 0.98217328 | 0.0331 | 0.1792 | 0.1076 | 0.1076 | TRUE | 0.5325 |
| Mir5134 | 0 | 0 | 0 | 0 | 0 | 1 | 0 | 0 | 0 | 0 | 0 | 1 | 0.18 | 0 | 0 | 1.859 | 0.894 | 0.82650734 | NA | 0 | 18.0643 | 10 | 10 | TRUE | 0.5383 |
| Mir5135 | 0 | 0 | 0 | 0 | 0 | 0 | 0 | 0 | 0 | 0 | 0 | 0 | 0 | 0 | 0 | NA | NA | NA | NA | NA | NA | NA | NA | NA | NA |
| Mir5136 | 0 | 0 | 0 | 0 | 0 | 0 | 0 | 0 | 0 | 0 | 0 | 0 | 0 | 0 | 0 | NA | NA | NA | NA | NA | NA | NA | NA | NA | NA |
| Mir532 | 0 | 0 | 0 | 0 | 0 | 0 | 0 | 0 | 0 | 0 | 0 | 0 | 0 | 0 | 0 | NA | NA | NA | NA | NA | NA | NA | NA | NA | NA |
| Mir539 | 0 | 0 | 0 | 0 | 0 | 0 | 0 | 0 | 0 | 0 | 0 | 0 | 0 | 0 | 0 | NA | NA | NA | NA | NA | NA | NA | NA | NA | NA |
| Mir540 | 0 | 0 | 0 | 0 | 0 | 0 | 0 | 0 | 0 | 0 | 0 | 0 | 0 | 0 | 0 | NA | NA | NA | NA | NA | NA | NA | NA | NA | NA |
| Mir541 | 0 | 0 | 0 | 0 | 0 | 0 | 0 | 0 | 0 | 0 | 0 | 0 | 0 | 0 | 0 | NA | NA | NA | NA | NA | NA | NA | NA | NA | NA |
| Mir542 | 0 | 0 | 0 | 0 | 0 | 0 | 0 | 0 | 0 | 0 | 0 | 0 | 0 | 0 | 0 | NA | NA | NA | NA | NA | NA | NA | NA | NA | NA |
| Mir543 | 0 | 0 | 0 | 0 | 0 | 0 | 0 | 0 | 0 | 0 | 0 | 0 | 0 | 0 | 0 | NA | NA | NA | NA | NA | NA | NA | NA | NA | NA |
| Mir544 | 0 | 0 | 0 | 0 | 0 | 0 | 0 | 0 | 0 | 0 | 0 | 0 | 0 | 0 | 0 | NA | NA | NA | NA | NA | NA | NA | NA | NA | NA |
| Mir546 | 0 | 0 | 0 | 0 | 0 | 0 | 0 | 0 | 0 | 0 | 0 | 0 | 0 | 0 | 0 | NA | NA | NA | NA | NA | NA | NA | NA | NA | NA |
| Mir547 | 0 | 0 | 0 | 0 | 0 | 0 | 0 | 0 | 0 | 0 | 0 | 0 | 0 | 0 | 0 | NA | NA | NA | NA | NA | NA | NA | NA | NA | NA |
| Mir551b | 0 | 0 | 0 | 0 | 0 | 0 | 0 | 0 | 0 | 0 | 0 | 0 | 0 | 0 | 0 | NA | NA | NA | NA | NA | NA | NA | NA | NA | NA |
| Mir568 | 1 | 2 | 1 | 0 | 0 | 2 | 1 | 2 | 1 | 0 | 0 | 2 | 1.04 | 1 | 1 | 0.512 | -0.967 | 0.63452993 | NA | 0 | 3.168 | 1.9006 | 1.9006 | TRUE | 0.9753 |
| Mir574 | 0 | 0 | 0 | 0 | 0 | 0 | 0 | 0 | 0 | 0 | 0 | 0 | 0 | 0 | 0 | NA | NA | NA | NA | NA | NA | NA | NA | NA | NA |
| Mir582 | 0 | 0 | 0 | 0 | 0 | 0 | 0 | 0 | 0 | 0 | 0 | 0 | 0 | 0 | 0 | NA | NA | NA | NA | NA | NA | NA | NA | NA | NA |
| Mir592 | 0 | 0 | 0 | 0 | 0 | 0 | 0 | 0 | 0 | 0 | 0 | 0 | 0 | 0 | 0 | NA | NA | NA | NA | NA | NA | NA | NA | NA | NA |
| Mir598 | 0 | 0 | 0 | 0 | 0 | 0 | 0 | 0 | 0 | 0 | 0 | 0 | 0 | 0 | 0 | NA | NA | NA | NA | NA | NA | NA | NA | NA | NA |
| Mir599 | 0 | 0 | 0 | 0 | 0 | 0 | 0 | 0 | 0 | 0 | 0 | 0 | 0 | 0 | 0 | NA | NA | NA | NA | NA | NA | NA | NA | NA | NA |
| Mir615 | 0 | 0 | 0 | 0 | 0 | 0 | 0 | 0 | 0 | 0 | 0 | 0 | 0 | 0 | 0 | NA | NA | NA | NA | NA | NA | NA | NA | NA | NA |
| Mir653 | 0 | 0 | 0 | 0 | 0 | 0 | 0 | 0 | 0 | 0 | 0 | 0 | 0 | 0 | 0 | NA | NA | NA | NA | NA | NA | NA | NA | NA | NA |
| Mir654 | 1 | 0 | 0 | 0 | 0 | 0 | 1 | 0 | 0 | 0 | 0 | 0 | 0.18 | 0 | 0 | 0.49 | -1.029 | 0.80087 | NA | 0 | 18.6165 | 10 | 10 | TRUE | 0.4614 |
| Mir664 | 0 | 0 | 0 | 0 | 0 | 0 | 0 | 0 | 0 | 0 | 0 | 0 | 0 | 0 | 0 | NA | NA | NA | NA | NA | NA | NA | NA | NA | NA |
| Mir665 | 0 | 0 | 0 | 0 | 0 | 0 | 0 | 0 | 0 | 0 | 0 | 0 | 0 | 0 | 0 | NA | NA | NA | NA | NA | NA | NA | NA | NA | NA |
| Mir666 | 0 | 0 | 0 | 0 | 0 | 0 | 0 | 0 | 0 | 0 | 0 | 0 | 0 | 0 | 0 | NA | NA | NA | NA | NA | NA | NA | NA | NA | NA |
| Mir667 | 0 | 0 | 0 | 0 | 0 | 0 | 0 | 0 | 0 | 0 | 0 | 0 | 0 | 0 | 0 | NA | NA | NA | NA | NA | NA | NA | NA | NA | NA |
| Mir668 | 0 | 0 | 0 | 0 | 0 | 0 | 0 | 0 | 0 | 0 | 0 | 0 | 0 | 0 | 0 | NA | NA | NA | NA | NA | NA | NA | NA | NA | NA |
| Mir669a-1 | 0 | 0 | 0 | 0 | 0 | 0 | 0 | 0 | 0 | 0 | 0 | 0 | 0 | 0 | 0 | NA | NA | NA | NA | NA | NA | NA | NA | NA | NA |
| Mir669a-10 | 0 | 0 | 0 | 0 | 0 | 0 | 0 | 0 | 0 | 0 | 0 | 0 | 0 | 0 | 0 | NA | NA | NA | NA | NA | NA | NA | NA | NA | NA |
| Mir669a-11 | 0 | 0 | 0 | 0 | 0 | 0 | 0 | 0 | 0 | 0 | 0 | 0 | 0 | 0 | 0 | NA | NA | NA | NA | NA | NA | NA | NA | NA | NA |
| Mir669a-12 | 0 | 0 | 0 | 0 | 0 | 0 | 0 | 0 | 0 | 0 | 0 | 0 | 0 | 0 | 0 | NA | NA | NA | NA | NA | NA | NA | NA | NA | NA |
| Mir669a-2 | 0 | 0 | 0 | 0 | 0 | 0 | 0 | 0 | 0 | 0 | 0 | 0 | 0 | 0 | 0 | NA | NA | NA | NA | NA | NA | NA | NA | NA | NA |
| Mir669a-3 | 0 | 0 | 0 | 0 | 0 | 0 | 0 | 0 | 0 | 0 | 0 | 0 | 0 | 0 | 0 | NA | NA | NA | NA | NA | NA | NA | NA | NA | NA |
| Mir669a-4 | 0 | 0 | 0 | 0 | 0 | 0 | 0 | 0 | 0 | 0 | 0 | 0 | 0 | 0 | 0 | NA | NA | NA | NA | NA | NA | NA | NA | NA | NA |
| Mir669a-5 | 0 | 0 | 0 | 0 | 0 | 0 | 0 | 0 | 0 | 0 | 0 | 0 | 0 | 0 | 0 | NA | NA | NA | NA | NA | NA | NA | NA | NA | NA |
| Mir669a-6 | 0 | 0 | 0 | 0 | 0 | 0 | 0 | 0 | 0 | 0 | 0 | 0 | 0 | 0 | 0 | NA | NA | NA | NA | NA | NA | NA | NA | NA | NA |
| Mir669a-7 | 0 | 0 | 0 | 0 | 0 | 0 | 0 | 0 | 0 | 0 | 0 | 0 | 0 | 0 | 0 | NA | NA | NA | NA | NA | NA | NA | NA | NA | NA |
| Mir669a-8 | 0 | 0 | 0 | 0 | 0 | 0 | 0 | 0 | 0 | 0 | 0 | 0 | 0 | 0 | 0 | NA | NA | NA | NA | NA | NA | NA | NA | NA | NA |
| Mir669a-9 | 0 | 0 | 0 | 0 | 0 | 0 | 0 | 0 | 0 | 0 | 0 | 0 | 0 | 0 | 0 | NA | NA | NA | NA | NA | NA | NA | NA | NA | NA |
| Mir669b | 0 | 0 | 0 | 0 | 0 | 0 | 0 | 0 | 0 | 0 | 0 | 0 | 0 | 0 | 0 | NA | NA | NA | NA | NA | NA | NA | NA | NA | NA |
| Mir669c | 0 | 0 | 0 | 0 | 0 | 0 | 0 | 0 | 0 | 0 | 0 | 0 | 0 | 0 | 0 | NA | NA | NA | NA | NA | NA | NA | NA | NA | NA |
| Mir669e | 0 | 0 | 0 | 0 | 0 | 0 | 0 | 0 | 0 | 0 | 0 | 0 | 0 | 0 | 0 | NA | NA | NA | NA | NA | NA | NA | NA | NA | NA |
| Mir669g | 0 | 0 | 0 | 0 | 0 | 0 | 0 | 0 | 0 | 0 | 0 | 0 | 0 | 0 | 0 | NA | NA | NA | NA | NA | NA | NA | NA | NA | NA |
| Mir669h | 0 | 0 | 0 | 0 | 0 | 0 | 0 | 0 | 0 | 0 | 0 | 0 | 0 | 0 | 0 | NA | NA | NA | NA | NA | NA | NA | NA | NA | NA |
| Mir669i | 0 | 0 | 0 | 1 | 2 | 0 | 0 | 0 | 0 | 1 | 2 | 0 | 0.49 | 0 | 1 | 5.204 | 2.38 | 0.47267317 | NA | 0 | 6.7348 | 6.4099 | 6.4099 | TRUE | 0.1222 |
| Mir669j | 0 | 0 | 0 | 0 | 0 | 0 | 0 | 0 | 0 | 0 | 0 | 0 | 0 | 0 | 0 | NA | NA | NA | NA | NA | NA | NA | NA | NA | NA |
| Mir669k | 0 | 0 | 0 | 0 | 0 | 0 | 0 | 0 | 0 | 0 | 0 | 0 | 0 | 0 | 0 | NA | NA | NA | NA | NA | NA | NA | NA | NA | NA |
| Mir669m-1 | 0 | 0 | 0 | 0 | 0 | 0 | 0 | 0 | 0 | 0 | 0 | 0 | 0 | 0 | 0 | NA | NA | NA | NA | NA | NA | NA | NA | NA | NA |
| Mir669m-2 | 0 | 0 | 0 | 0 | 0 | 0 | 0 | 0 | 0 | 0 | 0 | 0 | 0 | 0 | 0 | NA | NA | NA | NA | NA | NA | NA | NA | NA | NA |
| Mir669p-1 | 0 | 0 | 0 | 0 | 0 | 0 | 0 | 0 | 0 | 0 | 0 | 0 | 0 | 0 | 0 | NA | NA | NA | NA | NA | NA | NA | NA | NA | NA |
| Mir669p-2 | 0 | 0 | 0 | 0 | 0 | 0 | 0 | 0 | 0 | 0 | 0 | 0 | 0 | 0 | 0 | NA | NA | NA | NA | NA | NA | NA | NA | NA | NA |
| Mir670 | 0 | 0 | 0 | 0 | 0 | 0 | 0 | 0 | 0 | 0 | 0 | 0 | 0 | 0 | 0 | NA | NA | NA | NA | NA | NA | NA | NA | NA | NA |
| Mir671 | 0 | 0 | 0 | 0 | 0 | 0 | 0 | 0 | 0 | 0 | 0 | 0 | 0 | 0 | 0 | NA | NA | NA | NA | NA | NA | NA | NA | NA | NA |
| Mir672 | 0 | 0 | 0 | 0 | 0 | 0 | 0 | 0 | 0 | 0 | 0 | 0 | 0 | 0 | 0 | NA | NA | NA | NA | NA | NA | NA | NA | NA | NA |
| Mir673 | 0 | 0 | 0 | 0 | 0 | 0 | 0 | 0 | 0 | 0 | 0 | 0 | 0 | 0 | 0 | NA | NA | NA | NA | NA | NA | NA | NA | NA | NA |
| Mir674 | 0 | 0 | 1 | 0 | 0 | 0 | 0 | 0 | 1 | 0 | 0 | 0 | 0.17 | 0 | 0 | 0.49 | -1.029 | 0.80087 | NA | 0 | 19.6928 | 10 | 10 | TRUE | 0.4098 |
| Mir675 | 0 | 0 | 0 | 0 | 0 | 0 | 0 | 0 | 0 | 0 | 0 | 0 | 0 | 0 | 0 | NA | NA | NA | NA | NA | NA | NA | NA | NA | NA |
| Mir676 | 0 | 0 | 0 | 0 | 0 | 0 | 0 | 0 | 0 | 0 | 0 | 0 | 0 | 0 | 0 | NA | NA | NA | NA | NA | NA | NA | NA | NA | NA |
| Mir677 | 4 | 3 | 3 | 8 | 3 | 3 | 4 | 3 | 3 | 6 | 3 | 3 | 3.84 | 3 | 4 | 1.274 | 0.349 | 0.71192047 | NA | 0 | 0.8642 | 0.3821 | 0.3821 | TRUE | 0.3963 |
| Mir678 | 0 | 0 | 0 | 0 | 0 | 0 | 0 | 0 | 0 | 0 | 0 | 0 | 0 | 0 | 0 | NA | NA | NA | NA | NA | NA | NA | NA | NA | NA |
| Mir679 | 0 | 0 | 0 | 0 | 0 | 1 | 0 | 0 | 0 | 0 | 0 | 1 | 0.18 | 0 | 0 | 1.859 | 0.894 | 0.82650734 | NA | 0 | 18.0643 | 10 | 10 | TRUE | 0.5383 |
| Mir680-2 | 1 | 2 | 0 | 0 | 3 | 1 | 1 | 2 | 0 | 0 | 3 | 1 | 1.23 | 1 | 1 | 1.375 | 0.459 | 0.8150902 | NA | 0 | 2.6846 | 1.9283 | 1.9283 | TRUE | 0.3847 |
| Mir680-3 | 0 | 0 | 0 | 0 | 0 | 0 | 0 | 0 | 0 | 0 | 0 | 0 | 0 | 0 | 0 | NA | NA | NA | NA | NA | NA | NA | NA | NA | NA |
| Mir681 | 0 | 0 | 0 | 0 | 0 | 0 | 0 | 0 | 0 | 0 | 0 | 0 | 0 | 0 | 0 | NA | NA | NA | NA | NA | NA | NA | NA | NA | NA |
| Mir682 | 32 | 49 | 49 | 58 | 47 | 30 | 34 | 49 | 49 | 46 | 51 | 33 | 43.52 | 44 | 43 | 0.977 | -0.033 | 0.91828817 | 0.95991788 | 0.0177 | 0.0826 | 0.0526 | 0.0526 | TRUE | 0.3033 |
| Mir683-1 | 0 | 0 | 0 | 0 | 0 | 0 | 0 | 0 | 0 | 0 | 0 | 0 | 0 | 0 | 0 | NA | NA | NA | NA | NA | NA | NA | NA | NA | NA |
| Mir683-2 | 0 | 0 | 0 | 0 | 0 | 0 | 0 | 0 | 0 | 0 | 0 | 0 | 0 | 0 | 0 | NA | NA | NA | NA | NA | NA | NA | NA | NA | NA |
| Mir684-1 | 3 | 5 | 4 | 10 | 5 | 1 | 3 | 5 | 4 | 8 | 5 | 1 | 4.42 | 4 | 5 | 1.199 | 0.262 | 0.78716365 | NA | 0 | 0.7519 | 0.4515 | 0.4515 | TRUE | 0.3284 |

| | | | | | | | | | | | | | | | | | | | | | | | | | |
|---|---|---|---|---|---|---|---|---|---|---|---|---|---|---|---|---|---|---|---|---|---|---|---|---|---|
| Mir684-2 | 0 | 0 | 0 | 0 | 0 | 0 | 0 | 0 | 0 | 0 | 0 | 0 | 0 | 0 | 0 | NA | NA | NA | NA | NA | NA | NA | NA | NA | NA |
| Mir686 | 0 | 0 | 0 | 0 | 0 | 0 | 0 | 0 | 0 | 0 | 0 | 0 | 0 | 0 | 0 | NA | NA | NA | NA | NA | NA | NA | NA | NA | NA |
| Mir687 | 0 | 0 | 0 | 0 | 0 | 0 | 0 | 0 | 0 | 0 | 0 | 0 | 0 | 0 | 0 | NA | NA | NA | NA | NA | NA | NA | NA | NA | NA |
| Mir688 | 0 | 0 | 0 | 1 | 1 | 0 | 0 | 0 | 0 | 1 | 1 | 0 | 0.31 | 0 | 1 | 3.32 | 1.731 | 0.65383052 | NA | 0 | 10.6224 | 8.9541 | 8.9541 | TRUE | 0.202 |
| Mir690 | 0 | 0 | 0 | 0 | 0 | 0 | 0 | 0 | 0 | 0 | 0 | 0 | 0 | 0 | 0 | NA | NA | NA | NA | NA | NA | NA | NA | NA | NA |
| Mir691 | 0 | 0 | 0 | 0 | 0 | 0 | 0 | 0 | 0 | 0 | 0 | 0 | 0 | 0 | 0 | NA | NA | NA | NA | NA | NA | NA | NA | NA | NA |
| Mir692-1 | 0 | 0 | 0 | 0 | 0 | 0 | 0 | 0 | 0 | 0 | 0 | 0 | 0 | 0 | 0 | NA | NA | NA | NA | NA | NA | NA | NA | NA | NA |
| Mir693 | 0 | 0 | 0 | 0 | 0 | 0 | 0 | 0 | 0 | 0 | 0 | 0 | 0 | 0 | 0 | NA | NA | NA | NA | NA | NA | NA | NA | NA | NA |
| Mir694 | 0 | 0 | 0 | 0 | 0 | 0 | 0 | 0 | 0 | 0 | 0 | 0 | 0 | 0 | 0 | NA | NA | NA | NA | NA | NA | NA | NA | NA | NA |
| Mir695 | 0 | 0 | 0 | 0 | 0 | 0 | 0 | 0 | 0 | 0 | 0 | 0 | 0 | 0 | 0 | NA | NA | NA | NA | NA | NA | NA | NA | NA | NA |
| Mir697 | 2 | 0 | 0 | 3 | 0 | 1 | 2 | 0 | 0 | 2 | 0 | 1 | 0.93 | 1 | 1 | 1.686 | 0.753 | 0.75180119 | NA | 0 | 3.551 | 2.9406 | 2.9406 | TRUE | 0.4643 |
| Mir698 | 0 | 0 | 0 | 0 | 0 | 0 | 0 | 0 | 0 | 0 | 0 | 0 | 0 | 0 | 0 | NA | NA | NA | NA | NA | NA | NA | NA | NA | NA |
| Mir7-1 | 2 | 2 | 1 | 3 | 0 | 1 | 2 | 2 | 1 | 2 | 0 | 1 | 1.43 | 2 | 1 | 0.705 | -0.504 | 0.75253972 | NA | 0 | 2.311 | 1.1355 | 1.1355 | TRUE | 0.3584 |
| Mir7-2 | 0 | 0 | 0 | 0 | 0 | 0 | 0 | 0 | 0 | 0 | 0 | 0 | 0 | 0 | 0 | NA | NA | NA | NA | NA | NA | NA | NA | NA | NA |
| Mir700 | 0 | 0 | 0 | 0 | 0 | 0 | 0 | 0 | 0 | 0 | 0 | 0 | 0 | 0 | 0 | NA | NA | NA | NA | NA | NA | NA | NA | NA | NA |
| Mir701 | 0 | 0 | 0 | 0 | 0 | 0 | 0 | 0 | 0 | 0 | 0 | 0 | 0 | 0 | 0 | NA | NA | NA | NA | NA | NA | NA | NA | NA | NA |
| Mir702 | 0 | 0 | 0 | 0 | 0 | 0 | 0 | 0 | 0 | 0 | 0 | 0 | 0 | 0 | 0 | NA | NA | NA | NA | NA | NA | NA | NA | NA | NA |
| Mir703 | 273 | 370 | 338 | 373 | 266 | 289 | 290 | 370 | 339 | 293 | 286 | 316 | 315.78 | 333 | 298 | 0.896 | -0.159 | 0.25964631 | 0.47879594 | 0.0056 | 0.0173 | 0.0111 | 0.0111 | TRUE | 0.1434 |
| Mir704 | 0 | 0 | 0 | 1 | 0 | 0 | 0 | 0 | 0 | 1 | 0 | 0 | 0.13 | 0 | 0 | 1.859 | 0.894 | 0.82650734 | NA | 0 | 25.1515 | 10 | 10 | TRUE | 0.2563 |
| Mir705 | 0 | 0 | 0 | 0 | 0 | 0 | 0 | 0 | 0 | 0 | 0 | 0 | 0 | 0 | 0 | NA | NA | NA | NA | NA | NA | NA | NA | NA | NA |
| Mir706 | 0 | 0 | 0 | 0 | 0 | 0 | 0 | 0 | 0 | 0 | 0 | 0 | 0 | 0 | 0 | NA | NA | NA | NA | NA | NA | NA | NA | NA | NA |
| Mir707 | 4 | 3 | 4 | 3 | 1 | 2 | 4 | 3 | 4 | 2 | 1 | 2 | 2.81 | 4 | 2 | 0.507 | -0.981 | 0.37464557 | NA | 0 | 1.1776 | 0.4876 | 0.4876 | TRUE | 0.105 |
| Mir708 | 0 | 0 | 0 | 0 | 0 | 0 | 0 | 0 | 0 | 0 | 0 | 0 | 0 | 0 | 0 | NA | NA | NA | NA | NA | NA | NA | NA | NA | NA |
| Mir709 | 0 | 0 | 0 | 0 | 1 | 0 | 0 | 0 | 0 | 0 | 1 | 0 | 0.18 | 0 | 0 | 1.859 | 0.894 | 0.82650734 | NA | 0 | 18.3785 | 10 | 10 | TRUE | 0.5204 |
| Mir710 | 0 | 0 | 0 | 0 | 0 | 0 | 0 | 0 | 0 | 0 | 0 | 0 | 0 | 0 | 0 | NA | NA | NA | NA | NA | NA | NA | NA | NA | NA |
| Mir711 | 0 | 2 | 0 | 0 | 0 | 0 | 0 | 2 | 0 | 0 | 0 | 0 | 0.33 | 1 | 0 | 0.268 | -1.897 | 0.63866967 | NA | 0 | 9.903 | 10 | 10 | TRUE | 0.9739 |
| Mir713 | 0 | 0 | 0 | 0 | 0 | 0 | 0 | 0 | 0 | 0 | 0 | 0 | 0 | 0 | 0 | NA | NA | NA | NA | NA | NA | NA | NA | NA | NA |
| Mir717 | 0 | 0 | 0 | 0 | 1 | 0 | 0 | 0 | 0 | 0 | 1 | 0 | 0.18 | 0 | 0 | 1.859 | 0.894 | 0.82650734 | NA | 0 | 18.3785 | 10 | 10 | TRUE | 0.5204 |
| Mir718 | 0 | 0 | 0 | 0 | 0 | 0 | 0 | 0 | 0 | 0 | 0 | 0 | 0 | 0 | 0 | NA | NA | NA | NA | NA | NA | NA | NA | NA | NA |
| Mir719 | 0 | 0 | 0 | 0 | 0 | 0 | 0 | 0 | 0 | 0 | 0 | 0 | 0 | 0 | 0 | NA | NA | NA | NA | NA | NA | NA | NA | NA | NA |
| Mir721 | 0 | 0 | 0 | 0 | 0 | 0 | 0 | 0 | 0 | 0 | 0 | 0 | 0 | 0 | 0 | NA | NA | NA | NA | NA | NA | NA | NA | NA | NA |
| Mir741 | 0 | 0 | 0 | 0 | 0 | 0 | 0 | 0 | 0 | 0 | 0 | 0 | 0 | 0 | 0 | NA | NA | NA | NA | NA | NA | NA | NA | NA | NA |
| Mir742 | 0 | 0 | 0 | 0 | 0 | 0 | 0 | 0 | 0 | 0 | 0 | 0 | 0 | 0 | 0 | NA | NA | NA | NA | NA | NA | NA | NA | NA | NA |
| Mir743 | 0 | 0 | 0 | 0 | 0 | 0 | 0 | 0 | 0 | 0 | 0 | 0 | 0 | 0 | 0 | NA | NA | NA | NA | NA | NA | NA | NA | NA | NA |
| Mir743b | 0 | 0 | 0 | 0 | 0 | 0 | 0 | 0 | 0 | 0 | 0 | 0 | 0 | 0 | 0 | NA | NA | NA | NA | NA | NA | NA | NA | NA | NA |
| Mir744 | 0 | 0 | 0 | 0 | 0 | 0 | 0 | 0 | 0 | 0 | 0 | 0 | 0 | 0 | 0 | NA | NA | NA | NA | NA | NA | NA | NA | NA | NA |
| Mir758 | 0 | 0 | 0 | 0 | 0 | 0 | 0 | 0 | 0 | 0 | 0 | 0 | 0 | 0 | 0 | NA | NA | NA | NA | NA | NA | NA | NA | NA | NA |
| Mir759 | 0 | 0 | 0 | 0 | 0 | 0 | 0 | 0 | 0 | 0 | 0 | 0 | 0 | 0 | 0 | NA | NA | NA | NA | NA | NA | NA | NA | NA | NA |
| Mir760 | 0 | 0 | 0 | 0 | 0 | 0 | 0 | 0 | 0 | 0 | 0 | 0 | 0 | 0 | 0 | NA | NA | NA | NA | NA | NA | NA | NA | NA | NA |
| Mir761 | 2 | 0 | 0 | 2 | 1 | 0 | 2 | 0 | 0 | 2 | 1 | 0 | 0.8 | 1 | 1 | 1.29 | 0.367 | 0.88218031 | NA | 0 | 4.1487 | 3.1517 | 3.1517 | TRUE | 0.9743 |
| Mir762 | 0 | 2 | 1 | 1 | 1 | 1 | 0 | 2 | 1 | 1 | 1 | 1 | 0.99 | 1 | 1 | 0.968 | -0.047 | 0.97991429 | NA | 0 | 3.3244 | 1.4549 | 1.4549 | TRUE | 0.1811 |
| Mir764 | 0 | 0 | 0 | 0 | 0 | 0 | 0 | 0 | 0 | 0 | 0 | 0 | 0 | 0 | 0 | NA | NA | NA | NA | NA | NA | NA | NA | NA | NA |
| Mir767 | 0 | 0 | 0 | 0 | 0 | 0 | 0 | 0 | 0 | 0 | 0 | 0 | 0 | 0 | 0 | NA | NA | NA | NA | NA | NA | NA | NA | NA | NA |
| Mir770 | 0 | 0 | 0 | 0 | 0 | 0 | 0 | 0 | 0 | 0 | 0 | 0 | 0 | 0 | 0 | NA | NA | NA | NA | NA | NA | NA | NA | NA | NA |
| Mir7b | 0 | 0 | 0 | 0 | 0 | 0 | 0 | 0 | 0 | 0 | 0 | 0 | 0 | 0 | 0 | NA | NA | NA | NA | NA | NA | NA | NA | NA | NA |
| Mir802 | 0 | 0 | 0 | 0 | 0 | 0 | 0 | 0 | 0 | 0 | 0 | 0 | 0 | 0 | 0 | NA | NA | NA | NA | NA | NA | NA | NA | NA | NA |
| Mir804 | 0 | 0 | 0 | 0 | 0 | 0 | 0 | 0 | 0 | 0 | 0 | 0 | 0 | 0 | 0 | NA | NA | NA | NA | NA | NA | NA | NA | NA | NA |
| Mir871 | 0 | 0 | 0 | 0 | 0 | 0 | 0 | 0 | 0 | 0 | 0 | 0 | 0 | 0 | 0 | NA | NA | NA | NA | NA | NA | NA | NA | NA | NA |
| Mir872 | 0 | 0 | 0 | 0 | 0 | 0 | 0 | 0 | 0 | 0 | 0 | 0 | 0 | 0 | 0 | NA | NA | NA | NA | NA | NA | NA | NA | NA | NA |
| Mir873 | 0 | 0 | 0 | 0 | 0 | 0 | 0 | 0 | 0 | 0 | 0 | 0 | 0 | 0 | 0 | NA | NA | NA | NA | NA | NA | NA | NA | NA | NA |
| Mir874 | 0 | 0 | 0 | 0 | 0 | 0 | 0 | 0 | 0 | 0 | 0 | 0 | 0 | 0 | 0 | NA | NA | NA | NA | NA | NA | NA | NA | NA | NA |
| Mir875 | 0 | 0 | 0 | 0 | 0 | 0 | 0 | 0 | 0 | 0 | 0 | 0 | 0 | 0 | 0 | NA | NA | NA | NA | NA | NA | NA | NA | NA | NA |
| Mir876 | 0 | 0 | 0 | 0 | 0 | 0 | 0 | 0 | 0 | 0 | 0 | 0 | 0 | 0 | 0 | NA | NA | NA | NA | NA | NA | NA | NA | NA | NA |
| Mir877 | 0 | 0 | 0 | 1 | 0 | 0 | 0 | 0 | 0 | 1 | 0 | 0 | 0.13 | 0 | 0 | 1.859 | 0.894 | 0.82650734 | NA | 0 | 25.1515 | 10 | 10 | TRUE | 0.2563 |
| Mir878 | 0 | 0 | 0 | 0 | 0 | 0 | 0 | 0 | 0 | 0 | 0 | 0 | 0 | 0 | 0 | NA | NA | NA | NA | NA | NA | NA | NA | NA | NA |
| Mir879 | 0 | 0 | 0 | 0 | 0 | 0 | 0 | 0 | 0 | 0 | 0 | 0 | 0 | 0 | 0 | NA | NA | NA | NA | NA | NA | NA | NA | NA | NA |
| Mir880 | 0 | 0 | 0 | 0 | 0 | 0 | 0 | 0 | 0 | 0 | 0 | 0 | 0 | 0 | 0 | NA | NA | NA | NA | NA | NA | NA | NA | NA | NA |
| Mir881 | 0 | 0 | 0 | 0 | 0 | 0 | 0 | 0 | 0 | 0 | 0 | 0 | 0 | 0 | 0 | NA | NA | NA | NA | NA | NA | NA | NA | NA | NA |
| Mir882 | 0 | 0 | 0 | 0 | 0 | 0 | 0 | 0 | 0 | 0 | 0 | 0 | 0 | 0 | 0 | NA | NA | NA | NA | NA | NA | NA | NA | NA | NA |
| Mir883a | 0 | 0 | 0 | 0 | 0 | 0 | 0 | 0 | 0 | 0 | 0 | 0 | 0 | 0 | 0 | NA | NA | NA | NA | NA | NA | NA | NA | NA | NA |
| Mir883b | 0 | 0 | 0 | 0 | 0 | 0 | 0 | 0 | 0 | 0 | 0 | 0 | 0 | 0 | 0 | NA | NA | NA | NA | NA | NA | NA | NA | NA | NA |
| Mir9-1 | 0 | 0 | 0 | 0 | 0 | 0 | 0 | 0 | 0 | 0 | 0 | 0 | 0 | 0 | 0 | NA | NA | NA | NA | NA | NA | NA | NA | NA | NA |
| Mir9-2 | 0 | 0 | 0 | 0 | 0 | 0 | 0 | 0 | 0 | 0 | 0 | 0 | 0 | 0 | 0 | NA | NA | NA | NA | NA | NA | NA | NA | NA | NA |
| Mir9-3 | 0 | 0 | 0 | 0 | 0 | 0 | 0 | 0 | 0 | 0 | 0 | 0 | 0 | 0 | 0 | NA | NA | NA | NA | NA | NA | NA | NA | NA | NA |
| Mir92-1 | 0 | 0 | 0 | 0 | 0 | 0 | 0 | 0 | 0 | 0 | 0 | 0 | 0 | 0 | 0 | NA | NA | NA | NA | NA | NA | NA | NA | NA | NA |
| Mir92-2 | 1 | 3 | 0 | 0 | 1 | 1 | 1 | 3 | 0 | 0 | 1 | 1 | 1.04 | 1 | 1 | 0.511 | -0.969 | 0.64094994 | NA | 0 | 3.1803 | 2.0313 | 2.0313 | TRUE | 0.2916 |
| Mir92b | 0 | 0 | 0 | 0 | 0 | 0 | 0 | 0 | 0 | 0 | 0 | 0 | 0 | 0 | 0 | NA | NA | NA | NA | NA | NA | NA | NA | NA | NA |
| Mir93 | 0 | 1 | 0 | 0 | 0 | 0 | 0 | 1 | 0 | 0 | 0 | 0 | 0.17 | 0 | 0 | 0.49 | -1.029 | 0.80087 | NA | 0 | 19.7991 | 10 | 10 | TRUE | 0.405 |
| Mir96 | 0 | 0 | 0 | 0 | 0 | 0 | 0 | 0 | 0 | 0 | 0 | 0 | 0 | 0 | 0 | NA | NA | NA | NA | NA | NA | NA | NA | NA | NA |

| | | | | | | | | | | | | | | | | | | | | | | | | | |
|---|---|---|---|---|---|---|---|---|---|---|---|---|---|---|---|---|---|---|---|---|---|---|---|---|---|
| Mir98 | 0 | 0 | 0 | 0 | 0 | 0 | 0 | 0 | 0 | 0 | 0 | 0 | 0 | 0 | 0 | NA | NA | NA | NA | NA | NA | NA | NA | NA | NA |
| Mir99a | 0 | 0 | 0 | 0 | 0 | 0 | 0 | 0 | 0 | 0 | 0 | 0 | 0 | 0 | 0 | NA | NA | NA | NA | NA | NA | NA | NA | NA | NA |
| Mir99b | 0 | 0 | 0 | 0 | 0 | 0 | 0 | 0 | 0 | 0 | 0 | 0 | 0 | 0 | 0 | NA | NA | NA | NA | NA | NA | NA | NA | NA | NA |
| Mira | 5 | 5 | 7 | 19 | 11 | 4 | 5 | 5 | 7 | 15 | 12 | 4 | 8.08 | 6 | 10 | 1.819 | 0.863 | 0.24064544 | 0.45703258 | 0 | 0.4146 | 0.2552 | 0.2552 | TRUE | 0.4281 |
| Mirg | 565 | 464 | 397 | 524 | 443 | 270 | 600 | 463 | 399 | 412 | 477 | 296 | 441.05 | 487 | 395 | 0.81 | -0.303 | 0.15252764 | 0.34375257 | 0.0459 | 0.0144 | 0.0301 | 0.0301 | TRUE | 0.5241 |
| Mirlet7a-1 | 0 | 0 | 0 | 0 | 0 | 0 | 0 | 0 | 0 | 0 | 0 | 0 | 0 | 0 | 0 | NA | NA | NA | NA | NA | NA | NA | NA | NA | NA |
| Mirlet7a-2 | 0 | 0 | 0 | 0 | 0 | 0 | 0 | 0 | 0 | 0 | 0 | 0 | 0 | 0 | 0 | NA | NA | NA | NA | NA | NA | NA | NA | NA | NA |
| Mirlet7b | 0 | 0 | 0 | 0 | 0 | 0 | 0 | 0 | 0 | 0 | 0 | 0 | 0 | 0 | 0 | NA | NA | NA | NA | NA | NA | NA | NA | NA | NA |
| Mirlet7c-1 | 0 | 0 | 0 | 0 | 0 | 0 | 0 | 0 | 0 | 0 | 0 | 0 | 0 | 0 | 0 | NA | NA | NA | NA | NA | NA | NA | NA | NA | NA |
| Mirlet7c-2 | 0 | 0 | 0 | 0 | 0 | 0 | 0 | 0 | 0 | 0 | 0 | 0 | 0 | 0 | 0 | NA | NA | NA | NA | NA | NA | NA | NA | NA | NA |
| Mirlet7d | 5 | 5 | 3 | 4 | 3 | 0 | 5 | 5 | 3 | 3 | 3 | 0 | 3.28 | 4 | 2 | 0.49 | -1.028 | 0.36556716 | NA | 0 | 1.0108 | 0.5902 | 0.5902 | TRUE | 0.6245 |
| Mirlet7e | 0 | 1 | 0 | 0 | 0 | 0 | 0 | 1 | 0 | 0 | 0 | 0 | 0.17 | 0 | 0 | 0.49 | -1.029 | 0.80087 | NA | 0 | 19.7991 | 10 | 10 | TRUE | 0.405 |
| Mirlet7f-1 | 0 | 1 | 0 | 0 | 0 | 0 | 0 | 1 | 0 | 0 | 0 | 0 | 0.17 | 0 | 0 | 0.49 | -1.029 | 0.80087 | NA | 0 | 19.7991 | 10 | 10 | TRUE | 0.405 |
| Mirlet7f-2 | 0 | 0 | 0 | 0 | 0 | 0 | 0 | 0 | 0 | 0 | 0 | 0 | 0 | 0 | 0 | NA | NA | NA | NA | NA | NA | NA | NA | NA | NA |
| Mirlet7g | 0 | 1 | 0 | 0 | 0 | 0 | 0 | 1 | 0 | 0 | 0 | 0 | 0.17 | 0 | 0 | 0.49 | -1.029 | 0.80087 | NA | 0 | 19.7991 | 10 | 10 | TRUE | 0.405 |
| Mirlet7i | 0 | 0 | 0 | 0 | 0 | 0 | 0 | 0 | 0 | 0 | 0 | 0 | 0 | 0 | 0 | NA | NA | NA | NA | NA | NA | NA | NA | NA | NA |
| Mis12 | 2996 | 2582 | 2504 | 3778 | 2665 | 2457 | 3182 | 2579 | 2514 | 2970 | 2867 | 2690 | 2800.36 | 2758 | 2842 | 1.031 | 0.044 | 0.6990843 | 0.8354264 | 0.0094 | 0.0081 | 0.0088 | 0.0088 | TRUE | 0.219 |
| Mis18a | 1825 | 1775 | 1774 | 2174 | 1607 | 1596 | 1938 | 1773 | 1781 | 1709 | 1729 | 1747 | 1779.6 | 1831 | 1728 | 0.944 | -0.083 | 0.27704201 | 0.49805749 | 8.00E-04 | 0.0088 | 0.0036 | 0.0036 | TRUE | 0.0318 |
| Mis18bp1 | 5046 | 4711 | 4481 | 6084 | 4219 | 4104 | 5360 | 4705 | 4499 | 4783 | 4539 | 4492 | 4729.78 | 4855 | 4605 | 0.949 | -0.076 | 0.40269706 | 0.61555903 | 0.0045 | 0.0076 | 0.0057 | 0.0057 | TRUE | 0.1008 |
| Mitd1 | 947 | 877 | 937 | 1223 | 913 | 859 | 1006 | 876 | 941 | 961 | 982 | 940 | 951.11 | 941 | 961 | 1.022 | 0.031 | 0.7332966 | 0.85696162 | 0.0015 | 0.0104 | 0.0051 | 0.0051 | TRUE | 0.0436 |
| Mitf | 1504 | 1551 | 1475 | 1559 | 1358 | 1537 | 1598 | 1549 | 1481 | 1226 | 1461 | 1682 | 1499.46 | 1543 | 1456 | 0.943 | -0.084 | 0.50812351 | 0.70231565 | 0.0126 | 0.0091 | 0.011 | 0.011 | TRUE | 0.2153 |
| Mixl1 | 31 | 35 | 23 | 79 | 47 | 47 | 33 | 35 | 23 | 62 | 51 | 51 | 42.52 | 30 | 55 | 1.812 | 0.858 | 0.00666468 | 0.03880224 | 0 | 0.0844 | 0.0462 | 0.0462 | TRUE | 0.2947 |
| Mki67 | 13228 | 13919 | 16426 | 18208 | 12896 | 13431 | 14050 | 13901 | 16493 | 14314 | 13875 | 14702 | 14556.05 | 14815 | 14297 | 0.965 | -0.051 | 0.57020842 | 0.74900702 | 0.0046 | 0.0071 | 0.0058 | 0.0058 | TRUE | 0.1202 |
| Mki67ip | 3404 | 2982 | 2842 | 4694 | 3173 | 2662 | 3616 | 2978 | 2854 | 3690 | 3414 | 2914 | 3244.24 | 3149 | 3339 | 1.061 | 0.085 | 0.50931928 | 0.70294037 | 0.013 | 0.0079 | 0.0117 | 0.0117 | TRUE | 0.2038 |
| Mkks | 1828 | 1603 | 1512 | 2213 | 1548 | 1373 | 1942 | 1601 | 1518 | 1740 | 1666 | 1503 | 1661.5 | 1687 | 1636 | 0.97 | -0.043 | 0.71931754 | 0.8479218 | 0.0107 | 0.0089 | 0.0099 | 0.0099 | TRUE | 0.21 |
| Mkl1 | 2327 | 1851 | 1717 | 2734 | 2000 | 1757 | 2472 | 1849 | 1724 | 2149 | 2152 | 1923 | 2044.8 | 2015 | 2075 | 1.03 | 0.043 | 0.76744498 | 0.87860653 | 0.0202 | 0.0085 | 0.0146 | 0.0146 | TRUE | 0.4754 |
| Mkl2 | 1969 | 1487 | 1505 | 2462 | 1707 | 1625 | 2091 | 1485 | 1511 | 1935 | 1837 | 1779 | 1773.09 | 1696 | 1850 | 1.091 | 0.126 | 0.37966996 | 0.59453481 | 0.0194 | 0.0088 | 0.0143 | 0.0143 | TRUE | 0.5015 |
| Mkln1 | 1065 | 1668 | 1705 | 1917 | 1374 | 1581 | 1131 | 1666 | 1712 | 1507 | 1478 | 1731 | 1537.51 | 1503 | 1572 | 1.046 | 0.064 | 0.69435238 | 0.8325472 | 0.0283 | 0.009 | 0.0186 | 0.0186 | TRUE | 0.5627 |
| Mknk1 | 1285 | 1385 | 1311 | 1408 | 1043 | 1071 | 1365 | 1383 | 1316 | 1107 | 1122 | 1172 | 1244.32 | 1355 | 1134 | 0.836 | -0.258 | 0.00092156 | 0.00836781 | 0 | 0.0096 | 0.0035 | 0.0035 | TRUE | 0.0104 |
| Mknk2 | 4493 | 3466 | 3331 | 5271 | 3963 | 3305 | 4772 | 3462 | 3345 | 4144 | 4264 | 3618 | 3934 | 3860 | 4009 | 1.039 | 0.055 | 0.71191574 | 0.84325407 | 0.0232 | 0.0077 | 0.0156 | 0.0156 | TRUE | 0.5206 |
| Mkrn1 | 67510 | 66805 | 67321 | 66866 | 48520 | 50888 | 71708 | 66719 | 67597 | 52565 | 52205 | 55705 | 61082.99 | 68675 | 53492 | 0.779 | -0.36 | 3.78E-08 | 1.56E-06 | 0.0014 | 0.007 | 0.0031 | 0.0031 | TRUE | 0.0183 |
| Mkrn2 | 3226 | 4091 | 4024 | 4182 | 3124 | 3420 | 3427 | 4086 | 4041 | 3288 | 3361 | 3744 | 3657.56 | 3851 | 3464 | 0.899 | -0.153 | 0.13469351 | 0.31740995 | 0.0067 | 0.0078 | 0.0073 | 0.0073 | TRUE | 0.1129 |
| Mkrn3 | 145 | 170 | 202 | 370 | 274 | 281 | 154 | 170 | 203 | 291 | 295 | 308 | 236.65 | 176 | 298 | 1.694 | 0.761 | 7.38E-07 | 2.12E-05 | 0.0032 | 0.0208 | 0.0124 | 0.0124 | TRUE | 0.1982 |
| Mks1 | 1151 | 1497 | 1564 | 1639 | 1314 | 1403 | 1223 | 1495 | 1570 | 1288 | 1414 | 1536 | 1421.02 | 1429 | 1413 | 0.988 | -0.018 | 0.88553048 | 0.94387096 | 0.0117 | 0.0092 | 0.0106 | 0.0106 | TRUE | 0.1921 |
| Mkx | 241 | 307 | 279 | 394 | 290 | 282 | 256 | 307 | 280 | 310 | 312 | 309 | 295.53 | 281 | 310 | 1.103 | 0.142 | 0.27380689 | 0.49433176 | 0 | 0.0181 | 0.0087 | 0.0087 | TRUE | 0.072 |
| Mlana | 37 | 21 | 17 | 40 | 36 | 23 | 39 | 21 | 17 | 31 | 39 | 25 | 28.78 | 26 | 32 | 1.236 | 0.306 | 0.48196602 | 0.68083046 | 0.0792 | 0.1214 | 0.1014 | 0.1014 | TRUE | 0.9927 |
| Mlc1 | 7 | 5 | 4 | 7 | 3 | 3 | 7 | 5 | 4 | 6 | 3 | 3 | 4.74 | 5 | 4 | 0.745 | -0.425 | 0.62108072 | 0.78565991 | 0 | 0.7014 | 0.319 | 0.319 | TRUE | 0.2376 |
| Mlec | 10489 | 11259 | 10700 | 14502 | 10652 | 10302 | 11141 | 11244 | 10744 | 11400 | 11461 | 11277 | 11211.33 | 11043 | 11379 | 1.03 | 0.043 | 0.43584915 | 0.64407502 | 2.00E-04 | 0.0072 | 0.0021 | 0.0021 | TRUE | 0.0069 |
| Mlf1 | 267 | 306 | 244 | 476 | 348 | 341 | 284 | 306 | 245 | 374 | 374 | 373 | 326.02 | 278 | 374 | 1.345 | 0.428 | 0.00116069 | 0.00998171 | 0.0019 | 0.017 | 0.0093 | 0.0093 | TRUE | 0.1227 |
| Mlf1ip | 1816 | 1969 | 1859 | 2330 | 1734 | 1625 | 1929 | 1966 | 1867 | 1832 | 1866 | 1779 | 1873.03 | 1921 | 1826 | 0.95 | -0.073 | 0.29951276 | 0.51972817 | 0 | 0.0087 | 0.0031 | 0.0031 | TRUE | 0.0074 |
| Mlf2 | 14084 | 14456 | 13783 | 18762 | 13293 | 13467 | 14960 | 14437 | 13840 | 14749 | 14302 | 14742 | 14505.01 | 14412 | 14598 | 1.013 | 0.019 | 0.7652863 | 0.87700158 | 7.00E-04 | 0.0071 | 0.0027 | 0.0027 | TRUE | 0.0148 |
| Mlh1 | 1743 | 2150 | 2141 | 2719 | 1923 | 1778 | 1851 | 2147 | 2150 | 2137 | 2069 | 1946 | 2050.2 | 2049 | 2051 | 1.001 | 0.001 | 0.98896609 | 0.99419805 | 0.0042 | 0.0085 | 0.0059 | 0.0059 | TRUE | 0.0862 |
| Mlh3 | 3256 | 2908 | 3055 | 4471 | 3181 | 2772 | 3458 | 2904 | 3068 | 3515 | 3423 | 3034 | 3233.66 | 3143 | 3324 | 1.058 | 0.081 | 0.42776716 | 0.6378693 | 0.0062 | 0.0079 | 0.0072 | 0.0072 | TRUE | 0.0932 |
| Mlip | 0 | 0 | 2 | 2 | 1 | 0 | 0 | 0 | 2 | 2 | 1 | 0 | 0.78 | 1 | 1 | 1.34 | 0.422 | 0.86518368 | NA | 0 | 4.252 | 3.1559 | 3.1559 | TRUE | 0.9739 |
| Mlkl | 49 | 50 | 40 | 80 | 35 | 43 | 52 | 50 | 40 | 63 | 38 | 47 | 48.29 | 47 | 49 | 1.047 | 0.067 | 0.83133918 | 0.91442092 | 0.0247 | 0.0751 | 0.0503 | 0.0503 | TRUE | 0.4166 |
| Mll1 | 4331 | 3434 | 4279 | 5120 | 3793 | 3472 | 4600 | 3430 | 4297 | 4025 | 4081 | 3801 | 4038.84 | 4109 | 3969 | 0.966 | -0.05 | 0.67671658 | 0.82025622 | 0.0119 | 0.0077 | 0.0101 | 0.0101 | TRUE | 0.2548 |
| Mll2 | 5000 | 4229 | 4998 | 5573 | 3979 | 3301 | 5311 | 4224 | 5019 | 4381 | 4281 | 3613 | 4471.44 | 4851 | 4092 | 0.844 | -0.245 | 0.04078811 | 0.14287252 | 0.0116 | 0.0076 | 0.0101 | 0.0101 | TRUE | 0.1561 |
| Mll3 | 4806 | 4044 | 4916 | 6040 | 3984 | 3802 | 5105 | 4039 | 4936 | 4748 | 4287 | 4162 | 4546.06 | 4693 | 4399 | 0.938 | -0.093 | 0.40985394 | 0.62158382 | 0.0092 | 0.0076 | 0.009 | 0.009 | TRUE | 0.1814 |
| Mll5 | 1012 | 1180 | 1259 | 1626 | 1274 | 1154 | 1075 | 1178 | 1264 | 1278 | 1371 | 1263 | 1238.3 | 1172 | 1304 | 1.112 | 0.153 | 0.11294285 | 0.28302522 | 0.003 | 0.0096 | 0.0059 | 0.0059 | TRUE | 0.0632 |
| Mllt1 | 1611 | 1432 | 1499 | 2054 | 1506 | 1297 | 1711 | 1430 | 1505 | 1615 | 1620 | 1420 | 1550.22 | 1549 | 1552 | 1.002 | 0.003 | 0.97488557 | 0.98689956 | 0.0064 | 0.009 | 0.0075 | 0.0075 | TRUE | 0.101 |
| Mllt10 | 3712 | 4746 | 4910 | 6132 | 4734 | 4721 | 3943 | 4740 | 4930 | 4820 | 5093 | 5168 | 4782.45 | 4538 | 5027 | 1.108 | 0.148 | 0.15265362 | 0.3439013 | 0.007 | 0.0076 | 0.0075 | 0.0075 | TRUE | 0.1599 |
| Mllt11 | 388 | 474 | 457 | 437 | 328 | 301 | 412 | 473 | 459 | 344 | 353 | 329 | 395.05 | 448 | 342 | 0.763 | -0.39 | 0.00085874 | 0.007902 | 0 | 0.0152 | 0.0073 | 0.0073 | TRUE | 0.0566 |
| Mllt3 | 1233 | 1316 | 1320 | 1749 | 1271 | 1244 | 1310 | 1314 | 1325 | 1375 | 1368 | 1362 | 1342.26 | 1316 | 1368 | 1.039 | 0.056 | 0.43511672 | 0.643475 | 0 | 0.0094 | 0.0029 | 0.0029 | TRUE | 4.00E-04 |
| Mllt4 | 10046 | 11356 | 12616 | 13668 | 10479 | 11200 | 10671 | 11341 | 12668 | 10745 | 11275 | 12260 | 11493.23 | 11560 | 11427 | 0.988 | -0.017 | 0.85918475 | 0.93059974 | 0.0049 | 0.0072 | 0.0064 | 0.0064 | TRUE | 0.0859 |
| Mllt6 | 6759 | 6308 | 6095 | 9904 | 8484 | 6018 | 7179 | 6300 | 6120 | 7786 | 9128 | 6588 | 7183.47 | 6533 | 7834 | 1.199 | 0.262 | 0.0464201 | 0.15587882 | 0.0167 | 0.0074 | 0.0123 | 0.0123 | TRUE | 0.1801 |
| Mlph | 56 | 72 | 59 | 84 | 65 | 71 | 59 | 72 | 59 | 66 | 70 | 78 | 67.39 | 63 | 71 | 1.117 | 0.16 | 0.50575936 | 0.70029387 | 0 | 0.0558 | 0.0268 | 0.0268 | TRUE | 0.1175 |
| Mlst8 | 1044 | 1097 | 1108 | 1370 | 1005 | 1047 | 1109 | 1096 | 1113 | 1077 | 1081 | 1146 | 1103.58 | 1106 | 1101 | 0.996 | -0.006 | 0.93745103 | 0.96865027 | 0 | 0.0099 | 0.0036 | 0.0036 | TRUE | 0.0147 |
| Mlx | 2468 | 2314 | 2306 | 3136 | 2047 | 1945 | 2621 | 2311 | 2315 | 2465 | 2202 | 2129 | 2340.79 | 2416 | 2265 | 0.938 | -0.092 | 0.34833425 | 0.56702733 | 0.0051 | 0.0083 | 0.0065 | 0.0065 | TRUE | 0.073 |
| Mlxip | 4428 | 3923 | 4214 | 4260 | 3031 | 3312 | 4703 | 3918 | 4231 | 3349 | 3261 | 3626 | 3848.02 | 4284 | 3412 | 0.796 | -0.329 | 0.00063222 | 0.00624379 | 0.0054 | 0.0078 | 0.0064 | 0.0064 | TRUE | 0.0891 |
| Mlxipl | 239 | 336 | 318 | 298 | 216 | 233 | 254 | 336 | 319 | 234 | 232 | 255 | 271.74 | 303 | 240 | 0.793 | -0.334 | 0.03079394 | 0.11742794 | 0.0086 | 0.019 | 0.0135 | 0.0135 | TRUE | 0.224 |
| Mlycd | 443 | 835 | 825 | 631 | 437 | 491 | 471 | 834 | 828 | 496 | 470 | 537 | 606.09 | 711 | 501 | 0.705 | -0.505 | 0.01650985 | 0.07607166 | 0.0502 | 0.0123 | 0.0303 | 0.0303 | TRUE | 1.0309 |
| Mmaa | 134 | 158 | 145 | 264 | 162 | 155 | 142 | 158 | 146 | 208 | 174 | 170 | 166.21 | 149 | 184 | 1.241 | 0.312 | 0.06420663 | 0.19500617 | 0 | 0.0267 | 0.0144 | 0.0144 | TRUE | 0.1476 |
| Mmab | 848 | 1212 | 1095 | 916 | 817 | 842 | 901 | 1210 | 1099 | 720 | 879 | 922 | 955.25 | 1070 | 840 | 0.784 | -0.351 | 0.01732089 | 0.07827651 | 0.0171 | 0.0104 | 0.0146 | 0.0146 | TRUE | 0.2288 |
| Mmachc | 1159 | 1338 | 1366 | 1537 | 1128 | 1108 | 1231 | 1336 | 1372 | 1208 | 1214 | 1213 | 1262.29 | 1313 | 1212 | 0.923 | -0.116 | 0.1583331 | 0.35216891 | 8.00E-04 | 0.0095 | 0.0041 | 0.0041 | TRUE | 0.0355 |
| Mmadhc | 2547 | 2766 | 2722 | 3519 | 2410 | 2461 | 2705 | 2762 | 2733 | 2766 | 2593 | 2694 | 2709.05 | 2733 | 2684 | 0.982 | -0.026 | 0.69788475 | 0.83456201 | 3.00E-04 | 0.0081 | 0.0028 | 0.0028 | TRUE | 0.0107 |
| Mmd | 768 | 1019 | 1033 | 1439 | 1021 | 1153 | 816 | 1018 | 1037 | 1131 | 1099 | 1262 | 1060.43 | 957 | 1164 | 1.216 | 0.282 | 0.02288933 | 0.09571438 | 0.0095 | 0.01 | 0.0101 | 0.0101 | TRUE | 0.1976 |
| Mmd2 | 37 | 24 | 27 | 19 | 16 | 11 | 39 | 24 | 27 | 15 | 17 | 12 | 22.43 | 30 | 15 | 0.49 | -1.028 | 0.01793534 | 0.0803169 | 0 | 0.1538 | 0.0859 | 0.0859 | TRUE | 0.4582 |
| Mme | 889 | 928 | 735 | 3557 | 2845 | 2511 | 944 | 927 | 738 | 2796 | 3061 | 2749 | 1869.18 | 870 | 2869 | 3.299 | 1.722 | 3.87E-49 | 5.08E-46 | 0.0092 | 0.0087 | 0.0091 | 0.0091 | TRUE | 0.2093 |
| Mmel1 | 121 | 116 | 107 | 79 | 81 | 85 | 129 | 116 | 107 | 62 | 87 | 93 | 99.02 | 117 | 81 | 0.684 | -0.548 | 0.01779492 | 0.07980139 | 0.0152 | 0.0402 | 0.0281 | 0.0281 | TRUE | 0.4337 |

| | | | | | | | | | | | | | | | | | | | | | | | | | |
|---|---|---|---|---|---|---|---|---|---|---|---|---|---|---|---|---|---|---|---|---|---|---|---|---|---|
| Mmgt1 | 1315 | 1616 | 1621 | 1701 | 1285 | 1425 | 1397 | 1614 | 1628 | 1337 | 1383 | 1560 | 1486.33 | 1546 | 1427 | 0.922 | -0.117 | 0.27001141 | 0.49018427 | 0.0059 | 0.0091 | 0.0075 | 0.0075 | TRUE | 0.0857 |
| Mmgt2 | 186 | 297 | 283 | 401 | 302 | 333 | 198 | 297 | 284 | 315 | 325 | 365 | 297.17 | 260 | 335 | 1.289 | 0.366 | 0.04207148 | 0.1461245 | 0.0219 | 0.018 | 0.0199 | 0.0199 | TRUE | 0.48 |
| Mmp10 | 0 | 0 | 0 | 0 | 0 | 0 | 0 | 0 | 0 | 0 | 0 | 0 | 0 | 0 | 0 | NA | NA | NA | NA | NA | NA | NA | NA | NA | NA |
| Mmp11 | 1679 | 2466 | 2560 | 2144 | 1756 | 1809 | 1783 | 2463 | 2571 | 1685 | 1889 | 1980 | 2061.96 | 2272 | 1851 | 0.815 | -0.296 | 0.04650769 | 0.15606794 | 0.0202 | 0.0085 | 0.0154 | 0.0154 | TRUE | 0.4283 |
| Mmp12 | 1 | 0 | 0 | 0 | 0 | 0 | 1 | 0 | 0 | 0 | 0 | 0 | 0.18 | 0 | 0 | 0.49 | -1.029 | 0.80087 | NA | 0 | 18.6165 | 10 | 10 | TRUE | 0.4614 |
| Mmp13 | 0 | 0 | 0 | 0 | 0 | 0 | 0 | 0 | 0 | 0 | 0 | 0 | 0 | 0 | 0 | NA | NA | NA | NA | NA | NA | NA | NA | NA | NA |
| Mmp14 | 577 | 591 | 631 | 671 | 503 | 417 | 613 | 590 | 634 | 527 | 541 | 456 | 560.31 | 612 | 508 | 0.831 | -0.267 | 0.01648859 | 0.07601969 | 0.0025 | 0.0128 | 0.0071 | 0.0071 | TRUE | 0.0899 |
| Mmp15 | 1174 | 1616 | 1691 | 1430 | 1056 | 1090 | 1247 | 1614 | 1698 | 1124 | 1136 | 1193 | 1335.4 | 1520 | 1151 | 0.757 | -0.401 | 0.00200628 | 0.01541308 | 0.0131 | 0.0094 | 0.0114 | 0.0114 | TRUE | 0.2958 |
| Mmp16 | 3 | 3 | 2 | 9 | 5 | 4 | 3 | 3 | 2 | 7 | 5 | 4 | 4.17 | 3 | 5 | 2.078 | 1.055 | 0.25399642 | NA | 0 | 0.7968 | 0.3427 | 0.3427 | TRUE | 0.1498 |
| Mmp17 | 1663 | 2072 | 1954 | 1642 | 1393 | 1526 | 1766 | 2069 | 1962 | 1291 | 1499 | 1670 | 1709.63 | 1932 | 1487 | 0.769 | -0.38 | 0.00171081 | 0.01359343 | 0.0104 | 0.0088 | 0.01 | 0.01 | TRUE | 0.1629 |
| Mmp19 | 33 | 15 | 11 | 45 | 36 | 29 | 35 | 15 | 11 | 35 | 39 | 32 | 27.82 | 20 | 35 | 1.74 | 0.799 | 0.09494005 | 0.25246913 | 0.127 | 0.1253 | 0.1261 | 0.1261 | TRUE | 2.1069 |
| Mmp1a | 0 | 0 | 0 | 0 | 0 | 0 | 0 | 0 | 0 | 0 | 0 | 0 | 0 | 0 | 0 | NA | NA | NA | NA | NA | NA | NA | NA | NA | NA |
| Mmp1b | 0 | 1 | 0 | 1 | 1 | 0 | 0 | 1 | 0 | 1 | 1 | 0 | 0.48 | 0 | 1 | 1.712 | 0.776 | 0.78614772 | NA | 0 | 6.9165 | 4.1067 | 4.1067 | TRUE | 0.405 |
| Mmp2 | 2697 | 3406 | 3261 | 2654 | 1955 | 2465 | 2865 | 3402 | 3274 | 2086 | 2103 | 2698 | 2738.14 | 3180 | 2296 | 0.722 | -0.471 | 0.0002528 | 0.00294853 | 0.0148 | 0.0081 | 0.0115 | 0.0115 | TRUE | 0.2835 |
| Mmp20 | 3 | 1 | 1 | 11 | 2 | 3 | 3 | 1 | 1 | 9 | 2 | 3 | 3.21 | 2 | 5 | 2.795 | 1.483 | 0.21309868 | NA | 0 | 1.0325 | 0.6223 | 0.6223 | TRUE | 1.3145 |
| Mmp21 | 8 | 3 | 5 | 4 | 5 | 4 | 8 | 3 | 5 | 3 | 5 | 4 | 4.9 | 5 | 4 | 0.776 | -0.367 | 0.67342198 | 0.81805326 | 0 | 0.6789 | 0.3376 | 0.3376 | TRUE | 0.3265 |
| Mmp23 | 10 | 37 | 36 | 38 | 41 | 35 | 11 | 37 | 36 | 30 | 44 | 38 | 32.67 | 28 | 37 | 1.333 | 0.415 | 0.3853643 | 0.59920343 | 0.1654 | 0.1077 | 0.1333 | 0.1333 | TRUE | 1.8201 |
| Mmp24 | 66 | 52 | 49 | 91 | 82 | 42 | 70 | 52 | 49 | 72 | 88 | 46 | 62.83 | 57 | 69 | 1.204 | 0.268 | 0.39765404 | 0.61114531 | 0.0531 | 0.0593 | 0.0564 | 0.0564 | TRUE | 0.2717 |
| Mmp25 | 745 | 643 | 604 | 387 | 271 | 275 | 791 | 642 | 606 | 304 | 292 | 301 | 489.47 | 680 | 299 | 0.44 | -1.185 | 8.21E-18 | 1.62E-15 | 0.0094 | 0.0136 | 0.0113 | 0.0113 | TRUE | 0.2409 |
| Mmp27 | 0 | 0 | 0 | 0 | 0 | 0 | 0 | 0 | 0 | 0 | 0 | 0 | 0 | 0 | 0 | NA | NA | NA | NA | NA | NA | NA | NA | NA | NA |
| Mmp28 | 141 | 106 | 87 | 221 | 137 | 129 | 150 | 106 | 87 | 174 | 147 | 141 | 134.22 | 114 | 154 | 1.353 | 0.436 | 0.06603801 | 0.19823509 | 0.0342 | 0.0314 | 0.0328 | 0.0328 | TRUE | 0.7263 |
| Mmp3 | 0 | 0 | 0 | 1 | 0 | 0 | 0 | 0 | 0 | 1 | 0 | 0 | 0.13 | 0 | 0 | 1.859 | 0.894 | 0.82650734 | NA | 0 | 25.1515 | 10 | 10 | TRUE | 0.2563 |
| Mmp7 | 0 | 0 | 0 | 0 | 0 | 1 | 0 | 0 | 0 | 0 | 0 | 1 | 0.18 | 0 | 0 | 1.859 | 0.894 | 0.82650734 | NA | 0 | 18.0643 | 10 | 10 | TRUE | 0.5383 |
| Mmp8 | 0 | 0 | 0 | 0 | 0 | 0 | 0 | 0 | 0 | 0 | 0 | 0 | 0 | 0 | 0 | NA | NA | NA | NA | NA | NA | NA | NA | NA | NA |
| Mmp9 | 138 | 132 | 130 | 170 | 115 | 118 | 147 | 132 | 131 | 134 | 124 | 129 | 132.58 | 137 | 129 | 0.947 | -0.079 | 0.64802522 | 0.80243071 | 0 | 0.0318 | 0.0138 | 0.0138 | TRUE | 0.0439 |
| Mmrn1 | 0 | 0 | 0 | 1 | 0 | 0 | 0 | 0 | 0 | 1 | 0 | 0 | 0.13 | 0 | 0 | 1.859 | 0.894 | 0.82650734 | NA | 0 | 25.1515 | 10 | 10 | TRUE | 0.2563 |
| Mmrn2 | 1281 | 1154 | 965 | 1962 | 1445 | 1461 | 1361 | 1153 | 969 | 1542 | 1555 | 1599 | 1363.09 | 1161 | 1565 | 1.349 | 0.432 | 0.00094571 | 0.00852331 | 0.0133 | 0.0093 | 0.0115 | 0.0115 | TRUE | 0.2713 |
| Mms19 | 7399 | 8554 | 8984 | 8307 | 6616 | 7251 | 7859 | 8543 | 9021 | 6530 | 7118 | 7937 | 7834.83 | 8474 | 7195 | 0.849 | -0.236 | 0.01881016 | 0.08321847 | 0.007 | 0.0073 | 0.0072 | 0.0072 | TRUE | 0.0993 |
| Mms22l | 4515 | 4363 | 4619 | 6052 | 4488 | 4345 | 4796 | 4357 | 4638 | 4758 | 4829 | 4756 | 4688.96 | 4597 | 4781 | 1.04 | 0.057 | 0.40617155 | 0.61887077 | 9.00E-04 | 0.0076 | 0.0031 | 0.0031 | TRUE | 0.0254 |
| Mn1 | 83 | 106 | 113 | 151 | 129 | 131 | 88 | 106 | 113 | 119 | 139 | 143 | 118.06 | 102 | 134 | 1.298 | 0.377 | 0.05821407 | 0.18235677 | 0 | 0.0348 | 0.0199 | 0.0199 | TRUE | 0.1435 |
| Mnat1 | 1857 | 2678 | 2632 | 2953 | 2306 | 2290 | 1972 | 2675 | 2643 | 2321 | 2481 | 2507 | 2433.18 | 2430 | 2436 | 1.002 | 0.003 | 0.97889436 | 0.98860997 | 0.0147 | 0.0083 | 0.0117 | 0.0117 | TRUE | 0.3282 |
| Mnd1 | 951 | 1164 | 1238 | 1185 | 854 | 984 | 1010 | 1162 | 1243 | 932 | 919 | 1077 | 1057.21 | 1138 | 976 | 0.856 | -0.224 | 0.05899243 | 0.18387917 | 0.008 | 0.01 | 0.0091 | 0.0091 | TRUE | 0.1161 |
| Mnda | 0 | 0 | 0 | 0 | 0 | 0 | 0 | 0 | 0 | 0 | 0 | 0 | 0 | 0 | 0 | NA | NA | NA | NA | NA | NA | NA | NA | NA | NA |
| Mndal | 1 | 0 | 0 | 0 | 0 | 0 | 1 | 0 | 0 | 0 | 0 | 0 | 0.18 | 0 | 0 | 0.49 | -1.029 | 0.80087 | NA | 0 | 18.6165 | 10 | 10 | TRUE | 0.4614 |
| Mnf1 | 1630 | 2138 | 2098 | 2274 | 1733 | 1893 | 1731 | 2135 | 2107 | 1788 | 1865 | 2072 | 1949.6 | 1991 | 1908 | 0.958 | -0.062 | 0.5862 | 0.76113826 | 0.0091 | 0.0086 | 0.0089 | 0.0089 | TRUE | 0.157 |
| Mns1 | 549 | 534 | 613 | 1054 | 865 | 695 | 583 | 533 | 616 | 829 | 931 | 761 | 708.67 | 577 | 840 | 1.455 | 0.541 | 4.90E-06 | 0.00010847 | 0.0063 | 0.0116 | 0.0086 | 0.0086 | TRUE | 0.1037 |
| Mnt | 2252 | 2721 | 2927 | 2897 | 2431 | 2374 | 2392 | 2717 | 2939 | 2277 | 2616 | 2599 | 2590.04 | 2683 | 2497 | 0.93 | -0.104 | 0.33596637 | 0.55602254 | 0.0076 | 0.0082 | 0.0081 | 0.0081 | TRUE | 0.1089 |
| Mnx1 | 48 | 45 | 52 | 72 | 66 | 54 | 51 | 45 | 52 | 57 | 71 | 59 | 55.81 | 49 | 62 | 1.257 | 0.33 | 0.20905797 | 0.41885953 | 0 | 0.0659 | 0.0315 | 0.0315 | TRUE | 0.1277 |
| Moap1 | 6 | 10 | 6 | 10 | 8 | 10 | 6 | 10 | 6 | 8 | 9 | 11 | 8.3 | 7 | 9 | 1.216 | 0.282 | 0.66558112 | 0.81276361 | 0 | 0.4038 | 0.185 | 0.185 | TRUE | 0.2493 |
| Mob1a | 2102 | 1912 | 1903 | 3188 | 2157 | 2046 | 2233 | 1910 | 1911 | 2506 | 2321 | 2240 | 2186.61 | 2018 | 2356 | 1.168 | 0.224 | 0.02317547 | 0.0965516 | 0.0052 | 0.0084 | 0.0066 | 0.0066 | TRUE | 0.1048 |
| Mob1b | 592 | 540 | 473 | 903 | 600 | 448 | 629 | 539 | 475 | 710 | 646 | 490 | 581.48 | 548 | 615 | 1.125 | 0.17 | 0.31932541 | 0.54057938 | 0.0259 | 0.0126 | 0.0193 | 0.0193 | TRUE | 0.3671 |
| Mob2 | 869 | 813 | 882 | 1073 | 747 | 807 | 923 | 812 | 886 | 844 | 804 | 883 | 858.54 | 874 | 844 | 0.966 | -0.05 | 0.604904 | 0.77559108 | 0.002 | 0.0107 | 0.0056 | 0.0056 | TRUE | 0.0454 |
| Mob3a | 943 | 993 | 1096 | 1115 | 780 | 803 | 1002 | 992 | 1100 | 877 | 839 | 879 | 948.1 | 1031 | 865 | 0.839 | -0.254 | 0.00470869 | 0.02995781 | 0.0011 | 0.0104 | 0.0047 | 0.0047 | TRUE | 0.0414 |
| Mob3b | 679 | 556 | 531 | 982 | 688 | 669 | 721 | 555 | 534 | 772 | 740 | 732 | 675.87 | 603 | 748 | 1.241 | 0.311 | 0.02310334 | 0.09632192 | 0.0126 | 0.0118 | 0.012 | 0.012 | TRUE | 0.3399 |
| Mob3c | 199 | 290 | 291 | 307 | 267 | 289 | 211 | 290 | 292 | 241 | 287 | 316 | 273.03 | 264 | 281 | 1.062 | 0.087 | 0.63295925 | 0.79357325 | 0.0216 | 0.019 | 0.0203 | 0.0203 | TRUE | 0.3401 |
| Mob4 | 1981 | 2292 | 2184 | 2798 | 1957 | 2019 | 2104 | 2289 | 2193 | 2200 | 2106 | 2210 | 2183.58 | 2195 | 2172 | 0.989 | -0.016 | 0.83243476 | 0.91498787 | 8.00E-04 | 0.0084 | 0.0034 | 0.0034 | TRUE | 0.0169 |
| Mobp | 192 | 135 | 90 | 334 | 233 | 275 | 204 | 135 | 90 | 263 | 251 | 301 | 207.24 | 143 | 272 | 1.899 | 0.925 | 0.00057483 | 0.00579681 | 0.0767 | 0.0228 | 0.0467 | 0.0467 | TRUE | 0.702 |
| Mocos | 125 | 139 | 147 | 167 | 108 | 159 | 133 | 139 | 148 | 131 | 116 | 174 | 140.12 | 140 | 140 | 1.003 | 0.005 | 0.9805689 | 0.98941416 | 0.0155 | 0.0304 | 0.023 | 0.023 | TRUE | 0.4322 |
| Mocs1 | 436 | 559 | 604 | 541 | 444 | 466 | 463 | 558 | 606 | 425 | 478 | 510 | 506.83 | 542 | 471 | 0.866 | -0.207 | 0.14407451 | 0.33120207 | 0.0116 | 0.0134 | 0.0125 | 0.0125 | TRUE | 0.1908 |
| Mocs2 | 1018 | 968 | 898 | 1334 | 912 | 822 | 1081 | 967 | 902 | 1049 | 981 | 900 | 979.91 | 983 | 977 | 0.994 | -0.008 | 0.94119855 | 0.97017081 | 0.0061 | 0.0103 | 0.0079 | 0.0079 | TRUE | 0.0904 |
| Mocs3 | 322 | 322 | 301 | 567 | 433 | 271 | 342 | 322 | 302 | 446 | 466 | 297 | 362.35 | 322 | 403 | 1.253 | 0.325 | 0.08066744 | 0.2258631 | 0.0283 | 0.016 | 0.0221 | 0.0221 | TRUE | 0.6 |
| Mog | 2 | 1 | 3 | 6 | 7 | 3 | 2 | 1 | 3 | 5 | 8 | 3 | 3.61 | 2 | 5 | 2.529 | 1.338 | 0.20107542 | NA | 0 | 0.9192 | 0.4451 | 0.4451 | TRUE | 0.2938 |
| Mogat1 | 3 | 1 | 2 | 0 | 0 | 0 | 3 | 1 | 2 | 0 | 0 | 0 | 1.03 | 2 | 0 | 0.087 | -3.522 | 0.10550605 | NA | 0 | 3.1985 | 2.1648 | 2.1648 | TRUE | 0.0737 |
| Mogat2 | 145 | 146 | 147 | 134 | 62 | 88 | 154 | 146 | 148 | 105 | 67 | 96 | 119.3 | 149 | 89 | 0.603 | -0.73 | 0.00078307 | 0.00736471 | 0.0137 | 0.0345 | 0.0251 | 0.0251 | TRUE | 0.4542 |
| Mogs | 1938 | 2185 | 2152 | 1996 | 1547 | 1511 | 2058 | 2182 | 2161 | 1569 | 1664 | 1654 | 1881.52 | 2134 | 1629 | 0.763 | -0.39 | 1.17E-07 | 4.08E-06 | 5.00E-04 | 0.0087 | 0.0034 | 0.0034 | TRUE | 0.013 |
| Mon1a | 264 | 246 | 253 | 382 | 229 | 230 | 280 | 246 | 254 | 300 | 246 | 252 | 263.1 | 260 | 266 | 1.027 | 0.039 | 0.79345481 | 0.89356439 | 0.0053 | 0.0194 | 0.0119 | 0.0119 | TRUE | 0.1499 |
| Mon1b | 1670 | 1687 | 1679 | 2003 | 1443 | 1405 | 1774 | 1685 | 1686 | 1575 | 1553 | 1538 | 1634.95 | 1715 | 1555 | 0.907 | -0.141 | 0.04949783 | 0.16318557 | 0 | 0.0089 | 0.0031 | 0.0031 | TRUE | 0.0109 |
| Mon2 | 6098 | 5508 | 5892 | 6601 | 4713 | 4751 | 6477 | 5501 | 5916 | 5189 | 5071 | 5201 | 5559.17 | 5965 | 5154 | 0.864 | -0.211 | 0.01144797 | 0.05818803 | 0.0032 | 0.0075 | 0.0048 | 0.0048 | TRUE | 0.0688 |
| Morc1 | 7009 | 6977 | 7151 | 7765 | 6172 | 6599 | 7445 | 6968 | 7180 | 6104 | 6641 | 7224 | 6926.95 | 7198 | 6656 | 0.925 | -0.113 | 0.19018316 | 0.39517265 | 0.0039 | 0.0074 | 0.0052 | 0.0052 | TRUE | 0.0679 |
| Morc2a | 5427 | 4458 | 4593 | 7268 | 5033 | 4146 | 5764 | 4452 | 4612 | 5714 | 5415 | 4538 | 5082.62 | 4943 | 5222 | 1.057 | 0.08 | 0.54849004 | 0.73326245 | 0.0144 | 0.0076 | 0.0125 | 0.0125 | TRUE | 0.2575 |
| Morc2b | 39 | 33 | 12 | 34 | 33 | 22 | 41 | 33 | 12 | 27 | 36 | 24 | 28.79 | 29 | 29 | 0.999 | -0.001 | 0.99785302 | 0.99860587 | 0.153 | 0.1213 | 0.1361 | 0.1361 | TRUE | 0.722 |
| Morc3 | 8855 | 8465 | 8540 | 11185 | 8430 | 8372 | 9406 | 8454 | 8575 | 8793 | 9070 | 9164 | 8910.35 | 8812 | 9009 | 1.022 | 0.032 | 0.65720551 | 0.80696343 | 0.0016 | 0.0073 | 0.0036 | 0.0036 | TRUE | 0.0424 |
| Morc4 | 556 | 666 | 733 | 1006 | 682 | 733 | 591 | 665 | 736 | 791 | 734 | 802 | 719.79 | 664 | 776 | 1.168 | 0.224 | 0.05148792 | 0.16728458 | 0.0055 | 0.0115 | 0.0082 | 0.0082 | TRUE | 0.1092 |
| Morf4l1 | 12498 | 13488 | 13056 | 17374 | 12139 | 12582 | 13275 | 13471 | 13110 | 13658 | 13061 | 13773 | 13391.19 | 13285 | 13497 | 1.016 | 0.023 | 0.68992103 | 0.82966922 | 4.00E-04 | 0.0071 | 0.0023 | 0.0023 | TRUE | 0.0097 |
| Morf4l2 | 16318 | 16910 | 16591 | 23146 | 16002 | 15452 | 17333 | 16888 | 16659 | 18196 | 17217 | 16915 | 17201.21 | 16960 | 17443 | 1.029 | 0.041 | 0.51267946 | 0.70600768 | 9.00E-04 | 0.0071 | 0.0027 | 0.0027 | TRUE | 0.0175 |
| Morn1 | 60 | 133 | 143 | 94 | 98 | 132 | 64 | 133 | 144 | 74 | 105 | 144 | 110.66 | 114 | 108 | 0.948 | -0.077 | 0.82693634 | 0.91237775 | 0.1316 | 0.0367 | 0.0805 | 0.0805 | TRUE | 0.4305 |
| Morn2 | 171 | 213 | 191 | 261 | 176 | 166 | 182 | 213 | 192 | 205 | 189 | 182 | 193.73 | 196 | 192 | 0.985 | -0.022 | 0.8856261 | 0.94387096 | 0 | 0.0239 | 0.0118 | 0.0118 | TRUE | 0.0657 |
| Morn3 | 1 | 1 | 5 | 6 | 5 | 7 | 1 | 1 | 5 | 5 | 5 | 8 | 4.14 | 2 | 6 | 2.482 | 1.311 | 0.20227293 | NA | 0 | 0.8026 | 0.4637 | 0.4637 | TRUE | 1.0269 |
| Morn4 | 657 | 877 | 854 | 1011 | 671 | 820 | 698 | 876 | 858 | 795 | 722 | 898 | 807.59 | 811 | 805 | 0.993 | -0.011 | 0.93614635 | 0.96816629 | 0.0122 | 0.011 | 0.0116 | 0.0116 | TRUE | 0.1743 |

| | | | | | | | | | | | | | | | | | | | | | | | | |
|---|---|---|---|---|---|---|---|---|---|---|---|---|---|---|---|---|---|---|---|---|---|---|---|---|
| Morn5 | 8 | 11 | 8 | 3 | 6 | 16 | 8 | 11 | 8 | 2 | 6 | 18 | 8.97 | 9 | 9 | 0.938 | -0.093 | 0.90781774 | 0.95354464 | 0.3272 | 0.374 | 0.3506 | 0.3506 | TRUE | 0.8625 |
| Mos | 6 | 7 | 13 | 8 | 8 | 8 | 6 | 7 | 13 | 6 | 9 | 9 | 8.35 | 9 | 8 | 0.887 | -0.174 | 0.79535581 | 0.8946193 | 0 | 0.4017 | 0.2032 | 0.2032 | TRUE | 0.5681 |
| Mospd1 | 399 | 544 | 534 | 573 | 369 | 364 | 424 | 543 | 536 | 450 | 397 | 398 | 458.2 | 501 | 415 | 0.83 | -0.269 | 0.05406193 | 0.17281409 | 0.01 | 0.0141 | 0.0119 | 0.0119 | TRUE | 0.2104 |
| Mospd2 | 878 | 1024 | 1002 | 1035 | 766 | 902 | 933 | 1023 | 1006 | 814 | 824 | 987 | 931.09 | 987 | 875 | 0.886 | -0.175 | 0.11622394 | 0.2880396 | 0.0058 | 0.0104 | 0.0079 | 0.0079 | TRUE | 0.1481 |
| Mospd3 | 1324 | 876 | 755 | 1581 | 1086 | 1021 | 1406 | 875 | 758 | 1243 | 1168 | 1118 | 1094.71 | 1013 | 1176 | 1.162 | 0.216 | 0.29852685 | 0.51905693 | 0.054 | 0.0099 | 0.0302 | 0.0302 | TRUE | 1.3752 |
| Mospd4 | 2 | 0 | 1 | 1 | 1 | 4 | 2 | 0 | 1 | 1 | 1 | 4 | 1.56 | 1 | 2 | 1.975 | 0.982 | 0.55545411 | NA | 0 | 2.1167 | 1.2638 | 1.2638 | TRUE | 0.6255 |
| Mov10 | 4857 | 6642 | 6927 | 7840 | 5956 | 7365 | 5159 | 6633 | 6955 | 6163 | 6408 | 8062 | 6563.58 | 6249 | 6878 | 1.1 | 0.138 | 0.34641975 | 0.56554521 | 0.021 | 0.0074 | 0.0153 | 0.0153 | TRUE | 0.284 |
| Mov10l1 | 833 | 1472 | 1367 | 1668 | 1333 | 1400 | 885 | 1470 | 1373 | 1311 | 1434 | 1533 | 1334.25 | 1243 | 1426 | 1.147 | 0.198 | 0.2741128 | 0.49465156 | 0.0376 | 0.0094 | 0.0229 | 0.0229 | TRUE | 0.7594 |
| Moxd1 | 478 | 406 | 394 | 389 | 295 | 320 | 508 | 405 | 396 | 306 | 317 | 350 | 380.38 | 436 | 324 | 0.743 | -0.428 | 0.00320358 | 0.02231751 | 0.0099 | 0.0156 | 0.0125 | 0.0125 | TRUE | 0.2358 |
| Moxd2 | 0 | 0 | 0 | 0 | 0 | 0 | 0 | 0 | 0 | 0 | 0 | 0 | 0 | 0 | 0 | NA | NA | NA | NA | NA | NA | NA | NA | NA | NA |
| Mpc1 | 350 | 537 | 582 | 576 | 409 | 474 | 372 | 536 | 584 | 453 | 440 | 519 | 484.03 | 497 | 471 | 0.945 | -0.081 | 0.65125729 | 0.80449923 | 0.029 | 0.0137 | 0.0213 | 0.0213 | TRUE | 0.5659 |
| Mpc2 | 230 | 375 | 336 | 582 | 328 | 394 | 244 | 375 | 337 | 458 | 353 | 431 | 366.32 | 319 | 414 | 1.299 | 0.378 | 0.04386046 | 0.15007283 | 0.0293 | 0.0159 | 0.0225 | 0.0225 | TRUE | 0.4691 |
| Mpdu1 | 2014 | 2428 | 2537 | 4347 | 3121 | 2715 | 2139 | 2425 | 2547 | 3417 | 3358 | 2972 | 2809.8 | 2370 | 3249 | 1.371 | 0.455 | 7.92E-06 | 0.00016243 | 0.0061 | 0.0081 | 0.0071 | 0.0071 | TRUE | 0.088 |
| Mpdz | 904 | 1033 | 1194 | 1332 | 937 | 1029 | 960 | 1032 | 1199 | 1047 | 1008 | 1126 | 1062.07 | 1064 | 1060 | 0.997 | -0.005 | 0.96751419 | 0.98366749 | 0.0071 | 0.01 | 0.0084 | 0.0084 | TRUE | 0.1483 |
| Mpeg1 | 4 | 4 | 1 | 5 | 6 | 2 | 4 | 4 | 1 | 4 | 6 | 2 | 3.64 | 3 | 4 | 1.362 | 0.446 | 0.66782174 | NA | 0 | 0.9127 | 0.4942 | 0.4942 | TRUE | 0.332 |
| Mpg | 2065 | 2572 | 2505 | 2767 | 2206 | 2349 | 2193 | 2569 | 2515 | 2175 | 2374 | 2571 | 2399.57 | 2426 | 2373 | 0.978 | -0.032 | 0.75533434 | 0.87077942 | 0.0067 | 0.0083 | 0.0073 | 0.0073 | TRUE | 0.0849 |
| Mphosph10 | 5149 | 4718 | 4495 | 5919 | 4137 | 3817 | 5469 | 4712 | 4513 | 4653 | 4451 | 4178 | 4662.84 | 4898 | 4427 | 0.904 | -0.145 | 0.14300535 | 0.32931528 | 0.0064 | 0.0076 | 0.0069 | 0.0069 | TRUE | 0.1266 |
| Mphosph6 | 3150 | 2801 | 2615 | 3786 | 2750 | 2690 | 3346 | 2797 | 2626 | 2976 | 2959 | 2945 | 2941.45 | 2923 | 2960 | 1.013 | 0.018 | 0.86335048 | 0.9324082 | 0.0077 | 0.008 | 0.0078 | 0.0078 | TRUE | 0.1942 |
| Mphosph8 | 9286 | 10185 | 10111 | 9933 | 7566 | 7729 | 9863 | 10172 | 10153 | 7809 | 8141 | 8461 | 9099.57 | 10063 | 8137 | 0.808 | -0.307 | 1.33E-06 | 3.53E-05 | 8.00E-04 | 0.0073 | 0.0028 | 0.0028 | TRUE | 0.0153 |
| Mphosph9 | 1548 | 1486 | 1613 | 2107 | 1544 | 1280 | 1644 | 1484 | 1620 | 1656 | 1661 | 1401 | 1577.79 | 1583 | 1573 | 0.994 | -0.008 | 0.93710918 | 0.96855085 | 0.0055 | 0.009 | 0.0069 | 0.0069 | TRUE | 0.1087 |
| Mpi | 1774 | 1810 | 1716 | 2085 | 1505 | 1580 | 1884 | 1808 | 1723 | 1639 | 1619 | 1730 | 1733.82 | 1805 | 1663 | 0.921 | -0.119 | 0.13079327 | 0.31137681 | 0.001 | 0.0088 | 0.0039 | 0.0039 | TRUE | 0.0191 |
| Mpl | 0 | 1 | 1 | 1 | 4 | 1 | 0 | 1 | 1 | 1 | 4 | 1 | 1.36 | 1 | 2 | 3.006 | 1.588 | 0.37276278 | NA | 0 | 2.4212 | 1.2881 | 1.2881 | TRUE | 0.7858 |
| Mplkip | 874 | 1049 | 1079 | 1101 | 807 | 915 | 928 | 1048 | 1083 | 866 | 868 | 1002 | 965.81 | 1020 | 912 | 0.893 | -0.163 | 0.13852955 | 0.3230815 | 0.0053 | 0.0103 | 0.0077 | 0.0077 | TRUE | 0.0872 |
| Mpnd | 2862 | 3926 | 3918 | 3480 | 2618 | 2832 | 3040 | 3921 | 3934 | 2736 | 2817 | 3100 | 3257.92 | 3632 | 2884 | 0.794 | -0.333 | 0.00634214 | 0.03751995 | 0.0117 | 0.0079 | 0.0104 | 0.0104 | TRUE | 0.2468 |
| Mpo | 4 | 1 | 1 | 1 | 3 | 3 | 4 | 1 | 1 | 1 | 3 | 3 | 2.26 | 2 | 2 | 1.148 | 0.199 | 0.88025097 | NA | 0 | 1.4658 | 0.8089 | 0.8089 | TRUE | 0.7313 |
| Mpp1 | 897 | 879 | 799 | 1615 | 1082 | 1115 | 953 | 878 | 802 | 1270 | 1164 | 1221 | 1047.87 | 878 | 1218 | 1.389 | 0.474 | 2.11E-06 | 5.26E-05 | 0.0031 | 0.01 | 0.0062 | 0.0062 | TRUE | 0.0672 |
| Mpp2 | 1033 | 1187 | 1192 | 1269 | 887 | 979 | 1097 | 1185 | 1197 | 998 | 954 | 1072 | 1083.87 | 1160 | 1008 | 0.869 | -0.203 | 0.02592723 | 0.10426422 | 0.0018 | 0.0099 | 0.0051 | 0.0051 | TRUE | 0.0355 |
| Mpp3 | 358 | 544 | 506 | 412 | 410 | 420 | 380 | 543 | 508 | 324 | 441 | 460 | 442.73 | 477 | 408 | 0.854 | -0.229 | 0.22580532 | 0.43859625 | 0.0325 | 0.0143 | 0.0234 | 0.0234 | TRUE | 0.3908 |
| Mpp4 | 2 | 2 | 2 | 6 | 5 | 4 | 2 | 2 | 2 | 5 | 5 | 4 | 3.43 | 2 | 5 | 2.361 | 1.239 | 0.22147716 | NA | 0 | 0.9662 | 0.3891 | 0.3891 | TRUE | 0.0178 |
| Mpp5 | 2657 | 2602 | 2603 | 3710 | 2630 | 2729 | 2822 | 2599 | 2614 | 2917 | 2830 | 2987 | 2794.68 | 2678 | 2911 | 1.087 | 0.121 | 0.09848878 | 0.25854945 | 9.00E-04 | 0.0081 | 0.0035 | 0.0035 | TRUE | 0.0267 |
| Mpp6 | 3688 | 4858 | 4850 | 5700 | 4178 | 4335 | 3917 | 4852 | 4870 | 4481 | 4495 | 4745 | 4560.08 | 4546 | 4574 | 1.006 | 0.009 | 0.93548423 | 0.96807672 | 0.0078 | 0.0076 | 0.0077 | 0.0077 | TRUE | 0.1782 |
| Mpp7 | 1840 | 1702 | 1608 | 2649 | 1857 | 1681 | 1954 | 1700 | 1615 | 2082 | 1998 | 1840 | 1864.9 | 1756 | 1973 | 1.124 | 0.169 | 0.10523666 | 0.27039689 | 0.0062 | 0.0087 | 0.0073 | 0.0073 | TRUE | 0.1172 |
| Mppe1 | 135 | 212 | 172 | 378 | 235 | 221 | 143 | 212 | 173 | 297 | 253 | 242 | 219.96 | 176 | 264 | 1.502 | 0.587 | 0.00173537 | 0.01372132 | 0.0192 | 0.0219 | 0.0206 | 0.0206 | TRUE | 0.3396 |
| Mpped1 | 91 | 74 | 80 | 116 | 68 | 66 | 97 | 74 | 80 | 91 | 73 | 72 | 81.25 | 84 | 79 | 0.949 | -0.076 | 0.7470563 | 0.8655848 | 0.0075 | 0.0475 | 0.0276 | 0.0276 | TRUE | 0.1933 |
| Mpped2 | 204 | 185 | 180 | 432 | 272 | 255 | 217 | 185 | 181 | 340 | 293 | 279 | 248.93 | 194 | 304 | 1.57 | 0.651 | 2.64E-05 | 0.00045224 | 0.0067 | 0.0201 | 0.0131 | 0.0131 | TRUE | 0.1267 |
| Mprip | 11039 | 11553 | 11857 | 14918 | 10666 | 10053 | 11725 | 11538 | 11906 | 11727 | 11476 | 11005 | 11562.84 | 11723 | 11403 | 0.973 | -0.04 | 0.50352252 | 0.69859971 | 5.00E-04 | 0.0072 | 0.0025 | 0.0025 | TRUE | 0.0114 |
| Mpst | 1169 | 1708 | 1741 | 1363 | 1086 | 1366 | 1242 | 1706 | 1748 | 1071 | 1168 | 1495 | 1405.15 | 1565 | 1245 | 0.795 | -0.331 | 0.052652 | 0.16996057 | 0.0309 | 0.0092 | 0.0203 | 0.0203 | TRUE | 0.3933 |
| Mptx1 | 0 | 0 | 0 | 0 | 0 | 0 | 0 | 0 | 0 | 0 | 0 | 0 | 0 | 0 | 0 | NA | NA | NA | NA | NA | NA | NA | NA | NA | NA |
| Mptx2 | 0 | 0 | 0 | 0 | 0 | 0 | 0 | 0 | 0 | 0 | 0 | 0 | 0 | 0 | 0 | NA | NA | NA | NA | NA | NA | NA | NA | NA | NA |
| Mpv17 | 1823 | 1894 | 1824 | 2133 | 1460 | 1569 | 1936 | 1892 | 1831 | 1677 | 1571 | 1718 | 1770.76 | 1886 | 1655 | 0.878 | -0.188 | 0.01440577 | 0.06872589 | 8.00E-04 | 0.0088 | 0.0037 | 0.0037 | TRUE | 0.0234 |
| Mpv17l | 735 | 779 | 692 | 945 | 625 | 666 | 781 | 778 | 695 | 743 | 672 | 729 | 732.99 | 751 | 715 | 0.952 | -0.07 | 0.48774566 | 0.68458483 | 0.0022 | 0.0114 | 0.0061 | 0.0061 | TRUE | 0.0512 |
| Mpv17l2 | 1206 | 1514 | 1524 | 1600 | 1263 | 1190 | 1281 | 1512 | 1530 | 1258 | 1359 | 1303 | 1373.77 | 1441 | 1307 | 0.906 | -0.142 | 0.1603934 | 0.3548001 | 0.0048 | 0.0093 | 0.0067 | 0.0067 | TRUE | 0.1132 |
| Mpz | 7 | 14 | 15 | 18 | 12 | 12 | 7 | 14 | 15 | 14 | 13 | 13 | 12.78 | 12 | 13 | 1.101 | 0.139 | 0.79754555 | 0.89574083 | 0 | 0.2647 | 0.1328 | 0.1328 | TRUE | 0.436 |
| Mpzl1 | 2322 | 2590 | 2441 | 3328 | 2464 | 2560 | 2466 | 2587 | 2451 | 2616 | 2651 | 2802 | 2595.62 | 2501 | 2690 | 1.075 | 0.104 | 0.14204961 | 0.32778442 | 7.00E-04 | 0.0082 | 0.0033 | 0.0033 | TRUE | 0.0161 |
| Mpzl2 | 985 | 1220 | 1338 | 1647 | 1276 | 1445 | 1046 | 1218 | 1343 | 1295 | 1373 | 1582 | 1309.6 | 1202 | 1417 | 1.177 | 0.235 | 0.06628391 | 0.19871315 | 0.012 | 0.0094 | 0.011 | 0.011 | TRUE | 0.1547 |
| Mpzl3 | 51 | 39 | 42 | 70 | 48 | 57 | 54 | 39 | 42 | 55 | 52 | 62 | 50.73 | 45 | 56 | 1.25 | 0.322 | 0.25290583 | 0.47086616 | 0 | 0.0718 | 0.0372 | 0.0372 | TRUE | 0.2359 |
| Mr1 | 10 | 8 | 2 | 6 | 12 | 5 | 11 | 8 | 2 | 5 | 13 | 5 | 7.29 | 7 | 8 | 1.113 | 0.154 | 0.85442395 | 0.92752251 | 0.2734 | 0.459 | 0.3693 | 0.3693 | TRUE | 0.6872 |
| Mrap | 28 | 27 | 16 | 36 | 20 | 14 | 30 | 27 | 16 | 28 | 22 | 15 | 22.99 | 24 | 22 | 0.905 | -0.144 | 0.74964473 | 0.8673408 | 0.0512 | 0.1502 | 0.1031 | 0.1031 | TRUE | 0.2944 |
| Mrap2 | 3 | 2 | 1 | 2 | 0 | 3 | 3 | 2 | 1 | 2 | 0 | 3 | 1.84 | 2 | 2 | 0.785 | -0.35 | 0.81220708 | NA | 0 | 1.7967 | 1.0093 | 1.0093 | TRUE | 0.2272 |
| Mras | 1634 | 1185 | 1088 | 1484 | 1092 | 1012 | 1736 | 1183 | 1092 | 1167 | 1175 | 1108 | 1243.48 | 1337 | 1150 | 0.86 | -0.217 | 0.20390802 | 0.41206317 | 0.0332 | 0.0096 | 0.0203 | 0.0203 | TRUE | 0.8152 |
| Mrc1 | 78 | 58 | 50 | 131 | 101 | 64 | 83 | 58 | 50 | 103 | 109 | 70 | 78.78 | 64 | 94 | 1.48 | 0.566 | 0.05134864 | 0.16701811 | 0.0461 | 0.0487 | 0.0475 | 0.0475 | TRUE | 0.5965 |
| Mrc2 | 364 | 378 | 432 | 610 | 354 | 367 | 387 | 378 | 434 | 480 | 381 | 402 | 410.01 | 400 | 421 | 1.056 | 0.079 | 0.56806087 | 0.74740657 | 0.0082 | 0.0149 | 0.0113 | 0.0113 | TRUE | 0.1804 |
| Mre11a | 3767 | 4048 | 4183 | 5409 | 4000 | 4025 | 4001 | 4043 | 4200 | 4252 | 4304 | 4406 | 4201.01 | 4081 | 4321 | 1.058 | 0.082 | 0.18604526 | 0.38980244 | 3.00E-04 | 0.0077 | 0.0025 | 0.0025 | TRUE | 0.0079 |
| Mreg | 13568 | 10739 | 9894 | 25099 | 16585 | 12245 | 14412 | 10725 | 9935 | 19731 | 17844 | 13404 | 14341.79 | 11691 | 16993 | 1.454 | 0.54 | 0.00208851 | 0.01594383 | 0.039 | 0.0071 | 0.0221 | 0.0221 | TRUE | 0.5067 |
| Mrfap1 | 10160 | 10621 | 9768 | 15248 | 11469 | 10221 | 10792 | 10607 | 9808 | 11987 | 12340 | 11188 | 11120.39 | 10402 | 11838 | 1.138 | 0.187 | 0.01415836 | 0.06791533 | 0.0024 | 0.0072 | 0.0041 | 0.0041 | TRUE | 0.0305 |
| Mrgbp | 598 | 648 | 591 | 725 | 570 | 524 | 635 | 647 | 593 | 570 | 613 | 574 | 605.43 | 625 | 586 | 0.936 | -0.095 | 0.33422905 | 0.5548783 | 0 | 0.0123 | 0.0054 | 0.0054 | TRUE | 0.0235 |
| Mrgpra1 | 0 | 0 | 0 | 0 | 0 | 0 | 0 | 0 | 0 | 0 | 0 | 0 | 0 | 0 | 0 | NA | NA | NA | NA | NA | NA | NA | NA | NA | NA |
| Mrgpra2a | 0 | 0 | 0 | 0 | 0 | 0 | 0 | 0 | 0 | 0 | 0 | 0 | 0 | 0 | 0 | NA | NA | NA | NA | NA | NA | NA | NA | NA | NA |
| Mrgpra2b | 0 | 0 | 0 | 0 | 0 | 0 | 0 | 0 | 0 | 0 | 0 | 0 | 0 | 0 | 0 | NA | NA | NA | NA | NA | NA | NA | NA | NA | NA |
| Mrgpra3 | 0 | 0 | 1 | 0 | 0 | 0 | 0 | 0 | 1 | 0 | 0 | 0 | 0.17 | 0 | 0 | 0.49 | -1.029 | 0.80087 | NA | 0 | 19.6928 | 10 | 10 | TRUE | 0.4098 |
| Mrgpra4 | 0 | 0 | 0 | 0 | 0 | 0 | 0 | 0 | 0 | 0 | 0 | 0 | 0 | 0 | 0 | NA | NA | NA | NA | NA | NA | NA | NA | NA | NA |
| Mrgpra6 | 0 | 0 | 0 | 0 | 0 | 0 | 0 | 0 | 0 | 0 | 0 | 0 | 0 | 0 | 0 | NA | NA | NA | NA | NA | NA | NA | NA | NA | NA |
| Mrgpra9 | 0 | 0 | 0 | 0 | 0 | 0 | 0 | 0 | 0 | 0 | 0 | 0 | 0 | 0 | 0 | NA | NA | NA | NA | NA | NA | NA | NA | NA | NA |
| Mrgprb1 | 0 | 0 | 0 | 1 | 0 | 0 | 0 | 0 | 0 | 1 | 0 | 0 | 0.13 | 0 | 0 | 1.859 | 0.894 | 0.82650734 | NA | 0 | 25.1515 | 10 | 10 | TRUE | 0.2563 |
| Mrgprb2 | 0 | 0 | 0 | 0 | 0 | 0 | 0 | 0 | 0 | 0 | 0 | 0 | 0 | 0 | 0 | NA | NA | NA | NA | NA | NA | NA | NA | NA | NA |
| Mrgprb3 | 0 | 0 | 0 | 0 | 0 | 0 | 0 | 0 | 0 | 0 | 0 | 0 | 0 | 0 | 0 | NA | NA | NA | NA | NA | NA | NA | NA | NA | NA |
| Mrgprb4 | 0 | 0 | 0 | 0 | 0 | 0 | 0 | 0 | 0 | 0 | 0 | 0 | 0 | 0 | 0 | NA | NA | NA | NA | NA | NA | NA | NA | NA | NA |
| Mrgprb5 | 0 | 0 | 0 | 0 | 0 | 0 | 0 | 0 | 0 | 0 | 0 | 0 | 0 | 0 | 0 | NA | NA | NA | NA | NA | NA | NA | NA | NA | NA |
| Mrgprb8 | 0 | 0 | 0 | 0 | 0 | 0 | 0 | 0 | 0 | 0 | 0 | 0 | 0 | 0 | 0 | NA | NA | NA | NA | NA | NA | NA | NA | NA | NA |

| | | | | | | | | | | | | | | | | | | | | | | | | | |
|---|---|---|---|---|---|---|---|---|---|---|---|---|---|---|---|---|---|---|---|---|---|---|---|---|---|
| Mrgprd | 0 | 0 | 0 | 0 | 0 | 0 | 0 | 0 | 0 | 0 | 0 | 0 | 0 | 0 | 0 | NA | NA | NA | NA | NA | NA | NA | NA | NA | NA |
| Mrgpre | 15 | 19 | 13 | 12 | 11 | 12 | 16 | 19 | 13 | 9 | 12 | 13 | 13.73 | 16 | 11 | 0.711 | -0.491 | 0.33744233 | 0.55746731 | 0 | 0.2469 | 0.1143 | 0.1143 | TRUE | 0.1297 |
| Mrgprf | 33 | 26 | 22 | 44 | 16 | 18 | 35 | 26 | 22 | 35 | 17 | 20 | 25.77 | 28 | 24 | 0.873 | -0.196 | 0.65624112 | 0.80670515 | 0.0648 | 0.1347 | 0.1004 | 0.1004 | TRUE | 1.0864 |
| Mrgprg | 0 | 0 | 0 | 0 | 0 | 0 | 0 | 0 | 0 | 0 | 0 | 0 | 0 | 0 | 0 | NA | NA | NA | NA | NA | NA | NA | NA | NA | NA |
| Mrgprh | 0 | 0 | 0 | 0 | 0 | 0 | 0 | 0 | 0 | 0 | 0 | 0 | 0 | 0 | 0 | NA | NA | NA | NA | NA | NA | NA | NA | NA | NA |
| Mrgprx1 | 0 | 0 | 0 | 0 | 0 | 0 | 0 | 0 | 0 | 0 | 0 | 0 | 0 | 0 | 0 | NA | NA | NA | NA | NA | NA | NA | NA | NA | NA |
| Mrgprx2 | 0 | 0 | 0 | 0 | 0 | 0 | 0 | 0 | 0 | 0 | 0 | 0 | 0 | 0 | 0 | NA | NA | NA | NA | NA | NA | NA | NA | NA | NA |
| Mri1 | 1684 | 1876 | 1869 | 2116 | 1665 | 1811 | 1789 | 1874 | 1877 | 1663 | 1791 | 1982 | 1829.38 | 1847 | 1812 | 0.981 | -0.028 | 0.76440437 | 0.87630366 | 0.0038 | 0.0087 | 0.0057 | 0.0057 | TRUE | 0.0811 |
| Mrm1 | 1427 | 1769 | 1859 | 2264 | 1721 | 1623 | 1516 | 1767 | 1867 | 1780 | 1852 | 1777 | 1759.53 | 1717 | 1803 | 1.05 | 0.07 | 0.48494711 | 0.68236871 | 0.0053 | 0.0088 | 0.0068 | 0.0068 | TRUE | 0.1257 |
| Mro | 8 | 13 | 9 | 8 | 5 | 10 | 8 | 13 | 9 | 6 | 5 | 11 | 8.86 | 10 | 7 | 0.734 | -0.446 | 0.49478398 | 0.69076234 | 0 | 0.3789 | 0.1917 | 0.1917 | TRUE | 0.3941 |
| Mroh1 | 2299 | 2483 | 2552 | 2458 | 1983 | 1991 | 2442 | 2480 | 2562 | 1932 | 2134 | 2179 | 2288.26 | 2495 | 2082 | 0.834 | -0.262 | 0.00117761 | 0.01007776 | 0.0019 | 0.0083 | 0.0043 | 0.0043 | TRUE | 0.0489 |
| Mroh2a | 1 | 0 | 1 | 0 | 1 | 1 | 1 | 0 | 1 | 0 | 1 | 1 | 0.71 | 1 | 1 | 1.014 | 0.02 | 0.99311765 | NA | 0 | 4.6723 | 2.3783 | 2.3783 | TRUE | 0.3742 |
| Mroh2b | 0 | 0 | 0 | 12 | 9 | 7 | 0 | 0 | 0 | 9 | 10 | 8 | 4.46 | 0 | 9 | 47.579 | 5.572 | 0.0001186 | NA | 0 | 0.745 | 0.4556 | 0.4556 | TRUE | 0.07 |
| Mroh4 | 15 | 19 | 13 | 13 | 14 | 10 | 16 | 19 | 13 | 10 | 15 | 11 | 14.03 | 16 | 12 | 0.75 | -0.414 | 0.41684124 | 0.62807995 | 0 | 0.2417 | 0.1149 | 0.1149 | TRUE | 0.1738 |
| Mroh5 | 0 | 0 | 0 | 0 | 0 | 0 | 0 | 0 | 0 | 0 | 0 | 0 | 0 | 0 | 0 | NA | NA | NA | NA | NA | NA | NA | NA | NA | NA |
| Mroh8 | 34 | 44 | 43 | 25 | 23 | 21 | 36 | 44 | 43 | 20 | 25 | 23 | 31.77 | 41 | 23 | 0.543 | -0.881 | 0.0104666 | 0.05457162 | 0 | 0.1106 | 0.051 | 0.051 | TRUE | 0.0869 |
| Mroh9 | 0 | 0 | 0 | 0 | 0 | 0 | 0 | 0 | 0 | 0 | 0 | 0 | 0 | 0 | 0 | NA | NA | NA | NA | NA | NA | NA | NA | NA | NA |
| Mrp63 | 948 | 1443 | 1462 | 1636 | 1219 | 1232 | 1007 | 1441 | 1468 | 1286 | 1312 | 1349 | 1310.39 | 1305 | 1316 | 1.007 | 0.011 | 0.9433658 | 0.9717062 | 0.0209 | 0.0094 | 0.0153 | 0.0153 | TRUE | 0.4791 |
| Mrpl1 | 2700 | 2701 | 2809 | 3825 | 2782 | 2646 | 2868 | 2698 | 2821 | 3007 | 2993 | 2896 | 2880.43 | 2796 | 2965 | 1.061 | 0.086 | 0.20000671 | 0.40765056 | 3.00E-04 | 0.008 | 0.0029 | 0.0029 | TRUE | 0.0114 |
| Mrpl10 | 5270 | 5381 | 5128 | 6924 | 4889 | 4749 | 5598 | 5374 | 5149 | 5443 | 5260 | 5199 | 5337.11 | 5374 | 5301 | 0.987 | -0.019 | 0.77243118 | 0.88115034 | 8.00E-04 | 0.0075 | 0.0031 | 0.0031 | TRUE | 0.0163 |
| Mrpl11 | 2288 | 2296 | 2211 | 3143 | 2264 | 2136 | 2430 | 2293 | 2220 | 2471 | 2436 | 2338 | 2364.71 | 2314 | 2415 | 1.044 | 0.062 | 0.40635479 | 0.61899051 | 0.001 | 0.0083 | 0.0036 | 0.0036 | TRUE | 0.0231 |
| Mrpl12 | 4353 | 4512 | 4443 | 6613 | 4499 | 4074 | 4624 | 4506 | 4461 | 5199 | 4841 | 4460 | 4681.66 | 4530 | 4833 | 1.067 | 0.094 | 0.25193789 | 0.46974399 | 0.0028 | 0.0076 | 0.0046 | 0.0046 | TRUE | 0.0555 |
| Mrpl13 | 1740 | 1956 | 1888 | 2955 | 1886 | 1769 | 1848 | 1953 | 1896 | 2323 | 2029 | 1936 | 1997.68 | 1899 | 2096 | 1.105 | 0.144 | 0.13431342 | 0.31691521 | 0.0043 | 0.0086 | 0.0061 | 0.0061 | TRUE | 0.1101 |
| Mrpl14 | 710 | 954 | 893 | 1108 | 821 | 918 | 754 | 953 | 897 | 871 | 883 | 1005 | 893.81 | 868 | 920 | 1.059 | 0.083 | 0.50208367 | 0.6972672 | 0.0091 | 0.0106 | 0.0098 | 0.0098 | TRUE | 0.1554 |
| Mrpl15 | 7753 | 8967 | 9046 | 10382 | 7384 | 7398 | 8235 | 8955 | 9083 | 8162 | 7945 | 8098 | 8413.02 | 8758 | 8068 | 0.921 | -0.118 | 0.0843774 | 0.23298325 | 0.0013 | 0.0073 | 0.0033 | 0.0033 | TRUE | 0.0333 |
| Mrpl16 | 1720 | 1524 | 1463 | 2431 | 1631 | 1500 | 1827 | 1522 | 1469 | 1911 | 1755 | 1642 | 1687.65 | 1606 | 1769 | 1.102 | 0.141 | 0.2246716 | 0.43704002 | 0.0093 | 0.0089 | 0.0091 | 0.0091 | TRUE | 0.1742 |
| Mrpl17 | 3399 | 3591 | 3452 | 5445 | 3773 | 3434 | 3610 | 3586 | 3466 | 4280 | 4060 | 3759 | 3793.65 | 3554 | 4033 | 1.135 | 0.183 | 0.01906834 | 0.08386666 | 0.0019 | 0.0078 | 0.0041 | 0.0041 | TRUE | 0.0429 |
| Mrpl18 | 3306 | 3192 | 3254 | 4526 | 3118 | 3155 | 3512 | 3188 | 3267 | 3558 | 3355 | 3454 | 3388.87 | 3322 | 3456 | 1.04 | 0.057 | 0.4385253 | 0.64591156 | 0.0013 | 0.0079 | 0.0036 | 0.0036 | TRUE | 0.0301 |
| Mrpl19 | 2068 | 2160 | 2172 | 3045 | 2107 | 2057 | 2197 | 2157 | 2181 | 2394 | 2267 | 2252 | 2241.19 | 2178 | 2304 | 1.058 | 0.082 | 0.23269503 | 0.44696944 | 0 | 0.0084 | 0.0029 | 0.0029 | TRUE | 0.0144 |
| Mrpl2 | 4242 | 4574 | 4620 | 5442 | 3966 | 3991 | 4506 | 4568 | 4639 | 4278 | 4267 | 4369 | 4437.81 | 4571 | 4305 | 0.942 | -0.087 | 0.13618132 | 0.3196208 | 0 | 0.0076 | 0.0022 | 0.0022 | TRUE | 0.0021 |
| Mrpl20 | 2717 | 2035 | 1921 | 3538 | 2277 | 1899 | 2886 | 2032 | 1929 | 2781 | 2450 | 2079 | 2359.53 | 2282 | 2437 | 1.068 | 0.095 | 0.58265175 | 0.75868406 | 0.0347 | 0.0083 | 0.0211 | 0.0211 | TRUE | 0.6476 |
| Mrpl21 | 1666 | 2059 | 2057 | 2293 | 1623 | 1692 | 1770 | 2056 | 2065 | 1803 | 1746 | 1852 | 1882.06 | 1964 | 1800 | 0.917 | -0.126 | 0.17456222 | 0.37396928 | 0.0033 | 0.0087 | 0.0056 | 0.0056 | TRUE | 0.0901 |
| Mrpl22 | 2111 | 2346 | 2277 | 2935 | 2147 | 2137 | 2242 | 2343 | 2286 | 2307 | 2310 | 2339 | 2304.69 | 2290 | 2319 | 1.012 | 0.018 | 0.78763824 | 0.89063883 | 0 | 0.0083 | 0.0026 | 0.0026 | TRUE | 0.0049 |
| Mrpl23 | 782 | 1023 | 1098 | 896 | 678 | 765 | 831 | 1022 | 1103 | 704 | 729 | 837 | 871.01 | 985 | 757 | 0.768 | -0.381 | 0.00522513 | 0.03226731 | 0.0127 | 0.0107 | 0.0122 | 0.0122 | TRUE | 0.2233 |
| Mrpl24 | 4326 | 5737 | 5730 | 5914 | 4467 | 4781 | 4595 | 5730 | 5754 | 4649 | 4806 | 5234 | 5127.83 | 5360 | 4896 | 0.913 | -0.131 | 0.2416667 | 0.45839653 | 0.0092 | 0.0075 | 0.0088 | 0.0088 | TRUE | 0.1896 |
| Mrpl27 | 1383 | 1660 | 1559 | 1851 | 1297 | 1389 | 1469 | 1658 | 1565 | 1455 | 1395 | 1520 | 1510.56 | 1564 | 1457 | 0.931 | -0.103 | 0.23687415 | 0.45221065 | 0.002 | 0.0091 | 0.0048 | 0.0048 | TRUE | 0.0339 |
| Mrpl28 | 4867 | 5477 | 5185 | 5676 | 4167 | 4317 | 5170 | 5470 | 5206 | 4462 | 4483 | 4726 | 4919.49 | 5282 | 4557 | 0.863 | -0.213 | 0.00122322 | 0.01039472 | 7.00E-04 | 0.0076 | 0.0029 | 0.0029 | TRUE | 0.0127 |
| Mrpl3 | 2609 | 2633 | 2562 | 3858 | 2542 | 2454 | 2771 | 2630 | 2573 | 3033 | 2735 | 2686 | 2737.92 | 2658 | 2818 | 1.061 | 0.085 | 0.30469177 | 0.52525943 | 0.0024 | 0.0081 | 0.0046 | 0.0046 | TRUE | 0.0548 |
| Mrpl30 | 2166 | 2645 | 2675 | 2999 | 2114 | 2312 | 2301 | 2642 | 2686 | 2358 | 2275 | 2531 | 2465.2 | 2543 | 2388 | 0.939 | -0.091 | 0.33533421 | 0.55561415 | 0.0043 | 0.0082 | 0.006 | 0.006 | TRUE | 0.0839 |
| Mrpl32 | 1399 | 1794 | 1741 | 1859 | 1349 | 1451 | 1486 | 1792 | 1748 | 1461 | 1451 | 1588 | 1587.83 | 1675 | 1500 | 0.895 | -0.16 | 0.12189163 | 0.29685061 | 0.0052 | 0.009 | 0.007 | 0.007 | TRUE | 0.1174 |
| Mrpl33 | 1285 | 1428 | 1265 | 2006 | 1369 | 1233 | 1365 | 1426 | 1270 | 1577 | 1473 | 1350 | 1410.15 | 1354 | 1467 | 1.084 | 0.117 | 0.23228373 | 0.44654725 | 0.0041 | 0.0092 | 0.0062 | 0.0062 | TRUE | 0.0582 |
| Mrpl34 | 1969 | 1580 | 1471 | 2453 | 1689 | 1527 | 2091 | 1578 | 1477 | 1928 | 1817 | 1672 | 1760.6 | 1715 | 1806 | 1.053 | 0.075 | 0.60372887 | 0.77458812 | 0.0194 | 0.0088 | 0.0143 | 0.0143 | TRUE | 0.4428 |
| Mrpl35 | 2499 | 2778 | 2695 | 3369 | 2330 | 2360 | 2654 | 2774 | 2706 | 2648 | 2507 | 2583 | 2645.6 | 2711 | 2579 | 0.952 | -0.072 | 0.28420833 | 0.50581513 | 3.00E-04 | 0.0081 | 0.0028 | 0.0028 | TRUE | 0.0074 |
| Mrpl36 | 1351 | 1147 | 1192 | 1887 | 1267 | 1200 | 1435 | 1146 | 1197 | 1483 | 1363 | 1314 | 1322.94 | 1259 | 1387 | 1.102 | 0.14 | 0.22394983 | 0.4361201 | 0.0084 | 0.0094 | 0.0088 | 0.0088 | TRUE | 0.1786 |
| Mrpl37 | 3087 | 2833 | 2731 | 4343 | 2958 | 2669 | 3279 | 2829 | 2742 | 3414 | 3183 | 2922 | 3061.48 | 2950 | 3173 | 1.076 | 0.106 | 0.31409672 | 0.53452022 | 0.0073 | 0.008 | 0.0076 | 0.0076 | TRUE | 0.1152 |
| Mrpl38 | 4239 | 5104 | 5154 | 5616 | 4204 | 4173 | 4503 | 5097 | 5175 | 4415 | 4523 | 4568 | 4713.54 | 4925 | 4502 | 0.914 | -0.13 | 0.11007762 | 0.27845135 | 0.0026 | 0.0076 | 0.0045 | 0.0045 | TRUE | 0.0685 |
| Mrpl39 | 3987 | 4279 | 4330 | 5177 | 3613 | 3568 | 4235 | 4273 | 4348 | 4070 | 3887 | 3906 | 4119.83 | 4285 | 3954 | 0.923 | -0.116 | 0.05913011 | 0.18413436 | 2.00E-04 | 0.0077 | 0.0025 | 0.0025 | TRUE | 0.0081 |
| Mrpl4 | 2936 | 2999 | 2874 | 3855 | 2775 | 2733 | 3119 | 2995 | 2886 | 3030 | 2986 | 2992 | 3001.23 | 3000 | 3003 | 1.001 | 0.002 | 0.98094266 | 0.98961586 | 4.00E-04 | 0.008 | 0.0029 | 0.0029 | TRUE | 0.0145 |
| Mrpl40 | 2171 | 2487 | 2608 | 3537 | 2501 | 2565 | 2306 | 2484 | 2619 | 2781 | 2691 | 2808 | 2614.62 | 2470 | 2760 | 1.117 | 0.16 | 0.04297542 | 0.14812247 | 0.0017 | 0.0082 | 0.0041 | 0.0041 | TRUE | 0.0405 |
| Mrpl41 | 543 | 634 | 614 | 876 | 574 | 586 | 577 | 633 | 617 | 689 | 618 | 641 | 629.03 | 609 | 649 | 1.067 | 0.094 | 0.35327726 | 0.57170972 | 0.0013 | 0.0121 | 0.0058 | 0.0058 | TRUE | 0.0349 |
| Mrpl42 | 5190 | 5466 | 5329 | 6134 | 4274 | 4667 | 5513 | 5459 | 5351 | 4822 | 4599 | 5109 | 5141.99 | 5441 | 4843 | 0.89 | -0.168 | 0.016592 | 0.07624971 | 0.0012 | 0.0075 | 0.0033 | 0.0033 | TRUE | 0.0277 |
| Mrpl43 | 2118 | 2183 | 2089 | 3101 | 2076 | 1917 | 2250 | 2180 | 2098 | 2438 | 2234 | 2098 | 2216.22 | 2176 | 2257 | 1.038 | 0.054 | 0.54051163 | 0.72639311 | 0.0028 | 0.0084 | 0.0051 | 0.0051 | TRUE | 0.0607 |
| Mrpl44 | 2098 | 1978 | 1946 | 2996 | 2168 | 2015 | 2228 | 1975 | 1954 | 2355 | 2333 | 2206 | 2175.25 | 2052 | 2298 | 1.12 | 0.163 | 0.059292 | 0.18444842 | 0.0027 | 0.0084 | 0.0049 | 0.0049 | TRUE | 0.0677 |
| Mrpl45 | 8573 | 8265 | 8228 | 10558 | 7846 | 7877 | 9106 | 8254 | 8262 | 8300 | 8442 | 8623 | 8497.74 | 8541 | 8455 | 0.99 | -0.015 | 0.83698417 | 0.91717601 | 0.0014 | 0.0073 | 0.0035 | 0.0035 | TRUE | 0.0409 |
| Mrpl46 | 1382 | 1248 | 1139 | 2288 | 1336 | 1276 | 1468 | 1246 | 1144 | 1799 | 1437 | 1397 | 1415.15 | 1286 | 1544 | 1.202 | 0.265 | 0.05857937 | 0.18288804 | 0.0175 | 0.0092 | 0.0135 | 0.0135 | TRUE | 0.253 |
| Mrpl47 | 2400 | 2689 | 2635 | 2864 | 2065 | 2195 | 2549 | 2686 | 2646 | 2251 | 2222 | 2403 | 2459.43 | 2627 | 2292 | 0.872 | -0.197 | 0.00648548 | 0.03813907 | 6.00E-04 | 0.0082 | 0.0034 | 0.0034 | TRUE | 0.0213 |
| Mrpl48 | 1138 | 1580 | 1490 | 1725 | 1275 | 1320 | 1209 | 1578 | 1496 | 1356 | 1372 | 1445 | 1409.28 | 1428 | 1391 | 0.974 | -0.038 | 0.7471495 | 0.8655848 | 0.0095 | 0.0092 | 0.0094 | 0.0094 | TRUE | 0.2156 |
| Mrpl49 | 2665 | 2461 | 2398 | 3394 | 2509 | 2439 | 2831 | 2458 | 2408 | 2668 | 2700 | 2670 | 2622.31 | 2566 | 2679 | 1.044 | 0.063 | 0.48049894 | 0.67974072 | 0.0033 | 0.0082 | 0.0053 | 0.0053 | TRUE | 0.0989 |
| Mrpl50 | 2320 | 2242 | 2232 | 3052 | 2244 | 2170 | 2464 | 2239 | 2241 | 2399 | 2414 | 2375 | 2355.6 | 2315 | 2396 | 1.035 | 0.05 | 0.50482122 | 0.69965204 | 0.001 | 0.0083 | 0.0036 | 0.0036 | TRUE | 0.0385 |
| Mrpl51 | 2615 | 3147 | 2953 | 4286 | 3021 | 2797 | 2778 | 3143 | 2965 | 3369 | 3250 | 3062 | 3094.52 | 2962 | 3227 | 1.09 | 0.124 | 0.13612379 | 0.31954629 | 0.0026 | 0.008 | 0.0047 | 0.0047 | TRUE | 0.0359 |
| Mrpl52 | 1314 | 1693 | 1562 | 1902 | 1367 | 1408 | 1396 | 1691 | 1568 | 1495 | 1471 | 1541 | 1527.04 | 1552 | 1502 | 0.968 | -0.047 | 0.63036713 | 0.79245579 | 0.0042 | 0.0091 | 0.0062 | 0.0062 | TRUE | 0.0927 |
| Mrpl53 | 800 | 1009 | 967 | 1023 | 729 | 789 | 850 | 1008 | 971 | 804 | 784 | 864 | 880.11 | 943 | 817 | 0.867 | -0.207 | 0.04939093 | 0.1629694 | 0.004 | 0.0106 | 0.0068 | 0.0068 | TRUE | 0.0883 |
| Mrpl54 | 620 | 863 | 861 | 1015 | 779 | 806 | 659 | 862 | 865 | 798 | 838 | 882 | 817.22 | 795 | 839 | 1.055 | 0.077 | 0.55875208 | 0.74101748 | 0.0116 | 0.0109 | 0.0114 | 0.0114 | TRUE | 0.266 |
| Mrpl55 | 995 | 1057 | 1060 | 1701 | 1077 | 1046 | 1057 | 1056 | 1064 | 1337 | 1159 | 1145 | 1136.31 | 1059 | 1214 | 1.148 | 0.199 | 0.03953116 | 0.13942429 | 0.0032 | 0.0098 | 0.0058 | 0.0058 | TRUE | 0.0977 |
| Mrpl9 | 3062 | 3543 | 3555 | 4053 | 2828 | 3008 | 3252 | 3538 | 3570 | 3186 | 3043 | 3293 | 3313.67 | 3453 | 3174 | 0.919 | -0.122 | 0.11142319 | 0.28068465 | 0.0016 | 0.0079 | 0.0039 | 0.0039 | TRUE | 0.0314 |
| Mrps10 | 2271 | 2581 | 2524 | 3026 | 2230 | 2279 | 2412 | 2578 | 2534 | 2379 | 2399 | 2495 | 2466.18 | 2508 | 2424 | 0.966 | -0.049 | 0.47912779 | 0.67843674 | 5.00E-04 | 0.0082 | 0.0031 | 0.0031 | TRUE | 0.0135 |
| Mrps11 | 1155 | 983 | 927 | 1649 | 1069 | 900 | 1227 | 982 | 931 | 1296 | 1150 | 985 | 1095.17 | 1047 | 1144 | 1.094 | 0.13 | 0.37985504 | 0.59453654 | 0.0179 | 0.0099 | 0.0148 | 0.0148 | TRUE | 0.2711 |
| Mrps12 | 1762 | 1578 | 1555 | 2022 | 1593 | 1453 | 1872 | 1576 | 1561 | 1590 | 1714 | 1591 | 1650.49 | 1670 | 1632 | 0.977 | -0.033 | 0.7442793 | 0.86347991 | 0.0056 | 0.0089 | 0.007 | 0.007 | TRUE | 0.1346 |
| Mrps14 | 2440 | 2526 | 2321 | 3434 | 2253 | 2328 | 2592 | 2523 | 2331 | 2700 | 2424 | 2548 | 2519.5 | 2482 | 2557 | 1.031 | 0.044 | 0.59952809 | 0.77164657 | 0.0026 | 0.0082 | 0.0047 | 0.0047 | TRUE | 0.0344 |

| | | | | | | | | | | | | | | | | | | | | | | | | | |
|---|---|---|---|---|---|---|---|---|---|---|---|---|---|---|---|---|---|---|---|---|---|---|---|---|---|
| Mrps15 | 1388 | 1639 | 1659 | 2357 | 1761 | 1620 | 1474 | 1637 | 1666 | 1853 | 1895 | 1773 | 1716.33 | 1592 | 1840 | 1.156 | 0.209 | 0.0141743 | 0.0679328 | 0.0018 | 0.0088 | 0.0046 | 0.0046 | TRUE | 0.0504 |
| Mrps16 | 1557 | 1974 | 1984 | 2293 | 1682 | 1627 | 1654 | 1971 | 1992 | 1803 | 1810 | 1781 | 1835.12 | 1872 | 1798 | 0.96 | -0.059 | 0.54743199 | 0.73246078 | 0.0048 | 0.0087 | 0.0064 | 0.0064 | TRUE | 0.1256 |
| Mrps17 | 2249 | 2343 | 2304 | 3011 | 1966 | 1886 | 2389 | 2340 | 2313 | 2367 | 2115 | 2065 | 2264.85 | 2347 | 2182 | 0.93 | -0.104 | 0.21495376 | 0.42675287 | 0.0022 | 0.0084 | 0.0046 | 0.0046 | TRUE | 0.0677 |
| Mrps18a | 1631 | 1883 | 1813 | 2394 | 1740 | 1821 | 1732 | 1881 | 1820 | 1882 | 1872 | 1993 | 1863.48 | 1811 | 1916 | 1.057 | 0.081 | 0.29365591 | 0.51450133 | 9.00E-04 | 0.0087 | 0.0037 | 0.0037 | TRUE | 0.0174 |
| Mrps18b | 2722 | 2535 | 2538 | 3494 | 2471 | 2272 | 2891 | 2532 | 2548 | 2747 | 2659 | 2487 | 2643.97 | 2657 | 2631 | 0.991 | -0.014 | 0.87816797 | 0.93976544 | 0.0037 | 0.0081 | 0.0054 | 0.0054 | TRUE | 0.0719 |
| Mrps18c | 1042 | 1126 | 1019 | 1620 | 1178 | 1127 | 1107 | 1125 | 1023 | 1274 | 1267 | 1234 | 1171.53 | 1085 | 1258 | 1.16 | 0.214 | 0.0093588 | 0.05003628 | 4.00E-04 | 0.0097 | 0.004 | 0.004 | TRUE | 0.0297 |
| Mrps2 | 3643 | 3958 | 3870 | 5273 | 3783 | 3902 | 3870 | 3953 | 3886 | 4145 | 4070 | 4271 | 4032.52 | 3903 | 4162 | 1.066 | 0.093 | 0.12710104 | 0.30619768 | 0 | 0.0077 | 0.0024 | 0.0024 | TRUE | 0.0063 |
| Mrps21 | 838 | 842 | 865 | 1138 | 811 | 789 | 890 | 841 | 869 | 895 | 873 | 864 | 871.74 | 867 | 877 | 1.013 | 0.018 | 0.82805914 | 0.91291407 | 0 | 0.0107 | 0.0039 | 0.0039 | TRUE | 0.008 |
| Mrps22 | 1882 | 1888 | 1904 | 2620 | 1915 | 1884 | 1999 | 1886 | 1912 | 2060 | 2060 | 2062 | 1996.47 | 1932 | 2061 | 1.067 | 0.093 | 0.17387351 | 0.3731541 | 0 | 0.0086 | 0.0029 | 0.0029 | TRUE | 0.011 |
| Mrps23 | 3132 | 3457 | 3263 | 4278 | 3098 | 2908 | 3327 | 3453 | 3276 | 3363 | 3333 | 3183 | 3322.54 | 3352 | 3293 | 0.983 | -0.025 | 0.70473713 | 0.83847762 | 5.00E-04 | 0.0079 | 0.0029 | 0.0029 | TRUE | 0.0103 |
| Mrps24 | 2703 | 3449 | 3338 | 3588 | 2756 | 3043 | 2871 | 3445 | 3352 | 2821 | 2965 | 3331 | 3130.71 | 3223 | 3039 | 0.943 | -0.085 | 0.42884193 | 0.63860634 | 0.0074 | 0.008 | 0.008 | 0.008 | TRUE | 0.1104 |
| Mrps25 | 4122 | 3731 | 3605 | 4962 | 3489 | 3174 | 4378 | 3726 | 3620 | 3901 | 3754 | 3474 | 3808.9 | 3908 | 3710 | 0.95 | -0.075 | 0.46244577 | 0.66603001 | 0.0065 | 0.0078 | 0.0072 | 0.0072 | TRUE | 0.1346 |
| Mrps26 | 1572 | 1576 | 1458 | 2236 | 1534 | 1430 | 1670 | 1574 | 1464 | 1758 | 1650 | 1565 | 1613.55 | 1569 | 1658 | 1.057 | 0.08 | 0.38257678 | 0.59692727 | 0.0029 | 0.0089 | 0.0055 | 0.0055 | TRUE | 0.0415 |
| Mrps27 | 4551 | 4246 | 4514 | 5678 | 4106 | 4005 | 4834 | 4241 | 4533 | 4464 | 4418 | 4384 | 4478.75 | 4536 | 4422 | 0.975 | -0.036 | 0.63065889 | 0.79261236 | 0.0018 | 0.0076 | 0.0039 | 0.0039 | TRUE | 0.0402 |
| Mrps28 | 1345 | 1488 | 1536 | 1708 | 1280 | 1294 | 1429 | 1486 | 1542 | 1343 | 1377 | 1416 | 1432.23 | 1486 | 1379 | 0.928 | -0.109 | 0.16181105 | 0.35694259 | 4.00E-04 | 0.0092 | 0.0036 | 0.0036 | TRUE | 0.0135 |
| Mrps30 | 3641 | 4160 | 4332 | 5342 | 3848 | 4074 | 3867 | 4155 | 4350 | 4199 | 4140 | 4460 | 4195.18 | 4124 | 4266 | 1.034 | 0.049 | 0.53396449 | 0.72173954 | 0.0021 | 0.0077 | 0.0042 | 0.0042 | TRUE | 0.036 |
| Mrps31 | 3775 | 4356 | 4525 | 4816 | 3570 | 3737 | 4010 | 4350 | 4544 | 3786 | 3841 | 4091 | 4103.58 | 4301 | 3906 | 0.908 | -0.139 | 0.08382561 | 0.23198764 | 0.0025 | 0.0077 | 0.0044 | 0.0044 | TRUE | 0.0427 |
| Mrps33 | 541 | 588 | 692 | 785 | 534 | 511 | 575 | 587 | 695 | 617 | 575 | 559 | 601.29 | 619 | 584 | 0.944 | -0.083 | 0.48003693 | 0.67933086 | 0.0053 | 0.0124 | 0.0084 | 0.0084 | TRUE | 0.1359 |
| Mrps34 | 965 | 1070 | 1041 | 1315 | 1003 | 882 | 1025 | 1069 | 1045 | 1034 | 1079 | 965 | 1036.22 | 1046 | 1026 | 0.981 | -0.028 | 0.74692867 | 0.8655848 | 6.00E-04 | 0.0101 | 0.0044 | 0.0044 | TRUE | 0.0308 |
| Mrps35 | 780 | 1011 | 1045 | 1298 | 947 | 1089 | 828 | 1010 | 1049 | 1020 | 1019 | 1192 | 1019.81 | 962 | 1077 | 1.118 | 0.161 | 0.20173885 | 0.40927453 | 0.0103 | 0.0101 | 0.0105 | 0.0105 | TRUE | 0.1761 |
| Mrps36 | 610 | 677 | 683 | 744 | 574 | 633 | 648 | 676 | 686 | 585 | 618 | 693 | 650.87 | 670 | 632 | 0.941 | -0.087 | 0.41510679 | 0.62663326 | 0.0027 | 0.012 | 0.0067 | 0.0067 | TRUE | 0.0834 |
| Mrps5 | 5403 | 5354 | 5332 | 6366 | 4658 | 4621 | 5739 | 5347 | 5354 | 5004 | 5012 | 5058 | 5252.42 | 5480 | 5025 | 0.917 | -0.125 | 0.05180804 | 0.16792362 | 6.00E-04 | 0.0075 | 0.0028 | 0.0028 | TRUE | 0.0208 |
| Mrps6 | 962 | 736 | 670 | 986 | 707 | 616 | 1022 | 735 | 673 | 775 | 761 | 674 | 773.29 | 810 | 737 | 0.91 | -0.136 | 0.41752968 | 0.62838739 | 0.0266 | 0.0112 | 0.0189 | 0.0189 | TRUE | 0.6205 |
| Mrps7 | 3103 | 2986 | 2921 | 4484 | 3130 | 2886 | 3296 | 2982 | 2933 | 3525 | 3368 | 3159 | 3210.48 | 3070 | 3351 | 1.092 | 0.127 | 0.13943859 | 0.32412923 | 0.0031 | 0.0079 | 0.005 | 0.005 | TRUE | 0.0501 |
| Mrps9 | 3266 | 3718 | 3606 | 4976 | 3591 | 3582 | 3469 | 3713 | 3621 | 3912 | 3864 | 3921 | 3749.93 | 3601 | 3899 | 1.083 | 0.115 | 0.07323186 | 0.21204568 | 3.00E-04 | 0.0078 | 0.0027 | 0.0027 | TRUE | 0.0124 |
| Mrrf | 1282 | 1418 | 1469 | 1603 | 1185 | 1190 | 1362 | 1416 | 1475 | 1260 | 1275 | 1303 | 1348.45 | 1418 | 1279 | 0.902 | -0.149 | 0.05491423 | 0.17478428 | 0 | 0.0093 | 0.0036 | 0.0036 | TRUE | 0.0151 |
| Mrs2 | 1017 | 1357 | 1327 | 1519 | 1203 | 1193 | 1080 | 1355 | 1332 | 1194 | 1294 | 1306 | 1260.39 | 1256 | 1265 | 1.006 | 0.009 | 0.93702435 | 0.96852664 | 0.0081 | 0.0095 | 0.0087 | 0.0087 | TRUE | 0.1791 |
| Mrto4 | 4731 | 4668 | 4578 | 6498 | 4362 | 3917 | 5025 | 4662 | 4597 | 5108 | 4693 | 4288 | 4728.86 | 4761 | 4696 | 0.987 | -0.019 | 0.83374979 | 0.91554004 | 0.0048 | 0.0076 | 0.0059 | 0.0059 | TRUE | 0.0722 |
| Mrvi1 | 30 | 20 | 21 | 31 | 15 | 18 | 32 | 20 | 21 | 24 | 16 | 20 | 22.19 | 24 | 20 | 0.834 | -0.262 | 0.54142292 | 0.72706022 | 0 | 0.1554 | 0.0876 | 0.0876 | TRUE | 0.4288 |
| Ms4a1 | 0 | 0 | 0 | 1 | 0 | 1 | 0 | 0 | 0 | 1 | 0 | 1 | 0.31 | 0 | 1 | 3.348 | 1.743 | 0.65064889 | NA | 0 | 10.5167 | 8.8968 | 8.8968 | TRUE | 0.2082 |
| Ms4a10 | 5 | 0 | 1 | 4 | 3 | 5 | 5 | 0 | 1 | 3 | 3 | 5 | 3.03 | 2 | 4 | 1.887 | 0.916 | 0.4747559 | NA | 0.3788 | 1.0953 | 0.8142 | 0.8142 | TRUE | 1.5934 |
| Ms4a13 | 3 | 0 | 2 | 1 | 0 | 0 | 3 | 0 | 2 | 1 | 0 | 0 | 1 | 2 | 0 | 0.202 | -2.305 | 0.32045215 | NA | 0 | 3.3119 | 2.584 | 2.584 | TRUE | 0.1559 |
| Ms4a15 | 44 | 78 | 58 | 68 | 67 | 77 | 47 | 78 | 58 | 53 | 72 | 84 | 65.45 | 61 | 70 | 1.14 | 0.189 | 0.53295695 | 0.72090883 | 0.0438 | 0.0572 | 0.0506 | 0.0506 | TRUE | 0.4024 |
| Ms4a18 | 0 | 0 | 0 | 0 | 0 | 0 | 0 | 0 | 0 | 0 | 0 | 0 | 0 | 0 | 0 | NA | NA | NA | NA | NA | NA | NA | NA | NA | NA |
| Ms4a2 | 0 | 0 | 0 | 0 | 0 | 0 | 0 | 0 | 0 | 0 | 0 | 0 | 0 | 0 | 0 | NA | NA | NA | NA | NA | NA | NA | NA | NA | NA |
| Ms4a3 | 0 | 0 | 0 | 0 | 0 | 0 | 0 | 0 | 0 | 0 | 0 | 0 | 0 | 0 | 0 | NA | NA | NA | NA | NA | NA | NA | NA | NA | NA |
| Ms4a4b | 0 | 0 | 0 | 0 | 0 | 0 | 0 | 0 | 0 | 0 | 0 | 0 | 0 | 0 | 0 | NA | NA | NA | NA | NA | NA | NA | NA | NA | NA |
| Ms4a4c | 0 | 0 | 0 | 0 | 0 | 0 | 0 | 0 | 0 | 0 | 0 | 0 | 0 | 0 | 0 | NA | NA | NA | NA | NA | NA | NA | NA | NA | NA |
| Ms4a4d | 44 | 20 | 11 | 44 | 27 | 19 | 47 | 20 | 11 | 35 | 29 | 21 | 27.03 | 26 | 28 | 1.093 | 0.129 | 0.81472706 | 0.9053671 | 0.2365 | 0.1288 | 0.1803 | 0.1803 | TRUE | 1.0655 |
| Ms4a5 | 2 | 1 | 1 | 3 | 3 | 0 | 2 | 1 | 1 | 2 | 3 | 0 | 1.62 | 1 | 2 | 1.374 | 0.458 | 0.76781712 | NA | 0 | 2.0419 | 1.1031 | 1.1031 | TRUE | 0.5605 |
| Ms4a6b | 0 | 0 | 0 | 0 | 0 | 0 | 0 | 0 | 0 | 0 | 0 | 0 | 0 | 0 | 0 | NA | NA | NA | NA | NA | NA | NA | NA | NA | NA |
| Ms4a6c | 0 | 0 | 0 | 0 | 0 | 0 | 0 | 0 | 0 | 0 | 0 | 0 | 0 | 0 | 0 | NA | NA | NA | NA | NA | NA | NA | NA | NA | NA |
| Ms4a6d | 9 | 12 | 11 | 8 | 8 | 15 | 10 | 12 | 11 | 6 | 9 | 16 | 10.65 | 11 | 10 | 0.946 | -0.081 | 0.89668245 | 0.95009983 | 0 | 0.3162 | 0.1831 | 0.1831 | TRUE | 0.8437 |
| Ms4a7 | 1 | 3 | 1 | 0 | 0 | 1 | 1 | 3 | 1 | 0 | 0 | 1 | 1.03 | 2 | 0 | 0.206 | -2.276 | 0.26774836 | NA | 0 | 3.2173 | 1.7372 | 1.7372 | TRUE | 0.5383 |
| Ms4a8a | 0 | 0 | 0 | 1 | 0 | 0 | 0 | 0 | 0 | 1 | 0 | 0 | 0.13 | 0 | 0 | 1.859 | 0.894 | 0.82650734 | NA | 0 | 25.1515 | 10 | 10 | TRUE | 0.2563 |
| Msantd1 | 2 | 4 | 0 | 5 | 0 | 2 | 2 | 4 | 0 | 4 | 0 | 2 | 2.04 | 2 | 2 | 1.023 | 0.033 | 0.98399467 | NA | 1.1005 | 1.6219 | 1.4316 | 1.4316 | TRUE | 0.2203 |
| Msantd2 | 1478 | 1329 | 1379 | 1584 | 1307 | 1158 | 1570 | 1327 | 1385 | 1245 | 1406 | 1268 | 1366.82 | 1427 | 1306 | 0.915 | -0.128 | 0.21528291 | 0.42717126 | 0.0051 | 0.0093 | 0.007 | 0.007 | TRUE | 0.0916 |
| Msantd3 | 1571 | 1514 | 1574 | 1381 | 1066 | 1057 | 1669 | 1512 | 1580 | 1086 | 1147 | 1157 | 1358.47 | 1587 | 1130 | 0.712 | -0.491 | 3.66E-09 | 1.91E-07 | 0.0012 | 0.0093 | 0.0042 | 0.0042 | TRUE | 0.0243 |
| Msantd4 | 2135 | 2089 | 1998 | 3201 | 2114 | 2141 | 2268 | 2086 | 2006 | 2516 | 2275 | 2344 | 2249.14 | 2120 | 2378 | 1.122 | 0.167 | 0.05461048 | 0.17410453 | 0.0027 | 0.0084 | 0.005 | 0.005 | TRUE | 0.0448 |
| Msc | 426 | 189 | 163 | 433 | 335 | 224 | 452 | 189 | 164 | 340 | 360 | 245 | 291.82 | 268 | 315 | 1.176 | 0.234 | 0.49925036 | 0.6949847 | 0.1712 | 0.0182 | 0.0833 | 0.0833 | TRUE | 4.0176 |
| Msgn1 | 6 | 1 | 1 | 3 | 4 | 0 | 6 | 1 | 1 | 2 | 4 | 0 | 2.51 | 3 | 2 | 0.803 | -0.316 | 0.82727599 | NA | 0.8574 | 1.3214 | 1.1086 | 1.1086 | TRUE | 0.5101 |
| Msh2 | 17605 | 18521 | 19215 | 19899 | 15215 | 16533 | 18700 | 18497 | 19294 | 15643 | 16370 | 18098 | 17767 | 18830 | 16704 | 0.887 | -0.173 | 0.02827101 | 0.11053305 | 0.003 | 0.0071 | 0.0044 | 0.0044 | TRUE | 0.0652 |
| Msh3 | 3082 | 2371 | 2500 | 3942 | 2763 | 2475 | 3274 | 2368 | 2510 | 3099 | 2973 | 2709 | 2822.14 | 2717 | 2927 | 1.077 | 0.108 | 0.42717673 | 0.63729014 | 0.0152 | 0.0081 | 0.0129 | 0.0129 | TRUE | 0.3889 |
| Msh4 | 23 | 22 | 27 | 39 | 34 | 33 | 24 | 22 | 27 | 31 | 37 | 36 | 29.48 | 24 | 35 | 1.4 | 0.486 | 0.16501446 | 0.36167622 | 0 | 0.1187 | 0.0532 | 0.0532 | TRUE | 0.0783 |
| Msh5 | 70 | 141 | 133 | 103 | 123 | 106 | 74 | 141 | 134 | 81 | 132 | 116 | 113.01 | 116 | 110 | 0.94 | -0.089 | 0.76634132 | 0.87779299 | 0.0779 | 0.0361 | 0.056 | 0.056 | TRUE | 0.984 |
| Msh6 | 35453 | 35746 | 37335 | 45245 | 31754 | 31150 | 37657 | 35700 | 37488 | 35568 | 34165 | 34099 | 35779.58 | 36948 | 34611 | 0.937 | -0.094 | 0.10939988 | 0.27744894 | 7.00E-04 | 0.007 | 0.0025 | 0.0025 | TRUE | 0.0107 |
| Msi1 | 675 | 1052 | 1039 | 925 | 762 | 933 | 717 | 1051 | 1043 | 727 | 820 | 1021 | 896.54 | 937 | 856 | 0.913 | -0.132 | 0.47288228 | 0.67393434 | 0.0363 | 0.0106 | 0.0231 | 0.0231 | TRUE | 0.5015 |
| Msi2 | 5628 | 6467 | 6901 | 6070 | 4793 | 4755 | 5978 | 6459 | 6929 | 4772 | 5157 | 5205 | 5749.96 | 6455 | 5045 | 0.781 | -0.356 | 3.16E-05 | 0.00052596 | 0.0036 | 0.0075 | 0.0051 | 0.0051 | TRUE | 0.051 |
| Msl1 | 2808 | 2869 | 2871 | 4046 | 2934 | 2972 | 2983 | 2865 | 2883 | 3181 | 3157 | 3253 | 3053.57 | 2910 | 3197 | 1.099 | 0.136 | 0.03145759 | 0.11915204 | 0 | 0.008 | 0.0025 | 0.0025 | TRUE | 0.0057 |
| Msl2 | 1929 | 2133 | 2201 | 2806 | 2038 | 2040 | 2049 | 2130 | 2210 | 2206 | 2193 | 2233 | 2170.16 | 2130 | 2211 | 1.038 | 0.054 | 0.44361607 | 0.65031254 | 3.00E-04 | 0.0084 | 0.0031 | 0.0031 | TRUE | 0.0133 |
| Msl3 | 266 | 321 | 364 | 409 | 319 | 309 | 283 | 321 | 365 | 322 | 343 | 338 | 328.6 | 323 | 334 | 1.034 | 0.048 | 0.72839749 | 0.85395791 | 0.0056 | 0.0169 | 0.0109 | 0.0109 | TRUE | 0.1514 |
| Msl3l2 | 167 | 270 | 222 | 447 | 365 | 242 | 177 | 270 | 223 | 351 | 393 | 265 | 279.83 | 223 | 336 | 1.506 | 0.591 | 0.00465971 | 0.02970615 | 0.0371 | 0.0187 | 0.0277 | 0.0277 | TRUE | 0.3097 |
| Msln | 19 | 7 | 6 | 33 | 23 | 26 | 20 | 7 | 6 | 26 | 25 | 28 | 18.72 | 11 | 26 | 2.398 | 1.262 | 0.02188938 | 0.09290615 | 0.1086 | 0.1828 | 0.1533 | 0.1533 | TRUE | 1.858 |
| Mslnl | 7 | 7 | 5 | 20 | 16 | 19 | 7 | 7 | 5 | 16 | 17 | 21 | 12.2 | 6 | 18 | 2.756 | 1.463 | 0.0102665 | 0.05379508 | 0 | 0.277 | 0.1277 | 0.1277 | TRUE | 0.0986 |
| Msmb | 0 | 1 | 0 | 0 | 0 | 0 | 0 | 1 | 0 | 0 | 0 | 0 | 0.17 | 0 | 0 | 0.49 | -1.029 | 0.80087 | NA | 0 | 19.7991 | 10 | 10 | TRUE | 0.405 |
| Msmp | 1 | 0 | 1 | 0 | 0 | 0 | 1 | 0 | 1 | 0 | 0 | 0 | 0.34 | 1 | 0 | 0.261 | -1.939 | 0.60367425 | NA | 0 | 9.5732 | 8.3134 | 8.3134 | TRUE | 0.2529 |
| Msn | 7233 | 7260 | 7241 | 8347 | 5990 | 6044 | 7683 | 7251 | 7271 | 6562 | 6445 | 6616 | 6971.14 | 7402 | 6541 | 0.884 | -0.178 | 0.00341032 | 0.02338589 | 4.00E-04 | 0.0074 | 0.0025 | 0.0025 | TRUE | 0.0135 |
| Msr1 | 0 | 0 | 0 | 0 | 1 | 0 | 0 | 0 | 0 | 0 | 1 | 0 | 0.18 | 0 | 0 | 1.859 | 0.894 | 0.82650734 | NA | 0 | 18.3785 | 10 | 10 | TRUE | 0.5204 |
| Msra | 228 | 329 | 320 | 371 | 319 | 264 | 242 | 329 | 321 | 292 | 343 | 289 | 302.66 | 297 | 308 | 1.034 | 0.048 | 0.76901261 | 0.87928573 | 0.0149 | 0.0178 | 0.0163 | 0.0163 | TRUE | 0.294 |

| | | | | | | | | | | | | | | | | | | | | | | | | | |
|---|---|---|---|---|---|---|---|---|---|---|---|---|---|---|---|---|---|---|---|---|---|---|---|---|---|
| Msrb1 | 947 | 916 | 919 | 1172 | 806 | 805 | 1006 | 915 | 923 | 921 | 867 | 881 | 918.87 | 948 | 890 | 0.94 | -0.09 | 0.31668474 | 0.53753247 | 9.00E-04 | 0.0105 | 0.0047 | 0.0047 | TRUE | 0.034 |
| Msrb2 | 423 | 280 | 253 | 476 | 349 | 340 | 449 | 280 | 254 | 374 | 376 | 372 | 350.81 | 328 | 374 | 1.142 | 0.192 | 0.36976941 | 0.58696949 | 0.0449 | 0.0163 | 0.03 | 0.03 | TRUE | 1.1923 |
| Msrb3 | 329 | 285 | 265 | 527 | 312 | 277 | 349 | 285 | 266 | 414 | 336 | 303 | 325.56 | 300 | 351 | 1.173 | 0.231 | 0.18462073 | 0.38767649 | 0.0203 | 0.017 | 0.0187 | 0.0187 | TRUE | 0.2941 |
| Mss51 | 70 | 64 | 60 | 65 | 44 | 55 | 74 | 64 | 60 | 51 | 47 | 60 | 59.53 | 66 | 53 | 0.799 | -0.325 | 0.20773736 | 0.4170613 | 0 | 0.0622 | 0.0309 | 0.0309 | TRUE | 0.1115 |
| Mst1 | 15 | 43 | 30 | 49 | 27 | 45 | 16 | 43 | 30 | 39 | 29 | 49 | 34.31 | 30 | 39 | 1.309 | 0.388 | 0.36959464 | 0.5868566 | 0.1081 | 0.1029 | 0.1053 | 0.1053 | TRUE | 0.2718 |
| Mst1r | 47 | 32 | 30 | 34 | 16 | 26 | 50 | 32 | 30 | 27 | 17 | 28 | 30.73 | 37 | 24 | 0.65 | -0.622 | 0.11571591 | 0.28727709 | 0.0401 | 0.1141 | 0.0788 | 0.0788 | TRUE | 0.618 |
| Mstn | 0 | 0 | 0 | 0 | 0 | 0 | 0 | 0 | 0 | 0 | 0 | 0 | 0 | 0 | 0 | NA | NA | NA | NA | NA | NA | NA | NA | NA | NA |
| Msto1 | 4214 | 4476 | 4574 | 5313 | 3946 | 3995 | 4476 | 4470 | 4593 | 4177 | 4246 | 4373 | 4389.08 | 4513 | 4265 | 0.945 | -0.082 | 0.17698311 | 0.37747649 | 2.00E-04 | 0.0077 | 0.0024 | 0.0024 | TRUE | 0.0059 |
| Msx1 | 19 | 20 | 19 | 36 | 34 | 28 | 20 | 20 | 19 | 28 | 37 | 31 | 25.79 | 20 | 32 | 1.607 | 0.684 | 0.06818846 | 0.20230782 | 0 | 0.1346 | 0.0602 | 0.0602 | TRUE | 0.1114 |
| Msx1as | 0 | 1 | 0 | 0 | 0 | 0 | 0 | 1 | 0 | 0 | 0 | 0 | 0.17 | 0 | 0 | 0.49 | -1.029 | 0.80087 | NA | 0 | 19.7991 | 10 | 10 | TRUE | 0.405 |
| Msx2 | 215 | 146 | 126 | 180 | 101 | 131 | 228 | 146 | 127 | 142 | 109 | 143 | 149.04 | 167 | 131 | 0.788 | -0.344 | 0.18289273 | 0.3850219 | 0.0534 | 0.029 | 0.0414 | 0.0414 | TRUE | 1.0933 |
| Msx3 | 14 | 17 | 11 | 32 | 22 | 21 | 15 | 17 | 11 | 25 | 24 | 23 | 19.12 | 14 | 24 | 1.678 | 0.746 | 0.08833906 | 0.24101084 | 0 | 0.1792 | 0.0821 | 0.0821 | TRUE | 0.1786 |
| Mt1 | 2738 | 4534 | 4692 | 4608 | 4190 | 4682 | 2908 | 4528 | 4711 | 3622 | 4508 | 5125 | 4233.91 | 4049 | 4418 | 1.091 | 0.126 | 0.51246108 | 0.70577508 | 0.0481 | 0.0077 | 0.0263 | 0.0263 | TRUE | 0.6865 |
| Mt2 | 2414 | 4913 | 5071 | 6260 | 5052 | 5525 | 2564 | 4907 | 5092 | 4921 | 5436 | 6048 | 4827.89 | 4188 | 5468 | 1.306 | 0.385 | 0.08356187 | 0.23158281 | 0.0719 | 0.0076 | 0.0354 | 0.0354 | TRUE | 1.3996 |
| Mt3 | 394 | 391 | 348 | 306 | 223 | 235 | 418 | 390 | 349 | 241 | 240 | 257 | 316.03 | 386 | 246 | 0.637 | -0.652 | 7.46E-07 | 2.13E-05 | 0 | 0.0173 | 0.0092 | 0.0092 | TRUE | 0.0796 |
| Mt4 | 19 | 19 | 15 | 19 | 20 | 22 | 20 | 19 | 15 | 15 | 22 | 24 | 19.13 | 18 | 20 | 1.107 | 0.146 | 0.74364106 | 0.86319155 | 0 | 0.1792 | 0.0912 | 0.0912 | TRUE | 0.3489 |
| Mta1 | 4850 | 6353 | 6737 | 6478 | 5123 | 5605 | 5152 | 6345 | 6765 | 5092 | 5512 | 6136 | 5833.52 | 6087 | 5580 | 0.917 | -0.126 | 0.31385596 | 0.53428346 | 0.0136 | 0.0075 | 0.0111 | 0.0111 | TRUE | 0.2203 |
| Mta2 | 5279 | 5302 | 5276 | 7416 | 5317 | 5025 | 5607 | 5295 | 5298 | 5830 | 5721 | 5501 | 5541.89 | 5400 | 5684 | 1.053 | 0.074 | 0.25672393 | 0.47534648 | 7.00E-04 | 0.0075 | 0.0029 | 0.0029 | TRUE | 0.0137 |
| Mta3 | 2757 | 3425 | 3479 | 5006 | 3582 | 3786 | 2928 | 3421 | 3493 | 3935 | 3854 | 4144 | 3629.33 | 3281 | 3978 | 1.212 | 0.278 | 0.00304298 | 0.02146414 | 0.0047 | 0.0078 | 0.0061 | 0.0061 | TRUE | 0.107 |
| Mtag2 | 2 | 3 | 2 | 4 | 5 | 2 | 2 | 3 | 2 | 3 | 5 | 2 | 2.97 | 2 | 3 | 1.496 | 0.581 | 0.58940641 | NA | 0 | 1.1148 | 0.4826 | 0.4826 | TRUE | 0.2675 |
| Mtap | 5411 | 5646 | 5448 | 7270 | 5417 | 5122 | 5747 | 5639 | 5470 | 5715 | 5828 | 5607 | 5667.8 | 5619 | 5717 | 1.017 | 0.025 | 0.67865502 | 0.82146945 | 2.00E-04 | 0.0075 | 0.0024 | 0.0024 | TRUE | 0.0065 |
| Mtap7d3 | 558 | 460 | 489 | 343 | 262 | 241 | 593 | 459 | 491 | 270 | 282 | 264 | 393.08 | 514 | 272 | 0.529 | -0.92 | 4.41E-11 | 3.46E-09 | 0.008 | 0.0153 | 0.0112 | 0.0112 | TRUE | 0.2057 |
| Mtbp | 1243 | 1217 | 1170 | 1839 | 1278 | 1238 | 1320 | 1215 | 1175 | 1446 | 1375 | 1355 | 1314.41 | 1237 | 1392 | 1.126 | 0.172 | 0.04732842 | 0.15788153 | 0.0016 | 0.0094 | 0.0046 | 0.0046 | TRUE | 0.0415 |
| Mtch1 | 2155 | 2144 | 2109 | 3189 | 2226 | 2081 | 2289 | 2141 | 2118 | 2507 | 2395 | 2278 | 2287.97 | 2183 | 2393 | 1.097 | 0.134 | 0.0897882 | 0.24374174 | 0.0016 | 0.0083 | 0.004 | 0.004 | TRUE | 0.0219 |
| Mtch2 | 8369 | 9461 | 9442 | 10670 | 7507 | 7714 | 8889 | 9449 | 9481 | 8388 | 8077 | 8444 | 8788.01 | 9273 | 8303 | 0.895 | -0.159 | 0.01192429 | 0.05993305 | 8.00E-04 | 0.0073 | 0.0028 | 0.0028 | TRUE | 0.016 |
| Mtcp1 | 207 | 232 | 241 | 236 | 214 | 219 | 220 | 232 | 242 | 186 | 230 | 240 | 224.84 | 231 | 219 | 0.941 | -0.087 | 0.58133545 | 0.75775394 | 0.0066 | 0.0216 | 0.0137 | 0.0137 | TRUE | 0.2075 |
| Mtdh | 8530 | 9280 | 9085 | 9462 | 7006 | 7392 | 9060 | 9268 | 9122 | 7438 | 7538 | 8092 | 8419.78 | 9150 | 7689 | 0.84 | -0.251 | 0.00011256 | 0.00151219 | 9.00E-04 | 0.0073 | 0.0029 | 0.0029 | TRUE | 0.0255 |
| Mterf | 119 | 129 | 138 | 200 | 140 | 136 | 126 | 129 | 139 | 157 | 151 | 149 | 141.75 | 131 | 152 | 1.161 | 0.215 | 0.19291 | 0.39845161 | 0 | 0.0301 | 0.0126 | 0.0126 | TRUE | 0.0244 |
| Mterfd1 | 911 | 901 | 896 | 1162 | 803 | 845 | 968 | 900 | 900 | 913 | 864 | 925 | 911.6 | 923 | 901 | 0.977 | -0.034 | 0.7023212 | 0.83715683 | 0 | 0.0105 | 0.0044 | 0.0044 | TRUE | 0.0218 |
| Mterfd2 | 882 | 710 | 734 | 1220 | 819 | 698 | 937 | 709 | 737 | 959 | 881 | 764 | 831.21 | 794 | 868 | 1.094 | 0.13 | 0.37220359 | 0.58880591 | 0.0169 | 0.0109 | 0.014 | 0.014 | TRUE | 0.2913 |
| Mterfd3 | 297 | 263 | 270 | 382 | 254 | 259 | 315 | 263 | 271 | 300 | 273 | 284 | 284.39 | 283 | 286 | 1.011 | 0.016 | 0.90653375 | 0.95352883 | 0.0025 | 0.0185 | 0.01 | 0.01 | TRUE | 0.1112 |
| Mtf1 | 2066 | 1935 | 1701 | 2343 | 1681 | 1682 | 2194 | 1933 | 1708 | 1842 | 1809 | 1841 | 1887.78 | 1945 | 1831 | 0.941 | -0.087 | 0.42031476 | 0.63075209 | 0.0071 | 0.0086 | 0.0079 | 0.0079 | TRUE | 0.1518 |
| Mtf2 | 13250 | 11924 | 11489 | 20561 | 14747 | 13672 | 14074 | 11909 | 11536 | 16163 | 15867 | 14966 | 14085.85 | 12506 | 15665 | 1.253 | 0.325 | 0.00084433 | 0.00780581 | 0.0067 | 0.0071 | 0.0068 | 0.0068 | TRUE | 0.1469 |
| Mtfmt | 802 | 815 | 744 | 969 | 696 | 637 | 852 | 814 | 747 | 762 | 749 | 697 | 770.13 | 804 | 736 | 0.916 | -0.127 | 0.20237752 | 0.40995781 | 0.002 | 0.0112 | 0.0058 | 0.0058 | TRUE | 0.0461 |
| Mtfp1 | 351 | 350 | 325 | 469 | 330 | 301 | 373 | 350 | 326 | 369 | 355 | 329 | 350.32 | 350 | 351 | 1.006 | 0.009 | 0.94332883 | 0.9717062 | 0 | 0.0163 | 0.008 | 0.008 | TRUE | 0.0386 |
| Mtfr1 | 3626 | 4024 | 3937 | 4991 | 3448 | 3584 | 3851 | 4019 | 3953 | 3924 | 3710 | 3923 | 3896.67 | 3941 | 3852 | 0.978 | -0.033 | 0.61386325 | 0.78155079 | 5.00E-04 | 0.0077 | 0.0028 | 0.0028 | TRUE | 0.0126 |
| Mtfr1l | 3735 | 4104 | 4184 | 4001 | 3065 | 3156 | 3967 | 4099 | 4201 | 3145 | 3298 | 3455 | 3694.15 | 4089 | 3299 | 0.807 | -0.31 | 1.59E-05 | 0.00029362 | 0.0011 | 0.0078 | 0.0034 | 0.0034 | TRUE | 0.0206 |
| Mtfr2 | 1477 | 1385 | 1310 | 1768 | 1299 | 1334 | 1569 | 1383 | 1315 | 1390 | 1398 | 1460 | 1419.2 | 1422 | 1416 | 0.995 | -0.007 | 0.94495694 | 0.97246347 | 0.0038 | 0.0092 | 0.006 | 0.006 | TRUE | 0.097 |
| Mtg1 | 1233 | 1399 | 1425 | 1546 | 1119 | 1069 | 1310 | 1397 | 1431 | 1215 | 1204 | 1170 | 1287.87 | 1379 | 1196 | 0.868 | -0.205 | 0.01056471 | 0.05497411 | 5.00E-04 | 0.0095 | 0.0038 | 0.0038 | TRUE | 0.0232 |
| Mthfd1 | 15205 | 15179 | 15031 | 20474 | 14719 | 14330 | 16150 | 15159 | 15093 | 16095 | 15837 | 15686 | 15670.12 | 15467 | 15873 | 1.026 | 0.037 | 0.5385037 | 0.72493004 | 7.00E-04 | 0.0071 | 0.0026 | 0.0026 | TRUE | 0.0182 |
| Mthfd1l | 5502 | 6490 | 6660 | 7602 | 5769 | 6162 | 5844 | 6482 | 6687 | 5976 | 6207 | 6745 | 6323.59 | 6338 | 6309 | 0.995 | -0.007 | 0.93917664 | 0.96945089 | 0.0042 | 0.0074 | 0.0054 | 0.0054 | TRUE | 0.0566 |
| Mthfd2 | 4818 | 3871 | 3666 | 5756 | 3920 | 3457 | 5118 | 3866 | 3681 | 4525 | 4218 | 3784 | 4198.57 | 4222 | 4176 | 0.989 | -0.015 | 0.91253033 | 0.95675253 | 0.0199 | 0.0077 | 0.0141 | 0.0141 | TRUE | 0.4195 |
| Mthfd2l | 80 | 59 | 65 | 94 | 71 | 73 | 85 | 59 | 65 | 74 | 76 | 80 | 73.23 | 70 | 77 | 1.101 | 0.139 | 0.56935174 | 0.74821263 | 0 | 0.0519 | 0.029 | 0.029 | TRUE | 0.3215 |
| Mthfr | 737 | 524 | 504 | 1068 | 727 | 547 | 783 | 523 | 506 | 840 | 782 | 599 | 672.13 | 604 | 740 | 1.227 | 0.295 | 0.13946732 | 0.32412923 | 0.0451 | 0.0118 | 0.0271 | 0.0271 | TRUE | 0.785 |
| Mthfs | 158 | 207 | 199 | 191 | 130 | 150 | 168 | 207 | 200 | 150 | 140 | 164 | 171.43 | 192 | 151 | 0.79 | -0.34 | 0.04363085 | 0.14952757 | 0.0032 | 0.0261 | 0.0146 | 0.0146 | TRUE | 0.124 |
| Mthfsd | 290 | 325 | 356 | 586 | 413 | 385 | 308 | 325 | 357 | 461 | 444 | 421 | 386.09 | 330 | 442 | 1.341 | 0.423 | 0.00040432 | 0.00437375 | 0 | 0.0154 | 0.0076 | 0.0076 | TRUE | 0.0603 |
| Mtif2 | 3986 | 4021 | 4062 | 5232 | 3795 | 3650 | 4234 | 4016 | 4079 | 4113 | 4083 | 3995 | 4086.67 | 4110 | 4064 | 0.989 | -0.016 | 0.7985316 | 0.89619481 | 2.00E-04 | 0.0077 | 0.0025 | 0.0025 | TRUE | 0.0085 |
| Mtif3 | 760 | 765 | 750 | 1133 | 706 | 685 | 807 | 764 | 753 | 891 | 760 | 750 | 787.41 | 775 | 800 | 1.034 | 0.049 | 0.65146245 | 0.80454668 | 0.0044 | 0.0111 | 0.0072 | 0.0072 | TRUE | 0.1205 |
| Mtl5 | 296 | 435 | 451 | 524 | 402 | 444 | 314 | 434 | 453 | 412 | 433 | 486 | 422.03 | 400 | 444 | 1.106 | 0.145 | 0.38092732 | 0.59526263 | 0.0198 | 0.0147 | 0.0173 | 0.0173 | TRUE | 0.4043 |
| Mtm1 | 615 | 662 | 645 | 731 | 582 | 670 | 653 | 661 | 648 | 575 | 626 | 733 | 649.38 | 654 | 645 | 0.984 | -0.023 | 0.84760289 | 0.9238415 | 0.0064 | 0.012 | 0.0088 | 0.0088 | TRUE | 0.1692 |
| Mtmr1 | 629 | 581 | 726 | 1076 | 748 | 728 | 668 | 580 | 729 | 846 | 805 | 797 | 737.49 | 659 | 816 | 1.239 | 0.309 | 0.0070202 | 0.0401751 | 0.0054 | 0.0114 | 0.0081 | 0.0081 | TRUE | 0.1301 |
| Mtmr10 | 1335 | 1011 | 993 | 1694 | 1125 | 1045 | 1418 | 1010 | 997 | 1332 | 1210 | 1144 | 1185.14 | 1142 | 1229 | 1.077 | 0.107 | 0.48789256 | 0.68461223 | 0.0226 | 0.0097 | 0.0163 | 0.0163 | TRUE | 0.5363 |
| Mtmr11 | 122 | 84 | 71 | 85 | 69 | 81 | 130 | 84 | 71 | 67 | 74 | 89 | 85.75 | 95 | 77 | 0.805 | -0.312 | 0.27473667 | 0.49532999 | 0.0489 | 0.0453 | 0.0471 | 0.0471 | TRUE | 0.9718 |
| Mtmr12 | 2838 | 2650 | 2594 | 4044 | 2966 | 2855 | 3014 | 2647 | 2605 | 3179 | 3191 | 3125 | 2960.21 | 2755 | 3165 | 1.149 | 0.2 | 0.01801029 | 0.08058392 | 0.0029 | 0.008 | 0.0048 | 0.0048 | TRUE | 0.082 |
| Mtmr14 | 4071 | 4716 | 4982 | 5251 | 3876 | 4401 | 4324 | 4710 | 5002 | 4128 | 4170 | 4818 | 4525.39 | 4679 | 4372 | 0.934 | -0.098 | 0.31932575 | 0.54057938 | 0.0062 | 0.0076 | 0.0068 | 0.0068 | TRUE | 0.0965 |
| Mtmr2 | 3916 | 4139 | 4179 | 4844 | 3476 | 3500 | 4159 | 4134 | 4196 | 3808 | 3740 | 3831 | 3978.09 | 4163 | 3793 | 0.911 | -0.134 | 0.02071221 | 0.08908451 | 0 | 0.0077 | 0.0022 | 0.0022 | TRUE | 0.0018 |
| Mtmr3 | 3280 | 4365 | 4663 | 4803 | 3559 | 4029 | 3484 | 4359 | 4682 | 3776 | 3829 | 4410 | 4090.14 | 4175 | 4005 | 0.959 | -0.06 | 0.63904338 | 0.79663233 | 0.015 | 0.0077 | 0.0117 | 0.0117 | TRUE | 0.2551 |
| Mtmr4 | 4034 | 3982 | 3873 | 4915 | 3667 | 3434 | 4285 | 3977 | 3889 | 3864 | 3945 | 3759 | 3953.15 | 4050 | 3856 | 0.952 | -0.071 | 0.32671195 | 0.54780107 | 0.0012 | 0.0077 | 0.0035 | 0.0035 | TRUE | 0.0312 |
| Mtmr6 | 1133 | 1191 | 1298 | 1646 | 1141 | 1163 | 1203 | 1189 | 1303 | 1294 | 1228 | 1273 | 1248.49 | 1232 | 1265 | 1.027 | 0.038 | 0.64321094 | 0.79917464 | 8.00E-04 | 0.0095 | 0.0041 | 0.0041 | TRUE | 0.0308 |
| Mtmr7 | 4993 | 5362 | 5398 | 7531 | 5167 | 5114 | 5303 | 5355 | 5420 | 5920 | 5559 | 5598 | 5526.07 | 5359 | 5692 | 1.062 | 0.087 | 0.1611528 | 0.35598067 | 5.00E-04 | 0.0075 | 0.0026 | 0.0026 | TRUE | 0.0151 |
| Mtmr9 | 2892 | 3046 | 3119 | 4023 | 2812 | 2907 | 3072 | 3042 | 3132 | 3163 | 3026 | 3182 | 3102.66 | 3082 | 3124 | 1.014 | 0.019 | 0.76135862 | 0.87423727 | 0 | 0.008 | 0.0026 | 0.0026 | TRUE | 0.009 |
| Mtnr1a | 9 | 5 | 1 | 8 | 3 | 6 | 10 | 5 | 1 | 6 | 3 | 7 | 5.27 | 5 | 5 | 1.044 | 0.062 | 0.94812343 | 0.97406219 | 0.2536 | 0.6316 | 0.4582 | 0.4582 | TRUE | 0.2248 |
| Mtnr1b | 0 | 0 | 0 | 0 | 0 | 0 | 0 | 0 | 0 | 0 | 0 | 0 | 0 | 0 | 0 | NA | NA | NA | NA | NA | NA | NA | NA | NA | NA |
| Mto1 | 1819 | 1494 | 1543 | 1941 | 1374 | 1309 | 1932 | 1492 | 1549 | 1526 | 1478 | 1433 | 1568.44 | 1658 | 1479 | 0.893 | -0.164 | 0.1662689 | 0.36351681 | 0.0099 | 0.009 | 0.0095 | 0.0095 | TRUE | 0.252 |
| Mtor | 8259 | 7538 | 8266 | 9422 | 6858 | 6817 | 8773 | 7528 | 8300 | 7407 | 7379 | 7462 | 7808.1 | 8200 | 7416 | 0.904 | -0.145 | 0.06893595 | 0.20364446 | 0.0028 | 0.0073 | 0.0045 | 0.0045 | TRUE | 0.0628 |
| Mtpap | 2170 | 2197 | 2268 | 3041 | 2115 | 2193 | 2305 | 2194 | 2277 | 2391 | 2276 | 2401 | 2307.2 | 2259 | 2356 | 1.043 | 0.061 | 0.38035662 | 0.59482144 | 3.00E-04 | 0.0083 | 0.003 | 0.003 | TRUE | 0.0105 |
| Mtpn | 6500 | 7979 | 8329 | 9562 | 6798 | 7558 | 6904 | 7969 | 8363 | 7517 | 7314 | 8273 | 7723.43 | 7745 | 7701 | 0.994 | -0.008 | 0.93359291 | 0.96713012 | 0.0069 | 0.0073 | 0.007 | 0.007 | TRUE | 0.1101 |
| Mtr | 2929 | 2570 | 2819 | 4222 | 2940 | 2631 | 3111 | 2567 | 2831 | 3319 | 3163 | 2880 | 2978.45 | 2836 | 3121 | 1.101 | 0.139 | 0.17910249 | 0.38024408 | 0.0066 | 0.008 | 0.0073 | 0.0073 | TRUE | 0.0872 |
| Mtrf1 | 535 | 643 | 653 | 773 | 530 | 602 | 568 | 642 | 656 | 608 | 570 | 659 | 617.17 | 622 | 612 | 0.984 | -0.024 | 0.83441027 | 0.91571799 | 0.0037 | 0.0122 | 0.0075 | 0.0075 | TRUE | 0.0667 |

| | | | | | | | | | | | | | | | | | | | | | | | | | |
|---|---|---|---|---|---|---|---|---|---|---|---|---|---|---|---|---|---|---|---|---|---|---|---|---|---|
| Mtrf1l | 136 | 137 | 127 | 190 | 139 | 119 | 144 | 137 | 128 | 149 | 150 | 130 | 139.66 | 136 | 143 | 1.052 | 0.073 | 0.67145413 | 0.81667744 | 0 | 0.0305 | 0.0142 | 0.0142 | TRUE | 0.0604 |
| Mtrr | 2197 | 2460 | 2322 | 3114 | 2322 | 2149 | 2334 | 2457 | 2332 | 2448 | 2498 | 2352 | 2403.45 | 2374 | 2433 | 1.025 | 0.035 | 0.61354095 | 0.78135644 | 5.00E-04 | 0.0083 | 0.0031 | 0.0031 | TRUE | 0.0113 |
| Mtss1 | 6215 | 5432 | 5578 | 4805 | 3787 | 3493 | 6601 | 5425 | 5601 | 3777 | 4075 | 3824 | 4883.81 | 5876 | 3892 | 0.662 | -0.594 | 1.78E-09 | 1.02E-07 | 0.0064 | 0.0076 | 0.0068 | 0.0068 | TRUE | 0.1422 |
| Mtss1l | 1473 | 1444 | 1467 | 1759 | 1394 | 1238 | 1565 | 1442 | 1473 | 1383 | 1500 | 1355 | 1452.93 | 1493 | 1413 | 0.946 | -0.08 | 0.34453066 | 0.5641524 | 0.0016 | 0.0092 | 0.0045 | 0.0045 | TRUE | 0.0345 |
| Mttp | 244 | 325 | 307 | 423 | 296 | 279 | 259 | 325 | 308 | 333 | 318 | 305 | 308.07 | 297 | 319 | 1.073 | 0.101 | 0.46389917 | 0.66720935 | 0.004 | 0.0176 | 0.0105 | 0.0105 | TRUE | 0.1403 |
| Mtus1 | 2687 | 2800 | 3042 | 3520 | 2668 | 2685 | 2854 | 2796 | 3054 | 2767 | 2871 | 2939 | 2880.31 | 2901 | 2859 | 0.985 | -0.022 | 0.76678598 | 0.87807656 | 0.0012 | 0.008 | 0.0036 | 0.0036 | TRUE | 0.0258 |
| Mtus2 | 795 | 694 | 595 | 947 | 665 | 523 | 844 | 693 | 597 | 744 | 715 | 573 | 694.57 | 711 | 677 | 0.953 | -0.069 | 0.66906043 | 0.81504759 | 0.0233 | 0.0116 | 0.0176 | 0.0176 | TRUE | 0.3138 |
| Mtx1 | 1987 | 1843 | 1769 | 3198 | 2212 | 1935 | 2111 | 1841 | 1776 | 2514 | 2380 | 2118 | 2123.26 | 1909 | 2337 | 1.225 | 0.293 | 0.00661607 | 0.03861402 | 0.0073 | 0.0085 | 0.0079 | 0.0079 | TRUE | 0.1027 |
| Mtx2 | 2291 | 2596 | 2370 | 3379 | 2398 | 2302 | 2433 | 2593 | 2380 | 2656 | 2580 | 2520 | 2527.02 | 2469 | 2585 | 1.048 | 0.067 | 0.35920392 | 0.57780359 | 0.001 | 0.0082 | 0.0035 | 0.0035 | TRUE | 0.0235 |
| Mtx3 | 358 | 344 | 316 | 605 | 550 | 459 | 380 | 344 | 317 | 476 | 592 | 502 | 435.16 | 347 | 523 | 1.506 | 0.591 | 1.80E-05 | 0.00032495 | 0.0086 | 0.0145 | 0.0113 | 0.0113 | TRUE | 0.152 |
| Muc1 | 37 | 14 | 18 | 34 | 17 | 25 | 39 | 14 | 18 | 27 | 18 | 27 | 23.96 | 24 | 24 | 1.02 | 0.028 | 0.95439786 | 0.97676022 | 0.129 | 0.1444 | 0.1369 | 0.1369 | TRUE | 1.8758 |
| Muc13 | 6 | 6 | 3 | 14 | 6 | 5 | 6 | 6 | 3 | 11 | 6 | 5 | 6.39 | 5 | 7 | 1.516 | 0.601 | 0.4383538 | 0.64591156 | 0 | 0.5228 | 0.2699 | 0.2699 | TRUE | 0.531 |
| Muc15 | 0 | 0 | 0 | 0 | 0 | 0 | 0 | 0 | 0 | 0 | 0 | 0 | 0 | 0 | 0 | NA | NA | NA | NA | NA | NA | NA | NA | NA | NA |
| Muc19 | 0 | 1 | 0 | 0 | 0 | 1 | 0 | 1 | 0 | 0 | 0 | 1 | 0.35 | 0 | 0 | 0.954 | -0.067 | 0.98567705 | NA | 0 | 9.4494 | 8.1588 | 8.1588 | TRUE | 0.5383 |
| Muc2 | 113 | 111 | 124 | 116 | 122 | 109 | 120 | 111 | 125 | 91 | 131 | 119 | 116.19 | 119 | 114 | 0.956 | -0.064 | 0.76005438 | 0.87360281 | 0.0117 | 0.0353 | 0.0233 | 0.0233 | TRUE | 0.3402 |
| Muc20 | 15 | 3 | 3 | 5 | 11 | 5 | 16 | 3 | 3 | 4 | 12 | 5 | 7.2 | 7 | 7 | 0.964 | -0.054 | 0.95375802 | 0.97651612 | 0.485 | 0.4647 | 0.4751 | 0.4751 | TRUE | 2.8044 |
| Muc4 | 31 | 24 | 29 | 46 | 33 | 34 | 33 | 24 | 29 | 36 | 36 | 37 | 32.48 | 29 | 36 | 1.267 | 0.342 | 0.30646851 | 0.52694007 | 0 | 0.1083 | 0.0492 | 0.0492 | TRUE | 0.1355 |
| Muc5ac | 3 | 2 | 0 | 4 | 1 | 1 | 3 | 2 | 0 | 3 | 1 | 1 | 1.75 | 2 | 2 | 1.058 | 0.082 | 0.95715805 | NA | 0 | 1.8896 | 1.1069 | 1.1069 | TRUE | 0.3974 |
| Muc5b | 0 | 1 | 0 | 2 | 0 | 0 | 0 | 1 | 0 | 2 | 0 | 0 | 0.43 | 0 | 1 | 1.493 | 0.578 | 0.86962266 | NA | 0 | 7.6954 | 7.0545 | 7.0545 | TRUE | 0.9718 |
| Muc6 | 0 | 0 | 0 | 1 | 0 | 0 | 0 | 0 | 0 | 1 | 0 | 0 | 0.13 | 0 | 0 | 1.859 | 0.894 | 0.82650734 | NA | 0 | 25.1515 | 10 | 10 | TRUE | 0.2563 |
| Mucl1 | 0 | 0 | 0 | 0 | 0 | 0 | 0 | 0 | 0 | 0 | 0 | 0 | 0 | 0 | 0 | NA | NA | NA | NA | NA | NA | NA | NA | NA | NA |
| Mug-ps1 | 0 | 0 | 0 | 0 | 0 | 0 | 0 | 0 | 0 | 0 | 0 | 0 | 0 | 0 | 0 | NA | NA | NA | NA | NA | NA | NA | NA | NA | NA |
| Mug1 | 0 | 0 | 0 | 0 | 0 | 0 | 0 | 0 | 0 | 0 | 0 | 0 | 0 | 0 | 0 | NA | NA | NA | NA | NA | NA | NA | NA | NA | NA |
| Mug2 | 0 | 0 | 0 | 0 | 0 | 0 | 0 | 0 | 0 | 0 | 0 | 0 | 0 | 0 | 0 | NA | NA | NA | NA | NA | NA | NA | NA | NA | NA |
| Mul1 | 1304 | 1216 | 1085 | 1855 | 1285 | 1097 | 1385 | 1214 | 1089 | 1458 | 1383 | 1201 | 1288.44 | 1229 | 1347 | 1.096 | 0.133 | 0.28843653 | 0.50948829 | 0.0109 | 0.0095 | 0.0105 | 0.0105 | TRUE | 0.1462 |
| Mum1 | 3167 | 2978 | 2819 | 4171 | 3185 | 3028 | 3364 | 2974 | 2831 | 3279 | 3427 | 3315 | 3198.17 | 3056 | 3340 | 1.093 | 0.128 | 0.15186785 | 0.34295352 | 0.0039 | 0.0079 | 0.0055 | 0.0055 | TRUE | 0.094 |
| Mum1l1 | 1205 | 936 | 998 | 1423 | 990 | 838 | 1280 | 935 | 1002 | 1119 | 1065 | 917 | 1052.99 | 1072 | 1034 | 0.965 | -0.052 | 0.72232515 | 0.84975528 | 0.0182 | 0.01 | 0.0143 | 0.0143 | TRUE | 0.3424 |
| Mup1 | 0 | 0 | 0 | 0 | 0 | 0 | 0 | 0 | 0 | 0 | 0 | 0 | 0 | 0 | 0 | NA | NA | NA | NA | NA | NA | NA | NA | NA | NA |
| Mup10 | 0 | 0 | 0 | 0 | 0 | 0 | 0 | 0 | 0 | 0 | 0 | 0 | 0 | 0 | 0 | NA | NA | NA | NA | NA | NA | NA | NA | NA | NA |
| Mup11 | 0 | 0 | 0 | 0 | 0 | 0 | 0 | 0 | 0 | 0 | 0 | 0 | 0 | 0 | 0 | NA | NA | NA | NA | NA | NA | NA | NA | NA | NA |
| Mup12 | 0 | 0 | 0 | 0 | 0 | 0 | 0 | 0 | 0 | 0 | 0 | 0 | 0 | 0 | 0 | NA | NA | NA | NA | NA | NA | NA | NA | NA | NA |
| Mup13 | 0 | 0 | 0 | 0 | 0 | 0 | 0 | 0 | 0 | 0 | 0 | 0 | 0 | 0 | 0 | NA | NA | NA | NA | NA | NA | NA | NA | NA | NA |
| Mup14 | 0 | 0 | 0 | 0 | 0 | 0 | 0 | 0 | 0 | 0 | 0 | 0 | 0 | 0 | 0 | NA | NA | NA | NA | NA | NA | NA | NA | NA | NA |
| Mup15 | 0 | 0 | 0 | 0 | 0 | 0 | 0 | 0 | 0 | 0 | 0 | 0 | 0 | 0 | 0 | NA | NA | NA | NA | NA | NA | NA | NA | NA | NA |
| Mup16 | 0 | 0 | 0 | 0 | 0 | 0 | 0 | 0 | 0 | 0 | 0 | 0 | 0 | 0 | 0 | NA | NA | NA | NA | NA | NA | NA | NA | NA | NA |
| Mup17 | 0 | 0 | 0 | 0 | 0 | 0 | 0 | 0 | 0 | 0 | 0 | 0 | 0 | 0 | 0 | NA | NA | NA | NA | NA | NA | NA | NA | NA | NA |
| Mup19 | 0 | 0 | 0 | 0 | 0 | 0 | 0 | 0 | 0 | 0 | 0 | 0 | 0 | 0 | 0 | NA | NA | NA | NA | NA | NA | NA | NA | NA | NA |
| Mup2 | 0 | 0 | 0 | 0 | 0 | 0 | 0 | 0 | 0 | 0 | 0 | 0 | 0 | 0 | 0 | NA | NA | NA | NA | NA | NA | NA | NA | NA | NA |
| Mup20 | 1 | 0 | 0 | 0 | 0 | 0 | 1 | 0 | 0 | 0 | 0 | 0 | 0.18 | 0 | 0 | 0.49 | -1.029 | 0.80087 | NA | 0 | 18.6165 | 10 | 10 | TRUE | 0.4614 |
| Mup21 | 0 | 0 | 0 | 0 | 0 | 0 | 0 | 0 | 0 | 0 | 0 | 0 | 0 | 0 | 0 | NA | NA | NA | NA | NA | NA | NA | NA | NA | NA |
| Mup3 | 5 | 2 | 0 | 17 | 13 | 6 | 5 | 2 | 0 | 13 | 14 | 7 | 6.87 | 2 | 11 | 4.712 | 2.236 | 0.01964208 | 0.08576826 | 0.2104 | 0.4864 | 0.4068 | 0.4068 | TRUE | 1.0879 |
| Mup4 | 0 | 0 | 0 | 0 | 0 | 0 | 0 | 0 | 0 | 0 | 0 | 0 | 0 | 0 | 0 | NA | NA | NA | NA | NA | NA | NA | NA | NA | NA |
| Mup5 | 0 | 0 | 0 | 0 | 0 | 0 | 0 | 0 | 0 | 0 | 0 | 0 | 0 | 0 | 0 | NA | NA | NA | NA | NA | NA | NA | NA | NA | NA |
| Mup6 | 51 | 26 | 35 | 60 | 22 | 37 | 54 | 26 | 35 | 47 | 24 | 41 | 37.77 | 38 | 37 | 0.973 | -0.039 | 0.92409274 | 0.96274037 | 0.0994 | 0.0941 | 0.0965 | 0.0965 | TRUE | 0.515 |
| Mup7 | 0 | 0 | 0 | 0 | 0 | 0 | 0 | 0 | 0 | 0 | 0 | 0 | 0 | 0 | 0 | NA | NA | NA | NA | NA | NA | NA | NA | NA | NA |
| Mup8 | 0 | 0 | 0 | 0 | 0 | 0 | 0 | 0 | 0 | 0 | 0 | 0 | 0 | 0 | 0 | NA | NA | NA | NA | NA | NA | NA | NA | NA | NA |
| Mup9 | 0 | 0 | 0 | 0 | 1 | 0 | 0 | 0 | 0 | 0 | 1 | 0 | 0.18 | 0 | 0 | 1.859 | 0.894 | 0.82650734 | NA | 0 | 18.3785 | 10 | 10 | TRUE | 0.5204 |
| Murc | 99 | 51 | 47 | 67 | 43 | 36 | 105 | 51 | 47 | 53 | 46 | 39 | 56.94 | 68 | 46 | 0.684 | -0.548 | 0.14151988 | 0.32701377 | 0.0998 | 0.0648 | 0.0821 | 0.0821 | TRUE | 2.0446 |
| Mus81 | 226 | 301 | 306 | 471 | 333 | 249 | 240 | 301 | 307 | 370 | 358 | 273 | 308.17 | 283 | 334 | 1.182 | 0.241 | 0.16374016 | 0.35961366 | 0.0191 | 0.0176 | 0.0184 | 0.0184 | TRUE | 0.2862 |
| Musk | 0 | 0 | 0 | 0 | 1 | 0 | 0 | 0 | 0 | 0 | 1 | 0 | 0.18 | 0 | 0 | 1.859 | 0.894 | 0.82650734 | NA | 0 | 18.3785 | 10 | 10 | TRUE | 0.5204 |
| Mustn1 | 3 | 4 | 1 | 3 | 3 | 5 | 3 | 4 | 1 | 2 | 3 | 5 | 3.21 | 3 | 3 | 1.334 | 0.416 | 0.69910826 | NA | 0 | 1.034 | 0.5115 | 0.5115 | TRUE | 0.3716 |
| Mut | 4400 | 4033 | 4117 | 5073 | 3556 | 3709 | 4674 | 4028 | 4134 | 3988 | 3826 | 4060 | 4118.23 | 4279 | 3958 | 0.925 | -0.112 | 0.18902382 | 0.39422915 | 0.0034 | 0.0077 | 0.005 | 0.005 | TRUE | 0.0793 |
| Mutyh | 536 | 701 | 687 | 744 | 606 | 610 | 569 | 700 | 690 | 585 | 652 | 668 | 643.98 | 653 | 635 | 0.971 | -0.043 | 0.72801481 | 0.85382581 | 0.0074 | 0.012 | 0.0094 | 0.0094 | TRUE | 0.1472 |
| Mvb12a | 1707 | 2305 | 2294 | 2315 | 1697 | 2075 | 1813 | 2302 | 2303 | 1820 | 1826 | 2271 | 2055.96 | 2139 | 1972 | 0.921 | -0.118 | 0.38778663 | 0.60163001 | 0.0154 | 0.0085 | 0.013 | 0.013 | TRUE | 0.2152 |
| Mvb12b | 461 | 770 | 804 | 581 | 501 | 617 | 490 | 769 | 807 | 457 | 539 | 675 | 622.86 | 689 | 557 | 0.808 | -0.308 | 0.14974507 | 0.34036305 | 0.0529 | 0.0122 | 0.0313 | 0.0313 | TRUE | 0.752 |
| Mvd | 4555 | 3928 | 3844 | 5100 | 3777 | 3498 | 4838 | 3923 | 3860 | 4009 | 4064 | 3829 | 4087.18 | 4207 | 3967 | 0.943 | -0.084 | 0.43206114 | 0.64176836 | 0.0083 | 0.0077 | 0.0081 | 0.0081 | TRUE | 0.2095 |
| Mvk | 1241 | 1194 | 1217 | 1629 | 1247 | 1158 | 1318 | 1192 | 1222 | 1281 | 1342 | 1268 | 1270.42 | 1244 | 1297 | 1.042 | 0.06 | 0.47662283 | 0.67645532 | 0.001 | 0.0095 | 0.0043 | 0.0043 | TRUE | 0.0322 |
| Mvp | 1334 | 1347 | 1368 | 1763 | 1221 | 1347 | 1417 | 1345 | 1374 | 1386 | 1314 | 1475 | 1385 | 1379 | 1392 | 1.009 | 0.013 | 0.87364055 | 0.93766987 | 0.0012 | 0.0093 | 0.0043 | 0.0043 | TRUE | 0.0319 |
| Mx1 | 17 | 11 | 8 | 16 | 10 | 9 | 18 | 11 | 8 | 13 | 11 | 10 | 11.71 | 12 | 11 | 0.903 | -0.148 | 0.79857481 | 0.89619481 | 0 | 0.2882 | 0.1559 | 0.1559 | TRUE | 0.606 |
| Mx2 | 5 | 6 | 7 | 13 | 13 | 4 | 5 | 6 | 7 | 10 | 14 | 4 | 7.82 | 6 | 9 | 1.563 | 0.644 | 0.37357936 | 0.58997634 | 0 | 0.4282 | 0.2428 | 0.2428 | TRUE | 0.2236 |
| Mxd1 | 367 | 357 | 311 | 399 | 308 | 277 | 390 | 357 | 312 | 314 | 331 | 303 | 334.48 | 353 | 316 | 0.896 | -0.158 | 0.23879226 | 0.45463924 | 0.0042 | 0.0168 | 0.01 | 0.01 | TRUE | 0.116 |
| Mxd3 | 274 | 352 | 357 | 419 | 310 | 340 | 291 | 352 | 358 | 329 | 334 | 372 | 339.36 | 334 | 345 | 1.032 | 0.046 | 0.73875715 | 0.86021984 | 0.0055 | 0.0166 | 0.0107 | 0.0107 | TRUE | 0.1404 |
| Mxd4 | 296 | 304 | 270 | 344 | 229 | 213 | 314 | 304 | 271 | 270 | 246 | 233 | 273.18 | 296 | 250 | 0.846 | -0.242 | 0.0810987 | 0.22667707 | 0.0026 | 0.019 | 0.0102 | 0.0102 | TRUE | 0.063 |
| Mxi1 | 2453 | 2939 | 2929 | 2878 | 2246 | 2550 | 2606 | 2935 | 2941 | 2262 | 2417 | 2791 | 2658.69 | 2827 | 2490 | 0.88 | -0.184 | 0.08704718 | 0.23864211 | 0.0077 | 0.0081 | 0.0079 | 0.0079 | TRUE | 0.1354 |
| Mxra7 | 1469 | 1687 | 1707 | 1180 | 919 | 1177 | 1560 | 1685 | 1714 | 928 | 989 | 1288 | 1360.67 | 1653 | 1068 | 0.646 | -0.631 | 3.89E-06 | 8.95E-05 | 0.0164 | 0.0093 | 0.0127 | 0.0127 | TRUE | 0.3867 |
| Mxra8 | 351 | 369 | 370 | 352 | 261 | 304 | 373 | 369 | 372 | 277 | 281 | 333 | 333.86 | 371 | 297 | 0.798 | -0.325 | 0.01152377 | 0.05846025 | 0.0017 | 0.0168 | 0.0089 | 0.0089 | TRUE | 0.1233 |
| Myadm | 3706 | 4110 | 4466 | 6225 | 4309 | 4539 | 3936 | 4105 | 4484 | 4894 | 4636 | 4969 | 4503.99 | 4175 | 4833 | 1.158 | 0.211 | 0.00869059 | 0.04733008 | 0.0026 | 0.0076 | 0.0044 | 0.0044 | TRUE | 0.0511 |
| Myadml2 | 6 | 2 | 4 | 8 | 8 | 1 | 6 | 2 | 4 | 6 | 9 | 1 | 4.73 | 4 | 5 | 1.304 | 0.383 | 0.70071535 | 0.83629834 | 0.2597 | 0.7035 | 0.4989 | 0.4989 | TRUE | 0.4628 |
| Myb | 203 | 380 | 443 | 178 | 170 | 229 | 216 | 380 | 445 | 140 | 183 | 251 | 268.91 | 347 | 191 | 0.55 | -0.863 | 0.00310143 | 0.02177905 | 0.1036 | 0.0192 | 0.0572 | 0.0572 | TRUE | 0.4465 |

| | | | | | | | | | | | | | | | | | | | | | | | | | |
|---|---|---|---|---|---|---|---|---|---|---|---|---|---|---|---|---|---|---|---|---|---|---|---|---|---|
| Mybbp1a | 49215 | 46715 | 48560 | 67624 | 48486 | 44563 | 52275 | 46655 | 48759 | 53161 | 52168 | 48781 | 50299.82 | 49230 | 51370 | 1.043 | 0.061 | 0.41754374 | 0.62838739 | 0.0026 | 0.007 | 0.0041 | 0.0041 | TRUE | 0.0358 |
| Mybl1 | 932 | 1256 | 1304 | 1207 | 969 | 1079 | 990 | 1254 | 1309 | 949 | 1043 | 1181 | 1121.04 | 1184 | 1058 | 0.892 | -0.165 | 0.23756951 | 0.45313174 | 0.0152 | 0.0098 | 0.0132 | 0.0132 | TRUE | 0.2468 |
| Mybl2 | 20962 | 24781 | 24827 | 22322 | 17930 | 19155 | 22265 | 24749 | 24929 | 17548 | 19292 | 20968 | 21625.14 | 23981 | 19269 | 0.803 | -0.316 | 0.0007992 | 0.00748511 | 0.0058 | 0.0071 | 0.0063 | 0.0063 | TRUE | 0.0749 |
| Mybpc1 | 40 | 35 | 20 | 80 | 58 | 53 | 42 | 35 | 20 | 63 | 62 | 58 | 46.81 | 32 | 61 | 1.885 | 0.914 | 0.00602366 | 0.0360551 | 0.0244 | 0.0773 | 0.0561 | 0.0561 | TRUE | 0.7766 |
| Mybpc2 | 65 | 61 | 62 | 45 | 22 | 35 | 69 | 61 | 62 | 35 | 24 | 38 | 48.26 | 64 | 32 | 0.508 | -0.976 | 0.00111247 | 0.00966718 | 0 | 0.0752 | 0.0416 | 0.0416 | TRUE | 0.3528 |
| Mybpc3 | 128 | 83 | 69 | 77 | 74 | 55 | 136 | 83 | 69 | 61 | 80 | 60 | 81.42 | 96 | 67 | 0.695 | -0.525 | 0.08967065 | 0.24349956 | 0.0654 | 0.0474 | 0.0563 | 0.0563 | TRUE | 1.1563 |
| Mybph | 81 | 59 | 45 | 38 | 31 | 52 | 86 | 59 | 45 | 30 | 33 | 57 | 51.72 | 63 | 40 | 0.628 | -0.67 | 0.07456959 | 0.21465806 | 0.0931 | 0.0706 | 0.0814 | 0.0814 | TRUE | 0.4336 |
| Mybphl | 1 | 2 | 2 | 5 | 4 | 2 | 1 | 2 | 2 | 4 | 4 | 2 | 2.58 | 2 | 3 | 2.06 | 1.043 | 0.37154418 | NA | 0 | 1.2829 | 0.5399 | 0.5399 | TRUE | 0.1367 |
| Myc | 968 | 476 | 408 | 644 | 431 | 235 | 1028 | 475 | 410 | 506 | 464 | 257 | 523.41 | 638 | 409 | 0.642 | -0.64 | 0.2172372 | 0.42935653 | 0.1919 | 0.0132 | 0.0821 | 0.1919 | TRUE | 3.3729 |
| Mycbp | 964 | 973 | 951 | 1501 | 1100 | 975 | 1024 | 972 | 955 | 1180 | 1184 | 1067 | 1063.56 | 984 | 1144 | 1.164 | 0.219 | 0.01457876 | 0.06932051 | 0.0014 | 0.01 | 0.0048 | 0.0048 | TRUE | 0.04 |
| Mycbp2 | 6549 | 5698 | 6979 | 8505 | 5838 | 5478 | 6956 | 5691 | 7008 | 6686 | 6281 | 5997 | 6436.39 | 6552 | 6321 | 0.965 | -0.051 | 0.62542526 | 0.78901854 | 0.0068 | 0.0074 | 0.0078 | 0.0078 | TRUE | 0.1613 |
| Mycbpap | 12 | 8 | 8 | 16 | 17 | 7 | 13 | 8 | 8 | 13 | 18 | 8 | 11.22 | 10 | 13 | 1.343 | 0.425 | 0.48438376 | 0.68193347 | 0 | 0.3006 | 0.175 | 0.175 | TRUE | 0.1646 |
| Mycl1 | 836 | 730 | 672 | 1012 | 803 | 742 | 888 | 729 | 675 | 796 | 864 | 812 | 793.93 | 764 | 824 | 1.079 | 0.109 | 0.38952439 | 0.60361355 | 0.009 | 0.0111 | 0.0103 | 0.0103 | TRUE | 0.238 |
| Mycn | 6128 | 7055 | 7248 | 5414 | 4575 | 5089 | 6509 | 7046 | 7278 | 4256 | 4922 | 5571 | 5930.31 | 6944 | 4916 | 0.708 | -0.499 | 1.15E-05 | 0.00022411 | 0.0103 | 0.0075 | 0.0091 | 0.0091 | TRUE | 0.1693 |
| Mycs | 0 | 0 | 0 | 1 | 0 | 1 | 0 | 0 | 0 | 1 | 0 | 1 | 0.31 | 0 | 1 | 3.348 | 1.743 | 0.65064889 | NA | 0 | 10.5167 | 8.8968 | 8.8968 | TRUE | 0.2082 |
| Myct1 | 1 | 0 | 0 | 0 | 0 | 0 | 1 | 0 | 0 | 0 | 0 | 0 | 0.18 | 0 | 0 | 0.49 | -1.029 | 0.80087 | NA | 0 | 18.6165 | 10 | 10 | TRUE | 0.4614 |
| Myd88 | 365 | 338 | 285 | 506 | 338 | 329 | 388 | 338 | 286 | 398 | 364 | 360 | 355.5 | 337 | 374 | 1.111 | 0.152 | 0.30413942 | 0.5250533 | 0.0098 | 0.0162 | 0.0128 | 0.0128 | TRUE | 0.1995 |
| Myef2 | 8114 | 7648 | 7449 | 11200 | 7854 | 7447 | 8619 | 7638 | 7480 | 8805 | 8450 | 8152 | 8190.52 | 7912 | 8469 | 1.071 | 0.098 | 0.24099241 | 0.45755776 | 0.0036 | 0.0073 | 0.0049 | 0.0049 | TRUE | 0.0744 |
| Myeov2 | 892 | 932 | 852 | 1305 | 857 | 843 | 947 | 931 | 855 | 1026 | 922 | 923 | 934.09 | 911 | 957 | 1.051 | 0.072 | 0.45452346 | 0.65959798 | 0.0024 | 0.0104 | 0.0057 | 0.0057 | TRUE | 0.0491 |
| Myf5 | 1 | 2 | 2 | 0 | 0 | 1 | 1 | 2 | 2 | 0 | 0 | 1 | 1.03 | 2 | 0 | 0.206 | -2.277 | 0.2572845 | NA | 0 | 3.2145 | 1.6104 | 1.6104 | TRUE | 0.5383 |
| Myf6 | 1 | 1 | 3 | 0 | 0 | 0 | 1 | 1 | 3 | 0 | 0 | 0 | 0.85 | 2 | 0 | 0.106 | -3.239 | 0.16662365 | NA | 0 | 3.9032 | 2.6513 | 2.6513 | TRUE | 0.362 |
| Myg1 | 1685 | 1479 | 1524 | 2313 | 1631 | 1506 | 1790 | 1477 | 1530 | 1818 | 1755 | 1649 | 1669.8 | 1599 | 1741 | 1.089 | 0.123 | 0.23442648 | 0.44932517 | 0.0058 | 0.0089 | 0.0071 | 0.0071 | TRUE | 0.1308 |
| Myh1 | 1 | 2 | 0 | 3 | 1 | 2 | 1 | 2 | 0 | 2 | 1 | 2 | 1.45 | 1 | 2 | 1.852 | 0.889 | 0.57938865 | NA | 0 | 2.2833 | 1.0953 | 1.0953 | TRUE | 0.3328 |
| Myh10 | 9862 | 10081 | 11725 | 15665 | 10654 | 11167 | 10475 | 10068 | 11773 | 12315 | 11463 | 12224 | 11386.33 | 10772 | 12001 | 1.114 | 0.156 | 0.06881373 | 0.20343583 | 0.004 | 0.0072 | 0.0052 | 0.0052 | TRUE | 0.0808 |
| Myh11 | 53 | 36 | 24 | 88 | 61 | 41 | 56 | 36 | 24 | 69 | 66 | 45 | 49.34 | 39 | 60 | 1.552 | 0.634 | 0.08967601 | 0.24349956 | 0.0859 | 0.0737 | 0.0793 | 0.0793 | TRUE | 0.6084 |
| Myh13 | 1110 | 904 | 862 | 560 | 489 | 357 | 1179 | 903 | 866 | 440 | 526 | 391 | 717.42 | 983 | 452 | 0.46 | -1.119 | 9.35E-12 | 8.29E-10 | 0.0242 | 0.0115 | 0.0178 | 0.0178 | TRUE | 0.3647 |
| Myh14 | 1953 | 1910 | 1977 | 1823 | 1427 | 1546 | 2074 | 1908 | 1985 | 1433 | 1535 | 1692 | 1771.31 | 1989 | 1553 | 0.781 | -0.358 | 0.00013608 | 0.00176644 | 0.0036 | 0.0088 | 0.0058 | 0.0058 | TRUE | 0.0733 |
| Myh15 | 1 | 0 | 0 | 1 | 2 | 0 | 1 | 0 | 0 | 1 | 2 | 0 | 0.67 | 0 | 1 | 2.668 | 1.416 | 0.58602256 | NA | 0 | 4.9484 | 3.3683 | 3.3683 | TRUE | 0.3203 |
| Myh2 | 1 | 1 | 0 | 0 | 1 | 5 | 1 | 1 | 0 | 0 | 1 | 5 | 1.44 | 1 | 2 | 3.123 | 1.643 | 0.4121021 | NA | 1.4301 | 2.3027 | 1.9161 | 1.9161 | TRUE | 0.9096 |
| Myh3 | 1846 | 1576 | 1572 | 1779 | 1379 | 1273 | 1961 | 1574 | 1578 | 1399 | 1484 | 1393 | 1564.82 | 1704 | 1425 | 0.836 | -0.258 | 0.02222262 | 0.0937905 | 0.0082 | 0.009 | 0.0085 | 0.0085 | TRUE | 0.2084 |
| Myh4 | 16 | 2 | 3 | 17 | 25 | 12 | 17 | 2 | 3 | 13 | 27 | 13 | 12.57 | 7 | 18 | 2.437 | 1.285 | 0.11592624 | 0.28752771 | 0.5249 | 0.2691 | 0.3836 | 0.3836 | TRUE | 3.5122 |
| Myh6 | 532 | 401 | 430 | 465 | 391 | 352 | 565 | 400 | 432 | 366 | 421 | 385 | 428.15 | 466 | 391 | 0.838 | -0.255 | 0.11172613 | 0.28113131 | 0.0166 | 0.0146 | 0.0161 | 0.0161 | TRUE | 0.4017 |
| Myh7 | 79 | 63 | 66 | 159 | 126 | 103 | 84 | 63 | 66 | 125 | 136 | 113 | 97.74 | 71 | 125 | 1.755 | 0.811 | 0.00019111 | 0.00234386 | 3.00E-04 | 0.0406 | 0.0229 | 0.0229 | TRUE | 0.2215 |
| Myh7b | 632 | 923 | 988 | 868 | 799 | 876 | 671 | 922 | 992 | 682 | 860 | 959 | 847.68 | 862 | 834 | 0.966 | -0.049 | 0.78346717 | 0.88827864 | 0.034 | 0.0108 | 0.0222 | 0.0222 | TRUE | 0.443 |
| Myh8 | 12 | 3 | 2 | 12 | 4 | 5 | 13 | 3 | 2 | 9 | 4 | 5 | 6.16 | 6 | 6 | 1.102 | 0.14 | 0.87967514 | 0.94060925 | 0.3751 | 0.5417 | 0.4576 | 0.4576 | TRUE | 2.2676 |
| Myh9 | 21659 | 25651 | 28140 | 29104 | 22084 | 22874 | 23006 | 25618 | 28256 | 22879 | 23761 | 25039 | 24759.77 | 25627 | 23893 | 0.932 | -0.101 | 0.29095587 | 0.51170493 | 0.0062 | 0.007 | 0.0066 | 0.0066 | TRUE | 0.0986 |
| Myl1 | 0 | 0 | 0 | 0 | 1 | 0 | 0 | 0 | 0 | 0 | 1 | 0 | 0.18 | 0 | 0 | 1.859 | 0.894 | 0.82650734 | NA | 0 | 18.3785 | 10 | 10 | TRUE | 0.5204 |
| Myl10 | 7 | 3 | 3 | 4 | 3 | 3 | 7 | 3 | 3 | 3 | 3 | 3 | 3.85 | 4 | 3 | 0.721 | -0.472 | 0.6240923 | NA | 0 | 0.8626 | 0.4026 | 0.4026 | TRUE | 0.583 |
| Myl12a | 4694 | 4432 | 4202 | 7839 | 5102 | 4760 | 4986 | 4426 | 4219 | 6162 | 5489 | 5211 | 5082.3 | 4544 | 5621 | 1.237 | 0.307 | 0.00272816 | 0.01968366 | 0.0073 | 0.0076 | 0.0074 | 0.0074 | TRUE | 0.0881 |
| Myl12b | 3424 | 3516 | 3424 | 4801 | 3395 | 3251 | 3637 | 3511 | 3438 | 3774 | 3653 | 3559 | 3595.35 | 3529 | 3662 | 1.038 | 0.054 | 0.41705644 | 0.62807995 | 6.00E-04 | 0.0078 | 0.0029 | 0.0029 | TRUE | 0.0089 |
| Myl2 | 22 | 28 | 11 | 10 | 5 | 6 | 23 | 28 | 11 | 8 | 5 | 7 | 13.7 | 21 | 7 | 0.321 | -1.641 | 0.00673807 | 0.03902025 | 0 | 0.2474 | 0.1666 | 0.1666 | TRUE | 0.4048 |
| Myl3 | 34 | 106 | 98 | 12 | 7 | 24 | 36 | 106 | 98 | 9 | 8 | 26 | 47.27 | 80 | 14 | 0.178 | -2.492 | 6.29E-06 | 0.00013447 | 0.3176 | 0.0766 | 0.1787 | 0.1787 | TRUE | 2.1433 |
| Myl4 | 286 | 394 | 394 | 290 | 218 | 208 | 304 | 393 | 396 | 228 | 235 | 228 | 297.19 | 364 | 230 | 0.631 | -0.664 | 1.15E-05 | 0.00022411 | 0.0087 | 0.018 | 0.013 | 0.013 | TRUE | 0.2392 |
| Myl6 | 6104 | 6551 | 6704 | 7910 | 5423 | 5205 | 6484 | 6543 | 6732 | 6218 | 5835 | 5698 | 6251.39 | 6586 | 5917 | 0.899 | -0.154 | 0.02175164 | 0.09244594 | 0.001 | 0.0074 | 0.0031 | 0.0031 | TRUE | 0.0245 |
| Myl6b | 403 | 368 | 350 | 369 | 258 | 264 | 428 | 368 | 351 | 290 | 278 | 289 | 333.95 | 382 | 286 | 0.748 | -0.42 | 0.00141164 | 0.01167507 | 0.0033 | 0.0168 | 0.0094 | 0.0094 | TRUE | 0.1243 |
| Myl7 | 30 | 37 | 43 | 32 | 48 | 28 | 32 | 37 | 43 | 25 | 52 | 31 | 36.57 | 37 | 36 | 0.951 | -0.072 | 0.85050908 | 0.92562639 | 0.0572 | 0.097 | 0.0775 | 0.0775 | TRUE | 1.0418 |
| Myl9 | 52 | 22 | 23 | 90 | 49 | 31 | 55 | 22 | 23 | 71 | 53 | 34 | 42.95 | 33 | 53 | 1.581 | 0.661 | 0.14964394 | 0.34028031 | 0.1753 | 0.0836 | 0.1269 | 0.1269 | TRUE | 0.4321 |
| Mylip | 361 | 506 | 437 | 491 | 389 | 393 | 383 | 505 | 439 | 386 | 419 | 430 | 427.05 | 442 | 412 | 0.929 | -0.107 | 0.44035818 | 0.64746865 | 0.0088 | 0.0146 | 0.0115 | 0.0115 | TRUE | 0.1788 |
| Mylk | 108 | 83 | 75 | 121 | 83 | 65 | 115 | 83 | 75 | 95 | 89 | 71 | 88.08 | 91 | 85 | 0.941 | -0.088 | 0.72693883 | 0.852905 | 0.0249 | 0.0443 | 0.0348 | 0.0348 | TRUE | 0.4906 |
| Mylk2 | 42 | 41 | 32 | 28 | 16 | 13 | 45 | 41 | 32 | 22 | 17 | 14 | 28.52 | 39 | 18 | 0.46 | -1.121 | 0.00348917 | 0.02379211 | 0 | 0.1224 | 0.066 | 0.066 | TRUE | 0.2654 |
| Mylk3 | 4 | 8 | 4 | 21 | 9 | 5 | 4 | 8 | 4 | 17 | 10 | 5 | 7.99 | 5 | 11 | 1.974 | 0.981 | 0.19389397 | 0.39959437 | 0.1252 | 0.4194 | 0.271 | 0.271 | TRUE | 0.2363 |
| Mylk4 | 0 | 0 | 0 | 2 | 0 | 0 | 0 | 0 | 0 | 2 | 0 | 0 | 0.26 | 0 | 1 | 2.886 | 1.529 | 0.70676491 | NA | 0 | 12.5792 | 10 | 10 | TRUE | 0.9669 |
| Mylpf | 2330 | 1591 | 1375 | 4491 | 2616 | 2079 | 2475 | 1589 | 1381 | 3530 | 2815 | 2276 | 2344.23 | 1815 | 2874 | 1.584 | 0.664 | 0.00297215 | 0.02107661 | 0.0746 | 0.0083 | 0.0355 | 0.0355 | TRUE | 0.4593 |
| Mynn | 1664 | 1875 | 1920 | 2233 | 1620 | 1684 | 1767 | 1873 | 1928 | 1755 | 1743 | 1843 | 1818.29 | 1856 | 1780 | 0.959 | -0.06 | 0.43253222 | 0.64193794 | 9.00E-04 | 0.0087 | 0.0037 | 0.0037 | TRUE | 0.0209 |
| Myo10 | 9636 | 9858 | 10869 | 12073 | 9566 | 9694 | 10235 | 9845 | 10914 | 9491 | 10292 | 10612 | 10231.49 | 10331 | 10132 | 0.981 | -0.028 | 0.72072511 | 0.84874759 | 0.0024 | 0.0072 | 0.0044 | 0.0044 | TRUE | 0.0376 |
| Myo15 | 29 | 35 | 27 | 43 | 34 | 29 | 31 | 35 | 27 | 34 | 37 | 32 | 32.5 | 31 | 34 | 1.099 | 0.137 | 0.68035998 | 0.82277546 | 0 | 0.1083 | 0.0486 | 0.0486 | TRUE | 0.088 |
| Myo16 | 21 | 16 | 18 | 22 | 9 | 26 | 22 | 16 | 18 | 17 | 10 | 28 | 18.63 | 19 | 18 | 0.982 | -0.026 | 0.95954399 | 0.97922535 | 0.0842 | 0.1837 | 0.1373 | 0.1373 | TRUE | 0.3012 |
| Myo18a | 14760 | 15076 | 16155 | 11850 | 9451 | 10268 | 15678 | 15057 | 16221 | 9316 | 10169 | 11240 | 12946.63 | 15652 | 10242 | 0.654 | -0.612 | 1.44E-11 | 1.23E-09 | 0.005 | 0.0072 | 0.0058 | 0.0058 | TRUE | 0.0889 |
| Myo18b | 292 | 352 | 364 | 199 | 194 | 198 | 310 | 352 | 365 | 156 | 209 | 217 | 268.18 | 342 | 194 | 0.564 | -0.827 | 1.08E-06 | 2.95E-05 | 0.0142 | 0.0192 | 0.0167 | 0.0167 | TRUE | 0.3371 |
| Myo19 | 1730 | 1360 | 1306 | 2029 | 1581 | 1384 | 1838 | 1358 | 1311 | 1595 | 1701 | 1515 | 1553.05 | 1502 | 1604 | 1.068 | 0.094 | 0.50847274 | 0.70249055 | 0.0186 | 0.009 | 0.0141 | 0.0141 | TRUE | 0.4578 |
| Myo1a | 250 | 194 | 185 | 164 | 156 | 140 | 266 | 194 | 186 | 129 | 168 | 153 | 182.51 | 215 | 150 | 0.696 | -0.523 | 0.01051033 | 0.05472727 | 0.0238 | 0.025 | 0.0244 | 0.0244 | TRUE | 0.4584 |
| Myo1b | 5964 | 6498 | 6686 | 7029 | 5125 | 5171 | 6335 | 6490 | 6713 | 5526 | 5514 | 5660 | 6039.7 | 6513 | 5567 | 0.855 | -0.227 | 0.00017829 | 0.0022107 | 3.00E-04 | 0.0074 | 0.0025 | 0.0025 | TRUE | 0.0089 |
| Myo1c | 5905 | 6082 | 6098 | 7275 | 5495 | 5368 | 6272 | 6074 | 6123 | 5719 | 5912 | 5876 | 5996.13 | 6156 | 5836 | 0.948 | -0.077 | 0.18007185 | 0.38147659 | 1.00E-04 | 0.0075 | 0.0022 | 0.0022 | TRUE | 0.0038 |
| Myo1d | 961 | 730 | 728 | 1217 | 826 | 717 | 1021 | 729 | 731 | 957 | 889 | 785 | 851.85 | 827 | 877 | 1.061 | 0.086 | 0.58941993 | 0.76400346 | 0.023 | 0.0108 | 0.017 | 0.017 | TRUE | 0.498 |
| Myo1e | 3683 | 3860 | 3792 | 4366 | 3085 | 3217 | 3912 | 3855 | 3808 | 3432 | 3319 | 3522 | 3641.26 | 3858 | 3424 | 0.888 | -0.172 | 0.00651321 | 0.03815987 | 2.00E-04 | 0.0078 | 0.0026 | 0.0026 | TRUE | 0.0086 |
| Myo1f | 181 | 193 | 179 | 307 | 266 | 221 | 192 | 193 | 180 | 241 | 286 | 242 | 222.37 | 188 | 256 | 1.361 | 0.444 | 0.00277417 | 0.01994266 | 0 | 0.0217 | 0.0113 | 0.0113 | TRUE | 0.1115 |
| Myo1g | 334 | 336 | 326 | 360 | 242 | 226 | 355 | 336 | 327 | 283 | 260 | 247 | 301.41 | 339 | 263 | 0.779 | -0.36 | 0.00461342 | 0.02947057 | 0 | 0.0178 | 0.0083 | 0.0083 | TRUE | 0.0501 |
| Myo1h | 70 | 64 | 68 | 66 | 75 | 64 | 74 | 64 | 68 | 52 | 81 | 70 | 68.2 | 69 | 68 | 0.975 | -0.037 | 0.88822266 | 0.94527988 | 0.015 | 0.0552 | 0.0352 | 0.0352 | TRUE | 0.4201 |
| Myo3a | 4 | 7 | 7 | 7 | 3 | 10 | 4 | 7 | 7 | 6 | 3 | 11 | 6.32 | 6 | 7 | 1.067 | 0.094 | 0.90584974 | 0.953297 | 0 | 0.5278 | 0.2982 | 0.2982 | TRUE | 0.587 |

| | | | | | | | | | | | | | | | | | | | | | | | | | |
|---|---|---|---|---|---|---|---|---|---|---|---|---|---|---|---|---|---|---|---|---|---|---|---|---|---|
| Myo3b | 7 | 7 | 11 | 3 | 5 | 4 | 7 | 7 | 11 | 2 | 5 | 4 | 6.26 | 8 | 4 | 0.465 | -1.104 | 0.15882266 | 0.35281001 | 0 | 0.5328 | 0.2589 | 0.2589 | TRUE | 0.314 |
| Myo5a | 5241 | 5224 | 5832 | 7268 | 5114 | 4751 | 5567 | 5217 | 5856 | 5714 | 5502 | 5201 | 5509.44 | 5547 | 5472 | 0.987 | -0.019 | 0.80832278 | 0.90136219 | 0.0026 | 0.0075 | 0.0043 | 0.0043 | TRUE | 0.033 |
| Myo5b | 722 | 742 | 688 | 717 | 539 | 583 | 767 | 741 | 691 | 564 | 580 | 638 | 663.42 | 733 | 594 | 0.81 | -0.305 | 0.00324149 | 0.02248229 | 0.002 | 0.0119 | 0.0062 | 0.0062 | TRUE | 0.0491 |
| Myo5c | 885 | 801 | 797 | 1195 | 888 | 783 | 940 | 800 | 800 | 939 | 955 | 857 | 882.04 | 847 | 917 | 1.084 | 0.116 | 0.28257562 | 0.5039899 | 0.0045 | 0.0106 | 0.0073 | 0.0073 | TRUE | 0.1097 |
| Myo6 | 1119 | 1506 | 1686 | 1843 | 1360 | 1428 | 1189 | 1504 | 1693 | 1449 | 1463 | 1563 | 1476.8 | 1462 | 1492 | 1.02 | 0.029 | 0.83318274 | 0.91523598 | 0.0159 | 0.0091 | 0.0127 | 0.0127 | TRUE | 0.321 |
| Myo7a | 229 | 190 | 196 | 244 | 171 | 174 | 243 | 190 | 197 | 192 | 184 | 190 | 199.34 | 210 | 189 | 0.9 | -0.152 | 0.34780201 | 0.56662263 | 0.0047 | 0.0234 | 0.0138 | 0.0138 | TRUE | 0.2077 |
| Myo7b | 9 | 13 | 8 | 11 | 12 | 13 | 10 | 13 | 8 | 9 | 13 | 14 | 11.06 | 10 | 12 | 1.158 | 0.211 | 0.71560387 | 0.84567137 | 0 | 0.3048 | 0.1504 | 0.1504 | TRUE | 0.2921 |
| Myo9a | 2231 | 1806 | 1957 | 3537 | 2587 | 2081 | 2370 | 1804 | 1965 | 2781 | 2783 | 2278 | 2330.06 | 2046 | 2614 | 1.278 | 0.354 | 0.00784438 | 0.04365617 | 0.0157 | 0.0083 | 0.0123 | 0.0123 | TRUE | 0.2312 |
| Myo9b | 5715 | 5819 | 5913 | 8901 | 6797 | 5819 | 6070 | 5811 | 5937 | 6997 | 7313 | 6370 | 6416.56 | 5939 | 6893 | 1.161 | 0.215 | 0.00603124 | 0.03608679 | 0.0024 | 0.0074 | 0.0043 | 0.0043 | TRUE | 0.0536 |
| Myoc | 0 | 0 | 1 | 2 | 0 | 1 | 0 | 0 | 1 | 2 | 0 | 1 | 0.61 | 0 | 1 | 2.474 | 1.307 | 0.62050174 | NA | 0 | 5.3914 | 3.4771 | 3.4771 | TRUE | 0.4098 |
| Myocd | 10 | 4 | 5 | 10 | 3 | 3 | 11 | 4 | 5 | 8 | 3 | 3 | 5.67 | 7 | 5 | 0.753 | -0.41 | 0.63333177 | 0.79371543 | 0.1236 | 0.5881 | 0.3545 | 0.3545 | TRUE | 0.7889 |
| Myod1 | 10 | 14 | 13 | 7 | 8 | 6 | 11 | 14 | 13 | 6 | 9 | 7 | 9.72 | 13 | 7 | 0.543 | -0.881 | 0.14813047 | 0.33781287 | 0 | 0.3457 | 0.1552 | 0.1552 | TRUE | 0.1178 |
| Myof | 3112 | 2781 | 2908 | 3593 | 2726 | 2652 | 3305 | 2777 | 2920 | 2825 | 2933 | 2903 | 2943.9 | 3001 | 2887 | 0.962 | -0.056 | 0.53399064 | 0.72173954 | 0.0037 | 0.008 | 0.0055 | 0.0055 | TRUE | 0.0955 |
| Myog | 4 | 9 | 1 | 9 | 6 | 8 | 4 | 9 | 1 | 7 | 6 | 9 | 6.09 | 5 | 7 | 1.561 | 0.642 | 0.46196964 | 0.66583069 | 0.1333 | 0.548 | 0.3756 | 0.3756 | TRUE | 0.4839 |
| Myom1 | 109 | 89 | 60 | 148 | 127 | 76 | 116 | 89 | 60 | 116 | 137 | 83 | 100.18 | 88 | 112 | 1.271 | 0.346 | 0.24592844 | 0.46297247 | 0.0699 | 0.0398 | 0.054 | 0.054 | TRUE | 0.2555 |
| Myom2 | 991 | 1089 | 966 | 540 | 537 | 654 | 1053 | 1088 | 970 | 425 | 578 | 716 | 804.73 | 1037 | 573 | 0.551 | -0.859 | 1.71E-06 | 4.37E-05 | 0.0321 | 0.011 | 0.0219 | 0.0219 | TRUE | 0.4247 |
| Myom3 | 63 | 59 | 61 | 137 | 97 | 80 | 67 | 59 | 61 | 108 | 104 | 88 | 81.12 | 62 | 100 | 1.607 | 0.684 | 0.00225216 | 0.01689067 | 0 | 0.0475 | 0.0231 | 0.0231 | TRUE | 0.1083 |
| Myot | 39 | 13 | 7 | 51 | 28 | 29 | 41 | 13 | 7 | 40 | 30 | 32 | 27.23 | 20 | 34 | 1.669 | 0.739 | 0.21739324 | 0.4294633 | 0.3294 | 0.1279 | 0.2194 | 0.2194 | TRUE | 3.4757 |
| Myoz1 | 2 | 1 | 2 | 1 | 1 | 1 | 2 | 1 | 2 | 1 | 1 | 1 | 1.35 | 2 | 1 | 0.571 | -0.809 | 0.60062408 | NA | 0 | 2.4509 | 0.9209 | 0.9209 | TRUE | 0.1051 |
| Myoz2 | 1 | 1 | 4 | 2 | 1 | 1 | 1 | 1 | 4 | 2 | 1 | 1 | 1.64 | 2 | 1 | 0.622 | -0.685 | 0.63947687 | NA | 0 | 2.0198 | 0.9042 | 0.9042 | TRUE | 0.6782 |
| Myoz3 | 53 | 57 | 58 | 57 | 67 | 52 | 56 | 57 | 58 | 45 | 72 | 57 | 57.55 | 57 | 58 | 1.007 | 0.01 | 0.97293057 | 0.98561635 | 0.0138 | 0.0642 | 0.0392 | 0.0392 | TRUE | 0.2467 |
| Mypn | 428 | 374 | 341 | 914 | 604 | 522 | 455 | 374 | 342 | 719 | 650 | 571 | 518.39 | 390 | 647 | 1.66 | 0.731 | 1.08E-06 | 2.95E-05 | 0.0144 | 0.0133 | 0.0141 | 0.0141 | TRUE | 0.2381 |
| Mypop | 98 | 86 | 78 | 193 | 138 | 110 | 104 | 86 | 78 | 152 | 148 | 120 | 114.82 | 89 | 140 | 1.573 | 0.653 | 0.0018543 | 0.01448539 | 0.0087 | 0.0356 | 0.0225 | 0.0225 | TRUE | 0.1919 |
| Myrf | 4798 | 4306 | 4897 | 3308 | 2554 | 2554 | 5096 | 4300 | 4917 | 2600 | 2748 | 2796 | 3743.01 | 4771 | 2715 | 0.569 | -0.814 | 5.89E-19 | 1.31E-16 | 0.0041 | 0.0078 | 0.0057 | 0.0057 | TRUE | 0.0909 |
| Myrfl | 22 | 34 | 22 | 20 | 15 | 30 | 23 | 34 | 22 | 16 | 16 | 33 | 24.02 | 26 | 22 | 0.807 | -0.31 | 0.50259272 | 0.69778982 | 0.0765 | 0.1441 | 0.1115 | 0.1115 | TRUE | 1.1068 |
| Myrip | 150 | 181 | 127 | 233 | 167 | 146 | 159 | 181 | 128 | 183 | 180 | 160 | 165.05 | 156 | 174 | 1.119 | 0.162 | 0.38418158 | 0.59833236 | 0.0112 | 0.0269 | 0.019 | 0.019 | TRUE | 0.2694 |
| Mysm1 | 3202 | 2526 | 2580 | 3541 | 2630 | 2310 | 3401 | 2523 | 2591 | 2784 | 2830 | 2529 | 2776.08 | 2838 | 2714 | 0.956 | -0.064 | 0.6244837 | 0.78834114 | 0.014 | 0.0081 | 0.012 | 0.012 | TRUE | 0.3651 |
| Myt1 | 3 | 2 | 4 | 5 | 4 | 2 | 3 | 2 | 4 | 4 | 4 | 2 | 3.27 | 3 | 3 | 1.141 | 0.19 | 0.85178614 | NA | 0 | 1.0142 | 0.4369 | 0.4369 | TRUE | 0.1356 |
| Myt1l | 14 | 14 | 12 | 48 | 46 | 25 | 15 | 14 | 12 | 38 | 49 | 27 | 25.92 | 14 | 38 | 2.804 | 1.488 | 0.00048913 | 0.00508543 | 0 | 0.134 | 0.0809 | 0.0809 | TRUE | 0.099 |
| Myzap | 160 | 125 | 91 | 212 | 163 | 168 | 170 | 125 | 91 | 167 | 175 | 184 | 152.02 | 129 | 175 | 1.363 | 0.446 | 0.05925514 | 0.1843701 | 0.038 | 0.0286 | 0.0336 | 0.0336 | TRUE | 0.3805 |
| Mzb1 | 52 | 46 | 48 | 39 | 32 | 37 | 55 | 46 | 48 | 31 | 34 | 41 | 42.49 | 50 | 35 | 0.703 | -0.508 | 0.0902527 | 0.24458348 | 0 | 0.0844 | 0.0405 | 0.0405 | TRUE | 0.1186 |
| Mzf1 | 60 | 65 | 74 | 104 | 92 | 64 | 64 | 65 | 74 | 82 | 99 | 70 | 75.63 | 68 | 84 | 1.235 | 0.304 | 0.20765462 | 0.41694826 | 0 | 0.0505 | 0.0286 | 0.0286 | TRUE | 0.2283 |
| Mzt1 | 2988 | 4134 | 4098 | 3957 | 2998 | 3414 | 3174 | 4129 | 4115 | 3111 | 3226 | 3737 | 3581.8 | 3806 | 3358 | 0.882 | -0.181 | 0.16423017 | 0.36033051 | 0.0149 | 0.0078 | 0.0119 | 0.0119 | TRUE | 0.2565 |
| Mzt2 | 462 | 502 | 478 | 723 | 491 | 522 | 491 | 501 | 480 | 568 | 528 | 571 | 523.35 | 491 | 556 | 1.134 | 0.181 | 0.07081048 | 0.20727967 | 0 | 0.0132 | 0.0053 | 0.0053 | TRUE | 0.0216 |
| N28178 | 2 | 1 | 2 | 2 | 2 | 2 | 2 | 1 | 2 | 2 | 2 | 2 | 1.84 | 2 | 2 | 1.145 | 0.195 | 0.88196197 | NA | 0 | 1.7966 | 0.6956 | 0.6956 | TRUE | 0.1053 |
| N4bp1 | 17775 | 17377 | 17422 | 18929 | 14025 | 15349 | 18880 | 17355 | 17494 | 14880 | 15090 | 16802 | 16750.12 | 17910 | 15591 | 0.87 | -0.2 | 0.0131276 | 0.0641972 | 0.0033 | 0.0071 | 0.0046 | 0.0046 | TRUE | 0.0564 |
| N4bp2 | 3371 | 3187 | 3469 | 5028 | 3545 | 3250 | 3581 | 3183 | 3483 | 3953 | 3814 | 3558 | 3595.2 | 3416 | 3775 | 1.106 | 0.145 | 0.08446973 | 0.23319738 | 0.0028 | 0.0078 | 0.0048 | 0.0048 | TRUE | 0.0432 |
| N4bp2l1 | 54 | 46 | 51 | 43 | 32 | 32 | 57 | 46 | 51 | 34 | 34 | 35 | 42.96 | 51 | 34 | 0.668 | -0.581 | 0.04639844 | 0.15587882 | 0 | 0.0836 | 0.0373 | 0.0373 | TRUE | 0.079 |
| N4bp2l2 | 3364 | 3424 | 3571 | 4318 | 3225 | 3350 | 3573 | 3420 | 3586 | 3394 | 3470 | 3667 | 3518.31 | 3526 | 3510 | 0.995 | -0.007 | 0.92177372 | 0.96126008 | 9.00E-04 | 0.0078 | 0.0032 | 0.0032 | TRUE | 0.0184 |
| N4bp3 | 2997 | 3451 | 3478 | 3199 | 2802 | 2983 | 3183 | 3447 | 3492 | 2515 | 3015 | 3265 | 3152.85 | 3374 | 2932 | 0.868 | -0.204 | 0.07264738 | 0.21105073 | 0.0092 | 0.0079 | 0.009 | 0.009 | TRUE | 0.1898 |
| N6amt1 | 485 | 503 | 518 | 723 | 598 | 510 | 515 | 502 | 520 | 568 | 643 | 558 | 551.28 | 512 | 590 | 1.15 | 0.202 | 0.05857116 | 0.18288804 | 0.0015 | 0.0129 | 0.0064 | 0.0064 | TRUE | 0.0718 |
| N6amt2 | 746 | 706 | 687 | 1060 | 671 | 567 | 792 | 705 | 690 | 833 | 722 | 621 | 727.2 | 729 | 725 | 0.997 | -0.005 | 0.97086733 | 0.98475753 | 0.0123 | 0.0114 | 0.0119 | 0.0119 | TRUE | 0.2058 |
| Naa10 | 2447 | 2571 | 2588 | 3468 | 2286 | 2237 | 2599 | 2568 | 2599 | 2726 | 2460 | 2449 | 2566.68 | 2589 | 2545 | 0.984 | -0.024 | 0.76023507 | 0.87368315 | 0.0016 | 0.0082 | 0.0039 | 0.0039 | TRUE | 0.0481 |
| Naa11 | 551 | 694 | 677 | 1157 | 888 | 939 | 585 | 693 | 680 | 910 | 955 | 1028 | 808.5 | 653 | 964 | 1.476 | 0.561 | 3.04E-07 | 9.62E-06 | 0.0046 | 0.011 | 0.0074 | 0.0074 | TRUE | 0.0955 |
| Naa15 | 9369 | 10636 | 10842 | 14804 | 10656 | 10418 | 9952 | 10622 | 10887 | 11638 | 11465 | 11404 | 10994.57 | 10487 | 11502 | 1.097 | 0.133 | 0.03917657 | 0.13860316 | 0.001 | 0.0072 | 0.0029 | 0.0029 | TRUE | 0.0243 |
| Naa16 | 1799 | 2105 | 2103 | 2186 | 1760 | 1985 | 1911 | 2102 | 2112 | 1718 | 1894 | 2173 | 1984.96 | 2042 | 1928 | 0.944 | -0.083 | 0.44987275 | 0.65547305 | 0.0081 | 0.0086 | 0.0083 | 0.0083 | TRUE | 0.1485 |
| Naa20 | 2131 | 2330 | 2120 | 2880 | 2093 | 2013 | 2263 | 2327 | 2129 | 2264 | 2252 | 2204 | 2239.78 | 2240 | 2240 | 1 | 0 | 0.99683567 | 0.99828918 | 7.00E-04 | 0.0084 | 0.0033 | 0.0033 | TRUE | 0.0229 |
| Naa25 | 7648 | 7420 | 7683 | 8549 | 6041 | 6202 | 8124 | 7410 | 7715 | 6721 | 6500 | 6789 | 7209.64 | 7750 | 6670 | 0.861 | -0.216 | 0.00136603 | 0.01136938 | 0.0011 | 0.0074 | 0.0032 | 0.0032 | TRUE | 0.0217 |
| Naa30 | 3464 | 3810 | 3909 | 4538 | 3358 | 3478 | 3679 | 3805 | 3925 | 3567 | 3613 | 3807 | 3732.86 | 3803 | 3662 | 0.963 | -0.055 | 0.42471393 | 0.63482617 | 7.00E-04 | 0.0078 | 0.0031 | 0.0031 | TRUE | 0.0144 |
| Naa35 | 3768 | 3547 | 3533 | 5408 | 3973 | 3796 | 4002 | 3542 | 3548 | 4251 | 4275 | 4155 | 3962.26 | 3697 | 4227 | 1.143 | 0.193 | 0.01442999 | 0.06882064 | 0.0023 | 0.0077 | 0.0042 | 0.0042 | TRUE | 0.0631 |
| Naa38 | 2026 | 2165 | 2073 | 2988 | 2217 | 2262 | 2152 | 2162 | 2082 | 2349 | 2385 | 2476 | 2267.68 | 2132 | 2403 | 1.127 | 0.173 | 0.01128862 | 0.05758217 | 0 | 0.0084 | 0.0029 | 0.0029 | TRUE | 0.0084 |
| Naa40 | 3194 | 3330 | 3287 | 5132 | 3848 | 3483 | 3393 | 3326 | 3300 | 4034 | 4140 | 3813 | 3667.68 | 3340 | 3996 | 1.197 | 0.259 | 0.00012698 | 0.00167739 | 7.00E-04 | 0.0078 | 0.003 | 0.003 | TRUE | 0.0193 |
| Naa50 | 14396 | 15935 | 15134 | 20334 | 14610 | 14734 | 15291 | 15914 | 15196 | 15985 | 15719 | 16129 | 15705.8 | 15467 | 15944 | 1.031 | 0.044 | 0.43235394 | 0.64190268 | 3.00E-04 | 0.0071 | 0.0022 | 0.0022 | TRUE | 0.0078 |
| Naa60 | 1632 | 1516 | 1576 | 2098 | 1396 | 1415 | 1733 | 1514 | 1582 | 1649 | 1502 | 1549 | 1588.37 | 1610 | 1567 | 0.974 | -0.038 | 0.67313896 | 0.81799428 | 0.0029 | 0.009 | 0.0053 | 0.0053 | TRUE | 0.054 |
| Naaa | 455 | 395 | 382 | 527 | 359 | 345 | 483 | 394 | 384 | 414 | 386 | 378 | 406.59 | 420 | 393 | 0.936 | -0.096 | 0.4764929 | 0.67643978 | 0.0069 | 0.015 | 0.0106 | 0.0106 | TRUE | 0.1957 |
| Naalad2 | 11 | 7 | 5 | 66 | 55 | 36 | 12 | 7 | 5 | 52 | 59 | 39 | 29.03 | 8 | 50 | 6.392 | 2.676 | 8.00E-09 | 3.88E-07 | 0.0217 | 0.1204 | 0.0803 | 0.0803 | TRUE | 0.3701 |
| Naaladl1 | 49 | 48 | 50 | 42 | 44 | 37 | 52 | 48 | 50 | 33 | 47 | 41 | 45.18 | 50 | 40 | 0.799 | -0.323 | 0.27073719 | 0.49074002 | 0 | 0.0798 | 0.0397 | 0.0397 | TRUE | 0.2249 |
| Nab1 | 132 | 151 | 168 | 187 | 134 | 158 | 140 | 151 | 169 | 147 | 144 | 173 | 153.97 | 153 | 155 | 1.007 | 0.011 | 0.95093968 | 0.97524031 | 0 | 0.0283 | 0.0155 | 0.0155 | TRUE | 0.107 |
| Nab2 | 188 | 247 | 219 | 393 | 293 | 261 | 200 | 247 | 220 | 309 | 315 | 286 | 262.7 | 222 | 303 | 1.366 | 0.449 | 0.00156646 | 0.01268283 | 0.002 | 0.0194 | 0.0106 | 0.0106 | TRUE | 0.1034 |
| Nabp1 | 5110 | 4354 | 4139 | 5746 | 4210 | 4203 | 5428 | 4348 | 4156 | 4517 | 4530 | 4601 | 4596.62 | 4644 | 4549 | 0.98 | -0.03 | 0.79409784 | 0.8940332 | 0.0088 | 0.0076 | 0.0091 | 0.0091 | TRUE | 0.2652 |
| Nabp2 | 677 | 747 | 747 | 829 | 633 | 637 | 719 | 746 | 750 | 652 | 681 | 697 | 707.54 | 738 | 677 | 0.915 | -0.127 | 0.15730519 | 0.35047569 | 0 | 0.0116 | 0.0044 | 0.0044 | TRUE | 0.0131 |
| Naca | 20581 | 24237 | 24629 | 29176 | 20361 | 21563 | 21861 | 24206 | 24730 | 22936 | 21907 | 23604 | 23207.28 | 23599 | 22816 | 0.967 | -0.049 | 0.5291935 | 0.71760295 | 0.0026 | 0.007 | 0.0043 | 0.0043 | TRUE | 0.0508 |
| Nacad | 70 | 63 | 83 | 148 | 66 | 85 | 74 | 63 | 83 | 116 | 71 | 93 | 83.5 | 73 | 93 | 1.278 | 0.354 | 0.18048318 | 0.38188633 | 0.0299 | 0.0464 | 0.0381 | 0.0381 | TRUE | 0.168 |
| Nacc1 | 14462 | 16199 | 17505 | 19610 | 14670 | 14530 | 15361 | 16178 | 17577 | 15416 | 15784 | 15905 | 16036.9 | 16372 | 15702 | 0.959 | -0.06 | 0.4209085 | 0.63116133 | 0.0024 | 0.0071 | 0.004 | 0.004 | TRUE | 0.0507 |
| Nacc2 | 738 | 751 | 757 | 842 | 632 | 667 | 784 | 750 | 760 | 662 | 680 | 730 | 727.68 | 765 | 691 | 0.902 | -0.148 | 0.10985307 | 0.27819566 | 0 | 0.0114 | 0.0048 | 0.0048 | TRUE | 0.029 |
| Nadk | 9542 | 9577 | 9692 | 11510 | 8049 | 8351 | 10135 | 9565 | 9732 | 9048 | 8660 | 9141 | 9380.28 | 9811 | 8950 | 0.912 | -0.132 | 0.03370128 | 0.12474457 | 7.00E-04 | 0.0073 | 0.0027 | 0.0027 | TRUE | 0.0102 |
| Nadkd1 | 6009 | 8220 | 8242 | 7986 | 6343 | 7258 | 6383 | 8209 | 8276 | 6278 | 6825 | 7945 | 7319.25 | 7623 | 7016 | 0.92 | -0.12 | 0.36995387 | 0.58707239 | 0.0159 | 0.0074 | 0.0127 | 0.0127 | TRUE | 0.2471 |
| Nadsyn1 | 489 | 712 | 691 | 823 | 620 | 590 | 519 | 711 | 694 | 647 | 667 | 646 | 647.37 | 641 | 653 | 1.018 | 0.026 | 0.8542101 | 0.92748187 | 0.013 | 0.012 | 0.0125 | 0.0125 | TRUE | 0.3241 |
| Nae1 | 4596 | 5408 | 5409 | 6583 | 4759 | 4986 | 4882 | 5401 | 5431 | 5175 | 5120 | 5458 | 5244.56 | 5238 | 5251 | 1.002 | 0.003 | 0.96440411 | 0.98213662 | 0.0022 | 0.0075 | 0.004 | 0.004 | TRUE | 0.0431 |

| | | | | | | | | | | | | | | | | | | | | | | | | | |
|---|---|---|---|---|---|---|---|---|---|---|---|---|---|---|---|---|---|---|---|---|---|---|---|---|---|
| Naf1 | 4414 | 4125 | 4304 | 5371 | 4144 | 4243 | 4688 | 4120 | 4322 | 4222 | 4459 | 4645 | 4409.23 | 4377 | 4442 | 1.015 | 0.021 | 0.7992429 | 0.89649805 | 0.0031 | 0.0077 | 0.0047 | 0.0047 | TRUE | 0.0472 |
| Naga | 615 | 1027 | 1135 | 716 | 602 | 836 | 653 | 1026 | 1140 | 563 | 648 | 915 | 824.05 | 940 | 709 | 0.753 | -0.408 | 0.07927331 | 0.22326762 | 0.0685 | 0.0109 | 0.0378 | 0.0378 | TRUE | 0.8453 |
| Nagk | 461 | 555 | 589 | 866 | 652 | 602 | 490 | 554 | 591 | 681 | 702 | 659 | 612.77 | 545 | 681 | 1.248 | 0.319 | 0.00378399 | 0.0253253 | 0.0029 | 0.0123 | 0.0071 | 0.0071 | TRUE | 0.0917 |
| Naglu | 230 | 486 | 429 | 273 | 178 | 184 | 244 | 485 | 431 | 215 | 192 | 201 | 294.66 | 387 | 203 | 0.524 | -0.933 | 0.00011667 | 0.00155805 | 0.0609 | 0.0181 | 0.0385 | 0.0385 | TRUE | 0.7307 |
| Nagpa | 446 | 375 | 363 | 795 | 534 | 464 | 474 | 375 | 364 | 625 | 575 | 508 | 486.7 | 404 | 569 | 1.41 | 0.496 | 0.00080857 | 0.00755496 | 0.0133 | 0.0137 | 0.0137 | 0.0137 | TRUE | 0.2586 |
| Nags | 166 | 268 | 320 | 141 | 146 | 177 | 176 | 268 | 321 | 111 | 157 | 194 | 204.5 | 255 | 154 | 0.601 | -0.735 | 0.00711901 | 0.04059327 | 0.0774 | 0.023 | 0.0485 | 0.0485 | TRUE | 0.2549 |
| Naif1 | 151 | 154 | 144 | 246 | 164 | 153 | 160 | 154 | 145 | 193 | 176 | 167 | 166.02 | 153 | 179 | 1.175 | 0.232 | 0.1470305 | 0.3361306 | 0 | 0.0267 | 0.0124 | 0.0124 | TRUE | 0.0568 |
| Naip1 | 2 | 1 | 0 | 1 | 4 | 1 | 2 | 1 | 0 | 1 | 4 | 1 | 1.55 | 1 | 2 | 1.964 | 0.974 | 0.55852582 | NA | 0 | 2.1306 | 1.2611 | 1.2611 | TRUE | 0.6129 |
| Naip2 | 0 | 0 | 0 | 2 | 2 | 0 | 0 | 0 | 0 | 2 | 2 | 0 | 0.62 | 0 | 1 | 6.637 | 2.73 | 0.366597 | NA | 0 | 5.3147 | 5.1972 | 5.1972 | TRUE | 0.3481 |
| Naip5 | 10 | 11 | 16 | 46 | 25 | 8 | 11 | 11 | 16 | 36 | 27 | 9 | 18.25 | 13 | 24 | 1.923 | 0.943 | 0.1136595 | 0.28427491 | 0.2051 | 0.1874 | 0.1953 | 0.1953 | TRUE | 0.2841 |
| Naip6 | 8 | 9 | 4 | 13 | 5 | 2 | 8 | 9 | 4 | 10 | 5 | 2 | 6.55 | 7 | 6 | 0.848 | -0.237 | 0.77531012 | 0.88309317 | 0.1656 | 0.51 | 0.3452 | 0.3452 | TRUE | 0.4321 |
| Naip7 | 0 | 0 | 0 | 0 | 0 | 2 | 0 | 0 | 0 | 0 | 0 | 2 | 0.36 | 0 | 1 | 3.832 | 1.938 | 0.63022034 | NA | 10 | 9.0356 | 10 | 10 | TRUE | 0.9744 |
| Nalcn | 29 | 31 | 43 | 54 | 36 | 33 | 31 | 31 | 43 | 42 | 39 | 36 | 37.04 | 35 | 39 | 1.12 | 0.164 | 0.61253063 | 0.7806103 | 0 | 0.0958 | 0.0486 | 0.0486 | TRUE | 0.3012 |
| Nampt | 10944 | 13224 | 13163 | 13183 | 10207 | 12010 | 11624 | 13207 | 13217 | 10363 | 10982 | 13147 | 12090.14 | 12683 | 11497 | 0.906 | -0.142 | 0.20687473 | 0.41598394 | 0.0106 | 0.0072 | 0.009 | 0.009 | TRUE | 0.1924 |
| Nanog | 13640 | 17061 | 15668 | 31506 | 21763 | 20265 | 14488 | 17039 | 15732 | 24768 | 23416 | 22183 | 19604.3 | 15753 | 23456 | 1.489 | 0.574 | 9.76E-11 | 7.16E-09 | 0.0047 | 0.0071 | 0.0056 | 0.0056 | TRUE | 0.0624 |
| Nanos1 | 416 | 650 | 640 | 688 | 470 | 461 | 442 | 649 | 643 | 541 | 506 | 505 | 547.47 | 578 | 517 | 0.895 | -0.16 | 0.32604364 | 0.54717603 | 0.0216 | 0.0129 | 0.0174 | 0.0174 | TRUE | 0.4943 |
| Nanos2 | 0 | 1 | 0 | 0 | 1 | 1 | 0 | 1 | 0 | 0 | 1 | 1 | 0.53 | 0 | 1 | 1.924 | 0.944 | 0.7364225 | NA | 0 | 6.2438 | 3.9727 | 3.9727 | TRUE | 0.405 |
| Nanos3 | 15 | 12 | 17 | 62 | 43 | 16 | 16 | 12 | 17 | 49 | 46 | 18 | 26.25 | 15 | 38 | 2.516 | 1.331 | 0.00744006 | 0.0420289 | 0.1301 | 0.1324 | 0.1312 | 0.1312 | TRUE | 1.5013 |
| Nanp | 11 | 11 | 12 | 19 | 9 | 9 | 12 | 11 | 12 | 15 | 10 | 10 | 11.53 | 12 | 12 | 1.005 | 0.007 | 0.98934094 | 0.99437047 | 0 | 0.2926 | 0.1338 | 0.1338 | TRUE | 0.3309 |
| Nans | 4561 | 4761 | 4665 | 6295 | 4371 | 4465 | 4845 | 4755 | 4684 | 4949 | 4703 | 4888 | 4803.8 | 4761 | 4847 | 1.018 | 0.026 | 0.67261163 | 0.81756762 | 3.00E-04 | 0.0076 | 0.0025 | 0.0025 | TRUE | 0.0081 |
| Nap1l1 | 33612 | 29806 | 29628 | 47298 | 32752 | 28799 | 35702 | 29768 | 29750 | 37182 | 35239 | 31525 | 33194.2 | 31740 | 34649 | 1.092 | 0.127 | 0.23640218 | 0.45162196 | 0.0092 | 0.007 | 0.0082 | 0.0082 | TRUE | 0.146 |
| Nap1l2 | 44 | 12 | 11 | 66 | 49 | 27 | 47 | 12 | 11 | 52 | 53 | 30 | 33.99 | 23 | 45 | 1.933 | 0.951 | 0.09921131 | 0.25984818 | 0.3301 | 0.1038 | 0.2067 | 0.2067 | TRUE | 4.4541 |
| Nap1l3 | 105 | 106 | 115 | 173 | 137 | 112 | 112 | 106 | 115 | 136 | 147 | 123 | 123.14 | 111 | 135 | 1.22 | 0.287 | 0.11474901 | 0.28595694 | 0 | 0.0337 | 0.0157 | 0.0157 | TRUE | 0.0646 |
| Nap1l4 | 7399 | 7793 | 7569 | 10774 | 7760 | 7260 | 7859 | 7783 | 7600 | 8470 | 8349 | 7947 | 8001.37 | 7747 | 8255 | 1.066 | 0.092 | 0.13365926 | 0.31591096 | 5.00E-04 | 0.0073 | 0.0026 | 0.0026 | TRUE | 0.013 |
| Nap1l5 | 74 | 78 | 64 | 119 | 75 | 54 | 79 | 78 | 64 | 94 | 81 | 59 | 75.69 | 74 | 78 | 1.063 | 0.089 | 0.7325581 | 0.85654284 | 0.0198 | 0.0504 | 0.0354 | 0.0354 | TRUE | 0.3541 |
| Napa | 2945 | 2820 | 2671 | 4344 | 2908 | 2672 | 3128 | 2816 | 2682 | 3415 | 3129 | 2925 | 3015.85 | 2875 | 3156 | 1.098 | 0.135 | 0.17143436 | 0.37034626 | 0.0057 | 0.008 | 0.0067 | 0.0067 | TRUE | 0.0716 |
| Napb | 150 | 176 | 153 | 223 | 177 | 171 | 159 | 176 | 154 | 175 | 190 | 187 | 173.61 | 163 | 184 | 1.129 | 0.175 | 0.25935321 | 0.47842325 | 0 | 0.0259 | 0.0117 | 0.0117 | TRUE | 0.0512 |
| Napepld | 1306 | 1534 | 1455 | 1329 | 1051 | 1115 | 1387 | 1532 | 1461 | 1045 | 1131 | 1221 | 1296.05 | 1460 | 1132 | 0.774 | -0.369 | 0.00012478 | 0.00165103 | 0.0035 | 0.0094 | 0.0059 | 0.0059 | TRUE | 0.0563 |
| Napg | 2154 | 2302 | 2201 | 2860 | 1979 | 1975 | 2288 | 2299 | 2210 | 2248 | 2129 | 2162 | 2222.76 | 2266 | 2180 | 0.963 | -0.055 | 0.42292391 | 0.63322156 | 0 | 0.0084 | 0.003 | 0.003 | TRUE | 0.0094 |
| Naprt1 | 272 | 344 | 382 | 313 | 244 | 235 | 289 | 344 | 384 | 246 | 263 | 257 | 296.98 | 339 | 255 | 0.753 | -0.41 | 0.00588535 | 0.03540185 | 0.0078 | 0.018 | 0.0125 | 0.0125 | TRUE | 0.186 |
| Napsa | 16 | 16 | 13 | 11 | 3 | 12 | 17 | 16 | 13 | 9 | 3 | 13 | 11.84 | 15 | 8 | 0.544 | -0.879 | 0.15604816 | 0.3487095 | 0.0142 | 0.2852 | 0.1848 | 0.1848 | TRUE | 0.3866 |
| Narf | 2544 | 2944 | 3132 | 3239 | 2373 | 2460 | 2702 | 2940 | 3145 | 2546 | 2553 | 2693 | 2763.26 | 2929 | 2597 | 0.887 | -0.174 | 0.04254134 | 0.14725943 | 0.0028 | 0.0081 | 0.0049 | 0.0049 | TRUE | 0.0556 |
| Narfl | 520 | 494 | 464 | 870 | 642 | 514 | 552 | 493 | 466 | 684 | 691 | 563 | 574.82 | 504 | 646 | 1.283 | 0.36 | 0.00559606 | 0.03399887 | 0.0085 | 0.0126 | 0.0104 | 0.0104 | TRUE | 0.1465 |
| Narg2 | 1396 | 1623 | 1617 | 1948 | 1448 | 1472 | 1483 | 1621 | 1624 | 1531 | 1558 | 1611 | 1571.33 | 1576 | 1567 | 0.994 | -0.009 | 0.91159097 | 0.95633982 | 0.001 | 0.009 | 0.004 | 0.004 | TRUE | 0.0319 |
| Nars | 9023 | 8003 | 7709 | 14059 | 9543 | 8245 | 9584 | 7993 | 7741 | 11052 | 10268 | 9025 | 9277.09 | 8439 | 10115 | 1.199 | 0.262 | 0.02570614 | 0.1036672 | 0.012 | 0.0073 | 0.0098 | 0.0098 | TRUE | 0.1719 |
| Nars2 | 1229 | 1481 | 1354 | 1638 | 1196 | 1193 | 1305 | 1479 | 1360 | 1288 | 1287 | 1306 | 1337.41 | 1381 | 1294 | 0.936 | -0.095 | 0.26053847 | 0.47988003 | 0.0014 | 0.0094 | 0.0044 | 0.0044 | TRUE | 0.0462 |
| Nasp | 35408 | 30254 | 30865 | 43924 | 31839 | 28936 | 37610 | 30215 | 30992 | 34530 | 34257 | 31675 | 33212.94 | 32939 | 33487 | 1.017 | 0.024 | 0.81913037 | 0.90763834 | 0.0069 | 0.007 | 0.0078 | 0.0078 | TRUE | 0.1883 |
| Nat1 | 25 | 43 | 45 | 59 | 44 | 52 | 27 | 43 | 45 | 46 | 47 | 57 | 44.22 | 38 | 50 | 1.308 | 0.387 | 0.22823792 | 0.44159023 | 0.0141 | 0.0814 | 0.0513 | 0.0513 | TRUE | 0.5072 |
| Nat10 | 6190 | 6325 | 6341 | 7781 | 6001 | 5554 | 6575 | 6317 | 6367 | 6117 | 6457 | 6080 | 6318.66 | 6420 | 6218 | 0.969 | -0.046 | 0.4623081 | 0.66599473 | 5.00E-04 | 0.0074 | 0.0027 | 0.0027 | TRUE | 0.0137 |
| Nat14 | 20 | 35 | 28 | 27 | 21 | 20 | 21 | 35 | 28 | 21 | 23 | 22 | 25 | 28 | 22 | 0.777 | -0.364 | 0.35362136 | 0.57203241 | 0 | 0.1387 | 0.0704 | 0.0704 | TRUE | 0.1501 |
| Nat2 | 245 | 299 | 311 | 406 | 273 | 340 | 260 | 299 | 312 | 319 | 294 | 372 | 309.37 | 290 | 328 | 1.13 | 0.176 | 0.2368859 | 0.45221065 | 0.0082 | 0.0176 | 0.0127 | 0.0127 | TRUE | 0.1494 |
| Nat3 | 3 | 1 | 2 | 1 | 0 | 5 | 3 | 1 | 2 | 1 | 0 | 5 | 2.08 | 2 | 2 | 0.993 | -0.011 | 0.99445792 | NA | 0.7389 | 1.5942 | 1.189 | 1.189 | TRUE | 1.7201 |
| Nat6 | 223 | 239 | 224 | 288 | 202 | 193 | 237 | 239 | 225 | 226 | 217 | 211 | 225.91 | 234 | 218 | 0.936 | -0.095 | 0.48344158 | 0.68146464 | 0 | 0.0215 | 0.0088 | 0.0088 | TRUE | 0.0125 |
| Nat8 | 0 | 0 | 0 | 0 | 1 | 2 | 0 | 0 | 0 | 0 | 1 | 2 | 0.54 | 0 | 1 | 5.713 | 2.514 | 0.43612448 | NA | 0 | 6.0606 | 6.0547 | 6.0547 | TRUE | 0.0576 |
| Nat8l | 311 | 481 | 487 | 527 | 372 | 342 | 330 | 480 | 489 | 414 | 400 | 374 | 414.77 | 433 | 396 | 0.915 | -0.128 | 0.45033477 | 0.6557127 | 0.022 | 0.0148 | 0.0185 | 0.0185 | TRUE | 0.4956 |
| Nat9 | 552 | 719 | 762 | 812 | 678 | 630 | 586 | 718 | 765 | 638 | 729 | 690 | 687.83 | 690 | 686 | 0.993 | -0.01 | 0.93827273 | 0.96911836 | 0.0101 | 0.0117 | 0.0108 | 0.0108 | TRUE | 0.2022 |
| Nav1 | 3809 | 3362 | 3560 | 3336 | 2614 | 2578 | 4046 | 3358 | 3575 | 2623 | 2813 | 2822 | 3205.86 | 3660 | 2753 | 0.752 | -0.411 | 1.62E-05 | 0.0002984 | 0.005 | 0.0079 | 0.0062 | 0.0062 | TRUE | 0.1037 |
| Nav2 | 1586 | 1541 | 1988 | 1794 | 1293 | 1502 | 1685 | 1539 | 1996 | 1410 | 1391 | 1644 | 1610.91 | 1740 | 1482 | 0.851 | -0.232 | 0.06675498 | 0.1993297 | 0.0126 | 0.0089 | 0.0109 | 0.0109 | TRUE | 0.2007 |
| Nav3 | 4 | 0 | 1 | 7 | 5 | 3 | 4 | 0 | 1 | 6 | 5 | 3 | 3.24 | 2 | 5 | 2.745 | 1.457 | 0.23494755 | NA | 0 | 1.0248 | 0.6852 | 0.6852 | TRUE | 1.1755 |
| Nbas | 1804 | 1798 | 2073 | 2844 | 2096 | 2030 | 1916 | 1796 | 2082 | 2236 | 2255 | 2222 | 2084.4 | 1931 | 2238 | 1.159 | 0.213 | 0.01123982 | 0.05738895 | 0.0021 | 0.0085 | 0.0046 | 0.0046 | TRUE | 0.0563 |
| Nbea | 661 | 615 | 674 | 1101 | 691 | 625 | 702 | 614 | 677 | 866 | 743 | 684 | 714.37 | 664 | 764 | 1.153 | 0.205 | 0.09998514 | 0.26113025 | 0.0084 | 0.0115 | 0.0098 | 0.0098 | TRUE | 0.1632 |
| Nbeal1 | 1796 | 1620 | 1970 | 2178 | 1579 | 1512 | 1908 | 1618 | 1978 | 1712 | 1699 | 1655 | 1761.64 | 1835 | 1689 | 0.921 | -0.119 | 0.23452298 | 0.44941796 | 0.0047 | 0.0088 | 0.0067 | 0.0067 | TRUE | 0.1293 |
| Nbeal2 | 1935 | 1810 | 1901 | 2198 | 1681 | 1491 | 2055 | 1808 | 1909 | 1728 | 1809 | 1632 | 1823.41 | 1924 | 1723 | 0.896 | -0.159 | 0.07243021 | 0.2105692 | 0.0026 | 0.0087 | 0.0051 | 0.0051 | TRUE | 0.043 |
| Nbl1 | 88 | 73 | 51 | 92 | 58 | 48 | 93 | 73 | 51 | 72 | 62 | 53 | 67.48 | 72 | 62 | 0.865 | -0.209 | 0.4837661 | 0.68148223 | 0.0424 | 0.0557 | 0.0492 | 0.0492 | TRUE | 0.1722 |
| Nbn | 1417 | 1456 | 1487 | 1983 | 1521 | 1470 | 1505 | 1454 | 1493 | 1559 | 1637 | 1609 | 1542.81 | 1484 | 1602 | 1.079 | 0.109 | 0.13011583 | 0.31070082 | 0 | 0.009 | 0.0031 | 0.0031 | TRUE | 0.0068 |
| Nbr1 | 11123 | 11041 | 11312 | 15962 | 11524 | 11205 | 11815 | 11027 | 11358 | 12548 | 12399 | 12266 | 11902.1 | 11400 | 12404 | 1.088 | 0.122 | 0.04111509 | 0.14362506 | 5.00E-04 | 0.0072 | 0.0025 | 0.0025 | TRUE | 0.0124 |
| Ncald | 33 | 27 | 25 | 51 | 24 | 36 | 35 | 27 | 25 | 40 | 26 | 39 | 32.07 | 29 | 35 | 1.216 | 0.282 | 0.43848346 | 0.64591156 | 0 | 0.1096 | 0.0638 | 0.0638 | TRUE | 0.359 |
| Ncam1 | 731 | 960 | 959 | 1090 | 804 | 911 | 776 | 959 | 963 | 857 | 865 | 997 | 902.88 | 899 | 906 | 1.007 | 0.01 | 0.93650255 | 0.96825513 | 0.0098 | 0.0106 | 0.0101 | 0.0101 | TRUE | 0.1694 |
| Ncam2 | 3 | 0 | 3 | 2 | 1 | 1 | 3 | 0 | 3 | 2 | 1 | 1 | 1.66 | 2 | 1 | 0.613 | -0.706 | 0.64899478 | NA | 0 | 1.9952 | 1.0994 | 1.0994 | TRUE | 0.7241 |
| Ncan | 791 | 626 | 610 | 355 | 243 | 257 | 840 | 625 | 613 | 279 | 261 | 281 | 483.29 | 693 | 274 | 0.396 | -1.337 | 1.09E-17 | 2.13E-15 | 0.0162 | 0.0137 | 0.015 | 0.015 | TRUE | 0.4086 |
| Ncapd2 | 22935 | 28350 | 29529 | 29092 | 21764 | 25014 | 24361 | 28313 | 29650 | 22870 | 23417 | 27382 | 25998.83 | 27441 | 24556 | 0.895 | -0.16 | 0.14582085 | 0.33433441 | 0.01 | 0.007 | 0.0087 | 0.0087 | TRUE | 0.124 |
| Ncapd3 | 7008 | 6623 | 7286 | 7814 | 5654 | 5961 | 7444 | 6614 | 7316 | 6143 | 6083 | 6525 | 6687.58 | 7125 | 6250 | 0.877 | -0.189 | 0.01595816 | 0.07424548 | 0.0025 | 0.0074 | 0.0043 | 0.0043 | TRUE | 0.0479 |
| Ncapg | 8285 | 9540 | 10499 | 13326 | 9557 | 10084 | 8800 | 9528 | 10542 | 10476 | 10283 | 11039 | 10111.17 | 9623 | 10599 | 1.101 | 0.139 | 0.11891572 | 0.29200355 | 0.0047 | 0.0072 | 0.0056 | 0.0056 | TRUE | 0.0852 |
| Ncapg2 | 4295 | 4197 | 4281 | 6512 | 4530 | 4227 | 4562 | 4192 | 4299 | 5119 | 4874 | 4627 | 4612.09 | 4351 | 4873 | 1.12 | 0.164 | 0.03113854 | 0.11834144 | 0.0019 | 0.0076 | 0.004 | 0.004 | TRUE | 0.024 |
| Ncaph | 5332 | 5455 | 5459 | 6704 | 4845 | 5076 | 5664 | 5448 | 5481 | 5270 | 5213 | 5556 | 5438.74 | 5531 | 5346 | 0.967 | -0.049 | 0.44137519 | 0.6484368 | 5.00E-04 | 0.0075 | 0.0027 | 0.0027 | TRUE | 0.0143 |
| Ncaph2 | 7503 | 9724 | 9820 | 12829 | 9712 | 10351 | 7970 | 9711 | 9860 | 10085 | 10450 | 11331 | 9901.12 | 9180 | 10622 | 1.157 | 0.21 | 0.04726186 | 0.15785128 | 0.0082 | 0.0072 | 0.008 | 0.008 | TRUE | 0.1625 |
| Ncbp1 | 5684 | 5566 | 5522 | 8465 | 6222 | 6040 | 6037 | 5559 | 5545 | 6655 | 6694 | 6612 | 6183.61 | 5714 | 6654 | 1.165 | 0.22 | 0.00101829 | 0.00903796 | 0.001 | 0.0074 | 0.0031 | 0.0031 | TRUE | 0.0299 |
| Ncbp2 | 4957 | 4486 | 4529 | 5515 | 3997 | 4016 | 5265 | 4480 | 4548 | 4335 | 4301 | 4396 | 4554.19 | 4764 | 4344 | 0.912 | -0.133 | 0.12826415 | 0.30801003 | 0.0039 | 0.0076 | 0.0053 | 0.0053 | TRUE | 0.1029 |
| Nccrp1 | 327 | 384 | 390 | 147 | 108 | 149 | 347 | 384 | 392 | 116 | 116 | 163 | 252.88 | 374 | 132 | 0.35 | -1.514 | 3.55E-18 | 7.19E-16 | 0.0128 | 0.0199 | 0.0168 | 0.0168 | TRUE | 0.4321 |

| | | | | | | | | | | | | | | | | | | | | | | | | | |
|---|---|---|---|---|---|---|---|---|---|---|---|---|---|---|---|---|---|---|---|---|---|---|---|---|---|
| Ncdn | 1811 | 1748 | 1524 | 2604 | 1915 | 1494 | 1924 | 1746 | 1530 | 2047 | 2060 | 1635 | 1823.75 | 1733 | 1914 | 1.105 | 0.144 | 0.27001781 | 0.49018427 | 0.0136 | 0.0087 | 0.0117 | 0.0117 | TRUE | 0.1951 |
| Nceh1 | 1880 | 2000 | 1974 | 2385 | 1665 | 1647 | 1997 | 1997 | 1982 | 1875 | 1791 | 1803 | 1907.61 | 1992 | 1823 | 0.916 | -0.127 | 0.0610982 | 0.18838455 | 0 | 0.0086 | 0.0028 | 0.0028 | TRUE | 0.0077 |
| Ncf1 | 22 | 15 | 12 | 20 | 16 | 17 | 23 | 15 | 12 | 16 | 17 | 19 | 16.99 | 17 | 17 | 1.022 | 0.032 | 0.94712201 | 0.97354069 | 0 | 0.2008 | 0.1072 | 0.1072 | TRUE | 0.5494 |
| Ncf2 | 53 | 51 | 61 | 54 | 51 | 41 | 56 | 51 | 61 | 42 | 55 | 45 | 51.78 | 56 | 47 | 0.841 | -0.25 | 0.36020999 | 0.57859578 | 0 | 0.0705 | 0.0344 | 0.0344 | TRUE | 0.146 |
| Ncf4 | 0 | 0 | 1 | 0 | 1 | 2 | 0 | 0 | 1 | 0 | 1 | 2 | 0.71 | 0 | 1 | 2.906 | 1.539 | 0.55074121 | NA | 0 | 4.6368 | 3.3331 | 3.3331 | TRUE | 0.1901 |
| Nck1 | 1362 | 1446 | 1377 | 1800 | 1268 | 1239 | 1447 | 1444 | 1383 | 1415 | 1364 | 1356 | 1401.51 | 1425 | 1378 | 0.968 | -0.047 | 0.53254589 | 0.7204764 | 0 | 0.0093 | 0.0033 | 0.0033 | TRUE | 0.008 |
| Nck2 | 534 | 670 | 638 | 845 | 686 | 674 | 567 | 669 | 641 | 664 | 738 | 738 | 669.52 | 626 | 713 | 1.139 | 0.187 | 0.0909359 | 0.24596829 | 0.0039 | 0.0118 | 0.0073 | 0.0073 | TRUE | 0.0779 |
| Nckap1 | 8965 | 9739 | 10436 | 11179 | 8247 | 8700 | 9522 | 9726 | 10479 | 8788 | 8873 | 9524 | 9485.42 | 9909 | 9062 | 0.914 | -0.129 | 0.08195404 | 0.22845159 | 0.0021 | 0.0073 | 0.0039 | 0.0039 | TRUE | 0.0309 |
| Nckap1l | 1145 | 773 | 785 | 1531 | 1023 | 835 | 1216 | 772 | 788 | 1204 | 1101 | 914 | 999.12 | 925 | 1073 | 1.16 | 0.214 | 0.26497328 | 0.48470871 | 0.0396 | 0.0102 | 0.0256 | 0.0256 | TRUE | 0.8983 |
| Nckap5 | 41 | 17 | 13 | 75 | 45 | 31 | 44 | 17 | 13 | 59 | 48 | 34 | 35.82 | 25 | 47 | 1.934 | 0.951 | 0.05323895 | 0.17132927 | 0.1963 | 0.0989 | 0.1433 | 0.1433 | TRUE | 1.1706 |
| Nckap5l | 1011 | 671 | 591 | 902 | 686 | 556 | 1074 | 670 | 593 | 709 | 738 | 609 | 732.2 | 779 | 685 | 0.88 | -0.185 | 0.38664416 | 0.60061236 | 0.0567 | 0.0114 | 0.0314 | 0.0314 | TRUE | 1.2959 |
| Nckipsd | 689 | 843 | 812 | 1020 | 803 | 801 | 732 | 842 | 815 | 802 | 864 | 877 | 821.95 | 796 | 848 | 1.063 | 0.088 | 0.37985087 | 0.59453654 | 0.0026 | 0.0109 | 0.006 | 0.006 | TRUE | 0.0591 |
| Ncl | 127228 | 114466 | 114329 | 154973 | 112136 | 101933 | 135139 | 114318 | 114798 | 121828 | 120651 | 111582 | 119719.33 | 121418 | 118020 | 0.972 | -0.041 | 0.65966805 | 0.80828811 | 0.0051 | 0.0069 | 0.0062 | 0.0062 | TRUE | 0.1197 |
| Ncln | 3728 | 3523 | 3415 | 5159 | 3848 | 3284 | 3960 | 3518 | 3429 | 4056 | 4140 | 3595 | 3782.99 | 3636 | 3930 | 1.081 | 0.113 | 0.24291005 | 0.4595404 | 0.0054 | 0.0078 | 0.0064 | 0.0064 | TRUE | 0.0738 |
| Ncmap | 65 | 75 | 49 | 118 | 70 | 47 | 69 | 75 | 49 | 93 | 75 | 51 | 68.78 | 64 | 73 | 1.143 | 0.193 | 0.52886865 | 0.71740917 | 0.0516 | 0.0548 | 0.0533 | 0.0533 | TRUE | 0.2466 |
| Ncoa1 | 2157 | 1971 | 2030 | 2376 | 1731 | 1687 | 2291 | 1968 | 2038 | 1868 | 1862 | 1847 | 1979.14 | 2099 | 1859 | 0.886 | -0.175 | 0.04428343 | 0.15104634 | 0.0025 | 0.0086 | 0.005 | 0.005 | TRUE | 0.077 |
| Ncoa2 | 2312 | 2397 | 2654 | 3619 | 2670 | 2496 | 2456 | 2394 | 2665 | 2845 | 2873 | 2732 | 2660.76 | 2505 | 2817 | 1.125 | 0.169 | 0.02756259 | 0.10873367 | 0.0014 | 0.0081 | 0.0039 | 0.0039 | TRUE | 0.038 |
| Ncoa3 | 10426 | 9759 | 10507 | 15501 | 10530 | 9903 | 11074 | 9746 | 10550 | 12186 | 11330 | 10840 | 10954.41 | 10457 | 11452 | 1.095 | 0.131 | 0.11839991 | 0.29119031 | 0.0037 | 0.0072 | 0.005 | 0.005 | TRUE | 0.0432 |
| Ncoa4 | 2179 | 1630 | 1658 | 2706 | 1810 | 1574 | 2314 | 1628 | 1665 | 2127 | 1947 | 1723 | 1900.81 | 1869 | 1932 | 1.034 | 0.049 | 0.75324627 | 0.86983454 | 0.0249 | 0.0086 | 0.0169 | 0.0169 | TRUE | 0.5245 |
| Ncoa5 | 2376 | 1779 | 1710 | 3259 | 2179 | 1604 | 2524 | 1777 | 1717 | 2562 | 2344 | 1756 | 2113.29 | 2006 | 2221 | 1.108 | 0.147 | 0.4244077 | 0.63456714 | 0.0415 | 0.0085 | 0.0241 | 0.0241 | TRUE | 0.6165 |
| Ncoa6 | 1797 | 2038 | 2206 | 2939 | 2264 | 2396 | 1909 | 2035 | 2215 | 2310 | 2436 | 2623 | 2254.72 | 2053 | 2456 | 1.196 | 0.258 | 0.00617278 | 0.03678015 | 0.0042 | 0.0084 | 0.0059 | 0.0059 | TRUE | 0.0577 |
| Ncoa7 | 364 | 303 | 253 | 529 | 377 | 277 | 387 | 303 | 254 | 416 | 406 | 303 | 344.66 | 315 | 375 | 1.194 | 0.256 | 0.19308393 | 0.39859714 | 0.0322 | 0.0165 | 0.025 | 0.025 | TRUE | 0.4542 |
| Ncor1 | 11475 | 10515 | 11509 | 18123 | 12708 | 11759 | 12188 | 10501 | 11556 | 14247 | 13673 | 12872 | 12506.36 | 11415 | 13597 | 1.191 | 0.252 | 0.00318117 | 0.02221848 | 0.004 | 0.0072 | 0.0052 | 0.0052 | TRUE | 0.06 |
| Ncor2 | 8707 | 8220 | 9084 | 8076 | 6098 | 6587 | 9248 | 8209 | 9121 | 6349 | 6561 | 7211 | 7783.23 | 8859 | 6707 | 0.757 | -0.402 | 4.11E-06 | 9.34E-05 | 0.0041 | 0.0073 | 0.0054 | 0.0054 | TRUE | 0.0526 |
| Ncr1 | 0 | 0 | 0 | 0 | 0 | 0 | 0 | 0 | 0 | 0 | 0 | 0 | 0 | 0 | 0 | NA | NA | NA | NA | NA | NA | NA | NA | NA | NA |
| Ncrna00085 | 30 | 25 | 28 | 46 | 46 | 34 | 32 | 25 | 28 | 36 | 49 | 37 | 34.64 | 28 | 41 | 1.443 | 0.529 | 0.11693059 | 0.28910947 | 0 | 0.102 | 0.0519 | 0.0519 | TRUE | 0.2407 |
| Ncrna00086 | 486 | 508 | 553 | 740 | 522 | 532 | 516 | 507 | 555 | 582 | 562 | 582 | 550.76 | 526 | 575 | 1.093 | 0.129 | 0.19506716 | 0.40091206 | 0 | 0.0129 | 0.0053 | 0.0053 | TRUE | 0.0273 |
| Ncs1 | 555 | 713 | 714 | 898 | 716 | 655 | 590 | 712 | 717 | 706 | 770 | 717 | 701.97 | 673 | 731 | 1.086 | 0.119 | 0.30431349 | 0.52512118 | 0.0054 | 0.0116 | 0.0082 | 0.0082 | TRUE | 0.1374 |
| Ncstn | 1634 | 1597 | 1564 | 2376 | 1751 | 1598 | 1736 | 1595 | 1570 | 1868 | 1884 | 1749 | 1733.67 | 1634 | 1834 | 1.123 | 0.167 | 0.04289848 | 0.14795428 | 0.0014 | 0.0088 | 0.0043 | 0.0043 | TRUE | 0.0358 |
| Nctc1 | 0 | 0 | 0 | 0 | 0 | 0 | 0 | 0 | 0 | 0 | 0 | 0 | 0 | 0 | 0 | NA | NA | NA | NA | NA | NA | NA | NA | NA | NA |
| Ndc80 | 1999 | 2535 | 2712 | 3159 | 2183 | 2329 | 2123 | 2532 | 2723 | 2483 | 2349 | 2549 | 2459.96 | 2459 | 2460 | 1 | 0.001 | 0.99611554 | 0.99814068 | 0.0084 | 0.0082 | 0.0084 | 0.0084 | TRUE | 0.1728 |
| Nde1 | 3149 | 3948 | 3914 | 4195 | 3122 | 3314 | 3345 | 3943 | 3930 | 3298 | 3359 | 3628 | 3583.72 | 3739 | 3428 | 0.917 | -0.126 | 0.18916211 | 0.39431214 | 0.0051 | 0.0078 | 0.0063 | 0.0063 | TRUE | 0.1034 |
| Ndel1 | 2490 | 3436 | 3554 | 3367 | 2470 | 2907 | 2645 | 3432 | 3569 | 2647 | 2658 | 3182 | 3021.93 | 3215 | 2829 | 0.88 | -0.185 | 0.17841404 | 0.37934029 | 0.0174 | 0.008 | 0.0133 | 0.0133 | TRUE | 0.2921 |
| Ndfip1 | 2111 | 2962 | 2943 | 3258 | 2532 | 2700 | 2242 | 2958 | 2955 | 2561 | 2724 | 2956 | 2732.76 | 2718 | 2747 | 1.01 | 0.015 | 0.90966455 | 0.95489676 | 0.0145 | 0.0081 | 0.0115 | 0.0115 | TRUE | 0.2845 |
| Ndfip2 | 1753 | 1946 | 1848 | 2498 | 1693 | 1753 | 1862 | 1943 | 1856 | 1964 | 1822 | 1919 | 1894.22 | 1887 | 1902 | 1.008 | 0.012 | 0.8762823 | 0.9390993 | 6.00E-04 | 0.0086 | 0.0034 | 0.0034 | TRUE | 0.0162 |
| Ndn | 222 | 272 | 237 | 725 | 490 | 407 | 236 | 272 | 238 | 570 | 527 | 446 | 381.35 | 249 | 514 | 2.072 | 1.051 | 3.24E-13 | 3.55E-11 | 0.009 | 0.0155 | 0.012 | 0.012 | TRUE | 0.1573 |
| Ndnf | 23 | 23 | 41 | 44 | 32 | 19 | 24 | 23 | 41 | 35 | 34 | 21 | 29.73 | 29 | 30 | 1.018 | 0.026 | 0.95005629 | 0.97481103 | 0.0579 | 0.1177 | 0.0891 | 0.0891 | TRUE | 0.7917 |
| Ndnl2 | 782 | 822 | 791 | 1295 | 968 | 916 | 831 | 821 | 794 | 1018 | 1042 | 1003 | 918.01 | 815 | 1021 | 1.252 | 0.324 | 7.35E-05 | 0.00107706 | 0 | 0.0105 | 0.0037 | 0.0037 | TRUE | 0.0061 |
| Ndor1 | 1591 | 1447 | 1251 | 2267 | 1798 | 1444 | 1690 | 1445 | 1256 | 1782 | 1935 | 1581 | 1614.76 | 1464 | 1766 | 1.207 | 0.271 | 0.04356104 | 0.14942251 | 0.0152 | 0.0089 | 0.0124 | 0.0124 | TRUE | 0.2195 |
| Ndp | 0 | 0 | 0 | 0 | 1 | 0 | 0 | 0 | 0 | 0 | 1 | 0 | 0.18 | 0 | 0 | 1.859 | 0.894 | 0.82650734 | NA | 0 | 18.3785 | 10 | 10 | TRUE | 0.5204 |
| Ndrg1 | 2588 | 4520 | 5200 | 4105 | 3475 | 4610 | 2749 | 4514 | 5221 | 3227 | 3739 | 5046 | 4082.79 | 4161 | 4004 | 0.962 | -0.056 | 0.80793678 | 0.90112858 | 0.0779 | 0.0077 | 0.0378 | 0.0378 | TRUE | 0.7056 |
| Ndrg2 | 2754 | 4483 | 5019 | 2416 | 1944 | 2943 | 2925 | 4477 | 5040 | 1899 | 2092 | 3222 | 3275.75 | 4147 | 2404 | 0.58 | -0.787 | 0.00065341 | 0.00642491 | 0.0785 | 0.0079 | 0.0381 | 0.0381 | TRUE | 0.7188 |
| Ndrg3 | 3404 | 3755 | 3813 | 3442 | 2423 | 2792 | 3616 | 3750 | 3829 | 2706 | 2607 | 3056 | 3260.59 | 3732 | 2790 | 0.747 | -0.42 | 1.53E-06 | 3.96E-05 | 0.0035 | 0.0079 | 0.0052 | 0.0052 | TRUE | 0.084 |
| Ndrg4 | 3243 | 2738 | 2594 | 3209 | 2291 | 2092 | 3445 | 2734 | 2605 | 2523 | 2465 | 2290 | 2676.9 | 2928 | 2426 | 0.829 | -0.271 | 0.02696823 | 0.10720009 | 0.0126 | 0.0081 | 0.0104 | 0.0104 | TRUE | 0.2891 |
| Ndst1 | 1226 | 1168 | 1159 | 1453 | 1108 | 1090 | 1302 | 1166 | 1164 | 1142 | 1192 | 1193 | 1193.34 | 1211 | 1176 | 0.971 | -0.043 | 0.62779353 | 0.7906068 | 0.0016 | 0.0097 | 0.0047 | 0.0047 | TRUE | 0.052 |
| Ndst2 | 567 | 627 | 712 | 637 | 472 | 536 | 602 | 626 | 715 | 501 | 508 | 587 | 589.78 | 648 | 532 | 0.82 | -0.287 | 0.01902948 | 0.08376584 | 0.0063 | 0.0125 | 0.009 | 0.009 | TRUE | 0.097 |
| Ndst3 | 9 | 1 | 4 | 5 | 3 | 5 | 10 | 1 | 4 | 4 | 3 | 5 | 4.53 | 5 | 4 | 0.869 | -0.202 | 0.8403604 | NA | 0.2563 | 0.7334 | 0.5019 | 0.5019 | TRUE | 0.389 |
| Ndst4 | 0 | 0 | 0 | 0 | 3 | 0 | 0 | 0 | 0 | 0 | 3 | 0 | 0.54 | 0 | 1 | 5.683 | 2.507 | 0.52963929 | NA | 10 | 6.1308 | 10 | 10 | TRUE | 1.4428 |
| Ndufa1 | 1517 | 1775 | 1884 | 1994 | 1352 | 1430 | 1611 | 1773 | 1892 | 1568 | 1455 | 1565 | 1643.89 | 1759 | 1529 | 0.87 | -0.202 | 0.03072789 | 0.11725441 | 0.0032 | 0.0089 | 0.0057 | 0.0057 | TRUE | 0.0645 |
| Ndufa10 | 3704 | 3868 | 4151 | 5501 | 4011 | 3649 | 3934 | 3863 | 4168 | 4324 | 4316 | 3994 | 4099.97 | 3988 | 4211 | 1.056 | 0.079 | 0.28436124 | 0.5058795 | 0.0014 | 0.0077 | 0.0037 | 0.0037 | TRUE | 0.0246 |
| Ndufa11 | 2843 | 3030 | 3132 | 4020 | 2721 | 2739 | 3020 | 3026 | 3145 | 3160 | 2928 | 2998 | 3046.14 | 3064 | 3029 | 0.989 | -0.016 | 0.81791337 | 0.90711833 | 8.00E-04 | 0.008 | 0.0031 | 0.0031 | TRUE | 0.0179 |
| Ndufa12 | 2388 | 2515 | 2467 | 3096 | 2133 | 2228 | 2536 | 2512 | 2477 | 2434 | 2295 | 2439 | 2448.84 | 2508 | 2389 | 0.953 | -0.07 | 0.30296288 | 0.52371006 | 2.00E-04 | 0.0082 | 0.0029 | 0.0029 | TRUE | 0.0142 |
| Ndufa13 | 1774 | 2352 | 2392 | 3009 | 2125 | 2212 | 1884 | 2349 | 2402 | 2365 | 2286 | 2421 | 2284.71 | 2212 | 2357 | 1.066 | 0.092 | 0.40871272 | 0.62120237 | 0.0086 | 0.0083 | 0.0085 | 0.0085 | TRUE | 0.2025 |
| Ndufa2 | 1506 | 1760 | 1789 | 2394 | 1642 | 1579 | 1600 | 1758 | 1796 | 1882 | 1767 | 1728 | 1755.14 | 1718 | 1792 | 1.044 | 0.062 | 0.47248657 | 0.67362214 | 0.0023 | 0.0088 | 0.0048 | 0.0048 | TRUE | 0.0435 |
| Ndufa3 | 1676 | 1713 | 1682 | 1903 | 1356 | 1444 | 1780 | 1711 | 1689 | 1496 | 1459 | 1581 | 1619.26 | 1727 | 1512 | 0.876 | -0.192 | 0.01228224 | 0.06126358 | 5.00E-04 | 0.0089 | 0.0036 | 0.0036 | TRUE | 0.0186 |
| Ndufa4 | 5003 | 5462 | 5506 | 7263 | 4882 | 5037 | 5314 | 5455 | 5529 | 5710 | 5253 | 5514 | 5462.29 | 5433 | 5492 | 1.011 | 0.016 | 0.80960763 | 0.90202964 | 9.00E-04 | 0.0075 | 0.003 | 0.003 | TRUE | 0.0177 |
| Ndufa4l2 | 823 | 2224 | 2615 | 1168 | 935 | 1480 | 874 | 2221 | 2626 | 918 | 1006 | 1620 | 1544.22 | 1907 | 1181 | 0.619 | -0.691 | 0.18205649 | 0.38387633 | 0.1925 | 0.009 | 0.08 | 0.1925 | TRUE | 0.7858 |
| Ndufa5 | 1175 | 1132 | 1121 | 1781 | 1244 | 1170 | 1248 | 1131 | 1126 | 1400 | 1338 | 1281 | 1253.92 | 1168 | 1340 | 1.148 | 0.199 | 0.02512569 | 0.10216097 | 0.0019 | 0.0095 | 0.0049 | 0.0049 | TRUE | 0.0427 |
| Ndufa6 | 2630 | 3581 | 3564 | 3495 | 2668 | 3140 | 2794 | 3576 | 3579 | 2747 | 2871 | 3437 | 3167.31 | 3316 | 3018 | 0.91 | -0.136 | 0.30873149 | 0.529319 | 0.0159 | 0.0079 | 0.0126 | 0.0126 | TRUE | 0.2308 |
| Ndufa7 | 4481 | 6424 | 6386 | 6572 | 4906 | 5517 | 4760 | 6416 | 6412 | 5166 | 5279 | 6039 | 5678.62 | 5863 | 5495 | 0.937 | -0.094 | 0.48613555 | 0.68299749 | 0.0152 | 0.0075 | 0.0129 | 0.0129 | TRUE | 0.3301 |
| Ndufa8 | 2056 | 2464 | 2456 | 3164 | 2346 | 2316 | 2184 | 2461 | 2466 | 2487 | 2524 | 2535 | 2442.9 | 2370 | 2515 | 1.061 | 0.085 | 0.2888585 | 0.50977692 | 0.0018 | 0.0083 | 0.0043 | 0.0043 | TRUE | 0.0571 |
| Ndufa9 | 6996 | 8155 | 8310 | 9526 | 6759 | 6848 | 7431 | 8144 | 8344 | 7489 | 7272 | 7496 | 7696.11 | 7973 | 7419 | 0.93 | -0.104 | 0.15370817 | 0.34542149 | 0.0018 | 0.0073 | 0.0037 | 0.0037 | TRUE | 0.0431 |
| Ndufab1 | 1626 | 1875 | 2017 | 2298 | 1543 | 1654 | 1727 | 1873 | 2025 | 1807 | 1660 | 1811 | 1817.03 | 1875 | 1759 | 0.938 | -0.092 | 0.32548477 | 0.54670349 | 0.0035 | 0.0087 | 0.0057 | 0.0057 | TRUE | 0.0594 |
| Ndufaf1 | 1302 | 1297 | 1195 | 1636 | 1169 | 1223 | 1383 | 1295 | 1200 | 1286 | 1258 | 1339 | 1293.47 | 1293 | 1294 | 1.001 | 0.002 | 0.98445588 | 0.99187671 | 0.0022 | 0.0095 | 0.0051 | 0.0051 | TRUE | 0.0474 |
| Ndufaf2 | 936 | 860 | 795 | 1260 | 839 | 803 | 994 | 859 | 798 | 991 | 903 | 879 | 903.93 | 884 | 924 | 1.047 | 0.066 | 0.57102173 | 0.74982555 | 0.0072 | 0.0105 | 0.0086 | 0.0086 | TRUE | 0.1415 |
| Ndufaf3 | 514 | 726 | 674 | 717 | 565 | 555 | 546 | 725 | 677 | 564 | 608 | 608 | 621.15 | 649 | 593 | 0.912 | -0.132 | 0.31721691 | 0.53814619 | 0.01 | 0.0122 | 0.011 | 0.011 | TRUE | 0.2267 |
| Ndufaf4 | 1412 | 1231 | 1088 | 1928 | 1345 | 1216 | 1500 | 1229 | 1092 | 1516 | 1447 | 1331 | 1352.59 | 1274 | 1431 | 1.124 | 0.169 | 0.20353899 | 0.41140804 | 0.0142 | 0.0093 | 0.012 | 0.012 | TRUE | 0.2882 |
| Ndufaf5 | 618 | 863 | 797 | 899 | 601 | 627 | 656 | 862 | 800 | 707 | 647 | 686 | 726.38 | 773 | 680 | 0.88 | -0.185 | 0.14660318 | 0.33539677 | 0.0093 | 0.0114 | 0.0103 | 0.0103 | TRUE | 0.2047 |
| Ndufaf6 | 821 | 1200 | 1139 | 1097 | 840 | 886 | 872 | 1198 | 1144 | 862 | 904 | 970 | 991.7 | 1071 | 912 | 0.851 | -0.233 | 0.09118243 | 0.24633938 | 0.0149 | 0.0102 | 0.0127 | 0.0127 | TRUE | 0.3157 |

| | | | | | | | | | | | | | | | | | | | | | | | | | |
|---|---|---|---|---|---|---|---|---|---|---|---|---|---|---|---|---|---|---|---|---|---|---|---|---|---|
| Ndufaf7 | 1562 | 2107 | 2094 | 1847 | 1344 | 1535 | 1659 | 2104 | 2103 | 1452 | 1446 | 1680 | 1740.72 | 1955 | 1526 | 0.78 | -0.358 | 0.00415494 | 0.02721418 | 0.0116 | 0.0088 | 0.0107 | 0.0107 | TRUE | 0.2118 |
| Ndufb10 | 2592 | 2795 | 2612 | 3402 | 2317 | 2324 | 2753 | 2791 | 2623 | 2674 | 2493 | 2544 | 2646.43 | 2722 | 2570 | 0.945 | -0.082 | 0.25125845 | 0.46875424 | 9.00E-04 | 0.0081 | 0.0033 | 0.0033 | TRUE | 0.0155 |
| Ndufb11 | 2603 | 3175 | 3299 | 3463 | 2411 | 2634 | 2765 | 3171 | 3313 | 2722 | 2594 | 2883 | 2908.01 | 3083 | 2733 | 0.887 | -0.174 | 0.07382554 | 0.21338322 | 0.0051 | 0.008 | 0.0065 | 0.0065 | TRUE | 0.0986 |
| Ndufb2 | 975 | 1417 | 1333 | 1515 | 1082 | 1078 | 1036 | 1415 | 1338 | 1191 | 1164 | 1180 | 1220.74 | 1263 | 1178 | 0.933 | -0.1 | 0.43750148 | 0.64548863 | 0.0126 | 0.0096 | 0.0112 | 0.0112 | TRUE | 0.2969 |
| Ndufb3 | 1793 | 2088 | 2140 | 2356 | 1679 | 1627 | 1904 | 2085 | 2149 | 1852 | 1807 | 1781 | 1929.7 | 2046 | 1813 | 0.886 | -0.174 | 0.03147688 | 0.1191965 | 0.0016 | 0.0086 | 0.0042 | 0.0042 | TRUE | 0.0442 |
| Ndufb4 | 610 | 738 | 771 | 684 | 554 | 568 | 648 | 737 | 774 | 538 | 596 | 622 | 652.45 | 720 | 585 | 0.812 | -0.301 | 0.01007487 | 0.05301996 | 0.0055 | 0.012 | 0.0083 | 0.0083 | TRUE | 0.0894 |
| Ndufb5 | 3379 | 4352 | 4392 | 4495 | 3359 | 3586 | 3589 | 4346 | 4410 | 3534 | 3614 | 3925 | 3903.11 | 4115 | 3691 | 0.897 | -0.157 | 0.13533199 | 0.31824306 | 0.0073 | 0.0077 | 0.0077 | 0.0077 | TRUE | 0.152 |
| Ndufb6 | 2695 | 3391 | 3157 | 3783 | 2562 | 2766 | 2863 | 3387 | 3170 | 2974 | 2757 | 3028 | 3029.57 | 3140 | 2920 | 0.93 | -0.105 | 0.25632746 | 0.47500842 | 0.0042 | 0.008 | 0.0058 | 0.0058 | TRUE | 0.0723 |
| Ndufb7 | 2894 | 3559 | 3499 | 4325 | 3054 | 3062 | 3074 | 3554 | 3513 | 3400 | 3286 | 3352 | 3363.24 | 3380 | 3346 | 0.99 | -0.015 | 0.85992873 | 0.9309173 | 0.003 | 0.0079 | 0.0048 | 0.0048 | TRUE | 0.0763 |
| Ndufb8 | 4470 | 6303 | 6359 | 6380 | 4729 | 5276 | 4748 | 6295 | 6385 | 5015 | 5088 | 5775 | 5551.15 | 5809 | 5293 | 0.911 | -0.134 | 0.30457827 | 0.52512118 | 0.0154 | 0.0075 | 0.0122 | 0.0122 | TRUE | 0.3113 |
| Ndufb9 | 3886 | 5139 | 5006 | 5494 | 3950 | 4274 | 4128 | 5132 | 5027 | 4319 | 4250 | 4679 | 4589 | 4762 | 4416 | 0.927 | -0.109 | 0.30448138 | 0.52512118 | 0.0078 | 0.0076 | 0.0079 | 0.0079 | TRUE | 0.1653 |
| Ndufc1 | 1859 | 1841 | 1951 | 2511 | 1698 | 1754 | 1975 | 1839 | 1959 | 1974 | 1827 | 1920 | 1915.52 | 1924 | 1907 | 0.992 | -0.012 | 0.8743391 | 0.93803661 | 0.001 | 0.0086 | 0.0037 | 0.0037 | TRUE | 0.0183 |
| Ndufc2 | 2583 | 2911 | 2805 | 3568 | 2594 | 2597 | 2744 | 2907 | 2817 | 2805 | 2791 | 2843 | 2817.68 | 2823 | 2813 | 0.997 | -0.005 | 0.93786847 | 0.96893419 | 0 | 0.0081 | 0.0027 | 0.0027 | TRUE | 0.0084 |
| Ndufs1 | 6773 | 7026 | 6973 | 9780 | 6879 | 6619 | 7194 | 7017 | 7002 | 7688 | 7401 | 7246 | 7257.98 | 7071 | 7445 | 1.053 | 0.075 | 0.21404559 | 0.4257159 | 4.00E-04 | 0.0074 | 0.0025 | 0.0025 | TRUE | 0.0101 |
| Ndufs2 | 6077 | 8131 | 8188 | 9164 | 6740 | 7450 | 6455 | 8121 | 8222 | 7204 | 7252 | 8155 | 7568 | 7599 | 7537 | 0.992 | -0.012 | 0.91781321 | 0.95980308 | 0.0113 | 0.0073 | 0.0096 | 0.0096 | TRUE | 0.2117 |
| Ndufs3 | 2177 | 2488 | 2565 | 3028 | 2103 | 2188 | 2312 | 2485 | 2576 | 2380 | 2263 | 2395 | 2401.81 | 2458 | 2346 | 0.955 | -0.067 | 0.39024259 | 0.60395507 | 0.0013 | 0.0083 | 0.004 | 0.004 | TRUE | 0.0322 |
| Ndufs4 | 2295 | 3008 | 2926 | 3253 | 2352 | 2537 | 2438 | 3004 | 2938 | 2557 | 2531 | 2777 | 2707.47 | 2793 | 2622 | 0.938 | -0.092 | 0.38443745 | 0.5985945 | 0.0069 | 0.0081 | 0.0077 | 0.0077 | TRUE | 0.1502 |
| Ndufs5 | 0 | 0 | 0 | 0 | 0 | 0 | 0 | 0 | 0 | 0 | 0 | 0 | 0 | 0 | 0 | NA | NA | NA | NA | NA | NA | NA | NA | NA | NA |
| Ndufs6 | 2690 | 3082 | 3089 | 3720 | 2568 | 2679 | 2857 | 3078 | 3102 | 2924 | 2763 | 2933 | 2942.82 | 3012 | 2873 | 0.954 | -0.068 | 0.3554857 | 0.57387109 | 0.001 | 0.008 | 0.0036 | 0.0036 | TRUE | 0.0245 |
| Ndufs7 | 1739 | 2077 | 2187 | 2637 | 1878 | 1954 | 1847 | 2074 | 2196 | 2073 | 2021 | 2139 | 2058.33 | 2039 | 2078 | 1.019 | 0.027 | 0.7714787 | 0.88094525 | 0.0037 | 0.0085 | 0.0056 | 0.0056 | TRUE | 0.0818 |
| Ndufs8 | 1902 | 2270 | 2341 | 2780 | 1993 | 2128 | 2020 | 2267 | 2351 | 2185 | 2144 | 2329 | 2216.19 | 2213 | 2219 | 1.003 | 0.004 | 0.96271654 | 0.98130897 | 0.0035 | 0.0084 | 0.0054 | 0.0054 | TRUE | 0.0698 |
| Ndufv1 | 6754 | 7953 | 8053 | 8908 | 6644 | 6686 | 7174 | 7943 | 8086 | 7003 | 7149 | 7319 | 7445.49 | 7734 | 7157 | 0.925 | -0.112 | 0.13582896 | 0.31903692 | 0.0021 | 0.0073 | 0.0039 | 0.0039 | TRUE | 0.049 |
| Ndufv2 | 4635 | 6199 | 6123 | 6560 | 4771 | 5255 | 4923 | 6191 | 6148 | 5157 | 5133 | 5752 | 5550.83 | 5754 | 5347 | 0.929 | -0.106 | 0.34802349 | 0.56682051 | 0.0097 | 0.0075 | 0.009 | 0.009 | TRUE | 0.1944 |
| Ndufv3 | 3197 | 3743 | 3871 | 3807 | 3004 | 3263 | 3396 | 3738 | 3887 | 2993 | 3232 | 3572 | 3469.6 | 3674 | 3266 | 0.889 | -0.17 | 0.08513638 | 0.2346141 | 0.0059 | 0.0079 | 0.0068 | 0.0068 | TRUE | 0.0817 |
| Neat1 | 4711 | 4512 | 4009 | 5620 | 5931 | 4195 | 5004 | 4506 | 4025 | 4418 | 6381 | 4592 | 4821.17 | 4512 | 5130 | 1.137 | 0.185 | 0.23222801 | 0.4465074 | 0.028 | 0.0076 | 0.0171 | 0.0171 | TRUE | 0.5541 |
| Neb | 354 | 245 | 282 | 338 | 271 | 219 | 376 | 245 | 283 | 266 | 292 | 240 | 283.48 | 301 | 266 | 0.882 | -0.18 | 0.33834793 | 0.5583911 | 0.0246 | 0.0185 | 0.022 | 0.022 | TRUE | 0.528 |
| Nebl | 38 | 57 | 51 | 32 | 26 | 27 | 40 | 57 | 51 | 25 | 28 | 30 | 38.53 | 49 | 28 | 0.554 | -0.852 | 0.00815068 | 0.04507453 | 0 | 0.0924 | 0.0467 | 0.0467 | TRUE | 0.2051 |
| Necab1 | 78 | 28 | 33 | 152 | 122 | 87 | 83 | 28 | 33 | 119 | 131 | 95 | 81.66 | 48 | 115 | 2.411 | 1.269 | 0.00131623 | 0.01104818 | 0.1633 | 0.0472 | 0.0976 | 0.0976 | TRUE | 3.1777 |
| Necab2 | 945 | 1026 | 1000 | 1205 | 887 | 810 | 1004 | 1025 | 1004 | 947 | 954 | 887 | 970.14 | 1011 | 929 | 0.92 | -0.121 | 0.14553182 | 0.33391445 | 0 | 0.0103 | 0.0039 | 0.0039 | TRUE | 0.0187 |
| Necab3 | 681 | 771 | 697 | 566 | 467 | 502 | 723 | 770 | 700 | 445 | 502 | 550 | 615.02 | 731 | 499 | 0.681 | -0.554 | 2.20E-06 | 5.44E-05 | 0.0049 | 0.0123 | 0.0082 | 0.0082 | TRUE | 0.1104 |
| Necap1 | 6639 | 7487 | 7164 | 9471 | 6727 | 7180 | 7052 | 7477 | 7193 | 7445 | 7238 | 7860 | 7377.56 | 7241 | 7514 | 1.038 | 0.053 | 0.42903181 | 0.6387285 | 0.0011 | 0.0073 | 0.0031 | 0.0031 | TRUE | 0.0196 |
| Necap2 | 2522 | 2250 | 2078 | 2706 | 2037 | 2001 | 2679 | 2247 | 2087 | 2127 | 2192 | 2190 | 2253.63 | 2338 | 2170 | 0.928 | -0.107 | 0.32659668 | 0.5476962 | 0.0081 | 0.0084 | 0.0082 | 0.0082 | TRUE | 0.1973 |
| Nedd1 | 1994 | 2353 | 2380 | 3060 | 2258 | 2407 | 2118 | 2350 | 2390 | 2406 | 2429 | 2635 | 2387.93 | 2286 | 2490 | 1.089 | 0.123 | 0.15491306 | 0.34705807 | 0.0027 | 0.0083 | 0.0049 | 0.0049 | TRUE | 0.0498 |
| Nedd4 | 76010 | 92386 | 101004 | 95375 | 69566 | 77682 | 80736 | 92267 | 101419 | 74976 | 74849 | 85035 | 84880.29 | 91474 | 78287 | 0.856 | -0.225 | 0.03583992 | 0.12997637 | 0.0092 | 0.0069 | 0.0082 | 0.0082 | TRUE | 0.1291 |
| Nedd4l | 2366 | 2802 | 2763 | 3375 | 2560 | 2694 | 2513 | 2798 | 2774 | 2653 | 2754 | 2949 | 2740.4 | 2695 | 2785 | 1.033 | 0.047 | 0.5798393 | 0.75667925 | 0.0025 | 0.0081 | 0.0048 | 0.0048 | TRUE | 0.0423 |
| Nedd8 | 3973 | 3549 | 3301 | 4947 | 3273 | 3035 | 4220 | 3544 | 3315 | 3889 | 3522 | 3322 | 3635.3 | 3693 | 3578 | 0.969 | -0.045 | 0.69682598 | 0.83386531 | 0.0108 | 0.0078 | 0.0095 | 0.0095 | TRUE | 0.1894 |
| Nedd9 | 94 | 44 | 52 | 180 | 102 | 80 | 100 | 44 | 52 | 142 | 110 | 88 | 89.14 | 65 | 113 | 1.737 | 0.796 | 0.02133703 | 0.09106953 | 0.1092 | 0.0439 | 0.0741 | 0.0741 | TRUE | 0.7944 |
| Nefh | 1747 | 1402 | 1405 | 3419 | 2356 | 1448 | 1856 | 1400 | 1411 | 2688 | 2535 | 1585 | 1912.38 | 1556 | 2269 | 1.459 | 0.545 | 0.00643252 | 0.03791247 | 0.0478 | 0.0086 | 0.0283 | 0.0283 | TRUE | 0.8404 |
| Nefl | 535 | 651 | 788 | 944 | 712 | 671 | 568 | 650 | 791 | 742 | 766 | 735 | 708.72 | 670 | 748 | 1.116 | 0.158 | 0.24993643 | 0.46722738 | 0.0123 | 0.0116 | 0.0121 | 0.0121 | TRUE | 0.2964 |
| Nefm | 175 | 117 | 132 | 153 | 136 | 92 | 186 | 117 | 133 | 120 | 146 | 101 | 133.76 | 145 | 122 | 0.845 | -0.244 | 0.31666167 | 0.53753247 | 0.0389 | 0.0315 | 0.0352 | 0.0352 | TRUE | 0.6234 |
| Negr1 | 3 | 2 | 1 | 7 | 3 | 3 | 3 | 2 | 1 | 6 | 3 | 3 | 3.03 | 2 | 4 | 1.977 | 0.983 | 0.36959404 | NA | 0 | 1.0928 | 0.4958 | 0.4958 | TRUE | 0.2338 |
| Neil1 | 181 | 227 | 258 | 280 | 218 | 212 | 192 | 227 | 259 | 220 | 235 | 232 | 227.46 | 226 | 229 | 1.011 | 0.016 | 0.91970031 | 0.96064694 | 0.0069 | 0.0214 | 0.0139 | 0.0139 | TRUE | 0.185 |
| Neil2 | 447 | 587 | 636 | 399 | 376 | 485 | 475 | 586 | 639 | 314 | 405 | 531 | 491.46 | 567 | 417 | 0.733 | -0.448 | 0.02811357 | 0.11006223 | 0.0421 | 0.0136 | 0.0279 | 0.0279 | TRUE | 0.4038 |
| Neil3 | 605 | 718 | 714 | 964 | 707 | 723 | 643 | 717 | 717 | 758 | 761 | 791 | 731.09 | 692 | 770 | 1.112 | 0.153 | 0.11021851 | 0.27862899 | 6.00E-04 | 0.0114 | 0.0052 | 0.0052 | TRUE | 0.0459 |
| Nek1 | 4048 | 3829 | 4070 | 4710 | 3417 | 3535 | 4300 | 3824 | 4087 | 3703 | 3676 | 3870 | 3909.86 | 4070 | 3750 | 0.921 | -0.118 | 0.11914587 | 0.29243211 | 0.0017 | 0.0077 | 0.0039 | 0.0039 | TRUE | 0.0341 |
| Nek10 | 25 | 36 | 35 | 85 | 65 | 57 | 27 | 36 | 35 | 67 | 70 | 62 | 49.47 | 33 | 66 | 2.037 | 1.026 | 0.00030791 | 0.00348874 | 0 | 0.0735 | 0.0352 | 0.0352 | TRUE | 0.1715 |
| Nek11 | 2 | 5 | 5 | 1 | 4 | 9 | 2 | 5 | 5 | 1 | 4 | 10 | 4.51 | 4 | 5 | 1.199 | 0.262 | 0.80562888 | NA | 0.4451 | 0.7368 | 0.5893 | 0.5893 | TRUE | 0.2993 |
| Nek2 | 6967 | 7489 | 7258 | 8855 | 6306 | 6581 | 7400 | 7479 | 7288 | 6961 | 6785 | 7204 | 7186.21 | 7389 | 6983 | 0.945 | -0.082 | 0.17194632 | 0.37078718 | 3.00E-04 | 0.0074 | 0.0024 | 0.0024 | TRUE | 0.0092 |
| Nek3 | 1582 | 1485 | 1385 | 2302 | 1701 | 1617 | 1680 | 1483 | 1391 | 1810 | 1830 | 1770 | 1660.67 | 1518 | 1803 | 1.188 | 0.249 | 0.00950028 | 0.0506211 | 0.004 | 0.0089 | 0.006 | 0.006 | TRUE | 0.105 |
| Nek4 | 1041 | 1027 | 1126 | 1712 | 1284 | 1208 | 1106 | 1026 | 1131 | 1346 | 1382 | 1322 | 1218.62 | 1088 | 1350 | 1.242 | 0.312 | 0.000156 | 0.00196673 | 5.00E-04 | 0.0096 | 0.0041 | 0.0041 | TRUE | 0.0296 |
| Nek5 | 13 | 18 | 26 | 15 | 8 | 15 | 14 | 18 | 26 | 12 | 9 | 16 | 15.79 | 19 | 12 | 0.634 | -0.658 | 0.20741012 | 0.41661646 | 0.0331 | 0.2156 | 0.1302 | 0.1302 | TRUE | 0.3562 |
| Nek6 | 1998 | 2307 | 2297 | 2101 | 1481 | 1658 | 2122 | 2304 | 2306 | 1652 | 1593 | 1815 | 1965.45 | 2244 | 1687 | 0.751 | -0.413 | 2.56E-06 | 6.24E-05 | 0.0026 | 0.0086 | 0.005 | 0.005 | TRUE | 0.0525 |
| Nek7 | 1112 | 1556 | 1641 | 1779 | 1282 | 1419 | 1181 | 1554 | 1648 | 1399 | 1379 | 1553 | 1452.34 | 1461 | 1444 | 0.988 | -0.018 | 0.89844519 | 0.95064253 | 0.0166 | 0.0092 | 0.0131 | 0.0131 | TRUE | 0.337 |
| Nek8 | 783 | 1014 | 935 | 955 | 848 | 806 | 832 | 1013 | 939 | 751 | 912 | 882 | 888.11 | 928 | 848 | 0.913 | -0.131 | 0.28781254 | 0.5090214 | 0.0092 | 0.0106 | 0.0098 | 0.0098 | TRUE | 0.1245 |
| Nek9 | 4704 | 6509 | 6778 | 6651 | 4804 | 4995 | 4996 | 6501 | 6806 | 5228 | 5169 | 5468 | 5694.67 | 6101 | 5288 | 0.867 | -0.206 | 0.09556308 | 0.2535275 | 0.0128 | 0.0075 | 0.0109 | 0.0109 | TRUE | 0.3057 |
| Nelfa | 3856 | 3873 | 3944 | 5567 | 4310 | 4111 | 4096 | 3868 | 3960 | 4376 | 4637 | 4500 | 4239.62 | 3975 | 4504 | 1.133 | 0.18 | 0.00577965 | 0.03491054 | 6.00E-04 | 0.0077 | 0.0028 | 0.0028 | TRUE | 0.0086 |
| Nelfb | 4025 | 4396 | 4323 | 5575 | 4069 | 4222 | 4275 | 4390 | 4341 | 4383 | 4378 | 4622 | 4398.1 | 4335 | 4461 | 1.029 | 0.041 | 0.51323392 | 0.70632889 | 3.00E-04 | 0.0077 | 0.0026 | 0.0026 | TRUE | 0.012 |
| Nelfcd | 3907 | 3861 | 3716 | 4673 | 3584 | 3344 | 4150 | 3856 | 3731 | 3674 | 3856 | 3661 | 3821.24 | 3912 | 3730 | 0.953 | -0.069 | 0.35646638 | 0.57504222 | 0.0015 | 0.0078 | 0.0038 | 0.0038 | TRUE | 0.0342 |
| Nelfe | 2403 | 2834 | 2717 | 3757 | 2772 | 2631 | 2552 | 2830 | 2728 | 2953 | 2983 | 2880 | 2821.15 | 2703 | 2939 | 1.087 | 0.12 | 0.10205906 | 0.2652291 | 0.001 | 0.0081 | 0.0035 | 0.0035 | TRUE | 0.0289 |
| Nell1 | 11 | 29 | 14 | 15 | 11 | 15 | 12 | 29 | 14 | 12 | 12 | 16 | 15.79 | 18 | 13 | 0.727 | -0.459 | 0.40274682 | 0.61557543 | 0.0895 | 0.2155 | 0.1523 | 0.1523 | TRUE | 1.3722 |
| Nell2 | 1088 | 1229 | 1251 | 1006 | 727 | 667 | 1156 | 1227 | 1256 | 791 | 782 | 730 | 990.4 | 1213 | 768 | 0.633 | -0.659 | 8.01E-14 | 9.64E-12 | 8.00E-04 | 0.0102 | 0.0046 | 0.0046 | TRUE | 0.0212 |
| Nemf | 4489 | 4148 | 4259 | 5426 | 3991 | 3859 | 4768 | 4143 | 4276 | 4265 | 4294 | 4224 | 4328.52 | 4396 | 4261 | 0.97 | -0.045 | 0.57641665 | 0.75418558 | 0.0024 | 0.0077 | 0.0044 | 0.0044 | TRUE | 0.0667 |
| Nenf | 271 | 436 | 435 | 401 | 341 | 356 | 288 | 435 | 437 | 315 | 367 | 390 | 371.98 | 387 | 357 | 0.922 | -0.117 | 0.53526952 | 0.72254971 | 0.0301 | 0.0158 | 0.0229 | 0.0229 | TRUE | 0.5702 |
| Neo1 | 4679 | 5049 | 5576 | 5854 | 4081 | 4218 | 4970 | 5042 | 5599 | 4602 | 4391 | 4617 | 4870.24 | 5204 | 4537 | 0.872 | -0.198 | 0.01121544 | 0.05732309 | 0.0023 | 0.0076 | 0.0042 | 0.0042 | TRUE | 0.0538 |
| Nepn | 0 | 1 | 1 | 1 | 0 | 1 | 0 | 1 | 1 | 1 | 0 | 1 | 0.65 | 1 | 1 | 0.94 | -0.089 | 0.96942491 | NA | 0 | 5.0967 | 2.3884 | 2.3884 | TRUE | 0.2246 |
| Nes | 920 | 912 | 974 | 1892 | 1276 | 1073 | 977 | 911 | 978 | 1487 | 1373 | 1175 | 1150.14 | 955 | 1345 | 1.409 | 0.495 | 1.24E-05 | 0.00023988 | 0.0073 | 0.0098 | 0.0083 | 0.0083 | TRUE | 0.1466 |
| Nespas | 96 | 73 | 81 | 147 | 124 | 93 | 102 | 73 | 81 | 116 | 133 | 102 | 101.16 | 85 | 117 | 1.371 | 0.455 | 0.04366828 | 0.14956267 | 0.0124 | 0.0395 | 0.0265 | 0.0265 | TRUE | 0.2664 |
| Net1 | 3456 | 3150 | 3108 | 4882 | 3544 | 3612 | 3671 | 3146 | 3121 | 3838 | 3813 | 3954 | 3590.41 | 3313 | 3868 | 1.168 | 0.224 | 0.01275317 | 0.06295555 | 0.0042 | 0.0078 | 0.0055 | 0.0055 | TRUE | 0.1086 |
| Neto1 | 0 | 0 | 0 | 0 | 3 | 0 | 0 | 0 | 0 | 0 | 3 | 0 | 0.54 | 0 | 1 | 5.683 | 2.507 | 0.52963929 | NA | 10 | 6.1308 | 10 | 10 | TRUE | 1.4428 |

| | | | | | | | | | | | | | | | | | | | | | | | | | |
|---|---|---|---|---|---|---|---|---|---|---|---|---|---|---|---|---|---|---|---|---|---|---|---|---|---|
| Neto2 | 240 | 128 | 131 | 652 | 403 | 224 | 255 | 128 | 132 | 513 | 434 | 245 | 284.28 | 172 | 397 | 2.321 | 1.215 | 0.00017644 | 0.00219462 | 0.1426 | 0.0185 | 0.0714 | 0.0714 | TRUE | 0.555 |
| Neu1 | 826 | 1369 | 1417 | 855 | 712 | 838 | 877 | 1367 | 1423 | 672 | 766 | 917 | 1003.82 | 1222 | 785 | 0.642 | -0.64 | 0.00108788 | 0.00950437 | 0.0448 | 0.0102 | 0.0266 | 0.0266 | TRUE | 0.7301 |
| Neu2 | 0 | 3 | 0 | 2 | 3 | 2 | 0 | 3 | 0 | 2 | 3 | 2 | 1.66 | 1 | 2 | 2.293 | 1.197 | 0.48153092 | NA | 0.154 | 1.9864 | 1.3667 | 1.3667 | TRUE | 1.4418 |
| Neu3 | 389 | 493 | 479 | 598 | 378 | 455 | 413 | 492 | 481 | 470 | 407 | 498 | 460.23 | 462 | 458 | 0.992 | -0.012 | 0.92657002 | 0.96376135 | 0.0073 | 0.0141 | 0.0104 | 0.0104 | TRUE | 0.1093 |
| Neu4 | 0 | 0 | 1 | 1 | 0 | 0 | 0 | 0 | 1 | 1 | 0 | 0 | 0.3 | 0 | 0 | 0.954 | -0.067 | 0.98640617 | NA | 0 | 11.0483 | 9.2783 | 9.2783 | TRUE | 0.4098 |
| Neurl1a | 1417 | 1939 | 1917 | 1613 | 1300 | 1623 | 1505 | 1936 | 1925 | 1268 | 1399 | 1777 | 1634.97 | 1789 | 1481 | 0.827 | -0.273 | 0.07920371 | 0.22317416 | 0.0244 | 0.0089 | 0.0169 | 0.0169 | TRUE | 0.3659 |
| Neurl1b | 282 | 415 | 367 | 314 | 240 | 294 | 300 | 414 | 369 | 247 | 258 | 322 | 318.23 | 361 | 276 | 0.762 | -0.392 | 0.02491882 | 0.10145059 | 0.0203 | 0.0173 | 0.0188 | 0.0188 | TRUE | 0.2505 |
| Neurl2 | 63 | 105 | 81 | 82 | 69 | 71 | 67 | 105 | 81 | 64 | 74 | 78 | 78.26 | 84 | 72 | 0.851 | -0.232 | 0.36183889 | 0.58006064 | 0.0185 | 0.049 | 0.0339 | 0.0339 | TRUE | 0.3241 |
| Neurl3 | 26 | 19 | 13 | 32 | 32 | 24 | 28 | 19 | 13 | 25 | 34 | 26 | 24.25 | 20 | 28 | 1.439 | 0.525 | 0.2238094 | 0.43595429 | 0.0291 | 0.1428 | 0.0912 | 0.0912 | TRUE | 0.4196 |
| Neurl4 | 4204 | 4282 | 4357 | 5545 | 4199 | 3816 | 4465 | 4276 | 4375 | 4359 | 4518 | 4177 | 4361.81 | 4372 | 4351 | 0.995 | -0.007 | 0.91851078 | 0.96002333 | 7.00E-04 | 0.0077 | 0.003 | 0.003 | TRUE | 0.0147 |
| Neurod1 | 108 | 117 | 140 | 104 | 84 | 78 | 115 | 117 | 141 | 82 | 90 | 85 | 104.94 | 124 | 86 | 0.69 | -0.534 | 0.00761778 | 0.0428181 | 0 | 0.0383 | 0.0191 | 0.0191 | TRUE | 0.1389 |
| Neurod2 | 1 | 0 | 0 | 2 | 1 | 0 | 1 | 0 | 0 | 2 | 1 | 0 | 0.62 | 0 | 1 | 2.462 | 1.3 | 0.62119797 | NA | 0 | 5.3343 | 3.4416 | 3.4416 | TRUE | 0.4614 |
| Neurod4 | 3 | 2 | 0 | 3 | 0 | 0 | 3 | 2 | 0 | 2 | 0 | 0 | 1.26 | 2 | 1 | 0.487 | -1.038 | 0.63055079 | NA | 2.113 | 2.6277 | 2.4767 | 2.4767 | TRUE | 0.7145 |
| Neurod6 | 0 | 0 | 0 | 0 | 0 | 0 | 0 | 0 | 0 | 0 | 0 | 0 | 0 | 0 | 0 | NA | NA | NA | NA | NA | NA | NA | NA | NA | NA |
| Neurog1 | 2 | 3 | 2 | 50 | 36 | 23 | 2 | 3 | 2 | 39 | 39 | 25 | 18.39 | 2 | 34 | 14.498 | 3.858 | 2.03E-08 | 9.05E-07 | 0 | 0.186 | 0.1122 | 0.1122 | TRUE | 0.3657 |
| Neurog2 | 0 | 3 | 1 | 2 | 4 | 2 | 0 | 3 | 1 | 2 | 4 | 2 | 2.01 | 1 | 3 | 1.974 | 0.981 | 0.4975069 | NA | 0 | 1.6452 | 0.9447 | 0.9447 | TRUE | 0.7354 |
| Neurog3 | 26 | 70 | 94 | 34 | 53 | 66 | 28 | 70 | 94 | 27 | 57 | 72 | 57.99 | 64 | 52 | 0.808 | -0.308 | 0.5307789 | 0.71904843 | 0.2658 | 0.0637 | 0.1568 | 0.1568 | TRUE | 0.3319 |
| Nexn | 129 | 189 | 210 | 164 | 136 | 136 | 137 | 189 | 211 | 129 | 146 | 149 | 160.13 | 179 | 141 | 0.788 | -0.344 | 0.09562405 | 0.25355123 | 0.0212 | 0.0275 | 0.0243 | 0.0243 | TRUE | 0.4429 |
| Nf1 | 2816 | 2942 | 3449 | 3394 | 2343 | 2318 | 2991 | 2938 | 3463 | 2668 | 2521 | 2537 | 2853.15 | 3131 | 2575 | 0.823 | -0.281 | 0.00227474 | 0.01703568 | 0.0042 | 0.0081 | 0.0058 | 0.0058 | TRUE | 0.1049 |
| Nf2 | 5500 | 5135 | 5230 | 7254 | 5024 | 4787 | 5842 | 5128 | 5251 | 5703 | 5406 | 5240 | 5428.33 | 5407 | 5450 | 1.008 | 0.012 | 0.88912447 | 0.9459843 | 0.0032 | 0.0075 | 0.0048 | 0.0048 | TRUE | 0.0602 |
| Nfam1 | 181 | 146 | 131 | 354 | 287 | 283 | 192 | 146 | 132 | 278 | 309 | 310 | 227.75 | 157 | 299 | 1.91 | 0.934 | 1.48E-07 | 5.02E-06 | 0.0139 | 0.0214 | 0.0178 | 0.0178 | TRUE | 0.4138 |
| Nfasc | 416 | 518 | 606 | 567 | 420 | 480 | 442 | 517 | 608 | 446 | 452 | 525 | 498.46 | 522 | 474 | 0.907 | -0.141 | 0.34630788 | 0.56542111 | 0.0141 | 0.0135 | 0.0142 | 0.0142 | TRUE | 0.2171 |
| Nfat5 | 5068 | 2802 | 3352 | 5629 | 3942 | 3312 | 5383 | 2798 | 3366 | 4425 | 4241 | 3626 | 3973.2 | 3849 | 4097 | 1.065 | 0.09 | 0.67007763 | 0.81584564 | 0.0629 | 0.0077 | 0.0322 | 0.0322 | TRUE | 1.4307 |
| Nfatc1 | 147 | 203 | 202 | 215 | 191 | 168 | 156 | 203 | 203 | 169 | 206 | 184 | 186.69 | 187 | 186 | 0.991 | -0.013 | 0.9427985 | 0.97150254 | 0.0102 | 0.0246 | 0.0173 | 0.0173 | TRUE | 0.224 |
| Nfatc2 | 447 | 432 | 430 | 530 | 335 | 344 | 475 | 431 | 432 | 417 | 360 | 377 | 415.27 | 446 | 385 | 0.864 | -0.21 | 0.07716705 | 0.21945161 | 0.0021 | 0.0148 | 0.0078 | 0.0078 | TRUE | 0.0651 |
| Nfatc2ip | 4059 | 3168 | 3003 | 5401 | 3749 | 3356 | 4311 | 3164 | 3015 | 4246 | 4034 | 3674 | 3740.64 | 3497 | 3985 | 1.14 | 0.189 | 0.19393903 | 0.39960772 | 0.0216 | 0.0078 | 0.0149 | 0.0149 | TRUE | 0.5045 |
| Nfatc3 | 10257 | 11069 | 12233 | 10977 | 7925 | 8657 | 10895 | 11055 | 12283 | 8629 | 8527 | 9476 | 10144.2 | 11411 | 8877 | 0.778 | -0.362 | 1.76E-05 | 0.00031913 | 0.0038 | 0.0072 | 0.005 | 0.005 | TRUE | 0.0547 |
| Nfatc4 | 643 | 1131 | 1208 | 635 | 565 | 770 | 683 | 1130 | 1213 | 499 | 608 | 843 | 829.24 | 1009 | 650 | 0.644 | -0.635 | 0.00861493 | 0.04702949 | 0.0745 | 0.0109 | 0.0409 | 0.0409 | TRUE | 0.9504 |
| Nfe2 | 82 | 51 | 41 | 74 | 44 | 38 | 87 | 51 | 41 | 58 | 47 | 42 | 54.39 | 60 | 49 | 0.826 | -0.276 | 0.43368401 | 0.6424232 | 0.0747 | 0.0675 | 0.071 | 0.071 | TRUE | 1.2017 |
| Nfe2l1 | 16074 | 16557 | 16344 | 17467 | 12830 | 12770 | 17073 | 16536 | 16411 | 13731 | 13804 | 13979 | 15255.73 | 16673 | 13838 | 0.83 | -0.269 | 6.29E-07 | 1.83E-05 | 2.00E-04 | 0.0071 | 0.002 | 0.002 | TRUE | 0.0054 |
| Nfe2l2 | 2359 | 2990 | 3222 | 3078 | 2223 | 2319 | 2506 | 2986 | 3235 | 2420 | 2392 | 2539 | 2679.51 | 2909 | 2450 | 0.842 | -0.248 | 0.02427732 | 0.09963462 | 0.0081 | 0.0081 | 0.0084 | 0.0084 | TRUE | 0.1783 |
| Nfe2l3 | 229 | 271 | 284 | 215 | 169 | 214 | 243 | 271 | 285 | 169 | 182 | 234 | 230.69 | 266 | 195 | 0.73 | -0.455 | 0.00854258 | 0.04678251 | 0.0131 | 0.0212 | 0.0171 | 0.0171 | TRUE | 0.3284 |
| Nfia | 22 | 39 | 31 | 53 | 46 | 48 | 23 | 39 | 31 | 42 | 49 | 53 | 39.52 | 31 | 48 | 1.529 | 0.612 | 0.06388297 | 0.19428502 | 0 | 0.0903 | 0.0521 | 0.0521 | TRUE | 0.275 |
| Nfib | 273 | 247 | 231 | 535 | 353 | 310 | 290 | 247 | 232 | 421 | 380 | 339 | 318.05 | 256 | 380 | 1.487 | 0.572 | 0.00014555 | 0.00186037 | 0.0093 | 0.0173 | 0.0131 | 0.0131 | TRUE | 0.1464 |
| Nfic | 1045 | 1412 | 1536 | 1392 | 1173 | 1351 | 1110 | 1410 | 1542 | 1094 | 1262 | 1479 | 1316.28 | 1354 | 1278 | 0.943 | -0.084 | 0.59266339 | 0.7659991 | 0.0245 | 0.0094 | 0.0171 | 0.0171 | TRUE | 0.2984 |
| Nfil3 | 302 | 447 | 472 | 429 | 361 | 397 | 321 | 446 | 474 | 337 | 388 | 435 | 400.23 | 414 | 387 | 0.933 | -0.1 | 0.57819314 | 0.75546867 | 0.0264 | 0.0151 | 0.0208 | 0.0208 | TRUE | 0.4423 |
| Nfix | 90 | 125 | 110 | 180 | 166 | 133 | 96 | 125 | 110 | 142 | 179 | 146 | 132.76 | 110 | 156 | 1.403 | 0.489 | 0.01374409 | 0.06630601 | 0.0094 | 0.0317 | 0.0206 | 0.0206 | TRUE | 0.1785 |
| Nfkb1 | 1849 | 1879 | 1995 | 2800 | 1861 | 1802 | 1964 | 1877 | 2003 | 2201 | 2002 | 1973 | 2003.29 | 1948 | 2059 | 1.058 | 0.081 | 0.32483147 | 0.54594394 | 0.0019 | 0.0085 | 0.0044 | 0.0044 | TRUE | 0.0453 |
| Nfkb2 | 710 | 890 | 992 | 946 | 719 | 831 | 754 | 889 | 996 | 744 | 774 | 910 | 844.33 | 880 | 809 | 0.919 | -0.122 | 0.3769176 | 0.59206639 | 0.0136 | 0.0108 | 0.0126 | 0.0126 | TRUE | 0.1846 |
| Nfkbia | 904 | 815 | 806 | 742 | 526 | 517 | 960 | 814 | 809 | 583 | 566 | 566 | 716.44 | 861 | 572 | 0.664 | -0.59 | 6.41E-08 | 2.46E-06 | 0.004 | 0.0115 | 0.0071 | 0.0071 | TRUE | 0.1197 |
| Nfkbib | 2070 | 1883 | 2020 | 2301 | 1726 | 1638 | 2199 | 1881 | 2028 | 1809 | 1857 | 1793 | 1927.76 | 2036 | 1820 | 0.894 | -0.162 | 0.06364969 | 0.19379978 | 0.0024 | 0.0086 | 0.005 | 0.005 | TRUE | 0.059 |
| Nfkbid | 309 | 191 | 170 | 243 | 173 | 194 | 328 | 191 | 171 | 191 | 186 | 212 | 213.2 | 230 | 196 | 0.855 | -0.226 | 0.37139102 | 0.58805132 | 0.0616 | 0.0224 | 0.0412 | 0.0412 | TRUE | 1.5338 |
| Nfkbie | 132 | 133 | 109 | 195 | 152 | 124 | 140 | 133 | 109 | 153 | 164 | 136 | 139.18 | 127 | 151 | 1.185 | 0.244 | 0.18941489 | 0.39459142 | 0.0041 | 0.0306 | 0.0177 | 0.0177 | TRUE | 0.1575 |
| Nfkbil1 | 302 | 371 | 409 | 453 | 334 | 362 | 321 | 371 | 411 | 356 | 359 | 396 | 368.95 | 368 | 370 | 1.008 | 0.011 | 0.9373234 | 0.96864531 | 0.0065 | 0.0158 | 0.0108 | 0.0108 | TRUE | 0.1391 |
| Nfkbiz | 114 | 110 | 125 | 174 | 222 | 157 | 121 | 110 | 126 | 137 | 239 | 172 | 150.66 | 119 | 183 | 1.531 | 0.614 | 0.00953182 | 0.05072061 | 0.038 | 0.0288 | 0.0335 | 0.0335 | TRUE | 0.3265 |
| Nfrkb | 2430 | 2205 | 2275 | 2761 | 2126 | 1963 | 2581 | 2202 | 2284 | 2170 | 2287 | 2149 | 2279.05 | 2356 | 2202 | 0.935 | -0.097 | 0.2822772 | 0.50368548 | 0.0035 | 0.0083 | 0.0055 | 0.0055 | TRUE | 0.0845 |
| Nfs1 | 2704 | 3231 | 3154 | 3483 | 2435 | 2623 | 2872 | 3227 | 3167 | 2738 | 2620 | 2871 | 2915.86 | 3089 | 2743 | 0.888 | -0.171 | 0.03883538 | 0.13776689 | 0.0024 | 0.008 | 0.0046 | 0.0046 | TRUE | 0.0455 |
| Nfu1 | 2626 | 2894 | 2974 | 3552 | 2615 | 2461 | 2789 | 2890 | 2986 | 2792 | 2814 | 2694 | 2827.6 | 2888 | 2767 | 0.958 | -0.062 | 0.36189368 | 0.58006064 | 5.00E-04 | 0.0081 | 0.003 | 0.003 | TRUE | 0.0109 |
| Nfx1 | 7407 | 7658 | 7722 | 9380 | 6885 | 6777 | 7868 | 7648 | 7754 | 7374 | 7408 | 7418 | 7578.25 | 7757 | 7400 | 0.954 | -0.068 | 0.20847265 | 0.41795228 | 0 | 0.0073 | 0.002 | 0.002 | TRUE | 0.0019 |
| Nfxl1 | 2116 | 2666 | 2678 | 3032 | 2255 | 2276 | 2248 | 2663 | 2689 | 2384 | 2426 | 2491 | 2483.39 | 2533 | 2434 | 0.961 | -0.058 | 0.54275304 | 0.7283165 | 0.0042 | 0.0082 | 0.0062 | 0.0062 | TRUE | 0.1175 |
| Nfya | 2098 | 2592 | 2630 | 3533 | 2834 | 3050 | 2228 | 2589 | 2641 | 2777 | 3049 | 3339 | 2770.53 | 2486 | 3055 | 1.228 | 0.297 | 0.00614209 | 0.03662499 | 0.0078 | 0.0081 | 0.0081 | 0.0081 | TRUE | 0.0993 |
| Nfyb | 4168 | 5044 | 4851 | 6278 | 4776 | 4650 | 4427 | 5037 | 4871 | 4935 | 5139 | 5090 | 4916.61 | 4778 | 5055 | 1.058 | 0.081 | 0.29569617 | 0.5165093 | 0.0021 | 0.0076 | 0.0041 | 0.0041 | TRUE | 0.0503 |
| Nfyc | 2031 | 1226 | 1152 | 2432 | 1746 | 1270 | 2157 | 1224 | 1157 | 1912 | 1879 | 1390 | 1619.85 | 1513 | 1727 | 1.142 | 0.191 | 0.4093261 | 0.62130006 | 0.0758 | 0.0089 | 0.0382 | 0.0382 | TRUE | 1.6714 |
| Ngb | 49 | 65 | 39 | 42 | 46 | 33 | 52 | 65 | 39 | 33 | 49 | 36 | 45.79 | 52 | 39 | 0.756 | -0.403 | 0.22368877 | 0.43578437 | 0.034 | 0.0788 | 0.057 | 0.057 | TRUE | 0.1463 |
| Ngdn | 1978 | 2101 | 2186 | 3198 | 2145 | 2068 | 2101 | 2098 | 2195 | 2514 | 2308 | 2264 | 2246.65 | 2131 | 2362 | 1.109 | 0.149 | 0.05722559 | 0.18026002 | 0.0016 | 0.0084 | 0.004 | 0.004 | TRUE | 0.0393 |
| Ngef | 101 | 173 | 166 | 183 | 146 | 146 | 107 | 173 | 167 | 144 | 157 | 160 | 151.25 | 149 | 154 | 1.029 | 0.041 | 0.84999691 | 0.92526408 | 0.0265 | 0.0287 | 0.0276 | 0.0276 | TRUE | 0.6158 |
| Ngf | 18 | 29 | 17 | 23 | 20 | 21 | 19 | 29 | 17 | 18 | 22 | 23 | 21.29 | 22 | 21 | 0.955 | -0.067 | 0.8770232 | 0.93932608 | 0 | 0.1616 | 0.0866 | 0.0866 | TRUE | 0.4894 |
| Ngfr | 1733 | 1544 | 1584 | 1600 | 1258 | 1057 | 1841 | 1542 | 1591 | 1258 | 1354 | 1157 | 1456.94 | 1658 | 1256 | 0.758 | -0.4 | 0.00024054 | 0.00282666 | 0.0067 | 0.0092 | 0.0078 | 0.0078 | TRUE | 0.1122 |
| Ngfrap1 | 8011 | 7381 | 7119 | 12215 | 8290 | 7415 | 8509 | 7371 | 7148 | 9602 | 8920 | 8117 | 8277.95 | 7676 | 8880 | 1.157 | 0.21 | 0.04198352 | 0.14594756 | 0.0078 | 0.0073 | 0.0076 | 0.0076 | TRUE | 0.1099 |
| Ngly1 | 527 | 538 | 561 | 810 | 522 | 542 | 560 | 537 | 563 | 637 | 562 | 593 | 575.35 | 553 | 597 | 1.081 | 0.112 | 0.27414084 | 0.49465156 | 0 | 0.0126 | 0.0058 | 0.0058 | TRUE | 0.0415 |
| Ngp | 0 | 0 | 0 | 0 | 1 | 1 | 0 | 0 | 0 | 0 | 1 | 1 | 0.36 | 0 | 1 | 3.794 | 1.924 | 0.60247382 | NA | 0 | 9.1135 | 8.1425 | 8.1425 | TRUE | 0.3482 |
| Ngrn | 1853 | 1925 | 1963 | 2583 | 1945 | 1796 | 1968 | 1923 | 1971 | 2031 | 2093 | 1966 | 1991.84 | 1954 | 2030 | 1.039 | 0.055 | 0.42849936 | 0.63836934 | 0 | 0.0086 | 0.003 | 0.003 | TRUE | 0.0089 |
| Nhej1 | 210 | 174 | 176 | 259 | 192 | 180 | 223 | 174 | 177 | 204 | 207 | 197 | 196.8 | 191 | 203 | 1.06 | 0.084 | 0.60680973 | 0.77683466 | 0.0047 | 0.0236 | 0.0141 | 0.0141 | TRUE | 0.2267 |
| Nhlh1 | 0 | 0 | 2 | 0 | 0 | 0 | 0 | 0 | 2 | 0 | 0 | 0 | 0.33 | 1 | 0 | 0.267 | -1.904 | 0.63742563 | NA | 0.0046 | 9.8498 | 10 | 10 | TRUE | 0.974 |
| Nhlh2 | 0 | 2 | 1 | 0 | 1 | 1 | 0 | 2 | 1 | 0 | 1 | 1 | 0.86 | 1 | 1 | 0.688 | -0.539 | 0.8028361 | NA | 0 | 3.8287 | 2.1472 | 2.1472 | TRUE | 0.1651 |
| Nhlrc1 | 270 | 229 | 232 | 422 | 326 | 298 | 287 | 229 | 233 | 332 | 351 | 326 | 292.86 | 250 | 336 | 1.348 | 0.431 | 0.00242919 | 0.01795506 | 0.0039 | 0.0182 | 0.0111 | 0.0111 | TRUE | 0.1877 |
| Nhlrc2 | 1296 | 1307 | 1394 | 1863 | 1343 | 1303 | 1377 | 1305 | 1400 | 1465 | 1445 | 1426 | 1402.91 | 1361 | 1445 | 1.063 | 0.088 | 0.24619297 | 0.46320719 | 0 | 0.0093 | 0.0034 | 0.0034 | TRUE | 0.0153 |
| Nhlrc3 | 320 | 469 | 445 | 370 | 264 | 316 | 340 | 468 | 447 | 291 | 284 | 346 | 362.66 | 418 | 307 | 0.733 | -0.449 | 0.0064925 | 0.0381519 | 0.0175 | 0.016 | 0.0168 | 0.0168 | TRUE | 0.3083 |
| Nhlrc4 | 2 | 2 | 2 | 7 | 2 | 1 | 2 | 2 | 2 | 6 | 2 | 1 | 2.48 | 2 | 3 | 1.473 | 0.558 | 0.64583302 | NA | 0 | 1.3354 | 0.6515 | 0.6515 | TRUE | 0.9836 |

| | | | | | | | | | | | | | | | | | | | | | | | | | |
|---|---|---|---|---|---|---|---|---|---|---|---|---|---|---|---|---|---|---|---|---|---|---|---|---|---|
| Nhp2 | 8896 | 9282 | 9182 | 12660 | 8822 | 8293 | 9449 | 9270 | 9220 | 9952 | 9492 | 9078 | 9410.18 | 9313 | 9507 | 1.021 | 0.03 | 0.64498698 | 0.80042827 | 0.001 | 0.0073 | 0.003 | 0.003 | TRUE | 0.0206 |
| Nhp2l1 | 5208 | 5203 | 5188 | 7279 | 5178 | 4795 | 5532 | 5196 | 5209 | 5722 | 5571 | 5249 | 5413.28 | 5312 | 5514 | 1.038 | 0.054 | 0.44531824 | 0.65195946 | 0.0013 | 0.0075 | 0.0034 | 0.0034 | TRUE | 0.0215 |
| Nhs | 268 | 243 | 320 | 229 | 137 | 167 | 285 | 243 | 321 | 180 | 147 | 183 | 226.48 | 283 | 170 | 0.602 | -0.732 | 2.23E-05 | 0.00039156 | 0.0124 | 0.0214 | 0.0168 | 0.0168 | TRUE | 0.1452 |
| Nhsl1 | 1266 | 991 | 1114 | 1334 | 950 | 1011 | 1345 | 990 | 1119 | 1049 | 1022 | 1107 | 1105.09 | 1151 | 1059 | 0.92 | -0.12 | 0.35135421 | 0.57017785 | 0.0117 | 0.0099 | 0.011 | 0.011 | TRUE | 0.2589 |
| Nhsl2 | 157 | 154 | 156 | 194 | 123 | 143 | 167 | 154 | 157 | 153 | 132 | 157 | 153.1 | 159 | 147 | 0.926 | -0.11 | 0.50504233 | 0.69971255 | 0 | 0.0284 | 0.0133 | 0.0133 | TRUE | 0.0766 |
| Niacr1 | 0 | 0 | 0 | 0 | 0 | 0 | 0 | 0 | 0 | 0 | 0 | 0 | 0 | 0 | 0 | NA | NA | NA | NA | NA | NA | NA | NA | NA | NA |
| Nicn1 | 591 | 840 | 864 | 975 | 660 | 784 | 628 | 839 | 868 | 766 | 710 | 858 | 778.17 | 778 | 778 | 1 | 0 | 0.99827908 | 0.99885317 | 0.0183 | 0.0111 | 0.0148 | 0.0148 | TRUE | 0.3371 |
| Nid1 | 4378 | 5992 | 6689 | 6731 | 5033 | 6553 | 4650 | 5984 | 6716 | 5291 | 5415 | 7173 | 5871.8 | 5783 | 5960 | 1.03 | 0.043 | 0.79267667 | 0.89313448 | 0.0316 | 0.0075 | 0.0192 | 0.0192 | TRUE | 0.3865 |
| Nid2 | 42381 | 51510 | 56419 | 37220 | 29319 | 32217 | 45016 | 51443 | 56651 | 29259 | 31545 | 35267 | 41530.29 | 51037 | 32024 | 0.627 | -0.672 | 3.04E-09 | 1.64E-07 | 0.011 | 0.007 | 0.0092 | 0.0092 | TRUE | 0.1304 |
| Nif3l1 | 1939 | 1638 | 1507 | 2441 | 1666 | 1480 | 2060 | 1636 | 1513 | 1919 | 1793 | 1620 | 1756.69 | 1736 | 1777 | 1.024 | 0.034 | 0.79968807 | 0.89674997 | 0.0161 | 0.0088 | 0.0127 | 0.0127 | TRUE | 0.3198 |
| Nim1 | 116 | 178 | 180 | 147 | 88 | 102 | 123 | 178 | 181 | 116 | 95 | 112 | 133.94 | 161 | 108 | 0.669 | -0.58 | 0.00753046 | 0.04235751 | 0.021 | 0.0315 | 0.0262 | 0.0262 | TRUE | 0.4314 |
| Nin | 949 | 778 | 826 | 2287 | 1668 | 1332 | 1008 | 777 | 829 | 1798 | 1795 | 1458 | 1277.5 | 871 | 1684 | 1.933 | 0.951 | 3.54E-12 | 3.32E-10 | 0.0151 | 0.0095 | 0.0126 | 0.0126 | TRUE | 0.2227 |
| Ninj1 | 800 | 1220 | 1121 | 747 | 625 | 765 | 850 | 1218 | 1126 | 587 | 672 | 837 | 881.81 | 1065 | 699 | 0.656 | -0.609 | 0.00055881 | 0.00568252 | 0.031 | 0.0106 | 0.0212 | 0.0212 | TRUE | 0.3718 |
| Ninj2 | 17 | 21 | 15 | 66 | 19 | 14 | 18 | 21 | 15 | 52 | 20 | 15 | 23.62 | 18 | 29 | 1.641 | 0.715 | 0.19268904 | 0.39842218 | 0.197 | 0.1464 | 0.1722 | 0.1722 | TRUE | 2.8415 |
| Ninl | 2097 | 2163 | 2123 | 3004 | 2415 | 2252 | 2227 | 2160 | 2132 | 2362 | 2598 | 2465 | 2324.06 | 2173 | 2475 | 1.138 | 0.187 | 0.01198593 | 0.06007058 | 0.001 | 0.0083 | 0.0036 | 0.0036 | TRUE | 0.0231 |
| Nip7 | 5182 | 4860 | 4447 | 6552 | 4494 | 4078 | 5504 | 4854 | 4465 | 5151 | 4835 | 4464 | 4878.86 | 4941 | 4817 | 0.975 | -0.036 | 0.72915539 | 0.85433232 | 0.0079 | 0.0076 | 0.0078 | 0.0078 | TRUE | 0.121 |
| Nipa1 | 249 | 260 | 224 | 324 | 206 | 184 | 264 | 260 | 225 | 255 | 222 | 201 | 237.8 | 250 | 226 | 0.908 | -0.139 | 0.37480769 | 0.59085248 | 0.007 | 0.0208 | 0.0135 | 0.0135 | TRUE | 0.146 |
| Nipa2 | 2078 | 1913 | 1712 | 2868 | 1980 | 1861 | 2207 | 1911 | 1719 | 2255 | 2130 | 2037 | 2043.15 | 1946 | 2141 | 1.101 | 0.138 | 0.21771921 | 0.42970873 | 0.0086 | 0.0085 | 0.0086 | 0.0086 | TRUE | 0.1669 |
| Nipal1 | 102 | 73 | 86 | 89 | 60 | 49 | 108 | 73 | 86 | 70 | 65 | 54 | 75.96 | 89 | 63 | 0.707 | -0.501 | 0.0507175 | 0.16568278 | 0.017 | 0.0503 | 0.0338 | 0.0338 | TRUE | 0.2734 |
| Nipal2 | 4 | 7 | 12 | 25 | 14 | 26 | 4 | 7 | 12 | 20 | 15 | 28 | 14.41 | 8 | 21 | 2.697 | 1.432 | 0.01522512 | 0.07151 | 0.0671 | 0.2355 | 0.1619 | 0.1619 | TRUE | 0.4266 |
| Nipal3 | 1520 | 1448 | 1534 | 1413 | 1020 | 1084 | 1615 | 1446 | 1540 | 1111 | 1097 | 1187 | 1332.63 | 1534 | 1132 | 0.738 | -0.439 | 4.04E-07 | 1.24E-05 | 0.0017 | 0.0094 | 0.0046 | 0.0046 | TRUE | 0.0301 |
| Nipal4 | 174 | 183 | 149 | 150 | 93 | 113 | 185 | 183 | 150 | 118 | 100 | 124 | 143.15 | 173 | 114 | 0.662 | -0.596 | 0.00137685 | 0.01143536 | 0.0054 | 0.0299 | 0.0178 | 0.0178 | TRUE | 0.1436 |
| Nipbl | 7486 | 7241 | 7869 | 9090 | 6546 | 6370 | 7951 | 7232 | 7901 | 7146 | 7043 | 6973 | 7374.39 | 7695 | 7054 | 0.917 | -0.125 | 0.06849853 | 0.20277005 | 0.0012 | 0.0073 | 0.0033 | 0.0033 | TRUE | 0.0339 |
| Nipsnap1 | 3706 | 4142 | 4244 | 3955 | 2905 | 3268 | 3936 | 4137 | 4261 | 3109 | 3126 | 3577 | 3691.09 | 4111 | 3271 | 0.795 | -0.331 | 0.00015891 | 0.00200181 | 0.0037 | 0.0078 | 0.0052 | 0.0052 | TRUE | 0.0813 |
| Nipsnap3a | 1 | 1 | 0 | 0 | 0 | 2 | 1 | 1 | 0 | 0 | 0 | 2 | 0.71 | 1 | 1 | 1.031 | 0.044 | 0.9865942 | NA | 0 | 4.6577 | 3.4964 | 3.4964 | TRUE | 0.9748 |
| Nipsnap3b | 1609 | 1866 | 1850 | 2058 | 1504 | 1609 | 1709 | 1864 | 1858 | 1618 | 1618 | 1761 | 1737.93 | 1810 | 1666 | 0.92 | -0.121 | 0.14793186 | 0.33750633 | 0.0018 | 0.0088 | 0.0044 | 0.0044 | TRUE | 0.03 |
| Nisch | 15170 | 13962 | 15169 | 20519 | 15332 | 15058 | 16113 | 13944 | 15231 | 16130 | 16496 | 16483 | 15733.09 | 15096 | 16370 | 1.084 | 0.117 | 0.12830497 | 0.30801003 | 0.0026 | 0.0071 | 0.0042 | 0.0042 | TRUE | 0.0545 |
| Nit1 | 843 | 917 | 949 | 1380 | 931 | 893 | 895 | 916 | 953 | 1085 | 1002 | 978 | 971.37 | 921 | 1022 | 1.11 | 0.15 | 0.0933226 | 0.24976093 | 0.0012 | 0.0103 | 0.0047 | 0.0047 | TRUE | 0.0367 |
| Nit2 | 4171 | 4857 | 4891 | 4461 | 3152 | 3549 | 4430 | 4851 | 4911 | 3507 | 3391 | 3885 | 4162.56 | 4731 | 3594 | 0.76 | -0.397 | 5.92E-06 | 0.00012784 | 0.0037 | 0.0077 | 0.0053 | 0.0053 | TRUE | 0.0604 |
| Nkain1 | 3611 | 4170 | 3873 | 4749 | 3426 | 3592 | 3836 | 4165 | 3889 | 3733 | 3686 | 3932 | 3873.42 | 3963 | 3784 | 0.955 | -0.067 | 0.35223878 | 0.57073249 | 0.0012 | 0.0078 | 0.0035 | 0.0035 | TRUE | 0.0241 |
| Nkain2 | 0 | 0 | 1 | 0 | 0 | 0 | 0 | 0 | 1 | 0 | 0 | 0 | 0.17 | 0 | 0 | 0.49 | -1.029 | 0.80087 | NA | 0 | 19.6928 | 10 | 10 | TRUE | 0.4098 |
| Nkain3 | 0 | 0 | 1 | 1 | 1 | 0 | 0 | 0 | 1 | 1 | 1 | 0 | 0.48 | 0 | 1 | 1.712 | 0.776 | 0.78603958 | NA | 0 | 6.9035 | 4.1008 | 4.1008 | TRUE | 0.4098 |
| Nkain4 | 9 | 15 | 12 | 62 | 37 | 61 | 10 | 15 | 12 | 49 | 40 | 67 | 31.99 | 12 | 52 | 4.228 | 2.08 | 5.35E-07 | 1.59E-05 | 0 | 0.1099 | 0.0728 | 0.0728 | TRUE | 0.2046 |
| Nkap | 1397 | 1596 | 1749 | 2143 | 1501 | 1644 | 1484 | 1594 | 1756 | 1685 | 1615 | 1800 | 1655.54 | 1611 | 1700 | 1.055 | 0.077 | 0.4307441 | 0.64055214 | 0.0042 | 0.0089 | 0.0062 | 0.0062 | TRUE | 0.0746 |
| Nkapl | 14 | 16 | 16 | 51 | 22 | 15 | 15 | 16 | 16 | 40 | 24 | 16 | 21.18 | 16 | 27 | 1.73 | 0.79 | 0.1033331 | 0.26757014 | 0.0783 | 0.1624 | 0.1189 | 0.1189 | TRUE | 0.434 |
| Nkd1 | 360 | 376 | 366 | 492 | 358 | 380 | 382 | 376 | 368 | 387 | 385 | 416 | 385.55 | 375 | 396 | 1.055 | 0.077 | 0.48581954 | 0.6827493 | 0 | 0.0154 | 0.0062 | 0.0062 | TRUE | 0.0216 |
| Nkd2 | 464 | 520 | 478 | 632 | 520 | 422 | 493 | 519 | 480 | 497 | 559 | 462 | 501.73 | 497 | 506 | 1.017 | 0.025 | 0.8336008 | 0.91548463 | 0.0033 | 0.0135 | 0.0079 | 0.0079 | TRUE | 0.096 |
| Nkg7 | 3 | 2 | 1 | 2 | 3 | 6 | 3 | 2 | 1 | 2 | 3 | 7 | 2.93 | 2 | 4 | 1.81 | 0.856 | 0.46398711 | NA | 0 | 1.1328 | 0.6038 | 0.6038 | TRUE | 0.3402 |
| Nkiras1 | 398 | 459 | 439 | 574 | 455 | 426 | 423 | 458 | 441 | 451 | 490 | 466 | 454.84 | 441 | 469 | 1.063 | 0.088 | 0.40736097 | 0.62022369 | 0 | 0.0141 | 0.006 | 0.006 | TRUE | 0.0169 |
| Nkiras2 | 2356 | 2806 | 2747 | 3542 | 2431 | 2440 | 2502 | 2802 | 2758 | 2784 | 2616 | 2671 | 2689.03 | 2687 | 2690 | 1.001 | 0.002 | 0.98350516 | 0.99129879 | 0.002 | 0.0081 | 0.0042 | 0.0042 | TRUE | 0.0439 |
| Nkpd1 | 10 | 10 | 4 | 8 | 4 | 5 | 11 | 10 | 4 | 6 | 4 | 5 | 6.78 | 8 | 5 | 0.658 | -0.603 | 0.42500121 | 0.63512594 | 0 | 0.4927 | 0.2591 | 0.2591 | TRUE | 0.6065 |
| Nkrf | 1218 | 1072 | 943 | 1717 | 1093 | 921 | 1294 | 1071 | 947 | 1350 | 1176 | 1008 | 1140.86 | 1104 | 1178 | 1.068 | 0.095 | 0.5349626 | 0.72238272 | 0.0221 | 0.0098 | 0.0161 | 0.0161 | TRUE | 0.2508 |
| Nktr | 2506 | 2120 | 2164 | 4050 | 2993 | 2367 | 2662 | 2117 | 2173 | 3184 | 3220 | 2591 | 2657.85 | 2317 | 2998 | 1.294 | 0.372 | 0.00413628 | 0.02712572 | 0.0147 | 0.0081 | 0.0118 | 0.0118 | TRUE | 0.2046 |
| Nkx1-2 | 7 | 4 | 4 | 2 | 3 | 2 | 7 | 4 | 4 | 2 | 3 | 2 | 3.74 | 5 | 2 | 0.446 | -1.165 | 0.24336102 | NA | 0 | 0.8879 | 0.4063 | 0.4063 | TRUE | 0.2973 |
| Nkx2-1 | 0 | 0 | 0 | 4 | 0 | 0 | 0 | 0 | 0 | 3 | 0 | 0 | 0.52 | 0 | 1 | 5.676 | 2.505 | 0.52992595 | NA | 10 | 6.293 | 10 | 10 | TRUE | 1.8971 |
| Nkx2-2 | 6 | 3 | 3 | 8 | 9 | 5 | 6 | 3 | 3 | 6 | 10 | 5 | 5.64 | 4 | 7 | 1.735 | 0.795 | 0.33256747 | 0.55303898 | 0 | 0.5912 | 0.2916 | 0.2916 | TRUE | 0.3687 |
| Nkx2-2as | 0 | 1 | 0 | 1 | 0 | 0 | 0 | 1 | 0 | 1 | 0 | 0 | 0.3 | 0 | 0 | 0.954 | -0.067 | 0.98642001 | NA | 0 | 11.0817 | 9.3014 | 9.3014 | TRUE | 0.405 |
| Nkx2-3 | 5 | 7 | 11 | 17 | 13 | 7 | 5 | 7 | 11 | 13 | 14 | 8 | 9.73 | 8 | 12 | 1.504 | 0.589 | 0.35710822 | 0.57566545 | 0 | 0.3456 | 0.1874 | 0.1874 | TRUE | 0.3899 |
| Nkx2-4 | 66 | 128 | 119 | 106 | 77 | 81 | 70 | 128 | 119 | 83 | 83 | 89 | 95.38 | 106 | 85 | 0.801 | -0.319 | 0.23145492 | 0.44525843 | 0.0399 | 0.0414 | 0.0407 | 0.0407 | TRUE | 0.845 |
| Nkx2-5 | 20 | 24 | 30 | 29 | 16 | 18 | 21 | 24 | 30 | 23 | 17 | 20 | 22.51 | 25 | 20 | 0.797 | -0.328 | 0.41875789 | 0.62931391 | 0 | 0.1533 | 0.0739 | 0.0739 | TRUE | 0.1868 |
| Nkx2-6 | 0 | 0 | 0 | 1 | 0 | 0 | 0 | 0 | 0 | 1 | 0 | 0 | 0.13 | 0 | 0 | 1.859 | 0.894 | 0.82650734 | NA | 0 | 25.1515 | 10 | 10 | TRUE | 0.2563 |
| Nkx2-9 | 38 | 86 | 64 | 112 | 98 | 102 | 40 | 86 | 64 | 88 | 105 | 112 | 82.61 | 63 | 102 | 1.595 | 0.674 | 0.02378171 | 0.09818983 | 0.0564 | 0.0468 | 0.0512 | 0.0512 | TRUE | 0.3405 |
| Nkx3-1 | 171 | 212 | 178 | 222 | 160 | 181 | 182 | 212 | 179 | 175 | 172 | 198 | 186.15 | 191 | 182 | 0.951 | -0.073 | 0.64930003 | 0.8033859 | 0 | 0.0246 | 0.0131 | 0.0131 | TRUE | 0.1015 |
| Nkx3-2 | 0 | 0 | 0 | 0 | 0 | 1 | 0 | 0 | 0 | 0 | 0 | 1 | 0.18 | 0 | 0 | 1.859 | 0.894 | 0.82650734 | NA | 0 | 18.0643 | 10 | 10 | TRUE | 0.5383 |
| Nkx6-1 | 2 | 5 | 8 | 9 | 8 | 6 | 2 | 5 | 8 | 7 | 9 | 7 | 6.23 | 5 | 8 | 1.46 | 0.546 | 0.48906398 | 0.68540227 | 0 | 0.5354 | 0.2823 | 0.2823 | TRUE | 0.2756 |
| Nkx6-2 | 231 | 348 | 363 | 289 | 215 | 214 | 245 | 348 | 364 | 227 | 231 | 234 | 275.03 | 319 | 231 | 0.723 | -0.468 | 0.008483 | 0.04650467 | 0.0192 | 0.0189 | 0.0191 | 0.0191 | TRUE | 0.4593 |
| Nkx6-3 | 12 | 14 | 19 | 11 | 13 | 10 | 13 | 14 | 19 | 9 | 14 | 11 | 13.23 | 15 | 11 | 0.725 | -0.463 | 0.3832548 | 0.59760644 | 0 | 0.2559 | 0.126 | 0.126 | TRUE | 0.2212 |
| Nle1 | 2293 | 2578 | 2686 | 3426 | 2627 | 2514 | 2436 | 2575 | 2697 | 2693 | 2826 | 2752 | 2663.16 | 2569 | 2757 | 1.073 | 0.102 | 0.17278133 | 0.37156189 | 0.001 | 0.0081 | 0.0036 | 0.0036 | TRUE | 0.025 |
| Nlgn1 | 0 | 1 | 1 | 2 | 5 | 2 | 0 | 1 | 1 | 2 | 5 | 2 | 1.86 | 1 | 3 | 4.454 | 2.155 | 0.17742104 | NA | 0 | 1.7806 | 0.9213 | 0.9213 | TRUE | 0.5567 |
| Nlgn2 | 936 | 1434 | 1492 | 1427 | 1185 | 1270 | 994 | 1432 | 1498 | 1122 | 1275 | 1390 | 1285.24 | 1308 | 1262 | 0.964 | -0.052 | 0.75255858 | 0.86948897 | 0.0289 | 0.0095 | 0.0191 | 0.0191 | TRUE | 0.5282 |
| Nlgn3 | 34 | 38 | 38 | 29 | 21 | 31 | 36 | 38 | 38 | 23 | 23 | 34 | 31.92 | 37 | 27 | 0.702 | -0.51 | 0.14426492 | 0.33139329 | 0 | 0.1101 | 0.0557 | 0.0557 | TRUE | 0.3558 |
| Nlk | 4998 | 4757 | 4861 | 5837 | 4424 | 4184 | 5309 | 4751 | 4881 | 4589 | 4760 | 4580 | 4811.53 | 4980 | 4643 | 0.932 | -0.101 | 0.17036197 | 0.36901257 | 0.0016 | 0.0076 | 0.0037 | 0.0037 | TRUE | 0.0405 |
| Nln | 4150 | 4924 | 5069 | 5211 | 3875 | 4186 | 4408 | 4918 | 5090 | 4096 | 4169 | 4582 | 4543.91 | 4805 | 4282 | 0.891 | -0.166 | 0.06485576 | 0.19618431 | 0.0043 | 0.0076 | 0.0056 | 0.0056 | TRUE | 0.0636 |
| Nlrc3 | 7 | 13 | 5 | 32 | 8 | 17 | 7 | 13 | 5 | 25 | 9 | 19 | 12.97 | 8 | 18 | 2.075 | 1.053 | 0.10429131 | 0.26872775 | 0.1657 | 0.2609 | 0.2149 | 0.2149 | TRUE | 0.1905 |
| Nlrc4 | 77 | 39 | 29 | 71 | 58 | 38 | 82 | 39 | 29 | 56 | 62 | 42 | 51.61 | 50 | 53 | 1.07 | 0.097 | 0.81527305 | 0.90584619 | 0.1427 | 0.0707 | 0.105 | 0.105 | TRUE | 2.0007 |
| Nlrc5 | 119 | 107 | 104 | 95 | 87 | 76 | 126 | 107 | 104 | 75 | 94 | 83 | 98.2 | 112 | 84 | 0.742 | -0.43 | 0.04139391 | 0.14434331 | 0 | 0.0405 | 0.0216 | 0.0216 | TRUE | 0.1134 |
| Nlrp10 | 1 | 0 | 1 | 3 | 4 | 4 | 1 | 0 | 1 | 2 | 4 | 4 | 2.18 | 1 | 3 | 5.332 | 2.415 | 0.10604235 | NA | 0 | 1.515 | 0.7268 | 0.7268 | TRUE | 0.2573 |
| Nlrp12 | 11 | 7 | 9 | 20 | 16 | 22 | 12 | 7 | 9 | 16 | 17 | 24 | 14.12 | 9 | 19 | 2.051 | 1.036 | 0.05137963 | 0.16708444 | 0 | 0.2402 | 0.1226 | 0.1226 | TRUE | 0.2693 |
| Nlrp14 | 807 | 1242 | 1201 | 1548 | 1238 | 1230 | 857 | 1240 | 1206 | 1217 | 1332 | 1346 | 1199.81 | 1101 | 1298 | 1.179 | 0.237 | 0.11301328 | 0.28312717 | 0.0206 | 0.0096 | 0.0153 | 0.0153 | TRUE | 0.4482 |

| | | | | | | | | | | | | | | | | | | | | | | | | | |
|---|---|---|---|---|---|---|---|---|---|---|---|---|---|---|---|---|---|---|---|---|---|---|---|---|---|
| Nlrp1a | 0 | 0 | 1 | 1 | 1 | 1 | 0 | 0 | 1 | 1 | 1 | 1 | 0.66 | 0 | 1 | 2.676 | 1.42 | 0.53608725 | NA | 0 | 4.9975 | 2.297 | 2.297 | TRUE | 0.4098 |
| Nlrp1b | 0 | 0 | 0 | 0 | 1 | 0 | 0 | 0 | 0 | 0 | 1 | 0 | 0.18 | 0 | 0 | 1.859 | 0.894 | 0.82650734 | NA | 0 | 18.3785 | 10 | 10 | TRUE | 0.5204 |
| Nlrp1c-ps | 0 | 0 | 0 | 0 | 0 | 0 | 0 | 0 | 0 | 0 | 0 | 0 | 0 | 0 | 0 | NA | NA | NA | NA | NA | NA | NA | NA | NA | NA |
| Nlrp2 | 0 | 0 | 1 | 3 | 3 | 1 | 0 | 0 | 1 | 2 | 3 | 1 | 1.28 | 0 | 2 | 6.1 | 2.609 | 0.16990347 | NA | 0 | 2.579 | 1.3839 | 1.3839 | TRUE | 0.393 |
| Nlrp3 | 142 | 159 | 137 | 139 | 107 | 173 | 151 | 159 | 138 | 109 | 115 | 189 | 143.49 | 149 | 138 | 0.922 | -0.117 | 0.63242267 | 0.79314332 | 0.0434 | 0.0299 | 0.0365 | 0.0365 | TRUE | 1.078 |
| Nlrp4a | 38 | 29 | 31 | 105 | 69 | 79 | 40 | 29 | 31 | 83 | 74 | 86 | 57.29 | 33 | 81 | 2.427 | 1.279 | 3.09E-06 | 7.34E-05 | 0 | 0.0644 | 0.0329 | 0.0329 | TRUE | 0.2172 |
| Nlrp4b | 0 | 0 | 0 | 0 | 0 | 0 | 0 | 0 | 0 | 0 | 0 | 0 | 0 | 0 | 0 | NA | NA | NA | NA | NA | NA | NA | NA | NA | NA |
| Nlrp4c | 477 | 557 | 458 | 2484 | 1640 | 1209 | 507 | 556 | 460 | 1953 | 1765 | 1323 | 1093.92 | 508 | 1680 | 3.312 | 1.728 | 1.87E-27 | 9.53E-25 | 0.0237 | 0.0099 | 0.017 | 0.017 | TRUE | 0.2804 |
| Nlrp4e | 5 | 1 | 1 | 14 | 9 | 7 | 5 | 1 | 1 | 11 | 10 | 8 | 5.94 | 2 | 10 | 3.942 | 1.979 | 0.03200528 | 0.12050304 | 0 | 0.5611 | 0.35 | 0.35 | TRUE | 1.0862 |
| Nlrp4f | 1300 | 1037 | 880 | 1902 | 1249 | 1022 | 1381 | 1036 | 884 | 1495 | 1344 | 1119 | 1209.65 | 1100 | 1319 | 1.2 | 0.263 | 0.14049382 | 0.32571932 | 0.0359 | 0.0096 | 0.0221 | 0.0221 | TRUE | 0.5957 |
| Nlrp4g | 27 | 10 | 9 | 7 | 4 | 7 | 29 | 10 | 9 | 6 | 4 | 8 | 10.86 | 16 | 6 | 0.366 | -1.449 | 0.04413729 | 0.15067822 | 0.2084 | 0.3102 | 0.2571 | 0.2571 | TRUE | 2.2592 |
| Nlrp5 | 5 | 1 | 0 | 6 | 6 | 5 | 5 | 1 | 0 | 5 | 6 | 5 | 3.83 | 2 | 5 | 2.664 | 1.413 | 0.22643243 | NA | 0.1589 | 0.868 | 0.6493 | 0.6493 | TRUE | 1.5947 |
| Nlrp5-ps | 28 | 14 | 21 | 70 | 54 | 34 | 30 | 14 | 21 | 55 | 58 | 37 | 35.86 | 22 | 50 | 2.332 | 1.222 | 0.00187911 | 0.01464937 | 0.051 | 0.0988 | 0.0778 | 0.0778 | TRUE | 0.5198 |
| Nlrp6 | 9 | 8 | 16 | 22 | 11 | 22 | 10 | 8 | 16 | 17 | 12 | 24 | 14.47 | 11 | 18 | 1.58 | 0.66 | 0.23538179 | 0.45048436 | 0.0533 | 0.2346 | 0.1495 | 0.1495 | TRUE | 0.2287 |
| Nlrp9a | 1 | 0 | 0 | 0 | 0 | 0 | 1 | 0 | 0 | 0 | 0 | 0 | 0.18 | 0 | 0 | 0.49 | -1.029 | 0.80087 | NA | 0 | 18.6165 | 10 | 10 | TRUE | 0.4614 |
| Nlrp9b | 126 | 106 | 85 | 236 | 179 | 192 | 134 | 106 | 85 | 186 | 193 | 210 | 152.22 | 108 | 196 | 1.81 | 0.856 | 2.67E-05 | 0.00045547 | 0.0166 | 0.0285 | 0.0227 | 0.0227 | TRUE | 0.4126 |
| Nlrp9c | 0 | 0 | 0 | 1 | 1 | 1 | 0 | 0 | 0 | 1 | 1 | 1 | 0.49 | 0 | 1 | 5.224 | 2.385 | 0.38465 | NA | 0 | 6.6923 | 3.9249 | 3.9249 | TRUE | 0.07 |
| Nlrx1 | 786 | 741 | 695 | 828 | 597 | 601 | 835 | 740 | 698 | 651 | 642 | 658 | 703.98 | 758 | 650 | 0.859 | -0.22 | 0.03765576 | 0.13479496 | 0.003 | 0.0116 | 0.0066 | 0.0066 | TRUE | 0.0937 |
| Nmb | 151 | 132 | 137 | 187 | 134 | 138 | 160 | 132 | 138 | 147 | 144 | 151 | 145.34 | 143 | 147 | 1.03 | 0.042 | 0.80494354 | 0.89931073 | 0 | 0.0296 | 0.0142 | 0.0142 | TRUE | 0.1118 |
| Nmbr | 9 | 4 | 4 | 4 | 6 | 1 | 10 | 4 | 4 | 3 | 6 | 1 | 4.71 | 6 | 3 | 0.609 | -0.716 | 0.45925402 | 0.66378277 | 0.1598 | 0.7062 | 0.4488 | 0.4488 | TRUE | 0.4869 |
| Nmd3 | 4933 | 5036 | 4847 | 6794 | 4609 | 4466 | 5240 | 5029 | 4867 | 5341 | 4959 | 4889 | 5054.13 | 5045 | 5063 | 1.004 | 0.005 | 0.94022051 | 0.96979647 | 0.0016 | 0.0076 | 0.0036 | 0.0036 | TRUE | 0.0284 |
| Nme1 | 8764 | 8315 | 8165 | 11067 | 7587 | 7280 | 9309 | 8304 | 8199 | 8700 | 8163 | 7969 | 8440.66 | 8604 | 8277 | 0.962 | -0.056 | 0.50166403 | 0.69705269 | 0.0032 | 0.0073 | 0.0048 | 0.0048 | TRUE | 0.0627 |
| Nme2 | 19069 | 23851 | 24010 | 29057 | 20569 | 21324 | 20255 | 23820 | 24109 | 22842 | 22131 | 23342 | 22749.87 | 22728 | 22772 | 1.002 | 0.003 | 0.97530328 | 0.98697924 | 0.0038 | 0.007 | 0.0057 | 0.0057 | TRUE | 0.1108 |
| Nme3 | 509 | 661 | 654 | 685 | 503 | 552 | 541 | 660 | 657 | 538 | 541 | 604 | 590.24 | 619 | 561 | 0.906 | -0.143 | 0.24642614 | 0.46346686 | 0.0067 | 0.0125 | 0.0093 | 0.0093 | TRUE | 0.1436 |
| Nme4 | 1264 | 1967 | 1971 | 1765 | 1349 | 1529 | 1343 | 1964 | 1979 | 1388 | 1451 | 1674 | 1633.14 | 1762 | 1504 | 0.853 | -0.229 | 0.15679714 | 0.34972176 | 0.0253 | 0.0089 | 0.0182 | 0.0182 | TRUE | 0.5226 |
| Nme5 | 7 | 9 | 5 | 19 | 17 | 14 | 7 | 9 | 5 | 15 | 18 | 15 | 11.67 | 7 | 16 | 2.259 | 1.176 | 0.04018287 | 0.14137057 | 0 | 0.2893 | 0.1349 | 0.1349 | TRUE | 0.1872 |
| Nme6 | 953 | 1099 | 1039 | 1462 | 1019 | 1044 | 1012 | 1098 | 1043 | 1149 | 1096 | 1143 | 1090.27 | 1051 | 1129 | 1.075 | 0.104 | 0.20609241 | 0.41502674 | 0 | 0.0099 | 0.004 | 0.004 | TRUE | 0.018 |
| Nme7 | 1501 | 1456 | 1358 | 1943 | 1327 | 1496 | 1594 | 1454 | 1364 | 1527 | 1428 | 1638 | 1500.81 | 1471 | 1531 | 1.041 | 0.058 | 0.56060295 | 0.74238705 | 0.0047 | 0.0091 | 0.0065 | 0.0065 | TRUE | 0.0649 |
| Nme8 | 5 | 3 | 9 | 2 | 3 | 2 | 5 | 3 | 9 | 2 | 3 | 2 | 4.06 | 6 | 2 | 0.396 | -1.336 | 0.18215799 | NA | 0 | 0.8192 | 0.4216 | 0.4216 | TRUE | 0.2028 |
| Nme9 | 0 | 0 | 0 | 0 | 0 | 1 | 0 | 0 | 0 | 0 | 0 | 1 | 0.18 | 0 | 0 | 1.859 | 0.894 | 0.82650734 | NA | 0 | 18.0643 | 10 | 10 | TRUE | 0.5383 |
| Nmi | 36 | 41 | 64 | 102 | 76 | 50 | 38 | 41 | 64 | 80 | 82 | 55 | 60.02 | 48 | 72 | 1.513 | 0.597 | 0.05825609 | 0.18235677 | 0.0459 | 0.0618 | 0.0542 | 0.0542 | TRUE | 0.7289 |
| Nmnat1 | 315 | 279 | 267 | 347 | 241 | 232 | 335 | 279 | 268 | 273 | 259 | 254 | 277.89 | 294 | 262 | 0.893 | -0.163 | 0.25640731 | 0.47510057 | 0.0045 | 0.0188 | 0.0112 | 0.0112 | TRUE | 0.1646 |
| Nmnat2 | 1738 | 2425 | 2409 | 2180 | 1658 | 1949 | 1846 | 2422 | 2419 | 1714 | 1784 | 2133 | 2052.99 | 2229 | 1877 | 0.842 | -0.249 | 0.07533824 | 0.21627863 | 0.0175 | 0.0085 | 0.0136 | 0.0136 | TRUE | 0.273 |
| Nmnat3 | 163 | 194 | 204 | 152 | 112 | 129 | 173 | 194 | 205 | 119 | 121 | 141 | 158.82 | 191 | 127 | 0.665 | -0.589 | 0.00057906 | 0.00582826 | 0 | 0.0276 | 0.0146 | 0.0146 | TRUE | 0.0924 |
| Nmral1 | 630 | 660 | 587 | 992 | 681 | 663 | 669 | 659 | 589 | 780 | 733 | 726 | 692.67 | 639 | 746 | 1.168 | 0.225 | 0.02606173 | 0.10453855 | 0.0016 | 0.0117 | 0.0059 | 0.0059 | TRUE | 0.055 |
| Nmrk1 | 450 | 451 | 395 | 453 | 367 | 336 | 478 | 450 | 397 | 356 | 395 | 368 | 407.3 | 442 | 373 | 0.844 | -0.245 | 0.04931682 | 0.16282706 | 0.0036 | 0.015 | 0.0087 | 0.0087 | TRUE | 0.0924 |
| Nmrk2 | 1 | 1 | 0 | 1 | 1 | 0 | 1 | 1 | 0 | 1 | 1 | 0 | 0.65 | 1 | 1 | 0.919 | -0.122 | 0.95812821 | NA | 0 | 5.0456 | 2.3857 | 2.3857 | TRUE | 0.2521 |
| Nms | 0 | 1 | 0 | 0 | 1 | 0 | 0 | 1 | 0 | 0 | 1 | 0 | 0.35 | 0 | 0 | 0.954 | -0.067 | 0.98571955 | NA | 0 | 9.5346 | 8.2194 | 8.2194 | TRUE | 0.5204 |
| Nmt1 | 11377 | 12307 | 12527 | 15335 | 11053 | 10933 | 12084 | 12291 | 12578 | 12055 | 11892 | 11968 | 12144.89 | 12318 | 11972 | 0.972 | -0.041 | 0.44632711 | 0.65236459 | 1.00E-04 | 0.0072 | 0.002 | 0.002 | TRUE | 0.0042 |
| Nmt2 | 3868 | 4056 | 4073 | 5918 | 4375 | 4250 | 4108 | 4051 | 4090 | 4652 | 4707 | 4652 | 4376.8 | 4083 | 4670 | 1.144 | 0.194 | 0.00061048 | 0.00609394 | 0 | 0.0077 | 0.0021 | 0.0021 | TRUE | 6.00E-04 |
| Nmu | 7 | 4 | 11 | 5 | 5 | 3 | 7 | 4 | 11 | 4 | 5 | 3 | 5.84 | 7 | 4 | 0.559 | -0.839 | 0.30450583 | 0.52512118 | 0 | 0.5705 | 0.296 | 0.296 | TRUE | 0.1275 |
| Nmur1 | 6 | 1 | 1 | 3 | 0 | 0 | 6 | 1 | 1 | 2 | 0 | 0 | 1.79 | 3 | 1 | 0.308 | -1.7 | 0.3487977 | NA | 1.2944 | 1.8484 | 1.6182 | 1.6182 | TRUE | 1.4642 |
| Nmur2 | 9 | 2 | 1 | 5 | 2 | 4 | 10 | 2 | 1 | 4 | 2 | 4 | 3.84 | 4 | 3 | 0.842 | -0.249 | 0.8238551 | NA | 0.4228 | 0.8655 | 0.6376 | 0.6376 | TRUE | 2.0957 |
| Nnat | 904 | 811 | 833 | 708 | 491 | 555 | 960 | 810 | 836 | 557 | 528 | 608 | 716.5 | 869 | 564 | 0.649 | -0.623 | 5.09E-08 | 2.02E-06 | 0.0052 | 0.0115 | 0.008 | 0.008 | TRUE | 0.1001 |
| Nnmt | 1 | 0 | 1 | 10 | 6 | 5 | 1 | 0 | 1 | 8 | 6 | 5 | 3.64 | 1 | 6 | 9.714 | 3.28 | 0.01373206 | NA | 0 | 0.9113 | 0.4549 | 0.4549 | TRUE | 0.2581 |
| Nnt | 1614 | 1955 | 2064 | 2097 | 1397 | 1483 | 1714 | 1952 | 2072 | 1648 | 1503 | 1623 | 1752.38 | 1913 | 1591 | 0.832 | -0.265 | 0.0084198 | 0.04627077 | 0.0049 | 0.0088 | 0.0067 | 0.0067 | TRUE | 0.0995 |
| Noa1 | 2424 | 2174 | 2232 | 2466 | 1884 | 1803 | 2575 | 2171 | 2241 | 1939 | 2027 | 1974 | 2154.4 | 2329 | 1980 | 0.85 | -0.234 | 0.01148756 | 0.05837041 | 0.0038 | 0.0084 | 0.0057 | 0.0057 | TRUE | 0.1029 |
| Nob1 | 4029 | 4572 | 4670 | 5564 | 3949 | 4107 | 4280 | 4566 | 4689 | 4374 | 4249 | 4496 | 4442.23 | 4512 | 4373 | 0.969 | -0.045 | 0.52332787 | 0.71320549 | 0.0011 | 0.0076 | 0.0034 | 0.0034 | TRUE | 0.0246 |
| Nobox | 16 | 14 | 15 | 22 | 14 | 22 | 17 | 14 | 15 | 17 | 15 | 24 | 17.08 | 15 | 19 | 1.223 | 0.29 | 0.53237538 | 0.72039355 | 0 | 0.1998 | 0.0962 | 0.0962 | TRUE | 0.2847 |
| Noc2l | 12685 | 14586 | 14405 | 14912 | 11370 | 12230 | 13474 | 14567 | 14464 | 11723 | 12233 | 13388 | 13308.12 | 14168 | 12448 | 0.878 | -0.187 | 0.02002559 | 0.08693741 | 0.0031 | 0.0072 | 0.0046 | 0.0046 | TRUE | 0.0533 |
| Noc3l | 3944 | 3272 | 3142 | 5216 | 3738 | 3191 | 4189 | 3268 | 3155 | 4100 | 4022 | 3493 | 3704.54 | 3537 | 3872 | 1.095 | 0.131 | 0.31681987 | 0.53770396 | 0.0147 | 0.0078 | 0.012 | 0.012 | TRUE | 0.3158 |
| Noc4l | 3855 | 3405 | 3498 | 5020 | 3619 | 3340 | 4095 | 3401 | 3512 | 3946 | 3894 | 3656 | 3750.66 | 3669 | 3832 | 1.045 | 0.063 | 0.51363976 | 0.70657132 | 0.0049 | 0.0078 | 0.0064 | 0.0064 | TRUE | 0.125 |
| Nod1 | 49 | 56 | 57 | 87 | 67 | 49 | 52 | 56 | 57 | 68 | 72 | 54 | 59.89 | 55 | 65 | 1.177 | 0.235 | 0.35968015 | 0.57800887 | 0 | 0.0619 | 0.0308 | 0.0308 | TRUE | 0.1828 |
| Nod2 | 148 | 196 | 213 | 141 | 116 | 138 | 157 | 196 | 214 | 111 | 125 | 151 | 158.92 | 189 | 129 | 0.679 | -0.558 | 0.00571426 | 0.03458377 | 0.0181 | 0.0276 | 0.0229 | 0.0229 | TRUE | 0.2299 |
| Nodal | 2507 | 2315 | 2331 | 3059 | 1975 | 1630 | 2663 | 2312 | 2341 | 2405 | 2125 | 1784 | 2271.58 | 2439 | 2105 | 0.864 | -0.211 | 0.09718867 | 0.25620253 | 0.0134 | 0.0084 | 0.0113 | 0.0113 | TRUE | 0.214 |
| Nog | 41 | 55 | 58 | 53 | 31 | 30 | 44 | 55 | 58 | 42 | 33 | 33 | 44.1 | 52 | 36 | 0.692 | -0.531 | 0.07864118 | 0.22208213 | 0 | 0.0816 | 0.0424 | 0.0424 | TRUE | 0.1695 |
| Nol10 | 5167 | 4721 | 4558 | 6740 | 4679 | 4123 | 5488 | 4715 | 4577 | 5298 | 5034 | 4513 | 4937.66 | 4927 | 4948 | 1.005 | 0.007 | 0.94848656 | 0.9742329 | 0.0079 | 0.0076 | 0.0078 | 0.0078 | TRUE | 0.1211 |
| Nol11 | 11213 | 10933 | 11042 | 12814 | 9185 | 9298 | 11910 | 10919 | 11087 | 10073 | 9882 | 10178 | 10675.06 | 11305 | 10044 | 0.889 | -0.171 | 0.00918119 | 0.04934573 | 0.0011 | 0.0072 | 0.003 | 0.003 | TRUE | 0.0267 |
| Nol12 | 1082 | 1084 | 1101 | 1598 | 1165 | 1005 | 1149 | 1083 | 1106 | 1256 | 1253 | 1100 | 1157.87 | 1113 | 1203 | 1.082 | 0.114 | 0.21839628 | 0.43051243 | 0.0024 | 0.0097 | 0.0053 | 0.0053 | TRUE | 0.0663 |
| Nol3 | 173 | 281 | 274 | 282 | 223 | 220 | 184 | 281 | 275 | 222 | 240 | 241 | 240.33 | 247 | 234 | 0.948 | -0.077 | 0.68737809 | 0.82764541 | 0.0233 | 0.0206 | 0.022 | 0.022 | TRUE | 0.5442 |
| Nol4 | 13 | 9 | 10 | 34 | 25 | 27 | 14 | 9 | 10 | 27 | 27 | 30 | 19.34 | 11 | 28 | 2.537 | 1.343 | 0.00290843 | 0.02071322 | 0 | 0.1773 | 0.0822 | 0.0822 | TRUE | 0.1839 |
| Nol6 | 13984 | 11234 | 11154 | 15588 | 11037 | 9514 | 14853 | 11219 | 11200 | 12254 | 11875 | 10415 | 11969.42 | 12424 | 11515 | 0.927 | -0.11 | 0.40581659 | 0.61852911 | 0.0166 | 0.0072 | 0.0124 | 0.0124 | TRUE | 0.3576 |
| Nol7 | 2538 | 2678 | 2712 | 3582 | 2572 | 2536 | 2696 | 2675 | 2723 | 2816 | 2767 | 2776 | 2742.12 | 2698 | 2786 | 1.033 | 0.047 | 0.44528107 | 0.65195946 | 0 | 0.0081 | 0.0023 | 0.0023 | TRUE | 0.0011 |
| Nol8 | 4856 | 4520 | 4724 | 5399 | 3934 | 3593 | 5158 | 4514 | 4743 | 4244 | 4233 | 3933 | 4470.93 | 4805 | 4137 | 0.861 | -0.216 | 0.00890893 | 0.04829904 | 0.0029 | 0.0076 | 0.0047 | 0.0047 | TRUE | 0.0502 |
| Nol9 | 13894 | 12010 | 12196 | 13712 | 10094 | 9846 | 14758 | 11994 | 12246 | 10779 | 10861 | 10778 | 11902.71 | 12999 | 10806 | 0.831 | -0.267 | 0.0065037 | 0.03815231 | 0.0068 | 0.0072 | 0.0068 | 0.0068 | TRUE | 0.1711 |
| Nolc1 | 15091 | 12676 | 13086 | 15907 | 12048 | 10676 | 16029 | 12660 | 13140 | 12505 | 12963 | 11687 | 13163.83 | 13943 | 12385 | 0.888 | -0.171 | 0.11708762 | 0.28936163 | 0.0098 | 0.0072 | 0.0085 | 0.0085 | TRUE | 0.2095 |
| Nom1 | 2688 | 2094 | 2130 | 3371 | 2366 | 2045 | 2855 | 2091 | 2139 | 2650 | 2546 | 2239 | 2419.91 | 2362 | 2478 | 1.05 | 0.07 | 0.61881853 | 0.78456639 | 0.0188 | 0.0083 | 0.0138 | 0.0138 | TRUE | 0.4043 |
| Nomo1 | 11349 | 11532 | 11549 | 14386 | 9937 | 10176 | 12055 | 11517 | 11596 | 11309 | 10692 | 11139 | 11384.69 | 11723 | 11047 | 0.942 | -0.086 | 0.15712559 | 0.35017448 | 6.00E-04 | 0.0072 | 0.0025 | 0.0025 | TRUE | 0.0096 |
| Nono | 15186 | 16333 | 16917 | 21391 | 15360 | 14948 | 16130 | 16312 | 16986 | 16816 | 16526 | 16363 | 16522.31 | 16476 | 16568 | 1.006 | 0.008 | 0.88630644 | 0.94413246 | 3.00E-04 | 0.0071 | 0.0023 | 0.0023 | TRUE | 0.009 |

| | | | | | | | | | | | | | | | | | | | | | | | | | |
|---|---|---|---|---|---|---|---|---|---|---|---|---|---|---|---|---|---|---|---|---|---|---|---|---|---|
| Nop10 | 5685 | 4786 | 4647 | 5737 | 3986 | 4112 | 6038 | 4780 | 4666 | 4510 | 4289 | 4501 | 4797.38 | 5161 | 4433 | 0.859 | -0.219 | 0.0559787 | 0.17746656 | 0.011 | 0.0076 | 0.0093 | 0.0093 | TRUE | 0.2691 |
| Nop14 | 5340 | 5144 | 5084 | 7086 | 4870 | 4761 | 5672 | 5137 | 5105 | 5570 | 5240 | 5212 | 5322.7 | 5305 | 5341 | 1.007 | 0.01 | 0.8959625 | 0.9496715 | 0.0023 | 0.0075 | 0.0041 | 0.0041 | TRUE | 0.0446 |
| Nop16 | 3312 | 2340 | 1952 | 3225 | 2298 | 2028 | 3518 | 2337 | 1960 | 2535 | 2473 | 2220 | 2507.11 | 2605 | 2409 | 0.925 | -0.112 | 0.56088388 | 0.74256648 | 0.0452 | 0.0082 | 0.0265 | 0.0265 | TRUE | 0.9981 |
| Nop2 | 13099 | 10712 | 10210 | 16777 | 11559 | 9612 | 13913 | 10698 | 10252 | 13189 | 12437 | 10522 | 11835.15 | 11621 | 12049 | 1.037 | 0.052 | 0.70957428 | 0.84171221 | 0.0171 | 0.0072 | 0.0141 | 0.0141 | TRUE | 0.3639 |
| Nop56 | 11582 | 9599 | 9331 | 15814 | 11417 | 9247 | 12302 | 9587 | 9369 | 12432 | 12284 | 10122 | 11016.01 | 10419 | 11613 | 1.115 | 0.157 | 0.24369793 | 0.46027726 | 0.0177 | 0.0072 | 0.0129 | 0.0129 | TRUE | 0.3053 |
| Nop58 | 13326 | 12850 | 12492 | 17650 | 12499 | 11168 | 14155 | 12833 | 12543 | 13875 | 13448 | 12225 | 13179.93 | 13177 | 13183 | 1.001 | 0.001 | 0.9929004 | 0.99616826 | 0.0033 | 0.0072 | 0.0053 | 0.0053 | TRUE | 0.0515 |
| Nop9 | 4319 | 4400 | 4423 | 5837 | 4175 | 3998 | 4588 | 4394 | 4441 | 4589 | 4492 | 4376 | 4480.02 | 4474 | 4486 | 1.003 | 0.004 | 0.94865176 | 0.9742329 | 3.00E-04 | 0.0076 | 0.0025 | 0.0025 | TRUE | 0.0059 |
| Nos1 | 477 | 378 | 403 | 451 | 354 | 293 | 507 | 378 | 405 | 355 | 381 | 321 | 390.83 | 430 | 352 | 0.82 | -0.286 | 0.06105601 | 0.1883018 | 0.0133 | 0.0153 | 0.0142 | 0.0142 | TRUE | 0.2827 |
| Nos1ap | 72 | 111 | 130 | 109 | 91 | 99 | 76 | 111 | 131 | 86 | 98 | 108 | 101.64 | 106 | 97 | 0.915 | -0.128 | 0.6106289 | 0.77944634 | 0.0323 | 0.0393 | 0.0359 | 0.0359 | TRUE | 0.3747 |
| Nos2 | 47 | 44 | 35 | 39 | 40 | 42 | 50 | 44 | 35 | 31 | 43 | 46 | 41.45 | 43 | 40 | 0.921 | -0.118 | 0.71581063 | 0.8456753 | 0 | 0.0864 | 0.0521 | 0.0521 | TRUE | 0.3637 |
| Nos3 | 503 | 619 | 668 | 523 | 440 | 474 | 534 | 618 | 671 | 411 | 473 | 519 | 537.77 | 608 | 468 | 0.768 | -0.381 | 0.00653657 | 0.03822575 | 0.0115 | 0.013 | 0.0122 | 0.0122 | TRUE | 0.1364 |
| Nosip | 1693 | 1742 | 1837 | 2525 | 1838 | 1740 | 1798 | 1740 | 1845 | 1985 | 1978 | 1905 | 1874.97 | 1794 | 1956 | 1.09 | 0.125 | 0.0793383 | 0.22337087 | 0 | 0.0087 | 0.0031 | 0.0031 | TRUE | 0.0086 |
| Nostrin | 16 | 9 | 3 | 31 | 21 | 13 | 17 | 9 | 3 | 24 | 23 | 14 | 15.03 | 10 | 20 | 2.129 | 1.09 | 0.0877595 | 0.24001852 | 0.2037 | 0.2261 | 0.2167 | 0.2167 | TRUE | 0.5864 |
| Notch1 | 4043 | 4279 | 4709 | 4047 | 3047 | 3485 | 4294 | 4273 | 4728 | 3181 | 3278 | 3815 | 3928.48 | 4432 | 3425 | 0.772 | -0.372 | 0.00016837 | 0.0021048 | 0.0063 | 0.0077 | 0.0068 | 0.0068 | TRUE | 0.1202 |
| Notch2 | 2522 | 2314 | 2410 | 3823 | 2684 | 2384 | 2679 | 2311 | 2420 | 3005 | 2888 | 2610 | 2652.09 | 2470 | 2834 | 1.148 | 0.199 | 0.03891492 | 0.13798699 | 0.0049 | 0.0081 | 0.0063 | 0.0063 | TRUE | 0.0662 |
| Notch3 | 3748 | 4874 | 4832 | 4969 | 3619 | 3706 | 3981 | 4868 | 4852 | 3906 | 3894 | 4057 | 4259.57 | 4567 | 3952 | 0.865 | -0.209 | 0.03695998 | 0.13293794 | 0.0061 | 0.0077 | 0.007 | 0.007 | TRUE | 0.1532 |
| Notch4 | 2378 | 3420 | 4037 | 2767 | 2238 | 2263 | 2526 | 3416 | 4054 | 2175 | 2408 | 2477 | 2842.56 | 3332 | 2353 | 0.706 | -0.502 | 0.00206806 | 0.01581069 | 0.0282 | 0.0081 | 0.0188 | 0.0188 | TRUE | 0.2134 |
| Noto | 1 | 1 | 0 | 3 | 4 | 0 | 1 | 1 | 0 | 2 | 4 | 0 | 1.45 | 1 | 2 | 3.256 | 1.703 | 0.3638759 | NA | 0 | 2.2729 | 1.5688 | 1.5688 | TRUE | 0.1032 |
| Notum | 802 | 524 | 522 | 453 | 357 | 268 | 852 | 523 | 524 | 356 | 384 | 293 | 488.82 | 633 | 344 | 0.545 | -0.877 | 3.82E-05 | 0.00061465 | 0.0505 | 0.0136 | 0.0304 | 0.0304 | TRUE | 1.0728 |
| Nov | 10 | 14 | 15 | 43 | 19 | 21 | 11 | 14 | 15 | 34 | 20 | 23 | 19.48 | 13 | 26 | 1.961 | 0.972 | 0.03603247 | 0.13040481 | 0 | 0.176 | 0.0975 | 0.0975 | TRUE | 0.5439 |
| Nova1 | 106 | 91 | 89 | 227 | 151 | 96 | 113 | 91 | 89 | 178 | 162 | 105 | 123.14 | 98 | 148 | 1.529 | 0.613 | 0.01388926 | 0.06684817 | 0.0388 | 0.0337 | 0.0362 | 0.0362 | TRUE | 0.6682 |
| Nova2 | 802 | 964 | 943 | 909 | 789 | 801 | 852 | 963 | 947 | 715 | 849 | 877 | 866.97 | 921 | 814 | 0.882 | -0.181 | 0.12048903 | 0.29467406 | 0.007 | 0.0107 | 0.0086 | 0.0086 | TRUE | 0.139 |
| Nox1 | 2 | 0 | 0 | 3 | 2 | 2 | 2 | 0 | 0 | 2 | 2 | 2 | 1.47 | 1 | 2 | 3.226 | 1.69 | 0.33871732 | NA | 0 | 2.247 | 1.2922 | 1.2922 | TRUE | 0.9745 |
| Nox3 | 0 | 0 | 0 | 0 | 0 | 0 | 0 | 0 | 0 | 0 | 0 | 0 | 0 | 0 | 0 | NA | NA | NA | NA | NA | NA | NA | NA | NA | NA |
| Nox4 | 184 | 234 | 197 | 357 | 237 | 245 | 195 | 234 | 198 | 281 | 255 | 268 | 238.46 | 209 | 268 | 1.283 | 0.36 | 0.01317273 | 0.06434197 | 0 | 0.0207 | 0.0109 | 0.0109 | TRUE | 0.1178 |
| Noxa1 | 11 | 8 | 6 | 15 | 10 | 8 | 12 | 8 | 6 | 12 | 11 | 9 | 9.5 | 9 | 11 | 1.228 | 0.296 | 0.63146384 | 0.79271062 | 0 | 0.3536 | 0.1687 | 0.1687 | TRUE | 0.2989 |
| Noxo1 | 44 | 79 | 87 | 124 | 106 | 105 | 47 | 79 | 87 | 97 | 114 | 115 | 89.91 | 71 | 109 | 1.527 | 0.611 | 0.02355379 | 0.09763215 | 0.0374 | 0.0435 | 0.0407 | 0.0407 | TRUE | 0.7861 |
| Noxred1 | 246 | 272 | 285 | 377 | 271 | 255 | 261 | 272 | 286 | 296 | 292 | 279 | 281.03 | 273 | 289 | 1.059 | 0.083 | 0.50747275 | 0.70172367 | 0 | 0.0186 | 0.0077 | 0.0077 | TRUE | 0.0199 |
| Npas1 | 13 | 27 | 24 | 11 | 7 | 6 | 14 | 27 | 24 | 9 | 8 | 7 | 14.6 | 22 | 8 | 0.353 | -1.504 | 0.00633001 | 0.03747634 | 0 | 0.2325 | 0.1313 | 0.1313 | TRUE | 0.5425 |
| Npas2 | 18 | 27 | 24 | 40 | 32 | 33 | 19 | 27 | 24 | 31 | 34 | 36 | 28.7 | 23 | 34 | 1.447 | 0.533 | 0.13693565 | 0.320866 | 0 | 0.1217 | 0.0564 | 0.0564 | TRUE | 0.1439 |
| Npas3 | 33 | 31 | 26 | 26 | 20 | 13 | 35 | 31 | 26 | 20 | 22 | 14 | 24.72 | 31 | 19 | 0.613 | -0.705 | 0.07490674 | 0.21543201 | 0 | 0.1402 | 0.0704 | 0.0704 | TRUE | 0.2056 |
| Npas4 | 0 | 0 | 0 | 3 | 0 | 0 | 0 | 0 | 0 | 2 | 0 | 0 | 0.39 | 0 | 1 | 4.278 | 2.097 | 0.60136958 | NA | 10 | 8.3884 | 10 | 10 | TRUE | 1.4411 |
| Npat | 3288 | 3321 | 3429 | 4400 | 3261 | 3184 | 3492 | 3317 | 3443 | 3459 | 3509 | 3485 | 3450.86 | 3417 | 3484 | 1.02 | 0.028 | 0.65329271 | 0.80540538 | 0 | 0.0079 | 0.0025 | 0.0025 | TRUE | 0.0081 |
| Npb | 8 | 8 | 9 | 14 | 2 | 12 | 8 | 8 | 9 | 11 | 2 | 13 | 8.64 | 8 | 9 | 1.037 | 0.052 | 0.94423734 | 0.97228643 | 0.1504 | 0.3884 | 0.2851 | 0.2851 | TRUE | 1.2541 |
| Npbwr1 | 2 | 0 | 0 | 3 | 3 | 0 | 2 | 0 | 0 | 2 | 3 | 0 | 1.29 | 1 | 2 | 2.686 | 1.425 | 0.50800302 | NA | 1.8421 | 2.5705 | 2.3476 | 2.3476 | TRUE | 0.9073 |
| Npc1 | 3916 | 3194 | 3426 | 3423 | 2474 | 2275 | 4159 | 3190 | 3440 | 2691 | 2662 | 2490 | 3105.42 | 3596 | 2614 | 0.727 | -0.46 | 6.19E-05 | 0.00093196 | 0.0102 | 0.008 | 0.0092 | 0.0092 | TRUE | 0.2278 |
| Npc1l1 | 50 | 34 | 21 | 69 | 23 | 41 | 53 | 34 | 21 | 54 | 25 | 45 | 38.67 | 36 | 41 | 1.153 | 0.206 | 0.64591092 | 0.80114085 | 0.1474 | 0.0921 | 0.1184 | 0.1184 | TRUE | 0.3641 |
| Npc2 | 4074 | 5292 | 5391 | 5374 | 3979 | 4816 | 4327 | 5285 | 5413 | 4225 | 4281 | 5272 | 4800.54 | 5008 | 4593 | 0.917 | -0.125 | 0.32672867 | 0.54780107 | 0.0142 | 0.0076 | 0.0116 | 0.0116 | TRUE | 0.2036 |
| Npcd | 2 | 12 | 9 | 1 | 3 | 5 | 2 | 12 | 9 | 1 | 3 | 5 | 5.44 | 8 | 3 | 0.396 | -1.336 | 0.19002972 | 0.39514156 | 0.425 | 0.6126 | 0.5237 | 0.5237 | TRUE | 0.3616 |
| Npdc1 | 986 | 1440 | 1452 | 1102 | 852 | 1030 | 1047 | 1438 | 1458 | 866 | 917 | 1127 | 1142.32 | 1314 | 970 | 0.737 | -0.439 | 0.00634749 | 0.03753754 | 0.0244 | 0.0098 | 0.0178 | 0.0178 | TRUE | 0.3789 |
| Npepl1 | 1263 | 1515 | 1722 | 1981 | 1516 | 1518 | 1342 | 1513 | 1729 | 1557 | 1631 | 1662 | 1572.29 | 1528 | 1617 | 1.058 | 0.081 | 0.46944571 | 0.67151259 | 0.0079 | 0.009 | 0.0084 | 0.0084 | TRUE | 0.1599 |
| Npepps | 18386 | 19383 | 19602 | 22597 | 17256 | 18107 | 19529 | 19358 | 19682 | 17764 | 18566 | 19821 | 19120.16 | 19523 | 18717 | 0.959 | -0.061 | 0.36912114 | 0.5865181 | 0.0015 | 0.0071 | 0.0033 | 0.0033 | TRUE | 0.0325 |
| Npff | 5 | 2 | 0 | 18 | 11 | 1 | 5 | 2 | 0 | 14 | 12 | 1 | 5.73 | 2 | 9 | 3.762 | 1.912 | 0.1156334 | 0.28713974 | 1.1294 | 0.5817 | 0.7984 | 0.7984 | TRUE | 1.0137 |
| Npffr1 | 5 | 3 | 2 | 2 | 2 | 0 | 5 | 3 | 2 | 2 | 2 | 0 | 2.34 | 3 | 1 | 0.368 | -1.441 | 0.28086624 | NA | 0 | 1.4149 | 0.7522 | 0.7522 | TRUE | 0.3673 |
| Npffr2 | 0 | 2 | 1 | 3 | 2 | 0 | 0 | 2 | 1 | 2 | 2 | 0 | 1.25 | 1 | 1 | 1.517 | 0.602 | 0.74659055 | NA | 0 | 2.6383 | 1.6713 | 1.6713 | TRUE | 0.2931 |
| Nphp1 | 680 | 842 | 810 | 1181 | 853 | 845 | 722 | 841 | 813 | 928 | 918 | 925 | 857.95 | 792 | 924 | 1.166 | 0.221 | 0.0216212 | 0.0919659 | 0.0017 | 0.0107 | 0.0055 | 0.0055 | TRUE | 0.07 |
| Nphp3 | 2126 | 2163 | 2314 | 2423 | 1896 | 2151 | 2258 | 2160 | 2324 | 1905 | 2040 | 2355 | 2173.54 | 2247 | 2100 | 0.934 | -0.099 | 0.3336879 | 0.5542839 | 0.006 | 0.0084 | 0.0071 | 0.0071 | TRUE | 0.136 |
| Nphp4 | 728 | 693 | 713 | 927 | 759 | 733 | 773 | 692 | 716 | 729 | 817 | 802 | 754.84 | 727 | 783 | 1.075 | 0.105 | 0.29968333 | 0.51980705 | 0.0023 | 0.0113 | 0.006 | 0.006 | TRUE | 0.045 |
| Nphs1 | 2561 | 3888 | 4450 | 1441 | 1345 | 1596 | 2720 | 3883 | 4468 | 1133 | 1447 | 1747 | 2566.42 | 3690 | 1442 | 0.391 | -1.356 | 1.77E-11 | 1.50E-09 | 0.051 | 0.0082 | 0.0288 | 0.0288 | TRUE | 0.2444 |
| Nphs1as | 66 | 77 | 73 | 50 | 55 | 46 | 70 | 77 | 73 | 39 | 59 | 50 | 61.52 | 73 | 49 | 0.67 | -0.578 | 0.02899487 | 0.11230529 | 0 | 0.0605 | 0.0336 | 0.0336 | TRUE | 0.3175 |
| Nphs2 | 5 | 5 | 3 | 8 | 3 | 10 | 5 | 5 | 3 | 6 | 3 | 11 | 5.63 | 4 | 7 | 1.533 | 0.617 | 0.46507714 | 0.66788976 | 0 | 0.5921 | 0.3265 | 0.3265 | TRUE | 0.2222 |
| Npl | 1433 | 1433 | 1350 | 1832 | 1187 | 1297 | 1522 | 1431 | 1356 | 1440 | 1277 | 1420 | 1407.65 | 1436 | 1379 | 0.961 | -0.058 | 0.53433892 | 0.72176462 | 0.0031 | 0.0092 | 0.0055 | 0.0055 | TRUE | 0.0497 |
| Nploc4 | 9145 | 8323 | 8144 | 11778 | 8111 | 7674 | 9714 | 8312 | 8177 | 9259 | 8727 | 8400 | 8764.93 | 8734 | 8795 | 1.007 | 0.01 | 0.91365316 | 0.95742061 | 0.0046 | 0.0073 | 0.0063 | 0.0063 | TRUE | 0.1174 |
| Npm1 | 181581 | 197527 | 199271 | 255733 | 183020 | 179515 | 192871 | 197272 | 200089 | 201037 | 196918 | 196507 | 197449.19 | 196744 | 198154 | 1.007 | 0.01 | 0.84182127 | 0.920175 | 2.00E-04 | 0.0069 | 0.0019 | 0.0019 | TRUE | 0.0036 |
| Npm2 | 12 | 16 | 11 | 37 | 27 | 19 | 13 | 16 | 11 | 29 | 29 | 21 | 19.78 | 13 | 26 | 1.991 | 0.993 | 0.02587417 | 0.10412075 | 0 | 0.1734 | 0.0862 | 0.0862 | TRUE | 0.1939 |
| Npm3 | 4525 | 5772 | 5825 | 5542 | 4148 | 4351 | 4806 | 5765 | 5849 | 4357 | 4463 | 4763 | 5000.39 | 5473 | 4528 | 0.827 | -0.274 | 0.00636023 | 0.03758472 | 0.0064 | 0.0076 | 0.0071 | 0.0071 | TRUE | 0.1384 |
| Npm3-ps1 | 359 | 550 | 549 | 541 | 369 | 369 | 381 | 549 | 551 | 425 | 397 | 404 | 451.35 | 494 | 409 | 0.828 | -0.273 | 0.09521721 | 0.25297254 | 0.0198 | 0.0142 | 0.0171 | 0.0171 | TRUE | 0.4602 |
| Npnt | 497 | 691 | 675 | 740 | 571 | 634 | 528 | 690 | 678 | 582 | 614 | 694 | 630.98 | 632 | 630 | 0.996 | -0.006 | 0.96509345 | 0.98247298 | 0.0132 | 0.0121 | 0.0127 | 0.0127 | TRUE | 0.2425 |
| Nppa | 0 | 0 | 1 | 0 | 0 | 0 | 0 | 0 | 1 | 0 | 0 | 0 | 0.17 | 0 | 0 | 0.49 | -1.029 | 0.80087 | NA | 0 | 19.6928 | 10 | 10 | TRUE | 0.4098 |
| Nppb | 24 | 22 | 17 | 28 | 23 | 19 | 25 | 22 | 17 | 22 | 25 | 21 | 22.02 | 21 | 23 | 1.048 | 0.067 | 0.86904725 | 0.93551388 | 0 | 0.1565 | 0.0735 | 0.0735 | TRUE | 0.1848 |
| Nppc | 32 | 37 | 31 | 66 | 32 | 32 | 34 | 37 | 31 | 52 | 34 | 35 | 37.24 | 34 | 40 | 1.2 | 0.262 | 0.43459524 | 0.64317865 | 0 | 0.0954 | 0.0543 | 0.0543 | TRUE | 0.5288 |
| Npr1 | 549 | 428 | 376 | 1155 | 692 | 593 | 583 | 427 | 378 | 908 | 745 | 649 | 614.96 | 463 | 767 | 1.66 | 0.731 | 0.00013018 | 0.00170531 | 0.0388 | 0.0123 | 0.0246 | 0.0246 | TRUE | 0.509 |
| Npr2 | 117 | 139 | 137 | 182 | 152 | 132 | 124 | 139 | 138 | 143 | 164 | 144 | 141.96 | 134 | 150 | 1.124 | 0.168 | 0.32490157 | 0.54594394 | 0 | 0.0301 | 0.014 | 0.014 | TRUE | 0.0595 |
| Npr3 | 279 | 311 | 349 | 252 | 171 | 172 | 296 | 311 | 350 | 198 | 184 | 188 | 254.62 | 319 | 190 | 0.596 | -0.746 | 1.11E-07 | 3.90E-06 | 0 | 0.0198 | 0.01 | 0.01 | TRUE | 0.0837 |
| Nprl2 | 1079 | 1076 | 1099 | 1150 | 815 | 970 | 1146 | 1075 | 1104 | 904 | 877 | 1062 | 1027.83 | 1108 | 948 | 0.855 | -0.227 | 0.03139346 | 0.11908073 | 0.0046 | 0.0101 | 0.007 | 0.007 | TRUE | 0.1303 |
| Nprl3 | 2030 | 2037 | 2138 | 2837 | 2054 | 1761 | 2156 | 2034 | 2147 | 2230 | 2210 | 1928 | 2117.54 | 2112 | 2123 | 1.005 | 0.008 | 0.93193404 | 0.96628017 | 0.0033 | 0.0085 | 0.0053 | 0.0053 | TRUE | 0.0774 |
| Nps | 6 | 0 | 4 | 5 | 2 | 4 | 6 | 0 | 4 | 4 | 2 | 4 | 3.48 | 3 | 3 | 1.015 | 0.022 | 0.98529002 | NA | 0.3701 | 0.9549 | 0.7254 | 0.7254 | TRUE | 0.404 |
| Npsr1 | 2 | 2 | 2 | 2 | 1 | 2 | 2 | 2 | 2 | 2 | 1 | 2 | 1.83 | 2 | 2 | 0.788 | -0.344 | 0.7929913 | NA | 0 | 1.8092 | 0.6873 | 0.6873 | TRUE | 0.0607 |
| Nptn | 4462 | 4053 | 3909 | 5213 | 3686 | 3628 | 4739 | 4048 | 3925 | 4098 | 3966 | 3971 | 4124.6 | 4237 | 4012 | 0.947 | -0.079 | 0.4031266 | 0.6160365 | 0.0052 | 0.0077 | 0.0061 | 0.0061 | TRUE | 0.1306 |

| | | | | | | | | | | | | | | | | | | | | | | | | | |
|---|---|---|---|---|---|---|---|---|---|---|---|---|---|---|---|---|---|---|---|---|---|---|---|---|---|
| Nptx1 | 140 | 148 | 125 | 185 | 142 | 124 | 149 | 148 | 126 | 145 | 153 | 136 | 142.66 | 141 | 145 | 1.029 | 0.041 | 0.8123741 | 0.9037712 | 0 | 0.03 | 0.0146 | 0.0146 | TRUE | 0.093 |
| Nptx2 | 3171 | 3035 | 2891 | 2383 | 1866 | 1681 | 3368 | 3031 | 2903 | 1873 | 2008 | 1840 | 2503.88 | 3101 | 1907 | 0.615 | -0.701 | 6.46E-15 | 8.93E-13 | 0.0035 | 0.0082 | 0.0054 | 0.0054 | TRUE | 0.069 |
| Nptxr | 81 | 95 | 94 | 100 | 70 | 63 | 86 | 95 | 94 | 79 | 75 | 69 | 83.03 | 92 | 74 | 0.811 | -0.302 | 0.15529367 | 0.34751585 | 0 | 0.0466 | 0.0204 | 0.0204 | TRUE | 0.0344 |
| Npvf | 0 | 0 | 0 | 1 | 0 | 0 | 0 | 0 | 0 | 1 | 0 | 0 | 0.13 | 0 | 0 | 1.859 | 0.894 | 0.82650734 | NA | 0 | 25.1515 | 10 | 10 | TRUE | 0.2563 |
| Npw | 78 | 73 | 66 | 66 | 49 | 49 | 83 | 73 | 66 | 52 | 53 | 54 | 63.38 | 74 | 53 | 0.713 | -0.489 | 0.04583292 | 0.15485794 | 0 | 0.0589 | 0.027 | 0.027 | TRUE | 0.0972 |
| Npy | 4 | 16 | 20 | 16 | 5 | 9 | 4 | 16 | 20 | 13 | 5 | 10 | 11.35 | 13 | 9 | 0.695 | -0.525 | 0.45945684 | 0.66378277 | 0.2474 | 0.2971 | 0.2734 | 0.2734 | TRUE | 0.672 |
| Npy1r | 31 | 18 | 13 | 12 | 1 | 4 | 33 | 18 | 13 | 9 | 1 | 4 | 13.14 | 21 | 5 | 0.242 | -2.047 | 0.00593446 | 0.03562937 | 0.3147 | 0.2576 | 0.2801 | 0.2801 | TRUE | 0.8469 |
| Npy2r | 0 | 0 | 0 | 0 | 0 | 1 | 0 | 0 | 0 | 0 | 0 | 1 | 0.18 | 0 | 0 | 1.859 | 0.894 | 0.82650734 | NA | 0 | 18.0643 | 10 | 10 | TRUE | 0.5383 |
| Npy5r | 3 | 5 | 4 | 3 | 2 | 2 | 3 | 5 | 4 | 2 | 2 | 2 | 3.15 | 4 | 2 | 0.55 | -0.862 | 0.40317971 | NA | 0 | 1.053 | 0.424 | 0.424 | TRUE | 0.0684 |
| Npy6r | 0 | 0 | 0 | 0 | 0 | 0 | 0 | 0 | 0 | 0 | 0 | 0 | 0 | 0 | 0 | NA | NA | NA | NA | NA | NA | NA | NA | NA | NA |
| Nqo1 | 1018 | 835 | 772 | 1250 | 887 | 693 | 1081 | 834 | 775 | 983 | 954 | 759 | 897.67 | 897 | 899 | 1.003 | 0.004 | 0.98001665 | 0.98917334 | 0.0241 | 0.0106 | 0.0175 | 0.0175 | TRUE | 0.3846 |
| Nqo2 | 1221 | 1662 | 1729 | 1782 | 1301 | 1576 | 1297 | 1660 | 1736 | 1401 | 1400 | 1725 | 1536.45 | 1564 | 1509 | 0.964 | -0.053 | 0.71040548 | 0.84213468 | 0.0186 | 0.009 | 0.014 | 0.014 | TRUE | 0.2688 |
| Nr0b1 | 1601 | 1012 | 1064 | 1275 | 882 | 772 | 1701 | 1011 | 1068 | 1002 | 949 | 845 | 1095.99 | 1260 | 932 | 0.74 | -0.434 | 0.02649245 | 0.10579589 | 0.0484 | 0.0099 | 0.0266 | 0.0266 | TRUE | 1.1225 |
| Nr0b2 | 2 | 8 | 6 | 6 | 3 | 5 | 2 | 8 | 6 | 5 | 3 | 5 | 4.93 | 5 | 4 | 0.829 | -0.27 | 0.75935885 | 0.87312151 | 0 | 0.6757 | 0.3568 | 0.3568 | TRUE | 0.4222 |
| Nr1d1 | 472 | 620 | 692 | 537 | 399 | 523 | 501 | 619 | 695 | 422 | 429 | 573 | 539.89 | 605 | 475 | 0.783 | -0.352 | 0.04193383 | 0.14580696 | 0.0261 | 0.013 | 0.0198 | 0.0198 | TRUE | 0.3738 |
| Nr1d2 | 1431 | 1366 | 1365 | 1651 | 1331 | 1374 | 1520 | 1364 | 1371 | 1298 | 1432 | 1504 | 1414.81 | 1418 | 1411 | 0.994 | -0.008 | 0.93377234 | 0.96713012 | 0.004 | 0.0092 | 0.0061 | 0.0061 | TRUE | 0.0611 |
| Nr1h2 | 2096 | 1999 | 1978 | 3188 | 2567 | 2366 | 2226 | 1996 | 1986 | 2506 | 2762 | 2590 | 2344.49 | 2069 | 2619 | 1.265 | 0.34 | 8.30E-05 | 0.00118821 | 0.0027 | 0.0083 | 0.0049 | 0.0049 | TRUE | 0.0529 |
| Nr1h3 | 13 | 11 | 8 | 28 | 24 | 12 | 14 | 11 | 8 | 22 | 26 | 13 | 15.63 | 11 | 20 | 1.867 | 0.9 | 0.08673478 | 0.23803391 | 0 | 0.2176 | 0.1283 | 0.1283 | TRUE | 0.4013 |
| Nr1h4 | 2 | 0 | 0 | 2 | 0 | 0 | 2 | 0 | 0 | 2 | 0 | 0 | 0.62 | 1 | 1 | 0.783 | -0.353 | 0.91186553 | NA | 0 | 5.3542 | 5.6997 | 5.6997 | TRUE | 0.9742 |
| Nr1h5 | 2 | 0 | 1 | 3 | 0 | 2 | 2 | 0 | 1 | 2 | 0 | 2 | 1.28 | 1 | 1 | 1.487 | 0.572 | 0.75839437 | NA | 0 | 2.582 | 1.6821 | 1.6821 | TRUE | 0.3029 |
| Nr1i2 | 19 | 21 | 14 | 34 | 22 | 15 | 20 | 21 | 14 | 27 | 24 | 16 | 20.34 | 18 | 22 | 1.22 | 0.286 | 0.51714938 | 0.70874821 | 0 | 0.1689 | 0.0913 | 0.0913 | TRUE | 0.2816 |
| Nr1i3 | 0 | 2 | 2 | 2 | 3 | 0 | 0 | 2 | 2 | 2 | 3 | 0 | 1.47 | 1 | 2 | 1.192 | 0.253 | 0.88546044 | NA | 0 | 2.2517 | 1.5446 | 1.5446 | TRUE | 0.1569 |
| Nr2c1 | 433 | 510 | 495 | 573 | 427 | 399 | 460 | 509 | 497 | 450 | 459 | 437 | 468.82 | 489 | 449 | 0.918 | -0.123 | 0.24422203 | 0.46080511 | 0 | 0.0139 | 0.0059 | 0.0059 | TRUE | 0.0306 |
| Nr2c2 | 3136 | 3039 | 3164 | 3868 | 2914 | 2648 | 3331 | 3035 | 3177 | 3041 | 3135 | 2899 | 3102.95 | 3181 | 3025 | 0.951 | -0.072 | 0.33660143 | 0.55668597 | 0.0014 | 0.008 | 0.0038 | 0.0038 | TRUE | 0.0206 |
| Nr2c2ap | 1088 | 1101 | 1032 | 1536 | 1056 | 964 | 1156 | 1100 | 1036 | 1207 | 1136 | 1055 | 1115.06 | 1097 | 1133 | 1.034 | 0.048 | 0.61692602 | 0.7832683 | 0.0029 | 0.0099 | 0.0057 | 0.0057 | TRUE | 0.0429 |
| Nr2e1 | 0 | 0 | 1 | 1 | 0 | 0 | 0 | 0 | 1 | 1 | 0 | 0 | 0.3 | 0 | 0 | 0.954 | -0.067 | 0.98640617 | NA | 0 | 11.0483 | 9.2783 | 9.2783 | TRUE | 0.4098 |
| Nr2e3 | 3 | 7 | 14 | 16 | 16 | 12 | 3 | 7 | 14 | 13 | 17 | 13 | 11.19 | 8 | 14 | 1.759 | 0.815 | 0.21472206 | 0.42657719 | 0.0929 | 0.3012 | 0.2135 | 0.2135 | TRUE | 0.7554 |
| Nr2f1 | 7 | 13 | 13 | 38 | 34 | 20 | 7 | 13 | 13 | 30 | 37 | 22 | 20.3 | 11 | 30 | 2.633 | 1.397 | 0.00323728 | 0.02247287 | 0 | 0.1692 | 0.1 | 0.1 | TRUE | 0.1516 |
| Nr2f2 | 1 | 0 | 0 | 11 | 6 | 6 | 1 | 0 | 0 | 9 | 6 | 7 | 3.79 | 0 | 7 | 19.831 | 4.31 | 0.00328725 | NA | 0 | 0.8764 | 0.4815 | 0.4815 | TRUE | 0.4614 |
| Nr2f6 | 1295 | 1637 | 1722 | 1633 | 1349 | 1279 | 1376 | 1635 | 1729 | 1284 | 1451 | 1400 | 1479.12 | 1580 | 1378 | 0.872 | -0.198 | 0.08212292 | 0.22884146 | 0.0076 | 0.0091 | 0.0086 | 0.0086 | TRUE | 0.1537 |
| Nr3c1 | 1008 | 1364 | 1406 | 1877 | 1343 | 1360 | 1071 | 1362 | 1412 | 1476 | 1445 | 1489 | 1375.66 | 1282 | 1470 | 1.147 | 0.197 | 0.10191515 | 0.26494243 | 0.0094 | 0.0093 | 0.0098 | 0.0098 | TRUE | 0.2478 |
| Nr3c2 | 160 | 135 | 153 | 271 | 202 | 154 | 170 | 135 | 154 | 213 | 217 | 169 | 176.23 | 153 | 200 | 1.31 | 0.389 | 0.03183962 | 0.12006324 | 0.0103 | 0.0256 | 0.0179 | 0.0179 | TRUE | 0.1937 |
| Nr4a1 | 1101 | 1699 | 1830 | 1497 | 1234 | 1444 | 1169 | 1697 | 1838 | 1177 | 1328 | 1581 | 1464.83 | 1568 | 1362 | 0.868 | -0.204 | 0.25911719 | 0.47804378 | 0.0354 | 0.0092 | 0.0229 | 0.0229 | TRUE | 0.5946 |
| Nr4a2 | 60 | 35 | 32 | 57 | 42 | 30 | 64 | 35 | 32 | 45 | 45 | 33 | 42.28 | 44 | 41 | 0.943 | -0.084 | 0.82082403 | 0.90868509 | 0.0654 | 0.0848 | 0.0754 | 0.0754 | TRUE | 1.2337 |
| Nr4a3 | 0 | 1 | 1 | 2 | 1 | 1 | 0 | 1 | 1 | 2 | 1 | 1 | 0.96 | 1 | 1 | 1.868 | 0.902 | 0.63074385 | NA | 0 | 3.4472 | 1.3898 | 1.3898 | TRUE | 0.2229 |
| Nr5a1 | 2 | 1 | 2 | 2 | 1 | 3 | 2 | 1 | 2 | 2 | 1 | 3 | 1.84 | 2 | 2 | 1.147 | 0.198 | 0.88279883 | NA | 0 | 1.7936 | 0.7536 | 0.7536 | TRUE | 0.262 |
| Nr5a2 | 3344 | 3045 | 3068 | 3893 | 2754 | 2883 | 3552 | 3041 | 3081 | 3060 | 2963 | 3156 | 3142.16 | 3225 | 3060 | 0.949 | -0.076 | 0.39991787 | 0.61294737 | 0.0039 | 0.008 | 0.0055 | 0.0055 | TRUE | 0.0957 |
| Nr6a1 | 5247 | 4607 | 4718 | 6288 | 4382 | 4250 | 5573 | 4601 | 4737 | 4943 | 4715 | 4652 | 4870.31 | 4970 | 4770 | 0.96 | -0.059 | 0.53882238 | 0.72512264 | 0.0054 | 0.0076 | 0.0065 | 0.0065 | TRUE | 0.137 |
| Nradd | 946 | 1334 | 1280 | 945 | 746 | 978 | 1005 | 1332 | 1285 | 743 | 803 | 1071 | 1039.74 | 1207 | 872 | 0.721 | -0.471 | 0.00530125 | 0.03266069 | 0.0293 | 0.0101 | 0.0196 | 0.0196 | TRUE | 0.4699 |
| Nrap | 23 | 28 | 22 | 31 | 37 | 23 | 24 | 28 | 22 | 24 | 40 | 25 | 27.31 | 25 | 30 | 1.193 | 0.254 | 0.51510935 | 0.70749402 | 0 | 0.1275 | 0.073 | 0.073 | TRUE | 0.5473 |
| Nrarp | 313 | 433 | 432 | 400 | 313 | 312 | 332 | 432 | 434 | 314 | 337 | 342 | 365.24 | 399 | 331 | 0.827 | -0.274 | 0.06002097 | 0.18579996 | 0.0096 | 0.0159 | 0.0126 | 0.0126 | TRUE | 0.246 |
| Nras | 7312 | 8176 | 8264 | 9806 | 6958 | 7036 | 7767 | 8165 | 8298 | 7709 | 7486 | 7702 | 7854.52 | 8077 | 7632 | 0.945 | -0.082 | 0.18486387 | 0.38808372 | 5.00E-04 | 0.0073 | 0.0026 | 0.0026 | TRUE | 0.0137 |
| Nrbf2 | 803 | 789 | 714 | 989 | 743 | 679 | 853 | 788 | 717 | 777 | 799 | 743 | 779.67 | 786 | 773 | 0.984 | -0.023 | 0.82725516 | 0.912538 | 0.0031 | 0.0111 | 0.0065 | 0.0065 | TRUE | 0.0701 |
| Nrbp1 | 2108 | 2452 | 2396 | 2911 | 2199 | 2383 | 2239 | 2449 | 2406 | 2288 | 2366 | 2609 | 2392.78 | 2365 | 2421 | 1.023 | 0.033 | 0.7019756 | 0.83705948 | 0.003 | 0.0083 | 0.005 | 0.005 | TRUE | 0.0554 |
| Nrbp2 | 259 | 327 | 345 | 351 | 307 | 438 | 275 | 327 | 346 | 276 | 330 | 479 | 338.97 | 316 | 362 | 1.142 | 0.192 | 0.37151955 | 0.58813674 | 0.0447 | 0.0166 | 0.0303 | 0.0303 | TRUE | 0.7514 |
| Nrcam | 135 | 216 | 292 | 113 | 90 | 81 | 143 | 216 | 293 | 89 | 97 | 89 | 154.44 | 217 | 92 | 0.42 | -1.251 | 3.08E-06 | 7.32E-05 | 0.0603 | 0.0282 | 0.0441 | 0.0441 | TRUE | 0.102 |
| Nrd1 | 7662 | 7865 | 8164 | 11038 | 7872 | 7820 | 8138 | 7855 | 8198 | 8677 | 8470 | 8560 | 8316.33 | 8064 | 8569 | 1.063 | 0.088 | 0.12143703 | 0.29634919 | 2.00E-04 | 0.0073 | 0.0022 | 0.0022 | TRUE | 0.0063 |
| Nrde2 | 2158 | 1812 | 1640 | 2929 | 2039 | 1855 | 2292 | 1810 | 1647 | 2303 | 2194 | 2031 | 2045.92 | 1916 | 2176 | 1.136 | 0.184 | 0.17261031 | 0.37130521 | 0.0166 | 0.0085 | 0.0126 | 0.0126 | TRUE | 0.3555 |
| Nrep | 275 | 388 | 396 | 402 | 324 | 366 | 292 | 387 | 398 | 316 | 349 | 401 | 357.08 | 359 | 355 | 0.987 | -0.019 | 0.90821856 | 0.95368996 | 0.0182 | 0.0161 | 0.0172 | 0.0172 | TRUE | 0.3017 |
| Nrf1 | 2420 | 2307 | 2321 | 3845 | 2683 | 2345 | 2570 | 2304 | 2331 | 3023 | 2887 | 2567 | 2613.56 | 2402 | 2826 | 1.177 | 0.235 | 0.01417492 | 0.0679328 | 0.0047 | 0.0082 | 0.0062 | 0.0062 | TRUE | 0.0774 |
| Nrg1 | 33 | 36 | 25 | 73 | 49 | 45 | 35 | 36 | 25 | 57 | 53 | 49 | 42.58 | 32 | 53 | 1.664 | 0.734 | 0.01658746 | 0.07624971 | 0 | 0.0843 | 0.0426 | 0.0426 | TRUE | 0.2485 |
| Nrg2 | 189 | 194 | 196 | 324 | 234 | 232 | 201 | 194 | 197 | 255 | 252 | 254 | 225.29 | 197 | 254 | 1.286 | 0.363 | 0.00648922 | 0.03814685 | 0 | 0.0215 | 0.0083 | 0.0083 | TRUE | 0.0028 |
| Nrg3 | 0 | 1 | 0 | 1 | 0 | 0 | 0 | 1 | 0 | 1 | 0 | 0 | 0.3 | 0 | 0 | 0.954 | -0.067 | 0.98642001 | NA | 0 | 11.0817 | 9.3014 | 9.3014 | TRUE | 0.405 |
| Nrg4 | 95 | 97 | 78 | 177 | 107 | 110 | 101 | 97 | 78 | 139 | 115 | 120 | 108.46 | 92 | 125 | 1.362 | 0.446 | 0.03057337 | 0.11684108 | 0 | 0.0373 | 0.0212 | 0.0212 | TRUE | 0.1643 |
| Nrgn | 180 | 184 | 176 | 280 | 230 | 186 | 191 | 184 | 177 | 220 | 247 | 204 | 203.81 | 184 | 224 | 1.217 | 0.283 | 0.06226309 | 0.19097888 | 0 | 0.0231 | 0.0116 | 0.0116 | TRUE | 0.0899 |
| Nrip1 | 467 | 572 | 621 | 733 | 544 | 576 | 496 | 571 | 624 | 576 | 585 | 631 | 580.48 | 564 | 597 | 1.059 | 0.082 | 0.49647205 | 0.69197217 | 0.0058 | 0.0126 | 0.0089 | 0.0089 | TRUE | 0.1277 |
| Nrip2 | 16 | 27 | 32 | 22 | 11 | 16 | 17 | 27 | 32 | 17 | 12 | 18 | 20.46 | 25 | 16 | 0.614 | -0.703 | 0.12600663 | 0.30425238 | 0 | 0.168 | 0.1006 | 0.1006 | TRUE | 0.3268 |
| Nrip3 | 262 | 190 | 196 | 227 | 155 | 128 | 278 | 190 | 197 | 178 | 167 | 140 | 191.7 | 222 | 162 | 0.732 | -0.45 | 0.02861146 | 0.11136721 | 0.0261 | 0.0241 | 0.0251 | 0.0251 | TRUE | 0.545 |
| Nrk | 544 | 493 | 512 | 686 | 514 | 618 | 578 | 492 | 514 | 539 | 553 | 676 | 558.85 | 528 | 589 | 1.115 | 0.158 | 0.23818148 | 0.45396957 | 0.0091 | 0.0128 | 0.0111 | 0.0111 | TRUE | 0.1926 |
| Nrl | 0 | 3 | 0 | 1 | 0 | 1 | 0 | 3 | 0 | 1 | 0 | 1 | 0.81 | 1 | 1 | 0.629 | -0.669 | 0.79043222 | NA | 1.8364 | 4.06 | 3.2898 | 3.2898 | TRUE | 1.442 |
| Nrm | 868 | 1229 | 1262 | 1332 | 1043 | 1181 | 922 | 1227 | 1267 | 1047 | 1122 | 1293 | 1146.45 | 1139 | 1154 | 1.013 | 0.018 | 0.90294431 | 0.95205059 | 0.0196 | 0.0098 | 0.0149 | 0.0149 | TRUE | 0.3314 |
| Nrn1 | 5 | 4 | 6 | 11 | 12 | 9 | 5 | 4 | 6 | 9 | 13 | 10 | 7.79 | 5 | 11 | 2.038 | 1.027 | 0.13501543 | 0.31796579 | 0 | 0.4298 | 0.1936 | 0.1936 | TRUE | 0.1417 |
| Nrn1l | 82 | 96 | 123 | 51 | 49 | 54 | 87 | 96 | 124 | 40 | 53 | 59 | 76.4 | 102 | 51 | 0.492 | -1.024 | 0.00012749 | 0.00168131 | 0.0229 | 0.05 | 0.0368 | 0.0368 | TRUE | 0.3288 |
| Nron | 0 | 0 | 0 | 1 | 0 | 0 | 0 | 0 | 0 | 1 | 0 | 0 | 0.13 | 0 | 0 | 1.859 | 0.894 | 0.82650734 | NA | 0 | 25.1515 | 10 | 10 | TRUE | 0.2563 |
| Nrp | 27 | 29 | 39 | 44 | 42 | 30 | 29 | 29 | 39 | 35 | 45 | 33 | 34.9 | 32 | 38 | 1.16 | 0.214 | 0.5296129 | 0.71789091 | 0 | 0.1013 | 0.0546 | 0.0546 | TRUE | 0.2418 |
| Nrp1 | 43 | 26 | 24 | 83 | 56 | 30 | 46 | 26 | 24 | 65 | 60 | 33 | 42.35 | 32 | 53 | 1.665 | 0.735 | 0.06852214 | 0.20280187 | 0.1003 | 0.0847 | 0.0921 | 0.0921 | TRUE | 0.9449 |
| Nrp2 | 950 | 895 | 1019 | 1006 | 761 | 1065 | 1009 | 894 | 1023 | 791 | 819 | 1166 | 950.26 | 975 | 925 | 0.948 | -0.077 | 0.63339584 | 0.79373265 | 0.0254 | 0.0104 | 0.018 | 0.018 | TRUE | 0.6138 |
| Nrsn1 | 51 | 62 | 75 | 71 | 45 | 57 | 54 | 62 | 75 | 56 | 48 | 62 | 59.67 | 64 | 55 | 0.87 | -0.201 | 0.45075389 | 0.65626238 | 0 | 0.0621 | 0.0346 | 0.0346 | TRUE | 0.2188 |
| Nrsn2 | 172 | 158 | 161 | 212 | 153 | 162 | 183 | 158 | 162 | 167 | 165 | 177 | 168.46 | 168 | 170 | 1.013 | 0.018 | 0.90745192 | 0.95354464 | 0 | 0.0265 | 0.0121 | 0.0121 | TRUE | 0.0669 |

| | | | | | | | | | | | | | | | | | | | | | | | | | |
|---|---|---|---|---|---|---|---|---|---|---|---|---|---|---|---|---|---|---|---|---|---|---|---|---|---|
| Nrtn | 521 | 701 | 697 | 650 | 550 | 523 | 553 | 700 | 700 | 511 | 592 | 573 | 604.77 | 651 | 559 | 0.856 | -0.224 | 0.09396285 | 0.25088575 | 0.0102 | 0.0124 | 0.0112 | 0.0112 | TRUE | 0.2018 |
| Nrxn1 | 14 | 10 | 10 | 10 | 10 | 15 | 15 | 10 | 10 | 8 | 11 | 16 | 11.66 | 12 | 12 | 0.993 | -0.01 | 0.98699092 | 0.99290468 | 0 | 0.2895 | 0.1583 | 0.1583 | TRUE | 0.4593 |
| Nrxn2 | 472 | 449 | 415 | 491 | 363 | 304 | 501 | 448 | 417 | 386 | 391 | 333 | 412.63 | 455 | 370 | 0.813 | -0.299 | 0.02252633 | 0.09459141 | 0.0056 | 0.0149 | 0.0099 | 0.0099 | TRUE | 0.0892 |
| Nrxn3 | 3 | 4 | 1 | 9 | 7 | 8 | 3 | 4 | 1 | 7 | 8 | 9 | 5.26 | 3 | 8 | 2.85 | 1.511 | 0.08609554 | 0.23673272 | 0 | 0.6334 | 0.3079 | 0.3079 | TRUE | 0.373 |
| Nsa2 | 4038 | 5083 | 5024 | 5848 | 4378 | 4280 | 4289 | 5076 | 5045 | 4597 | 4710 | 4685 | 4733.83 | 4803 | 4664 | 0.971 | -0.043 | 0.6348066 | 0.79482512 | 0.0037 | 0.0076 | 0.0056 | 0.0056 | TRUE | 0.1067 |
| Nsd1 | 11555 | 9458 | 10824 | 14756 | 10320 | 9597 | 12273 | 9446 | 10868 | 11600 | 11104 | 10505 | 10966.13 | 10862 | 11070 | 1.019 | 0.027 | 0.80327101 | 0.89842886 | 0.0077 | 0.0072 | 0.0086 | 0.0086 | TRUE | 0.1591 |
| Nsdhl | 3040 | 2441 | 2442 | 3224 | 2162 | 2104 | 3229 | 2438 | 2452 | 2534 | 2326 | 2303 | 2547.11 | 2706 | 2388 | 0.883 | -0.18 | 0.16077628 | 0.35539782 | 0.0148 | 0.0082 | 0.0115 | 0.0115 | TRUE | 0.346 |
| Nsf | 9303 | 8196 | 8457 | 14573 | 9873 | 8960 | 9881 | 8185 | 8492 | 11456 | 10623 | 9808 | 9740.93 | 8853 | 10629 | 1.201 | 0.264 | 0.01095327 | 0.05632704 | 0.0081 | 0.0072 | 0.0077 | 0.0077 | TRUE | 0.1261 |
| Nsfl1c | 4532 | 4374 | 4344 | 6589 | 4756 | 4113 | 4814 | 4368 | 4362 | 5180 | 5117 | 4502 | 4723.87 | 4515 | 4933 | 1.093 | 0.128 | 0.15343556 | 0.34502219 | 0.0042 | 0.0076 | 0.0056 | 0.0056 | TRUE | 0.0708 |
| Nsg1 | 22 | 37 | 54 | 67 | 45 | 39 | 23 | 37 | 54 | 53 | 48 | 43 | 43.05 | 38 | 48 | 1.256 | 0.329 | 0.36736311 | 0.58475554 | 0.0591 | 0.0834 | 0.0722 | 0.0722 | TRUE | 0.3199 |
| Nsg2 | 47 | 99 | 117 | 101 | 100 | 96 | 50 | 99 | 117 | 79 | 108 | 105 | 93.06 | 89 | 97 | 1.093 | 0.128 | 0.69308557 | 0.83159723 | 0.0908 | 0.0423 | 0.0649 | 0.0649 | TRUE | 0.8599 |
| Nsl1 | 1467 | 1789 | 1805 | 1859 | 1413 | 1551 | 1558 | 1787 | 1812 | 1461 | 1520 | 1698 | 1639.47 | 1719 | 1560 | 0.907 | -0.141 | 0.17250702 | 0.37119832 | 0.0055 | 0.0089 | 0.0071 | 0.0071 | TRUE | 0.0806 |
| Nsmaf | 954 | 1100 | 1137 | 1341 | 936 | 981 | 1013 | 1099 | 1142 | 1054 | 1007 | 1074 | 1064.78 | 1085 | 1045 | 0.964 | -0.054 | 0.5482139 | 0.73305889 | 0.0014 | 0.01 | 0.0048 | 0.0048 | TRUE | 0.0392 |
| Nsmce1 | 907 | 1368 | 1373 | 1830 | 1269 | 1538 | 963 | 1366 | 1379 | 1439 | 1365 | 1684 | 1365.97 | 1236 | 1496 | 1.21 | 0.275 | 0.08158581 | 0.22771085 | 0.0234 | 0.0093 | 0.0172 | 0.0172 | TRUE | 0.4457 |
| Nsmce2 | 1361 | 1433 | 1419 | 2097 | 1486 | 1515 | 1446 | 1431 | 1425 | 1648 | 1599 | 1658 | 1534.56 | 1434 | 1635 | 1.141 | 0.19 | 0.00698636 | 0.04001046 | 0 | 0.0091 | 0.0029 | 0.0029 | TRUE | 0.0045 |
| Nsmce4a | 3255 | 4236 | 4422 | 5485 | 4000 | 4341 | 3457 | 4231 | 4440 | 4312 | 4304 | 4752 | 4249.27 | 4043 | 4456 | 1.102 | 0.14 | 0.21434495 | 0.42607047 | 0.0095 | 0.0077 | 0.0089 | 0.0089 | TRUE | 0.195 |
| Nsmf | 468 | 497 | 471 | 539 | 409 | 390 | 497 | 496 | 473 | 424 | 440 | 427 | 459.51 | 489 | 430 | 0.88 | -0.185 | 0.07090315 | 0.20739697 | 0 | 0.0141 | 0.0053 | 0.0053 | TRUE | 0.0094 |
| Nsun2 | 27111 | 28023 | 28084 | 31463 | 23296 | 24548 | 28797 | 27987 | 28199 | 24734 | 25065 | 26872 | 26942.21 | 28328 | 25557 | 0.902 | -0.149 | 0.01895428 | 0.08357464 | 0.0011 | 0.007 | 0.0029 | 0.0029 | TRUE | 0.0248 |
| Nsun3 | 179 | 154 | 147 | 298 | 182 | 135 | 190 | 154 | 148 | 234 | 196 | 148 | 178.23 | 164 | 193 | 1.18 | 0.239 | 0.26733569 | 0.48702174 | 0.0304 | 0.0254 | 0.0279 | 0.0279 | TRUE | 0.2116 |
| Nsun4 | 1117 | 913 | 887 | 1276 | 990 | 947 | 1186 | 912 | 891 | 1003 | 1065 | 1037 | 1015.64 | 996 | 1035 | 1.039 | 0.055 | 0.67422661 | 0.81838909 | 0.0124 | 0.0101 | 0.0114 | 0.0114 | TRUE | 0.3317 |
| Nsun5 | 1320 | 1302 | 1298 | 1598 | 1108 | 997 | 1402 | 1300 | 1303 | 1256 | 1192 | 1091 | 1257.58 | 1335 | 1180 | 0.885 | -0.177 | 0.05578698 | 0.17697081 | 0.0026 | 0.0095 | 0.0054 | 0.0054 | TRUE | 0.0512 |
| Nsun6 | 510 | 585 | 629 | 619 | 524 | 542 | 542 | 584 | 632 | 487 | 564 | 593 | 566.87 | 586 | 548 | 0.933 | -0.1 | 0.42094779 | 0.63116133 | 0.0065 | 0.0127 | 0.0093 | 0.0093 | TRUE | 0.1174 |
| Nsun7 | 12 | 10 | 7 | 9 | 5 | 6 | 13 | 10 | 7 | 7 | 5 | 7 | 8.13 | 10 | 6 | 0.644 | -0.635 | 0.33905429 | 0.55910074 | 0 | 0.4121 | 0.1903 | 0.1903 | TRUE | 0.1498 |
| Nt5c | 1101 | 1281 | 1193 | 1428 | 954 | 1039 | 1169 | 1279 | 1198 | 1123 | 1026 | 1137 | 1155.51 | 1215 | 1095 | 0.901 | -0.15 | 0.09374047 | 0.25046464 | 0.0017 | 0.0098 | 0.0049 | 0.0049 | TRUE | 0.0356 |
| Nt5c1a | 1 | 1 | 1 | 5 | 1 | 1 | 1 | 1 | 1 | 4 | 1 | 1 | 1.53 | 1 | 2 | 2.056 | 1.04 | 0.49635165 | NA | 0 | 2.1634 | 0.9503 | 0.9503 | TRUE | 0.7898 |
| Nt5c1b | 0 | 1 | 0 | 0 | 0 | 0 | 0 | 1 | 0 | 0 | 0 | 0 | 0.17 | 0 | 0 | 0.49 | -1.029 | 0.80087 | NA | 0 | 19.7991 | 10 | 10 | TRUE | 0.405 |
| Nt5c2 | 2372 | 1961 | 1998 | 2909 | 1995 | 1704 | 2519 | 1958 | 2006 | 2287 | 2146 | 1865 | 2130.46 | 2161 | 2099 | 0.972 | -0.041 | 0.75269809 | 0.86948897 | 0.0147 | 0.0084 | 0.0118 | 0.0118 | TRUE | 0.2539 |
| Nt5c3 | 730 | 840 | 864 | 918 | 718 | 716 | 775 | 839 | 868 | 722 | 773 | 784 | 793.3 | 827 | 760 | 0.917 | -0.125 | 0.19105225 | 0.39611884 | 0.0014 | 0.0111 | 0.0054 | 0.0054 | TRUE | 0.0355 |
| Nt5c3l | 1731 | 1637 | 1568 | 2362 | 1595 | 1579 | 1839 | 1635 | 1574 | 1857 | 1716 | 1728 | 1724.89 | 1683 | 1767 | 1.051 | 0.072 | 0.44353929 | 0.65031254 | 0.0035 | 0.0088 | 0.0057 | 0.0057 | TRUE | 0.079 |
| Nt5dc1 | 597 | 660 | 658 | 907 | 591 | 570 | 634 | 659 | 661 | 713 | 636 | 624 | 654.47 | 651 | 658 | 1.011 | 0.016 | 0.87317314 | 0.93761484 | 0.0017 | 0.0119 | 0.0059 | 0.0059 | TRUE | 0.067 |
| Nt5dc2 | 6269 | 7188 | 7151 | 7713 | 5816 | 6308 | 6659 | 7179 | 7180 | 6063 | 6258 | 6905 | 6707.33 | 7006 | 6409 | 0.915 | -0.129 | 0.11439363 | 0.28525158 | 0.0031 | 0.0074 | 0.0047 | 0.0047 | TRUE | 0.0559 |
| Nt5dc3 | 1185 | 1348 | 1193 | 1314 | 937 | 816 | 1259 | 1346 | 1198 | 1033 | 1008 | 893 | 1122.87 | 1268 | 978 | 0.772 | -0.373 | 0.00018255 | 0.00225293 | 0.0037 | 0.0098 | 0.0062 | 0.0062 | TRUE | 0.0678 |
| Nt5e | 40 | 17 | 15 | 93 | 42 | 28 | 42 | 17 | 15 | 73 | 45 | 31 | 37.25 | 25 | 50 | 2.016 | 1.011 | 0.04611771 | 0.15538742 | 0.2222 | 0.0954 | 0.1548 | 0.1548 | TRUE | 0.5773 |
| Nt5m | 197 | 192 | 194 | 293 | 194 | 195 | 209 | 192 | 195 | 230 | 209 | 213 | 208.05 | 199 | 217 | 1.098 | 0.134 | 0.34736058 | 0.56626058 | 0 | 0.0227 | 0.0099 | 0.0099 | TRUE | 0.0318 |
| Ntan1 | 1852 | 2353 | 2319 | 2114 | 1500 | 1844 | 1967 | 2350 | 2329 | 1662 | 1614 | 2019 | 1989.99 | 2215 | 1765 | 0.796 | -0.329 | 0.00747431 | 0.04212792 | 0.0118 | 0.0086 | 0.0104 | 0.0104 | TRUE | 0.1898 |
| Ntf3 | 5 | 5 | 8 | 10 | 3 | 12 | 5 | 5 | 8 | 8 | 3 | 13 | 7.09 | 6 | 8 | 1.318 | 0.398 | 0.60961525 | 0.77891178 | 0.0954 | 0.4713 | 0.2933 | 0.2933 | TRUE | 0.1608 |
| Ntf5 | 40 | 63 | 62 | 88 | 68 | 67 | 42 | 63 | 62 | 69 | 73 | 73 | 63.89 | 56 | 72 | 1.283 | 0.359 | 0.1678528 | 0.36558665 | 0 | 0.0585 | 0.033 | 0.033 | TRUE | 0.3592 |
| Nthl1 | 848 | 985 | 975 | 1152 | 806 | 845 | 901 | 984 | 979 | 906 | 867 | 925 | 926.88 | 955 | 899 | 0.942 | -0.086 | 0.33011364 | 0.55093867 | 6.00E-04 | 0.0105 | 0.0046 | 0.0046 | TRUE | 0.0287 |
| Ntm | 21 | 13 | 13 | 22 | 11 | 10 | 22 | 13 | 13 | 17 | 12 | 11 | 14.74 | 16 | 13 | 0.84 | -0.251 | 0.6322256 | 0.79302228 | 0 | 0.2305 | 0.1301 | 0.1301 | TRUE | 0.5177 |
| Ntmt1 | 1174 | 1184 | 1109 | 1459 | 1055 | 1066 | 1247 | 1182 | 1114 | 1147 | 1135 | 1167 | 1165.33 | 1181 | 1150 | 0.974 | -0.039 | 0.64659106 | 0.80159447 | 8.00E-04 | 0.0097 | 0.0043 | 0.0043 | TRUE | 0.0299 |
| Ntn1 | 1816 | 1752 | 2110 | 539 | 416 | 461 | 1929 | 1750 | 2119 | 424 | 448 | 505 | 1195.54 | 1933 | 459 | 0.237 | -2.077 | 7.66E-71 | 2.01E-67 | 0.0075 | 0.0097 | 0.0085 | 0.0085 | TRUE | 0.0788 |
| Ntn3 | 26 | 29 | 25 | 45 | 23 | 14 | 28 | 29 | 25 | 35 | 25 | 15 | 26.19 | 27 | 25 | 0.936 | -0.096 | 0.820943 | 0.90868924 | 0.0465 | 0.1327 | 0.0915 | 0.0915 | TRUE | 0.2496 |
| Ntn4 | 15 | 13 | 11 | 23 | 18 | 19 | 16 | 13 | 11 | 18 | 19 | 21 | 16.37 | 13 | 19 | 1.456 | 0.542 | 0.24414323 | 0.46071154 | 0 | 0.2082 | 0.0925 | 0.0925 | TRUE | 0.1185 |
| Ntn5 | 16 | 35 | 32 | 17 | 20 | 14 | 17 | 35 | 32 | 13 | 22 | 15 | 22.38 | 28 | 17 | 0.592 | -0.757 | 0.1031988 | 0.26735607 | 0.0581 | 0.1541 | 0.1079 | 0.1079 | TRUE | 0.7363 |
| Ntng1 | 0 | 0 | 0 | 0 | 1 | 0 | 0 | 0 | 0 | 0 | 1 | 0 | 0.18 | 0 | 0 | 1.859 | 0.894 | 0.82650734 | NA | 0 | 18.3785 | 10 | 10 | TRUE | 0.5204 |
| Ntng2 | 21 | 46 | 22 | 30 | 25 | 30 | 22 | 46 | 22 | 24 | 27 | 33 | 28.94 | 30 | 28 | 0.917 | -0.125 | 0.77140239 | 0.88080416 | 0.0762 | 0.1207 | 0.0988 | 0.0988 | TRUE | 1.4211 |
| Ntpcr | 1840 | 1982 | 1830 | 1839 | 1361 | 1322 | 1954 | 1979 | 1838 | 1446 | 1464 | 1447 | 1688.09 | 1924 | 1452 | 0.755 | -0.406 | 3.75E-08 | 1.55E-06 | 0 | 0.0089 | 0.0033 | 0.0033 | TRUE | 0.0187 |
| Ntrk1 | 240 | 314 | 293 | 256 | 256 | 199 | 255 | 314 | 294 | 201 | 275 | 218 | 259.54 | 288 | 231 | 0.803 | -0.317 | 0.06420332 | 0.19500617 | 0.015 | 0.0196 | 0.0173 | 0.0173 | TRUE | 0.3006 |
| Ntrk2 | 167 | 198 | 187 | 276 | 232 | 207 | 177 | 198 | 188 | 217 | 250 | 227 | 209.35 | 188 | 231 | 1.229 | 0.297 | 0.04304563 | 0.14822702 | 0 | 0.0226 | 0.0107 | 0.0107 | TRUE | 0.0537 |
| Ntrk3 | 45 | 51 | 64 | 30 | 27 | 35 | 48 | 51 | 64 | 24 | 29 | 38 | 42.32 | 54 | 30 | 0.552 | -0.856 | 0.0092146 | 0.04943996 | 0.0151 | 0.0847 | 0.0522 | 0.0522 | TRUE | 0.3414 |
| Nts | 1 | 0 | 0 | 0 | 1 | 1 | 1 | 0 | 0 | 0 | 1 | 1 | 0.54 | 0 | 1 | 1.923 | 0.944 | 0.73523723 | NA | 0 | 6.1214 | 3.9144 | 3.9144 | TRUE | 0.4614 |
| Ntsr1 | 5 | 4 | 4 | 3 | 1 | 4 | 5 | 4 | 4 | 2 | 1 | 4 | 3.52 | 4 | 2 | 0.584 | -0.777 | 0.44378496 | NA | 0 | 0.9422 | 0.4385 | 0.4385 | TRUE | 0.3484 |
| Ntsr2 | 1 | 1 | 0 | 0 | 0 | 0 | 1 | 1 | 0 | 0 | 0 | 0 | 0.34 | 1 | 0 | 0.261 | -1.936 | 0.60454921 | NA | 0 | 9.5983 | 8.3269 | 8.3269 | TRUE | 0.2507 |
| Nuak1 | 697 | 805 | 836 | 1107 | 874 | 906 | 740 | 804 | 839 | 870 | 940 | 992 | 864.35 | 794 | 934 | 1.174 | 0.232 | 0.02237164 | 0.09421777 | 0.0031 | 0.0107 | 0.0062 | 0.0062 | TRUE | 0.0443 |
| Nuak2 | 405 | 462 | 577 | 436 | 339 | 436 | 430 | 461 | 579 | 343 | 365 | 477 | 442.62 | 490 | 395 | 0.804 | -0.315 | 0.07448886 | 0.21446484 | 0.0261 | 0.0143 | 0.0202 | 0.0202 | TRUE | 0.3788 |
| Nub1 | 4536 | 5617 | 5801 | 6499 | 4900 | 5154 | 4818 | 5610 | 5825 | 5109 | 5272 | 5642 | 5379.26 | 5418 | 5341 | 0.986 | -0.021 | 0.83074835 | 0.91425507 | 0.006 | 0.0075 | 0.0066 | 0.0066 | TRUE | 0.1141 |
| Nubp1 | 2513 | 2812 | 2715 | 3514 | 2643 | 2522 | 2669 | 2808 | 2726 | 2762 | 2844 | 2761 | 2761.77 | 2734 | 2789 | 1.02 | 0.028 | 0.66491706 | 0.81250574 | 0 | 0.0081 | 0.0027 | 0.0027 | TRUE | 0.0068 |
| Nubp2 | 3156 | 3001 | 2845 | 3945 | 2990 | 2684 | 3352 | 2997 | 2857 | 3101 | 3217 | 2938 | 3077.07 | 3069 | 3085 | 1.006 | 0.008 | 0.92898265 | 0.9650846 | 0.0041 | 0.008 | 0.0056 | 0.0056 | TRUE | 0.0792 |
| Nubpl | 1463 | 2181 | 2345 | 1978 | 1476 | 1754 | 1554 | 2178 | 2355 | 1555 | 1588 | 1920 | 1858.31 | 2029 | 1688 | 0.832 | -0.266 | 0.10592975 | 0.27152998 | 0.0278 | 0.0087 | 0.019 | 0.019 | TRUE | 0.5065 |
| Nucb1 | 2475 | 3017 | 3121 | 3402 | 2550 | 2651 | 2629 | 3013 | 3134 | 2674 | 2744 | 2902 | 2849.3 | 2925 | 2773 | 0.948 | -0.077 | 0.41040926 | 0.62180353 | 0.0044 | 0.0081 | 0.006 | 0.006 | TRUE | 0.0951 |
| Nucb2 | 579 | 608 | 584 | 724 | 495 | 497 | 615 | 607 | 586 | 569 | 533 | 544 | 575.73 | 603 | 549 | 0.911 | -0.134 | 0.16146835 | 0.356478 | 0 | 0.0126 | 0.0049 | 0.0049 | TRUE | 0.0134 |
| Nucks1 | 18555 | 17979 | 17183 | 23753 | 16583 | 14816 | 19709 | 17956 | 17254 | 18673 | 17842 | 16218 | 17941.92 | 18306 | 17578 | 0.96 | -0.058 | 0.51324304 | 0.70632889 | 0.0044 | 0.0071 | 0.0057 | 0.0057 | TRUE | 0.0559 |
| Nudc | 15103 | 15099 | 15170 | 20096 | 15601 | 14141 | 16042 | 15080 | 15232 | 15798 | 16786 | 15480 | 15736.17 | 15451 | 16021 | 1.037 | 0.052 | 0.43621031 | 0.64448796 | 0.0014 | 0.0071 | 0.0032 | 0.0032 | TRUE | 0.0213 |
| Nudcd1 | 4121 | 4135 | 4067 | 4923 | 3626 | 3584 | 4377 | 4130 | 4084 | 3870 | 3901 | 3923 | 4047.55 | 4197 | 3898 | 0.929 | -0.106 | 0.09888528 | 0.25937447 | 5.00E-04 | 0.0077 | 0.0028 | 0.0028 | TRUE | 0.0172 |
| Nudcd2 | 6963 | 7847 | 7690 | 9741 | 6807 | 6763 | 7396 | 7837 | 7722 | 7658 | 7324 | 7403 | 7556.51 | 7652 | 7462 | 0.975 | -0.036 | 0.55775818 | 0.7401735 | 5.00E-04 | 0.0073 | 0.0026 | 0.0026 | TRUE | 0.0104 |
| Nudcd3 | 4705 | 4550 | 4444 | 6458 | 4753 | 4364 | 4998 | 4544 | 4462 | 5077 | 5114 | 4777 | 4828.62 | 4668 | 4989 | 1.069 | 0.096 | 0.21797723 | 0.42995709 | 0.0023 | 0.0076 | 0.0042 | 0.0042 | TRUE | 0.0464 |
| Nudt1 | 945 | 928 | 935 | 1504 | 992 | 811 | 1004 | 927 | 939 | 1182 | 1067 | 888 | 1001.14 | 957 | 1046 | 1.095 | 0.131 | 0.29666282 | 0.51741303 | 0.0104 | 0.0102 | 0.0103 | 0.0103 | TRUE | 0.2075 |
| Nudt10 | 40 | 51 | 45 | 145 | 77 | 71 | 42 | 51 | 45 | 114 | 83 | 78 | 68.86 | 46 | 92 | 1.991 | 0.994 | 0.00023497 | 0.00277393 | 0.0192 | 0.0547 | 0.0362 | 0.0362 | TRUE | 0.4819 |
| Nudt11 | 250 | 239 | 280 | 552 | 368 | 353 | 266 | 239 | 281 | 434 | 396 | 386 | 333.61 | 262 | 405 | 1.551 | 0.633 | 1.07E-06 | 2.93E-05 | 0.0023 | 0.0168 | 0.009 | 0.009 | TRUE | 0.0674 |

| | | | | | | | | | | | | | | | | | | | | | | | | |
|---|---|---|---|---|---|---|---|---|---|---|---|---|---|---|---|---|---|---|---|---|---|---|---|---|
| Nudt12 | 514 | 552 | 548 | 652 | 437 | 432 | 546 | 551 | 550 | 513 | 470 | 473 | 517.19 | 549 | 485 | 0.885 | -0.176 | 0.07994688 | 0.22447967 | 0 | 0.0133 | 0.0054 | 0.0054 | TRUE | 0.0302 |
| Nudt13 | 180 | 187 | 214 | 247 | 202 | 154 | 191 | 187 | 215 | 194 | 217 | 169 | 195.49 | 198 | 193 | 0.979 | -0.031 | 0.84996937 | 0.92526408 | 0.005 | 0.0238 | 0.0143 | 0.0143 | TRUE | 0.1277 |
| Nudt14 | 583 | 939 | 922 | 846 | 667 | 808 | 619 | 938 | 926 | 665 | 718 | 884 | 791.67 | 828 | 756 | 0.912 | -0.132 | 0.47187848 | 0.67314138 | 0.0359 | 0.0111 | 0.0232 | 0.0232 | TRUE | 0.5743 |
| Nudt15 | 65 | 48 | 41 | 79 | 66 | 60 | 69 | 48 | 41 | 62 | 71 | 66 | 59.49 | 53 | 66 | 1.257 | 0.33 | 0.24620602 | 0.46320719 | 0.0175 | 0.0623 | 0.0414 | 0.0414 | TRUE | 0.5922 |
| Nudt16 | 243 | 370 | 422 | 455 | 338 | 390 | 258 | 370 | 424 | 358 | 364 | 427 | 366.61 | 351 | 383 | 1.091 | 0.125 | 0.51628538 | 0.70829955 | 0.0327 | 0.0159 | 0.0242 | 0.0242 | TRUE | 0.5766 |
| Nudt16l1 | 2943 | 2985 | 2946 | 3696 | 2672 | 2547 | 3126 | 2981 | 2958 | 2906 | 2875 | 2788 | 2938.96 | 3022 | 2856 | 0.945 | -0.081 | 0.22304191 | 0.43505698 | 3.00E-04 | 0.008 | 0.0028 | 0.0028 | TRUE | 0.011 |
| Nudt17 | 426 | 618 | 623 | 474 | 381 | 446 | 452 | 617 | 626 | 373 | 410 | 488 | 494.34 | 565 | 424 | 0.748 | -0.418 | 0.01354494 | 0.06561143 | 0.0235 | 0.0136 | 0.0186 | 0.0186 | TRUE | 0.354 |
| Nudt18 | 357 | 391 | 355 | 517 | 340 | 399 | 379 | 390 | 356 | 406 | 366 | 437 | 389.19 | 375 | 403 | 1.074 | 0.103 | 0.40320888 | 0.61609882 | 0.0022 | 0.0154 | 0.0083 | 0.0083 | TRUE | 0.0718 |
| Nudt19 | 1019 | 981 | 971 | 1534 | 1028 | 959 | 1082 | 980 | 975 | 1206 | 1106 | 1050 | 1066.47 | 1012 | 1121 | 1.108 | 0.148 | 0.13264112 | 0.31402218 | 0.0034 | 0.01 | 0.0061 | 0.0061 | TRUE | 0.0547 |
| Nudt2 | 613 | 748 | 798 | 833 | 573 | 644 | 651 | 747 | 801 | 655 | 617 | 705 | 695.96 | 733 | 659 | 0.898 | -0.155 | 0.18956477 | 0.3947026 | 0.0062 | 0.0116 | 0.0086 | 0.0086 | TRUE | 0.1125 |
| Nudt21 | 4523 | 5251 | 4915 | 5577 | 3789 | 4269 | 4804 | 5244 | 4935 | 4384 | 4077 | 4673 | 4686.27 | 4994 | 4378 | 0.877 | -0.19 | 0.02193459 | 0.09302296 | 0.003 | 0.0076 | 0.0047 | 0.0047 | TRUE | 0.0437 |
| Nudt22 | 273 | 283 | 242 | 366 | 295 | 263 | 290 | 283 | 243 | 288 | 317 | 288 | 284.77 | 272 | 298 | 1.094 | 0.13 | 0.34181613 | 0.56204134 | 0.0022 | 0.0185 | 0.01 | 0.01 | TRUE | 0.0972 |
| Nudt3 | 3241 | 4433 | 4469 | 4517 | 3379 | 3700 | 3443 | 4427 | 4487 | 3551 | 3636 | 4050 | 3932.31 | 4119 | 3746 | 0.909 | -0.137 | 0.26323781 | 0.48264194 | 0.0123 | 0.0077 | 0.0106 | 0.0106 | TRUE | 0.251 |
| Nudt4 | 14031 | 12442 | 11592 | 23901 | 15632 | 13785 | 14903 | 12426 | 11640 | 18789 | 16819 | 15090 | 14944.49 | 12990 | 16899 | 1.301 | 0.38 | 0.00216726 | 0.01641606 | 0.0145 | 0.0071 | 0.011 | 0.011 | TRUE | 0.203 |
| Nudt5 | 1735 | 2121 | 2059 | 2864 | 2005 | 1639 | 1843 | 2118 | 2067 | 2251 | 2157 | 1794 | 2038.57 | 2009 | 2067 | 1.029 | 0.042 | 0.71600735 | 0.84571767 | 0.0093 | 0.0085 | 0.009 | 0.009 | TRUE | 0.1611 |
| Nudt6 | 87 | 128 | 111 | 143 | 101 | 76 | 92 | 128 | 111 | 112 | 109 | 83 | 106 | 110 | 101 | 0.919 | -0.121 | 0.59089816 | 0.76478301 | 0.0159 | 0.038 | 0.0272 | 0.0272 | TRUE | 0.2314 |
| Nudt7 | 109 | 115 | 107 | 136 | 95 | 103 | 116 | 115 | 107 | 107 | 102 | 113 | 109.99 | 113 | 107 | 0.952 | -0.071 | 0.70133466 | 0.83650923 | 0 | 0.0369 | 0.0155 | 0.0155 | TRUE | 0.0179 |
| Nudt8 | 432 | 558 | 546 | 646 | 430 | 462 | 459 | 557 | 548 | 508 | 463 | 506 | 506.77 | 521 | 492 | 0.944 | -0.083 | 0.49377915 | 0.69010942 | 0.0049 | 0.0134 | 0.0088 | 0.0088 | TRUE | 0.1275 |
| Nudt9 | 1271 | 1425 | 1344 | 2164 | 1613 | 1579 | 1350 | 1423 | 1350 | 1701 | 1735 | 1728 | 1547.97 | 1374 | 1721 | 1.253 | 0.325 | 7.48E-06 | 0.00015498 | 0 | 0.009 | 0.0031 | 0.0031 | TRUE | 0.0117 |
| Nuf2 | 2190 | 2146 | 2065 | 3237 | 2299 | 2270 | 2326 | 2143 | 2073 | 2545 | 2474 | 2485 | 2341 | 2181 | 2501 | 1.147 | 0.198 | 0.01048495 | 0.05463353 | 0.0013 | 0.0083 | 0.0039 | 0.0039 | TRUE | 0.0409 |
| Nufip1 | 3221 | 3790 | 3897 | 4497 | 3175 | 3393 | 3421 | 3785 | 3913 | 3535 | 3416 | 3714 | 3630.81 | 3706 | 3555 | 0.959 | -0.06 | 0.47161234 | 0.67308567 | 0.0028 | 0.0078 | 0.0048 | 0.0048 | TRUE | 0.055 |
| Nufip2 | 12861 | 10341 | 10249 | 13192 | 10190 | 8918 | 13661 | 10328 | 10291 | 10371 | 10964 | 9762 | 10895.98 | 11427 | 10366 | 0.907 | -0.141 | 0.26734937 | 0.48702174 | 0.0159 | 0.0072 | 0.0115 | 0.0115 | TRUE | 0.3575 |
| Nuggc | 12 | 24 | 28 | 57 | 52 | 48 | 13 | 24 | 28 | 45 | 56 | 53 | 36.36 | 22 | 51 | 2.35 | 1.233 | 0.0009377 | 0.00847047 | 0.0184 | 0.0975 | 0.0669 | 0.0669 | TRUE | 0.6927 |
| Numa1 | 9937 | 10819 | 12011 | 12469 | 9419 | 9418 | 10555 | 10805 | 12060 | 9802 | 10134 | 10309 | 10611.02 | 11140 | 10082 | 0.905 | -0.144 | 0.0665259 | 0.19895694 | 0.0028 | 0.0072 | 0.0044 | 0.0044 | TRUE | 0.0638 |
| Numb | 1447 | 1391 | 1347 | 1922 | 1446 | 1282 | 1537 | 1389 | 1353 | 1511 | 1556 | 1403 | 1458.13 | 1426 | 1490 | 1.045 | 0.064 | 0.48888602 | 0.68533555 | 0.0027 | 0.0092 | 0.0054 | 0.0054 | TRUE | 0.0552 |
| Numbl | 924 | 1152 | 1101 | 1429 | 1087 | 1045 | 981 | 1151 | 1106 | 1123 | 1170 | 1144 | 1112.39 | 1079 | 1146 | 1.061 | 0.085 | 0.36566879 | 0.58308847 | 0.0025 | 0.0099 | 0.0055 | 0.0055 | TRUE | 0.0745 |
| Nup107 | 7944 | 7782 | 8034 | 10766 | 7805 | 7574 | 8438 | 7772 | 8067 | 8463 | 8398 | 8291 | 8238.15 | 8092 | 8384 | 1.036 | 0.051 | 0.41769691 | 0.62855792 | 7.00E-04 | 0.0073 | 0.0028 | 0.0028 | TRUE | 0.017 |
| Nup133 | 12682 | 13755 | 14569 | 15894 | 11601 | 12415 | 13471 | 13737 | 14629 | 12495 | 12482 | 13590 | 13400.56 | 13946 | 12856 | 0.922 | -0.118 | 0.10702628 | 0.27341772 | 0.0021 | 0.0071 | 0.0038 | 0.0038 | TRUE | 0.0305 |
| Nup153 | 12201 | 10280 | 10737 | 14819 | 10902 | 9924 | 12960 | 10267 | 10781 | 11650 | 11730 | 10863 | 11375.04 | 11336 | 11414 | 1.007 | 0.01 | 0.92422343 | 0.96274037 | 0.007 | 0.0072 | 0.0079 | 0.0079 | TRUE | 0.1918 |
| Nup155 | 7277 | 6789 | 7368 | 9123 | 6609 | 6525 | 7729 | 6780 | 7398 | 7172 | 7111 | 7143 | 7222.21 | 7302 | 7142 | 0.978 | -0.032 | 0.66754939 | 0.8140242 | 0.0018 | 0.0074 | 0.0039 | 0.0039 | TRUE | 0.0478 |
| Nup160 | 13568 | 13074 | 13825 | 15465 | 11089 | 11551 | 14412 | 13057 | 13882 | 12157 | 11931 | 12644 | 13013.89 | 13784 | 12244 | 0.888 | -0.171 | 0.01374719 | 0.06630601 | 0.0016 | 0.0072 | 0.0034 | 0.0034 | TRUE | 0.026 |
| Nup188 | 15579 | 17995 | 19979 | 19714 | 14988 | 16413 | 16548 | 17972 | 20061 | 15498 | 16126 | 17967 | 17361.8 | 18194 | 16530 | 0.909 | -0.138 | 0.17280031 | 0.37156189 | 0.0076 | 0.0071 | 0.0074 | 0.0074 | TRUE | 0.0986 |
| Nup205 | 13256 | 12498 | 13858 | 17456 | 12263 | 12239 | 14080 | 12482 | 13915 | 13723 | 13194 | 13398 | 13465.21 | 13492 | 13438 | 0.996 | -0.006 | 0.93882538 | 0.96930832 | 0.0018 | 0.0071 | 0.0039 | 0.0039 | TRUE | 0.0525 |
| Nup210 | 6200 | 7015 | 7753 | 7820 | 6017 | 6372 | 6585 | 7006 | 7785 | 6147 | 6474 | 6975 | 6828.8 | 7125 | 6532 | 0.917 | -0.126 | 0.18141922 | 0.38324992 | 0.0055 | 0.0074 | 0.0062 | 0.0062 | TRUE | 0.08 |
| Nup210l | 52 | 44 | 39 | 77 | 90 | 119 | 55 | 44 | 39 | 61 | 97 | 130 | 70.99 | 46 | 96 | 2.07 | 1.05 | 0.00184834 | 0.01445319 | 0.0781 | 0.0533 | 0.0657 | 0.0657 | TRUE | 0.1185 |
| Nup214 | 5491 | 4548 | 4849 | 7637 | 5303 | 5002 | 5832 | 4542 | 4869 | 6004 | 5706 | 5475 | 5404.71 | 5081 | 5728 | 1.128 | 0.173 | 0.11533429 | 0.28675672 | 0.0095 | 0.0075 | 0.0085 | 0.0085 | TRUE | 0.2037 |
| Nup35 | 994 | 987 | 942 | 1404 | 919 | 727 | 1056 | 986 | 946 | 1104 | 989 | 796 | 979.29 | 996 | 963 | 0.968 | -0.047 | 0.73047803 | 0.8551885 | 0.014 | 0.0103 | 0.0123 | 0.0123 | TRUE | 0.2726 |
| Nup37 | 2521 | 3167 | 2985 | 4183 | 2884 | 2922 | 2678 | 3163 | 2997 | 3288 | 3103 | 3199 | 3071.31 | 2946 | 3197 | 1.085 | 0.118 | 0.18173142 | 0.38360117 | 0.0034 | 0.008 | 0.0053 | 0.0053 | TRUE | 0.0768 |
| Nup43 | 4192 | 4455 | 4532 | 5393 | 3873 | 4086 | 4453 | 4449 | 4551 | 4240 | 4167 | 4473 | 4388.66 | 4484 | 4293 | 0.957 | -0.063 | 0.32796785 | 0.54906156 | 4.00E-04 | 0.0077 | 0.0028 | 0.0028 | TRUE | 0.0161 |
| Nup50 | 6606 | 6436 | 6400 | 9606 | 6815 | 6321 | 7017 | 6428 | 6426 | 7551 | 7333 | 6919 | 6945.67 | 6624 | 7268 | 1.097 | 0.134 | 0.07437418 | 0.21421286 | 0.0021 | 0.0074 | 0.0039 | 0.0039 | TRUE | 0.0329 |
| Nup54 | 4087 | 4138 | 4308 | 5865 | 4355 | 4292 | 4341 | 4133 | 4326 | 4611 | 4686 | 4698 | 4465.67 | 4267 | 4665 | 1.093 | 0.129 | 0.03444803 | 0.12649925 | 2.00E-04 | 0.0076 | 0.0024 | 0.0024 | TRUE | 0.0092 |
| Nup62 | 7736 | 7427 | 7164 | 10988 | 7737 | 6984 | 8217 | 7417 | 7193 | 8638 | 8325 | 7645 | 7905.89 | 7609 | 8203 | 1.078 | 0.109 | 0.21586542 | 0.42782475 | 0.0042 | 0.0073 | 0.0054 | 0.0054 | TRUE | 0.0595 |
| Nup62-il4i1 | 77 | 76 | 79 | 90 | 84 | 85 | 82 | 76 | 79 | 71 | 90 | 93 | 81.86 | 79 | 85 | 1.067 | 0.093 | 0.68033612 | 0.82277546 | 0 | 0.0471 | 0.0245 | 0.0245 | TRUE | 0.2232 |
| Nup62cl | 379 | 442 | 410 | 1054 | 686 | 632 | 403 | 441 | 412 | 829 | 738 | 692 | 585.69 | 419 | 753 | 1.801 | 0.849 | 2.43E-13 | 2.75E-11 | 0.0043 | 0.0125 | 0.0078 | 0.0078 | TRUE | 0.0949 |
| Nup85 | 12518 | 11296 | 11370 | 15187 | 10905 | 10622 | 13296 | 11281 | 11417 | 11939 | 11733 | 11627 | 11882.3 | 11998 | 11766 | 0.981 | -0.028 | 0.74690815 | 0.8655848 | 0.0037 | 0.0072 | 0.0054 | 0.0054 | TRUE | 0.1095 |
| Nup88 | 7204 | 6939 | 6809 | 10244 | 7189 | 7147 | 7652 | 6930 | 6837 | 8053 | 7735 | 7824 | 7505.06 | 7140 | 7871 | 1.103 | 0.141 | 0.0566158 | 0.17876766 | 0.002 | 0.0073 | 0.0038 | 0.0038 | TRUE | 0.048 |
| Nup93 | 15719 | 14317 | 14349 | 19736 | 14168 | 13530 | 16696 | 14299 | 14408 | 15515 | 15244 | 14811 | 15162.05 | 15134 | 15190 | 1.004 | 0.005 | 0.94999059 | 0.97481103 | 0.0033 | 0.0071 | 0.0052 | 0.0052 | TRUE | 0.0997 |
| Nup98 | 7585 | 6478 | 6890 | 8381 | 5860 | 5789 | 8057 | 6470 | 6918 | 6588 | 6305 | 6337 | 6779.17 | 7148 | 6410 | 0.897 | -0.157 | 0.1118144 | 0.28113131 | 0.0066 | 0.0074 | 0.0069 | 0.0069 | TRUE | 0.1508 |
| Nupl1 | 4126 | 4014 | 4132 | 5310 | 3928 | 3899 | 4383 | 4009 | 4149 | 4174 | 4226 | 4268 | 4201.5 | 4180 | 4223 | 1.01 | 0.015 | 0.82802063 | 0.91291407 | 8.00E-04 | 0.0077 | 0.003 | 0.003 | TRUE | 0.0218 |
| Nupl2 | 401 | 438 | 381 | 651 | 445 | 362 | 426 | 437 | 383 | 512 | 479 | 396 | 438.79 | 415 | 462 | 1.115 | 0.157 | 0.25449629 | 0.47255892 | 0.0089 | 0.0144 | 0.0114 | 0.0114 | TRUE | 0.178 |
| Nupr1 | 94 | 85 | 79 | 145 | 105 | 90 | 100 | 85 | 79 | 114 | 113 | 99 | 98.26 | 88 | 109 | 1.236 | 0.306 | 0.14015841 | 0.32504722 | 0 | 0.0404 | 0.0206 | 0.0206 | TRUE | 0.1283 |
| Nupr1l | 22 | 35 | 38 | 25 | 9 | 17 | 23 | 35 | 38 | 20 | 10 | 19 | 24.07 | 32 | 16 | 0.501 | -0.998 | 0.02524897 | 0.10247729 | 0.0388 | 0.1438 | 0.0972 | 0.0972 | TRUE | 0.5198 |
| Nus1 | 4847 | 5327 | 5272 | 6230 | 4660 | 4800 | 5148 | 5320 | 5294 | 4898 | 5014 | 5254 | 5154.65 | 5254 | 5055 | 0.962 | -0.056 | 0.38124999 | 0.59555331 | 5.00E-04 | 0.0075 | 0.0028 | 0.0028 | TRUE | 0.0144 |
| Nusap1 | 3513 | 5493 | 5933 | 5816 | 4313 | 5105 | 3731 | 5486 | 5957 | 4572 | 4641 | 5588 | 4995.92 | 5058 | 4934 | 0.975 | -0.036 | 0.83289824 | 0.91523598 | 0.0355 | 0.0076 | 0.021 | 0.021 | TRUE | 0.6414 |
| Nutf2 | 1444 | 1137 | 1042 | 1384 | 894 | 878 | 1534 | 1136 | 1046 | 1088 | 962 | 961 | 1121.1 | 1239 | 1004 | 0.811 | -0.302 | 0.05094234 | 0.16617633 | 0.0233 | 0.0098 | 0.0164 | 0.0164 | TRUE | 0.5208 |
| Nutf2-ps1 | 0 | 0 | 0 | 0 | 0 | 0 | 0 | 0 | 0 | 0 | 0 | 0 | 0 | 0 | 0 | NA | NA | NA | NA | NA | NA | NA | NA | NA | NA |
| Nvl | 4928 | 5424 | 5496 | 6722 | 4833 | 4988 | 5234 | 5417 | 5519 | 5284 | 5200 | 5460 | 5352.41 | 5390 | 5315 | 0.986 | -0.02 | 0.7439186 | 0.86335793 | 4.00E-04 | 0.0075 | 0.0026 | 0.0026 | TRUE | 0.0078 |
| Nwd1 | 4 | 4 | 4 | 2 | 1 | 4 | 4 | 4 | 4 | 2 | 1 | 4 | 3.21 | 4 | 2 | 0.563 | -0.829 | 0.43767381 | NA | 0 | 1.0318 | 0.4855 | 0.4855 | TRUE | 0.533 |
| Nxf1 | 4774 | 2969 | 2926 | 6308 | 4408 | 3589 | 5071 | 2965 | 2938 | 4959 | 4743 | 3929 | 4100.72 | 3658 | 4544 | 1.242 | 0.313 | 0.12710349 | 0.30619768 | 0.0564 | 0.0077 | 0.0301 | 0.0301 | TRUE | 1.3881 |
| Nxf2 | 22 | 28 | 29 | 48 | 36 | 33 | 23 | 28 | 29 | 38 | 39 | 36 | 32.17 | 27 | 38 | 1.398 | 0.484 | 0.14560893 | 0.33399418 | 0 | 0.1093 | 0.0476 | 0.0476 | TRUE | 0.0764 |
| Nxf3 | 120 | 135 | 142 | 598 | 428 | 420 | 127 | 135 | 143 | 470 | 461 | 460 | 299.2 | 135 | 464 | 3.433 | 1.78 | 6.07E-42 | 6.84E-39 | 0 | 0.0179 | 0.0076 | 0.0076 | TRUE | 0.0252 |
| Nxf7 | 71 | 74 | 50 | 134 | 104 | 63 | 75 | 74 | 50 | 105 | 112 | 69 | 80.95 | 66 | 95 | 1.438 | 0.525 | 0.06523243 | 0.19690761 | 0.0433 | 0.0476 | 0.0455 | 0.0455 | TRUE | 0.5432 |
| Nxn | 4935 | 5141 | 5339 | 4731 | 3520 | 3754 | 5242 | 5134 | 5361 | 3719 | 3787 | 4109 | 4558.82 | 5246 | 3872 | 0.738 | -0.439 | 1.04E-09 | 6.33E-08 | 0.0013 | 0.0076 | 0.0035 | 0.0035 | TRUE | 0.0348 |
| Nxnl1 | 1 | 2 | 3 | 1 | 1 | 2 | 1 | 2 | 3 | 1 | 1 | 2 | 1.69 | 2 | 1 | 0.65 | -0.621 | 0.66095833 | NA | 0 | 1.9595 | 0.8241 | 0.8241 | TRUE | 0.1674 |
| Nxnl2 | 48 | 74 | 88 | 41 | 42 | 53 | 51 | 74 | 88 | 32 | 45 | 58 | 58.11 | 71 | 45 | 0.629 | -0.668 | 0.04598114 | 0.15519249 | 0.0617 | 0.0636 | 0.0626 | 0.0626 | TRUE | 0.2592 |
| Nxpe2 | 2 | 0 | 1 | 1 | 0 | 1 | 2 | 0 | 1 | 1 | 0 | 1 | 0.83 | 1 | 1 | 0.611 | -0.711 | 0.74305956 | NA | 0 | 3.953 | 2.1268 | 2.1268 | TRUE | 0.1454 |
| Nxpe3 | 643 | 634 | 693 | 655 | 458 | 520 | 683 | 633 | 696 | 515 | 493 | 569 | 598.15 | 671 | 526 | 0.783 | -0.352 | 0.00102054 | 0.00905284 | 0.002 | 0.0124 | 0.0066 | 0.0066 | TRUE | 0.0591 |
| Nxpe4 | 34 | 21 | 21 | 46 | 37 | 25 | 36 | 21 | 21 | 36 | 40 | 27 | 30.25 | 26 | 34 | 1.328 | 0.409 | 0.29501948 | 0.51594065 | 0.0298 | 0.1158 | 0.076 | 0.076 | TRUE | 0.6849 |
| Nxpe5 | 0 | 0 | 0 | 0 | 1 | 0 | 0 | 0 | 0 | 0 | 1 | 0 | 0.18 | 0 | 0 | 1.859 | 0.894 | 0.82650734 | NA | 0 | 18.3785 | 10 | 10 | TRUE | 0.5204 |

| | | | | | | | | | | | | | | | | | | | | | | | | | |
|---|---|---|---|---|---|---|---|---|---|---|---|---|---|---|---|---|---|---|---|---|---|---|---|---|---|
| Nxph1 | 0 | 0 | 0 | 0 | 0 | 0 | 0 | 0 | 0 | 0 | 0 | 0 | 0 | 0 | 0 | NA | NA | NA | NA | NA | NA | NA | NA | NA | NA |
| Nxph2 | 0 | 0 | 0 | 0 | 0 | 0 | 0 | 0 | 0 | 0 | 0 | 0 | 0 | 0 | 0 | NA | NA | NA | NA | NA | NA | NA | NA | NA | NA |
| Nxph3 | 157 | 125 | 137 | 125 | 105 | 88 | 167 | 125 | 138 | 98 | 113 | 96 | 122.79 | 143 | 102 | 0.716 | -0.481 | 0.0162189 | 0.07508204 | 0.0076 | 0.0337 | 0.0205 | 0.0205 | TRUE | 0.2153 |
| Nxph4 | 453 | 1042 | 1194 | 599 | 535 | 811 | 481 | 1041 | 1199 | 471 | 576 | 888 | 775.83 | 907 | 645 | 0.711 | -0.493 | 0.2902423 | 0.51138358 | 0.1551 | 0.0111 | 0.0698 | 0.1551 | TRUE | 0.9747 |
| Nxt1 | 1494 | 1632 | 1589 | 2078 | 1571 | 1531 | 1587 | 1630 | 1596 | 1634 | 1690 | 1676 | 1635.35 | 1604 | 1667 | 1.039 | 0.055 | 0.43332934 | 0.64227707 | 0 | 0.0089 | 0.0029 | 0.0029 | TRUE | 0.0038 |
| Nxt2 | 2251 | 3413 | 3388 | 2771 | 2075 | 2632 | 2391 | 3409 | 3402 | 2178 | 2233 | 2881 | 2748.92 | 3067 | 2431 | 0.792 | -0.336 | 0.04363907 | 0.14952757 | 0.0306 | 0.0081 | 0.0196 | 0.0196 | TRUE | 0.4514 |
| Nyap1 | 716 | 808 | 877 | 725 | 479 | 631 | 761 | 807 | 881 | 570 | 515 | 691 | 704.02 | 816 | 592 | 0.725 | -0.464 | 0.0006209 | 0.00616676 | 0.0121 | 0.0116 | 0.0118 | 0.0118 | TRUE | 0.2443 |
| Nyap2 | 0 | 0 | 0 | 1 | 0 | 0 | 0 | 0 | 0 | 1 | 0 | 0 | 0.13 | 0 | 0 | 1.859 | 0.894 | 0.82650734 | NA | 0 | 25.1515 | 10 | 10 | TRUE | 0.2563 |
| Nynrin | 1249 | 1102 | 1131 | 1218 | 934 | 836 | 1327 | 1101 | 1136 | 957 | 1005 | 915 | 1073.41 | 1188 | 959 | 0.808 | -0.308 | 0.00392016 | 0.02594267 | 0.0049 | 0.01 | 0.0073 | 0.0073 | TRUE | 0.1252 |
| Nyx | 13 | 6 | 4 | 12 | 11 | 6 | 14 | 6 | 4 | 9 | 12 | 7 | 8.61 | 8 | 9 | 1.176 | 0.233 | 0.74550839 | 0.86463404 | 0.1064 | 0.3896 | 0.2554 | 0.2554 | TRUE | 1.153 |
| Oacyl | 54 | 62 | 68 | 59 | 55 | 51 | 57 | 62 | 68 | 46 | 59 | 56 | 58.16 | 62 | 54 | 0.855 | -0.225 | 0.38391813 | 0.59804511 | 0 | 0.0636 | 0.031 | 0.031 | TRUE | 0.1421 |
| Oaf | 183 | 266 | 234 | 140 | 97 | 120 | 194 | 266 | 235 | 110 | 104 | 131 | 173.46 | 232 | 115 | 0.496 | -1.01 | 1.24E-07 | 4.30E-06 | 0.014 | 0.0259 | 0.0199 | 0.0199 | TRUE | 0.1537 |
| Oard1 | 2094 | 2085 | 1950 | 3198 | 2190 | 2266 | 2224 | 2082 | 1958 | 2514 | 2356 | 2480 | 2269.22 | 2088 | 2450 | 1.174 | 0.231 | 0.00495649 | 0.03110777 | 0.0019 | 0.0084 | 0.0044 | 0.0044 | TRUE | 0.0391 |
| Oas1a | 156 | 188 | 144 | 290 | 205 | 232 | 166 | 188 | 145 | 228 | 221 | 254 | 200.09 | 166 | 234 | 1.409 | 0.495 | 0.00258302 | 0.01884424 | 0.0051 | 0.0234 | 0.0143 | 0.0143 | TRUE | 0.1415 |
| Oas1b | 20 | 18 | 24 | 33 | 29 | 28 | 21 | 18 | 24 | 26 | 31 | 31 | 25.19 | 21 | 29 | 1.381 | 0.466 | 0.22004972 | 0.43222937 | 0 | 0.1377 | 0.063 | 0.063 | TRUE | 0.0961 |
| Oas1c | 412 | 442 | 408 | 380 | 306 | 301 | 438 | 441 | 410 | 299 | 329 | 329 | 374.36 | 430 | 319 | 0.741 | -0.432 | 0.0002097 | 0.0025293 | 0 | 0.0157 | 0.007 | 0.007 | TRUE | 0.0386 |
| Oas1d | 48 | 32 | 26 | 30 | 24 | 26 | 51 | 32 | 26 | 24 | 26 | 28 | 31.15 | 36 | 26 | 0.713 | -0.488 | 0.21134362 | 0.42189873 | 0.0404 | 0.1127 | 0.0767 | 0.0767 | TRUE | 0.8715 |
| Oas1e | 3 | 1 | 1 | 3 | 3 | 1 | 3 | 1 | 1 | 2 | 3 | 1 | 1.98 | 2 | 2 | 1.301 | 0.38 | 0.77577421 | NA | 0 | 1.6721 | 0.7646 | 0.7646 | TRUE | 0.4027 |
| Oas1f | 3 | 4 | 3 | 11 | 9 | 11 | 3 | 4 | 3 | 9 | 10 | 12 | 6.76 | 3 | 10 | 2.962 | 1.567 | 0.03931745 | 0.13899559 | 0 | 0.4942 | 0.2178 | 0.2178 | TRUE | 0.0896 |
| Oas1g | 68 | 75 | 90 | 147 | 121 | 127 | 72 | 75 | 90 | 116 | 130 | 139 | 103.71 | 79 | 128 | 1.614 | 0.691 | 0.00085682 | 0.00789353 | 0 | 0.0387 | 0.0207 | 0.0207 | TRUE | 0.143 |
| Oas1h | 8 | 8 | 5 | 6 | 5 | 2 | 8 | 8 | 5 | 5 | 5 | 2 | 5.63 | 7 | 4 | 0.577 | -0.794 | 0.32708369 | 0.54824943 | 0 | 0.5918 | 0.2839 | 0.2839 | TRUE | 0.2442 |
| Oas2 | 102 | 93 | 100 | 98 | 98 | 85 | 108 | 93 | 100 | 77 | 105 | 93 | 96.19 | 100 | 92 | 0.909 | -0.137 | 0.52710844 | 0.71594509 | 0 | 0.0412 | 0.0236 | 0.0236 | TRUE | 0.2162 |
| Oas3 | 309 | 364 | 337 | 339 | 271 | 343 | 328 | 364 | 338 | 266 | 292 | 375 | 327.28 | 343 | 311 | 0.904 | -0.146 | 0.36450767 | 0.5818841 | 0.0144 | 0.017 | 0.0156 | 0.0156 | TRUE | 0.365 |
| Oasl1 | 87 | 79 | 56 | 72 | 66 | 58 | 92 | 79 | 56 | 57 | 71 | 63 | 69.77 | 76 | 64 | 0.838 | -0.255 | 0.35014242 | 0.56908968 | 0.0237 | 0.0541 | 0.0392 | 0.0392 | TRUE | 0.3335 |
| Oasl2 | 5 | 11 | 9 | 21 | 15 | 11 | 5 | 11 | 9 | 17 | 16 | 12 | 11.67 | 8 | 15 | 1.765 | 0.819 | 0.15414141 | 0.34607548 | 0 | 0.2892 | 0.1453 | 0.1453 | TRUE | 0.3048 |
| Oat | 4019 | 4333 | 4252 | 4792 | 3263 | 3408 | 4269 | 4327 | 4269 | 3767 | 3511 | 3731 | 3979.04 | 4288 | 3670 | 0.856 | -0.225 | 0.00053329 | 0.00544824 | 4.00E-04 | 0.0077 | 0.0028 | 0.0028 | TRUE | 0.0173 |
| Oaz1 | 6 | 5 | 7 | 6 | 8 | 2 | 6 | 5 | 7 | 5 | 9 | 2 | 5.65 | 6 | 5 | 0.842 | -0.248 | 0.76397378 | 0.8758737 | 0 | 0.5898 | 0.3149 | 0.3149 | TRUE | 0.3539 |
| Oaz1-ps | 0 | 0 | 0 | 0 | 0 | 0 | 0 | 0 | 0 | 0 | 0 | 0 | 0 | 0 | 0 | NA | NA | NA | NA | NA | NA | NA | NA | NA | NA |
| Oaz2 | 2910 | 2556 | 2272 | 3989 | 2665 | 2434 | 3091 | 2553 | 2281 | 3136 | 2867 | 2664 | 2765.43 | 2642 | 2889 | 1.094 | 0.13 | 0.3162579 | 0.53698133 | 0.0138 | 0.0081 | 0.0117 | 0.0117 | TRUE | 0.2682 |
| Oaz3 | 2 | 2 | 6 | 3 | 0 | 1 | 2 | 2 | 6 | 2 | 0 | 1 | 2.27 | 3 | 1 | 0.358 | -1.483 | 0.29063803 | NA | 0 | 1.4604 | 0.8667 | 0.8667 | TRUE | 0.6814 |
| Obfc1 | 613 | 717 | 673 | 897 | 613 | 641 | 651 | 716 | 676 | 705 | 660 | 702 | 684.89 | 681 | 689 | 1.012 | 0.017 | 0.85971976 | 0.93089328 | 0 | 0.0117 | 0.0051 | 0.0051 | TRUE | 0.0241 |
| Obox1 | 1 | 0 | 0 | 0 | 1 | 0 | 1 | 0 | 0 | 0 | 1 | 0 | 0.36 | 0 | 0 | 0.954 | -0.067 | 0.98557668 | NA | 0 | 9.2518 | 8.0179 | 8.0179 | TRUE | 0.5204 |
| Obox2 | 0 | 3 | 0 | 3 | 1 | 1 | 0 | 3 | 0 | 2 | 1 | 1 | 1.25 | 1 | 1 | 1.523 | 0.607 | 0.75059079 | NA | 0.5179 | 2.6337 | 1.797 | 1.797 | TRUE | 1.4418 |
| Obox3 | 0 | 1 | 0 | 2 | 5 | 1 | 0 | 1 | 0 | 2 | 5 | 1 | 1.51 | 0 | 3 | 7.272 | 2.862 | 0.12996003 | NA | 0 | 2.1922 | 1.3908 | 1.3908 | TRUE | 0.8468 |
| Obox5 | 2 | 0 | 0 | 1 | 0 | 1 | 2 | 0 | 0 | 1 | 0 | 1 | 0.67 | 1 | 1 | 0.902 | -0.15 | 0.95485009 | NA | 0 | 4.9422 | 3.5197 | 3.5197 | TRUE | 0.9743 |
| Obox6 | 83 | 42 | 35 | 105 | 100 | 102 | 88 | 42 | 35 | 83 | 108 | 112 | 77.84 | 55 | 101 | 1.827 | 0.869 | 0.01570481 | 0.07336973 | 0.1151 | 0.0492 | 0.0791 | 0.0791 | TRUE | 2.2722 |
| Obp1a | 0 | 0 | 0 | 0 | 0 | 0 | 0 | 0 | 0 | 0 | 0 | 0 | 0 | 0 | 0 | NA | NA | NA | NA | NA | NA | NA | NA | NA | NA |
| Obp2a | 0 | 0 | 0 | 0 | 0 | 0 | 0 | 0 | 0 | 0 | 0 | 0 | 0 | 0 | 0 | NA | NA | NA | NA | NA | NA | NA | NA | NA | NA |
| Obp2b | 0 | 0 | 0 | 0 | 0 | 0 | 0 | 0 | 0 | 0 | 0 | 0 | 0 | 0 | 0 | NA | NA | NA | NA | NA | NA | NA | NA | NA | NA |
| Obscn | 67 | 47 | 59 | 105 | 82 | 62 | 71 | 47 | 59 | 83 | 88 | 68 | 69.33 | 59 | 80 | 1.349 | 0.432 | 0.0998136 | 0.26089825 | 0.0124 | 0.0544 | 0.0349 | 0.0349 | TRUE | 0.2542 |
| Obsl1 | 830 | 872 | 896 | 1050 | 747 | 743 | 882 | 871 | 900 | 825 | 804 | 813 | 849.11 | 884 | 814 | 0.921 | -0.118 | 0.15044951 | 0.34117752 | 0 | 0.0108 | 0.0037 | 0.0037 | TRUE | 0.0029 |
| Oc90 | 8 | 0 | 1 | 8 | 4 | 5 | 8 | 0 | 1 | 6 | 4 | 5 | 4.26 | 3 | 5 | 1.713 | 0.776 | 0.52233226 | NA | 0.8475 | 0.78 | 0.8071 | 0.8071 | TRUE | 2.8278 |
| Oca2 | 0 | 2 | 0 | 0 | 3 | 1 | 0 | 2 | 0 | 0 | 3 | 1 | 1.05 | 1 | 1 | 2.106 | 1.075 | 0.64801158 | NA | 2.359 | 3.1346 | 2.8847 | 2.8847 | TRUE | 0.5428 |
| Ocel1 | 390 | 348 | 351 | 502 | 382 | 334 | 414 | 348 | 352 | 395 | 411 | 366 | 380.92 | 371 | 391 | 1.052 | 0.073 | 0.56778337 | 0.74733351 | 0.0037 | 0.0156 | 0.0092 | 0.0092 | TRUE | 0.1154 |
| Ociad1 | 4592 | 5871 | 5971 | 6590 | 4939 | 5195 | 4878 | 5863 | 5996 | 5181 | 5314 | 5687 | 5486.3 | 5579 | 5394 | 0.967 | -0.049 | 0.63234947 | 0.79311458 | 0.006 | 0.0075 | 0.0073 | 0.0073 | TRUE | 0.1473 |
| Ociad2 | 128 | 133 | 157 | 202 | 164 | 130 | 136 | 133 | 158 | 159 | 176 | 142 | 150.66 | 142 | 159 | 1.12 | 0.163 | 0.35718862 | 0.57567739 | 0.0032 | 0.0288 | 0.016 | 0.016 | TRUE | 0.0953 |
| Ocln | 520 | 705 | 700 | 777 | 581 | 602 | 552 | 704 | 703 | 611 | 625 | 659 | 642.37 | 653 | 632 | 0.966 | -0.049 | 0.69733652 | 0.83408871 | 0.0085 | 0.012 | 0.0101 | 0.0101 | TRUE | 0.2131 |
| Ocm | 1 | 0 | 0 | 4 | 5 | 0 | 1 | 0 | 0 | 3 | 5 | 0 | 1.6 | 0 | 3 | 7.786 | 2.961 | 0.15364717 | NA | 0 | 2.0689 | 1.9311 | 1.9311 | TRUE | 0.2166 |
| Ocrl | 2245 | 2337 | 2369 | 3164 | 2266 | 2391 | 2385 | 2334 | 2379 | 2487 | 2438 | 2617 | 2440 | 2366 | 2514 | 1.063 | 0.088 | 0.20144053 | 0.40905369 | 3.00E-04 | 0.0083 | 0.003 | 0.003 | TRUE | 0.0153 |
| Ocstamp | 7 | 18 | 10 | 16 | 14 | 9 | 7 | 18 | 10 | 13 | 15 | 10 | 12.16 | 12 | 13 | 1.055 | 0.078 | 0.89427066 | 0.94899775 | 0.0331 | 0.2779 | 0.164 | 0.164 | TRUE | 0.8192 |
| Odam | 0 | 0 | 0 | 0 | 0 | 0 | 0 | 0 | 0 | 0 | 0 | 0 | 0 | 0 | 0 | NA | NA | NA | NA | NA | NA | NA | NA | NA | NA |
| Odc1 | 36128 | 31928 | 31236 | 48438 | 33294 | 28670 | 38374 | 31887 | 31364 | 38078 | 35822 | 31384 | 34484.94 | 33875 | 35095 | 1.036 | 0.051 | 0.65412375 | 0.80568839 | 0.0094 | 0.007 | 0.0093 | 0.0093 | TRUE | 0.1652 |
| Odf1 | 5 | 1 | 6 | 5 | 2 | 3 | 5 | 1 | 6 | 4 | 2 | 3 | 3.62 | 4 | 3 | 0.769 | -0.38 | 0.71693526 | NA | 0 | 0.9178 | 0.5104 | 0.5104 | TRUE | 0.7684 |
| Odf2 | 10594 | 13132 | 13699 | 14211 | 11096 | 11319 | 11253 | 13115 | 13755 | 11172 | 11939 | 12390 | 12270.6 | 12708 | 11834 | 0.931 | -0.103 | 0.29503352 | 0.51594065 | 0.0065 | 0.0072 | 0.0069 | 0.0069 | TRUE | 0.1226 |
| Odf2l | 778 | 844 | 815 | 1180 | 872 | 862 | 826 | 843 | 818 | 928 | 938 | 944 | 882.84 | 829 | 937 | 1.129 | 0.175 | 0.03051449 | 0.11670091 | 0 | 0.0106 | 0.0036 | 0.0036 | TRUE | 0.0025 |
| Odf3 | 3 | 0 | 1 | 1 | 2 | 6 | 3 | 0 | 1 | 1 | 2 | 7 | 2.28 | 1 | 3 | 2.25 | 1.17 | 0.43884315 | NA | 0.754 | 1.4501 | 1.1198 | 1.1198 | TRUE | 0.6838 |
| Odf3b | 7 | 14 | 19 | 20 | 8 | 14 | 7 | 14 | 19 | 16 | 9 | 15 | 13.36 | 13 | 13 | 0.982 | -0.026 | 0.96422648 | 0.98203715 | 0.0678 | 0.2535 | 0.1673 | 0.1673 | TRUE | 0.2473 |
| Odf3l1 | 3 | 10 | 7 | 9 | 6 | 5 | 3 | 10 | 7 | 7 | 6 | 5 | 6.53 | 7 | 6 | 0.941 | -0.087 | 0.90892951 | 0.95437296 | 0 | 0.5111 | 0.2653 | 0.2653 | TRUE | 0.2412 |
| Odf3l2 | 0 | 1 | 0 | 1 | 3 | 1 | 0 | 1 | 0 | 1 | 3 | 1 | 1.02 | 0 | 2 | 4.595 | 2.2 | 0.28830267 | NA | 0 | 3.2435 | 1.8018 | 1.8018 | TRUE | 0.4455 |
| Odf4 | 0 | 1 | 2 | 1 | 0 | 0 | 0 | 1 | 2 | 1 | 0 | 0 | 0.63 | 1 | 0 | 0.347 | -1.527 | 0.56139244 | NA | 0 | 5.2182 | 3.4771 | 3.4771 | TRUE | 0.1019 |
| Ofcc1 | 0 | 0 | 1 | 0 | 0 | 1 | 0 | 0 | 1 | 0 | 0 | 1 | 0.35 | 0 | 0 | 0.954 | -0.067 | 0.98566485 | NA | 0 | 9.4252 | 8.1416 | 8.1416 | TRUE | 0.5383 |
| Ofd1 | 2177 | 2027 | 2008 | 2826 | 2053 | 1831 | 2312 | 2024 | 2016 | 2222 | 2209 | 2004 | 2131.3 | 2117 | 2145 | 1.013 | 0.019 | 0.8393345 | 0.91841152 | 0.0042 | 0.0084 | 0.0059 | 0.0059 | TRUE | 0.0781 |
| Ogdh | 11790 | 10093 | 10212 | 16822 | 11156 | 9789 | 12523 | 10080 | 10254 | 13224 | 12003 | 10716 | 11466.64 | 10952 | 11981 | 1.094 | 0.13 | 0.28003216 | 0.50109681 | 0.0128 | 0.0072 | 0.0103 | 0.0103 | TRUE | 0.1923 |
| Ogdhl | 5385 | 7144 | 7104 | 7967 | 6383 | 6831 | 5720 | 7135 | 7133 | 6263 | 6868 | 7478 | 6766.02 | 6663 | 6870 | 1.031 | 0.044 | 0.70849845 | 0.84126423 | 0.0117 | 0.0074 | 0.0098 | 0.0098 | TRUE | 0.1868 |
| Ogfod1 | 4744 | 4245 | 4394 | 6139 | 4260 | 3783 | 5039 | 4240 | 4412 | 4826 | 4583 | 4141 | 4540.19 | 4564 | 4517 | 0.99 | -0.014 | 0.8870329 | 0.94442566 | 0.0069 | 0.0076 | 0.0072 | 0.0072 | TRUE | 0.1011 |
| Ogfod2 | 275 | 287 | 315 | 342 | 325 | 287 | 292 | 287 | 316 | 269 | 350 | 314 | 304.62 | 298 | 311 | 1.039 | 0.055 | 0.70541136 | 0.83898258 | 0.0073 | 0.0177 | 0.0121 | 0.0121 | TRUE | 0.1693 |
| Ogfod3 | 530 | 742 | 786 | 832 | 594 | 676 | 563 | 741 | 789 | 654 | 639 | 740 | 687.73 | 698 | 678 | 0.971 | -0.043 | 0.7723482 | 0.88115034 | 0.017 | 0.0117 | 0.0145 | 0.0145 | TRUE | 0.3356 |
| Ogfr | 1524 | 1530 | 1444 | 1766 | 1294 | 1232 | 1619 | 1528 | 1450 | 1388 | 1392 | 1349 | 1454.31 | 1532 | 1376 | 0.899 | -0.154 | 0.05710783 | 0.18003291 | 0.001 | 0.0092 | 0.0041 | 0.0041 | TRUE | 0.0292 |
| Ogfrl1 | 684 | 845 | 898 | 1147 | 892 | 927 | 727 | 844 | 902 | 902 | 960 | 1015 | 891.38 | 824 | 959 | 1.162 | 0.217 | 0.0577033 | 0.18143868 | 0.0065 | 0.0106 | 0.0083 | 0.0083 | TRUE | 0.1264 |

| | | | | | | | | | | | | | | | | | | | | | | | | | |
|---|---|---|---|---|---|---|---|---|---|---|---|---|---|---|---|---|---|---|---|---|---|---|---|---|---|
| Ogg1 | 852 | 973 | 920 | 1344 | 869 | 807 | 905 | 972 | 924 | 1057 | 935 | 883 | 945.9 | 934 | 958 | 1.028 | 0.04 | 0.70062505 | 0.83629834 | 0.004 | 0.0104 | 0.0067 | 0.0067 | TRUE | 0.0989 |
| Ogn | 1 | 0 | 0 | 1 | 1 | 2 | 1 | 0 | 0 | 1 | 1 | 2 | 0.85 | 0 | 1 | 3.644 | 1.866 | 0.38437313 | NA | 0 | 3.8725 | 1.9476 | 1.9476 | TRUE | 0.4614 |
| Ogt | 9524 | 8893 | 8927 | 11608 | 9318 | 8282 | 10116 | 8882 | 8964 | 9125 | 10026 | 9066 | 9363.03 | 9321 | 9406 | 1.009 | 0.013 | 0.88001567 | 0.94067144 | 0.0034 | 0.0073 | 0.0053 | 0.0053 | TRUE | 0.0681 |
| Oip5 | 1809 | 2421 | 2429 | 2299 | 1804 | 2183 | 1921 | 2418 | 2439 | 1807 | 1941 | 2390 | 2152.71 | 2259 | 2046 | 0.905 | -0.144 | 0.30942602 | 0.52983032 | 0.0179 | 0.0084 | 0.014 | 0.014 | TRUE | 0.2606 |
| Oit1 | 4 | 4 | 2 | 12 | 7 | 7 | 4 | 4 | 2 | 9 | 8 | 8 | 5.81 | 3 | 8 | 2.42 | 1.275 | 0.11339465 | 0.28383206 | 0 | 0.5736 | 0.2582 | 0.2582 | TRUE | 0.1928 |
| Oit3 | 0 | 0 | 1 | 4 | 1 | 0 | 0 | 0 | 1 | 3 | 1 | 0 | 0.87 | 0 | 1 | 3.94 | 1.978 | 0.41505635 | NA | 0 | 3.7903 | 2.9051 | 2.9051 | TRUE | 0.2422 |
| Ola1 | 6444 | 6733 | 6784 | 8229 | 5870 | 6180 | 6845 | 6724 | 6812 | 6469 | 6316 | 6765 | 6655.09 | 6794 | 6517 | 0.959 | -0.06 | 0.32625024 | 0.54734802 | 4.00E-04 | 0.0074 | 0.0026 | 0.0026 | TRUE | 0.0134 |
| Olah | 2 | 1 | 0 | 8 | 6 | 6 | 2 | 1 | 0 | 6 | 6 | 7 | 3.74 | 1 | 6 | 6.264 | 2.647 | 0.02683929 | NA | 0 | 0.8879 | 0.457 | 0.457 | TRUE | 0.3775 |
| Olfm1 | 567 | 712 | 621 | 897 | 644 | 628 | 602 | 711 | 624 | 705 | 693 | 687 | 670.4 | 646 | 695 | 1.077 | 0.107 | 0.30935021 | 0.52975803 | 0.0024 | 0.0118 | 0.0064 | 0.0064 | TRUE | 0.093 |
| Olfm2 | 352 | 399 | 384 | 313 | 224 | 218 | 374 | 398 | 386 | 246 | 241 | 239 | 313.94 | 386 | 242 | 0.627 | -0.673 | 1.50E-08 | 6.88E-07 | 0 | 0.0174 | 0.0069 | 0.0069 | TRUE | 0.0092 |
| Olfm3 | 0 | 0 | 0 | 0 | 0 | 0 | 0 | 0 | 0 | 0 | 0 | 0 | 0 | 0 | 0 | NA | NA | NA | NA | NA | NA | NA | NA | NA | NA |
| Olfm4 | 0 | 1 | 0 | 4 | 0 | 2 | 0 | 1 | 0 | 3 | 0 | 2 | 1.06 | 0 | 2 | 4.926 | 2.3 | 0.31109267 | NA | 0 | 3.1284 | 2.445 | 2.445 | TRUE | 0.3249 |
| Olfml1 | 78 | 78 | 75 | 54 | 52 | 64 | 83 | 78 | 75 | 42 | 56 | 70 | 67.42 | 79 | 56 | 0.708 | -0.499 | 0.06231759 | 0.19104136 | 0.0147 | 0.0558 | 0.0363 | 0.0363 | TRUE | 0.2988 |
| Olfml2a | 38 | 33 | 22 | 41 | 23 | 23 | 40 | 33 | 22 | 32 | 25 | 25 | 29.59 | 32 | 27 | 0.868 | -0.205 | 0.5946375 | 0.7677918 | 0.0217 | 0.1182 | 0.073 | 0.073 | TRUE | 0.2795 |
| Olfml2b | 190 | 225 | 227 | 200 | 143 | 128 | 202 | 225 | 228 | 157 | 154 | 140 | 184.28 | 218 | 150 | 0.69 | -0.535 | 0.00052107 | 0.00534696 | 0 | 0.0248 | 0.0115 | 0.0115 | TRUE | 0.0463 |
| Olfml3 | 2 | 0 | 3 | 1 | 1 | 2 | 2 | 0 | 3 | 1 | 1 | 2 | 1.53 | 2 | 1 | 0.776 | -0.365 | 0.81938143 | NA | 0 | 2.1583 | 1.1789 | 1.1789 | TRUE | 0.6096 |
| Olfr1 | 0 | 0 | 0 | 0 | 0 | 0 | 0 | 0 | 0 | 0 | 0 | 0 | 0 | 0 | 0 | NA | NA | NA | NA | NA | NA | NA | NA | NA | NA |
| Olfr10 | 0 | 0 | 0 | 0 | 0 | 0 | 0 | 0 | 0 | 0 | 0 | 0 | 0 | 0 | 0 | NA | NA | NA | NA | NA | NA | NA | NA | NA | NA |
| Olfr100 | 0 | 0 | 0 | 0 | 0 | 0 | 0 | 0 | 0 | 0 | 0 | 0 | 0 | 0 | 0 | NA | NA | NA | NA | NA | NA | NA | NA | NA | NA |
| Olfr1000 | 0 | 0 | 0 | 0 | 0 | 0 | 0 | 0 | 0 | 0 | 0 | 0 | 0 | 0 | 0 | NA | NA | NA | NA | NA | NA | NA | NA | NA | NA |
| Olfr1002 | 0 | 0 | 0 | 0 | 0 | 0 | 0 | 0 | 0 | 0 | 0 | 0 | 0 | 0 | 0 | NA | NA | NA | NA | NA | NA | NA | NA | NA | NA |
| Olfr1006 | 0 | 0 | 0 | 0 | 0 | 0 | 0 | 0 | 0 | 0 | 0 | 0 | 0 | 0 | 0 | NA | NA | NA | NA | NA | NA | NA | NA | NA | NA |
| Olfr1008 | 0 | 0 | 0 | 0 | 0 | 0 | 0 | 0 | 0 | 0 | 0 | 0 | 0 | 0 | 0 | NA | NA | NA | NA | NA | NA | NA | NA | NA | NA |
| Olfr1009 | 0 | 0 | 0 | 0 | 0 | 0 | 0 | 0 | 0 | 0 | 0 | 0 | 0 | 0 | 0 | NA | NA | NA | NA | NA | NA | NA | NA | NA | NA |
| Olfr101 | 0 | 0 | 0 | 0 | 0 | 0 | 0 | 0 | 0 | 0 | 0 | 0 | 0 | 0 | 0 | NA | NA | NA | NA | NA | NA | NA | NA | NA | NA |
| Olfr1010 | 0 | 0 | 0 | 0 | 0 | 0 | 0 | 0 | 0 | 0 | 0 | 0 | 0 | 0 | 0 | NA | NA | NA | NA | NA | NA | NA | NA | NA | NA |
| Olfr1012 | 0 | 0 | 0 | 0 | 0 | 0 | 0 | 0 | 0 | 0 | 0 | 0 | 0 | 0 | 0 | NA | NA | NA | NA | NA | NA | NA | NA | NA | NA |
| Olfr1013 | 0 | 0 | 0 | 0 | 0 | 0 | 0 | 0 | 0 | 0 | 0 | 0 | 0 | 0 | 0 | NA | NA | NA | NA | NA | NA | NA | NA | NA | NA |
| Olfr1014 | 0 | 0 | 0 | 0 | 0 | 0 | 0 | 0 | 0 | 0 | 0 | 0 | 0 | 0 | 0 | NA | NA | NA | NA | NA | NA | NA | NA | NA | NA |
| Olfr1015 | 0 | 0 | 0 | 0 | 0 | 0 | 0 | 0 | 0 | 0 | 0 | 0 | 0 | 0 | 0 | NA | NA | NA | NA | NA | NA | NA | NA | NA | NA |
| Olfr1016 | 0 | 0 | 0 | 0 | 0 | 0 | 0 | 0 | 0 | 0 | 0 | 0 | 0 | 0 | 0 | NA | NA | NA | NA | NA | NA | NA | NA | NA | NA |
| Olfr1018 | 0 | 0 | 0 | 0 | 0 | 0 | 0 | 0 | 0 | 0 | 0 | 0 | 0 | 0 | 0 | NA | NA | NA | NA | NA | NA | NA | NA | NA | NA |
| Olfr1019 | 0 | 0 | 0 | 0 | 0 | 0 | 0 | 0 | 0 | 0 | 0 | 0 | 0 | 0 | 0 | NA | NA | NA | NA | NA | NA | NA | NA | NA | NA |
| Olfr102 | 0 | 0 | 0 | 0 | 0 | 0 | 0 | 0 | 0 | 0 | 0 | 0 | 0 | 0 | 0 | NA | NA | NA | NA | NA | NA | NA | NA | NA | NA |
| Olfr1020 | 0 | 0 | 0 | 0 | 0 | 0 | 0 | 0 | 0 | 0 | 0 | 0 | 0 | 0 | 0 | NA | NA | NA | NA | NA | NA | NA | NA | NA | NA |
| Olfr1022 | 0 | 0 | 0 | 0 | 0 | 0 | 0 | 0 | 0 | 0 | 0 | 0 | 0 | 0 | 0 | NA | NA | NA | NA | NA | NA | NA | NA | NA | NA |
| Olfr1023 | 0 | 0 | 0 | 0 | 0 | 0 | 0 | 0 | 0 | 0 | 0 | 0 | 0 | 0 | 0 | NA | NA | NA | NA | NA | NA | NA | NA | NA | NA |
| Olfr1024 | 0 | 0 | 0 | 0 | 0 | 0 | 0 | 0 | 0 | 0 | 0 | 0 | 0 | 0 | 0 | NA | NA | NA | NA | NA | NA | NA | NA | NA | NA |
| Olfr1026 | 0 | 0 | 0 | 0 | 0 | 0 | 0 | 0 | 0 | 0 | 0 | 0 | 0 | 0 | 0 | NA | NA | NA | NA | NA | NA | NA | NA | NA | NA |
| Olfr1028 | 0 | 0 | 0 | 0 | 0 | 0 | 0 | 0 | 0 | 0 | 0 | 0 | 0 | 0 | 0 | NA | NA | NA | NA | NA | NA | NA | NA | NA | NA |
| Olfr1029 | 0 | 0 | 0 | 0 | 0 | 0 | 0 | 0 | 0 | 0 | 0 | 0 | 0 | 0 | 0 | NA | NA | NA | NA | NA | NA | NA | NA | NA | NA |
| Olfr103 | 0 | 0 | 0 | 0 | 0 | 0 | 0 | 0 | 0 | 0 | 0 | 0 | 0 | 0 | 0 | NA | NA | NA | NA | NA | NA | NA | NA | NA | NA |
| Olfr1030 | 0 | 0 | 0 | 0 | 0 | 0 | 0 | 0 | 0 | 0 | 0 | 0 | 0 | 0 | 0 | NA | NA | NA | NA | NA | NA | NA | NA | NA | NA |
| Olfr1031 | 0 | 0 | 0 | 0 | 0 | 0 | 0 | 0 | 0 | 0 | 0 | 0 | 0 | 0 | 0 | NA | NA | NA | NA | NA | NA | NA | NA | NA | NA |
| Olfr1032 | 0 | 0 | 0 | 0 | 0 | 0 | 0 | 0 | 0 | 0 | 0 | 0 | 0 | 0 | 0 | NA | NA | NA | NA | NA | NA | NA | NA | NA | NA |
| Olfr1033 | 0 | 0 | 0 | 0 | 0 | 0 | 0 | 0 | 0 | 0 | 0 | 0 | 0 | 0 | 0 | NA | NA | NA | NA | NA | NA | NA | NA | NA | NA |
| Olfr1034 | 0 | 0 | 0 | 0 | 0 | 0 | 0 | 0 | 0 | 0 | 0 | 0 | 0 | 0 | 0 | NA | NA | NA | NA | NA | NA | NA | NA | NA | NA |
| Olfr1036 | 0 | 0 | 0 | 0 | 0 | 0 | 0 | 0 | 0 | 0 | 0 | 0 | 0 | 0 | 0 | NA | NA | NA | NA | NA | NA | NA | NA | NA | NA |
| Olfr1037 | 0 | 0 | 0 | 0 | 0 | 0 | 0 | 0 | 0 | 0 | 0 | 0 | 0 | 0 | 0 | NA | NA | NA | NA | NA | NA | NA | NA | NA | NA |
| Olfr1038-ps | 0 | 0 | 0 | 0 | 0 | 0 | 0 | 0 | 0 | 0 | 0 | 0 | 0 | 0 | 0 | NA | NA | NA | NA | NA | NA | NA | NA | NA | NA |
| Olfr1039 | 0 | 0 | 0 | 0 | 0 | 0 | 0 | 0 | 0 | 0 | 0 | 0 | 0 | 0 | 0 | NA | NA | NA | NA | NA | NA | NA | NA | NA | NA |
| Olfr1040 | 0 | 0 | 0 | 0 | 0 | 0 | 0 | 0 | 0 | 0 | 0 | 0 | 0 | 0 | 0 | NA | NA | NA | NA | NA | NA | NA | NA | NA | NA |
| Olfr1042 | 0 | 0 | 0 | 0 | 0 | 0 | 0 | 0 | 0 | 0 | 0 | 0 | 0 | 0 | 0 | NA | NA | NA | NA | NA | NA | NA | NA | NA | NA |
| Olfr1043 | 1 | 0 | 0 | 0 | 0 | 0 | 1 | 0 | 0 | 0 | 0 | 0 | 0.18 | 0 | 0 | 0.49 | -1.029 | 0.80087 | NA | 0 | 18.6165 | 10 | 10 | TRUE | 0.4614 |
| Olfr1044 | 0 | 0 | 0 | 0 | 0 | 0 | 0 | 0 | 0 | 0 | 0 | 0 | 0 | 0 | 0 | NA | NA | NA | NA | NA | NA | NA | NA | NA | NA |
| Olfr1045 | 0 | 0 | 0 | 0 | 0 | 0 | 0 | 0 | 0 | 0 | 0 | 0 | 0 | 0 | 0 | NA | NA | NA | NA | NA | NA | NA | NA | NA | NA |
| Olfr1046 | 0 | 0 | 0 | 0 | 0 | 0 | 0 | 0 | 0 | 0 | 0 | 0 | 0 | 0 | 0 | NA | NA | NA | NA | NA | NA | NA | NA | NA | NA |
| Olfr1047 | 0 | 0 | 0 | 0 | 0 | 0 | 0 | 0 | 0 | 0 | 0 | 0 | 0 | 0 | 0 | NA | NA | NA | NA | NA | NA | NA | NA | NA | NA |
| Olfr1048 | 0 | 0 | 0 | 0 | 0 | 0 | 0 | 0 | 0 | 0 | 0 | 0 | 0 | 0 | 0 | NA | NA | NA | NA | NA | NA | NA | NA | NA | NA |
| Olfr1049 | 0 | 0 | 0 | 0 | 0 | 0 | 0 | 0 | 0 | 0 | 0 | 0 | 0 | 0 | 0 | NA | NA | NA | NA | NA | NA | NA | NA | NA | NA |
| Olfr1051 | 0 | 0 | 0 | 0 | 0 | 0 | 0 | 0 | 0 | 0 | 0 | 0 | 0 | 0 | 0 | NA | NA | NA | NA | NA | NA | NA | NA | NA | NA |
| Olfr1052 | 0 | 0 | 0 | 0 | 0 | 0 | 0 | 0 | 0 | 0 | 0 | 0 | 0 | 0 | 0 | NA | NA | NA | NA | NA | NA | NA | NA | NA | NA |
| Olfr1053 | 0 | 0 | 0 | 0 | 0 | 0 | 0 | 0 | 0 | 0 | 0 | 0 | 0 | 0 | 0 | NA | NA | NA | NA | NA | NA | NA | NA | NA | NA |
| Olfr1054 | 0 | 0 | 0 | 0 | 0 | 0 | 0 | 0 | 0 | 0 | 0 | 0 | 0 | 0 | 0 | NA | NA | NA | NA | NA | NA | NA | NA | NA | NA |
| Olfr1055 | 0 | 0 | 0 | 0 | 0 | 0 | 0 | 0 | 0 | 0 | 0 | 0 | 0 | 0 | 0 | NA | NA | NA | NA | NA | NA | NA | NA | NA | NA |
| Olfr1056 | 0 | 0 | 0 | 0 | 0 | 0 | 0 | 0 | 0 | 0 | 0 | 0 | 0 | 0 | 0 | NA | NA | NA | NA | NA | NA | NA | NA | NA | NA |
| Olfr1057 | 0 | 0 | 0 | 0 | 0 | 0 | 0 | 0 | 0 | 0 | 0 | 0 | 0 | 0 | 0 | NA | NA | NA | NA | NA | NA | NA | NA | NA | NA |
| Olfr1058 | 0 | 0 | 0 | 0 | 0 | 0 | 0 | 0 | 0 | 0 | 0 | 0 | 0 | 0 | 0 | NA | NA | NA | NA | NA | NA | NA | NA | NA | NA |
| Olfr1061 | 0 | 0 | 0 | 0 | 0 | 0 | 0 | 0 | 0 | 0 | 0 | 0 | 0 | 0 | 0 | NA | NA | NA | NA | NA | NA | NA | NA | NA | NA |

| | | | | | | | | | | | | | | | | | | | | | | | | | |
|---|---|---|---|---|---|---|---|---|---|---|---|---|---|---|---|---|---|---|---|---|---|---|---|---|---|
| Olfr1062 | 0 | 0 | 0 | 0 | 0 | 0 | 0 | 0 | 0 | 0 | 0 | 0 | 0 | 0 | 0 | NA | NA | NA | NA | NA | NA | NA | NA | NA | NA |
| Olfr1065 | 0 | 0 | 0 | 0 | 0 | 0 | 0 | 0 | 0 | 0 | 0 | 0 | 0 | 0 | 0 | NA | NA | NA | NA | NA | NA | NA | NA | NA | NA |
| Olfr1066 | 0 | 0 | 0 | 0 | 0 | 0 | 0 | 0 | 0 | 0 | 0 | 0 | 0 | 0 | 0 | NA | NA | NA | NA | NA | NA | NA | NA | NA | NA |
| Olfr107 | 0 | 0 | 0 | 0 | 0 | 0 | 0 | 0 | 0 | 0 | 0 | 0 | 0 | 0 | 0 | NA | NA | NA | NA | NA | NA | NA | NA | NA | NA |
| Olfr1076 | 0 | 0 | 0 | 0 | 0 | 0 | 0 | 0 | 0 | 0 | 0 | 0 | 0 | 0 | 0 | NA | NA | NA | NA | NA | NA | NA | NA | NA | NA |
| Olfr1077-ps1 | 0 | 0 | 0 | 0 | 0 | 0 | 0 | 0 | 0 | 0 | 0 | 0 | 0 | 0 | 0 | NA | NA | NA | NA | NA | NA | NA | NA | NA | NA |
| Olfr1079 | 0 | 0 | 0 | 0 | 0 | 0 | 0 | 0 | 0 | 0 | 0 | 0 | 0 | 0 | 0 | NA | NA | NA | NA | NA | NA | NA | NA | NA | NA |
| Olfr108 | 0 | 0 | 0 | 0 | 0 | 0 | 0 | 0 | 0 | 0 | 0 | 0 | 0 | 0 | 0 | NA | NA | NA | NA | NA | NA | NA | NA | NA | NA |
| Olfr1080 | 0 | 0 | 0 | 0 | 0 | 0 | 0 | 0 | 0 | 0 | 0 | 0 | 0 | 0 | 0 | NA | NA | NA | NA | NA | NA | NA | NA | NA | NA |
| Olfr1082 | 0 | 0 | 0 | 0 | 0 | 0 | 0 | 0 | 0 | 0 | 0 | 0 | 0 | 0 | 0 | NA | NA | NA | NA | NA | NA | NA | NA | NA | NA |
| Olfr1084 | 0 | 0 | 0 | 0 | 0 | 0 | 0 | 0 | 0 | 0 | 0 | 0 | 0 | 0 | 0 | NA | NA | NA | NA | NA | NA | NA | NA | NA | NA |
| Olfr1085 | 0 | 0 | 0 | 0 | 0 | 0 | 0 | 0 | 0 | 0 | 0 | 0 | 0 | 0 | 0 | NA | NA | NA | NA | NA | NA | NA | NA | NA | NA |
| Olfr1086 | 0 | 0 | 0 | 0 | 0 | 0 | 0 | 0 | 0 | 0 | 0 | 0 | 0 | 0 | 0 | NA | NA | NA | NA | NA | NA | NA | NA | NA | NA |
| Olfr1087 | 0 | 0 | 0 | 0 | 0 | 0 | 0 | 0 | 0 | 0 | 0 | 0 | 0 | 0 | 0 | NA | NA | NA | NA | NA | NA | NA | NA | NA | NA |
| Olfr1089 | 0 | 0 | 0 | 0 | 0 | 0 | 0 | 0 | 0 | 0 | 0 | 0 | 0 | 0 | 0 | NA | NA | NA | NA | NA | NA | NA | NA | NA | NA |
| Olfr109 | 0 | 0 | 0 | 0 | 0 | 0 | 0 | 0 | 0 | 0 | 0 | 0 | 0 | 0 | 0 | NA | NA | NA | NA | NA | NA | NA | NA | NA | NA |
| Olfr1090 | 0 | 0 | 0 | 0 | 0 | 0 | 0 | 0 | 0 | 0 | 0 | 0 | 0 | 0 | 0 | NA | NA | NA | NA | NA | NA | NA | NA | NA | NA |
| Olfr1093 | 0 | 0 | 0 | 0 | 0 | 0 | 0 | 0 | 0 | 0 | 0 | 0 | 0 | 0 | 0 | NA | NA | NA | NA | NA | NA | NA | NA | NA | NA |
| Olfr1094 | 0 | 0 | 0 | 0 | 0 | 0 | 0 | 0 | 0 | 0 | 0 | 0 | 0 | 0 | 0 | NA | NA | NA | NA | NA | NA | NA | NA | NA | NA |
| Olfr1095 | 0 | 0 | 0 | 0 | 0 | 0 | 0 | 0 | 0 | 0 | 0 | 0 | 0 | 0 | 0 | NA | NA | NA | NA | NA | NA | NA | NA | NA | NA |
| Olfr1097 | 0 | 0 | 0 | 0 | 0 | 0 | 0 | 0 | 0 | 0 | 0 | 0 | 0 | 0 | 0 | NA | NA | NA | NA | NA | NA | NA | NA | NA | NA |
| Olfr1098 | 0 | 0 | 0 | 0 | 0 | 0 | 0 | 0 | 0 | 0 | 0 | 0 | 0 | 0 | 0 | NA | NA | NA | NA | NA | NA | NA | NA | NA | NA |
| Olfr1099 | 0 | 0 | 0 | 0 | 0 | 0 | 0 | 0 | 0 | 0 | 0 | 0 | 0 | 0 | 0 | NA | NA | NA | NA | NA | NA | NA | NA | NA | NA |
| Olfr11 | 0 | 0 | 0 | 0 | 0 | 0 | 0 | 0 | 0 | 0 | 0 | 0 | 0 | 0 | 0 | NA | NA | NA | NA | NA | NA | NA | NA | NA | NA |
| Olfr110 | 0 | 0 | 0 | 0 | 0 | 0 | 0 | 0 | 0 | 0 | 0 | 0 | 0 | 0 | 0 | NA | NA | NA | NA | NA | NA | NA | NA | NA | NA |
| Olfr1100 | 0 | 0 | 0 | 0 | 0 | 0 | 0 | 0 | 0 | 0 | 0 | 0 | 0 | 0 | 0 | NA | NA | NA | NA | NA | NA | NA | NA | NA | NA |
| Olfr1101 | 0 | 0 | 0 | 0 | 0 | 0 | 0 | 0 | 0 | 0 | 0 | 0 | 0 | 0 | 0 | NA | NA | NA | NA | NA | NA | NA | NA | NA | NA |
| Olfr1102 | 0 | 0 | 0 | 0 | 0 | 0 | 0 | 0 | 0 | 0 | 0 | 0 | 0 | 0 | 0 | NA | NA | NA | NA | NA | NA | NA | NA | NA | NA |
| Olfr1104 | 0 | 0 | 0 | 0 | 0 | 0 | 0 | 0 | 0 | 0 | 0 | 0 | 0 | 0 | 0 | NA | NA | NA | NA | NA | NA | NA | NA | NA | NA |
| Olfr1105 | 0 | 0 | 0 | 0 | 0 | 0 | 0 | 0 | 0 | 0 | 0 | 0 | 0 | 0 | 0 | NA | NA | NA | NA | NA | NA | NA | NA | NA | NA |
| Olfr1106 | 0 | 0 | 0 | 0 | 0 | 0 | 0 | 0 | 0 | 0 | 0 | 0 | 0 | 0 | 0 | NA | NA | NA | NA | NA | NA | NA | NA | NA | NA |
| Olfr1107 | 0 | 0 | 0 | 0 | 0 | 0 | 0 | 0 | 0 | 0 | 0 | 0 | 0 | 0 | 0 | NA | NA | NA | NA | NA | NA | NA | NA | NA | NA |
| Olfr1109 | 0 | 0 | 0 | 0 | 0 | 0 | 0 | 0 | 0 | 0 | 0 | 0 | 0 | 0 | 0 | NA | NA | NA | NA | NA | NA | NA | NA | NA | NA |
| Olfr111 | 0 | 0 | 0 | 0 | 0 | 0 | 0 | 0 | 0 | 0 | 0 | 0 | 0 | 0 | 0 | NA | NA | NA | NA | NA | NA | NA | NA | NA | NA |
| Olfr1110 | 0 | 0 | 0 | 0 | 0 | 0 | 0 | 0 | 0 | 0 | 0 | 0 | 0 | 0 | 0 | NA | NA | NA | NA | NA | NA | NA | NA | NA | NA |
| Olfr1111 | 0 | 0 | 0 | 0 | 0 | 0 | 0 | 0 | 0 | 0 | 0 | 0 | 0 | 0 | 0 | NA | NA | NA | NA | NA | NA | NA | NA | NA | NA |
| Olfr1112 | 0 | 0 | 0 | 0 | 0 | 0 | 0 | 0 | 0 | 0 | 0 | 0 | 0 | 0 | 0 | NA | NA | NA | NA | NA | NA | NA | NA | NA | NA |
| Olfr1113 | 0 | 0 | 0 | 0 | 0 | 0 | 0 | 0 | 0 | 0 | 0 | 0 | 0 | 0 | 0 | NA | NA | NA | NA | NA | NA | NA | NA | NA | NA |
| Olfr1115 | 0 | 0 | 0 | 0 | 0 | 0 | 0 | 0 | 0 | 0 | 0 | 0 | 0 | 0 | 0 | NA | NA | NA | NA | NA | NA | NA | NA | NA | NA |
| Olfr1116-ps | 0 | 0 | 0 | 0 | 0 | 0 | 0 | 0 | 0 | 0 | 0 | 0 | 0 | 0 | 0 | NA | NA | NA | NA | NA | NA | NA | NA | NA | NA |
| Olfr1118 | 0 | 0 | 0 | 0 | 0 | 0 | 0 | 0 | 0 | 0 | 0 | 0 | 0 | 0 | 0 | NA | NA | NA | NA | NA | NA | NA | NA | NA | NA |
| Olfr112 | 0 | 0 | 0 | 0 | 0 | 0 | 0 | 0 | 0 | 0 | 0 | 0 | 0 | 0 | 0 | NA | NA | NA | NA | NA | NA | NA | NA | NA | NA |
| Olfr1120 | 0 | 0 | 0 | 0 | 0 | 0 | 0 | 0 | 0 | 0 | 0 | 0 | 0 | 0 | 0 | NA | NA | NA | NA | NA | NA | NA | NA | NA | NA |
| Olfr1121 | 0 | 0 | 0 | 0 | 0 | 0 | 0 | 0 | 0 | 0 | 0 | 0 | 0 | 0 | 0 | NA | NA | NA | NA | NA | NA | NA | NA | NA | NA |
| Olfr1122 | 0 | 0 | 0 | 0 | 0 | 0 | 0 | 0 | 0 | 0 | 0 | 0 | 0 | 0 | 0 | NA | NA | NA | NA | NA | NA | NA | NA | NA | NA |
| Olfr1123 | 0 | 0 | 0 | 0 | 0 | 0 | 0 | 0 | 0 | 0 | 0 | 0 | 0 | 0 | 0 | NA | NA | NA | NA | NA | NA | NA | NA | NA | NA |
| Olfr1124 | 0 | 0 | 0 | 0 | 0 | 0 | 0 | 0 | 0 | 0 | 0 | 0 | 0 | 0 | 0 | NA | NA | NA | NA | NA | NA | NA | NA | NA | NA |
| Olfr1126 | 0 | 0 | 0 | 0 | 0 | 0 | 0 | 0 | 0 | 0 | 0 | 0 | 0 | 0 | 0 | NA | NA | NA | NA | NA | NA | NA | NA | NA | NA |
| Olfr1128 | 0 | 0 | 0 | 0 | 0 | 0 | 0 | 0 | 0 | 0 | 0 | 0 | 0 | 0 | 0 | NA | NA | NA | NA | NA | NA | NA | NA | NA | NA |
| Olfr1129 | 0 | 0 | 0 | 0 | 0 | 0 | 0 | 0 | 0 | 0 | 0 | 0 | 0 | 0 | 0 | NA | NA | NA | NA | NA | NA | NA | NA | NA | NA |
| Olfr113 | 0 | 0 | 0 | 0 | 0 | 0 | 0 | 0 | 0 | 0 | 0 | 0 | 0 | 0 | 0 | NA | NA | NA | NA | NA | NA | NA | NA | NA | NA |
| Olfr1130 | 0 | 0 | 0 | 0 | 0 | 0 | 0 | 0 | 0 | 0 | 0 | 0 | 0 | 0 | 0 | NA | NA | NA | NA | NA | NA | NA | NA | NA | NA |
| Olfr1131 | 0 | 0 | 0 | 0 | 0 | 0 | 0 | 0 | 0 | 0 | 0 | 0 | 0 | 0 | 0 | NA | NA | NA | NA | NA | NA | NA | NA | NA | NA |
| Olfr1132 | 0 | 0 | 0 | 0 | 0 | 0 | 0 | 0 | 0 | 0 | 0 | 0 | 0 | 0 | 0 | NA | NA | NA | NA | NA | NA | NA | NA | NA | NA |
| Olfr1133 | 0 | 0 | 0 | 0 | 0 | 0 | 0 | 0 | 0 | 0 | 0 | 0 | 0 | 0 | 0 | NA | NA | NA | NA | NA | NA | NA | NA | NA | NA |
| Olfr1134 | 0 | 0 | 0 | 0 | 0 | 0 | 0 | 0 | 0 | 0 | 0 | 0 | 0 | 0 | 0 | NA | NA | NA | NA | NA | NA | NA | NA | NA | NA |
| Olfr1135 | 0 | 0 | 0 | 0 | 0 | 0 | 0 | 0 | 0 | 0 | 0 | 0 | 0 | 0 | 0 | NA | NA | NA | NA | NA | NA | NA | NA | NA | NA |
| Olfr1136 | 0 | 0 | 0 | 0 | 0 | 0 | 0 | 0 | 0 | 0 | 0 | 0 | 0 | 0 | 0 | NA | NA | NA | NA | NA | NA | NA | NA | NA | NA |
| Olfr1137 | 0 | 0 | 0 | 0 | 0 | 0 | 0 | 0 | 0 | 0 | 0 | 0 | 0 | 0 | 0 | NA | NA | NA | NA | NA | NA | NA | NA | NA | NA |
| Olfr1138 | 0 | 0 | 0 | 0 | 0 | 0 | 0 | 0 | 0 | 0 | 0 | 0 | 0 | 0 | 0 | NA | NA | NA | NA | NA | NA | NA | NA | NA | NA |
| Olfr114 | 0 | 0 | 0 | 0 | 0 | 0 | 0 | 0 | 0 | 0 | 0 | 0 | 0 | 0 | 0 | NA | NA | NA | NA | NA | NA | NA | NA | NA | NA |
| Olfr1140 | 0 | 0 | 0 | 0 | 0 | 0 | 0 | 0 | 0 | 0 | 0 | 0 | 0 | 0 | 0 | NA | NA | NA | NA | NA | NA | NA | NA | NA | NA |
| Olfr1141 | 0 | 0 | 0 | 0 | 0 | 0 | 0 | 0 | 0 | 0 | 0 | 0 | 0 | 0 | 0 | NA | NA | NA | NA | NA | NA | NA | NA | NA | NA |
| Olfr1143 | 0 | 0 | 0 | 0 | 0 | 0 | 0 | 0 | 0 | 0 | 0 | 0 | 0 | 0 | 0 | NA | NA | NA | NA | NA | NA | NA | NA | NA | NA |
| Olfr1145 | 0 | 0 | 0 | 0 | 0 | 0 | 0 | 0 | 0 | 0 | 0 | 0 | 0 | 0 | 0 | NA | NA | NA | NA | NA | NA | NA | NA | NA | NA |
| Olfr1148 | 0 | 0 | 0 | 0 | 0 | 0 | 0 | 0 | 0 | 0 | 0 | 0 | 0 | 0 | 0 | NA | NA | NA | NA | NA | NA | NA | NA | NA | NA |
| Olfr115 | 0 | 0 | 0 | 0 | 0 | 0 | 0 | 0 | 0 | 0 | 0 | 0 | 0 | 0 | 0 | NA | NA | NA | NA | NA | NA | NA | NA | NA | NA |
| Olfr1151 | 0 | 0 | 0 | 0 | 0 | 0 | 0 | 0 | 0 | 0 | 0 | 0 | 0 | 0 | 0 | NA | NA | NA | NA | NA | NA | NA | NA | NA | NA |
| Olfr1152 | 0 | 0 | 0 | 0 | 0 | 0 | 0 | 0 | 0 | 0 | 0 | 0 | 0 | 0 | 0 | NA | NA | NA | NA | NA | NA | NA | NA | NA | NA |

| | | | | | | | | | | | | | | | | | | | | | | | | | |
|---|---|---|---|---|---|---|---|---|---|---|---|---|---|---|---|---|---|---|---|---|---|---|---|---|---|
| Olfr1153 | 0 | 0 | 0 | 0 | 0 | 0 | 0 | 0 | 0 | 0 | 0 | 0 | 0 | 0 | 0 | NA | NA | NA | NA | NA | NA | NA | NA | NA | NA |
| Olfr1154 | 0 | 0 | 0 | 0 | 0 | 0 | 0 | 0 | 0 | 0 | 0 | 0 | 0 | 0 | 0 | NA | NA | NA | NA | NA | NA | NA | NA | NA | NA |
| Olfr1155 | 0 | 0 | 0 | 0 | 0 | 0 | 0 | 0 | 0 | 0 | 0 | 0 | 0 | 0 | 0 | NA | NA | NA | NA | NA | NA | NA | NA | NA | NA |
| Olfr1156 | 0 | 0 | 0 | 0 | 0 | 0 | 0 | 0 | 0 | 0 | 0 | 0 | 0 | 0 | 0 | NA | NA | NA | NA | NA | NA | NA | NA | NA | NA |
| Olfr1157 | 0 | 0 | 0 | 0 | 0 | 0 | 0 | 0 | 0 | 0 | 0 | 0 | 0 | 0 | 0 | NA | NA | NA | NA | NA | NA | NA | NA | NA | NA |
| Olfr1158 | 0 | 0 | 0 | 0 | 0 | 0 | 0 | 0 | 0 | 0 | 0 | 0 | 0 | 0 | 0 | NA | NA | NA | NA | NA | NA | NA | NA | NA | NA |
| Olfr116 | 0 | 0 | 0 | 0 | 0 | 0 | 0 | 0 | 0 | 0 | 0 | 0 | 0 | 0 | 0 | NA | NA | NA | NA | NA | NA | NA | NA | NA | NA |
| Olfr1160 | 0 | 0 | 0 | 0 | 0 | 0 | 0 | 0 | 0 | 0 | 0 | 0 | 0 | 0 | 0 | NA | NA | NA | NA | NA | NA | NA | NA | NA | NA |
| Olfr1161 | 0 | 0 | 0 | 0 | 0 | 0 | 0 | 0 | 0 | 0 | 0 | 0 | 0 | 0 | 0 | NA | NA | NA | NA | NA | NA | NA | NA | NA | NA |
| Olfr1162 | 0 | 0 | 0 | 0 | 0 | 0 | 0 | 0 | 0 | 0 | 0 | 0 | 0 | 0 | 0 | NA | NA | NA | NA | NA | NA | NA | NA | NA | NA |
| Olfr1163 | 0 | 0 | 0 | 0 | 0 | 0 | 0 | 0 | 0 | 0 | 0 | 0 | 0 | 0 | 0 | NA | NA | NA | NA | NA | NA | NA | NA | NA | NA |
| Olfr1164 | 0 | 0 | 0 | 0 | 0 | 0 | 0 | 0 | 0 | 0 | 0 | 0 | 0 | 0 | 0 | NA | NA | NA | NA | NA | NA | NA | NA | NA | NA |
| Olfr1166 | 0 | 0 | 0 | 0 | 0 | 0 | 0 | 0 | 0 | 0 | 0 | 0 | 0 | 0 | 0 | NA | NA | NA | NA | NA | NA | NA | NA | NA | NA |
| Olfr1167 | 0 | 0 | 0 | 0 | 0 | 0 | 0 | 0 | 0 | 0 | 0 | 0 | 0 | 0 | 0 | NA | NA | NA | NA | NA | NA | NA | NA | NA | NA |
| Olfr1168 | 0 | 0 | 0 | 0 | 0 | 0 | 0 | 0 | 0 | 0 | 0 | 0 | 0 | 0 | 0 | NA | NA | NA | NA | NA | NA | NA | NA | NA | NA |
| Olfr117 | 0 | 0 | 0 | 0 | 0 | 0 | 0 | 0 | 0 | 0 | 0 | 0 | 0 | 0 | 0 | NA | NA | NA | NA | NA | NA | NA | NA | NA | NA |
| Olfr1170 | 0 | 0 | 0 | 0 | 0 | 0 | 0 | 0 | 0 | 0 | 0 | 0 | 0 | 0 | 0 | NA | NA | NA | NA | NA | NA | NA | NA | NA | NA |
| Olfr1173 | 0 | 0 | 0 | 0 | 0 | 0 | 0 | 0 | 0 | 0 | 0 | 0 | 0 | 0 | 0 | NA | NA | NA | NA | NA | NA | NA | NA | NA | NA |
| Olfr1176 | 0 | 0 | 0 | 0 | 0 | 0 | 0 | 0 | 0 | 0 | 0 | 0 | 0 | 0 | 0 | NA | NA | NA | NA | NA | NA | NA | NA | NA | NA |
| Olfr1178 | 0 | 0 | 0 | 0 | 0 | 0 | 0 | 0 | 0 | 0 | 0 | 0 | 0 | 0 | 0 | NA | NA | NA | NA | NA | NA | NA | NA | NA | NA |
| Olfr1179 | 0 | 0 | 0 | 0 | 0 | 0 | 0 | 0 | 0 | 0 | 0 | 0 | 0 | 0 | 0 | NA | NA | NA | NA | NA | NA | NA | NA | NA | NA |
| Olfr118 | 1 | 0 | 0 | 0 | 1 | 1 | 1 | 0 | 0 | 0 | 1 | 1 | 0.54 | 0 | 1 | 1.923 | 0.944 | 0.73523723 | NA | 0 | 6.1214 | 3.9144 | 3.9144 | TRUE | 0.4614 |
| Olfr1180 | 0 | 0 | 0 | 0 | 0 | 0 | 0 | 0 | 0 | 0 | 0 | 0 | 0 | 0 | 0 | NA | NA | NA | NA | NA | NA | NA | NA | NA | NA |
| Olfr1181 | 0 | 0 | 0 | 0 | 0 | 0 | 0 | 0 | 0 | 0 | 0 | 0 | 0 | 0 | 0 | NA | NA | NA | NA | NA | NA | NA | NA | NA | NA |
| Olfr1182 | 0 | 0 | 0 | 0 | 0 | 0 | 0 | 0 | 0 | 0 | 0 | 0 | 0 | 0 | 0 | NA | NA | NA | NA | NA | NA | NA | NA | NA | NA |
| Olfr1183 | 0 | 0 | 0 | 0 | 0 | 0 | 0 | 0 | 0 | 0 | 0 | 0 | 0 | 0 | 0 | NA | NA | NA | NA | NA | NA | NA | NA | NA | NA |
| Olfr1184 | 0 | 0 | 0 | 0 | 0 | 0 | 0 | 0 | 0 | 0 | 0 | 0 | 0 | 0 | 0 | NA | NA | NA | NA | NA | NA | NA | NA | NA | NA |
| Olfr1186 | 0 | 0 | 0 | 0 | 0 | 0 | 0 | 0 | 0 | 0 | 0 | 0 | 0 | 0 | 0 | NA | NA | NA | NA | NA | NA | NA | NA | NA | NA |
| Olfr1188 | 0 | 0 | 0 | 0 | 0 | 0 | 0 | 0 | 0 | 0 | 0 | 0 | 0 | 0 | 0 | NA | NA | NA | NA | NA | NA | NA | NA | NA | NA |
| Olfr1189 | 0 | 0 | 0 | 0 | 0 | 0 | 0 | 0 | 0 | 0 | 0 | 0 | 0 | 0 | 0 | NA | NA | NA | NA | NA | NA | NA | NA | NA | NA |
| Olfr119 | 0 | 0 | 0 | 0 | 0 | 0 | 0 | 0 | 0 | 0 | 0 | 0 | 0 | 0 | 0 | NA | NA | NA | NA | NA | NA | NA | NA | NA | NA |
| Olfr1193 | 0 | 0 | 0 | 0 | 0 | 0 | 0 | 0 | 0 | 0 | 0 | 0 | 0 | 0 | 0 | NA | NA | NA | NA | NA | NA | NA | NA | NA | NA |
| Olfr1195 | 0 | 0 | 0 | 0 | 0 | 0 | 0 | 0 | 0 | 0 | 0 | 0 | 0 | 0 | 0 | NA | NA | NA | NA | NA | NA | NA | NA | NA | NA |
| Olfr1196 | 0 | 0 | 0 | 1 | 1 | 0 | 0 | 0 | 0 | 1 | 1 | 0 | 0.31 | 0 | 1 | 3.32 | 1.731 | 0.65383052 | NA | 0 | 10.6224 | 8.9541 | 8.9541 | TRUE | 0.202 |
| Olfr1197 | 0 | 0 | 1 | 0 | 0 | 0 | 0 | 0 | 1 | 0 | 0 | 0 | 0.17 | 0 | 0 | 0.49 | -1.029 | 0.80087 | NA | 0 | 19.6928 | 10 | 10 | TRUE | 0.4098 |
| Olfr1198 | 0 | 0 | 0 | 0 | 0 | 0 | 0 | 0 | 0 | 0 | 0 | 0 | 0 | 0 | 0 | NA | NA | NA | NA | NA | NA | NA | NA | NA | NA |
| Olfr1199 | 0 | 0 | 0 | 0 | 0 | 0 | 0 | 0 | 0 | 0 | 0 | 0 | 0 | 0 | 0 | NA | NA | NA | NA | NA | NA | NA | NA | NA | NA |
| Olfr12 | 1 | 0 | 0 | 0 | 0 | 0 | 1 | 0 | 0 | 0 | 0 | 0 | 0.18 | 0 | 0 | 0.49 | -1.029 | 0.80087 | NA | 0 | 18.6165 | 10 | 10 | TRUE | 0.4614 |
| Olfr120 | 0 | 0 | 0 | 0 | 0 | 0 | 0 | 0 | 0 | 0 | 0 | 0 | 0 | 0 | 0 | NA | NA | NA | NA | NA | NA | NA | NA | NA | NA |
| Olfr1200 | 0 | 0 | 0 | 0 | 0 | 0 | 0 | 0 | 0 | 0 | 0 | 0 | 0 | 0 | 0 | NA | NA | NA | NA | NA | NA | NA | NA | NA | NA |
| Olfr1201 | 0 | 0 | 0 | 0 | 0 | 0 | 0 | 0 | 0 | 0 | 0 | 0 | 0 | 0 | 0 | NA | NA | NA | NA | NA | NA | NA | NA | NA | NA |
| Olfr1202 | 0 | 0 | 0 | 0 | 0 | 0 | 0 | 0 | 0 | 0 | 0 | 0 | 0 | 0 | 0 | NA | NA | NA | NA | NA | NA | NA | NA | NA | NA |
| Olfr1204 | 0 | 0 | 0 | 0 | 0 | 0 | 0 | 0 | 0 | 0 | 0 | 0 | 0 | 0 | 0 | NA | NA | NA | NA | NA | NA | NA | NA | NA | NA |
| Olfr1205 | 0 | 0 | 0 | 0 | 0 | 0 | 0 | 0 | 0 | 0 | 0 | 0 | 0 | 0 | 0 | NA | NA | NA | NA | NA | NA | NA | NA | NA | NA |
| Olfr1206 | 0 | 0 | 0 | 0 | 0 | 0 | 0 | 0 | 0 | 0 | 0 | 0 | 0 | 0 | 0 | NA | NA | NA | NA | NA | NA | NA | NA | NA | NA |
| Olfr1208 | 0 | 0 | 0 | 0 | 0 | 0 | 0 | 0 | 0 | 0 | 0 | 0 | 0 | 0 | 0 | NA | NA | NA | NA | NA | NA | NA | NA | NA | NA |
| Olfr1209 | 0 | 0 | 0 | 0 | 0 | 0 | 0 | 0 | 0 | 0 | 0 | 0 | 0 | 0 | 0 | NA | NA | NA | NA | NA | NA | NA | NA | NA | NA |
| Olfr121 | 0 | 1 | 0 | 1 | 0 | 0 | 0 | 1 | 0 | 1 | 0 | 0 | 0.3 | 0 | 0 | 0.954 | -0.067 | 0.98642001 | NA | 0 | 11.0817 | 9.3014 | 9.3014 | TRUE | 0.405 |
| Olfr1211 | 0 | 0 | 0 | 0 | 0 | 0 | 0 | 0 | 0 | 0 | 0 | 0 | 0 | 0 | 0 | NA | NA | NA | NA | NA | NA | NA | NA | NA | NA |
| Olfr1212 | 0 | 0 | 0 | 0 | 0 | 0 | 0 | 0 | 0 | 0 | 0 | 0 | 0 | 0 | 0 | NA | NA | NA | NA | NA | NA | NA | NA | NA | NA |
| Olfr1213 | 0 | 0 | 0 | 0 | 0 | 0 | 0 | 0 | 0 | 0 | 0 | 0 | 0 | 0 | 0 | NA | NA | NA | NA | NA | NA | NA | NA | NA | NA |
| Olfr1214 | 0 | 0 | 0 | 0 | 0 | 0 | 0 | 0 | 0 | 0 | 0 | 0 | 0 | 0 | 0 | NA | NA | NA | NA | NA | NA | NA | NA | NA | NA |
| Olfr1215 | 0 | 0 | 0 | 0 | 0 | 0 | 0 | 0 | 0 | 0 | 0 | 0 | 0 | 0 | 0 | NA | NA | NA | NA | NA | NA | NA | NA | NA | NA |
| Olfr1216 | 0 | 0 | 0 | 0 | 0 | 0 | 0 | 0 | 0 | 0 | 0 | 0 | 0 | 0 | 0 | NA | NA | NA | NA | NA | NA | NA | NA | NA | NA |
| Olfr1217 | 0 | 0 | 0 | 0 | 0 | 0 | 0 | 0 | 0 | 0 | 0 | 0 | 0 | 0 | 0 | NA | NA | NA | NA | NA | NA | NA | NA | NA | NA |
| Olfr1218 | 0 | 0 | 0 | 0 | 0 | 0 | 0 | 0 | 0 | 0 | 0 | 0 | 0 | 0 | 0 | NA | NA | NA | NA | NA | NA | NA | NA | NA | NA |
| Olfr1219 | 0 | 0 | 0 | 0 | 0 | 0 | 0 | 0 | 0 | 0 | 0 | 0 | 0 | 0 | 0 | NA | NA | NA | NA | NA | NA | NA | NA | NA | NA |
| Olfr122 | 0 | 0 | 0 | 0 | 0 | 0 | 0 | 0 | 0 | 0 | 0 | 0 | 0 | 0 | 0 | NA | NA | NA | NA | NA | NA | NA | NA | NA | NA |
| Olfr1220 | 0 | 0 | 0 | 0 | 0 | 0 | 0 | 0 | 0 | 0 | 0 | 0 | 0 | 0 | 0 | NA | NA | NA | NA | NA | NA | NA | NA | NA | NA |
| Olfr1221 | 0 | 0 | 0 | 0 | 0 | 0 | 0 | 0 | 0 | 0 | 0 | 0 | 0 | 0 | 0 | NA | NA | NA | NA | NA | NA | NA | NA | NA | NA |
| Olfr1222 | 0 | 0 | 0 | 0 | 0 | 0 | 0 | 0 | 0 | 0 | 0 | 0 | 0 | 0 | 0 | NA | NA | NA | NA | NA | NA | NA | NA | NA | NA |
| Olfr1223 | 0 | 0 | 0 | 0 | 0 | 0 | 0 | 0 | 0 | 0 | 0 | 0 | 0 | 0 | 0 | NA | NA | NA | NA | NA | NA | NA | NA | NA | NA |
| Olfr1225 | 0 | 0 | 0 | 0 | 0 | 0 | 0 | 0 | 0 | 0 | 0 | 0 | 0 | 0 | 0 | NA | NA | NA | NA | NA | NA | NA | NA | NA | NA |
| Olfr1226 | 0 | 0 | 0 | 0 | 0 | 0 | 0 | 0 | 0 | 0 | 0 | 0 | 0 | 0 | 0 | NA | NA | NA | NA | NA | NA | NA | NA | NA | NA |
| Olfr1228 | 0 | 0 | 0 | 0 | 0 | 0 | 0 | 0 | 0 | 0 | 0 | 0 | 0 | 0 | 0 | NA | NA | NA | NA | NA | NA | NA | NA | NA | NA |
| Olfr1229 | 0 | 0 | 0 | 0 | 0 | 0 | 0 | 0 | 0 | 0 | 0 | 0 | 0 | 0 | 0 | NA | NA | NA | NA | NA | NA | NA | NA | NA | NA |
| Olfr123 | 0 | 0 | 0 | 0 | 0 | 0 | 0 | 0 | 0 | 0 | 0 | 0 | 0 | 0 | 0 | NA | NA | NA | NA | NA | NA | NA | NA | NA | NA |
| Olfr1230 | 0 | 0 | 0 | 0 | 0 | 0 | 0 | 0 | 0 | 0 | 0 | 0 | 0 | 0 | 0 | NA | NA | NA | NA | NA | NA | NA | NA | NA | NA |
| Olfr1231 | 0 | 0 | 0 | 0 | 0 | 0 | 0 | 0 | 0 | 0 | 0 | 0 | 0 | 0 | 0 | NA | NA | NA | NA | NA | NA | NA | NA | NA | NA |

| | | | | | | | | | | | | | | | | | | | | | | | | | |
|---|---|---|---|---|---|---|---|---|---|---|---|---|---|---|---|---|---|---|---|---|---|---|---|---|---|
| Olfr1232 | 0 | 0 | 0 | 0 | 0 | 0 | 0 | 0 | 0 | 0 | 0 | 0 | 0 | 0 | 0 | NA | NA | NA | NA | NA | NA | NA | NA | NA | NA |
| Olfr1233 | 0 | 0 | 0 | 0 | 0 | 0 | 0 | 0 | 0 | 0 | 0 | 0 | 0 | 0 | 0 | NA | NA | NA | NA | NA | NA | NA | NA | NA | NA |
| Olfr1234 | 0 | 0 | 0 | 0 | 0 | 0 | 0 | 0 | 0 | 0 | 0 | 0 | 0 | 0 | 0 | NA | NA | NA | NA | NA | NA | NA | NA | NA | NA |
| Olfr1238 | 0 | 0 | 0 | 0 | 0 | 0 | 0 | 0 | 0 | 0 | 0 | 0 | 0 | 0 | 0 | NA | NA | NA | NA | NA | NA | NA | NA | NA | NA |
| Olfr1239 | 0 | 0 | 0 | 0 | 0 | 0 | 0 | 0 | 0 | 0 | 0 | 0 | 0 | 0 | 0 | NA | NA | NA | NA | NA | NA | NA | NA | NA | NA |
| Olfr124 | 0 | 0 | 0 | 0 | 0 | 0 | 0 | 0 | 0 | 0 | 0 | 0 | 0 | 0 | 0 | NA | NA | NA | NA | NA | NA | NA | NA | NA | NA |
| Olfr1240 | 0 | 0 | 0 | 0 | 0 | 0 | 0 | 0 | 0 | 0 | 0 | 0 | 0 | 0 | 0 | NA | NA | NA | NA | NA | NA | NA | NA | NA | NA |
| Olfr1241 | 0 | 0 | 0 | 0 | 0 | 0 | 0 | 0 | 0 | 0 | 0 | 0 | 0 | 0 | 0 | NA | NA | NA | NA | NA | NA | NA | NA | NA | NA |
| Olfr1242 | 0 | 0 | 0 | 0 | 0 | 0 | 0 | 0 | 0 | 0 | 0 | 0 | 0 | 0 | 0 | NA | NA | NA | NA | NA | NA | NA | NA | NA | NA |
| Olfr1243 | 0 | 0 | 0 | 0 | 0 | 0 | 0 | 0 | 0 | 0 | 0 | 0 | 0 | 0 | 0 | NA | NA | NA | NA | NA | NA | NA | NA | NA | NA |
| Olfr1245 | 0 | 0 | 0 | 0 | 0 | 0 | 0 | 0 | 0 | 0 | 0 | 0 | 0 | 0 | 0 | NA | NA | NA | NA | NA | NA | NA | NA | NA | NA |
| Olfr1246 | 0 | 0 | 0 | 0 | 0 | 0 | 0 | 0 | 0 | 0 | 0 | 0 | 0 | 0 | 0 | NA | NA | NA | NA | NA | NA | NA | NA | NA | NA |
| Olfr1247 | 0 | 0 | 0 | 0 | 0 | 0 | 0 | 0 | 0 | 0 | 0 | 0 | 0 | 0 | 0 | NA | NA | NA | NA | NA | NA | NA | NA | NA | NA |
| Olfr1248 | 0 | 0 | 0 | 0 | 0 | 0 | 0 | 0 | 0 | 0 | 0 | 0 | 0 | 0 | 0 | NA | NA | NA | NA | NA | NA | NA | NA | NA | NA |
| Olfr1249 | 0 | 0 | 0 | 0 | 0 | 0 | 0 | 0 | 0 | 0 | 0 | 0 | 0 | 0 | 0 | NA | NA | NA | NA | NA | NA | NA | NA | NA | NA |
| Olfr125 | 0 | 0 | 0 | 0 | 0 | 0 | 0 | 0 | 0 | 0 | 0 | 0 | 0 | 0 | 0 | NA | NA | NA | NA | NA | NA | NA | NA | NA | NA |
| Olfr1250 | 0 | 0 | 0 | 0 | 0 | 0 | 0 | 0 | 0 | 0 | 0 | 0 | 0 | 0 | 0 | NA | NA | NA | NA | NA | NA | NA | NA | NA | NA |
| Olfr1251 | 0 | 0 | 0 | 0 | 0 | 0 | 0 | 0 | 0 | 0 | 0 | 0 | 0 | 0 | 0 | NA | NA | NA | NA | NA | NA | NA | NA | NA | NA |
| Olfr1252 | 0 | 0 | 0 | 0 | 0 | 0 | 0 | 0 | 0 | 0 | 0 | 0 | 0 | 0 | 0 | NA | NA | NA | NA | NA | NA | NA | NA | NA | NA |
| Olfr1253 | 0 | 0 | 0 | 0 | 0 | 0 | 0 | 0 | 0 | 0 | 0 | 0 | 0 | 0 | 0 | NA | NA | NA | NA | NA | NA | NA | NA | NA | NA |
| Olfr1254 | 0 | 0 | 0 | 0 | 0 | 0 | 0 | 0 | 0 | 0 | 0 | 0 | 0 | 0 | 0 | NA | NA | NA | NA | NA | NA | NA | NA | NA | NA |
| Olfr1255 | 0 | 0 | 0 | 0 | 0 | 0 | 0 | 0 | 0 | 0 | 0 | 0 | 0 | 0 | 0 | NA | NA | NA | NA | NA | NA | NA | NA | NA | NA |
| Olfr1256 | 0 | 0 | 0 | 0 | 0 | 0 | 0 | 0 | 0 | 0 | 0 | 0 | 0 | 0 | 0 | NA | NA | NA | NA | NA | NA | NA | NA | NA | NA |
| Olfr1257 | 0 | 0 | 0 | 0 | 0 | 0 | 0 | 0 | 0 | 0 | 0 | 0 | 0 | 0 | 0 | NA | NA | NA | NA | NA | NA | NA | NA | NA | NA |
| Olfr1258 | 0 | 0 | 0 | 0 | 0 | 0 | 0 | 0 | 0 | 0 | 0 | 0 | 0 | 0 | 0 | NA | NA | NA | NA | NA | NA | NA | NA | NA | NA |
| Olfr1259 | 0 | 0 | 0 | 2 | 4 | 4 | 0 | 0 | 0 | 2 | 4 | 4 | 1.71 | 0 | 3 | 18.017 | 4.171 | 0.02508981 | NA | 0 | 1.9345 | 1.354 | 1.354 | TRUE | 0.4246 |
| Olfr126 | 0 | 0 | 0 | 0 | 0 | 0 | 0 | 0 | 0 | 0 | 0 | 0 | 0 | 0 | 0 | NA | NA | NA | NA | NA | NA | NA | NA | NA | NA |
| Olfr1260 | 0 | 0 | 0 | 0 | 0 | 0 | 0 | 0 | 0 | 0 | 0 | 0 | 0 | 0 | 0 | NA | NA | NA | NA | NA | NA | NA | NA | NA | NA |
| Olfr1261 | 0 | 0 | 0 | 0 | 0 | 0 | 0 | 0 | 0 | 0 | 0 | 0 | 0 | 0 | 0 | NA | NA | NA | NA | NA | NA | NA | NA | NA | NA |
| Olfr1262 | 0 | 0 | 0 | 0 | 0 | 0 | 0 | 0 | 0 | 0 | 0 | 0 | 0 | 0 | 0 | NA | NA | NA | NA | NA | NA | NA | NA | NA | NA |
| Olfr1263 | 0 | 0 | 0 | 0 | 0 | 0 | 0 | 0 | 0 | 0 | 0 | 0 | 0 | 0 | 0 | NA | NA | NA | NA | NA | NA | NA | NA | NA | NA |
| Olfr1264 | 0 | 0 | 0 | 0 | 0 | 0 | 0 | 0 | 0 | 0 | 0 | 0 | 0 | 0 | 0 | NA | NA | NA | NA | NA | NA | NA | NA | NA | NA |
| Olfr1265 | 0 | 0 | 0 | 0 | 0 | 0 | 0 | 0 | 0 | 0 | 0 | 0 | 0 | 0 | 0 | NA | NA | NA | NA | NA | NA | NA | NA | NA | NA |
| Olfr1269 | 0 | 0 | 0 | 0 | 0 | 0 | 0 | 0 | 0 | 0 | 0 | 0 | 0 | 0 | 0 | NA | NA | NA | NA | NA | NA | NA | NA | NA | NA |
| Olfr127 | 0 | 0 | 0 | 0 | 0 | 0 | 0 | 0 | 0 | 0 | 0 | 0 | 0 | 0 | 0 | NA | NA | NA | NA | NA | NA | NA | NA | NA | NA |
| Olfr1270 | 0 | 0 | 0 | 0 | 0 | 0 | 0 | 0 | 0 | 0 | 0 | 0 | 0 | 0 | 0 | NA | NA | NA | NA | NA | NA | NA | NA | NA | NA |
| Olfr1271 | 0 | 0 | 0 | 0 | 0 | 0 | 0 | 0 | 0 | 0 | 0 | 0 | 0 | 0 | 0 | NA | NA | NA | NA | NA | NA | NA | NA | NA | NA |
| Olfr1272 | 0 | 0 | 0 | 0 | 0 | 0 | 0 | 0 | 0 | 0 | 0 | 0 | 0 | 0 | 0 | NA | NA | NA | NA | NA | NA | NA | NA | NA | NA |
| Olfr1273-ps | 0 | 0 | 0 | 0 | 0 | 0 | 0 | 0 | 0 | 0 | 0 | 0 | 0 | 0 | 0 | NA | NA | NA | NA | NA | NA | NA | NA | NA | NA |
| Olfr1274-ps | 0 | 0 | 0 | 0 | 0 | 0 | 0 | 0 | 0 | 0 | 0 | 0 | 0 | 0 | 0 | NA | NA | NA | NA | NA | NA | NA | NA | NA | NA |
| Olfr1275 | 0 | 0 | 0 | 0 | 0 | 0 | 0 | 0 | 0 | 0 | 0 | 0 | 0 | 0 | 0 | NA | NA | NA | NA | NA | NA | NA | NA | NA | NA |
| Olfr1276 | 0 | 0 | 0 | 0 | 0 | 0 | 0 | 0 | 0 | 0 | 0 | 0 | 0 | 0 | 0 | NA | NA | NA | NA | NA | NA | NA | NA | NA | NA |
| Olfr1277 | 0 | 0 | 0 | 0 | 0 | 0 | 0 | 0 | 0 | 0 | 0 | 0 | 0 | 0 | 0 | NA | NA | NA | NA | NA | NA | NA | NA | NA | NA |
| Olfr1278 | 0 | 0 | 0 | 0 | 0 | 0 | 0 | 0 | 0 | 0 | 0 | 0 | 0 | 0 | 0 | NA | NA | NA | NA | NA | NA | NA | NA | NA | NA |
| Olfr1279 | 0 | 0 | 0 | 0 | 0 | 0 | 0 | 0 | 0 | 0 | 0 | 0 | 0 | 0 | 0 | NA | NA | NA | NA | NA | NA | NA | NA | NA | NA |
| Olfr128 | 0 | 0 | 0 | 0 | 0 | 0 | 0 | 0 | 0 | 0 | 0 | 0 | 0 | 0 | 0 | NA | NA | NA | NA | NA | NA | NA | NA | NA | NA |
| Olfr1280 | 0 | 0 | 0 | 0 | 0 | 0 | 0 | 0 | 0 | 0 | 0 | 0 | 0 | 0 | 0 | NA | NA | NA | NA | NA | NA | NA | NA | NA | NA |
| Olfr1281 | 0 | 0 | 0 | 0 | 0 | 0 | 0 | 0 | 0 | 0 | 0 | 0 | 0 | 0 | 0 | NA | NA | NA | NA | NA | NA | NA | NA | NA | NA |
| Olfr1282 | 0 | 0 | 0 | 0 | 0 | 0 | 0 | 0 | 0 | 0 | 0 | 0 | 0 | 0 | 0 | NA | NA | NA | NA | NA | NA | NA | NA | NA | NA |
| Olfr1283 | 0 | 0 | 0 | 0 | 0 | 0 | 0 | 0 | 0 | 0 | 0 | 0 | 0 | 0 | 0 | NA | NA | NA | NA | NA | NA | NA | NA | NA | NA |
| Olfr1284 | 0 | 0 | 0 | 0 | 0 | 0 | 0 | 0 | 0 | 0 | 0 | 0 | 0 | 0 | 0 | NA | NA | NA | NA | NA | NA | NA | NA | NA | NA |
| Olfr1286 | 0 | 0 | 0 | 0 | 0 | 0 | 0 | 0 | 0 | 0 | 0 | 0 | 0 | 0 | 0 | NA | NA | NA | NA | NA | NA | NA | NA | NA | NA |
| Olfr1287 | 0 | 0 | 0 | 0 | 0 | 0 | 0 | 0 | 0 | 0 | 0 | 0 | 0 | 0 | 0 | NA | NA | NA | NA | NA | NA | NA | NA | NA | NA |
| Olfr1288 | 0 | 0 | 0 | 0 | 0 | 0 | 0 | 0 | 0 | 0 | 0 | 0 | 0 | 0 | 0 | NA | NA | NA | NA | NA | NA | NA | NA | NA | NA |
| Olfr1289 | 0 | 0 | 0 | 0 | 0 | 0 | 0 | 0 | 0 | 0 | 0 | 0 | 0 | 0 | 0 | NA | NA | NA | NA | NA | NA | NA | NA | NA | NA |
| Olfr129 | 0 | 0 | 0 | 0 | 0 | 0 | 0 | 0 | 0 | 0 | 0 | 0 | 0 | 0 | 0 | NA | NA | NA | NA | NA | NA | NA | NA | NA | NA |
| Olfr1290 | 0 | 0 | 0 | 0 | 0 | 0 | 0 | 0 | 0 | 0 | 0 | 0 | 0 | 0 | 0 | NA | NA | NA | NA | NA | NA | NA | NA | NA | NA |
| Olfr1294 | 0 | 0 | 0 | 0 | 0 | 0 | 0 | 0 | 0 | 0 | 0 | 0 | 0 | 0 | 0 | NA | NA | NA | NA | NA | NA | NA | NA | NA | NA |
| Olfr1295 | 0 | 0 | 0 | 0 | 0 | 0 | 0 | 0 | 0 | 0 | 0 | 0 | 0 | 0 | 0 | NA | NA | NA | NA | NA | NA | NA | NA | NA | NA |
| Olfr1297 | 0 | 0 | 0 | 0 | 0 | 0 | 0 | 0 | 0 | 0 | 0 | 0 | 0 | 0 | 0 | NA | NA | NA | NA | NA | NA | NA | NA | NA | NA |
| Olfr1298 | 0 | 0 | 0 | 0 | 0 | 0 | 0 | 0 | 0 | 0 | 0 | 0 | 0 | 0 | 0 | NA | NA | NA | NA | NA | NA | NA | NA | NA | NA |
| Olfr1299 | 0 | 0 | 0 | 0 | 0 | 0 | 0 | 0 | 0 | 0 | 0 | 0 | 0 | 0 | 0 | NA | NA | NA | NA | NA | NA | NA | NA | NA | NA |
| Olfr13 | 0 | 0 | 0 | 0 | 0 | 0 | 0 | 0 | 0 | 0 | 0 | 0 | 0 | 0 | 0 | NA | NA | NA | NA | NA | NA | NA | NA | NA | NA |
| Olfr130 | 0 | 0 | 0 | 0 | 0 | 0 | 0 | 0 | 0 | 0 | 0 | 0 | 0 | 0 | 0 | NA | NA | NA | NA | NA | NA | NA | NA | NA | NA |
| Olfr1300-ps1 | 0 | 0 | 0 | 0 | 0 | 0 | 0 | 0 | 0 | 0 | 0 | 0 | 0 | 0 | 0 | NA | NA | NA | NA | NA | NA | NA | NA | NA | NA |
| Olfr1301 | 0 | 0 | 0 | 0 | 0 | 0 | 0 | 0 | 0 | 0 | 0 | 0 | 0 | 0 | 0 | NA | NA | NA | NA | NA | NA | NA | NA | NA | NA |
| Olfr1302 | 0 | 0 | 0 | 0 | 0 | 0 | 0 | 0 | 0 | 0 | 0 | 0 | 0 | 0 | 0 | NA | NA | NA | NA | NA | NA | NA | NA | NA | NA |
| Olfr1303 | 0 | 0 | 0 | 0 | 0 | 0 | 0 | 0 | 0 | 0 | 0 | 0 | 0 | 0 | 0 | NA | NA | NA | NA | NA | NA | NA | NA | NA | NA |
| Olfr1305 | 0 | 0 | 0 | 0 | 0 | 0 | 0 | 0 | 0 | 0 | 0 | 0 | 0 | 0 | 0 | NA | NA | NA | NA | NA | NA | NA | NA | NA | NA |

| | | | | | | | | | | | | | | | | | | | | | | | | | |
|---|---|---|---|---|---|---|---|---|---|---|---|---|---|---|---|---|---|---|---|---|---|---|---|---|---|
| Olfr1306 | 0 | 0 | 0 | 0 | 0 | 0 | 0 | 0 | 0 | 0 | 0 | 0 | 0 | 0 | 0 | NA | NA | NA | NA | NA | NA | NA | NA | NA | NA |
| Olfr1307 | 0 | 0 | 0 | 0 | 0 | 0 | 0 | 0 | 0 | 0 | 0 | 0 | 0 | 0 | 0 | NA | NA | NA | NA | NA | NA | NA | NA | NA | NA |
| Olfr1308 | 0 | 0 | 0 | 0 | 0 | 0 | 0 | 0 | 0 | 0 | 0 | 0 | 0 | 0 | 0 | NA | NA | NA | NA | NA | NA | NA | NA | NA | NA |
| Olfr1309 | 0 | 0 | 0 | 0 | 0 | 0 | 0 | 0 | 0 | 0 | 0 | 0 | 0 | 0 | 0 | NA | NA | NA | NA | NA | NA | NA | NA | NA | NA |
| Olfr131 | 0 | 0 | 0 | 0 | 0 | 0 | 0 | 0 | 0 | 0 | 0 | 0 | 0 | 0 | 0 | NA | NA | NA | NA | NA | NA | NA | NA | NA | NA |
| Olfr1310 | 0 | 0 | 0 | 0 | 0 | 0 | 0 | 0 | 0 | 0 | 0 | 0 | 0 | 0 | 0 | NA | NA | NA | NA | NA | NA | NA | NA | NA | NA |
| Olfr1311 | 0 | 0 | 0 | 0 | 0 | 0 | 0 | 0 | 0 | 0 | 0 | 0 | 0 | 0 | 0 | NA | NA | NA | NA | NA | NA | NA | NA | NA | NA |
| Olfr1312 | 0 | 0 | 0 | 0 | 0 | 0 | 0 | 0 | 0 | 0 | 0 | 0 | 0 | 0 | 0 | NA | NA | NA | NA | NA | NA | NA | NA | NA | NA |
| Olfr1313 | 0 | 0 | 0 | 0 | 0 | 0 | 0 | 0 | 0 | 0 | 0 | 0 | 0 | 0 | 0 | NA | NA | NA | NA | NA | NA | NA | NA | NA | NA |
| Olfr1314 | 0 | 0 | 0 | 0 | 0 | 0 | 0 | 0 | 0 | 0 | 0 | 0 | 0 | 0 | 0 | NA | NA | NA | NA | NA | NA | NA | NA | NA | NA |
| Olfr1316 | 0 | 0 | 0 | 0 | 0 | 0 | 0 | 0 | 0 | 0 | 0 | 0 | 0 | 0 | 0 | NA | NA | NA | NA | NA | NA | NA | NA | NA | NA |
| Olfr1317 | 0 | 0 | 0 | 0 | 0 | 0 | 0 | 0 | 0 | 0 | 0 | 0 | 0 | 0 | 0 | NA | NA | NA | NA | NA | NA | NA | NA | NA | NA |
| Olfr1318 | 0 | 0 | 0 | 0 | 0 | 0 | 0 | 0 | 0 | 0 | 0 | 0 | 0 | 0 | 0 | NA | NA | NA | NA | NA | NA | NA | NA | NA | NA |
| Olfr132 | 0 | 0 | 0 | 0 | 0 | 0 | 0 | 0 | 0 | 0 | 0 | 0 | 0 | 0 | 0 | NA | NA | NA | NA | NA | NA | NA | NA | NA | NA |
| Olfr1320 | 0 | 0 | 0 | 0 | 0 | 0 | 0 | 0 | 0 | 0 | 0 | 0 | 0 | 0 | 0 | NA | NA | NA | NA | NA | NA | NA | NA | NA | NA |
| Olfr1321 | 0 | 0 | 0 | 0 | 0 | 0 | 0 | 0 | 0 | 0 | 0 | 0 | 0 | 0 | 0 | NA | NA | NA | NA | NA | NA | NA | NA | NA | NA |
| Olfr1322 | 0 | 0 | 0 | 0 | 0 | 0 | 0 | 0 | 0 | 0 | 0 | 0 | 0 | 0 | 0 | NA | NA | NA | NA | NA | NA | NA | NA | NA | NA |
| Olfr1323 | 0 | 0 | 0 | 0 | 0 | 0 | 0 | 0 | 0 | 0 | 0 | 0 | 0 | 0 | 0 | NA | NA | NA | NA | NA | NA | NA | NA | NA | NA |
| Olfr1324 | 0 | 0 | 0 | 0 | 0 | 0 | 0 | 0 | 0 | 0 | 0 | 0 | 0 | 0 | 0 | NA | NA | NA | NA | NA | NA | NA | NA | NA | NA |
| Olfr1325 | 0 | 0 | 0 | 0 | 0 | 0 | 0 | 0 | 0 | 0 | 0 | 0 | 0 | 0 | 0 | NA | NA | NA | NA | NA | NA | NA | NA | NA | NA |
| Olfr1328 | 1 | 0 | 1 | 0 | 0 | 0 | 1 | 0 | 1 | 0 | 0 | 0 | 0.34 | 1 | 0 | 0.261 | -1.939 | 0.60367425 | NA | 0 | 9.5732 | 8.3134 | 8.3134 | TRUE | 0.2529 |
| Olfr1329 | 0 | 0 | 0 | 0 | 0 | 0 | 0 | 0 | 0 | 0 | 0 | 0 | 0 | 0 | 0 | NA | NA | NA | NA | NA | NA | NA | NA | NA | NA |
| Olfr133 | 0 | 0 | 0 | 0 | 0 | 0 | 0 | 0 | 0 | 0 | 0 | 0 | 0 | 0 | 0 | NA | NA | NA | NA | NA | NA | NA | NA | NA | NA |
| Olfr1330 | 0 | 0 | 0 | 0 | 0 | 0 | 0 | 0 | 0 | 0 | 0 | 0 | 0 | 0 | 0 | NA | NA | NA | NA | NA | NA | NA | NA | NA | NA |
| Olfr1331 | 0 | 0 | 0 | 0 | 0 | 0 | 0 | 0 | 0 | 0 | 0 | 0 | 0 | 0 | 0 | NA | NA | NA | NA | NA | NA | NA | NA | NA | NA |
| Olfr1333 | 0 | 0 | 0 | 0 | 0 | 0 | 0 | 0 | 0 | 0 | 0 | 0 | 0 | 0 | 0 | NA | NA | NA | NA | NA | NA | NA | NA | NA | NA |
| Olfr1335 | 0 | 0 | 0 | 0 | 0 | 0 | 0 | 0 | 0 | 0 | 0 | 0 | 0 | 0 | 0 | NA | NA | NA | NA | NA | NA | NA | NA | NA | NA |
| Olfr1336 | 1 | 0 | 2 | 1 | 0 | 0 | 1 | 0 | 2 | 1 | 0 | 0 | 0.64 | 1 | 0 | 0.341 | -1.551 | 0.55438633 | NA | 0 | 5.1324 | 3.4582 | 3.4582 | TRUE | 0.1148 |
| Olfr1337 | 0 | 0 | 0 | 0 | 0 | 0 | 0 | 0 | 0 | 0 | 0 | 0 | 0 | 0 | 0 | NA | NA | NA | NA | NA | NA | NA | NA | NA | NA |
| Olfr1338 | 0 | 0 | 0 | 0 | 0 | 0 | 0 | 0 | 0 | 0 | 0 | 0 | 0 | 0 | 0 | NA | NA | NA | NA | NA | NA | NA | NA | NA | NA |
| Olfr1339 | 0 | 0 | 0 | 0 | 0 | 0 | 0 | 0 | 0 | 0 | 0 | 0 | 0 | 0 | 0 | NA | NA | NA | NA | NA | NA | NA | NA | NA | NA |
| Olfr134 | 0 | 0 | 0 | 0 | 0 | 0 | 0 | 0 | 0 | 0 | 0 | 0 | 0 | 0 | 0 | NA | NA | NA | NA | NA | NA | NA | NA | NA | NA |
| Olfr1340 | 0 | 0 | 0 | 0 | 0 | 0 | 0 | 0 | 0 | 0 | 0 | 0 | 0 | 0 | 0 | NA | NA | NA | NA | NA | NA | NA | NA | NA | NA |
| Olfr1341 | 0 | 0 | 0 | 0 | 0 | 0 | 0 | 0 | 0 | 0 | 0 | 0 | 0 | 0 | 0 | NA | NA | NA | NA | NA | NA | NA | NA | NA | NA |
| Olfr1342 | 6 | 2 | 1 | 2 | 2 | 0 | 6 | 2 | 1 | 2 | 2 | 0 | 2.18 | 3 | 1 | 0.405 | -1.303 | 0.37158009 | NA | 0 | 1.516 | 0.9818 | 0.9818 | TRUE | 1.0618 |
| Olfr1344 | 37 | 52 | 37 | 25 | 13 | 25 | 39 | 52 | 37 | 20 | 14 | 27 | 31.57 | 43 | 20 | 0.474 | -1.077 | 0.00532079 | 0.03274268 | 0.0213 | 0.1113 | 0.0717 | 0.0717 | TRUE | 0.4423 |
| Olfr1346 | 0 | 0 | 0 | 0 | 0 | 1 | 0 | 0 | 0 | 0 | 0 | 1 | 0.18 | 0 | 0 | 1.859 | 0.894 | 0.82650734 | NA | 0 | 18.0643 | 10 | 10 | TRUE | 0.5383 |
| Olfr1347 | 0 | 0 | 0 | 1 | 1 | 0 | 0 | 0 | 0 | 1 | 1 | 0 | 0.31 | 0 | 1 | 3.32 | 1.731 | 0.65383052 | NA | 0 | 10.6224 | 8.9541 | 8.9541 | TRUE | 0.202 |
| Olfr1348 | 0 | 0 | 0 | 1 | 0 | 0 | 0 | 0 | 0 | 1 | 0 | 0 | 0.13 | 0 | 0 | 1.859 | 0.894 | 0.82650734 | NA | 0 | 25.1515 | 10 | 10 | TRUE | 0.2563 |
| Olfr1349 | 8 | 2 | 2 | 6 | 6 | 1 | 8 | 2 | 2 | 5 | 6 | 1 | 4.13 | 4 | 4 | 0.993 | -0.011 | 0.99212061 | NA | 0.3893 | 0.8049 | 0.5994 | 0.5994 | TRUE | 1.0766 |
| Olfr135 | 0 | 0 | 0 | 0 | 0 | 0 | 0 | 0 | 0 | 0 | 0 | 0 | 0 | 0 | 0 | NA | NA | NA | NA | NA | NA | NA | NA | NA | NA |
| Olfr1350 | 0 | 0 | 0 | 0 | 1 | 0 | 0 | 0 | 0 | 0 | 1 | 0 | 0.18 | 0 | 0 | 1.859 | 0.894 | 0.82650734 | NA | 0 | 18.3785 | 10 | 10 | TRUE | 0.5204 |
| Olfr1351 | 0 | 0 | 0 | 0 | 0 | 0 | 0 | 0 | 0 | 0 | 0 | 0 | 0 | 0 | 0 | NA | NA | NA | NA | NA | NA | NA | NA | NA | NA |
| Olfr1352 | 0 | 0 | 0 | 0 | 0 | 0 | 0 | 0 | 0 | 0 | 0 | 0 | 0 | 0 | 0 | NA | NA | NA | NA | NA | NA | NA | NA | NA | NA |
| Olfr1353 | 0 | 0 | 0 | 0 | 0 | 0 | 0 | 0 | 0 | 0 | 0 | 0 | 0 | 0 | 0 | NA | NA | NA | NA | NA | NA | NA | NA | NA | NA |
| Olfr1354 | 0 | 0 | 0 | 0 | 0 | 0 | 0 | 0 | 0 | 0 | 0 | 0 | 0 | 0 | 0 | NA | NA | NA | NA | NA | NA | NA | NA | NA | NA |
| Olfr1355 | 0 | 0 | 0 | 0 | 0 | 0 | 0 | 0 | 0 | 0 | 0 | 0 | 0 | 0 | 0 | NA | NA | NA | NA | NA | NA | NA | NA | NA | NA |
| Olfr1356 | 0 | 0 | 0 | 0 | 0 | 0 | 0 | 0 | 0 | 0 | 0 | 0 | 0 | 0 | 0 | NA | NA | NA | NA | NA | NA | NA | NA | NA | NA |
| Olfr1357 | 0 | 0 | 0 | 0 | 0 | 0 | 0 | 0 | 0 | 0 | 0 | 0 | 0 | 0 | 0 | NA | NA | NA | NA | NA | NA | NA | NA | NA | NA |
| Olfr1359 | 0 | 0 | 0 | 0 | 0 | 0 | 0 | 0 | 0 | 0 | 0 | 0 | 0 | 0 | 0 | NA | NA | NA | NA | NA | NA | NA | NA | NA | NA |
| Olfr136 | 0 | 0 | 0 | 0 | 0 | 0 | 0 | 0 | 0 | 0 | 0 | 0 | 0 | 0 | 0 | NA | NA | NA | NA | NA | NA | NA | NA | NA | NA |
| Olfr1360 | 0 | 0 | 0 | 0 | 0 | 0 | 0 | 0 | 0 | 0 | 0 | 0 | 0 | 0 | 0 | NA | NA | NA | NA | NA | NA | NA | NA | NA | NA |
| Olfr1361 | 0 | 0 | 0 | 0 | 0 | 0 | 0 | 0 | 0 | 0 | 0 | 0 | 0 | 0 | 0 | NA | NA | NA | NA | NA | NA | NA | NA | NA | NA |
| Olfr1362 | 0 | 0 | 0 | 0 | 0 | 0 | 0 | 0 | 0 | 0 | 0 | 0 | 0 | 0 | 0 | NA | NA | NA | NA | NA | NA | NA | NA | NA | NA |
| Olfr1364 | 0 | 0 | 0 | 0 | 0 | 0 | 0 | 0 | 0 | 0 | 0 | 0 | 0 | 0 | 0 | NA | NA | NA | NA | NA | NA | NA | NA | NA | NA |
| Olfr1366 | 0 | 0 | 0 | 0 | 0 | 0 | 0 | 0 | 0 | 0 | 0 | 0 | 0 | 0 | 0 | NA | NA | NA | NA | NA | NA | NA | NA | NA | NA |
| Olfr1367 | 0 | 0 | 0 | 0 | 0 | 0 | 0 | 0 | 0 | 0 | 0 | 0 | 0 | 0 | 0 | NA | NA | NA | NA | NA | NA | NA | NA | NA | NA |
| Olfr1368 | 0 | 0 | 0 | 0 | 0 | 0 | 0 | 0 | 0 | 0 | 0 | 0 | 0 | 0 | 0 | NA | NA | NA | NA | NA | NA | NA | NA | NA | NA |
| Olfr137 | 0 | 0 | 0 | 0 | 0 | 0 | 0 | 0 | 0 | 0 | 0 | 0 | 0 | 0 | 0 | NA | NA | NA | NA | NA | NA | NA | NA | NA | NA |
| Olfr1370 | 0 | 0 | 0 | 0 | 0 | 0 | 0 | 0 | 0 | 0 | 0 | 0 | 0 | 0 | 0 | NA | NA | NA | NA | NA | NA | NA | NA | NA | NA |
| Olfr1371 | 0 | 0 | 1 | 0 | 0 | 2 | 0 | 0 | 1 | 0 | 0 | 2 | 0.53 | 0 | 1 | 1.955 | 0.967 | 0.77212926 | NA | 0 | 6.1967 | 6.3475 | 6.3475 | TRUE | 0.9745 |
| Olfr1372-ps1 | 45 | 87 | 69 | 89 | 62 | 64 | 48 | 87 | 69 | 70 | 67 | 70 | 68.45 | 68 | 69 | 1.012 | 0.018 | 0.94977863 | 0.97481103 | 0.0272 | 0.055 | 0.0418 | 0.0418 | TRUE | 0.2401 |
| Olfr1373 | 0 | 0 | 0 | 0 | 0 | 0 | 0 | 0 | 0 | 0 | 0 | 0 | 0 | 0 | 0 | NA | NA | NA | NA | NA | NA | NA | NA | NA | NA |
| Olfr1377 | 0 | 0 | 0 | 0 | 0 | 0 | 0 | 0 | 0 | 0 | 0 | 0 | 0 | 0 | 0 | NA | NA | NA | NA | NA | NA | NA | NA | NA | NA |
| Olfr1378 | 0 | 0 | 0 | 0 | 0 | 0 | 0 | 0 | 0 | 0 | 0 | 0 | 0 | 0 | 0 | NA | NA | NA | NA | NA | NA | NA | NA | NA | NA |
| Olfr138 | 0 | 0 | 0 | 0 | 0 | 0 | 0 | 0 | 0 | 0 | 0 | 0 | 0 | 0 | 0 | NA | NA | NA | NA | NA | NA | NA | NA | NA | NA |
| Olfr1380 | 0 | 0 | 0 | 0 | 0 | 0 | 0 | 0 | 0 | 0 | 0 | 0 | 0 | 0 | 0 | NA | NA | NA | NA | NA | NA | NA | NA | NA | NA |
| Olfr1381 | 0 | 0 | 0 | 0 | 0 | 0 | 0 | 0 | 0 | 0 | 0 | 0 | 0 | 0 | 0 | NA | NA | NA | NA | NA | NA | NA | NA | NA | NA |
| Olfr1382 | 0 | 0 | 0 | 0 | 0 | 0 | 0 | 0 | 0 | 0 | 0 | 0 | 0 | 0 | 0 | NA | NA | NA | NA | NA | NA | NA | NA | NA | NA |

| | | | | | | | | | | | | | | | | | | | | | | | | | |
|---|---|---|---|---|---|---|---|---|---|---|---|---|---|---|---|---|---|---|---|---|---|---|---|---|---|
| Olfr1383 | 0 | 0 | 0 | 0 | 0 | 0 | 0 | 0 | 0 | 0 | 0 | 0 | 0 | 0 | 0 | NA | NA | NA | NA | NA | NA | NA | NA | NA | NA |
| Olfr1384 | 0 | 0 | 0 | 0 | 0 | 0 | 0 | 0 | 0 | 0 | 0 | 0 | 0 | 0 | 0 | NA | NA | NA | NA | NA | NA | NA | NA | NA | NA |
| Olfr1385 | 0 | 0 | 0 | 0 | 0 | 0 | 0 | 0 | 0 | 0 | 0 | 0 | 0 | 0 | 0 | NA | NA | NA | NA | NA | NA | NA | NA | NA | NA |
| Olfr1386 | 0 | 0 | 0 | 0 | 0 | 0 | 0 | 0 | 0 | 0 | 0 | 0 | 0 | 0 | 0 | NA | NA | NA | NA | NA | NA | NA | NA | NA | NA |
| Olfr1387 | 0 | 0 | 0 | 0 | 0 | 0 | 0 | 0 | 0 | 0 | 0 | 0 | 0 | 0 | 0 | NA | NA | NA | NA | NA | NA | NA | NA | NA | NA |
| Olfr1388 | 0 | 0 | 0 | 0 | 0 | 0 | 0 | 0 | 0 | 0 | 0 | 0 | 0 | 0 | 0 | NA | NA | NA | NA | NA | NA | NA | NA | NA | NA |
| Olfr1389 | 0 | 0 | 0 | 1 | 0 | 0 | 0 | 0 | 0 | 1 | 0 | 0 | 0.13 | 0 | 0 | 1.859 | 0.894 | 0.82650734 | NA | | 0 | 25.1515 | 10 | 10 | TRUE | 0.2563 |
| Olfr139 | 0 | 0 | 0 | 0 | 0 | 0 | 0 | 0 | 0 | 0 | 0 | 0 | 0 | 0 | 0 | NA | NA | NA | NA | NA | NA | NA | NA | NA | NA |
| Olfr1390 | 0 | 0 | 0 | 0 | 0 | 0 | 0 | 0 | 0 | 0 | 0 | 0 | 0 | 0 | 0 | NA | NA | NA | NA | NA | NA | NA | NA | NA | NA |
| Olfr1391 | 0 | 0 | 0 | 0 | 0 | 0 | 0 | 0 | 0 | 0 | 0 | 0 | 0 | 0 | 0 | NA | NA | NA | NA | NA | NA | NA | NA | NA | NA |
| Olfr1392 | 0 | 0 | 0 | 0 | 1 | 0 | 0 | 0 | 0 | 0 | 1 | 0 | 0.18 | 0 | 0 | 1.859 | 0.894 | 0.82650734 | NA | | 0 | 18.3785 | 10 | 10 | TRUE | 0.5204 |
| Olfr1393 | 0 | 0 | 1 | 3 | 1 | 1 | 0 | 0 | 1 | 2 | 1 | 1 | 0.92 | 0 | 1 | 4.189 | 2.067 | 0.32121847 | NA | | 0 | 3.5794 | 1.8085 | 1.8085 | TRUE | 0.4098 |
| Olfr1394 | 0 | 0 | 0 | 0 | 0 | 0 | 0 | 0 | 0 | 0 | 0 | 0 | 0 | 0 | 0 | NA | NA | NA | NA | NA | NA | NA | NA | NA | NA |
| Olfr1395 | 0 | 0 | 0 | 0 | 0 | 0 | 0 | 0 | 0 | 0 | 0 | 0 | 0 | 0 | 0 | NA | NA | NA | NA | NA | NA | NA | NA | NA | NA |
| Olfr1396 | 0 | 0 | 0 | 0 | 0 | 0 | 0 | 0 | 0 | 0 | 0 | 0 | 0 | 0 | 0 | NA | NA | NA | NA | NA | NA | NA | NA | NA | NA |
| Olfr140 | 0 | 0 | 0 | 0 | 0 | 0 | 0 | 0 | 0 | 0 | 0 | 0 | 0 | 0 | 0 | NA | NA | NA | NA | NA | NA | NA | NA | NA | NA |
| Olfr1402 | 0 | 0 | 0 | 0 | 0 | 0 | 0 | 0 | 0 | 0 | 0 | 0 | 0 | 0 | 0 | NA | NA | NA | NA | NA | NA | NA | NA | NA | NA |
| Olfr1404 | 0 | 0 | 0 | 0 | 0 | 0 | 0 | 0 | 0 | 0 | 0 | 0 | 0 | 0 | 0 | NA | NA | NA | NA | NA | NA | NA | NA | NA | NA |
| Olfr1406 | 1 | 0 | 0 | 0 | 1 | 0 | 1 | 0 | 0 | 0 | 1 | 0 | 0.36 | 0 | 0 | 0.954 | -0.067 | 0.98557668 | NA | | 0 | 9.2518 | 8.0179 | 8.0179 | TRUE | 0.5204 |
| Olfr1408 | 0 | 0 | 0 | 0 | 0 | 0 | 0 | 0 | 0 | 0 | 0 | 0 | 0 | 0 | 0 | NA | NA | NA | NA | NA | NA | NA | NA | NA | NA |
| Olfr141 | 0 | 0 | 0 | 0 | 0 | 0 | 0 | 0 | 0 | 0 | 0 | 0 | 0 | 0 | 0 | NA | NA | NA | NA | NA | NA | NA | NA | NA | NA |
| Olfr1410 | 0 | 0 | 0 | 0 | 0 | 0 | 0 | 0 | 0 | 0 | 0 | 0 | 0 | 0 | 0 | NA | NA | NA | NA | NA | NA | NA | NA | NA | NA |
| Olfr1411 | 0 | 0 | 0 | 0 | 0 | 0 | 0 | 0 | 0 | 0 | 0 | 0 | 0 | 0 | 0 | NA | NA | NA | NA | NA | NA | NA | NA | NA | NA |
| Olfr1412 | 0 | 0 | 0 | 0 | 0 | 0 | 0 | 0 | 0 | 0 | 0 | 0 | 0 | 0 | 0 | NA | NA | NA | NA | NA | NA | NA | NA | NA | NA |
| Olfr1413 | 0 | 0 | 0 | 0 | 0 | 0 | 0 | 0 | 0 | 0 | 0 | 0 | 0 | 0 | 0 | NA | NA | NA | NA | NA | NA | NA | NA | NA | NA |
| Olfr1414 | 0 | 0 | 0 | 0 | 0 | 0 | 0 | 0 | 0 | 0 | 0 | 0 | 0 | 0 | 0 | NA | NA | NA | NA | NA | NA | NA | NA | NA | NA |
| Olfr1415 | 0 | 0 | 0 | 0 | 0 | 0 | 0 | 0 | 0 | 0 | 0 | 0 | 0 | 0 | 0 | NA | NA | NA | NA | NA | NA | NA | NA | NA | NA |
| Olfr1416 | 0 | 0 | 0 | 0 | 0 | 0 | 0 | 0 | 0 | 0 | 0 | 0 | 0 | 0 | 0 | NA | NA | NA | NA | NA | NA | NA | NA | NA | NA |
| Olfr1417 | 0 | 0 | 0 | 0 | 0 | 0 | 0 | 0 | 0 | 0 | 0 | 0 | 0 | 0 | 0 | NA | NA | NA | NA | NA | NA | NA | NA | NA | NA |
| Olfr1418 | 0 | 0 | 0 | 0 | 0 | 0 | 0 | 0 | 0 | 0 | 0 | 0 | 0 | 0 | 0 | NA | NA | NA | NA | NA | NA | NA | NA | NA | NA |
| Olfr1419 | 0 | 0 | 0 | 0 | 1 | 0 | 0 | 0 | 0 | 0 | 1 | 0 | 0.18 | 0 | 0 | 1.859 | 0.894 | 0.82650734 | NA | | 0 | 18.3785 | 10 | 10 | TRUE | 0.5204 |
| Olfr142 | 0 | 0 | 0 | 0 | 0 | 0 | 0 | 0 | 0 | 0 | 0 | 0 | 0 | 0 | 0 | NA | NA | NA | NA | NA | NA | NA | NA | NA | NA |
| Olfr1420 | 0 | 0 | 0 | 0 | 0 | 0 | 0 | 0 | 0 | 0 | 0 | 0 | 0 | 0 | 0 | NA | NA | NA | NA | NA | NA | NA | NA | NA | NA |
| Olfr1423 | 0 | 0 | 0 | 0 | 0 | 0 | 0 | 0 | 0 | 0 | 0 | 0 | 0 | 0 | 0 | NA | NA | NA | NA | NA | NA | NA | NA | NA | NA |
| Olfr1424 | 0 | 0 | 0 | 0 | 0 | 0 | 0 | 0 | 0 | 0 | 0 | 0 | 0 | 0 | 0 | NA | NA | NA | NA | NA | NA | NA | NA | NA | NA |
| Olfr1425 | 0 | 0 | 0 | 0 | 0 | 0 | 0 | 0 | 0 | 0 | 0 | 0 | 0 | 0 | 0 | NA | NA | NA | NA | NA | NA | NA | NA | NA | NA |
| Olfr1426 | 0 | 0 | 0 | 0 | 0 | 0 | 0 | 0 | 0 | 0 | 0 | 0 | 0 | 0 | 0 | NA | NA | NA | NA | NA | NA | NA | NA | NA | NA |
| Olfr1427 | 0 | 0 | 0 | 0 | 0 | 0 | 0 | 0 | 0 | 0 | 0 | 0 | 0 | 0 | 0 | NA | NA | NA | NA | NA | NA | NA | NA | NA | NA |
| Olfr1428 | 1 | 0 | 0 | 0 | 0 | 0 | 1 | 0 | 0 | 0 | 0 | 0 | 0.18 | 0 | 0 | 0.49 | -1.029 | 0.80087 | NA | | 0 | 18.6165 | 10 | 10 | TRUE | 0.4614 |
| Olfr143 | 0 | 0 | 0 | 0 | 0 | 0 | 0 | 0 | 0 | 0 | 0 | 0 | 0 | 0 | 0 | NA | NA | NA | NA | NA | NA | NA | NA | NA | NA |
| Olfr1431 | 0 | 0 | 0 | 0 | 0 | 0 | 0 | 0 | 0 | 0 | 0 | 0 | 0 | 0 | 0 | NA | NA | NA | NA | NA | NA | NA | NA | NA | NA |
| Olfr1433 | 0 | 0 | 0 | 0 | 0 | 0 | 0 | 0 | 0 | 0 | 0 | 0 | 0 | 0 | 0 | NA | NA | NA | NA | NA | NA | NA | NA | NA | NA |
| Olfr1434 | 0 | 0 | 0 | 0 | 0 | 0 | 0 | 0 | 0 | 0 | 0 | 0 | 0 | 0 | 0 | NA | NA | NA | NA | NA | NA | NA | NA | NA | NA |
| Olfr1436 | 0 | 0 | 0 | 0 | 0 | 0 | 0 | 0 | 0 | 0 | 0 | 0 | 0 | 0 | 0 | NA | NA | NA | NA | NA | NA | NA | NA | NA | NA |
| Olfr1437 | 0 | 0 | 0 | 0 | 0 | 0 | 0 | 0 | 0 | 0 | 0 | 0 | 0 | 0 | 0 | NA | NA | NA | NA | NA | NA | NA | NA | NA | NA |
| Olfr1440 | 0 | 0 | 0 | 0 | 0 | 0 | 0 | 0 | 0 | 0 | 0 | 0 | 0 | 0 | 0 | NA | NA | NA | NA | NA | NA | NA | NA | NA | NA |
| Olfr1441 | 0 | 0 | 0 | 0 | 0 | 0 | 0 | 0 | 0 | 0 | 0 | 0 | 0 | 0 | 0 | NA | NA | NA | NA | NA | NA | NA | NA | NA | NA |
| Olfr1442 | 0 | 0 | 0 | 0 | 0 | 0 | 0 | 0 | 0 | 0 | 0 | 0 | 0 | 0 | 0 | NA | NA | NA | NA | NA | NA | NA | NA | NA | NA |
| Olfr1443 | 0 | 0 | 0 | 0 | 0 | 0 | 0 | 0 | 0 | 0 | 0 | 0 | 0 | 0 | 0 | NA | NA | NA | NA | NA | NA | NA | NA | NA | NA |
| Olfr1444 | 0 | 0 | 0 | 0 | 0 | 0 | 0 | 0 | 0 | 0 | 0 | 0 | 0 | 0 | 0 | NA | NA | NA | NA | NA | NA | NA | NA | NA | NA |
| Olfr1445 | 0 | 0 | 0 | 0 | 0 | 0 | 0 | 0 | 0 | 0 | 0 | 0 | 0 | 0 | 0 | NA | NA | NA | NA | NA | NA | NA | NA | NA | NA |
| Olfr1446 | 0 | 0 | 0 | 0 | 0 | 0 | 0 | 0 | 0 | 0 | 0 | 0 | 0 | 0 | 0 | NA | NA | NA | NA | NA | NA | NA | NA | NA | NA |
| Olfr1447 | 0 | 0 | 0 | 0 | 0 | 0 | 0 | 0 | 0 | 0 | 0 | 0 | 0 | 0 | 0 | NA | NA | NA | NA | NA | NA | NA | NA | NA | NA |
| Olfr1448 | 0 | 0 | 0 | 0 | 0 | 0 | 0 | 0 | 0 | 0 | 0 | 0 | 0 | 0 | 0 | NA | NA | NA | NA | NA | NA | NA | NA | NA | NA |
| Olfr1449 | 0 | 0 | 0 | 0 | 0 | 0 | 0 | 0 | 0 | 0 | 0 | 0 | 0 | 0 | 0 | NA | NA | NA | NA | NA | NA | NA | NA | NA | NA |
| Olfr145 | 0 | 0 | 0 | 0 | 0 | 0 | 0 | 0 | 0 | 0 | 0 | 0 | 0 | 0 | 0 | NA | NA | NA | NA | NA | NA | NA | NA | NA | NA |
| Olfr1450 | 0 | 0 | 0 | 0 | 0 | 0 | 0 | 0 | 0 | 0 | 0 | 0 | 0 | 0 | 0 | NA | NA | NA | NA | NA | NA | NA | NA | NA | NA |
| Olfr1451 | 0 | 0 | 0 | 0 | 0 | 0 | 0 | 0 | 0 | 0 | 0 | 0 | 0 | 0 | 0 | NA | NA | NA | NA | NA | NA | NA | NA | NA | NA |
| Olfr1453 | 0 | 0 | 0 | 0 | 0 | 0 | 0 | 0 | 0 | 0 | 0 | 0 | 0 | 0 | 0 | NA | NA | NA | NA | NA | NA | NA | NA | NA | NA |
| Olfr1454 | 0 | 0 | 0 | 0 | 0 | 0 | 0 | 0 | 0 | 0 | 0 | 0 | 0 | 0 | 0 | NA | NA | NA | NA | NA | NA | NA | NA | NA | NA |
| Olfr1457 | 0 | 0 | 0 | 0 | 0 | 0 | 0 | 0 | 0 | 0 | 0 | 0 | 0 | 0 | 0 | NA | NA | NA | NA | NA | NA | NA | NA | NA | NA |
| Olfr1459 | 0 | 0 | 0 | 0 | 0 | 0 | 0 | 0 | 0 | 0 | 0 | 0 | 0 | 0 | 0 | NA | NA | NA | NA | NA | NA | NA | NA | NA | NA |
| Olfr146 | 0 | 0 | 0 | 0 | 0 | 0 | 0 | 0 | 0 | 0 | 0 | 0 | 0 | 0 | 0 | NA | NA | NA | NA | NA | NA | NA | NA | NA | NA |
| Olfr1461 | 0 | 0 | 0 | 0 | 0 | 0 | 0 | 0 | 0 | 0 | 0 | 0 | 0 | 0 | 0 | NA | NA | NA | NA | NA | NA | NA | NA | NA | NA |
| Olfr1462 | 0 | 0 | 0 | 0 | 0 | 0 | 0 | 0 | 0 | 0 | 0 | 0 | 0 | 0 | 0 | NA | NA | NA | NA | NA | NA | NA | NA | NA | NA |
| Olfr1463 | 0 | 0 | 0 | 0 | 0 | 0 | 0 | 0 | 0 | 0 | 0 | 0 | 0 | 0 | 0 | NA | NA | NA | NA | NA | NA | NA | NA | NA | NA |
| Olfr1465 | 0 | 0 | 0 | 0 | 0 | 0 | 0 | 0 | 0 | 0 | 0 | 0 | 0 | 0 | 0 | NA | NA | NA | NA | NA | NA | NA | NA | NA | NA |
| Olfr1466 | 0 | 0 | 0 | 0 | 0 | 0 | 0 | 0 | 0 | 0 | 0 | 0 | 0 | 0 | 0 | NA | NA | NA | NA | NA | NA | NA | NA | NA | NA |
| Olfr1467 | 0 | 0 | 0 | 0 | 0 | 0 | 0 | 0 | 0 | 0 | 0 | 0 | 0 | 0 | 0 | NA | NA | NA | NA | NA | NA | NA | NA | NA | NA |

| | | | | | | | | | | | | | | | | | | | | | | | | | |
|---|---|---|---|---|---|---|---|---|---|---|---|---|---|---|---|---|---|---|---|---|---|---|---|---|---|
| Olfr1469 | 0 | 0 | 0 | 0 | 0 | 0 | 0 | 0 | 0 | 0 | 0 | 0 | 0 | 0 | 0 | NA | NA | NA | NA | NA | NA | NA | NA | NA | NA |
| Olfr147 | 5 | 0 | 1 | 0 | 3 | 0 | 5 | 0 | 1 | 0 | 3 | 0 | 1.59 | 2 | 1 | 0.501 | -0.997 | 0.64668547 | NA | 4.0871 | 2.0783 | 2.7004 | 2.7004 | TRUE | 1.2714 |
| Olfr1471 | 0 | 0 | 0 | 0 | 0 | 0 | 0 | 0 | 0 | 0 | 0 | 0 | 0 | 0 | 0 | NA | NA | NA | NA | NA | NA | NA | NA | NA | NA |
| Olfr1472 | 0 | 0 | 0 | 0 | 0 | 0 | 0 | 0 | 0 | 0 | 0 | 0 | 0 | 0 | 0 | NA | NA | NA | NA | NA | NA | NA | NA | NA | NA |
| Olfr1474 | 0 | 0 | 0 | 0 | 0 | 0 | 0 | 0 | 0 | 0 | 0 | 0 | 0 | 0 | 0 | NA | NA | NA | NA | NA | NA | NA | NA | NA | NA |
| Olfr1475 | 0 | 0 | 0 | 0 | 0 | 0 | 0 | 0 | 0 | 0 | 0 | 0 | 0 | 0 | 0 | NA | NA | NA | NA | NA | NA | NA | NA | NA | NA |
| Olfr1477 | 0 | 0 | 0 | 0 | 0 | 0 | 0 | 0 | 0 | 0 | 0 | 0 | 0 | 0 | 0 | NA | NA | NA | NA | NA | NA | NA | NA | NA | NA |
| Olfr148 | 0 | 0 | 0 | 0 | 0 | 0 | 0 | 0 | 0 | 0 | 0 | 0 | 0 | 0 | 0 | NA | NA | NA | NA | NA | NA | NA | NA | NA | NA |
| Olfr1480 | 0 | 0 | 0 | 0 | 0 | 0 | 0 | 0 | 0 | 0 | 0 | 0 | 0 | 0 | 0 | NA | NA | NA | NA | NA | NA | NA | NA | NA | NA |
| Olfr1484 | 0 | 0 | 0 | 0 | 0 | 0 | 0 | 0 | 0 | 0 | 0 | 0 | 0 | 0 | 0 | NA | NA | NA | NA | NA | NA | NA | NA | NA | NA |
| Olfr1487 | 0 | 0 | 0 | 0 | 0 | 0 | 0 | 0 | 0 | 0 | 0 | 0 | 0 | 0 | 0 | NA | NA | NA | NA | NA | NA | NA | NA | NA | NA |
| Olfr1489 | 0 | 0 | 0 | 0 | 0 | 0 | 0 | 0 | 0 | 0 | 0 | 0 | 0 | 0 | 0 | NA | NA | NA | NA | NA | NA | NA | NA | NA | NA |
| Olfr149 | 0 | 0 | 0 | 0 | 0 | 0 | 0 | 0 | 0 | 0 | 0 | 0 | 0 | 0 | 0 | NA | NA | NA | NA | NA | NA | NA | NA | NA | NA |
| Olfr1490 | 0 | 0 | 0 | 0 | 0 | 0 | 0 | 0 | 0 | 0 | 0 | 0 | 0 | 0 | 0 | NA | NA | NA | NA | NA | NA | NA | NA | NA | NA |
| Olfr1491 | 0 | 0 | 0 | 0 | 0 | 0 | 0 | 0 | 0 | 0 | 0 | 0 | 0 | 0 | 0 | NA | NA | NA | NA | NA | NA | NA | NA | NA | NA |
| Olfr1494 | 0 | 0 | 0 | 0 | 0 | 0 | 0 | 0 | 0 | 0 | 0 | 0 | 0 | 0 | 0 | NA | NA | NA | NA | NA | NA | NA | NA | NA | NA |
| Olfr1495 | 0 | 0 | 0 | 0 | 0 | 0 | 0 | 0 | 0 | 0 | 0 | 0 | 0 | 0 | 0 | NA | NA | NA | NA | NA | NA | NA | NA | NA | NA |
| Olfr1496 | 0 | 0 | 0 | 0 | 0 | 0 | 0 | 0 | 0 | 0 | 0 | 0 | 0 | 0 | 0 | NA | NA | NA | NA | NA | NA | NA | NA | NA | NA |
| Olfr1497 | 0 | 0 | 0 | 0 | 0 | 0 | 0 | 0 | 0 | 0 | 0 | 0 | 0 | 0 | 0 | NA | NA | NA | NA | NA | NA | NA | NA | NA | NA |
| Olfr1499 | 0 | 0 | 0 | 0 | 0 | 0 | 0 | 0 | 0 | 0 | 0 | 0 | 0 | 0 | 0 | NA | NA | NA | NA | NA | NA | NA | NA | NA | NA |
| Olfr15 | 0 | 0 | 0 | 0 | 0 | 0 | 0 | 0 | 0 | 0 | 0 | 0 | 0 | 0 | 0 | NA | NA | NA | NA | NA | NA | NA | NA | NA | NA |
| Olfr150 | 0 | 0 | 0 | 0 | 0 | 0 | 0 | 0 | 0 | 0 | 0 | 0 | 0 | 0 | 0 | NA | NA | NA | NA | NA | NA | NA | NA | NA | NA |
| Olfr1500 | 0 | 0 | 0 | 0 | 0 | 0 | 0 | 0 | 0 | 0 | 0 | 0 | 0 | 0 | 0 | NA | NA | NA | NA | NA | NA | NA | NA | NA | NA |
| Olfr1501 | 0 | 0 | 0 | 0 | 0 | 0 | 0 | 0 | 0 | 0 | 0 | 0 | 0 | 0 | 0 | NA | NA | NA | NA | NA | NA | NA | NA | NA | NA |
| Olfr1502 | 0 | 0 | 0 | 0 | 0 | 0 | 0 | 0 | 0 | 0 | 0 | 0 | 0 | 0 | 0 | NA | NA | NA | NA | NA | NA | NA | NA | NA | NA |
| Olfr1504 | 0 | 1 | 0 | 0 | 0 | 0 | 0 | 1 | 0 | 0 | 0 | 0 | 0.17 | 0 | 0 | 0.49 | -1.029 | 0.80087 | NA | 0 | 19.7991 | 10 | 10 | TRUE | 0.405 |
| Olfr1505 | 0 | 0 | 0 | 0 | 0 | 0 | 0 | 0 | 0 | 0 | 0 | 0 | 0 | 0 | 0 | NA | NA | NA | NA | NA | NA | NA | NA | NA | NA |
| Olfr1506 | 0 | 0 | 0 | 0 | 0 | 0 | 0 | 0 | 0 | 0 | 0 | 0 | 0 | 0 | 0 | NA | NA | NA | NA | NA | NA | NA | NA | NA | NA |
| Olfr1507 | 0 | 0 | 0 | 0 | 0 | 0 | 0 | 0 | 0 | 0 | 0 | 0 | 0 | 0 | 0 | NA | NA | NA | NA | NA | NA | NA | NA | NA | NA |
| Olfr1508 | 0 | 0 | 0 | 0 | 0 | 0 | 0 | 0 | 0 | 0 | 0 | 0 | 0 | 0 | 0 | NA | NA | NA | NA | NA | NA | NA | NA | NA | NA |
| Olfr1509 | 0 | 0 | 0 | 0 | 0 | 0 | 0 | 0 | 0 | 0 | 0 | 0 | 0 | 0 | 0 | NA | NA | NA | NA | NA | NA | NA | NA | NA | NA |
| Olfr151 | 0 | 0 | 0 | 0 | 0 | 0 | 0 | 0 | 0 | 0 | 0 | 0 | 0 | 0 | 0 | NA | NA | NA | NA | NA | NA | NA | NA | NA | NA |
| Olfr1510 | 0 | 0 | 0 | 0 | 0 | 0 | 0 | 0 | 0 | 0 | 0 | 0 | 0 | 0 | 0 | NA | NA | NA | NA | NA | NA | NA | NA | NA | NA |
| Olfr1511 | 0 | 0 | 0 | 0 | 0 | 0 | 0 | 0 | 0 | 0 | 0 | 0 | 0 | 0 | 0 | NA | NA | NA | NA | NA | NA | NA | NA | NA | NA |
| Olfr1512 | 0 | 0 | 0 | 0 | 0 | 0 | 0 | 0 | 0 | 0 | 0 | 0 | 0 | 0 | 0 | NA | NA | NA | NA | NA | NA | NA | NA | NA | NA |
| Olfr1513 | 0 | 0 | 0 | 0 | 0 | 0 | 0 | 0 | 0 | 0 | 0 | 0 | 0 | 0 | 0 | NA | NA | NA | NA | NA | NA | NA | NA | NA | NA |
| Olfr152 | 0 | 0 | 0 | 0 | 0 | 0 | 0 | 0 | 0 | 0 | 0 | 0 | 0 | 0 | 0 | NA | NA | NA | NA | NA | NA | NA | NA | NA | NA |
| Olfr153 | 6 | 2 | 0 | 9 | 5 | 4 | 6 | 2 | 0 | 7 | 5 | 4 | 4.2 | 3 | 5 | 2.042 | 1.03 | 0.35436184 | NA | 0.3024 | 0.7912 | 0.6197 | 0.6197 | TRUE | 1.0214 |
| Olfr1532-ps1 | 0 | 0 | 0 | 0 | 0 | 0 | 0 | 0 | 0 | 0 | 0 | 0 | 0 | 0 | 0 | NA | NA | NA | NA | NA | NA | NA | NA | NA | NA |
| Olfr1535 | 0 | 0 | 0 | 0 | 0 | 0 | 0 | 0 | 0 | 0 | 0 | 0 | 0 | 0 | 0 | NA | NA | NA | NA | NA | NA | NA | NA | NA | NA |
| Olfr1537 | 0 | 0 | 0 | 0 | 0 | 0 | 0 | 0 | 0 | 0 | 0 | 0 | 0 | 0 | 0 | NA | NA | NA | NA | NA | NA | NA | NA | NA | NA |
| Olfr154 | 0 | 0 | 0 | 0 | 0 | 0 | 0 | 0 | 0 | 0 | 0 | 0 | 0 | 0 | 0 | NA | NA | NA | NA | NA | NA | NA | NA | NA | NA |
| Olfr155 | 0 | 0 | 0 | 0 | 0 | 0 | 0 | 0 | 0 | 0 | 0 | 0 | 0 | 0 | 0 | NA | NA | NA | NA | NA | NA | NA | NA | NA | NA |
| Olfr156 | 1 | 0 | 0 | 0 | 0 | 0 | 1 | 0 | 0 | 0 | 0 | 0 | 0.18 | 0 | 0 | 0.49 | -1.029 | 0.80087 | NA | 0 | 18.6165 | 10 | 10 | TRUE | 0.4614 |
| Olfr157 | 3 | 0 | 0 | 1 | 1 | 0 | 3 | 0 | 0 | 1 | 1 | 0 | 0.84 | 1 | 1 | 0.596 | -0.747 | 0.76594026 | NA | 2.0314 | 3.9222 | 3.2678 | 3.2678 | TRUE | 1.4428 |
| Olfr159 | 0 | 0 | 0 | 0 | 0 | 0 | 0 | 0 | 0 | 0 | 0 | 0 | 0 | 0 | 0 | NA | NA | NA | NA | NA | NA | NA | NA | NA | NA |
| Olfr16 | 0 | 0 | 0 | 0 | 0 | 0 | 0 | 0 | 0 | 0 | 0 | 0 | 0 | 0 | 0 | NA | NA | NA | NA | NA | NA | NA | NA | NA | NA |
| Olfr160 | 0 | 1 | 0 | 0 | 0 | 0 | 0 | 1 | 0 | 0 | 0 | 0 | 0.17 | 0 | 0 | 0.49 | -1.029 | 0.80087 | NA | 0 | 19.7991 | 10 | 10 | TRUE | 0.405 |
| Olfr161 | 0 | 1 | 0 | 0 | 0 | 2 | 0 | 1 | 0 | 0 | 0 | 2 | 0.53 | 0 | 1 | 1.955 | 0.967 | 0.77223757 | NA | 0 | 6.2072 | 6.3559 | 6.3559 | TRUE | 0.9745 |
| Olfr164 | 0 | 0 | 0 | 0 | 0 | 0 | 0 | 0 | 0 | 0 | 0 | 0 | 0 | 0 | 0 | NA | NA | NA | NA | NA | NA | NA | NA | NA | NA |
| Olfr165 | 0 | 0 | 0 | 0 | 0 | 0 | 0 | 0 | 0 | 0 | 0 | 0 | 0 | 0 | 0 | NA | NA | NA | NA | NA | NA | NA | NA | NA | NA |
| Olfr166 | 0 | 0 | 0 | 0 | 0 | 0 | 0 | 0 | 0 | 0 | 0 | 0 | 0 | 0 | 0 | NA | NA | NA | NA | NA | NA | NA | NA | NA | NA |
| Olfr167 | 0 | 0 | 0 | 0 | 0 | 0 | 0 | 0 | 0 | 0 | 0 | 0 | 0 | 0 | 0 | NA | NA | NA | NA | NA | NA | NA | NA | NA | NA |
| Olfr168 | 0 | 0 | 0 | 0 | 0 | 0 | 0 | 0 | 0 | 0 | 0 | 0 | 0 | 0 | 0 | NA | NA | NA | NA | NA | NA | NA | NA | NA | NA |
| Olfr169 | 0 | 0 | 0 | 0 | 0 | 0 | 0 | 0 | 0 | 0 | 0 | 0 | 0 | 0 | 0 | NA | NA | NA | NA | NA | NA | NA | NA | NA | NA |
| Olfr17 | 0 | 0 | 0 | 0 | 0 | 0 | 0 | 0 | 0 | 0 | 0 | 0 | 0 | 0 | 0 | NA | NA | NA | NA | NA | NA | NA | NA | NA | NA |
| Olfr170 | 0 | 0 | 0 | 0 | 0 | 0 | 0 | 0 | 0 | 0 | 0 | 0 | 0 | 0 | 0 | NA | NA | NA | NA | NA | NA | NA | NA | NA | NA |
| Olfr171 | 0 | 0 | 0 | 0 | 0 | 0 | 0 | 0 | 0 | 0 | 0 | 0 | 0 | 0 | 0 | NA | NA | NA | NA | NA | NA | NA | NA | NA | NA |
| Olfr172 | 0 | 0 | 0 | 0 | 0 | 0 | 0 | 0 | 0 | 0 | 0 | 0 | 0 | 0 | 0 | NA | NA | NA | NA | NA | NA | NA | NA | NA | NA |
| Olfr173 | 0 | 0 | 0 | 0 | 0 | 0 | 0 | 0 | 0 | 0 | 0 | 0 | 0 | 0 | 0 | NA | NA | NA | NA | NA | NA | NA | NA | NA | NA |
| Olfr175-ps1 | 0 | 0 | 0 | 0 | 0 | 0 | 0 | 0 | 0 | 0 | 0 | 0 | 0 | 0 | 0 | NA | NA | NA | NA | NA | NA | NA | NA | NA | NA |
| Olfr176 | 0 | 0 | 0 | 0 | 0 | 0 | 0 | 0 | 0 | 0 | 0 | 0 | 0 | 0 | 0 | NA | NA | NA | NA | NA | NA | NA | NA | NA | NA |
| Olfr177 | 0 | 0 | 0 | 0 | 0 | 0 | 0 | 0 | 0 | 0 | 0 | 0 | 0 | 0 | 0 | NA | NA | NA | NA | NA | NA | NA | NA | NA | NA |
| Olfr178 | 0 | 0 | 0 | 0 | 0 | 0 | 0 | 0 | 0 | 0 | 0 | 0 | 0 | 0 | 0 | NA | NA | NA | NA | NA | NA | NA | NA | NA | NA |
| Olfr179 | 0 | 0 | 0 | 1 | 0 | 0 | 0 | 0 | 0 | 1 | 0 | 0 | 0.13 | 0 | 0 | 1.859 | 0.894 | 0.82650734 | NA | 0 | 25.1515 | 10 | 10 | TRUE | 0.2563 |
| Olfr18 | 0 | 0 | 0 | 3 | 0 | 0 | 0 | 0 | 0 | 2 | 0 | 0 | 0.39 | 0 | 1 | 4.278 | 2.097 | 0.60136958 | NA | 10 | 8.3884 | 10 | 10 | TRUE | 1.4411 |
| Olfr180 | 0 | 0 | 0 | 0 | 0 | 0 | 0 | 0 | 0 | 0 | 0 | 0 | 0 | 0 | 0 | NA | NA | NA | NA | NA | NA | NA | NA | NA | NA |
| Olfr181 | 0 | 0 | 0 | 0 | 0 | 0 | 0 | 0 | 0 | 0 | 0 | 0 | 0 | 0 | 0 | NA | NA | NA | NA | NA | NA | NA | NA | NA | NA |
| Olfr183 | 0 | 0 | 0 | 0 | 0 | 0 | 0 | 0 | 0 | 0 | 0 | 0 | 0 | 0 | 0 | NA | NA | NA | NA | NA | NA | NA | NA | NA | NA |

| | | | | | | | | | | | | | | | | | | | | | | | | | |
|---|---|---|---|---|---|---|---|---|---|---|---|---|---|---|---|---|---|---|---|---|---|---|---|---|---|
| Olfr186 | 0 | 0 | 0 | 0 | 0 | 0 | 0 | 0 | 0 | 0 | 0 | 0 | 0 | 0 | 0 | NA | NA | NA | NA | NA | NA | NA | NA | NA | NA |
| Olfr187 | 0 | 0 | 0 | 0 | 0 | 0 | 0 | 0 | 0 | 0 | 0 | 0 | 0 | 0 | 0 | NA | NA | NA | NA | NA | NA | NA | NA | NA | NA |
| Olfr19 | 0 | 0 | 0 | 0 | 0 | 0 | 0 | 0 | 0 | 0 | 0 | 0 | 0 | 0 | 0 | NA | NA | NA | NA | NA | NA | NA | NA | NA | NA |
| Olfr190 | 0 | 0 | 0 | 0 | 0 | 0 | 0 | 0 | 0 | 0 | 0 | 0 | 0 | 0 | 0 | NA | NA | NA | NA | NA | NA | NA | NA | NA | NA |
| Olfr191 | 0 | 0 | 0 | 0 | 0 | 0 | 0 | 0 | 0 | 0 | 0 | 0 | 0 | 0 | 0 | NA | NA | NA | NA | NA | NA | NA | NA | NA | NA |
| Olfr192 | 0 | 0 | 0 | 0 | 0 | 0 | 0 | 0 | 0 | 0 | 0 | 0 | 0 | 0 | 0 | NA | NA | NA | NA | NA | NA | NA | NA | NA | NA |
| Olfr193 | 0 | 0 | 0 | 0 | 0 | 0 | 0 | 0 | 0 | 0 | 0 | 0 | 0 | 0 | 0 | NA | NA | NA | NA | NA | NA | NA | NA | NA | NA |
| Olfr194 | 0 | 0 | 0 | 0 | 0 | 0 | 0 | 0 | 0 | 0 | 0 | 0 | 0 | 0 | 0 | NA | NA | NA | NA | NA | NA | NA | NA | NA | NA |
| Olfr195 | 0 | 0 | 0 | 0 | 0 | 0 | 0 | 0 | 0 | 0 | 0 | 0 | 0 | 0 | 0 | NA | NA | NA | NA | NA | NA | NA | NA | NA | NA |
| Olfr196 | 0 | 0 | 0 | 0 | 0 | 0 | 0 | 0 | 0 | 0 | 0 | 0 | 0 | 0 | 0 | NA | NA | NA | NA | NA | NA | NA | NA | NA | NA |
| Olfr197 | 0 | 0 | 0 | 0 | 0 | 0 | 0 | 0 | 0 | 0 | 0 | 0 | 0 | 0 | 0 | NA | NA | NA | NA | NA | NA | NA | NA | NA | NA |
| Olfr198 | 0 | 0 | 0 | 0 | 0 | 0 | 0 | 0 | 0 | 0 | 0 | 0 | 0 | 0 | 0 | NA | NA | NA | NA | NA | NA | NA | NA | NA | NA |
| Olfr199 | 0 | 0 | 0 | 0 | 0 | 0 | 0 | 0 | 0 | 0 | 0 | 0 | 0 | 0 | 0 | NA | NA | NA | NA | NA | NA | NA | NA | NA | NA |
| Olfr2 | 0 | 0 | 0 | 0 | 0 | 0 | 0 | 0 | 0 | 0 | 0 | 0 | 0 | 0 | 0 | NA | NA | NA | NA | NA | NA | NA | NA | NA | NA |
| Olfr20 | 1 | 0 | 1 | 0 | 1 | 0 | 1 | 0 | 1 | 0 | 1 | 0 | 0.52 | 1 | 0 | 0.509 | -0.975 | 0.72714826 | NA | 0 | 6.2976 | 3.8857 | 3.8857 | TRUE | 0.5204 |
| Olfr201 | 0 | 0 | 0 | 0 | 0 | 0 | 0 | 0 | 0 | 0 | 0 | 0 | 0 | 0 | 0 | NA | NA | NA | NA | NA | NA | NA | NA | NA | NA |
| Olfr202 | 0 | 0 | 0 | 0 | 0 | 0 | 0 | 0 | 0 | 0 | 0 | 0 | 0 | 0 | 0 | NA | NA | NA | NA | NA | NA | NA | NA | NA | NA |
| Olfr203 | 0 | 0 | 0 | 0 | 0 | 0 | 0 | 0 | 0 | 0 | 0 | 0 | 0 | 0 | 0 | NA | NA | NA | NA | NA | NA | NA | NA | NA | NA |
| Olfr204 | 0 | 0 | 0 | 0 | 0 | 0 | 0 | 0 | 0 | 0 | 0 | 0 | 0 | 0 | 0 | NA | NA | NA | NA | NA | NA | NA | NA | NA | NA |
| Olfr205 | 0 | 0 | 0 | 0 | 0 | 0 | 0 | 0 | 0 | 0 | 0 | 0 | 0 | 0 | 0 | NA | NA | NA | NA | NA | NA | NA | NA | NA | NA |
| Olfr206 | 0 | 0 | 0 | 0 | 0 | 0 | 0 | 0 | 0 | 0 | 0 | 0 | 0 | 0 | 0 | NA | NA | NA | NA | NA | NA | NA | NA | NA | NA |
| Olfr207 | 0 | 0 | 0 | 0 | 0 | 0 | 0 | 0 | 0 | 0 | 0 | 0 | 0 | 0 | 0 | NA | NA | NA | NA | NA | NA | NA | NA | NA | NA |
| Olfr208 | 0 | 0 | 0 | 0 | 0 | 0 | 0 | 0 | 0 | 0 | 0 | 0 | 0 | 0 | 0 | NA | NA | NA | NA | NA | NA | NA | NA | NA | NA |
| Olfr209 | 0 | 0 | 0 | 0 | 0 | 0 | 0 | 0 | 0 | 0 | 0 | 0 | 0 | 0 | 0 | NA | NA | NA | NA | NA | NA | NA | NA | NA | NA |
| Olfr211 | 0 | 0 | 0 | 0 | 0 | 0 | 0 | 0 | 0 | 0 | 0 | 0 | 0 | 0 | 0 | NA | NA | NA | NA | NA | NA | NA | NA | NA | NA |
| Olfr212 | 0 | 0 | 0 | 0 | 0 | 0 | 0 | 0 | 0 | 0 | 0 | 0 | 0 | 0 | 0 | NA | NA | NA | NA | NA | NA | NA | NA | NA | NA |
| Olfr213 | 0 | 0 | 0 | 0 | 0 | 0 | 0 | 0 | 0 | 0 | 0 | 0 | 0 | 0 | 0 | NA | NA | NA | NA | NA | NA | NA | NA | NA | NA |
| Olfr214 | 0 | 0 | 0 | 0 | 0 | 0 | 0 | 0 | 0 | 0 | 0 | 0 | 0 | 0 | 0 | NA | NA | NA | NA | NA | NA | NA | NA | NA | NA |
| Olfr215 | 0 | 0 | 0 | 0 | 0 | 0 | 0 | 0 | 0 | 0 | 0 | 0 | 0 | 0 | 0 | NA | NA | NA | NA | NA | NA | NA | NA | NA | NA |
| Olfr218 | 0 | 0 | 0 | 0 | 0 | 0 | 0 | 0 | 0 | 0 | 0 | 0 | 0 | 0 | 0 | NA | NA | NA | NA | NA | NA | NA | NA | NA | NA |
| Olfr220 | 0 | 0 | 0 | 0 | 0 | 0 | 0 | 0 | 0 | 0 | 0 | 0 | 0 | 0 | 0 | NA | NA | NA | NA | NA | NA | NA | NA | NA | NA |
| Olfr221 | 0 | 0 | 0 | 0 | 0 | 0 | 0 | 0 | 0 | 0 | 0 | 0 | 0 | 0 | 0 | NA | NA | NA | NA | NA | NA | NA | NA | NA | NA |
| Olfr222 | 0 | 1 | 0 | 0 | 0 | 0 | 0 | 1 | 0 | 0 | 0 | 0 | 0.17 | 0 | 0 | 0.49 | -1.029 | 0.80087 | NA | 0 | 19.7991 | 10 | 10 | TRUE | 0.405 |
| Olfr223 | 0 | 0 | 0 | 0 | 0 | 0 | 0 | 0 | 0 | 0 | 0 | 0 | 0 | 0 | 0 | NA | NA | NA | NA | NA | NA | NA | NA | NA | NA |
| Olfr224 | 0 | 0 | 0 | 0 | 0 | 0 | 0 | 0 | 0 | 0 | 0 | 0 | 0 | 0 | 0 | NA | NA | NA | NA | NA | NA | NA | NA | NA | NA |
| Olfr225 | 0 | 0 | 0 | 0 | 0 | 0 | 0 | 0 | 0 | 0 | 0 | 0 | 0 | 0 | 0 | NA | NA | NA | NA | NA | NA | NA | NA | NA | NA |
| Olfr228 | 0 | 0 | 0 | 0 | 0 | 0 | 0 | 0 | 0 | 0 | 0 | 0 | 0 | 0 | 0 | NA | NA | NA | NA | NA | NA | NA | NA | NA | NA |
| Olfr229 | 0 | 0 | 0 | 0 | 0 | 0 | 0 | 0 | 0 | 0 | 0 | 0 | 0 | 0 | 0 | NA | NA | NA | NA | NA | NA | NA | NA | NA | NA |
| Olfr23 | 0 | 0 | 0 | 0 | 0 | 0 | 0 | 0 | 0 | 0 | 0 | 0 | 0 | 0 | 0 | NA | NA | NA | NA | NA | NA | NA | NA | NA | NA |
| Olfr231 | 0 | 0 | 0 | 0 | 0 | 0 | 0 | 0 | 0 | 0 | 0 | 0 | 0 | 0 | 0 | NA | NA | NA | NA | NA | NA | NA | NA | NA | NA |
| Olfr235 | 0 | 0 | 0 | 0 | 0 | 0 | 0 | 0 | 0 | 0 | 0 | 0 | 0 | 0 | 0 | NA | NA | NA | NA | NA | NA | NA | NA | NA | NA |
| Olfr237-ps1 | 0 | 0 | 0 | 0 | 0 | 0 | 0 | 0 | 0 | 0 | 0 | 0 | 0 | 0 | 0 | NA | NA | NA | NA | NA | NA | NA | NA | NA | NA |
| Olfr239 | 0 | 0 | 0 | 0 | 0 | 0 | 0 | 0 | 0 | 0 | 0 | 0 | 0 | 0 | 0 | NA | NA | NA | NA | NA | NA | NA | NA | NA | NA |
| Olfr24 | 1 | 0 | 0 | 0 | 0 | 1 | 1 | 0 | 0 | 0 | 0 | 1 | 0.36 | 0 | 0 | 0.954 | -0.067 | 0.98553519 | NA | 0 | 9.1716 | 7.9605 | 7.9605 | TRUE | 0.5383 |
| Olfr242 | 0 | 0 | 0 | 0 | 0 | 0 | 0 | 0 | 0 | 0 | 0 | 0 | 0 | 0 | 0 | NA | NA | NA | NA | NA | NA | NA | NA | NA | NA |
| Olfr243 | 0 | 0 | 0 | 0 | 0 | 0 | 0 | 0 | 0 | 0 | 0 | 0 | 0 | 0 | 0 | NA | NA | NA | NA | NA | NA | NA | NA | NA | NA |
| Olfr247 | 0 | 0 | 0 | 0 | 0 | 0 | 0 | 0 | 0 | 0 | 0 | 0 | 0 | 0 | 0 | NA | NA | NA | NA | NA | NA | NA | NA | NA | NA |
| Olfr248 | 0 | 0 | 0 | 0 | 0 | 0 | 0 | 0 | 0 | 0 | 0 | 0 | 0 | 0 | 0 | NA | NA | NA | NA | NA | NA | NA | NA | NA | NA |
| Olfr25 | 0 | 0 | 0 | 0 | 0 | 0 | 0 | 0 | 0 | 0 | 0 | 0 | 0 | 0 | 0 | NA | NA | NA | NA | NA | NA | NA | NA | NA | NA |
| Olfr250 | 0 | 0 | 0 | 0 | 0 | 0 | 0 | 0 | 0 | 0 | 0 | 0 | 0 | 0 | 0 | NA | NA | NA | NA | NA | NA | NA | NA | NA | NA |
| Olfr251 | 0 | 0 | 0 | 0 | 0 | 0 | 0 | 0 | 0 | 0 | 0 | 0 | 0 | 0 | 0 | NA | NA | NA | NA | NA | NA | NA | NA | NA | NA |
| Olfr257 | 0 | 0 | 0 | 0 | 0 | 0 | 0 | 0 | 0 | 0 | 0 | 0 | 0 | 0 | 0 | NA | NA | NA | NA | NA | NA | NA | NA | NA | NA |
| Olfr259 | 0 | 0 | 0 | 0 | 0 | 0 | 0 | 0 | 0 | 0 | 0 | 0 | 0 | 0 | 0 | NA | NA | NA | NA | NA | NA | NA | NA | NA | NA |
| Olfr26 | 0 | 0 | 0 | 0 | 1 | 0 | 0 | 0 | 0 | 0 | 1 | 0 | 0.18 | 0 | 0 | 1.859 | 0.894 | 0.82650734 | NA | 0 | 18.3785 | 10 | 10 | TRUE | 0.5204 |
| Olfr262 | 1 | 0 | 0 | 0 | 0 | 0 | 1 | 0 | 0 | 0 | 0 | 0 | 0.18 | 0 | 0 | 0.49 | -1.029 | 0.80087 | NA | 0 | 18.6165 | 10 | 10 | TRUE | 0.4614 |
| Olfr263 | 0 | 0 | 0 | 0 | 0 | 0 | 0 | 0 | 0 | 0 | 0 | 0 | 0 | 0 | 0 | NA | NA | NA | NA | NA | NA | NA | NA | NA | NA |
| Olfr266 | 0 | 0 | 0 | 0 | 0 | 0 | 0 | 0 | 0 | 0 | 0 | 0 | 0 | 0 | 0 | NA | NA | NA | NA | NA | NA | NA | NA | NA | NA |
| Olfr267 | 0 | 0 | 0 | 0 | 0 | 0 | 0 | 0 | 0 | 0 | 0 | 0 | 0 | 0 | 0 | NA | NA | NA | NA | NA | NA | NA | NA | NA | NA |
| Olfr27 | 0 | 0 | 0 | 0 | 0 | 0 | 0 | 0 | 0 | 0 | 0 | 0 | 0 | 0 | 0 | NA | NA | NA | NA | NA | NA | NA | NA | NA | NA |
| Olfr270 | 0 | 0 | 0 | 0 | 0 | 0 | 0 | 0 | 0 | 0 | 0 | 0 | 0 | 0 | 0 | NA | NA | NA | NA | NA | NA | NA | NA | NA | NA |
| Olfr272 | 0 | 0 | 0 | 0 | 0 | 0 | 0 | 0 | 0 | 0 | 0 | 0 | 0 | 0 | 0 | NA | NA | NA | NA | NA | NA | NA | NA | NA | NA |
| Olfr273 | 0 | 0 | 0 | 0 | 0 | 0 | 0 | 0 | 0 | 0 | 0 | 0 | 0 | 0 | 0 | NA | NA | NA | NA | NA | NA | NA | NA | NA | NA |
| Olfr275 | 0 | 0 | 0 | 0 | 0 | 0 | 0 | 0 | 0 | 0 | 0 | 0 | 0 | 0 | 0 | NA | NA | NA | NA | NA | NA | NA | NA | NA | NA |
| Olfr279 | 0 | 0 | 0 | 0 | 0 | 0 | 0 | 0 | 0 | 0 | 0 | 0 | 0 | 0 | 0 | NA | NA | NA | NA | NA | NA | NA | NA | NA | NA |
| Olfr281 | 0 | 0 | 0 | 0 | 0 | 0 | 0 | 0 | 0 | 0 | 0 | 0 | 0 | 0 | 0 | NA | NA | NA | NA | NA | NA | NA | NA | NA | NA |
| Olfr282 | 0 | 0 | 0 | 0 | 0 | 0 | 0 | 0 | 0 | 0 | 0 | 0 | 0 | 0 | 0 | NA | NA | NA | NA | NA | NA | NA | NA | NA | NA |
| Olfr283 | 0 | 0 | 0 | 0 | 0 | 0 | 0 | 0 | 0 | 0 | 0 | 0 | 0 | 0 | 0 | NA | NA | NA | NA | NA | NA | NA | NA | NA | NA |
| Olfr284 | 0 | 0 | 0 | 0 | 0 | 0 | 0 | 0 | 0 | 0 | 0 | 0 | 0 | 0 | 0 | NA | NA | NA | NA | NA | NA | NA | NA | NA | NA |
| Olfr285 | 0 | 0 | 0 | 0 | 0 | 0 | 0 | 0 | 0 | 0 | 0 | 0 | 0 | 0 | 0 | NA | NA | NA | NA | NA | NA | NA | NA | NA | NA |

| | | | | | | | | | | | | | | | | | | | | | | | | | |
|---|---|---|---|---|---|---|---|---|---|---|---|---|---|---|---|---|---|---|---|---|---|---|---|---|---|
| Olfr286 | 0 | 0 | 0 | 0 | 0 | 1 | 0 | 0 | 0 | 0 | 0 | 1 | 0.18 | 0 | 0 | 1.859 | 0.894 | 0.82650734 | NA | 0 | 18.0643 | 10 | 10 | TRUE | 0.5383 |
| Olfr287 | 5 | 2 | 3 | 5 | 2 | 7 | 5 | 2 | 3 | 4 | 2 | 8 | 4.01 | 3 | 5 | 1.33 | 0.411 | 0.67787078 | NA | 0 | 0.8283 | 0.4492 | 0.4492 | TRUE | 0.4135 |
| Olfr288 | 1 | 0 | 0 | 0 | 0 | 1 | 1 | 0 | 0 | 0 | 0 | 1 | 0.36 | 0 | 0 | 0.954 | -0.067 | 0.98553519 | NA | 0 | 9.1716 | 7.9605 | 7.9605 | TRUE | 0.5383 |
| Olfr29-ps1 | 0 | 0 | 0 | 0 | 0 | 0 | 0 | 0 | 0 | 0 | 0 | 0 | 0 | 0 | 0 | NA | NA | NA | NA | NA | NA | NA | NA | NA | NA |
| Olfr290 | 0 | 0 | 0 | 0 | 0 | 0 | 0 | 0 | 0 | 0 | 0 | 0 | 0 | 0 | 0 | NA | NA | NA | NA | NA | NA | NA | NA | NA | NA |
| Olfr291 | 0 | 0 | 0 | 0 | 0 | 0 | 0 | 0 | 0 | 0 | 0 | 0 | 0 | 0 | 0 | NA | NA | NA | NA | NA | NA | NA | NA | NA | NA |
| Olfr292 | 0 | 0 | 0 | 0 | 0 | 0 | 0 | 0 | 0 | 0 | 0 | 0 | 0 | 0 | 0 | NA | NA | NA | NA | NA | NA | NA | NA | NA | NA |
| Olfr293 | 0 | 1 | 0 | 2 | 0 | 0 | 0 | 1 | 0 | 2 | 0 | 0 | 0.43 | 0 | 1 | 1.493 | 0.578 | 0.86962266 | NA | 0 | 7.6954 | 7.0545 | 7.0545 | TRUE | 0.9718 |
| Olfr294 | 0 | 0 | 0 | 0 | 0 | 0 | 0 | 0 | 0 | 0 | 0 | 0 | 0 | 0 | 0 | NA | NA | NA | NA | NA | NA | NA | NA | NA | NA |
| Olfr295 | 0 | 0 | 0 | 0 | 0 | 0 | 0 | 0 | 0 | 0 | 0 | 0 | 0 | 0 | 0 | NA | NA | NA | NA | NA | NA | NA | NA | NA | NA |
| Olfr297 | 0 | 0 | 0 | 0 | 0 | 0 | 0 | 0 | 0 | 0 | 0 | 0 | 0 | 0 | 0 | NA | NA | NA | NA | NA | NA | NA | NA | NA | NA |
| Olfr298 | 0 | 0 | 0 | 0 | 0 | 0 | 0 | 0 | 0 | 0 | 0 | 0 | 0 | 0 | 0 | NA | NA | NA | NA | NA | NA | NA | NA | NA | NA |
| Olfr299 | 0 | 0 | 0 | 0 | 0 | 0 | 0 | 0 | 0 | 0 | 0 | 0 | 0 | 0 | 0 | NA | NA | NA | NA | NA | NA | NA | NA | NA | NA |
| Olfr3 | 0 | 0 | 0 | 0 | 0 | 0 | 0 | 0 | 0 | 0 | 0 | 0 | 0 | 0 | 0 | NA | NA | NA | NA | NA | NA | NA | NA | NA | NA |
| Olfr30 | 0 | 0 | 0 | 0 | 0 | 0 | 0 | 0 | 0 | 0 | 0 | 0 | 0 | 0 | 0 | NA | NA | NA | NA | NA | NA | NA | NA | NA | NA |
| Olfr301 | 0 | 0 | 0 | 0 | 0 | 0 | 0 | 0 | 0 | 0 | 0 | 0 | 0 | 0 | 0 | NA | NA | NA | NA | NA | NA | NA | NA | NA | NA |
| Olfr303 | 0 | 0 | 0 | 0 | 0 | 0 | 0 | 0 | 0 | 0 | 0 | 0 | 0 | 0 | 0 | NA | NA | NA | NA | NA | NA | NA | NA | NA | NA |
| Olfr304 | 0 | 0 | 0 | 0 | 0 | 0 | 0 | 0 | 0 | 0 | 0 | 0 | 0 | 0 | 0 | NA | NA | NA | NA | NA | NA | NA | NA | NA | NA |
| Olfr305 | 0 | 0 | 0 | 0 | 0 | 0 | 0 | 0 | 0 | 0 | 0 | 0 | 0 | 0 | 0 | NA | NA | NA | NA | NA | NA | NA | NA | NA | NA |
| Olfr307 | 6 | 2 | 7 | 3 | 6 | 6 | 6 | 2 | 7 | 2 | 6 | 7 | 5.13 | 5 | 5 | 0.982 | -0.026 | 0.97677363 | 0.98735331 | 0 | 0.6491 | 0.3925 | 0.3925 | TRUE | 0.6192 |
| Olfr308 | 0 | 0 | 0 | 0 | 0 | 0 | 0 | 0 | 0 | 0 | 0 | 0 | 0 | 0 | 0 | NA | NA | NA | NA | NA | NA | NA | NA | NA | NA |
| Olfr309 | 0 | 0 | 0 | 0 | 0 | 0 | 0 | 0 | 0 | 0 | 0 | 0 | 0 | 0 | 0 | NA | NA | NA | NA | NA | NA | NA | NA | NA | NA |
| Olfr31 | 1 | 0 | 0 | 1 | 0 | 0 | 1 | 0 | 0 | 1 | 0 | 0 | 0.31 | 0 | 0 | 0.954 | -0.067 | 0.98625928 | NA | 0 | 10.7014 | 9.0384 | 9.0384 | TRUE | 0.4614 |
| Olfr310 | 0 | 0 | 0 | 0 | 0 | 0 | 0 | 0 | 0 | 0 | 0 | 0 | 0 | 0 | 0 | NA | NA | NA | NA | NA | NA | NA | NA | NA | NA |
| Olfr311 | 0 | 0 | 0 | 0 | 0 | 0 | 0 | 0 | 0 | 0 | 0 | 0 | 0 | 0 | 0 | NA | NA | NA | NA | NA | NA | NA | NA | NA | NA |
| Olfr312 | 0 | 0 | 0 | 0 | 0 | 0 | 0 | 0 | 0 | 0 | 0 | 0 | 0 | 0 | 0 | NA | NA | NA | NA | NA | NA | NA | NA | NA | NA |
| Olfr313 | 0 | 0 | 0 | 0 | 0 | 0 | 0 | 0 | 0 | 0 | 0 | 0 | 0 | 0 | 0 | NA | NA | NA | NA | NA | NA | NA | NA | NA | NA |
| Olfr314 | 0 | 0 | 0 | 0 | 0 | 0 | 0 | 0 | 0 | 0 | 0 | 0 | 0 | 0 | 0 | NA | NA | NA | NA | NA | NA | NA | NA | NA | NA |
| Olfr315 | 0 | 0 | 0 | 0 | 0 | 0 | 0 | 0 | 0 | 0 | 0 | 0 | 0 | 0 | 0 | NA | NA | NA | NA | NA | NA | NA | NA | NA | NA |
| Olfr316 | 1 | 1 | 1 | 0 | 2 | 1 | 1 | 1 | 1 | 0 | 2 | 1 | 1.05 | 1 | 1 | 1.017 | 0.024 | 0.98943646 | NA | 0 | 3.1387 | 1.4741 | 1.4741 | TRUE | 0.2366 |
| Olfr317 | 0 | 0 | 0 | 0 | 0 | 0 | 0 | 0 | 0 | 0 | 0 | 0 | 0 | 0 | 0 | NA | NA | NA | NA | NA | NA | NA | NA | NA | NA |
| Olfr318 | 0 | 0 | 0 | 0 | 0 | 0 | 0 | 0 | 0 | 0 | 0 | 0 | 0 | 0 | 0 | NA | NA | NA | NA | NA | NA | NA | NA | NA | NA |
| Olfr319 | 0 | 0 | 0 | 0 | 0 | 0 | 0 | 0 | 0 | 0 | 0 | 0 | 0 | 0 | 0 | NA | NA | NA | NA | NA | NA | NA | NA | NA | NA |
| Olfr32 | 0 | 0 | 0 | 0 | 0 | 0 | 0 | 0 | 0 | 0 | 0 | 0 | 0 | 0 | 0 | NA | NA | NA | NA | NA | NA | NA | NA | NA | NA |
| Olfr320 | 0 | 0 | 0 | 0 | 0 | 0 | 0 | 0 | 0 | 0 | 0 | 0 | 0 | 0 | 0 | NA | NA | NA | NA | NA | NA | NA | NA | NA | NA |
| Olfr323 | 0 | 0 | 0 | 2 | 0 | 0 | 0 | 0 | 0 | 2 | 0 | 0 | 0.26 | 0 | 1 | 2.886 | 1.529 | 0.70676491 | NA | 0 | 12.5792 | 10 | 10 | TRUE | 0.9669 |
| Olfr324 | 0 | 0 | 0 | 0 | 0 | 0 | 0 | 0 | 0 | 0 | 0 | 0 | 0 | 0 | 0 | NA | NA | NA | NA | NA | NA | NA | NA | NA | NA |
| Olfr325 | 0 | 0 | 0 | 0 | 0 | 0 | 0 | 0 | 0 | 0 | 0 | 0 | 0 | 0 | 0 | NA | NA | NA | NA | NA | NA | NA | NA | NA | NA |
| Olfr328 | 0 | 0 | 0 | 3 | 0 | 1 | 0 | 0 | 0 | 2 | 0 | 1 | 0.58 | 0 | 1 | 6.21 | 2.635 | 0.40034089 | NA | 0 | 5.7314 | 5.6518 | 5.6518 | TRUE | 0.0653 |
| Olfr329-ps | 5 | 2 | 0 | 15 | 13 | 5 | 5 | 2 | 0 | 12 | 14 | 5 | 6.43 | 2 | 10 | 4.335 | 2.116 | 0.03286466 | 0.1224525 | 0.2978 | 0.5195 | 0.4499 | 0.4499 | TRUE | 1 |
| Olfr33 | 0 | 0 | 0 | 0 | 0 | 0 | 0 | 0 | 0 | 0 | 0 | 0 | 0 | 0 | 0 | NA | NA | NA | NA | NA | NA | NA | NA | NA | NA |
| Olfr330 | 0 | 0 | 0 | 0 | 0 | 1 | 0 | 0 | 0 | 0 | 0 | 1 | 0.18 | 0 | 0 | 1.859 | 0.894 | 0.82650734 | NA | 0 | 18.0643 | 10 | 10 | TRUE | 0.5383 |
| Olfr331 | 0 | 0 | 0 | 0 | 0 | 0 | 0 | 0 | 0 | 0 | 0 | 0 | 0 | 0 | 0 | NA | NA | NA | NA | NA | NA | NA | NA | NA | NA |
| Olfr332 | 0 | 0 | 0 | 0 | 0 | 0 | 0 | 0 | 0 | 0 | 0 | 0 | 0 | 0 | 0 | NA | NA | NA | NA | NA | NA | NA | NA | NA | NA |
| Olfr338 | 0 | 0 | 0 | 0 | 0 | 0 | 0 | 0 | 0 | 0 | 0 | 0 | 0 | 0 | 0 | NA | NA | NA | NA | NA | NA | NA | NA | NA | NA |
| Olfr339 | 0 | 0 | 0 | 0 | 0 | 0 | 0 | 0 | 0 | 0 | 0 | 0 | 0 | 0 | 0 | NA | NA | NA | NA | NA | NA | NA | NA | NA | NA |
| Olfr340 | 0 | 1 | 1 | 1 | 1 | 2 | 0 | 1 | 1 | 1 | 1 | 2 | 1.01 | 1 | 1 | 1.97 | 0.978 | 0.60105973 | NA | 0 | 3.2719 | 1.3982 | 1.3982 | TRUE | 0.2229 |
| Olfr341 | 0 | 0 | 0 | 1 | 0 | 0 | 0 | 0 | 0 | 1 | 0 | 0 | 0.13 | 0 | 0 | 1.859 | 0.894 | 0.82650734 | NA | 0 | 25.1515 | 10 | 10 | TRUE | 0.2563 |
| Olfr342 | 0 | 0 | 0 | 0 | 0 | 0 | 0 | 0 | 0 | 0 | 0 | 0 | 0 | 0 | 0 | NA | NA | NA | NA | NA | NA | NA | NA | NA | NA |
| Olfr344 | 0 | 0 | 0 | 0 | 0 | 0 | 0 | 0 | 0 | 0 | 0 | 0 | 0 | 0 | 0 | NA | NA | NA | NA | NA | NA | NA | NA | NA | NA |
| Olfr345 | 0 | 0 | 0 | 0 | 0 | 0 | 0 | 0 | 0 | 0 | 0 | 0 | 0 | 0 | 0 | NA | NA | NA | NA | NA | NA | NA | NA | NA | NA |
| Olfr346 | 0 | 0 | 0 | 0 | 0 | 0 | 0 | 0 | 0 | 0 | 0 | 0 | 0 | 0 | 0 | NA | NA | NA | NA | NA | NA | NA | NA | NA | NA |
| Olfr347 | 0 | 0 | 0 | 0 | 0 | 0 | 0 | 0 | 0 | 0 | 0 | 0 | 0 | 0 | 0 | NA | NA | NA | NA | NA | NA | NA | NA | NA | NA |
| Olfr348 | 0 | 0 | 0 | 0 | 0 | 0 | 0 | 0 | 0 | 0 | 0 | 0 | 0 | 0 | 0 | NA | NA | NA | NA | NA | NA | NA | NA | NA | NA |
| Olfr350 | 0 | 0 | 0 | 0 | 0 | 0 | 0 | 0 | 0 | 0 | 0 | 0 | 0 | 0 | 0 | NA | NA | NA | NA | NA | NA | NA | NA | NA | NA |
| Olfr351 | 0 | 0 | 0 | 0 | 0 | 0 | 0 | 0 | 0 | 0 | 0 | 0 | 0 | 0 | 0 | NA | NA | NA | NA | NA | NA | NA | NA | NA | NA |
| Olfr352 | 0 | 0 | 0 | 0 | 0 | 0 | 0 | 0 | 0 | 0 | 0 | 0 | 0 | 0 | 0 | NA | NA | NA | NA | NA | NA | NA | NA | NA | NA |
| Olfr353 | 0 | 0 | 0 | 0 | 0 | 0 | 0 | 0 | 0 | 0 | 0 | 0 | 0 | 0 | 0 | NA | NA | NA | NA | NA | NA | NA | NA | NA | NA |
| Olfr354 | 0 | 0 | 0 | 0 | 0 | 0 | 0 | 0 | 0 | 0 | 0 | 0 | 0 | 0 | 0 | NA | NA | NA | NA | NA | NA | NA | NA | NA | NA |
| Olfr355 | 0 | 0 | 0 | 0 | 0 | 0 | 0 | 0 | 0 | 0 | 0 | 0 | 0 | 0 | 0 | NA | NA | NA | NA | NA | NA | NA | NA | NA | NA |
| Olfr356 | 0 | 0 | 0 | 0 | 0 | 0 | 0 | 0 | 0 | 0 | 0 | 0 | 0 | 0 | 0 | NA | NA | NA | NA | NA | NA | NA | NA | NA | NA |
| Olfr357 | 0 | 0 | 0 | 0 | 0 | 0 | 0 | 0 | 0 | 0 | 0 | 0 | 0 | 0 | 0 | NA | NA | NA | NA | NA | NA | NA | NA | NA | NA |
| Olfr358 | 0 | 0 | 0 | 0 | 0 | 0 | 0 | 0 | 0 | 0 | 0 | 0 | 0 | 0 | 0 | NA | NA | NA | NA | NA | NA | NA | NA | NA | NA |
| Olfr360 | 0 | 0 | 0 | 0 | 0 | 0 | 0 | 0 | 0 | 0 | 0 | 0 | 0 | 0 | 0 | NA | NA | NA | NA | NA | NA | NA | NA | NA | NA |
| Olfr361 | 0 | 0 | 0 | 0 | 0 | 0 | 0 | 0 | 0 | 0 | 0 | 0 | 0 | 0 | 0 | NA | NA | NA | NA | NA | NA | NA | NA | NA | NA |
| Olfr362 | 0 | 0 | 0 | 0 | 0 | 0 | 0 | 0 | 0 | 0 | 0 | 0 | 0 | 0 | 0 | NA | NA | NA | NA | NA | NA | NA | NA | NA | NA |
| Olfr365 | 0 | 0 | 0 | 0 | 0 | 0 | 0 | 0 | 0 | 0 | 0 | 0 | 0 | 0 | 0 | NA | NA | NA | NA | NA | NA | NA | NA | NA | NA |
| Olfr366 | 0 | 0 | 0 | 0 | 0 | 0 | 0 | 0 | 0 | 0 | 0 | 0 | 0 | 0 | 0 | NA | NA | NA | NA | NA | NA | NA | NA | NA | NA |
| Olfr367-ps | 0 | 0 | 0 | 0 | 0 | 0 | 0 | 0 | 0 | 0 | 0 | 0 | 0 | 0 | 0 | NA | NA | NA | NA | NA | NA | NA | NA | NA | NA |

| | | | | | | | | | | | | | | | | | | | | | | | | | |
|---|---|---|---|---|---|---|---|---|---|---|---|---|---|---|---|---|---|---|---|---|---|---|---|---|---|
| Olfr368 | 0 | 0 | 0 | 0 | 0 | 0 | 0 | 0 | 0 | 0 | 0 | 0 | 0 | 0 | 0 | NA | NA | NA | NA | NA | NA | NA | NA | NA | NA |
| Olfr370 | 0 | 0 | 0 | 0 | 0 | 0 | 0 | 0 | 0 | 0 | 0 | 0 | 0 | 0 | 0 | NA | NA | NA | NA | NA | NA | NA | NA | NA | NA |
| Olfr371 | 0 | 0 | 0 | 0 | 0 | 0 | 0 | 0 | 0 | 0 | 0 | 0 | 0 | 0 | 0 | NA | NA | NA | NA | NA | NA | NA | NA | NA | NA |
| Olfr372 | 0 | 0 | 0 | 0 | 0 | 0 | 0 | 0 | 0 | 0 | 0 | 0 | 0 | 0 | 0 | NA | NA | NA | NA | NA | NA | NA | NA | NA | NA |
| Olfr373 | 0 | 0 | 0 | 0 | 0 | 0 | 0 | 0 | 0 | 0 | 0 | 0 | 0 | 0 | 0 | NA | NA | NA | NA | NA | NA | NA | NA | NA | NA |
| Olfr374 | 0 | 0 | 0 | 0 | 0 | 0 | 0 | 0 | 0 | 0 | 0 | 0 | 0 | 0 | 0 | NA | NA | NA | NA | NA | NA | NA | NA | NA | NA |
| Olfr376 | 12 | 8 | 1 | 62 | 38 | 22 | 13 | 8 | 1 | 49 | 41 | 24 | 22.57 | 7 | 38 | 5.278 | 2.4 | 0.00017145 | 0.00213938 | 0.3427 | 0.1528 | 0.2102 | 0.2102 | TRUE | 0.8234 |
| Olfr378 | 0 | 0 | 0 | 0 | 0 | 0 | 0 | 0 | 0 | 0 | 0 | 0 | 0 | 0 | 0 | NA | NA | NA | NA | NA | NA | NA | NA | NA | NA |
| Olfr38 | 0 | 0 | 0 | 0 | 0 | 0 | 0 | 0 | 0 | 0 | 0 | 0 | 0 | 0 | 0 | NA | NA | NA | NA | NA | NA | NA | NA | NA | NA |
| Olfr380 | 0 | 0 | 0 | 0 | 0 | 0 | 0 | 0 | 0 | 0 | 0 | 0 | 0 | 0 | 0 | NA | NA | NA | NA | NA | NA | NA | NA | NA | NA |
| Olfr381 | 0 | 0 | 0 | 0 | 0 | 0 | 0 | 0 | 0 | 0 | 0 | 0 | 0 | 0 | 0 | NA | NA | NA | NA | NA | NA | NA | NA | NA | NA |
| Olfr382 | 0 | 0 | 0 | 0 | 0 | 0 | 0 | 0 | 0 | 0 | 0 | 0 | 0 | 0 | 0 | NA | NA | NA | NA | NA | NA | NA | NA | NA | NA |
| Olfr384 | 0 | 0 | 0 | 0 | 0 | 0 | 0 | 0 | 0 | 0 | 0 | 0 | 0 | 0 | 0 | NA | NA | NA | NA | NA | NA | NA | NA | NA | NA |
| Olfr385 | 0 | 0 | 0 | 0 | 0 | 0 | 0 | 0 | 0 | 0 | 0 | 0 | 0 | 0 | 0 | NA | NA | NA | NA | NA | NA | NA | NA | NA | NA |
| Olfr389 | 0 | 0 | 0 | 0 | 0 | 0 | 0 | 0 | 0 | 0 | 0 | 0 | 0 | 0 | 0 | NA | NA | NA | NA | NA | NA | NA | NA | NA | NA |
| Olfr39 | 0 | 0 | 0 | 0 | 0 | 0 | 0 | 0 | 0 | 0 | 0 | 0 | 0 | 0 | 0 | NA | NA | NA | NA | NA | NA | NA | NA | NA | NA |
| Olfr390 | 0 | 0 | 0 | 0 | 0 | 0 | 0 | 0 | 0 | 0 | 0 | 0 | 0 | 0 | 0 | NA | NA | NA | NA | NA | NA | NA | NA | NA | NA |
| Olfr391-ps | 0 | 0 | 0 | 0 | 0 | 0 | 0 | 0 | 0 | 0 | 0 | 0 | 0 | 0 | 0 | NA | NA | NA | NA | NA | NA | NA | NA | NA | NA |
| Olfr392 | 0 | 0 | 0 | 0 | 0 | 0 | 0 | 0 | 0 | 0 | 0 | 0 | 0 | 0 | 0 | NA | NA | NA | NA | NA | NA | NA | NA | NA | NA |
| Olfr393 | 0 | 0 | 0 | 0 | 0 | 0 | 0 | 0 | 0 | 0 | 0 | 0 | 0 | 0 | 0 | NA | NA | NA | NA | NA | NA | NA | NA | NA | NA |
| Olfr394 | 0 | 0 | 0 | 0 | 0 | 0 | 0 | 0 | 0 | 0 | 0 | 0 | 0 | 0 | 0 | NA | NA | NA | NA | NA | NA | NA | NA | NA | NA |
| Olfr395 | 0 | 0 | 0 | 0 | 0 | 0 | 0 | 0 | 0 | 0 | 0 | 0 | 0 | 0 | 0 | NA | NA | NA | NA | NA | NA | NA | NA | NA | NA |
| Olfr397 | 0 | 0 | 0 | 0 | 0 | 0 | 0 | 0 | 0 | 0 | 0 | 0 | 0 | 0 | 0 | NA | NA | NA | NA | NA | NA | NA | NA | NA | NA |
| Olfr398 | 0 | 0 | 0 | 0 | 0 | 0 | 0 | 0 | 0 | 0 | 0 | 0 | 0 | 0 | 0 | NA | NA | NA | NA | NA | NA | NA | NA | NA | NA |
| Olfr399 | 0 | 0 | 0 | 0 | 0 | 0 | 0 | 0 | 0 | 0 | 0 | 0 | 0 | 0 | 0 | NA | NA | NA | NA | NA | NA | NA | NA | NA | NA |
| Olfr401 | 0 | 0 | 0 | 0 | 0 | 0 | 0 | 0 | 0 | 0 | 0 | 0 | 0 | 0 | 0 | NA | NA | NA | NA | NA | NA | NA | NA | NA | NA |
| Olfr402 | 0 | 0 | 0 | 0 | 0 | 0 | 0 | 0 | 0 | 0 | 0 | 0 | 0 | 0 | 0 | NA | NA | NA | NA | NA | NA | NA | NA | NA | NA |
| Olfr403 | 0 | 0 | 0 | 0 | 0 | 0 | 0 | 0 | 0 | 0 | 0 | 0 | 0 | 0 | 0 | NA | NA | NA | NA | NA | NA | NA | NA | NA | NA |
| Olfr406 | 0 | 0 | 0 | 0 | 0 | 0 | 0 | 0 | 0 | 0 | 0 | 0 | 0 | 0 | 0 | NA | NA | NA | NA | NA | NA | NA | NA | NA | NA |
| Olfr410 | 0 | 0 | 0 | 0 | 0 | 0 | 0 | 0 | 0 | 0 | 0 | 0 | 0 | 0 | 0 | NA | NA | NA | NA | NA | NA | NA | NA | NA | NA |
| Olfr411 | 0 | 0 | 0 | 0 | 0 | 0 | 0 | 0 | 0 | 0 | 0 | 0 | 0 | 0 | 0 | NA | NA | NA | NA | NA | NA | NA | NA | NA | NA |
| Olfr412 | 0 | 0 | 0 | 0 | 0 | 0 | 0 | 0 | 0 | 0 | 0 | 0 | 0 | 0 | 0 | NA | NA | NA | NA | NA | NA | NA | NA | NA | NA |
| Olfr414 | 0 | 0 | 0 | 0 | 0 | 0 | 0 | 0 | 0 | 0 | 0 | 0 | 0 | 0 | 0 | NA | NA | NA | NA | NA | NA | NA | NA | NA | NA |
| Olfr417 | 0 | 0 | 0 | 0 | 0 | 0 | 0 | 0 | 0 | 0 | 0 | 0 | 0 | 0 | 0 | NA | NA | NA | NA | NA | NA | NA | NA | NA | NA |
| Olfr418-ps1 | 0 | 0 | 0 | 0 | 0 | 0 | 0 | 0 | 0 | 0 | 0 | 0 | 0 | 0 | 0 | NA | NA | NA | NA | NA | NA | NA | NA | NA | NA |
| Olfr419 | 0 | 0 | 0 | 0 | 0 | 0 | 0 | 0 | 0 | 0 | 0 | 0 | 0 | 0 | 0 | NA | NA | NA | NA | NA | NA | NA | NA | NA | NA |
| Olfr420 | 0 | 0 | 0 | 0 | 0 | 0 | 0 | 0 | 0 | 0 | 0 | 0 | 0 | 0 | 0 | NA | NA | NA | NA | NA | NA | NA | NA | NA | NA |
| Olfr421-ps1 | 0 | 0 | 0 | 0 | 0 | 0 | 0 | 0 | 0 | 0 | 0 | 0 | 0 | 0 | 0 | NA | NA | NA | NA | NA | NA | NA | NA | NA | NA |
| Olfr424 | 0 | 0 | 0 | 0 | 0 | 0 | 0 | 0 | 0 | 0 | 0 | 0 | 0 | 0 | 0 | NA | NA | NA | NA | NA | NA | NA | NA | NA | NA |
| Olfr426 | 0 | 0 | 0 | 0 | 0 | 0 | 0 | 0 | 0 | 0 | 0 | 0 | 0 | 0 | 0 | NA | NA | NA | NA | NA | NA | NA | NA | NA | NA |
| Olfr427 | 0 | 0 | 0 | 0 | 0 | 0 | 0 | 0 | 0 | 0 | 0 | 0 | 0 | 0 | 0 | NA | NA | NA | NA | NA | NA | NA | NA | NA | NA |
| Olfr429 | 0 | 0 | 0 | 0 | 0 | 0 | 0 | 0 | 0 | 0 | 0 | 0 | 0 | 0 | 0 | NA | NA | NA | NA | NA | NA | NA | NA | NA | NA |
| Olfr43 | 0 | 0 | 0 | 0 | 0 | 0 | 0 | 0 | 0 | 0 | 0 | 0 | 0 | 0 | 0 | NA | NA | NA | NA | NA | NA | NA | NA | NA | NA |
| Olfr430 | 0 | 0 | 0 | 0 | 0 | 0 | 0 | 0 | 0 | 0 | 0 | 0 | 0 | 0 | 0 | NA | NA | NA | NA | NA | NA | NA | NA | NA | NA |
| Olfr432 | 0 | 0 | 0 | 0 | 0 | 0 | 0 | 0 | 0 | 0 | 0 | 0 | 0 | 0 | 0 | NA | NA | NA | NA | NA | NA | NA | NA | NA | NA |
| Olfr433 | 0 | 0 | 0 | 0 | 0 | 0 | 0 | 0 | 0 | 0 | 0 | 0 | 0 | 0 | 0 | NA | NA | NA | NA | NA | NA | NA | NA | NA | NA |
| Olfr434 | 0 | 0 | 0 | 0 | 0 | 0 | 0 | 0 | 0 | 0 | 0 | 0 | 0 | 0 | 0 | NA | NA | NA | NA | NA | NA | NA | NA | NA | NA |
| Olfr435 | 0 | 0 | 0 | 0 | 0 | 0 | 0 | 0 | 0 | 0 | 0 | 0 | 0 | 0 | 0 | NA | NA | NA | NA | NA | NA | NA | NA | NA | NA |
| Olfr437 | 0 | 0 | 0 | 0 | 0 | 0 | 0 | 0 | 0 | 0 | 0 | 0 | 0 | 0 | 0 | NA | NA | NA | NA | NA | NA | NA | NA | NA | NA |
| Olfr44 | 0 | 0 | 0 | 0 | 0 | 0 | 0 | 0 | 0 | 0 | 0 | 0 | 0 | 0 | 0 | NA | NA | NA | NA | NA | NA | NA | NA | NA | NA |
| Olfr441 | 0 | 0 | 0 | 0 | 0 | 0 | 0 | 0 | 0 | 0 | 0 | 0 | 0 | 0 | 0 | NA | NA | NA | NA | NA | NA | NA | NA | NA | NA |
| Olfr444 | 0 | 0 | 0 | 0 | 0 | 0 | 0 | 0 | 0 | 0 | 0 | 0 | 0 | 0 | 0 | NA | NA | NA | NA | NA | NA | NA | NA | NA | NA |
| Olfr446 | 0 | 0 | 0 | 0 | 0 | 0 | 0 | 0 | 0 | 0 | 0 | 0 | 0 | 0 | 0 | NA | NA | NA | NA | NA | NA | NA | NA | NA | NA |
| Olfr447 | 0 | 0 | 0 | 0 | 0 | 0 | 0 | 0 | 0 | 0 | 0 | 0 | 0 | 0 | 0 | NA | NA | NA | NA | NA | NA | NA | NA | NA | NA |
| Olfr448 | 0 | 0 | 0 | 0 | 0 | 0 | 0 | 0 | 0 | 0 | 0 | 0 | 0 | 0 | 0 | NA | NA | NA | NA | NA | NA | NA | NA | NA | NA |
| Olfr449 | 0 | 0 | 0 | 0 | 0 | 0 | 0 | 0 | 0 | 0 | 0 | 0 | 0 | 0 | 0 | NA | NA | NA | NA | NA | NA | NA | NA | NA | NA |
| Olfr45 | 0 | 0 | 0 | 0 | 0 | 0 | 0 | 0 | 0 | 0 | 0 | 0 | 0 | 0 | 0 | NA | NA | NA | NA | NA | NA | NA | NA | NA | NA |
| Olfr450 | 0 | 0 | 0 | 0 | 0 | 0 | 0 | 0 | 0 | 0 | 0 | 0 | 0 | 0 | 0 | NA | NA | NA | NA | NA | NA | NA | NA | NA | NA |
| Olfr452 | 0 | 0 | 0 | 0 | 0 | 0 | 0 | 0 | 0 | 0 | 0 | 0 | 0 | 0 | 0 | NA | NA | NA | NA | NA | NA | NA | NA | NA | NA |
| Olfr453 | 0 | 0 | 0 | 0 | 0 | 0 | 0 | 0 | 0 | 0 | 0 | 0 | 0 | 0 | 0 | NA | NA | NA | NA | NA | NA | NA | NA | NA | NA |
| Olfr455 | 1 | 0 | 2 | 3 | 1 | 0 | 1 | 0 | 2 | 2 | 1 | 0 | 1.08 | 1 | 1 | 1.158 | 0.212 | 0.91438814 | NA | 0 | 3.0457 | 1.8907 | 1.8907 | TRUE | 0.2388 |
| Olfr456 | 0 | 0 | 0 | 0 | 0 | 0 | 0 | 0 | 0 | 0 | 0 | 0 | 0 | 0 | 0 | NA | NA | NA | NA | NA | NA | NA | NA | NA | NA |
| Olfr457 | 0 | 0 | 0 | 0 | 0 | 0 | 0 | 0 | 0 | 0 | 0 | 0 | 0 | 0 | 0 | NA | NA | NA | NA | NA | NA | NA | NA | NA | NA |
| Olfr458 | 0 | 0 | 0 | 0 | 0 | 0 | 0 | 0 | 0 | 0 | 0 | 0 | 0 | 0 | 0 | NA | NA | NA | NA | NA | NA | NA | NA | NA | NA |
| Olfr459 | 1 | 0 | 0 | 1 | 0 | 1 | 1 | 0 | 0 | 1 | 0 | 1 | 0.49 | 0 | 1 | 1.724 | 0.785 | 0.78200058 | NA | 0 | 6.7235 | 4.0295 | 4.0295 | TRUE | 0.4614 |
| Olfr46 | 0 | 0 | 0 | 0 | 0 | 0 | 0 | 0 | 0 | 0 | 0 | 0 | 0 | 0 | 0 | NA | NA | NA | NA | NA | NA | NA | NA | NA | NA |
| Olfr460 | 1 | 0 | 0 | 0 | 0 | 0 | 1 | 0 | 0 | 0 | 0 | 0 | 0.18 | 0 | 0 | 0.49 | -1.029 | 0.80087 | NA | 0 | 18.6165 | 10 | 10 | TRUE | 0.4614 |
| Olfr461 | 0 | 0 | 0 | 0 | 0 | 0 | 0 | 0 | 0 | 0 | 0 | 0 | 0 | 0 | 0 | NA | NA | NA | NA | NA | NA | NA | NA | NA | NA |
| Olfr462 | 0 | 0 | 0 | 0 | 0 | 0 | 0 | 0 | 0 | 0 | 0 | 0 | 0 | 0 | 0 | NA | NA | NA | NA | NA | NA | NA | NA | NA | NA |

| | | | | | | | | | | | | | | | | | | | | | | | | | |
|---|---|---|---|---|---|---|---|---|---|---|---|---|---|---|---|---|---|---|---|---|---|---|---|---|---|
| Olfr463 | 0 | 0 | 0 | 0 | 0 | 0 | 0 | 0 | 0 | 0 | 0 | 0 | 0 | 0 | 0 | NA | NA | NA | NA | NA | NA | NA | NA | NA | NA |
| Olfr464 | 3 | 0 | 0 | 1 | 0 | 0 | 3 | 0 | 0 | 1 | 0 | 0 | 0.66 | 1 | 0 | 0.331 | -1.597 | 0.61164788 | NA | 8.2589 | 4.9826 | 5.6428 | 5.6428 | TRUE | 1.4426 |
| Olfr466 | 0 | 0 | 0 | 0 | 0 | 0 | 0 | 0 | 0 | 0 | 0 | 0 | 0 | 0 | 0 | NA | NA | NA | NA | NA | NA | NA | NA | NA | NA |
| Olfr467 | 0 | 0 | 0 | 0 | 0 | 0 | 0 | 0 | 0 | 0 | 0 | 0 | 0 | 0 | 0 | NA | NA | NA | NA | NA | NA | NA | NA | NA | NA |
| Olfr469 | 0 | 0 | 0 | 0 | 0 | 0 | 0 | 0 | 0 | 0 | 0 | 0 | 0 | 0 | 0 | NA | NA | NA | NA | NA | NA | NA | NA | NA | NA |
| Olfr47 | 0 | 0 | 0 | 0 | 0 | 0 | 0 | 0 | 0 | 0 | 0 | 0 | 0 | 0 | 0 | NA | NA | NA | NA | NA | NA | NA | NA | NA | NA |
| Olfr470 | 0 | 0 | 0 | 0 | 0 | 0 | 0 | 0 | 0 | 0 | 0 | 0 | 0 | 0 | 0 | NA | NA | NA | NA | NA | NA | NA | NA | NA | NA |
| Olfr472 | 0 | 0 | 0 | 0 | 0 | 0 | 0 | 0 | 0 | 0 | 0 | 0 | 0 | 0 | 0 | NA | NA | NA | NA | NA | NA | NA | NA | NA | NA |
| Olfr473 | 0 | 0 | 0 | 0 | 0 | 0 | 0 | 0 | 0 | 0 | 0 | 0 | 0 | 0 | 0 | NA | NA | NA | NA | NA | NA | NA | NA | NA | NA |
| Olfr474 | 0 | 0 | 0 | 0 | 0 | 0 | 0 | 0 | 0 | 0 | 0 | 0 | 0 | 0 | 0 | NA | NA | NA | NA | NA | NA | NA | NA | NA | NA |
| Olfr476 | 0 | 0 | 0 | 0 | 0 | 0 | 0 | 0 | 0 | 0 | 0 | 0 | 0 | 0 | 0 | NA | NA | NA | NA | NA | NA | NA | NA | NA | NA |
| Olfr477 | 0 | 0 | 0 | 0 | 0 | 0 | 0 | 0 | 0 | 0 | 0 | 0 | 0 | 0 | 0 | NA | NA | NA | NA | NA | NA | NA | NA | NA | NA |
| Olfr478 | 0 | 0 | 0 | 0 | 0 | 0 | 0 | 0 | 0 | 0 | 0 | 0 | 0 | 0 | 0 | NA | NA | NA | NA | NA | NA | NA | NA | NA | NA |
| Olfr479 | 0 | 0 | 0 | 0 | 0 | 0 | 0 | 0 | 0 | 0 | 0 | 0 | 0 | 0 | 0 | NA | NA | NA | NA | NA | NA | NA | NA | NA | NA |
| Olfr48 | 0 | 0 | 0 | 0 | 0 | 0 | 0 | 0 | 0 | 0 | 0 | 0 | 0 | 0 | 0 | NA | NA | NA | NA | NA | NA | NA | NA | NA | NA |
| Olfr480 | 0 | 0 | 0 | 0 | 0 | 0 | 0 | 0 | 0 | 0 | 0 | 0 | 0 | 0 | 0 | NA | NA | NA | NA | NA | NA | NA | NA | NA | NA |
| Olfr481 | 0 | 0 | 0 | 0 | 0 | 0 | 0 | 0 | 0 | 0 | 0 | 0 | 0 | 0 | 0 | NA | NA | NA | NA | NA | NA | NA | NA | NA | NA |
| Olfr482 | 0 | 0 | 0 | 0 | 0 | 0 | 0 | 0 | 0 | 0 | 0 | 0 | 0 | 0 | 0 | NA | NA | NA | NA | NA | NA | NA | NA | NA | NA |
| Olfr483 | 0 | 0 | 0 | 0 | 0 | 0 | 0 | 0 | 0 | 0 | 0 | 0 | 0 | 0 | 0 | NA | NA | NA | NA | NA | NA | NA | NA | NA | NA |
| Olfr484 | 0 | 0 | 0 | 0 | 0 | 0 | 0 | 0 | 0 | 0 | 0 | 0 | 0 | 0 | 0 | NA | NA | NA | NA | NA | NA | NA | NA | NA | NA |
| Olfr485 | 0 | 0 | 0 | 0 | 0 | 0 | 0 | 0 | 0 | 0 | 0 | 0 | 0 | 0 | 0 | NA | NA | NA | NA | NA | NA | NA | NA | NA | NA |
| Olfr486 | 0 | 0 | 0 | 0 | 0 | 0 | 0 | 0 | 0 | 0 | 0 | 0 | 0 | 0 | 0 | NA | NA | NA | NA | NA | NA | NA | NA | NA | NA |
| Olfr487 | 0 | 0 | 0 | 0 | 0 | 0 | 0 | 0 | 0 | 0 | 0 | 0 | 0 | 0 | 0 | NA | NA | NA | NA | NA | NA | NA | NA | NA | NA |
| Olfr488 | 0 | 0 | 0 | 0 | 0 | 0 | 0 | 0 | 0 | 0 | 0 | 0 | 0 | 0 | 0 | NA | NA | NA | NA | NA | NA | NA | NA | NA | NA |
| Olfr49 | 0 | 0 | 0 | 0 | 0 | 0 | 0 | 0 | 0 | 0 | 0 | 0 | 0 | 0 | 0 | NA | NA | NA | NA | NA | NA | NA | NA | NA | NA |
| Olfr490 | 0 | 0 | 0 | 0 | 0 | 0 | 0 | 0 | 0 | 0 | 0 | 0 | 0 | 0 | 0 | NA | NA | NA | NA | NA | NA | NA | NA | NA | NA |
| Olfr491 | 0 | 0 | 0 | 0 | 0 | 0 | 0 | 0 | 0 | 0 | 0 | 0 | 0 | 0 | 0 | NA | NA | NA | NA | NA | NA | NA | NA | NA | NA |
| Olfr492 | 0 | 0 | 0 | 0 | 0 | 0 | 0 | 0 | 0 | 0 | 0 | 0 | 0 | 0 | 0 | NA | NA | NA | NA | NA | NA | NA | NA | NA | NA |
| Olfr493 | 0 | 0 | 0 | 0 | 0 | 0 | 0 | 0 | 0 | 0 | 0 | 0 | 0 | 0 | 0 | NA | NA | NA | NA | NA | NA | NA | NA | NA | NA |
| Olfr494 | 0 | 0 | 0 | 0 | 0 | 0 | 0 | 0 | 0 | 0 | 0 | 0 | 0 | 0 | 0 | NA | NA | NA | NA | NA | NA | NA | NA | NA | NA |
| Olfr495 | 0 | 0 | 0 | 0 | 0 | 0 | 0 | 0 | 0 | 0 | 0 | 0 | 0 | 0 | 0 | NA | NA | NA | NA | NA | NA | NA | NA | NA | NA |
| Olfr497 | 0 | 0 | 0 | 0 | 0 | 0 | 0 | 0 | 0 | 0 | 0 | 0 | 0 | 0 | 0 | NA | NA | NA | NA | NA | NA | NA | NA | NA | NA |
| Olfr498 | 0 | 0 | 0 | 0 | 0 | 0 | 0 | 0 | 0 | 0 | 0 | 0 | 0 | 0 | 0 | NA | NA | NA | NA | NA | NA | NA | NA | NA | NA |
| Olfr5 | 0 | 0 | 0 | 1 | 0 | 0 | 0 | 0 | 0 | 1 | 0 | 0 | 0.13 | 0 | 0 | 1.859 | 0.894 | 0.82650734 | NA | 0 | 25.1515 | 10 | 10 | TRUE | 0.2563 |
| Olfr50 | 0 | 0 | 0 | 0 | 0 | 0 | 0 | 0 | 0 | 0 | 0 | 0 | 0 | 0 | 0 | NA | NA | NA | NA | NA | NA | NA | NA | NA | NA |
| Olfr502 | 0 | 0 | 0 | 0 | 0 | 0 | 0 | 0 | 0 | 0 | 0 | 0 | 0 | 0 | 0 | NA | NA | NA | NA | NA | NA | NA | NA | NA | NA |
| Olfr503 | 4 | 3 | 0 | 7 | 5 | 4 | 4 | 3 | 0 | 6 | 5 | 4 | 3.75 | 2 | 5 | 2.124 | 1.087 | 0.31824535 | NA | 0 | 0.8852 | 0.546 | 0.546 | TRUE | 0.8304 |
| Olfr504 | 0 | 0 | 0 | 1 | 0 | 0 | 0 | 0 | 0 | 1 | 0 | 0 | 0.13 | 0 | 0 | 1.859 | 0.894 | 0.82650734 | NA | 0 | 25.1515 | 10 | 10 | TRUE | 0.2563 |
| Olfr506 | 0 | 0 | 0 | 0 | 0 | 0 | 0 | 0 | 0 | 0 | 0 | 0 | 0 | 0 | 0 | NA | NA | NA | NA | NA | NA | NA | NA | NA | NA |
| Olfr507 | 0 | 0 | 0 | 0 | 0 | 0 | 0 | 0 | 0 | 0 | 0 | 0 | 0 | 0 | 0 | NA | NA | NA | NA | NA | NA | NA | NA | NA | NA |
| Olfr508 | 0 | 0 | 0 | 0 | 0 | 0 | 0 | 0 | 0 | 0 | 0 | 0 | 0 | 0 | 0 | NA | NA | NA | NA | NA | NA | NA | NA | NA | NA |
| Olfr509 | 0 | 0 | 0 | 0 | 0 | 0 | 0 | 0 | 0 | 0 | 0 | 0 | 0 | 0 | 0 | NA | NA | NA | NA | NA | NA | NA | NA | NA | NA |
| Olfr51 | 0 | 0 | 0 | 0 | 0 | 0 | 0 | 0 | 0 | 0 | 0 | 0 | 0 | 0 | 0 | NA | NA | NA | NA | NA | NA | NA | NA | NA | NA |
| Olfr510 | 0 | 0 | 0 | 0 | 0 | 0 | 0 | 0 | 0 | 0 | 0 | 0 | 0 | 0 | 0 | NA | NA | NA | NA | NA | NA | NA | NA | NA | NA |
| Olfr512 | 0 | 0 | 0 | 0 | 0 | 0 | 0 | 0 | 0 | 0 | 0 | 0 | 0 | 0 | 0 | NA | NA | NA | NA | NA | NA | NA | NA | NA | NA |
| Olfr513 | 0 | 0 | 0 | 0 | 0 | 0 | 0 | 0 | 0 | 0 | 0 | 0 | 0 | 0 | 0 | NA | NA | NA | NA | NA | NA | NA | NA | NA | NA |
| Olfr514 | 0 | 0 | 0 | 0 | 0 | 0 | 0 | 0 | 0 | 0 | 0 | 0 | 0 | 0 | 0 | NA | NA | NA | NA | NA | NA | NA | NA | NA | NA |
| Olfr516 | 0 | 0 | 0 | 0 | 0 | 0 | 0 | 0 | 0 | 0 | 0 | 0 | 0 | 0 | 0 | NA | NA | NA | NA | NA | NA | NA | NA | NA | NA |
| Olfr517 | 0 | 0 | 0 | 0 | 0 | 0 | 0 | 0 | 0 | 0 | 0 | 0 | 0 | 0 | 0 | NA | NA | NA | NA | NA | NA | NA | NA | NA | NA |
| Olfr518 | 0 | 0 | 0 | 0 | 0 | 0 | 0 | 0 | 0 | 0 | 0 | 0 | 0 | 0 | 0 | NA | NA | NA | NA | NA | NA | NA | NA | NA | NA |
| Olfr519 | 0 | 0 | 0 | 0 | 0 | 0 | 0 | 0 | 0 | 0 | 0 | 0 | 0 | 0 | 0 | NA | NA | NA | NA | NA | NA | NA | NA | NA | NA |
| Olfr52 | 0 | 0 | 0 | 0 | 0 | 0 | 0 | 0 | 0 | 0 | 0 | 0 | 0 | 0 | 0 | NA | NA | NA | NA | NA | NA | NA | NA | NA | NA |
| Olfr520 | 0 | 0 | 0 | 0 | 0 | 0 | 0 | 0 | 0 | 0 | 0 | 0 | 0 | 0 | 0 | NA | NA | NA | NA | NA | NA | NA | NA | NA | NA |
| Olfr521 | 0 | 0 | 0 | 0 | 0 | 0 | 0 | 0 | 0 | 0 | 0 | 0 | 0 | 0 | 0 | NA | NA | NA | NA | NA | NA | NA | NA | NA | NA |
| Olfr522 | 0 | 0 | 0 | 0 | 0 | 0 | 0 | 0 | 0 | 0 | 0 | 0 | 0 | 0 | 0 | NA | NA | NA | NA | NA | NA | NA | NA | NA | NA |
| Olfr523 | 0 | 0 | 0 | 0 | 0 | 0 | 0 | 0 | 0 | 0 | 0 | 0 | 0 | 0 | 0 | NA | NA | NA | NA | NA | NA | NA | NA | NA | NA |
| Olfr524 | 0 | 0 | 0 | 0 | 0 | 0 | 0 | 0 | 0 | 0 | 0 | 0 | 0 | 0 | 0 | NA | NA | NA | NA | NA | NA | NA | NA | NA | NA |
| Olfr525 | 0 | 0 | 0 | 0 | 0 | 0 | 0 | 0 | 0 | 0 | 0 | 0 | 0 | 0 | 0 | NA | NA | NA | NA | NA | NA | NA | NA | NA | NA |
| Olfr527 | 0 | 0 | 0 | 0 | 0 | 0 | 0 | 0 | 0 | 0 | 0 | 0 | 0 | 0 | 0 | NA | NA | NA | NA | NA | NA | NA | NA | NA | NA |
| Olfr53 | 0 | 0 | 0 | 0 | 0 | 0 | 0 | 0 | 0 | 0 | 0 | 0 | 0 | 0 | 0 | NA | NA | NA | NA | NA | NA | NA | NA | NA | NA |
| Olfr530 | 0 | 0 | 0 | 0 | 0 | 0 | 0 | 0 | 0 | 0 | 0 | 0 | 0 | 0 | 0 | NA | NA | NA | NA | NA | NA | NA | NA | NA | NA |
| Olfr531 | 0 | 0 | 0 | 0 | 0 | 0 | 0 | 0 | 0 | 0 | 0 | 0 | 0 | 0 | 0 | NA | NA | NA | NA | NA | NA | NA | NA | NA | NA |
| Olfr532 | 0 | 0 | 0 | 0 | 0 | 0 | 0 | 0 | 0 | 0 | 0 | 0 | 0 | 0 | 0 | NA | NA | NA | NA | NA | NA | NA | NA | NA | NA |
| Olfr533 | 0 | 0 | 0 | 0 | 0 | 0 | 0 | 0 | 0 | 0 | 0 | 0 | 0 | 0 | 0 | NA | NA | NA | NA | NA | NA | NA | NA | NA | NA |
| Olfr535 | 0 | 0 | 0 | 0 | 0 | 0 | 0 | 0 | 0 | 0 | 0 | 0 | 0 | 0 | 0 | NA | NA | NA | NA | NA | NA | NA | NA | NA | NA |
| Olfr536 | 0 | 0 | 1 | 0 | 0 | 0 | 0 | 0 | 1 | 0 | 0 | 0 | 0.17 | 0 | 0 | 0.49 | -1.029 | 0.80087 | NA | 0 | 19.6928 | 10 | 10 | TRUE | 0.4098 |
| Olfr538 | 0 | 0 | 0 | 0 | 0 | 0 | 0 | 0 | 0 | 0 | 0 | 0 | 0 | 0 | 0 | NA | NA | NA | NA | NA | NA | NA | NA | NA | NA |
| Olfr539 | 0 | 0 | 0 | 0 | 0 | 0 | 0 | 0 | 0 | 0 | 0 | 0 | 0 | 0 | 0 | NA | NA | NA | NA | NA | NA | NA | NA | NA | NA |
| Olfr54 | 0 | 0 | 0 | 0 | 0 | 0 | 0 | 0 | 0 | 0 | 0 | 0 | 0 | 0 | 0 | NA | NA | NA | NA | NA | NA | NA | NA | NA | NA |

| | | | | | | | | | | | | | | | | | | | | | | | | | |
|---|---|---|---|---|---|---|---|---|---|---|---|---|---|---|---|---|---|---|---|---|---|---|---|---|---|
| Olfr541 | 0 | 0 | 0 | 0 | 0 | 0 | 0 | 0 | 0 | 0 | 0 | 0 | 0 | 0 | 0 | NA | NA | NA | NA | NA | NA | NA | NA | NA | NA |
| Olfr543 | 0 | 0 | 0 | 0 | 0 | 0 | 0 | 0 | 0 | 0 | 0 | 0 | 0 | 0 | 0 | NA | NA | NA | NA | NA | NA | NA | NA | NA | NA |
| Olfr544 | 0 | 0 | 0 | 0 | 0 | 0 | 0 | 0 | 0 | 0 | 0 | 0 | 0 | 0 | 0 | NA | NA | NA | NA | NA | NA | NA | NA | NA | NA |
| Olfr545 | 0 | 0 | 0 | 0 | 0 | 0 | 0 | 0 | 0 | 0 | 0 | 0 | 0 | 0 | 0 | NA | NA | NA | NA | NA | NA | NA | NA | NA | NA |
| Olfr547 | 0 | 0 | 0 | 0 | 0 | 0 | 0 | 0 | 0 | 0 | 0 | 0 | 0 | 0 | 0 | NA | NA | NA | NA | NA | NA | NA | NA | NA | NA |
| Olfr549 | 0 | 0 | 0 | 0 | 0 | 0 | 0 | 0 | 0 | 0 | 0 | 0 | 0 | 0 | 0 | NA | NA | NA | NA | NA | NA | NA | NA | NA | NA |
| Olfr55 | 0 | 0 | 0 | 1 | 0 | 0 | 0 | 0 | 0 | 1 | 0 | 0 | 0.13 | 0 | 0 | 1.859 | 0.894 | 0.82650734 | NA | 0 | 25.1515 | 10 | 10 | TRUE | 0.2563 |
| Olfr550 | 0 | 0 | 0 | 0 | 0 | 0 | 0 | 0 | 0 | 0 | 0 | 0 | 0 | 0 | 0 | NA | NA | NA | NA | NA | NA | NA | NA | NA | NA |
| Olfr551 | 0 | 0 | 0 | 0 | 0 | 0 | 0 | 0 | 0 | 0 | 0 | 0 | 0 | 0 | 0 | NA | NA | NA | NA | NA | NA | NA | NA | NA | NA |
| Olfr552 | 0 | 0 | 0 | 0 | 0 | 0 | 0 | 0 | 0 | 0 | 0 | 0 | 0 | 0 | 0 | NA | NA | NA | NA | NA | NA | NA | NA | NA | NA |
| Olfr553 | 0 | 0 | 0 | 0 | 0 | 0 | 0 | 0 | 0 | 0 | 0 | 0 | 0 | 0 | 0 | NA | NA | NA | NA | NA | NA | NA | NA | NA | NA |
| Olfr554 | 0 | 0 | 0 | 0 | 0 | 0 | 0 | 0 | 0 | 0 | 0 | 0 | 0 | 0 | 0 | NA | NA | NA | NA | NA | NA | NA | NA | NA | NA |
| Olfr555 | 0 | 0 | 0 | 0 | 0 | 0 | 0 | 0 | 0 | 0 | 0 | 0 | 0 | 0 | 0 | NA | NA | NA | NA | NA | NA | NA | NA | NA | NA |
| Olfr556 | 0 | 0 | 0 | 0 | 0 | 0 | 0 | 0 | 0 | 0 | 0 | 0 | 0 | 0 | 0 | NA | NA | NA | NA | NA | NA | NA | NA | NA | NA |
| Olfr557 | 0 | 0 | 0 | 0 | 0 | 1 | 0 | 0 | 0 | 0 | 0 | 1 | 0.18 | 0 | 0 | 1.859 | 0.894 | 0.82650734 | NA | 0 | 18.0643 | 10 | 10 | TRUE | 0.5383 |
| Olfr558 | 0 | 0 | 0 | 1 | 0 | 0 | 0 | 0 | 0 | 1 | 0 | 0 | 0.13 | 0 | 0 | 1.859 | 0.894 | 0.82650734 | NA | 0 | 25.1515 | 10 | 10 | TRUE | 0.2563 |
| Olfr559 | 0 | 0 | 0 | 0 | 0 | 0 | 0 | 0 | 0 | 0 | 0 | 0 | 0 | 0 | 0 | NA | NA | NA | NA | NA | NA | NA | NA | NA | NA |
| Olfr56 | 0 | 0 | 0 | 0 | 0 | 0 | 0 | 0 | 0 | 0 | 0 | 0 | 0 | 0 | 0 | NA | NA | NA | NA | NA | NA | NA | NA | NA | NA |
| Olfr560 | 0 | 0 | 0 | 0 | 0 | 0 | 0 | 0 | 0 | 0 | 0 | 0 | 0 | 0 | 0 | NA | NA | NA | NA | NA | NA | NA | NA | NA | NA |
| Olfr561 | 0 | 0 | 0 | 0 | 0 | 0 | 0 | 0 | 0 | 0 | 0 | 0 | 0 | 0 | 0 | NA | NA | NA | NA | NA | NA | NA | NA | NA | NA |
| Olfr564 | 0 | 0 | 0 | 0 | 0 | 0 | 0 | 0 | 0 | 0 | 0 | 0 | 0 | 0 | 0 | NA | NA | NA | NA | NA | NA | NA | NA | NA | NA |
| Olfr566 | 0 | 1 | 0 | 0 | 0 | 0 | 0 | 1 | 0 | 0 | 0 | 0 | 0.17 | 0 | 0 | 0.49 | -1.029 | 0.80087 | NA | 0 | 19.7991 | 10 | 10 | TRUE | 0.405 |
| Olfr568 | 0 | 0 | 0 | 0 | 0 | 0 | 0 | 0 | 0 | 0 | 0 | 0 | 0 | 0 | 0 | NA | NA | NA | NA | NA | NA | NA | NA | NA | NA |
| Olfr569 | 0 | 0 | 0 | 0 | 0 | 0 | 0 | 0 | 0 | 0 | 0 | 0 | 0 | 0 | 0 | NA | NA | NA | NA | NA | NA | NA | NA | NA | NA |
| Olfr57 | 0 | 0 | 0 | 0 | 0 | 0 | 0 | 0 | 0 | 0 | 0 | 0 | 0 | 0 | 0 | NA | NA | NA | NA | NA | NA | NA | NA | NA | NA |
| Olfr570 | 0 | 0 | 0 | 1 | 0 | 0 | 0 | 0 | 0 | 1 | 0 | 0 | 0.13 | 0 | 0 | 1.859 | 0.894 | 0.82650734 | NA | 0 | 25.1515 | 10 | 10 | TRUE | 0.2563 |
| Olfr571 | 0 | 0 | 0 | 0 | 0 | 0 | 0 | 0 | 0 | 0 | 0 | 0 | 0 | 0 | 0 | NA | NA | NA | NA | NA | NA | NA | NA | NA | NA |
| Olfr572 | 0 | 0 | 0 | 0 | 0 | 0 | 0 | 0 | 0 | 0 | 0 | 0 | 0 | 0 | 0 | NA | NA | NA | NA | NA | NA | NA | NA | NA | NA |
| Olfr574 | 0 | 0 | 0 | 0 | 0 | 0 | 0 | 0 | 0 | 0 | 0 | 0 | 0 | 0 | 0 | NA | NA | NA | NA | NA | NA | NA | NA | NA | NA |
| Olfr575 | 0 | 0 | 0 | 0 | 0 | 0 | 0 | 0 | 0 | 0 | 0 | 0 | 0 | 0 | 0 | NA | NA | NA | NA | NA | NA | NA | NA | NA | NA |
| Olfr576 | 0 | 0 | 0 | 0 | 0 | 0 | 0 | 0 | 0 | 0 | 0 | 0 | 0 | 0 | 0 | NA | NA | NA | NA | NA | NA | NA | NA | NA | NA |
| Olfr577 | 0 | 0 | 0 | 0 | 0 | 0 | 0 | 0 | 0 | 0 | 0 | 0 | 0 | 0 | 0 | NA | NA | NA | NA | NA | NA | NA | NA | NA | NA |
| Olfr578 | 0 | 0 | 0 | 0 | 0 | 0 | 0 | 0 | 0 | 0 | 0 | 0 | 0 | 0 | 0 | NA | NA | NA | NA | NA | NA | NA | NA | NA | NA |
| Olfr58 | 0 | 2 | 1 | 0 | 0 | 0 | 0 | 2 | 1 | 0 | 0 | 0 | 0.5 | 1 | 0 | 0.179 | -2.485 | 0.44922046 | NA | 0 | 6.5925 | 6.2606 | 6.2606 | TRUE | 0.0526 |
| Olfr582 | 0 | 0 | 0 | 0 | 0 | 0 | 0 | 0 | 0 | 0 | 0 | 0 | 0 | 0 | 0 | NA | NA | NA | NA | NA | NA | NA | NA | NA | NA |
| Olfr583 | 0 | 0 | 0 | 0 | 0 | 0 | 0 | 0 | 0 | 0 | 0 | 0 | 0 | 0 | 0 | NA | NA | NA | NA | NA | NA | NA | NA | NA | NA |
| Olfr584 | 0 | 0 | 0 | 0 | 0 | 0 | 0 | 0 | 0 | 0 | 0 | 0 | 0 | 0 | 0 | NA | NA | NA | NA | NA | NA | NA | NA | NA | NA |
| Olfr585 | 0 | 0 | 0 | 0 | 0 | 0 | 0 | 0 | 0 | 0 | 0 | 0 | 0 | 0 | 0 | NA | NA | NA | NA | NA | NA | NA | NA | NA | NA |
| Olfr586 | 0 | 0 | 0 | 0 | 0 | 0 | 0 | 0 | 0 | 0 | 0 | 0 | 0 | 0 | 0 | NA | NA | NA | NA | NA | NA | NA | NA | NA | NA |
| Olfr589 | 0 | 0 | 0 | 0 | 0 | 0 | 0 | 0 | 0 | 0 | 0 | 0 | 0 | 0 | 0 | NA | NA | NA | NA | NA | NA | NA | NA | NA | NA |
| Olfr59 | 0 | 0 | 0 | 0 | 0 | 0 | 0 | 0 | 0 | 0 | 0 | 0 | 0 | 0 | 0 | NA | NA | NA | NA | NA | NA | NA | NA | NA | NA |
| Olfr591 | 0 | 0 | 0 | 0 | 0 | 0 | 0 | 0 | 0 | 0 | 0 | 0 | 0 | 0 | 0 | NA | NA | NA | NA | NA | NA | NA | NA | NA | NA |
| Olfr592 | 0 | 0 | 0 | 0 | 0 | 0 | 0 | 0 | 0 | 0 | 0 | 0 | 0 | 0 | 0 | NA | NA | NA | NA | NA | NA | NA | NA | NA | NA |
| Olfr593 | 0 | 0 | 0 | 0 | 0 | 0 | 0 | 0 | 0 | 0 | 0 | 0 | 0 | 0 | 0 | NA | NA | NA | NA | NA | NA | NA | NA | NA | NA |
| Olfr594 | 0 | 0 | 0 | 0 | 0 | 0 | 0 | 0 | 0 | 0 | 0 | 0 | 0 | 0 | 0 | NA | NA | NA | NA | NA | NA | NA | NA | NA | NA |
| Olfr596 | 0 | 0 | 0 | 0 | 0 | 0 | 0 | 0 | 0 | 0 | 0 | 0 | 0 | 0 | 0 | NA | NA | NA | NA | NA | NA | NA | NA | NA | NA |
| Olfr597 | 0 | 0 | 0 | 0 | 0 | 0 | 0 | 0 | 0 | 0 | 0 | 0 | 0 | 0 | 0 | NA | NA | NA | NA | NA | NA | NA | NA | NA | NA |
| Olfr598 | 0 | 0 | 0 | 0 | 0 | 0 | 0 | 0 | 0 | 0 | 0 | 0 | 0 | 0 | 0 | NA | NA | NA | NA | NA | NA | NA | NA | NA | NA |
| Olfr599 | 0 | 0 | 0 | 0 | 0 | 0 | 0 | 0 | 0 | 0 | 0 | 0 | 0 | 0 | 0 | NA | NA | NA | NA | NA | NA | NA | NA | NA | NA |
| Olfr6 | 0 | 0 | 0 | 0 | 0 | 0 | 0 | 0 | 0 | 0 | 0 | 0 | 0 | 0 | 0 | NA | NA | NA | NA | NA | NA | NA | NA | NA | NA |
| Olfr60 | 0 | 0 | 0 | 0 | 0 | 0 | 0 | 0 | 0 | 0 | 0 | 0 | 0 | 0 | 0 | NA | NA | NA | NA | NA | NA | NA | NA | NA | NA |
| Olfr600 | 0 | 0 | 0 | 0 | 0 | 0 | 0 | 0 | 0 | 0 | 0 | 0 | 0 | 0 | 0 | NA | NA | NA | NA | NA | NA | NA | NA | NA | NA |
| Olfr601 | 0 | 0 | 0 | 0 | 0 | 0 | 0 | 0 | 0 | 0 | 0 | 0 | 0 | 0 | 0 | NA | NA | NA | NA | NA | NA | NA | NA | NA | NA |
| Olfr603 | 0 | 0 | 0 | 0 | 0 | 0 | 0 | 0 | 0 | 0 | 0 | 0 | 0 | 0 | 0 | NA | NA | NA | NA | NA | NA | NA | NA | NA | NA |
| Olfr605 | 0 | 0 | 0 | 0 | 0 | 0 | 0 | 0 | 0 | 0 | 0 | 0 | 0 | 0 | 0 | NA | NA | NA | NA | NA | NA | NA | NA | NA | NA |
| Olfr606 | 0 | 0 | 0 | 0 | 0 | 0 | 0 | 0 | 0 | 0 | 0 | 0 | 0 | 0 | 0 | NA | NA | NA | NA | NA | NA | NA | NA | NA | NA |
| Olfr608 | 0 | 0 | 0 | 0 | 0 | 0 | 0 | 0 | 0 | 0 | 0 | 0 | 0 | 0 | 0 | NA | NA | NA | NA | NA | NA | NA | NA | NA | NA |
| Olfr609 | 0 | 0 | 0 | 0 | 0 | 0 | 0 | 0 | 0 | 0 | 0 | 0 | 0 | 0 | 0 | NA | NA | NA | NA | NA | NA | NA | NA | NA | NA |
| Olfr61 | 0 | 0 | 0 | 0 | 0 | 0 | 0 | 0 | 0 | 0 | 0 | 0 | 0 | 0 | 0 | NA | NA | NA | NA | NA | NA | NA | NA | NA | NA |
| Olfr610 | 0 | 0 | 0 | 0 | 0 | 0 | 0 | 0 | 0 | 0 | 0 | 0 | 0 | 0 | 0 | NA | NA | NA | NA | NA | NA | NA | NA | NA | NA |
| Olfr611 | 0 | 0 | 0 | 0 | 0 | 0 | 0 | 0 | 0 | 0 | 0 | 0 | 0 | 0 | 0 | NA | NA | NA | NA | NA | NA | NA | NA | NA | NA |
| Olfr612 | 0 | 0 | 0 | 0 | 0 | 0 | 0 | 0 | 0 | 0 | 0 | 0 | 0 | 0 | 0 | NA | NA | NA | NA | NA | NA | NA | NA | NA | NA |
| Olfr613 | 0 | 0 | 0 | 0 | 0 | 0 | 0 | 0 | 0 | 0 | 0 | 0 | 0 | 0 | 0 | NA | NA | NA | NA | NA | NA | NA | NA | NA | NA |
| Olfr615 | 0 | 0 | 0 | 0 | 0 | 0 | 0 | 0 | 0 | 0 | 0 | 0 | 0 | 0 | 0 | NA | NA | NA | NA | NA | NA | NA | NA | NA | NA |
| Olfr616 | 0 | 0 | 0 | 0 | 0 | 0 | 0 | 0 | 0 | 0 | 0 | 0 | 0 | 0 | 0 | NA | NA | NA | NA | NA | NA | NA | NA | NA | NA |
| Olfr617 | 0 | 0 | 0 | 0 | 0 | 0 | 0 | 0 | 0 | 0 | 0 | 0 | 0 | 0 | 0 | NA | NA | NA | NA | NA | NA | NA | NA | NA | NA |
| Olfr618 | 0 | 0 | 0 | 0 | 0 | 0 | 0 | 0 | 0 | 0 | 0 | 0 | 0 | 0 | 0 | NA | NA | NA | NA | NA | NA | NA | NA | NA | NA |
| Olfr619 | 0 | 0 | 0 | 0 | 0 | 0 | 0 | 0 | 0 | 0 | 0 | 0 | 0 | 0 | 0 | NA | NA | NA | NA | NA | NA | NA | NA | NA | NA |
| Olfr62 | 0 | 0 | 0 | 0 | 0 | 0 | 0 | 0 | 0 | 0 | 0 | 0 | 0 | 0 | 0 | NA | NA | NA | NA | NA | NA | NA | NA | NA | NA |

| | | | | | | | | | | | | | | | | | | | | | | | | | |
|---|---|---|---|---|---|---|---|---|---|---|---|---|---|---|---|---|---|---|---|---|---|---|---|---|---|
| Olfr620 | 0 | 0 | 0 | 0 | 0 | 0 | 0 | 0 | 0 | 0 | 0 | 0 | 0 | 0 | 0 | NA | NA | NA | NA | NA | NA | NA | NA | NA | NA |
| Olfr622 | 0 | 0 | 0 | 0 | 0 | 0 | 0 | 0 | 0 | 0 | 0 | 0 | 0 | 0 | 0 | NA | NA | NA | NA | NA | NA | NA | NA | NA | NA |
| Olfr623 | 0 | 0 | 0 | 0 | 0 | 0 | 0 | 0 | 0 | 0 | 0 | 0 | 0 | 0 | 0 | NA | NA | NA | NA | NA | NA | NA | NA | NA | NA |
| Olfr624 | 0 | 0 | 0 | 0 | 0 | 0 | 0 | 0 | 0 | 0 | 0 | 0 | 0 | 0 | 0 | NA | NA | NA | NA | NA | NA | NA | NA | NA | NA |
| Olfr628 | 0 | 0 | 0 | 0 | 0 | 0 | 0 | 0 | 0 | 0 | 0 | 0 | 0 | 0 | 0 | NA | NA | NA | NA | NA | NA | NA | NA | NA | NA |
| Olfr629 | 0 | 0 | 0 | 0 | 0 | 0 | 0 | 0 | 0 | 0 | 0 | 0 | 0 | 0 | 0 | NA | NA | NA | NA | NA | NA | NA | NA | NA | NA |
| Olfr63 | 0 | 0 | 0 | 0 | 0 | 0 | 0 | 0 | 0 | 0 | 0 | 0 | 0 | 0 | 0 | NA | NA | NA | NA | NA | NA | NA | NA | NA | NA |
| Olfr630 | 0 | 0 | 0 | 0 | 0 | 0 | 0 | 0 | 0 | 0 | 0 | 0 | 0 | 0 | 0 | NA | NA | NA | NA | NA | NA | NA | NA | NA | NA |
| Olfr631 | 0 | 0 | 0 | 0 | 0 | 0 | 0 | 0 | 0 | 0 | 0 | 0 | 0 | 0 | 0 | NA | NA | NA | NA | NA | NA | NA | NA | NA | NA |
| Olfr632 | 0 | 0 | 0 | 0 | 0 | 0 | 0 | 0 | 0 | 0 | 0 | 0 | 0 | 0 | 0 | NA | NA | NA | NA | NA | NA | NA | NA | NA | NA |
| Olfr633 | 0 | 0 | 0 | 0 | 0 | 0 | 0 | 0 | 0 | 0 | 0 | 0 | 0 | 0 | 0 | NA | NA | NA | NA | NA | NA | NA | NA | NA | NA |
| Olfr635 | 0 | 0 | 0 | 0 | 0 | 0 | 0 | 0 | 0 | 0 | 0 | 0 | 0 | 0 | 0 | NA | NA | NA | NA | NA | NA | NA | NA | NA | NA |
| Olfr638 | 0 | 0 | 0 | 0 | 0 | 0 | 0 | 0 | 0 | 0 | 0 | 0 | 0 | 0 | 0 | NA | NA | NA | NA | NA | NA | NA | NA | NA | NA |
| Olfr639 | 0 | 0 | 0 | 0 | 0 | 0 | 0 | 0 | 0 | 0 | 0 | 0 | 0 | 0 | 0 | NA | NA | NA | NA | NA | NA | NA | NA | NA | NA |
| Olfr64 | 0 | 0 | 0 | 0 | 0 | 0 | 0 | 0 | 0 | 0 | 0 | 0 | 0 | 0 | 0 | NA | NA | NA | NA | NA | NA | NA | NA | NA | NA |
| Olfr640 | 0 | 0 | 0 | 0 | 0 | 0 | 0 | 0 | 0 | 0 | 0 | 0 | 0 | 0 | 0 | NA | NA | NA | NA | NA | NA | NA | NA | NA | NA |
| Olfr641 | 0 | 0 | 0 | 0 | 0 | 0 | 0 | 0 | 0 | 0 | 0 | 0 | 0 | 0 | 0 | NA | NA | NA | NA | NA | NA | NA | NA | NA | NA |
| Olfr642 | 0 | 0 | 0 | 0 | 0 | 0 | 0 | 0 | 0 | 0 | 0 | 0 | 0 | 0 | 0 | NA | NA | NA | NA | NA | NA | NA | NA | NA | NA |
| Olfr643 | 0 | 0 | 0 | 0 | 0 | 0 | 0 | 0 | 0 | 0 | 0 | 0 | 0 | 0 | 0 | NA | NA | NA | NA | NA | NA | NA | NA | NA | NA |
| Olfr644 | 0 | 0 | 0 | 0 | 0 | 0 | 0 | 0 | 0 | 0 | 0 | 0 | 0 | 0 | 0 | NA | NA | NA | NA | NA | NA | NA | NA | NA | NA |
| Olfr645 | 0 | 0 | 0 | 0 | 0 | 0 | 0 | 0 | 0 | 0 | 0 | 0 | 0 | 0 | 0 | NA | NA | NA | NA | NA | NA | NA | NA | NA | NA |
| Olfr646 | 0 | 0 | 0 | 0 | 0 | 0 | 0 | 0 | 0 | 0 | 0 | 0 | 0 | 0 | 0 | NA | NA | NA | NA | NA | NA | NA | NA | NA | NA |
| Olfr648 | 0 | 0 | 0 | 0 | 0 | 0 | 0 | 0 | 0 | 0 | 0 | 0 | 0 | 0 | 0 | NA | NA | NA | NA | NA | NA | NA | NA | NA | NA |
| Olfr649 | 0 | 0 | 0 | 0 | 0 | 0 | 0 | 0 | 0 | 0 | 0 | 0 | 0 | 0 | 0 | NA | NA | NA | NA | NA | NA | NA | NA | NA | NA |
| Olfr65 | 0 | 0 | 0 | 0 | 0 | 0 | 0 | 0 | 0 | 0 | 0 | 0 | 0 | 0 | 0 | NA | NA | NA | NA | NA | NA | NA | NA | NA | NA |
| Olfr651 | 0 | 0 | 0 | 0 | 0 | 0 | 0 | 0 | 0 | 0 | 0 | 0 | 0 | 0 | 0 | NA | NA | NA | NA | NA | NA | NA | NA | NA | NA |
| Olfr652 | 0 | 0 | 0 | 0 | 0 | 0 | 0 | 0 | 0 | 0 | 0 | 0 | 0 | 0 | 0 | NA | NA | NA | NA | NA | NA | NA | NA | NA | NA |
| Olfr653 | 0 | 0 | 0 | 0 | 0 | 0 | 0 | 0 | 0 | 0 | 0 | 0 | 0 | 0 | 0 | NA | NA | NA | NA | NA | NA | NA | NA | NA | NA |
| Olfr654 | 0 | 0 | 0 | 0 | 0 | 0 | 0 | 0 | 0 | 0 | 0 | 0 | 0 | 0 | 0 | NA | NA | NA | NA | NA | NA | NA | NA | NA | NA |
| Olfr655 | 0 | 0 | 0 | 0 | 0 | 0 | 0 | 0 | 0 | 0 | 0 | 0 | 0 | 0 | 0 | NA | NA | NA | NA | NA | NA | NA | NA | NA | NA |
| Olfr656 | 0 | 0 | 0 | 0 | 0 | 0 | 0 | 0 | 0 | 0 | 0 | 0 | 0 | 0 | 0 | NA | NA | NA | NA | NA | NA | NA | NA | NA | NA |
| Olfr657 | 0 | 0 | 0 | 0 | 0 | 0 | 0 | 0 | 0 | 0 | 0 | 0 | 0 | 0 | 0 | NA | NA | NA | NA | NA | NA | NA | NA | NA | NA |
| Olfr658 | 2 | 3 | 3 | 6 | 6 | 3 | 2 | 3 | 3 | 5 | 6 | 3 | 3.76 | 3 | 5 | 1.774 | 0.827 | 0.39055709 | NA | 0 | 0.882 | 0.3786 | 0.3786 | TRUE | 0.1537 |
| Olfr659 | 0 | 0 | 0 | 0 | 0 | 0 | 0 | 0 | 0 | 0 | 0 | 0 | 0 | 0 | 0 | NA | NA | NA | NA | NA | NA | NA | NA | NA | NA |
| Olfr66 | 0 | 0 | 0 | 0 | 0 | 0 | 0 | 0 | 0 | 0 | 0 | 0 | 0 | 0 | 0 | NA | NA | NA | NA | NA | NA | NA | NA | NA | NA |
| Olfr661 | 0 | 0 | 0 | 0 | 0 | 0 | 0 | 0 | 0 | 0 | 0 | 0 | 0 | 0 | 0 | NA | NA | NA | NA | NA | NA | NA | NA | NA | NA |
| Olfr663 | 0 | 0 | 0 | 0 | 0 | 0 | 0 | 0 | 0 | 0 | 0 | 0 | 0 | 0 | 0 | NA | NA | NA | NA | NA | NA | NA | NA | NA | NA |
| Olfr665 | 0 | 0 | 0 | 0 | 0 | 0 | 0 | 0 | 0 | 0 | 0 | 0 | 0 | 0 | 0 | NA | NA | NA | NA | NA | NA | NA | NA | NA | NA |
| Olfr666 | 0 | 0 | 0 | 0 | 0 | 0 | 0 | 0 | 0 | 0 | 0 | 0 | 0 | 0 | 0 | NA | NA | NA | NA | NA | NA | NA | NA | NA | NA |
| Olfr667 | 0 | 0 | 0 | 0 | 0 | 0 | 0 | 0 | 0 | 0 | 0 | 0 | 0 | 0 | 0 | NA | NA | NA | NA | NA | NA | NA | NA | NA | NA |
| Olfr668 | 0 | 0 | 0 | 0 | 0 | 0 | 0 | 0 | 0 | 0 | 0 | 0 | 0 | 0 | 0 | NA | NA | NA | NA | NA | NA | NA | NA | NA | NA |
| Olfr669 | 0 | 0 | 0 | 0 | 0 | 0 | 0 | 0 | 0 | 0 | 0 | 0 | 0 | 0 | 0 | NA | NA | NA | NA | NA | NA | NA | NA | NA | NA |
| Olfr67 | 0 | 0 | 0 | 0 | 0 | 0 | 0 | 0 | 0 | 0 | 0 | 0 | 0 | 0 | 0 | NA | NA | NA | NA | NA | NA | NA | NA | NA | NA |
| Olfr670 | 0 | 0 | 0 | 0 | 0 | 0 | 0 | 0 | 0 | 0 | 0 | 0 | 0 | 0 | 0 | NA | NA | NA | NA | NA | NA | NA | NA | NA | NA |
| Olfr671 | 0 | 0 | 0 | 0 | 0 | 0 | 0 | 0 | 0 | 0 | 0 | 0 | 0 | 0 | 0 | NA | NA | NA | NA | NA | NA | NA | NA | NA | NA |
| Olfr672 | 0 | 0 | 0 | 0 | 0 | 0 | 0 | 0 | 0 | 0 | 0 | 0 | 0 | 0 | 0 | NA | NA | NA | NA | NA | NA | NA | NA | NA | NA |
| Olfr675 | 0 | 0 | 0 | 0 | 0 | 0 | 0 | 0 | 0 | 0 | 0 | 0 | 0 | 0 | 0 | NA | NA | NA | NA | NA | NA | NA | NA | NA | NA |
| Olfr676 | 0 | 0 | 0 | 0 | 0 | 0 | 0 | 0 | 0 | 0 | 0 | 0 | 0 | 0 | 0 | NA | NA | NA | NA | NA | NA | NA | NA | NA | NA |
| Olfr677 | 0 | 0 | 0 | 0 | 0 | 0 | 0 | 0 | 0 | 0 | 0 | 0 | 0 | 0 | 0 | NA | NA | NA | NA | NA | NA | NA | NA | NA | NA |
| Olfr678 | 0 | 0 | 0 | 0 | 0 | 0 | 0 | 0 | 0 | 0 | 0 | 0 | 0 | 0 | 0 | NA | NA | NA | NA | NA | NA | NA | NA | NA | NA |
| Olfr679 | 0 | 0 | 0 | 0 | 0 | 0 | 0 | 0 | 0 | 0 | 0 | 0 | 0 | 0 | 0 | NA | NA | NA | NA | NA | NA | NA | NA | NA | NA |
| Olfr68 | 0 | 0 | 0 | 0 | 0 | 0 | 0 | 0 | 0 | 0 | 0 | 0 | 0 | 0 | 0 | NA | NA | NA | NA | NA | NA | NA | NA | NA | NA |
| Olfr681 | 0 | 0 | 0 | 0 | 0 | 0 | 0 | 0 | 0 | 0 | 0 | 0 | 0 | 0 | 0 | NA | NA | NA | NA | NA | NA | NA | NA | NA | NA |
| Olfr683 | 0 | 0 | 0 | 0 | 0 | 0 | 0 | 0 | 0 | 0 | 0 | 0 | 0 | 0 | 0 | NA | NA | NA | NA | NA | NA | NA | NA | NA | NA |
| Olfr684 | 0 | 0 | 0 | 0 | 0 | 0 | 0 | 0 | 0 | 0 | 0 | 0 | 0 | 0 | 0 | NA | NA | NA | NA | NA | NA | NA | NA | NA | NA |
| Olfr685 | 0 | 0 | 0 | 0 | 0 | 0 | 0 | 0 | 0 | 0 | 0 | 0 | 0 | 0 | 0 | NA | NA | NA | NA | NA | NA | NA | NA | NA | NA |
| Olfr686 | 0 | 0 | 0 | 0 | 0 | 0 | 0 | 0 | 0 | 0 | 0 | 0 | 0 | 0 | 0 | NA | NA | NA | NA | NA | NA | NA | NA | NA | NA |
| Olfr688 | 0 | 0 | 0 | 0 | 0 | 0 | 0 | 0 | 0 | 0 | 0 | 0 | 0 | 0 | 0 | NA | NA | NA | NA | NA | NA | NA | NA | NA | NA |
| Olfr689 | 0 | 0 | 0 | 0 | 0 | 0 | 0 | 0 | 0 | 0 | 0 | 0 | 0 | 0 | 0 | NA | NA | NA | NA | NA | NA | NA | NA | NA | NA |
| Olfr69 | 0 | 0 | 0 | 0 | 0 | 0 | 0 | 0 | 0 | 0 | 0 | 0 | 0 | 0 | 0 | NA | NA | NA | NA | NA | NA | NA | NA | NA | NA |
| Olfr690 | 1 | 0 | 0 | 1 | 0 | 0 | 1 | 0 | 0 | 1 | 0 | 0 | 0.31 | 0 | 0 | 0.954 | -0.067 | 0.98625928 | NA | 0 | 10.7014 | 9.0384 | 9.0384 | TRUE | 0.4614 |
| Olfr691 | 0 | 0 | 0 | 0 | 2 | 0 | 0 | 0 | 0 | 0 | 2 | 0 | 0.36 | 0 | 1 | 3.775 | 1.916 | 0.63423067 | NA | 10 | 9.1927 | 10 | 10 | TRUE | 0.9743 |
| Olfr692 | 0 | 0 | 0 | 0 | 0 | 0 | 0 | 0 | 0 | 0 | 0 | 0 | 0 | 0 | 0 | NA | NA | NA | NA | NA | NA | NA | NA | NA | NA |
| Olfr693 | 0 | 0 | 0 | 0 | 0 | 0 | 0 | 0 | 0 | 0 | 0 | 0 | 0 | 0 | 0 | NA | NA | NA | NA | NA | NA | NA | NA | NA | NA |
| Olfr694 | 0 | 0 | 0 | 0 | 0 | 0 | 0 | 0 | 0 | 0 | 0 | 0 | 0 | 0 | 0 | NA | NA | NA | NA | NA | NA | NA | NA | NA | NA |
| Olfr695 | 0 | 0 | 0 | 0 | 0 | 0 | 0 | 0 | 0 | 0 | 0 | 0 | 0 | 0 | 0 | NA | NA | NA | NA | NA | NA | NA | NA | NA | NA |
| Olfr697 | 0 | 0 | 0 | 1 | 0 | 1 | 0 | 0 | 0 | 1 | 0 | 1 | 0.31 | 0 | 1 | 3.348 | 1.743 | 0.65064889 | NA | 0 | 10.5167 | 8.8968 | 8.8968 | TRUE | 0.2082 |
| Olfr698 | 0 | 0 | 0 | 0 | 0 | 0 | 0 | 0 | 0 | 0 | 0 | 0 | 0 | 0 | 0 | NA | NA | NA | NA | NA | NA | NA | NA | NA | NA |
| Olfr699 | 0 | 0 | 0 | 2 | 1 | 1 | 0 | 0 | 0 | 2 | 1 | 1 | 0.62 | 0 | 1 | 6.676 | 2.739 | 0.27800695 | NA | 0 | 5.2881 | 3.2034 | 3.2034 | TRUE | 0.07 |

| | | | | | | | | | | | | | | | | | | | | | | | | | |
|---|---|---|---|---|---|---|---|---|---|---|---|---|---|---|---|---|---|---|---|---|---|---|---|---|---|
| Olfr70 | 0 | 0 | 0 | 0 | 0 | 0 | 0 | 0 | 0 | 0 | 0 | 0 | 0 | 0 | 0 | NA | NA | NA | NA | NA | NA | NA | NA | NA | NA |
| Olfr700 | 0 | 0 | 0 | 0 | 0 | 0 | 0 | 0 | 0 | 0 | 0 | 0 | 0 | 0 | 0 | NA | NA | NA | NA | NA | NA | NA | NA | NA | NA |
| Olfr701 | 0 | 9 | 3 | 2 | 5 | 3 | 0 | 9 | 3 | 2 | 5 | 3 | 3.71 | 4 | 3 | 0.837 | -0.256 | 0.83805087 | NA | 0.8158 | 0.8958 | 0.861 | 0.861 | TRUE | 0.4365 |
| Olfr702 | 0 | 0 | 0 | 0 | 0 | 0 | 0 | 0 | 0 | 0 | 0 | 0 | 0 | 0 | 0 | NA | NA | NA | NA | NA | NA | NA | NA | NA | NA |
| Olfr703 | 0 | 0 | 0 | 0 | 0 | 0 | 0 | 0 | 0 | 0 | 0 | 0 | 0 | 0 | 0 | NA | NA | NA | NA | NA | NA | NA | NA | NA | NA |
| Olfr704 | 0 | 0 | 0 | 0 | 0 | 0 | 0 | 0 | 0 | 0 | 0 | 0 | 0 | 0 | 0 | NA | NA | NA | NA | NA | NA | NA | NA | NA | NA |
| Olfr705 | 0 | 1 | 0 | 0 | 0 | 0 | 0 | 1 | 0 | 0 | 0 | 0 | 0.17 | 0 | 0 | 0.49 | -1.029 | 0.80087 | NA | 0 | 19.7991 | 10 | 10 | TRUE | 0.405 |
| Olfr706 | 0 | 0 | 0 | 0 | 0 | 0 | 0 | 0 | 0 | 0 | 0 | 0 | 0 | 0 | 0 | NA | NA | NA | NA | NA | NA | NA | NA | NA | NA |
| Olfr707 | 0 | 0 | 0 | 0 | 0 | 0 | 0 | 0 | 0 | 0 | 0 | 0 | 0 | 0 | 0 | NA | NA | NA | NA | NA | NA | NA | NA | NA | NA |
| Olfr71 | 0 | 0 | 0 | 0 | 0 | 0 | 0 | 0 | 0 | 0 | 0 | 0 | 0 | 0 | 0 | NA | NA | NA | NA | NA | NA | NA | NA | NA | NA |
| Olfr710 | 0 | 0 | 0 | 0 | 0 | 0 | 0 | 0 | 0 | 0 | 0 | 0 | 0 | 0 | 0 | NA | NA | NA | NA | NA | NA | NA | NA | NA | NA |
| Olfr711 | 0 | 0 | 0 | 0 | 0 | 0 | 0 | 0 | 0 | 0 | 0 | 0 | 0 | 0 | 0 | NA | NA | NA | NA | NA | NA | NA | NA | NA | NA |
| Olfr713 | 0 | 0 | 0 | 0 | 0 | 0 | 0 | 0 | 0 | 0 | 0 | 0 | 0 | 0 | 0 | NA | NA | NA | NA | NA | NA | NA | NA | NA | NA |
| Olfr714 | 0 | 0 | 0 | 0 | 0 | 0 | 0 | 0 | 0 | 0 | 0 | 0 | 0 | 0 | 0 | NA | NA | NA | NA | NA | NA | NA | NA | NA | NA |
| Olfr715 | 0 | 0 | 0 | 0 | 0 | 0 | 0 | 0 | 0 | 0 | 0 | 0 | 0 | 0 | 0 | NA | NA | NA | NA | NA | NA | NA | NA | NA | NA |
| Olfr716 | 0 | 0 | 0 | 0 | 0 | 0 | 0 | 0 | 0 | 0 | 0 | 0 | 0 | 0 | 0 | NA | NA | NA | NA | NA | NA | NA | NA | NA | NA |
| Olfr720 | 0 | 0 | 0 | 0 | 0 | 0 | 0 | 0 | 0 | 0 | 0 | 0 | 0 | 0 | 0 | NA | NA | NA | NA | NA | NA | NA | NA | NA | NA |
| Olfr722 | 0 | 0 | 0 | 0 | 0 | 0 | 0 | 0 | 0 | 0 | 0 | 0 | 0 | 0 | 0 | NA | NA | NA | NA | NA | NA | NA | NA | NA | NA |
| Olfr723 | 0 | 0 | 0 | 0 | 0 | 0 | 0 | 0 | 0 | 0 | 0 | 0 | 0 | 0 | 0 | NA | NA | NA | NA | NA | NA | NA | NA | NA | NA |
| Olfr724 | 0 | 0 | 0 | 0 | 0 | 0 | 0 | 0 | 0 | 0 | 0 | 0 | 0 | 0 | 0 | NA | NA | NA | NA | NA | NA | NA | NA | NA | NA |
| Olfr725 | 0 | 0 | 0 | 0 | 0 | 0 | 0 | 0 | 0 | 0 | 0 | 0 | 0 | 0 | 0 | NA | NA | NA | NA | NA | NA | NA | NA | NA | NA |
| Olfr726 | 0 | 0 | 0 | 0 | 0 | 0 | 0 | 0 | 0 | 0 | 0 | 0 | 0 | 0 | 0 | NA | NA | NA | NA | NA | NA | NA | NA | NA | NA |
| Olfr727 | 0 | 0 | 0 | 0 | 0 | 0 | 0 | 0 | 0 | 0 | 0 | 0 | 0 | 0 | 0 | NA | NA | NA | NA | NA | NA | NA | NA | NA | NA |
| Olfr728 | 0 | 0 | 0 | 0 | 0 | 0 | 0 | 0 | 0 | 0 | 0 | 0 | 0 | 0 | 0 | NA | NA | NA | NA | NA | NA | NA | NA | NA | NA |
| Olfr729 | 0 | 0 | 0 | 0 | 0 | 0 | 0 | 0 | 0 | 0 | 0 | 0 | 0 | 0 | 0 | NA | NA | NA | NA | NA | NA | NA | NA | NA | NA |
| Olfr73 | 0 | 0 | 0 | 0 | 0 | 0 | 0 | 0 | 0 | 0 | 0 | 0 | 0 | 0 | 0 | NA | NA | NA | NA | NA | NA | NA | NA | NA | NA |
| Olfr730 | 0 | 0 | 0 | 0 | 0 | 0 | 0 | 0 | 0 | 0 | 0 | 0 | 0 | 0 | 0 | NA | NA | NA | NA | NA | NA | NA | NA | NA | NA |
| Olfr731 | 0 | 0 | 0 | 1 | 0 | 0 | 0 | 0 | 0 | 1 | 0 | 0 | 0.13 | 0 | 0 | 1.859 | 0.894 | 0.82650734 | NA | 0 | 25.1515 | 10 | 10 | TRUE | 0.2563 |
| Olfr732 | 0 | 0 | 0 | 0 | 0 | 0 | 0 | 0 | 0 | 0 | 0 | 0 | 0 | 0 | 0 | NA | NA | NA | NA | NA | NA | NA | NA | NA | NA |
| Olfr733 | 0 | 0 | 0 | 0 | 0 | 0 | 0 | 0 | 0 | 0 | 0 | 0 | 0 | 0 | 0 | NA | NA | NA | NA | NA | NA | NA | NA | NA | NA |
| Olfr734 | 0 | 0 | 0 | 0 | 0 | 0 | 0 | 0 | 0 | 0 | 0 | 0 | 0 | 0 | 0 | NA | NA | NA | NA | NA | NA | NA | NA | NA | NA |
| Olfr735 | 0 | 0 | 0 | 0 | 0 | 0 | 0 | 0 | 0 | 0 | 0 | 0 | 0 | 0 | 0 | NA | NA | NA | NA | NA | NA | NA | NA | NA | NA |
| Olfr736 | 0 | 0 | 0 | 0 | 0 | 0 | 0 | 0 | 0 | 0 | 0 | 0 | 0 | 0 | 0 | NA | NA | NA | NA | NA | NA | NA | NA | NA | NA |
| Olfr738 | 0 | 0 | 0 | 0 | 0 | 0 | 0 | 0 | 0 | 0 | 0 | 0 | 0 | 0 | 0 | NA | NA | NA | NA | NA | NA | NA | NA | NA | NA |
| Olfr739 | 0 | 0 | 0 | 0 | 0 | 0 | 0 | 0 | 0 | 0 | 0 | 0 | 0 | 0 | 0 | NA | NA | NA | NA | NA | NA | NA | NA | NA | NA |
| Olfr74 | 0 | 0 | 0 | 0 | 0 | 0 | 0 | 0 | 0 | 0 | 0 | 0 | 0 | 0 | 0 | NA | NA | NA | NA | NA | NA | NA | NA | NA | NA |
| Olfr740 | 0 | 0 | 0 | 0 | 0 | 0 | 0 | 0 | 0 | 0 | 0 | 0 | 0 | 0 | 0 | NA | NA | NA | NA | NA | NA | NA | NA | NA | NA |
| Olfr741 | 0 | 0 | 0 | 0 | 0 | 0 | 0 | 0 | 0 | 0 | 0 | 0 | 0 | 0 | 0 | NA | NA | NA | NA | NA | NA | NA | NA | NA | NA |
| Olfr742 | 0 | 0 | 0 | 0 | 0 | 0 | 0 | 0 | 0 | 0 | 0 | 0 | 0 | 0 | 0 | NA | NA | NA | NA | NA | NA | NA | NA | NA | NA |
| Olfr743 | 0 | 0 | 0 | 0 | 0 | 0 | 0 | 0 | 0 | 0 | 0 | 0 | 0 | 0 | 0 | NA | NA | NA | NA | NA | NA | NA | NA | NA | NA |
| Olfr744 | 0 | 0 | 0 | 0 | 0 | 0 | 0 | 0 | 0 | 0 | 0 | 0 | 0 | 0 | 0 | NA | NA | NA | NA | NA | NA | NA | NA | NA | NA |
| Olfr745 | 0 | 0 | 0 | 0 | 0 | 0 | 0 | 0 | 0 | 0 | 0 | 0 | 0 | 0 | 0 | NA | NA | NA | NA | NA | NA | NA | NA | NA | NA |
| Olfr746 | 0 | 0 | 0 | 0 | 0 | 0 | 0 | 0 | 0 | 0 | 0 | 0 | 0 | 0 | 0 | NA | NA | NA | NA | NA | NA | NA | NA | NA | NA |
| Olfr747 | 0 | 0 | 0 | 0 | 0 | 0 | 0 | 0 | 0 | 0 | 0 | 0 | 0 | 0 | 0 | NA | NA | NA | NA | NA | NA | NA | NA | NA | NA |
| Olfr748 | 0 | 0 | 0 | 0 | 0 | 0 | 0 | 0 | 0 | 0 | 0 | 0 | 0 | 0 | 0 | NA | NA | NA | NA | NA | NA | NA | NA | NA | NA |
| Olfr749 | 0 | 0 | 0 | 0 | 0 | 0 | 0 | 0 | 0 | 0 | 0 | 0 | 0 | 0 | 0 | NA | NA | NA | NA | NA | NA | NA | NA | NA | NA |
| Olfr75-ps1 | 0 | 0 | 0 | 0 | 0 | 0 | 0 | 0 | 0 | 0 | 0 | 0 | 0 | 0 | 0 | NA | NA | NA | NA | NA | NA | NA | NA | NA | NA |
| Olfr750 | 0 | 0 | 0 | 0 | 0 | 0 | 0 | 0 | 0 | 0 | 0 | 0 | 0 | 0 | 0 | NA | NA | NA | NA | NA | NA | NA | NA | NA | NA |
| Olfr76 | 1 | 0 | 0 | 0 | 0 | 1 | 1 | 0 | 0 | 0 | 0 | 1 | 0.36 | 0 | 0 | 0.954 | -0.067 | 0.98553519 | NA | 0 | 9.1716 | 7.9605 | 7.9605 | TRUE | 0.5383 |
| Olfr761 | 0 | 0 | 0 | 0 | 0 | 0 | 0 | 0 | 0 | 0 | 0 | 0 | 0 | 0 | 0 | NA | NA | NA | NA | NA | NA | NA | NA | NA | NA |
| Olfr763 | 0 | 0 | 0 | 0 | 0 | 0 | 0 | 0 | 0 | 0 | 0 | 0 | 0 | 0 | 0 | NA | NA | NA | NA | NA | NA | NA | NA | NA | NA |
| Olfr765 | 0 | 0 | 0 | 0 | 0 | 0 | 0 | 0 | 0 | 0 | 0 | 0 | 0 | 0 | 0 | NA | NA | NA | NA | NA | NA | NA | NA | NA | NA |
| Olfr767 | 0 | 0 | 0 | 0 | 0 | 0 | 0 | 0 | 0 | 0 | 0 | 0 | 0 | 0 | 0 | NA | NA | NA | NA | NA | NA | NA | NA | NA | NA |
| Olfr768 | 0 | 0 | 0 | 0 | 0 | 0 | 0 | 0 | 0 | 0 | 0 | 0 | 0 | 0 | 0 | NA | NA | NA | NA | NA | NA | NA | NA | NA | NA |
| Olfr769 | 0 | 0 | 0 | 0 | 0 | 0 | 0 | 0 | 0 | 0 | 0 | 0 | 0 | 0 | 0 | NA | NA | NA | NA | NA | NA | NA | NA | NA | NA |
| Olfr77 | 0 | 0 | 0 | 0 | 0 | 0 | 0 | 0 | 0 | 0 | 0 | 0 | 0 | 0 | 0 | NA | NA | NA | NA | NA | NA | NA | NA | NA | NA |
| Olfr770 | 0 | 0 | 0 | 0 | 0 | 0 | 0 | 0 | 0 | 0 | 0 | 0 | 0 | 0 | 0 | NA | NA | NA | NA | NA | NA | NA | NA | NA | NA |
| Olfr771 | 0 | 0 | 0 | 0 | 0 | 0 | 0 | 0 | 0 | 0 | 0 | 0 | 0 | 0 | 0 | NA | NA | NA | NA | NA | NA | NA | NA | NA | NA |
| Olfr772 | 0 | 0 | 0 | 1 | 0 | 1 | 0 | 0 | 0 | 1 | 0 | 1 | 0.31 | 0 | 1 | 3.348 | 1.743 | 0.65064889 | NA | 0 | 10.5167 | 8.8968 | 8.8968 | TRUE | 0.2082 |
| Olfr773 | 0 | 0 | 0 | 0 | 0 | 0 | 0 | 0 | 0 | 0 | 0 | 0 | 0 | 0 | 0 | NA | NA | NA | NA | NA | NA | NA | NA | NA | NA |
| Olfr774 | 0 | 0 | 0 | 0 | 0 | 0 | 0 | 0 | 0 | 0 | 0 | 0 | 0 | 0 | 0 | NA | NA | NA | NA | NA | NA | NA | NA | NA | NA |
| Olfr775 | 0 | 0 | 0 | 0 | 0 | 0 | 0 | 0 | 0 | 0 | 0 | 0 | 0 | 0 | 0 | NA | NA | NA | NA | NA | NA | NA | NA | NA | NA |
| Olfr776 | 0 | 0 | 0 | 0 | 0 | 0 | 0 | 0 | 0 | 0 | 0 | 0 | 0 | 0 | 0 | NA | NA | NA | NA | NA | NA | NA | NA | NA | NA |
| Olfr777 | 0 | 0 | 0 | 0 | 0 | 0 | 0 | 0 | 0 | 0 | 0 | 0 | 0 | 0 | 0 | NA | NA | NA | NA | NA | NA | NA | NA | NA | NA |
| Olfr78 | 0 | 0 | 0 | 0 | 0 | 0 | 0 | 0 | 0 | 0 | 0 | 0 | 0 | 0 | 0 | NA | NA | NA | NA | NA | NA | NA | NA | NA | NA |
| Olfr780 | 0 | 0 | 0 | 0 | 0 | 0 | 0 | 0 | 0 | 0 | 0 | 0 | 0 | 0 | 0 | NA | NA | NA | NA | NA | NA | NA | NA | NA | NA |
| Olfr781 | 0 | 0 | 0 | 0 | 0 | 0 | 0 | 0 | 0 | 0 | 0 | 0 | 0 | 0 | 0 | NA | NA | NA | NA | NA | NA | NA | NA | NA | NA |
| Olfr782 | 0 | 0 | 0 | 0 | 0 | 0 | 0 | 0 | 0 | 0 | 0 | 0 | 0 | 0 | 0 | NA | NA | NA | NA | NA | NA | NA | NA | NA | NA |
| Olfr784 | 0 | 0 | 0 | 0 | 0 | 0 | 0 | 0 | 0 | 0 | 0 | 0 | 0 | 0 | 0 | NA | NA | NA | NA | NA | NA | NA | NA | NA | NA |

| | | | | | | | | | | | | | | | | | | | | | | | | | |
|---|---|---|---|---|---|---|---|---|---|---|---|---|---|---|---|---|---|---|---|---|---|---|---|---|---|
| Olfr786 | 0 | 0 | 0 | 0 | 0 | 0 | 0 | 0 | 0 | 0 | 0 | 0 | 0 | 0 | 0 | NA | NA | NA | NA | NA | NA | NA | NA | NA | NA |
| Olfr787 | 1 | 1 | 0 | 2 | 3 | 0 | 1 | 1 | 0 | 2 | 3 | 0 | 1.14 | 1 | 2 | 2.342 | 1.228 | 0.53526505 | NA | 0 | 2.8879 | 1.7729 | 1.7729 | TRUE | 0.1007 |
| Olfr788 | 0 | 0 | 0 | 0 | 1 | 0 | 0 | 0 | 0 | 0 | 1 | 0 | 0.18 | 0 | 0 | 1.859 | 0.894 | 0.82650734 | NA | 0 | 18.3785 | 10 | 10 | TRUE | 0.5204 |
| Olfr790 | 0 | 0 | 0 | 0 | 0 | 0 | 0 | 0 | 0 | 0 | 0 | 0 | 0 | 0 | 0 | NA | NA | NA | NA | NA | NA | NA | NA | NA | NA |
| Olfr791 | 0 | 0 | 0 | 0 | 0 | 0 | 0 | 0 | 0 | 0 | 0 | 0 | 0 | 0 | 0 | NA | NA | NA | NA | NA | NA | NA | NA | NA | NA |
| Olfr792 | 0 | 0 | 0 | 0 | 0 | 0 | 0 | 0 | 0 | 0 | 0 | 0 | 0 | 0 | 0 | NA | NA | NA | NA | NA | NA | NA | NA | NA | NA |
| Olfr794 | 0 | 0 | 0 | 0 | 0 | 0 | 0 | 0 | 0 | 0 | 0 | 0 | 0 | 0 | 0 | NA | NA | NA | NA | NA | NA | NA | NA | NA | NA |
| Olfr796 | 0 | 0 | 0 | 0 | 0 | 0 | 0 | 0 | 0 | 0 | 0 | 0 | 0 | 0 | 0 | NA | NA | NA | NA | NA | NA | NA | NA | NA | NA |
| Olfr798 | 0 | 0 | 0 | 0 | 0 | 0 | 0 | 0 | 0 | 0 | 0 | 0 | 0 | 0 | 0 | NA | NA | NA | NA | NA | NA | NA | NA | NA | NA |
| Olfr799 | 0 | 0 | 0 | 0 | 0 | 0 | 0 | 0 | 0 | 0 | 0 | 0 | 0 | 0 | 0 | NA | NA | NA | NA | NA | NA | NA | NA | NA | NA |
| Olfr8 | 0 | 0 | 0 | 0 | 0 | 0 | 0 | 0 | 0 | 0 | 0 | 0 | 0 | 0 | 0 | NA | NA | NA | NA | NA | NA | NA | NA | NA | NA |
| Olfr800 | 0 | 0 | 0 | 0 | 0 | 0 | 0 | 0 | 0 | 0 | 0 | 0 | 0 | 0 | 0 | NA | NA | NA | NA | NA | NA | NA | NA | NA | NA |
| Olfr801 | 0 | 0 | 0 | 0 | 0 | 0 | 0 | 0 | 0 | 0 | 0 | 0 | 0 | 0 | 0 | NA | NA | NA | NA | NA | NA | NA | NA | NA | NA |
| Olfr802 | 0 | 0 | 0 | 0 | 0 | 0 | 0 | 0 | 0 | 0 | 0 | 0 | 0 | 0 | 0 | NA | NA | NA | NA | NA | NA | NA | NA | NA | NA |
| Olfr803 | 0 | 0 | 0 | 0 | 0 | 0 | 0 | 0 | 0 | 0 | 0 | 0 | 0 | 0 | 0 | NA | NA | NA | NA | NA | NA | NA | NA | NA | NA |
| Olfr804 | 0 | 0 | 0 | 0 | 0 | 0 | 0 | 0 | 0 | 0 | 0 | 0 | 0 | 0 | 0 | NA | NA | NA | NA | NA | NA | NA | NA | NA | NA |
| Olfr805 | 0 | 0 | 0 | 0 | 0 | 0 | 0 | 0 | 0 | 0 | 0 | 0 | 0 | 0 | 0 | NA | NA | NA | NA | NA | NA | NA | NA | NA | NA |
| Olfr806 | 0 | 0 | 0 | 0 | 0 | 0 | 0 | 0 | 0 | 0 | 0 | 0 | 0 | 0 | 0 | NA | NA | NA | NA | NA | NA | NA | NA | NA | NA |
| Olfr807 | 0 | 0 | 0 | 0 | 0 | 0 | 0 | 0 | 0 | 0 | 0 | 0 | 0 | 0 | 0 | NA | NA | NA | NA | NA | NA | NA | NA | NA | NA |
| Olfr808 | 0 | 0 | 0 | 0 | 0 | 0 | 0 | 0 | 0 | 0 | 0 | 0 | 0 | 0 | 0 | NA | NA | NA | NA | NA | NA | NA | NA | NA | NA |
| Olfr809 | 0 | 0 | 0 | 0 | 0 | 0 | 0 | 0 | 0 | 0 | 0 | 0 | 0 | 0 | 0 | NA | NA | NA | NA | NA | NA | NA | NA | NA | NA |
| Olfr810 | 0 | 0 | 0 | 0 | 0 | 0 | 0 | 0 | 0 | 0 | 0 | 0 | 0 | 0 | 0 | NA | NA | NA | NA | NA | NA | NA | NA | NA | NA |
| Olfr811 | 0 | 0 | 0 | 0 | 0 | 0 | 0 | 0 | 0 | 0 | 0 | 0 | 0 | 0 | 0 | NA | NA | NA | NA | NA | NA | NA | NA | NA | NA |
| Olfr812 | 0 | 0 | 0 | 0 | 0 | 0 | 0 | 0 | 0 | 0 | 0 | 0 | 0 | 0 | 0 | NA | NA | NA | NA | NA | NA | NA | NA | NA | NA |
| Olfr813 | 0 | 0 | 0 | 0 | 0 | 0 | 0 | 0 | 0 | 0 | 0 | 0 | 0 | 0 | 0 | NA | NA | NA | NA | NA | NA | NA | NA | NA | NA |
| Olfr814 | 0 | 0 | 0 | 0 | 0 | 0 | 0 | 0 | 0 | 0 | 0 | 0 | 0 | 0 | 0 | NA | NA | NA | NA | NA | NA | NA | NA | NA | NA |
| Olfr815 | 4 | 2 | 1 | 18 | 13 | 5 | 4 | 2 | 1 | 14 | 14 | 5 | 6.81 | 2 | 11 | 4.703 | 2.234 | 0.01297404 | 0.06378636 | 0 | 0.4907 | 0.326 | 0.326 | TRUE | 0.6903 |
| Olfr816 | 0 | 0 | 0 | 0 | 0 | 0 | 0 | 0 | 0 | 0 | 0 | 0 | 0 | 0 | 0 | NA | NA | NA | NA | NA | NA | NA | NA | NA | NA |
| Olfr818 | 0 | 0 | 0 | 0 | 0 | 0 | 0 | 0 | 0 | 0 | 0 | 0 | 0 | 0 | 0 | NA | NA | NA | NA | NA | NA | NA | NA | NA | NA |
| Olfr819 | 0 | 0 | 0 | 0 | 0 | 1 | 0 | 0 | 0 | 0 | 0 | 1 | 0.18 | 0 | 0 | 1.859 | 0.894 | 0.82650734 | NA | 0 | 18.0643 | 10 | 10 | TRUE | 0.5383 |
| Olfr820 | 0 | 0 | 0 | 0 | 0 | 0 | 0 | 0 | 0 | 0 | 0 | 0 | 0 | 0 | 0 | NA | NA | NA | NA | NA | NA | NA | NA | NA | NA |
| Olfr821 | 0 | 0 | 0 | 0 | 0 | 0 | 0 | 0 | 0 | 0 | 0 | 0 | 0 | 0 | 0 | NA | NA | NA | NA | NA | NA | NA | NA | NA | NA |
| Olfr822 | 0 | 0 | 0 | 0 | 0 | 0 | 0 | 0 | 0 | 0 | 0 | 0 | 0 | 0 | 0 | NA | NA | NA | NA | NA | NA | NA | NA | NA | NA |
| Olfr823 | 0 | 0 | 0 | 0 | 0 | 0 | 0 | 0 | 0 | 0 | 0 | 0 | 0 | 0 | 0 | NA | NA | NA | NA | NA | NA | NA | NA | NA | NA |
| Olfr824 | 0 | 0 | 0 | 0 | 0 | 0 | 0 | 0 | 0 | 0 | 0 | 0 | 0 | 0 | 0 | NA | NA | NA | NA | NA | NA | NA | NA | NA | NA |
| Olfr825 | 0 | 0 | 0 | 0 | 0 | 0 | 0 | 0 | 0 | 0 | 0 | 0 | 0 | 0 | 0 | NA | NA | NA | NA | NA | NA | NA | NA | NA | NA |
| Olfr826 | 0 | 0 | 0 | 0 | 0 | 0 | 0 | 0 | 0 | 0 | 0 | 0 | 0 | 0 | 0 | NA | NA | NA | NA | NA | NA | NA | NA | NA | NA |
| Olfr827 | 0 | 0 | 0 | 0 | 0 | 0 | 0 | 0 | 0 | 0 | 0 | 0 | 0 | 0 | 0 | NA | NA | NA | NA | NA | NA | NA | NA | NA | NA |
| Olfr828 | 0 | 0 | 0 | 0 | 0 | 0 | 0 | 0 | 0 | 0 | 0 | 0 | 0 | 0 | 0 | NA | NA | NA | NA | NA | NA | NA | NA | NA | NA |
| Olfr829 | 0 | 0 | 0 | 0 | 0 | 0 | 0 | 0 | 0 | 0 | 0 | 0 | 0 | 0 | 0 | NA | NA | NA | NA | NA | NA | NA | NA | NA | NA |
| Olfr830 | 0 | 0 | 0 | 0 | 0 | 0 | 0 | 0 | 0 | 0 | 0 | 0 | 0 | 0 | 0 | NA | NA | NA | NA | NA | NA | NA | NA | NA | NA |
| Olfr832 | 0 | 0 | 0 | 0 | 0 | 0 | 0 | 0 | 0 | 0 | 0 | 0 | 0 | 0 | 0 | NA | NA | NA | NA | NA | NA | NA | NA | NA | NA |
| Olfr834 | 0 | 0 | 0 | 0 | 0 | 0 | 0 | 0 | 0 | 0 | 0 | 0 | 0 | 0 | 0 | NA | NA | NA | NA | NA | NA | NA | NA | NA | NA |
| Olfr835 | 0 | 0 | 0 | 0 | 0 | 0 | 0 | 0 | 0 | 0 | 0 | 0 | 0 | 0 | 0 | NA | NA | NA | NA | NA | NA | NA | NA | NA | NA |
| Olfr836 | 0 | 0 | 0 | 1 | 0 | 1 | 0 | 0 | 0 | 1 | 0 | 1 | 0.31 | 0 | 1 | 3.348 | 1.743 | 0.65064889 | NA | 0 | 10.5167 | 8.8968 | 8.8968 | TRUE | 0.2082 |
| Olfr837 | 0 | 0 | 0 | 0 | 1 | 0 | 0 | 0 | 0 | 0 | 1 | 0 | 0.18 | 0 | 0 | 1.859 | 0.894 | 0.82650734 | NA | 0 | 18.3785 | 10 | 10 | TRUE | 0.5204 |
| Olfr843 | 0 | 0 | 0 | 0 | 0 | 0 | 0 | 0 | 0 | 0 | 0 | 0 | 0 | 0 | 0 | NA | NA | NA | NA | NA | NA | NA | NA | NA | NA |
| Olfr845 | 0 | 0 | 0 | 0 | 0 | 0 | 0 | 0 | 0 | 0 | 0 | 0 | 0 | 0 | 0 | NA | NA | NA | NA | NA | NA | NA | NA | NA | NA |
| Olfr846 | 1 | 0 | 0 | 2 | 0 | 0 | 1 | 0 | 0 | 2 | 0 | 0 | 0.44 | 0 | 1 | 1.494 | 0.579 | 0.86844164 | NA | 0 | 7.5101 | 6.9157 | 6.9157 | TRUE | 0.9721 |
| Olfr847 | 0 | 0 | 0 | 0 | 0 | 0 | 0 | 0 | 0 | 0 | 0 | 0 | 0 | 0 | 0 | NA | NA | NA | NA | NA | NA | NA | NA | NA | NA |
| Olfr849 | 0 | 0 | 0 | 0 | 0 | 0 | 0 | 0 | 0 | 0 | 0 | 0 | 0 | 0 | 0 | NA | NA | NA | NA | NA | NA | NA | NA | NA | NA |
| Olfr850 | 0 | 0 | 0 | 0 | 0 | 0 | 0 | 0 | 0 | 0 | 0 | 0 | 0 | 0 | 0 | NA | NA | NA | NA | NA | NA | NA | NA | NA | NA |
| Olfr851 | 0 | 0 | 0 | 0 | 0 | 0 | 0 | 0 | 0 | 0 | 0 | 0 | 0 | 0 | 0 | NA | NA | NA | NA | NA | NA | NA | NA | NA | NA |
| Olfr853 | 0 | 0 | 0 | 0 | 0 | 0 | 0 | 0 | 0 | 0 | 0 | 0 | 0 | 0 | 0 | NA | NA | NA | NA | NA | NA | NA | NA | NA | NA |
| Olfr854 | 0 | 0 | 0 | 0 | 0 | 0 | 0 | 0 | 0 | 0 | 0 | 0 | 0 | 0 | 0 | NA | NA | NA | NA | NA | NA | NA | NA | NA | NA |
| Olfr855 | 0 | 0 | 0 | 0 | 0 | 0 | 0 | 0 | 0 | 0 | 0 | 0 | 0 | 0 | 0 | NA | NA | NA | NA | NA | NA | NA | NA | NA | NA |
| Olfr856-ps1 | 0 | 0 | 0 | 0 | 0 | 0 | 0 | 0 | 0 | 0 | 0 | 0 | 0 | 0 | 0 | NA | NA | NA | NA | NA | NA | NA | NA | NA | NA |
| Olfr857 | 0 | 1 | 0 | 0 | 1 | 1 | 0 | 1 | 0 | 0 | 1 | 1 | 0.53 | 0 | 1 | 1.924 | 0.944 | 0.7364225 | NA | 0 | 6.2438 | 3.9727 | 3.9727 | TRUE | 0.405 |
| Olfr859 | 1 | 0 | 1 | 2 | 3 | 0 | 1 | 0 | 1 | 2 | 3 | 0 | 1.14 | 1 | 2 | 2.338 | 1.225 | 0.53592605 | NA | 0 | 2.8857 | 1.7735 | 1.7735 | TRUE | 0.1009 |
| Olfr860 | 0 | 0 | 0 | 0 | 0 | 0 | 0 | 0 | 0 | 0 | 0 | 0 | 0 | 0 | 0 | NA | NA | NA | NA | NA | NA | NA | NA | NA | NA |
| Olfr862 | 0 | 0 | 0 | 0 | 0 | 0 | 0 | 0 | 0 | 0 | 0 | 0 | 0 | 0 | 0 | NA | NA | NA | NA | NA | NA | NA | NA | NA | NA |
| Olfr866 | 0 | 0 | 0 | 0 | 0 | 0 | 0 | 0 | 0 | 0 | 0 | 0 | 0 | 0 | 0 | NA | NA | NA | NA | NA | NA | NA | NA | NA | NA |
| Olfr867 | 0 | 0 | 0 | 0 | 0 | 0 | 0 | 0 | 0 | 0 | 0 | 0 | 0 | 0 | 0 | NA | NA | NA | NA | NA | NA | NA | NA | NA | NA |
| Olfr868 | 0 | 0 | 0 | 0 | 0 | 0 | 0 | 0 | 0 | 0 | 0 | 0 | 0 | 0 | 0 | NA | NA | NA | NA | NA | NA | NA | NA | NA | NA |
| Olfr869 | 0 | 0 | 0 | 0 | 0 | 0 | 0 | 0 | 0 | 0 | 0 | 0 | 0 | 0 | 0 | NA | NA | NA | NA | NA | NA | NA | NA | NA | NA |
| Olfr870 | 0 | 0 | 0 | 0 | 0 | 0 | 0 | 0 | 0 | 0 | 0 | 0 | 0 | 0 | 0 | NA | NA | NA | NA | NA | NA | NA | NA | NA | NA |
| Olfr871 | 0 | 0 | 0 | 0 | 0 | 0 | 0 | 0 | 0 | 0 | 0 | 0 | 0 | 0 | 0 | NA | NA | NA | NA | NA | NA | NA | NA | NA | NA |
| Olfr872 | 0 | 0 | 0 | 0 | 0 | 0 | 0 | 0 | 0 | 0 | 0 | 0 | 0 | 0 | 0 | NA | NA | NA | NA | NA | NA | NA | NA | NA | NA |
| Olfr873 | 0 | 0 | 0 | 0 | 0 | 0 | 0 | 0 | 0 | 0 | 0 | 0 | 0 | 0 | 0 | NA | NA | NA | NA | NA | NA | NA | NA | NA | NA |

| | | | | | | | | | | | | | | | | | | | | | | | | | | |
|---|---|---|---|---|---|---|---|---|---|---|---|---|---|---|---|---|---|---|---|---|---|---|---|---|---|---|
| Olfr874 | 0 | 0 | 0 | 0 | 0 | 0 | 0 | 0 | 0 | 0 | 0 | 0 | 0 | 0 | 0 | NA | NA | NA | NA | NA | NA | NA | NA | NA | NA | |
| Olfr875 | 0 | 0 | 0 | 0 | 0 | 0 | 0 | 0 | 0 | 0 | 0 | 0 | 0 | 0 | 0 | NA | NA | NA | NA | NA | NA | NA | NA | NA | NA | |
| Olfr876 | 0 | 0 | 0 | 0 | 1 | 0 | 0 | 0 | 0 | 0 | 1 | 0 | 0.18 | 0 | 0 | 1.859 | 0.894 | 0.82650734 | NA | | 0 | 18.3785 | 10 | 10 | TRUE | 0.5204 |
| Olfr877 | 0 | 0 | 0 | 0 | 0 | 0 | 0 | 0 | 0 | 0 | 0 | 0 | 0 | 0 | 0 | NA | NA | NA | NA | NA | NA | NA | NA | NA | NA | |
| Olfr878 | 0 | 0 | 0 | 0 | 0 | 0 | 0 | 0 | 0 | 0 | 0 | 0 | 0 | 0 | 0 | NA | NA | NA | NA | NA | NA | NA | NA | NA | NA | |
| Olfr881 | 0 | 0 | 0 | 1 | 2 | 1 | 0 | 0 | 0 | 1 | 2 | 1 | 0.67 | 0 | 1 | 7.1 | 2.828 | 0.25693906 | NA | | 0 | 4.9086 | 3.1164 | 3.1164 | TRUE | 0.1607 |
| Olfr883 | 0 | 0 | 0 | 0 | 0 | 0 | 0 | 0 | 0 | 0 | 0 | 0 | 0 | 0 | 0 | NA | NA | NA | NA | NA | NA | NA | NA | NA | NA | |
| Olfr884 | 0 | 0 | 0 | 0 | 0 | 0 | 0 | 0 | 0 | 0 | 0 | 0 | 0 | 0 | 0 | NA | NA | NA | NA | NA | NA | NA | NA | NA | NA | |
| Olfr885 | 0 | 0 | 0 | 0 | 0 | 0 | 0 | 0 | 0 | 0 | 0 | 0 | 0 | 0 | 0 | NA | NA | NA | NA | NA | NA | NA | NA | NA | NA | |
| Olfr887 | 0 | 0 | 0 | 0 | 0 | 0 | 0 | 0 | 0 | 0 | 0 | 0 | 0 | 0 | 0 | NA | NA | NA | NA | NA | NA | NA | NA | NA | NA | |
| Olfr888 | 0 | 0 | 0 | 0 | 0 | 0 | 0 | 0 | 0 | 0 | 0 | 0 | 0 | 0 | 0 | NA | NA | NA | NA | NA | NA | NA | NA | NA | NA | |
| Olfr889 | 0 | 0 | 0 | 0 | 0 | 0 | 0 | 0 | 0 | 0 | 0 | 0 | 0 | 0 | 0 | NA | NA | NA | NA | NA | NA | NA | NA | NA | NA | |
| Olfr890 | 0 | 0 | 0 | 0 | 0 | 0 | 0 | 0 | 0 | 0 | 0 | 0 | 0 | 0 | 0 | NA | NA | NA | NA | NA | NA | NA | NA | NA | NA | |
| Olfr891 | 0 | 0 | 0 | 0 | 0 | 0 | 0 | 0 | 0 | 0 | 0 | 0 | 0 | 0 | 0 | NA | NA | NA | NA | NA | NA | NA | NA | NA | NA | |
| Olfr893 | 0 | 0 | 0 | 0 | 0 | 0 | 0 | 0 | 0 | 0 | 0 | 0 | 0 | 0 | 0 | NA | NA | NA | NA | NA | NA | NA | NA | NA | NA | |
| Olfr894 | 0 | 0 | 0 | 0 | 0 | 0 | 0 | 0 | 0 | 0 | 0 | 0 | 0 | 0 | 0 | NA | NA | NA | NA | NA | NA | NA | NA | NA | NA | |
| Olfr895 | 0 | 0 | 0 | 0 | 0 | 0 | 0 | 0 | 0 | 0 | 0 | 0 | 0 | 0 | 0 | NA | NA | NA | NA | NA | NA | NA | NA | NA | NA | |
| Olfr898 | 0 | 0 | 0 | 0 | 0 | 0 | 0 | 0 | 0 | 0 | 0 | 0 | 0 | 0 | 0 | NA | NA | NA | NA | NA | NA | NA | NA | NA | NA | |
| Olfr899 | 0 | 0 | 0 | 0 | 0 | 0 | 0 | 0 | 0 | 0 | 0 | 0 | 0 | 0 | 0 | NA | NA | NA | NA | NA | NA | NA | NA | NA | NA | |
| Olfr9 | 0 | 0 | 0 | 0 | 0 | 0 | 0 | 0 | 0 | 0 | 0 | 0 | 0 | 0 | 0 | NA | NA | NA | NA | NA | NA | NA | NA | NA | NA | |
| Olfr90 | 0 | 0 | 0 | 0 | 0 | 0 | 0 | 0 | 0 | 0 | 0 | 0 | 0 | 0 | 0 | NA | NA | NA | NA | NA | NA | NA | NA | NA | NA | |
| Olfr900 | 0 | 0 | 0 | 0 | 0 | 0 | 0 | 0 | 0 | 0 | 0 | 0 | 0 | 0 | 0 | NA | NA | NA | NA | NA | NA | NA | NA | NA | NA | |
| Olfr901 | 3 | 0 | 0 | 4 | 3 | 3 | 3 | 0 | 0 | 3 | 3 | 3 | 2.14 | 1 | 3 | 3.086 | 1.626 | 0.29318364 | NA | | 0 | 1.546 | 1.0818 | 1.0818 | TRUE | 1.4433 |
| Olfr902 | 0 | 0 | 0 | 0 | 0 | 0 | 0 | 0 | 0 | 0 | 0 | 0 | 0 | 0 | 0 | NA | NA | NA | NA | NA | NA | NA | NA | NA | NA | |
| Olfr904 | 0 | 0 | 0 | 0 | 0 | 0 | 0 | 0 | 0 | 0 | 0 | 0 | 0 | 0 | 0 | NA | NA | NA | NA | NA | NA | NA | NA | NA | NA | |
| Olfr905 | 0 | 0 | 0 | 0 | 0 | 0 | 0 | 0 | 0 | 0 | 0 | 0 | 0 | 0 | 0 | NA | NA | NA | NA | NA | NA | NA | NA | NA | NA | |
| Olfr906 | 0 | 0 | 0 | 0 | 0 | 0 | 0 | 0 | 0 | 0 | 0 | 0 | 0 | 0 | 0 | NA | NA | NA | NA | NA | NA | NA | NA | NA | NA | |
| Olfr907 | 0 | 0 | 0 | 0 | 0 | 0 | 0 | 0 | 0 | 0 | 0 | 0 | 0 | 0 | 0 | NA | NA | NA | NA | NA | NA | NA | NA | NA | NA | |
| Olfr908 | 0 | 0 | 0 | 0 | 0 | 0 | 0 | 0 | 0 | 0 | 0 | 0 | 0 | 0 | 0 | NA | NA | NA | NA | NA | NA | NA | NA | NA | NA | |
| Olfr91 | 0 | 0 | 0 | 0 | 0 | 0 | 0 | 0 | 0 | 0 | 0 | 0 | 0 | 0 | 0 | NA | NA | NA | NA | NA | NA | NA | NA | NA | NA | |
| Olfr910 | 0 | 0 | 0 | 0 | 0 | 0 | 0 | 0 | 0 | 0 | 0 | 0 | 0 | 0 | 0 | NA | NA | NA | NA | NA | NA | NA | NA | NA | NA | |
| Olfr911-ps1 | 0 | 0 | 0 | 0 | 0 | 0 | 0 | 0 | 0 | 0 | 0 | 0 | 0 | 0 | 0 | NA | NA | NA | NA | NA | NA | NA | NA | NA | NA | |
| Olfr912 | 0 | 0 | 0 | 0 | 0 | 0 | 0 | 0 | 0 | 0 | 0 | 0 | 0 | 0 | 0 | NA | NA | NA | NA | NA | NA | NA | NA | NA | NA | |
| Olfr913 | 0 | 0 | 0 | 0 | 0 | 0 | 0 | 0 | 0 | 0 | 0 | 0 | 0 | 0 | 0 | NA | NA | NA | NA | NA | NA | NA | NA | NA | NA | |
| Olfr914 | 0 | 0 | 0 | 0 | 0 | 0 | 0 | 0 | 0 | 0 | 0 | 0 | 0 | 0 | 0 | NA | NA | NA | NA | NA | NA | NA | NA | NA | NA | |
| Olfr915 | 0 | 0 | 0 | 1 | 0 | 0 | 0 | 0 | 0 | 1 | 0 | 0 | 0.13 | 0 | 0 | 1.859 | 0.894 | 0.82650734 | NA | | 0 | 25.1515 | 10 | 10 | TRUE | 0.2563 |
| Olfr916 | 0 | 0 | 0 | 0 | 0 | 0 | 0 | 0 | 0 | 0 | 0 | 0 | 0 | 0 | 0 | NA | NA | NA | NA | NA | NA | NA | NA | NA | NA | |
| Olfr917 | 0 | 0 | 0 | 0 | 0 | 0 | 0 | 0 | 0 | 0 | 0 | 0 | 0 | 0 | 0 | NA | NA | NA | NA | NA | NA | NA | NA | NA | NA | |
| Olfr918 | 0 | 0 | 0 | 0 | 0 | 0 | 0 | 0 | 0 | 0 | 0 | 0 | 0 | 0 | 0 | NA | NA | NA | NA | NA | NA | NA | NA | NA | NA | |
| Olfr919 | 0 | 0 | 0 | 0 | 0 | 0 | 0 | 0 | 0 | 0 | 0 | 0 | 0 | 0 | 0 | NA | NA | NA | NA | NA | NA | NA | NA | NA | NA | |
| Olfr92 | 0 | 0 | 0 | 0 | 0 | 0 | 0 | 0 | 0 | 0 | 0 | 0 | 0 | 0 | 0 | NA | NA | NA | NA | NA | NA | NA | NA | NA | NA | |
| Olfr920 | 3 | 6 | 5 | 5 | 0 | 4 | 3 | 6 | 5 | 4 | 0 | 4 | 3.75 | 5 | 3 | 0.593 | -0.754 | 0.4906895 | NA | | 0 | 0.8851 | 0.5818 | 0.5818 | TRUE | 0.8247 |
| Olfr921 | 0 | 0 | 0 | 1 | 0 | 0 | 0 | 0 | 0 | 1 | 0 | 0 | 0.13 | 0 | 0 | 1.859 | 0.894 | 0.82650734 | NA | | 0 | 25.1515 | 10 | 10 | TRUE | 0.2563 |
| Olfr922 | 0 | 0 | 0 | 0 | 0 | 0 | 0 | 0 | 0 | 0 | 0 | 0 | 0 | 0 | 0 | NA | NA | NA | NA | NA | NA | NA | NA | NA | NA | |
| Olfr923 | 0 | 0 | 0 | 0 | 0 | 0 | 0 | 0 | 0 | 0 | 0 | 0 | 0 | 0 | 0 | NA | NA | NA | NA | NA | NA | NA | NA | NA | NA | |
| Olfr924 | 0 | 0 | 0 | 0 | 0 | 0 | 0 | 0 | 0 | 0 | 0 | 0 | 0 | 0 | 0 | NA | NA | NA | NA | NA | NA | NA | NA | NA | NA | |
| Olfr926 | 1 | 0 | 0 | 0 | 0 | 0 | 1 | 0 | 0 | 0 | 0 | 0 | 0.18 | 0 | 0 | 0.49 | -1.029 | 0.80087 | NA | | 0 | 18.6165 | 10 | 10 | TRUE | 0.4614 |
| Olfr93 | 0 | 0 | 0 | 0 | 0 | 0 | 0 | 0 | 0 | 0 | 0 | 0 | 0 | 0 | 0 | NA | NA | NA | NA | NA | NA | NA | NA | NA | NA | |
| Olfr930 | 0 | 0 | 0 | 0 | 0 | 0 | 0 | 0 | 0 | 0 | 0 | 0 | 0 | 0 | 0 | NA | NA | NA | NA | NA | NA | NA | NA | NA | NA | |
| Olfr933 | 0 | 0 | 0 | 0 | 0 | 0 | 0 | 0 | 0 | 0 | 0 | 0 | 0 | 0 | 0 | NA | NA | NA | NA | NA | NA | NA | NA | NA | NA | |
| Olfr934 | 0 | 0 | 0 | 0 | 0 | 0 | 0 | 0 | 0 | 0 | 0 | 0 | 0 | 0 | 0 | NA | NA | NA | NA | NA | NA | NA | NA | NA | NA | |
| Olfr935 | 0 | 0 | 0 | 0 | 0 | 0 | 0 | 0 | 0 | 0 | 0 | 0 | 0 | 0 | 0 | NA | NA | NA | NA | NA | NA | NA | NA | NA | NA | |
| Olfr936 | 0 | 0 | 0 | 0 | 0 | 0 | 0 | 0 | 0 | 0 | 0 | 0 | 0 | 0 | 0 | NA | NA | NA | NA | NA | NA | NA | NA | NA | NA | |
| Olfr937 | 0 | 0 | 0 | 0 | 0 | 0 | 0 | 0 | 0 | 0 | 0 | 0 | 0 | 0 | 0 | NA | NA | NA | NA | NA | NA | NA | NA | NA | NA | |
| Olfr938 | 0 | 0 | 0 | 0 | 0 | 0 | 0 | 0 | 0 | 0 | 0 | 0 | 0 | 0 | 0 | NA | NA | NA | NA | NA | NA | NA | NA | NA | NA | |
| Olfr94 | 0 | 0 | 0 | 0 | 0 | 0 | 0 | 0 | 0 | 0 | 0 | 0 | 0 | 0 | 0 | NA | NA | NA | NA | NA | NA | NA | NA | NA | NA | |
| Olfr943 | 0 | 0 | 0 | 0 | 0 | 0 | 0 | 0 | 0 | 0 | 0 | 0 | 0 | 0 | 0 | NA | NA | NA | NA | NA | NA | NA | NA | NA | NA | |
| Olfr944 | 0 | 0 | 0 | 0 | 0 | 0 | 0 | 0 | 0 | 0 | 0 | 0 | 0 | 0 | 0 | NA | NA | NA | NA | NA | NA | NA | NA | NA | NA | |
| Olfr945 | 0 | 0 | 0 | 0 | 0 | 0 | 0 | 0 | 0 | 0 | 0 | 0 | 0 | 0 | 0 | NA | NA | NA | NA | NA | NA | NA | NA | NA | NA | |
| Olfr947-ps1 | 0 | 0 | 0 | 0 | 0 | 0 | 0 | 0 | 0 | 0 | 0 | 0 | 0 | 0 | 0 | NA | NA | NA | NA | NA | NA | NA | NA | NA | NA | |
| Olfr948 | 0 | 0 | 0 | 0 | 0 | 0 | 0 | 0 | 0 | 0 | 0 | 0 | 0 | 0 | 0 | NA | NA | NA | NA | NA | NA | NA | NA | NA | NA | |
| Olfr95 | 0 | 0 | 0 | 0 | 0 | 0 | 0 | 0 | 0 | 0 | 0 | 0 | 0 | 0 | 0 | NA | NA | NA | NA | NA | NA | NA | NA | NA | NA | |
| Olfr951 | 0 | 0 | 0 | 0 | 0 | 0 | 0 | 0 | 0 | 0 | 0 | 0 | 0 | 0 | 0 | NA | NA | NA | NA | NA | NA | NA | NA | NA | NA | |
| Olfr952 | 0 | 0 | 0 | 0 | 0 | 0 | 0 | 0 | 0 | 0 | 0 | 0 | 0 | 0 | 0 | NA | NA | NA | NA | NA | NA | NA | NA | NA | NA | |
| Olfr954 | 0 | 0 | 0 | 0 | 0 | 0 | 0 | 0 | 0 | 0 | 0 | 0 | 0 | 0 | 0 | NA | NA | NA | NA | NA | NA | NA | NA | NA | NA | |
| Olfr955 | 0 | 0 | 0 | 0 | 0 | 0 | 0 | 0 | 0 | 0 | 0 | 0 | 0 | 0 | 0 | NA | NA | NA | NA | NA | NA | NA | NA | NA | NA | |
| Olfr957 | 0 | 0 | 0 | 0 | 0 | 0 | 0 | 0 | 0 | 0 | 0 | 0 | 0 | 0 | 0 | NA | NA | NA | NA | NA | NA | NA | NA | NA | NA | |
| Olfr958 | 0 | 0 | 0 | 0 | 0 | 0 | 0 | 0 | 0 | 0 | 0 | 0 | 0 | 0 | 0 | NA | NA | NA | NA | NA | NA | NA | NA | NA | NA | |
| Olfr959 | 0 | 0 | 0 | 0 | 0 | 0 | 0 | 0 | 0 | 0 | 0 | 0 | 0 | 0 | 0 | NA | NA | NA | NA | NA | NA | NA | NA | NA | NA | |

| | | | | | | | | | | | | | | | | | | | | | | | | | |
|---|---|---|---|---|---|---|---|---|---|---|---|---|---|---|---|---|---|---|---|---|---|---|---|---|---|
| Olfr96 | 0 | 0 | 0 | 0 | 0 | 0 | 0 | 0 | 0 | 0 | 0 | 0 | 0 | 0 | 0 | NA | NA | NA | NA | NA | NA | NA | NA | NA | NA |
| Olfr960 | 0 | 0 | 0 | 1 | 2 | 1 | 0 | 0 | 0 | 1 | 2 | 1 | 0.67 | 0 | 1 | 7.1 | 2.828 | 0.25693906 | NA | 0 | 4.9086 | 3.1164 | 3.1164 | TRUE | 0.1607 |
| Olfr961 | 0 | 0 | 0 | 0 | 0 | 0 | 0 | 0 | 0 | 0 | 0 | 0 | 0 | 0 | 0 | NA | NA | NA | NA | NA | NA | NA | NA | NA | NA |
| Olfr963 | 0 | 0 | 0 | 0 | 0 | 0 | 0 | 0 | 0 | 0 | 0 | 0 | 0 | 0 | 0 | NA | NA | NA | NA | NA | NA | NA | NA | NA | NA |
| Olfr965 | 0 | 0 | 0 | 0 | 0 | 0 | 0 | 0 | 0 | 0 | 0 | 0 | 0 | 0 | 0 | NA | NA | NA | NA | NA | NA | NA | NA | NA | NA |
| Olfr967 | 0 | 0 | 0 | 0 | 0 | 0 | 0 | 0 | 0 | 0 | 0 | 0 | 0 | 0 | 0 | NA | NA | NA | NA | NA | NA | NA | NA | NA | NA |
| Olfr968 | 0 | 1 | 0 | 0 | 1 | 0 | 0 | 1 | 0 | 0 | 1 | 0 | 0.35 | 0 | 0 | 0.954 | -0.067 | 0.98571955 | NA | 0 | 9.5346 | 8.2194 | 8.2194 | TRUE | 0.5204 |
| Olfr969 | 0 | 0 | 0 | 0 | 0 | 0 | 0 | 0 | 0 | 0 | 0 | 0 | 0 | 0 | 0 | NA | NA | NA | NA | NA | NA | NA | NA | NA | NA |
| Olfr97 | 0 | 0 | 0 | 0 | 0 | 0 | 0 | 0 | 0 | 0 | 0 | 0 | 0 | 0 | 0 | NA | NA | NA | NA | NA | NA | NA | NA | NA | NA |
| Olfr970 | 0 | 0 | 0 | 0 | 0 | 0 | 0 | 0 | 0 | 0 | 0 | 0 | 0 | 0 | 0 | NA | NA | NA | NA | NA | NA | NA | NA | NA | NA |
| Olfr971 | 0 | 0 | 0 | 0 | 0 | 0 | 0 | 0 | 0 | 0 | 0 | 0 | 0 | 0 | 0 | NA | NA | NA | NA | NA | NA | NA | NA | NA | NA |
| Olfr972 | 0 | 0 | 0 | 0 | 0 | 0 | 0 | 0 | 0 | 0 | 0 | 0 | 0 | 0 | 0 | NA | NA | NA | NA | NA | NA | NA | NA | NA | NA |
| Olfr974 | 0 | 0 | 0 | 0 | 0 | 0 | 0 | 0 | 0 | 0 | 0 | 0 | 0 | 0 | 0 | NA | NA | NA | NA | NA | NA | NA | NA | NA | NA |
| Olfr975 | 0 | 0 | 0 | 2 | 0 | 0 | 0 | 0 | 0 | 2 | 0 | 0 | 0.26 | 0 | 1 | 2.886 | 1.529 | 0.70676491 | NA | 0 | 12.5792 | 10 | 10 | TRUE | 0.9669 |
| Olfr976 | 4 | 0 | 0 | 0 | 1 | 1 | 4 | 0 | 0 | 0 | 1 | 1 | 1.07 | 1 | 1 | 0.498 | -1.006 | 0.67631032 | NA | 3.1821 | 3.0862 | 3.1199 | 3.1199 | TRUE | 1.8994 |
| Olfr978 | 0 | 0 | 0 | 0 | 0 | 0 | 0 | 0 | 0 | 0 | 0 | 0 | 0 | 0 | 0 | NA | NA | NA | NA | NA | NA | NA | NA | NA | NA |
| Olfr979 | 0 | 0 | 0 | 1 | 0 | 0 | 0 | 0 | 0 | 1 | 0 | 0 | 0.13 | 0 | 0 | 1.859 | 0.894 | 0.82650734 | NA | 0 | 25.1515 | 10 | 10 | TRUE | 0.2563 |
| Olfr98 | 0 | 0 | 0 | 0 | 0 | 0 | 0 | 0 | 0 | 0 | 0 | 0 | 0 | 0 | 0 | NA | NA | NA | NA | NA | NA | NA | NA | NA | NA |
| Olfr980 | 0 | 0 | 0 | 0 | 0 | 0 | 0 | 0 | 0 | 0 | 0 | 0 | 0 | 0 | 0 | NA | NA | NA | NA | NA | NA | NA | NA | NA | NA |
| Olfr981 | 0 | 0 | 0 | 0 | 0 | 0 | 0 | 0 | 0 | 0 | 0 | 0 | 0 | 0 | 0 | NA | NA | NA | NA | NA | NA | NA | NA | NA | NA |
| Olfr982 | 0 | 0 | 0 | 0 | 0 | 0 | 0 | 0 | 0 | 0 | 0 | 0 | 0 | 0 | 0 | NA | NA | NA | NA | NA | NA | NA | NA | NA | NA |
| Olfr983 | 1 | 0 | 0 | 0 | 0 | 0 | 1 | 0 | 0 | 0 | 0 | 0 | 0.18 | 0 | 0 | 0.49 | -1.029 | 0.80087 | NA | 0 | 18.6165 | 10 | 10 | TRUE | 0.4614 |
| Olfr984 | 0 | 0 | 0 | 0 | 1 | 0 | 0 | 0 | 0 | 0 | 1 | 0 | 0.18 | 0 | 0 | 1.859 | 0.894 | 0.82650734 | NA | 0 | 18.3785 | 10 | 10 | TRUE | 0.5204 |
| Olfr985 | 92 | 92 | 100 | 92 | 75 | 77 | 98 | 92 | 100 | 72 | 81 | 84 | 87.89 | 97 | 79 | 0.815 | -0.294 | 0.15678869 | 0.34972176 | 0 | 0.0444 | 0.0197 | 0.0197 | TRUE | 0.06 |
| Olfr986 | 0 | 0 | 0 | 0 | 0 | 0 | 0 | 0 | 0 | 0 | 0 | 0 | 0 | 0 | 0 | NA | NA | NA | NA | NA | NA | NA | NA | NA | NA |
| Olfr987 | 0 | 0 | 0 | 0 | 0 | 0 | 0 | 0 | 0 | 0 | 0 | 0 | 0 | 0 | 0 | NA | NA | NA | NA | NA | NA | NA | NA | NA | NA |
| Olfr988 | 0 | 0 | 0 | 0 | 0 | 0 | 0 | 0 | 0 | 0 | 0 | 0 | 0 | 0 | 0 | NA | NA | NA | NA | NA | NA | NA | NA | NA | NA |
| Olfr99 | 0 | 0 | 0 | 0 | 0 | 0 | 0 | 0 | 0 | 0 | 0 | 0 | 0 | 0 | 0 | NA | NA | NA | NA | NA | NA | NA | NA | NA | NA |
| Olfr992 | 0 | 0 | 0 | 0 | 0 | 0 | 0 | 0 | 0 | 0 | 0 | 0 | 0 | 0 | 0 | NA | NA | NA | NA | NA | NA | NA | NA | NA | NA |
| Olfr993 | 0 | 0 | 0 | 0 | 0 | 0 | 0 | 0 | 0 | 0 | 0 | 0 | 0 | 0 | 0 | NA | NA | NA | NA | NA | NA | NA | NA | NA | NA |
| Olfr994 | 0 | 0 | 0 | 0 | 0 | 0 | 0 | 0 | 0 | 0 | 0 | 0 | 0 | 0 | 0 | NA | NA | NA | NA | NA | NA | NA | NA | NA | NA |
| Olfr995 | 0 | 0 | 0 | 0 | 0 | 0 | 0 | 0 | 0 | 0 | 0 | 0 | 0 | 0 | 0 | NA | NA | NA | NA | NA | NA | NA | NA | NA | NA |
| Olfr996 | 0 | 0 | 0 | 0 | 0 | 0 | 0 | 0 | 0 | 0 | 0 | 0 | 0 | 0 | 0 | NA | NA | NA | NA | NA | NA | NA | NA | NA | NA |
| Olfr998 | 0 | 0 | 0 | 0 | 0 | 0 | 0 | 0 | 0 | 0 | 0 | 0 | 0 | 0 | 0 | NA | NA | NA | NA | NA | NA | NA | NA | NA | NA |
| Olig1 | 46 | 75 | 95 | 112 | 102 | 85 | 49 | 75 | 95 | 88 | 110 | 93 | 85 | 73 | 97 | 1.323 | 0.404 | 0.15168877 | 0.3428955 | 0.0445 | 0.0457 | 0.0451 | 0.0451 | TRUE | 0.3348 |
| Olig2 | 5 | 0 | 5 | 5 | 4 | 3 | 5 | 0 | 5 | 4 | 4 | 3 | 3.64 | 3 | 4 | 1.12 | 0.164 | 0.88492335 | NA | 0 | 0.9116 | 0.6457 | 0.6457 | TRUE | 1.1487 |
| Olig3 | 8 | 4 | 4 | 22 | 10 | 7 | 8 | 4 | 4 | 17 | 11 | 8 | 8.7 | 5 | 12 | 2.201 | 1.138 | 0.11139455 | 0.28068465 | 0 | 0.3854 | 0.2343 | 0.2343 | TRUE | 0.3784 |
| Olr1 | 0 | 0 | 0 | 8 | 5 | 1 | 0 | 0 | 0 | 6 | 5 | 1 | 2.13 | 0 | 4 | 22.825 | 4.513 | 0.01381775 | NA | 0.6399 | 1.5556 | 1.3066 | 1.3066 | TRUE | 0.681 |
| Oma1 | 1177 | 1277 | 1250 | 1502 | 1048 | 1034 | 1250 | 1275 | 1255 | 1181 | 1128 | 1132 | 1203.48 | 1260 | 1147 | 0.91 | -0.135 | 0.07437406 | 0.21421286 | 0 | 0.0096 | 0.0033 | 0.0033 | TRUE | 0.0084 |
| Omd | 0 | 0 | 0 | 0 | 0 | 0 | 0 | 0 | 0 | 0 | 0 | 0 | 0 | 0 | 0 | NA | NA | NA | NA | NA | NA | NA | NA | NA | NA |
| Omg | 0 | 0 | 0 | 0 | 0 | 0 | 0 | 0 | 0 | 0 | 0 | 0 | 0 | 0 | 0 | NA | NA | NA | NA | NA | NA | NA | NA | NA | NA |
| Omp | 25 | 16 | 15 | 31 | 16 | 18 | 27 | 16 | 15 | 24 | 17 | 20 | 19.81 | 19 | 20 | 1.074 | 0.103 | 0.82146826 | 0.90875327 | 0 | 0.1732 | 0.099 | 0.099 | TRUE | 0.5716 |
| Omt2a | 1 | 0 | 0 | 1 | 0 | 0 | 1 | 0 | 0 | 1 | 0 | 0 | 0.31 | 0 | 0 | 0.954 | -0.067 | 0.98625928 | NA | 0 | 10.7014 | 9.0384 | 9.0384 | TRUE | 0.4614 |
| Omt2b | 1 | 4 | 1 | 2 | 1 | 1 | 1 | 4 | 1 | 2 | 1 | 1 | 1.63 | 2 | 1 | 0.623 | -0.683 | 0.6406703 | NA | 0 | 2.0231 | 0.9035 | 0.9035 | TRUE | 0.673 |
| Onecut1 | 11 | 16 | 13 | 35 | 37 | 25 | 12 | 16 | 13 | 28 | 40 | 27 | 22.57 | 14 | 32 | 2.314 | 1.211 | 0.00491612 | 0.03097762 | 0 | 0.1529 | 0.0804 | 0.0804 | TRUE | 0.3379 |
| Onecut2 | 34 | 33 | 35 | 48 | 43 | 33 | 36 | 33 | 35 | 38 | 46 | 36 | 37.39 | 35 | 40 | 1.151 | 0.203 | 0.51572374 | 0.7079811 | 0 | 0.095 | 0.0431 | 0.0431 | TRUE | 0.1299 |
| Onecut3 | 3 | 10 | 10 | 8 | 8 | 4 | 3 | 10 | 10 | 6 | 9 | 4 | 7.08 | 8 | 6 | 0.827 | -0.274 | 0.72019347 | 0.84830169 | 0 | 0.4721 | 0.2791 | 0.2791 | TRUE | 0.7194 |
| Ooep | 1222 | 1451 | 1501 | 2632 | 1888 | 1988 | 1298 | 1449 | 1507 | 2069 | 2031 | 2176 | 1755.15 | 1418 | 2092 | 1.475 | 0.561 | 3.71E-10 | 2.51E-08 | 0.0027 | 0.0088 | 0.0052 | 0.0052 | TRUE | 0.0657 |
| Oog1 | 0 | 0 | 0 | 0 | 0 | 0 | 0 | 0 | 0 | 0 | 0 | 0 | 0 | 0 | 0 | NA | NA | NA | NA | NA | NA | NA | NA | NA | NA |
| Oog2 | 0 | 0 | 0 | 0 | 0 | 0 | 0 | 0 | 0 | 0 | 0 | 0 | 0 | 0 | 0 | NA | NA | NA | NA | NA | NA | NA | NA | NA | NA |
| Oog3 | 0 | 0 | 0 | 0 | 0 | 0 | 0 | 0 | 0 | 0 | 0 | 0 | 0 | 0 | 0 | NA | NA | NA | NA | NA | NA | NA | NA | NA | NA |
| Oog4 | 3 | 2 | 0 | 0 | 6 | 1 | 3 | 2 | 0 | 0 | 6 | 1 | 2.12 | 2 | 2 | 1.436 | 0.522 | 0.76620406 | NA | 1.9683 | 1.5592 | 1.7243 | 1.7243 | TRUE | 1.5566 |
| Oosp1 | 0 | 0 | 0 | 0 | 0 | 0 | 0 | 0 | 0 | 0 | 0 | 0 | 0 | 0 | 0 | NA | NA | NA | NA | NA | NA | NA | NA | NA | NA |
| Opa1 | 6128 | 6429 | 6685 | 7573 | 5376 | 5765 | 6509 | 6421 | 6712 | 5953 | 5784 | 6311 | 6281.73 | 6547 | 6016 | 0.919 | -0.122 | 0.06929064 | 0.20428374 | 0.001 | 0.0074 | 0.0031 | 0.0031 | TRUE | 0.0222 |
| Opa3 | 713 | 703 | 687 | 913 | 691 | 565 | 757 | 702 | 690 | 718 | 743 | 618 | 704.82 | 716 | 693 | 0.968 | -0.046 | 0.67724922 | 0.82056964 | 0.0043 | 0.0116 | 0.0075 | 0.0075 | TRUE | 0.1023 |
| Opalin | 2 | 1 | 0 | 0 | 0 | 1 | 2 | 1 | 0 | 0 | 0 | 1 | 0.7 | 1 | 0 | 0.337 | -1.569 | 0.54084423 | NA | 0 | 4.6935 | 3.253 | 3.253 | TRUE | 0.2932 |
| Opcml | 0 | 0 | 1 | 0 | 1 | 0 | 0 | 0 | 1 | 0 | 1 | 0 | 0.35 | 0 | 0 | 0.954 | -0.067 | 0.98570726 | NA | 0 | 9.5099 | 8.2019 | 8.2019 | TRUE | 0.5204 |
| Ophn1 | 213 | 268 | 272 | 252 | 168 | 150 | 226 | 268 | 273 | 198 | 181 | 164 | 218.35 | 256 | 181 | 0.71 | -0.495 | 0.00173911 | 0.01374214 | 0.0051 | 0.022 | 0.0133 | 0.0133 | TRUE | 0.1114 |
| Oplah | 330 | 372 | 363 | 471 | 398 | 426 | 351 | 372 | 364 | 370 | 428 | 466 | 391.89 | 362 | 421 | 1.161 | 0.215 | 0.09962478 | 0.26057745 | 0.0053 | 0.0153 | 0.0098 | 0.0098 | TRUE | 0.1387 |
| Opn1mw | 0 | 0 | 0 | 0 | 0 | 0 | 0 | 0 | 0 | 0 | 0 | 0 | 0 | 0 | 0 | NA | NA | NA | NA | NA | NA | NA | NA | NA | NA |
| Opn1sw | 0 | 0 | 2 | 1 | 0 | 0 | 0 | 0 | 2 | 1 | 0 | 0 | 0.47 | 1 | 0 | 0.52 | -0.943 | 0.78484416 | NA | 0 | 7.0808 | 6.8441 | 6.8441 | TRUE | 0.9739 |
| Opn3 | 98 | 106 | 120 | 116 | 67 | 72 | 104 | 106 | 120 | 91 | 72 | 79 | 95.42 | 110 | 81 | 0.736 | -0.442 | 0.03567241 | 0.1296073 | 0 | 0.0414 | 0.0213 | 0.0213 | TRUE | 0.1362 |
| Opn4 | 17 | 31 | 23 | 37 | 23 | 30 | 18 | 31 | 23 | 29 | 25 | 33 | 26.46 | 24 | 29 | 1.2 | 0.263 | 0.50206802 | 0.6972672 | 0 | 0.1314 | 0.0721 | 0.0721 | TRUE | 0.3815 |
| Opn5 | 0 | 0 | 0 | 0 | 0 | 0 | 0 | 0 | 0 | 0 | 0 | 0 | 0 | 0 | 0 | NA | NA | NA | NA | NA | NA | NA | NA | NA | NA |
| Oprd1 | 52 | 55 | 78 | 45 | 36 | 28 | 55 | 55 | 78 | 35 | 39 | 31 | 48.87 | 63 | 35 | 0.556 | -0.847 | 0.00510919 | 0.03185068 | 0 | 0.0743 | 0.0438 | 0.0438 | TRUE | 0.4089 |
| Oprk1 | 107 | 100 | 75 | 113 | 81 | 88 | 114 | 100 | 75 | 89 | 87 | 96 | 93.52 | 96 | 91 | 0.943 | -0.085 | 0.71220855 | 0.84337812 | 0.0116 | 0.0421 | 0.0272 | 0.0272 | TRUE | 0.3537 |
| Oprl1 | 162 | 155 | 137 | 235 | 180 | 100 | 172 | 155 | 138 | 185 | 194 | 109 | 158.72 | 155 | 163 | 1.053 | 0.075 | 0.75761638 | 0.8719099 | 0.0449 | 0.0277 | 0.0359 | 0.0359 | TRUE | 0.8381 |
| Oprm1 | 0 | 0 | 0 | 0 | 0 | 0 | 0 | 0 | 0 | 0 | 0 | 0 | 0 | 0 | 0 | NA | NA | NA | NA | NA | NA | NA | NA | NA | NA |
| Optc | 7 | 15 | 18 | 15 | 7 | 14 | 7 | 15 | 18 | 12 | 8 | 15 | 12.52 | 13 | 12 | 0.854 | -0.228 | 0.70003623 | 0.8359939 | 0.0603 | 0.27 | 0.1733 | 0.1733 | TRUE | 0.4615 |

| | | | | | | | | | | | | | | | | | | | | | | | | | |
|---|---|---|---|---|---|---|---|---|---|---|---|---|---|---|---|---|---|---|---|---|---|---|---|---|---|
| Optn | 429 | 564 | 659 | 477 | 384 | 452 | 456 | 563 | 662 | 375 | 413 | 495 | 493.93 | 560 | 428 | 0.762 | -0.392 | 0.02336911 | 0.09709633 | 0.0241 | 0.0136 | 0.0195 | 0.0195 | TRUE | 0.1608 |
| Orai1 | 514 | 585 | 621 | 715 | 521 | 536 | 546 | 584 | 624 | 562 | 561 | 587 | 577.19 | 585 | 570 | 0.974 | -0.038 | 0.71205352 | 0.84325786 | 0 | 0.0126 | 0.0059 | 0.0059 | TRUE | 0.0399 |
| Orai2 | 101 | 98 | 111 | 94 | 60 | 66 | 107 | 98 | 111 | 74 | 65 | 72 | 87.88 | 105 | 70 | 0.667 | -0.584 | 0.00537893 | 0.03301033 | 0 | 0.0444 | 0.02 | 0.02 | TRUE | 0.0429 |
| Orai3 | 505 | 803 | 825 | 643 | 459 | 533 | 536 | 802 | 828 | 505 | 494 | 583 | 624.92 | 722 | 527 | 0.73 | -0.454 | 0.01069612 | 0.05538383 | 0.0302 | 0.0122 | 0.0212 | 0.0212 | TRUE | 0.5969 |
| Oraov1 | 583 | 453 | 521 | 934 | 676 | 551 | 619 | 452 | 523 | 734 | 727 | 603 | 609.92 | 531 | 688 | 1.296 | 0.374 | 0.01232572 | 0.06138491 | 0.0156 | 0.0123 | 0.0144 | 0.0144 | TRUE | 0.2418 |
| Orc1 | 4571 | 4239 | 4344 | 5794 | 4119 | 4117 | 4855 | 4234 | 4362 | 4555 | 4432 | 4507 | 4490.64 | 4484 | 4498 | 1.003 | 0.005 | 0.95201023 | 0.97545023 | 0.0025 | 0.0076 | 0.0043 | 0.0043 | TRUE | 0.0639 |
| Orc2 | 7393 | 7009 | 6716 | 9169 | 6476 | 6319 | 7853 | 7000 | 6744 | 7208 | 6968 | 6917 | 7114.84 | 7199 | 7031 | 0.977 | -0.034 | 0.68168065 | 0.82355168 | 0.0029 | 0.0074 | 0.0048 | 0.0048 | TRUE | 0.077 |
| Orc3 | 3723 | 4101 | 4271 | 5563 | 4043 | 4130 | 3954 | 4096 | 4289 | 4373 | 4350 | 4521 | 4263.81 | 4113 | 4415 | 1.073 | 0.102 | 0.12894152 | 0.30890254 | 7.00E-04 | 0.0077 | 0.003 | 0.003 | TRUE | 0.017 |
| Orc4 | 2594 | 2775 | 2669 | 3482 | 2434 | 2594 | 2755 | 2771 | 2680 | 2737 | 2619 | 2840 | 2733.72 | 2735 | 2732 | 0.999 | -0.002 | 0.97787984 | 0.98799114 | 6.00E-04 | 0.0081 | 0.0031 | 0.0031 | TRUE | 0.0156 |
| Orc5 | 3850 | 3678 | 3739 | 4516 | 3324 | 3239 | 4089 | 3673 | 3754 | 3550 | 3576 | 3546 | 3698.19 | 3839 | 3557 | 0.927 | -0.11 | 0.13152569 | 0.31236609 | 0.0013 | 0.0078 | 0.0035 | 0.0035 | TRUE | 0.0395 |
| Orc6 | 4322 | 4252 | 4169 | 7008 | 4724 | 4384 | 4591 | 4247 | 4186 | 5509 | 5083 | 4799 | 4735.7 | 4341 | 5130 | 1.182 | 0.241 | 0.00450677 | 0.02890639 | 0.0034 | 0.0076 | 0.005 | 0.005 | TRUE | 0.0513 |
| ORF63 | 3 | 1 | 0 | 2 | 6 | 5 | 3 | 1 | 0 | 2 | 6 | 5 | 2.95 | 1 | 4 | 3.217 | 1.686 | 0.20505366 | NA | 0.3514 | 1.1245 | 0.7942 | 0.7942 | TRUE | 0.7869 |
| Orm1 | 0 | 0 | 0 | 0 | 0 | 0 | 0 | 0 | 0 | 0 | 0 | 0 | 0 | 0 | 0 | NA | NA | NA | NA | NA | NA | NA | NA | NA | NA |
| Orm2 | 0 | 0 | 0 | 0 | 0 | 0 | 0 | 0 | 0 | 0 | 0 | 0 | 0 | 0 | 0 | NA | NA | NA | NA | NA | NA | NA | NA | NA | NA |
| Orm3 | 0 | 0 | 0 | 0 | 0 | 0 | 0 | 0 | 0 | 0 | 0 | 0 | 0 | 0 | 0 | NA | NA | NA | NA | NA | NA | NA | NA | NA | NA |
| Ormdl1 | 871 | 1062 | 997 | 1368 | 892 | 979 | 925 | 1061 | 1001 | 1075 | 960 | 1072 | 1015.62 | 996 | 1036 | 1.04 | 0.057 | 0.5659042 | 0.74602482 | 0.0034 | 0.0101 | 0.0062 | 0.0062 | TRUE | 0.0484 |
| Ormdl2 | 2314 | 2345 | 2469 | 3007 | 2054 | 2087 | 2458 | 2342 | 2479 | 2364 | 2210 | 2285 | 2356.23 | 2426 | 2286 | 0.943 | -0.085 | 0.23263504 | 0.44690862 | 6.00E-04 | 0.0083 | 0.0033 | 0.0033 | TRUE | 0.0113 |
| Ormdl3 | 1178 | 1201 | 1124 | 1609 | 1095 | 1081 | 1251 | 1199 | 1129 | 1265 | 1178 | 1183 | 1200.94 | 1193 | 1209 | 1.014 | 0.02 | 0.81632186 | 0.90643681 | 0.0014 | 0.0096 | 0.0046 | 0.0046 | TRUE | 0.0268 |
| Os9 | 1065 | 1205 | 1139 | 1441 | 1186 | 1140 | 1131 | 1203 | 1144 | 1133 | 1276 | 1248 | 1189.19 | 1159 | 1219 | 1.05 | 0.071 | 0.42783422 | 0.63790314 | 0.0019 | 0.0097 | 0.0049 | 0.0049 | TRUE | 0.0475 |
| Osbp | 3307 | 3517 | 3553 | 3944 | 2928 | 3055 | 3513 | 3512 | 3568 | 3100 | 3150 | 3344 | 3364.61 | 3531 | 3198 | 0.906 | -0.143 | 0.0320532 | 0.1206259 | 5.00E-04 | 0.0079 | 0.0029 | 0.0029 | TRUE | 0.0192 |
| Osbp2 | 196 | 188 | 170 | 150 | 101 | 137 | 208 | 188 | 171 | 118 | 109 | 150 | 157.2 | 189 | 126 | 0.663 | -0.592 | 0.00190559 | 0.01481267 | 0.0108 | 0.0279 | 0.0196 | 0.0196 | TRUE | 0.2806 |
| Osbpl10 | 1113 | 1329 | 1388 | 1409 | 1128 | 1228 | 1182 | 1327 | 1394 | 1108 | 1214 | 1344 | 1261.45 | 1301 | 1222 | 0.938 | -0.092 | 0.41312036 | 0.62447137 | 0.0069 | 0.0095 | 0.0083 | 0.0083 | TRUE | 0.0919 |
| Osbpl11 | 2632 | 2230 | 2201 | 3397 | 2360 | 2184 | 2796 | 2227 | 2210 | 2670 | 2539 | 2391 | 2472.2 | 2411 | 2533 | 1.051 | 0.072 | 0.5356977 | 0.72271961 | 0.0102 | 0.0082 | 0.0093 | 0.0093 | TRUE | 0.2358 |
| Osbpl1a | 3842 | 4985 | 5082 | 5071 | 3904 | 4829 | 4081 | 4979 | 5103 | 3986 | 4200 | 5286 | 4605.88 | 4721 | 4491 | 0.951 | -0.072 | 0.59810501 | 0.77063216 | 0.0185 | 0.0076 | 0.0134 | 0.0134 | TRUE | 0.2918 |
| Osbpl2 | 1075 | 1189 | 1300 | 1338 | 956 | 1103 | 1142 | 1187 | 1305 | 1052 | 1029 | 1207 | 1153.75 | 1211 | 1096 | 0.904 | -0.146 | 0.16778957 | 0.36558665 | 0.0051 | 0.0098 | 0.0072 | 0.0072 | TRUE | 0.0933 |
| Osbpl3 | 825 | 1009 | 1026 | 1412 | 977 | 1122 | 876 | 1008 | 1030 | 1110 | 1051 | 1228 | 1050.6 | 971 | 1130 | 1.163 | 0.217 | 0.04669485 | 0.15657569 | 0.0054 | 0.01 | 0.0076 | 0.0076 | TRUE | 0.087 |
| Osbpl5 | 104 | 193 | 170 | 125 | 87 | 121 | 110 | 193 | 171 | 98 | 94 | 132 | 133.04 | 158 | 108 | 0.682 | -0.551 | 0.03378843 | 0.12497916 | 0.0505 | 0.0317 | 0.0408 | 0.0408 | TRUE | 0.6095 |
| Osbpl6 | 197 | 332 | 316 | 153 | 110 | 153 | 209 | 332 | 317 | 120 | 118 | 167 | 210.7 | 286 | 135 | 0.472 | -1.083 | 3.03E-06 | 7.23E-05 | 0.0449 | 0.0225 | 0.0334 | 0.0334 | TRUE | 0.6153 |
| Osbpl7 | 861 | 937 | 996 | 1080 | 910 | 890 | 915 | 936 | 1000 | 849 | 979 | 974 | 942.13 | 950 | 934 | 0.982 | -0.027 | 0.79181773 | 0.89301263 | 0.0034 | 0.0104 | 0.0063 | 0.0063 | TRUE | 0.0787 |
| Osbpl8 | 1487 | 1686 | 1764 | 2329 | 1622 | 1596 | 1579 | 1684 | 1771 | 1831 | 1745 | 1747 | 1726.27 | 1678 | 1774 | 1.058 | 0.081 | 0.31965672 | 0.54077291 | 0.0014 | 0.0088 | 0.0042 | 0.0042 | TRUE | 0.0318 |
| Osbpl9 | 2354 | 2704 | 2720 | 3598 | 2559 | 2580 | 2500 | 2701 | 2731 | 2828 | 2753 | 2824 | 2723.01 | 2644 | 2802 | 1.06 | 0.084 | 0.243477 | 0.46008258 | 8.00E-04 | 0.0081 | 0.0033 | 0.0033 | TRUE | 0.0273 |
| Oscar | 7 | 4 | 7 | 6 | 5 | 5 | 7 | 4 | 7 | 5 | 5 | 5 | 5.67 | 6 | 5 | 0.842 | -0.249 | 0.74855873 | 0.8665787 | 0 | 0.5878 | 0.2571 | 0.2571 | TRUE | 0.2302 |
| Oscp1 | 32 | 43 | 49 | 76 | 67 | 47 | 34 | 43 | 49 | 60 | 72 | 51 | 51.57 | 42 | 61 | 1.451 | 0.537 | 0.06743667 | 0.200795 | 0.0093 | 0.0708 | 0.0419 | 0.0419 | TRUE | 0.2073 |
| Osgep | 2108 | 2617 | 2719 | 2464 | 1763 | 1999 | 2239 | 2614 | 2730 | 1937 | 1897 | 2188 | 2267.49 | 2528 | 2007 | 0.794 | -0.333 | 0.00220529 | 0.01662318 | 0.0076 | 0.0084 | 0.0081 | 0.0081 | TRUE | 0.1206 |
| Osgepl1 | 689 | 656 | 646 | 904 | 623 | 626 | 732 | 655 | 649 | 711 | 670 | 685 | 683.64 | 679 | 689 | 1.016 | 0.023 | 0.81756531 | 0.90711028 | 0.0012 | 0.0117 | 0.0056 | 0.0056 | TRUE | 0.0552 |
| Osgin1 | 15 | 5 | 5 | 4 | 3 | 4 | 16 | 5 | 5 | 3 | 3 | 4 | 6.12 | 9 | 3 | 0.414 | -1.274 | 0.14382474 | 0.33076755 | 0 | 0.5455 | 0.351 | 0.351 | TRUE | 1.6258 |
| Osgin2 | 825 | 900 | 823 | 1221 | 884 | 853 | 876 | 899 | 826 | 960 | 951 | 934 | 907.71 | 867 | 948 | 1.094 | 0.129 | 0.12828248 | 0.30801003 | 0 | 0.0105 | 0.0041 | 0.0041 | TRUE | 0.0203 |
| Osm | 12 | 26 | 22 | 24 | 15 | 21 | 13 | 26 | 22 | 19 | 16 | 23 | 19.8 | 20 | 19 | 0.95 | -0.073 | 0.87339849 | 0.93762803 | 0 | 0.1733 | 0.1028 | 0.1028 | TRUE | 0.5499 |
| Osmr | 9 | 9 | 5 | 15 | 9 | 12 | 10 | 9 | 5 | 12 | 10 | 13 | 9.7 | 8 | 12 | 1.472 | 0.558 | 0.36435277 | 0.58180878 | 0 | 0.3467 | 0.1647 | 0.1647 | TRUE | 0.2934 |
| Osr1 | 0 | 0 | 0 | 1 | 0 | 0 | 0 | 0 | 0 | 1 | 0 | 0 | 0.13 | 0 | 0 | 1.859 | 0.894 | 0.82650734 | NA | 0 | 25.1515 | 10 | 10 | TRUE | 0.2563 |
| Osr2 | 12 | 2 | 5 | 21 | 18 | 16 | 13 | 2 | 5 | 17 | 19 | 18 | 12.19 | 7 | 18 | 2.719 | 1.443 | 0.03300891 | 0.12282999 | 0.1252 | 0.2771 | 0.2243 | 0.2243 | TRUE | 1.5908 |
| Ost4 | 2880 | 2672 | 2410 | 3881 | 2680 | 2534 | 3059 | 2669 | 2420 | 3051 | 2884 | 2774 | 2809.3 | 2716 | 2903 | 1.069 | 0.097 | 0.3663797 | 0.5838679 | 0.0072 | 0.0081 | 0.0079 | 0.0079 | TRUE | 0.148 |
| Ostc | 4285 | 4978 | 4603 | 5552 | 3967 | 3934 | 4551 | 4972 | 4622 | 4365 | 4268 | 4306 | 4514.01 | 4715 | 4313 | 0.915 | -0.128 | 0.0595795 | 0.18494151 | 9.00E-04 | 0.0076 | 0.0031 | 0.0031 | TRUE | 0.0277 |
| Ostf1 | 1574 | 2053 | 2015 | 2137 | 1490 | 1798 | 1672 | 2050 | 2023 | 1680 | 1603 | 1968 | 1832.8 | 1915 | 1750 | 0.914 | -0.13 | 0.28815862 | 0.50913413 | 0.0113 | 0.0087 | 0.0103 | 0.0103 | TRUE | 0.1489 |
| Ostm1 | 1540 | 1610 | 1603 | 1908 | 1390 | 1440 | 1636 | 1608 | 1610 | 1500 | 1496 | 1576 | 1570.84 | 1618 | 1524 | 0.942 | -0.087 | 0.22841666 | 0.44176426 | 0 | 0.009 | 0.0031 | 0.0031 | TRUE | 0.0107 |
| Ostn | 0 | 0 | 0 | 1 | 0 | 0 | 0 | 0 | 0 | 1 | 0 | 0 | 0.13 | 0 | 0 | 1.859 | 0.894 | 0.82650734 | NA | 0 | 25.1515 | 10 | 10 | TRUE | 0.2563 |
| Otc | 1 | 1 | 1 | 1 | 2 | 1 | 1 | 1 | 1 | 1 | 2 | 1 | 1.18 | 1 | 1 | 1.293 | 0.371 | 0.81961764 | NA | 0 | 2.7919 | 1.039 | 1.039 | TRUE | 0.1601 |
| Otoa | 0 | 0 | 1 | 1 | 2 | 3 | 0 | 0 | 1 | 1 | 2 | 3 | 1.2 | 0 | 2 | 5.579 | 2.48 | 0.20607493 | NA | 0 | 2.7424 | 1.532 | 1.532 | TRUE | 0.307 |
| Otof | 20 | 13 | 17 | 14 | 18 | 10 | 21 | 13 | 17 | 11 | 19 | 11 | 15.44 | 17 | 14 | 0.801 | -0.32 | 0.53427171 | 0.72175874 | 0 | 0.2203 | 0.1253 | 0.1253 | TRUE | 0.3857 |
| Otog | 38 | 23 | 32 | 29 | 22 | 17 | 40 | 23 | 32 | 23 | 24 | 19 | 26.76 | 32 | 22 | 0.684 | -0.547 | 0.16678873 | 0.36429745 | 0 | 0.13 | 0.0743 | 0.0743 | TRUE | 0.1558 |
| Otogl | 3 | 2 | 0 | 1 | 2 | 0 | 3 | 2 | 0 | 1 | 2 | 0 | 1.35 | 2 | 1 | 0.566 | -0.821 | 0.65814323 | NA | 0 | 2.4406 | 1.6831 | 1.6831 | TRUE | 0.2666 |
| Otol1 | 3 | 1 | 4 | 12 | 7 | 4 | 3 | 1 | 4 | 9 | 8 | 4 | 4.92 | 3 | 7 | 2.633 | 1.397 | 0.13189515 | 0.31290508 | 0 | 0.6759 | 0.364 | 0.364 | TRUE | 0.3298 |
| Otop1 | 46 | 67 | 47 | 72 | 50 | 69 | 49 | 67 | 47 | 57 | 54 | 76 | 58.15 | 54 | 62 | 1.137 | 0.185 | 0.5187668 | 0.71026891 | 0.019 | 0.0636 | 0.042 | 0.042 | TRUE | 0.3473 |
| Otop2 | 18 | 14 | 12 | 13 | 11 | 12 | 19 | 14 | 12 | 10 | 12 | 13 | 13.39 | 15 | 12 | 0.776 | -0.365 | 0.48224425 | 0.68089732 | 0 | 0.2529 | 0.1188 | 0.1188 | TRUE | 0.2449 |
| Otop3 | 1 | 1 | 0 | 1 | 1 | 0 | 1 | 1 | 0 | 1 | 1 | 0 | 0.65 | 1 | 1 | 0.919 | -0.122 | 0.95812821 | NA | 0 | 5.0456 | 2.3857 | 2.3857 | TRUE | 0.2521 |
| Otor | 0 | 0 | 0 | 0 | 0 | 1 | 0 | 0 | 0 | 0 | 0 | 1 | 0.18 | 0 | 0 | 1.859 | 0.894 | 0.82650734 | NA | 0 | 18.0643 | 10 | 10 | TRUE | 0.5383 |
| Otos | 5 | 4 | 2 | 1 | 3 | 3 | 5 | 4 | 2 | 1 | 3 | 3 | 3.1 | 4 | 2 | 0.627 | -0.674 | 0.54453076 | NA | 0 | 1.069 | 0.5495 | 0.5495 | TRUE | 0.5278 |
| Otp | 1 | 0 | 0 | 0 | 1 | 1 | 1 | 0 | 0 | 0 | 1 | 1 | 0.54 | 0 | 1 | 1.923 | 0.944 | 0.73523723 | NA | 0 | 6.1214 | 3.9144 | 3.9144 | TRUE | 0.4614 |
| Ott | 0 | 0 | 0 | 0 | 0 | 0 | 0 | 0 | 0 | 0 | 0 | 0 | 0 | 0 | 0 | NA | NA | NA | NA | NA | NA | NA | NA | NA | NA |
| OTTMUSG0C | 0 | 0 | 0 | 0 | 0 | 0 | 0 | 0 | 0 | 0 | 0 | 0 | 0 | 0 | 0 | NA | NA | NA | NA | NA | NA | NA | NA | NA | NA |
| Otub1 | 2795 | 2858 | 2852 | 3746 | 2787 | 2909 | 2969 | 2854 | 2864 | 2945 | 2999 | 3184 | 2969.1 | 2896 | 3043 | 1.05 | 0.071 | 0.30891304 | 0.52946929 | 8.00E-04 | 0.008 | 0.0032 | 0.0032 | TRUE | 0.02 |
| Otub2 | 304 | 352 | 310 | 352 | 238 | 255 | 323 | 352 | 311 | 277 | 256 | 279 | 299.61 | 329 | 271 | 0.824 | -0.279 | 0.02799075 | 0.10984575 | 0 | 0.0179 | 0.0083 | 0.0083 | TRUE | 0.043 |
| Otud1 | 414 | 400 | 381 | 579 | 474 | 373 | 440 | 399 | 383 | 455 | 510 | 408 | 432.54 | 407 | 458 | 1.125 | 0.169 | 0.19645421 | 0.40292288 | 0.0062 | 0.0145 | 0.0101 | 0.0101 | TRUE | 0.1112 |
| Otud3 | 773 | 754 | 685 | 909 | 694 | 630 | 821 | 753 | 688 | 715 | 747 | 690 | 735.47 | 754 | 717 | 0.951 | -0.072 | 0.49608226 | 0.69196835 | 0.0033 | 0.0114 | 0.0067 | 0.0067 | TRUE | 0.0713 |
| Otud4 | 17329 | 15976 | 16392 | 21782 | 15769 | 14719 | 18406 | 15955 | 16459 | 17123 | 16966 | 16112 | 16837.2 | 16940 | 16734 | 0.988 | -0.018 | 0.82811775 | 0.91291488 | 0.0028 | 0.0071 | 0.0047 | 0.0047 | TRUE | 0.0701 |
| Otud5 | 3627 | 3596 | 3440 | 4595 | 3555 | 3234 | 3853 | 3591 | 3454 | 3612 | 3825 | 3540 | 3645.88 | 3633 | 3659 | 1.007 | 0.01 | 0.89294754 | 0.94826074 | 0.002 | 0.0078 | 0.0041 | 0.0041 | TRUE | 0.034 |
| Otud6a | 67 | 98 | 48 | 249 | 189 | 155 | 71 | 98 | 48 | 196 | 203 | 170 | 131 | 72 | 190 | 2.619 | 1.389 | 7.92E-08 | 2.95E-06 | 0.049 | 0.0321 | 0.0386 | 0.0386 | TRUE | 0.68 |
| Otud6b | 3224 | 2828 | 2897 | 4594 | 3114 | 2941 | 3424 | 2824 | 2909 | 3611 | 3350 | 3219 | 3223.17 | 3052 | 3393 | 1.112 | 0.153 | 0.13548396 | 0.31845798 | 0.0069 | 0.0079 | 0.0073 | 0.0073 | TRUE | 0.1377 |
| Otud7a | 103 | 128 | 131 | 174 | 110 | 139 | 109 | 128 | 132 | 137 | 118 | 152 | 129.35 | 123 | 136 | 1.104 | 0.142 | 0.45565664 | 0.66053461 | 0.004 | 0.0324 | 0.0185 | 0.0185 | TRUE | 0.122 |

| | | | | | | | | | | | | | | | | | | | | | | | | |
|---|---|---|---|---|---|---|---|---|---|---|---|---|---|---|---|---|---|---|---|---|---|---|---|---|
| Otud7b | 2943 | 2620 | 2687 | 3744 | 2657 | 2360 | 3126 | 2617 | 2698 | 2943 | 2859 | 2583 | 2804.34 | 2814 | 2795 | 0.994 | -0.009 | 0.93034174 | 0.96566962 | 0.0065 | 0.0081 | 0.0072 | 0.0072 | TRUE | 0.1143 |
| Otx1 | 15 | 16 | 13 | 37 | 33 | 26 | 16 | 16 | 13 | 29 | 36 | 28 | 23 | 15 | 31 | 2.067 | 1.047 | 0.00975775 | 0.05171343 | 0 | 0.1501 | 0.0685 | 0.0685 | TRUE | 0.1017 |
| Otx2 | 1735 | 2249 | 2640 | 2534 | 1916 | 2003 | 1843 | 2246 | 2651 | 1992 | 2061 | 2193 | 2164.32 | 2247 | 2082 | 0.926 | -0.11 | 0.42040591 | 0.63082884 | 0.0161 | 0.0084 | 0.013 | 0.013 | TRUE | 0.1714 |
| Otx2os1 | 1 | 4 | 0 | 1 | 2 | 5 | 1 | 4 | 0 | 1 | 2 | 5 | 2.24 | 2 | 3 | 1.627 | 0.702 | 0.6395225 | NA | 0.7605 | 1.4745 | 1.1393 | 1.1393 | TRUE | 0.785 |
| Ovca2 | 522 | 534 | 562 | 775 | 533 | 383 | 554 | 533 | 564 | 609 | 573 | 419 | 542.34 | 550 | 534 | 0.971 | -0.042 | 0.7855808 | 0.88984346 | 0.0176 | 0.013 | 0.0154 | 0.0154 | TRUE | 0.4068 |
| Ovch2 | 0 | 0 | 0 | 0 | 0 | 0 | 0 | 0 | 0 | 0 | 0 | 0 | 0 | 0 | 0 | NA | NA | NA | NA | NA | NA | NA | NA | NA | NA |
| Ovgp1 | 215 | 190 | 139 | 147 | 126 | 127 | 228 | 190 | 140 | 116 | 136 | 139 | 157.97 | 186 | 130 | 0.698 | -0.518 | 0.01916385 | 0.08408354 | 0.0296 | 0.0278 | 0.0287 | 0.0287 | TRUE | 0.1928 |
| Ovol1 | 35 | 54 | 39 | 44 | 47 | 40 | 37 | 54 | 39 | 35 | 51 | 44 | 43.2 | 43 | 43 | 0.983 | -0.025 | 0.93792761 | 0.96893419 | 0.0182 | 0.0832 | 0.0514 | 0.0514 | TRUE | 0.3501 |
| Ovol2 | 168 | 177 | 192 | 185 | 164 | 168 | 178 | 177 | 193 | 145 | 176 | 184 | 175.63 | 183 | 168 | 0.919 | -0.122 | 0.46526851 | 0.66796058 | 0 | 0.0257 | 0.0144 | 0.0144 | TRUE | 0.1693 |
| Oxa1l | 4220 | 5140 | 5319 | 5452 | 4109 | 4581 | 4482 | 5133 | 5341 | 4286 | 4421 | 5015 | 4779.7 | 4985 | 4574 | 0.917 | -0.125 | 0.22796972 | 0.44159023 | 0.0073 | 0.0076 | 0.0075 | 0.0075 | TRUE | 0.0949 |
| Oxct1 | 2755 | 3639 | 3870 | 4754 | 3431 | 3500 | 2926 | 3634 | 3886 | 3737 | 3692 | 3831 | 3617.76 | 3482 | 3753 | 1.078 | 0.108 | 0.34976426 | 0.56865085 | 0.0091 | 0.0078 | 0.0093 | 0.0093 | TRUE | 0.2368 |
| Oxct2a | 8 | 4 | 1 | 11 | 4 | 7 | 8 | 4 | 1 | 9 | 4 | 8 | 5.69 | 4 | 7 | 1.547 | 0.629 | 0.48939006 | 0.68561557 | 0.1867 | 0.5864 | 0.4119 | 0.4119 | TRUE | 0.4719 |
| Oxct2b | 11 | 8 | 4 | 19 | 17 | 11 | 12 | 8 | 4 | 15 | 18 | 12 | 11.49 | 8 | 15 | 1.918 | 0.94 | 0.12345277 | 0.29946124 | 0 | 0.2935 | 0.1709 | 0.1709 | TRUE | 0.3625 |
| Oxgr1 | 1 | 2 | 0 | 3 | 1 | 2 | 1 | 2 | 0 | 2 | 1 | 2 | 1.45 | 1 | 2 | 1.852 | 0.889 | 0.57938865 | NA | 0 | 2.2833 | 1.0953 | 1.0953 | TRUE | 0.3328 |
| Oxld1 | 81 | 163 | 210 | 148 | 130 | 148 | 86 | 163 | 211 | 116 | 140 | 162 | 146.32 | 153 | 139 | 0.907 | -0.14 | 0.6525509 | 0.8050714 | 0.1017 | 0.0294 | 0.0629 | 0.0629 | TRUE | 0.3255 |
| Oxnad1 | 668 | 677 | 667 | 1214 | 813 | 707 | 710 | 676 | 670 | 954 | 875 | 774 | 776.4 | 685 | 868 | 1.268 | 0.343 | 0.00204074 | 0.01563975 | 0.005 | 0.0111 | 0.0076 | 0.0076 | TRUE | 0.1054 |
| Oxr1 | 4335 | 4515 | 4797 | 5513 | 4139 | 4452 | 4605 | 4509 | 4817 | 4334 | 4453 | 4873 | 4598.5 | 4644 | 4553 | 0.98 | -0.029 | 0.71319957 | 0.84398136 | 0.0023 | 0.0076 | 0.0041 | 0.0041 | TRUE | 0.0458 |
| Oxsm | 514 | 604 | 662 | 993 | 771 | 738 | 546 | 603 | 665 | 781 | 830 | 808 | 705.32 | 605 | 806 | 1.332 | 0.414 | 0.00013832 | 0.0017896 | 0.0031 | 0.0116 | 0.007 | 0.007 | TRUE | 0.0888 |
| Oxsr1 | 7514 | 8209 | 8287 | 8913 | 6932 | 7224 | 7981 | 8198 | 8321 | 7007 | 7458 | 7908 | 7812.26 | 8167 | 7458 | 0.913 | -0.131 | 0.07293387 | 0.21153236 | 0.0019 | 0.0073 | 0.0037 | 0.0037 | TRUE | 0.0344 |
| Oxt | 11 | 8 | 3 | 5 | 3 | 3 | 12 | 8 | 3 | 4 | 3 | 3 | 5.52 | 8 | 3 | 0.464 | -1.106 | 0.20548576 | 0.41412906 | 0 | 0.6036 | 0.3432 | 0.3432 | TRUE | 0.1578 |
| Oxtr | 15 | 11 | 3 | 22 | 3 | 9 | 16 | 11 | 3 | 17 | 3 | 10 | 10.05 | 10 | 10 | 1.031 | 0.045 | 0.95674648 | 0.97766996 | 0.4491 | 0.3347 | 0.3901 | 0.3901 | TRUE | 0.2208 |
| P2rx1 | 57 | 20 | 21 | 43 | 30 | 25 | 61 | 20 | 21 | 34 | 32 | 27 | 32.51 | 34 | 31 | 0.924 | -0.114 | 0.81612401 | 0.90637793 | 0.1806 | 0.1082 | 0.1436 | 0.1436 | TRUE | 3.2514 |
| P2rx2 | 42 | 70 | 78 | 92 | 56 | 73 | 45 | 70 | 78 | 72 | 60 | 80 | 67.55 | 64 | 71 | 1.101 | 0.138 | 0.6312065 | 0.79265149 | 0.0327 | 0.0557 | 0.0449 | 0.0449 | TRUE | 0.613 |
| P2rx3 | 18 | 32 | 34 | 60 | 36 | 26 | 19 | 32 | 34 | 47 | 39 | 28 | 33.26 | 28 | 38 | 1.348 | 0.431 | 0.26371986 | 0.48297603 | 0.0427 | 0.1059 | 0.0763 | 0.0763 | TRUE | 0.2848 |
| P2rx4 | 259 | 373 | 378 | 340 | 257 | 287 | 275 | 373 | 380 | 267 | 277 | 314 | 314.19 | 343 | 286 | 0.834 | -0.262 | 0.11469901 | 0.28592269 | 0.0161 | 0.0174 | 0.0167 | 0.0167 | TRUE | 0.3336 |
| P2rx5 | 307 | 349 | 334 | 344 | 271 | 299 | 326 | 349 | 335 | 270 | 292 | 327 | 316.55 | 337 | 296 | 0.878 | -0.188 | 0.15172887 | 0.3428955 | 0.0021 | 0.0173 | 0.0092 | 0.0092 | TRUE | 0.0923 |
| P2rx6 | 7 | 10 | 11 | 8 | 8 | 9 | 7 | 10 | 11 | 6 | 9 | 10 | 8.87 | 9 | 8 | 0.859 | -0.22 | 0.72705569 | 0.85295465 | 0 | 0.3783 | 0.1716 | 0.1716 | TRUE | 0.1723 |
| P2rx7 | 469 | 666 | 641 | 533 | 364 | 556 | 498 | 665 | 644 | 419 | 392 | 609 | 537.7 | 602 | 473 | 0.785 | -0.35 | 0.07244157 | 0.2105692 | 0.0397 | 0.013 | 0.0255 | 0.0255 | TRUE | 0.7216 |
| P2ry1 | 56 | 46 | 42 | 110 | 89 | 61 | 59 | 46 | 42 | 86 | 96 | 67 | 66.1 | 49 | 83 | 1.692 | 0.758 | 0.00547931 | 0.03345709 | 0.0161 | 0.0567 | 0.0375 | 0.0375 | TRUE | 0.2628 |
| P2ry10 | 0 | 0 | 0 | 0 | 0 | 0 | 0 | 0 | 0 | 0 | 0 | 0 | 0 | 0 | 0 | NA | NA | NA | NA | NA | NA | NA | NA | NA | NA |
| P2ry12 | 0 | 0 | 0 | 0 | 0 | 0 | 0 | 0 | 0 | 0 | 0 | 0 | 0 | 0 | 0 | NA | NA | NA | NA | NA | NA | NA | NA | NA | NA |
| P2ry13 | 0 | 0 | 0 | 0 | 0 | 0 | 0 | 0 | 0 | 0 | 0 | 0 | 0 | 0 | 0 | NA | NA | NA | NA | NA | NA | NA | NA | NA | NA |
| P2ry14 | 0 | 0 | 0 | 0 | 0 | 1 | 0 | 0 | 0 | 0 | 0 | 1 | 0.18 | 0 | 0 | 1.859 | 0.894 | 0.82650734 | NA | 0 | 18.0643 | 10 | 10 | TRUE | 0.5383 |
| P2ry2 | 200 | 288 | 269 | 233 | 153 | 162 | 212 | 288 | 270 | 183 | 165 | 177 | 215.88 | 257 | 175 | 0.682 | -0.552 | 0.00110387 | 0.00960301 | 0.01 | 0.0222 | 0.0158 | 0.0158 | TRUE | 0.2505 |
| P2ry4 | 4 | 3 | 4 | 6 | 3 | 5 | 4 | 3 | 4 | 5 | 3 | 5 | 4.11 | 4 | 4 | 1.195 | 0.257 | 0.77494701 | NA | 0 | 0.8079 | 0.3373 | 0.3373 | TRUE | 0.0969 |
| P2ry6 | 24 | 19 | 20 | 50 | 26 | 30 | 25 | 19 | 20 | 39 | 28 | 33 | 27.45 | 21 | 33 | 1.562 | 0.644 | 0.08757917 | 0.23964239 | 0 | 0.1269 | 0.064 | 0.064 | TRUE | 0.1972 |
| P4ha1 | 3232 | 5139 | 5384 | 4603 | 3578 | 4482 | 3433 | 5132 | 5406 | 3619 | 3850 | 4906 | 4390.98 | 4657 | 4125 | 0.886 | -0.175 | 0.33596848 | 0.55602254 | 0.0377 | 0.0077 | 0.0237 | 0.0237 | TRUE | 0.6438 |
| P4ha2 | 1413 | 2222 | 2204 | 3165 | 2544 | 2450 | 1501 | 2219 | 2213 | 2488 | 2737 | 2682 | 2306.7 | 1978 | 2636 | 1.332 | 0.414 | 0.00694911 | 0.03986953 | 0.0224 | 0.0083 | 0.0165 | 0.0165 | TRUE | 0.5369 |
| P4ha3 | 3 | 10 | 4 | 14 | 7 | 4 | 3 | 10 | 4 | 11 | 8 | 4 | 6.68 | 6 | 8 | 1.346 | 0.429 | 0.59685586 | 0.76965176 | 0.1344 | 0.4998 | 0.3219 | 0.3219 | TRUE | 0.4329 |
| P4hb | 38911 | 38008 | 36866 | 43862 | 32683 | 32611 | 41330 | 37959 | 37017 | 34481 | 35165 | 35698 | 36941.71 | 38769 | 35115 | 0.906 | -0.143 | 0.04040109 | 0.14197994 | 0.0018 | 0.007 | 0.0035 | 0.0035 | TRUE | 0.0409 |
| P4htm | 162 | 190 | 170 | 248 | 210 | 178 | 172 | 190 | 171 | 195 | 226 | 195 | 191.38 | 178 | 205 | 1.154 | 0.207 | 0.18025576 | 0.3815587 | 0 | 0.0241 | 0.012 | 0.012 | TRUE | 0.0822 |
| Pa2g4 | 20723 | 20065 | 19884 | 29310 | 20838 | 18538 | 22011 | 20039 | 19966 | 23041 | 22420 | 20293 | 21295.1 | 20672 | 21918 | 1.06 | 0.085 | 0.30877002 | 0.529319 | 0.0037 | 0.0071 | 0.0049 | 0.0049 | TRUE | 0.0515 |
| Pabpc1 | 131670 | 134294 | 133869 | 156489 | 117323 | 114159 | 139857 | 134121 | 134419 | 123019 | 126232 | 124965 | 130435.46 | 136132 | 124739 | 0.916 | -0.126 | 0.01844862 | 0.08205628 | 4.00E-04 | 0.0069 | 0.0021 | 0.0021 | TRUE | 0.007 |
| Pabpc1l | 45 | 55 | 58 | 44 | 25 | 32 | 48 | 55 | 58 | 35 | 27 | 35 | 42.91 | 54 | 32 | 0.601 | -0.734 | 0.01444526 | 0.0688518 | 0 | 0.0837 | 0.0403 | 0.0403 | TRUE | 0.1303 |
| Pabpc2 | 0 | 0 | 0 | 0 | 0 | 0 | 0 | 0 | 0 | 0 | 0 | 0 | 0 | 0 | 0 | NA | NA | NA | NA | NA | NA | NA | NA | NA | NA |
| Pabpc4 | 14832 | 18120 | 18578 | 23509 | 16952 | 15631 | 15754 | 18097 | 18654 | 18481 | 18239 | 17111 | 17722.65 | 17502 | 17944 | 1.025 | 0.036 | 0.68589365 | 0.82648902 | 0.0038 | 0.0071 | 0.0057 | 0.0057 | TRUE | 0.0933 |
| Pabpc4l | 437 | 452 | 408 | 730 | 556 | 512 | 464 | 451 | 410 | 574 | 598 | 560 | 509.64 | 442 | 577 | 1.308 | 0.387 | 0.00028051 | 0.00322228 | 0 | 0.0134 | 0.0062 | 0.0062 | TRUE | 0.0469 |
| Pabpc5 | 4 | 3 | 5 | 6 | 3 | 2 | 4 | 3 | 5 | 5 | 3 | 2 | 3.73 | 4 | 3 | 0.842 | -0.249 | 0.79423372 | NA | 0 | 0.8894 | 0.3886 | 0.3886 | TRUE | 0.2301 |
| Pabpc6 | 24 | 20 | 14 | 77 | 57 | 14 | 25 | 20 | 14 | 61 | 61 | 15 | 32.78 | 20 | 46 | 2.316 | 1.212 | 0.02642384 | 0.10564178 | 0.2566 | 0.1074 | 0.1783 | 0.1783 | TRUE | 2.5447 |
| Pabpn1 | 11723 | 12220 | 11862 | 15934 | 12151 | 11509 | 12452 | 12204 | 11911 | 12526 | 13074 | 12598 | 12460.84 | 12189 | 12733 | 1.045 | 0.063 | 0.27669132 | 0.49765942 | 4.00E-04 | 0.0072 | 0.0023 | 0.0023 | TRUE | 0.0067 |
| Pabpn1l | 0 | 1 | 3 | 0 | 0 | 0 | 0 | 1 | 3 | 0 | 0 | 0 | 0.67 | 1 | 0 | 0.134 | -2.902 | 0.33534003 | NA | 6.5513 | 4.935 | 5.1586 | 5.1586 | TRUE | 0.154 |
| Pacrg | 11 | 5 | 16 | 27 | 18 | 10 | 12 | 5 | 16 | 21 | 19 | 11 | 14.05 | 11 | 17 | 1.585 | 0.664 | 0.26671524 | 0.4864284 | 0.1112 | 0.2414 | 0.1827 | 0.1827 | TRUE | 0.5112 |
| Pacrgl | 409 | 331 | 293 | 657 | 467 | 405 | 434 | 331 | 294 | 516 | 502 | 443 | 420.25 | 353 | 487 | 1.382 | 0.467 | 0.00516018 | 0.03198394 | 0.0207 | 0.0147 | 0.0176 | 0.0176 | TRUE | 0.4609 |
| Pacs1 | 2860 | 3476 | 3851 | 3550 | 2801 | 3171 | 3038 | 3472 | 3867 | 2791 | 3014 | 3471 | 3275.29 | 3459 | 3092 | 0.894 | -0.162 | 0.19022006 | 0.39517265 | 0.013 | 0.0079 | 0.0108 | 0.0108 | TRUE | 0.1397 |
| Pacs2 | 2122 | 2070 | 2246 | 2613 | 2016 | 2054 | 2254 | 2067 | 2255 | 2054 | 2169 | 2248 | 2174.69 | 2192 | 2157 | 0.984 | -0.024 | 0.76823151 | 0.87915741 | 0.0018 | 0.0084 | 0.0042 | 0.0042 | TRUE | 0.0301 |
| Pacsin1 | 1898 | 2161 | 2209 | 2025 | 1558 | 1644 | 2016 | 2158 | 2218 | 1592 | 1676 | 1800 | 1910.02 | 2131 | 1689 | 0.792 | -0.336 | 0.00010449 | 0.00142691 | 0.0023 | 0.0086 | 0.0049 | 0.0049 | TRUE | 0.0393 |
| Pacsin2 | 5075 | 4958 | 4883 | 6809 | 5129 | 4770 | 5391 | 4952 | 4903 | 5353 | 5518 | 5222 | 5222.98 | 5082 | 5364 | 1.056 | 0.078 | 0.27853954 | 0.49967307 | 0.0015 | 0.0075 | 0.0036 | 0.0036 | TRUE | 0.0344 |
| Pacsin3 | 635 | 624 | 708 | 833 | 623 | 527 | 674 | 623 | 711 | 655 | 670 | 577 | 651.77 | 669 | 634 | 0.948 | -0.078 | 0.48377951 | 0.68148223 | 0.0037 | 0.012 | 0.0073 | 0.0073 | TRUE | 0.0711 |
| Padi1 | 9 | 19 | 13 | 20 | 19 | 15 | 10 | 19 | 13 | 16 | 20 | 16 | 15.7 | 14 | 17 | 1.257 | 0.33 | 0.50865395 | 0.70252018 | 0 | 0.2168 | 0.1149 | 0.1149 | TRUE | 0.4553 |
| Padi2 | 746 | 1025 | 1011 | 639 | 521 | 701 | 792 | 1024 | 1015 | 502 | 561 | 767 | 776.91 | 944 | 610 | 0.646 | -0.631 | 0.00044795 | 0.0047737 | 0.0337 | 0.0111 | 0.022 | 0.022 | TRUE | 0.5947 |
| Padi3 | 572 | 765 | 753 | 482 | 417 | 517 | 608 | 764 | 756 | 379 | 449 | 566 | 586.86 | 709 | 465 | 0.654 | -0.614 | 0.00035535 | 0.00393583 | 0.0257 | 0.0125 | 0.0195 | 0.0195 | TRUE | 0.4219 |
| Padi4 | 214 | 206 | 155 | 212 | 180 | 170 | 227 | 206 | 156 | 167 | 194 | 186 | 189.18 | 196 | 182 | 0.927 | -0.109 | 0.56168215 | 0.74274244 | 0.0163 | 0.0243 | 0.0202 | 0.0202 | TRUE | 0.3565 |
| Padi6 | 47 | 35 | 33 | 38 | 22 | 16 | 50 | 35 | 33 | 30 | 24 | 18 | 31.51 | 39 | 24 | 0.609 | -0.715 | 0.06117931 | 0.1885346 | 0.0279 | 0.1114 | 0.0717 | 0.0717 | TRUE | 0.3682 |
| Paf1 | 3305 | 2843 | 2681 | 4059 | 3082 | 2539 | 3510 | 2839 | 2692 | 3191 | 3316 | 2779 | 3054.68 | 3014 | 3095 | 1.027 | 0.039 | 0.75923998 | 0.87304848 | 0.0139 | 0.008 | 0.0112 | 0.0112 | TRUE | 0.252 |
| Pafah1b1 | 12066 | 14191 | 14812 | 19260 | 13533 | 13776 | 12816 | 14173 | 14873 | 15141 | 14561 | 15080 | 14440.52 | 13954 | 14927 | 1.07 | 0.097 | 0.22008839 | 0.43222937 | 0.0029 | 0.0071 | 0.0045 | 0.0045 | TRUE | 0.0622 |
| Pafah1b2 | 2488 | 2814 | 2736 | 3523 | 2515 | 2558 | 2643 | 2810 | 2747 | 2770 | 2706 | 2800 | 2745.99 | 2733 | 2759 | 1.009 | 0.013 | 0.84363467 | 0.92132635 | 2.00E-04 | 0.0081 | 0.0028 | 0.0028 | TRUE | 0.0102 |
| Pafah1b3 | 994 | 1430 | 1420 | 1451 | 1161 | 1365 | 1056 | 1428 | 1426 | 1141 | 1249 | 1494 | 1298.97 | 1303 | 1295 | 0.993 | -0.011 | 0.94538564 | 0.97275189 | 0.0232 | 0.0094 | 0.0165 | 0.0165 | TRUE | 0.3305 |
| Pafah2 | 846 | 871 | 769 | 1160 | 821 | 743 | 899 | 870 | 772 | 912 | 883 | 813 | 858.2 | 847 | 869 | 1.028 | 0.039 | 0.70502846 | 0.83859041 | 0.0037 | 0.0107 | 0.0066 | 0.0066 | TRUE | 0.071 |
| Pag1 | 470 | 448 | 356 | 556 | 409 | 389 | 499 | 447 | 357 | 437 | 440 | 426 | 434.51 | 434 | 434 | 1 | 0 | 0.99728042 | 0.9983565 | 0.0118 | 0.0145 | 0.0131 | 0.0131 | TRUE | 0.2802 |

| | | | | | | | | | | | | | | | | | | | | | | | | | |
|---|---|---|---|---|---|---|---|---|---|---|---|---|---|---|---|---|---|---|---|---|---|---|---|---|---|
| Pagr1a | 1345 | 1477 | 1373 | 1939 | 1346 | 1356 | 1429 | 1475 | 1379 | 1524 | 1448 | 1484 | 1456.54 | 1428 | 1485 | 1.041 | 0.058 | 0.44420655 | 0.65087811 | 0 | 0.0092 | 0.0035 | 0.0035 | TRUE | 0.0108 |
| Pah | 0 | 0 | 1 | 1 | 0 | 0 | 0 | 0 | 1 | 1 | 0 | 0 | 0.3 | 0 | 0 | 0.954 | -0.067 | 0.98640617 | NA | 0 | 11.0483 | 9.2783 | 9.2783 | TRUE | 0.4098 |
| Paics | 23906 | 28507 | 28961 | 31057 | 23020 | 25771 | 25392 | 28470 | 29080 | 24415 | 24768 | 28210 | 26722.6 | 27647 | 25798 | 0.933 | -0.1 | 0.28547621 | 0.50675641 | 0.0058 | 0.007 | 0.0063 | 0.0063 | TRUE | 0.0819 |
| Paip1 | 1107 | 1718 | 1741 | 2295 | 1634 | 1766 | 1176 | 1716 | 1748 | 1804 | 1758 | 1933 | 1689.19 | 1547 | 1832 | 1.184 | 0.244 | 0.11415646 | 0.28497856 | 0.0219 | 0.0089 | 0.0166 | 0.0166 | TRUE | 0.5287 |
| Paip2 | 8030 | 8639 | 8437 | 10160 | 7060 | 7710 | 8529 | 8628 | 8472 | 7987 | 7596 | 8440 | 8275.28 | 8543 | 8008 | 0.937 | -0.093 | 0.16993514 | 0.3685667 | 0.0013 | 0.0073 | 0.0032 | 0.0032 | TRUE | 0.027 |
| Paip2b | 6886 | 7028 | 6710 | 9204 | 6484 | 6498 | 7314 | 7019 | 6738 | 7235 | 6976 | 7113 | 7065.93 | 7024 | 7108 | 1.012 | 0.017 | 0.78713113 | 0.89032073 | 7.00E-04 | 0.0074 | 0.0029 | 0.0029 | TRUE | 0.016 |
| Pak1 | 1818 | 1787 | 1767 | 1829 | 1355 | 1464 | 1931 | 1785 | 1774 | 1438 | 1458 | 1603 | 1664.71 | 1830 | 1500 | 0.819 | -0.288 | 0.00087412 | 0.00801547 | 0.002 | 0.0089 | 0.0048 | 0.0048 | TRUE | 0.0431 |
| Pak1ip1 | 4719 | 4877 | 4736 | 6710 | 4784 | 4486 | 5012 | 4871 | 4755 | 5275 | 5147 | 4911 | 4995.23 | 4879 | 5111 | 1.048 | 0.067 | 0.30927561 | 0.52974532 | 7.00E-04 | 0.0076 | 0.0029 | 0.0029 | TRUE | 0.0143 |
| Pak2 | 6482 | 6757 | 6833 | 7766 | 5816 | 5851 | 6885 | 6748 | 6861 | 6105 | 6258 | 6405 | 6543.65 | 6831 | 6256 | 0.916 | -0.127 | 0.02790664 | 0.10967118 | 2.00E-04 | 0.0074 | 0.0023 | 0.0023 | TRUE | 0.0055 |
| Pak3 | 206 | 219 | 207 | 289 | 167 | 172 | 219 | 219 | 208 | 227 | 180 | 188 | 206.75 | 215 | 198 | 0.926 | -0.111 | 0.48187838 | 0.68077623 | 0.0043 | 0.0228 | 0.0131 | 0.0131 | TRUE | 0.1887 |
| Pak4 | 5449 | 6377 | 6540 | 6327 | 4919 | 5237 | 5788 | 6369 | 6567 | 4974 | 5293 | 5733 | 5787.08 | 6241 | 5333 | 0.854 | -0.227 | 0.01144756 | 0.05818803 | 0.0044 | 0.0075 | 0.0056 | 0.0056 | TRUE | 0.0524 |
| Pak6 | 205 | 207 | 228 | 182 | 183 | 150 | 218 | 207 | 229 | 143 | 197 | 164 | 192.93 | 218 | 168 | 0.769 | -0.38 | 0.02770861 | 0.10920046 | 0.0086 | 0.024 | 0.0162 | 0.0162 | TRUE | 0.2385 |
| Pak7 | 37 | 32 | 22 | 32 | 25 | 27 | 39 | 32 | 22 | 25 | 27 | 30 | 29.16 | 31 | 27 | 0.873 | -0.196 | 0.60351287 | 0.77438672 | 0 | 0.1199 | 0.0679 | 0.0679 | TRUE | 0.2486 |
| Palb2 | 1863 | 1975 | 1958 | 2534 | 1878 | 1798 | 1979 | 1972 | 1966 | 1992 | 2021 | 1968 | 1983.03 | 1972 | 1994 | 1.011 | 0.015 | 0.81360979 | 0.90463543 | 0 | 0.0086 | 0.0026 | 0.0026 | TRUE | 0.0016 |
| Pald1 | 7907 | 10525 | 10527 | 7872 | 6195 | 6940 | 8399 | 10511 | 10570 | 6188 | 6665 | 7597 | 8321.83 | 9827 | 6817 | 0.694 | -0.528 | 1.75E-05 | 0.00031853 | 0.0133 | 0.0073 | 0.0108 | 0.0108 | TRUE | 0.1974 |
| Palld | 1755 | 1581 | 1679 | 2875 | 2048 | 2019 | 1864 | 1579 | 1686 | 2260 | 2204 | 2210 | 1967.12 | 1710 | 2225 | 1.302 | 0.38 | 1.79E-05 | 0.00032376 | 0.0029 | 0.0086 | 0.0051 | 0.0051 | TRUE | 0.0751 |
| Palm | 2004 | 2798 | 2961 | 1733 | 1500 | 1838 | 2129 | 2794 | 2973 | 1362 | 1614 | 2012 | 2147.4 | 2632 | 1663 | 0.631 | -0.663 | 0.00010989 | 0.0014853 | 0.033 | 0.0084 | 0.0207 | 0.0207 | TRUE | 0.4069 |
| Palm2 | 207 | 137 | 148 | 150 | 122 | 144 | 220 | 137 | 149 | 118 | 131 | 158 | 152.02 | 169 | 136 | 0.804 | -0.315 | 0.18078973 | 0.38227853 | 0.038 | 0.0286 | 0.0333 | 0.0333 | TRUE | 0.752 |
| Palm3 | 3603 | 3564 | 3602 | 3462 | 2649 | 3010 | 3827 | 3559 | 3617 | 2722 | 2850 | 3295 | 3311.64 | 3668 | 2956 | 0.806 | -0.312 | 0.00123444 | 0.01046751 | 0.0055 | 0.0079 | 0.0064 | 0.0064 | TRUE | 0.1221 |
| Palmd | 4 | 3 | 0 | 5 | 3 | 1 | 4 | 3 | 0 | 4 | 3 | 1 | 2.58 | 2 | 3 | 1.16 | 0.214 | 0.87137045 | NA | 0 | 1.2823 | 0.8707 | 0.8707 | TRUE | 0.7432 |
| Pam | 687 | 743 | 787 | 984 | 678 | 727 | 730 | 742 | 790 | 774 | 729 | 796 | 760.14 | 754 | 766 | 1.016 | 0.023 | 0.80184073 | 0.89762453 | 0 | 0.0112 | 0.0049 | 0.0049 | TRUE | 0.021 |
| Pam16 | 812 | 930 | 904 | 1023 | 695 | 722 | 862 | 929 | 908 | 804 | 748 | 790 | 840.22 | 900 | 781 | 0.868 | -0.204 | 0.0219037 | 0.09291694 | 0 | 0.0108 | 0.0045 | 0.0045 | TRUE | 0.016 |
| Pamr1 | 1 | 4 | 6 | 2 | 4 | 9 | 1 | 4 | 6 | 2 | 4 | 10 | 4.47 | 4 | 5 | 1.386 | 0.471 | 0.65674036 | NA | 0.4146 | 0.7442 | 0.5777 | 0.5777 | TRUE | 0.3953 |
| Pan2 | 3535 | 3873 | 3750 | 4993 | 3841 | 3779 | 3755 | 3868 | 3765 | 3925 | 4133 | 4137 | 3930.45 | 3796 | 4065 | 1.07 | 0.098 | 0.1215684 | 0.29652827 | 4.00E-04 | 0.0077 | 0.0026 | 0.0026 | TRUE | 0.0112 |
| Pan3 | 2024 | 1742 | 1724 | 1941 | 1582 | 1503 | 2150 | 1740 | 1731 | 1526 | 1702 | 1645 | 1748.99 | 1874 | 1624 | 0.867 | -0.206 | 0.0709801 | 0.20748529 | 0.0088 | 0.0088 | 0.0088 | 0.0088 | TRUE | 0.2006 |
| Pank1 | 746 | 643 | 619 | 878 | 619 | 594 | 792 | 642 | 622 | 690 | 666 | 650 | 677.09 | 685 | 669 | 0.977 | -0.034 | 0.78369668 | 0.88834728 | 0.0078 | 0.0118 | 0.0096 | 0.0096 | TRUE | 0.2191 |
| Pank2 | 4046 | 4160 | 4130 | 4434 | 3317 | 3425 | 4298 | 4155 | 4147 | 3486 | 3569 | 3749 | 3900.48 | 4200 | 3601 | 0.857 | -0.222 | 0.0008137 | 0.00758041 | 5.00E-04 | 0.0077 | 0.0029 | 0.0029 | TRUE | 0.0156 |
| Pank3 | 7037 | 6926 | 6847 | 9293 | 6675 | 6276 | 7475 | 6917 | 6875 | 7305 | 7182 | 6870 | 7104.02 | 7089 | 7119 | 1.004 | 0.006 | 0.927521 | 0.96439193 | 0.0014 | 0.0074 | 0.0034 | 0.0034 | TRUE | 0.0276 |
| Pank4 | 1195 | 1517 | 1455 | 1526 | 1257 | 1403 | 1269 | 1515 | 1461 | 1200 | 1352 | 1536 | 1388.87 | 1415 | 1363 | 0.962 | -0.056 | 0.6518129 | 0.80491647 | 0.0112 | 0.0093 | 0.0103 | 0.0103 | TRUE | 0.1484 |
| Panx1 | 491 | 540 | 562 | 674 | 476 | 496 | 522 | 539 | 564 | 530 | 512 | 543 | 535.01 | 542 | 528 | 0.975 | -0.036 | 0.71469939 | 0.8452151 | 0 | 0.0131 | 0.0053 | 0.0053 | TRUE | 0.0156 |
| Panx2 | 37 | 51 | 26 | 37 | 30 | 33 | 39 | 51 | 26 | 29 | 32 | 36 | 35.64 | 39 | 32 | 0.835 | -0.26 | 0.47598401 | 0.67596072 | 0.0345 | 0.0993 | 0.0678 | 0.0678 | TRUE | 0.1879 |
| Panx3 | 1 | 0 | 1 | 1 | 1 | 0 | 1 | 0 | 1 | 1 | 1 | 0 | 0.65 | 1 | 1 | 0.917 | -0.125 | 0.95730863 | NA | 0 | 5.0387 | 2.3854 | 2.3854 | TRUE | 0.2549 |
| Paox | 964 | 1199 | 1319 | 1114 | 893 | 949 | 1024 | 1197 | 1324 | 876 | 961 | 1039 | 1070.2 | 1182 | 959 | 0.81 | -0.304 | 0.01624469 | 0.07514487 | 0.0103 | 0.01 | 0.0106 | 0.0106 | TRUE | 0.1633 |
| Papd4 | 1593 | 1699 | 1580 | 2154 | 1504 | 1544 | 1692 | 1697 | 1586 | 1693 | 1618 | 1690 | 1662.83 | 1658 | 1667 | 1.006 | 0.008 | 0.91528962 | 0.95830774 | 4.00E-04 | 0.0089 | 0.0035 | 0.0035 | TRUE | 0.0174 |
| Papd5 | 7965 | 6929 | 6771 | 9038 | 6592 | 6192 | 8460 | 6920 | 6799 | 7105 | 7093 | 6778 | 7192.46 | 7393 | 6992 | 0.946 | -0.08 | 0.43747786 | 0.64548863 | 0.0077 | 0.0074 | 0.0076 | 0.0076 | TRUE | 0.1945 |
| Papd7 | 2941 | 3176 | 3302 | 3439 | 2641 | 2768 | 3124 | 3172 | 3316 | 2703 | 2842 | 3030 | 3031.06 | 3204 | 2858 | 0.892 | -0.165 | 0.03268416 | 0.12195283 | 0.0016 | 0.008 | 0.004 | 0.004 | TRUE | 0.0334 |
| Papln | 115 | 100 | 101 | 158 | 130 | 128 | 122 | 100 | 101 | 124 | 140 | 140 | 121.27 | 108 | 135 | 1.248 | 0.319 | 0.09135726 | 0.24661325 | 0 | 0.0341 | 0.0174 | 0.0174 | TRUE | 0.1314 |
| Papola | 19414 | 16491 | 15597 | 22396 | 15819 | 15053 | 20621 | 16470 | 15661 | 17606 | 17020 | 16478 | 17309.33 | 17584 | 17035 | 0.969 | -0.046 | 0.69170491 | 0.83095304 | 0.0101 | 0.0071 | 0.0095 | 0.0095 | TRUE | 0.2792 |
| Papolb | 37 | 21 | 17 | 71 | 30 | 33 | 39 | 21 | 17 | 56 | 32 | 36 | 33.59 | 26 | 41 | 1.621 | 0.697 | 0.10992594 | 0.27829083 | 0.1053 | 0.105 | 0.1053 | 0.1053 | TRUE | 1.2613 |
| Papolg | 2448 | 2686 | 2669 | 3038 | 2210 | 2338 | 2600 | 2683 | 2680 | 2388 | 2378 | 2559 | 2548.01 | 2654 | 2442 | 0.92 | -0.121 | 0.08598511 | 0.23655278 | 6.00E-04 | 0.0082 | 0.0032 | 0.0032 | TRUE | 0.0212 |
| Pappa | 0 | 0 | 1 | 5 | 2 | 0 | 0 | 0 | 1 | 4 | 2 | 0 | 1.18 | 0 | 2 | 5.636 | 2.495 | 0.26121578 | NA | 0 | 2.7962 | 2.315 | 2.315 | TRUE | 0.1749 |
| Pappa2 | 2 | 10 | 7 | 26 | 11 | 19 | 2 | 10 | 7 | 20 | 12 | 21 | 12.04 | 6 | 18 | 2.765 | 1.467 | 0.0258175 | 0.1039555 | 0.0842 | 0.2806 | 0.2051 | 0.2051 | TRUE | 0.7884 |
| Papss1 | 1702 | 2162 | 2278 | 2283 | 1651 | 2009 | 1808 | 2159 | 2287 | 1795 | 1776 | 2199 | 2004.11 | 2085 | 1923 | 0.922 | -0.117 | 0.3671448 | 0.58453273 | 0.0133 | 0.0085 | 0.0116 | 0.0116 | TRUE | 0.1892 |
| Papss2 | 39 | 34 | 24 | 101 | 79 | 44 | 41 | 34 | 24 | 79 | 85 | 48 | 52.01 | 33 | 71 | 2.145 | 1.101 | 0.00151368 | 0.01235125 | 0.0581 | 0.0702 | 0.0645 | 0.0645 | TRUE | 0.6739 |
| Paqr3 | 387 | 370 | 356 | 531 | 402 | 372 | 411 | 370 | 357 | 417 | 433 | 407 | 399.2 | 379 | 419 | 1.105 | 0.144 | 0.2130593 | 0.42424003 | 0 | 0.0152 | 0.0071 | 0.0071 | TRUE | 0.0606 |
| Paqr4 | 109 | 120 | 128 | 178 | 101 | 90 | 116 | 120 | 129 | 140 | 109 | 99 | 118.54 | 122 | 116 | 0.959 | -0.061 | 0.76979745 | 0.87970853 | 0.0112 | 0.0347 | 0.0227 | 0.0227 | TRUE | 0.3667 |
| Paqr5 | 171 | 173 | 197 | 159 | 121 | 140 | 182 | 173 | 198 | 125 | 130 | 153 | 160.11 | 184 | 136 | 0.737 | -0.44 | 0.00987594 | 0.05221089 | 0 | 0.0275 | 0.0145 | 0.0145 | TRUE | 0.1205 |
| Paqr6 | 94 | 104 | 94 | 57 | 50 | 69 | 100 | 104 | 94 | 45 | 54 | 76 | 78.71 | 99 | 58 | 0.579 | -0.787 | 0.00237624 | 0.01763671 | 0.0177 | 0.0488 | 0.0348 | 0.0348 | TRUE | 0.5739 |
| Paqr7 | 381 | 364 | 327 | 240 | 200 | 219 | 405 | 364 | 328 | 189 | 215 | 240 | 290.02 | 366 | 215 | 0.585 | -0.773 | 5.39E-07 | 1.60E-05 | 0.009 | 0.0183 | 0.0135 | 0.0135 | TRUE | 0.132 |
| Paqr8 | 449 | 406 | 410 | 422 | 294 | 300 | 477 | 405 | 412 | 332 | 316 | 328 | 378.42 | 431 | 325 | 0.755 | -0.405 | 0.00098975 | 0.00884441 | 0.0022 | 0.0156 | 0.0082 | 0.0082 | TRUE | 0.0977 |
| Paqr9 | 82 | 89 | 105 | 201 | 159 | 110 | 87 | 89 | 105 | 158 | 171 | 120 | 121.82 | 94 | 150 | 1.599 | 0.677 | 0.0015175 | 0.01236349 | 0.0143 | 0.0339 | 0.0241 | 0.0241 | TRUE | 0.2947 |
| Pard3 | 5215 | 6613 | 7076 | 7829 | 5702 | 6077 | 5539 | 6604 | 7105 | 6155 | 6135 | 6652 | 6365.09 | 6416 | 6314 | 0.984 | -0.023 | 0.82919537 | 0.91333678 | 0.0089 | 0.0074 | 0.0083 | 0.0083 | TRUE | 0.1743 |
| Pard3b | 13 | 10 | 10 | 20 | 30 | 19 | 14 | 10 | 10 | 16 | 32 | 21 | 17.11 | 11 | 23 | 2.021 | 1.015 | 0.04832354 | 0.16025697 | 0 | 0.1995 | 0.1235 | 0.1235 | TRUE | 0.3719 |
| Pard6a | 616 | 501 | 498 | 809 | 544 | 479 | 654 | 500 | 500 | 636 | 585 | 524 | 566.72 | 551 | 582 | 1.057 | 0.079 | 0.59190067 | 0.76551516 | 0.0152 | 0.0127 | 0.014 | 0.014 | TRUE | 0.3091 |
| Pard6b | 1718 | 1605 | 1577 | 1995 | 1436 | 1486 | 1825 | 1603 | 1583 | 1568 | 1545 | 1627 | 1625.21 | 1670 | 1580 | 0.946 | -0.08 | 0.37340545 | 0.58981981 | 0.0027 | 0.0089 | 0.0052 | 0.0052 | TRUE | 0.0786 |
| Pard6g | 744 | 874 | 936 | 1087 | 932 | 958 | 790 | 873 | 940 | 855 | 1003 | 1049 | 918.16 | 868 | 969 | 1.115 | 0.157 | 0.19275324 | 0.39842218 | 0.0085 | 0.0105 | 0.0094 | 0.0094 | TRUE | 0.1304 |
| Parg | 4622 | 4365 | 4528 | 5694 | 4226 | 3951 | 4909 | 4359 | 4547 | 4476 | 4547 | 4325 | 4527.24 | 4605 | 4449 | 0.966 | -0.049 | 0.51286662 | 0.70608697 | 0.0018 | 0.0076 | 0.0039 | 0.0039 | TRUE | 0.0406 |
| Park2 | 14 | 10 | 14 | 21 | 15 | 22 | 15 | 10 | 14 | 17 | 16 | 24 | 15.94 | 13 | 19 | 1.453 | 0.539 | 0.27096052 | 0.49096002 | 0 | 0.2136 | 0.1072 | 0.1072 | TRUE | 0.2772 |
| Park7 | 9011 | 10139 | 9792 | 12198 | 8537 | 8363 | 9571 | 10126 | 9832 | 9589 | 9185 | 9155 | 9576.4 | 9843 | 9310 | 0.946 | -0.08 | 0.18860788 | 0.39374235 | 6.00E-04 | 0.0072 | 0.0026 | 0.0026 | TRUE | 0.0085 |
| Parl | 4164 | 5248 | 5042 | 5604 | 4191 | 4197 | 4423 | 5241 | 5063 | 4405 | 4509 | 4594 | 4705.96 | 4909 | 4503 | 0.917 | -0.125 | 0.15525478 | 0.34749626 | 0.0037 | 0.0076 | 0.0053 | 0.0053 | TRUE | 0.0913 |
| Parm1 | 226 | 250 | 254 | 624 | 442 | 444 | 240 | 250 | 255 | 491 | 476 | 486 | 366.15 | 248 | 484 | 1.95 | 0.963 | 1.83E-17 | 3.39E-15 | 0 | 0.0159 | 0.0062 | 0.0062 | TRUE | 0.0091 |
| Parn | 3150 | 3996 | 4198 | 4056 | 3071 | 3484 | 3346 | 3991 | 4215 | 3189 | 3304 | 3814 | 3643.07 | 3851 | 3436 | 0.892 | -0.165 | 0.16286255 | 0.35825218 | 0.0111 | 0.0078 | 0.0098 | 0.0098 | TRUE | 0.1598 |
| Parp1 | 39323 | 45730 | 47631 | 50727 | 39132 | 42065 | 41768 | 45671 | 47827 | 39878 | 42104 | 46047 | 43882.25 | 45089 | 42676 | 0.946 | -0.079 | 0.37467735 | 0.59076389 | 0.0049 | 0.007 | 0.0057 | 0.0057 | TRUE | 0.0584 |
| Parp10 | 253 | 295 | 306 | 227 | 167 | 180 | 269 | 295 | 307 | 178 | 180 | 197 | 237.63 | 290 | 185 | 0.637 | -0.652 | 3.62E-06 | 8.37E-05 | 0 | 0.0208 | 0.0099 | 0.0099 | TRUE | 0.0465 |
| Parp11 | 954 | 873 | 871 | 1123 | 849 | 741 | 1013 | 872 | 875 | 883 | 913 | 811 | 894.53 | 920 | 869 | 0.945 | -0.081 | 0.44567884 | 0.65230081 | 0.0044 | 0.0106 | 0.007 | 0.007 | TRUE | 0.0935 |
| Parp12 | 1547 | 1818 | 1794 | 2343 | 1693 | 1418 | 1643 | 1816 | 1801 | 1842 | 1822 | 1552 | 1745.98 | 1753 | 1739 | 0.992 | -0.012 | 0.90796979 | 0.95356627 | 0.0054 | 0.0088 | 0.0068 | 0.0068 | TRUE | 0.1051 |
| Parp14 | 376 | 265 | 369 | 376 | 277 | 251 | 399 | 265 | 371 | 296 | 298 | 275 | 317.15 | 345 | 290 | 0.84 | -0.251 | 0.15306357 | 0.34455792 | 0.0209 | 0.0173 | 0.0191 | 0.0191 | TRUE | 0.4741 |
| Parp16 | 431 | 440 | 456 | 462 | 354 | 369 | 458 | 439 | 458 | 363 | 381 | 404 | 417.18 | 452 | 383 | 0.846 | -0.241 | 0.02769656 | 0.10918024 | 0 | 0.0148 | 0.0063 | 0.0063 | TRUE | 0.0268 |
| Parp2 | 2115 | 2440 | 2607 | 3343 | 2486 | 2514 | 2247 | 2437 | 2618 | 2628 | 2675 | 2752 | 2559.3 | 2434 | 2685 | 1.103 | 0.141 | 0.09566995 | 0.25355511 | 0.0026 | 0.0082 | 0.0048 | 0.0048 | TRUE | 0.0547 |

| | | | | | | | | | | | | | | | | | | | | | | | | | |
|---|---|---|---|---|---|---|---|---|---|---|---|---|---|---|---|---|---|---|---|---|---|---|---|---|---|
| Parp3 | 29 | 28 | 33 | 47 | 29 | 38 | 31 | 28 | 33 | 37 | 31 | 42 | 33.61 | 31 | 37 | 1.194 | 0.256 | 0.43802543 | 0.64579729 | 0 | 0.1049 | 0.0487 | 0.0487 | TRUE | 0.1092 |
| Parp4 | 626 | 511 | 564 | 912 | 621 | 544 | 665 | 510 | 566 | 717 | 668 | 595 | 620.36 | 580 | 660 | 1.139 | 0.188 | 0.17043384 | 0.36901257 | 0.0109 | 0.0122 | 0.0119 | 0.0119 | TRUE | 0.1887 |
| Parp6 | 663 | 625 | 649 | 657 | 552 | 566 | 704 | 624 | 652 | 516 | 594 | 620 | 618.34 | 660 | 577 | 0.872 | -0.197 | 0.08893972 | 0.24223057 | 0.0048 | 0.0122 | 0.0081 | 0.0081 | TRUE | 0.1026 |
| Parp8 | 20 | 26 | 21 | 80 | 66 | 40 | 21 | 26 | 21 | 63 | 71 | 44 | 41 | 23 | 59 | 2.605 | 1.381 | 5.87E-05 | 0.00088799 | 0 | 0.0873 | 0.0545 | 0.0545 | TRUE | 0.3296 |
| Parp9 | 512 | 433 | 446 | 735 | 513 | 448 | 544 | 432 | 448 | 578 | 552 | 490 | 507.38 | 475 | 540 | 1.139 | 0.188 | 0.16278952 | 0.35814149 | 0.0092 | 0.0134 | 0.0111 | 0.0111 | TRUE | 0.1873 |
| Parpbp | 1337 | 1644 | 1740 | 2391 | 1690 | 1565 | 1420 | 1642 | 1747 | 1880 | 1818 | 1713 | 1703.37 | 1603 | 1804 | 1.125 | 0.17 | 0.10088873 | 0.26296762 | 0.0056 | 0.0088 | 0.0072 | 0.0072 | TRUE | 0.1196 |
| Pars2 | 447 | 526 | 472 | 636 | 510 | 488 | 475 | 525 | 474 | 500 | 549 | 534 | 509.49 | 491 | 528 | 1.072 | 0.101 | 0.34996388 | 0.56885811 | 0 | 0.0134 | 0.0064 | 0.0064 | TRUE | 0.0429 |
| Parva | 449 | 496 | 482 | 713 | 458 | 454 | 477 | 495 | 484 | 561 | 493 | 497 | 501.09 | 485 | 517 | 1.067 | 0.093 | 0.39164188 | 0.60534847 | 0 | 0.0135 | 0.0065 | 0.0065 | TRUE | 0.0675 |
| Parvb | 5364 | 7662 | 8278 | 6129 | 4838 | 5834 | 5698 | 7652 | 8312 | 4818 | 5205 | 6386 | 6345.23 | 7221 | 5470 | 0.757 | -0.401 | 0.01217144 | 0.06078781 | 0.0286 | 0.0074 | 0.0183 | 0.0183 | TRUE | 0.4154 |
| Parvg | 11 | 14 | 8 | 10 | 15 | 6 | 12 | 14 | 8 | 8 | 16 | 7 | 10.71 | 11 | 10 | 0.901 | -0.151 | 0.8106713 | 0.90276832 | 0.0576 | 0.3145 | 0.1919 | 0.1919 | TRUE | 0.8509 |
| Pask | 2664 | 3024 | 3105 | 4316 | 3234 | 2954 | 2830 | 3020 | 3118 | 3393 | 3480 | 3234 | 3178.93 | 2989 | 3369 | 1.127 | 0.172 | 0.02235962 | 0.09421777 | 0.0014 | 0.0079 | 0.0038 | 0.0038 | TRUE | 0.0264 |
| Pate2 | 12 | 12 | 10 | 24 | 11 | 9 | 13 | 12 | 10 | 19 | 12 | 10 | 12.55 | 12 | 14 | 1.184 | 0.243 | 0.65924381 | 0.80795659 | 0 | 0.2693 | 0.1397 | 0.1397 | TRUE | 0.6276 |
| Pate4 | 1 | 0 | 0 | 0 | 0 | 0 | 1 | 0 | 0 | 0 | 0 | 0 | 0.18 | 0 | 0 | 0.49 | -1.029 | 0.80087 | NA | 0 | 18.6165 | 10 | 10 | TRUE | 0.4614 |
| Patl1 | 3735 | 4055 | 4363 | 4587 | 3355 | 3917 | 3967 | 4050 | 4381 | 3606 | 3610 | 4288 | 3983.57 | 4133 | 3835 | 0.928 | -0.108 | 0.27300831 | 0.49328527 | 0.0063 | 0.0077 | 0.0068 | 0.0068 | TRUE | 0.1296 |
| Patl2 | 92 | 152 | 202 | 98 | 71 | 88 | 98 | 152 | 203 | 77 | 76 | 96 | 117.02 | 151 | 83 | 0.551 | -0.86 | 0.00281113 | 0.02012581 | 0.0666 | 0.0351 | 0.0505 | 0.0505 | TRUE | 0.1194 |
| Patz1 | 1734 | 1889 | 1884 | 2587 | 1891 | 1800 | 1842 | 1887 | 1892 | 2034 | 2035 | 1970 | 1943.13 | 1874 | 2013 | 1.075 | 0.104 | 0.12290651 | 0.29859542 | 0 | 0.0086 | 0.0027 | 0.0027 | TRUE | 0.0041 |
| Pawr | 2171 | 2545 | 2506 | 3398 | 2478 | 2383 | 2306 | 2542 | 2516 | 2671 | 2666 | 2609 | 2551.66 | 2455 | 2649 | 1.079 | 0.11 | 0.13859811 | 0.3230815 | 9.00E-04 | 0.0082 | 0.0036 | 0.0036 | TRUE | 0.0339 |
| Pax1 | 4 | 4 | 4 | 8 | 9 | 4 | 4 | 4 | 4 | 6 | 10 | 4 | 5.44 | 4 | 7 | 1.658 | 0.729 | 0.37075948 | 0.58760362 | 0 | 0.613 | 0.2809 | 0.2809 | TRUE | 0.2862 |
| Pax2 | 7 | 2 | 0 | 19 | 5 | 8 | 7 | 2 | 0 | 15 | 5 | 9 | 6.42 | 3 | 10 | 3.138 | 1.65 | 0.11196014 | 0.28141396 | 0.6384 | 0.5202 | 0.5623 | 0.5623 | TRUE | 0.9351 |
| Pax3 | 4 | 3 | 0 | 4 | 3 | 3 | 4 | 3 | 0 | 3 | 3 | 3 | 2.82 | 2 | 3 | 1.339 | 0.421 | 0.72931662 | NA | 0 | 1.1765 | 0.7019 | 0.7019 | TRUE | 0.8145 |
| Pax4 | 0 | 0 | 0 | 0 | 0 | 0 | 0 | 0 | 0 | 0 | 0 | 0 | 0 | 0 | 0 | NA | NA | NA | NA | NA | NA | NA | NA | NA | NA |
| Pax5 | 0 | 1 | 3 | 3 | 0 | 2 | 0 | 1 | 3 | 2 | 0 | 2 | 1.43 | 1 | 1 | 1.144 | 0.194 | 0.91426024 | NA | 0 | 2.3164 | 1.6486 | 1.6486 | TRUE | 0.5574 |
| Pax6 | 98 | 109 | 150 | 149 | 103 | 117 | 104 | 109 | 151 | 117 | 111 | 128 | 119.93 | 121 | 119 | 0.978 | -0.032 | 0.88242829 | 0.94171859 | 0.0154 | 0.0344 | 0.0249 | 0.0249 | TRUE | 0.4589 |
| Pax6os1 | 0 | 0 | 0 | 0 | 0 | 0 | 0 | 0 | 0 | 0 | 0 | 0 | 0 | 0 | 0 | NA | NA | NA | NA | NA | NA | NA | NA | NA | NA |
| Pax7 | 13 | 9 | 16 | 6 | 4 | 7 | 14 | 9 | 16 | 5 | 4 | 8 | 9.26 | 13 | 6 | 0.426 | -1.232 | 0.06178653 | 0.18984945 | 0 | 0.3628 | 0.1856 | 0.1856 | TRUE | 0.3033 |
| Pax8 | 215 | 240 | 269 | 140 | 124 | 172 | 228 | 240 | 270 | 110 | 133 | 188 | 194.99 | 246 | 144 | 0.582 | -0.781 | 0.00033456 | 0.00373 | 0.0345 | 0.0238 | 0.0287 | 0.0287 | TRUE | 0.7464 |
| Pax9 | 8 | 12 | 13 | 17 | 10 | 12 | 8 | 12 | 13 | 13 | 11 | 13 | 11.8 | 11 | 12 | 1.112 | 0.153 | 0.7782069 | 0.88511424 | 0 | 0.2861 | 0.1285 | 0.1285 | TRUE | 0.1567 |
| Paxbp1 | 1725 | 1602 | 1606 | 2185 | 1829 | 1672 | 1832 | 1600 | 1613 | 1718 | 1968 | 1830 | 1760.1 | 1682 | 1839 | 1.093 | 0.128 | 0.19015285 | 0.39517265 | 0.0045 | 0.0088 | 0.0063 | 0.0063 | TRUE | 0.0739 |
| Paxip1 | 4294 | 3779 | 3624 | 5885 | 4325 | 4055 | 4561 | 3774 | 3639 | 4626 | 4653 | 4439 | 4282.1 | 3991 | 4573 | 1.146 | 0.196 | 0.05948765 | 0.18472911 | 0.0078 | 0.0077 | 0.0076 | 0.0076 | TRUE | 0.1895 |
| Pbdc1 | 3089 | 2969 | 2981 | 3968 | 2798 | 2764 | 3281 | 2965 | 2993 | 3119 | 3010 | 3026 | 3065.82 | 3080 | 3052 | 0.991 | -0.013 | 0.86450357 | 0.93294221 | 0.0014 | 0.008 | 0.0037 | 0.0037 | TRUE | 0.0396 |
| Pbk | 2132 | 2474 | 2525 | 3073 | 2207 | 2490 | 2265 | 2471 | 2535 | 2416 | 2375 | 2726 | 2464.46 | 2424 | 2506 | 1.033 | 0.047 | 0.60850673 | 0.77824912 | 0.0037 | 0.0082 | 0.0058 | 0.0058 | TRUE | 0.0712 |
| Pbld1 | 14 | 27 | 17 | 20 | 18 | 18 | 15 | 27 | 17 | 16 | 19 | 20 | 18.95 | 20 | 18 | 0.924 | -0.115 | 0.8029081 | 0.8981819 | 0 | 0.1808 | 0.0988 | 0.0988 | TRUE | 0.577 |
| Pbld2 | 34 | 37 | 28 | 38 | 31 | 35 | 36 | 37 | 28 | 30 | 33 | 38 | 33.79 | 34 | 34 | 0.999 | -0.001 | 0.99752608 | 0.99847569 | 0 | 0.1044 | 0.0506 | 0.0506 | TRUE | 0.15 |
| Pbp2 | 4 | 1 | 2 | 9 | 3 | 6 | 4 | 1 | 2 | 7 | 3 | 7 | 4.02 | 2 | 6 | 2.36 | 1.238 | 0.22081093 | NA | 0 | 0.8262 | 0.4382 | 0.4382 | TRUE | 0.4433 |
| Pbrm1 | 8213 | 7657 | 8138 | 9912 | 7051 | 7065 | 8724 | 7647 | 8171 | 7792 | 7586 | 7734 | 7942.4 | 8181 | 7704 | 0.942 | -0.086 | 0.24724925 | 0.46398502 | 0.0021 | 0.0073 | 0.0039 | 0.0039 | TRUE | 0.0411 |
| Pbsn | 0 | 0 | 0 | 0 | 0 | 0 | 0 | 0 | 0 | 0 | 0 | 0 | 0 | 0 | 0 | NA | NA | NA | NA | NA | NA | NA | NA | NA | NA |
| Pbx1 | 381 | 341 | 355 | 558 | 364 | 445 | 405 | 341 | 356 | 439 | 392 | 487 | 403.19 | 367 | 439 | 1.196 | 0.258 | 0.05827651 | 0.18236848 | 0.0073 | 0.0151 | 0.0109 | 0.0109 | TRUE | 0.1017 |
| Pbx2 | 2224 | 2433 | 2409 | 3089 | 2160 | 2127 | 2362 | 2430 | 2419 | 2428 | 2324 | 2328 | 2381.95 | 2404 | 2360 | 0.982 | -0.026 | 0.69659848 | 0.83378245 | 0 | 0.0083 | 0.0027 | 0.0027 | TRUE | 0.0079 |
| Pbx3 | 272 | 375 | 381 | 663 | 478 | 439 | 289 | 375 | 383 | 521 | 514 | 481 | 427.01 | 349 | 505 | 1.449 | 0.535 | 0.00014084 | 0.00181033 | 0.0091 | 0.0146 | 0.0118 | 0.0118 | TRUE | 0.2535 |
| Pbx4 | 245 | 289 | 350 | 242 | 194 | 250 | 260 | 289 | 351 | 190 | 209 | 274 | 262.16 | 300 | 224 | 0.745 | -0.425 | 0.02705372 | 0.10723237 | 0.0253 | 0.0195 | 0.0227 | 0.0227 | TRUE | 0.4035 |
| Pbxip1 | 692 | 607 | 553 | 891 | 696 | 560 | 735 | 606 | 555 | 700 | 749 | 613 | 659.8 | 632 | 687 | 1.088 | 0.122 | 0.38973276 | 0.6037429 | 0.0138 | 0.0119 | 0.0129 | 0.0129 | TRUE | 0.2373 |
| Pcbd1 | 158 | 252 | 237 | 228 | 173 | 217 | 168 | 252 | 238 | 179 | 186 | 238 | 210.06 | 219 | 201 | 0.915 | -0.129 | 0.53410045 | 0.72173954 | 0.0297 | 0.0226 | 0.0261 | 0.0261 | TRUE | 0.4554 |
| Pcbd2 | 90 | 156 | 171 | 264 | 175 | 135 | 96 | 156 | 172 | 208 | 188 | 148 | 161.12 | 141 | 181 | 1.286 | 0.363 | 0.1482887 | 0.33797824 | 0.0521 | 0.0274 | 0.0391 | 0.0391 | TRUE | 0.8123 |
| Pcbp1 | 14367 | 13758 | 13362 | 27150 | 18853 | 16005 | 15260 | 13740 | 13417 | 21343 | 20285 | 17520 | 16927.54 | 14139 | 19716 | 1.395 | 0.48 | 2.13E-06 | 5.30E-05 | 0.0073 | 0.0071 | 0.0073 | 0.0073 | TRUE | 0.1161 |
| Pcbp2 | 37543 | 38476 | 38408 | 47576 | 32862 | 33911 | 39877 | 38426 | 38566 | 37401 | 35357 | 37121 | 37791.38 | 38956 | 36626 | 0.94 | -0.089 | 0.12666846 | 0.30533622 | 6.00E-04 | 0.007 | 0.0024 | 0.0024 | TRUE | 0.0112 |
| Pcbp3 | 1785 | 1990 | 1955 | 1964 | 1468 | 1555 | 1896 | 1987 | 1963 | 1544 | 1579 | 1702 | 1778.68 | 1949 | 1608 | 0.825 | -0.278 | 0.00038098 | 0.00416698 | 0.001 | 0.0088 | 0.0038 | 0.0038 | TRUE | 0.031 |
| Pcbp4 | 2326 | 2539 | 2792 | 2889 | 2130 | 2230 | 2471 | 2536 | 2803 | 2271 | 2292 | 2441 | 2468.96 | 2603 | 2335 | 0.897 | -0.158 | 0.06174996 | 0.18977404 | 0.0025 | 0.0082 | 0.0047 | 0.0047 | TRUE | 0.0549 |
| Pcca | 1246 | 1427 | 1455 | 1347 | 965 | 1055 | 1323 | 1425 | 1461 | 1059 | 1038 | 1155 | 1243.61 | 1403 | 1084 | 0.772 | -0.373 | 3.34E-05 | 0.00055063 | 0.002 | 0.0096 | 0.005 | 0.005 | TRUE | 0.0382 |
| Pccb | 1117 | 1209 | 1124 | 1611 | 1156 | 1096 | 1186 | 1207 | 1129 | 1266 | 1244 | 1200 | 1205.41 | 1174 | 1237 | 1.054 | 0.076 | 0.34276405 | 0.56283962 | 0 | 0.0096 | 0.0037 | 0.0037 | TRUE | 0.0139 |
| Pcdh1 | 642 | 775 | 938 | 807 | 607 | 690 | 682 | 774 | 942 | 634 | 653 | 755 | 740.1 | 799 | 681 | 0.851 | -0.232 | 0.11179835 | 0.28113131 | 0.0161 | 0.0114 | 0.014 | 0.014 | TRUE | 0.2893 |
| Pcdh10 | 11 | 7 | 12 | 18 | 6 | 11 | 12 | 7 | 12 | 14 | 6 | 12 | 10.56 | 10 | 11 | 1.074 | 0.104 | 0.86606209 | 0.93370586 | 0 | 0.3188 | 0.1778 | 0.1778 | TRUE | 0.4342 |
| Pcdh11x | 0 | 0 | 0 | 0 | 0 | 0 | 0 | 0 | 0 | 0 | 0 | 0 | 0 | 0 | 0 | NA | NA | NA | NA | NA | NA | NA | NA | NA | NA |
| Pcdh12 | 2 | 4 | 0 | 9 | 1 | 2 | 2 | 4 | 0 | 7 | 1 | 2 | 2.74 | 2 | 3 | 1.73 | 0.791 | 0.56658186 | NA | 0.7109 | 1.2078 | 0.9819 | 0.9819 | TRUE | 0.5141 |
| Pcdh15 | 1 | 1 | 1 | 0 | 3 | 3 | 1 | 1 | 1 | 0 | 3 | 3 | 1.6 | 1 | 2 | 2.064 | 1.046 | 0.52629343 | NA | 0 | 2.0709 | 1.2302 | 1.2302 | TRUE | 1.0179 |
| Pcdh17 | 7 | 3 | 1 | 20 | 11 | 5 | 7 | 3 | 1 | 16 | 12 | 5 | 7.41 | 4 | 11 | 2.937 | 1.554 | 0.07503898 | 0.21569433 | 0.2622 | 0.4514 | 0.3701 | 0.3701 | TRUE | 0.4784 |
| Pcdh18 | 11 | 1 | 1 | 13 | 6 | 8 | 12 | 1 | 1 | 10 | 6 | 9 | 6.52 | 5 | 8 | 1.881 | 0.912 | 0.36829768 | 0.58564866 | 0.6395 | 0.5122 | 0.5691 | 0.5691 | TRUE | 3.3454 |
| Pcdh19 | 65 | 53 | 56 | 249 | 165 | 97 | 69 | 53 | 56 | 196 | 178 | 106 | 109.61 | 59 | 160 | 2.701 | 1.433 | 3.59E-07 | 1.11E-05 | 0.0542 | 0.037 | 0.0455 | 0.0455 | TRUE | 0.7698 |
| Pcdh20 | 1 | 1 | 1 | 7 | 6 | 1 | 1 | 1 | 1 | 6 | 6 | 1 | 2.69 | 1 | 4 | 4.294 | 2.102 | 0.11540464 | NA | 0 | 1.2333 | 0.6733 | 0.6733 | TRUE | 0.6888 |
| Pcdh7 | 0 | 0 | 0 | 2 | 3 | 0 | 0 | 0 | 0 | 2 | 3 | 0 | 0.8 | 0 | 2 | 8.522 | 3.091 | 0.27029335 | NA | 5.3026 | 4.1249 | 4.3597 | 4.3597 | TRUE | 0.0518 |
| Pcdh8 | 79 | 48 | 23 | 99 | 63 | 43 | 84 | 48 | 23 | 78 | 68 | 47 | 57.94 | 52 | 64 | 1.249 | 0.32 | 0.47017576 | 0.67200834 | 0.1957 | 0.0638 | 0.1243 | 0.1243 | TRUE | 0.3858 |
| Pcdh9 | 9 | 10 | 3 | 19 | 9 | 5 | 10 | 10 | 3 | 15 | 10 | 5 | 8.78 | 8 | 10 | 1.354 | 0.437 | 0.55402097 | 0.73738555 | 0.1598 | 0.3823 | 0.2772 | 0.2772 | TRUE | 0.2779 |
| Pcdha1 | 0 | 0 | 1 | 0 | 2 | 0 | 0 | 0 | 1 | 0 | 2 | 0 | 0.53 | 0 | 1 | 1.927 | 0.947 | 0.77736506 | NA | 0 | 6.2701 | 6.3814 | 6.3814 | TRUE | 0.9744 |
| Pcdha10 | 0 | 0 | 0 | 2 | 4 | 0 | 0 | 0 | 0 | 2 | 4 | 0 | 0.98 | 0 | 2 | 10.41 | 3.38 | 0.20365372 | NA | 6.3568 | 3.3709 | 3.8435 | 3.8435 | TRUE | 0.1105 |
| Pcdha11 | 1 | 0 | 0 | 0 | 0 | 0 | 1 | 0 | 0 | 0 | 0 | 0 | 0.18 | 0 | 0 | 0.49 | -1.029 | 0.80087 | NA | 0 | 18.6165 | 10 | 10 | TRUE | 0.4614 |
| Pcdha12 | 0 | 0 | 0 | 0 | 0 | 1 | 0 | 0 | 0 | 0 | 0 | 1 | 0.18 | 0 | 0 | 1.859 | 0.894 | 0.82650734 | NA | 0 | 18.0643 | 10 | 10 | TRUE | 0.5383 |
| Pcdha2 | 0 | 1 | 0 | 0 | 0 | 0 | 0 | 1 | 0 | 0 | 0 | 0 | 0.17 | 0 | 0 | 0.49 | -1.029 | 0.80087 | NA | 0 | 19.7991 | 10 | 10 | TRUE | 0.405 |
| Pcdha3 | 0 | 1 | 0 | 3 | 0 | 0 | 0 | 1 | 0 | 2 | 0 | 0 | 0.56 | 0 | 1 | 2.222 | 1.152 | 0.72351576 | NA | 0 | 5.895 | 6.0404 | 6.0404 | TRUE | 1.4404 |
| Pcdha4 | 0 | 0 | 0 | 0 | 0 | 0 | 0 | 0 | 0 | 0 | 0 | 0 | 0 | 0 | 0 | NA | NA | NA | NA | NA | NA | NA | NA | NA | NA |
| Pcdha4-g | 0 | 0 | 0 | 0 | 0 | 0 | 0 | 0 | 0 | 0 | 0 | 0 | 0 | 0 | 0 | NA | NA | NA | NA | NA | NA | NA | NA | NA | NA |

| | | | | | | | | | | | | | | | | | | | | | | | | |
|---|---|---|---|---|---|---|---|---|---|---|---|---|---|---|---|---|---|---|---|---|---|---|---|---|
| Pcdha5 | 1 | 0 | 0 | 0 | 1 | 0 | 1 | 0 | 0 | 0 | 1 | 0 | 0.36 | 0 | 0 | 0.954 | -0.067 | 0.98557668 | NA | 0 | 9.2518 | 8.0179 | 8.0179 | TRUE | 0.5204 |
| Pcdha6 | 2 | 1 | 0 | 6 | 4 | 2 | 2 | 1 | 0 | 5 | 4 | 2 | 2.39 | 1 | 4 | 3.658 | 1.871 | 0.17324448 | NA | 0 | 1.386 | 0.734 | 0.734 | TRUE | 0.3784 |
| Pcdha7 | 3 | 0 | 0 | 5 | 2 | 2 | 3 | 0 | 0 | 4 | 2 | 2 | 1.91 | 1 | 3 | 2.667 | 1.415 | 0.38664796 | NA | 0.2095 | 1.732 | 1.2602 | 1.2602 | TRUE | 1.4432 |
| Pcdha8 | 0 | 0 | 0 | 0 | 1 | 0 | 0 | 0 | 0 | 0 | 1 | 0 | 0.18 | 0 | 0 | 1.859 | 0.894 | 0.82650734 | NA | 0 | 18.3785 | 10 | 10 | TRUE | 0.5204 |
| Pcdha9 | 0 | 0 | 0 | 1 | 0 | 0 | 0 | 0 | 0 | 1 | 0 | 0 | 0.13 | 0 | 0 | 1.859 | 0.894 | 0.82650734 | NA | 0 | 25.1515 | 10 | 10 | TRUE | 0.2563 |
| Pcdhac1 | 2 | 0 | 0 | 0 | 1 | 0 | 2 | 0 | 0 | 0 | 1 | 0 | 0.53 | 1 | 0 | 0.497 | -1.009 | 0.76131406 | NA | 0 | 6.1834 | 6.228 | 6.228 | TRUE | 0.9742 |
| Pcdhac2 | 8 | 16 | 18 | 41 | 34 | 27 | 8 | 16 | 18 | 32 | 37 | 30 | 23.49 | 14 | 33 | 2.303 | 1.203 | 0.00538455 | 0.0330191 | 0 | 0.1472 | 0.0839 | 0.0839 | TRUE | 0.5125 |
| Pcdhb1 | 2 | 1 | 0 | 1 | 0 | 0 | 2 | 1 | 0 | 1 | 0 | 0 | 0.65 | 1 | 0 | 0.337 | -1.57 | 0.54866551 | NA | 0 | 5.0634 | 3.4431 | 3.4431 | TRUE | 0.1066 |
| Pcdhb10 | 0 | 0 | 0 | 0 | 0 | 0 | 0 | 0 | 0 | 0 | 0 | 0 | 0 | 0 | 0 | NA | NA | NA | NA | NA | NA | NA | NA | NA | NA |
| Pcdhb11 | 0 | 2 | 0 | 3 | 7 | 0 | 0 | 2 | 0 | 2 | 8 | 0 | 1.98 | 1 | 3 | 4.89 | 2.29 | 0.2532114 | NA | 2.818 | 1.6697 | 1.9851 | 1.9851 | TRUE | 0.3837 |
| Pcdhb12 | 1 | 0 | 1 | 0 | 0 | 0 | 1 | 0 | 1 | 0 | 0 | 0 | 0.34 | 1 | 0 | 0.261 | -1.939 | 0.60367425 | NA | 0 | 9.5732 | 8.3134 | 8.3134 | TRUE | 0.2529 |
| Pcdhb13 | 0 | 0 | 0 | 0 | 0 | 0 | 0 | 0 | 0 | 0 | 0 | 0 | 0 | 0 | 0 | NA | NA | NA | NA | NA | NA | NA | NA | NA | NA |
| Pcdhb14 | 1 | 1 | 0 | 5 | 4 | 2 | 1 | 1 | 0 | 4 | 4 | 2 | 2.08 | 1 | 3 | 5.111 | 2.354 | 0.11942017 | NA | 0 | 1.5902 | 0.758 | 0.758 | TRUE | 0.2538 |
| Pcdhb15 | 1 | 0 | 0 | 0 | 0 | 0 | 1 | 0 | 0 | 0 | 0 | 0 | 0.18 | 0 | 0 | 0.49 | -1.029 | 0.80087 | NA | 0 | 18.6165 | 10 | 10 | TRUE | 0.4614 |
| Pcdhb16 | 1 | 0 | 0 | 0 | 0 | 1 | 1 | 0 | 0 | 0 | 0 | 1 | 0.36 | 0 | 0 | 0.954 | -0.067 | 0.98553519 | NA | 0 | 9.1716 | 7.9605 | 7.9605 | TRUE | 0.5383 |
| Pcdhb17 | 1 | 0 | 0 | 1 | 0 | 0 | 1 | 0 | 0 | 1 | 0 | 0 | 0.31 | 0 | 0 | 0.954 | -0.067 | 0.98625928 | NA | 0 | 10.7014 | 9.0384 | 9.0384 | TRUE | 0.4614 |
| Pcdhb18 | 3 | 2 | 6 | 9 | 1 | 3 | 3 | 2 | 6 | 7 | 1 | 3 | 3.77 | 4 | 4 | 1.047 | 0.067 | 0.95018571 | NA | 0.2403 | 0.8799 | 0.5594 | 0.5594 | TRUE | 0.438 |
| Pcdhb19 | 1 | 1 | 1 | 1 | 2 | 1 | 1 | 1 | 1 | 1 | 2 | 1 | 1.18 | 1 | 1 | 1.293 | 0.371 | 0.81961764 | NA | 0 | 2.7919 | 1.039 | 1.039 | TRUE | 0.1601 |
| Pcdhb2 | 1 | 3 | 1 | 6 | 5 | 3 | 1 | 3 | 1 | 5 | 5 | 3 | 3.07 | 2 | 4 | 2.637 | 1.399 | 0.21251428 | NA | 0 | 1.0787 | 0.4967 | 0.4967 | TRUE | 0.3597 |
| Pcdhb20 | 0 | 2 | 1 | 1 | 1 | 0 | 0 | 2 | 1 | 1 | 1 | 0 | 0.81 | 1 | 1 | 0.625 | -0.678 | 0.7553135 | NA | 0 | 4.0711 | 2.1302 | 2.1302 | TRUE | 0.1304 |
| Pcdhb21 | 0 | 0 | 0 | 1 | 1 | 0 | 0 | 0 | 0 | 1 | 1 | 0 | 0.31 | 0 | 1 | 3.32 | 1.731 | 0.65383052 | NA | 0 | 10.6224 | 8.9541 | 8.9541 | TRUE | 0.202 |
| Pcdhb22 | 4 | 6 | 5 | 8 | 7 | 4 | 4 | 6 | 5 | 6 | 8 | 4 | 5.58 | 5 | 6 | 1.193 | 0.255 | 0.74397018 | 0.86335793 | 0 | 0.5976 | 0.2581 | 0.2581 | TRUE | 0.122 |
| Pcdhb3 | 2 | 1 | 0 | 5 | 4 | 1 | 2 | 1 | 0 | 4 | 4 | 1 | 2.08 | 1 | 3 | 3.042 | 1.605 | 0.275918 | NA | 0 | 1.5943 | 0.9116 | 0.9116 | TRUE | 0.3849 |
| Pcdhb4 | 1 | 2 | 0 | 1 | 2 | 0 | 1 | 2 | 0 | 1 | 2 | 0 | 1 | 1 | 1 | 0.955 | -0.066 | 0.9741481 | NA | 0 | 3.3027 | 1.9774 | 1.9774 | TRUE | 0.2713 |
| Pcdhb5 | 0 | 0 | 0 | 2 | 0 | 1 | 0 | 0 | 0 | 2 | 0 | 1 | 0.44 | 0 | 1 | 4.782 | 2.258 | 0.50653358 | NA | 0 | 7.4188 | 6.7775 | 6.7775 | TRUE | 0.2804 |
| Pcdhb6 | 0 | 0 | 0 | 1 | 0 | 1 | 0 | 0 | 0 | 1 | 0 | 1 | 0.31 | 0 | 1 | 3.348 | 1.743 | 0.65064889 | NA | 0 | 10.5167 | 8.8968 | 8.8968 | TRUE | 0.2082 |
| Pcdhb7 | 0 | 2 | 0 | 3 | 4 | 1 | 0 | 2 | 0 | 2 | 4 | 1 | 1.63 | 1 | 2 | 3.833 | 1.938 | 0.27490309 | NA | 0 | 2.0334 | 1.3239 | 1.3239 | TRUE | 0.7059 |
| Pcdhb8 | 1 | 0 | 0 | 4 | 1 | 2 | 1 | 0 | 0 | 3 | 1 | 2 | 1.25 | 0 | 2 | 5.907 | 2.562 | 0.18154603 | NA | 0 | 2.6524 | 1.424 | 1.424 | TRUE | 0.4115 |
| Pcdhb9 | 0 | 0 | 0 | 0 | 0 | 0 | 0 | 0 | 0 | 0 | 0 | 0 | 0 | 0 | 0 | NA | NA | NA | NA | NA | NA | NA | NA | NA | NA |
| Pcdhga1 | 94 | 36 | 55 | 168 | 107 | 94 | 100 | 36 | 55 | 132 | 115 | 103 | 90.19 | 64 | 117 | 1.839 | 0.879 | 0.01210037 | 0.06049035 | 0.1166 | 0.0434 | 0.0762 | 0.0762 | TRUE | 1.2216 |
| Pcdhga10 | 70 | 26 | 25 | 98 | 67 | 35 | 74 | 26 | 25 | 77 | 72 | 38 | 52.14 | 42 | 62 | 1.501 | 0.586 | 0.22305502 | 0.43505698 | 0.2361 | 0.0701 | 0.1468 | 0.1468 | TRUE | 3.467 |
| Pcdhga11 | 199 | 164 | 178 | 229 | 153 | 133 | 211 | 164 | 179 | 180 | 165 | 146 | 174.02 | 185 | 164 | 0.888 | -0.171 | 0.33623678 | 0.55634982 | 0.0086 | 0.0258 | 0.017 | 0.017 | TRUE | 0.1706 |
| Pcdhga12 | 278 | 181 | 186 | 313 | 182 | 179 | 295 | 181 | 187 | 246 | 196 | 196 | 216.77 | 221 | 213 | 0.965 | -0.051 | 0.82121793 | 0.90875327 | 0.0435 | 0.0221 | 0.0326 | 0.0326 | TRUE | 0.9445 |
| Pcdhga2 | 27 | 17 | 24 | 46 | 49 | 39 | 29 | 17 | 24 | 36 | 53 | 43 | 33.55 | 23 | 44 | 1.881 | 0.912 | 0.01309993 | 0.06414534 | 0.0176 | 0.1051 | 0.0641 | 0.0641 | TRUE | 0.3039 |
| Pcdhga3 | 32 | 24 | 22 | 62 | 51 | 34 | 34 | 24 | 22 | 49 | 55 | 37 | 36.81 | 27 | 47 | 1.765 | 0.82 | 0.01892179 | 0.08350154 | 0.0142 | 0.0964 | 0.0584 | 0.0584 | TRUE | 0.3479 |
| Pcdhga4 | 13 | 12 | 21 | 45 | 26 | 28 | 14 | 12 | 21 | 35 | 28 | 31 | 23.48 | 16 | 31 | 2.011 | 1.008 | 0.01655415 | 0.07619066 | 0 | 0.1472 | 0.0793 | 0.0793 | TRUE | 0.4434 |
| Pcdhga5 | 35 | 30 | 34 | 79 | 36 | 35 | 37 | 30 | 34 | 62 | 39 | 38 | 40.07 | 34 | 46 | 1.388 | 0.473 | 0.16849558 | 0.36640749 | 0.0309 | 0.0891 | 0.0593 | 0.0593 | TRUE | 0.7851 |
| Pcdhga6 | 5 | 2 | 2 | 15 | 11 | 3 | 5 | 2 | 2 | 12 | 12 | 3 | 6.04 | 3 | 9 | 2.93 | 1.551 | 0.0896689 | 0.24349956 | 0.191 | 0.5525 | 0.3822 | 0.3822 | TRUE | 0.8978 |
| Pcdhga7 | 23 | 25 | 16 | 61 | 52 | 40 | 24 | 25 | 16 | 48 | 56 | 44 | 35.53 | 22 | 49 | 2.257 | 1.174 | 0.00073174 | 0.00699527 | 0 | 0.0996 | 0.0538 | 0.0538 | TRUE | 0.3064 |
| Pcdhga8 | 17 | 7 | 4 | 28 | 18 | 7 | 18 | 7 | 4 | 22 | 19 | 8 | 13.02 | 10 | 16 | 1.706 | 0.771 | 0.27908729 | 0.50014371 | 0.3081 | 0.26 | 0.2827 | 0.2827 | TRUE | 1.8604 |
| Pcdhga9 | 56 | 31 | 32 | 46 | 33 | 29 | 59 | 31 | 32 | 36 | 36 | 32 | 37.66 | 41 | 35 | 0.847 | -0.24 | 0.51881434 | 0.71026891 | 0.0506 | 0.0944 | 0.0727 | 0.0727 | TRUE | 1.1731 |
| Pcdhgb1 | 13 | 5 | 5 | 8 | 7 | 6 | 14 | 5 | 5 | 6 | 8 | 7 | 7.37 | 8 | 7 | 0.858 | -0.221 | 0.76650522 | 0.87786206 | 0.0772 | 0.454 | 0.264 | 0.264 | TRUE | 1.1699 |
| Pcdhgb2 | 24 | 24 | 15 | 57 | 27 | 19 | 25 | 24 | 15 | 45 | 29 | 21 | 26.53 | 21 | 32 | 1.483 | 0.569 | 0.20435313 | 0.41273627 | 0.0794 | 0.1311 | 0.1056 | 0.1056 | TRUE | 0.3682 |
| Pcdhgb4 | 50 | 32 | 18 | 54 | 49 | 25 | 53 | 32 | 18 | 42 | 53 | 27 | 37.61 | 34 | 41 | 1.192 | 0.253 | 0.5787503 | 0.75588355 | 0.1543 | 0.0945 | 0.1227 | 0.1227 | TRUE | 0.3926 |
| Pcdhgb5 | 31 | 22 | 18 | 54 | 35 | 22 | 33 | 22 | 18 | 42 | 38 | 24 | 29.53 | 24 | 35 | 1.439 | 0.526 | 0.20187537 | 0.40940958 | 0.0524 | 0.1185 | 0.0871 | 0.0871 | TRUE | 0.5552 |
| Pcdhgb6 | 13 | 6 | 11 | 31 | 21 | 16 | 14 | 6 | 11 | 24 | 23 | 18 | 15.89 | 10 | 22 | 2.106 | 1.074 | 0.03798376 | 0.13547712 | 0 | 0.2143 | 0.1206 | 0.1206 | TRUE | 0.4802 |
| Pcdhgb7 | 48 | 28 | 31 | 45 | 45 | 36 | 51 | 28 | 31 | 35 | 48 | 39 | 38.88 | 37 | 41 | 1.118 | 0.161 | 0.6539249 | 0.80556928 | 0.0393 | 0.0916 | 0.0665 | 0.0665 | TRUE | 0.8187 |
| Pcdhgb8 | 50 | 72 | 95 | 87 | 61 | 76 | 53 | 72 | 95 | 68 | 66 | 83 | 72.94 | 73 | 72 | 0.983 | -0.025 | 0.9295596 | 0.9654152 | 0.0354 | 0.0521 | 0.044 | 0.044 | TRUE | 0.2458 |
| Pcdhgc3 | 281 | 338 | 406 | 396 | 241 | 259 | 298 | 338 | 408 | 311 | 259 | 284 | 316.3 | 348 | 285 | 0.819 | -0.287 | 0.07390656 | 0.21345069 | 0.0137 | 0.0173 | 0.0154 | 0.0154 | TRUE | 0.258 |
| Pcdhgc4 | 5 | 3 | 6 | 4 | 3 | 4 | 5 | 3 | 6 | 3 | 3 | 4 | 4.18 | 5 | 3 | 0.746 | -0.422 | 0.64063683 | NA | 0 | 0.795 | 0.3454 | 0.3454 | TRUE | 0.212 |
| Pcdhgc5 | 13 | 21 | 24 | 25 | 7 | 17 | 14 | 21 | 24 | 20 | 8 | 19 | 17.45 | 20 | 16 | 0.783 | -0.352 | 0.50490846 | 0.69967429 | 0.0799 | 0.1957 | 0.1436 | 0.1436 | TRUE | 0.8393 |
| Pced1a | 515 | 753 | 815 | 905 | 757 | 719 | 547 | 752 | 818 | 711 | 814 | 787 | 738.4 | 706 | 771 | 1.091 | 0.126 | 0.42893403 | 0.63864321 | 0.0224 | 0.0114 | 0.017 | 0.017 | TRUE | 0.4555 |
| Pced1b | 1402 | 1360 | 1491 | 1718 | 1418 | 1474 | 1489 | 1358 | 1497 | 1351 | 1526 | 1614 | 1472.38 | 1448 | 1497 | 1.033 | 0.046 | 0.64741933 | 0.80206565 | 0.005 | 0.0091 | 0.0067 | 0.0067 | TRUE | 0.0899 |
| Pcf11 | 3433 | 2894 | 2438 | 3526 | 2315 | 1897 | 3646 | 2890 | 2448 | 2772 | 2491 | 2077 | 2720.66 | 2995 | 2447 | 0.817 | -0.291 | 0.07530164 | 0.2162129 | 0.0301 | 0.0081 | 0.019 | 0.019 | TRUE | 0.3291 |
| Pcgf1 | 368 | 522 | 522 | 468 | 383 | 389 | 391 | 521 | 524 | 368 | 412 | 426 | 440.36 | 479 | 402 | 0.838 | -0.255 | 0.09275607 | 0.24873893 | 0.0142 | 0.0144 | 0.0143 | 0.0143 | TRUE | 0.2976 |
| Pcgf2 | 4813 | 4861 | 4805 | 5510 | 4130 | 3942 | 5112 | 4855 | 4825 | 4332 | 4444 | 4315 | 4647 | 4931 | 4364 | 0.885 | -0.176 | 0.00503677 | 0.03151273 | 4.00E-04 | 0.0076 | 0.0026 | 0.0026 | TRUE | 0.0126 |
| Pcgf3 | 3775 | 3732 | 3711 | 5221 | 3721 | 3585 | 4010 | 3727 | 3726 | 4104 | 4004 | 3924 | 3915.9 | 3821 | 4011 | 1.05 | 0.07 | 0.30559381 | 0.52646921 | 8.00E-04 | 0.0077 | 0.0031 | 0.0031 | TRUE | 0.0226 |
| Pcgf5 | 938 | 1330 | 1340 | 1486 | 1156 | 1282 | 996 | 1328 | 1346 | 1168 | 1244 | 1403 | 1247.57 | 1223 | 1272 | 1.039 | 0.055 | 0.6976442 | 0.8343376 | 0.0173 | 0.0095 | 0.0136 | 0.0136 | TRUE | 0.3152 |
| Pcgf6 | 3205 | 2668 | 2808 | 4100 | 2888 | 2657 | 3404 | 2665 | 2820 | 3223 | 3107 | 2909 | 3021.21 | 2963 | 3080 | 1.04 | 0.056 | 0.61682577 | 0.7832683 | 0.0094 | 0.008 | 0.0088 | 0.0088 | TRUE | 0.2061 |
| Pcid2 | 4030 | 4101 | 4055 | 5624 | 3991 | 3894 | 4281 | 4096 | 4072 | 4421 | 4294 | 4263 | 4237.62 | 4150 | 4326 | 1.043 | 0.06 | 0.33440233 | 0.5548783 | 3.00E-04 | 0.0077 | 0.0026 | 0.0026 | TRUE | 0.0093 |
| Pcif1 | 2158 | 2543 | 2720 | 2629 | 1883 | 2177 | 2292 | 2540 | 2731 | 2067 | 2026 | 2383 | 2339.81 | 2521 | 2159 | 0.856 | -0.224 | 0.03525311 | 0.12870648 | 0.0073 | 0.0083 | 0.0078 | 0.0078 | TRUE | 0.0995 |
| Pck1 | 6 | 1 | 0 | 5 | 4 | 1 | 6 | 1 | 0 | 4 | 4 | 1 | 2.78 | 2 | 3 | 1.286 | 0.363 | 0.79655092 | NA | 0.8929 | 1.1905 | 1.0586 | 1.0586 | TRUE | 2.0129 |
| Pck2 | 550 | 744 | 770 | 867 | 722 | 645 | 584 | 743 | 773 | 682 | 777 | 706 | 710.81 | 700 | 722 | 1.03 | 0.042 | 0.75573383 | 0.87098531 | 0.0118 | 0.0115 | 0.0117 | 0.0117 | TRUE | 0.2464 |
| Pclo | 108 | 43 | 47 | 174 | 124 | 80 | 115 | 43 | 47 | 137 | 133 | 88 | 93.77 | 68 | 119 | 1.752 | 0.809 | 0.03738297 | 0.13412289 | 0.1647 | 0.042 | 0.0972 | 0.0972 | TRUE | 3.1141 |
| Pcm1 | 12819 | 13268 | 14689 | 18940 | 13771 | 13102 | 13616 | 13251 | 14749 | 14889 | 14817 | 14342 | 14277.39 | 13872 | 14683 | 1.058 | 0.082 | 0.24135345 | 0.45807782 | 0.0017 | 0.0071 | 0.0035 | 0.0035 | TRUE | 0.0374 |
| Pcmt1 | 2124 | 2341 | 2365 | 2516 | 1793 | 1846 | 2256 | 2338 | 2375 | 1978 | 1929 | 2021 | 2149.42 | 2323 | 1976 | 0.851 | -0.234 | 0.00069566 | 0.00673539 | 0 | 0.0084 | 0.0029 | 0.0029 | TRUE | 0.0076 |
| Pcmtd1 | 2018 | 2326 | 2464 | 3305 | 2395 | 2580 | 2143 | 2323 | 2474 | 2598 | 2577 | 2824 | 2489.97 | 2313 | 2666 | 1.152 | 0.204 | 0.02127554 | 0.09083859 | 0.0033 | 0.0082 | 0.0053 | 0.0053 | TRUE | 0.0499 |
| Pcmtd2 | 1541 | 1767 | 1787 | 1869 | 1380 | 1568 | 1637 | 1765 | 1794 | 1469 | 1485 | 1716 | 1644.39 | 1732 | 1557 | 0.898 | -0.155 | 0.11130648 | 0.28056988 | 0.0042 | 0.0089 | 0.0062 | 0.0062 | TRUE | 0.096 |
| Pcna | 8548 | 8415 | 8332 | 12399 | 8505 | 8027 | 9079 | 8404 | 8366 | 9747 | 9151 | 8787 | 8922.44 | 8616 | 9228 | 1.071 | 0.099 | 0.19054078 | 0.39568257 | 0.0023 | 0.0073 | 0.004 | 0.004 | TRUE | 0.0297 |

| | | | | | | | | | | | | | | | | | | | | | | | | | |
|---|---|---|---|---|---|---|---|---|---|---|---|---|---|---|---|---|---|---|---|---|---|---|---|---|---|
| Pcnp | 2296 | 2583 | 2484 | 2773 | 2017 | 2422 | 2439 | 2580 | 2494 | 2180 | 2170 | 2651 | 2418.99 | 2504 | 2334 | 0.932 | -0.102 | 0.3226374 | 0.543774 | 0.0066 | 0.0083 | 0.0073 | 0.0073 | TRUE | 0.1705 |
| Pcnt | 7517 | 6382 | 7127 | 9915 | 7153 | 6605 | 7984 | 6374 | 7156 | 7794 | 7696 | 7230 | 7372.53 | 7171 | 7573 | 1.056 | 0.079 | 0.43225588 | 0.64183202 | 0.0062 | 0.0074 | 0.0071 | 0.0071 | TRUE | 0.1199 |
| Pcnx | 2080 | 2183 | 2506 | 3351 | 2487 | 2642 | 2209 | 2180 | 2516 | 2634 | 2676 | 2892 | 2518.01 | 2302 | 2734 | 1.187 | 0.248 | 0.00685982 | 0.0395587 | 0.004 | 0.0082 | 0.0057 | 0.0057 | TRUE | 0.0805 |
| Pcnxl2 | 129 | 107 | 102 | 125 | 94 | 83 | 137 | 107 | 102 | 98 | 101 | 91 | 106.09 | 115 | 97 | 0.84 | -0.252 | 0.22609647 | 0.43894554 | 0.0061 | 0.038 | 0.0218 | 0.0218 | TRUE | 0.264 |
| Pcnxl3 | 3490 | 3102 | 3200 | 4275 | 3292 | 2921 | 3707 | 3098 | 3213 | 3361 | 3542 | 3197 | 3353.05 | 3339 | 3367 | 1.008 | 0.012 | 0.90224615 | 0.95205059 | 0.0055 | 0.0079 | 0.0065 | 0.0065 | TRUE | 0.1125 |
| Pcnxl4 | 1272 | 1153 | 1246 | 1752 | 1196 | 1161 | 1351 | 1152 | 1251 | 1377 | 1287 | 1271 | 1281.45 | 1251 | 1312 | 1.049 | 0.069 | 0.4694211 | 0.67151259 | 0.0033 | 0.0095 | 0.0058 | 0.0058 | TRUE | 0.0584 |
| Pcolce | 2564 | 2970 | 3366 | 2848 | 2286 | 2768 | 2723 | 2966 | 3380 | 2239 | 2460 | 3030 | 2799.65 | 3023 | 2576 | 0.852 | -0.231 | 0.09096702 | 0.24600797 | 0.018 | 0.0081 | 0.0132 | 0.0132 | TRUE | 0.2876 |
| Pcolce2 | 1923 | 1824 | 1652 | 1878 | 1315 | 1313 | 2043 | 1822 | 1659 | 1476 | 1415 | 1437 | 1641.91 | 1841 | 1443 | 0.784 | -0.351 | 0.00048836 | 0.00508071 | 0.0048 | 0.0089 | 0.0067 | 0.0067 | TRUE | 0.1105 |
| Pcp2 | 0 | 0 | 0 | 2 | 1 | 2 | 0 | 0 | 0 | 2 | 1 | 2 | 0.81 | 0 | 2 | 8.576 | 3.1 | 0.18121756 | NA | 0 | 4.093 | 2.5705 | 2.5705 | TRUE | 0.07 |
| Pcp4 | 2 | 4 | 2 | 4 | 3 | 3 | 2 | 4 | 2 | 3 | 3 | 3 | 2.96 | 3 | 3 | 1.184 | 0.244 | 0.81667751 | NA | 0 | 1.1184 | 0.4587 | 0.4587 | TRUE | 0.2084 |
| Pcp4l1 | 25 | 19 | 16 | 30 | 23 | 22 | 27 | 19 | 16 | 24 | 25 | 24 | 22.33 | 21 | 24 | 1.177 | 0.235 | 0.56608449 | 0.74613777 | 0 | 0.1544 | 0.0761 | 0.0761 | TRUE | 0.3472 |
| Pcsk1 | 18 | 31 | 25 | 109 | 63 | 55 | 19 | 31 | 25 | 86 | 68 | 60 | 48.14 | 25 | 71 | 2.849 | 1.511 | 3.50E-06 | 8.15E-05 | 0.0198 | 0.0753 | 0.0492 | 0.0492 | TRUE | 0.3138 |
| Pcsk1n | 900 | 1224 | 1139 | 1406 | 1271 | 1663 | 956 | 1222 | 1144 | 1105 | 1368 | 1820 | 1269.21 | 1107 | 1431 | 1.291 | 0.369 | 0.04454196 | 0.15176405 | 0.0384 | 0.0095 | 0.0235 | 0.0235 | TRUE | 0.3189 |
| Pcsk2 | 1 | 0 | 0 | 1 | 4 | 2 | 1 | 0 | 0 | 1 | 4 | 2 | 1.39 | 0 | 2 | 6.532 | 2.708 | 0.15780005 | NA | 0 | 2.3766 | 1.4429 | 1.4429 | TRUE | 0.3186 |
| Pcsk4 | 60 | 105 | 112 | 65 | 63 | 75 | 64 | 105 | 112 | 51 | 68 | 82 | 80.34 | 94 | 67 | 0.71 | -0.493 | 0.10389339 | 0.26817827 | 0.0594 | 0.0479 | 0.0535 | 0.0535 | TRUE | 0.597 |
| Pcsk5 | 11 | 9 | 8 | 39 | 33 | 26 | 12 | 9 | 8 | 31 | 36 | 28 | 20.56 | 10 | 32 | 3.302 | 1.724 | 0.00014668 | 0.00187171 | 0 | 0.1672 | 0.0795 | 0.0795 | TRUE | 0.119 |
| Pcsk6 | 1579 | 1887 | 1920 | 2612 | 1805 | 1778 | 1677 | 1885 | 1928 | 2053 | 1942 | 1946 | 1905.22 | 1830 | 1980 | 1.083 | 0.114 | 0.18986209 | 0.39508927 | 0.0024 | 0.0086 | 0.005 | 0.005 | TRUE | 0.0641 |
| Pcsk7 | 520 | 360 | 330 | 662 | 446 | 359 | 552 | 360 | 331 | 520 | 480 | 393 | 439.41 | 414 | 464 | 1.122 | 0.166 | 0.43023632 | 0.64003841 | 0.0465 | 0.0144 | 0.0298 | 0.0298 | TRUE | 0.9739 |
| Pcsk9 | 157 | 159 | 191 | 82 | 68 | 76 | 167 | 159 | 192 | 64 | 73 | 83 | 123.03 | 173 | 73 | 0.424 | -1.236 | 6.71E-10 | 4.28E-08 | 0 | 0.0337 | 0.0193 | 0.0193 | TRUE | 0.125 |
| Pctp | 1270 | 1226 | 1136 | 1273 | 863 | 788 | 1349 | 1224 | 1141 | 1001 | 929 | 863 | 1084.32 | 1238 | 931 | 0.753 | -0.41 | 0.00012952 | 0.00170093 | 0.0054 | 0.0099 | 0.0073 | 0.0073 | TRUE | 0.0734 |
| Pcx | 2847 | 3233 | 3409 | 2962 | 2230 | 2530 | 3024 | 3229 | 3423 | 2328 | 2399 | 2769 | 2862.19 | 3225 | 2499 | 0.774 | -0.369 | 0.00020828 | 0.00252229 | 0.0059 | 0.0081 | 0.0068 | 0.0068 | TRUE | 0.1081 |
| Pcyox1 | 3463 | 3382 | 3365 | 4083 | 3023 | 2989 | 3678 | 3378 | 3379 | 3210 | 3253 | 3272 | 3361.5 | 3478 | 3245 | 0.933 | -0.1 | 0.15485318 | 0.34702251 | 8.00E-04 | 0.0079 | 0.0033 | 0.0033 | TRUE | 0.0307 |
| Pcyox1l | 1002 | 1097 | 1068 | 1054 | 845 | 802 | 1064 | 1096 | 1072 | 829 | 909 | 878 | 974.65 | 1077 | 872 | 0.808 | -0.307 | 0.00029454 | 0.0033661 | 0 | 0.0103 | 0.0041 | 0.0041 | TRUE | 0.0237 |
| Pcyt1a | 1454 | 1548 | 1508 | 2428 | 1652 | 1440 | 1544 | 1546 | 1514 | 1909 | 1777 | 1576 | 1644.51 | 1535 | 1754 | 1.144 | 0.194 | 0.04409479 | 0.1505983 | 0.0042 | 0.0089 | 0.0061 | 0.0061 | TRUE | 0.0944 |
| Pcyt1b | 2764 | 2405 | 2550 | 2516 | 1898 | 2020 | 2936 | 2402 | 2560 | 1978 | 2042 | 2211 | 2354.91 | 2633 | 2077 | 0.789 | -0.342 | 0.00094306 | 0.00850777 | 0.0065 | 0.0083 | 0.0073 | 0.0073 | TRUE | 0.1228 |
| Pcyt2 | 2179 | 1796 | 1684 | 2422 | 1929 | 1708 | 2314 | 1794 | 1691 | 1904 | 2075 | 1870 | 1941.37 | 1933 | 1950 | 1.009 | 0.013 | 0.92482325 | 0.96292 | 0.0152 | 0.0086 | 0.0122 | 0.0122 | TRUE | 0.3596 |
| Pdap1 | 7850 | 7157 | 7246 | 10034 | 7145 | 6374 | 8338 | 7148 | 7276 | 7888 | 7688 | 6977 | 7552.41 | 7587 | 7518 | 0.991 | -0.013 | 0.88944455 | 0.94613336 | 0.0045 | 0.0073 | 0.0062 | 0.0062 | TRUE | 0.0914 |
| Pdc | 0 | 2 | 0 | 7 | 1 | 4 | 0 | 2 | 0 | 6 | 1 | 4 | 2.16 | 1 | 4 | 5.468 | 2.451 | 0.14267827 | NA | 0 | 1.5327 | 1.1245 | 1.1245 | TRUE | 0.9428 |
| Pdcd1 | 12 | 12 | 4 | 16 | 11 | 12 | 13 | 12 | 4 | 13 | 12 | 13 | 11.05 | 10 | 13 | 1.309 | 0.389 | 0.52998318 | 0.7181798 | 0 | 0.3051 | 0.1833 | 0.1833 | TRUE | 0.8808 |
| Pdcd10 | 1767 | 1842 | 1711 | 2454 | 1689 | 1708 | 1877 | 1840 | 1718 | 1929 | 1817 | 1870 | 1841.77 | 1812 | 1872 | 1.034 | 0.048 | 0.53392911 | 0.72173954 | 0.001 | 0.0087 | 0.0038 | 0.0038 | TRUE | 0.0247 |
| Pdcd11 | 12012 | 8891 | 9201 | 13699 | 9610 | 7947 | 12759 | 8880 | 9239 | 10769 | 10340 | 8699 | 10114.21 | 10293 | 9936 | 0.965 | -0.051 | 0.74065668 | 0.86141414 | 0.0229 | 0.0072 | 0.0168 | 0.0168 | TRUE | 0.5369 |
| Pdcd1lg2 | 0 | 1 | 1 | 0 | 0 | 0 | 0 | 1 | 1 | 0 | 0 | 0 | 0.33 | 1 | 0 | 0.268 | -1.901 | 0.61401143 | NA | 0 | 9.8764 | 8.4772 | 8.4772 | TRUE | 0.2278 |
| Pdcd2 | 1623 | 2071 | 2131 | 2513 | 1869 | 1874 | 1724 | 2068 | 2140 | 1976 | 2011 | 2051 | 1994.97 | 1977 | 2013 | 1.018 | 0.025 | 0.80799901 | 0.90112858 | 0.0062 | 0.0086 | 0.0072 | 0.0072 | TRUE | 0.1515 |
| Pdcd2l | 1589 | 1886 | 1895 | 2089 | 1665 | 1687 | 1688 | 1884 | 1903 | 1642 | 1791 | 1847 | 1792.41 | 1825 | 1760 | 0.964 | -0.053 | 0.56361043 | 0.74449445 | 0.0035 | 0.0087 | 0.0055 | 0.0055 | TRUE | 0.0518 |
| Pdcd4 | 2245 | 2859 | 2799 | 2673 | 2150 | 2535 | 2385 | 2855 | 2810 | 2101 | 2313 | 2775 | 2539.99 | 2683 | 2396 | 0.893 | -0.164 | 0.20418248 | 0.41249726 | 0.0144 | 0.0082 | 0.0116 | 0.0116 | TRUE | 0.2309 |
| Pdcd5 | 1852 | 1929 | 1982 | 2780 | 2060 | 1814 | 1967 | 1927 | 1990 | 2185 | 2216 | 1986 | 2045.23 | 1961 | 2129 | 1.086 | 0.119 | 0.13093867 | 0.31162492 | 0.0014 | 0.0085 | 0.004 | 0.004 | TRUE | 0.0414 |
| Pdcd6 | 2246 | 2506 | 2548 | 3244 | 2317 | 2237 | 2386 | 2503 | 2558 | 2550 | 2493 | 2449 | 2489.79 | 2482 | 2497 | 1.006 | 0.009 | 0.89744505 | 0.95022175 | 4.00E-04 | 0.0082 | 0.0031 | 0.0031 | TRUE | 0.014 |
| Pdcd6ip | 8142 | 8657 | 8978 | 10361 | 7630 | 8261 | 8648 | 8646 | 9015 | 8145 | 8209 | 9043 | 8617.72 | 8770 | 8466 | 0.965 | -0.051 | 0.48215461 | 0.68089732 | 0.0019 | 0.0073 | 0.0037 | 0.0037 | TRUE | 0.0433 |
| Pdcd7 | 1407 | 1910 | 1892 | 2535 | 1985 | 1911 | 1494 | 1908 | 1900 | 1993 | 2136 | 2092 | 1920.37 | 1767 | 2074 | 1.173 | 0.23 | 0.04623914 | 0.15563032 | 0.0088 | 0.0086 | 0.0091 | 0.0091 | TRUE | 0.2194 |
| Pdcl | 3054 | 3167 | 3003 | 4547 | 3319 | 3333 | 3244 | 3163 | 3015 | 3574 | 3571 | 3648 | 3369.36 | 3141 | 3598 | 1.146 | 0.196 | 0.00296691 | 0.02105946 | 4.00E-04 | 0.0079 | 0.0028 | 0.0028 | TRUE | 0.0148 |
| Pdcl2 | 144 | 159 | 170 | 179 | 139 | 184 | 153 | 159 | 171 | 141 | 150 | 201 | 162.36 | 161 | 164 | 1.015 | 0.022 | 0.90949969 | 0.95489676 | 0.0147 | 0.0272 | 0.0208 | 0.0208 | TRUE | 0.4152 |
| Pdcl3 | 2715 | 2824 | 2610 | 3274 | 2382 | 2461 | 2884 | 2820 | 2621 | 2574 | 2563 | 2694 | 2692.58 | 2775 | 2610 | 0.941 | -0.088 | 0.23640572 | 0.45162196 | 0.0011 | 0.0081 | 0.0036 | 0.0036 | TRUE | 0.0287 |
| Pddc1 | 1243 | 1615 | 1726 | 1725 | 1303 | 1352 | 1320 | 1613 | 1733 | 1356 | 1402 | 1480 | 1484.05 | 1555 | 1413 | 0.908 | -0.14 | 0.24259359 | 0.4591056 | 0.0093 | 0.0091 | 0.0096 | 0.0096 | TRUE | 0.21 |
| Pde10a | 58 | 104 | 91 | 166 | 86 | 107 | 62 | 104 | 91 | 130 | 93 | 117 | 99.5 | 86 | 113 | 1.326 | 0.407 | 0.11874263 | 0.29185114 | 0.0375 | 0.04 | 0.0388 | 0.0388 | TRUE | 0.5563 |
| Pde11a | 14 | 19 | 31 | 32 | 20 | 24 | 15 | 19 | 31 | 25 | 22 | 26 | 22.99 | 22 | 24 | 1.122 | 0.166 | 0.704259 | 0.83814573 | 0.0285 | 0.1502 | 0.0935 | 0.0935 | TRUE | 0.83 |
| Pde12 | 1789 | 2100 | 2064 | 2383 | 1695 | 1822 | 1900 | 2097 | 2072 | 1873 | 1824 | 1994 | 1960.25 | 2023 | 1897 | 0.937 | -0.093 | 0.25943852 | 0.4784696 | 0.0017 | 0.0086 | 0.0044 | 0.0044 | TRUE | 0.0341 |
| Pde1a | 0 | 0 | 2 | 1 | 2 | 0 | 0 | 0 | 2 | 1 | 2 | 0 | 0.82 | 1 | 1 | 1.45 | 0.536 | 0.82805985 | NA | 0 | 4.0032 | 3.1289 | 3.1289 | TRUE | 0.6991 |
| Pde1b | 573 | 835 | 909 | 620 | 559 | 663 | 609 | 834 | 913 | 487 | 601 | 726 | 694.98 | 785 | 605 | 0.769 | -0.378 | 0.04915551 | 0.16239647 | 0.0374 | 0.0116 | 0.0252 | 0.0252 | TRUE | 0.3445 |
| Pde1c | 21 | 20 | 25 | 49 | 33 | 31 | 22 | 20 | 25 | 39 | 36 | 34 | 29.22 | 22 | 36 | 1.606 | 0.684 | 0.05178051 | 0.16790341 | 0 | 0.1196 | 0.0528 | 0.0528 | TRUE | 0.0623 |
| Pde2a | 1289 | 1181 | 1151 | 976 | 702 | 714 | 1369 | 1179 | 1156 | 767 | 755 | 782 | 1001.42 | 1235 | 768 | 0.622 | -0.685 | 1.72E-11 | 1.46E-09 | 0.0038 | 0.0102 | 0.0064 | 0.0064 | TRUE | 0.1082 |
| Pde3a | 27 | 25 | 29 | 37 | 34 | 18 | 29 | 25 | 29 | 29 | 37 | 20 | 28.02 | 28 | 29 | 1.033 | 0.047 | 0.90254193 | 0.95205059 | 0 | 0.1245 | 0.071 | 0.071 | TRUE | 0.2228 |
| Pde3b | 1028 | 1286 | 1333 | 1632 | 1168 | 1155 | 1092 | 1284 | 1338 | 1283 | 1257 | 1264 | 1253.12 | 1238 | 1268 | 1.024 | 0.034 | 0.73996192 | 0.86079654 | 0.0048 | 0.0095 | 0.0068 | 0.0068 | TRUE | 0.1275 |
| Pde4a | 471 | 633 | 634 | 853 | 685 | 677 | 500 | 632 | 637 | 671 | 737 | 741 | 652.96 | 590 | 716 | 1.213 | 0.279 | 0.02927138 | 0.11304316 | 0.0088 | 0.0119 | 0.0102 | 0.0102 | TRUE | 0.2049 |
| Pde4b | 9 | 4 | 8 | 5 | 13 | 8 | 10 | 4 | 8 | 4 | 14 | 9 | 8.04 | 7 | 9 | 1.219 | 0.286 | 0.70696387 | 0.84000559 | 0.1613 | 0.4165 | 0.289 | 0.289 | TRUE | 0.1825 |
| Pde4c | 325 | 482 | 424 | 291 | 305 | 292 | 345 | 481 | 426 | 229 | 328 | 320 | 354.81 | 417 | 292 | 0.698 | -0.519 | 0.00638222 | 0.03768643 | 0.0302 | 0.0162 | 0.0232 | 0.0232 | TRUE | 0.3644 |
| Pde4d | 631 | 543 | 547 | 586 | 398 | 378 | 670 | 542 | 549 | 461 | 428 | 414 | 510.74 | 587 | 434 | 0.741 | -0.433 | 0.0007472 | 0.00709506 | 0.0071 | 0.0134 | 0.0099 | 0.0099 | TRUE | 0.1781 |
| Pde4dip | 2392 | 2277 | 2280 | 2447 | 1790 | 1894 | 2541 | 2274 | 2289 | 1924 | 1926 | 2073 | 2171.17 | 2368 | 1974 | 0.834 | -0.263 | 0.00185954 | 0.01451196 | 0.0022 | 0.0084 | 0.0047 | 0.0047 | TRUE | 0.0491 |
| Pde5a | 230 | 287 | 287 | 455 | 301 | 300 | 244 | 287 | 288 | 358 | 324 | 328 | 304.84 | 273 | 337 | 1.234 | 0.303 | 0.02300935 | 0.09608248 | 0.0021 | 0.0177 | 0.0095 | 0.0095 | TRUE | 0.0935 |
| Pde6a | 9 | 13 | 10 | 21 | 16 | 19 | 10 | 13 | 10 | 17 | 17 | 21 | 14.52 | 11 | 18 | 1.665 | 0.736 | 0.13952025 | 0.32412923 | 0 | 0.2338 | 0.1048 | 0.1048 | TRUE | 0.1092 |
| Pde6b | 4 | 1 | 2 | 3 | 3 | 3 | 4 | 1 | 2 | 2 | 3 | 3 | 2.69 | 2 | 3 | 1.221 | 0.288 | 0.80126674 | NA | 0 | 1.2327 | 0.563 | 0.563 | TRUE | 0.4439 |
| Pde6c | 0 | 0 | 0 | 1 | 0 | 0 | 0 | 0 | 0 | 1 | 0 | 0 | 0.13 | 0 | 0 | 1.859 | 0.894 | 0.82650734 | NA | 0 | 25.1515 | 10 | 10 | TRUE | 0.2563 |
| Pde6d | 1176 | 1335 | 1283 | 1592 | 1151 | 1084 | 1249 | 1333 | 1288 | 1252 | 1238 | 1187 | 1257.86 | 1290 | 1226 | 0.95 | -0.074 | 0.34684846 | 0.56592295 | 0 | 0.0095 | 0.0036 | 0.0036 | TRUE | 0.0102 |
| Pde6g | 3 | 10 | 4 | 3 | 1 | 6 | 3 | 10 | 4 | 2 | 1 | 7 | 4.53 | 6 | 3 | 0.573 | -0.804 | 0.43065885 | NA | 0.2818 | 0.7338 | 0.5124 | 0.5124 | TRUE | 0.9677 |
| Pde6h | 0 | 3 | 0 | 1 | 0 | 0 | 0 | 3 | 0 | 1 | 0 | 0 | 0.63 | 1 | 0 | 0.348 | -1.521 | 0.63222046 | NA | 7.8645 | 5.2331 | 5.7681 | 5.7681 | TRUE | 1.4421 |
| Pde7a | 1733 | 1807 | 2083 | 2335 | 1671 | 1664 | 1841 | 1805 | 2092 | 1836 | 1798 | 1822 | 1865.33 | 1913 | 1819 | 0.951 | -0.073 | 0.40895331 | 0.62126578 | 0.0026 | 0.0087 | 0.005 | 0.005 | TRUE | 0.0814 |
| Pde7b | 3 | 5 | 1 | 0 | 0 | 0 | 3 | 5 | 1 | 0 | 0 | 0 | 1.53 | 3 | 0 | 0.059 | -4.094 | 0.03836229 | NA | 0 | 2.1592 | 1.6496 | 1.6496 | TRUE | 0.0677 |
| Pde8a | 1422 | 1891 | 2024 | 2169 | 1588 | 1831 | 1510 | 1889 | 2032 | 1705 | 1709 | 2004 | 1808.21 | 1810 | 1806 | 0.997 | -0.004 | 0.97539619 | 0.98697924 | 0.0154 | 0.0087 | 0.0123 | 0.0123 | TRUE | 0.2532 |
| Pde8b | 31 | 35 | 23 | 35 | 31 | 30 | 33 | 35 | 23 | 28 | 33 | 33 | 30.78 | 30 | 31 | 1.026 | 0.037 | 0.91703314 | 0.95930534 | 0 | 0.1139 | 0.0587 | 0.0587 | TRUE | 0.2943 |
| Pde9a | 477 | 485 | 502 | 589 | 382 | 436 | 507 | 484 | 504 | 463 | 411 | 477 | 474.4 | 498 | 450 | 0.904 | -0.145 | 0.19113892 | 0.39617243 | 7.00E-04 | 0.0138 | 0.0067 | 0.0067 | TRUE | 0.0655 |

| | | | | | | | | | | | | | | | | | | | | | | | | |
|---|---|---|---|---|---|---|---|---|---|---|---|---|---|---|---|---|---|---|---|---|---|---|---|---|
| Pdf | 729 | 752 | 835 | 1176 | 847 | 762 | 774 | 751 | 838 | 924 | 911 | 834 | 838.95 | 788 | 890 | 1.13 | 0.177 | 0.06933726 | 0.20437064 | 0.0019 | 0.0108 | 0.0056 | 0.0056 | TRUE | 0.0375 |
| Pdgfa | 485 | 897 | 797 | 747 | 694 | 684 | 515 | 896 | 800 | 587 | 747 | 749 | 715.66 | 737 | 694 | 0.941 | -0.088 | 0.66683433 | 0.81378141 | 0.0478 | 0.0115 | 0.0286 | 0.0286 | TRUE | 0.8184 |
| Pdgfb | 284 | 607 | 662 | 295 | 297 | 358 | 302 | 606 | 665 | 232 | 320 | 392 | 419.32 | 524 | 315 | 0.599 | -0.74 | 0.01058154 | 0.05499872 | 0.1134 | 0.0148 | 0.0578 | 0.0578 | TRUE | 1.1179 |
| Pdgfc | 59 | 34 | 35 | 82 | 63 | 50 | 63 | 34 | 35 | 64 | 68 | 55 | 53.12 | 44 | 62 | 1.424 | 0.51 | 0.1176167 | 0.29016904 | 0.0431 | 0.0689 | 0.0569 | 0.0569 | TRUE | 1.0563 |
| Pdgfd | 9 | 6 | 4 | 22 | 10 | 14 | 10 | 6 | 4 | 17 | 11 | 15 | 10.49 | 7 | 14 | 2.238 | 1.162 | 0.06520584 | 0.19686571 | 0 | 0.3209 | 0.1741 | 0.1741 | TRUE | 0.395 |
| Pdgfra | 344 | 308 | 350 | 531 | 359 | 318 | 365 | 308 | 351 | 417 | 386 | 348 | 362.7 | 341 | 384 | 1.127 | 0.172 | 0.20114347 | 0.40877592 | 0.0054 | 0.016 | 0.0103 | 0.0103 | TRUE | 0.0863 |
| Pdgfrb | 1205 | 1256 | 1292 | 1007 | 796 | 909 | 1280 | 1254 | 1297 | 792 | 856 | 995 | 1079.12 | 1277 | 881 | 0.689 | -0.538 | 7.73E-07 | 2.20E-05 | 0.0057 | 0.01 | 0.0076 | 0.0076 | TRUE | 0.1516 |
| Pdgfrl | 271 | 221 | 197 | 489 | 344 | 248 | 288 | 221 | 198 | 384 | 370 | 271 | 288.73 | 236 | 342 | 1.455 | 0.541 | 0.0068197 | 0.03938497 | 0.0324 | 0.0183 | 0.0253 | 0.0253 | TRUE | 0.4152 |
| Pdha1 | 10769 | 11957 | 12004 | 15289 | 11149 | 10859 | 11439 | 11942 | 12053 | 12019 | 11996 | 11887 | 11889.16 | 11811 | 11967 | 1.013 | 0.019 | 0.73768311 | 0.85954034 | 3.00E-04 | 0.0072 | 0.0022 | 0.0022 | TRUE | 0.0093 |
| Pdha2 | 139 | 66 | 58 | 257 | 194 | 135 | 148 | 66 | 58 | 202 | 209 | 148 | 138.39 | 91 | 186 | 2.06 | 1.042 | 0.00247763 | 0.01823482 | 0.1362 | 0.0307 | 0.0772 | 0.0772 | TRUE | 2.8676 |
| Pdhb | 5135 | 5040 | 4810 | 6235 | 4476 | 4370 | 5454 | 5033 | 4830 | 4901 | 4816 | 4784 | 4969.76 | 5106 | 4834 | 0.947 | -0.079 | 0.28823586 | 0.50919086 | 0.0017 | 0.0076 | 0.0038 | 0.0038 | TRUE | 0.0434 |
| Pdhx | 2362 | 2260 | 2187 | 3194 | 2120 | 2125 | 2509 | 2257 | 2196 | 2511 | 2281 | 2326 | 2346.66 | 2321 | 2373 | 1.023 | 0.033 | 0.71235856 | 0.84341563 | 0.0034 | 0.0083 | 0.0053 | 0.0053 | TRUE | 0.0608 |
| Pdia2 | 10 | 18 | 25 | 10 | 10 | 10 | 11 | 18 | 25 | 8 | 11 | 11 | 13.88 | 18 | 10 | 0.545 | -0.877 | 0.11823486 | 0.29101127 | 0 | 0.2443 | 0.1484 | 0.1484 | TRUE | 0.1163 |
| Pdia3 | 16588 | 16967 | 16283 | 23000 | 15508 | 15335 | 17619 | 16945 | 16350 | 18081 | 16686 | 16787 | 17077.89 | 16971 | 17185 | 1.013 | 0.018 | 0.79356602 | 0.89360725 | 0.0014 | 0.0071 | 0.0034 | 0.0034 | TRUE | 0.0256 |
| Pdia4 | 8962 | 7965 | 7696 | 9985 | 6927 | 6752 | 9519 | 7955 | 7728 | 7849 | 7453 | 7391 | 7982.52 | 8401 | 7564 | 0.901 | -0.151 | 0.13041679 | 0.31094991 | 0.0071 | 0.0073 | 0.0071 | 0.0071 | TRUE | 0.1656 |
| Pdia5 | 1698 | 1898 | 1897 | 1756 | 1262 | 1424 | 1804 | 1896 | 1905 | 1380 | 1358 | 1559 | 1650.16 | 1868 | 1432 | 0.766 | -0.384 | 1.49E-05 | 0.00027752 | 0.0025 | 0.0089 | 0.005 | 0.005 | TRUE | 0.0706 |
| Pdia6 | 20971 | 22290 | 20937 | 22638 | 16489 | 17285 | 22275 | 22261 | 21023 | 17796 | 17741 | 18921 | 20002.94 | 21853 | 18153 | 0.831 | -0.268 | 3.45E-05 | 0.00056367 | 0.0011 | 0.0071 | 0.003 | 0.003 | TRUE | 0.0167 |
| Pdik1l | 529 | 664 | 658 | 671 | 577 | 596 | 562 | 663 | 661 | 527 | 621 | 652 | 614.41 | 629 | 600 | 0.953 | -0.069 | 0.59131353 | 0.76506948 | 0.0089 | 0.0123 | 0.0104 | 0.0104 | TRUE | 0.1375 |
| Pdilt | 0 | 0 | 0 | 2 | 1 | 1 | 0 | 0 | 0 | 2 | 1 | 1 | 0.62 | 0 | 1 | 6.676 | 2.739 | 0.27800695 | NA | 0 | 5.2881 | 3.2034 | 3.2034 | TRUE | 0.07 |
| Pdk1 | 18839 | 23528 | 23568 | 21379 | 16562 | 17535 | 20010 | 23498 | 23665 | 16806 | 17820 | 19195 | 20165.62 | 22391 | 17940 | 0.801 | -0.32 | 0.00101539 | 0.00901726 | 0.0064 | 0.0071 | 0.0068 | 0.0068 | TRUE | 0.1058 |
| Pdk2 | 594 | 851 | 885 | 582 | 470 | 628 | 631 | 850 | 889 | 458 | 506 | 687 | 670.02 | 790 | 550 | 0.696 | -0.524 | 0.00581469 | 0.03504365 | 0.0374 | 0.0118 | 0.0244 | 0.0244 | TRUE | 0.5539 |
| Pdk3 | 2031 | 2688 | 2902 | 2901 | 2152 | 2589 | 2157 | 2685 | 2914 | 2281 | 2315 | 2834 | 2530.96 | 2585 | 2477 | 0.958 | -0.062 | 0.65621583 | 0.80670515 | 0.0188 | 0.0082 | 0.0138 | 0.0138 | TRUE | 0.254 |
| Pdk4 | 467 | 841 | 865 | 603 | 451 | 665 | 496 | 840 | 869 | 474 | 485 | 728 | 648.62 | 735 | 562 | 0.765 | -0.387 | 0.10529912 | 0.27039689 | 0.07 | 0.012 | 0.0396 | 0.0396 | TRUE | 0.9538 |
| Pdlim1 | 2304 | 2669 | 2763 | 3551 | 2560 | 2565 | 2447 | 2666 | 2774 | 2792 | 2754 | 2808 | 2706.81 | 2629 | 2785 | 1.059 | 0.083 | 0.28673664 | 0.50796477 | 0.0015 | 0.0081 | 0.004 | 0.004 | TRUE | 0.0442 |
| Pdlim2 | 136 | 165 | 158 | 167 | 132 | 132 | 144 | 165 | 159 | 131 | 142 | 144 | 147.62 | 156 | 139 | 0.891 | -0.167 | 0.314719 | 0.53540589 | 0 | 0.0292 | 0.0131 | 0.0131 | TRUE | 0.043 |
| Pdlim3 | 148 | 97 | 70 | 186 | 168 | 110 | 157 | 97 | 70 | 146 | 181 | 120 | 128.63 | 108 | 149 | 1.381 | 0.466 | 0.12936506 | 0.30962138 | 0.0928 | 0.0325 | 0.0599 | 0.0599 | TRUE | 0.8633 |
| Pdlim4 | 273 | 402 | 342 | 236 | 170 | 246 | 290 | 401 | 343 | 186 | 183 | 269 | 278.76 | 345 | 213 | 0.615 | -0.702 | 0.0005457 | 0.00556714 | 0.0341 | 0.0187 | 0.0259 | 0.0259 | TRUE | 0.3442 |
| Pdlim5 | 3103 | 3756 | 4033 | 4557 | 3254 | 3275 | 3296 | 3751 | 4050 | 3582 | 3501 | 3585 | 3627.52 | 3699 | 3556 | 0.961 | -0.057 | 0.5489746 | 0.73376503 | 0.0046 | 0.0078 | 0.0062 | 0.0062 | TRUE | 0.1103 |
| Pdlim7 | 1501 | 1193 | 1153 | 1680 | 1151 | 1002 | 1594 | 1191 | 1158 | 1321 | 1238 | 1097 | 1266.58 | 1314 | 1219 | 0.928 | -0.108 | 0.46178872 | 0.66571988 | 0.0188 | 0.0095 | 0.0148 | 0.0148 | TRUE | 0.4158 |
| Pdp1 | 809 | 932 | 1013 | 972 | 759 | 867 | 859 | 931 | 1017 | 764 | 817 | 949 | 889.51 | 936 | 843 | 0.9 | -0.152 | 0.21136837 | 0.42189873 | 0.0083 | 0.0106 | 0.0095 | 0.0095 | TRUE | 0.1422 |
| Pdp2 | 2246 | 2393 | 2397 | 2449 | 1882 | 1899 | 2386 | 2390 | 2407 | 1925 | 2025 | 2079 | 2201.88 | 2394 | 2010 | 0.839 | -0.253 | 0.00029085 | 0.00332771 | 3.00E-04 | 0.0084 | 0.0031 | 0.0031 | TRUE | 0.0169 |
| Pdpk1 | 7464 | 6923 | 7283 | 8992 | 6474 | 6289 | 7928 | 6914 | 7313 | 7069 | 6966 | 6884 | 7178.96 | 7385 | 6973 | 0.944 | -0.083 | 0.27900858 | 0.50005946 | 0.0023 | 0.0074 | 0.0041 | 0.0041 | TRUE | 0.0505 |
| Pdpn | 1617 | 2354 | 2269 | 1179 | 888 | 1179 | 1718 | 2351 | 2278 | 927 | 955 | 1291 | 1586.61 | 2116 | 1058 | 0.5 | -1.001 | 2.55E-09 | 1.39E-07 | 0.0306 | 0.009 | 0.0196 | 0.0196 | TRUE | 0.442 |
| Pdpr | 1645 | 1777 | 1943 | 2352 | 1633 | 1641 | 1747 | 1775 | 1951 | 1849 | 1757 | 1796 | 1812.54 | 1824 | 1801 | 0.987 | -0.018 | 0.82088375 | 0.90868743 | 0.0016 | 0.0087 | 0.0042 | 0.0042 | TRUE | 0.0446 |
| Pdrg1 | 981 | 1005 | 916 | 1289 | 912 | 776 | 1042 | 1004 | 920 | 1013 | 981 | 849 | 968.25 | 989 | 948 | 0.96 | -0.059 | 0.58701116 | 0.76162718 | 0.0053 | 0.0103 | 0.0074 | 0.0074 | TRUE | 0.0973 |
| Pds5a | 8151 | 6216 | 6994 | 10254 | 7174 | 6556 | 8658 | 6208 | 7023 | 8061 | 7719 | 7177 | 7474.12 | 7296 | 7652 | 1.049 | 0.069 | 0.59375371 | 0.76728255 | 0.0131 | 0.0073 | 0.0119 | 0.0119 | TRUE | 0.3252 |
| Pds5b | 5299 | 5045 | 5515 | 7294 | 5152 | 4997 | 5628 | 5038 | 5538 | 5734 | 5543 | 5470 | 5491.97 | 5401 | 5582 | 1.034 | 0.048 | 0.52037651 | 0.71109881 | 0.0017 | 0.0075 | 0.0038 | 0.0038 | TRUE | 0.0422 |
| Pdss1 | 673 | 563 | 504 | 1003 | 642 | 515 | 715 | 562 | 506 | 788 | 691 | 564 | 637.69 | 594 | 681 | 1.147 | 0.198 | 0.25388218 | 0.47188737 | 0.0278 | 0.0121 | 0.0202 | 0.0202 | TRUE | 0.3191 |
| Pdss2 | 586 | 475 | 492 | 737 | 462 | 438 | 622 | 474 | 494 | 579 | 497 | 479 | 524.46 | 530 | 518 | 0.98 | -0.029 | 0.84255 | 0.92065253 | 0.0143 | 0.0132 | 0.0138 | 0.0138 | TRUE | 0.2686 |
| Pdx1 | 1 | 2 | 1 | 9 | 3 | 1 | 1 | 2 | 1 | 7 | 3 | 1 | 2.58 | 1 | 4 | 2.864 | 1.518 | 0.2431418 | NA | 0 | 1.2854 | 0.7174 | 0.7174 | TRUE | 0.2311 |
| Pdxdc1 | 4620 | 5137 | 5151 | 6841 | 4826 | 4667 | 4907 | 5130 | 5172 | 5378 | 5192 | 5109 | 5148.15 | 5070 | 5226 | 1.031 | 0.044 | 0.48730307 | 0.68433301 | 5.00E-04 | 0.0075 | 0.0027 | 0.0027 | TRUE | 0.0096 |
| Pdxk | 2026 | 1980 | 1976 | 2330 | 1679 | 1507 | 2152 | 1977 | 1984 | 1832 | 1807 | 1650 | 1900.22 | 2038 | 1763 | 0.865 | -0.209 | 0.01368514 | 0.06610782 | 0.0019 | 0.0086 | 0.0046 | 0.0046 | TRUE | 0.0375 |
| Pdxk-ps | 7 | 11 | 6 | 5 | 2 | 3 | 7 | 11 | 6 | 4 | 2 | 3 | 5.64 | 8 | 3 | 0.388 | -1.365 | 0.09989109 | 0.26096381 | 0 | 0.5915 | 0.2794 | 0.2794 | TRUE | 0.2822 |
| Pdxp | 1019 | 1552 | 1654 | 1367 | 1027 | 1120 | 1082 | 1550 | 1661 | 1075 | 1105 | 1226 | 1283.13 | 1431 | 1135 | 0.793 | -0.335 | 0.03898979 | 0.13815928 | 0.0268 | 0.0095 | 0.0182 | 0.0182 | TRUE | 0.5454 |
| Pdyn | 32 | 22 | 22 | 24 | 15 | 10 | 34 | 22 | 22 | 19 | 16 | 11 | 20.67 | 26 | 15 | 0.596 | -0.747 | 0.09980688 | 0.26089825 | 0 | 0.1663 | 0.0974 | 0.0974 | TRUE | 0.4318 |
| Pdzd11 | 648 | 798 | 732 | 1180 | 766 | 731 | 688 | 797 | 735 | 928 | 824 | 800 | 795.38 | 740 | 851 | 1.151 | 0.202 | 0.06462645 | 0.19582871 | 0.0047 | 0.011 | 0.0074 | 0.0074 | TRUE | 0.0769 |
| Pdzd2 | 283 | 235 | 220 | 348 | 243 | 228 | 301 | 235 | 221 | 274 | 261 | 250 | 256.8 | 252 | 262 | 1.039 | 0.056 | 0.7305809 | 0.85524545 | 0.0106 | 0.0197 | 0.015 | 0.015 | TRUE | 0.3122 |
| Pdzd3 | 25 | 28 | 27 | 37 | 27 | 23 | 27 | 28 | 27 | 29 | 29 | 25 | 27.49 | 27 | 28 | 1.023 | 0.032 | 0.92720119 | 0.96418653 | 0 | 0.1267 | 0.0536 | 0.0536 | TRUE | 0.0392 |
| Pdzd4 | 5433 | 4944 | 5105 | 4337 | 3249 | 3538 | 5771 | 4938 | 5126 | 3409 | 3496 | 3873 | 4435.4 | 5278 | 3593 | 0.681 | -0.555 | 5.61E-09 | 2.82E-07 | 0.0054 | 0.0076 | 0.0063 | 0.0063 | TRUE | 0.0812 |
| Pdzd7 | 86 | 102 | 85 | 125 | 72 | 71 | 91 | 102 | 85 | 98 | 77 | 78 | 88.67 | 93 | 84 | 0.915 | -0.129 | 0.5615813 | 0.74269575 | 0 | 0.0441 | 0.0241 | 0.0241 | TRUE | 0.214 |
| Pdzd8 | 2272 | 2250 | 2291 | 4087 | 2911 | 2539 | 2413 | 2247 | 2300 | 3213 | 3132 | 2779 | 2680.84 | 2320 | 3041 | 1.311 | 0.391 | 7.54E-06 | 0.0001558 | 0.003 | 0.0081 | 0.0051 | 0.0051 | TRUE | 0.0686 |
| Pdzd9 | 0 | 0 | 3 | 1 | 0 | 0 | 0 | 0 | 3 | 1 | 0 | 0 | 0.63 | 1 | 0 | 0.347 | -1.528 | 0.63042337 | NA | 7.8967 | 5.2108 | 5.757 | 5.757 | TRUE | 1.4422 |
| Pdzk1 | 589 | 840 | 821 | 244 | 207 | 305 | 626 | 839 | 824 | 192 | 223 | 334 | 506.22 | 763 | 250 | 0.326 | -1.617 | 5.17E-14 | 6.36E-12 | 0.0531 | 0.0134 | 0.0306 | 0.0306 | TRUE | 0.9639 |
| Pdzk1ip1 | 12 | 25 | 14 | 17 | 10 | 10 | 13 | 25 | 14 | 13 | 11 | 11 | 14.47 | 17 | 12 | 0.68 | -0.556 | 0.29484311 | 0.51583645 | 0 | 0.2345 | 0.132 | 0.132 | TRUE | 0.7675 |
| Pdzrn3 | 307 | 190 | 194 | 425 | 284 | 269 | 326 | 190 | 195 | 334 | 306 | 294 | 274.13 | 237 | 311 | 1.317 | 0.397 | 0.07417756 | 0.21399826 | 0.0459 | 0.0189 | 0.0319 | 0.0319 | TRUE | 1.192 |
| Pdzrn4 | 3 | 11 | 4 | 4 | 4 | 4 | 3 | 11 | 4 | 3 | 4 | 4 | 5 | 6 | 4 | 0.643 | -0.638 | 0.47914702 | 0.67843674 | 0 | 0.6655 | 0.3768 | 0.3768 | TRUE | 1.2102 |
| Pea15a | 1023 | 939 | 939 | 1016 | 689 | 663 | 1087 | 938 | 943 | 799 | 741 | 726 | 872.17 | 989 | 755 | 0.765 | -0.387 | 0.00020568 | 0.00250114 | 0.0038 | 0.0107 | 0.0067 | 0.0067 | TRUE | 0.0883 |
| Pea15b | 0 | 0 | 0 | 0 | 0 | 0 | 0 | 0 | 0 | 0 | 0 | 0 | 0 | 0 | 0 | NA | NA | NA | NA | NA | NA | NA | NA | NA | NA |
| Pear1 | 119 | 125 | 108 | 138 | 112 | 86 | 126 | 125 | 108 | 108 | 121 | 94 | 113.8 | 120 | 108 | 0.899 | -0.153 | 0.43492972 | 0.6433672 | 0 | 0.0359 | 0.019 | 0.019 | TRUE | 0.1109 |
| Pebp1 | 10033 | 11729 | 12143 | 10396 | 7825 | 8218 | 10657 | 11714 | 12193 | 8173 | 8419 | 8996 | 10025.19 | 11521 | 8529 | 0.74 | -0.434 | 1.63E-07 | 5.49E-06 | 0.0034 | 0.0072 | 0.0048 | 0.0048 | TRUE | 0.0526 |
| Pebp4 | 0 | 0 | 0 | 0 | 0 | 0 | 0 | 0 | 0 | 0 | 0 | 0 | 0 | 0 | 0 | NA | NA | NA | NA | NA | NA | NA | NA | NA | NA |
| Pecam1 | 6325 | 7418 | 7966 | 8723 | 6314 | 6404 | 6718 | 7408 | 7999 | 6857 | 6793 | 7010 | 7131.07 | 7375 | 6887 | 0.934 | -0.099 | 0.24711823 | 0.46388344 | 0.0037 | 0.0074 | 0.0051 | 0.0051 | TRUE | 0.074 |
| Pecr | 1590 | 1865 | 1838 | 1921 | 1524 | 1560 | 1689 | 1863 | 1846 | 1510 | 1640 | 1708 | 1709.09 | 1799 | 1619 | 0.899 | -0.153 | 0.08600716 | 0.2365722 | 0.0028 | 0.0088 | 0.0051 | 0.0051 | TRUE | 0.043 |
| Pef1 | 2124 | 2722 | 2606 | 2673 | 1908 | 2150 | 2256 | 2718 | 2617 | 2101 | 2053 | 2354 | 2349.83 | 2530 | 2169 | 0.857 | -0.223 | 0.034352 | 0.1262348 | 0.0067 | 0.0083 | 0.0076 | 0.0076 | TRUE | 0.1088 |
| Peg10 | 17777 | 10220 | 11375 | 25579 | 15712 | 12531 | 18882 | 10207 | 11422 | 20108 | 16905 | 13717 | 15206.89 | 13504 | 16910 | 1.252 | 0.325 | 0.14314978 | 0.32950355 | 0.0708 | 0.0071 | 0.0354 | 0.0354 | TRUE | 0.6635 |
| Peg12 | 94 | 83 | 74 | 542 | 385 | 280 | 100 | 83 | 74 | 426 | 414 | 307 | 233.98 | 86 | 382 | 4.473 | 2.161 | 3.74E-27 | 1.84E-24 | 0.0225 | 0.021 | 0.0216 | 0.0216 | TRUE | 0.3393 |
| Peg13 | 253 | 188 | 173 | 458 | 289 | 185 | 269 | 188 | 174 | 360 | 311 | 203 | 250.62 | 210 | 291 | 1.39 | 0.475 | 0.05914198 | 0.18413436 | 0.0654 | 0.02 | 0.0415 | 0.0415 | TRUE | 0.4418 |
| Peg3 | 1975 | 1610 | 2121 | 2969 | 2232 | 2410 | 2098 | 1608 | 2130 | 2334 | 2401 | 2638 | 2201.51 | 1945 | 2458 | 1.263 | 0.337 | 0.00796739 | 0.04418483 | 0.0127 | 0.0084 | 0.0112 | 0.0112 | TRUE | 0.2785 |
| Peg3as | 0 | 0 | 0 | 0 | 0 | 0 | 0 | 0 | 0 | 0 | 0 | 0 | 0 | 0 | 0 | NA | NA | NA | NA | NA | NA | NA | NA | NA | NA |

| | | | | | | | | | | | | | | | | | | | | | | | | | |
|---|---|---|---|---|---|---|---|---|---|---|---|---|---|---|---|---|---|---|---|---|---|---|---|---|---|
| Peli1 | 1860 | 1958 | 1868 | 2673 | 1950 | 1882 | 1976 | 1955 | 1876 | 2101 | 2098 | 2060 | 2011.05 | 1936 | 2086 | 1.078 | 0.109 | 0.11130165 | 0.28056988 | 0 | 0.0085 | 0.0029 | 0.0029 | TRUE | 0.0089 |
| Peli2 | 541 | 671 | 678 | 681 | 523 | 494 | 575 | 670 | 681 | 535 | 563 | 541 | 594.06 | 642 | 546 | 0.85 | -0.234 | 0.03442534 | 0.1264454 | 0.003 | 0.0124 | 0.0071 | 0.0071 | TRUE | 0.0979 |
| Peli3 | 90 | 109 | 79 | 109 | 78 | 71 | 96 | 109 | 79 | 86 | 84 | 78 | 88.52 | 95 | 83 | 0.873 | -0.196 | 0.37564994 | 0.59147378 | 0 | 0.0441 | 0.024 | 0.024 | TRUE | 0.1951 |
| Pelo | 1370 | 1185 | 1176 | 2368 | 1684 | 1367 | 1455 | 1183 | 1181 | 1862 | 1812 | 1496 | 1498.22 | 1273 | 1723 | 1.354 | 0.438 | 0.00075923 | 0.00717474 | 0.0132 | 0.0091 | 0.0115 | 0.0115 | TRUE | 0.1873 |
| Pelp1 | 14623 | 11923 | 12170 | 19389 | 14122 | 11894 | 15532 | 11908 | 12220 | 15242 | 15194 | 13020 | 13852.69 | 13220 | 14485 | 1.096 | 0.132 | 0.29372871 | 0.51457172 | 0.0142 | 0.0071 | 0.0113 | 0.0113 | TRUE | 0.2862 |
| Pemt | 149 | 207 | 202 | 259 | 207 | 185 | 158 | 207 | 203 | 204 | 223 | 203 | 199.44 | 189 | 210 | 1.106 | 0.145 | 0.3815022 | 0.59580679 | 0.006 | 0.0234 | 0.0148 | 0.0148 | TRUE | 0.218 |
| Penk | 52 | 62 | 61 | 115 | 89 | 61 | 55 | 62 | 61 | 90 | 96 | 67 | 71.89 | 59 | 84 | 1.42 | 0.506 | 0.04241708 | 0.14693657 | 0 | 0.0527 | 0.0305 | 0.0305 | TRUE | 0.293 |
| Peo1 | 2592 | 2394 | 2312 | 3498 | 2426 | 2156 | 2753 | 2391 | 2321 | 2750 | 2610 | 2360 | 2530.95 | 2488 | 2573 | 1.035 | 0.049 | 0.63677557 | 0.79562895 | 0.0068 | 0.0082 | 0.0074 | 0.0074 | TRUE | 0.1047 |
| Pepd | 2932 | 2578 | 2498 | 3071 | 2355 | 2204 | 3114 | 2575 | 2508 | 2414 | 2534 | 2413 | 2592.98 | 2732 | 2454 | 0.898 | -0.155 | 0.139621 | 0.32417231 | 0.0072 | 0.0082 | 0.0076 | 0.0076 | TRUE | 0.1811 |
| Per1 | 2314 | 2279 | 2362 | 2298 | 1730 | 1858 | 2458 | 2276 | 2372 | 1807 | 1861 | 2034 | 2134.56 | 2369 | 1901 | 0.802 | -0.318 | 0.00012722 | 0.00167911 | 0.002 | 0.0084 | 0.0045 | 0.0045 | TRUE | 0.0452 |
| Per2 | 42 | 36 | 46 | 42 | 26 | 25 | 45 | 36 | 46 | 33 | 28 | 27 | 35.85 | 42 | 29 | 0.701 | -0.512 | 0.11341045 | 0.28383206 | 0 | 0.0988 | 0.0469 | 0.0469 | TRUE | 0.1322 |
| Per3 | 832 | 984 | 1138 | 821 | 696 | 770 | 884 | 983 | 1143 | 645 | 749 | 843 | 874.38 | 1003 | 746 | 0.742 | -0.43 | 0.00264309 | 0.01919286 | 0.0155 | 0.0107 | 0.0136 | 0.0136 | TRUE | 0.1772 |
| Perp | 1153 | 1636 | 1636 | 2274 | 1622 | 1775 | 1225 | 1634 | 1643 | 1788 | 1745 | 1943 | 1662.85 | 1501 | 1825 | 1.216 | 0.282 | 0.0315994 | 0.11948077 | 0.0134 | 0.0089 | 0.0118 | 0.0118 | TRUE | 0.3103 |
| Pes1 | 12327 | 12161 | 12228 | 16608 | 11982 | 11525 | 13093 | 12145 | 12278 | 13056 | 12892 | 12616 | 12680.11 | 12505 | 12855 | 1.028 | 0.04 | 0.52745672 | 0.71617144 | 9.00E-04 | 0.0072 | 0.0028 | 0.0028 | TRUE | 0.0206 |
| Pet100 | 516 | 476 | 455 | 993 | 653 | 529 | 548 | 475 | 457 | 781 | 703 | 579 | 590.44 | 493 | 688 | 1.395 | 0.481 | 0.00094957 | 0.00854828 | 0.0144 | 0.0125 | 0.0135 | 0.0135 | TRUE | 0.2225 |
| Pet112 | 800 | 854 | 805 | 1382 | 973 | 941 | 850 | 853 | 808 | 1086 | 1047 | 1030 | 945.72 | 837 | 1054 | 1.261 | 0.334 | 6.13E-05 | 0.00092586 | 0 | 0.0104 | 0.0039 | 0.0039 | TRUE | 0.0107 |
| Pet117 | 1 | 2 | 0 | 1 | 2 | 2 | 1 | 2 | 0 | 1 | 2 | 2 | 1.36 | 1 | 2 | 1.644 | 0.717 | 0.66514503 | NA | 0 | 2.4213 | 1.185 | 1.185 | TRUE | 0.3149 |
| Pet2 | 5 | 6 | 3 | 80 | 70 | 36 | 5 | 6 | 3 | 63 | 75 | 39 | 31.99 | 5 | 59 | 12.437 | 3.637 | 1.06E-11 | 9.30E-10 | 0 | 0.1099 | 0.0908 | 0.0908 | TRUE | 0.138 |
| Pex1 | 1375 | 1305 | 1370 | 1761 | 1234 | 1239 | 1460 | 1303 | 1376 | 1384 | 1328 | 1356 | 1367.96 | 1380 | 1356 | 0.983 | -0.024 | 0.76939288 | 0.87946546 | 0.0011 | 0.0093 | 0.0042 | 0.0042 | TRUE | 0.0313 |
| Pex10 | 682 | 810 | 853 | 967 | 709 | 729 | 724 | 809 | 857 | 760 | 763 | 798 | 785.15 | 797 | 774 | 0.971 | -0.043 | 0.6721329 | 0.81721246 | 0.0026 | 0.0111 | 0.0062 | 0.0062 | TRUE | 0.074 |
| Pex11a | 251 | 231 | 214 | 287 | 198 | 195 | 267 | 231 | 215 | 226 | 213 | 213 | 227.38 | 238 | 217 | 0.917 | -0.125 | 0.40170779 | 0.61434455 | 0.0026 | 0.0214 | 0.0115 | 0.0115 | TRUE | 0.1264 |
| Pex11b | 694 | 784 | 765 | 1066 | 779 | 773 | 737 | 783 | 768 | 838 | 838 | 846 | 801.77 | 763 | 841 | 1.102 | 0.14 | 0.09935629 | 0.26013402 | 0 | 0.011 | 0.004 | 0.004 | TRUE | 0.0101 |
| Pex11c | 149 | 184 | 207 | 271 | 178 | 225 | 158 | 184 | 208 | 213 | 192 | 246 | 200.12 | 183 | 217 | 1.182 | 0.242 | 0.17145059 | 0.37034626 | 0.0116 | 0.0234 | 0.0174 | 0.0174 | TRUE | 0.151 |
| Pex12 | 685 | 714 | 695 | 1073 | 774 | 703 | 728 | 713 | 698 | 844 | 833 | 770 | 764.06 | 713 | 816 | 1.145 | 0.195 | 0.03255066 | 0.121608 | 0 | 0.0112 | 0.0047 | 0.0047 | TRUE | 0.028 |
| Pex13 | 1629 | 1806 | 1793 | 2475 | 1719 | 1708 | 1730 | 1804 | 1800 | 1946 | 1850 | 1870 | 1833.2 | 1778 | 1889 | 1.062 | 0.087 | 0.21928699 | 0.43140756 | 0 | 0.0087 | 0.0031 | 0.0031 | TRUE | 0.0088 |
| Pex14 | 2047 | 2030 | 2133 | 2637 | 1940 | 1893 | 2174 | 2027 | 2142 | 2073 | 2087 | 2072 | 2095.99 | 2114 | 2077 | 0.983 | -0.025 | 0.71599479 | 0.84571767 | 0 | 0.0085 | 0.003 | 0.003 | TRUE | 0.0158 |
| Pex16 | 462 | 572 | 604 | 895 | 587 | 568 | 491 | 571 | 606 | 704 | 632 | 622 | 604.23 | 556 | 653 | 1.174 | 0.231 | 0.05802729 | 0.1819496 | 0.0064 | 0.0124 | 0.009 | 0.009 | TRUE | 0.1229 |
| Pex19 | 3815 | 4114 | 4067 | 4554 | 3581 | 3918 | 4052 | 4109 | 4084 | 3580 | 3853 | 4289 | 3994.4 | 4082 | 3907 | 0.957 | -0.064 | 0.47203589 | 0.67320282 | 0.0035 | 0.0077 | 0.0054 | 0.0054 | TRUE | 0.0887 |
| Pex2 | 236 | 235 | 254 | 468 | 292 | 289 | 251 | 235 | 255 | 368 | 314 | 316 | 289.81 | 247 | 333 | 1.352 | 0.435 | 0.00132597 | 0.01110045 | 0 | 0.0183 | 0.0097 | 0.0097 | TRUE | 0.1027 |
| Pex26 | 983 | 1022 | 957 | 1392 | 1034 | 989 | 1044 | 1021 | 961 | 1094 | 1113 | 1083 | 1052.52 | 1009 | 1097 | 1.087 | 0.121 | 0.14090444 | 0.3262876 | 0 | 0.01 | 0.0039 | 0.0039 | TRUE | 0.0204 |
| Pex3 | 766 | 736 | 713 | 1314 | 931 | 845 | 814 | 735 | 716 | 1033 | 1002 | 925 | 870.71 | 755 | 987 | 1.308 | 0.388 | 0.00010524 | 0.00143467 | 0.0027 | 0.0107 | 0.006 | 0.006 | TRUE | 0.0545 |
| Pex5 | 4479 | 4596 | 4772 | 6088 | 4313 | 4519 | 4757 | 4590 | 4792 | 4786 | 4641 | 4947 | 4752.06 | 4713 | 4791 | 1.017 | 0.024 | 0.71168345 | 0.84307281 | 5.00E-04 | 0.0076 | 0.0027 | 0.0027 | TRUE | 0.0097 |
| Pex5l | 16 | 22 | 24 | 18 | 13 | 23 | 17 | 22 | 24 | 14 | 14 | 25 | 19.4 | 21 | 18 | 0.837 | -0.256 | 0.58118253 | 0.75767985 | 0.0176 | 0.1768 | 0.103 | 0.103 | TRUE | 0.6219 |
| Pex6 | 999 | 943 | 914 | 1384 | 1098 | 804 | 1061 | 942 | 918 | 1088 | 1181 | 880 | 1011.69 | 974 | 1050 | 1.079 | 0.11 | 0.41062856 | 0.62196522 | 0.0123 | 0.0102 | 0.0118 | 0.0118 | TRUE | 0.2357 |
| Pex7 | 1655 | 1918 | 1890 | 2194 | 1564 | 1630 | 1758 | 1916 | 1898 | 1725 | 1683 | 1784 | 1793.83 | 1857 | 1731 | 0.932 | -0.102 | 0.18845021 | 0.39351738 | 9.00E-04 | 0.0087 | 0.0038 | 0.0038 | TRUE | 0.0262 |
| Pf4 | 1 | 1 | 1 | 1 | 1 | 0 | 1 | 1 | 1 | 1 | 1 | 0 | 0.82 | 1 | 1 | 0.618 | -0.695 | 0.72575676 | NA | 0 | 4.0188 | 1.5537 | 1.5537 | TRUE | 0.1881 |
| Pfas | 13123 | 11569 | 11434 | 15982 | 12188 | 10649 | 13939 | 11554 | 11481 | 12564 | 13114 | 11657 | 12384.72 | 12325 | 12445 | 1.01 | 0.014 | 0.89153694 | 0.94746449 | 0.0065 | 0.0072 | 0.0076 | 0.0076 | TRUE | 0.1604 |
| Pfdn1 | 3986 | 4370 | 4416 | 5660 | 4157 | 4008 | 4234 | 4364 | 4434 | 4449 | 4473 | 4387 | 4390.31 | 4344 | 4436 | 1.021 | 0.03 | 0.61198752 | 0.78028154 | 0 | 0.0077 | 0.0024 | 0.0024 | TRUE | 0.006 |
| Pfdn2 | 3791 | 4355 | 4316 | 4900 | 3640 | 3752 | 4027 | 4349 | 4334 | 3852 | 3916 | 4107 | 4097.56 | 4237 | 3958 | 0.934 | -0.098 | 0.16708059 | 0.3645851 | 0.0011 | 0.0077 | 0.0034 | 0.0034 | TRUE | 0.0228 |
| Pfdn4 | 2106 | 2237 | 2109 | 2534 | 1849 | 1919 | 2237 | 2234 | 2118 | 1992 | 1989 | 2101 | 2111.8 | 2196 | 2027 | 0.923 | -0.116 | 0.1073892 | 0.27411271 | 5.00E-04 | 0.0085 | 0.0032 | 0.0032 | TRUE | 0.0119 |
| Pfdn5 | 2401 | 3261 | 3337 | 3469 | 2463 | 2699 | 2550 | 3257 | 3351 | 2727 | 2650 | 2954 | 2914.89 | 3053 | 2777 | 0.91 | -0.137 | 0.26052839 | 0.47988003 | 0.0114 | 0.008 | 0.0103 | 0.0103 | TRUE | 0.2513 |
| Pfkfb1 | 464 | 624 | 612 | 642 | 403 | 497 | 493 | 623 | 615 | 505 | 434 | 544 | 535.48 | 577 | 494 | 0.856 | -0.224 | 0.11885603 | 0.29199879 | 0.0129 | 0.0131 | 0.0129 | 0.0129 | TRUE | 0.189 |
| Pfkfb2 | 707 | 789 | 778 | 908 | 698 | 665 | 751 | 788 | 781 | 714 | 751 | 728 | 752.15 | 773 | 731 | 0.944 | -0.082 | 0.34612968 | 0.56538943 | 0 | 0.0113 | 0.0042 | 0.0042 | TRUE | 0.0075 |
| Pfkfb3 | 1066 | 1777 | 1926 | 1894 | 1612 | 1838 | 1132 | 1775 | 1934 | 1489 | 1734 | 2012 | 1679.37 | 1614 | 1745 | 1.081 | 0.112 | 0.56896087 | 0.7479289 | 0.049 | 0.0089 | 0.0274 | 0.0274 | TRUE | 0.7634 |
| Pfkfb4 | 206 | 159 | 155 | 199 | 147 | 145 | 219 | 159 | 156 | 156 | 158 | 159 | 167.76 | 178 | 158 | 0.888 | -0.171 | 0.36712064 | 0.58453273 | 0.0135 | 0.0265 | 0.0199 | 0.0199 | TRUE | 0.4322 |
| Pfkl | 21836 | 29656 | 31054 | 23247 | 18748 | 23350 | 23194 | 29618 | 31181 | 18275 | 20172 | 25560 | 24666.64 | 27998 | 21336 | 0.762 | -0.392 | 0.01076684 | 0.05569518 | 0.0271 | 0.007 | 0.017 | 0.017 | TRUE | 0.3671 |
| Pfkm | 2532 | 3157 | 3133 | 3176 | 2391 | 2768 | 2689 | 3153 | 3146 | 2497 | 2573 | 3030 | 2847.92 | 2996 | 2700 | 0.901 | -0.151 | 0.17870652 | 0.37980855 | 0.0091 | 0.0081 | 0.0087 | 0.0087 | TRUE | 0.1385 |
| Pfkp | 18083 | 21151 | 20554 | 22067 | 18269 | 21322 | 19207 | 21124 | 20638 | 17347 | 19656 | 23340 | 20218.9 | 20323 | 20114 | 0.99 | -0.015 | 0.89886042 | 0.95089051 | 0.0099 | 0.0071 | 0.01 | 0.01 | TRUE | 0.2407 |
| Pfn1 | 15026 | 12169 | 11702 | 28925 | 20196 | 16844 | 15960 | 12153 | 11750 | 22739 | 21730 | 18438 | 17128.37 | 13288 | 20969 | 1.578 | 0.658 | 2.31E-06 | 5.70E-05 | 0.0209 | 0.0071 | 0.0139 | 0.0139 | TRUE | 0.3784 |
| Pfn2 | 890 | 1361 | 1359 | 1426 | 996 | 1037 | 945 | 1359 | 1365 | 1121 | 1072 | 1135 | 1166.16 | 1223 | 1109 | 0.907 | -0.141 | 0.34699394 | 0.5660139 | 0.0207 | 0.0097 | 0.0154 | 0.0154 | TRUE | 0.4719 |
| Pfn3 | 0 | 0 | 0 | 0 | 0 | 0 | 0 | 0 | 0 | 0 | 0 | 0 | 0 | 0 | 0 | NA | NA | NA | NA | NA | NA | NA | NA | NA | NA |
| Pfn4 | 96 | 105 | 103 | 180 | 125 | 75 | 102 | 105 | 103 | 142 | 134 | 82 | 111.39 | 103 | 119 | 1.159 | 0.212 | 0.38796927 | 0.6017358 | 0.0327 | 0.0365 | 0.0346 | 0.0346 | TRUE | 0.7247 |
| Pfpl | 0 | 0 | 0 | 0 | 0 | 0 | 0 | 0 | 0 | 0 | 0 | 0 | 0 | 0 | 0 | NA | NA | NA | NA | NA | NA | NA | NA | NA | NA |
| Pga5 | 146 | 189 | 140 | 84 | 78 | 100 | 155 | 189 | 141 | 66 | 84 | 109 | 123.97 | 162 | 86 | 0.532 | -0.911 | 0.00016571 | 0.00207432 | 0.033 | 0.0335 | 0.0333 | 0.0333 | TRUE | 0.5043 |
| Pgam1 | 12151 | 13364 | 13856 | 15647 | 11224 | 13045 | 12907 | 13347 | 13913 | 12300 | 12076 | 14280 | 13137.12 | 13389 | 12885 | 0.962 | -0.055 | 0.53871529 | 0.7250911 | 0.0044 | 0.0072 | 0.0058 | 0.0058 | TRUE | 0.1094 |
| Pgam2 | 85 | 141 | 126 | 116 | 73 | 112 | 90 | 141 | 127 | 91 | 79 | 123 | 108.33 | 119 | 98 | 0.815 | -0.294 | 0.25607743 | 0.47476821 | 0.0411 | 0.0373 | 0.0391 | 0.0391 | TRUE | 0.473 |
| Pgam5 | 3856 | 4538 | 4473 | 4924 | 3610 | 3859 | 4096 | 4532 | 4491 | 3871 | 3884 | 4224 | 4183.09 | 4373 | 3993 | 0.913 | -0.131 | 0.10076873 | 0.26271835 | 0.0024 | 0.0077 | 0.0044 | 0.0044 | TRUE | 0.0374 |
| Pgap1 | 1737 | 1591 | 1649 | 2229 | 1602 | 1484 | 1845 | 1589 | 1656 | 1752 | 1724 | 1624 | 1698.35 | 1697 | 1700 | 1.003 | 0.004 | 0.96726295 | 0.98366569 | 0.0032 | 0.0088 | 0.0054 | 0.0054 | TRUE | 0.0704 |
| Pgap2 | 679 | 864 | 1024 | 967 | 737 | 804 | 721 | 863 | 1028 | 760 | 793 | 880 | 840.93 | 871 | 811 | 0.931 | -0.104 | 0.47869411 | 0.67816074 | 0.0173 | 0.0108 | 0.0142 | 0.0142 | TRUE | 0.2117 |
| Pgap3 | 176 | 229 | 228 | 338 | 223 | 238 | 187 | 229 | 229 | 266 | 240 | 261 | 235.13 | 215 | 256 | 1.189 | 0.25 | 0.0948073 | 0.25228627 | 0.0028 | 0.0209 | 0.0118 | 0.0118 | TRUE | 0.1391 |
| Pgbd1 | 173 | 266 | 338 | 289 | 228 | 225 | 184 | 266 | 339 | 227 | 245 | 246 | 251.27 | 263 | 239 | 0.91 | -0.136 | 0.53792637 | 0.72421466 | 0.0427 | 0.02 | 0.0311 | 0.0311 | TRUE | 0.1924 |
| Pgbd5 | 302 | 322 | 295 | 412 | 286 | 309 | 321 | 322 | 296 | 324 | 308 | 338 | 318.07 | 313 | 323 | 1.033 | 0.047 | 0.69621713 | 0.83365646 | 0 | 0.0173 | 0.0075 | 0.0075 | TRUE | 0.0247 |
| Pgc | 284 | 279 | 271 | 223 | 195 | 241 | 302 | 279 | 272 | 175 | 210 | 264 | 250.22 | 284 | 216 | 0.758 | -0.399 | 0.02580859 | 0.10394612 | 0.018 | 0.0201 | 0.019 | 0.019 | TRUE | 0.4011 |
| Pgd | 11083 | 8712 | 8561 | 10797 | 7891 | 7227 | 11772 | 8701 | 8596 | 8488 | 8490 | 7911 | 8993.01 | 9690 | 8296 | 0.856 | -0.224 | 0.08986968 | 0.24390184 | 0.018 | 0.0073 | 0.0124 | 0.0124 | TRUE | 0.4317 |
| Pgf | 89 | 106 | 93 | 102 | 70 | 79 | 95 | 106 | 93 | 80 | 75 | 86 | 89.29 | 98 | 80 | 0.823 | -0.281 | 0.17514117 | 0.37485771 | 0 | 0.0438 | 0.0196 | 0.0196 | TRUE | 0.0495 |
| Pggt1b | 726 | 728 | 765 | 1276 | 924 | 757 | 771 | 727 | 768 | 1003 | 994 | 829 | 848.71 | 755 | 942 | 1.248 | 0.32 | 0.00369108 | 0.02484276 | 0.0051 | 0.0108 | 0.0075 | 0.0075 | TRUE | 0.1299 |
| Pgk1 | 23900 | 37372 | 39355 | 30766 | 23238 | 30164 | 25386 | 37324 | 39517 | 24186 | 25003 | 33019 | 30739.01 | 34076 | 27403 | 0.804 | -0.314 | 0.08099747 | 0.22654586 | 0.0407 | 0.007 | 0.0234 | 0.0234 | TRUE | 0.6091 |
| Pgk2 | 1 | 0 | 1 | 0 | 0 | 1 | 1 | 0 | 1 | 0 | 0 | 1 | 0.53 | 1 | 0 | 0.509 | -0.975 | 0.72674596 | NA | 0 | 6.2603 | 3.8683 | 3.8683 | TRUE | 0.5383 |
| Pgls | 2487 | 3310 | 3348 | 3748 | 2744 | 2832 | 2642 | 3306 | 3362 | 2946 | 2952 | 3100 | 3051.32 | 3103 | 2999 | 0.966 | -0.049 | 0.6564214 | 0.80670515 | 0.0088 | 0.008 | 0.0085 | 0.0085 | TRUE | 0.2052 |

| | | | | | | | | | | | | | | | | | | | | | | | | |
|---|---|---|---|---|---|---|---|---|---|---|---|---|---|---|---|---|---|---|---|---|---|---|---|---|
| Pglyrp1 | 5 | 22 | 8 | 12 | 13 | 15 | 5 | 22 | 8 | 9 | 14 | 16 | 12.53 | 12 | 13 | 1.117 | 0.159 | 0.81237255 | 0.9037712 | 0.2151 | 0.2699 | 0.2431 | 0.2431 | TRUE | 2.2561 |
| Pglyrp2 | 0 | 0 | 0 | 0 | 0 | 0 | 0 | 0 | 0 | 0 | 0 | 0 | 0 | 0 | 0 | NA | NA | NA | NA | NA | NA | NA | NA | NA | NA |
| Pglyrp3 | 0 | 0 | 0 | 1 | 0 | 1 | 0 | 0 | 0 | 1 | 0 | 1 | 0.31 | 0 | 1 | 3.348 | 1.743 | 0.65064889 | NA | 0 | 10.5167 | 8.8968 | 8.8968 | TRUE | 0.2082 |
| Pglyrp4 | 0 | 0 | 0 | 0 | 0 | 0 | 0 | 0 | 0 | 0 | 0 | 0 | 0 | 0 | 0 | NA | NA | NA | NA | NA | NA | NA | NA | NA | NA |
| Pgm1 | 2813 | 3554 | 3572 | 3561 | 2797 | 3275 | 2988 | 3549 | 3587 | 2799 | 3009 | 3585 | 3252.96 | 3375 | 3131 | 0.928 | -0.109 | 0.37985954 | 0.59453654 | 0.0119 | 0.0079 | 0.0107 | 0.0107 | TRUE | 0.1949 |
| Pgm2 | 3862 | 6266 | 6779 | 5275 | 4061 | 5106 | 4102 | 6258 | 6807 | 4147 | 4369 | 5589 | 5212.06 | 5722 | 4702 | 0.822 | -0.284 | 0.13530042 | 0.31824306 | 0.0446 | 0.0075 | 0.0258 | 0.0258 | TRUE | 0.7478 |
| Pgm2l1 | 1841 | 1312 | 1220 | 2405 | 1616 | 1494 | 1955 | 1310 | 1225 | 1891 | 1739 | 1635 | 1625.92 | 1497 | 1755 | 1.173 | 0.23 | 0.18929482 | 0.39444549 | 0.0336 | 0.0089 | 0.0215 | 0.0215 | TRUE | 0.8636 |
| Pgm3 | 3002 | 3055 | 3120 | 3627 | 2689 | 2808 | 3189 | 3051 | 3133 | 2851 | 2893 | 3074 | 3031.8 | 3124 | 2939 | 0.941 | -0.088 | 0.20248995 | 0.4100233 | 7.00E-04 | 0.008 | 0.0031 | 0.0031 | TRUE | 0.0193 |
| Pgm5 | 15 | 10 | 10 | 27 | 17 | 15 | 16 | 10 | 10 | 21 | 18 | 16 | 15.32 | 12 | 18 | 1.567 | 0.648 | 0.19092278 | 0.39611884 | 0 | 0.222 | 0.1083 | 0.1083 | TRUE | 0.3116 |
| Pgp | 1837 | 2372 | 2327 | 2887 | 2332 | 2129 | 1951 | 2369 | 2337 | 2270 | 2509 | 2331 | 2294.31 | 2219 | 2370 | 1.068 | 0.094 | 0.36310785 | 0.58070747 | 0.0061 | 0.0083 | 0.0073 | 0.0073 | TRUE | 0.1345 |
| Pgpep1 | 2058 | 3046 | 3082 | 2436 | 1943 | 2294 | 2186 | 3042 | 3095 | 1915 | 2091 | 2511 | 2473.23 | 2774 | 2172 | 0.783 | -0.353 | 0.02638509 | 0.10556712 | 0.0262 | 0.0082 | 0.0179 | 0.0179 | TRUE | 0.4171 |
| Pgpep1l | 18 | 21 | 17 | 39 | 37 | 29 | 19 | 21 | 17 | 31 | 40 | 32 | 26.56 | 19 | 34 | 1.782 | 0.834 | 0.0269498 | 0.10720009 | 0 | 0.1309 | 0.0612 | 0.0612 | TRUE | 0.1457 |
| Pgr | 39 | 42 | 70 | 85 | 64 | 51 | 41 | 42 | 70 | 67 | 69 | 56 | 57.53 | 51 | 64 | 1.247 | 0.318 | 0.29798818 | 0.51872997 | 0.0338 | 0.0642 | 0.0497 | 0.0497 | TRUE | 0.8747 |
| Pgr15l | 75 | 53 | 45 | 101 | 70 | 43 | 80 | 53 | 45 | 79 | 75 | 47 | 63.26 | 59 | 67 | 1.141 | 0.19 | 0.56200708 | 0.74295871 | 0.0647 | 0.059 | 0.0617 | 0.0617 | TRUE | 0.7626 |
| Pgrmc1 | 3908 | 4340 | 4509 | 5536 | 3822 | 3592 | 4151 | 4334 | 4528 | 4352 | 4112 | 3932 | 4234.85 | 4338 | 4132 | 0.953 | -0.07 | 0.36294328 | 0.5806209 | 0.0019 | 0.0077 | 0.004 | 0.004 | TRUE | 0.0267 |
| Pgrmc2 | 3065 | 3521 | 3381 | 3972 | 3096 | 3187 | 3256 | 3516 | 3395 | 3122 | 3331 | 3489 | 3351.53 | 3389 | 3314 | 0.977 | -0.033 | 0.67326794 | 0.81803018 | 0.002 | 0.0079 | 0.0041 | 0.0041 | TRUE | 0.0316 |
| Pgs1 | 5702 | 5380 | 5449 | 7210 | 5389 | 4908 | 6057 | 5373 | 5471 | 5668 | 5798 | 5373 | 5623.28 | 5634 | 5613 | 0.996 | -0.005 | 0.94828037 | 0.97415997 | 0.0027 | 0.0075 | 0.0044 | 0.0044 | TRUE | 0.0525 |
| Phactr1 | 135 | 133 | 131 | 217 | 125 | 165 | 143 | 133 | 132 | 171 | 134 | 181 | 148.91 | 136 | 162 | 1.193 | 0.255 | 0.1660642 | 0.36326832 | 0.0061 | 0.029 | 0.0176 | 0.0176 | TRUE | 0.222 |
| Phactr2 | 202 | 232 | 236 | 363 | 256 | 238 | 215 | 232 | 237 | 285 | 275 | 261 | 250.76 | 228 | 274 | 1.203 | 0.267 | 0.04504815 | 0.1529778 | 0 | 0.02 | 0.0088 | 0.0088 | TRUE | 0.0277 |
| Phactr3 | 257 | 207 | 182 | 246 | 138 | 158 | 273 | 207 | 183 | 193 | 148 | 173 | 196.21 | 221 | 171 | 0.78 | -0.359 | 0.07728771 | 0.21955724 | 0.0256 | 0.0237 | 0.0246 | 0.0246 | TRUE | 0.4646 |
| Phactr4 | 2740 | 2958 | 2956 | 3039 | 2279 | 2522 | 2910 | 2954 | 2968 | 2389 | 2452 | 2761 | 2739.08 | 2944 | 2534 | 0.86 | -0.217 | 0.00944806 | 0.05041095 | 0.0026 | 0.0081 | 0.0047 | 0.0047 | TRUE | 0.0739 |
| Phax | 1818 | 1784 | 1917 | 2676 | 1847 | 1787 | 1931 | 1782 | 1925 | 2104 | 1987 | 1956 | 1947.45 | 1879 | 2016 | 1.073 | 0.102 | 0.19373579 | 0.39947733 | 0.0013 | 0.0086 | 0.0039 | 0.0039 | TRUE | 0.025 |
| Phb | 12872 | 11642 | 11379 | 13848 | 9552 | 8827 | 13672 | 11627 | 11426 | 10886 | 10277 | 9663 | 11258.52 | 12242 | 10275 | 0.839 | -0.252 | 0.01026489 | 0.05379508 | 0.0068 | 0.0072 | 0.0069 | 0.0069 | TRUE | 0.1277 |
| Phb2 | 20284 | 21007 | 20686 | 26605 | 19468 | 18256 | 21545 | 20980 | 20771 | 20915 | 20946 | 19984 | 20856.87 | 21099 | 20615 | 0.977 | -0.033 | 0.55873886 | 0.74101748 | 5.00E-04 | 0.0071 | 0.0023 | 0.0023 | TRUE | 0.0088 |
| Phc1 | 35271 | 38199 | 42104 | 41823 | 30582 | 30777 | 37464 | 38150 | 42277 | 32878 | 32904 | 33690 | 36227.19 | 39297 | 33157 | 0.844 | -0.245 | 0.0007761 | 0.00731219 | 0.0022 | 0.007 | 0.0038 | 0.0038 | TRUE | 0.0539 |
| Phc2 | 1700 | 1516 | 1437 | 2304 | 1666 | 1567 | 1806 | 1514 | 1443 | 1811 | 1793 | 1715 | 1680.28 | 1588 | 1773 | 1.117 | 0.16 | 0.13577063 | 0.31898918 | 0.0068 | 0.0089 | 0.0077 | 0.0077 | TRUE | 0.1736 |
| Phc3 | 1115 | 1170 | 1280 | 1284 | 986 | 1102 | 1184 | 1168 | 1285 | 1009 | 1061 | 1206 | 1152.44 | 1212 | 1092 | 0.9 | -0.152 | 0.14033038 | 0.32538823 | 0.0044 | 0.0098 | 0.0068 | 0.0068 | TRUE | 0.0993 |
| Phex | 1 | 0 | 1 | 2 | 3 | 3 | 1 | 0 | 1 | 2 | 3 | 3 | 1.69 | 1 | 3 | 3.892 | 1.96 | 0.21961789 | NA | 0 | 1.9543 | 0.9073 | 0.9073 | TRUE | 0.2569 |
| Phf1 | 402 | 456 | 403 | 495 | 421 | 369 | 427 | 455 | 405 | 389 | 453 | 404 | 422.18 | 429 | 415 | 0.967 | -0.049 | 0.68487327 | 0.82582731 | 0.0027 | 0.0147 | 0.0081 | 0.0081 | TRUE | 0.0716 |
| Phf10 | 2371 | 2762 | 2774 | 3353 | 2503 | 2524 | 2518 | 2758 | 2785 | 2636 | 2693 | 2763 | 2692.35 | 2687 | 2697 | 1.003 | 0.005 | 0.94936418 | 0.9746726 | 0.0014 | 0.0081 | 0.0038 | 0.0038 | TRUE | 0.0366 |
| Phf11a | 20 | 5 | 4 | 48 | 30 | 15 | 21 | 5 | 4 | 38 | 32 | 16 | 19.45 | 10 | 29 | 2.885 | 1.529 | 0.02487332 | 0.10140264 | 0.3824 | 0.1763 | 0.2667 | 0.2667 | TRUE | 2.137 |
| Phf11b | 21 | 10 | 19 | 38 | 17 | 14 | 22 | 10 | 19 | 30 | 18 | 15 | 19.14 | 17 | 21 | 1.252 | 0.324 | 0.52627261 | 0.71548759 | 0.0894 | 0.179 | 0.1357 | 0.1357 | TRUE | 0.852 |
| Phf11c | 56 | 46 | 30 | 74 | 56 | 39 | 59 | 46 | 30 | 58 | 60 | 43 | 49.44 | 45 | 54 | 1.193 | 0.255 | 0.45021917 | 0.65567698 | 0.0482 | 0.0735 | 0.0616 | 0.0616 | TRUE | 0.2694 |
| Phf11d | 2165 | 2656 | 2523 | 3455 | 2572 | 2674 | 2300 | 2653 | 2533 | 2716 | 2767 | 2927 | 2649.34 | 2495 | 2803 | 1.123 | 0.168 | 0.05078044 | 0.16585402 | 0.0028 | 0.0081 | 0.0049 | 0.0049 | TRUE | 0.0568 |
| Phf12 | 5276 | 4200 | 4032 | 6373 | 4854 | 4039 | 5604 | 4195 | 4049 | 5010 | 5223 | 4421 | 4750.18 | 4616 | 4885 | 1.058 | 0.082 | 0.55896084 | 0.74102381 | 0.0169 | 0.0076 | 0.0139 | 0.0139 | TRUE | 0.4272 |
| Phf13 | 2962 | 3265 | 3341 | 4570 | 3557 | 3701 | 3146 | 3261 | 3355 | 3593 | 3827 | 4051 | 3538.78 | 3254 | 3824 | 1.175 | 0.232 | 0.00294448 | 0.02092851 | 0.0019 | 0.0078 | 0.0041 | 0.0041 | TRUE | 0.0345 |
| Phf14 | 1336 | 1496 | 1563 | 2701 | 1815 | 1693 | 1419 | 1494 | 1569 | 2123 | 1953 | 1853 | 1735.33 | 1494 | 1976 | 1.324 | 0.404 | 8.13E-06 | 0.00016559 | 0.0031 | 0.0088 | 0.0053 | 0.0053 | TRUE | 0.052 |
| Phf15 | 1121 | 1475 | 1580 | 2053 | 1634 | 1456 | 1191 | 1473 | 1586 | 1614 | 1758 | 1594 | 1536.02 | 1417 | 1655 | 1.168 | 0.224 | 0.06992556 | 0.2055668 | 0.0108 | 0.009 | 0.0104 | 0.0104 | TRUE | 0.2336 |
| Phf16 | 1872 | 1999 | 2075 | 2516 | 1766 | 1892 | 1988 | 1996 | 2084 | 1978 | 1900 | 2071 | 2002.9 | 2023 | 1983 | 0.98 | -0.029 | 0.69931819 | 0.83551598 | 7.00E-04 | 0.0085 | 0.0035 | 0.0035 | TRUE | 0.0178 |
| Phf17 | 13992 | 11898 | 11826 | 19127 | 13153 | 11341 | 14862 | 11883 | 11875 | 15036 | 14152 | 12415 | 13370.27 | 12873 | 13868 | 1.077 | 0.107 | 0.37408206 | 0.59035644 | 0.0122 | 0.0071 | 0.0105 | 0.0105 | TRUE | 0.2233 |
| Phf19 | 510 | 854 | 863 | 724 | 620 | 644 | 542 | 853 | 867 | 569 | 667 | 705 | 700.39 | 754 | 647 | 0.858 | -0.222 | 0.24113387 | 0.45777122 | 0.0378 | 0.0116 | 0.0243 | 0.0243 | TRUE | 0.7146 |
| Phf2 | 329 | 507 | 523 | 676 | 469 | 501 | 349 | 506 | 525 | 531 | 505 | 548 | 494.23 | 460 | 528 | 1.147 | 0.198 | 0.23534461 | 0.45046786 | 0.0223 | 0.0136 | 0.0179 | 0.0179 | TRUE | 0.5113 |
| Phf20 | 6534 | 7281 | 7836 | 8100 | 5894 | 6792 | 6940 | 7272 | 7868 | 6368 | 6342 | 7435 | 7037.35 | 7360 | 6715 | 0.912 | -0.133 | 0.16949873 | 0.36794046 | 0.0061 | 0.0074 | 0.0066 | 0.0066 | TRUE | 0.1072 |
| Phf20l1 | 502 | 512 | 527 | 893 | 740 | 613 | 533 | 511 | 529 | 702 | 796 | 671 | 623.82 | 524 | 723 | 1.378 | 0.463 | 2.03E-05 | 0.00036149 | 0.0029 | 0.0122 | 0.0068 | 0.0068 | TRUE | 0.0904 |
| Phf21a | 2023 | 2453 | 2665 | 2577 | 1912 | 1948 | 2149 | 2450 | 2676 | 2026 | 2057 | 2132 | 2248.33 | 2425 | 2072 | 0.854 | -0.227 | 0.02477895 | 0.10114224 | 0.0056 | 0.0084 | 0.0069 | 0.0069 | TRUE | 0.1199 |
| Phf21b | 250 | 220 | 234 | 282 | 235 | 217 | 266 | 220 | 235 | 222 | 253 | 238 | 238.72 | 240 | 238 | 0.987 | -0.018 | 0.90146224 | 0.95205059 | 0.0027 | 0.0207 | 0.0113 | 0.0113 | TRUE | 0.0941 |
| Phf23 | 3883 | 3821 | 3885 | 5883 | 4229 | 3763 | 4124 | 3816 | 3901 | 4625 | 4550 | 4119 | 4189.26 | 3947 | 4431 | 1.123 | 0.167 | 0.03627576 | 0.13110479 | 0.0024 | 0.0077 | 0.0044 | 0.0044 | TRUE | 0.046 |
| Phf3 | 10067 | 8467 | 8982 | 12476 | 8790 | 8248 | 10693 | 8456 | 9019 | 9808 | 9457 | 9029 | 9410.29 | 9389 | 9431 | 1.005 | 0.007 | 0.94995993 | 0.97481103 | 0.0067 | 0.0073 | 0.0078 | 0.0078 | TRUE | 0.1801 |
| Phf5a | 3821 | 3224 | 2986 | 5300 | 3342 | 2973 | 4059 | 3220 | 2998 | 4166 | 3596 | 3254 | 3548.89 | 3426 | 3672 | 1.072 | 0.101 | 0.47785453 | 0.67733636 | 0.0178 | 0.0078 | 0.0142 | 0.0142 | TRUE | 0.3175 |
| Phf6 | 1977 | 1938 | 1930 | 2782 | 1733 | 1828 | 2100 | 1935 | 1938 | 2187 | 1865 | 2001 | 2004.33 | 1991 | 2018 | 1.014 | 0.02 | 0.82824035 | 0.9129575 | 0.0039 | 0.0085 | 0.0057 | 0.0057 | TRUE | 0.0665 |
| Phf7 | 398 | 531 | 537 | 758 | 494 | 449 | 423 | 530 | 539 | 596 | 532 | 492 | 518.53 | 497 | 540 | 1.086 | 0.119 | 0.39891128 | 0.61202615 | 0.0116 | 0.0133 | 0.0124 | 0.0124 | TRUE | 0.1993 |
| Phf8 | 2565 | 2818 | 2975 | 3325 | 2253 | 2477 | 2724 | 2814 | 2987 | 2614 | 2424 | 2711 | 2712.58 | 2842 | 2583 | 0.909 | -0.138 | 0.09168593 | 0.24710705 | 0.0021 | 0.0081 | 0.0044 | 0.0044 | TRUE | 0.0347 |
| Phgdh | 4032 | 4999 | 5083 | 5999 | 4823 | 4952 | 4283 | 4993 | 5104 | 4716 | 5189 | 5421 | 4950.84 | 4793 | 5109 | 1.066 | 0.092 | 0.36285099 | 0.58058374 | 0.0064 | 0.0076 | 0.0071 | 0.0071 | TRUE | 0.1056 |
| Phgr1 | 0 | 0 | 0 | 0 | 0 | 0 | 0 | 0 | 0 | 0 | 0 | 0 | 0 | 0 | 0 | NA | NA | NA | NA | NA | NA | NA | NA | NA | NA |
| Phip | 3407 | 3458 | 3949 | 5377 | 3942 | 3711 | 3619 | 3454 | 3965 | 4227 | 4241 | 4062 | 3928.03 | 3679 | 4177 | 1.135 | 0.183 | 0.02249975 | 0.09450494 | 0.0024 | 0.0077 | 0.0044 | 0.0044 | TRUE | 0.0564 |
| Phka1 | 1538 | 1726 | 1644 | 1819 | 1307 | 1426 | 1634 | 1724 | 1651 | 1430 | 1406 | 1561 | 1567.56 | 1670 | 1466 | 0.878 | -0.188 | 0.02141208 | 0.09133658 | 0.0013 | 0.009 | 0.0042 | 0.0042 | TRUE | 0.0382 |
| Phka2 | 867 | 1093 | 1109 | 1416 | 1077 | 990 | 921 | 1092 | 1114 | 1113 | 1159 | 1084 | 1080.28 | 1042 | 1119 | 1.073 | 0.102 | 0.33137929 | 0.55213544 | 0.0048 | 0.01 | 0.007 | 0.007 | TRUE | 0.1228 |
| Phkb | 2129 | 2272 | 2502 | 3180 | 2189 | 2017 | 2261 | 2269 | 2512 | 2500 | 2355 | 2208 | 2350.96 | 2347 | 2354 | 1.003 | 0.005 | 0.95699758 | 0.97770215 | 0.0033 | 0.0083 | 0.0052 | 0.0052 | TRUE | 0.0457 |
| Phkg1 | 168 | 200 | 185 | 251 | 170 | 162 | 178 | 200 | 186 | 197 | 183 | 177 | 186.92 | 188 | 186 | 0.99 | -0.014 | 0.926609 | 0.96376135 | 0 | 0.0245 | 0.011 | 0.011 | TRUE | 0.0343 |
| Phkg2 | 1388 | 1603 | 1625 | 1519 | 1217 | 1227 | 1474 | 1601 | 1632 | 1194 | 1309 | 1343 | 1425.6 | 1569 | 1282 | 0.816 | -0.293 | 0.0012597 | 0.01064378 | 0.0027 | 0.0092 | 0.0052 | 0.0052 | TRUE | 0.0448 |
| Phlda1 | 150 | 149 | 163 | 328 | 239 | 176 | 159 | 149 | 164 | 258 | 257 | 193 | 196.58 | 157 | 236 | 1.503 | 0.588 | 0.00080068 | 0.00749454 | 0.0105 | 0.0237 | 0.0168 | 0.0168 | TRUE | 0.2748 |
| Phlda2 | 1089 | 1324 | 1472 | 1141 | 812 | 859 | 1157 | 1322 | 1478 | 897 | 874 | 940 | 1111.33 | 1319 | 904 | 0.685 | -0.546 | 1.80E-06 | 4.58E-05 | 0.0075 | 0.0099 | 0.0085 | 0.0085 | TRUE | 0.1386 |
| Phlda3 | 1691 | 2421 | 2200 | 2614 | 1799 | 2018 | 1796 | 2418 | 2209 | 2055 | 1936 | 2209 | 2103.77 | 2141 | 2067 | 0.965 | -0.051 | 0.68329726 | 0.82451561 | 0.013 | 0.0085 | 0.0109 | 0.0109 | TRUE | 0.2398 |
| Phldb1 | 1966 | 1710 | 1759 | 2156 | 1642 | 1494 | 2088 | 1708 | 1766 | 1695 | 1767 | 1635 | 1776.54 | 1854 | 1699 | 0.917 | -0.126 | 0.22552232 | 0.43826245 | 0.006 | 0.0088 | 0.0072 | 0.0072 | TRUE | 0.1471 |
| Phldb2 | 216 | 235 | 223 | 503 | 326 | 270 | 229 | 235 | 224 | 395 | 351 | 296 | 288.3 | 229 | 347 | 1.518 | 0.602 | 7.84E-05 | 0.00113688 | 0.0087 | 0.0183 | 0.0131 | 0.0131 | TRUE | 0.1691 |
| Phldb3 | 203 | 264 | 269 | 128 | 127 | 148 | 216 | 264 | 270 | 101 | 137 | 162 | 191.44 | 250 | 133 | 0.53 | -0.917 | 1.36E-05 | 0.00025939 | 0.0286 | 0.0241 | 0.0263 | 0.0263 | TRUE | 0.5133 |
| Phlpp1 | 810 | 756 | 812 | 1192 | 906 | 842 | 860 | 755 | 815 | 937 | 975 | 922 | 877.38 | 810 | 945 | 1.166 | 0.221 | 0.01766101 | 0.07933035 | 0.0012 | 0.0107 | 0.0051 | 0.0051 | TRUE | 0.0423 |
| Phlpp2 | 668 | 899 | 944 | 1486 | 1080 | 1224 | 710 | 898 | 948 | 1168 | 1162 | 1340 | 1037.55 | 852 | 1223 | 1.435 | 0.521 | 9.59E-05 | 0.00133489 | 0.0128 | 0.0101 | 0.0119 | 0.0119 | TRUE | 0.2524 |
| Phospho1 | 1467 | 1508 | 1479 | 1778 | 1156 | 1312 | 1558 | 1506 | 1485 | 1398 | 1244 | 1436 | 1437.84 | 1516 | 1359 | 0.897 | -0.157 | 0.07801115 | 0.22101584 | 0.0021 | 0.0092 | 0.005 | 0.005 | TRUE | 0.0652 |

| | | | | | | | | | | | | | | | | | | | | | | | | | |
|---|---|---|---|---|---|---|---|---|---|---|---|---|---|---|---|---|---|---|---|---|---|---|---|---|---|
| Phospho2 | 745 | 664 | 603 | 930 | 615 | 514 | 791 | 663 | 605 | 731 | 662 | 563 | 669.23 | 686 | 652 | 0.951 | -0.073 | 0.62188051 | 0.78629946 | 0.0164 | 0.0118 | 0.0143 | 0.0143 | TRUE | 0.2091 |
| Phox2a | 129 | 187 | 185 | 173 | 145 | 148 | 137 | 187 | 186 | 136 | 156 | 162 | 160.59 | 170 | 151 | 0.888 | -0.171 | 0.37006369 | 0.58718757 | 0.0129 | 0.0274 | 0.0201 | 0.0201 | TRUE | 0.2996 |
| Phox2b | 0 | 0 | 0 | 1 | 2 | 0 | 0 | 0 | 0 | 1 | 2 | 0 | 0.49 | 0 | 1 | 5.204 | 2.38 | 0.47267317 | NA | 0 | 6.7348 | 6.4099 | 6.4099 | TRUE | 0.1222 |
| Phpt1 | 580 | 940 | 883 | 963 | 706 | 757 | 616 | 939 | 887 | 757 | 760 | 829 | 797.8 | 814 | 782 | 0.96 | -0.059 | 0.71870951 | 0.84765105 | 0.0247 | 0.011 | 0.0179 | 0.0179 | TRUE | 0.5344 |
| Phrf1 | 6915 | 6731 | 6977 | 9398 | 7043 | 6462 | 7345 | 6722 | 7006 | 7388 | 7578 | 7074 | 7185.4 | 7024 | 7347 | 1.046 | 0.065 | 0.35307692 | 0.57159736 | 0.0014 | 0.0074 | 0.0034 | 0.0034 | TRUE | 0.0194 |
| Phtf1 | 963 | 1197 | 1187 | 1752 | 1338 | 1226 | 1023 | 1195 | 1192 | 1377 | 1440 | 1342 | 1261.52 | 1137 | 1386 | 1.219 | 0.286 | 0.00323477 | 0.02247131 | 0.0032 | 0.0095 | 0.006 | 0.006 | TRUE | 0.0914 |
| Phtf2 | 3720 | 3089 | 3011 | 4334 | 3097 | 2730 | 3951 | 3085 | 3023 | 3407 | 3332 | 2988 | 3297.89 | 3353 | 3242 | 0.967 | -0.048 | 0.70044766 | 0.83619323 | 0.0118 | 0.0079 | 0.0109 | 0.0109 | TRUE | 0.2956 |
| Phxr4 | 0 | 0 | 0 | 0 | 0 | 0 | 0 | 0 | 0 | 0 | 0 | 0 | 0 | 0 | 0 | NA | NA | NA | NA | NA | NA | NA | NA | NA | NA |
| Phyh | 2150 | 2449 | 2456 | 2446 | 1893 | 2221 | 2284 | 2446 | 2466 | 1923 | 2037 | 2431 | 2264.41 | 2399 | 2130 | 0.888 | -0.172 | 0.11529704 | 0.28673367 | 0.008 | 0.0084 | 0.0082 | 0.0082 | TRUE | 0.1842 |
| Phyhd1 | 37 | 48 | 45 | 52 | 37 | 47 | 39 | 48 | 45 | 41 | 40 | 51 | 44.09 | 44 | 44 | 0.994 | -0.009 | 0.97651364 | 0.98729439 | 0 | 0.0816 | 0.0396 | 0.0396 | TRUE | 0.1566 |
| Phyhip | 531 | 641 | 713 | 586 | 494 | 568 | 564 | 640 | 716 | 461 | 532 | 622 | 589.01 | 640 | 538 | 0.839 | -0.253 | 0.09271733 | 0.24873893 | 0.0157 | 0.0125 | 0.0147 | 0.0147 | TRUE | 0.2164 |
| Phyhipl | 505 | 437 | 430 | 581 | 418 | 400 | 536 | 436 | 432 | 457 | 450 | 438 | 458.16 | 468 | 448 | 0.958 | -0.062 | 0.62683899 | 0.78984618 | 0.0057 | 0.0141 | 0.0095 | 0.0095 | TRUE | 0.187 |
| Pi15 | 1 | 4 | 0 | 1 | 2 | 0 | 1 | 4 | 0 | 1 | 2 | 0 | 1.33 | 2 | 1 | 0.577 | -0.794 | 0.67935808 | NA | 0.9878 | 2.4793 | 1.8554 | 1.8554 | TRUE | 0.6208 |
| Pi16 | 18 | 19 | 10 | 14 | 5 | 17 | 19 | 19 | 10 | 11 | 5 | 19 | 13.86 | 16 | 12 | 0.726 | -0.463 | 0.44696319 | 0.65297364 | 0.1317 | 0.2447 | 0.193 | 0.193 | TRUE | 0.3513 |
| Pi4k2a | 3201 | 3061 | 3030 | 3976 | 3009 | 2810 | 3400 | 3057 | 3042 | 3126 | 3237 | 3076 | 3156.44 | 3166 | 3146 | 0.994 | -0.009 | 0.90772439 | 0.95354464 | 0.002 | 0.0079 | 0.0041 | 0.0041 | TRUE | 0.0504 |
| Pi4k2b | 3308 | 3615 | 3560 | 3776 | 2821 | 2916 | 3514 | 3610 | 3575 | 2968 | 3035 | 3192 | 3315.71 | 3566 | 3065 | 0.859 | -0.219 | 0.00098442 | 0.00880175 | 5.00E-04 | 0.0079 | 0.0029 | 0.0029 | TRUE | 0.0158 |
| Pi4ka | 7296 | 8073 | 8940 | 8183 | 6024 | 6387 | 7750 | 8063 | 8977 | 6433 | 6481 | 6992 | 7449.13 | 8263 | 6635 | 0.803 | -0.317 | 0.00021976 | 0.00262185 | 0.0039 | 0.0073 | 0.0052 | 0.0052 | TRUE | 0.0697 |
| Pi4kb | 1720 | 1417 | 1388 | 2361 | 1619 | 1357 | 1827 | 1415 | 1394 | 1856 | 1742 | 1485 | 1619.87 | 1545 | 1694 | 1.097 | 0.134 | 0.33930745 | 0.55942318 | 0.0165 | 0.0089 | 0.0135 | 0.0135 | TRUE | 0.3058 |
| Pianp | 448 | 645 | 684 | 978 | 847 | 980 | 476 | 644 | 687 | 769 | 911 | 1073 | 759.96 | 602 | 918 | 1.522 | 0.606 | 0.00055296 | 0.00563025 | 0.0288 | 0.0112 | 0.0208 | 0.0208 | TRUE | 0.2281 |
| Pias1 | 2130 | 2052 | 2136 | 2654 | 1848 | 1941 | 2262 | 2049 | 2145 | 2086 | 1988 | 2125 | 2109.33 | 2152 | 2066 | 0.96 | -0.058 | 0.45365082 | 0.65902006 | 0.0013 | 0.0085 | 0.0039 | 0.0039 | TRUE | 0.0242 |
| Pias2 | 8216 | 8435 | 8590 | 10974 | 7805 | 8108 | 8727 | 8424 | 8625 | 8627 | 8398 | 8875 | 8612.72 | 8592 | 8633 | 1.005 | 0.007 | 0.90682779 | 0.95352883 | 3.00E-04 | 0.0073 | 0.0024 | 0.0024 | TRUE | 0.0073 |
| Pias3 | 875 | 1047 | 1007 | 1003 | 755 | 877 | 929 | 1046 | 1011 | 788 | 812 | 960 | 924.5 | 995 | 853 | 0.857 | -0.223 | 0.04846989 | 0.16053487 | 0.0061 | 0.0105 | 0.0081 | 0.0081 | TRUE | 0.1399 |
| Pias4 | 2753 | 3080 | 3042 | 3312 | 2612 | 2720 | 2924 | 3076 | 3054 | 2604 | 2810 | 2977 | 2907.69 | 3018 | 2797 | 0.926 | -0.111 | 0.17222436 | 0.37111375 | 0.0021 | 0.008 | 0.0044 | 0.0044 | TRUE | 0.0453 |
| Pibf1 | 435 | 443 | 481 | 800 | 613 | 598 | 462 | 442 | 483 | 629 | 660 | 655 | 555.08 | 462 | 648 | 1.4 | 0.485 | 9.95E-07 | 2.74E-05 | 0 | 0.0128 | 0.0052 | 0.0052 | TRUE | 0.0176 |
| Picalm | 9560 | 9516 | 9529 | 12272 | 8850 | 9002 | 10154 | 9504 | 9568 | 9647 | 9522 | 9854 | 9708.28 | 9742 | 9674 | 0.993 | -0.01 | 0.87131916 | 0.93670819 | 6.00E-04 | 0.0072 | 0.0027 | 0.0027 | TRUE | 0.0167 |
| Pick1 | 2074 | 1989 | 1968 | 2818 | 2039 | 1833 | 2203 | 1986 | 1976 | 2215 | 2194 | 2007 | 2096.85 | 2055 | 2139 | 1.041 | 0.058 | 0.50588298 | 0.70029387 | 0.0027 | 0.0085 | 0.005 | 0.005 | TRUE | 0.0477 |
| Pid1 | 269 | 294 | 316 | 534 | 358 | 367 | 286 | 294 | 317 | 420 | 385 | 402 | 350.56 | 299 | 402 | 1.347 | 0.43 | 0.00030163 | 0.00342743 | 0 | 0.0163 | 0.0073 | 0.0073 | TRUE | 0.0329 |
| Piezo1 | 9696 | 9054 | 9901 | 12595 | 8620 | 7644 | 10299 | 9042 | 9942 | 9901 | 9275 | 8368 | 9471.03 | 9761 | 9181 | 0.941 | -0.088 | 0.3504519 | 0.56929934 | 0.0055 | 0.0073 | 0.0063 | 0.0063 | TRUE | 0.0733 |
| Piezo2 | 99 | 92 | 90 | 154 | 126 | 95 | 105 | 92 | 90 | 121 | 136 | 104 | 108 | 96 | 120 | 1.256 | 0.329 | 0.10585091 | 0.27146026 | 0 | 0.0374 | 0.0204 | 0.0204 | TRUE | 0.1309 |
| Pif1 | 1983 | 2723 | 2874 | 2140 | 1694 | 2020 | 2106 | 2719 | 2886 | 1682 | 1823 | 2211 | 2237.95 | 2570 | 1905 | 0.741 | -0.433 | 0.00402574 | 0.02649604 | 0.0224 | 0.0084 | 0.0159 | 0.0159 | TRUE | 0.3023 |
| Pifo | 26 | 16 | 27 | 30 | 22 | 24 | 28 | 16 | 27 | 24 | 24 | 26 | 24.04 | 24 | 25 | 1.04 | 0.056 | 0.88977903 | 0.94623031 | 0 | 0.144 | 0.0771 | 0.0771 | TRUE | 0.4766 |
| Piga | 1810 | 2129 | 2135 | 2391 | 1952 | 2001 | 1923 | 2126 | 2144 | 1880 | 2100 | 2190 | 2060.47 | 2064 | 2057 | 0.996 | -0.006 | 0.94734155 | 0.97367016 | 0.0045 | 0.0085 | 0.0061 | 0.0061 | TRUE | 0.07 |
| Pigb | 385 | 362 | 398 | 531 | 429 | 407 | 409 | 362 | 400 | 417 | 462 | 446 | 415.77 | 390 | 442 | 1.131 | 0.177 | 0.12554613 | 0.30336443 | 0 | 0.0148 | 0.0072 | 0.0072 | TRUE | 0.0473 |
| Pigc | 187 | 181 | 174 | 232 | 150 | 162 | 199 | 181 | 175 | 182 | 161 | 177 | 179.2 | 185 | 173 | 0.942 | -0.086 | 0.58147274 | 0.75780764 | 0 | 0.0253 | 0.0118 | 0.0118 | TRUE | 0.046 |
| Pigf | 1150 | 1216 | 1146 | 1388 | 918 | 1011 | 1222 | 1214 | 1151 | 1091 | 988 | 1107 | 1128.7 | 1196 | 1062 | 0.889 | -0.17 | 0.05429413 | 0.17320531 | 0.0014 | 0.0098 | 0.0048 | 0.0048 | TRUE | 0.0437 |
| Pigg | 1215 | 1392 | 1595 | 1866 | 1384 | 1470 | 1291 | 1390 | 1602 | 1467 | 1489 | 1609 | 1474.57 | 1428 | 1522 | 1.066 | 0.092 | 0.3945404 | 0.60815981 | 0.0063 | 0.0091 | 0.0077 | 0.0077 | TRUE | 0.1371 |
| Pigh | 649 | 764 | 782 | 1056 | 815 | 848 | 689 | 763 | 785 | 830 | 877 | 928 | 812.15 | 746 | 878 | 1.176 | 0.234 | 0.02025909 | 0.08766156 | 0.0026 | 0.011 | 0.0061 | 0.0061 | TRUE | 0.0515 |
| Pigk | 1205 | 1203 | 1253 | 1802 | 1181 | 1264 | 1280 | 1201 | 1258 | 1417 | 1271 | 1384 | 1301.74 | 1246 | 1357 | 1.089 | 0.123 | 0.14826091 | 0.33796373 | 0.0014 | 0.0094 | 0.0045 | 0.0045 | TRUE | 0.0369 |
| Pigl | 1134 | 993 | 992 | 2134 | 1397 | 927 | 1205 | 992 | 996 | 1678 | 1503 | 1015 | 1231.29 | 1064 | 1399 | 1.315 | 0.395 | 0.03136745 | 0.11901069 | 0.0355 | 0.0096 | 0.0235 | 0.0235 | TRUE | 0.6874 |
| Pigm | 1537 | 1751 | 1863 | 2182 | 1711 | 1655 | 1633 | 1749 | 1871 | 1715 | 1841 | 1812 | 1769.98 | 1751 | 1789 | 1.022 | 0.031 | 0.72316901 | 0.85005881 | 0.0024 | 0.0088 | 0.0048 | 0.0048 | TRUE | 0.0434 |
| Pign | 1338 | 1615 | 1721 | 2231 | 1571 | 1513 | 1421 | 1613 | 1728 | 1754 | 1690 | 1656 | 1643.75 | 1587 | 1700 | 1.071 | 0.099 | 0.31334323 | 0.53381393 | 0.0043 | 0.0089 | 0.0064 | 0.0064 | TRUE | 0.1007 |
| Pigo | 2535 | 2730 | 2836 | 3133 | 2365 | 2414 | 2693 | 2726 | 2848 | 2463 | 2545 | 2643 | 2652.79 | 2756 | 2550 | 0.925 | -0.112 | 0.11063321 | 0.27922979 | 7.00E-04 | 0.0081 | 0.0032 | 0.0032 | TRUE | 0.0122 |
| Pigp | 835 | 1133 | 1058 | 1182 | 789 | 893 | 887 | 1132 | 1062 | 929 | 849 | 978 | 972.74 | 1027 | 919 | 0.894 | -0.161 | 0.18691116 | 0.39108023 | 0.0092 | 0.0103 | 0.0097 | 0.0097 | TRUE | 0.1692 |
| Pigq | 2976 | 3204 | 3284 | 4387 | 3267 | 3125 | 3161 | 3200 | 3297 | 3449 | 3515 | 3421 | 3340.5 | 3219 | 3462 | 1.075 | 0.104 | 0.09189288 | 0.24742196 | 0 | 0.0079 | 0.0025 | 0.0025 | TRUE | 0.0055 |
| Pigr | 1 | 0 | 0 | 2 | 0 | 0 | 1 | 0 | 0 | 2 | 0 | 0 | 0.44 | 0 | 1 | 1.494 | 0.579 | 0.86844164 | NA | 0 | 7.5101 | 6.9157 | 6.9157 | TRUE | 0.9721 |
| Pigs | 5586 | 6015 | 6292 | 6469 | 4864 | 4910 | 5933 | 6007 | 6318 | 5085 | 5233 | 5375 | 5658.66 | 6086 | 5231 | 0.859 | -0.219 | 0.00075874 | 0.00717434 | 7.00E-04 | 0.0075 | 0.0029 | 0.0029 | TRUE | 0.0135 |
| Pigt | 5598 | 6066 | 6120 | 6273 | 4793 | 4764 | 5946 | 6058 | 6145 | 4931 | 5157 | 5215 | 5575.44 | 6050 | 5101 | 0.843 | -0.246 | 5.76E-05 | 0.00087583 | 4.00E-04 | 0.0075 | 0.0025 | 0.0025 | TRUE | 0.0105 |
| Pigu | 1347 | 1576 | 1569 | 1810 | 1338 | 1308 | 1431 | 1574 | 1575 | 1423 | 1440 | 1432 | 1479.08 | 1527 | 1432 | 0.937 | -0.093 | 0.24259306 | 0.4591056 | 8.00E-04 | 0.0091 | 0.0039 | 0.0039 | TRUE | 0.0362 |
| Pigv | 785 | 1101 | 1041 | 1039 | 841 | 867 | 834 | 1100 | 1045 | 817 | 905 | 949 | 941.56 | 993 | 890 | 0.896 | -0.159 | 0.22826406 | 0.44159023 | 0.0124 | 0.0104 | 0.0115 | 0.0115 | TRUE | 0.2335 |
| Pigw | 289 | 398 | 445 | 433 | 342 | 355 | 307 | 397 | 447 | 340 | 368 | 389 | 374.71 | 384 | 366 | 0.951 | -0.072 | 0.65910802 | 0.80794771 | 0.0173 | 0.0157 | 0.0165 | 0.0165 | TRUE | 0.3487 |
| Pigx | 1370 | 1696 | 1760 | 1722 | 1225 | 1485 | 1455 | 1694 | 1767 | 1354 | 1318 | 1626 | 1535.59 | 1639 | 1433 | 0.874 | -0.195 | 0.11123627 | 0.28048257 | 0.0109 | 0.009 | 0.0101 | 0.0101 | TRUE | 0.166 |
| Pigyl | 656 | 1075 | 915 | 710 | 515 | 684 | 697 | 1074 | 919 | 558 | 554 | 749 | 758.36 | 897 | 620 | 0.691 | -0.532 | 0.0043266 | 0.02802021 | 0.0364 | 0.0112 | 0.0237 | 0.0237 | TRUE | 0.2185 |
| Pigz | 0 | 2 | 3 | 2 | 2 | 2 | 0 | 2 | 3 | 2 | 2 | 2 | 1.82 | 2 | 2 | 1.163 | 0.218 | 0.87942537 | NA | 0 | 1.8165 | 0.9287 | 0.9287 | TRUE | 0.5077 |
| Pih1d1 | 932 | 1118 | 1125 | 1439 | 1126 | 1039 | 990 | 1117 | 1130 | 1131 | 1212 | 1137 | 1119.37 | 1079 | 1160 | 1.075 | 0.104 | 0.26517372 | 0.48484318 | 0.0023 | 0.0098 | 0.0054 | 0.0054 | TRUE | 0.0616 |
| Pih1d2 | 34 | 31 | 24 | 54 | 36 | 28 | 36 | 31 | 24 | 42 | 39 | 31 | 33.83 | 30 | 37 | 1.234 | 0.303 | 0.38126438 | 0.59555331 | 0 | 0.1043 | 0.0567 | 0.0567 | TRUE | 0.2205 |
| Pih1d3 | 0 | 0 | 0 | 0 | 0 | 0 | 0 | 0 | 0 | 0 | 0 | 0 | 0 | 0 | 0 | NA | NA | NA | NA | NA | NA | NA | NA | NA | NA |
| Pik3ap1 | 111 | 65 | 66 | 99 | 81 | 75 | 118 | 65 | 66 | 78 | 87 | 82 | 82.69 | 83 | 82 | 0.993 | -0.011 | 0.97007904 | 0.98456155 | 0.0496 | 0.0467 | 0.048 | 0.048 | TRUE | 1.2466 |
| Pik3c2a | 4372 | 4476 | 4447 | 5425 | 3792 | 3854 | 4644 | 4470 | 4465 | 4265 | 4080 | 4219 | 4357.13 | 4526 | 4188 | 0.925 | -0.112 | 0.07101997 | 0.20748529 | 3.00E-04 | 0.0077 | 0.0025 | 0.0025 | TRUE | 0.0062 |
| Pik3c2b | 1376 | 1575 | 1575 | 1894 | 1447 | 1457 | 1462 | 1573 | 1581 | 1489 | 1557 | 1595 | 1542.78 | 1539 | 1547 | 1.005 | 0.007 | 0.93288375 | 0.96671764 | 9.00E-04 | 0.009 | 0.0039 | 0.0039 | TRUE | 0.023 |
| Pik3c2g | 13 | 3 | 7 | 9 | 13 | 14 | 14 | 3 | 7 | 7 | 14 | 15 | 10.04 | 8 | 12 | 1.519 | 0.603 | 0.40412817 | 0.61693403 | 0.1983 | 0.3351 | 0.2708 | 0.2708 | TRUE | 0.5597 |
| Pik3c3 | 2325 | 2139 | 2309 | 3002 | 2229 | 2314 | 2470 | 2136 | 2318 | 2360 | 2398 | 2533 | 2369.25 | 2308 | 2430 | 1.053 | 0.074 | 0.38820002 | 0.60203448 | 0.0027 | 0.0083 | 0.0049 | 0.0049 | TRUE | 0.0515 |
| Pik3ca | 2170 | 2192 | 2261 | 2986 | 2138 | 2087 | 2305 | 2189 | 2270 | 2347 | 2300 | 2285 | 2282.77 | 2255 | 2311 | 1.025 | 0.036 | 0.59235116 | 0.76572104 | 0 | 0.0083 | 0.0028 | 0.0028 | TRUE | 0.0079 |
| Pik3cb | 3631 | 3293 | 3185 | 4569 | 3430 | 3287 | 3857 | 3289 | 3198 | 3592 | 3690 | 3598 | 3537.33 | 3448 | 3627 | 1.052 | 0.073 | 0.43788499 | 0.64575242 | 0.0047 | 0.0078 | 0.0061 | 0.0061 | TRUE | 0.1306 |
| Pik3cd | 3008 | 3332 | 3281 | 2611 | 2102 | 2285 | 3195 | 3328 | 3294 | 2053 | 2262 | 2501 | 2772.11 | 3272 | 2272 | 0.694 | -0.528 | 2.18E-08 | 9.53E-07 | 0.0045 | 0.0081 | 0.006 | 0.006 | TRUE | 0.0945 |
| Pik3cg | 4 | 0 | 0 | 10 | 4 | 4 | 4 | 0 | 0 | 8 | 4 | 4 | 3.47 | 1 | 5 | 3.994 | 1.998 | 0.14196903 | NA | 0.6268 | 0.9576 | 0.8783 | 0.8783 | TRUE | 1.9004 |
| Pik3ip1 | 141 | 286 | 341 | 219 | 175 | 267 | 150 | 286 | 342 | 172 | 188 | 292 | 238.42 | 259 | 217 | 0.838 | -0.256 | 0.41020368 | 0.62167122 | 0.1204 | 0.0207 | 0.0653 | 0.0653 | TRUE | 0.577 |
| Pik3r1 | 732 | 771 | 713 | 988 | 742 | 630 | 778 | 770 | 716 | 777 | 798 | 690 | 754.69 | 755 | 755 | 1.001 | 0.002 | 0.98621564 | 0.99257183 | 0.0025 | 0.0113 | 0.0062 | 0.0062 | TRUE | 0.0658 |
| Pik3r2 | 2220 | 1835 | 1689 | 3468 | 2580 | 1965 | 2358 | 1833 | 1696 | 2726 | 2776 | 2151 | 2256.63 | 1962 | 2551 | 1.3 | 0.379 | 0.01336208 | 0.06494505 | 0.0245 | 0.0084 | 0.0165 | 0.0165 | TRUE | 0.376 |
| Pik3r3 | 456 | 278 | 232 | 636 | 394 | 286 | 484 | 278 | 233 | 500 | 424 | 313 | 371.99 | 332 | 412 | 1.245 | 0.316 | 0.25315808 | 0.47107236 | 0.0982 | 0.0158 | 0.0525 | 0.0525 | TRUE | 0.9256 |

| | | | | | | | | | | | | | | | | | | | | | | | | | |
|---|---|---|---|---|---|---|---|---|---|---|---|---|---|---|---|---|---|---|---|---|---|---|---|---|---|
| Pik3r4 | 2907 | 2358 | 2399 | 3935 | 2793 | 2467 | 3088 | 2355 | 2409 | 3093 | 3005 | 2701 | 2775.09 | 2617 | 2933 | 1.121 | 0.165 | 0.19188511 | 0.39737553 | 0.0141 | 0.0081 | 0.0111 | 0.0111 | TRUE | 0.2997 |
| Pik3r5 | 764 | 673 | 661 | 689 | 480 | 501 | 812 | 672 | 664 | 542 | 516 | 548 | 625.64 | 716 | 535 | 0.749 | -0.418 | 0.00042193 | 0.00452389 | 0.0057 | 0.0122 | 0.0085 | 0.0085 | TRUE | 0.1609 |
| Pik3r6 | 28 | 34 | 32 | 33 | 31 | 25 | 30 | 34 | 32 | 26 | 33 | 27 | 30.42 | 32 | 29 | 0.9 | -0.152 | 0.65828932 | 0.80743861 | 0 | 0.1152 | 0.0518 | 0.0518 | TRUE | 0.1138 |
| Pikfyve | 5393 | 5450 | 6149 | 6036 | 4451 | 4759 | 5728 | 5443 | 6174 | 4745 | 4789 | 5209 | 5348.17 | 5782 | 4914 | 0.85 | -0.235 | 0.00459785 | 0.02938301 | 0.0031 | 0.0075 | 0.0048 | 0.0048 | TRUE | 0.043 |
| Pilra | 0 | 0 | 0 | 0 | 0 | 0 | 0 | 0 | 0 | 0 | 0 | 0 | 0 | 0 | 0 | NA | NA | NA | NA | NA | NA | NA | NA | NA | NA |
| Pilrb1 | 0 | 0 | 0 | 0 | 0 | 0 | 0 | 0 | 0 | 0 | 0 | 0 | 0 | 0 | 0 | NA | NA | NA | NA | NA | NA | NA | NA | NA | NA |
| Pilrb2 | 0 | 0 | 0 | 0 | 0 | 0 | 0 | 0 | 0 | 0 | 0 | 0 | 0 | 0 | 0 | NA | NA | NA | NA | NA | NA | NA | NA | NA | NA |
| Pim1 | 3622 | 5828 | 6416 | 6068 | 4586 | 5355 | 3847 | 5820 | 6442 | 4770 | 4934 | 5862 | 5279.39 | 5370 | 5189 | 0.966 | -0.05 | 0.78285198 | 0.88783644 | 0.041 | 0.0075 | 0.0232 | 0.0232 | TRUE | 0.7499 |
| Pim2 | 571 | 799 | 770 | 887 | 606 | 771 | 607 | 798 | 773 | 697 | 652 | 844 | 728.49 | 726 | 731 | 1.006 | 0.009 | 0.95103503 | 0.97527468 | 0.0184 | 0.0114 | 0.0151 | 0.0151 | TRUE | 0.2437 |
| Pim3 | 11243 | 16356 | 16834 | 10047 | 8323 | 10243 | 11942 | 16335 | 16903 | 7898 | 8955 | 11213 | 12207.64 | 15060 | 9355 | 0.621 | -0.687 | 3.47E-05 | 0.00056583 | 0.0325 | 0.0072 | 0.0198 | 0.0198 | TRUE | 0.4011 |
| Pin1 | 3166 | 2680 | 2616 | 4784 | 3434 | 2699 | 3363 | 2677 | 2627 | 3761 | 3695 | 2954 | 3179.37 | 2889 | 3470 | 1.202 | 0.265 | 0.05366954 | 0.17222299 | 0.018 | 0.0079 | 0.0133 | 0.0133 | TRUE | 0.2501 |
| Pin1-ps1 | 0 | 1 | 0 | 1 | 1 | 1 | 0 | 1 | 0 | 1 | 1 | 1 | 0.66 | 0 | 1 | 2.676 | 1.42 | 0.53620179 | NA | 0 | 5.0043 | 2.2992 | 2.2992 | TRUE | 0.405 |
| Pin4 | 480 | 468 | 460 | 717 | 487 | 415 | 510 | 467 | 462 | 564 | 524 | 454 | 496.84 | 480 | 514 | 1.074 | 0.102 | 0.40940792 | 0.62130006 | 0.0055 | 0.0135 | 0.0091 | 0.0091 | TRUE | 0.1186 |
| Pinc | 0 | 0 | 0 | 0 | 0 | 0 | 0 | 0 | 0 | 0 | 0 | 0 | 0 | 0 | 0 | NA | NA | NA | NA | NA | NA | NA | NA | NA | NA |
| Pink1 | 1157 | 1456 | 1422 | 1400 | 1099 | 1200 | 1229 | 1454 | 1428 | 1101 | 1182 | 1314 | 1284.59 | 1370 | 1199 | 0.874 | -0.194 | 0.08321433 | 0.23091204 | 0.0069 | 0.0095 | 0.0083 | 0.0083 | TRUE | 0.0975 |
| Pinlyp | 2 | 1 | 6 | 2 | 0 | 2 | 2 | 1 | 6 | 2 | 0 | 2 | 2.15 | 3 | 1 | 0.415 | -1.27 | 0.38306827 | NA | 0 | 1.5381 | 0.9745 | 0.9745 | TRUE | 0.8485 |
| Pinx1 | 802 | 585 | 569 | 997 | 681 | 567 | 852 | 584 | 571 | 784 | 733 | 621 | 690.77 | 669 | 713 | 1.066 | 0.092 | 0.60696916 | 0.77691265 | 0.0315 | 0.0117 | 0.0215 | 0.0215 | TRUE | 0.671 |
| Pion | 78 | 35 | 40 | 49 | 43 | 47 | 83 | 35 | 40 | 39 | 46 | 51 | 49.03 | 53 | 45 | 0.861 | -0.215 | 0.58057929 | 0.75708116 | 0.104 | 0.0741 | 0.0887 | 0.0887 | TRUE | 2.0416 |
| Pip | 0 | 0 | 0 | 0 | 0 | 0 | 0 | 0 | 0 | 0 | 0 | 0 | 0 | 0 | 0 | NA | NA | NA | NA | NA | NA | NA | NA | NA | NA |
| Pip4k2a | 1155 | 1196 | 1163 | 1379 | 1161 | 1114 | 1227 | 1194 | 1168 | 1084 | 1249 | 1219 | 1190.29 | 1196 | 1184 | 0.989 | -0.016 | 0.85970067 | 0.93089328 | 0.0025 | 0.0097 | 0.0053 | 0.0053 | TRUE | 0.068 |
| Pip4k2b | 7413 | 6815 | 6956 | 8604 | 6015 | 5896 | 7874 | 6806 | 6985 | 6764 | 6472 | 6454 | 6892.39 | 7222 | 6563 | 0.909 | -0.138 | 0.09517454 | 0.25292214 | 0.0033 | 0.0074 | 0.0048 | 0.0048 | TRUE | 0.0761 |
| Pip4k2c | 2600 | 3152 | 3123 | 3818 | 2951 | 3111 | 2762 | 3148 | 3136 | 3001 | 3175 | 3405 | 3104.57 | 3015 | 3194 | 1.059 | 0.083 | 0.37031257 | 0.58728707 | 0.0042 | 0.008 | 0.0058 | 0.0058 | TRUE | 0.0655 |
| Pip5k1a | 1201 | 1214 | 1106 | 1779 | 1217 | 1040 | 1276 | 1212 | 1111 | 1399 | 1309 | 1138 | 1240.84 | 1200 | 1282 | 1.07 | 0.097 | 0.38228541 | 0.59655738 | 0.0065 | 0.0096 | 0.0081 | 0.0081 | TRUE | 0.1146 |
| Pip5k1b | 422 | 395 | 368 | 428 | 304 | 311 | 448 | 394 | 370 | 336 | 327 | 340 | 369.37 | 404 | 334 | 0.829 | -0.271 | 0.03072135 | 0.11725441 | 0.0027 | 0.0158 | 0.0086 | 0.0086 | TRUE | 0.1042 |
| Pip5k1c | 2564 | 2708 | 2625 | 3048 | 2318 | 2000 | 2723 | 2705 | 2636 | 2396 | 2494 | 2189 | 2523.86 | 2688 | 2360 | 0.878 | -0.188 | 0.01868402 | 0.08287906 | 0.0017 | 0.0082 | 0.0042 | 0.0042 | TRUE | 0.0476 |
| Pip5kl1 | 103 | 104 | 105 | 103 | 74 | 92 | 109 | 104 | 105 | 81 | 80 | 101 | 96.67 | 106 | 87 | 0.818 | -0.29 | 0.15865004 | 0.3526252 | 0 | 0.041 | 0.0202 | 0.0202 | TRUE | 0.1673 |
| Pipox | 4292 | 5323 | 5645 | 4776 | 3661 | 4366 | 4559 | 5316 | 5668 | 3755 | 3939 | 4779 | 4669.33 | 5181 | 4158 | 0.802 | -0.318 | 0.01151611 | 0.05844019 | 0.0143 | 0.0076 | 0.0112 | 0.0112 | TRUE | 0.208 |
| Pir | 1324 | 1298 | 1400 | 1594 | 1079 | 1044 | 1406 | 1296 | 1406 | 1253 | 1161 | 1143 | 1277.54 | 1369 | 1186 | 0.867 | -0.207 | 0.01719832 | 0.07785645 | 0.0016 | 0.0095 | 0.0046 | 0.0046 | TRUE | 0.0307 |
| Pira1 | 0 | 0 | 0 | 0 | 0 | 0 | 0 | 0 | 0 | 0 | 0 | 0 | 0 | 0 | 0 | NA | NA | NA | NA | NA | NA | NA | NA | NA | NA |
| Pira11 | 0 | 0 | 0 | 0 | 0 | 0 | 0 | 0 | 0 | 0 | 0 | 0 | 0 | 0 | 0 | NA | NA | NA | NA | NA | NA | NA | NA | NA | NA |
| Pira2 | 0 | 0 | 0 | 0 | 0 | 0 | 0 | 0 | 0 | 0 | 0 | 0 | 0 | 0 | 0 | NA | NA | NA | NA | NA | NA | NA | NA | NA | NA |
| Pira4 | 0 | 0 | 0 | 0 | 0 | 0 | 0 | 0 | 0 | 0 | 0 | 0 | 0 | 0 | 0 | NA | NA | NA | NA | NA | NA | NA | NA | NA | NA |
| Pira6 | 0 | 0 | 0 | 0 | 0 | 0 | 0 | 0 | 0 | 0 | 0 | 0 | 0 | 0 | 0 | NA | NA | NA | NA | NA | NA | NA | NA | NA | NA |
| Pira7 | 0 | 0 | 0 | 0 | 0 | 0 | 0 | 0 | 0 | 0 | 0 | 0 | 0 | 0 | 0 | NA | NA | NA | NA | NA | NA | NA | NA | NA | NA |
| Pirt | 41 | 33 | 54 | 88 | 65 | 41 | 44 | 33 | 54 | 69 | 70 | 45 | 52.45 | 44 | 61 | 1.412 | 0.498 | 0.12224028 | 0.297435 | 0.0391 | 0.0697 | 0.0551 | 0.0551 | TRUE | 0.3186 |
| Pisd | 7408 | 7256 | 7186 | 6748 | 5163 | 5589 | 7869 | 7247 | 7216 | 5305 | 5555 | 6118 | 6551.43 | 7444 | 5659 | 0.76 | -0.396 | 3.45E-06 | 8.05E-05 | 0.0037 | 0.0074 | 0.0051 | 0.0051 | TRUE | 0.0612 |
| Pisd-ps1 | 123 | 182 | 157 | 196 | 206 | 169 | 131 | 182 | 158 | 154 | 222 | 185 | 171.8 | 157 | 187 | 1.188 | 0.249 | 0.23129839 | 0.44504552 | 0.0244 | 0.0261 | 0.0253 | 0.0253 | TRUE | 0.1973 |
| Pisd-ps2 | 26 | 18 | 22 | 47 | 41 | 44 | 28 | 18 | 22 | 37 | 44 | 48 | 32.82 | 23 | 43 | 1.905 | 0.93 | 0.00872865 | 0.04751707 | 0 | 0.1073 | 0.0565 | 0.0565 | TRUE | 0.2132 |
| Pisd-ps3 | 5058 | 6441 | 5702 | 7514 | 7014 | 6385 | 5372 | 6433 | 5725 | 5907 | 7547 | 6989 | 6328.92 | 5843 | 6814 | 1.166 | 0.221 | 0.06018175 | 0.18615151 | 0.0115 | 0.0074 | 0.0098 | 0.0098 | TRUE | 0.1663 |
| Pithd1 | 1354 | 1382 | 1332 | 1823 | 1345 | 1186 | 1438 | 1380 | 1337 | 1433 | 1447 | 1298 | 1389.06 | 1385 | 1393 | 1.006 | 0.009 | 0.92019287 | 0.96082634 | 0.0017 | 0.0093 | 0.0046 | 0.0046 | TRUE | 0.0416 |
| Pitpna | 4294 | 4030 | 4040 | 6602 | 4654 | 4135 | 4561 | 4025 | 4057 | 5190 | 5007 | 4526 | 4561.03 | 4214 | 4908 | 1.165 | 0.22 | 0.01652745 | 0.07613054 | 0.0047 | 0.0076 | 0.0059 | 0.0059 | TRUE | 0.063 |
| Pitpnb | 8240 | 8136 | 7856 | 10221 | 7239 | 6987 | 8752 | 8125 | 7888 | 8035 | 7789 | 7648 | 8039.69 | 8255 | 7824 | 0.948 | -0.077 | 0.27612304 | 0.49686396 | 0.0016 | 0.0073 | 0.0035 | 0.0035 | TRUE | 0.0338 |
| Pitpnc1 | 5109 | 4318 | 4677 | 4030 | 2766 | 3112 | 5427 | 4312 | 4696 | 3168 | 2976 | 3407 | 3997.66 | 4812 | 3184 | 0.662 | -0.596 | 5.18E-08 | 2.04E-06 | 0.0088 | 0.0077 | 0.0084 | 0.0084 | TRUE | 0.1521 |
| Pitpnm1 | 524 | 562 | 631 | 378 | 338 | 367 | 557 | 561 | 634 | 297 | 364 | 402 | 469 | 584 | 354 | 0.605 | -0.726 | 4.93E-07 | 1.48E-05 | 0.0117 | 0.0139 | 0.0127 | 0.0127 | TRUE | 0.2401 |
| Pitpnm2 | 793 | 866 | 861 | 1034 | 821 | 794 | 842 | 865 | 865 | 813 | 883 | 869 | 856.18 | 857 | 855 | 0.996 | -0.005 | 0.95192758 | 0.97542891 | 0 | 0.0108 | 0.0043 | 0.0043 | TRUE | 0.0234 |
| Pitpnm3 | 45 | 80 | 88 | 115 | 112 | 175 | 48 | 80 | 88 | 90 | 121 | 192 | 103.09 | 72 | 134 | 1.855 | 0.891 | 0.00881872 | 0.04789559 | 0.1108 | 0.0389 | 0.0727 | 0.0727 | TRUE | 0.3986 |
| Pitrm1 | 9744 | 9723 | 10207 | 11804 | 8625 | 8482 | 10350 | 9710 | 10249 | 9279 | 9280 | 9285 | 9692.24 | 10103 | 9281 | 0.919 | -0.122 | 0.03917134 | 0.13860316 | 4.00E-04 | 0.0072 | 0.0024 | 0.0024 | TRUE | 0.0141 |
| Pitx1 | 9 | 14 | 11 | 52 | 44 | 31 | 10 | 14 | 11 | 41 | 47 | 34 | 26.12 | 12 | 41 | 3.526 | 1.818 | 1.29E-05 | 0.00024697 | 0 | 0.133 | 0.069 | 0.069 | TRUE | 0.0983 |
| Pitx2 | 65 | 158 | 182 | 493 | 327 | 337 | 69 | 158 | 183 | 388 | 352 | 369 | 252.98 | 137 | 370 | 2.703 | 1.435 | 9.34E-07 | 2.60E-05 | 0.1082 | 0.0199 | 0.0566 | 0.0566 | TRUE | 1.9048 |
| Pitx3 | 37 | 18 | 41 | 46 | 29 | 29 | 39 | 18 | 41 | 36 | 31 | 32 | 32.93 | 33 | 33 | 1.011 | 0.015 | 0.97002474 | 0.98456155 | 0.061 | 0.107 | 0.0855 | 0.0855 | TRUE | 1.0979 |
| Piwil1 | 237 | 150 | 161 | 240 | 175 | 135 | 252 | 150 | 162 | 189 | 188 | 148 | 181.32 | 188 | 175 | 0.934 | -0.099 | 0.67512522 | 0.8190765 | 0.0446 | 0.0251 | 0.0346 | 0.0346 | TRUE | 0.9511 |
| Piwil2 | 826 | 1223 | 1366 | 2018 | 1597 | 1552 | 877 | 1221 | 1372 | 1586 | 1718 | 1699 | 1412.33 | 1157 | 1668 | 1.441 | 0.527 | 0.0008821 | 0.00806993 | 0.0238 | 0.0092 | 0.0174 | 0.0174 | TRUE | 0.5335 |
| Piwil4 | 400 | 480 | 443 | 435 | 319 | 300 | 425 | 479 | 445 | 342 | 343 | 328 | 393.78 | 450 | 338 | 0.751 | -0.412 | 0.00029513 | 0.0033682 | 0 | 0.0153 | 0.0068 | 0.0068 | TRUE | 0.0388 |
| Pja1 | 2218 | 2085 | 2067 | 2659 | 2011 | 1997 | 2356 | 2082 | 2075 | 2090 | 2164 | 2186 | 2158.96 | 2171 | 2147 | 0.989 | -0.017 | 0.84496588 | 0.9220148 | 0.0024 | 0.0084 | 0.0047 | 0.0047 | TRUE | 0.0668 |
| Pja2 | 3485 | 3581 | 3784 | 4163 | 2992 | 3235 | 3702 | 3576 | 3800 | 3273 | 3219 | 3541 | 3518.44 | 3693 | 3344 | 0.906 | -0.143 | 0.05292102 | 0.17065434 | 0.0014 | 0.0078 | 0.0037 | 0.0037 | TRUE | 0.0319 |
| Pkd1 | 1651 | 1971 | 2046 | 2014 | 1529 | 1598 | 1754 | 1968 | 2054 | 1583 | 1645 | 1749 | 1792.36 | 1925 | 1659 | 0.861 | -0.216 | 0.02280953 | 0.09545021 | 0.0037 | 0.0087 | 0.0059 | 0.0059 | TRUE | 0.0734 |
| Pkd1l2 | 28 | 31 | 29 | 21 | 33 | 19 | 30 | 31 | 29 | 17 | 36 | 21 | 27.11 | 30 | 25 | 0.802 | -0.318 | 0.44371843 | 0.65040214 | 0.0403 | 0.1284 | 0.0865 | 0.0865 | TRUE | 0.9498 |
| Pkd1l3 | 24 | 11 | 11 | 27 | 16 | 14 | 25 | 11 | 11 | 21 | 17 | 15 | 16.88 | 16 | 18 | 1.142 | 0.191 | 0.71719807 | 0.84674362 | 0.0756 | 0.2021 | 0.1415 | 0.1415 | TRUE | 1.2909 |
| Pkd2 | 571 | 390 | 350 | 848 | 562 | 358 | 607 | 389 | 351 | 667 | 605 | 392 | 501.77 | 449 | 555 | 1.236 | 0.306 | 0.21921133 | 0.43131916 | 0.0743 | 0.0135 | 0.0425 | 0.0425 | TRUE | 1.085 |
| Pkd2l1 | 324 | 366 | 302 | 384 | 405 | 350 | 344 | 366 | 303 | 302 | 436 | 383 | 355.61 | 338 | 374 | 1.104 | 0.142 | 0.39882875 | 0.61201859 | 0.0191 | 0.0162 | 0.0177 | 0.0177 | TRUE | 0.3245 |
| Pkd2l2 | 9 | 9 | 9 | 10 | 7 | 0 | 10 | 9 | 9 | 8 | 8 | 0 | 7.16 | 9 | 5 | 0.568 | -0.815 | 0.36014352 | 0.57854793 | 0.3029 | 0.4668 | 0.4223 | 0.4223 | TRUE | 1.4011 |
| Pkdcc | 66 | 116 | 121 | 119 | 110 | 97 | 70 | 116 | 121 | 94 | 118 | 106 | 104.26 | 102 | 106 | 1.031 | 0.044 | 0.86498033 | 0.93316559 | 0.0383 | 0.0385 | 0.0384 | 0.0384 | TRUE | 0.7355 |
| Pkdrej | 1 | 4 | 0 | 9 | 8 | 4 | 1 | 4 | 0 | 7 | 9 | 4 | 4.19 | 2 | 7 | 3.954 | 1.983 | 0.0796287 | NA | 0 | 0.7938 | 0.548 | 0.548 | TRUE | 1.1249 |
| Pkhd1 | 35 | 23 | 21 | 69 | 59 | 29 | 37 | 23 | 21 | 54 | 63 | 32 | 38.45 | 27 | 50 | 1.848 | 0.886 | 0.02677169 | 0.10670786 | 0.0799 | 0.0926 | 0.0866 | 0.0866 | TRUE | 0.4195 |
| Pkhd1l1 | 6 | 3 | 2 | 2 | 6 | 1 | 6 | 3 | 2 | 2 | 6 | 1 | 3.42 | 4 | 3 | 0.792 | -0.336 | 0.76747289 | NA | 0.2976 | 0.9711 | 0.6299 | 0.6299 | TRUE | 1.1755 |
| Pkia | 164 | 135 | 123 | 214 | 172 | 105 | 174 | 135 | 124 | 168 | 185 | 115 | 150.13 | 144 | 156 | 1.085 | 0.118 | 0.61902115 | 0.78469067 | 0.0387 | 0.0288 | 0.0336 | 0.0336 | TRUE | 0.5394 |
| Pkib | 11 | 8 | 10 | 25 | 11 | 13 | 12 | 8 | 10 | 20 | 12 | 14 | 12.57 | 10 | 15 | 1.556 | 0.638 | 0.24458313 | 0.4611329 | 0 | 0.2689 | 0.1331 | 0.1331 | TRUE | 0.3621 |
| Pkig | 470 | 362 | 380 | 742 | 532 | 461 | 499 | 362 | 382 | 583 | 572 | 505 | 483.78 | 414 | 553 | 1.338 | 0.42 | 0.00605685 | 0.03622624 | 0.0159 | 0.0137 | 0.0148 | 0.0148 | TRUE | 0.3706 |
| Pklr | 30 | 25 | 27 | 36 | 27 | 26 | 32 | 25 | 27 | 28 | 29 | 28 | 28.29 | 28 | 28 | 1.023 | 0.033 | 0.92570876 | 0.96333323 | 0 | 0.1233 | 0.0538 | 0.0538 | TRUE | 0.0914 |

| | | | | | | | | | | | | | | | | | | | | | | | | | |
|---|---|---|---|---|---|---|---|---|---|---|---|---|---|---|---|---|---|---|---|---|---|---|---|---|---|
| Pkm | 160893 | 194794 | 199608 | 192524 | 148271 | 165549 | 170897 | 194542 | 200428 | 151347 | 159530 | 181219 | 176327.34 | 188622 | 164032 | 0.87 | -0.202 | 0.04782765 | 0.15907585 | 0.0079 | 0.0069 | 0.0075 | 0.0075 | TRUE | 0.1029 |
| Pkmyt1 | 1377 | 1482 | 1383 | 1760 | 1400 | 1209 | 1463 | 1480 | 1389 | 1384 | 1506 | 1323 | 1424.12 | 1444 | 1404 | 0.973 | -0.04 | 0.64636727 | 0.80145477 | 0.002 | 0.0092 | 0.0048 | 0.0048 | TRUE | 0.0475 |
| Pkn1 | 2193 | 3082 | 3105 | 2843 | 2089 | 2344 | 2329 | 3078 | 3118 | 2235 | 2248 | 2566 | 2595.6 | 2842 | 2350 | 0.827 | -0.275 | 0.03780485 | 0.13505279 | 0.0148 | 0.0082 | 0.0122 | 0.0122 | TRUE | 0.3015 |
| Pkn2 | 8335 | 9058 | 9656 | 11376 | 8173 | 8199 | 8853 | 9046 | 9696 | 8943 | 8794 | 8975 | 9051.14 | 9198 | 8904 | 0.968 | -0.047 | 0.47544798 | 0.67567262 | 0.001 | 0.0073 | 0.003 | 0.003 | TRUE | 0.0273 |
| Pkn3 | 1454 | 1746 | 1828 | 1738 | 1407 | 1399 | 1544 | 1744 | 1836 | 1366 | 1514 | 1531 | 1589.2 | 1708 | 1470 | 0.86 | -0.217 | 0.03221028 | 0.12092848 | 0.0047 | 0.009 | 0.0068 | 0.0068 | TRUE | 0.0843 |
| Pknox1 | 1515 | 1755 | 1801 | 2107 | 1611 | 1627 | 1609 | 1753 | 1808 | 1656 | 1733 | 1781 | 1723.5 | 1723 | 1723 | 0.999 | -0.001 | 0.99263818 | 0.99616826 | 0.0019 | 0.0088 | 0.0045 | 0.0045 | TRUE | 0.0404 |
| Pknox2 | 90 | 45 | 38 | 111 | 57 | 52 | 96 | 45 | 38 | 87 | 61 | 57 | 64.03 | 60 | 68 | 1.156 | 0.21 | 0.5910336 | 0.76481919 | 0.1335 | 0.0584 | 0.0941 | 0.0941 | TRUE | 2.3654 |
| Pkp1 | 12 | 12 | 8 | 19 | 6 | 8 | 13 | 12 | 8 | 15 | 6 | 9 | 10.49 | 11 | 10 | 0.937 | -0.094 | 0.87991523 | 0.94062783 | 0 | 0.3211 | 0.1819 | 0.1819 | TRUE | 0.7868 |
| Pkp2 | 2893 | 3056 | 3057 | 4954 | 4212 | 4243 | 3073 | 3052 | 3070 | 3894 | 4532 | 4645 | 3710.9 | 3065 | 4357 | 1.421 | 0.507 | 2.31E-08 | 1.00E-06 | 0.0039 | 0.0078 | 0.0057 | 0.0057 | TRUE | 0.1061 |
| Pkp3 | 356 | 421 | 514 | 550 | 415 | 414 | 378 | 420 | 516 | 432 | 447 | 453 | 441.13 | 438 | 444 | 1.012 | 0.018 | 0.90081275 | 0.95187048 | 0.0106 | 0.0144 | 0.0124 | 0.0124 | TRUE | 0.2801 |
| Pkp4 | 6906 | 5624 | 5662 | 8818 | 6367 | 5841 | 7335 | 5617 | 5685 | 6932 | 6851 | 6394 | 6468.97 | 6212 | 6726 | 1.083 | 0.115 | 0.3383268 | 0.5583911 | 0.0116 | 0.0074 | 0.0102 | 0.0102 | TRUE | 0.3048 |
| Pla1a | 337 | 321 | 309 | 187 | 136 | 173 | 358 | 321 | 310 | 147 | 146 | 189 | 245.25 | 330 | 161 | 0.487 | -1.038 | 1.18E-10 | 8.58E-09 | 0.0077 | 0.0203 | 0.0141 | 0.0141 | TRUE | 0.2434 |
| Pla2g10 | 1244 | 1373 | 1313 | 1057 | 823 | 886 | 1321 | 1371 | 1318 | 831 | 885 | 970 | 1116.21 | 1337 | 895 | 0.669 | -0.58 | 4.35E-10 | 2.88E-08 | 0.0019 | 0.0099 | 0.0053 | 0.0053 | TRUE | 0.0626 |
| Pla2g12a | 1555 | 1822 | 1827 | 2251 | 1704 | 1645 | 1652 | 1820 | 1835 | 1770 | 1833 | 1801 | 1784.92 | 1769 | 1801 | 1.018 | 0.026 | 0.74567841 | 0.86476763 | 0.0012 | 0.0087 | 0.004 | 0.004 | TRUE | 0.0402 |
| Pla2g12b | 27 | 38 | 29 | 36 | 32 | 28 | 29 | 38 | 29 | 28 | 34 | 31 | 31.52 | 32 | 31 | 0.971 | -0.042 | 0.90183087 | 0.95205059 | 0 | 0.1114 | 0.0531 | 0.0531 | TRUE | 0.1893 |
| Pla2g15 | 2150 | 2858 | 2834 | 3136 | 2378 | 2669 | 2284 | 2854 | 2846 | 2465 | 2559 | 2922 | 2654.85 | 2661 | 2649 | 0.995 | -0.007 | 0.95065499 | 0.97507515 | 0.0116 | 0.0081 | 0.01 | 0.01 | TRUE | 0.1865 |
| Pla2g16 | 698 | 753 | 793 | 704 | 523 | 519 | 741 | 752 | 796 | 553 | 563 | 568 | 662.33 | 763 | 561 | 0.735 | -0.444 | 1.46E-06 | 3.82E-05 | 0 | 0.0119 | 0.0046 | 0.0046 | TRUE | 0.017 |
| Pla2g1b | 1223 | 629 | 634 | 832 | 575 | 422 | 1299 | 628 | 637 | 654 | 619 | 462 | 716.42 | 855 | 578 | 0.677 | -0.563 | 0.03830306 | 0.13633535 | 0.1048 | 0.0115 | 0.0517 | 0.0517 | TRUE | 2.4579 |
| Pla2g2a | 0 | 0 | 0 | 0 | 0 | 1 | 0 | 0 | 0 | 0 | 0 | 1 | 0.18 | 0 | 0 | 1.859 | 0.894 | 0.82650734 | NA | 0 | 18.0643 | 10 | 10 | TRUE | 0.5383 |
| Pla2g2c | 17 | 60 | 65 | 58 | 34 | 38 | 18 | 60 | 65 | 46 | 37 | 42 | 44.5 | 48 | 42 | 0.864 | -0.211 | 0.64375042 | 0.79971894 | 0.1815 | 0.0809 | 0.1274 | 0.1274 | TRUE | 2.3096 |
| Pla2g2d | 2 | 1 | 0 | 7 | 0 | 3 | 2 | 1 | 0 | 6 | 0 | 3 | 1.98 | 1 | 3 | 2.883 | 1.527 | 0.3694468 | NA | 0 | 1.6666 | 1.4281 | 1.4281 | TRUE | 0.1504 |
| Pla2g2e | 11 | 10 | 9 | 15 | 9 | 10 | 12 | 10 | 9 | 12 | 10 | 11 | 10.52 | 10 | 11 | 1.061 | 0.085 | 0.88009871 | 0.94069646 | 0 | 0.32 | 0.1358 | 0.1358 | TRUE | 0.0514 |
| Pla2g2f | 20 | 8 | 15 | 14 | 9 | 12 | 21 | 8 | 15 | 11 | 10 | 13 | 13.02 | 15 | 11 | 0.765 | -0.387 | 0.50033926 | 0.69619328 | 0 | 0.2599 | 0.1593 | 0.1593 | TRUE | 0.1764 |
| Pla2g3 | 6 | 4 | 3 | 5 | 1 | 6 | 6 | 4 | 3 | 4 | 1 | 7 | 4.16 | 4 | 4 | 0.867 | -0.206 | 0.8338976 | NA | 0 | 0.799 | 0.4558 | 0.4558 | TRUE | 0.2117 |
| Pla2g4a | 239 | 213 | 172 | 425 | 274 | 311 | 254 | 213 | 173 | 334 | 295 | 340 | 268.11 | 213 | 323 | 1.518 | 0.602 | 0.00049074 | 0.00509874 | 0.0152 | 0.0192 | 0.0176 | 0.0176 | TRUE | 0.2916 |
| Pla2g4b | 427 | 481 | 436 | 415 | 358 | 383 | 454 | 480 | 438 | 326 | 385 | 419 | 417.07 | 457 | 377 | 0.822 | -0.283 | 0.0342299 | 0.12607987 | 0.0068 | 0.0148 | 0.0104 | 0.0104 | TRUE | 0.1684 |
| Pla2g4c | 43 | 25 | 11 | 51 | 41 | 21 | 46 | 25 | 11 | 40 | 44 | 23 | 31.48 | 27 | 36 | 1.318 | 0.398 | 0.44987926 | 0.65547305 | 0.2328 | 0.1116 | 0.1678 | 0.1678 | TRUE | 0.4223 |
| Pla2g4d | 1 | 3 | 1 | 4 | 5 | 0 | 1 | 3 | 1 | 3 | 5 | 0 | 2.26 | 2 | 3 | 1.686 | 0.753 | 0.60055629 | NA | 0 | 1.4618 | 1.0185 | 1.0185 | TRUE | 0.1835 |
| Pla2g4e | 17 | 24 | 16 | 23 | 13 | 21 | 18 | 24 | 16 | 18 | 14 | 23 | 18.86 | 19 | 18 | 0.946 | -0.08 | 0.86016538 | 0.9309173 | 0 | 0.1816 | 0.0951 | 0.0951 | TRUE | 0.2263 |
| Pla2g4f | 59 | 81 | 65 | 116 | 101 | 71 | 63 | 81 | 65 | 91 | 109 | 78 | 81.07 | 70 | 93 | 1.328 | 0.409 | 0.09145899 | 0.24674841 | 0.0108 | 0.0475 | 0.0298 | 0.0298 | TRUE | 0.1956 |
| Pla2g5 | 289 | 475 | 458 | 362 | 277 | 323 | 307 | 474 | 460 | 285 | 298 | 354 | 362.9 | 414 | 312 | 0.753 | -0.409 | 0.03322544 | 0.12342876 | 0.0317 | 0.016 | 0.0238 | 0.0238 | TRUE | 0.584 |
| Pla2g6 | 4381 | 5012 | 5028 | 4201 | 3437 | 3597 | 4653 | 5006 | 5049 | 3302 | 3698 | 3937 | 4274.26 | 4903 | 3646 | 0.743 | -0.428 | 3.20E-06 | 7.58E-05 | 0.0045 | 0.0077 | 0.0058 | 0.0058 | TRUE | 0.0836 |
| Pla2g7 | 91 | 84 | 86 | 70 | 46 | 86 | 97 | 84 | 86 | 55 | 49 | 94 | 77.59 | 89 | 66 | 0.741 | -0.432 | 0.13923395 | 0.32391948 | 0.0474 | 0.0494 | 0.0484 | 0.0484 | TRUE | 1.1516 |
| Pla2r1 | 9 | 9 | 7 | 24 | 21 | 17 | 10 | 9 | 7 | 19 | 23 | 19 | 14.27 | 9 | 20 | 2.347 | 1.231 | 0.01669205 | 0.07648271 | 0 | 0.2377 | 0.1062 | 0.1062 | TRUE | 0.0737 |
| Plaa | 9815 | 9940 | 10009 | 12392 | 8923 | 8767 | 10425 | 9927 | 10050 | 9742 | 9601 | 9597 | 9890.27 | 10134 | 9647 | 0.952 | -0.071 | 0.20825359 | 0.41767238 | 2.00E-04 | 0.0072 | 0.0022 | 0.0022 | TRUE | 0.0077 |
| Plac1 | 13 | 1 | 2 | 16 | 13 | 4 | 14 | 1 | 2 | 13 | 14 | 4 | 7.96 | 6 | 10 | 1.86 | 0.895 | 0.36746609 | 0.58475554 | 0.7425 | 0.4208 | 0.572 | 0.572 | TRUE | 3.4954 |
| Plac1l | 0 | 0 | 0 | 0 | 0 | 0 | 0 | 0 | 0 | 0 | 0 | 0 | 0 | 0 | 0 | NA | NA | NA | NA | NA | NA | NA | NA | NA | NA |
| Plac8 | 547 | 562 | 642 | 470 | 324 | 412 | 581 | 561 | 645 | 369 | 349 | 451 | 492.67 | 596 | 390 | 0.653 | -0.614 | 6.53E-06 | 0.00013904 | 0.0091 | 0.0136 | 0.0112 | 0.0112 | TRUE | 0.2115 |
| Plac8l1 | 1 | 1 | 0 | 1 | 0 | 0 | 1 | 1 | 0 | 1 | 0 | 0 | 0.47 | 1 | 0 | 0.51 | -0.973 | 0.73452857 | NA | 0 | 6.9499 | 4.1835 | 4.1835 | TRUE | 0.2562 |
| Plac9a | 0 | 0 | 0 | 0 | 0 | 0 | 0 | 0 | 0 | 0 | 0 | 0 | 0 | 0 | 0 | NA | NA | NA | NA | NA | NA | NA | NA | NA | NA |
| Plac9b | 0 | 0 | 0 | 0 | 0 | 0 | 0 | 0 | 0 | 0 | 0 | 0 | 0 | 0 | 0 | NA | NA | NA | NA | NA | NA | NA | NA | NA | NA |
| Plag1 | 478 | 487 | 570 | 493 | 383 | 351 | 508 | 486 | 572 | 388 | 412 | 384 | 458.38 | 522 | 395 | 0.755 | -0.405 | 0.00052255 | 0.00535872 | 0.0025 | 0.0141 | 0.0076 | 0.0076 | TRUE | 0.0831 |
| Plagl1 | 700 | 424 | 368 | 1122 | 729 | 520 | 744 | 423 | 370 | 882 | 784 | 569 | 628.68 | 512 | 745 | 1.456 | 0.542 | 0.03755879 | 0.13461602 | 0.0985 | 0.0121 | 0.0474 | 0.0474 | TRUE | 1.5377 |
| Plagl2 | 1201 | 925 | 890 | 1368 | 935 | 815 | 1276 | 924 | 894 | 1075 | 1006 | 892 | 1011.12 | 1031 | 991 | 0.962 | -0.056 | 0.72318597 | 0.85005881 | 0.0233 | 0.0102 | 0.0169 | 0.0169 | TRUE | 0.5136 |
| Plat | 53 | 54 | 50 | 119 | 61 | 78 | 56 | 54 | 50 | 94 | 66 | 85 | 67.5 | 53 | 82 | 1.531 | 0.615 | 0.01603923 | 0.07451272 | 0 | 0.0557 | 0.0314 | 0.0314 | TRUE | 0.2628 |
| Plau | 149 | 103 | 89 | 174 | 157 | 112 | 158 | 103 | 89 | 137 | 169 | 123 | 129.8 | 117 | 143 | 1.223 | 0.29 | 0.26335569 | 0.48264194 | 0.0493 | 0.0323 | 0.0407 | 0.0407 | TRUE | 0.956 |
| Plaur | 335 | 402 | 459 | 225 | 163 | 177 | 356 | 401 | 461 | 177 | 175 | 194 | 294.03 | 406 | 182 | 0.448 | -1.16 | 1.79E-14 | 2.28E-12 | 0.0078 | 0.0181 | 0.0125 | 0.0125 | TRUE | 0.1415 |
| Plb1 | 166 | 225 | 191 | 227 | 162 | 182 | 176 | 225 | 192 | 178 | 174 | 199 | 190.8 | 198 | 184 | 0.93 | -0.105 | 0.52337448 | 0.71320739 | 0.005 | 0.0242 | 0.0143 | 0.0143 | TRUE | 0.1569 |
| Plbd1 | 131 | 148 | 140 | 151 | 122 | 147 | 139 | 148 | 141 | 119 | 131 | 161 | 139.74 | 143 | 137 | 0.957 | -0.063 | 0.7345114 | 0.85755524 | 0.0061 | 0.0305 | 0.0181 | 0.0181 | TRUE | 0.2365 |
| Plbd2 | 3281 | 3674 | 3582 | 4267 | 3362 | 3243 | 3485 | 3669 | 3597 | 3354 | 3617 | 3550 | 3545.44 | 3584 | 3507 | 0.978 | -0.032 | 0.6454323 | 0.80077998 | 9.00E-04 | 0.0078 | 0.0031 | 0.0031 | TRUE | 0.018 |
| Plcb1 | 68 | 67 | 95 | 111 | 80 | 69 | 72 | 67 | 95 | 87 | 86 | 76 | 80.57 | 78 | 83 | 1.063 | 0.088 | 0.71391618 | 0.84470263 | 0.0082 | 0.0478 | 0.0286 | 0.0286 | TRUE | 0.3463 |
| Plcb2 | 19 | 11 | 13 | 12 | 10 | 8 | 20 | 11 | 13 | 9 | 11 | 9 | 12.2 | 15 | 10 | 0.656 | -0.609 | 0.27487624 | 0.49548018 | 0 | 0.277 | 0.1386 | 0.1386 | TRUE | 0.4516 |
| Plcb3 | 8047 | 7674 | 7667 | 8167 | 6283 | 6341 | 8547 | 7664 | 7698 | 6420 | 6760 | 6941 | 7338.58 | 7970 | 6707 | 0.841 | -0.249 | 0.00142205 | 0.01175504 | 0.0025 | 0.0074 | 0.0043 | 0.0043 | TRUE | 0.049 |
| Plcb4 | 1257 | 1281 | 1368 | 1587 | 1093 | 1093 | 1335 | 1279 | 1374 | 1248 | 1176 | 1196 | 1268.03 | 1329 | 1207 | 0.908 | -0.139 | 0.08123874 | 0.22701938 | 0 | 0.0095 | 0.0038 | 0.0038 | TRUE | 0.0131 |
| Plcd1 | 1866 | 2321 | 2286 | 1879 | 1496 | 1489 | 1982 | 2318 | 2295 | 1477 | 1610 | 1630 | 1885.35 | 2198 | 1572 | 0.715 | -0.485 | 6.56E-07 | 1.90E-05 | 0.0042 | 0.0087 | 0.0063 | 0.0063 | TRUE | 0.0895 |
| Plcd3 | 1337 | 1584 | 1704 | 1571 | 1218 | 1366 | 1420 | 1582 | 1711 | 1235 | 1310 | 1495 | 1458.98 | 1571 | 1347 | 0.857 | -0.223 | 0.05149421 | 0.16728458 | 0.0082 | 0.0092 | 0.0088 | 0.0088 | TRUE | 0.1111 |
| Plcd4 | 26 | 41 | 30 | 60 | 34 | 59 | 28 | 41 | 30 | 47 | 37 | 65 | 41.17 | 33 | 50 | 1.499 | 0.584 | 0.09649807 | 0.25497864 | 0.0385 | 0.0869 | 0.0636 | 0.0636 | TRUE | 0.2543 |
| Plce1 | 1448 | 1183 | 1310 | 1601 | 1070 | 901 | 1538 | 1181 | 1315 | 1259 | 1151 | 986 | 1238.5 | 1345 | 1132 | 0.842 | -0.247 | 0.07126081 | 0.20801879 | 0.0151 | 0.0096 | 0.0127 | 0.0127 | TRUE | 0.1891 |
| Plcg1 | 8820 | 9004 | 9627 | 8965 | 7066 | 7383 | 9368 | 8992 | 9667 | 7048 | 7603 | 8082 | 8459.89 | 9342 | 7578 | 0.811 | -0.302 | 0.00014519 | 0.00185867 | 0.0028 | 0.0073 | 0.0044 | 0.0044 | TRUE | 0.046 |
| Plcg2 | 6598 | 6925 | 7214 | 4504 | 3253 | 3670 | 7008 | 6916 | 7244 | 3541 | 3500 | 4017 | 5371.01 | 7056 | 3686 | 0.522 | -0.937 | 2.19E-30 | 1.28E-27 | 0.003 | 0.0075 | 0.0046 | 0.0046 | TRUE | 0.0747 |
| Plch1 | 159 | 106 | 134 | 332 | 215 | 183 | 169 | 106 | 135 | 261 | 231 | 200 | 183.66 | 137 | 231 | 1.697 | 0.763 | 0.00031088 | 0.00350733 | 0.0281 | 0.0248 | 0.0264 | 0.0264 | TRUE | 0.3794 |
| Plch2 | 65 | 76 | 79 | 72 | 59 | 44 | 69 | 76 | 79 | 57 | 63 | 48 | 65.42 | 75 | 56 | 0.75 | -0.414 | 0.09105987 | 0.24609259 | 0 | 0.0573 | 0.0278 | 0.0278 | TRUE | 0.1159 |
| Plcl1 | 8 | 4 | 10 | 14 | 29 | 10 | 8 | 4 | 10 | 11 | 31 | 11 | 12.61 | 7 | 18 | 2.346 | 1.23 | 0.07443062 | 0.21433626 | 0.2259 | 0.2681 | 0.2468 | 0.2468 | TRUE | 2.1382 |
| Plcl2 | 405 | 497 | 536 | 573 | 453 | 430 | 430 | 496 | 538 | 450 | 487 | 471 | 478.88 | 488 | 469 | 0.96 | -0.058 | 0.63841524 | 0.79637583 | 0.005 | 0.0138 | 0.009 | 0.009 | TRUE | 0.1247 |
| Plcxd1 | 25 | 40 | 40 | 19 | 28 | 26 | 27 | 40 | 40 | 15 | 30 | 28 | 30.03 | 36 | 24 | 0.68 | -0.557 | 0.1777097 | 0.37833021 | 0.0573 | 0.1166 | 0.0886 | 0.0886 | TRUE | 0.8632 |
| Plcxd2 | 619 | 600 | 653 | 685 | 554 | 471 | 657 | 599 | 656 | 538 | 596 | 516 | 593.76 | 637 | 550 | 0.863 | -0.213 | 0.04877872 | 0.16135445 | 0.0022 | 0.0125 | 0.0067 | 0.0067 | TRUE | 0.0607 |
| Plcxd3 | 1 | 1 | 4 | 5 | 0 | 1 | 1 | 1 | 4 | 4 | 0 | 1 | 1.85 | 2 | 2 | 0.861 | -0.216 | 0.89009247 | NA | 0.6267 | 1.7873 | 1.2288 | 1.2288 | TRUE | 1.0685 |
| Plcz1 | 0 | 0 | 0 | 2 | 0 | 0 | 0 | 0 | 0 | 2 | 0 | 0 | 0.26 | 0 | 1 | 2.886 | 1.529 | 0.70676491 | NA | 0 | 12.5792 | 10 | 10 | TRUE | 0.9669 |
| Pld1 | 240 | 239 | 285 | 386 | 265 | 310 | 255 | 239 | 286 | 303 | 285 | 339 | 284.62 | 260 | 309 | 1.189 | 0.25 | 0.07971318 | 0.22402643 | 0.0044 | 0.0185 | 0.0111 | 0.0111 | TRUE | 0.0878 |

| | | | | | | | | | | | | | | | | | | | | | | | | | |
|---|---|---|---|---|---|---|---|---|---|---|---|---|---|---|---|---|---|---|---|---|---|---|---|---|---|
| Pld2 | 808 | 958 | 981 | 1079 | 776 | 764 | 858 | 957 | 985 | 848 | 835 | 836 | 886.58 | 933 | 840 | 0.9 | -0.152 | 0.10441814 | 0.26901061 | 0.0015 | 0.0106 | 0.0052 | 0.0052 | TRUE | 0.0586 |
| Pld3 | 2965 | 3392 | 3495 | 3146 | 2291 | 2538 | 3149 | 3388 | 3509 | 2473 | 2465 | 2778 | 2960.45 | 3349 | 2572 | 0.768 | -0.381 | 1.41E-05 | 0.00026534 | 0.0034 | 0.008 | 0.0052 | 0.0052 | TRUE | 0.0591 |
| Pld4 | 100 | 87 | 97 | 58 | 47 | 55 | 106 | 87 | 97 | 46 | 51 | 60 | 74.48 | 97 | 52 | 0.536 | -0.901 | 0.00018169 | 0.00224397 | 0 | 0.0511 | 0.0271 | 0.0271 | TRUE | 0.1458 |
| Pld5 | 0 | 0 | 1 | 1 | 1 | 3 | 0 | 0 | 1 | 1 | 1 | 3 | 1.03 | 0 | 2 | 4.619 | 2.208 | 0.2863257 | NA | 0 | 3.2209 | 1.7997 | 1.7997 | TRUE | 0.4588 |
| Pld6 | 1027 | 1347 | 1341 | 1093 | 879 | 1010 | 1091 | 1345 | 1347 | 859 | 946 | 1106 | 1115.53 | 1261 | 970 | 0.769 | -0.38 | 0.00500639 | 0.03137101 | 0.0137 | 0.0099 | 0.0123 | 0.0123 | TRUE | 0.1773 |
| Pldi | 2 | 3 | 0 | 2 | 0 | 0 | 2 | 3 | 0 | 2 | 0 | 0 | 1.12 | 2 | 1 | 0.334 | -1.582 | 0.47988215 | NA | 0 | 2.9604 | 2.4562 | 2.4562 | TRUE | 0.7978 |
| Plec | 6586 | 6541 | 7610 | 6756 | 5779 | 5946 | 6995 | 6533 | 7641 | 5311 | 6218 | 6509 | 6534.5 | 7056 | 6013 | 0.852 | -0.231 | 0.03025238 | 0.11592337 | 0.0083 | 0.0074 | 0.0081 | 0.0081 | TRUE | 0.1278 |
| Plek | 0 | 1 | 0 | 5 | 4 | 2 | 0 | 1 | 0 | 4 | 4 | 2 | 1.9 | 0 | 3 | 9.53 | 3.253 | 0.05554491 | NA | 0 | 1.7374 | 0.9392 | 0.9392 | TRUE | 0.405 |
| Plek2 | 274 | 362 | 322 | 406 | 357 | 363 | 291 | 362 | 323 | 319 | 384 | 397 | 346.09 | 325 | 367 | 1.125 | 0.17 | 0.25393796 | 0.47188737 | 0.01 | 0.0164 | 0.013 | 0.013 | TRUE | 0.1565 |
| Plekha1 | 3066 | 3041 | 2860 | 3655 | 2672 | 2742 | 3257 | 3037 | 2872 | 2873 | 2875 | 3002 | 2985.86 | 3055 | 2917 | 0.955 | -0.067 | 0.39427289 | 0.60787039 | 0.0018 | 0.008 | 0.0041 | 0.0041 | TRUE | 0.0403 |
| Plekha2 | 5820 | 9269 | 10011 | 8413 | 7061 | 7789 | 6182 | 9257 | 10052 | 6614 | 7597 | 8526 | 8038.02 | 8497 | 7579 | 0.892 | -0.165 | 0.35221082 | 0.57073249 | 0.0353 | 0.0073 | 0.0226 | 0.0226 | TRUE | 0.6936 |
| Plekha3 | 1301 | 1691 | 1678 | 2173 | 1624 | 1570 | 1382 | 1689 | 1685 | 1708 | 1747 | 1719 | 1654.96 | 1585 | 1725 | 1.088 | 0.121 | 0.24064682 | 0.45703258 | 0.0053 | 0.0089 | 0.0071 | 0.0071 | TRUE | 0.1511 |
| Plekha4 | 364 | 429 | 429 | 547 | 381 | 432 | 387 | 428 | 431 | 430 | 410 | 473 | 426.45 | 415 | 438 | 1.053 | 0.074 | 0.52898446 | 0.71750455 | 0.0019 | 0.0146 | 0.0077 | 0.0077 | TRUE | 0.0552 |
| Plekha5 | 3522 | 3647 | 3838 | 5084 | 3877 | 3742 | 3741 | 3642 | 3854 | 3997 | 4171 | 4096 | 3916.88 | 3746 | 4088 | 1.091 | 0.126 | 0.04855519 | 0.16075543 | 4.00E-04 | 0.0077 | 0.0027 | 0.0027 | TRUE | 0.0078 |
| Plekha6 | 437 | 409 | 365 | 452 | 362 | 353 | 464 | 408 | 366 | 355 | 389 | 386 | 395.06 | 413 | 377 | 0.912 | -0.133 | 0.31800221 | 0.53887305 | 0.0059 | 0.0152 | 0.0102 | 0.0102 | TRUE | 0.134 |
| Plekha7 | 3619 | 4267 | 4603 | 4488 | 3539 | 3476 | 3844 | 4261 | 4622 | 3528 | 3808 | 3805 | 3978.05 | 4242 | 3714 | 0.875 | -0.192 | 0.03948057 | 0.13932675 | 0.0048 | 0.0077 | 0.006 | 0.006 | TRUE | 0.082 |
| Plekha8 | 3074 | 2971 | 2935 | 4330 | 3010 | 2850 | 3265 | 2967 | 2947 | 3404 | 3239 | 3120 | 3156.93 | 3060 | 3254 | 1.064 | 0.089 | 0.26611306 | 0.48577954 | 0.0022 | 0.0079 | 0.0043 | 0.0043 | TRUE | 0.0418 |
| Plekhb1 | 918 | 1152 | 1051 | 1083 | 678 | 741 | 975 | 1151 | 1055 | 851 | 729 | 811 | 928.82 | 1060 | 797 | 0.753 | -0.41 | 0.00018542 | 0.00228478 | 0.0054 | 0.0105 | 0.0076 | 0.0076 | TRUE | 0.0663 |
| Plekhb2 | 1395 | 1556 | 1376 | 1683 | 1388 | 1415 | 1482 | 1554 | 1382 | 1323 | 1493 | 1549 | 1463.79 | 1473 | 1455 | 0.987 | -0.018 | 0.85271175 | 0.92636518 | 0.0045 | 0.0092 | 0.0064 | 0.0064 | TRUE | 0.0778 |
| Plekhd1 | 630 | 636 | 596 | 810 | 665 | 584 | 669 | 635 | 598 | 637 | 715 | 639 | 649.06 | 634 | 664 | 1.046 | 0.065 | 0.5355742 | 0.72271961 | 0.0022 | 0.012 | 0.0064 | 0.0064 | TRUE | 0.0537 |
| Plekhf1 | 142 | 159 | 120 | 163 | 106 | 125 | 151 | 159 | 120 | 128 | 114 | 137 | 134.86 | 143 | 126 | 0.882 | -0.182 | 0.34466656 | 0.5641524 | 0.007 | 0.0313 | 0.0192 | 0.0192 | TRUE | 0.2046 |
| Plekhf2 | 4078 | 3844 | 3607 | 5822 | 4078 | 3788 | 4332 | 3839 | 3622 | 4577 | 4388 | 4147 | 4150.57 | 3931 | 4371 | 1.112 | 0.153 | 0.10393108 | 0.26817827 | 0.0051 | 0.0077 | 0.0062 | 0.0062 | TRUE | 0.0966 |
| Plekhg1 | 288 | 276 | 237 | 626 | 460 | 380 | 306 | 276 | 238 | 492 | 495 | 416 | 370.42 | 273 | 468 | 1.714 | 0.778 | 9.46E-08 | 3.42E-06 | 0.0092 | 0.0158 | 0.0124 | 0.0124 | TRUE | 0.1427 |
| Plekhg2 | 1156 | 1220 | 1147 | 1548 | 1234 | 1092 | 1228 | 1218 | 1152 | 1217 | 1328 | 1195 | 1223 | 1199 | 1247 | 1.039 | 0.055 | 0.5205626 | 0.71109881 | 0.0013 | 0.0096 | 0.0046 | 0.0046 | TRUE | 0.0381 |
| Plekhg3 | 1324 | 1711 | 1744 | 2005 | 1501 | 1490 | 1406 | 1709 | 1751 | 1576 | 1615 | 1631 | 1614.75 | 1622 | 1607 | 0.991 | -0.014 | 0.89800326 | 0.95044288 | 0.0065 | 0.0089 | 0.0075 | 0.0075 | TRUE | 0.1626 |
| Plekhg4 | 278 | 318 | 272 | 331 | 290 | 276 | 295 | 318 | 273 | 260 | 312 | 302 | 293.39 | 295 | 291 | 0.984 | -0.023 | 0.87034007 | 0.93630208 | 0.0045 | 0.0181 | 0.0108 | 0.0108 | TRUE | 0.1065 |
| Plekhg5 | 2372 | 3195 | 3221 | 2461 | 2168 | 2370 | 2519 | 3191 | 3234 | 1935 | 2333 | 2594 | 2634.37 | 2981 | 2287 | 0.767 | -0.383 | 0.00725866 | 0.04125534 | 0.019 | 0.0082 | 0.0143 | 0.0143 | TRUE | 0.2229 |
| Plekhg6 | 415 | 419 | 373 | 268 | 191 | 229 | 441 | 418 | 375 | 211 | 206 | 251 | 316.78 | 411 | 223 | 0.54 | -0.89 | 4.05E-10 | 2.73E-08 | 0.0054 | 0.0173 | 0.0111 | 0.0111 | TRUE | 0.1315 |
| Plekhh1 | 1752 | 2686 | 2748 | 2969 | 2165 | 2503 | 1861 | 2683 | 2759 | 2334 | 2329 | 2740 | 2451.01 | 2434 | 2468 | 1.014 | 0.019 | 0.90218845 | 0.95205059 | 0.0268 | 0.0082 | 0.0175 | 0.0175 | TRUE | 0.5138 |
| Plekhh2 | 889 | 764 | 767 | 1156 | 766 | 653 | 944 | 763 | 770 | 909 | 824 | 715 | 820.86 | 826 | 816 | 0.989 | -0.015 | 0.90991984 | 0.95498953 | 0.0134 | 0.0109 | 0.0122 | 0.0122 | TRUE | 0.1863 |
| Plekhh3 | 132 | 265 | 285 | 251 | 206 | 264 | 140 | 265 | 286 | 197 | 222 | 289 | 233.16 | 230 | 236 | 1.022 | 0.032 | 0.90672803 | 0.95352883 | 0.0828 | 0.021 | 0.0494 | 0.0494 | TRUE | 1.2814 |
| Plekhj1 | 1299 | 1392 | 1295 | 1892 | 1413 | 1346 | 1380 | 1390 | 1300 | 1487 | 1520 | 1473 | 1425.22 | 1357 | 1493 | 1.101 | 0.139 | 0.06642004 | 0.19885665 | 0 | 0.0092 | 0.0034 | 0.0034 | TRUE | 0.016 |
| Plekhm1 | 3876 | 4006 | 3977 | 4708 | 3557 | 3614 | 4117 | 4001 | 3993 | 3701 | 3827 | 3956 | 3932.57 | 4037 | 3828 | 0.948 | -0.077 | 0.23239435 | 0.44654725 | 5.00E-04 | 0.0077 | 0.0027 | 0.0027 | TRUE | 0.0105 |
| Plekhm2 | 483 | 639 | 571 | 803 | 508 | 546 | 513 | 638 | 573 | 631 | 547 | 598 | 583.34 | 575 | 592 | 1.03 | 0.043 | 0.7293903 | 0.85448591 | 0.0068 | 0.0126 | 0.0094 | 0.0094 | TRUE | 0.109 |
| Plekhm3 | 1521 | 1380 | 1410 | 1589 | 1208 | 1037 | 1616 | 1378 | 1416 | 1249 | 1300 | 1135 | 1348.94 | 1470 | 1228 | 0.836 | -0.259 | 0.01238479 | 0.06156096 | 0.005 | 0.0093 | 0.007 | 0.007 | TRUE | 0.0902 |
| Plekhn1 | 80 | 141 | 122 | 102 | 105 | 147 | 85 | 141 | 123 | 80 | 113 | 161 | 117.06 | 116 | 118 | 1.011 | 0.016 | 0.95675792 | 0.97766996 | 0.0829 | 0.035 | 0.058 | 0.058 | TRUE | 0.3091 |
| Plekho1 | 135 | 175 | 159 | 195 | 146 | 119 | 143 | 175 | 160 | 153 | 157 | 130 | 153.08 | 159 | 147 | 0.923 | -0.116 | 0.51068601 | 0.70422593 | 0 | 0.0284 | 0.0157 | 0.0157 | TRUE | 0.0968 |
| Plekho2 | 737 | 615 | 623 | 994 | 680 | 665 | 783 | 614 | 626 | 781 | 732 | 728 | 710.6 | 674 | 747 | 1.109 | 0.149 | 0.23344038 | 0.44812824 | 0.0085 | 0.0115 | 0.0099 | 0.0099 | TRUE | 0.2331 |
| Plekhs1 | 43 | 31 | 17 | 47 | 32 | 36 | 46 | 31 | 17 | 37 | 34 | 39 | 34.08 | 31 | 37 | 1.185 | 0.245 | 0.54952361 | 0.73404177 | 0.0762 | 0.1036 | 0.0909 | 0.0909 | TRUE | 0.2318 |
| Plg | 0 | 0 | 1 | 7 | 0 | 2 | 0 | 0 | 1 | 6 | 0 | 2 | 1.45 | 0 | 3 | 7.129 | 2.834 | 0.18715184 | NA | 1.8007 | 2.2799 | 2.1377 | 2.1377 | TRUE | 0.1701 |
| Plgrkt | 314 | 348 | 317 | 437 | 303 | 304 | 334 | 348 | 318 | 344 | 326 | 333 | 333.62 | 333 | 334 | 1.004 | 0.005 | 0.96273794 | 0.98130897 | 0 | 0.0168 | 0.0069 | 0.0069 | TRUE | 0.0175 |
| Plin1 | 2 | 0 | 1 | 0 | 2 | 0 | 2 | 0 | 1 | 0 | 2 | 0 | 0.88 | 1 | 1 | 0.671 | -0.576 | 0.81366899 | NA | 0 | 3.7504 | 3.1118 | 3.1118 | TRUE | 0.5044 |
| Plin2 | 1264 | 1857 | 1934 | 2232 | 1626 | 1803 | 1343 | 1855 | 1942 | 1755 | 1749 | 1974 | 1769.48 | 1713 | 1826 | 1.066 | 0.092 | 0.53406232 | 0.72173954 | 0.021 | 0.0088 | 0.0151 | 0.0151 | TRUE | 0.4309 |
| Plin3 | 3545 | 4406 | 4641 | 4666 | 3599 | 3899 | 3765 | 4400 | 4660 | 3668 | 3872 | 4268 | 4105.7 | 4275 | 3936 | 0.92 | -0.12 | 0.2710308 | 0.49100595 | 0.0084 | 0.0077 | 0.0083 | 0.0083 | TRUE | 0.1323 |
| Plin4 | 10 | 14 | 16 | 6 | 11 | 13 | 11 | 14 | 16 | 5 | 12 | 14 | 11.91 | 14 | 10 | 0.741 | -0.433 | 0.47985126 | 0.67912904 | 0.0687 | 0.2835 | 0.1839 | 0.1839 | TRUE | 1.0339 |
| Plin5 | 244 | 242 | 278 | 256 | 206 | 185 | 259 | 242 | 279 | 201 | 222 | 203 | 234.23 | 260 | 209 | 0.801 | -0.32 | 0.02319315 | 0.0965516 | 0 | 0.021 | 0.01 | 0.01 | TRUE | 0.0468 |
| Plk1 | 7623 | 9141 | 9197 | 9913 | 7599 | 7992 | 8097 | 9129 | 9235 | 7793 | 8176 | 8749 | 8529.72 | 8820 | 8239 | 0.934 | -0.099 | 0.25785346 | 0.47660433 | 0.0041 | 0.0073 | 0.0054 | 0.0054 | TRUE | 0.0628 |
| Plk1s1 | 294 | 423 | 451 | 631 | 495 | 472 | 312 | 422 | 453 | 496 | 533 | 517 | 455.48 | 396 | 515 | 1.3 | 0.378 | 0.01494839 | 0.07064591 | 0.0162 | 0.0141 | 0.0152 | 0.0152 | TRUE | 0.3889 |
| Plk2 | 2191 | 2151 | 1766 | 2593 | 1795 | 1851 | 2327 | 2148 | 1773 | 2038 | 1931 | 2026 | 2040.77 | 2083 | 1998 | 0.96 | -0.059 | 0.60660641 | 0.77676352 | 0.0095 | 0.0085 | 0.0091 | 0.0091 | TRUE | 0.2048 |
| Plk3 | 640 | 515 | 537 | 1184 | 709 | 570 | 680 | 514 | 539 | 931 | 763 | 624 | 675.15 | 578 | 773 | 1.339 | 0.421 | 0.01664479 | 0.07641362 | 0.0294 | 0.0118 | 0.0208 | 0.0208 | TRUE | 0.1849 |
| Plk4 | 3300 | 3487 | 3655 | 4513 | 3221 | 3531 | 3505 | 3482 | 3670 | 3548 | 3466 | 3865 | 3589.38 | 3552 | 3626 | 1.021 | 0.029 | 0.69977807 | 0.83581217 | 0.0016 | 0.0078 | 0.0039 | 0.0039 | TRUE | 0.0401 |
| Plk5 | 595 | 769 | 880 | 490 | 441 | 569 | 632 | 768 | 884 | 385 | 474 | 623 | 627.69 | 761 | 494 | 0.648 | -0.626 | 0.001407 | 0.0116489 | 0.0407 | 0.0122 | 0.026 | 0.026 | TRUE | 0.3642 |
| Pllp | 148 | 125 | 75 | 187 | 125 | 95 | 157 | 125 | 75 | 147 | 134 | 104 | 123.81 | 119 | 128 | 1.082 | 0.113 | 0.69350329 | 0.83197184 | 0.0718 | 0.0335 | 0.0516 | 0.0516 | TRUE | 0.3636 |
| Pln | 0 | 0 | 0 | 0 | 0 | 0 | 0 | 0 | 0 | 0 | 0 | 0 | 0 | 0 | 0 | NA | NA | NA | NA | NA | NA | NA | NA | NA | NA |
| Plod1 | 3174 | 3150 | 3172 | 3218 | 2279 | 2431 | 3371 | 3146 | 3185 | 2530 | 2452 | 2661 | 2890.87 | 3234 | 2548 | 0.788 | -0.344 | 2.79E-06 | 6.72E-05 | 0.001 | 0.008 | 0.0035 | 0.0035 | TRUE | 0.0181 |
| Plod2 | 5147 | 5984 | 5820 | 7819 | 5599 | 6229 | 5467 | 5976 | 5844 | 6147 | 6024 | 6819 | 6046.11 | 5762 | 6330 | 1.098 | 0.135 | 0.09943768 | 0.26028966 | 0.0031 | 0.0074 | 0.0047 | 0.0047 | TRUE | 0.0554 |
| Plod3 | 1469 | 1591 | 1546 | 1979 | 1532 | 1518 | 1560 | 1589 | 1552 | 1556 | 1648 | 1662 | 1594.57 | 1567 | 1622 | 1.034 | 0.049 | 0.50991171 | 0.70343107 | 0 | 0.009 | 0.0033 | 0.0033 | TRUE | 0.0159 |
| Plp1 | 158 | 208 | 192 | 144 | 138 | 157 | 168 | 208 | 193 | 113 | 148 | 172 | 166.98 | 190 | 144 | 0.759 | -0.399 | 0.05106624 | 0.16640842 | 0.0213 | 0.0266 | 0.024 | 0.024 | TRUE | 0.4088 |
| Plp2 | 2221 | 3163 | 3370 | 3081 | 2389 | 2921 | 2359 | 3159 | 3384 | 2422 | 2570 | 3197 | 2848.63 | 2967 | 2730 | 0.92 | -0.121 | 0.4494367 | 0.65500976 | 0.0281 | 0.0081 | 0.018 | 0.018 | TRUE | 0.39 |
| Plrg1 | 4382 | 4713 | 4803 | 6278 | 4663 | 4588 | 4654 | 4707 | 4823 | 4935 | 5017 | 5022 | 4859.79 | 4728 | 4991 | 1.056 | 0.078 | 0.17731309 | 0.37791649 | 0 | 0.0076 | 0.0022 | 0.0022 | TRUE | 0.0037 |
| Pls1 | 1973 | 2180 | 2112 | 2171 | 1695 | 1720 | 2096 | 2177 | 2121 | 1707 | 1824 | 1883 | 1967.79 | 2131 | 1805 | 0.846 | -0.241 | 0.00153643 | 0.01248462 | 0.001 | 0.0086 | 0.0037 | 0.0037 | TRUE | 0.028 |
| Pls3 | 1009 | 773 | 714 | 1121 | 694 | 669 | 1072 | 772 | 717 | 881 | 747 | 732 | 820.16 | 854 | 787 | 0.923 | -0.116 | 0.49164954 | 0.68798656 | 0.0265 | 0.0109 | 0.0193 | 0.0193 | TRUE | 0.593 |
| Plscr1 | 879 | 991 | 944 | 1620 | 1150 | 1223 | 934 | 990 | 948 | 1274 | 1237 | 1339 | 1120.14 | 957 | 1283 | 1.34 | 0.423 | 3.17E-07 | 9.96E-06 | 0 | 0.0098 | 0.004 | 0.004 | TRUE | 0.0167 |
| Plscr2 | 2 | 1 | 1 | 11 | 3 | 4 | 2 | 1 | 1 | 9 | 3 | 4 | 3.4 | 1 | 5 | 4.028 | 2.01 | 0.08221417 | NA | 0 | 0.9768 | 0.5013 | 0.5013 | TRUE | 0.745 |
| Plscr3 | 843 | 807 | 803 | 970 | 809 | 751 | 895 | 806 | 806 | 763 | 870 | 822 | 827.12 | 836 | 818 | 0.978 | -0.032 | 0.7539389 | 0.87020836 | 0.0029 | 0.0109 | 0.0062 | 0.0062 | TRUE | 0.0458 |
| Plscr4 | 17 | 17 | 20 | 35 | 15 | 18 | 18 | 17 | 20 | 28 | 16 | 20 | 19.75 | 18 | 21 | 1.162 | 0.216 | 0.62444371 | 0.78834114 | 0 | 0.1737 | 0.0897 | 0.0897 | TRUE | 0.4699 |
| Plscr5 | 0 | 0 | 0 | 0 | 0 | 0 | 0 | 0 | 0 | 0 | 0 | 0 | 0 | 0 | 0 | NA | NA | NA | NA | NA | NA | NA | NA | NA | NA |
| Pltp | 2144 | 3914 | 4147 | 2312 | 1945 | 2228 | 2277 | 3909 | 4164 | 1818 | 2093 | 2439 | 2783.23 | 3450 | 2117 | 0.613 | -0.705 | 0.00081789 | 0.00761498 | 0.0543 | 0.0081 | 0.0316 | 0.0316 | TRUE | 1.0745 |
| Plvap | 196 | 167 | 128 | 273 | 139 | 109 | 208 | 167 | 129 | 215 | 150 | 119 | 164.5 | 168 | 161 | 0.965 | -0.052 | 0.84674493 | 0.92344497 | 0.0665 | 0.0269 | 0.0459 | 0.0459 | TRUE | 0.3838 |

| | | | | | | | | | | | | | | | | | | | | | | | | | |
|---|---|---|---|---|---|---|---|---|---|---|---|---|---|---|---|---|---|---|---|---|---|---|---|---|---|
| Plxdc1 | 1746 | 2389 | 2608 | 982 | 834 | 1011 | 1855 | 2386 | 2619 | 772 | 897 | 1107 | 1605.86 | 2287 | 925 | 0.404 | -1.306 | 9.70E-15 | 1.31E-12 | 0.0302 | 0.009 | 0.0198 | 0.0198 | TRUE | 0.3316 |
| Plxdc2 | 92 | 104 | 93 | 189 | 141 | 115 | 98 | 104 | 93 | 149 | 152 | 126 | 120.19 | 98 | 142 | 1.447 | 0.533 | 0.00448383 | 0.02877093 | 0 | 0.0343 | 0.0167 | 0.0167 | TRUE | 0.0976 |
| Plxna1 | 9759 | 10250 | 10642 | 12768 | 9354 | 10077 | 10366 | 10237 | 10686 | 10037 | 10064 | 11031 | 10403.44 | 10430 | 10377 | 0.995 | -0.007 | 0.91629632 | 0.95891617 | 0.0013 | 0.0072 | 0.0034 | 0.0034 | TRUE | 0.037 |
| Plxna2 | 505 | 437 | 477 | 757 | 522 | 516 | 536 | 436 | 479 | 595 | 562 | 565 | 528.9 | 484 | 574 | 1.187 | 0.248 | 0.03403687 | 0.12550866 | 0.0036 | 0.0131 | 0.0079 | 0.0079 | TRUE | 0.1037 |
| Plxna3 | 734 | 694 | 736 | 943 | 864 | 762 | 780 | 693 | 739 | 741 | 930 | 834 | 786.14 | 737 | 835 | 1.131 | 0.178 | 0.13260953 | 0.31402218 | 0.0071 | 0.0111 | 0.0088 | 0.0088 | TRUE | 0.1185 |
| Plxna4 | 43 | 36 | 44 | 35 | 22 | 20 | 46 | 36 | 44 | 28 | 24 | 22 | 33.15 | 42 | 25 | 0.585 | -0.774 | 0.02215534 | 0.09360675 | 0 | 0.1063 | 0.0504 | 0.0504 | TRUE | 0.1204 |
| Plxnb1 | 1049 | 1008 | 1045 | 1116 | 829 | 798 | 1114 | 1007 | 1049 | 877 | 892 | 874 | 968.84 | 1057 | 881 | 0.834 | -0.262 | 0.00226052 | 0.01694532 | 0 | 0.0103 | 0.0043 | 0.0043 | TRUE | 0.0269 |
| Plxnb2 | 4733 | 7445 | 8156 | 5714 | 4709 | 5546 | 5027 | 7435 | 8189 | 4492 | 5067 | 6071 | 6046.94 | 6884 | 5210 | 0.757 | -0.402 | 0.02786112 | 0.10959798 | 0.0403 | 0.0074 | 0.0239 | 0.0239 | TRUE | 0.6792 |
| Plxnb3 | 36 | 52 | 65 | 46 | 39 | 42 | 38 | 52 | 65 | 36 | 42 | 46 | 46.59 | 52 | 41 | 0.794 | -0.333 | 0.29390988 | 0.51480884 | 0.0227 | 0.0776 | 0.0508 | 0.0508 | TRUE | 0.1528 |
| Plxnc1 | 113 | 81 | 81 | 121 | 96 | 83 | 120 | 81 | 81 | 95 | 103 | 91 | 95.25 | 94 | 96 | 1.026 | 0.037 | 0.87531741 | 0.93851164 | 0.0177 | 0.0415 | 0.0298 | 0.0298 | TRUE | 0.5514 |
| Plxnd1 | 3063 | 3785 | 3746 | 3418 | 2697 | 3110 | 3253 | 3780 | 3761 | 2687 | 2902 | 3404 | 3298.02 | 3598 | 2998 | 0.833 | -0.264 | 0.02296837 | 0.09593674 | 0.0105 | 0.0079 | 0.0094 | 0.0094 | TRUE | 0.1708 |
| Pm20d1 | 495 | 700 | 717 | 742 | 515 | 554 | 526 | 699 | 720 | 583 | 554 | 606 | 614.78 | 648 | 581 | 0.896 | -0.158 | 0.26885596 | 0.48886282 | 0.0138 | 0.0123 | 0.0131 | 0.0131 | TRUE | 0.3199 |
| Pm20d2 | 491 | 471 | 520 | 585 | 423 | 406 | 522 | 470 | 522 | 460 | 455 | 444 | 478.91 | 505 | 453 | 0.898 | -0.155 | 0.14466213 | 0.33211224 | 0 | 0.0138 | 0.006 | 0.006 | TRUE | 0.0415 |
| Pmaip1 | 1988 | 2696 | 2751 | 2747 | 2047 | 2146 | 2112 | 2693 | 2762 | 2159 | 2202 | 2349 | 2379.58 | 2522 | 2237 | 0.887 | -0.173 | 0.14705635 | 0.33614098 | 0.0104 | 0.0083 | 0.0099 | 0.0099 | TRUE | 0.2454 |
| Pmch | 0 | 0 | 0 | 0 | 0 | 0 | 0 | 0 | 0 | 0 | 0 | 0 | 0 | 0 | 0 | NA | NA | NA | NA | NA | NA | NA | NA | NA | NA |
| Pmel | 554 | 394 | 371 | 383 | 247 | 267 | 588 | 393 | 373 | 301 | 266 | 292 | 368.93 | 451 | 286 | 0.635 | -0.655 | 0.00072483 | 0.00697075 | 0.0326 | 0.0158 | 0.0242 | 0.0242 | TRUE | 0.8128 |
| Pmepa1 | 1483 | 1543 | 1440 | 1806 | 1406 | 1093 | 1575 | 1541 | 1446 | 1420 | 1513 | 1196 | 1448.52 | 1521 | 1376 | 0.905 | -0.143 | 0.19499152 | 0.40087767 | 0.0069 | 0.0092 | 0.0081 | 0.0081 | TRUE | 0.1548 |
| Pmf1 | 1733 | 2015 | 2073 | 3120 | 2186 | 2074 | 1841 | 2012 | 2082 | 2453 | 2352 | 2270 | 2168.28 | 1978 | 2358 | 1.192 | 0.254 | 0.0023453 | 0.01746462 | 0.002 | 0.0084 | 0.0046 | 0.0046 | TRUE | 0.0445 |
| Pmfbp1 | 20 | 34 | 34 | 53 | 30 | 30 | 21 | 34 | 34 | 42 | 32 | 33 | 32.69 | 30 | 36 | 1.199 | 0.262 | 0.46547291 | 0.6680031 | 0 | 0.1077 | 0.0623 | 0.0623 | TRUE | 0.4003 |
| Pml | 8520 | 10773 | 11373 | 12730 | 9153 | 9119 | 9050 | 10759 | 11420 | 10007 | 9848 | 9982 | 10177.68 | 10410 | 9946 | 0.955 | -0.066 | 0.51173672 | 0.70520852 | 0.0063 | 0.0072 | 0.0071 | 0.0071 | TRUE | 0.1595 |
| Pmm1 | 9933 | 10224 | 9948 | 10612 | 7865 | 8748 | 10551 | 10211 | 9989 | 8342 | 8462 | 9576 | 9521.81 | 10250 | 8793 | 0.858 | -0.221 | 0.00638043 | 0.03768643 | 0.0032 | 0.0072 | 0.0046 | 0.0046 | TRUE | 0.0739 |
| Pmm2 | 361 | 334 | 286 | 652 | 499 | 309 | 383 | 334 | 287 | 513 | 537 | 338 | 398.65 | 335 | 463 | 1.384 | 0.468 | 0.01950269 | 0.08531106 | 0.0381 | 0.0152 | 0.0264 | 0.0264 | TRUE | 0.632 |
| Pmp2 | 0 | 0 | 0 | 0 | 0 | 0 | 0 | 0 | 0 | 0 | 0 | 0 | 0 | 0 | 0 | NA | NA | NA | NA | NA | NA | NA | NA | NA | NA |
| Pmp22 | 173 | 342 | 360 | 144 | 125 | 166 | 184 | 342 | 361 | 113 | 134 | 182 | 219.37 | 296 | 143 | 0.483 | -1.051 | 0.00020077 | 0.00244904 | 0.0878 | 0.0219 | 0.0524 | 0.0524 | TRUE | 1.2273 |
| Pmpca | 7302 | 7726 | 7608 | 11401 | 7881 | 7680 | 7756 | 7716 | 7639 | 8963 | 8479 | 8407 | 8160.05 | 7704 | 8616 | 1.119 | 0.162 | 0.00752772 | 0.0423572 | 5.00E-04 | 0.0073 | 0.0025 | 0.0025 | TRUE | 0.0152 |
| Pmpcb | 6451 | 6876 | 6949 | 7449 | 5233 | 5572 | 6852 | 6867 | 6978 | 5856 | 5630 | 6099 | 6380.4 | 6899 | 5862 | 0.85 | -0.235 | 0.00020976 | 0.0025293 | 6.00E-04 | 0.0074 | 0.0027 | 0.0027 | TRUE | 0.0151 |
| Pms1 | 1649 | 1538 | 1620 | 2278 | 1606 | 1551 | 1752 | 1536 | 1627 | 1791 | 1728 | 1698 | 1688.46 | 1638 | 1739 | 1.062 | 0.087 | 0.30354748 | 0.52431834 | 0.0019 | 0.0089 | 0.0045 | 0.0045 | TRUE | 0.0441 |
| Pms2 | 1045 | 1126 | 1147 | 1843 | 1224 | 1075 | 1110 | 1125 | 1152 | 1449 | 1317 | 1177 | 1221.46 | 1129 | 1314 | 1.165 | 0.221 | 0.03222876 | 0.12094983 | 0.0049 | 0.0096 | 0.0068 | 0.0068 | TRUE | 0.1 |
| Pmvk | 2084 | 2187 | 2081 | 2282 | 1689 | 1757 | 2214 | 2184 | 2090 | 1794 | 1817 | 1923 | 2003.63 | 2163 | 1845 | 0.853 | -0.23 | 0.00184473 | 0.01444641 | 6.00E-04 | 0.0085 | 0.0034 | 0.0034 | TRUE | 0.0165 |
| Pnck | 143 | 209 | 184 | 211 | 173 | 153 | 152 | 209 | 185 | 166 | 186 | 167 | 177.48 | 182 | 173 | 0.951 | -0.073 | 0.68146171 | 0.82347641 | 0.0088 | 0.0255 | 0.017 | 0.017 | TRUE | 0.2189 |
| Pnkd | 1929 | 2232 | 2218 | 2306 | 1825 | 1947 | 2049 | 2229 | 2227 | 1813 | 1964 | 2131 | 2068.81 | 2168 | 1969 | 0.907 | -0.14 | 0.13178427 | 0.31279177 | 0.0037 | 0.0085 | 0.0057 | 0.0057 | TRUE | 0.0626 |
| Pnkp | 999 | 1238 | 1254 | 1544 | 1164 | 1057 | 1061 | 1236 | 1259 | 1214 | 1252 | 1157 | 1196.65 | 1185 | 1208 | 1.019 | 0.027 | 0.79196991 | 0.89301263 | 0.0042 | 0.0097 | 0.0064 | 0.0064 | TRUE | 0.1005 |
| Pnldc1 | 488 | 542 | 584 | 777 | 595 | 550 | 518 | 541 | 586 | 611 | 640 | 602 | 583.18 | 548 | 618 | 1.125 | 0.17 | 0.09539563 | 0.25325364 | 0 | 0.0126 | 0.0058 | 0.0058 | TRUE | 0.0424 |
| Pnlip | 36 | 27 | 19 | 16 | 21 | 23 | 38 | 27 | 19 | 13 | 23 | 25 | 24.11 | 28 | 20 | 0.709 | -0.497 | 0.28599705 | 0.50728131 | 0.0816 | 0.1436 | 0.1135 | 0.1135 | TRUE | 0.2649 |
| Pnliprp1 | 1 | 0 | 0 | 0 | 0 | 0 | 1 | 0 | 0 | 0 | 0 | 0 | 0.18 | 0 | 0 | 0.49 | -1.029 | 0.80087 | NA | 0 | 18.6165 | 10 | 10 | TRUE | 0.4614 |
| Pnliprp2 | 131 | 176 | 126 | 133 | 101 | 116 | 139 | 176 | 127 | 105 | 109 | 127 | 130.27 | 147 | 114 | 0.769 | -0.38 | 0.06446747 | 0.19553479 | 0.0133 | 0.0322 | 0.0226 | 0.0226 | TRUE | 0.3054 |
| Pnma1 | 22 | 36 | 43 | 34 | 20 | 31 | 23 | 36 | 43 | 27 | 22 | 34 | 30.78 | 34 | 28 | 0.799 | -0.323 | 0.40915143 | 0.62130006 | 0.0367 | 0.1139 | 0.0775 | 0.0775 | TRUE | 0.3329 |
| Pnma2 | 914 | 988 | 1015 | 943 | 781 | 757 | 971 | 987 | 1019 | 741 | 840 | 829 | 897.83 | 992 | 803 | 0.808 | -0.307 | 0.00106825 | 0.00937591 | 0.0016 | 0.0106 | 0.0052 | 0.0052 | TRUE | 0.057 |
| Pnma3 | 51 | 63 | 51 | 60 | 60 | 44 | 54 | 63 | 51 | 47 | 65 | 48 | 54.7 | 56 | 53 | 0.946 | -0.08 | 0.7726341 | 0.88119062 | 0 | 0.0671 | 0.0366 | 0.0366 | TRUE | 0.2717 |
| Pnma5 | 1201 | 1530 | 1249 | 3342 | 2686 | 2437 | 1276 | 1528 | 1254 | 2627 | 2890 | 2668 | 2040.45 | 1353 | 2728 | 2.016 | 1.012 | 7.90E-22 | 2.44E-19 | 0.0069 | 0.0085 | 0.0074 | 0.0074 | TRUE | 0.1551 |
| Pnmal1 | 19 | 24 | 9 | 31 | 27 | 35 | 20 | 24 | 9 | 24 | 29 | 38 | 24.15 | 18 | 30 | 1.717 | 0.78 | 0.09619981 | 0.25449008 | 0.0761 | 0.1433 | 0.1136 | 0.1136 | TRUE | 0.9418 |
| Pnmal2 | 76 | 82 | 65 | 177 | 122 | 116 | 81 | 82 | 65 | 139 | 131 | 127 | 104.21 | 76 | 132 | 1.747 | 0.805 | 6.51E-05 | 0.00097405 | 0 | 0.0385 | 0.0189 | 0.0189 | TRUE | 0.1409 |
| Pnmt | 3 | 10 | 8 | 9 | 4 | 10 | 3 | 10 | 8 | 7 | 4 | 11 | 7.26 | 7 | 7 | 1.046 | 0.065 | 0.93211792 | 0.96628017 | 0.0772 | 0.461 | 0.2851 | 0.2851 | TRUE | 0.3581 |
| Pnn | 7711 | 7915 | 8218 | 10088 | 8047 | 8104 | 8190 | 7905 | 8252 | 7930 | 8658 | 8871 | 8301.09 | 8116 | 8486 | 1.045 | 0.064 | 0.37737817 | 0.59235754 | 0.0018 | 0.0073 | 0.0037 | 0.0037 | TRUE | 0.0404 |
| Pno1 | 5853 | 5186 | 4884 | 7809 | 5287 | 4669 | 6217 | 5179 | 4904 | 6139 | 5688 | 5111 | 5539.76 | 5433 | 5646 | 1.039 | 0.056 | 0.63795554 | 0.79611447 | 0.01 | 0.0075 | 0.0099 | 0.0099 | TRUE | 0.1939 |
| Pnoc | 12 | 16 | 18 | 20 | 10 | 15 | 13 | 16 | 18 | 16 | 11 | 16 | 14.95 | 16 | 14 | 0.919 | -0.122 | 0.80434381 | 0.8990865 | 0 | 0.2273 | 0.1084 | 0.1084 | TRUE | 0.1896 |
| Pnp | 4275 | 3212 | 3228 | 3609 | 2659 | 2342 | 4541 | 3208 | 3241 | 2837 | 2861 | 2564 | 3208.61 | 3663 | 2754 | 0.752 | -0.411 | 0.00479652 | 0.03038179 | 0.0228 | 0.0079 | 0.015 | 0.015 | TRUE | 0.5338 |
| Pnp2 | 33 | 11 | 21 | 19 | 16 | 15 | 35 | 11 | 21 | 15 | 17 | 16 | 19.28 | 22 | 16 | 0.724 | -0.466 | 0.37559111 | 0.59147378 | 0.1139 | 0.1778 | 0.1462 | 0.1462 | TRUE | 0.2159 |
| Pnpla1 | 11 | 10 | 12 | 20 | 9 | 12 | 12 | 10 | 12 | 16 | 10 | 13 | 12.04 | 11 | 13 | 1.153 | 0.205 | 0.70481647 | 0.83847762 | 0 | 0.2804 | 0.1289 | 0.1289 | TRUE | 0.1983 |
| Pnpla2 | 2249 | 2677 | 2784 | 2192 | 1651 | 1954 | 2389 | 2674 | 2795 | 1723 | 1776 | 2139 | 2249.39 | 2619 | 1879 | 0.717 | -0.48 | 3.13E-05 | 0.00052247 | 0.0097 | 0.0084 | 0.0091 | 0.0091 | TRUE | 0.1754 |
| Pnpla3 | 520 | 548 | 509 | 588 | 406 | 413 | 552 | 547 | 511 | 462 | 437 | 452 | 493.65 | 537 | 450 | 0.84 | -0.252 | 0.01396237 | 0.06715906 | 0 | 0.0136 | 0.0055 | 0.0055 | TRUE | 0.0208 |
| Pnpla5 | 2 | 3 | 0 | 1 | 4 | 2 | 2 | 3 | 0 | 1 | 4 | 2 | 2.07 | 2 | 2 | 1.395 | 0.481 | 0.74131205 | NA | 0 | 1.601 | 1.0229 | 1.0229 | TRUE | 0.3632 |
| Pnpla6 | 7439 | 6576 | 6363 | 9660 | 6860 | 6253 | 7902 | 6568 | 6389 | 7594 | 7381 | 6845 | 7112.99 | 6953 | 7273 | 1.046 | 0.065 | 0.53383538 | 0.72173954 | 0.007 | 0.0074 | 0.0078 | 0.0078 | TRUE | 0.1738 |
| Pnpla7 | 704 | 796 | 826 | 916 | 694 | 646 | 748 | 795 | 829 | 720 | 747 | 707 | 757.68 | 791 | 725 | 0.916 | -0.126 | 0.1724679 | 0.37119832 | 0 | 0.0113 | 0.0049 | 0.0049 | TRUE | 0.0265 |
| Pnpla8 | 2584 | 2751 | 2715 | 3489 | 2464 | 2545 | 2745 | 2747 | 2726 | 2743 | 2651 | 2786 | 2733.01 | 2739 | 2727 | 0.995 | -0.007 | 0.91653659 | 0.95897673 | 0 | 0.0081 | 0.0026 | 0.0026 | TRUE | 0.007 |
| Pnpo | 2426 | 2479 | 2453 | 2997 | 2196 | 2212 | 2577 | 2476 | 2463 | 2356 | 2363 | 2421 | 2442.64 | 2505 | 2380 | 0.95 | -0.074 | 0.26041535 | 0.47988003 | 0 | 0.0083 | 0.0027 | 0.0027 | TRUE | 0.0075 |
| Pnpt1 | 5721 | 6409 | 6336 | 7402 | 5457 | 5932 | 6077 | 6401 | 6362 | 5819 | 5871 | 6494 | 6170.54 | 6280 | 6061 | 0.965 | -0.051 | 0.49576495 | 0.69173643 | 0.002 | 0.0074 | 0.0039 | 0.0039 | TRUE | 0.0473 |
| Pnrc1 | 2332 | 2201 | 2238 | 2802 | 2198 | 2398 | 2477 | 2198 | 2247 | 2203 | 2365 | 2625 | 2352.49 | 2307 | 2398 | 1.039 | 0.055 | 0.58145456 | 0.75780764 | 0.0049 | 0.0083 | 0.0067 | 0.0067 | TRUE | 0.0833 |
| Pnrc2 | 2379 | 2481 | 2550 | 3445 | 2478 | 2576 | 2527 | 2478 | 2560 | 2708 | 2666 | 2820 | 2626.56 | 2522 | 2731 | 1.083 | 0.115 | 0.08262123 | 0.2296215 | 0 | 0.0082 | 0.0028 | 0.0028 | TRUE | 0.0096 |
| Poc1a | 1028 | 1129 | 1161 | 1442 | 1052 | 1002 | 1092 | 1128 | 1166 | 1134 | 1132 | 1097 | 1124.59 | 1129 | 1121 | 0.993 | -0.01 | 0.903183 | 0.95212067 | 0 | 0.0098 | 0.0036 | 0.0036 | TRUE | 0.01 |
| Poc1b | 991 | 1142 | 1132 | 1713 | 1193 | 1282 | 1053 | 1141 | 1137 | 1347 | 1284 | 1403 | 1227.23 | 1110 | 1345 | 1.211 | 0.276 | 0.00118582 | 0.01013147 | 0.0011 | 0.0096 | 0.0044 | 0.0044 | TRUE | 0.0242 |
| Poc5 | 1351 | 1419 | 1421 | 1790 | 1361 | 1195 | 1435 | 1417 | 1427 | 1407 | 1464 | 1308 | 1409.77 | 1426 | 1393 | 0.977 | -0.034 | 0.6784896 | 0.82135505 | 8.00E-04 | 0.0092 | 0.004 | 0.004 | TRUE | 0.0334 |
| Podn | 17 | 26 | 18 | 38 | 28 | 34 | 18 | 26 | 18 | 30 | 30 | 37 | 26.55 | 21 | 32 | 1.559 | 0.641 | 0.09402319 | 0.25099515 | 0 | 0.131 | 0.0658 | 0.0658 | TRUE | 0.2773 |
| Podnl1 | 22 | 15 | 22 | 29 | 17 | 28 | 23 | 15 | 22 | 23 | 18 | 31 | 22.03 | 20 | 24 | 1.186 | 0.246 | 0.56920819 | 0.74819165 | 0 | 0.1564 | 0.0889 | 0.0889 | TRUE | 0.3285 |
| Podxl | 1725 | 2858 | 3000 | 2838 | 2223 | 2741 | 1832 | 2854 | 3012 | 2231 | 2392 | 3000 | 2553.69 | 2566 | 2541 | 0.99 | -0.015 | 0.93895684 | 0.96938059 | 0.0464 | 0.0082 | 0.0259 | 0.0259 | TRUE | 0.7586 |
| Podxl2 | 162 | 194 | 207 | 219 | 155 | 174 | 172 | 194 | 208 | 172 | 167 | 190 | 183.85 | 191 | 176 | 0.921 | -0.118 | 0.45938156 | 0.66378277 | 0 | 0.0248 | 0.0129 | 0.0129 | TRUE | 0.0815 |
| Pof1b | 11 | 22 | 16 | 29 | 32 | 34 | 12 | 22 | 16 | 23 | 34 | 37 | 24.03 | 17 | 31 | 1.881 | 0.912 | 0.03831082 | 0.13633535 | 0.0372 | 0.144 | 0.0931 | 0.0931 | TRUE | 0.4676 |
| Pofut1 | 1272 | 1265 | 1191 | 2322 | 1540 | 1220 | 1351 | 1263 | 1196 | 1825 | 1657 | 1335 | 1438.02 | 1270 | 1606 | 1.265 | 0.34 | 0.00956041 | 0.0508556 | 0.0128 | 0.0092 | 0.0117 | 0.0117 | TRUE | 0.2604 |
| Pofut2 | 1425 | 1367 | 1368 | 1870 | 1267 | 1176 | 1514 | 1365 | 1374 | 1470 | 1363 | 1287 | 1395.51 | 1418 | 1373 | 0.97 | -0.044 | 0.63895122 | 0.7965804 | 0.0032 | 0.0093 | 0.0056 | 0.0056 | TRUE | 0.0466 |

| | | | | | | | | | | | | | | | | | | | | | | | | | |
|---|---|---|---|---|---|---|---|---|---|---|---|---|---|---|---|---|---|---|---|---|---|---|---|---|---|
| Pogk | 244 | 234 | 226 | 440 | 334 | 266 | 259 | 234 | 227 | 346 | 359 | 291 | 286.04 | 240 | 332 | 1.386 | 0.471 | 0.00109812 | 0.00957007 | 0.0051 | 0.0184 | 0.0114 | 0.0114 | TRUE | 0.1276 |
| Poglut1 | 1485 | 1639 | 1697 | 2824 | 2092 | 1861 | 1577 | 1637 | 1704 | 2220 | 2251 | 2037 | 1904.37 | 1639 | 2169 | 1.323 | 0.404 | 6.81E-07 | 1.96E-05 | 0.0017 | 0.0086 | 0.0042 | 0.0042 | TRUE | 0.0339 |
| Pogz | 5434 | 5498 | 5682 | 6783 | 4877 | 4787 | 5772 | 5491 | 5705 | 5332 | 5247 | 5240 | 5464.64 | 5656 | 5273 | 0.932 | -0.101 | 0.08856912 | 0.2414297 | 2.00E-04 | 0.0075 | 0.0023 | 0.0023 | TRUE | 0.008 |
| Pola1 | 4363 | 4641 | 4999 | 6302 | 4786 | 4596 | 4634 | 4635 | 5020 | 4954 | 5149 | 5031 | 4903.91 | 4763 | 5045 | 1.059 | 0.083 | 0.22619833 | 0.43904737 | 0.001 | 0.0076 | 0.0032 | 0.0032 | TRUE | 0.0271 |
| Pola2 | 2570 | 2566 | 2563 | 4276 | 3000 | 2847 | 2730 | 2563 | 2574 | 3361 | 3228 | 3116 | 2928.63 | 2622 | 3235 | 1.234 | 0.304 | 2.52E-05 | 0.00043613 | 0.001 | 0.008 | 0.0034 | 0.0034 | TRUE | 0.0156 |
| Polb | 2088 | 2539 | 2433 | 2791 | 2061 | 2024 | 2218 | 2536 | 2443 | 2194 | 2218 | 2216 | 2303.95 | 2399 | 2209 | 0.921 | -0.119 | 0.14067876 | 0.32593224 | 0.0017 | 0.0083 | 0.0043 | 0.0043 | TRUE | 0.0526 |
| Pold1 | 8683 | 10614 | 11091 | 10719 | 8304 | 8829 | 9223 | 10600 | 11137 | 8426 | 8935 | 9665 | 9664.25 | 10320 | 9009 | 0.873 | -0.196 | 0.04938448 | 0.1629694 | 0.0068 | 0.0072 | 0.0071 | 0.0071 | TRUE | 0.1056 |
| Pold2 | 6903 | 8860 | 8914 | 8201 | 6384 | 6927 | 7332 | 8849 | 8951 | 6447 | 6869 | 7583 | 7671.64 | 8377 | 6966 | 0.831 | -0.266 | 0.01484366 | 0.07030455 | 0.0091 | 0.0073 | 0.0085 | 0.0085 | TRUE | 0.1453 |
| Pold3 | 4124 | 4133 | 4383 | 5673 | 4208 | 4119 | 4380 | 4128 | 4401 | 4460 | 4528 | 4509 | 4400.86 | 4303 | 4499 | 1.045 | 0.064 | 0.31210439 | 0.53255705 | 4.00E-04 | 0.0077 | 0.0027 | 0.0027 | TRUE | 0.0155 |
| Pold4 | 216 | 359 | 391 | 306 | 217 | 271 | 229 | 359 | 393 | 241 | 233 | 297 | 291.88 | 327 | 257 | 0.785 | -0.349 | 0.10804101 | 0.27511586 | 0.0437 | 0.0182 | 0.0306 | 0.0306 | TRUE | 0.7667 |
| Poldip2 | 6063 | 7085 | 7045 | 7669 | 5587 | 5724 | 6440 | 7076 | 7074 | 6029 | 6011 | 6266 | 6482.6 | 6863 | 6102 | 0.889 | -0.17 | 0.01717262 | 0.07778476 | 0.0015 | 0.0074 | 0.0035 | 0.0035 | TRUE | 0.0355 |
| Poldip3 | 7218 | 7624 | 7518 | 12364 | 9043 | 8412 | 7667 | 7614 | 7549 | 9720 | 9730 | 9208 | 8581.23 | 7610 | 9553 | 1.255 | 0.328 | 2.70E-08 | 1.16E-06 | 3.00E-04 | 0.0073 | 0.0024 | 0.0024 | TRUE | 0.0121 |
| Pole | 5056 | 4391 | 4728 | 7594 | 5491 | 4748 | 5370 | 4385 | 4747 | 5970 | 5908 | 5197 | 5263.05 | 4834 | 5692 | 1.178 | 0.236 | 0.0244723 | 0.10014975 | 0.0079 | 0.0075 | 0.0077 | 0.0077 | TRUE | 0.1145 |
| Pole2 | 1900 | 1763 | 1770 | 2430 | 1745 | 1797 | 2018 | 1761 | 1777 | 1910 | 1878 | 1967 | 1885.17 | 1852 | 1918 | 1.036 | 0.051 | 0.55957334 | 0.74136428 | 0.0024 | 0.0087 | 0.0049 | 0.0049 | TRUE | 0.0741 |
| Pole3 | 574 | 600 | 605 | 1064 | 803 | 760 | 610 | 599 | 607 | 836 | 864 | 832 | 724.79 | 605 | 844 | 1.394 | 0.479 | 3.64E-08 | 1.52E-06 | 0 | 0.0114 | 0.004 | 0.004 | TRUE | 0.0049 |
| Pole4 | 2489 | 2544 | 2315 | 3788 | 2550 | 2406 | 2644 | 2541 | 2325 | 2978 | 2744 | 2634 | 2644.03 | 2503 | 2785 | 1.113 | 0.155 | 0.08514653 | 0.2346141 | 0.0035 | 0.0081 | 0.0054 | 0.0054 | TRUE | 0.0472 |
| Polg | 6610 | 7331 | 7130 | 8163 | 6275 | 5912 | 7021 | 7322 | 7159 | 6417 | 6752 | 6472 | 6857.01 | 7167 | 6547 | 0.913 | -0.131 | 0.03067242 | 0.11710614 | 4.00E-04 | 0.0074 | 0.0025 | 0.0025 | TRUE | 0.0091 |
| Polg2 | 422 | 628 | 579 | 512 | 442 | 438 | 448 | 627 | 581 | 402 | 476 | 479 | 502.39 | 552 | 452 | 0.818 | -0.29 | 0.06475654 | 0.19603458 | 0.0179 | 0.0135 | 0.0158 | 0.0158 | TRUE | 0.3158 |
| Polh | 1154 | 1359 | 1386 | 1208 | 932 | 1151 | 1226 | 1357 | 1392 | 950 | 1003 | 1260 | 1197.84 | 1325 | 1071 | 0.807 | -0.309 | 0.01758089 | 0.07913406 | 0.0129 | 0.0097 | 0.0113 | 0.0113 | TRUE | 0.284 |
| Poli | 1968 | 2507 | 2584 | 3053 | 2374 | 2454 | 2090 | 2504 | 2595 | 2400 | 2554 | 2686 | 2471.56 | 2396 | 2547 | 1.062 | 0.087 | 0.41645564 | 0.62782817 | 0.0068 | 0.0082 | 0.0079 | 0.0079 | TRUE | 0.1507 |
| Polk | 1170 | 1606 | 1531 | 1498 | 1119 | 1406 | 1243 | 1604 | 1537 | 1178 | 1204 | 1539 | 1384.11 | 1461 | 1307 | 0.894 | -0.162 | 0.26683701 | 0.48648172 | 0.0192 | 0.0093 | 0.0146 | 0.0146 | TRUE | 0.2888 |
| Poll | 1090 | 1280 | 1285 | 1574 | 1160 | 1170 | 1158 | 1278 | 1290 | 1237 | 1248 | 1281 | 1248.76 | 1242 | 1255 | 1.01 | 0.015 | 0.86218887 | 0.93192349 | 0.0011 | 0.0095 | 0.0043 | 0.0043 | TRUE | 0.0421 |
| Polm | 420 | 464 | 445 | 569 | 452 | 476 | 446 | 463 | 447 | 447 | 486 | 521 | 468.5 | 452 | 485 | 1.07 | 0.098 | 0.37896926 | 0.59408638 | 0 | 0.0139 | 0.0068 | 0.0068 | TRUE | 0.0569 |
| Poln | 75 | 85 | 59 | 80 | 53 | 45 | 80 | 85 | 59 | 63 | 57 | 49 | 65.49 | 75 | 56 | 0.759 | -0.397 | 0.13215022 | 0.31323614 | 0.0115 | 0.0572 | 0.0347 | 0.0347 | TRUE | 0.2994 |
| Polq | 1297 | 1011 | 1041 | 1494 | 1101 | 935 | 1378 | 1010 | 1045 | 1174 | 1185 | 1024 | 1135.87 | 1144 | 1128 | 0.986 | -0.02 | 0.88595137 | 0.94396169 | 0.0172 | 0.0098 | 0.0137 | 0.0137 | TRUE | 0.3806 |
| Polr1a | 9421 | 7818 | 8432 | 11386 | 8109 | 7146 | 10007 | 7808 | 8467 | 8951 | 8725 | 7822 | 8629.88 | 8761 | 8499 | 0.97 | -0.043 | 0.70042681 | 0.83619323 | 0.0089 | 0.0073 | 0.0091 | 0.0091 | TRUE | 0.1891 |
| Polr1b | 7264 | 5322 | 5201 | 7851 | 5826 | 5073 | 7716 | 5315 | 5222 | 6172 | 6268 | 5553 | 6041.1 | 6084 | 5998 | 0.986 | -0.021 | 0.89465107 | 0.9490474 | 0.0234 | 0.0074 | 0.0172 | 0.0172 | TRUE | 0.6708 |
| Polr1c | 5948 | 6322 | 6307 | 8006 | 6041 | 6005 | 6318 | 6314 | 6333 | 6294 | 6500 | 6573 | 6388.57 | 6322 | 6456 | 1.021 | 0.03 | 0.59745635 | 0.77004845 | 1.00E-04 | 0.0074 | 0.0022 | 0.0022 | TRUE | 0.006 |
| Polr1d | 5139 | 5509 | 5556 | 6135 | 4235 | 4755 | 5459 | 5502 | 5579 | 4823 | 4557 | 5205 | 5187.3 | 5513 | 4862 | 0.882 | -0.182 | 0.01683515 | 0.07691888 | 0.002 | 0.0075 | 0.004 | 0.004 | TRUE | 0.0461 |
| Polr1e | 2321 | 2098 | 1941 | 2593 | 1848 | 1575 | 2465 | 2095 | 1949 | 2038 | 1988 | 1724 | 2043.4 | 2170 | 1917 | 0.884 | -0.178 | 0.1361095 | 0.31954629 | 0.0107 | 0.0085 | 0.0098 | 0.0098 | TRUE | 0.1714 |
| Polr2a | 43379 | 30988 | 32152 | 51257 | 35125 | 29007 | 46076 | 30948 | 32284 | 40294 | 37792 | 31753 | 36524.58 | 36436 | 36613 | 1.005 | 0.007 | 0.96559199 | 0.98266337 | 0.0268 | 0.007 | 0.0191 | 0.0191 | TRUE | 0.6558 |
| Polr2b | 17933 | 15143 | 15286 | 20436 | 14509 | 13671 | 19048 | 15123 | 15349 | 16065 | 15611 | 14965 | 16026.88 | 16507 | 15547 | 0.942 | -0.086 | 0.42130659 | 0.63145942 | 0.0091 | 0.0071 | 0.0082 | 0.0082 | TRUE | 0.2218 |
| Polr2c | 5036 | 6057 | 5910 | 6855 | 4908 | 5202 | 5349 | 6049 | 5934 | 5389 | 5281 | 5694 | 5616.09 | 5777 | 5455 | 0.944 | -0.083 | 0.29891134 | 0.51926751 | 0.0026 | 0.0075 | 0.0044 | 0.0044 | TRUE | 0.0512 |
| Polr2d | 2088 | 1675 | 1640 | 2759 | 1943 | 1704 | 2218 | 1673 | 1647 | 2169 | 2091 | 1865 | 1943.69 | 1846 | 2042 | 1.107 | 0.146 | 0.28594259 | 0.50724177 | 0.0162 | 0.0086 | 0.013 | 0.013 | TRUE | 0.375 |
| Polr2e | 8364 | 9401 | 9439 | 11486 | 8246 | 8396 | 8884 | 9389 | 9478 | 9029 | 8872 | 9191 | 9140.5 | 9250 | 9031 | 0.976 | -0.035 | 0.57221502 | 0.75089236 | 6.00E-04 | 0.0073 | 0.0026 | 0.0026 | TRUE | 0.0146 |
| Polr2f | 2883 | 2898 | 2855 | 3902 | 2712 | 2567 | 3062 | 2894 | 2867 | 3067 | 2918 | 2810 | 2936.44 | 2941 | 2932 | 0.997 | -0.004 | 0.95836088 | 0.97866252 | 0.0013 | 0.008 | 0.0036 | 0.0036 | TRUE | 0.0203 |
| Polr2g | 2762 | 3040 | 2936 | 4067 | 2696 | 2547 | 2934 | 3036 | 2948 | 3197 | 2901 | 2788 | 2967.31 | 2973 | 2962 | 0.997 | -0.004 | 0.95714451 | 0.97778892 | 0.0024 | 0.008 | 0.0044 | 0.0044 | TRUE | 0.0595 |
| Polr2h | 2931 | 2802 | 2800 | 3460 | 2503 | 2471 | 3113 | 2798 | 2811 | 2720 | 2693 | 2705 | 2806.85 | 2907 | 2706 | 0.931 | -0.103 | 0.17336353 | 0.37246828 | 0.0014 | 0.0081 | 0.0038 | 0.0038 | TRUE | 0.0463 |
| Polr2i | 1196 | 1089 | 1112 | 1546 | 1056 | 984 | 1270 | 1088 | 1117 | 1215 | 1136 | 1077 | 1150.53 | 1158 | 1143 | 0.988 | -0.018 | 0.86233586 | 0.93194197 | 0.0045 | 0.0098 | 0.0067 | 0.0067 | TRUE | 0.0856 |
| Polr2j | 1667 | 1553 | 1537 | 2483 | 1716 | 1510 | 1771 | 1551 | 1543 | 1952 | 1846 | 1653 | 1719.36 | 1622 | 1817 | 1.121 | 0.165 | 0.1119625 | 0.28141396 | 0.0058 | 0.0088 | 0.0072 | 0.0072 | TRUE | 0.0776 |
| Polr2k | 305 | 298 | 300 | 364 | 281 | 304 | 324 | 298 | 301 | 286 | 302 | 333 | 307.35 | 308 | 307 | 0.997 | -0.005 | 0.96966577 | 0.98448447 | 0 | 0.0176 | 0.0087 | 0.0087 | TRUE | 0.0596 |
| Polr2l | 872 | 634 | 630 | 1231 | 794 | 624 | 926 | 633 | 633 | 968 | 854 | 668 | 782.84 | 731 | 835 | 1.144 | 0.194 | 0.30422591 | 0.52512118 | 0.0362 | 0.0111 | 0.0244 | 0.0244 | TRUE | 0.6237 |
| Polr2m | 9454 | 8965 | 9079 | 16183 | 11175 | 10534 | 10042 | 8953 | 9116 | 12722 | 12024 | 11531 | 10731.34 | 9370 | 12092 | 1.291 | 0.368 | 4.28E-06 | 9.67E-05 | 0.003 | 0.0072 | 0.0045 | 0.0045 | TRUE | 0.0479 |
| Polr3a | 2826 | 2734 | 2871 | 3981 | 2802 | 2696 | 3002 | 2730 | 2883 | 3130 | 3015 | 2951 | 2951.75 | 2872 | 3032 | 1.056 | 0.079 | 0.28436293 | 0.5058795 | 0.0011 | 0.008 | 0.0036 | 0.0036 | TRUE | 0.0225 |
| Polr3b | 6624 | 6197 | 6586 | 7797 | 5833 | 5527 | 7036 | 6189 | 6613 | 6129 | 6276 | 6050 | 6382.23 | 6613 | 6152 | 0.93 | -0.104 | 0.16516927 | 0.36181246 | 0.002 | 0.0074 | 0.0039 | 0.0039 | TRUE | 0.0383 |
| Polr3c | 2419 | 2122 | 2150 | 2978 | 2165 | 1932 | 2569 | 2119 | 2159 | 2341 | 2329 | 2115 | 2272.14 | 2282 | 2262 | 0.991 | -0.013 | 0.90419348 | 0.95248677 | 0.0069 | 0.0084 | 0.0075 | 0.0075 | TRUE | 0.1461 |
| Polr3d | 1742 | 1831 | 1869 | 2467 | 1784 | 1685 | 1850 | 1829 | 1877 | 1939 | 1919 | 1844 | 1876.49 | 1852 | 1901 | 1.027 | 0.038 | 0.57893007 | 0.75593055 | 0 | 0.0087 | 0.0029 | 0.0029 | TRUE | 0.0081 |
| Polr3e | 3916 | 3659 | 3854 | 4981 | 3631 | 3325 | 4159 | 3654 | 3870 | 3916 | 3907 | 3640 | 3857.62 | 3894 | 3821 | 0.981 | -0.027 | 0.73796903 | 0.85973853 | 0.0027 | 0.0078 | 0.0045 | 0.0045 | TRUE | 0.043 |
| Polr3f | 1293 | 1312 | 1299 | 2289 | 1689 | 1456 | 1373 | 1310 | 1304 | 1799 | 1817 | 1594 | 1533.09 | 1329 | 1737 | 1.307 | 0.386 | 1.28E-05 | 0.00024586 | 0.0024 | 0.0091 | 0.005 | 0.005 | TRUE | 0.0617 |
| Polr3g | 5868 | 5502 | 5396 | 6603 | 4476 | 4300 | 6233 | 5495 | 5418 | 5191 | 4816 | 4707 | 5309.93 | 5715 | 4905 | 0.858 | -0.22 | 0.01263816 | 0.06252481 | 0.0042 | 0.0075 | 0.0054 | 0.0054 | TRUE | 0.0764 |
| Polr3gl | 311 | 362 | 316 | 310 | 201 | 219 | 330 | 362 | 317 | 244 | 216 | 240 | 284.81 | 336 | 233 | 0.694 | -0.527 | 7.61E-05 | 0.00111048 | 0 | 0.0185 | 0.0092 | 0.0092 | TRUE | 0.0491 |
| Polr3h | 1688 | 1463 | 1327 | 1945 | 1307 | 1130 | 1793 | 1461 | 1332 | 1529 | 1406 | 1237 | 1459.79 | 1529 | 1391 | 0.91 | -0.136 | 0.32753795 | 0.54874958 | 0.0166 | 0.0092 | 0.0131 | 0.0131 | TRUE | 0.2746 |
| Polr3k | 1430 | 1448 | 1476 | 1881 | 1356 | 1303 | 1519 | 1446 | 1482 | 1479 | 1459 | 1426 | 1468.52 | 1482 | 1455 | 0.982 | -0.027 | 0.71477802 | 0.8452151 | 0 | 0.0091 | 0.0031 | 0.0031 | TRUE | 0.0056 |
| Polrmt | 3328 | 3174 | 3071 | 3860 | 2904 | 2721 | 3535 | 3170 | 3084 | 3034 | 3125 | 2979 | 3154.33 | 3263 | 3046 | 0.934 | -0.099 | 0.22829868 | 0.44159023 | 0.0025 | 0.0079 | 0.0046 | 0.0046 | TRUE | 0.0645 |
| Pom121 | 6260 | 6167 | 6392 | 8350 | 6051 | 5907 | 6649 | 6159 | 6418 | 6564 | 6511 | 6466 | 6461.21 | 6409 | 6514 | 1.016 | 0.024 | 0.70705943 | 0.84005585 | 5.00E-04 | 0.0074 | 0.0027 | 0.0027 | TRUE | 0.0142 |
| Pom121l12 | 0 | 0 | 0 | 0 | 0 | 0 | 0 | 0 | 0 | 0 | 0 | 0 | 0 | 0 | 0 | NA | NA | NA | NA | NA | NA | NA | NA | NA | NA |
| Pom121l2 | 22 | 13 | 18 | 16 | 11 | 11 | 23 | 13 | 18 | 13 | 12 | 12 | 15.15 | 18 | 12 | 0.672 | -0.573 | 0.25119403 | 0.46875424 | 0 | 0.2244 | 0.1114 | 0.1114 | TRUE | 0.1388 |
| Pomc | 27 | 33 | 31 | 68 | 52 | 44 | 29 | 33 | 31 | 53 | 56 | 48 | 41.72 | 31 | 52 | 1.699 | 0.764 | 0.00999648 | 0.05266013 | 0 | 0.0859 | 0.0377 | 0.0377 | TRUE | 0.04 |
| Pomgnt1 | 913 | 1060 | 1034 | 1841 | 1225 | 1156 | 970 | 1059 | 1038 | 1447 | 1318 | 1265 | 1182.89 | 1022 | 1343 | 1.315 | 0.396 | 2.55E-05 | 0.0004397 | 0.0028 | 0.0097 | 0.0055 | 0.0055 | TRUE | 0.0562 |
| Pomp | 4260 | 4267 | 4197 | 5140 | 3489 | 3647 | 4525 | 4261 | 4214 | 4041 | 3754 | 3992 | 4131.24 | 4333 | 3929 | 0.907 | -0.141 | 0.04771602 | 0.15877152 | 0.0011 | 0.0077 | 0.0034 | 0.0034 | TRUE | 0.0184 |
| Pomt1 | 1179 | 1461 | 1581 | 1534 | 1251 | 1269 | 1252 | 1459 | 1587 | 1206 | 1346 | 1389 | 1373.32 | 1433 | 1314 | 0.916 | -0.127 | 0.28105369 | 0.50229789 | 0.0085 | 0.0093 | 0.0092 | 0.0092 | TRUE | 0.1458 |
| Pomt2 | 1480 | 1697 | 1667 | 1862 | 1341 | 1328 | 1572 | 1695 | 1674 | 1464 | 1443 | 1454 | 1550.16 | 1647 | 1454 | 0.883 | -0.18 | 0.0158224 | 0.07378795 | 0 | 0.009 | 0.0034 | 0.0034 | TRUE | 0.0189 |
| Pon1 | 5 | 0 | 2 | 8 | 2 | 1 | 5 | 0 | 2 | 6 | 2 | 1 | 2.81 | 2 | 3 | 1.343 | 0.426 | 0.75733898 | NA | 0.7964 | 1.1797 | 1.0062 | 1.0062 | TRUE | 0.4629 |
| Pon2 | 116 | 114 | 109 | 196 | 141 | 126 | 123 | 114 | 109 | 154 | 152 | 138 | 131.7 | 115 | 148 | 1.283 | 0.36 | 0.03938109 | 0.13910805 | 0 | 0.0319 | 0.0142 | 0.0142 | TRUE | 0.0349 |
| Pon3 | 80 | 114 | 124 | 131 | 87 | 79 | 85 | 114 | 125 | 103 | 94 | 86 | 101.07 | 108 | 94 | 0.877 | -0.189 | 0.3993597 | 0.6124758 | 0.0131 | 0.0395 | 0.0265 | 0.0265 | TRUE | 0.3325 |
| Pop1 | 1931 | 1048 | 1024 | 1960 | 1213 | 894 | 2051 | 1047 | 1028 | 1541 | 1305 | 979 | 1325.07 | 1375 | 1275 | 0.927 | -0.109 | 0.68025086 | 0.82277546 | 0.0972 | 0.0094 | 0.0494 | 0.0494 | TRUE | 1.402 |
| Pop4 | 2418 | 2161 | 2239 | 3301 | 2410 | 2126 | 2568 | 2158 | 2248 | 2595 | 2593 | 2327 | 2415 | 2325 | 2505 | 1.078 | 0.108 | 0.27728173 | 0.49821001 | 0.0056 | 0.0083 | 0.0067 | 0.0067 | TRUE | 0.1014 |
| Pop5 | 1993 | 2012 | 1850 | 2414 | 1645 | 1664 | 2117 | 2009 | 1858 | 1898 | 1770 | 1822 | 1912.17 | 1995 | 1830 | 0.918 | -0.124 | 0.14422346 | 0.33139329 | 0.0023 | 0.0086 | 0.0047 | 0.0047 | TRUE | 0.0437 |
| Pop7 | 587 | 518 | 472 | 930 | 599 | 535 | 623 | 517 | 474 | 731 | 644 | 586 | 596 | 538 | 654 | 1.217 | 0.283 | 0.05132141 | 0.16701811 | 0.0139 | 0.0124 | 0.0135 | 0.0135 | TRUE | 0.2231 |

| | | | | | | | | | | | | | | | | | | | | | | | | | |
|---|---|---|---|---|---|---|---|---|---|---|---|---|---|---|---|---|---|---|---|---|---|---|---|---|---|
| Popdc2 | 13 | 15 | 11 | 22 | 16 | 15 | 14 | 15 | 11 | 17 | 17 | 16 | 15.13 | 13 | 17 | 1.28 | 0.356 | 0.45553468 | 0.66053461 | 0 | 0.2247 | 0.0968 | 0.0968 | TRUE | 0.0929 |
| Popdc3 | 214 | 141 | 149 | 161 | 129 | 137 | 227 | 141 | 150 | 127 | 139 | 150 | 155.51 | 173 | 139 | 0.802 | -0.319 | 0.16068708 | 0.35525043 | 0.0334 | 0.0281 | 0.0307 | 0.0307 | TRUE | 0.8134 |
| Por | 3472 | 3104 | 3212 | 4229 | 3151 | 3065 | 3688 | 3100 | 3225 | 3325 | 3390 | 3355 | 3347.16 | 3338 | 3357 | 1.006 | 0.008 | 0.92626426 | 0.9637204 | 0.0039 | 0.0079 | 0.0054 | 0.0054 | TRUE | 0.1022 |
| Porcn | 576 | 815 | 902 | 789 | 559 | 497 | 612 | 814 | 906 | 620 | 601 | 544 | 682.87 | 777 | 588 | 0.758 | -0.401 | 0.01106729 | 0.05674686 | 0.0208 | 0.0117 | 0.0164 | 0.0164 | TRUE | 0.4088 |
| Postn | 0 | 0 | 0 | 2 | 8 | 6 | 0 | 0 | 0 | 2 | 9 | 7 | 2.79 | 0 | 6 | 29.401 | 4.878 | 0.00452682 | NA | 0.685 | 1.1872 | 1.0387 | 1.0387 | TRUE | 0.2333 |
| Pot1a | 2454 | 2585 | 2768 | 3607 | 2592 | 2675 | 2607 | 2582 | 2779 | 2836 | 2789 | 2928 | 2753.37 | 2656 | 2851 | 1.073 | 0.102 | 0.14836213 | 0.33809673 | 7.00E-04 | 0.0081 | 0.0032 | 0.0032 | TRUE | 0.0201 |
| Pot1b | 371 | 343 | 326 | 514 | 401 | 335 | 394 | 343 | 327 | 404 | 431 | 367 | 377.7 | 355 | 401 | 1.131 | 0.177 | 0.18113634 | 0.38280307 | 0.0051 | 0.0156 | 0.01 | 0.01 | TRUE | 0.1068 |
| Poteg | 1 | 2 | 0 | 0 | 2 | 0 | 1 | 2 | 0 | 0 | 2 | 0 | 0.87 | 1 | 1 | 0.682 | -0.552 | 0.82171562 | NA | 0 | 3.7998 | 3.1174 | 3.1174 | TRUE | 0.5602 |
| Pou1f1 | 0 | 0 | 0 | 0 | 0 | 0 | 0 | 0 | 0 | 0 | 0 | 0 | 0 | 0 | 0 | NA | NA | NA | NA | NA | NA | NA | NA | NA | NA |
| Pou2af1 | 1 | 0 | 0 | 1 | 3 | 1 | 1 | 0 | 0 | 1 | 3 | 1 | 1.03 | 0 | 2 | 4.594 | 2.2 | 0.2873376 | NA | 0 | 3.2102 | 1.7891 | 1.7891 | TRUE | 0.4614 |
| Pou2f1 | 3632 | 3741 | 4483 | 5132 | 3794 | 3754 | 3858 | 3736 | 4501 | 4034 | 4082 | 4109 | 4053.54 | 4032 | 4075 | 1.011 | 0.015 | 0.8684324 | 0.93526226 | 0.0048 | 0.0077 | 0.006 | 0.006 | TRUE | 0.1265 |
| Pou2f2 | 67 | 95 | 101 | 83 | 60 | 55 | 71 | 95 | 101 | 65 | 65 | 60 | 76.24 | 89 | 63 | 0.71 | -0.493 | 0.04130867 | 0.14410979 | 0 | 0.0501 | 0.0287 | 0.0287 | TRUE | 0.2901 |
| Pou2f3 | 733 | 894 | 815 | 450 | 353 | 413 | 779 | 893 | 818 | 354 | 380 | 452 | 612.57 | 830 | 395 | 0.475 | -1.074 | 6.58E-17 | 1.12E-14 | 0.008 | 0.0123 | 0.01 | 0.01 | TRUE | 0.1799 |
| Pou3f1 | 346 | 383 | 399 | 407 | 343 | 310 | 368 | 383 | 401 | 320 | 369 | 339 | 363.17 | 384 | 343 | 0.892 | -0.165 | 0.17165531 | 0.37055804 | 0 | 0.016 | 0.0077 | 0.0077 | TRUE | 0.051 |
| Pou3f2 | 7 | 10 | 7 | 20 | 16 | 10 | 7 | 10 | 7 | 16 | 17 | 11 | 11.39 | 8 | 15 | 1.798 | 0.847 | 0.13922053 | 0.32391948 | 0 | 0.2962 | 0.1403 | 0.1403 | TRUE | 0.1959 |
| Pou3f3 | 44 | 72 | 54 | 75 | 75 | 75 | 47 | 72 | 54 | 59 | 81 | 82 | 65.77 | 58 | 74 | 1.275 | 0.35 | 0.21449669 | 0.42623685 | 0.0261 | 0.057 | 0.0418 | 0.0418 | TRUE | 0.4027 |
| Pou3f4 | 6 | 4 | 3 | 4 | 5 | 1 | 6 | 4 | 3 | 3 | 5 | 1 | 3.83 | 4 | 3 | 0.721 | -0.471 | 0.63794661 | NA | 0 | 0.8664 | 0.4552 | 0.4552 | TRUE | 0.2413 |
| Pou4f1 | 251 | 217 | 215 | 444 | 379 | 265 | 267 | 217 | 216 | 349 | 408 | 290 | 291.02 | 233 | 349 | 1.498 | 0.583 | 0.00081208 | 0.00758002 | 0.0183 | 0.0182 | 0.0183 | 0.0183 | TRUE | 0.1253 |
| Pou4f2 | 411 | 297 | 336 | 294 | 207 | 206 | 437 | 297 | 337 | 231 | 223 | 225 | 291.65 | 357 | 226 | 0.635 | -0.655 | 0.00014992 | 0.00190069 | 0.0176 | 0.0182 | 0.0179 | 0.0179 | TRUE | 0.4335 |
| Pou4f3 | 45 | 33 | 31 | 36 | 23 | 18 | 48 | 33 | 31 | 28 | 25 | 20 | 30.77 | 37 | 24 | 0.656 | -0.609 | 0.10267064 | 0.26642339 | 0 | 0.114 | 0.0665 | 0.0665 | TRUE | 0.4304 |
| Pou5f1 | 33282 | 36480 | 37153 | 43595 | 31651 | 31531 | 35351 | 36433 | 37306 | 34271 | 34055 | 34516 | 35321.82 | 36363 | 34281 | 0.943 | -0.085 | 0.11844228 | 0.29124909 | 3.00E-04 | 0.007 | 0.0021 | 0.0021 | TRUE | 0.0073 |
| Pou5f2 | 0 | 0 | 2 | 0 | 2 | 5 | 0 | 0 | 2 | 0 | 2 | 5 | 1.61 | 1 | 2 | 3.712 | 1.892 | 0.37262395 | NA | 2.8423 | 2.0588 | 2.2927 | 2.2927 | TRUE | 0.3288 |
| Pou6f1 | 427 | 412 | 419 | 1399 | 1088 | 909 | 454 | 411 | 421 | 1100 | 1171 | 995 | 758.53 | 429 | 1089 | 2.541 | 1.345 | 2.42E-35 | 1.91E-32 | 0.0036 | 0.0112 | 0.0068 | 0.0068 | TRUE | 0.0659 |
| Pou6f2 | 6 | 10 | 6 | 11 | 10 | 6 | 6 | 10 | 6 | 9 | 11 | 7 | 8.06 | 7 | 9 | 1.159 | 0.213 | 0.74889836 | 0.86684463 | 0 | 0.4156 | 0.1952 | 0.1952 | TRUE | 0.2492 |
| Pp2d1 | 4 | 0 | 0 | 1 | 0 | 0 | 4 | 0 | 0 | 1 | 0 | 0 | 0.84 | 1 | 0 | 0.248 | -2.013 | 0.4979877 | NA | 9.4889 | 3.9329 | 4.9952 | 4.9952 | TRUE | 1.8992 |
| Ppa1 | 9096 | 7275 | 6827 | 12507 | 8430 | 6974 | 9662 | 7266 | 6855 | 9832 | 9070 | 7634 | 8386.42 | 7928 | 8845 | 1.116 | 0.158 | 0.29491314 | 0.51584451 | 0.0236 | 0.0073 | 0.0163 | 0.0163 | TRUE | 0.447 |
| Ppa2 | 2442 | 2859 | 2818 | 3611 | 2476 | 2592 | 2594 | 2855 | 2830 | 2839 | 2664 | 2837 | 2769.8 | 2760 | 2780 | 1.007 | 0.011 | 0.88967051 | 0.94620362 | 0.0017 | 0.0081 | 0.004 | 0.004 | TRUE | 0.0334 |
| Ppan | 4093 | 3985 | 3915 | 5284 | 3998 | 3598 | 4347 | 3980 | 3931 | 4154 | 4302 | 3939 | 4108.74 | 4086 | 4132 | 1.011 | 0.016 | 0.8377591 | 0.91757893 | 0.0022 | 0.0077 | 0.0042 | 0.0042 | TRUE | 0.038 |
| Ppap2a | 890 | 874 | 860 | 1327 | 902 | 957 | 945 | 873 | 864 | 1043 | 970 | 1048 | 957.17 | 894 | 1020 | 1.142 | 0.192 | 0.03292259 | 0.12263908 | 0.001 | 0.0103 | 0.0048 | 0.0048 | TRUE | 0.0299 |
| Ppap2b | 501 | 601 | 529 | 945 | 677 | 702 | 532 | 600 | 531 | 743 | 728 | 768 | 650.55 | 554 | 746 | 1.346 | 0.429 | 2.15E-05 | 0.00037855 | 9.00E-04 | 0.012 | 0.0058 | 0.0058 | TRUE | 0.0613 |
| Ppap2c | 1358 | 1754 | 1767 | 1787 | 1369 | 1406 | 1442 | 1752 | 1774 | 1405 | 1473 | 1539 | 1564.21 | 1656 | 1472 | 0.888 | -0.171 | 0.11561675 | 0.28713974 | 0.0063 | 0.009 | 0.0078 | 0.0078 | TRUE | 0.1531 |
| Ppapdc1a | 4 | 2 | 4 | 1 | 1 | 2 | 4 | 2 | 4 | 1 | 1 | 2 | 2.39 | 3 | 1 | 0.385 | -1.376 | 0.27309813 | NA | 0 | 1.3878 | 0.6166 | 0.6166 | TRUE | 0.1972 |
| Ppapdc1b | 779 | 895 | 872 | 873 | 662 | 775 | 827 | 894 | 876 | 686 | 712 | 848 | 807.3 | 866 | 749 | 0.864 | -0.211 | 0.06488592 | 0.1962003 | 0.0057 | 0.011 | 0.0082 | 0.0082 | TRUE | 0.1577 |
| Ppapdc2 | 59 | 141 | 166 | 173 | 155 | 150 | 63 | 141 | 167 | 136 | 167 | 164 | 139.52 | 124 | 156 | 1.259 | 0.332 | 0.30343994 | 0.5242845 | 0.1134 | 0.0305 | 0.0677 | 0.0677 | TRUE | 1.2614 |
| Ppapdc3 | 33 | 23 | 35 | 41 | 29 | 22 | 35 | 23 | 35 | 32 | 31 | 24 | 30.11 | 31 | 29 | 0.944 | -0.083 | 0.82130163 | 0.90875327 | 0 | 0.1163 | 0.0643 | 0.0643 | TRUE | 0.3555 |
| Ppara | 131 | 127 | 115 | 79 | 73 | 84 | 139 | 127 | 115 | 62 | 79 | 92 | 102.34 | 127 | 78 | 0.606 | -0.723 | 0.00127723 | 0.01075522 | 0.0116 | 0.0391 | 0.026 | 0.026 | TRUE | 0.3202 |
| Ppard | 1843 | 1964 | 1908 | 2038 | 1574 | 1662 | 1958 | 1961 | 1916 | 1602 | 1694 | 1819 | 1824.98 | 1945 | 1705 | 0.876 | -0.191 | 0.01871826 | 0.08297482 | 0.0016 | 0.0087 | 0.0042 | 0.0042 | TRUE | 0.0414 |
| Pparg | 2 | 6 | 8 | 14 | 2 | 6 | 2 | 6 | 8 | 11 | 2 | 7 | 5.98 | 5 | 7 | 1.239 | 0.309 | 0.73281994 | 0.8565538 | 0.2744 | 0.5579 | 0.4212 | 0.4212 | TRUE | 0.1633 |
| Ppargc1a | 11 | 16 | 7 | 11 | 16 | 5 | 12 | 16 | 7 | 9 | 17 | 5 | 11 | 12 | 10 | 0.899 | -0.153 | 0.81740568 | 0.90708637 | 0.1394 | 0.3063 | 0.2262 | 0.2262 | TRUE | 0.4705 |
| Ppargc1b | 207 | 379 | 415 | 260 | 226 | 266 | 220 | 379 | 417 | 204 | 243 | 291 | 292.3 | 339 | 246 | 0.726 | -0.461 | 0.06284728 | 0.19213555 | 0.0665 | 0.0182 | 0.0408 | 0.0408 | TRUE | 1.0621 |
| Ppat | 8146 | 7499 | 7542 | 10353 | 7444 | 7191 | 8652 | 7489 | 7573 | 8139 | 8009 | 7872 | 7955.74 | 7905 | 8007 | 1.013 | 0.019 | 0.8212685 | 0.90875327 | 0.0027 | 0.0073 | 0.0047 | 0.0047 | TRUE | 0.0835 |
| Ppbp | 0 | 0 | 0 | 0 | 0 | 0 | 0 | 0 | 0 | 0 | 0 | 0 | 0 | 0 | 0 | NA | NA | NA | NA | NA | NA | NA | NA | NA | NA |
| Ppcdc | 497 | 611 | 495 | 719 | 560 | 401 | 528 | 610 | 497 | 565 | 603 | 439 | 540.31 | 545 | 536 | 0.983 | -0.024 | 0.87357236 | 0.93766049 | 0.017 | 0.013 | 0.0151 | 0.0151 | TRUE | 0.2851 |
| Ppcs | 757 | 744 | 667 | 1089 | 763 | 650 | 804 | 743 | 670 | 856 | 821 | 712 | 767.57 | 739 | 796 | 1.079 | 0.109 | 0.3629917 | 0.58063946 | 0.0074 | 0.0112 | 0.0091 | 0.0091 | TRUE | 0.1014 |
| Ppdpf | 1922 | 1909 | 1937 | 2660 | 1863 | 1736 | 2042 | 1907 | 1945 | 2091 | 2004 | 1900 | 1981.48 | 1965 | 1998 | 1.018 | 0.026 | 0.74084523 | 0.86150638 | 0.0013 | 0.0086 | 0.0039 | 0.0039 | TRUE | 0.0223 |
| Ppef1 | 25 | 29 | 16 | 32 | 27 | 24 | 27 | 29 | 16 | 25 | 29 | 26 | 25.34 | 24 | 27 | 1.123 | 0.167 | 0.67596406 | 0.8197851 | 0 | 0.1369 | 0.0753 | 0.0753 | TRUE | 0.4922 |
| Ppef2 | 181 | 183 | 153 | 247 | 163 | 182 | 192 | 183 | 154 | 194 | 175 | 199 | 182.9 | 176 | 189 | 1.077 | 0.107 | 0.51331104 | 0.70632889 | 0.0026 | 0.0249 | 0.0139 | 0.0139 | TRUE | 0.1353 |
| Ppfia1 | 4454 | 3993 | 3923 | 5918 | 4382 | 4004 | 4731 | 3988 | 3939 | 4652 | 4715 | 4383 | 4401.32 | 4219 | 4583 | 1.086 | 0.12 | 0.21777199 | 0.42971348 | 0.0059 | 0.0077 | 0.0066 | 0.0066 | TRUE | 0.1368 |
| Ppfia2 | 2 | 0 | 0 | 0 | 0 | 0 | 2 | 0 | 0 | 0 | 0 | 0 | 0.35 | 1 | 0 | 0.254 | -1.975 | 0.62446185 | NA | 10 | 9.3117 | 10 | 10 | TRUE | 0.9741 |
| Ppfia3 | 268 | 299 | 293 | 417 | 363 | 349 | 285 | 299 | 294 | 328 | 391 | 382 | 329.65 | 293 | 367 | 1.251 | 0.323 | 0.01279792 | 0.06315668 | 0 | 0.0169 | 0.009 | 0.009 | TRUE | 0.1059 |
| Ppfia4 | 1046 | 1199 | 1147 | 1233 | 1210 | 1356 | 1111 | 1197 | 1152 | 969 | 1302 | 1484 | 1202.62 | 1153 | 1252 | 1.084 | 0.117 | 0.45271242 | 0.65809138 | 0.023 | 0.0096 | 0.0165 | 0.0165 | TRUE | 0.4132 |
| Ppfibp1 | 1620 | 1484 | 1672 | 2194 | 1518 | 1478 | 1721 | 1482 | 1679 | 1725 | 1633 | 1618 | 1642.93 | 1627 | 1659 | 1.02 | 0.028 | 0.7560677 | 0.87112296 | 0.0031 | 0.0089 | 0.0054 | 0.0054 | TRUE | 0.0736 |
| Ppfibp2 | 991 | 1224 | 1310 | 974 | 753 | 935 | 1053 | 1222 | 1315 | 766 | 810 | 1024 | 1031.63 | 1197 | 867 | 0.723 | -0.468 | 0.00113757 | 0.00984185 | 0.0174 | 0.0101 | 0.0139 | 0.0139 | TRUE | 0.2972 |
| Pphln1 | 1704 | 1812 | 1769 | 2998 | 2178 | 2035 | 1810 | 1810 | 1776 | 2357 | 2343 | 2228 | 2053.95 | 1799 | 2309 | 1.284 | 0.361 | 1.69E-07 | 5.63E-06 | 0 | 0.0085 | 0.0029 | 0.0029 | TRUE | 0.0114 |
| Ppia | 33380 | 38234 | 38352 | 46827 | 31662 | 32997 | 35455 | 38185 | 38509 | 36812 | 34066 | 36120 | 36524.67 | 37383 | 35666 | 0.954 | -0.068 | 0.33063366 | 0.55147568 | 0.0018 | 0.007 | 0.0035 | 0.0035 | TRUE | 0.0249 |
| Ppib | 5229 | 6174 | 6052 | 8266 | 5680 | 5541 | 5554 | 6166 | 6077 | 6498 | 6111 | 6065 | 6078.65 | 5932 | 6225 | 1.049 | 0.07 | 0.35772538 | 0.57624805 | 0.002 | 0.0074 | 0.004 | 0.004 | TRUE | 0.0379 |
| Ppic | 1078 | 1561 | 1577 | 1532 | 1032 | 1043 | 1145 | 1559 | 1583 | 1204 | 1110 | 1142 | 1290.65 | 1429 | 1152 | 0.806 | -0.311 | 0.02471757 | 0.10097007 | 0.0161 | 0.0095 | 0.013 | 0.013 | TRUE | 0.3629 |
| Ppid | 9071 | 8915 | 8996 | 12486 | 9168 | 9045 | 9635 | 8903 | 9033 | 9816 | 9864 | 9901 | 9525.39 | 9190 | 9860 | 1.073 | 0.102 | 0.1057605 | 0.27131663 | 8.00E-04 | 0.0072 | 0.0027 | 0.0027 | TRUE | 0.0218 |
| Ppie | 2418 | 2592 | 2539 | 3070 | 2213 | 2301 | 2568 | 2589 | 2549 | 2413 | 2381 | 2519 | 2503.28 | 2569 | 2438 | 0.949 | -0.076 | 0.24992613 | 0.46722738 | 0 | 0.0082 | 0.0027 | 0.0027 | TRUE | 0.0101 |
| Ppif | 3890 | 3717 | 3707 | 4397 | 3213 | 3047 | 4132 | 3712 | 3722 | 3457 | 3457 | 3335 | 3635.88 | 3855 | 3416 | 0.886 | -0.174 | 0.02241693 | 0.09428263 | 0.0017 | 0.0078 | 0.0039 | 0.0039 | TRUE | 0.0478 |
| Ppifos | 0 | 0 | 0 | 0 | 0 | 0 | 0 | 0 | 0 | 0 | 0 | 0 | 0 | 0 | 0 | NA | NA | NA | NA | NA | NA | NA | NA | NA | NA |
| Ppig | 8022 | 8135 | 7624 | 10825 | 7977 | 7420 | 8521 | 8124 | 7655 | 8510 | 8583 | 8122 | 8252.58 | 8100 | 8405 | 1.038 | 0.053 | 0.45665268 | 0.66124918 | 0.0017 | 0.0073 | 0.0036 | 0.0036 | TRUE | 0.0282 |
| Ppih | 1493 | 1482 | 1339 | 2217 | 1485 | 1290 | 1586 | 1480 | 1344 | 1743 | 1598 | 1412 | 1527.19 | 1470 | 1584 | 1.078 | 0.109 | 0.33717688 | 0.55726226 | 0.0079 | 0.0091 | 0.0086 | 0.0086 | TRUE | 0.1084 |
| Ppil1 | 4754 | 4271 | 4163 | 6019 | 4346 | 4115 | 5050 | 4265 | 4180 | 4732 | 4676 | 4505 | 4567.9 | 4498 | 4638 | 1.031 | 0.044 | 0.6448618 | 0.80034312 | 0.0047 | 0.0076 | 0.0064 | 0.0064 | TRUE | 0.1398 |
| Ppil2 | 2545 | 2505 | 2501 | 4208 | 2941 | 2585 | 2703 | 2502 | 2511 | 3308 | 3164 | 2830 | 2836.38 | 2572 | 3101 | 1.206 | 0.27 | 0.00261966 | 0.01904901 | 0.0036 | 0.0081 | 0.0055 | 0.0055 | TRUE | 0.0707 |
| Ppil3 | 962 | 1044 | 942 | 1261 | 826 | 809 | 1022 | 1043 | 946 | 991 | 889 | 886 | 962.66 | 1004 | 922 | 0.92 | -0.12 | 0.21102388 | 0.42172688 | 0.0025 | 0.0103 | 0.0056 | 0.0056 | TRUE | 0.0536 |
| Ppil4 | 3317 | 3019 | 2801 | 4143 | 3108 | 2711 | 3523 | 3015 | 2813 | 3257 | 3344 | 2968 | 3153.23 | 3117 | 3190 | 1.023 | 0.033 | 0.75764172 | 0.8719099 | 0.0083 | 0.0079 | 0.0082 | 0.0082 | TRUE | 0.1577 |
| Ppil6 | 20 | 27 | 33 | 21 | 9 | 18 | 21 | 27 | 33 | 17 | 10 | 20 | 21.21 | 27 | 16 | 0.565 | -0.824 | 0.0701485 | 0.20603021 | 0.0176 | 0.1623 | 0.0984 | 0.0984 | TRUE | 0.2752 |
| Ppip5k1 | 904 | 1059 | 971 | 1026 | 798 | 816 | 960 | 1058 | 975 | 807 | 859 | 893 | 925.2 | 998 | 853 | 0.854 | -0.228 | 0.01477036 | 0.07004152 | 0.0017 | 0.0105 | 0.0052 | 0.0052 | TRUE | 0.0332 |

| | | | | | | | | | | | | | | | | | | | | | | | | | |
|---|---|---|---|---|---|---|---|---|---|---|---|---|---|---|---|---|---|---|---|---|---|---|---|---|---|
| Ppip5k2 | 6849 | 5110 | 5186 | 7233 | 5172 | 4606 | 7275 | 5103 | 5207 | 5686 | 5565 | 5042 | 5646.38 | 5862 | 5431 | 0.927 | -0.11 | 0.44915878 | 0.65472572 | 0.0219 | 0.0075 | 0.015 | 0.015 | TRUE | 0.5421 |
| Ppl | 515 | 566 | 615 | 827 | 672 | 679 | 547 | 565 | 618 | 650 | 723 | 743 | 641.04 | 577 | 705 | 1.222 | 0.289 | 0.00775162 | 0.04336828 | 0.0031 | 0.012 | 0.0069 | 0.0069 | TRUE | 0.0582 |
| Ppm1a | 5440 | 6109 | 6188 | 7697 | 5818 | 6091 | 5778 | 6101 | 6213 | 6051 | 6260 | 6668 | 6178.48 | 6031 | 6326 | 1.049 | 0.069 | 0.34708763 | 0.56607908 | 0.0017 | 0.0074 | 0.0037 | 0.0037 | TRUE | 0.0272 |
| Ppm1b | 7309 | 8733 | 8410 | 8988 | 6802 | 7427 | 7763 | 8722 | 8445 | 7066 | 7319 | 8130 | 7907.32 | 8310 | 7505 | 0.903 | -0.147 | 0.0947469 | 0.25228627 | 0.0043 | 0.0073 | 0.0055 | 0.0055 | TRUE | 0.0648 |
| Ppm1d | 1872 | 1598 | 1475 | 2681 | 1708 | 1521 | 1988 | 1596 | 1481 | 2108 | 1838 | 1665 | 1779.28 | 1688 | 1870 | 1.108 | 0.148 | 0.29341122 | 0.51437903 | 0.0178 | 0.0088 | 0.0138 | 0.0138 | TRUE | 0.2908 |
| Ppm1e | 131 | 138 | 137 | 229 | 160 | 122 | 139 | 138 | 138 | 180 | 172 | 134 | 150.04 | 138 | 162 | 1.176 | 0.233 | 0.20253667 | 0.41006526 | 0.0064 | 0.0289 | 0.0174 | 0.0174 | TRUE | 0.2391 |
| Ppm1f | 4200 | 3811 | 3631 | 5256 | 3506 | 3496 | 4461 | 3806 | 3646 | 4132 | 3772 | 3827 | 3940.69 | 3971 | 3910 | 0.985 | -0.022 | 0.82918004 | 0.91333678 | 0.0067 | 0.0077 | 0.0071 | 0.0071 | TRUE | 0.1417 |
| Ppm1g | 13248 | 11595 | 11751 | 15562 | 11048 | 9829 | 14072 | 11580 | 11799 | 12234 | 11887 | 10759 | 12055.16 | 12484 | 11627 | 0.931 | -0.102 | 0.32344764 | 0.5447321 | 0.0082 | 0.0072 | 0.0077 | 0.0077 | TRUE | 0.1513 |
| Ppm1h | 695 | 570 | 569 | 572 | 453 | 456 | 738 | 569 | 571 | 450 | 487 | 499 | 552.51 | 626 | 479 | 0.764 | -0.388 | 0.00519271 | 0.03211744 | 0.0107 | 0.0129 | 0.0121 | 0.0121 | TRUE | 0.2862 |
| Ppm1j | 1424 | 2070 | 2157 | 1266 | 1010 | 1344 | 1513 | 2067 | 2166 | 995 | 1087 | 1471 | 1549.81 | 1915 | 1184 | 0.618 | -0.694 | 0.00014027 | 0.00180749 | 0.0386 | 0.009 | 0.0233 | 0.0233 | TRUE | 0.5366 |
| Ppm1k | 565 | 655 | 694 | 658 | 484 | 538 | 600 | 654 | 697 | 517 | 521 | 589 | 596.35 | 650 | 542 | 0.833 | -0.264 | 0.01980992 | 0.08626231 | 0.0037 | 0.0124 | 0.0076 | 0.0076 | TRUE | 0.0647 |
| Ppm1l | 298 | 283 | 312 | 525 | 394 | 355 | 317 | 283 | 313 | 413 | 424 | 389 | 356.28 | 304 | 409 | 1.344 | 0.426 | 0.00036136 | 0.00399399 | 0 | 0.0162 | 0.0074 | 0.0074 | TRUE | 0.0436 |
| Ppm1m | 2095 | 2817 | 2907 | 2410 | 1925 | 1985 | 2225 | 2813 | 2919 | 1895 | 2071 | 2173 | 2349.36 | 2652 | 2046 | 0.771 | -0.375 | 0.00244634 | 0.01803939 | 0.0116 | 0.0083 | 0.0106 | 0.0106 | TRUE | 0.2404 |
| Ppm1n | 69 | 79 | 116 | 112 | 84 | 81 | 73 | 79 | 116 | 88 | 90 | 89 | 89.29 | 89 | 89 | 0.993 | -0.01 | 0.96746063 | 0.98366749 | 0.0213 | 0.0438 | 0.0327 | 0.0327 | TRUE | 0.6629 |
| Ppme1 | 2616 | 3162 | 3487 | 3741 | 2875 | 3305 | 2779 | 3158 | 3501 | 2941 | 3093 | 3618 | 3181.66 | 3146 | 3217 | 1.022 | 0.032 | 0.79282895 | 0.89317844 | 0.0122 | 0.0079 | 0.0103 | 0.0103 | TRUE | 0.1438 |
| Ppox | 360 | 481 | 467 | 536 | 498 | 473 | 382 | 480 | 469 | 421 | 536 | 518 | 467.77 | 444 | 492 | 1.105 | 0.144 | 0.33368125 | 0.5542839 | 0.0138 | 0.0139 | 0.0139 | 0.0139 | TRUE | 0.1897 |
| Ppp1ca | 10184 | 11223 | 11141 | 16607 | 11626 | 11319 | 10817 | 11209 | 11187 | 13055 | 12509 | 12390 | 11861.14 | 11071 | 12651 | 1.143 | 0.193 | 0.0010624 | 0.00932968 | 5.00E-04 | 0.0072 | 0.0024 | 0.0024 | TRUE | 0.0096 |
| Ppp1cb | 9739 | 10798 | 10869 | 15206 | 10751 | 11003 | 10345 | 10784 | 10914 | 11954 | 11567 | 12045 | 11267.99 | 10681 | 11855 | 1.11 | 0.151 | 0.01117875 | 0.05718822 | 5.00E-04 | 0.0072 | 0.0024 | 0.0024 | TRUE | 0.0093 |
| Ppp1cc | 7229 | 8306 | 8006 | 9909 | 7458 | 7275 | 7678 | 8295 | 8039 | 7790 | 8024 | 7964 | 7965.05 | 8004 | 7926 | 0.99 | -0.014 | 0.81980694 | 0.90781402 | 6.00E-04 | 0.0073 | 0.0027 | 0.0027 | TRUE | 0.0154 |
| Ppp1r10 | 4057 | 2750 | 2067 | 3735 | 2403 | 1577 | 4309 | 2746 | 2075 | 2936 | 2585 | 1726 | 2729.85 | 3043 | 2416 | 0.794 | -0.333 | 0.38040679 | 0.59482144 | 0.1035 | 0.0081 | 0.0473 | 0.1035 | TRUE | 0.5261 |
| Ppp1r11 | 774 | 736 | 653 | 1393 | 954 | 657 | 822 | 735 | 656 | 1095 | 1026 | 719 | 842.26 | 738 | 947 | 1.285 | 0.362 | 0.03557994 | 0.12950116 | 0.0295 | 0.0108 | 0.0201 | 0.0201 | TRUE | 0.524 |
| Ppp1r12a | 3922 | 3673 | 3768 | 5623 | 4005 | 3533 | 4166 | 3668 | 3783 | 4420 | 4309 | 3867 | 4035.75 | 3872 | 4199 | 1.085 | 0.117 | 0.1976131 | 0.40433284 | 0.0043 | 0.0077 | 0.0057 | 0.0057 | TRUE | 0.0579 |
| Ppp1r12b | 284 | 313 | 355 | 282 | 249 | 276 | 302 | 313 | 356 | 222 | 268 | 302 | 293.74 | 324 | 264 | 0.813 | -0.299 | 0.06299565 | 0.1924239 | 0.0126 | 0.0181 | 0.0152 | 0.0152 | TRUE | 0.2337 |
| Ppp1r12c | 3429 | 3464 | 3422 | 4332 | 3345 | 3214 | 3642 | 3460 | 3436 | 3405 | 3599 | 3518 | 3510.08 | 3513 | 3507 | 0.998 | -0.002 | 0.9722484 | 0.985305 | 6.00E-04 | 0.0078 | 0.003 | 0.003 | TRUE | 0.0126 |
| Ppp1r13b | 1295 | 1292 | 1400 | 1576 | 1165 | 1075 | 1376 | 1290 | 1406 | 1239 | 1253 | 1177 | 1290.13 | 1357 | 1223 | 0.901 | -0.15 | 0.06750246 | 0.20087746 | 7.00E-04 | 0.0095 | 0.0041 | 0.0041 | TRUE | 0.0225 |
| Ppp1r13l | 861 | 989 | 947 | 1021 | 809 | 966 | 915 | 988 | 951 | 803 | 870 | 1057 | 930.61 | 951 | 910 | 0.956 | -0.065 | 0.60072541 | 0.77266232 | 0.0099 | 0.0104 | 0.0101 | 0.0101 | TRUE | 0.2373 |
| Ppp1r14a | 394 | 527 | 512 | 739 | 556 | 525 | 418 | 526 | 514 | 581 | 598 | 575 | 535.46 | 486 | 585 | 1.201 | 0.265 | 0.03104308 | 0.1180925 | 0.0055 | 0.0131 | 0.009 | 0.009 | TRUE | 0.1714 |
| Ppp1r14b | 4185 | 4280 | 4113 | 5809 | 4186 | 3684 | 4445 | 4274 | 4130 | 4567 | 4504 | 4033 | 4325.46 | 4283 | 4368 | 1.02 | 0.029 | 0.72455435 | 0.850916 | 0.0027 | 0.0077 | 0.0045 | 0.0045 | TRUE | 0.0545 |
| Ppp1r14c | 184 | 181 | 154 | 240 | 191 | 177 | 195 | 181 | 155 | 189 | 206 | 194 | 186.46 | 177 | 196 | 1.107 | 0.147 | 0.36101832 | 0.57936315 | 0 | 0.0246 | 0.0132 | 0.0132 | TRUE | 0.1308 |
| Ppp1r14d | 47 | 53 | 54 | 66 | 53 | 59 | 50 | 53 | 54 | 52 | 57 | 65 | 55.09 | 52 | 58 | 1.1 | 0.137 | 0.59718443 | 0.77001249 | 0 | 0.0667 | 0.0303 | 0.0303 | TRUE | 0.0861 |
| Ppp1r15a | 781 | 617 | 609 | 1036 | 750 | 599 | 830 | 616 | 612 | 814 | 807 | 656 | 722.39 | 686 | 759 | 1.108 | 0.148 | 0.3484619 | 0.56705955 | 0.0213 | 0.0115 | 0.0165 | 0.0165 | TRUE | 0.3959 |
| Ppp1r15b | 4312 | 4677 | 4245 | 6660 | 4894 | 4695 | 4580 | 4671 | 4262 | 5236 | 5266 | 5139 | 4859.02 | 4504 | 5214 | 1.157 | 0.211 | 0.00195919 | 0.01514725 | 9.00E-04 | 0.0076 | 0.0031 | 0.0031 | TRUE | 0.027 |
| Ppp1r16a | 1141 | 1098 | 1086 | 1600 | 1301 | 1090 | 1212 | 1097 | 1090 | 1258 | 1400 | 1193 | 1208.29 | 1133 | 1284 | 1.133 | 0.18 | 0.07360062 | 0.21283994 | 0.004 | 0.0096 | 0.0065 | 0.0065 | TRUE | 0.0741 |
| Ppp1r16b | 435 | 337 | 273 | 428 | 276 | 270 | 462 | 337 | 274 | 336 | 297 | 296 | 333.62 | 358 | 310 | 0.867 | -0.205 | 0.30364247 | 0.524425 | 0.0341 | 0.0168 | 0.0257 | 0.0257 | TRUE | 0.4467 |
| Ppp1r17 | 0 | 0 | 0 | 0 | 0 | 0 | 0 | 0 | 0 | 0 | 0 | 0 | 0 | 0 | 0 | NA | NA | NA | NA | NA | NA | NA | NA | NA | NA |
| Ppp1r18 | 725 | 688 | 737 | 1106 | 783 | 747 | 770 | 687 | 740 | 869 | 842 | 818 | 787.81 | 732 | 843 | 1.152 | 0.204 | 0.02894476 | 0.11224628 | 7.00E-04 | 0.0111 | 0.005 | 0.005 | TRUE | 0.0348 |
| Ppp1r1a | 258 | 378 | 427 | 308 | 266 | 314 | 274 | 378 | 429 | 242 | 286 | 344 | 325.39 | 360 | 291 | 0.805 | -0.312 | 0.12744973 | 0.30689116 | 0.0376 | 0.017 | 0.0272 | 0.0272 | TRUE | 0.4224 |
| Ppp1r1b | 185 | 225 | 273 | 233 | 179 | 165 | 197 | 225 | 274 | 183 | 193 | 181 | 208.62 | 232 | 186 | 0.799 | -0.323 | 0.05998119 | 0.18578558 | 0.0106 | 0.0227 | 0.0164 | 0.0164 | TRUE | 0.2822 |
| Ppp1r1c | 0 | 0 | 0 | 0 | 0 | 0 | 0 | 0 | 0 | 0 | 0 | 0 | 0 | 0 | 0 | NA | NA | NA | NA | NA | NA | NA | NA | NA | NA |
| Ppp1r2 | 4973 | 4815 | 4513 | 6386 | 4549 | 4269 | 5282 | 4809 | 4532 | 5020 | 4894 | 4673 | 4868.37 | 4874 | 4862 | 0.998 | -0.003 | 0.97036353 | 0.9846576 | 0.0034 | 0.0076 | 0.005 | 0.005 | TRUE | 0.0652 |
| Ppp1r2-ps3 | 2 | 0 | 1 | 2 | 0 | 0 | 2 | 0 | 1 | 2 | 0 | 0 | 0.78 | 1 | 1 | 0.543 | -0.88 | 0.72386184 | NA | 0 | 4.212 | 3.1127 | 3.1127 | TRUE | 0.402 |
| Ppp1r2-ps7 | 1 | 0 | 0 | 1 | 1 | 0 | 1 | 0 | 0 | 1 | 1 | 0 | 0.49 | 0 | 1 | 1.712 | 0.776 | 0.78488768 | NA | 0 | 6.7665 | 4.0387 | 4.0387 | TRUE | 0.4614 |
| Ppp1r2-ps9 | 0 | 0 | 0 | 0 | 0 | 0 | 0 | 0 | 0 | 0 | 0 | 0 | 0 | 0 | 0 | NA | NA | NA | NA | NA | NA | NA | NA | NA | NA |
| Ppp1r21 | 1759 | 1705 | 1623 | 2194 | 1699 | 1602 | 1868 | 1703 | 1630 | 1725 | 1828 | 1754 | 1751.21 | 1734 | 1769 | 1.02 | 0.029 | 0.73755985 | 0.85946021 | 0.0023 | 0.0088 | 0.0048 | 0.0048 | TRUE | 0.0556 |
| Ppp1r26 | 269 | 318 | 364 | 449 | 388 | 316 | 286 | 318 | 365 | 353 | 417 | 346 | 347.53 | 323 | 372 | 1.151 | 0.203 | 0.17245065 | 0.37119832 | 0.01 | 0.0164 | 0.013 | 0.013 | TRUE | 0.151 |
| Ppp1r27 | 0 | 0 | 1 | 1 | 0 | 0 | 0 | 0 | 1 | 1 | 0 | 0 | 0.3 | 0 | 0 | 0.954 | -0.067 | 0.98640617 | NA | 0 | 11.0483 | 9.2783 | 9.2783 | TRUE | 0.4098 |
| Ppp1r32 | 5 | 9 | 7 | 12 | 5 | 4 | 5 | 9 | 7 | 9 | 5 | 4 | 6.75 | 7 | 6 | 0.915 | -0.128 | 0.86287453 | 0.93221844 | 0 | 0.4947 | 0.2473 | 0.2473 | TRUE | 0.5465 |
| Ppp1r35 | 812 | 1056 | 1020 | 1124 | 874 | 871 | 862 | 1055 | 1024 | 884 | 940 | 953 | 953.12 | 980 | 926 | 0.943 | -0.084 | 0.44292762 | 0.64974531 | 0.0055 | 0.0104 | 0.0076 | 0.0076 | TRUE | 0.1314 |
| Ppp1r36 | 49 | 67 | 53 | 53 | 33 | 58 | 52 | 67 | 53 | 42 | 36 | 63 | 52.14 | 57 | 47 | 0.813 | -0.298 | 0.34583388 | 0.56517324 | 0.0331 | 0.0701 | 0.0526 | 0.0526 | TRUE | 0.7152 |
| Ppp1r37 | 1076 | 1133 | 1115 | 1769 | 1398 | 1146 | 1143 | 1132 | 1120 | 1391 | 1504 | 1254 | 1257.22 | 1132 | 1383 | 1.223 | 0.29 | 0.00257496 | 0.01881938 | 0.0034 | 0.0095 | 0.0059 | 0.0059 | TRUE | 0.0778 |
| Ppp1r3a | 0 | 0 | 0 | 0 | 0 | 1 | 0 | 0 | 0 | 0 | 0 | 1 | 0.18 | 0 | 0 | 1.859 | 0.894 | 0.82650734 | NA | 0 | 18.0643 | 10 | 10 | TRUE | 0.5383 |
| Ppp1r3b | 593 | 1111 | 1161 | 957 | 877 | 1121 | 630 | 1110 | 1166 | 752 | 944 | 1227 | 971.37 | 969 | 974 | 1.005 | 0.008 | 0.97453796 | 0.98673767 | 0.0814 | 0.0103 | 0.0415 | 0.0415 | TRUE | 0.5781 |
| Ppp1r3c | 124 | 232 | 244 | 174 | 184 | 195 | 132 | 232 | 245 | 137 | 198 | 213 | 192.77 | 203 | 183 | 0.898 | -0.155 | 0.5678033 | 0.74733351 | 0.0748 | 0.024 | 0.0479 | 0.0479 | TRUE | 1.0136 |
| Ppp1r3d | 25 | 45 | 27 | 36 | 45 | 18 | 27 | 45 | 27 | 28 | 48 | 20 | 32.5 | 33 | 32 | 0.975 | -0.037 | 0.93393484 | 0.9672349 | 0.1135 | 0.1083 | 0.1106 | 0.1106 | TRUE | 0.6425 |
| Ppp1r3e | 3 | 6 | 11 | 12 | 6 | 9 | 3 | 6 | 11 | 9 | 6 | 10 | 7.66 | 7 | 8 | 1.272 | 0.347 | 0.63522285 | 0.79491607 | 0 | 0.4369 | 0.253 | 0.253 | TRUE | 0.5212 |
| Ppp1r3f | 210 | 211 | 183 | 181 | 127 | 124 | 223 | 211 | 184 | 142 | 137 | 136 | 172.03 | 206 | 138 | 0.673 | -0.572 | 0.00040581 | 0.00438329 | 0 | 0.0261 | 0.0129 | 0.0129 | TRUE | 0.0965 |
| Ppp1r3g | 25 | 54 | 74 | 90 | 98 | 83 | 27 | 54 | 74 | 71 | 105 | 91 | 70.31 | 52 | 89 | 1.718 | 0.781 | 0.03404029 | 0.12550866 | 0.1177 | 0.0538 | 0.0824 | 0.0824 | TRUE | 0.4916 |
| Ppp1r42 | 1 | 2 | 1 | 2 | 2 | 2 | 1 | 2 | 1 | 2 | 2 | 2 | 1.66 | 1 | 2 | 1.439 | 0.525 | 0.70499898 | NA | 0 | 1.9881 | 0.7617 | 0.7617 | TRUE | 0.1097 |
| Ppp1r7 | 2679 | 2961 | 2993 | 4482 | 3369 | 2770 | 2846 | 2957 | 3005 | 3523 | 3625 | 3032 | 3164.75 | 2936 | 3393 | 1.156 | 0.209 | 0.02436125 | 0.09979885 | 0.0043 | 0.0079 | 0.0059 | 0.0059 | TRUE | 0.1047 |
| Ppp1r8 | 2969 | 2912 | 2798 | 4846 | 3346 | 3120 | 3154 | 2908 | 2809 | 3810 | 3600 | 3415 | 3282.72 | 2957 | 3608 | 1.221 | 0.288 | 0.00062545 | 0.00620419 | 0.0028 | 0.0079 | 0.0048 | 0.0048 | TRUE | 0.0409 |
| Ppp1r9a | 1562 | 1565 | 1722 | 2159 | 1652 | 1553 | 1659 | 1563 | 1729 | 1697 | 1777 | 1700 | 1687.64 | 1650 | 1725 | 1.045 | 0.063 | 0.42154205 | 0.63163227 | 0.001 | 0.0089 | 0.0039 | 0.0039 | TRUE | 0.026 |
| Ppp1r9b | 1685 | 1639 | 1611 | 2558 | 2021 | 1715 | 1790 | 1637 | 1618 | 2011 | 2174 | 1877 | 1851.16 | 1682 | 2021 | 1.202 | 0.265 | 0.00418761 | 0.02738268 | 0.0035 | 0.0087 | 0.0056 | 0.0056 | TRUE | 0.0526 |
| Ppp2ca | 19605 | 20743 | 20038 | 29123 | 20257 | 18947 | 20824 | 20716 | 20120 | 22894 | 21795 | 20740 | 21181.74 | 20553 | 21810 | 1.061 | 0.086 | 0.19597001 | 0.40229584 | 0.0013 | 0.0071 | 0.0031 | 0.0031 | TRUE | 0.0232 |
| Ppp2cb | 4895 | 5389 | 4999 | 7474 | 5218 | 5079 | 5199 | 5382 | 5020 | 5875 | 5614 | 5560 | 5441.73 | 5200 | 5683 | 1.093 | 0.128 | 0.0521798 | 0.16888567 | 8.00E-04 | 0.0075 | 0.003 | 0.003 | TRUE | 0.0114 |
| Ppp2r1a | 11244 | 11437 | 11507 | 16813 | 11967 | 11633 | 11943 | 11422 | 11554 | 13217 | 12876 | 12734 | 12291.09 | 11640 | 12942 | 1.112 | 0.153 | 0.00726865 | 0.04129725 | 3.00E-04 | 0.0072 | 0.0023 | 0.0023 | TRUE | 0.0063 |
| Ppp2r1b | 5111 | 4322 | 4543 | 6736 | 4898 | 4455 | 5429 | 4316 | 4562 | 5295 | 5270 | 4877 | 4958.14 | 4769 | 5147 | 1.08 | 0.11 | 0.29522483 | 0.51607626 | 0.008 | 0.0076 | 0.0078 | 0.0078 | TRUE | 0.1783 |
| Ppp2r2a | 8879 | 8578 | 8132 | 10311 | 7721 | 7660 | 9431 | 8567 | 8165 | 8106 | 8307 | 8385 | 8493.58 | 8721 | 8266 | 0.948 | -0.077 | 0.32578036 | 0.54702521 | 0.0027 | 0.0073 | 0.0043 | 0.0043 | TRUE | 0.0619 |
| Ppp2r2b | 18 | 14 | 10 | 34 | 32 | 29 | 19 | 14 | 10 | 27 | 34 | 32 | 22.67 | 14 | 31 | 2.151 | 1.105 | 0.01042661 | 0.05441711 | 0 | 0.1522 | 0.0827 | 0.0827 | TRUE | 0.3522 |
| Ppp2r2c | 1292 | 1536 | 1459 | 1786 | 1321 | 1494 | 1372 | 1534 | 1465 | 1404 | 1421 | 1635 | 1472.01 | 1457 | 1487 | 1.02 | 0.028 | 0.77595794 | 0.88357581 | 0.0045 | 0.0091 | 0.0064 | 0.0064 | TRUE | 0.0911 |

| | | | | | | | | | | | | | | | | | | | | | | | | | |
|---|---|---|---|---|---|---|---|---|---|---|---|---|---|---|---|---|---|---|---|---|---|---|---|---|---|
| Ppp2r2d | 4372 | 3941 | 4020 | 5445 | 3715 | 3658 | 4644 | 3936 | 4037 | 4280 | 3997 | 4004 | 4149.68 | 4206 | 4094 | 0.974 | -0.038 | 0.67443586 | 0.81856117 | 0.004 | 0.0077 | 0.0058 | 0.0058 | TRUE | 0.1011 |
| Ppp2r3a | 794 | 738 | 728 | 1237 | 793 | 778 | 843 | 737 | 731 | 972 | 853 | 852 | 831.45 | 770 | 892 | 1.16 | 0.214 | 0.05224914 | 0.16904071 | 0.0051 | 0.0109 | 0.0076 | 0.0076 | TRUE | 0.0807 |
| Ppp2r3c | 897 | 835 | 846 | 1388 | 970 | 904 | 953 | 834 | 849 | 1091 | 1044 | 990 | 960.09 | 879 | 1042 | 1.186 | 0.247 | 0.01184145 | 0.05966881 | 0.0027 | 0.0103 | 0.0059 | 0.0059 | TRUE | 0.0642 |
| Ppp2r3d | 89 | 93 | 92 | 130 | 113 | 97 | 95 | 93 | 92 | 102 | 122 | 106 | 101.63 | 93 | 110 | 1.177 | 0.235 | 0.2311898 | 0.44494515 | 0 | 0.0393 | 0.0178 | 0.0178 | TRUE | 0.0824 |
| Ppp2r4 | 4146 | 4678 | 4772 | 7010 | 4861 | 4458 | 4404 | 4672 | 4792 | 5511 | 5230 | 4880 | 4914.69 | 4623 | 5207 | 1.127 | 0.172 | 0.03073356 | 0.11725441 | 0.0025 | 0.0076 | 0.0044 | 0.0044 | TRUE | 0.0367 |
| Ppp2r5a | 4881 | 5584 | 5704 | 6814 | 4878 | 5035 | 5184 | 5577 | 5727 | 5357 | 5248 | 5512 | 5434.23 | 5496 | 5372 | 0.977 | -0.033 | 0.64142374 | 0.79835032 | 0.0013 | 0.0075 | 0.0034 | 0.0034 | TRUE | 0.03 |
| Ppp2r5b | 1107 | 891 | 849 | 1258 | 895 | 741 | 1176 | 890 | 852 | 989 | 963 | 811 | 946.87 | 973 | 921 | 0.948 | -0.078 | 0.60796643 | 0.77781039 | 0.0202 | 0.0104 | 0.0155 | 0.0155 | TRUE | 0.3971 |
| Ppp2r5c | 9675 | 13523 | 14011 | 10108 | 7630 | 8885 | 10277 | 13506 | 14069 | 7946 | 8209 | 9726 | 10622.03 | 12617 | 8627 | 0.684 | -0.549 | 7.85E-05 | 0.00113688 | 0.0192 | 0.0072 | 0.0138 | 0.0138 | TRUE | 0.3218 |
| Ppp2r5d | 5201 | 5380 | 5347 | 6030 | 4409 | 4596 | 5524 | 5373 | 5369 | 4740 | 4744 | 5031 | 5130.26 | 5422 | 4838 | 0.892 | -0.165 | 0.0091461 | 0.04921608 | 4.00E-04 | 0.0075 | 0.0027 | 0.0027 | TRUE | 0.0147 |
| Ppp2r5e | 2712 | 2803 | 2936 | 3776 | 2752 | 2732 | 2881 | 2799 | 2948 | 2968 | 2961 | 2991 | 2924.67 | 2876 | 2973 | 1.034 | 0.048 | 0.44869764 | 0.65453747 | 0 | 0.008 | 0.0025 | 0.0025 | TRUE | 0.0066 |
| Ppp3ca | 801 | 1016 | 1044 | 1443 | 1062 | 1082 | 851 | 1015 | 1048 | 1134 | 1143 | 1184 | 1062.54 | 971 | 1154 | 1.187 | 0.248 | 0.02039456 | 0.08812687 | 0.0048 | 0.01 | 0.0073 | 0.0073 | TRUE | 0.1396 |
| Ppp3cb | 4135 | 4825 | 4753 | 5366 | 3657 | 3837 | 4392 | 4819 | 4773 | 4218 | 3935 | 4200 | 4389.44 | 4661 | 4118 | 0.883 | -0.179 | 0.01711093 | 0.07766148 | 0.0017 | 0.0077 | 0.0038 | 0.0038 | TRUE | 0.031 |
| Ppp3cc | 538 | 662 | 702 | 731 | 539 | 557 | 571 | 661 | 705 | 575 | 580 | 610 | 616.96 | 646 | 588 | 0.91 | -0.136 | 0.23682398 | 0.45220191 | 0.0046 | 0.0122 | 0.0079 | 0.0079 | TRUE | 0.1185 |
| Ppp3r1 | 4836 | 4581 | 4610 | 7103 | 4963 | 4713 | 5137 | 4575 | 4629 | 5584 | 5340 | 5159 | 5070.59 | 4780 | 5361 | 1.122 | 0.166 | 0.03782382 | 0.13508997 | 0.0026 | 0.0076 | 0.0044 | 0.0044 | TRUE | 0.0518 |
| Ppp3r2 | 0 | 0 | 0 | 0 | 0 | 0 | 0 | 0 | 0 | 0 | 0 | 0 | 0 | 0 | 0 | NA | NA | NA | NA | NA | NA | NA | NA | NA | NA |
| Ppp4c | 5568 | 5524 | 5386 | 7881 | 5541 | 5396 | 5914 | 5517 | 5408 | 6195 | 5962 | 5907 | 5817.19 | 5613 | 6021 | 1.073 | 0.102 | 0.14208127 | 0.3277973 | 0.0012 | 0.0075 | 0.0033 | 0.0033 | TRUE | 0.0268 |
| Ppp4r1 | 4636 | 3969 | 4284 | 5735 | 4054 | 3731 | 4924 | 3964 | 4302 | 4508 | 4362 | 4084 | 4357.36 | 4397 | 4318 | 0.982 | -0.026 | 0.80192998 | 0.89766073 | 0.0071 | 0.0077 | 0.0073 | 0.0073 | TRUE | 0.1341 |
| Ppp4r1l-ps | 270 | 355 | 323 | 283 | 263 | 295 | 287 | 355 | 324 | 222 | 283 | 323 | 299 | 322 | 276 | 0.855 | -0.226 | 0.19747502 | 0.40422792 | 0.0198 | 0.0179 | 0.0189 | 0.0189 | TRUE | 0.3432 |
| Ppp4r2 | 5579 | 5693 | 5747 | 7496 | 5573 | 5543 | 5926 | 5686 | 5771 | 5893 | 5996 | 6068 | 5889.8 | 5794 | 5986 | 1.033 | 0.047 | 0.42097164 | 0.63116133 | 2.00E-04 | 0.0075 | 0.0023 | 0.0023 | TRUE | 0.0048 |
| Ppp4r4 | 214 | 228 | 222 | 208 | 162 | 183 | 227 | 228 | 223 | 164 | 174 | 200 | 202.68 | 226 | 179 | 0.791 | -0.338 | 0.02549832 | 0.10306497 | 0 | 0.0232 | 0.0115 | 0.0115 | TRUE | 0.1092 |
| Ppp5c | 6131 | 6413 | 6261 | 6874 | 5107 | 5244 | 6512 | 6405 | 6287 | 5404 | 5495 | 5740 | 5973.77 | 6401 | 5546 | 0.866 | -0.207 | 0.00078724 | 0.00738191 | 4.00E-04 | 0.0075 | 0.0026 | 0.0026 | TRUE | 0.0114 |
| Ppp6c | 1710 | 1718 | 1792 | 2317 | 1598 | 1656 | 1816 | 1716 | 1799 | 1821 | 1719 | 1813 | 1780.83 | 1777 | 1784 | 1.004 | 0.006 | 0.9302365 | 0.96562394 | 4.00E-04 | 0.0088 | 0.0034 | 0.0034 | TRUE | 0.0122 |
| Ppp6r1 | 6308 | 5025 | 5074 | 8607 | 6108 | 4912 | 6700 | 5019 | 5095 | 6766 | 6572 | 5377 | 5921.42 | 5605 | 6238 | 1.113 | 0.155 | 0.27677006 | 0.49774428 | 0.02 | 0.0075 | 0.0144 | 0.0144 | TRUE | 0.3565 |
| Ppp6r2 | 549 | 829 | 893 | 927 | 674 | 775 | 583 | 828 | 897 | 729 | 725 | 848 | 768.33 | 769 | 767 | 0.997 | -0.004 | 0.97944207 | 0.98867206 | 0.0274 | 0.0112 | 0.0193 | 0.0193 | TRUE | 0.5288 |
| Ppp6r3 | 15451 | 16394 | 16879 | 19769 | 14236 | 14780 | 16412 | 16373 | 16948 | 15541 | 15317 | 16179 | 16128.29 | 16578 | 15679 | 0.946 | -0.08 | 0.16688127 | 0.3643482 | 5.00E-04 | 0.0071 | 0.0024 | 0.0024 | TRUE | 0.0095 |
| Pprc1 | 15479 | 13953 | 14557 | 16463 | 12371 | 12428 | 16441 | 13935 | 14617 | 12942 | 13310 | 13604 | 14141.66 | 14998 | 13285 | 0.886 | -0.175 | 0.03882714 | 0.13776689 | 0.0039 | 0.0071 | 0.0051 | 0.0051 | TRUE | 0.0867 |
| Ppt1 | 1070 | 1325 | 1422 | 1806 | 1410 | 1418 | 1137 | 1323 | 1428 | 1420 | 1517 | 1552 | 1396.11 | 1296 | 1496 | 1.154 | 0.207 | 0.05887311 | 0.18363368 | 0.0065 | 0.0093 | 0.0079 | 0.0079 | TRUE | 0.1383 |
| Ppt2 | 417 | 599 | 602 | 563 | 401 | 527 | 443 | 598 | 604 | 443 | 431 | 577 | 516.09 | 548 | 484 | 0.881 | -0.183 | 0.2899493 | 0.51113003 | 0.025 | 0.0133 | 0.0197 | 0.0197 | TRUE | 0.3301 |
| Pptc7 | 1966 | 1956 | 1787 | 2217 | 1516 | 1419 | 2088 | 1953 | 1794 | 1743 | 1631 | 1553 | 1793.89 | 1945 | 1642 | 0.845 | -0.243 | 0.01018887 | 0.05349496 | 0.0037 | 0.0087 | 0.0059 | 0.0059 | TRUE | 0.0558 |
| Ppwd1 | 2337 | 2566 | 2735 | 3498 | 2611 | 2575 | 2482 | 2563 | 2746 | 2750 | 2809 | 2819 | 2694.85 | 2597 | 2793 | 1.075 | 0.104 | 0.15372264 | 0.34542149 | 9.00E-04 | 0.0081 | 0.0035 | 0.0035 | TRUE | 0.0306 |
| Ppy | 0 | 3 | 2 | 2 | 2 | 1 | 0 | 3 | 2 | 2 | 2 | 1 | 1.64 | 2 | 2 | 0.956 | -0.065 | 0.96556663 | NA | 0 | 2.0192 | 1.0439 | 1.0439 | TRUE | 0.4768 |
| Ppyr1 | 53 | 37 | 33 | 54 | 44 | 27 | 56 | 37 | 33 | 42 | 47 | 30 | 40.96 | 42 | 40 | 0.948 | -0.077 | 0.82834478 | 0.91297372 | 0.0427 | 0.0873 | 0.066 | 0.066 | TRUE | 0.6452 |
| Pqbp1 | 1539 | 1570 | 1535 | 1890 | 1419 | 1366 | 1635 | 1568 | 1541 | 1486 | 1527 | 1495 | 1541.97 | 1581 | 1503 | 0.95 | -0.074 | 0.31080948 | 0.53139156 | 0 | 0.009 | 0.0032 | 0.0032 | TRUE | 0.0104 |
| Pqlc1 | 788 | 698 | 655 | 645 | 491 | 496 | 837 | 697 | 658 | 507 | 528 | 543 | 628.34 | 731 | 526 | 0.72 | -0.474 | 0.00013547 | 0.00176289 | 0.0074 | 0.0121 | 0.0095 | 0.0095 | TRUE | 0.191 |
| Pqlc2 | 344 | 356 | 344 | 553 | 380 | 331 | 365 | 356 | 345 | 435 | 409 | 362 | 378.71 | 355 | 402 | 1.133 | 0.181 | 0.14303339 | 0.32933177 | 0.0024 | 0.0156 | 0.0083 | 0.0083 | TRUE | 0.0841 |
| Pqlc3 | 547 | 742 | 730 | 895 | 630 | 812 | 581 | 741 | 733 | 704 | 678 | 889 | 720.89 | 685 | 757 | 1.104 | 0.142 | 0.34802751 | 0.56682051 | 0.018 | 0.0115 | 0.0152 | 0.0152 | TRUE | 0.2732 |
| Pradc1 | 459 | 344 | 350 | 599 | 391 | 328 | 488 | 344 | 351 | 471 | 421 | 359 | 405.53 | 394 | 417 | 1.06 | 0.084 | 0.64169023 | 0.79835032 | 0.0264 | 0.015 | 0.0208 | 0.0208 | TRUE | 0.4911 |
| Praf2 | 1188 | 1385 | 1366 | 1832 | 1275 | 1428 | 1262 | 1383 | 1372 | 1440 | 1372 | 1563 | 1398.64 | 1339 | 1458 | 1.089 | 0.123 | 0.17822892 | 0.37899779 | 0.0024 | 0.0093 | 0.0053 | 0.0053 | TRUE | 0.0466 |
| Pram1 | 14 | 17 | 20 | 18 | 18 | 22 | 15 | 17 | 20 | 14 | 19 | 24 | 18.26 | 17 | 19 | 1.097 | 0.134 | 0.77085855 | 0.88050269 | 0 | 0.1874 | 0.0967 | 0.0967 | TRUE | 0.231 |
| Prame | 20 | 22 | 24 | 115 | 94 | 69 | 21 | 22 | 24 | 90 | 101 | 76 | 55.73 | 22 | 89 | 3.968 | 1.988 | 2.05E-11 | 1.72E-09 | 0 | 0.066 | 0.0352 | 0.0352 | TRUE | 0.1173 |
| Pramef12 | 1378 | 1490 | 1477 | 2070 | 1442 | 1643 | 1464 | 1488 | 1483 | 1627 | 1552 | 1799 | 1568.69 | 1478 | 1659 | 1.122 | 0.166 | 0.05714505 | 0.18011426 | 0.002 | 0.009 | 0.0049 | 0.0049 | TRUE | 0.064 |
| Pramef17 | 11 | 3 | 0 | 12 | 10 | 5 | 12 | 3 | 0 | 9 | 11 | 5 | 6.72 | 5 | 8 | 1.764 | 0.819 | 0.43310739 | 0.6421895 | 0.8351 | 0.4968 | 0.6235 | 0.6235 | TRUE | 1.526 |
| Pramef6 | 15 | 4 | 6 | 22 | 17 | 8 | 16 | 4 | 6 | 17 | 18 | 9 | 11.72 | 9 | 15 | 1.725 | 0.787 | 0.2530625 | 0.47095006 | 0.2035 | 0.2881 | 0.249 | 0.249 | TRUE | 1.6228 |
| Pramef8 | 1744 | 1295 | 1037 | 2613 | 1888 | 1641 | 1852 | 1293 | 1041 | 2054 | 2031 | 1796 | 1678.14 | 1395 | 1960 | 1.405 | 0.491 | 0.00992193 | 0.0523899 | 0.046 | 0.0089 | 0.0255 | 0.0255 | TRUE | 0.8688 |
| Pramel1 | 407 | 313 | 278 | 543 | 385 | 320 | 432 | 313 | 279 | 427 | 414 | 350 | 369.24 | 341 | 397 | 1.165 | 0.22 | 0.23661849 | 0.45191896 | 0.0281 | 0.0158 | 0.0223 | 0.0223 | TRUE | 0.6154 |
| Pramel3 | 9 | 9 | 11 | 191 | 138 | 95 | 10 | 9 | 11 | 150 | 148 | 104 | 72.04 | 10 | 134 | 13.629 | 3.769 | 3.15E-25 | 1.34E-22 | 0 | 0.0526 | 0.0397 | 0.0397 | TRUE | 0.3867 |
| Pramel4 | 0 | 0 | 0 | 0 | 0 | 0 | 0 | 0 | 0 | 0 | 0 | 0 | 0 | 0 | 0 | NA | NA | NA | NA | NA | NA | NA | NA | NA | NA |
| Pramel5 | 3 | 2 | 1 | 2 | 6 | 1 | 3 | 2 | 1 | 2 | 6 | 1 | 2.55 | 2 | 3 | 1.46 | 0.546 | 0.66780778 | NA | 0.2746 | 1.298 | 0.7545 | 0.7545 | TRUE | 1.1785 |
| Pramel6 | 30 | 13 | 9 | 176 | 109 | 59 | 32 | 13 | 9 | 138 | 117 | 65 | 62.35 | 18 | 107 | 5.984 | 2.581 | 3.91E-08 | 1.60E-06 | 0.2268 | 0.0597 | 0.1258 | 0.1258 | TRUE | 0.8099 |
| Pramel7 | 23 | 12 | 13 | 357 | 229 | 94 | 24 | 12 | 13 | 281 | 246 | 103 | 113.23 | 16 | 210 | 12.802 | 3.678 | 1.19E-17 | 2.28E-15 | 0.1889 | 0.036 | 0.0998 | 0.0998 | TRUE | 0.5291 |
| Prap1 | 0 | 0 | 0 | 0 | 0 | 0 | 0 | 0 | 0 | 0 | 0 | 0 | 0 | 0 | 0 | NA | NA | NA | NA | NA | NA | NA | NA | NA | NA |
| Prb1 | 0 | 0 | 0 | 0 | 0 | 0 | 0 | 0 | 0 | 0 | 0 | 0 | 0 | 0 | 0 | NA | NA | NA | NA | NA | NA | NA | NA | NA | NA |
| Prc1 | 7695 | 10487 | 10610 | 10572 | 7938 | 8717 | 8173 | 10473 | 10654 | 8311 | 8541 | 9542 | 9282.38 | 9767 | 8798 | 0.901 | -0.151 | 0.21433177 | 0.42607047 | 0.0126 | 0.0073 | 0.0105 | 0.0105 | TRUE | 0.2487 |
| Prcc | 4741 | 4332 | 4203 | 6168 | 4372 | 4012 | 5036 | 4326 | 4220 | 4849 | 4704 | 4392 | 4587.83 | 4527 | 4648 | 1.027 | 0.038 | 0.6909455 | 0.830485 | 0.0057 | 0.0076 | 0.0065 | 0.0065 | TRUE | 0.1173 |
| Prcp | 1354 | 1404 | 1355 | 1606 | 1062 | 1123 | 1438 | 1402 | 1361 | 1263 | 1143 | 1229 | 1305.9 | 1400 | 1212 | 0.866 | -0.208 | 0.01188056 | 0.05975174 | 9.00E-04 | 0.0094 | 0.0042 | 0.0042 | TRUE | 0.0294 |
| Prdm1 | 662 | 918 | 923 | 620 | 543 | 658 | 703 | 917 | 927 | 487 | 584 | 720 | 723.11 | 849 | 597 | 0.702 | -0.51 | 0.00336399 | 0.0231285 | 0.0281 | 0.0115 | 0.0203 | 0.0203 | TRUE | 0.3801 |
| Prdm10 | 645 | 547 | 604 | 748 | 546 | 591 | 685 | 546 | 606 | 588 | 587 | 647 | 610.05 | 612 | 607 | 0.991 | -0.012 | 0.91836707 | 0.95993674 | 0.0063 | 0.0123 | 0.009 | 0.009 | TRUE | 0.1251 |
| Prdm11 | 99 | 89 | 80 | 95 | 60 | 42 | 105 | 89 | 80 | 75 | 65 | 46 | 76.6 | 91 | 62 | 0.68 | -0.556 | 0.03552486 | 0.12942897 | 0.0225 | 0.0499 | 0.037 | 0.037 | TRUE | 0.4121 |
| Prdm12 | 5 | 3 | 4 | 5 | 5 | 1 | 5 | 3 | 4 | 4 | 5 | 1 | 3.79 | 4 | 3 | 0.852 | -0.231 | 0.81577265 | NA | 0 | 0.8766 | 0.445 | 0.445 | TRUE | 0.4574 |
| Prdm13 | 2 | 8 | 3 | 26 | 12 | 7 | 2 | 8 | 3 | 20 | 13 | 8 | 9.02 | 4 | 14 | 3.141 | 1.651 | 0.03431057 | 0.12614807 | 0.1871 | 0.372 | 0.2871 | 0.2871 | TRUE | 0.3166 |
| Prdm14 | 1053 | 1738 | 1750 | 2506 | 1853 | 2075 | 1118 | 1736 | 1757 | 1970 | 1994 | 2271 | 1807.76 | 1537 | 2078 | 1.352 | 0.435 | 0.01129362 | 0.05758907 | 0.0302 | 0.0087 | 0.0207 | 0.0207 | TRUE | 0.6826 |
| Prdm15 | 1158 | 1250 | 1332 | 1190 | 977 | 1033 | 1230 | 1248 | 1337 | 935 | 1051 | 1131 | 1155.55 | 1272 | 1039 | 0.816 | -0.294 | 0.00441289 | 0.02846493 | 0.0047 | 0.0098 | 0.0068 | 0.0068 | TRUE | 0.0941 |
| Prdm16 | 267 | 302 | 264 | 472 | 420 | 374 | 284 | 302 | 265 | 371 | 452 | 409 | 347.11 | 284 | 411 | 1.447 | 0.533 | 7.25E-05 | 0.0010658 | 0.0047 | 0.0164 | 0.01 | 0.01 | TRUE | 0.0885 |
| Prdm2 | 5742 | 5483 | 5623 | 7566 | 5619 | 5403 | 6099 | 5476 | 5646 | 5948 | 6046 | 5914 | 5854.83 | 5740 | 5969 | 1.04 | 0.056 | 0.42388947 | 0.63416469 | 0.0014 | 0.0075 | 0.0034 | 0.0034 | TRUE | 0.0364 |
| Prdm4 | 3067 | 3307 | 3523 | 4079 | 3099 | 3225 | 3258 | 3303 | 3537 | 3207 | 3334 | 3530 | 3361.51 | 3366 | 3357 | 0.997 | -0.004 | 0.95500455 | 0.97712175 | 0.0019 | 0.0079 | 0.004 | 0.004 | TRUE | 0.0247 |
| Prdm5 | 2114 | 2517 | 2516 | 1913 | 1401 | 1898 | 2245 | 2514 | 2526 | 1504 | 1507 | 2078 | 2062.4 | 2428 | 1696 | 0.698 | -0.518 | 0.00027112 | 0.00313263 | 0.0203 | 0.0085 | 0.0141 | 0.0141 | TRUE | 0.4647 |
| Prdm6 | 16 | 18 | 12 | 39 | 23 | 31 | 17 | 18 | 12 | 31 | 25 | 34 | 22.73 | 16 | 30 | 1.902 | 0.928 | 0.02580632 | 0.10394612 | 0 | 0.1519 | 0.0758 | 0.0758 | TRUE | 0.1949 |
| Prdm8 | 7 | 4 | 4 | 11 | 10 | 11 | 7 | 4 | 4 | 9 | 11 | 12 | 7.82 | 5 | 11 | 2.033 | 1.023 | 0.14152197 | 0.32701377 | 0 | 0.4284 | 0.2026 | 0.2026 | TRUE | 0.2959 |
| Prdm9 | 54 | 56 | 54 | 108 | 65 | 54 | 57 | 56 | 54 | 85 | 70 | 59 | 63.58 | 56 | 71 | 1.285 | 0.362 | 0.16085256 | 0.35546681 | 0 | 0.0587 | 0.0321 | 0.0321 | TRUE | 0.2624 |

| | | | | | | | | | | | | | | | | | | | | | | | | | |
|---|---|---|---|---|---|---|---|---|---|---|---|---|---|---|---|---|---|---|---|---|---|---|---|---|---|
| Prdx1 | 45249 | 43474 | 42137 | 55456 | 37178 | 37320 | 48062 | 43418 | 42310 | 43595 | 40001 | 40853 | 43039.89 | 44597 | 41483 | 0.93 | -0.104 | 0.19221436 | 0.39784841 | 0.0033 | 0.007 | 0.0046 | 0.0046 | TRUE | 0.0566 |
| Prdx2 | 8093 | 9244 | 9106 | 11326 | 8437 | 8598 | 8596 | 9232 | 9143 | 8904 | 9078 | 9412 | 9060.8 | 8990 | 9131 | 1.016 | 0.022 | 0.73395633 | 0.85735379 | 9.00E-04 | 0.0073 | 0.003 | 0.003 | TRUE | 0.018 |
| Prdx3 | 2844 | 3178 | 3105 | 3805 | 2744 | 2771 | 3021 | 3174 | 3118 | 2991 | 2952 | 3033 | 3048.22 | 3104 | 2992 | 0.964 | -0.053 | 0.40324551 | 0.61609882 | 0 | 0.008 | 0.0026 | 0.0026 | TRUE | 0.0067 |
| Prdx4 | 2783 | 3377 | 3472 | 4318 | 2828 | 3111 | 2956 | 3373 | 3486 | 3394 | 3043 | 3405 | 3276.27 | 3272 | 3281 | 1.003 | 0.004 | 0.96571303 | 0.98272313 | 0.0054 | 0.0079 | 0.0064 | 0.0064 | TRUE | 0.0864 |
| Prdx5 | 1756 | 1938 | 1916 | 2141 | 1493 | 1536 | 1865 | 1935 | 1924 | 1683 | 1606 | 1681 | 1782.57 | 1908 | 1657 | 0.868 | -0.203 | 0.00397714 | 0.02623285 | 0 | 0.0088 | 0.003 | 0.003 | TRUE | 0.0085 |
| Prdx6 | 9334 | 8360 | 7856 | 12772 | 8880 | 8289 | 9914 | 8349 | 7888 | 10040 | 9554 | 9074 | 9136.69 | 8717 | 9556 | 1.096 | 0.133 | 0.20783761 | 0.41715638 | 0.0085 | 0.0073 | 0.0079 | 0.0079 | TRUE | 0.1762 |
| Prdx6b | 0 | 0 | 0 | 0 | 0 | 0 | 0 | 0 | 0 | 0 | 0 | 0 | 0 | 0 | 0 | NA | NA | NA | NA | NA | NA | NA | NA | NA | NA |
| Preb | 3330 | 2952 | 2713 | 4115 | 3036 | 2774 | 3537 | 2948 | 2724 | 3235 | 3267 | 3037 | 3124.57 | 3070 | 3180 | 1.036 | 0.051 | 0.65200032 | 0.80502185 | 0.0095 | 0.008 | 0.0088 | 0.0088 | TRUE | 0.2151 |
| Prelid1 | 6688 | 7995 | 8138 | 10585 | 7847 | 7848 | 7104 | 7985 | 8171 | 8321 | 8443 | 8591 | 8102.46 | 7753 | 8452 | 1.09 | 0.124 | 0.11553815 | 0.28713974 | 0.0026 | 0.0073 | 0.0044 | 0.0044 | TRUE | 0.0655 |
| Prelid2 | 1700 | 2520 | 2594 | 2387 | 1882 | 2339 | 1806 | 2517 | 2605 | 1876 | 2025 | 2560 | 2231.48 | 2309 | 2154 | 0.932 | -0.101 | 0.54987733 | 0.73422496 | 0.0323 | 0.0084 | 0.0201 | 0.0201 | TRUE | 0.4399 |
| Prelp | 4 | 1 | 2 | 5 | 3 | 1 | 4 | 1 | 2 | 4 | 3 | 1 | 2.58 | 2 | 3 | 1.163 | 0.218 | 0.85603528 | NA | 0 | 1.2815 | 0.655 | 0.655 | TRUE | 0.4443 |
| Prep | 4756 | 4979 | 5022 | 6070 | 4310 | 4349 | 5052 | 4973 | 5043 | 4772 | 4637 | 4761 | 4872.77 | 5023 | 4723 | 0.941 | -0.088 | 0.12226227 | 0.29744262 | 0 | 0.0076 | 0.0022 | 0.0022 | TRUE | 0.0031 |
| Prepl | 2449 | 2534 | 2677 | 3398 | 2505 | 2260 | 2601 | 2531 | 2688 | 2671 | 2695 | 2474 | 2610.06 | 2607 | 2613 | 1.003 | 0.004 | 0.95594833 | 0.97741618 | 0.0011 | 0.0082 | 0.0036 | 0.0036 | TRUE | 0.0261 |
| Prex1 | 1999 | 2366 | 2592 | 1673 | 1437 | 1785 | 2123 | 2363 | 2603 | 1315 | 1546 | 1954 | 1984.02 | 2363 | 1605 | 0.679 | -0.559 | 0.00028473 | 0.00326842 | 0.0247 | 0.0086 | 0.0166 | 0.0166 | TRUE | 0.4354 |
| Prex2 | 1388 | 924 | 883 | 2042 | 1522 | 979 | 1474 | 923 | 887 | 1605 | 1638 | 1072 | 1266.37 | 1095 | 1438 | 1.314 | 0.394 | 0.07741918 | 0.21985151 | 0.0685 | 0.0095 | 0.0352 | 0.0352 | TRUE | 1.1008 |
| Prf1 | 6 | 7 | 5 | 10 | 1 | 2 | 6 | 7 | 5 | 8 | 1 | 2 | 4.92 | 6 | 4 | 0.631 | -0.665 | 0.48751017 | 0.68450195 | 0.1894 | 0.6767 | 0.4508 | 0.4508 | TRUE | 1.9143 |
| Prg2 | 1 | 0 | 1 | 1 | 0 | 1 | 1 | 0 | 1 | 1 | 0 | 1 | 0.66 | 1 | 1 | 0.922 | -0.117 | 0.95998349 | NA | 0 | 5.0149 | 2.384 | 2.384 | TRUE | 0.2549 |
| Prg3 | 0 | 0 | 0 | 0 | 0 | 0 | 0 | 0 | 0 | 0 | 0 | 0 | 0 | 0 | 0 | NA | NA | NA | NA | NA | NA | NA | NA | NA | NA |
| Prg4 | 1 | 0 | 1 | 0 | 2 | 1 | 1 | 0 | 1 | 0 | 2 | 1 | 0.89 | 1 | 1 | 1.53 | 0.614 | 0.77623094 | NA | 0 | 3.7275 | 2.1477 | 2.1477 | TRUE | 0.1683 |
| Prh1 | 0 | 0 | 0 | 0 | 0 | 0 | 0 | 0 | 0 | 0 | 0 | 0 | 0 | 0 | 0 | NA | NA | NA | NA | NA | NA | NA | NA | NA | NA |
| Prhoxnb | 1 | 0 | 0 | 0 | 1 | 0 | 1 | 0 | 0 | 0 | 1 | 0 | 0.36 | 0 | 0 | 0.954 | -0.067 | 0.98557668 | NA | 0 | 9.2518 | 8.0179 | 8.0179 | TRUE | 0.5204 |
| Prickle1 | 378 | 332 | 305 | 589 | 474 | 398 | 402 | 332 | 306 | 463 | 510 | 436 | 408 | 347 | 470 | 1.356 | 0.439 | 0.0022162 | 0.01666687 | 0.0096 | 0.015 | 0.0123 | 0.0123 | TRUE | 0.2185 |
| Prickle2 | 757 | 575 | 578 | 675 | 500 | 493 | 804 | 574 | 580 | 531 | 538 | 540 | 594.49 | 653 | 536 | 0.821 | -0.284 | 0.06295012 | 0.19237538 | 0.0176 | 0.0124 | 0.0151 | 0.0151 | TRUE | 0.4815 |
| Prickle3 | 115 | 143 | 147 | 129 | 97 | 134 | 122 | 143 | 148 | 101 | 104 | 147 | 127.5 | 138 | 117 | 0.851 | -0.233 | 0.27874267 | 0.49992374 | 0.0182 | 0.0327 | 0.0256 | 0.0256 | TRUE | 0.4608 |
| Prim1 | 2756 | 3338 | 3326 | 4299 | 3000 | 3044 | 2927 | 3334 | 3340 | 3380 | 3228 | 3332 | 3256.7 | 3200 | 3313 | 1.035 | 0.05 | 0.54634518 | 0.73136617 | 0.0024 | 0.0079 | 0.0047 | 0.0047 | TRUE | 0.0674 |
| Prim2 | 3090 | 3522 | 3631 | 3729 | 2764 | 2835 | 3282 | 3517 | 3646 | 2931 | 2974 | 3103 | 3242.36 | 3482 | 3003 | 0.862 | -0.214 | 0.00445889 | 0.02868092 | 0.0014 | 0.0079 | 0.0038 | 0.0038 | TRUE | 0.0305 |
| Prima1 | 43 | 46 | 55 | 74 | 52 | 62 | 46 | 46 | 55 | 58 | 56 | 68 | 54.81 | 49 | 61 | 1.237 | 0.307 | 0.24662933 | 0.46362786 | 0 | 0.067 | 0.032 | 0.032 | TRUE | 0.1026 |
| Prkaa1 | 3171 | 3507 | 3586 | 3687 | 2730 | 2976 | 3368 | 3502 | 3601 | 2898 | 2937 | 3258 | 3260.8 | 3490 | 3031 | 0.868 | -0.204 | 0.01092954 | 0.05627155 | 0.0022 | 0.0079 | 0.0043 | 0.0043 | TRUE | 0.0516 |
| Prkaa2 | 4364 | 5228 | 5644 | 4158 | 3076 | 3440 | 4635 | 5221 | 5667 | 3269 | 3310 | 3766 | 4311.28 | 5174 | 3448 | 0.666 | -0.586 | 3.07E-08 | 1.31E-06 | 0.0078 | 0.0077 | 0.0078 | 0.0078 | TRUE | 0.1012 |
| Prkab1 | 1106 | 1283 | 1302 | 1644 | 1204 | 1152 | 1175 | 1281 | 1307 | 1292 | 1295 | 1261 | 1268.72 | 1254 | 1283 | 1.023 | 0.032 | 0.69723442 | 0.83403726 | 8.00E-04 | 0.0095 | 0.0042 | 0.0042 | TRUE | 0.0368 |
| Prkab2 | 859 | 949 | 1025 | 1740 | 1335 | 1274 | 912 | 948 | 1029 | 1368 | 1436 | 1395 | 1181.37 | 963 | 1400 | 1.452 | 0.539 | 5.41E-10 | 3.53E-08 | 0.0011 | 0.0097 | 0.0045 | 0.0045 | TRUE | 0.043 |
| Prkaca | 5186 | 5501 | 5605 | 8081 | 5699 | 5639 | 5508 | 5494 | 5628 | 6353 | 6132 | 6173 | 5881.26 | 5543 | 6219 | 1.122 | 0.166 | 0.00383591 | 0.02551668 | 1.00E-04 | 0.0075 | 0.0022 | 0.0022 | TRUE | 0.0044 |
| Prkacb | 2734 | 3061 | 3147 | 3569 | 2795 | 2829 | 2904 | 3057 | 3160 | 2806 | 3007 | 3097 | 3005.11 | 3040 | 2970 | 0.976 | -0.035 | 0.65761691 | 0.80725118 | 0.0019 | 0.008 | 0.004 | 0.004 | TRUE | 0.0289 |
| Prkag1 | 1015 | 1207 | 1217 | 1657 | 1111 | 1128 | 1078 | 1205 | 1222 | 1303 | 1195 | 1235 | 1206.38 | 1168 | 1244 | 1.065 | 0.091 | 0.32253558 | 0.54376341 | 0.0024 | 0.0096 | 0.0053 | 0.0053 | TRUE | 0.0545 |
| Prkag2 | 671 | 791 | 823 | 981 | 685 | 690 | 713 | 790 | 826 | 771 | 737 | 755 | 765.43 | 776 | 754 | 0.972 | -0.041 | 0.677453 | 0.82064429 | 0.0017 | 0.0112 | 0.0057 | 0.0057 | TRUE | 0.0604 |
| Prkag3 | 22 | 13 | 16 | 16 | 10 | 16 | 23 | 13 | 16 | 13 | 11 | 18 | 15.54 | 17 | 14 | 0.778 | -0.362 | 0.4770371 | 0.67668578 | 0 | 0.2188 | 0.121 | 0.121 | TRUE | 0.4196 |
| Prkar1a | 10687 | 11913 | 11661 | 15028 | 10909 | 10864 | 11351 | 11898 | 11709 | 11814 | 11737 | 11892 | 11733.6 | 11653 | 11814 | 1.014 | 0.02 | 0.71889631 | 0.84776027 | 2.00E-04 | 0.0072 | 0.0021 | 0.0021 | TRUE | 0.0062 |
| Prkar1b | 153 | 327 | 324 | 183 | 151 | 227 | 163 | 327 | 325 | 144 | 162 | 248 | 228.21 | 272 | 185 | 0.68 | -0.557 | 0.06434915 | 0.19526081 | 0.1088 | 0.0213 | 0.0609 | 0.0609 | TRUE | 1.3713 |
| Prkar2a | 4060 | 4040 | 4108 | 7267 | 5110 | 5093 | 4312 | 4035 | 4125 | 5713 | 5498 | 5575 | 4876.33 | 4157 | 5595 | 1.346 | 0.429 | 1.92E-11 | 1.62E-09 | 5.00E-04 | 0.0076 | 0.0027 | 0.0027 | TRUE | 0.0129 |
| Prkar2b | 399 | 525 | 496 | 693 | 488 | 390 | 424 | 524 | 498 | 545 | 525 | 427 | 490.49 | 482 | 499 | 1.036 | 0.051 | 0.72290697 | 0.85005881 | 0.0122 | 0.0136 | 0.0129 | 0.0129 | TRUE | 0.1825 |
| Prkca | 883 | 736 | 765 | 1033 | 719 | 596 | 938 | 735 | 768 | 812 | 774 | 652 | 779.86 | 814 | 746 | 0.918 | -0.124 | 0.37204283 | 0.58861064 | 0.0133 | 0.0111 | 0.0125 | 0.0125 | TRUE | 0.2109 |
| Prkcb | 2210 | 2968 | 3039 | 1634 | 1405 | 1609 | 2347 | 2964 | 3051 | 1285 | 1512 | 1761 | 2153.43 | 2787 | 1519 | 0.545 | -0.877 | 3.23E-09 | 1.73E-07 | 0.021 | 0.0084 | 0.0153 | 0.0153 | TRUE | 0.2341 |
| Prkcd | 1587 | 1573 | 1488 | 2406 | 1682 | 1791 | 1686 | 1571 | 1494 | 1891 | 1810 | 1961 | 1735.4 | 1584 | 1887 | 1.192 | 0.253 | 0.00276751 | 0.01990388 | 0.0018 | 0.0088 | 0.0046 | 0.0046 | TRUE | 0.0381 |
| Prkcdbp | 9 | 13 | 7 | 38 | 24 | 18 | 10 | 13 | 7 | 30 | 26 | 20 | 17.5 | 10 | 25 | 2.562 | 1.357 | 0.00625135 | 0.03716408 | 0 | 0.1952 | 0.1064 | 0.1064 | TRUE | 0.2501 |
| Prkce | 54 | 52 | 65 | 217 | 116 | 131 | 57 | 52 | 65 | 171 | 125 | 143 | 102.23 | 58 | 146 | 2.523 | 1.335 | 5.00E-09 | 2.54E-07 | 0.0124 | 0.0391 | 0.0254 | 0.0254 | TRUE | 0.1705 |
| Prkcg | 625 | 649 | 666 | 539 | 456 | 434 | 664 | 648 | 669 | 424 | 491 | 475 | 561.7 | 660 | 463 | 0.7 | -0.515 | 1.12E-06 | 3.05E-05 | 0.0011 | 0.0128 | 0.0062 | 0.0062 | TRUE | 0.069 |
| Prkch | 125 | 134 | 141 | 435 | 311 | 240 | 133 | 134 | 142 | 342 | 335 | 263 | 224.58 | 136 | 313 | 2.305 | 1.205 | 4.86E-13 | 5.14E-11 | 0.0088 | 0.0216 | 0.0147 | 0.0147 | TRUE | 0.2186 |
| Prkci | 12899 | 13598 | 13976 | 14269 | 10416 | 11561 | 13701 | 13580 | 14033 | 11217 | 11207 | 12655 | 12732.39 | 13771 | 11693 | 0.849 | -0.236 | 0.00198951 | 0.01530149 | 0.0026 | 0.0072 | 0.0041 | 0.0041 | TRUE | 0.0632 |
| Prkcq | 18 | 11 | 15 | 27 | 24 | 8 | 19 | 11 | 15 | 21 | 26 | 9 | 16.83 | 15 | 19 | 1.243 | 0.314 | 0.56821043 | 0.74742773 | 0.1067 | 0.2027 | 0.1577 | 0.1577 | TRUE | 0.6605 |
| Prkcsh | 5948 | 6718 | 6759 | 7137 | 5533 | 5744 | 6318 | 6709 | 6787 | 5611 | 5953 | 6288 | 6277.56 | 6605 | 5951 | 0.901 | -0.151 | 0.04718983 | 0.15772971 | 0.0021 | 0.0074 | 0.004 | 0.004 | TRUE | 0.0307 |
| Prkcz | 1727 | 2152 | 2163 | 2020 | 1365 | 1789 | 1834 | 2149 | 2172 | 1588 | 1469 | 1958 | 1861.74 | 2052 | 1672 | 0.814 | -0.296 | 0.02499044 | 0.10165562 | 0.0151 | 0.0087 | 0.012 | 0.012 | TRUE | 0.2695 |
| Prkd1 | 268 | 315 | 351 | 406 | 313 | 292 | 285 | 315 | 352 | 319 | 337 | 320 | 321.21 | 317 | 325 | 1.024 | 0.034 | 0.79519587 | 0.8946193 | 0.0028 | 0.0172 | 0.0096 | 0.0096 | TRUE | 0.1069 |
| Prkd2 | 324 | 402 | 369 | 442 | 333 | 338 | 344 | 401 | 371 | 347 | 358 | 370 | 365.31 | 372 | 358 | 0.963 | -0.055 | 0.64710962 | 0.80180791 | 0 | 0.0159 | 0.0077 | 0.0077 | TRUE | 0.055 |
| Prkd3 | 4045 | 3679 | 3757 | 4531 | 3512 | 3390 | 4297 | 3674 | 3772 | 3562 | 3779 | 3711 | 3799.11 | 3914 | 3684 | 0.941 | -0.088 | 0.31707826 | 0.53796883 | 0.0037 | 0.0078 | 0.0053 | 0.0053 | TRUE | 0.0886 |
| Prkdc | 2428 | 2411 | 2788 | 3223 | 2296 | 1943 | 2579 | 2408 | 2799 | 2534 | 2470 | 2127 | 2486.21 | 2595 | 2377 | 0.916 | -0.126 | 0.22478982 | 0.43721605 | 0.0065 | 0.0082 | 0.0074 | 0.0074 | TRUE | 0.102 |
| Prkg1 | 8 | 11 | 10 | 19 | 15 | 8 | 8 | 11 | 10 | 15 | 16 | 9 | 11.56 | 10 | 13 | 1.354 | 0.438 | 0.44015336 | 0.64740266 | 0 | 0.2919 | 0.1431 | 0.1431 | TRUE | 0.3359 |
| Prkg2 | 58 | 94 | 69 | 89 | 73 | 52 | 62 | 94 | 69 | 70 | 79 | 57 | 71.7 | 75 | 69 | 0.914 | -0.13 | 0.62904127 | 0.79135669 | 0.0227 | 0.0529 | 0.0382 | 0.0382 | TRUE | 0.4511 |
| Prkra | 967 | 1387 | 1299 | 1062 | 775 | 1077 | 1027 | 1385 | 1304 | 835 | 834 | 1179 | 1094.05 | 1239 | 949 | 0.766 | -0.385 | 0.02556705 | 0.10326339 | 0.0321 | 0.0099 | 0.0205 | 0.0205 | TRUE | 0.5315 |
| Prkrip1 | 1000 | 1158 | 1117 | 1436 | 1009 | 907 | 1062 | 1157 | 1122 | 1129 | 1086 | 993 | 1091.27 | 1114 | 1069 | 0.961 | -0.057 | 0.53449575 | 0.72187592 | 0.0021 | 0.0099 | 0.0053 | 0.0053 | TRUE | 0.046 |
| Prkrir | 2316 | 2314 | 2348 | 3212 | 2266 | 2186 | 2460 | 2311 | 2358 | 2525 | 2438 | 2393 | 2414.11 | 2376 | 2452 | 1.032 | 0.046 | 0.51146266 | 0.70495404 | 5.00E-04 | 0.0083 | 0.0031 | 0.0031 | TRUE | 0.0115 |
| Prkx | 684 | 593 | 619 | 1104 | 758 | 747 | 727 | 592 | 622 | 868 | 816 | 818 | 740.24 | 647 | 834 | 1.29 | 0.368 | 0.00102174 | 0.00905841 | 0.0045 | 0.0114 | 0.0077 | 0.0077 | TRUE | 0.1362 |
| Prl | 0 | 0 | 0 | 0 | 0 | 0 | 0 | 0 | 0 | 0 | 0 | 0 | 0 | 0 | 0 | NA | NA | NA | NA | NA | NA | NA | NA | NA | NA |
| Prl2a1 | 0 | 0 | 0 | 0 | 0 | 0 | 0 | 0 | 0 | 0 | 0 | 0 | 0 | 0 | 0 | NA | NA | NA | NA | NA | NA | NA | NA | NA | NA |
| Prl2b1 | 0 | 0 | 0 | 1 | 0 | 0 | 0 | 0 | 0 | 1 | 0 | 0 | 0.13 | 0 | 0 | 1.859 | 0.894 | 0.82650734 | NA | 0 | 25.1515 | 10 | 10 | TRUE | 0.2563 |
| Prl2c1 | 0 | 0 | 0 | 0 | 0 | 0 | 0 | 0 | 0 | 0 | 0 | 0 | 0 | 0 | 0 | NA | NA | NA | NA | NA | NA | NA | NA | NA | NA |
| Prl2c2 | 0 | 0 | 0 | 0 | 0 | 0 | 0 | 0 | 0 | 0 | 0 | 0 | 0 | 0 | 0 | NA | NA | NA | NA | NA | NA | NA | NA | NA | NA |
| Prl2c3 | 0 | 0 | 0 | 0 | 0 | 0 | 0 | 0 | 0 | 0 | 0 | 0 | 0 | 0 | 0 | NA | NA | NA | NA | NA | NA | NA | NA | NA | NA |
| Prl2c4 | 0 | 0 | 0 | 0 | 0 | 0 | 0 | 0 | 0 | 0 | 0 | 0 | 0 | 0 | 0 | NA | NA | NA | NA | NA | NA | NA | NA | NA | NA |
| Prl2c5 | 0 | 0 | 0 | 0 | 0 | 0 | 0 | 0 | 0 | 0 | 0 | 0 | 0 | 0 | 0 | NA | NA | NA | NA | NA | NA | NA | NA | NA | NA |

| | | | | | | | | | | | | | | | | | | | | | | | | | |
|---|---|---|---|---|---|---|---|---|---|---|---|---|---|---|---|---|---|---|---|---|---|---|---|---|---|
| Prl3a1 | 0 | 0 | 0 | 0 | 0 | 0 | 0 | 0 | 0 | 0 | 0 | 0 | 0 | 0 | 0 | NA | NA | NA | NA | NA | NA | NA | NA | NA | NA |
| Prl3b1 | 0 | 0 | 0 | 0 | 0 | 0 | 0 | 0 | 0 | 0 | 0 | 0 | 0 | 0 | 0 | NA | NA | NA | NA | NA | NA | NA | NA | NA | NA |
| Prl3c1 | 0 | 0 | 0 | 0 | 0 | 0 | 0 | 0 | 0 | 0 | 0 | 0 | 0 | 0 | 0 | NA | NA | NA | NA | NA | NA | NA | NA | NA | NA |
| Prl3d1 | 0 | 0 | 0 | 0 | 0 | 0 | 0 | 0 | 0 | 0 | 0 | 0 | 0 | 0 | 0 | NA | NA | NA | NA | NA | NA | NA | NA | NA | NA |
| Prl3d2 | 0 | 0 | 0 | 0 | 0 | 0 | 0 | 0 | 0 | 0 | 0 | 0 | 0 | 0 | 0 | NA | NA | NA | NA | NA | NA | NA | NA | NA | NA |
| Prl3d3 | 0 | 0 | 0 | 0 | 0 | 0 | 0 | 0 | 0 | 0 | 0 | 0 | 0 | 0 | 0 | NA | NA | NA | NA | NA | NA | NA | NA | NA | NA |
| Prl4a1 | 0 | 0 | 0 | 0 | 0 | 0 | 0 | 0 | 0 | 0 | 0 | 0 | 0 | 0 | 0 | NA | NA | NA | NA | NA | NA | NA | NA | NA | NA |
| Prl5a1 | 0 | 0 | 0 | 0 | 0 | 0 | 0 | 0 | 0 | 0 | 0 | 0 | 0 | 0 | 0 | NA | NA | NA | NA | NA | NA | NA | NA | NA | NA |
| Prl6a1 | 0 | 0 | 0 | 0 | 0 | 0 | 0 | 0 | 0 | 0 | 0 | 0 | 0 | 0 | 0 | NA | NA | NA | NA | NA | NA | NA | NA | NA | NA |
| Prl7a1 | 0 | 0 | 0 | 0 | 0 | 0 | 0 | 0 | 0 | 0 | 0 | 0 | 0 | 0 | 0 | NA | NA | NA | NA | NA | NA | NA | NA | NA | NA |
| Prl7a2 | 0 | 0 | 0 | 0 | 0 | 0 | 0 | 0 | 0 | 0 | 0 | 0 | 0 | 0 | 0 | NA | NA | NA | NA | NA | NA | NA | NA | NA | NA |
| Prl7b1 | 0 | 0 | 0 | 0 | 0 | 0 | 0 | 0 | 0 | 0 | 0 | 0 | 0 | 0 | 0 | NA | NA | NA | NA | NA | NA | NA | NA | NA | NA |
| Prl7c1 | 0 | 0 | 0 | 0 | 0 | 0 | 0 | 0 | 0 | 0 | 0 | 0 | 0 | 0 | 0 | NA | NA | NA | NA | NA | NA | NA | NA | NA | NA |
| Prl7d1 | 0 | 0 | 0 | 0 | 0 | 0 | 0 | 0 | 0 | 0 | 0 | 0 | 0 | 0 | 0 | NA | NA | NA | NA | NA | NA | NA | NA | NA | NA |
| Prl8a1 | 0 | 0 | 0 | 0 | 0 | 0 | 0 | 0 | 0 | 0 | 0 | 0 | 0 | 0 | 0 | NA | NA | NA | NA | NA | NA | NA | NA | NA | NA |
| Prl8a2 | 0 | 0 | 0 | 0 | 0 | 0 | 0 | 0 | 0 | 0 | 0 | 0 | 0 | 0 | 0 | NA | NA | NA | NA | NA | NA | NA | NA | NA | NA |
| Prl8a6 | 0 | 0 | 0 | 0 | 0 | 0 | 0 | 0 | 0 | 0 | 0 | 0 | 0 | 0 | 0 | NA | NA | NA | NA | NA | NA | NA | NA | NA | NA |
| Prl8a8 | 0 | 0 | 0 | 0 | 0 | 0 | 0 | 0 | 0 | 0 | 0 | 0 | 0 | 0 | 0 | NA | NA | NA | NA | NA | NA | NA | NA | NA | NA |
| Prl8a9 | 0 | 0 | 0 | 4 | 1 | 0 | 0 | 0 | 0 | 3 | 1 | 0 | 0.7 | 0 | 1 | 7.607 | 2.927 | 0.32216877 | NA | 6.5513 | 4.6905 | 4.9568 | 4.9568 | TRUE | 0.1456 |
| Prlh | 0 | 0 | 0 | 0 | 0 | 0 | 0 | 0 | 0 | 0 | 0 | 0 | 0 | 0 | 0 | NA | NA | NA | NA | NA | NA | NA | NA | NA | NA |
| Prlhr | 3 | 1 | 6 | 3 | 3 | 2 | 3 | 1 | 6 | 2 | 3 | 2 | 3 | 3 | 2 | 0.759 | -0.398 | 0.72328949 | NA | 0 | 1.1059 | 0.5703 | 0.5703 | TRUE | 0.2148 |
| Prlr | 22 | 20 | 21 | 11 | 19 | 10 | 23 | 20 | 21 | 9 | 20 | 11 | 17.41 | 21 | 13 | 0.613 | -0.705 | 0.15985699 | 0.35445491 | 0.0258 | 0.1961 | 0.1205 | 0.1205 | TRUE | 0.8735 |
| Prm1 | 0 | 0 | 0 | 0 | 0 | 0 | 0 | 0 | 0 | 0 | 0 | 0 | 0 | 0 | 0 | NA | NA | NA | NA | NA | NA | NA | NA | NA | NA |
| Prm2 | 1 | 0 | 0 | 0 | 0 | 0 | 1 | 0 | 0 | 0 | 0 | 0 | 0.18 | 0 | 0 | 0.49 | -1.029 | 0.80087 | NA | 0 | 18.6165 | 10 | 10 | TRUE | 0.4614 |
| Prm3 | 1 | 0 | 0 | 0 | 0 | 0 | 1 | 0 | 0 | 0 | 0 | 0 | 0.18 | 0 | 0 | 0.49 | -1.029 | 0.80087 | NA | 0 | 18.6165 | 10 | 10 | TRUE | 0.4614 |
| Prmt1 | 10985 | 10073 | 10032 | 15377 | 10484 | 9541 | 11668 | 10060 | 10073 | 12088 | 11280 | 10444 | 10935.61 | 10600 | 11271 | 1.063 | 0.089 | 0.35886825 | 0.57738141 | 0.0062 | 0.0072 | 0.0066 | 0.0066 | TRUE | 0.0948 |
| Prmt10 | 2690 | 2613 | 2699 | 3787 | 2785 | 2492 | 2857 | 2610 | 2710 | 2977 | 2996 | 2728 | 2813.06 | 2726 | 2900 | 1.064 | 0.09 | 0.25741877 | 0.47607984 | 0.0018 | 0.0081 | 0.0042 | 0.0042 | TRUE | 0.0326 |
| Prmt2 | 1178 | 940 | 921 | 1141 | 819 | 862 | 1251 | 939 | 925 | 897 | 881 | 944 | 972.76 | 1038 | 907 | 0.874 | -0.194 | 0.15767267 | 0.35109605 | 0.0149 | 0.0103 | 0.0126 | 0.0126 | TRUE | 0.3837 |
| Prmt3 | 4207 | 4604 | 4733 | 5352 | 4176 | 4219 | 4469 | 4598 | 4752 | 4207 | 4493 | 4618 | 4522.98 | 4606 | 4439 | 0.963 | -0.054 | 0.45413655 | 0.65923992 | 0.0013 | 0.0076 | 0.0035 | 0.0035 | TRUE | 0.0259 |
| Prmt5 | 11663 | 11806 | 11668 | 13841 | 10342 | 10655 | 12388 | 11791 | 11716 | 10881 | 11127 | 11664 | 11594.41 | 11965 | 11224 | 0.938 | -0.092 | 0.15132024 | 0.34239864 | 0.001 | 0.0072 | 0.0029 | 0.0029 | TRUE | 0.0143 |
| Prmt6 | 967 | 822 | 757 | 1851 | 1240 | 925 | 1027 | 821 | 760 | 1455 | 1334 | 1013 | 1068.33 | 869 | 1267 | 1.459 | 0.545 | 0.00116459 | 0.01000428 | 0.0278 | 0.01 | 0.0193 | 0.0193 | TRUE | 0.3694 |
| Prmt7 | 11895 | 15422 | 15796 | 13821 | 10578 | 11817 | 12635 | 15402 | 15861 | 10865 | 11381 | 12936 | 13179.89 | 14633 | 11727 | 0.801 | -0.319 | 0.00578283 | 0.03491054 | 0.0112 | 0.0072 | 0.0096 | 0.0096 | TRUE | 0.1744 |
| Prmt8 | 496 | 890 | 1015 | 759 | 579 | 748 | 527 | 889 | 1019 | 597 | 623 | 819 | 745.55 | 812 | 680 | 0.837 | -0.257 | 0.26744029 | 0.48710169 | 0.0693 | 0.0113 | 0.0375 | 0.0375 | TRUE | 0.7398 |
| Prnd | 2 | 3 | 5 | 3 | 5 | 0 | 2 | 3 | 5 | 2 | 5 | 0 | 2.98 | 3 | 2 | 0.761 | -0.394 | 0.74977235 | NA | 0.2358 | 1.1125 | 0.7608 | 0.7608 | TRUE | 0.3059 |
| Prnp | 644 | 711 | 777 | 785 | 525 | 626 | 684 | 710 | 780 | 617 | 565 | 685 | 673.59 | 725 | 622 | 0.858 | -0.22 | 0.05700244 | 0.17977254 | 0.0052 | 0.0118 | 0.0082 | 0.0082 | TRUE | 0.0887 |
| Prob1 | 85 | 55 | 53 | 117 | 101 | 76 | 90 | 55 | 53 | 92 | 109 | 83 | 80.38 | 66 | 95 | 1.433 | 0.519 | 0.06410969 | 0.19484295 | 0.0388 | 0.0479 | 0.0436 | 0.0436 | TRUE | 0.8847 |
| Proc | 20 | 18 | 12 | 32 | 21 | 17 | 21 | 18 | 12 | 25 | 23 | 19 | 19.6 | 17 | 22 | 1.303 | 0.382 | 0.3921452 | 0.60582958 | 0 | 0.1749 | 0.0915 | 0.0915 | TRUE | 0.3316 |
| Proca1 | 81 | 78 | 90 | 132 | 121 | 119 | 86 | 78 | 90 | 104 | 130 | 130 | 103.09 | 85 | 121 | 1.426 | 0.512 | 0.01382306 | 0.06659051 | 0 | 0.0389 | 0.0211 | 0.0211 | TRUE | 0.1816 |
| Procr | 215 | 273 | 291 | 260 | 170 | 221 | 228 | 273 | 292 | 204 | 183 | 242 | 237.07 | 264 | 210 | 0.792 | -0.336 | 0.04955935 | 0.16329979 | 0.0129 | 0.0208 | 0.0168 | 0.0168 | TRUE | 0.1879 |
| Prodh | 784 | 868 | 892 | 1157 | 973 | 737 | 833 | 867 | 896 | 910 | 1047 | 807 | 893.08 | 865 | 921 | 1.065 | 0.09 | 0.44895683 | 0.65455233 | 0.0075 | 0.0106 | 0.0091 | 0.0091 | TRUE | 0.1668 |
| Prodh2 | 6 | 7 | 7 | 8 | 12 | 7 | 6 | 7 | 7 | 6 | 13 | 8 | 7.88 | 7 | 9 | 1.303 | 0.382 | 0.57568043 | 0.7538092 | 0 | 0.4252 | 0.205 | 0.205 | TRUE | 0.4534 |
| Prok1 | 0 | 0 | 0 | 1 | 0 | 0 | 0 | 0 | 0 | 1 | 0 | 0 | 0.13 | 0 | 0 | 1.859 | 0.894 | 0.82650734 | NA | 0 | 25.1515 | 10 | 10 | TRUE | 0.2563 |
| Prok2 | 51 | 86 | 69 | 63 | 40 | 48 | 54 | 86 | 69 | 50 | 43 | 53 | 59.08 | 70 | 49 | 0.693 | -0.53 | 0.06073092 | 0.18759264 | 0.0168 | 0.0627 | 0.0401 | 0.0401 | TRUE | 0.1525 |
| Prokr1 | 94 | 83 | 54 | 60 | 48 | 49 | 100 | 83 | 54 | 47 | 52 | 54 | 64.9 | 79 | 51 | 0.643 | -0.638 | 0.0314351 | 0.11909762 | 0.0368 | 0.0577 | 0.0472 | 0.0472 | TRUE | 0.2708 |
| Prokr2 | 39 | 17 | 18 | 39 | 23 | 15 | 41 | 17 | 18 | 31 | 25 | 16 | 24.72 | 25 | 24 | 0.948 | -0.077 | 0.87791945 | 0.93970448 | 0.1382 | 0.1402 | 0.1393 | 0.1393 | TRUE | 1.2369 |
| Prol1 | 0 | 0 | 0 | 0 | 0 | 0 | 0 | 0 | 0 | 0 | 0 | 0 | 0 | 0 | 0 | NA | NA | NA | NA | NA | NA | NA | NA | NA | NA |
| Prom1 | 693 | 1344 | 1391 | 666 | 528 | 691 | 736 | 1342 | 1397 | 524 | 568 | 756 | 887.19 | 1158 | 616 | 0.531 | -0.912 | 0.00013018 | 0.00170531 | 0.0762 | 0.0106 | 0.0397 | 0.0397 | TRUE | 1.2165 |
| Prom2 | 30 | 38 | 42 | 47 | 40 | 60 | 32 | 38 | 42 | 37 | 43 | 66 | 42.94 | 37 | 49 | 1.291 | 0.368 | 0.28147207 | 0.50281771 | 0.0374 | 0.0836 | 0.0605 | 0.0605 | TRUE | 0.739 |
| Prop1 | 1 | 0 | 0 | 1 | 0 | 0 | 1 | 0 | 0 | 1 | 0 | 0 | 0.31 | 0 | 0 | 0.954 | -0.067 | 0.98625928 | NA | 0 | 10.7014 | 9.0384 | 9.0384 | TRUE | 0.4614 |
| Prorsd1 | 547 | 628 | 553 | 705 | 518 | 485 | 581 | 627 | 555 | 554 | 557 | 531 | 567.66 | 588 | 547 | 0.932 | -0.102 | 0.31728357 | 0.53815218 | 0 | 0.0127 | 0.0058 | 0.0058 | TRUE | 0.0407 |
| Pros1 | 1512 | 1394 | 1264 | 2505 | 1611 | 1524 | 1606 | 1392 | 1269 | 1969 | 1733 | 1668 | 1606.37 | 1422 | 1790 | 1.259 | 0.333 | 0.00535559 | 0.0329055 | 0.0101 | 0.009 | 0.0097 | 0.0097 | TRUE | 0.1528 |
| Prosapip1 | 485 | 414 | 438 | 606 | 399 | 296 | 515 | 413 | 440 | 476 | 429 | 324 | 433.02 | 456 | 410 | 0.901 | -0.151 | 0.37860373 | 0.59369041 | 0.0231 | 0.0145 | 0.0189 | 0.0189 | TRUE | 0.3818 |
| Prosc | 4471 | 4551 | 4392 | 5356 | 3936 | 3728 | 4749 | 4545 | 4410 | 4210 | 4235 | 4081 | 4371.73 | 4568 | 4175 | 0.914 | -0.129 | 0.0486709 | 0.16103156 | 6.00E-04 | 0.0077 | 0.0029 | 0.0029 | TRUE | 0.0146 |
| Proser1 | 3057 | 2613 | 2606 | 2798 | 2025 | 1975 | 3247 | 2610 | 2617 | 2200 | 2179 | 2162 | 2502.28 | 2825 | 2180 | 0.772 | -0.373 | 0.00052514 | 0.00538179 | 0.0078 | 0.0082 | 0.0079 | 0.0079 | TRUE | 0.2076 |
| Proser2 | 461 | 676 | 668 | 813 | 650 | 747 | 490 | 675 | 671 | 639 | 699 | 818 | 665.29 | 612 | 719 | 1.173 | 0.231 | 0.15190647 | 0.34295352 | 0.0222 | 0.0119 | 0.0171 | 0.0171 | TRUE | 0.3568 |
| Prox1 | 30 | 26 | 35 | 76 | 34 | 45 | 32 | 26 | 35 | 60 | 37 | 49 | 39.76 | 31 | 49 | 1.576 | 0.656 | 0.05253536 | 0.16968825 | 0.0217 | 0.0898 | 0.0561 | 0.0561 | TRUE | 0.1616 |
| Prox2 | 13 | 9 | 14 | 7 | 12 | 6 | 14 | 9 | 14 | 6 | 13 | 7 | 10.31 | 12 | 9 | 0.67 | -0.579 | 0.35959687 | 0.57800887 | 0 | 0.3266 | 0.1865 | 0.1865 | TRUE | 0.6631 |
| Proz | 0 | 4 | 1 | 4 | 2 | 2 | 0 | 4 | 1 | 3 | 2 | 2 | 2.08 | 2 | 2 | 1.497 | 0.582 | 0.68084648 | NA | 0 | 1.5902 | 0.9441 | 0.9441 | TRUE | 1.1416 |
| Prp2 | 0 | 0 | 0 | 0 | 0 | 0 | 0 | 0 | 0 | 0 | 0 | 0 | 0 | 0 | 0 | NA | NA | NA | NA | NA | NA | NA | NA | NA | NA |
| Prpf18 | 2984 | 2877 | 2597 | 3701 | 2633 | 2648 | 3170 | 2873 | 2608 | 2909 | 2833 | 2899 | 2881.92 | 2884 | 2880 | 0.999 | -0.001 | 0.98844843 | 0.99386769 | 0.0045 | 0.008 | 0.0059 | 0.0059 | TRUE | 0.0912 |
| Prpf19 | 8677 | 7472 | 7526 | 12978 | 8967 | 7859 | 9217 | 7462 | 7557 | 10202 | 9648 | 8603 | 8781.48 | 8079 | 9484 | 1.174 | 0.232 | 0.04199454 | 0.14595369 | 0.011 | 0.0073 | 0.0092 | 0.0092 | TRUE | 0.1853 |
| Prpf3 | 3706 | 3704 | 3762 | 4368 | 3279 | 3258 | 3936 | 3699 | 3777 | 3434 | 3528 | 3566 | 3656.88 | 3804 | 3509 | 0.922 | -0.117 | 0.07234339 | 0.21036134 | 4.00E-04 | 0.0078 | 0.0028 | 0.0028 | TRUE | 0.0112 |
| Prpf31 | 3612 | 2876 | 2870 | 5521 | 3903 | 2913 | 3837 | 2872 | 2882 | 4340 | 4199 | 3189 | 3553.16 | 3197 | 3909 | 1.223 | 0.291 | 0.06545653 | 0.19715481 | 0.0275 | 0.0078 | 0.0177 | 0.0177 | TRUE | 0.3721 |
| Prpf38a | 3780 | 4049 | 3886 | 5453 | 3754 | 3632 | 4015 | 4044 | 3902 | 4287 | 4039 | 3976 | 4043.72 | 3987 | 4101 | 1.029 | 0.041 | 0.54002298 | 0.72598384 | 8.00E-04 | 0.0077 | 0.003 | 0.003 | TRUE | 0.0195 |
| Prpf38b | 5631 | 5072 | 5186 | 6641 | 5174 | 4978 | 5981 | 5065 | 5207 | 5221 | 5567 | 5449 | 5415.1 | 5418 | 5412 | 0.999 | -0.002 | 0.98586267 | 0.99253326 | 0.0044 | 0.0075 | 0.0056 | 0.0056 | TRUE | 0.1007 |
| Prpf39 | 1121 | 933 | 943 | 1849 | 1466 | 1371 | 1191 | 932 | 947 | 1454 | 1577 | 1501 | 1266.83 | 1023 | 1511 | 1.477 | 0.562 | 2.63E-06 | 6.37E-05 | 0.0097 | 0.0095 | 0.0095 | 0.0095 | TRUE | 0.2444 |
| Prpf4 | 3397 | 3347 | 3333 | 4602 | 3294 | 3145 | 3608 | 3343 | 3347 | 3618 | 3544 | 3443 | 3483.69 | 3433 | 3535 | 1.03 | 0.043 | 0.54277982 | 0.7283165 | 9.00E-04 | 0.0078 | 0.0033 | 0.0033 | TRUE | 0.0243 |
| Prpf40a | 7973 | 8227 | 8314 | 11577 | 8090 | 7742 | 8469 | 8216 | 8348 | 9101 | 8704 | 8475 | 8552.23 | 8344 | 8760 | 1.05 | 0.07 | 0.25420817 | 0.47219071 | 6.00E-04 | 0.0073 | 0.0026 | 0.0026 | TRUE | 0.0143 |
| Prpf40b | 2121 | 2378 | 2377 | 2298 | 1842 | 1850 | 2253 | 2375 | 2387 | 1807 | 1982 | 2025 | 2138.01 | 2338 | 1938 | 0.828 | -0.272 | 0.00081269 | 0.00758002 | 0.0019 | 0.0084 | 0.0043 | 0.0043 | TRUE | 0.0436 |

| | | | | | | | | | | | | | | | | | | | | | | | | | |
|---|---|---|---|---|---|---|---|---|---|---|---|---|---|---|---|---|---|---|---|---|---|---|---|---|---|
| Prpf4b | 3872 | 3177 | 3272 | 4979 | 3846 | 3473 | 4113 | 3173 | 3285 | 3914 | 4138 | 3802 | 3737.5 | 3524 | 3951 | 1.121 | 0.165 | 0.15429637 | 0.34626671 | 0.0111 | 0.0078 | 0.0094 | 0.0094 | TRUE | 0.2598 |
| Prpf6 | 6268 | 6258 | 6543 | 7813 | 5762 | 5496 | 6658 | 6250 | 6570 | 6142 | 6200 | 6016 | 6305.88 | 6493 | 6119 | 0.943 | -0.085 | 0.16681862 | 0.36431228 | 4.00E-04 | 0.0074 | 0.0026 | 0.0026 | TRUE | 0.0131 |
| Prpf8 | 56153 | 47380 | 54643 | 69038 | 48051 | 47554 | 59644 | 47319 | 54867 | 54272 | 51700 | 52055 | 53309.68 | 53943 | 52676 | 0.977 | -0.034 | 0.72920503 | 0.85433232 | 0.0059 | 0.007 | 0.0071 | 0.0071 | TRUE | 0.1413 |
| Prph | 2062 | 2942 | 3106 | 1785 | 1532 | 1626 | 2190 | 2938 | 3119 | 1403 | 1648 | 1780 | 2179.77 | 2749 | 1610 | 0.586 | -0.772 | 4.32E-07 | 1.32E-05 | 0.0223 | 0.0084 | 0.0163 | 0.0163 | TRUE | 0.3833 |
| Prph2 | 45 | 33 | 21 | 42 | 24 | 24 | 48 | 33 | 21 | 33 | 26 | 26 | 31.16 | 34 | 28 | 0.841 | -0.249 | 0.54064961 | 0.72651663 | 0.0603 | 0.1126 | 0.0872 | 0.0872 | TRUE | 0.21 |
| Prpmp5 | 0 | 0 | 0 | 0 | 0 | 0 | 0 | 0 | 0 | 0 | 0 | 0 | 0 | 0 | 0 | NA | NA | NA | NA | NA | NA | NA | NA | NA | NA |
| Prps1 | 9525 | 12042 | 11727 | 12906 | 10303 | 11810 | 10117 | 12026 | 11775 | 10146 | 11085 | 12928 | 11346.3 | 11306 | 11386 | 1.007 | 0.01 | 0.93186335 | 0.96628017 | 0.0095 | 0.0072 | 0.0098 | 0.0098 | TRUE | 0.1714 |
| Prps1l1 | 0 | 0 | 0 | 0 | 0 | 0 | 0 | 0 | 0 | 0 | 0 | 0 | 0 | 0 | 0 | NA | NA | NA | NA | NA | NA | NA | NA | NA | NA |
| Prps1l3 | 4824 | 4730 | 4725 | 6814 | 4911 | 4725 | 5124 | 4724 | 4744 | 5357 | 5284 | 5172 | 5067.51 | 4864 | 5271 | 1.084 | 0.116 | 0.08668494 | 0.23797996 | 9.00E-04 | 0.0076 | 0.0031 | 0.0031 | TRUE | 0.0266 |
| Prps2 | 4925 | 5101 | 5227 | 6125 | 4366 | 4565 | 5231 | 5094 | 5248 | 4815 | 4698 | 4997 | 5013.96 | 5191 | 4837 | 0.932 | -0.102 | 0.1002778 | 0.26180673 | 3.00E-04 | 0.0076 | 0.0026 | 0.0026 | TRUE | 0.0101 |
| Prpsap1 | 9393 | 7994 | 7805 | 9814 | 7268 | 6904 | 9977 | 7984 | 7837 | 7715 | 7820 | 7558 | 8148.36 | 8599 | 7698 | 0.895 | -0.16 | 0.14235295 | 0.32814832 | 0.0096 | 0.0073 | 0.0084 | 0.0084 | TRUE | 0.2398 |
| Prpsap2 | 2845 | 3666 | 3755 | 3998 | 2825 | 3182 | 3022 | 3661 | 3770 | 3143 | 3040 | 3483 | 3353.2 | 3484 | 3222 | 0.924 | -0.113 | 0.30961556 | 0.52992476 | 0.0089 | 0.0079 | 0.0087 | 0.0087 | TRUE | 0.1638 |
| Prr11 | 2422 | 3668 | 4018 | 3343 | 2456 | 3006 | 2573 | 3663 | 4034 | 2628 | 2643 | 3291 | 3138.57 | 3423 | 2854 | 0.833 | -0.263 | 0.12499735 | 0.30223182 | 0.0329 | 0.008 | 0.0208 | 0.0208 | TRUE | 0.5742 |
| Prr12 | 3677 | 3353 | 3358 | 4663 | 3525 | 3093 | 3906 | 3349 | 3372 | 3666 | 3793 | 3386 | 3578.37 | 3542 | 3615 | 1.021 | 0.03 | 0.75625061 | 0.87119893 | 0.0052 | 0.0078 | 0.0062 | 0.0062 | TRUE | 0.0979 |
| Prr13 | 1124 | 1895 | 2121 | 1590 | 1396 | 1735 | 1194 | 1893 | 2130 | 1250 | 1502 | 1899 | 1644.55 | 1739 | 1550 | 0.891 | -0.166 | 0.44692464 | 0.65297364 | 0.0645 | 0.0089 | 0.0338 | 0.0338 | TRUE | 0.7675 |
| Prr14 | 860 | 801 | 773 | 1683 | 1087 | 1050 | 913 | 800 | 776 | 1323 | 1170 | 1149 | 1021.93 | 830 | 1214 | 1.465 | 0.55 | 5.18E-07 | 1.54E-05 | 0.0054 | 0.0101 | 0.0076 | 0.0076 | TRUE | 0.0917 |
| Prr14l | 3864 | 3878 | 4250 | 5846 | 4062 | 4303 | 4104 | 3873 | 4267 | 4596 | 4370 | 4710 | 4320.19 | 4081 | 4559 | 1.117 | 0.16 | 0.03116628 | 0.11838982 | 0.0015 | 0.0077 | 0.0037 | 0.0037 | TRUE | 0.0244 |
| Prr15 | 11 | 21 | 16 | 23 | 15 | 13 | 12 | 21 | 16 | 18 | 16 | 14 | 16.2 | 16 | 16 | 0.997 | -0.004 | 0.99269345 | 0.99616826 | 0 | 0.2103 | 0.1057 | 0.1057 | TRUE | 0.2132 |
| Prr15l | 95 | 196 | 222 | 183 | 138 | 194 | 101 | 196 | 223 | 144 | 148 | 212 | 170.71 | 173 | 168 | 0.969 | -0.045 | 0.88068911 | 0.94100864 | 0.0974 | 0.0262 | 0.059 | 0.059 | TRUE | 1.2515 |
| Prr16 | 10 | 9 | 11 | 19 | 22 | 12 | 11 | 9 | 11 | 15 | 24 | 13 | 13.73 | 10 | 17 | 1.682 | 0.75 | 0.16003374 | 0.35450171 | 0 | 0.2468 | 0.1271 | 0.1271 | TRUE | 0.4909 |
| Prr18 | 29 | 51 | 51 | 33 | 41 | 26 | 31 | 51 | 51 | 26 | 44 | 28 | 38.58 | 44 | 33 | 0.736 | -0.443 | 0.23472995 | 0.44950957 | 0.0535 | 0.0923 | 0.0737 | 0.0737 | TRUE | 0.6003 |
| Prr19 | 94 | 73 | 72 | 87 | 48 | 41 | 100 | 73 | 72 | 68 | 52 | 45 | 68.33 | 82 | 55 | 0.679 | -0.559 | 0.04582312 | 0.15485794 | 0.0269 | 0.0551 | 0.0413 | 0.0413 | TRUE | 0.4312 |
| Prr22 | 22 | 6 | 2 | 20 | 11 | 10 | 23 | 6 | 2 | 16 | 12 | 11 | 11.65 | 10 | 13 | 1.238 | 0.308 | 0.7057252 | 0.83911335 | 0.5074 | 0.2898 | 0.3927 | 0.3927 | TRUE | 2.2463 |
| Prr23a | 66 | 71 | 64 | 120 | 73 | 57 | 70 | 71 | 64 | 94 | 79 | 62 | 73.42 | 68 | 78 | 1.153 | 0.206 | 0.41165319 | 0.62279432 | 0.0116 | 0.0518 | 0.0316 | 0.0316 | TRUE | 0.1777 |
| Prr24 | 145 | 215 | 227 | 266 | 220 | 200 | 154 | 215 | 228 | 209 | 237 | 219 | 210.24 | 199 | 222 | 1.112 | 0.153 | 0.40857838 | 0.62117776 | 0.0173 | 0.0226 | 0.02 | 0.02 | TRUE | 0.4169 |
| Prr3 | 751 | 711 | 739 | 984 | 790 | 647 | 798 | 710 | 742 | 774 | 850 | 708 | 763.6 | 750 | 777 | 1.037 | 0.052 | 0.63615321 | 0.79536224 | 0.0044 | 0.0112 | 0.0074 | 0.0074 | TRUE | 0.0772 |
| Prr5 | 1318 | 1098 | 1011 | 1483 | 1114 | 992 | 1400 | 1097 | 1015 | 1166 | 1199 | 1086 | 1160.33 | 1171 | 1150 | 0.983 | -0.025 | 0.85438968 | 0.92752251 | 0.0147 | 0.0097 | 0.0124 | 0.0124 | TRUE | 0.3513 |
| Prr5l | 87 | 88 | 74 | 164 | 85 | 78 | 92 | 88 | 74 | 129 | 91 | 85 | 93.39 | 85 | 102 | 1.209 | 0.274 | 0.26560454 | 0.48518818 | 0.0238 | 0.0422 | 0.0328 | 0.0328 | TRUE | 0.5708 |
| Prr7 | 195 | 246 | 253 | 325 | 310 | 269 | 207 | 246 | 254 | 255 | 334 | 294 | 265.06 | 236 | 294 | 1.246 | 0.318 | 0.05010198 | 0.16442122 | 0.0113 | 0.0193 | 0.0151 | 0.0151 | TRUE | 0.1597 |
| Prr9 | 0 | 0 | 0 | 0 | 0 | 0 | 0 | 0 | 0 | 0 | 0 | 0 | 0 | 0 | 0 | NA | NA | NA | NA | NA | NA | NA | NA | NA | NA |
| Prrc1 | 5704 | 7654 | 7688 | 9214 | 7461 | 8483 | 6059 | 7644 | 7720 | 7243 | 8028 | 9286 | 7663.2 | 7141 | 8186 | 1.146 | 0.197 | 0.13484824 | 0.31767385 | 0.0164 | 0.0073 | 0.0124 | 0.0124 | TRUE | 0.2144 |
| Prrc2a | 23233 | 24380 | 25184 | 29576 | 22535 | 22831 | 24678 | 24349 | 25287 | 23250 | 24246 | 24992 | 24467.03 | 24771 | 24163 | 0.975 | -0.036 | 0.55306346 | 0.73679508 | 8.00E-04 | 0.007 | 0.0026 | 0.0026 | TRUE | 0.0134 |
| Prrc2b | 16408 | 22663 | 25854 | 20654 | 15681 | 17390 | 17428 | 22634 | 25960 | 16237 | 16872 | 19036 | 19694.42 | 22007 | 17382 | 0.79 | -0.34 | 0.01977387 | 0.08617669 | 0.0226 | 0.0071 | 0.0153 | 0.0153 | TRUE | 0.279 |
| Prrc2c | 26518 | 22872 | 25943 | 34938 | 24291 | 22266 | 28167 | 22842 | 26050 | 27466 | 26136 | 24374 | 25838.93 | 25686 | 25992 | 1.012 | 0.017 | 0.86431934 | 0.93287105 | 0.0057 | 0.007 | 0.0072 | 0.0072 | TRUE | 0.1148 |
| Prrg1 | 139 | 139 | 143 | 197 | 111 | 122 | 148 | 139 | 144 | 155 | 119 | 134 | 139.65 | 144 | 136 | 0.953 | -0.07 | 0.69856221 | 0.83505557 | 0 | 0.0305 | 0.0164 | 0.0164 | TRUE | 0.1676 |
| Prrg2 | 86 | 113 | 130 | 143 | 86 | 103 | 91 | 113 | 131 | 112 | 93 | 113 | 108.74 | 112 | 106 | 0.95 | -0.074 | 0.73249296 | 0.85653017 | 0.0122 | 0.0372 | 0.025 | 0.025 | TRUE | 0.2244 |
| Prrg3 | 316 | 500 | 565 | 711 | 522 | 627 | 336 | 499 | 567 | 559 | 562 | 686 | 534.87 | 467 | 602 | 1.287 | 0.364 | 0.06347708 | 0.19340955 | 0.0398 | 0.0131 | 0.0259 | 0.0259 | TRUE | 0.7024 |
| Prrg4 | 912 | 943 | 989 | 925 | 715 | 735 | 969 | 942 | 993 | 727 | 769 | 805 | 867.43 | 968 | 767 | 0.792 | -0.337 | 0.00016141 | 0.00202688 | 0 | 0.0107 | 0.0046 | 0.0046 | TRUE | 0.0258 |
| Prrt1 | 1 | 0 | 1 | 1 | 2 | 1 | 1 | 0 | 1 | 1 | 2 | 1 | 1.02 | 1 | 1 | 1.936 | 0.953 | 0.61010723 | NA | 0 | 3.2479 | 1.4013 | 1.4013 | TRUE | 0.256 |
| Prrt2 | 35 | 69 | 47 | 52 | 46 | 47 | 37 | 69 | 47 | 41 | 49 | 51 | 49.18 | 51 | 47 | 0.92 | -0.12 | 0.71236141 | 0.84341563 | 0.0369 | 0.0739 | 0.0558 | 0.0558 | TRUE | 0.5442 |
| Prrt3 | 70 | 36 | 30 | 156 | 91 | 44 | 74 | 36 | 30 | 123 | 98 | 48 | 68.19 | 47 | 90 | 1.923 | 0.943 | 0.03175351 | 0.11986925 | 0.2022 | 0.0552 | 0.1227 | 0.1227 | TRUE | 0.4688 |
| Prrt4 | 66 | 98 | 76 | 151 | 104 | 109 | 70 | 98 | 76 | 119 | 112 | 119 | 99.03 | 81 | 117 | 1.432 | 0.518 | 0.01567218 | 0.07326072 | 0.0011 | 0.0402 | 0.0226 | 0.0226 | TRUE | 0.2936 |
| Prrx1 | 2 | 1 | 1 | 1 | 2 | 1 | 2 | 1 | 1 | 1 | 2 | 1 | 1.36 | 1 | 1 | 0.964 | -0.053 | 0.97239284 | NA | 0 | 2.4293 | 0.9602 | 0.9602 | TRUE | 0.16 |
| Prrx2 | 165 | 184 | 146 | 140 | 100 | 116 | 175 | 184 | 147 | 110 | 108 | 127 | 141.71 | 169 | 115 | 0.681 | -0.555 | 0.00256579 | 0.01876978 | 0 | 0.0302 | 0.0171 | 0.0171 | TRUE | 0.1392 |
| Prrxl1 | 0 | 0 | 0 | 2 | 0 | 0 | 0 | 0 | 0 | 2 | 0 | 0 | 0.26 | 0 | 1 | 2.886 | 1.529 | 0.70676491 | NA | 0 | 12.5792 | 10 | 10 | TRUE | 0.9669 |
| Prss1 | 0 | 0 | 0 | 0 | 0 | 0 | 0 | 0 | 0 | 0 | 0 | 0 | 0 | 0 | 0 | NA | NA | NA | NA | NA | NA | NA | NA | NA | NA |
| Prss12 | 16 | 16 | 6 | 26 | 12 | 12 | 17 | 16 | 6 | 20 | 13 | 13 | 14.25 | 13 | 15 | 1.205 | 0.269 | 0.64047118 | 0.79767036 | 0.0864 | 0.2381 | 0.1681 | 0.1681 | TRUE | 0.9271 |
| Prss16 | 178 | 240 | 218 | 276 | 244 | 225 | 189 | 240 | 219 | 217 | 263 | 246 | 228.91 | 216 | 242 | 1.117 | 0.16 | 0.31763265 | 0.53856183 | 0.0074 | 0.0213 | 0.0141 | 0.0141 | TRUE | 0.1274 |
| Prss2 | 0 | 0 | 0 | 0 | 0 | 0 | 0 | 0 | 0 | 0 | 0 | 0 | 0 | 0 | 0 | NA | NA | NA | NA | NA | NA | NA | NA | NA | NA |
| Prss21 | 2 | 0 | 0 | 0 | 1 | 0 | 2 | 0 | 0 | 0 | 1 | 0 | 0.53 | 1 | 0 | 0.497 | -1.009 | 0.76131406 | NA | 0 | 6.1834 | 6.228 | 6.228 | TRUE | 0.9742 |
| Prss22 | 6 | 3 | 8 | 3 | 3 | 1 | 6 | 3 | 8 | 2 | 3 | 1 | 4.01 | 6 | 2 | 0.386 | -1.373 | 0.17192055 | NA | 0 | 0.8277 | 0.4225 | 0.4225 | TRUE | 0.326 |
| Prss23 | 43 | 33 | 33 | 54 | 48 | 36 | 46 | 33 | 33 | 42 | 52 | 39 | 40.88 | 37 | 44 | 1.195 | 0.257 | 0.41631374 | 0.62772693 | 0 | 0.0875 | 0.0472 | 0.0472 | TRUE | 0.2761 |
| Prss27 | 2 | 17 | 6 | 3 | 3 | 9 | 2 | 17 | 6 | 2 | 3 | 10 | 6.76 | 8 | 5 | 0.603 | -0.73 | 0.45648692 | 0.66122104 | 0.575 | 0.4942 | 0.5356 | 0.5356 | TRUE | 0.6067 |
| Prss28 | 0 | 0 | 0 | 0 | 0 | 0 | 0 | 0 | 0 | 0 | 0 | 0 | 0 | 0 | 0 | NA | NA | NA | NA | NA | NA | NA | NA | NA | NA |
| Prss29 | 0 | 0 | 0 | 0 | 0 | 0 | 0 | 0 | 0 | 0 | 0 | 0 | 0 | 0 | 0 | NA | NA | NA | NA | NA | NA | NA | NA | NA | NA |
| Prss3 | 0 | 0 | 0 | 0 | 0 | 0 | 0 | 0 | 0 | 0 | 0 | 0 | 0 | 0 | 0 | NA | NA | NA | NA | NA | NA | NA | NA | NA | NA |
| Prss30 | 0 | 0 | 0 | 0 | 0 | 0 | 0 | 0 | 0 | 0 | 0 | 0 | 0 | 0 | 0 | NA | NA | NA | NA | NA | NA | NA | NA | NA | NA |
| Prss32 | 9 | 2 | 2 | 10 | 11 | 5 | 10 | 2 | 2 | 8 | 12 | 5 | 6.46 | 5 | 8 | 1.869 | 0.902 | 0.30927477 | 0.52974532 | 0.2498 | 0.5172 | 0.3948 | 0.3948 | TRUE | 1.2915 |
| Prss33 | 24 | 23 | 23 | 23 | 12 | 14 | 25 | 23 | 23 | 18 | 13 | 15 | 19.65 | 24 | 15 | 0.653 | -0.615 | 0.1484699 | 0.33827816 | 0 | 0.1746 | 0.078 | 0.078 | TRUE | 0.1311 |
| Prss34 | 2 | 1 | 0 | 0 | 0 | 0 | 2 | 1 | 0 | 0 | 0 | 0 | 0.52 | 1 | 0 | 0.173 | -2.534 | 0.43568356 | NA | 0 | 6.3362 | 6.1226 | 6.1226 | TRUE | 0.0614 |
| Prss35 | 326 | 293 | 287 | 437 | 316 | 318 | 346 | 293 | 288 | 344 | 340 | 348 | 326.45 | 309 | 344 | 1.113 | 0.155 | 0.23126882 | 0.44504293 | 0.0018 | 0.017 | 0.009 | 0.009 | TRUE | 0.1242 |
| Prss36 | 597 | 813 | 805 | 733 | 553 | 571 | 634 | 812 | 808 | 576 | 595 | 625 | 675.11 | 751 | 599 | 0.796 | -0.329 | 0.01022998 | 0.05367149 | 0.0091 | 0.0118 | 0.0103 | 0.0103 | TRUE | 0.2196 |
| Prss37 | 0 | 0 | 0 | 0 | 0 | 0 | 0 | 0 | 0 | 0 | 0 | 0 | 0 | 0 | 0 | NA | NA | NA | NA | NA | NA | NA | NA | NA | NA |
| Prss38 | 114 | 205 | 182 | 227 | 163 | 213 | 121 | 205 | 183 | 178 | 175 | 233 | 182.59 | 170 | 195 | 1.152 | 0.204 | 0.3799222 | 0.59457561 | 0.0422 | 0.0249 | 0.0333 | 0.0333 | TRUE | 0.6567 |
| Prss39 | 1 | 8 | 1 | 5 | 2 | 0 | 1 | 8 | 1 | 4 | 2 | 0 | 2.69 | 3 | 2 | 0.621 | -0.688 | 0.63655519 | NA | 1.0392 | 1.2317 | 1.141 | 1.141 | TRUE | 0.8768 |
| Prss40 | 0 | 0 | 1 | 0 | 0 | 1 | 0 | 0 | 1 | 0 | 0 | 1 | 0.35 | 0 | 0 | 0.954 | -0.067 | 0.98566485 | NA | 0 | 9.4252 | 8.1416 | 8.1416 | TRUE | 0.5383 |
| Prss41 | 95 | 149 | 172 | 128 | 119 | 131 | 101 | 149 | 173 | 101 | 128 | 143 | 132.41 | 141 | 124 | 0.877 | -0.189 | 0.45187771 | 0.65723121 | 0.0446 | 0.0318 | 0.0381 | 0.0381 | TRUE | 0.4495 |
| Prss42 | 61 | 81 | 66 | 129 | 142 | 182 | 65 | 81 | 66 | 101 | 153 | 199 | 110.9 | 71 | 151 | 2.129 | 1.09 | 0.00012618 | 0.00166814 | 0.0589 | 0.0366 | 0.0478 | 0.0478 | TRUE | 0.1184 |
| Prss43 | 0 | 0 | 2 | 3 | 2 | 1 | 0 | 0 | 2 | 2 | 2 | 1 | 1.27 | 1 | 2 | 2.788 | 1.479 | 0.42986696 | NA | 0 | 2.6033 | 1.5139 | 1.5139 | TRUE | 0.9738 |

| | | | | | | | | | | | | | | | | | | | | | | | | |
|---|---|---|---|---|---|---|---|---|---|---|---|---|---|---|---|---|---|---|---|---|---|---|---|---|
| Prss44 | 26 | 26 | 25 | 13 | 19 | 18 | 28 | 26 | 25 | 10 | 20 | 20 | 21.51 | 26 | 17 | 0.63 | -0.667 | 0.13201324 | 0.31299142 | 0 | 0.1601 | 0.0921 | 0.0921 | TRUE | 0.7334 |
| Prss45 | 7 | 10 | 8 | 9 | 11 | 15 | 7 | 10 | 8 | 7 | 12 | 16 | 10.13 | 8 | 12 | 1.367 | 0.451 | 0.47033706 | 0.67209973 | 0 | 0.3321 | 0.1795 | 0.1795 | TRUE | 0.1692 |
| Prss46 | 2 | 0 | 0 | 0 | 1 | 0 | 2 | 0 | 0 | 0 | 1 | 0 | 0.53 | 1 | 0 | 0.497 | -1.009 | 0.76131406 | NA | 0 | 6.1834 | 6.228 | 6.228 | TRUE | 0.9742 |
| Prss48 | 8 | 10 | 14 | 11 | 16 | 14 | 8 | 10 | 14 | 9 | 17 | 15 | 12.29 | 11 | 14 | 1.248 | 0.32 | 0.57694146 | 0.75452079 | 0 | 0.275 | 0.1539 | 0.1539 | TRUE | 0.574 |
| Prss50 | 336 | 557 | 578 | 409 | 395 | 444 | 357 | 556 | 580 | 322 | 425 | 486 | 454.35 | 498 | 411 | 0.824 | -0.28 | 0.20237963 | 0.40995781 | 0.0529 | 0.0142 | 0.0325 | 0.0325 | TRUE | 0.7113 |
| Prss51 | 0 | 0 | 0 | 0 | 0 | 0 | 0 | 0 | 0 | 0 | 0 | 0 | 0 | 0 | 0 | NA | NA | NA | NA | NA | NA | NA | NA | NA | NA |
| Prss52 | 0 | 0 | 0 | 0 | 0 | 0 | 0 | 0 | 0 | 0 | 0 | 0 | 0 | 0 | 0 | NA | NA | NA | NA | NA | NA | NA | NA | NA | NA |
| Prss53 | 16 | 24 | 21 | 17 | 14 | 19 | 17 | 24 | 21 | 13 | 15 | 21 | 18.55 | 21 | 16 | 0.786 | -0.348 | 0.44161121 | 0.6486159 | 0 | 0.1845 | 0.0924 | 0.0924 | TRUE | 0.2466 |
| Prss54 | 0 | 1 | 0 | 1 | 0 | 0 | 0 | 1 | 0 | 1 | 0 | 0 | 0.3 | 0 | 0 | 0.954 | -0.067 | 0.98642001 | NA | 0 | 11.0817 | 9.3014 | 9.3014 | TRUE | 0.405 |
| Prss55 | 1 | 0 | 0 | 0 | 0 | 0 | 1 | 0 | 0 | 0 | 0 | 0 | 0.18 | 0 | 0 | 0.49 | -1.029 | 0.80087 | NA | 0 | 18.6165 | 10 | 10 | TRUE | 0.4614 |
| Prss56 | 32 | 22 | 22 | 24 | 27 | 18 | 34 | 22 | 22 | 19 | 29 | 20 | 24.28 | 26 | 23 | 0.863 | -0.212 | 0.61280654 | 0.78070959 | 0.0217 | 0.1426 | 0.085 | 0.085 | TRUE | 0.4315 |
| Prss57 | 1 | 1 | 1 | 0 | 1 | 0 | 1 | 1 | 1 | 0 | 1 | 0 | 0.69 | 1 | 0 | 0.342 | -1.548 | 0.49536145 | NA | 0 | 4.7804 | 2.2174 | 2.2174 | TRUE | 0.5204 |
| Prss58 | 0 | 0 | 0 | 0 | 0 | 0 | 0 | 0 | 0 | 0 | 0 | 0 | 0 | 0 | 0 | NA | NA | NA | NA | NA | NA | NA | NA | NA | NA |
| Prss8 | 160 | 262 | 246 | 156 | 133 | 171 | 170 | 262 | 247 | 123 | 143 | 187 | 188.59 | 226 | 151 | 0.665 | -0.589 | 0.01156363 | 0.0585871 | 0.0435 | 0.0244 | 0.0337 | 0.0337 | TRUE | 0.516 |
| Prtg | 1097 | 1147 | 1221 | 1563 | 928 | 895 | 1165 | 1146 | 1226 | 1229 | 998 | 980 | 1123.94 | 1179 | 1069 | 0.908 | -0.139 | 0.22904249 | 0.44227427 | 0.0077 | 0.0098 | 0.0088 | 0.0088 | TRUE | 0.209 |
| Prtn3 | 19 | 24 | 46 | 13 | 10 | 28 | 20 | 24 | 46 | 10 | 11 | 31 | 23.66 | 30 | 17 | 0.565 | -0.824 | 0.15255433 | 0.34375257 | 0.244 | 0.1461 | 0.1932 | 0.1932 | TRUE | 2.184 |
| Prune | 3097 | 3396 | 3379 | 3928 | 2845 | 3070 | 3290 | 3392 | 3393 | 3088 | 3061 | 3361 | 3263.93 | 3358 | 3170 | 0.944 | -0.084 | 0.24797868 | 0.4646691 | 0.0011 | 0.0079 | 0.0035 | 0.0035 | TRUE | 0.0333 |
| Prune2 | 1803 | 1803 | 2086 | 1964 | 1402 | 1352 | 1915 | 1801 | 2095 | 1544 | 1508 | 1480 | 1723.79 | 1937 | 1511 | 0.78 | -0.358 | 4.81E-05 | 0.0007528 | 0.0027 | 0.0088 | 0.005 | 0.005 | TRUE | 0.0616 |
| Prx | 857 | 870 | 849 | 933 | 660 | 647 | 910 | 869 | 852 | 733 | 710 | 708 | 797.24 | 877 | 717 | 0.818 | -0.289 | 0.00086585 | 0.00795348 | 0 | 0.011 | 0.0042 | 0.0042 | TRUE | 0.0128 |
| Psap | 13681 | 15661 | 15691 | 18845 | 13470 | 13838 | 14532 | 15641 | 15755 | 14814 | 14493 | 15148 | 15063.84 | 15309 | 14818 | 0.968 | -0.047 | 0.47052105 | 0.6721364 | 0.0011 | 0.0071 | 0.003 | 0.003 | TRUE | 0.0241 |
| Psapl1 | 0 | 1 | 0 | 0 | 1 | 1 | 0 | 1 | 0 | 0 | 1 | 1 | 0.53 | 0 | 1 | 1.924 | 0.944 | 0.7364225 | NA | 0 | 6.2438 | 3.9727 | 3.9727 | TRUE | 0.405 |
| Psat1 | 9422 | 10275 | 9964 | 13004 | 9195 | 9139 | 10008 | 10262 | 10005 | 10223 | 9893 | 10004 | 10065.75 | 10092 | 10040 | 0.995 | -0.007 | 0.89453783 | 0.9490474 | 2.00E-04 | 0.0072 | 0.0021 | 0.0021 | TRUE | 0.0031 |
| Psca | 2 | 2 | 4 | 3 | 0 | 2 | 2 | 2 | 4 | 2 | 0 | 2 | 2.11 | 3 | 1 | 0.571 | -0.807 | 0.55159311 | NA | 0 | 1.5651 | 0.8223 | 0.8223 | TRUE | 0.4697 |
| Psd | 131 | 167 | 126 | 122 | 95 | 113 | 139 | 167 | 127 | 96 | 102 | 124 | 125.71 | 144 | 107 | 0.741 | -0.432 | 0.03533172 | 0.12893379 | 0.0114 | 0.0331 | 0.0222 | 0.0222 | TRUE | 0.1985 |
| Psd2 | 142 | 183 | 159 | 154 | 116 | 136 | 151 | 183 | 160 | 121 | 125 | 149 | 148 | 165 | 132 | 0.798 | -0.326 | 0.07146323 | 0.20833976 | 0.0044 | 0.0292 | 0.0167 | 0.0167 | TRUE | 0.1308 |
| Psd3 | 288 | 228 | 246 | 392 | 245 | 234 | 306 | 228 | 247 | 308 | 264 | 256 | 268.09 | 260 | 276 | 1.064 | 0.089 | 0.59146242 | 0.76507383 | 0.0132 | 0.0192 | 0.0161 | 0.0161 | TRUE | 0.2607 |
| Psd4 | 293 | 234 | 213 | 475 | 350 | 334 | 311 | 234 | 214 | 373 | 377 | 366 | 312.4 | 253 | 372 | 1.472 | 0.557 | 0.00072773 | 0.00698182 | 0.0158 | 0.0174 | 0.0163 | 0.0163 | TRUE | 0.4477 |
| Psen1 | 2301 | 2547 | 2709 | 3026 | 2294 | 2197 | 2444 | 2544 | 2720 | 2379 | 2468 | 2405 | 2493.31 | 2569 | 2417 | 0.941 | -0.088 | 0.24184026 | 0.45850647 | 0.0013 | 0.0082 | 0.0037 | 0.0037 | TRUE | 0.032 |
| Psen2 | 249 | 325 | 355 | 309 | 236 | 220 | 264 | 325 | 356 | 243 | 254 | 241 | 280.53 | 315 | 246 | 0.779 | -0.359 | 0.01913306 | 0.08403416 | 0.0086 | 0.0186 | 0.0133 | 0.0133 | TRUE | 0.2216 |
| Psenen | 584 | 710 | 695 | 729 | 525 | 573 | 620 | 709 | 698 | 573 | 565 | 627 | 632.07 | 676 | 588 | 0.87 | -0.201 | 0.06101832 | 0.18822235 | 0.0026 | 0.0121 | 0.0067 | 0.0067 | TRUE | 0.0602 |
| Psg-ps1 | 0 | 0 | 0 | 0 | 0 | 0 | 0 | 0 | 0 | 0 | 0 | 0 | 0 | 0 | 0 | NA | NA | NA | NA | NA | NA | NA | NA | NA | NA |
| Psg16 | 5 | 6 | 11 | 9 | 7 | 14 | 5 | 6 | 11 | 7 | 8 | 15 | 8.71 | 7 | 10 | 1.324 | 0.405 | 0.56158949 | 0.74269575 | 0.0756 | 0.385 | 0.2321 | 0.2321 | TRUE | 0.7057 |
| Psg17 | 0 | 0 | 0 | 2 | 0 | 1 | 0 | 0 | 0 | 2 | 0 | 1 | 0.44 | 0 | 1 | 4.782 | 2.258 | 0.50653358 | NA | 0 | 7.4188 | 6.7775 | 6.7775 | TRUE | 0.2804 |
| Psg18 | 0 | 0 | 0 | 0 | 0 | 0 | 0 | 0 | 0 | 0 | 0 | 0 | 0 | 0 | 0 | NA | NA | NA | NA | NA | NA | NA | NA | NA | NA |
| Psg19 | 0 | 0 | 0 | 0 | 0 | 0 | 0 | 0 | 0 | 0 | 0 | 0 | 0 | 0 | 0 | NA | NA | NA | NA | NA | NA | NA | NA | NA | NA |
| Psg20 | 0 | 0 | 0 | 0 | 0 | 0 | 0 | 0 | 0 | 0 | 0 | 0 | 0 | 0 | 0 | NA | NA | NA | NA | NA | NA | NA | NA | NA | NA |
| Psg21 | 0 | 0 | 0 | 0 | 0 | 0 | 0 | 0 | 0 | 0 | 0 | 0 | 0 | 0 | 0 | NA | NA | NA | NA | NA | NA | NA | NA | NA | NA |
| Psg22 | 0 | 0 | 0 | 0 | 0 | 0 | 0 | 0 | 0 | 0 | 0 | 0 | 0 | 0 | 0 | NA | NA | NA | NA | NA | NA | NA | NA | NA | NA |
| Psg23 | 0 | 0 | 0 | 0 | 0 | 0 | 0 | 0 | 0 | 0 | 0 | 0 | 0 | 0 | 0 | NA | NA | NA | NA | NA | NA | NA | NA | NA | NA |
| Psg25 | 0 | 0 | 0 | 0 | 0 | 0 | 0 | 0 | 0 | 0 | 0 | 0 | 0 | 0 | 0 | NA | NA | NA | NA | NA | NA | NA | NA | NA | NA |
| Psg26 | 0 | 0 | 0 | 0 | 0 | 0 | 0 | 0 | 0 | 0 | 0 | 0 | 0 | 0 | 0 | NA | NA | NA | NA | NA | NA | NA | NA | NA | NA |
| Psg27 | 0 | 0 | 0 | 0 | 0 | 0 | 0 | 0 | 0 | 0 | 0 | 0 | 0 | 0 | 0 | NA | NA | NA | NA | NA | NA | NA | NA | NA | NA |
| Psg28 | 0 | 0 | 0 | 0 | 0 | 0 | 0 | 0 | 0 | 0 | 0 | 0 | 0 | 0 | 0 | NA | NA | NA | NA | NA | NA | NA | NA | NA | NA |
| Psg29 | 0 | 1 | 0 | 0 | 0 | 0 | 0 | 1 | 0 | 0 | 0 | 0 | 0.17 | 0 | 0 | 0.49 | -1.029 | 0.80087 | NA | 0 | 19.7991 | 10 | 10 | TRUE | 0.405 |
| Psip1 | 12151 | 11156 | 11049 | 15096 | 10955 | 10522 | 12907 | 11142 | 11094 | 11867 | 11787 | 11518 | 11719.11 | 11714 | 11724 | 1.001 | 0.001 | 0.98771175 | 0.99338029 | 0.0031 | 0.0072 | 0.005 | 0.005 | TRUE | 0.0969 |
| Pskh1 | 1135 | 1435 | 1426 | 1793 | 1330 | 1300 | 1206 | 1433 | 1432 | 1410 | 1431 | 1423 | 1389.02 | 1357 | 1421 | 1.047 | 0.066 | 0.49781392 | 0.69335228 | 0.004 | 0.0093 | 0.0062 | 0.0062 | TRUE | 0.1138 |
| Psma1 | 7550 | 7893 | 7795 | 12138 | 8036 | 7690 | 8019 | 7883 | 7827 | 9542 | 8646 | 8418 | 8389.22 | 7910 | 8869 | 1.121 | 0.165 | 0.02722381 | 0.10774752 | 0.0021 | 0.0073 | 0.0039 | 0.0039 | TRUE | 0.0542 |
| Psma2 | 8103 | 7814 | 7585 | 11531 | 7680 | 7770 | 8607 | 7804 | 7616 | 9065 | 8263 | 8505 | 8310.06 | 8009 | 8611 | 1.075 | 0.105 | 0.19627319 | 0.4026562 | 0.0031 | 0.0073 | 0.0046 | 0.0046 | TRUE | 0.052 |
| Psma3 | 6495 | 7595 | 7263 | 9778 | 6699 | 6718 | 6899 | 7585 | 7293 | 7687 | 7208 | 7354 | 7337.53 | 7259 | 7416 | 1.022 | 0.031 | 0.65978227 | 0.80836526 | 0.0013 | 0.0074 | 0.0034 | 0.0034 | TRUE | 0.023 |
| Psma4 | 9562 | 9544 | 9279 | 12947 | 8995 | 8265 | 10157 | 9532 | 9317 | 10178 | 9678 | 9047 | 9651.44 | 9669 | 9634 | 0.997 | -0.005 | 0.95027388 | 0.97481103 | 0.0021 | 0.0072 | 0.0042 | 0.0042 | TRUE | 0.0347 |
| Psma5 | 5550 | 5445 | 5150 | 7844 | 5420 | 5112 | 5895 | 5438 | 5171 | 6166 | 5832 | 5596 | 5683 | 5501 | 5865 | 1.066 | 0.093 | 0.263588 | 0.48279061 | 0.0031 | 0.0075 | 0.0048 | 0.0048 | TRUE | 0.0477 |
| Psma6 | 7693 | 8345 | 8039 | 11461 | 7769 | 7166 | 8171 | 8334 | 8072 | 9010 | 8359 | 7844 | 8298.43 | 8192 | 8404 | 1.026 | 0.037 | 0.62819355 | 0.79087692 | 0.0022 | 0.0073 | 0.0041 | 0.0041 | TRUE | 0.0488 |
| Psma7 | 5520 | 6327 | 6146 | 8955 | 6177 | 6104 | 5863 | 6319 | 6171 | 7040 | 6646 | 6682 | 6453.48 | 6118 | 6789 | 1.11 | 0.15 | 0.0251681 | 0.10230464 | 0.001 | 0.0074 | 0.0031 | 0.0031 | TRUE | 0.0161 |
| Psma8 | 679 | 669 | 672 | 1214 | 899 | 806 | 721 | 668 | 675 | 954 | 967 | 882 | 811.34 | 688 | 934 | 1.359 | 0.443 | 1.98E-06 | 5.00E-05 | 8.00E-04 | 0.011 | 0.005 | 0.005 | TRUE | 0.0279 |
| Psmb1 | 7318 | 9095 | 8983 | 11593 | 8067 | 8618 | 7773 | 9083 | 9020 | 9114 | 8680 | 9434 | 8850.5 | 8625 | 9076 | 1.052 | 0.073 | 0.40932066 | 0.62130006 | 0.0042 | 0.0073 | 0.0056 | 0.0056 | TRUE | 0.0912 |
| Psmb10 | 1276 | 1066 | 1158 | 1350 | 898 | 873 | 1355 | 1065 | 1163 | 1061 | 966 | 956 | 1094.3 | 1194 | 994 | 0.833 | -0.263 | 0.02545058 | 0.10300401 | 0.008 | 0.0099 | 0.0091 | 0.0091 | TRUE | 0.1664 |
| Psmb11 | 2 | 0 | 0 | 0 | 1 | 0 | 2 | 0 | 0 | 0 | 1 | 0 | 0.53 | 1 | 0 | 0.497 | -1.009 | 0.76131406 | NA | 0 | 6.1834 | 6.228 | 6.228 | TRUE | 0.9742 |
| Psmb2 | 5875 | 5389 | 5281 | 7471 | 5232 | 4839 | 6240 | 5382 | 5303 | 5873 | 5629 | 5297 | 5620.75 | 5642 | 5600 | 0.993 | -0.01 | 0.91117376 | 0.95603547 | 0.0053 | 0.0075 | 0.0061 | 0.0061 | TRUE | 0.1049 |
| Psmb3 | 9133 | 9543 | 8928 | 12446 | 8629 | 8357 | 9701 | 9531 | 8965 | 9784 | 9284 | 9148 | 9402.1 | 9399 | 9405 | 1.001 | 0.001 | 0.98543963 | 0.99223403 | 0.0014 | 0.0073 | 0.0033 | 0.0033 | TRUE | 0.02 |
| Psmb4 | 9382 | 9972 | 9614 | 12906 | 9356 | 8782 | 9965 | 9959 | 9653 | 10146 | 10066 | 9613 | 9900.56 | 9859 | 9942 | 1.008 | 0.012 | 0.8373961 | 0.91730875 | 4.00E-04 | 0.0072 | 0.0024 | 0.0024 | TRUE | 0.0102 |
| Psmb5 | 3146 | 3182 | 2959 | 4233 | 2964 | 2753 | 3342 | 3178 | 2971 | 3328 | 3189 | 3014 | 3170.16 | 3164 | 3177 | 1.005 | 0.007 | 0.93647058 | 0.96825513 | 0.0026 | 0.0079 | 0.0046 | 0.0046 | TRUE | 0.0344 |
| Psmb6 | 14137 | 14518 | 14626 | 19573 | 13453 | 13106 | 15016 | 14499 | 14686 | 15387 | 14475 | 14347 | 14734.87 | 14734 | 14736 | 1 | 0 | 0.9949801 | 0.99757333 | 7.00E-04 | 0.0071 | 0.0027 | 0.0027 | TRUE | 0.0183 |
| Psmb7 | 9653 | 10364 | 10626 | 14352 | 9721 | 10006 | 10253 | 10351 | 10670 | 11282 | 10459 | 10953 | 10661.37 | 10425 | 10898 | 1.046 | 0.064 | 0.30779634 | 0.52830165 | 8.00E-04 | 0.0072 | 0.0028 | 0.0028 | TRUE | 0.0152 |
| Psmb8 | 4 | 7 | 8 | 8 | 3 | 2 | 4 | 7 | 8 | 6 | 3 | 2 | 5.16 | 6 | 4 | 0.623 | -0.683 | 0.427471 | 0.63765927 | 0 | 0.645 | 0.3324 | 0.3324 | TRUE | 0.6127 |
| Psmb9 | 121 | 157 | 180 | 182 | 124 | 109 | 129 | 157 | 181 | 143 | 133 | 119 | 143.64 | 156 | 132 | 0.851 | -0.233 | 0.23456814 | 0.44941796 | 0.0119 | 0.0298 | 0.0208 | 0.0208 | TRUE | 0.1956 |
| Psmc1 | 7749 | 7267 | 6791 | 11791 | 8036 | 7257 | 8231 | 7258 | 6819 | 9269 | 8646 | 7944 | 8027.77 | 7436 | 8620 | 1.159 | 0.213 | 0.03627508 | 0.13110479 | 0.0075 | 0.0073 | 0.0074 | 0.0074 | TRUE | 0.1067 |
| Psmc2 | 7183 | 6552 | 6253 | 9119 | 6462 | 5775 | 7630 | 6544 | 6279 | 7169 | 6953 | 6322 | 6815.81 | 6818 | 6815 | 1 | 0 | 0.99686513 | 0.99828918 | 0.0073 | 0.0074 | 0.0074 | 0.0074 | TRUE | 0.1325 |
| Psmc3 | 8149 | 6842 | 6858 | 12909 | 8999 | 7266 | 8656 | 6833 | 6886 | 10148 | 9682 | 7954 | 8359.87 | 7458 | 9261 | 1.242 | 0.313 | 0.01840527 | 0.08197227 | 0.0173 | 0.0073 | 0.0125 | 0.0125 | TRUE | 0.2407 |
| Psmc3ip | 774 | 948 | 1021 | 1304 | 948 | 999 | 822 | 947 | 1025 | 1025 | 1020 | 1094 | 988.79 | 931 | 1046 | 1.123 | 0.167 | 0.12837032 | 0.30807314 | 0.0053 | 0.0102 | 0.0077 | 0.0077 | TRUE | 0.1247 |

| | | | | | | | | | | | | | | | | | | | | | | | | | |
|---|---|---|---|---|---|---|---|---|---|---|---|---|---|---|---|---|---|---|---|---|---|---|---|---|---|
| Psmc4 | 6837 | 5287 | 5260 | 10395 | 7024 | 5554 | 7262 | 5280 | 5282 | 8172 | 7557 | 6080 | 6605.45 | 5941 | 7270 | 1.224 | 0.291 | 0.06547388 | 0.19715481 | 0.0292 | 0.0074 | 0.0179 | 0.0179 | TRUE | 0.4611 |
| Psmc5 | 10275 | 9195 | 8943 | 14947 | 10455 | 9135 | 10914 | 9183 | 8980 | 11750 | 11249 | 10000 | 10345.92 | 9692 | 11000 | 1.135 | 0.183 | 0.09076565 | 0.24567338 | 0.0092 | 0.0072 | 0.0083 | 0.0083 | TRUE | 0.1485 |
| Psmc6 | 6262 | 7518 | 7592 | 9660 | 6869 | 7096 | 6651 | 7508 | 7623 | 7594 | 7391 | 7768 | 7422.51 | 7261 | 7584 | 1.044 | 0.063 | 0.43370482 | 0.6424232 | 0.0027 | 0.0073 | 0.0045 | 0.0045 | TRUE | 0.0658 |
| Psmd1 | 18874 | 17919 | 17818 | 25677 | 17771 | 17413 | 20048 | 17896 | 17891 | 20185 | 19121 | 19061 | 19033.59 | 18612 | 19456 | 1.045 | 0.064 | 0.40256496 | 0.61541674 | 0.0027 | 0.0071 | 0.0042 | 0.0042 | TRUE | 0.0557 |
| Psmd10 | 1276 | 1532 | 1518 | 1979 | 1411 | 1433 | 1355 | 1530 | 1524 | 1556 | 1518 | 1569 | 1508.69 | 1470 | 1548 | 1.053 | 0.074 | 0.38446913 | 0.5985945 | 0.0017 | 0.0091 | 0.0046 | 0.0046 | TRUE | 0.0556 |
| Psmd11 | 11962 | 12721 | 12551 | 15497 | 11306 | 11241 | 12706 | 12705 | 12603 | 12183 | 12165 | 12305 | 12444.17 | 12671 | 12218 | 0.964 | -0.053 | 0.30060474 | 0.52100417 | 0 | 0.0072 | 0.0018 | 0.0018 | TRUE | 5.00E-04 |
| Psmd12 | 12326 | 10951 | 10541 | 16731 | 11728 | 10600 | 13092 | 10937 | 10584 | 13153 | 12619 | 11603 | 11998.02 | 11538 | 12458 | 1.08 | 0.111 | 0.29524195 | 0.51607626 | 0.0085 | 0.0072 | 0.008 | 0.008 | TRUE | 0.1697 |
| Psmd13 | 9125 | 8676 | 8248 | 11900 | 8231 | 7224 | 9692 | 8665 | 8282 | 9355 | 8856 | 7908 | 8792.95 | 8880 | 8706 | 0.981 | -0.028 | 0.77634883 | 0.8837657 | 0.0056 | 0.0073 | 0.007 | 0.007 | TRUE | 0.0785 |
| Psmd14 | 5842 | 5823 | 5603 | 9362 | 6362 | 5760 | 6205 | 5815 | 5626 | 7360 | 6845 | 6305 | 6359.46 | 5882 | 6837 | 1.163 | 0.217 | 0.01288774 | 0.06342133 | 0.004 | 0.0074 | 0.0053 | 0.0053 | TRUE | 0.0564 |
| Psmd2 | 30188 | 27713 | 26871 | 37215 | 25603 | 24383 | 32065 | 27677 | 26981 | 29256 | 27547 | 26691 | 28369.56 | 28908 | 27831 | 0.963 | -0.055 | 0.55219157 | 0.73619319 | 0.005 | 0.007 | 0.0061 | 0.0061 | TRUE | 0.1117 |
| Psmd3 | 25829 | 22714 | 22635 | 32133 | 22957 | 21191 | 27435 | 22685 | 22728 | 25260 | 24700 | 23197 | 24334.2 | 24283 | 24386 | 1.004 | 0.006 | 0.95018936 | 0.97481103 | 0.0058 | 0.007 | 0.007 | 0.007 | TRUE | 0.1578 |
| Psmd4 | 8106 | 8665 | 8618 | 11331 | 8351 | 7707 | 8610 | 8654 | 8653 | 8908 | 8985 | 8437 | 8707.74 | 8639 | 8777 | 1.016 | 0.023 | 0.70116831 | 0.83650923 | 4.00E-04 | 0.0073 | 0.0024 | 0.0024 | TRUE | 0.0139 |
| Psmd5 | 5040 | 4265 | 4117 | 6437 | 4618 | 4287 | 5353 | 4259 | 4134 | 5060 | 4969 | 4693 | 4744.75 | 4582 | 4907 | 1.071 | 0.099 | 0.39013203 | 0.60390248 | 0.0098 | 0.0076 | 0.0094 | 0.0094 | TRUE | 0.2638 |
| Psmd6 | 4273 | 4109 | 4117 | 6847 | 4946 | 4175 | 4539 | 4104 | 4134 | 5383 | 5322 | 4570 | 4675.11 | 4259 | 5092 | 1.196 | 0.258 | 0.00673923 | 0.03902025 | 0.0053 | 0.0076 | 0.0063 | 0.0063 | TRUE | 0.0974 |
| Psmd7 | 14313 | 12197 | 10866 | 17488 | 12570 | 11483 | 15203 | 12181 | 10911 | 13748 | 13525 | 12570 | 13022.83 | 12765 | 13281 | 1.04 | 0.057 | 0.65343085 | 0.80540538 | 0.0128 | 0.0072 | 0.0116 | 0.0116 | TRUE | 0.3412 |
| Psmd8 | 10786 | 11243 | 10928 | 13472 | 9458 | 9451 | 11457 | 11228 | 10973 | 10591 | 10176 | 10346 | 10795.08 | 11219 | 10371 | 0.924 | -0.113 | 0.04747936 | 0.1581843 | 3.00E-04 | 0.0072 | 0.0023 | 0.0023 | TRUE | 0.0045 |
| Psmd9 | 906 | 871 | 873 | 1628 | 1107 | 1032 | 962 | 870 | 877 | 1280 | 1191 | 1130 | 1051.56 | 903 | 1200 | 1.331 | 0.412 | 1.71E-05 | 0.00031231 | 0.0027 | 0.01 | 0.0056 | 0.0056 | TRUE | 0.0416 |
| Psme1 | 2017 | 2422 | 2478 | 2377 | 1840 | 1990 | 2142 | 2419 | 2488 | 1869 | 1980 | 2178 | 2179.36 | 2350 | 2009 | 0.854 | -0.227 | 0.02359864 | 0.09771535 | 0.0054 | 0.0084 | 0.0068 | 0.0068 | TRUE | 0.072 |
| Psme2 | 1288 | 1399 | 1420 | 1290 | 1011 | 1036 | 1368 | 1397 | 1426 | 1014 | 1088 | 1134 | 1237.84 | 1397 | 1079 | 0.771 | -0.375 | 8.29E-06 | 0.00016864 | 0.001 | 0.0096 | 0.0043 | 0.0043 | TRUE | 0.0342 |
| Psme3 | 13571 | 12466 | 12414 | 17904 | 12752 | 11729 | 14415 | 12450 | 12465 | 14075 | 13720 | 12839 | 13327.33 | 13110 | 13545 | 1.033 | 0.047 | 0.59457048 | 0.7677918 | 0.004 | 0.0072 | 0.0056 | 0.0056 | TRUE | 0.0926 |
| Psme4 | 30375 | 28098 | 30298 | 34284 | 24518 | 25783 | 32264 | 28062 | 30422 | 26951 | 26380 | 28224 | 28717.09 | 30249 | 27185 | 0.899 | -0.154 | 0.04972258 | 0.16368701 | 0.003 | 0.007 | 0.0044 | 0.0044 | TRUE | 0.049 |
| Psmf1 | 1805 | 1783 | 1822 | 2661 | 1895 | 1723 | 1917 | 1781 | 1829 | 2092 | 2039 | 1886 | 1924.05 | 1842 | 2006 | 1.089 | 0.123 | 0.12783986 | 0.30739486 | 0.0016 | 0.0086 | 0.0042 | 0.0042 | TRUE | 0.0326 |
| Psmg1 | 3652 | 4321 | 4152 | 4934 | 3599 | 3780 | 3879 | 4315 | 4169 | 3879 | 3872 | 4138 | 4042.06 | 4121 | 3963 | 0.961 | -0.057 | 0.45687742 | 0.66151388 | 0.0018 | 0.0077 | 0.004 | 0.004 | TRUE | 0.0321 |
| Psmg2 | 1890 | 2162 | 2200 | 3173 | 2141 | 2191 | 2008 | 2159 | 2209 | 2494 | 2304 | 2398 | 2262.02 | 2125 | 2399 | 1.129 | 0.175 | 0.02630834 | 0.10529217 | 0.0016 | 0.0084 | 0.004 | 0.004 | TRUE | 0.0283 |
| Psmg3 | 808 | 822 | 824 | 1065 | 776 | 674 | 858 | 821 | 827 | 837 | 835 | 738 | 819.42 | 835 | 803 | 0.962 | -0.055 | 0.56449788 | 0.74512706 | 0.0015 | 0.0109 | 0.0054 | 0.0054 | TRUE | 0.059 |
| Psmg4 | 590 | 724 | 660 | 946 | 720 | 669 | 627 | 723 | 663 | 744 | 775 | 732 | 710.52 | 671 | 750 | 1.118 | 0.161 | 0.10673253 | 0.27288789 | 0.0014 | 0.0115 | 0.0057 | 0.0057 | TRUE | 0.0549 |
| Psors1c2 | 92 | 72 | 60 | 92 | 64 | 59 | 98 | 72 | 60 | 72 | 69 | 65 | 72.61 | 77 | 69 | 0.898 | -0.155 | 0.55424063 | 0.7374912 | 0.019 | 0.0523 | 0.0359 | 0.0359 | TRUE | 0.5232 |
| Pspc1 | 4147 | 5849 | 5809 | 5567 | 4417 | 4690 | 4405 | 5841 | 5833 | 4376 | 4752 | 5134 | 5056.97 | 5360 | 4754 | 0.887 | -0.173 | 0.18152788 | 0.38337678 | 0.0148 | 0.0076 | 0.0119 | 0.0119 | TRUE | 0.2959 |
| Psph | 1213 | 1216 | 1129 | 1573 | 1109 | 975 | 1288 | 1214 | 1134 | 1237 | 1193 | 1067 | 1188.93 | 1212 | 1166 | 0.963 | -0.055 | 0.58226754 | 0.7585095 | 0.0041 | 0.0097 | 0.0064 | 0.0064 | TRUE | 0.0646 |
| Pspn | 1 | 0 | 0 | 0 | 0 | 0 | 1 | 0 | 0 | 0 | 0 | 0 | 0.18 | 0 | 0 | 0.49 | -1.029 | 0.80087 | NA | 0 | 18.6165 | 10 | 10 | TRUE | 0.4614 |
| Psrc1 | 3864 | 5573 | 5628 | 5792 | 4354 | 4885 | 4104 | 5566 | 5651 | 4553 | 4685 | 5347 | 4984.4 | 5107 | 4862 | 0.952 | -0.071 | 0.6065318 | 0.77676352 | 0.0191 | 0.0076 | 0.0136 | 0.0136 | TRUE | 0.3593 |
| Pstk | 1380 | 1762 | 1603 | 2074 | 1444 | 1454 | 1466 | 1760 | 1610 | 1630 | 1554 | 1592 | 1601.8 | 1612 | 1592 | 0.988 | -0.018 | 0.85150667 | 0.92588509 | 0.0038 | 0.009 | 0.0059 | 0.0059 | TRUE | 0.0779 |
| Pstpip1 | 136 | 142 | 138 | 233 | 178 | 182 | 144 | 142 | 139 | 183 | 192 | 199 | 166.46 | 142 | 191 | 1.349 | 0.432 | 0.00512604 | 0.03191786 | 0 | 0.0267 | 0.011 | 0.011 | TRUE | 0.0164 |
| Pstpip2 | 587 | 578 | 591 | 594 | 482 | 526 | 623 | 577 | 593 | 467 | 519 | 576 | 559.25 | 598 | 521 | 0.869 | -0.203 | 0.08360355 | 0.23165762 | 0.0045 | 0.0128 | 0.0082 | 0.0082 | TRUE | 0.1007 |
| Ptafr | 19 | 9 | 18 | 20 | 18 | 7 | 20 | 9 | 18 | 16 | 19 | 8 | 15 | 16 | 14 | 0.91 | -0.136 | 0.81195678 | 0.90349812 | 0.1032 | 0.2265 | 0.1695 | 0.1695 | TRUE | 0.5937 |
| Ptar1 | 258 | 272 | 304 | 319 | 265 | 238 | 274 | 272 | 305 | 251 | 285 | 261 | 274.56 | 284 | 266 | 0.934 | -0.098 | 0.4651033 | 0.66788976 | 0 | 0.0189 | 0.0093 | 0.0093 | TRUE | 0.0502 |
| Ptbp1 | 34176 | 33442 | 34903 | 42581 | 31398 | 31039 | 36301 | 33399 | 35046 | 33474 | 33782 | 33977 | 34329.88 | 34915 | 33744 | 0.966 | -0.049 | 0.41952004 | 0.63003905 | 9.00E-04 | 0.007 | 0.0026 | 0.0026 | TRUE | 0.0177 |
| Ptbp2 | 868 | 1187 | 1324 | 1860 | 1454 | 1523 | 922 | 1185 | 1329 | 1462 | 1564 | 1667 | 1355.11 | 1145 | 1564 | 1.365 | 0.449 | 0.00165591 | 0.01324391 | 0.0173 | 0.0093 | 0.0139 | 0.0139 | TRUE | 0.3482 |
| Ptbp3 | 6615 | 7448 | 7849 | 10235 | 7968 | 7950 | 7026 | 7438 | 7881 | 8046 | 8573 | 8703 | 7944.58 | 7448 | 8441 | 1.133 | 0.18 | 0.01851041 | 0.08223835 | 0.0023 | 0.0073 | 0.0041 | 0.0041 | TRUE | 0.0315 |
| Ptcd1 | 964 | 736 | 765 | 1252 | 767 | 693 | 1024 | 735 | 768 | 984 | 825 | 759 | 849.2 | 842 | 856 | 1.017 | 0.025 | 0.87751904 | 0.93954572 | 0.0244 | 0.0108 | 0.0177 | 0.0177 | TRUE | 0.4214 |
| Ptcd2 | 2874 | 3048 | 3079 | 3243 | 2362 | 2721 | 3053 | 3044 | 3092 | 2549 | 2541 | 2979 | 2876.29 | 3063 | 2690 | 0.878 | -0.188 | 0.03617441 | 0.13082125 | 0.0038 | 0.008 | 0.0054 | 0.0054 | TRUE | 0.1063 |
| Ptcd3 | 7026 | 6399 | 6283 | 8986 | 6352 | 5870 | 7463 | 6391 | 6309 | 7064 | 6834 | 6426 | 6747.75 | 6721 | 6775 | 1.008 | 0.012 | 0.90040842 | 0.95169273 | 0.0054 | 0.0074 | 0.0062 | 0.0062 | TRUE | 0.1137 |
| Ptch1 | 2338 | 2881 | 3335 | 2168 | 1693 | 2194 | 2483 | 2877 | 3349 | 1704 | 1822 | 2402 | 2439.48 | 2903 | 1976 | 0.68 | -0.556 | 0.00048228 | 0.00503741 | 0.0284 | 0.0083 | 0.0178 | 0.0178 | TRUE | 0.3261 |
| Ptch2 | 809 | 652 | 630 | 779 | 613 | 576 | 859 | 651 | 633 | 612 | 660 | 631 | 674.25 | 714 | 634 | 0.888 | -0.172 | 0.22545177 | 0.43817934 | 0.014 | 0.0118 | 0.013 | 0.013 | TRUE | 0.371 |
| Ptchd1 | 60 | 91 | 104 | 90 | 54 | 55 | 64 | 91 | 104 | 71 | 58 | 60 | 74.68 | 86 | 63 | 0.732 | -0.451 | 0.09232028 | 0.24796925 | 0.0247 | 0.051 | 0.038 | 0.038 | TRUE | 0.3852 |
| Ptchd2 | 99 | 69 | 104 | 95 | 71 | 67 | 105 | 69 | 104 | 75 | 76 | 73 | 83.82 | 93 | 75 | 0.806 | -0.31 | 0.20781918 | 0.41715638 | 0.0169 | 0.0462 | 0.0317 | 0.0317 | TRUE | 0.494 |
| Ptchd3 | 57 | 97 | 84 | 108 | 80 | 90 | 61 | 97 | 84 | 85 | 86 | 99 | 85.21 | 81 | 90 | 1.111 | 0.152 | 0.53657814 | 0.72331397 | 0.0169 | 0.0456 | 0.032 | 0.032 | TRUE | 0.4304 |
| Ptchd4 | 1 | 0 | 3 | 6 | 4 | 1 | 1 | 0 | 3 | 5 | 4 | 1 | 2.36 | 1 | 3 | 2.506 | 1.326 | 0.34733382 | NA | 0.2223 | 1.3999 | 0.9144 | 0.9144 | TRUE | 0.7228 |
| Ptcra | 5 | 8 | 6 | 10 | 7 | 5 | 5 | 8 | 6 | 8 | 8 | 5 | 6.7 | 6 | 7 | 1.084 | 0.117 | 0.86983288 | 0.9361313 | 0 | 0.4987 | 0.2164 | 0.2164 | TRUE | 0.1115 |
| Ptdss1 | 1735 | 1660 | 1584 | 2888 | 1927 | 1805 | 1843 | 1658 | 1591 | 2270 | 2073 | 1976 | 1901.79 | 1697 | 2106 | 1.242 | 0.313 | 0.00143573 | 0.01184044 | 0.0047 | 0.0086 | 0.0064 | 0.0064 | TRUE | 0.0679 |
| Ptdss2 | 3129 | 3641 | 3814 | 5010 | 3550 | 3779 | 3324 | 3636 | 3830 | 3938 | 3820 | 4137 | 3780.71 | 3597 | 3965 | 1.102 | 0.141 | 0.09403565 | 0.25099515 | 0.0029 | 0.0078 | 0.0048 | 0.0048 | TRUE | 0.0535 |
| Pten | 5615 | 5480 | 5557 | 6838 | 5019 | 4823 | 5964 | 5473 | 5580 | 5375 | 5400 | 5280 | 5512 | 5672 | 5352 | 0.944 | -0.084 | 0.2073374 | 0.41661082 | 8.00E-04 | 0.0075 | 0.003 | 0.003 | TRUE | 0.0246 |
| Pter | 808 | 755 | 795 | 1163 | 755 | 693 | 858 | 754 | 798 | 914 | 812 | 759 | 815.95 | 803 | 828 | 1.033 | 0.046 | 0.67939723 | 0.82193713 | 0.0055 | 0.0109 | 0.0078 | 0.0078 | TRUE | 0.1007 |
| Ptf1a | 5 | 7 | 0 | 2 | 2 | 0 | 5 | 7 | 0 | 2 | 2 | 0 | 2.67 | 4 | 1 | 0.307 | -1.704 | 0.27365008 | NA | 1.2072 | 1.2403 | 1.2337 | 1.2337 | TRUE | 0.5565 |
| Ptgdr | 0 | 1 | 1 | 2 | 0 | 0 | 0 | 1 | 1 | 2 | 0 | 0 | 0.6 | 1 | 1 | 0.834 | -0.262 | 0.92267544 | NA | 0 | 5.536 | 3.6136 | 3.6136 | TRUE | 0.9722 |
| Ptgds | 61 | 39 | 30 | 42 | 22 | 23 | 65 | 39 | 30 | 33 | 24 | 25 | 35.96 | 45 | 27 | 0.617 | -0.698 | 0.08106778 | 0.22666213 | 0.0733 | 0.0985 | 0.0859 | 0.0859 | TRUE | 0.641 |
| Ptger1 | 1 | 5 | 3 | 7 | 2 | 2 | 1 | 5 | 3 | 6 | 2 | 2 | 3.15 | 3 | 3 | 1.106 | 0.146 | 0.89549017 | NA | 0 | 1.0521 | 0.5718 | 0.5718 | TRUE | 0.2517 |
| Ptger2 | 30 | 38 | 42 | 30 | 16 | 15 | 32 | 38 | 42 | 24 | 17 | 16 | 28.2 | 37 | 19 | 0.516 | -0.954 | 0.0113758 | 0.0579332 | 0 | 0.1237 | 0.0634 | 0.0634 | TRUE | 0.2779 |
| Ptger3 | 7 | 8 | 6 | 27 | 20 | 5 | 7 | 8 | 6 | 21 | 22 | 5 | 11.61 | 7 | 16 | 2.265 | 1.18 | 0.08475077 | 0.23379598 | 0.1786 | 0.2906 | 0.2357 | 0.2357 | TRUE | 1.458 |
| Ptger4 | 81 | 108 | 103 | 121 | 76 | 101 | 86 | 108 | 103 | 95 | 82 | 111 | 97.46 | 99 | 96 | 0.966 | -0.051 | 0.81748197 | 0.90708637 | 0.0064 | 0.0407 | 0.0243 | 0.0243 | TRUE | 0.1625 |
| Ptges | 413 | 313 | 280 | 433 | 304 | 281 | 439 | 313 | 281 | 340 | 327 | 308 | 334.58 | 344 | 325 | 0.946 | -0.081 | 0.66158994 | 0.80957379 | 0.026 | 0.0167 | 0.0214 | 0.0214 | TRUE | 0.6544 |
| Ptges2 | 990 | 1149 | 1182 | 1277 | 1007 | 939 | 1052 | 1148 | 1187 | 1004 | 1083 | 1028 | 1083.53 | 1129 | 1038 | 0.919 | -0.121 | 0.18189745 | 0.38376383 | 0.0018 | 0.0099 | 0.005 | 0.005 | TRUE | 0.0424 |
| Ptges3 | 16256 | 18386 | 17768 | 22952 | 16567 | 16497 | 17267 | 18362 | 17841 | 18043 | 17825 | 18059 | 17899.44 | 17823 | 17976 | 1.009 | 0.012 | 0.82939104 | 0.91345091 | 4.00E-04 | 0.0071 | 0.0023 | 0.0023 | TRUE | 0.0091 |
| Ptges3l | 48 | 50 | 57 | 82 | 36 | 45 | 51 | 50 | 57 | 64 | 39 | 49 | 51.77 | 53 | 51 | 0.972 | -0.041 | 0.88965193 | 0.94620362 | 0.018 | 0.0705 | 0.0447 | 0.0447 | TRUE | 0.3521 |
| Ptgfr | 15 | 11 | 7 | 17 | 11 | 5 | 16 | 11 | 7 | 13 | 12 | 5 | 10.77 | 11 | 10 | 0.916 | -0.127 | 0.84196182 | 0.92021591 | 0.0726 | 0.3128 | 0.2006 | 0.2006 | TRUE | 0.2674 |
| Ptgfrn | 5410 | 6837 | 7069 | 6225 | 4971 | 5590 | 5746 | 6828 | 7098 | 4894 | 5348 | 6119 | 6005.63 | 6557 | 5454 | 0.831 | -0.266 | 0.02620325 | 0.10497272 | 0.0121 | 0.0075 | 0.0102 | 0.0102 | TRUE | 0.1428 |
| Ptgir | 44 | 45 | 55 | 37 | 27 | 30 | 47 | 45 | 55 | 29 | 29 | 33 | 39.65 | 49 | 30 | 0.618 | -0.695 | 0.02319516 | 0.0965516 | 0 | 0.09 | 0.0409 | 0.0409 | TRUE | 0.102 |
| Ptgis | 541 | 649 | 616 | 788 | 551 | 493 | 575 | 648 | 619 | 619 | 593 | 540 | 598.88 | 614 | 584 | 0.952 | -0.07 | 0.51819492 | 0.70964076 | 0.0026 | 0.0124 | 0.0069 | 0.0069 | TRUE | 0.0509 |
| Ptgr1 | 1513 | 1319 | 1256 | 2376 | 1645 | 1510 | 1607 | 1317 | 1261 | 1868 | 1770 | 1653 | 1579.37 | 1395 | 1764 | 1.265 | 0.339 | 0.00399069 | 0.02630219 | 0.0098 | 0.009 | 0.0093 | 0.0093 | TRUE | 0.2117 |

| | | | | | | | | | | | | | | | | | | | | | | | | | |
|---|---|---|---|---|---|---|---|---|---|---|---|---|---|---|---|---|---|---|---|---|---|---|---|---|---|
| Ptgr2 | 1489 | 1707 | 1680 | 1978 | 1445 | 1421 | 1582 | 1705 | 1687 | 1555 | 1555 | 1556 | 1606.41 | 1658 | 1555 | 0.938 | -0.092 | 0.21178407 | 0.42244319 | 0 | 0.009 | 0.0033 | 0.0033 | TRUE | 0.0194 |
| Ptgs1 | 158 | 109 | 102 | 167 | 137 | 85 | 168 | 109 | 102 | 131 | 147 | 93 | 125.14 | 126 | 124 | 0.983 | -0.025 | 0.92522174 | 0.9630806 | 0.0558 | 0.0332 | 0.0441 | 0.0441 | TRUE | 0.8302 |
| Ptgs2 | 9 | 13 | 10 | 35 | 17 | 16 | 10 | 13 | 10 | 28 | 18 | 18 | 15.98 | 11 | 21 | 1.959 | 0.97 | 0.05244253 | 0.16942309 | 0 | 0.213 | 0.1104 | 0.1104 | TRUE | 0.4682 |
| Pth | 0 | 0 | 0 | 0 | 0 | 0 | 0 | 0 | 0 | 0 | 0 | 0 | 0 | 0 | 0 | NA | NA | NA | NA | NA | NA | NA | NA | NA | NA |
| Pth1r | 101 | 126 | 111 | 124 | 96 | 73 | 107 | 126 | 111 | 97 | 103 | 80 | 104.21 | 115 | 93 | 0.816 | -0.294 | 0.15116573 | 0.34221126 | 0 | 0.0385 | 0.0206 | 0.0206 | TRUE | 0.1459 |
| Pth2 | 1 | 2 | 1 | 1 | 1 | 5 | 1 | 2 | 1 | 1 | 1 | 5 | 1.9 | 1 | 2 | 1.76 | 0.815 | 0.57579653 | NA | 0 | 1.741 | 0.9488 | 0.9488 | TRUE | 1.1714 |
| Pth2r | 2 | 0 | 1 | 0 | 0 | 0 | 2 | 0 | 1 | 0 | 0 | 0 | 0.52 | 1 | 0 | 0.172 | -2.536 | 0.43509252 | NA | 0 | 6.3253 | 6.1167 | 6.1167 | TRUE | 0.0605 |
| Pthlh | 0 | 0 | 0 | 1 | 0 | 0 | 0 | 0 | 0 | 1 | 0 | 0 | 0.13 | 0 | 0 | 1.859 | 0.894 | 0.82650734 | NA | 0 | 25.1515 | 10 | 10 | TRUE | 0.2563 |
| Ptk2 | 5745 | 5877 | 5800 | 6799 | 4878 | 4885 | 6102 | 5869 | 5824 | 5345 | 5248 | 5347 | 5622.68 | 5932 | 5313 | 0.896 | -0.159 | 0.0070785 | 0.04042075 | 2.00E-04 | 0.0075 | 0.0023 | 0.0023 | TRUE | 0.0077 |
| Ptk2b | 730 | 811 | 894 | 760 | 596 | 690 | 775 | 810 | 898 | 597 | 641 | 755 | 746.17 | 828 | 664 | 0.802 | -0.319 | 0.01079887 | 0.05571471 | 0.0084 | 0.0113 | 0.0099 | 0.0099 | TRUE | 0.1665 |
| Ptk6 | 18 | 23 | 33 | 13 | 15 | 10 | 19 | 23 | 33 | 10 | 16 | 11 | 18.76 | 25 | 12 | 0.492 | -1.025 | 0.03294703 | 0.12267246 | 0 | 0.1826 | 0.1064 | 0.1064 | TRUE | 0.4842 |
| Ptk7 | 5481 | 6077 | 6263 | 5474 | 3938 | 4289 | 5822 | 6069 | 6289 | 4303 | 4237 | 4695 | 5235.82 | 6060 | 4412 | 0.728 | -0.458 | 1.62E-09 | 9.40E-08 | 0.002 | 0.0075 | 0.004 | 0.004 | TRUE | 0.0381 |
| Ptma | 55884 | 53575 | 52741 | 76995 | 57652 | 52588 | 59359 | 53506 | 52958 | 60527 | 62030 | 57566 | 57657.56 | 55274 | 60041 | 1.086 | 0.119 | 0.11734063 | 0.28984926 | 0.0027 | 0.007 | 0.0042 | 0.0042 | TRUE | 0.0512 |
| Ptms | 6793 | 7495 | 7351 | 9209 | 7156 | 7069 | 7215 | 7485 | 7381 | 7239 | 7699 | 7738 | 7459.8 | 7360 | 7559 | 1.027 | 0.038 | 0.54651564 | 0.73153226 | 6.00E-04 | 0.0073 | 0.0027 | 0.0027 | TRUE | 0.0169 |
| Ptn | 5 | 5 | 2 | 9 | 4 | 4 | 5 | 5 | 2 | 7 | 4 | 4 | 4.68 | 4 | 5 | 1.299 | 0.377 | 0.66858367 | 0.81492803 | 0 | 0.7111 | 0.3431 | 0.3431 | TRUE | 0.3464 |
| Ptov1 | 4166 | 4679 | 5064 | 5664 | 4208 | 4183 | 4425 | 4673 | 5085 | 4453 | 4528 | 4579 | 4623.65 | 4728 | 4520 | 0.956 | -0.065 | 0.40624119 | 0.61887724 | 0.0023 | 0.0076 | 0.0042 | 0.0042 | TRUE | 0.0532 |
| Ptp4a1 | 300 | 362 | 322 | 374 | 248 | 245 | 319 | 362 | 323 | 294 | 267 | 268 | 305.42 | 335 | 276 | 0.828 | -0.273 | 0.03426646 | 0.12613824 | 0 | 0.0177 | 0.0087 | 0.0087 | TRUE | 0.0572 |
| Ptp4a2 | 10586 | 12112 | 11449 | 14547 | 10457 | 10177 | 11244 | 12096 | 11496 | 11436 | 11251 | 11140 | 11443.95 | 11612 | 11276 | 0.971 | -0.042 | 0.48870023 | 0.68519691 | 7.00E-04 | 0.0072 | 0.0026 | 0.0026 | TRUE | 0.0163 |
| Ptp4a3 | 3760 | 4633 | 4341 | 4227 | 3131 | 3558 | 3994 | 4627 | 4359 | 3323 | 3369 | 3895 | 3927.69 | 4327 | 3529 | 0.815 | -0.294 | 0.00319404 | 0.02229044 | 0.0064 | 0.0077 | 0.0069 | 0.0069 | TRUE | 0.0997 |
| Ptpdc1 | 429 | 438 | 425 | 678 | 519 | 404 | 456 | 437 | 427 | 533 | 558 | 442 | 475.58 | 440 | 511 | 1.163 | 0.218 | 0.08543042 | 0.23527302 | 0.0058 | 0.0138 | 0.0095 | 0.0095 | TRUE | 0.1577 |
| Ptpla | 242 | 314 | 290 | 372 | 283 | 267 | 257 | 314 | 291 | 292 | 304 | 292 | 291.84 | 287 | 296 | 1.031 | 0.044 | 0.74325152 | 0.86303195 | 0.0014 | 0.0182 | 0.0094 | 0.0094 | TRUE | 0.0939 |
| Ptplad1 | 1699 | 1708 | 1610 | 2462 | 1733 | 1618 | 1805 | 1706 | 1617 | 1935 | 1865 | 1771 | 1783.04 | 1709 | 1857 | 1.087 | 0.121 | 0.14959334 | 0.34028031 | 0.0019 | 0.0088 | 0.0045 | 0.0045 | TRUE | 0.0288 |
| Ptplad2 | 109 | 113 | 127 | 156 | 107 | 114 | 116 | 113 | 128 | 123 | 115 | 125 | 119.78 | 119 | 121 | 1.018 | 0.026 | 0.88434424 | 0.94318889 | 0 | 0.0344 | 0.015 | 0.015 | TRUE | 0.0429 |
| Ptplb | 1804 | 1869 | 1829 | 2203 | 1407 | 1416 | 1916 | 1867 | 1837 | 1732 | 1514 | 1550 | 1735.83 | 1873 | 1599 | 0.854 | -0.227 | 0.00837629 | 0.04608178 | 0.0023 | 0.0088 | 0.0048 | 0.0048 | TRUE | 0.0657 |
| Ptpmt1 | 657 | 884 | 871 | 1185 | 750 | 757 | 698 | 883 | 875 | 932 | 807 | 829 | 837.07 | 819 | 856 | 1.046 | 0.065 | 0.60984071 | 0.77891178 | 0.0103 | 0.0108 | 0.0105 | 0.0105 | TRUE | 0.1962 |
| Ptpn1 | 3132 | 2683 | 2700 | 3184 | 2278 | 2222 | 3327 | 2680 | 2711 | 2503 | 2451 | 2432 | 2683.95 | 2906 | 2462 | 0.847 | -0.239 | 0.02383163 | 0.09831873 | 0.0074 | 0.0081 | 0.0077 | 0.0077 | TRUE | 0.1947 |
| Ptpn11 | 6192 | 6004 | 5950 | 8212 | 5837 | 5397 | 6577 | 5996 | 5974 | 6456 | 6280 | 5908 | 6198.57 | 6182 | 6215 | 1.005 | 0.008 | 0.92004506 | 0.96079922 | 0.0024 | 0.0074 | 0.0041 | 0.0041 | TRUE | 0.0379 |
| Ptpn12 | 2114 | 1853 | 1909 | 3040 | 2178 | 2048 | 2245 | 1851 | 1917 | 2390 | 2343 | 2242 | 2164.66 | 2004 | 2325 | 1.16 | 0.215 | 0.03047124 | 0.11659204 | 0.0054 | 0.0084 | 0.0066 | 0.0066 | TRUE | 0.1336 |
| Ptpn13 | 3773 | 4242 | 4581 | 4144 | 2979 | 3314 | 4008 | 4237 | 4600 | 3258 | 3205 | 3628 | 3822.42 | 4282 | 3364 | 0.785 | -0.348 | 0.00013125 | 0.001715 | 0.0044 | 0.0078 | 0.0057 | 0.0057 | TRUE | 0.057 |
| Ptpn14 | 1381 | 2068 | 1995 | 3251 | 2237 | 2228 | 1467 | 2065 | 2003 | 2556 | 2407 | 2439 | 2156.14 | 1845 | 2467 | 1.337 | 0.419 | 0.0021208 | 0.01613567 | 0.0157 | 0.0084 | 0.0129 | 0.0129 | TRUE | 0.3877 |
| Ptpn18 | 105 | 139 | 132 | 114 | 101 | 95 | 112 | 139 | 133 | 90 | 109 | 104 | 114.2 | 128 | 101 | 0.786 | -0.348 | 0.08025723 | 0.22499414 | 0 | 0.0358 | 0.0196 | 0.0196 | TRUE | 0.122 |
| Ptpn2 | 2656 | 2788 | 2803 | 3756 | 2545 | 2529 | 2821 | 2784 | 2815 | 2953 | 2738 | 2768 | 2813.23 | 2807 | 2820 | 1.005 | 0.007 | 0.91353702 | 0.95736251 | 6.00E-04 | 0.0081 | 0.003 | 0.003 | TRUE | 0.0211 |
| Ptpn20 | 90 | 94 | 100 | 200 | 167 | 111 | 96 | 94 | 100 | 157 | 180 | 122 | 124.72 | 97 | 153 | 1.583 | 0.662 | 0.00159204 | 0.01284385 | 0.0134 | 0.0333 | 0.0232 | 0.0232 | TRUE | 0.231 |
| Ptpn21 | 840 | 977 | 1095 | 1285 | 912 | 949 | 892 | 976 | 1099 | 1010 | 981 | 1039 | 999.62 | 989 | 1010 | 1.021 | 0.03 | 0.77847042 | 0.88535012 | 0.0048 | 0.0102 | 0.0071 | 0.0071 | TRUE | 0.1137 |
| Ptpn22 | 2 | 2 | 0 | 12 | 4 | 1 | 2 | 2 | 0 | 9 | 4 | 1 | 3.16 | 1 | 5 | 3.68 | 1.88 | 0.16234048 | NA | 0.5626 | 1.0498 | 0.8336 | 0.8336 | TRUE | 0.1592 |
| Ptpn23 | 6395 | 5489 | 5317 | 7462 | 5582 | 4979 | 6793 | 5482 | 5339 | 5866 | 6006 | 5450 | 5822.6 | 5871 | 5774 | 0.984 | -0.024 | 0.83125357 | 0.91439242 | 0.0085 | 0.0075 | 0.0089 | 0.0089 | TRUE | 0.2298 |
| Ptpn3 | 3277 | 3470 | 3636 | 3760 | 2864 | 3027 | 3481 | 3466 | 3651 | 2956 | 3081 | 3314 | 3324.67 | 3533 | 3117 | 0.882 | -0.181 | 0.01821506 | 0.08131559 | 0.0017 | 0.0079 | 0.0039 | 0.0039 | TRUE | 0.0369 |
| Ptpn4 | 780 | 607 | 621 | 1097 | 752 | 627 | 828 | 606 | 624 | 862 | 809 | 686 | 736.02 | 686 | 786 | 1.147 | 0.198 | 0.20500676 | 0.41342113 | 0.0208 | 0.0114 | 0.0162 | 0.0162 | TRUE | 0.3878 |
| Ptpn5 | 25 | 35 | 37 | 24 | 18 | 16 | 27 | 35 | 37 | 19 | 19 | 18 | 25.73 | 33 | 19 | 0.565 | -0.824 | 0.03113777 | 0.11834144 | 0 | 0.1349 | 0.0636 | 0.0636 | TRUE | 0.1894 |
| Ptpn6 | 2004 | 2233 | 2314 | 2627 | 2172 | 2343 | 2129 | 2230 | 2324 | 2065 | 2337 | 2565 | 2274.85 | 2228 | 2322 | 1.042 | 0.059 | 0.56625861 | 0.74618019 | 0.0064 | 0.0084 | 0.0072 | 0.0072 | TRUE | 0.1154 |
| Ptpn7 | 205 | 169 | 118 | 227 | 153 | 134 | 218 | 169 | 118 | 178 | 165 | 147 | 165.79 | 168 | 163 | 0.972 | -0.041 | 0.86352894 | 0.9324082 | 0.0435 | 0.0268 | 0.0349 | 0.0349 | TRUE | 0.1851 |
| Ptpn9 | 382 | 452 | 461 | 847 | 585 | 526 | 406 | 451 | 463 | 666 | 629 | 576 | 531.85 | 440 | 624 | 1.419 | 0.505 | 1.13E-05 | 0.00022161 | 0.0032 | 0.0131 | 0.0076 | 0.0076 | TRUE | 0.0531 |
| Ptpra | 2041 | 2735 | 2770 | 3384 | 2442 | 2570 | 2168 | 2731 | 2781 | 2660 | 2627 | 2813 | 2630.28 | 2560 | 2700 | 1.055 | 0.077 | 0.5011306 | 0.69669611 | 0.0096 | 0.0082 | 0.009 | 0.009 | TRUE | 0.2175 |
| Ptprb | 32 | 23 | 19 | 28 | 16 | 25 | 34 | 23 | 19 | 22 | 17 | 27 | 23.77 | 25 | 22 | 0.877 | -0.189 | 0.65752347 | 0.80725118 | 0.0268 | 0.1455 | 0.0894 | 0.0894 | TRUE | 0.5212 |
| Ptprc | 19 | 14 | 9 | 30 | 18 | 19 | 20 | 14 | 9 | 24 | 19 | 21 | 17.82 | 14 | 21 | 1.485 | 0.571 | 0.23589368 | 0.45108089 | 0 | 0.1917 | 0.1084 | 0.1084 | TRUE | 0.3421 |
| Ptprcap | 28 | 21 | 26 | 32 | 24 | 21 | 30 | 21 | 26 | 25 | 26 | 23 | 25.13 | 26 | 25 | 0.965 | -0.051 | 0.89200664 | 0.94755432 | 0 | 0.138 | 0.0634 | 0.0634 | TRUE | 0.1562 |
| Ptprd | 59 | 44 | 47 | 132 | 65 | 84 | 63 | 44 | 47 | 104 | 70 | 92 | 69.91 | 51 | 89 | 1.736 | 0.796 | 0.00387221 | 0.02571475 | 0.0237 | 0.054 | 0.0392 | 0.0392 | TRUE | 0.2914 |
| Ptpre | 152 | 146 | 127 | 191 | 133 | 111 | 161 | 146 | 128 | 150 | 143 | 122 | 141.59 | 145 | 138 | 0.957 | -0.063 | 0.7339444 | 0.85735379 | 0.0056 | 0.0302 | 0.0179 | 0.0179 | TRUE | 0.1153 |
| Ptprf | 11998 | 13408 | 14398 | 16950 | 12361 | 12974 | 12744 | 13391 | 14457 | 13325 | 13300 | 14202 | 13569.72 | 13531 | 13609 | 1.006 | 0.008 | 0.91493987 | 0.95806876 | 0.0021 | 0.0071 | 0.0042 | 0.0042 | TRUE | 0.0439 |
| Ptprg | 2449 | 2517 | 2653 | 3410 | 2398 | 2450 | 2601 | 2514 | 2664 | 2681 | 2580 | 2682 | 2620.27 | 2593 | 2648 | 1.021 | 0.03 | 0.65304517 | 0.80536621 | 3.00E-04 | 0.0082 | 0.0029 | 0.0029 | TRUE | 0.0087 |
| Ptprh | 1 | 0 | 1 | 1 | 1 | 1 | 1 | 0 | 1 | 1 | 1 | 1 | 0.84 | 1 | 1 | 1.427 | 0.513 | 0.79501633 | NA | 0 | 3.9421 | 1.5671 | 1.5671 | TRUE | 0.2557 |
| Ptprj | 654 | 883 | 886 | 1046 | 777 | 803 | 695 | 882 | 890 | 822 | 836 | 879 | 833.91 | 822 | 846 | 1.028 | 0.04 | 0.74467022 | 0.8638525 | 0.0088 | 0.0109 | 0.0097 | 0.0097 | TRUE | 0.217 |
| Ptprk | 2379 | 2054 | 2219 | 3143 | 2110 | 2085 | 2527 | 2051 | 2228 | 2471 | 2270 | 2282 | 2304.96 | 2269 | 2341 | 1.032 | 0.046 | 0.65319706 | 0.80540538 | 0.0062 | 0.0083 | 0.0071 | 0.0071 | TRUE | 0.1197 |
| Ptprm | 109 | 108 | 121 | 212 | 143 | 131 | 116 | 108 | 121 | 167 | 154 | 143 | 134.84 | 115 | 155 | 1.348 | 0.43 | 0.01418176 | 0.06793806 | 0 | 0.0313 | 0.0146 | 0.0146 | TRUE | 0.0531 |
| Ptprn | 253 | 239 | 229 | 297 | 232 | 198 | 269 | 239 | 230 | 233 | 250 | 217 | 239.53 | 246 | 233 | 0.95 | -0.074 | 0.60437264 | 0.77516185 | 0 | 0.0207 | 0.0107 | 0.0107 | TRUE | 0.073 |
| Ptprn2 | 209 | 296 | 306 | 194 | 144 | 167 | 222 | 296 | 307 | 153 | 155 | 183 | 219.19 | 275 | 164 | 0.593 | -0.754 | 3.40E-05 | 0.00055846 | 0.0162 | 0.0219 | 0.019 | 0.019 | TRUE | 0.3148 |
| Ptpro | 82 | 42 | 35 | 113 | 115 | 104 | 87 | 42 | 35 | 89 | 124 | 114 | 81.77 | 55 | 109 | 1.989 | 0.992 | 0.00515213 | 0.03198394 | 0.1127 | 0.0472 | 0.0767 | 0.0767 | TRUE | 2.1991 |
| Ptprq | 6 | 6 | 7 | 36 | 48 | 33 | 6 | 6 | 7 | 28 | 52 | 36 | 22.58 | 6 | 39 | 5.952 | 2.573 | 5.47E-07 | 1.62E-05 | 0 | 0.1528 | 0.0985 | 0.0985 | TRUE | 0.1847 |
| Ptprr | 15 | 7 | 4 | 17 | 14 | 11 | 16 | 7 | 4 | 13 | 15 | 12 | 11.23 | 9 | 13 | 1.51 | 0.595 | 0.36051271 | 0.57896411 | 0.0986 | 0.3001 | 0.2112 | 0.2112 | TRUE | 1.2954 |
| Ptprs | 4814 | 6751 | 7084 | 4948 | 4077 | 4448 | 5113 | 6742 | 7113 | 3890 | 4387 | 4869 | 5352.34 | 6323 | 4382 | 0.693 | -0.529 | 0.00021592 | 0.0025858 | 0.0201 | 0.0075 | 0.0146 | 0.0146 | TRUE | 0.3415 |
| Ptprt | 32 | 59 | 46 | 77 | 38 | 54 | 34 | 59 | 46 | 61 | 41 | 59 | 49.94 | 46 | 54 | 1.156 | 0.209 | 0.52002418 | 0.71102318 | 0.037 | 0.0729 | 0.0559 | 0.0559 | TRUE | 0.2227 |
| Ptpru | 663 | 687 | 698 | 847 | 584 | 637 | 704 | 686 | 701 | 666 | 628 | 697 | 680.45 | 697 | 664 | 0.952 | -0.07 | 0.4518311 | 0.65723121 | 0 | 0.0117 | 0.0048 | 0.0048 | TRUE | 0.0246 |
| Ptprv | 224 | 287 | 265 | 392 | 307 | 303 | 238 | 287 | 266 | 308 | 330 | 332 | 293.47 | 264 | 323 | 1.225 | 0.293 | 0.02813351 | 0.11007735 | 0 | 0.0181 | 0.0093 | 0.0093 | TRUE | 0.0795 |
| Ptprz1 | 183 | 120 | 101 | 193 | 132 | 105 | 194 | 120 | 101 | 152 | 142 | 115 | 137.39 | 138 | 136 | 0.986 | -0.02 | 0.94012738 | 0.96979647 | 0.0613 | 0.0309 | 0.0455 | 0.0455 | TRUE | 1.273 |
| Ptrf | 2869 | 3212 | 3289 | 2968 | 2181 | 2210 | 3047 | 3208 | 3303 | 2333 | 2347 | 2419 | 2776.13 | 3186 | 2366 | 0.743 | -0.429 | 8.40E-10 | 5.28E-08 | 7.00E-04 | 0.0081 | 0.0032 | 0.0032 | TRUE | 0.0175 |
| Ptrh1 | 288 | 441 | 474 | 426 | 356 | 357 | 306 | 440 | 476 | 335 | 383 | 391 | 388.5 | 407 | 370 | 0.906 | -0.143 | 0.43217232 | 0.64176836 | 0.0272 | 0.0154 | 0.0213 | 0.0213 | TRUE | 0.5436 |
| Ptrh2 | 1276 | 1305 | 1258 | 2109 | 1409 | 1224 | 1355 | 1303 | 1263 | 1658 | 1516 | 1340 | 1405.93 | 1307 | 1505 | 1.152 | 0.204 | 0.05091526 | 0.16612235 | 0.0052 | 0.0092 | 0.0072 | 0.0072 | TRUE | 0.1099 |
| Ptrhd1 | 374 | 355 | 325 | 425 | 321 | 331 | 397 | 355 | 326 | 334 | 345 | 362 | 353.32 | 359 | 347 | 0.966 | -0.05 | 0.69584927 | 0.83345494 | 0.0029 | 0.0162 | 0.009 | 0.009 | TRUE | 0.0961 |
| Pts | 496 | 636 | 653 | 663 | 575 | 536 | 527 | 635 | 656 | 521 | 619 | 587 | 590.72 | 606 | 576 | 0.948 | -0.077 | 0.55667778 | 0.73929959 | 0.0089 | 0.0125 | 0.0105 | 0.0105 | TRUE | 0.152 |

| | | | | | | | | | | | | | | | | | | | | | | | | | |
|---|---|---|---|---|---|---|---|---|---|---|---|---|---|---|---|---|---|---|---|---|---|---|---|---|---|
| Pttg1 | 4221 | 5710 | 5448 | 7231 | 5341 | 5935 | 4483 | 5703 | 5470 | 5684 | 5747 | 6497 | 5597.38 | 5219 | 5976 | 1.145 | 0.195 | 0.08789059 | 0.24028609 | 0.0103 | 0.0075 | 0.0093 | 0.0093 | TRUE | 0.185 |
| Pttg1ip | 3078 | 3938 | 4208 | 4253 | 3162 | 3521 | 3269 | 3933 | 4225 | 3343 | 3402 | 3854 | 3671.23 | 3809 | 3533 | 0.927 | -0.109 | 0.35638788 | 0.57497439 | 0.0109 | 0.0078 | 0.0098 | 0.0098 | TRUE | 0.1867 |
| Ptx3 | 2 | 2 | 0 | 19 | 4 | 7 | 2 | 2 | 0 | 15 | 4 | 8 | 5.17 | 1 | 9 | 6.647 | 2.733 | 0.01473424 | 0.06989126 | 0 | 0.644 | 0.4769 | 0.4769 | TRUE | 0.273 |
| Ptx4 | 0 | 0 | 0 | 0 | 0 | 0 | 0 | 0 | 0 | 0 | 0 | 0 | 0 | 0 | 0 | NA | NA | NA | NA | NA | NA | NA | NA | NA | NA |
| Puf60 | 8624 | 7329 | 7260 | 10847 | 7866 | 7519 | 9160 | 7320 | 7290 | 8527 | 8463 | 8231 | 8165.11 | 7923 | 8407 | 1.061 | 0.086 | 0.42413378 | 0.63431045 | 0.008 | 0.0073 | 0.0081 | 0.0081 | TRUE | 0.2276 |
| Pum1 | 9686 | 8883 | 8942 | 12755 | 9331 | 8931 | 10288 | 8872 | 8979 | 10027 | 10040 | 9776 | 9663.57 | 9380 | 9948 | 1.061 | 0.085 | 0.30410894 | 0.5250533 | 0.0035 | 0.0072 | 0.0048 | 0.0048 | TRUE | 0.0877 |
| Pum2 | 11639 | 11870 | 12723 | 15739 | 11654 | 12027 | 12363 | 11855 | 12775 | 12373 | 12539 | 13165 | 12511.63 | 12331 | 12692 | 1.029 | 0.042 | 0.5256684 | 0.71503596 | 0.0011 | 0.0072 | 0.003 | 0.003 | TRUE | 0.014 |
| Pura | 752 | 775 | 782 | 1464 | 1013 | 855 | 799 | 774 | 785 | 1151 | 1090 | 936 | 922.45 | 786 | 1059 | 1.349 | 0.432 | 6.22E-05 | 0.00093449 | 0.005 | 0.0105 | 0.0073 | 0.0073 | TRUE | 0.1221 |
| Purb | 3511 | 3468 | 3878 | 5382 | 4087 | 3452 | 3729 | 3464 | 3894 | 4231 | 4397 | 3779 | 3915.63 | 3696 | 4136 | 1.119 | 0.163 | 0.07587816 | 0.21720036 | 0.0043 | 0.0077 | 0.0058 | 0.0058 | TRUE | 0.0689 |
| Purg | 310 | 270 | 290 | 455 | 360 | 304 | 329 | 270 | 291 | 358 | 387 | 333 | 327.99 | 297 | 359 | 1.212 | 0.277 | 0.04305275 | 0.14822702 | 0.0045 | 0.0169 | 0.0104 | 0.0104 | TRUE | 0.1028 |
| Pus1 | 2938 | 3611 | 3425 | 3840 | 3009 | 3016 | 3121 | 3606 | 3439 | 3019 | 3237 | 3301 | 3287.29 | 3389 | 3186 | 0.94 | -0.09 | 0.3034614 | 0.5242845 | 0.0033 | 0.0079 | 0.0052 | 0.0052 | TRUE | 0.0581 |
| Pus10 | 2052 | 1867 | 1922 | 2799 | 1975 | 1860 | 2180 | 1865 | 1930 | 2200 | 2125 | 2036 | 2055.91 | 1992 | 2120 | 1.065 | 0.091 | 0.31825506 | 0.53921139 | 0.0035 | 0.0085 | 0.0055 | 0.0055 | TRUE | 0.0824 |
| Pus3 | 920 | 817 | 718 | 1045 | 727 | 667 | 977 | 816 | 721 | 821 | 782 | 730 | 807.99 | 838 | 778 | 0.929 | -0.106 | 0.42836346 | 0.63827572 | 0.0118 | 0.011 | 0.0117 | 0.0117 | TRUE | 0.2498 |
| Pus7 | 2804 | 2742 | 2805 | 4277 | 3112 | 3093 | 2978 | 2738 | 2817 | 3362 | 3348 | 3386 | 3104.94 | 2844 | 3365 | 1.183 | 0.243 | 0.00036315 | 0.00400812 | 5.00E-04 | 0.008 | 0.003 | 0.003 | TRUE | 0.0205 |
| Pus7l | 535 | 489 | 505 | 887 | 643 | 565 | 568 | 488 | 507 | 697 | 692 | 618 | 595.22 | 521 | 669 | 1.285 | 0.362 | 0.00131995 | 0.01106764 | 0.0035 | 0.0124 | 0.0075 | 0.0075 | TRUE | 0.0721 |
| Pusl1 | 754 | 656 | 646 | 991 | 750 | 683 | 801 | 655 | 649 | 779 | 807 | 748 | 739.72 | 702 | 778 | 1.109 | 0.15 | 0.20313866 | 0.41080946 | 0.0062 | 0.0114 | 0.0086 | 0.0086 | TRUE | 0.1801 |
| Pvalb | 1 | 0 | 1 | 1 | 1 | 1 | 1 | 0 | 1 | 1 | 1 | 1 | 0.84 | 1 | 1 | 1.427 | 0.513 | 0.79501633 | NA | 0 | 3.9421 | 1.5671 | 1.5671 | TRUE | 0.2557 |
| Pvr | 906 | 1397 | 1640 | 1343 | 1013 | 1235 | 962 | 1395 | 1647 | 1056 | 1090 | 1352 | 1250.31 | 1335 | 1166 | 0.873 | -0.196 | 0.309262 | 0.52974532 | 0.0421 | 0.0095 | 0.0259 | 0.0259 | TRUE | 0.3531 |
| Pvrl1 | 1666 | 1560 | 1482 | 2224 | 1595 | 1468 | 1770 | 1558 | 1488 | 1748 | 1716 | 1607 | 1647.85 | 1605 | 1690 | 1.054 | 0.075 | 0.438653 | 0.64603933 | 0.0042 | 0.0089 | 0.0062 | 0.0062 | TRUE | 0.0964 |
| Pvrl2 | 4309 | 5487 | 5876 | 5495 | 3989 | 4557 | 4577 | 5480 | 5900 | 4320 | 4292 | 4988 | 4926.16 | 5319 | 4533 | 0.852 | -0.231 | 0.05153641 | 0.1673183 | 0.0116 | 0.0076 | 0.0099 | 0.0099 | TRUE | 0.1814 |
| Pvrl3 | 1623 | 1845 | 1928 | 1964 | 1427 | 1574 | 1724 | 1843 | 1936 | 1544 | 1535 | 1723 | 1717.46 | 1834 | 1601 | 0.872 | -0.197 | 0.03061947 | 0.1169606 | 0.003 | 0.0088 | 0.0054 | 0.0054 | TRUE | 0.0531 |
| Pvrl4 | 210 | 236 | 236 | 259 | 196 | 222 | 223 | 236 | 237 | 204 | 211 | 243 | 225.54 | 232 | 219 | 0.943 | -0.085 | 0.55612183 | 0.73889464 | 0 | 0.0215 | 0.0107 | 0.0107 | TRUE | 0.0966 |
| Pvt1 | 467 | 521 | 540 | 413 | 342 | 399 | 496 | 520 | 542 | 325 | 368 | 437 | 448 | 519 | 377 | 0.723 | -0.469 | 0.00077927 | 0.00733767 | 0.0095 | 0.0143 | 0.0117 | 0.0117 | TRUE | 0.2236 |
| Pwp1 | 5961 | 6125 | 6120 | 7851 | 5718 | 5613 | 6332 | 6117 | 6145 | 6172 | 6152 | 6144 | 6177.04 | 6198 | 6156 | 0.993 | -0.01 | 0.8625257 | 0.93208328 | 0 | 0.0074 | 0.0021 | 0.0021 | TRUE | 0.0043 |
| Pwp2 | 5151 | 4599 | 4338 | 6545 | 4629 | 4093 | 5471 | 4593 | 4356 | 5145 | 4981 | 4480 | 4837.71 | 4807 | 4869 | 1.013 | 0.019 | 0.8660358 | 0.93370586 | 0.0096 | 0.0076 | 0.0087 | 0.0087 | TRUE | 0.1781 |
| Pwwp2a | 3624 | 3381 | 3747 | 3870 | 3094 | 3125 | 3849 | 3377 | 3762 | 3042 | 3329 | 3421 | 3463.4 | 3663 | 3264 | 0.891 | -0.167 | 0.06244524 | 0.19131436 | 0.0038 | 0.0079 | 0.0055 | 0.0055 | TRUE | 0.0569 |
| Pwwp2b | 478 | 595 | 696 | 356 | 298 | 359 | 508 | 594 | 699 | 280 | 321 | 393 | 465.71 | 600 | 331 | 0.55 | -0.862 | 7.62E-07 | 2.18E-05 | 0.0242 | 0.014 | 0.0195 | 0.0195 | TRUE | 0.1823 |
| Pxdc1 | 47 | 136 | 145 | 101 | 82 | 139 | 50 | 136 | 146 | 79 | 88 | 152 | 108.52 | 111 | 106 | 0.962 | -0.056 | 0.89082348 | 0.94702533 | 0.1977 | 0.0373 | 0.109 | 0.109 | TRUE | 2.2591 |
| Pxdn | 2813 | 2796 | 3096 | 2598 | 2075 | 2078 | 2988 | 2792 | 3109 | 2042 | 2233 | 2275 | 2573.1 | 2963 | 2183 | 0.736 | -0.441 | 1.86E-07 | 6.14E-06 | 0.0025 | 0.0082 | 0.0048 | 0.0048 | TRUE | 0.0394 |
| Pxk | 979 | 1014 | 1070 | 1222 | 829 | 946 | 1040 | 1013 | 1074 | 961 | 892 | 1036 | 1002.52 | 1042 | 963 | 0.924 | -0.115 | 0.22256103 | 0.4346499 | 0.002 | 0.0102 | 0.0054 | 0.0054 | TRUE | 0.0505 |
| Pxmp2 | 618 | 727 | 627 | 912 | 716 | 627 | 656 | 726 | 630 | 717 | 770 | 686 | 697.62 | 671 | 724 | 1.08 | 0.111 | 0.29395366 | 0.51480884 | 0.0029 | 0.0116 | 0.0066 | 0.0066 | TRUE | 0.062 |
| Pxmp4 | 458 | 561 | 586 | 538 | 415 | 420 | 486 | 560 | 588 | 423 | 447 | 460 | 494.06 | 545 | 443 | 0.812 | -0.301 | 0.01135971 | 0.05786997 | 0.0038 | 0.0136 | 0.0081 | 0.0081 | TRUE | 0.1024 |
| Pxn | 4329 | 3845 | 4006 | 4674 | 3578 | 3482 | 4598 | 3840 | 4022 | 3674 | 3850 | 3812 | 3966.05 | 4153 | 3779 | 0.91 | -0.137 | 0.13434536 | 0.31691521 | 0.0045 | 0.0077 | 0.0057 | 0.0057 | TRUE | 0.1066 |
| Pxt1 | 82 | 77 | 60 | 123 | 84 | 92 | 87 | 77 | 60 | 97 | 90 | 101 | 85.34 | 75 | 96 | 1.285 | 0.362 | 0.11438375 | 0.28525158 | 1.00E-04 | 0.0455 | 0.0259 | 0.0259 | TRUE | 0.2649 |
| Pycard | 814 | 1595 | 1760 | 1090 | 950 | 1220 | 865 | 1593 | 1767 | 857 | 1022 | 1335 | 1239.88 | 1408 | 1071 | 0.76 | -0.395 | 0.11534267 | 0.28675672 | 0.0911 | 0.0096 | 0.0445 | 0.0445 | TRUE | 1.3701 |
| Pycr1 | 900 | 1485 | 1433 | 1181 | 931 | 1231 | 956 | 1483 | 1439 | 928 | 1002 | 1348 | 1192.59 | 1293 | 1093 | 0.845 | -0.244 | 0.21660988 | 0.42865383 | 0.0453 | 0.0097 | 0.0272 | 0.0272 | TRUE | 0.6223 |
| Pycr2 | 3471 | 4201 | 4273 | 5264 | 3573 | 3486 | 3687 | 4196 | 4291 | 4138 | 3844 | 3816 | 3995.23 | 4058 | 3933 | 0.969 | -0.045 | 0.61429707 | 0.78173136 | 0.0035 | 0.0077 | 0.0055 | 0.0055 | TRUE | 0.0776 |
| Pycrl | 1990 | 1750 | 1806 | 2084 | 1555 | 1345 | 2114 | 1748 | 1813 | 1638 | 1673 | 1472 | 1743.09 | 1892 | 1594 | 0.843 | -0.246 | 0.02169677 | 0.09226246 | 0.0067 | 0.0088 | 0.0077 | 0.0077 | TRUE | 0.1272 |
| Pydc3 | 0 | 0 | 0 | 0 | 0 | 0 | 0 | 0 | 0 | 0 | 0 | 0 | 0 | 0 | 0 | NA | NA | NA | NA | NA | NA | NA | NA | NA | NA |
| Pydc4 | 0 | 0 | 0 | 0 | 0 | 0 | 0 | 0 | 0 | 0 | 0 | 0 | 0 | 0 | 0 | NA | NA | NA | NA | NA | NA | NA | NA | NA | NA |
| Pygb | 3833 | 4065 | 4117 | 3822 | 2922 | 3350 | 4071 | 4060 | 4134 | 3005 | 3144 | 3667 | 3680.09 | 4088 | 3272 | 0.8 | -0.322 | 0.00072748 | 0.00698182 | 0.0053 | 0.0078 | 0.0063 | 0.0063 | TRUE | 0.1353 |
| Pygl | 418 | 799 | 874 | 674 | 647 | 918 | 444 | 798 | 878 | 530 | 696 | 1005 | 725.07 | 707 | 744 | 1.052 | 0.073 | 0.79292908 | 0.89322745 | 0.1104 | 0.0114 | 0.0536 | 0.0536 | TRUE | 0.4813 |
| Pygm | 522 | 670 | 616 | 257 | 219 | 302 | 554 | 669 | 619 | 202 | 236 | 331 | 435.06 | 614 | 256 | 0.416 | -1.267 | 2.15E-11 | 1.79E-09 | 0.0341 | 0.0145 | 0.023 | 0.023 | TRUE | 0.7144 |
| Pygo1 | 78 | 112 | 129 | 170 | 126 | 126 | 83 | 112 | 130 | 134 | 136 | 138 | 121.9 | 108 | 136 | 1.254 | 0.326 | 0.12576574 | 0.30376374 | 0.0136 | 0.0339 | 0.0244 | 0.0244 | TRUE | 0.4045 |
| Pygo2 | 2199 | 1997 | 1785 | 2447 | 1737 | 1553 | 2336 | 1994 | 1792 | 1924 | 1869 | 1700 | 1935.84 | 2041 | 1831 | 0.897 | -0.156 | 0.18891498 | 0.39412263 | 0.0103 | 0.0086 | 0.0096 | 0.0096 | TRUE | 0.1929 |
| Pyhin1 | 0 | 0 | 0 | 0 | 0 | 0 | 0 | 0 | 0 | 0 | 0 | 0 | 0 | 0 | 0 | NA | NA | NA | NA | NA | NA | NA | NA | NA | NA |
| Pyroxd1 | 168 | 119 | 121 | 234 | 154 | 130 | 178 | 119 | 121 | 184 | 166 | 142 | 151.79 | 139 | 164 | 1.179 | 0.237 | 0.28189277 | 0.50334119 | 0.0272 | 0.0286 | 0.0284 | 0.0284 | TRUE | 0.6064 |
| Pyroxd2 | 10 | 15 | 13 | 11 | 16 | 7 | 11 | 15 | 13 | 9 | 17 | 8 | 12.03 | 13 | 11 | 0.86 | -0.218 | 0.70779547 | 0.84080366 | 0 | 0.2808 | 0.1604 | 0.1604 | TRUE | 0.7791 |
| Pyurf | 345 | 337 | 317 | 615 | 415 | 478 | 366 | 337 | 318 | 483 | 447 | 523 | 412.42 | 340 | 484 | 1.423 | 0.509 | 3.76E-05 | 0.00060826 | 0.003 | 0.0149 | 0.0085 | 0.0085 | TRUE | 0.0548 |
| Pyy | 12 | 19 | 21 | 16 | 12 | 7 | 13 | 19 | 21 | 13 | 13 | 8 | 14.33 | 18 | 11 | 0.631 | -0.665 | 0.20461921 | 0.4130621 | 0 | 0.2369 | 0.1249 | 0.1249 | TRUE | 0.2802 |
| Pzp | 53 | 45 | 56 | 183 | 107 | 126 | 56 | 45 | 56 | 144 | 115 | 138 | 92.4 | 52 | 132 | 2.528 | 1.338 | 4.03E-09 | 2.08E-07 | 0 | 0.0426 | 0.0238 | 0.0238 | TRUE | 0.1328 |
| Qars | 8222 | 9620 | 10071 | 10616 | 7806 | 8558 | 8733 | 9608 | 10112 | 8345 | 8399 | 9368 | 9094.24 | 9484 | 8704 | 0.918 | -0.124 | 0.16801295 | 0.36575573 | 0.0047 | 0.0073 | 0.0057 | 0.0057 | TRUE | 0.0586 |
| Qdpr | 6801 | 7929 | 7935 | 7280 | 5391 | 5984 | 7224 | 7919 | 7968 | 5723 | 5800 | 6550 | 6864 | 7704 | 6024 | 0.782 | -0.355 | 4.87E-05 | 0.0007602 | 0.0041 | 0.0074 | 0.0054 | 0.0054 | TRUE | 0.0709 |
| Qk | 9322 | 11219 | 11056 | 13452 | 10308 | 10440 | 9902 | 11205 | 11101 | 10575 | 11091 | 11428 | 10883.57 | 10736 | 11031 | 1.027 | 0.039 | 0.62493423 | 0.78868818 | 0.0025 | 0.0072 | 0.0045 | 0.0045 | TRUE | 0.0564 |
| Qpct | 313 | 393 | 357 | 340 | 304 | 339 | 332 | 392 | 358 | 267 | 327 | 371 | 341.48 | 361 | 322 | 0.888 | -0.171 | 0.2823782 | 0.50369474 | 0.0141 | 0.0166 | 0.0153 | 0.0153 | TRUE | 0.2638 |
| Qpctl | 172 | 225 | 226 | 393 | 296 | 273 | 183 | 225 | 227 | 309 | 318 | 299 | 260.1 | 212 | 309 | 1.459 | 0.545 | 0.00015009 | 0.00190136 | 0.0015 | 0.0196 | 0.0109 | 0.0109 | TRUE | 0.1519 |
| Qprt | 700 | 776 | 825 | 737 | 533 | 609 | 744 | 775 | 828 | 579 | 573 | 667 | 694.4 | 782 | 606 | 0.774 | -0.369 | 0.00066342 | 0.00649503 | 0.0034 | 0.0116 | 0.007 | 0.007 | TRUE | 0.0864 |
| Qrfp | 2 | 3 | 2 | 1 | 2 | 0 | 2 | 3 | 2 | 1 | 2 | 0 | 1.68 | 2 | 1 | 0.409 | -1.291 | 0.39443879 | NA | 0 | 1.9705 | 0.9403 | 0.9403 | TRUE | 0.435 |
| Qrfpr | 2 | 2 | 0 | 3 | 1 | 2 | 2 | 2 | 0 | 2 | 1 | 2 | 1.62 | 1 | 2 | 1.382 | 0.466 | 0.7580283 | NA | 0 | 2.0352 | 1.0191 | 1.0191 | TRUE | 0.4916 |
| Qrich1 | 5642 | 5145 | 5097 | 7340 | 5084 | 4687 | 5993 | 5138 | 5118 | 5770 | 5470 | 5131 | 5436.66 | 5416 | 5457 | 1.008 | 0.011 | 0.90747835 | 0.95354464 | 0.0056 | 0.0075 | 0.0064 | 0.0064 | TRUE | 0.1056 |
| Qrich2 | 30 | 42 | 28 | 37 | 37 | 38 | 32 | 42 | 28 | 29 | 40 | 42 | 35.4 | 34 | 37 | 1.076 | 0.106 | 0.76100773 | 0.87423727 | 0 | 0.1 | 0.0584 | 0.0584 | TRUE | 0.3003 |
| Qrsl1 | 1241 | 1280 | 1285 | 1461 | 1093 | 1145 | 1318 | 1278 | 1290 | 1149 | 1176 | 1253 | 1244.12 | 1295 | 1193 | 0.92 | -0.12 | 0.13243743 | 0.31372831 | 0 | 0.0096 | 0.0038 | 0.0038 | TRUE | 0.0235 |
| Qser1 | 5264 | 5261 | 5867 | 5271 | 3927 | 4094 | 5591 | 5254 | 5891 | 4144 | 4225 | 4482 | 4931.16 | 5579 | 4284 | 0.768 | -0.381 | 8.97E-07 | 2.50E-05 | 0.0022 | 0.0076 | 0.0041 | 0.0041 | TRUE | 0.0317 |
| Qsox1 | 962 | 1100 | 1035 | 1288 | 896 | 872 | 1022 | 1099 | 1039 | 1013 | 964 | 955 | 1015.12 | 1053 | 977 | 0.928 | -0.107 | 0.20179599 | 0.40930123 | 0 | 0.0101 | 0.0041 | 0.0041 | TRUE | 0.017 |
| Qsox2 | 1219 | 1735 | 1702 | 1638 | 1219 | 1292 | 1295 | 1733 | 1709 | 1288 | 1312 | 1414 | 1458.35 | 1579 | 1338 | 0.847 | -0.24 | 0.06599283 | 0.19817615 | 0.0124 | 0.0092 | 0.0115 | 0.0115 | TRUE | 0.2977 |
| Qtrt1 | 924 | 1072 | 1159 | 1309 | 1031 | 1061 | 981 | 1071 | 1164 | 1029 | 1109 | 1161 | 1085.93 | 1072 | 1100 | 1.025 | 0.036 | 0.72981927 | 0.85492495 | 0.0046 | 0.0099 | 0.0068 | 0.0068 | TRUE | 0.0672 |
| Qtrtd1 | 1942 | 1574 | 1439 | 2389 | 1746 | 1450 | 2063 | 1572 | 1445 | 1878 | 1879 | 1587 | 1737.25 | 1693 | 1781 | 1.052 | 0.074 | 0.62059264 | 0.78523764 | 0.0216 | 0.0088 | 0.0154 | 0.0154 | TRUE | 0.4391 |
| R3hcc1 | 784 | 814 | 772 | 1686 | 1161 | 881 | 833 | 813 | 775 | 1325 | 1249 | 964 | 993.3 | 807 | 1179 | 1.463 | 0.549 | 4.84E-05 | 0.00075619 | 0.0137 | 0.0102 | 0.0121 | 0.0121 | TRUE | 0.3031 |
| R3hcc1l | 1132 | 1085 | 1192 | 1393 | 1030 | 1025 | 1202 | 1084 | 1197 | 1095 | 1108 | 1122 | 1134.7 | 1161 | 1108 | 0.955 | -0.067 | 0.43242556 | 0.64190268 | 9.00E-04 | 0.0098 | 0.0044 | 0.0044 | TRUE | 0.0409 |

| | | | | | | | | | | | | | | | | | | | | | | | | | |
|---|---|---|---|---|---|---|---|---|---|---|---|---|---|---|---|---|---|---|---|---|---|---|---|---|---|
| R3hdm1 | 5899 | 6891 | 7215 | 6345 | 5009 | 5538 | 6266 | 6882 | 7245 | 4988 | 5389 | 6062 | 6138.67 | 6798 | 5480 | 0.806 | -0.311 | 0.00237154 | 0.01761012 | 0.0073 | 0.0074 | 0.0074 | 0.0074 | TRUE | 0.1053 |
| R3hdm2 | 4048 | 5039 | 5136 | 5615 | 4225 | 4611 | 4300 | 5032 | 5157 | 4414 | 4546 | 5047 | 4749.44 | 4830 | 4669 | 0.967 | -0.049 | 0.62987032 | 0.79195749 | 0.0071 | 0.0076 | 0.0073 | 0.0073 | TRUE | 0.1122 |
| R3hdm4 | 1653 | 1950 | 1967 | 2027 | 1481 | 1626 | 1756 | 1947 | 1975 | 1593 | 1593 | 1780 | 1774.2 | 1893 | 1655 | 0.874 | -0.194 | 0.03564837 | 0.12960084 | 0.0033 | 0.0088 | 0.0056 | 0.0056 | TRUE | 0.0515 |
| R3hdml | 70 | 40 | 42 | 32 | 21 | 12 | 74 | 40 | 42 | 25 | 23 | 13 | 36.23 | 52 | 20 | 0.393 | -1.347 | 0.00126728 | 0.01069033 | 0.0857 | 0.0978 | 0.0921 | 0.0921 | TRUE | 1.1199 |
| R74862 | 58 | 74 | 88 | 71 | 37 | 39 | 62 | 74 | 88 | 56 | 40 | 43 | 60.36 | 75 | 46 | 0.622 | -0.685 | 0.01515544 | 0.0712676 | 0.0173 | 0.0615 | 0.04 | 0.04 | TRUE | 0.2068 |
| Rab1 | 14443 | 15895 | 15463 | 19035 | 13338 | 13898 | 15341 | 15874 | 15526 | 14964 | 14351 | 15214 | 15211.7 | 15580 | 14843 | 0.953 | -0.07 | 0.23099444 | 0.44489489 | 5.00E-04 | 0.0071 | 0.0024 | 0.0024 | TRUE | 0.0102 |
| Rab10 | 5011 | 5800 | 6100 | 7814 | 5570 | 5490 | 5323 | 5793 | 6125 | 6143 | 5993 | 6010 | 5897.59 | 5747 | 6049 | 1.053 | 0.074 | 0.34135881 | 0.56159399 | 0.0023 | 0.0075 | 0.0042 | 0.0042 | TRUE | 0.0507 |
| Rab11a | 2350 | 2850 | 2819 | 3371 | 2359 | 2607 | 2496 | 2846 | 2831 | 2650 | 2538 | 2854 | 2702.49 | 2724 | 2681 | 0.984 | -0.024 | 0.79609888 | 0.89531728 | 0.0041 | 0.0081 | 0.0056 | 0.0056 | TRUE | 0.065 |
| Rab11b | 2069 | 2029 | 1949 | 2875 | 2062 | 1968 | 2198 | 2026 | 1957 | 2260 | 2219 | 2154 | 2135.67 | 2060 | 2211 | 1.074 | 0.102 | 0.19765684 | 0.40433284 | 0.0014 | 0.0084 | 0.0041 | 0.0041 | TRUE | 0.0409 |
| Rab11fip1 | 4340 | 3690 | 3849 | 4083 | 3183 | 3393 | 4610 | 3685 | 3865 | 3210 | 3425 | 3714 | 3751.42 | 4053 | 3450 | 0.851 | -0.233 | 0.03736418 | 0.13408597 | 0.0096 | 0.0078 | 0.0088 | 0.0088 | TRUE | 0.1754 |
| Rab11fip2 | 321 | 551 | 545 | 693 | 472 | 569 | 341 | 550 | 547 | 545 | 508 | 623 | 519 | 479 | 559 | 1.164 | 0.219 | 0.2588882 | 0.4778449 | 0.038 | 0.0133 | 0.0252 | 0.0252 | TRUE | 0.7386 |
| Rab11fip3 | 1922 | 1879 | 1828 | 2443 | 1864 | 1779 | 2042 | 1877 | 1836 | 1920 | 2006 | 1947 | 1937.84 | 1918 | 1958 | 1.021 | 0.03 | 0.70663888 | 0.8398044 | 0.0013 | 0.0086 | 0.004 | 0.004 | TRUE | 0.0383 |
| Rab11fip4 | 2386 | 2453 | 2475 | 2526 | 1997 | 1935 | 2534 | 2450 | 2485 | 1986 | 2149 | 2118 | 2286.98 | 2490 | 2084 | 0.837 | -0.257 | 0.0003264 | 0.00364845 | 6.00E-04 | 0.0083 | 0.0033 | 0.0033 | TRUE | 0.0213 |
| Rab11fip5 | 1894 | 2036 | 2145 | 1867 | 1427 | 1519 | 2012 | 2033 | 2154 | 1468 | 1535 | 1663 | 1810.8 | 2066 | 1555 | 0.752 | -0.411 | 1.20E-06 | 3.23E-05 | 0.0019 | 0.0087 | 0.0046 | 0.0046 | TRUE | 0.0438 |
| Rab12 | 1053 | 1092 | 973 | 1924 | 1431 | 1247 | 1118 | 1091 | 977 | 1512 | 1540 | 1365 | 1267.21 | 1062 | 1472 | 1.387 | 0.472 | 1.45E-06 | 3.80E-05 | 0.0037 | 0.0095 | 0.0061 | 0.0061 | TRUE | 0.0588 |
| Rab13 | 123 | 119 | 114 | 234 | 140 | 128 | 131 | 119 | 114 | 184 | 151 | 140 | 139.78 | 121 | 158 | 1.31 | 0.39 | 0.03914442 | 0.13855158 | 0.0076 | 0.0305 | 0.0185 | 0.0185 | TRUE | 0.2298 |
| Rab14 | 5120 | 6283 | 6445 | 8051 | 5812 | 6082 | 5438 | 6275 | 6471 | 6329 | 6253 | 6658 | 6237.47 | 6061 | 6413 | 1.058 | 0.081 | 0.3671675 | 0.58453273 | 0.0044 | 0.0074 | 0.0057 | 0.0057 | TRUE | 0.0985 |
| Rab15 | 505 | 526 | 543 | 814 | 633 | 593 | 536 | 525 | 545 | 640 | 681 | 649 | 596.18 | 535 | 657 | 1.225 | 0.293 | 0.00191937 | 0.01490514 | 0 | 0.0124 | 0.0047 | 0.0047 | TRUE | 0.0122 |
| Rab17 | 29 | 45 | 32 | 42 | 48 | 33 | 31 | 45 | 32 | 33 | 52 | 36 | 38.11 | 36 | 40 | 1.113 | 0.154 | 0.65776517 | 0.80727354 | 0.027 | 0.0933 | 0.061 | 0.061 | TRUE | 0.4442 |
| Rab18 | 5374 | 6036 | 5924 | 6817 | 4948 | 5614 | 5708 | 6028 | 5948 | 5359 | 5324 | 6145 | 5752.13 | 5895 | 5609 | 0.951 | -0.072 | 0.39604633 | 0.60952797 | 0.0035 | 0.0075 | 0.005 | 0.005 | TRUE | 0.0849 |
| Rab19 | 9 | 9 | 8 | 19 | 6 | 14 | 10 | 9 | 8 | 15 | 6 | 15 | 10.55 | 9 | 12 | 1.391 | 0.476 | 0.43849264 | 0.64591156 | 0 | 0.3192 | 0.175 | 0.175 | TRUE | 0.6282 |
| Rab1b | 4157 | 3559 | 3436 | 6064 | 4142 | 3725 | 4415 | 3554 | 3450 | 4767 | 4457 | 4078 | 4120.19 | 3806 | 4434 | 1.165 | 0.22 | 0.06588818 | 0.19794066 | 0.0124 | 0.0077 | 0.0101 | 0.0101 | TRUE | 0.2379 |
| Rab20 | 78 | 83 | 72 | 87 | 87 | 65 | 83 | 83 | 72 | 68 | 83 | 71 | 76.74 | 79 | 74 | 0.932 | -0.102 | 0.65231342 | 0.80502827 | 0 | 0.0498 | 0.0237 | 0.0237 | TRUE | 0.0934 |
| Rab21 | 1427 | 1511 | 1413 | 1850 | 1296 | 1152 | 1516 | 1509 | 1419 | 1454 | 1394 | 1261 | 1425.56 | 1481 | 1370 | 0.926 | -0.111 | 0.21833263 | 0.43044278 | 0.0026 | 0.0092 | 0.0052 | 0.0052 | TRUE | 0.0576 |
| Rab22a | 832 | 1036 | 1064 | 1169 | 851 | 955 | 884 | 1035 | 1068 | 919 | 916 | 1045 | 977.79 | 996 | 960 | 0.964 | -0.054 | 0.63787219 | 0.79611447 | 0.0068 | 0.0103 | 0.0083 | 0.0083 | TRUE | 0.1147 |
| Rab23 | 2038 | 1940 | 1750 | 2418 | 1692 | 1667 | 2165 | 1937 | 1757 | 1901 | 1820 | 1825 | 1900.92 | 1953 | 1849 | 0.947 | -0.079 | 0.4287927 | 0.63860634 | 0.005 | 0.0086 | 0.0066 | 0.0066 | TRUE | 0.1083 |
| Rab24 | 828 | 1101 | 1121 | 1450 | 1155 | 1032 | 879 | 1100 | 1126 | 1140 | 1243 | 1130 | 1102.82 | 1035 | 1171 | 1.131 | 0.177 | 0.14135677 | 0.32680725 | 0.0086 | 0.0099 | 0.0096 | 0.0096 | TRUE | 0.2052 |
| Rab25 | 34 | 36 | 51 | 35 | 38 | 53 | 36 | 36 | 51 | 28 | 41 | 58 | 41.62 | 41 | 42 | 1.016 | 0.023 | 0.95122641 | 0.97528069 | 0.0675 | 0.0861 | 0.0769 | 0.0769 | TRUE | 0.2528 |
| Rab26 | 122 | 113 | 113 | 154 | 117 | 126 | 130 | 113 | 113 | 121 | 126 | 138 | 123.46 | 119 | 128 | 1.08 | 0.111 | 0.54195126 | 0.7276458 | 0 | 0.0336 | 0.0157 | 0.0157 | TRUE | 0.0644 |
| Rab27a | 1675 | 1802 | 1829 | 2482 | 1944 | 2052 | 1779 | 1800 | 1837 | 1951 | 2092 | 2246 | 1950.72 | 1805 | 2096 | 1.161 | 0.215 | 0.01059036 | 0.05499872 | 0.0022 | 0.0086 | 0.0046 | 0.0046 | TRUE | 0.0473 |
| Rab27b | 5 | 3 | 1 | 6 | 4 | 7 | 5 | 3 | 1 | 5 | 4 | 8 | 4.33 | 3 | 6 | 1.793 | 0.842 | 0.38144592 | NA | 0 | 0.7673 | 0.4134 | 0.4134 | TRUE | 0.3459 |
| Rab28 | 1012 | 1133 | 1108 | 1446 | 1026 | 1119 | 1075 | 1132 | 1113 | 1137 | 1104 | 1225 | 1130.76 | 1107 | 1155 | 1.044 | 0.062 | 0.4686541 | 0.6710497 | 8.00E-04 | 0.0098 | 0.0043 | 0.0043 | TRUE | 0.0325 |
| Rab2a | 3727 | 4284 | 4473 | 5472 | 3840 | 4115 | 3959 | 4278 | 4491 | 4302 | 4132 | 4505 | 4277.72 | 4243 | 4313 | 1.016 | 0.023 | 0.77281675 | 0.88133519 | 0.0027 | 0.0077 | 0.0045 | 0.0045 | TRUE | 0.0417 |
| Rab2b | 740 | 1076 | 1081 | 1111 | 883 | 994 | 786 | 1075 | 1085 | 873 | 950 | 1088 | 976.26 | 982 | 970 | 0.987 | -0.018 | 0.90499538 | 0.95269332 | 0.0212 | 0.0103 | 0.0159 | 0.0159 | TRUE | 0.3626 |
| Rab30 | 259 | 301 | 292 | 289 | 202 | 215 | 275 | 301 | 293 | 227 | 217 | 235 | 258.13 | 290 | 226 | 0.782 | -0.354 | 0.00653629 | 0.03822575 | 0 | 0.0197 | 0.0083 | 0.0083 | TRUE | 0.0214 |
| Rab31 | 2411 | 2066 | 2049 | 2295 | 1677 | 1749 | 2561 | 2063 | 2057 | 1804 | 1804 | 1915 | 2034.12 | 2227 | 1841 | 0.827 | -0.275 | 0.01310803 | 0.06416507 | 0.0083 | 0.0085 | 0.0083 | 0.0083 | TRUE | 0.2076 |
| Rab32 | 66 | 72 | 84 | 124 | 109 | 103 | 70 | 72 | 84 | 97 | 117 | 113 | 92.31 | 75 | 109 | 1.441 | 0.528 | 0.01347754 | 0.06536524 | 0 | 0.0426 | 0.0216 | 0.0216 | TRUE | 0.0981 |
| Rab33a | 114 | 96 | 126 | 116 | 92 | 86 | 121 | 96 | 127 | 91 | 99 | 94 | 104.63 | 115 | 95 | 0.827 | -0.274 | 0.182194 | 0.38410413 | 0 | 0.0384 | 0.0208 | 0.0208 | TRUE | 0.2054 |
| Rab33b | 292 | 319 | 284 | 503 | 316 | 313 | 310 | 319 | 285 | 395 | 340 | 343 | 331.99 | 305 | 359 | 1.182 | 0.242 | 0.0637882 | 0.19410909 | 0.003 | 0.0168 | 0.0092 | 0.0092 | TRUE | 0.0934 |
| Rab34 | 902 | 1211 | 1289 | 1389 | 996 | 1138 | 958 | 1209 | 1294 | 1092 | 1072 | 1246 | 1145.18 | 1154 | 1137 | 0.984 | -0.023 | 0.86609627 | 0.93370586 | 0.0145 | 0.0098 | 0.0123 | 0.0123 | TRUE | 0.2632 |
| Rab35 | 3732 | 3246 | 3063 | 4750 | 3403 | 3131 | 3964 | 3242 | 3076 | 3734 | 3661 | 3427 | 3517.38 | 3427 | 3607 | 1.053 | 0.074 | 0.51156015 | 0.7050268 | 0.0085 | 0.0078 | 0.009 | 0.009 | TRUE | 0.2281 |
| Rab36 | 7 | 10 | 6 | 19 | 8 | 6 | 7 | 10 | 6 | 15 | 9 | 7 | 8.93 | 8 | 10 | 1.305 | 0.384 | 0.56513645 | 0.74551564 | 0 | 0.376 | 0.2073 | 0.2073 | TRUE | 0.7872 |
| Rab37 | 44 | 64 | 68 | 69 | 49 | 46 | 47 | 64 | 68 | 54 | 53 | 50 | 56.04 | 60 | 52 | 0.879 | -0.186 | 0.4943357 | 0.69052608 | 0 | 0.0657 | 0.0354 | 0.0354 | TRUE | 0.2992 |
| Rab38 | 36 | 37 | 22 | 117 | 60 | 42 | 38 | 37 | 22 | 92 | 65 | 46 | 49.96 | 32 | 68 | 2.096 | 1.068 | 0.00422922 | 0.02756336 | 0.0823 | 0.0728 | 0.0774 | 0.0774 | TRUE | 0.1836 |
| Rab39 | 88 | 168 | 154 | 209 | 182 | 191 | 93 | 168 | 155 | 164 | 196 | 209 | 164.18 | 139 | 190 | 1.365 | 0.449 | 0.06251382 | 0.19148727 | 0.0451 | 0.027 | 0.0355 | 0.0355 | TRUE | 0.8276 |
| Rab39b | 35 | 33 | 48 | 60 | 39 | 52 | 37 | 33 | 48 | 47 | 42 | 57 | 44.06 | 39 | 49 | 1.232 | 0.301 | 0.33129428 | 0.55212268 | 0 | 0.0817 | 0.0464 | 0.0464 | TRUE | 0.2848 |
| Rab3a | 278 | 491 | 436 | 351 | 287 | 332 | 295 | 490 | 438 | 276 | 309 | 363 | 361.93 | 408 | 316 | 0.774 | -0.37 | 0.07269495 | 0.21109955 | 0.0403 | 0.016 | 0.0279 | 0.0279 | TRUE | 0.663 |
| Rab3b | 516 | 408 | 359 | 477 | 350 | 328 | 548 | 407 | 360 | 375 | 377 | 359 | 404.44 | 438 | 370 | 0.845 | -0.244 | 0.15339368 | 0.34502219 | 0.0214 | 0.015 | 0.0185 | 0.0185 | TRUE | 0.5479 |
| Rab3c | 6 | 1 | 1 | 5 | 4 | 2 | 6 | 1 | 1 | 4 | 4 | 2 | 3.13 | 3 | 3 | 1.262 | 0.335 | 0.77444636 | NA | 0.2411 | 1.0583 | 0.662 | 0.662 | TRUE | 1.4611 |
| Rab3d | 815 | 957 | 994 | 835 | 617 | 680 | 866 | 956 | 998 | 656 | 664 | 744 | 814.03 | 940 | 688 | 0.731 | -0.451 | 2.08E-05 | 0.00036771 | 0.0038 | 0.011 | 0.0068 | 0.0068 | TRUE | 0.0591 |
| Rab3gap1 | 3679 | 3275 | 3200 | 4586 | 3342 | 3196 | 3908 | 3271 | 3213 | 3605 | 3596 | 3499 | 3515.19 | 3464 | 3567 | 1.03 | 0.042 | 0.66536784 | 0.8127356 | 0.0058 | 0.0078 | 0.0066 | 0.0066 | TRUE | 0.1525 |
| Rab3gap2 | 5026 | 5308 | 5825 | 6054 | 4436 | 4618 | 5339 | 5301 | 5849 | 4759 | 4773 | 5055 | 5179.29 | 5496 | 4862 | 0.885 | -0.177 | 0.0180393 | 0.08066795 | 0.0018 | 0.0075 | 0.0038 | 0.0038 | TRUE | 0.0385 |
| Rab3il1 | 152 | 494 | 533 | 199 | 163 | 226 | 161 | 493 | 535 | 156 | 175 | 247 | 294.87 | 396 | 193 | 0.486 | -1.04 | 0.05414915 | 0.17296651 | 0.2063 | 0.0181 | 0.0966 | 0.2063 | TRUE | 3.0873 |
| Rab3ip | 4202 | 4426 | 4436 | 4735 | 3551 | 3672 | 4463 | 4420 | 4454 | 3722 | 3821 | 4020 | 4150.05 | 4446 | 3854 | 0.867 | -0.206 | 0.00146029 | 0.01200817 | 4.00E-04 | 0.0077 | 0.0028 | 0.0028 | TRUE | 0.0171 |
| Rab40b | 22 | 9 | 15 | 22 | 14 | 17 | 23 | 9 | 15 | 17 | 15 | 19 | 16.4 | 16 | 17 | 1.079 | 0.109 | 0.83228324 | 0.91488509 | 0.0415 | 0.2078 | 0.131 | 0.131 | TRUE | 0.298 |
| Rab40c | 1777 | 1933 | 1794 | 2125 | 1511 | 1594 | 1887 | 1931 | 1801 | 1671 | 1626 | 1745 | 1776.75 | 1873 | 1681 | 0.897 | -0.157 | 0.03864022 | 0.1372598 | 6.00E-04 | 0.0088 | 0.0036 | 0.0036 | TRUE | 0.0136 |
| Rab42 | 5 | 12 | 13 | 10 | 13 | 3 | 5 | 12 | 13 | 8 | 14 | 3 | 9.25 | 10 | 8 | 0.825 | -0.277 | 0.70633551 | 0.8396577 | 0.183 | 0.3632 | 0.2799 | 0.2799 | TRUE | 0.3066 |
| Rab43 | 237 | 219 | 221 | 459 | 341 | 219 | 252 | 219 | 222 | 361 | 367 | 240 | 276.64 | 231 | 323 | 1.4 | 0.485 | 0.0116922 | 0.05908087 | 0.0271 | 0.0188 | 0.023 | 0.023 | TRUE | 0.5595 |
| Rab44 | 51 | 82 | 72 | 62 | 60 | 63 | 54 | 82 | 72 | 49 | 65 | 69 | 65.1 | 69 | 61 | 0.868 | -0.203 | 0.4662051 | 0.66857491 | 0.0231 | 0.0575 | 0.0407 | 0.0407 | TRUE | 0.3244 |
| Rab4a | 1591 | 1568 | 1500 | 1974 | 1268 | 1283 | 1690 | 1566 | 1506 | 1552 | 1364 | 1404 | 1513.77 | 1587 | 1440 | 0.908 | -0.139 | 0.13788234 | 0.32222259 | 0.0031 | 0.0091 | 0.0057 | 0.0057 | TRUE | 0.0567 |
| Rab4b | 1078 | 1360 | 1382 | 1269 | 975 | 1072 | 1145 | 1358 | 1388 | 998 | 1049 | 1173 | 1185.17 | 1297 | 1073 | 0.827 | -0.274 | 0.01754854 | 0.07905614 | 0.0075 | 0.0097 | 0.0088 | 0.0088 | TRUE | 0.1256 |
| Rab5a | 3510 | 3411 | 3267 | 4892 | 3504 | 3391 | 3728 | 3407 | 3280 | 3846 | 3770 | 3712 | 3623.84 | 3472 | 3776 | 1.088 | 0.121 | 0.11828323 | 0.2910395 | 0.002 | 0.0078 | 0.0041 | 0.0041 | TRUE | 0.0507 |
| Rab5b | 2821 | 3335 | 3281 | 3231 | 2395 | 2821 | 2996 | 3331 | 3294 | 2540 | 2577 | 3088 | 2971.07 | 3207 | 2735 | 0.852 | -0.23 | 0.0282916 | 0.11058257 | 0.0074 | 0.008 | 0.0076 | 0.0076 | TRUE | 0.1541 |
| Rab5c | 6999 | 6899 | 6957 | 9064 | 6422 | 6461 | 7434 | 6890 | 6986 | 7125 | 6910 | 7073 | 7069.58 | 7103 | 7036 | 0.991 | -0.014 | 0.83215543 | 0.91487212 | 7.00E-04 | 0.0074 | 0.0028 | 0.0028 | TRUE | 0.0202 |
| Rab6a | 5620 | 6775 | 6801 | 8394 | 5916 | 6393 | 5969 | 6766 | 6829 | 6599 | 6365 | 6998 | 6587.79 | 6521 | 6654 | 1.02 | 0.029 | 0.73535002 | 0.85804993 | 0.0038 | 0.0074 | 0.0051 | 0.0051 | TRUE | 0.0668 |
| Rab6b | 1009 | 1166 | 1211 | 1256 | 983 | 1299 | 1072 | 1164 | 1216 | 987 | 1058 | 1422 | 1153.2 | 1151 | 1156 | 1.003 | 0.005 | 0.9741627 | 0.98667442 | 0.0204 | 0.0098 | 0.0153 | 0.0153 | TRUE | 0.4849 |
| Rab7 | 1816 | 1902 | 2003 | 2825 | 1936 | 2021 | 1929 | 1900 | 2011 | 2221 | 2083 | 2212 | 2059.3 | 1947 | 2172 | 1.116 | 0.158 | 0.03004428 | 0.11540632 | 6.00E-04 | 0.0085 | 0.0034 | 0.0034 | TRUE | 0.0154 |
| Rab7l1 | 170 | 231 | 250 | 345 | 241 | 228 | 181 | 231 | 251 | 271 | 259 | 250 | 240.4 | 221 | 260 | 1.178 | 0.236 | 0.14921505 | 0.33959881 | 0.0097 | 0.0206 | 0.0151 | 0.0151 | TRUE | 0.2744 |
| Rab8a | 3338 | 2990 | 3011 | 4810 | 3467 | 2969 | 3546 | 2986 | 3023 | 3781 | 3730 | 3250 | 3386.1 | 3185 | 3587 | 1.127 | 0.172 | 0.10519651 | 0.27035349 | 0.0077 | 0.0079 | 0.0078 | 0.0078 | TRUE | 0.119 |

| | | | | | | | | | | | | | | | | | | | | | | | | | |
|---|---|---|---|---|---|---|---|---|---|---|---|---|---|---|---|---|---|---|---|---|---|---|---|---|---|
| Rab8b | 1447 | 1542 | 1572 | 1825 | 1361 | 1310 | 1537 | 1540 | 1578 | 1435 | 1464 | 1434 | 1498.08 | 1552 | 1444 | 0.931 | -0.104 | 0.14066501 | 0.32593224 | 0 | 0.0091 | 0.0029 | 0.0029 | TRUE | 0.0027 |
| Rab9 | 2645 | 2889 | 2963 | 2760 | 1948 | 2422 | 2809 | 2885 | 2975 | 2170 | 2096 | 2651 | 2597.8 | 2890 | 2306 | 0.798 | -0.326 | 0.00269595 | 0.01950482 | 0.0084 | 0.0082 | 0.0081 | 0.0081 | TRUE | 0.207 |
| Rab9b | 7 | 16 | 23 | 25 | 17 | 15 | 7 | 16 | 23 | 20 | 18 | 16 | 16.81 | 15 | 18 | 1.168 | 0.224 | 0.67123036 | 0.81655574 | 0.0656 | 0.2029 | 0.1414 | 0.1414 | TRUE | 0.2662 |
| Rabac1 | 2374 | 3146 | 3018 | 3155 | 2312 | 2747 | 2522 | 3142 | 3030 | 2480 | 2488 | 3007 | 2778.12 | 2898 | 2658 | 0.917 | -0.125 | 0.30975914 | 0.53005547 | 0.0123 | 0.0081 | 0.0106 | 0.0106 | TRUE | 0.1592 |
| Rabep1 | 8565 | 8800 | 9104 | 10617 | 7658 | 7181 | 9098 | 8789 | 9141 | 8346 | 8240 | 7861 | 8579.01 | 9009 | 8149 | 0.905 | -0.145 | 0.01815597 | 0.08114364 | 5.00E-04 | 0.0073 | 0.0026 | 0.0026 | TRUE | 0.0116 |
| Rabep2 | 1062 | 1100 | 985 | 1502 | 1141 | 1093 | 1128 | 1099 | 989 | 1181 | 1228 | 1196 | 1136.75 | 1072 | 1202 | 1.121 | 0.165 | 0.06507167 | 0.19649825 | 0.0016 | 0.0098 | 0.0049 | 0.0049 | TRUE | 0.0548 |
| Rabepk | 1117 | 1432 | 1313 | 1491 | 1111 | 1171 | 1186 | 1430 | 1318 | 1172 | 1195 | 1282 | 1264.05 | 1311 | 1216 | 0.927 | -0.11 | 0.28136148 | 0.50267709 | 0.0043 | 0.0095 | 0.0067 | 0.0067 | TRUE | 0.0834 |
| Rabgap1 | 3610 | 3768 | 3823 | 4538 | 3198 | 3307 | 3834 | 3763 | 3839 | 3567 | 3441 | 3620 | 3677.43 | 3812 | 3543 | 0.929 | -0.106 | 0.08792805 | 0.24030519 | 0 | 0.0078 | 0.0025 | 0.0025 | TRUE | 0.0076 |
| Rabgap1l | 740 | 1014 | 1044 | 965 | 720 | 825 | 786 | 1013 | 1048 | 759 | 775 | 903 | 880.56 | 949 | 812 | 0.855 | -0.226 | 0.10853332 | 0.27600573 | 0.0143 | 0.0106 | 0.0131 | 0.0131 | TRUE | 0.2681 |
| Rabgef1 | 965 | 1125 | 1010 | 1422 | 1032 | 1054 | 1025 | 1124 | 1014 | 1118 | 1110 | 1154 | 1090.78 | 1054 | 1127 | 1.069 | 0.096 | 0.26243751 | 0.48186844 | 8.00E-04 | 0.0099 | 0.0044 | 0.0044 | TRUE | 0.0398 |
| Rabggta | 2301 | 2361 | 2307 | 2642 | 2009 | 2023 | 2444 | 2358 | 2316 | 2077 | 2162 | 2214 | 2261.91 | 2373 | 2151 | 0.906 | -0.142 | 0.04439617 | 0.15134402 | 5.00E-04 | 0.0084 | 0.0032 | 0.0032 | TRUE | 0.0113 |
| Rabggtb | 4822 | 5595 | 5520 | 6853 | 4934 | 4813 | 5122 | 5588 | 5543 | 5387 | 5309 | 5269 | 5369.47 | 5418 | 5322 | 0.982 | -0.026 | 0.7030979 | 0.83762068 | 9.00E-04 | 0.0075 | 0.0031 | 0.0031 | TRUE | 0.0277 |
| Rabif | 2073 | 2232 | 2141 | 2691 | 1907 | 1967 | 2202 | 2229 | 2150 | 2115 | 2052 | 2153 | 2150.21 | 2194 | 2107 | 0.96 | -0.058 | 0.38945684 | 0.60356817 | 0 | 0.0084 | 0.0028 | 0.0028 | TRUE | 0.0062 |
| Rabl2 | 199 | 148 | 182 | 342 | 258 | 229 | 211 | 148 | 183 | 269 | 278 | 251 | 223.18 | 181 | 266 | 1.473 | 0.558 | 0.00105857 | 0.00930642 | 0.0106 | 0.0217 | 0.0163 | 0.0163 | TRUE | 0.273 |
| Rabl3 | 618 | 625 | 578 | 887 | 631 | 649 | 656 | 624 | 580 | 697 | 679 | 710 | 657.94 | 620 | 695 | 1.122 | 0.166 | 0.09007863 | 0.24423588 | 0 | 0.0119 | 0.0053 | 0.0053 | TRUE | 0.0374 |
| Rabl5 | 168 | 209 | 222 | 318 | 231 | 239 | 178 | 209 | 223 | 250 | 249 | 262 | 228.37 | 203 | 254 | 1.245 | 0.316 | 0.03240441 | 0.12140286 | 0 | 0.0213 | 0.0112 | 0.0112 | TRUE | 0.1228 |
| Rabl6 | 2887 | 4186 | 4353 | 4625 | 3543 | 3937 | 3067 | 4181 | 4371 | 3636 | 3812 | 4310 | 3895.92 | 3873 | 3919 | 1.012 | 0.017 | 0.90666182 | 0.95352883 | 0.0214 | 0.0077 | 0.0148 | 0.0148 | TRUE | 0.4031 |
| Rac1 | 8127 | 9869 | 10272 | 13712 | 9924 | 9963 | 8632 | 9856 | 10314 | 10779 | 10678 | 10906 | 10194.29 | 9601 | 10788 | 1.124 | 0.168 | 0.05222641 | 0.16900183 | 0.004 | 0.0072 | 0.0053 | 0.0053 | TRUE | 0.0951 |
| Rac2 | 115 | 117 | 90 | 47 | 30 | 49 | 122 | 117 | 90 | 37 | 32 | 54 | 75.37 | 110 | 41 | 0.371 | -1.429 | 2.89E-07 | 9.17E-06 | 0.0245 | 0.0506 | 0.0392 | 0.0392 | TRUE | 0.512 |
| Rac3 | 585 | 752 | 686 | 561 | 393 | 483 | 621 | 751 | 689 | 441 | 423 | 529 | 575.63 | 687 | 464 | 0.675 | -0.567 | 2.03E-05 | 0.00036149 | 0.0092 | 0.0126 | 0.011 | 0.011 | TRUE | 0.1675 |
| Racgap1 | 7193 | 7867 | 7761 | 8752 | 6434 | 6752 | 7640 | 7857 | 7793 | 6880 | 6923 | 7391 | 7413.97 | 7763 | 7065 | 0.91 | -0.136 | 0.03169033 | 0.11971732 | 7.00E-04 | 0.0073 | 0.0028 | 0.0028 | TRUE | 0.0199 |
| Rad1 | 1696 | 1785 | 1698 | 2160 | 1572 | 1567 | 1801 | 1783 | 1705 | 1698 | 1691 | 1715 | 1732.31 | 1763 | 1701 | 0.965 | -0.051 | 0.46663151 | 0.66912557 | 0 | 0.0088 | 0.003 | 0.003 | TRUE | 0.01 |
| Rad17 | 2574 | 2497 | 2473 | 3169 | 2162 | 2162 | 2734 | 2494 | 2483 | 2491 | 2326 | 2367 | 2482.5 | 2570 | 2395 | 0.932 | -0.101 | 0.19833401 | 0.4053588 | 0.0016 | 0.0082 | 0.0041 | 0.0041 | TRUE | 0.0375 |
| Rad18 | 2923 | 3211 | 3284 | 3744 | 2709 | 2831 | 3105 | 3207 | 3297 | 2943 | 2915 | 3099 | 3094.34 | 3203 | 2986 | 0.932 | -0.102 | 0.13900284 | 0.32369007 | 6.00E-04 | 0.008 | 0.0031 | 0.0031 | TRUE | 0.0132 |
| Rad21 | 19446 | 20207 | 20075 | 29230 | 20070 | 20434 | 20655 | 20181 | 20157 | 22978 | 21594 | 22368 | 21322.34 | 20331 | 22313 | 1.098 | 0.134 | 0.0198882 | 0.0864838 | 5.00E-04 | 0.0071 | 0.0024 | 0.0024 | TRUE | 0.0097 |
| Rad21l | 6 | 6 | 4 | 13 | 13 | 12 | 6 | 6 | 4 | 10 | 14 | 13 | 8.95 | 5 | 12 | 2.27 | 1.182 | 0.0684264 | 0.2027087 | 0 | 0.3748 | 0.1703 | 0.1703 | TRUE | 0.1253 |
| Rad23a | 4671 | 4966 | 5136 | 7314 | 5416 | 4914 | 4961 | 4960 | 5157 | 5750 | 5827 | 5379 | 5339.04 | 5026 | 5652 | 1.125 | 0.169 | 0.01167864 | 0.05903701 | 9.00E-04 | 0.0075 | 0.0031 | 0.0031 | TRUE | 0.0216 |
| Rad23b | 16995 | 16757 | 16644 | 23833 | 16591 | 15827 | 18052 | 16735 | 16712 | 18736 | 17851 | 17325 | 17568.51 | 17166 | 17971 | 1.047 | 0.066 | 0.34236702 | 0.56253908 | 0.0017 | 0.0071 | 0.0034 | 0.0034 | TRUE | 0.0249 |
| Rad50 | 12355 | 11796 | 12507 | 14326 | 10950 | 11065 | 13123 | 11781 | 12558 | 11262 | 11782 | 12112 | 12103.03 | 12487 | 11719 | 0.938 | -0.092 | 0.20939099 | 0.41942021 | 0.002 | 0.0072 | 0.0038 | 0.0038 | TRUE | 0.03 |
| Rad51 | 5092 | 5155 | 4781 | 6926 | 4881 | 4404 | 5409 | 5148 | 4801 | 5445 | 5252 | 4821 | 5145.8 | 5119 | 5173 | 1.011 | 0.015 | 0.85744674 | 0.92971607 | 0.0035 | 0.0075 | 0.005 | 0.005 | TRUE | 0.043 |
| Rad51ap1 | 1775 | 2017 | 2106 | 2732 | 2008 | 2137 | 1885 | 2014 | 2115 | 2148 | 2160 | 2339 | 2110.31 | 2005 | 2216 | 1.105 | 0.144 | 0.08693077 | 0.23841657 | 0.0021 | 0.0085 | 0.0046 | 0.0046 | TRUE | 0.0327 |
| Rad51ap2 | 19 | 12 | 12 | 23 | 16 | 14 | 20 | 12 | 12 | 18 | 17 | 15 | 15.81 | 15 | 17 | 1.151 | 0.203 | 0.67695038 | 0.82041351 | 0 | 0.2153 | 0.1079 | 0.1079 | TRUE | 0.4511 |
| Rad51b | 384 | 572 | 601 | 522 | 391 | 456 | 408 | 571 | 603 | 410 | 421 | 499 | 485.47 | 527 | 443 | 0.839 | -0.252 | 0.1441228 | 0.33125982 | 0.0251 | 0.0137 | 0.0194 | 0.0194 | TRUE | 0.4572 |
| Rad51c | 1331 | 1605 | 1620 | 2049 | 1552 | 1799 | 1414 | 1603 | 1627 | 1611 | 1670 | 1969 | 1648.87 | 1548 | 1750 | 1.13 | 0.176 | 0.11665734 | 0.28879604 | 0.008 | 0.0089 | 0.0085 | 0.0085 | TRUE | 0.1444 |
| Rad51d | 1484 | 2061 | 2007 | 2306 | 1766 | 1984 | 1576 | 2058 | 2015 | 1813 | 1900 | 2172 | 1922.43 | 1883 | 1962 | 1.041 | 0.058 | 0.6567259 | 0.80670515 | 0.0145 | 0.0086 | 0.0118 | 0.0118 | TRUE | 0.2452 |
| Rad52 | 593 | 632 | 656 | 828 | 599 | 550 | 630 | 631 | 659 | 651 | 644 | 602 | 636.2 | 640 | 632 | 0.989 | -0.016 | 0.86707 | 0.93417775 | 0 | 0.0121 | 0.0049 | 0.0049 | TRUE | 0.0203 |
| Rad54b | 2236 | 2657 | 2683 | 2610 | 2137 | 2244 | 2375 | 2654 | 2694 | 2052 | 2299 | 2456 | 2421.68 | 2574 | 2269 | 0.881 | -0.183 | 0.07037166 | 0.20653235 | 0.0057 | 0.0083 | 0.007 | 0.007 | TRUE | 0.0864 |
| Rad54l | 2708 | 3046 | 2893 | 4077 | 3072 | 2914 | 2876 | 3042 | 2905 | 3205 | 3305 | 3190 | 3087.24 | 2941 | 3233 | 1.099 | 0.136 | 0.03743734 | 0.13422636 | 3.00E-04 | 0.008 | 0.0028 | 0.0028 | TRUE | 0.011 |
| Rad54l2 | 6061 | 5410 | 6053 | 7513 | 5101 | 4671 | 6438 | 5403 | 6078 | 5906 | 5488 | 5113 | 5737.73 | 5973 | 5502 | 0.921 | -0.118 | 0.23092349 | 0.44486689 | 0.0062 | 0.0075 | 0.0068 | 0.0068 | TRUE | 0.085 |
| Rad9a | 1706 | 1809 | 1747 | 2402 | 1844 | 1787 | 1812 | 1807 | 1754 | 1888 | 1984 | 1956 | 1866.89 | 1791 | 1943 | 1.084 | 0.117 | 0.09340679 | 0.24989155 | 0 | 0.0087 | 0.003 | 0.003 | TRUE | 0.0075 |
| Rad9b | 273 | 382 | 414 | 577 | 417 | 425 | 290 | 382 | 416 | 454 | 449 | 465 | 409.11 | 363 | 456 | 1.257 | 0.33 | 0.03121792 | 0.11852889 | 0.0138 | 0.015 | 0.0144 | 0.0144 | TRUE | 0.3461 |
| Radil | 726 | 737 | 772 | 659 | 540 | 548 | 771 | 736 | 775 | 518 | 581 | 600 | 663.55 | 761 | 566 | 0.743 | -0.429 | 3.09E-05 | 0.00051633 | 0.0058 | 0.0119 | 0.0061 | 0.0061 | TRUE | 0.0691 |
| Rae1 | 6534 | 6936 | 6643 | 7611 | 5847 | 5691 | 6940 | 6927 | 6670 | 5983 | 6291 | 6230 | 6506.91 | 6846 | 6168 | 0.901 | -0.151 | 0.01335218 | 0.06494505 | 4.00E-04 | 0.0074 | 0.0025 | 0.0025 | TRUE | 0.0085 |
| Raet1a | 10 | 20 | 23 | 10 | 15 | 13 | 11 | 20 | 23 | 8 | 16 | 14 | 15.32 | 18 | 13 | 0.701 | -0.513 | 0.35305334 | 0.57159736 | 0.0759 | 0.2219 | 0.1522 | 0.1522 | TRUE | 0.615 |
| Raet1b | 0 | 0 | 0 | 0 | 0 | 0 | 0 | 0 | 0 | 0 | 0 | 0 | 0 | 0 | 0 | NA | NA | NA | NA | NA | NA | NA | NA | NA | NA |
| Raet1c | 0 | 1 | 0 | 0 | 1 | 0 | 0 | 1 | 0 | 0 | 1 | 0 | 0.35 | 0 | 0 | 0.954 | -0.067 | 0.98571955 | NA | 0 | 9.5346 | 8.2194 | 8.2194 | TRUE | 0.5204 |
| Raet1d | 8 | 11 | 9 | 18 | 4 | 15 | 8 | 11 | 9 | 14 | 4 | 16 | 10.57 | 9 | 11 | 1.229 | 0.297 | 0.6527873 | 0.80520057 | 0.1 | 0.3187 | 0.2183 | 0.2183 | TRUE | 1.0747 |
| Raet1e | 6 | 10 | 7 | 12 | 9 | 6 | 6 | 10 | 7 | 9 | 10 | 7 | 8.18 | 8 | 9 | 1.101 | 0.139 | 0.83125529 | 0.91439242 | 0 | 0.4097 | 0.1854 | 0.1854 | TRUE | 0.177 |
| Raf1 | 6420 | 7182 | 7541 | 11197 | 8277 | 7668 | 6819 | 7173 | 7572 | 8802 | 8906 | 8394 | 7944.24 | 7188 | 8701 | 1.21 | 0.276 | 0.00012132 | 0.0016107 | 0.0017 | 0.0073 | 0.0036 | 0.0036 | TRUE | 0.0267 |
| Rag1 | 1 | 0 | 0 | 0 | 0 | 0 | 1 | 0 | 0 | 0 | 0 | 0 | 0.18 | 0 | 0 | 0.49 | -1.029 | 0.80087 | NA | 0 | 18.6165 | 10 | 10 | TRUE | 0.4614 |
| Rag2 | 0 | 0 | 0 | 0 | 0 | 0 | 0 | 0 | 0 | 0 | 0 | 0 | 0 | 0 | 0 | NA | NA | NA | NA | NA | NA | NA | NA | NA | NA |
| Rai1 | 617 | 597 | 542 | 925 | 683 | 545 | 655 | 596 | 544 | 727 | 735 | 597 | 642.41 | 598 | 686 | 1.148 | 0.199 | 0.12603606 | 0.30427348 | 0.0093 | 0.012 | 0.0106 | 0.0106 | TRUE | 0.1513 |
| Rai14 | 6452 | 6644 | 6783 | 6306 | 4548 | 4750 | 6853 | 6635 | 6811 | 4957 | 4893 | 5200 | 5891.62 | 6766 | 5017 | 0.741 | -0.432 | 2.92E-12 | 2.83E-10 | 4.00E-04 | 0.0075 | 0.0026 | 0.0026 | TRUE | 0.0123 |
| Rai2 | 10 | 11 | 8 | 6 | 4 | 2 | 11 | 11 | 8 | 5 | 4 | 2 | 6.81 | 10 | 4 | 0.384 | -1.38 | 0.06675417 | 0.1993297 | 0 | 0.4908 | 0.2281 | 0.2281 | TRUE | 0.1807 |
| Rala | 2505 | 2512 | 2523 | 3553 | 2514 | 2548 | 2661 | 2509 | 2533 | 2793 | 2705 | 2789 | 2665.01 | 2568 | 2762 | 1.076 | 0.106 | 0.11483333 | 0.28611259 | 2.00E-04 | 0.0081 | 0.0029 | 0.0029 | TRUE | 0.0122 |
| Ralb | 1761 | 2175 | 2120 | 2445 | 1863 | 1937 | 1870 | 2172 | 2129 | 1922 | 2004 | 2120 | 2036.38 | 2057 | 2015 | 0.979 | -0.03 | 0.74576296 | 0.8648021 | 0.0039 | 0.0085 | 0.0057 | 0.0057 | TRUE | 0.0759 |
| Ralbp1 | 6812 | 7660 | 7631 | 8007 | 6139 | 6474 | 7236 | 7650 | 7662 | 6294 | 6605 | 7087 | 7089.08 | 7516 | 6662 | 0.886 | -0.174 | 0.02073844 | 0.08914875 | 0.0021 | 0.0074 | 0.004 | 0.004 | TRUE | 0.0379 |
| Ralgapa1 | 1949 | 2180 | 2358 | 2556 | 1854 | 1893 | 2070 | 2177 | 2368 | 2009 | 1995 | 2072 | 2115.23 | 2205 | 2025 | 0.918 | -0.123 | 0.13611382 | 0.31954629 | 0.0018 | 0.0085 | 0.0044 | 0.0044 | TRUE | 0.0505 |
| Ralgapa2 | 2303 | 2237 | 2367 | 2874 | 2131 | 2021 | 2446 | 2234 | 2377 | 2259 | 2293 | 2212 | 2303.58 | 2352 | 2255 | 0.959 | -0.061 | 0.40416891 | 0.61693403 | 8.00E-04 | 0.0083 | 0.0034 | 0.0034 | TRUE | 0.0235 |
| Ralgapb | 6837 | 6667 | 6841 | 7757 | 5491 | 5650 | 7262 | 6658 | 6869 | 6098 | 5908 | 6185 | 6496.72 | 6930 | 6064 | 0.875 | -0.193 | 0.00416883 | 0.0272825 | 0.001 | 0.0074 | 0.0031 | 0.0031 | TRUE | 0.0214 |
| Ralgds | 1543 | 1735 | 1682 | 2065 | 1557 | 1673 | 1639 | 1733 | 1689 | 1623 | 1675 | 1831 | 1698.42 | 1687 | 1710 | 1.013 | 0.019 | 0.82349452 | 0.9101083 | 0.0018 | 0.0088 | 0.0044 | 0.0044 | TRUE | 0.0462 |
| Ralgps1 | 619 | 702 | 740 | 789 | 531 | 561 | 657 | 701 | 743 | 620 | 571 | 614 | 651.22 | 700 | 602 | 0.86 | -0.218 | 0.03097341 | 0.11791279 | 0.0013 | 0.012 | 0.0058 | 0.0058 | TRUE | 0.0338 |
| Ralgps2 | 3264 | 3907 | 3990 | 3743 | 2965 | 3298 | 3467 | 3902 | 4006 | 2942 | 3190 | 3610 | 3519.68 | 3792 | 3247 | 0.856 | -0.224 | 0.03558572 | 0.12950116 | 0.0078 | 0.0078 | 0.0079 | 0.0079 | TRUE | 0.1158 |
| Raly | 12887 | 13405 | 13193 | 18687 | 13828 | 13072 | 13688 | 13388 | 13247 | 14690 | 14878 | 14309 | 14033.47 | 13441 | 14626 | 1.088 | 0.122 | 0.02715856 | 0.10754325 | 2.00E-04 | 0.0071 | 0.0021 | 0.0021 | TRUE | 0.0044 |
| Ralyl | 1 | 1 | 0 | 1 | 0 | 0 | 1 | 1 | 0 | 1 | 0 | 0 | 0.47 | 1 | 0 | 0.51 | -0.973 | 0.73452857 | NA | 0 | 6.9499 | 4.1835 | 4.1835 | TRUE | 0.2562 |
| Ramp1 | 4 | 3 | 0 | 10 | 13 | 2 | 4 | 3 | 0 | 8 | 14 | 2 | 5.21 | 2 | 8 | 3.338 | 1.739 | 0.12175232 | 0.29675128 | 0.6321 | 0.6388 | 0.6361 | 0.6361 | TRUE | 0.144 |
| Ramp2 | 55 | 82 | 90 | 52 | 40 | 50 | 58 | 82 | 90 | 41 | 43 | 55 | 61.56 | 77 | 46 | 0.598 | -0.742 | 0.00950413 | 0.0506245 | 0.0222 | 0.0604 | 0.0418 | 0.0418 | TRUE | 0.3976 |
| Ramp3 | 1011 | 973 | 852 | 1033 | 789 | 799 | 1074 | 972 | 855 | 812 | 849 | 875 | 906.12 | 967 | 845 | 0.874 | -0.195 | 0.08349481 | 0.23154066 | 0.0061 | 0.0105 | 0.008 | 0.008 | TRUE | 0.1216 |
| Ran | 29015 | 30481 | 30102 | 40535 | 27997 | 27598 | 30819 | 30442 | 30226 | 31865 | 30123 | 30210 | 30614.19 | 30496 | 30733 | 1.008 | 0.011 | 0.84307707 | 0.92105202 | 4.00E-04 | 0.007 | 0.0023 | 0.0023 | TRUE | 0.0128 |

| | | | | | | | | | | | | | | | | | | | | | | | | |
|---|---|---|---|---|---|---|---|---|---|---|---|---|---|---|---|---|---|---|---|---|---|---|---|---|
| Ranbp1 | 9912 | 9958 | 9957 | 15300 | 10948 | 9622 | 10528 | 9945 | 9998 | 12028 | 11779 | 10533 | 10801.86 | 10157 | 11447 | 1.127 | 0.173 | 0.02942905 | 0.11340213 | 0.0028 | 0.0072 | 0.0044 | 0.0044 | TRUE | 0.0595 |
| Ranbp10 | 3121 | 2247 | 2226 | 3796 | 2542 | 2034 | 3315 | 2244 | 2235 | 2984 | 2735 | 2227 | 2623.33 | 2598 | 2649 | 1.02 | 0.028 | 0.87273149 | 0.93739587 | 0.037 | 0.0082 | 0.022 | 0.022 | TRUE | 0.7063 |
| Ranbp17 | 6171 | 5655 | 5743 | 7805 | 5661 | 5476 | 6555 | 5648 | 5767 | 6136 | 6091 | 5994 | 6031.65 | 5990 | 6074 | 1.014 | 0.02 | 0.80629653 | 0.90019698 | 0.0032 | 0.0074 | 0.0047 | 0.0047 | TRUE | 0.083 |
| Ranbp2 | 15857 | 12917 | 14279 | 20669 | 14105 | 12727 | 16843 | 12900 | 14338 | 16248 | 15176 | 13932 | 14906.17 | 14694 | 15119 | 1.029 | 0.041 | 0.72451358 | 0.850916 | 0.0098 | 0.0071 | 0.0098 | 0.0098 | TRUE | 0.2002 |
| Ranbp3 | 3618 | 3768 | 3638 | 5181 | 3867 | 3897 | 3843 | 3763 | 3653 | 4073 | 4161 | 4266 | 3959.74 | 3753 | 4167 | 1.11 | 0.151 | 0.01756278 | 0.07908012 | 3.00E-04 | 0.0077 | 0.0026 | 0.0026 | TRUE | 0.0066 |
| Ranbp3l | 0 | 0 | 0 | 1 | 0 | 0 | 0 | 0 | 0 | 1 | 0 | 0 | 0.13 | 0 | 0 | 1.859 | 0.894 | 0.82650734 | NA | 0 | 25.1515 | 10 | 10 | TRUE | 0.2563 |
| Ranbp6 | 479 | 427 | 418 | 684 | 520 | 412 | 509 | 426 | 420 | 538 | 559 | 451 | 483.86 | 452 | 516 | 1.144 | 0.194 | 0.16094243 | 0.35561557 | 0.01 | 0.0137 | 0.0117 | 0.0117 | TRUE | 0.1409 |
| Ranbp9 | 2266 | 2490 | 2468 | 3244 | 2506 | 2425 | 2407 | 2487 | 2478 | 2550 | 2696 | 2655 | 2545.47 | 2457 | 2634 | 1.071 | 0.099 | 0.13713028 | 0.32113122 | 0 | 0.0082 | 0.0028 | 0.0028 | TRUE | 0.0096 |
| Rangap1 | 15966 | 15971 | 15866 | 19405 | 13877 | 13478 | 16959 | 15950 | 15931 | 15255 | 14931 | 14754 | 15629.92 | 16280 | 14980 | 0.92 | -0.12 | 0.0475532 | 0.15835777 | 7.00E-04 | 0.0071 | 0.0026 | 0.0026 | TRUE | 0.0162 |
| Rangrf | 2588 | 2903 | 2806 | 3019 | 2220 | 2447 | 2749 | 2899 | 2818 | 2373 | 2389 | 2679 | 2651.03 | 2822 | 2480 | 0.879 | -0.187 | 0.02229763 | 0.09405678 | 0.0022 | 0.0081 | 0.0044 | 0.0044 | TRUE | 0.0589 |
| Rap1a | 2005 | 2489 | 2389 | 2623 | 1884 | 2191 | 2130 | 2486 | 2399 | 2062 | 2027 | 2398 | 2250.29 | 2338 | 2162 | 0.925 | -0.113 | 0.2848263 | 0.50626428 | 0.007 | 0.0084 | 0.0076 | 0.0076 | TRUE | 0.1095 |
| Rap1b | 6992 | 7022 | 6468 | 9486 | 6561 | 6100 | 7427 | 7013 | 6495 | 7457 | 7059 | 6677 | 7021.34 | 6978 | 7064 | 1.013 | 0.018 | 0.83103192 | 0.91427424 | 0.0036 | 0.0074 | 0.005 | 0.005 | TRUE | 0.0449 |
| Rap1gap | 1567 | 1437 | 1465 | 1461 | 1090 | 1095 | 1664 | 1435 | 1471 | 1149 | 1173 | 1199 | 1348.42 | 1523 | 1174 | 0.77 | -0.377 | 4.28E-05 | 0.0006781 | 0.0028 | 0.0093 | 0.0054 | 0.0054 | TRUE | 0.0785 |
| Rap1gap2 | 1476 | 1684 | 1638 | 1062 | 848 | 950 | 1568 | 1682 | 1645 | 835 | 912 | 1040 | 1280.25 | 1632 | 929 | 0.569 | -0.814 | 1.82E-14 | 2.30E-12 | 0.0056 | 0.0095 | 0.0073 | 0.0073 | TRUE | 0.1293 |
| Rap1gds1 | 3278 | 3889 | 4386 | 5145 | 3731 | 3999 | 3482 | 3884 | 4404 | 4045 | 4014 | 4378 | 4034.38 | 3923 | 4146 | 1.056 | 0.079 | 0.45223024 | 0.65768329 | 0.0069 | 0.0077 | 0.0078 | 0.0078 | TRUE | 0.1399 |
| Rap2a | 759 | 955 | 980 | 1290 | 861 | 944 | 806 | 954 | 984 | 1014 | 926 | 1033 | 952.97 | 915 | 991 | 1.084 | 0.116 | 0.29935179 | 0.51963201 | 0.0061 | 0.0104 | 0.0079 | 0.0079 | TRUE | 0.1274 |
| Rap2b | 1027 | 1053 | 1012 | 1760 | 1214 | 1184 | 1091 | 1052 | 1016 | 1384 | 1306 | 1296 | 1190.75 | 1053 | 1329 | 1.263 | 0.337 | 3.82E-05 | 0.00061465 | 0 | 0.0097 | 0.004 | 0.004 | TRUE | 0.0163 |
| Rap2c | 1334 | 1506 | 1463 | 2171 | 1555 | 1513 | 1417 | 1504 | 1469 | 1707 | 1673 | 1656 | 1571 | 1463 | 1679 | 1.147 | 0.198 | 0.0062844 | 0.03729025 | 0 | 0.009 | 0.0032 | 0.0032 | TRUE | 0.0092 |
| Rapgef1 | 7351 | 6775 | 7092 | 8025 | 5511 | 5737 | 7808 | 6766 | 7121 | 6309 | 5929 | 6280 | 6702.27 | 7232 | 6173 | 0.854 | -0.228 | 0.00514264 | 0.03195814 | 0.003 | 0.0074 | 0.0046 | 0.0046 | TRUE | 0.0592 |
| Rapgef2 | 1287 | 1165 | 1216 | 1631 | 1152 | 1151 | 1367 | 1163 | 1221 | 1282 | 1239 | 1260 | 1255.52 | 1250 | 1260 | 1.008 | 0.012 | 0.89602216 | 0.9496715 | 0.0028 | 0.0095 | 0.0055 | 0.0055 | TRUE | 0.0793 |
| Rapgef3 | 369 | 565 | 572 | 299 | 259 | 303 | 392 | 564 | 574 | 235 | 279 | 332 | 395.99 | 510 | 282 | 0.551 | -0.86 | 1.02E-05 | 0.00020168 | 0.0343 | 0.0152 | 0.0246 | 0.0246 | TRUE | 0.4769 |
| Rapgef4 | 57 | 50 | 55 | 49 | 37 | 40 | 61 | 50 | 55 | 39 | 40 | 44 | 47.97 | 55 | 41 | 0.736 | -0.443 | 0.10934241 | 0.27739239 | 0 | 0.0756 | 0.0339 | 0.0339 | TRUE | 0.0596 |
| Rapgef5 | 1321 | 1285 | 1318 | 1561 | 1151 | 1232 | 1403 | 1283 | 1323 | 1227 | 1238 | 1349 | 1304.01 | 1336 | 1271 | 0.951 | -0.073 | 0.40063113 | 0.61357733 | 0.0016 | 0.0094 | 0.0046 | 0.0046 | TRUE | 0.0334 |
| Rapgef6 | 4960 | 4303 | 4261 | 5397 | 3637 | 3452 | 5268 | 4297 | 4278 | 4243 | 3913 | 3779 | 4296.5 | 4614 | 3978 | 0.862 | -0.214 | 0.04954717 | 0.16329979 | 0.009 | 0.0077 | 0.0083 | 0.0083 | TRUE | 0.1868 |
| Rapgefl1 | 353 | 372 | 357 | 378 | 274 | 271 | 375 | 372 | 358 | 297 | 295 | 297 | 332.26 | 368 | 296 | 0.804 | -0.314 | 0.00598425 | 0.03587208 | 0 | 0.0168 | 0.0064 | 0.0064 | TRUE | 0.0063 |
| Raph1 | 2885 | 3134 | 3619 | 3451 | 2842 | 2734 | 3064 | 3130 | 3634 | 2713 | 3058 | 2993 | 3098.62 | 3276 | 2921 | 0.891 | -0.166 | 0.09639423 | 0.25485389 | 0.006 | 0.008 | 0.0069 | 0.0069 | TRUE | 0.111 |
| Rapsn | 29 | 38 | 29 | 75 | 52 | 45 | 31 | 38 | 29 | 59 | 56 | 49 | 43.67 | 33 | 55 | 1.681 | 0.749 | 0.01194914 | 0.05999451 | 0 | 0.0823 | 0.0394 | 0.0394 | TRUE | 0.1432 |
| Rara | 4070 | 5405 | 5473 | 3263 | 2674 | 2894 | 4323 | 5398 | 5495 | 2565 | 2877 | 3168 | 3971.11 | 5072 | 2870 | 0.566 | -0.822 | 6.26E-11 | 4.77E-09 | 0.0133 | 0.0077 | 0.0111 | 0.0111 | TRUE | 0.2033 |
| Rarb | 20 | 12 | 10 | 32 | 17 | 8 | 21 | 12 | 10 | 25 | 18 | 9 | 15.91 | 14 | 17 | 1.223 | 0.291 | 0.61626707 | 0.78303103 | 0.143 | 0.2139 | 0.1795 | 0.1795 | TRUE | 0.2366 |
| Rarg | 7253 | 5868 | 6582 | 8645 | 5641 | 4366 | 7704 | 5860 | 6609 | 6796 | 6069 | 4779 | 6303.01 | 6724 | 5881 | 0.875 | -0.193 | 0.2008299 | 0.40850494 | 0.0243 | 0.0074 | 0.0162 | 0.0162 | TRUE | 0.3275 |
| Rarres1 | 4 | 5 | 5 | 3 | 3 | 1 | 4 | 5 | 5 | 2 | 3 | 1 | 3.49 | 5 | 2 | 0.47 | -1.088 | 0.28142399 | NA | 0 | 0.9507 | 0.4108 | 0.4108 | TRUE | 0.1639 |
| Rarres2 | 2 | 0 | 1 | 1 | 2 | 3 | 2 | 0 | 1 | 1 | 2 | 3 | 1.56 | 1 | 2 | 1.969 | 0.977 | 0.54366999 | NA | 0 | 2.1209 | 1.1308 | 1.1308 | TRUE | 0.2762 |
| Rars | 10295 | 10034 | 9998 | 12915 | 9105 | 8667 | 10935 | 10021 | 10039 | 10153 | 9796 | 9487 | 10071.96 | 10332 | 9812 | 0.95 | -0.074 | 0.2959551 | 0.51663572 | 0.0017 | 0.0072 | 0.0035 | 0.0035 | TRUE | 0.0319 |
| Rars2 | 1710 | 1801 | 1748 | 2425 | 1696 | 1525 | 1816 | 1799 | 1755 | 1906 | 1825 | 1669 | 1795.11 | 1790 | 1800 | 1.006 | 0.009 | 0.91246386 | 0.95675253 | 0.0019 | 0.0087 | 0.0044 | 0.0044 | TRUE | 0.0484 |
| Rasa1 | 2609 | 2813 | 2936 | 4222 | 3034 | 2998 | 2771 | 2809 | 2948 | 3319 | 3264 | 3282 | 3065.64 | 2843 | 3288 | 1.157 | 0.21 | 0.00122623 | 0.01041468 | 2.00E-04 | 0.008 | 0.0027 | 0.0027 | TRUE | 0.0127 |
| Rasa2 | 1271 | 1273 | 1248 | 1641 | 1292 | 1274 | 1350 | 1271 | 1253 | 1290 | 1390 | 1395 | 1324.87 | 1291 | 1358 | 1.051 | 0.072 | 0.38667372 | 0.60061236 | 0.0011 | 0.0094 | 0.0042 | 0.0042 | TRUE | 0.0241 |
| Rasa3 | 1351 | 1575 | 1570 | 2209 | 1524 | 1540 | 1435 | 1573 | 1576 | 1737 | 1640 | 1686 | 1607.74 | 1528 | 1688 | 1.104 | 0.143 | 0.07592326 | 0.21724631 | 0.0012 | 0.009 | 0.0041 | 0.0041 | TRUE | 0.034 |
| Rasa4 | 881 | 1036 | 1076 | 728 | 670 | 730 | 936 | 1035 | 1080 | 572 | 721 | 799 | 857.19 | 1017 | 697 | 0.684 | -0.547 | 0.00011595 | 0.00154978 | 0.0145 | 0.0107 | 0.0133 | 0.0133 | TRUE | 0.3002 |
| Rasal1 | 564 | 641 | 621 | 653 | 566 | 606 | 599 | 640 | 624 | 513 | 609 | 663 | 608.08 | 621 | 595 | 0.956 | -0.064 | 0.6080251 | 0.77782236 | 0.0075 | 0.0123 | 0.0096 | 0.0096 | TRUE | 0.1776 |
| Rasal2 | 1805 | 1830 | 2097 | 2156 | 1597 | 1585 | 1917 | 1828 | 2106 | 1695 | 1718 | 1735 | 1833.11 | 1950 | 1716 | 0.88 | -0.185 | 0.02936233 | 0.11328342 | 0.0022 | 0.0087 | 0.0046 | 0.0046 | TRUE | 0.059 |
| Rasal3 | 56 | 64 | 69 | 71 | 57 | 41 | 59 | 64 | 69 | 56 | 61 | 45 | 59.12 | 64 | 54 | 0.842 | -0.248 | 0.33795176 | 0.55812432 | 0 | 0.0626 | 0.0314 | 0.0314 | TRUE | 0.1661 |
| Rasd1 | 132 | 183 | 210 | 260 | 178 | 229 | 140 | 183 | 211 | 204 | 192 | 251 | 196.74 | 178 | 216 | 1.209 | 0.274 | 0.17426886 | 0.37354831 | 0.0248 | 0.0236 | 0.0241 | 0.0241 | TRUE | 0.3635 |
| Rasd2 | 392 | 447 | 430 | 575 | 427 | 395 | 416 | 446 | 432 | 452 | 459 | 432 | 439.73 | 431 | 448 | 1.038 | 0.054 | 0.60838594 | 0.77822084 | 0 | 0.0144 | 0.0057 | 0.0057 | TRUE | 0.0108 |
| Rasef | 26 | 44 | 20 | 60 | 41 | 42 | 28 | 44 | 20 | 47 | 44 | 46 | 38.15 | 31 | 46 | 1.498 | 0.583 | 0.1115151 | 0.28087132 | 0.0393 | 0.0933 | 0.0692 | 0.0692 | TRUE | 0.8459 |
| Rasgef1a | 171 | 283 | 296 | 212 | 209 | 181 | 182 | 283 | 297 | 167 | 225 | 198 | 225.19 | 254 | 197 | 0.772 | -0.373 | 0.09353571 | 0.25004155 | 0.0411 | 0.0215 | 0.0311 | 0.0311 | TRUE | 0.6826 |
| Rasgef1b | 109 | 148 | 137 | 186 | 157 | 134 | 116 | 148 | 138 | 146 | 169 | 147 | 143.83 | 134 | 154 | 1.149 | 0.2 | 0.2706988 | 0.49074002 | 0.0032 | 0.0298 | 0.0168 | 0.0168 | TRUE | 0.1389 |
| Rasgef1c | 87 | 74 | 72 | 170 | 118 | 98 | 92 | 74 | 72 | 134 | 127 | 107 | 101.08 | 79 | 123 | 1.547 | 0.63 | 0.00337106 | 0.02316704 | 0.0049 | 0.0395 | 0.0229 | 0.0229 | TRUE | 0.1818 |
| Rasgrf1 | 5 | 4 | 3 | 16 | 10 | 7 | 5 | 4 | 3 | 13 | 11 | 8 | 7.22 | 4 | 11 | 2.542 | 1.346 | 0.06730606 | 0.20047213 | 0 | 0.4632 | 0.2183 | 0.2183 | TRUE | 0.1677 |
| Rasgrf2 | 29 | 32 | 28 | 58 | 48 | 39 | 31 | 32 | 28 | 46 | 52 | 43 | 38.47 | 30 | 47 | 1.539 | 0.622 | 0.04350155 | 0.14928338 | 0 | 0.0925 | 0.0411 | 0.0411 | TRUE | 0.0648 |
| Rasgrp1 | 230 | 370 | 471 | 202 | 126 | 227 | 244 | 370 | 473 | 159 | 136 | 248 | 271.6 | 362 | 181 | 0.499 | -1.004 | 0.00043402 | 0.0046409 | 0.098 | 0.019 | 0.0545 | 0.0545 | TRUE | 0.1734 |
| Rasgrp2 | 796 | 1032 | 1054 | 961 | 788 | 908 | 845 | 1031 | 1058 | 755 | 848 | 994 | 921.96 | 978 | 866 | 0.884 | -0.178 | 0.20748156 | 0.41670688 | 0.015 | 0.0105 | 0.0133 | 0.0133 | TRUE | 0.199 |
| Rasgrp3 | 123 | 86 | 53 | 66 | 57 | 43 | 131 | 86 | 53 | 52 | 61 | 47 | 71.67 | 90 | 53 | 0.595 | -0.75 | 0.03209573 | 0.12067454 | 0.0948 | 0.0529 | 0.0734 | 0.0734 | TRUE | 0.1818 |
| Rasgrp4 | 399 | 819 | 840 | 675 | 642 | 606 | 424 | 818 | 843 | 531 | 691 | 663 | 661.66 | 695 | 628 | 0.903 | -0.147 | 0.54251107 | 0.72821145 | 0.0758 | 0.0119 | 0.0405 | 0.0405 | TRUE | 1.3727 |
| Rasip1 | 469 | 406 | 389 | 461 | 325 | 299 | 498 | 405 | 391 | 362 | 350 | 327 | 388.94 | 431 | 346 | 0.804 | -0.314 | 0.02364949 | 0.09783886 | 0.0078 | 0.0154 | 0.0113 | 0.0113 | TRUE | 0.2102 |
| Rasl10a | 100 | 103 | 95 | 141 | 113 | 122 | 106 | 103 | 95 | 111 | 122 | 134 | 111.74 | 101 | 122 | 1.199 | 0.261 | 0.17143615 | 0.37034626 | 0 | 0.0364 | 0.0173 | 0.0173 | TRUE | 0.0722 |
| Rasl10b | 147 | 261 | 239 | 292 | 178 | 170 | 156 | 261 | 240 | 230 | 192 | 186 | 210.66 | 219 | 203 | 0.926 | -0.112 | 0.60922645 | 0.77872758 | 0.037 | 0.0225 | 0.0296 | 0.0296 | TRUE | 0.6831 |
| Rasl11a | 286 | 253 | 208 | 329 | 214 | 179 | 304 | 253 | 209 | 259 | 230 | 196 | 241.69 | 255 | 228 | 0.897 | -0.156 | 0.41006718 | 0.62158382 | 0.0229 | 0.0205 | 0.0217 | 0.0217 | TRUE | 0.2025 |
| Rasl11b | 147 | 196 | 259 | 217 | 174 | 207 | 156 | 196 | 260 | 171 | 187 | 227 | 199.39 | 204 | 195 | 0.953 | -0.07 | 0.75266174 | 0.86948897 | 0.038 | 0.0234 | 0.0306 | 0.0306 | TRUE | 0.3522 |
| Rasl12 | 31 | 27 | 33 | 54 | 36 | 53 | 33 | 27 | 33 | 42 | 39 | 58 | 38.7 | 31 | 46 | 1.493 | 0.578 | 0.07917709 | 0.22317416 | 0 | 0.092 | 0.051 | 0.051 | TRUE | 0.3557 |
| Rasl2-9 | 16 | 12 | 11 | 23 | 14 | 13 | 17 | 12 | 11 | 18 | 15 | 14 | 14.57 | 13 | 16 | 1.193 | 0.254 | 0.60916867 | 0.77871682 | 0 | 0.2331 | 0.1093 | 0.1093 | TRUE | 0.2319 |
| Rassf1 | 399 | 499 | 658 | 534 | 416 | 474 | 424 | 498 | 661 | 420 | 448 | 519 | 494.85 | 528 | 462 | 0.875 | -0.193 | 0.28588699 | 0.50720017 | 0.0294 | 0.0136 | 0.0216 | 0.0216 | TRUE | 0.5166 |
| Rassf10 | 184 | 160 | 144 | 184 | 166 | 131 | 195 | 160 | 145 | 145 | 179 | 143 | 161.08 | 167 | 156 | 0.933 | -0.1 | 0.60182893 | 0.77334576 | 0.0135 | 0.0274 | 0.0204 | 0.0204 | TRUE | 0.2402 |
| Rassf2 | 644 | 690 | 677 | 705 | 577 | 569 | 684 | 689 | 680 | 554 | 621 | 623 | 641.8 | 684 | 599 | 0.874 | -0.194 | 0.04999092 | 0.16415923 | 0 | 0.012 | 0.0055 | 0.0055 | TRUE | 0.054 |
| Rassf3 | 2506 | 3334 | 3202 | 3290 | 2561 | 2607 | 2662 | 3330 | 3215 | 2586 | 2755 | 2854 | 2900.37 | 3069 | 2732 | 0.89 | -0.168 | 0.11774916 | 0.29026879 | 0.0075 | 0.008 | 0.008 | 0.008 | TRUE | 0.1632 |
| Rassf4 | 594 | 716 | 753 | 567 | 460 | 624 | 631 | 715 | 756 | 446 | 495 | 683 | 620.97 | 701 | 541 | 0.771 | -0.375 | 0.03021754 | 0.115818 | 0.0285 | 0.0122 | 0.0199 | 0.0199 | TRUE | 0.6106 |
| Rassf5 | 279 | 329 | 312 | 396 | 294 | 307 | 296 | 329 | 313 | 311 | 316 | 336 | 316.98 | 313 | 321 | 1.026 | 0.037 | 0.76242016 | 0.87472228 | 0 | 0.0173 | 0.0075 | 0.0075 | TRUE | 0.0234 |
| Rassf6 | 262 | 421 | 408 | 280 | 223 | 291 | 278 | 420 | 410 | 220 | 240 | 319 | 314.5 | 369 | 260 | 0.701 | -0.513 | 0.01510789 | 0.07112887 | 0.0409 | 0.0174 | 0.0288 | 0.0288 | TRUE | 0.5304 |
| Rassf7 | 958 | 1044 | 1055 | 1176 | 899 | 922 | 1018 | 1043 | 1059 | 924 | 967 | 1009 | 1003.43 | 1040 | 967 | 0.929 | -0.106 | 0.20593034 | 0.41485928 | 0 | 0.0102 | 0.0041 | 0.0041 | TRUE | 0.0185 |
| Rassf8 | 101 | 121 | 118 | 218 | 136 | 160 | 107 | 121 | 118 | 171 | 146 | 175 | 139.91 | 115 | 164 | 1.423 | 0.509 | 0.00402682 | 0.02649604 | 0 | 0.0305 | 0.0152 | 0.0152 | TRUE | 0.0928 |

| | | | | | | | | | | | | | | | | | | | | | | | | | |
|---|---|---|---|---|---|---|---|---|---|---|---|---|---|---|---|---|---|---|---|---|---|---|---|---|---|
| Rassf9 | 1 | 3 | 0 | 10 | 10 | 5 | 1 | 3 | 0 | 8 | 11 | 5 | 4.69 | 1 | 8 | 5.914 | 2.564 | 0.02051484 | 0.08835612 | 0 | 0.709 | 0.4457 | 0.4457 | TRUE | 0.534 |
| Raver1 | 2288 | 1835 | 1820 | 3186 | 2349 | 2007 | 2430 | 1833 | 1827 | 2505 | 2527 | 2197 | 2219.88 | 2030 | 2410 | 1.187 | 0.248 | 0.06534411 | 0.19701918 | 0.0168 | 0.0084 | 0.0126 | 0.0126 | TRUE | 0.3591 |
| Raver1-fdx1l | 1 | 2 | 0 | 0 | 0 | 0 | 1 | 2 | 0 | 0 | 0 | 0 | 0.51 | 1 | 0 | 0.176 | -2.508 | 0.44268408 | NA | 0 | 6.4675 | 6.1932 | 6.1932 | TRUE | 0.064 |
| Raver2 | 2778 | 3361 | 3648 | 3399 | 2664 | 2867 | 2951 | 3357 | 3663 | 2672 | 2866 | 3138 | 3107.85 | 3324 | 2892 | 0.87 | -0.201 | 0.06826197 | 0.20245789 | 0.0086 | 0.008 | 0.0084 | 0.0084 | TRUE | 0.1168 |
| Rax | 42 | 56 | 68 | 50 | 32 | 57 | 45 | 56 | 68 | 39 | 34 | 62 | 50.82 | 56 | 45 | 0.802 | -0.318 | 0.34392534 | 0.56374875 | 0.0499 | 0.0717 | 0.0613 | 0.0613 | TRUE | 0.4847 |
| Rb1 | 1737 | 2185 | 2241 | 2412 | 1693 | 1967 | 1845 | 2182 | 2250 | 1896 | 1822 | 2153 | 2024.71 | 2092 | 1957 | 0.935 | -0.097 | 0.39114492 | 0.60481742 | 0.0084 | 0.0085 | 0.0087 | 0.0087 | TRUE | 0.1292 |
| Rb1cc1 | 2271 | 2229 | 2418 | 3392 | 2500 | 2373 | 2412 | 2226 | 2428 | 2667 | 2690 | 2598 | 2503.37 | 2355 | 2652 | 1.126 | 0.171 | 0.01908869 | 0.08390936 | 9.00E-04 | 0.0082 | 0.0034 | 0.0034 | TRUE | 0.028 |
| Rbak | 425 | 414 | 429 | 555 | 435 | 357 | 451 | 413 | 431 | 436 | 468 | 391 | 431.8 | 432 | 432 | 1 | 0 | 0.99835565 | 0.9988623 | 0.0024 | 0.0145 | 0.008 | 0.008 | TRUE | 0.0758 |
| Rbbp4 | 15983 | 17624 | 16698 | 22266 | 15767 | 15687 | 16977 | 17601 | 16767 | 17504 | 16964 | 17172 | 17164.09 | 17115 | 17213 | 1.006 | 0.008 | 0.88228846 | 0.94169383 | 3.00E-04 | 0.0071 | 0.0022 | 0.0022 | TRUE | 0.0076 |
| Rbbp5 | 4927 | 4549 | 4474 | 6523 | 4748 | 4350 | 5233 | 4543 | 4492 | 5128 | 5109 | 4762 | 4877.84 | 4756 | 5000 | 1.051 | 0.072 | 0.41992776 | 0.63035126 | 0.0041 | 0.0076 | 0.0056 | 0.0056 | TRUE | 0.0937 |
| Rbbp6 | 5481 | 5427 | 5320 | 7621 | 5602 | 5032 | 5822 | 5420 | 5342 | 5991 | 6027 | 5508 | 5685.06 | 5528 | 5842 | 1.057 | 0.08 | 0.29229506 | 0.51317582 | 0.0021 | 0.0075 | 0.004 | 0.004 | TRUE | 0.0303 |
| Rbbp7 | 12999 | 15060 | 15389 | 18523 | 13260 | 14199 | 13807 | 15041 | 15452 | 14561 | 14267 | 15543 | 14778.55 | 14767 | 14790 | 1.002 | 0.002 | 0.97665457 | 0.98733516 | 0.0021 | 0.0071 | 0.0042 | 0.0042 | TRUE | 0.0395 |
| Rbbp8 | 2146 | 2169 | 2068 | 3151 | 2116 | 1927 | 2279 | 2166 | 2076 | 2477 | 2277 | 2109 | 2230.88 | 2174 | 2288 | 1.053 | 0.075 | 0.41808767 | 0.62897231 | 0.0037 | 0.0084 | 0.0057 | 0.0057 | TRUE | 0.0645 |
| Rbbp8nl | 7 | 5 | 2 | 7 | 2 | 10 | 7 | 5 | 2 | 6 | 2 | 11 | 5.51 | 5 | 6 | 1.287 | 0.364 | 0.69474557 | 0.83270212 | 0.253 | 0.6052 | 0.4352 | 0.4352 | TRUE | 0.2807 |
| Rbbp9 | 1341 | 1571 | 1578 | 1868 | 1285 | 1473 | 1424 | 1569 | 1584 | 1468 | 1383 | 1612 | 1506.89 | 1526 | 1488 | 0.975 | -0.037 | 0.70186913 | 0.83704195 | 0.004 | 0.0091 | 0.006 | 0.006 | TRUE | 0.0634 |
| Rbck1 | 1779 | 1930 | 1877 | 2801 | 2053 | 1934 | 1890 | 1928 | 1885 | 2202 | 2209 | 2117 | 2038.29 | 1901 | 2176 | 1.145 | 0.195 | 0.0037313 | 0.02507456 | 0 | 0.0085 | 0.0028 | 0.0028 | TRUE | 0.0067 |
| Rbfa | 2097 | 2141 | 2161 | 2696 | 1969 | 1894 | 2227 | 2138 | 2170 | 2119 | 2119 | 2073 | 2141.11 | 2178 | 2104 | 0.966 | -0.05 | 0.44895502 | 0.65455233 | 0 | 0.0084 | 0.0027 | 0.0027 | TRUE | 0.0046 |
| Rbfox1 | 4 | 7 | 1 | 7 | 3 | 6 | 4 | 7 | 1 | 6 | 3 | 7 | 4.59 | 4 | 5 | 1.252 | 0.324 | 0.73526501 | NA | 0 | 0.7246 | 0.4402 | 0.4402 | TRUE | 0.2854 |
| Rbfox2 | 4987 | 5343 | 5772 | 7220 | 5311 | 5565 | 5297 | 5336 | 5796 | 5676 | 5714 | 6092 | 5651.79 | 5476 | 5827 | 1.064 | 0.089 | 0.22654358 | 0.43938089 | 0.0018 | 0.0075 | 0.0038 | 0.0038 | TRUE | 0.0318 |
| Rbfox3 | 137 | 152 | 169 | 129 | 101 | 112 | 146 | 152 | 170 | 101 | 109 | 123 | 133.28 | 156 | 111 | 0.71 | -0.495 | 0.0064752 | 0.03809281 | 0 | 0.0316 | 0.0161 | 0.0161 | TRUE | 0.0827 |
| Rbks | 39 | 79 | 74 | 94 | 48 | 57 | 41 | 79 | 74 | 74 | 52 | 62 | 63.76 | 65 | 63 | 0.968 | -0.047 | 0.88220789 | 0.94169383 | 0.0556 | 0.0586 | 0.0572 | 0.0572 | TRUE | 0.8257 |
| Rbl1 | 1714 | 2021 | 2002 | 2232 | 1673 | 1821 | 1821 | 2018 | 2010 | 1755 | 1800 | 1993 | 1899.54 | 1950 | 1849 | 0.948 | -0.077 | 0.39787786 | 0.61124685 | 0.0032 | 0.0086 | 0.0055 | 0.0055 | TRUE | 0.0557 |
| Rbl2 | 596 | 856 | 866 | 749 | 682 | 725 | 633 | 855 | 870 | 589 | 734 | 794 | 745.62 | 786 | 706 | 0.896 | -0.158 | 0.34181988 | 0.56204134 | 0.0256 | 0.0113 | 0.0185 | 0.0185 | TRUE | 0.3417 |
| Rbm10 | 2598 | 3027 | 3080 | 4264 | 3153 | 2774 | 2760 | 3023 | 3093 | 3352 | 3392 | 3037 | 3109.38 | 2959 | 3260 | 1.102 | 0.14 | 0.10381583 | 0.26807506 | 0.0031 | 0.008 | 0.005 | 0.005 | TRUE | 0.0435 |
| Rbm11 | 105 | 122 | 92 | 113 | 85 | 87 | 112 | 122 | 92 | 89 | 91 | 95 | 100.21 | 109 | 92 | 0.845 | -0.243 | 0.23921861 | 0.45517625 | 0 | 0.0398 | 0.0207 | 0.0207 | TRUE | 0.1712 |
| Rbm12 | 3333 | 3159 | 3221 | 5605 | 3977 | 3512 | 3540 | 3155 | 3234 | 4406 | 4279 | 3844 | 3743.17 | 3310 | 4176 | 1.262 | 0.336 | 0.00018103 | 0.00223763 | 0.004 | 0.0078 | 0.0055 | 0.0055 | TRUE | 0.0587 |
| Rbm12b1 | 161 | 134 | 121 | 278 | 202 | 157 | 171 | 134 | 121 | 219 | 217 | 172 | 172.35 | 142 | 203 | 1.429 | 0.515 | 0.00907367 | 0.04892646 | 0.018 | 0.026 | 0.0221 | 0.0221 | TRUE | 0.324 |
| Rbm12b2 | 323 | 279 | 280 | 573 | 448 | 352 | 343 | 279 | 281 | 450 | 482 | 385 | 370.11 | 301 | 439 | 1.461 | 0.547 | 0.00022262 | 0.00264796 | 0.0104 | 0.0158 | 0.013 | 0.013 | TRUE | 0.1676 |
| Rbm14 | 5786 | 5995 | 6117 | 7979 | 5855 | 5717 | 6146 | 5987 | 6142 | 6272 | 6300 | 6258 | 6184.23 | 6092 | 6277 | 1.03 | 0.043 | 0.43371305 | 0.6424232 | 0 | 0.0074 | 0.002 | 0.002 | TRUE | 0.0027 |
| Rbm15 | 1811 | 1501 | 1441 | 1895 | 1447 | 1254 | 1924 | 1499 | 1447 | 1490 | 1557 | 1373 | 1548.14 | 1623 | 1473 | 0.908 | -0.14 | 0.28356348 | 0.50512347 | 0.014 | 0.009 | 0.0116 | 0.0116 | TRUE | 0.3154 |
| Rbm15b | 1333 | 1322 | 1330 | 2377 | 1763 | 1420 | 1416 | 1320 | 1335 | 1869 | 1897 | 1554 | 1565.26 | 1357 | 1773 | 1.307 | 0.387 | 0.00023856 | 0.00280793 | 0.0061 | 0.009 | 0.0073 | 0.0073 | TRUE | 0.1393 |
| Rbm17 | 5932 | 6640 | 6547 | 8937 | 6678 | 6224 | 6301 | 6631 | 6574 | 7026 | 7185 | 6813 | 6755 | 6502 | 7008 | 1.078 | 0.108 | 0.08146812 | 0.22753944 | 5.00E-04 | 0.0074 | 0.0026 | 0.0026 | TRUE | 0.0089 |
| Rbm18 | 1794 | 1856 | 1924 | 2389 | 1687 | 1651 | 1906 | 1854 | 1932 | 1878 | 1815 | 1807 | 1865.25 | 1897 | 1833 | 0.967 | -0.049 | 0.48348786 | 0.68146464 | 0 | 0.0087 | 0.0029 | 0.0029 | TRUE | 0.0056 |
| Rbm19 | 4923 | 3644 | 3526 | 6188 | 4520 | 3657 | 5229 | 3639 | 3540 | 4865 | 4863 | 4003 | 4356.63 | 4136 | 4577 | 1.107 | 0.146 | 0.36740982 | 0.58475554 | 0.027 | 0.0077 | 0.0187 | 0.0187 | TRUE | 0.65 |
| Rbm20 | 434 | 346 | 392 | 287 | 215 | 214 | 461 | 346 | 394 | 226 | 231 | 234 | 315.22 | 400 | 230 | 0.576 | -0.796 | 9.63E-08 | 3.47E-06 | 0.0087 | 0.0174 | 0.0126 | 0.0126 | TRUE | 0.1845 |
| Rbm22 | 2581 | 2748 | 2851 | 3602 | 2615 | 2810 | 2741 | 2744 | 2863 | 2832 | 2814 | 3076 | 2844.97 | 2783 | 2907 | 1.044 | 0.063 | 0.39618669 | 0.60964502 | 0.0011 | 0.0081 | 0.0036 | 0.0036 | TRUE | 0.031 |
| Rbm24 | 62 | 90 | 65 | 65 | 45 | 51 | 66 | 90 | 65 | 51 | 48 | 56 | 62.72 | 74 | 52 | 0.702 | -0.511 | 0.05232594 | 0.16921976 | 0 | 0.0594 | 0.0335 | 0.0335 | TRUE | 0.3411 |
| Rbm25 | 10523 | 8105 | 7935 | 13825 | 10455 | 8324 | 11177 | 8095 | 7968 | 10868 | 11249 | 9112 | 9744.73 | 9080 | 10410 | 1.147 | 0.197 | 0.18842207 | 0.39351072 | 0.0249 | 0.0072 | 0.0161 | 0.0161 | TRUE | 0.4987 |
| Rbm26 | 6090 | 5266 | 5190 | 7004 | 5268 | 4952 | 6469 | 5259 | 5211 | 5506 | 5668 | 5421 | 5588.99 | 5646 | 5532 | 0.98 | -0.03 | 0.77626155 | 0.88373755 | 0.0066 | 0.0075 | 0.0076 | 0.0076 | TRUE | 0.1977 |
| Rbm27 | 5016 | 3914 | 4122 | 6476 | 4688 | 4007 | 5328 | 3909 | 4139 | 5091 | 5044 | 4386 | 4649.49 | 4459 | 4840 | 1.086 | 0.119 | 0.37328306 | 0.58968554 | 0.0154 | 0.0076 | 0.0126 | 0.0126 | TRUE | 0.3541 |
| Rbm28 | 7663 | 6285 | 6289 | 8888 | 6497 | 5698 | 8139 | 6277 | 6315 | 6987 | 6990 | 6237 | 6824.32 | 6910 | 6738 | 0.975 | -0.036 | 0.76654301 | 0.87786206 | 0.0114 | 0.0074 | 0.0106 | 0.0106 | TRUE | 0.2953 |
| Rbm3 | 4389 | 4436 | 4665 | 8540 | 6016 | 5292 | 4662 | 4430 | 4684 | 6713 | 6473 | 5793 | 5459.26 | 4592 | 6326 | 1.378 | 0.463 | 2.65E-08 | 1.14E-06 | 0.003 | 0.0075 | 0.0048 | 0.0048 | TRUE | 0.0662 |
| Rbm31y | 0 | 0 | 0 | 0 | 0 | 0 | 0 | 0 | 0 | 0 | 0 | 0 | 0 | 0 | 0 | NA | NA | NA | NA | NA | NA | NA | NA | NA | NA |
| Rbm33 | 7828 | 6152 | 6321 | 7121 | 5317 | 5428 | 8315 | 6144 | 6347 | 5598 | 5721 | 5942 | 6344.38 | 6935 | 5754 | 0.83 | -0.269 | 0.03209673 | 0.12067454 | 0.0152 | 0.0074 | 0.0112 | 0.0112 | TRUE | 0.3694 |
| Rbm34 | 3050 | 2758 | 2596 | 4026 | 2846 | 2585 | 3240 | 2754 | 2607 | 3165 | 3062 | 2830 | 2942.91 | 2867 | 3019 | 1.053 | 0.075 | 0.48222059 | 0.68089732 | 0.0069 | 0.008 | 0.0079 | 0.0079 | TRUE | 0.1568 |
| Rbm38 | 4522 | 4236 | 4193 | 5617 | 4272 | 3908 | 4803 | 4231 | 4210 | 4416 | 4596 | 4278 | 4422.31 | 4415 | 4430 | 1.004 | 0.005 | 0.95145794 | 0.97533692 | 0.0032 | 0.0076 | 0.0049 | 0.0049 | TRUE | 0.0721 |
| Rbm39 | 13283 | 13831 | 13719 | 17136 | 13295 | 12372 | 14109 | 13813 | 13775 | 13471 | 14305 | 13543 | 13836.01 | 13899 | 13773 | 0.991 | -0.013 | 0.82285888 | 0.90959702 | 5.00E-04 | 0.0071 | 0.0024 | 0.0024 | TRUE | 0.0139 |
| Rbm4 | 604 | 598 | 516 | 898 | 656 | 617 | 642 | 597 | 518 | 706 | 706 | 675 | 640.68 | 586 | 696 | 1.189 | 0.249 | 0.02808627 | 0.11006223 | 0.0042 | 0.012 | 0.0077 | 0.0077 | TRUE | 0.1196 |
| Rbm41 | 779 | 765 | 819 | 1228 | 953 | 878 | 827 | 764 | 822 | 965 | 1025 | 961 | 894.27 | 804 | 984 | 1.223 | 0.29 | 0.00106953 | 0.0093819 | 0 | 0.0106 | 0.0045 | 0.0045 | TRUE | 0.0232 |
| Rbm42 | 2408 | 2246 | 2304 | 3836 | 2764 | 2346 | 2558 | 2243 | 2313 | 3016 | 2974 | 2568 | 2611.97 | 2371 | 2853 | 1.203 | 0.267 | 0.00750705 | 0.04227102 | 0.0056 | 0.0082 | 0.0068 | 0.0068 | TRUE | 0.0918 |
| Rbm43 | 304 | 271 | 251 | 479 | 361 | 326 | 323 | 271 | 252 | 377 | 388 | 357 | 327.9 | 282 | 374 | 1.328 | 0.409 | 0.00332983 | 0.02297381 | 0.0051 | 0.017 | 0.0109 | 0.0109 | TRUE | 0.1801 |
| Rbm44 | 745 | 968 | 930 | 1226 | 902 | 1019 | 791 | 967 | 934 | 964 | 970 | 1115 | 956.94 | 897 | 1016 | 1.132 | 0.179 | 0.12937526 | 0.30962138 | 0.0074 | 0.0103 | 0.009 | 0.009 | TRUE | 0.1264 |
| Rbm45 | 1282 | 1281 | 1213 | 1869 | 1438 | 1188 | 1362 | 1279 | 1218 | 1469 | 1547 | 1300 | 1362.66 | 1286 | 1439 | 1.119 | 0.162 | 0.10784917 | 0.27490073 | 0.0043 | 0.0093 | 0.0066 | 0.0066 | TRUE | 0.0839 |
| Rbm46 | 18 | 25 | 15 | 39 | 29 | 34 | 19 | 25 | 15 | 31 | 31 | 37 | 26.37 | 20 | 33 | 1.67 | 0.74 | 0.05678616 | 0.17926968 | 0 | 0.1318 | 0.068 | 0.068 | TRUE | 0.298 |
| Rbm47 | 1606 | 1504 | 1425 | 1812 | 1379 | 1449 | 1706 | 1502 | 1431 | 1424 | 1484 | 1586 | 1522.18 | 1546 | 1498 | 0.969 | -0.046 | 0.64759098 | 0.80215228 | 0.005 | 0.0091 | 0.0067 | 0.0067 | TRUE | 0.0979 |
| Rbm48 | 616 | 676 | 644 | 811 | 626 | 599 | 654 | 675 | 647 | 638 | 674 | 656 | 657.14 | 659 | 656 | 0.995 | -0.008 | 0.93108031 | 0.96611833 | 0 | 0.0119 | 0.0045 | 0.0045 | TRUE | 0.0073 |
| Rbm4b | 212 | 225 | 261 | 292 | 239 | 226 | 225 | 225 | 262 | 230 | 257 | 247 | 241.01 | 237 | 245 | 1.029 | 0.041 | 0.77337927 | 0.88178542 | 0 | 0.0206 | 0.0107 | 0.0107 | TRUE | 0.0926 |
| Rbm5 | 4114 | 4377 | 3888 | 5862 | 4274 | 3808 | 4370 | 4371 | 3904 | 4608 | 4599 | 4168 | 4336.73 | 4215 | 4458 | 1.058 | 0.081 | 0.34185027 | 0.56204134 | 0.0033 | 0.0077 | 0.005 | 0.005 | TRUE | 0.0508 |
| Rbm6 | 3737 | 3852 | 3690 | 5316 | 3802 | 3526 | 3969 | 3847 | 3705 | 4179 | 4091 | 3860 | 3941.84 | 3840 | 4043 | 1.053 | 0.075 | 0.2927293 | 0.5137339 | 0.001 | 0.0077 | 0.0034 | 0.0034 | TRUE | 0.0191 |
| Rbm7 | 2543 | 2854 | 2835 | 3297 | 2382 | 2569 | 2701 | 2850 | 2847 | 2592 | 2563 | 2812 | 2727.5 | 2799 | 2656 | 0.948 | -0.076 | 0.31077594 | 0.53139156 | 0.0012 | 0.0081 | 0.0037 | 0.0037 | TRUE | 0.0319 |
| Rbm8a | 8084 | 7729 | 7325 | 11871 | 8536 | 7606 | 8587 | 7719 | 7355 | 9332 | 9184 | 8326 | 8417.16 | 7887 | 8947 | 1.135 | 0.182 | 0.04501709 | 0.15295338 | 0.0049 | 0.0073 | 0.0058 | 0.0058 | TRUE | 0.0735 |
| Rbms1 | 847 | 1145 | 1177 | 1780 | 1174 | 1359 | 900 | 1144 | 1182 | 1399 | 1263 | 1488 | 1229.18 | 1075 | 1383 | 1.287 | 0.364 | 0.00534192 | 0.03284711 | 0.0122 | 0.0096 | 0.0114 | 0.0114 | TRUE | 0.2425 |
| Rbms2 | 2917 | 2780 | 2713 | 4257 | 2919 | 3038 | 3098 | 2776 | 2724 | 3347 | 3141 | 3326 | 3068.61 | 2866 | 3271 | 1.141 | 0.191 | 0.02153067 | 0.09168102 | 0.0026 | 0.008 | 0.0046 | 0.0046 | TRUE | 0.0607 |
| Rbms3 | 4 | 4 | 2 | 7 | 10 | 4 | 4 | 4 | 2 | 6 | 11 | 4 | 5.15 | 3 | 7 | 2.006 | 1.005 | 0.25374155 | 0.47171285 | 0 | 0.6468 | 0.3372 | 0.3372 | TRUE | 0.5881 |
| Rbmx | 282 | 296 | 292 | 401 | 390 | 324 | 300 | 296 | 293 | 315 | 420 | 355 | 329.65 | 296 | 363 | 1.224 | 0.291 | 0.04580539 | 0.15485794 | 0.0083 | 0.0169 | 0.0122 | 0.0122 | TRUE | 0.211 |
| Rbmx2 | 1043 | 1119 | 1181 | 1540 | 1142 | 1121 | 1108 | 1118 | 1186 | 1211 | 1229 | 1227 | 1179.62 | 1137 | 1222 | 1.075 | 0.104 | 0.18505162 | 0.38837447 | 0 | 0.0097 | 0.0036 | 0.0036 | TRUE | 0.0169 |
| Rbmxl1 | 4572 | 3691 | 3721 | 6340 | 4587 | 3879 | 4856 | 3686 | 3736 | 4984 | 4935 | 4246 | 4407.38 | 4093 | 4722 | 1.154 | 0.206 | 0.11498917 | 0.28628401 | 0.0165 | 0.0077 | 0.0121 | 0.0121 | TRUE | 0.3238 |
| Rbmxl2 | 4537 | 4920 | 5072 | 6182 | 4829 | 4399 | 4819 | 4914 | 5093 | 4860 | 5196 | 4815 | 4949.41 | 4942 | 4957 | 1.003 | 0.004 | 0.95082309 | 0.97518415 | 0.001 | 0.0076 | 0.0032 | 0.0032 | TRUE | 0.0215 |
| Rbmy | 0 | 0 | 0 | 0 | 0 | 0 | 0 | 0 | 0 | 0 | 0 | 0 | 0 | 0 | 0 | NA | NA | NA | NA | NA | NA | NA | NA | NA | NA |

| | | | | | | | | | | | | | | | | | | | | | | | | |
|---|---|---|---|---|---|---|---|---|---|---|---|---|---|---|---|---|---|---|---|---|---|---|---|---|
| Rbp1 | 588 | 703 | 673 | 375 | 299 | 267 | 625 | 702 | 676 | 295 | 322 | 292 | 485.2 | 668 | 303 | 0.453 | -1.141 | 3.42E-24 | 1.31E-21 | 0 | 0.0137 | 0.0067 | 0.0067 | TRUE | 0.037 |
| Rbp2 | 0 | 3 | 0 | 2 | 3 | 0 | 0 | 3 | 0 | 2 | 3 | 0 | 1.3 | 1 | 2 | 1.593 | 0.672 | 0.75578505 | NA | 2.5633 | 2.5423 | 2.5497 | 2.5497 | TRUE | 0.4642 |
| Rbp3 | 2 | 0 | 0 | 4 | 2 | 0 | 2 | 0 | 0 | 3 | 2 | 0 | 1.24 | 1 | 2 | 2.57 | 1.362 | 0.53209554 | NA | 1.8421 | 2.6706 | 2.4182 | 2.4182 | TRUE | 0.7508 |
| Rbp4 | 320 | 372 | 270 | 394 | 303 | 275 | 340 | 372 | 271 | 310 | 326 | 301 | 319.88 | 328 | 312 | 0.953 | -0.069 | 0.65224631 | 0.80502827 | 0.0105 | 0.0172 | 0.0137 | 0.0137 | TRUE | 0.2597 |
| Rbp7 | 62 | 53 | 58 | 57 | 32 | 41 | 66 | 53 | 58 | 45 | 34 | 45 | 50.19 | 59 | 41 | 0.704 | -0.507 | 0.07137914 | 0.20821006 | 0 | 0.0725 | 0.0364 | 0.0364 | TRUE | 0.1531 |
| Rbpj | 4265 | 4396 | 4509 | 7581 | 5256 | 5262 | 4530 | 4390 | 4528 | 5960 | 5655 | 5760 | 5137.14 | 4483 | 5792 | 1.292 | 0.37 | 1.49E-09 | 8.65E-08 | 4.00E-04 | 0.0075 | 0.0025 | 0.0025 | TRUE | 0.008 |
| Rbpjl | 1 | 1 | 0 | 2 | 1 | 0 | 1 | 1 | 0 | 2 | 1 | 0 | 0.78 | 1 | 1 | 1.327 | 0.408 | 0.85171912 | NA | 0 | 4.2045 | 2.1498 | 2.1498 | TRUE | 0.2789 |
| Rbpms | 7536 | 8809 | 9190 | 9143 | 7167 | 7554 | 8005 | 8798 | 9228 | 7188 | 7711 | 8269 | 8199.62 | 8677 | 7723 | 0.89 | -0.168 | 0.06435956 | 0.19526081 | 0.0048 | 0.0073 | 0.0058 | 0.0058 | TRUE | 0.056 |
| Rbpms2 | 6882 | 7920 | 8183 | 9373 | 6810 | 7102 | 7310 | 7910 | 8217 | 7368 | 7327 | 7774 | 7651 | 7812 | 7490 | 0.959 | -0.061 | 0.41737383 | 0.62831155 | 0.0021 | 0.0073 | 0.0039 | 0.0039 | TRUE | 0.0386 |
| Rbx1 | 3078 | 3448 | 3260 | 4188 | 2935 | 2800 | 3269 | 3444 | 3273 | 3292 | 3158 | 3065 | 3250.25 | 3329 | 3172 | 0.953 | -0.069 | 0.31762817 | 0.53856183 | 8.00E-04 | 0.0079 | 0.0032 | 0.0032 | TRUE | 0.0137 |
| Rc3h1 | 2131 | 1878 | 2043 | 2837 | 1953 | 1831 | 2263 | 1876 | 2051 | 2230 | 2101 | 2004 | 2087.72 | 2063 | 2112 | 1.024 | 0.034 | 0.73031078 | 0.85505616 | 0.0054 | 0.0085 | 0.0066 | 0.0066 | TRUE | 0.0868 |
| Rc3h2 | 2444 | 1930 | 1881 | 3570 | 2331 | 2007 | 2596 | 1928 | 1889 | 2806 | 2508 | 2197 | 2320.61 | 2138 | 2504 | 1.172 | 0.229 | 0.1286854 | 0.30845382 | 0.0241 | 0.0083 | 0.0159 | 0.0159 | TRUE | 0.4258 |
| Rcan1 | 1341 | 1226 | 1159 | 2163 | 1585 | 1450 | 1424 | 1224 | 1164 | 1700 | 1705 | 1587 | 1467.59 | 1271 | 1664 | 1.31 | 0.39 | 0.00017234 | 0.00214869 | 0.0055 | 0.0091 | 0.0071 | 0.0071 | TRUE | 0.1335 |
| Rcan2 | 104 | 87 | 79 | 137 | 85 | 91 | 110 | 87 | 79 | 108 | 91 | 100 | 95.91 | 92 | 100 | 1.084 | 0.116 | 0.59982407 | 0.77190154 | 0.0073 | 0.0413 | 0.0247 | 0.0247 | TRUE | 0.2818 |
| Rcan3 | 1156 | 1805 | 1816 | 1833 | 1369 | 1587 | 1228 | 1803 | 1823 | 1441 | 1473 | 1737 | 1584.19 | 1618 | 1550 | 0.958 | -0.062 | 0.70422106 | 0.83814573 | 0.0285 | 0.009 | 0.0187 | 0.0187 | TRUE | 0.5354 |
| Rcbtb1 | 2736 | 3677 | 3761 | 4340 | 3246 | 3466 | 2906 | 3672 | 3776 | 3412 | 3492 | 3794 | 3508.86 | 3451 | 3566 | 1.033 | 0.047 | 0.69181108 | 0.83095304 | 0.0113 | 0.0078 | 0.0097 | 0.0097 | TRUE | 0.232 |
| Rcbtb2 | 192 | 309 | 309 | 552 | 378 | 361 | 204 | 309 | 310 | 434 | 407 | 395 | 343.1 | 274 | 412 | 1.502 | 0.587 | 0.00103231 | 0.00913161 | 0.0236 | 0.0165 | 0.02 | 0.02 | TRUE | 0.5577 |
| Rcc1 | 7360 | 7463 | 6930 | 9398 | 6652 | 6178 | 7818 | 7453 | 6958 | 7388 | 7157 | 6763 | 7256.22 | 7410 | 7103 | 0.959 | -0.061 | 0.43532132 | 0.64353622 | 0.0025 | 0.0074 | 0.0042 | 0.0042 | TRUE | 0.0347 |
| Rcc2 | 21414 | 20372 | 19947 | 33511 | 25232 | 22048 | 22745 | 20346 | 20029 | 26344 | 27148 | 24135 | 23457.81 | 21040 | 25876 | 1.23 | 0.299 | 0.00051344 | 0.00528397 | 0.0042 | 0.007 | 0.0053 | 0.0053 | TRUE | 0.0615 |
| Rccd1 | 745 | 561 | 471 | 768 | 670 | 461 | 791 | 560 | 473 | 604 | 721 | 505 | 608.96 | 608 | 610 | 1.003 | 0.004 | 0.98390539 | 0.99158695 | 0.0487 | 0.0123 | 0.0295 | 0.0295 | TRUE | 0.6272 |
| Rce1 | 1174 | 1290 | 1197 | 1671 | 1249 | 1131 | 1247 | 1288 | 1202 | 1314 | 1344 | 1238 | 1272.13 | 1246 | 1299 | 1.043 | 0.06 | 0.46147333 | 0.66558634 | 6.00E-04 | 0.0095 | 0.004 | 0.004 | TRUE | 0.0194 |
| Rchy1 | 2457 | 2461 | 2498 | 2757 | 2087 | 2223 | 2610 | 2458 | 2508 | 2167 | 2245 | 2433 | 2403.68 | 2525 | 2282 | 0.903 | -0.147 | 0.06546804 | 0.19715481 | 0.0016 | 0.0083 | 0.0042 | 0.0042 | TRUE | 0.0405 |
| Rcl1 | 4901 | 4116 | 4092 | 5292 | 3810 | 3606 | 5206 | 4111 | 4109 | 4160 | 4099 | 3947 | 4272.01 | 4475 | 4069 | 0.909 | -0.137 | 0.2211196 | 0.43313412 | 0.01 | 0.0077 | 0.0088 | 0.0088 | TRUE | 0.2482 |
| Rcn1 | 3956 | 4620 | 4204 | 4882 | 3499 | 3883 | 4202 | 4614 | 4221 | 3838 | 3765 | 4251 | 4148.4 | 4346 | 3951 | 0.909 | -0.138 | 0.10415112 | 0.26850868 | 0.0033 | 0.0077 | 0.0049 | 0.0049 | TRUE | 0.0532 |
| Rcn2 | 1357 | 1528 | 1523 | 2254 | 1508 | 1536 | 1441 | 1526 | 1529 | 1772 | 1623 | 1681 | 1595.41 | 1499 | 1692 | 1.129 | 0.176 | 0.02688298 | 0.10704328 | 0.001 | 0.009 | 0.0039 | 0.0039 | TRUE | 0.0212 |
| Rcn3 | 2173 | 2670 | 2823 | 1622 | 1255 | 1664 | 2308 | 2667 | 2835 | 1275 | 1350 | 1822 | 2042.69 | 2603 | 1482 | 0.569 | -0.813 | 8.86E-08 | 3.24E-06 | 0.0245 | 0.0085 | 0.0161 | 0.0161 | TRUE | 0.4809 |
| Rcor1 | 1281 | 1339 | 1333 | 1866 | 1356 | 1273 | 1361 | 1337 | 1338 | 1467 | 1459 | 1393 | 1392.63 | 1345 | 1440 | 1.071 | 0.098 | 0.18133726 | 0.38312809 | 0 | 0.0093 | 0.0032 | 0.0032 | TRUE | 0.0093 |
| Rcor2 | 4002 | 4167 | 4151 | 6049 | 4551 | 4180 | 4251 | 4162 | 4168 | 4755 | 4897 | 4576 | 4468 | 4194 | 4743 | 1.131 | 0.178 | 0.00494134 | 0.03107894 | 4.00E-04 | 0.0076 | 0.0027 | 0.0027 | TRUE | 0.0114 |
| Rcor3 | 86 | 143 | 119 | 168 | 129 | 128 | 91 | 143 | 119 | 132 | 139 | 140 | 127.44 | 118 | 137 | 1.16 | 0.214 | 0.3120689 | 0.53255705 | 0.0152 | 0.0328 | 0.0243 | 0.0243 | TRUE | 0.27 |
| Rcsd1 | 86 | 116 | 96 | 151 | 132 | 110 | 91 | 116 | 96 | 119 | 142 | 120 | 114.12 | 101 | 127 | 1.252 | 0.325 | 0.10556455 | 0.27090208 | 0.004 | 0.0358 | 0.0201 | 0.0201 | TRUE | 0.1583 |
| Rcvrn | 1 | 0 | 0 | 0 | 1 | 2 | 1 | 0 | 0 | 0 | 1 | 2 | 0.72 | 0 | 1 | 2.905 | 1.539 | 0.54944501 | NA | 0 | 4.5747 | 3.3003 | 3.3003 | TRUE | 0.2242 |
| Rd3 | 0 | 0 | 2 | 1 | 1 | 0 | 0 | 0 | 2 | 1 | 1 | 0 | 0.65 | 1 | 1 | 0.932 | -0.102 | 0.96941655 | NA | 0 | 5.1142 | 3.5427 | 3.5427 | TRUE | 0.9739 |
| Rdh1 | 1 | 1 | 2 | 2 | 1 | 2 | 1 | 1 | 2 | 2 | 1 | 2 | 1.48 | 1 | 2 | 1.185 | 0.244 | 0.86702896 | NA | 0 | 2.2262 | 0.8554 | 0.8554 | TRUE | 0.1118 |
| Rdh10 | 185 | 211 | 204 | 269 | 196 | 164 | 197 | 211 | 205 | 211 | 211 | 180 | 202.32 | 204 | 201 | 0.985 | -0.022 | 0.88377055 | 0.94289583 | 0 | 0.0232 | 0.0112 | 0.0112 | TRUE | 0.0879 |
| Rdh11 | 2992 | 3506 | 3583 | 2712 | 2075 | 2611 | 3178 | 3501 | 3598 | 2132 | 2233 | 2858 | 2916.65 | 3426 | 2408 | 0.702 | -0.51 | 6.82E-05 | 0.00101165 | 0.0149 | 0.008 | 0.0114 | 0.0114 | TRUE | 0.324 |
| Rdh12 | 73 | 82 | 91 | 116 | 71 | 62 | 78 | 82 | 91 | 91 | 76 | 68 | 81.04 | 84 | 78 | 0.944 | -0.084 | 0.71615452 | 0.84576481 | 0 | 0.0476 | 0.0259 | 0.0259 | TRUE | 0.2075 |
| Rdh13 | 1465 | 1393 | 1417 | 1717 | 1224 | 1322 | 1556 | 1391 | 1423 | 1350 | 1317 | 1447 | 1413.99 | 1457 | 1371 | 0.941 | -0.087 | 0.32492872 | 0.54594394 | 0.002 | 0.0092 | 0.005 | 0.005 | TRUE | 0.0427 |
| Rdh14 | 303 | 410 | 406 | 468 | 371 | 380 | 322 | 409 | 408 | 368 | 399 | 416 | 387 | 380 | 394 | 1.037 | 0.052 | 0.71041097 | 0.84213468 | 0.0083 | 0.0154 | 0.0116 | 0.0116 | TRUE | 0.2015 |
| Rdh16 | 6 | 1 | 1 | 5 | 4 | 2 | 6 | 1 | 1 | 4 | 4 | 2 | 3.13 | 3 | 3 | 1.262 | 0.335 | 0.77444636 | NA | 0.2411 | 1.0583 | 0.662 | 0.662 | TRUE | 1.4611 |
| Rdh18-ps | 3 | 0 | 0 | 1 | 1 | 1 | 3 | 0 | 0 | 1 | 1 | 1 | 1.02 | 1 | 1 | 0.933 | -0.1 | 0.9616121 | NA | 0.6399 | 3.2245 | 2.1155 | 2.1155 | TRUE | 1.443 |
| Rdh19 | 0 | 0 | 0 | 0 | 0 | 0 | 0 | 0 | 0 | 0 | 0 | 0 | 0 | 0 | 0 | NA | NA | NA | NA | NA | NA | NA | NA | NA | NA |
| Rdh5 | 5 | 4 | 4 | 18 | 11 | 6 | 5 | 4 | 4 | 14 | 12 | 7 | 7.65 | 4 | 11 | 2.472 | 1.306 | 0.07297851 | 0.21158403 | 0 | 0.4378 | 0.2222 | 0.2222 | TRUE | 0.3607 |
| Rdh7 | 0 | 1 | 1 | 0 | 1 | 0 | 0 | 1 | 1 | 0 | 1 | 0 | 0.51 | 1 | 0 | 0.52 | -0.943 | 0.73680291 | NA | 0 | 6.4273 | 3.9121 | 3.9121 | TRUE | 0.5204 |
| Rdh8 | 0 | 0 | 0 | 0 | 0 | 0 | 0 | 0 | 0 | 0 | 0 | 0 | 0 | 0 | 0 | NA | NA | NA | NA | NA | NA | NA | NA | NA | NA |
| Rdh9 | 2 | 3 | 4 | 1 | 2 | 2 | 2 | 3 | 4 | 1 | 2 | 2 | 2.38 | 3 | 2 | 0.544 | -0.878 | 0.46899487 | NA | 0 | 1.3927 | 0.5995 | 0.5995 | TRUE | 0.2496 |
| Rdm1 | 1984 | 2929 | 3049 | 3206 | 2555 | 2683 | 2107 | 2925 | 3062 | 2520 | 2749 | 2937 | 2716.73 | 2698 | 2735 | 1.014 | 0.019 | 0.89536462 | 0.94945374 | 0.0223 | 0.0081 | 0.0154 | 0.0154 | TRUE | 0.4444 |
| Rdx | 9777 | 10764 | 11271 | 15675 | 11352 | 11192 | 10385 | 10750 | 11317 | 12322 | 12214 | 12251 | 11540.03 | 10817 | 12262 | 1.134 | 0.181 | 0.00397851 | 0.02623285 | 8.00E-04 | 0.0072 | 0.0028 | 0.0028 | TRUE | 0.02 |
| Rec8 | 1134 | 1133 | 1128 | 1519 | 1051 | 1172 | 1205 | 1132 | 1133 | 1194 | 1131 | 1283 | 1179.42 | 1157 | 1203 | 1.04 | 0.057 | 0.52470671 | 0.71437805 | 0.0017 | 0.0097 | 0.0049 | 0.0049 | TRUE | 0.0398 |
| Reck | 615 | 668 | 692 | 821 | 624 | 640 | 653 | 667 | 695 | 645 | 671 | 701 | 672.1 | 672 | 672 | 1 | 0 | 0.99936177 | 0.99950602 | 0 | 0.0118 | 0.0048 | 0.0048 | TRUE | 0.0157 |
| Recql | 218 | 205 | 275 | 588 | 362 | 311 | 232 | 205 | 276 | 462 | 389 | 340 | 317.43 | 238 | 397 | 1.677 | 0.746 | 2.05E-05 | 0.0003637 | 0.0201 | 0.0173 | 0.0187 | 0.0187 | TRUE | 0.2141 |
| Recql4 | 1079 | 761 | 754 | 1385 | 1039 | 732 | 1146 | 760 | 757 | 1089 | 1118 | 801 | 945.19 | 888 | 1003 | 1.13 | 0.176 | 0.36676544 | 0.58430551 | 0.0444 | 0.0104 | 0.0265 | 0.0265 | TRUE | 0.7701 |
| Recql5 | 2313 | 2131 | 2156 | 3027 | 2148 | 2005 | 2457 | 2128 | 2165 | 2380 | 2311 | 2195 | 2272.57 | 2250 | 2295 | 1.021 | 0.029 | 0.74413363 | 0.86342055 | 0.0035 | 0.0084 | 0.0054 | 0.0054 | TRUE | 0.0781 |
| Reep1 | 1004 | 1048 | 1064 | 1803 | 1095 | 1128 | 1066 | 1047 | 1068 | 1417 | 1178 | 1235 | 1168.62 | 1060 | 1277 | 1.205 | 0.27 | 0.00732963 | 0.04152402 | 0.0042 | 0.0097 | 0.0064 | 0.0064 | TRUE | 0.1142 |
| Reep2 | 589 | 845 | 862 | 585 | 413 | 537 | 626 | 844 | 866 | 460 | 444 | 588 | 637.86 | 779 | 497 | 0.638 | -0.648 | 0.00012865 | 0.00169226 | 0.0246 | 0.0121 | 0.019 | 0.019 | TRUE | 0.348 |
| Reep3 | 628 | 720 | 680 | 1152 | 795 | 688 | 667 | 719 | 683 | 906 | 855 | 753 | 763.84 | 690 | 838 | 1.217 | 0.283 | 0.00873178 | 0.04751707 | 0.004 | 0.0112 | 0.0071 | 0.0071 | TRUE | 0.0923 |
| Reep4 | 1807 | 2359 | 2405 | 2261 | 1684 | 1950 | 1919 | 2356 | 2415 | 1777 | 1812 | 2135 | 2069.01 | 2230 | 1908 | 0.855 | -0.226 | 0.06897623 | 0.2037253 | 0.012 | 0.0085 | 0.0106 | 0.0106 | TRUE | 0.1795 |
| Reep5 | 2006 | 2674 | 2645 | 2472 | 1815 | 2030 | 2131 | 2671 | 2656 | 1943 | 1953 | 2222 | 2262.57 | 2486 | 2039 | 0.82 | -0.286 | 0.01528646 | 0.07171268 | 0.01 | 0.0084 | 0.0096 | 0.0096 | TRUE | 0.1888 |
| Reep6 | 1172 | 1077 | 1106 | 1318 | 1106 | 964 | 1245 | 1076 | 1111 | 1036 | 1190 | 1055 | 1118.73 | 1144 | 1094 | 0.956 | -0.065 | 0.53518418 | 0.72254971 | 0.0045 | 0.0098 | 0.007 | 0.007 | TRUE | 0.0714 |
| Reg1 | 1 | 0 | 0 | 4 | 0 | 4 | 1 | 0 | 0 | 3 | 0 | 4 | 1.43 | 0 | 2 | 6.877 | 2.782 | 0.18881057 | NA | 0 | 2.3093 | 2.0342 | 2.0342 | TRUE | 0.3649 |
| Reg2 | 0 | 0 | 0 | 0 | 0 | 0 | 0 | 0 | 0 | 0 | 0 | 0 | 0 | 0 | 0 | NA | NA | NA | NA | NA | NA | NA | NA | NA | NA |
| Reg3a | 0 | 0 | 0 | 0 | 0 | 0 | 0 | 0 | 0 | 0 | 0 | 0 | 0 | 0 | 0 | NA | NA | NA | NA | NA | NA | NA | NA | NA | NA |
| Reg3b | 0 | 0 | 0 | 0 | 0 | 0 | 0 | 0 | 0 | 0 | 0 | 0 | 0 | 0 | 0 | NA | NA | NA | NA | NA | NA | NA | NA | NA | NA |
| Reg3d | 0 | 0 | 0 | 0 | 0 | 0 | 0 | 0 | 0 | 0 | 0 | 0 | 0 | 0 | 0 | NA | NA | NA | NA | NA | NA | NA | NA | NA | NA |
| Reg3g | 0 | 0 | 0 | 0 | 0 | 0 | 0 | 0 | 0 | 0 | 0 | 0 | 0 | 0 | 0 | NA | NA | NA | NA | NA | NA | NA | NA | NA | NA |
| Reg4 | 3 | 0 | 0 | 3 | 0 | 1 | 3 | 0 | 0 | 2 | 0 | 1 | 1.11 | 1 | 1 | 1.121 | 0.164 | 0.94281481 | NA | 2.645 | 2.984 | 2.8773 | 2.8773 | TRUE | 0.7206 |
| Rel | 67 | 57 | 81 | 98 | 89 | 71 | 71 | 57 | 81 | 77 | 96 | 78 | 76.66 | 70 | 84 | 1.194 | 0.256 | 0.29886526 | 0.51926751 | 0.0096 | 0.0499 | 0.0305 | 0.0305 | TRUE | 0.2381 |
| Rela | 644 | 543 | 529 | 922 | 675 | 579 | 684 | 542 | 531 | 725 | 726 | 634 | 640.4 | 586 | 695 | 1.187 | 0.248 | 0.06727409 | 0.20042442 | 0.0111 | 0.012 | 0.0116 | 0.0116 | TRUE | 0.2509 |
| Relb | 572 | 777 | 827 | 677 | 482 | 555 | 608 | 776 | 830 | 532 | 519 | 608 | 645.38 | 738 | 553 | 0.748 | -0.418 | 0.00409753 | 0.02691639 | 0.0152 | 0.012 | 0.0137 | 0.0137 | TRUE | 0.2814 |
| Rell1 | 1952 | 2332 | 2212 | 2514 | 2027 | 2434 | 2073 | 2329 | 2221 | 1976 | 2181 | 2664 | 2240.85 | 2208 | 2274 | 1.029 | 0.042 | 0.73979602 | 0.86066702 | 0.013 | 0.0084 | 0.0109 | 0.0109 | TRUE | 0.2729 |

| | | | | | | | | | | | | | | | | | | | | | | | | | |
|---|---|---|---|---|---|---|---|---|---|---|---|---|---|---|---|---|---|---|---|---|---|---|---|---|---|
| Rell2 | 98 | 119 | 108 | 128 | 108 | 126 | 104 | 119 | 108 | 101 | 116 | 138 | 114.36 | 110 | 118 | 1.065 | 0.091 | 0.65550762 | 0.80619668 | 0.0075 | 0.0357 | 0.0214 | 0.0214 | TRUE | 0.2103 |
| Reln | 262 | 219 | 243 | 168 | 121 | 116 | 278 | 219 | 244 | 132 | 130 | 127 | 188.37 | 247 | 130 | 0.526 | -0.927 | 1.82E-08 | 8.21E-07 | 0 | 0.0244 | 0.0137 | 0.0137 | TRUE | 0.1348 |
| Relt | 435 | 542 | 509 | 619 | 445 | 400 | 462 | 541 | 511 | 487 | 479 | 438 | 486.28 | 505 | 468 | 0.927 | -0.109 | 0.34516821 | 0.56455387 | 0.0026 | 0.0137 | 0.0076 | 0.0076 | TRUE | 0.0634 |
| Rem1 | 28 | 26 | 21 | 22 | 17 | 15 | 30 | 26 | 21 | 17 | 18 | 16 | 21.47 | 26 | 17 | 0.678 | -0.561 | 0.17181745 | 0.37064898 | 0 | 0.1604 | 0.0732 | 0.0732 | TRUE | 0.1473 |
| Rem2 | 158 | 82 | 84 | 115 | 75 | 59 | 168 | 82 | 84 | 90 | 81 | 65 | 94.96 | 111 | 79 | 0.708 | -0.498 | 0.12728747 | 0.30654725 | 0.094 | 0.0416 | 0.066 | 0.066 | TRUE | 1.9416 |
| Ren1 | 0 | 0 | 0 | 0 | 0 | 0 | 0 | 0 | 0 | 0 | 0 | 0 | 0 | 0 | 0 | NA | NA | NA | NA | NA | NA | NA | NA | NA | NA |
| Ren2 | 0 | 0 | 0 | 0 | 1 | 1 | 0 | 0 | 0 | 0 | 1 | 1 | 0.36 | 0 | 1 | 3.794 | 1.924 | 0.60247382 | NA | 0 | 9.1135 | 8.1425 | 8.1425 | TRUE | 0.3482 |
| Renbp | 911 | 1263 | 1284 | 1237 | 970 | 1149 | 968 | 1261 | 1289 | 972 | 1044 | 1258 | 1132.02 | 1173 | 1091 | 0.93 | -0.105 | 0.48239016 | 0.68092108 | 0.0203 | 0.0098 | 0.0152 | 0.0152 | TRUE | 0.2797 |
| Rep15 | 0 | 1 | 0 | 1 | 0 | 0 | 0 | 1 | 0 | 1 | 0 | 0 | 0.3 | 0 | 0 | 0.954 | -0.067 | 0.98642001 | NA | 0 | 11.0817 | 9.3014 | 9.3014 | TRUE | 0.405 |
| Repin1 | 388 | 626 | 617 | 543 | 384 | 445 | 412 | 625 | 620 | 427 | 413 | 487 | 497.33 | 552 | 442 | 0.8 | -0.321 | 0.07254943 | 0.21080501 | 0.0285 | 0.0135 | 0.021 | 0.021 | TRUE | 0.5738 |
| Reps1 | 3427 | 4082 | 4126 | 4737 | 3458 | 3738 | 3640 | 4077 | 4143 | 3724 | 3721 | 4092 | 3899.34 | 3953 | 3846 | 0.972 | -0.04 | 0.64433688 | 0.79996825 | 0.0031 | 0.0077 | 0.0052 | 0.0052 | TRUE | 0.0583 |
| Reps2 | 277 | 282 | 322 | 426 | 363 | 435 | 294 | 282 | 323 | 335 | 391 | 476 | 350.14 | 300 | 401 | 1.333 | 0.415 | 0.01079588 | 0.05571471 | 0.0161 | 0.0163 | 0.0162 | 0.0162 | TRUE | 0.2596 |
| Rer1 | 3838 | 4507 | 4330 | 5158 | 3842 | 3929 | 4077 | 4501 | 4348 | 4055 | 4134 | 4301 | 4235.85 | 4309 | 4163 | 0.966 | -0.05 | 0.49265554 | 0.68896642 | 0.0013 | 0.0077 | 0.0036 | 0.0036 | TRUE | 0.0269 |
| Rere | 5437 | 5869 | 6541 | 6595 | 5031 | 5092 | 5775 | 5861 | 6568 | 5184 | 5413 | 5574 | 5729.31 | 6068 | 5390 | 0.888 | -0.171 | 0.03596911 | 0.13026862 | 0.003 | 0.0075 | 0.0046 | 0.0046 | TRUE | 0.0634 |
| Rerg | 276 | 243 | 206 | 175 | 126 | 142 | 293 | 243 | 207 | 138 | 136 | 155 | 195.21 | 248 | 143 | 0.577 | -0.794 | 1.56E-05 | 0.00028877 | 0.0143 | 0.0238 | 0.0188 | 0.0188 | TRUE | 0.1861 |
| Rergl | 0 | 0 | 0 | 0 | 0 | 0 | 0 | 0 | 0 | 0 | 0 | 0 | 0 | 0 | 0 | NA | NA | NA | NA | NA | NA | NA | NA | NA | NA |
| Resp18 | 3 | 4 | 0 | 2 | 3 | 3 | 3 | 4 | 0 | 2 | 3 | 3 | 2.54 | 2 | 3 | 1.112 | 0.153 | 0.90565314 | NA | 0 | 1.3018 | 0.81 | 0.81 | TRUE | 0.8245 |
| Rest | 6740 | 6979 | 6918 | 8660 | 6518 | 5989 | 7159 | 6970 | 6946 | 6808 | 7013 | 6556 | 6908.69 | 7025 | 6792 | 0.967 | -0.049 | 0.43099736 | 0.64068713 | 5.00E-04 | 0.0074 | 0.0026 | 0.0026 | TRUE | 0.0112 |
| Ret | 804 | 789 | 711 | 954 | 765 | 694 | 854 | 788 | 714 | 750 | 823 | 760 | 781.44 | 785 | 778 | 0.99 | -0.015 | 0.89161965 | 0.94748855 | 0.004 | 0.0111 | 0.007 | 0.007 | TRUE | 0.0751 |
| Retn | 4 | 3 | 2 | 2 | 2 | 3 | 4 | 3 | 2 | 2 | 2 | 3 | 2.71 | 3 | 2 | 0.747 | -0.421 | 0.70648686 | NA | 0 | 1.2225 | 0.5205 | 0.5205 | TRUE | 0.1372 |
| Retnla | 0 | 0 | 0 | 0 | 0 | 0 | 0 | 0 | 0 | 0 | 0 | 0 | 0 | 0 | 0 | NA | NA | NA | NA | NA | NA | NA | NA | NA | NA |
| Retnlb | 0 | 0 | 0 | 0 | 0 | 0 | 0 | 0 | 0 | 0 | 0 | 0 | 0 | 0 | 0 | NA | NA | NA | NA | NA | NA | NA | NA | NA | NA |
| Retnlg | 0 | 0 | 0 | 0 | 0 | 0 | 0 | 0 | 0 | 0 | 0 | 0 | 0 | 0 | 0 | NA | NA | NA | NA | NA | NA | NA | NA | NA | NA |
| Retsat | 798 | 1032 | 994 | 1095 | 803 | 927 | 848 | 1031 | 998 | 861 | 864 | 1015 | 935.98 | 959 | 913 | 0.952 | -0.072 | 0.55398969 | 0.73738555 | 0.0087 | 0.0104 | 0.0095 | 0.0095 | TRUE | 0.1221 |
| Rev1 | 2936 | 3347 | 3281 | 3284 | 2726 | 2869 | 3119 | 3343 | 3294 | 2582 | 2933 | 3141 | 3068.48 | 3252 | 2885 | 0.887 | -0.174 | 0.07109071 | 0.20759909 | 0.005 | 0.008 | 0.0063 | 0.0063 | TRUE | 0.1042 |
| Rev3l | 4506 | 3815 | 4425 | 5025 | 3560 | 3435 | 4786 | 3810 | 4443 | 3950 | 3830 | 3760 | 4096.69 | 4346 | 3847 | 0.885 | -0.176 | 0.08221598 | 0.22889833 | 0.0064 | 0.0077 | 0.0071 | 0.0071 | TRUE | 0.142 |
| Rex2 | 1105 | 856 | 808 | 1520 | 1141 | 990 | 1174 | 855 | 811 | 1195 | 1228 | 1084 | 1057.7 | 947 | 1169 | 1.235 | 0.305 | 0.04625043 | 0.1556351 | 0.0225 | 0.01 | 0.0159 | 0.0159 | TRUE | 0.5237 |
| Rexo1 | 5113 | 4740 | 4826 | 5653 | 4526 | 4291 | 5431 | 4734 | 4846 | 4444 | 4870 | 4697 | 4836.9 | 5004 | 4670 | 0.933 | -0.1 | 0.24462966 | 0.4611329 | 0.0035 | 0.0076 | 0.0051 | 0.0051 | TRUE | 0.0679 |
| Rexo2 | 1828 | 1508 | 1437 | 2351 | 1697 | 1363 | 1942 | 1506 | 1443 | 1848 | 1826 | 1492 | 1676.11 | 1630 | 1722 | 1.057 | 0.08 | 0.58080194 | 0.75724627 | 0.0195 | 0.0089 | 0.0144 | 0.0144 | TRUE | 0.3363 |
| Rexo4 | 3000 | 2993 | 2871 | 3306 | 2463 | 2324 | 3187 | 2989 | 2883 | 2599 | 2650 | 2544 | 2808.57 | 3020 | 2598 | 0.86 | -0.217 | 0.00317475 | 0.02218527 | 0.001 | 0.0081 | 0.0035 | 0.0035 | TRUE | 0.0283 |
| Rfc1 | 12507 | 12510 | 12964 | 15424 | 11075 | 11296 | 13285 | 12494 | 13017 | 12125 | 11916 | 12365 | 12533.69 | 12932 | 12135 | 0.938 | -0.092 | 0.12338529 | 0.2994547 | 5.00E-04 | 0.0072 | 0.0025 | 0.0025 | TRUE | 0.0107 |
| Rfc2 | 4395 | 4691 | 4714 | 6018 | 4335 | 4094 | 4668 | 4685 | 4733 | 4731 | 4664 | 4482 | 4660.53 | 4695 | 4626 | 0.985 | -0.021 | 0.7237332 | 0.85044852 | 2.00E-04 | 0.0076 | 0.0024 | 0.0024 | TRUE | 0.009 |
| Rfc3 | 1700 | 1863 | 1835 | 2853 | 2103 | 1990 | 1806 | 1861 | 1843 | 2243 | 2263 | 2178 | 2032.12 | 1837 | 2228 | 1.213 | 0.279 | 3.20E-05 | 0.0005319 | 0 | 0.0085 | 0.0027 | 0.0027 | TRUE | 0.0045 |
| Rfc4 | 2279 | 2853 | 2788 | 3265 | 2293 | 2570 | 2421 | 2849 | 2799 | 2567 | 2467 | 2813 | 2652.76 | 2690 | 2616 | 0.972 | -0.041 | 0.68274924 | 0.82414825 | 0.0058 | 0.0081 | 0.0067 | 0.0067 | TRUE | 0.0927 |
| Rfc5 | 8372 | 7875 | 8227 | 12197 | 8574 | 8227 | 8893 | 7865 | 8261 | 9588 | 9225 | 9006 | 8806.22 | 8340 | 9273 | 1.112 | 0.153 | 0.04263299 | 0.14742545 | 0.0023 | 0.0073 | 0.004 | 0.004 | TRUE | 0.0411 |
| Rfesd | 296 | 306 | 301 | 409 | 282 | 268 | 314 | 306 | 302 | 322 | 303 | 293 | 306.76 | 307 | 306 | 0.998 | -0.004 | 0.97620108 | 0.98709087 | 0 | 0.0176 | 0.0072 | 0.0072 | TRUE | 0.0237 |
| Rffl | 1447 | 1684 | 1690 | 2687 | 1982 | 1927 | 1537 | 1682 | 1697 | 2112 | 2133 | 2109 | 1878.33 | 1639 | 2118 | 1.292 | 0.37 | 1.37E-06 | 3.62E-05 | 7.00E-04 | 0.0087 | 0.0037 | 0.0037 | TRUE | 0.0353 |
| Rfk | 1673 | 1899 | 1800 | 2984 | 2150 | 2104 | 1777 | 1897 | 1807 | 2346 | 2313 | 2303 | 2073.86 | 1827 | 2321 | 1.27 | 0.345 | 6.36E-07 | 1.85E-05 | 0 | 0.0085 | 0.003 | 0.003 | TRUE | 0.0134 |
| Rfng | 1158 | 1050 | 1034 | 1247 | 928 | 909 | 1230 | 1049 | 1038 | 980 | 998 | 995 | 1048.45 | 1106 | 991 | 0.897 | -0.157 | 0.11785145 | 0.29041792 | 0.0038 | 0.01 | 0.0063 | 0.0063 | TRUE | 0.1153 |
| Rfpl3s | 0 | 0 | 0 | 1 | 0 | 0 | 0 | 0 | 0 | 1 | 0 | 0 | 0.13 | 0 | 0 | 1.859 | 0.894 | 0.82650734 | NA | 0 | 25.1515 | 10 | 10 | TRUE | 0.2563 |
| Rfpl4 | 1 | 0 | 0 | 0 | 0 | 0 | 1 | 0 | 0 | 0 | 0 | 0 | 0.18 | 0 | 0 | 0.49 | -1.029 | 0.80087 | NA | 0 | 18.6165 | 10 | 10 | TRUE | 0.4614 |
| Rfpl4b | 4 | 3 | 1 | 12 | 7 | 2 | 4 | 3 | 1 | 9 | 8 | 2 | 4.57 | 3 | 6 | 2.362 | 1.24 | 0.2191293 | NA | 0 | 0.7282 | 0.4715 | 0.4715 | TRUE | 0.5104 |
| Rft1 | 599 | 587 | 531 | 796 | 584 | 495 | 636 | 586 | 533 | 626 | 628 | 542 | 591.94 | 585 | 599 | 1.024 | 0.034 | 0.77450175 | 0.8825015 | 0.0055 | 0.0125 | 0.0086 | 0.0086 | TRUE | 0.0791 |
| Rftn1 | 566 | 686 | 687 | 873 | 594 | 622 | 601 | 685 | 690 | 686 | 639 | 681 | 663.73 | 659 | 669 | 1.015 | 0.022 | 0.83211557 | 0.91487212 | 0.002 | 0.0119 | 0.0063 | 0.0063 | TRUE | 0.0681 |
| Rftn2 | 9 | 32 | 21 | 24 | 12 | 16 | 10 | 32 | 21 | 19 | 13 | 18 | 18.65 | 21 | 17 | 0.788 | -0.343 | 0.52640362 | 0.71554235 | 0.1285 | 0.1836 | 0.1571 | 0.1571 | TRUE | 0.1572 |
| Rfwd2 | 1230 | 1306 | 1380 | 1754 | 1206 | 1271 | 1306 | 1304 | 1386 | 1379 | 1298 | 1391 | 1344.03 | 1332 | 1356 | 1.018 | 0.026 | 0.74667965 | 0.86556561 | 5.00E-04 | 0.0094 | 0.0038 | 0.0038 | TRUE | 0.0167 |
| Rfwd3 | 12064 | 11733 | 12240 | 15352 | 11011 | 11273 | 12814 | 11718 | 12290 | 12069 | 11847 | 12340 | 12179.66 | 12274 | 12085 | 0.985 | -0.022 | 0.73223496 | 0.85630933 | 0.001 | 0.0072 | 0.003 | 0.003 | TRUE | 0.0192 |
| Rfx1 | 2491 | 2355 | 2505 | 3113 | 2531 | 2408 | 2646 | 2352 | 2515 | 2447 | 2723 | 2636 | 2553.24 | 2504 | 2602 | 1.039 | 0.055 | 0.52112272 | 0.7115529 | 0.0026 | 0.0082 | 0.0049 | 0.0049 | TRUE | 0.0344 |
| Rfx2 | 3647 | 3316 | 3332 | 3177 | 2589 | 2828 | 3874 | 3312 | 3346 | 2498 | 2786 | 3096 | 3151.66 | 3511 | 2793 | 0.795 | -0.33 | 0.00320544 | 0.02232063 | 0.0093 | 0.0079 | 0.0087 | 0.0087 | TRUE | 0.1091 |
| Rfx3 | 288 | 389 | 486 | 465 | 344 | 401 | 306 | 388 | 488 | 366 | 370 | 439 | 392.84 | 394 | 392 | 0.992 | -0.011 | 0.95190056 | 0.97542891 | 0.0295 | 0.0153 | 0.0224 | 0.0224 | TRUE | 0.2356 |
| Rfx4 | 49 | 50 | 34 | 29 | 19 | 35 | 52 | 50 | 34 | 23 | 20 | 38 | 36.28 | 45 | 27 | 0.596 | -0.747 | 0.04946728 | 0.16315305 | 0.0484 | 0.0977 | 0.0749 | 0.0749 | TRUE | 0.7556 |
| Rfx5 | 828 | 775 | 726 | 1028 | 777 | 826 | 879 | 774 | 729 | 808 | 836 | 904 | 821.8 | 794 | 849 | 1.069 | 0.097 | 0.38186454 | 0.59601856 | 0.005 | 0.0109 | 0.0076 | 0.0076 | TRUE | 0.1042 |
| Rfx6 | 2 | 3 | 1 | 4 | 3 | 1 | 2 | 3 | 1 | 3 | 3 | 1 | 2.27 | 2 | 2 | 1.233 | 0.302 | 0.80710066 | NA | 0 | 1.4612 | 0.6583 | 0.6583 | TRUE | 0.2315 |
| Rfx7 | 4688 | 4002 | 4151 | 6126 | 4400 | 4039 | 4979 | 3997 | 4168 | 4816 | 4734 | 4421 | 4519.27 | 4381 | 4657 | 1.063 | 0.088 | 0.39891024 | 0.61202615 | 0.0071 | 0.0076 | 0.0077 | 0.0077 | TRUE | 0.1734 |
| Rfx8 | 14 | 40 | 29 | 23 | 23 | 19 | 15 | 40 | 29 | 18 | 25 | 21 | 24.59 | 28 | 21 | 0.753 | -0.409 | 0.37960094 | 0.59448576 | 0.0873 | 0.1409 | 0.1149 | 0.1149 | TRUE | 0.2067 |
| Rfxank | 388 | 372 | 398 | 599 | 477 | 424 | 412 | 372 | 400 | 471 | 513 | 464 | 438.59 | 395 | 483 | 1.224 | 0.291 | 0.00946627 | 0.05049106 | 0 | 0.0144 | 0.0068 | 0.0068 | TRUE | 0.0345 |
| Rfxap | 786 | 819 | 815 | 1025 | 824 | 838 | 835 | 818 | 818 | 806 | 887 | 917 | 846.81 | 824 | 870 | 1.055 | 0.077 | 0.41239223 | 0.62355002 | 0.0014 | 0.0108 | 0.0051 | 0.0051 | TRUE | 0.0518 |
| Rgag1 | 2 | 1 | 0 | 2 | 2 | 0 | 2 | 1 | 0 | 2 | 2 | 0 | 1.14 | 1 | 1 | 1.213 | 0.279 | 0.8846965 | NA | 0 | 2.8937 | 1.7759 | 1.7759 | TRUE | 0.2368 |
| Rgag4 | 19 | 25 | 19 | 38 | 21 | 29 | 20 | 25 | 19 | 30 | 23 | 32 | 24.74 | 21 | 28 | 1.313 | 0.393 | 0.31324199 | 0.53375678 | 0 | 0.1401 | 0.0681 | 0.0681 | TRUE | 0.1736 |
| Rgcc | 67 | 182 | 119 | 37 | 47 | 90 | 71 | 182 | 119 | 29 | 51 | 99 | 91.77 | 124 | 60 | 0.475 | -1.073 | 0.01933555 | 0.08468765 | 0.2602 | 0.0428 | 0.1392 | 0.1392 | TRUE | 0.2855 |
| Rgl1 | 397 | 405 | 458 | 427 | 333 | 314 | 422 | 404 | 460 | 336 | 358 | 344 | 387.29 | 429 | 346 | 0.806 | -0.311 | 0.00747566 | 0.04212792 | 0 | 0.0154 | 0.0071 | 0.0071 | TRUE | 0.0472 |
| Rgl2 | 1688 | 1548 | 1466 | 1868 | 1496 | 1221 | 1793 | 1546 | 1472 | 1468 | 1610 | 1337 | 1537.61 | 1604 | 1472 | 0.918 | -0.124 | 0.28433079 | 0.5058795 | 0.0088 | 0.009 | 0.009 | 0.009 | TRUE | 0.1282 |
| Rgl3 | 80 | 201 | 210 | 71 | 68 | 80 | 85 | 201 | 211 | 56 | 73 | 88 | 118.85 | 166 | 72 | 0.434 | -1.203 | 0.00061181 | 0.00609706 | 0.1301 | 0.0346 | 0.079 | 0.079 | TRUE | 1.9053 |
| Rgma | 123 | 112 | 89 | 123 | 118 | 101 | 131 | 112 | 89 | 97 | 127 | 111 | 111.01 | 111 | 112 | 1.005 | 0.007 | 0.97619959 | 0.98709087 | 0.0181 | 0.0366 | 0.0274 | 0.0274 | TRUE | 0.2369 |
| Rgmb | 232 | 263 | 273 | 359 | 280 | 275 | 246 | 263 | 274 | 282 | 301 | 301 | 277.95 | 261 | 295 | 1.127 | 0.173 | 0.17458435 | 0.37396928 | 0 | 0.0188 | 0.0081 | 0.0081 | TRUE | 0.0264 |
| Rgn | 7 | 9 | 9 | 8 | 11 | 9 | 7 | 9 | 9 | 6 | 12 | 10 | 8.91 | 8 | 9 | 1.084 | 0.116 | 0.85473135 | 0.92766466 | 0 | 0.3768 | 0.1765 | 0.1765 | TRUE | 0.3517 |
| Rgp1 | 1596 | 1981 | 2180 | 2177 | 1651 | 1746 | 1695 | 1978 | 2189 | 1711 | 1776 | 1911 | 1876.94 | 1954 | 1799 | 0.921 | -0.119 | 0.29842521 | 0.51905693 | 0.0088 | 0.0087 | 0.009 | 0.009 | TRUE | 0.162 |
| Rgr | 0 | 0 | 0 | 0 | 0 | 0 | 0 | 0 | 0 | 0 | 0 | 0 | 0 | 0 | 0 | NA | NA | NA | NA | NA | NA | NA | NA | NA | NA |
| Rgs1 | 0 | 0 | 0 | 0 | 0 | 0 | 0 | 0 | 0 | 0 | 0 | 0 | 0 | 0 | 0 | NA | NA | NA | NA | NA | NA | NA | NA | NA | NA |

| | | | | | | | | | | | | | | | | | | | | | | | | | |
|---|---|---|---|---|---|---|---|---|---|---|---|---|---|---|---|---|---|---|---|---|---|---|---|---|---|
| Rgs10 | 288 | 313 | 312 | 334 | 233 | 236 | 306 | 313 | 313 | 263 | 251 | 258 | 283.9 | 311 | 257 | 0.829 | -0.271 | 0.02573637 | 0.10376127 | 0 | 0.0185 | 0.0071 | 0.0071 | TRUE | 0.0054 |
| Rgs11 | 193 | 258 | 234 | 268 | 247 | 236 | 205 | 258 | 235 | 211 | 266 | 258 | 238.73 | 233 | 245 | 1.05 | 0.07 | 0.67222524 | 0.81726174 | 0.0105 | 0.0207 | 0.0154 | 0.0154 | TRUE | 0.178 |
| Rgs12 | 2347 | 2655 | 2788 | 2661 | 2045 | 2162 | 2493 | 2652 | 2799 | 2092 | 2200 | 2367 | 2433.79 | 2648 | 2220 | 0.838 | -0.255 | 0.00357902 | 0.02423711 | 0.003 | 0.0083 | 0.0051 | 0.0051 | TRUE | 0.0406 |
| Rgs13 | 0 | 0 | 0 | 0 | 0 | 0 | 0 | 0 | 0 | 0 | 0 | 0 | 0 | 0 | 0 | NA | NA | NA | NA | NA | NA | NA | NA | NA | NA |
| Rgs14 | 51 | 80 | 65 | 82 | 66 | 81 | 54 | 80 | 65 | 64 | 71 | 89 | 70.58 | 66 | 75 | 1.119 | 0.162 | 0.5413694 | 0.72705025 | 0.0181 | 0.0536 | 0.0363 | 0.0363 | TRUE | 0.2793 |
| Rgs16 | 314 | 262 | 249 | 174 | 131 | 146 | 334 | 262 | 250 | 137 | 141 | 160 | 213.79 | 282 | 146 | 0.517 | -0.952 | 6.19E-08 | 2.39E-06 | 0.0124 | 0.0223 | 0.0171 | 0.0171 | TRUE | 0.2878 |
| Rgs17 | 300 | 321 | 317 | 173 | 135 | 150 | 319 | 321 | 318 | 136 | 145 | 164 | 233.83 | 319 | 148 | 0.463 | -1.109 | 1.01E-14 | 1.35E-12 | 0 | 0.021 | 0.0099 | 0.0099 | TRUE | 0.0872 |
| Rgs18 | 0 | 0 | 0 | 0 | 0 | 0 | 0 | 0 | 0 | 0 | 0 | 0 | 0 | 0 | 0 | NA | NA | NA | NA | NA | NA | NA | NA | NA | NA |
| Rgs19 | 1558 | 1702 | 1670 | 2070 | 1472 | 1373 | 1655 | 1700 | 1677 | 1627 | 1584 | 1503 | 1624.26 | 1677 | 1571 | 0.937 | -0.093 | 0.21087264 | 0.42147804 | 0 | 0.0089 | 0.0034 | 0.0034 | TRUE | 0.0173 |
| Rgs2 | 41 | 40 | 27 | 53 | 42 | 31 | 44 | 40 | 27 | 42 | 45 | 34 | 38.57 | 37 | 40 | 1.095 | 0.131 | 0.69555843 | 0.83329644 | 0 | 0.0923 | 0.0543 | 0.0543 | TRUE | 0.3927 |
| Rgs20 | 61 | 61 | 85 | 69 | 38 | 43 | 65 | 61 | 85 | 54 | 41 | 47 | 58.88 | 70 | 47 | 0.677 | -0.563 | 0.0424548 | 0.14703492 | 0.0115 | 0.0629 | 0.0379 | 0.0379 | TRUE | 0.3156 |
| Rgs22 | 150 | 244 | 250 | 182 | 162 | 181 | 159 | 244 | 251 | 143 | 174 | 198 | 194.92 | 218 | 172 | 0.785 | -0.348 | 0.12007807 | 0.2939912 | 0.0384 | 0.0238 | 0.031 | 0.031 | TRUE | 0.6016 |
| Rgs3 | 671 | 881 | 973 | 530 | 497 | 577 | 713 | 880 | 977 | 417 | 535 | 632 | 692.1 | 857 | 528 | 0.615 | -0.702 | 9.80E-05 | 0.00135768 | 0.0307 | 0.0117 | 0.0218 | 0.0218 | TRUE | 0.4078 |
| Rgs4 | 4 | 4 | 2 | 4 | 3 | 3 | 4 | 4 | 2 | 3 | 3 | 3 | 3.32 | 3 | 3 | 0.943 | -0.085 | 0.93187542 | NA | 0 | 0.9998 | 0.4156 | 0.4156 | TRUE | 0.1924 |
| Rgs5 | 5 | 2 | 2 | 31 | 17 | 12 | 5 | 2 | 2 | 24 | 18 | 13 | 10.85 | 3 | 18 | 6.085 | 2.605 | 0.00037114 | 0.00408773 | 0 | 0.3105 | 0.1939 | 0.1939 | TRUE | 0.2632 |
| Rgs6 | 2 | 4 | 2 | 7 | 3 | 6 | 2 | 4 | 2 | 6 | 3 | 7 | 3.9 | 3 | 5 | 1.882 | 0.912 | 0.34537168 | NA | 0 | 0.8507 | 0.3896 | 0.3896 | TRUE | 0.2086 |
| Rgs7 | 14 | 14 | 17 | 26 | 19 | 15 | 15 | 14 | 17 | 20 | 20 | 16 | 17.2 | 15 | 19 | 1.251 | 0.323 | 0.47183231 | 0.67314138 | 0 | 0.1984 | 0.0868 | 0.0868 | TRUE | 0.0724 |
| Rgs7bp | 2 | 5 | 3 | 19 | 7 | 12 | 2 | 5 | 3 | 15 | 8 | 13 | 7.62 | 3 | 12 | 3.522 | 1.816 | 0.01879438 | 0.083172 | 0 | 0.4391 | 0.2392 | 0.2392 | TRUE | 0.3749 |
| Rgs8 | 5 | 5 | 3 | 10 | 2 | 3 | 5 | 5 | 3 | 8 | 2 | 3 | 4.44 | 4 | 4 | 1.028 | 0.039 | 0.96651687 | NA | 0 | 0.7496 | 0.4084 | 0.4084 | TRUE | 1.0727 |
| Rgs9 | 145 | 242 | 225 | 339 | 339 | 312 | 154 | 242 | 226 | 266 | 365 | 342 | 265.73 | 207 | 324 | 1.561 | 0.642 | 0.0021094 | 0.01606442 | 0.0362 | 0.0193 | 0.0275 | 0.0275 | TRUE | 0.5436 |
| Rgs9bp | 205 | 417 | 449 | 389 | 311 | 354 | 218 | 416 | 451 | 306 | 335 | 388 | 352.16 | 362 | 343 | 0.946 | -0.08 | 0.75252404 | 0.86948897 | 0.0753 | 0.0163 | 0.0432 | 0.0432 | TRUE | 1.3786 |
| Rgsl1 | 1 | 1 | 0 | 1 | 0 | 0 | 1 | 1 | 0 | 1 | 0 | 0 | 0.47 | 1 | 0 | 0.51 | -0.973 | 0.73452857 | NA | 0 | 6.9499 | 4.1835 | 4.1835 | TRUE | 0.2562 |
| Rhag | 0 | 0 | 0 | 0 | 0 | 0 | 0 | 0 | 0 | 0 | 0 | 0 | 0 | 0 | 0 | NA | NA | NA | NA | NA | NA | NA | NA | NA | NA |
| Rhbdd1 | 863 | 983 | 1044 | 1101 | 751 | 752 | 917 | 982 | 1048 | 866 | 808 | 823 | 907.23 | 982 | 832 | 0.848 | -0.238 | 0.01200625 | 0.06015328 | 0.0019 | 0.0105 | 0.0054 | 0.0054 | TRUE | 0.0413 |
| Rhbdd2 | 866 | 1014 | 1082 | 1324 | 1030 | 983 | 920 | 1013 | 1086 | 1041 | 1108 | 1076 | 1040.68 | 1006 | 1075 | 1.068 | 0.094 | 0.33215728 | 0.55255612 | 0.0029 | 0.0101 | 0.0059 | 0.0059 | TRUE | 0.067 |
| Rhbdd3 | 1131 | 1309 | 1258 | 1522 | 1138 | 1064 | 1201 | 1307 | 1263 | 1196 | 1224 | 1165 | 1226.23 | 1257 | 1195 | 0.951 | -0.073 | 0.36375967 | 0.58120643 | 0 | 0.0096 | 0.0039 | 0.0039 | TRUE | 0.0181 |
| Rhbdf1 | 507 | 698 | 765 | 801 | 653 | 752 | 539 | 697 | 768 | 630 | 703 | 823 | 693.2 | 668 | 719 | 1.074 | 0.104 | 0.52705512 | 0.71593432 | 0.0238 | 0.0117 | 0.0178 | 0.0178 | TRUE | 0.3371 |
| Rhbdf2 | 4807 | 5209 | 5127 | 4601 | 3705 | 4292 | 5106 | 5202 | 5148 | 3617 | 3986 | 4698 | 4626.3 | 5152 | 4100 | 0.796 | -0.33 | 0.00225062 | 0.01688715 | 0.0087 | 0.0076 | 0.0082 | 0.0082 | TRUE | 0.1978 |
| Rhbdl1 | 42 | 43 | 33 | 42 | 39 | 39 | 45 | 43 | 33 | 33 | 42 | 43 | 39.73 | 40 | 39 | 0.97 | -0.044 | 0.88878345 | 0.94574906 | 0 | 0.0898 | 0.0464 | 0.0464 | TRUE | 0.1805 |
| Rhbdl2 | 137 | 88 | 68 | 103 | 54 | 42 | 146 | 88 | 68 | 81 | 58 | 46 | 81.12 | 101 | 62 | 0.618 | -0.695 | 0.04622968 | 0.15563032 | 0.104 | 0.0475 | 0.0747 | 0.0747 | TRUE | 0.6365 |
| Rhbdl3 | 79 | 111 | 92 | 117 | 85 | 76 | 84 | 111 | 92 | 92 | 91 | 83 | 92.3 | 96 | 89 | 0.929 | -0.106 | 0.62014234 | 0.78498274 | 0 | 0.0426 | 0.0223 | 0.0223 | TRUE | 0.1865 |
| Rhbg | 47 | 37 | 26 | 67 | 33 | 34 | 50 | 37 | 26 | 53 | 36 | 37 | 39.73 | 38 | 42 | 1.119 | 0.163 | 0.65573017 | 0.80634463 | 0.0499 | 0.0898 | 0.0705 | 0.0705 | TRUE | 0.2543 |
| Rhcg | 10 | 12 | 8 | 25 | 12 | 8 | 11 | 12 | 8 | 20 | 13 | 9 | 11.99 | 10 | 14 | 1.368 | 0.452 | 0.43878258 | 0.6460857 | 0 | 0.2816 | 0.1602 | 0.1602 | TRUE | 0.3244 |
| Rhd | 424 | 405 | 373 | 403 | 269 | 322 | 450 | 404 | 375 | 317 | 289 | 352 | 364.68 | 410 | 319 | 0.78 | -0.359 | 0.00873396 | 0.04751707 | 0.006 | 0.0159 | 0.0107 | 0.0107 | TRUE | 0.0877 |
| Rheb | 3718 | 4101 | 3886 | 4715 | 3357 | 3278 | 3949 | 4096 | 3902 | 3707 | 3612 | 3588 | 3808.93 | 3982 | 3636 | 0.913 | -0.131 | 0.03565945 | 0.12960084 | 2.00E-04 | 0.0078 | 0.0025 | 0.0025 | TRUE | 0.0076 |
| Rhebl1 | 1528 | 1495 | 1504 | 1830 | 1358 | 1367 | 1623 | 1493 | 1510 | 1439 | 1461 | 1496 | 1503.73 | 1542 | 1465 | 0.95 | -0.074 | 0.34262084 | 0.56278024 | 6.00E-04 | 0.0091 | 0.0037 | 0.0037 | TRUE | 0.0252 |
| Rhno1 | 1604 | 1545 | 1447 | 1885 | 1504 | 1388 | 1704 | 1543 | 1453 | 1482 | 1618 | 1519 | 1553.19 | 1567 | 1540 | 0.983 | -0.025 | 0.79013291 | 0.8919706 | 0.0036 | 0.009 | 0.0058 | 0.0058 | TRUE | 0.0704 |
| Rho | 11 | 12 | 9 | 24 | 24 | 20 | 12 | 12 | 9 | 19 | 26 | 22 | 16.55 | 11 | 22 | 2.027 | 1.019 | 0.03296898 | 0.12272521 | 0 | 0.206 | 0.0957 | 0.0957 | TRUE | 0.1203 |
| Rhoa | 14966 | 16521 | 15775 | 22700 | 15843 | 15840 | 15897 | 16500 | 15840 | 17845 | 17046 | 17339 | 16744.4 | 16079 | 17410 | 1.083 | 0.115 | 0.04523985 | 0.1534128 | 4.00E-04 | 0.0071 | 0.0023 | 0.0023 | TRUE | 0.0064 |
| Rhob | 1685 | 1529 | 955 | 3186 | 1937 | 1168 | 1790 | 1527 | 959 | 2505 | 2084 | 1279 | 1690.49 | 1425 | 1956 | 1.372 | 0.457 | 0.20793252 | 0.4172612 | 0.0942 | 0.0089 | 0.0484 | 0.0942 | TRUE | 0.3561 |
| Rhobtb1 | 545 | 634 | 613 | 813 | 563 | 483 | 579 | 633 | 616 | 639 | 606 | 529 | 600.2 | 609 | 591 | 0.972 | -0.041 | 0.71737886 | 0.8467669 | 0.004 | 0.0124 | 0.0077 | 0.0077 | TRUE | 0.0987 |
| Rhobtb2 | 1123 | 1370 | 1342 | 1462 | 1091 | 1198 | 1193 | 1368 | 1348 | 1149 | 1174 | 1311 | 1257.19 | 1303 | 1211 | 0.929 | -0.106 | 0.29435692 | 0.51515728 | 0.0041 | 0.0095 | 0.0066 | 0.0066 | TRUE | 0.0652 |
| Rhobtb3 | 860 | 1224 | 1298 | 1605 | 1152 | 1291 | 913 | 1222 | 1303 | 1262 | 1239 | 1413 | 1225.6 | 1146 | 1305 | 1.138 | 0.186 | 0.19977587 | 0.40732488 | 0.0175 | 0.0096 | 0.0144 | 0.0144 | TRUE | 0.3772 |
| Rhoc | 1368 | 1547 | 1655 | 2094 | 1454 | 1441 | 1453 | 1545 | 1662 | 1646 | 1564 | 1577 | 1574.63 | 1553 | 1596 | 1.028 | 0.039 | 0.64546766 | 0.80077998 | 0.002 | 0.009 | 0.0047 | 0.0047 | TRUE | 0.045 |
| Rhod | 382 | 420 | 429 | 464 | 309 | 345 | 406 | 419 | 431 | 365 | 332 | 378 | 388.48 | 419 | 358 | 0.856 | -0.224 | 0.05064473 | 0.16557427 | 0 | 0.0154 | 0.0069 | 0.0069 | TRUE | 0.0434 |
| Rhof | 277 | 179 | 188 | 219 | 159 | 148 | 294 | 179 | 189 | 172 | 171 | 162 | 194.5 | 221 | 168 | 0.764 | -0.387 | 0.07803939 | 0.22101584 | 0.0347 | 0.0238 | 0.0296 | 0.0296 | TRUE | 0.929 |
| Rhog | 376 | 482 | 497 | 517 | 386 | 447 | 399 | 481 | 499 | 406 | 415 | 489 | 448.47 | 460 | 437 | 0.949 | -0.076 | 0.58686669 | 0.76150238 | 0.0098 | 0.0142 | 0.0119 | 0.0119 | TRUE | 0.1525 |
| Rhoh | 27 | 6 | 5 | 40 | 31 | 23 | 29 | 6 | 5 | 31 | 33 | 25 | 21.61 | 13 | 30 | 2.281 | 1.19 | 0.06797064 | 0.20177545 | 0.3687 | 0.1593 | 0.2509 | 0.2509 | TRUE | 4.2413 |
| Rhoj | 5 | 1 | 0 | 4 | 7 | 1 | 5 | 1 | 0 | 3 | 8 | 1 | 3.01 | 2 | 4 | 1.878 | 0.909 | 0.51539632 | NA | 0.9546 | 1.1002 | 1.0368 | 1.0368 | TRUE | 0.7823 |
| Rhoq | 696 | 828 | 812 | 998 | 676 | 771 | 739 | 827 | 815 | 785 | 727 | 844 | 789.57 | 794 | 785 | 0.989 | -0.016 | 0.87714903 | 0.93932608 | 0.0031 | 0.0111 | 0.0065 | 0.0065 | TRUE | 0.0489 |
| Rhot1 | 7564 | 8641 | 9016 | 9112 | 6682 | 7279 | 8034 | 8630 | 9053 | 7163 | 7189 | 7968 | 8006.29 | 8572 | 7440 | 0.868 | -0.205 | 0.01434412 | 0.0684732 | 0.0035 | 0.0073 | 0.0049 | 0.0049 | TRUE | 0.0469 |
| Rhot2 | 1901 | 1691 | 1640 | 2609 | 1871 | 1601 | 2019 | 1689 | 1647 | 2051 | 2013 | 1753 | 1861.89 | 1785 | 1939 | 1.087 | 0.12 | 0.29772348 | 0.51843801 | 0.0092 | 0.0087 | 0.009 | 0.009 | TRUE | 0.1588 |
| Rhou | 1645 | 2361 | 2313 | 1934 | 1405 | 1751 | 1747 | 2358 | 2322 | 1520 | 1512 | 1917 | 1896.09 | 2142 | 1650 | 0.77 | -0.378 | 0.01188064 | 0.05975174 | 0.0218 | 0.0086 | 0.0157 | 0.0157 | TRUE | 0.3148 |
| Rhov | 179 | 231 | 235 | 205 | 143 | 186 | 190 | 231 | 236 | 161 | 154 | 204 | 195.9 | 219 | 173 | 0.788 | -0.344 | 0.05529795 | 0.17583856 | 0.0125 | 0.0237 | 0.0181 | 0.0181 | TRUE | 0.2483 |
| Rhox1 | 128 | 141 | 119 | 372 | 262 | 215 | 136 | 141 | 119 | 292 | 282 | 235 | 200.99 | 132 | 270 | 2.047 | 1.034 | 5.52E-10 | 3.59E-08 | 0.006 | 0.0233 | 0.0143 | 0.0143 | TRUE | 0.1366 |
| Rhox10 | 209 | 178 | 171 | 421 | 241 | 204 | 222 | 178 | 172 | 331 | 259 | 223 | 230.84 | 191 | 271 | 1.429 | 0.515 | 0.00929133 | 0.04974297 | 0.0262 | 0.0212 | 0.0237 | 0.0237 | TRUE | 0.386 |
| Rhox11 | 1 | 0 | 1 | 5 | 1 | 0 | 1 | 0 | 1 | 4 | 1 | 0 | 1.18 | 1 | 2 | 2.516 | 1.331 | 0.50897544 | NA | 0 | 2.8016 | 1.8981 | 1.8981 | TRUE | 0.5172 |
| Rhox12 | 1 | 0 | 0 | 3 | 1 | 0 | 1 | 0 | 0 | 2 | 1 | 0 | 0.75 | 0 | 1 | 3.204 | 1.68 | 0.5025945 | NA | 0 | 4.4029 | 3.1058 | 3.1058 | TRUE | 0.2352 |
| Rhox13 | 314 | 378 | 314 | 1910 | 1338 | 1107 | 334 | 378 | 315 | 1501 | 1440 | 1212 | 863.2 | 342 | 1384 | 4.048 | 2.017 | 8.38E-56 | 1.89E-52 | 0.0094 | 0.0107 | 0.01 | 0.01 | TRUE | 0.1419 |
| Rhox2a | 9 | 26 | 25 | 90 | 60 | 46 | 10 | 26 | 25 | 71 | 65 | 50 | 41.05 | 20 | 62 | 3.058 | 1.613 | 5.37E-05 | 0.00082297 | 0.0725 | 0.0872 | 0.0818 | 0.0818 | TRUE | 1.102 |
| Rhox2b | 0 | 5 | 2 | 13 | 9 | 8 | 0 | 5 | 2 | 10 | 10 | 9 | 5.94 | 2 | 10 | 4.055 | 2.02 | 0.03283331 | 0.12236459 | 0 | 0.5612 | 0.3778 | 0.3778 | TRUE | 0.9805 |
| Rhox2c | 0 | 4 | 3 | 10 | 7 | 4 | 0 | 4 | 3 | 8 | 8 | 4 | 4.46 | 2 | 7 | 2.809 | 1.49 | 0.14909206 | NA | 0.0041 | 0.7451 | 0.4783 | 0.4783 | TRUE | 0.7365 |
| Rhox2d | 3 | 2 | 2 | 13 | 7 | 9 | 3 | 2 | 2 | 10 | 8 | 10 | 5.8 | 2 | 9 | 3.866 | 1.951 | 0.022403 | 0.0942498 | 0 | 0.575 | 0.259 | 0.259 | TRUE | 0.0824 |
| Rhox2e | 0 | 0 | 0 | 0 | 1 | 0 | 0 | 0 | 0 | 0 | 1 | 0 | 0.18 | 0 | 0 | 1.859 | 0.894 | 0.82650734 | NA | 0 | 18.3785 | 10 | 10 | TRUE | 0.5204 |
| Rhox2f | 0 | 2 | 0 | 2 | 3 | 4 | 0 | 2 | 0 | 2 | 3 | 4 | 1.86 | 1 | 3 | 4.484 | 2.165 | 0.19762345 | NA | 0 | 1.7756 | 1.1155 | 1.1155 | TRUE | 0.8596 |
| Rhox2g | 1 | 3 | 4 | 6 | 5 | 2 | 1 | 3 | 4 | 5 | 5 | 2 | 3.39 | 3 | 4 | 1.523 | 0.607 | 0.56262255 | NA | 0 | 0.9777 | 0.4843 | 0.4843 | TRUE | 0.3089 |
| Rhox2h | 0 | 1 | 1 | 6 | 3 | 2 | 0 | 1 | 1 | 5 | 3 | 2 | 2.02 | 1 | 3 | 5.052 | 2.337 | 0.1257367 | NA | 0 | 1.6356 | 0.7804 | 0.7804 | TRUE | 0.2221 |
| Rhox3a | 0 | 0 | 0 | 0 | 0 | 0 | 0 | 0 | 0 | 0 | 0 | 0 | 0 | 0 | 0 | NA | NA | NA | NA | NA | NA | NA | NA | NA | NA |
| Rhox3c | 0 | 0 | 0 | 0 | 1 | 0 | 0 | 0 | 0 | 0 | 1 | 0 | 0.18 | 0 | 0 | 1.859 | 0.894 | 0.82650734 | NA | 0 | 18.3785 | 10 | 10 | TRUE | 0.5204 |

| | | | | | | | | | | | | | | | | | | | | | | | | | |
|---|---|---|---|---|---|---|---|---|---|---|---|---|---|---|---|---|---|---|---|---|---|---|---|---|---|
| Rhox3e | 0 | 0 | 0 | 0 | 0 | 0 | 0 | 0 | 0 | 0 | 0 | 0 | 0 | 0 | 0 | NA | NA | NA | NA | NA | NA | NA | NA | NA | NA |
| Rhox3f | 0 | 0 | 0 | 0 | 0 | 0 | 0 | 0 | 0 | 0 | 0 | 0 | 0 | 0 | 0 | NA | NA | NA | NA | NA | NA | NA | NA | NA | NA |
| Rhox3g-ps | 0 | 0 | 0 | 0 | 0 | 0 | 0 | 0 | 0 | 0 | 0 | 0 | 0 | 0 | 0 | NA | NA | NA | NA | NA | NA | NA | NA | NA | NA |
| Rhox3h | 0 | 0 | 0 | 0 | 0 | 0 | 0 | 0 | 0 | 0 | 0 | 0 | 0 | 0 | 0 | NA | NA | NA | NA | NA | NA | NA | NA | NA | NA |
| Rhox4a | 1 | 1 | 0 | 4 | 3 | 2 | 1 | 1 | 0 | 3 | 3 | 2 | 1.77 | 1 | 3 | 4.194 | 2.068 | 0.18542675 | NA | 0 | 1.8677 | 0.8392 | 0.8392 | TRUE | 0.2537 |
| Rhox4b | 0 | 0 | 0 | 0 | 2 | 2 | 0 | 0 | 0 | 0 | 2 | 2 | 0.72 | 0 | 1 | 7.604 | 2.927 | 0.31659722 | NA | 0 | 4.5602 | 4.8112 | 4.8112 | TRUE | 0.6686 |
| Rhox4c | 0 | 0 | 0 | 0 | 1 | 0 | 0 | 0 | 0 | 0 | 1 | 0 | 0.18 | 0 | 0 | 1.859 | 0.894 | 0.82650734 | NA | 0 | 18.3785 | 10 | 10 | TRUE | 0.5204 |
| Rhox4d | 0 | 0 | 0 | 0 | 2 | 0 | 0 | 0 | 0 | 0 | 2 | 0 | 0.36 | 0 | 1 | 3.775 | 1.916 | 0.63423067 | NA | 10 | 9.1927 | 10 | 10 | TRUE | 0.9743 |
| Rhox4e | 0 | 0 | 1 | 2 | 2 | 5 | 0 | 0 | 1 | 2 | 2 | 5 | 1.7 | 0 | 3 | 8.281 | 3.05 | 0.0895272 | NA | 0 | 1.9445 | 1.1633 | 1.1633 | TRUE | 0.5785 |
| Rhox4f | 0 | 0 | 0 | 1 | 0 | 0 | 0 | 0 | 0 | 1 | 0 | 0 | 0.13 | 0 | 0 | 1.859 | 0.894 | 0.82650734 | NA | 0 | 25.1515 | 10 | 10 | TRUE | 0.2563 |
| Rhox4g | 0 | 2 | 2 | 5 | 6 | 8 | 0 | 2 | 2 | 4 | 6 | 9 | 3.86 | 1 | 6 | 4.677 | 2.225 | 0.05219481 | NA | 0 | 0.8608 | 0.4913 | 0.4913 | TRUE | 0.2854 |
| Rhox5 | 633 | 863 | 696 | 3103 | 2627 | 2379 | 672 | 862 | 699 | 2439 | 2826 | 2604 | 1683.85 | 744 | 2623 | 3.522 | 1.817 | 1.08E-50 | 1.71E-47 | 0.011 | 0.0089 | 0.0097 | 0.0097 | TRUE | 0.2265 |
| Rhox6 | 86 | 123 | 113 | 249 | 212 | 245 | 91 | 123 | 113 | 196 | 228 | 268 | 169.95 | 109 | 231 | 2.104 | 1.073 | 7.91E-08 | 2.95E-06 | 0.0178 | 0.0263 | 0.022 | 0.022 | TRUE | 0.1363 |
| Rhox7 | 15 | 29 | 20 | 23 | 31 | 29 | 16 | 29 | 20 | 18 | 33 | 32 | 24.69 | 22 | 28 | 1.266 | 0.34 | 0.44539605 | 0.65201285 | 0.0617 | 0.1403 | 0.1018 | 0.1018 | TRUE | 0.7107 |
| Rhox8 | 0 | 0 | 1 | 0 | 0 | 0 | 0 | 0 | 1 | 0 | 0 | 0 | 0.17 | 0 | 0 | 0.49 | -1.029 | 0.80087 | NA | 0 | 19.6928 | 10 | 10 | TRUE | 0.4098 |
| Rhox9 | 76 | 124 | 104 | 207 | 228 | 245 | 81 | 124 | 104 | 163 | 245 | 268 | 164.2 | 103 | 225 | 2.18 | 1.124 | 7.50E-06 | 0.0001552 | 0.0502 | 0.027 | 0.0383 | 0.0383 | TRUE | 0.6846 |
| Rhpn1 | 255 | 228 | 214 | 238 | 214 | 200 | 271 | 228 | 215 | 187 | 230 | 219 | 224.95 | 238 | 212 | 0.89 | -0.168 | 0.30587394 | 0.52665845 | 0.009 | 0.0215 | 0.015 | 0.015 | TRUE | 0.1615 |
| Rhpn2 | 2189 | 2296 | 2370 | 2639 | 1929 | 2231 | 2325 | 2293 | 2380 | 2075 | 2075 | 2442 | 2265.02 | 2333 | 2197 | 0.942 | -0.087 | 0.35245816 | 0.57102929 | 0.0042 | 0.0084 | 0.0058 | 0.0058 | TRUE | 0.1141 |
| Rian | 15885 | 18367 | 18334 | 14512 | 11545 | 10883 | 16873 | 18343 | 18409 | 11408 | 12422 | 11913 | 14894.72 | 17875 | 11914 | 0.666 | -0.585 | 7.66E-16 | 1.17E-13 | 0.002 | 0.0071 | 0.0037 | 0.0037 | TRUE | 0.0294 |
| Ribc1 | 261 | 315 | 339 | 488 | 333 | 294 | 277 | 315 | 340 | 384 | 358 | 322 | 332.66 | 311 | 355 | 1.143 | 0.192 | 0.16979056 | 0.36840511 | 0.0061 | 0.0168 | 0.0111 | 0.0111 | TRUE | 0.0994 |
| Ribc2 | 1 | 1 | 1 | 7 | 5 | 1 | 1 | 1 | 1 | 6 | 5 | 1 | 2.51 | 1 | 4 | 3.953 | 1.983 | 0.14093046 | NA | 0 | 1.321 | 0.6859 | 0.6859 | TRUE | 0.6032 |
| Ric3 | 778 | 765 | 824 | 1349 | 989 | 854 | 826 | 764 | 827 | 1060 | 1064 | 935 | 912.86 | 806 | 1020 | 1.266 | 0.341 | 0.00054439 | 0.0055574 | 0.0026 | 0.0105 | 0.0059 | 0.0059 | TRUE | 0.0623 |
| Ric8 | 1687 | 1409 | 1289 | 2245 | 1636 | 1495 | 1792 | 1407 | 1294 | 1765 | 1760 | 1637 | 1609.16 | 1498 | 1721 | 1.149 | 0.201 | 0.13033383 | 0.31086811 | 0.0151 | 0.009 | 0.012 | 0.012 | TRUE | 0.3545 |
| Ric8b | 1467 | 1275 | 1426 | 1421 | 1029 | 1100 | 1558 | 1273 | 1432 | 1117 | 1107 | 1204 | 1281.96 | 1421 | 1143 | 0.804 | -0.315 | 0.00267097 | 0.01935073 | 0.0049 | 0.0095 | 0.0071 | 0.0071 | TRUE | 0.0997 |
| Rictor | 3601 | 3490 | 3869 | 4388 | 3183 | 3279 | 3825 | 3485 | 3885 | 3450 | 3425 | 3589 | 3609.81 | 3732 | 3488 | 0.935 | -0.098 | 0.19773905 | 0.40436123 | 0.0016 | 0.0078 | 0.0039 | 0.0039 | TRUE | 0.0406 |
| Rif1 | 23577 | 22701 | 25078 | 30930 | 22176 | 22179 | 25043 | 22672 | 25181 | 24315 | 23860 | 24278 | 24224.79 | 24299 | 24151 | 0.994 | -0.009 | 0.8997099 | 0.95140627 | 0.0014 | 0.007 | 0.0034 | 0.0034 | TRUE | 0.042 |
| Riiad1 | 1 | 2 | 1 | 1 | 2 | 2 | 1 | 2 | 1 | 1 | 2 | 2 | 1.53 | 1 | 2 | 1.23 | 0.299 | 0.83816522 | NA | 0 | 2.1576 | 0.8707 | 0.8707 | TRUE | 0.2434 |
| Rilp | 8 | 12 | 13 | 22 | 12 | 11 | 8 | 12 | 13 | 17 | 13 | 12 | 12.63 | 11 | 14 | 1.268 | 0.343 | 0.52280069 | 0.71285662 | 0 | 0.2677 | 0.1271 | 0.1271 | TRUE | 0.216 |
| Rilpl1 | 1141 | 1120 | 1119 | 1126 | 818 | 819 | 1212 | 1119 | 1124 | 885 | 880 | 897 | 1019.32 | 1152 | 887 | 0.771 | -0.376 | 6.66E-06 | 0.00014118 | 0 | 0.0101 | 0.004 | 0.004 | TRUE | 0.0252 |
| Rilpl2 | 483 | 476 | 466 | 448 | 335 | 333 | 513 | 475 | 468 | 352 | 360 | 365 | 422.25 | 485 | 359 | 0.739 | -0.436 | 6.11E-05 | 0.00092334 | 0 | 0.0147 | 0.0061 | 0.0061 | TRUE | 0.0284 |
| Rimbp2 | 185 | 181 | 158 | 286 | 249 | 325 | 197 | 181 | 159 | 225 | 268 | 356 | 230.74 | 179 | 283 | 1.58 | 0.66 | 0.00126749 | 0.01069033 | 0.0298 | 0.0212 | 0.0256 | 0.0256 | TRUE | 0.3525 |
| Rimbp3 | 66 | 63 | 72 | 187 | 164 | 83 | 70 | 63 | 72 | 147 | 176 | 91 | 103.27 | 68 | 138 | 2.021 | 1.015 | 0.00026865 | 0.00310638 | 0.0511 | 0.0388 | 0.0449 | 0.0449 | TRUE | 0.2605 |
| Rimkla | 267 | 258 | 208 | 523 | 350 | 311 | 284 | 258 | 209 | 411 | 377 | 340 | 313.05 | 250 | 376 | 1.507 | 0.592 | 0.00019613 | 0.00239799 | 0.0124 | 0.0174 | 0.0148 | 0.0148 | TRUE | 0.2316 |
| Rimklb | 1843 | 2362 | 2374 | 3120 | 2551 | 2577 | 1958 | 2359 | 2384 | 2453 | 2745 | 2821 | 2453.11 | 2234 | 2673 | 1.196 | 0.258 | 0.01841924 | 0.08197227 | 0.0078 | 0.0082 | 0.0082 | 0.0082 | TRUE | 0.141 |
| Rims1 | 209 | 237 | 221 | 349 | 270 | 243 | 222 | 237 | 222 | 274 | 291 | 266 | 251.91 | 227 | 277 | 1.22 | 0.287 | 0.02856837 | 0.11125441 | 0 | 0.02 | 0.0084 | 0.0084 | TRUE | 0.0195 |
| Rims2 | 436 | 467 | 479 | 830 | 572 | 529 | 463 | 466 | 481 | 652 | 615 | 579 | 542.91 | 470 | 615 | 1.311 | 0.391 | 0.00015192 | 0.00192308 | 0 | 0.013 | 0.0058 | 0.0058 | TRUE | 0.0338 |
| Rims3 | 364 | 447 | 414 | 479 | 355 | 365 | 387 | 446 | 416 | 377 | 382 | 400 | 401.14 | 416 | 386 | 0.926 | -0.11 | 0.34013669 | 0.56027529 | 0 | 0.0151 | 0.0071 | 0.0071 | TRUE | 0.0465 |
| Rims4 | 147 | 134 | 138 | 137 | 114 | 94 | 156 | 134 | 139 | 108 | 123 | 103 | 126.96 | 143 | 111 | 0.777 | -0.363 | 0.04781028 | 0.15905163 | 0 | 0.0329 | 0.0163 | 0.0163 | TRUE | 0.078 |
| Rin1 | 152 | 310 | 261 | 261 | 270 | 365 | 161 | 310 | 262 | 205 | 291 | 400 | 271.39 | 244 | 299 | 1.218 | 0.285 | 0.3249263 | 0.54594394 | 0.1023 | 0.019 | 0.0566 | 0.0566 | TRUE | 0.2141 |
| Rin2 | 688 | 712 | 743 | 841 | 609 | 647 | 731 | 711 | 746 | 661 | 655 | 708 | 702.09 | 729 | 675 | 0.925 | -0.113 | 0.2196447 | 0.43181262 | 0 | 0.0116 | 0.0046 | 0.0046 | TRUE | 0.0214 |
| Rin3 | 156 | 117 | 84 | 345 | 224 | 191 | 166 | 117 | 84 | 271 | 241 | 209 | 181.37 | 122 | 240 | 1.971 | 0.979 | 8.78E-05 | 0.00124893 | 0.0552 | 0.0251 | 0.0387 | 0.0387 | TRUE | 0.6841 |
| Ring1 | 164 | 212 | 172 | 400 | 311 | 216 | 174 | 212 | 173 | 314 | 335 | 236 | 240.69 | 186 | 295 | 1.586 | 0.666 | 0.00029474 | 0.0033661 | 0.0193 | 0.0206 | 0.02 | 0.02 | TRUE | 0.3318 |
| Rinl | 659 | 561 | 596 | 1757 | 1290 | 1079 | 700 | 560 | 598 | 1381 | 1388 | 1181 | 968.17 | 619 | 1317 | 2.127 | 1.089 | 1.07E-18 | 2.26E-16 | 0.009 | 0.0103 | 0.0098 | 0.0098 | TRUE | 0.1507 |
| Rint1 | 996 | 939 | 889 | 1215 | 872 | 770 | 1058 | 938 | 893 | 955 | 938 | 843 | 937.43 | 963 | 912 | 0.948 | -0.077 | 0.47654805 | 0.67645532 | 0.005 | 0.0104 | 0.0073 | 0.0073 | TRUE | 0.0887 |
| Riok1 | 1416 | 1260 | 1249 | 2001 | 1392 | 1175 | 1504 | 1258 | 1254 | 1573 | 1498 | 1286 | 1395.58 | 1339 | 1452 | 1.086 | 0.118 | 0.32597384 | 0.5471171 | 0.0099 | 0.0093 | 0.0098 | 0.0098 | TRUE | 0.1396 |
| Riok2 | 4377 | 3385 | 3343 | 6543 | 4202 | 3411 | 4649 | 3381 | 3357 | 5144 | 4521 | 3734 | 4130.84 | 3796 | 4466 | 1.177 | 0.235 | 0.14780257 | 0.33726015 | 0.0311 | 0.0077 | 0.0188 | 0.0188 | TRUE | 0.409 |
| Riok3 | 4012 | 5091 | 5437 | 6076 | 4416 | 4330 | 4261 | 5084 | 5459 | 4776 | 4751 | 4740 | 4845.48 | 4935 | 4756 | 0.964 | -0.053 | 0.6096431 | 0.77891178 | 0.0066 | 0.0076 | 0.0077 | 0.0077 | TRUE | 0.1735 |
| Ripk1 | 276 | 248 | 238 | 454 | 262 | 212 | 293 | 248 | 239 | 357 | 282 | 232 | 275.11 | 260 | 290 | 1.121 | 0.164 | 0.38873481 | 0.60274532 | 0.0261 | 0.0189 | 0.0226 | 0.0226 | TRUE | 0.3051 |
| Ripk2 | 804 | 866 | 882 | 1015 | 754 | 788 | 854 | 865 | 886 | 798 | 811 | 863 | 846.04 | 868 | 824 | 0.948 | -0.077 | 0.37598132 | 0.59147378 | 0 | 0.0108 | 0.0042 | 0.0042 | TRUE | 0.0197 |
| Ripk3 | 776 | 695 | 722 | 724 | 539 | 540 | 824 | 694 | 725 | 569 | 580 | 591 | 663.92 | 748 | 580 | 0.776 | -0.366 | 0.00061463 | 0.00611605 | 0.003 | 0.0119 | 0.0067 | 0.0067 | TRUE | 0.0941 |
| Ripk4 | 1292 | 1569 | 1628 | 1603 | 1178 | 1204 | 1372 | 1567 | 1635 | 1260 | 1267 | 1318 | 1403.26 | 1525 | 1282 | 0.84 | -0.251 | 0.00905783 | 0.0488745 | 0.0037 | 0.0093 | 0.0059 | 0.0059 | TRUE | 0.0916 |
| Ripply1 | 133 | 159 | 187 | 78 | 56 | 106 | 141 | 159 | 188 | 61 | 60 | 116 | 120.91 | 163 | 79 | 0.484 | -1.046 | 0.0002686 | 0.00310638 | 0.0689 | 0.0342 | 0.05 | 0.05 | TRUE | 1.4853 |
| Ripply2 | 0 | 2 | 4 | 3 | 1 | 1 | 0 | 2 | 4 | 2 | 1 | 1 | 1.76 | 2 | 1 | 0.765 | -0.387 | 0.79956823 | NA | 0 | 1.8818 | 1.1013 | 1.1013 | TRUE | 0.1454 |
| Ripply3 | 106 | 127 | 130 | 167 | 106 | 87 | 113 | 127 | 131 | 131 | 114 | 95 | 118.42 | 124 | 113 | 0.925 | -0.113 | 0.57680394 | 0.75440349 | 0.0075 | 0.0347 | 0.0212 | 0.0212 | TRUE | 0.2052 |
| Rit1 | 365 | 362 | 332 | 537 | 316 | 317 | 388 | 362 | 333 | 422 | 340 | 347 | 365.29 | 361 | 370 | 1.028 | 0.039 | 0.78032352 | 0.88629118 | 0.0079 | 0.0159 | 0.0116 | 0.0116 | TRUE | 0.185 |
| Rit2 | 0 | 0 | 0 | 0 | 1 | 0 | 0 | 0 | 0 | 0 | 1 | 0 | 0.18 | 0 | 0 | 1.859 | 0.894 | 0.82650734 | NA | 0 | 18.3785 | 10 | 10 | TRUE | 0.5204 |
| Rlbp1 | 130 | 120 | 117 | 279 | 220 | 197 | 138 | 120 | 117 | 219 | 237 | 216 | 174.52 | 125 | 224 | 1.79 | 0.84 | 1.64E-07 | 5.52E-06 | 0 | 0.0258 | 0.0123 | 0.0123 | TRUE | 0.0816 |
| Rlf | 4766 | 5090 | 5683 | 6209 | 4801 | 5409 | 5062 | 5083 | 5706 | 4881 | 5166 | 5921 | 5303.28 | 5284 | 5323 | 1.007 | 0.01 | 0.92080218 | 0.96114949 | 0.0071 | 0.0075 | 0.0073 | 0.0073 | TRUE | 0.1178 |
| Rlim | 4554 | 4421 | 5356 | 12752 | 8372 | 9366 | 4837 | 4415 | 5378 | 10025 | 9008 | 10253 | 7319.23 | 4877 | 9762 | 2.002 | 1.001 | 3.01E-23 | 1.01E-20 | 0.007 | 0.0074 | 0.0072 | 0.0072 | TRUE | 0.0986 |
| Rln1 | 89 | 116 | 101 | 126 | 88 | 66 | 95 | 116 | 101 | 99 | 95 | 72 | 96.3 | 104 | 89 | 0.856 | -0.224 | 0.31144315 | 0.53199265 | 0.0076 | 0.0411 | 0.025 | 0.025 | TRUE | 0.2372 |
| Rln3 | 55 | 76 | 98 | 101 | 94 | 85 | 58 | 76 | 98 | 79 | 101 | 93 | 84.38 | 77 | 91 | 1.171 | 0.227 | 0.38326079 | 0.59760644 | 0.0274 | 0.0459 | 0.037 | 0.037 | TRUE | 0.3042 |
| Rltpr | 784 | 816 | 831 | 939 | 757 | 854 | 833 | 815 | 834 | 738 | 814 | 935 | 828.27 | 827 | 829 | 1.001 | 0.001 | 0.99297969 | 0.99616826 | 0.0061 | 0.0109 | 0.0081 | 0.0081 | TRUE | 0.1471 |
| Rmdn1 | 78 | 148 | 146 | 167 | 125 | 123 | 83 | 148 | 147 | 131 | 134 | 135 | 129.61 | 126 | 133 | 1.06 | 0.084 | 0.73438151 | 0.85753055 | 0.0404 | 0.0323 | 0.0362 | 0.0362 | TRUE | 0.8924 |
| Rmdn2 | 78 | 40 | 44 | 138 | 105 | 68 | 83 | 40 | 44 | 108 | 113 | 74 | 77.15 | 56 | 98 | 1.778 | 0.831 | 0.01339318 | 0.06505611 | 0.0875 | 0.0496 | 0.0671 | 0.0671 | TRUE | 1.5071 |
| Rmdn3 | 2363 | 2380 | 2408 | 2949 | 2169 | 2013 | 2510 | 2377 | 2418 | 2318 | 2334 | 2204 | 2360.04 | 2435 | 2285 | 0.939 | -0.091 | 0.19285883 | 0.39842218 | 4.00E-04 | 0.0083 | 0.0031 | 0.0031 | TRUE | 0.0116 |
| Rmi1 | 829 | 757 | 716 | 1200 | 841 | 736 | 881 | 756 | 719 | 943 | 905 | 806 | 834.9 | 785 | 885 | 1.128 | 0.174 | 0.14606358 | 0.33475656 | 0.0072 | 0.0108 | 0.0091 | 0.0091 | TRUE | 0.1332 |
| Rmi2 | 261 | 406 | 445 | 382 | 330 | 332 | 277 | 405 | 447 | 300 | 355 | 363 | 358.05 | 376 | 339 | 0.9 | -0.151 | 0.43408226 | 0.6426683 | 0.0325 | 0.0161 | 0.0242 | 0.0242 | TRUE | 0.6058 |
| Rmnd1 | 8 | 7 | 6 | 4 | 4 | 9 | 8 | 7 | 6 | 3 | 4 | 10 | 6.47 | 7 | 6 | 0.79 | -0.341 | 0.66332185 | 0.81128824 | 0 | 0.5162 | 0.2839 | 0.2839 | TRUE | 0.8101 |
| Rmnd5a | 8105 | 8384 | 8537 | 11508 | 8143 | 8456 | 8609 | 8373 | 8572 | 9047 | 8761 | 9256 | 8769.77 | 8518 | 9021 | 1.059 | 0.083 | 0.15643906 | 0.34938501 | 3.00E-04 | 0.0073 | 0.0023 | 0.0023 | TRUE | 0.0077 |

| | | | | | | | | | | | | | | | | | | | | | | | | | |
|---|---|---|---|---|---|---|---|---|---|---|---|---|---|---|---|---|---|---|---|---|---|---|---|---|---|
| Rmnd5b | 2400 | 2419 | 2555 | 3184 | 2460 | 2417 | 2549 | 2416 | 2565 | 2503 | 2647 | 2646 | 2554.37 | 2510 | 2599 | 1.035 | 0.049 | 0.4859164 | 0.68281125 | 7.00E-04 | 0.0082 | 0.0032 | 0.0032 | TRUE | 0.0131 |
| Rmrp | 15 | 15 | 15 | 8 | 8 | 5 | 16 | 15 | 15 | 6 | 9 | 5 | 11.06 | 15 | 7 | 0.442 | -1.179 | 0.04098283 | 0.14332177 | 0 | 0.3048 | 0.1345 | 0.1345 | TRUE | 0.1258 |
| Rmst | 1 | 0 | 0 | 1 | 0 | 0 | 1 | 0 | 0 | 1 | 0 | 0 | 0.31 | 0 | 0 | 0.954 | -0.067 | 0.98625928 | NA | 0 | 10.7014 | 9.0384 | 9.0384 | TRUE | 0.4614 |
| Rn4.5s | 0 | 0 | 0 | 0 | 0 | 0 | 0 | 0 | 0 | 0 | 0 | 0 | 0 | 0 | 0 | NA | NA | NA | NA | NA | NA | NA | NA | NA | NA |
| Rn45s | 111021 | 104296 | 88204 | 106700 | 89088 | 75211 | 117924 | 104161 | 88566 | 83879 | 95853 | 82330 | 95452.34 | 103550 | 87354 | 0.844 | -0.245 | 0.04329458 | 0.14876736 | 0.0136 | 0.0069 | 0.0106 | 0.0106 | TRUE | 0.1852 |
| Rnase1 | 19 | 13 | 12 | 7 | 7 | 7 | 20 | 13 | 12 | 6 | 8 | 8 | 10.99 | 15 | 7 | 0.454 | -1.14 | 0.05701798 | 0.1797856 | 0 | 0.3068 | 0.153 | 0.153 | TRUE | 0.3865 |
| Rnase10 | 1 | 0 | 0 | 1 | 0 | 0 | 1 | 0 | 0 | 1 | 0 | 0 | 0.31 | 0 | 0 | 0.954 | -0.067 | 0.98625928 | NA | 0 | 10.7014 | 9.0384 | 9.0384 | TRUE | 0.4614 |
| Rnase11 | 0 | 0 | 0 | 0 | 0 | 0 | 0 | 0 | 0 | 0 | 0 | 0 | 0 | 0 | 0 | NA | NA | NA | NA | NA | NA | NA | NA | NA | NA |
| Rnase12 | 2 | 5 | 1 | 2 | 0 | 9 | 2 | 5 | 1 | 2 | 0 | 10 | 3.26 | 3 | 4 | 1.386 | 0.471 | 0.73894687 | NA | 1.2416 | 1.0182 | 1.1228 | 1.1228 | TRUE | 2.0554 |
| Rnase13 | 0 | 0 | 0 | 0 | 0 | 0 | 0 | 0 | 0 | 0 | 0 | 0 | 0 | 0 | 0 | NA | NA | NA | NA | NA | NA | NA | NA | NA | NA |
| Rnase4 | 182 | 168 | 209 | 124 | 100 | 110 | 193 | 168 | 210 | 97 | 108 | 120 | 149.41 | 190 | 108 | 0.568 | -0.816 | 9.23E-06 | 0.00018601 | 0.0058 | 0.029 | 0.0172 | 0.0172 | TRUE | 0.1172 |
| Rnase6 | 4 | 3 | 2 | 8 | 6 | 6 | 4 | 3 | 2 | 6 | 6 | 7 | 4.76 | 3 | 6 | 2.093 | 1.066 | 0.21938961 | 0.43155562 | 0 | 0.6989 | 0.3004 | 0.3004 | TRUE | 0.1366 |
| Rnase9 | 0 | 0 | 0 | 0 | 0 | 0 | 0 | 0 | 0 | 0 | 0 | 0 | 0 | 0 | 0 | NA | NA | NA | NA | NA | NA | NA | NA | NA | NA |
| Rnaseh1 | 851 | 848 | 775 | 1145 | 852 | 797 | 904 | 847 | 778 | 900 | 917 | 872 | 869.71 | 843 | 896 | 1.064 | 0.089 | 0.35392768 | 0.57229316 | 0.0018 | 0.0107 | 0.0055 | 0.0055 | TRUE | 0.0538 |
| Rnaseh2a | 4451 | 5042 | 4805 | 5685 | 4209 | 4242 | 4728 | 5035 | 4825 | 4469 | 4529 | 4644 | 4704.87 | 4863 | 4547 | 0.935 | -0.097 | 0.12660794 | 0.30523699 | 5.00E-04 | 0.0076 | 0.0027 | 0.0027 | TRUE | 0.0118 |
| Rnaseh2b | 1208 | 1429 | 1370 | 2034 | 1489 | 1564 | 1283 | 1427 | 1376 | 1599 | 1602 | 1712 | 1499.83 | 1362 | 1638 | 1.202 | 0.265 | 0.0015406 | 0.01250558 | 0.0015 | 0.0091 | 0.0044 | 0.0044 | TRUE | 0.0306 |
| Rnaseh2c | 899 | 977 | 983 | 1098 | 817 | 852 | 955 | 976 | 987 | 863 | 879 | 933 | 932.09 | 973 | 892 | 0.916 | -0.127 | 0.13290064 | 0.31444779 | 0 | 0.0104 | 0.004 | 0.004 | TRUE | 0.019 |
| Rnasek | 2035 | 2050 | 1965 | 2674 | 1786 | 1912 | 2162 | 2047 | 1973 | 2102 | 1922 | 2093 | 2049.78 | 2061 | 2039 | 0.99 | -0.015 | 0.85624208 | 0.928793 | 0.0018 | 0.0085 | 0.0043 | 0.0043 | TRUE | 0.0303 |
| Rnasel | 49 | 77 | 91 | 46 | 36 | 53 | 52 | 77 | 91 | 36 | 39 | 58 | 58.87 | 73 | 44 | 0.599 | -0.739 | 0.02408923 | 0.09901922 | 0.0553 | 0.0629 | 0.0592 | 0.0592 | TRUE | 0.2698 |
| Rnaset2a | 0 | 0 | 0 | 0 | 0 | 0 | 0 | 0 | 0 | 0 | 0 | 0 | 0 | 0 | 0 | NA | NA | NA | NA | NA | NA | NA | NA | NA | NA |
| Rnaset2b | 2 | 4 | 2 | 1 | 0 | 4 | 2 | 4 | 2 | 1 | 0 | 4 | 2.22 | 3 | 2 | 0.619 | -0.691 | 0.62749021 | NA | 0.3134 | 1.494 | 0.9833 | 0.9833 | TRUE | 1.2842 |
| Rnd1 | 512 | 316 | 302 | 561 | 390 | 348 | 544 | 316 | 303 | 441 | 420 | 381 | 400.71 | 388 | 414 | 1.069 | 0.096 | 0.66955274 | 0.8154066 | 0.0554 | 0.0151 | 0.0341 | 0.0341 | TRUE | 1.4242 |
| Rnd2 | 928 | 914 | 936 | 1074 | 732 | 732 | 986 | 913 | 940 | 844 | 788 | 801 | 878.59 | 946 | 811 | 0.858 | -0.221 | 0.01221189 | 0.06097053 | 0 | 0.0107 | 0.0044 | 0.0044 | TRUE | 0.016 |
| Rnd3 | 639 | 555 | 517 | 884 | 619 | 510 | 679 | 554 | 519 | 695 | 666 | 558 | 611.89 | 584 | 640 | 1.097 | 0.133 | 0.35934542 | 0.57787504 | 0.0139 | 0.0123 | 0.0136 | 0.0136 | TRUE | 0.2348 |
| Rnf10 | 22990 | 24972 | 25414 | 28569 | 20987 | 21099 | 24419 | 24940 | 25518 | 22459 | 22581 | 23096 | 23835.52 | 24959 | 22712 | 0.91 | -0.136 | 0.01243343 | 0.06171896 | 3.00E-04 | 0.007 | 0.0021 | 0.0021 | TRUE | 0.0047 |
| Rnf103 | 276 | 387 | 336 | 588 | 390 | 357 | 293 | 387 | 337 | 462 | 420 | 391 | 381.61 | 339 | 424 | 1.252 | 0.325 | 0.02611896 | 0.10468825 | 0.0101 | 0.0155 | 0.0127 | 0.0127 | TRUE | 0.1713 |
| Rnf11 | 979 | 1325 | 1221 | 2057 | 1450 | 1508 | 1040 | 1323 | 1226 | 1617 | 1560 | 1651 | 1402.85 | 1196 | 1609 | 1.345 | 0.427 | 9.39E-05 | 0.00131066 | 0.0066 | 0.0093 | 0.0079 | 0.0079 | TRUE | 0.1563 |
| Rnf111 | 3160 | 3078 | 3013 | 3964 | 2819 | 2646 | 3356 | 3074 | 3025 | 3116 | 3033 | 2896 | 3083.6 | 3152 | 3015 | 0.957 | -0.063 | 0.41715433 | 0.62816098 | 0.0018 | 0.008 | 0.0041 | 0.0041 | TRUE | 0.039 |
| Rnf112 | 25 | 20 | 16 | 22 | 20 | 15 | 27 | 20 | 16 | 17 | 22 | 16 | 19.64 | 21 | 18 | 0.882 | -0.181 | 0.68165768 | 0.82355168 | 0 | 0.1747 | 0.0889 | 0.0889 | TRUE | 0.3031 |
| Rnf113a1 | 121 | 131 | 141 | 259 | 180 | 169 | 129 | 131 | 142 | 204 | 194 | 185 | 163.87 | 134 | 194 | 1.454 | 0.54 | 0.00064975 | 0.00639294 | 0 | 0.027 | 0.0117 | 0.0117 | TRUE | 0.0279 |
| Rnf113a2 | 551 | 542 | 552 | 837 | 632 | 555 | 585 | 541 | 554 | 658 | 680 | 608 | 604.39 | 560 | 649 | 1.158 | 0.212 | 0.03640979 | 0.13148414 | 0 | 0.0124 | 0.0057 | 0.0057 | TRUE | 0.0346 |
| Rnf114 | 3677 | 4627 | 4761 | 4931 | 3741 | 4178 | 3906 | 4621 | 4781 | 3876 | 4025 | 4573 | 4297.02 | 4436 | 4158 | 0.937 | -0.094 | 0.39789953 | 0.61124685 | 0.0086 | 0.0077 | 0.0086 | 0.0086 | TRUE | 0.1329 |
| Rnf115 | 3043 | 3515 | 3519 | 3661 | 2628 | 2920 | 3232 | 3510 | 3533 | 2878 | 2828 | 3196 | 3196.35 | 3425 | 2967 | 0.866 | -0.208 | 0.01473062 | 0.06989126 | 0.003 | 0.0079 | 0.0049 | 0.0049 | TRUE | 0.0549 |
| Rnf121 | 745 | 815 | 857 | 1060 | 871 | 845 | 791 | 814 | 861 | 833 | 937 | 925 | 860.2 | 822 | 898 | 1.092 | 0.126 | 0.18912282 | 0.39431214 | 0.002 | 0.0107 | 0.0055 | 0.0055 | TRUE | 0.0501 |
| Rnf122 | 325 | 388 | 378 | 388 | 340 | 340 | 345 | 387 | 380 | 305 | 366 | 372 | 359.21 | 371 | 348 | 0.935 | -0.097 | 0.47008424 | 0.67198975 | 0.0054 | 0.0161 | 0.0102 | 0.0102 | TRUE | 0.1403 |
| Rnf123 | 618 | 875 | 822 | 917 | 709 | 773 | 656 | 874 | 825 | 721 | 763 | 846 | 780.93 | 785 | 777 | 0.988 | -0.017 | 0.8987047 | 0.95078955 | 0.0131 | 0.0111 | 0.0122 | 0.0122 | TRUE | 0.2429 |
| Rnf125 | 543 | 603 | 570 | 755 | 604 | 591 | 577 | 602 | 572 | 594 | 650 | 647 | 606.94 | 584 | 630 | 1.078 | 0.108 | 0.2711506 | 0.49106313 | 0 | 0.0123 | 0.0053 | 0.0053 | TRUE | 0.0322 |
| Rnf126 | 4162 | 4471 | 4653 | 5161 | 4050 | 4226 | 4421 | 4465 | 4672 | 4057 | 4358 | 4626 | 4433.14 | 4519 | 4347 | 0.961 | -0.057 | 0.47183291 | 0.67314138 | 0.0022 | 0.0076 | 0.0042 | 0.0042 | TRUE | 0.0419 |
| Rnf128 | 494 | 696 | 682 | 772 | 594 | 662 | 525 | 695 | 685 | 607 | 639 | 725 | 645.88 | 635 | 657 | 1.033 | 0.047 | 0.74136652 | 0.86165152 | 0.0145 | 0.012 | 0.0133 | 0.0133 | TRUE | 0.2696 |
| Rnf13 | 1330 | 1621 | 1719 | 1742 | 1233 | 1376 | 1413 | 1619 | 1726 | 1369 | 1327 | 1506 | 1493.33 | 1586 | 1401 | 0.883 | -0.18 | 0.09551092 | 0.25348616 | 0.0063 | 0.0091 | 0.0077 | 0.0077 | TRUE | 0.1096 |
| Rnf130 | 903 | 1463 | 1559 | 1656 | 1254 | 1375 | 959 | 1461 | 1565 | 1302 | 1349 | 1505 | 1356.98 | 1328 | 1385 | 1.042 | 0.06 | 0.73465577 | 0.85766031 | 0.035 | 0.0093 | 0.0218 | 0.0218 | TRUE | 0.7094 |
| Rnf133 | 0 | 0 | 0 | 0 | 0 | 0 | 0 | 0 | 0 | 0 | 0 | 0 | 0 | 0 | 0 | NA | NA | NA | NA | NA | NA | NA | NA | NA | NA |
| Rnf135 | 225 | 275 | 221 | 329 | 243 | 234 | 239 | 275 | 222 | 259 | 261 | 256 | 251.96 | 245 | 259 | 1.055 | 0.077 | 0.58543047 | 0.76057738 | 0.0017 | 0.02 | 0.0105 | 0.0105 | TRUE | 0.124 |
| Rnf138 | 3965 | 4918 | 4969 | 4868 | 3774 | 4111 | 4212 | 4912 | 4989 | 3827 | 4061 | 4500 | 4416.69 | 4704 | 4129 | 0.878 | -0.189 | 0.06923486 | 0.20425473 | 0.0072 | 0.0076 | 0.0075 | 0.0075 | TRUE | 0.1021 |
| Rnf139 | 1506 | 1511 | 1657 | 2099 | 1471 | 1498 | 1600 | 1509 | 1664 | 1650 | 1583 | 1640 | 1607.51 | 1591 | 1624 | 1.021 | 0.03 | 0.69922239 | 0.83549402 | 8.00E-04 | 0.009 | 0.0038 | 0.0038 | TRUE | 0.0245 |
| Rnf14 | 3144 | 3428 | 3567 | 4186 | 2873 | 2997 | 3339 | 3424 | 3582 | 3291 | 3091 | 3281 | 3334.54 | 3448 | 3221 | 0.934 | -0.098 | 0.16257619 | 0.35788551 | 8.00E-04 | 0.0079 | 0.0033 | 0.0033 | TRUE | 0.015 |
| Rnf141 | 510 | 628 | 611 | 729 | 576 | 566 | 542 | 627 | 614 | 573 | 620 | 620 | 599.13 | 594 | 604 | 1.015 | 0.022 | 0.83801683 | 0.91779748 | 0.0024 | 0.0124 | 0.0067 | 0.0067 | TRUE | 0.0693 |
| Rnf144a | 656 | 896 | 877 | 1045 | 733 | 781 | 697 | 895 | 881 | 821 | 789 | 855 | 822.89 | 824 | 822 | 0.997 | -0.005 | 0.96961279 | 0.98448447 | 0.0089 | 0.0109 | 0.0098 | 0.0098 | TRUE | 0.2156 |
| Rnf144b | 250 | 196 | 216 | 222 | 149 | 147 | 266 | 196 | 217 | 175 | 160 | 161 | 195.65 | 226 | 165 | 0.733 | -0.449 | 0.00927075 | 0.04966378 | 0.0094 | 0.0237 | 0.0162 | 0.0162 | TRUE | 0.2542 |
| Rnf145 | 6785 | 7256 | 7117 | 9286 | 6957 | 7416 | 7207 | 7247 | 7146 | 7300 | 7485 | 8118 | 7417.15 | 7200 | 7634 | 1.06 | 0.084 | 0.22667011 | 0.43951819 | 0.0014 | 0.0073 | 0.0034 | 0.0034 | TRUE | 0.0374 |
| Rnf146 | 1437 | 1501 | 1586 | 1921 | 1449 | 1355 | 1526 | 1499 | 1593 | 1510 | 1559 | 1483 | 1528.39 | 1539 | 1517 | 0.986 | -0.021 | 0.78130806 | 0.88682105 | 0 | 0.0091 | 0.0034 | 0.0034 | TRUE | 0.0111 |
| Rnf148 | 0 | 0 | 0 | 0 | 0 | 0 | 0 | 0 | 0 | 0 | 0 | 0 | 0 | 0 | 0 | NA | NA | NA | NA | NA | NA | NA | NA | NA | NA |
| Rnf149 | 1873 | 1771 | 1778 | 2825 | 1903 | 1616 | 1989 | 1769 | 1785 | 2221 | 2048 | 1769 | 1930.12 | 1848 | 2013 | 1.09 | 0.124 | 0.26428477 | 0.48381628 | 0.0078 | 0.0086 | 0.0084 | 0.0084 | TRUE | 0.1349 |
| Rnf150 | 4 | 5 | 4 | 11 | 9 | 8 | 4 | 5 | 4 | 9 | 10 | 9 | 6.72 | 4 | 9 | 2.039 | 1.028 | 0.1549681 | 0.34713207 | 0 | 0.4968 | 0.2068 | 0.2068 | TRUE | 0.0241 |
| Rnf151 | 7 | 4 | 2 | 3 | 6 | 1 | 7 | 4 | 2 | 2 | 6 | 1 | 3.89 | 4 | 3 | 0.734 | -0.447 | 0.67508354 | NA | 0.2445 | 0.8536 | 0.5561 | 0.5561 | TRUE | 0.5487 |
| Rnf152 | 89 | 110 | 124 | 111 | 89 | 118 | 95 | 110 | 125 | 87 | 96 | 129 | 106.85 | 110 | 104 | 0.945 | -0.082 | 0.72395783 | 0.8506491 | 0.0217 | 0.0377 | 0.0298 | 0.0298 | TRUE | 0.427 |
| Rnf157 | 571 | 894 | 1001 | 1144 | 751 | 712 | 607 | 893 | 1005 | 899 | 808 | 779 | 831.87 | 835 | 829 | 0.993 | -0.01 | 0.95592301 | 0.97741618 | 0.0345 | 0.0109 | 0.0224 | 0.0224 | TRUE | 0.6778 |
| Rnf165 | 1280 | 1274 | 1353 | 1287 | 926 | 1011 | 1360 | 1272 | 1359 | 1012 | 996 | 1107 | 1184.21 | 1330 | 1038 | 0.78 | -0.358 | 4.23E-05 | 0.00067081 | 0.0013 | 0.0097 | 0.0047 | 0.0047 | TRUE | 0.0388 |
| Rnf166 | 723 | 743 | 733 | 726 | 636 | 615 | 768 | 742 | 736 | 571 | 684 | 673 | 695.71 | 749 | 643 | 0.857 | -0.223 | 0.04169226 | 0.14515382 | 0.0039 | 0.0116 | 0.0072 | 0.0072 | TRUE | 0.1189 |
| Rnf167 | 3896 | 4547 | 4227 | 5298 | 3971 | 4075 | 4138 | 4541 | 4244 | 4165 | 4273 | 4461 | 4303.65 | 4308 | 4300 | 0.998 | -0.003 | 0.9648577 | 0.9823598 | 0.0015 | 0.0077 | 0.0036 | 0.0036 | TRUE | 0.0273 |
| Rnf168 | 2144 | 2002 | 1986 | 3514 | 2626 | 2181 | 2277 | 1999 | 1994 | 2762 | 2825 | 2387 | 2374.36 | 2090 | 2658 | 1.272 | 0.347 | 0.00078636 | 0.00738191 | 0.0063 | 0.0083 | 0.0073 | 0.0073 | TRUE | 0.0958 |
| Rnf169 | 2419 | 2737 | 2738 | 3008 | 2204 | 2281 | 2569 | 2733 | 2749 | 2365 | 2371 | 2497 | 2547.51 | 2684 | 2411 | 0.898 | -0.155 | 0.03030075 | 0.1160805 | 8.00E-04 | 0.0082 | 0.0033 | 0.0033 | TRUE | 0.0169 |
| Rnf17 | 713 | 1305 | 1318 | 3383 | 2490 | 2594 | 757 | 1303 | 1323 | 2659 | 2679 | 2840 | 1927.02 | 1128 | 2726 | 2.416 | 1.273 | 2.82E-11 | 2.30E-09 | 0.0459 | 0.0086 | 0.0257 | 0.0257 | TRUE | 0.9874 |
| Rnf170 | 900 | 1117 | 1148 | 1278 | 1019 | 1045 | 956 | 1116 | 1153 | 1005 | 1096 | 1144 | 1078.2 | 1075 | 1082 | 1.005 | 0.008 | 0.94349641 | 0.97171381 | 0.0061 | 0.01 | 0.0078 | 0.0078 | TRUE | 0.1112 |
| Rnf180 | 30 | 36 | 37 | 57 | 41 | 40 | 32 | 36 | 37 | 45 | 44 | 44 | 39.61 | 35 | 44 | 1.264 | 0.338 | 0.25614793 | 0.47484311 | 0 | 0.0901 | 0.0382 | 0.0382 | TRUE | 0.0417 |
| Rnf181 | 4452 | 4998 | 4794 | 5930 | 4310 | 4505 | 4729 | 4992 | 4814 | 4662 | 4637 | 4931 | 4794.08 | 4845 | 4743 | 0.979 | -0.031 | 0.64111058 | 0.79813687 | 7.00E-04 | 0.0076 | 0.0029 | 0.0029 | TRUE | 0.0145 |
| Rnf182 | 58 | 44 | 44 | 79 | 68 | 60 | 62 | 44 | 44 | 62 | 73 | 66 | 58.45 | 50 | 67 | 1.341 | 0.424 | 0.11664046 | 0.28879604 | 0 | 0.0633 | 0.035 | 0.035 | TRUE | 0.3321 |
| Rnf183 | 1 | 5 | 8 | 0 | 4 | 3 | 1 | 5 | 8 | 0 | 4 | 3 | 3.61 | 5 | 2 | 0.52 | -0.945 | 0.46133458 | NA | 0.8189 | 0.9188 | 0.8767 | 0.8767 | TRUE | 0.2871 |
| Rnf185 | 1407 | 1397 | 1344 | 2121 | 1388 | 1314 | 1494 | 1395 | 1350 | 1667 | 1493 | 1438 | 1473.06 | 1413 | 1533 | 1.086 | 0.119 | 0.21283057 | 0.42394087 | 0.0034 | 0.0091 | 0.0059 | 0.0059 | TRUE | 0.0723 |

| | | | | | | | | | | | | | | | | | | | | | | | | | |
|---|---|---|---|---|---|---|---|---|---|---|---|---|---|---|---|---|---|---|---|---|---|---|---|---|---|
| Rnf186 | 4 | 3 | 3 | 4 | 1 | 2 | 4 | 3 | 3 | 3 | 1 | 2 | 2.78 | 3 | 2 | 0.64 | -0.643 | 0.56128261 | NA | 0 | 1.1929 | 0.5106 | 0.5106 | TRUE | 0.2201 |
| Rnf187 | 8497 | 9646 | 9619 | 14835 | 10490 | 9922 | 9025 | 9634 | 9658 | 11662 | 11287 | 10861 | 10354.54 | 9439 | 11270 | 1.194 | 0.256 | 0.00013642 | 0.00176945 | 0.0012 | 0.0072 | 0.0031 | 0.0031 | TRUE | 0.0179 |
| Rnf19a | 1384 | 2023 | 2191 | 2688 | 2036 | 2301 | 1470 | 2020 | 2200 | 2113 | 2191 | 2519 | 2085.49 | 1897 | 2274 | 1.199 | 0.261 | 0.09300861 | 0.24915172 | 0.0239 | 0.0085 | 0.017 | 0.017 | TRUE | 0.4673 |
| Rnf19b | 3891 | 3870 | 3881 | 4961 | 3565 | 3276 | 4133 | 3865 | 3897 | 3900 | 3836 | 3586 | 3869.44 | 3965 | 3774 | 0.952 | -0.071 | 0.32983521 | 0.55084296 | 0.0013 | 0.0078 | 0.0035 | 0.0035 | TRUE | 0.0229 |
| Rnf2 | 4922 | 5427 | 5472 | 7352 | 5258 | 5002 | 5228 | 5420 | 5494 | 5780 | 5657 | 5475 | 5509.14 | 5381 | 5637 | 1.048 | 0.067 | 0.28187852 | 0.50334119 | 4.00E-04 | 0.0075 | 0.0026 | 0.0026 | TRUE | 0.0077 |
| Rnf20 | 2654 | 2098 | 2194 | 3238 | 2412 | 2155 | 2819 | 2095 | 2203 | 2545 | 2595 | 2359 | 2436.15 | 2372 | 2500 | 1.054 | 0.076 | 0.55067687 | 0.73491924 | 0.0137 | 0.0083 | 0.0112 | 0.0112 | TRUE | 0.3281 |
| Rnf207 | 90 | 114 | 152 | 105 | 128 | 124 | 96 | 114 | 153 | 83 | 138 | 136 | 119.68 | 121 | 119 | 0.978 | -0.032 | 0.90757403 | 0.95354464 | 0.0596 | 0.0344 | 0.0467 | 0.0467 | TRUE | 0.7754 |
| Rnf208 | 344 | 503 | 464 | 483 | 370 | 406 | 365 | 502 | 466 | 380 | 398 | 444 | 425.98 | 444 | 407 | 0.915 | -0.128 | 0.40134013 | 0.6141287 | 0.0143 | 0.0146 | 0.0144 | 0.0144 | TRUE | 0.2787 |
| Rnf214 | 1475 | 1559 | 1803 | 1895 | 1483 | 1627 | 1567 | 1557 | 1810 | 1490 | 1596 | 1781 | 1633.4 | 1645 | 1622 | 0.986 | -0.021 | 0.84749583 | 0.92381672 | 0.0072 | 0.0089 | 0.0079 | 0.0079 | TRUE | 0.0938 |
| Rnf215 | 211 | 238 | 255 | 380 | 268 | 271 | 224 | 238 | 256 | 299 | 288 | 297 | 266.93 | 239 | 295 | 1.231 | 0.3 | 0.02097786 | 0.08988517 | 0 | 0.0192 | 0.0084 | 0.0084 | TRUE | 0.0416 |
| Rnf216 | 3951 | 4204 | 4344 | 4575 | 3589 | 3833 | 4197 | 4199 | 4362 | 3597 | 3862 | 4196 | 4068.49 | 4253 | 3885 | 0.913 | -0.131 | 0.11424598 | 0.28501859 | 0.0029 | 0.0077 | 0.0047 | 0.0047 | TRUE | 0.0597 |
| Rnf217 | 75 | 94 | 95 | 299 | 202 | 169 | 80 | 94 | 95 | 235 | 217 | 185 | 151.05 | 90 | 212 | 2.374 | 1.247 | 3.13E-11 | 2.53E-09 | 0 | 0.0287 | 0.0174 | 0.0174 | TRUE | 0.1367 |
| Rnf219 | 382 | 387 | 422 | 842 | 591 | 505 | 406 | 387 | 424 | 662 | 636 | 553 | 511.1 | 406 | 617 | 1.524 | 0.608 | 2.76E-07 | 8.82E-06 | 0.0039 | 0.0133 | 0.008 | 0.008 | TRUE | 0.0954 |
| Rnf220 | 6930 | 7330 | 7085 | 8018 | 6167 | 6169 | 7361 | 7321 | 7114 | 6303 | 6635 | 6753 | 6914.48 | 7265 | 6564 | 0.903 | -0.147 | 0.01950493 | 0.08531106 | 6.00E-04 | 0.0074 | 0.0027 | 0.0027 | TRUE | 0.0149 |
| Rnf222 | 11 | 10 | 15 | 11 | 12 | 6 | 12 | 10 | 15 | 9 | 13 | 7 | 10.81 | 12 | 10 | 0.764 | -0.389 | 0.50955036 | 0.70311708 | 0 | 0.3117 | 0.1568 | 0.1568 | TRUE | 0.324 |
| Rnf223 | 5 | 1 | 4 | 6 | 5 | 1 | 5 | 1 | 4 | 5 | 5 | 1 | 3.59 | 3 | 4 | 1.099 | 0.136 | 0.90101 | NA | 0 | 0.9256 | 0.5776 | 0.5776 | TRUE | 0.5772 |
| Rnf24 | 1200 | 995 | 1058 | 1272 | 899 | 842 | 1275 | 994 | 1062 | 1000 | 967 | 922 | 1036.6 | 1110 | 963 | 0.868 | -0.204 | 0.08428074 | 0.23276049 | 0.0081 | 0.0101 | 0.0091 | 0.0091 | TRUE | 0.2001 |
| Rnf25 | 751 | 713 | 698 | 966 | 696 | 603 | 798 | 712 | 701 | 759 | 749 | 660 | 729.83 | 737 | 723 | 0.982 | -0.026 | 0.80997838 | 0.90218778 | 0.004 | 0.0114 | 0.0072 | 0.0072 | TRUE | 0.0668 |
| Rnf26 | 1406 | 1822 | 1816 | 2272 | 1670 | 1718 | 1493 | 1820 | 1823 | 1786 | 1797 | 1881 | 1766.67 | 1712 | 1821 | 1.063 | 0.089 | 0.39506539 | 0.60855272 | 0.0055 | 0.0088 | 0.0072 | 0.0072 | TRUE | 0.1502 |
| Rnf31 | 2443 | 3225 | 3421 | 2938 | 2377 | 2680 | 2595 | 3221 | 3435 | 2310 | 2558 | 2934 | 2841.93 | 3084 | 2601 | 0.843 | -0.247 | 0.06982269 | 0.20540329 | 0.0168 | 0.0081 | 0.013 | 0.013 | TRUE | 0.2332 |
| Rnf32 | 133 | 158 | 148 | 151 | 113 | 121 | 141 | 158 | 149 | 119 | 122 | 132 | 136.74 | 149 | 124 | 0.831 | -0.267 | 0.11743701 | 0.28999788 | 0 | 0.031 | 0.0137 | 0.0137 | TRUE | 0.0326 |
| Rnf34 | 2381 | 2625 | 2439 | 3017 | 2253 | 2204 | 2529 | 2622 | 2449 | 2372 | 2424 | 2413 | 2468.02 | 2533 | 2403 | 0.948 | -0.076 | 0.25895031 | 0.4778477 | 3.00E-04 | 0.0082 | 0.0029 | 0.0029 | TRUE | 0.0114 |
| Rnf38 | 3102 | 3167 | 3434 | 3872 | 2836 | 3028 | 3295 | 3163 | 3448 | 3044 | 3051 | 3315 | 3219.29 | 3302 | 3137 | 0.95 | -0.074 | 0.33354984 | 0.55422968 | 0.0017 | 0.0079 | 0.004 | 0.004 | TRUE | 0.0297 |
| Rnf39 | 77 | 52 | 46 | 81 | 67 | 30 | 82 | 52 | 46 | 64 | 72 | 33 | 58.09 | 60 | 56 | 0.941 | -0.088 | 0.8137105 | 0.90468363 | 0.1049 | 0.0636 | 0.0831 | 0.0831 | TRUE | 1.0567 |
| Rnf4 | 11115 | 11872 | 11057 | 16723 | 12087 | 11346 | 11806 | 11857 | 11102 | 13146 | 13005 | 12420 | 12222.72 | 11588 | 12857 | 1.11 | 0.15 | 0.02004195 | 0.08696054 | 0.001 | 0.0072 | 0.0029 | 0.0029 | TRUE | 0.0165 |
| Rnf40 | 4589 | 4163 | 4090 | 6182 | 4564 | 4013 | 4874 | 4158 | 4107 | 4860 | 4911 | 4393 | 4550.33 | 4380 | 4721 | 1.078 | 0.109 | 0.27192575 | 0.49187272 | 0.0063 | 0.0076 | 0.0068 | 0.0068 | TRUE | 0.1188 |
| Rnf41 | 1088 | 1100 | 1051 | 1660 | 1231 | 1064 | 1156 | 1099 | 1055 | 1305 | 1324 | 1165 | 1183.95 | 1103 | 1265 | 1.147 | 0.198 | 0.03400761 | 0.12550866 | 0.0026 | 0.0097 | 0.0054 | 0.0054 | TRUE | 0.0564 |
| Rnf43 | 28 | 66 | 67 | 56 | 31 | 46 | 30 | 66 | 67 | 44 | 33 | 50 | 48.44 | 54 | 42 | 0.783 | -0.353 | 0.35331313 | 0.57170972 | 0.0924 | 0.0749 | 0.0832 | 0.0832 | TRUE | 1.2757 |
| Rnf44 | 12335 | 11064 | 11377 | 14558 | 11221 | 10411 | 13102 | 11050 | 11424 | 11444 | 12073 | 11396 | 11748.22 | 11859 | 11638 | 0.981 | -0.027 | 0.75968094 | 0.87323722 | 0.0039 | 0.0072 | 0.0056 | 0.0056 | TRUE | 0.1028 |
| Rnf5 | 1973 | 1903 | 1844 | 2019 | 1517 | 1524 | 2096 | 1901 | 1852 | 1587 | 1632 | 1668 | 1789.24 | 1950 | 1629 | 0.836 | -0.259 | 0.00197848 | 0.01525612 | 0.002 | 0.0087 | 0.0045 | 0.0045 | TRUE | 0.052 |
| Rnf6 | 2673 | 2914 | 2866 | 3311 | 2420 | 2371 | 2839 | 2910 | 2878 | 2603 | 2604 | 2595 | 2738.21 | 2876 | 2601 | 0.904 | -0.145 | 0.01792842 | 0.08030875 | 0 | 0.0081 | 0.0023 | 0.0023 | TRUE | 0.0015 |
| Rnf7 | 3004 | 3155 | 3050 | 4949 | 3361 | 3221 | 3191 | 3151 | 3063 | 3891 | 3616 | 3526 | 3406.14 | 3135 | 3678 | 1.174 | 0.231 | 0.00150104 | 0.01226033 | 0.0013 | 0.0079 | 0.0035 | 0.0035 | TRUE | 0.0317 |
| Rnf8 | 872 | 824 | 743 | 1573 | 1113 | 990 | 926 | 823 | 746 | 1237 | 1198 | 1084 | 1002.17 | 832 | 1173 | 1.411 | 0.497 | 1.41E-05 | 0.00026534 | 0.0067 | 0.0102 | 0.0084 | 0.0084 | TRUE | 0.1167 |
| Rnft1 | 1920 | 2266 | 2250 | 2778 | 1951 | 2079 | 2039 | 2263 | 2259 | 2184 | 2099 | 2276 | 2186.75 | 2187 | 2186 | 0.999 | -0.001 | 0.9915605 | 0.99560047 | 0.0021 | 0.0084 | 0.0044 | 0.0044 | TRUE | 0.0422 |
| Rnft2 | 426 | 533 | 575 | 401 | 326 | 424 | 452 | 532 | 577 | 315 | 351 | 464 | 448.71 | 520 | 377 | 0.722 | -0.47 | 0.00725252 | 0.04123533 | 0.0253 | 0.0142 | 0.0198 | 0.0198 | TRUE | 0.4686 |
| Rngtt | 2893 | 2922 | 2885 | 3555 | 2491 | 2543 | 3073 | 2918 | 2897 | 2795 | 2680 | 2784 | 2857.75 | 2963 | 2753 | 0.929 | -0.106 | 0.11804519 | 0.29058978 | 4.00E-04 | 0.0081 | 0.0029 | 0.0029 | TRUE | 0.0128 |
| Rnh1 | 1898 | 2182 | 2288 | 2623 | 1971 | 2130 | 2016 | 2179 | 2297 | 2062 | 2121 | 2332 | 2167.81 | 2164 | 2172 | 1.003 | 0.004 | 0.9644985 | 0.98213662 | 0.0038 | 0.0084 | 0.0056 | 0.0056 | TRUE | 0.0501 |
| Rnls | 75 | 84 | 84 | 133 | 70 | 75 | 80 | 84 | 84 | 105 | 75 | 82 | 84.98 | 83 | 87 | 1.063 | 0.088 | 0.69938778 | 0.83553583 | 0 | 0.0457 | 0.0258 | 0.0258 | TRUE | 0.3135 |
| Rnmt | 5024 | 5590 | 5538 | 6628 | 4857 | 4948 | 5336 | 5583 | 5561 | 5210 | 5226 | 5416 | 5388.75 | 5493 | 5284 | 0.962 | -0.056 | 0.35750219 | 0.57600619 | 3.00E-04 | 0.0075 | 0.0025 | 0.0025 | TRUE | 0.0076 |
| Rnmtl1 | 1100 | 885 | 838 | 1145 | 816 | 675 | 1168 | 884 | 841 | 900 | 878 | 739 | 901.78 | 964 | 839 | 0.871 | -0.2 | 0.19104267 | 0.39611884 | 0.0207 | 0.0106 | 0.0158 | 0.0158 | TRUE | 0.4066 |
| Rnpc3 | 1040 | 1168 | 1174 | 1509 | 1186 | 1169 | 1105 | 1166 | 1179 | 1186 | 1276 | 1280 | 1198.66 | 1150 | 1247 | 1.084 | 0.116 | 0.16309605 | 0.35861577 | 8.00E-04 | 0.0097 | 0.0041 | 0.0041 | TRUE | 0.0229 |
| Rnpep | 1511 | 1842 | 1861 | 2045 | 1558 | 1475 | 1605 | 1840 | 1869 | 1608 | 1676 | 1615 | 1701.96 | 1771 | 1633 | 0.922 | -0.118 | 0.19451244 | 0.40034585 | 0.0031 | 0.0088 | 0.0053 | 0.0053 | TRUE | 0.0809 |
| Rnpepl1 | 687 | 938 | 992 | 1500 | 1118 | 1080 | 730 | 937 | 996 | 1179 | 1203 | 1182 | 1037.81 | 888 | 1188 | 1.338 | 0.42 | 0.00111053 | 0.00965559 | 0.011 | 0.0101 | 0.011 | 0.011 | TRUE | 0.2865 |
| Rnps1 | 4557 | 5256 | 5403 | 6093 | 4446 | 4984 | 4840 | 5249 | 5425 | 4790 | 4784 | 5456 | 5090.66 | 5171 | 5010 | 0.969 | -0.046 | 0.60674234 | 0.77681144 | 0.0038 | 0.0076 | 0.0056 | 0.0056 | TRUE | 0.0737 |
| Rnu11 | 0 | 1 | 0 | 0 | 0 | 0 | 0 | 1 | 0 | 0 | 0 | 0 | 0.17 | 0 | 0 | 0.49 | -1.029 | 0.80087 | NA | 0 | 19.7991 | 10 | 10 | TRUE | 0.405 |
| Rnu12 | 1 | 0 | 1 | 0 | 0 | 0 | 1 | 0 | 1 | 0 | 0 | 0 | 0.34 | 1 | 0 | 0.261 | -1.939 | 0.60367425 | NA | 0 | 9.5732 | 8.3134 | 8.3134 | TRUE | 0.2529 |
| Rnu6 | 0 | 0 | 0 | 0 | 0 | 0 | 0 | 0 | 0 | 0 | 0 | 0 | 0 | 0 | 0 | NA | NA | NA | NA | NA | NA | NA | NA | NA | NA |
| Rnu7 | 0 | 0 | 0 | 0 | 0 | 0 | 0 | 0 | 0 | 0 | 0 | 0 | 0 | 0 | 0 | NA | NA | NA | NA | NA | NA | NA | NA | NA | NA |
| Rnu73b | 2 | 2 | 1 | 0 | 6 | 3 | 2 | 2 | 1 | 0 | 6 | 3 | 2.48 | 2 | 3 | 1.859 | 0.895 | 0.53509956 | NA | 0 | 1.3366 | 1.0455 | 1.0455 | TRUE | 0.1293 |
| Robo1 | 2138 | 2310 | 2257 | 2320 | 1728 | 1812 | 2271 | 2307 | 2266 | 1824 | 1859 | 1984 | 2085.13 | 2281 | 1889 | 0.827 | -0.273 | 0.0001594 | 0.00200645 | 5.00E-04 | 0.0085 | 0.0033 | 0.0033 | TRUE | 0.023 |
| Robo2 | 6 | 2 | 7 | 15 | 8 | 7 | 6 | 2 | 7 | 12 | 9 | 8 | 7.24 | 5 | 10 | 1.841 | 0.881 | 0.24191917 | 0.45854454 | 0 | 0.4617 | 0.2566 | 0.2566 | TRUE | 0.6052 |
| Robo3 | 39 | 41 | 35 | 78 | 54 | 51 | 41 | 41 | 35 | 61 | 58 | 56 | 48.79 | 39 | 58 | 1.495 | 0.58 | 0.03410133 | 0.12563605 | 0 | 0.0744 | 0.0326 | 0.0326 | TRUE | 0.0609 |
| Robo4 | 1092 | 685 | 732 | 1164 | 751 | 618 | 1160 | 684 | 735 | 915 | 808 | 676 | 829.77 | 860 | 800 | 0.931 | -0.103 | 0.62271837 | 0.78704337 | 0.0524 | 0.0109 | 0.0303 | 0.0303 | TRUE | 1.1081 |
| Rock1 | 3956 | 3837 | 4074 | 6103 | 4273 | 3990 | 4202 | 3832 | 4091 | 4798 | 4597 | 4368 | 4314.6 | 4042 | 4588 | 1.135 | 0.183 | 0.01638214 | 0.07566008 | 0.0019 | 0.0077 | 0.004 | 0.004 | TRUE | 0.0251 |
| Rock2 | 5339 | 5075 | 5188 | 6811 | 4890 | 4700 | 5671 | 5068 | 5209 | 5354 | 5261 | 5145 | 5284.87 | 5316 | 5253 | 0.988 | -0.017 | 0.81810876 | 0.90726388 | 0.0014 | 0.0075 | 0.0037 | 0.0037 | TRUE | 0.0415 |
| Rogdi | 1591 | 1542 | 1556 | 2096 | 1642 | 1342 | 1690 | 1540 | 1562 | 1648 | 1767 | 1469 | 1612.63 | 1597 | 1628 | 1.019 | 0.028 | 0.78219857 | 0.88739843 | 0.0048 | 0.0089 | 0.0065 | 0.0065 | TRUE | 0.0863 |
| Rom1 | 128 | 138 | 123 | 130 | 71 | 104 | 136 | 138 | 124 | 102 | 76 | 114 | 114.95 | 133 | 97 | 0.737 | -0.44 | 0.0372698 | 0.1338691 | 0.0089 | 0.0356 | 0.0233 | 0.0233 | TRUE | 0.3273 |
| Romo1 | 817 | 1060 | 1033 | 1112 | 768 | 818 | 868 | 1059 | 1037 | 874 | 826 | 895 | 926.6 | 988 | 865 | 0.876 | -0.191 | 0.08248015 | 0.22939111 | 0.0056 | 0.0105 | 0.0077 | 0.0077 | TRUE | 0.1342 |
| Ropn1 | 1 | 1 | 1 | 2 | 1 | 3 | 1 | 1 | 1 | 2 | 1 | 3 | 1.5 | 1 | 2 | 1.919 | 0.94 | 0.53201045 | NA | 0 | 2.2039 | 0.8817 | 0.8817 | TRUE | 0.2633 |
| Ropn1l | 42 | 73 | 72 | 39 | 47 | 52 | 45 | 73 | 72 | 31 | 51 | 57 | 54.66 | 63 | 46 | 0.721 | -0.472 | 0.17146008 | 0.37034626 | 0.0672 | 0.0672 | 0.0672 | 0.0672 | TRUE | 0.7819 |
| Ror1 | 25 | 22 | 24 | 25 | 23 | 19 | 27 | 22 | 24 | 20 | 25 | 21 | 22.97 | 24 | 22 | 0.895 | -0.16 | 0.68212117 | 0.82364225 | 0 | 0.1503 | 0.0665 | 0.0665 | TRUE | 0.0791 |
| Ror2 | 1484 | 1962 | 1825 | 1546 | 1179 | 1364 | 1576 | 1959 | 1832 | 1215 | 1269 | 1493 | 1557.54 | 1789 | 1326 | 0.74 | -0.434 | 0.00044366 | 0.00473112 | 0.0111 | 0.009 | 0.0103 | 0.0103 | TRUE | 0.1459 |
| Rora | 21 | 57 | 65 | 58 | 52 | 53 | 22 | 57 | 65 | 46 | 56 | 58 | 50.68 | 48 | 53 | 1.099 | 0.137 | 0.73206155 | 0.85627965 | 0.1228 | 0.0719 | 0.0952 | 0.0952 | TRUE | 1.724 |
| Rorb | 2 | 1 | 0 | 2 | 1 | 0 | 2 | 1 | 0 | 2 | 1 | 0 | 0.96 | 1 | 1 | 0.879 | -0.185 | 0.92762908 | NA | 0 | 3.4319 | 1.9663 | 1.9663 | TRUE | 0.1856 |
| Rorc | 21 | 18 | 15 | 5 | 6 | 5 | 22 | 18 | 15 | 4 | 6 | 5 | 11.87 | 18 | 5 | 0.282 | -1.824 | 0.00244517 | 0.01803939 | 0 | 0.2845 | 0.1399 | 0.1399 | TRUE | 0.1664 |
| Ros1 | 1 | 0 | 0 | 1 | 0 | 0 | 1 | 0 | 0 | 1 | 0 | 0 | 0.31 | 0 | 0 | 0.954 | -0.067 | 0.98625928 | NA | 0 | 10.7014 | 9.0384 | 9.0384 | TRUE | 0.4614 |
| Rp1 | 0 | 0 | 1 | 2 | 0 | 1 | 0 | 0 | 1 | 2 | 0 | 1 | 0.61 | 0 | 1 | 2.474 | 1.307 | 0.62050174 | NA | 0 | 5.3914 | 3.4771 | 3.4771 | TRUE | 0.4098 |
| Rp1l1 | 0 | 0 | 0 | 0 | 0 | 0 | 0 | 0 | 0 | 0 | 0 | 0 | 0 | 0 | 0 | NA | NA | NA | NA | NA | NA | NA | NA | NA | NA |
| Rp2h | 1318 | 1615 | 1611 | 1822 | 1340 | 1524 | 1400 | 1613 | 1618 | 1432 | 1442 | 1668 | 1528.8 | 1544 | 1514 | 0.98 | -0.029 | 0.7882694 | 0.8909931 | 0.0064 | 0.0091 | 0.0075 | 0.0075 | TRUE | 0.0947 |

| | | | | | | | | | | | | | | | | | | | | | | | | | |
|---|---|---|---|---|---|---|---|---|---|---|---|---|---|---|---|---|---|---|---|---|---|---|---|---|---|
| Rp9 | 1318 | 1714 | 1786 | 1899 | 1493 | 1583 | 1400 | 1712 | 1793 | 1493 | 1606 | 1733 | 1622.86 | 1635 | 1611 | 0.985 | -0.023 | 0.8516168 | 0.925941 | 0.0106 | 0.0089 | 0.0098 | 0.0098 | TRUE | 0.1902 |
| Rpa1 | 29455 | 27313 | 27669 | 38561 | 27656 | 25939 | 31286 | 27278 | 27783 | 30314 | 29756 | 28394 | 29135.14 | 28782 | 29488 | 1.025 | 0.035 | 0.66405673 | 0.81189943 | 0.0028 | 0.007 | 0.0046 | 0.0046 | TRUE | 0.0709 |
| Rpa2 | 4408 | 4100 | 4007 | 6354 | 4631 | 4175 | 4682 | 4095 | 4023 | 4995 | 4983 | 4570 | 4558.02 | 4267 | 4849 | 1.137 | 0.185 | 0.04158121 | 0.14488911 | 0.0045 | 0.0076 | 0.0057 | 0.0057 | TRUE | 0.0882 |
| Rpa3 | 1620 | 1715 | 1572 | 2263 | 1569 | 1502 | 1721 | 1713 | 1578 | 1779 | 1688 | 1644 | 1687.21 | 1671 | 1704 | 1.02 | 0.029 | 0.71910842 | 0.84792001 | 0.0014 | 0.0089 | 0.0042 | 0.0042 | TRUE | 0.0282 |
| Rpain | 809 | 940 | 856 | 1011 | 697 | 812 | 859 | 939 | 860 | 795 | 750 | 889 | 848.53 | 886 | 811 | 0.915 | -0.128 | 0.22281272 | 0.4348722 | 0.0034 | 0.0108 | 0.0067 | 0.0067 | TRUE | 0.0811 |
| Rpap1 | 7774 | 8175 | 8608 | 10891 | 8407 | 8618 | 8257 | 8164 | 8643 | 8562 | 9045 | 9434 | 8684.33 | 8355 | 9014 | 1.079 | 0.109 | 0.11786489 | 0.29041792 | 0.0015 | 0.0073 | 0.0034 | 0.0034 | TRUE | 0.0237 |
| Rpap2 | 1639 | 1831 | 1750 | 2189 | 1591 | 1548 | 1741 | 1829 | 1757 | 1721 | 1712 | 1695 | 1742.32 | 1776 | 1709 | 0.963 | -0.055 | 0.42932125 | 0.63907651 | 0 | 0.0088 | 0.0029 | 0.0029 | TRUE | 0.0083 |
| Rpap3 | 2803 | 2691 | 2513 | 3592 | 2492 | 2491 | 2977 | 2688 | 2523 | 2824 | 2681 | 2727 | 2736.65 | 2729 | 2744 | 1.006 | 0.008 | 0.92677017 | 0.96386542 | 0.0035 | 0.0081 | 0.0052 | 0.0052 | TRUE | 0.0764 |
| Rpe | 1777 | 1841 | 1850 | 2986 | 2153 | 2026 | 1887 | 1839 | 1858 | 2347 | 2316 | 2218 | 2077.56 | 1861 | 2294 | 1.233 | 0.302 | 1.11E-05 | 0.00021805 | 0 | 0.0085 | 0.0029 | 0.0029 | TRUE | 0.0101 |
| Rpe65 | 0 | 0 | 0 | 1 | 0 | 0 | 0 | 0 | 0 | 1 | 0 | 0 | 0.13 | 0 | 0 | 1.859 | 0.894 | 0.82650734 | NA | 0 | 25.1515 | 10 | 10 | TRUE | 0.2563 |
| Rpf1 | 2412 | 2478 | 2358 | 3230 | 2277 | 2337 | 2562 | 2475 | 2368 | 2539 | 2450 | 2558 | 2491.96 | 2468 | 2516 | 1.019 | 0.028 | 0.69468077 | 0.83270212 | 6.00E-04 | 0.0082 | 0.0032 | 0.0032 | TRUE | 0.0154 |
| Rpf2 | 2250 | 2466 | 2367 | 3134 | 2154 | 2155 | 2390 | 2463 | 2377 | 2464 | 2318 | 2359 | 2394.95 | 2410 | 2380 | 0.988 | -0.017 | 0.80026956 | 0.89686328 | 3.00E-04 | 0.0083 | 0.003 | 0.003 | TRUE | 0.0118 |
| Rpgr | 1737 | 1990 | 2054 | 1956 | 1560 | 1582 | 1845 | 1987 | 2062 | 1538 | 1678 | 1732 | 1807.12 | 1965 | 1649 | 0.839 | -0.254 | 0.00443859 | 0.02857366 | 0.003 | 0.0087 | 0.0052 | 0.0052 | TRUE | 0.0434 |
| Rpgrip1 | 97 | 76 | 63 | 149 | 110 | 77 | 103 | 76 | 63 | 117 | 118 | 84 | 93.66 | 81 | 106 | 1.325 | 0.406 | 0.12339311 | 0.2994547 | 0.0358 | 0.0421 | 0.0391 | 0.0391 | TRUE | 0.5342 |
| Rpgrip1l | 653 | 543 | 602 | 1044 | 736 | 613 | 694 | 542 | 604 | 821 | 792 | 671 | 687.33 | 613 | 761 | 1.242 | 0.313 | 0.02053243 | 0.08840772 | 0.0109 | 0.0117 | 0.0117 | 0.0117 | TRUE | 0.1527 |
| Rph3a | 113 | 69 | 56 | 67 | 59 | 41 | 120 | 69 | 56 | 53 | 63 | 45 | 67.7 | 82 | 54 | 0.658 | -0.605 | 0.07634494 | 0.21799969 | 0.0818 | 0.0556 | 0.0685 | 0.0685 | TRUE | 1.1759 |
| Rph3al | 135 | 188 | 153 | 168 | 108 | 139 | 143 | 188 | 154 | 132 | 116 | 152 | 147.53 | 162 | 133 | 0.825 | -0.277 | 0.15464875 | 0.34671216 | 0.0118 | 0.0292 | 0.0205 | 0.0205 | TRUE | 0.2134 |
| Rpia | 391 | 373 | 393 | 498 | 374 | 350 | 415 | 373 | 395 | 391 | 402 | 383 | 393.24 | 394 | 392 | 0.996 | -0.006 | 0.95576116 | 0.97741618 | 0 | 0.0153 | 0.0065 | 0.0065 | TRUE | 0.0266 |
| Rpl10 | 32314 | 37971 | 38549 | 41871 | 29487 | 31424 | 34323 | 37922 | 38707 | 32916 | 31726 | 34399 | 34998.81 | 36984 | 33014 | 0.893 | -0.164 | 0.03392574 | 0.12537411 | 0.0028 | 0.007 | 0.0043 | 0.0043 | TRUE | 0.0485 |
| Rpl10a | 24349 | 34840 | 35979 | 33271 | 25274 | 28632 | 25863 | 34795 | 36127 | 26155 | 27193 | 31342 | 30245.87 | 32262 | 28230 | 0.875 | -0.193 | 0.16813781 | 0.36589195 | 0.0196 | 0.007 | 0.014 | 0.014 | TRUE | 0.3684 |
| Rpl10l | 1877 | 1198 | 1027 | 5225 | 3624 | 2800 | 1994 | 1196 | 1031 | 4107 | 3899 | 3065 | 2548.85 | 1407 | 3690 | 2.623 | 1.391 | 5.89E-10 | 3.81E-08 | 0.0773 | 0.0082 | 0.0359 | 0.0359 | TRUE | 1.5982 |
| Rpl11 | 24069 | 32119 | 32673 | 34768 | 25190 | 26314 | 25566 | 32078 | 32807 | 27332 | 27103 | 28805 | 28948.3 | 30150 | 27747 | 0.92 | -0.12 | 0.27275019 | 0.49295859 | 0.0092 | 0.007 | 0.0086 | 0.0086 | TRUE | 0.2165 |
| Rpl12 | 44289 | 57333 | 59561 | 60755 | 44222 | 47697 | 47043 | 57259 | 59806 | 47761 | 47580 | 52212 | 51943.35 | 54703 | 49184 | 0.899 | -0.153 | 0.15414534 | 0.34607548 | 0.009 | 0.007 | 0.0083 | 0.0083 | TRUE | 0.1837 |
| Rpl13 | 46339 | 61151 | 63258 | 64917 | 48832 | 51787 | 49220 | 61072 | 63518 | 51033 | 52540 | 56689 | 55678.65 | 57937 | 53421 | 0.922 | -0.117 | 0.29609844 | 0.51682876 | 0.0102 | 0.007 | 0.009 | 0.009 | TRUE | 0.2121 |
| Rpl13a | 65430 | 83855 | 83791 | 93118 | 65215 | 68300 | 69498 | 83747 | 84135 | 73202 | 70167 | 74765 | 75919.08 | 79127 | 72711 | 0.919 | -0.122 | 0.20165125 | 0.40921241 | 0.0061 | 0.0069 | 0.0066 | 0.0066 | TRUE | 0.1388 |
| Rpl14 | 29381 | 36594 | 37059 | 41330 | 29823 | 31685 | 31208 | 36547 | 37211 | 32490 | 32088 | 34684 | 34038.01 | 34989 | 33087 | 0.946 | -0.081 | 0.37975892 | 0.59453654 | 0.0053 | 0.007 | 0.006 | 0.006 | TRUE | 0.1094 |
| Rpl15 | 18219 | 21920 | 21712 | 25646 | 17814 | 18849 | 19352 | 21892 | 21801 | 20161 | 19167 | 20633 | 20500.91 | 21015 | 19987 | 0.951 | -0.072 | 0.36120184 | 0.57953971 | 0.003 | 0.0071 | 0.0045 | 0.0045 | TRUE | 0.0586 |
| Rpl17 | 30687 | 43127 | 44123 | 45742 | 32611 | 36688 | 32595 | 43071 | 44304 | 35959 | 35087 | 40161 | 38529.58 | 39990 | 37069 | 0.927 | -0.109 | 0.39729269 | 0.61079056 | 0.0148 | 0.007 | 0.012 | 0.012 | TRUE | 0.3204 |
| Rpl18 | 30677 | 38527 | 38403 | 41511 | 30581 | 31028 | 32584 | 38477 | 38561 | 32633 | 32903 | 33965 | 34853.89 | 36541 | 33167 | 0.908 | -0.14 | 0.11402408 | 0.28478082 | 0.0046 | 0.007 | 0.0056 | 0.0056 | TRUE | 0.1098 |
| Rpl18a | 35540 | 47437 | 48232 | 50932 | 36899 | 40672 | 37750 | 47376 | 48430 | 40039 | 39701 | 44522 | 42969.54 | 44519 | 41421 | 0.93 | -0.104 | 0.36224106 | 0.58014481 | 0.0106 | 0.007 | 0.0094 | 0.0094 | TRUE | 0.2166 |
| Rpl19 | 47382 | 60616 | 62202 | 68932 | 48554 | 51225 | 50328 | 60538 | 62457 | 54189 | 52241 | 56074 | 55971.18 | 57774 | 54168 | 0.938 | -0.093 | 0.3505898 | 0.56934743 | 0.007 | 0.007 | 0.0071 | 0.0071 | TRUE | 0.1556 |
| Rpl21 | 31936 | 36487 | 37121 | 43231 | 31357 | 31708 | 33922 | 36440 | 37273 | 33985 | 33738 | 34709 | 35011.23 | 35878 | 34144 | 0.952 | -0.071 | 0.27211351 | 0.49206309 | 0.0012 | 0.007 | 0.003 | 0.003 | TRUE | 0.0279 |
| Rpl22 | 17354 | 25697 | 27092 | 24635 | 18579 | 21991 | 18433 | 25664 | 27203 | 19366 | 19990 | 24073 | 22454.77 | 23767 | 21143 | 0.89 | -0.169 | 0.2777996 | 0.49891315 | 0.0266 | 0.007 | 0.0174 | 0.0174 | TRUE | 0.4716 |
| Rpl22l1 | 8820 | 12169 | 11860 | 11715 | 8633 | 10160 | 9368 | 12153 | 11909 | 9209 | 9289 | 11122 | 10508.36 | 11143 | 9873 | 0.886 | -0.175 | 0.17413658 | 0.37335066 | 0.0154 | 0.0072 | 0.0118 | 0.0118 | TRUE | 0.2372 |
| Rpl23 | 43092 | 53509 | 54164 | 60844 | 41934 | 44435 | 45771 | 53440 | 54386 | 47831 | 45118 | 48641 | 49197.98 | 51199 | 47197 | 0.922 | -0.117 | 0.19350986 | 0.39916787 | 0.005 | 0.007 | 0.0059 | 0.0059 | TRUE | 0.1053 |
| Rpl23a | 34979 | 42467 | 44403 | 44690 | 33591 | 36641 | 37154 | 42412 | 44585 | 35132 | 36142 | 40109 | 39255.72 | 41384 | 37128 | 0.897 | -0.157 | 0.10952039 | 0.27762038 | 0.0067 | 0.007 | 0.0069 | 0.0069 | TRUE | 0.0979 |
| Rpl24 | 14507 | 18336 | 18752 | 20431 | 14671 | 15859 | 15409 | 18312 | 18829 | 16061 | 15785 | 17360 | 16959.47 | 17517 | 16402 | 0.936 | -0.095 | 0.33880623 | 0.55879656 | 0.0068 | 0.0071 | 0.007 | 0.007 | TRUE | 0.1355 |
| Rpl26 | 2 | 1 | 1 | 4 | 1 | 3 | 2 | 1 | 1 | 3 | 1 | 3 | 1.94 | 1 | 2 | 1.843 | 0.882 | 0.51275444 | NA | 0 | 1.7063 | 0.7426 | 0.7426 | TRUE | 0.2473 |
| Rpl27 | 14866 | 18845 | 18567 | 20722 | 14341 | 14839 | 15790 | 18821 | 18643 | 16290 | 15430 | 16244 | 16869.65 | 17751 | 15988 | 0.901 | -0.151 | 0.09736772 | 0.2563981 | 0.005 | 0.0071 | 0.0059 | 0.0059 | TRUE | 0.1142 |
| Rpl27a | 23681 | 30310 | 31166 | 31746 | 23326 | 25307 | 25153 | 30271 | 31294 | 24956 | 25097 | 27702 | 27412.38 | 28906 | 25918 | 0.897 | -0.157 | 0.13260259 | 0.31402218 | 0.0082 | 0.007 | 0.0079 | 0.0079 | TRUE | 0.1578 |
| Rpl28 | 19491 | 27470 | 28062 | 32337 | 23529 | 25181 | 20703 | 27435 | 28177 | 25421 | 25316 | 27565 | 25769.3 | 25438 | 26101 | 1.026 | 0.037 | 0.76882005 | 0.87928573 | 0.0113 | 0.007 | 0.0114 | 0.0114 | TRUE | 0.3245 |
| Rpl29 | 1753 | 1921 | 2045 | 2380 | 1597 | 1587 | 1862 | 1919 | 2053 | 1871 | 1718 | 1737 | 1860.06 | 1945 | 1775 | 0.914 | -0.131 | 0.11424184 | 0.28501859 | 0.0019 | 0.0087 | 0.0044 | 0.0044 | TRUE | 0.029 |
| Rpl3 | 49663 | 66782 | 69191 | 68903 | 50219 | 56780 | 52751 | 66696 | 69475 | 54166 | 54033 | 62155 | 59879.18 | 62974 | 56785 | 0.902 | -0.149 | 0.22225916 | 0.43444058 | 0.0135 | 0.007 | 0.0108 | 0.0108 | TRUE | 0.2469 |
| Rpl30 | 136 | 92 | 79 | 184 | 124 | 102 | 144 | 92 | 79 | 145 | 133 | 112 | 117.56 | 105 | 130 | 1.239 | 0.309 | 0.24198637 | 0.4585617 | 0.0474 | 0.0349 | 0.0415 | 0.0415 | TRUE | 1.0355 |
| Rpl31 | 15134 | 18572 | 18981 | 21311 | 14666 | 15300 | 16075 | 18548 | 19059 | 16753 | 15780 | 16748 | 17160.49 | 17894 | 16427 | 0.918 | -0.123 | 0.16254441 | 0.35788551 | 0.0045 | 0.0071 | 0.0056 | 0.0056 | TRUE | 0.0967 |
| Rpl31-ps12 | 0 | 2 | 1 | 0 | 1 | 1 | 0 | 2 | 1 | 0 | 1 | 1 | 0.86 | 1 | 1 | 0.688 | -0.539 | 0.8028361 | NA | 0 | 3.8287 | 2.1472 | 2.1472 | TRUE | 0.1651 |
| Rpl32 | 47926 | 63312 | 64216 | 65126 | 46253 | 50894 | 50906 | 63230 | 64480 | 51197 | 49765 | 55711 | 55881.6 | 59539 | 52224 | 0.877 | -0.189 | 0.0861871 | 0.23687768 | 0.0097 | 0.007 | 0.0087 | 0.0087 | TRUE | 0.197 |
| Rpl34 | 182 | 170 | 166 | 280 | 222 | 167 | 193 | 170 | 167 | 220 | 239 | 183 | 195.26 | 177 | 214 | 1.213 | 0.279 | 0.09830698 | 0.25811514 | 0.0071 | 0.0238 | 0.0153 | 0.0153 | TRUE | 0.1687 |
| Rpl34-ps1 | 0 | 0 | 2 | 2 | 0 | 2 | 0 | 0 | 2 | 2 | 0 | 2 | 0.96 | 1 | 1 | 1.87 | 0.903 | 0.69743206 | NA | 0 | 3.4328 | 2.7436 | 2.7436 | TRUE | 0.9739 |
| Rpl35 | 12830 | 15969 | 15901 | 17723 | 13175 | 13476 | 13628 | 15948 | 15966 | 13932 | 14175 | 14752 | 14733.65 | 15181 | 14286 | 0.941 | -0.088 | 0.31614153 | 0.53684153 | 0.0043 | 0.0071 | 0.0054 | 0.0054 | TRUE | 0.0979 |
| Rpl35a | 12741 | 17462 | 17619 | 18461 | 12749 | 14223 | 13533 | 17439 | 17691 | 14513 | 13717 | 15569 | 15410.5 | 16221 | 14600 | 0.9 | -0.152 | 0.20524428 | 0.41379429 | 0.0123 | 0.0071 | 0.0103 | 0.0103 | TRUE | 0.257 |
| Rpl36 | 18454 | 26369 | 27335 | 27187 | 19879 | 21670 | 19601 | 26335 | 27447 | 21372 | 21389 | 23721 | 23310.95 | 24461 | 22161 | 0.906 | -0.143 | 0.28545712 | 0.50675641 | 0.0169 | 0.007 | 0.0128 | 0.0128 | TRUE | 0.3696 |
| Rpl36a | 12000 | 14787 | 15665 | 17134 | 11851 | 12811 | 12746 | 14768 | 15729 | 13469 | 12751 | 14024 | 13914.56 | 14414 | 13415 | 0.931 | -0.104 | 0.29271237 | 0.5137339 | 0.0066 | 0.0071 | 0.0069 | 0.0069 | TRUE | 0.1253 |
| Rpl36al | 5984 | 6743 | 6833 | 7930 | 5607 | 5772 | 6356 | 6734 | 6861 | 6234 | 6033 | 6318 | 6422.75 | 6650 | 6195 | 0.932 | -0.102 | 0.11887676 | 0.29199879 | 8.00E-04 | 0.0074 | 0.0029 | 0.0029 | TRUE | 0.0183 |
| Rpl37 | 15560 | 19352 | 20338 | 21659 | 15688 | 16861 | 16527 | 19327 | 20422 | 17027 | 16879 | 18457 | 18106.49 | 18759 | 17454 | 0.93 | -0.104 | 0.29584359 | 0.51661261 | 0.0069 | 0.0071 | 0.0071 | 0.0071 | TRUE | 0.1324 |
| Rpl37a | 14412 | 17280 | 17383 | 21256 | 14832 | 15635 | 15308 | 17258 | 17454 | 16710 | 15958 | 17115 | 16633.87 | 16673 | 16594 | 0.995 | -0.007 | 0.93161643 | 0.96628017 | 0.0025 | 0.0071 | 0.0046 | 0.0046 | TRUE | 0.0627 |
| Rpl38 | 13477 | 17214 | 17448 | 18640 | 13801 | 14680 | 14315 | 17192 | 17520 | 14653 | 14849 | 16070 | 15766.38 | 16342 | 15191 | 0.929 | -0.105 | 0.29341518 | 0.51437903 | 0.007 | 0.0071 | 0.0072 | 0.0072 | TRUE | 0.144 |
| Rpl39 | 8544 | 11801 | 11976 | 13114 | 8460 | 9985 | 9075 | 11786 | 12025 | 10309 | 9102 | 10930 | 10537.99 | 10962 | 10114 | 0.923 | -0.116 | 0.36895214 | 0.58632924 | 0.0146 | 0.0072 | 0.012 | 0.012 | TRUE | 0.277 |
| Rpl39l | 2070 | 2254 | 2095 | 4235 | 3338 | 3080 | 2199 | 2251 | 2104 | 3329 | 3591 | 3372 | 2807.61 | 2185 | 3431 | 1.57 | 0.651 | 6.00E-19 | 1.31E-16 | 0.0011 | 0.0081 | 0.0035 | 0.0035 | TRUE | 0.0203 |
| Rpl3l | 87 | 119 | 74 | 113 | 93 | 83 | 92 | 119 | 74 | 89 | 100 | 91 | 94.22 | 95 | 93 | 0.978 | -0.032 | 0.89447826 | 0.9490474 | 0.0191 | 0.0419 | 0.0306 | 0.0306 | TRUE | 0.3109 |
| Rpl4 | 112503 | 142366 | 146574 | 153276 | 111773 | 118605 | 119498 | 142182 | 147176 | 120494 | 120261 | 129832 | 129907.06 | 136285 | 123529 | 0.906 | -0.142 | 0.15044692 | 0.34117752 | 0.0068 | 0.0069 | 0.007 | 0.007 | TRUE | 0.1422 |
| Rpl41 | 23413 | 30592 | 30061 | 36649 | 24943 | 25324 | 24869 | 30553 | 30184 | 28811 | 26837 | 27721 | 28162.41 | 28535 | 27790 | 0.974 | -0.038 | 0.70045714 | 0.83619323 | 0.0058 | 0.007 | 0.0071 | 0.0071 | TRUE | 0.1546 |
| Rpl5 | 36143 | 45386 | 45783 | 51241 | 34903 | 37817 | 38390 | 45327 | 45971 | 40282 | 37553 | 41397 | 41486.74 | 43229 | 39744 | 0.919 | -0.121 | 0.20155925 | 0.40908409 | 0.0059 | 0.007 | 0.0065 | 0.0065 | TRUE | 0.1174 |
| Rpl6 | 42319 | 49172 | 50996 | 57028 | 41278 | 43592 | 44950 | 49109 | 51205 | 44831 | 44413 | 47718 | 47037.66 | 48421 | 45654 | 0.943 | -0.085 | 0.2749222 | 0.49548018 | 0.0029 | 0.007 | 0.0043 | 0.0043 | TRUE | 0.0481 |
| Rpl7 | 55368 | 68781 | 70182 | 81032 | 56820 | 60369 | 58811 | 68692 | 70470 | 63701 | 61135 | 66083 | 64815.36 | 65991 | 63640 | 0.964 | -0.052 | 0.56742392 | 0.74721633 | 0.0047 | 0.007 | 0.006 | 0.006 | TRUE | 0.1109 |
| Rpl7a | 58916 | 70795 | 71650 | 79992 | 56882 | 59342 | 62579 | 70704 | 71944 | 62883 | 61202 | 64959 | 65711.85 | 68409 | 63015 | 0.921 | -0.119 | 0.1376044 | 0.32173028 | 0.0032 | 0.007 | 0.0046 | 0.0046 | TRUE | 0.068 |
| Rpl7l1 | 9699 | 9093 | 8385 | 13258 | 9296 | 8219 | 10302 | 9081 | 8419 | 10422 | 10002 | 8997 | 9537.34 | 9267 | 9807 | 1.058 | 0.082 | 0.43073133 | 0.64055214 | 0.0072 | 0.0072 | 0.0077 | 0.0077 | TRUE | 0.1164 |
| Rpl8 | 71139 | 88353 | 89938 | 91402 | 67345 | 71631 | 75562 | 88239 | 90307 | 71853 | 72459 | 78411 | 79471.99 | 84703 | 74241 | 0.876 | -0.19 | 0.04089783 | 0.14311971 | 0.0056 | 0.0069 | 0.0062 | 0.0062 | TRUE | 0.1091 |
| Rpl9 | 35297 | 49661 | 48917 | 51288 | 34531 | 38704 | 37492 | 49597 | 49118 | 40319 | 37153 | 42368 | 42674.3 | 45402 | 39947 | 0.88 | -0.185 | 0.13477974 | 0.31755991 | 0.0135 | 0.007 | 0.011 | 0.011 | TRUE | 0.2844 |

| | | | | | | | | | | | | | | | | | | | | | | | | | |
|---|---|---|---|---|---|---|---|---|---|---|---|---|---|---|---|---|---|---|---|---|---|---|---|---|---|
| Rplp0 | 31241 | 38356 | 40039 | 52543 | 37314 | 37818 | 33183 | 38306 | 40203 | 41305 | 40148 | 41398 | 39090.64 | 37231 | 40950 | 1.1 | 0.137 | 0.12556195 | 0.30336443 | 0.0049 | 0.007 | 0.0058 | 0.0058 | TRUE | 0.1107 |
| Rplp1 | 43070 | 59188 | 60918 | 64074 | 45607 | 49935 | 45748 | 59112 | 61168 | 50370 | 49070 | 54662 | 53354.94 | 55343 | 51367 | 0.928 | -0.108 | 0.37717069 | 0.59226764 | 0.0127 | 0.007 | 0.0107 | 0.0107 | TRUE | 0.2816 |
| Rplp2 | 27280 | 34662 | 36033 | 35704 | 25878 | 27744 | 28976 | 34617 | 36181 | 28068 | 27843 | 30370 | 31009.23 | 33258 | 28760 | 0.865 | -0.21 | 0.04059141 | 0.14242663 | 0.0076 | 0.007 | 0.0075 | 0.0075 | TRUE | 0.1552 |
| Rplp2-ps1 | 0 | 0 | 0 | 0 | 0 | 0 | 0 | 0 | 0 | 0 | 0 | 0 | 0 | 0 | 0 | NA | NA | NA | NA | NA | NA | NA | NA | NA | NA |
| Rpn1 | 13588 | 16193 | 15847 | 15910 | 11771 | 12602 | 14433 | 16172 | 15912 | 12507 | 12665 | 13795 | 14247.33 | 15506 | 12989 | 0.838 | -0.256 | 0.00154562 | 0.01253985 | 0.0032 | 0.0071 | 0.0046 | 0.0046 | TRUE | 0.0448 |
| Rpn2 | 13125 | 16152 | 16368 | 15897 | 11798 | 12807 | 13941 | 16131 | 16435 | 12497 | 12694 | 14019 | 14286.27 | 15502 | 13070 | 0.843 | -0.246 | 0.00917549 | 0.04934053 | 0.0058 | 0.0071 | 0.0064 | 0.0064 | TRUE | 0.0949 |
| Rpp14 | 1020 | 1038 | 1014 | 1730 | 1134 | 1033 | 1083 | 1037 | 1018 | 1360 | 1220 | 1131 | 1141.52 | 1046 | 1237 | 1.184 | 0.244 | 0.01575848 | 0.07356015 | 0.0042 | 0.0098 | 0.0064 | 0.0064 | TRUE | 0.0931 |
| Rpp21 | 585 | 610 | 543 | 850 | 601 | 608 | 621 | 609 | 545 | 668 | 647 | 666 | 626.03 | 592 | 660 | 1.116 | 0.158 | 0.11685639 | 0.28898118 | 7.00E-04 | 0.0122 | 0.0057 | 0.0057 | TRUE | 0.0562 |
| Rpp25 | 1220 | 957 | 1011 | 2195 | 1576 | 1283 | 1296 | 956 | 1015 | 1726 | 1696 | 1404 | 1348.74 | 1089 | 1609 | 1.478 | 0.564 | 9.24E-05 | 0.00129627 | 0.0188 | 0.0093 | 0.0142 | 0.0142 | TRUE | 0.3299 |
| Rpp25l | 706 | 698 | 644 | 1083 | 765 | 712 | 750 | 697 | 647 | 851 | 823 | 779 | 757.92 | 698 | 818 | 1.173 | 0.23 | 0.02307511 | 0.09626652 | 0.0023 | 0.0112 | 0.0061 | 0.0061 | TRUE | 0.0498 |
| Rpp30 | 1582 | 1718 | 1772 | 2246 | 1662 | 1737 | 1680 | 1716 | 1779 | 1766 | 1788 | 1901 | 1771.78 | 1725 | 1818 | 1.054 | 0.075 | 0.31957012 | 0.5406844 | 7.00E-04 | 0.0088 | 0.0036 | 0.0036 | TRUE | 0.019 |
| Rpp38 | 318 | 375 | 362 | 649 | 451 | 383 | 338 | 375 | 363 | 510 | 485 | 419 | 415.08 | 359 | 471 | 1.317 | 0.397 | 0.00171528 | 0.01362203 | 0.0044 | 0.0148 | 0.0091 | 0.0091 | TRUE | 0.1074 |
| Rpp40 | 1941 | 1793 | 1876 | 2430 | 1701 | 1676 | 2062 | 1791 | 1884 | 1910 | 1830 | 1835 | 1885.19 | 1912 | 1858 | 0.972 | -0.04 | 0.63567044 | 0.79519306 | 0.0021 | 0.0087 | 0.0047 | 0.0047 | TRUE | 0.0564 |
| Rpph1 | 43 | 19 | 19 | 33 | 18 | 17 | 46 | 19 | 19 | 26 | 19 | 19 | 24.61 | 28 | 21 | 0.77 | -0.377 | 0.43972956 | 0.64702067 | 0.1199 | 0.1408 | 0.1303 | 0.1303 | TRUE | 1.9376 |
| Rprd1a | 2824 | 2919 | 3109 | 2799 | 2053 | 2547 | 3000 | 2915 | 3122 | 2200 | 2209 | 2788 | 2705.66 | 3012 | 2399 | 0.796 | -0.329 | 0.00376804 | 0.02525689 | 0.0099 | 0.0081 | 0.0089 | 0.0089 | TRUE | 0.2427 |
| Rprd1b | 1394 | 1245 | 1278 | 2812 | 1829 | 1594 | 1481 | 1243 | 1283 | 2211 | 1968 | 1745 | 1655.11 | 1336 | 1975 | 1.479 | 0.565 | 3.15E-06 | 7.48E-05 | 0.0105 | 0.0089 | 0.01 | 0.01 | TRUE | 0.1338 |
| Rprd2 | 2265 | 2160 | 2240 | 3088 | 2346 | 2022 | 2406 | 2157 | 2249 | 2428 | 2524 | 2213 | 2329.56 | 2271 | 2388 | 1.052 | 0.073 | 0.40875248 | 0.62120294 | 0.0031 | 0.0083 | 0.0052 | 0.0052 | TRUE | 0.0491 |
| Rprl1 | 0 | 0 | 0 | 0 | 0 | 0 | 0 | 0 | 0 | 0 | 0 | 0 | 0 | 0 | 0 | NA | NA | NA | NA | NA | NA | NA | NA | NA | NA |
| Rprl2 | 0 | 0 | 0 | 0 | 0 | 0 | 0 | 0 | 0 | 0 | 0 | 0 | 0 | 0 | 0 | NA | NA | NA | NA | NA | NA | NA | NA | NA | NA |
| Rprl3 | 2 | 2 | 0 | 2 | 2 | 0 | 2 | 2 | 0 | 2 | 2 | 0 | 1.31 | 1 | 1 | 0.916 | -0.126 | 0.94438853 | NA | 0 | 2.5263 | 1.606 | 1.606 | TRUE | 0.4904 |
| Rprm | 66 | 100 | 63 | 119 | 89 | 82 | 70 | 100 | 63 | 94 | 96 | 90 | 85.38 | 78 | 93 | 1.196 | 0.258 | 0.29342304 | 0.51437903 | 0.0166 | 0.0455 | 0.0317 | 0.0317 | TRUE | 0.5788 |
| Rprml | 47 | 75 | 61 | 84 | 49 | 55 | 50 | 75 | 61 | 66 | 53 | 60 | 60.84 | 62 | 60 | 0.964 | -0.053 | 0.84440732 | 0.92161735 | 0.0093 | 0.0611 | 0.0366 | 0.0366 | TRUE | 0.2295 |
| Rps10 | 21343 | 30240 | 31338 | 32133 | 22854 | 24612 | 22670 | 30201 | 31467 | 25260 | 24589 | 26942 | 26854.89 | 28113 | 25597 | 0.911 | -0.135 | 0.29591636 | 0.51662528 | 0.0154 | 0.007 | 0.012 | 0.012 | TRUE | 0.351 |
| Rps11 | 34118 | 40421 | 41509 | 49575 | 33440 | 36330 | 36239 | 40369 | 41679 | 38972 | 35979 | 39769 | 38834.64 | 39429 | 38240 | 0.97 | -0.044 | 0.60056775 | 0.77260641 | 0.0035 | 0.007 | 0.0051 | 0.0051 | TRUE | 0.0613 |
| Rps12 | 36259 | 43695 | 47040 | 51781 | 36455 | 39102 | 38513 | 43639 | 47233 | 40706 | 39223 | 42803 | 42019.66 | 43128 | 40911 | 0.949 | -0.076 | 0.42329586 | 0.6335416 | 0.0059 | 0.007 | 0.0065 | 0.0065 | TRUE | 0.1073 |
| Rps13 | 15091 | 18387 | 19094 | 21784 | 15189 | 16157 | 16029 | 18363 | 19172 | 17125 | 16342 | 17686 | 17453.11 | 17855 | 17051 | 0.955 | -0.066 | 0.4615047 | 0.66558634 | 0.0047 | 0.0071 | 0.0058 | 0.0058 | TRUE | 0.0978 |
| Rps14 | 27933 | 44479 | 42424 | 44944 | 32552 | 34140 | 29670 | 44422 | 42598 | 35331 | 35024 | 37372 | 37402.76 | 38897 | 35909 | 0.923 | -0.115 | 0.43327212 | 0.64227707 | 0.0208 | 0.007 | 0.0156 | 0.0156 | TRUE | 0.5271 |
| Rps15 | 28890 | 40656 | 42714 | 42474 | 30705 | 34083 | 30686 | 40604 | 42889 | 33390 | 33037 | 37309 | 36319.13 | 38060 | 34579 | 0.909 | -0.138 | 0.29733358 | 0.51819934 | 0.0168 | 0.007 | 0.0127 | 0.0127 | TRUE | 0.3516 |
| Rps15a | 21964 | 26850 | 27929 | 29582 | 20234 | 22508 | 23330 | 26815 | 28044 | 23255 | 21771 | 24639 | 24642.13 | 26063 | 23222 | 0.891 | -0.167 | 0.08394878 | 0.2321814 | 0.0062 | 0.007 | 0.0067 | 0.0067 | TRUE | 0.103 |
| Rps15a-ps4 | 337 | 477 | 541 | 512 | 376 | 437 | 358 | 476 | 543 | 402 | 405 | 478 | 443.83 | 459 | 428 | 0.932 | -0.102 | 0.55848927 | 0.74083202 | 0.0244 | 0.0143 | 0.0194 | 0.0194 | TRUE | 0.3915 |
| Rps15a-ps6 | 835 | 1139 | 1150 | 1130 | 914 | 930 | 887 | 1138 | 1155 | 888 | 983 | 1018 | 1011.49 | 1060 | 963 | 0.908 | -0.139 | 0.28523416 | 0.5065548 | 0.0121 | 0.0102 | 0.0112 | 0.0112 | TRUE | 0.2423 |
| Rps16 | 25052 | 32403 | 32440 | 36544 | 25856 | 27637 | 26610 | 32361 | 32573 | 28728 | 27819 | 30253 | 29724.09 | 30515 | 28933 | 0.948 | -0.077 | 0.44286743 | 0.64974531 | 0.0067 | 0.007 | 0.0072 | 0.0072 | TRUE | 0.1534 |
| Rps17 | 23367 | 29686 | 31123 | 32077 | 22959 | 24303 | 24820 | 29648 | 31251 | 25216 | 24702 | 26603 | 27040.12 | 28573 | 25507 | 0.893 | -0.164 | 0.10924867 | 0.27719918 | 0.0076 | 0.007 | 0.0075 | 0.0075 | TRUE | 0.1615 |
| Rps18 | 37157 | 53300 | 53202 | 54092 | 38629 | 42289 | 39467 | 53231 | 53420 | 42523 | 41562 | 46292 | 46082.7 | 48706 | 43459 | 0.892 | -0.164 | 0.20015988 | 0.40776564 | 0.0151 | 0.007 | 0.0119 | 0.0119 | TRUE | 0.3371 |
| Rps19 | 22007 | 29672 | 29454 | 30527 | 22081 | 24138 | 23375 | 29634 | 29575 | 23998 | 23758 | 26423 | 26127.09 | 27528 | 24726 | 0.898 | -0.155 | 0.1692544 | 0.36769703 | 0.0103 | 0.007 | 0.0091 | 0.0091 | TRUE | 0.2131 |
| Rps19-ps3 | 9 | 9 | 8 | 17 | 16 | 7 | 10 | 9 | 8 | 13 | 17 | 8 | 10.8 | 9 | 13 | 1.443 | 0.529 | 0.37641595 | 0.59184749 | 0 | 0.3118 | 0.1618 | 0.1618 | TRUE | 0.2213 |
| Rps19bp1 | 1338 | 1292 | 1292 | 1479 | 1131 | 1042 | 1421 | 1290 | 1297 | 1163 | 1217 | 1141 | 1254.84 | 1336 | 1174 | 0.878 | -0.187 | 0.02781003 | 0.10951806 | 0.0012 | 0.0095 | 0.0044 | 0.0044 | TRUE | 0.0369 |
| Rps2 | 58902 | 76031 | 76997 | 83437 | 62851 | 64018 | 62564 | 75933 | 77313 | 65592 | 67624 | 70078 | 69850.59 | 71937 | 67765 | 0.942 | -0.086 | 0.38638918 | 0.60046608 | 0.0069 | 0.007 | 0.0071 | 0.0071 | TRUE | 0.1591 |
| Rps20 | 39065 | 48487 | 49631 | 57615 | 41905 | 42921 | 41494 | 48424 | 49835 | 45292 | 45087 | 46984 | 46186.08 | 46584 | 45788 | 0.983 | -0.025 | 0.78076484 | 0.88649531 | 0.004 | 0.007 | 0.0057 | 0.0057 | TRUE | 0.1119 |
| Rps21 | 15256 | 19816 | 19913 | 21094 | 15031 | 16364 | 16205 | 19790 | 19995 | 16582 | 16172 | 17913 | 17776.27 | 18663 | 16889 | 0.905 | -0.144 | 0.16583936 | 0.36290577 | 0.0079 | 0.0071 | 0.0077 | 0.0077 | TRUE | 0.1624 |
| Rps23 | 23184 | 30788 | 31346 | 34684 | 24442 | 26282 | 24626 | 30748 | 31475 | 27266 | 26298 | 28770 | 28197.03 | 28950 | 27445 | 0.948 | -0.077 | 0.48429448 | 0.68186865 | 0.0087 | 0.007 | 0.0087 | 0.0087 | TRUE | 0.2089 |
| Rps24 | 31326 | 39881 | 40863 | 44792 | 32158 | 33742 | 33274 | 39830 | 41031 | 35212 | 34600 | 36936 | 36813.66 | 38045 | 35583 | 0.935 | -0.097 | 0.32451141 | 0.54588245 | 0.0066 | 0.007 | 0.0069 | 0.0069 | TRUE | 0.1473 |
| Rps25 | 22051 | 25957 | 26138 | 29699 | 20538 | 22446 | 23422 | 25923 | 26245 | 23347 | 22098 | 24571 | 24267.7 | 25197 | 23339 | 0.926 | -0.111 | 0.16931515 | 0.36777835 | 0.0032 | 0.007 | 0.0046 | 0.0046 | TRUE | 0.0465 |
| Rps26 | 27068 | 32037 | 33465 | 35831 | 26059 | 28088 | 28751 | 31996 | 33602 | 28168 | 28038 | 30747 | 30216.86 | 31450 | 28984 | 0.922 | -0.118 | 0.17570595 | 0.37581152 | 0.0044 | 0.007 | 0.0054 | 0.0054 | TRUE | 0.069 |
| Rps27 | 51 | 93 | 111 | 78 | 51 | 52 | 54 | 93 | 111 | 61 | 55 | 57 | 71.94 | 86 | 58 | 0.67 | -0.578 | 0.0599208 | 0.18563561 | 0.0548 | 0.0527 | 0.0536 | 0.0536 | TRUE | 0.3808 |
| Rps27a | 39028 | 50363 | 51330 | 55091 | 38708 | 41204 | 41455 | 50298 | 51541 | 43308 | 41647 | 45104 | 45558.88 | 47765 | 43353 | 0.908 | -0.14 | 0.16855063 | 0.36640749 | 0.0074 | 0.007 | 0.0074 | 0.0074 | TRUE | 0.1635 |
| Rps27l | 7865 | 9957 | 9551 | 11198 | 7717 | 8110 | 8354 | 9944 | 9590 | 8803 | 8303 | 8878 | 8978.68 | 9296 | 8661 | 0.932 | -0.102 | 0.25497036 | 0.47316067 | 0.0045 | 0.0073 | 0.0057 | 0.0057 | TRUE | 0.0959 |
| Rps28 | 16034 | 18728 | 19262 | 21039 | 14415 | 15419 | 17031 | 18704 | 19341 | 16539 | 15510 | 16879 | 17333.87 | 18359 | 16309 | 0.888 | -0.171 | 0.03141833 | 0.11909665 | 0.003 | 0.0071 | 0.0045 | 0.0045 | TRUE | 0.0489 |
| Rps29 | 13212 | 15589 | 16562 | 18246 | 12228 | 13441 | 14033 | 15569 | 16630 | 14344 | 13157 | 14713 | 14740.97 | 15411 | 14071 | 0.913 | -0.131 | 0.15219213 | 0.34327323 | 0.0052 | 0.0071 | 0.006 | 0.006 | TRUE | 0.0747 |
| Rps3 | 58491 | 81151 | 79480 | 80523 | 58918 | 62037 | 62128 | 81046 | 79806 | 63301 | 63392 | 67909 | 69597.1 | 74327 | 64867 | 0.873 | -0.196 | 0.08790987 | 0.24029714 | 0.0109 | 0.007 | 0.0095 | 0.0095 | TRUE | 0.2524 |
| Rps3a1 | 12421 | 14965 | 15324 | 16679 | 11755 | 12744 | 13193 | 14946 | 15387 | 13112 | 12648 | 13950 | 13872.6 | 14509 | 13237 | 0.912 | -0.132 | 0.13063933 | 0.31124736 | 0.0043 | 0.0071 | 0.0055 | 0.0055 | TRUE | 0.0769 |
| Rps4x | 49542 | 62304 | 62944 | 66720 | 46568 | 51287 | 52622 | 62224 | 63202 | 52450 | 50104 | 56142 | 56124.06 | 59349 | 52899 | 0.891 | -0.166 | 0.08753502 | 0.23956314 | 0.0065 | 0.007 | 0.0068 | 0.0068 | TRUE | 0.1204 |
| Rps4y2 | 8209 | 8272 | 8130 | 12213 | 8895 | 8250 | 8719 | 8261 | 8163 | 9601 | 9570 | 9031 | 8891.07 | 8381 | 9401 | 1.122 | 0.166 | 0.01198403 | 0.06007058 | 0.001 | 0.0073 | 0.003 | 0.003 | TRUE | 0.0152 |
| Rps5 | 41582 | 50180 | 51506 | 57279 | 41281 | 42965 | 44167 | 50115 | 51717 | 45028 | 44416 | 47032 | 47079.36 | 48666 | 45492 | 0.935 | -0.097 | 0.24336861 | 0.46008258 | 0.0038 | 0.007 | 0.005 | 0.005 | TRUE | 0.0801 |
| Rps6 | 22255 | 28112 | 29299 | 32127 | 22614 | 24153 | 23639 | 28076 | 29419 | 25256 | 24331 | 26439 | 26193.33 | 27045 | 25342 | 0.937 | -0.094 | 0.34921533 | 0.56805117 | 0.0071 | 0.007 | 0.0072 | 0.0072 | TRUE | 0.1485 |
| Rps6ka1 | 7494 | 8623 | 8730 | 8761 | 6873 | 7459 | 7960 | 8612 | 8766 | 6887 | 7395 | 8165 | 7964.14 | 8446 | 7482 | 0.886 | -0.175 | 0.0534578 | 0.17175218 | 0.0048 | 0.0073 | 0.0058 | 0.0058 | TRUE | 0.0777 |
| Rps6ka2 | 1539 | 1876 | 1990 | 1805 | 1240 | 1359 | 1635 | 1874 | 1998 | 1419 | 1334 | 1488 | 1624.53 | 1836 | 1414 | 0.77 | -0.377 | 0.00031309 | 0.00352715 | 0.0056 | 0.0089 | 0.0073 | 0.0073 | TRUE | 0.1102 |
| Rps6ka3 | 794 | 942 | 975 | 1366 | 940 | 919 | 843 | 941 | 979 | 1074 | 1011 | 1006 | 975.73 | 921 | 1030 | 1.119 | 0.162 | 0.09264532 | 0.24858829 | 0.0025 | 0.0103 | 0.0057 | 0.0057 | TRUE | 0.0644 |
| Rps6ka4 | 469 | 672 | 759 | 732 | 567 | 596 | 498 | 671 | 762 | 575 | 610 | 652 | 628.22 | 644 | 612 | 0.951 | -0.073 | 0.65671198 | 0.80670515 | 0.0232 | 0.0121 | 0.0178 | 0.0178 | TRUE | 0.4461 |
| Rps6ka5 | 1164 | 1129 | 1170 | 1039 | 705 | 859 | 1236 | 1128 | 1175 | 817 | 759 | 940 | 1009.06 | 1180 | 839 | 0.711 | -0.493 | 6.92E-06 | 0.00014547 | 0.0057 | 0.0102 | 0.0076 | 0.0076 | TRUE | 0.1308 |
| Rps6ka6 | 4111 | 4846 | 5288 | 6884 | 4751 | 5223 | 4367 | 4840 | 5310 | 5412 | 5112 | 5717 | 5126.15 | 4839 | 5414 | 1.119 | 0.162 | 0.09760369 | 0.25673873 | 0.006 | 0.0075 | 0.0067 | 0.0067 | TRUE | 0.0886 |
| Rps6kb1 | 4838 | 4308 | 4308 | 6049 | 4430 | 4125 | 5139 | 4302 | 4326 | 4755 | 4766 | 4515 | 4634.01 | 4589 | 4679 | 1.02 | 0.028 | 0.76637139 | 0.87779299 | 0.0054 | 0.0076 | 0.0063 | 0.0063 | TRUE | 0.1337 |
| Rps6kb2 | 321 | 306 | 295 | 456 | 385 | 292 | 341 | 306 | 296 | 358 | 414 | 320 | 339.19 | 314 | 364 | 1.159 | 0.213 | 0.14158758 | 0.3270532 | 0.0081 | 0.0166 | 0.0121 | 0.0121 | TRUE | 0.1602 |
| Rps6kc1 | 1637 | 1555 | 1640 | 2205 | 1619 | 1442 | 1739 | 1553 | 1647 | 1733 | 1742 | 1578 | 1665.39 | 1646 | 1684 | 1.024 | 0.034 | 0.69850619 | 0.83505557 | 0.0025 | 0.0089 | 0.0049 | 0.0049 | TRUE | 0.0361 |
| Rps6kl1 | 346 | 275 | 260 | 456 | 294 | 285 | 368 | 275 | 261 | 358 | 316 | 312 | 315 | 301 | 329 | 1.095 | 0.131 | 0.43759333 | 0.64550463 | 0.0171 | 0.0174 | 0.0172 | 0.0172 | TRUE | 0.4173 |
| Rps7 | 18884 | 23180 | 23940 | 27468 | 19576 | 20500 | 20058 | 23150 | 24038 | 21593 | 21063 | 22440 | 22057.12 | 22415 | 21699 | 0.968 | -0.047 | 0.60083667 | 0.77266232 | 0.0043 | 0.0071 | 0.0057 | 0.0057 | TRUE | 0.1035 |
| Rps8 | 41246 | 53349 | 53987 | 59255 | 44431 | 45771 | 43811 | 53280 | 54209 | 46582 | 47805 | 50104 | 49298.26 | 50433 | 48164 | 0.955 | -0.066 | 0.50885204 | 0.70257778 | 0.0066 | 0.007 | 0.0073 | 0.0073 | TRUE | 0.1616 |
| Rps9 | 37283 | 46272 | 46313 | 52396 | 37759 | 39259 | 39601 | 46212 | 46503 | 41190 | 40626 | 42975 | 42851.28 | 44105 | 41597 | 0.943 | -0.084 | 0.33033712 | 0.55109763 | 0.0043 | 0.007 | 0.0054 | 0.0054 | TRUE | 0.0977 |
| Rpsa | 67715 | 88954 | 91526 | 98681 | 71024 | 76188 | 71925 | 88839 | 91902 | 77575 | 76417 | 83400 | 81676.45 | 84222 | 79131 | 0.94 | -0.09 | 0.40778151 | 0.62062543 | 0.0089 | 0.0069 | 0.0085 | 0.0085 | TRUE | 0.1998 |

| | | | | | | | | | | | | | | | | | | | | | | | | | |
|---|---|---|---|---|---|---|---|---|---|---|---|---|---|---|---|---|---|---|---|---|---|---|---|---|---|
| Rptn | 0 | 0 | 0 | 0 | 0 | 0 | 0 | 0 | 0 | 0 | 0 | 0 | 0 | 0 | 0 | NA | NA | NA | NA | NA | NA | NA | NA | NA | NA |
| Rptor | 5937 | 5809 | 6157 | 8691 | 6416 | 5884 | 6306 | 5801 | 6182 | 6832 | 6903 | 6441 | 6411.05 | 6096 | 6725 | 1.103 | 0.142 | 0.04439949 | 0.15134402 | 0.0014 | 0.0074 | 0.0034 | 0.0034 | TRUE | 0.0219 |
| Rpusd1 | 414 | 379 | 380 | 585 | 464 | 397 | 440 | 379 | 382 | 460 | 499 | 435 | 432.25 | 400 | 465 | 1.162 | 0.217 | 0.07776468 | 0.22059811 | 0.0035 | 0.0145 | 0.0085 | 0.0085 | TRUE | 0.0864 |
| Rpusd2 | 1227 | 1190 | 1229 | 1574 | 1257 | 1098 | 1303 | 1188 | 1234 | 1237 | 1352 | 1202 | 1252.92 | 1242 | 1264 | 1.018 | 0.025 | 0.78002154 | 0.88617899 | 0.0021 | 0.0095 | 0.0051 | 0.0051 | TRUE | 0.0443 |
| Rpusd3 | 818 | 810 | 830 | 1378 | 964 | 825 | 869 | 809 | 833 | 1083 | 1037 | 903 | 922.47 | 837 | 1008 | 1.205 | 0.27 | 0.0098814 | 0.05221089 | 0.0041 | 0.0105 | 0.0068 | 0.0068 | TRUE | 0.0976 |
| Rpusd4 | 1533 | 1648 | 1640 | 1792 | 1367 | 1369 | 1628 | 1646 | 1647 | 1409 | 1471 | 1499 | 1549.84 | 1640 | 1460 | 0.889 | -0.17 | 0.01934099 | 0.08468796 | 0 | 0.009 | 0.0031 | 0.0031 | TRUE | 0.0116 |
| Rqcd1 | 5907 | 6193 | 6277 | 8316 | 5958 | 5839 | 6274 | 6185 | 6303 | 6537 | 6410 | 6392 | 6350.26 | 6254 | 6446 | 1.031 | 0.044 | 0.42607457 | 0.63621327 | 0 | 0.0074 | 0.002 | 0.002 | TRUE | 0.0019 |
| Rrad | 27 | 32 | 43 | 65 | 34 | 30 | 29 | 32 | 43 | 51 | 37 | 33 | 37.39 | 35 | 40 | 1.169 | 0.225 | 0.52060981 | 0.71109881 | 0.0277 | 0.095 | 0.0619 | 0.0619 | TRUE | 0.486 |
| Rraga | 2675 | 2340 | 2135 | 4247 | 2918 | 2560 | 2841 | 2337 | 2144 | 3339 | 3140 | 2802 | 2767.11 | 2441 | 3094 | 1.268 | 0.342 | 0.00712746 | 0.04062676 | 0.0142 | 0.0081 | 0.0113 | 0.0113 | TRUE | 0.2496 |
| Rragb | 579 | 697 | 721 | 530 | 419 | 421 | 615 | 696 | 724 | 417 | 451 | 461 | 560.56 | 678 | 443 | 0.652 | -0.618 | 5.83E-08 | 2.27E-06 | 0.0035 | 0.0128 | 0.0075 | 0.0075 | TRUE | 0.0781 |
| Rragc | 1705 | 2016 | 2045 | 2216 | 1658 | 1751 | 1811 | 2013 | 2053 | 1742 | 1784 | 1917 | 1886.75 | 1959 | 1814 | 0.925 | -0.112 | 0.20820534 | 0.41762872 | 0.0027 | 0.0086 | 0.0051 | 0.0051 | TRUE | 0.0528 |
| Rragd | 1252 | 1466 | 1618 | 1361 | 1185 | 1227 | 1330 | 1464 | 1625 | 1070 | 1275 | 1343 | 1351.11 | 1473 | 1229 | 0.834 | -0.262 | 0.03490291 | 0.12790164 | 0.0106 | 0.0093 | 0.0104 | 0.0104 | TRUE | 0.1577 |
| Rras | 995 | 1501 | 1474 | 1074 | 849 | 1119 | 1057 | 1499 | 1480 | 844 | 913 | 1225 | 1169.78 | 1345 | 994 | 0.738 | -0.437 | 0.0163359 | 0.07546861 | 0.0367 | 0.0097 | 0.023 | 0.023 | TRUE | 0.4902 |
| Rras2 | 3945 | 3887 | 3775 | 4933 | 3663 | 3485 | 4190 | 3882 | 3790 | 3878 | 3941 | 3815 | 3916.12 | 3954 | 3878 | 0.981 | -0.028 | 0.69579652 | 0.83345494 | 0.0011 | 0.0077 | 0.0034 | 0.0034 | TRUE | 0.0331 |
| Rrbp1 | 10356 | 7911 | 8090 | 11649 | 8398 | 7503 | 11000 | 7901 | 8123 | 9158 | 9036 | 8213 | 8905.06 | 9008 | 8802 | 0.977 | -0.033 | 0.80797943 | 0.90112858 | 0.0165 | 0.0073 | 0.0134 | 0.0134 | TRUE | 0.457 |
| Rreb1 | 2334 | 2088 | 2411 | 3191 | 2310 | 2011 | 2479 | 2085 | 2421 | 2509 | 2485 | 2201 | 2363.44 | 2328 | 2398 | 1.03 | 0.043 | 0.6748379 | 0.81886009 | 0.0065 | 0.0083 | 0.0072 | 0.0072 | TRUE | 0.1013 |
| Rrh | 0 | 1 | 0 | 0 | 1 | 0 | 0 | 1 | 0 | 0 | 1 | 0 | 0.35 | 0 | 0 | 0.954 | -0.067 | 0.98571955 | NA | 0 | 9.5346 | 8.2194 | 8.2194 | TRUE | 0.5204 |
| Rrm1 | 9657 | 10079 | 10645 | 14846 | 10690 | 10971 | 10257 | 10066 | 10689 | 11671 | 11502 | 12009 | 11032.36 | 10337 | 11727 | 1.134 | 0.182 | 0.00260139 | 0.01893359 | 6.00E-04 | 0.0072 | 0.0025 | 0.0025 | TRUE | 0.0108 |
| Rrm2 | 11467 | 13482 | 13757 | 18955 | 13220 | 14296 | 12180 | 13465 | 13813 | 14901 | 14224 | 15649 | 14038.69 | 13153 | 14925 | 1.135 | 0.182 | 0.02413481 | 0.09915453 | 0.0032 | 0.0071 | 0.0046 | 0.0046 | TRUE | 0.0511 |
| Rrm2b | 701 | 1000 | 1044 | 1194 | 854 | 984 | 745 | 999 | 1048 | 939 | 919 | 1077 | 954.37 | 931 | 978 | 1.051 | 0.071 | 0.63104545 | 0.79265149 | 0.019 | 0.0104 | 0.0148 | 0.0148 | TRUE | 0.3628 |
| Rrn3 | 15883 | 15765 | 15779 | 16784 | 12129 | 12223 | 16871 | 15745 | 15844 | 13194 | 13050 | 13380 | 14680.55 | 16153 | 13208 | 0.818 | -0.29 | 1.94E-06 | 4.90E-05 | 7.00E-04 | 0.0071 | 0.0026 | 0.0026 | TRUE | 0.0185 |
| Rrnad1 | 353 | 362 | 385 | 447 | 397 | 366 | 375 | 362 | 387 | 351 | 427 | 401 | 383.71 | 375 | 393 | 1.047 | 0.067 | 0.59446522 | 0.7677918 | 0.0034 | 0.0155 | 0.0088 | 0.0088 | TRUE | 0.1055 |
| Rrp1 | 7964 | 8014 | 8121 | 9306 | 6809 | 6236 | 8459 | 8004 | 8154 | 7316 | 7326 | 6826 | 7680.86 | 8206 | 7156 | 0.872 | -0.197 | 0.0029217 | 0.0207947 | 0.001 | 0.0073 | 0.003 | 0.003 | TRUE | 0.0197 |
| Rrp12 | 7471 | 5920 | 6278 | 8832 | 6340 | 5304 | 7936 | 5912 | 6304 | 6943 | 6821 | 5806 | 6620.37 | 6717 | 6523 | 0.971 | -0.042 | 0.74999081 | 0.86759976 | 0.0142 | 0.0074 | 0.0124 | 0.0124 | TRUE | 0.3071 |
| Rrp15 | 3658 | 3286 | 3121 | 4624 | 3477 | 3072 | 3885 | 3282 | 3134 | 3635 | 3741 | 3363 | 3506.64 | 3434 | 3580 | 1.043 | 0.06 | 0.5682479 | 0.74742773 | 0.0077 | 0.0078 | 0.0078 | 0.0078 | TRUE | 0.1608 |
| Rrp1b | 5914 | 6089 | 6067 | 8629 | 6418 | 5484 | 6282 | 6081 | 6092 | 6783 | 6905 | 6003 | 6357.78 | 6152 | 6564 | 1.067 | 0.094 | 0.24291075 | 0.4595404 | 0.0027 | 0.0074 | 0.0045 | 0.0045 | TRUE | 0.0679 |
| Rrp36 | 1678 | 1675 | 1560 | 1919 | 1408 | 1315 | 1782 | 1673 | 1566 | 1509 | 1515 | 1439 | 1580.76 | 1674 | 1488 | 0.889 | -0.17 | 0.04575432 | 0.15478347 | 0.0019 | 0.009 | 0.0046 | 0.0046 | TRUE | 0.0386 |
| Rrp7a | 3696 | 3241 | 3257 | 5503 | 3981 | 3188 | 3926 | 3237 | 3270 | 4326 | 4283 | 3490 | 3755.35 | 3478 | 4033 | 1.16 | 0.214 | 0.08080342 | 0.22612512 | 0.0127 | 0.0078 | 0.0106 | 0.0106 | TRUE | 0.1685 |
| Rrp8 | 940 | 979 | 965 | 1895 | 1292 | 1120 | 998 | 978 | 969 | 1490 | 1390 | 1226 | 1175.16 | 982 | 1369 | 1.395 | 0.481 | 2.10E-06 | 5.26E-05 | 0.0044 | 0.0097 | 0.0065 | 0.0065 | TRUE | 0.0992 |
| Rrp9 | 5038 | 5458 | 5468 | 6758 | 5052 | 4835 | 5351 | 5451 | 5490 | 5313 | 5436 | 5293 | 5388.93 | 5431 | 5347 | 0.984 | -0.023 | 0.69194079 | 0.83104554 | 0 | 0.0075 | 0.0022 | 0.0022 | TRUE | 0.0025 |
| Rrs1 | 2586 | 2656 | 2660 | 3978 | 2948 | 2501 | 2747 | 2653 | 2671 | 3127 | 3172 | 2738 | 2851.18 | 2690 | 3012 | 1.12 | 0.164 | 0.05604082 | 0.17757093 | 0.0028 | 0.0081 | 0.0049 | 0.0049 | TRUE | 0.0765 |
| Rs1 | 0 | 0 | 0 | 0 | 0 | 0 | 0 | 0 | 0 | 0 | 0 | 0 | 0 | 0 | 0 | NA | NA | NA | NA | NA | NA | NA | NA | NA | NA |
| Rsad1 | 336 | 395 | 362 | 628 | 415 | 378 | 357 | 394 | 363 | 494 | 447 | 414 | 411.47 | 371 | 452 | 1.217 | 0.283 | 0.02181511 | 0.09266577 | 0.0034 | 0.0149 | 0.0085 | 0.0085 | TRUE | 0.0821 |
| Rsad2 | 28 | 51 | 29 | 26 | 29 | 25 | 30 | 51 | 29 | 20 | 31 | 27 | 31.47 | 37 | 26 | 0.714 | -0.487 | 0.22006136 | 0.43222937 | 0.0499 | 0.1116 | 0.0809 | 0.0809 | TRUE | 0.8587 |
| Rsbn1 | 2209 | 2540 | 2708 | 2692 | 2179 | 2461 | 2346 | 2537 | 2719 | 2116 | 2344 | 2694 | 2459.47 | 2534 | 2385 | 0.941 | -0.088 | 0.44052846 | 0.64757144 | 0.0086 | 0.0082 | 0.0091 | 0.0091 | TRUE | 0.1556 |
| Rsbn1l | 2868 | 2956 | 3102 | 2870 | 2296 | 2678 | 3046 | 2952 | 3115 | 2256 | 2470 | 2931 | 2795.21 | 3038 | 2552 | 0.84 | -0.252 | 0.0235375 | 0.09759028 | 0.009 | 0.0081 | 0.0085 | 0.0085 | TRUE | 0.204 |
| Rsc1a1 | 0 | 0 | 3 | 0 | 0 | 2 | 0 | 0 | 3 | 0 | 0 | 2 | 0.87 | 1 | 1 | 0.707 | -0.5 | 0.86485107 | NA | 10 | 3.807 | 5.0241 | 5.0241 | TRUE | 1.4421 |
| Rsf1 | 3357 | 3040 | 3398 | 4568 | 3287 | 3225 | 3566 | 3036 | 3412 | 3591 | 3537 | 3530 | 3445.27 | 3338 | 3553 | 1.064 | 0.09 | 0.28798487 | 0.50903254 | 0.0029 | 0.0079 | 0.0049 | 0.0049 | TRUE | 0.0762 |
| Rsg1 | 48 | 35 | 44 | 48 | 40 | 28 | 51 | 35 | 44 | 38 | 43 | 31 | 40.26 | 43 | 37 | 0.858 | -0.221 | 0.49214785 | 0.6883982 | 0 | 0.0887 | 0.0495 | 0.0495 | TRUE | 0.2251 |
| Rsl1 | 157 | 205 | 186 | 236 | 200 | 175 | 167 | 205 | 187 | 186 | 215 | 192 | 191.76 | 186 | 198 | 1.059 | 0.082 | 0.60872705 | 0.77840466 | 0.0032 | 0.0241 | 0.0133 | 0.0133 | TRUE | 0.0874 |
| Rsl1d1 | 12906 | 14036 | 14358 | 16927 | 12170 | 11983 | 13708 | 14018 | 14417 | 13307 | 13094 | 13117 | 13610.24 | 14048 | 13173 | 0.938 | -0.093 | 0.09504296 | 0.25270021 | 2.00E-04 | 0.0071 | 0.0022 | 0.0022 | TRUE | 0.0065 |
| Rsl24d1 | 3877 | 4152 | 3912 | 5361 | 3964 | 3669 | 4118 | 4147 | 3928 | 4214 | 4265 | 4016 | 4114.75 | 4064 | 4165 | 1.025 | 0.036 | 0.59182209 | 0.7654763 | 6.00E-04 | 0.0077 | 0.0029 | 0.0029 | TRUE | 0.0118 |
| Rslcan18 | 126 | 116 | 134 | 127 | 94 | 95 | 134 | 116 | 135 | 100 | 101 | 104 | 114.87 | 128 | 102 | 0.793 | -0.334 | 0.07141419 | 0.20823529 | 0 | 0.0356 | 0.0159 | 0.0159 | TRUE | 0.0721 |
| Rsph1 | 138 | 160 | 178 | 170 | 138 | 179 | 147 | 160 | 179 | 134 | 148 | 196 | 160.53 | 162 | 159 | 0.981 | -0.027 | 0.89361782 | 0.94852549 | 0.019 | 0.0274 | 0.0231 | 0.0231 | TRUE | 0.4182 |
| Rsph3a | 107 | 132 | 153 | 118 | 92 | 114 | 114 | 132 | 154 | 93 | 99 | 125 | 119.28 | 133 | 106 | 0.79 | -0.341 | 0.11386277 | 0.28460279 | 0.0153 | 0.0345 | 0.0249 | 0.0249 | TRUE | 0.2445 |
| Rsph3b | 151 | 223 | 230 | 197 | 166 | 178 | 160 | 223 | 231 | 155 | 179 | 195 | 190.39 | 205 | 176 | 0.858 | -0.222 | 0.25777759 | 0.47651994 | 0.0206 | 0.0242 | 0.0224 | 0.0224 | TRUE | 0.3852 |
| Rsph4a | 16 | 3 | 2 | 18 | 5 | 4 | 17 | 3 | 2 | 14 | 5 | 4 | 7.65 | 7 | 8 | 1.108 | 0.148 | 0.87870195 | 0.94003168 | 0.6593 | 0.4375 | 0.5515 | 0.5515 | TRUE | 3.5159 |
| Rsph6a | 10 | 4 | 7 | 11 | 6 | 2 | 11 | 4 | 7 | 9 | 6 | 2 | 6.49 | 7 | 6 | 0.816 | -0.294 | 0.72080684 | 0.84878046 | 0.1241 | 0.5146 | 0.3318 | 0.3318 | TRUE | 0.2855 |
| Rsph9 | 21 | 31 | 32 | 21 | 27 | 41 | 22 | 31 | 32 | 17 | 29 | 45 | 29.31 | 28 | 30 | 1.046 | 0.065 | 0.88501158 | 0.94366363 | 0.1066 | 0.1193 | 0.113 | 0.113 | TRUE | 0.2418 |
| Rspo1 | 200 | 168 | 159 | 172 | 131 | 118 | 212 | 168 | 160 | 135 | 141 | 129 | 157.53 | 180 | 135 | 0.752 | -0.412 | 0.0235678 | 0.09763248 | 0.0077 | 0.0278 | 0.0174 | 0.0174 | TRUE | 0.2639 |
| Rspo2 | 3 | 1 | 1 | 1 | 0 | 2 | 3 | 1 | 1 | 1 | 0 | 2 | 1.36 | 2 | 1 | 0.569 | -0.813 | 0.63050956 | NA | 0 | 2.4279 | 1.2637 | 1.2637 | TRUE | 0.4463 |
| Rspo3 | 14 | 2 | 8 | 12 | 15 | 5 | 15 | 2 | 8 | 9 | 16 | 5 | 9.32 | 8 | 10 | 1.25 | 0.322 | 0.69067033 | 0.83034167 | 0.3673 | 0.3602 | 0.3637 | 0.3637 | TRUE | 0.2679 |
| Rspo4 | 75 | 118 | 128 | 100 | 62 | 85 | 80 | 118 | 129 | 79 | 67 | 93 | 94.07 | 109 | 80 | 0.73 | -0.454 | 0.07997408 | 0.22447967 | 0.0331 | 0.0419 | 0.0376 | 0.0376 | TRUE | 0.5317 |
| Rspry1 | 1986 | 2136 | 2175 | 2958 | 2125 | 2108 | 2109 | 2133 | 2184 | 2325 | 2286 | 2308 | 2224.32 | 2142 | 2306 | 1.077 | 0.107 | 0.09990486 | 0.26096381 | 0 | 0.0084 | 0.0026 | 0.0026 | TRUE | 0.0035 |
| Rsrc1 | 1010 | 1512 | 1485 | 2029 | 1536 | 1565 | 1073 | 1510 | 1491 | 1595 | 1653 | 1713 | 1505.79 | 1358 | 1654 | 1.217 | 0.284 | 0.04372727 | 0.14971024 | 0.0162 | 0.0091 | 0.0136 | 0.0136 | TRUE | 0.4045 |
| Rsrc2 | 5988 | 6210 | 6231 | 7496 | 5784 | 6233 | 6360 | 6202 | 6257 | 5893 | 6223 | 6823 | 6292.98 | 6273 | 6313 | 1.006 | 0.009 | 0.91118821 | 0.95603547 | 0.0027 | 0.0074 | 0.0044 | 0.0044 | TRUE | 0.0609 |
| Rsu1 | 2061 | 1688 | 1799 | 2958 | 2036 | 1714 | 2189 | 1686 | 1806 | 2325 | 2191 | 1876 | 2012.26 | 1894 | 2131 | 1.126 | 0.171 | 0.19287049 | 0.39842218 | 0.0147 | 0.0085 | 0.0119 | 0.0119 | TRUE | 0.2246 |
| Rtbdn | 21 | 21 | 11 | 22 | 12 | 22 | 22 | 21 | 11 | 17 | 13 | 24 | 18.1 | 18 | 18 | 0.999 | -0.002 | 0.99690014 | 0.99828918 | 0.0507 | 0.1889 | 0.1247 | 0.1247 | TRUE | 0.479 |
| Rtca | 1919 | 2054 | 2031 | 2767 | 2068 | 1945 | 2038 | 2051 | 2039 | 2175 | 2225 | 2129 | 2109.73 | 2043 | 2176 | 1.065 | 0.091 | 0.16642143 | 0.36370349 | 0 | 0.0085 | 0.0027 | 0.0027 | TRUE | 0.0045 |
| Rtdr1 | 2 | 1 | 0 | 2 | 5 | 2 | 2 | 1 | 0 | 2 | 5 | 2 | 2.04 | 1 | 3 | 2.92 | 1.546 | 0.29602447 | NA | 0 | 1.6186 | 0.9193 | 0.9193 | TRUE | 0.5567 |
| Rtel1 | 3787 | 2650 | 2468 | 5082 | 3647 | 2464 | 4022 | 2647 | 2478 | 3995 | 3924 | 2697 | 3293.9 | 3049 | 3539 | 1.161 | 0.215 | 0.29766019 | 0.5184073 | 0.0524 | 0.0079 | 0.0305 | 0.0305 | TRUE | 0.947 |
| Rtf1 | 4642 | 4227 | 4398 | 5485 | 4066 | 4097 | 4931 | 4222 | 4416 | 4312 | 4375 | 4485 | 4456.61 | 4523 | 4391 | 0.971 | -0.043 | 0.60905493 | 0.77863454 | 0.0029 | 0.0076 | 0.0048 | 0.0048 | TRUE | 0.0757 |
| Rtfdc1 | 2350 | 2257 | 2207 | 3512 | 2626 | 2328 | 2496 | 2254 | 2216 | 2761 | 2825 | 2548 | 2516.82 | 2322 | 2711 | 1.168 | 0.224 | 0.00996086 | 0.05252512 | 0.003 | 0.0082 | 0.005 | 0.005 | TRUE | 0.0519 |
| Rtkn | 318 | 421 | 429 | 471 | 344 | 360 | 338 | 420 | 431 | 370 | 370 | 394 | 387.24 | 396 | 378 | 0.953 | -0.069 | 0.61011134 | 0.77891178 | 0.0066 | 0.0154 | 0.0107 | 0.0107 | TRUE | 0.1901 |
| Rtkn2 | 1504 | 1765 | 1776 | 1896 | 1480 | 1462 | 1598 | 1763 | 1783 | 1490 | 1592 | 1600 | 1637.8 | 1715 | 1561 | 0.91 | -0.136 | 0.10965069 | 0.27781698 | 0.002 | 0.0089 | 0.0046 | 0.0046 | TRUE | 0.0428 |
| Rtl1 | 0 | 0 | 1 | 1 | 0 | 0 | 0 | 0 | 1 | 1 | 0 | 0 | 0.3 | 0 | 0 | 0.954 | -0.067 | 0.98640617 | NA | 0 | 11.0483 | 9.2783 | 9.2783 | TRUE | 0.4098 |
| Rtn1 | 699 | 1319 | 1375 | 855 | 866 | 892 | 742 | 1317 | 1381 | 672 | 932 | 976 | 1003.46 | 1147 | 860 | 0.749 | -0.416 | 0.07384009 | 0.21338322 | 0.0728 | 0.0102 | 0.038 | 0.038 | TRUE | 1.1377 |
| Rtn2 | 740 | 729 | 822 | 930 | 702 | 662 | 786 | 728 | 825 | 731 | 755 | 725 | 758.42 | 780 | 737 | 0.945 | -0.082 | 0.39065659 | 0.60450043 | 9.00E-04 | 0.0112 | 0.0052 | 0.0052 | TRUE | 0.0403 |
| Rtn3 | 6724 | 6713 | 7209 | 9315 | 6308 | 5787 | 7142 | 6704 | 7239 | 7323 | 6787 | 6335 | 6921.59 | 7028 | 6815 | 0.97 | -0.044 | 0.59393829 | 0.76745819 | 0.0029 | 0.0074 | 0.0048 | 0.0048 | TRUE | 0.0522 |

| | | | | | | | | | | | | | | | | | | | | | | | | | |
|---|---|---|---|---|---|---|---|---|---|---|---|---|---|---|---|---|---|---|---|---|---|---|---|---|---|
| Rtn4 | 31077 | 29636 | 29360 | 36394 | 26677 | 26732 | 33009 | 29598 | 29481 | 28610 | 28703 | 29262 | 29777.14 | 30696 | 28858 | 0.94 | -0.089 | 0.21844985 | 0.43051243 | 0.0022 | 0.007 | 0.0037 | 0.0037 | TRUE | 0.0532 |
| Rtn4ip1 | 1323 | 1820 | 1762 | 1682 | 1256 | 1334 | 1405 | 1818 | 1769 | 1322 | 1351 | 1460 | 1521.01 | 1664 | 1378 | 0.828 | -0.273 | 0.02370972 | 0.09797529 | 0.0096 | 0.0091 | 0.0098 | 0.0098 | TRUE | 0.2226 |
| Rtn4r | 64 | 104 | 116 | 73 | 51 | 85 | 68 | 104 | 116 | 57 | 55 | 93 | 82.27 | 96 | 68 | 0.709 | -0.497 | 0.11013029 | 0.27853993 | 0.0682 | 0.0469 | 0.0571 | 0.0571 | TRUE | 0.8501 |
| Rtn4rl1 | 70 | 112 | 122 | 218 | 151 | 135 | 74 | 112 | 123 | 171 | 162 | 148 | 131.72 | 103 | 160 | 1.56 | 0.641 | 0.00410832 | 0.02697604 | 0.0234 | 0.0319 | 0.028 | 0.028 | TRUE | 0.5662 |
| Rtn4rl2 | 57 | 83 | 59 | 43 | 27 | 32 | 61 | 83 | 59 | 34 | 29 | 35 | 50.09 | 68 | 33 | 0.483 | -1.049 | 0.00042055 | 0.00451217 | 0 | 0.0727 | 0.0413 | 0.0413 | TRUE | 0.3545 |
| Rtp1 | 0 | 0 | 1 | 0 | 0 | 0 | 0 | 0 | 1 | 0 | 0 | 0 | 0.17 | 0 | 0 | 0.49 | -1.029 | 0.80087 | NA | 0 | 19.6928 | 10 | 10 | TRUE | 0.4098 |
| Rtp2 | 1 | 0 | 0 | 2 | 0 | 1 | 1 | 0 | 0 | 2 | 0 | 1 | 0.62 | 0 | 1 | 2.474 | 1.307 | 0.61898859 | NA | 0 | 5.3076 | 3.4358 | 3.4358 | TRUE | 0.4614 |
| Rtp3 | 103 | 95 | 109 | 48 | 32 | 51 | 109 | 95 | 109 | 38 | 34 | 56 | 73.62 | 104 | 43 | 0.406 | -1.301 | 8.43E-07 | 2.37E-05 | 0.0082 | 0.0517 | 0.0339 | 0.0339 | TRUE | 0.5178 |
| Rtp4 | 4 | 4 | 2 | 4 | 3 | 6 | 4 | 4 | 2 | 3 | 3 | 7 | 3.87 | 3 | 4 | 1.25 | 0.322 | 0.73750276 | NA | 0 | 0.8592 | 0.4023 | 0.4023 | TRUE | 0.339 |
| Rttn | 1540 | 1529 | 1701 | 2154 | 1530 | 1590 | 1636 | 1527 | 1708 | 1693 | 1646 | 1741 | 1658.46 | 1624 | 1693 | 1.043 | 0.061 | 0.45515111 | 0.66016583 | 0.0013 | 0.0089 | 0.0042 | 0.0042 | TRUE | 0.0328 |
| Rubie | 0 | 1 | 0 | 4 | 3 | 1 | 0 | 1 | 0 | 3 | 3 | 1 | 1.41 | 0 | 2 | 6.85 | 2.776 | 0.13541812 | NA | 0 | 2.3418 | 1.2958 | 1.2958 | TRUE | 0.405 |
| Rufy1 | 939 | 1030 | 993 | 1083 | 834 | 854 | 997 | 1029 | 997 | 851 | 897 | 935 | 951.11 | 1008 | 894 | 0.887 | -0.174 | 0.04254763 | 0.14725943 | 0 | 0.0104 | 0.0042 | 0.0042 | TRUE | 0.0223 |
| Rufy2 | 1518 | 1622 | 1510 | 1906 | 1503 | 1522 | 1612 | 1620 | 1516 | 1498 | 1617 | 1666 | 1588.34 | 1583 | 1594 | 1.006 | 0.009 | 0.91432373 | 0.95780402 | 0.0016 | 0.009 | 0.0044 | 0.0044 | TRUE | 0.0341 |
| Rufy3 | 409 | 598 | 640 | 691 | 541 | 532 | 434 | 597 | 643 | 543 | 582 | 582 | 563.66 | 558 | 569 | 1.019 | 0.027 | 0.86331078 | 0.9324082 | 0.0194 | 0.0127 | 0.0162 | 0.0162 | TRUE | 0.4372 |
| Rufy4 | 28 | 24 | 17 | 51 | 48 | 28 | 30 | 24 | 17 | 40 | 52 | 31 | 32.19 | 24 | 41 | 1.732 | 0.793 | 0.04078677 | 0.14287252 | 0.0345 | 0.1092 | 0.0745 | 0.0745 | TRUE | 0.1538 |
| Rundc1 | 783 | 776 | 732 | 1145 | 839 | 724 | 832 | 775 | 735 | 900 | 903 | 793 | 822.84 | 781 | 865 | 1.109 | 0.15 | 0.14949782 | 0.3401442 | 0.0033 | 0.0109 | 0.0065 | 0.0065 | TRUE | 0.0628 |
| Rundc3a | 553 | 797 | 825 | 1005 | 948 | 904 | 587 | 796 | 828 | 790 | 1020 | 990 | 835.23 | 737 | 933 | 1.264 | 0.338 | 0.03820343 | 0.13604526 | 0.025 | 0.0108 | 0.018 | 0.018 | TRUE | 0.3726 |
| Rundc3b | 358 | 404 | 364 | 684 | 531 | 432 | 380 | 403 | 365 | 538 | 571 | 473 | 455.19 | 383 | 527 | 1.377 | 0.462 | 0.00014861 | 0.00188841 | 0.0037 | 0.0141 | 0.0084 | 0.0084 | TRUE | 0.0919 |
| Runx1 | 75 | 74 | 94 | 144 | 85 | 92 | 80 | 74 | 94 | 113 | 91 | 101 | 92.22 | 83 | 102 | 1.235 | 0.305 | 0.1621445 | 0.35742865 | 0 | 0.0426 | 0.0233 | 0.0233 | TRUE | 0.1463 |
| Runx1t1 | 186 | 113 | 99 | 126 | 116 | 89 | 198 | 113 | 99 | 99 | 125 | 97 | 121.85 | 137 | 107 | 0.784 | -0.351 | 0.22604249 | 0.43889477 | 0.0737 | 0.0339 | 0.0523 | 0.0523 | TRUE | 1.5581 |
| Runx2 | 0 | 3 | 4 | 5 | 8 | 3 | 0 | 3 | 4 | 4 | 9 | 3 | 3.81 | 2 | 5 | 2.226 | 1.154 | 0.30645934 | NA | 0.192 | 0.8725 | 0.6081 | 0.6081 | TRUE | 0.7394 |
| Runx3 | 257 | 297 | 283 | 231 | 196 | 213 | 273 | 297 | 284 | 182 | 211 | 233 | 246.57 | 285 | 209 | 0.73 | -0.454 | 0.00240495 | 0.01780795 | 0.0042 | 0.0203 | 0.012 | 0.012 | TRUE | 0.1525 |
| Rusc1 | 333 | 365 | 373 | 589 | 381 | 386 | 354 | 365 | 375 | 463 | 410 | 423 | 398.04 | 365 | 432 | 1.188 | 0.248 | 0.03057326 | 0.11684108 | 0 | 0.0152 | 0.0069 | 0.0069 | TRUE | 0.049 |
| Rusc2 | 1276 | 1641 | 1612 | 1573 | 1184 | 1331 | 1355 | 1639 | 1619 | 1237 | 1274 | 1457 | 1430.05 | 1538 | 1323 | 0.859 | -0.218 | 0.05836737 | 0.18250795 | 0.0082 | 0.0092 | 0.0089 | 0.0089 | TRUE | 0.1291 |
| Ruvbl1 | 4095 | 3950 | 4041 | 5317 | 3876 | 4001 | 4350 | 3945 | 4058 | 4180 | 4170 | 4380 | 4180.33 | 4118 | 4243 | 1.031 | 0.043 | 0.54804999 | 0.732964 | 0.0013 | 0.0077 | 0.0035 | 0.0035 | TRUE | 0.0296 |
| Ruvbl2 | 12144 | 12778 | 13325 | 13846 | 10627 | 10804 | 12899 | 12762 | 13380 | 10885 | 11434 | 11827 | 12197.6 | 13014 | 11382 | 0.875 | -0.193 | 0.00293242 | 0.0208616 | 0.0011 | 0.0072 | 0.003 | 0.003 | TRUE | 0.018 |
| Rwdd1 | 1895 | 2197 | 2117 | 3140 | 2058 | 1990 | 2013 | 2194 | 2126 | 2468 | 2214 | 2178 | 2198.96 | 2111 | 2287 | 1.084 | 0.116 | 0.1798355 | 0.38118068 | 0.0026 | 0.0084 | 0.005 | 0.005 | TRUE | 0.0593 |
| Rwdd2a | 425 | 452 | 426 | 541 | 387 | 399 | 451 | 451 | 428 | 425 | 416 | 437 | 434.84 | 443 | 426 | 0.961 | -0.058 | 0.58162503 | 0.75788087 | 0 | 0.0145 | 0.0056 | 0.0056 | TRUE | 0.0113 |
| Rwdd2b | 365 | 347 | 353 | 643 | 471 | 454 | 388 | 347 | 354 | 505 | 507 | 497 | 432.99 | 363 | 503 | 1.387 | 0.472 | 1.41E-05 | 0.00026534 | 0 | 0.0145 | 0.0061 | 0.0061 | TRUE | 0.0403 |
| Rwdd3 | 257 | 306 | 292 | 366 | 288 | 250 | 273 | 306 | 293 | 288 | 310 | 274 | 290.51 | 291 | 291 | 0.999 | -0.002 | 0.98990897 | 0.99457546 | 0 | 0.0182 | 0.0086 | 0.0086 | TRUE | 0.0367 |
| Rwdd4a | 1956 | 2064 | 2002 | 2741 | 1902 | 1849 | 2078 | 2061 | 2010 | 2155 | 2046 | 2024 | 2062.4 | 2050 | 2075 | 1.013 | 0.019 | 0.79187569 | 0.89301263 | 0 | 0.0085 | 0.0031 | 0.0031 | TRUE | 0.0141 |
| Rxfp1 | 1 | 0 | 0 | 1 | 1 | 0 | 1 | 0 | 0 | 1 | 1 | 0 | 0.49 | 0 | 1 | 1.712 | 0.776 | 0.78488768 | NA | 0 | 6.7665 | 4.0387 | 4.0387 | TRUE | 0.4614 |
| Rxfp2 | 7 | 6 | 6 | 12 | 11 | 3 | 7 | 6 | 6 | 9 | 12 | 3 | 7.33 | 6 | 8 | 1.271 | 0.346 | 0.64173759 | 0.79835032 | 0 | 0.4561 | 0.2606 | 0.2606 | TRUE | 0.5856 |
| Rxfp3 | 0 | 1 | 0 | 1 | 0 | 0 | 0 | 1 | 0 | 1 | 0 | 0 | 0.3 | 0 | 0 | 0.954 | -0.067 | 0.98642001 | NA | 0 | 11.0817 | 9.3014 | 9.3014 | TRUE | 0.405 |
| Rxfp4 | 1 | 2 | 0 | 1 | 1 | 0 | 1 | 2 | 0 | 1 | 1 | 0 | 0.82 | 1 | 1 | 0.617 | -0.697 | 0.74855989 | NA | 0 | 4.0232 | 2.1283 | 2.1283 | TRUE | 0.1568 |
| Rxra | 816 | 809 | 882 | 1789 | 1353 | 1011 | 867 | 808 | 886 | 1406 | 1456 | 1107 | 1088.19 | 854 | 1323 | 1.551 | 0.633 | 5.97E-07 | 1.75E-05 | 0.0112 | 0.0099 | 0.0106 | 0.0106 | TRUE | 0.243 |
| Rxrb | 1622 | 2155 | 2078 | 2326 | 1829 | 1889 | 1723 | 2152 | 2087 | 1829 | 1968 | 2068 | 1970.97 | 1987 | 1955 | 0.983 | -0.025 | 0.82641248 | 0.91211327 | 0.0084 | 0.0086 | 0.0085 | 0.0085 | TRUE | 0.1633 |
| Rxrg | 120 | 76 | 70 | 58 | 47 | 32 | 127 | 76 | 70 | 46 | 51 | 35 | 67.47 | 91 | 44 | 0.481 | -1.057 | 0.0010422 | 0.00920877 | 0.0604 | 0.0557 | 0.058 | 0.058 | TRUE | 1.1416 |
| Rybp | 8149 | 8016 | 7929 | 10723 | 7572 | 7213 | 8656 | 8006 | 7962 | 8430 | 8147 | 7896 | 8182.54 | 8208 | 8158 | 0.994 | -0.009 | 0.90187978 | 0.95205059 | 0.0012 | 0.0073 | 0.0034 | 0.0034 | TRUE | 0.0278 |
| Ryk | 3484 | 4267 | 4093 | 4974 | 3603 | 3528 | 3701 | 4261 | 4110 | 3910 | 3877 | 3862 | 3953.44 | 4024 | 3883 | 0.965 | -0.052 | 0.51745483 | 0.70899987 | 0.0022 | 0.0077 | 0.0043 | 0.0043 | TRUE | 0.06 |
| Ryr1 | 1479 | 1472 | 1701 | 1294 | 1007 | 1013 | 1571 | 1470 | 1708 | 1017 | 1083 | 1109 | 1326.44 | 1583 | 1070 | 0.675 | -0.566 | 2.08E-09 | 1.18E-07 | 0.0032 | 0.0094 | 0.0057 | 0.0057 | TRUE | 0.0577 |
| Ryr2 | 590 | 711 | 841 | 765 | 554 | 669 | 627 | 710 | 844 | 601 | 596 | 732 | 685.17 | 727 | 643 | 0.884 | -0.178 | 0.2287531 | 0.44219804 | 0.0164 | 0.0117 | 0.0143 | 0.0143 | TRUE | 0.2361 |
| Ryr3 | 376 | 229 | 201 | 180 | 148 | 94 | 399 | 229 | 202 | 142 | 159 | 103 | 205.59 | 277 | 135 | 0.487 | -1.038 | 0.00027598 | 0.00317715 | 0.0896 | 0.0229 | 0.0532 | 0.0532 | TRUE | 1.6814 |
| S100a1 | 75 | 149 | 143 | 72 | 54 | 92 | 80 | 149 | 144 | 57 | 58 | 101 | 97.91 | 124 | 72 | 0.576 | -0.795 | 0.01619049 | 0.07497251 | 0.0986 | 0.0406 | 0.0679 | 0.0679 | TRUE | 1.0876 |
| S100a10 | 1464 | 1676 | 1690 | 2451 | 1709 | 1869 | 1555 | 1674 | 1697 | 1927 | 1839 | 2046 | 1789.55 | 1642 | 1937 | 1.18 | 0.238 | 0.00445806 | 0.02868092 | 0.0017 | 0.0087 | 0.0045 | 0.0045 | TRUE | 0.0286 |
| S100a11 | 1644 | 1422 | 1367 | 1789 | 1357 | 1250 | 1746 | 1420 | 1373 | 1406 | 1460 | 1368 | 1462.29 | 1513 | 1411 | 0.933 | -0.1 | 0.38411196 | 0.59828302 | 0.0085 | 0.0092 | 0.0088 | 0.0088 | TRUE | 0.2184 |
| S100a13 | 39 | 75 | 66 | 53 | 37 | 54 | 41 | 75 | 66 | 42 | 40 | 59 | 53.86 | 61 | 47 | 0.766 | -0.384 | 0.2434626 | 0.46008258 | 0.0498 | 0.0681 | 0.0593 | 0.0593 | TRUE | 0.6587 |
| S100a14 | 26 | 17 | 21 | 17 | 17 | 18 | 28 | 17 | 21 | 13 | 18 | 20 | 19.51 | 22 | 17 | 0.776 | -0.365 | 0.41634869 | 0.62772693 | 0 | 0.1758 | 0.0934 | 0.0934 | TRUE | 0.2865 |
| S100a16 | 21 | 22 | 24 | 47 | 17 | 24 | 22 | 22 | 24 | 37 | 18 | 26 | 24.98 | 23 | 27 | 1.205 | 0.269 | 0.51971363 | 0.71085972 | 1.00E-04 | 0.1388 | 0.0855 | 0.0855 | TRUE | 0.2568 |
| S100a2 | 0 | 1 | 0 | 0 | 0 | 0 | 0 | 1 | 0 | 0 | 0 | 0 | 0.17 | 0 | 0 | 0.49 | -1.029 | 0.80087 | NA | 0 | 19.7991 | 10 | 10 | TRUE | 0.405 |
| S100a3 | 4 | 5 | 7 | 11 | 3 | 1 | 4 | 5 | 7 | 9 | 3 | 1 | 4.87 | 5 | 4 | 0.824 | -0.28 | 0.77015544 | 0.88001822 | 0.2149 | 0.6829 | 0.458 | 0.458 | TRUE | 1.0348 |
| S100a4 | 9 | 1 | 0 | 4 | 0 | 6 | 10 | 1 | 0 | 3 | 0 | 7 | 3.38 | 4 | 3 | 0.923 | -0.115 | 0.94361784 | NA | 2.6516 | 0.982 | 1.6113 | 1.6113 | TRUE | 0.6251 |
| S100a5 | 1 | 0 | 0 | 0 | 0 | 0 | 1 | 0 | 0 | 0 | 0 | 0 | 0.18 | 0 | 0 | 0.49 | -1.029 | 0.80087 | NA | 0 | 18.6165 | 10 | 10 | TRUE | 0.4614 |
| S100a6 | 382 | 253 | 268 | 1001 | 673 | 753 | 406 | 253 | 269 | 787 | 724 | 824 | 543.8 | 309 | 778 | 2.52 | 1.333 | 3.98E-13 | 4.27E-11 | 0.0323 | 0.013 | 0.022 | 0.022 | TRUE | 0.8394 |
| S100a7a | 0 | 0 | 0 | 0 | 0 | 0 | 0 | 0 | 0 | 0 | 0 | 0 | 0 | 0 | 0 | NA | NA | NA | NA | NA | NA | NA | NA | NA | NA |
| S100a8 | 0 | 0 | 0 | 0 | 0 | 0 | 0 | 0 | 0 | 0 | 0 | 0 | 0 | 0 | 0 | NA | NA | NA | NA | NA | NA | NA | NA | NA | NA |
| S100a9 | 0 | 0 | 0 | 0 | 0 | 0 | 0 | 0 | 0 | 0 | 0 | 0 | 0 | 0 | 0 | NA | NA | NA | NA | NA | NA | NA | NA | NA | NA |
| S100b | 0 | 0 | 1 | 0 | 2 | 0 | 0 | 0 | 1 | 0 | 2 | 0 | 0.53 | 0 | 1 | 1.927 | 0.947 | 0.77736506 | NA | 0 | 6.2701 | 6.3814 | 6.3814 | TRUE | 0.9744 |
| S100g | 1 | 1 | 0 | 2 | 0 | 0 | 1 | 1 | 0 | 2 | 0 | 0 | 0.61 | 1 | 1 | 0.818 | -0.291 | 0.91402744 | NA | 0 | 5.4476 | 3.5995 | 3.5995 | TRUE | 0.9722 |
| S100pbp | 1309 | 1444 | 1520 | 1965 | 1415 | 1492 | 1390 | 1442 | 1526 | 1545 | 1522 | 1633 | 1509.86 | 1453 | 1567 | 1.078 | 0.108 | 0.18221323 | 0.38410413 | 0.0011 | 0.0091 | 0.0041 | 0.0041 | TRUE | 0.0235 |
| S100z | 0 | 1 | 0 | 1 | 0 | 0 | 0 | 1 | 0 | 1 | 0 | 0 | 0.3 | 0 | 0 | 0.954 | -0.067 | 0.98642001 | NA | 0 | 11.0817 | 9.3014 | 9.3014 | TRUE | 0.405 |
| S1pr1 | 27 | 22 | 21 | 58 | 46 | 29 | 29 | 22 | 21 | 46 | 49 | 32 | 33.09 | 24 | 42 | 1.775 | 0.828 | 0.02207662 | 0.09337449 | 0 | 0.1064 | 0.0614 | 0.0614 | TRUE | 0.3374 |
| S1pr2 | 2055 | 2394 | 2653 | 2838 | 2201 | 2094 | 2183 | 2391 | 2664 | 2231 | 2368 | 2292 | 2354.82 | 2413 | 2297 | 0.952 | -0.071 | 0.46221509 | 0.66594103 | 0.0048 | 0.0083 | 0.0063 | 0.0063 | TRUE | 0.1007 |
| S1pr3 | 257 | 201 | 196 | 162 | 133 | 157 | 273 | 201 | 197 | 127 | 143 | 172 | 185.47 | 224 | 147 | 0.658 | -0.604 | 0.00304084 | 0.02146414 | 0.0239 | 0.0247 | 0.0243 | 0.0243 | TRUE | 0.4085 |
| S1pr4 | 100 | 86 | 103 | 59 | 53 | 66 | 106 | 86 | 103 | 46 | 57 | 72 | 78.53 | 98 | 58 | 0.59 | -0.761 | 0.00278452 | 0.01998971 | 0.0154 | 0.0489 | 0.0331 | 0.0331 | TRUE | 0.3495 |
| S1pr5 | 18 | 32 | 34 | 34 | 23 | 27 | 19 | 32 | 34 | 27 | 25 | 30 | 27.71 | 28 | 27 | 0.948 | -0.077 | 0.84312851 | 0.92105202 | 0 | 0.1258 | 0.0717 | 0.0717 | TRUE | 0.5094 |
| Saa1 | 1 | 2 | 3 | 1 | 0 | 0 | 1 | 2 | 3 | 1 | 0 | 0 | 1.14 | 2 | 0 | 0.172 | -2.538 | 0.19527801 | NA | 0 | 2.8892 | 1.5165 | 1.5165 | TRUE | 0.2141 |
| Saa2 | 2 | 1 | 0 | 0 | 0 | 0 | 2 | 1 | 0 | 0 | 0 | 0 | 0.52 | 1 | 0 | 0.173 | -2.534 | 0.43568356 | NA | 0 | 6.3362 | 6.1226 | 6.1226 | TRUE | 0.0614 |
| Saa3 | 10 | 11 | 6 | 13 | 9 | 10 | 11 | 11 | 6 | 10 | 10 | 11 | 9.75 | 9 | 10 | 1.117 | 0.16 | 0.79113513 | 0.89260914 | 0 | 0.3449 | 0.1595 | 0.1595 | TRUE | 0.3039 |

| | | | | | | | | | | | | | | | | | | | | | | | | | |
|---|---|---|---|---|---|---|---|---|---|---|---|---|---|---|---|---|---|---|---|---|---|---|---|---|---|
| Saa4 | 0 | 0 | 0 | 0 | 0 | 0 | 0 | 0 | 0 | 0 | 0 | 0 | 0 | 0 | 0 | NA | NA | NA | NA | NA | NA | NA | NA | NA | NA |
| Saal1 | 1263 | 1538 | 1512 | 2343 | 1641 | 1700 | 1342 | 1536 | 1518 | 1842 | 1766 | 1861 | 1644.03 | 1465 | 1823 | 1.244 | 0.315 | 0.00033464 | 0.00373 | 0.0022 | 0.0089 | 0.0049 | 0.0049 | TRUE | 0.0653 |
| Sac3d1 | 1081 | 1027 | 1018 | 1346 | 983 | 891 | 1148 | 1026 | 1022 | 1058 | 1058 | 975 | 1047.86 | 1065 | 1030 | 0.968 | -0.047 | 0.61634564 | 0.78306778 | 0.0023 | 0.01 | 0.0054 | 0.0054 | TRUE | 0.055 |
| Sacm1l | 2168 | 2275 | 2204 | 3083 | 2204 | 2153 | 2303 | 2272 | 2213 | 2424 | 2371 | 2357 | 2323.28 | 2263 | 2384 | 1.054 | 0.076 | 0.24695454 | 0.46379674 | 0 | 0.0083 | 0.0027 | 0.0027 | TRUE | 0.0045 |
| Sacs | 292 | 283 | 349 | 343 | 221 | 252 | 310 | 283 | 350 | 270 | 238 | 276 | 287.75 | 314 | 261 | 0.831 | -0.267 | 0.06495848 | 0.19626919 | 0.0055 | 0.0184 | 0.0116 | 0.0116 | TRUE | 0.1148 |
| Sae1 | 16451 | 15518 | 14879 | 19475 | 13824 | 13881 | 17474 | 15498 | 14940 | 15310 | 14874 | 15195 | 15548.39 | 15971 | 15126 | 0.947 | -0.078 | 0.33876269 | 0.55879656 | 0.0035 | 0.0071 | 0.0048 | 0.0048 | TRUE | 0.0829 |
| Safb | 6600 | 5975 | 5846 | 9238 | 6772 | 6129 | 7010 | 5967 | 5870 | 7262 | 7286 | 6709 | 6684.21 | 6282 | 7086 | 1.128 | 0.174 | 0.06909688 | 0.20400524 | 0.0059 | 0.0074 | 0.0064 | 0.0064 | TRUE | 0.1252 |
| Safb2 | 3328 | 2800 | 2658 | 3955 | 3211 | 2556 | 3535 | 2796 | 2669 | 3109 | 3455 | 2798 | 3060.35 | 3000 | 3121 | 1.04 | 0.057 | 0.67087073 | 0.81643311 | 0.0167 | 0.008 | 0.0126 | 0.0126 | TRUE | 0.2951 |
| Sag | 0 | 3 | 1 | 5 | 0 | 2 | 0 | 3 | 1 | 4 | 0 | 2 | 1.69 | 1 | 2 | 1.552 | 0.634 | 0.71642786 | NA | 0.9655 | 1.9601 | 1.5805 | 1.5805 | TRUE | 0.2482 |
| Sall1 | 9037 | 9445 | 9841 | 12466 | 9696 | 9066 | 9599 | 9433 | 9881 | 9800 | 10432 | 9924 | 9844.89 | 9638 | 10052 | 1.043 | 0.061 | 0.32783055 | 0.54900652 | 7.00E-04 | 0.0072 | 0.0027 | 0.0027 | TRUE | 0.0134 |
| Sall2 | 320 | 207 | 209 | 354 | 281 | 202 | 340 | 207 | 210 | 278 | 302 | 221 | 259.71 | 252 | 267 | 1.061 | 0.086 | 0.70953063 | 0.84171221 | 0.0499 | 0.0196 | 0.0342 | 0.0342 | TRUE | 1.0239 |
| Sall3 | 95 | 75 | 85 | 158 | 125 | 109 | 101 | 75 | 85 | 124 | 134 | 119 | 106.53 | 87 | 126 | 1.449 | 0.535 | 0.0089772 | 0.04858902 | 0 | 0.0378 | 0.0204 | 0.0204 | TRUE | 0.1798 |
| Sall4 | 8176 | 7803 | 7885 | 11543 | 8219 | 7196 | 8684 | 7793 | 7917 | 9074 | 8843 | 7877 | 8364.86 | 8131 | 8598 | 1.058 | 0.081 | 0.36063388 | 0.57904078 | 0.0043 | 0.0073 | 0.0055 | 0.0055 | TRUE | 0.0656 |
| Samd1 | 3921 | 4574 | 4431 | 6456 | 4719 | 4292 | 4165 | 4568 | 4449 | 5075 | 5077 | 4698 | 4672.15 | 4394 | 4950 | 1.127 | 0.172 | 0.02213168 | 0.09357813 | 0.0018 | 0.0076 | 0.0039 | 0.0039 | TRUE | 0.0253 |
| Samd10 | 393 | 523 | 577 | 575 | 425 | 494 | 417 | 522 | 579 | 452 | 457 | 541 | 494.86 | 506 | 483 | 0.953 | -0.069 | 0.65527165 | 0.80609504 | 0.0166 | 0.0136 | 0.0152 | 0.0152 | TRUE | 0.2733 |
| Samd11 | 0 | 1 | 2 | 2 | 2 | 3 | 0 | 1 | 2 | 2 | 2 | 3 | 1.67 | 1 | 2 | 2.283 | 1.191 | 0.43440582 | NA | 0 | 1.9806 | 0.9566 | 0.9566 | TRUE | 0.3292 |
| Samd12 | 45 | 46 | 54 | 64 | 37 | 40 | 48 | 46 | 54 | 50 | 40 | 44 | 46.98 | 49 | 45 | 0.909 | -0.137 | 0.62711748 | 0.78999388 | 0 | 0.077 | 0.0361 | 0.0361 | TRUE | 0.1129 |
| Samd14 | 116 | 133 | 117 | 162 | 130 | 97 | 123 | 133 | 117 | 127 | 140 | 106 | 124.49 | 124 | 124 | 1.001 | 0.001 | 0.99659545 | 0.99828918 | 0 | 0.0334 | 0.0183 | 0.0183 | TRUE | 0.1567 |
| Samd3 | 0 | 0 | 0 | 0 | 0 | 0 | 0 | 0 | 0 | 0 | 0 | 0 | 0 | 0 | 0 | NA | NA | NA | NA | NA | NA | NA | NA | NA | NA |
| Samd4 | 61 | 42 | 43 | 133 | 72 | 61 | 65 | 42 | 43 | 105 | 77 | 67 | 66.45 | 50 | 83 | 1.671 | 0.741 | 0.01415482 | 0.06791533 | 0.0424 | 0.0565 | 0.0496 | 0.0496 | TRUE | 0.5338 |
| Samd4b | 7352 | 7124 | 7370 | 9251 | 6955 | 6503 | 7809 | 7115 | 7400 | 7272 | 7483 | 7119 | 7366.38 | 7441 | 7291 | 0.98 | -0.029 | 0.66730936 | 0.81392029 | 0.0011 | 0.0074 | 0.0032 | 0.0032 | TRUE | 0.0228 |
| Samd5 | 11 | 9 | 5 | 12 | 9 | 4 | 12 | 9 | 5 | 9 | 10 | 4 | 8.2 | 9 | 8 | 0.924 | -0.114 | 0.87093209 | 0.93648357 | 0 | 0.4088 | 0.2324 | 0.2324 | TRUE | 0.2986 |
| Samd7 | 24 | 23 | 14 | 72 | 56 | 55 | 25 | 23 | 14 | 57 | 60 | 60 | 39.93 | 21 | 59 | 2.835 | 1.503 | 8.85E-06 | 0.00017939 | 0 | 0.0894 | 0.0496 | 0.0496 | TRUE | 0.4538 |
| Samd8 | 3219 | 3279 | 3568 | 4076 | 2800 | 2854 | 3419 | 3275 | 3583 | 3204 | 3013 | 3124 | 3269.6 | 3426 | 3114 | 0.909 | -0.137 | 0.05802645 | 0.1819496 | 0.001 | 0.0079 | 0.0035 | 0.0035 | TRUE | 0.0196 |
| Samd9l | 15 | 5 | 4 | 20 | 9 | 4 | 16 | 5 | 4 | 16 | 10 | 4 | 9.12 | 8 | 10 | 1.216 | 0.282 | 0.7264378 | 0.85254683 | 0.3442 | 0.3681 | 0.3569 | 0.3569 | TRUE | 0.4438 |
| Samhd1 | 4673 | 4468 | 4708 | 5352 | 3777 | 3810 | 4964 | 4462 | 4727 | 4207 | 4064 | 4171 | 4432.48 | 4718 | 4147 | 0.879 | -0.186 | 0.00944328 | 0.05040251 | 0.0012 | 0.0076 | 0.0035 | 0.0035 | TRUE | 0.0274 |
| Samm50 | 11131 | 12214 | 11953 | 12247 | 8897 | 9561 | 11823 | 12198 | 12002 | 9628 | 9573 | 10466 | 10948.28 | 12008 | 9889 | 0.823 | -0.28 | 2.89E-05 | 0.00048799 | 0.0013 | 0.0072 | 0.0031 | 0.0031 | TRUE | 0.0317 |
| Samsn1 | 0 | 0 | 0 | 0 | 0 | 0 | 0 | 0 | 0 | 0 | 0 | 0 | 0 | 0 | 0 | NA | NA | NA | NA | NA | NA | NA | NA | NA | NA |
| Samt1 | 0 | 0 | 0 | 0 | 0 | 0 | 0 | 0 | 0 | 0 | 0 | 0 | 0 | 0 | 0 | NA | NA | NA | NA | NA | NA | NA | NA | NA | NA |
| Samt2 | 0 | 0 | 0 | 0 | 0 | 0 | 0 | 0 | 0 | 0 | 0 | 0 | 0 | 0 | 0 | NA | NA | NA | NA | NA | NA | NA | NA | NA | NA |
| Samt3 | 0 | 0 | 0 | 0 | 0 | 0 | 0 | 0 | 0 | 0 | 0 | 0 | 0 | 0 | 0 | NA | NA | NA | NA | NA | NA | NA | NA | NA | NA |
| Samt4 | 0 | 0 | 0 | 0 | 0 | 0 | 0 | 0 | 0 | 0 | 0 | 0 | 0 | 0 | 0 | NA | NA | NA | NA | NA | NA | NA | NA | NA | NA |
| Sap130 | 7529 | 7825 | 8126 | 9845 | 7048 | 7202 | 7997 | 7815 | 8159 | 7739 | 7583 | 7884 | 7862.95 | 7990 | 7735 | 0.968 | -0.047 | 0.41979669 | 0.63033203 | 2.00E-04 | 0.0073 | 0.0023 | 0.0023 | TRUE | 0.0045 |
| Sap18 | 556 | 490 | 479 | 684 | 431 | 475 | 591 | 489 | 481 | 538 | 464 | 520 | 513.72 | 520 | 507 | 0.976 | -0.035 | 0.78742837 | 0.89059306 | 0.0076 | 0.0133 | 0.0102 | 0.0102 | TRUE | 0.1618 |
| Sap25 | 1 | 0 | 0 | 0 | 0 | 0 | 1 | 0 | 0 | 0 | 0 | 0 | 0.18 | 0 | 0 | 0.49 | -1.029 | 0.80087 | NA | 0 | 18.6165 | 10 | 10 | TRUE | 0.4614 |
| Sap30 | 2848 | 3680 | 3728 | 4342 | 3429 | 3164 | 3025 | 3675 | 3743 | 3413 | 3689 | 3463 | 3501.64 | 3481 | 3522 | 1.012 | 0.017 | 0.87389068 | 0.9378107 | 0.0073 | 0.0078 | 0.0075 | 0.0075 | TRUE | 0.1595 |
| Sap30bp | 2450 | 2110 | 1957 | 3376 | 2360 | 1994 | 2602 | 2107 | 1965 | 2654 | 2539 | 2183 | 2341.76 | 2225 | 2459 | 1.105 | 0.145 | 0.27405812 | 0.49464606 | 0.0151 | 0.0083 | 0.0122 | 0.0122 | TRUE | 0.2663 |
| Sap30l | 1746 | 2012 | 1987 | 2274 | 1725 | 1787 | 1855 | 2009 | 1995 | 1788 | 1856 | 1956 | 1909.82 | 1953 | 1867 | 0.955 | -0.066 | 0.40845047 | 0.62109317 | 0.0015 | 0.0086 | 0.0041 | 0.0041 | TRUE | 0.0234 |
| Sapcd1 | 39 | 72 | 52 | 28 | 28 | 43 | 41 | 72 | 52 | 22 | 30 | 47 | 44.13 | 55 | 33 | 0.593 | -0.754 | 0.05079574 | 0.16586965 | 0.0852 | 0.0816 | 0.0832 | 0.0832 | TRUE | 0.5086 |
| Sapcd2 | 733 | 1161 | 1109 | 1529 | 1135 | 1152 | 779 | 1160 | 1114 | 1202 | 1221 | 1261 | 1122.64 | 1018 | 1228 | 1.207 | 0.271 | 0.07635241 | 0.21799969 | 0.0219 | 0.0098 | 0.016 | 0.016 | TRUE | 0.5014 |
| Sar1a | 6466 | 6303 | 5944 | 8144 | 5619 | 5552 | 6868 | 6295 | 5968 | 6402 | 6046 | 6078 | 6276.12 | 6377 | 6175 | 0.969 | -0.046 | 0.56585354 | 0.7460204 | 0.0027 | 0.0074 | 0.0045 | 0.0045 | TRUE | 0.0553 |
| Sar1b | 3704 | 4486 | 3953 | 4939 | 3543 | 3304 | 3934 | 4480 | 3969 | 3883 | 3812 | 3617 | 3949.2 | 4128 | 3771 | 0.914 | -0.13 | 0.11949988 | 0.29294718 | 0.003 | 0.0077 | 0.0048 | 0.0048 | TRUE | 0.0679 |
| Sardh | 17 | 10 | 7 | 25 | 11 | 10 | 18 | 10 | 7 | 20 | 12 | 11 | 12.92 | 12 | 14 | 1.228 | 0.296 | 0.61963839 | 0.78485124 | 0.0901 | 0.2619 | 0.1785 | 0.1785 | TRUE | 0.8423 |
| Sarm1 | 775 | 1012 | 1018 | 1078 | 708 | 655 | 823 | 1011 | 1022 | 847 | 762 | 717 | 863.71 | 952 | 775 | 0.815 | -0.295 | 0.01873839 | 0.08301738 | 0.0097 | 0.0107 | 0.0102 | 0.0102 | TRUE | 0.1663 |
| Sarnp | 4135 | 4148 | 4182 | 5334 | 3864 | 3905 | 4392 | 4143 | 4199 | 4193 | 4157 | 4275 | 4226.53 | 4245 | 4208 | 0.991 | -0.012 | 0.8440919 | 0.92161735 | 3.00E-04 | 0.0077 | 0.0026 | 0.0026 | TRUE | 0.0112 |
| Sars | 8966 | 8821 | 8990 | 11136 | 8053 | 7861 | 9523 | 8810 | 9027 | 8754 | 8665 | 8605 | 8897.31 | 9120 | 8675 | 0.951 | -0.072 | 0.24698002 | 0.46379674 | 7.00E-04 | 0.0073 | 0.0027 | 0.0027 | TRUE | 0.0183 |
| Sars2 | 1023 | 1154 | 1180 | 1771 | 1262 | 1080 | 1087 | 1153 | 1185 | 1392 | 1358 | 1182 | 1226.04 | 1142 | 1311 | 1.149 | 0.201 | 0.04323155 | 0.14864827 | 0.004 | 0.0096 | 0.0063 | 0.0063 | TRUE | 0.0874 |
| Sart1 | 5555 | 4878 | 4889 | 7467 | 5576 | 4580 | 5900 | 4872 | 4909 | 5870 | 5999 | 5014 | 5427.35 | 5227 | 5628 | 1.077 | 0.107 | 0.34911534 | 0.56797812 | 0.01 | 0.0075 | 0.0092 | 0.0092 | TRUE | 0.1547 |
| Sart3 | 7631 | 6136 | 6046 | 9440 | 7291 | 6292 | 8105 | 6128 | 6071 | 7421 | 7845 | 6888 | 7076.27 | 6768 | 7385 | 1.091 | 0.126 | 0.32994782 | 0.55091436 | 0.0147 | 0.0074 | 0.0119 | 0.0119 | TRUE | 0.3645 |
| Sash1 | 2403 | 2163 | 2337 | 1790 | 1347 | 1446 | 2552 | 2160 | 2347 | 1407 | 1449 | 1583 | 1916.42 | 2353 | 1480 | 0.629 | -0.67 | 8.74E-12 | 7.78E-10 | 0.0046 | 0.0086 | 0.0064 | 0.0064 | TRUE | 0.0664 |
| Sash3 | 6 | 9 | 1 | 7 | 5 | 11 | 6 | 9 | 1 | 6 | 5 | 12 | 6.55 | 5 | 8 | 1.391 | 0.476 | 0.5920317 | 0.76555069 | 0.2887 | 0.51 | 0.4104 | 0.4104 | TRUE | 0.7069 |
| Sass6 | 733 | 802 | 856 | 1097 | 813 | 765 | 779 | 801 | 860 | 862 | 875 | 837 | 835.6 | 813 | 858 | 1.056 | 0.078 | 0.38390409 | 0.59804511 | 0 | 0.0108 | 0.0046 | 0.0046 | TRUE | 0.0298 |
| Sat1 | 2352 | 2297 | 2131 | 2207 | 1498 | 1807 | 2498 | 2294 | 2140 | 1735 | 1612 | 1978 | 2042.8 | 2311 | 1775 | 0.768 | -0.381 | 0.00050005 | 0.00517509 | 0.0078 | 0.0085 | 0.0081 | 0.0081 | TRUE | 0.1196 |
| Sat2 | 94 | 115 | 110 | 149 | 118 | 106 | 100 | 115 | 110 | 117 | 127 | 116 | 114.21 | 108 | 120 | 1.106 | 0.145 | 0.43139099 | 0.64121182 | 0 | 0.0357 | 0.0158 | 0.0158 | TRUE | 0.046 |
| Satb1 | 1562 | 1960 | 1856 | 1919 | 1388 | 1360 | 1659 | 1957 | 1864 | 1509 | 1493 | 1489 | 1661.82 | 1827 | 1497 | 0.819 | -0.287 | 0.00151618 | 0.01236349 | 0.003 | 0.0089 | 0.0053 | 0.0053 | TRUE | 0.0775 |
| Satb2 | 185 | 166 | 135 | 471 | 282 | 224 | 197 | 166 | 136 | 370 | 303 | 245 | 236.12 | 166 | 306 | 1.851 | 0.888 | 2.42E-05 | 0.00041936 | 0.0337 | 0.0209 | 0.0272 | 0.0272 | TRUE | 0.1359 |
| Satl1 | 0 | 0 | 0 | 1 | 0 | 0 | 0 | 0 | 0 | 1 | 0 | 0 | 0.13 | 0 | 0 | 1.859 | 0.894 | 0.82650734 | NA | 0 | 25.1515 | 10 | 10 | TRUE | 0.2563 |
| Sav1 | 2413 | 2372 | 2284 | 3370 | 2347 | 2315 | 2563 | 2369 | 2293 | 2649 | 2525 | 2534 | 2488.99 | 2408 | 2569 | 1.067 | 0.094 | 0.22738142 | 0.44072728 | 0.0014 | 0.0082 | 0.004 | 0.004 | TRUE | 0.0381 |
| Saysd1 | 524 | 331 | 286 | 514 | 351 | 327 | 557 | 331 | 287 | 404 | 378 | 358 | 385.67 | 392 | 380 | 0.971 | -0.042 | 0.85873238 | 0.9304418 | 0.0618 | 0.0154 | 0.0372 | 0.0372 | TRUE | 1.56 |
| Sbds | 1520 | 1510 | 1531 | 2419 | 1670 | 1579 | 1615 | 1508 | 1537 | 1902 | 1797 | 1728 | 1681.13 | 1553 | 1809 | 1.165 | 0.221 | 0.00588266 | 0.03539922 | 0.0013 | 0.0089 | 0.004 | 0.004 | TRUE | 0.0249 |
| Sbf1 | 6253 | 6253 | 6230 | 8101 | 6123 | 5747 | 6642 | 6245 | 6256 | 6368 | 6588 | 6291 | 6398.27 | 6381 | 6416 | 1.005 | 0.008 | 0.90167113 | 0.95205059 | 6.00E-04 | 0.0074 | 0.0028 | 0.0028 | TRUE | 0.0156 |
| Sbf2 | 7852 | 8869 | 9875 | 7612 | 5667 | 6508 | 8340 | 8858 | 9916 | 5984 | 6097 | 7124 | 7719.77 | 9038 | 6402 | 0.708 | -0.498 | 2.59E-06 | 6.29E-05 | 0.0086 | 0.0073 | 0.0079 | 0.0079 | TRUE | 0.1186 |
| Sbk1 | 2440 | 2549 | 2635 | 2942 | 2201 | 2485 | 2592 | 2546 | 2646 | 2313 | 2368 | 2720 | 2530.73 | 2595 | 2467 | 0.95 | -0.073 | 0.41459329 | 0.62619037 | 0.0036 | 0.0082 | 0.0054 | 0.0054 | TRUE | 0.0971 |
| Sbk2 | 18 | 17 | 21 | 23 | 21 | 18 | 19 | 17 | 21 | 18 | 23 | 20 | 19.59 | 19 | 20 | 1.052 | 0.073 | 0.86286628 | 0.93221844 | 0 | 0.175 | 0.0769 | 0.0769 | TRUE | 0.06 |
| Sbno1 | 12790 | 12823 | 13497 | 15326 | 11548 | 11597 | 13585 | 12806 | 13552 | 12048 | 12425 | 12695 | 12851.98 | 13314 | 12389 | 0.93 | -0.104 | 0.09414645 | 0.25112088 | 8.00E-04 | 0.0072 | 0.0027 | 0.0027 | TRUE | 0.0136 |
| Sbno2 | 3349 | 3718 | 4290 | 3739 | 2939 | 3396 | 3557 | 3713 | 4308 | 2939 | 3162 | 3717 | 3566.17 | 3859 | 3273 | 0.848 | -0.238 | 0.04821628 | 0.16003096 | 0.0123 | 0.0078 | 0.0102 | 0.0102 | TRUE | 0.1713 |
| Sbp | 0 | 0 | 1 | 0 | 2 | 1 | 0 | 0 | 1 | 0 | 2 | 1 | 0.71 | 0 | 1 | 2.893 | 1.532 | 0.55266254 | NA | 0 | 4.6572 | 3.3372 | 3.3372 | TRUE | 0.1942 |
| Sbpl | 1 | 0 | 0 | 0 | 0 | 1 | 1 | 0 | 0 | 0 | 0 | 1 | 0.36 | 0 | 0 | 0.954 | -0.067 | 0.98553519 | NA | 0 | 9.1716 | 7.9605 | 7.9605 | TRUE | 0.5383 |
| Sbsn | 71 | 54 | 54 | 95 | 63 | 60 | 75 | 54 | 54 | 75 | 68 | 66 | 65.29 | 61 | 70 | 1.138 | 0.187 | 0.46729893 | 0.66963926 | 0 | 0.0574 | 0.0323 | 0.0323 | TRUE | 0.3496 |

| | | | | | | | | | | | | | | | | | | | | | | | | |
|---|---|---|---|---|---|---|---|---|---|---|---|---|---|---|---|---|---|---|---|---|---|---|---|---|
| Sbspon | 2 | 1 | 5 | 4 | 4 | 6 | 2 | 1 | 5 | 3 | 4 | 7 | 3.69 | 3 | 5 | 1.7 | 0.766 | 0.46093186 | NA | 0 | 0.8989 | 0.4834 | 0.4834 | TRUE | 0.6671 |
| Sc4mol | 11254 | 10862 | 10507 | 10759 | 8127 | 9241 | 11954 | 10848 | 10550 | 8458 | 8744 | 10116 | 10111.6 | 11117 | 9106 | 0.819 | -0.288 | 0.00343432 | 0.02348923 | 0.0068 | 0.0072 | 0.0069 | 0.0069 | TRUE | 0.1148 |
| Sc5d | 1036 | 1117 | 1098 | 1478 | 1025 | 1045 | 1100 | 1116 | 1103 | 1162 | 1103 | 1144 | 1121.19 | 1106 | 1136 | 1.028 | 0.039 | 0.61045104 | 0.77928238 | 0 | 0.0098 | 0.0034 | 0.0034 | TRUE | 0.0078 |
| Scaf1 | 6010 | 5543 | 5308 | 8289 | 6345 | 5472 | 6384 | 5536 | 5330 | 6516 | 6827 | 5990 | 6097.05 | 5750 | 6444 | 1.121 | 0.165 | 0.09700996 | 0.2559453 | 0.0066 | 0.0074 | 0.0069 | 0.0069 | TRUE | 0.1133 |
| Scaf11 | 7689 | 7875 | 8542 | 10866 | 7747 | 8020 | 8167 | 7865 | 8577 | 8542 | 8335 | 8779 | 8377.57 | 8203 | 8552 | 1.043 | 0.06 | 0.3680143 | 0.58529007 | 0.0011 | 0.0073 | 0.0031 | 0.0031 | TRUE | 0.0195 |
| Scaf4 | 4735 | 3987 | 3992 | 4719 | 3466 | 3217 | 5029 | 3982 | 4008 | 3710 | 3729 | 3522 | 3996.68 | 4340 | 3654 | 0.842 | -0.248 | 0.02538438 | 0.10284163 | 0.0096 | 0.0077 | 0.0086 | 0.0086 | TRUE | 0.235 |
| Scaf8 | 2962 | 3434 | 3632 | 5222 | 3815 | 3558 | 3146 | 3430 | 3647 | 4105 | 4105 | 3895 | 3721.21 | 3408 | 4035 | 1.184 | 0.244 | 0.00329213 | 0.02274555 | 0.0028 | 0.0078 | 0.0047 | 0.0047 | TRUE | 0.0546 |
| Scai | 1024 | 905 | 962 | 1014 | 843 | 883 | 1088 | 904 | 966 | 797 | 907 | 967 | 938.03 | 986 | 890 | 0.902 | -0.149 | 0.21275253 | 0.42383894 | 0.0076 | 0.0104 | 0.0092 | 0.0092 | TRUE | 0.1028 |
| Scamp1 | 2113 | 2488 | 2421 | 3258 | 2427 | 2380 | 2244 | 2485 | 2431 | 2561 | 2611 | 2605 | 2489.65 | 2387 | 2592 | 1.086 | 0.119 | 0.10806952 | 0.27513681 | 8.00E-04 | 0.0082 | 0.0035 | 0.0035 | TRUE | 0.0328 |
| Scamp2 | 1471 | 1422 | 1510 | 1792 | 1279 | 1420 | 1562 | 1420 | 1516 | 1409 | 1376 | 1554 | 1473.02 | 1499 | 1446 | 0.964 | -0.052 | 0.55932224 | 0.74125763 | 0.0023 | 0.0091 | 0.0051 | 0.0051 | TRUE | 0.0505 |
| Scamp3 | 3099 | 3322 | 3496 | 3456 | 2647 | 2952 | 3292 | 3318 | 3510 | 2717 | 2848 | 3231 | 3152.67 | 3373 | 2932 | 0.869 | -0.203 | 0.02706645 | 0.10723237 | 0.0043 | 0.0079 | 0.0058 | 0.0058 | TRUE | 0.0965 |
| Scamp4 | 2906 | 2611 | 2478 | 3624 | 2383 | 2271 | 3087 | 2608 | 2488 | 2849 | 2564 | 2486 | 2680.22 | 2728 | 2633 | 0.966 | -0.05 | 0.65040367 | 0.80412061 | 0.0087 | 0.0081 | 0.0085 | 0.0085 | TRUE | 0.1608 |
| Scamp5 | 2270 | 2442 | 2201 | 2824 | 1872 | 1835 | 2411 | 2439 | 2210 | 2220 | 2014 | 2009 | 2217.15 | 2353 | 2081 | 0.885 | -0.177 | 0.03893855 | 0.13803973 | 0.0024 | 0.0084 | 0.0048 | 0.0048 | TRUE | 0.0422 |
| Scand1 | 579 | 852 | 869 | 926 | 631 | 753 | 615 | 851 | 873 | 728 | 679 | 824 | 761.6 | 780 | 744 | 0.954 | -0.069 | 0.66305737 | 0.81107023 | 0.0215 | 0.0112 | 0.0165 | 0.0165 | TRUE | 0.4023 |
| Scap | 3071 | 2995 | 2888 | 3859 | 2730 | 2631 | 3262 | 2991 | 2900 | 3034 | 2937 | 2880 | 3000.66 | 3051 | 2950 | 0.967 | -0.048 | 0.53950566 | 0.72541204 | 0.0018 | 0.008 | 0.0041 | 0.0041 | TRUE | 0.0443 |
| Scaper | 996 | 1313 | 1357 | 1644 | 1205 | 1238 | 1058 | 1311 | 1363 | 1292 | 1297 | 1355 | 1279.31 | 1244 | 1315 | 1.056 | 0.079 | 0.49448688 | 0.69052608 | 0.0085 | 0.0095 | 0.0089 | 0.0089 | TRUE | 0.2044 |
| Scara3 | 24 | 34 | 39 | 68 | 46 | 36 | 25 | 34 | 39 | 53 | 49 | 39 | 40.16 | 33 | 47 | 1.447 | 0.533 | 0.10156903 | 0.26421685 | 0 | 0.0889 | 0.0506 | 0.0506 | TRUE | 0.2565 |
| Scara5 | 44 | 44 | 50 | 87 | 57 | 72 | 47 | 44 | 50 | 68 | 61 | 79 | 58.24 | 47 | 69 | 1.479 | 0.565 | 0.02917074 | 0.11279256 | 0 | 0.0635 | 0.0304 | 0.0304 | TRUE | 0.1135 |
| Scarb1 | 4493 | 5713 | 5900 | 5148 | 3985 | 4699 | 4772 | 5706 | 5924 | 4047 | 4288 | 5144 | 4980.1 | 5467 | 4493 | 0.822 | -0.284 | 0.02406573 | 0.09897384 | 0.0142 | 0.0076 | 0.0112 | 0.0112 | TRUE | 0.1954 |
| Scarb2 | 1510 | 2410 | 2594 | 1917 | 1415 | 1815 | 1604 | 2407 | 2605 | 1507 | 1522 | 1987 | 1938.61 | 2205 | 1672 | 0.758 | -0.4 | 0.03410144 | 0.12563605 | 0.0414 | 0.0086 | 0.0251 | 0.0251 | TRUE | 0.688 |
| Scarf1 | 104 | 113 | 118 | 220 | 149 | 144 | 110 | 113 | 118 | 173 | 160 | 158 | 138.78 | 114 | 164 | 1.439 | 0.525 | 0.00187993 | 0.01464937 | 0 | 0.0306 | 0.0131 | 0.0131 | TRUE | 0.0288 |
| Scarf2 | 35 | 69 | 67 | 108 | 82 | 65 | 37 | 69 | 67 | 85 | 88 | 71 | 69.61 | 58 | 81 | 1.408 | 0.493 | 0.09301327 | 0.24915172 | 0.039 | 0.0542 | 0.0474 | 0.0474 | TRUE | 0.8058 |
| Scarna10 | 5 | 2 | 3 | 2 | 2 | 4 | 5 | 2 | 3 | 2 | 2 | 4 | 3.07 | 3 | 3 | 0.773 | -0.372 | 0.73291708 | NA | 0 | 1.0798 | 0.5208 | 0.5208 | TRUE | 0.3207 |
| Scarna13 | 1 | 4 | 2 | 5 | 1 | 2 | 1 | 4 | 2 | 4 | 1 | 2 | 2.38 | 2 | 2 | 1.039 | 0.055 | 0.96483074 | NA | 0 | 1.393 | 0.688 | 0.688 | TRUE | 0.4372 |
| Scarna17 | 7 | 4 | 5 | 3 | 8 | 3 | 7 | 4 | 5 | 2 | 9 | 3 | 5.12 | 5 | 5 | 0.853 | -0.23 | 0.79668107 | 0.89536272 | 0.0954 | 0.6508 | 0.3749 | 0.3749 | TRUE | 0.923 |
| Scarna2 | 0 | 0 | 0 | 0 | 1 | 0 | 0 | 0 | 0 | 0 | 1 | 0 | 0.18 | 0 | 0 | 1.859 | 0.894 | 0.82650734 | NA | 0 | 18.3785 | 10 | 10 | TRUE | 0.5204 |
| Scarna3a | 0 | 0 | 0 | 0 | 0 | 0 | 0 | 0 | 0 | 0 | 0 | 0 | 0 | 0 | 0 | NA | NA | NA | NA | NA | NA | NA | NA | NA | NA |
| Scarna3b | 0 | 0 | 0 | 0 | 0 | 0 | 0 | 0 | 0 | 0 | 0 | 0 | 0 | 0 | 0 | NA | NA | NA | NA | NA | NA | NA | NA | NA | NA |
| Scarna6 | 3 | 2 | 1 | 2 | 3 | 1 | 3 | 2 | 1 | 2 | 3 | 1 | 2.01 | 2 | 2 | 0.95 | -0.075 | 0.954367 | NA | 0 | 1.6428 | 0.728 | 0.728 | TRUE | 0.251 |
| Scarna8 | 0 | 1 | 1 | 4 | 0 | 1 | 0 | 1 | 1 | 3 | 0 | 1 | 1.04 | 1 | 1 | 2.166 | 1.115 | 0.5872675 | NA | 0 | 3.1736 | 1.9471 | 1.9471 | TRUE | 0.3035 |
| Scarna9 | 0 | 0 | 0 | 1 | 0 | 0 | 0 | 0 | 0 | 1 | 0 | 0 | 0.13 | 0 | 0 | 1.859 | 0.894 | 0.82650734 | NA | 0 | 25.1515 | 10 | 10 | TRUE | 0.2563 |
| Sccpdh | 1771 | 2418 | 2277 | 2722 | 1958 | 2166 | 1881 | 2415 | 2286 | 2140 | 2107 | 2371 | 2199.98 | 2194 | 2206 | 1.005 | 0.007 | 0.95014531 | 0.97481103 | 0.01 | 0.0084 | 0.0093 | 0.0093 | TRUE | 0.1881 |
| Scd1 | 11487 | 13902 | 14579 | 11183 | 8344 | 9832 | 12201 | 13884 | 14639 | 8791 | 8978 | 10763 | 11542.61 | 13575 | 9511 | 0.701 | -0.513 | 5.25E-06 | 0.00011557 | 0.0106 | 0.0072 | 0.0091 | 0.0091 | TRUE | 0.1619 |
| Scd2 | 48963 | 43192 | 43272 | 49436 | 35328 | 34349 | 52007 | 43136 | 43450 | 38863 | 38011 | 37600 | 42177.85 | 46198 | 38158 | 0.826 | -0.276 | 0.00319642 | 0.02229698 | 0.006 | 0.007 | 0.0063 | 0.0063 | TRUE | 0.1482 |
| Scd3 | 159 | 163 | 177 | 131 | 114 | 118 | 169 | 163 | 178 | 103 | 123 | 129 | 144.04 | 170 | 118 | 0.693 | -0.529 | 0.00273557 | 0.01972812 | 0 | 0.0298 | 0.0153 | 0.0153 | TRUE | 0.1455 |
| Scd4 | 3 | 3 | 0 | 1 | 1 | 0 | 3 | 3 | 0 | 1 | 1 | 0 | 1.34 | 2 | 1 | 0.307 | -1.705 | 0.37239074 | NA | 0 | 2.464 | 1.6107 | 1.6107 | TRUE | 0.7155 |
| Scel | 5 | 7 | 10 | 11 | 5 | 6 | 5 | 7 | 10 | 9 | 5 | 7 | 7.16 | 7 | 7 | 0.93 | -0.105 | 0.88143269 | 0.94135674 | 0 | 0.4673 | 0.2206 | 0.2206 | TRUE | 0.2598 |
| Scfd1 | 1661 | 1763 | 1659 | 2422 | 1671 | 1576 | 1764 | 1761 | 1666 | 1904 | 1798 | 1725 | 1769.65 | 1730 | 1809 | 1.046 | 0.065 | 0.41177761 | 0.62279981 | 0.0013 | 0.0088 | 0.004 | 0.004 | TRUE | 0.0261 |
| Scfd2 | 479 | 295 | 328 | 868 | 628 | 507 | 509 | 295 | 329 | 682 | 676 | 555 | 507.63 | 378 | 638 | 1.691 | 0.757 | 0.00026411 | 0.0030652 | 0.0464 | 0.0134 | 0.0289 | 0.0289 | TRUE | 1.0552 |
| Scg2 | 0 | 1 | 0 | 0 | 0 | 0 | 0 | 1 | 0 | 0 | 0 | 0 | 0.17 | 0 | 0 | 0.49 | -1.029 | 0.80087 | NA | 0 | 19.7991 | 10 | 10 | TRUE | 0.405 |
| Scg3 | 6 | 6 | 8 | 21 | 13 | 9 | 6 | 6 | 8 | 17 | 14 | 10 | 10.12 | 7 | 14 | 1.992 | 0.995 | 0.10339785 | 0.26760005 | 0 | 0.3323 | 0.1576 | 0.1576 | TRUE | 0.2156 |
| Scg5 | 12 | 30 | 24 | 11 | 6 | 14 | 13 | 30 | 24 | 9 | 6 | 15 | 16.21 | 22 | 10 | 0.452 | -1.147 | 0.04443822 | 0.15144329 | 0.1101 | 0.2102 | 0.1633 | 0.1633 | TRUE | 0.2978 |
| Scgb1a1 | 0 | 0 | 0 | 0 | 0 | 0 | 0 | 0 | 0 | 0 | 0 | 0 | 0 | 0 | 0 | NA | NA | NA | NA | NA | NA | NA | NA | NA | NA |
| Scgb1b2 | 0 | 0 | 0 | 0 | 0 | 0 | 0 | 0 | 0 | 0 | 0 | 0 | 0 | 0 | 0 | NA | NA | NA | NA | NA | NA | NA | NA | NA | NA |
| Scgb1b20 | 0 | 0 | 0 | 0 | 0 | 0 | 0 | 0 | 0 | 0 | 0 | 0 | 0 | 0 | 0 | NA | NA | NA | NA | NA | NA | NA | NA | NA | NA |
| Scgb1b24 | 0 | 0 | 0 | 0 | 0 | 0 | 0 | 0 | 0 | 0 | 0 | 0 | 0 | 0 | 0 | NA | NA | NA | NA | NA | NA | NA | NA | NA | NA |
| Scgb1b27 | 0 | 0 | 0 | 0 | 0 | 0 | 0 | 0 | 0 | 0 | 0 | 0 | 0 | 0 | 0 | NA | NA | NA | NA | NA | NA | NA | NA | NA | NA |
| Scgb1b29 | 0 | 0 | 0 | 0 | 0 | 0 | 0 | 0 | 0 | 0 | 0 | 0 | 0 | 0 | 0 | NA | NA | NA | NA | NA | NA | NA | NA | NA | NA |
| Scgb1b3 | 0 | 0 | 0 | 0 | 0 | 0 | 0 | 0 | 0 | 0 | 0 | 0 | 0 | 0 | 0 | NA | NA | NA | NA | NA | NA | NA | NA | NA | NA |
| Scgb1b30 | 0 | 0 | 0 | 0 | 0 | 0 | 0 | 0 | 0 | 0 | 0 | 0 | 0 | 0 | 0 | NA | NA | NA | NA | NA | NA | NA | NA | NA | NA |
| Scgb1b7 | 0 | 0 | 0 | 0 | 0 | 0 | 0 | 0 | 0 | 0 | 0 | 0 | 0 | 0 | 0 | NA | NA | NA | NA | NA | NA | NA | NA | NA | NA |
| Scgb1c1 | 0 | 1 | 1 | 1 | 1 | 1 | 0 | 1 | 1 | 1 | 1 | 1 | 0.83 | 1 | 1 | 1.45 | 0.536 | 0.78654731 | NA | 0 | 3.9925 | 1.5656 | 1.5656 | TRUE | 0.2233 |
| Scgb2b19 | 0 | 0 | 0 | 0 | 0 | 0 | 0 | 0 | 0 | 0 | 0 | 0 | 0 | 0 | 0 | NA | NA | NA | NA | NA | NA | NA | NA | NA | NA |
| Scgb2b2 | 0 | 0 | 0 | 0 | 0 | 0 | 0 | 0 | 0 | 0 | 0 | 0 | 0 | 0 | 0 | NA | NA | NA | NA | NA | NA | NA | NA | NA | NA |
| Scgb2b20 | 0 | 0 | 0 | 0 | 0 | 0 | 0 | 0 | 0 | 0 | 0 | 0 | 0 | 0 | 0 | NA | NA | NA | NA | NA | NA | NA | NA | NA | NA |
| Scgb2b23-ps | 0 | 0 | 0 | 0 | 0 | 0 | 0 | 0 | 0 | 0 | 0 | 0 | 0 | 0 | 0 | NA | NA | NA | NA | NA | NA | NA | NA | NA | NA |
| Scgb2b24 | 0 | 0 | 0 | 0 | 0 | 0 | 0 | 0 | 0 | 0 | 0 | 0 | 0 | 0 | 0 | NA | NA | NA | NA | NA | NA | NA | NA | NA | NA |
| Scgb2b26 | 0 | 0 | 0 | 0 | 0 | 0 | 0 | 0 | 0 | 0 | 0 | 0 | 0 | 0 | 0 | NA | NA | NA | NA | NA | NA | NA | NA | NA | NA |
| Scgb2b27 | 0 | 0 | 0 | 0 | 0 | 0 | 0 | 0 | 0 | 0 | 0 | 0 | 0 | 0 | 0 | NA | NA | NA | NA | NA | NA | NA | NA | NA | NA |
| Scgb2b3 | 0 | 0 | 0 | 0 | 0 | 0 | 0 | 0 | 0 | 0 | 0 | 0 | 0 | 0 | 0 | NA | NA | NA | NA | NA | NA | NA | NA | NA | NA |
| Scgb2b7 | 0 | 0 | 0 | 0 | 0 | 0 | 0 | 0 | 0 | 0 | 0 | 0 | 0 | 0 | 0 | NA | NA | NA | NA | NA | NA | NA | NA | NA | NA |
| Scgb3a1 | 4 | 5 | 8 | 8 | 4 | 6 | 4 | 5 | 8 | 6 | 4 | 7 | 5.74 | 6 | 6 | 0.995 | -0.007 | 0.9928239 | 0.99616826 | 0 | 0.5809 | 0.2639 | 0.2639 | TRUE | 0.2734 |
| Scgb3a2 | 0 | 0 | 0 | 0 | 0 | 0 | 0 | 0 | 0 | 0 | 0 | 0 | 0 | 0 | 0 | NA | NA | NA | NA | NA | NA | NA | NA | NA | NA |
| Scgn | 5 | 4 | 4 | 3 | 1 | 2 | 5 | 4 | 4 | 2 | 1 | 2 | 3.16 | 4 | 2 | 0.429 | -1.222 | 0.25001192 | NA | 0 | 1.0502 | 0.4423 | 0.4423 | TRUE | 0.1047 |
| Schip1 | 150 | 150 | 209 | 143 | 106 | 144 | 159 | 150 | 210 | 112 | 114 | 158 | 150.51 | 173 | 128 | 0.738 | -0.439 | 0.04796649 | 0.15936591 | 0.0285 | 0.0288 | 0.0287 | 0.0287 | TRUE | 0.4056 |
| Scimp | 0 | 0 | 0 | 0 | 0 | 1 | 0 | 0 | 0 | 0 | 0 | 1 | 0.18 | 0 | 0 | 1.859 | 0.894 | 0.82650734 | NA | 0 | 18.0643 | 10 | 10 | TRUE | 0.5383 |
| Scin | 11 | 2 | 5 | 31 | 12 | 14 | 12 | 2 | 5 | 24 | 13 | 15 | 11.88 | 6 | 17 | 2.854 | 1.513 | 0.03195783 | 0.12038187 | 0.1969 | 0.2841 | 0.2482 | 0.2482 | TRUE | 1.3043 |
| Sclt1 | 403 | 316 | 389 | 673 | 471 | 438 | 428 | 316 | 391 | 529 | 507 | 479 | 441.59 | 378 | 505 | 1.338 | 0.42 | 0.00307604 | 0.02165876 | 0.0101 | 0.0144 | 0.0122 | 0.0122 | TRUE | 0.241 |
| Scly | 513 | 484 | 483 | 637 | 456 | 380 | 545 | 483 | 485 | 501 | 491 | 416 | 486.77 | 504 | 469 | 0.932 | -0.102 | 0.40993521 | 0.62158382 | 0.0052 | 0.0137 | 0.009 | 0.009 | TRUE | 0.1114 |

| | | | | | | | | | | | | | | | | | | | | | | | | |
|---|---|---|---|---|---|---|---|---|---|---|---|---|---|---|---|---|---|---|---|---|---|---|---|---|
| Scmh1 | 927 | 964 | 998 | 1239 | 830 | 866 | 985 | 963 | 1002 | 974 | 893 | 948 | 960.75 | 983 | 938 | 0.955 | -0.066 | 0.43523538 | 0.64352914 | 0 | 0.0103 | 0.0041 | 0.0041 | TRUE | 0.0211 |
| Scml2 | 861 | 1307 | 1208 | 3631 | 2812 | 2511 | 915 | 1305 | 1213 | 2854 | 3026 | 2749 | 2010.24 | 1144 | 2876 | 2.513 | 1.329 | 2.70E-22 | 8.50E-20 | 0.0162 | 0.0085 | 0.0129 | 0.0129 | TRUE | 0.3682 |
| Scml4 | 18 | 14 | 13 | 20 | 23 | 13 | 19 | 14 | 13 | 16 | 25 | 14 | 16.81 | 15 | 18 | 1.182 | 0.242 | 0.62005764 | 0.78498274 | 0 | 0.2029 | 0.1109 | 0.1109 | TRUE | 0.4671 |
| Scn10a | 0 | 0 | 0 | 3 | 3 | 0 | 0 | 0 | 0 | 2 | 3 | 0 | 0.93 | 0 | 2 | 9.951 | 3.315 | 0.21293448 | NA | 0 | 3.5454 | 3.8418 | 3.8418 | TRUE | 0.2828 |
| Scn11a | 19 | 24 | 12 | 31 | 15 | 18 | 20 | 24 | 12 | 24 | 16 | 20 | 19.4 | 19 | 20 | 1.079 | 0.11 | 0.81390925 | 0.90478612 | 0 | 0.1767 | 0.1056 | 0.1056 | TRUE | 0.5153 |
| Scn1a | 11 | 13 | 8 | 14 | 5 | 7 | 12 | 13 | 8 | 11 | 5 | 8 | 9.46 | 11 | 8 | 0.747 | -0.421 | 0.5059487 | 0.70029387 | 0 | 0.3552 | 0.1816 | 0.1816 | TRUE | 0.4024 |
| Scn1b | 262 | 576 | 615 | 474 | 382 | 518 | 278 | 575 | 618 | 373 | 411 | 567 | 470.29 | 490 | 450 | 0.917 | -0.125 | 0.65681489 | 0.80670515 | 0.1074 | 0.0139 | 0.0546 | 0.0546 | TRUE | 1.6599 |
| Scn2a1 | 22 | 26 | 32 | 46 | 34 | 26 | 23 | 26 | 32 | 36 | 37 | 28 | 30.45 | 27 | 34 | 1.245 | 0.316 | 0.37096037 | 0.58772345 | 0 | 0.1151 | 0.0564 | 0.0564 | TRUE | 0.1637 |
| Scn2b | 222 | 189 | 225 | 305 | 208 | 230 | 236 | 189 | 226 | 240 | 224 | 252 | 227.64 | 217 | 239 | 1.1 | 0.138 | 0.36483183 | 0.5822837 | 0.0035 | 0.0214 | 0.0123 | 0.0123 | TRUE | 0.1417 |
| Scn3a | 82 | 36 | 25 | 149 | 80 | 51 | 87 | 36 | 25 | 117 | 86 | 56 | 67.86 | 49 | 86 | 1.757 | 0.813 | 0.08163635 | 0.22776729 | 0.2452 | 0.0554 | 0.141 | 0.141 | TRUE | 0.53 |
| Scn3b | 297 | 267 | 232 | 355 | 283 | 288 | 315 | 267 | 233 | 279 | 304 | 315 | 285.65 | 272 | 299 | 1.102 | 0.14 | 0.36752991 | 0.58475554 | 0.0097 | 0.0184 | 0.0139 | 0.0139 | TRUE | 0.2201 |
| Scn4a | 50 | 25 | 17 | 73 | 52 | 18 | 53 | 25 | 17 | 57 | 56 | 20 | 38.03 | 32 | 44 | 1.406 | 0.492 | 0.36048625 | 0.57896411 | 0.2807 | 0.0935 | 0.1811 | 0.1811 | TRUE | 1.7877 |
| Scn4b | 19 | 37 | 31 | 27 | 20 | 16 | 20 | 37 | 31 | 21 | 22 | 18 | 24.75 | 29 | 20 | 0.683 | -0.55 | 0.18163487 | 0.38350003 | 0 | 0.14 | 0.0805 | 0.0805 | TRUE | 0.3353 |
| Scn5a | 73 | 111 | 82 | 122 | 94 | 72 | 78 | 111 | 82 | 96 | 101 | 79 | 91.1 | 90 | 92 | 1.019 | 0.028 | 0.90675713 | 0.95352883 | 0.0157 | 0.0431 | 0.0297 | 0.0297 | TRUE | 0.3844 |
| Scn7a | 2 | 2 | 1 | 4 | 4 | 1 | 2 | 2 | 1 | 3 | 4 | 1 | 2.28 | 2 | 3 | 1.678 | 0.747 | 0.5511416 | NA | 0 | 1.453 | 0.6611 | 0.6611 | TRUE | 0.2802 |
| Scn8a | 277 | 441 | 470 | 485 | 374 | 411 | 294 | 440 | 472 | 381 | 402 | 450 | 406.69 | 402 | 411 | 1.021 | 0.03 | 0.87291028 | 0.93752406 | 0.0313 | 0.015 | 0.0231 | 0.0231 | TRUE | 0.6306 |
| Scn9a | 35 | 35 | 38 | 37 | 29 | 27 | 37 | 35 | 38 | 29 | 31 | 30 | 33.36 | 37 | 30 | 0.814 | -0.297 | 0.35472303 | 0.57316787 | 0 | 0.1057 | 0.0442 | 0.0442 | TRUE | 0.0137 |
| Scnm1 | 616 | 635 | 603 | 907 | 618 | 658 | 654 | 634 | 605 | 713 | 665 | 720 | 665.36 | 631 | 699 | 1.108 | 0.149 | 0.11978314 | 0.29349382 | 0 | 0.0119 | 0.0051 | 0.0051 | TRUE | 0.0214 |
| Scnn1a | 63 | 118 | 105 | 45 | 45 | 67 | 67 | 118 | 105 | 35 | 48 | 73 | 74.56 | 97 | 52 | 0.537 | -0.897 | 0.00899507 | 0.04865233 | 0.0917 | 0.0511 | 0.0703 | 0.0703 | TRUE | 0.8606 |
| Scnn1b | 4 | 6 | 3 | 15 | 11 | 12 | 4 | 6 | 3 | 12 | 12 | 13 | 8.34 | 4 | 12 | 2.768 | 1.469 | 0.0321315 | 0.12074768 | 0 | 0.4021 | 0.1832 | 0.1832 | TRUE | 0.1826 |
| Scnn1g | 1 | 2 | 3 | 0 | 0 | 3 | 1 | 2 | 3 | 0 | 0 | 3 | 1.56 | 2 | 1 | 0.517 | -0.952 | 0.59684197 | NA | 0.7602 | 2.1197 | 1.6142 | 1.6142 | TRUE | 1.3778 |
| Sco1 | 1981 | 2343 | 2239 | 2363 | 1822 | 1837 | 2104 | 2340 | 2248 | 1858 | 1960 | 2011 | 2086.87 | 2231 | 1943 | 0.87 | -0.2 | 0.01340972 | 0.06511642 | 0.0018 | 0.0085 | 0.0042 | 0.0042 | TRUE | 0.0297 |
| Sco2 | 77 | 125 | 129 | 140 | 118 | 93 | 82 | 125 | 130 | 110 | 127 | 102 | 112.5 | 112 | 113 | 1.006 | 0.009 | 0.96944438 | 0.98448447 | 0.0263 | 0.0362 | 0.0314 | 0.0314 | TRUE | 0.5468 |
| Scoc | 1285 | 1500 | 1493 | 1759 | 1182 | 1234 | 1365 | 1498 | 1499 | 1383 | 1272 | 1351 | 1394.57 | 1454 | 1335 | 0.919 | -0.123 | 0.152087 | 0.3431743 | 0.0016 | 0.0093 | 0.0046 | 0.0046 | TRUE | 0.0344 |
| Scp2 | 2325 | 2649 | 2674 | 3327 | 2458 | 2615 | 2470 | 2646 | 2685 | 2615 | 2645 | 2863 | 2653.79 | 2600 | 2708 | 1.041 | 0.058 | 0.46237905 | 0.66599473 | 0.0017 | 0.0081 | 0.0041 | 0.0041 | TRUE | 0.0302 |
| Scp2d1 | 1 | 0 | 0 | 0 | 0 | 0 | 1 | 0 | 0 | 0 | 0 | 0 | 0.18 | 0 | 0 | 0.49 | -1.029 | 0.80087 | NA | 0 | 18.6165 | 10 | 10 | TRUE | 0.4614 |
| Scpep1 | 12042 | 16355 | 16382 | 14927 | 10945 | 13293 | 12791 | 16334 | 16449 | 11734 | 11776 | 14551 | 13939.29 | 15191 | 12687 | 0.835 | -0.26 | 0.05041509 | 0.16500246 | 0.0174 | 0.0071 | 0.0127 | 0.0127 | TRUE | 0.2336 |
| Scrg1 | 0 | 0 | 0 | 0 | 0 | 0 | 0 | 0 | 0 | 0 | 0 | 0 | 0 | 0 | 0 | NA | NA | NA | NA | NA | NA | NA | NA | NA | NA |
| Scrib | 8525 | 8231 | 8094 | 10044 | 7568 | 7076 | 9055 | 8220 | 8127 | 7896 | 8143 | 7746 | 8197.83 | 8467 | 7928 | 0.936 | -0.095 | 0.19654409 | 0.40305465 | 0.002 | 0.0073 | 0.0038 | 0.0038 | TRUE | 0.0449 |
| Scrn1 | 351 | 490 | 507 | 385 | 311 | 303 | 373 | 489 | 509 | 303 | 335 | 332 | 390.04 | 457 | 323 | 0.705 | -0.503 | 0.0009952 | 0.00888208 | 0.0132 | 0.0153 | 0.0142 | 0.0142 | TRUE | 0.2992 |
| Scrn2 | 542 | 657 | 722 | 774 | 571 | 560 | 576 | 656 | 725 | 608 | 614 | 613 | 632.11 | 652 | 612 | 0.938 | -0.093 | 0.42493646 | 0.63508934 | 0.005 | 0.0121 | 0.0081 | 0.0081 | TRUE | 0.1232 |
| Scrn3 | 850 | 1172 | 1279 | 999 | 768 | 850 | 903 | 1170 | 1284 | 785 | 826 | 930 | 983.28 | 1119 | 847 | 0.757 | -0.403 | 0.00665667 | 0.03878427 | 0.0176 | 0.0103 | 0.0148 | 0.0148 | TRUE | 0.3412 |
| Scrt1 | 2 | 5 | 1 | 5 | 2 | 4 | 2 | 5 | 1 | 4 | 2 | 4 | 3.1 | 3 | 3 | 1.29 | 0.367 | 0.73890628 | NA | 0 | 1.0706 | 0.548 | 0.548 | TRUE | 0.6608 |
| Scrt2 | 2 | 7 | 6 | 2 | 4 | 4 | 2 | 7 | 6 | 2 | 4 | 4 | 4.23 | 5 | 3 | 0.659 | -0.601 | 0.53632157 | NA | 0 | 0.7853 | 0.4338 | 0.4338 | TRUE | 0.4937 |
| Sct | 60 | 87 | 72 | 106 | 55 | 66 | 64 | 87 | 72 | 83 | 59 | 72 | 72.94 | 74 | 71 | 0.967 | -0.048 | 0.85043219 | 0.92561014 | 0.0136 | 0.0521 | 0.0332 | 0.0332 | TRUE | 0.2087 |
| Sctr | 5 | 0 | 3 | 5 | 2 | 6 | 5 | 0 | 3 | 4 | 2 | 7 | 3.5 | 3 | 4 | 1.526 | 0.61 | 0.60807046 | NA | 0.3534 | 0.9494 | 0.7169 | 0.7169 | TRUE | 0.3973 |
| Scube1 | 109 | 118 | 113 | 141 | 115 | 101 | 116 | 118 | 113 | 111 | 124 | 111 | 115.37 | 116 | 115 | 0.993 | -0.01 | 0.95615991 | 0.97741618 | 0 | 0.0355 | 0.0151 | 0.0151 | TRUE | 0.0419 |
| Scube2 | 132 | 131 | 122 | 99 | 76 | 85 | 140 | 131 | 123 | 78 | 82 | 93 | 107.7 | 131 | 84 | 0.64 | -0.644 | 0.00099306 | 0.00886889 | 0 | 0.0375 | 0.0178 | 0.0178 | TRUE | 0.0761 |
| Scube3 | 400 | 394 | 460 | 649 | 478 | 404 | 425 | 393 | 462 | 510 | 514 | 442 | 457.83 | 427 | 489 | 1.147 | 0.198 | 0.11037787 | 0.27886389 | 0.0044 | 0.0141 | 0.0088 | 0.0088 | TRUE | 0.0789 |
| Scx | 15 | 35 | 35 | 22 | 24 | 23 | 16 | 35 | 35 | 17 | 26 | 25 | 25.72 | 29 | 23 | 0.786 | -0.347 | 0.43926373 | 0.6465162 | 0.0733 | 0.135 | 0.1055 | 0.1055 | TRUE | 0.9486 |
| Scyl1 | 1728 | 1426 | 1343 | 2167 | 1585 | 1480 | 1835 | 1424 | 1349 | 1704 | 1705 | 1620 | 1606.18 | 1536 | 1676 | 1.092 | 0.127 | 0.32832297 | 0.54936451 | 0.0127 | 0.009 | 0.0114 | 0.0114 | TRUE | 0.3495 |
| Scyl2 | 2833 | 2882 | 2852 | 3866 | 2677 | 2764 | 3009 | 2878 | 2864 | 3039 | 2880 | 3026 | 2949.37 | 2917 | 2982 | 1.022 | 0.032 | 0.63594872 | 0.79536224 | 5.00E-04 | 0.008 | 0.0029 | 0.0029 | TRUE | 0.0107 |
| Scyl3 | 1105 | 1146 | 1019 | 1551 | 1042 | 875 | 1174 | 1145 | 1023 | 1219 | 1121 | 958 | 1106.61 | 1114 | 1099 | 0.988 | -0.017 | 0.88596658 | 0.94396169 | 0.0091 | 0.0099 | 0.0095 | 0.0095 | TRUE | 0.1508 |
| Sdad1 | 7934 | 7540 | 7923 | 10557 | 7843 | 7754 | 8427 | 7530 | 7956 | 8299 | 8439 | 8488 | 8189.79 | 7971 | 8409 | 1.055 | 0.077 | 0.2706708 | 0.49074002 | 0.0015 | 0.0073 | 0.0034 | 0.0034 | TRUE | 0.0306 |
| Sdc1 | 1491 | 2142 | 2237 | 2438 | 1911 | 2087 | 1584 | 2139 | 2246 | 1917 | 2056 | 2285 | 2037.73 | 1990 | 2086 | 1.048 | 0.067 | 0.64259202 | 0.79888727 | 0.0205 | 0.0085 | 0.0147 | 0.0147 | TRUE | 0.3846 |
| Sdc2 | 295 | 371 | 344 | 624 | 442 | 451 | 313 | 371 | 345 | 491 | 476 | 494 | 414.85 | 343 | 487 | 1.418 | 0.504 | 1.44E-05 | 0.00026945 | 0 | 0.0148 | 0.0072 | 0.0072 | TRUE | 0.0646 |
| Sdc3 | 805 | 1062 | 913 | 689 | 566 | 620 | 855 | 1061 | 917 | 542 | 609 | 679 | 776.96 | 944 | 610 | 0.645 | -0.633 | 1.54E-06 | 3.99E-05 | 0.0105 | 0.0111 | 0.0112 | 0.0112 | TRUE | 0.1391 |
| Sdc4 | 3176 | 4352 | 3761 | 4484 | 3459 | 4100 | 3373 | 4346 | 3776 | 3525 | 3722 | 4488 | 3871.84 | 3832 | 3912 | 1.02 | 0.029 | 0.8236212 | 0.91017184 | 0.0159 | 0.0078 | 0.0121 | 0.0121 | TRUE | 0.202 |
| Sdcbp | 2501 | 2601 | 2561 | 3830 | 2498 | 2686 | 2657 | 2598 | 2572 | 3011 | 2688 | 2940 | 2744.08 | 2609 | 2880 | 1.104 | 0.143 | 0.06197598 | 0.19032032 | 0.0016 | 0.0081 | 0.0039 | 0.0039 | TRUE | 0.041 |
| Sdcbp2 | 131 | 253 | 258 | 176 | 161 | 149 | 139 | 253 | 259 | 138 | 173 | 163 | 187.59 | 217 | 158 | 0.728 | -0.459 | 0.06924613 | 0.20425473 | 0.0578 | 0.0245 | 0.0405 | 0.0405 | TRUE | 1.0685 |
| Sdccag3 | 2763 | 2802 | 2569 | 3236 | 2294 | 2213 | 2935 | 2798 | 2580 | 2544 | 2468 | 2422 | 2624.55 | 2771 | 2478 | 0.895 | -0.161 | 0.04553225 | 0.15417008 | 0.0018 | 0.0082 | 0.0043 | 0.0043 | TRUE | 0.0443 |
| Sdccag8 | 164 | 131 | 181 | 234 | 170 | 157 | 174 | 131 | 182 | 184 | 183 | 172 | 170.92 | 162 | 180 | 1.108 | 0.148 | 0.41409252 | 0.62570102 | 0.0092 | 0.0262 | 0.0177 | 0.0177 | TRUE | 0.3072 |
| Sde2 | 3809 | 3680 | 3378 | 5114 | 3732 | 3267 | 4046 | 3675 | 3392 | 4020 | 4015 | 3576 | 3787.47 | 3704 | 3870 | 1.045 | 0.064 | 0.51456897 | 0.70725471 | 0.0051 | 0.0078 | 0.0066 | 0.0066 | TRUE | 0.079 |
| Sdf2 | 1723 | 2304 | 2336 | 2110 | 1631 | 1913 | 1830 | 2301 | 2346 | 1659 | 1755 | 2094 | 1997.4 | 2159 | 1836 | 0.85 | -0.235 | 0.08219148 | 0.22887185 | 0.0158 | 0.0086 | 0.0126 | 0.0126 | TRUE | 0.2144 |
| Sdf2l1 | 2275 | 2749 | 2685 | 2374 | 1716 | 1947 | 2416 | 2745 | 2696 | 1866 | 1846 | 2131 | 2283.63 | 2619 | 1948 | 0.743 | -0.428 | 1.15E-05 | 0.0002238 | 0.005 | 0.0083 | 0.0064 | 0.0064 | TRUE | 0.0812 |
| Sdf4 | 4211 | 4576 | 4450 | 6771 | 4729 | 4396 | 4473 | 4570 | 4468 | 5323 | 5088 | 4812 | 4789.04 | 4504 | 5074 | 1.127 | 0.173 | 0.01302744 | 0.06391556 | 0.001 | 0.0076 | 0.0033 | 0.0033 | TRUE | 0.0249 |
| Sdha | 17682 | 19379 | 19672 | 20610 | 14770 | 15708 | 18781 | 19354 | 19753 | 16202 | 15892 | 17195 | 17862.77 | 19296 | 16430 | 0.851 | -0.232 | 0.00030871 | 0.00349026 | 0.0011 | 0.0071 | 0.0029 | 0.0029 | TRUE | 0.0203 |
| Sdhaf1 | 348 | 446 | 399 | 547 | 369 | 368 | 370 | 445 | 401 | 430 | 397 | 403 | 407.59 | 405 | 410 | 1.012 | 0.018 | 0.88501303 | 0.94366363 | 0.0028 | 0.015 | 0.0084 | 0.0084 | TRUE | 0.087 |
| Sdhaf2 | 1117 | 1053 | 968 | 1749 | 1220 | 1063 | 1186 | 1052 | 972 | 1375 | 1313 | 1164 | 1176.88 | 1070 | 1284 | 1.201 | 0.264 | 0.02153097 | 0.09168102 | 0.0074 | 0.0097 | 0.0086 | 0.0086 | TRUE | 0.1079 |
| Sdhb | 4069 | 5273 | 5237 | 5447 | 3962 | 4310 | 4322 | 5266 | 5259 | 4282 | 4263 | 4718 | 4684.92 | 4949 | 4421 | 0.893 | -0.163 | 0.11940994 | 0.29280684 | 0.0074 | 0.0076 | 0.0077 | 0.0077 | TRUE | 0.1495 |
| Sdhc | 5975 | 7634 | 7709 | 7299 | 5353 | 6014 | 6347 | 7624 | 7741 | 5738 | 5759 | 6583 | 6632 | 7237 | 6027 | 0.833 | -0.264 | 0.01450885 | 0.06905059 | 0.0087 | 0.0074 | 0.0083 | 0.0083 | TRUE | 0.1413 |
| Sdhd | 5613 | 6145 | 6033 | 7371 | 5064 | 5415 | 5962 | 6137 | 6058 | 5795 | 5449 | 5928 | 5887.91 | 6052 | 5724 | 0.946 | -0.081 | 0.22032822 | 0.43242065 | 8.00E-04 | 0.0075 | 0.0029 | 0.0029 | TRUE | 0.0214 |
| Sdk1 | 40 | 72 | 64 | 85 | 38 | 46 | 42 | 72 | 64 | 67 | 41 | 50 | 56.12 | 59 | 53 | 0.889 | -0.169 | 0.60078008 | 0.77266232 | 0.0487 | 0.0656 | 0.0574 | 0.0574 | TRUE | 0.5242 |
| Sdk2 | 673 | 866 | 1026 | 1157 | 861 | 817 | 715 | 865 | 1030 | 910 | 926 | 894 | 890.03 | 870 | 910 | 1.046 | 0.065 | 0.64621921 | 0.80133418 | 0.0154 | 0.0106 | 0.0132 | 0.0132 | TRUE | 0.2192 |
| Sdpr | 7 | 3 | 0 | 13 | 4 | 4 | 7 | 3 | 0 | 10 | 4 | 4 | 4.89 | 3 | 6 | 1.847 | 0.885 | 0.42094835 | 0.63116133 | 0.5947 | 0.6808 | 0.647 | 0.647 | TRUE | 0.5595 |
| Sdr16c5 | 0 | 0 | 0 | 0 | 0 | 0 | 0 | 0 | 0 | 0 | 0 | 0 | 0 | 0 | 0 | NA | NA | NA | NA | NA | NA | NA | NA | NA | NA |
| Sdr16c6 | 0 | 0 | 0 | 0 | 0 | 0 | 0 | 0 | 0 | 0 | 0 | 0 | 0 | 0 | 0 | NA | NA | NA | NA | NA | NA | NA | NA | NA | NA |
| Sdr39u1 | 563 | 811 | 733 | 951 | 756 | 720 | 598 | 810 | 736 | 748 | 813 | 788 | 748.86 | 715 | 783 | 1.095 | 0.131 | 0.32127207 | 0.54237525 | 0.0111 | 0.0113 | 0.0112 | 0.0112 | TRUE | 0.2396 |
| Sdr42e1 | 94 | 106 | 94 | 114 | 73 | 89 | 100 | 106 | 94 | 90 | 79 | 97 | 94.28 | 100 | 89 | 0.885 | -0.176 | 0.3919725 | 0.60563505 | 0 | 0.0418 | 0.0198 | 0.0198 | TRUE | 0.0857 |

| | | | | | | | | | | | | | | | | | | | | | | | | | |
|---|---|---|---|---|---|---|---|---|---|---|---|---|---|---|---|---|---|---|---|---|---|---|---|---|---|
| Sdr9c7 | 5 | 1 | 0 | 4 | 1 | 2 | 5 | 1 | 0 | 3 | 1 | 2 | 2.12 | 2 | 2 | 1.039 | 0.055 | 0.97042714 | NA | 0.628 | 1.561 | 1.131 | 1.131 | TRUE | 1.5964 |
| Sds | 9 | 9 | 7 | 8 | 2 | 3 | 10 | 9 | 7 | 6 | 2 | 3 | 6.22 | 9 | 4 | 0.473 | -1.081 | 0.17177684 | 0.37062439 | 0 | 0.5368 | 0.27 | 0.27 | TRUE | 0.6143 |
| Sdsl | 7 | 3 | 3 | 4 | 0 | 0 | 7 | 3 | 3 | 3 | 0 | 0 | 2.76 | 4 | 1 | 0.258 | -1.957 | 0.17857981 | NA | 0.5488 | 1.1985 | 0.9872 | 0.9872 | TRUE | 1.8893 |
| Sebox | 0 | 1 | 0 | 2 | 0 | 3 | 0 | 1 | 0 | 2 | 0 | 3 | 0.98 | 0 | 2 | 4.416 | 2.143 | 0.35716898 | NA | 0 | 3.383 | 2.5996 | 2.5996 | TRUE | 0.1602 |
| Sec1 | 26 | 17 | 12 | 48 | 32 | 17 | 28 | 17 | 12 | 38 | 34 | 19 | 24.57 | 19 | 30 | 1.616 | 0.692 | 0.14996762 | 0.34065871 | 0.1037 | 0.141 | 0.1234 | 0.1234 | TRUE | 0.815 |
| Sec11a | 1538 | 1891 | 1795 | 2135 | 1543 | 1630 | 1634 | 1889 | 1802 | 1678 | 1660 | 1784 | 1741.23 | 1775 | 1707 | 0.962 | -0.056 | 0.52863414 | 0.71718529 | 0.0026 | 0.0088 | 0.0052 | 0.0052 | TRUE | 0.0582 |
| Sec11c | 684 | 757 | 706 | 708 | 459 | 493 | 727 | 756 | 709 | 557 | 494 | 540 | 630.26 | 731 | 530 | 0.726 | -0.461 | 4.09E-06 | 9.32E-05 | 0 | 0.0121 | 0.0056 | 0.0056 | TRUE | 0.041 |
| Sec13 | 6577 | 6210 | 5956 | 8324 | 5529 | 5620 | 6986 | 6202 | 5980 | 6544 | 5949 | 6152 | 6302.15 | 6389 | 6215 | 0.973 | -0.04 | 0.65490777 | 0.80596172 | 0.0038 | 0.0074 | 0.0055 | 0.0055 | TRUE | 0.0813 |
| Sec14l1 | 7826 | 8307 | 8715 | 9326 | 7246 | 7988 | 8313 | 8296 | 8751 | 7331 | 7796 | 8744 | 8205.23 | 8453 | 7957 | 0.941 | -0.088 | 0.32090958 | 0.54202034 | 0.0044 | 0.0073 | 0.0055 | 0.0055 | TRUE | 0.0913 |
| Sec14l2 | 269 | 263 | 239 | 457 | 341 | 328 | 286 | 263 | 240 | 359 | 367 | 359 | 312.26 | 263 | 362 | 1.377 | 0.462 | 0.00027407 | 0.00315746 | 0 | 0.0175 | 0.0083 | 0.0083 | TRUE | 0.0644 |
| Sec14l3 | 5 | 9 | 7 | 4 | 5 | 4 | 5 | 9 | 7 | 3 | 5 | 4 | 5.71 | 7 | 4 | 0.596 | -0.747 | 0.34435111 | 0.56409489 | 0 | 0.5844 | 0.2628 | 0.2628 | TRUE | 0.1451 |
| Sec14l4 | 197 | 306 | 298 | 217 | 212 | 228 | 209 | 306 | 299 | 171 | 228 | 250 | 243.72 | 271 | 216 | 0.793 | -0.334 | 0.11578618 | 0.28736108 | 0.0364 | 0.0204 | 0.0283 | 0.0283 | TRUE | 0.4448 |
| Sec14l5 | 195 | 208 | 175 | 115 | 119 | 151 | 207 | 208 | 176 | 90 | 128 | 165 | 162.38 | 197 | 128 | 0.646 | -0.63 | 0.00773393 | 0.04330961 | 0.0411 | 0.0272 | 0.0339 | 0.0339 | TRUE | 0.315 |
| Sec16a | 6325 | 5479 | 6511 | 7238 | 5196 | 4930 | 6718 | 5472 | 6538 | 5690 | 5591 | 5397 | 5900.85 | 6243 | 5559 | 0.891 | -0.167 | 0.0880102 | 0.24041601 | 0.0059 | 0.0075 | 0.0067 | 0.0067 | TRUE | 0.1424 |
| Sec16b | 69 | 50 | 39 | 69 | 57 | 56 | 73 | 50 | 39 | 54 | 61 | 61 | 56.54 | 54 | 59 | 1.089 | 0.123 | 0.68796127 | 0.82822114 | 0.033 | 0.0652 | 0.0498 | 0.0498 | TRUE | 0.6133 |
| Sec22a | 952 | 935 | 853 | 1175 | 773 | 791 | 1011 | 934 | 857 | 924 | 832 | 866 | 903.79 | 934 | 874 | 0.937 | -0.094 | 0.3616446 | 0.58001409 | 0.0038 | 0.0105 | 0.0066 | 0.0066 | TRUE | 0.0626 |
| Sec22b | 2181 | 2210 | 2183 | 2913 | 1975 | 2100 | 2317 | 2207 | 2192 | 2290 | 2125 | 2299 | 2238.24 | 2239 | 2238 | 1 | 0 | 0.99974677 | 0.99974677 | 9.00E-04 | 0.0084 | 0.0036 | 0.0036 | TRUE | 0.0233 |
| Sec22c | 819 | 923 | 884 | 1033 | 770 | 776 | 870 | 922 | 888 | 812 | 828 | 849 | 861.56 | 893 | 830 | 0.929 | -0.107 | 0.2066284 | 0.41562851 | 0 | 0.0107 | 0.004 | 0.004 | TRUE | 0.0094 |
| Sec23a | 5199 | 5957 | 5932 | 6263 | 4642 | 5116 | 5522 | 5949 | 5956 | 4923 | 4995 | 5600 | 5491.03 | 5809 | 5173 | 0.89 | -0.168 | 0.04387432 | 0.15007283 | 0.0032 | 0.0075 | 0.0048 | 0.0048 | TRUE | 0.0635 |
| Sec23b | 4771 | 4598 | 4719 | 6871 | 4601 | 4539 | 5068 | 4592 | 4738 | 5401 | 4950 | 4969 | 4953.09 | 4799 | 5107 | 1.064 | 0.09 | 0.24703558 | 0.46383871 | 0.0022 | 0.0076 | 0.0042 | 0.0042 | TRUE | 0.0313 |
| Sec23ip | 4159 | 3492 | 3345 | 5113 | 3545 | 3173 | 4418 | 3487 | 3359 | 4019 | 3814 | 3473 | 3761.8 | 3755 | 3769 | 1.004 | 0.006 | 0.96253674 | 0.98130897 | 0.0137 | 0.0078 | 0.011 | 0.011 | TRUE | 0.29 |
| Sec24a | 3928 | 3852 | 3857 | 4130 | 3061 | 3083 | 4172 | 3847 | 3873 | 3247 | 3293 | 3375 | 3634.51 | 3964 | 3305 | 0.834 | -0.262 | 0.00016473 | 0.00206365 | 8.00E-04 | 0.0078 | 0.0032 | 0.0032 | TRUE | 0.0256 |
| Sec24b | 3582 | 3554 | 3664 | 4949 | 3621 | 3455 | 3805 | 3549 | 3679 | 3891 | 3896 | 3782 | 3766.95 | 3678 | 3856 | 1.049 | 0.069 | 0.2923137 | 0.51317582 | 5.00E-04 | 0.0078 | 0.0028 | 0.0028 | TRUE | 0.0113 |
| Sec24c | 8165 | 7741 | 8270 | 10104 | 7220 | 7455 | 8673 | 7731 | 8304 | 7943 | 7768 | 8161 | 8096.59 | 8236 | 7957 | 0.966 | -0.05 | 0.49451019 | 0.69052608 | 0.0018 | 0.0073 | 0.0037 | 0.0037 | TRUE | 0.0351 |
| Sec24d | 3055 | 2376 | 2384 | 3276 | 2329 | 2108 | 3245 | 2373 | 2394 | 2575 | 2506 | 2308 | 2566.73 | 2671 | 2463 | 0.922 | -0.116 | 0.39556195 | 0.60896077 | 0.0178 | 0.0082 | 0.0131 | 0.0131 | TRUE | 0.429 |
| Sec31a | 12438 | 11413 | 11625 | 13967 | 10229 | 10014 | 13211 | 11398 | 11673 | 10980 | 11006 | 10962 | 11538.3 | 12094 | 10983 | 0.908 | -0.139 | 0.08312692 | 0.2307423 | 0.0031 | 0.0072 | 0.0046 | 0.0046 | TRUE | 0.0798 |
| Sec31b | 267 | 396 | 409 | 306 | 289 | 300 | 284 | 395 | 411 | 241 | 311 | 328 | 328.28 | 363 | 293 | 0.805 | -0.313 | 0.10364287 | 0.26782464 | 0.0301 | 0.0169 | 0.0235 | 0.0235 | TRUE | 0.4179 |
| Sec61a1 | 13158 | 13646 | 13464 | 15140 | 11100 | 10796 | 13976 | 13628 | 13519 | 11902 | 11943 | 11818 | 12797.75 | 13708 | 11888 | 0.867 | -0.206 | 0.00010322 | 0.00141301 | 1.00E-04 | 0.0072 | 0.0019 | 0.0019 | TRUE | 0.0036 |
| Sec61a2 | 3051 | 3863 | 4064 | 3603 | 2686 | 2983 | 3241 | 3858 | 4081 | 2832 | 2890 | 3265 | 3361.19 | 3727 | 2996 | 0.804 | -0.315 | 0.00540185 | 0.03309943 | 0.0094 | 0.0079 | 0.009 | 0.009 | TRUE | 0.158 |
| Sec61b | 3014 | 3050 | 2892 | 4139 | 2686 | 2710 | 3201 | 3046 | 2904 | 3254 | 2890 | 2967 | 3043.6 | 3050 | 3037 | 0.996 | -0.006 | 0.94549115 | 0.97275189 | 0.0029 | 0.008 | 0.0047 | 0.0047 | TRUE | 0.0481 |
| Sec61g | 1205 | 1403 | 1329 | 1899 | 1165 | 1271 | 1280 | 1401 | 1334 | 1493 | 1253 | 1391 | 1358.87 | 1338 | 1379 | 1.031 | 0.044 | 0.65355702 | 0.80540538 | 0.0042 | 0.0093 | 0.0063 | 0.0063 | TRUE | 0.0762 |
| Sec62 | 5074 | 5492 | 5382 | 6775 | 4851 | 4857 | 5389 | 5485 | 5404 | 5326 | 5219 | 5317 | 5356.77 | 5426 | 5287 | 0.974 | -0.037 | 0.50440149 | 0.69937761 | 0 | 0.0075 | 0.0021 | 0.0021 | TRUE | 0.0015 |
| Sec63 | 3597 | 3172 | 3217 | 4483 | 3102 | 3034 | 3821 | 3168 | 3230 | 3524 | 3338 | 3321 | 3400.28 | 3406 | 3394 | 0.997 | -0.005 | 0.96171262 | 0.98074043 | 0.0056 | 0.0079 | 0.0065 | 0.0065 | TRUE | 0.1375 |
| Secisbp2 | 1877 | 1948 | 2020 | 2202 | 1714 | 1855 | 1994 | 1945 | 2028 | 1731 | 1844 | 2031 | 1928.88 | 1989 | 1869 | 0.939 | -0.091 | 0.30274491 | 0.52360605 | 0.0027 | 0.0086 | 0.0051 | 0.0051 | TRUE | 0.0693 |
| Secisbp2l | 2996 | 2712 | 2817 | 3740 | 2697 | 2570 | 3182 | 2708 | 2829 | 2940 | 2902 | 2813 | 2895.75 | 2906 | 2885 | 0.993 | -0.01 | 0.90627904 | 0.95350897 | 0.0034 | 0.008 | 0.0052 | 0.0052 | TRUE | 0.0835 |
| Sectm1a | 0 | 1 | 2 | 1 | 3 | 1 | 0 | 1 | 2 | 1 | 3 | 1 | 1.35 | 1 | 2 | 1.657 | 0.729 | 0.66738955 | NA | 0 | 2.4426 | 1.2739 | 1.2739 | TRUE | 0.299 |
| Sectm1b | 0 | 0 | 0 | 0 | 0 | 0 | 0 | 0 | 0 | 0 | 0 | 0 | 0 | 0 | 0 | NA | NA | NA | NA | NA | NA | NA | NA | NA | NA |
| Seh1l | 7034 | 8764 | 9388 | 9616 | 7184 | 7895 | 7471 | 8753 | 9427 | 7559 | 7730 | 8642 | 8263.63 | 8550 | 7977 | 0.933 | -0.1 | 0.35940888 | 0.57789754 | 0.0091 | 0.0073 | 0.0085 | 0.0085 | TRUE | 0.1487 |
| Sel1l | 4160 | 4059 | 4010 | 5265 | 3850 | 3653 | 4419 | 4054 | 4026 | 4139 | 4142 | 3999 | 4129.83 | 4166 | 4093 | 0.983 | -0.025 | 0.72438885 | 0.85090198 | 0.0011 | 0.0077 | 0.0035 | 0.0035 | TRUE | 0.0341 |
| Sel1l2 | 0 | 0 | 0 | 1 | 1 | 0 | 0 | 0 | 0 | 1 | 1 | 0 | 0.31 | 0 | 1 | 3.32 | 1.731 | 0.65383052 | NA | 0 | 10.6224 | 8.9541 | 8.9541 | TRUE | 0.202 |
| Sel1l3 | 54 | 98 | 111 | 87 | 71 | 85 | 57 | 98 | 111 | 68 | 76 | 93 | 84.09 | 89 | 79 | 0.888 | -0.171 | 0.56207097 | 0.74295871 | 0.0561 | 0.0461 | 0.0509 | 0.0509 | TRUE | 0.8773 |
| Sele | 0 | 1 | 0 | 1 | 1 | 1 | 0 | 1 | 0 | 1 | 1 | 1 | 0.66 | 0 | 1 | 2.676 | 1.42 | 0.53620179 | NA | 0 | 5.0043 | 2.2992 | 2.2992 | TRUE | 0.405 |
| Selenbp1 | 254 | 407 | 391 | 336 | 273 | 308 | 270 | 406 | 393 | 264 | 294 | 337 | 327.32 | 356 | 298 | 0.836 | -0.259 | 0.17237297 | 0.37119832 | 0.0288 | 0.017 | 0.0229 | 0.0229 | TRUE | 0.5114 |
| Selenbp2 | 15 | 12 | 32 | 21 | 11 | 24 | 16 | 12 | 32 | 17 | 12 | 26 | 19.11 | 20 | 18 | 0.906 | -0.142 | 0.79735635 | 0.89565589 | 0.1568 | 0.1793 | 0.1683 | 0.1683 | TRUE | 1.1516 |
| Selk | 764 | 742 | 727 | 1141 | 785 | 695 | 812 | 741 | 730 | 897 | 845 | 761 | 797.48 | 761 | 834 | 1.098 | 0.135 | 0.20627059 | 0.41518859 | 0.0039 | 0.011 | 0.0069 | 0.0069 | TRUE | 0.0696 |
| Sell | 2 | 0 | 1 | 0 | 1 | 0 | 2 | 0 | 1 | 0 | 1 | 0 | 0.7 | 1 | 0 | 0.337 | -1.571 | 0.54072415 | NA | 0 | 4.7083 | 3.2627 | 3.2627 | TRUE | 0.278 |
| Selm | 572 | 586 | 506 | 677 | 464 | 480 | 608 | 585 | 508 | 532 | 499 | 525 | 542.96 | 567 | 519 | 0.916 | -0.127 | 0.26513719 | 0.48483985 | 0.0031 | 0.013 | 0.0074 | 0.0074 | TRUE | 0.0971 |
| Selp | 1 | 3 | 0 | 5 | 5 | 7 | 1 | 3 | 0 | 4 | 5 | 8 | 3.51 | 1 | 6 | 4.135 | 2.048 | 0.08707257 | NA | 0 | 0.9468 | 0.569 | 0.569 | TRUE | 0.7013 |
| Selplg | 54 | 75 | 74 | 41 | 29 | 39 | 57 | 75 | 74 | 32 | 31 | 43 | 52.11 | 69 | 35 | 0.511 | -0.968 | 0.00085604 | 0.00789098 | 0 | 0.0701 | 0.0395 | 0.0395 | TRUE | 0.2156 |
| Selrc1 | 1094 | 1004 | 891 | 1466 | 961 | 820 | 1162 | 1003 | 895 | 1152 | 1034 | 898 | 1023.91 | 1020 | 1028 | 1.009 | 0.013 | 0.92471614 | 0.96292 | 0.0154 | 0.0101 | 0.0129 | 0.0129 | TRUE | 0.1773 |
| Sema3a | 3 | 4 | 4 | 27 | 24 | 14 | 3 | 4 | 4 | 21 | 26 | 15 | 12.26 | 4 | 21 | 5.565 | 2.476 | 0.00013723 | 0.00177697 | 0 | 0.2756 | 0.1439 | 0.1439 | TRUE | 0.0955 |
| Sema3b | 99 | 126 | 105 | 135 | 106 | 95 | 105 | 126 | 105 | 106 | 114 | 104 | 110.1 | 112 | 108 | 0.963 | -0.055 | 0.77500025 | 0.8829953 | 0 | 0.0368 | 0.0176 | 0.0176 | TRUE | 0.1151 |
| Sema3c | 40 | 59 | 47 | 117 | 85 | 64 | 42 | 59 | 47 | 92 | 91 | 70 | 67.02 | 49 | 84 | 1.708 | 0.772 | 0.00334318 | 0.02305505 | 0.0093 | 0.0561 | 0.0339 | 0.0339 | TRUE | 0.2246 |
| Sema3d | 20 | 11 | 8 | 33 | 20 | 21 | 21 | 11 | 8 | 26 | 22 | 23 | 18.45 | 13 | 24 | 1.762 | 0.817 | 0.10366912 | 0.26782464 | 0.0373 | 0.1854 | 0.1228 | 0.1228 | TRUE | 1.0591 |
| Sema3e | 174 | 176 | 197 | 341 | 221 | 225 | 185 | 176 | 198 | 268 | 238 | 246 | 218.42 | 186 | 251 | 1.35 | 0.433 | 0.00270696 | 0.01955757 | 0 | 0.022 | 0.0103 | 0.0103 | TRUE | 0.0439 |
| Sema3f | 314 | 285 | 337 | 239 | 227 | 229 | 334 | 285 | 338 | 188 | 244 | 251 | 273.22 | 319 | 228 | 0.711 | -0.492 | 0.00290894 | 0.02071322 | 0.013 | 0.019 | 0.0159 | 0.0159 | TRUE | 0.2765 |
| Sema3g | 20 | 18 | 23 | 32 | 29 | 9 | 21 | 18 | 23 | 25 | 31 | 10 | 21.42 | 21 | 22 | 1.067 | 0.094 | 0.85099398 | 0.92587032 | 0.1018 | 0.1607 | 0.1337 | 0.1337 | TRUE | 0.6619 |
| Sema4a | 795 | 691 | 695 | 658 | 495 | 527 | 844 | 690 | 698 | 517 | 533 | 577 | 643.19 | 744 | 542 | 0.728 | -0.457 | 0.00017658 | 0.00219462 | 0.0068 | 0.012 | 0.0091 | 0.0091 | TRUE | 0.1637 |
| Sema4b | 2327 | 2388 | 2454 | 3064 | 2434 | 2362 | 2472 | 2385 | 2464 | 2409 | 2619 | 2586 | 2488.96 | 2440 | 2538 | 1.039 | 0.056 | 0.44095306 | 0.648035 | 9.00E-04 | 0.0082 | 0.0034 | 0.0034 | TRUE | 0.0246 |
| Sema4c | 201 | 369 | 375 | 309 | 257 | 336 | 213 | 369 | 377 | 243 | 277 | 368 | 307.63 | 320 | 296 | 0.924 | -0.114 | 0.64220626 | 0.79855543 | 0.0664 | 0.0176 | 0.0404 | 0.0404 | TRUE | 0.9494 |
| Sema4d | 2475 | 2636 | 2573 | 3097 | 2333 | 2337 | 2629 | 2633 | 2584 | 2435 | 2510 | 2558 | 2558.01 | 2615 | 2501 | 0.956 | -0.065 | 0.31785349 | 0.53882043 | 0 | 0.0082 | 0.0026 | 0.0026 | TRUE | 0.0067 |
| Sema4f | 439 | 510 | 507 | 571 | 464 | 441 | 466 | 509 | 509 | 449 | 499 | 483 | 485.93 | 495 | 477 | 0.962 | -0.056 | 0.60720374 | 0.77714983 | 0 | 0.0137 | 0.0064 | 0.0064 | TRUE | 0.0329 |
| Sema4g | 1732 | 1748 | 1779 | 1366 | 1174 | 1292 | 1840 | 1746 | 1786 | 1074 | 1263 | 1414 | 1520.5 | 1791 | 1250 | 0.698 | -0.52 | 7.70E-06 | 0.00015872 | 0.0086 | 0.0091 | 0.009 | 0.009 | TRUE | 0.1875 |
| Sema5a | 201 | 238 | 277 | 299 | 222 | 280 | 213 | 238 | 278 | 235 | 239 | 307 | 251.62 | 243 | 260 | 1.068 | 0.094 | 0.58950684 | 0.76400346 | 0.0162 | 0.02 | 0.0181 | 0.0181 | TRUE | 0.2655 |
| Sema5b | 827 | 1217 | 1304 | 1063 | 852 | 868 | 878 | 1215 | 1309 | 836 | 917 | 950 | 1017.62 | 1134 | 901 | 0.794 | -0.334 | 0.03282276 | 0.12235419 | 0.0228 | 0.0101 | 0.0166 | 0.0166 | TRUE | 0.4653 |
| Sema6a | 61 | 75 | 77 | 71 | 46 | 33 | 65 | 75 | 77 | 56 | 49 | 36 | 59.74 | 72 | 47 | 0.656 | -0.608 | 0.02613019 | 0.10470666 | 0.0061 | 0.062 | 0.0365 | 0.0365 | TRUE | 0.3135 |
| Sema6b | 483 | 596 | 489 | 341 | 263 | 297 | 513 | 595 | 491 | 268 | 283 | 325 | 412.57 | 533 | 292 | 0.547 | -0.871 | 2.62E-10 | 1.82E-08 | 0.0078 | 0.0149 | 0.0111 | 0.0111 | TRUE | 0.1222 |
| Sema6c | 1090 | 1375 | 1374 | 868 | 688 | 766 | 1158 | 1373 | 1380 | 682 | 740 | 839 | 1028.62 | 1304 | 754 | 0.577 | -0.792 | 7.49E-11 | 5.63E-09 | 0.0086 | 0.0101 | 0.0096 | 0.0096 | TRUE | 0.1145 |
| Sema6d | 10 | 15 | 6 | 14 | 15 | 10 | 11 | 15 | 6 | 11 | 16 | 11 | 11.62 | 11 | 13 | 1.199 | 0.262 | 0.65872082 | 0.80762941 | 0 | 0.2904 | 0.1667 | 0.1667 | TRUE | 0.2355 |

| | | | | | | | | | | | | | | | | | | | | | | | | | |
|---|---|---|---|---|---|---|---|---|---|---|---|---|---|---|---|---|---|---|---|---|---|---|---|---|---|
| Sema7a | 952 | 1090 | 1059 | 671 | 535 | 547 | 1011 | 1089 | 1063 | 527 | 576 | 599 | 810.84 | 1054 | 567 | 0.537 | -0.897 | 1.87E-20 | 4.91E-18 | 0.0013 | 0.011 | 0.0054 | 0.0054 | TRUE | 0.047 |
| Senp1 | 2797 | 2791 | 2926 | 4102 | 3037 | 2937 | 2971 | 2787 | 2938 | 3225 | 3268 | 3215 | 3067.27 | 2899 | 3236 | 1.116 | 0.159 | 0.01524188 | 0.0715674 | 2.00E-04 | 0.008 | 0.0028 | 0.0028 | TRUE | 0.0137 |
| Senp2 | 5384 | 5793 | 5773 | 7082 | 5278 | 5070 | 5719 | 5786 | 5797 | 5567 | 5679 | 5550 | 5682.84 | 5767 | 5599 | 0.971 | -0.043 | 0.44038592 | 0.64746865 | 0 | 0.0075 | 0.002 | 0.002 | TRUE | 0.0019 |
| Senp3 | 11916 | 12223 | 11679 | 17890 | 12152 | 10958 | 12657 | 12207 | 11727 | 14064 | 13075 | 11995 | 12620.81 | 12197 | 13045 | 1.07 | 0.097 | 0.24950115 | 0.466779 | 0.0038 | 0.0072 | 0.005 | 0.005 | TRUE | 0.0605 |
| Senp5 | 1265 | 1360 | 1323 | 1753 | 1189 | 1088 | 1344 | 1358 | 1328 | 1378 | 1279 | 1191 | 1313.11 | 1343 | 1283 | 0.956 | -0.065 | 0.46185055 | 0.66571988 | 0.002 | 0.0094 | 0.0049 | 0.0049 | TRUE | 0.0522 |
| Senp6 | 3562 | 3921 | 4018 | 5436 | 3879 | 3850 | 3783 | 3916 | 4034 | 4273 | 4174 | 4214 | 4065.88 | 3911 | 4220 | 1.079 | 0.11 | 0.08187526 | 0.22835308 | 3.00E-04 | 0.0077 | 0.0026 | 0.0026 | TRUE | 0.0099 |
| Senp7 | 380 | 418 | 399 | 330 | 271 | 291 | 404 | 417 | 401 | 259 | 292 | 319 | 348.55 | 407 | 290 | 0.709 | -0.495 | 0.00011008 | 0.00148641 | 0.0022 | 0.0164 | 0.0089 | 0.0089 | TRUE | 0.1028 |
| Senp8 | 629 | 728 | 781 | 965 | 686 | 611 | 668 | 727 | 784 | 759 | 738 | 669 | 724.15 | 726 | 722 | 0.994 | -0.008 | 0.93946261 | 0.96945854 | 0.0039 | 0.0115 | 0.0072 | 0.0072 | TRUE | 0.0579 |
| Sephs1 | 4212 | 3791 | 3719 | 5288 | 3795 | 3549 | 4474 | 3786 | 3734 | 4157 | 4083 | 3885 | 4019.9 | 3998 | 4042 | 1.011 | 0.016 | 0.86796162 | 0.93494677 | 0.0055 | 0.0077 | 0.0064 | 0.0064 | TRUE | 0.1317 |
| Sephs2 | 6674 | 5640 | 5248 | 8183 | 5894 | 4742 | 7089 | 5633 | 5270 | 6433 | 6342 | 5191 | 5992.75 | 5997 | 5989 | 0.999 | -0.002 | 0.98913823 | 0.99430772 | 0.0191 | 0.0075 | 0.0136 | 0.0136 | TRUE | 0.3093 |
| Sepn1 | 559 | 803 | 805 | 596 | 451 | 466 | 594 | 802 | 808 | 469 | 485 | 510 | 611.32 | 735 | 488 | 0.664 | -0.592 | 4.49E-05 | 0.00070855 | 0.0144 | 0.0123 | 0.0134 | 0.0134 | TRUE | 0.3314 |
| Sepp1 | 396 | 497 | 484 | 483 | 372 | 436 | 421 | 496 | 486 | 380 | 400 | 477 | 443.36 | 468 | 419 | 0.894 | -0.161 | 0.24220517 | 0.45880608 | 0.0089 | 0.0143 | 0.0114 | 0.0114 | TRUE | 0.168 |
| Sepsecs | 863 | 764 | 766 | 1103 | 827 | 785 | 917 | 763 | 769 | 867 | 890 | 859 | 844.17 | 816 | 872 | 1.069 | 0.096 | 0.37167907 | 0.58818853 | 0.0041 | 0.0108 | 0.0071 | 0.0071 | TRUE | 0.1366 |
| Sep-01 | 3116 | 2282 | 2075 | 3624 | 2720 | 2450 | 3310 | 2279 | 2084 | 2849 | 2927 | 2682 | 2688.28 | 2558 | 2819 | 1.102 | 0.141 | 0.39750888 | 0.61106336 | 0.0315 | 0.0081 | 0.0196 | 0.0196 | TRUE | 0.8024 |
| Sep-02 | 11063 | 14026 | 13933 | 16354 | 11740 | 12180 | 11751 | 14008 | 13990 | 12856 | 12632 | 13333 | 13094.94 | 13250 | 12940 | 0.977 | -0.034 | 0.71033855 | 0.84213468 | 0.0043 | 0.0072 | 0.006 | 0.006 | TRUE | 0.1197 |
| Sep-03 | 1165 | 1454 | 1562 | 1184 | 997 | 1136 | 1237 | 1452 | 1568 | 931 | 1073 | 1244 | 1250.83 | 1419 | 1083 | 0.762 | -0.393 | 0.00515313 | 0.03198394 | 0.0162 | 0.0095 | 0.0134 | 0.0134 | TRUE | 0.2029 |
| Sep-04 | 41 | 27 | 22 | 104 | 84 | 61 | 44 | 27 | 22 | 82 | 90 | 67 | 55.25 | 31 | 80 | 2.589 | 1.373 | 2.69E-05 | 0.00045797 | 0.0374 | 0.0665 | 0.0543 | 0.0543 | TRUE | 0.8415 |
| Sep-05 | 149 | 130 | 124 | 170 | 108 | 101 | 158 | 130 | 125 | 134 | 116 | 111 | 128.83 | 138 | 120 | 0.877 | -0.189 | 0.32925943 | 0.5502542 | 0.0063 | 0.0325 | 0.0192 | 0.0192 | TRUE | 0.1768 |
| Sep-06 | 1871 | 2229 | 2248 | 1590 | 1159 | 1401 | 1987 | 2226 | 2257 | 1250 | 1247 | 1534 | 1750.21 | 2157 | 1344 | 0.622 | -0.684 | 2.16E-09 | 1.21E-07 | 0.0089 | 0.0088 | 0.0088 | 0.0088 | TRUE | 0.1826 |
| Sep-07 | 5247 | 5570 | 5633 | 10076 | 6893 | 6772 | 5573 | 5563 | 5656 | 7921 | 7416 | 7413 | 6590.43 | 5597 | 7583 | 1.355 | 0.438 | 3.21E-12 | 3.07E-10 | 6.00E-04 | 0.0074 | 0.0027 | 0.0027 | TRUE | 0.0187 |
| Sep-08 | 3259 | 3906 | 3921 | 5035 | 3876 | 3587 | 3462 | 3901 | 3937 | 3958 | 4170 | 3927 | 3892.45 | 3767 | 4018 | 1.067 | 0.093 | 0.25675099 | 0.47534648 | 0.0026 | 0.0077 | 0.0046 | 0.0046 | TRUE | 0.0609 |
| Sep-09 | 7805 | 8806 | 9121 | 9250 | 7228 | 7869 | 8290 | 8795 | 9158 | 7272 | 7777 | 8614 | 8317.62 | 8748 | 7888 | 0.901 | -0.15 | 0.09726869 | 0.25625993 | 0.0048 | 0.0073 | 0.0057 | 0.0057 | TRUE | 0.0792 |
| Sep-10 | 1757 | 2338 | 2441 | 2634 | 1916 | 2008 | 1866 | 2335 | 2451 | 2071 | 2061 | 2198 | 2163.74 | 2217 | 2110 | 0.951 | -0.072 | 0.54133163 | 0.72705025 | 0.0104 | 0.0084 | 0.0095 | 0.0095 | TRUE | 0.2319 |
| Sep-11 | 14430 | 11869 | 11569 | 19191 | 13649 | 11966 | 15327 | 11854 | 11616 | 15086 | 14685 | 13099 | 13611.33 | 12932 | 14290 | 1.105 | 0.144 | 0.25212338 | 0.47003427 | 0.0146 | 0.0071 | 0.0113 | 0.0113 | TRUE | 0.3207 |
| Sep-12 | 2 | 4 | 1 | 9 | 4 | 6 | 2 | 4 | 1 | 7 | 4 | 7 | 4.18 | 2 | 6 | 2.529 | 1.338 | 0.17115682 | NA | 0 | 0.7954 | 0.3931 | 0.3931 | TRUE | 0.3855 |
| Sep-14 | 6 | 7 | 2 | 0 | 2 | 1 | 6 | 7 | 2 | 0 | 2 | 1 | 3.1 | 5 | 1 | 0.199 | -2.332 | 0.08022092 | NA | 0 | 1.0686 | 0.7027 | 0.7027 | TRUE | 0.591 |
| Sep-15 | 7432 | 8140 | 7710 | 9916 | 6864 | 6683 | 7894 | 8129 | 7742 | 7795 | 7385 | 7316 | 7710.21 | 7922 | 7499 | 0.947 | -0.079 | 0.21306145 | 0.42424003 | 7.00E-04 | 0.0073 | 0.0028 | 0.0028 | TRUE | 0.0147 |
| Sepw1 | 1659 | 1829 | 1704 | 2692 | 1630 | 1658 | 1762 | 1827 | 1711 | 2116 | 1754 | 1815 | 1830.79 | 1767 | 1895 | 1.074 | 0.102 | 0.30436626 | 0.52512118 | 0.005 | 0.0087 | 0.0066 | 0.0066 | TRUE | 0.1282 |
| Serac1 | 1506 | 1452 | 1488 | 1752 | 1420 | 1380 | 1600 | 1450 | 1494 | 1377 | 1528 | 1511 | 1493.27 | 1515 | 1472 | 0.971 | -0.042 | 0.6309086 | 0.79265149 | 0.0023 | 0.0091 | 0.0049 | 0.0049 | TRUE | 0.0392 |
| Serbp1 | 56969 | 61712 | 60583 | 72390 | 53871 | 53346 | 60511 | 61632 | 60832 | 56907 | 57962 | 58396 | 59373.34 | 60992 | 57755 | 0.947 | -0.079 | 0.11600759 | 0.28765413 | 1.00E-04 | 0.007 | 0.0018 | 0.0018 | TRUE | 0.002 |
| Serf1 | 88 | 107 | 111 | 189 | 147 | 131 | 93 | 107 | 111 | 149 | 158 | 143 | 126.99 | 104 | 150 | 1.442 | 0.528 | 0.00335822 | 0.0231247 | 0 | 0.0328 | 0.0152 | 0.0152 | TRUE | 0.0743 |
| Serf2 | 3306 | 4054 | 4127 | 4310 | 3070 | 3242 | 3512 | 4049 | 4144 | 3388 | 3303 | 3549 | 3657.41 | 3902 | 3413 | 0.875 | -0.193 | 0.03366854 | 0.12468189 | 0.0041 | 0.0078 | 0.0057 | 0.0057 | TRUE | 0.0928 |
| Sergef | 319 | 340 | 323 | 441 | 339 | 339 | 339 | 340 | 324 | 347 | 365 | 371 | 347.54 | 334 | 361 | 1.078 | 0.109 | 0.34131835 | 0.56159399 | 0 | 0.0164 | 0.0065 | 0.0065 | TRUE | 0.0146 |
| Serhl | 294 | 453 | 447 | 397 | 261 | 328 | 312 | 452 | 449 | 312 | 281 | 359 | 360.91 | 404 | 317 | 0.784 | -0.351 | 0.05329903 | 0.17145098 | 0.0259 | 0.016 | 0.021 | 0.021 | TRUE | 0.455 |
| Serinc1 | 3885 | 4105 | 4197 | 5695 | 4048 | 4045 | 4127 | 4100 | 4214 | 4477 | 4355 | 4428 | 4283.46 | 4147 | 4420 | 1.066 | 0.092 | 0.11555189 | 0.28713974 | 0 | 0.0077 | 0.0022 | 0.0022 | TRUE | 0.0025 |
| Serinc2 | 566 | 812 | 844 | 677 | 522 | 608 | 601 | 811 | 847 | 532 | 562 | 666 | 669.83 | 753 | 587 | 0.778 | -0.363 | 0.02404306 | 0.09890765 | 0.0221 | 0.0118 | 0.0171 | 0.0171 | TRUE | 0.3676 |
| Serinc3 | 6180 | 7814 | 7909 | 8643 | 6440 | 7064 | 6564 | 7804 | 7941 | 6794 | 6929 | 7733 | 7294.3 | 7436 | 7152 | 0.962 | -0.056 | 0.5850434 | 0.76019975 | 0.0066 | 0.0074 | 0.0076 | 0.0076 | TRUE | 0.1284 |
| Serinc4 | 7 | 3 | 3 | 6 | 2 | 3 | 7 | 3 | 3 | 5 | 2 | 3 | 3.93 | 4 | 3 | 0.771 | -0.375 | 0.69950286 | NA | 0 | 0.8446 | 0.424 | 0.424 | TRUE | 0.5831 |
| Serinc5 | 1451 | 2092 | 2061 | 1920 | 1424 | 1798 | 1541 | 2089 | 2069 | 1509 | 1532 | 1968 | 1784.94 | 1900 | 1670 | 0.878 | -0.187 | 0.23116399 | 0.44494515 | 0.0245 | 0.0087 | 0.017 | 0.017 | TRUE | 0.3292 |
| Serp1 | 3236 | 3565 | 3460 | 4879 | 3403 | 3380 | 3437 | 3560 | 3474 | 3835 | 3661 | 3700 | 3611.44 | 3490 | 3732 | 1.069 | 0.097 | 0.12270709 | 0.29824877 | 2.00E-04 | 0.0078 | 0.0026 | 0.0026 | TRUE | 0.0073 |
| Serp2 | 145 | 154 | 116 | 165 | 129 | 140 | 154 | 154 | 116 | 130 | 139 | 153 | 141.01 | 141 | 141 | 0.992 | -0.011 | 0.95329444 | 0.97644888 | 0.0086 | 0.0303 | 0.0195 | 0.0195 | TRUE | 0.2492 |
| Serpina10 | 0 | 0 | 0 | 0 | 0 | 0 | 0 | 0 | 0 | 0 | 0 | 0 | 0 | 0 | 0 | NA | NA | NA | NA | NA | NA | NA | NA | NA | NA |
| Serpina11 | 0 | 0 | 0 | 0 | 0 | 0 | 0 | 0 | 0 | 0 | 0 | 0 | 0 | 0 | 0 | NA | NA | NA | NA | NA | NA | NA | NA | NA | NA |
| Serpina12 | 5 | 8 | 8 | 12 | 6 | 5 | 5 | 8 | 8 | 9 | 6 | 5 | 7.12 | 7 | 7 | 1.013 | 0.018 | 0.97923911 | 0.98867206 | 0 | 0.4699 | 0.2165 | 0.2165 | TRUE | 0.2794 |
| Serpina1a | 0 | 0 | 0 | 0 | 0 | 0 | 0 | 0 | 0 | 0 | 0 | 0 | 0 | 0 | 0 | NA | NA | NA | NA | NA | NA | NA | NA | NA | NA |
| Serpina1b | 0 | 0 | 0 | 1 | 0 | 1 | 0 | 0 | 0 | 1 | 0 | 1 | 0.31 | 0 | 1 | 3.348 | 1.743 | 0.65064889 | NA | 0 | 10.5167 | 8.8968 | 8.8968 | TRUE | 0.2082 |
| Serpina1c | 0 | 0 | 0 | 0 | 0 | 0 | 0 | 0 | 0 | 0 | 0 | 0 | 0 | 0 | 0 | NA | NA | NA | NA | NA | NA | NA | NA | NA | NA |
| Serpina1d | 0 | 0 | 0 | 0 | 0 | 0 | 0 | 0 | 0 | 0 | 0 | 0 | 0 | 0 | 0 | NA | NA | NA | NA | NA | NA | NA | NA | NA | NA |
| Serpina1e | 0 | 0 | 0 | 0 | 0 | 0 | 0 | 0 | 0 | 0 | 0 | 0 | 0 | 0 | 0 | NA | NA | NA | NA | NA | NA | NA | NA | NA | NA |
| Serpina1f | 0 | 0 | 0 | 0 | 0 | 0 | 0 | 0 | 0 | 0 | 0 | 0 | 0 | 0 | 0 | NA | NA | NA | NA | NA | NA | NA | NA | NA | NA |
| Serpina3a | 0 | 0 | 0 | 0 | 0 | 0 | 0 | 0 | 0 | 0 | 0 | 0 | 0 | 0 | 0 | NA | NA | NA | NA | NA | NA | NA | NA | NA | NA |
| Serpina3b | 0 | 0 | 0 | 0 | 0 | 0 | 0 | 0 | 0 | 0 | 0 | 0 | 0 | 0 | 0 | NA | NA | NA | NA | NA | NA | NA | NA | NA | NA |
| Serpina3c | 0 | 0 | 0 | 0 | 0 | 0 | 0 | 0 | 0 | 0 | 0 | 0 | 0 | 0 | 0 | NA | NA | NA | NA | NA | NA | NA | NA | NA | NA |
| Serpina3f | 0 | 0 | 0 | 0 | 0 | 0 | 0 | 0 | 0 | 0 | 0 | 0 | 0 | 0 | 0 | NA | NA | NA | NA | NA | NA | NA | NA | NA | NA |
| Serpina3g | 0 | 0 | 0 | 0 | 0 | 1 | 0 | 0 | 0 | 0 | 0 | 1 | 0.18 | 0 | 0 | 1.859 | 0.894 | 0.82650734 | NA | 0 | 18.0643 | 10 | 10 | TRUE | 0.5383 |
| Serpina3h | 4 | 5 | 3 | 1 | 5 | 3 | 4 | 5 | 3 | 1 | 5 | 3 | 3.62 | 4 | 3 | 0.748 | -0.419 | 0.68967606 | NA | 0 | 0.9176 | 0.5074 | 0.5074 | TRUE | 0.3656 |
| Serpina3i | 8 | 22 | 23 | 11 | 7 | 9 | 8 | 22 | 23 | 9 | 8 | 10 | 13.27 | 18 | 9 | 0.485 | -1.045 | 0.07953551 | 0.22376634 | 0 | 0.2552 | 0.1713 | 0.1713 | TRUE | 1.0213 |
| Serpina3j | 0 | 0 | 0 | 0 | 0 | 0 | 0 | 0 | 0 | 0 | 0 | 0 | 0 | 0 | 0 | NA | NA | NA | NA | NA | NA | NA | NA | NA | NA |
| Serpina3k | 1 | 2 | 1 | 6 | 6 | 3 | 1 | 2 | 1 | 5 | 6 | 3 | 3.09 | 1 | 5 | 3.547 | 1.826 | 0.113491 | NA | 0 | 1.0742 | 0.4823 | 0.4823 | TRUE | 0.1207 |
| Serpina3m | 23 | 21 | 7 | 65 | 67 | 59 | 24 | 21 | 7 | 51 | 72 | 65 | 40.03 | 17 | 63 | 3.582 | 1.841 | 1.82E-05 | 0.00032682 | 0.1078 | 0.0892 | 0.0957 | 0.0957 | TRUE | 1.2971 |
| Serpina3n | 0 | 0 | 0 | 2 | 2 | 2 | 0 | 0 | 0 | 2 | 2 | 2 | 0.99 | 0 | 2 | 10.45 | 3.385 | 0.11522384 | NA | 0 | 3.3496 | 2.0792 | 2.0792 | TRUE | 0.07 |
| Serpina4-ps1 | 0 | 0 | 0 | 0 | 0 | 0 | 0 | 0 | 0 | 0 | 0 | 0 | 0 | 0 | 0 | NA | NA | NA | NA | NA | NA | NA | NA | NA | NA |
| Serpina5 | 0 | 0 | 0 | 0 | 0 | 0 | 0 | 0 | 0 | 0 | 0 | 0 | 0 | 0 | 0 | NA | NA | NA | NA | NA | NA | NA | NA | NA | NA |
| Serpina6 | 0 | 0 | 0 | 0 | 0 | 0 | 0 | 0 | 0 | 0 | 0 | 0 | 0 | 0 | 0 | NA | NA | NA | NA | NA | NA | NA | NA | NA | NA |
| Serpina7 | 8 | 1 | 2 | 9 | 13 | 9 | 8 | 1 | 2 | 7 | 14 | 10 | 7.07 | 4 | 10 | 2.697 | 1.431 | 0.10832747 | 0.27561556 | 0.272 | 0.4729 | 0.3903 | 0.3903 | TRUE | 1.2284 |
| Serpina9 | 0 | 0 | 0 | 3 | 1 | 0 | 0 | 0 | 0 | 2 | 1 | 0 | 0.57 | 0 | 1 | 6.182 | 2.628 | 0.40213576 | NA | 0 | 5.7626 | 5.6688 | 5.6688 | TRUE | 0.0684 |
| Serpinb10 | 1 | 0 | 1 | 2 | 0 | 0 | 1 | 0 | 1 | 2 | 0 | 0 | 0.61 | 1 | 1 | 0.816 | -0.293 | 0.91321207 | NA | 0 | 5.4395 | 3.5982 | 3.5982 | TRUE | 0.9722 |
| Serpinb11 | 0 | 0 | 0 | 0 | 0 | 0 | 0 | 0 | 0 | 0 | 0 | 0 | 0 | 0 | 0 | NA | NA | NA | NA | NA | NA | NA | NA | NA | NA |
| Serpinb12 | 0 | 0 | 0 | 2 | 0 | 0 | 0 | 0 | 0 | 2 | 0 | 0 | 0.26 | 0 | 1 | 2.886 | 1.529 | 0.70676491 | NA | 0 | 12.5792 | 10 | 10 | TRUE | 0.9669 |

| | | | | | | | | | | | | | | | | | | | | | | | | | |
|---|---|---|---|---|---|---|---|---|---|---|---|---|---|---|---|---|---|---|---|---|---|---|---|---|---|
| Serpinb13 | 0 | 0 | 0 | 0 | 0 | 0 | 0 | 0 | 0 | 0 | 0 | 0 | 0 | 0 | 0 | NA | NA | NA | NA | NA | NA | NA | NA | NA | NA |
| Serpinb1a | 108 | 165 | 184 | 252 | 188 | 230 | 115 | 165 | 185 | 198 | 202 | 252 | 186.07 | 155 | 217 | 1.401 | 0.487 | 0.02348599 | 0.09742795 | 0.0311 | 0.0246 | 0.0277 | 0.0277 | TRUE | 0.5305 |
| Serpinb1b | 0 | 0 | 0 | 0 | 0 | 0 | 0 | 0 | 0 | 0 | 0 | 0 | 0 | 0 | 0 | NA | NA | NA | NA | NA | NA | NA | NA | NA | NA |
| Serpinb1c | 1 | 3 | 7 | 5 | 2 | 9 | 1 | 3 | 7 | 4 | 2 | 10 | 4.5 | 4 | 5 | 1.42 | 0.506 | 0.62927523 | NA | 0.3782 | 0.7384 | 0.5603 | 0.5603 | TRUE | 0.7403 |
| Serpinb2 | 0 | 0 | 0 | 1 | 1 | 0 | 0 | 0 | 0 | 1 | 1 | 0 | 0.31 | 0 | 1 | 3.32 | 1.731 | 0.65383052 | NA | 0 | 10.6224 | 8.9541 | 8.9541 | TRUE | 0.202 |
| Serpinb3a | 0 | 0 | 0 | 0 | 0 | 0 | 0 | 0 | 0 | 0 | 0 | 0 | 0 | 0 | 0 | NA | NA | NA | NA | NA | NA | NA | NA | NA | NA |
| Serpinb3b | 0 | 0 | 0 | 0 | 0 | 0 | 0 | 0 | 0 | 0 | 0 | 0 | 0 | 0 | 0 | NA | NA | NA | NA | NA | NA | NA | NA | NA | NA |
| Serpinb3c | 0 | 0 | 0 | 0 | 0 | 0 | 0 | 0 | 0 | 0 | 0 | 0 | 0 | 0 | 0 | NA | NA | NA | NA | NA | NA | NA | NA | NA | NA |
| Serpinb3d | 0 | 0 | 0 | 0 | 0 | 0 | 0 | 0 | 0 | 0 | 0 | 0 | 0 | 0 | 0 | NA | NA | NA | NA | NA | NA | NA | NA | NA | NA |
| Serpinb5 | 48 | 74 | 60 | 42 | 61 | 80 | 51 | 74 | 60 | 33 | 66 | 88 | 61.89 | 62 | 62 | 0.998 | -0.004 | 0.99252622 | 0.99616826 | 0.1145 | 0.0601 | 0.0864 | 0.0864 | TRUE | 0.3402 |
| Serpinb6a | 1428 | 1631 | 1521 | 3655 | 2539 | 2514 | 1517 | 1629 | 1527 | 2873 | 2732 | 2752 | 2171.66 | 1558 | 2786 | 1.789 | 0.839 | 3.05E-30 | 1.72E-27 | 6.00E-04 | 0.0084 | 0.0034 | 0.0034 | TRUE | 0.0194 |
| Serpinb6b | 32 | 26 | 22 | 72 | 49 | 42 | 34 | 26 | 22 | 57 | 53 | 46 | 39.56 | 27 | 52 | 1.902 | 0.928 | 0.00429122 | 0.02787527 | 0 | 0.0902 | 0.0479 | 0.0479 | TRUE | 0.2763 |
| Serpinb6c | 85 | 80 | 53 | 121 | 96 | 113 | 90 | 80 | 53 | 95 | 103 | 124 | 90.92 | 74 | 107 | 1.44 | 0.526 | 0.04272954 | 0.1475652 | 0.0303 | 0.0431 | 0.0371 | 0.0371 | TRUE | 0.573 |
| Serpinb6d | 0 | 0 | 0 | 0 | 1 | 0 | 0 | 0 | 0 | 0 | 1 | 0 | 0.18 | 0 | 0 | 1.859 | 0.894 | 0.82650734 | NA | 0 | 18.3785 | 10 | 10 | TRUE | 0.5204 |
| Serpinb6e | 0 | 0 | 0 | 0 | 0 | 0 | 0 | 0 | 0 | 0 | 0 | 0 | 0 | 0 | 0 | NA | NA | NA | NA | NA | NA | NA | NA | NA | NA |
| Serpinb7 | 0 | 0 | 0 | 0 | 0 | 0 | 0 | 0 | 0 | 0 | 0 | 0 | 0 | 0 | 0 | NA | NA | NA | NA | NA | NA | NA | NA | NA | NA |
| Serpinb8 | 9 | 8 | 8 | 15 | 17 | 16 | 10 | 8 | 8 | 12 | 18 | 18 | 12.2 | 9 | 16 | 1.847 | 0.885 | 0.11040846 | 0.27888569 | 0 | 0.277 | 0.1303 | 0.1303 | TRUE | 0.2987 |
| Serpinb9 | 26 | 21 | 16 | 52 | 30 | 25 | 28 | 21 | 16 | 41 | 32 | 27 | 27.53 | 22 | 33 | 1.568 | 0.649 | 0.10234066 | 0.26578575 | 0 | 0.1266 | 0.0755 | 0.0755 | TRUE | 0.3283 |
| Serpinb9b | 143 | 61 | 47 | 143 | 94 | 60 | 152 | 61 | 47 | 112 | 101 | 66 | 89.87 | 87 | 93 | 1.077 | 0.107 | 0.80225854 | 0.89777365 | 0.2149 | 0.0436 | 0.1214 | 0.1214 | TRUE | 4.0592 |
| Serpinb9c | 0 | 0 | 0 | 0 | 0 | 0 | 0 | 0 | 0 | 0 | 0 | 0 | 0 | 0 | 0 | NA | NA | NA | NA | NA | NA | NA | NA | NA | NA |
| Serpinb9d | 0 | 0 | 0 | 0 | 0 | 0 | 0 | 0 | 0 | 0 | 0 | 0 | 0 | 0 | 0 | NA | NA | NA | NA | NA | NA | NA | NA | NA | NA |
| Serpinb9e | 0 | 0 | 1 | 0 | 2 | 0 | 0 | 0 | 1 | 0 | 2 | 0 | 0.53 | 0 | 1 | 1.927 | 0.947 | 0.77736506 | NA | 0 | 6.2701 | 6.3814 | 6.3814 | TRUE | 0.9744 |
| Serpinb9f | 0 | 0 | 0 | 0 | 0 | 0 | 0 | 0 | 0 | 0 | 0 | 0 | 0 | 0 | 0 | NA | NA | NA | NA | NA | NA | NA | NA | NA | NA |
| Serpinb9g | 0 | 0 | 0 | 0 | 0 | 0 | 0 | 0 | 0 | 0 | 0 | 0 | 0 | 0 | 0 | NA | NA | NA | NA | NA | NA | NA | NA | NA | NA |
| Serpinc1 | 0 | 3 | 0 | 3 | 0 | 3 | 0 | 3 | 0 | 2 | 0 | 3 | 1.44 | 1 | 2 | 1.878 | 0.909 | 0.66361726 | NA | 2.5914 | 2.2951 | 2.3798 | 2.3798 | TRUE | 1.3048 |
| Serpind1 | 0 | 0 | 0 | 0 | 0 | 0 | 0 | 0 | 0 | 0 | 0 | 0 | 0 | 0 | 0 | NA | NA | NA | NA | NA | NA | NA | NA | NA | NA |
| Serpine1 | 516 | 493 | 423 | 1042 | 825 | 735 | 548 | 492 | 425 | 819 | 888 | 805 | 662.76 | 488 | 837 | 1.714 | 0.778 | 2.95E-10 | 2.03E-08 | 0.007 | 0.0119 | 0.0093 | 0.0093 | TRUE | 0.1516 |
| Serpine2 | 1559 | 1787 | 1693 | 2400 | 1498 | 1771 | 1656 | 1785 | 1700 | 1887 | 1612 | 1939 | 1762.94 | 1714 | 1813 | 1.058 | 0.081 | 0.41306776 | 0.62445171 | 0.0046 | 0.0088 | 0.0065 | 0.0065 | TRUE | 0.1121 |
| Serpine3 | 1 | 6 | 2 | 6 | 4 | 6 | 1 | 6 | 2 | 5 | 4 | 7 | 4.11 | 3 | 5 | 1.704 | 0.769 | 0.43941548 | NA | 0.0087 | 0.8088 | 0.4507 | 0.4507 | TRUE | 0.9793 |
| Serpinf1 | 20 | 17 | 13 | 24 | 9 | 12 | 21 | 17 | 13 | 19 | 10 | 13 | 15.49 | 17 | 14 | 0.825 | -0.278 | 0.58871929 | 0.76346659 | 0 | 0.2195 | 0.1253 | 0.1253 | TRUE | 0.4121 |
| Serpinf2 | 18 | 10 | 13 | 11 | 9 | 5 | 19 | 10 | 13 | 9 | 10 | 5 | 10.99 | 14 | 8 | 0.569 | -0.815 | 0.17642298 | 0.3767831 | 0 | 0.3066 | 0.164 | 0.164 | TRUE | 0.3371 |
| Serping1 | 159 | 150 | 125 | 331 | 241 | 251 | 169 | 150 | 126 | 260 | 259 | 275 | 206.41 | 148 | 265 | 1.789 | 0.839 | 2.22E-07 | 7.19E-06 | 0.0026 | 0.0229 | 0.0136 | 0.0136 | TRUE | 0.1863 |
| Serpinh1 | 1529 | 2090 | 2254 | 2851 | 1769 | 1870 | 1624 | 2087 | 2263 | 2241 | 1903 | 2047 | 2027.7 | 1991 | 2064 | 1.036 | 0.052 | 0.70718766 | 0.84014491 | 0.0173 | 0.0085 | 0.0132 | 0.0132 | TRUE | 0.3144 |
| Serpini1 | 332 | 287 | 284 | 379 | 215 | 234 | 353 | 287 | 285 | 298 | 231 | 256 | 284.98 | 308 | 262 | 0.852 | -0.23 | 0.15381647 | 0.3455831 | 0.0124 | 0.0185 | 0.0153 | 0.0153 | TRUE | 0.1789 |
| Serpini2 | 0 | 1 | 0 | 0 | 0 | 0 | 0 | 1 | 0 | 0 | 0 | 0 | 0.17 | 0 | 0 | 0.49 | -1.029 | 0.80087 | NA | 0 | 19.7991 | 10 | 10 | TRUE | 0.405 |
| Sertad1 | 353 | 306 | 292 | 504 | 355 | 332 | 375 | 306 | 293 | 396 | 382 | 363 | 352.56 | 325 | 380 | 1.174 | 0.232 | 0.09634282 | 0.25478184 | 0.0065 | 0.0162 | 0.0111 | 0.0111 | TRUE | 0.2071 |
| Sertad2 | 1890 | 1820 | 2020 | 2369 | 1696 | 1785 | 2008 | 1818 | 2028 | 1862 | 1825 | 1954 | 1915.76 | 1951 | 1880 | 0.964 | -0.053 | 0.51817434 | 0.70964076 | 0.0019 | 0.0086 | 0.0044 | 0.0044 | TRUE | 0.0435 |
| Sertad3 | 402 | 497 | 485 | 603 | 442 | 454 | 427 | 496 | 487 | 474 | 476 | 497 | 476.15 | 470 | 482 | 1.025 | 0.035 | 0.75227128 | 0.86942757 | 0.0013 | 0.0138 | 0.0069 | 0.0069 | TRUE | 0.0739 |
| Sertad4 | 203 | 360 | 381 | 273 | 236 | 252 | 216 | 360 | 383 | 215 | 254 | 276 | 283.68 | 320 | 248 | 0.776 | -0.366 | 0.10736642 | 0.27409893 | 0.0502 | 0.0185 | 0.0338 | 0.0338 | TRUE | 0.9077 |
| Sertm1 | 59 | 87 | 60 | 134 | 82 | 90 | 63 | 87 | 60 | 105 | 88 | 99 | 83.65 | 70 | 97 | 1.394 | 0.479 | 0.0463388 | 0.15580006 | 0.0106 | 0.0463 | 0.0294 | 0.0294 | TRUE | 0.4088 |
| Sesn1 | 2257 | 2170 | 2002 | 2650 | 1834 | 1653 | 2397 | 2167 | 2010 | 2083 | 1973 | 1809 | 2073.45 | 2191 | 1955 | 0.893 | -0.164 | 0.10793798 | 0.27502387 | 0.0057 | 0.0085 | 0.007 | 0.007 | TRUE | 0.0814 |
| Sesn2 | 2178 | 2367 | 2066 | 3102 | 2231 | 2045 | 2313 | 2364 | 2074 | 2439 | 2400 | 2239 | 2304.9 | 2250 | 2359 | 1.049 | 0.068 | 0.43178457 | 0.64149457 | 0.0027 | 0.0083 | 0.005 | 0.005 | TRUE | 0.0569 |
| Sesn3 | 713 | 727 | 717 | 986 | 676 | 710 | 757 | 726 | 720 | 775 | 727 | 777 | 747.17 | 734 | 760 | 1.035 | 0.05 | 0.57761285 | 0.75499296 | 0 | 0.0113 | 0.0045 | 0.0045 | TRUE | 0.0162 |
| Sestd1 | 1700 | 1767 | 1985 | 2122 | 1635 | 1752 | 1806 | 1765 | 1993 | 1668 | 1759 | 1918 | 1818.12 | 1855 | 1782 | 0.96 | -0.059 | 0.5356958 | 0.72271961 | 0.0037 | 0.0087 | 0.0059 | 0.0059 | TRUE | 0.0537 |
| Set | 37766 | 37482 | 37661 | 54069 | 39734 | 35584 | 40114 | 37434 | 37816 | 42505 | 42751 | 38952 | 39928.64 | 38455 | 41403 | 1.077 | 0.107 | 0.13510764 | 0.31796579 | 0.002 | 0.007 | 0.0036 | 0.0036 | TRUE | 0.0328 |
| Setbp1 | 88 | 119 | 117 | 210 | 169 | 147 | 93 | 119 | 117 | 165 | 182 | 161 | 139.61 | 110 | 169 | 1.537 | 0.62 | 0.00069289 | 0.00671321 | 0 | 0.0305 | 0.0166 | 0.0166 | TRUE | 0.1666 |
| Setd1a | 5803 | 4777 | 5036 | 6510 | 4871 | 4583 | 6164 | 4771 | 5057 | 5118 | 5241 | 5017 | 5227.78 | 5331 | 5125 | 0.962 | -0.057 | 0.60638445 | 0.77676352 | 0.0084 | 0.0075 | 0.0085 | 0.0085 | TRUE | 0.2279 |
| Setd1b | 758 | 405 | 385 | 1126 | 739 | 449 | 805 | 404 | 387 | 885 | 795 | 492 | 628 | 532 | 724 | 1.362 | 0.445 | 0.13264071 | 0.31402218 | 0.1264 | 0.0121 | 0.0616 | 0.0616 | TRUE | 2.3308 |
| Setd2 | 6366 | 4805 | 5430 | 8127 | 5705 | 5600 | 6762 | 4799 | 5452 | 6389 | 6138 | 6130 | 5945 | 5671 | 6219 | 1.097 | 0.133 | 0.29786148 | 0.51858608 | 0.0139 | 0.0075 | 0.0116 | 0.0116 | TRUE | 0.3451 |
| Setd3 | 5317 | 6333 | 6444 | 7032 | 5196 | 5560 | 5648 | 6325 | 6470 | 5528 | 5591 | 6086 | 5941.29 | 6148 | 5735 | 0.933 | -0.1 | 0.24387738 | 0.46043022 | 0.0037 | 0.0075 | 0.0052 | 0.0052 | TRUE | 0.0617 |
| Setd4 | 719 | 695 | 660 | 777 | 570 | 521 | 764 | 694 | 663 | 611 | 613 | 570 | 652.49 | 707 | 598 | 0.847 | -0.24 | 0.020799 | 0.08931169 | 0.002 | 0.012 | 0.0062 | 0.0062 | TRUE | 0.0581 |
| Setd5 | 16652 | 14845 | 15522 | 22581 | 16494 | 15433 | 17687 | 14826 | 15586 | 17751 | 17747 | 16894 | 16748.45 | 16033 | 17464 | 1.089 | 0.123 | 0.15969434 | 0.3541976 | 0.0046 | 0.0071 | 0.0055 | 0.0055 | TRUE | 0.0996 |
| Setd6 | 4847 | 5936 | 5958 | 4934 | 3890 | 4465 | 5148 | 5928 | 5982 | 3879 | 4185 | 4888 | 5001.82 | 5686 | 4317 | 0.759 | -0.398 | 0.00045808 | 0.00485536 | 0.0102 | 0.0076 | 0.0091 | 0.0091 | TRUE | 0.1625 |
| Setd7 | 870 | 1098 | 1196 | 1952 | 1233 | 1186 | 924 | 1097 | 1201 | 1535 | 1327 | 1298 | 1230.17 | 1074 | 1387 | 1.292 | 0.369 | 0.00378622 | 0.0253253 | 0.0116 | 0.0096 | 0.0109 | 0.0109 | TRUE | 0.1773 |
| Setd8 | 6015 | 6675 | 6720 | 7510 | 5467 | 5655 | 6389 | 6666 | 6748 | 5904 | 5882 | 6190 | 6296.53 | 6601 | 5992 | 0.908 | -0.14 | 0.02701696 | 0.10722484 | 6.00E-04 | 0.0074 | 0.0027 | 0.0027 | TRUE | 0.0101 |
| Setdb1 | 5573 | 4912 | 5135 | 7550 | 5721 | 5106 | 5920 | 4906 | 5156 | 5935 | 6155 | 5589 | 5610.21 | 5327 | 5893 | 1.106 | 0.146 | 0.12841755 | 0.30813961 | 0.0058 | 0.0075 | 0.0065 | 0.0065 | TRUE | 0.1153 |
| Setdb2 | 786 | 743 | 757 | 1114 | 786 | 776 | 835 | 742 | 760 | 876 | 846 | 849 | 817.98 | 779 | 857 | 1.101 | 0.139 | 0.13516777 | 0.31804657 | 8.00E-04 | 0.0109 | 0.005 | 0.005 | TRUE | 0.0463 |
| Setmar | 364 | 333 | 347 | 456 | 335 | 337 | 387 | 333 | 348 | 358 | 360 | 369 | 359.24 | 356 | 362 | 1.019 | 0.027 | 0.81979479 | 0.90781402 | 0 | 0.0161 | 0.0075 | 0.0075 | TRUE | 0.0645 |
| Setx | 16742 | 14434 | 16528 | 19405 | 13734 | 13569 | 17783 | 14415 | 16596 | 15255 | 14777 | 14853 | 15613.2 | 16265 | 14962 | 0.92 | -0.12 | 0.1991056 | 0.40635268 | 0.0056 | 0.0071 | 0.0063 | 0.0063 | TRUE | 0.121 |
| Sez6 | 70 | 65 | 89 | 83 | 65 | 67 | 74 | 65 | 89 | 65 | 70 | 73 | 72.86 | 76 | 69 | 0.91 | -0.136 | 0.57013328 | 0.74900702 | 0 | 0.0521 | 0.0277 | 0.0277 | TRUE | 0.2123 |
| Sez6l | 150 | 119 | 121 | 99 | 69 | 78 | 159 | 119 | 121 | 78 | 74 | 85 | 106.19 | 133 | 79 | 0.594 | -0.751 | 0.0004829 | 0.00503754 | 0.0094 | 0.0379 | 0.0233 | 0.0233 | TRUE | 0.2977 |
| Sez6l2 | 42 | 42 | 32 | 62 | 31 | 26 | 45 | 42 | 32 | 49 | 33 | 28 | 38.21 | 40 | 37 | 0.942 | -0.087 | 0.80425966 | 0.89905616 | 0.0307 | 0.0931 | 0.0626 | 0.0626 | TRUE | 0.6641 |
| Sf1 | 18321 | 17012 | 16621 | 22564 | 16827 | 15228 | 19460 | 16990 | 16689 | 17738 | 18105 | 16669 | 17608.62 | 17713 | 17504 | 0.988 | -0.017 | 0.84489837 | 0.92200492 | 0.0037 | 0.0071 | 0.0054 | 0.0054 | TRUE | 0.091 |
| Sf3a1 | 7793 | 7592 | 7892 | 11778 | 8854 | 7952 | 8278 | 7582 | 7924 | 9259 | 9526 | 8705 | 8545.69 | 7928 | 9163 | 1.156 | 0.209 | 0.00407732 | 0.02679476 | 0.0018 | 0.0073 | 0.0037 | 0.0037 | TRUE | 0.0233 |
| Sf3a2 | 3558 | 3127 | 3046 | 4863 | 3459 | 3096 | 3779 | 3123 | 3059 | 3823 | 3722 | 3389 | 3482.39 | 3320 | 3645 | 1.098 | 0.135 | 0.21601541 | 0.42801446 | 0.0087 | 0.0078 | 0.0083 | 0.0083 | TRUE | 0.1775 |
| Sf3a3 | 4297 | 4082 | 3979 | 6540 | 4727 | 4095 | 4564 | 4077 | 3995 | 5141 | 5086 | 4483 | 4557.68 | 4212 | 4903 | 1.164 | 0.22 | 0.01977128 | 0.08617669 | 0.0051 | 0.0076 | 0.0062 | 0.0062 | TRUE | 0.0684 |
| Sf3b1 | 23684 | 28280 | 29012 | 32093 | 23941 | 26318 | 25157 | 28243 | 29131 | 25229 | 25759 | 28809 | 27054.74 | 27510 | 26599 | 0.967 | -0.049 | 0.59740802 | 0.77004845 | 0.0047 | 0.007 | 0.0061 | 0.0061 | TRUE | 0.0685 |
| Sf3b2 | 17824 | 18294 | 18785 | 23194 | 17639 | 17287 | 18932 | 18270 | 18862 | 18233 | 18978 | 18923 | 18699.98 | 18688 | 18711 | 1.001 | 0.002 | 0.97533203 | 0.98697924 | 3.00E-04 | 0.0071 | 0.0022 | 0.0022 | TRUE | 0.0062 |
| Sf3b3 | 21003 | 18872 | 19810 | 35519 | 25060 | 21435 | 22309 | 18848 | 19891 | 27922 | 26963 | 23464 | 23232.85 | 20349 | 26116 | 1.283 | 0.36 | 0.00043939 | 0.00468881 | 0.0078 | 0.007 | 0.0075 | 0.0075 | TRUE | 0.0966 |

| | | | | | | | | | | | | | | | | | | | | | | | | | |
|---|---|---|---|---|---|---|---|---|---|---|---|---|---|---|---|---|---|---|---|---|---|---|---|---|---|
| Sf3b4 | 3193 | 2709 | 2713 | 4870 | 3396 | 2647 | 3392 | 2706 | 2724 | 3828 | 3654 | 2898 | 3200.17 | 2941 | 3460 | 1.177 | 0.235 | 0.09117095 | 0.24633938 | 0.0186 | 0.0079 | 0.0137 | 0.0137 | TRUE | 0.2452 |
| Sf3b5 | 3112 | 3226 | 3137 | 4370 | 3239 | 3255 | 3305 | 3222 | 3150 | 3435 | 3485 | 3563 | 3360.11 | 3226 | 3494 | 1.083 | 0.115 | 0.06840374 | 0.20267966 | 2.00E-04 | 0.0079 | 0.0026 | 0.0026 | TRUE | 0.0057 |
| Sfi1 | 289 | 285 | 235 | 290 | 288 | 247 | 307 | 285 | 236 | 228 | 310 | 270 | 272.63 | 276 | 269 | 0.974 | -0.037 | 0.83009513 | 0.9138554 | 0.0172 | 0.019 | 0.0181 | 0.0181 | TRUE | 0.2151 |
| Sfmbt1 | 1390 | 1377 | 1367 | 2382 | 1609 | 1478 | 1476 | 1375 | 1373 | 1873 | 1731 | 1618 | 1574.32 | 1408 | 1741 | 1.237 | 0.307 | 0.00075195 | 0.00713022 | 0.003 | 0.009 | 0.0053 | 0.0053 | TRUE | 0.0543 |
| Sfmbt2 | 264 | 309 | 317 | 1687 | 1332 | 1205 | 280 | 309 | 318 | 1326 | 1433 | 1319 | 830.95 | 302 | 1359 | 4.491 | 2.167 | 6.32E-99 | 1.99E-95 | 0.0013 | 0.0109 | 0.0055 | 0.0055 | TRUE | 0.045 |
| Sfn | 655 | 897 | 904 | 2435 | 1708 | 1524 | 696 | 896 | 908 | 1914 | 1838 | 1668 | 1319.91 | 833 | 1807 | 2.169 | 1.117 | 1.43E-18 | 2.97E-16 | 0.0113 | 0.0094 | 0.0107 | 0.0107 | TRUE | 0.2458 |
| Sfpi1 | 16 | 16 | 12 | 30 | 24 | 24 | 17 | 16 | 12 | 24 | 26 | 26 | 20.12 | 15 | 25 | 1.679 | 0.747 | 0.07845833 | 0.22169438 | 0 | 0.1707 | 0.0765 | 0.0765 | TRUE | 0.1372 |
| Sfpq | 22374 | 19927 | 18605 | 31502 | 24120 | 19215 | 23765 | 19901 | 18681 | 24764 | 25952 | 21034 | 22349.61 | 20782 | 23917 | 1.151 | 0.203 | 0.09737904 | 0.2563981 | 0.0138 | 0.0071 | 0.0107 | 0.0107 | TRUE | 0.1928 |
| Sfr1 | 8379 | 7969 | 7676 | 10760 | 7366 | 7173 | 8900 | 7959 | 7708 | 8459 | 7925 | 7852 | 8133.7 | 8189 | 8079 | 0.987 | -0.019 | 0.81817893 | 0.90726388 | 0.003 | 0.0073 | 0.005 | 0.005 | TRUE | 0.0704 |
| Sfrp1 | 3523 | 3865 | 3842 | 3446 | 2797 | 3210 | 3742 | 3860 | 3858 | 2709 | 3009 | 3514 | 3448.68 | 3820 | 3077 | 0.805 | -0.313 | 0.00397131 | 0.02621826 | 0.0084 | 0.0079 | 0.0082 | 0.0082 | TRUE | 0.1868 |
| Sfrp2 | 933 | 1248 | 1214 | 1136 | 915 | 924 | 991 | 1246 | 1219 | 893 | 984 | 1011 | 1057.56 | 1152 | 963 | 0.835 | -0.26 | 0.03035822 | 0.11627243 | 0.009 | 0.01 | 0.0094 | 0.0094 | TRUE | 0.1785 |
| Sfrp4 | 282 | 267 | 328 | 308 | 233 | 208 | 300 | 267 | 329 | 242 | 251 | 228 | 269.34 | 299 | 240 | 0.805 | -0.313 | 0.02730069 | 0.10797053 | 0.0034 | 0.0191 | 0.0108 | 0.0108 | TRUE | 0.0995 |
| Sfrp5 | 167 | 215 | 189 | 231 | 136 | 172 | 177 | 215 | 190 | 182 | 146 | 188 | 183.01 | 194 | 172 | 0.888 | -0.172 | 0.32248048 | 0.54376341 | 0.0075 | 0.0249 | 0.0162 | 0.0162 | TRUE | 0.1752 |
| Sfrs18 | 3817 | 3913 | 3504 | 4549 | 3780 | 3517 | 4054 | 3908 | 3518 | 3576 | 4067 | 3850 | 3828.95 | 3827 | 3831 | 1.001 | 0.001 | 0.98938979 | 0.99437047 | 0.0045 | 0.0078 | 0.0058 | 0.0058 | TRUE | 0.0606 |
| Sfswap | 2151 | 1976 | 2006 | 2901 | 2240 | 1873 | 2285 | 1973 | 2014 | 2281 | 2410 | 2050 | 2168.89 | 2091 | 2247 | 1.075 | 0.104 | 0.30581004 | 0.52665845 | 0.0059 | 0.0084 | 0.007 | 0.007 | TRUE | 0.0795 |
| Sft2d1 | 730 | 919 | 775 | 1070 | 714 | 705 | 775 | 918 | 778 | 841 | 768 | 772 | 808.75 | 824 | 794 | 0.964 | -0.053 | 0.63209011 | 0.79293961 | 0.0049 | 0.011 | 0.0075 | 0.0075 | TRUE | 0.1188 |
| Sft2d2 | 1912 | 2119 | 2013 | 2088 | 1554 | 1724 | 2031 | 2116 | 2021 | 1641 | 1672 | 1887 | 1894.84 | 2056 | 1733 | 0.843 | -0.247 | 0.00457278 | 0.02924649 | 0.0025 | 0.0086 | 0.0049 | 0.0049 | TRUE | 0.0719 |
| Sft2d3 | 149 | 207 | 239 | 266 | 226 | 221 | 158 | 207 | 240 | 209 | 243 | 242 | 216.53 | 202 | 231 | 1.145 | 0.195 | 0.30103424 | 0.52157663 | 0.0197 | 0.0221 | 0.0209 | 0.0209 | TRUE | 0.3528 |
| Sfta2 | 0 | 1 | 0 | 1 | 2 | 0 | 0 | 1 | 0 | 1 | 2 | 0 | 0.66 | 0 | 1 | 2.668 | 1.416 | 0.58756221 | NA | 0 | 5.028 | 3.4088 | 3.4088 | TRUE | 0.2713 |
| Sftpa1 | 0 | 0 | 0 | 0 | 0 | 0 | 0 | 0 | 0 | 0 | 0 | 0 | 0 | 0 | 0 | NA | NA | NA | NA | NA | NA | NA | NA | NA | NA |
| Sftpb | 0 | 0 | 0 | 2 | 0 | 0 | 0 | 0 | 0 | 2 | 0 | 0 | 0.26 | 0 | 1 | 2.886 | 1.529 | 0.70676491 | NA | 0 | 12.5792 | 10 | 10 | TRUE | 0.9669 |
| Sftpc | 0 | 1 | 2 | 0 | 0 | 0 | 0 | 1 | 2 | 0 | 0 | 0 | 0.5 | 1 | 0 | 0.178 | -2.487 | 0.44860443 | NA | 0 | 6.5806 | 6.2542 | 6.2542 | TRUE | 0.0529 |
| Sftpd | 7 | 7 | 4 | 8 | 2 | 3 | 7 | 7 | 4 | 6 | 2 | 3 | 5.03 | 6 | 4 | 0.653 | -0.614 | 0.48036036 | 0.67960563 | 0 | 0.6621 | 0.3403 | 0.3403 | TRUE | 0.6099 |
| Sfxn1 | 3976 | 4636 | 4649 | 4473 | 3297 | 3648 | 4223 | 4630 | 4668 | 3516 | 3547 | 3993 | 4096.39 | 4507 | 3685 | 0.818 | -0.291 | 0.00090926 | 0.00827041 | 0.0037 | 0.0077 | 0.0053 | 0.0053 | TRUE | 0.0646 |
| Sfxn2 | 603 | 624 | 605 | 866 | 651 | 632 | 640 | 623 | 607 | 681 | 700 | 692 | 657.37 | 623 | 691 | 1.108 | 0.147 | 0.10196355 | 0.26502456 | 0 | 0.0119 | 0.0043 | 0.0043 | TRUE | 0.0064 |
| Sfxn3 | 448 | 606 | 592 | 502 | 441 | 480 | 476 | 605 | 594 | 395 | 474 | 525 | 511.68 | 558 | 465 | 0.83 | -0.268 | 0.08348522 | 0.23154066 | 0.0171 | 0.0133 | 0.0153 | 0.0153 | TRUE | 0.211 |
| Sfxn4 | 19 | 27 | 23 | 46 | 40 | 17 | 20 | 27 | 23 | 36 | 43 | 19 | 28.01 | 23 | 33 | 1.396 | 0.481 | 0.259439 | 0.4784696 | 0.0612 | 0.1245 | 0.0944 | 0.0944 | TRUE | 0.5465 |
| Sfxn5 | 234 | 218 | 237 | 232 | 165 | 189 | 249 | 218 | 238 | 182 | 178 | 207 | 211.84 | 235 | 189 | 0.804 | -0.315 | 0.03496535 | 0.12807094 | 0 | 0.0225 | 0.0113 | 0.0113 | TRUE | 0.0709 |
| Sgca | 7 | 6 | 6 | 5 | 2 | 0 | 7 | 6 | 6 | 4 | 2 | 0 | 4.26 | 6 | 2 | 0.329 | -1.604 | 0.12234083 | NA | 0 | 0.781 | 0.4694 | 0.4694 | TRUE | 0.6722 |
| Sgcb | 990 | 1029 | 990 | 1432 | 1203 | 1222 | 1052 | 1028 | 994 | 1126 | 1294 | 1338 | 1138.51 | 1025 | 1253 | 1.221 | 0.288 | 0.00382355 | 0.02546666 | 0.0039 | 0.0098 | 0.0063 | 0.0063 | TRUE | 0.097 |
| Sgcd | 21 | 28 | 28 | 31 | 20 | 39 | 22 | 28 | 28 | 24 | 22 | 43 | 27.83 | 26 | 30 | 1.123 | 0.167 | 0.68522054 | 0.82611973 | 0.0442 | 0.1253 | 0.0859 | 0.0859 | TRUE | 0.939 |
| Sgce | 337 | 630 | 640 | 580 | 329 | 462 | 358 | 629 | 643 | 456 | 354 | 506 | 490.91 | 543 | 439 | 0.807 | -0.309 | 0.18391767 | 0.38664225 | 0.0642 | 0.0136 | 0.0369 | 0.0369 | TRUE | 1.0373 |
| Sgcg | 1 | 1 | 4 | 3 | 3 | 2 | 1 | 1 | 4 | 2 | 3 | 2 | 2.31 | 2 | 2 | 1.271 | 0.346 | 0.78043577 | NA | 0 | 1.4338 | 0.6713 | 0.6713 | TRUE | 0.6786 |
| Sgcz | 3 | 0 | 0 | 17 | 8 | 2 | 3 | 0 | 0 | 13 | 9 | 2 | 4.56 | 1 | 8 | 7.818 | 2.967 | 0.03140457 | NA | 1.0313 | 0.7297 | 0.8182 | 0.8182 | TRUE | 0.5068 |
| Sgip1 | 13 | 13 | 16 | 10 | 10 | 9 | 14 | 13 | 16 | 8 | 11 | 10 | 11.89 | 14 | 10 | 0.659 | -0.602 | 0.26653392 | 0.48621015 | 0 | 0.284 | 0.1241 | 0.1241 | TRUE | 0.1 |
| Sgk1 | 14126 | 11261 | 8214 | 14489 | 9938 | 9605 | 15004 | 11246 | 8248 | 11390 | 10693 | 10514 | 11182.58 | 11499 | 10866 | 0.945 | -0.082 | 0.65812655 | 0.80743861 | 0.0382 | 0.0072 | 0.0245 | 0.0245 | TRUE | 0.2374 |
| Sgk2 | 76 | 54 | 61 | 134 | 75 | 67 | 81 | 54 | 61 | 105 | 81 | 73 | 75.88 | 65 | 86 | 1.333 | 0.414 | 0.12438877 | 0.30099106 | 0.0272 | 0.0503 | 0.0389 | 0.0389 | TRUE | 0.3776 |
| Sgk3 | 983 | 1357 | 1331 | 1205 | 909 | 1073 | 1044 | 1355 | 1336 | 947 | 978 | 1175 | 1139.28 | 1245 | 1033 | 0.829 | -0.27 | 0.05357618 | 0.1719584 | 0.0156 | 0.0098 | 0.0133 | 0.0133 | TRUE | 0.2389 |
| Sgms1 | 1494 | 1548 | 1566 | 1780 | 1285 | 1428 | 1587 | 1546 | 1572 | 1399 | 1383 | 1563 | 1508.4 | 1568 | 1448 | 0.923 | -0.116 | 0.17226739 | 0.37111375 | 0.0015 | 0.0091 | 0.0045 | 0.0045 | TRUE | 0.0569 |
| Sgms2 | 312 | 353 | 347 | 410 | 321 | 330 | 331 | 353 | 348 | 322 | 345 | 361 | 343.55 | 344 | 343 | 0.995 | -0.008 | 0.94782895 | 0.97406219 | 0 | 0.0165 | 0.0072 | 0.0072 | TRUE | 0.0342 |
| Sgol1 | 2491 | 3085 | 2989 | 3760 | 2675 | 2827 | 2646 | 3081 | 3001 | 2956 | 2878 | 3095 | 2942.79 | 2909 | 2976 | 1.023 | 0.033 | 0.71240065 | 0.84341563 | 0.0036 | 0.008 | 0.0053 | 0.0053 | TRUE | 0.076 |
| Sgol2 | 6562 | 8062 | 8451 | 10163 | 7560 | 8132 | 6970 | 8052 | 8486 | 7989 | 8134 | 8902 | 8088.75 | 7836 | 8342 | 1.064 | 0.09 | 0.36224081 | 0.58014481 | 0.0062 | 0.0073 | 0.0069 | 0.0069 | TRUE | 0.114 |
| Sgpl1 | 4490 | 5577 | 5695 | 6183 | 4433 | 4938 | 4769 | 5570 | 5718 | 4861 | 4770 | 5405 | 5182.17 | 5352 | 5012 | 0.936 | -0.095 | 0.34482409 | 0.56427418 | 0.0066 | 0.0075 | 0.0071 | 0.0071 | TRUE | 0.1107 |
| Sgpp1 | 981 | 1147 | 1047 | 1843 | 1347 | 1421 | 1042 | 1146 | 1051 | 1449 | 1449 | 1556 | 1282.07 | 1080 | 1485 | 1.374 | 0.459 | 9.45E-08 | 3.42E-06 | 0.0014 | 0.0095 | 0.0045 | 0.0045 | TRUE | 0.0343 |
| Sgpp2 | 128 | 288 | 269 | 175 | 178 | 233 | 136 | 288 | 270 | 138 | 192 | 255 | 212.97 | 231 | 195 | 0.84 | -0.252 | 0.4182583 | 0.62904262 | 0.1172 | 0.0224 | 0.0651 | 0.0651 | TRUE | 0.485 |
| Sgsh | 190 | 302 | 293 | 375 | 275 | 237 | 202 | 302 | 294 | 295 | 296 | 259 | 274.62 | 266 | 283 | 1.066 | 0.092 | 0.61660408 | 0.78320685 | 0.0223 | 0.0189 | 0.0206 | 0.0206 | TRUE | 0.4911 |
| Sgsm1 | 1416 | 1606 | 1595 | 1473 | 1146 | 1260 | 1504 | 1604 | 1602 | 1158 | 1233 | 1379 | 1413.29 | 1570 | 1257 | 0.8 | -0.322 | 0.00085997 | 0.00790869 | 0.0036 | 0.0092 | 0.006 | 0.006 | TRUE | 0.0868 |
| Sgsm2 | 2712 | 3186 | 3380 | 3552 | 2731 | 2718 | 2881 | 3182 | 3394 | 2792 | 2938 | 2975 | 3027.06 | 3152 | 2902 | 0.92 | -0.12 | 0.17494764 | 0.37459602 | 0.0034 | 0.008 | 0.0053 | 0.0053 | TRUE | 0.0688 |
| Sgsm3 | 2497 | 2414 | 2366 | 2707 | 2099 | 1882 | 2652 | 2411 | 2376 | 2128 | 2258 | 2060 | 2314.24 | 2480 | 2149 | 0.867 | -0.206 | 0.01384866 | 0.06669345 | 0.0023 | 0.0083 | 0.0046 | 0.0046 | TRUE | 0.0448 |
| Sgta | 4766 | 4741 | 4645 | 6433 | 4542 | 4270 | 5062 | 4735 | 4664 | 5057 | 4887 | 4674 | 4846.58 | 4820 | 4873 | 1.011 | 0.016 | 0.82536522 | 0.91142649 | 0.0015 | 0.0076 | 0.0036 | 0.0036 | TRUE | 0.0234 |
| Sgtb | 913 | 1151 | 1205 | 913 | 730 | 868 | 970 | 1150 | 1210 | 718 | 785 | 950 | 963.76 | 1110 | 818 | 0.736 | -0.442 | 0.0016859 | 0.01343606 | 0.0154 | 0.0103 | 0.0132 | 0.0132 | TRUE | 0.237 |
| Sh2b1 | 2337 | 2266 | 2250 | 3099 | 2295 | 2194 | 2482 | 2263 | 2259 | 2436 | 2469 | 2402 | 2385.29 | 2335 | 2436 | 1.043 | 0.061 | 0.41486282 | 0.62638488 | 0.001 | 0.0083 | 0.0036 | 0.0036 | TRUE | 0.0368 |
| Sh2b2 | 350 | 376 | 422 | 331 | 307 | 312 | 372 | 376 | 424 | 260 | 330 | 342 | 350.51 | 391 | 311 | 0.793 | -0.334 | 0.02562427 | 0.10346799 | 0.0106 | 0.0163 | 0.0133 | 0.0133 | TRUE | 0.2435 |
| Sh2b3 | 1136 | 1306 | 1442 | 1332 | 1128 | 1100 | 1207 | 1304 | 1448 | 1047 | 1214 | 1204 | 1237.29 | 1320 | 1155 | 0.874 | -0.194 | 0.07967687 | 0.22396429 | 0.0063 | 0.0096 | 0.008 | 0.008 | TRUE | 0.0869 |
| Sh2d1a | 1 | 0 | 1 | 9 | 1 | 1 | 1 | 0 | 1 | 7 | 1 | 1 | 1.89 | 1 | 3 | 4.629 | 2.211 | 0.19448619 | NA | 0 | 1.7543 | 1.1914 | 1.1914 | TRUE | 2.1567 |
| Sh2d1b1 | 0 | 0 | 0 | 0 | 0 | 0 | 0 | 0 | 0 | 0 | 0 | 0 | 0 | 0 | 0 | NA | NA | NA | NA | NA | NA | NA | NA | NA | NA |
| Sh2d1b2 | 0 | 0 | 0 | 0 | 0 | 0 | 0 | 0 | 0 | 0 | 0 | 0 | 0 | 0 | 0 | NA | NA | NA | NA | NA | NA | NA | NA | NA | NA |
| Sh2d2a | 3 | 5 | 3 | 5 | 3 | 3 | 3 | 5 | 3 | 4 | 3 | 3 | 3.61 | 4 | 3 | 0.937 | -0.094 | 0.92161984 | NA | 0 | 0.9206 | 0.3778 | 0.3778 | TRUE | 0.1409 |
| Sh2d3c | 459 | 422 | 391 | 511 | 311 | 340 | 488 | 421 | 393 | 402 | 335 | 372 | 401.68 | 434 | 370 | 0.853 | -0.229 | 0.09789546 | 0.25720594 | 0.008 | 0.0151 | 0.0113 | 0.0113 | TRUE | 0.1344 |
| Sh2d4a | 141 | 169 | 137 | 207 | 164 | 145 | 150 | 169 | 138 | 163 | 176 | 159 | 159 | 152 | 166 | 1.091 | 0.126 | 0.45395964 | 0.65915199 | 0 | 0.0276 | 0.0139 | 0.0139 | TRUE | 0.0988 |
| Sh2d4b | 17 | 5 | 9 | 21 | 23 | 20 | 18 | 5 | 9 | 17 | 25 | 22 | 15.87 | 11 | 21 | 1.968 | 0.977 | 0.08877045 | 0.24189486 | 0.1021 | 0.2145 | 0.1661 | 0.1661 | TRUE | 1.0456 |
| Sh2d5 | 288 | 489 | 572 | 287 | 277 | 324 | 306 | 488 | 574 | 226 | 298 | 355 | 374.49 | 456 | 293 | 0.64 | -0.643 | 0.009744 | 0.05165792 | 0.0722 | 0.0157 | 0.0418 | 0.0418 | TRUE | 0.3787 |
| Sh2d7 | 1 | 0 | 0 | 5 | 4 | 0 | 1 | 0 | 0 | 4 | 4 | 0 | 1.55 | 0 | 3 | 7.559 | 2.918 | 0.16002525 | NA | 0 | 2.1331 | 1.9316 | 1.9316 | TRUE | 0.831 |
| Sh3bgr | 16 | 8 | 11 | 19 | 11 | 11 | 17 | 8 | 11 | 15 | 12 | 12 | 12.47 | 12 | 13 | 1.087 | 0.12 | 0.82913248 | 0.91333678 | 0 | 0.271 | 0.1424 | 0.1424 | TRUE | 0.4893 |
| Sh3bgrl | 1198 | 1430 | 1377 | 2151 | 1433 | 1351 | 1272 | 1428 | 1383 | 1691 | 1542 | 1479 | 1465.82 | 1361 | 1571 | 1.155 | 0.207 | 0.02812267 | 0.11006223 | 0.0035 | 0.0092 | 0.0057 | 0.0057 | TRUE | 0.0554 |
| Sh3bgrl2 | 3780 | 3406 | 3333 | 5140 | 3644 | 3356 | 4015 | 3402 | 3347 | 4041 | 3921 | 3674 | 3733.06 | 3588 | 3879 | 1.081 | 0.113 | 0.25336948 | 0.47129891 | 0.0061 | 0.0078 | 0.0068 | 0.0068 | TRUE | 0.1318 |
| Sh3bgrl3 | 1624 | 1717 | 1701 | 2173 | 1529 | 1529 | 1725 | 1715 | 1708 | 1708 | 1645 | 1674 | 1695.8 | 1716 | 1676 | 0.977 | -0.034 | 0.62298344 | 0.78725221 | 0 | 0.0088 | 0.0028 | 0.0028 | TRUE | 0.0036 |
| Sh3bp1 | 761 | 833 | 783 | 952 | 820 | 823 | 808 | 832 | 786 | 748 | 882 | 901 | 826.34 | 809 | 844 | 1.042 | 0.059 | 0.58586351 | 0.76088926 | 0.0045 | 0.0109 | 0.0072 | 0.0072 | TRUE | 0.121 |
| Sh3bp2 | 288 | 240 | 244 | 300 | 270 | 220 | 306 | 240 | 245 | 236 | 291 | 241 | 259.63 | 264 | 256 | 0.97 | -0.045 | 0.78674209 | 0.89028751 | 0.0122 | 0.0196 | 0.0157 | 0.0157 | TRUE | 0.2187 |

| | | | | | | | | | | | | | | | | | | | | | | | | | |
|---|---|---|---|---|---|---|---|---|---|---|---|---|---|---|---|---|---|---|---|---|---|---|---|---|---|
| Sh3bp4 | 1534 | 1596 | 1500 | 2456 | 1829 | 1813 | 1629 | 1594 | 1506 | 1931 | 1968 | 1985 | 1768.78 | 1576 | 1961 | 1.244 | 0.315 | 1.82E-05 | 0.00032759 | 3.00E-04 | 0.0088 | 0.0033 | 0.0033 | TRUE | 0.0184 |
| Sh3bp5 | 415 | 484 | 450 | 529 | 405 | 423 | 441 | 483 | 452 | 416 | 436 | 463 | 448.45 | 459 | 438 | 0.954 | -0.068 | 0.5373464 | 0.72405189 | 0 | 0.0142 | 0.0066 | 0.0066 | TRUE | 0.0281 |
| Sh3bp5l | 2438 | 2206 | 2019 | 3424 | 2434 | 2201 | 2590 | 2203 | 2027 | 2692 | 2619 | 2409 | 2423.31 | 2273 | 2573 | 1.132 | 0.179 | 0.11052164 | 0.27912687 | 0.009 | 0.0083 | 0.0087 | 0.0087 | TRUE | 0.179 |
| Sh3d19 | 2158 | 2736 | 2789 | 3193 | 2300 | 2469 | 2292 | 2732 | 2800 | 2510 | 2475 | 2703 | 2585.42 | 2608 | 2563 | 0.982 | -0.026 | 0.80049192 | 0.89700593 | 0.0065 | 0.0082 | 0.0072 | 0.0072 | TRUE | 0.1361 |
| Sh3d21 | 119 | 109 | 99 | 167 | 117 | 108 | 126 | 109 | 99 | 131 | 126 | 118 | 118.34 | 111 | 125 | 1.125 | 0.169 | 0.37397372 | 0.59024452 | 0 | 0.0347 | 0.0176 | 0.0176 | TRUE | 0.1324 |
| Sh3gl1 | 2989 | 2886 | 2750 | 4367 | 3129 | 2708 | 3175 | 2882 | 2761 | 3433 | 3367 | 2964 | 3097.06 | 2939 | 3255 | 1.108 | 0.147 | 0.12700673 | 0.30605808 | 0.0052 | 0.008 | 0.0064 | 0.0064 | TRUE | 0.0736 |
| Sh3gl2 | 1484 | 1668 | 1707 | 1620 | 1283 | 1496 | 1576 | 1666 | 1714 | 1274 | 1380 | 1638 | 1541.28 | 1652 | 1431 | 0.865 | -0.209 | 0.06746559 | 0.20084321 | 0.0085 | 0.009 | 0.0088 | 0.0088 | TRUE | 0.1923 |
| Sh3gl3 | 878 | 711 | 686 | 882 | 593 | 598 | 933 | 710 | 689 | 693 | 638 | 655 | 719.58 | 777 | 662 | 0.853 | -0.23 | 0.10128128 | 0.26359873 | 0.0139 | 0.0115 | 0.0128 | 0.0128 | TRUE | 0.3615 |
| Sh3glb1 | 3068 | 3255 | 3164 | 4499 | 3281 | 3114 | 3259 | 3251 | 3177 | 3537 | 3530 | 3409 | 3360.37 | 3229 | 3492 | 1.082 | 0.113 | 0.06669597 | 0.19928507 | 0 | 0.0079 | 0.0024 | 0.0024 | TRUE | 0.0052 |
| Sh3glb2 | 1157 | 1034 | 957 | 1356 | 1066 | 908 | 1229 | 1033 | 961 | 1066 | 1147 | 994 | 1071.57 | 1074 | 1069 | 0.995 | -0.007 | 0.95648937 | 0.97766996 | 0.0097 | 0.01 | 0.0098 | 0.0098 | TRUE | 0.1893 |
| Sh3kbp1 | 401 | 333 | 297 | 485 | 357 | 301 | 426 | 333 | 298 | 381 | 384 | 329 | 358.6 | 352 | 365 | 1.037 | 0.053 | 0.74815657 | 0.86643111 | 0.0176 | 0.0161 | 0.0169 | 0.0169 | TRUE | 0.3797 |
| Sh3pxd2a | 2803 | 2935 | 3040 | 2803 | 2073 | 2202 | 2977 | 2931 | 3052 | 2203 | 2230 | 2410 | 2634.22 | 2987 | 2281 | 0.763 | -0.389 | 1.01E-07 | 3.60E-06 | 8.00E-04 | 0.0082 | 0.0035 | 0.0035 | TRUE | 0.0293 |
| Sh3pxd2b | 1635 | 1528 | 1552 | 1953 | 1301 | 1330 | 1737 | 1526 | 1558 | 1535 | 1400 | 1456 | 1535.34 | 1607 | 1464 | 0.911 | -0.134 | 0.14098551 | 0.32640025 | 0.0026 | 0.009 | 0.0053 | 0.0053 | TRUE | 0.0598 |
| Sh3rf1 | 1475 | 1463 | 1423 | 2171 | 1484 | 1436 | 1567 | 1461 | 1429 | 1707 | 1597 | 1572 | 1555.33 | 1486 | 1625 | 1.095 | 0.131 | 0.11583661 | 0.287441 | 0.0015 | 0.009 | 0.0043 | 0.0043 | TRUE | 0.0273 |
| Sh3rf2 | 13 | 10 | 7 | 49 | 23 | 35 | 14 | 10 | 7 | 39 | 25 | 38 | 22.07 | 10 | 34 | 3.316 | 1.73 | 0.00025017 | 0.00292927 | 0 | 0.1562 | 0.0965 | 0.0965 | TRUE | 0.3687 |
| Sh3rf3 | 111 | 126 | 115 | 128 | 104 | 85 | 118 | 126 | 115 | 101 | 112 | 93 | 110.8 | 120 | 102 | 0.85 | -0.234 | 0.21664822 | 0.42867592 | 0 | 0.0366 | 0.0168 | 0.0168 | TRUE | 0.0676 |
| Sh3tc1 | 319 | 431 | 456 | 584 | 416 | 463 | 339 | 430 | 458 | 459 | 448 | 507 | 440.11 | 409 | 471 | 1.151 | 0.203 | 0.16331682 | 0.35895107 | 0.0116 | 0.0144 | 0.0129 | 0.0129 | TRUE | 0.2574 |
| Sh3tc2 | 153 | 129 | 132 | 175 | 133 | 105 | 163 | 129 | 133 | 138 | 143 | 115 | 136.58 | 142 | 132 | 0.935 | -0.096 | 0.614874 | 0.78208006 | 0.0069 | 0.031 | 0.0191 | 0.0191 | TRUE | 0.1763 |
| Sh3yl1 | 190 | 161 | 139 | 352 | 245 | 214 | 202 | 161 | 140 | 277 | 264 | 234 | 212.79 | 168 | 258 | 1.546 | 0.629 | 0.00050187 | 0.00517876 | 0.014 | 0.0224 | 0.0186 | 0.0186 | TRUE | 0.3393 |
| Shank1 | 81 | 60 | 47 | 77 | 76 | 40 | 86 | 60 | 47 | 61 | 82 | 44 | 63.21 | 64 | 62 | 0.965 | -0.052 | 0.87769687 | 0.93959377 | 0.0753 | 0.059 | 0.0668 | 0.0668 | TRUE | 0.2932 |
| Shank2 | 64 | 53 | 50 | 93 | 64 | 47 | 68 | 53 | 50 | 73 | 69 | 51 | 60.76 | 57 | 64 | 1.136 | 0.185 | 0.50299248 | 0.69812793 | 0.0133 | 0.0611 | 0.0382 | 0.0382 | TRUE | 0.2604 |
| Shank3 | 583 | 668 | 663 | 617 | 501 | 467 | 619 | 667 | 666 | 485 | 539 | 511 | 581.23 | 651 | 512 | 0.785 | -0.349 | 0.0006111 | 0.00609632 | 0 | 0.0126 | 0.0057 | 0.0057 | TRUE | 0.026 |
| Sharpin | 931 | 1107 | 1125 | 1332 | 1049 | 983 | 989 | 1106 | 1130 | 1047 | 1129 | 1076 | 1079.32 | 1075 | 1084 | 1.008 | 0.011 | 0.903518 | 0.95228213 | 0.0023 | 0.01 | 0.0054 | 0.0054 | TRUE | 0.0579 |
| Shb | 612 | 686 | 670 | 810 | 667 | 702 | 650 | 685 | 673 | 637 | 718 | 768 | 688.46 | 669 | 708 | 1.055 | 0.078 | 0.47483706 | 0.67524193 | 0.0037 | 0.0117 | 0.007 | 0.007 | TRUE | 0.0948 |
| Shbg | 3 | 5 | 2 | 6 | 7 | 2 | 3 | 5 | 2 | 5 | 8 | 2 | 4.1 | 3 | 5 | 1.416 | 0.502 | 0.60448457 | NA | 0 | 0.8096 | 0.4258 | 0.4258 | TRUE | 0.1489 |
| Shc1 | 3203 | 3238 | 3315 | 4642 | 3288 | 2960 | 3402 | 3234 | 3329 | 3649 | 3538 | 3240 | 3398.6 | 3322 | 3476 | 1.047 | 0.066 | 0.39352201 | 0.6070053 | 0.0017 | 0.0079 | 0.004 | 0.004 | TRUE | 0.0425 |
| Shc2 | 384 | 421 | 426 | 448 | 347 | 274 | 408 | 420 | 428 | 352 | 373 | 300 | 380.26 | 419 | 342 | 0.817 | -0.291 | 0.0218376 | 0.09273631 | 0.0029 | 0.0156 | 0.009 | 0.009 | TRUE | 0.125 |
| Shc3 | 14 | 7 | 7 | 25 | 15 | 11 | 15 | 7 | 7 | 20 | 16 | 12 | 12.79 | 10 | 16 | 1.674 | 0.744 | 0.19953925 | 0.40713235 | 0.0331 | 0.2645 | 0.1584 | 0.1584 | TRUE | 0.5907 |
| Shc4 | 23 | 20 | 18 | 30 | 19 | 12 | 24 | 20 | 18 | 24 | 20 | 13 | 19.94 | 21 | 19 | 0.924 | -0.113 | 0.79900029 | 0.89640568 | 0 | 0.1721 | 0.0927 | 0.0927 | TRUE | 0.3583 |
| Shcbp1 | 2619 | 2528 | 2668 | 4306 | 3158 | 2943 | 2782 | 2525 | 2679 | 3385 | 3398 | 3222 | 2998.33 | 2662 | 3335 | 1.253 | 0.325 | 1.05E-05 | 0.00020784 | 0.0011 | 0.008 | 0.0036 | 0.0036 | TRUE | 0.0247 |
| Shcbp1l | 134 | 164 | 102 | 141 | 130 | 97 | 142 | 164 | 102 | 111 | 140 | 106 | 127.57 | 136 | 119 | 0.873 | -0.197 | 0.40299632 | 0.61589709 | 0.0311 | 0.0327 | 0.032 | 0.032 | TRUE | 0.405 |
| Shd | 16 | 10 | 14 | 13 | 9 | 8 | 17 | 10 | 14 | 10 | 10 | 9 | 11.62 | 14 | 10 | 0.702 | -0.511 | 0.35930499 | 0.57787504 | 0 | 0.2905 | 0.1353 | 0.1353 | TRUE | 0.234 |
| She | 206 | 238 | 238 | 304 | 254 | 222 | 219 | 238 | 239 | 239 | 273 | 243 | 241.79 | 232 | 252 | 1.084 | 0.116 | 0.40144984 | 0.6141287 | 0 | 0.0205 | 0.0097 | 0.0097 | TRUE | 0.0607 |
| Shf | 273 | 323 | 288 | 290 | 245 | 207 | 290 | 323 | 289 | 228 | 264 | 227 | 269.99 | 301 | 240 | 0.795 | -0.331 | 0.01680148 | 0.0768318 | 0.0017 | 0.0191 | 0.01 | 0.01 | TRUE | 0.0839 |
| Shfm1 | 2741 | 3209 | 3341 | 4005 | 2782 | 3017 | 2911 | 3205 | 3355 | 3148 | 2993 | 3303 | 3152.54 | 3157 | 3148 | 0.997 | -0.004 | 0.96127115 | 0.98035361 | 0.0034 | 0.0079 | 0.0052 | 0.0052 | TRUE | 0.0561 |
| Shh | 2 | 0 | 2 | 5 | 0 | 1 | 2 | 0 | 2 | 4 | 0 | 1 | 1.53 | 1 | 2 | 1.263 | 0.337 | 0.85161451 | NA | 0.9578 | 2.1654 | 1.6822 | 1.6822 | TRUE | 0.7889 |
| Shisa2 | 373 | 454 | 416 | 432 | 353 | 323 | 396 | 453 | 418 | 340 | 380 | 354 | 390.05 | 422 | 358 | 0.845 | -0.242 | 0.04281113 | 0.14775471 | 0.0014 | 0.0153 | 0.0077 | 0.0077 | TRUE | 0.0477 |
| Shisa3 | 17 | 36 | 23 | 18 | 22 | 7 | 18 | 36 | 23 | 14 | 24 | 8 | 20.43 | 26 | 15 | 0.588 | -0.765 | 0.15200748 | 0.34313181 | 0.136 | 0.1681 | 0.1534 | 0.1534 | TRUE | 0.493 |
| Shisa4 | 115 | 223 | 264 | 206 | 163 | 186 | 122 | 223 | 265 | 162 | 175 | 204 | 191.81 | 203 | 180 | 0.885 | -0.176 | 0.51883136 | 0.71026891 | 0.0759 | 0.0241 | 0.0483 | 0.0483 | TRUE | 0.5971 |
| Shisa5 | 1182 | 1660 | 1611 | 1900 | 1406 | 1455 | 1255 | 1658 | 1618 | 1494 | 1513 | 1593 | 1521.68 | 1510 | 1533 | 1.015 | 0.021 | 0.8645816 | 0.93296258 | 0.0113 | 0.0091 | 0.0103 | 0.0103 | TRUE | 0.2615 |
| Shisa6 | 23 | 23 | 20 | 33 | 22 | 12 | 24 | 23 | 20 | 26 | 24 | 13 | 21.71 | 22 | 21 | 0.939 | -0.091 | 0.8342877 | 0.91571799 | 0 | 0.1587 | 0.091 | 0.091 | TRUE | 0.5386 |
| Shisa7 | 199 | 238 | 273 | 505 | 334 | 295 | 211 | 238 | 274 | 397 | 359 | 323 | 300.41 | 241 | 360 | 1.494 | 0.579 | 0.00018649 | 0.00229435 | 0.0101 | 0.0179 | 0.0139 | 0.0139 | TRUE | 0.1595 |
| Shisa9 | 1 | 1 | 4 | 3 | 2 | 3 | 1 | 1 | 4 | 2 | 2 | 3 | 2.31 | 2 | 2 | 1.273 | 0.348 | 0.77920636 | NA | 0 | 1.4319 | 0.6721 | 0.6721 | TRUE | 0.6786 |
| Shkbp1 | 1483 | 1512 | 1581 | 2047 | 1447 | 1435 | 1575 | 1510 | 1587 | 1609 | 1557 | 1571 | 1568.28 | 1557 | 1579 | 1.014 | 0.02 | 0.77950526 | 0.88603691 | 0 | 0.009 | 0.0031 | 0.0031 | TRUE | 0.0086 |
| Shmt1 | 10267 | 9651 | 9523 | 13097 | 9379 | 9042 | 10905 | 9639 | 9562 | 10296 | 10091 | 9898 | 10065.16 | 10035 | 10095 | 1.006 | 0.009 | 0.91250896 | 0.95675253 | 0.0023 | 0.0072 | 0.0044 | 0.0044 | TRUE | 0.0702 |
| Shmt2 | 8726 | 9417 | 9782 | 10071 | 7769 | 8022 | 9269 | 9405 | 9822 | 7917 | 8359 | 8781 | 8925.48 | 9499 | 8352 | 0.879 | -0.186 | 0.00891137 | 0.04829904 | 0.0016 | 0.0073 | 0.0035 | 0.0035 | TRUE | 0.0256 |
| Shoc2 | 2686 | 3087 | 3115 | 4176 | 3113 | 3140 | 2853 | 3083 | 3128 | 3283 | 3349 | 3437 | 3188.88 | 3021 | 3356 | 1.111 | 0.151 | 0.03638174 | 0.13142759 | 0.001 | 0.0079 | 0.0035 | 0.0035 | TRUE | 0.0287 |
| Shox2 | 3 | 3 | 3 | 10 | 3 | 5 | 3 | 3 | 3 | 8 | 3 | 5 | 4.29 | 3 | 5 | 1.828 | 0.87 | 0.34357079 | NA | 0 | 0.7743 | 0.3541 | 0.3541 | TRUE | 0.1508 |
| Shpk | 171 | 197 | 205 | 220 | 150 | 162 | 182 | 197 | 206 | 173 | 161 | 177 | 182.65 | 195 | 170 | 0.876 | -0.191 | 0.20761389 | 0.41691957 | 0 | 0.0249 | 0.0111 | 0.0111 | TRUE | 0.0369 |
| Shprh | 2440 | 2360 | 2419 | 3214 | 2295 | 2266 | 2592 | 2357 | 2429 | 2527 | 2469 | 2480 | 2475.66 | 2459 | 2492 | 1.014 | 0.02 | 0.78748983 | 0.89059871 | 8.00E-04 | 0.0082 | 0.0034 | 0.0034 | TRUE | 0.0268 |
| Shq1 | 2431 | 2394 | 2331 | 3365 | 2475 | 2357 | 2582 | 2391 | 2341 | 2645 | 2663 | 2580 | 2533.66 | 2438 | 2629 | 1.079 | 0.109 | 0.13930995 | 0.32392695 | 9.00E-04 | 0.0082 | 0.0036 | 0.0036 | TRUE | 0.0324 |
| Shroom1 | 31 | 18 | 14 | 59 | 49 | 24 | 33 | 18 | 14 | 46 | 53 | 26 | 31.72 | 22 | 42 | 1.94 | 0.956 | 0.03569877 | 0.12964334 | 0.118 | 0.1108 | 0.114 | 0.114 | TRUE | 1.0191 |
| Shroom2 | 2445 | 2436 | 2668 | 2704 | 1925 | 1861 | 2597 | 2433 | 2679 | 2126 | 2071 | 2037 | 2323.81 | 2570 | 2078 | 0.809 | -0.306 | 4.23E-05 | 0.00067081 | 0.0011 | 0.0083 | 0.0036 | 0.0036 | TRUE | 0.0264 |
| Shroom3 | 2736 | 2255 | 2317 | 2870 | 2131 | 2030 | 2906 | 2252 | 2327 | 2256 | 2293 | 2222 | 2375.98 | 2495 | 2257 | 0.905 | -0.144 | 0.20702655 | 0.41621705 | 0.0098 | 0.0083 | 0.009 | 0.009 | TRUE | 0.2516 |
| Shroom4 | 327 | 253 | 287 | 479 | 315 | 298 | 347 | 253 | 288 | 377 | 339 | 326 | 321.64 | 296 | 347 | 1.175 | 0.233 | 0.13712859 | 0.32113122 | 0.0122 | 0.0171 | 0.0146 | 0.0146 | TRUE | 0.2565 |
| Siae | 786 | 890 | 1037 | 1294 | 907 | 1093 | 835 | 889 | 1041 | 1017 | 976 | 1196 | 992.43 | 922 | 1063 | 1.153 | 0.205 | 0.10951869 | 0.27762038 | 0.0113 | 0.0102 | 0.0108 | 0.0108 | TRUE | 0.1537 |
| Siah1a | 1064 | 1149 | 1113 | 1412 | 1054 | 937 | 1130 | 1148 | 1118 | 1110 | 1134 | 1026 | 1110.83 | 1132 | 1090 | 0.963 | -0.054 | 0.51988614 | 0.71100654 | 4.00E-04 | 0.0099 | 0.0041 | 0.0041 | TRUE | 0.0309 |
| Siah1b | 1711 | 1637 | 1415 | 2196 | 1438 | 1332 | 1817 | 1635 | 1421 | 1726 | 1547 | 1458 | 1600.78 | 1624 | 1577 | 0.972 | -0.041 | 0.73136161 | 0.85590527 | 0.0106 | 0.009 | 0.0099 | 0.0099 | TRUE | 0.145 |
| Siah2 | 519 | 681 | 591 | 834 | 673 | 623 | 551 | 680 | 593 | 656 | 724 | 682 | 647.75 | 608 | 687 | 1.129 | 0.175 | 0.13457486 | 0.3173142 | 0.0053 | 0.012 | 0.0083 | 0.0083 | TRUE | 0.1258 |
| Siah3 | 0 | 0 | 0 | 0 | 0 | 0 | 0 | 0 | 0 | 0 | 0 | 0 | 0 | 0 | 0 | NA | NA | NA | NA | NA | NA | NA | NA | NA | NA |
| Sidt1 | 7 | 5 | 3 | 10 | 4 | 7 | 7 | 5 | 3 | 8 | 4 | 8 | 5.88 | 5 | 7 | 1.297 | 0.375 | 0.63758169 | 0.79603827 | 0 | 0.5674 | 0.2831 | 0.2831 | TRUE | 0.2731 |
| Sidt2 | 2014 | 2297 | 2418 | 3335 | 2305 | 2394 | 2139 | 2294 | 2428 | 2622 | 2480 | 2621 | 2430.59 | 2287 | 2574 | 1.126 | 0.171 | 0.0355202 | 0.12942897 | 0.0019 | 0.0083 | 0.0043 | 0.0043 | TRUE | 0.0386 |
| Sigirr | 451 | 631 | 629 | 697 | 503 | 599 | 479 | 630 | 632 | 548 | 541 | 656 | 580.94 | 580 | 582 | 1.001 | 0.002 | 0.99045968 | 0.99500191 | 0.0161 | 0.0126 | 0.0144 | 0.0144 | TRUE | 0.2714 |
| Siglec1 | 22 | 17 | 9 | 36 | 14 | 21 | 23 | 17 | 9 | 28 | 15 | 23 | 19.29 | 16 | 22 | 1.356 | 0.439 | 0.39228282 | 0.60598282 | 0.0906 | 0.1777 | 0.1371 | 0.1371 | TRUE | 0.3287 |
| Siglec15 | 1 | 0 | 1 | 1 | 3 | 2 | 1 | 0 | 1 | 1 | 3 | 2 | 1.38 | 1 | 2 | 2.966 | 1.569 | 0.36347209 | NA | 0 | 2.3972 | 1.1594 | 1.1594 | TRUE | 0.2032 |
| Siglec5 | 182 | 168 | 174 | 223 | 183 | 181 | 193 | 168 | 175 | 175 | 197 | 198 | 184.36 | 179 | 190 | 1.062 | 0.087 | 0.57507936 | 0.75313869 | 0 | 0.0248 | 0.012 | 0.012 | TRUE | 0.055 |
| Siglece | 10 | 10 | 5 | 7 | 3 | 5 | 11 | 10 | 5 | 6 | 3 | 5 | 6.64 | 9 | 5 | 0.559 | -0.839 | 0.26816527 | 0.4879603 | 0 | 0.5031 | 0.2518 | 0.2518 | TRUE | 0.4072 |
| Siglecg | 29 | 38 | 25 | 44 | 22 | 40 | 31 | 38 | 25 | 35 | 24 | 44 | 32.65 | 31 | 34 | 1.087 | 0.12 | 0.75053336 | 0.86797274 | 0.0329 | 0.1078 | 0.0726 | 0.0726 | TRUE | 0.2017 |
| Siglech | 0 | 0 | 0 | 0 | 0 | 0 | 0 | 0 | 0 | 0 | 0 | 0 | 0 | 0 | 0 | NA | NA | NA | NA | NA | NA | NA | NA | NA | NA |

| | | | | | | | | | | | | | | | | | | | | | | | | | |
|---|---|---|---|---|---|---|---|---|---|---|---|---|---|---|---|---|---|---|---|---|---|---|---|---|---|
| Sigmar1 | 4187 | 3655 | 3481 | 5150 | 3550 | 3232 | 4447 | 3650 | 3495 | 4049 | 3820 | 3538 | 3833.16 | 3864 | 3802 | 0.984 | -0.023 | 0.84090137 | 0.9194881 | 0.0104 | 0.0078 | 0.0092 | 0.0092 | TRUE | 0.2118 |
| Sik1 | 6928 | 8161 | 7623 | 10583 | 7973 | 7575 | 7359 | 8150 | 7654 | 8320 | 8578 | 8292 | 8058.92 | 7721 | 8397 | 1.087 | 0.121 | 0.07899415 | 0.22291925 | 0.0013 | 0.0073 | 0.0033 | 0.0033 | TRUE | 0.0289 |
| Sik2 | 807 | 755 | 904 | 1143 | 813 | 801 | 857 | 754 | 908 | 899 | 875 | 877 | 861.5 | 840 | 884 | 1.052 | 0.074 | 0.47289065 | 0.67393434 | 0.0033 | 0.0107 | 0.0064 | 0.0064 | TRUE | 0.0952 |
| Sik3 | 2067 | 2210 | 2373 | 2698 | 1956 | 1991 | 2196 | 2207 | 2383 | 2121 | 2105 | 2179 | 2198.39 | 2262 | 2135 | 0.944 | -0.083 | 0.25595893 | 0.47460431 | 8.00E-04 | 0.0084 | 0.0034 | 0.0034 | TRUE | 0.0266 |
| Sike1 | 1584 | 1859 | 1928 | 2363 | 1773 | 1784 | 1682 | 1857 | 1936 | 1858 | 1908 | 1953 | 1865.52 | 1825 | 1906 | 1.044 | 0.062 | 0.46733232 | 0.66963926 | 0.002 | 0.0087 | 0.0047 | 0.0047 | TRUE | 0.0561 |
| Sil1 | 1153 | 1646 | 1524 | 1285 | 1016 | 1134 | 1225 | 1644 | 1530 | 1010 | 1093 | 1241 | 1290.58 | 1466 | 1115 | 0.76 | -0.396 | 0.004162 | 0.02724909 | 0.0153 | 0.0095 | 0.013 | 0.013 | TRUE | 0.2494 |
| Sim1 | 4 | 4 | 4 | 17 | 4 | 5 | 4 | 4 | 4 | 13 | 4 | 5 | 5.9 | 4 | 7 | 1.928 | 0.947 | 0.26164027 | 0.48118655 | 0 | 0.5653 | 0.3258 | 0.3258 | TRUE | 1.4607 |
| Sim2 | 4 | 8 | 4 | 9 | 7 | 6 | 4 | 8 | 4 | 7 | 8 | 7 | 6.24 | 5 | 7 | 1.3 | 0.379 | 0.61430247 | 0.78173136 | 0 | 0.535 | 0.2438 | 0.2438 | TRUE | 0.3795 |
| Simc1 | 1751 | 1556 | 1702 | 1841 | 1379 | 1382 | 1860 | 1554 | 1709 | 1447 | 1484 | 1513 | 1594.44 | 1708 | 1481 | 0.867 | -0.205 | 0.02956597 | 0.1138185 | 0.0033 | 0.009 | 0.0058 | 0.0058 | TRUE | 0.0748 |
| Sin3a | 13379 | 13425 | 14409 | 16609 | 12302 | 12839 | 14211 | 13408 | 14468 | 13057 | 13236 | 14054 | 13738.98 | 14029 | 13449 | 0.959 | -0.061 | 0.37168588 | 0.58818853 | 0.0014 | 0.0071 | 0.0033 | 0.0033 | TRUE | 0.0189 |
| Sin3b | 4596 | 4949 | 5142 | 7154 | 5385 | 5425 | 4882 | 4943 | 5163 | 5624 | 5794 | 5939 | 5390.64 | 4996 | 5786 | 1.158 | 0.211 | 0.00092894 | 0.00841058 | 5.00E-04 | 0.0075 | 0.0028 | 0.0028 | TRUE | 0.0105 |
| Sipa1 | 1519 | 1482 | 1430 | 1443 | 1123 | 1220 | 1613 | 1480 | 1436 | 1134 | 1208 | 1335 | 1367.92 | 1510 | 1226 | 0.811 | -0.301 | 0.00258125 | 0.01884424 | 0.0042 | 0.0093 | 0.0065 | 0.0065 | TRUE | 0.0728 |
| Sipa1l1 | 3298 | 3686 | 3994 | 4925 | 3828 | 4199 | 3503 | 3681 | 4010 | 3872 | 4119 | 4596 | 3963.58 | 3731 | 4196 | 1.124 | 0.169 | 0.08476068 | 0.23379598 | 0.0059 | 0.0077 | 0.0066 | 0.0066 | TRUE | 0.0849 |
| Sipa1l2 | 5878 | 6966 | 7389 | 7065 | 5308 | 5950 | 6243 | 6957 | 7419 | 5554 | 5711 | 6513 | 6399.68 | 6873 | 5926 | 0.862 | -0.214 | 0.0356054 | 0.12953124 | 0.0072 | 0.0074 | 0.0073 | 0.0073 | TRUE | 0.0915 |
| Sipa1l3 | 2730 | 2577 | 2605 | 3464 | 2622 | 2337 | 2900 | 2574 | 2616 | 2723 | 2821 | 2558 | 2698.59 | 2697 | 2701 | 1.002 | 0.003 | 0.97525762 | 0.98697924 | 0.0029 | 0.0081 | 0.0049 | 0.0049 | TRUE | 0.0527 |
| Sirpa | 391 | 517 | 502 | 696 | 549 | 633 | 415 | 516 | 504 | 547 | 591 | 693 | 544.41 | 478 | 610 | 1.273 | 0.348 | 0.01413418 | 0.0678554 | 0.0124 | 0.013 | 0.0126 | 0.0126 | TRUE | 0.1639 |
| Sirpb1a | 0 | 0 | 0 | 0 | 0 | 0 | 0 | 0 | 0 | 0 | 0 | 0 | 0 | 0 | 0 | NA | NA | NA | NA | NA | NA | NA | NA | NA | NA |
| Sirpb1b | 0 | 0 | 0 | 0 | 0 | 0 | 0 | 0 | 0 | 0 | 0 | 0 | 0 | 0 | 0 | NA | NA | NA | NA | NA | NA | NA | NA | NA | NA |
| Sirt1 | 5479 | 4518 | 4124 | 7489 | 5308 | 4389 | 5820 | 4512 | 4141 | 5887 | 5711 | 4804 | 5145.93 | 4824 | 5467 | 1.134 | 0.181 | 0.20929768 | 0.41928656 | 0.0213 | 0.0075 | 0.0148 | 0.0148 | TRUE | 0.3968 |
| Sirt2 | 826 | 876 | 887 | 1237 | 879 | 848 | 877 | 875 | 891 | 972 | 946 | 928 | 914.89 | 881 | 949 | 1.078 | 0.108 | 0.18366376 | 0.38643874 | 0 | 0.0105 | 0.0036 | 0.0036 | TRUE | 0.0059 |
| Sirt3 | 618 | 984 | 1031 | 882 | 657 | 807 | 656 | 983 | 1035 | 693 | 707 | 883 | 826.34 | 891 | 761 | 0.853 | -0.229 | 0.21760347 | 0.42962457 | 0.0372 | 0.0109 | 0.0236 | 0.0236 | TRUE | 0.6314 |
| Sirt4 | 293 | 364 | 376 | 411 | 341 | 359 | 311 | 364 | 378 | 323 | 367 | 393 | 355.88 | 351 | 361 | 1.026 | 0.038 | 0.78829049 | 0.8909931 | 0.0073 | 0.0162 | 0.0114 | 0.0114 | TRUE | 0.1098 |
| Sirt5 | 344 | 436 | 462 | 428 | 327 | 334 | 365 | 435 | 464 | 336 | 352 | 366 | 386.44 | 421 | 351 | 0.832 | -0.265 | 0.04732633 | 0.15788153 | 0.0059 | 0.0154 | 0.0103 | 0.0103 | TRUE | 0.1553 |
| Sirt6 | 625 | 651 | 726 | 1112 | 891 | 785 | 664 | 650 | 729 | 874 | 959 | 859 | 789.19 | 681 | 897 | 1.317 | 0.397 | 7.70E-05 | 0.00111998 | 0.0023 | 0.0111 | 0.006 | 0.006 | TRUE | 0.0451 |
| Sirt7 | 847 | 752 | 679 | 991 | 824 | 726 | 900 | 751 | 682 | 779 | 887 | 795 | 798.8 | 778 | 820 | 1.055 | 0.077 | 0.55698541 | 0.73964586 | 0.0111 | 0.011 | 0.0111 | 0.0111 | TRUE | 0.223 |
| Sis | 5 | 3 | 5 | 4 | 0 | 6 | 5 | 3 | 5 | 3 | 0 | 7 | 3.84 | 4 | 3 | 0.727 | -0.459 | 0.68247063 | NA | 0.2548 | 0.8648 | 0.6409 | 0.6409 | TRUE | 0.2578 |
| Sit1 | 16 | 20 | 14 | 22 | 15 | 16 | 17 | 20 | 14 | 17 | 16 | 18 | 17 | 17 | 17 | 0.999 | -0.001 | 0.99746978 | 0.99847569 | 0 | 0.2007 | 0.0888 | 0.0888 | TRUE | 0.1174 |
| Siva1 | 2487 | 3216 | 3328 | 3654 | 2640 | 2801 | 2642 | 3212 | 3342 | 2872 | 2840 | 3066 | 2995.71 | 3065 | 2926 | 0.955 | -0.067 | 0.53585424 | 0.72271961 | 0.0082 | 0.008 | 0.0081 | 0.0081 | TRUE | 0.1771 |
| Six1 | 432 | 801 | 841 | 1004 | 752 | 718 | 459 | 800 | 844 | 789 | 809 | 786 | 747.94 | 701 | 795 | 1.133 | 0.18 | 0.3835424 | 0.59786822 | 0.0505 | 0.0113 | 0.0296 | 0.0296 | TRUE | 1.0756 |
| Six2 | 1 | 20 | 14 | 36 | 27 | 24 | 1 | 20 | 14 | 28 | 29 | 26 | 19.79 | 12 | 28 | 2.373 | 1.247 | 0.06605885 | 0.19823509 | 0.4613 | 0.1734 | 0.2704 | 0.2704 | TRUE | 1.3693 |
| Six3 | 1 | 3 | 2 | 12 | 9 | 6 | 1 | 3 | 2 | 9 | 10 | 7 | 5.29 | 2 | 9 | 4.222 | 2.078 | 0.02370427 | 0.09797529 | 0 | 0.6294 | 0.3002 | 0.3002 | TRUE | 0.1623 |
| Six3os1 | 3 | 1 | 4 | 7 | 5 | 5 | 3 | 1 | 4 | 6 | 5 | 5 | 4.09 | 3 | 5 | 1.999 | 0.999 | 0.29499413 | NA | 0 | 0.8118 | 0.3793 | 0.3793 | TRUE | 0.3794 |
| Six4 | 972 | 1405 | 1523 | 1471 | 1190 | 1213 | 1032 | 1403 | 1529 | 1156 | 1280 | 1328 | 1288.24 | 1321 | 1255 | 0.949 | -0.076 | 0.61978464 | 0.78489204 | 0.0221 | 0.0095 | 0.016 | 0.016 | TRUE | 0.439 |
| Six5 | 236 | 355 | 337 | 326 | 279 | 277 | 251 | 355 | 338 | 256 | 300 | 303 | 300.55 | 315 | 286 | 0.909 | -0.138 | 0.42202436 | 0.63217478 | 0.0179 | 0.0179 | 0.0179 | 0.0179 | TRUE | 0.3538 |
| Six6 | 10 | 7 | 16 | 10 | 5 | 5 | 11 | 7 | 16 | 8 | 5 | 5 | 8.73 | 11 | 6 | 0.562 | -0.831 | 0.21891178 | 0.4310996 | 0 | 0.3842 | 0.2063 | 0.2063 | TRUE | 0.2148 |
| Ska1 | 1020 | 1067 | 1012 | 1631 | 1117 | 1075 | 1083 | 1066 | 1016 | 1282 | 1202 | 1177 | 1137.66 | 1055 | 1220 | 1.158 | 0.211 | 0.01216808 | 0.06078781 | 8.00E-04 | 0.0098 | 0.0042 | 0.0042 | TRUE | 0.0245 |
| Ska2 | 1472 | 1967 | 2058 | 2134 | 1483 | 1701 | 1564 | 1964 | 2066 | 1678 | 1596 | 1862 | 1788.27 | 1865 | 1712 | 0.918 | -0.124 | 0.33135968 | 0.55213544 | 0.0125 | 0.0087 | 0.0112 | 0.0112 | TRUE | 0.2408 |
| Ska3 | 2797 | 2706 | 2749 | 3492 | 2397 | 2324 | 2971 | 2703 | 2760 | 2745 | 2579 | 2544 | 2716.97 | 2811 | 2623 | 0.933 | -0.099 | 0.20155846 | 0.40908409 | 0.0015 | 0.0081 | 0.004 | 0.004 | TRUE | 0.0299 |
| Skap1 | 200 | 246 | 274 | 168 | 130 | 148 | 212 | 246 | 275 | 132 | 140 | 162 | 194.53 | 244 | 145 | 0.59 | -0.762 | 1.17E-05 | 0.00022765 | 0.0091 | 0.0238 | 0.0163 | 0.0163 | TRUE | 0.1384 |
| Skap2 | 2393 | 2508 | 2440 | 3017 | 2203 | 2273 | 2542 | 2505 | 2450 | 2372 | 2370 | 2488 | 2454.46 | 2499 | 2410 | 0.964 | -0.052 | 0.43345817 | 0.64234492 | 0 | 0.0082 | 0.0028 | 0.0028 | TRUE | 0.0096 |
| Ski | 7267 | 8130 | 8870 | 9719 | 7156 | 6639 | 7719 | 8120 | 8906 | 7640 | 7699 | 7267 | 7891.99 | 8248 | 7535 | 0.914 | -0.13 | 0.10578885 | 0.27134521 | 0.003 | 0.0073 | 0.0046 | 0.0046 | TRUE | 0.0595 |
| Skida1 | 100 | 119 | 132 | 241 | 227 | 171 | 106 | 119 | 133 | 189 | 244 | 187 | 163.08 | 119 | 207 | 1.732 | 0.793 | 3.19E-05 | 0.0005297 | 0.0123 | 0.0271 | 0.0195 | 0.0195 | TRUE | 0.2646 |
| Skil | 10791 | 12350 | 13435 | 10798 | 8272 | 9426 | 11462 | 12334 | 13490 | 8489 | 8900 | 10318 | 10832.18 | 12429 | 9236 | 0.743 | -0.429 | 4.94E-05 | 0.00076831 | 0.0086 | 0.0072 | 0.0079 | 0.0079 | TRUE | 0.1283 |
| Skint1 | 0 | 0 | 0 | 0 | 0 | 0 | 0 | 0 | 0 | 0 | 0 | 0 | 0 | 0 | 0 | NA | NA | NA | NA | NA | NA | NA | NA | NA | NA |
| Skint10 | 0 | 0 | 0 | 0 | 0 | 0 | 0 | 0 | 0 | 0 | 0 | 0 | 0 | 0 | 0 | NA | NA | NA | NA | NA | NA | NA | NA | NA | NA |
| Skint11 | 0 | 0 | 0 | 0 | 0 | 0 | 0 | 0 | 0 | 0 | 0 | 0 | 0 | 0 | 0 | NA | NA | NA | NA | NA | NA | NA | NA | NA | NA |
| Skint2 | 2 | 0 | 0 | 3 | 0 | 0 | 2 | 0 | 0 | 2 | 0 | 0 | 0.75 | 1 | 1 | 1.163 | 0.218 | 0.94242245 | NA | 8.7098 | 4.4164 | 5.2331 | 5.2331 | TRUE | 1.4402 |
| Skint3 | 0 | 0 | 0 | 0 | 0 | 0 | 0 | 0 | 0 | 0 | 0 | 0 | 0 | 0 | 0 | NA | NA | NA | NA | NA | NA | NA | NA | NA | NA |
| Skint4 | 0 | 0 | 0 | 0 | 0 | 0 | 0 | 0 | 0 | 0 | 0 | 0 | 0 | 0 | 0 | NA | NA | NA | NA | NA | NA | NA | NA | NA | NA |
| Skint5 | 0 | 0 | 1 | 1 | 0 | 0 | 0 | 0 | 1 | 1 | 0 | 0 | 0.3 | 0 | 0 | 0.954 | -0.067 | 0.98640617 | NA | 0 | 11.0483 | 9.2783 | 9.2783 | TRUE | 0.4098 |
| Skint6 | 0 | 0 | 0 | 0 | 0 | 0 | 0 | 0 | 0 | 0 | 0 | 0 | 0 | 0 | 0 | NA | NA | NA | NA | NA | NA | NA | NA | NA | NA |
| Skint7 | 0 | 0 | 0 | 0 | 0 | 0 | 0 | 0 | 0 | 0 | 0 | 0 | 0 | 0 | 0 | NA | NA | NA | NA | NA | NA | NA | NA | NA | NA |
| Skint8 | 0 | 0 | 0 | 0 | 0 | 0 | 0 | 0 | 0 | 0 | 0 | 0 | 0 | 0 | 0 | NA | NA | NA | NA | NA | NA | NA | NA | NA | NA |
| Skint9 | 0 | 0 | 0 | 0 | 0 | 0 | 0 | 0 | 0 | 0 | 0 | 0 | 0 | 0 | 0 | NA | NA | NA | NA | NA | NA | NA | NA | NA | NA |
| Skiv2l | 1619 | 1368 | 1409 | 2043 | 1643 | 1311 | 1720 | 1366 | 1415 | 1606 | 1768 | 1435 | 1551.6 | 1500 | 1603 | 1.069 | 0.096 | 0.44940051 | 0.65500976 | 0.0114 | 0.009 | 0.0109 | 0.0109 | TRUE | 0.1967 |
| Skiv2l2 | 6559 | 6030 | 6296 | 8103 | 5725 | 5994 | 6967 | 6022 | 6322 | 6370 | 6160 | 6561 | 6400.32 | 6437 | 6364 | 0.989 | -0.016 | 0.84068758 | 0.91938181 | 0.0026 | 0.0074 | 0.0047 | 0.0047 | TRUE | 0.0632 |
| Skor1 | 1 | 3 | 7 | 7 | 4 | 0 | 1 | 3 | 7 | 6 | 4 | 0 | 3.48 | 4 | 3 | 0.899 | -0.154 | 0.90273933 | NA | 0.7391 | 0.953 | 0.864 | 0.864 | TRUE | 0.5278 |
| Skor2 | 0 | 0 | 1 | 0 | 0 | 0 | 0 | 0 | 1 | 0 | 0 | 0 | 0.17 | 0 | 0 | 0.49 | -1.029 | 0.80087 | NA | 0 | 19.6928 | 10 | 10 | TRUE | 0.4098 |
| Skp1a | 14432 | 14585 | 14145 | 23970 | 17250 | 17051 | 15329 | 14566 | 14203 | 18843 | 18560 | 18665 | 16694.48 | 14699 | 18689 | 1.271 | 0.347 | 9.79E-09 | 4.62E-07 | 7.00E-04 | 0.0071 | 0.0026 | 0.0026 | TRUE | 0.0172 |
| Skp2 | 2632 | 2365 | 2332 | 3474 | 2400 | 2300 | 2796 | 2362 | 2342 | 2731 | 2582 | 2518 | 2555.02 | 2500 | 2610 | 1.045 | 0.063 | 0.52271378 | 0.71279973 | 0.0049 | 0.0082 | 0.0066 | 0.0066 | TRUE | 0.1297 |
| Sla | 1 | 0 | 0 | 0 | 0 | 0 | 1 | 0 | 0 | 0 | 0 | 0 | 0.18 | 0 | 0 | 0.49 | -1.029 | 0.80087 | NA | 0 | 18.6165 | 10 | 10 | TRUE | 0.4614 |
| Sla2 | 1 | 5 | 5 | 4 | 3 | 4 | 1 | 5 | 5 | 3 | 3 | 4 | 3.64 | 4 | 3 | 0.961 | -0.058 | 0.95416026 | NA | 0 | 0.9125 | 0.4604 | 0.4604 | TRUE | 0.5687 |
| Slain1 | 1146 | 1319 | 1272 | 1598 | 1254 | 1366 | 1217 | 1317 | 1277 | 1256 | 1349 | 1495 | 1318.75 | 1270 | 1367 | 1.075 | 0.104 | 0.28866013 | 0.50971201 | 0.004 | 0.0094 | 0.0062 | 0.0062 | TRUE | 0.0809 |
| Slain2 | 1391 | 1827 | 1926 | 2165 | 1612 | 1695 | 1477 | 1825 | 1934 | 1702 | 1734 | 1855 | 1754.64 | 1745 | 1764 | 1.01 | 0.015 | 0.9010404 | 0.95196343 | 0.0102 | 0.0088 | 0.0095 | 0.0095 | TRUE | 0.2169 |
| Slamf1 | 0 | 0 | 0 | 1 | 0 | 0 | 0 | 0 | 0 | 1 | 0 | 0 | 0.13 | 0 | 0 | 1.859 | 0.894 | 0.82650734 | NA | 0 | 25.1515 | 10 | 10 | TRUE | 0.2563 |
| Slamf6 | 0 | 0 | 1 | 12 | 6 | 3 | 0 | 0 | 1 | 9 | 6 | 3 | 3.36 | 0 | 6 | 17.64 | 4.141 | 0.00713869 | NA | 0 | 0.9866 | 0.6312 | 0.6312 | TRUE | 0.282 |
| Slamf7 | 4 | 1 | 0 | 5 | 2 | 5 | 4 | 1 | 0 | 4 | 2 | 5 | 2.8 | 2 | 4 | 2.228 | 1.156 | 0.37727631 | NA | 0.2107 | 1.1833 | 0.8119 | 0.8119 | TRUE | 0.8794 |
| Slamf8 | 0 | 0 | 0 | 0 | 1 | 1 | 0 | 0 | 0 | 0 | 1 | 1 | 0.36 | 0 | 1 | 3.794 | 1.924 | 0.60247382 | NA | 0 | 9.1135 | 8.1425 | 8.1425 | TRUE | 0.3482 |
| Slamf9 | 67 | 55 | 40 | 86 | 74 | 38 | 71 | 55 | 40 | 68 | 80 | 42 | 59.18 | 55 | 63 | 1.139 | 0.188 | 0.58325552 | 0.75875085 | 0.0729 | 0.0626 | 0.0675 | 0.0675 | TRUE | 0.3266 |

| | | | | | | | | | | | | | | | | | | | | | | | | |
|---|---|---|---|---|---|---|---|---|---|---|---|---|---|---|---|---|---|---|---|---|---|---|---|---|
| Slbp | 3706 | 3658 | 3590 | 7229 | 5059 | 4539 | 3936 | 3653 | 3605 | 5683 | 5443 | 4969 | 4548.19 | 3731 | 5365 | 1.438 | 0.524 | 4.60E-10 | 3.01E-08 | 0.0031 | 0.0076 | 0.0049 | 0.0049 | TRUE | 0.0508 |
| Slc10a1 | 0 | 0 | 0 | 2 | 0 | 0 | 0 | 0 | 0 | 2 | 0 | 0 | 0.26 | 0 | 1 | 2.886 | 1.529 | 0.70676491 | NA | 0 | 12.5792 | 10 | 10 | TRUE | 0.9669 |
| Slc10a2 | 0 | 0 | 0 | 0 | 0 | 0 | 0 | 0 | 0 | 0 | 0 | 0 | 0 | 0 | 0 | NA | NA | NA | NA | NA | NA | NA | NA | NA | NA |
| Slc10a3 | 82 | 67 | 58 | 149 | 109 | 52 | 87 | 67 | 58 | 117 | 117 | 57 | 83.93 | 71 | 97 | 1.378 | 0.463 | 0.15756151 | 0.35089807 | 0.0856 | 0.0462 | 0.0649 | 0.0649 | TRUE | 1.2269 |
| Slc10a4 | 9 | 6 | 1 | 18 | 12 | 12 | 10 | 6 | 1 | 14 | 13 | 13 | 9.46 | 6 | 13 | 2.445 | 1.29 | 0.08058827 | 0.22576168 | 0 | 0.3552 | 0.263 | 0.263 | TRUE | 0.6909 |
| Slc10a5 | 2 | 2 | 1 | 2 | 2 | 0 | 2 | 2 | 1 | 2 | 2 | 0 | 1.47 | 2 | 1 | 0.738 | -0.439 | 0.77862381 | NA | 0 | 2.2404 | 1.0667 | 1.0667 | TRUE | 0.374 |
| Slc10a6 | 42 | 48 | 46 | 69 | 45 | 50 | 45 | 48 | 46 | 54 | 48 | 55 | 49.36 | 46 | 52 | 1.135 | 0.183 | 0.49363859 | 0.68997411 | 0 | 0.0737 | 0.0313 | 0.0313 | TRUE | 0.0353 |
| Slc10a7 | 1310 | 1160 | 1096 | 1913 | 1328 | 1158 | 1391 | 1159 | 1100 | 1504 | 1429 | 1268 | 1308.46 | 1217 | 1400 | 1.151 | 0.203 | 0.09793092 | 0.25725624 | 0.0106 | 0.0094 | 0.0101 | 0.0101 | TRUE | 0.1885 |
| Slc11a1 | 165 | 163 | 187 | 179 | 127 | 175 | 175 | 163 | 188 | 141 | 137 | 192 | 165.79 | 175 | 157 | 0.889 | -0.169 | 0.37692992 | 0.59206639 | 0.0139 | 0.0268 | 0.0203 | 0.0203 | TRUE | 0.396 |
| Slc11a2 | 2794 | 2510 | 2336 | 3534 | 2392 | 2420 | 2968 | 2507 | 2346 | 2778 | 2574 | 2649 | 2636.82 | 2607 | 2667 | 1.023 | 0.033 | 0.75587422 | 0.87101981 | 0.0078 | 0.0082 | 0.0079 | 0.0079 | TRUE | 0.1777 |
| Slc12a1 | 2 | 2 | 1 | 1 | 3 | 4 | 2 | 2 | 1 | 1 | 3 | 4 | 2.25 | 2 | 3 | 1.599 | 0.677 | 0.60032778 | NA | 0 | 1.4691 | 0.7239 | 0.7239 | TRUE | 0.5789 |
| Slc12a2 | 4508 | 3688 | 3668 | 5813 | 3938 | 3458 | 4788 | 3683 | 3683 | 4570 | 4237 | 3785 | 4124.45 | 4051 | 4197 | 1.036 | 0.051 | 0.69470488 | 0.83270212 | 0.0159 | 0.0077 | 0.0121 | 0.0121 | TRUE | 0.3078 |
| Slc12a3 | 60 | 72 | 73 | 77 | 60 | 48 | 64 | 72 | 73 | 61 | 65 | 53 | 64.43 | 70 | 60 | 0.851 | -0.233 | 0.33584019 | 0.55598526 | 0 | 0.058 | 0.0268 | 0.0268 | TRUE | 0.0758 |
| Slc12a4 | 2901 | 3215 | 3465 | 3496 | 2505 | 2600 | 3081 | 3211 | 3479 | 2748 | 2695 | 2846 | 3010.18 | 3257 | 2763 | 0.848 | -0.237 | 0.00254796 | 0.01864798 | 0.0018 | 0.008 | 0.0041 | 0.0041 | TRUE | 0.0433 |
| Slc12a5 | 202 | 179 | 178 | 135 | 134 | 104 | 215 | 179 | 179 | 106 | 144 | 114 | 156.03 | 191 | 121 | 0.635 | -0.656 | 0.00062912 | 0.00622877 | 0.0114 | 0.028 | 0.0198 | 0.0198 | TRUE | 0.2676 |
| Slc12a6 | 3228 | 3502 | 3634 | 4003 | 2823 | 3030 | 3429 | 3497 | 3649 | 3147 | 3037 | 3317 | 3346.02 | 3525 | 3167 | 0.898 | -0.155 | 0.03197552 | 0.12041974 | 0.001 | 0.0079 | 0.0034 | 0.0034 | TRUE | 0.0205 |
| Slc12a7 | 6595 | 9591 | 9167 | 9488 | 7429 | 8699 | 7005 | 9579 | 9205 | 7459 | 7993 | 9522 | 8460.43 | 8596 | 8325 | 0.968 | -0.046 | 0.74642855 | 0.86538305 | 0.0218 | 0.0073 | 0.0147 | 0.0147 | TRUE | 0.3201 |
| Slc12a8 | 594 | 885 | 911 | 540 | 502 | 670 | 631 | 884 | 915 | 425 | 540 | 733 | 687.93 | 810 | 566 | 0.698 | -0.519 | 0.01617694 | 0.07497251 | 0.0543 | 0.0117 | 0.0321 | 0.0321 | TRUE | 0.5648 |
| Slc12a9 | 675 | 713 | 659 | 841 | 681 | 647 | 717 | 712 | 662 | 661 | 733 | 708 | 698.81 | 697 | 701 | 1.004 | 0.006 | 0.94892059 | 0.9743734 | 0.0011 | 0.0116 | 0.0054 | 0.0054 | TRUE | 0.0304 |
| Slc13a1 | 0 | 0 | 0 | 0 | 0 | 0 | 0 | 0 | 0 | 0 | 0 | 0 | 0 | 0 | 0 | NA | NA | NA | NA | NA | NA | NA | NA | NA | NA |
| Slc13a2 | 6 | 11 | 9 | 11 | 10 | 10 | 6 | 11 | 9 | 9 | 11 | 11 | 9.46 | 9 | 10 | 1.141 | 0.19 | 0.7554608 | 0.87079791 | 0 | 0.3552 | 0.161 | 0.161 | TRUE | 0.174 |
| Slc13a3 | 35 | 27 | 20 | 17 | 15 | 13 | 37 | 27 | 20 | 13 | 16 | 14 | 21.33 | 28 | 14 | 0.518 | -0.948 | 0.03366378 | 0.12468189 | 0 | 0.1614 | 0.0918 | 0.0918 | TRUE | 0.1921 |
| Slc13a4 | 171 | 170 | 150 | 191 | 131 | 128 | 182 | 170 | 151 | 150 | 141 | 140 | 155.54 | 168 | 144 | 0.861 | -0.216 | 0.19310682 | 0.39859714 | 0 | 0.0281 | 0.0135 | 0.0135 | TRUE | 0.0816 |
| Slc13a5 | 554 | 609 | 571 | 515 | 390 | 417 | 588 | 608 | 573 | 405 | 420 | 456 | 508.49 | 590 | 427 | 0.723 | -0.469 | 9.03E-06 | 0.00018228 | 0 | 0.0134 | 0.006 | 0.006 | TRUE | 0.0414 |
| Slc14a1 | 0 | 0 | 0 | 2 | 0 | 0 | 0 | 0 | 0 | 2 | 0 | 0 | 0.26 | 0 | 1 | 2.886 | 1.529 | 0.70676491 | NA | 0 | 12.5792 | 10 | 10 | TRUE | 0.9669 |
| Slc14a2 | 1 | 1 | 2 | 2 | 4 | 1 | 1 | 1 | 2 | 2 | 4 | 1 | 1.84 | 1 | 2 | 1.693 | 0.759 | 0.58635164 | NA | 0 | 1.7974 | 0.8144 | 0.8144 | TRUE | 0.5247 |
| Slc15a1 | 863 | 1517 | 1602 | 1264 | 983 | 1222 | 917 | 1515 | 1609 | 994 | 1058 | 1338 | 1238.21 | 1347 | 1130 | 0.838 | -0.254 | 0.22344611 | 0.43546176 | 0.0556 | 0.0096 | 0.0306 | 0.0306 | TRUE | 0.9365 |
| Slc15a2 | 364 | 329 | 337 | 313 | 239 | 329 | 387 | 329 | 338 | 246 | 257 | 360 | 319.49 | 351 | 288 | 0.818 | -0.29 | 0.10499988 | 0.27022343 | 0.0231 | 0.0172 | 0.0199 | 0.0199 | TRUE | 0.5359 |
| Slc15a3 | 0 | 7 | 5 | 3 | 3 | 3 | 0 | 7 | 5 | 2 | 3 | 3 | 3.48 | 4 | 3 | 0.73 | -0.455 | 0.70160098 | NA | 0 | 0.9535 | 0.7213 | 0.7213 | TRUE | 0.5827 |
| Slc15a4 | 1272 | 1240 | 1286 | 1751 | 1310 | 1225 | 1351 | 1238 | 1291 | 1376 | 1409 | 1341 | 1334.62 | 1293 | 1375 | 1.064 | 0.089 | 0.26304301 | 0.48235255 | 4.00E-04 | 0.0094 | 0.0038 | 0.0038 | TRUE | 0.018 |
| Slc15a5 | 0 | 4 | 0 | 0 | 0 | 1 | 0 | 4 | 0 | 0 | 0 | 1 | 0.85 | 1 | 0 | 0.261 | -1.935 | 0.51113784 | NA | 9.1219 | 3.8907 | 4.8708 | 4.8708 | TRUE | 1.8984 |
| Slc16a1 | 9961 | 10367 | 9622 | 13302 | 9623 | 9358 | 10580 | 10354 | 9662 | 10457 | 10354 | 10244 | 10275 | 10199 | 10352 | 1.015 | 0.022 | 0.74108165 | 0.86159073 | 9.00E-04 | 0.0072 | 0.003 | 0.003 | TRUE | 0.0259 |
| Slc16a10 | 1019 | 1017 | 1017 | 1143 | 914 | 815 | 1082 | 1016 | 1021 | 899 | 983 | 892 | 982.22 | 1040 | 925 | 0.889 | -0.17 | 0.05719505 | 0.18023584 | 0.001 | 0.0103 | 0.0047 | 0.0047 | TRUE | 0.036 |
| Slc16a11 | 43 | 88 | 60 | 30 | 29 | 43 | 46 | 88 | 60 | 24 | 31 | 47 | 49.28 | 65 | 34 | 0.52 | -0.942 | 0.01363368 | 0.06589961 | 0.0923 | 0.0738 | 0.0827 | 0.0827 | TRUE | 0.2944 |
| Slc16a12 | 5 | 6 | 5 | 10 | 7 | 6 | 5 | 6 | 5 | 8 | 8 | 7 | 6.38 | 5 | 8 | 1.349 | 0.432 | 0.54911624 | 0.73382997 | 0 | 0.5232 | 0.2145 | 0.2145 | TRUE | 0.0202 |
| Slc16a13 | 220 | 199 | 207 | 393 | 254 | 196 | 234 | 199 | 208 | 309 | 273 | 215 | 239.51 | 214 | 266 | 1.249 | 0.32 | 0.07191849 | 0.20935741 | 0.0167 | 0.0207 | 0.0186 | 0.0186 | TRUE | 0.3028 |
| Slc16a14 | 49 | 76 | 80 | 140 | 103 | 77 | 52 | 76 | 80 | 110 | 111 | 84 | 85.57 | 69 | 102 | 1.466 | 0.552 | 0.03018356 | 0.11577216 | 0.0222 | 0.0454 | 0.0346 | 0.0346 | TRUE | 0.4188 |
| Slc16a2 | 47 | 47 | 38 | 77 | 55 | 39 | 50 | 47 | 38 | 61 | 59 | 43 | 49.57 | 45 | 54 | 1.208 | 0.273 | 0.35221897 | 0.57073249 | 0 | 0.0734 | 0.0418 | 0.0418 | TRUE | 0.2678 |
| Slc16a3 | 8821 | 13430 | 14181 | 15116 | 11977 | 14334 | 9369 | 13413 | 14239 | 11883 | 12887 | 15691 | 12913.62 | 12340 | 13487 | 1.093 | 0.128 | 0.44614262 | 0.65236459 | 0.0279 | 0.0072 | 0.0203 | 0.0203 | TRUE | 0.5421 |
| Slc16a4 | 4 | 7 | 5 | 16 | 7 | 5 | 4 | 7 | 5 | 13 | 8 | 5 | 6.97 | 5 | 9 | 1.595 | 0.674 | 0.36553559 | 0.58306561 | 0 | 0.4793 | 0.2488 | 0.2488 | TRUE | 0.6675 |
| Slc16a5 | 8 | 3 | 6 | 24 | 16 | 7 | 8 | 3 | 6 | 19 | 17 | 8 | 10.21 | 6 | 15 | 2.526 | 1.337 | 0.05405039 | 0.17281409 | 0.1043 | 0.3296 | 0.2259 | 0.2259 | TRUE | 0.7115 |
| Slc16a6 | 873 | 1423 | 1513 | 2460 | 1842 | 1579 | 927 | 1421 | 1519 | 1934 | 1982 | 1728 | 1585.31 | 1289 | 1881 | 1.459 | 0.545 | 0.00192501 | 0.01494159 | 0.0353 | 0.009 | 0.0216 | 0.0216 | TRUE | 0.7227 |
| Slc16a7 | 26 | 31 | 20 | 34 | 31 | 23 | 28 | 31 | 20 | 27 | 33 | 25 | 27.32 | 26 | 28 | 1.082 | 0.114 | 0.76328992 | 0.87540784 | 0 | 0.1275 | 0.0664 | 0.0664 | TRUE | 0.2653 |
| Slc16a8 | 5 | 4 | 4 | 1 | 1 | 3 | 5 | 4 | 4 | 1 | 1 | 3 | 3.08 | 4 | 2 | 0.374 | -1.419 | 0.20969649 | NA | 0 | 1.0772 | 0.5135 | 0.5135 | TRUE | 0.454 |
| Slc16a9 | 236 | 248 | 238 | 275 | 197 | 205 | 251 | 248 | 239 | 216 | 212 | 224 | 231.65 | 246 | 217 | 0.885 | -0.176 | 0.18393208 | 0.38664225 | 0 | 0.0211 | 0.0084 | 0.0084 | TRUE | 0.0079 |
| Slc17a1 | 2 | 0 | 0 | 0 | 0 | 0 | 2 | 0 | 0 | 0 | 0 | 0 | 0.35 | 1 | 0 | 0.254 | -1.975 | 0.62446185 | NA | 10 | 9.3117 | 10 | 10 | TRUE | 0.9741 |
| Slc17a2 | 0 | 0 | 0 | 0 | 0 | 0 | 0 | 0 | 0 | 0 | 0 | 0 | 0 | 0 | 0 | NA | NA | NA | NA | NA | NA | NA | NA | NA | NA |
| Slc17a3 | 0 | 0 | 0 | 0 | 0 | 0 | 0 | 0 | 0 | 0 | 0 | 0 | 0 | 0 | 0 | NA | NA | NA | NA | NA | NA | NA | NA | NA | NA |
| Slc17a4 | 0 | 0 | 0 | 0 | 0 | 0 | 0 | 0 | 0 | 0 | 0 | 0 | 0 | 0 | 0 | NA | NA | NA | NA | NA | NA | NA | NA | NA | NA |
| Slc17a5 | 544 | 765 | 737 | 691 | 512 | 581 | 578 | 764 | 740 | 543 | 551 | 636 | 635.32 | 694 | 577 | 0.83 | -0.269 | 0.05788634 | 0.18183298 | 0.0135 | 0.0121 | 0.0129 | 0.0129 | TRUE | 0.2517 |
| Slc17a6 | 0 | 0 | 0 | 0 | 0 | 0 | 0 | 0 | 0 | 0 | 0 | 0 | 0 | 0 | 0 | NA | NA | NA | NA | NA | NA | NA | NA | NA | NA |
| Slc17a7 | 792 | 913 | 903 | 885 | 780 | 713 | 841 | 912 | 907 | 696 | 839 | 780 | 829.2 | 887 | 772 | 0.869 | -0.202 | 0.05914435 | 0.18413436 | 0.0043 | 0.0109 | 0.0071 | 0.0071 | TRUE | 0.092 |
| Slc17a8 | 1 | 0 | 0 | 1 | 0 | 0 | 1 | 0 | 0 | 1 | 0 | 0 | 0.31 | 0 | 0 | 0.954 | -0.067 | 0.98625928 | NA | 0 | 10.7014 | 9.0384 | 9.0384 | TRUE | 0.4614 |
| Slc17a9 | 1127 | 1457 | 1518 | 812 | 905 | 973 | 1197 | 1455 | 1524 | 638 | 974 | 1065 | 1142.26 | 1392 | 892 | 0.64 | -0.643 | 0.00070099 | 0.00677869 | 0.042 | 0.0098 | 0.0251 | 0.0251 | TRUE | 0.7516 |
| Slc18a1 | 34 | 17 | 20 | 88 | 44 | 20 | 36 | 17 | 20 | 69 | 47 | 22 | 35.26 | 24 | 46 | 1.908 | 0.932 | 0.06276991 | 0.19201067 | 0.2022 | 0.1003 | 0.1493 | 0.1493 | TRUE | 0.1311 |
| Slc18a2 | 28 | 30 | 13 | 29 | 21 | 15 | 30 | 30 | 13 | 23 | 23 | 16 | 22.43 | 24 | 21 | 0.854 | -0.229 | 0.62477955 | 0.78857419 | 0.0664 | 0.1538 | 0.1126 | 0.1126 | TRUE | 0.9885 |
| Slc18a3 | 33 | 13 | 21 | 46 | 45 | 19 | 35 | 13 | 21 | 36 | 48 | 21 | 29.08 | 23 | 35 | 1.53 | 0.613 | 0.2123202 | 0.42329847 | 0.1583 | 0.1202 | 0.1381 | 0.1381 | TRUE | 0.2713 |
| Slc18b1 | 155 | 138 | 114 | 192 | 140 | 129 | 165 | 138 | 114 | 151 | 151 | 141 | 143.28 | 139 | 148 | 1.064 | 0.089 | 0.64354152 | 0.79952239 | 0.0092 | 0.0299 | 0.0197 | 0.0197 | TRUE | 0.2309 |
| Slc19a1 | 1165 | 1142 | 1238 | 1286 | 954 | 991 | 1237 | 1141 | 1243 | 1011 | 1026 | 1085 | 1123.87 | 1207 | 1041 | 0.862 | -0.215 | 0.01249625 | 0.06197826 | 0.001 | 0.0098 | 0.0044 | 0.0044 | TRUE | 0.028 |
| Slc19a2 | 4545 | 5334 | 5413 | 5450 | 4096 | 4775 | 4828 | 5327 | 5435 | 4284 | 4407 | 5227 | 4918.05 | 5197 | 4639 | 0.893 | -0.164 | 0.11544231 | 0.2869592 | 0.0077 | 0.0076 | 0.0076 | 0.0076 | TRUE | 0.1492 |
| Slc19a3 | 238 | 337 | 354 | 270 | 161 | 245 | 253 | 337 | 355 | 212 | 173 | 268 | 266.41 | 315 | 218 | 0.691 | -0.533 | 0.00995958 | 0.05252512 | 0.035 | 0.0193 | 0.0269 | 0.0269 | TRUE | 0.4322 |
| Slc1a1 | 614 | 894 | 819 | 1035 | 760 | 855 | 652 | 893 | 822 | 814 | 818 | 936 | 822.44 | 789 | 856 | 1.084 | 0.116 | 0.40788631 | 0.62066049 | 0.0147 | 0.0109 | 0.0129 | 0.0129 | TRUE | 0.272 |
| Slc1a2 | 11 | 9 | 14 | 11 | 14 | 14 | 12 | 9 | 14 | 9 | 15 | 15 | 12.29 | 12 | 13 | 1.11 | 0.151 | 0.78706098 | 0.89032073 | 0 | 0.2749 | 0.1433 | 0.1433 | TRUE | 0.4587 |
| Slc1a3 | 1302 | 1098 | 1171 | 1136 | 824 | 1007 | 1383 | 1097 | 1176 | 893 | 887 | 1102 | 1089.55 | 1219 | 961 | 0.788 | -0.344 | 0.01061475 | 0.05507099 | 0.0141 | 0.0099 | 0.0121 | 0.0121 | TRUE | 0.1967 |
| Slc1a4 | 575 | 698 | 647 | 728 | 472 | 609 | 611 | 697 | 650 | 572 | 508 | 667 | 617.38 | 653 | 582 | 0.892 | -0.165 | 0.20837568 | 0.41781099 | 0.0092 | 0.0122 | 0.0108 | 0.0108 | TRUE | 0.1834 |
| Slc1a5 | 2038 | 2474 | 2426 | 2223 | 1677 | 1994 | 2165 | 2471 | 2436 | 1748 | 1804 | 2183 | 2134.35 | 2357 | 1912 | 0.81 | -0.303 | 0.00818113 | 0.04521123 | 0.0095 | 0.0084 | 0.009 | 0.009 | TRUE | 0.1853 |
| Slc1a6 | 198 | 211 | 188 | 151 | 138 | 156 | 210 | 211 | 189 | 119 | 148 | 171 | 174.63 | 203 | 146 | 0.714 | -0.485 | 0.00892249 | 0.04834268 | 0.012 | 0.0258 | 0.0189 | 0.0189 | TRUE | 0.307 |
| Slc1a7 | 7 | 11 | 11 | 13 | 9 | 7 | 7 | 11 | 11 | 10 | 10 | 8 | 9.51 | 10 | 9 | 0.938 | -0.092 | 0.87916286 | 0.9402059 | 0 | 0.3535 | 0.1572 | 0.1572 | TRUE | 0.1468 |
| Slc20a1 | 7347 | 6756 | 6432 | 7065 | 5347 | 5295 | 7804 | 6747 | 6458 | 5554 | 5753 | 5796 | 6352.12 | 7003 | 5701 | 0.814 | -0.297 | 0.00132363 | 0.01108676 | 0.0051 | 0.0074 | 0.006 | 0.006 | TRUE | 0.122 |

| | | | | | | | | | | | | | | | | | | | | | | | | | |
|---|---|---|---|---|---|---|---|---|---|---|---|---|---|---|---|---|---|---|---|---|---|---|---|---|---|
| Slc20a2 | 6725 | 6570 | 6774 | 7801 | 5466 | 5651 | 7143 | 6562 | 6802 | 6133 | 5881 | 6186 | 6451 | 6836 | 6067 | 0.888 | -0.172 | 0.01078722 | 0.05571471 | 0.001 | 0.0074 | 0.0031 | 0.0031 | TRUE | 0.0189 |
| Slc22a1 | 4 | 3 | 6 | 13 | 3 | 3 | 4 | 3 | 6 | 10 | 3 | 3 | 5 | 4 | 5 | 1.294 | 0.372 | 0.68360576 | 0.82477223 | 0.1543 | 0.6658 | 0.3984 | 0.3984 | TRUE | 1.4789 |
| Slc22a12 | 0 | 0 | 0 | 0 | 0 | 0 | 0 | 0 | 0 | 0 | 0 | 0 | 0 | 0 | 0 | NA | NA | NA | NA | NA | NA | NA | NA | NA | NA |
| Slc22a13 | 5 | 1 | 0 | 3 | 1 | 3 | 5 | 1 | 0 | 2 | 1 | 3 | 2.17 | 2 | 2 | 1.075 | 0.105 | 0.94347717 | NA | 0.6139 | 1.5242 | 1.1098 | 1.1098 | TRUE | 1.5964 |
| Slc22a13b-ps | 13 | 10 | 7 | 20 | 13 | 7 | 14 | 10 | 7 | 16 | 14 | 8 | 11.37 | 10 | 13 | 1.226 | 0.294 | 0.62339707 | 0.7875856 | 0 | 0.2968 | 0.1704 | 0.1704 | TRUE | 0.3875 |
| Slc22a14 | 16 | 13 | 10 | 21 | 15 | 14 | 17 | 13 | 10 | 17 | 16 | 15 | 14.67 | 13 | 16 | 1.203 | 0.267 | 0.59022708 | 0.76449681 | 0 | 0.2315 | 0.1076 | 0.1076 | TRUE | 0.232 |
| Slc22a15 | 22 | 35 | 36 | 36 | 25 | 24 | 23 | 35 | 36 | 28 | 27 | 26 | 29.32 | 31 | 27 | 0.862 | -0.214 | 0.55565671 | 0.73856545 | 0 | 0.1193 | 0.0606 | 0.0606 | TRUE | 0.3328 |
| Slc22a16 | 7 | 0 | 1 | 4 | 3 | 5 | 7 | 0 | 1 | 3 | 3 | 5 | 3.38 | 3 | 4 | 1.41 | 0.496 | 0.70167756 | NA | 0.7857 | 0.9814 | 0.8987 | 0.8987 | TRUE | 2.4221 |
| Slc22a17 | 246 | 291 | 247 | 240 | 257 | 218 | 261 | 291 | 248 | 189 | 277 | 239 | 250.63 | 267 | 235 | 0.877 | -0.19 | 0.28625816 | 0.5074591 | 0.0178 | 0.02 | 0.0189 | 0.0189 | TRUE | 0.3271 |
| Slc22a18 | 81 | 252 | 237 | 83 | 79 | 131 | 86 | 252 | 238 | 65 | 85 | 143 | 144.89 | 192 | 98 | 0.508 | -0.976 | 0.01707798 | 0.07760122 | 0.2208 | 0.0296 | 0.113 | 0.113 | TRUE | 2.4994 |
| Slc22a19 | 0 | 0 | 0 | 0 | 0 | 0 | 0 | 0 | 0 | 0 | 0 | 0 | 0 | 0 | 0 | NA | NA | NA | NA | NA | NA | NA | NA | NA | NA |
| Slc22a2 | 0 | 0 | 0 | 1 | 0 | 0 | 0 | 0 | 0 | 1 | 0 | 0 | 0.13 | 0 | 0 | 1.859 | 0.894 | 0.82650734 | NA | 0 | 25.1515 | 10 | 10 | TRUE | 0.2563 |
| Slc22a20 | 3 | 6 | 5 | 7 | 10 | 4 | 3 | 6 | 5 | 6 | 11 | 4 | 5.81 | 5 | 7 | 1.441 | 0.527 | 0.51671095 | 0.70855113 | 0 | 0.5743 | 0.297 | 0.297 | TRUE | 0.5863 |
| Slc22a21 | 71 | 72 | 77 | 150 | 99 | 82 | 75 | 72 | 77 | 118 | 107 | 90 | 89.81 | 75 | 105 | 1.404 | 0.49 | 0.02404336 | 0.09890765 | 0 | 0.0436 | 0.0225 | 0.0225 | TRUE | 0.149 |
| Slc22a22 | 0 | 0 | 0 | 0 | 0 | 0 | 0 | 0 | 0 | 0 | 0 | 0 | 0 | 0 | 0 | NA | NA | NA | NA | NA | NA | NA | NA | NA | NA |
| Slc22a23 | 231 | 244 | 223 | 449 | 292 | 282 | 245 | 244 | 224 | 353 | 314 | 309 | 281.47 | 238 | 325 | 1.372 | 0.456 | 0.00062732 | 0.00621789 | 0 | 0.0186 | 0.0092 | 0.0092 | TRUE | 0.0671 |
| Slc22a26 | 0 | 0 | 0 | 0 | 0 | 0 | 0 | 0 | 0 | 0 | 0 | 0 | 0 | 0 | 0 | NA | NA | NA | NA | NA | NA | NA | NA | NA | NA |
| Slc22a27 | 0 | 0 | 0 | 0 | 1 | 0 | 0 | 0 | 0 | 0 | 1 | 0 | 0.18 | 0 | 0 | 1.859 | 0.894 | 0.82650734 | NA | 0 | 18.3785 | 10 | 10 | TRUE | 0.5204 |
| Slc22a28 | 0 | 0 | 0 | 0 | 0 | 1 | 0 | 0 | 0 | 0 | 0 | 1 | 0.18 | 0 | 0 | 1.859 | 0.894 | 0.82650734 | NA | 0 | 18.0643 | 10 | 10 | TRUE | 0.5383 |
| Slc22a29 | 0 | 0 | 0 | 0 | 1 | 0 | 0 | 0 | 0 | 0 | 1 | 0 | 0.18 | 0 | 0 | 1.859 | 0.894 | 0.82650734 | NA | 0 | 18.3785 | 10 | 10 | TRUE | 0.5204 |
| Slc22a3 | 0 | 3 | 1 | 3 | 2 | 0 | 0 | 3 | 1 | 2 | 2 | 0 | 1.42 | 1 | 1 | 1.14 | 0.189 | 0.91654559 | NA | 0 | 2.3295 | 1.6462 | 1.6462 | TRUE | 0.544 |
| Slc22a30 | 0 | 0 | 1 | 0 | 0 | 1 | 0 | 0 | 1 | 0 | 0 | 1 | 0.35 | 0 | 0 | 0.954 | -0.067 | 0.98566485 | NA | 0 | 9.4252 | 8.1416 | 8.1416 | TRUE | 0.5383 |
| Slc22a4 | 1 | 3 | 1 | 4 | 0 | 1 | 1 | 3 | 1 | 3 | 0 | 1 | 1.55 | 2 | 1 | 0.872 | -0.197 | 0.90232219 | NA | 0 | 2.132 | 1.2281 | 1.2281 | TRUE | 0.6134 |
| Slc22a5 | 490 | 730 | 797 | 799 | 660 | 695 | 520 | 729 | 800 | 628 | 710 | 761 | 691.47 | 683 | 700 | 1.023 | 0.033 | 0.84799634 | 0.92391533 | 0.0274 | 0.0117 | 0.0196 | 0.0196 | TRUE | 0.5105 |
| Slc22a6 | 3 | 1 | 0 | 2 | 1 | 1 | 3 | 1 | 0 | 2 | 1 | 1 | 1.32 | 1 | 1 | 0.912 | -0.133 | 0.93698999 | NA | 0 | 2.5002 | 1.297 | 1.297 | TRUE | 0.5271 |
| Slc22a7 | 313 | 395 | 368 | 259 | 219 | 263 | 332 | 394 | 370 | 204 | 236 | 288 | 303.93 | 365 | 243 | 0.661 | -0.598 | 0.00030372 | 0.00344875 | 0.015 | 0.0177 | 0.0163 | 0.0163 | TRUE | 0.2964 |
| Slc22a8 | 0 | 0 | 1 | 0 | 0 | 0 | 0 | 0 | 1 | 0 | 0 | 0 | 0.17 | 0 | 0 | 0.49 | -1.029 | 0.80087 | NA | 0 | 19.6928 | 10 | 10 | TRUE | 0.4098 |
| Slc23a1 | 24 | 17 | 22 | 33 | 28 | 28 | 25 | 17 | 22 | 26 | 30 | 31 | 25.21 | 21 | 29 | 1.34 | 0.423 | 0.2717135 | 0.4917344 | 0 | 0.1376 | 0.0658 | 0.0658 | TRUE | 0.1945 |
| Slc23a2 | 2768 | 2595 | 2613 | 3696 | 2680 | 2369 | 2940 | 2592 | 2624 | 2906 | 2884 | 2593 | 2756.29 | 2719 | 2794 | 1.028 | 0.04 | 0.66011296 | 0.80864478 | 0.0041 | 0.0081 | 0.0056 | 0.0056 | TRUE | 0.0616 |
| Slc23a3 | 4 | 2 | 0 | 5 | 1 | 1 | 4 | 2 | 0 | 4 | 1 | 1 | 2.06 | 2 | 2 | 1.011 | 0.016 | 0.99131012 | NA | 0.5186 | 1.6078 | 1.1078 | 1.1078 | TRUE | 0.2082 |
| Slc24a1 | 446 | 445 | 419 | 395 | 282 | 305 | 474 | 444 | 421 | 311 | 303 | 334 | 381.11 | 446 | 316 | 0.708 | -0.499 | 2.25E-05 | 0.00039343 | 0 | 0.0155 | 0.0073 | 0.0073 | TRUE | 0.0331 |
| Slc24a2 | 14 | 7 | 14 | 60 | 23 | 21 | 15 | 7 | 14 | 47 | 25 | 23 | 21.8 | 12 | 32 | 2.672 | 1.418 | 0.00630136 | 0.03736281 | 0.113 | 0.158 | 0.1361 | 0.1361 | TRUE | 1.4255 |
| Slc24a3 | 8 | 2 | 2 | 10 | 6 | 2 | 8 | 2 | 2 | 8 | 6 | 2 | 4.83 | 4 | 5 | 1.345 | 0.428 | 0.66720589 | 0.81391997 | 0.3102 | 0.6883 | 0.5026 | 0.5026 | TRUE | 1.3652 |
| Slc24a4 | 0 | 0 | 1 | 2 | 1 | 2 | 0 | 0 | 1 | 2 | 1 | 2 | 0.97 | 0 | 2 | 4.402 | 2.138 | 0.2928876 | NA | 0 | 3.3907 | 1.6736 | 1.6736 | TRUE | 0.4098 |
| Slc24a5 | 6 | 12 | 22 | 31 | 20 | 21 | 6 | 12 | 22 | 24 | 22 | 23 | 18.22 | 13 | 23 | 1.7 | 0.765 | 0.14747445 | 0.33665755 | 0.079 | 0.1877 | 0.142 | 0.142 | TRUE | 0.7142 |
| Slc24a6 | 2033 | 2176 | 2116 | 2040 | 1648 | 1644 | 2159 | 2173 | 2125 | 1604 | 1773 | 1800 | 1938.96 | 2152 | 1726 | 0.801 | -0.32 | 6.37E-05 | 0.00095529 | 0.0015 | 0.0086 | 0.0041 | 0.0041 | TRUE | 0.0474 |
| Slc25a1 | 5341 | 6340 | 6453 | 6208 | 4840 | 5365 | 5673 | 6332 | 6479 | 4880 | 5208 | 5873 | 5740.84 | 6161 | 5320 | 0.863 | -0.212 | 0.03404299 | 0.12550866 | 0.0067 | 0.0075 | 0.007 | 0.007 | TRUE | 0.1005 |
| Slc25a10 | 389 | 266 | 236 | 676 | 476 | 304 | 413 | 266 | 237 | 531 | 512 | 333 | 382.03 | 305 | 459 | 1.505 | 0.59 | 0.01744334 | 0.0786721 | 0.0724 | 0.0155 | 0.0416 | 0.0416 | TRUE | 1.0757 |
| Slc25a11 | 3736 | 3648 | 3668 | 5648 | 4020 | 3728 | 3968 | 3643 | 3683 | 4440 | 4325 | 4081 | 4023.47 | 3765 | 4282 | 1.138 | 0.186 | 0.01330744 | 0.06481934 | 0.0017 | 0.0077 | 0.0038 | 0.0038 | TRUE | 0.0271 |
| Slc25a12 | 1624 | 1665 | 1740 | 2308 | 1853 | 1774 | 1725 | 1663 | 1747 | 1814 | 1994 | 1942 | 1814.16 | 1712 | 1917 | 1.119 | 0.162 | 0.03662683 | 0.13204078 | 0.0011 | 0.0087 | 0.0038 | 0.0038 | TRUE | 0.0271 |
| Slc25a13 | 9513 | 10877 | 10611 | 11032 | 8088 | 8517 | 10104 | 10863 | 10655 | 8672 | 8702 | 9323 | 9719.98 | 10541 | 8899 | 0.844 | -0.244 | 0.00037272 | 0.00409652 | 0.0014 | 0.0072 | 0.0033 | 0.0033 | TRUE | 0.0211 |
| Slc25a14 | 195 | 186 | 193 | 323 | 214 | 217 | 207 | 186 | 194 | 254 | 230 | 238 | 218.06 | 196 | 241 | 1.233 | 0.302 | 0.03293029 | 0.12263908 | 0 | 0.022 | 0.0098 | 0.0098 | TRUE | 0.0285 |
| Slc25a15 | 4314 | 4226 | 4197 | 5796 | 4160 | 3908 | 4582 | 4221 | 4214 | 4556 | 4476 | 4278 | 4387.87 | 4339 | 4437 | 1.023 | 0.032 | 0.6533394 | 0.80540538 | 0.0013 | 0.0077 | 0.0035 | 0.0035 | TRUE | 0.0292 |
| Slc25a16 | 911 | 993 | 923 | 1497 | 970 | 920 | 968 | 992 | 927 | 1177 | 1044 | 1007 | 1018.95 | 962 | 1076 | 1.12 | 0.163 | 0.09743656 | 0.2564613 | 0.0032 | 0.0101 | 0.006 | 0.006 | TRUE | 0.0831 |
| Slc25a17 | 2910 | 3515 | 3455 | 3638 | 2583 | 2893 | 3091 | 3510 | 3469 | 2860 | 2779 | 3167 | 3146.08 | 3357 | 2935 | 0.874 | -0.194 | 0.03573917 | 0.12973029 | 0.0043 | 0.008 | 0.0058 | 0.0058 | TRUE | 0.0582 |
| Slc25a18 | 7 | 6 | 3 | 8 | 3 | 2 | 7 | 6 | 3 | 6 | 3 | 2 | 4.69 | 5 | 4 | 0.732 | -0.45 | 0.62004921 | 0.78498274 | 0 | 0.7092 | 0.3788 | 0.3788 | TRUE | 0.6104 |
| Slc25a19 | 1316 | 1593 | 1612 | 1339 | 1023 | 1261 | 1398 | 1591 | 1619 | 1053 | 1101 | 1380 | 1356.84 | 1536 | 1178 | 0.766 | -0.384 | 0.00304245 | 0.02146414 | 0.0132 | 0.0093 | 0.0114 | 0.0114 | TRUE | 0.2695 |
| Slc25a2 | 9 | 11 | 5 | 16 | 8 | 8 | 10 | 11 | 5 | 13 | 9 | 9 | 9.25 | 9 | 10 | 1.183 | 0.242 | 0.70495215 | 0.83856289 | 0 | 0.363 | 0.1869 | 0.1869 | TRUE | 0.4055 |
| Slc25a20 | 1610 | 1775 | 1504 | 3683 | 2549 | 2353 | 1710 | 1773 | 1510 | 2895 | 2743 | 2576 | 2201.09 | 1664 | 2738 | 1.646 | 0.719 | 7.04E-14 | 8.55E-12 | 0.0044 | 0.0084 | 0.0062 | 0.0062 | TRUE | 0.0794 |
| Slc25a21 | 1 | 0 | 3 | 2 | 0 | 1 | 1 | 0 | 3 | 2 | 0 | 1 | 1.12 | 1 | 1 | 0.672 | -0.572 | 0.77114751 | NA | 0 | 2.939 | 1.8802 | 1.8802 | TRUE | 0.3351 |
| Slc25a22 | 731 | 767 | 699 | 1009 | 746 | 713 | 776 | 766 | 702 | 793 | 803 | 780 | 770.11 | 748 | 792 | 1.059 | 0.083 | 0.36538909 | 0.58293715 | 0 | 0.0112 | 0.0047 | 0.0047 | TRUE | 0.0348 |
| Slc25a23 | 781 | 1031 | 1112 | 1026 | 861 | 980 | 830 | 1030 | 1117 | 807 | 926 | 1073 | 963.58 | 992 | 935 | 0.942 | -0.087 | 0.5693981 | 0.74821263 | 0.0206 | 0.0103 | 0.0156 | 0.0156 | TRUE | 0.2439 |
| Slc25a24 | 194 | 217 | 237 | 617 | 399 | 373 | 206 | 217 | 238 | 485 | 429 | 408 | 330.57 | 220 | 441 | 2.005 | 1.003 | 2.02E-13 | 2.33E-11 | 0.0043 | 0.0169 | 0.01 | 0.01 | TRUE | 0.093 |
| Slc25a25 | 1442 | 1703 | 1700 | 1670 | 1214 | 1155 | 1532 | 1701 | 1707 | 1313 | 1306 | 1264 | 1470.46 | 1647 | 1294 | 0.786 | -0.347 | 3.32E-05 | 0.00054919 | 0.0015 | 0.0091 | 0.0043 | 0.0043 | TRUE | 0.0446 |
| Slc25a26 | 858 | 782 | 686 | 1133 | 789 | 730 | 911 | 781 | 689 | 891 | 849 | 799 | 819.97 | 794 | 846 | 1.067 | 0.094 | 0.46096621 | 0.66511381 | 0.01 | 0.0109 | 0.0104 | 0.0104 | TRUE | 0.1985 |
| Slc25a27 | 89 | 98 | 75 | 84 | 87 | 81 | 95 | 98 | 75 | 66 | 94 | 89 | 86 | 89 | 83 | 0.922 | -0.117 | 0.63069269 | 0.79261236 | 0.0161 | 0.0452 | 0.0308 | 0.0308 | TRUE | 0.3315 |
| Slc25a28 | 686 | 796 | 752 | 1004 | 865 | 765 | 729 | 795 | 755 | 789 | 931 | 837 | 806.01 | 760 | 852 | 1.121 | 0.165 | 0.11364262 | 0.28427491 | 0.0034 | 0.011 | 0.0066 | 0.0066 | TRUE | 0.0764 |
| Slc25a29 | 79 | 95 | 94 | 84 | 78 | 65 | 84 | 95 | 94 | 66 | 84 | 71 | 82.38 | 91 | 74 | 0.806 | -0.311 | 0.16034969 | 0.3548001 | 0 | 0.0469 | 0.0232 | 0.0232 | TRUE | 0.1308 |
| Slc25a3 | 24808 | 23461 | 23028 | 31333 | 22026 | 20958 | 26350 | 23431 | 23123 | 24632 | 23699 | 22942 | 24029.29 | 24301 | 23758 | 0.978 | -0.033 | 0.68371071 | 0.82480383 | 0.0028 | 0.007 | 0.0046 | 0.0046 | TRUE | 0.0666 |
| Slc25a30 | 1566 | 2181 | 2304 | 1740 | 1461 | 1990 | 1663 | 2178 | 2313 | 1368 | 1572 | 2178 | 1878.86 | 2051 | 1706 | 0.831 | -0.267 | 0.15516676 | 0.34747838 | 0.0431 | 0.0087 | 0.0249 | 0.0249 | TRUE | 0.7068 |
| Slc25a31 | 89 | 128 | 112 | 368 | 273 | 225 | 95 | 128 | 112 | 289 | 294 | 246 | 194.03 | 112 | 276 | 2.476 | 1.308 | 1.40E-13 | 1.65E-11 | 0.0076 | 0.0239 | 0.0162 | 0.0162 | TRUE | 0.1745 |
| Slc25a32 | 1418 | 1218 | 1144 | 1440 | 1031 | 1007 | 1506 | 1216 | 1149 | 1132 | 1109 | 1102 | 1202.49 | 1290 | 1114 | 0.864 | -0.211 | 0.08125924 | 0.22703646 | 0.0097 | 0.0096 | 0.0097 | 0.0097 | TRUE | 0.2561 |
| Slc25a33 | 1293 | 1668 | 1720 | 1371 | 1056 | 1085 | 1373 | 1666 | 1727 | 1078 | 1136 | 1188 | 1361.33 | 1589 | 1134 | 0.713 | -0.488 | 1.76E-05 | 0.0003193 | 0.0079 | 0.0093 | 0.0085 | 0.0085 | TRUE | 0.1689 |
| Slc25a34 | 34 | 30 | 26 | 42 | 19 | 25 | 36 | 30 | 26 | 33 | 20 | 27 | 28.83 | 31 | 27 | 0.885 | -0.177 | 0.63858516 | 0.79637583 | 0 | 0.1212 | 0.0671 | 0.0671 | TRUE | 0.2781 |
| Slc25a35 | 111 | 187 | 189 | 210 | 131 | 179 | 118 | 187 | 190 | 165 | 141 | 196 | 166.07 | 165 | 167 | 1.014 | 0.02 | 0.93159934 | 0.96628017 | 0.0403 | 0.0267 | 0.0333 | 0.0333 | TRUE | 0.649 |
| Slc25a36 | 5748 | 8146 | 8259 | 5288 | 4116 | 5026 | 6105 | 8135 | 8293 | 4157 | 4429 | 5502 | 6103.52 | 7511 | 4696 | 0.625 | -0.678 | 7.06E-06 | 0.00014753 | 0.0244 | 0.0074 | 0.0162 | 0.0162 | TRUE | 0.3271 |
| Slc25a37 | 1458 | 1369 | 1334 | 1325 | 1182 | 976 | 1549 | 1367 | 1339 | 1042 | 1272 | 1068 | 1272.85 | 1418 | 1127 | 0.794 | -0.332 | 0.00399975 | 0.02634386 | 0.0082 | 0.0095 | 0.0088 | 0.0088 | TRUE | 0.1497 |
| Slc25a38 | 767 | 748 | 683 | 976 | 766 | 728 | 815 | 747 | 686 | 767 | 824 | 797 | 772.64 | 749 | 796 | 1.062 | 0.087 | 0.39993922 | 0.61294737 | 0.003 | 0.0112 | 0.0064 | 0.0064 | TRUE | 0.0688 |
| Slc25a39 | 9277 | 12193 | 11858 | 14557 | 10536 | 11295 | 9854 | 12177 | 11907 | 11444 | 11336 | 12364 | 11513.6 | 11313 | 11715 | 1.035 | 0.05 | 0.62297448 | 0.78725221 | 0.0061 | 0.0072 | 0.0075 | 0.0075 | TRUE | 0.1554 |

| | | | | | | | | | | | | | | | | | | | | | | | | | |
|---|---|---|---|---|---|---|---|---|---|---|---|---|---|---|---|---|---|---|---|---|---|---|---|---|---|
| Slc25a4 | 5786 | 8623 | 8297 | 19520 | 13541 | 13608 | 6146 | 8612 | 8331 | 15345 | 14569 | 14896 | 11316.53 | 7696 | 14937 | 1.941 | 0.957 | 2.74E-13 | 3.07E-11 | 0.0156 | 0.0072 | 0.0123 | 0.0123 | TRUE | 0.379 |
| Slc25a40 | 1313 | 1501 | 1298 | 1750 | 1237 | 1233 | 1395 | 1499 | 1303 | 1376 | 1331 | 1350 | 1375.57 | 1399 | 1352 | 0.967 | -0.049 | 0.5747258 | 0.75283624 | 0.0019 | 0.0093 | 0.0047 | 0.0047 | TRUE | 0.0474 |
| Slc25a41 | 42 | 32 | 22 | 62 | 46 | 30 | 45 | 32 | 22 | 49 | 49 | 33 | 38.29 | 33 | 44 | 1.335 | 0.417 | 0.26520049 | 0.48484318 | 0.0528 | 0.0929 | 0.0742 | 0.0742 | TRUE | 0.3335 |
| Slc25a42 | 593 | 872 | 883 | 708 | 567 | 612 | 630 | 871 | 887 | 557 | 610 | 670 | 703.99 | 796 | 612 | 0.768 | -0.38 | 0.01529537 | 0.07173315 | 0.0206 | 0.0116 | 0.0163 | 0.0163 | TRUE | 0.393 |
| Slc25a43 | 40 | 62 | 65 | 68 | 63 | 53 | 42 | 62 | 65 | 53 | 68 | 58 | 58.16 | 56 | 60 | 1.052 | 0.073 | 0.79651604 | 0.89536272 | 0.0136 | 0.0636 | 0.04 | 0.04 | TRUE | 0.389 |
| Slc25a44 | 836 | 658 | 634 | 871 | 549 | 542 | 888 | 657 | 637 | 685 | 591 | 593 | 675.07 | 727 | 623 | 0.858 | -0.222 | 0.15323622 | 0.34475084 | 0.0199 | 0.0118 | 0.0159 | 0.0159 | TRUE | 0.4407 |
| Slc25a45 | 4 | 5 | 4 | 15 | 8 | 7 | 4 | 5 | 4 | 12 | 9 | 8 | 6.89 | 4 | 10 | 2.135 | 1.094 | 0.1332244 | 0.31511926 | 0 | 0.4853 | 0.2152 | 0.2152 | TRUE | 0.2166 |
| Slc25a46 | 2490 | 2443 | 2391 | 3284 | 2269 | 2381 | 2645 | 2440 | 2401 | 2582 | 2441 | 2606 | 2519.13 | 2495 | 2543 | 1.019 | 0.028 | 0.72000799 | 0.8481969 | 0.0015 | 0.0082 | 0.0039 | 0.0039 | TRUE | 0.0333 |
| Slc25a47 | 15 | 18 | 15 | 33 | 16 | 16 | 16 | 18 | 15 | 26 | 17 | 18 | 18.27 | 16 | 20 | 1.251 | 0.323 | 0.47432581 | 0.67494286 | 0 | 0.1872 | 0.092 | 0.092 | TRUE | 0.4014 |
| Slc25a48 | 6 | 15 | 11 | 12 | 13 | 15 | 6 | 15 | 11 | 9 | 14 | 16 | 12.04 | 11 | 13 | 1.214 | 0.28 | 0.63369668 | 0.7939204 | 0 | 0.2805 | 0.1643 | 0.1643 | TRUE | 0.2651 |
| Slc25a5 | 25523 | 27173 | 27329 | 37570 | 25821 | 25981 | 27110 | 27138 | 27441 | 29535 | 27782 | 28440 | 27907.62 | 27230 | 28586 | 1.05 | 0.07 | 0.21295263 | 0.42413043 | 4.00E-04 | 0.007 | 0.0023 | 0.0023 | TRUE | 0.0104 |
| Slc25a51 | 1575 | 1710 | 1723 | 2393 | 1692 | 1795 | 1673 | 1708 | 1730 | 1881 | 1820 | 1965 | 1796.23 | 1704 | 1889 | 1.109 | 0.149 | 0.04167632 | 0.14513557 | 3.00E-04 | 0.0087 | 0.0033 | 0.0033 | TRUE | 0.0146 |
| Slc25a53 | 57 | 64 | 78 | 79 | 57 | 54 | 61 | 64 | 78 | 62 | 61 | 59 | 64.22 | 68 | 61 | 0.9 | -0.151 | 0.53752038 | 0.72416259 | 0 | 0.0582 | 0.0279 | 0.0279 | TRUE | 0.1726 |
| Slc25a54 | 0 | 0 | 0 | 0 | 0 | 0 | 0 | 0 | 0 | 0 | 0 | 0 | 0 | 0 | 0 | NA | NA | NA | NA | NA | NA | NA | NA | NA | NA |
| Slc26a1 | 0 | 0 | 0 | 5 | 2 | 0 | 0 | 0 | 0 | 4 | 2 | 0 | 1.01 | 0 | 2 | 10.917 | 3.448 | 0.18671364 | NA | 5.9256 | 3.2567 | 3.6764 | 3.6764 | TRUE | 0.0627 |
| Slc26a10 | 160 | 186 | 197 | 132 | 112 | 135 | 170 | 186 | 198 | 104 | 121 | 148 | 154.26 | 185 | 124 | 0.669 | -0.581 | 0.0024465 | 0.01803939 | 0.011 | 0.0283 | 0.0198 | 0.0198 | TRUE | 0.2809 |
| Slc26a11 | 323 | 430 | 412 | 435 | 306 | 339 | 343 | 429 | 414 | 342 | 329 | 371 | 371.42 | 395 | 347 | 0.878 | -0.188 | 0.16671454 | 0.36418584 | 0.0061 | 0.0158 | 0.0106 | 0.0106 | TRUE | 0.1525 |
| Slc26a2 | 1768 | 1531 | 1575 | 2345 | 1632 | 1521 | 1878 | 1529 | 1581 | 1843 | 1756 | 1665 | 1708.8 | 1663 | 1755 | 1.056 | 0.078 | 0.46404048 | 0.66733357 | 0.0062 | 0.0088 | 0.0077 | 0.0077 | TRUE | 0.1541 |
| Slc26a3 | 1 | 3 | 1 | 2 | 2 | 1 | 1 | 3 | 1 | 2 | 2 | 1 | 1.65 | 2 | 2 | 0.948 | -0.076 | 0.95689918 | NA | 0 | 2.0073 | 0.8346 | 0.8346 | TRUE | 0.3591 |
| Slc26a4 | 22 | 10 | 13 | 27 | 6 | 9 | 23 | 10 | 13 | 21 | 6 | 10 | 13.99 | 15 | 12 | 0.826 | -0.277 | 0.66585996 | 0.81297171 | 0.2056 | 0.2424 | 0.2241 | 0.2241 | TRUE | 1.6741 |
| Slc26a5 | 10 | 3 | 6 | 9 | 4 | 6 | 11 | 3 | 6 | 7 | 4 | 7 | 6.26 | 7 | 6 | 0.923 | -0.115 | 0.88576355 | 0.94393667 | 0 | 0.5328 | 0.3008 | 0.3008 | TRUE | 0.3011 |
| Slc26a6 | 321 | 240 | 250 | 309 | 209 | 259 | 341 | 240 | 251 | 243 | 225 | 284 | 263.83 | 277 | 251 | 0.903 | -0.147 | 0.42715904 | 0.63729014 | 0.0215 | 0.0194 | 0.0207 | 0.0207 | TRUE | 0.4501 |
| Slc26a7 | 2 | 0 | 2 | 4 | 0 | 3 | 2 | 0 | 2 | 3 | 0 | 3 | 1.76 | 1 | 2 | 1.579 | 0.659 | 0.69335891 | NA | 0 | 1.8786 | 1.4129 | 1.4129 | TRUE | 0.667 |
| Slc26a9 | 19 | 20 | 19 | 27 | 24 | 21 | 20 | 20 | 19 | 21 | 26 | 23 | 21.54 | 20 | 23 | 1.178 | 0.237 | 0.55326921 | 0.73682023 | 0 | 0.1598 | 0.0681 | 0.0681 | TRUE | 0.0489 |
| Slc27a1 | 806 | 1193 | 1173 | 1216 | 1114 | 1203 | 856 | 1191 | 1178 | 956 | 1199 | 1317 | 1116.13 | 1075 | 1157 | 1.075 | 0.105 | 0.53284027 | 0.72081283 | 0.029 | 0.0099 | 0.0194 | 0.0194 | TRUE | 0.3788 |
| Slc27a2 | 462 | 486 | 518 | 512 | 460 | 411 | 491 | 485 | 520 | 402 | 495 | 450 | 473.93 | 499 | 449 | 0.898 | -0.154 | 0.20167437 | 0.40921241 | 0.0041 | 0.0139 | 0.0084 | 0.0084 | TRUE | 0.1015 |
| Slc27a3 | 704 | 1022 | 990 | 1202 | 929 | 861 | 748 | 1021 | 994 | 945 | 1000 | 943 | 941.58 | 921 | 963 | 1.045 | 0.063 | 0.64155441 | 0.79835032 | 0.0136 | 0.0104 | 0.0121 | 0.0121 | TRUE | 0.3206 |
| Slc27a4 | 1531 | 1599 | 1543 | 2463 | 1710 | 1570 | 1626 | 1597 | 1549 | 1936 | 1840 | 1719 | 1711.19 | 1591 | 1832 | 1.152 | 0.204 | 0.01266935 | 0.06265943 | 0.0016 | 0.0088 | 0.0043 | 0.0043 | TRUE | 0.035 |
| Slc27a5 | 33 | 27 | 31 | 37 | 21 | 31 | 35 | 27 | 31 | 29 | 23 | 34 | 29.79 | 31 | 29 | 0.92 | -0.12 | 0.73884083 | 0.86025377 | 0 | 0.1175 | 0.0594 | 0.0594 | TRUE | 0.1955 |
| Slc27a6 | 6 | 3 | 2 | 9 | 3 | 3 | 6 | 3 | 2 | 7 | 3 | 3 | 4.16 | 4 | 4 | 1.224 | 0.291 | 0.76295442 | NA | 0 | 0.7987 | 0.4307 | 0.4307 | TRUE | 0.58 |
| Slc28a1 | 343 | 331 | 415 | 285 | 212 | 241 | 364 | 331 | 417 | 224 | 228 | 264 | 304.59 | 371 | 239 | 0.643 | -0.637 | 1.82E-05 | 0.00032682 | 0.0079 | 0.0177 | 0.0125 | 0.0125 | TRUE | 0.1371 |
| Slc28a2 | 2 | 0 | 2 | 6 | 2 | 2 | 2 | 0 | 2 | 5 | 2 | 2 | 2.2 | 1 | 3 | 2.241 | 1.164 | 0.39474831 | NA | 0 | 1.5055 | 0.8159 | 0.8159 | TRUE | 0.498 |
| Slc28a3 | 38 | 40 | 31 | 73 | 52 | 57 | 40 | 40 | 31 | 57 | 56 | 62 | 47.86 | 37 | 58 | 1.577 | 0.657 | 0.02003893 | 0.08696054 | 0 | 0.0757 | 0.0354 | 0.0354 | TRUE | 0.148 |
| Slc29a1 | 12823 | 14534 | 14746 | 15542 | 11964 | 11616 | 13620 | 14515 | 14807 | 12218 | 12873 | 12716 | 13458.01 | 14314 | 12602 | 0.88 | -0.184 | 0.00543975 | 0.03325416 | 0.0012 | 0.0071 | 0.0031 | 0.0031 | TRUE | 0.022 |
| Slc29a2 | 683 | 728 | 839 | 929 | 669 | 737 | 725 | 727 | 842 | 730 | 720 | 807 | 758.64 | 765 | 752 | 0.983 | -0.025 | 0.81858751 | 0.90729179 | 0.0043 | 0.0112 | 0.0073 | 0.0073 | TRUE | 0.0933 |
| Slc29a3 | 1117 | 914 | 985 | 1276 | 969 | 915 | 1186 | 913 | 989 | 1003 | 1043 | 1002 | 1022.6 | 1029 | 1016 | 0.987 | -0.019 | 0.87259227 | 0.93737812 | 0.0084 | 0.0101 | 0.0092 | 0.0092 | TRUE | 0.2119 |
| Slc29a4 | 1907 | 2017 | 2014 | 1564 | 1209 | 1172 | 2026 | 2014 | 2022 | 1229 | 1301 | 1283 | 1645.91 | 2021 | 1271 | 0.629 | -0.67 | 5.16E-21 | 1.54E-18 | 0 | 0.0089 | 0.003 | 0.003 | TRUE | 0.0103 |
| Slc2a1 | 33148 | 41724 | 42661 | 61481 | 43709 | 44289 | 35209 | 41670 | 42836 | 48332 | 47028 | 48481 | 43926.06 | 39905 | 47947 | 1.202 | 0.265 | 0.00421885 | 0.02754129 | 0.0054 | 0.007 | 0.0062 | 0.0062 | TRUE | 0.1297 |
| Slc2a10 | 11 | 8 | 6 | 3 | 1 | 4 | 12 | 8 | 6 | 2 | 1 | 4 | 5.59 | 9 | 2 | 0.302 | -1.726 | 0.05238602 | 0.16927521 | 0 | 0.5968 | 0.3222 | 0.3222 | TRUE | 0.3202 |
| Slc2a12 | 32 | 45 | 47 | 57 | 42 | 45 | 34 | 45 | 47 | 45 | 45 | 49 | 44.23 | 42 | 46 | 1.101 | 0.139 | 0.63653205 | 0.79545072 | 0 | 0.0814 | 0.0396 | 0.0396 | TRUE | 0.2071 |
| Slc2a13 | 2 | 7 | 3 | 5 | 7 | 2 | 2 | 7 | 3 | 4 | 8 | 2 | 4.3 | 4 | 5 | 1.118 | 0.16 | 0.86963846 | NA | 0.1179 | 0.7737 | 0.4554 | 0.4554 | TRUE | 0.5717 |
| Slc2a2 | 0 | 0 | 0 | 3 | 4 | 0 | 0 | 0 | 0 | 2 | 4 | 0 | 1.11 | 0 | 2 | 11.838 | 3.565 | 0.16030473 | NA | 6.0598 | 2.9739 | 3.4262 | 3.4262 | TRUE | 0.0624 |
| Slc2a3 | 93243 | 135981 | 142683 | 165404 | 125212 | 139259 | 99041 | 135805 | 143269 | 130028 | 134720 | 152441 | 132550.63 | 126038 | 139063 | 1.103 | 0.142 | 0.32382167 | 0.54518737 | 0.0188 | 0.0069 | 0.0149 | 0.0149 | TRUE | 0.43 |
| Slc2a4 | 135 | 224 | 260 | 141 | 127 | 175 | 143 | 224 | 261 | 111 | 137 | 192 | 177.87 | 209 | 147 | 0.696 | -0.522 | 0.06035627 | 0.18661813 | 0.0773 | 0.0254 | 0.0499 | 0.0499 | TRUE | 0.4146 |
| Slc2a4rg-ps | 130 | 105 | 115 | 183 | 166 | 130 | 138 | 105 | 115 | 144 | 179 | 142 | 137.2 | 119 | 155 | 1.296 | 0.374 | 0.05889038 | 0.18363368 | 0.0106 | 0.0309 | 0.0208 | 0.0208 | TRUE | 0.1843 |
| Slc2a5 | 488 | 499 | 495 | 493 | 355 | 411 | 518 | 498 | 497 | 388 | 382 | 450 | 455.53 | 504 | 407 | 0.805 | -0.314 | 0.0066901 | 0.03889283 | 0.0017 | 0.0141 | 0.0074 | 0.0074 | TRUE | 0.0974 |
| Slc2a6 | 32 | 51 | 43 | 50 | 36 | 35 | 34 | 51 | 43 | 39 | 39 | 38 | 40.74 | 43 | 39 | 0.907 | -0.14 | 0.65060397 | 0.80417914 | 0 | 0.0878 | 0.0444 | 0.0444 | TRUE | 0.2068 |
| Slc2a7 | 55 | 66 | 49 | 75 | 44 | 39 | 58 | 66 | 49 | 59 | 47 | 43 | 53.75 | 58 | 50 | 0.864 | -0.21 | 0.455823 | 0.66071504 | 0 | 0.0682 | 0.0387 | 0.0387 | TRUE | 0.2488 |
| Slc2a8 | 1047 | 1414 | 1406 | 1332 | 1198 | 1214 | 1112 | 1412 | 1412 | 1047 | 1289 | 1329 | 1266.84 | 1312 | 1222 | 0.93 | -0.104 | 0.45483164 | 0.65982383 | 0.0166 | 0.0095 | 0.0133 | 0.0133 | TRUE | 0.2128 |
| Slc2a9 | 25 | 19 | 16 | 58 | 41 | 37 | 27 | 19 | 16 | 46 | 44 | 41 | 31.97 | 21 | 44 | 2.123 | 1.086 | 0.00249427 | 0.01834013 | 0 | 0.11 | 0.0567 | 0.0567 | TRUE | 0.3464 |
| Slc30a1 | 1568 | 2123 | 1910 | 2628 | 2113 | 1777 | 1665 | 2120 | 1918 | 2066 | 2273 | 1945 | 1998.03 | 1901 | 2095 | 1.102 | 0.14 | 0.22999708 | 0.44367832 | 0.0096 | 0.0086 | 0.0093 | 0.0093 | TRUE | 0.1418 |
| Slc30a10 | 65 | 98 | 93 | 86 | 86 | 96 | 69 | 98 | 93 | 68 | 93 | 105 | 87.59 | 87 | 89 | 1.012 | 0.017 | 0.94677406 | 0.97350026 | 0.031 | 0.0445 | 0.0378 | 0.0378 | TRUE | 0.449 |
| Slc30a2 | 1105 | 1250 | 1304 | 1850 | 1374 | 1234 | 1174 | 1248 | 1309 | 1454 | 1478 | 1351 | 1335.82 | 1244 | 1428 | 1.148 | 0.199 | 0.02283843 | 0.0955458 | 0.0018 | 0.0094 | 0.0048 | 0.0048 | TRUE | 0.029 |
| Slc30a3 | 223 | 282 | 285 | 147 | 125 | 165 | 237 | 282 | 286 | 116 | 134 | 181 | 205.89 | 268 | 144 | 0.532 | -0.909 | 5.82E-06 | 0.00012576 | 0.0243 | 0.0229 | 0.0237 | 0.0237 | TRUE | 0.5219 |
| Slc30a4 | 672 | 636 | 650 | 1114 | 842 | 689 | 714 | 635 | 653 | 876 | 906 | 754 | 756.25 | 667 | 845 | 1.268 | 0.342 | 0.00239775 | 0.01777963 | 0.0052 | 0.0113 | 0.0078 | 0.0078 | TRUE | 0.1033 |
| Slc30a5 | 3284 | 3334 | 3134 | 4303 | 3075 | 2997 | 3488 | 3330 | 3147 | 3383 | 3309 | 3281 | 3322.77 | 3322 | 3324 | 1.001 | 0.001 | 0.98499595 | 0.99199132 | 0.0011 | 0.0079 | 0.0034 | 0.0034 | TRUE | 0.0257 |
| Slc30a6 | 753 | 791 | 705 | 936 | 634 | 632 | 800 | 790 | 708 | 736 | 682 | 692 | 734.58 | 766 | 703 | 0.919 | -0.122 | 0.22184875 | 0.43411891 | 0.0018 | 0.0114 | 0.0058 | 0.0058 | TRUE | 0.0522 |
| Slc30a7 | 1357 | 1511 | 1581 | 1945 | 1372 | 1501 | 1441 | 1509 | 1587 | 1529 | 1476 | 1643 | 1531.03 | 1512 | 1549 | 1.024 | 0.034 | 0.69100506 | 0.83049088 | 0.0019 | 0.0091 | 0.0047 | 0.0047 | TRUE | 0.033 |
| Slc30a8 | 0 | 0 | 0 | 0 | 0 | 1 | 0 | 0 | 0 | 0 | 0 | 1 | 0.18 | 0 | 0 | 1.859 | 0.894 | 0.82650734 | NA | 0 | 18.0643 | 10 | 10 | TRUE | 0.5383 |
| Slc30a9 | 3729 | 4652 | 4615 | 4694 | 3430 | 3436 | 3961 | 4646 | 4634 | 3690 | 3690 | 3761 | 4063.76 | 4414 | 3714 | 0.841 | -0.249 | 0.00473254 | 0.03009746 | 0.0036 | 0.0077 | 0.0054 | 0.0054 | TRUE | 0.0979 |
| Slc31a1 | 3903 | 5088 | 5044 | 4556 | 3440 | 4121 | 4146 | 5081 | 5065 | 3582 | 3701 | 4511 | 4347.62 | 4764 | 3931 | 0.825 | -0.278 | 0.02791845 | 0.10967118 | 0.0142 | 0.0077 | 0.0113 | 0.0113 | TRUE | 0.2022 |
| Slc31a2 | 562 | 644 | 641 | 684 | 432 | 525 | 597 | 643 | 644 | 538 | 465 | 575 | 576.83 | 628 | 526 | 0.837 | -0.256 | 0.02904194 | 0.11243236 | 0.0044 | 0.0126 | 0.0082 | 0.0082 | TRUE | 0.1162 |
| Slc32a1 | 108 | 113 | 126 | 107 | 76 | 87 | 115 | 113 | 127 | 84 | 82 | 95 | 102.53 | 118 | 87 | 0.736 | -0.442 | 0.02441996 | 0.09996147 | 0 | 0.039 | 0.0178 | 0.0178 | TRUE | 0.0599 |
| Slc33a1 | 656 | 708 | 625 | 1180 | 831 | 724 | 697 | 707 | 628 | 928 | 894 | 793 | 774.28 | 677 | 872 | 1.288 | 0.365 | 0.00075226 | 0.00713022 | 0.0042 | 0.0112 | 0.0072 | 0.0072 | TRUE | 0.0736 |
| Slc34a1 | 1 | 0 | 0 | 0 | 0 | 0 | 1 | 0 | 0 | 0 | 0 | 0 | 0.18 | 0 | 0 | 0.49 | -1.029 | 0.80087 | NA | 0 | 18.6165 | 10 | 10 | TRUE | 0.4614 |
| Slc34a2 | 28 | 20 | 7 | 89 | 50 | 23 | 30 | 20 | 7 | 70 | 54 | 25 | 34.28 | 19 | 50 | 2.642 | 1.402 | 0.01165179 | 0.05895801 | 0.2894 | 0.103 | 0.1856 | 0.1856 | TRUE | 0.3114 |
| Slc34a3 | 21 | 46 | 50 | 63 | 46 | 32 | 22 | 46 | 50 | 50 | 49 | 35 | 42.08 | 39 | 45 | 1.132 | 0.179 | 0.63668971 | 0.7955847 | 0.0738 | 0.0852 | 0.0799 | 0.0799 | TRUE | 1.0472 |
| Slc35a1 | 1078 | 1085 | 1047 | 1437 | 1047 | 925 | 1145 | 1084 | 1051 | 1130 | 1127 | 1013 | 1091.44 | 1093 | 1090 | 0.997 | -0.004 | 0.96646158 | 0.98304089 | 0.0019 | 0.0099 | 0.0051 | 0.0051 | TRUE | 0.0449 |
| Slc35a2 | 519 | 732 | 814 | 974 | 692 | 717 | 551 | 731 | 817 | 766 | 745 | 785 | 732.46 | 700 | 765 | 1.093 | 0.128 | 0.39764498 | 0.61114531 | 0.0186 | 0.0114 | 0.0151 | 0.0151 | TRUE | 0.4055 |

| | | | | | | | | | | | | | | | | | | | | | | | | | |
|---|---|---|---|---|---|---|---|---|---|---|---|---|---|---|---|---|---|---|---|---|---|---|---|---|---|
| Slc35a3 | 1734 | 2077 | 2166 | 2250 | 1572 | 1770 | 1842 | 2074 | 2175 | 1769 | 1691 | 1938 | 1914.79 | 2030 | 1799 | 0.886 | -0.175 | 0.08215198 | 0.22887185 | 0.0052 | 0.0086 | 0.0068 | 0.0068 | TRUE | 0.0795 |
| Slc35a4 | 2412 | 2077 | 1983 | 3139 | 2199 | 1822 | 2562 | 2074 | 1991 | 2468 | 2366 | 1994 | 2242.59 | 2209 | 2276 | 1.031 | 0.044 | 0.73938884 | 0.86065684 | 0.0151 | 0.0084 | 0.012 | 0.012 | TRUE | 0.236 |
| Slc35a5 | 576 | 594 | 577 | 809 | 627 | 582 | 612 | 593 | 579 | 636 | 675 | 637 | 622.02 | 595 | 649 | 1.091 | 0.126 | 0.18056221 | 0.38193351 | 0 | 0.0122 | 0.0047 | 0.0047 | TRUE | 0.0135 |
| Slc35b1 | 2579 | 2206 | 1994 | 3603 | 2515 | 2017 | 2739 | 2203 | 2002 | 2832 | 2706 | 2208 | 2448.5 | 2315 | 2582 | 1.116 | 0.158 | 0.27807781 | 0.49924225 | 0.0201 | 0.0082 | 0.0149 | 0.0149 | TRUE | 0.3113 |
| Slc35b2 | 1244 | 1600 | 1470 | 1582 | 1117 | 1231 | 1321 | 1598 | 1476 | 1244 | 1202 | 1348 | 1364.72 | 1465 | 1265 | 0.863 | -0.213 | 0.04083926 | 0.1429782 | 0.0051 | 0.0093 | 0.0071 | 0.0071 | TRUE | 0.0883 |
| Slc35b3 | 256 | 212 | 201 | 338 | 228 | 214 | 272 | 212 | 202 | 266 | 245 | 234 | 238.46 | 229 | 248 | 1.09 | 0.124 | 0.45418197 | 0.65924517 | 0.0107 | 0.0207 | 0.0156 | 0.0156 | TRUE | 0.3012 |
| Slc35b4 | 1661 | 1560 | 1496 | 1918 | 1436 | 1288 | 1764 | 1558 | 1502 | 1508 | 1545 | 1410 | 1547.86 | 1608 | 1488 | 0.925 | -0.112 | 0.24570496 | 0.4627727 | 0.0038 | 0.009 | 0.0061 | 0.0061 | TRUE | 0.0866 |
| Slc35c1 | 598 | 690 | 645 | 763 | 545 | 618 | 635 | 689 | 648 | 600 | 586 | 676 | 639.11 | 657 | 621 | 0.944 | -0.084 | 0.42850724 | 0.63836934 | 0.0022 | 0.0121 | 0.0065 | 0.0065 | TRUE | 0.0702 |
| Slc35c2 | 1735 | 1687 | 1620 | 1837 | 1330 | 1328 | 1843 | 1685 | 1627 | 1444 | 1431 | 1454 | 1580.53 | 1718 | 1443 | 0.84 | -0.252 | 0.00242937 | 0.01795506 | 0.0016 | 0.009 | 0.0043 | 0.0043 | TRUE | 0.0485 |
| Slc35d1 | 237 | 316 | 293 | 445 | 360 | 292 | 252 | 316 | 294 | 350 | 387 | 320 | 319.72 | 287 | 352 | 1.226 | 0.294 | 0.04404678 | 0.15049954 | 0.0075 | 0.0172 | 0.0122 | 0.0122 | TRUE | 0.1294 |
| Slc35d2 | 607 | 812 | 832 | 815 | 548 | 659 | 645 | 811 | 835 | 641 | 590 | 721 | 707.13 | 764 | 651 | 0.851 | -0.232 | 0.09506651 | 0.2527202 | 0.0133 | 0.0116 | 0.0125 | 0.0125 | TRUE | 0.2189 |
| Slc35d3 | 27 | 34 | 33 | 63 | 28 | 37 | 29 | 34 | 33 | 50 | 30 | 41 | 35.99 | 32 | 40 | 1.263 | 0.337 | 0.32501747 | 0.54603489 | 0 | 0.0984 | 0.0562 | 0.0562 | TRUE | 0.1837 |
| Slc35e1 | 4382 | 4641 | 4449 | 6284 | 4496 | 4045 | 4654 | 4635 | 4467 | 4940 | 4837 | 4428 | 4660.34 | 4585 | 4735 | 1.033 | 0.047 | 0.52766388 | 0.71626773 | 0.0016 | 0.0076 | 0.0037 | 0.0037 | TRUE | 0.0391 |
| Slc35e2 | 1790 | 1605 | 1460 | 2255 | 1715 | 1423 | 1901 | 1603 | 1466 | 1773 | 1845 | 1558 | 1690.98 | 1657 | 1725 | 1.042 | 0.059 | 0.6362091 | 0.79536224 | 0.012 | 0.0089 | 0.0106 | 0.0106 | TRUE | 0.2007 |
| Slc35e3 | 753 | 662 | 568 | 987 | 659 | 594 | 800 | 661 | 570 | 776 | 709 | 650 | 694.41 | 677 | 712 | 1.052 | 0.074 | 0.6187565 | 0.78455081 | 0.0168 | 0.0116 | 0.0143 | 0.0143 | TRUE | 0.2954 |
| Slc35e4 | 248 | 225 | 225 | 233 | 188 | 191 | 263 | 225 | 226 | 183 | 202 | 209 | 218.1 | 238 | 198 | 0.831 | -0.267 | 0.0762738 | 0.21787537 | 0 | 0.022 | 0.0117 | 0.0117 | TRUE | 0.0952 |
| Slc35f1 | 45 | 49 | 46 | 85 | 50 | 58 | 48 | 49 | 46 | 67 | 54 | 63 | 54.51 | 48 | 61 | 1.292 | 0.37 | 0.15715173 | 0.35018326 | 0 | 0.0673 | 0.0305 | 0.0305 | TRUE | 0.097 |
| Slc35f2 | 3967 | 4344 | 4635 | 4201 | 3123 | 3599 | 4214 | 4338 | 4654 | 3302 | 3360 | 3940 | 3968.07 | 4402 | 3534 | 0.803 | -0.317 | 0.00113599 | 0.00983397 | 0.0059 | 0.0077 | 0.0066 | 0.0066 | TRUE | 0.1221 |
| Slc35f3 | 92 | 129 | 90 | 200 | 151 | 141 | 98 | 129 | 90 | 157 | 162 | 154 | 131.83 | 106 | 158 | 1.495 | 0.58 | 0.00342718 | 0.02345689 | 0.0072 | 0.0319 | 0.0204 | 0.0204 | TRUE | 0.3684 |
| Slc35f4 | 38 | 11 | 7 | 10 | 9 | 6 | 40 | 11 | 7 | 8 | 10 | 7 | 13.75 | 19 | 8 | 0.414 | -1.272 | 0.08987811 | 0.24390184 | 0.3837 | 0.2465 | 0.3179 | 0.3179 | TRUE | 2.8666 |
| Slc35f5 | 1284 | 1427 | 1368 | 1410 | 1096 | 1276 | 1364 | 1425 | 1374 | 1108 | 1179 | 1397 | 1307.84 | 1388 | 1228 | 0.884 | -0.177 | 0.10528307 | 0.27039689 | 0.0066 | 0.0094 | 0.0079 | 0.0079 | TRUE | 0.1721 |
| Slc35f6 | 920 | 906 | 833 | 1168 | 841 | 776 | 977 | 905 | 836 | 918 | 905 | 849 | 898.49 | 906 | 891 | 0.984 | -0.023 | 0.8126773 | 0.90398098 | 0.0027 | 0.0106 | 0.006 | 0.006 | TRUE | 0.0557 |
| Slc35g1 | 3379 | 3173 | 2902 | 4526 | 3305 | 3236 | 3589 | 3169 | 2914 | 3558 | 3556 | 3542 | 3388.03 | 3224 | 3552 | 1.102 | 0.14 | 0.14209676 | 0.3277973 | 0.0052 | 0.0079 | 0.0063 | 0.0063 | TRUE | 0.1191 |
| Slc35g2 | 52 | 64 | 65 | 82 | 51 | 52 | 55 | 64 | 65 | 64 | 55 | 57 | 60.11 | 61 | 59 | 0.959 | -0.061 | 0.80726238 | 0.90068918 | 0 | 0.0617 | 0.0284 | 0.0284 | TRUE | 0.0711 |
| Slc35g3 | 0 | 2 | 2 | 1 | 1 | 2 | 0 | 2 | 2 | 1 | 1 | 2 | 1.34 | 1 | 1 | 0.986 | -0.02 | 0.99025923 | NA | 0 | 2.4603 | 1.2132 | 1.2132 | TRUE | 0.4385 |
| Slc36a1 | 1322 | 1543 | 1547 | 1921 | 1494 | 1480 | 1404 | 1541 | 1553 | 1510 | 1607 | 1620 | 1539.37 | 1499 | 1579 | 1.052 | 0.074 | 0.38075894 | 0.5952354 | 0.0016 | 0.009 | 0.0044 | 0.0044 | TRUE | 0.037 |
| Slc36a2 | 13 | 14 | 21 | 22 | 20 | 16 | 14 | 14 | 21 | 17 | 22 | 18 | 17.53 | 16 | 19 | 1.148 | 0.199 | 0.66469631 | 0.81242949 | 0 | 0.1948 | 0.0949 | 0.0949 | TRUE | 0.3211 |
| Slc36a3 | 7 | 25 | 20 | 15 | 8 | 10 | 7 | 25 | 20 | 12 | 9 | 11 | 13.97 | 17 | 11 | 0.598 | -0.741 | 0.21608732 | 0.42810315 | 0.1199 | 0.2427 | 0.1832 | 0.1832 | TRUE | 0.5124 |
| Slc36a4 | 201 | 178 | 170 | 298 | 222 | 184 | 213 | 178 | 171 | 234 | 239 | 201 | 206.08 | 187 | 225 | 1.203 | 0.266 | 0.10365378 | 0.26782464 | 0.006 | 0.0229 | 0.0144 | 0.0144 | TRUE | 0.1588 |
| Slc37a1 | 563 | 652 | 682 | 585 | 458 | 533 | 598 | 651 | 685 | 460 | 493 | 583 | 578.35 | 645 | 512 | 0.793 | -0.335 | 0.00901727 | 0.04872229 | 0.0075 | 0.0126 | 0.0101 | 0.0101 | TRUE | 0.1718 |
| Slc37a2 | 143 | 127 | 109 | 335 | 223 | 150 | 152 | 127 | 109 | 263 | 240 | 164 | 175.94 | 129 | 222 | 1.725 | 0.786 | 0.00059099 | 0.00592187 | 0.0377 | 0.0256 | 0.0316 | 0.0316 | TRUE | 0.5654 |
| Slc37a3 | 1620 | 1350 | 1394 | 2455 | 1632 | 1416 | 1721 | 1348 | 1400 | 1930 | 1756 | 1550 | 1617.43 | 1490 | 1745 | 1.172 | 0.23 | 0.07973655 | 0.22405219 | 0.0142 | 0.0089 | 0.0117 | 0.0117 | TRUE | 0.2214 |
| Slc37a4 | 311 | 514 | 562 | 583 | 432 | 537 | 330 | 513 | 564 | 458 | 465 | 588 | 486.49 | 469 | 504 | 1.072 | 0.1 | 0.62830089 | 0.79087692 | 0.0456 | 0.0137 | 0.0289 | 0.0289 | TRUE | 0.7754 |
| Slc38a1 | 4674 | 3960 | 4192 | 5805 | 4111 | 3670 | 4965 | 3955 | 4209 | 4563 | 4423 | 4017 | 4355.45 | 4376 | 4334 | 0.991 | -0.013 | 0.90213512 | 0.95205059 | 0.0089 | 0.0077 | 0.0084 | 0.0084 | TRUE | 0.1683 |
| Slc38a10 | 7219 | 7545 | 7473 | 10189 | 7355 | 6961 | 7668 | 7535 | 7504 | 8010 | 7914 | 7620 | 7708.34 | 7569 | 7848 | 1.037 | 0.052 | 0.36556254 | 0.58306561 | 2.00E-04 | 0.0073 | 0.0023 | 0.0023 | TRUE | 0.0078 |
| Slc38a11 | 4 | 0 | 1 | 9 | 3 | 3 | 4 | 0 | 1 | 7 | 3 | 3 | 3.14 | 2 | 4 | 2.653 | 1.408 | 0.26692463 | NA | 0.2948 | 1.0561 | 0.7564 | 0.7564 | TRUE | 1.1758 |
| Slc38a2 | 25662 | 16482 | 15916 | 16365 | 12198 | 12042 | 27258 | 16461 | 15981 | 12865 | 13124 | 13182 | 16478.45 | 19900 | 13057 | 0.656 | -0.608 | 0.00118482 | 0.01012844 | 0.0513 | 0.0071 | 0.0253 | 0.0253 | TRUE | 1.2798 |
| Slc38a3 | 400 | 490 | 447 | 333 | 282 | 345 | 425 | 489 | 449 | 262 | 303 | 378 | 384.32 | 454 | 314 | 0.69 | -0.536 | 0.0008703 | 0.00798511 | 0.016 | 0.0155 | 0.016 | 0.016 | TRUE | 0.3499 |
| Slc38a4 | 22368 | 27686 | 28071 | 25534 | 18385 | 21423 | 23759 | 27650 | 28186 | 20073 | 19781 | 23451 | 23816.68 | 26532 | 21102 | 0.795 | -0.33 | 0.0018025 | 0.014172 | 0.0088 | 0.007 | 0.008 | 0.008 | TRUE | 0.116 |
| Slc38a5 | 250 | 175 | 168 | 102 | 65 | 86 | 266 | 175 | 169 | 80 | 70 | 94 | 142.21 | 203 | 81 | 0.401 | -1.318 | 6.80E-08 | 2.58E-06 | 0.0388 | 0.0301 | 0.0344 | 0.0344 | TRUE | 0.784 |
| Slc38a6 | 180 | 174 | 170 | 286 | 216 | 167 | 191 | 174 | 171 | 225 | 232 | 183 | 195.95 | 179 | 213 | 1.197 | 0.26 | 0.11218579 | 0.28161615 | 0.0048 | 0.0237 | 0.0141 | 0.0141 | TRUE | 0.164 |
| Slc38a7 | 994 | 1130 | 1151 | 1713 | 1266 | 1218 | 1056 | 1129 | 1156 | 1347 | 1362 | 1333 | 1230.35 | 1114 | 1347 | 1.21 | 0.275 | 0.0005881 | 0.00589675 | 0 | 0.0096 | 0.0038 | 0.0038 | TRUE | 0.0243 |
| Slc38a8 | 218 | 235 | 192 | 168 | 107 | 103 | 232 | 235 | 193 | 132 | 115 | 113 | 169.83 | 220 | 120 | 0.548 | -0.867 | 5.50E-07 | 1.62E-05 | 0 | 0.0263 | 0.0152 | 0.0152 | TRUE | 0.1267 |
| Slc38a9 | 1543 | 1506 | 1591 | 1681 | 1331 | 1154 | 1639 | 1504 | 1598 | 1321 | 1432 | 1263 | 1459.55 | 1580 | 1339 | 0.847 | -0.239 | 0.00675772 | 0.03908423 | 0.002 | 0.0092 | 0.0049 | 0.0049 | TRUE | 0.0436 |
| Slc39a1 | 1573 | 1658 | 1696 | 2181 | 1584 | 1550 | 1671 | 1656 | 1703 | 1715 | 1704 | 1697 | 1690.86 | 1677 | 1705 | 1.017 | 0.025 | 0.7177401 | 0.8469816 | 0 | 0.0089 | 0.0027 | 0.0027 | TRUE | 0.0023 |
| Slc39a10 | 6005 | 5233 | 5460 | 6985 | 4763 | 4954 | 6378 | 5226 | 5482 | 5491 | 5125 | 5423 | 5520.95 | 5695 | 5346 | 0.939 | -0.091 | 0.34676208 | 0.56586979 | 0.006 | 0.0075 | 0.0066 | 0.0066 | TRUE | 0.1339 |
| Slc39a11 | 1721 | 2462 | 2355 | 2060 | 1531 | 1773 | 1828 | 2459 | 2365 | 1619 | 1647 | 1941 | 1976.5 | 2217 | 1736 | 0.783 | -0.354 | 0.00998763 | 0.0526311 | 0.0163 | 0.0086 | 0.0131 | 0.0131 | TRUE | 0.285 |
| Slc39a12 | 0 | 1 | 1 | 0 | 0 | 0 | 0 | 1 | 1 | 0 | 0 | 0 | 0.33 | 1 | 0 | 0.268 | -1.901 | 0.61401143 | NA | 0 | 9.8764 | 8.4772 | 8.4772 | TRUE | 0.2278 |
| Slc39a13 | 278 | 370 | 361 | 354 | 264 | 287 | 295 | 370 | 362 | 278 | 284 | 314 | 317.3 | 342 | 292 | 0.852 | -0.232 | 0.10527261 | 0.27039689 | 0.0065 | 0.0173 | 0.0116 | 0.0116 | TRUE | 0.1634 |
| Slc39a14 | 3138 | 2761 | 3090 | 3602 | 2779 | 2733 | 3333 | 2757 | 3103 | 2832 | 2990 | 2992 | 3001.1 | 3064 | 2938 | 0.959 | -0.061 | 0.51514464 | 0.70749402 | 0.0043 | 0.008 | 0.006 | 0.006 | TRUE | 0.0934 |
| Slc39a2 | 13 | 15 | 12 | 27 | 15 | 16 | 14 | 15 | 12 | 21 | 16 | 18 | 15.95 | 14 | 18 | 1.352 | 0.435 | 0.3541998 | 0.57261574 | 0 | 0.2134 | 0.0948 | 0.0948 | TRUE | 0.124 |
| Slc39a3 | 164 | 176 | 149 | 237 | 159 | 142 | 174 | 176 | 150 | 186 | 171 | 155 | 168.73 | 167 | 171 | 1.03 | 0.042 | 0.80119311 | 0.89741637 | 0 | 0.0264 | 0.0142 | 0.0142 | TRUE | 0.0846 |
| Slc39a4 | 956 | 1385 | 1612 | 1119 | 928 | 1164 | 1015 | 1383 | 1619 | 880 | 998 | 1274 | 1194.93 | 1339 | 1051 | 0.784 | -0.351 | 0.06867904 | 0.2032281 | 0.043 | 0.0097 | 0.0259 | 0.0259 | TRUE | 0.2764 |
| Slc39a5 | 5 | 15 | 9 | 7 | 5 | 7 | 5 | 15 | 9 | 6 | 5 | 8 | 7.98 | 10 | 6 | 0.627 | -0.673 | 0.34578683 | 0.56517324 | 0 | 0.4198 | 0.2356 | 0.2356 | TRUE | 0.2514 |
| Slc39a6 | 1884 | 1968 | 1867 | 2378 | 1820 | 1882 | 2001 | 1965 | 1875 | 1869 | 1958 | 2060 | 1954.84 | 1947 | 1962 | 1.007 | 0.011 | 0.89136039 | 0.94740454 | 0.0013 | 0.0086 | 0.0039 | 0.0039 | TRUE | 0.0228 |
| Slc39a7 | 4498 | 4114 | 3768 | 6201 | 4313 | 3878 | 4778 | 4109 | 3783 | 4875 | 4641 | 4245 | 4405.03 | 4223 | 4587 | 1.086 | 0.119 | 0.27941365 | 0.5005155 | 0.0092 | 0.0077 | 0.0086 | 0.0086 | TRUE | 0.1604 |
| Slc39a8 | 76 | 122 | 101 | 102 | 63 | 74 | 81 | 122 | 101 | 80 | 68 | 81 | 88.83 | 101 | 76 | 0.754 | -0.408 | 0.08813396 | 0.24065941 | 0.0155 | 0.044 | 0.0299 | 0.0299 | TRUE | 0.1442 |
| Slc39a9 | 2141 | 1934 | 1931 | 2732 | 1892 | 1803 | 2274 | 1932 | 1939 | 2148 | 2036 | 1974 | 2050.26 | 2048 | 2053 | 1.003 | 0.004 | 0.9700379 | 0.98456155 | 0.0048 | 0.0085 | 0.0063 | 0.0063 | TRUE | 0.1123 |
| Slc3a1 | 5 | 0 | 1 | 3 | 0 | 3 | 5 | 0 | 1 | 2 | 0 | 3 | 1.99 | 2 | 2 | 0.905 | -0.145 | 0.93260302 | NA | 1.5237 | 1.66 | 1.6082 | 1.6082 | TRUE | 1.5948 |
| Slc3a2 | 9542 | 10861 | 11102 | 12662 | 9381 | 9314 | 10135 | 10847 | 11148 | 9954 | 10093 | 10196 | 10395.46 | 10710 | 10081 | 0.941 | -0.087 | 0.18630095 | 0.3900622 | 0.0011 | 0.0072 | 0.0031 | 0.0031 | TRUE | 0.0269 |
| Slc40a1 | 168 | 167 | 138 | 368 | 255 | 278 | 178 | 167 | 139 | 289 | 274 | 304 | 225.29 | 161 | 289 | 1.795 | 0.844 | 5.17E-08 | 2.04E-06 | 0.0022 | 0.0215 | 0.0125 | 0.0125 | TRUE | 0.1613 |
| Slc41a1 | 2197 | 2411 | 2371 | 2836 | 2084 | 2125 | 2334 | 2408 | 2381 | 2229 | 2242 | 2326 | 2320.01 | 2374 | 2266 | 0.954 | -0.068 | 0.30603326 | 0.52665845 | 0 | 0.0083 | 0.0027 | 0.0027 | TRUE | 0.0065 |
| Slc41a2 | 220 | 216 | 209 | 200 | 144 | 162 | 234 | 216 | 210 | 157 | 155 | 177 | 191.46 | 220 | 163 | 0.742 | -0.431 | 0.00453039 | 0.02903428 | 0 | 0.0241 | 0.0113 | 0.0113 | TRUE | 0.058 |
| Slc41a3 | 227 | 304 | 309 | 282 | 219 | 275 | 241 | 304 | 310 | 222 | 236 | 301 | 268.89 | 285 | 253 | 0.885 | -0.177 | 0.31952205 | 0.5406844 | 0.0189 | 0.0192 | 0.0191 | 0.0191 | TRUE | 0.3051 |
| Slc43a1 | 422 | 575 | 583 | 473 | 333 | 364 | 448 | 574 | 585 | 372 | 358 | 398 | 456.08 | 536 | 376 | 0.701 | -0.512 | 0.00027872 | 0.00320405 | 0.0102 | 0.0141 | 0.012 | 0.012 | TRUE | 0.2378 |
| Slc43a2 | 1386 | 2170 | 2348 | 2518 | 1827 | 1834 | 1472 | 2167 | 2358 | 1979 | 1966 | 2008 | 1991.64 | 1999 | 1984 | 0.992 | -0.011 | 0.94698457 | 0.97351104 | 0.0293 | 0.0086 | 0.0188 | 0.0188 | TRUE | 0.6418 |
| Slc43a3 | 244 | 279 | 196 | 681 | 426 | 332 | 259 | 279 | 197 | 535 | 458 | 363 | 348.62 | 245 | 452 | 1.851 | 0.888 | 4.72E-06 | 0.0001051 | 0.0317 | 0.0164 | 0.024 | 0.024 | TRUE | 0.1456 |
| Slc44a1 | 3393 | 3789 | 3756 | 4064 | 2965 | 3222 | 3604 | 3784 | 3771 | 3195 | 3190 | 3527 | 3511.91 | 3720 | 3304 | 0.888 | -0.171 | 0.02387119 | 0.09843043 | 0.0016 | 0.0078 | 0.0039 | 0.0039 | TRUE | 0.042 |
| Slc44a2 | 2234 | 3214 | 3464 | 2847 | 2353 | 2498 | 2373 | 3210 | 3478 | 2238 | 2532 | 2734 | 2760.87 | 3020 | 2501 | 0.828 | -0.272 | 0.07277464 | 0.2112648 | 0.0226 | 0.0081 | 0.0163 | 0.0163 | TRUE | 0.4266 |

| | | | | | | | | | | | | | | | | | | | | | | | | | |
|---|---|---|---|---|---|---|---|---|---|---|---|---|---|---|---|---|---|---|---|---|---|---|---|---|---|
| Slc44a3 | 149 | 227 | 201 | 165 | 121 | 168 | 158 | 227 | 202 | 130 | 130 | 184 | 171.77 | 196 | 148 | 0.754 | -0.408 | 0.0615824 | 0.18940672 | 0.0307 | 0.0261 | 0.0283 | 0.0283 | TRUE | 0.46 |
| Slc44a4 | 335 | 423 | 467 | 412 | 284 | 364 | 356 | 422 | 469 | 324 | 306 | 398 | 379.18 | 416 | 343 | 0.823 | -0.281 | 0.08148807 | 0.22755486 | 0.0157 | 0.0156 | 0.0161 | 0.0161 | TRUE | 0.2258 |
| Slc44a5 | 8 | 9 | 3 | 11 | 6 | 6 | 8 | 9 | 3 | 9 | 6 | 7 | 7.03 | 7 | 7 | 1.066 | 0.093 | 0.90046351 | 0.95169273 | 0 | 0.4757 | 0.254 | 0.254 | TRUE | 0.6328 |
| Slc45a1 | 29 | 21 | 29 | 22 | 9 | 20 | 31 | 21 | 29 | 17 | 10 | 22 | 21.63 | 27 | 16 | 0.606 | -0.724 | 0.11331199 | 0.28367568 | 0.0251 | 0.1592 | 0.1016 | 0.1016 | TRUE | 0.5151 |
| Slc45a2 | 1 | 0 | 1 | 3 | 1 | 0 | 1 | 0 | 1 | 2 | 1 | 0 | 0.92 | 1 | 1 | 1.726 | 0.787 | 0.70823394 | NA | 0 | 3.6005 | 2.0291 | 2.0291 | TRUE | 0.1755 |
| Slc45a3 | 415 | 599 | 539 | 565 | 393 | 420 | 441 | 598 | 541 | 444 | 423 | 460 | 484.5 | 527 | 442 | 0.839 | -0.253 | 0.07156428 | 0.20855724 | 0.0107 | 0.0137 | 0.0121 | 0.0121 | TRUE | 0.2361 |
| Slc45a4 | 3931 | 4428 | 4428 | 4519 | 3462 | 3758 | 4175 | 4422 | 4446 | 3552 | 3725 | 4114 | 4072.5 | 4348 | 3797 | 0.873 | -0.196 | 0.01972054 | 0.08601559 | 0.0031 | 0.0077 | 0.0048 | 0.0048 | TRUE | 0.0646 |
| Slc46a1 | 720 | 526 | 495 | 1189 | 784 | 618 | 765 | 525 | 497 | 935 | 844 | 676 | 706.98 | 596 | 818 | 1.375 | 0.459 | 0.01685764 | 0.07697705 | 0.0411 | 0.0116 | 0.0252 | 0.0252 | TRUE | 0.7215 |
| Slc46a2 | 3 | 1 | 0 | 0 | 1 | 1 | 3 | 1 | 0 | 0 | 1 | 1 | 1.06 | 1 | 1 | 0.499 | -1.002 | 0.62935735 | NA | 0 | 3.1169 | 2.0316 | 2.0316 | TRUE | 0.3789 |
| Slc46a3 | 127 | 146 | 164 | 82 | 45 | 65 | 135 | 146 | 165 | 64 | 48 | 71 | 104.9 | 149 | 61 | 0.414 | -1.273 | 1.10E-08 | 5.12E-07 | 0.0068 | 0.0383 | 0.0244 | 0.0244 | TRUE | 0.2715 |
| Slc47a1 | 528 | 419 | 409 | 412 | 240 | 254 | 561 | 418 | 411 | 324 | 258 | 278 | 375.02 | 463 | 287 | 0.62 | -0.689 | 5.22E-05 | 0.0008042 | 0.0193 | 0.0157 | 0.0181 | 0.0181 | TRUE | 0.3914 |
| Slc47a2 | 433 | 480 | 500 | 440 | 451 | 439 | 460 | 479 | 502 | 346 | 485 | 481 | 458.84 | 480 | 437 | 0.907 | -0.14 | 0.3710548 | 0.58781402 | 0.0167 | 0.0141 | 0.0155 | 0.0155 | TRUE | 0.4039 |
| Slc48a1 | 884 | 1129 | 1003 | 1041 | 735 | 854 | 939 | 1128 | 1007 | 818 | 791 | 935 | 936.27 | 1025 | 848 | 0.827 | -0.274 | 0.01783207 | 0.07989985 | 0.0068 | 0.0104 | 0.0085 | 0.0085 | TRUE | 0.0937 |
| Slc4a1 | 61 | 84 | 91 | 114 | 82 | 97 | 65 | 84 | 91 | 90 | 88 | 106 | 87.35 | 80 | 95 | 1.18 | 0.238 | 0.30147254 | 0.52210661 | 0.007 | 0.0446 | 0.0268 | 0.0268 | TRUE | 0.251 |
| Slc4a10 | 2 | 2 | 6 | 3 | 3 | 3 | 2 | 2 | 6 | 2 | 3 | 3 | 3.17 | 3 | 3 | 0.865 | -0.21 | 0.84349319 | NA | 0 | 1.0463 | 0.497 | 0.497 | TRUE | 0.6823 |
| Slc4a11 | 762 | 844 | 859 | 1144 | 864 | 960 | 809 | 843 | 863 | 899 | 930 | 1051 | 899.1 | 838 | 960 | 1.144 | 0.194 | 0.05213364 | 0.1687709 | 0.0029 | 0.0106 | 0.0061 | 0.0061 | TRUE | 0.081 |
| Slc4a1ap | 1904 | 1802 | 1822 | 2603 | 1876 | 1809 | 2022 | 1800 | 1829 | 2046 | 2018 | 1980 | 1949.42 | 1884 | 2015 | 1.07 | 0.098 | 0.22658953 | 0.43941596 | 0.0014 | 0.0086 | 0.0042 | 0.0042 | TRUE | 0.0498 |
| Slc4a2 | 729 | 500 | 486 | 941 | 679 | 497 | 774 | 499 | 488 | 740 | 731 | 544 | 629.34 | 587 | 672 | 1.144 | 0.195 | 0.34429139 | 0.56408957 | 0.0476 | 0.0121 | 0.0289 | 0.0289 | TRUE | 0.9091 |
| Slc4a3 | 316 | 288 | 281 | 344 | 266 | 237 | 336 | 288 | 282 | 270 | 286 | 259 | 286.91 | 302 | 272 | 0.902 | -0.149 | 0.27333155 | 0.49381273 | 0.0023 | 0.0184 | 0.0099 | 0.0099 | TRUE | 0.1073 |
| Slc4a4 | 191 | 185 | 193 | 434 | 279 | 226 | 203 | 185 | 194 | 341 | 300 | 247 | 245.03 | 194 | 296 | 1.533 | 0.617 | 0.00020062 | 0.00244904 | 0.0113 | 0.0203 | 0.0155 | 0.0155 | TRUE | 0.1779 |
| Slc4a5 | 817 | 636 | 608 | 1470 | 1077 | 918 | 868 | 635 | 610 | 1156 | 1159 | 1005 | 905.46 | 704 | 1107 | 1.572 | 0.652 | 2.07E-05 | 0.00036678 | 0.0217 | 0.0105 | 0.0157 | 0.0157 | TRUE | 0.4842 |
| Slc4a7 | 6851 | 7166 | 7598 | 8134 | 5935 | 6278 | 7277 | 7157 | 7629 | 6394 | 6386 | 6872 | 6952.53 | 7354 | 6551 | 0.891 | -0.167 | 0.0149689 | 0.0706643 | 0.0012 | 0.0074 | 0.0033 | 0.0033 | TRUE | 0.0224 |
| Slc4a8 | 1365 | 1271 | 1235 | 1448 | 1170 | 1175 | 1450 | 1269 | 1240 | 1138 | 1259 | 1286 | 1273.78 | 1320 | 1228 | 0.93 | -0.105 | 0.3068943 | 0.52738472 | 0.0046 | 0.0095 | 0.0068 | 0.0068 | TRUE | 0.089 |
| Slc4a9 | 31 | 26 | 27 | 40 | 37 | 42 | 33 | 26 | 27 | 31 | 40 | 46 | 33.87 | 29 | 39 | 1.356 | 0.439 | 0.20113401 | 0.40877592 | 0 | 0.1042 | 0.0545 | 0.0545 | TRUE | 0.2562 |
| Slc50a1 | 279 | 385 | 330 | 494 | 393 | 378 | 296 | 385 | 331 | 388 | 423 | 414 | 372.86 | 337 | 408 | 1.208 | 0.273 | 0.04604732 | 0.15528248 | 0.0064 | 0.0157 | 0.0108 | 0.0108 | TRUE | 0.1702 |
| Slc51a | 24 | 13 | 21 | 56 | 59 | 36 | 25 | 13 | 21 | 44 | 63 | 39 | 34.41 | 20 | 49 | 2.468 | 1.303 | 0.00099566 | 0.00888208 | 0.0464 | 0.1026 | 0.0771 | 0.0771 | TRUE | 0.5189 |
| Slc51b | 0 | 0 | 0 | 0 | 0 | 0 | 0 | 0 | 0 | 0 | 0 | 0 | 0 | 0 | 0 | NA | NA | NA | NA | NA | NA | NA | NA | NA | NA |
| Slc52a2 | 888 | 844 | 834 | 1216 | 833 | 816 | 943 | 843 | 837 | 956 | 896 | 893 | 894.83 | 874 | 915 | 1.048 | 0.067 | 0.48241264 | 0.68092108 | 0.0019 | 0.0106 | 0.0054 | 0.0054 | TRUE | 0.0558 |
| Slc52a3 | 275 | 462 | 483 | 633 | 526 | 556 | 292 | 461 | 485 | 498 | 566 | 609 | 485.11 | 413 | 558 | 1.349 | 0.431 | 0.02785606 | 0.10959798 | 0.0385 | 0.0137 | 0.0256 | 0.0256 | TRUE | 0.7499 |
| Slc5a1 | 55 | 22 | 18 | 111 | 72 | 55 | 58 | 22 | 18 | 87 | 77 | 60 | 53.9 | 33 | 75 | 2.296 | 1.199 | 0.00667339 | 0.03883861 | 0.1834 | 0.068 | 0.1186 | 0.1186 | TRUE | 2.8023 |
| Slc5a10 | 4 | 8 | 2 | 7 | 2 | 8 | 4 | 8 | 2 | 6 | 2 | 9 | 5.11 | 5 | 6 | 1.15 | 0.201 | 0.82941482 | 0.91345091 | 0.1928 | 0.6516 | 0.4323 | 0.4323 | TRUE | 0.2094 |
| Slc5a11 | 186 | 166 | 190 | 143 | 108 | 111 | 198 | 166 | 191 | 112 | 116 | 122 | 150.71 | 185 | 117 | 0.631 | -0.664 | 9.35E-05 | 0.00130672 | 0 | 0.0288 | 0.0139 | 0.0139 | TRUE | 0.0871 |
| Slc5a12 | 0 | 0 | 0 | 1 | 0 | 0 | 0 | 0 | 0 | 1 | 0 | 0 | 0.13 | 0 | 0 | 1.859 | 0.894 | 0.82650734 | NA | 0 | 25.1515 | 10 | 10 | TRUE | 0.2563 |
| Slc5a2 | 86 | 56 | 68 | 58 | 40 | 41 | 91 | 56 | 68 | 46 | 43 | 45 | 58.18 | 72 | 45 | 0.621 | -0.688 | 0.01667513 | 0.07645354 | 0.0198 | 0.0635 | 0.0414 | 0.0414 | TRUE | 0.3061 |
| Slc5a3 | 1165 | 1063 | 1206 | 1208 | 979 | 1100 | 1237 | 1062 | 1211 | 950 | 1053 | 1204 | 1119.52 | 1170 | 1069 | 0.913 | -0.132 | 0.2783402 | 0.49954274 | 0.0091 | 0.0098 | 0.0097 | 0.0097 | TRUE | 0.1458 |
| Slc5a4a | 11 | 7 | 6 | 9 | 12 | 8 | 12 | 7 | 6 | 7 | 13 | 9 | 8.91 | 8 | 10 | 1.157 | 0.21 | 0.74909714 | 0.86688385 | 0 | 0.3768 | 0.1977 | 0.1977 | TRUE | 0.3853 |
| Slc5a4b | 166 | 141 | 111 | 420 | 297 | 279 | 176 | 141 | 111 | 330 | 320 | 305 | 230.62 | 143 | 318 | 2.232 | 1.159 | 3.47E-10 | 2.36E-08 | 0.0173 | 0.0212 | 0.0194 | 0.0194 | TRUE | 0.4297 |
| Slc5a5 | 228 | 284 | 294 | 176 | 140 | 198 | 242 | 284 | 295 | 138 | 151 | 217 | 221.12 | 274 | 169 | 0.614 | -0.705 | 0.00053277 | 0.00544824 | 0.0293 | 0.0218 | 0.0251 | 0.0251 | TRUE | 0.652 |
| Slc5a6 | 671 | 531 | 549 | 1054 | 672 | 539 | 713 | 530 | 551 | 829 | 723 | 590 | 655.99 | 598 | 714 | 1.195 | 0.257 | 0.12769183 | 0.30730165 | 0.0256 | 0.0119 | 0.019 | 0.019 | TRUE | 0.2589 |
| Slc5a7 | 2 | 0 | 0 | 6 | 0 | 0 | 2 | 0 | 0 | 5 | 0 | 0 | 1.14 | 1 | 2 | 2.298 | 1.201 | 0.66193504 | NA | 10 | 2.8963 | 4.4004 | 4.4004 | TRUE | 2.7727 |
| Slc5a8 | 3 | 3 | 2 | 4 | 6 | 5 | 3 | 3 | 2 | 3 | 6 | 5 | 3.88 | 3 | 5 | 1.819 | 0.863 | 0.36935532 | NA | 0 | 0.8566 | 0.3797 | 0.3797 | TRUE | 0.2914 |
| Slc5a9 | 1 | 1 | 3 | 3 | 3 | 8 | 1 | 1 | 3 | 2 | 3 | 9 | 3.24 | 2 | 5 | 2.777 | 1.474 | 0.21158218 | NA | 0 | 1.0249 | 0.598 | 0.598 | TRUE | 0.9573 |
| Slc6a1 | 207 | 140 | 112 | 140 | 109 | 82 | 220 | 140 | 112 | 110 | 117 | 90 | 131.54 | 157 | 106 | 0.673 | -0.572 | 0.03813749 | 0.13584113 | 0.0626 | 0.0319 | 0.0469 | 0.0469 | TRUE | 0.6743 |
| Slc6a11 | 13 | 9 | 17 | 41 | 26 | 20 | 14 | 9 | 17 | 32 | 28 | 22 | 20.33 | 13 | 27 | 2.071 | 1.051 | 0.02196488 | 0.09310133 | 0 | 0.169 | 0.0953 | 0.0953 | TRUE | 0.3462 |
| Slc6a12 | 89 | 49 | 28 | 58 | 44 | 42 | 95 | 49 | 28 | 46 | 47 | 46 | 51.75 | 57 | 46 | 0.811 | -0.303 | 0.47923927 | 0.67850644 | 0.1575 | 0.0706 | 0.1123 | 0.1123 | TRUE | 0.484 |
| Slc6a13 | 59 | 57 | 45 | 110 | 67 | 82 | 63 | 57 | 45 | 86 | 72 | 90 | 68.85 | 55 | 83 | 1.51 | 0.595 | 0.01823731 | 0.08139186 | 0 | 0.0548 | 0.0305 | 0.0305 | TRUE | 0.2031 |
| Slc6a14 | 12 | 4 | 6 | 70 | 36 | 29 | 13 | 4 | 6 | 55 | 39 | 32 | 24.71 | 8 | 42 | 5.583 | 2.481 | 2.49E-06 | 6.10E-05 | 0.0896 | 0.1402 | 0.1206 | 0.1206 | TRUE | 0.6529 |
| Slc6a15 | 442 | 537 | 564 | 576 | 383 | 424 | 469 | 536 | 566 | 453 | 412 | 464 | 483.52 | 524 | 443 | 0.845 | -0.242 | 0.04642622 | 0.15587882 | 0.0044 | 0.0137 | 0.0086 | 0.0086 | TRUE | 0.0959 |
| Slc6a17 | 24 | 41 | 25 | 68 | 42 | 35 | 25 | 41 | 25 | 53 | 45 | 38 | 38.08 | 30 | 45 | 1.501 | 0.586 | 0.09440893 | 0.25177842 | 0.0247 | 0.0934 | 0.0611 | 0.0611 | TRUE | 0.6044 |
| Slc6a18 | 0 | 0 | 0 | 0 | 0 | 0 | 0 | 0 | 0 | 0 | 0 | 0 | 0 | 0 | 0 | NA | NA | NA | NA | NA | NA | NA | NA | NA | NA |
| Slc6a19 | 4 | 5 | 10 | 18 | 11 | 7 | 4 | 5 | 10 | 14 | 12 | 8 | 8.82 | 6 | 11 | 1.755 | 0.811 | 0.23599167 | 0.45112853 | 0 | 0.3803 | 0.2152 | 0.2152 | TRUE | 0.6053 |
| Slc6a2 | 25 | 40 | 30 | 41 | 37 | 46 | 27 | 40 | 30 | 32 | 40 | 50 | 36.5 | 32 | 41 | 1.257 | 0.33 | 0.34686642 | 0.56592295 | 0.0224 | 0.0972 | 0.0607 | 0.0607 | TRUE | 0.2371 |
| Slc6a20a | 1 | 0 | 0 | 4 | 0 | 2 | 1 | 0 | 0 | 3 | 0 | 2 | 1.07 | 0 | 2 | 4.927 | 2.301 | 0.30986572 | NA | 0 | 3.0974 | 2.4266 | 2.4266 | TRUE | 0.3769 |
| Slc6a20b | 0 | 0 | 0 | 0 | 0 | 0 | 0 | 0 | 0 | 0 | 0 | 0 | 0 | 0 | 0 | NA | NA | NA | NA | NA | NA | NA | NA | NA | NA |
| Slc6a3 | 8 | 23 | 16 | 25 | 12 | 15 | 8 | 23 | 16 | 20 | 13 | 16 | 16.09 | 16 | 16 | 1.034 | 0.048 | 0.92846796 | 0.96476961 | 0.0647 | 0.2117 | 0.143 | 0.143 | TRUE | 0.2082 |
| Slc6a4 | 1 | 1 | 1 | 7 | 1 | 2 | 1 | 1 | 1 | 6 | 1 | 2 | 1.97 | 1 | 3 | 2.937 | 1.554 | 0.27822843 | NA | 0 | 1.6774 | 0.8172 | 0.8172 | TRUE | 0.7265 |
| Slc6a5 | 3 | 1 | 3 | 2 | 1 | 4 | 3 | 1 | 3 | 2 | 1 | 4 | 2.37 | 2 | 2 | 0.965 | -0.052 | 0.96676233 | NA | 0 | 1.3965 | 0.6985 | 0.6985 | TRUE | 0.5381 |
| Slc6a6 | 12664 | 11669 | 11934 | 12294 | 9267 | 10048 | 13451 | 11654 | 11983 | 9665 | 9971 | 10999 | 11287.13 | 12363 | 10212 | 0.826 | -0.276 | 0.00253511 | 0.0185798 | 0.0052 | 0.0072 | 0.0059 | 0.0059 | TRUE | 0.0725 |
| Slc6a7 | 106 | 175 | 168 | 67 | 84 | 83 | 113 | 175 | 169 | 53 | 90 | 91 | 114.99 | 152 | 78 | 0.509 | -0.974 | 0.00059924 | 0.00600079 | 0.0626 | 0.0356 | 0.0484 | 0.0484 | TRUE | 0.8327 |
| Slc6a8 | 3564 | 3756 | 3760 | 4703 | 3377 | 3262 | 3786 | 3751 | 3775 | 3697 | 3633 | 3571 | 3702.26 | 3771 | 3634 | 0.964 | -0.053 | 0.37078621 | 0.58760362 | 0 | 0.0078 | 0.0023 | 0.0023 | TRUE | 0.0029 |
| Slc6a9 | 78 | 66 | 69 | 108 | 84 | 69 | 83 | 66 | 69 | 85 | 90 | 76 | 78.14 | 73 | 84 | 1.152 | 0.204 | 0.36983435 | 0.58700084 | 0 | 0.0491 | 0.0246 | 0.0246 | TRUE | 0.1327 |
| Slc7a1 | 7035 | 6461 | 6570 | 7680 | 5482 | 5065 | 7472 | 6453 | 6597 | 6037 | 5898 | 5544 | 6333.7 | 6841 | 5826 | 0.852 | -0.231 | 0.00773248 | 0.04330961 | 0.004 | 0.0074 | 0.0053 | 0.0053 | TRUE | 0.0796 |
| Slc7a10 | 2 | 2 | 2 | 4 | 3 | 3 | 2 | 2 | 2 | 3 | 3 | 3 | 2.63 | 2 | 3 | 1.574 | 0.655 | 0.55481241 | NA | 0 | 1.259 | 0.4839 | 0.4839 | TRUE | 0.001 |
| Slc7a11 | 14 | 10 | 11 | 17 | 20 | 9 | 15 | 10 | 11 | 13 | 22 | 10 | 13.44 | 12 | 15 | 1.245 | 0.316 | 0.56772758 | 0.74733351 | 0 | 0.252 | 0.1448 | 0.1448 | TRUE | 0.5139 |
| Slc7a12 | 0 | 0 | 0 | 0 | 0 | 0 | 0 | 0 | 0 | 0 | 0 | 0 | 0 | 0 | 0 | NA | NA | NA | NA | NA | NA | NA | NA | NA | NA |
| Slc7a13 | 1 | 0 | 0 | 3 | 4 | 1 | 1 | 0 | 0 | 2 | 4 | 1 | 1.47 | 0 | 2 | 7.058 | 2.819 | 0.12907767 | NA | 0 | 2.2483 | 1.2968 | 1.2968 | TRUE | 0.364 |
| Slc7a14 | 21 | 21 | 30 | 51 | 33 | 24 | 22 | 21 | 30 | 40 | 36 | 26 | 29.21 | 24 | 34 | 1.395 | 0.481 | 0.20228805 | 0.40993025 | 0 | 0.1197 | 0.0672 | 0.0672 | TRUE | 0.2591 |
| Slc7a15 | 1 | 1 | 0 | 4 | 3 | 6 | 1 | 1 | 0 | 3 | 3 | 7 | 2.5 | 1 | 4 | 6.271 | 2.649 | 0.06981422 | NA | 0 | 1.3246 | 0.6754 | 0.6754 | TRUE | 0.342 |
| Slc7a2 | 412 | 454 | 429 | 872 | 556 | 510 | 438 | 453 | 431 | 685 | 598 | 558 | 527.3 | 441 | 614 | 1.396 | 0.482 | 5.58E-05 | 0.00084977 | 0.0048 | 0.0132 | 0.0083 | 0.0083 | TRUE | 0.1267 |

| | | | | | | | | | | | | | | | | | | | | | | | | | |
|---|---|---|---|---|---|---|---|---|---|---|---|---|---|---|---|---|---|---|---|---|---|---|---|---|---|
| Slc7a3 | 2854 | 2006 | 2085 | 3698 | 2650 | 2230 | 3031 | 2003 | 2094 | 2907 | 2851 | 2441 | 2554.64 | 2376 | 2733 | 1.151 | 0.202 | 0.22029169 | 0.43242065 | 0.0293 | 0.0082 | 0.0192 | 0.0192 | TRUE | 0.7047 |
| Slc7a4 | 263 | 319 | 298 | 288 | 231 | 236 | 279 | 319 | 299 | 226 | 249 | 258 | 271.74 | 299 | 244 | 0.815 | -0.295 | 0.02914994 | 0.11273977 | 0 | 0.019 | 0.0095 | 0.0095 | TRUE | 0.0503 |
| Slc7a5 | 18746 | 16479 | 15608 | 22321 | 16721 | 14600 | 19912 | 16458 | 15672 | 17547 | 17991 | 15982 | 17260.19 | 17347 | 17173 | 0.99 | -0.014 | 0.89576636 | 0.94959182 | 0.0083 | 0.0071 | 0.0087 | 0.0087 | TRUE | 0.2045 |
| Slc7a6 | 5090 | 4773 | 4652 | 6491 | 4349 | 4195 | 5406 | 4767 | 4671 | 5103 | 4679 | 4592 | 4869.75 | 4948 | 4791 | 0.969 | -0.046 | 0.61215484 | 0.78032054 | 0.0041 | 0.0076 | 0.0057 | 0.0057 | TRUE | 0.0799 |
| Slc7a6os | 2863 | 2885 | 2704 | 3721 | 2821 | 2798 | 3041 | 2881 | 2715 | 2925 | 3035 | 3063 | 2943.44 | 2879 | 3008 | 1.045 | 0.063 | 0.40821803 | 0.62087325 | 0.0014 | 0.008 | 0.0038 | 0.0038 | TRUE | 0.0302 |
| Slc7a7 | 4793 | 3881 | 4372 | 2850 | 2027 | 1839 | 5091 | 3876 | 4390 | 2240 | 2181 | 2013 | 3298.57 | 4452 | 2145 | 0.482 | -1.053 | 1.68E-19 | 3.93E-17 | 0.0104 | 0.0079 | 0.0095 | 0.0095 | TRUE | 0.1559 |
| Slc7a8 | 128 | 85 | 100 | 115 | 77 | 76 | 136 | 85 | 100 | 90 | 83 | 83 | 96.28 | 107 | 85 | 0.801 | -0.321 | 0.18227066 | 0.38412252 | 0.0214 | 0.0411 | 0.0312 | 0.0312 | TRUE | 0.5154 |
| Slc7a9 | 1 | 1 | 2 | 5 | 4 | 2 | 1 | 1 | 2 | 4 | 4 | 2 | 2.42 | 1 | 3 | 2.569 | 1.361 | 0.27059457 | NA | 0 | 1.3708 | 0.5848 | 0.5848 | TRUE | 0.1371 |
| Slc8a1 | 15 | 11 | 15 | 43 | 22 | 35 | 16 | 11 | 15 | 34 | 24 | 38 | 22.96 | 14 | 32 | 2.288 | 1.194 | 0.00580983 | 0.0350278 | 0 | 0.1504 | 0.0832 | 0.0832 | TRUE | 0.3259 |
| Slc8a2 | 310 | 351 | 347 | 348 | 272 | 278 | 329 | 351 | 348 | 274 | 293 | 304 | 316.46 | 343 | 290 | 0.845 | -0.243 | 0.04545191 | 0.15393334 | 0 | 0.0173 | 0.0075 | 0.0075 | TRUE | 0.0307 |
| Slc8a3 | 42 | 41 | 31 | 53 | 39 | 35 | 45 | 41 | 31 | 42 | 42 | 38 | 39.77 | 39 | 41 | 1.047 | 0.066 | 0.83010233 | 0.9138554 | 0 | 0.0897 | 0.044 | 0.044 | TRUE | 0.2282 |
| Slc9a1 | 2802 | 2418 | 2465 | 3350 | 2438 | 2190 | 2976 | 2415 | 2475 | 2634 | 2623 | 2397 | 2586.69 | 2622 | 2551 | 0.973 | -0.039 | 0.71523294 | 0.84540034 | 0.0076 | 0.0082 | 0.0079 | 0.0079 | TRUE | 0.169 |
| Slc9a2 | 4 | 3 | 3 | 8 | 5 | 8 | 4 | 3 | 3 | 6 | 5 | 9 | 5.11 | 3 | 7 | 1.99 | 0.993 | 0.23602544 | 0.45112853 | 0 | 0.6511 | 0.284 | 0.284 | TRUE | 0.1454 |
| Slc9a3 | 0 | 1 | 0 | 0 | 0 | 0 | 0 | 1 | 0 | 0 | 0 | 0 | 0.17 | 0 | 0 | 0.49 | -1.029 | 0.80087 | NA | 0 | 19.7991 | 10 | 10 | TRUE | 0.405 |
| Slc9a3r1 | 4945 | 4088 | 4044 | 5756 | 4691 | 4039 | 5252 | 4083 | 4061 | 4525 | 5047 | 4421 | 4564.88 | 4465 | 4664 | 1.045 | 0.063 | 0.60966485 | 0.77891178 | 0.0114 | 0.0076 | 0.0108 | 0.0108 | TRUE | 0.2895 |
| Slc9a3r2 | 1008 | 823 | 774 | 765 | 596 | 607 | 1071 | 822 | 777 | 601 | 641 | 664 | 762.81 | 890 | 635 | 0.714 | -0.486 | 0.00067979 | 0.00662235 | 0.015 | 0.0112 | 0.0134 | 0.0134 | TRUE | 0.3746 |
| Slc9a4 | 0 | 0 | 0 | 1 | 1 | 0 | 0 | 0 | 0 | 1 | 1 | 0 | 0.31 | 0 | 1 | 3.32 | 1.731 | 0.65383052 | NA | 0 | 10.6224 | 8.9541 | 8.9541 | TRUE | 0.202 |
| Slc9a5 | 1668 | 2338 | 2423 | 1466 | 1203 | 1386 | 1772 | 2335 | 2433 | 1152 | 1294 | 1517 | 1750.61 | 2180 | 1321 | 0.606 | -0.723 | 2.13E-06 | 5.31E-05 | 0.0222 | 0.0088 | 0.0162 | 0.0162 | TRUE | 0.3243 |
| Slc9a6 | 1318 | 1313 | 1380 | 1766 | 1245 | 1137 | 1400 | 1311 | 1386 | 1388 | 1340 | 1245 | 1344.9 | 1366 | 1324 | 0.97 | -0.043 | 0.60998375 | 0.77891178 | 0.0014 | 0.0094 | 0.0045 | 0.0045 | TRUE | 0.0328 |
| Slc9a7 | 54 | 68 | 69 | 112 | 66 | 57 | 57 | 68 | 69 | 88 | 71 | 62 | 69.33 | 65 | 74 | 1.144 | 0.194 | 0.44210789 | 0.64905093 | 0 | 0.0544 | 0.0317 | 0.0317 | TRUE | 0.2886 |
| Slc9a8 | 991 | 1068 | 1063 | 1290 | 969 | 837 | 1053 | 1067 | 1067 | 1014 | 1043 | 916 | 1026.59 | 1062 | 991 | 0.933 | -0.099 | 0.26497245 | 0.48470871 | 0.0011 | 0.0101 | 0.0048 | 0.0048 | TRUE | 0.0505 |
| Slc9a9 | 15 | 9 | 7 | 18 | 11 | 9 | 16 | 9 | 7 | 14 | 12 | 10 | 11.3 | 11 | 12 | 1.133 | 0.18 | 0.76109529 | 0.87423727 | 0 | 0.2985 | 0.1645 | 0.1645 | TRUE | 0.6535 |
| Slc9b1 | 1224 | 1143 | 1135 | 1568 | 1130 | 1045 | 1300 | 1142 | 1140 | 1233 | 1216 | 1144 | 1195.61 | 1194 | 1198 | 1.004 | 0.005 | 0.95440348 | 0.97676022 | 0.0028 | 0.0097 | 0.0056 | 0.0056 | TRUE | 0.0724 |
| Slc9b2 | 191 | 201 | 188 | 183 | 131 | 145 | 203 | 201 | 189 | 144 | 141 | 159 | 172.65 | 198 | 148 | 0.748 | -0.419 | 0.0067237 | 0.03901628 | 0 | 0.026 | 0.0113 | 0.0113 | TRUE | 0.0407 |
| Slc9c1 | 0 | 0 | 0 | 0 | 0 | 0 | 0 | 0 | 0 | 0 | 0 | 0 | 0 | 0 | 0 | NA | NA | NA | NA | NA | NA | NA | NA | NA | NA |
| Slco1a1 | 0 | 0 | 0 | 0 | 0 | 0 | 0 | 0 | 0 | 0 | 0 | 0 | 0 | 0 | 0 | NA | NA | NA | NA | NA | NA | NA | NA | NA | NA |
| Slco1a4 | 0 | 1 | 0 | 0 | 2 | 0 | 0 | 1 | 0 | 0 | 2 | 0 | 0.53 | 0 | 1 | 1.927 | 0.947 | 0.77747358 | NA | 0 | 6.2809 | 6.39 | 6.39 | TRUE | 0.9744 |
| Slco1a5 | 1 | 4 | 0 | 6 | 2 | 0 | 1 | 4 | 0 | 5 | 2 | 0 | 1.99 | 2 | 2 | 1.387 | 0.472 | 0.78215875 | NA | 1.4581 | 1.6644 | 1.585 | 1.585 | TRUE | 0.2848 |
| Slco1a6 | 0 | 0 | 0 | 0 | 0 | 0 | 0 | 0 | 0 | 0 | 0 | 0 | 0 | 0 | 0 | NA | NA | NA | NA | NA | NA | NA | NA | NA | NA |
| Slco1b2 | 1 | 0 | 0 | 1 | 1 | 0 | 1 | 0 | 0 | 1 | 1 | 0 | 0.49 | 0 | 1 | 1.712 | 0.776 | 0.78488768 | NA | 0 | 6.7665 | 4.0387 | 4.0387 | TRUE | 0.4614 |
| Slco1c1 | 0 | 0 | 0 | 0 | 0 | 0 | 0 | 0 | 0 | 0 | 0 | 0 | 0 | 0 | 0 | NA | NA | NA | NA | NA | NA | NA | NA | NA | NA |
| Slco2a1 | 35 | 30 | 32 | 86 | 69 | 59 | 37 | 30 | 32 | 68 | 74 | 65 | 50.95 | 33 | 69 | 2.081 | 1.057 | 0.00013685 | 0.0017735 | 0 | 0.0716 | 0.0328 | 0.0328 | TRUE | 0.078 |
| Slco2b1 | 8 | 10 | 10 | 8 | 3 | 7 | 8 | 10 | 10 | 6 | 3 | 8 | 7.62 | 9 | 6 | 0.604 | -0.727 | 0.29323206 | 0.51433007 | 0 | 0.4394 | 0.2065 | 0.2065 | TRUE | 0.2903 |
| Slco3a1 | 173 | 158 | 161 | 288 | 178 | 165 | 184 | 158 | 162 | 226 | 192 | 181 | 183.63 | 168 | 200 | 1.194 | 0.256 | 0.12873111 | 0.30851649 | 0.006 | 0.0248 | 0.015 | 0.015 | TRUE | 0.1618 |
| Slco4a1 | 1499 | 1956 | 2154 | 778 | 677 | 1063 | 1592 | 1953 | 2163 | 612 | 728 | 1164 | 1368.69 | 1903 | 835 | 0.438 | -1.191 | 8.66E-08 | 3.18E-06 | 0.071 | 0.0093 | 0.0348 | 0.0348 | TRUE | 1.206 |
| Slco4c1 | 436 | 377 | 412 | 482 | 284 | 384 | 463 | 377 | 414 | 379 | 306 | 420 | 393.02 | 418 | 368 | 0.882 | -0.181 | 0.24930897 | 0.46649675 | 0.0152 | 0.0153 | 0.0152 | 0.0152 | TRUE | 0.2471 |
| Slco5a1 | 306 | 305 | 335 | 412 | 298 | 329 | 325 | 305 | 336 | 324 | 321 | 360 | 328.44 | 322 | 335 | 1.039 | 0.055 | 0.65546584 | 0.80619668 | 0 | 0.0169 | 0.008 | 0.008 | TRUE | 0.0477 |
| Slco6b1 | 0 | 0 | 0 | 0 | 0 | 0 | 0 | 0 | 0 | 0 | 0 | 0 | 0 | 0 | 0 | NA | NA | NA | NA | NA | NA | NA | NA | NA | NA |
| Slco6c1 | 0 | 0 | 0 | 0 | 0 | 0 | 0 | 0 | 0 | 0 | 0 | 0 | 0 | 0 | 0 | NA | NA | NA | NA | NA | NA | NA | NA | NA | NA |
| Slco6d1 | 0 | 0 | 0 | 0 | 0 | 0 | 0 | 0 | 0 | 0 | 0 | 0 | 0 | 0 | 0 | NA | NA | NA | NA | NA | NA | NA | NA | NA | NA |
| Slfn1 | 0 | 0 | 0 | 0 | 0 | 0 | 0 | 0 | 0 | 0 | 0 | 0 | 0 | 0 | 0 | NA | NA | NA | NA | NA | NA | NA | NA | NA | NA |
| Slfn10-ps | 63 | 46 | 66 | 143 | 99 | 80 | 67 | 46 | 66 | 112 | 107 | 88 | 80.94 | 60 | 102 | 1.717 | 0.78 | 0.00182126 | 0.01428393 | 0.0134 | 0.0476 | 0.0317 | 0.0317 | TRUE | 0.3547 |
| Slfn14-ps | 1 | 1 | 0 | 0 | 0 | 0 | 0 | 1 | 1 | 0 | 0 | 0 | 0.34 | 1 | 0 | 0.261 | -1.936 | 0.60454921 | NA | 0 | 9.5983 | 8.3269 | 8.3269 | TRUE | 0.2507 |
| Slfn2 | 0 | 0 | 0 | 0 | 0 | 0 | 0 | 0 | 0 | 0 | 0 | 0 | 0 | 0 | 0 | NA | NA | NA | NA | NA | NA | NA | NA | NA | NA |
| Slfn3 | 29 | 35 | 26 | 196 | 134 | 117 | 31 | 35 | 26 | 154 | 144 | 128 | 86.37 | 31 | 142 | 4.65 | 2.217 | 4.41E-19 | 9.93E-17 | 0 | 0.045 | 0.0243 | 0.0243 | TRUE | 0.1143 |
| Slfn4 | 9 | 9 | 2 | 30 | 21 | 14 | 10 | 9 | 2 | 24 | 23 | 15 | 13.68 | 7 | 21 | 3.012 | 1.59 | 0.0110611 | 0.05674686 | 0.0584 | 0.2478 | 0.1838 | 0.1838 | TRUE | 1.0149 |
| Slfn5 | 120 | 145 | 152 | 79 | 57 | 66 | 127 | 145 | 153 | 62 | 61 | 72 | 103.43 | 142 | 65 | 0.459 | -1.123 | 4.79E-08 | 1.91E-06 | 0 | 0.0388 | 0.0194 | 0.0194 | TRUE | 0.0781 |
| Slfn8 | 44 | 40 | 50 | 126 | 85 | 79 | 47 | 40 | 50 | 99 | 91 | 86 | 68.98 | 46 | 92 | 2.029 | 1.02 | 2.59E-05 | 0.00044542 | 0 | 0.0547 | 0.0259 | 0.0259 | TRUE | 0.0955 |
| Slfn9 | 382 | 389 | 381 | 1360 | 1032 | 958 | 406 | 388 | 383 | 1069 | 1110 | 1049 | 734.16 | 392 | 1076 | 2.744 | 1.456 | 4.51E-55 | 8.89E-52 | 0 | 0.0114 | 0.0045 | 0.0045 | TRUE | 0.0102 |
| Slfnl1 | 0 | 1 | 0 | 3 | 1 | 0 | 0 | 1 | 0 | 2 | 1 | 0 | 0.74 | 0 | 1 | 3.203 | 1.679 | 0.50423271 | NA | 0 | 4.4659 | 3.139 | 3.139 | TRUE | 0.1967 |
| Slirp | 1783 | 1835 | 1765 | 2502 | 1700 | 1609 | 1894 | 1833 | 1772 | 1967 | 1829 | 1761 | 1842.67 | 1833 | 1852 | 1.011 | 0.016 | 0.84027356 | 0.91912023 | 0.0017 | 0.0087 | 0.0042 | 0.0042 | TRUE | 0.0362 |
| Slit1 | 38 | 50 | 35 | 37 | 34 | 36 | 40 | 50 | 35 | 29 | 37 | 39 | 38.42 | 42 | 35 | 0.832 | -0.265 | 0.415039 | 0.62659089 | 0 | 0.0927 | 0.0502 | 0.0502 | TRUE | 0.2238 |
| Slit2 | 226 | 273 | 254 | 451 | 353 | 378 | 240 | 273 | 255 | 355 | 380 | 414 | 319.31 | 256 | 383 | 1.492 | 0.578 | 1.08E-05 | 0.00021254 | 0.0026 | 0.0172 | 0.0091 | 0.0091 | TRUE | 0.0574 |
| Slit3 | 316 | 335 | 413 | 485 | 308 | 352 | 336 | 335 | 415 | 381 | 331 | 385 | 363.81 | 362 | 366 | 1.013 | 0.018 | 0.89990734 | 0.95148767 | 0.0084 | 0.016 | 0.0119 | 0.0119 | TRUE | 0.1893 |
| Slitrk1 | 0 | 0 | 0 | 0 | 0 | 0 | 0 | 0 | 0 | 0 | 0 | 0 | 0 | 0 | 0 | NA | NA | NA | NA | NA | NA | NA | NA | NA | NA |
| Slitrk2 | 1 | 0 | 1 | 1 | 2 | 1 | 1 | 0 | 1 | 1 | 2 | 1 | 1.02 | 1 | 1 | 1.936 | 0.953 | 0.61010723 | NA | 0 | 3.2479 | 1.4013 | 1.4013 | TRUE | 0.256 |
| Slitrk3 | 0 | 0 | 0 | 1 | 0 | 0 | 0 | 0 | 0 | 1 | 0 | 0 | 0.13 | 0 | 0 | 1.859 | 0.894 | 0.82650734 | NA | 0 | 25.1515 | 10 | 10 | TRUE | 0.2563 |
| Slitrk4 | 49 | 45 | 58 | 80 | 43 | 47 | 52 | 45 | 58 | 63 | 46 | 51 | 52.64 | 52 | 53 | 1.041 | 0.058 | 0.83660983 | 0.9170844 | 0 | 0.0695 | 0.0376 | 0.0376 | TRUE | 0.2217 |
| Slitrk5 | 40 | 36 | 31 | 65 | 51 | 31 | 42 | 36 | 31 | 51 | 55 | 34 | 41.58 | 36 | 47 | 1.282 | 0.359 | 0.27681919 | 0.49777588 | 0.0178 | 0.0861 | 0.0539 | 0.0539 | TRUE | 0.4186 |
| Slitrk6 | 0 | 0 | 0 | 0 | 0 | 1 | 0 | 0 | 0 | 0 | 0 | 1 | 0.18 | 0 | 0 | 1.859 | 0.894 | 0.82650734 | NA | 0 | 18.0643 | 10 | 10 | TRUE | 0.5383 |
| Slk | 4587 | 5661 | 6139 | 6594 | 5173 | 5243 | 4872 | 5654 | 6164 | 5184 | 5566 | 5739 | 5529.82 | 5563 | 5496 | 0.988 | -0.018 | 0.86651146 | 0.93389495 | 0.0081 | 0.0075 | 0.0079 | 0.0079 | TRUE | 0.1439 |
| Slmap | 4705 | 4288 | 4417 | 6020 | 4314 | 4220 | 4998 | 4282 | 4435 | 4732 | 4642 | 4619 | 4618.11 | 4572 | 4664 | 1.02 | 0.029 | 0.72642912 | 0.85254683 | 0.0032 | 0.0076 | 0.0048 | 0.0048 | TRUE | 0.0807 |
| Slmo1 | 164 | 161 | 174 | 280 | 204 | 185 | 174 | 161 | 175 | 220 | 219 | 203 | 191.97 | 170 | 214 | 1.261 | 0.335 | 0.02292282 | 0.09578079 | 0 | 0.0241 | 0.0103 | 0.0103 | TRUE | 0.0237 |
| Slmo2 | 2373 | 2114 | 2009 | 3683 | 2552 | 2131 | 2521 | 2111 | 2017 | 2895 | 2746 | 2333 | 2437.14 | 2216 | 2658 | 1.2 | 0.263 | 0.03474261 | 0.12746232 | 0.0127 | 0.0083 | 0.0107 | 0.0107 | TRUE | 0.1743 |
| Sln | 0 | 0 | 0 | 0 | 0 | 0 | 0 | 0 | 0 | 0 | 0 | 0 | 0 | 0 | 0 | NA | NA | NA | NA | NA | NA | NA | NA | NA | NA |
| Slpi | 2 | 3 | 1 | 2 | 2 | 1 | 2 | 3 | 1 | 2 | 2 | 1 | 1.82 | 2 | 2 | 0.787 | -0.346 | 0.79634939 | NA | 0 | 1.8132 | 0.743 | 0.743 | TRUE | 0.1846 |
| Sltm | 5481 | 4586 | 4385 | 5897 | 4224 | 3671 | 5822 | 4580 | 4403 | 4636 | 4545 | 4018 | 4667.31 | 4935 | 4400 | 0.892 | -0.165 | 0.19062426 | 0.39573321 | 0.0148 | 0.0076 | 0.0113 | 0.0113 | TRUE | 0.301 |
| Slu7 | 2893 | 2889 | 2866 | 4227 | 3065 | 2843 | 3073 | 2885 | 2878 | 3323 | 3298 | 3112 | 3094.79 | 2945 | 3244 | 1.102 | 0.14 | 0.05034837 | 0.16493186 | 8.00E-04 | 0.008 | 0.0034 | 0.0034 | TRUE | 0.0174 |
| Slurp1 | 0 | 0 | 0 | 0 | 0 | 0 | 0 | 0 | 0 | 0 | 0 | 0 | 0 | 0 | 0 | NA | NA | NA | NA | NA | NA | NA | NA | NA | NA |

| | | | | | | | | | | | | | | | | | | | | | | | | | |
|---|---|---|---|---|---|---|---|---|---|---|---|---|---|---|---|---|---|---|---|---|---|---|---|---|---|
| Slx | 0 | 0 | 0 | 0 | 0 | 0 | 0 | 0 | 0 | 0 | 0 | 0 | 0 | 0 | 0 | NA | NA | NA | NA | NA | NA | NA | NA | NA | NA |
| Slx1b | 811 | 955 | 930 | 994 | 689 | 754 | 861 | 954 | 934 | 781 | 741 | 825 | 849.52 | 916 | 782 | 0.854 | -0.228 | 0.01698194 | 0.07733081 | 0.0015 | 0.0108 | 0.0054 | 0.0054 | TRUE | 0.0325 |
| Slx4 | 1851 | 1435 | 1520 | 2438 | 1745 | 1590 | 1966 | 1433 | 1526 | 1917 | 1878 | 1741 | 1743.34 | 1642 | 1845 | 1.124 | 0.169 | 0.20293455 | 0.41057686 | 0.0153 | 0.0088 | 0.0121 | 0.0121 | TRUE | 0.3593 |
| Slx4ip | 471 | 502 | 552 | 654 | 504 | 532 | 500 | 501 | 554 | 514 | 542 | 582 | 532.44 | 518 | 546 | 1.052 | 0.073 | 0.50702199 | 0.70128482 | 0.0019 | 0.0131 | 0.0068 | 0.0068 | TRUE | 0.0424 |
| Slxl1 | 0 | 0 | 0 | 0 | 0 | 0 | 0 | 0 | 0 | 0 | 0 | 0 | 0 | 0 | 0 | NA | NA | NA | NA | NA | NA | NA | NA | NA | NA |
| Sly | 0 | 0 | 1 | 0 | 0 | 0 | 0 | 0 | 1 | 0 | 0 | 0 | 0.17 | 0 | 0 | 0.49 | -1.029 | 0.80087 | NA | 0 | 19.6928 | 10 | 10 | TRUE | 0.4098 |
| Smad1 | 643 | 720 | 685 | 1195 | 848 | 788 | 683 | 719 | 688 | 939 | 912 | 863 | 800.71 | 697 | 905 | 1.3 | 0.378 | 2.58E-05 | 0.00044452 | 0 | 0.011 | 0.0045 | 0.0045 | TRUE | 0.0196 |
| Smad2 | 4204 | 4446 | 4699 | 5377 | 3985 | 4221 | 4465 | 4440 | 4718 | 4227 | 4288 | 4621 | 4459.85 | 4541 | 4379 | 0.964 | -0.053 | 0.46444942 | 0.6677075 | 0.0014 | 0.0076 | 0.0036 | 0.0036 | TRUE | 0.0284 |
| Smad3 | 686 | 824 | 907 | 1096 | 831 | 838 | 729 | 823 | 911 | 862 | 894 | 917 | 855.89 | 821 | 891 | 1.085 | 0.117 | 0.29018774 | 0.51137888 | 0.0054 | 0.0108 | 0.0077 | 0.0077 | TRUE | 0.1137 |
| Smad4 | 4536 | 4839 | 4773 | 6473 | 4707 | 4978 | 4818 | 4833 | 4793 | 5089 | 5064 | 5449 | 5007.6 | 4815 | 5201 | 1.08 | 0.111 | 0.08560329 | 0.2356668 | 6.00E-04 | 0.0076 | 0.0028 | 0.0028 | TRUE | 0.0211 |
| Smad5 | 3188 | 3235 | 3310 | 4108 | 2975 | 2998 | 3386 | 3231 | 3324 | 3229 | 3201 | 3282 | 3275.45 | 3314 | 3237 | 0.977 | -0.034 | 0.59107139 | 0.76481919 | 0 | 0.0079 | 0.0025 | 0.0025 | TRUE | 0.0058 |
| Smad6 | 110 | 220 | 209 | 373 | 274 | 313 | 117 | 220 | 210 | 293 | 295 | 343 | 246.18 | 182 | 310 | 1.701 | 0.766 | 0.00107145 | 0.00938986 | 0.0528 | 0.0203 | 0.0351 | 0.0351 | TRUE | 1.044 |
| Smad7 | 3031 | 3789 | 3578 | 4856 | 3781 | 3265 | 3219 | 3784 | 3593 | 3817 | 4068 | 3574 | 3675.97 | 3532 | 3820 | 1.081 | 0.113 | 0.23356878 | 0.44826562 | 0.005 | 0.0078 | 0.0062 | 0.0062 | TRUE | 0.0728 |
| Smad9 | 85 | 94 | 73 | 163 | 122 | 113 | 90 | 94 | 73 | 128 | 131 | 124 | 106.76 | 86 | 128 | 1.489 | 0.574 | 0.00381577 | 0.02543232 | 0 | 0.0378 | 0.0186 | 0.0186 | TRUE | 0.1559 |
| Smagp | 1165 | 1754 | 1903 | 1155 | 935 | 1117 | 1237 | 1752 | 1911 | 908 | 1006 | 1223 | 1339.45 | 1633 | 1046 | 0.64 | -0.645 | 0.00030937 | 0.00349383 | 0.0339 | 0.0094 | 0.0222 | 0.0222 | TRUE | 0.5412 |
| Smap1 | 4325 | 4547 | 4457 | 5451 | 3999 | 4081 | 4594 | 4541 | 4475 | 4285 | 4303 | 4467 | 4444.24 | 4537 | 4352 | 0.959 | -0.06 | 0.31544248 | 0.53617717 | 1.00E-04 | 0.0076 | 0.0024 | 0.0024 | TRUE | 0.0065 |
| Smap2 | 2142 | 1677 | 1529 | 2244 | 1665 | 1533 | 2275 | 1675 | 1535 | 1764 | 1791 | 1678 | 1786.48 | 1828 | 1744 | 0.954 | -0.067 | 0.65128083 | 0.80449923 | 0.0199 | 0.0087 | 0.0155 | 0.0155 | TRUE | 0.5511 |
| Smarca1 | 1158 | 1150 | 1114 | 829 | 595 | 570 | 1230 | 1149 | 1119 | 652 | 640 | 624 | 902.15 | 1166 | 639 | 0.548 | -0.867 | 2.08E-22 | 6.70E-20 | 0 | 0.0106 | 0.0045 | 0.0045 | TRUE | 0.0277 |
| Smarca2 | 80 | 123 | 117 | 214 | 167 | 143 | 85 | 123 | 117 | 168 | 180 | 157 | 138.29 | 108 | 168 | 1.549 | 0.631 | 0.00158342 | 0.01279389 | 0.0099 | 0.0307 | 0.0212 | 0.0212 | TRUE | 0.3474 |
| Smarca4 | 17560 | 17398 | 18526 | 19912 | 14693 | 14942 | 18652 | 17376 | 18602 | 15653 | 15809 | 16356 | 17074.64 | 18210 | 15939 | 0.875 | -0.192 | 0.00244535 | 0.01803939 | 0.001 | 0.0071 | 0.0028 | 0.0028 | TRUE | 0.0197 |
| Smarca5 | 23233 | 21297 | 20950 | 32788 | 23258 | 21203 | 24678 | 21270 | 21036 | 25775 | 25024 | 23210 | 23498.77 | 22328 | 24670 | 1.105 | 0.144 | 0.11766958 | 0.29020869 | 0.0055 | 0.007 | 0.0061 | 0.0061 | TRUE | 0.1037 |
| Smarca5-ps | 74 | 35 | 40 | 120 | 97 | 39 | 79 | 35 | 40 | 94 | 104 | 43 | 65.85 | 51 | 80 | 1.576 | 0.656 | 0.1228303 | 0.29845625 | 0.1806 | 0.0569 | 0.1142 | 0.1142 | TRUE | 1.755 |
| Smarcad1 | 5364 | 3779 | 3857 | 7997 | 5304 | 4370 | 5698 | 3774 | 3873 | 6287 | 5707 | 4784 | 5020.25 | 4448 | 5593 | 1.257 | 0.33 | 0.05636701 | 0.1782676 | 0.0385 | 0.0076 | 0.0214 | 0.0214 | TRUE | 0.735 |
| Smarcal1 | 2189 | 1995 | 1995 | 2778 | 2037 | 1840 | 2325 | 1992 | 2003 | 2184 | 2192 | 2014 | 2118.4 | 2107 | 2130 | 1.011 | 0.016 | 0.86551662 | 0.93364615 | 0.0045 | 0.0085 | 0.0061 | 0.0061 | TRUE | 0.099 |
| Smarcb1 | 2691 | 2633 | 2645 | 4726 | 3330 | 2963 | 2858 | 2630 | 2656 | 3715 | 3583 | 3243 | 3114.22 | 2715 | 3514 | 1.295 | 0.373 | 1.38E-05 | 0.00026151 | 0.003 | 0.008 | 0.005 | 0.005 | TRUE | 0.0549 |
| Smarcc1 | 21725 | 20546 | 21844 | 24687 | 18397 | 18293 | 23076 | 20519 | 21934 | 19407 | 19794 | 20025 | 20792.42 | 21843 | 19742 | 0.904 | -0.146 | 0.03779605 | 0.13505193 | 0.0018 | 0.0071 | 0.0035 | 0.0035 | TRUE | 0.0344 |
| Smarcc2 | 1526 | 1526 | 1695 | 2419 | 1827 | 1746 | 1621 | 1524 | 1702 | 1902 | 1966 | 1911 | 1770.92 | 1616 | 1926 | 1.192 | 0.254 | 0.00127492 | 0.01074725 | 0.001 | 0.0088 | 0.0039 | 0.0039 | TRUE | 0.0298 |
| Smarcd1 | 7882 | 8086 | 8545 | 9286 | 6891 | 6930 | 8372 | 8076 | 8580 | 7300 | 7414 | 7586 | 7887.99 | 8343 | 7433 | 0.891 | -0.167 | 0.00606963 | 0.03626145 | 5.00E-04 | 0.0073 | 0.0025 | 0.0025 | TRUE | 0.0096 |
| Smarcd2 | 4162 | 4878 | 4724 | 6779 | 5188 | 5289 | 4421 | 4872 | 4743 | 5329 | 5582 | 5790 | 5122.77 | 4679 | 5567 | 1.19 | 0.25 | 0.000827 | 0.00769075 | 0.0018 | 0.0075 | 0.0038 | 0.0038 | TRUE | 0.0283 |
| Smarcd3 | 402 | 502 | 559 | 271 | 219 | 267 | 427 | 501 | 561 | 213 | 236 | 292 | 371.76 | 496 | 247 | 0.496 | -1.012 | 2.13E-09 | 1.20E-07 | 0.0193 | 0.0158 | 0.0176 | 0.0176 | TRUE | 0.2287 |
| Smarce1 | 4011 | 4150 | 4306 | 5108 | 3928 | 3742 | 4260 | 4145 | 4324 | 4016 | 4226 | 4096 | 4177.79 | 4243 | 4113 | 0.969 | -0.045 | 0.47097539 | 0.67235914 | 3.00E-04 | 0.0077 | 0.0026 | 0.0026 | TRUE | 0.0071 |
| Smc1a | 8601 | 9192 | 9908 | 12485 | 9107 | 9486 | 9136 | 9180 | 9949 | 9815 | 9799 | 10384 | 9710.3 | 9422 | 9999 | 1.061 | 0.086 | 0.22116431 | 0.43313412 | 0.0016 | 0.0072 | 0.0034 | 0.0034 | TRUE | 0.0293 |
| Smc1b | 615 | 1601 | 1764 | 2119 | 1571 | 1800 | 653 | 1599 | 1771 | 1666 | 1690 | 1970 | 1558.31 | 1341 | 1775 | 1.324 | 0.405 | 0.34488988 | 0.56427418 | 0.1316 | 0.009 | 0.0586 | 0.1316 | TRUE | 2.4177 |
| Smc2 | 12135 | 11188 | 11438 | 16852 | 12115 | 11336 | 12890 | 11174 | 11485 | 13248 | 13035 | 12409 | 12373.3 | 11850 | 12897 | 1.089 | 0.122 | 0.13538292 | 0.31831536 | 0.0034 | 0.0072 | 0.0048 | 0.0048 | TRUE | 0.072 |
| Smc3 | 16695 | 15766 | 17046 | 19759 | 14853 | 15614 | 17733 | 15746 | 17116 | 15533 | 15981 | 17092 | 16533.42 | 16865 | 16202 | 0.961 | -0.058 | 0.46372387 | 0.66707888 | 0.0029 | 0.0071 | 0.0044 | 0.0044 | TRUE | 0.0412 |
| Smc4 | 12458 | 15709 | 15881 | 16989 | 12374 | 14027 | 13233 | 15689 | 15946 | 13355 | 13314 | 15355 | 14481.89 | 14956 | 14008 | 0.937 | -0.095 | 0.36942419 | 0.58676318 | 0.0081 | 0.0071 | 0.0079 | 0.0079 | TRUE | 0.1242 |
| Smc5 | 5259 | 5550 | 6065 | 7689 | 5865 | 5919 | 5586 | 5543 | 6090 | 6044 | 6310 | 6479 | 6008.81 | 5740 | 6278 | 1.094 | 0.129 | 0.07903216 | 0.22294674 | 0.0018 | 0.0075 | 0.0037 | 0.0037 | TRUE | 0.0348 |
| Smc6 | 5597 | 6371 | 6264 | 9029 | 6235 | 6340 | 5945 | 6363 | 6290 | 7098 | 6708 | 6940 | 6557.33 | 6199 | 6915 | 1.116 | 0.158 | 0.01589085 | 0.07401963 | 8.00E-04 | 0.0074 | 0.0029 | 0.0029 | TRUE | 0.0157 |
| Smchd1 | 5818 | 6437 | 7190 | 8120 | 5770 | 6116 | 6180 | 6429 | 7220 | 6383 | 6208 | 6695 | 6519.06 | 6610 | 6429 | 0.973 | -0.04 | 0.6420397 | 0.79842557 | 0.0034 | 0.0074 | 0.0052 | 0.0052 | TRUE | 0.0796 |
| Smcp | 0 | 0 | 0 | 0 | 0 | 0 | 0 | 0 | 0 | 0 | 0 | 0 | 0 | 0 | 0 | NA | NA | NA | NA | NA | NA | NA | NA | NA | NA |
| Smcr7 | 786 | 802 | 740 | 1225 | 849 | 759 | 835 | 801 | 743 | 963 | 913 | 831 | 847.7 | 793 | 902 | 1.139 | 0.188 | 0.06780428 | 0.20154715 | 0.0034 | 0.0108 | 0.0065 | 0.0065 | TRUE | 0.0568 |
| Smcr7l | 3410 | 3215 | 3088 | 4502 | 3162 | 2954 | 3622 | 3211 | 3101 | 3539 | 3402 | 3234 | 3351.4 | 3311 | 3392 | 1.025 | 0.035 | 0.69667621 | 0.8338122 | 0.0041 | 0.0079 | 0.0055 | 0.0055 | TRUE | 0.0818 |
| Smcr8 | 3802 | 3075 | 3085 | 4730 | 3167 | 2841 | 4038 | 3071 | 3098 | 3718 | 3407 | 3110 | 3407.14 | 3402 | 3412 | 1.003 | 0.004 | 0.97287169 | 0.98561635 | 0.0161 | 0.0079 | 0.0123 | 0.0123 | TRUE | 0.3248 |
| Smek1 | 5232 | 4019 | 3690 | 6015 | 4308 | 3750 | 5557 | 4014 | 3705 | 4729 | 4635 | 4105 | 4457.48 | 4425 | 4490 | 1.015 | 0.021 | 0.892042 | 0.94755432 | 0.0262 | 0.0076 | 0.017 | 0.017 | TRUE | 0.6094 |
| Smek2 | 6282 | 5633 | 5391 | 8070 | 5474 | 5306 | 6673 | 5626 | 5413 | 6344 | 5890 | 5808 | 5958.9 | 5904 | 6014 | 1.019 | 0.027 | 0.79187396 | 0.89301263 | 0.0073 | 0.0075 | 0.0073 | 0.0073 | TRUE | 0.1581 |
| Smg1 | 12639 | 10889 | 13176 | 14484 | 10373 | 10155 | 13425 | 10875 | 13230 | 11386 | 11161 | 11116 | 11865.51 | 12510 | 11221 | 0.897 | -0.157 | 0.11179014 | 0.28113131 | 0.0064 | 0.0072 | 0.0069 | 0.0069 | TRUE | 0.1599 |
| Smg5 | 10689 | 9302 | 9298 | 13558 | 9288 | 8583 | 11354 | 9290 | 9336 | 10658 | 9993 | 9395 | 10004.46 | 9993 | 10015 | 1.002 | 0.003 | 0.97442778 | 0.98671997 | 0.007 | 0.0072 | 0.008 | 0.008 | TRUE | 0.1732 |
| Smg6 | 6279 | 6237 | 6721 | 8039 | 5978 | 5846 | 6669 | 6229 | 6749 | 6320 | 6432 | 6399 | 6466.32 | 6549 | 6384 | 0.975 | -0.037 | 0.56797162 | 0.74740657 | 7.00E-04 | 0.0074 | 0.0029 | 0.0029 | TRUE | 0.0223 |
| Smg7 | 6770 | 5868 | 6136 | 7833 | 6095 | 5747 | 7191 | 5860 | 6161 | 6158 | 6558 | 6291 | 6369.85 | 6404 | 6336 | 0.989 | -0.016 | 0.87269106 | 0.93739587 | 0.0051 | 0.0074 | 0.0066 | 0.0066 | TRUE | 0.1408 |
| Smg8 | 2641 | 2579 | 2497 | 3801 | 2587 | 2370 | 2805 | 2576 | 2507 | 2988 | 2783 | 2594 | 2708.99 | 2629 | 2788 | 1.061 | 0.086 | 0.34330837 | 0.5633815 | 0.0037 | 0.0081 | 0.0055 | 0.0055 | TRUE | 0.0481 |
| Smg9 | 769 | 823 | 969 | 885 | 699 | 794 | 817 | 822 | 973 | 696 | 752 | 869 | 821.45 | 871 | 772 | 0.886 | -0.175 | 0.17206685 | 0.37089495 | 0.0099 | 0.0109 | 0.0106 | 0.0106 | TRUE | 0.1418 |
| Smgc | 0 | 0 | 0 | 0 | 0 | 0 | 0 | 0 | 0 | 0 | 0 | 0 | 0 | 0 | 0 | NA | NA | NA | NA | NA | NA | NA | NA | NA | NA |
| Smim1 | 124 | 102 | 83 | 121 | 74 | 76 | 132 | 102 | 83 | 95 | 80 | 83 | 95.81 | 106 | 86 | 0.817 | -0.291 | 0.22725271 | 0.44059153 | 0.0216 | 0.0413 | 0.0314 | 0.0314 | TRUE | 0.2986 |
| Smim11 | 374 | 261 | 232 | 413 | 278 | 262 | 397 | 261 | 233 | 325 | 299 | 287 | 300.24 | 297 | 304 | 1.024 | 0.034 | 0.87245562 | 0.93737812 | 0.0393 | 0.0179 | 0.0284 | 0.0284 | TRUE | 0.9788 |
| Smim12 | 1029 | 1123 | 1110 | 1393 | 1003 | 1007 | 1093 | 1122 | 1115 | 1095 | 1079 | 1102 | 1100.94 | 1110 | 1092 | 0.984 | -0.023 | 0.762124 | 0.87457944 | 0 | 0.0099 | 0.0032 | 0.0032 | TRUE | 0.0021 |
| Smim13 | 801 | 844 | 888 | 1110 | 805 | 777 | 851 | 843 | 892 | 873 | 866 | 851 | 862.44 | 862 | 863 | 1.002 | 0.003 | 0.97608954 | 0.98709087 | 0 | 0.0107 | 0.0039 | 0.0039 | TRUE | 0.011 |
| Smim14 | 1311 | 1819 | 1692 | 1894 | 1363 | 1358 | 1393 | 1817 | 1699 | 1489 | 1467 | 1487 | 1558.35 | 1636 | 1481 | 0.905 | -0.144 | 0.20893165 | 0.41871284 | 0.008 | 0.009 | 0.0088 | 0.0088 | TRUE | 0.2038 |
| Smim15 | 1784 | 2019 | 2005 | 2665 | 1863 | 1875 | 1895 | 2016 | 2013 | 2095 | 2004 | 2052 | 2012.75 | 1975 | 2050 | 1.039 | 0.055 | 0.44596844 | 0.65236459 | 4.00E-04 | 0.0085 | 0.0032 | 0.0032 | TRUE | 0.0151 |
| Smim18 | 0 | 0 | 0 | 0 | 0 | 0 | 0 | 0 | 0 | 0 | 0 | 0 | 0 | 0 | 0 | NA | NA | NA | NA | NA | NA | NA | NA | NA | NA |
| Smim3 | 166 | 210 | 226 | 367 | 282 | 244 | 176 | 210 | 227 | 289 | 303 | 267 | 245.33 | 204 | 286 | 1.401 | 0.486 | 0.0014237 | 0.01176249 | 0.0046 | 0.0203 | 0.0125 | 0.0125 | TRUE | 0.1538 |
| Smim4 | 99 | 126 | 127 | 136 | 93 | 102 | 105 | 126 | 128 | 107 | 100 | 112 | 112.86 | 120 | 106 | 0.888 | -0.171 | 0.37126475 | 0.58797215 | 0 | 0.0361 | 0.0175 | 0.0175 | TRUE | 0.109 |
| Smim5 | 0 | 0 | 0 | 0 | 0 | 0 | 0 | 0 | 0 | 0 | 0 | 0 | 0 | 0 | 0 | NA | NA | NA | NA | NA | NA | NA | NA | NA | NA |
| Smim6 | 0 | 0 | 0 | 0 | 0 | 0 | 0 | 0 | 0 | 0 | 0 | 0 | 0 | 0 | 0 | NA | NA | NA | NA | NA | NA | NA | NA | NA | NA |
| Smim7 | 2489 | 2836 | 2882 | 3388 | 2343 | 2516 | 2644 | 2832 | 2894 | 2663 | 2521 | 2754 | 2718.07 | 2790 | 2646 | 0.948 | -0.076 | 0.32582411 | 0.54704044 | 0.0015 | 0.0081 | 0.004 | 0.004 | TRUE | 0.0254 |
| Smim8 | 320 | 436 | 403 | 686 | 437 | 380 | 340 | 435 | 405 | 539 | 470 | 416 | 434.24 | 393 | 475 | 1.21 | 0.275 | 0.07042634 | 0.20661537 | 0.0141 | 0.0145 | 0.0143 | 0.0143 | TRUE | 0.1676 |
| Smim9 | 1 | 0 | 0 | 0 | 1 | 1 | 1 | 0 | 0 | 0 | 1 | 1 | 0.54 | 0 | 1 | 1.923 | 0.944 | 0.73523723 | NA | 0 | 6.1214 | 3.9144 | 3.9144 | TRUE | 0.4614 |
| Smlr1 | 0 | 0 | 0 | 0 | 0 | 0 | 0 | 0 | 0 | 0 | 0 | 0 | 0 | 0 | 0 | NA | NA | NA | NA | NA | NA | NA | NA | NA | NA |
| Smn1 | 2847 | 3033 | 3019 | 4082 | 2941 | 2695 | 3024 | 3029 | 3031 | 3209 | 3164 | 2950 | 3067.98 | 3028 | 3108 | 1.027 | 0.038 | 0.58235629 | 0.75852039 | 7.00E-04 | 0.008 | 0.0031 | 0.0031 | TRUE | 0.0237 |
| Smndc1 | 4099 | 3990 | 3968 | 5869 | 4261 | 3842 | 4354 | 3985 | 3984 | 4614 | 4585 | 4206 | 4287.83 | 4108 | 4468 | 1.088 | 0.122 | 0.124201 | 0.30072124 | 0.0023 | 0.0077 | 0.0043 | 0.0043 | TRUE | 0.0334 |

| | | | | | | | | | | | | | | | | | | | | | | | | |
|---|---|---|---|---|---|---|---|---|---|---|---|---|---|---|---|---|---|---|---|---|---|---|---|---|
| Smo | 2132 | 2442 | 2506 | 3095 | 2144 | 2135 | 2265 | 2439 | 2516 | 2433 | 2307 | 2337 | 2382.77 | 2407 | 2359 | 0.98 | -0.029 | 0.71149208 | 0.84297274 | 0.0014 | 0.0083 | 0.0039 | 0.0039 | TRUE | 0.0321 |
| Smoc1 | 1148 | 1244 | 1140 | 1958 | 1473 | 1388 | 1219 | 1242 | 1145 | 1539 | 1585 | 1519 | 1374.99 | 1202 | 1548 | 1.288 | 0.365 | 3.35E-06 | 7.85E-05 | 3.00E-04 | 0.0093 | 0.0037 | 0.0037 | TRUE | 0.0211 |
| Smoc2 | 20 | 10 | 12 | 10 | 8 | 9 | 21 | 10 | 12 | 8 | 9 | 10 | 11.6 | 14 | 9 | 0.607 | -0.72 | 0.22234367 | 0.43444058 | 0 | 0.2909 | 0.1595 | 0.1595 | TRUE | 0.7309 |
| Smok2a | 3 | 0 | 0 | 0 | 0 | 0 | 3 | 0 | 0 | 0 | 0 | 0 | 0.53 | 1 | 0 | 0.169 | -2.562 | 0.52101829 | NA | 10 | 6.2101 | 10 | 10 | TRUE | 1.4425 |
| Smok2b | 0 | 1 | 0 | 0 | 0 | 0 | 0 | 1 | 0 | 0 | 0 | 0 | 0.17 | 0 | 0 | 0.49 | -1.029 | 0.80087 | NA | 0 | 19.7991 | 10 | 10 | TRUE | 0.405 |
| Smok3a | 0 | 0 | 0 | 0 | 0 | 0 | 0 | 0 | 0 | 0 | 0 | 0 | 0 | 0 | 0 | NA | NA | NA | NA | NA | NA | NA | NA | NA | NA |
| Smok3b | 0 | 0 | 0 | 0 | 0 | 0 | 0 | 0 | 0 | 0 | 0 | 0 | 0 | 0 | 0 | NA | NA | NA | NA | NA | NA | NA | NA | NA | NA |
| Smok4a | 1 | 1 | 0 | 2 | 0 | 0 | 1 | 1 | 0 | 2 | 0 | 0 | 0.61 | 1 | 1 | 0.818 | -0.291 | 0.91402744 | NA | 0 | 5.4476 | 3.5995 | 3.5995 | TRUE | 0.9722 |
| Smox | 456 | 472 | 442 | 670 | 529 | 502 | 484 | 471 | 444 | 527 | 569 | 550 | 507.49 | 466 | 549 | 1.175 | 0.232 | 0.02472969 | 0.10099345 | 0 | 0.0134 | 0.0057 | 0.0057 | TRUE | 0.0211 |
| Smpd1 | 475 | 730 | 752 | 709 | 483 | 570 | 505 | 729 | 755 | 557 | 520 | 624 | 614.95 | 663 | 567 | 0.855 | -0.226 | 0.18418434 | 0.38701777 | 0.0262 | 0.0123 | 0.0193 | 0.0193 | TRUE | 0.5125 |
| Smpd2 | 603 | 550 | 568 | 798 | 594 | 570 | 640 | 549 | 570 | 627 | 639 | 624 | 608.42 | 586 | 630 | 1.074 | 0.103 | 0.32124032 | 0.54237525 | 0.0015 | 0.0123 | 0.0062 | 0.0062 | TRUE | 0.0748 |
| Smpd3 | 120 | 96 | 87 | 85 | 69 | 67 | 127 | 96 | 87 | 67 | 74 | 73 | 87.52 | 103 | 71 | 0.689 | -0.537 | 0.02332959 | 0.09700879 | 0.013 | 0.0445 | 0.0286 | 0.0286 | TRUE | 0.394 |
| Smpd4 | 5961 | 5557 | 5671 | 8397 | 5964 | 5778 | 6332 | 5550 | 5694 | 6601 | 6417 | 6325 | 6153.11 | 5859 | 6448 | 1.101 | 0.138 | 0.07715241 | 0.21944956 | 0.0025 | 0.0074 | 0.0043 | 0.0043 | TRUE | 0.0608 |
| Smpd5 | 7 | 6 | 2 | 3 | 3 | 2 | 7 | 6 | 2 | 2 | 3 | 2 | 3.87 | 5 | 2 | 0.503 | -0.991 | 0.32283509 | NA | 0 | 0.8585 | 0.4361 | 0.4361 | TRUE | 0.596 |
| Smpdl3a | 33 | 36 | 26 | 77 | 53 | 81 | 35 | 36 | 26 | 61 | 57 | 89 | 50.56 | 32 | 69 | 2.118 | 1.083 | 0.00069614 | 0.00673586 | 0.0284 | 0.0721 | 0.0505 | 0.0505 | TRUE | 0.5501 |
| Smpdl3b | 394 | 275 | 262 | 628 | 370 | 311 | 418 | 275 | 263 | 494 | 398 | 340 | 364.74 | 319 | 411 | 1.291 | 0.369 | 0.090788 | 0.24567338 | 0.0505 | 0.0159 | 0.0315 | 0.0315 | TRUE | 0.7105 |
| Smpx | 0 | 0 | 0 | 0 | 0 | 0 | 0 | 0 | 0 | 0 | 0 | 0 | 0 | 0 | 0 | NA | NA | NA | NA | NA | NA | NA | NA | NA | NA |
| Smr2 | 0 | 0 | 0 | 0 | 0 | 0 | 0 | 0 | 0 | 0 | 0 | 0 | 0 | 0 | 0 | NA | NA | NA | NA | NA | NA | NA | NA | NA | NA |
| Smr3a | 7 | 3 | 2 | 0 | 3 | 0 | 7 | 3 | 2 | 0 | 3 | 0 | 2.61 | 4 | 1 | 0.249 | -2.008 | 0.19583664 | NA | 0.8732 | 1.2685 | 1.1387 | 1.1387 | TRUE | 1.4448 |
| Sms | 1842 | 1738 | 1742 | 2958 | 2172 | 2041 | 1957 | 1736 | 1749 | 2325 | 2337 | 2234 | 2056.32 | 1814 | 2299 | 1.268 | 0.342 | 3.47E-05 | 0.00056583 | 0.0019 | 0.0085 | 0.0044 | 0.0044 | TRUE | 0.057 |
| Smtn | 1188 | 1214 | 1203 | 1375 | 1085 | 1120 | 1262 | 1212 | 1208 | 1081 | 1167 | 1226 | 1192.76 | 1227 | 1158 | 0.943 | -0.085 | 0.32990208 | 0.55089631 | 0.0016 | 0.0097 | 0.0047 | 0.0047 | TRUE | 0.0423 |
| Smtnl1 | 94 | 58 | 47 | 149 | 127 | 95 | 100 | 58 | 47 | 117 | 137 | 104 | 93.79 | 68 | 119 | 1.749 | 0.806 | 0.00775416 | 0.04336828 | 0.0692 | 0.042 | 0.0545 | 0.0545 | TRUE | 1.4296 |
| Smtnl2 | 6438 | 7392 | 7573 | 7817 | 6542 | 6765 | 6838 | 7382 | 7604 | 6145 | 7039 | 7405 | 7069.02 | 7275 | 6863 | 0.943 | -0.084 | 0.37932139 | 0.59440208 | 0.0055 | 0.0074 | 0.0065 | 0.0065 | TRUE | 0.1028 |
| Smu1 | 6426 | 7161 | 7051 | 9279 | 6751 | 6670 | 6826 | 7152 | 7080 | 7294 | 7264 | 7301 | 7152.78 | 7019 | 7286 | 1.038 | 0.054 | 0.34361346 | 0.56358897 | 1.00E-04 | 0.0074 | 0.0022 | 0.0022 | TRUE | 0.0071 |
| Smug1 | 1074 | 1427 | 1423 | 1261 | 1036 | 1186 | 1141 | 1425 | 1429 | 991 | 1115 | 1298 | 1233.17 | 1332 | 1135 | 0.851 | -0.232 | 0.0960532 | 0.25427176 | 0.0157 | 0.0096 | 0.0132 | 0.0132 | TRUE | 0.19 |
| Smurf1 | 2255 | 2506 | 2566 | 3525 | 2520 | 2548 | 2395 | 2503 | 2577 | 2771 | 2711 | 2789 | 2624.36 | 2492 | 2757 | 1.107 | 0.146 | 0.03217329 | 0.12083124 | 3.00E-04 | 0.0082 | 0.003 | 0.003 | TRUE | 0.0138 |
| Smurf2 | 5964 | 6458 | 6697 | 7490 | 5541 | 5697 | 6335 | 6450 | 6724 | 5888 | 5962 | 6236 | 6265.84 | 6503 | 6029 | 0.927 | -0.109 | 0.08907237 | 0.24246624 | 7.00E-04 | 0.0074 | 0.0028 | 0.0028 | TRUE | 0.011 |
| Smyd1 | 69 | 32 | 28 | 174 | 109 | 75 | 73 | 32 | 28 | 137 | 117 | 82 | 78.25 | 44 | 112 | 2.532 | 1.34 | 0.00055608 | 0.00565836 | 0.1482 | 0.049 | 0.0927 | 0.0927 | TRUE | 1.1405 |
| Smyd2 | 1681 | 1730 | 1777 | 2712 | 1865 | 1695 | 1786 | 1728 | 1784 | 2132 | 2007 | 1855 | 1881.94 | 1766 | 1998 | 1.132 | 0.179 | 0.03317404 | 0.12331393 | 0.0022 | 0.0087 | 0.0046 | 0.0046 | TRUE | 0.0469 |
| Smyd3 | 998 | 1366 | 1394 | 1195 | 875 | 1052 | 1060 | 1364 | 1400 | 939 | 941 | 1152 | 1142.74 | 1275 | 1011 | 0.792 | -0.336 | 0.01874741 | 0.083034 | 0.0167 | 0.0098 | 0.0138 | 0.0138 | TRUE | 0.2597 |
| Smyd4 | 1762 | 1567 | 1586 | 2276 | 1597 | 1571 | 1872 | 1565 | 1593 | 1789 | 1718 | 1720 | 1709.37 | 1677 | 1742 | 1.04 | 0.056 | 0.56440888 | 0.74511232 | 0.0046 | 0.0088 | 0.0063 | 0.0063 | TRUE | 0.1248 |
| Smyd5 | 5381 | 4806 | 4799 | 6932 | 5067 | 4677 | 5716 | 4800 | 4819 | 5449 | 5452 | 5120 | 5225.83 | 5112 | 5340 | 1.045 | 0.063 | 0.50416994 | 0.69924096 | 0.0049 | 0.0075 | 0.0063 | 0.0063 | TRUE | 0.1302 |
| Snai1 | 99 | 79 | 68 | 116 | 75 | 45 | 105 | 79 | 68 | 91 | 81 | 49 | 78.91 | 84 | 74 | 0.882 | -0.182 | 0.54769519 | 0.73261374 | 0.0589 | 0.0487 | 0.053 | 0.053 | TRUE | 0.7352 |
| Snai2 | 33 | 35 | 32 | 75 | 49 | 44 | 35 | 35 | 32 | 59 | 53 | 48 | 43.66 | 34 | 53 | 1.571 | 0.652 | 0.02518049 | 0.10230464 | 0 | 0.0824 | 0.037 | 0.037 | TRUE | 0.0825 |
| Snai3 | 213 | 284 | 323 | 300 | 266 | 263 | 226 | 284 | 324 | 236 | 286 | 288 | 274.02 | 278 | 270 | 0.968 | -0.046 | 0.79256037 | 0.89313105 | 0.0188 | 0.0189 | 0.0189 | 0.0189 | TRUE | 0.2862 |
| Snap23 | 2109 | 2140 | 2165 | 2599 | 1743 | 1875 | 2240 | 2137 | 2174 | 2043 | 1875 | 2052 | 2087.04 | 2184 | 1990 | 0.912 | -0.133 | 0.08017446 | 0.22480268 | 0.001 | 0.0085 | 0.0037 | 0.0037 | TRUE | 0.0305 |
| Snap25 | 9 | 13 | 12 | 15 | 14 | 16 | 10 | 13 | 12 | 12 | 15 | 18 | 13.16 | 12 | 15 | 1.271 | 0.346 | 0.50760253 | 0.70184159 | 0 | 0.2572 | 0.1185 | 0.1185 | TRUE | 0.1797 |
| Snap29 | 1192 | 1207 | 1294 | 1800 | 1208 | 1163 | 1266 | 1205 | 1299 | 1415 | 1300 | 1273 | 1293.12 | 1257 | 1329 | 1.059 | 0.082 | 0.34325437 | 0.5633815 | 0.0017 | 0.0095 | 0.0046 | 0.0046 | TRUE | 0.0394 |
| Snap47 | 4670 | 4929 | 4892 | 6506 | 4607 | 4545 | 4960 | 4923 | 4912 | 5115 | 4957 | 4975 | 4973.61 | 4932 | 5016 | 1.017 | 0.025 | 0.66697008 | 0.81379599 | 0 | 0.0076 | 0.0021 | 0.0021 | TRUE | 0.0037 |
| Snap91 | 172 | 288 | 271 | 372 | 311 | 323 | 183 | 288 | 272 | 292 | 335 | 354 | 287.18 | 248 | 327 | 1.318 | 0.399 | 0.03954127 | 0.13942429 | 0.0287 | 0.0184 | 0.0234 | 0.0234 | TRUE | 0.5759 |
| Snapc1 | 1187 | 1227 | 1201 | 1630 | 1203 | 1257 | 1261 | 1225 | 1206 | 1281 | 1294 | 1376 | 1273.98 | 1231 | 1317 | 1.07 | 0.097 | 0.21623884 | 0.42827661 | 0 | 0.0095 | 0.0037 | 0.0037 | TRUE | 0.018 |
| Snapc2 | 1392 | 1392 | 1290 | 2060 | 1501 | 1409 | 1479 | 1390 | 1295 | 1619 | 1615 | 1542 | 1490.14 | 1388 | 1592 | 1.148 | 0.199 | 0.02049528 | 0.08835387 | 0.0018 | 0.0091 | 0.0046 | 0.0046 | TRUE | 0.0411 |
| Snapc3 | 1213 | 1143 | 1198 | 1418 | 1054 | 1052 | 1288 | 1142 | 1203 | 1115 | 1134 | 1152 | 1172.2 | 1211 | 1134 | 0.936 | -0.096 | 0.26592612 | 0.48555067 | 0.0011 | 0.0097 | 0.0045 | 0.0045 | TRUE | 0.0373 |
| Snapc4 | 2172 | 1918 | 1790 | 2305 | 1992 | 1590 | 2307 | 1916 | 1797 | 1812 | 2143 | 1741 | 1952.62 | 2007 | 1899 | 0.946 | -0.08 | 0.53766622 | 0.72419528 | 0.0127 | 0.0086 | 0.0116 | 0.0116 | TRUE | 0.2071 |
| Snapc5 | 720 | 597 | 548 | 925 | 608 | 536 | 765 | 596 | 550 | 727 | 654 | 587 | 646.55 | 637 | 656 | 1.031 | 0.044 | 0.77519389 | 0.88305321 | 0.0192 | 0.012 | 0.0157 | 0.0157 | TRUE | 0.3604 |
| Snapin | 1959 | 1913 | 1841 | 2366 | 1636 | 1743 | 2081 | 1911 | 1849 | 1860 | 1760 | 1908 | 1894.68 | 1947 | 1843 | 0.947 | -0.079 | 0.35038887 | 0.5692556 | 0.002 | 0.0086 | 0.0046 | 0.0046 | TRUE | 0.0438 |
| Snca | 38 | 57 | 58 | 99 | 54 | 46 | 40 | 57 | 58 | 78 | 58 | 50 | 56.97 | 52 | 62 | 1.204 | 0.268 | 0.37123237 | 0.58797215 | 0.0288 | 0.0647 | 0.0471 | 0.0471 | TRUE | 0.475 |
| Sncaip | 25 | 22 | 19 | 47 | 25 | 19 | 27 | 22 | 19 | 37 | 27 | 21 | 25.37 | 23 | 28 | 1.266 | 0.34 | 0.40649232 | 0.6191402 | 0 | 0.1367 | 0.0812 | 0.0812 | TRUE | 0.3524 |
| Sncb | 76 | 132 | 132 | 120 | 100 | 135 | 81 | 132 | 133 | 94 | 108 | 148 | 115.8 | 115 | 117 | 1.009 | 0.013 | 0.96197283 | 0.98087895 | 0.0552 | 0.0354 | 0.0449 | 0.0449 | TRUE | 0.6719 |
| Sncg | 167 | 160 | 132 | 268 | 174 | 190 | 177 | 160 | 133 | 211 | 187 | 208 | 179.27 | 157 | 202 | 1.292 | 0.37 | 0.03175615 | 0.11986925 | 0.0057 | 0.0253 | 0.0157 | 0.0157 | TRUE | 0.1907 |
| Snd1 | 16721 | 17734 | 18082 | 19188 | 14090 | 14373 | 17761 | 17711 | 18156 | 15084 | 15160 | 15734 | 16600.93 | 17876 | 15326 | 0.857 | -0.222 | 5.61E-05 | 0.00085456 | 3.00E-04 | 0.0071 | 0.0021 | 0.0021 | TRUE | 0.0066 |
| Sned1 | 236 | 150 | 156 | 249 | 206 | 148 | 251 | 150 | 157 | 196 | 222 | 162 | 189.42 | 186 | 193 | 1.041 | 0.058 | 0.80780507 | 0.90112858 | 0.0487 | 0.0243 | 0.0361 | 0.0361 | TRUE | 1.0003 |
| Snf8 | 2623 | 2645 | 2554 | 4152 | 2817 | 2513 | 2786 | 2642 | 2564 | 3264 | 3031 | 2751 | 2839.66 | 2664 | 3015 | 1.132 | 0.179 | 0.05065279 | 0.16557427 | 0.004 | 0.0081 | 0.0057 | 0.0057 | TRUE | 0.0713 |
| Snhg1 | 2459 | 3251 | 3751 | 3202 | 2344 | 3045 | 2612 | 3247 | 3766 | 2517 | 2522 | 3333 | 2999.58 | 3208 | 2791 | 0.87 | -0.202 | 0.21435894 | 0.42607047 | 0.0299 | 0.008 | 0.0187 | 0.0187 | TRUE | 0.2308 |
| Snhg10 | 37 | 43 | 43 | 48 | 37 | 52 | 39 | 43 | 43 | 38 | 40 | 57 | 43.31 | 42 | 45 | 1.066 | 0.092 | 0.76750836 | 0.87860653 | 0 | 0.083 | 0.0459 | 0.0459 | TRUE | 0.4147 |
| Snhg11 | 1695 | 1966 | 1688 | 2042 | 2028 | 1905 | 1800 | 1963 | 1695 | 1605 | 2182 | 2085 | 1888.56 | 1819 | 1957 | 1.075 | 0.104 | 0.43517699 | 0.6435038 | 0.0156 | 0.0086 | 0.0124 | 0.0124 | TRUE | 0.3034 |
| Snhg12 | 906 | 1023 | 1188 | 950 | 700 | 875 | 962 | 1022 | 1193 | 747 | 753 | 958 | 939.11 | 1059 | 819 | 0.773 | -0.372 | 0.0077959 | 0.04354001 | 0.0155 | 0.0104 | 0.013 | 0.013 | TRUE | 0.2576 |
| Snhg3 | 803 | 916 | 869 | 1095 | 768 | 907 | 853 | 915 | 873 | 861 | 826 | 993 | 886.72 | 880 | 893 | 1.014 | 0.021 | 0.84574257 | 0.92273463 | 0.0041 | 0.0106 | 0.0069 | 0.0069 | TRUE | 0.1108 |
| Snhg4 | 1106 | 1191 | 1143 | 1079 | 947 | 1032 | 1175 | 1189 | 1148 | 848 | 1019 | 1130 | 1084.79 | 1171 | 999 | 0.852 | -0.231 | 0.05793985 | 0.18189239 | 0.0088 | 0.0099 | 0.0098 | 0.0098 | TRUE | 0.2137 |
| Snhg5 | 1314 | 1472 | 1457 | 1957 | 1574 | 1230 | 1396 | 1470 | 1463 | 1538 | 1694 | 1346 | 1484.53 | 1443 | 1526 | 1.058 | 0.081 | 0.44571683 | 0.65230081 | 0.0059 | 0.0091 | 0.0074 | 0.0074 | TRUE | 0.1257 |
| Snhg6 | 915 | 1526 | 1585 | 1111 | 914 | 1219 | 972 | 1524 | 1592 | 873 | 983 | 1334 | 1213.1 | 1363 | 1063 | 0.78 | -0.358 | 0.09167454 | 0.24710705 | 0.0555 | 0.0096 | 0.0316 | 0.0316 | TRUE | 0.7548 |
| Snhg7 | 620 | 812 | 839 | 520 | 453 | 660 | 659 | 811 | 842 | 409 | 487 | 722 | 655.1 | 771 | 539 | 0.699 | -0.517 | 0.01314443 | 0.06424219 | 0.0515 | 0.0119 | 0.0297 | 0.0297 | TRUE | 1.0244 |
| Snhg8 | 1347 | 1494 | 1441 | 1347 | 1047 | 1097 | 1431 | 1492 | 1447 | 1059 | 1127 | 1201 | 1292.67 | 1457 | 1129 | 0.774 | -0.369 | 1.63E-05 | 0.00030078 | 0.0014 | 0.0095 | 0.0045 | 0.0045 | TRUE | 0.0373 |
| Snhg9 | 10 | 8 | 5 | 12 | 7 | 4 | 11 | 8 | 5 | 9 | 8 | 4 | 7.5 | 8 | 7 | 0.917 | -0.125 | 0.86128529 | 0.93144485 | 0 | 0.4464 | 0.235 | 0.235 | TRUE | 0.2014 |
| Snip1 | 1610 | 1718 | 1588 | 2226 | 1673 | 1472 | 1710 | 1716 | 1595 | 1750 | 1800 | 1611 | 1696.95 | 1674 | 1720 | 1.028 | 0.04 | 0.63124878 | 0.79265149 | 0.0018 | 0.0088 | 0.0045 | 0.0045 | TRUE | 0.0364 |
| Snn | 1324 | 1437 | 1410 | 1637 | 1159 | 1278 | 1406 | 1435 | 1416 | 1287 | 1247 | 1399 | 1365.02 | 1419 | 1311 | 0.923 | -0.115 | 0.16393297 | 0.35985398 | 9.00E-04 | 0.0093 | 0.0042 | 0.0042 | TRUE | 0.0404 |
| Snora15 | 5 | 6 | 2 | 2 | 3 | 7 | 5 | 6 | 2 | 2 | 3 | 8 | 4.3 | 4 | 4 | 0.916 | -0.127 | 0.89951097 | NA | 0.2044 | 0.7738 | 0.4891 | 0.4891 | TRUE | 0.8053 |
| Snora16a | 54 | 59 | 45 | 54 | 43 | 45 | 57 | 59 | 45 | 42 | 46 | 49 | 49.91 | 54 | 46 | 0.852 | -0.231 | 0.40573276 | 0.61850512 | 0 | 0.0729 | 0.0354 | 0.0354 | TRUE | 0.1659 |
| Snora17 | 28 | 26 | 18 | 36 | 23 | 16 | 30 | 26 | 18 | 28 | 25 | 18 | 24.06 | 25 | 24 | 0.965 | -0.052 | 0.90088386 | 0.95188185 | 0 | 0.1438 | 0.0842 | 0.0842 | TRUE | 0.3272 |

| | | | | | | | | | | | | | | | | | | | | | | | | | |
|---|---|---|---|---|---|---|---|---|---|---|---|---|---|---|---|---|---|---|---|---|---|---|---|---|---|
| Snora19 | 0 | 1 | 0 | 0 | 0 | 0 | 0 | 1 | 0 | 0 | 0 | 0 | 0.17 | 0 | 0 | 0.49 | -1.029 | 0.80087 | NA | 0 | 19.7991 | 10 | 10 | TRUE | 0.405 |
| Snora20 | 0 | 0 | 0 | 0 | 0 | 1 | 0 | 0 | 0 | 0 | 0 | 1 | 0.18 | 0 | 0 | 1.859 | 0.894 | 0.82650734 | NA | 0 | 18.0643 | 10 | 10 | TRUE | 0.5383 |
| Snora21 | 5 | 6 | 6 | 4 | 1 | 4 | 5 | 6 | 6 | 3 | 1 | 4 | 4.32 | 6 | 3 | 0.498 | -1.007 | 0.27727036 | NA | 0 | 0.7693 | 0.3608 | 0.3608 | TRUE | 0.3223 |
| Snora23 | 11 | 8 | 7 | 5 | 9 | 7 | 12 | 8 | 7 | 4 | 10 | 8 | 8 | 9 | 7 | 0.784 | -0.351 | 0.61534599 | 0.78236493 | 0 | 0.4189 | 0.2235 | 0.2235 | TRUE | 0.5475 |
| Snora24 | 0 | 0 | 0 | 0 | 0 | 0 | 0 | 0 | 0 | 0 | 0 | 0 | 0 | 0 | 0 | NA | NA | NA | NA | NA | NA | NA | NA | NA | NA |
| Snora26 | 0 | 0 | 0 | 0 | 0 | 0 | 0 | 0 | 0 | 0 | 0 | 0 | 0 | 0 | 0 | NA | NA | NA | NA | NA | NA | NA | NA | NA | NA |
| Snora28 | 21 | 28 | 18 | 29 | 23 | 22 | 22 | 28 | 18 | 23 | 25 | 24 | 23.33 | 23 | 24 | 1.046 | 0.065 | 0.8703906 | 0.93630208 | 0 | 0.1481 | 0.0706 | 0.0706 | TRUE | 0.2348 |
| Snora2b | 6 | 3 | 3 | 2 | 4 | 0 | 6 | 3 | 3 | 2 | 4 | 0 | 3.04 | 4 | 2 | 0.473 | -1.08 | 0.37881237 | NA | 0.1414 | 1.0896 | 0.7103 | 0.7103 | TRUE | 0.6593 |
| Snora3 | 12 | 19 | 16 | 24 | 12 | 11 | 13 | 19 | 16 | 19 | 13 | 12 | 15.27 | 16 | 15 | 0.926 | -0.111 | 0.82345554 | 0.9101083 | 0 | 0.2227 | 0.1124 | 0.1124 | TRUE | 0.3606 |
| Snora30 | 7 | 8 | 11 | 1 | 4 | 4 | 7 | 8 | 11 | 1 | 4 | 4 | 5.99 | 9 | 3 | 0.343 | -1.543 | 0.07576339 | 0.21702511 | 0 | 0.5569 | 0.3255 | 0.3255 | TRUE | 0.8478 |
| Snora31 | 24 | 13 | 12 | 19 | 19 | 11 | 25 | 13 | 12 | 15 | 20 | 12 | 16.32 | 17 | 16 | 0.94 | -0.089 | 0.86614897 | 0.93370586 | 0.0645 | 0.2087 | 0.1391 | 0.1391 | TRUE | 0.9524 |
| Snora33 | 19 | 21 | 15 | 22 | 15 | 19 | 20 | 21 | 15 | 17 | 16 | 21 | 18.41 | 19 | 18 | 0.963 | -0.054 | 0.90230657 | 0.95205059 | 0 | 0.1859 | 0.0857 | 0.0857 | TRUE | 0.1557 |
| Snora34 | 10 | 7 | 9 | 13 | 5 | 5 | 11 | 7 | 9 | 10 | 5 | 5 | 7.95 | 9 | 7 | 0.807 | -0.309 | 0.6496723 | 0.80359435 | 0 | 0.4211 | 0.2087 | 0.2087 | TRUE | 0.5406 |
| Snora35 | 0 | 0 | 0 | 0 | 0 | 0 | 0 | 0 | 0 | 0 | 0 | 0 | 0 | 0 | 0 | NA | NA | NA | NA | NA | NA | NA | NA | NA | NA |
| Snora36b | 0 | 0 | 0 | 0 | 0 | 0 | 0 | 0 | 0 | 0 | 0 | 0 | 0 | 0 | 0 | NA | NA | NA | NA | NA | NA | NA | NA | NA | NA |
| Snora41 | 14 | 13 | 12 | 8 | 8 | 7 | 15 | 13 | 12 | 6 | 9 | 8 | 10.41 | 13 | 8 | 0.561 | -0.834 | 0.15185257 | 0.34295352 | 0 | 0.3234 | 0.1401 | 0.1401 | TRUE | 0.0777 |
| Snora43 | 16 | 35 | 30 | 29 | 21 | 22 | 17 | 35 | 30 | 23 | 23 | 24 | 25.26 | 27 | 23 | 0.844 | -0.245 | 0.55553884 | 0.73847101 | 0 | 0.1373 | 0.0844 | 0.0844 | TRUE | 0.6722 |
| Snora44 | 14 | 13 | 17 | 25 | 12 | 15 | 15 | 13 | 17 | 20 | 13 | 16 | 15.65 | 15 | 16 | 1.099 | 0.136 | 0.77694591 | 0.88406254 | 0 | 0.2174 | 0.1021 | 0.1021 | TRUE | 0.1885 |
| Snora47 | 0 | 0 | 0 | 1 | 0 | 0 | 0 | 0 | 0 | 1 | 0 | 0 | 0.13 | 0 | 0 | 1.859 | 0.894 | 0.82650734 | NA | 0 | 25.1515 | 10 | 10 | TRUE | 0.2563 |
| Snora52 | 15 | 23 | 16 | 23 | 21 | 14 | 16 | 23 | 16 | 18 | 23 | 15 | 18.49 | 18 | 19 | 1.017 | 0.024 | 0.95732482 | 0.97784646 | 0 | 0.185 | 0.0925 | 0.0925 | TRUE | 0.257 |
| Snora5c | 11 | 9 | 15 | 26 | 15 | 17 | 12 | 9 | 15 | 20 | 16 | 19 | 15.15 | 12 | 18 | 1.55 | 0.632 | 0.20084144 | 0.40850494 | 0 | 0.2243 | 0.1068 | 0.1068 | TRUE | 0.2143 |
| Snora61 | 66 | 51 | 61 | 90 | 72 | 72 | 70 | 51 | 61 | 71 | 77 | 79 | 68.22 | 61 | 76 | 1.244 | 0.315 | 0.1970165 | 0.40371603 | 0 | 0.0552 | 0.0281 | 0.0281 | TRUE | 0.1745 |
| Snora62 | 3 | 3 | 3 | 4 | 3 | 2 | 3 | 3 | 3 | 3 | 3 | 2 | 2.96 | 3 | 3 | 0.936 | -0.095 | 0.92704662 | NA | 0 | 1.1201 | 0.439 | 0.439 | TRUE | 0.0451 |
| Snora64 | 20 | 26 | 40 | 48 | 26 | 27 | 21 | 26 | 40 | 38 | 28 | 30 | 30.44 | 29 | 32 | 1.095 | 0.131 | 0.73732396 | 0.85938343 | 0.0344 | 0.1151 | 0.0764 | 0.0764 | TRUE | 0.7236 |
| Snora65 | 166 | 244 | 255 | 175 | 114 | 177 | 176 | 244 | 256 | 138 | 123 | 194 | 188.34 | 225 | 152 | 0.67 | -0.578 | 0.01224169 | 0.06109999 | 0.0405 | 0.0244 | 0.0328 | 0.0328 | TRUE | 0.611 |
| Snora68 | 35 | 24 | 33 | 52 | 32 | 33 | 37 | 24 | 33 | 41 | 34 | 36 | 34.29 | 31 | 37 | 1.188 | 0.248 | 0.4644572 | 0.6677075 | 0 | 0.103 | 0.0536 | 0.0536 | TRUE | 0.2962 |
| Snora69 | 4 | 4 | 6 | 2 | 3 | 5 | 4 | 4 | 6 | 2 | 3 | 5 | 4.09 | 5 | 3 | 0.701 | -0.512 | 0.58994004 | NA | 0 | 0.8123 | 0.396 | 0.396 | TRUE | 0.3859 |
| Snora70 | 14 | 36 | 13 | 31 | 22 | 15 | 15 | 36 | 13 | 24 | 24 | 16 | 21.39 | 21 | 21 | 1.011 | 0.016 | 0.97589084 | 0.98709087 | 0.1358 | 0.1609 | 0.1487 | 0.1487 | TRUE | 2.0531 |
| Snora74a | 14 | 14 | 10 | 9 | 16 | 5 | 15 | 14 | 10 | 7 | 17 | 5 | 11.44 | 13 | 10 | 0.759 | -0.398 | 0.53239331 | 0.72039355 | 0.099 | 0.2948 | 0.2031 | 0.2031 | TRUE | 1.3267 |
| Snora75 | 8 | 10 | 3 | 7 | 6 | 4 | 8 | 10 | 3 | 6 | 6 | 4 | 6.31 | 7 | 5 | 0.761 | -0.393 | 0.61872106 | 0.78455081 | 0 | 0.5294 | 0.2893 | 0.2893 | TRUE | 0.7059 |
| Snora78 | 12 | 15 | 13 | 11 | 10 | 3 | 13 | 15 | 13 | 9 | 11 | 3 | 10.58 | 14 | 8 | 0.56 | -0.835 | 0.1799372 | 0.38129377 | 0 | 0.3183 | 0.1781 | 0.1781 | TRUE | 0.6003 |
| Snora7a | 47 | 45 | 43 | 43 | 30 | 38 | 50 | 45 | 43 | 34 | 32 | 42 | 40.95 | 46 | 36 | 0.778 | -0.362 | 0.22926547 | 0.44264598 | 0 | 0.0873 | 0.0404 | 0.0404 | TRUE | 0.1273 |
| Snora81 | 51 | 63 | 53 | 44 | 40 | 49 | 54 | 63 | 53 | 35 | 43 | 54 | 50.26 | 57 | 44 | 0.764 | -0.388 | 0.18553105 | 0.3890589 | 0 | 0.0724 | 0.0416 | 0.0416 | TRUE | 0.3083 |
| Snord100 | 14 | 22 | 7 | 33 | 13 | 9 | 15 | 22 | 7 | 26 | 14 | 10 | 15.61 | 15 | 17 | 1.149 | 0.201 | 0.7449006 | 0.86405621 | 0.2015 | 0.218 | 0.2099 | 0.2099 | TRUE | 0.2992 |
| Snord104 | 115 | 108 | 76 | 135 | 94 | 61 | 122 | 108 | 76 | 106 | 101 | 67 | 96.73 | 102 | 91 | 0.898 | -0.155 | 0.57273759 | 0.7512479 | 0.0469 | 0.041 | 0.0438 | 0.0438 | TRUE | 0.5093 |
| Snord11 | 0 | 0 | 0 | 1 | 0 | 0 | 0 | 0 | 0 | 1 | 0 | 0 | 0.13 | 0 | 0 | 1.859 | 0.894 | 0.82650734 | NA | 0 | 25.1515 | 10 | 10 | TRUE | 0.2563 |
| Snord110 | 3 | 0 | 1 | 1 | 0 | 0 | 3 | 0 | 1 | 1 | 0 | 0 | 0.83 | 1 | 0 | 0.251 | -1.993 | 0.42105822 | NA | 0 | 3.9787 | 3.0538 | 3.0538 | TRUE | 0.2422 |
| Snord111 | 3 | 2 | 1 | 1 | 1 | 1 | 3 | 2 | 1 | 1 | 1 | 1 | 1.52 | 2 | 1 | 0.474 | -1.077 | 0.47297327 | NA | 0 | 2.1684 | 0.8746 | 0.8746 | TRUE | 0.1519 |
| Snord116 | 0 | 0 | 0 | 0 | 0 | 0 | 0 | 0 | 0 | 0 | 0 | 0 | 0 | 0 | 0 | NA | NA | NA | NA | NA | NA | NA | NA | NA | NA |
| Snord116l1 | 0 | 0 | 0 | 0 | 0 | 0 | 0 | 0 | 0 | 0 | 0 | 0 | 0 | 0 | 0 | NA | NA | NA | NA | NA | NA | NA | NA | NA | NA |
| Snord116l2 | 0 | 0 | 0 | 0 | 0 | 0 | 0 | 0 | 0 | 0 | 0 | 0 | 0 | 0 | 0 | NA | NA | NA | NA | NA | NA | NA | NA | NA | NA |
| Snord118 | 0 | 0 | 0 | 0 | 0 | 0 | 0 | 0 | 0 | 0 | 0 | 0 | 0 | 0 | 0 | NA | NA | NA | NA | NA | NA | NA | NA | NA | NA |
| Snord12 | 63 | 50 | 42 | 69 | 45 | 32 | 67 | 50 | 42 | 54 | 48 | 35 | 49.45 | 53 | 46 | 0.872 | -0.198 | 0.53573586 | 0.72271961 | 0.0314 | 0.0735 | 0.0532 | 0.0532 | TRUE | 0.4264 |
| Snord123 | 1 | 0 | 0 | 0 | 0 | 0 | 1 | 0 | 0 | 0 | 0 | 0 | 0.18 | 0 | 0 | 0.49 | -1.029 | 0.80087 | NA | 0 | 18.6165 | 10 | 10 | TRUE | 0.4614 |
| Snord14a | 2 | 1 | 4 | 1 | 1 | 0 | 2 | 1 | 4 | 1 | 1 | 0 | 1.5 | 2 | 1 | 0.266 | -1.911 | 0.25814152 | NA | 0 | 2.2028 | 1.0752 | 1.0752 | TRUE | 0.1979 |
| Snord14c | 0 | 0 | 0 | 0 | 0 | 0 | 0 | 0 | 0 | 0 | 0 | 0 | 0 | 0 | 0 | NA | NA | NA | NA | NA | NA | NA | NA | NA | NA |
| Snord14d | 0 | 0 | 0 | 0 | 0 | 0 | 0 | 0 | 0 | 0 | 0 | 0 | 0 | 0 | 0 | NA | NA | NA | NA | NA | NA | NA | NA | NA | NA |
| Snord15a | 1 | 1 | 0 | 4 | 1 | 0 | 1 | 1 | 0 | 3 | 1 | 0 | 1.05 | 1 | 1 | 2.125 | 1.088 | 0.59608725 | NA | 0 | 3.1538 | 1.9521 | 1.9521 | TRUE | 0.3239 |
| Snord15b | 11 | 25 | 19 | 16 | 13 | 9 | 12 | 25 | 19 | 13 | 14 | 10 | 15.36 | 19 | 12 | 0.653 | -0.614 | 0.23785863 | 0.45362833 | 0 | 0.2214 | 0.1277 | 0.1277 | TRUE | 0.174 |
| Snord16a | 8 | 8 | 6 | 9 | 9 | 5 | 8 | 8 | 6 | 7 | 10 | 5 | 7.46 | 7 | 7 | 0.987 | -0.018 | 0.97844233 | 0.98828002 | 0 | 0.4487 | 0.2021 | 0.2021 | TRUE | 0.178 |
| Snord17 | 13 | 5 | 2 | 15 | 14 | 6 | 14 | 5 | 2 | 12 | 15 | 7 | 9.04 | 7 | 11 | 1.618 | 0.694 | 0.38521645 | 0.59920343 | 0.3093 | 0.3714 | 0.3428 | 0.3428 | TRUE | 1.1841 |
| Snord19 | 14 | 7 | 9 | 6 | 11 | 12 | 15 | 7 | 9 | 5 | 12 | 13 | 10.1 | 10 | 10 | 0.947 | -0.079 | 0.90691962 | 0.95353218 | 0.1134 | 0.3332 | 0.2253 | 0.2253 | TRUE | 0.9366 |
| Snord1a | 0 | 0 | 0 | 0 | 0 | 0 | 0 | 0 | 0 | 0 | 0 | 0 | 0 | 0 | 0 | NA | NA | NA | NA | NA | NA | NA | NA | NA | NA |
| Snord1b | 4 | 3 | 9 | 10 | 3 | 6 | 4 | 3 | 9 | 8 | 3 | 7 | 5.66 | 5 | 6 | 1.095 | 0.131 | 0.87713997 | 0.93932608 | 0 | 0.5893 | 0.3411 | 0.3411 | TRUE | 0.7426 |
| Snord1c | 3 | 6 | 1 | 7 | 3 | 4 | 3 | 6 | 1 | 6 | 3 | 4 | 3.88 | 3 | 4 | 1.297 | 0.375 | 0.70843124 | NA | 0 | 0.8555 | 0.4646 | 0.4646 | TRUE | 0.3268 |
| Snord2 | 33 | 27 | 29 | 29 | 24 | 21 | 35 | 27 | 29 | 23 | 26 | 23 | 27.12 | 30 | 24 | 0.785 | -0.349 | 0.33661501 | 0.55668597 | 0 | 0.1284 | 0.0577 | 0.0577 | TRUE | 0.1169 |
| Snord22 | 81 | 114 | 63 | 115 | 105 | 94 | 86 | 114 | 63 | 90 | 113 | 103 | 94.9 | 88 | 102 | 1.161 | 0.215 | 0.41283465 | 0.62415914 | 0.0366 | 0.0416 | 0.0392 | 0.0392 | TRUE | 0.2837 |
| Snord23 | 1 | 1 | 1 | 1 | 0 | 0 | 1 | 1 | 1 | 1 | 0 | 0 | 0.64 | 1 | 0 | 0.342 | -1.549 | 0.50187412 | NA | 0 | 5.1396 | 2.332 | 2.332 | TRUE | 0.2563 |
| Snord32a | 16 | 20 | 17 | 19 | 21 | 14 | 17 | 20 | 17 | 15 | 23 | 15 | 17.82 | 18 | 18 | 0.972 | -0.041 | 0.92832471 | 0.96476961 | 0 | 0.1918 | 0.0917 | 0.0917 | TRUE | 0.2883 |
| Snord33 | 2 | 11 | 1 | 13 | 6 | 1 | 2 | 11 | 1 | 10 | 6 | 1 | 5.31 | 5 | 6 | 1.273 | 0.348 | 0.76022993 | 0.87368315 | 0.8694 | 0.6269 | 0.7479 | 0.7479 | TRUE | 0.6385 |
| Snord34 | 7 | 11 | 5 | 16 | 11 | 7 | 7 | 11 | 5 | 13 | 12 | 8 | 9.25 | 8 | 11 | 1.376 | 0.461 | 0.47524997 | 0.67546567 | 0 | 0.3629 | 0.1892 | 0.1892 | TRUE | 0.3735 |
| Snord35a | 7 | 6 | 10 | 11 | 3 | 15 | 7 | 6 | 10 | 9 | 3 | 16 | 8.63 | 8 | 9 | 1.201 | 0.265 | 0.72456264 | 0.850916 | 0.1806 | 0.3888 | 0.2891 | 0.2891 | TRUE | 0.2445 |
| Snord35b | 9 | 4 | 6 | 9 | 5 | 7 | 10 | 4 | 6 | 7 | 5 | 8 | 6.62 | 7 | 7 | 1.033 | 0.047 | 0.94939736 | 0.9746726 | 0 | 0.5048 | 0.2447 | 0.2447 | TRUE | 0.3936 |
| Snord37 | 6 | 5 | 4 | 11 | 7 | 1 | 6 | 5 | 4 | 9 | 8 | 1 | 5.44 | 5 | 6 | 1.143 | 0.193 | 0.83175153 | 0.91463474 | 9.00E-04 | 0.6122 | 0.4091 | 0.4091 | TRUE | 1.0213 |
| Snord38a | 2 | 1 | 2 | 1 | 1 | 3 | 2 | 1 | 2 | 1 | 1 | 3 | 1.71 | 2 | 2 | 0.98 | -0.029 | 0.9836917 | NA | 0 | 1.9302 | 0.8567 | 0.8567 | TRUE | 0.4562 |
| Snord42a | 5 | 3 | 1 | 3 | 2 | 1 | 5 | 3 | 1 | 2 | 2 | 1 | 2.49 | 3 | 2 | 0.613 | -0.707 | 0.56417711 | NA | 0 | 1.3321 | 0.6607 | 0.6607 | TRUE | 0.1508 |
| Snord42b | 17 | 14 | 34 | 37 | 20 | 34 | 18 | 14 | 34 | 29 | 22 | 37 | 25.67 | 22 | 29 | 1.325 | 0.406 | 0.38531859 | 0.59920343 | 0.0989 | 0.1353 | 0.118 | 0.118 | TRUE | 0.6148 |
| Snord43 | 6 | 5 | 4 | 4 | 3 | 4 | 6 | 5 | 4 | 3 | 3 | 4 | 4.36 | 5 | 3 | 0.696 | -0.524 | 0.55115217 | NA | 0 | 0.7633 | 0.3195 | 0.3195 | TRUE | 0.0883 |
| Snord45b | 1 | 6 | 3 | 13 | 2 | 0 | 1 | 6 | 3 | 10 | 2 | 0 | 3.74 | 3 | 4 | 1.26 | 0.333 | 0.80394272 | NA | 1.2002 | 0.8878 | 1.0299 | 1.0299 | TRUE | 1.3547 |

| | | | | | | | | | | | | | | | | | | | | | | | | | |
|---|---|---|---|---|---|---|---|---|---|---|---|---|---|---|---|---|---|---|---|---|---|---|---|---|---|
| Snord45c | 0 | 0 | 0 | 0 | 0 | 0 | 0 | 0 | 0 | 0 | 0 | 0 | 0 | 0 | 0 | NA | NA | NA | NA | NA | NA | NA | NA | NA | NA |
| Snord47 | 45 | 37 | 24 | 43 | 23 | 17 | 48 | 37 | 24 | 34 | 25 | 19 | 31 | 36 | 26 | 0.717 | -0.48 | 0.24861046 | 0.46552109 | 0.0694 | 0.1132 | 0.092 | 0.092 | TRUE | 0.1729 |
| Snord49a | 23 | 24 | 17 | 28 | 29 | 17 | 24 | 24 | 17 | 22 | 31 | 19 | 22.88 | 22 | 24 | 1.095 | 0.131 | 0.75526209 | 0.87077942 | 0 | 0.1509 | 0.0842 | 0.0842 | TRUE | 0.3882 |
| Snord49b | 24 | 17 | 20 | 28 | 24 | 18 | 25 | 17 | 20 | 22 | 26 | 20 | 21.68 | 21 | 23 | 1.081 | 0.112 | 0.78586171 | 0.89005269 | 0 | 0.1588 | 0.077 | 0.077 | TRUE | 0.2018 |
| Snord4a | 3 | 7 | 7 | 15 | 5 | 5 | 3 | 7 | 7 | 12 | 5 | 5 | 6.64 | 6 | 7 | 1.336 | 0.417 | 0.59217083 | 0.76555069 | 0 | 0.5029 | 0.2852 | 0.2852 | TRUE | 0.8556 |
| Snord52 | 21 | 17 | 14 | 15 | 14 | 13 | 22 | 17 | 14 | 12 | 15 | 14 | 15.74 | 18 | 14 | 0.767 | -0.383 | 0.4282404 | 0.63819387 | 0 | 0.2162 | 0.1036 | 0.1036 | TRUE | 0.2406 |
| Snord53 | 0 | 5 | 5 | 10 | 10 | 2 | 0 | 5 | 5 | 8 | 11 | 2 | 5.14 | 3 | 7 | 2.071 | 1.051 | 0.33363808 | 0.5542839 | 0.5948 | 0.6482 | 0.6285 | 0.6285 | TRUE | 0.4742 |
| Snord55 | 6 | 8 | 4 | 13 | 13 | 10 | 6 | 8 | 4 | 10 | 14 | 11 | 8.92 | 6 | 12 | 1.905 | 0.93 | 0.15183624 | 0.34295352 | 0 | 0.3761 | 0.1781 | 0.1781 | TRUE | 0.2216 |
| Snord57 | 9 | 10 | 7 | 8 | 10 | 11 | 10 | 10 | 7 | 6 | 11 | 12 | 9.28 | 9 | 10 | 1.08 | 0.111 | 0.85992875 | 0.9309173 | 0 | 0.362 | 0.1774 | 0.1774 | TRUE | 0.4193 |
| Snord58b | 1 | 5 | 2 | 5 | 5 | 2 | 1 | 5 | 2 | 4 | 5 | 2 | 3.26 | 3 | 4 | 1.421 | 0.507 | 0.64130125 | NA | 0 | 1.0173 | 0.5362 | 0.5362 | TRUE | 0.5895 |
| Snord61 | 0 | 0 | 0 | 0 | 0 | 0 | 0 | 0 | 0 | 0 | 0 | 0 | 0 | 0 | 0 | NA | NA | NA | NA | NA | NA | NA | NA | NA | NA |
| Snord64 | 3 | 0 | 6 | 2 | 1 | 4 | 3 | 0 | 6 | 2 | 1 | 4 | 2.71 | 3 | 2 | 0.754 | -0.407 | 0.76714027 | NA | 0.6743 | 1.2242 | 0.9803 | 0.9803 | TRUE | 0.1724 |
| Snord65 | 22 | 19 | 13 | 9 | 15 | 8 | 23 | 19 | 13 | 7 | 16 | 9 | 14.56 | 18 | 11 | 0.571 | -0.81 | 0.14722084 | 0.3363323 | 0.0566 | 0.2331 | 0.1507 | 0.1507 | TRUE | 0.4529 |
| Snord66 | 2 | 0 | 0 | 0 | 1 | 0 | 2 | 0 | 0 | 0 | 1 | 0 | 0.53 | 1 | 0 | 0.497 | -1.009 | 0.76131406 | NA | 0 | 6.1834 | 6.228 | 6.228 | TRUE | 0.9742 |
| Snord67 | 1 | 2 | 0 | 1 | 0 | 1 | 1 | 2 | 0 | 1 | 0 | 1 | 0.82 | 1 | 1 | 0.62 | -0.689 | 0.75103894 | NA | 0 | 4.008 | 2.1286 | 2.1286 | TRUE | 0.1579 |
| Snord68 | 7 | 12 | 3 | 11 | 11 | 11 | 7 | 12 | 3 | 9 | 12 | 12 | 9.16 | 7 | 11 | 1.441 | 0.527 | 0.44576669 | 0.65231325 | 0.0542 | 0.3666 | 0.23 | 0.23 | TRUE | 0.2764 |
| Snord69 | 9 | 4 | 4 | 9 | 5 | 7 | 10 | 4 | 4 | 7 | 5 | 8 | 6.28 | 6 | 7 | 1.152 | 0.205 | 0.7914037 | 0.89278443 | 0 | 0.5314 | 0.2714 | 0.2714 | TRUE | 0.6705 |
| Snord7 | 0 | 0 | 0 | 0 | 0 | 0 | 0 | 0 | 0 | 0 | 0 | 0 | 0 | 0 | 0 | NA | NA | NA | NA | NA | NA | NA | NA | NA | NA |
| Snord70 | 1 | 1 | 1 | 2 | 0 | 0 | 1 | 1 | 1 | 2 | 0 | 0 | 0.77 | 1 | 1 | 0.562 | -0.832 | 0.70497892 | NA | 0 | 4.2695 | 2.1321 | 2.1321 | TRUE | 0.9716 |
| Snord71 | 2 | 0 | 3 | 1 | 1 | 2 | 2 | 0 | 3 | 1 | 1 | 2 | 1.53 | 2 | 1 | 0.776 | -0.365 | 0.81938143 | NA | 0 | 2.1583 | 1.1789 | 1.1789 | TRUE | 0.6096 |
| Snord72 | 3 | 15 | 6 | 15 | 5 | 4 | 3 | 15 | 6 | 12 | 5 | 4 | 7.62 | 8 | 7 | 0.904 | -0.146 | 0.86093385 | 0.93125634 | 0.3025 | 0.439 | 0.3698 | 0.3698 | TRUE | 1.0004 |
| Snord73a | 6 | 3 | 1 | 5 | 4 | 3 | 6 | 3 | 1 | 4 | 4 | 3 | 3.65 | 3 | 4 | 1.118 | 0.162 | 0.87519227 | NA | 0 | 0.9098 | 0.4862 | 0.4862 | TRUE | 0.4132 |
| Snord8 | 6 | 0 | 1 | 1 | 2 | 3 | 6 | 0 | 1 | 1 | 2 | 3 | 2.27 | 2 | 2 | 0.834 | -0.261 | 0.86354448 | NA | 0.953 | 1.4604 | 1.2172 | 1.2172 | TRUE | 1.8998 |
| Snord82 | 0 | 0 | 0 | 0 | 0 | 0 | 0 | 0 | 0 | 0 | 0 | 0 | 0 | 0 | 0 | NA | NA | NA | NA | NA | NA | NA | NA | NA | NA |
| Snord83b | 8 | 11 | 7 | 10 | 4 | 9 | 8 | 11 | 7 | 8 | 4 | 10 | 8.09 | 9 | 7 | 0.832 | -0.266 | 0.69491868 | 0.83281701 | 0 | 0.4142 | 0.2064 | 0.2064 | TRUE | 0.3208 |
| Snord85 | 1 | 0 | 2 | 0 | 0 | 0 | 1 | 0 | 2 | 0 | 0 | 0 | 0.51 | 1 | 0 | 0.175 | -2.513 | 0.44147732 | NA | 0 | 6.4448 | 6.181 | 6.181 | TRUE | 0.063 |
| Snord87 | 8 | 6 | 1 | 5 | 1 | 2 | 8 | 6 | 1 | 4 | 1 | 2 | 3.78 | 5 | 2 | 0.478 | -1.065 | 0.34368638 | NA | 0 | 0.8781 | 0.6142 | 0.6142 | TRUE | 0.2967 |
| Snord88a | 6 | 5 | 5 | 5 | 3 | 1 | 6 | 5 | 5 | 4 | 3 | 1 | 4.11 | 5 | 3 | 0.516 | -0.955 | 0.31171171 | NA | 0 | 0.8091 | 0.3755 | 0.3755 | TRUE | 0.2919 |
| Snord88c | 3 | 0 | 0 | 3 | 5 | 7 | 3 | 0 | 0 | 2 | 5 | 8 | 3.1 | 1 | 5 | 4.884 | 2.288 | 0.11207826 | NA | 0.5174 | 1.0704 | 0.9071 | 0.9071 | TRUE | 0.8479 |
| Snord89 | 2 | 3 | 5 | 5 | 3 | 4 | 2 | 3 | 5 | 4 | 3 | 4 | 3.61 | 3 | 4 | 1.135 | 0.183 | 0.85020794 | NA | 0 | 0.9187 | 0.3968 | 0.3968 | TRUE | 0.2618 |
| Snord90 | 0 | 0 | 0 | 0 | 0 | 0 | 0 | 0 | 0 | 0 | 0 | 0 | 0 | 0 | 0 | NA | NA | NA | NA | NA | NA | NA | NA | NA | NA |
| Snord91a | 1 | 1 | 1 | 0 | 2 | 0 | 1 | 1 | 1 | 0 | 2 | 0 | 0.87 | 1 | 1 | 0.675 | -0.567 | 0.79277114 | NA | 0 | 3.7959 | 2.1461 | 2.1461 | TRUE | 0.975 |
| Snord92 | 1 | 2 | 0 | 0 | 0 | 0 | 1 | 2 | 0 | 0 | 0 | 0 | 0.51 | 1 | 0 | 0.176 | -2.508 | 0.44268408 | NA | 0 | 6.4675 | 6.1932 | 6.1932 | TRUE | 0.064 |
| Snord93 | 0 | 0 | 0 | 0 | 0 | 0 | 0 | 0 | 0 | 0 | 0 | 0 | 0 | 0 | 0 | NA | NA | NA | NA | NA | NA | NA | NA | NA | NA |
| Snord95 | 5 | 11 | 4 | 9 | 11 | 8 | 5 | 11 | 4 | 7 | 12 | 9 | 8 | 7 | 9 | 1.349 | 0.432 | 0.54403441 | 0.72913925 | 0 | 0.4189 | 0.2372 | 0.2372 | TRUE | 0.7815 |
| Snord96a | 94 | 101 | 101 | 148 | 89 | 65 | 100 | 101 | 101 | 116 | 96 | 71 | 97.56 | 101 | 94 | 0.943 | -0.084 | 0.72292872 | 0.85005881 | 0.0194 | 0.0407 | 0.0302 | 0.0302 | TRUE | 0.2424 |
| Snord98 | 0 | 1 | 0 | 0 | 0 | 0 | 0 | 1 | 0 | 0 | 0 | 0 | 0.17 | 0 | 0 | 0.49 | -1.029 | 0.80087 | NA | 0 | 19.7991 | 10 | 10 | TRUE | 0.405 |
| Snord99 | 43 | 28 | 31 | 26 | 27 | 20 | 46 | 28 | 31 | 20 | 29 | 22 | 29.36 | 35 | 24 | 0.679 | -0.558 | 0.14821216 | 0.33790144 | 0.0217 | 0.1191 | 0.0722 | 0.0722 | TRUE | 0.5006 |
| Snph | 40 | 40 | 61 | 40 | 27 | 32 | 42 | 40 | 61 | 31 | 29 | 35 | 39.87 | 48 | 32 | 0.664 | -0.591 | 0.07227089 | 0.21018927 | 0 | 0.0895 | 0.0518 | 0.0518 | TRUE | 0.4808 |
| Snrk | 434 | 491 | 509 | 743 | 530 | 477 | 461 | 490 | 511 | 584 | 570 | 522 | 523.15 | 487 | 559 | 1.147 | 0.198 | 0.06530971 | 0.19695312 | 0.0011 | 0.0132 | 0.0064 | 0.0064 | TRUE | 0.0378 |
| Snrnp200 | 19575 | 19272 | 21179 | 24727 | 17965 | 18427 | 20792 | 19247 | 21266 | 19438 | 19329 | 20171 | 20040.68 | 20435 | 19646 | 0.961 | -0.057 | 0.40619771 | 0.61887077 | 0.0015 | 0.0071 | 0.0033 | 0.0033 | TRUE | 0.0317 |
| Snrnp25 | 2107 | 1990 | 1960 | 2655 | 1942 | 1823 | 2238 | 1987 | 1968 | 2087 | 2089 | 1996 | 2060.95 | 2064 | 2057 | 0.997 | -0.005 | 0.95750215 | 0.97796425 | 0.0024 | 0.0085 | 0.0047 | 0.0047 | TRUE | 0.0652 |
| Snrnp27 | 2071 | 2440 | 2366 | 2742 | 1892 | 1886 | 2200 | 2437 | 2376 | 2156 | 2036 | 2065 | 2211.35 | 2338 | 2086 | 0.892 | -0.164 | 0.03489505 | 0.12790164 | 0.0014 | 0.0084 | 0.0039 | 0.0039 | TRUE | 0.032 |
| Snrnp35 | 174 | 204 | 256 | 443 | 301 | 265 | 185 | 204 | 257 | 348 | 324 | 290 | 267.97 | 215 | 321 | 1.492 | 0.577 | 0.00063187 | 0.00624379 | 0.0135 | 0.0192 | 0.0166 | 0.0166 | TRUE | 0.3177 |
| Snrnp40 | 6205 | 6282 | 5974 | 8731 | 6302 | 5872 | 6591 | 6274 | 5999 | 6864 | 6781 | 6428 | 6489.21 | 6288 | 6691 | 1.064 | 0.09 | 0.20655999 | 0.41554389 | 0.0015 | 0.0074 | 0.0035 | 0.0035 | TRUE | 0.0217 |
| Snrnp48 | 1457 | 1918 | 1905 | 1611 | 1374 | 1472 | 1548 | 1916 | 1913 | 1266 | 1478 | 1611 | 1622.01 | 1792 | 1452 | 0.81 | -0.305 | 0.01978878 | 0.08621787 | 0.0132 | 0.0089 | 0.0117 | 0.0117 | TRUE | 0.1715 |
| Snrnp70 | 18879 | 16859 | 16598 | 23967 | 17210 | 16255 | 20053 | 16837 | 16666 | 18841 | 18517 | 17794 | 18117.96 | 17852 | 18384 | 1.03 | 0.042 | 0.65279026 | 0.80520057 | 0.005 | 0.0071 | 0.0063 | 0.0063 | TRUE | 0.1422 |
| Snrpa | 6496 | 7448 | 7653 | 9361 | 6921 | 7215 | 6900 | 7438 | 7684 | 7359 | 7447 | 7898 | 7454.35 | 7341 | 7568 | 1.031 | 0.044 | 0.55820947 | 0.74059171 | 0.0019 | 0.0073 | 0.0039 | 0.0039 | TRUE | 0.0337 |
| Snrpa1 | 5458 | 5664 | 5554 | 7438 | 5056 | 5099 | 5797 | 5657 | 5577 | 5847 | 5440 | 5582 | 5649.94 | 5677 | 5623 | 0.991 | -0.013 | 0.83452149 | 0.91571799 | 7.00E-04 | 0.0075 | 0.0028 | 0.0028 | TRUE | 0.015 |
| Snrpb | 9274 | 8725 | 8994 | 12894 | 9195 | 8463 | 9851 | 8714 | 9031 | 10136 | 9893 | 9264 | 9481.48 | 9199 | 9764 | 1.062 | 0.086 | 0.2796015 | 0.5007807 | 0.0029 | 0.0073 | 0.0045 | 0.0045 | TRUE | 0.047 |
| Snrpb2 | 3700 | 3534 | 3399 | 5102 | 3506 | 3278 | 3930 | 3529 | 3413 | 4011 | 3772 | 3588 | 3707.29 | 3624 | 3790 | 1.046 | 0.065 | 0.46476709 | 0.66784862 | 0.0038 | 0.0078 | 0.0055 | 0.0055 | TRUE | 0.0662 |
| Snrpc | 4086 | 4330 | 4499 | 5524 | 3711 | 3805 | 4340 | 4324 | 4517 | 4343 | 3993 | 4165 | 4280.41 | 4394 | 4167 | 0.949 | -0.076 | 0.26543 | 0.48505136 | 8.00E-04 | 0.0077 | 0.0031 | 0.0031 | TRUE | 0.0168 |
| Snrpd1 | 4079 | 4633 | 4509 | 6468 | 4391 | 4352 | 4333 | 4627 | 4528 | 5085 | 4724 | 4764 | 4676.7 | 4496 | 4858 | 1.081 | 0.112 | 0.10787222 | 0.27490073 | 0.0011 | 0.0076 | 0.0033 | 0.0033 | TRUE | 0.0206 |
| Snrpd2 | 6972 | 7913 | 7887 | 9268 | 6300 | 6882 | 7405 | 7903 | 7919 | 7286 | 6778 | 7533 | 7470.88 | 7742 | 7199 | 0.93 | -0.105 | 0.15681158 | 0.34972176 | 0.0019 | 0.0073 | 0.0038 | 0.0038 | TRUE | 0.0317 |
| Snrpd3 | 5847 | 5282 | 5050 | 7889 | 5444 | 4995 | 6211 | 5275 | 5071 | 6202 | 5857 | 5468 | 5680.57 | 5519 | 5842 | 1.059 | 0.082 | 0.42686438 | 0.63700492 | 0.007 | 0.0075 | 0.0076 | 0.0076 | TRUE | 0.1465 |
| Snrpe | 3632 | 4435 | 4342 | 5552 | 3913 | 3966 | 3858 | 4429 | 4360 | 4365 | 4210 | 4341 | 4260.51 | 4216 | 4305 | 1.021 | 0.03 | 0.70907671 | 0.84150527 | 0.0027 | 0.0077 | 0.0045 | 0.0045 | TRUE | 0.067 |
| Snrpf | 3598 | 3472 | 3353 | 4324 | 3148 | 3114 | 3822 | 3468 | 3367 | 3399 | 3387 | 3409 | 3475.17 | 3552 | 3398 | 0.957 | -0.064 | 0.41038239 | 0.62180353 | 0.0018 | 0.0079 | 0.004 | 0.004 | TRUE | 0.0535 |
| Snrpg | 2997 | 3090 | 3033 | 3959 | 2733 | 2680 | 3183 | 3086 | 3045 | 3112 | 2941 | 2934 | 3050.21 | 3105 | 2996 | 0.965 | -0.051 | 0.44885391 | 0.65455233 | 5.00E-04 | 0.008 | 0.003 | 0.003 | TRUE | 0.0144 |
| Snrpn | 0 | 0 | 0 | 0 | 0 | 0 | 0 | 0 | 0 | 0 | 0 | 0 | 0 | 0 | 0 | NA | NA | NA | NA | NA | NA | NA | NA | NA | NA |
| Snta1 | 972 | 1012 | 1057 | 1038 | 742 | 727 | 1032 | 1011 | 1061 | 816 | 798 | 796 | 919.1 | 1035 | 803 | 0.777 | -0.365 | 7.92E-06 | 0.00016243 | 0 | 0.0105 | 0.0037 | 0.0037 | TRUE | 0.0061 |
| Sntb1 | 97 | 69 | 62 | 173 | 105 | 64 | 103 | 69 | 62 | 136 | 113 | 70 | 92.2 | 78 | 106 | 1.369 | 0.453 | 0.15025884 | 0.34099028 | 0.0797 | 0.0426 | 0.0604 | 0.0604 | TRUE | 0.3398 |
| Sntb2 | 5134 | 4950 | 5309 | 5774 | 4338 | 4297 | 5453 | 4944 | 5331 | 4539 | 4667 | 4704 | 4939.64 | 5243 | 4637 | 0.884 | -0.177 | 0.01169574 | 0.05908087 | 0.0012 | 0.0076 | 0.0034 | 0.0034 | TRUE | 0.0304 |
| Sntg1 | 1 | 1 | 0 | 0 | 2 | 5 | 1 | 1 | 0 | 0 | 2 | 5 | 1.61 | 1 | 2 | 3.64 | 1.864 | 0.32814916 | NA | 0 | 2.0476 | 1.6752 | 1.6752 | TRUE | 0.1497 |
| Sntg2 | 24 | 31 | 38 | 18 | 23 | 27 | 25 | 31 | 38 | 14 | 25 | 30 | 27.18 | 31 | 23 | 0.713 | -0.487 | 0.24774444 | 0.46439568 | 0.0481 | 0.1281 | 0.0902 | 0.0902 | TRUE | 0.7349 |
| Sntn | 0 | 0 | 0 | 0 | 0 | 0 | 0 | 0 | 0 | 0 | 0 | 0 | 0 | 0 | 0 | NA | NA | NA | NA | NA | NA | NA | NA | NA | NA |
| Snupn | 1203 | 1190 | 1107 | 1410 | 977 | 1018 | 1278 | 1188 | 1112 | 1108 | 1051 | 1114 | 1141.97 | 1193 | 1091 | 0.915 | -0.127 | 0.16289126 | 0.35826537 | 0.0021 | 0.0098 | 0.0051 | 0.0051 | TRUE | 0.0465 |
| Snurf | 0 | 0 | 0 | 0 | 0 | 0 | 0 | 0 | 0 | 0 | 0 | 0 | 0 | 0 | 0 | NA | NA | NA | NA | NA | NA | NA | NA | NA | NA |
| Snw1 | 4697 | 5007 | 4889 | 6833 | 4866 | 4650 | 4989 | 5001 | 4909 | 5372 | 5236 | 5090 | 5099.32 | 4966 | 5233 | 1.054 | 0.076 | 0.20735448 | 0.41661082 | 2.00E-04 | 0.0075 | 0.0024 | 0.0024 | TRUE | 0.0069 |
| Snx1 | 1815 | 1777 | 1823 | 2864 | 1973 | 1855 | 1928 | 1775 | 1830 | 2251 | 2123 | 2031 | 1989.65 | 1844 | 2135 | 1.158 | 0.212 | 0.00907883 | 0.04893757 | 0.0018 | 0.0086 | 0.0043 | 0.0043 | TRUE | 0.0282 |

| | | | | | | | | | | | | | | | | | | | | | | | | | |
|---|---|---|---|---|---|---|---|---|---|---|---|---|---|---|---|---|---|---|---|---|---|---|---|---|---|
| Snx10 | 3020 | 3734 | 3826 | 3035 | 2312 | 2770 | 3208 | 3729 | 3842 | 2386 | 2488 | 3032 | 3114.05 | 3593 | 2635 | 0.733 | -0.448 | 0.00025671 | 0.00299033 | 0.0126 | 0.008 | 0.0105 | 0.0105 | TRUE | 0.2101 |
| Snx11 | 1361 | 1306 | 1155 | 1548 | 1240 | 1027 | 1446 | 1304 | 1160 | 1217 | 1334 | 1124 | 1264.16 | 1303 | 1225 | 0.94 | -0.089 | 0.44860293 | 0.65445985 | 0.0082 | 0.0095 | 0.0091 | 0.0091 | TRUE | 0.1115 |
| Snx12 | 1472 | 1574 | 1636 | 1927 | 1436 | 1250 | 1564 | 1572 | 1643 | 1515 | 1545 | 1368 | 1534.41 | 1593 | 1476 | 0.927 | -0.109 | 0.19774526 | 0.40436123 | 0.0017 | 0.0091 | 0.0045 | 0.0045 | TRUE | 0.0481 |
| Snx13 | 1638 | 2064 | 2287 | 2569 | 1921 | 2014 | 1740 | 2061 | 2296 | 2020 | 2067 | 2205 | 2064.77 | 2032 | 2097 | 1.031 | 0.045 | 0.70362065 | 0.83769895 | 0.0101 | 0.0085 | 0.0094 | 0.0094 | TRUE | 0.1914 |
| Snx14 | 1878 | 2044 | 1986 | 2515 | 1810 | 1776 | 1995 | 2041 | 1994 | 1977 | 1947 | 1944 | 1983.16 | 2010 | 1956 | 0.973 | -0.039 | 0.55355592 | 0.73700313 | 0 | 0.0086 | 0.0026 | 0.0026 | TRUE | 0.0022 |
| Snx15 | 422 | 565 | 607 | 599 | 432 | 438 | 448 | 564 | 609 | 471 | 465 | 479 | 506.19 | 540 | 472 | 0.872 | -0.198 | 0.15451238 | 0.34655422 | 0.0106 | 0.0134 | 0.0119 | 0.0119 | TRUE | 0.2595 |
| Snx16 | 756 | 992 | 901 | 1282 | 890 | 857 | 803 | 991 | 905 | 1008 | 958 | 938 | 933.66 | 900 | 968 | 1.076 | 0.106 | 0.32637708 | 0.54744437 | 0.005 | 0.0104 | 0.0073 | 0.0073 | TRUE | 0.1042 |
| Snx17 | 2276 | 2260 | 2286 | 3026 | 2148 | 1986 | 2418 | 2257 | 2295 | 2379 | 2311 | 2174 | 2305.65 | 2323 | 2288 | 0.985 | -0.021 | 0.77963608 | 0.88603691 | 0.0013 | 0.0083 | 0.0038 | 0.0038 | TRUE | 0.0228 |
| Snx18 | 2848 | 3096 | 3015 | 4417 | 3403 | 3463 | 3025 | 3092 | 3027 | 3472 | 3661 | 3791 | 3344.83 | 3048 | 3641 | 1.194 | 0.256 | 0.00020669 | 0.00250763 | 8.00E-04 | 0.0079 | 0.0031 | 0.0031 | TRUE | 0.0205 |
| Snx19 | 1028 | 911 | 921 | 1756 | 1214 | 1053 | 1092 | 910 | 925 | 1380 | 1306 | 1153 | 1127.64 | 976 | 1280 | 1.313 | 0.393 | 0.00083132 | 0.00772182 | 0.0081 | 0.0098 | 0.0091 | 0.0091 | TRUE | 0.1295 |
| Snx2 | 6974 | 7774 | 7739 | 8805 | 6441 | 6709 | 7408 | 7764 | 7771 | 6922 | 6930 | 7344 | 7356.39 | 7648 | 7065 | 0.924 | -0.114 | 0.07388435 | 0.21342564 | 7.00E-04 | 0.0074 | 0.0028 | 0.0028 | TRUE | 0.0145 |
| Snx20 | 48 | 59 | 35 | 93 | 65 | 43 | 51 | 59 | 35 | 73 | 70 | 47 | 55.86 | 48 | 63 | 1.316 | 0.396 | 0.2111967 | 0.42189873 | 0.041 | 0.0659 | 0.054 | 0.054 | TRUE | 0.4644 |
| Snx21 | 362 | 477 | 478 | 288 | 263 | 347 | 385 | 476 | 480 | 226 | 283 | 380 | 371.68 | 447 | 296 | 0.661 | -0.597 | 0.00311563 | 0.021869 | 0.0374 | 0.0158 | 0.0266 | 0.0266 | TRUE | 0.616 |
| Snx22 | 83 | 83 | 72 | 128 | 91 | 92 | 88 | 83 | 72 | 101 | 98 | 101 | 90.43 | 81 | 100 | 1.231 | 0.3 | 0.14639583 | 0.33511683 | 0 | 0.0433 | 0.0195 | 0.0195 | TRUE | 0.0849 |
| Snx24 | 645 | 493 | 447 | 512 | 348 | 346 | 685 | 492 | 449 | 402 | 374 | 379 | 463.66 | 542 | 385 | 0.711 | -0.491 | 0.00456567 | 0.02922472 | 0.0243 | 0.014 | 0.0194 | 0.0194 | TRUE | 0.6203 |
| Snx25 | 1361 | 1364 | 1263 | 2362 | 1695 | 1477 | 1446 | 1362 | 1268 | 1857 | 1824 | 1617 | 1562.23 | 1359 | 1766 | 1.3 | 0.379 | 0.00010067 | 0.00138664 | 0.0039 | 0.009 | 0.0062 | 0.0062 | TRUE | 0.0652 |
| Snx27 | 6914 | 8146 | 8552 | 8999 | 6559 | 6939 | 7344 | 8135 | 8587 | 7074 | 7057 | 7596 | 7632.28 | 8022 | 7242 | 0.903 | -0.148 | 0.08483119 | 0.23394955 | 0.0038 | 0.0073 | 0.0052 | 0.0052 | TRUE | 0.0667 |
| Snx29 | 638 | 757 | 830 | 697 | 522 | 537 | 678 | 756 | 833 | 548 | 562 | 588 | 660.75 | 756 | 566 | 0.748 | -0.419 | 0.0002327 | 0.00275127 | 0.0047 | 0.0119 | 0.0078 | 0.0078 | TRUE | 0.0959 |
| Snx3 | 5686 | 6455 | 6525 | 9136 | 6573 | 6894 | 6040 | 6447 | 6552 | 7182 | 7072 | 7547 | 6806.45 | 6346 | 7267 | 1.145 | 0.195 | 0.00476081 | 0.03017986 | 0.0013 | 0.0074 | 0.0033 | 0.0033 | TRUE | 0.0217 |
| Snx30 | 2516 | 2602 | 2708 | 2586 | 2016 | 1998 | 2672 | 2599 | 2719 | 2033 | 2169 | 2187 | 2396.55 | 2663 | 2130 | 0.799 | -0.323 | 6.04E-06 | 0.00012999 | 7.00E-04 | 0.0083 | 0.0033 | 0.0033 | TRUE | 0.0197 |
| Snx31 | 157 | 97 | 103 | 140 | 115 | 107 | 167 | 97 | 103 | 110 | 124 | 117 | 119.66 | 122 | 117 | 0.956 | -0.065 | 0.79667875 | 0.89536272 | 0.0393 | 0.0344 | 0.0368 | 0.0368 | TRUE | 1.0099 |
| Snx32 | 72 | 74 | 77 | 99 | 76 | 58 | 76 | 74 | 77 | 78 | 82 | 63 | 75.13 | 76 | 74 | 0.982 | -0.027 | 0.90725577 | 0.95354464 | 0 | 0.0508 | 0.0241 | 0.0241 | TRUE | 0.1383 |
| Snx33 | 42 | 43 | 36 | 94 | 67 | 44 | 45 | 43 | 36 | 74 | 72 | 48 | 52.98 | 41 | 65 | 1.577 | 0.657 | 0.02690343 | 0.10709767 | 0.0176 | 0.0691 | 0.0436 | 0.0436 | TRUE | 0.4167 |
| Snx4 | 5090 | 5176 | 5143 | 6839 | 4823 | 4815 | 5406 | 5169 | 5164 | 5376 | 5189 | 5271 | 5262.7 | 5246 | 5279 | 1.006 | 0.009 | 0.88149256 | 0.94135694 | 3.00E-04 | 0.0075 | 0.0025 | 0.0025 | TRUE | 0.0086 |
| Snx5 | 22189 | 21494 | 21698 | 39149 | 26266 | 21698 | 23569 | 21466 | 21787 | 30776 | 28261 | 23752 | 24935.05 | 22274 | 27596 | 1.239 | 0.309 | 0.00477818 | 0.03027778 | 0.0095 | 0.007 | 0.0086 | 0.0086 | TRUE | 0.1817 |
| Snx6 | 3868 | 4399 | 4272 | 5938 | 4282 | 4332 | 4108 | 4393 | 4290 | 4668 | 4607 | 4742 | 4468.1 | 4264 | 4672 | 1.096 | 0.132 | 0.03758253 | 0.13462451 | 3.00E-04 | 0.0076 | 0.0027 | 0.0027 | TRUE | 0.0123 |
| Snx7 | 294 | 335 | 325 | 734 | 502 | 423 | 312 | 335 | 326 | 577 | 540 | 463 | 425.56 | 324 | 527 | 1.626 | 0.701 | 5.10E-08 | 2.02E-06 | 0.0053 | 0.0146 | 0.0094 | 0.0094 | TRUE | 0.1287 |
| Snx8 | 1912 | 2359 | 2556 | 2060 | 1633 | 1680 | 2031 | 2356 | 2566 | 1619 | 1757 | 1839 | 2028.13 | 2318 | 1738 | 0.75 | -0.416 | 0.00020552 | 0.00250114 | 0.0081 | 0.0085 | 0.0085 | 0.0085 | TRUE | 0.1417 |
| Snx9 | 362 | 548 | 629 | 858 | 584 | 556 | 385 | 547 | 632 | 674 | 628 | 609 | 579.14 | 521 | 637 | 1.223 | 0.29 | 0.10454868 | 0.26925895 | 0.0302 | 0.0126 | 0.0213 | 0.0213 | TRUE | 0.5896 |
| Soat1 | 1886 | 2398 | 2720 | 1964 | 1536 | 1931 | 2003 | 2395 | 2731 | 1544 | 1653 | 2114 | 2073.28 | 2376 | 1770 | 0.744 | -0.426 | 0.00623983 | 0.03710957 | 0.0256 | 0.0085 | 0.017 | 0.017 | TRUE | 0.2623 |
| Soat2 | 129 | 143 | 157 | 117 | 81 | 94 | 137 | 143 | 158 | 92 | 87 | 103 | 119.92 | 146 | 94 | 0.644 | -0.635 | 0.00065499 | 0.00643468 | 0 | 0.0344 | 0.0164 | 0.0164 | TRUE | 0.0608 |
| Sobp | 502 | 409 | 428 | 656 | 535 | 345 | 533 | 408 | 430 | 516 | 576 | 378 | 473.4 | 457 | 490 | 1.072 | 0.1 | 0.58479388 | 0.75998583 | 0.0307 | 0.0139 | 0.0222 | 0.0222 | TRUE | 0.4577 |
| Socs1 | 229 | 262 | 243 | 278 | 241 | 213 | 243 | 262 | 244 | 219 | 259 | 233 | 243.32 | 250 | 237 | 0.947 | -0.078 | 0.57644328 | 0.75418558 | 0 | 0.0204 | 0.0101 | 0.0101 | TRUE | 0.0743 |
| Socs2 | 2054 | 1841 | 1785 | 2763 | 2024 | 1774 | 2182 | 1839 | 1792 | 2172 | 2178 | 1942 | 2017.39 | 1938 | 2097 | 1.083 | 0.115 | 0.28663453 | 0.50796477 | 0.0073 | 0.0085 | 0.0079 | 0.0079 | TRUE | 0.1465 |
| Socs3 | 15927 | 24997 | 26816 | 20841 | 17741 | 22349 | 16917 | 24965 | 26926 | 16384 | 19088 | 24464 | 21457.4 | 22936 | 19979 | 0.871 | -0.199 | 0.29863531 | 0.51910525 | 0.0448 | 0.0071 | 0.0264 | 0.0264 | TRUE | 0.6448 |
| Socs4 | 470 | 535 | 575 | 703 | 533 | 562 | 499 | 534 | 577 | 553 | 573 | 615 | 558.7 | 537 | 580 | 1.08 | 0.11 | 0.31453456 | 0.53514985 | 0.0023 | 0.0128 | 0.0069 | 0.0069 | TRUE | 0.0507 |
| Socs5 | 682 | 763 | 740 | 854 | 588 | 602 | 724 | 762 | 743 | 671 | 633 | 659 | 698.74 | 743 | 654 | 0.881 | -0.183 | 0.04235008 | 0.14676895 | 0 | 0.0116 | 0.0044 | 0.0044 | TRUE | 0.0098 |
| Socs6 | 1529 | 1569 | 1589 | 2036 | 1481 | 1613 | 1624 | 1567 | 1596 | 1601 | 1593 | 1766 | 1624.38 | 1596 | 1653 | 1.036 | 0.051 | 0.53080655 | 0.71904843 | 0.0012 | 0.0089 | 0.0041 | 0.0041 | TRUE | 0.0421 |
| Socs7 | 2365 | 2648 | 2659 | 3087 | 2382 | 2333 | 2512 | 2645 | 2670 | 2427 | 2563 | 2554 | 2561.67 | 2609 | 2515 | 0.963 | -0.054 | 0.44427191 | 0.65087811 | 6.00E-04 | 0.0082 | 0.0032 | 0.0032 | TRUE | 0.0127 |
| Sod1 | 19877 | 22364 | 22685 | 25300 | 17599 | 18192 | 21113 | 22335 | 22778 | 19889 | 18935 | 19914 | 20827.41 | 22075 | 19579 | 0.887 | -0.173 | 0.00710957 | 0.04056038 | 0.0011 | 0.0071 | 0.0029 | 0.0029 | TRUE | 0.0178 |
| Sod2 | 6049 | 6624 | 6511 | 6176 | 4918 | 5059 | 6425 | 6615 | 6538 | 4855 | 5291 | 5538 | 5877.12 | 6526 | 5228 | 0.801 | -0.32 | 2.46E-05 | 0.00042718 | 0.002 | 0.0075 | 0.004 | 0.004 | TRUE | 0.048 |
| Sod3 | 1 | 0 | 0 | 2 | 4 | 0 | 1 | 0 | 0 | 2 | 4 | 0 | 1.16 | 0 | 2 | 5.341 | 2.417 | 0.28284963 | NA | 0 | 2.8559 | 2.3991 | 2.3991 | TRUE | 0.2338 |
| Soga2 | 5503 | 5912 | 5840 | 6199 | 4569 | 4383 | 5845 | 5904 | 5864 | 4873 | 4916 | 4798 | 5366.75 | 5871 | 4862 | 0.828 | -0.272 | 1.02E-06 | 2.81E-05 | 0 | 0.0075 | 0.002 | 0.002 | TRUE | 0.0016 |
| Soga3 | 247 | 286 | 320 | 161 | 151 | 129 | 262 | 286 | 321 | 127 | 162 | 141 | 216.59 | 290 | 143 | 0.493 | -1.021 | 1.01E-09 | 6.14E-08 | 0.0079 | 0.0221 | 0.0149 | 0.0149 | TRUE | 0.1396 |
| Sohlh1 | 4 | 4 | 6 | 14 | 8 | 10 | 4 | 4 | 6 | 11 | 9 | 11 | 7.47 | 5 | 10 | 2.145 | 1.101 | 0.11492076 | 0.28620404 | 0 | 0.4478 | 0.1965 | 0.1965 | TRUE | 0.1073 |
| Sohlh2 | 6159 | 7102 | 6664 | 11088 | 8594 | 8511 | 6542 | 7093 | 6691 | 8717 | 9247 | 9317 | 7934.32 | 6775 | 9094 | 1.342 | 0.424 | 8.65E-10 | 5.42E-08 | 0.0013 | 0.0073 | 0.0033 | 0.0033 | TRUE | 0.0205 |
| Solh | 1011 | 947 | 995 | 1317 | 945 | 937 | 1074 | 946 | 999 | 1035 | 1017 | 1026 | 1016.08 | 1006 | 1026 | 1.02 | 0.029 | 0.74715137 | 0.8655848 | 0.001 | 0.0101 | 0.0047 | 0.0047 | TRUE | 0.041 |
| Son | 14668 | 12807 | 13323 | 19288 | 13575 | 12265 | 15580 | 12790 | 13378 | 15163 | 14606 | 13426 | 14157.12 | 13916 | 14398 | 1.035 | 0.049 | 0.62423807 | 0.78833236 | 0.0062 | 0.0071 | 0.0072 | 0.0072 | TRUE | 0.1337 |
| Sorbs1 | 1945 | 2108 | 2231 | 2513 | 1923 | 2108 | 2066 | 2105 | 2240 | 1976 | 2069 | 2308 | 2127.24 | 2137 | 2118 | 0.99 | -0.014 | 0.87557839 | 0.93865053 | 0.0037 | 0.0085 | 0.0055 | 0.0055 | TRUE | 0.0744 |
| Sorbs2 | 127 | 107 | 91 | 181 | 119 | 108 | 135 | 107 | 91 | 142 | 128 | 118 | 120.28 | 111 | 129 | 1.171 | 0.227 | 0.28794213 | 0.50903254 | 0.0144 | 0.0343 | 0.0246 | 0.0246 | TRUE | 0.3459 |
| Sorbs3 | 395 | 537 | 579 | 594 | 406 | 474 | 420 | 536 | 581 | 467 | 437 | 519 | 493.32 | 512 | 474 | 0.925 | -0.113 | 0.45805492 | 0.66261049 | 0.0155 | 0.0136 | 0.0146 | 0.0146 | TRUE | 0.291 |
| Sorcs1 | 8 | 4 | 8 | 7 | 10 | 7 | 8 | 4 | 8 | 6 | 11 | 8 | 7.41 | 7 | 8 | 1.155 | 0.208 | 0.77174358 | 0.88094525 | 0 | 0.4516 | 0.2347 | 0.2347 | TRUE | 0.3526 |
| Sorcs2 | 193 | 192 | 173 | 278 | 174 | 188 | 205 | 192 | 174 | 219 | 187 | 206 | 197 | 190 | 204 | 1.075 | 0.104 | 0.50285086 | 0.69802532 | 0 | 0.0236 | 0.0123 | 0.0123 | TRUE | 0.062 |
| Sorcs3 | 18 | 32 | 31 | 16 | 14 | 20 | 19 | 32 | 31 | 13 | 15 | 22 | 21.96 | 27 | 17 | 0.596 | -0.748 | 0.09718402 | 0.25620253 | 0.0331 | 0.1569 | 0.0979 | 0.0979 | TRUE | 0.4292 |
| Sord | 2223 | 2490 | 2392 | 3336 | 2387 | 2362 | 2361 | 2487 | 2402 | 2623 | 2568 | 2586 | 2504.36 | 2417 | 2592 | 1.073 | 0.101 | 0.1217031 | 0.29667717 | 0 | 0.0082 | 0.0027 | 0.0027 | TRUE | 0.0078 |
| Sorl1 | 4263 | 5308 | 6162 | 4990 | 3741 | 4457 | 4528 | 5301 | 6187 | 3923 | 4025 | 4879 | 4807.2 | 5339 | 4276 | 0.801 | -0.321 | 0.02056931 | 0.08854233 | 0.0193 | 0.0076 | 0.0136 | 0.0136 | TRUE | 0.1796 |
| Sort1 | 2301 | 2686 | 2846 | 2586 | 1889 | 1755 | 2444 | 2683 | 2858 | 2033 | 2032 | 1921 | 2328.46 | 2662 | 1995 | 0.75 | -0.415 | 2.43E-06 | 5.98E-05 | 0.0029 | 0.0083 | 0.0052 | 0.0052 | TRUE | 0.0617 |
| Sos1 | 3790 | 3494 | 3536 | 4535 | 3312 | 3165 | 4026 | 3489 | 3551 | 3565 | 3564 | 3465 | 3609.8 | 3689 | 3531 | 0.957 | -0.063 | 0.45187604 | 0.65723121 | 0.0029 | 0.0078 | 0.0047 | 0.0047 | TRUE | 0.0776 |
| Sos2 | 1032 | 1286 | 1275 | 1574 | 1158 | 1249 | 1096 | 1284 | 1280 | 1237 | 1246 | 1367 | 1251.88 | 1220 | 1283 | 1.051 | 0.072 | 0.48066334 | 0.6799123 | 0.0043 | 0.0095 | 0.0067 | 0.0067 | TRUE | 0.0944 |
| Sost | 8 | 17 | 11 | 34 | 17 | 9 | 8 | 17 | 11 | 27 | 18 | 10 | 15.23 | 12 | 18 | 1.517 | 0.601 | 0.29719021 | 0.51810368 | 0.1172 | 0.2232 | 0.1712 | 0.1712 | TRUE | 0.1355 |
| Sostdc1 | 5 | 3 | 1 | 1 | 3 | 7 | 5 | 3 | 1 | 1 | 3 | 8 | 3.5 | 3 | 4 | 1.23 | 0.299 | 0.80015174 | NA | 0.4866 | 0.9487 | 0.7119 | 0.7119 | TRUE | 0.3652 |
| Sowaha | 81 | 192 | 235 | 142 | 123 | 164 | 86 | 192 | 236 | 112 | 132 | 180 | 156.21 | 171 | 141 | 0.822 | -0.282 | 0.41569701 | 0.62722408 | 0.1442 | 0.028 | 0.0802 | 0.0802 | TRUE | 0.5665 |
| Sowahb | 334 | 379 | 330 | 391 | 286 | 323 | 355 | 379 | 331 | 307 | 308 | 354 | 338.88 | 355 | 323 | 0.909 | -0.138 | 0.28366902 | 0.50519735 | 0.0025 | 0.0166 | 0.009 | 0.009 | TRUE | 0.076 |
| Sowahc | 342 | 429 | 437 | 555 | 422 | 397 | 363 | 428 | 439 | 436 | 454 | 435 | 425.9 | 410 | 442 | 1.076 | 0.106 | 0.38320862 | 0.59760644 | 0.0027 | 0.0146 | 0.0082 | 0.0082 | TRUE | 0.114 |
| Sowahd | 0 | 0 | 0 | 0 | 0 | 1 | 0 | 0 | 0 | 0 | 0 | 1 | 0.18 | 0 | 0 | 1.859 | 0.894 | 0.82650734 | NA | 0 | 18.0643 | 10 | 10 | TRUE | 0.5383 |
| Sox1 | 127 | 214 | 171 | 233 | 185 | 135 | 135 | 214 | 172 | 183 | 199 | 148 | 175.05 | 174 | 177 | 1.019 | 0.027 | 0.90291975 | 0.95205059 | 0.0312 | 0.0257 | 0.0284 | 0.0284 | TRUE | 0.2241 |
| Sox10 | 61 | 66 | 45 | 42 | 39 | 32 | 65 | 66 | 45 | 33 | 42 | 35 | 47.65 | 59 | 37 | 0.623 | -0.682 | 0.02499496 | 0.10165562 | 0 | 0.076 | 0.0446 | 0.0446 | TRUE | 0.3474 |
| Sox11 | 1184 | 1087 | 1253 | 852 | 671 | 662 | 1258 | 1086 | 1258 | 670 | 722 | 725 | 952.96 | 1201 | 706 | 0.587 | -0.768 | 6.96E-14 | 8.51E-12 | 0.0038 | 0.0104 | 0.0065 | 0.0065 | TRUE | 0.0843 |
| Sox12 | 2004 | 2620 | 2958 | 3486 | 2866 | 2827 | 2129 | 2617 | 2970 | 2740 | 3084 | 3095 | 2772.34 | 2572 | 2973 | 1.156 | 0.209 | 0.11409487 | 0.28491202 | 0.0152 | 0.0081 | 0.0122 | 0.0122 | TRUE | 0.2751 |

| | | | | | | | | | | | | | | | | | | | | | | | | | |
|---|---|---|---|---|---|---|---|---|---|---|---|---|---|---|---|---|---|---|---|---|---|---|---|---|---|
| Sox13 | 1371 | 1416 | 1642 | 1761 | 1363 | 1389 | 1456 | 1414 | 1649 | 1384 | 1467 | 1520 | 1481.75 | 1506 | 1457 | 0.967 | -0.049 | 0.61076593 | 0.77955817 | 0.0038 | 0.0091 | 0.006 | 0.006 | TRUE | 0.0825 |
| Sox14 | 1 | 2 | 0 | 0 | 0 | 4 | 1 | 2 | 0 | 0 | 0 | 4 | 1.24 | 1 | 1 | 1.4 | 0.486 | 0.83205655 | NA | 3.4897 | 2.6644 | 2.9352 | 2.9352 | TRUE | 1.3676 |
| Sox15 | 897 | 1425 | 1466 | 1456 | 1156 | 1430 | 953 | 1423 | 1472 | 1145 | 1244 | 1565 | 1300.28 | 1283 | 1318 | 1.027 | 0.038 | 0.8365035 | 0.91703157 | 0.04 | 0.0094 | 0.024 | 0.024 | TRUE | 0.6065 |
| Sox17 | 24 | 41 | 39 | 88 | 55 | 39 | 25 | 41 | 39 | 69 | 59 | 43 | 46.11 | 35 | 57 | 1.625 | 0.701 | 0.03728386 | 0.13387059 | 0.0366 | 0.0784 | 0.0585 | 0.0585 | TRUE | 0.2686 |
| Sox18 | 39 | 73 | 80 | 68 | 51 | 39 | 41 | 73 | 80 | 53 | 55 | 43 | 57.61 | 65 | 50 | 0.776 | -0.366 | 0.25581715 | 0.47448015 | 0.0496 | 0.0641 | 0.0571 | 0.0571 | TRUE | 0.86 |
| Sox2 | 35158 | 32750 | 32196 | 38188 | 29022 | 29122 | 37344 | 32708 | 32328 | 30020 | 31226 | 31879 | 32584.14 | 34127 | 31042 | 0.91 | -0.137 | 0.0979977 | 0.25738881 | 0.0038 | 0.007 | 0.0049 | 0.0049 | TRUE | 0.0832 |
| Sox21 | 87 | 89 | 78 | 299 | 219 | 198 | 92 | 89 | 78 | 235 | 236 | 217 | 157.84 | 86 | 229 | 2.651 | 1.407 | 1.72E-16 | 2.83E-14 | 0 | 0.0278 | 0.013 | 0.013 | TRUE | 0.0659 |
| Sox2ot | 41 | 29 | 28 | 58 | 42 | 38 | 44 | 29 | 28 | 46 | 45 | 42 | 38.83 | 34 | 44 | 1.32 | 0.401 | 0.21876466 | 0.43086366 | 0 | 0.0917 | 0.0501 | 0.0501 | TRUE | 0.4604 |
| Sox3 | 334 | 375 | 332 | 459 | 362 | 254 | 355 | 375 | 333 | 361 | 389 | 278 | 348.5 | 354 | 343 | 0.969 | -0.046 | 0.76967658 | 0.87966225 | 0.0133 | 0.0164 | 0.0148 | 0.0148 | TRUE | 0.3013 |
| Sox30 | 13 | 16 | 14 | 30 | 24 | 23 | 14 | 16 | 14 | 24 | 26 | 25 | 19.74 | 15 | 25 | 1.697 | 0.763 | 0.07074133 | 0.20719261 | 0 | 0.1738 | 0.0738 | 0.0738 | TRUE | 0.0304 |
| Sox4 | 1973 | 1781 | 1860 | 1610 | 1239 | 1159 | 2096 | 1779 | 1868 | 1266 | 1333 | 1269 | 1601.58 | 1914 | 1289 | 0.674 | -0.57 | 1.07E-09 | 6.46E-08 | 0.0035 | 0.009 | 0.0057 | 0.0057 | TRUE | 0.0831 |
| Sox5 | 7 | 18 | 9 | 25 | 10 | 9 | 7 | 18 | 9 | 20 | 11 | 10 | 12.45 | 11 | 14 | 1.181 | 0.241 | 0.69642072 | 0.83375949 | 0.1141 | 0.2715 | 0.1935 | 0.1935 | TRUE | 0.9468 |
| Sox6 | 19 | 22 | 27 | 43 | 28 | 15 | 20 | 22 | 27 | 34 | 30 | 16 | 24.94 | 23 | 27 | 1.169 | 0.225 | 0.59817975 | 0.77063216 | 0.0379 | 0.139 | 0.0911 | 0.0911 | TRUE | 0.6703 |
| Sox7 | 26 | 25 | 21 | 69 | 56 | 25 | 28 | 25 | 21 | 54 | 60 | 27 | 35.92 | 25 | 47 | 1.934 | 0.951 | 0.01728593 | 0.07818575 | 0.0686 | 0.0986 | 0.0839 | 0.0839 | TRUE | 1.0186 |
| Sox8 | 76 | 79 | 63 | 114 | 89 | 62 | 81 | 79 | 63 | 90 | 96 | 68 | 79.35 | 74 | 85 | 1.139 | 0.188 | 0.44217268 | 0.64905093 | 0.0107 | 0.0484 | 0.0304 | 0.0304 | TRUE | 0.2599 |
| Sox9 | 6 | 7 | 8 | 16 | 10 | 12 | 6 | 7 | 8 | 13 | 11 | 13 | 9.65 | 7 | 12 | 1.705 | 0.77 | 0.1997529 | 0.40732488 | 0 | 0.3485 | 0.1479 | 0.1479 | TRUE | 0.0367 |
| Sp1 | 12406 | 13584 | 15355 | 15732 | 11580 | 12556 | 13177 | 13566 | 15418 | 12367 | 12459 | 13745 | 13455.5 | 14054 | 12857 | 0.915 | -0.129 | 0.16020916 | 0.35474763 | 0.0053 | 0.0071 | 0.006 | 0.006 | TRUE | 0.0882 |
| Sp100 | 1 | 0 | 1 | 5 | 6 | 0 | 1 | 0 | 1 | 4 | 6 | 0 | 2.08 | 1 | 3 | 5.076 | 2.344 | 0.18308624 | NA | 0 | 1.5943 | 1.3459 | 1.3459 | TRUE | 0.1264 |
| Sp110 | 46 | 37 | 27 | 153 | 139 | 64 | 49 | 37 | 27 | 120 | 150 | 70 | 75.47 | 38 | 113 | 3.017 | 1.593 | 6.93E-06 | 0.00014559 | 0.0976 | 0.0506 | 0.0726 | 0.0726 | TRUE | 0.1763 |
| Sp140 | 33 | 37 | 28 | 102 | 55 | 43 | 35 | 37 | 28 | 80 | 59 | 47 | 47.76 | 33 | 62 | 1.875 | 0.907 | 0.00558184 | 0.03393859 | 0.033 | 0.0759 | 0.0541 | 0.0541 | TRUE | 0.2656 |
| Sp2 | 3010 | 2996 | 3015 | 3516 | 2487 | 2603 | 3197 | 2992 | 3027 | 2764 | 2676 | 2849 | 2917.65 | 3072 | 2763 | 0.899 | -0.153 | 0.02934957 | 0.11326193 | 6.00E-04 | 0.008 | 0.0032 | 0.0032 | TRUE | 0.0153 |
| Sp3 | 5305 | 5843 | 5731 | 7458 | 5384 | 5586 | 5635 | 5835 | 5755 | 5863 | 5793 | 6115 | 5832.56 | 5742 | 5924 | 1.032 | 0.045 | 0.46106949 | 0.66520197 | 3.00E-04 | 0.0075 | 0.0025 | 0.0025 | TRUE | 0.0096 |
| Sp4 | 1224 | 1218 | 1198 | 1591 | 1265 | 1292 | 1300 | 1216 | 1203 | 1251 | 1361 | 1414 | 1290.92 | 1240 | 1342 | 1.082 | 0.113 | 0.20618248 | 0.41515512 | 0.0022 | 0.0095 | 0.005 | 0.005 | TRUE | 0.044 |
| Sp5 | 34 | 102 | 121 | 82 | 59 | 78 | 36 | 102 | 121 | 64 | 63 | 85 | 78.8 | 86 | 71 | 0.82 | -0.286 | 0.48214055 | 0.68089732 | 0.1749 | 0.0487 | 0.1065 | 0.1065 | TRUE | 1.0022 |
| Sp6 | 144 | 204 | 228 | 236 | 215 | 263 | 153 | 204 | 229 | 186 | 231 | 288 | 215.06 | 195 | 235 | 1.199 | 0.262 | 0.23803625 | 0.45383564 | 0.0399 | 0.0222 | 0.0309 | 0.0309 | TRUE | 0.2065 |
| Sp7 | 10 | 9 | 10 | 7 | 7 | 3 | 11 | 9 | 10 | 6 | 8 | 3 | 7.66 | 10 | 6 | 0.552 | -0.858 | 0.21373378 | 0.42520297 | 0 | 0.4369 | 0.2023 | 0.2023 | TRUE | 0.2202 |
| Sp8 | 12 | 8 | 7 | 19 | 21 | 6 | 13 | 8 | 7 | 15 | 23 | 7 | 11.98 | 9 | 15 | 1.594 | 0.673 | 0.29762397 | 0.5184073 | 0.1372 | 0.282 | 0.2121 | 0.2121 | TRUE | 0.132 |
| Sp9 | 12 | 11 | 10 | 18 | 19 | 8 | 13 | 11 | 10 | 14 | 20 | 9 | 12.85 | 11 | 14 | 1.285 | 0.362 | 0.51772788 | 0.70925078 | 0 | 0.2632 | 0.146 | 0.146 | TRUE | 0.1705 |
| Spa17 | 36 | 46 | 39 | 82 | 43 | 39 | 38 | 46 | 39 | 64 | 46 | 43 | 46.13 | 41 | 51 | 1.253 | 0.326 | 0.29025914 | 0.51138358 | 0 | 0.0783 | 0.0464 | 0.0464 | TRUE | 0.4807 |
| Spaca1 | 23 | 19 | 9 | 44 | 34 | 27 | 24 | 19 | 9 | 35 | 37 | 30 | 25.53 | 17 | 34 | 1.928 | 0.947 | 0.03242138 | 0.12143197 | 0.0376 | 0.136 | 0.0973 | 0.0973 | TRUE | 0.7897 |
| Spaca3 | 0 | 0 | 1 | 0 | 1 | 1 | 0 | 0 | 1 | 0 | 1 | 1 | 0.53 | 0 | 1 | 1.924 | 0.944 | 0.73632075 | NA | 0 | 6.2332 | 3.9677 | 3.9677 | TRUE | 0.4098 |
| Spaca4 | 0 | 0 | 0 | 0 | 0 | 0 | 0 | 0 | 0 | 0 | 0 | 0 | 0 | 0 | 0 | NA | NA | NA | NA | NA | NA | NA | NA | NA | NA |
| Spaca5 | 0 | 0 | 0 | 0 | 0 | 0 | 0 | 0 | 0 | 0 | 0 | 0 | 0 | 0 | 0 | NA | NA | NA | NA | NA | NA | NA | NA | NA | NA |
| Spaca7 | 0 | 0 | 0 | 0 | 1 | 0 | 0 | 0 | 0 | 0 | 1 | 0 | 0.18 | 0 | 0 | 1.859 | 0.894 | 0.82650734 | NA | 0 | 18.3785 | 10 | 10 | TRUE | 0.5204 |
| Spag1 | 1520 | 1430 | 1434 | 1438 | 1215 | 1245 | 1615 | 1428 | 1440 | 1130 | 1307 | 1363 | 1380.52 | 1494 | 1267 | 0.847 | -0.239 | 0.02628964 | 0.10526533 | 0.0059 | 0.0093 | 0.0076 | 0.0076 | TRUE | 0.1093 |
| Spag11a | 0 | 0 | 0 | 0 | 0 | 0 | 0 | 0 | 0 | 0 | 0 | 0 | 0 | 0 | 0 | NA | NA | NA | NA | NA | NA | NA | NA | NA | NA |
| Spag11b | 0 | 1 | 0 | 1 | 1 | 0 | 0 | 1 | 0 | 1 | 1 | 0 | 0.48 | 0 | 1 | 1.712 | 0.776 | 0.78614772 | NA | 0 | 6.9165 | 4.1067 | 4.1067 | TRUE | 0.405 |
| Spag16 | 391 | 509 | 509 | 710 | 587 | 586 | 415 | 508 | 511 | 558 | 632 | 641 | 544.32 | 478 | 610 | 1.274 | 0.35 | 0.00654264 | 0.03824704 | 0.0076 | 0.013 | 0.01 | 0.01 | TRUE | 0.1526 |
| Spag17 | 2 | 0 | 0 | 3 | 0 | 0 | 2 | 0 | 0 | 2 | 0 | 0 | 0.75 | 1 | 1 | 1.163 | 0.218 | 0.94242245 | NA | 8.7098 | 4.4164 | 5.2331 | 5.2331 | TRUE | 1.4402 |
| Spag4 | 121 | 162 | 143 | 117 | 108 | 104 | 129 | 162 | 144 | 92 | 116 | 114 | 125.99 | 145 | 107 | 0.738 | -0.438 | 0.02700733 | 0.10721359 | 0.0074 | 0.0331 | 0.0201 | 0.0201 | TRUE | 0.1746 |
| Spag5 | 9492 | 9607 | 9922 | 10017 | 7361 | 7328 | 10082 | 9595 | 9963 | 7875 | 7920 | 8022 | 8909.29 | 9880 | 7939 | 0.804 | -0.316 | 3.22E-08 | 1.36E-06 | 2.00E-04 | 0.0073 | 0.0022 | 0.0022 | TRUE | 0.0078 |
| Spag6 | 30 | 29 | 27 | 29 | 22 | 24 | 32 | 29 | 27 | 23 | 24 | 26 | 26.78 | 29 | 24 | 0.825 | -0.277 | 0.44214955 | 0.64905093 | 0 | 0.1299 | 0.0559 | 0.0559 | TRUE | 0.0367 |
| Spag7 | 3662 | 4432 | 4876 | 5031 | 3789 | 4130 | 3890 | 4426 | 4896 | 3955 | 4077 | 4521 | 4294.11 | 4404 | 4184 | 0.95 | -0.074 | 0.49645048 | 0.69197217 | 0.008 | 0.0077 | 0.0083 | 0.0083 | TRUE | 0.1269 |
| Spag8 | 46 | 37 | 43 | 62 | 46 | 39 | 49 | 37 | 43 | 49 | 49 | 43 | 44.99 | 43 | 47 | 1.095 | 0.131 | 0.64909523 | 0.80332157 | 0 | 0.0801 | 0.0379 | 0.0379 | TRUE | 0.1176 |
| Spag9 | 5712 | 7143 | 7531 | 6621 | 5016 | 5820 | 6067 | 7134 | 7562 | 5205 | 5397 | 6371 | 6289.26 | 6921 | 5658 | 0.817 | -0.291 | 0.01433925 | 0.0684732 | 0.012 | 0.0074 | 0.01 | 0.01 | TRUE | 0.1481 |
| Spam1 | 0 | 0 | 0 | 0 | 0 | 0 | 0 | 0 | 0 | 0 | 0 | 0 | 0 | 0 | 0 | NA | NA | NA | NA | NA | NA | NA | NA | NA | NA |
| Sparc | 8331 | 8523 | 8237 | 17251 | 11444 | 11160 | 8849 | 8512 | 8271 | 13561 | 12313 | 12216 | 10620.43 | 8544 | 12697 | 1.486 | 0.572 | 1.71E-14 | 2.19E-12 | 0.0022 | 0.0072 | 0.0039 | 0.0039 | TRUE | 0.0435 |
| Sparcl1 | 1 | 1 | 7 | 2 | 7 | 6 | 1 | 1 | 7 | 2 | 8 | 7 | 4.13 | 3 | 6 | 1.691 | 0.758 | 0.51042902 | NA | 0.5808 | 0.8052 | 0.6928 | 0.6928 | TRUE | 1.7617 |
| Spast | 3443 | 3523 | 3491 | 4438 | 3142 | 3071 | 3657 | 3518 | 3505 | 3489 | 3381 | 3362 | 3485.33 | 3560 | 3411 | 0.958 | -0.062 | 0.3291329 | 0.55024931 | 2.00E-04 | 0.0078 | 0.0026 | 0.0026 | TRUE | 0.0069 |
| Spata1 | 27 | 35 | 33 | 45 | 33 | 38 | 29 | 35 | 33 | 35 | 36 | 42 | 34.87 | 32 | 38 | 1.159 | 0.212 | 0.50831128 | 0.70245207 | 0 | 0.1014 | 0.0454 | 0.0454 | TRUE | 0.0625 |
| Spata13 | 2744 | 2119 | 2399 | 2098 | 1679 | 1672 | 2915 | 2116 | 2409 | 1649 | 1807 | 1830 | 2120.96 | 2480 | 1762 | 0.71 | -0.493 | 0.00013903 | 0.00179543 | 0.0141 | 0.0085 | 0.0116 | 0.0116 | TRUE | 0.2847 |
| Spata16 | 4 | 6 | 3 | 23 | 13 | 13 | 4 | 6 | 3 | 18 | 14 | 14 | 9.93 | 4 | 15 | 3.508 | 1.811 | 0.00580203 | 0.03499415 | 0 | 0.3388 | 0.1634 | 0.1634 | TRUE | 0.1828 |
| Spata17 | 13 | 9 | 11 | 31 | 15 | 9 | 14 | 9 | 11 | 24 | 16 | 10 | 14.03 | 11 | 17 | 1.51 | 0.595 | 0.29119215 | 0.51183356 | 0.0673 | 0.2417 | 0.1546 | 0.1546 | TRUE | 0.2046 |
| Spata18 | 13 | 12 | 8 | 16 | 16 | 23 | 14 | 12 | 8 | 13 | 17 | 25 | 14.8 | 11 | 18 | 1.612 | 0.689 | 0.20098111 | 0.4086996 | 0 | 0.2295 | 0.1368 | 0.1368 | TRUE | 0.2734 |
| Spata19 | 0 | 0 | 1 | 0 | 0 | 0 | 0 | 0 | 1 | 0 | 0 | 0 | 0.17 | 0 | 0 | 0.49 | -1.029 | 0.80087 | NA | 0 | 19.6928 | 10 | 10 | TRUE | 0.4098 |
| Spata2 | 1888 | 1575 | 1431 | 2311 | 1619 | 1450 | 2005 | 1573 | 1437 | 1817 | 1742 | 1587 | 1693.53 | 1672 | 1715 | 1.027 | 0.038 | 0.78408089 | 0.88852736 | 0.0168 | 0.0088 | 0.0131 | 0.0131 | TRUE | 0.3671 |
| Spata20 | 20 | 16 | 11 | 25 | 13 | 6 | 21 | 16 | 11 | 20 | 14 | 7 | 14.75 | 16 | 14 | 0.847 | -0.24 | 0.6779851 | 0.82103663 | 0.1084 | 0.2303 | 0.1729 | 0.1729 | TRUE | 0.2988 |
| Spata21 | 81 | 86 | 76 | 50 | 55 | 58 | 86 | 86 | 76 | 39 | 59 | 63 | 68.37 | 83 | 54 | 0.646 | -0.63 | 0.01915043 | 0.08408354 | 0.0159 | 0.0551 | 0.0367 | 0.0367 | TRUE | 0.5529 |
| Spata22 | 20 | 19 | 16 | 63 | 61 | 43 | 21 | 19 | 16 | 50 | 66 | 47 | 36.42 | 19 | 54 | 2.88 | 1.526 | 1.44E-05 | 0.00026894 | 0 | 0.0974 | 0.0529 | 0.0529 | TRUE | 0.2783 |
| Spata24 | 213 | 348 | 335 | 375 | 281 | 279 | 226 | 348 | 336 | 295 | 302 | 305 | 302.12 | 303 | 301 | 0.991 | -0.014 | 0.94044945 | 0.96984234 | 0.0232 | 0.0178 | 0.0205 | 0.0205 | TRUE | 0.5535 |
| Spata25 | 0 | 0 | 1 | 0 | 0 | 0 | 0 | 0 | 1 | 0 | 0 | 0 | 0.17 | 0 | 0 | 0.49 | -1.029 | 0.80087 | NA | 0 | 19.6928 | 10 | 10 | TRUE | 0.4098 |
| Spata2l | 22 | 18 | 13 | 29 | 21 | 17 | 23 | 18 | 13 | 23 | 23 | 19 | 19.73 | 18 | 22 | 1.182 | 0.241 | 0.58483059 | 0.75998583 | 0 | 0.1739 | 0.0896 | 0.0896 | TRUE | 0.2288 |
| Spata3 | 0 | 4 | 4 | 1 | 1 | 0 | 0 | 4 | 4 | 1 | 1 | 0 | 1.65 | 3 | 1 | 0.235 | -2.086 | 0.25415282 | NA | 0 | 2.0089 | 1.4502 | 1.4502 | TRUE | 0.8458 |
| Spata31 | 1 | 0 | 0 | 0 | 0 | 0 | 1 | 0 | 0 | 0 | 0 | 0 | 0.18 | 0 | 0 | 0.49 | -1.029 | 0.80087 | NA | 0 | 18.6165 | 10 | 10 | TRUE | 0.4614 |
| Spata31d1a | 154 | 116 | 145 | 234 | 187 | 125 | 164 | 116 | 146 | 184 | 201 | 137 | 157.83 | 142 | 174 | 1.23 | 0.299 | 0.16855674 | 0.36640749 | 0.0274 | 0.0278 | 0.0276 | 0.0276 | TRUE | 0.3589 |
| Spata31d1b | 224 | 155 | 172 | 150 | 109 | 89 | 238 | 155 | 173 | 118 | 117 | 97 | 149.68 | 189 | 111 | 0.59 | -0.762 | 0.00051503 | 0.0052919 | 0.0262 | 0.0289 | 0.0275 | 0.0275 | TRUE | 0.5629 |
| Spata31d1c | 0 | 1 | 4 | 3 | 4 | 2 | 0 | 1 | 4 | 2 | 4 | 2 | 2.31 | 2 | 3 | 1.743 | 0.802 | 0.5601043 | NA | 0.1191 | 1.4324 | 0.8942 | 0.8942 | TRUE | 1.1464 |
| Spata31d1d | 6 | 1 | 2 | 4 | 5 | 4 | 6 | 1 | 2 | 3 | 5 | 4 | 3.71 | 3 | 4 | 1.375 | 0.459 | 0.66146163 | NA | 0.0954 | 0.894 | 0.5139 | 0.5139 | TRUE | 1.058 |
| Spata32 | 1 | 1 | 3 | 5 | 4 | 6 | 1 | 1 | 3 | 4 | 4 | 7 | 3.31 | 2 | 5 | 2.888 | 1.53 | 0.16483769 | NA | 0 | 1.0014 | 0.4737 | 0.4737 | TRUE | 0.3635 |

| | | | | | | | | | | | | | | | | | | | | | | | | | |
|---|---|---|---|---|---|---|---|---|---|---|---|---|---|---|---|---|---|---|---|---|---|---|---|---|---|
| Spata4 | 2 | 3 | 3 | 2 | 3 | 3 | 2 | 3 | 3 | 2 | 3 | 3 | 2.7 | 3 | 3 | 0.974 | -0.039 | 0.97215961 | NA | 0 | 1.2258 | 0.5058 | 0.5058 | TRUE | 0.2183 |
| Spata5 | 3414 | 3496 | 3488 | 4552 | 3276 | 3078 | 3626 | 3491 | 3502 | 3578 | 3525 | 3369 | 3515.44 | 3540 | 3491 | 0.986 | -0.02 | 0.76229615 | 0.87464971 | 4.00E-04 | 0.0078 | 0.0028 | 0.0028 | TRUE | 0.0112 |
| Spata5l1 | 227 | 342 | 348 | 343 | 297 | 268 | 241 | 342 | 349 | 270 | 320 | 293 | 302.44 | 311 | 294 | 0.945 | -0.081 | 0.64472791 | 0.80030289 | 0.0206 | 0.0178 | 0.0192 | 0.0192 | TRUE | 0.4302 |
| Spata6 | 776 | 832 | 881 | 1162 | 802 | 912 | 824 | 831 | 885 | 913 | 863 | 998 | 885.75 | 847 | 925 | 1.092 | 0.127 | 0.19248878 | 0.39815515 | 0.0023 | 0.0106 | 0.0057 | 0.0057 | TRUE | 0.0558 |
| Spata7 | 521 | 559 | 539 | 729 | 586 | 568 | 553 | 558 | 541 | 573 | 630 | 622 | 579.71 | 551 | 608 | 1.103 | 0.141 | 0.15234623 | 0.34350317 | 0 | 0.0126 | 0.0053 | 0.0053 | TRUE | 0.0323 |
| Spata9 | 1 | 0 | 0 | 1 | 0 | 0 | 1 | 0 | 0 | 1 | 0 | 0 | 0.31 | 0 | 0 | 0.954 | -0.067 | 0.98625928 | NA | 0 | 10.7014 | 9.0384 | 9.0384 | TRUE | 0.4614 |
| Spatc1 | 13 | 23 | 18 | 5 | 7 | 14 | 14 | 23 | 18 | 4 | 8 | 15 | 13.61 | 18 | 9 | 0.477 | -1.068 | 0.08725955 | 0.23905813 | 0.1326 | 0.249 | 0.1973 | 0.1973 | TRUE | 0.7803 |
| Spatc1l | 2 | 1 | 0 | 1 | 0 | 0 | 2 | 1 | 0 | 1 | 0 | 0 | 0.65 | 1 | 0 | 0.337 | -1.57 | 0.54866551 | NA | 0 | 5.0634 | 3.4431 | 3.4431 | TRUE | 0.1066 |
| Spats1 | 78 | 105 | 130 | 184 | 131 | 130 | 83 | 105 | 131 | 145 | 141 | 142 | 124.36 | 106 | 143 | 1.343 | 0.426 | 0.04514418 | 0.15318718 | 0.0141 | 0.0334 | 0.0243 | 0.0243 | TRUE | 0.304 |
| Spats2 | 2008 | 2335 | 2313 | 3022 | 2236 | 2341 | 2133 | 2332 | 2322 | 2376 | 2406 | 2563 | 2355.23 | 2262 | 2448 | 1.082 | 0.113 | 0.15085114 | 0.34184256 | 0.0014 | 0.0083 | 0.004 | 0.004 | TRUE | 0.0303 |
| Spats2l | 56 | 75 | 71 | 158 | 126 | 129 | 59 | 75 | 71 | 124 | 136 | 141 | 101.11 | 68 | 134 | 1.943 | 0.959 | 3.36E-06 | 7.85E-05 | 0 | 0.0395 | 0.0195 | 0.0195 | TRUE | 0.1162 |
| Spc24 | 1998 | 2957 | 3100 | 2996 | 2342 | 2424 | 2122 | 2953 | 3113 | 2355 | 2520 | 2653 | 2619.44 | 2729 | 2509 | 0.919 | -0.122 | 0.40890358 | 0.62126578 | 0.0195 | 0.0082 | 0.0152 | 0.0152 | TRUE | 0.4589 |
| Spc25 | 3409 | 4069 | 4122 | 4914 | 3455 | 3599 | 3621 | 4064 | 4139 | 3863 | 3717 | 3940 | 3890.61 | 3941 | 3840 | 0.974 | -0.038 | 0.64579045 | 0.80105443 | 0.0024 | 0.0077 | 0.0046 | 0.0046 | TRUE | 0.0613 |
| Spcs1 | 942 | 1264 | 1237 | 1710 | 1197 | 1320 | 1001 | 1262 | 1242 | 1344 | 1288 | 1445 | 1263.69 | 1168 | 1359 | 1.163 | 0.218 | 0.06419529 | 0.19500617 | 0.0083 | 0.0095 | 0.0092 | 0.0092 | TRUE | 0.1883 |
| Spcs2 | 5448 | 5634 | 5238 | 7235 | 5198 | 5157 | 5787 | 5627 | 5260 | 5688 | 5593 | 5645 | 5599.74 | 5558 | 5642 | 1.015 | 0.022 | 0.74703374 | 0.8655848 | 8.00E-04 | 0.0075 | 0.0031 | 0.0031 | TRUE | 0.0269 |
| Spcs3 | 10615 | 13378 | 13168 | 14455 | 10555 | 11616 | 11275 | 13361 | 13222 | 11363 | 11357 | 12716 | 12215.54 | 12619 | 11812 | 0.936 | -0.096 | 0.32705998 | 0.54824943 | 0.0063 | 0.0072 | 0.0068 | 0.0068 | TRUE | 0.1061 |
| Spdef | 17 | 46 | 58 | 32 | 35 | 34 | 18 | 46 | 58 | 25 | 38 | 37 | 37.04 | 41 | 33 | 0.813 | -0.299 | 0.5137787 | 0.70663057 | 0.1546 | 0.0958 | 0.1237 | 0.1237 | TRUE | 0.5141 |
| Spdl1 | 3747 | 3573 | 3293 | 4854 | 3428 | 3048 | 3980 | 3568 | 3307 | 3816 | 3688 | 3337 | 3615.93 | 3618 | 3614 | 0.999 | -0.001 | 0.98841993 | 0.99386769 | 0.0065 | 0.0078 | 0.007 | 0.007 | TRUE | 0.0929 |
| Spdya | 30 | 24 | 12 | 47 | 23 | 33 | 32 | 24 | 12 | 37 | 25 | 36 | 27.62 | 23 | 33 | 1.448 | 0.534 | 0.2314655 | 0.44525843 | 0.0816 | 0.1262 | 0.1061 | 0.1061 | TRUE | 0.4706 |
| Spdyb | 6 | 1 | 3 | 4 | 6 | 2 | 6 | 1 | 3 | 3 | 6 | 2 | 3.7 | 3 | 4 | 1.136 | 0.184 | 0.86349981 | NA | 0.1774 | 0.8984 | 0.5496 | 0.5496 | TRUE | 0.4116 |
| Specc1 | 1655 | 1771 | 1892 | 2640 | 1900 | 1759 | 1758 | 1769 | 1900 | 2075 | 2044 | 1926 | 1911.92 | 1809 | 2015 | 1.114 | 0.156 | 0.04588629 | 0.15497186 | 0.0012 | 0.0086 | 0.0039 | 0.0039 | TRUE | 0.0234 |
| Specc1l | 4483 | 4142 | 3941 | 5623 | 4113 | 3882 | 4762 | 4137 | 3957 | 4420 | 4425 | 4249 | 4325.12 | 4285 | 4365 | 1.019 | 0.027 | 0.77078614 | 0.88050269 | 0.0047 | 0.0077 | 0.0059 | 0.0059 | TRUE | 0.1151 |
| Speer1-ps1 | 0 | 0 | 0 | 0 | 0 | 0 | 0 | 0 | 0 | 0 | 0 | 0 | 0 | 0 | 0 | NA | NA | NA | NA | NA | NA | NA | NA | NA | NA |
| Speer2 | 0 | 0 | 0 | 0 | 0 | 0 | 0 | 0 | 0 | 0 | 0 | 0 | 0 | 0 | 0 | NA | NA | NA | NA | NA | NA | NA | NA | NA | NA |
| Speer3 | 1 | 0 | 0 | 0 | 0 | 0 | 1 | 0 | 0 | 0 | 0 | 0 | 0.18 | 0 | 0 | 0.49 | -1.029 | 0.80087 | NA | 0 | 18.6165 | 10 | 10 | TRUE | 0.4614 |
| Speer4a | 1 | 1 | 0 | 2 | 0 | 1 | 1 | 1 | 0 | 2 | 0 | 1 | 0.79 | 1 | 1 | 1.332 | 0.414 | 0.8494818 | NA | 0 | 4.1879 | 2.1489 | 2.1489 | TRUE | 0.2882 |
| Speer4b | 6 | 2 | 2 | 10 | 4 | 6 | 6 | 2 | 2 | 8 | 4 | 7 | 4.85 | 3 | 6 | 1.832 | 0.874 | 0.33880218 | 0.55879656 | 0 | 0.6859 | 0.3745 | 0.3745 | TRUE | 0.7593 |
| Speer4d | 0 | 0 | 0 | 0 | 1 | 0 | 0 | 0 | 0 | 0 | 1 | 0 | 0.18 | 0 | 0 | 1.859 | 0.894 | 0.82650734 | NA | 0 | 18.3785 | 10 | 10 | TRUE | 0.5204 |
| Speer4e | 0 | 1 | 0 | 0 | 0 | 0 | 0 | 1 | 0 | 0 | 0 | 0 | 0.17 | 0 | 0 | 0.49 | -1.029 | 0.80087 | NA | 0 | 19.7991 | 10 | 10 | TRUE | 0.405 |
| Speer4f | 1 | 0 | 0 | 1 | 2 | 0 | 1 | 0 | 0 | 1 | 2 | 0 | 0.67 | 0 | 1 | 2.668 | 1.416 | 0.58602256 | NA | 0 | 4.9484 | 3.3683 | 3.3683 | TRUE | 0.3203 |
| Speer5-ps1 | 22 | 20 | 13 | 20 | 22 | 25 | 23 | 20 | 13 | 16 | 24 | 27 | 20.53 | 19 | 22 | 1.174 | 0.232 | 0.61440286 | 0.78179603 | 0.0331 | 0.1674 | 0.1034 | 0.1034 | TRUE | 0.4619 |
| Speer6-ps1 | 0 | 0 | 0 | 0 | 0 | 0 | 0 | 0 | 0 | 0 | 0 | 0 | 0 | 0 | 0 | NA | NA | NA | NA | NA | NA | NA | NA | NA | NA |
| Speer7-ps1 | 5 | 3 | 1 | 15 | 14 | 4 | 5 | 3 | 1 | 12 | 15 | 4 | 6.76 | 3 | 10 | 3.388 | 1.76 | 0.04949537 | 0.16318557 | 0.2057 | 0.4944 | 0.3662 | 0.3662 | TRUE | 0.2792 |
| Speer8-ps1 | 4 | 0 | 0 | 9 | 13 | 2 | 4 | 0 | 0 | 7 | 14 | 2 | 4.58 | 1 | 8 | 5.566 | 2.477 | 0.07237022 | NA | 1.4424 | 0.7257 | 0.9388 | 0.9388 | TRUE | 0.5331 |
| Speer9-ps1 | 1 | 0 | 2 | 4 | 2 | 1 | 1 | 0 | 2 | 3 | 2 | 1 | 1.58 | 1 | 2 | 2.116 | 1.081 | 0.49148447 | NA | 0 | 2.0961 | 1.0551 | 1.0551 | TRUE | 0.3329 |
| Spef1 | 50 | 53 | 28 | 88 | 48 | 69 | 53 | 53 | 28 | 69 | 52 | 76 | 55.09 | 45 | 66 | 1.466 | 0.552 | 0.09645557 | 0.25495179 | 0.0534 | 0.0667 | 0.0606 | 0.0606 | TRUE | 0.8328 |
| Spef2 | 11 | 5 | 7 | 21 | 17 | 11 | 12 | 5 | 7 | 17 | 18 | 12 | 11.76 | 8 | 16 | 1.989 | 0.992 | 0.09453047 | 0.25193271 | 0 | 0.2871 | 0.1574 | 0.1574 | TRUE | 0.4865 |
| Speg | 578 | 514 | 580 | 404 | 321 | 331 | 614 | 513 | 582 | 318 | 345 | 362 | 455.83 | 570 | 342 | 0.599 | -0.74 | 2.65E-09 | 1.45E-07 | 0.0045 | 0.0141 | 0.0088 | 0.0088 | TRUE | 0.0887 |
| Spem1 | 4 | 7 | 5 | 3 | 2 | 3 | 4 | 7 | 5 | 2 | 2 | 3 | 4.01 | 5 | 2 | 0.476 | -1.071 | 0.25441546 | NA | 0 | 0.8287 | 0.3543 | 0.3543 | TRUE | 0.1408 |
| Spen | 5104 | 3640 | 4135 | 6141 | 4165 | 3591 | 5421 | 3635 | 4152 | 4828 | 4481 | 3931 | 4408.07 | 4403 | 4413 | 1.003 | 0.004 | 0.98095703 | 0.98961586 | 0.026 | 0.0077 | 0.0169 | 0.0169 | TRUE | 0.4984 |
| Spert | 0 | 0 | 0 | 0 | 0 | 0 | 0 | 0 | 0 | 0 | 0 | 0 | 0 | 0 | 0 | NA | NA | NA | NA | NA | NA | NA | NA | NA | NA |
| Spesp1 | 204 | 175 | 210 | 257 | 222 | 254 | 217 | 175 | 211 | 202 | 239 | 278 | 220.21 | 201 | 240 | 1.19 | 0.251 | 0.15999941 | 0.35447545 | 0.0153 | 0.0219 | 0.0185 | 0.0185 | TRUE | 0.1726 |
| Spg11 | 1677 | 2234 | 2499 | 2196 | 1769 | 2086 | 1781 | 2231 | 2509 | 1726 | 1903 | 2283 | 2072.46 | 2174 | 1971 | 0.906 | -0.142 | 0.35588035 | 0.57433183 | 0.0225 | 0.0085 | 0.0166 | 0.0166 | TRUE | 0.3017 |
| Spg20 | 1531 | 1584 | 1439 | 2570 | 1800 | 1731 | 1626 | 1582 | 1445 | 2020 | 1937 | 1895 | 1750.82 | 1551 | 1951 | 1.258 | 0.331 | 7.04E-05 | 0.00103709 | 0.0018 | 0.0088 | 0.0044 | 0.0044 | TRUE | 0.0432 |
| Spg21 | 2931 | 3609 | 3562 | 4403 | 3355 | 3664 | 3113 | 3604 | 3577 | 3461 | 3610 | 4011 | 3562.68 | 3431 | 3694 | 1.076 | 0.106 | 0.28232286 | 0.50369474 | 0.0058 | 0.0078 | 0.0067 | 0.0067 | TRUE | 0.0798 |
| Spg7 | 4637 | 5034 | 5126 | 6004 | 4298 | 4159 | 4925 | 5028 | 5147 | 4720 | 4624 | 4553 | 4832.8 | 5033 | 4632 | 0.92 | -0.12 | 0.04696496 | 0.15716769 | 2.00E-04 | 0.0076 | 0.0024 | 0.0024 | TRUE | 0.0048 |
| Sphk1 | 177 | 265 | 231 | 235 | 154 | 184 | 188 | 265 | 232 | 185 | 166 | 201 | 206.08 | 228 | 184 | 0.805 | -0.312 | 0.08386374 | 0.23205244 | 0.0144 | 0.0229 | 0.0186 | 0.0186 | TRUE | 0.2253 |
| Sphk2 | 1012 | 792 | 801 | 1458 | 1062 | 782 | 1075 | 791 | 804 | 1146 | 1143 | 856 | 969.17 | 890 | 1048 | 1.179 | 0.237 | 0.15458081 | 0.34660912 | 0.0273 | 0.0103 | 0.0189 | 0.0189 | TRUE | 0.392 |
| Sphkap | 8 | 3 | 5 | 6 | 2 | 4 | 8 | 3 | 5 | 5 | 2 | 4 | 4.63 | 5 | 4 | 0.691 | -0.534 | 0.55589726 | 0.73882292 | 0 | 0.7189 | 0.3697 | 0.3697 | TRUE | 0.2995 |
| Spib | 10 | 31 | 14 | 9 | 8 | 7 | 11 | 31 | 14 | 7 | 9 | 8 | 13.16 | 19 | 8 | 0.417 | -1.261 | 0.04389369 | 0.15010653 | 0.1323 | 0.2572 | 0.1918 | 0.1918 | TRUE | 1.4387 |
| Spic | 146 | 117 | 99 | 103 | 81 | 94 | 155 | 117 | 99 | 81 | 87 | 103 | 107.06 | 124 | 90 | 0.728 | -0.457 | 0.05522658 | 0.17564703 | 0.0252 | 0.0377 | 0.0314 | 0.0314 | TRUE | 0.4462 |
| Spice1 | 2507 | 2609 | 2628 | 2889 | 2357 | 2333 | 2663 | 2606 | 2639 | 2271 | 2536 | 2554 | 2544.7 | 2636 | 2454 | 0.93 | -0.104 | 0.18808091 | 0.39305845 | 0.0019 | 0.0082 | 0.0041 | 0.0041 | TRUE | 0.0525 |
| Spin1 | 7466 | 7122 | 6962 | 7899 | 5735 | 6039 | 7930 | 7113 | 6991 | 6210 | 6171 | 6611 | 6837.39 | 7345 | 6331 | 0.862 | -0.214 | 0.00781387 | 0.04357862 | 0.0029 | 0.0074 | 0.0045 | 0.0045 | TRUE | 0.0593 |
| Spin2 | 26 | 31 | 25 | 72 | 44 | 37 | 28 | 31 | 25 | 57 | 47 | 41 | 38.02 | 28 | 48 | 1.736 | 0.795 | 0.0144698 | 0.0689271 | 0 | 0.0936 | 0.0478 | 0.0478 | TRUE | 0.2156 |
| Spin4 | 201 | 223 | 212 | 236 | 216 | 194 | 213 | 223 | 213 | 186 | 232 | 212 | 213.23 | 216 | 210 | 0.968 | -0.047 | 0.75584901 | 0.87101981 | 0 | 0.0224 | 0.012 | 0.012 | TRUE | 0.1251 |
| Spink10 | 15 | 14 | 6 | 9 | 8 | 3 | 16 | 14 | 6 | 7 | 9 | 3 | 9.15 | 12 | 6 | 0.532 | -0.911 | 0.19713467 | 0.40376033 | 0.0954 | 0.3669 | 0.2403 | 0.2403 | TRUE | 0.752 |
| Spink11 | 0 | 0 | 0 | 0 | 0 | 0 | 0 | 0 | 0 | 0 | 0 | 0 | 0 | 0 | 0 | NA | NA | NA | NA | NA | NA | NA | NA | NA | NA |
| Spink12 | 0 | 0 | 0 | 0 | 0 | 0 | 0 | 0 | 0 | 0 | 0 | 0 | 0 | 0 | 0 | NA | NA | NA | NA | NA | NA | NA | NA | NA | NA |
| Spink13 | 2 | 0 | 0 | 0 | 0 | 0 | 2 | 0 | 0 | 0 | 0 | 0 | 0.35 | 1 | 0 | 0.254 | -1.975 | 0.62446185 | NA | 10 | 9.3117 | 10 | 10 | TRUE | 0.9741 |
| Spink14 | 0 | 0 | 0 | 0 | 0 | 0 | 0 | 0 | 0 | 0 | 0 | 0 | 0 | 0 | 0 | NA | NA | NA | NA | NA | NA | NA | NA | NA | NA |
| Spink2 | 123 | 117 | 113 | 116 | 84 | 90 | 131 | 117 | 113 | 91 | 90 | 99 | 106.84 | 120 | 93 | 0.776 | -0.367 | 0.05502242 | 0.17503298 | 0 | 0.0377 | 0.0168 | 0.0168 | TRUE | 0.0559 |
| Spink3 | 8 | 2 | 2 | 4 | 2 | 2 | 8 | 2 | 2 | 3 | 2 | 2 | 3.33 | 4 | 2 | 0.609 | -0.715 | 0.51942763 | NA | 0 | 0.9958 | 0.5712 | 0.5712 | TRUE | 1.3659 |
| Spink4 | 0 | 0 | 0 | 0 | 0 | 0 | 0 | 0 | 0 | 0 | 0 | 0 | 0 | 0 | 0 | NA | NA | NA | NA | NA | NA | NA | NA | NA | NA |
| Spink5 | 0 | 1 | 0 | 1 | 2 | 2 | 0 | 1 | 0 | 1 | 2 | 2 | 1.02 | 0 | 2 | 4.603 | 2.203 | 0.27756188 | NA | 0 | 3.2336 | 1.6747 | 1.6747 | TRUE | 0.405 |
| Spink6 | 0 | 0 | 0 | 0 | 0 | 0 | 0 | 0 | 0 | 0 | 0 | 0 | 0 | 0 | 0 | NA | NA | NA | NA | NA | NA | NA | NA | NA | NA |
| Spink7 | 0 | 0 | 0 | 0 | 1 | 0 | 0 | 0 | 0 | 0 | 1 | 0 | 0.18 | 0 | 0 | 1.859 | 0.894 | 0.82650734 | NA | 0 | 18.3785 | 10 | 10 | TRUE | 0.5204 |
| Spink8 | 0 | 0 | 0 | 0 | 0 | 0 | 0 | 0 | 0 | 0 | 0 | 0 | 0 | 0 | 0 | NA | NA | NA | NA | NA | NA | NA | NA | NA | NA |
| Spinkl | 0 | 0 | 0 | 0 | 0 | 0 | 0 | 0 | 0 | 0 | 0 | 0 | 0 | 0 | 0 | NA | NA | NA | NA | NA | NA | NA | NA | NA | NA |
| Spint1 | 1897 | 2287 | 2415 | 3011 | 2212 | 2245 | 2015 | 2284 | 2425 | 2367 | 2380 | 2458 | 2321.4 | 2241 | 2402 | 1.071 | 0.099 | 0.28502795 | 0.50627409 | 0.0039 | 0.0083 | 0.0058 | 0.0058 | TRUE | 0.0942 |

| | | | | | | | | | | | | | | | | | | | | | | | | | |
|---|---|---|---|---|---|---|---|---|---|---|---|---|---|---|---|---|---|---|---|---|---|---|---|---|---|
| Spint2 | 7592 | 9869 | 9639 | 8561 | 6442 | 7189 | 8064 | 9856 | 9679 | 6730 | 6931 | 7869 | 8188.26 | 9200 | 7177 | 0.78 | -0.358 | 0.00104594 | 0.0092314 | 0.0092 | 0.0073 | 0.0085 | 0.0085 | TRUE | 0.1424 |
| Spint3 | 0 | 0 | 0 | 0 | 0 | 0 | 0 | 0 | 0 | 0 | 0 | 0 | 0 | 0 | 0 | NA | NA | NA | NA | NA | NA | NA | NA | NA | NA |
| Spint4 | 0 | 0 | 0 | 0 | 0 | 0 | 0 | 0 | 0 | 0 | 0 | 0 | 0 | 0 | 0 | NA | NA | NA | NA | NA | NA | NA | NA | NA | NA |
| Spint5 | 2 | 1 | 0 | 12 | 5 | 2 | 2 | 1 | 0 | 9 | 5 | 2 | 3.35 | 1 | 5 | 5.587 | 2.482 | 0.05958178 | NA | 0 | 0.9891 | 0.6695 | 0.6695 | TRUE | 0.1572 |
| Spire1 | 3125 | 3969 | 4189 | 4037 | 3072 | 3328 | 3319 | 3964 | 4206 | 3174 | 3305 | 3643 | 3601.88 | 3830 | 3374 | 0.881 | -0.183 | 0.1039724 | 0.26817827 | 0.0093 | 0.0078 | 0.0089 | 0.0089 | TRUE | 0.1652 |
| Spire2 | 1812 | 1930 | 1917 | 2058 | 1646 | 1638 | 1925 | 1928 | 1925 | 1618 | 1771 | 1793 | 1826.49 | 1926 | 1727 | 0.896 | -0.158 | 0.04224129 | 0.14655295 | 0.0011 | 0.0087 | 0.0038 | 0.0038 | TRUE | 0.0383 |
| Spn | 1497 | 1171 | 1150 | 948 | 703 | 742 | 1590 | 1169 | 1155 | 745 | 756 | 812 | 1038.03 | 1305 | 771 | 0.591 | -0.759 | 1.41E-07 | 4.81E-06 | 0.0175 | 0.0101 | 0.0139 | 0.0139 | TRUE | 0.4386 |
| Spn-ps | 0 | 0 | 0 | 0 | 0 | 0 | 0 | 0 | 0 | 0 | 0 | 0 | 0 | 0 | 0 | NA | NA | NA | NA | NA | NA | NA | NA | NA | NA |
| Spns1 | 2466 | 2156 | 2056 | 3358 | 2453 | 2063 | 2619 | 2153 | 2064 | 2640 | 2639 | 2258 | 2395.72 | 2279 | 2512 | 1.103 | 0.141 | 0.24441087 | 0.4610511 | 0.0118 | 0.0083 | 0.0102 | 0.0102 | TRUE | 0.2064 |
| Spns2 | 264 | 272 | 267 | 306 | 235 | 220 | 280 | 272 | 268 | 241 | 253 | 241 | 259.06 | 273 | 245 | 0.895 | -0.16 | 0.20641822 | 0.41536466 | 0 | 0.0196 | 0.0077 | 0.0077 | TRUE | 0.009 |
| Spns3 | 18 | 20 | 21 | 29 | 21 | 16 | 19 | 20 | 21 | 23 | 23 | 18 | 20.51 | 20 | 21 | 1.049 | 0.069 | 0.86696404 | 0.93412739 | 0 | 0.1675 | 0.0735 | 0.0735 | TRUE | 0.1051 |
| Spo11 | 0 | 0 | 0 | 3 | 1 | 1 | 0 | 0 | 0 | 2 | 1 | 1 | 0.75 | 0 | 1 | 8.123 | 3.022 | 0.20670853 | NA | 0 | 4.3714 | 2.8067 | 2.8067 | TRUE | 0.217 |
| Spock1 | 254 | 231 | 196 | 207 | 184 | 179 | 270 | 231 | 197 | 163 | 198 | 196 | 208.99 | 233 | 186 | 0.796 | -0.328 | 0.06633223 | 0.19871336 | 0.014 | 0.0227 | 0.0182 | 0.0182 | TRUE | 0.18 |
| Spock2 | 876 | 926 | 973 | 902 | 692 | 739 | 930 | 925 | 977 | 709 | 745 | 809 | 849.14 | 944 | 754 | 0.798 | -0.326 | 0.00058624 | 0.00588558 | 0.0014 | 0.0108 | 0.0053 | 0.0053 | TRUE | 0.0473 |
| Spock3 | 41 | 44 | 42 | 65 | 53 | 59 | 44 | 44 | 42 | 51 | 57 | 65 | 50.4 | 43 | 58 | 1.326 | 0.408 | 0.13395172 | 0.31645992 | 0 | 0.0723 | 0.033 | 0.033 | TRUE | 0.0951 |
| Spon1 | 31 | 22 | 21 | 18 | 30 | 22 | 33 | 22 | 21 | 14 | 32 | 24 | 24.42 | 25 | 23 | 0.919 | -0.122 | 0.78934684 | 0.891612 | 0.0748 | 0.1418 | 0.109 | 0.109 | TRUE | 0.2891 |
| Spon2 | 117 | 140 | 97 | 119 | 80 | 76 | 124 | 140 | 97 | 94 | 86 | 83 | 104.05 | 120 | 88 | 0.729 | -0.457 | 0.03641407 | 0.13148414 | 0.0108 | 0.0386 | 0.0245 | 0.0245 | TRUE | 0.2874 |
| Spop | 4870 | 4084 | 4305 | 5033 | 3439 | 3366 | 5173 | 4079 | 4323 | 3957 | 3700 | 3685 | 4152.59 | 4525 | 3781 | 0.836 | -0.259 | 0.01602882 | 0.07448633 | 0.0085 | 0.0077 | 0.0081 | 0.0081 | TRUE | 0.191 |
| Spopl | 602 | 557 | 556 | 843 | 537 | 544 | 639 | 556 | 558 | 663 | 578 | 595 | 598.33 | 584 | 612 | 1.049 | 0.068 | 0.55083259 | 0.73506486 | 0.0043 | 0.0124 | 0.0078 | 0.0078 | TRUE | 0.0782 |
| Spp1 | 4827 | 4327 | 4214 | 11754 | 7920 | 7394 | 5127 | 4321 | 4231 | 9240 | 8521 | 8094 | 6589.21 | 4560 | 8618 | 1.89 | 0.919 | 6.97E-19 | 1.51E-16 | 0.0079 | 0.0074 | 0.0076 | 0.0076 | TRUE | 0.1441 |
| Spp2 | 0 | 0 | 0 | 0 | 0 | 0 | 0 | 0 | 0 | 0 | 0 | 0 | 0 | 0 | 0 | NA | NA | NA | NA | NA | NA | NA | NA | NA | NA |
| Sppl2a | 1703 | 1736 | 1718 | 1972 | 1393 | 1461 | 1809 | 1734 | 1725 | 1550 | 1499 | 1599 | 1652.67 | 1756 | 1549 | 0.882 | -0.18 | 0.01468087 | 0.06976401 | 0 | 0.0089 | 0.0033 | 0.0033 | TRUE | 0.0095 |
| Sppl2b | 255 | 257 | 246 | 564 | 394 | 294 | 271 | 257 | 247 | 443 | 424 | 322 | 327.28 | 258 | 396 | 1.538 | 0.621 | 9.28E-05 | 0.00129819 | 0.0133 | 0.017 | 0.015 | 0.015 | TRUE | 0.3053 |
| Sppl2c | 0 | 0 | 0 | 0 | 0 | 0 | 0 | 0 | 0 | 0 | 0 | 0 | 0 | 0 | 0 | NA | NA | NA | NA | NA | NA | NA | NA | NA | NA |
| Sppl3 | 3482 | 3346 | 3383 | 4421 | 3223 | 3089 | 3698 | 3342 | 3397 | 3475 | 3468 | 3381 | 3460.28 | 3479 | 3441 | 0.989 | -0.015 | 0.83306545 | 0.91523598 | 0.0013 | 0.0079 | 0.0035 | 0.0035 | TRUE | 0.0369 |
| Spr | 2419 | 2057 | 1931 | 3066 | 2020 | 1854 | 2569 | 2054 | 1939 | 2410 | 2173 | 2029 | 2195.97 | 2187 | 2204 | 1.008 | 0.012 | 0.92798838 | 0.96471962 | 0.0146 | 0.0084 | 0.0117 | 0.0117 | TRUE | 0.2819 |
| Spred1 | 2196 | 2508 | 2732 | 3368 | 2443 | 2333 | 2333 | 2505 | 2743 | 2648 | 2629 | 2554 | 2568.42 | 2527 | 2610 | 1.033 | 0.047 | 0.58941109 | 0.76400346 | 0.0031 | 0.0082 | 0.005 | 0.005 | TRUE | 0.0681 |
| Spred2 | 2837 | 2485 | 2689 | 3612 | 2526 | 2326 | 3013 | 2482 | 2700 | 2839 | 2718 | 2546 | 2716.45 | 2732 | 2701 | 0.989 | -0.016 | 0.87486961 | 0.93825131 | 0.0059 | 0.0081 | 0.0068 | 0.0068 | TRUE | 0.0985 |
| Spred3 | 204 | 128 | 131 | 172 | 139 | 77 | 217 | 128 | 132 | 135 | 150 | 84 | 140.85 | 159 | 123 | 0.777 | -0.364 | 0.21149823 | 0.42203316 | 0.0782 | 0.0303 | 0.0539 | 0.0539 | TRUE | 1.0699 |
| Sprn | 1 | 0 | 1 | 1 | 4 | 2 | 1 | 0 | 1 | 1 | 4 | 2 | 1.56 | 1 | 2 | 3.482 | 1.8 | 0.28863368 | NA | 0 | 2.122 | 1.1217 | 1.1217 | TRUE | 0.1582 |
| Sprr1a | 0 | 0 | 0 | 0 | 0 | 1 | 0 | 0 | 0 | 0 | 0 | 1 | 0.18 | 0 | 0 | 1.859 | 0.894 | 0.82650734 | NA | 0 | 18.0643 | 10 | 10 | TRUE | 0.5383 |
| Sprr1b | 0 | 0 | 0 | 0 | 0 | 0 | 0 | 0 | 0 | 0 | 0 | 0 | 0 | 0 | 0 | NA | NA | NA | NA | NA | NA | NA | NA | NA | NA |
| Sprr2a1 | 0 | 0 | 0 | 0 | 0 | 0 | 0 | 0 | 0 | 0 | 0 | 0 | 0 | 0 | 0 | NA | NA | NA | NA | NA | NA | NA | NA | NA | NA |
| Sprr2a2 | 0 | 0 | 0 | 0 | 0 | 0 | 0 | 0 | 0 | 0 | 0 | 0 | 0 | 0 | 0 | NA | NA | NA | NA | NA | NA | NA | NA | NA | NA |
| Sprr2b | 0 | 0 | 0 | 0 | 0 | 0 | 0 | 0 | 0 | 0 | 0 | 0 | 0 | 0 | 0 | NA | NA | NA | NA | NA | NA | NA | NA | NA | NA |
| Sprr2d | 0 | 0 | 0 | 0 | 0 | 0 | 0 | 0 | 0 | 0 | 0 | 0 | 0 | 0 | 0 | NA | NA | NA | NA | NA | NA | NA | NA | NA | NA |
| Sprr2e | 0 | 1 | 1 | 3 | 2 | 1 | 0 | 1 | 1 | 2 | 2 | 1 | 1.27 | 1 | 2 | 2.793 | 1.482 | 0.39066658 | NA | 0 | 2.6051 | 1.1315 | 1.1315 | TRUE | 0.2222 |
| Sprr2f | 0 | 0 | 0 | 0 | 0 | 0 | 0 | 0 | 0 | 0 | 0 | 0 | 0 | 0 | 0 | NA | NA | NA | NA | NA | NA | NA | NA | NA | NA |
| Sprr2g | 0 | 0 | 0 | 0 | 0 | 0 | 0 | 0 | 0 | 0 | 0 | 0 | 0 | 0 | 0 | NA | NA | NA | NA | NA | NA | NA | NA | NA | NA |
| Sprr2h | 0 | 0 | 0 | 0 | 0 | 0 | 0 | 0 | 0 | 0 | 0 | 0 | 0 | 0 | 0 | NA | NA | NA | NA | NA | NA | NA | NA | NA | NA |
| Sprr2i | 0 | 0 | 0 | 0 | 0 | 0 | 0 | 0 | 0 | 0 | 0 | 0 | 0 | 0 | 0 | NA | NA | NA | NA | NA | NA | NA | NA | NA | NA |
| Sprr2j-ps | 0 | 0 | 0 | 0 | 0 | 0 | 0 | 0 | 0 | 0 | 0 | 0 | 0 | 0 | 0 | NA | NA | NA | NA | NA | NA | NA | NA | NA | NA |
| Sprr2k | 0 | 0 | 0 | 2 | 0 | 0 | 0 | 0 | 0 | 2 | 0 | 0 | 0.26 | 0 | 1 | 2.886 | 1.529 | 0.70676491 | NA | 0 | 12.5792 | 10 | 10 | TRUE | 0.9669 |
| Sprr3 | 0 | 0 | 0 | 0 | 0 | 0 | 0 | 0 | 0 | 0 | 0 | 0 | 0 | 0 | 0 | NA | NA | NA | NA | NA | NA | NA | NA | NA | NA |
| Sprr4 | 0 | 0 | 0 | 0 | 0 | 0 | 0 | 0 | 0 | 0 | 0 | 0 | 0 | 0 | 0 | NA | NA | NA | NA | NA | NA | NA | NA | NA | NA |
| Sprtn | 1343 | 1128 | 1070 | 1751 | 1216 | 1021 | 1427 | 1127 | 1074 | 1376 | 1308 | 1118 | 1238.32 | 1209 | 1267 | 1.049 | 0.069 | 0.6222358 | 0.78656528 | 0.0167 | 0.0096 | 0.0133 | 0.0133 | TRUE | 0.2958 |
| Spry1 | 1015 | 1351 | 1364 | 983 | 788 | 823 | 1078 | 1349 | 1370 | 773 | 848 | 901 | 1053.08 | 1266 | 841 | 0.663 | -0.592 | 2.69E-06 | 6.49E-05 | 0.0108 | 0.01 | 0.0105 | 0.0105 | TRUE | 0.201 |
| Spry2 | 4715 | 4046 | 4367 | 5162 | 3738 | 3510 | 5008 | 4041 | 4385 | 4058 | 4022 | 3842 | 4225.99 | 4478 | 3974 | 0.888 | -0.172 | 0.08046137 | 0.22552634 | 0.0061 | 0.0077 | 0.0067 | 0.0067 | TRUE | 0.1305 |
| Spry3 | 12 | 5 | 1 | 18 | 12 | 10 | 13 | 5 | 1 | 14 | 13 | 11 | 9.46 | 6 | 13 | 2.046 | 1.033 | 0.19328137 | 0.39886684 | 0.2989 | 0.3552 | 0.3333 | 0.3333 | TRUE | 0.9468 |
| Spry4 | 12729 | 12740 | 12343 | 11079 | 8648 | 6365 | 13520 | 12724 | 12394 | 8709 | 9305 | 6967 | 10603.22 | 12879 | 8327 | 0.647 | -0.629 | 8.89E-08 | 3.24E-06 | 0.0115 | 0.0072 | 0.0099 | 0.0099 | TRUE | 0.2487 |
| Spryd3 | 1224 | 1317 | 1293 | 1468 | 1122 | 1108 | 1300 | 1315 | 1298 | 1154 | 1207 | 1213 | 1247.97 | 1304 | 1191 | 0.913 | -0.132 | 0.07957697 | 0.22381823 | 0 | 0.0095 | 0.0033 | 0.0033 | TRUE | 0.0094 |
| Spryd4 | 1211 | 1137 | 1003 | 1537 | 1121 | 1037 | 1286 | 1136 | 1007 | 1208 | 1206 | 1135 | 1163.08 | 1143 | 1183 | 1.036 | 0.051 | 0.65364459 | 0.80540538 | 0.0072 | 0.0097 | 0.0083 | 0.0083 | TRUE | 0.1435 |
| Spryd7 | 1375 | 1509 | 1484 | 1630 | 1201 | 1353 | 1460 | 1507 | 1490 | 1281 | 1292 | 1481 | 1418.71 | 1486 | 1351 | 0.909 | -0.138 | 0.13072458 | 0.31130728 | 0.0025 | 0.0092 | 0.0053 | 0.0053 | TRUE | 0.0834 |
| Spsb1 | 747 | 1100 | 1086 | 985 | 782 | 801 | 793 | 1099 | 1090 | 774 | 841 | 877 | 912.5 | 994 | 831 | 0.835 | -0.26 | 0.07381053 | 0.21338322 | 0.0174 | 0.0105 | 0.0141 | 0.0141 | TRUE | 0.371 |
| Spsb2 | 121 | 138 | 129 | 202 | 151 | 136 | 129 | 138 | 130 | 159 | 162 | 149 | 144.34 | 132 | 157 | 1.188 | 0.249 | 0.13049492 | 0.31106435 | 0 | 0.0297 | 0.0125 | 0.0125 | TRUE | 0.0189 |
| Spsb3 | 883 | 839 | 764 | 1120 | 839 | 708 | 938 | 838 | 767 | 880 | 903 | 775 | 850.19 | 848 | 853 | 1.007 | 0.01 | 0.93460751 | 0.9675739 | 0.0071 | 0.0108 | 0.0087 | 0.0087 | TRUE | 0.1026 |
| Spsb4 | 88 | 71 | 62 | 119 | 69 | 61 | 93 | 71 | 62 | 94 | 74 | 67 | 76.87 | 75 | 78 | 1.042 | 0.059 | 0.82406223 | 0.91048052 | 0.0248 | 0.0498 | 0.0374 | 0.0374 | TRUE | 0.3873 |
| Spt1 | 0 | 0 | 0 | 0 | 0 | 0 | 0 | 0 | 0 | 0 | 0 | 0 | 0 | 0 | 0 | NA | NA | NA | NA | NA | NA | NA | NA | NA | NA |
| Spta1 | 299 | 114 | 137 | 117 | 71 | 25 | 318 | 114 | 138 | 92 | 76 | 27 | 127.46 | 190 | 65 | 0.346 | -1.533 | 0.00103017 | 0.00911777 | 0.305 | 0.0328 | 0.147 | 0.147 | TRUE | 2.4947 |
| Sptan1 | 18210 | 16330 | 18833 | 30334 | 20677 | 18858 | 19342 | 16309 | 18910 | 23846 | 22247 | 20643 | 20216.32 | 18187 | 22245 | 1.223 | 0.291 | 0.0030328 | 0.02142112 | 0.0066 | 0.0071 | 0.0069 | 0.0069 | TRUE | 0.0999 |
| Sptb | 1545 | 1832 | 2142 | 1970 | 1525 | 1708 | 1641 | 1830 | 2151 | 1549 | 1641 | 1870 | 1780.11 | 1874 | 1687 | 0.899 | -0.153 | 0.23364322 | 0.44829938 | 0.0134 | 0.0088 | 0.0113 | 0.0113 | TRUE | 0.2021 |
| Sptbn1 | 20996 | 19465 | 22217 | 27995 | 19320 | 18652 | 22301 | 19440 | 22308 | 22007 | 20787 | 20418 | 21210.28 | 21350 | 21071 | 0.987 | -0.019 | 0.82129627 | 0.90875327 | 0.0031 | 0.0071 | 0.005 | 0.005 | TRUE | 0.075 |
| Sptbn2 | 1626 | 1642 | 1689 | 2223 | 1736 | 1734 | 1727 | 1640 | 1696 | 1748 | 1868 | 1898 | 1762.74 | 1688 | 1838 | 1.088 | 0.122 | 0.11029571 | 0.27877947 | 8.00E-04 | 0.0088 | 0.0036 | 0.0036 | TRUE | 0.023 |
| Sptbn4 | 1211 | 1779 | 1925 | 1411 | 1231 | 1395 | 1286 | 1777 | 1933 | 1109 | 1324 | 1527 | 1492.77 | 1665 | 1320 | 0.792 | -0.336 | 0.05405869 | 0.17281409 | 0.032 | 0.0091 | 0.0212 | 0.0212 | TRUE | 0.4773 |
| Sptlc1 | 2273 | 2310 | 2185 | 3171 | 2192 | 2028 | 2414 | 2307 | 2194 | 2493 | 2358 | 2220 | 2331.09 | 2305 | 2357 | 1.023 | 0.033 | 0.69286416 | 0.83159723 | 0.0024 | 0.0083 | 0.0046 | 0.0046 | TRUE | 0.0313 |
| Sptlc2 | 3057 | 3047 | 3226 | 4476 | 3053 | 3348 | 3247 | 3043 | 3239 | 3519 | 3285 | 3665 | 3332.97 | 3176 | 3490 | 1.099 | 0.136 | 0.07911369 | 0.22309694 | 0.0017 | 0.0079 | 0.004 | 0.004 | TRUE | 0.0316 |
| Sptlc3 | 1 | 0 | 0 | 0 | 0 | 0 | 1 | 0 | 0 | 0 | 0 | 0 | 0.18 | 0 | 0 | 0.49 | -1.029 | 0.80087 | NA | 0 | 18.6165 | 10 | 10 | TRUE | 0.4614 |
| Sptssa | 1164 | 1131 | 1052 | 1606 | 1118 | 1164 | 1236 | 1130 | 1056 | 1263 | 1203 | 1274 | 1193.64 | 1141 | 1247 | 1.093 | 0.128 | 0.17039291 | 0.36901257 | 0.0027 | 0.0097 | 0.0055 | 0.0055 | TRUE | 0.064 |
| Sptssb | 1 | 0 | 0 | 0 | 0 | 0 | 1 | 0 | 0 | 0 | 0 | 0 | 0.18 | 0 | 0 | 0.49 | -1.029 | 0.80087 | NA | 0 | 18.6165 | 10 | 10 | TRUE | 0.4614 |

| | | | | | | | | | | | | | | | | | | | | | | | | |
|---|---|---|---|---|---|---|---|---|---|---|---|---|---|---|---|---|---|---|---|---|---|---|---|---|
| Spty2d1 | 1419 | 1394 | 1352 | 1687 | 1205 | 1175 | 1507 | 1392 | 1358 | 1326 | 1297 | 1286 | 1360.98 | 1419 | 1303 | 0.919 | -0.122 | 0.13417329 | 0.31679168 | 9.00E-04 | 0.0093 | 0.0041 | 0.0041 | TRUE | 0.0354 |
| Spz1 | 24 | 11 | 10 | 57 | 34 | 15 | 25 | 11 | 10 | 45 | 37 | 16 | 24.05 | 15 | 33 | 2.122 | 1.086 | 0.05033913 | 0.16493186 | 0.2072 | 0.1439 | 0.1738 | 0.1738 | TRUE | 0.5828 |
| Sqle | 8274 | 7446 | 7257 | 8580 | 6414 | 6424 | 8788 | 7436 | 7287 | 6745 | 6901 | 7032 | 7364.95 | 7837 | 6893 | 0.879 | -0.185 | 0.04826253 | 0.16015076 | 0.0056 | 0.0074 | 0.0062 | 0.0062 | TRUE | 0.1375 |
| Sqrdl | 25 | 22 | 32 | 41 | 40 | 33 | 27 | 22 | 32 | 32 | 43 | 36 | 32.01 | 27 | 37 | 1.375 | 0.46 | 0.19113292 | 0.39617243 | 0 | 0.1098 | 0.0569 | 0.0569 | TRUE | 0.1872 |
| Sqstm1 | 13233 | 13008 | 13265 | 14468 | 10323 | 9973 | 14056 | 12991 | 13319 | 11374 | 11107 | 10917 | 12294 | 13455 | 11133 | 0.827 | -0.273 | 1.71E-05 | 0.00031187 | 9.00E-04 | 0.0072 | 0.0028 | 0.0028 | TRUE | 0.0186 |
| Sra1 | 1510 | 1935 | 1994 | 2535 | 1790 | 1876 | 1604 | 1933 | 2002 | 1993 | 1926 | 2054 | 1918.48 | 1846 | 1991 | 1.078 | 0.109 | 0.30617671 | 0.52666802 | 0.0064 | 0.0086 | 0.0076 | 0.0076 | TRUE | 0.1587 |
| Srbd1 | 3164 | 3605 | 3866 | 3979 | 3024 | 3144 | 3361 | 3600 | 3882 | 3128 | 3254 | 3442 | 3444.36 | 3614 | 3275 | 0.906 | -0.143 | 0.0990028 | 0.2595656 | 0.0033 | 0.0079 | 0.0051 | 0.0051 | TRUE | 0.051 |
| Src | 2667 | 3578 | 3821 | 4332 | 3409 | 3725 | 2833 | 3573 | 3837 | 3405 | 3668 | 4078 | 3565.64 | 3414 | 3717 | 1.088 | 0.122 | 0.35304346 | 0.57159736 | 0.014 | 0.0078 | 0.0122 | 0.0122 | TRUE | 0.2695 |
| Srcin1 | 628 | 845 | 864 | 678 | 595 | 595 | 667 | 844 | 868 | 533 | 640 | 651 | 700.5 | 793 | 608 | 0.766 | -0.385 | 0.00711067 | 0.04056038 | 0.0148 | 0.0116 | 0.0133 | 0.0133 | TRUE | 0.2274 |
| Srcrb4d | 340 | 315 | 328 | 366 | 298 | 254 | 361 | 315 | 329 | 288 | 321 | 278 | 315.25 | 335 | 296 | 0.882 | -0.182 | 0.16504764 | 0.36167622 | 0.0018 | 0.0174 | 0.0091 | 0.0091 | TRUE | 0.0602 |
| Srd5a1 | 520 | 286 | 273 | 491 | 316 | 217 | 552 | 286 | 274 | 386 | 340 | 238 | 345.93 | 371 | 321 | 0.868 | -0.205 | 0.47638182 | 0.67634297 | 0.1054 | 0.0164 | 0.0566 | 0.0566 | TRUE | 2.0984 |
| Srd5a2 | 2 | 4 | 0 | 2 | 2 | 0 | 2 | 4 | 0 | 2 | 2 | 0 | 1.64 | 2 | 1 | 0.615 | -0.7 | 0.68430013 | NA | 0 | 2.015 | 1.4989 | 1.4989 | TRUE | 0.1492 |
| Srd5a3 | 512 | 548 | 514 | 805 | 584 | 501 | 544 | 547 | 516 | 633 | 628 | 548 | 569.47 | 536 | 603 | 1.127 | 0.173 | 0.10939387 | 0.27744894 | 0.002 | 0.0127 | 0.0066 | 0.0066 | TRUE | 0.0723 |
| Srebf1 | 9064 | 9327 | 9473 | 9546 | 7109 | 7322 | 9628 | 9315 | 9512 | 7504 | 7649 | 8015 | 8603.78 | 9485 | 7723 | 0.814 | -0.297 | 1.20E-06 | 3.23E-05 | 5.00E-04 | 0.0073 | 0.0026 | 0.0026 | TRUE | 0.0134 |
| Srebf2 | 14900 | 17101 | 18279 | 18338 | 13363 | 15185 | 15826 | 17079 | 18354 | 14416 | 14378 | 16622 | 16112.57 | 17086 | 15139 | 0.886 | -0.175 | 0.06793139 | 0.20177545 | 0.0062 | 0.0071 | 0.0065 | 0.0065 | TRUE | 0.0898 |
| Srek1 | 2784 | 2586 | 2625 | 3780 | 2906 | 2738 | 2957 | 2583 | 2636 | 2972 | 3127 | 2997 | 2878.49 | 2725 | 3032 | 1.113 | 0.154 | 0.06538785 | 0.19707572 | 0.0026 | 0.008 | 0.0047 | 0.0047 | TRUE | 0.0671 |
| Srek1ip1 | 303 | 466 | 442 | 620 | 426 | 522 | 322 | 465 | 444 | 487 | 458 | 571 | 458.03 | 410 | 505 | 1.231 | 0.3 | 0.07711025 | 0.21943021 | 0.023 | 0.0141 | 0.0186 | 0.0186 | TRUE | 0.4074 |
| Srf | 2144 | 1987 | 1894 | 2925 | 2319 | 1800 | 2277 | 1984 | 1902 | 2299 | 2495 | 1970 | 2154.73 | 2054 | 2255 | 1.098 | 0.135 | 0.26127271 | 0.48067109 | 0.0108 | 0.0084 | 0.0099 | 0.0099 | TRUE | 0.1464 |
| Srfbp1 | 2605 | 2672 | 2714 | 3466 | 2434 | 2325 | 2767 | 2669 | 2725 | 2725 | 2619 | 2545 | 2674.88 | 2720 | 2630 | 0.967 | -0.048 | 0.47778304 | 0.67729589 | 4.00E-04 | 0.0081 | 0.003 | 0.003 | TRUE | 0.0125 |
| Srgap1 | 550 | 604 | 581 | 754 | 528 | 532 | 584 | 603 | 583 | 593 | 568 | 582 | 585.67 | 590 | 581 | 0.985 | -0.022 | 0.81391742 | 0.90478612 | 0 | 0.0125 | 0.0046 | 0.0046 | TRUE | 0.0044 |
| Srgap2 | 3536 | 3892 | 3957 | 4701 | 3423 | 3597 | 3756 | 3887 | 3973 | 3696 | 3683 | 3937 | 3822.01 | 3872 | 3772 | 0.974 | -0.038 | 0.57747599 | 0.75490686 | 7.00E-04 | 0.0078 | 0.0031 | 0.0031 | TRUE | 0.0178 |
| Srgap3 | 944 | 851 | 855 | 1183 | 844 | 895 | 1003 | 850 | 859 | 930 | 908 | 980 | 921.48 | 904 | 939 | 1.04 | 0.056 | 0.59084103 | 0.76478301 | 0.0039 | 0.0105 | 0.0067 | 0.0067 | TRUE | 0.1087 |
| Srgn | 32 | 47 | 31 | 61 | 30 | 31 | 34 | 47 | 31 | 48 | 32 | 34 | 37.7 | 37 | 38 | 1.026 | 0.037 | 0.91410494 | 0.95770315 | 0.0259 | 0.0943 | 0.0606 | 0.0606 | TRUE | 0.4385 |
| Sri | 955 | 1271 | 1225 | 1804 | 1285 | 1201 | 1014 | 1269 | 1230 | 1418 | 1383 | 1315 | 1271.53 | 1171 | 1372 | 1.171 | 0.228 | 0.0394871 | 0.13932675 | 0.0064 | 0.0095 | 0.008 | 0.008 | TRUE | 0.1637 |
| Srl | 4424 | 3557 | 3567 | 3912 | 2766 | 2732 | 4699 | 3552 | 3582 | 3075 | 2976 | 2991 | 3479.18 | 3944 | 3014 | 0.764 | -0.388 | 0.0016361 | 0.01311872 | 0.0135 | 0.0079 | 0.0106 | 0.0106 | TRUE | 0.3407 |
| Srm | 10329 | 9878 | 9841 | 9637 | 7314 | 6878 | 10971 | 9865 | 9881 | 7576 | 7869 | 7529 | 8948.7 | 10239 | 7658 | 0.748 | -0.419 | 1.40E-08 | 6.46E-07 | 0.002 | 0.0073 | 0.0038 | 0.0038 | TRUE | 0.0477 |
| Srms | 54 | 63 | 62 | 93 | 51 | 47 | 57 | 63 | 62 | 73 | 55 | 51 | 60.33 | 61 | 60 | 0.99 | -0.014 | 0.95687651 | 0.97770215 | 0 | 0.0615 | 0.034 | 0.034 | TRUE | 0.3675 |
| Srp14 | 3430 | 3464 | 3238 | 4417 | 3066 | 2891 | 3643 | 3460 | 3251 | 3472 | 3299 | 3165 | 3381.64 | 3451 | 3312 | 0.96 | -0.059 | 0.46468681 | 0.66783795 | 0.0023 | 0.0079 | 0.0044 | 0.0044 | TRUE | 0.0313 |
| Srp19 | 2251 | 1906 | 1899 | 2772 | 1947 | 1802 | 2391 | 1904 | 1907 | 2179 | 2095 | 1973 | 2074.64 | 2067 | 2082 | 1.008 | 0.011 | 0.92342505 | 0.96235363 | 0.0095 | 0.0085 | 0.0091 | 0.0091 | TRUE | 0.2268 |
| Srp54a | 336 | 296 | 377 | 436 | 286 | 272 | 357 | 296 | 379 | 343 | 308 | 298 | 329.88 | 344 | 316 | 0.921 | -0.118 | 0.41693976 | 0.62807995 | 0.0081 | 0.0169 | 0.0122 | 0.0122 | TRUE | 0.1721 |
| Srp54b | 0 | 0 | 0 | 0 | 0 | 0 | 0 | 0 | 0 | 0 | 0 | 0 | 0 | 0 | 0 | NA | NA | NA | NA | NA | NA | NA | NA | NA | NA |
| Srp54c | 0 | 0 | 0 | 0 | 0 | 0 | 0 | 0 | 0 | 0 | 0 | 0 | 0 | 0 | 0 | NA | NA | NA | NA | NA | NA | NA | NA | NA | NA |
| Srp68 | 11389 | 13399 | 13499 | 13520 | 10293 | 10736 | 12097 | 13382 | 13554 | 10628 | 11075 | 11752 | 12081.42 | 13011 | 11152 | 0.857 | -0.223 | 0.00553723 | 0.03370634 | 0.003 | 0.0072 | 0.0046 | 0.0046 | TRUE | 0.0461 |
| Srp72 | 12836 | 13804 | 14004 | 16486 | 12040 | 11562 | 13634 | 13786 | 14061 | 12960 | 12954 | 12656 | 13342.08 | 13827 | 12857 | 0.93 | -0.105 | 0.0500267 | 0.16424252 | 1.00E-04 | 0.0072 | 0.002 | 0.002 | TRUE | 0.0027 |
| Srp9 | 3294 | 3150 | 2919 | 3588 | 2570 | 2472 | 3499 | 3146 | 2931 | 2821 | 2765 | 2706 | 2977.91 | 3192 | 2764 | 0.866 | -0.207 | 0.02091047 | 0.0896683 | 0.0037 | 0.008 | 0.0055 | 0.0055 | TRUE | 0.0858 |
| Srpk1 | 7894 | 7471 | 7216 | 9372 | 7070 | 6329 | 8385 | 7461 | 7246 | 7368 | 7607 | 6928 | 7499.05 | 7697 | 7301 | 0.949 | -0.076 | 0.37811717 | 0.59334036 | 0.004 | 0.0073 | 0.0053 | 0.0053 | TRUE | 0.0745 |
| Srpk2 | 4021 | 4627 | 4825 | 4896 | 3496 | 3760 | 4271 | 4621 | 4845 | 3849 | 3761 | 4116 | 4243.85 | 4579 | 3909 | 0.854 | -0.229 | 0.00538413 | 0.0330191 | 0.0028 | 0.0077 | 0.0046 | 0.0046 | TRUE | 0.0421 |
| Srpk3 | 7 | 8 | 4 | 4 | 6 | 3 | 7 | 8 | 4 | 3 | 6 | 3 | 5.39 | 6 | 4 | 0.656 | -0.609 | 0.46450514 | 0.66771555 | 0 | 0.6184 | 0.3054 | 0.3054 | TRUE | 0.3181 |
| Srpr | 6442 | 7081 | 7152 | 8408 | 6237 | 6366 | 6843 | 7072 | 7181 | 6610 | 6711 | 6969 | 6897.45 | 7032 | 6763 | 0.962 | -0.056 | 0.35782709 | 0.57635303 | 5.00E-04 | 0.0074 | 0.0026 | 0.0026 | TRUE | 0.0086 |
| Srprb | 2792 | 3001 | 2872 | 3286 | 2443 | 2367 | 2966 | 2997 | 2884 | 2583 | 2629 | 2591 | 2774.88 | 2949 | 2601 | 0.882 | -0.181 | 0.00392134 | 0.02594267 | 0 | 0.0081 | 0.0025 | 0.0025 | TRUE | 0.0045 |
| Srpx | 1 | 3 | 4 | 4 | 1 | 0 | 1 | 3 | 4 | 3 | 1 | 0 | 2.05 | 3 | 1 | 0.549 | -0.866 | 0.54913516 | NA | 0 | 1.6146 | 0.986 | 0.986 | TRUE | 0.9235 |
| Srpx2 | 0 | 0 | 0 | 1 | 0 | 0 | 0 | 0 | 0 | 1 | 0 | 0 | 0.13 | 0 | 0 | 1.859 | 0.894 | 0.82650734 | NA | 0 | 25.1515 | 10 | 10 | TRUE | 0.2563 |
| Srr | 222 | 278 | 265 | 400 | 229 | 249 | 236 | 278 | 266 | 314 | 246 | 273 | 268.82 | 260 | 278 | 1.072 | 0.1 | 0.51391885 | 0.70667203 | 0.0078 | 0.0192 | 0.0131 | 0.0131 | TRUE | 0.1583 |
| Srrd | 495 | 286 | 308 | 572 | 360 | 283 | 526 | 286 | 309 | 450 | 387 | 310 | 377.91 | 374 | 382 | 1.025 | 0.036 | 0.88556571 | 0.94387096 | 0.0712 | 0.0156 | 0.0413 | 0.0413 | TRUE | 1.1167 |
| Srrm1 | 14458 | 14959 | 15933 | 20529 | 15683 | 15762 | 15357 | 14940 | 15998 | 16138 | 16874 | 17254 | 16093.55 | 15432 | 16755 | 1.086 | 0.119 | 0.06607767 | 0.19824557 | 0.0011 | 0.0071 | 0.0029 | 0.0029 | TRUE | 0.0128 |
| Srrm2 | 31832 | 28556 | 31774 | 37901 | 27741 | 27063 | 33811 | 28519 | 31904 | 29795 | 29848 | 29625 | 30583.65 | 31411 | 29756 | 0.947 | -0.078 | 0.34446438 | 0.5641524 | 0.0036 | 0.007 | 0.0049 | 0.0049 | TRUE | 0.0794 |
| Srrm3 | 57 | 40 | 43 | 39 | 31 | 30 | 61 | 40 | 43 | 31 | 33 | 33 | 40.09 | 48 | 32 | 0.674 | -0.569 | 0.07713169 | 0.21943021 | 0 | 0.0891 | 0.049 | 0.049 | TRUE | 0.4172 |
| Srrm4 | 39 | 33 | 30 | 56 | 39 | 39 | 41 | 33 | 30 | 44 | 42 | 43 | 38.86 | 35 | 43 | 1.234 | 0.304 | 0.3261783 | 0.54728555 | 0 | 0.0917 | 0.0429 | 0.0429 | TRUE | 0.1877 |
| Srrt | 9302 | 6032 | 5722 | 11506 | 8428 | 6489 | 9880 | 6024 | 5745 | 9045 | 9068 | 7103 | 7811.07 | 7216 | 8405 | 1.165 | 0.22 | 0.27420981 | 0.49471942 | 0.0502 | 0.0073 | 0.0291 | 0.0291 | TRUE | 1.2724 |
| Srsf1 | 20060 | 19132 | 17575 | 28709 | 22479 | 18994 | 21307 | 19107 | 17647 | 22569 | 24186 | 20792 | 20934.74 | 19354 | 22516 | 1.163 | 0.218 | 0.02940577 | 0.11339555 | 0.0073 | 0.0071 | 0.0072 | 0.0072 | TRUE | 0.0953 |
| Srsf10 | 6786 | 6407 | 6079 | 10049 | 7574 | 6511 | 7208 | 6399 | 6104 | 7900 | 8149 | 7127 | 7147.8 | 6570 | 7725 | 1.176 | 0.234 | 0.01508058 | 0.07108517 | 0.006 | 0.0074 | 0.0065 | 0.0065 | TRUE | 0.0879 |
| Srsf11 | 8441 | 8521 | 8856 | 10678 | 8209 | 7915 | 8966 | 8510 | 8892 | 8394 | 8832 | 8664 | 8709.83 | 8789 | 8630 | 0.982 | -0.026 | 0.66484159 | 0.81250574 | 5.00E-04 | 0.0073 | 0.0026 | 0.0026 | TRUE | 0.0095 |
| Srsf12 | 324 | 292 | 307 | 252 | 215 | 221 | 344 | 292 | 308 | 198 | 231 | 242 | 269.23 | 315 | 224 | 0.709 | -0.496 | 0.00078597 | 0.00738191 | 0.0054 | 0.0191 | 0.0119 | 0.0119 | TRUE | 0.1206 |
| Srsf2 | 19640 | 15964 | 13864 | 27544 | 20597 | 15641 | 20861 | 15943 | 13921 | 21653 | 22161 | 17122 | 18610.17 | 16908 | 20312 | 1.201 | 0.265 | 0.1018441 | 0.26480137 | 0.0315 | 0.0071 | 0.0188 | 0.0188 | TRUE | 0.5115 |
| Srsf3 | 13098 | 12035 | 10822 | 21140 | 15392 | 12442 | 13912 | 12019 | 10866 | 16619 | 16561 | 13620 | 13932.91 | 12266 | 15600 | 1.272 | 0.347 | 0.00485963 | 0.03068296 | 0.0138 | 0.0071 | 0.0109 | 0.0109 | TRUE | 0.1685 |
| Srsf4 | 4370 | 4190 | 4183 | 5664 | 4181 | 3996 | 4642 | 4185 | 4200 | 4453 | 4498 | 4374 | 4391.97 | 4342 | 4442 | 1.023 | 0.033 | 0.65478246 | 0.80595188 | 0.0014 | 0.0077 | 0.0037 | 0.0037 | TRUE | 0.0442 |
| Srsf5 | 8298 | 7748 | 6506 | 11633 | 8888 | 6779 | 8814 | 7738 | 6533 | 9145 | 9563 | 7421 | 8202.2 | 7695 | 8710 | 1.132 | 0.179 | 0.19898076 | 0.4062556 | 0.0193 | 0.0073 | 0.0138 | 0.0138 | TRUE | 0.2132 |
| Srsf6 | 15664 | 11775 | 10191 | 19221 | 14172 | 11306 | 16638 | 11760 | 10233 | 15110 | 15248 | 12376 | 13560.84 | 12877 | 14245 | 1.106 | 0.146 | 0.40465018 | 0.61732751 | 0.033 | 0.0071 | 0.022 | 0.022 | TRUE | 0.7981 |
| Srsf7 | 11170 | 12661 | 11838 | 16652 | 12761 | 12196 | 11865 | 12645 | 11887 | 13090 | 13730 | 13350 | 12761.13 | 12132 | 13390 | 1.104 | 0.142 | 0.02318373 | 0.0965516 | 8.00E-04 | 0.0072 | 0.0028 | 0.0028 | TRUE | 0.0167 |
| Srsf9 | 3433 | 3244 | 3005 | 4851 | 3537 | 3145 | 3646 | 3240 | 3017 | 3813 | 3806 | 3443 | 3494.23 | 3301 | 3687 | 1.117 | 0.16 | 0.10515832 | 0.27035167 | 0.0059 | 0.0078 | 0.0067 | 0.0067 | TRUE | 0.1016 |
| Srxn1 | 1174 | 1009 | 935 | 1678 | 1125 | 934 | 1247 | 1008 | 939 | 1319 | 1210 | 1022 | 1124.25 | 1065 | 1184 | 1.113 | 0.155 | 0.28698868 | 0.50821107 | 0.0179 | 0.0098 | 0.0143 | 0.0143 | TRUE | 0.2682 |
| Sry | 0 | 0 | 0 | 0 | 0 | 0 | 0 | 0 | 0 | 0 | 0 | 0 | 0 | 0 | 0 | NA | NA | NA | NA | NA | NA | NA | NA | NA | NA |
| Ss18 | 4554 | 4672 | 4705 | 6853 | 4951 | 4887 | 4837 | 4666 | 4724 | 5387 | 5327 | 5350 | 5048.55 | 4742 | 5355 | 1.129 | 0.175 | 0.00220701 | 0.01662318 | 0 | 0.0076 | 0.0022 | 0.0022 | TRUE | 0.0037 |
| Ss18l1 | 417 | 623 | 701 | 644 | 517 | 614 | 443 | 622 | 704 | 506 | 556 | 672 | 583.94 | 590 | 578 | 0.979 | -0.03 | 0.8737612 | 0.93773555 | 0.0361 | 0.0125 | 0.024 | 0.024 | TRUE | 0.5537 |
| Ssb | 14064 | 14060 | 13769 | 18357 | 13088 | 12925 | 14938 | 14042 | 13826 | 14431 | 14082 | 14148 | 14244.5 | 14269 | 14220 | 0.997 | -0.005 | 0.93856937 | 0.96917088 | 7.00E-04 | 0.0071 | 0.0027 | 0.0027 | TRUE | 0.0206 |
| Ssbp1 | 2472 | 2601 | 2490 | 3741 | 2569 | 2416 | 2626 | 2598 | 2500 | 2941 | 2764 | 2645 | 2678.87 | 2575 | 2783 | 1.082 | 0.113 | 0.13556187 | 0.31859362 | 0.0014 | 0.0081 | 0.0038 | 0.0038 | TRUE | 0.0304 |
| Ssbp2 | 56 | 92 | 73 | 152 | 112 | 113 | 59 | 92 | 73 | 119 | 121 | 124 | 98.06 | 75 | 121 | 1.616 | 0.692 | 0.00203454 | 0.01559978 | 0.0062 | 0.0405 | 0.0254 | 0.0254 | TRUE | 0.362 |
| Ssbp3 | 3102 | 3169 | 3114 | 3895 | 2895 | 2803 | 3295 | 3165 | 3127 | 3062 | 3115 | 3068 | 3138.61 | 3196 | 3082 | 0.964 | -0.052 | 0.40928558 | 0.62130006 | 0 | 0.008 | 0.0026 | 0.0026 | TRUE | 0.0089 |
| Ssbp4 | 2581 | 3256 | 3378 | 4801 | 3716 | 3635 | 2741 | 3252 | 3392 | 3774 | 3998 | 3979 | 3522.76 | 3128 | 3917 | 1.252 | 0.324 | 0.0012371 | 0.01048441 | 0.0061 | 0.0078 | 0.007 | 0.007 | TRUE | 0.1419 |

| | | | | | | | | | | | | | | | | | | | | | | | | | |
|---|---|---|---|---|---|---|---|---|---|---|---|---|---|---|---|---|---|---|---|---|---|---|---|---|---|
| Ssc5d | 53 | 76 | 61 | 58 | 56 | 44 | 56 | 76 | 61 | 46 | 60 | 48 | 57.91 | 64 | 51 | 0.792 | -0.336 | 0.2186701 | 0.43078502 | 0 | 0.0638 | 0.0362 | 0.0362 | TRUE | 0.2126 |
| Ssfa2 | 1695 | 2155 | 2154 | 2496 | 1803 | 2081 | 1800 | 2152 | 2163 | 1962 | 1940 | 2278 | 2049.25 | 2038 | 2060 | 1.01 | 0.015 | 0.8973584 | 0.95019381 | 0.0089 | 0.0085 | 0.0087 | 0.0087 | TRUE | 0.1259 |
| Ssh1 | 942 | 1033 | 1016 | 1409 | 1004 | 957 | 1001 | 1032 | 1020 | 1108 | 1080 | 1048 | 1047.98 | 1018 | 1079 | 1.061 | 0.085 | 0.28503397 | 0.50627409 | 0 | 0.01 | 0.0036 | 0.0036 | TRUE | 0.0075 |
| Ssh2 | 3026 | 2372 | 2366 | 4234 | 2780 | 2087 | 3214 | 2369 | 2376 | 3328 | 2991 | 2285 | 2760.48 | 2653 | 2868 | 1.081 | 0.113 | 0.50588919 | 0.70029387 | 0.0335 | 0.0081 | 0.0204 | 0.0204 | TRUE | 0.4151 |
| Ssh3 | 248 | 315 | 263 | 229 | 226 | 235 | 263 | 315 | 264 | 180 | 243 | 257 | 253.75 | 281 | 227 | 0.805 | -0.314 | 0.08241354 | 0.22928761 | 0.0191 | 0.0199 | 0.0195 | 0.0195 | TRUE | 0.3837 |
| Ssna1 | 1226 | 1307 | 1277 | 1758 | 1252 | 1215 | 1302 | 1305 | 1282 | 1382 | 1347 | 1330 | 1324.81 | 1296 | 1353 | 1.044 | 0.062 | 0.39511518 | 0.60856999 | 0 | 0.0094 | 0.0031 | 0.0031 | TRUE | 0.0044 |
| Sspn | 8 | 13 | 12 | 5 | 7 | 15 | 8 | 13 | 12 | 4 | 8 | 16 | 10.24 | 11 | 9 | 0.815 | -0.296 | 0.66948136 | 0.8154066 | 0.1608 | 0.3288 | 0.2471 | 0.2471 | TRUE | 0.8173 |
| Sspo | 26 | 18 | 13 | 9 | 12 | 6 | 28 | 18 | 13 | 7 | 13 | 7 | 14.2 | 20 | 9 | 0.45 | -1.152 | 0.04596375 | 0.15519235 | 0.0709 | 0.2389 | 0.1582 | 0.1582 | TRUE | 0.2769 |
| Ssr1 | 9665 | 10734 | 10475 | 13088 | 9274 | 9734 | 10266 | 10720 | 10518 | 10289 | 9978 | 10655 | 10404.42 | 10501 | 10307 | 0.982 | -0.027 | 0.65996189 | 0.80852252 | 6.00E-04 | 0.0072 | 0.0026 | 0.0026 | TRUE | 0.0106 |
| Ssr2 | 6269 | 7125 | 6955 | 8319 | 5683 | 6054 | 6659 | 7116 | 6984 | 6540 | 6115 | 6627 | 6673.25 | 6920 | 6427 | 0.929 | -0.106 | 0.12471347 | 0.30163791 | 0.0013 | 0.0074 | 0.0033 | 0.0033 | TRUE | 0.022 |
| Ssr3 | 14538 | 15979 | 15664 | 17983 | 12731 | 13295 | 15442 | 15958 | 15728 | 14137 | 13698 | 14553 | 14919.45 | 15709 | 14129 | 0.899 | -0.153 | 0.00878265 | 0.04774903 | 5.00E-04 | 0.0071 | 0.0024 | 0.0024 | TRUE | 0.0087 |
| Ssr4 | 1340 | 1845 | 1655 | 1916 | 1426 | 1476 | 1423 | 1843 | 1662 | 1506 | 1534 | 1616 | 1597.32 | 1643 | 1552 | 0.945 | -0.082 | 0.46732792 | 0.66963926 | 0.0078 | 0.009 | 0.0086 | 0.0086 | TRUE | 0.1639 |
| Ssrp1 | 15287 | 14303 | 14768 | 21552 | 15752 | 15152 | 16237 | 14285 | 14829 | 16942 | 16948 | 16586 | 15971.26 | 15117 | 16825 | 1.113 | 0.155 | 0.03580087 | 0.12989448 | 0.0022 | 0.0071 | 0.0038 | 0.0038 | TRUE | 0.0514 |
| Sssca1 | 879 | 620 | 561 | 1341 | 859 | 672 | 934 | 619 | 563 | 1054 | 924 | 736 | 805.03 | 705 | 905 | 1.284 | 0.36 | 0.08350262 | 0.23154066 | 0.0538 | 0.011 | 0.03 | 0.03 | TRUE | 0.7341 |
| Sst | 0 | 2 | 0 | 0 | 1 | 1 | 0 | 2 | 0 | 0 | 1 | 1 | 0.69 | 1 | 1 | 1.047 | 0.067 | 0.97977773 | NA | 0 | 4.7494 | 3.5066 | 3.5066 | TRUE | 0.9738 |
| Sstr1 | 0 | 0 | 0 | 0 | 0 | 0 | 0 | 0 | 0 | 0 | 0 | 0 | 0 | 0 | 0 | NA | NA | NA | NA | NA | NA | NA | NA | NA | NA |
| Sstr2 | 50 | 62 | 65 | 31 | 38 | 31 | 53 | 62 | 65 | 24 | 41 | 34 | 46.58 | 60 | 33 | 0.544 | -0.878 | 0.00501334 | 0.03138958 | 0.0093 | 0.0776 | 0.0472 | 0.0472 | TRUE | 0.4484 |
| Sstr3 | 1 | 0 | 1 | 0 | 1 | 0 | 1 | 0 | 1 | 0 | 1 | 0 | 0.52 | 1 | 0 | 0.509 | -0.975 | 0.72714826 | NA | 0 | 6.2976 | 3.8857 | 3.8857 | TRUE | 0.5204 |
| Sstr4 | 0 | 2 | 0 | 8 | 0 | 2 | 0 | 2 | 0 | 6 | 0 | 2 | 1.75 | 1 | 3 | 4.278 | 2.097 | 0.30758354 | NA | 2.6626 | 1.8938 | 2.113 | 2.113 | TRUE | 0.3606 |
| Sstr5 | 27 | 30 | 25 | 30 | 15 | 20 | 29 | 30 | 25 | 24 | 16 | 22 | 24.23 | 28 | 21 | 0.74 | -0.434 | 0.26586954 | 0.48550358 | 0 | 0.1429 | 0.0675 | 0.0675 | TRUE | 0.1802 |
| Ssty1 | 0 | 0 | 0 | 0 | 0 | 0 | 0 | 0 | 0 | 0 | 0 | 0 | 0 | 0 | 0 | NA | NA | NA | NA | NA | NA | NA | NA | NA | NA |
| Ssty2 | 0 | 0 | 0 | 0 | 0 | 0 | 0 | 0 | 0 | 0 | 0 | 0 | 0 | 0 | 0 | NA | NA | NA | NA | NA | NA | NA | NA | NA | NA |
| Ssu2 | 17 | 29 | 32 | 26 | 6 | 20 | 18 | 29 | 32 | 20 | 6 | 22 | 21.32 | 26 | 16 | 0.62 | -0.69 | 0.19334571 | 0.39893361 | 0.1433 | 0.1614 | 0.1536 | 0.1536 | TRUE | 1.2332 |
| Ssu72 | 3144 | 3480 | 3345 | 5169 | 3484 | 3380 | 3339 | 3476 | 3359 | 4063 | 3749 | 3700 | 3614.28 | 3391 | 3837 | 1.132 | 0.179 | 0.01351199 | 0.06549202 | 0.0011 | 0.0078 | 0.0035 | 0.0035 | TRUE | 0.0328 |
| Ssx2ip | 1911 | 2065 | 1960 | 2532 | 1907 | 1937 | 2030 | 2062 | 1968 | 1990 | 2052 | 2120 | 2037.14 | 2020 | 2054 | 1.016 | 0.024 | 0.73974451 | 0.86066702 | 3.00E-04 | 0.0085 | 0.0031 | 0.0031 | TRUE | 0.0096 |
| Ssx9 | 0 | 0 | 0 | 0 | 0 | 0 | 0 | 0 | 0 | 0 | 0 | 0 | 0 | 0 | 0 | NA | NA | NA | NA | NA | NA | NA | NA | NA | NA |
| Ssxb1 | 0 | 0 | 0 | 0 | 0 | 0 | 0 | 0 | 0 | 0 | 0 | 0 | 0 | 0 | 0 | NA | NA | NA | NA | NA | NA | NA | NA | NA | NA |
| Ssxb10 | 0 | 0 | 0 | 0 | 0 | 0 | 0 | 0 | 0 | 0 | 0 | 0 | 0 | 0 | 0 | NA | NA | NA | NA | NA | NA | NA | NA | NA | NA |
| Ssxb2 | 0 | 0 | 0 | 0 | 0 | 0 | 0 | 0 | 0 | 0 | 0 | 0 | 0 | 0 | 0 | NA | NA | NA | NA | NA | NA | NA | NA | NA | NA |
| Ssxb3 | 0 | 0 | 0 | 0 | 0 | 0 | 0 | 0 | 0 | 0 | 0 | 0 | 0 | 0 | 0 | NA | NA | NA | NA | NA | NA | NA | NA | NA | NA |
| Ssxb5 | 0 | 0 | 0 | 0 | 0 | 0 | 0 | 0 | 0 | 0 | 0 | 0 | 0 | 0 | 0 | NA | NA | NA | NA | NA | NA | NA | NA | NA | NA |
| Ssxb6 | 0 | 0 | 0 | 0 | 0 | 1 | 0 | 0 | 0 | 0 | 0 | 1 | 0.18 | 0 | 0 | 1.859 | 0.894 | 0.82650734 | NA | 0 | 18.0643 | 10 | 10 | TRUE | 0.5383 |
| Ssxb8 | 0 | 0 | 0 | 0 | 0 | 0 | 0 | 0 | 0 | 0 | 0 | 0 | 0 | 0 | 0 | NA | NA | NA | NA | NA | NA | NA | NA | NA | NA |
| Ssxb9 | 0 | 0 | 0 | 1 | 0 | 0 | 0 | 0 | 0 | 1 | 0 | 0 | 0.13 | 0 | 0 | 1.859 | 0.894 | 0.82650734 | NA | 0 | 25.1515 | 10 | 10 | TRUE | 0.2563 |
| St13 | 15028 | 17725 | 17688 | 21319 | 15615 | 15997 | 15962 | 17702 | 17761 | 16759 | 16801 | 17511 | 17082.74 | 17142 | 17024 | 0.993 | -0.01 | 0.8900411 | 0.94632117 | 0.0016 | 0.0071 | 0.0037 | 0.0037 | TRUE | 0.0443 |
| St14 | 2129 | 3697 | 3776 | 4012 | 3130 | 4035 | 2261 | 3692 | 3792 | 3154 | 3368 | 4417 | 3447.28 | 3248 | 3646 | 1.122 | 0.166 | 0.41033474 | 0.62180353 | 0.055 | 0.0079 | 0.0291 | 0.0291 | TRUE | 0.858 |
| St18 | 0 | 0 | 0 | 1 | 1 | 0 | 0 | 0 | 0 | 1 | 1 | 0 | 0.31 | 0 | 1 | 3.32 | 1.731 | 0.65383052 | NA | 0 | 10.6224 | 8.9541 | 8.9541 | TRUE | 0.202 |
| St3gal1 | 330 | 368 | 413 | 360 | 302 | 279 | 351 | 368 | 415 | 283 | 325 | 305 | 341.01 | 378 | 304 | 0.805 | -0.314 | 0.01709395 | 0.07762908 | 0.0035 | 0.0166 | 0.0095 | 0.0095 | TRUE | 0.085 |
| St3gal2 | 1258 | 1381 | 1442 | 1941 | 1301 | 1329 | 1336 | 1379 | 1448 | 1526 | 1400 | 1455 | 1423.97 | 1388 | 1460 | 1.053 | 0.074 | 0.36588923 | 0.58332204 | 0.0011 | 0.0092 | 0.0041 | 0.0041 | TRUE | 0.0193 |
| St3gal3 | 440 | 412 | 475 | 587 | 489 | 438 | 467 | 411 | 477 | 461 | 526 | 479 | 470.47 | 452 | 489 | 1.081 | 0.112 | 0.34431209 | 0.56408957 | 0.0033 | 0.0139 | 0.0081 | 0.0081 | TRUE | 0.0716 |
| St3gal4 | 503 | 955 | 899 | 674 | 589 | 660 | 534 | 954 | 903 | 530 | 634 | 722 | 712.8 | 797 | 629 | 0.788 | -0.344 | 0.11496709 | 0.28627423 | 0.0575 | 0.0115 | 0.0328 | 0.0328 | TRUE | 0.9835 |
| St3gal5 | 591 | 692 | 652 | 915 | 618 | 659 | 628 | 691 | 655 | 719 | 665 | 721 | 679.86 | 658 | 702 | 1.067 | 0.094 | 0.33465178 | 0.55489149 | 0 | 0.0117 | 0.0053 | 0.0053 | TRUE | 0.0245 |
| St3gal6 | 43 | 48 | 46 | 96 | 63 | 74 | 46 | 48 | 46 | 75 | 68 | 81 | 60.68 | 47 | 75 | 1.604 | 0.682 | 0.0061116 | 0.03647076 | 0 | 0.0612 | 0.0271 | 0.0271 | TRUE | 0.0564 |
| St5 | 1173 | 1220 | 1193 | 1899 | 1401 | 1332 | 1246 | 1218 | 1198 | 1493 | 1507 | 1458 | 1353.43 | 1221 | 1486 | 1.218 | 0.284 | 0.00010841 | 0.0014689 | 0 | 0.0093 | 0.0031 | 0.0031 | TRUE | 0.0039 |
| St6gal1 | 774 | 719 | 714 | 981 | 741 | 730 | 822 | 718 | 717 | 771 | 797 | 799 | 770.78 | 752 | 789 | 1.049 | 0.069 | 0.48881446 | 0.68529614 | 0.0019 | 0.0112 | 0.0058 | 0.0058 | TRUE | 0.0773 |
| St6gal2 | 16 | 9 | 4 | 16 | 13 | 7 | 17 | 9 | 4 | 13 | 14 | 8 | 10.7 | 10 | 12 | 1.149 | 0.201 | 0.76879137 | 0.87928573 | 0.1586 | 0.3147 | 0.2416 | 0.2416 | TRUE | 0.4031 |
| St6galnac1 | 1 | 0 | 2 | 4 | 1 | 3 | 1 | 0 | 2 | 3 | 1 | 3 | 1.76 | 1 | 2 | 2.464 | 1.301 | 0.393501 | NA | 0 | 1.8761 | 0.9814 | 0.9814 | TRUE | 0.3771 |
| St6galnac2 | 141 | 235 | 252 | 508 | 474 | 357 | 150 | 235 | 253 | 399 | 510 | 391 | 322.94 | 213 | 433 | 2.037 | 1.026 | 1.50E-06 | 3.89E-05 | 0.0431 | 0.0171 | 0.0292 | 0.0292 | TRUE | 0.7211 |
| St6galnac3 | 363 | 438 | 429 | 318 | 242 | 306 | 386 | 437 | 431 | 250 | 260 | 335 | 349.85 | 418 | 282 | 0.672 | -0.573 | 0.00017707 | 0.00219907 | 0.0114 | 0.0163 | 0.0138 | 0.0138 | TRUE | 0.3002 |
| St6galnac4 | 675 | 699 | 711 | 755 | 548 | 569 | 717 | 698 | 714 | 594 | 590 | 623 | 655.83 | 710 | 602 | 0.848 | -0.238 | 0.00867246 | 0.04724767 | 0 | 0.0119 | 0.0044 | 0.0044 | TRUE | 0.0104 |
| St6galnac5 | 3 | 5 | 3 | 4 | 5 | 8 | 3 | 5 | 3 | 3 | 5 | 9 | 4.75 | 4 | 6 | 1.517 | 0.601 | 0.50110886 | 0.69669611 | 0 | 0.7011 | 0.3519 | 0.3519 | TRUE | 0.2378 |
| St6galnac6 | 2300 | 2808 | 2908 | 1662 | 1329 | 1757 | 2443 | 2804 | 2920 | 1307 | 1430 | 1923 | 2137.85 | 2722 | 1553 | 0.57 | -0.811 | 1.35E-07 | 4.62E-06 | 0.0255 | 0.0084 | 0.0165 | 0.0165 | TRUE | 0.5221 |
| St7 | 136 | 178 | 207 | 222 | 163 | 171 | 144 | 178 | 208 | 175 | 175 | 187 | 177.86 | 177 | 179 | 1.012 | 0.017 | 0.92632521 | 0.9637204 | 0.011 | 0.0254 | 0.0182 | 0.0182 | TRUE | 0.2101 |
| St7l | 209 | 332 | 366 | 349 | 265 | 335 | 222 | 332 | 368 | 274 | 285 | 367 | 307.88 | 307 | 309 | 1.004 | 0.005 | 0.97944853 | 0.98867206 | 0.0424 | 0.0176 | 0.0296 | 0.0296 | TRUE | 0.6584 |
| St8sia1 | 441 | 435 | 388 | 656 | 496 | 507 | 468 | 434 | 390 | 516 | 534 | 555 | 482.8 | 431 | 535 | 1.241 | 0.311 | 0.00723281 | 0.0411529 | 0.0024 | 0.0137 | 0.0076 | 0.0076 | TRUE | 0.0812 |
| St8sia2 | 0 | 2 | 2 | 10 | 5 | 4 | 0 | 2 | 2 | 8 | 5 | 4 | 3.6 | 1 | 6 | 4.386 | 2.133 | 0.06454419 | NA | 0 | 0.921 | 0.499 | 0.499 | TRUE | 0.4329 |
| St8sia3 | 20 | 26 | 13 | 26 | 26 | 13 | 21 | 26 | 13 | 20 | 28 | 14 | 20.48 | 20 | 21 | 1.04 | 0.056 | 0.90675559 | 0.95352883 | 0.0575 | 0.1677 | 0.1161 | 0.1161 | TRUE | 0.2462 |
| St8sia4 | 13 | 11 | 4 | 17 | 13 | 8 | 14 | 11 | 4 | 13 | 14 | 9 | 10.82 | 10 | 12 | 1.261 | 0.335 | 0.60248661 | 0.77387557 | 0.0668 | 0.3114 | 0.2042 | 0.2042 | TRUE | 0.8457 |
| St8sia5 | 174 | 123 | 112 | 162 | 106 | 145 | 185 | 123 | 112 | 127 | 114 | 159 | 136.71 | 140 | 133 | 0.952 | -0.07 | 0.7767055 | 0.88398031 | 0.0429 | 0.031 | 0.0368 | 0.0368 | TRUE | 0.802 |
| St8sia6 | 490 | 349 | 337 | 740 | 500 | 517 | 520 | 349 | 338 | 582 | 538 | 566 | 482.17 | 402 | 562 | 1.397 | 0.483 | 0.00673086 | 0.03902025 | 0.0279 | 0.0137 | 0.0207 | 0.0207 | TRUE | 0.7537 |
| Stab1 | 26 | 28 | 19 | 49 | 29 | 23 | 28 | 28 | 19 | 39 | 31 | 25 | 28.26 | 25 | 32 | 1.281 | 0.357 | 0.3505004 | 0.56931295 | 0 | 0.1235 | 0.0697 | 0.0697 | TRUE | 0.227 |
| Stab2 | 17 | 22 | 30 | 73 | 71 | 73 | 18 | 22 | 30 | 57 | 76 | 80 | 47.31 | 23 | 71 | 3.027 | 1.598 | 1.47E-06 | 3.84E-05 | 0.0204 | 0.0765 | 0.0507 | 0.0507 | TRUE | 0.3753 |
| Stac | 0 | 1 | 2 | 3 | 2 | 4 | 0 | 1 | 2 | 2 | 2 | 4 | 1.98 | 1 | 3 | 2.907 | 1.539 | 0.28722988 | NA | 0 | 1.6686 | 0.8371 | 0.8371 | TRUE | 0.358 |
| Stac2 | 2162 | 3123 | 3040 | 2144 | 1657 | 2141 | 2296 | 3119 | 3052 | 1685 | 1783 | 2344 | 2379.97 | 2822 | 1937 | 0.686 | -0.544 | 0.00085414 | 0.00787808 | 0.0293 | 0.0083 | 0.0187 | 0.0187 | TRUE | 0.4061 |
| Stac3 | 15 | 15 | 7 | 21 | 8 | 14 | 16 | 15 | 7 | 17 | 9 | 15 | 13.06 | 13 | 14 | 1.075 | 0.104 | 0.85769012 | 0.92990093 | 0.0539 | 0.2591 | 0.1652 | 0.1652 | TRUE | 0.6239 |
| Stag1 | 4924 | 5327 | 5415 | 7641 | 5294 | 5265 | 5230 | 5320 | 5437 | 6007 | 5696 | 5763 | 5575.61 | 5329 | 5822 | 1.093 | 0.128 | 0.03728719 | 0.13387059 | 4.00E-04 | 0.0075 | 0.0025 | 0.0025 | TRUE | 0.0095 |
| Stag2 | 3689 | 4459 | 5136 | 5585 | 3908 | 4162 | 3918 | 4453 | 5157 | 4390 | 4205 | 4556 | 4446.65 | 4509 | 4384 | 0.972 | -0.041 | 0.71732967 | 0.8467669 | 0.01 | 0.0076 | 0.009 | 0.009 | TRUE | 0.1922 |
| Stag3 | 2439 | 3730 | 4046 | 3637 | 3135 | 3243 | 2591 | 3725 | 4063 | 2859 | 3373 | 3550 | 3360.1 | 3460 | 3261 | 0.942 | -0.086 | 0.61000442 | 0.77891178 | 0.0329 | 0.0079 | 0.0201 | 0.0201 | TRUE | 0.5864 |
| Stam | 3011 | 3136 | 3185 | 4218 | 3032 | 3428 | 3198 | 3132 | 3198 | 3316 | 3262 | 3752 | 3309.81 | 3176 | 3443 | 1.084 | 0.116 | 0.15995153 | 0.35445491 | 0.0027 | 0.0079 | 0.0046 | 0.0046 | TRUE | 0.0743 |

| | | | | | | | | | | | | | | | | | | | | | | | | | |
|---|---|---|---|---|---|---|---|---|---|---|---|---|---|---|---|---|---|---|---|---|---|---|---|---|---|
| Stam2 | 3102 | 3074 | 3104 | 4136 | 2895 | 2951 | 3295 | 3070 | 3117 | 3251 | 3115 | 3230 | 3179.7 | 3161 | 3199 | 1.012 | 0.018 | 0.79534091 | 0.8946193 | 6.00E-04 | 0.0079 | 0.003 | 0.003 | TRUE | 0.0167 |
| Stambp | 131 | 178 | 217 | 240 | 209 | 198 | 139 | 178 | 218 | 189 | 225 | 217 | 194.18 | 178 | 210 | 1.176 | 0.233 | 0.24174602 | 0.45849186 | 0.0229 | 0.0239 | 0.0234 | 0.0234 | TRUE | 0.2277 |
| Stambpl1 | 467 | 569 | 617 | 394 | 324 | 331 | 496 | 568 | 620 | 310 | 349 | 362 | 450.75 | 561 | 340 | 0.605 | -0.726 | 6.24E-08 | 2.40E-06 | 0.0076 | 0.0142 | 0.0106 | 0.0106 | TRUE | 0.1201 |
| Stap1 | 0 | 2 | 2 | 1 | 0 | 2 | 0 | 2 | 2 | 1 | 0 | 2 | 1.16 | 1 | 1 | 0.731 | -0.452 | 0.81362651 | NA | 0 | 2.8384 | 1.7737 | 1.7737 | TRUE | 0.4402 |
| Stap2 | 327 | 276 | 258 | 281 | 220 | 209 | 347 | 276 | 259 | 221 | 237 | 229 | 261.4 | 294 | 229 | 0.778 | -0.362 | 0.02263612 | 0.0949083 | 0.0097 | 0.0195 | 0.0143 | 0.0143 | TRUE | 0.281 |
| Star | 93 | 105 | 118 | 113 | 101 | 72 | 99 | 105 | 118 | 89 | 109 | 79 | 99.74 | 107 | 92 | 0.857 | -0.223 | 0.30172826 | 0.52237237 | 0.0057 | 0.0399 | 0.0236 | 0.0236 | TRUE | 0.2235 |
| Stard10 | 350 | 351 | 368 | 524 | 380 | 372 | 372 | 351 | 370 | 412 | 409 | 407 | 386.64 | 364 | 409 | 1.125 | 0.17 | 0.11681387 | 0.28895749 | 0 | 0.0154 | 0.0059 | 0.0059 | TRUE | 0.012 |
| Stard13 | 316 | 276 | 306 | 352 | 238 | 276 | 336 | 276 | 307 | 277 | 256 | 302 | 292.24 | 306 | 278 | 0.909 | -0.137 | 0.33205307 | 0.55255612 | 0.0045 | 0.0182 | 0.011 | 0.011 | TRUE | 0.0864 |
| Stard3 | 1283 | 1213 | 1095 | 1424 | 1019 | 1078 | 1363 | 1211 | 1099 | 1119 | 1096 | 1180 | 1178.26 | 1224 | 1132 | 0.924 | -0.113 | 0.28817147 | 0.50913413 | 0.0053 | 0.0097 | 0.0074 | 0.0074 | TRUE | 0.1164 |
| Stard3nl | 2329 | 2365 | 2160 | 2991 | 2156 | 2132 | 2474 | 2362 | 2169 | 2351 | 2320 | 2334 | 2334.91 | 2335 | 2335 | 1 | 0 | 0.99714987 | 0.99828918 | 0.0018 | 0.0083 | 0.0042 | 0.0042 | TRUE | 0.0469 |
| Stard4 | 3387 | 4045 | 4218 | 4979 | 3738 | 4030 | 3598 | 4040 | 4235 | 3914 | 4022 | 4411 | 4036.69 | 3958 | 4116 | 1.04 | 0.056 | 0.55245624 | 0.7362971 | 0.0045 | 0.0077 | 0.0062 | 0.0062 | TRUE | 0.0769 |
| Stard5 | 733 | 821 | 791 | 930 | 658 | 646 | 779 | 820 | 794 | 731 | 708 | 707 | 756.5 | 798 | 715 | 0.897 | -0.156 | 0.07187445 | 0.20926783 | 0 | 0.0113 | 0.0041 | 0.0041 | TRUE | 0.0072 |
| Stard6 | 12 | 4 | 7 | 17 | 6 | 5 | 13 | 4 | 7 | 13 | 6 | 5 | 8.18 | 8 | 8 | 1.087 | 0.12 | 0.87316763 | 0.93761484 | 0.1632 | 0.4098 | 0.287 | 0.287 | TRUE | 0.525 |
| Stard7 | 4488 | 4988 | 4853 | 5310 | 3911 | 3904 | 4767 | 4982 | 4873 | 4174 | 4208 | 4274 | 4546.23 | 4874 | 4219 | 0.865 | -0.208 | 0.00046076 | 0.00487722 | 0 | 0.0076 | 0.0023 | 0.0023 | TRUE | 0.0046 |
| Stard8 | 577 | 549 | 580 | 495 | 379 | 427 | 613 | 548 | 582 | 389 | 408 | 467 | 501.31 | 581 | 421 | 0.724 | -0.466 | 9.40E-05 | 0.00131109 | 0.0039 | 0.0135 | 0.0082 | 0.0082 | TRUE | 0.1034 |
| Stat1 | 762 | 867 | 904 | 920 | 663 | 656 | 809 | 866 | 908 | 723 | 713 | 718 | 789.61 | 861 | 718 | 0.834 | -0.262 | 0.0043364 | 0.02805322 | 0 | 0.0111 | 0.0048 | 0.0048 | TRUE | 0.0324 |
| Stat2 | 4220 | 4219 | 4173 | 4326 | 3420 | 3496 | 4482 | 4214 | 4190 | 3401 | 3680 | 3827 | 3965.58 | 4295 | 3636 | 0.846 | -0.241 | 0.00216806 | 0.01641606 | 0.0021 | 0.0077 | 0.0042 | 0.0042 | TRUE | 0.0395 |
| Stat3 | 8135 | 9189 | 9777 | 8270 | 6614 | 7335 | 8641 | 9177 | 9817 | 6501 | 7116 | 8029 | 8213.65 | 9212 | 7215 | 0.783 | -0.353 | 0.0005753 | 0.00579789 | 0.0075 | 0.0073 | 0.0074 | 0.0074 | TRUE | 0.1187 |
| Stat4 | 728 | 590 | 615 | 594 | 413 | 458 | 773 | 589 | 618 | 467 | 444 | 501 | 565.45 | 660 | 471 | 0.714 | -0.487 | 0.00041099 | 0.00442462 | 0.0111 | 0.0127 | 0.0119 | 0.0119 | TRUE | 0.2644 |
| Stat5a | 415 | 739 | 763 | 670 | 567 | 596 | 441 | 738 | 766 | 527 | 610 | 652 | 622.36 | 648 | 596 | 0.919 | -0.122 | 0.5554719 | 0.73847101 | 0.0481 | 0.0122 | 0.0292 | 0.0292 | TRUE | 0.9203 |
| Stat5b | 1058 | 1638 | 1704 | 1580 | 1244 | 1369 | 1124 | 1636 | 1711 | 1242 | 1338 | 1499 | 1424.96 | 1490 | 1360 | 0.912 | -0.133 | 0.42142196 | 0.63151231 | 0.0289 | 0.0092 | 0.019 | 0.019 | TRUE | 0.556 |
| Stat6 | 1740 | 2033 | 2204 | 2265 | 1748 | 1811 | 1848 | 2030 | 2213 | 1781 | 1881 | 1982 | 1955.89 | 2030 | 1881 | 0.926 | -0.111 | 0.25862091 | 0.47748367 | 0.0047 | 0.0086 | 0.0064 | 0.0064 | TRUE | 0.0748 |
| Stau1 | 3175 | 3379 | 3446 | 4831 | 3429 | 3327 | 3372 | 3375 | 3460 | 3798 | 3689 | 3642 | 3556.04 | 3402 | 3710 | 1.091 | 0.125 | 0.04238839 | 0.14686943 | 0 | 0.0078 | 0.0025 | 0.0025 | TRUE | 0.0054 |
| Stau2 | 340 | 456 | 414 | 616 | 455 | 435 | 361 | 455 | 416 | 484 | 490 | 476 | 447.04 | 411 | 483 | 1.176 | 0.234 | 0.05875686 | 0.18332606 | 0.004 | 0.0143 | 0.0088 | 0.0088 | TRUE | 0.1272 |
| Stbd1 | 132 | 206 | 204 | 167 | 121 | 147 | 140 | 206 | 205 | 131 | 130 | 161 | 162.19 | 184 | 141 | 0.765 | -0.386 | 0.06622482 | 0.19857374 | 0.0241 | 0.0272 | 0.0256 | 0.0256 | TRUE | 0.4535 |
| Stc1 | 154 | 354 | 354 | 360 | 330 | 354 | 164 | 354 | 355 | 283 | 355 | 388 | 316.36 | 291 | 342 | 1.174 | 0.231 | 0.40516313 | 0.6178914 | 0.0958 | 0.0173 | 0.0523 | 0.0523 | TRUE | 1.6393 |
| Stc2 | 374 | 523 | 623 | 519 | 473 | 513 | 397 | 522 | 626 | 408 | 509 | 562 | 503.94 | 515 | 493 | 0.955 | -0.066 | 0.73152142 | 0.85590756 | 0.0364 | 0.0134 | 0.0247 | 0.0247 | TRUE | 0.2281 |
| Steap1 | 610 | 584 | 469 | 1018 | 646 | 586 | 648 | 583 | 471 | 800 | 695 | 641 | 639.82 | 567 | 712 | 1.257 | 0.33 | 0.02903567 | 0.11243236 | 0.0165 | 0.0121 | 0.0149 | 0.0149 | TRUE | 0.2596 |
| Steap2 | 1162 | 1127 | 1196 | 2195 | 1477 | 1442 | 1234 | 1126 | 1201 | 1726 | 1589 | 1578 | 1408.98 | 1187 | 1631 | 1.375 | 0.46 | 8.54E-08 | 3.14E-06 | 0.0017 | 0.0092 | 0.0046 | 0.0046 | TRUE | 0.0318 |
| Steap3 | 3257 | 2796 | 2354 | 4101 | 2595 | 2292 | 3460 | 2792 | 2364 | 3224 | 2792 | 2509 | 2856.74 | 2872 | 2842 | 0.99 | -0.015 | 0.92421839 | 0.96274037 | 0.0258 | 0.0081 | 0.017 | 0.017 | TRUE | 0.3414 |
| Steap4 | 97 | 27 | 30 | 61 | 35 | 15 | 103 | 27 | 30 | 48 | 38 | 16 | 43.69 | 53 | 34 | 0.641 | -0.641 | 0.27366271 | 0.49433176 | 0.3833 | 0.0823 | 0.223 | 0.223 | TRUE | 2.942 |
| Stfa1 | 0 | 0 | 0 | 0 | 0 | 0 | 0 | 0 | 0 | 0 | 0 | 0 | 0 | 0 | 0 | NA | NA | NA | NA | NA | NA | NA | NA | NA | NA |
| Stfa2 | 0 | 0 | 0 | 0 | 0 | 0 | 0 | 0 | 0 | 0 | 0 | 0 | 0 | 0 | 0 | NA | NA | NA | NA | NA | NA | NA | NA | NA | NA |
| Stfa2l1 | 0 | 0 | 0 | 0 | 0 | 0 | 0 | 0 | 0 | 0 | 0 | 0 | 0 | 0 | 0 | NA | NA | NA | NA | NA | NA | NA | NA | NA | NA |
| Stfa3 | 0 | 0 | 0 | 0 | 0 | 0 | 0 | 0 | 0 | 0 | 0 | 0 | 0 | 0 | 0 | NA | NA | NA | NA | NA | NA | NA | NA | NA | NA |
| Stil | 1218 | 1396 | 1450 | 1653 | 1189 | 1310 | 1294 | 1394 | 1456 | 1299 | 1279 | 1434 | 1359.44 | 1381 | 1337 | 0.968 | -0.047 | 0.61134716 | 0.77987759 | 0.0029 | 0.0093 | 0.0055 | 0.0055 | TRUE | 0.047 |
| Stim1 | 2369 | 2716 | 2691 | 2988 | 2144 | 2284 | 2516 | 2712 | 2702 | 2349 | 2307 | 2500 | 2514.46 | 2643 | 2385 | 0.902 | -0.149 | 0.05098362 | 0.16620787 | 0.0012 | 0.0082 | 0.0038 | 0.0038 | TRUE | 0.0214 |
| Stim2 | 997 | 867 | 825 | 1532 | 1049 | 948 | 1059 | 866 | 828 | 1204 | 1129 | 1038 | 1020.67 | 918 | 1124 | 1.225 | 0.293 | 0.0190397 | 0.08378742 | 0.0104 | 0.0101 | 0.0103 | 0.0103 | TRUE | 0.2151 |
| Stip1 | 27411 | 27212 | 26831 | 37136 | 26839 | 25992 | 29115 | 27177 | 26941 | 29193 | 28877 | 28452 | 28292.7 | 27744 | 28841 | 1.04 | 0.056 | 0.36750949 | 0.58475554 | 0.001 | 0.007 | 0.0027 | 0.0027 | TRUE | 0.0229 |
| Stk10 | 1440 | 1900 | 1989 | 2618 | 2034 | 2075 | 1530 | 1898 | 1997 | 2058 | 2188 | 2271 | 1990.36 | 1808 | 2172 | 1.201 | 0.264 | 0.02522816 | 0.10244557 | 0.0097 | 0.0086 | 0.0096 | 0.0096 | TRUE | 0.2187 |
| Stk11 | 2578 | 2937 | 2965 | 3620 | 2688 | 2663 | 2738 | 2933 | 2977 | 2846 | 2892 | 2915 | 2883.6 | 2883 | 2884 | 1 | 0 | 0.99569902 | 0.99791351 | 7.00E-04 | 0.008 | 0.0032 | 0.0032 | TRUE | 0.0233 |
| Stk11ip | 1430 | 1251 | 1209 | 2096 | 1609 | 1377 | 1519 | 1249 | 1214 | 1648 | 1731 | 1507 | 1478.08 | 1327 | 1629 | 1.227 | 0.296 | 0.01151081 | 0.05843773 | 0.0093 | 0.0091 | 0.0092 | 0.0092 | TRUE | 0.1908 |
| Stk16 | 1117 | 911 | 873 | 1493 | 1056 | 865 | 1186 | 910 | 877 | 1174 | 1136 | 947 | 1038.27 | 991 | 1086 | 1.096 | 0.133 | 0.37187601 | 0.58840574 | 0.0194 | 0.0101 | 0.0149 | 0.0149 | TRUE | 0.3546 |
| Stk17b | 1004 | 1373 | 1345 | 1594 | 1111 | 1187 | 1066 | 1371 | 1351 | 1253 | 1195 | 1299 | 1256 | 1263 | 1249 | 0.989 | -0.016 | 0.89512579 | 0.94935942 | 0.0096 | 0.0095 | 0.0096 | 0.0096 | TRUE | 0.2211 |
| Stk19 | 352 | 367 | 342 | 582 | 428 | 314 | 374 | 367 | 343 | 458 | 461 | 344 | 390.93 | 361 | 421 | 1.166 | 0.222 | 0.13761048 | 0.32173028 | 0.0118 | 0.0153 | 0.0135 | 0.0135 | TRUE | 0.2881 |
| Stk24 | 4481 | 4871 | 5103 | 5491 | 4202 | 4426 | 4760 | 4865 | 5124 | 4317 | 4521 | 4845 | 4738.49 | 4916 | 4561 | 0.927 | -0.109 | 0.15990435 | 0.35445491 | 0.0021 | 0.0076 | 0.0041 | 0.0041 | TRUE | 0.0361 |
| Stk25 | 2903 | 3372 | 3162 | 4117 | 3204 | 3006 | 3083 | 3368 | 3175 | 3236 | 3447 | 3291 | 3266.74 | 3209 | 3325 | 1.036 | 0.051 | 0.4849525 | 0.68236871 | 0.0011 | 0.0079 | 0.0035 | 0.0035 | TRUE | 0.0228 |
| Stk3 | 647 | 831 | 866 | 1273 | 802 | 796 | 687 | 830 | 870 | 1001 | 863 | 871 | 853.62 | 796 | 912 | 1.147 | 0.197 | 0.11681416 | 0.28895749 | 0.0091 | 0.0108 | 0.0102 | 0.0102 | TRUE | 0.1674 |
| Stk30 | 315 | 476 | 488 | 624 | 516 | 571 | 335 | 475 | 490 | 491 | 555 | 625 | 495.12 | 433 | 557 | 1.283 | 0.36 | 0.03896316 | 0.13809593 | 0.0261 | 0.0136 | 0.0198 | 0.0198 | TRUE | 0.4563 |
| Stk31 | 625 | 623 | 586 | 1449 | 1038 | 872 | 664 | 622 | 588 | 1139 | 1117 | 955 | 847.49 | 625 | 1070 | 1.714 | 0.778 | 3.19E-12 | 3.06E-10 | 0.0054 | 0.0108 | 0.0077 | 0.0077 | TRUE | 0.1055 |
| Stk32a | 249 | 224 | 219 | 260 | 201 | 181 | 264 | 224 | 220 | 204 | 216 | 198 | 221.15 | 236 | 206 | 0.874 | -0.194 | 0.19337351 | 0.39893879 | 0 | 0.0218 | 0.0115 | 0.0115 | TRUE | 0.1213 |
| Stk32b | 65 | 46 | 52 | 39 | 39 | 34 | 69 | 46 | 52 | 31 | 42 | 37 | 46.17 | 56 | 37 | 0.654 | -0.613 | 0.04979029 | 0.16379495 | 0.0176 | 0.0783 | 0.0477 | 0.0477 | TRUE | 0.3591 |
| Stk32c | 182 | 170 | 176 | 210 | 142 | 133 | 193 | 170 | 177 | 165 | 153 | 146 | 167.21 | 180 | 155 | 0.861 | -0.216 | 0.17682878 | 0.37734319 | 0 | 0.0266 | 0.0124 | 0.0124 | TRUE | 0.0446 |
| Stk33 | 12 | 9 | 17 | 23 | 23 | 23 | 13 | 9 | 17 | 18 | 25 | 25 | 17.8 | 13 | 23 | 1.742 | 0.801 | 0.09482587 | 0.25229312 | 0 | 0.192 | 0.1043 | 0.1043 | TRUE | 0.3309 |
| Stk35 | 1772 | 1782 | 1683 | 3425 | 2375 | 2043 | 1882 | 1780 | 1690 | 2692 | 2555 | 2236 | 2139.33 | 1784 | 2494 | 1.399 | 0.485 | 1.27E-06 | 3.39E-05 | 0.0051 | 0.0084 | 0.0067 | 0.0067 | TRUE | 0.099 |
| Stk36 | 175 | 162 | 170 | 254 | 197 | 177 | 186 | 162 | 171 | 200 | 212 | 194 | 187.29 | 173 | 202 | 1.168 | 0.224 | 0.13755096 | 0.3217144 | 0 | 0.0245 | 0.0111 | 0.0111 | TRUE | 0.0461 |
| Stk38 | 6183 | 5686 | 5616 | 6136 | 4856 | 4683 | 6567 | 5679 | 5639 | 4824 | 5225 | 5126 | 5509.97 | 5962 | 5058 | 0.848 | -0.237 | 0.008201 | 0.045272 | 0.0045 | 0.0075 | 0.0056 | 0.0056 | TRUE | 0.0962 |
| Stk38l | 596 | 578 | 526 | 817 | 623 | 525 | 633 | 577 | 528 | 642 | 670 | 575 | 604.29 | 579 | 629 | 1.086 | 0.12 | 0.31395606 | 0.53438031 | 0.0054 | 0.0124 | 0.0085 | 0.0085 | TRUE | 0.0761 |
| Stk39 | 184 | 514 | 567 | 426 | 316 | 404 | 195 | 513 | 569 | 335 | 340 | 442 | 399.21 | 426 | 372 | 0.873 | -0.195 | 0.54849369 | 0.73326245 | 0.1548 | 0.0152 | 0.0738 | 0.0738 | TRUE | 2.5804 |
| Stk4 | 5904 | 5142 | 5098 | 7149 | 5196 | 4775 | 6271 | 5135 | 5119 | 5620 | 5591 | 5227 | 5493.82 | 5508 | 5479 | 0.995 | -0.007 | 0.94212906 | 0.97094617 | 0.0075 | 0.0075 | 0.0075 | 0.0075 | TRUE | 0.1787 |
| Stk40 | 3257 | 2896 | 2855 | 3613 | 2661 | 2487 | 3460 | 2892 | 2867 | 2840 | 2863 | 2722 | 2940.71 | 3073 | 2808 | 0.914 | -0.129 | 0.18946663 | 0.39464709 | 0.0058 | 0.008 | 0.0067 | 0.0067 | TRUE | 0.1469 |
| Stmn1 | 6478 | 5880 | 5829 | 6375 | 4645 | 4764 | 6881 | 5872 | 5853 | 5012 | 4998 | 5215 | 5638.39 | 6202 | 5075 | 0.818 | -0.289 | 0.00125995 | 0.01064378 | 0.0045 | 0.0075 | 0.0056 | 0.0056 | TRUE | 0.1117 |
| Stmn1-rs1 | 0 | 0 | 1 | 1 | 0 | 2 | 0 | 0 | 1 | 1 | 0 | 2 | 0.66 | 0 | 1 | 2.693 | 1.429 | 0.58329919 | NA | 0 | 4.974 | 3.3955 | 3.3955 | TRUE | 0.2781 |
| Stmn2 | 15125 | 17819 | 17681 | 16437 | 11906 | 13265 | 16065 | 17796 | 17754 | 12921 | 12810 | 14521 | 15311.21 | 17205 | 13417 | 0.78 | -0.359 | 2.76E-05 | 0.00046847 | 0.0041 | 0.0071 | 0.0052 | 0.0052 | TRUE | 0.0632 |
| Stmn3 | 71 | 95 | 108 | 117 | 86 | 75 | 75 | 95 | 108 | 92 | 93 | 82 | 90.89 | 93 | 89 | 0.957 | -0.064 | 0.77648131 | 0.88385271 | 0.0067 | 0.0431 | 0.0256 | 0.0256 | TRUE | 0.2549 |
| Stmn4 | 0 | 0 | 0 | 1 | 0 | 1 | 0 | 0 | 0 | 1 | 0 | 1 | 0.31 | 0 | 1 | 3.348 | 1.743 | 0.65064889 | NA | 0 | 10.5167 | 8.8968 | 8.8968 | TRUE | 0.2082 |
| Stmnd1 | 2 | 0 | 1 | 2 | 0 | 0 | 2 | 0 | 1 | 2 | 0 | 0 | 0.78 | 1 | 1 | 0.543 | -0.88 | 0.72386184 | NA | 0 | 4.212 | 3.1127 | 3.1127 | TRUE | 0.402 |
| Stom | 1309 | 1721 | 1895 | 1247 | 1016 | 1234 | 1390 | 1719 | 1903 | 980 | 1093 | 1351 | 1406.03 | 1671 | 1141 | 0.683 | -0.551 | 0.00049485 | 0.00512798 | 0.0248 | 0.0092 | 0.0173 | 0.0173 | TRUE | 0.3077 |
| Stoml1 | 1685 | 2621 | 2643 | 2388 | 1934 | 1952 | 1790 | 2618 | 2654 | 1877 | 2081 | 2137 | 2192.69 | 2354 | 2032 | 0.863 | -0.213 | 0.17183574 | 0.37064898 | 0.0231 | 0.0084 | 0.017 | 0.017 | TRUE | 0.5316 |

| | | | | | | | | | | | | | | | | | | | | | | | | | |
|---|---|---|---|---|---|---|---|---|---|---|---|---|---|---|---|---|---|---|---|---|---|---|---|---|---|
| Stoml2 | 4215 | 4662 | 4595 | 6387 | 4619 | 4431 | 4477 | 4656 | 4614 | 5021 | 4970 | 4850 | 4764.68 | 4582 | 4947 | 1.08 | 0.111 | 0.06435565 | 0.19526081 | 2.00E-04 | 0.0076 | 0.0024 | 0.0024 | TRUE | 0.0049 |
| Stoml3 | 1 | 2 | 1 | 1 | 2 | 1 | 1 | 2 | 1 | 1 | 2 | 1 | 1.35 | 1 | 1 | 0.972 | -0.041 | 0.97884658 | NA | 0 | 2.4483 | 0.9574 | 0.9574 | TRUE | 0.16 |
| Ston1 | 408 | 474 | 512 | 670 | 499 | 532 | 433 | 473 | 514 | 527 | 537 | 582 | 511.14 | 473 | 549 | 1.157 | 0.21 | 0.06762805 | 0.20113702 | 0.0029 | 0.0133 | 0.0076 | 0.0076 | TRUE | 0.065 |
| Ston2 | 2912 | 2576 | 2725 | 4304 | 2954 | 3081 | 3093 | 2573 | 2736 | 3383 | 3178 | 3373 | 3056.06 | 2801 | 3311 | 1.183 | 0.242 | 0.00965489 | 0.05127171 | 0.0046 | 0.008 | 0.006 | 0.006 | TRUE | 0.101 |
| Stox1 | 36 | 36 | 25 | 72 | 56 | 36 | 38 | 36 | 25 | 57 | 60 | 39 | 42.59 | 33 | 52 | 1.579 | 0.659 | 0.04790535 | 0.15919999 | 0.0219 | 0.0843 | 0.0552 | 0.0552 | TRUE | 0.3496 |
| Stox2 | 1327 | 1140 | 1256 | 2107 | 1452 | 1365 | 1410 | 1139 | 1261 | 1656 | 1562 | 1494 | 1420.34 | 1270 | 1571 | 1.238 | 0.308 | 0.00389251 | 0.02580609 | 0.006 | 0.0092 | 0.0075 | 0.0075 | TRUE | 0.1109 |
| Stpg1 | 47 | 62 | 49 | 123 | 62 | 53 | 50 | 62 | 49 | 97 | 67 | 58 | 63.74 | 54 | 74 | 1.384 | 0.469 | 0.11003324 | 0.2784095 | 0.0338 | 0.0586 | 0.046 | 0.046 | TRUE | 0.7349 |
| Stpg2 | 0 | 1 | 1 | 1 | 0 | 2 | 0 | 1 | 1 | 1 | 0 | 2 | 0.83 | 1 | 1 | 1.46 | 0.546 | 0.80141093 | NA | 0 | 3.9775 | 2.1434 | 2.1434 | TRUE | 0.302 |
| Stra13 | 1543 | 1905 | 1836 | 1898 | 1269 | 1457 | 1639 | 1903 | 1844 | 1492 | 1365 | 1595 | 1639.56 | 1795 | 1484 | 0.827 | -0.275 | 0.00693217 | 0.03984485 | 0.005 | 0.0089 | 0.0068 | 0.0068 | TRUE | 0.0696 |
| Stra6 | 102 | 88 | 91 | 110 | 62 | 54 | 108 | 88 | 91 | 86 | 67 | 59 | 83.32 | 96 | 71 | 0.744 | -0.426 | 0.07935489 | 0.22337772 | 0.0136 | 0.0464 | 0.0304 | 0.0304 | TRUE | 0.3799 |
| Stra8 | 856 | 690 | 587 | 3620 | 2194 | 1639 | 909 | 689 | 589 | 2846 | 2361 | 1794 | 1531.38 | 729 | 2334 | 3.202 | 1.679 | 3.33E-17 | 5.90E-15 | 0.0493 | 0.0091 | 0.0276 | 0.0276 | TRUE | 0.111 |
| Strada | 2101 | 1973 | 2068 | 2687 | 1972 | 1818 | 2232 | 1970 | 2076 | 2112 | 2122 | 1990 | 2083.79 | 2093 | 2075 | 0.992 | -0.012 | 0.88406248 | 0.94309013 | 0.0021 | 0.0085 | 0.0045 | 0.0045 | TRUE | 0.0406 |
| Stradb | 668 | 839 | 813 | 935 | 635 | 702 | 710 | 838 | 816 | 735 | 683 | 768 | 758.41 | 788 | 729 | 0.925 | -0.113 | 0.30201545 | 0.52259973 | 0.0042 | 0.0112 | 0.0073 | 0.0073 | TRUE | 0.0892 |
| Strap | 17040 | 17441 | 16609 | 21358 | 15014 | 14896 | 18099 | 17418 | 16677 | 16790 | 16154 | 16306 | 16907.55 | 17398 | 16417 | 0.944 | -0.084 | 0.18512284 | 0.38842056 | 0.001 | 0.0071 | 0.0028 | 0.0028 | TRUE | 0.0161 |
| Strbp | 6986 | 8104 | 8408 | 8732 | 6329 | 7081 | 7420 | 8094 | 8443 | 6864 | 6810 | 7751 | 7563.62 | 7986 | 7142 | 0.894 | -0.161 | 0.0728176 | 0.21131669 | 0.0047 | 0.0073 | 0.0057 | 0.0057 | TRUE | 0.0678 |
| Strc | 4 | 6 | 1 | 3 | 5 | 2 | 4 | 6 | 1 | 2 | 5 | 2 | 3.53 | 4 | 3 | 0.875 | -0.193 | 0.85724276 | NA | 0 | 0.9405 | 0.5456 | 0.5456 | TRUE | 0.421 |
| Strip1 | 3522 | 3147 | 3190 | 4636 | 3344 | 3146 | 3741 | 3143 | 3203 | 3644 | 3598 | 3444 | 3462.2 | 3362 | 3562 | 1.06 | 0.084 | 0.37028009 | 0.58728707 | 0.0047 | 0.0079 | 0.006 | 0.006 | TRUE | 0.1178 |
| Strip2 | 165 | 185 | 147 | 283 | 208 | 171 | 175 | 185 | 148 | 222 | 224 | 187 | 190.18 | 169 | 211 | 1.25 | 0.322 | 0.05555766 | 0.17645611 | 0.0059 | 0.0242 | 0.015 | 0.015 | TRUE | 0.1341 |
| Strn | 1464 | 1321 | 1505 | 1827 | 1401 | 1285 | 1555 | 1319 | 1511 | 1436 | 1507 | 1407 | 1455.96 | 1462 | 1450 | 0.992 | -0.012 | 0.90391029 | 0.95248677 | 0.0037 | 0.0092 | 0.0059 | 0.0059 | TRUE | 0.0878 |
| Strn3 | 5446 | 5296 | 5315 | 6879 | 4903 | 4728 | 5785 | 5289 | 5337 | 5408 | 5275 | 5176 | 5378.2 | 5470 | 5286 | 0.967 | -0.049 | 0.48187706 | 0.68077623 | 0.0012 | 0.0075 | 0.0033 | 0.0033 | TRUE | 0.0308 |
| Strn4 | 3641 | 4360 | 4601 | 5790 | 4443 | 4285 | 3867 | 4354 | 4620 | 4552 | 4780 | 4691 | 4477.38 | 4280 | 4674 | 1.092 | 0.127 | 0.15350677 | 0.34513311 | 0.0039 | 0.0076 | 0.0055 | 0.0055 | TRUE | 0.0866 |
| Stt3a | 12155 | 14237 | 14193 | 13613 | 9887 | 11074 | 12911 | 14219 | 14251 | 10701 | 10638 | 12122 | 12473.69 | 13794 | 11154 | 0.809 | -0.307 | 0.00040408 | 0.00437375 | 0.0042 | 0.0072 | 0.0053 | 0.0053 | TRUE | 0.0704 |
| Stt3b | 8797 | 9969 | 10374 | 11086 | 8413 | 8952 | 9344 | 9956 | 10417 | 8715 | 9052 | 9799 | 9547.15 | 9906 | 9189 | 0.927 | -0.109 | 0.18078484 | 0.38227853 | 0.0031 | 0.0072 | 0.0046 | 0.0046 | TRUE | 0.0412 |
| Stub1 | 2813 | 2864 | 2805 | 4041 | 2908 | 2624 | 2988 | 2860 | 2817 | 3177 | 3129 | 2872 | 2973.77 | 2888 | 3059 | 1.06 | 0.084 | 0.2727026 | 0.49295859 | 0.0014 | 0.008 | 0.0038 | 0.0038 | TRUE | 0.0344 |
| Stx11 | 106 | 121 | 132 | 120 | 115 | 123 | 113 | 121 | 133 | 94 | 124 | 135 | 119.78 | 122 | 118 | 0.958 | -0.062 | 0.76873343 | 0.87928573 | 0.0128 | 0.0344 | 0.0234 | 0.0234 | TRUE | 0.3341 |
| Stx12 | 2895 | 3605 | 3757 | 3749 | 2749 | 3141 | 3075 | 3600 | 3772 | 2947 | 2958 | 3438 | 3298.5 | 3482 | 3114 | 0.894 | -0.162 | 0.14725757 | 0.3363323 | 0.009 | 0.0079 | 0.0087 | 0.0087 | TRUE | 0.1273 |
| Stx16 | 2400 | 2484 | 2471 | 2741 | 2108 | 2025 | 2549 | 2481 | 2481 | 2155 | 2268 | 2217 | 2358.45 | 2504 | 2213 | 0.884 | -0.178 | 0.00732771 | 0.04152402 | 0 | 0.0083 | 0.0028 | 0.0028 | TRUE | 0.0067 |
| Stx17 | 660 | 799 | 759 | 1154 | 844 | 771 | 701 | 798 | 762 | 907 | 908 | 844 | 820.06 | 754 | 886 | 1.176 | 0.234 | 0.01547064 | 0.07242591 | 0.0016 | 0.0109 | 0.0055 | 0.0055 | TRUE | 0.0439 |
| Stx18 | 488 | 519 | 434 | 603 | 497 | 407 | 518 | 518 | 436 | 474 | 535 | 446 | 487.79 | 491 | 485 | 0.988 | -0.018 | 0.88907967 | 0.9459843 | 0.007 | 0.0137 | 0.0101 | 0.0101 | TRUE | 0.1125 |
| Stx19 | 0 | 0 | 0 | 0 | 0 | 0 | 0 | 0 | 0 | 0 | 0 | 0 | 0 | 0 | 0 | NA | NA | NA | NA | NA | NA | NA | NA | NA | NA |
| Stx1a | 356 | 342 | 335 | 375 | 314 | 318 | 378 | 342 | 336 | 295 | 338 | 348 | 339.47 | 352 | 327 | 0.927 | -0.11 | 0.40055917 | 0.61353979 | 0.0033 | 0.0166 | 0.0094 | 0.0094 | TRUE | 0.0901 |
| Stx1b | 100 | 123 | 97 | 87 | 82 | 71 | 106 | 123 | 97 | 68 | 88 | 78 | 93.47 | 109 | 78 | 0.715 | -0.485 | 0.02735327 | 0.10812425 | 0 | 0.0422 | 0.0238 | 0.0238 | TRUE | 0.1278 |
| Stx2 | 412 | 373 | 344 | 438 | 356 | 372 | 438 | 373 | 345 | 344 | 383 | 407 | 381.69 | 385 | 378 | 0.981 | -0.028 | 0.84149191 | 0.9199425 | 0.0084 | 0.0155 | 0.0117 | 0.0117 | TRUE | 0.1615 |
| Stx3 | 1139 | 1288 | 1463 | 1043 | 802 | 1028 | 1210 | 1286 | 1469 | 820 | 863 | 1125 | 1128.88 | 1322 | 936 | 0.707 | -0.499 | 0.00062441 | 0.00619775 | 0.0193 | 0.0098 | 0.0144 | 0.0144 | TRUE | 0.3708 |
| Stx4a | 553 | 616 | 632 | 974 | 685 | 675 | 587 | 615 | 635 | 766 | 737 | 739 | 679.8 | 612 | 747 | 1.221 | 0.288 | 0.00173355 | 0.01372132 | 0 | 0.0117 | 0.0046 | 0.0046 | TRUE | 0.0148 |
| Stx5a | 893 | 853 | 794 | 1451 | 992 | 887 | 949 | 852 | 797 | 1141 | 1067 | 971 | 962.77 | 866 | 1060 | 1.225 | 0.293 | 0.00856309 | 0.04684396 | 0.0056 | 0.0103 | 0.0079 | 0.0079 | TRUE | 0.0824 |
| Stx6 | 1393 | 1563 | 1538 | 1816 | 1374 | 1342 | 1480 | 1561 | 1544 | 1428 | 1478 | 1469 | 1493.31 | 1528 | 1458 | 0.954 | -0.068 | 0.35184927 | 0.57063668 | 0 | 0.0091 | 0.0032 | 0.0032 | TRUE | 0.0093 |
| Stx7 | 1903 | 2222 | 2393 | 2615 | 1848 | 2112 | 2021 | 2219 | 2403 | 2056 | 1988 | 2312 | 2166.54 | 2214 | 2119 | 0.956 | -0.064 | 0.53436758 | 0.72176462 | 0.0057 | 0.0084 | 0.0072 | 0.0072 | TRUE | 0.0764 |
| Stx8 | 738 | 950 | 994 | 1346 | 935 | 928 | 784 | 949 | 998 | 1058 | 1006 | 1016 | 968.45 | 910 | 1027 | 1.128 | 0.174 | 0.13051539 | 0.31106435 | 0.007 | 0.0103 | 0.0085 | 0.0085 | TRUE | 0.1746 |
| Stxbp1 | 2194 | 2047 | 2096 | 1859 | 1384 | 1481 | 2330 | 2044 | 2105 | 1461 | 1489 | 1621 | 1841.84 | 2160 | 1524 | 0.705 | -0.504 | 3.63E-08 | 1.51E-06 | 0.0032 | 0.0087 | 0.0055 | 0.0055 | TRUE | 0.0576 |
| Stxbp2 | 3327 | 4299 | 4502 | 4820 | 3844 | 4435 | 3534 | 4293 | 4520 | 3789 | 4136 | 4855 | 4187.94 | 4116 | 4260 | 1.035 | 0.049 | 0.7067413 | 0.8398044 | 0.0159 | 0.0077 | 0.0121 | 0.0121 | TRUE | 0.1861 |
| Stxbp3a | 1578 | 1574 | 1550 | 1828 | 1332 | 1416 | 1676 | 1572 | 1556 | 1437 | 1433 | 1550 | 1537.44 | 1601 | 1473 | 0.92 | -0.121 | 0.13698063 | 0.32092372 | 0.0011 | 0.009 | 0.0041 | 0.0041 | TRUE | 0.0245 |
| Stxbp3b | 14 | 11 | 15 | 8 | 9 | 11 | 15 | 11 | 15 | 6 | 10 | 12 | 11.49 | 14 | 9 | 0.675 | -0.567 | 0.32210388 | 0.54330584 | 0 | 0.2937 | 0.146 | 0.146 | TRUE | 0.357 |
| Stxbp4 | 1388 | 1036 | 1119 | 1631 | 1146 | 999 | 1474 | 1035 | 1124 | 1282 | 1233 | 1094 | 1206.89 | 1211 | 1203 | 0.994 | -0.009 | 0.95365413 | 0.97650213 | 0.02 | 0.0096 | 0.015 | 0.015 | TRUE | 0.4336 |
| Stxbp5 | 695 | 776 | 870 | 927 | 690 | 828 | 738 | 775 | 874 | 729 | 742 | 906 | 794.05 | 796 | 792 | 0.995 | -0.007 | 0.95506794 | 0.97712322 | 0.01 | 0.0111 | 0.0105 | 0.0105 | TRUE | 0.1855 |
| Stxbp5l | 28 | 22 | 18 | 39 | 35 | 20 | 30 | 22 | 18 | 31 | 38 | 22 | 26.67 | 23 | 30 | 1.296 | 0.375 | 0.35850545 | 0.57703316 | 0.0255 | 0.1304 | 0.0816 | 0.0816 | TRUE | 0.1935 |
| Stxbp6 | 116 | 162 | 186 | 259 | 156 | 191 | 123 | 162 | 187 | 204 | 168 | 209 | 175.38 | 157 | 194 | 1.23 | 0.299 | 0.13859623 | 0.3230815 | 0.0215 | 0.0257 | 0.0236 | 0.0236 | TRUE | 0.372 |
| Styk1 | 74 | 50 | 46 | 41 | 32 | 24 | 79 | 50 | 46 | 32 | 34 | 26 | 44.61 | 58 | 31 | 0.534 | -0.906 | 0.0076832 | 0.04310903 | 0.0368 | 0.0808 | 0.0587 | 0.0587 | TRUE | 0.7817 |
| Styx | 575 | 679 | 663 | 828 | 590 | 648 | 611 | 678 | 666 | 651 | 635 | 709 | 658.27 | 652 | 665 | 1.02 | 0.028 | 0.78078116 | 0.88649531 | 0.0016 | 0.0119 | 0.006 | 0.006 | TRUE | 0.0389 |
| Styxl1 | 6 | 9 | 9 | 11 | 3 | 4 | 6 | 9 | 9 | 9 | 3 | 4 | 6.78 | 8 | 5 | 0.682 | -0.552 | 0.46304099 | 0.66646126 | 0 | 0.4931 | 0.2568 | 0.2568 | TRUE | 0.759 |
| Sub1 | 10425 | 10195 | 9678 | 13706 | 10464 | 9421 | 11073 | 10182 | 9718 | 10775 | 11259 | 10313 | 10553.12 | 10324 | 10782 | 1.044 | 0.063 | 0.43473249 | 0.64326868 | 0.003 | 0.0072 | 0.0045 | 0.0045 | TRUE | 0.0492 |
| Sucla2 | 3960 | 4242 | 4174 | 6053 | 4433 | 4467 | 4206 | 4237 | 4191 | 4758 | 4770 | 4890 | 4508.62 | 4211 | 4806 | 1.141 | 0.19 | 0.0009072 | 0.00826119 | 0 | 0.0076 | 0.0022 | 0.0022 | TRUE | 0.0028 |
| Suclg1 | 4342 | 4671 | 4638 | 6406 | 4382 | 4386 | 4612 | 4665 | 4657 | 5036 | 4715 | 4801 | 4747.63 | 4645 | 4851 | 1.045 | 0.063 | 0.31354783 | 0.5339318 | 4.00E-04 | 0.0076 | 0.0026 | 0.0026 | TRUE | 0.0138 |
| Suclg2 | 7226 | 9268 | 9560 | 11617 | 8528 | 9601 | 7675 | 9256 | 9599 | 9132 | 9176 | 10510 | 9224.73 | 8843 | 9606 | 1.086 | 0.119 | 0.28610557 | 0.50736877 | 0.0098 | 0.0073 | 0.0089 | 0.0089 | TRUE | 0.163 |
| Sucnr1 | 0 | 0 | 0 | 0 | 0 | 0 | 0 | 0 | 0 | 0 | 0 | 0 | 0 | 0 | 0 | NA | NA | NA | NA | NA | NA | NA | NA | NA | NA |
| Suco | 3556 | 4024 | 4264 | 4739 | 3424 | 3477 | 3777 | 4019 | 4282 | 3725 | 3684 | 3806 | 3882.16 | 4026 | 3738 | 0.929 | -0.107 | 0.1590652 | 0.35313872 | 0.0017 | 0.0078 | 0.0039 | 0.0039 | TRUE | 0.0376 |
| Suds3 | 4030 | 3773 | 3706 | 5335 | 3923 | 3573 | 4281 | 3768 | 3721 | 4194 | 4221 | 3911 | 4016 | 3923 | 4109 | 1.047 | 0.067 | 0.44139091 | 0.6484368 | 0.0035 | 0.0077 | 0.0052 | 0.0052 | TRUE | 0.0771 |
| Sufu | 846 | 647 | 658 | 1064 | 767 | 677 | 899 | 646 | 661 | 836 | 825 | 741 | 768.04 | 735 | 801 | 1.09 | 0.125 | 0.40493026 | 0.61765571 | 0.0182 | 0.0112 | 0.0148 | 0.0148 | TRUE | 0.4461 |
| Sugp1 | 2742 | 2465 | 2318 | 4194 | 3020 | 2385 | 2912 | 2462 | 2328 | 3297 | 3249 | 2611 | 2809.82 | 2567 | 3052 | 1.189 | 0.25 | 0.05312046 | 0.17112251 | 0.0145 | 0.0081 | 0.0117 | 0.0117 | TRUE | 0.1939 |
| Sugp2 | 3118 | 3889 | 3991 | 3271 | 2991 | 3340 | 3312 | 3884 | 4007 | 2571 | 3218 | 3656 | 3441.49 | 3734 | 3148 | 0.843 | -0.247 | 0.0830771 | 0.23068521 | 0.0195 | 0.0079 | 0.0143 | 0.0143 | TRUE | 0.3146 |
| Sugt1 | 4123 | 4199 | 4417 | 5997 | 4231 | 4408 | 4379 | 4194 | 4435 | 4714 | 4552 | 4825 | 4516.66 | 4336 | 4697 | 1.083 | 0.116 | 0.07622134 | 0.21782262 | 5.00E-04 | 0.0076 | 0.0028 | 0.0028 | TRUE | 0.0101 |
| Sulf1 | 1529 | 1276 | 1435 | 1612 | 1119 | 1140 | 1624 | 1274 | 1441 | 1267 | 1204 | 1248 | 1343.07 | 1446 | 1240 | 0.857 | -0.222 | 0.04394956 | 0.15023242 | 0.0065 | 0.0094 | 0.008 | 0.008 | TRUE | 0.1385 |
| Sulf2 | 5618 | 6452 | 7022 | 11504 | 8274 | 7868 | 5967 | 6444 | 7051 | 9044 | 8902 | 8613 | 7670.07 | 6487 | 8853 | 1.365 | 0.449 | 1.07E-07 | 3.80E-06 | 0.0036 | 0.0073 | 0.005 | 0.005 | TRUE | 0.0705 |
| Sult1a1 | 1 | 0 | 0 | 2 | 0 | 1 | 1 | 0 | 0 | 2 | 0 | 1 | 0.62 | 0 | 1 | 2.474 | 1.307 | 0.61898859 | NA | 0 | 5.3076 | 3.4358 | 3.4358 | TRUE | 0.4614 |
| Sult1b1 | 0 | 0 | 0 | 0 | 0 | 0 | 0 | 0 | 0 | 0 | 0 | 0 | 0 | 0 | 0 | NA | NA | NA | NA | NA | NA | NA | NA | NA | NA |
| Sult1c1 | 0 | 0 | 0 | 0 | 2 | 0 | 0 | 0 | 0 | 0 | 2 | 0 | 0.36 | 0 | 1 | 3.775 | 1.916 | 0.63423067 | NA | 10 | 9.1927 | 10 | 10 | TRUE | 0.9743 |
| Sult1c2 | 0 | 0 | 0 | 0 | 0 | 0 | 0 | 0 | 0 | 0 | 0 | 0 | 0 | 0 | 0 | NA | NA | NA | NA | NA | NA | NA | NA | NA | NA |
| Sult1d1 | 0 | 0 | 0 | 0 | 0 | 0 | 0 | 0 | 0 | 0 | 0 | 0 | 0 | 0 | 0 | NA | NA | NA | NA | NA | NA | NA | NA | NA | NA |
| Sult1e1 | 0 | 0 | 0 | 0 | 0 | 0 | 0 | 0 | 0 | 0 | 0 | 0 | 0 | 0 | 0 | NA | NA | NA | NA | NA | NA | NA | NA | NA | NA |

| | | | | | | | | | | | | | | | | | | | | | | | | | |
|---|---|---|---|---|---|---|---|---|---|---|---|---|---|---|---|---|---|---|---|---|---|---|---|---|---|
| Sult2a1 | 0 | 0 | 0 | 0 | 0 | 0 | 0 | 0 | 0 | 0 | 0 | 0 | 0 | 0 | 0 | NA | NA | NA | NA | NA | NA | NA | NA | NA | NA |
| Sult2a2 | 0 | 0 | 0 | 0 | 0 | 0 | 0 | 0 | 0 | 0 | 0 | 0 | 0 | 0 | 0 | NA | NA | NA | NA | NA | NA | NA | NA | NA | NA |
| Sult2a3 | 0 | 0 | 0 | 2 | 2 | 1 | 0 | 0 | 0 | 2 | 2 | 1 | 0.8 | 0 | 2 | 8.55 | 3.096 | 0.18206245 | NA | 0 | 4.1089 | 2.5737 | 2.5737 | TRUE | 0.07 |
| Sult2a4 | 1 | 2 | 3 | 5 | 2 | 3 | 1 | 2 | 3 | 4 | 2 | 3 | 2.57 | 2 | 3 | 1.551 | 0.633 | 0.58314076 | NA | 0 | 1.2873 | 0.5519 | 0.5519 | TRUE | 0.1658 |
| Sult2a5 | 0 | 0 | 0 | 5 | 2 | 1 | 0 | 0 | 0 | 4 | 2 | 1 | 1.2 | 0 | 2 | 12.884 | 3.688 | 0.07760175 | NA | 0 | 2.761 | 1.9438 | 1.9438 | TRUE | 0.1448 |
| Sult2a6 | 0 | 0 | 0 | 0 | 0 | 0 | 0 | 0 | 0 | 0 | 0 | 0 | 0 | 0 | 0 | NA | NA | NA | NA | NA | NA | NA | NA | NA | NA |
| Sult2a7 | 0 | 0 | 0 | 1 | 0 | 0 | 0 | 0 | 0 | 1 | 0 | 0 | 0.13 | 0 | 0 | 1.859 | 0.894 | 0.82650734 | NA | 0 | 25.1515 | 10 | 10 | TRUE | 0.2563 |
| Sult2b1 | 136 | 140 | 123 | 123 | 83 | 125 | 144 | 140 | 124 | 97 | 89 | 137 | 121.77 | 136 | 108 | 0.789 | -0.341 | 0.12352167 | 0.29955471 | 0.019 | 0.034 | 0.027 | 0.027 | TRUE | 0.5349 |
| Sult3a1 | 0 | 0 | 0 | 0 | 0 | 0 | 0 | 0 | 0 | 0 | 0 | 0 | 0 | 0 | 0 | NA | NA | NA | NA | NA | NA | NA | NA | NA | NA |
| Sult4a1 | 2077 | 1930 | 1938 | 2792 | 2024 | 2044 | 2206 | 1928 | 1946 | 2195 | 2178 | 2237 | 2114.94 | 2027 | 2203 | 1.087 | 0.121 | 0.15446002 | 0.346511 | 0.0023 | 0.0085 | 0.0047 | 0.0047 | TRUE | 0.0724 |
| Sult5a1 | 202 | 249 | 203 | 190 | 134 | 159 | 215 | 249 | 204 | 149 | 144 | 174 | 189.11 | 223 | 156 | 0.7 | -0.515 | 0.00196807 | 0.01519352 | 0.0051 | 0.0243 | 0.0145 | 0.0145 | TRUE | 0.1189 |
| Sult6b1 | 263 | 414 | 416 | 485 | 423 | 392 | 279 | 413 | 418 | 381 | 455 | 429 | 396 | 370 | 422 | 1.138 | 0.186 | 0.29734365 | 0.51819934 | 0.0257 | 0.0152 | 0.0205 | 0.0205 | TRUE | 0.5235 |
| Sumf1 | 1165 | 1096 | 1042 | 1745 | 1233 | 1172 | 1237 | 1095 | 1046 | 1372 | 1327 | 1283 | 1226.61 | 1126 | 1327 | 1.179 | 0.238 | 0.01413701 | 0.0678554 | 0.0032 | 0.0096 | 0.006 | 0.006 | TRUE | 0.0891 |
| Sumf2 | 1693 | 1686 | 1649 | 2001 | 1445 | 1413 | 1798 | 1684 | 1656 | 1573 | 1555 | 1547 | 1635.4 | 1713 | 1558 | 0.91 | -0.136 | 0.07046521 | 0.20666393 | 4.00E-04 | 0.0089 | 0.0035 | 0.0035 | TRUE | 0.023 |
| Sumo1 | 4276 | 4060 | 4092 | 6211 | 4277 | 4027 | 4542 | 4055 | 4109 | 4883 | 4602 | 4408 | 4433 | 4235 | 4631 | 1.094 | 0.129 | 0.11871072 | 0.29181819 | 0.003 | 0.0076 | 0.0047 | 0.0047 | TRUE | 0.0488 |
| Sumo2 | 2606 | 2808 | 2754 | 3549 | 2353 | 2366 | 2768 | 2804 | 2765 | 2790 | 2532 | 2590 | 2708.22 | 2779 | 2637 | 0.949 | -0.075 | 0.30281051 | 0.52361883 | 0.001 | 0.0081 | 0.0034 | 0.0034 | TRUE | 0.0318 |
| Sumo3 | 8616 | 7372 | 6856 | 7936 | 5516 | 5682 | 9152 | 7362 | 6884 | 6239 | 5935 | 6220 | 6965.29 | 7799 | 6131 | 0.786 | -0.347 | 0.00298124 | 0.02111368 | 0.0119 | 0.0074 | 0.0097 | 0.0097 | TRUE | 0.2807 |
| Sun1 | 9215 | 11956 | 11679 | 12105 | 9420 | 10739 | 9788 | 11941 | 11727 | 9516 | 10135 | 11756 | 10810.39 | 11152 | 10469 | 0.939 | -0.091 | 0.43419206 | 0.6427102 | 0.0105 | 0.0072 | 0.0097 | 0.0097 | TRUE | 0.1412 |
| Sun2 | 2242 | 2567 | 2533 | 2253 | 1851 | 1774 | 2381 | 2564 | 2543 | 1771 | 1992 | 1942 | 2198.85 | 2496 | 1902 | 0.761 | -0.394 | 2.28E-06 | 5.63E-05 | 0.0023 | 0.0084 | 0.0045 | 0.0045 | TRUE | 0.0446 |
| Sun3 | 13 | 6 | 5 | 13 | 5 | 8 | 14 | 6 | 5 | 10 | 5 | 9 | 8.2 | 8 | 8 | 0.994 | -0.009 | 0.99003184 | 0.99463546 | 0.0847 | 0.4088 | 0.2513 | 0.2513 | TRUE | 0.9888 |
| Sun5 | 0 | 0 | 0 | 0 | 0 | 0 | 0 | 0 | 0 | 0 | 0 | 0 | 0 | 0 | 0 | NA | NA | NA | NA | NA | NA | NA | NA | NA | NA |
| Suox | 480 | 446 | 411 | 494 | 358 | 328 | 510 | 445 | 413 | 388 | 385 | 359 | 416.76 | 456 | 377 | 0.829 | -0.271 | 0.03257633 | 0.121608 | 0.0045 | 0.0148 | 0.0091 | 0.0091 | TRUE | 0.1227 |
| Supt16 | 25952 | 24276 | 23967 | 31629 | 22839 | 21615 | 27566 | 24245 | 24065 | 24864 | 24573 | 23661 | 24829.05 | 25292 | 24366 | 0.963 | -0.054 | 0.50269067 | 0.69786439 | 0.0032 | 0.007 | 0.0046 | 0.0046 | TRUE | 0.0757 |
| Supt20 | 3590 | 3822 | 3953 | 4290 | 3140 | 3095 | 3813 | 3817 | 3969 | 3372 | 3378 | 3388 | 3623.06 | 3866 | 3379 | 0.874 | -0.194 | 0.00136528 | 0.01136918 | 0 | 0.0078 | 0.0024 | 0.0024 | TRUE | 0.0066 |
| Supt3 | 698 | 857 | 922 | 976 | 780 | 796 | 741 | 856 | 926 | 767 | 839 | 871 | 833.49 | 841 | 826 | 0.981 | -0.028 | 0.81337706 | 0.90450545 | 0.0072 | 0.0109 | 0.0088 | 0.0088 | TRUE | 0.1268 |
| Supt4a | 1712 | 1675 | 1606 | 2814 | 1986 | 1814 | 1818 | 1673 | 1613 | 2212 | 2137 | 1986 | 1906.42 | 1701 | 2112 | 1.242 | 0.312 | 0.00040202 | 0.0043638 | 0.0026 | 0.0086 | 0.0051 | 0.0051 | TRUE | 0.0436 |
| Supt5 | 13702 | 11846 | 12244 | 17495 | 12802 | 11857 | 14554 | 11831 | 12294 | 13753 | 13774 | 12979 | 13197.61 | 12893 | 13502 | 1.047 | 0.067 | 0.49651682 | 0.69197343 | 0.0059 | 0.0072 | 0.0069 | 0.0069 | TRUE | 0.1552 |
| Supt6 | 12358 | 9318 | 10256 | 17908 | 12221 | 9843 | 13126 | 9306 | 10298 | 14078 | 13149 | 10775 | 11788.68 | 10910 | 12667 | 1.161 | 0.216 | 0.15171952 | 0.3428955 | 0.0254 | 0.0072 | 0.0162 | 0.0162 | TRUE | 0.3858 |
| Supt7l | 1109 | 1084 | 1054 | 1557 | 1138 | 1014 | 1178 | 1083 | 1058 | 1224 | 1224 | 1110 | 1146.21 | 1106 | 1186 | 1.073 | 0.102 | 0.27159697 | 0.49163634 | 0.0022 | 0.0098 | 0.0053 | 0.0053 | TRUE | 0.0382 |
| Supv3l1 | 2864 | 2868 | 2965 | 3882 | 2934 | 2667 | 3042 | 2864 | 2977 | 3052 | 3157 | 2919 | 3001.92 | 2961 | 3043 | 1.028 | 0.039 | 0.57889195 | 0.75593055 | 7.00E-04 | 0.008 | 0.0033 | 0.0033 | TRUE | 0.015 |
| Surf1 | 844 | 1141 | 1092 | 1146 | 887 | 963 | 896 | 1140 | 1096 | 901 | 954 | 1054 | 1006.98 | 1044 | 970 | 0.928 | -0.108 | 0.38657049 | 0.60061236 | 0.0094 | 0.0102 | 0.0102 | 0.0102 | TRUE | 0.1822 |
| Surf2 | 655 | 778 | 754 | 1135 | 753 | 755 | 696 | 777 | 757 | 892 | 810 | 826 | 793.12 | 743 | 843 | 1.135 | 0.183 | 0.06112515 | 0.18840451 | 0.0018 | 0.0111 | 0.0056 | 0.0056 | TRUE | 0.0367 |
| Surf4 | 6301 | 6698 | 6514 | 8995 | 6140 | 6383 | 6693 | 6689 | 6541 | 7071 | 6606 | 6987 | 6764.58 | 6641 | 6888 | 1.037 | 0.053 | 0.39486134 | 0.60835724 | 5.00E-04 | 0.0074 | 0.0026 | 0.0026 | TRUE | 0.0156 |
| Surf6 | 2879 | 3001 | 2882 | 3857 | 2788 | 2610 | 3058 | 2997 | 2894 | 3032 | 3000 | 2857 | 2972.97 | 2983 | 2963 | 0.994 | -0.009 | 0.89153207 | 0.94746449 | 5.00E-04 | 0.008 | 0.003 | 0.003 | TRUE | 0.0118 |
| Susd1 | 712 | 1059 | 1140 | 733 | 621 | 671 | 756 | 1058 | 1145 | 576 | 668 | 735 | 822.91 | 986 | 660 | 0.668 | -0.582 | 0.00073119 | 0.00699527 | 0.0293 | 0.0109 | 0.0201 | 0.0201 | TRUE | 0.495 |
| Susd2 | 1534 | 2182 | 2154 | 1499 | 1322 | 1707 | 1629 | 2179 | 2163 | 1178 | 1422 | 1869 | 1740.13 | 1990 | 1490 | 0.748 | -0.419 | 0.02160488 | 0.09194449 | 0.0391 | 0.0088 | 0.0234 | 0.0234 | TRUE | 0.5951 |
| Susd3 | 54 | 60 | 52 | 57 | 40 | 62 | 57 | 60 | 52 | 45 | 43 | 68 | 54.2 | 56 | 52 | 0.914 | -0.129 | 0.65782396 | 0.80728288 | 0.0157 | 0.0677 | 0.0428 | 0.0428 | TRUE | 0.563 |
| Susd4 | 539 | 659 | 709 | 546 | 435 | 506 | 573 | 658 | 712 | 429 | 468 | 554 | 565.62 | 648 | 484 | 0.746 | -0.424 | 0.00281872 | 0.02016187 | 0.0119 | 0.0127 | 0.0127 | 0.0127 | TRUE | 0.1859 |
| Susd5 | 770 | 1075 | 1083 | 749 | 608 | 745 | 818 | 1074 | 1087 | 589 | 654 | 816 | 839.57 | 993 | 686 | 0.69 | -0.535 | 0.00107429 | 0.00940796 | 0.0243 | 0.0108 | 0.0181 | 0.0181 | TRUE | 0.3196 |
| Suv39h1 | 1284 | 1736 | 1792 | 1901 | 1334 | 1420 | 1364 | 1734 | 1799 | 1494 | 1435 | 1554 | 1563.51 | 1632 | 1494 | 0.916 | -0.127 | 0.29760124 | 0.5184073 | 0.0102 | 0.009 | 0.0101 | 0.0101 | TRUE | 0.2489 |
| Suv39h2 | 1799 | 2095 | 2073 | 2546 | 1874 | 1860 | 1911 | 2092 | 2082 | 2001 | 2016 | 2036 | 2023.08 | 2028 | 2018 | 0.995 | -0.008 | 0.9182263 | 0.95991788 | 8.00E-04 | 0.0085 | 0.0035 | 0.0035 | TRUE | 0.0308 |
| Suv420h1 | 1940 | 2094 | 2134 | 2977 | 2266 | 2079 | 2061 | 2091 | 2143 | 2340 | 2438 | 2276 | 2224.81 | 2098 | 2351 | 1.121 | 0.164 | 0.01895955 | 0.08357464 | 3.00E-04 | 0.0084 | 0.0031 | 0.0031 | TRUE | 0.0123 |
| Suv420h2 | 4851 | 5826 | 5562 | 6102 | 4789 | 4831 | 5153 | 5818 | 5585 | 4797 | 5153 | 5288 | 5298.97 | 5519 | 5079 | 0.92 | -0.12 | 0.14175039 | 0.32733339 | 0.0028 | 0.0075 | 0.0046 | 0.0046 | TRUE | 0.041 |
| Suz12 | 30589 | 33030 | 32365 | 38051 | 27405 | 28977 | 32491 | 32987 | 32498 | 29913 | 29486 | 31720 | 31515.81 | 32659 | 30373 | 0.93 | -0.105 | 0.07979714 | 0.22418248 | 8.00E-04 | 0.007 | 0.0025 | 0.0025 | TRUE | 0.0184 |
| Sv2a | 975 | 1124 | 1107 | 807 | 562 | 677 | 1036 | 1123 | 1112 | 634 | 605 | 741 | 874.98 | 1090 | 660 | 0.605 | -0.725 | 4.07E-11 | 3.21E-09 | 0.005 | 0.0107 | 0.0075 | 0.0075 | TRUE | 0.1325 |
| Sv2b | 21 | 20 | 8 | 20 | 15 | 19 | 22 | 20 | 8 | 16 | 16 | 21 | 17.16 | 17 | 18 | 1.045 | 0.064 | 0.90346868 | 0.95228213 | 0.0682 | 0.1989 | 0.1391 | 0.1391 | TRUE | 1.0254 |
| Sv2c | 49 | 54 | 65 | 39 | 36 | 47 | 52 | 54 | 65 | 31 | 39 | 51 | 48.68 | 57 | 40 | 0.699 | -0.516 | 0.09561327 | 0.25355123 | 0.0184 | 0.0746 | 0.048 | 0.048 | TRUE | 0.4288 |
| Sva | 0 | 0 | 0 | 3 | 2 | 1 | 0 | 0 | 0 | 2 | 2 | 1 | 0.93 | 0 | 2 | 9.999 | 3.322 | 0.13367799 | NA | 0 | 3.5336 | 2.2742 | 2.2742 | TRUE | 0.1005 |
| Sval1 | 0 | 0 | 0 | 0 | 0 | 0 | 0 | 0 | 0 | 0 | 0 | 0 | 0 | 0 | 0 | NA | NA | NA | NA | NA | NA | NA | NA | NA | NA |
| Sval2 | 0 | 0 | 0 | 0 | 0 | 0 | 0 | 0 | 0 | 0 | 0 | 0 | 0 | 0 | 0 | NA | NA | NA | NA | NA | NA | NA | NA | NA | NA |
| Sval3 | 0 | 0 | 0 | 0 | 0 | 0 | 0 | 0 | 0 | 0 | 0 | 0 | 0 | 0 | 0 | NA | NA | NA | NA | NA | NA | NA | NA | NA | NA |
| Svep1 | 5 | 2 | 1 | 5 | 2 | 2 | 5 | 2 | 1 | 4 | 2 | 2 | 2.76 | 3 | 3 | 1.018 | 0.026 | 0.98234539 | NA | 0 | 1.1988 | 0.6172 | 0.6172 | TRUE | 0.7386 |
| Svil | 861 | 832 | 815 | 1280 | 942 | 854 | 915 | 831 | 818 | 1006 | 1014 | 935 | 919.74 | 855 | 985 | 1.153 | 0.206 | 0.02847866 | 0.11100349 | 0.0016 | 0.0105 | 0.0053 | 0.0053 | TRUE | 0.0444 |
| Svip | 955 | 1084 | 1149 | 1058 | 849 | 958 | 1014 | 1083 | 1154 | 832 | 913 | 1049 | 1007.43 | 1084 | 931 | 0.858 | -0.22 | 0.05991885 | 0.18563561 | 0.0075 | 0.0102 | 0.0089 | 0.0089 | TRUE | 0.1444 |
| Svop | 49 | 97 | 89 | 58 | 37 | 49 | 52 | 97 | 89 | 46 | 40 | 54 | 62.89 | 79 | 47 | 0.582 | -0.78 | 0.01319485 | 0.06441016 | 0.0493 | 0.0593 | 0.0544 | 0.0544 | TRUE | 0.8269 |
| Svopl | 12 | 7 | 7 | 7 | 5 | 5 | 13 | 7 | 7 | 6 | 5 | 5 | 7.19 | 9 | 5 | 0.613 | -0.706 | 0.32129166 | 0.54237525 | 0 | 0.4653 | 0.2181 | 0.2181 | TRUE | 0.4288 |
| Svs1 | 0 | 0 | 0 | 0 | 0 | 0 | 0 | 0 | 0 | 0 | 0 | 0 | 0 | 0 | 0 | NA | NA | NA | NA | NA | NA | NA | NA | NA | NA |
| Svs2 | 0 | 0 | 0 | 0 | 0 | 0 | 0 | 0 | 0 | 0 | 0 | 0 | 0 | 0 | 0 | NA | NA | NA | NA | NA | NA | NA | NA | NA | NA |
| Svs3a | 0 | 0 | 0 | 0 | 0 | 0 | 0 | 0 | 0 | 0 | 0 | 0 | 0 | 0 | 0 | NA | NA | NA | NA | NA | NA | NA | NA | NA | NA |
| Svs3b | 0 | 0 | 0 | 0 | 0 | 0 | 0 | 0 | 0 | 0 | 0 | 0 | 0 | 0 | 0 | NA | NA | NA | NA | NA | NA | NA | NA | NA | NA |
| Svs4 | 0 | 0 | 0 | 0 | 0 | 0 | 0 | 0 | 0 | 0 | 0 | 0 | 0 | 0 | 0 | NA | NA | NA | NA | NA | NA | NA | NA | NA | NA |
| Svs5 | 1 | 0 | 1 | 6 | 3 | 2 | 1 | 0 | 1 | 5 | 3 | 2 | 2.03 | 1 | 3 | 4.999 | 2.322 | 0.1277917 | NA | 0 | 1.6271 | 0.7844 | 0.7844 | TRUE | 0.2571 |
| Svs6 | 0 | 0 | 0 | 0 | 0 | 0 | 0 | 0 | 0 | 0 | 0 | 0 | 0 | 0 | 0 | NA | NA | NA | NA | NA | NA | NA | NA | NA | NA |
| Swap70 | 3316 | 3418 | 3649 | 4289 | 3314 | 3076 | 3522 | 3414 | 3664 | 3372 | 3566 | 3367 | 3484.04 | 3533 | 3435 | 0.972 | -0.041 | 0.55512062 | 0.73835068 | 7.00E-04 | 0.0078 | 0.0032 | 0.0032 | TRUE | 0.0134 |
| Swi5 | 5075 | 6200 | 6159 | 7548 | 5127 | 5487 | 5391 | 6192 | 6184 | 5934 | 5516 | 6006 | 5870.53 | 5922 | 5819 | 0.983 | -0.025 | 0.77021801 | 0.88002596 | 0.004 | 0.0075 | 0.0053 | 0.0053 | TRUE | 0.0751 |
| Swsap1 | 547 | 592 | 617 | 711 | 480 | 437 | 581 | 591 | 620 | 559 | 516 | 478 | 557.59 | 597 | 518 | 0.869 | -0.203 | 0.05983166 | 0.18554167 | 0.0018 | 0.0128 | 0.0066 | 0.0066 | TRUE | 0.059 |
| Swt1 | 2425 | 2517 | 2426 | 3062 | 2180 | 2200 | 2576 | 2514 | 2436 | 2407 | 2346 | 2408 | 2447.73 | 2509 | 2387 | 0.952 | -0.071 | 0.28324763 | 0.50491557 | 0 | 0.0082 | 0.0028 | 0.0028 | TRUE | 0.0078 |
| Syap1 | 655 | 773 | 792 | 913 | 688 | 790 | 696 | 772 | 795 | 718 | 740 | 865 | 764.29 | 754 | 774 | 1.025 | 0.036 | 0.75762467 | 0.8719099 | 0.0062 | 0.0112 | 0.0084 | 0.0084 | TRUE | 0.1226 |
| Sybu | 12 | 12 | 13 | 30 | 25 | 19 | 13 | 12 | 13 | 24 | 27 | 21 | 18.18 | 13 | 24 | 1.887 | 0.916 | 0.04025972 | 0.14155593 | 0 | 0.1881 | 0.0826 | 0.0826 | TRUE | 0.0719 |

| | | | | | | | | | | | | | | | | | | | | | | | | | |
|---|---|---|---|---|---|---|---|---|---|---|---|---|---|---|---|---|---|---|---|---|---|---|---|---|---|
| Syce1 | 1044 | 994 | 929 | 1971 | 1524 | 1130 | 1109 | 993 | 933 | 1549 | 1640 | 1237 | 1243.43 | 1012 | 1475 | 1.459 | 0.545 | 4.02E-05 | 0.0006437 | 0.0129 | 0.0096 | 0.0119 | 0.0119 | TRUE | 0.2383 |
| Syce1l | 2 | 0 | 0 | 1 | 1 | 0 | 2 | 0 | 0 | 1 | 1 | 0 | 0.66 | 1 | 1 | 0.896 | -0.159 | 0.95209679 | NA | 0 | 4.9654 | 3.5226 | 3.5226 | TRUE | 0.9743 |
| Syce2 | 1501 | 2208 | 2562 | 2709 | 2180 | 2423 | 1594 | 2205 | 2573 | 2130 | 2346 | 2652 | 2249.92 | 2124 | 2376 | 1.118 | 0.161 | 0.34842916 | 0.56705955 | 0.0341 | 0.0084 | 0.0209 | 0.0209 | TRUE | 0.4279 |
| Syce3 | 64 | 94 | 120 | 58 | 56 | 74 | 68 | 94 | 120 | 46 | 60 | 81 | 78.2 | 94 | 62 | 0.657 | -0.606 | 0.05418477 | 0.17303143 | 0.0676 | 0.049 | 0.058 | 0.058 | TRUE | 0.1488 |
| Sycn | 1 | 1 | 0 | 0 | 0 | 0 | 1 | 1 | 0 | 0 | 0 | 0 | 0.34 | 1 | 0 | 0.261 | -1.936 | 0.60454921 | NA | 0 | 9.5983 | 8.3269 | 8.3269 | TRUE | 0.2507 |
| Sycp1 | 1109 | 1211 | 1028 | 3728 | 2778 | 1903 | 1178 | 1209 | 1032 | 2931 | 2989 | 2083 | 1903.73 | 1140 | 2668 | 2.341 | 1.227 | 5.51E-16 | 8.61E-14 | 0.0206 | 0.0086 | 0.0159 | 0.0159 | TRUE | 0.444 |
| Sycp1-ps1 | 0 | 4 | 0 | 1 | 2 | 1 | 0 | 4 | 0 | 1 | 2 | 1 | 1.34 | 1 | 1 | 0.992 | -0.012 | 0.99499044 | NA | 1.1741 | 2.4693 | 1.9197 | 1.9197 | TRUE | 1.8981 |
| Sycp2 | 49 | 83 | 67 | 147 | 106 | 85 | 52 | 83 | 67 | 116 | 114 | 93 | 87.48 | 67 | 108 | 1.596 | 0.675 | 0.00646929 | 0.03809248 | 0.0176 | 0.0446 | 0.0321 | 0.0321 | TRUE | 0.3032 |
| Sycp3 | 319 | 690 | 726 | 660 | 507 | 646 | 339 | 689 | 729 | 519 | 546 | 707 | 588.07 | 586 | 591 | 1.008 | 0.011 | 0.96692073 | 0.9834445 | 0.0926 | 0.0125 | 0.0476 | 0.0476 | TRUE | 1.5896 |
| Syde1 | 75 | 139 | 131 | 243 | 183 | 133 | 80 | 139 | 132 | 191 | 197 | 146 | 147.26 | 117 | 178 | 1.524 | 0.608 | 0.014288 | 0.06832942 | 0.0463 | 0.0293 | 0.0372 | 0.0372 | TRUE | 0.7611 |
| Syde2 | 2875 | 2828 | 2856 | 3477 | 2682 | 2653 | 3054 | 2824 | 2868 | 2733 | 2886 | 2904 | 2878.16 | 2915 | 2841 | 0.974 | -0.038 | 0.60508593 | 0.77576129 | 0.0011 | 0.008 | 0.0035 | 0.0035 | TRUE | 0.0209 |
| Syf2 | 1708 | 1819 | 1913 | 2381 | 1760 | 1766 | 1814 | 1817 | 1921 | 1872 | 1894 | 1933 | 1875.05 | 1851 | 1900 | 1.026 | 0.037 | 0.59930839 | 0.77143379 | 0 | 0.0087 | 0.0031 | 0.0031 | TRUE | 0.0134 |
| Syk | 1672 | 1962 | 2020 | 1724 | 1299 | 1332 | 1776 | 1959 | 2028 | 1355 | 1398 | 1458 | 1662.45 | 1921 | 1404 | 0.73 | -0.454 | 2.33E-07 | 7.51E-06 | 0.0025 | 0.0089 | 0.0049 | 0.0049 | TRUE | 0.0527 |
| Sympk | 11352 | 13218 | 13784 | 14566 | 11163 | 11374 | 12058 | 13201 | 13841 | 11451 | 12011 | 12451 | 12501.89 | 13033 | 11971 | 0.918 | -0.123 | 0.12922834 | 0.30947303 | 0.0032 | 0.0072 | 0.0046 | 0.0046 | TRUE | 0.0524 |
| Syn1 | 862 | 679 | 724 | 877 | 638 | 654 | 916 | 678 | 727 | 689 | 686 | 716 | 735.41 | 774 | 697 | 0.902 | -0.149 | 0.26247348 | 0.48186844 | 0.0114 | 0.0114 | 0.0115 | 0.0115 | TRUE | 0.3046 |
| Syn2 | 24 | 23 | 11 | 31 | 29 | 20 | 25 | 23 | 11 | 24 | 31 | 22 | 22.83 | 20 | 26 | 1.302 | 0.381 | 0.40141103 | 0.6141287 | 0.0449 | 0.1512 | 0.1038 | 0.1038 | TRUE | 0.8177 |
| Syn3 | 60 | 52 | 46 | 92 | 60 | 75 | 64 | 52 | 46 | 72 | 65 | 82 | 63.47 | 54 | 73 | 1.354 | 0.437 | 0.09210293 | 0.24762974 | 0 | 0.0588 | 0.0324 | 0.0324 | TRUE | 0.2038 |
| Syna | 53 | 51 | 43 | 55 | 38 | 33 | 56 | 51 | 43 | 43 | 41 | 36 | 45.11 | 50 | 40 | 0.803 | -0.317 | 0.27436529 | 0.49488215 | 0 | 0.0799 | 0.0382 | 0.0382 | TRUE | 0.1206 |
| Synb | 0 | 0 | 0 | 0 | 0 | 0 | 0 | 0 | 0 | 0 | 0 | 0 | 0 | 0 | 0 | NA | NA | NA | NA | NA | NA | NA | NA | NA | NA |
| Sync | 52 | 55 | 47 | 136 | 79 | 66 | 55 | 55 | 47 | 107 | 85 | 72 | 70.25 | 52 | 88 | 1.689 | 0.756 | 0.00377431 | 0.02528823 | 0.0149 | 0.0538 | 0.0338 | 0.0338 | TRUE | 0.2889 |
| Syncrip | 10446 | 10031 | 10708 | 15593 | 11077 | 9960 | 11095 | 10018 | 10752 | 12258 | 11918 | 10903 | 11157.41 | 10622 | 11693 | 1.101 | 0.139 | 0.08452959 | 0.23332178 | 0.003 | 0.0072 | 0.0046 | 0.0046 | TRUE | 0.0427 |
| Syndig1 | 83 | 75 | 50 | 97 | 62 | 55 | 88 | 75 | 50 | 76 | 67 | 60 | 69.41 | 71 | 68 | 0.957 | -0.064 | 0.82236015 | 0.90923689 | 0.0317 | 0.0544 | 0.0434 | 0.0434 | TRUE | 0.4378 |
| Syndig1l | 8 | 12 | 16 | 13 | 13 | 15 | 8 | 12 | 16 | 10 | 14 | 16 | 12.86 | 12 | 13 | 1.099 | 0.136 | 0.80364641 | 0.89855367 | 0 | 0.263 | 0.1386 | 0.1386 | TRUE | 0.2455 |
| Syne1 | 582 | 817 | 990 | 797 | 591 | 775 | 618 | 816 | 994 | 627 | 636 | 848 | 756.5 | 809 | 704 | 0.869 | -0.203 | 0.29336253 | 0.51437903 | 0.0403 | 0.0113 | 0.0257 | 0.0257 | TRUE | 0.1846 |
| Syne2 | 2351 | 2072 | 2391 | 2867 | 1944 | 1976 | 2497 | 2069 | 2401 | 2254 | 2092 | 2163 | 2245.97 | 2322 | 2170 | 0.934 | -0.098 | 0.31613771 | 0.53684153 | 0.0047 | 0.0084 | 0.0064 | 0.0064 | TRUE | 0.1103 |
| Syne3 | 300 | 430 | 458 | 344 | 254 | 312 | 319 | 429 | 460 | 270 | 273 | 342 | 348.87 | 403 | 295 | 0.732 | -0.451 | 0.01206225 | 0.06035719 | 0.0242 | 0.0163 | 0.0203 | 0.0203 | TRUE | 0.3808 |
| Syne4 | 556 | 605 | 673 | 523 | 414 | 510 | 591 | 604 | 676 | 411 | 445 | 558 | 547.57 | 624 | 471 | 0.755 | -0.406 | 0.00494796 | 0.03107894 | 0.0132 | 0.0129 | 0.0132 | 0.0132 | TRUE | 0.2972 |
| Syngr1 | 1069 | 1518 | 1571 | 1419 | 1009 | 1212 | 1135 | 1516 | 1577 | 1116 | 1086 | 1327 | 1292.8 | 1409 | 1176 | 0.834 | -0.262 | 0.07858807 | 0.22197188 | 0.0195 | 0.0095 | 0.0152 | 0.0152 | TRUE | 0.3474 |
| Syngr2 | 6362 | 7274 | 7452 | 6860 | 5281 | 5491 | 6758 | 7265 | 7483 | 5393 | 5682 | 6011 | 6431.73 | 7169 | 5695 | 0.794 | -0.332 | 2.71E-05 | 0.00046036 | 0.0026 | 0.0074 | 0.0044 | 0.0044 | TRUE | 0.0306 |
| Syngr3 | 414 | 571 | 606 | 494 | 399 | 501 | 440 | 570 | 608 | 388 | 429 | 548 | 497.43 | 539 | 455 | 0.843 | -0.247 | 0.16526943 | 0.3619816 | 0.0271 | 0.0135 | 0.0208 | 0.0208 | TRUE | 0.3681 |
| Syngr4 | 226 | 514 | 570 | 451 | 342 | 444 | 240 | 513 | 572 | 355 | 368 | 486 | 422.38 | 442 | 403 | 0.911 | -0.135 | 0.63647132 | 0.79543785 | 0.1102 | 0.0147 | 0.0563 | 0.0563 | TRUE | 1.8411 |
| Synj1 | 1550 | 1426 | 1515 | 2301 | 1679 | 1468 | 1646 | 1424 | 1521 | 1809 | 1807 | 1607 | 1635.68 | 1530 | 1741 | 1.138 | 0.186 | 0.0541255 | 0.1729472 | 0.0039 | 0.0089 | 0.0061 | 0.0061 | TRUE | 0.054 |
| Synj2 | 628 | 635 | 609 | 458 | 398 | 412 | 667 | 634 | 612 | 360 | 428 | 451 | 525.33 | 638 | 413 | 0.646 | -0.63 | 3.47E-07 | 1.08E-05 | 0.0055 | 0.0132 | 0.009 | 0.009 | TRUE | 0.1547 |
| Synj2bp | 1072 | 1472 | 1558 | 1599 | 1159 | 1210 | 1139 | 1470 | 1564 | 1257 | 1247 | 1325 | 1333.62 | 1391 | 1276 | 0.917 | -0.125 | 0.33999083 | 0.56014202 | 0.0123 | 0.0094 | 0.0116 | 0.0116 | TRUE | 0.302 |
| Synm | 792 | 867 | 966 | 1297 | 1035 | 1008 | 841 | 866 | 970 | 1020 | 1114 | 1103 | 985.62 | 892 | 1079 | 1.208 | 0.273 | 0.00569306 | 0.03450841 | 0.003 | 0.0102 | 0.006 | 0.006 | TRUE | 0.0691 |
| Synpo | 227 | 284 | 228 | 248 | 198 | 250 | 241 | 284 | 229 | 195 | 213 | 274 | 239.22 | 251 | 227 | 0.902 | -0.149 | 0.40839022 | 0.62107131 | 0.018 | 0.0207 | 0.0193 | 0.0193 | TRUE | 0.3467 |
| Synpo2 | 146 | 75 | 68 | 142 | 114 | 74 | 155 | 75 | 68 | 112 | 123 | 81 | 102.26 | 99 | 105 | 1.059 | 0.083 | 0.80865742 | 0.90160787 | 0.1163 | 0.0391 | 0.0753 | 0.0753 | TRUE | 2.3132 |
| Synpo2l | 2 | 3 | 1 | 9 | 4 | 0 | 2 | 3 | 1 | 7 | 4 | 0 | 2.92 | 2 | 4 | 1.896 | 0.923 | 0.48805883 | NA | 0 | 1.1362 | 0.8999 | 0.8999 | TRUE | 0.2003 |
| Synpr | 73 | 53 | 55 | 77 | 45 | 50 | 78 | 53 | 55 | 61 | 48 | 55 | 58.23 | 62 | 55 | 0.886 | -0.175 | 0.53088037 | 0.71908667 | 0.0131 | 0.0635 | 0.0389 | 0.0389 | TRUE | 0.415 |
| Synrg | 2077 | 2433 | 2702 | 3031 | 2162 | 2327 | 2206 | 2430 | 2713 | 2383 | 2326 | 2547 | 2434.21 | 2450 | 2419 | 0.987 | -0.019 | 0.85352352 | 0.92686401 | 0.006 | 0.0083 | 0.0069 | 0.0069 | TRUE | 0.1073 |
| Syp | 69 | 83 | 112 | 59 | 60 | 63 | 73 | 83 | 112 | 46 | 65 | 69 | 74.76 | 89 | 60 | 0.665 | -0.589 | 0.03552074 | 0.12942897 | 0.0342 | 0.051 | 0.0427 | 0.0427 | TRUE | 0.4796 |
| Sypl | 3641 | 4333 | 4477 | 4565 | 3380 | 3801 | 3867 | 4327 | 4495 | 3589 | 3637 | 4161 | 4012.71 | 4230 | 3796 | 0.897 | -0.157 | 0.11247291 | 0.28215727 | 0.0061 | 0.0077 | 0.0068 | 0.0068 | TRUE | 0.086 |
| Sypl2 | 3 | 2 | 4 | 16 | 7 | 9 | 3 | 2 | 4 | 13 | 8 | 10 | 6.53 | 3 | 10 | 3.284 | 1.715 | 0.03209546 | 0.12067454 | 0 | 0.5116 | 0.2463 | 0.2463 | TRUE | 0.1443 |
| Sys1 | 1317 | 1340 | 1330 | 1973 | 1401 | 1275 | 1399 | 1338 | 1335 | 1551 | 1507 | 1396 | 1421.12 | 1357 | 1485 | 1.094 | 0.13 | 0.11380295 | 0.28449835 | 0.0012 | 0.0092 | 0.0042 | 0.0042 | TRUE | 0.0328 |
| Syt1 | 73 | 47 | 41 | 51 | 44 | 33 | 78 | 47 | 41 | 40 | 47 | 36 | 48.2 | 55 | 41 | 0.747 | -0.421 | 0.21689432 | 0.42884167 | 0.0498 | 0.0753 | 0.0626 | 0.0626 | TRUE | 1.0284 |
| Syt10 | 0 | 3 | 2 | 4 | 0 | 1 | 0 | 3 | 2 | 3 | 0 | 1 | 1.54 | 2 | 1 | 0.873 | -0.195 | 0.91221133 | NA | 0.7693 | 2.1453 | 1.6212 | 1.6212 | TRUE | 0.6065 |
| Syt11 | 528 | 623 | 582 | 686 | 460 | 525 | 561 | 622 | 584 | 539 | 495 | 575 | 562.72 | 589 | 536 | 0.91 | -0.136 | 0.21408689 | 0.42574436 | 0.0021 | 0.0128 | 0.0068 | 0.0068 | TRUE | 0.0511 |
| Syt12 | 51 | 52 | 63 | 94 | 77 | 60 | 54 | 52 | 63 | 74 | 83 | 66 | 65.3 | 56 | 74 | 1.313 | 0.393 | 0.11202695 | 0.28149099 | 0 | 0.0574 | 0.0284 | 0.0284 | TRUE | 0.0951 |
| Syt13 | 224 | 328 | 395 | 158 | 112 | 150 | 238 | 328 | 397 | 124 | 121 | 164 | 228.51 | 321 | 136 | 0.424 | -1.238 | 4.92E-08 | 1.96E-06 | 0.043 | 0.0213 | 0.0319 | 0.0319 | TRUE | 0.1343 |
| Syt14 | 144 | 143 | 183 | 207 | 159 | 152 | 153 | 143 | 184 | 163 | 171 | 166 | 163.28 | 160 | 167 | 1.042 | 0.06 | 0.72649256 | 0.85254767 | 0 | 0.0271 | 0.0148 | 0.0148 | TRUE | 0.1827 |
| Syt15 | 18 | 16 | 12 | 14 | 16 | 10 | 19 | 16 | 12 | 11 | 17 | 11 | 14.39 | 16 | 13 | 0.827 | -0.274 | 0.59495657 | 0.76795606 | 0 | 0.2359 | 0.1211 | 0.1211 | TRUE | 0.3015 |
| Syt16 | 10 | 34 | 28 | 22 | 21 | 19 | 11 | 34 | 28 | 17 | 23 | 21 | 22.23 | 24 | 20 | 0.829 | -0.27 | 0.59214419 | 0.76555069 | 0.1178 | 0.1551 | 0.1375 | 0.1375 | TRUE | 0.8354 |
| Syt17 | 166 | 191 | 187 | 249 | 146 | 160 | 176 | 191 | 188 | 196 | 157 | 175 | 180.47 | 185 | 176 | 0.955 | -0.067 | 0.67686521 | 0.82037332 | 0 | 0.0252 | 0.0132 | 0.0132 | TRUE | 0.1123 |
| Syt2 | 25 | 13 | 9 | 18 | 8 | 7 | 27 | 13 | 9 | 14 | 9 | 8 | 13.17 | 16 | 10 | 0.637 | -0.652 | 0.29685411 | 0.51768942 | 0.1476 | 0.2571 | 0.202 | 0.202 | TRUE | 0.884 |
| Syt3 | 364 | 291 | 282 | 320 | 249 | 257 | 387 | 291 | 283 | 252 | 268 | 281 | 293.53 | 320 | 267 | 0.833 | -0.263 | 0.10878732 | 0.27629527 | 0.0139 | 0.0181 | 0.0159 | 0.0159 | TRUE | 0.3715 |
| Syt4 | 23 | 17 | 11 | 35 | 19 | 12 | 24 | 17 | 11 | 28 | 20 | 13 | 18.92 | 17 | 20 | 1.179 | 0.237 | 0.64262508 | 0.79888727 | 0.0861 | 0.181 | 0.1354 | 0.1354 | TRUE | 0.2005 |
| Syt5 | 171 | 201 | 194 | 124 | 121 | 162 | 182 | 201 | 195 | 97 | 130 | 177 | 163.7 | 193 | 135 | 0.698 | -0.519 | 0.02498112 | 0.10165177 | 0.0381 | 0.027 | 0.0324 | 0.0324 | TRUE | 0.4435 |
| Syt6 | 8 | 8 | 5 | 13 | 10 | 5 | 8 | 8 | 5 | 10 | 11 | 5 | 7.99 | 7 | 9 | 1.239 | 0.31 | 0.64957725 | 0.80354219 | 0 | 0.4191 | 0.2084 | 0.2084 | TRUE | 0.3114 |
| Syt7 | 812 | 812 | 719 | 596 | 498 | 530 | 862 | 811 | 722 | 469 | 536 | 580 | 663.32 | 798 | 528 | 0.66 | -0.599 | 2.06E-06 | 5.19E-05 | 0.0083 | 0.0119 | 0.0099 | 0.0099 | TRUE | 0.1194 |
| Syt8 | 0 | 0 | 0 | 1 | 1 | 0 | 0 | 0 | 0 | 1 | 1 | 0 | 0.31 | 0 | 1 | 3.32 | 1.731 | 0.65383052 | NA | 0 | 10.6224 | 8.9541 | 8.9541 | TRUE | 0.202 |
| Syt9 | 1996 | 2125 | 2207 | 2680 | 1956 | 1876 | 2120 | 2122 | 2216 | 2107 | 2105 | 2054 | 2120.56 | 2153 | 2089 | 0.97 | -0.044 | 0.51678639 | 0.70855117 | 0 | 0.0085 | 0.0028 | 0.0028 | TRUE | 0.008 |
| Sytl1 | 188 | 230 | 208 | 107 | 92 | 89 | 200 | 230 | 209 | 84 | 99 | 97 | 153.13 | 213 | 93 | 0.437 | -1.193 | 8.21E-12 | 7.36E-10 | 0 | 0.0284 | 0.0143 | 0.0143 | TRUE | 0.0859 |
| Sytl2 | 14 | 5 | 6 | 21 | 15 | 15 | 15 | 5 | 6 | 17 | 16 | 16 | 12.49 | 9 | 16 | 1.907 | 0.931 | 0.12160838 | 0.29653795 | 0.0217 | 0.2706 | 0.1713 | 0.1713 | TRUE | 1.1921 |
| Sytl3 | 174 | 205 | 208 | 78 | 75 | 91 | 185 | 205 | 209 | 61 | 81 | 100 | 140.01 | 200 | 81 | 0.4 | -1.321 | 8.19E-10 | 5.19E-08 | 0.0173 | 0.0304 | 0.0247 | 0.0247 | TRUE | 0.4656 |
| Sytl4 | 19 | 26 | 13 | 75 | 34 | 27 | 20 | 26 | 13 | 59 | 37 | 30 | 30.72 | 20 | 42 | 2.13 | 1.091 | 0.01327749 | 0.06473338 | 0.0898 | 0.1142 | 0.1023 | 0.1023 | TRUE | 0.7065 |
| Sytl5 | 33 | 29 | 29 | 57 | 38 | 34 | 35 | 29 | 29 | 45 | 41 | 37 | 36.01 | 31 | 41 | 1.326 | 0.407 | 0.20115972 | 0.40877592 | 0 | 0.0984 | 0.0447 | 0.0447 | TRUE | 0.0829 |
| Syvn1 | 1794 | 1692 | 1646 | 2004 | 1499 | 1460 | 1906 | 1690 | 1653 | 1575 | 1613 | 1598 | 1672.42 | 1750 | 1595 | 0.912 | -0.133 | 0.12912446 | 0.30927122 | 0.0022 | 0.0089 | 0.0049 | 0.0049 | TRUE | 0.0733 |
| Szrd1 | 3789 | 3549 | 3347 | 5142 | 3868 | 3346 | 4025 | 3544 | 3361 | 4042 | 4162 | 3663 | 3799.41 | 3643 | 3956 | 1.086 | 0.119 | 0.23198502 | 0.44609457 | 0.0062 | 0.0078 | 0.0069 | 0.0069 | TRUE | 0.1018 |
| Szt2 | 2174 | 1752 | 1931 | 2420 | 1930 | 1696 | 2309 | 1750 | 1939 | 1902 | 2077 | 1857 | 1972.23 | 1999 | 1945 | 0.973 | -0.039 | 0.7435275 | 0.86316176 | 0.0111 | 0.0086 | 0.01 | 0.01 | TRUE | 0.2219 |

| | | | | | | | | | | | | | | | | | | | | | | | | | |
|---|---|---|---|---|---|---|---|---|---|---|---|---|---|---|---|---|---|---|---|---|---|---|---|---|---|
| T | 109 | 126 | 109 | 60 | 50 | 48 | 116 | 126 | 109 | 47 | 54 | 53 | 84.09 | 117 | 51 | 0.436 | -1.199 | 5.50E-08 | 2.15E-06 | 0 | 0.0461 | 0.0212 | 0.0212 | TRUE | 0.046 |
| T2 | 15 | 9 | 9 | 9 | 1 | 6 | 16 | 9 | 9 | 7 | 1 | 7 | 8.11 | 11 | 5 | 0.442 | -1.178 | 0.12947182 | 0.30966954 | 0.1062 | 0.413 | 0.2922 | 0.2922 | TRUE | 0.835 |
| Taar1 | 0 | 0 | 0 | 0 | 0 | 0 | 0 | 0 | 0 | 0 | 0 | 0 | 0 | 0 | 0 | NA | NA | NA | NA | NA | NA | NA | NA | NA | NA |
| Taar2 | 0 | 0 | 0 | 0 | 0 | 2 | 0 | 0 | 0 | 0 | 0 | 2 | 0.36 | 0 | 1 | 3.832 | 1.938 | 0.63022034 | NA | 10 | 9.0356 | 10 | 10 | TRUE | 0.9744 |
| Taar3 | 0 | 0 | 0 | 0 | 0 | 0 | 0 | 0 | 0 | 0 | 0 | 0 | 0 | 0 | 0 | NA | NA | NA | NA | NA | NA | NA | NA | NA | NA |
| Taar4 | 0 | 0 | 0 | 0 | 0 | 0 | 0 | 0 | 0 | 0 | 0 | 0 | 0 | 0 | 0 | NA | NA | NA | NA | NA | NA | NA | NA | NA | NA |
| Taar5 | 0 | 0 | 0 | 0 | 0 | 0 | 0 | 0 | 0 | 0 | 0 | 0 | 0 | 0 | 0 | NA | NA | NA | NA | NA | NA | NA | NA | NA | NA |
| Taar6 | 0 | 0 | 0 | 0 | 0 | 0 | 0 | 0 | 0 | 0 | 0 | 0 | 0 | 0 | 0 | NA | NA | NA | NA | NA | NA | NA | NA | NA | NA |
| Taar7a | 0 | 0 | 0 | 0 | 0 | 0 | 0 | 0 | 0 | 0 | 0 | 0 | 0 | 0 | 0 | NA | NA | NA | NA | NA | NA | NA | NA | NA | NA |
| Taar7b | 0 | 0 | 0 | 0 | 0 | 0 | 0 | 0 | 0 | 0 | 0 | 0 | 0 | 0 | 0 | NA | NA | NA | NA | NA | NA | NA | NA | NA | NA |
| Taar7d | 0 | 0 | 0 | 0 | 0 | 0 | 0 | 0 | 0 | 0 | 0 | 0 | 0 | 0 | 0 | NA | NA | NA | NA | NA | NA | NA | NA | NA | NA |
| Taar7e | 0 | 0 | 0 | 0 | 0 | 0 | 0 | 0 | 0 | 0 | 0 | 0 | 0 | 0 | 0 | NA | NA | NA | NA | NA | NA | NA | NA | NA | NA |
| Taar7f | 0 | 0 | 0 | 0 | 0 | 0 | 0 | 0 | 0 | 0 | 0 | 0 | 0 | 0 | 0 | NA | NA | NA | NA | NA | NA | NA | NA | NA | NA |
| Taar8a | 0 | 0 | 0 | 0 | 0 | 0 | 0 | 0 | 0 | 0 | 0 | 0 | 0 | 0 | 0 | NA | NA | NA | NA | NA | NA | NA | NA | NA | NA |
| Taar8b | 0 | 0 | 0 | 0 | 0 | 0 | 0 | 0 | 0 | 0 | 0 | 0 | 0 | 0 | 0 | NA | NA | NA | NA | NA | NA | NA | NA | NA | NA |
| Taar8c | 0 | 0 | 0 | 0 | 0 | 0 | 0 | 0 | 0 | 0 | 0 | 0 | 0 | 0 | 0 | NA | NA | NA | NA | NA | NA | NA | NA | NA | NA |
| Taar9 | 0 | 0 | 0 | 0 | 0 | 0 | 0 | 0 | 0 | 0 | 0 | 0 | 0 | 0 | 0 | NA | NA | NA | NA | NA | NA | NA | NA | NA | NA |
| Tab1 | 902 | 1101 | 968 | 1072 | 876 | 803 | 958 | 1100 | 972 | 843 | 943 | 879 | 948.98 | 1010 | 888 | 0.879 | -0.187 | 0.06565867 | 0.1975522 | 0.0035 | 0.0104 | 0.0064 | 0.0064 | TRUE | 0.0723 |
| Tab2 | 4806 | 5039 | 5074 | 6794 | 4912 | 5148 | 5105 | 5032 | 5095 | 5341 | 5285 | 5635 | 5248.89 | 5077 | 5420 | 1.067 | 0.094 | 0.12932631 | 0.30962138 | 3.00E-04 | 0.0075 | 0.0026 | 0.0026 | TRUE | 0.0145 |
| Tab3 | 2809 | 2679 | 2789 | 3996 | 2853 | 2811 | 2984 | 2676 | 2800 | 3141 | 3070 | 3077 | 2957.95 | 2820 | 3096 | 1.098 | 0.135 | 0.06646341 | 0.1989088 | 0.0011 | 0.008 | 0.0036 | 0.0036 | TRUE | 0.0312 |
| Tac1 | 0 | 0 | 0 | 2 | 1 | 1 | 0 | 0 | 0 | 2 | 1 | 1 | 0.62 | 0 | 1 | 6.676 | 2.739 | 0.27800695 | NA | 0 | 5.2881 | 3.2034 | 3.2034 | TRUE | 0.07 |
| Tac2 | 6 | 9 | 9 | 4 | 5 | 4 | 6 | 9 | 9 | 3 | 5 | 4 | 6.22 | 8 | 4 | 0.521 | -0.941 | 0.21534264 | 0.42717126 | 0 | 0.5368 | 0.2383 | 0.2383 | TRUE | 0.1371 |
| Tac4 | 0 | 1 | 1 | 0 | 0 | 0 | 0 | 1 | 1 | 0 | 0 | 0 | 0.33 | 1 | 0 | 0.268 | -1.901 | 0.61401143 | NA | 0 | 9.8764 | 8.4772 | 8.4772 | TRUE | 0.2278 |
| Tacc1 | 6777 | 7856 | 8271 | 7827 | 5533 | 5921 | 7198 | 7846 | 8305 | 6153 | 5953 | 6481 | 6989.47 | 7783 | 6196 | 0.796 | -0.329 | 7.32E-05 | 0.00107483 | 0.0033 | 0.0074 | 0.0048 | 0.0048 | TRUE | 0.0527 |
| Tacc2 | 3547 | 3627 | 3736 | 3802 | 3064 | 3308 | 3768 | 3622 | 3751 | 2989 | 3297 | 3621 | 3507.97 | 3714 | 3302 | 0.889 | -0.17 | 0.06480549 | 0.19610748 | 0.0044 | 0.0078 | 0.0058 | 0.0058 | TRUE | 0.0867 |
| Tacc3 | 4638 | 5073 | 5070 | 7250 | 5211 | 4910 | 4926 | 5066 | 5091 | 5699 | 5607 | 5375 | 5294.08 | 5028 | 5560 | 1.106 | 0.145 | 0.01891127 | 0.0835003 | 4.00E-04 | 0.0075 | 0.0026 | 0.0026 | TRUE | 0.0103 |
| Taco1 | 801 | 779 | 753 | 1167 | 786 | 635 | 851 | 778 | 756 | 917 | 846 | 695 | 807.18 | 795 | 819 | 1.032 | 0.046 | 0.72330187 | 0.85006834 | 0.0106 | 0.011 | 0.0108 | 0.0108 | TRUE | 0.2071 |
| Tacr1 | 3 | 4 | 14 | 8 | 4 | 6 | 3 | 4 | 14 | 6 | 4 | 7 | 6.4 | 7 | 6 | 0.808 | -0.307 | 0.71931698 | 0.8479218 | 0.2246 | 0.5217 | 0.3695 | 0.3695 | TRUE | 2.0052 |
| Tacr2 | 14 | 20 | 18 | 35 | 23 | 21 | 15 | 20 | 18 | 28 | 25 | 23 | 21.36 | 18 | 25 | 1.425 | 0.511 | 0.21131415 | 0.42189873 | 0 | 0.1611 | 0.0722 | 0.0722 | TRUE | 0.0905 |
| Tacr3 | 14 | 7 | 5 | 38 | 31 | 11 | 15 | 7 | 5 | 30 | 33 | 12 | 17.02 | 9 | 25 | 2.823 | 1.497 | 0.01615552 | 0.07496338 | 0.2047 | 0.2004 | 0.2025 | 0.2025 | TRUE | 1.1794 |
| Tacstd2 | 0 | 1 | 2 | 12 | 15 | 10 | 0 | 1 | 2 | 9 | 16 | 11 | 6.59 | 1 | 12 | 11.916 | 3.575 | 0.00092719 | 0.00840436 | 0 | 0.507 | 0.2971 | 0.2971 | TRUE | 0.3562 |
| Tada1 | 194 | 258 | 275 | 489 | 363 | 359 | 206 | 258 | 276 | 384 | 391 | 393 | 317.97 | 247 | 389 | 1.577 | 0.657 | 5.54E-06 | 0.00012063 | 0.0059 | 0.0173 | 0.0117 | 0.0117 | TRUE | 0.2261 |
| Tada2a | 5147 | 4810 | 4712 | 6481 | 4727 | 4520 | 5467 | 4804 | 4731 | 5095 | 5086 | 4948 | 5021.8 | 5001 | 5043 | 1.009 | 0.012 | 0.88203669 | 0.94168298 | 0.0031 | 0.0076 | 0.0047 | 0.0047 | TRUE | 0.081 |
| Tada2b | 744 | 662 | 622 | 983 | 735 | 606 | 790 | 661 | 625 | 773 | 791 | 663 | 717.15 | 692 | 742 | 1.074 | 0.102 | 0.43463087 | 0.64317865 | 0.0106 | 0.0115 | 0.011 | 0.011 | TRUE | 0.1813 |
| Tada3 | 5004 | 4529 | 4417 | 5809 | 4357 | 3957 | 5315 | 4523 | 4435 | 4567 | 4688 | 4332 | 4643.24 | 4758 | 4529 | 0.952 | -0.071 | 0.4583365 | 0.66283544 | 0.0055 | 0.0076 | 0.0064 | 0.0064 | TRUE | 0.1278 |
| Taf1 | 7470 | 5953 | 6383 | 9780 | 6885 | 6346 | 7934 | 5945 | 6409 | 7688 | 7408 | 6947 | 7055.3 | 6763 | 7348 | 1.087 | 0.12 | 0.31791209 | 0.53886184 | 0.0117 | 0.0074 | 0.0102 | 0.0102 | TRUE | 0.2801 |
| Taf10 | 2722 | 3444 | 3563 | 4519 | 3283 | 3110 | 2891 | 3440 | 3578 | 3552 | 3532 | 3404 | 3399.6 | 3303 | 3496 | 1.059 | 0.082 | 0.41070482 | 0.62196522 | 0.0055 | 0.0079 | 0.0069 | 0.0069 | TRUE | 0.1441 |
| Taf11 | 1285 | 1237 | 1092 | 1819 | 1265 | 1143 | 1365 | 1235 | 1096 | 1430 | 1361 | 1251 | 1289.83 | 1232 | 1347 | 1.094 | 0.13 | 0.2469196 | 0.46379674 | 0.007 | 0.0095 | 0.0083 | 0.0083 | TRUE | 0.1116 |
| Taf12 | 687 | 727 | 676 | 1114 | 780 | 625 | 730 | 726 | 679 | 876 | 839 | 684 | 755.61 | 712 | 800 | 1.125 | 0.17 | 0.1636424 | 0.35946629 | 0.008 | 0.0113 | 0.0095 | 0.0095 | TRUE | 0.187 |
| Taf13 | 815 | 750 | 701 | 1113 | 789 | 679 | 866 | 749 | 704 | 875 | 849 | 743 | 797.62 | 773 | 822 | 1.065 | 0.091 | 0.45332608 | 0.65875499 | 0.0081 | 0.011 | 0.0094 | 0.0094 | TRUE | 0.1301 |
| Taf15 | 10409 | 10783 | 11040 | 14653 | 11056 | 10128 | 11056 | 10769 | 11085 | 11519 | 11896 | 11087 | 11235.32 | 10970 | 11501 | 1.048 | 0.068 | 0.26210623 | 0.48147442 | 6.00E-04 | 0.0072 | 0.0026 | 0.0026 | TRUE | 0.012 |
| Taf1a | 493 | 484 | 436 | 649 | 527 | 445 | 524 | 483 | 438 | 510 | 567 | 487 | 501.53 | 482 | 521 | 1.083 | 0.115 | 0.3481077 | 0.56689258 | 0.0049 | 0.0135 | 0.0088 | 0.0088 | TRUE | 0.0738 |
| Taf1b | 1445 | 1555 | 1541 | 1936 | 1410 | 1388 | 1535 | 1553 | 1547 | 1522 | 1517 | 1519 | 1532.26 | 1545 | 1519 | 0.983 | -0.024 | 0.72616242 | 0.85247738 | 0 | 0.0091 | 0.0028 | 0.0028 | TRUE | 4.00E-04 |
| Taf1c | 1859 | 1385 | 1263 | 2135 | 1593 | 1277 | 1975 | 1383 | 1268 | 1678 | 1714 | 1398 | 1569.37 | 1542 | 1597 | 1.036 | 0.051 | 0.76936454 | 0.87946546 | 0.0336 | 0.009 | 0.021 | 0.021 | TRUE | 0.7247 |
| Taf1d | 2508 | 2465 | 2245 | 3867 | 2826 | 2374 | 2664 | 2462 | 2254 | 3040 | 3041 | 2599 | 2676.54 | 2460 | 2893 | 1.176 | 0.234 | 0.02522031 | 0.10244003 | 0.0068 | 0.0081 | 0.0075 | 0.0075 | TRUE | 0.0956 |
| Taf2 | 6766 | 5860 | 5589 | 7992 | 5629 | 5145 | 7187 | 5852 | 5612 | 6283 | 6056 | 5632 | 6103.7 | 6217 | 5990 | 0.964 | -0.053 | 0.63707916 | 0.79588218 | 0.0091 | 0.0074 | 0.0091 | 0.0091 | TRUE | 0.227 |
| Taf3 | 1912 | 2002 | 1948 | 2626 | 1824 | 1782 | 2031 | 1999 | 1956 | 2064 | 1963 | 1951 | 1993.97 | 1995 | 1993 | 0.999 | -0.001 | 0.98530154 | 0.99215832 | 0 | 0.0086 | 0.0031 | 0.0031 | TRUE | 0.0124 |
| Taf4a | 2405 | 2581 | 2702 | 3665 | 2892 | 2672 | 2555 | 2578 | 2713 | 2881 | 3112 | 2925 | 2793.83 | 2615 | 2973 | 1.136 | 0.184 | 0.0110657 | 0.05674686 | 0.001 | 0.0081 | 0.0034 | 0.0034 | TRUE | 0.0202 |
| Taf4b | 3563 | 2944 | 2889 | 3724 | 2587 | 2459 | 3785 | 2940 | 2901 | 2928 | 2783 | 2692 | 3004.72 | 3209 | 2801 | 0.873 | -0.196 | 0.10656784 | 0.27266637 | 0.0125 | 0.008 | 0.0103 | 0.0103 | TRUE | 0.2994 |
| Taf5 | 1983 | 2394 | 2505 | 2567 | 1892 | 1977 | 2106 | 2391 | 2515 | 2018 | 2036 | 2164 | 2205.05 | 2337 | 2073 | 0.886 | -0.174 | 0.06553313 | 0.19721209 | 0.0041 | 0.0084 | 0.006 | 0.006 | TRUE | 0.0904 |
| Taf5l | 5986 | 6474 | 6449 | 7071 | 5334 | 5586 | 6358 | 6466 | 6475 | 5559 | 5739 | 6115 | 6118.63 | 6433 | 5804 | 0.902 | -0.149 | 0.0270781 | 0.10725158 | 0.001 | 0.0074 | 0.0031 | 0.0031 | TRUE | 0.0267 |
| Taf6 | 2613 | 3193 | 3357 | 3575 | 2668 | 2722 | 2775 | 3189 | 3371 | 2810 | 2871 | 2980 | 2999.3 | 3112 | 2887 | 0.928 | -0.109 | 0.25469101 | 0.47275352 | 0.0047 | 0.008 | 0.0062 | 0.0062 | TRUE | 0.1083 |
| Taf6l | 1064 | 755 | 687 | 1505 | 1115 | 791 | 1130 | 754 | 690 | 1183 | 1200 | 866 | 970.44 | 858 | 1083 | 1.263 | 0.337 | 0.09919227 | 0.25984818 | 0.052 | 0.0103 | 0.029 | 0.029 | TRUE | 0.9139 |
| Taf7 | 5230 | 4617 | 4478 | 7829 | 5637 | 4909 | 5555 | 4611 | 4496 | 6155 | 6065 | 5374 | 5375.98 | 4887 | 5865 | 1.2 | 0.263 | 0.01678762 | 0.07679071 | 0.0094 | 0.0075 | 0.0085 | 0.0085 | TRUE | 0.1739 |
| Taf7l | 149 | 167 | 192 | 549 | 360 | 290 | 158 | 167 | 193 | 432 | 387 | 317 | 275.7 | 173 | 379 | 2.198 | 1.136 | 2.19E-11 | 1.81E-09 | 0.0144 | 0.0189 | 0.0165 | 0.0165 | TRUE | 0.188 |
| Taf8 | 2528 | 2634 | 2594 | 2852 | 2186 | 2204 | 2685 | 2631 | 2605 | 2242 | 2352 | 2413 | 2487.85 | 2640 | 2336 | 0.884 | -0.178 | 0.01019623 | 0.05351581 | 4.00E-04 | 0.0082 | 0.003 | 0.003 | TRUE | 0.0153 |
| Taf9 | 5080 | 5916 | 5729 | 6955 | 4707 | 4824 | 5396 | 5908 | 5753 | 5467 | 5064 | 5281 | 5478.21 | 5686 | 5271 | 0.927 | -0.109 | 0.1323736 | 0.31362421 | 0.0015 | 0.0075 | 0.0036 | 0.0036 | TRUE | 0.0242 |
| Taf9b | 73 | 176 | 162 | 365 | 323 | 457 | 78 | 176 | 163 | 287 | 348 | 500 | 258.45 | 139 | 378 | 2.723 | 1.445 | 2.59E-06 | 6.29E-05 | 0.121 | 0.0197 | 0.0631 | 0.0631 | TRUE | 0.6248 |
| Tagap | 128 | 163 | 171 | 159 | 124 | 140 | 136 | 163 | 172 | 125 | 133 | 153 | 147.02 | 157 | 137 | 0.872 | -0.198 | 0.28277502 | 0.50428852 | 0.0059 | 0.0293 | 0.0176 | 0.0176 | TRUE | 0.1401 |
| Tagap1 | 156 | 207 | 208 | 210 | 163 | 175 | 166 | 207 | 209 | 165 | 175 | 192 | 185.55 | 194 | 177 | 0.913 | -0.132 | 0.43303168 | 0.64213762 | 0.0056 | 0.0247 | 0.0149 | 0.0149 | TRUE | 0.1709 |
| Tagln | 58 | 42 | 48 | 87 | 47 | 41 | 62 | 42 | 48 | 68 | 51 | 45 | 52.6 | 51 | 55 | 1.089 | 0.123 | 0.68793719 | 0.82822114 | 0.0266 | 0.0695 | 0.0484 | 0.0484 | TRUE | 0.4574 |
| Tagln2 | 9146 | 12469 | 13182 | 7909 | 6272 | 7751 | 9715 | 12453 | 13236 | 6217 | 6748 | 8485 | 9475.68 | 11801 | 7150 | 0.606 | -0.723 | 2.10E-06 | 5.26E-05 | 0.0259 | 0.0073 | 0.0166 | 0.0166 | TRUE | 0.3252 |
| Tagln3 | 245 | 379 | 389 | 294 | 234 | 274 | 260 | 379 | 391 | 231 | 252 | 300 | 302.03 | 343 | 261 | 0.759 | -0.398 | 0.04028878 | 0.1416168 | 0.0298 | 0.0178 | 0.0238 | 0.0238 | TRUE | 0.505 |
| Tal1 | 16 | 30 | 40 | 37 | 33 | 44 | 17 | 30 | 40 | 29 | 36 | 48 | 33.31 | 29 | 38 | 1.285 | 0.361 | 0.38663284 | 0.60061236 | 0.0827 | 0.1058 | 0.0949 | 0.0949 | TRUE | 0.3211 |
| Tal2 | 53 | 70 | 92 | 43 | 19 | 30 | 56 | 70 | 92 | 34 | 20 | 33 | 50.94 | 73 | 29 | 0.4 | -1.32 | 8.89E-05 | 0.00125979 | 0.0427 | 0.0716 | 0.0581 | 0.0581 | TRUE | 0.1928 |
| Taldo1 | 11171 | 12839 | 12386 | 12852 | 9411 | 9901 | 11866 | 12822 | 12437 | 10103 | 10126 | 10838 | 11365.32 | 12375 | 10356 | 0.837 | -0.257 | 0.00017792 | 0.00220779 | 0.0014 | 0.0072 | 0.0033 | 0.0033 | TRUE | 0.0202 |
| Tamm41 | 1937 | 1878 | 1887 | 2224 | 1595 | 1694 | 2057 | 1876 | 1895 | 1748 | 1716 | 1854 | 1857.76 | 1943 | 1773 | 0.913 | -0.132 | 0.10342886 | 0.26760005 | 0.0013 | 0.0087 | 0.0042 | 0.0042 | TRUE | 0.0322 |
| Tanc1 | 5345 | 5455 | 5857 | 5991 | 4488 | 4830 | 5677 | 5448 | 5881 | 4710 | 4829 | 5287 | 5305.33 | 5669 | 4942 | 0.872 | -0.198 | 0.01113443 | 0.05699845 | 0.0023 | 0.0075 | 0.0042 | 0.0042 | TRUE | 0.0454 |
| Tanc2 | 1095 | 994 | 1046 | 1785 | 1219 | 1120 | 1163 | 993 | 1050 | 1403 | 1312 | 1226 | 1191.15 | 1069 | 1314 | 1.23 | 0.299 | 0.00357235 | 0.02420237 | 0.0043 | 0.0097 | 0.0067 | 0.0067 | TRUE | 0.071 |

| | | | | | | | | | | | | | | | | | | | | | | | | | |
|---|---|---|---|---|---|---|---|---|---|---|---|---|---|---|---|---|---|---|---|---|---|---|---|---|---|
| Tango2 | 463 | 392 | 341 | 631 | 435 | 384 | 492 | 391 | 342 | 496 | 468 | 420 | 435.02 | 408 | 461 | 1.131 | 0.178 | 0.26727678 | 0.48702174 | 0.0173 | 0.0145 | 0.0162 | 0.0162 | TRUE | 0.3637 |
| Tango6 | 3703 | 3850 | 4040 | 4979 | 3645 | 3599 | 3933 | 3845 | 4057 | 3914 | 3922 | 3940 | 3935.07 | 3945 | 3925 | 0.995 | -0.007 | 0.90503449 | 0.95269332 | 0 | 0.0077 | 0.0024 | 0.0024 | TRUE | 0.0075 |
| Tank | 472 | 493 | 495 | 633 | 460 | 457 | 501 | 492 | 497 | 498 | 495 | 500 | 497.26 | 497 | 498 | 1.001 | 0.002 | 0.98308884 | 0.99106917 | 0 | 0.0135 | 0.0048 | 0.0048 | TRUE | 7.00E-04 |
| Taok1 | 8575 | 7227 | 8240 | 10949 | 7740 | 6934 | 9108 | 7218 | 8274 | 8607 | 8328 | 7590 | 8187.5 | 8200 | 8175 | 0.997 | -0.004 | 0.96873323 | 0.98433582 | 0.0087 | 0.0073 | 0.0081 | 0.0081 | TRUE | 0.1341 |
| Taok2 | 3372 | 3247 | 3129 | 5207 | 3871 | 3425 | 3582 | 3243 | 3142 | 4093 | 4165 | 3749 | 3662.3 | 3322 | 4002 | 1.205 | 0.269 | 0.00203293 | 0.01559505 | 0.0035 | 0.0078 | 0.0052 | 0.0052 | TRUE | 0.0567 |
| Taok3 | 961 | 1086 | 1225 | 1278 | 912 | 1007 | 1021 | 1085 | 1230 | 1005 | 981 | 1102 | 1070.6 | 1112 | 1029 | 0.925 | -0.112 | 0.29888741 | 0.51926751 | 0.0052 | 0.01 | 0.0074 | 0.0074 | TRUE | 0.1037 |
| Tap1 | 269 | 407 | 373 | 318 | 277 | 297 | 286 | 406 | 375 | 250 | 298 | 325 | 323.31 | 356 | 291 | 0.816 | -0.293 | 0.09989459 | 0.26096381 | 0.0222 | 0.0171 | 0.0197 | 0.0197 | TRUE | 0.3347 |
| Tap2 | 147 | 210 | 174 | 159 | 115 | 132 | 156 | 210 | 175 | 125 | 124 | 144 | 155.63 | 180 | 131 | 0.726 | -0.462 | 0.01311616 | 0.06418492 | 0.0093 | 0.0281 | 0.0185 | 0.0185 | TRUE | 0.2215 |
| Tapbp | 963 | 998 | 1009 | 1195 | 792 | 923 | 1023 | 997 | 1013 | 939 | 852 | 1010 | 972.44 | 1011 | 934 | 0.924 | -0.114 | 0.23799314 | 0.45382998 | 0.0024 | 0.0103 | 0.0057 | 0.0057 | TRUE | 0.068 |
| Tapbpl | 113 | 62 | 63 | 85 | 75 | 60 | 120 | 62 | 63 | 67 | 81 | 66 | 76.4 | 82 | 71 | 0.87 | -0.201 | 0.52600773 | 0.71529414 | 0.0695 | 0.05 | 0.0592 | 0.0592 | TRUE | 1.5467 |
| Tapt1 | 617 | 836 | 858 | 951 | 732 | 738 | 655 | 835 | 862 | 748 | 788 | 808 | 782.48 | 784 | 781 | 0.995 | -0.007 | 0.95917773 | 0.97898705 | 0.0102 | 0.0111 | 0.0106 | 0.0106 | TRUE | 0.2427 |
| Tarbp2 | 2808 | 2679 | 2651 | 3369 | 2549 | 2291 | 2983 | 2676 | 2662 | 2648 | 2743 | 2508 | 2703.15 | 2774 | 2633 | 0.95 | -0.075 | 0.37430862 | 0.5904186 | 0.0026 | 0.0081 | 0.0047 | 0.0047 | TRUE | 0.0527 |
| Tardbp | 19971 | 17267 | 17172 | 28669 | 21402 | 18738 | 21213 | 17245 | 17243 | 22537 | 23027 | 20512 | 20296.03 | 18567 | 22025 | 1.186 | 0.247 | 0.02152154 | 0.09168102 | 0.0094 | 0.0071 | 0.0082 | 0.0082 | TRUE | 0.1901 |
| Tarm1 | 19 | 17 | 19 | 12 | 13 | 9 | 20 | 17 | 19 | 9 | 14 | 10 | 14.92 | 19 | 11 | 0.588 | -0.766 | 0.1202697 | 0.2942289 | 0 | 0.2277 | 0.1038 | 0.1038 | TRUE | 0.1811 |
| Tars | 7973 | 7818 | 7788 | 10408 | 7619 | 7303 | 8469 | 7808 | 7820 | 8182 | 8198 | 7994 | 8078.4 | 8032 | 8125 | 1.012 | 0.017 | 0.80129138 | 0.89741637 | 9.00E-04 | 0.0073 | 0.003 | 0.003 | TRUE | 0.0276 |
| Tars2 | 3784 | 4089 | 4065 | 4910 | 3734 | 3695 | 4019 | 4084 | 4082 | 3860 | 4018 | 4045 | 4017.81 | 4062 | 3974 | 0.978 | -0.032 | 0.6010727 | 0.77268877 | 0 | 0.0077 | 0.0024 | 0.0024 | TRUE | 0.0079 |
| Tarsl2 | 1845 | 2117 | 2239 | 2395 | 1609 | 1721 | 1960 | 2114 | 2248 | 1883 | 1731 | 1884 | 1970 | 2107 | 1833 | 0.87 | -0.201 | 0.02308401 | 0.09626681 | 0.0027 | 0.0086 | 0.0052 | 0.0052 | TRUE | 0.0453 |
| Tas1r1 | 19 | 25 | 28 | 31 | 22 | 20 | 20 | 25 | 28 | 24 | 24 | 22 | 23.87 | 24 | 23 | 0.955 | -0.066 | 0.86394186 | 0.93278279 | 0 | 0.1449 | 0.0656 | 0.0656 | TRUE | 0.133 |
| Tas1r2 | 2 | 1 | 1 | 5 | 2 | 4 | 2 | 1 | 1 | 4 | 2 | 4 | 2.43 | 1 | 3 | 2.559 | 1.356 | 0.27294626 | NA | 0 | 1.3619 | 0.589 | 0.589 | TRUE | 0.1505 |
| Tas1r3 | 2 | 1 | 4 | 3 | 1 | 1 | 2 | 1 | 4 | 2 | 1 | 1 | 1.94 | 2 | 1 | 0.651 | -0.619 | 0.64546603 | NA | 0 | 1.7009 | 0.775 | 0.775 | TRUE | 0.3032 |
| Tas2r102 | 0 | 0 | 0 | 0 | 0 | 0 | 0 | 0 | 0 | 0 | 0 | 0 | 0 | 0 | 0 | NA | NA | NA | NA | NA | NA | NA | NA | NA | NA |
| Tas2r103 | 0 | 0 | 0 | 0 | 0 | 0 | 0 | 0 | 0 | 0 | 0 | 0 | 0 | 0 | 0 | NA | NA | NA | NA | NA | NA | NA | NA | NA | NA |
| Tas2r104 | 0 | 0 | 0 | 0 | 0 | 0 | 0 | 0 | 0 | 0 | 0 | 0 | 0 | 0 | 0 | NA | NA | NA | NA | NA | NA | NA | NA | NA | NA |
| Tas2r105 | 0 | 0 | 0 | 0 | 0 | 0 | 0 | 0 | 0 | 0 | 0 | 0 | 0 | 0 | 0 | NA | NA | NA | NA | NA | NA | NA | NA | NA | NA |
| Tas2r106 | 0 | 0 | 0 | 0 | 0 | 0 | 0 | 0 | 0 | 0 | 0 | 0 | 0 | 0 | 0 | NA | NA | NA | NA | NA | NA | NA | NA | NA | NA |
| Tas2r107 | 0 | 0 | 0 | 1 | 0 | 0 | 0 | 0 | 0 | 1 | 0 | 0 | 0.13 | 0 | 0 | 1.859 | 0.894 | 0.82650734 | NA | 0 | 25.1515 | 10 | 10 | TRUE | 0.2563 |
| Tas2r108 | 0 | 1 | 1 | 0 | 0 | 0 | 0 | 1 | 1 | 0 | 0 | 0 | 0.33 | 1 | 0 | 0.268 | -1.901 | 0.61401143 | NA | 0 | 9.8764 | 8.4772 | 8.4772 | TRUE | 0.2278 |
| Tas2r109 | 0 | 0 | 0 | 0 | 0 | 0 | 0 | 0 | 0 | 0 | 0 | 0 | 0 | 0 | 0 | NA | NA | NA | NA | NA | NA | NA | NA | NA | NA |
| Tas2r110 | 0 | 0 | 0 | 0 | 0 | 0 | 0 | 0 | 0 | 0 | 0 | 0 | 0 | 0 | 0 | NA | NA | NA | NA | NA | NA | NA | NA | NA | NA |
| Tas2r113 | 0 | 0 | 0 | 0 | 0 | 0 | 0 | 0 | 0 | 0 | 0 | 0 | 0 | 0 | 0 | NA | NA | NA | NA | NA | NA | NA | NA | NA | NA |
| Tas2r114 | 0 | 0 | 0 | 0 | 0 | 0 | 0 | 0 | 0 | 0 | 0 | 0 | 0 | 0 | 0 | NA | NA | NA | NA | NA | NA | NA | NA | NA | NA |
| Tas2r115 | 0 | 0 | 0 | 0 | 0 | 0 | 0 | 0 | 0 | 0 | 0 | 0 | 0 | 0 | 0 | NA | NA | NA | NA | NA | NA | NA | NA | NA | NA |
| Tas2r116 | 0 | 0 | 0 | 0 | 0 | 0 | 0 | 0 | 0 | 0 | 0 | 0 | 0 | 0 | 0 | NA | NA | NA | NA | NA | NA | NA | NA | NA | NA |
| Tas2r117 | 0 | 0 | 0 | 0 | 0 | 0 | 0 | 0 | 0 | 0 | 0 | 0 | 0 | 0 | 0 | NA | NA | NA | NA | NA | NA | NA | NA | NA | NA |
| Tas2r118 | 0 | 0 | 0 | 0 | 0 | 0 | 0 | 0 | 0 | 0 | 0 | 0 | 0 | 0 | 0 | NA | NA | NA | NA | NA | NA | NA | NA | NA | NA |
| Tas2r119 | 0 | 0 | 0 | 0 | 0 | 0 | 0 | 0 | 0 | 0 | 0 | 0 | 0 | 0 | 0 | NA | NA | NA | NA | NA | NA | NA | NA | NA | NA |
| Tas2r120 | 0 | 0 | 0 | 0 | 0 | 0 | 0 | 0 | 0 | 0 | 0 | 0 | 0 | 0 | 0 | NA | NA | NA | NA | NA | NA | NA | NA | NA | NA |
| Tas2r121 | 0 | 0 | 0 | 0 | 0 | 0 | 0 | 0 | 0 | 0 | 0 | 0 | 0 | 0 | 0 | NA | NA | NA | NA | NA | NA | NA | NA | NA | NA |
| Tas2r122 | 0 | 0 | 0 | 0 | 0 | 0 | 0 | 0 | 0 | 0 | 0 | 0 | 0 | 0 | 0 | NA | NA | NA | NA | NA | NA | NA | NA | NA | NA |
| Tas2r123 | 0 | 1 | 0 | 0 | 0 | 0 | 0 | 1 | 0 | 0 | 0 | 0 | 0.17 | 0 | 0 | 0.49 | -1.029 | 0.80087 | NA | 0 | 19.7991 | 10 | 10 | TRUE | 0.405 |
| Tas2r124 | 0 | 0 | 0 | 0 | 0 | 0 | 0 | 0 | 0 | 0 | 0 | 0 | 0 | 0 | 0 | NA | NA | NA | NA | NA | NA | NA | NA | NA | NA |
| Tas2r125 | 0 | 0 | 0 | 0 | 0 | 0 | 0 | 0 | 0 | 0 | 0 | 0 | 0 | 0 | 0 | NA | NA | NA | NA | NA | NA | NA | NA | NA | NA |
| Tas2r126 | 0 | 0 | 0 | 0 | 0 | 0 | 0 | 0 | 0 | 0 | 0 | 0 | 0 | 0 | 0 | NA | NA | NA | NA | NA | NA | NA | NA | NA | NA |
| Tas2r129 | 0 | 0 | 0 | 0 | 0 | 0 | 0 | 0 | 0 | 0 | 0 | 0 | 0 | 0 | 0 | NA | NA | NA | NA | NA | NA | NA | NA | NA | NA |
| Tas2r130 | 0 | 0 | 0 | 0 | 0 | 0 | 0 | 0 | 0 | 0 | 0 | 0 | 0 | 0 | 0 | NA | NA | NA | NA | NA | NA | NA | NA | NA | NA |
| Tas2r131 | 0 | 0 | 0 | 0 | 0 | 0 | 0 | 0 | 0 | 0 | 0 | 0 | 0 | 0 | 0 | NA | NA | NA | NA | NA | NA | NA | NA | NA | NA |
| Tas2r134 | 0 | 0 | 0 | 0 | 0 | 0 | 0 | 0 | 0 | 0 | 0 | 0 | 0 | 0 | 0 | NA | NA | NA | NA | NA | NA | NA | NA | NA | NA |
| Tas2r135 | 0 | 0 | 0 | 0 | 0 | 0 | 0 | 0 | 0 | 0 | 0 | 0 | 0 | 0 | 0 | NA | NA | NA | NA | NA | NA | NA | NA | NA | NA |
| Tas2r136 | 0 | 0 | 0 | 0 | 0 | 0 | 0 | 0 | 0 | 0 | 0 | 0 | 0 | 0 | 0 | NA | NA | NA | NA | NA | NA | NA | NA | NA | NA |
| Tas2r137 | 1 | 0 | 1 | 0 | 0 | 0 | 1 | 0 | 1 | 0 | 0 | 0 | 0.34 | 1 | 0 | 0.261 | -1.939 | 0.60367425 | NA | 0 | 9.5732 | 8.3134 | 8.3134 | TRUE | 0.2529 |
| Tas2r138 | 0 | 0 | 0 | 0 | 0 | 0 | 0 | 0 | 0 | 0 | 0 | 0 | 0 | 0 | 0 | NA | NA | NA | NA | NA | NA | NA | NA | NA | NA |
| Tas2r139 | 0 | 0 | 0 | 0 | 0 | 0 | 0 | 0 | 0 | 0 | 0 | 0 | 0 | 0 | 0 | NA | NA | NA | NA | NA | NA | NA | NA | NA | NA |
| Tas2r140 | 0 | 0 | 0 | 0 | 0 | 0 | 0 | 0 | 0 | 0 | 0 | 0 | 0 | 0 | 0 | NA | NA | NA | NA | NA | NA | NA | NA | NA | NA |
| Tas2r143 | 0 | 0 | 0 | 0 | 0 | 0 | 0 | 0 | 0 | 0 | 0 | 0 | 0 | 0 | 0 | NA | NA | NA | NA | NA | NA | NA | NA | NA | NA |
| Tas2r144 | 0 | 0 | 0 | 0 | 0 | 0 | 0 | 0 | 0 | 0 | 0 | 0 | 0 | 0 | 0 | NA | NA | NA | NA | NA | NA | NA | NA | NA | NA |
| Tasp1 | 757 | 750 | 716 | 932 | 651 | 628 | 804 | 749 | 719 | 733 | 700 | 687 | 732.1 | 757 | 707 | 0.934 | -0.098 | 0.30443918 | 0.52512118 | 9.00E-04 | 0.0114 | 0.0052 | 0.0052 | TRUE | 0.0342 |
| Tat | 9 | 3 | 5 | 2 | 3 | 3 | 10 | 3 | 5 | 2 | 3 | 3 | 4.28 | 6 | 3 | 0.452 | -1.146 | 0.24122387 | NA | 0 | 0.7772 | 0.4163 | 0.4163 | TRUE | 0.3678 |
| Tatdn1 | 417 | 455 | 467 | 521 | 324 | 390 | 443 | 454 | 469 | 410 | 349 | 427 | 425.22 | 455 | 395 | 0.868 | -0.204 | 0.09522902 | 0.25297254 | 0.003 | 0.0147 | 0.0084 | 0.0084 | TRUE | 0.1174 |
| Tatdn2 | 2372 | 1842 | 1756 | 2961 | 1983 | 1612 | 2519 | 1840 | 1763 | 2328 | 2134 | 1765 | 2058.03 | 2041 | 2076 | 1.017 | 0.025 | 0.87858024 | 0.93996524 | 0.0288 | 0.0085 | 0.0186 | 0.0186 | TRUE | 0.5088 |
| Tatdn3 | 218 | 294 | 252 | 350 | 259 | 245 | 232 | 294 | 253 | 275 | 279 | 268 | 266.7 | 260 | 274 | 1.056 | 0.078 | 0.58407777 | 0.75938295 | 0.0033 | 0.0193 | 0.011 | 0.011 | TRUE | 0.149 |
| Tax1bp1 | 7197 | 6453 | 6508 | 9150 | 6429 | 6130 | 7644 | 6445 | 6535 | 7193 | 6917 | 6710 | 6907.39 | 6875 | 6940 | 1.01 | 0.014 | 0.87980931 | 0.94062783 | 0.0041 | 0.0074 | 0.0059 | 0.0059 | TRUE | 0.117 |
| Tax1bp3 | 231 | 202 | 208 | 439 | 329 | 297 | 245 | 202 | 209 | 345 | 354 | 325 | 280.03 | 219 | 341 | 1.563 | 0.644 | 3.24E-06 | 7.64E-05 | 0.0012 | 0.0187 | 0.01 | 0.01 | TRUE | 0.1234 |
| Taz | 703 | 850 | 737 | 790 | 690 | 684 | 747 | 849 | 740 | 621 | 742 | 749 | 741.3 | 779 | 704 | 0.903 | -0.148 | 0.21903563 | 0.43123579 | 0.0072 | 0.0113 | 0.009 | 0.009 | TRUE | 0.1307 |
| Tbata | 1 | 1 | 1 | 0 | 1 | 1 | 1 | 1 | 1 | 0 | 1 | 1 | 0.87 | 1 | 1 | 0.672 | -0.573 | 0.77064782 | NA | 0 | 3.7824 | 1.5699 | 1.5699 | TRUE | 0.387 |
| Tbc1d1 | 5071 | 5531 | 5789 | 6054 | 4453 | 4709 | 5386 | 5524 | 5813 | 4759 | 4791 | 5155 | 5238 | 5574 | 4902 | 0.879 | -0.186 | 0.00993453 | 0.05243889 | 0.0015 | 0.0075 | 0.0036 | 0.0036 | TRUE | 0.0247 |
| Tbc1d10a | 1544 | 1649 | 1657 | 1831 | 1331 | 1333 | 1640 | 1647 | 1664 | 1439 | 1432 | 1459 | 1546.89 | 1650 | 1443 | 0.875 | -0.193 | 0.00516381 | 0.03198888 | 0 | 0.009 | 0.0028 | 0.0028 | TRUE | 0.001 |
| Tbc1d10b | 4341 | 4235 | 3855 | 6937 | 4815 | 4540 | 4611 | 4230 | 3871 | 5453 | 5181 | 4970 | 4719.16 | 4237 | 5201 | 1.228 | 0.296 | 0.00119177 | 0.01017686 | 0.0046 | 0.0076 | 0.0058 | 0.0058 | TRUE | 0.0724 |
| Tbc1d10c | 61 | 55 | 84 | 91 | 78 | 73 | 65 | 55 | 84 | 72 | 84 | 80 | 73.24 | 68 | 79 | 1.15 | 0.202 | 0.42691989 | 0.63702748 | 0.0123 | 0.0519 | 0.0328 | 0.0328 | TRUE | 0.3979 |
| Tbc1d12 | 427 | 361 | 374 | 708 | 478 | 322 | 454 | 361 | 376 | 557 | 514 | 352 | 435.5 | 397 | 474 | 1.199 | 0.261 | 0.17044063 | 0.36901257 | 0.0335 | 0.0145 | 0.0239 | 0.0239 | TRUE | 0.5805 |

| | | | | | | | | | | | | | | | | | | | | | | | | | |
|---|---|---|---|---|---|---|---|---|---|---|---|---|---|---|---|---|---|---|---|---|---|---|---|---|---|
| Tbc1d13 | 3576 | 3235 | 3358 | 4235 | 2965 | 2925 | 3798 | 3231 | 3372 | 3329 | 3190 | 3202 | 3353.7 | 3467 | 3240 | 0.935 | -0.097 | 0.26540876 | 0.48505136 | 0.0035 | 0.0079 | 0.0052 | 0.0052 | TRUE | 0.0848 |
| Tbc1d14 | 3578 | 3627 | 3836 | 4447 | 3301 | 3574 | 3800 | 3622 | 3852 | 3496 | 3552 | 3912 | 3705.73 | 3758 | 3653 | 0.972 | -0.041 | 0.59771652 | 0.77025778 | 0.0019 | 0.0078 | 0.0041 | 0.0041 | TRUE | 0.0465 |
| Tbc1d15 | 5506 | 6040 | 5972 | 5509 | 4181 | 4945 | 5848 | 6032 | 5997 | 4331 | 4498 | 5413 | 5353.23 | 5959 | 4747 | 0.796 | -0.328 | 0.00128326 | 0.0107976 | 0.0074 | 0.0075 | 0.0073 | 0.0073 | TRUE | 0.1829 |
| Tbc1d16 | 3698 | 3452 | 3422 | 3120 | 2457 | 2395 | 3928 | 3448 | 3436 | 2453 | 2644 | 2622 | 3088.25 | 3604 | 2573 | 0.714 | -0.487 | 2.77E-08 | 1.19E-06 | 0.0034 | 0.008 | 0.0052 | 0.0052 | TRUE | 0.0751 |
| Tbc1d17 | 1181 | 1373 | 1442 | 1668 | 1251 | 1253 | 1254 | 1371 | 1448 | 1311 | 1346 | 1372 | 1350.41 | 1358 | 1343 | 0.989 | -0.017 | 0.85080842 | 0.92576408 | 0.0021 | 0.0093 | 0.0049 | 0.0049 | TRUE | 0.053 |
| Tbc1d19 | 465 | 537 | 567 | 793 | 619 | 570 | 494 | 536 | 569 | 623 | 666 | 624 | 585.48 | 533 | 638 | 1.195 | 0.257 | 0.0142243 | 0.06808672 | 0.0012 | 0.0125 | 0.0062 | 0.0062 | TRUE | 0.0479 |
| Tbc1d2 | 212 | 290 | 292 | 338 | 271 | 287 | 225 | 290 | 293 | 266 | 292 | 314 | 279.91 | 269 | 291 | 1.076 | 0.106 | 0.50303462 | 0.69812793 | 0.0103 | 0.0187 | 0.0143 | 0.0143 | TRUE | 0.2271 |
| Tbc1d20 | 1881 | 1726 | 1806 | 2256 | 1685 | 1794 | 1998 | 1724 | 1813 | 1773 | 1813 | 1964 | 1847.57 | 1845 | 1850 | 1.002 | 0.004 | 0.96912008 | 0.984481 | 0.0037 | 0.0087 | 0.0057 | 0.0057 | TRUE | 0.0633 |
| Tbc1d21 | 3 | 3 | 2 | 1 | 0 | 1 | 3 | 3 | 2 | 1 | 0 | 1 | 1.68 | 3 | 1 | 0.233 | -2.101 | 0.18655936 | NA | 0 | 1.9695 | 0.8717 | 0.8717 | TRUE | 0.1903 |
| Tbc1d22a | 1192 | 1373 | 1445 | 1466 | 1152 | 1257 | 1266 | 1371 | 1451 | 1152 | 1239 | 1376 | 1309.37 | 1363 | 1256 | 0.921 | -0.119 | 0.25624721 | 0.4749437 | 0.0052 | 0.0094 | 0.0072 | 0.0072 | TRUE | 0.0836 |
| Tbc1d22b | 2183 | 2374 | 2408 | 2455 | 1916 | 2078 | 2319 | 2371 | 2418 | 1930 | 2061 | 2275 | 2228.95 | 2369 | 2089 | 0.881 | -0.183 | 0.03871525 | 0.13743347 | 0.003 | 0.0084 | 0.0052 | 0.0052 | TRUE | 0.0732 |
| Tbc1d23 | 2604 | 2470 | 2485 | 3210 | 2328 | 2297 | 2766 | 2467 | 2495 | 2523 | 2505 | 2514 | 2545.1 | 2576 | 2514 | 0.976 | -0.035 | 0.65488074 | 0.80596172 | 0.0015 | 0.0082 | 0.004 | 0.004 | TRUE | 0.0503 |
| Tbc1d24 | 1353 | 1562 | 1577 | 1514 | 1229 | 1206 | 1437 | 1560 | 1583 | 1190 | 1322 | 1320 | 1402.21 | 1527 | 1277 | 0.836 | -0.259 | 0.00403957 | 0.02656888 | 0.0025 | 0.0093 | 0.0051 | 0.0051 | TRUE | 0.0444 |
| Tbc1d25 | 952 | 668 | 592 | 993 | 695 | 574 | 1011 | 667 | 594 | 781 | 748 | 628 | 738.25 | 757 | 719 | 0.95 | -0.075 | 0.70928231 | 0.84168239 | 0.0451 | 0.0114 | 0.0274 | 0.0274 | TRUE | 1.0137 |
| Tbc1d2b | 983 | 818 | 875 | 1361 | 1044 | 937 | 1044 | 817 | 879 | 1070 | 1123 | 1026 | 993.09 | 913 | 1073 | 1.175 | 0.233 | 0.04714235 | 0.15766097 | 0.0078 | 0.0102 | 0.0089 | 0.0089 | TRUE | 0.1865 |
| Tbc1d30 | 67 | 82 | 76 | 87 | 68 | 76 | 71 | 82 | 76 | 68 | 73 | 83 | 75.69 | 76 | 75 | 0.976 | -0.035 | 0.87676133 | 0.93932608 | 0 | 0.0504 | 0.0236 | 0.0236 | TRUE | 0.0826 |
| Tbc1d4 | 315 | 266 | 265 | 363 | 320 | 272 | 335 | 266 | 266 | 285 | 344 | 298 | 298.96 | 289 | 309 | 1.07 | 0.097 | 0.53162689 | 0.71978876 | 0.0105 | 0.0179 | 0.014 | 0.014 | TRUE | 0.2146 |
| Tbc1d5 | 1091 | 1134 | 1143 | 1665 | 1170 | 1116 | 1159 | 1133 | 1148 | 1309 | 1259 | 1222 | 1204.74 | 1147 | 1263 | 1.103 | 0.141 | 0.06974678 | 0.20534744 | 0 | 0.0096 | 0.0035 | 0.0035 | TRUE | 0.0125 |
| Tbc1d7 | 472 | 494 | 532 | 759 | 554 | 522 | 501 | 493 | 534 | 597 | 596 | 571 | 548.84 | 509 | 588 | 1.154 | 0.207 | 0.03598093 | 0.13027805 | 0 | 0.0129 | 0.0052 | 0.0052 | TRUE | 0.0209 |
| Tbc1d8 | 753 | 862 | 854 | 1255 | 949 | 1017 | 800 | 861 | 858 | 987 | 1021 | 1113 | 939.85 | 840 | 1040 | 1.238 | 0.308 | 0.00103748 | 0.00917212 | 0.0018 | 0.0104 | 0.0053 | 0.0053 | TRUE | 0.0446 |
| Tbc1d8b | 238 | 201 | 186 | 247 | 173 | 165 | 253 | 201 | 187 | 194 | 186 | 181 | 200.21 | 214 | 187 | 0.878 | -0.188 | 0.2685954 | 0.48850169 | 0.0089 | 0.0234 | 0.0159 | 0.0159 | TRUE | 0.2814 |
| Tbc1d9 | 715 | 732 | 719 | 931 | 595 | 643 | 759 | 731 | 722 | 732 | 640 | 704 | 714.73 | 737 | 692 | 0.939 | -0.09 | 0.35969801 | 0.57800887 | 0.0013 | 0.0115 | 0.0056 | 0.0056 | TRUE | 0.05 |
| Tbc1d9b | 2105 | 2802 | 2881 | 3602 | 2579 | 2647 | 2236 | 2798 | 2893 | 2832 | 2775 | 2898 | 2738.52 | 2642 | 2835 | 1.073 | 0.101 | 0.36957669 | 0.5868566 | 0.0093 | 0.0081 | 0.0088 | 0.0088 | TRUE | 0.2192 |
| Tbca | 3112 | 3691 | 3560 | 4189 | 2979 | 3181 | 3305 | 3686 | 3575 | 3293 | 3205 | 3482 | 3424.46 | 3522 | 3327 | 0.944 | -0.083 | 0.2954437 | 0.51637168 | 0.002 | 0.0079 | 0.0042 | 0.0042 | TRUE | 0.0351 |
| Tbcb | 2595 | 2757 | 2583 | 3713 | 2587 | 2452 | 2756 | 2753 | 2594 | 2919 | 2783 | 2684 | 2748.3 | 2701 | 2795 | 1.035 | 0.05 | 0.49559824 | 0.69160992 | 0.0012 | 0.0081 | 0.0035 | 0.0035 | TRUE | 0.0185 |
| Tbcc | 500 | 600 | 547 | 1101 | 774 | 566 | 531 | 599 | 549 | 866 | 833 | 620 | 666.24 | 560 | 773 | 1.381 | 0.466 | 0.0017356 | 0.01372132 | 0.0167 | 0.0118 | 0.0144 | 0.0144 | TRUE | 0.3519 |
| Tbccd1 | 1005 | 901 | 884 | 1377 | 1015 | 925 | 1067 | 900 | 888 | 1082 | 1092 | 1013 | 1007.01 | 952 | 1062 | 1.117 | 0.16 | 0.13612931 | 0.31954629 | 0.0049 | 0.0102 | 0.0073 | 0.0073 | TRUE | 0.1345 |
| Tbcd | 7592 | 7671 | 7569 | 9557 | 6860 | 6494 | 8064 | 7661 | 7600 | 7513 | 7381 | 7109 | 7554.64 | 7775 | 7334 | 0.943 | -0.084 | 0.18678975 | 0.39087807 | 7.00E-04 | 0.0073 | 0.0028 | 0.0028 | TRUE | 0.0129 |
| Tbce | 677 | 773 | 774 | 1046 | 770 | 740 | 719 | 772 | 777 | 822 | 828 | 810 | 788.18 | 756 | 820 | 1.085 | 0.117 | 0.18201674 | 0.38384382 | 0 | 0.0111 | 0.0043 | 0.0043 | TRUE | 0.0215 |
| Tbcel | 1752 | 2295 | 2403 | 2105 | 1643 | 1936 | 1861 | 2292 | 2413 | 1655 | 1768 | 2119 | 2017.94 | 2189 | 1847 | 0.844 | -0.245 | 0.07308072 | 0.21172467 | 0.0166 | 0.0085 | 0.013 | 0.013 | TRUE | 0.2073 |
| Tbck | 571 | 571 | 555 | 704 | 516 | 489 | 607 | 570 | 557 | 553 | 555 | 535 | 562.99 | 578 | 548 | 0.949 | -0.076 | 0.43556511 | 0.64371561 | 0 | 0.0128 | 0.0051 | 0.0051 | TRUE | 0.0216 |
| Tbk1 | 2014 | 1903 | 1892 | 2609 | 1913 | 1928 | 2139 | 1901 | 1900 | 2051 | 2058 | 2110 | 2026.55 | 1980 | 2073 | 1.047 | 0.067 | 0.41935103 | 0.62990518 | 0.0018 | 0.0085 | 0.0044 | 0.0044 | TRUE | 0.0597 |
| Tbkbp1 | 1137 | 1359 | 1475 | 1392 | 1094 | 1181 | 1208 | 1357 | 1481 | 1094 | 1177 | 1293 | 1268.36 | 1349 | 1188 | 0.88 | -0.184 | 0.10671355 | 0.27288789 | 0.0075 | 0.0095 | 0.0086 | 0.0086 | TRUE | 0.1001 |
| Tbl1x | 1963 | 2304 | 2558 | 3126 | 2398 | 2486 | 2085 | 2301 | 2569 | 2457 | 2580 | 2721 | 2452.24 | 2318 | 2586 | 1.115 | 0.157 | 0.12322352 | 0.29918123 | 0.0061 | 0.0082 | 0.0071 | 0.0071 | TRUE | 0.1081 |
| Tbl1xr1 | 3125 | 3474 | 3754 | 3942 | 2941 | 3233 | 3319 | 3470 | 3769 | 3099 | 3164 | 3539 | 3393.41 | 3519 | 3267 | 0.928 | -0.108 | 0.23912239 | 0.45504806 | 0.0043 | 0.0079 | 0.0057 | 0.0057 | TRUE | 0.064 |
| Tbl2 | 4114 | 5853 | 6437 | 4780 | 4215 | 4994 | 4370 | 5845 | 6463 | 3758 | 4535 | 5467 | 5073.02 | 5559 | 4587 | 0.825 | -0.278 | 0.11004342 | 0.2784095 | 0.0358 | 0.0076 | 0.0216 | 0.0216 | TRUE | 0.3584 |
| Tbl3 | 5831 | 5878 | 5729 | 7524 | 5353 | 5197 | 6194 | 5870 | 5753 | 5915 | 5759 | 5689 | 5863.28 | 5939 | 5788 | 0.975 | -0.037 | 0.56802402 | 0.74740657 | 7.00E-04 | 0.0075 | 0.0028 | 0.0028 | TRUE | 0.0171 |
| Tbp | 1499 | 1669 | 1634 | 2139 | 1467 | 1470 | 1592 | 1667 | 1641 | 1682 | 1578 | 1609 | 1628.14 | 1633 | 1623 | 0.994 | -0.008 | 0.90977874 | 0.95494678 | 0 | 0.0089 | 0.0033 | 0.0033 | TRUE | 0.0124 |
| Tbpl1 | 1600 | 1597 | 1632 | 2316 | 1662 | 1595 | 1699 | 1595 | 1639 | 1821 | 1788 | 1746 | 1714.66 | 1644 | 1785 | 1.086 | 0.119 | 0.10100172 | 0.26316973 | 0 | 0.0088 | 0.0032 | 0.0032 | TRUE | 0.0103 |
| Tbpl2 | 0 | 0 | 0 | 0 | 0 | 0 | 0 | 0 | 0 | 0 | 0 | 0 | 0 | 0 | 0 | NA | NA | NA | NA | NA | NA | NA | NA | NA | NA |
| Tbr1 | 0 | 2 | 1 | 4 | 4 | 0 | 0 | 2 | 1 | 3 | 4 | 0 | 1.74 | 1 | 2 | 2.478 | 1.309 | 0.44823307 | NA | 0 | 1.8985 | 1.4453 | 1.4453 | TRUE | 0.4655 |
| Tbrg1 | 6873 | 9059 | 9351 | 12345 | 8909 | 9063 | 7300 | 9047 | 9389 | 9705 | 9586 | 9921 | 9158.02 | 8579 | 9737 | 1.135 | 0.183 | 0.09006788 | 0.24423588 | 0.0085 | 0.0073 | 0.0083 | 0.0083 | TRUE | 0.2075 |
| Tbrg3 | 23 | 34 | 37 | 34 | 20 | 19 | 24 | 34 | 37 | 27 | 22 | 21 | 27.43 | 32 | 23 | 0.726 | -0.461 | 0.22369454 | 0.43578437 | 0 | 0.127 | 0.0664 | 0.0664 | TRUE | 0.273 |
| Tbrg4 | 4718 | 4318 | 4190 | 6201 | 4402 | 4037 | 5011 | 4312 | 4207 | 4875 | 4736 | 4419 | 4593.52 | 4510 | 4677 | 1.037 | 0.053 | 0.58299417 | 0.75875085 | 0.0048 | 0.0076 | 0.0064 | 0.0064 | TRUE | 0.1149 |
| Tbx1 | 32 | 62 | 55 | 85 | 63 | 68 | 34 | 62 | 55 | 67 | 68 | 74 | 60.03 | 50 | 70 | 1.379 | 0.463 | 0.11214454 | 0.28161615 | 0.022 | 0.0618 | 0.0442 | 0.0442 | TRUE | 0.6396 |
| Tbx10 | 0 | 0 | 0 | 2 | 2 | 1 | 0 | 0 | 0 | 2 | 2 | 1 | 0.8 | 0 | 2 | 8.55 | 3.096 | 0.18206245 | NA | 0 | 4.1089 | 2.5737 | 2.5737 | TRUE | 0.07 |
| Tbx15 | 11 | 22 | 7 | 33 | 28 | 27 | 12 | 22 | 7 | 26 | 30 | 30 | 21.05 | 14 | 29 | 2.097 | 1.069 | 0.03121108 | 0.11852889 | 0.0619 | 0.1634 | 0.1225 | 0.1225 | TRUE | 1.1576 |
| Tbx18 | 7 | 11 | 9 | 26 | 16 | 28 | 7 | 11 | 9 | 20 | 17 | 31 | 15.96 | 9 | 23 | 2.475 | 1.307 | 0.01202547 | 0.0602013 | 0 | 0.2133 | 0.118 | 0.118 | TRUE | 0.4917 |
| Tbx19 | 2 | 0 | 1 | 1 | 1 | 1 | 2 | 0 | 1 | 1 | 1 | 1 | 1.01 | 1 | 1 | 0.945 | -0.082 | 0.96468043 | NA | 0 | 3.2552 | 1.4594 | 1.4594 | TRUE | 0.198 |
| Tbx2 | 9 | 30 | 22 | 45 | 29 | 29 | 10 | 30 | 22 | 35 | 31 | 32 | 26.66 | 21 | 33 | 1.593 | 0.672 | 0.13903739 | 0.32369396 | 0.0789 | 0.1305 | 0.109 | 0.109 | TRUE | 0.5479 |
| Tbx20 | 49 | 51 | 31 | 97 | 44 | 56 | 52 | 51 | 31 | 76 | 47 | 61 | 53.17 | 45 | 61 | 1.387 | 0.472 | 0.149946 | 0.34065871 | 0.0463 | 0.0689 | 0.0581 | 0.0581 | TRUE | 0.2411 |
| Tbx21 | 76 | 53 | 79 | 93 | 67 | 68 | 81 | 53 | 79 | 73 | 72 | 74 | 72.1 | 71 | 73 | 1.032 | 0.046 | 0.8579124 | 0.92996525 | 0.0109 | 0.0526 | 0.0329 | 0.0329 | TRUE | 0.4525 |
| Tbx22 | 1 | 0 | 0 | 3 | 0 | 0 | 1 | 0 | 0 | 2 | 0 | 0 | 0.57 | 0 | 1 | 2.223 | 1.153 | 0.72182708 | NA | 0 | 5.7857 | 5.9519 | 5.9519 | TRUE | 1.4404 |
| Tbx3 | 3191 | 3017 | 2471 | 7560 | 5735 | 4519 | 3389 | 3013 | 2481 | 5943 | 6171 | 4947 | 4324 | 2961 | 5687 | 1.921 | 0.942 | 8.04E-12 | 7.24E-10 | 0.018 | 0.0077 | 0.0134 | 0.0134 | TRUE | 0.2444 |
| Tbx4 | 107 | 154 | 159 | 152 | 93 | 114 | 114 | 154 | 160 | 119 | 100 | 125 | 128.58 | 143 | 115 | 0.806 | -0.311 | 0.13870159 | 0.32322717 | 0.015 | 0.0325 | 0.0239 | 0.0239 | TRUE | 0.318 |
| Tbx5 | 15 | 20 | 20 | 20 | 14 | 20 | 16 | 20 | 20 | 16 | 15 | 22 | 18.11 | 19 | 18 | 0.935 | -0.096 | 0.82966472 | 0.91359855 | 0 | 0.1888 | 0.0891 | 0.0891 | TRUE | 0.2135 |
| Tbx6 | 48 | 69 | 79 | 36 | 34 | 51 | 51 | 69 | 79 | 28 | 37 | 56 | 53.32 | 66 | 40 | 0.6 | -0.737 | 0.03209292 | 0.12067454 | 0.0614 | 0.0687 | 0.0652 | 0.0652 | TRUE | 0.6514 |
| Tbxa2r | 78 | 55 | 66 | 44 | 34 | 43 | 83 | 55 | 66 | 35 | 37 | 47 | 53.72 | 68 | 40 | 0.577 | -0.793 | 0.00767639 | 0.04309416 | 0.0181 | 0.0682 | 0.0437 | 0.0437 | TRUE | 0.2012 |
| Tbxas1 | 1 | 1 | 0 | 0 | 0 | 0 | 1 | 1 | 0 | 0 | 0 | 0 | 0.34 | 1 | 0 | 0.261 | -1.936 | 0.60454921 | NA | 0 | 9.5983 | 8.3269 | 8.3269 | TRUE | 0.2507 |
| Tc2n | 85 | 87 | 67 | 96 | 63 | 64 | 90 | 87 | 67 | 75 | 68 | 70 | 76.29 | 81 | 71 | 0.875 | -0.193 | 0.41109729 | 0.62224822 | 0 | 0.0501 | 0.0264 | 0.0264 | TRUE | 0.219 |
| Tcaim | 796 | 964 | 953 | 988 | 760 | 738 | 845 | 963 | 957 | 777 | 818 | 808 | 861.24 | 922 | 801 | 0.868 | -0.204 | 0.033809 | 0.1250259 | 0.0019 | 0.0107 | 0.0055 | 0.0055 | TRUE | 0.0619 |
| Tcam1 | 29 | 39 | 38 | 86 | 56 | 50 | 31 | 39 | 38 | 68 | 60 | 55 | 48.42 | 36 | 61 | 1.696 | 0.762 | 0.0075263 | 0.0423572 | 0 | 0.0749 | 0.0364 | 0.0364 | TRUE | 0.1092 |
| Tcap | 1 | 0 | 1 | 1 | 0 | 0 | 1 | 0 | 1 | 1 | 0 | 0 | 0.48 | 1 | 0 | 0.509 | -0.975 | 0.7336918 | NA | 0 | 6.9367 | 4.1806 | 4.1806 | TRUE | 0.2562 |
| Tcea1 | 5516 | 6189 | 6289 | 8292 | 6012 | 6184 | 5859 | 6181 | 6315 | 6519 | 6469 | 6769 | 6351.87 | 6118 | 6586 | 1.076 | 0.106 | 0.1046754 | 0.26954128 | 8.00E-04 | 0.0074 | 0.0029 | 0.0029 | TRUE | 0.0167 |
| Tcea2 | 263 | 377 | 381 | 344 | 280 | 286 | 279 | 377 | 383 | 270 | 301 | 313 | 320.53 | 346 | 295 | 0.85 | -0.234 | 0.15032279 | 0.34103726 | 0.0148 | 0.0172 | 0.0159 | 0.0159 | TRUE | 0.3218 |
| Tcea3 | 3000 | 4206 | 4388 | 3300 | 2755 | 3245 | 3187 | 4201 | 4406 | 2594 | 2964 | 3552 | 3483.95 | 3931 | 3037 | 0.772 | -0.373 | 0.01713816 | 0.07771795 | 0.026 | 0.0078 | 0.0173 | 0.0173 | TRUE | 0.3338 |
| Tceal1 | 69 | 51 | 67 | 110 | 71 | 79 | 73 | 51 | 67 | 86 | 76 | 86 | 73.47 | 64 | 83 | 1.305 | 0.384 | 0.11526133 | 0.28673367 | 0 | 0.0517 | 0.0289 | 0.0289 | TRUE | 0.277 |
| Tceal3 | 62 | 66 | 78 | 91 | 72 | 61 | 66 | 66 | 78 | 72 | 77 | 67 | 70.98 | 70 | 72 | 1.027 | 0.038 | 0.87034508 | 0.93630208 | 0 | 0.0533 | 0.0248 | 0.0248 | TRUE | 0.0973 |
| Tceal5 | 59 | 39 | 38 | 68 | 38 | 35 | 63 | 39 | 38 | 53 | 41 | 38 | 45.4 | 47 | 44 | 0.957 | -0.064 | 0.84808273 | 0.92394563 | 0.0372 | 0.0795 | 0.0587 | 0.0587 | TRUE | 0.7055 |

| | | | | | | | | | | | | | | | | | | | | | | | | |
|---|---|---|---|---|---|---|---|---|---|---|---|---|---|---|---|---|---|---|---|---|---|---|---|---|
| Tceal6 | 11 | 17 | 8 | 11 | 11 | 6 | 12 | 17 | 8 | 9 | 12 | 7 | 10.62 | 12 | 9 | 0.736 | -0.442 | 0.47088386 | 0.67235019 | 0 | 0.317 | 0.1749 | 0.1749 | TRUE | 0.2499 |
| Tceal7 | 15 | 38 | 26 | 47 | 21 | 29 | 16 | 38 | 26 | 37 | 23 | 32 | 28.55 | 27 | 31 | 1.144 | 0.195 | 0.6551476 | 0.80606817 | 0.0789 | 0.1223 | 0.1018 | 0.1018 | TRUE | 0.2562 |
| Tceal8 | 1511 | 1273 | 1215 | 2271 | 1528 | 1366 | 1605 | 1271 | 1220 | 1785 | 1644 | 1495 | 1503.49 | 1365 | 1641 | 1.203 | 0.267 | 0.04366408 | 0.14956267 | 0.0146 | 0.0091 | 0.0119 | 0.0119 | TRUE | 0.2824 |
| Tceanc | 477 | 688 | 716 | 627 | 462 | 609 | 507 | 687 | 719 | 493 | 497 | 667 | 594.89 | 638 | 552 | 0.865 | -0.209 | 0.24349037 | 0.46008258 | 0.0297 | 0.0124 | 0.0215 | 0.0215 | TRUE | 0.3811 |
| Tceanc2 | 292 | 285 | 355 | 327 | 236 | 256 | 310 | 285 | 356 | 257 | 254 | 280 | 290.41 | 317 | 264 | 0.831 | -0.267 | 0.06044547 | 0.18682068 | 0.0049 | 0.0182 | 0.0111 | 0.0111 | TRUE | 0.1349 |
| Tceb1 | 2583 | 2491 | 2393 | 4129 | 2699 | 2517 | 2744 | 2488 | 2403 | 3246 | 2904 | 2755 | 2756.55 | 2545 | 2968 | 1.167 | 0.223 | 0.02333593 | 0.09700955 | 0.0055 | 0.0081 | 0.0066 | 0.0066 | TRUE | 0.0821 |
| Tceb2 | 5639 | 6153 | 6283 | 7774 | 5821 | 5850 | 5990 | 6145 | 6309 | 6111 | 6263 | 6404 | 6203.59 | 6148 | 6259 | 1.018 | 0.026 | 0.67449637 | 0.81857162 | 4.00E-04 | 0.0074 | 0.0025 | 0.0025 | TRUE | 0.0064 |
| Tceb3 | 2275 | 2113 | 2152 | 2875 | 2053 | 1898 | 2416 | 2110 | 2161 | 2260 | 2209 | 2078 | 2205.7 | 2229 | 2182 | 0.979 | -0.03 | 0.73289332 | 0.8565538 | 0.0031 | 0.0084 | 0.0051 | 0.0051 | TRUE | 0.0652 |
| Tcerg1 | 9901 | 9077 | 9355 | 11741 | 8752 | 8479 | 10517 | 9065 | 9393 | 9230 | 9417 | 9282 | 9483.89 | 9658 | 9310 | 0.964 | -0.053 | 0.50676401 | 0.70098947 | 0.0029 | 0.0073 | 0.0045 | 0.0045 | TRUE | 0.0738 |
| Tcerg1l | 141 | 82 | 63 | 120 | 79 | 55 | 150 | 82 | 63 | 94 | 85 | 60 | 89.08 | 98 | 80 | 0.815 | -0.294 | 0.40143772 | 0.6141287 | 0.1146 | 0.0439 | 0.0774 | 0.0774 | TRUE | 1.1534 |
| Tcf12 | 1460 | 1676 | 1826 | 2130 | 1571 | 1523 | 1551 | 1674 | 1833 | 1674 | 1690 | 1667 | 1681.67 | 1686 | 1677 | 0.995 | -0.008 | 0.9314412 | 0.96623853 | 0.0029 | 0.0089 | 0.0052 | 0.0052 | TRUE | 0.0706 |
| Tcf15 | 2363 | 2123 | 2162 | 2637 | 1963 | 1482 | 2510 | 2120 | 2171 | 2073 | 2112 | 1622 | 2101.4 | 2267 | 1936 | 0.854 | -0.227 | 0.07820882 | 0.22129701 | 0.0135 | 0.0085 | 0.0115 | 0.0115 | TRUE | 0.241 |
| Tcf19 | 825 | 607 | 666 | 943 | 666 | 663 | 876 | 606 | 669 | 741 | 717 | 726 | 722.48 | 717 | 728 | 1.016 | 0.022 | 0.88022718 | 0.94070627 | 0.0173 | 0.0115 | 0.0145 | 0.0145 | TRUE | 0.4445 |
| Tcf20 | 10603 | 9977 | 10594 | 13634 | 9852 | 9653 | 11262 | 9964 | 10637 | 10718 | 10600 | 10567 | 10624.79 | 10621 | 10628 | 1.001 | 0.001 | 0.98852074 | 0.99387704 | 0.0014 | 0.0072 | 0.0036 | 0.0036 | TRUE | 0.0358 |
| Tcf21 | 0 | 0 | 0 | 1 | 2 | 0 | 0 | 0 | 0 | 1 | 2 | 0 | 0.49 | 0 | 1 | 5.204 | 2.38 | 0.47267317 | NA | 0 | 6.7348 | 6.4099 | 6.4099 | TRUE | 0.1222 |
| Tcf23 | 6 | 3 | 2 | 17 | 7 | 8 | 6 | 3 | 2 | 13 | 8 | 9 | 6.84 | 4 | 10 | 2.649 | 1.405 | 0.07812901 | 0.22111084 | 0 | 0.4887 | 0.2739 | 0.2739 | TRUE | 0.5356 |
| Tcf25 | 14407 | 15006 | 15891 | 20372 | 15072 | 14906 | 15303 | 14987 | 15956 | 16015 | 16217 | 16317 | 15799 | 15415 | 16183 | 1.05 | 0.07 | 0.22572645 | 0.43849705 | 5.00E-04 | 0.0071 | 0.0023 | 0.0023 | TRUE | 0.0115 |
| Tcf3 | 5536 | 6250 | 6455 | 7162 | 5608 | 5736 | 5880 | 6242 | 6482 | 5630 | 6034 | 6279 | 6091.11 | 6201 | 5981 | 0.964 | -0.053 | 0.50346387 | 0.69859971 | 0.0024 | 0.0074 | 0.0043 | 0.0043 | TRUE | 0.0324 |
| Tcf4 | 2836 | 2688 | 2665 | 3540 | 2820 | 2547 | 3012 | 2685 | 2676 | 2783 | 3034 | 2788 | 2829.65 | 2791 | 2868 | 1.028 | 0.039 | 0.64812103 | 0.80243071 | 0.0027 | 0.0081 | 0.005 | 0.005 | TRUE | 0.0583 |
| Tcf7 | 5593 | 6543 | 6792 | 5002 | 3810 | 4469 | 5941 | 6535 | 6820 | 3932 | 4099 | 4892 | 5369.79 | 6432 | 4308 | 0.67 | -0.579 | 1.18E-07 | 4.10E-06 | 0.0092 | 0.0075 | 0.0084 | 0.0084 | TRUE | 0.171 |
| Tcf7l1 | 2208 | 1361 | 1361 | 3969 | 2887 | 2559 | 2345 | 1359 | 1367 | 3120 | 3106 | 2801 | 2349.78 | 1690 | 3009 | 1.781 | 0.832 | 3.04E-05 | 0.0005101 | 0.0574 | 0.0083 | 0.0282 | 0.0282 | TRUE | 1.3848 |
| Tcf7l2 | 145 | 123 | 122 | 291 | 175 | 181 | 154 | 123 | 123 | 229 | 188 | 198 | 169.09 | 133 | 205 | 1.546 | 0.628 | 0.00046746 | 0.00493162 | 0.0079 | 0.0264 | 0.017 | 0.017 | TRUE | 0.1906 |
| Tcfl5 | 3679 | 3175 | 3115 | 4493 | 3136 | 2693 | 3908 | 3171 | 3128 | 3532 | 3374 | 2948 | 3343.42 | 3402 | 3285 | 0.966 | -0.05 | 0.67658772 | 0.82016306 | 0.012 | 0.0079 | 0.0101 | 0.0101 | TRUE | 0.2053 |
| Tchh | 58 | 64 | 69 | 143 | 101 | 46 | 62 | 64 | 69 | 112 | 109 | 50 | 77.71 | 65 | 90 | 1.399 | 0.484 | 0.13950192 | 0.32412923 | 0.08 | 0.0493 | 0.064 | 0.064 | TRUE | 1.385 |
| Tchhl1 | 0 | 0 | 0 | 1 | 0 | 0 | 0 | 0 | 0 | 1 | 0 | 0 | 0.13 | 0 | 0 | 1.859 | 0.894 | 0.82650734 | NA | 0 | 25.1515 | 10 | 10 | TRUE | 0.2563 |
| Tchp | 526 | 472 | 523 | 702 | 563 | 518 | 559 | 471 | 525 | 552 | 606 | 567 | 546.65 | 518 | 575 | 1.108 | 0.149 | 0.18723515 | 0.39165422 | 0.0028 | 0.0129 | 0.0073 | 0.0073 | TRUE | 0.0738 |
| Tcirg1 | 554 | 562 | 522 | 672 | 549 | 524 | 588 | 561 | 524 | 528 | 591 | 574 | 561.07 | 558 | 564 | 1.01 | 0.014 | 0.89322926 | 0.94834594 | 0.0017 | 0.0128 | 0.0065 | 0.0065 | TRUE | 0.0385 |
| Tcl1 | 3125 | 3964 | 3952 | 3080 | 2523 | 2995 | 3319 | 3959 | 3968 | 2421 | 2715 | 3278 | 3276.79 | 3749 | 2805 | 0.748 | -0.419 | 0.00167571 | 0.01336835 | 0.0165 | 0.0079 | 0.0125 | 0.0125 | TRUE | 0.2647 |
| Tcl1b1 | 0 | 0 | 0 | 0 | 0 | 0 | 0 | 0 | 0 | 0 | 0 | 0 | 0 | 0 | 0 | NA | NA | NA | NA | NA | NA | NA | NA | NA | NA |
| Tcl1b2 | 0 | 0 | 0 | 1 | 1 | 0 | 0 | 0 | 0 | 1 | 1 | 0 | 0.31 | 0 | 1 | 3.32 | 1.731 | 0.65383052 | NA | 0 | 10.6224 | 8.9541 | 8.9541 | TRUE | 0.202 |
| Tcl1b3 | 0 | 0 | 0 | 0 | 0 | 0 | 0 | 0 | 0 | 0 | 0 | 0 | 0 | 0 | 0 | NA | NA | NA | NA | NA | NA | NA | NA | NA | NA |
| Tcl1b4 | 0 | 0 | 0 | 0 | 0 | 0 | 0 | 0 | 0 | 0 | 0 | 0 | 0 | 0 | 0 | NA | NA | NA | NA | NA | NA | NA | NA | NA | NA |
| Tcl1b5 | 0 | 0 | 0 | 0 | 0 | 0 | 0 | 0 | 0 | 0 | 0 | 0 | 0 | 0 | 0 | NA | NA | NA | NA | NA | NA | NA | NA | NA | NA |
| Tcn2 | 1309 | 1726 | 1726 | 1402 | 1117 | 1231 | 1390 | 1724 | 1733 | 1102 | 1202 | 1348 | 1416.46 | 1616 | 1217 | 0.753 | -0.41 | 0.00117599 | 0.01006929 | 0.0114 | 0.0092 | 0.0108 | 0.0108 | TRUE | 0.179 |
| Tcof1 | 15125 | 12523 | 12847 | 16613 | 12478 | 10943 | 16065 | 12507 | 12900 | 13060 | 13426 | 11979 | 13322.71 | 13824 | 12822 | 0.928 | -0.109 | 0.34405909 | 0.56385079 | 0.0115 | 0.0072 | 0.0094 | 0.0094 | TRUE | 0.2459 |
| Tcp1 | 33388 | 37078 | 37070 | 48006 | 34067 | 34063 | 35464 | 37030 | 37222 | 37739 | 36654 | 37287 | 36899.36 | 36572 | 37227 | 1.018 | 0.026 | 0.64451941 | 0.80010705 | 4.00E-04 | 0.007 | 0.0022 | 0.0022 | TRUE | 0.0086 |
| Tcp10a | 0 | 0 | 0 | 0 | 0 | 1 | 0 | 0 | 0 | 0 | 0 | 1 | 0.18 | 0 | 0 | 1.859 | 0.894 | 0.82650734 | NA | 0 | 18.0643 | 10 | 10 | TRUE | 0.5383 |
| Tcp10b | 12 | 2 | 3 | 2 | 1 | 3 | 13 | 2 | 3 | 2 | 1 | 3 | 3.95 | 6 | 2 | 0.332 | -1.591 | 0.17136203 | NA | 0.4639 | 0.8414 | 0.6381 | 0.6381 | TRUE | 2.2679 |
| Tcp10c | 0 | 0 | 0 | 1 | 1 | 0 | 0 | 0 | 0 | 1 | 1 | 0 | 0.31 | 0 | 1 | 3.32 | 1.731 | 0.65383052 | NA | 0 | 10.6224 | 8.9541 | 8.9541 | TRUE | 0.202 |
| Tcp11 | 467 | 806 | 889 | 848 | 706 | 741 | 496 | 805 | 893 | 667 | 760 | 811 | 738.51 | 731 | 746 | 1.019 | 0.027 | 0.89387043 | 0.94872978 | 0.0489 | 0.0114 | 0.029 | 0.029 | TRUE | 0.9335 |
| Tcp11l1 | 144 | 235 | 245 | 263 | 198 | 197 | 153 | 235 | 246 | 207 | 213 | 216 | 211.51 | 211 | 212 | 1.002 | 0.002 | 0.99134862 | 0.99557064 | 0.0267 | 0.0225 | 0.0246 | 0.0246 | TRUE | 0.6285 |
| Tcp11l2 | 475 | 416 | 415 | 475 | 347 | 382 | 505 | 415 | 417 | 373 | 373 | 418 | 416.94 | 446 | 388 | 0.871 | -0.199 | 0.13094973 | 0.31162492 | 0.0061 | 0.0148 | 0.0101 | 0.0101 | TRUE | 0.154 |
| Tcstv1 | 38 | 23 | 9 | 61 | 35 | 24 | 40 | 23 | 9 | 48 | 38 | 26 | 30.71 | 24 | 37 | 1.556 | 0.638 | 0.22844476 | 0.44176445 | 0.2326 | 0.1142 | 0.168 | 0.168 | TRUE | 0.3926 |
| Tcstv3 | 80 | 44 | 29 | 138 | 105 | 43 | 85 | 44 | 29 | 108 | 113 | 47 | 71.09 | 53 | 89 | 1.705 | 0.77 | 0.08801433 | 0.24041601 | 0.2259 | 0.0532 | 0.1315 | 0.1315 | TRUE | 1.4832 |
| Tcta | 146 | 238 | 214 | 292 | 222 | 208 | 155 | 238 | 215 | 230 | 239 | 228 | 217.29 | 203 | 232 | 1.144 | 0.194 | 0.29417836 | 0.51501621 | 0.0181 | 0.0221 | 0.0201 | 0.0201 | TRUE | 0.451 |
| Tcte1 | 25 | 18 | 19 | 17 | 27 | 10 | 27 | 18 | 19 | 13 | 29 | 11 | 19.5 | 21 | 18 | 0.835 | -0.261 | 0.61345873 | 0.78135113 | 0.102 | 0.1759 | 0.1406 | 0.1406 | TRUE | 1.4598 |
| Tcte2 | 27 | 18 | 21 | 26 | 22 | 13 | 29 | 18 | 21 | 20 | 24 | 14 | 21.01 | 23 | 19 | 0.865 | -0.209 | 0.63367452 | 0.7939204 | 0 | 0.1637 | 0.0915 | 0.0915 | TRUE | 0.311 |
| Tcte3 | 0 | 0 | 0 | 0 | 0 | 0 | 0 | 0 | 0 | 0 | 0 | 0 | 0 | 0 | 0 | NA | NA | NA | NA | NA | NA | NA | NA | NA | NA |
| Tctex1d1 | 1 | 0 | 1 | 0 | 2 | 1 | 1 | 0 | 1 | 0 | 2 | 1 | 0.89 | 1 | 1 | 1.53 | 0.614 | 0.77623094 | NA | 0 | 3.7275 | 2.1477 | 2.1477 | TRUE | 0.1683 |
| Tctex1d2 | 699 | 961 | 911 | 763 | 579 | 666 | 742 | 960 | 915 | 600 | 623 | 729 | 761.46 | 872 | 651 | 0.745 | -0.425 | 0.00199932 | 0.01536709 | 0.0122 | 0.0112 | 0.0123 | 0.0123 | TRUE | 0.2008 |
| Tctex1d4 | 0 | 1 | 6 | 5 | 2 | 4 | 0 | 1 | 6 | 4 | 2 | 4 | 2.91 | 2 | 3 | 1.485 | 0.57 | 0.66523429 | NA | 0.5617 | 1.1374 | 0.895 | 0.895 | TRUE | 1.9748 |
| Tctn1 | 332 | 367 | 384 | 544 | 427 | 454 | 353 | 367 | 386 | 428 | 459 | 497 | 414.8 | 369 | 461 | 1.25 | 0.322 | 0.00627692 | 0.03727392 | 0.0019 | 0.0148 | 0.0076 | 0.0076 | TRUE | 0.0526 |
| Tctn2 | 794 | 723 | 755 | 993 | 701 | 702 | 843 | 722 | 758 | 781 | 754 | 768 | 771.14 | 774 | 768 | 0.992 | -0.012 | 0.90633354 | 0.95350897 | 0.002 | 0.0112 | 0.0058 | 0.0058 | TRUE | 0.0712 |
| Tctn3 | 426 | 519 | 505 | 706 | 526 | 464 | 452 | 518 | 507 | 555 | 566 | 508 | 517.79 | 492 | 543 | 1.102 | 0.141 | 0.20871289 | 0.41832757 | 0.002 | 0.0133 | 0.0071 | 0.0071 | TRUE | 0.0585 |
| Tdg | 38 | 27 | 28 | 41 | 37 | 24 | 40 | 27 | 28 | 32 | 40 | 26 | 32.29 | 32 | 33 | 1.032 | 0.045 | 0.90126069 | 0.95202489 | 0.0142 | 0.1089 | 0.0641 | 0.0641 | TRUE | 0.3651 |
| Tdgf1 | 51062 | 69975 | 71449 | 80521 | 59249 | 66219 | 54237 | 69885 | 71742 | 63299 | 63748 | 72487 | 65899.76 | 65288 | 66511 | 1.019 | 0.027 | 0.828746 | 0.91322447 | 0.0111 | 0.007 | 0.011 | 0.011 | TRUE | 0.2685 |
| Tdh | 15157 | 16501 | 17166 | 10969 | 9322 | 10584 | 16099 | 16480 | 17236 | 8623 | 10030 | 11586 | 13342.38 | 16605 | 10080 | 0.607 | -0.72 | 4.10E-10 | 2.74E-08 | 0.0113 | 0.0072 | 0.0095 | 0.0095 | TRUE | 0.2088 |
| Tdo2 | 1 | 0 | 0 | 0 | 1 | 0 | 1 | 0 | 0 | 0 | 1 | 0 | 0.36 | 0 | 0 | 0.954 | -0.067 | 0.98557668 | NA | 0 | 9.2518 | 8.0179 | 8.0179 | TRUE | 0.5204 |
| Tdp1 | 2467 | 2584 | 2676 | 3114 | 2202 | 2222 | 2620 | 2581 | 2687 | 2448 | 2369 | 2432 | 2522.93 | 2629 | 2416 | 0.919 | -0.122 | 0.06120179 | 0.18856703 | 0 | 0.0082 | 0.0026 | 0.0026 | TRUE | 0.0045 |
| Tdp2 | 2999 | 3211 | 3226 | 4157 | 3018 | 3010 | 3185 | 3207 | 3239 | 3268 | 3247 | 3295 | 3240.26 | 3210 | 3270 | 1.018 | 0.026 | 0.65595877 | 0.80656285 | 0 | 0.0079 | 0.0022 | 0.0022 | TRUE | 7.00E-04 |
| Tdpoz1 | 0 | 0 | 0 | 3 | 7 | 2 | 0 | 0 | 0 | 2 | 8 | 2 | 2.01 | 0 | 4 | 21.3 | 4.413 | 0.01566987 | NA | 0.4822 | 1.6433 | 1.2805 | 1.2805 | TRUE | 0.9168 |
| Tdpoz2 | 0 | 0 | 0 | 4 | 3 | 1 | 0 | 0 | 0 | 3 | 3 | 1 | 1.24 | 0 | 2 | 13.319 | 3.735 | 0.06833944 | NA | 0 | 2.6541 | 1.8305 | 1.8305 | TRUE | 0.2388 |
| Tdpoz3 | 69 | 17 | 6 | 108 | 86 | 22 | 73 | 17 | 6 | 85 | 93 | 24 | 49.63 | 32 | 67 | 2.096 | 1.068 | 0.31520743 | 0.53603999 | 0.791 | 0.0733 | 0.3838 | 0.791 | TRUE | 5.5007 |
| Tdpoz4 | 1 | 3 | 0 | 51 | 29 | 14 | 1 | 3 | 0 | 40 | 31 | 15 | 15.11 | 1 | 29 | 21.361 | 4.417 | 1.59E-06 | 4.10E-05 | 0.204 | 0.2249 | 0.2178 | 0.2178 | TRUE | 0.4095 |
| Tdpoz5 | 0 | 0 | 0 | 0 | 0 | 0 | 0 | 0 | 0 | 0 | 0 | 0 | 0 | 0 | 0 | NA | NA | NA | NA | NA | NA | NA | NA | NA | NA |
| Tdrd1 | 229 | 327 | 322 | 305 | 280 | 260 | 243 | 327 | 323 | 240 | 301 | 285 | 286.46 | 298 | 275 | 0.922 | -0.117 | 0.49342172 | 0.68979325 | 0.0169 | 0.0184 | 0.0177 | 0.0177 | TRUE | 0.286 |
| Tdrd12 | 420 | 519 | 505 | 1282 | 1054 | 1124 | 446 | 518 | 507 | 1008 | 1134 | 1230 | 807.29 | 490 | 1124 | 2.289 | 1.194 | 1.57E-23 | 5.52E-21 | 0.0071 | 0.011 | 0.0088 | 0.0088 | TRUE | 0.1006 |
| Tdrd3 | 694 | 785 | 774 | 1041 | 759 | 818 | 737 | 784 | 777 | 818 | 817 | 895 | 804.79 | 766 | 843 | 1.1 | 0.138 | 0.13625111 | 0.31973701 | 7.00E-04 | 0.011 | 0.0049 | 0.0049 | TRUE | 0.0337 |
| Tdrd5 | 846 | 971 | 1005 | 972 | 748 | 744 | 899 | 970 | 1009 | 764 | 805 | 814 | 876.8 | 959 | 794 | 0.827 | -0.273 | 0.00318232 | 0.02221848 | 0.0013 | 0.0107 | 0.005 | 0.005 | TRUE | 0.0361 |

| | | | | | | | | | | | | | | | | | | | | | | | | |
|---|---|---|---|---|---|---|---|---|---|---|---|---|---|---|---|---|---|---|---|---|---|---|---|---|
| Tdrd6 | 7 | 8 | 15 | 32 | 19 | 13 | 7 | 8 | 15 | 25 | 20 | 14 | 15.05 | 10 | 20 | 1.973 | 0.98 | 0.07270434 | 0.21109955 | 0.0416 | 0.2258 | 0.1405 | 0.1405 | TRUE | 0.3763 |
| Tdrd7 | 399 | 427 | 470 | 624 | 440 | 467 | 424 | 426 | 472 | 491 | 473 | 511 | 466.22 | 441 | 492 | 1.115 | 0.158 | 0.14809853 | 0.33778887 | 0 | 0.014 | 0.0064 | 0.0064 | TRUE | 0.0446 |
| Tdrd9 | 56 | 74 | 69 | 116 | 54 | 64 | 59 | 74 | 69 | 91 | 58 | 70 | 70.34 | 67 | 73 | 1.089 | 0.123 | 0.6450907 | 0.80043824 | 0.0202 | 0.0537 | 0.037 | 0.037 | TRUE | 0.431 |
| Tdrkh | 387 | 574 | 588 | 771 | 529 | 498 | 411 | 573 | 590 | 606 | 569 | 545 | 549.19 | 525 | 573 | 1.093 | 0.128 | 0.41331235 | 0.62458201 | 0.0185 | 0.0129 | 0.0158 | 0.0158 | TRUE | 0.4176 |
| Tead1 | 8932 | 8275 | 8877 | 14218 | 10059 | 9850 | 9487 | 8264 | 8913 | 11177 | 10823 | 10782 | 9907.91 | 8888 | 10927 | 1.23 | 0.298 | 9.66E-05 | 0.00134008 | 0.0024 | 0.0072 | 0.0041 | 0.0041 | TRUE | 0.0461 |
| Tead2 | 5941 | 6030 | 6275 | 6574 | 4581 | 4912 | 6310 | 6022 | 6301 | 5168 | 4929 | 5377 | 5684.53 | 6211 | 5158 | 0.83 | -0.268 | 8.15E-05 | 0.00117147 | 0.001 | 0.0075 | 0.0032 | 0.0032 | TRUE | 0.0182 |
| Tead3 | 208 | 247 | 271 | 229 | 233 | 219 | 221 | 247 | 272 | 180 | 251 | 240 | 235.03 | 247 | 224 | 0.903 | -0.148 | 0.40979981 | 0.62158382 | 0.017 | 0.0209 | 0.0189 | 0.0189 | TRUE | 0.3412 |
| Tead4 | 1024 | 1315 | 1266 | 1236 | 994 | 948 | 1088 | 1313 | 1271 | 972 | 1069 | 1038 | 1125.17 | 1224 | 1026 | 0.838 | -0.255 | 0.01605049 | 0.0745431 | 0.0053 | 0.0098 | 0.0072 | 0.0072 | TRUE | 0.1134 |
| Tec | 249 | 232 | 214 | 204 | 169 | 182 | 264 | 232 | 215 | 160 | 182 | 199 | 208.75 | 237 | 180 | 0.759 | -0.397 | 0.01548722 | 0.07248202 | 0.0069 | 0.0227 | 0.0145 | 0.0145 | TRUE | 0.1122 |
| Tecpr1 | 913 | 995 | 1040 | 1128 | 920 | 958 | 970 | 994 | 1044 | 887 | 990 | 1049 | 988.84 | 1003 | 975 | 0.971 | -0.042 | 0.67369197 | 0.818099 | 0.0034 | 0.0102 | 0.0062 | 0.0062 | TRUE | 0.0778 |
| Tecpr2 | 2207 | 1956 | 2002 | 3079 | 2059 | 1975 | 2344 | 1953 | 2010 | 2420 | 2215 | 2162 | 2184.28 | 2102 | 2266 | 1.078 | 0.109 | 0.28894187 | 0.50981931 | 0.0062 | 0.0084 | 0.0071 | 0.0071 | TRUE | 0.1219 |
| Tecr | 12956 | 15206 | 14860 | 14477 | 10713 | 11594 | 13762 | 15186 | 14921 | 11381 | 11527 | 12691 | 13244.6 | 14623 | 11866 | 0.811 | -0.302 | 0.00016085 | 0.0020215 | 0.0031 | 0.0072 | 0.0045 | 0.0045 | TRUE | 0.0452 |
| Tecrl | 0 | 0 | 0 | 0 | 0 | 0 | 0 | 0 | 0 | 0 | 0 | 0 | 0 | 0 | 0 | NA | NA | NA | NA | NA | NA | NA | NA | NA | NA |
| Tecta | 2 | 2 | 3 | 8 | 7 | 4 | 2 | 2 | 3 | 6 | 8 | 4 | 4.22 | 2 | 6 | 2.553 | 1.352 | 0.15095251 | NA | 0 | 0.7872 | 0.3438 | 0.3438 | TRUE | 0.1237 |
| Tectb | 1 | 0 | 0 | 0 | 0 | 0 | 1 | 0 | 0 | 0 | 0 | 0 | 0.18 | 0 | 0 | 0.49 | -1.029 | 0.80087 | NA | 0 | 18.6165 | 10 | 10 | TRUE | 0.4614 |
| Teddm1 | 0 | 0 | 0 | 0 | 0 | 0 | 0 | 0 | 0 | 0 | 0 | 0 | 0 | 0 | 0 | NA | NA | NA | NA | NA | NA | NA | NA | NA | NA |
| Tef | 1171 | 1671 | 1582 | 1490 | 1135 | 1387 | 1244 | 1669 | 1588 | 1171 | 1221 | 1518 | 1401.99 | 1500 | 1303 | 0.868 | -0.204 | 0.1724931 | 0.37119832 | 0.0206 | 0.0093 | 0.0154 | 0.0154 | TRUE | 0.2689 |
| Tefm | 1296 | 1597 | 1485 | 1730 | 1165 | 1124 | 1377 | 1595 | 1491 | 1360 | 1253 | 1230 | 1384.41 | 1488 | 1281 | 0.862 | -0.214 | 0.02408932 | 0.09901922 | 0.0035 | 0.0093 | 0.0058 | 0.0058 | TRUE | 0.051 |
| Tek | 560 | 491 | 462 | 623 | 424 | 375 | 595 | 490 | 464 | 490 | 456 | 410 | 484.26 | 516 | 452 | 0.877 | -0.189 | 0.17669148 | 0.37715933 | 0.0106 | 0.0137 | 0.012 | 0.012 | TRUE | 0.2049 |
| Tekt1 | 496 | 473 | 448 | 608 | 477 | 471 | 527 | 472 | 450 | 478 | 513 | 516 | 492.64 | 483 | 502 | 1.039 | 0.055 | 0.6250846 | 0.78876985 | 0.0021 | 0.0136 | 0.0072 | 0.0072 | TRUE | 0.0725 |
| Tekt2 | 40 | 30 | 23 | 54 | 28 | 45 | 42 | 30 | 23 | 42 | 30 | 49 | 36.23 | 32 | 40 | 1.279 | 0.355 | 0.34329674 | 0.5633815 | 0.0446 | 0.0978 | 0.0729 | 0.0729 | TRUE | 0.5246 |
| Tekt3 | 0 | 1 | 1 | 3 | 0 | 0 | 0 | 1 | 1 | 2 | 0 | 0 | 0.73 | 1 | 1 | 1.234 | 0.303 | 0.90578576 | NA | 0 | 4.5393 | 3.3736 | 3.3736 | TRUE | 1.4393 |
| Tekt4 | 0 | 0 | 2 | 1 | 0 | 0 | 0 | 0 | 2 | 1 | 0 | 0 | 0.47 | 1 | 0 | 0.52 | -0.943 | 0.78484416 | NA | 0 | 7.0808 | 6.8441 | 6.8441 | TRUE | 0.9739 |
| Tekt5 | 9 | 16 | 9 | 10 | 13 | 7 | 10 | 16 | 9 | 8 | 14 | 8 | 10.68 | 12 | 10 | 0.846 | -0.241 | 0.69401176 | 0.83228303 | 0 | 0.3153 | 0.1752 | 0.1752 | TRUE | 0.4451 |
| Telo2 | 1528 | 1225 | 1165 | 1692 | 1301 | 1075 | 1623 | 1223 | 1170 | 1330 | 1400 | 1177 | 1320.48 | 1339 | 1302 | 0.973 | -0.039 | 0.78686962 | 0.89032073 | 0.0193 | 0.0094 | 0.0145 | 0.0145 | TRUE | 0.4138 |
| Ten1 | 590 | 676 | 691 | 717 | 446 | 483 | 627 | 675 | 694 | 564 | 480 | 529 | 594.65 | 665 | 524 | 0.789 | -0.342 | 0.00191533 | 0.01488108 | 0.0029 | 0.0124 | 0.0071 | 0.0071 | TRUE | 0.0631 |
| Tenc1 | 758 | 607 | 626 | 399 | 369 | 413 | 805 | 606 | 629 | 314 | 397 | 452 | 533.78 | 680 | 388 | 0.569 | -0.814 | 3.27E-06 | 7.68E-05 | 0.026 | 0.0131 | 0.02 | 0.02 | TRUE | 0.3334 |
| Tenm1 | 13 | 3 | 2 | 7 | 5 | 3 | 14 | 3 | 2 | 6 | 5 | 3 | 5.5 | 6 | 5 | 0.762 | -0.393 | 0.68571772 | 0.8264033 | 0.3904 | 0.6063 | 0.494 | 0.494 | TRUE | 2.5799 |
| Tenm2 | 3 | 3 | 4 | 5 | 3 | 0 | 3 | 3 | 4 | 4 | 3 | 0 | 2.89 | 3 | 2 | 0.721 | -0.473 | 0.69118591 | NA | 0 | 1.1457 | 0.6709 | 0.6709 | TRUE | 0.7071 |
| Tenm3 | 4106 | 3092 | 3495 | 4621 | 3170 | 2796 | 4361 | 3088 | 3509 | 3633 | 3411 | 3061 | 3510.45 | 3653 | 3368 | 0.922 | -0.117 | 0.3997208 | 0.61279125 | 0.018 | 0.0078 | 0.0136 | 0.0136 | TRUE | 0.3498 |
| Tenm4 | 614 | 339 | 389 | 808 | 542 | 372 | 652 | 339 | 391 | 635 | 583 | 407 | 501.15 | 461 | 542 | 1.178 | 0.236 | 0.35581059 | 0.574278 | 0.0846 | 0.0135 | 0.0452 | 0.0452 | TRUE | 1.5353 |
| Tep1 | 1899 | 1934 | 2148 | 2184 | 1595 | 1767 | 2017 | 1932 | 2157 | 1717 | 1716 | 1934 | 1912.11 | 2035 | 1789 | 0.879 | -0.187 | 0.04044631 | 0.14207554 | 0.0032 | 0.0086 | 0.0054 | 0.0054 | TRUE | 0.0602 |
| Tepp | 33 | 39 | 41 | 48 | 38 | 32 | 35 | 39 | 41 | 38 | 41 | 35 | 38.14 | 38 | 38 | 0.986 | -0.02 | 0.94737144 | 0.97367016 | 0 | 0.0933 | 0.0407 | 0.0407 | TRUE | 0.0412 |
| Terc | 35 | 12 | 11 | 58 | 54 | 26 | 37 | 12 | 11 | 46 | 58 | 28 | 32.06 | 20 | 44 | 2.205 | 1.141 | 0.03344239 | 0.12401914 | 0.2443 | 0.1097 | 0.1707 | 0.1707 | TRUE | 0.8188 |
| Terf1 | 5125 | 6530 | 6425 | 6443 | 4812 | 5563 | 5444 | 6522 | 6451 | 5065 | 5177 | 6090 | 5791.43 | 6139 | 5444 | 0.887 | -0.174 | 0.12165099 | 0.29659599 | 0.0099 | 0.0075 | 0.0089 | 0.0089 | TRUE | 0.131 |
| Terf2 | 2288 | 2245 | 2144 | 2718 | 2161 | 1973 | 2430 | 2242 | 2153 | 2137 | 2325 | 2160 | 2241.12 | 2275 | 2207 | 0.97 | -0.044 | 0.60393017 | 0.77478336 | 0.0023 | 0.0084 | 0.0047 | 0.0047 | TRUE | 0.043 |
| Terf2ip | 1554 | 1448 | 1409 | 2372 | 1622 | 1492 | 1651 | 1446 | 1415 | 1865 | 1745 | 1633 | 1625.77 | 1504 | 1748 | 1.163 | 0.218 | 0.03021579 | 0.115818 | 0.0049 | 0.0089 | 0.0067 | 0.0067 | TRUE | 0.0875 |
| Tert | 1165 | 1170 | 1152 | 1231 | 987 | 1152 | 1237 | 1168 | 1157 | 968 | 1062 | 1261 | 1142.23 | 1187 | 1097 | 0.923 | -0.116 | 0.32866516 | 0.54970381 | 0.0085 | 0.0098 | 0.0093 | 0.0093 | TRUE | 0.2043 |
| Tes | 1208 | 1756 | 1679 | 1601 | 1171 | 1468 | 1283 | 1754 | 1686 | 1259 | 1260 | 1607 | 1474.7 | 1574 | 1375 | 0.873 | -0.196 | 0.20276917 | 0.41037796 | 0.0227 | 0.0091 | 0.0164 | 0.0164 | TRUE | 0.3148 |
| Tesc | 309 | 427 | 405 | 432 | 316 | 293 | 328 | 426 | 407 | 340 | 340 | 321 | 360.28 | 387 | 334 | 0.861 | -0.215 | 0.12395905 | 0.30031984 | 0.0073 | 0.016 | 0.0114 | 0.0114 | TRUE | 0.2014 |
| Tescl | 9 | 15 | 13 | 20 | 15 | 11 | 10 | 15 | 13 | 16 | 16 | 12 | 13.58 | 13 | 15 | 1.169 | 0.226 | 0.66069873 | 0.80885977 | 0 | 0.2495 | 0.1163 | 0.1163 | TRUE | 0.1659 |
| Tesk1 | 321 | 309 | 302 | 583 | 390 | 407 | 341 | 309 | 303 | 458 | 420 | 446 | 379.38 | 318 | 441 | 1.391 | 0.476 | 5.27E-05 | 0.00081071 | 0 | 0.0156 | 0.0073 | 0.0073 | TRUE | 0.0463 |
| Tesk2 | 1459 | 1144 | 1118 | 1577 | 1077 | 1077 | 1550 | 1143 | 1123 | 1240 | 1159 | 1179 | 1232.05 | 1272 | 1193 | 0.938 | -0.092 | 0.51194573 | 0.70537329 | 0.0162 | 0.0096 | 0.0134 | 0.0134 | TRUE | 0.4385 |
| Tespa1 | 1 | 0 | 0 | 0 | 0 | 0 | 1 | 0 | 0 | 0 | 0 | 0 | 0.18 | 0 | 0 | 0.49 | -1.029 | 0.80087 | NA | 0 | 18.6165 | 10 | 10 | TRUE | 0.4614 |
| Tet1 | 37951 | 37011 | 46601 | 44442 | 30885 | 29903 | 40311 | 36963 | 46792 | 34937 | 33230 | 32734 | 37494.49 | 41355 | 33634 | 0.813 | -0.298 | 0.00336079 | 0.0231247 | 0.0079 | 0.007 | 0.0074 | 0.0074 | TRUE | 0.162 |
| Tet2 | 3805 | 3433 | 3536 | 5424 | 3617 | 3233 | 4042 | 3429 | 3551 | 4264 | 3892 | 3539 | 3785.88 | 3674 | 3898 | 1.062 | 0.086 | 0.41617886 | 0.62771911 | 0.0074 | 0.0078 | 0.0078 | 0.0078 | TRUE | 0.0933 |
| Tet3 | 657 | 608 | 609 | 1175 | 871 | 785 | 698 | 607 | 612 | 924 | 937 | 859 | 772.78 | 639 | 907 | 1.42 | 0.506 | 8.45E-07 | 2.37E-05 | 0.0025 | 0.0112 | 0.0063 | 0.0063 | TRUE | 0.0762 |
| Tex10 | 7064 | 6410 | 6510 | 9634 | 6853 | 6135 | 7503 | 6402 | 6537 | 7573 | 7373 | 6716 | 7017.38 | 6814 | 7221 | 1.06 | 0.084 | 0.37452477 | 0.59064141 | 0.0054 | 0.0074 | 0.0063 | 0.0063 | TRUE | 0.0955 |
| Tex101 | 47 | 58 | 53 | 126 | 79 | 68 | 50 | 58 | 53 | 99 | 85 | 74 | 69.93 | 54 | 86 | 1.611 | 0.688 | 0.00511384 | 0.03186279 | 0 | 0.054 | 0.0284 | 0.0284 | TRUE | 0.1793 |
| Tex11 | 497 | 557 | 587 | 830 | 587 | 570 | 528 | 556 | 589 | 652 | 632 | 624 | 596.93 | 558 | 636 | 1.14 | 0.19 | 0.0546312 | 0.17410453 | 0 | 0.0124 | 0.0053 | 0.0053 | TRUE | 0.0287 |
| Tex12 | 53 | 53 | 43 | 126 | 63 | 50 | 56 | 53 | 43 | 99 | 68 | 55 | 62.33 | 51 | 74 | 1.465 | 0.551 | 0.07400492 | 0.21361742 | 0.0443 | 0.0598 | 0.0519 | 0.0519 | TRUE | 0.5334 |
| Tex13 | 123 | 170 | 156 | 421 | 423 | 302 | 131 | 170 | 157 | 331 | 455 | 331 | 262.29 | 153 | 372 | 2.439 | 1.286 | 1.18E-11 | 1.03E-09 | 0.023 | 0.0195 | 0.0212 | 0.0212 | TRUE | 0.4301 |
| Tex13a | 0 | 0 | 0 | 0 | 0 | 0 | 0 | 0 | 0 | 0 | 0 | 0 | 0 | 0 | 0 | NA | NA | NA | NA | NA | NA | NA | NA | NA | NA |
| Tex14 | 2322 | 3097 | 3563 | 4084 | 3096 | 3066 | 2466 | 3093 | 3578 | 3211 | 3331 | 3356 | 3172.48 | 3046 | 3299 | 1.083 | 0.115 | 0.39315819 | 0.60663283 | 0.0171 | 0.0079 | 0.0128 | 0.0128 | TRUE | 0.2844 |
| Tex15 | 1688 | 1302 | 1185 | 2328 | 1706 | 1319 | 1793 | 1300 | 1190 | 1830 | 1836 | 1444 | 1565.44 | 1428 | 1703 | 1.193 | 0.255 | 0.13509937 | 0.31796579 | 0.033 | 0.009 | 0.0204 | 0.0204 | TRUE | 0.6017 |
| Tex16 | 29 | 15 | 14 | 56 | 45 | 24 | 31 | 15 | 14 | 44 | 48 | 26 | 29.76 | 20 | 39 | 1.995 | 0.997 | 0.02829826 | 0.11058257 | 0.1027 | 0.1176 | 0.1108 | 0.1108 | TRUE | 1.1819 |
| Tex19.1 | 7405 | 4217 | 3206 | 11733 | 7286 | 4183 | 7865 | 4212 | 3219 | 9224 | 7839 | 4579 | 6156.32 | 5099 | 7214 | 1.415 | 0.501 | 0.28585077 | 0.50719297 | 0.1585 | 0.0074 | 0.068 | 0.1585 | TRUE | 1.07 |
| Tex19.2 | 259 | 203 | 157 | 506 | 379 | 221 | 275 | 203 | 158 | 398 | 408 | 242 | 280.49 | 212 | 349 | 1.651 | 0.723 | 0.00509238 | 0.03181566 | 0.0741 | 0.0186 | 0.0442 | 0.0442 | TRUE | 0.6704 |
| Tex2 | 4368 | 4391 | 4254 | 5419 | 3708 | 3903 | 4640 | 4385 | 4271 | 4260 | 3990 | 4272 | 4303.07 | 4432 | 4174 | 0.942 | -0.086 | 0.2308912 | 0.44485903 | 0.0013 | 0.0077 | 0.0035 | 0.0035 | TRUE | 0.0204 |
| Tex21 | 147 | 185 | 186 | 227 | 145 | 148 | 156 | 185 | 187 | 178 | 156 | 162 | 170.69 | 176 | 165 | 0.943 | -0.085 | 0.60457954 | 0.77530112 | 0 | 0.0262 | 0.0137 | 0.0137 | TRUE | 0.1013 |
| Tex22 | 1 | 1 | 0 | 1 | 0 | 2 | 1 | 1 | 0 | 1 | 0 | 2 | 0.84 | 1 | 1 | 1.436 | 0.522 | 0.80963223 | NA | 0 | 3.9317 | 2.1449 | 2.1449 | TRUE | 0.3069 |
| Tex24 | 5 | 1 | 2 | 0 | 0 | 1 | 5 | 1 | 2 | 0 | 0 | 1 | 1.57 | 3 | 0 | 0.126 | -2.983 | 0.11090515 | NA | 0 | 2.107 | 1.3392 | 1.3392 | TRUE | 0.5307 |
| Tex26 | 31 | 37 | 59 | 32 | 36 | 23 | 33 | 37 | 59 | 25 | 39 | 25 | 36.36 | 43 | 30 | 0.686 | -0.544 | 0.15683425 | 0.34972286 | 0.0573 | 0.0975 | 0.0778 | 0.0778 | TRUE | 0.84 |
| Tex261 | 1120 | 1206 | 1273 | 1853 | 1337 | 1275 | 1190 | 1204 | 1278 | 1457 | 1439 | 1396 | 1327.2 | 1224 | 1431 | 1.169 | 0.225 | 0.00391208 | 0.02590313 | 0 | 0.0094 | 0.0036 | 0.0036 | TRUE | 0.018 |
| Tex264 | 880 | 1228 | 1214 | 1813 | 1347 | 1490 | 935 | 1226 | 1219 | 1425 | 1449 | 1631 | 1314.28 | 1127 | 1502 | 1.332 | 0.414 | 0.00152595 | 0.01240578 | 0.0125 | 0.0094 | 0.0115 | 0.0115 | TRUE | 0.265 |
| Tex28 | 0 | 0 | 0 | 0 | 0 | 0 | 0 | 0 | 0 | 0 | 0 | 0 | 0 | 0 | 0 | NA | NA | NA | NA | NA | NA | NA | NA | NA | NA |
| Tex29 | 2 | 0 | 0 | 0 | 0 | 0 | 2 | 0 | 0 | 0 | 0 | 0 | 0.35 | 1 | 0 | 0.254 | -1.975 | 0.62446185 | NA | 10 | 9.3117 | 10 | 10 | TRUE | 0.9741 |
| Tex30 | 1326 | 1477 | 1565 | 1873 | 1315 | 1318 | 1408 | 1475 | 1571 | 1472 | 1415 | 1443 | 1464.16 | 1485 | 1443 | 0.972 | -0.041 | 0.61524027 | 0.78235666 | 0.001 | 0.0092 | 0.0041 | 0.0041 | TRUE | 0.0312 |
| Tex33 | 0 | 0 | 0 | 0 | 0 | 0 | 0 | 0 | 0 | 0 | 0 | 0 | 0 | 0 | 0 | NA | NA | NA | NA | NA | NA | NA | NA | NA | NA |

| | | | | | | | | | | | | | | | | | | | | | | | | | |
|---|---|---|---|---|---|---|---|---|---|---|---|---|---|---|---|---|---|---|---|---|---|---|---|---|---|
| Tex35 | 10 | 7 | 6 | 17 | 5 | 7 | 11 | 7 | 6 | 13 | 5 | 8 | 8.34 | 8 | 9 | 1.139 | 0.188 | 0.78631876 | 0.89023346 | 0 | 0.4019 | 0.2279 | 0.2279 | TRUE | 0.8142 |
| Tex36 | 0 | 0 | 3 | 0 | 0 | 2 | 0 | 0 | 3 | 0 | 0 | 2 | 0.87 | 1 | 1 | 0.707 | -0.5 | 0.86485107 | NA | 10 | 3.807 | 5.0241 | 5.0241 | TRUE | 1.4421 |
| Tex37 | 0 | 0 | 0 | 0 | 0 | 0 | 0 | 0 | 0 | 0 | 0 | 0 | 0 | 0 | 0 | NA | NA | NA | NA | NA | NA | NA | NA | NA | NA |
| Tex38 | 0 | 0 | 0 | 1 | 1 | 1 | 0 | 0 | 0 | 1 | 1 | 1 | 0.49 | 0 | 1 | 5.224 | 2.385 | 0.38465 | NA | 0 | 6.6923 | 3.9249 | 3.9249 | TRUE | 0.07 |
| Tex40 | 110 | 164 | 162 | 219 | 151 | 139 | 117 | 164 | 163 | 172 | 162 | 152 | 155.01 | 148 | 162 | 1.099 | 0.136 | 0.48019801 | 0.67946422 | 0.0118 | 0.0282 | 0.0201 | 0.0201 | TRUE | 0.3439 |
| Tex9 | 686 | 473 | 446 | 986 | 766 | 616 | 729 | 472 | 448 | 775 | 824 | 674 | 653.74 | 550 | 758 | 1.38 | 0.464 | 0.01639075 | 0.07567769 | 0.04 | 0.0119 | 0.0254 | 0.0254 | TRUE | 0.9471 |
| Tfam | 5739 | 5764 | 5457 | 6950 | 5007 | 5105 | 6096 | 5757 | 5479 | 5464 | 5387 | 5588 | 5628.47 | 5777 | 5480 | 0.949 | -0.076 | 0.2804236 | 0.50174033 | 0.0013 | 0.0075 | 0.0034 | 0.0034 | TRUE | 0.0283 |
| Tfap2a | 91 | 88 | 76 | 150 | 102 | 106 | 97 | 88 | 76 | 118 | 110 | 116 | 100.76 | 87 | 115 | 1.32 | 0.4 | 0.04626072 | 0.15563653 | 0 | 0.0396 | 0.0189 | 0.0189 | TRUE | 0.1091 |
| Tfap2b | 0 | 0 | 0 | 0 | 1 | 0 | 0 | 0 | 0 | 0 | 1 | 0 | 0.18 | 0 | 0 | 1.859 | 0.894 | 0.82650734 | NA | 0 | 18.3785 | 10 | 10 | TRUE | 0.5204 |
| Tfap2c | 2672 | 3622 | 3102 | 4239 | 3024 | 3147 | 2838 | 3617 | 3115 | 3332 | 3254 | 3445 | 3266.85 | 3190 | 3344 | 1.048 | 0.068 | 0.5219069 | 0.71194565 | 0.0076 | 0.0079 | 0.0077 | 0.0077 | TRUE | 0.1669 |
| Tfap2d | 0 | 0 | 0 | 0 | 0 | 0 | 0 | 0 | 0 | 0 | 0 | 0 | 0 | 0 | 0 | NA | NA | NA | NA | NA | NA | NA | NA | NA | NA |
| Tfap2e | 21 | 31 | 28 | 49 | 25 | 25 | 22 | 31 | 28 | 39 | 27 | 27 | 29.03 | 27 | 31 | 1.148 | 0.199 | 0.59139966 | 0.76507383 | 0 | 0.1204 | 0.0649 | 0.0649 | TRUE | 0.3642 |
| Tfap4 | 581 | 812 | 910 | 1045 | 923 | 833 | 617 | 811 | 914 | 821 | 993 | 912 | 844.71 | 781 | 909 | 1.163 | 0.218 | 0.16783955 | 0.36558665 | 0.0213 | 0.0108 | 0.0168 | 0.0168 | TRUE | 0.3963 |
| Tfb1m | 1353 | 1508 | 1535 | 1666 | 1256 | 1163 | 1437 | 1506 | 1541 | 1310 | 1351 | 1273 | 1403.1 | 1495 | 1311 | 0.877 | -0.189 | 0.0149499 | 0.07064591 | 0 | 0.0093 | 0.0036 | 0.0036 | TRUE | 0.0136 |
| Tfb2m | 562 | 469 | 475 | 655 | 498 | 512 | 597 | 468 | 477 | 515 | 536 | 560 | 525.58 | 514 | 537 | 1.045 | 0.063 | 0.63173811 | 0.79278407 | 0.0082 | 0.0132 | 0.0105 | 0.0105 | TRUE | 0.2303 |
| Tfcp2 | 1065 | 1071 | 1055 | 1406 | 1068 | 977 | 1131 | 1070 | 1059 | 1105 | 1149 | 1069 | 1097.34 | 1087 | 1108 | 1.02 | 0.028 | 0.73395833 | 0.85735379 | 0 | 0.0099 | 0.004 | 0.004 | TRUE | 0.0152 |
| Tfcp2l1 | 8143 | 9930 | 10069 | 10356 | 7970 | 9037 | 8649 | 9917 | 10110 | 8141 | 8575 | 9892 | 9214.26 | 9559 | 8869 | 0.928 | -0.108 | 0.30860032 | 0.52922087 | 0.0085 | 0.0073 | 0.008 | 0.008 | TRUE | 0.1242 |
| Tfdp1 | 7929 | 6982 | 7220 | 11830 | 8756 | 7994 | 8422 | 6973 | 7250 | 9300 | 9421 | 8751 | 8352.68 | 7548 | 9157 | 1.213 | 0.279 | 0.00315363 | 0.02206701 | 0.0058 | 0.0073 | 0.0063 | 0.0063 | TRUE | 0.1251 |
| Tfdp2 | 4851 | 5089 | 5166 | 5424 | 4156 | 4380 | 5153 | 5082 | 5187 | 4264 | 4472 | 4795 | 4825.4 | 5141 | 4510 | 0.877 | -0.189 | 0.00933106 | 0.04993873 | 0.0015 | 0.0076 | 0.0036 | 0.0036 | TRUE | 0.037 |
| Tfe3 | 3052 | 3271 | 3622 | 3190 | 2463 | 2835 | 3242 | 3267 | 3637 | 2508 | 2650 | 3103 | 3067.75 | 3382 | 2754 | 0.814 | -0.297 | 0.00558407 | 0.0339391 | 0.008 | 0.008 | 0.008 | 0.008 | TRUE | 0.1493 |
| Tfeb | 1052 | 1246 | 1275 | 1256 | 1034 | 1071 | 1117 | 1244 | 1280 | 987 | 1113 | 1172 | 1152.38 | 1214 | 1091 | 0.897 | -0.156 | 0.14262998 | 0.32854682 | 0.0055 | 0.0098 | 0.0073 | 0.0073 | TRUE | 0.0847 |
| Tfec | 0 | 0 | 0 | 6 | 3 | 0 | 0 | 0 | 0 | 5 | 3 | 0 | 1.32 | 0 | 3 | 14.225 | 3.83 | 0.11269191 | NA | 6.3954 | 2.495 | 3.0186 | 3.0186 | TRUE | 0.1638 |
| Tff1 | 0 | 0 | 0 | 0 | 0 | 0 | 0 | 0 | 0 | 0 | 0 | 0 | 0 | 0 | 0 | NA | NA | NA | NA | NA | NA | NA | NA | NA | NA |
| Tff2 | 4 | 1 | 0 | 1 | 2 | 1 | 4 | 1 | 0 | 1 | 2 | 1 | 1.55 | 2 | 1 | 0.764 | -0.389 | 0.81447138 | NA | 0 | 2.1369 | 1.3182 | 1.3182 | TRUE | 0.9423 |
| Tff3 | 1 | 1 | 0 | 1 | 0 | 0 | 1 | 1 | 0 | 1 | 0 | 0 | 0.47 | 1 | 0 | 0.51 | -0.973 | 0.73452857 | NA | 0 | 6.9499 | 4.1835 | 4.1835 | TRUE | 0.2562 |
| Tfg | 3550 | 3432 | 3494 | 5082 | 3522 | 3449 | 3771 | 3428 | 3508 | 3995 | 3789 | 3775 | 3711.11 | 3569 | 3853 | 1.08 | 0.111 | 0.13134966 | 0.31213602 | 0.0014 | 0.0078 | 0.0036 | 0.0036 | TRUE | 0.0297 |
| Tfip11 | 2412 | 2078 | 2008 | 3330 | 2436 | 2056 | 2562 | 2075 | 2016 | 2618 | 2621 | 2251 | 2357.15 | 2218 | 2497 | 1.126 | 0.171 | 0.16132898 | 0.35629025 | 0.0121 | 0.0083 | 0.0103 | 0.0103 | TRUE | 0.2227 |
| Tfpi | 3566 | 3785 | 4170 | 6203 | 4442 | 4870 | 3788 | 3780 | 4187 | 4876 | 4779 | 5331 | 4456.93 | 3918 | 4995 | 1.275 | 0.35 | 2.94E-05 | 0.00049402 | 0.0032 | 0.0076 | 0.0048 | 0.0048 | TRUE | 0.0439 |
| Tfpi2 | 6 | 3 | 0 | 15 | 3 | 1 | 6 | 3 | 0 | 12 | 3 | 1 | 4.25 | 3 | 5 | 1.762 | 0.818 | 0.52913646 | NA | 1.1904 | 0.7826 | 0.9622 | 0.9622 | TRUE | 0.5761 |
| Tfpt | 465 | 522 | 462 | 859 | 603 | 475 | 494 | 521 | 464 | 675 | 649 | 520 | 553.86 | 493 | 615 | 1.248 | 0.32 | 0.01696351 | 0.07731702 | 0.0097 | 0.0129 | 0.0111 | 0.0111 | TRUE | 0.2103 |
| Tfr2 | 325 | 454 | 475 | 445 | 338 | 375 | 345 | 453 | 477 | 350 | 364 | 410 | 399.93 | 425 | 375 | 0.88 | -0.185 | 0.24051035 | 0.4569331 | 0.0155 | 0.0151 | 0.0153 | 0.0153 | TRUE | 0.3102 |
| Tfrc | 32968 | 43481 | 43040 | 36581 | 28702 | 36821 | 35018 | 43425 | 43217 | 28757 | 30882 | 40306 | 36934.08 | 40553 | 33315 | 0.821 | -0.284 | 0.05205968 | 0.16863529 | 0.024 | 0.007 | 0.0153 | 0.0153 | TRUE | 0.4125 |
| Tg | 14 | 11 | 8 | 28 | 17 | 11 | 15 | 11 | 8 | 22 | 18 | 12 | 14.37 | 11 | 17 | 1.561 | 0.643 | 0.23107549 | 0.44492876 | 0 | 0.2361 | 0.1344 | 0.1344 | TRUE | 0.2701 |
| Tgds | 915 | 1056 | 1003 | 1170 | 878 | 883 | 972 | 1055 | 1007 | 920 | 945 | 967 | 977.44 | 1011 | 944 | 0.933 | -0.101 | 0.23239134 | 0.44654725 | 0 | 0.0103 | 0.0041 | 0.0041 | TRUE | 0.0169 |
| Tgfa | 396 | 472 | 474 | 359 | 246 | 291 | 421 | 471 | 476 | 282 | 265 | 319 | 372.23 | 456 | 289 | 0.632 | -0.662 | 3.06E-07 | 9.64E-06 | 0.0033 | 0.0158 | 0.0092 | 0.0092 | TRUE | 0.0889 |
| Tgfb1 | 1064 | 1340 | 1349 | 989 | 820 | 808 | 1130 | 1338 | 1355 | 777 | 882 | 884 | 1061.2 | 1274 | 848 | 0.665 | -0.589 | 1.79E-07 | 5.93E-06 | 0.0068 | 0.01 | 0.0082 | 0.0082 | TRUE | 0.1171 |
| Tgfb1i1 | 63 | 55 | 62 | 60 | 48 | 38 | 67 | 55 | 62 | 47 | 52 | 42 | 54.09 | 61 | 47 | 0.764 | -0.389 | 0.1424562 | 0.32833833 | 0 | 0.0678 | 0.032 | 0.032 | TRUE | 0.0736 |
| Tgfb2 | 11 | 6 | 3 | 17 | 16 | 9 | 12 | 6 | 3 | 13 | 17 | 10 | 10.19 | 7 | 13 | 1.965 | 0.975 | 0.15144941 | 0.34246023 | 0.0753 | 0.3303 | 0.2219 | 0.2219 | TRUE | 0.6007 |
| Tgfb3 | 4 | 6 | 6 | 7 | 10 | 10 | 4 | 6 | 6 | 6 | 11 | 11 | 7.25 | 5 | 9 | 1.647 | 0.72 | 0.31867344 | 0.53986224 | 0 | 0.4616 | 0.2269 | 0.2269 | TRUE | 0.4979 |
| Tgfbi | 999 | 1377 | 1377 | 813 | 739 | 1020 | 1061 | 1375 | 1383 | 639 | 795 | 1117 | 1061.63 | 1273 | 850 | 0.667 | -0.584 | 0.00380719 | 0.02538983 | 0.0499 | 0.01 | 0.0284 | 0.0284 | TRUE | 0.8284 |
| Tgfbr1 | 1842 | 2043 | 2083 | 3072 | 2246 | 2157 | 1957 | 2040 | 2092 | 2415 | 2417 | 2361 | 2213.52 | 2030 | 2398 | 1.181 | 0.24 | 0.00047616 | 0.00499686 | 0 | 0.0084 | 0.003 | 0.003 | TRUE | 0.0119 |
| Tgfbr2 | 92 | 129 | 112 | 200 | 123 | 154 | 98 | 129 | 112 | 157 | 132 | 169 | 132.86 | 113 | 153 | 1.351 | 0.434 | 0.02780862 | 0.10951806 | 0.0084 | 0.0317 | 0.0203 | 0.0203 | TRUE | 0.1506 |
| Tgfbr3 | 138 | 222 | 251 | 325 | 212 | 216 | 147 | 222 | 252 | 255 | 228 | 236 | 223.39 | 207 | 240 | 1.161 | 0.215 | 0.30700203 | 0.5274549 | 0.0336 | 0.0217 | 0.0274 | 0.0274 | TRUE | 0.6986 |
| Tgfbrap1 | 535 | 511 | 471 | 726 | 533 | 474 | 568 | 510 | 473 | 571 | 573 | 519 | 535.77 | 517 | 554 | 1.073 | 0.102 | 0.38383168 | 0.59804511 | 0.0038 | 0.0131 | 0.0079 | 0.0079 | TRUE | 0.0864 |
| Tgif1 | 6731 | 8547 | 7900 | 8728 | 6438 | 6530 | 7150 | 8536 | 7932 | 6861 | 6927 | 7148 | 7425.7 | 7873 | 6979 | 0.886 | -0.174 | 0.04466804 | 0.15211197 | 0.0039 | 0.0073 | 0.0053 | 0.0053 | TRUE | 0.0788 |
| Tgif2 | 2154 | 2557 | 2682 | 3001 | 2165 | 2295 | 2288 | 2554 | 2693 | 2359 | 2329 | 2512 | 2455.91 | 2512 | 2400 | 0.955 | -0.066 | 0.46894216 | 0.67134026 | 0.0036 | 0.0082 | 0.0055 | 0.0055 | TRUE | 0.0733 |
| Tgif2lx1 | 0 | 0 | 0 | 0 | 0 | 0 | 0 | 0 | 0 | 0 | 0 | 0 | 0 | 0 | 0 | NA | NA | NA | NA | NA | NA | NA | NA | NA | NA |
| Tgif2lx2 | 0 | 0 | 0 | 0 | 0 | 0 | 0 | 0 | 0 | 0 | 0 | 0 | 0 | 0 | 0 | NA | NA | NA | NA | NA | NA | NA | NA | NA | NA |
| Tgm1 | 563 | 423 | 389 | 519 | 361 | 354 | 598 | 422 | 391 | 408 | 388 | 388 | 432.5 | 470 | 395 | 0.84 | -0.252 | 0.1506364 | 0.34155222 | 0.0251 | 0.0145 | 0.0198 | 0.0198 | TRUE | 0.6517 |
| Tgm2 | 613 | 1126 | 1116 | 1004 | 741 | 1177 | 651 | 1125 | 1121 | 789 | 797 | 1288 | 961.87 | 966 | 958 | 0.992 | -0.012 | 0.96260965 | 0.98130897 | 0.0842 | 0.0103 | 0.0426 | 0.0426 | TRUE | 1.0797 |
| Tgm3 | 313 | 213 | 229 | 89 | 64 | 115 | 332 | 213 | 230 | 70 | 69 | 126 | 173.31 | 258 | 88 | 0.34 | -1.556 | 4.18E-08 | 1.70E-06 | 0.0803 | 0.0259 | 0.0504 | 0.0504 | TRUE | 1.285 |
| Tgm4 | 0 | 0 | 0 | 0 | 0 | 0 | 0 | 0 | 0 | 0 | 0 | 0 | 0 | 0 | 0 | NA | NA | NA | NA | NA | NA | NA | NA | NA | NA |
| Tgm5 | 3 | 3 | 3 | 7 | 3 | 4 | 3 | 3 | 3 | 6 | 3 | 4 | 3.72 | 3 | 4 | 1.441 | 0.527 | 0.57764252 | NA | 0 | 0.8931 | 0.3671 | 0.3671 | TRUE | 0.1226 |
| Tgm6 | 0 | 1 | 2 | 0 | 4 | 3 | 0 | 1 | 2 | 0 | 4 | 3 | 1.77 | 1 | 2 | 2.453 | 1.294 | 0.46497461 | NA | 0 | 1.8726 | 1.5581 | 1.5581 | TRUE | 0.9449 |
| Tgm7 | 0 | 0 | 0 | 0 | 0 | 1 | 0 | 0 | 0 | 0 | 0 | 1 | 0.18 | 0 | 0 | 1.859 | 0.894 | 0.82650734 | NA | 0 | 18.0643 | 10 | 10 | TRUE | 0.5383 |
| Tgoln1 | 1845 | 1832 | 1813 | 2371 | 1661 | 1708 | 1960 | 1830 | 1820 | 1864 | 1787 | 1870 | 1855.08 | 1870 | 1840 | 0.984 | -0.023 | 0.76151186 | 0.87425863 | 6.00E-04 | 0.0087 | 0.0035 | 0.0035 | TRUE | 0.0212 |
| Tgoln2 | 0 | 0 | 0 | 0 | 0 | 0 | 0 | 0 | 0 | 0 | 0 | 0 | 0 | 0 | 0 | NA | NA | NA | NA | NA | NA | NA | NA | NA | NA |
| Tgs1 | 2969 | 2844 | 2800 | 3871 | 2689 | 2732 | 3154 | 2840 | 2811 | 3043 | 2893 | 2991 | 2955.38 | 2935 | 2976 | 1.014 | 0.02 | 0.7989156 | 0.89640568 | 0.002 | 0.008 | 0.0042 | 0.0042 | TRUE | 0.0513 |
| Tgtp1 | 10 | 2 | 5 | 1 | 2 | 2 | 11 | 2 | 5 | 1 | 2 | 2 | 3.79 | 6 | 2 | 0.283 | -1.821 | 0.11041243 | NA | 0 | 0.8751 | 0.5563 | 0.5563 | TRUE | 0.2008 |
| Tgtp2 | 0 | 0 | 0 | 0 | 0 | 0 | 0 | 0 | 0 | 0 | 0 | 0 | 0 | 0 | 0 | NA | NA | NA | NA | NA | NA | NA | NA | NA | NA |
| Th | 4 | 1 | 2 | 0 | 0 | 1 | 4 | 1 | 2 | 0 | 0 | 1 | 1.39 | 2 | 0 | 0.145 | -2.787 | 0.14244306 | NA | 0 | 2.3741 | 1.3898 | 1.3898 | TRUE | 0.4898 |
| Tha1 | 37 | 39 | 59 | 59 | 41 | 37 | 39 | 39 | 59 | 46 | 44 | 41 | 44.75 | 46 | 44 | 0.954 | -0.068 | 0.82718791 | 0.91252764 | 0.0115 | 0.0805 | 0.0472 | 0.0472 | TRUE | 0.521 |
| Thada | 3254 | 2202 | 2320 | 4961 | 3216 | 2497 | 3456 | 2199 | 2330 | 3900 | 3460 | 2733 | 3013.09 | 2662 | 3364 | 1.264 | 0.338 | 0.07587966 | 0.21720036 | 0.0486 | 0.008 | 0.0259 | 0.0259 | TRUE | 0.7678 |
| Thap1 | 930 | 934 | 806 | 1354 | 943 | 818 | 988 | 933 | 809 | 1064 | 1015 | 895 | 950.73 | 910 | 991 | 1.091 | 0.125 | 0.29057536 | 0.5114334 | 0.008 | 0.0104 | 0.009 | 0.009 | TRUE | 0.1118 |
| Thap11 | 2387 | 2703 | 2619 | 3806 | 2788 | 2564 | 2535 | 2700 | 2630 | 2992 | 3000 | 2807 | 2777.18 | 2622 | 2933 | 1.119 | 0.162 | 0.0229127 | 0.09578033 | 8.00E-04 | 0.0081 | 0.0033 | 0.0033 | TRUE | 0.017 |
| Thap2 | 225 | 216 | 236 | 240 | 157 | 150 | 239 | 216 | 237 | 189 | 169 | 164 | 202.24 | 231 | 174 | 0.757 | -0.402 | 0.00713917 | 0.04067884 | 0 | 0.0232 | 0.0111 | 0.0111 | TRUE | 0.0644 |
| Thap3 | 294 | 479 | 481 | 318 | 275 | 334 | 312 | 478 | 483 | 250 | 296 | 366 | 364.19 | 424 | 304 | 0.714 | -0.486 | 0.02216389 | 0.0936178 | 0.0444 | 0.0159 | 0.0297 | 0.0297 | TRUE | 0.6147 |
| Thap4 | 1814 | 2246 | 2352 | 2991 | 2217 | 2275 | 1927 | 2243 | 2362 | 2351 | 2385 | 2490 | 2293.09 | 2177 | 2409 | 1.106 | 0.146 | 0.14132027 | 0.32680725 | 0.0051 | 0.0083 | 0.0066 | 0.0066 | TRUE | 0.1221 |
| Thap6 | 345 | 279 | 277 | 665 | 438 | 296 | 366 | 279 | 278 | 523 | 471 | 324 | 373.55 | 308 | 439 | 1.43 | 0.516 | 0.01202728 | 0.0602013 | 0.0402 | 0.0157 | 0.0277 | 0.0277 | TRUE | 0.6033 |

| | | | | | | | | | | | | | | | | | | | | | | | | | |
|---|---|---|---|---|---|---|---|---|---|---|---|---|---|---|---|---|---|---|---|---|---|---|---|---|---|
| Thap7 | 536 | 514 | 475 | 799 | 527 | 493 | 569 | 513 | 477 | 628 | 567 | 540 | 549.07 | 520 | 578 | 1.114 | 0.156 | 0.19498704 | 0.40087767 | 0.0052 | 0.0129 | 0.0086 | 0.0086 | TRUE | 0.0802 |
| Thbd | 10 | 20 | 18 | 36 | 32 | 35 | 11 | 20 | 18 | 28 | 34 | 38 | 24.95 | 16 | 33 | 2.06 | 1.043 | 0.01232313 | 0.06138491 | 0 | 0.1389 | 0.0791 | 0.0791 | TRUE | 0.4149 |
| Thbs1 | 73 | 41 | 35 | 189 | 117 | 90 | 78 | 41 | 35 | 149 | 126 | 99 | 87.77 | 51 | 125 | 2.439 | 1.286 | 0.00010424 | 0.00142465 | 0.0911 | 0.0444 | 0.0652 | 0.0652 | TRUE | 0.6769 |
| Thbs2 | 23 | 26 | 23 | 22 | 11 | 13 | 24 | 26 | 23 | 17 | 12 | 14 | 19.48 | 24 | 14 | 0.595 | -0.748 | 0.08311381 | 0.2307423 | 0 | 0.1761 | 0.0799 | 0.0799 | TRUE | 0.1666 |
| Thbs3 | 230 | 194 | 204 | 187 | 116 | 129 | 244 | 194 | 205 | 147 | 125 | 141 | 175.99 | 214 | 138 | 0.644 | -0.634 | 0.00023752 | 0.00280192 | 0.006 | 0.0256 | 0.0155 | 0.0155 | TRUE | 0.1622 |
| Thbs4 | 56 | 54 | 53 | 110 | 89 | 54 | 59 | 54 | 53 | 86 | 96 | 59 | 68 | 55 | 80 | 1.452 | 0.538 | 0.04692261 | 0.15705929 | 0.0191 | 0.0554 | 0.0376 | 0.0376 | TRUE | 0.4715 |
| Theg | 0 | 4 | 2 | 0 | 2 | 4 | 0 | 4 | 2 | 0 | 2 | 4 | 2.09 | 2 | 2 | 1.055 | 0.078 | 0.96290267 | NA | 1.4075 | 1.584 | 1.5175 | 1.5175 | TRUE | 0.2008 |
| Them4 | 289 | 253 | 233 | 400 | 261 | 217 | 307 | 253 | 234 | 314 | 281 | 238 | 271.07 | 265 | 278 | 1.052 | 0.074 | 0.66717737 | 0.81391997 | 0.0161 | 0.0191 | 0.0175 | 0.0175 | TRUE | 0.2179 |
| Them5 | 22 | 13 | 24 | 21 | 17 | 13 | 23 | 13 | 24 | 17 | 18 | 14 | 18.25 | 20 | 16 | 0.813 | -0.299 | 0.52297749 | 0.71298212 | 0 | 0.1875 | 0.103 | 0.103 | TRUE | 0.5348 |
| Them6 | 597 | 715 | 703 | 850 | 585 | 593 | 634 | 714 | 706 | 668 | 629 | 649 | 666.81 | 685 | 649 | 0.948 | -0.077 | 0.43828865 | 0.64591156 | 0.001 | 0.0118 | 0.0056 | 0.0056 | TRUE | 0.0488 |
| Themis | 29 | 20 | 25 | 29 | 29 | 28 | 31 | 20 | 25 | 23 | 31 | 31 | 26.76 | 25 | 28 | 1.11 | 0.15 | 0.69622894 | 0.83365646 | 0 | 0.13 | 0.0693 | 0.0693 | TRUE | 0.2206 |
| Themis2 | 102 | 78 | 60 | 154 | 82 | 92 | 108 | 78 | 60 | 121 | 88 | 101 | 92.75 | 82 | 103 | 1.263 | 0.337 | 0.2174546 | 0.4294633 | 0.0434 | 0.0424 | 0.0429 | 0.0429 | TRUE | 0.4973 |
| Themis3 | 7 | 6 | 4 | 31 | 17 | 13 | 7 | 6 | 4 | 24 | 18 | 14 | 12.39 | 6 | 19 | 3.296 | 1.721 | 0.00425358 | 0.02771064 | 0 | 0.2728 | 0.1474 | 0.1474 | TRUE | 0.1759 |
| Thg1l | 2436 | 2457 | 2391 | 3026 | 2191 | 2189 | 2587 | 2454 | 2401 | 2379 | 2357 | 2396 | 2429.08 | 2481 | 2377 | 0.958 | -0.061 | 0.37503281 | 0.59102546 | 4.00E-04 | 0.0083 | 0.003 | 0.003 | TRUE | 0.0171 |
| Thnsl1 | 890 | 981 | 960 | 1449 | 1024 | 1046 | 945 | 980 | 964 | 1139 | 1102 | 1145 | 1045.81 | 963 | 1129 | 1.172 | 0.229 | 0.00352743 | 0.02397008 | 0 | 0.0101 | 0.0035 | 0.0035 | TRUE | 0.0051 |
| Thnsl2 | 432 | 485 | 485 | 644 | 410 | 449 | 459 | 484 | 487 | 506 | 441 | 492 | 478.19 | 477 | 480 | 1.007 | 0.01 | 0.92555684 | 0.96330226 | 0 | 0.0138 | 0.0067 | 0.0067 | TRUE | 0.0563 |
| Thoc1 | 2146 | 1869 | 1811 | 3088 | 2077 | 1771 | 2279 | 1867 | 1818 | 2428 | 2235 | 1939 | 2094.23 | 1988 | 2201 | 1.107 | 0.147 | 0.25003902 | 0.46730839 | 0.0137 | 0.0085 | 0.0113 | 0.0113 | TRUE | 0.1983 |
| Thoc2 | 5321 | 4747 | 4841 | 6432 | 4817 | 4236 | 5652 | 4741 | 4861 | 5056 | 5183 | 4637 | 5021.61 | 5085 | 4959 | 0.975 | -0.036 | 0.71254123 | 0.84351875 | 0.0052 | 0.0076 | 0.0067 | 0.0067 | TRUE | 0.116 |
| Thoc3 | 4276 | 4168 | 4195 | 5959 | 4274 | 3877 | 4542 | 4163 | 4212 | 4684 | 4599 | 4244 | 4407.29 | 4306 | 4509 | 1.048 | 0.067 | 0.39067163 | 0.60450043 | 0.0021 | 0.0077 | 0.0042 | 0.0042 | TRUE | 0.0321 |
| Thoc5 | 2897 | 2974 | 3212 | 3955 | 2915 | 2619 | 3077 | 2970 | 3225 | 3109 | 3136 | 2867 | 3064.14 | 3091 | 3037 | 0.983 | -0.025 | 0.74675065 | 0.86556561 | 0.0017 | 0.008 | 0.0039 | 0.0039 | TRUE | 0.029 |
| Thoc6 | 2608 | 2647 | 2616 | 3225 | 2321 | 2239 | 2770 | 2644 | 2627 | 2535 | 2497 | 2451 | 2587.32 | 2680 | 2494 | 0.931 | -0.103 | 0.12125621 | 0.29612636 | 0 | 0.0082 | 0.0028 | 0.0028 | TRUE | 0.0104 |
| Thoc7 | 1702 | 2003 | 1996 | 2555 | 1793 | 1884 | 1808 | 2000 | 2004 | 2009 | 1929 | 2062 | 1968.74 | 1937 | 2000 | 1.032 | 0.046 | 0.57479419 | 0.75283624 | 0.0015 | 0.0086 | 0.0043 | 0.0043 | TRUE | 0.0412 |
| Thop1 | 8802 | 7640 | 7719 | 9606 | 7110 | 6435 | 9349 | 7630 | 7751 | 7551 | 7650 | 7044 | 7829.27 | 8243 | 7415 | 0.9 | -0.153 | 0.13387422 | 0.31632422 | 0.0075 | 0.0073 | 0.0073 | 0.0073 | TRUE | 0.1681 |
| Thpo | 175 | 152 | 167 | 160 | 121 | 151 | 186 | 152 | 168 | 126 | 130 | 165 | 154.44 | 169 | 140 | 0.831 | -0.266 | 0.15599425 | 0.34868782 | 0.0096 | 0.0282 | 0.0189 | 0.0189 | TRUE | 0.2412 |
| Thra | 370 | 752 | 844 | 595 | 467 | 534 | 393 | 751 | 847 | 468 | 502 | 585 | 591.04 | 664 | 518 | 0.78 | -0.358 | 0.15138559 | 0.34241409 | 0.0814 | 0.0125 | 0.0432 | 0.0432 | TRUE | 1.149 |
| Thrap3 | 9621 | 7223 | 7646 | 14722 | 10563 | 8174 | 10219 | 7214 | 7677 | 11573 | 11365 | 8948 | 9499.4 | 8370 | 10629 | 1.27 | 0.345 | 0.02646036 | 0.10571469 | 0.028 | 0.0073 | 0.0173 | 0.0173 | TRUE | 0.456 |
| Thrb | 52 | 43 | 60 | 84 | 67 | 57 | 55 | 43 | 60 | 66 | 72 | 62 | 59.82 | 53 | 67 | 1.266 | 0.34 | 0.19282585 | 0.39842218 | 0 | 0.062 | 0.0322 | 0.0322 | TRUE | 0.2249 |
| Thrsp | 67 | 75 | 63 | 132 | 62 | 76 | 71 | 75 | 63 | 104 | 67 | 83 | 77.17 | 70 | 85 | 1.219 | 0.286 | 0.26176075 | 0.48126204 | 0.018 | 0.0496 | 0.0337 | 0.0337 | TRUE | 0.2257 |
| Thsd1 | 855 | 818 | 854 | 801 | 652 | 590 | 908 | 817 | 858 | 630 | 702 | 646 | 759.94 | 861 | 659 | 0.765 | -0.387 | 8.89E-05 | 0.00125979 | 0.0017 | 0.0112 | 0.0057 | 0.0057 | TRUE | 0.0371 |
| Thsd4 | 91 | 106 | 78 | 142 | 94 | 87 | 97 | 106 | 78 | 112 | 101 | 95 | 98.14 | 94 | 103 | 1.1 | 0.137 | 0.52153514 | 0.71165526 | 0 | 0.0405 | 0.0229 | 0.0229 | TRUE | 0.1992 |
| Thsd7a | 7 | 3 | 2 | 11 | 7 | 6 | 7 | 3 | 2 | 9 | 8 | 7 | 5.86 | 4 | 8 | 1.849 | 0.887 | 0.28442919 | 0.50589188 | 0 | 0.5687 | 0.3067 | 0.3067 | TRUE | 0.7885 |
| Thsd7b | 7 | 4 | 4 | 9 | 9 | 5 | 7 | 4 | 4 | 7 | 10 | 5 | 6.28 | 5 | 7 | 1.443 | 0.529 | 0.48811304 | 0.68476003 | 0 | 0.5315 | 0.2545 | 0.2545 | TRUE | 0.2964 |
| Thtpa | 509 | 653 | 604 | 695 | 515 | 558 | 541 | 652 | 606 | 546 | 554 | 611 | 585.09 | 600 | 570 | 0.95 | -0.074 | 0.52496463 | 0.71463336 | 0.0045 | 0.0125 | 0.0081 | 0.0081 | TRUE | 0.0866 |
| Thumpd1 | 6015 | 6398 | 6271 | 8331 | 6556 | 6177 | 6389 | 6390 | 6297 | 6549 | 7054 | 6762 | 6573.37 | 6359 | 6788 | 1.067 | 0.094 | 0.13075981 | 0.31134416 | 5.00E-04 | 0.0074 | 0.0026 | 0.0026 | TRUE | 0.0143 |
| Thumpd2 | 835 | 777 | 693 | 731 | 539 | 617 | 887 | 776 | 696 | 575 | 580 | 675 | 698.12 | 786 | 610 | 0.775 | -0.367 | 0.00492802 | 0.03104024 | 0.0096 | 0.0116 | 0.0108 | 0.0108 | TRUE | 0.1481 |
| Thumpd3 | 4979 | 5339 | 5430 | 6279 | 4558 | 4520 | 5289 | 5332 | 5452 | 4936 | 4904 | 4948 | 5143.5 | 5358 | 4929 | 0.92 | -0.12 | 0.03346869 | 0.12408747 | 0 | 0.0075 | 0.0021 | 0.0021 | TRUE | 0.0029 |
| Thy1 | 825 | 937 | 897 | 479 | 355 | 458 | 876 | 936 | 901 | 377 | 382 | 501 | 662.11 | 904 | 420 | 0.463 | -1.109 | 2.30E-16 | 3.71E-14 | 0.0109 | 0.0119 | 0.0114 | 0.0114 | TRUE | 0.3259 |
| Thyn1 | 2244 | 2466 | 2498 | 2751 | 1977 | 1945 | 2384 | 2463 | 2508 | 2163 | 2127 | 2129 | 2295.58 | 2452 | 2140 | 0.873 | -0.196 | 0.00302739 | 0.02139248 | 0 | 0.0083 | 0.0027 | 0.0027 | TRUE | 0.0071 |
| Tia1 | 1090 | 1203 | 1253 | 1588 | 1225 | 1033 | 1158 | 1201 | 1258 | 1248 | 1318 | 1131 | 1219.09 | 1206 | 1232 | 1.022 | 0.032 | 0.73801673 | 0.85973853 | 0.0029 | 0.0096 | 0.0056 | 0.0056 | TRUE | 0.0609 |
| Tial1 | 3984 | 4770 | 4864 | 4844 | 3740 | 4167 | 4232 | 4764 | 4884 | 3808 | 4024 | 4561 | 4378.82 | 4627 | 4131 | 0.893 | -0.164 | 0.10784847 | 0.27490073 | 0.0069 | 0.0077 | 0.0073 | 0.0073 | TRUE | 0.1009 |
| Tiam1 | 3479 | 3603 | 3798 | 3936 | 2771 | 3081 | 3695 | 3598 | 3814 | 3094 | 2981 | 3373 | 3425.92 | 3702 | 3149 | 0.851 | -0.234 | 0.00295817 | 0.02100686 | 0.002 | 0.0079 | 0.0042 | 0.0042 | TRUE | 0.0462 |
| Tiam2 | 112 | 101 | 98 | 138 | 102 | 79 | 119 | 101 | 98 | 108 | 110 | 86 | 103.82 | 106 | 101 | 0.961 | -0.058 | 0.78013131 | 0.88621658 | 0 | 0.0386 | 0.0216 | 0.0216 | TRUE | 0.1563 |
| Ticam1 | 149 | 152 | 162 | 255 | 151 | 140 | 158 | 152 | 163 | 200 | 162 | 153 | 164.82 | 158 | 172 | 1.097 | 0.133 | 0.44733523 | 0.65333562 | 0.0065 | 0.0269 | 0.0161 | 0.0161 | TRUE | 0.2395 |
| Ticam2 | 69 | 103 | 112 | 82 | 65 | 69 | 73 | 103 | 112 | 64 | 70 | 76 | 83.09 | 96 | 70 | 0.725 | -0.465 | 0.0602726 | 0.18639597 | 0.0167 | 0.0466 | 0.0318 | 0.0318 | TRUE | 0.4118 |
| Ticrr | 4516 | 3801 | 4195 | 7165 | 4848 | 3923 | 4797 | 3796 | 4212 | 5633 | 5216 | 4294 | 4658.03 | 4268 | 5048 | 1.183 | 0.242 | 0.06483544 | 0.19616048 | 0.0158 | 0.0076 | 0.0122 | 0.0122 | TRUE | 0.2074 |
| Tie1 | 64 | 105 | 77 | 58 | 35 | 48 | 68 | 105 | 77 | 46 | 38 | 53 | 64.33 | 83 | 46 | 0.542 | -0.883 | 0.00198377 | 0.01527738 | 0.025 | 0.0581 | 0.0418 | 0.0418 | TRUE | 0.4794 |
| Tifa | 6 | 3 | 5 | 4 | 6 | 3 | 6 | 3 | 5 | 3 | 6 | 3 | 4.55 | 5 | 4 | 0.888 | -0.171 | 0.84786772 | NA | 0 | 0.7317 | 0.3495 | 0.3495 | TRUE | 0.3188 |
| Tifab | 12 | 14 | 10 | 50 | 32 | 35 | 13 | 14 | 10 | 39 | 34 | 38 | 24.8 | 12 | 37 | 3.053 | 1.61 | 7.52E-05 | 0.00109884 | 0 | 0.1397 | 0.0644 | 0.0644 | TRUE | 0.1021 |
| Tigd2 | 798 | 675 | 590 | 1052 | 806 | 738 | 848 | 674 | 592 | 827 | 867 | 808 | 769.37 | 705 | 834 | 1.184 | 0.244 | 0.08990543 | 0.24390214 | 0.0158 | 0.0112 | 0.0135 | 0.0135 | TRUE | 0.3703 |
| Tigd3 | 134 | 119 | 101 | 175 | 146 | 111 | 142 | 119 | 101 | 138 | 157 | 122 | 129.79 | 121 | 139 | 1.149 | 0.2 | 0.33457346 | 0.5548783 | 0.0138 | 0.0323 | 0.0232 | 0.0232 | TRUE | 0.2402 |
| Tigd4 | 0 | 0 | 0 | 0 | 0 | 0 | 0 | 0 | 0 | 0 | 0 | 0 | 0 | 0 | 0 | NA | NA | NA | NA | NA | NA | NA | NA | NA | NA |
| Tigd5 | 16 | 15 | 9 | 58 | 32 | 15 | 17 | 15 | 9 | 46 | 34 | 16 | 22.91 | 14 | 32 | 2.374 | 1.247 | 0.01672165 | 0.07653332 | 0.1353 | 0.1507 | 0.1432 | 0.1432 | TRUE | 0.1848 |
| Tigit | 1 | 1 | 2 | 2 | 0 | 0 | 1 | 1 | 2 | 2 | 0 | 0 | 0.94 | 1 | 1 | 0.425 | -1.233 | 0.55053282 | NA | 0 | 3.5108 | 1.8559 | 1.8559 | TRUE | 0.9714 |
| Timd2 | 2 | 2 | 1 | 38 | 18 | 13 | 2 | 2 | 1 | 30 | 19 | 14 | 11.43 | 2 | 21 | 12.526 | 3.647 | 1.64E-05 | 0.00030155 | 0 | 0.2951 | 0.1935 | 0.1935 | TRUE | 0.1801 |
| Timd4 | 1 | 1 | 2 | 2 | 3 | 1 | 1 | 1 | 2 | 2 | 3 | 1 | 1.66 | 1 | 2 | 1.436 | 0.522 | 0.71258777 | NA | 0 | 1.9908 | 0.8231 | 0.8231 | TRUE | 0.2519 |
| Timeless | 4184 | 4271 | 4436 | 5961 | 4530 | 4160 | 4444 | 4265 | 4454 | 4686 | 4874 | 4554 | 4546.28 | 4388 | 4705 | 1.072 | 0.101 | 0.1236413 | 0.29973419 | 5.00E-04 | 0.0076 | 0.0029 | 0.0029 | TRUE | 0.0119 |
| Timm10 | 564 | 657 | 604 | 952 | 608 | 544 | 599 | 656 | 606 | 748 | 654 | 595 | 643.29 | 620 | 666 | 1.075 | 0.104 | 0.38730956 | 0.60112639 | 0.0066 | 0.012 | 0.009 | 0.009 | TRUE | 0.1427 |
| Timm10b | 800 | 879 | 901 | 1058 | 814 | 767 | 850 | 878 | 905 | 832 | 876 | 840 | 863.24 | 878 | 849 | 0.967 | -0.048 | 0.5717009 | 0.75046747 | 0 | 0.0107 | 0.0041 | 0.0041 | TRUE | 0.009 |
| Timm13 | 2357 | 2611 | 2534 | 3735 | 2519 | 2349 | 2504 | 2608 | 2544 | 2936 | 2710 | 2571 | 2645.56 | 2552 | 2739 | 1.074 | 0.103 | 0.20141851 | 0.40905369 | 0.0019 | 0.0081 | 0.0043 | 0.0043 | TRUE | 0.0488 |
| Timm17a | 3818 | 3892 | 3668 | 5314 | 3516 | 3400 | 4055 | 3887 | 3683 | 4177 | 3783 | 3722 | 3884.62 | 3875 | 3894 | 1.005 | 0.008 | 0.92625342 | 0.9637204 | 0.0029 | 0.0078 | 0.0047 | 0.0047 | TRUE | 0.0499 |
| Timm17b | 1997 | 2425 | 2329 | 2595 | 1861 | 1929 | 2121 | 2422 | 2339 | 2040 | 2002 | 2112 | 2172.58 | 2294 | 2051 | 0.894 | -0.162 | 0.05235328 | 0.16923389 | 0.002 | 0.0084 | 0.0045 | 0.0045 | TRUE | 0.0523 |
| Timm21 | 626 | 553 | 555 | 570 | 400 | 345 | 665 | 552 | 557 | 448 | 430 | 378 | 505.1 | 591 | 419 | 0.709 | -0.496 | 0.00014369 | 0.00184099 | 0.0076 | 0.0134 | 0.0102 | 0.0102 | TRUE | 0.1376 |
| Timm22 | 1061 | 1038 | 992 | 1748 | 1182 | 988 | 1127 | 1037 | 996 | 1374 | 1272 | 1082 | 1147.85 | 1053 | 1243 | 1.181 | 0.24 | 0.04128588 | 0.14408646 | 0.0078 | 0.0098 | 0.0091 | 0.0091 | TRUE | 0.153 |
| Timm23 | 2733 | 3142 | 3020 | 3978 | 2810 | 2681 | 2903 | 3138 | 3032 | 3127 | 3023 | 2935 | 3026.44 | 3024 | 3028 | 1.002 | 0.002 | 0.97477609 | 0.98685208 | 9.00E-04 | 0.008 | 0.0033 | 0.0033 | TRUE | 0.0149 |
| Timm44 | 4250 | 4539 | 4674 | 5763 | 4227 | 4388 | 4514 | 4533 | 4693 | 4530 | 4548 | 4803 | 4603.72 | 4580 | 4627 | 1.01 | 0.014 | 0.82151941 | 0.90875327 | 4.00E-04 | 0.0076 | 0.0028 | 0.0028 | TRUE | 0.0134 |
| Timm50 | 4338 | 4576 | 4712 | 5518 | 4214 | 4285 | 4608 | 4570 | 4731 | 4338 | 4534 | 4691 | 4578.6 | 4636 | 4521 | 0.975 | -0.037 | 0.57580567 | 0.75380936 | 6.00E-04 | 0.0076 | 0.0029 | 0.0029 | TRUE | 0.0155 |
| Timm8a1 | 509 | 499 | 492 | 653 | 496 | 429 | 541 | 498 | 494 | 513 | 534 | 470 | 508.27 | 511 | 506 | 0.99 | -0.014 | 0.89430342 | 0.94899775 | 0.0011 | 0.0134 | 0.0066 | 0.0066 | TRUE | 0.0432 |
| Timm8a2 | 77 | 74 | 60 | 126 | 106 | 111 | 82 | 74 | 60 | 99 | 114 | 122 | 91.76 | 72 | 112 | 1.546 | 0.629 | 0.00498365 | 0.03125331 | 0 | 0.0428 | 0.0246 | 0.0246 | TRUE | 0.1852 |
| Timm8b | 1521 | 1698 | 1634 | 1886 | 1303 | 1379 | 1616 | 1696 | 1641 | 1483 | 1402 | 1510 | 1557.7 | 1651 | 1465 | 0.887 | -0.172 | 0.02323405 | 0.09666247 | 3.00E-04 | 0.009 | 0.0035 | 0.0035 | TRUE | 0.0165 |

| | | | | | | | | | | | | | | | | | | | | | | | | | |
|---|---|---|---|---|---|---|---|---|---|---|---|---|---|---|---|---|---|---|---|---|---|---|---|---|---|
| Timm9 | 670 | 791 | 832 | 951 | 690 | 731 | 712 | 790 | 835 | 748 | 742 | 800 | 771.21 | 779 | 763 | 0.979 | -0.03 | 0.76893933 | 0.87928573 | 0.0027 | 0.0112 | 0.0063 | 0.0063 | TRUE | 0.0673 |
| Timmdc1 | 1600 | 2055 | 1995 | 2092 | 1556 | 1561 | 1699 | 2052 | 2003 | 1645 | 1674 | 1709 | 1797.08 | 1918 | 1676 | 0.873 | -0.195 | 0.0464588 | 0.15593704 | 0.0043 | 0.0087 | 0.0064 | 0.0064 | TRUE | 0.12 |
| Timp1 | 513 | 823 | 884 | 699 | 515 | 686 | 545 | 822 | 888 | 549 | 554 | 751 | 684.83 | 752 | 618 | 0.822 | -0.283 | 0.16440492 | 0.36058884 | 0.0444 | 0.0117 | 0.0284 | 0.0284 | TRUE | 0.6827 |
| Timp2 | 818 | 1013 | 929 | 1175 | 841 | 792 | 869 | 1012 | 933 | 924 | 905 | 867 | 918.15 | 938 | 899 | 0.958 | -0.061 | 0.52638679 | 0.71554235 | 0.0023 | 0.0105 | 0.0057 | 0.0057 | TRUE | 0.0567 |
| Timp3 | 661 | 721 | 735 | 689 | 530 | 569 | 702 | 720 | 738 | 542 | 570 | 623 | 649.16 | 720 | 578 | 0.802 | -0.319 | 0.0015872 | 0.01281789 | 0.0012 | 0.012 | 0.0058 | 0.0058 | TRUE | 0.0528 |
| Timp4 | 8 | 3 | 7 | 1 | 2 | 6 | 8 | 3 | 7 | 1 | 2 | 7 | 4.67 | 6 | 3 | 0.498 | -1.006 | 0.32751155 | 0.54874958 | 0.2982 | 0.7121 | 0.5189 | 0.5189 | TRUE | 1.1268 |
| Tinag | 0 | 0 | 0 | 0 | 0 | 0 | 0 | 0 | 0 | 0 | 0 | 0 | 0 | 0 | 0 | NA | NA | NA | NA | NA | NA | NA | NA | NA | NA |
| Tinagl1 | 229 | 429 | 431 | 630 | 448 | 485 | 243 | 428 | 433 | 495 | 482 | 531 | 435.44 | 368 | 503 | 1.365 | 0.448 | 0.03401459 | 0.12550866 | 0.0474 | 0.0145 | 0.0299 | 0.0299 | TRUE | 1.0022 |
| Tinf2 | 2626 | 2748 | 2598 | 3405 | 2382 | 2276 | 2789 | 2744 | 2609 | 2677 | 2563 | 2491 | 2645.58 | 2714 | 2577 | 0.95 | -0.074 | 0.30455114 | 0.52512118 | 9.00E-04 | 0.0081 | 0.0034 | 0.0034 | TRUE | 0.0142 |
| Tiparp | 2769 | 3490 | 3749 | 3566 | 3016 | 3051 | 2941 | 3485 | 3764 | 2803 | 3245 | 3340 | 3263.2 | 3397 | 3129 | 0.921 | -0.119 | 0.3220181 | 0.54330584 | 0.0113 | 0.0079 | 0.0101 | 0.0101 | TRUE | 0.1673 |
| Tipin | 6358 | 5724 | 5504 | 8056 | 5480 | 5298 | 6753 | 5717 | 5527 | 6333 | 5896 | 5799 | 6004.19 | 5999 | 6009 | 1.002 | 0.003 | 0.97740149 | 0.98786135 | 0.0068 | 0.0075 | 0.0071 | 0.0071 | TRUE | 0.1475 |
| Tiprl | 2754 | 3260 | 3060 | 3265 | 2359 | 2568 | 2925 | 3256 | 3073 | 2567 | 2538 | 2811 | 2861.58 | 3085 | 2639 | 0.855 | -0.226 | 0.00673692 | 0.03902025 | 0.0025 | 0.0081 | 0.0046 | 0.0046 | TRUE | 0.0393 |
| Tirap | 608 | 671 | 634 | 870 | 625 | 599 | 646 | 670 | 637 | 684 | 672 | 656 | 660.77 | 651 | 671 | 1.031 | 0.044 | 0.62706967 | 0.78999388 | 0 | 0.0119 | 0.0044 | 0.0044 | TRUE | 0.008 |
| Tjap1 | 4078 | 4268 | 4138 | 4554 | 3639 | 3666 | 4332 | 4262 | 4155 | 3580 | 3915 | 4013 | 4042.9 | 4250 | 3836 | 0.902 | -0.148 | 0.04858244 | 0.16080635 | 0.0016 | 0.0077 | 0.0038 | 0.0038 | TRUE | 0.0421 |
| Tjp1 | 13841 | 12249 | 13175 | 17397 | 12850 | 12202 | 14702 | 12233 | 13229 | 13676 | 13826 | 13357 | 13503.81 | 13388 | 13620 | 1.017 | 0.025 | 0.77583506 | 0.88349968 | 0.0035 | 0.0071 | 0.0054 | 0.0054 | TRUE | 0.09 |
| Tjp2 | 6591 | 6028 | 6441 | 11249 | 7910 | 7323 | 7001 | 6020 | 6467 | 8843 | 8511 | 8016 | 7476.4 | 6496 | 8457 | 1.302 | 0.381 | 9.62E-06 | 0.00019301 | 0.0039 | 0.0073 | 0.0052 | 0.0052 | TRUE | 0.0563 |
| Tjp3 | 805 | 1637 | 1844 | 846 | 826 | 1273 | 855 | 1635 | 1852 | 665 | 889 | 1393 | 1214.8 | 1447 | 982 | 0.679 | -0.559 | 0.20009136 | 0.40767872 | 0.1364 | 0.0096 | 0.0637 | 0.1364 | TRUE | 0.9455 |
| Tk1 | 2909 | 3121 | 3191 | 4469 | 3246 | 3198 | 3090 | 3117 | 3204 | 3513 | 3492 | 3501 | 3319.56 | 3137 | 3502 | 1.116 | 0.159 | 0.0086175 | 0.04702949 | 0 | 0.0079 | 0.0023 | 0.0023 | TRUE | 0.0043 |
| Tk2 | 1011 | 1388 | 1271 | 1569 | 1200 | 1193 | 1074 | 1386 | 1276 | 1233 | 1291 | 1306 | 1261.13 | 1245 | 1277 | 1.025 | 0.035 | 0.75828759 | 0.87245014 | 0.0079 | 0.0095 | 0.0086 | 0.0086 | TRUE | 0.1735 |
| Tkt | 32189 | 35126 | 35814 | 36090 | 27228 | 28449 | 34190 | 35081 | 35961 | 28371 | 29296 | 31142 | 32340.13 | 35077 | 29603 | 0.844 | -0.245 | 0.00022619 | 0.00268234 | 0.0014 | 0.007 | 0.0031 | 0.0031 | TRUE | 0.0253 |
| Tktl1 | 75 | 51 | 61 | 314 | 170 | 143 | 80 | 51 | 61 | 247 | 183 | 157 | 129.69 | 64 | 196 | 3.07 | 1.618 | 4.10E-10 | 2.74E-08 | 0.0438 | 0.0323 | 0.0378 | 0.0378 | TRUE | 0.3273 |
| Tktl2 | 573 | 395 | 301 | 1921 | 1312 | 1098 | 609 | 394 | 302 | 1510 | 1412 | 1202 | 904.84 | 435 | 1375 | 3.161 | 1.661 | 1.60E-13 | 1.87E-11 | 0.0695 | 0.0105 | 0.035 | 0.035 | TRUE | 1.4027 |
| Tlcd1 | 371 | 293 | 342 | 715 | 518 | 360 | 394 | 293 | 343 | 562 | 557 | 394 | 423.93 | 343 | 504 | 1.471 | 0.557 | 0.0021855 | 0.01651636 | 0.028 | 0.0147 | 0.0214 | 0.0214 | TRUE | 0.4213 |
| Tlcd2 | 245 | 253 | 248 | 293 | 222 | 238 | 260 | 253 | 249 | 230 | 239 | 261 | 248.61 | 254 | 243 | 0.956 | -0.065 | 0.62676906 | 0.78982116 | 0 | 0.0202 | 0.0088 | 0.0088 | TRUE | 0.0418 |
| Tle1 | 2480 | 2365 | 2213 | 3115 | 2320 | 2203 | 2634 | 2362 | 2222 | 2449 | 2496 | 2412 | 2429.12 | 2406 | 2452 | 1.019 | 0.028 | 0.75684711 | 0.87155626 | 0.0034 | 0.0083 | 0.0053 | 0.0053 | TRUE | 0.0831 |
| Tle2 | 50 | 56 | 43 | 84 | 78 | 60 | 53 | 56 | 43 | 66 | 84 | 66 | 61.31 | 51 | 72 | 1.414 | 0.5 | 0.05747854 | 0.18084012 | 0 | 0.0606 | 0.033 | 0.033 | TRUE | 0.1867 |
| Tle3 | 6121 | 7191 | 6845 | 6808 | 5406 | 5330 | 6502 | 7182 | 6873 | 5352 | 5817 | 5835 | 6259.89 | 6852 | 5668 | 0.827 | -0.274 | 0.00034192 | 0.00379767 | 0.0022 | 0.0074 | 0.0041 | 0.0041 | TRUE | 0.0292 |
| Tle4 | 3251 | 2415 | 2383 | 4756 | 3081 | 2741 | 3453 | 2412 | 2393 | 3739 | 3315 | 3000 | 3052 | 2753 | 3351 | 1.218 | 0.284 | 0.07493548 | 0.21543609 | 0.0299 | 0.008 | 0.018 | 0.018 | TRUE | 0.6011 |
| Tle6 | 520 | 472 | 463 | 939 | 734 | 697 | 552 | 471 | 465 | 738 | 790 | 763 | 629.92 | 496 | 764 | 1.539 | 0.622 | 1.80E-08 | 8.14E-07 | 0.0029 | 0.0121 | 0.0071 | 0.0071 | TRUE | 0.1129 |
| Tlk1 | 4363 | 4082 | 4126 | 4994 | 3694 | 3602 | 4634 | 4077 | 4143 | 3926 | 3975 | 3943 | 4116.22 | 4285 | 3948 | 0.921 | -0.118 | 0.13182134 | 0.31279177 | 0.0022 | 0.0077 | 0.0042 | 0.0042 | TRUE | 0.0619 |
| Tlk2 | 5271 | 5690 | 5967 | 7867 | 5574 | 5643 | 5599 | 5683 | 5991 | 6184 | 5997 | 6177 | 5938.62 | 5758 | 6119 | 1.063 | 0.088 | 0.16375509 | 0.35961366 | 5.00E-04 | 0.0075 | 0.0027 | 0.0027 | TRUE | 0.0154 |
| Tll1 | 6 | 3 | 1 | 19 | 12 | 8 | 6 | 3 | 1 | 15 | 13 | 9 | 7.83 | 3 | 12 | 3.578 | 1.839 | 0.02188041 | 0.09289308 | 0.0176 | 0.4277 | 0.2749 | 0.2749 | TRUE | 0.7587 |
| Tll2 | 2 | 5 | 3 | 3 | 4 | 4 | 2 | 5 | 3 | 2 | 4 | 4 | 3.53 | 3 | 3 | 1.07 | 0.098 | 0.92133803 | NA | 0 | 0.9406 | 0.4238 | 0.4238 | TRUE | 0.2564 |
| Tln1 | 6335 | 5559 | 5986 | 10219 | 7000 | 5897 | 6729 | 5552 | 6011 | 8033 | 7532 | 6455 | 6718.57 | 6097 | 7340 | 1.204 | 0.268 | 0.01913195 | 0.08403416 | 0.0105 | 0.0074 | 0.0093 | 0.0093 | TRUE | 0.1356 |
| Tln2 | 3975 | 4283 | 4788 | 5133 | 3533 | 3576 | 4222 | 4277 | 4808 | 4035 | 3801 | 3914 | 4176.37 | 4436 | 3917 | 0.883 | -0.179 | 0.02844899 | 0.11092646 | 0.0027 | 0.0077 | 0.0046 | 0.0046 | TRUE | 0.0656 |
| Tlr1 | 1 | 0 | 0 | 2 | 0 | 0 | 1 | 0 | 0 | 2 | 0 | 0 | 0.44 | 0 | 1 | 1.494 | 0.579 | 0.86844164 | NA | 0 | 7.5101 | 6.9157 | 6.9157 | TRUE | 0.9721 |
| Tlr11 | 0 | 0 | 0 | 0 | 0 | 0 | 0 | 0 | 0 | 0 | 0 | 0 | 0 | 0 | 0 | NA | NA | NA | NA | NA | NA | NA | NA | NA | NA |
| Tlr12 | 0 | 0 | 0 | 0 | 0 | 1 | 0 | 0 | 0 | 0 | 0 | 1 | 0.18 | 0 | 0 | 1.859 | 0.894 | 0.82650734 | NA | 0 | 18.0643 | 10 | 10 | TRUE | 0.5383 |
| Tlr13 | 0 | 0 | 0 | 1 | 0 | 0 | 0 | 0 | 0 | 1 | 0 | 0 | 0.13 | 0 | 0 | 1.859 | 0.894 | 0.82650734 | NA | 0 | 25.1515 | 10 | 10 | TRUE | 0.2563 |
| Tlr2 | 192 | 236 | 167 | 344 | 255 | 217 | 204 | 236 | 168 | 270 | 274 | 238 | 231.61 | 203 | 261 | 1.289 | 0.366 | 0.03203062 | 0.12056968 | 0.0122 | 0.0211 | 0.0166 | 0.0166 | TRUE | 0.2479 |
| Tlr3 | 104 | 70 | 62 | 131 | 99 | 84 | 110 | 70 | 62 | 103 | 107 | 92 | 90.68 | 81 | 101 | 1.245 | 0.316 | 0.23607046 | 0.45112853 | 0.0374 | 0.0432 | 0.0402 | 0.0402 | TRUE | 0.9385 |
| Tlr4 | 0 | 0 | 0 | 0 | 0 | 2 | 0 | 0 | 0 | 0 | 0 | 2 | 0.36 | 0 | 1 | 3.832 | 1.938 | 0.63022034 | NA | 10 | 9.0356 | 10 | 10 | TRUE | 0.9744 |
| Tlr5 | 26 | 20 | 15 | 36 | 14 | 5 | 28 | 20 | 15 | 28 | 15 | 5 | 18.58 | 21 | 16 | 0.794 | -0.334 | 0.58638308 | 0.76131329 | 0.2542 | 0.1842 | 0.2169 | 0.2169 | TRUE | 0.3843 |
| Tlr6 | 32 | 23 | 40 | 33 | 10 | 22 | 34 | 23 | 40 | 26 | 11 | 24 | 26.32 | 32 | 20 | 0.631 | -0.664 | 0.14644598 | 0.33513436 | 0.0851 | 0.1321 | 0.1108 | 0.1108 | TRUE | 0.6244 |
| Tlr7 | 3 | 1 | 0 | 6 | 2 | 2 | 3 | 1 | 0 | 5 | 2 | 2 | 2.21 | 1 | 3 | 2.223 | 1.152 | 0.41215168 | NA | 0.0177 | 1.4995 | 0.8925 | 0.8925 | TRUE | 0.7665 |
| Tlr8 | 0 | 0 | 0 | 0 | 0 | 0 | 0 | 0 | 0 | 0 | 0 | 0 | 0 | 0 | 0 | NA | NA | NA | NA | NA | NA | NA | NA | NA | NA |
| Tlr9 | 1 | 0 | 0 | 0 | 0 | 0 | 1 | 0 | 0 | 0 | 0 | 0 | 0.18 | 0 | 0 | 0.49 | -1.029 | 0.80087 | NA | 0 | 18.6165 | 10 | 10 | TRUE | 0.4614 |
| Tlx1 | 13 | 20 | 10 | 24 | 21 | 23 | 14 | 20 | 10 | 19 | 23 | 25 | 18.41 | 15 | 22 | 1.512 | 0.597 | 0.20465342 | 0.41307829 | 0 | 0.1858 | 0.1024 | 0.1024 | TRUE | 0.4733 |
| Tlx2 | 6 | 8 | 7 | 10 | 7 | 3 | 6 | 8 | 7 | 8 | 8 | 3 | 6.68 | 7 | 6 | 0.883 | -0.18 | 0.80612457 | 0.90019698 | 0 | 0.5002 | 0.2392 | 0.2392 | TRUE | 0.3669 |
| Tlx3 | 75 | 100 | 112 | 135 | 127 | 96 | 80 | 100 | 112 | 106 | 137 | 105 | 106.64 | 97 | 116 | 1.188 | 0.248 | 0.27066287 | 0.49074002 | 0.0161 | 0.0378 | 0.0272 | 0.0272 | TRUE | 0.2395 |
| Tm2d1 | 417 | 691 | 660 | 772 | 556 | 684 | 443 | 690 | 663 | 607 | 598 | 749 | 624.93 | 599 | 651 | 1.087 | 0.121 | 0.51429153 | 0.70699957 | 0.0344 | 0.0122 | 0.023 | 0.023 | TRUE | 0.6049 |
| Tm2d2 | 944 | 956 | 981 | 1355 | 888 | 916 | 1003 | 955 | 985 | 1065 | 955 | 1003 | 994.3 | 981 | 1008 | 1.029 | 0.041 | 0.64297504 | 0.79907039 | 0.001 | 0.0102 | 0.0045 | 0.0045 | TRUE | 0.0309 |
| Tm2d3 | 995 | 1290 | 1199 | 1171 | 900 | 1000 | 1057 | 1288 | 1204 | 921 | 968 | 1095 | 1088.78 | 1183 | 995 | 0.84 | -0.252 | 0.03091085 | 0.11776 | 0.0076 | 0.0099 | 0.0089 | 0.0089 | TRUE | 0.1039 |
| Tm4sf1 | 2 | 2 | 0 | 2 | 3 | 2 | 2 | 2 | 0 | 2 | 3 | 2 | 1.85 | 1 | 2 | 1.685 | 0.753 | 0.60011443 | NA | 0 | 1.7859 | 0.9006 | 0.9006 | TRUE | 0.492 |
| Tm4sf19 | 0 | 1 | 0 | 0 | 3 | 2 | 0 | 1 | 0 | 0 | 3 | 2 | 1.07 | 0 | 2 | 4.846 | 2.277 | 0.32387131 | NA | 0 | 3.0878 | 2.5617 | 2.5617 | TRUE | 0.3017 |
| Tm4sf20 | 0 | 0 | 0 | 0 | 0 | 0 | 0 | 0 | 0 | 0 | 0 | 0 | 0 | 0 | 0 | NA | NA | NA | NA | NA | NA | NA | NA | NA | NA |
| Tm4sf4 | 0 | 0 | 0 | 0 | 0 | 0 | 0 | 0 | 0 | 0 | 0 | 0 | 0 | 0 | 0 | NA | NA | NA | NA | NA | NA | NA | NA | NA | NA |
| Tm4sf5 | 513 | 589 | 571 | 293 | 225 | 302 | 545 | 588 | 573 | 230 | 242 | 331 | 418.25 | 569 | 268 | 0.469 | -1.092 | 5.19E-12 | 4.82E-10 | 0.016 | 0.0148 | 0.0153 | 0.0153 | TRUE | 0.464 |
| Tm6sf1 | 587 | 694 | 647 | 599 | 423 | 469 | 623 | 693 | 650 | 471 | 455 | 513 | 567.61 | 655 | 480 | 0.731 | -0.451 | 2.25E-05 | 0.00039343 | 0.0014 | 0.0127 | 0.0064 | 0.0064 | TRUE | 0.0418 |
| Tm6sf2 | 59 | 51 | 45 | 46 | 34 | 37 | 63 | 51 | 45 | 36 | 37 | 41 | 45.34 | 53 | 38 | 0.713 | -0.489 | 0.09657678 | 0.25514389 | 0 | 0.0796 | 0.0397 | 0.0397 | TRUE | 0.209 |
| Tm7sf2 | 612 | 923 | 961 | 761 | 601 | 654 | 650 | 922 | 965 | 598 | 647 | 716 | 749.6 | 846 | 654 | 0.772 | -0.373 | 0.02371111 | 0.09797529 | 0.025 | 0.0113 | 0.0182 | 0.0182 | TRUE | 0.4846 |
| Tm7sf3 | 2638 | 3851 | 4017 | 3966 | 2784 | 3099 | 2802 | 3846 | 4033 | 3118 | 2995 | 3392 | 3364.51 | 3560 | 3168 | 0.89 | -0.168 | 0.23805365 | 0.45383564 | 0.019 | 0.0079 | 0.0144 | 0.0144 | TRUE | 0.4219 |
| Tm9sf1 | 822 | 1001 | 924 | 1740 | 1241 | 1107 | 873 | 1000 | 928 | 1368 | 1335 | 1212 | 1119.25 | 934 | 1305 | 1.398 | 0.484 | 8.51E-07 | 2.39E-05 | 0.0034 | 0.0098 | 0.006 | 0.006 | TRUE | 0.0464 |
| Tm9sf2 | 5308 | 6172 | 6076 | 8107 | 5651 | 5751 | 5638 | 6164 | 6101 | 6373 | 6080 | 6295 | 6108.6 | 5968 | 6249 | 1.047 | 0.067 | 0.33674459 | 0.55677852 | 0.0012 | 0.0074 | 0.0033 | 0.0033 | TRUE | 0.0284 |
| Tm9sf3 | 6599 | 7902 | 8038 | 9002 | 6401 | 6974 | 7009 | 7892 | 8071 | 7077 | 6887 | 7634 | 7428.33 | 7657 | 7199 | 0.94 | -0.089 | 0.30614367 | 0.52666802 | 0.004 | 0.0073 | 0.0053 | 0.0053 | TRUE | 0.0668 |
| Tm9sf4 | 3358 | 3768 | 3930 | 4322 | 3084 | 3289 | 3567 | 3763 | 3946 | 3398 | 3318 | 3600 | 3598.7 | 3759 | 3439 | 0.915 | -0.129 | 0.0934703 | 0.24990906 | 0.0018 | 0.0078 | 0.004 | 0.004 | TRUE | 0.0242 |
| Tma16 | 1851 | 1764 | 1780 | 2277 | 1595 | 1605 | 1966 | 1762 | 1787 | 1790 | 1716 | 1757 | 1796.36 | 1838 | 1754 | 0.955 | -0.067 | 0.40764456 | 0.62059556 | 0.0015 | 0.0087 | 0.0041 | 0.0041 | TRUE | 0.0444 |
| Tma7 | 1508 | 1536 | 1479 | 2516 | 1725 | 1550 | 1602 | 1534 | 1485 | 1978 | 1856 | 1697 | 1691.91 | 1540 | 1844 | 1.198 | 0.26 | 0.004213 | 0.02751447 | 0.0032 | 0.0089 | 0.0054 | 0.0054 | TRUE | 0.0583 |
| Tmbim1 | 887 | 1106 | 1071 | 888 | 764 | 795 | 942 | 1105 | 1075 | 698 | 822 | 870 | 918.74 | 1041 | 797 | 0.764 | -0.388 | 0.00147913 | 0.01211262 | 0.0089 | 0.0105 | 0.0096 | 0.0096 | TRUE | 0.1441 |

| | | | | | | | | | | | | | | | | | | | | | | | | | |
|---|---|---|---|---|---|---|---|---|---|---|---|---|---|---|---|---|---|---|---|---|---|---|---|---|---|
| Tmbim4 | 1377 | 1564 | 1370 | 2099 | 1458 | 1420 | 1463 | 1562 | 1376 | 1650 | 1569 | 1554 | 1528.9 | 1467 | 1591 | 1.085 | 0.118 | 0.16797966 | 0.36573373 | 0.0019 | 0.0091 | 0.0046 | 0.0046 | TRUE | 0.0391 |
| Tmbim6 | 9746 | 9792 | 9376 | 13710 | 9910 | 9770 | 10352 | 9779 | 9414 | 10778 | 10663 | 10695 | 10280.15 | 9848 | 10712 | 1.088 | 0.121 | 0.0626537 | 0.19176677 | 0.001 | 0.0072 | 0.003 | 0.003 | TRUE | 0.0244 |
| Tmc1 | 3 | 3 | 1 | 5 | 5 | 2 | 3 | 3 | 1 | 4 | 5 | 2 | 3.11 | 2 | 4 | 1.607 | 0.684 | 0.52741636 | NA | 0 | 1.0647 | 0.5043 | 0.5043 | TRUE | 0.2367 |
| Tmc2 | 4 | 1 | 4 | 5 | 2 | 3 | 4 | 1 | 4 | 4 | 2 | 3 | 3.11 | 3 | 3 | 1.024 | 0.035 | 0.97433855 | NA | 0 | 1.0679 | 0.5179 | 0.5179 | TRUE | 0.4778 |
| Tmc3 | 225 | 240 | 242 | 273 | 234 | 199 | 239 | 240 | 243 | 215 | 252 | 218 | 234.32 | 241 | 228 | 0.946 | -0.079 | 0.57124642 | 0.74993329 | 0 | 0.021 | 0.0099 | 0.0099 | TRUE | 0.0886 |
| Tmc4 | 118 | 109 | 133 | 131 | 121 | 112 | 125 | 109 | 134 | 103 | 130 | 123 | 120.59 | 123 | 119 | 0.964 | -0.053 | 0.78649498 | 0.89023346 | 0 | 0.0342 | 0.0196 | 0.0196 | TRUE | 0.1492 |
| Tmc5 | 55 | 50 | 47 | 60 | 33 | 33 | 58 | 50 | 47 | 47 | 36 | 36 | 45.72 | 52 | 40 | 0.77 | -0.377 | 0.20175688 | 0.40927453 | 0 | 0.079 | 0.0407 | 0.0407 | TRUE | 0.244 |
| Tmc6 | 1976 | 2658 | 2567 | 2372 | 1964 | 2045 | 2099 | 2655 | 2578 | 1865 | 2113 | 2239 | 2257.89 | 2444 | 2072 | 0.847 | -0.239 | 0.05037014 | 0.16493186 | 0.0111 | 0.0084 | 0.0103 | 0.0103 | TRUE | 0.1843 |
| Tmc7 | 1401 | 1394 | 1440 | 1316 | 951 | 1086 | 1488 | 1392 | 1446 | 1035 | 1023 | 1189 | 1262.13 | 1442 | 1082 | 0.75 | -0.415 | 1.29E-05 | 0.00024673 | 0.003 | 0.0095 | 0.0057 | 0.0057 | TRUE | 0.0874 |
| Tmc8 | 703 | 954 | 1111 | 657 | 548 | 614 | 747 | 953 | 1116 | 516 | 590 | 672 | 765.54 | 939 | 593 | 0.631 | -0.665 | 8.96E-05 | 0.00126807 | 0.026 | 0.0112 | 0.0194 | 0.0194 | TRUE | 0.1694 |
| Tmcc1 | 1061 | 893 | 929 | 1636 | 1114 | 1104 | 1127 | 892 | 933 | 1286 | 1199 | 1209 | 1107.47 | 984 | 1231 | 1.252 | 0.324 | 0.00463835 | 0.02958193 | 0.0075 | 0.0099 | 0.0085 | 0.0085 | TRUE | 0.1924 |
| Tmcc2 | 310 | 457 | 401 | 503 | 386 | 404 | 329 | 456 | 403 | 395 | 415 | 442 | 406.88 | 396 | 417 | 1.053 | 0.074 | 0.61714907 | 0.7832683 | 0.0121 | 0.015 | 0.0135 | 0.0135 | TRUE | 0.2483 |
| Tmcc3 | 273 | 289 | 306 | 399 | 283 | 246 | 290 | 289 | 307 | 314 | 304 | 269 | 295.55 | 295 | 296 | 1.004 | 0.005 | 0.9685946 | 0.98425836 | 0 | 0.0181 | 0.0087 | 0.0087 | TRUE | 0.0672 |
| Tmco1 | 1444 | 1673 | 1714 | 2210 | 1459 | 1524 | 1534 | 1671 | 1721 | 1737 | 1570 | 1668 | 1650.17 | 1642 | 1658 | 1.01 | 0.015 | 0.86411591 | 0.93284299 | 0.0025 | 0.0089 | 0.0049 | 0.0049 | TRUE | 0.0398 |
| Tmco2 | 0 | 0 | 0 | 0 | 0 | 0 | 0 | 0 | 0 | 0 | 0 | 0 | 0 | 0 | 0 | NA | NA | NA | NA | NA | NA | NA | NA | NA | NA |
| Tmco3 | 891 | 751 | 792 | 1156 | 803 | 790 | 946 | 750 | 795 | 909 | 864 | 865 | 854.87 | 830 | 879 | 1.059 | 0.083 | 0.47081627 | 0.67231455 | 0.0066 | 0.0108 | 0.0084 | 0.0084 | TRUE | 0.176 |
| Tmco4 | 303 | 354 | 396 | 456 | 335 | 302 | 322 | 354 | 398 | 358 | 360 | 331 | 353.75 | 358 | 350 | 0.978 | -0.032 | 0.80937087 | 0.90182956 | 0.0038 | 0.0162 | 0.0096 | 0.0096 | TRUE | 0.1093 |
| Tmco5 | 0 | 0 | 0 | 0 | 0 | 0 | 0 | 0 | 0 | 0 | 0 | 0 | 0 | 0 | 0 | NA | NA | NA | NA | NA | NA | NA | NA | NA | NA |
| Tmco5b | 0 | 0 | 0 | 1 | 0 | 0 | 0 | 0 | 0 | 1 | 0 | 0 | 0.13 | 0 | 0 | 1.859 | 0.894 | 0.82650734 | NA | 0 | 25.1515 | 10 | 10 | TRUE | 0.2563 |
| Tmco6 | 1289 | 1783 | 1811 | 1663 | 1300 | 1367 | 1369 | 1781 | 1818 | 1307 | 1399 | 1496 | 1528.45 | 1656 | 1401 | 0.845 | -0.242 | 0.0627926 | 0.19204283 | 0.0126 | 0.0091 | 0.0116 | 0.0116 | TRUE | 0.2764 |
| Tmed1 | 1358 | 1585 | 1531 | 1565 | 1151 | 1131 | 1442 | 1583 | 1537 | 1230 | 1238 | 1238 | 1378.24 | 1521 | 1235 | 0.812 | -0.3 | 0.00013436 | 0.00175137 | 5.00E-04 | 0.0093 | 0.0037 | 0.0037 | TRUE | 0.0244 |
| Tmed10 | 6525 | 7331 | 7090 | 7549 | 5461 | 5598 | 6931 | 7322 | 7119 | 5934 | 5876 | 6128 | 6551.56 | 7124 | 5979 | 0.839 | -0.253 | 3.43E-05 | 0.00056194 | 4.00E-04 | 0.0074 | 0.0025 | 0.0025 | TRUE | 0.0072 |
| Tmed11 | 0 | 0 | 0 | 0 | 0 | 0 | 0 | 0 | 0 | 0 | 0 | 0 | 0 | 0 | 0 | NA | NA | NA | NA | NA | NA | NA | NA | NA | NA |
| Tmed2 | 1359 | 1444 | 1414 | 1550 | 1253 | 1327 | 1443 | 1442 | 1420 | 1218 | 1348 | 1453 | 1387.45 | 1435 | 1340 | 0.933 | -0.101 | 0.2833773 | 0.50501998 | 0.0033 | 0.0093 | 0.0056 | 0.0056 | TRUE | 0.0776 |
| Tmed3 | 962 | 1253 | 1278 | 1359 | 1021 | 1071 | 1022 | 1251 | 1283 | 1068 | 1099 | 1172 | 1149.28 | 1185 | 1113 | 0.938 | -0.092 | 0.42472012 | 0.63482617 | 0.0078 | 0.0098 | 0.0087 | 0.0087 | TRUE | 0.1741 |
| Tmed4 | 1101 | 1248 | 1337 | 1602 | 1119 | 1167 | 1169 | 1246 | 1342 | 1259 | 1204 | 1277 | 1249.86 | 1252 | 1247 | 0.995 | -0.007 | 0.93920933 | 0.96945089 | 0.002 | 0.0095 | 0.005 | 0.005 | TRUE | 0.0471 |
| Tmed5 | 1266 | 1427 | 1422 | 1763 | 1275 | 1329 | 1345 | 1425 | 1428 | 1386 | 1372 | 1455 | 1401.71 | 1399 | 1404 | 1.003 | 0.005 | 0.95324462 | 0.97644888 | 0 | 0.0093 | 0.0036 | 0.0036 | TRUE | 0.0139 |
| Tmed6 | 18 | 15 | 15 | 11 | 10 | 20 | 19 | 15 | 15 | 9 | 11 | 22 | 15.08 | 16 | 14 | 0.829 | -0.27 | 0.62146435 | 0.78589927 | 0.0639 | 0.2254 | 0.1478 | 0.1478 | TRUE | 1.0811 |
| Tmed7 | 5401 | 6188 | 5991 | 6752 | 4891 | 5040 | 5737 | 6180 | 6016 | 5308 | 5262 | 5517 | 5669.97 | 5978 | 5362 | 0.897 | -0.157 | 0.01698414 | 0.07733081 | 8.00E-04 | 0.0075 | 0.0029 | 0.0029 | TRUE | 0.0151 |
| Tmed8 | 1762 | 1888 | 1997 | 2895 | 2097 | 1930 | 1872 | 1886 | 2005 | 2276 | 2256 | 2113 | 2067.85 | 1921 | 2215 | 1.153 | 0.206 | 0.00696394 | 0.0399256 | 0.001 | 0.0085 | 0.0037 | 0.0037 | TRUE | 0.0195 |
| Tmed9 | 4684 | 4573 | 4131 | 6099 | 4164 | 4156 | 4975 | 4567 | 4148 | 4795 | 4480 | 4549 | 4585.74 | 4563 | 4608 | 1.01 | 0.014 | 0.87453006 | 0.93811777 | 0.0045 | 0.0076 | 0.0057 | 0.0057 | TRUE | 0.0773 |
| Tmeff1 | 8219 | 8331 | 8407 | 6409 | 4863 | 5685 | 8730 | 8320 | 8442 | 5038 | 5232 | 6223 | 6997.58 | 8497 | 5498 | 0.647 | -0.628 | 2.00E-10 | 1.41E-08 | 0.0068 | 0.0074 | 0.0069 | 0.0069 | TRUE | 0.162 |
| Tmeff2 | 40 | 52 | 39 | 78 | 53 | 46 | 42 | 52 | 39 | 61 | 57 | 50 | 50.38 | 44 | 56 | 1.266 | 0.341 | 0.22231158 | 0.43444058 | 0 | 0.0723 | 0.036 | 0.036 | TRUE | 0.1681 |
| Tmem100 | 6 | 5 | 2 | 6 | 1 | 1 | 6 | 5 | 2 | 5 | 1 | 1 | 3.38 | 4 | 2 | 0.54 | -0.89 | 0.42920045 | NA | 0.1738 | 0.9825 | 0.5956 | 0.5956 | TRUE | 1.1145 |
| Tmem101 | 1374 | 1753 | 1704 | 1986 | 1453 | 1428 | 1459 | 1751 | 1711 | 1561 | 1563 | 1563 | 1601.49 | 1640 | 1562 | 0.952 | -0.07 | 0.46572101 | 0.66818446 | 0.0041 | 0.009 | 0.0061 | 0.0061 | TRUE | 0.1118 |
| Tmem102 | 211 | 211 | 197 | 233 | 203 | 167 | 224 | 211 | 198 | 183 | 218 | 183 | 202.84 | 211 | 195 | 0.922 | -0.116 | 0.4529339 | 0.6583424 | 0.0021 | 0.0231 | 0.0124 | 0.0124 | TRUE | 0.1181 |
| Tmem104 | 686 | 533 | 580 | 691 | 571 | 470 | 729 | 532 | 582 | 543 | 614 | 514 | 585.9 | 614 | 557 | 0.907 | -0.14 | 0.34456364 | 0.5641524 | 0.0155 | 0.0125 | 0.0142 | 0.0142 | TRUE | 0.3094 |
| Tmem106a | 145 | 134 | 137 | 186 | 141 | 117 | 154 | 134 | 138 | 146 | 152 | 128 | 141.9 | 142 | 142 | 1.003 | 0.004 | 0.97945732 | 0.98867206 | 0 | 0.0301 | 0.0148 | 0.0148 | TRUE | 0.0718 |
| Tmem106b | 1718 | 2520 | 2811 | 2268 | 1696 | 2123 | 1825 | 2517 | 2823 | 1783 | 1825 | 2324 | 2182.63 | 2388 | 1977 | 0.828 | -0.273 | 0.11589677 | 0.28752771 | 0.0338 | 0.0084 | 0.0212 | 0.0212 | TRUE | 0.5152 |
| Tmem106c | 697 | 1146 | 1179 | 975 | 766 | 970 | 740 | 1145 | 1184 | 766 | 824 | 1062 | 953.53 | 1023 | 884 | 0.864 | -0.211 | 0.28462054 | 0.5060919 | 0.0426 | 0.0104 | 0.0271 | 0.0271 | TRUE | 0.696 |
| Tmem107 | 604 | 838 | 815 | 954 | 672 | 761 | 642 | 837 | 818 | 750 | 723 | 833 | 767.14 | 766 | 769 | 1.003 | 0.005 | 0.9708965 | 0.98475753 | 0.0117 | 0.0112 | 0.0115 | 0.0115 | TRUE | 0.2368 |
| Tmem108 | 446 | 340 | 270 | 509 | 421 | 279 | 474 | 340 | 271 | 400 | 453 | 305 | 373.82 | 362 | 386 | 1.069 | 0.097 | 0.67148573 | 0.81667744 | 0.0558 | 0.0157 | 0.0346 | 0.0346 | TRUE | 0.5256 |
| Tmem109 | 773 | 741 | 758 | 1107 | 734 | 750 | 821 | 740 | 761 | 870 | 790 | 821 | 800.53 | 774 | 827 | 1.07 | 0.097 | 0.31147125 | 0.53199265 | 0.0015 | 0.011 | 0.0054 | 0.0054 | TRUE | 0.0332 |
| Tmem11 | 3353 | 3592 | 3659 | 4396 | 3183 | 3238 | 3561 | 3587 | 3674 | 3456 | 3425 | 3545 | 3541.31 | 3607 | 3475 | 0.963 | -0.054 | 0.37493423 | 0.59093261 | 0 | 0.0078 | 0.0024 | 0.0024 | TRUE | 0.0037 |
| Tmem110 | 1279 | 1411 | 1386 | 1676 | 1269 | 1307 | 1359 | 1409 | 1392 | 1318 | 1365 | 1431 | 1378.84 | 1387 | 1371 | 0.988 | -0.017 | 0.82638147 | 0.91211327 | 0 | 0.0093 | 0.0036 | 0.0036 | TRUE | 0.0172 |
| Tmem115 | 554 | 511 | 420 | 962 | 664 | 567 | 588 | 510 | 422 | 756 | 714 | 621 | 601.98 | 507 | 697 | 1.377 | 0.462 | 0.00220639 | 0.01662318 | 0.0159 | 0.0124 | 0.0147 | 0.0147 | TRUE | 0.252 |
| Tmem116 | 39 | 60 | 49 | 84 | 54 | 62 | 41 | 60 | 49 | 66 | 58 | 68 | 57.09 | 50 | 64 | 1.275 | 0.35 | 0.19328806 | 0.39886684 | 0 | 0.0646 | 0.0344 | 0.0344 | TRUE | 0.2359 |
| Tmem117 | 20 | 28 | 34 | 41 | 45 | 48 | 21 | 28 | 34 | 32 | 48 | 53 | 36.09 | 28 | 44 | 1.584 | 0.664 | 0.06995333 | 0.20561012 | 0.0348 | 0.0982 | 0.0672 | 0.0672 | TRUE | 0.5193 |
| Tmem119 | 27 | 20 | 21 | 28 | 25 | 17 | 29 | 20 | 21 | 22 | 27 | 19 | 22.88 | 23 | 23 | 0.969 | -0.045 | 0.91252404 | 0.95675253 | 0 | 0.1509 | 0.0765 | 0.0765 | TRUE | 0.2362 |
| Tmem120a | 800 | 1039 | 990 | 1275 | 821 | 992 | 850 | 1038 | 994 | 1002 | 883 | 1086 | 975.5 | 961 | 990 | 1.031 | 0.044 | 0.71999918 | 0.8481969 | 0.0096 | 0.0103 | 0.0099 | 0.0099 | TRUE | 0.1208 |
| Tmem120b | 501 | 429 | 438 | 597 | 405 | 344 | 532 | 428 | 440 | 469 | 436 | 377 | 447 | 467 | 427 | 0.917 | -0.125 | 0.38364659 | 0.59797155 | 0.0111 | 0.0143 | 0.0126 | 0.0126 | TRUE | 0.173 |
| Tmem121 | 610 | 680 | 619 | 748 | 592 | 600 | 648 | 679 | 622 | 588 | 637 | 657 | 638.39 | 650 | 627 | 0.964 | -0.052 | 0.6031047 | 0.77412127 | 0.0012 | 0.0121 | 0.0058 | 0.0058 | TRUE | 0.0372 |
| Tmem123 | 1390 | 1966 | 1854 | 2257 | 1713 | 1893 | 1476 | 1963 | 1862 | 1774 | 1843 | 2072 | 1831.84 | 1767 | 1896 | 1.073 | 0.101 | 0.43289768 | 0.64199926 | 0.0138 | 0.0087 | 0.0115 | 0.0115 | TRUE | 0.2495 |
| Tmem125 | 129 | 117 | 106 | 95 | 71 | 71 | 137 | 117 | 106 | 75 | 76 | 78 | 98.18 | 120 | 76 | 0.635 | -0.655 | 0.00162206 | 0.01304593 | 0 | 0.0405 | 0.0205 | 0.0205 | TRUE | 0.1508 |
| Tmem126a | 1012 | 915 | 876 | 1254 | 860 | 766 | 1075 | 914 | 880 | 986 | 925 | 839 | 936.32 | 956 | 917 | 0.96 | -0.059 | 0.61646561 | 0.78309404 | 0.008 | 0.0104 | 0.0091 | 0.0091 | TRUE | 0.1403 |
| Tmem126b | 337 | 359 | 402 | 644 | 481 | 450 | 358 | 359 | 404 | 506 | 518 | 493 | 439.42 | 374 | 506 | 1.354 | 0.437 | 8.63E-05 | 0.00123143 | 0 | 0.0144 | 0.0066 | 0.0066 | TRUE | 0.058 |
| Tmem127 | 818 | 749 | 770 | 1205 | 839 | 722 | 869 | 748 | 773 | 947 | 903 | 790 | 838.4 | 797 | 880 | 1.106 | 0.145 | 0.20396107 | 0.41210273 | 0.0063 | 0.0108 | 0.0082 | 0.0082 | TRUE | 0.0937 |
| Tmem128 | 780 | 834 | 791 | 1197 | 846 | 845 | 828 | 833 | 794 | 941 | 910 | 925 | 871.98 | 818 | 925 | 1.131 | 0.178 | 0.03249538 | 0.12155199 | 0 | 0.0107 | 0.0038 | 0.0038 | TRUE | 0.0081 |
| Tmem129 | 315 | 403 | 409 | 446 | 330 | 336 | 335 | 402 | 411 | 351 | 355 | 368 | 370.2 | 383 | 358 | 0.934 | -0.098 | 0.44740358 | 0.65334295 | 0.0037 | 0.0158 | 0.0093 | 0.0093 | TRUE | 0.1366 |
| Tmem130 | 283 | 302 | 301 | 194 | 147 | 200 | 301 | 302 | 302 | 153 | 158 | 219 | 239.01 | 302 | 177 | 0.583 | -0.777 | 7.73E-06 | 0.00015915 | 0.0137 | 0.0207 | 0.0173 | 0.0173 | TRUE | 0.4599 |
| Tmem131 | 12069 | 11887 | 13501 | 15454 | 11147 | 10609 | 12819 | 11872 | 13556 | 12149 | 11993 | 11613 | 12333.82 | 12749 | 11918 | 0.935 | -0.097 | 0.19847967 | 0.40560008 | 0.0023 | 0.0072 | 0.004 | 0.004 | TRUE | 0.0444 |
| Tmem132a | 1578 | 1446 | 1210 | 1496 | 1140 | 1070 | 1676 | 1444 | 1215 | 1176 | 1227 | 1171 | 1318.18 | 1445 | 1191 | 0.824 | -0.278 | 0.02995838 | 0.11516053 | 0.0119 | 0.0094 | 0.0111 | 0.0111 | TRUE | 0.1552 |
| Tmem132b | 60 | 60 | 34 | 58 | 53 | 53 | 64 | 60 | 34 | 46 | 57 | 58 | 53.07 | 53 | 54 | 1.015 | 0.022 | 0.94650964 | 0.97329183 | 0.0414 | 0.069 | 0.0558 | 0.0558 | TRUE | 0.7854 |
| Tmem132c | 245 | 207 | 184 | 187 | 115 | 132 | 260 | 207 | 185 | 147 | 124 | 144 | 177.83 | 217 | 138 | 0.639 | -0.647 | 0.00066817 | 0.00652939 | 0.0152 | 0.0254 | 0.0202 | 0.0202 | TRUE | 0.3251 |
| Tmem132d | 79 | 61 | 94 | 58 | 41 | 48 | 84 | 61 | 94 | 46 | 44 | 53 | 63.58 | 80 | 48 | 0.594 | -0.752 | 0.00635957 | 0.03758472 | 0.0172 | 0.0587 | 0.038 | 0.038 | TRUE | 0.4012 |
| Tmem132e | 79 | 101 | 86 | 148 | 94 | 74 | 84 | 101 | 86 | 116 | 101 | 81 | 94.94 | 90 | 99 | 1.106 | 0.145 | 0.52373254 | 0.71357201 | 0.0118 | 0.0416 | 0.0268 | 0.0268 | TRUE | 0.2409 |
| Tmem134 | 749 | 978 | 943 | 1145 | 906 | 888 | 796 | 977 | 947 | 900 | 975 | 972 | 927.69 | 907 | 949 | 1.046 | 0.065 | 0.55896274 | 0.74102381 | 0.0058 | 0.0105 | 0.0078 | 0.0078 | TRUE | 0.1354 |
| Tmem135 | 1782 | 1761 | 1838 | 2165 | 1514 | 1580 | 1893 | 1759 | 1846 | 1702 | 1629 | 1730 | 1759.59 | 1833 | 1687 | 0.921 | -0.119 | 0.11399101 | 0.28478082 | 6.00E-04 | 0.0088 | 0.0035 | 0.0035 | TRUE | 0.015 |
| Tmem136 | 458 | 603 | 653 | 552 | 437 | 499 | 486 | 602 | 656 | 434 | 470 | 546 | 532.46 | 581 | 483 | 0.83 | -0.269 | 0.07825693 | 0.22135372 | 0.0165 | 0.0131 | 0.0149 | 0.0149 | TRUE | 0.2381 |

| | | | | | | | | | | | | | | | | | | | | | | | | | |
|---|---|---|---|---|---|---|---|---|---|---|---|---|---|---|---|---|---|---|---|---|---|---|---|---|---|
| Tmem138 | 502 | 592 | 642 | 626 | 481 | 528 | 533 | 591 | 645 | 492 | 518 | 578 | 559.45 | 590 | 529 | 0.896 | -0.158 | 0.1976747 | 0.40433284 | 0.0061 | 0.0128 | 0.0091 | 0.0091 | TRUE | 0.0817 |
| Tmem139 | 6 | 6 | 3 | 15 | 6 | 10 | 6 | 6 | 3 | 12 | 6 | 11 | 7.43 | 5 | 10 | 1.914 | 0.937 | 0.19645428 | 0.40292288 | 0 | 0.4504 | 0.2292 | 0.2292 | TRUE | 0.2741 |
| Tmem140 | 3 | 7 | 5 | 14 | 8 | 5 | 3 | 7 | 5 | 11 | 9 | 5 | 6.71 | 5 | 8 | 1.663 | 0.734 | 0.33151067 | 0.55215482 | 0 | 0.4976 | 0.2529 | 0.2529 | TRUE | 0.1909 |
| Tmem141 | 288 | 297 | 289 | 292 | 206 | 173 | 306 | 297 | 290 | 230 | 222 | 189 | 255.55 | 298 | 214 | 0.72 | -0.475 | 0.00065923 | 0.00646605 | 0 | 0.0198 | 0.01 | 0.01 | TRUE | 0.103 |
| Tmem143 | 677 | 887 | 951 | 861 | 641 | 690 | 719 | 886 | 955 | 677 | 690 | 755 | 780.28 | 853 | 707 | 0.828 | -0.272 | 0.03775162 | 0.13498494 | 0.011 | 0.0111 | 0.0111 | 0.0111 | TRUE | 0.2238 |
| Tmem144 | 214 | 278 | 259 | 348 | 311 | 297 | 227 | 278 | 260 | 274 | 335 | 325 | 283.05 | 255 | 311 | 1.216 | 0.283 | 0.060846 | 0.18772753 | 0.0077 | 0.0185 | 0.0128 | 0.0128 | TRUE | 0.1343 |
| Tmem145 | 52 | 115 | 146 | 29 | 32 | 31 | 55 | 115 | 147 | 23 | 34 | 34 | 67.97 | 106 | 30 | 0.286 | -1.808 | 4.86E-06 | 0.0001078 | 0.131 | 0.0554 | 0.0916 | 0.0916 | TRUE | 0.1925 |
| Tmem147 | 1395 | 1693 | 1611 | 1891 | 1308 | 1414 | 1482 | 1691 | 1618 | 1487 | 1407 | 1548 | 1538.65 | 1597 | 1481 | 0.927 | -0.109 | 0.22440972 | 0.43681551 | 0.0027 | 0.009 | 0.0052 | 0.0052 | TRUE | 0.0476 |
| Tmem14a | 99 | 113 | 118 | 184 | 158 | 148 | 105 | 113 | 118 | 145 | 170 | 162 | 135.52 | 112 | 159 | 1.412 | 0.498 | 0.00486528 | 0.03070635 | 0 | 0.0312 | 0.0149 | 0.0149 | TRUE | 0.0716 |
| Tmem14c | 3804 | 3734 | 3505 | 4318 | 2924 | 3136 | 4041 | 3729 | 3519 | 3394 | 3146 | 3433 | 3543.74 | 3763 | 3324 | 0.884 | -0.179 | 0.03618072 | 0.13082125 | 0.0031 | 0.0078 | 0.0049 | 0.0049 | TRUE | 0.0505 |
| Tmem150a | 603 | 664 | 715 | 714 | 520 | 529 | 640 | 663 | 718 | 561 | 559 | 579 | 620.24 | 674 | 566 | 0.84 | -0.251 | 0.01094096 | 0.05630045 | 0 | 0.0122 | 0.0054 | 0.0054 | TRUE | 0.0387 |
| Tmem150b | 1 | 0 | 0 | 0 | 0 | 0 | 1 | 0 | 0 | 0 | 0 | 0 | 0.18 | 0 | 0 | 0.49 | -1.029 | 0.80087 | NA | 0 | 18.6165 | 10 | 10 | TRUE | 0.4614 |
| Tmem150c | 37 | 37 | 32 | 52 | 35 | 28 | 39 | 37 | 32 | 41 | 38 | 31 | 36.26 | 36 | 37 | 1.013 | 0.019 | 0.95365599 | 0.97650213 | 0 | 0.0978 | 0.0466 | 0.0466 | TRUE | 0.1294 |
| Tmem151a | 63 | 90 | 104 | 57 | 47 | 65 | 67 | 90 | 104 | 45 | 51 | 71 | 71.29 | 87 | 56 | 0.633 | -0.659 | 0.02357244 | 0.09763248 | 0.0392 | 0.0531 | 0.0464 | 0.0464 | TRUE | 0.3251 |
| Tmem151b | 424 | 532 | 429 | 352 | 271 | 255 | 450 | 531 | 431 | 277 | 292 | 279 | 376.64 | 471 | 283 | 0.6 | -0.738 | 2.14E-08 | 9.43E-07 | 0.005 | 0.0157 | 0.0097 | 0.0097 | TRUE | 0.1477 |
| Tmem154 | 25 | 11 | 14 | 4 | 5 | 4 | 27 | 11 | 14 | 3 | 5 | 4 | 10.75 | 17 | 4 | 0.247 | -2.018 | 0.00372761 | 0.02506049 | 0 | 0.3134 | 0.2041 | 0.2041 | TRUE | 0.816 |
| Tmem158 | 232 | 341 | 335 | 409 | 323 | 313 | 246 | 341 | 336 | 322 | 348 | 343 | 322.51 | 308 | 338 | 1.094 | 0.13 | 0.41437356 | 0.6260657 | 0.0131 | 0.0171 | 0.0151 | 0.0151 | TRUE | 0.3401 |
| Tmem159 | 300 | 317 | 317 | 427 | 324 | 332 | 319 | 317 | 318 | 336 | 349 | 363 | 333.54 | 318 | 349 | 1.097 | 0.134 | 0.24622934 | 0.46320719 | 0 | 0.0168 | 0.0066 | 0.0066 | TRUE | 0.0143 |
| Tmem160 | 1542 | 1975 | 1773 | 1724 | 1277 | 1226 | 1638 | 1972 | 1780 | 1355 | 1374 | 1342 | 1576.98 | 1797 | 1357 | 0.755 | -0.405 | 2.18E-05 | 0.00038307 | 0.0039 | 0.009 | 0.0059 | 0.0059 | TRUE | 0.0883 |
| Tmem161a | 2437 | 2911 | 3032 | 2852 | 2090 | 2303 | 2589 | 2907 | 3044 | 2242 | 2249 | 2521 | 2591.99 | 2847 | 2337 | 0.821 | -0.285 | 0.00346567 | 0.02365231 | 0.0051 | 0.0082 | 0.0065 | 0.0065 | TRUE | 0.0762 |
| Tmem161b | 1120 | 1373 | 1359 | 1537 | 1121 | 1140 | 1190 | 1371 | 1365 | 1208 | 1206 | 1248 | 1264.63 | 1309 | 1221 | 0.933 | -0.101 | 0.27444828 | 0.49488215 | 0.0026 | 0.0095 | 0.0053 | 0.0053 | TRUE | 0.0754 |
| Tmem163 | 49 | 71 | 59 | 60 | 42 | 69 | 52 | 71 | 59 | 47 | 45 | 76 | 58.35 | 61 | 56 | 0.916 | -0.126 | 0.68089836 | 0.82311098 | 0.0363 | 0.0634 | 0.0503 | 0.0503 | TRUE | 0.7511 |
| Tmem164 | 927 | 978 | 984 | 1412 | 989 | 970 | 985 | 977 | 988 | 1110 | 1064 | 1062 | 1030.89 | 983 | 1079 | 1.098 | 0.135 | 0.08685344 | 0.23829686 | 0 | 0.0101 | 0.0035 | 0.0035 | TRUE | 0.0081 |
| Tmem165 | 3703 | 3865 | 3815 | 4507 | 3332 | 3391 | 3933 | 3860 | 3831 | 3543 | 3585 | 3712 | 3744 | 3875 | 3613 | 0.932 | -0.101 | 0.10140982 | 0.26388975 | 0 | 0.0078 | 0.0025 | 0.0025 | TRUE | 0.0069 |
| Tmem167 | 3009 | 3102 | 3008 | 4059 | 2836 | 2802 | 3196 | 3098 | 3020 | 3191 | 3051 | 3067 | 3103.98 | 3105 | 3103 | 1 | 0 | 0.99674764 | 0.99828918 | 4.00E-04 | 0.008 | 0.0028 | 0.0028 | TRUE | 0.008 |
| Tmem167b | 1370 | 1405 | 1404 | 2069 | 1439 | 1437 | 1455 | 1403 | 1410 | 1626 | 1548 | 1573 | 1502.65 | 1423 | 1582 | 1.113 | 0.154 | 0.0343124 | 0.12614807 | 0 | 0.0091 | 0.0032 | 0.0032 | TRUE | 0.0074 |
| Tmem168 | 1640 | 1815 | 1769 | 2456 | 1700 | 1774 | 1742 | 1813 | 1776 | 1931 | 1829 | 1942 | 1838.77 | 1777 | 1901 | 1.07 | 0.097 | 0.17512986 | 0.37485771 | 0 | 0.0087 | 0.0032 | 0.0032 | TRUE | 0.0129 |
| Tmem169 | 39 | 21 | 23 | 36 | 30 | 23 | 41 | 21 | 23 | 28 | 32 | 25 | 28.54 | 28 | 28 | 1.006 | 0.008 | 0.9839366 | 0.99158695 | 0.0433 | 0.1223 | 0.0844 | 0.0844 | TRUE | 0.991 |
| Tmem17 | 144 | 168 | 151 | 220 | 147 | 140 | 153 | 168 | 152 | 173 | 158 | 153 | 159.45 | 158 | 161 | 1.028 | 0.039 | 0.80687995 | 0.90045359 | 0 | 0.0276 | 0.0125 | 0.0125 | TRUE | 0.045 |
| Tmem170 | 2369 | 2502 | 2432 | 2711 | 2045 | 2202 | 2516 | 2499 | 2442 | 2131 | 2200 | 2410 | 2366.49 | 2486 | 2247 | 0.904 | -0.146 | 0.06550632 | 0.19720665 | 0.0016 | 0.0083 | 0.0041 | 0.0041 | TRUE | 0.0485 |
| Tmem170b | 662 | 951 | 1109 | 1088 | 862 | 1009 | 703 | 950 | 1114 | 855 | 927 | 1105 | 942.29 | 922 | 962 | 1.043 | 0.061 | 0.73385627 | 0.85735379 | 0.0337 | 0.0104 | 0.0218 | 0.0218 | TRUE | 0.3418 |
| Tmem171 | 1 | 5 | 2 | 0 | 1 | 0 | 1 | 5 | 2 | 0 | 1 | 0 | 1.52 | 3 | 0 | 0.13 | -2.949 | 0.11519638 | NA | 0 | 2.1696 | 1.3355 | 1.3355 | TRUE | 0.5495 |
| Tmem173 | 134 | 132 | 112 | 136 | 91 | 85 | 142 | 132 | 112 | 107 | 98 | 93 | 114.08 | 129 | 99 | 0.773 | -0.371 | 0.05796085 | 0.1819221 | 0 | 0.0358 | 0.0188 | 0.0188 | TRUE | 0.1276 |
| Tmem174 | 1 | 0 | 0 | 0 | 1 | 2 | 1 | 0 | 0 | 0 | 1 | 2 | 0.72 | 0 | 1 | 2.905 | 1.539 | 0.54944501 | NA | 0 | 4.5747 | 3.3003 | 3.3003 | TRUE | 0.2242 |
| Tmem175 | 382 | 578 | 591 | 824 | 577 | 567 | 406 | 577 | 593 | 648 | 621 | 621 | 577.61 | 525 | 630 | 1.198 | 0.261 | 0.09663413 | 0.25520989 | 0.0193 | 0.0126 | 0.016 | 0.016 | TRUE | 0.4608 |
| Tmem176a | 14 | 33 | 21 | 18 | 13 | 14 | 15 | 33 | 21 | 14 | 14 | 15 | 18.73 | 23 | 14 | 0.629 | -0.67 | 0.16685282 | 0.36433653 | 0 | 0.1828 | 0.1131 | 0.1131 | TRUE | 0.3297 |
| Tmem176b | 8 | 18 | 26 | 9 | 10 | 8 | 8 | 18 | 26 | 7 | 11 | 9 | 13.2 | 17 | 9 | 0.501 | -0.998 | 0.10175672 | 0.2646464 | 0.1099 | 0.2566 | 0.1835 | 0.1835 | TRUE | 0.1292 |
| Tmem177 | 1390 | 1345 | 1309 | 1332 | 1050 | 1085 | 1476 | 1343 | 1314 | 1047 | 1130 | 1188 | 1249.77 | 1378 | 1122 | 0.813 | -0.298 | 0.0017278 | 0.01369388 | 0.0032 | 0.0095 | 0.0057 | 0.0057 | TRUE | 0.0467 |
| Tmem178 | 13 | 10 | 12 | 24 | 30 | 20 | 14 | 10 | 12 | 19 | 32 | 22 | 18.15 | 12 | 24 | 2.028 | 1.02 | 0.03361675 | 0.12454861 | 0 | 0.1884 | 0.103 | 0.103 | TRUE | 0.4697 |
| Tmem178b | 6 | 26 | 27 | 29 | 22 | 16 | 6 | 26 | 27 | 23 | 24 | 18 | 20.57 | 20 | 22 | 1.075 | 0.104 | 0.85237959 | 0.92628008 | 0.1886 | 0.167 | 0.1767 | 0.1767 | TRUE | 1.8196 |
| Tmem179 | 154 | 142 | 138 | 136 | 105 | 79 | 164 | 142 | 139 | 107 | 113 | 86 | 125.05 | 148 | 102 | 0.692 | -0.532 | 0.00626793 | 0.03723458 | 0.0032 | 0.0332 | 0.019 | 0.019 | TRUE | 0.1641 |
| Tmem179b | 653 | 738 | 822 | 965 | 708 | 663 | 694 | 737 | 825 | 759 | 762 | 726 | 750.36 | 752 | 749 | 0.996 | -0.006 | 0.95146677 | 0.97533692 | 0.0029 | 0.0113 | 0.0065 | 0.0065 | TRUE | 0.0864 |
| Tmem18 | 1132 | 1206 | 1074 | 1867 | 1341 | 1213 | 1202 | 1204 | 1078 | 1468 | 1443 | 1328 | 1287.26 | 1161 | 1413 | 1.217 | 0.283 | 0.00209165 | 0.01595234 | 0.0026 | 0.0095 | 0.0053 | 0.0053 | TRUE | 0.0473 |
| Tmem180 | 3173 | 4524 | 4580 | 2161 | 1622 | 2282 | 3370 | 4518 | 4599 | 1699 | 1745 | 2498 | 3071.54 | 4162 | 1981 | 0.476 | -1.072 | 1.40E-09 | 8.17E-08 | 0.0387 | 0.008 | 0.0222 | 0.0222 | TRUE | 0.6299 |
| Tmem181a | 615 | 792 | 745 | 893 | 753 | 764 | 653 | 791 | 748 | 702 | 810 | 836 | 756.8 | 731 | 783 | 1.07 | 0.097 | 0.42428964 | 0.63448338 | 0.0078 | 0.0113 | 0.0093 | 0.0093 | TRUE | 0.1012 |
| Tmem181b-p | 409 | 324 | 254 | 409 | 471 | 353 | 434 | 324 | 255 | 322 | 507 | 386 | 371.29 | 338 | 405 | 1.198 | 0.261 | 0.2604525 | 0.47988003 | 0.0565 | 0.0158 | 0.0359 | 0.0359 | TRUE | 0.4389 |
| Tmem181c-p | 13 | 8 | 8 | 20 | 10 | 13 | 14 | 8 | 8 | 16 | 11 | 14 | 11.76 | 10 | 14 | 1.376 | 0.461 | 0.41652586 | 0.62787403 | 0 | 0.2871 | 0.1445 | 0.1445 | TRUE | 0.3811 |
| Tmem182 | 0 | 0 | 0 | 0 | 0 | 0 | 0 | 0 | 0 | 0 | 0 | 0 | 0 | 0 | 0 | NA | NA | NA | NA | NA | NA | NA | NA | NA | NA |
| Tmem183a | 5596 | 5240 | 5020 | 6884 | 4832 | 4671 | 5944 | 5233 | 5041 | 5412 | 5199 | 5113 | 5323.59 | 5406 | 5241 | 0.97 | -0.044 | 0.61243036 | 0.78054559 | 0.0036 | 0.0075 | 0.0053 | 0.0053 | TRUE | 0.0923 |
| Tmem184a | 55 | 87 | 78 | 71 | 56 | 86 | 58 | 87 | 78 | 56 | 60 | 94 | 72.31 | 74 | 70 | 0.934 | -0.098 | 0.7425765 | 0.86256566 | 0.0475 | 0.0525 | 0.05 | 0.05 | TRUE | 0.7857 |
| Tmem184b | 4329 | 5458 | 5370 | 5405 | 4048 | 4539 | 4598 | 5451 | 5392 | 4249 | 4355 | 4969 | 4835.7 | 5147 | 4524 | 0.879 | -0.186 | 0.07598625 | 0.21734486 | 0.0077 | 0.0076 | 0.0077 | 0.0077 | TRUE | 0.106 |
| Tmem184c | 2921 | 2644 | 2480 | 3220 | 2315 | 2177 | 3103 | 2641 | 2490 | 2531 | 2491 | 2383 | 2606.43 | 2745 | 2468 | 0.9 | -0.153 | 0.13951232 | 0.32412923 | 0.0067 | 0.0082 | 0.0073 | 0.0073 | TRUE | 0.1579 |
| Tmem185b | 403 | 418 | 350 | 621 | 472 | 367 | 428 | 417 | 351 | 488 | 508 | 402 | 432.45 | 399 | 466 | 1.169 | 0.225 | 0.1167637 | 0.28895749 | 0.0107 | 0.0145 | 0.0125 | 0.0125 | TRUE | 0.1643 |
| Tmem186 | 929 | 1032 | 1137 | 1117 | 802 | 883 | 987 | 1031 | 1142 | 878 | 863 | 967 | 977.78 | 1053 | 903 | 0.857 | -0.223 | 0.02789795 | 0.10967118 | 0.0037 | 0.0103 | 0.0064 | 0.0064 | TRUE | 0.0649 |
| Tmem189 | 1423 | 1477 | 1465 | 2096 | 1518 | 1482 | 1511 | 1475 | 1471 | 1648 | 1633 | 1622 | 1560.14 | 1486 | 1634 | 1.1 | 0.138 | 0.04729566 | 0.15785128 | 0 | 0.009 | 0.0028 | 0.0028 | TRUE | 0.0027 |
| Tmem19 | 2248 | 2570 | 2431 | 2904 | 2166 | 2133 | 2388 | 2567 | 2441 | 2283 | 2330 | 2335 | 2390.62 | 2465 | 2316 | 0.939 | -0.09 | 0.1909 | 0.39611884 | 4.00E-04 | 0.0083 | 0.003 | 0.003 | TRUE | 0.0158 |
| Tmem190 | 0 | 0 | 0 | 0 | 0 | 0 | 0 | 0 | 0 | 0 | 0 | 0 | 0 | 0 | 0 | NA | NA | NA | NA | NA | NA | NA | NA | NA | NA |
| Tmem191c | 265 | 459 | 437 | 467 | 480 | 421 | 281 | 458 | 439 | 367 | 516 | 461 | 420.52 | 393 | 448 | 1.139 | 0.187 | 0.37414536 | 0.59039725 | 0.0459 | 0.0147 | 0.0296 | 0.0296 | TRUE | 0.7033 |
| Tmem192 | 1100 | 1207 | 1220 | 1687 | 1249 | 1312 | 1168 | 1205 | 1225 | 1326 | 1344 | 1436 | 1284.18 | 1199 | 1369 | 1.14 | 0.189 | 0.0179684 | 0.08041929 | 5.00E-04 | 0.0095 | 0.0038 | 0.0038 | TRUE | 0.022 |
| Tmem194 | 3992 | 4592 | 4857 | 4114 | 3346 | 3657 | 4240 | 4586 | 4877 | 3234 | 3600 | 4003 | 4090.1 | 4568 | 3612 | 0.791 | -0.339 | 0.00133417 | 0.011144 | 0.0077 | 0.0077 | 0.0078 | 0.0078 | TRUE | 0.1089 |
| Tmem194b | 667 | 1088 | 1065 | 943 | 800 | 942 | 708 | 1087 | 1069 | 741 | 861 | 1031 | 916.28 | 955 | 878 | 0.918 | -0.123 | 0.51628503 | 0.70829955 | 0.04 | 0.0105 | 0.0247 | 0.0247 | TRUE | 0.6059 |
| Tmem196 | 0 | 1 | 0 | 0 | 0 | 0 | 0 | 1 | 0 | 0 | 0 | 0 | 0.17 | 0 | 0 | 0.49 | -1.029 | 0.80087 | NA | 0 | 19.7991 | 10 | 10 | TRUE | 0.405 |
| Tmem198 | 14 | 24 | 22 | 19 | 25 | 11 | 15 | 24 | 22 | 15 | 27 | 12 | 19.13 | 20 | 18 | 0.879 | -0.187 | 0.70556295 | 0.83909959 | 0.0603 | 0.1791 | 0.1231 | 0.1231 | TRUE | 0.8989 |
| Tmem198b | 464 | 493 | 523 | 641 | 459 | 470 | 493 | 492 | 525 | 504 | 494 | 514 | 503.77 | 503 | 504 | 1.001 | 0.002 | 0.98728479 | 0.9930142 | 0 | 0.0134 | 0.0052 | 0.0052 | TRUE | 0.0167 |
| Tmem199 | 2046 | 2892 | 2795 | 2733 | 2063 | 2409 | 2173 | 2888 | 2806 | 2148 | 2220 | 2637 | 2478.85 | 2622 | 2335 | 0.89 | -0.168 | 0.22030938 | 0.43242065 | 0.0169 | 0.0082 | 0.0131 | 0.0131 | TRUE | 0.2722 |
| Tmem2 | 6097 | 6626 | 6851 | 6228 | 4820 | 5023 | 6476 | 6617 | 6879 | 4896 | 5186 | 5498 | 5925.52 | 6657 | 5193 | 0.78 | -0.359 | 1.58E-06 | 4.08E-05 | 0.0019 | 0.0075 | 0.0039 | 0.0039 | TRUE | 0.0322 |
| Tmem200a | 19 | 11 | 10 | 16 | 3 | 10 | 20 | 11 | 10 | 13 | 3 | 11 | 11.33 | 14 | 9 | 0.659 | -0.602 | 0.36977738 | 0.58696949 | 0.1543 | 0.2978 | 0.2328 | 0.2328 | TRUE | 0.9154 |
| Tmem200b | 16 | 18 | 24 | 45 | 31 | 25 | 17 | 18 | 24 | 35 | 33 | 27 | 25.86 | 20 | 32 | 1.631 | 0.706 | 0.06704018 | 0.19995418 | 0 | 0.1343 | 0.0659 | 0.0659 | TRUE | 0.2074 |
| Tmem200c | 8 | 15 | 16 | 31 | 13 | 12 | 8 | 15 | 16 | 24 | 14 | 13 | 15.17 | 13 | 17 | 1.315 | 0.395 | 0.46275425 | 0.66629195 | 0.0565 | 0.224 | 0.1421 | 0.1421 | TRUE | 0.8011 |

| | | | | | | | | | | | | | | | | | | | | | | | | | |
|---|---|---|---|---|---|---|---|---|---|---|---|---|---|---|---|---|---|---|---|---|---|---|---|---|---|
| Tmem201 | 5089 | 5448 | 5533 | 6048 | 4892 | 4872 | 5405 | 5441 | 5556 | 4754 | 5263 | 5333 | 5292.2 | 5467 | 5117 | 0.936 | -0.096 | 0.19721548 | 0.40385389 | 0.0017 | 0.0075 | 0.0038 | 0.0038 | TRUE | 0.0474 |
| Tmem202 | 1 | 4 | 1 | 1 | 0 | 0 | 1 | 4 | 1 | 1 | 0 | 0 | 1.14 | 2 | 0 | 0.172 | -2.536 | 0.20960755 | NA | 0 | 2.8937 | 1.6924 | 1.6924 | TRUE | 0.6721 |
| Tmem203 | 170 | 214 | 193 | 261 | 179 | 187 | 181 | 214 | 194 | 205 | 193 | 205 | 198.43 | 196 | 201 | 1.025 | 0.035 | 0.81459606 | 0.90528531 | 0 | 0.0235 | 0.011 | 0.011 | TRUE | 0.0679 |
| Tmem204 | 0 | 0 | 2 | 0 | 0 | 1 | 0 | 0 | 2 | 0 | 0 | 1 | 0.52 | 1 | 0 | 0.52 | -0.944 | 0.77774021 | NA | 0 | 6.3774 | 6.2947 | 6.2947 | TRUE | 0.9739 |
| Tmem205 | 373 | 755 | 820 | 519 | 398 | 448 | 396 | 754 | 823 | 408 | 428 | 490 | 550.04 | 658 | 442 | 0.672 | -0.574 | 0.01843428 | 0.08201564 | 0.0746 | 0.0129 | 0.0409 | 0.0409 | TRUE | 1.4228 |
| Tmem206 | 675 | 809 | 888 | 788 | 671 | 628 | 717 | 808 | 892 | 619 | 722 | 687 | 740.91 | 806 | 676 | 0.838 | -0.254 | 0.03661001 | 0.13201032 | 0.0077 | 0.0113 | 0.0093 | 0.0093 | TRUE | 0.1091 |
| Tmem207 | 0 | 0 | 0 | 0 | 0 | 0 | 0 | 0 | 0 | 0 | 0 | 0 | 0 | 0 | 0 | NA | NA | NA | NA | NA | NA | NA | NA | NA | NA |
| Tmem208 | 754 | 807 | 794 | 924 | 685 | 655 | 801 | 806 | 797 | 726 | 737 | 717 | 764.08 | 801 | 727 | 0.907 | -0.141 | 0.09470372 | 0.25226601 | 0 | 0.0112 | 0.0038 | 0.0038 | TRUE | 0.0017 |
| Tmem209 | 6113 | 7344 | 7619 | 7447 | 5553 | 6648 | 6493 | 7335 | 7650 | 5854 | 5975 | 7277 | 6764.02 | 7159 | 6369 | 0.889 | -0.169 | 0.13961916 | 0.32417231 | 0.0109 | 0.0074 | 0.0093 | 0.0093 | TRUE | 0.1897 |
| Tmem210 | 0 | 0 | 0 | 0 | 0 | 0 | 0 | 0 | 0 | 0 | 0 | 0 | 0 | 0 | 0 | NA | NA | NA | NA | NA | NA | NA | NA | NA | NA |
| Tmem211 | 27 | 85 | 53 | 12 | 14 | 23 | 29 | 85 | 53 | 9 | 15 | 25 | 36.08 | 56 | 16 | 0.294 | -1.768 | 0.00063204 | 0.00624379 | 0.2168 | 0.0982 | 0.1537 | 0.1537 | TRUE | 0.1113 |
| Tmem212 | 0 | 0 | 0 | 0 | 0 | 0 | 0 | 0 | 0 | 0 | 0 | 0 | 0 | 0 | 0 | NA | NA | NA | NA | NA | NA | NA | NA | NA | NA |
| Tmem213 | 31 | 77 | 64 | 36 | 44 | 53 | 33 | 77 | 64 | 28 | 47 | 58 | 51.29 | 58 | 44 | 0.761 | -0.393 | 0.33429555 | 0.5548783 | 0.1309 | 0.0711 | 0.0997 | 0.0997 | TRUE | 0.5674 |
| Tmem214 | 2086 | 2207 | 2321 | 3034 | 2312 | 2179 | 2216 | 2204 | 2331 | 2385 | 2488 | 2385 | 2334.72 | 2250 | 2419 | 1.075 | 0.104 | 0.13234935 | 0.31361387 | 3.00E-04 | 0.0083 | 0.003 | 0.003 | TRUE | 0.0119 |
| Tmem215 | 24 | 38 | 21 | 73 | 52 | 57 | 25 | 38 | 21 | 57 | 56 | 62 | 43.38 | 28 | 58 | 2.076 | 1.054 | 0.00125722 | 0.01063209 | 0 | 0.0829 | 0.0504 | 0.0504 | TRUE | 0.605 |
| Tmem216 | 246 | 211 | 233 | 427 | 317 | 278 | 261 | 211 | 234 | 336 | 341 | 304 | 281.17 | 235 | 327 | 1.391 | 0.477 | 0.00074508 | 0.00707917 | 0.0031 | 0.0186 | 0.0107 | 0.0107 | TRUE | 0.1016 |
| Tmem217 | 4 | 6 | 8 | 4 | 4 | 7 | 4 | 6 | 8 | 3 | 4 | 8 | 5.56 | 6 | 5 | 0.812 | -0.301 | 0.71263363 | 0.84356482 | 0 | 0.599 | 0.2997 | 0.2997 | TRUE | 0.3922 |
| Tmem218 | 228 | 266 | 307 | 358 | 266 | 243 | 242 | 266 | 308 | 281 | 286 | 266 | 274.95 | 272 | 278 | 1.021 | 0.03 | 0.83222387 | 0.91488359 | 0.0043 | 0.0189 | 0.0113 | 0.0113 | TRUE | 0.1522 |
| Tmem219 | 179 | 183 | 171 | 272 | 166 | 165 | 190 | 183 | 172 | 214 | 179 | 181 | 186.27 | 182 | 191 | 1.056 | 0.079 | 0.61964035 | 0.78485124 | 0 | 0.0246 | 0.0129 | 0.0129 | TRUE | 0.1276 |
| Tmem220 | 107 | 165 | 191 | 168 | 127 | 125 | 114 | 165 | 192 | 132 | 137 | 137 | 145.96 | 157 | 135 | 0.861 | -0.216 | 0.33178572 | 0.55225401 | 0.0278 | 0.0295 | 0.0287 | 0.0287 | TRUE | 0.4027 |
| Tmem221 | 24 | 40 | 41 | 28 | 29 | 27 | 25 | 40 | 41 | 22 | 31 | 30 | 31.56 | 35 | 28 | 0.769 | -0.378 | 0.30934734 | 0.52975803 | 0 | 0.1113 | 0.0675 | 0.0675 | TRUE | 0.4225 |
| Tmem222 | 1413 | 1591 | 1541 | 1722 | 1263 | 1451 | 1501 | 1589 | 1547 | 1354 | 1359 | 1588 | 1489.68 | 1546 | 1434 | 0.927 | -0.11 | 0.25503226 | 0.47321987 | 0.0038 | 0.0091 | 0.006 | 0.006 | TRUE | 0.1061 |
| Tmem223 | 318 | 504 | 518 | 584 | 387 | 464 | 338 | 503 | 520 | 459 | 416 | 508 | 457.44 | 454 | 461 | 1.016 | 0.022 | 0.90185513 | 0.95205059 | 0.0295 | 0.0141 | 0.0217 | 0.0217 | TRUE | 0.5759 |
| Tmem225 | 0 | 0 | 0 | 1 | 0 | 0 | 0 | 0 | 0 | 0 | 1 | 0 | 0.13 | 0 | 0 | 1.859 | 0.894 | 0.82650734 | NA | 0 | 25.1515 | 10 | 10 | TRUE | 0.2563 |
| Tmem229a | 12 | 17 | 19 | 45 | 30 | 32 | 13 | 17 | 19 | 35 | 32 | 35 | 25.25 | 16 | 34 | 2.102 | 1.072 | 0.00594318 | 0.03566813 | 0 | 0.1374 | 0.0638 | 0.0638 | TRUE | 0.1631 |
| Tmem229b | 504 | 525 | 508 | 663 | 419 | 439 | 535 | 524 | 510 | 521 | 451 | 481 | 503.72 | 523 | 484 | 0.927 | -0.109 | 0.31398062 | 0.53438031 | 0 | 0.0134 | 0.0065 | 0.0065 | TRUE | 0.0551 |
| Tmem230 | 1817 | 2105 | 2210 | 2201 | 1550 | 1847 | 1930 | 2102 | 2219 | 1730 | 1668 | 2022 | 1945.19 | 2084 | 1807 | 0.867 | -0.207 | 0.05419819 | 0.17303925 | 0.0071 | 0.0086 | 0.0078 | 0.0078 | TRUE | 0.13 |
| Tmem231 | 1676 | 2121 | 2140 | 3500 | 2707 | 2590 | 1780 | 2118 | 2149 | 2751 | 2913 | 2835 | 2424.4 | 2016 | 2833 | 1.405 | 0.491 | 4.98E-07 | 1.49E-05 | 0.0049 | 0.0083 | 0.0064 | 0.0064 | TRUE | 0.1259 |
| Tmem232 | 0 | 0 | 0 | 1 | 1 | 2 | 0 | 0 | 0 | 1 | 1 | 2 | 0.68 | 0 | 1 | 7.126 | 2.833 | 0.25569871 | NA | 0 | 4.8859 | 3.111 | 3.111 | TRUE | 0.1691 |
| Tmem233 | 7 | 4 | 4 | 5 | 2 | 3 | 7 | 4 | 4 | 4 | 2 | 3 | 4.14 | 5 | 3 | 0.616 | -0.7 | 0.44968967 | NA | 0 | 0.8035 | 0.3636 | 0.3636 | TRUE | 0.2971 |
| Tmem234 | 1505 | 1583 | 1590 | 2129 | 1632 | 1316 | 1599 | 1581 | 1597 | 1674 | 1756 | 1441 | 1607.7 | 1592 | 1624 | 1.02 | 0.029 | 0.77085829 | 0.88050269 | 0.0045 | 0.009 | 0.0063 | 0.0063 | TRUE | 0.1153 |
| Tmem235 | 15 | 48 | 47 | 34 | 31 | 40 | 16 | 48 | 47 | 27 | 33 | 44 | 35.82 | 37 | 35 | 0.929 | -0.106 | 0.81852881 | 0.9072905 | 0.157 | 0.0989 | 0.126 | 0.126 | TRUE | 1.7065 |
| Tmem236 | 4 | 2 | 2 | 8 | 0 | 2 | 4 | 2 | 2 | 6 | 0 | 2 | 2.79 | 3 | 3 | 1.066 | 0.092 | 0.94427509 | NA | 0.5333 | 1.1882 | 0.8921 | 0.8921 | TRUE | 0.477 |
| Tmem237 | 1183 | 1457 | 1449 | 1558 | 1176 | 1245 | 1257 | 1455 | 1455 | 1225 | 1265 | 1363 | 1336.59 | 1389 | 1284 | 0.924 | -0.114 | 0.25070116 | 0.46815755 | 0.0043 | 0.0094 | 0.0063 | 0.0063 | TRUE | 0.0831 |
| Tmem238 | 118 | 177 | 165 | 197 | 155 | 140 | 125 | 177 | 166 | 155 | 167 | 153 | 157.11 | 156 | 158 | 1.014 | 0.02 | 0.91678822 | 0.95917638 | 0.0097 | 0.0279 | 0.0189 | 0.0189 | TRUE | 0.3047 |
| Tmem239 | 0 | 0 | 0 | 0 | 0 | 0 | 0 | 0 | 0 | 0 | 0 | 0 | 0 | 0 | 0 | NA | NA | NA | NA | NA | NA | NA | NA | NA | NA |
| Tmem240 | 1 | 6 | 4 | 4 | 4 | 9 | 1 | 6 | 4 | 3 | 4 | 10 | 4.73 | 4 | 6 | 1.533 | 0.616 | 0.52844658 | 0.71714503 | 0.2116 | 0.7036 | 0.4655 | 0.4655 | TRUE | 0.5604 |
| Tmem241 | 53 | 73 | 94 | 126 | 82 | 80 | 56 | 73 | 94 | 99 | 88 | 88 | 83.07 | 74 | 92 | 1.23 | 0.299 | 0.23857837 | 0.45439156 | 0.0203 | 0.0466 | 0.0342 | 0.0342 | TRUE | 0.3199 |
| Tmem242 | 1862 | 1836 | 1703 | 2465 | 1702 | 1616 | 1978 | 1834 | 1710 | 1938 | 1831 | 1769 | 1843.23 | 1841 | 1846 | 1.004 | 0.005 | 0.95391731 | 0.97651612 | 0.0032 | 0.0087 | 0.0053 | 0.0053 | TRUE | 0.0513 |
| Tmem243 | 434 | 568 | 524 | 687 | 535 | 531 | 461 | 567 | 526 | 540 | 576 | 581 | 541.89 | 518 | 566 | 1.09 | 0.125 | 0.28907633 | 0.50996392 | 0.0042 | 0.013 | 0.0081 | 0.0081 | TRUE | 0.1076 |
| Tmem245 | 1407 | 1362 | 1563 | 1338 | 1137 | 1125 | 1494 | 1360 | 1569 | 1052 | 1223 | 1231 | 1321.8 | 1474 | 1169 | 0.792 | -0.337 | 0.00147197 | 0.012079 | 0.0058 | 0.0094 | 0.0073 | 0.0073 | TRUE | 0.0946 |
| Tmem246 | 480 | 504 | 521 | 508 | 305 | 325 | 510 | 503 | 523 | 399 | 328 | 356 | 436.6 | 512 | 361 | 0.707 | -0.5 | 2.90E-05 | 0.00048801 | 0.0025 | 0.0144 | 0.008 | 0.008 | TRUE | 0.1047 |
| Tmem247 | 0 | 0 | 0 | 0 | 0 | 2 | 0 | 0 | 0 | 0 | 0 | 2 | 0.36 | 0 | 1 | 3.832 | 1.938 | 0.63022034 | NA | 10 | 9.0356 | 10 | 10 | TRUE | 0.9744 |
| Tmem248 | 5893 | 5595 | 5401 | 6634 | 4832 | 4859 | 6259 | 5588 | 5423 | 5215 | 5199 | 5319 | 5500.56 | 5757 | 5244 | 0.911 | -0.134 | 0.09462841 | 0.25210798 | 0.0028 | 0.0075 | 0.0045 | 0.0045 | TRUE | 0.071 |
| Tmem25 | 59 | 84 | 104 | 110 | 95 | 98 | 63 | 84 | 104 | 86 | 102 | 107 | 91.16 | 84 | 98 | 1.174 | 0.232 | 0.35674079 | 0.57524954 | 0.0253 | 0.043 | 0.0345 | 0.0345 | TRUE | 0.2214 |
| Tmem251 | 261 | 250 | 201 | 349 | 222 | 186 | 277 | 250 | 202 | 274 | 239 | 204 | 240.93 | 243 | 239 | 0.987 | -0.019 | 0.9163817 | 0.95892234 | 0.0197 | 0.0206 | 0.0202 | 0.0202 | TRUE | 0.2434 |
| Tmem252 | 84 | 50 | 46 | 43 | 34 | 20 | 89 | 50 | 46 | 34 | 37 | 22 | 46.27 | 62 | 31 | 0.5 | -0.999 | 0.00860869 | 0.04701395 | 0.0825 | 0.0781 | 0.0801 | 0.0801 | TRUE | 1.2879 |
| Tmem253 | 21 | 20 | 17 | 47 | 43 | 28 | 22 | 20 | 17 | 37 | 46 | 31 | 28.87 | 20 | 38 | 1.92 | 0.941 | 0.01241883 | 0.06169127 | 0 | 0.121 | 0.0633 | 0.0633 | TRUE | 0.1883 |
| Tmem254a | 0 | 0 | 0 | 0 | 0 | 0 | 0 | 0 | 0 | 0 | 0 | 0 | 0 | 0 | 0 | NA | NA | NA | NA | NA | NA | NA | NA | NA | NA |
| Tmem254b | 0 | 0 | 0 | 0 | 0 | 0 | 0 | 0 | 0 | 0 | 0 | 0 | 0 | 0 | 0 | NA | NA | NA | NA | NA | NA | NA | NA | NA | NA |
| Tmem254c | 0 | 0 | 0 | 0 | 0 | 0 | 0 | 0 | 0 | 0 | 0 | 0 | 0 | 0 | 0 | NA | NA | NA | NA | NA | NA | NA | NA | NA | NA |
| Tmem255a | 6 | 0 | 2 | 4 | 0 | 1 | 6 | 0 | 2 | 3 | 0 | 1 | 2.1 | 3 | 1 | 0.527 | -0.923 | 0.58792439 | NA | 1.5751 | 1.5732 | 1.5738 | 1.5738 | TRUE | 0.3494 |
| Tmem255b | 6 | 5 | 5 | 0 | 2 | 2 | 6 | 5 | 5 | 0 | 2 | 2 | 3.45 | 5 | 1 | 0.249 | -2.007 | 0.08717635 | NA | 0 | 0.9605 | 0.5402 | 0.5402 | TRUE | 0.7727 |
| Tmem256 | 1348 | 2087 | 2089 | 1961 | 1373 | 1573 | 1432 | 2084 | 2098 | 1542 | 1477 | 1722 | 1725.74 | 1871 | 1580 | 0.844 | -0.244 | 0.11890678 | 0.29200355 | 0.023 | 0.0088 | 0.0171 | 0.0171 | TRUE | 0.509 |
| Tmem258 | 3632 | 3958 | 3941 | 4070 | 2849 | 3019 | 3858 | 3953 | 3957 | 3200 | 3065 | 3305 | 3556.26 | 3923 | 3190 | 0.813 | -0.298 | 6.23E-06 | 0.0001336 | 4.00E-04 | 0.0078 | 0.0029 | 0.0029 | TRUE | 0.0139 |
| Tmem259 | 5385 | 5235 | 5043 | 6281 | 4837 | 4515 | 5720 | 5228 | 5064 | 4938 | 5204 | 4942 | 5182.68 | 5337 | 5028 | 0.942 | -0.086 | 0.26543738 | 0.48505136 | 0.0022 | 0.0075 | 0.0041 | 0.0041 | TRUE | 0.0478 |
| Tmem26 | 21 | 8 | 7 | 15 | 16 | 8 | 22 | 8 | 7 | 12 | 17 | 9 | 12.51 | 12 | 13 | 1.015 | 0.021 | 0.97410898 | 0.98667442 | 0.1838 | 0.2701 | 0.2276 | 0.2276 | TRUE | 1.8701 |
| Tmem27 | 0 | 0 | 0 | 0 | 0 | 0 | 0 | 0 | 0 | 0 | 0 | 0 | 0 | 0 | 0 | NA | NA | NA | NA | NA | NA | NA | NA | NA | NA |
| Tmem28 | 147 | 158 | 145 | 122 | 102 | 78 | 156 | 158 | 146 | 96 | 110 | 85 | 125.09 | 153 | 97 | 0.633 | -0.659 | 0.00039692 | 0.00431149 | 0 | 0.0332 | 0.0166 | 0.0166 | TRUE | 0.1178 |
| Tmem29 | 466 | 421 | 405 | 613 | 446 | 407 | 495 | 420 | 407 | 482 | 480 | 446 | 454.9 | 441 | 469 | 1.066 | 0.092 | 0.45660127 | 0.66123545 | 0.0042 | 0.0141 | 0.0087 | 0.0087 | TRUE | 0.1331 |
| Tmem30a | 2868 | 3507 | 3604 | 4427 | 2946 | 3332 | 3046 | 3502 | 3619 | 3480 | 3170 | 3647 | 3410.81 | 3389 | 3432 | 1.013 | 0.018 | 0.85573391 | 0.92836946 | 0.0063 | 0.0079 | 0.0069 | 0.0069 | TRUE | 0.095 |
| Tmem30b | 233 | 307 | 338 | 415 | 345 | 312 | 247 | 307 | 339 | 326 | 371 | 342 | 322.08 | 298 | 346 | 1.161 | 0.215 | 0.16277963 | 0.35814149 | 0.0111 | 0.0171 | 0.014 | 0.014 | TRUE | 0.2436 |
| Tmem30c | 0 | 0 | 0 | 0 | 0 | 0 | 0 | 0 | 0 | 0 | 0 | 0 | 0 | 0 | 0 | NA | NA | NA | NA | NA | NA | NA | NA | NA | NA |
| Tmem33 | 1926 | 2011 | 1923 | 2703 | 1944 | 2038 | 2046 | 2008 | 1931 | 2125 | 2092 | 2231 | 2072.08 | 1995 | 2149 | 1.077 | 0.107 | 0.13809515 | 0.32245868 | 5.00E-04 | 0.0085 | 0.0033 | 0.0033 | TRUE | 0.0132 |
| Tmem35 | 42 | 33 | 25 | 95 | 70 | 57 | 45 | 33 | 25 | 75 | 75 | 62 | 52.51 | 34 | 71 | 2.076 | 1.054 | 0.00054765 | 0.00558339 | 0.0111 | 0.0696 | 0.0452 | 0.0452 | TRUE | 0.4802 |
| Tmem37 | 236 | 255 | 235 | 257 | 180 | 208 | 251 | 255 | 236 | 202 | 194 | 228 | 227.45 | 247 | 208 | 0.84 | -0.251 | 0.07723083 | 0.21951427 | 0 | 0.0214 | 0.0102 | 0.0102 | TRUE | 0.0723 |
| Tmem38a | 602 | 707 | 681 | 704 | 497 | 512 | 639 | 706 | 684 | 553 | 535 | 560 | 612.99 | 676 | 549 | 0.812 | -0.3 | 0.00205953 | 0.01576073 | 0 | 0.0123 | 0.0052 | 0.0052 | TRUE | 0.0268 |
| Tmem38b | 1779 | 2076 | 1958 | 1851 | 1286 | 1510 | 1890 | 2073 | 1966 | 1455 | 1384 | 1653 | 1736.78 | 1976 | 1497 | 0.757 | -0.401 | 4.12E-05 | 0.00065695 | 0.0045 | 0.0088 | 0.0063 | 0.0063 | TRUE | 0.0982 |
| Tmem39a | 3289 | 3583 | 3347 | 3724 | 2636 | 2776 | 3493 | 3578 | 3361 | 2928 | 2836 | 3039 | 3205.85 | 3477 | 2934 | 0.844 | -0.245 | 0.00040359 | 0.00437186 | 6.00E-04 | 0.0079 | 0.0031 | 0.0031 | TRUE | 0.0116 |

| | | | | | | | | | | | | | | | | | | | | | | | | |
|---|---|---|---|---|---|---|---|---|---|---|---|---|---|---|---|---|---|---|---|---|---|---|---|---|
| Tmem39b | 703 | 510 | 566 | 994 | 708 | 551 | 747 | 509 | 568 | 781 | 762 | 603 | 661.78 | 608 | 715 | 1.178 | 0.236 | 0.1709454 | 0.36958887 | 0.0277 | 0.0119 | 0.0199 | 0.0199 | TRUE | 0.4654 |
| Tmem40 | 283 | 232 | 198 | 387 | 265 | 274 | 301 | 232 | 199 | 304 | 285 | 300 | 270.07 | 244 | 296 | 1.218 | 0.284 | 0.10608533 | 0.27179626 | 0.0177 | 0.0191 | 0.0186 | 0.0186 | TRUE | 0.458 |
| Tmem41a | 1203 | 1095 | 1048 | 1430 | 994 | 921 | 1278 | 1094 | 1052 | 1124 | 1069 | 1008 | 1104.25 | 1141 | 1067 | 0.936 | -0.095 | 0.38050434 | 0.59489635 | 0.0057 | 0.0099 | 0.0076 | 0.0076 | TRUE | 0.1307 |
| Tmem41b | 1898 | 1755 | 1713 | 2842 | 1955 | 1817 | 2016 | 1753 | 1720 | 2234 | 2103 | 1989 | 1969.23 | 1830 | 2109 | 1.153 | 0.206 | 0.03602554 | 0.13040481 | 0.0049 | 0.0086 | 0.0064 | 0.0064 | TRUE | 0.0957 |
| Tmem42 | 23 | 22 | 19 | 48 | 37 | 34 | 24 | 22 | 19 | 38 | 40 | 37 | 30.04 | 22 | 38 | 1.754 | 0.81 | 0.01987196 | 0.08643699 | 0 | 0.1166 | 0.0511 | 0.0511 | TRUE | 0.0692 |
| Tmem43 | 1711 | 1983 | 1894 | 2112 | 1571 | 1742 | 1817 | 1980 | 1902 | 1660 | 1690 | 1907 | 1826.18 | 1900 | 1752 | 0.922 | -0.117 | 0.194766 | 0.40071236 | 0.003 | 0.0087 | 0.0054 | 0.0054 | TRUE | 0.0711 |
| Tmem44 | 376 | 391 | 352 | 466 | 390 | 394 | 399 | 390 | 353 | 366 | 420 | 431 | 393.43 | 381 | 406 | 1.063 | 0.088 | 0.48679935 | 0.68377024 | 0.0037 | 0.0153 | 0.0089 | 0.0089 | TRUE | 0.0885 |
| Tmem45a | 389 | 397 | 442 | 434 | 317 | 340 | 413 | 396 | 444 | 341 | 341 | 372 | 384.65 | 418 | 351 | 0.84 | -0.251 | 0.03160084 | 0.11948077 | 0 | 0.0155 | 0.0072 | 0.0072 | TRUE | 0.0345 |
| Tmem45b | 45 | 31 | 34 | 47 | 26 | 17 | 48 | 31 | 34 | 37 | 28 | 19 | 32.74 | 38 | 28 | 0.749 | -0.417 | 0.28792089 | 0.50903254 | 0.05 | 0.1075 | 0.0801 | 0.0801 | TRUE | 0.2765 |
| Tmem47 | 1145 | 1311 | 1539 | 1648 | 1172 | 1209 | 1216 | 1309 | 1545 | 1296 | 1261 | 1323 | 1325.13 | 1357 | 1293 | 0.953 | -0.07 | 0.53099801 | 0.71913278 | 0.0067 | 0.0094 | 0.0081 | 0.0081 | TRUE | 0.1774 |
| Tmem48 | 4258 | 4762 | 4768 | 5348 | 3815 | 3968 | 4523 | 4756 | 4788 | 4204 | 4105 | 4344 | 4453.11 | 4689 | 4218 | 0.899 | -0.153 | 0.01945989 | 0.0851613 | 5.00E-04 | 0.0076 | 0.0029 | 0.0029 | TRUE | 0.0116 |
| Tmem5 | 643 | 757 | 780 | 1155 | 833 | 755 | 683 | 756 | 783 | 908 | 896 | 826 | 808.82 | 741 | 877 | 1.184 | 0.244 | 0.01510616 | 0.07112887 | 0.0024 | 0.011 | 0.006 | 0.006 | TRUE | 0.0546 |
| Tmem50a | 2751 | 3312 | 3339 | 2815 | 2129 | 2393 | 2922 | 3308 | 3353 | 2213 | 2291 | 2620 | 2784.27 | 3194 | 2375 | 0.743 | -0.429 | 2.63E-05 | 0.00045186 | 0.0063 | 0.0081 | 0.0071 | 0.0071 | TRUE | 0.0984 |
| Tmem50b | 1191 | 1452 | 1363 | 1733 | 1231 | 1422 | 1265 | 1450 | 1369 | 1362 | 1324 | 1557 | 1387.87 | 1361 | 1414 | 1.039 | 0.055 | 0.59750526 | 0.77004845 | 0.0053 | 0.0093 | 0.007 | 0.007 | TRUE | 0.0918 |
| Tmem51 | 674 | 649 | 622 | 738 | 527 | 543 | 716 | 648 | 625 | 580 | 567 | 594 | 621.7 | 663 | 580 | 0.876 | -0.191 | 0.06140379 | 0.18904169 | 0.0012 | 0.0122 | 0.0059 | 0.0059 | TRUE | 0.0572 |
| Tmem51as1 | 28 | 45 | 27 | 35 | 43 | 32 | 30 | 45 | 27 | 28 | 46 | 35 | 35.1 | 34 | 36 | 1.061 | 0.085 | 0.8205346 | 0.90842845 | 0.0454 | 0.1008 | 0.0736 | 0.0736 | TRUE | 0.4207 |
| Tmem52 | 29 | 23 | 10 | 20 | 21 | 12 | 31 | 23 | 10 | 16 | 23 | 13 | 19.21 | 21 | 17 | 0.806 | -0.311 | 0.56206849 | 0.74295871 | 0.1277 | 0.1784 | 0.1541 | 0.1541 | TRUE | 0.314 |
| Tmem52b | 0 | 0 | 0 | 0 | 0 | 0 | 0 | 0 | 0 | 0 | 0 | 0 | 0 | 0 | 0 | NA | NA | NA | NA | NA | NA | NA | NA | NA | NA |
| Tmem53 | 57 | 79 | 63 | 100 | 81 | 89 | 61 | 79 | 63 | 79 | 87 | 97 | 77.65 | 68 | 88 | 1.293 | 0.371 | 0.11601369 | 0.28765413 | 0 | 0.0493 | 0.027 | 0.027 | TRUE | 0.1935 |
| Tmem54 | 48 | 30 | 34 | 53 | 33 | 26 | 51 | 30 | 34 | 42 | 36 | 28 | 36.79 | 38 | 35 | 0.925 | -0.112 | 0.75343957 | 0.86987269 | 0.0317 | 0.0965 | 0.0651 | 0.0651 | TRUE | 0.5949 |
| Tmem55a | 1151 | 1591 | 1659 | 1602 | 1193 | 1421 | 1223 | 1589 | 1666 | 1259 | 1284 | 1556 | 1429.3 | 1493 | 1366 | 0.915 | -0.128 | 0.37833181 | 0.5934918 | 0.0179 | 0.0092 | 0.0146 | 0.0146 | TRUE | 0.3006 |
| Tmem55b | 4045 | 4898 | 5058 | 4358 | 3269 | 3854 | 4297 | 4892 | 5079 | 3426 | 3517 | 4219 | 4238.15 | 4756 | 3721 | 0.782 | -0.355 | 0.00176796 | 0.01394209 | 0.0101 | 0.0077 | 0.009 | 0.009 | TRUE | 0.1665 |
| Tmem56 | 44 | 52 | 40 | 93 | 77 | 62 | 47 | 52 | 40 | 73 | 83 | 68 | 60.44 | 46 | 75 | 1.611 | 0.688 | 0.00767782 | 0.04309416 | 0 | 0.0614 | 0.0304 | 0.0304 | TRUE | 0.1076 |
| Tmem57 | 4150 | 3482 | 3745 | 4819 | 3576 | 3273 | 4408 | 3478 | 3760 | 3788 | 3848 | 3583 | 3810.77 | 3882 | 3740 | 0.963 | -0.054 | 0.61128242 | 0.77987759 | 0.007 | 0.0078 | 0.0078 | 0.0078 | TRUE | 0.1708 |
| Tmem59 | 2567 | 3512 | 3648 | 4220 | 2938 | 3359 | 2727 | 3507 | 3663 | 3317 | 3161 | 3677 | 3342.09 | 3299 | 3385 | 1.026 | 0.037 | 0.77421361 | 0.88241777 | 0.0149 | 0.0079 | 0.0117 | 0.0117 | TRUE | 0.2796 |
| Tmem59l | 320 | 442 | 506 | 346 | 248 | 253 | 340 | 441 | 508 | 272 | 267 | 277 | 350.86 | 430 | 272 | 0.632 | -0.661 | 8.46E-05 | 0.00120841 | 0.0184 | 0.0163 | 0.0174 | 0.0174 | TRUE | 0.2228 |
| Tmem60 | 991 | 1154 | 1148 | 1134 | 804 | 981 | 1053 | 1153 | 1153 | 891 | 865 | 1074 | 1031.37 | 1120 | 943 | 0.842 | -0.248 | 0.03038761 | 0.1163285 | 0.0071 | 0.0101 | 0.0085 | 0.0085 | TRUE | 0.1718 |
| Tmem62 | 70 | 69 | 76 | 150 | 89 | 79 | 74 | 69 | 76 | 118 | 96 | 86 | 86.62 | 73 | 100 | 1.374 | 0.459 | 0.04323643 | 0.14864827 | 0 | 0.0449 | 0.0253 | 0.0253 | TRUE | 0.2616 |
| Tmem63a | 487 | 568 | 601 | 521 | 416 | 486 | 517 | 567 | 603 | 410 | 448 | 532 | 512.86 | 562 | 463 | 0.821 | -0.284 | 0.03740291 | 0.13413339 | 0.0099 | 0.0133 | 0.0115 | 0.0115 | TRUE | 0.1952 |
| Tmem63b | 3231 | 4150 | 4132 | 4764 | 3640 | 3824 | 3432 | 4145 | 4149 | 3745 | 3916 | 4186 | 3928.83 | 3909 | 3949 | 1.01 | 0.015 | 0.88755206 | 0.94475744 | 0.0071 | 0.0077 | 0.0074 | 0.0074 | TRUE | 0.1383 |
| Tmem63c | 280 | 278 | 277 | 274 | 216 | 231 | 297 | 278 | 278 | 215 | 232 | 253 | 258.98 | 284 | 233 | 0.819 | -0.288 | 0.03428202 | 0.12614807 | 0 | 0.0196 | 0.0094 | 0.0094 | TRUE | 0.0571 |
| Tmem64 | 9084 | 8160 | 7172 | 15142 | 9728 | 8124 | 9649 | 8149 | 7201 | 11903 | 10467 | 8893 | 9377.15 | 8333 | 10421 | 1.251 | 0.323 | 0.0231609 | 0.0965364 | 0.0212 | 0.0073 | 0.0144 | 0.0144 | TRUE | 0.195 |
| Tmem65 | 1366 | 1811 | 1798 | 2035 | 1495 | 1650 | 1451 | 1809 | 1805 | 1600 | 1609 | 1806 | 1679.91 | 1688 | 1672 | 0.99 | -0.015 | 0.89699869 | 0.9501066 | 0.0095 | 0.0089 | 0.0092 | 0.0092 | TRUE | 0.182 |
| Tmem66 | 3435 | 4605 | 4844 | 4028 | 3089 | 3509 | 3649 | 4599 | 4864 | 3166 | 3324 | 3841 | 3907.12 | 4371 | 3444 | 0.788 | -0.344 | 0.00888166 | 0.04818309 | 0.0156 | 0.0077 | 0.0122 | 0.0122 | TRUE | 0.254 |
| Tmem67 | 279 | 226 | 277 | 434 | 331 | 270 | 296 | 226 | 278 | 341 | 356 | 296 | 298.84 | 267 | 331 | 1.242 | 0.313 | 0.04491128 | 0.15269255 | 0.0106 | 0.0179 | 0.0141 | 0.0141 | TRUE | 0.2036 |
| Tmem68 | 508 | 667 | 619 | 747 | 492 | 553 | 540 | 666 | 622 | 587 | 529 | 605 | 591.53 | 609 | 574 | 0.942 | -0.086 | 0.4829978 | 0.68129391 | 0.0063 | 0.0125 | 0.0091 | 0.0091 | TRUE | 0.1162 |
| Tmem69 | 789 | 662 | 674 | 954 | 673 | 621 | 838 | 661 | 677 | 750 | 724 | 680 | 721.64 | 725 | 718 | 0.991 | -0.013 | 0.91492184 | 0.95806876 | 0.0085 | 0.0115 | 0.0099 | 0.0099 | TRUE | 0.2176 |
| Tmem70 | 1161 | 1394 | 1355 | 1878 | 1393 | 1493 | 1233 | 1392 | 1361 | 1476 | 1499 | 1634 | 1432.57 | 1329 | 1536 | 1.156 | 0.209 | 0.02268482 | 0.09505447 | 0.0028 | 0.0092 | 0.0053 | 0.0053 | TRUE | 0.0472 |
| Tmem71 | 0 | 0 | 0 | 0 | 0 | 0 | 0 | 0 | 0 | 0 | 0 | 0 | 0 | 0 | 0 | NA | NA | NA | NA | NA | NA | NA | NA | NA | NA |
| Tmem72 | 5 | 3 | 0 | 2 | 1 | 0 | 5 | 3 | 0 | 2 | 1 | 0 | 1.83 | 3 | 1 | 0.332 | -1.592 | 0.35817115 | NA | 0 | 1.8112 | 1.4452 | 1.4452 | TRUE | 0.122 |
| Tmem74 | 55 | 50 | 35 | 47 | 48 | 38 | 58 | 50 | 35 | 37 | 52 | 42 | 45.61 | 48 | 44 | 0.904 | -0.145 | 0.65293265 | 0.80529041 | 0.0251 | 0.0791 | 0.0531 | 0.0531 | TRUE | 0.38 |
| Tmem74b | 23 | 53 | 52 | 69 | 51 | 67 | 24 | 53 | 52 | 54 | 55 | 73 | 52.01 | 43 | 61 | 1.402 | 0.487 | 0.16859112 | 0.36640749 | 0.0706 | 0.0703 | 0.0704 | 0.0704 | TRUE | 1.0808 |
| Tmem79 | 222 | 220 | 200 | 341 | 219 | 170 | 236 | 220 | 201 | 268 | 236 | 186 | 224.36 | 219 | 230 | 1.055 | 0.077 | 0.66677441 | 0.81377126 | 0.0157 | 0.0216 | 0.0185 | 0.0185 | TRUE | 0.3003 |
| Tmem8 | 1518 | 1426 | 1452 | 2298 | 1677 | 1554 | 1612 | 1424 | 1458 | 1807 | 1804 | 1701 | 1634.41 | 1498 | 1771 | 1.182 | 0.242 | 0.00495119 | 0.03108688 | 0.0019 | 0.0089 | 0.0047 | 0.0047 | TRUE | 0.0533 |
| Tmem80 | 5 | 20 | 8 | 33 | 14 | 16 | 5 | 20 | 8 | 26 | 15 | 18 | 15.31 | 11 | 20 | 1.762 | 0.817 | 0.18225562 | 0.38412252 | 0.1735 | 0.2221 | 0.1997 | 0.1997 | TRUE | 1.8414 |
| Tmem81 | 295 | 323 | 306 | 378 | 304 | 270 | 313 | 323 | 307 | 297 | 327 | 296 | 310.5 | 314 | 307 | 0.974 | -0.038 | 0.75808892 | 0.87229725 | 0 | 0.0175 | 0.0075 | 0.0075 | TRUE | 0.0373 |
| Tmem82 | 125 | 131 | 139 | 153 | 135 | 120 | 133 | 131 | 140 | 120 | 145 | 131 | 133.34 | 135 | 132 | 0.981 | -0.027 | 0.87721657 | 0.93933463 | 0 | 0.0316 | 0.0148 | 0.0148 | TRUE | 0.075 |
| Tmem86a | 296 | 366 | 349 | 287 | 236 | 272 | 314 | 366 | 350 | 226 | 254 | 298 | 301.27 | 343 | 259 | 0.752 | -0.412 | 0.00682235 | 0.03938586 | 0.0091 | 0.0178 | 0.0133 | 0.0133 | TRUE | 0.1881 |
| Tmem86b | 55 | 54 | 46 | 80 | 45 | 43 | 58 | 54 | 46 | 63 | 48 | 47 | 52.82 | 53 | 53 | 1.006 | 0.009 | 0.97407186 | 0.98667442 | 0 | 0.0693 | 0.0373 | 0.0373 | TRUE | 0.2642 |
| Tmem87a | 3322 | 3800 | 3613 | 4501 | 3470 | 3438 | 3529 | 3795 | 3628 | 3538 | 3734 | 3763 | 3664.46 | 3651 | 3678 | 1.007 | 0.01 | 0.88053586 | 0.94090865 | 0.001 | 0.0078 | 0.0032 | 0.0032 | TRUE | 0.0146 |
| Tmem87b | 1266 | 1770 | 1953 | 1778 | 1436 | 1667 | 1345 | 1768 | 1961 | 1398 | 1545 | 1825 | 1640.17 | 1691 | 1589 | 0.939 | -0.091 | 0.57192947 | 0.75070503 | 0.0268 | 0.0089 | 0.0179 | 0.0179 | TRUE | 0.3867 |
| Tmem88 | 329 | 254 | 259 | 271 | 207 | 151 | 349 | 254 | 260 | 213 | 223 | 165 | 244.04 | 288 | 200 | 0.698 | -0.519 | 0.0068654 | 0.03956199 | 0.0244 | 0.0204 | 0.0224 | 0.0224 | TRUE | 0.3933 |
| Tmem88b | 4 | 8 | 11 | 8 | 9 | 8 | 4 | 8 | 11 | 6 | 10 | 9 | 8 | 8 | 8 | 1.05 | 0.07 | 0.91973619 | 0.96064694 | 0 | 0.4186 | 0.2194 | 0.2194 | TRUE | 0.2433 |
| Tmem89 | 0 | 0 | 0 | 0 | 0 | 0 | 0 | 0 | 0 | 0 | 0 | 0 | 0 | 0 | 0 | NA | NA | NA | NA | NA | NA | NA | NA | NA | NA |
| Tmem8b | 398 | 597 | 606 | 454 | 401 | 493 | 423 | 596 | 608 | 357 | 431 | 540 | 492.58 | 542 | 443 | 0.814 | -0.296 | 0.13335752 | 0.31533955 | 0.0391 | 0.0136 | 0.026 | 0.026 | TRUE | 0.3903 |
| Tmem8c | 0 | 0 | 0 | 0 | 0 | 0 | 0 | 0 | 0 | 0 | 0 | 0 | 0 | 0 | 0 | NA | NA | NA | NA | NA | NA | NA | NA | NA | NA |
| Tmem9 | 1283 | 1685 | 1582 | 1864 | 1448 | 1578 | 1363 | 1683 | 1588 | 1465 | 1558 | 1727 | 1564.13 | 1545 | 1583 | 1.024 | 0.035 | 0.76083908 | 0.8741862 | 0.0086 | 0.009 | 0.0088 | 0.0088 | TRUE | 0.1275 |
| Tmem91 | 47 | 54 | 55 | 48 | 26 | 44 | 50 | 54 | 55 | 38 | 28 | 48 | 45.49 | 53 | 38 | 0.715 | -0.484 | 0.11676312 | 0.28895749 | 0.0025 | 0.0793 | 0.0461 | 0.0461 | TRUE | 0.2496 |
| Tmem92 | 210 | 121 | 98 | 560 | 430 | 192 | 223 | 121 | 98 | 440 | 463 | 210 | 259.23 | 147 | 371 | 2.52 | 1.333 | 0.00014107 | 0.00181187 | 0.1654 | 0.0196 | 0.0837 | 0.0837 | TRUE | 2.0866 |
| Tmem95 | 5 | 4 | 7 | 9 | 4 | 2 | 5 | 4 | 7 | 7 | 4 | 2 | 4.98 | 5 | 4 | 0.85 | -0.234 | 0.78676308 | 0.89028751 | 0 | 0.6679 | 0.3418 | 0.3418 | TRUE | 0.3657 |
| Tmem97 | 5549 | 6537 | 6289 | 6771 | 4938 | 5396 | 5894 | 6529 | 6315 | 5323 | 5313 | 5907 | 5880 | 6246 | 5514 | 0.883 | -0.18 | 0.02745388 | 0.10846756 | 0.003 | 0.0075 | 0.0046 | 0.0046 | TRUE | 0.0471 |
| Tmem98 | 632 | 926 | 917 | 801 | 639 | 724 | 671 | 925 | 921 | 630 | 688 | 793 | 771.1 | 839 | 704 | 0.837 | -0.256 | 0.10344645 | 0.26760005 | 0.0216 | 0.0112 | 0.0165 | 0.0165 | TRUE | 0.3617 |
| Tmem9b | 473 | 559 | 580 | 706 | 461 | 487 | 502 | 558 | 582 | 555 | 496 | 533 | 537.86 | 547 | 528 | 0.965 | -0.052 | 0.64435688 | 0.79996825 | 0.0026 | 0.013 | 0.0072 | 0.0072 | TRUE | 0.0606 |
| Tmf1 | 3393 | 3745 | 3785 | 4742 | 3509 | 3643 | 3604 | 3740 | 3801 | 3728 | 3775 | 3988 | 3772.63 | 3715 | 3830 | 1.031 | 0.044 | 0.5175115 | 0.70901593 | 6.00E-04 | 0.0078 | 0.003 | 0.003 | TRUE | 0.0156 |
| Tmie | 51 | 66 | 54 | 51 | 44 | 56 | 54 | 66 | 54 | 40 | 47 | 61 | 53.84 | 58 | 49 | 0.847 | -0.24 | 0.40440578 | 0.61721385 | 0.0113 | 0.0681 | 0.0407 | 0.0407 | TRUE | 0.3355 |
| Tmigd1 | 0 | 0 | 0 | 2 | 1 | 2 | 0 | 0 | 0 | 2 | 1 | 2 | 0.81 | 0 | 2 | 8.576 | 3.1 | 0.18121756 | NA | 0 | 4.093 | 2.5705 | 2.5705 | TRUE | 0.07 |
| Tmlhe | 87 | 141 | 118 | 227 | 133 | 141 | 92 | 141 | 118 | 178 | 143 | 154 | 137.93 | 117 | 158 | 1.355 | 0.438 | 0.03967782 | 0.1398745 | 0.0195 | 0.0308 | 0.0253 | 0.0253 | TRUE | 0.3006 |
| Tmod1 | 7 | 15 | 14 | 17 | 15 | 8 | 7 | 15 | 14 | 13 | 16 | 9 | 12.46 | 12 | 13 | 1.048 | 0.068 | 0.90462662 | 0.95255222 | 0 | 0.2714 | 0.1542 | 0.1542 | TRUE | 0.4366 |
| Tmod2 | 672 | 856 | 921 | 788 | 650 | 619 | 714 | 855 | 925 | 619 | 699 | 678 | 748.31 | 831 | 665 | 0.8 | -0.322 | 0.01102397 | 0.05657993 | 0.0094 | 0.0113 | 0.0102 | 0.0102 | TRUE | 0.1803 |

| | | | | | | | | | | | | | | | | | | | | | | | | | |
|---|---|---|---|---|---|---|---|---|---|---|---|---|---|---|---|---|---|---|---|---|---|---|---|---|---|
| Tmod3 | 4895 | 4260 | 4257 | 5487 | 3823 | 3885 | 5199 | 4254 | 4274 | 4313 | 4113 | 4253 | 4401.31 | 4576 | 4226 | 0.924 | -0.114 | 0.25709002 | 0.47575076 | 0.0069 | 0.0077 | 0.0071 | 0.0071 | TRUE | 0.1727 |
| Tmod4 | 10 | 21 | 21 | 29 | 21 | 9 | 11 | 21 | 21 | 23 | 23 | 10 | 17.99 | 18 | 19 | 1.054 | 0.076 | 0.88678018 | 0.94438861 | 0.0983 | 0.1901 | 0.1476 | 0.1476 | TRUE | 0.7837 |
| Tmpo | 11715 | 12213 | 12235 | 16647 | 12005 | 11573 | 12443 | 12197 | 12285 | 13087 | 12917 | 12668 | 12599.59 | 12308 | 12891 | 1.047 | 0.067 | 0.21022802 | 0.42072279 | 1.00E-04 | 0.0072 | 0.002 | 0.002 | TRUE | 0.0028 |
| Tmppe | 1231 | 1167 | 1198 | 2053 | 1608 | 1341 | 1308 | 1165 | 1203 | 1614 | 1730 | 1468 | 1414.65 | 1225 | 1604 | 1.309 | 0.389 | 9.04E-05 | 0.00127735 | 0.0041 | 0.0092 | 0.0064 | 0.0064 | TRUE | 0.0652 |
| Tmprss11a | 1 | 0 | 0 | 0 | 0 | 0 | 1 | 0 | 0 | 0 | 0 | 0 | 0.18 | 0 | 0 | 0.49 | -1.029 | 0.80087 | NA | 0 | 18.6165 | 10 | 10 | TRUE | 0.4614 |
| Tmprss11bnl | 0 | 0 | 0 | 3 | 0 | 0 | 0 | 0 | 0 | 2 | 0 | 0 | 0.39 | 0 | 1 | 4.278 | 2.097 | 0.60136958 | NA | 10 | 8.3884 | 10 | 10 | TRUE | 1.4411 |
| Tmprss11c | 0 | 0 | 0 | 0 | 0 | 0 | 0 | 0 | 0 | 0 | 0 | 0 | 0 | 0 | 0 | NA | NA | NA | NA | NA | NA | NA | NA | NA | NA |
| Tmprss11d | 24 | 12 | 9 | 35 | 32 | 11 | 25 | 12 | 9 | 28 | 34 | 12 | 20.08 | 15 | 25 | 1.6 | 0.678 | 0.24266597 | 0.45918748 | 0.2149 | 0.1709 | 0.1897 | 0.1897 | TRUE | 0.5943 |
| Tmprss11e | 3 | 3 | 3 | 7 | 4 | 1 | 3 | 3 | 3 | 6 | 4 | 1 | 3.35 | 3 | 4 | 1.21 | 0.275 | 0.79283248 | NA | 0 | 0.9905 | 0.4887 | 0.4887 | TRUE | 0.5122 |
| Tmprss11f | 1 | 0 | 0 | 0 | 1 | 2 | 1 | 0 | 0 | 0 | 1 | 2 | 0.72 | 0 | 1 | 2.905 | 1.539 | 0.54944501 | NA | 0 | 4.5747 | 3.3003 | 3.3003 | TRUE | 0.2242 |
| Tmprss11g | 0 | 0 | 0 | 0 | 0 | 0 | 0 | 0 | 0 | 0 | 0 | 0 | 0 | 0 | 0 | NA | NA | NA | NA | NA | NA | NA | NA | NA | NA |
| Tmprss12 | 34 | 42 | 36 | 41 | 25 | 30 | 36 | 42 | 36 | 32 | 27 | 33 | 34.36 | 38 | 31 | 0.807 | -0.31 | 0.33729223 | 0.5573856 | 0 | 0.1028 | 0.046 | 0.046 | TRUE | 0.0713 |
| Tmprss13 | 14 | 24 | 16 | 11 | 13 | 24 | 15 | 24 | 16 | 9 | 14 | 26 | 17.3 | 18 | 16 | 0.877 | -0.189 | 0.73402824 | 0.85737195 | 0.1291 | 0.1973 | 0.1638 | 0.1638 | TRUE | 0.6932 |
| Tmprss15 | 1 | 0 | 0 | 0 | 0 | 0 | 1 | 0 | 0 | 0 | 0 | 0 | 0.18 | 0 | 0 | 0.49 | -1.029 | 0.80087 | NA | 0 | 18.6165 | 10 | 10 | TRUE | 0.4614 |
| Tmprss2 | 18 | 20 | 14 | 31 | 22 | 20 | 19 | 20 | 14 | 24 | 24 | 22 | 20.51 | 18 | 23 | 1.319 | 0.399 | 0.33868284 | 0.55879656 | 0 | 0.1675 | 0.0757 | 0.0757 | TRUE | 0.1672 |
| Tmprss3 | 26 | 16 | 22 | 42 | 14 | 13 | 28 | 16 | 22 | 33 | 15 | 14 | 21.33 | 22 | 21 | 0.965 | -0.052 | 0.91827953 | 0.95991788 | 0.113 | 0.1613 | 0.1373 | 0.1373 | TRUE | 1.0453 |
| Tmprss4 | 1 | 0 | 1 | 5 | 2 | 3 | 1 | 0 | 1 | 4 | 2 | 3 | 1.91 | 1 | 3 | 4.601 | 2.202 | 0.15275523 | NA | 0 | 1.7359 | 0.8065 | 0.8065 | TRUE | 0.2571 |
| Tmprss5 | 11 | 8 | 7 | 9 | 6 | 13 | 12 | 8 | 7 | 7 | 6 | 14 | 9.08 | 9 | 9 | 1.032 | 0.046 | 0.94496418 | 0.97246347 | 0 | 0.3698 | 0.2053 | 0.2053 | TRUE | 0.6634 |
| Tmprss6 | 19 | 23 | 11 | 17 | 11 | 13 | 20 | 23 | 11 | 13 | 12 | 14 | 15.6 | 18 | 13 | 0.728 | -0.457 | 0.36862185 | 0.58596087 | 0 | 0.218 | 0.1211 | 0.1211 | TRUE | 0.5977 |
| Tmprss7 | 9 | 1 | 5 | 3 | 2 | 3 | 10 | 1 | 5 | 2 | 2 | 3 | 3.9 | 5 | 2 | 0.5 | -0.999 | 0.35693486 | NA | 0 | 0.8526 | 0.5599 | 0.5599 | TRUE | 0.1248 |
| Tmprss9 | 10 | 4 | 10 | 5 | 4 | 2 | 11 | 4 | 10 | 4 | 4 | 2 | 5.85 | 8 | 3 | 0.427 | -1.229 | 0.14460396 | 0.33207537 | 0 | 0.5704 | 0.3097 | 0.3097 | TRUE | 0.6182 |
| Tmsb10 | 14569 | 14942 | 15093 | 18536 | 13964 | 14439 | 15475 | 14923 | 15155 | 14572 | 15024 | 15806 | 15159.04 | 15184 | 15134 | 0.997 | -0.005 | 0.93803298 | 0.96893419 | 7.00E-04 | 0.0071 | 0.0028 | 0.0028 | TRUE | 0.0184 |
| Tmsb15a | 5 | 6 | 9 | 15 | 10 | 11 | 5 | 6 | 9 | 12 | 11 | 12 | 9.16 | 7 | 12 | 1.699 | 0.764 | 0.22117462 | 0.43313412 | 0 | 0.3667 | 0.1638 | 0.1638 | TRUE | 0.2215 |
| Tmsb15b1 | 49 | 64 | 64 | 89 | 58 | 58 | 52 | 64 | 64 | 70 | 62 | 63 | 62.68 | 60 | 65 | 1.089 | 0.122 | 0.6196232 | 0.78485124 | 0 | 0.0595 | 0.0278 | 0.0278 | TRUE | 0.1142 |
| Tmsb15b2 | 27 | 45 | 37 | 52 | 35 | 32 | 29 | 45 | 37 | 41 | 38 | 35 | 37.39 | 37 | 38 | 1.027 | 0.038 | 0.90797985 | 0.95356627 | 0 | 0.095 | 0.0503 | 0.0503 | TRUE | 0.2031 |
| Tmsb15l | 29 | 32 | 17 | 22 | 14 | 27 | 31 | 32 | 17 | 17 | 15 | 30 | 23.62 | 27 | 21 | 0.772 | -0.374 | 0.42048071 | 0.63088107 | 0.0747 | 0.1464 | 0.1122 | 0.1122 | TRUE | 0.7275 |
| Tmsb4x | 7268 | 6262 | 5993 | 10505 | 7079 | 6578 | 7720 | 6254 | 6018 | 8258 | 7617 | 7201 | 7177.81 | 6664 | 7692 | 1.154 | 0.207 | 0.07489598 | 0.21543201 | 0.0118 | 0.0074 | 0.0096 | 0.0096 | TRUE | 0.2344 |
| Tmtc1 | 207 | 199 | 240 | 284 | 186 | 224 | 220 | 199 | 241 | 223 | 200 | 245 | 221.36 | 220 | 223 | 1.014 | 0.019 | 0.90114215 | 0.95196343 | 0.0048 | 0.0218 | 0.0131 | 0.0131 | TRUE | 0.0828 |
| Tmtc2 | 151 | 164 | 156 | 160 | 141 | 139 | 160 | 164 | 157 | 126 | 152 | 152 | 151.74 | 160 | 143 | 0.89 | -0.168 | 0.32088493 | 0.54202034 | 0 | 0.0286 | 0.0141 | 0.0141 | TRUE | 0.1316 |
| Tmtc3 | 3505 | 2997 | 2924 | 4545 | 3112 | 2956 | 3723 | 2993 | 2936 | 3573 | 3348 | 3236 | 3301.52 | 3217 | 3386 | 1.053 | 0.074 | 0.51482748 | 0.70742803 | 0.0086 | 0.0079 | 0.009 | 0.009 | TRUE | 0.2293 |
| Tmtc4 | 957 | 1400 | 1425 | 1498 | 1098 | 1315 | 1017 | 1398 | 1431 | 1178 | 1181 | 1439 | 1274 | 1282 | 1266 | 0.987 | -0.019 | 0.90471859 | 0.95255222 | 0.0233 | 0.0095 | 0.0165 | 0.0165 | TRUE | 0.3927 |
| Tmub1 | 121 | 166 | 138 | 198 | 114 | 160 | 129 | 166 | 139 | 156 | 123 | 175 | 147.72 | 145 | 151 | 1.047 | 0.067 | 0.74318641 | 0.86301988 | 0.0171 | 0.0292 | 0.0232 | 0.0232 | TRUE | 0.2724 |
| Tmub2 | 994 | 1106 | 1029 | 1235 | 948 | 902 | 1056 | 1105 | 1033 | 971 | 1020 | 987 | 1028.64 | 1065 | 993 | 0.932 | -0.101 | 0.21582391 | 0.42779624 | 0 | 0.0101 | 0.0039 | 0.0039 | TRUE | 0.013 |
| Tmx1 | 2506 | 3293 | 3219 | 4571 | 3296 | 3336 | 2662 | 3289 | 3232 | 3593 | 3546 | 3652 | 3329.04 | 3061 | 3597 | 1.175 | 0.233 | 0.02113193 | 0.09040097 | 0.0061 | 0.0079 | 0.007 | 0.007 | TRUE | 0.1577 |
| Tmx2 | 3379 | 4657 | 4562 | 4645 | 3406 | 3902 | 3589 | 4651 | 4581 | 3652 | 3665 | 4271 | 4068.06 | 4274 | 3863 | 0.904 | -0.146 | 0.24481502 | 0.46142717 | 0.0135 | 0.0077 | 0.0111 | 0.0111 | TRUE | 0.2388 |
| Tmx3 | 3043 | 2923 | 2723 | 4596 | 3180 | 3043 | 3232 | 2919 | 2734 | 3613 | 3421 | 3331 | 3208.53 | 2962 | 3455 | 1.167 | 0.223 | 0.01371837 | 0.06622779 | 0.004 | 0.0079 | 0.0056 | 0.0056 | TRUE | 0.0772 |
| Tmx4 | 1216 | 1334 | 1253 | 1791 | 1350 | 1362 | 1292 | 1332 | 1258 | 1408 | 1453 | 1491 | 1372.24 | 1294 | 1451 | 1.12 | 0.164 | 0.03215568 | 0.12080975 | 0 | 0.0093 | 0.0035 | 0.0035 | TRUE | 0.0082 |
| Tnc | 0 | 4 | 1 | 7 | 5 | 2 | 0 | 4 | 1 | 6 | 5 | 2 | 3.01 | 2 | 4 | 2.608 | 1.383 | 0.27750555 | NA | 0.1741 | 1.1008 | 0.7533 | 0.7533 | TRUE | 1.1411 |
| Tnf | 0 | 0 | 0 | 0 | 0 | 1 | 0 | 0 | 0 | 0 | 0 | 1 | 0.18 | 0 | 0 | 1.859 | 0.894 | 0.82650734 | NA | 0 | 18.0643 | 10 | 10 | TRUE | 0.5383 |
| Tnfaip1 | 3581 | 3417 | 3228 | 4926 | 3332 | 3213 | 3804 | 3413 | 3241 | 3872 | 3585 | 3517 | 3572.02 | 3486 | 3658 | 1.05 | 0.07 | 0.44208995 | 0.64905093 | 0.0041 | 0.0078 | 0.0057 | 0.0057 | TRUE | 0.0772 |
| Tnfaip2 | 86 | 79 | 69 | 57 | 47 | 55 | 91 | 79 | 69 | 45 | 51 | 60 | 65.85 | 80 | 52 | 0.646 | -0.629 | 0.01483013 | 0.07026158 | 0 | 0.0569 | 0.0322 | 0.0322 | TRUE | 0.1575 |
| Tnfaip3 | 86 | 160 | 186 | 223 | 190 | 232 | 91 | 160 | 187 | 175 | 204 | 254 | 178.6 | 146 | 211 | 1.443 | 0.53 | 0.05193384 | 0.16826221 | 0.0737 | 0.0253 | 0.0477 | 0.0477 | TRUE | 0.9325 |
| Tnfaip6 | 28 | 25 | 19 | 36 | 14 | 27 | 30 | 25 | 19 | 28 | 15 | 30 | 24.45 | 25 | 24 | 0.994 | -0.009 | 0.98397967 | 0.99158695 | 0.0386 | 0.1416 | 0.0941 | 0.0941 | TRUE | 0.6182 |
| Tnfaip8 | 478 | 678 | 648 | 758 | 585 | 602 | 508 | 677 | 651 | 596 | 629 | 659 | 619.97 | 612 | 628 | 1.025 | 0.036 | 0.78918644 | 0.891612 | 0.0112 | 0.0122 | 0.0117 | 0.0117 | TRUE | 0.2587 |
| Tnfaip8l1 | 122 | 125 | 108 | 122 | 95 | 110 | 130 | 125 | 108 | 96 | 102 | 120 | 113.57 | 121 | 106 | 0.875 | -0.192 | 0.33128464 | 0.55212268 | 0 | 0.0359 | 0.0194 | 0.0194 | TRUE | 0.1317 |
| Tnfaip8l2 | 31 | 21 | 27 | 42 | 22 | 27 | 33 | 21 | 27 | 33 | 24 | 30 | 27.88 | 27 | 29 | 1.072 | 0.1 | 0.79192975 | 0.89301263 | 0 | 0.1251 | 0.0674 | 0.0674 | TRUE | 0.2088 |
| Tnfaip8l3 | 35 | 61 | 50 | 26 | 23 | 34 | 37 | 61 | 50 | 20 | 25 | 37 | 38.45 | 49 | 27 | 0.55 | -0.863 | 0.0204755 | 0.08834803 | 0.0484 | 0.0926 | 0.0717 | 0.0717 | TRUE | 0.2723 |
| Tnfrsf10b | 2525 | 3433 | 3445 | 3044 | 2365 | 2803 | 2682 | 3429 | 3459 | 2393 | 2545 | 3068 | 2929.26 | 3190 | 2669 | 0.836 | -0.258 | 0.0616183 | 0.18948018 | 0.0178 | 0.008 | 0.0134 | 0.0134 | TRUE | 0.2355 |
| Tnfrsf11a | 97 | 95 | 88 | 142 | 102 | 101 | 103 | 95 | 88 | 112 | 110 | 111 | 103.03 | 95 | 111 | 1.161 | 0.215 | 0.26284971 | 0.48227847 | 0 | 0.0389 | 0.0168 | 0.0168 | TRUE | 0.0459 |
| Tnfrsf11b | 0 | 1 | 1 | 1 | 1 | 1 | 0 | 1 | 1 | 1 | 1 | 1 | 0.83 | 1 | 1 | 1.45 | 0.536 | 0.78654731 | NA | 0 | 3.9925 | 1.5656 | 1.5656 | TRUE | 0.2233 |
| Tnfrsf12a | 283 | 358 | 369 | 383 | 269 | 301 | 301 | 358 | 371 | 301 | 289 | 329 | 324.78 | 343 | 306 | 0.893 | -0.163 | 0.23992214 | 0.45609524 | 0.005 | 0.017 | 0.0107 | 0.0107 | TRUE | 0.1311 |
| Tnfrsf13b | 14 | 3 | 9 | 12 | 14 | 8 | 15 | 3 | 9 | 9 | 15 | 9 | 10.03 | 9 | 11 | 1.237 | 0.307 | 0.66492833 | 0.81250574 | 0.17 | 0.3355 | 0.2597 | 0.2597 | TRUE | 0.2389 |
| Tnfrsf13c | 47 | 46 | 49 | 50 | 36 | 38 | 50 | 46 | 49 | 39 | 39 | 42 | 44.12 | 48 | 40 | 0.824 | -0.279 | 0.32153122 | 0.54266349 | 0 | 0.0816 | 0.0341 | 0.0341 | TRUE | 0.0155 |
| Tnfrsf14 | 0 | 6 | 4 | 0 | 1 | 3 | 0 | 6 | 4 | 0 | 1 | 3 | 2.39 | 3 | 1 | 0.419 | -1.256 | 0.44636694 | NA | 1.5904 | 1.3826 | 1.4591 | 1.4591 | TRUE | 0.3583 |
| Tnfrsf17 | 9 | 8 | 10 | 5 | 2 | 2 | 10 | 8 | 10 | 4 | 2 | 2 | 5.98 | 9 | 3 | 0.308 | -1.7 | 0.03739156 | 0.13412319 | 0 | 0.5581 | 0.2531 | 0.2531 | TRUE | 0.2281 |
| Tnfrsf18 | 2 | 2 | 1 | 2 | 2 | 5 | 2 | 2 | 1 | 2 | 2 | 5 | 2.39 | 2 | 3 | 1.766 | 0.82 | 0.51058162 | NA | 0 | 1.3869 | 0.6578 | 0.6578 | TRUE | 0.5763 |
| Tnfrsf19 | 1551 | 1327 | 1172 | 1254 | 889 | 841 | 1647 | 1325 | 1177 | 986 | 957 | 921 | 1168.74 | 1383 | 955 | 0.69 | -0.535 | 8.14E-05 | 0.00117147 | 0.0144 | 0.0097 | 0.0124 | 0.0124 | TRUE | 0.3351 |
| Tnfrsf1a | 513 | 559 | 493 | 595 | 448 | 484 | 545 | 558 | 495 | 468 | 482 | 530 | 512.96 | 533 | 493 | 0.925 | -0.113 | 0.31346602 | 0.5339318 | 0.0022 | 0.0133 | 0.0071 | 0.0071 | TRUE | 0.0483 |
| Tnfrsf1b | 1551 | 2095 | 2072 | 1681 | 1250 | 1595 | 1647 | 2092 | 2081 | 1321 | 1345 | 1746 | 1705.44 | 1940 | 1471 | 0.758 | -0.4 | 0.0065095 | 0.03815231 | 0.0208 | 0.0088 | 0.015 | 0.015 | TRUE | 0.3211 |
| Tnfrsf21 | 928 | 966 | 1005 | 1347 | 1040 | 907 | 986 | 965 | 1009 | 1059 | 1119 | 993 | 1021.72 | 987 | 1057 | 1.071 | 0.1 | 0.25888693 | 0.4778449 | 0.001 | 0.0101 | 0.0046 | 0.0046 | TRUE | 0.0324 |
| Tnfrsf22 | 1052 | 836 | 820 | 901 | 653 | 596 | 1117 | 835 | 823 | 708 | 703 | 652 | 806.5 | 925 | 688 | 0.744 | -0.427 | 0.00266915 | 0.01934642 | 0.0153 | 0.011 | 0.0133 | 0.0133 | TRUE | 0.3922 |
| Tnfrsf23 | 126 | 82 | 56 | 99 | 75 | 47 | 134 | 82 | 56 | 78 | 81 | 51 | 80.32 | 91 | 70 | 0.774 | -0.369 | 0.29636176 | 0.51705948 | 0.1093 | 0.0479 | 0.0772 | 0.0772 | TRUE | 0.4088 |
| Tnfrsf25 | 98 | 108 | 113 | 101 | 102 | 90 | 104 | 108 | 113 | 79 | 110 | 99 | 102.18 | 108 | 96 | 0.879 | -0.186 | 0.37722039 | 0.5922867 | 0 | 0.0391 | 0.0222 | 0.0222 | TRUE | 0.2482 |
| Tnfrsf26 | 15 | 18 | 18 | 11 | 6 | 9 | 16 | 18 | 18 | 9 | 6 | 10 | 12.82 | 17 | 8 | 0.481 | -1.057 | 0.04734647 | 0.15790833 | 0 | 0.2638 | 0.1169 | 0.1169 | TRUE | 0.1017 |
| Tnfrsf4 | 13 | 18 | 12 | 18 | 13 | 14 | 14 | 18 | 12 | 14 | 14 | 15 | 14.55 | 15 | 14 | 0.99 | -0.015 | 0.97611922 | 0.98709087 | 0 | 0.2333 | 0.1045 | 0.1045 | TRUE | 0.1858 |
| Tnfrsf8 | 73 | 67 | 62 | 64 | 59 | 49 | 78 | 67 | 62 | 50 | 63 | 54 | 62.36 | 69 | 56 | 0.808 | -0.308 | 0.22425729 | 0.43666493 | 0 | 0.0597 | 0.0301 | 0.0301 | TRUE | 0.1179 |
| Tnfrsf9 | 44 | 56 | 42 | 48 | 38 | 37 | 47 | 56 | 42 | 38 | 41 | 41 | 43.99 | 48 | 40 | 0.821 | -0.285 | 0.32809576 | 0.54915913 | 0 | 0.0818 | 0.0385 | 0.0385 | TRUE | 0.1569 |
| Tnfsf10 | 0 | 0 | 0 | 0 | 1 | 0 | 0 | 0 | 0 | 0 | 1 | 0 | 0.18 | 0 | 0 | 1.859 | 0.894 | 0.82650734 | NA | 0 | 18.3785 | 10 | 10 | TRUE | 0.5204 |
| Tnfsf11 | 80 | 34 | 19 | 46 | 35 | 18 | 85 | 34 | 19 | 36 | 38 | 20 | 38.59 | 46 | 31 | 0.681 | -0.555 | 0.31312234 | 0.5336682 | 0.3003 | 0.0923 | 0.1915 | 0.1915 | TRUE | 0.895 |

| | | | | | | | | | | | | | | | | | | | | | | | |
|---|---|---|---|---|---|---|---|---|---|---|---|---|---|---|---|---|---|---|---|---|---|---|---|
| Tnfsf12 | 334 | 501 | 531 | 435 | 332 | 396 | 355 | 500 | 533 | 342 | 357 | 433 | 420.16 | 463 | 377 | 0.815 | -0.296 | 0.10533452 | 0.27039976 | 0.0285 | 0.0147 | 0.0216 | 0.0216 | TRUE | 0.4807 |
| Tnfsf13 | 9 | 26 | 14 | 23 | 17 | 9 | 10 | 26 | 14 | 18 | 18 | 10 | 15.97 | 17 | 15 | 0.935 | -0.097 | 0.8634864 | 0.9324082 | 0.1159 | 0.2132 | 0.1664 | 0.1664 | TRUE | 0.7657 |
| Tnfsf13b | 6 | 10 | 5 | 13 | 15 | 8 | 6 | 10 | 5 | 10 | 16 | 9 | 9.42 | 7 | 12 | 1.635 | 0.709 | 0.2755475 | 0.49611133 | 0 | 0.3568 | 0.1913 | 0.1913 | TRUE | 0.3935 |
| Tnfsf14 | 0 | 0 | 0 | 0 | 0 | 0 | 0 | 0 | 0 | 0 | 0 | 0 | 0 | 0 | 0 | NA | NA | NA | NA | NA | NA | NA | NA | NA | NA |
| Tnfsf15 | 1 | 0 | 2 | 1 | 0 | 0 | 1 | 0 | 2 | 1 | 0 | 0 | 0.64 | 1 | 0 | 0.341 | -1.551 | 0.55438633 | NA | 0 | 5.1324 | 3.4582 | 3.4582 | TRUE | 0.1148 |
| Tnfsf18 | 5 | 5 | 2 | 11 | 3 | 6 | 5 | 5 | 2 | 9 | 3 | 7 | 5.13 | 4 | 6 | 1.52 | 0.604 | 0.49093074 | 0.68729309 | 0 | 0.6496 | 0.3512 | 0.3512 | TRUE | 0.2614 |
| Tnfsf4 | 3 | 4 | 2 | 1 | 5 | 3 | 3 | 4 | 2 | 1 | 5 | 3 | 3.11 | 3 | 3 | 1 | 0.001 | 0.99961605 | NA | 0 | 1.0674 | 0.5849 | 0.5849 | TRUE | 0.2784 |
| Tnfsf8 | 0 | 0 | 0 | 0 | 0 | 0 | 0 | 0 | 0 | 0 | 0 | 0 | 0 | 0 | 0 | NA | NA | NA | NA | NA | NA | NA | NA | NA | NA |
| Tnfsf9 | 43 | 73 | 68 | 65 | 44 | 48 | 46 | 73 | 68 | 51 | 47 | 53 | 56.31 | 62 | 50 | 0.807 | -0.309 | 0.27826355 | 0.49951885 | 0.0138 | 0.0654 | 0.0407 | 0.0407 | TRUE | 0.4611 |
| Tnik | 615 | 532 | 563 | 1022 | 655 | 715 | 653 | 531 | 565 | 803 | 705 | 783 | 673.45 | 583 | 764 | 1.311 | 0.39 | 0.00109907 | 0.0095718 | 0.0059 | 0.0118 | 0.0088 | 0.0088 | TRUE | 0.1282 |
| Tnip1 | 3397 | 4153 | 4269 | 3963 | 3032 | 3456 | 3608 | 4148 | 4287 | 3115 | 3262 | 3783 | 3700.53 | 4014 | 3387 | 0.843 | -0.246 | 0.02602657 | 0.10445065 | 0.0089 | 0.0078 | 0.0085 | 0.0085 | TRUE | 0.127 |
| Tnip2 | 492 | 574 | 483 | 860 | 723 | 674 | 523 | 573 | 485 | 676 | 778 | 738 | 628.77 | 527 | 731 | 1.385 | 0.47 | 4.08E-05 | 0.00065201 | 0.0044 | 0.0121 | 0.0078 | 0.0078 | TRUE | 0.0697 |
| Tnip3 | 2 | 3 | 0 | 0 | 3 | 2 | 2 | 3 | 0 | 0 | 3 | 2 | 1.76 | 2 | 2 | 1.03 | 0.043 | 0.98013422 | NA | 0 | 1.8827 | 1.5476 | 1.5476 | TRUE | 0.7078 |
| Tnk1 | 715 | 872 | 872 | 919 | 810 | 945 | 759 | 871 | 876 | 722 | 872 | 1034 | 855.72 | 835 | 876 | 1.047 | 0.067 | 0.65222591 | 0.80502827 | 0.0181 | 0.0108 | 0.0146 | 0.0146 | TRUE | 0.1978 |
| Tnk2 | 7707 | 6755 | 6717 | 7487 | 5840 | 5558 | 8186 | 6746 | 6745 | 5886 | 6283 | 6084 | 6655.06 | 7226 | 6084 | 0.842 | -0.248 | 0.01291059 | 0.06351397 | 0.0069 | 0.0074 | 0.007 | 0.007 | TRUE | 0.1649 |
| Tnks | 6647 | 7204 | 8681 | 8771 | 6785 | 6956 | 7060 | 7195 | 8717 | 6895 | 7300 | 7614 | 7463.56 | 7657 | 7270 | 0.949 | -0.075 | 0.47298675 | 0.67398619 | 0.0077 | 0.0073 | 0.0077 | 0.0077 | TRUE | 0.1789 |
| Tnks1bp1 | 1392 | 947 | 962 | 1371 | 1058 | 928 | 1479 | 946 | 966 | 1078 | 1138 | 1016 | 1103.71 | 1130 | 1077 | 0.954 | -0.069 | 0.69716628 | 0.83403726 | 0.0309 | 0.0099 | 0.0215 | 0.0215 | TRUE | 0.8699 |
| Tnks2 | 26859 | 30293 | 31581 | 32342 | 23683 | 26502 | 28529 | 30254 | 31711 | 25425 | 25481 | 29011 | 28401.73 | 30165 | 26639 | 0.883 | -0.179 | 0.03673624 | 0.13240494 | 0.0043 | 0.007 | 0.0053 | 0.0053 | TRUE | 0.0742 |
| Tnmd | 0 | 0 | 0 | 0 | 0 | 0 | 0 | 0 | 0 | 0 | 0 | 0 | 0 | 0 | 0 | NA | NA | NA | NA | NA | NA | NA | NA | NA | NA |
| Tnn | 5 | 11 | 4 | 7 | 10 | 9 | 5 | 11 | 4 | 6 | 11 | 10 | 7.74 | 7 | 9 | 1.268 | 0.343 | 0.64091923 | 0.79802464 | 0.0772 | 0.4327 | 0.2567 | 0.2567 | TRUE | 0.7812 |
| Tnnc1 | 15 | 25 | 16 | 16 | 11 | 11 | 16 | 25 | 16 | 13 | 12 | 12 | 15.57 | 19 | 12 | 0.64 | -0.643 | 0.18974006 | 0.39495581 | 0 | 0.2185 | 0.1065 | 0.1065 | TRUE | 0.4038 |
| Tnnc2 | 13 | 9 | 3 | 8 | 4 | 3 | 14 | 9 | 3 | 6 | 4 | 3 | 6.61 | 9 | 4 | 0.547 | -0.87 | 0.29807569 | 0.51872997 | 0.1799 | 0.505 | 0.34 | 0.34 | TRUE | 0.1518 |
| Tnni1 | 69 | 83 | 81 | 63 | 38 | 59 | 73 | 83 | 81 | 50 | 41 | 65 | 65.42 | 79 | 52 | 0.651 | -0.619 | 0.01855003 | 0.08239118 | 0.004 | 0.0573 | 0.0339 | 0.0339 | TRUE | 0.3599 |
| Tnni2 | 8 | 5 | 2 | 6 | 3 | 8 | 8 | 5 | 2 | 5 | 3 | 9 | 5.37 | 5 | 6 | 1.076 | 0.105 | 0.90625912 | 0.95350897 | 0.1324 | 0.6208 | 0.387 | 0.387 | TRUE | 0.238 |
| Tnni3 | 72 | 109 | 99 | 68 | 70 | 78 | 76 | 109 | 99 | 53 | 75 | 85 | 83.15 | 95 | 71 | 0.746 | -0.422 | 0.11507513 | 0.28645281 | 0.0321 | 0.0465 | 0.0394 | 0.0394 | TRUE | 0.4945 |
| Tnni3k | 0 | 0 | 0 | 0 | 0 | 0 | 0 | 0 | 0 | 0 | 0 | 0 | 0 | 0 | 0 | NA | NA | NA | NA | NA | NA | NA | NA | NA | NA |
| Tnnt1 | 256 | 377 | 382 | 344 | 266 | 271 | 272 | 377 | 384 | 270 | 286 | 297 | 314.21 | 344 | 284 | 0.826 | -0.277 | 0.09588825 | 0.25396298 | 0.0159 | 0.0174 | 0.0167 | 0.0167 | TRUE | 0.3794 |
| Tnnt2 | 12 | 20 | 17 | 14 | 17 | 2 | 13 | 20 | 17 | 11 | 18 | 2 | 13.55 | 17 | 10 | 0.633 | -0.659 | 0.33482922 | 0.55512735 | 0.2714 | 0.2501 | 0.2589 | 0.2589 | TRUE | 0.357 |
| Tnnt3 | 0 | 2 | 1 | 3 | 5 | 2 | 0 | 2 | 1 | 2 | 5 | 2 | 2.15 | 1 | 3 | 3.247 | 1.699 | 0.23448211 | NA | 0 | 1.5358 | 0.818 | 0.818 | TRUE | 0.3941 |
| Tnp1 | 5 | 1 | 4 | 5 | 3 | 0 | 5 | 1 | 4 | 4 | 3 | 0 | 2.91 | 3 | 2 | 0.712 | -0.491 | 0.70164021 | NA | 0.3787 | 1.1374 | 0.8337 | 0.8337 | TRUE | 0.6857 |
| Tnp2 | 0 | 1 | 2 | 3 | 2 | 2 | 0 | 1 | 2 | 2 | 2 | 2 | 1.62 | 1 | 2 | 2.211 | 1.145 | 0.45134287 | NA | 0 | 2.0433 | 0.9409 | 0.9409 | TRUE | 0.3211 |
| Tnpo1 | 18594 | 17764 | 18516 | 22731 | 16080 | 16062 | 19750 | 17741 | 18592 | 17869 | 17301 | 17582 | 18139.33 | 18694 | 17584 | 0.941 | -0.088 | 0.19498087 | 0.40087767 | 0.0015 | 0.0071 | 0.0033 | 0.0033 | TRUE | 0.0298 |
| Tnpo2 | 11631 | 13333 | 13669 | 18920 | 13459 | 13924 | 12354 | 13316 | 13725 | 14873 | 14481 | 15242 | 13998.6 | 13132 | 14865 | 1.132 | 0.179 | 0.01069562 | 0.05538383 | 0.0017 | 0.0071 | 0.0035 | 0.0035 | TRUE | 0.0328 |
| Tnpo3 | 13254 | 13290 | 14060 | 15273 | 10963 | 11126 | 14078 | 13273 | 14118 | 12006 | 11796 | 12179 | 12908.29 | 13823 | 11994 | 0.868 | -0.205 | 0.0006848 | 0.00665476 | 6.00E-04 | 0.0072 | 0.0025 | 0.0025 | TRUE | 0.0148 |
| Tnr | 0 | 0 | 1 | 2 | 1 | 0 | 0 | 0 | 1 | 2 | 1 | 0 | 0.61 | 0 | 1 | 2.461 | 1.299 | 0.62272026 | NA | 0 | 5.419 | 3.4832 | 3.4832 | TRUE | 0.4098 |
| Tnrc18 | 2494 | 3136 | 3412 | 2242 | 1864 | 1819 | 2649 | 3132 | 3426 | 1762 | 2006 | 1991 | 2494.37 | 3069 | 1920 | 0.625 | -0.678 | 8.91E-09 | 4.26E-07 | 0.0101 | 0.0082 | 0.0096 | 0.0096 | TRUE | 0.1738 |
| Tnrc6a | 4909 | 5350 | 5065 | 6372 | 4741 | 4618 | 5214 | 5343 | 5086 | 5009 | 5101 | 5055 | 5134.74 | 5214 | 5055 | 0.969 | -0.045 | 0.44807494 | 0.65381052 | 2.00E-04 | 0.0075 | 0.0023 | 0.0023 | TRUE | 0.0057 |
| Tnrc6b | 2232 | 2196 | 2703 | 3526 | 2333 | 2307 | 2371 | 2193 | 2714 | 2772 | 2510 | 2525 | 2514.24 | 2426 | 2602 | 1.073 | 0.102 | 0.33180059 | 0.55225401 | 0.0069 | 0.0082 | 0.0075 | 0.0075 | TRUE | 0.131 |
| Tnrc6c | 5671 | 5565 | 5820 | 5592 | 4404 | 4039 | 6024 | 5558 | 5844 | 4396 | 4738 | 4421 | 5163.51 | 5809 | 4518 | 0.778 | -0.362 | 4.49E-07 | 1.36E-05 | 0.0014 | 0.0075 | 0.0035 | 0.0035 | TRUE | 0.022 |
| Tns1 | 3474 | 3115 | 3214 | 4123 | 3139 | 2965 | 3690 | 3111 | 3227 | 3241 | 3377 | 3246 | 3315.4 | 3343 | 3288 | 0.984 | -0.024 | 0.79200232 | 0.89301263 | 0.004 | 0.0079 | 0.0055 | 0.0055 | TRUE | 0.1002 |
| Tns3 | 10895 | 6727 | 7202 | 10963 | 7654 | 6806 | 11572 | 6718 | 7232 | 8618 | 8235 | 7450 | 8304.34 | 8507 | 8101 | 0.952 | -0.07 | 0.70992122 | 0.84199711 | 0.0439 | 0.0073 | 0.0258 | 0.0258 | TRUE | 1.2129 |
| Tns4 | 68 | 74 | 83 | 54 | 50 | 61 | 72 | 74 | 83 | 42 | 54 | 67 | 65.42 | 76 | 54 | 0.705 | -0.505 | 0.05775916 | 0.18150576 | 0.011 | 0.0573 | 0.0354 | 0.0354 | TRUE | 0.3215 |
| Tnxb | 40 | 43 | 33 | 39 | 26 | 29 | 42 | 43 | 33 | 31 | 28 | 32 | 34.82 | 39 | 30 | 0.763 | -0.39 | 0.2299931 | 0.44367832 | 0 | 0.1015 | 0.047 | 0.047 | TRUE | 0.1508 |
| Tob1 | 2105 | 2438 | 2397 | 2373 | 1950 | 2067 | 2236 | 2435 | 2407 | 1865 | 2098 | 2263 | 2217.3 | 2359 | 2075 | 0.879 | -0.186 | 0.05930534 | 0.18445353 | 0.005 | 0.0084 | 0.0066 | 0.0066 | TRUE | 0.0964 |
| Tob2 | 1000 | 843 | 774 | 1331 | 980 | 849 | 1062 | 842 | 777 | 1046 | 1054 | 929 | 951.9 | 894 | 1010 | 1.131 | 0.177 | 0.1985662 | 0.4057235 | 0.0148 | 0.0104 | 0.0126 | 0.0126 | TRUE | 0.3224 |
| Toe1 | 962 | 720 | 746 | 1331 | 903 | 709 | 1022 | 719 | 749 | 1046 | 972 | 776 | 880.66 | 830 | 931 | 1.123 | 0.168 | 0.32793433 | 0.54906156 | 0.0294 | 0.0106 | 0.02 | 0.02 | TRUE | 0.4844 |
| Tollip | 3760 | 4952 | 4976 | 4546 | 3368 | 3808 | 3994 | 4946 | 4996 | 3574 | 3624 | 4168 | 4216.96 | 4645 | 3789 | 0.815 | -0.294 | 0.011952 | 0.05999451 | 0.0108 | 0.0077 | 0.0096 | 0.0096 | TRUE | 0.1832 |
| Tom1 | 860 | 985 | 992 | 1270 | 890 | 806 | 913 | 984 | 996 | 998 | 958 | 882 | 955.25 | 964 | 946 | 0.982 | -0.027 | 0.7786502 | 0.88542689 | 0.002 | 0.0104 | 0.0054 | 0.0054 | TRUE | 0.041 |
| Tom1l1 | 1297 | 1674 | 1810 | 2140 | 1637 | 1652 | 1378 | 1672 | 1817 | 1682 | 1761 | 1808 | 1686.48 | 1622 | 1750 | 1.079 | 0.109 | 0.35383141 | 0.57222912 | 0.0099 | 0.0089 | 0.0094 | 0.0094 | TRUE | 0.2092 |
| Tom1l2 | 1035 | 1158 | 1203 | 1307 | 945 | 1005 | 1099 | 1157 | 1208 | 1027 | 1017 | 1100 | 1101.36 | 1155 | 1048 | 0.907 | -0.14 | 0.10655403 | 0.27266637 | 0.0011 | 0.0099 | 0.0045 | 0.0045 | TRUE | 0.022 |
| Tomm20 | 15955 | 19260 | 19082 | 20097 | 14742 | 15379 | 16947 | 19235 | 19160 | 15799 | 15861 | 16835 | 17306.24 | 18447 | 16165 | 0.876 | -0.191 | 0.01697131 | 0.07731702 | 0.003 | 0.0071 | 0.0045 | 0.0045 | TRUE | 0.0619 |
| Tomm20l | 5 | 13 | 9 | 23 | 14 | 12 | 5 | 13 | 9 | 18 | 15 | 13 | 12.27 | 9 | 15 | 1.696 | 0.762 | 0.18382216 | 0.38659864 | 0 | 0.2754 | 0.1497 | 0.1497 | TRUE | 0.3122 |
| Tomm22 | 5129 | 5380 | 5078 | 6883 | 4946 | 4855 | 5448 | 5373 | 5099 | 5411 | 5322 | 5315 | 5327.81 | 5307 | 5349 | 1.008 | 0.012 | 0.85182917 | 0.92610806 | 5.00E-04 | 0.0075 | 0.0026 | 0.0026 | TRUE | 0.0143 |
| Tomm34 | 4095 | 4791 | 4678 | 5410 | 4182 | 4643 | 4350 | 4785 | 4697 | 4253 | 4500 | 5082 | 4611.1 | 4611 | 4612 | 1 | 0 | 0.99908755 | 0.99946777 | 0.0052 | 0.0076 | 0.0062 | 0.0062 | TRUE | 0.097 |
| Tomm40 | 7559 | 7014 | 6549 | 11147 | 7872 | 6602 | 8029 | 7005 | 6576 | 8763 | 8470 | 7227 | 7678.24 | 7203 | 8153 | 1.132 | 0.179 | 0.11206424 | 0.28149099 | 0.0102 | 0.0073 | 0.009 | 0.009 | TRUE | 0.1227 |
| Tomm40l | 1168 | 1364 | 1307 | 1693 | 1294 | 1175 | 1241 | 1362 | 1312 | 1331 | 1392 | 1286 | 1320.77 | 1305 | 1336 | 1.024 | 0.034 | 0.68363222 | 0.82477223 | 0.001 | 0.0094 | 0.0043 | 0.0043 | TRUE | 0.0223 |
| Tomm5 | 2455 | 2435 | 2164 | 3070 | 2076 | 2026 | 2608 | 2432 | 2173 | 2413 | 2234 | 2218 | 2346.2 | 2404 | 2288 | 0.952 | -0.071 | 0.4635321 | 0.66686857 | 0.0046 | 0.0083 | 0.0063 | 0.0063 | TRUE | 0.0859 |
| Tomm6 | 3768 | 3685 | 3582 | 4433 | 3307 | 3219 | 4002 | 3680 | 3597 | 3485 | 3558 | 3524 | 3640.99 | 3760 | 3522 | 0.937 | -0.094 | 0.19726524 | 0.40387778 | 0.0013 | 0.0078 | 0.0036 | 0.0036 | TRUE | 0.0386 |
| Tomm7 | 3612 | 4756 | 4598 | 5002 | 3713 | 3903 | 3837 | 4750 | 4617 | 3932 | 3995 | 4272 | 4233.82 | 4401 | 4066 | 0.924 | -0.114 | 0.26836226 | 0.48824657 | 0.0069 | 0.0077 | 0.0075 | 0.0075 | TRUE | 0.1531 |
| Tomm70a | 18630 | 18542 | 18557 | 24755 | 17918 | 16961 | 19788 | 18518 | 18633 | 19460 | 19279 | 18566 | 19040.87 | 18980 | 19102 | 1.006 | 0.009 | 0.88127776 | 0.94135674 | 8.00E-04 | 0.0071 | 0.0028 | 0.0028 | TRUE | 0.017 |
| Tomt | 0 | 1 | 0 | 1 | 0 | 0 | 0 | 1 | 0 | 1 | 0 | 0 | 0.3 | 0 | 0 | 0.954 | -0.067 | 0.98642001 | NA | 0 | 11.0817 | 9.3014 | 9.3014 | TRUE | 0.405 |
| Tonsl | 2413 | 2303 | 2520 | 3344 | 2642 | 2498 | 2563 | 2300 | 2530 | 2629 | 2843 | 2734 | 2599.88 | 2464 | 2735 | 1.11 | 0.15 | 0.0630026 | 0.1924239 | 0.0019 | 0.0082 | 0.0043 | 0.0043 | TRUE | 0.0414 |
| Top1 | 8735 | 8278 | 8082 | 11488 | 8091 | 7551 | 9278 | 8267 | 8115 | 9031 | 8705 | 8266 | 8610.46 | 8553 | 8667 | 1.013 | 0.019 | 0.81737303 | 0.90708637 | 0.0029 | 0.0073 | 0.0049 | 0.0049 | TRUE | 0.067 |
| Top1mt | 1010 | 1095 | 1099 | 1150 | 919 | 977 | 1073 | 1094 | 1104 | 904 | 989 | 1069 | 1038.7 | 1090 | 987 | 0.905 | -0.145 | 0.13271566 | 0.31409066 | 0.0027 | 0.0101 | 0.0057 | 0.0057 | TRUE | 0.0677 |
| Top2a | 65636 | 85674 | 93193 | 82905 | 61722 | 68699 | 69717 | 85563 | 93576 | 65173 | 66409 | 75202 | 75940.07 | 82952 | 68928 | 0.831 | -0.267 | 0.02972147 | 0.11430556 | 0.0138 | 0.0069 | 0.0109 | 0.0109 | TRUE | 0.2386 |
| Top2b | 4072 | 4509 | 5068 | 5799 | 4042 | 4357 | 4325 | 4503 | 5089 | 4559 | 4349 | 4769 | 4599.04 | 4639 | 4559 | 0.983 | -0.025 | 0.78064851 | 0.88648487 | 0.0045 | 0.0076 | 0.0057 | 0.0057 | TRUE | 0.0877 |
| Top3a | 2129 | 1701 | 1630 | 2599 | 1836 | 1628 | 2261 | 1699 | 1637 | 2043 | 1975 | 1782 | 1899.59 | 1866 | 1933 | 1.037 | 0.052 | 0.7090797 | 0.84150527 | 0.018 | 0.0086 | 0.0136 | 0.0136 | TRUE | 0.4154 |
| Top3b | 3042 | 2962 | 2904 | 5353 | 3830 | 3600 | 3231 | 2958 | 2916 | 4208 | 4121 | 3941 | 3562.49 | 3035 | 4090 | 1.348 | 0.431 | 1.71E-08 | 7.80E-07 | 0.0017 | 0.0078 | 0.0039 | 0.0039 | TRUE | 0.0387 |
| Topaz1 | 46 | 50 | 53 | 152 | 104 | 84 | 49 | 50 | 53 | 119 | 112 | 92 | 79.23 | 51 | 108 | 2.133 | 1.093 | 3.25E-06 | 7.65E-05 | 0 | 0.0485 | 0.0251 | 0.0251 | TRUE | 0.1575 |

| | | | | | | | | | | | | | | | | | | | | | | | | | |
|---|---|---|---|---|---|---|---|---|---|---|---|---|---|---|---|---|---|---|---|---|---|---|---|---|---|
| Topbp1 | 12564 | 13024 | 13266 | 17248 | 12899 | 13235 | 13345 | 13007 | 13320 | 13559 | 13879 | 14488 | 13599.7 | 13224 | 13975 | 1.057 | 0.08 | 0.18111975 | 0.38280307 | 6.00E-04 | 0.0071 | 0.0025 | 0.0025 | TRUE | 0.0126 |
| Topors | 3343 | 3053 | 3068 | 4875 | 3399 | 3122 | 3551 | 3049 | 3081 | 3832 | 3657 | 3418 | 3431.25 | 3227 | 3636 | 1.127 | 0.173 | 0.06786055 | 0.2016764 | 0.005 | 0.0079 | 0.0061 | 0.0061 | TRUE | 0.0936 |
| Tor1a | 1003 | 1180 | 1177 | 1462 | 1028 | 960 | 1065 | 1178 | 1182 | 1149 | 1106 | 1051 | 1121.99 | 1142 | 1102 | 0.966 | -0.05 | 0.57675191 | 0.75440349 | 0.0018 | 0.0098 | 0.005 | 0.005 | TRUE | 0.0409 |
| Tor1aip1 | 645 | 700 | 616 | 1112 | 754 | 606 | 685 | 699 | 619 | 874 | 811 | 663 | 725.25 | 668 | 783 | 1.174 | 0.232 | 0.07812378 | 0.22111084 | 0.0108 | 0.0114 | 0.0111 | 0.0111 | TRUE | 0.2094 |
| Tor1aip2 | 8042 | 8167 | 7650 | 10328 | 7472 | 7289 | 8542 | 8156 | 7681 | 8119 | 8039 | 7979 | 8086.22 | 8126 | 8046 | 0.99 | -0.014 | 0.83411111 | 0.91571799 | 0.0011 | 0.0073 | 0.0032 | 0.0032 | TRUE | 0.0281 |
| Tor1b | 1253 | 1205 | 1116 | 1644 | 1287 | 1036 | 1331 | 1203 | 1121 | 1292 | 1385 | 1134 | 1244.35 | 1218 | 1270 | 1.043 | 0.061 | 0.59460594 | 0.7677918 | 0.0079 | 0.0096 | 0.0086 | 0.0086 | TRUE | 0.1041 |
| Tor2a | 1344 | 1658 | 1586 | 1949 | 1377 | 1517 | 1428 | 1656 | 1593 | 1532 | 1482 | 1661 | 1558.38 | 1559 | 1558 | 0.999 | -0.001 | 0.9914597 | 0.99557064 | 0.0039 | 0.009 | 0.006 | 0.006 | TRUE | 0.0649 |
| Tor3a | 623 | 703 | 779 | 981 | 717 | 797 | 662 | 702 | 782 | 771 | 771 | 872 | 760.18 | 715 | 805 | 1.124 | 0.169 | 0.128092 | 0.30785004 | 0.0048 | 0.0112 | 0.0076 | 0.0076 | TRUE | 0.0791 |
| Tor4a | 219 | 279 | 281 | 354 | 317 | 266 | 233 | 279 | 282 | 278 | 341 | 291 | 283.99 | 265 | 303 | 1.145 | 0.196 | 0.1960749 | 0.40230395 | 0.0079 | 0.0185 | 0.0129 | 0.0129 | TRUE | 0.1307 |
| Tox | 0 | 1 | 2 | 2 | 1 | 2 | 0 | 1 | 2 | 2 | 1 | 2 | 1.31 | 1 | 2 | 1.592 | 0.671 | 0.6851841 | NA | 0 | 2.5267 | 1.1664 | 1.1664 | TRUE | 0.2612 |
| Tox2 | 60 | 57 | 48 | 70 | 45 | 48 | 64 | 57 | 48 | 55 | 48 | 53 | 54.14 | 56 | 52 | 0.927 | -0.11 | 0.67958871 | 0.82209488 | 0 | 0.0678 | 0.0326 | 0.0326 | TRUE | 0.134 |
| Tox3 | 55 | 80 | 80 | 117 | 117 | 89 | 58 | 80 | 80 | 92 | 126 | 97 | 88.99 | 73 | 105 | 1.436 | 0.522 | 0.03259413 | 0.12164567 | 0.0182 | 0.0439 | 0.0314 | 0.0314 | TRUE | 0.289 |
| Tox4 | 5385 | 5695 | 5807 | 7236 | 5398 | 5120 | 5720 | 5688 | 5831 | 5688 | 5808 | 5605 | 5723.21 | 5746 | 5700 | 0.992 | -0.012 | 0.84000499 | 0.91895392 | 0 | 0.0075 | 0.0022 | 0.0022 | TRUE | 0.0033 |
| Tpbg | 176 | 224 | 255 | 457 | 369 | 363 | 187 | 224 | 256 | 359 | 397 | 397 | 303.39 | 222 | 384 | 1.727 | 0.788 | 2.53E-07 | 8.14E-06 | 0.0084 | 0.0178 | 0.0133 | 0.0133 | TRUE | 0.2086 |
| Tpbpa | 0 | 0 | 0 | 0 | 0 | 0 | 0 | 0 | 0 | 0 | 0 | 0 | 0 | 0 | 0 | NA | NA | NA | NA | NA | NA | NA | NA | NA | NA |
| Tpbpb | 0 | 0 | 0 | 0 | 0 | 0 | 0 | 0 | 0 | 0 | 0 | 0 | 0 | 0 | 0 | NA | NA | NA | NA | NA | NA | NA | NA | NA | NA |
| Tpcn1 | 1472 | 1695 | 1754 | 2110 | 1484 | 1589 | 1564 | 1693 | 1761 | 1659 | 1597 | 1739 | 1668.73 | 1673 | 1665 | 0.995 | -0.007 | 0.93614947 | 0.96816629 | 0.0021 | 0.0089 | 0.0047 | 0.0047 | TRUE | 0.039 |
| Tpcn2 | 243 | 168 | 171 | 337 | 244 | 183 | 258 | 168 | 172 | 265 | 263 | 200 | 220.89 | 199 | 243 | 1.22 | 0.287 | 0.1856568 | 0.38923023 | 0.0374 | 0.0218 | 0.0294 | 0.0294 | TRUE | 0.7205 |
| Tpd52 | 8463 | 12738 | 12640 | 11552 | 9125 | 10771 | 8989 | 12722 | 12692 | 9081 | 9818 | 11791 | 10848.74 | 11468 | 10230 | 0.892 | -0.165 | 0.29110591 | 0.51183021 | 0.0261 | 0.0072 | 0.0175 | 0.0175 | TRUE | 0.4368 |
| Tpd52l1 | 803 | 1057 | 1010 | 1154 | 797 | 986 | 853 | 1056 | 1014 | 907 | 858 | 1079 | 961.12 | 974 | 948 | 0.972 | -0.04 | 0.75964124 | 0.87323722 | 0.0124 | 0.0103 | 0.0115 | 0.0115 | TRUE | 0.1728 |
| Tpd52l2 | 3902 | 3973 | 3937 | 5229 | 3753 | 3516 | 4145 | 3968 | 3953 | 4111 | 4038 | 3849 | 4010.52 | 4022 | 3999 | 0.995 | -0.008 | 0.90685598 | 0.95352883 | 7.00E-04 | 0.0077 | 0.0029 | 0.0029 | TRUE | 0.0131 |
| Tpgs1 | 366 | 563 | 632 | 601 | 482 | 430 | 389 | 562 | 635 | 472 | 519 | 471 | 507.9 | 529 | 487 | 0.921 | -0.118 | 0.5150062 | 0.70743379 | 0.0303 | 0.0134 | 0.0218 | 0.0218 | TRUE | 0.6218 |
| Tpgs2 | 1230 | 1343 | 1493 | 1778 | 1222 | 1246 | 1306 | 1341 | 1499 | 1398 | 1315 | 1364 | 1370.56 | 1382 | 1359 | 0.983 | -0.024 | 0.78696838 | 0.89032073 | 0.0025 | 0.0093 | 0.0051 | 0.0051 | TRUE | 0.0659 |
| Tph1 | 25 | 4 | 8 | 17 | 12 | 5 | 27 | 4 | 8 | 13 | 13 | 5 | 11.72 | 13 | 10 | 0.831 | -0.267 | 0.73476307 | 0.85771731 | 0.436 | 0.288 | 0.362 | 0.362 | TRUE | 1.7749 |
| Tph2 | 22 | 22 | 19 | 53 | 33 | 38 | 23 | 22 | 19 | 42 | 36 | 42 | 30.53 | 21 | 40 | 1.848 | 0.886 | 0.01100867 | 0.05654356 | 0 | 0.1148 | 0.0514 | 0.0514 | TRUE | 0.0574 |
| Tpi1 | 21091 | 34093 | 36234 | 35814 | 29175 | 36879 | 22402 | 34049 | 36383 | 28154 | 31391 | 40370 | 32124.78 | 30945 | 33305 | 1.076 | 0.106 | 0.57944117 | 0.75622227 | 0.0389 | 0.007 | 0.0263 | 0.0263 | TRUE | 0.7138 |
| Tpk1 | 348 | 368 | 386 | 695 | 522 | 458 | 370 | 368 | 388 | 546 | 562 | 501 | 455.68 | 375 | 536 | 1.431 | 0.518 | 2.10E-06 | 5.26E-05 | 0 | 0.0141 | 0.0063 | 0.0063 | TRUE | 0.0368 |
| Tpm1 | 2326 | 2767 | 2652 | 3614 | 2570 | 2502 | 2471 | 2763 | 2663 | 2841 | 2765 | 2739 | 2707 | 2632 | 2782 | 1.057 | 0.08 | 0.2928143 | 0.51382589 | 0.0012 | 0.0081 | 0.0038 | 0.0038 | TRUE | 0.0349 |
| Tpm2 | 526 | 579 | 572 | 504 | 399 | 390 | 559 | 578 | 574 | 396 | 429 | 427 | 493.95 | 570 | 417 | 0.731 | -0.453 | 9.42E-06 | 0.0001893 | 0 | 0.0136 | 0.0055 | 0.0055 | TRUE | 0.0248 |
| Tpm3 | 5581 | 4946 | 5023 | 8892 | 6234 | 5195 | 5928 | 4940 | 5044 | 6990 | 6707 | 5687 | 5882.6 | 5304 | 6461 | 1.219 | 0.285 | 0.01285314 | 0.06328543 | 0.0106 | 0.0075 | 0.0093 | 0.0093 | TRUE | 0.134 |
| Tpm4 | 2727 | 3515 | 3660 | 5328 | 3896 | 3948 | 2897 | 3510 | 3675 | 4188 | 4192 | 4322 | 3797.34 | 3361 | 4234 | 1.26 | 0.333 | 0.00149699 | 0.01223981 | 0.0072 | 0.0078 | 0.0077 | 0.0077 | TRUE | 0.1771 |
| Tpmt | 739 | 696 | 736 | 991 | 696 | 731 | 785 | 695 | 739 | 779 | 749 | 800 | 757.86 | 740 | 776 | 1.049 | 0.07 | 0.46616859 | 0.66857491 | 9.00E-04 | 0.0113 | 0.0053 | 0.0053 | TRUE | 0.0336 |
| Tpo | 11 | 18 | 25 | 28 | 19 | 30 | 12 | 18 | 25 | 22 | 20 | 33 | 21.68 | 18 | 25 | 1.366 | 0.45 | 0.33130161 | 0.55212268 | 0.0481 | 0.1589 | 0.1074 | 0.1074 | TRUE | 0.2906 |
| Tpp1 | 1864 | 2168 | 2404 | 2663 | 1870 | 2035 | 1980 | 2165 | 2414 | 2093 | 2012 | 2228 | 2148.67 | 2186 | 2111 | 0.965 | -0.051 | 0.61407767 | 0.78169758 | 0.0058 | 0.0084 | 0.0069 | 0.0069 | TRUE | 0.1004 |
| Tpp2 | 7234 | 7185 | 7359 | 8872 | 6428 | 6364 | 7684 | 7176 | 7389 | 6974 | 6916 | 6966 | 7184.29 | 7416 | 6952 | 0.937 | -0.093 | 0.12352928 | 0.29955471 | 4.00E-04 | 0.0074 | 0.0025 | 0.0025 | TRUE | 0.0121 |
| Tppp | 58 | 109 | 74 | 92 | 77 | 59 | 62 | 109 | 74 | 72 | 83 | 65 | 77.42 | 82 | 73 | 0.897 | -0.157 | 0.57512062 | 0.75313869 | 0.0381 | 0.0495 | 0.0439 | 0.0439 | TRUE | 0.7737 |
| Tppp2 | 1 | 0 | 0 | 0 | 0 | 0 | 1 | 0 | 0 | 0 | 0 | 0 | 0.18 | 0 | 0 | 0.49 | -1.029 | 0.80087 | NA | 0 | 18.6165 | 10 | 10 | TRUE | 0.4614 |
| Tppp3 | 811 | 922 | 932 | 561 | 408 | 435 | 861 | 921 | 936 | 441 | 439 | 476 | 679.04 | 906 | 452 | 0.498 | -1.004 | 9.21E-25 | 3.73E-22 | 0 | 0.0118 | 0.0052 | 0.0052 | TRUE | 0.0243 |
| Tpr | 15442 | 12190 | 13423 | 23355 | 15906 | 14074 | 16402 | 12174 | 13478 | 18360 | 17114 | 15406 | 15489.08 | 14018 | 16960 | 1.21 | 0.275 | 0.03040397 | 0.1163629 | 0.0157 | 0.0071 | 0.0116 | 0.0116 | TRUE | 0.2706 |
| Tpra1 | 440 | 584 | 537 | 565 | 483 | 517 | 467 | 583 | 539 | 444 | 520 | 566 | 519.93 | 530 | 510 | 0.96 | -0.059 | 0.67850843 | 0.82135505 | 0.0118 | 0.0132 | 0.0125 | 0.0125 | TRUE | 0.1548 |
| Tprg | 1 | 0 | 0 | 0 | 1 | 0 | 1 | 0 | 0 | 0 | 1 | 0 | 0.36 | 0 | 0 | 0.954 | -0.067 | 0.98557668 | NA | 0 | 9.2518 | 8.0179 | 8.0179 | TRUE | 0.5204 |
| Tprgl | 3386 | 3963 | 3711 | 4189 | 3051 | 3085 | 3597 | 3958 | 3726 | 3293 | 3283 | 3377 | 3538.9 | 3760 | 3318 | 0.882 | -0.181 | 0.01024647 | 0.05372582 | 9.00E-04 | 0.0078 | 0.0033 | 0.0033 | TRUE | 0.0258 |
| Tprkb | 2273 | 2150 | 2080 | 2600 | 1804 | 1748 | 2414 | 2147 | 2089 | 2044 | 1941 | 1913 | 2091.41 | 2217 | 1966 | 0.887 | -0.172 | 0.05210848 | 0.16873862 | 0.003 | 0.0085 | 0.0052 | 0.0052 | TRUE | 0.0734 |
| Tprn | 1088 | 1196 | 1118 | 1612 | 1282 | 1246 | 1156 | 1194 | 1123 | 1267 | 1379 | 1364 | 1247.2 | 1158 | 1337 | 1.154 | 0.207 | 0.01250093 | 0.06198093 | 9.00E-04 | 0.0095 | 0.0041 | 0.0041 | TRUE | 0.0259 |
| Tpsab1 | 0 | 0 | 0 | 0 | 0 | 0 | 0 | 0 | 0 | 0 | 0 | 0 | 0 | 0 | 0 | NA | NA | NA | NA | NA | NA | NA | NA | NA | NA |
| Tpsb2 | 0 | 0 | 0 | 0 | 1 | 0 | 0 | 0 | 0 | 0 | 1 | 0 | 0.18 | 0 | 0 | 1.859 | 0.894 | 0.82650734 | NA | 0 | 18.3785 | 10 | 10 | TRUE | 0.5204 |
| Tpsg1 | 0 | 0 | 3 | 0 | 0 | 1 | 0 | 0 | 3 | 0 | 0 | 1 | 0.68 | 1 | 0 | 0.347 | -1.528 | 0.62224563 | NA | 7.9002 | 4.8199 | 5.4225 | 5.4225 | TRUE | 1.4421 |
| Tpst1 | 3059 | 3045 | 3020 | 4349 | 3078 | 3077 | 3249 | 3041 | 3032 | 3419 | 3312 | 3368 | 3236.92 | 3107 | 3366 | 1.084 | 0.116 | 0.08672198 | 0.23803391 | 6.00E-04 | 0.0079 | 0.003 | 0.003 | TRUE | 0.0192 |
| Tpst2 | 1645 | 1437 | 1337 | 1641 | 1115 | 1126 | 1747 | 1435 | 1342 | 1290 | 1200 | 1233 | 1374.53 | 1508 | 1241 | 0.823 | -0.281 | 0.01813508 | 0.08107328 | 0.0095 | 0.0093 | 0.0095 | 0.0095 | TRUE | 0.2307 |
| Tpt1 | 45298 | 48947 | 49527 | 60213 | 41384 | 41084 | 48114 | 48884 | 49730 | 47335 | 44527 | 44973 | 47260.49 | 48909 | 45612 | 0.933 | -0.101 | 0.08352269 | 0.23155281 | 7.00E-04 | 0.007 | 0.0024 | 0.0024 | TRUE | 0.0134 |
| Tpte | 6 | 8 | 15 | 8 | 10 | 9 | 6 | 8 | 15 | 6 | 11 | 10 | 9.39 | 10 | 9 | 0.902 | -0.148 | 0.82185072 | 0.909056 | 0 | 0.3578 | 0.2052 | 0.2052 | TRUE | 0.7563 |
| Tpx2 | 11823 | 13728 | 13743 | 15474 | 11130 | 11458 | 12558 | 13710 | 13799 | 12164 | 11975 | 12543 | 12791.67 | 13356 | 12227 | 0.915 | -0.127 | 0.06529654 | 0.19695312 | 0.0015 | 0.0072 | 0.0034 | 0.0034 | TRUE | 0.0334 |
| Tra2a | 6025 | 6602 | 6573 | 7992 | 6167 | 6578 | 6400 | 6593 | 6600 | 6283 | 6635 | 7201 | 6618.62 | 6531 | 6706 | 1.027 | 0.038 | 0.623275 | 0.78749446 | 0.0021 | 0.0074 | 0.0041 | 0.0041 | TRUE | 0.0507 |
| Tra2b | 7531 | 6990 | 6955 | 11401 | 7985 | 7132 | 7999 | 6981 | 6984 | 8963 | 8591 | 7807 | 7887.47 | 7321 | 8454 | 1.155 | 0.208 | 0.02656692 | 0.1060525 | 0.0055 | 0.0073 | 0.0062 | 0.0062 | TRUE | 0.08 |
| Trabd | 1635 | 1765 | 1636 | 2209 | 1573 | 1546 | 1737 | 1763 | 1643 | 1737 | 1692 | 1692 | 1710.57 | 1714 | 1707 | 0.996 | -0.005 | 0.94101729 | 0.97010197 | 0 | 0.0088 | 0.0033 | 0.0033 | TRUE | 0.0161 |
| Trabd2b | 22 | 20 | 22 | 59 | 62 | 31 | 23 | 20 | 22 | 46 | 67 | 34 | 35.41 | 22 | 49 | 2.247 | 1.168 | 0.00223617 | 0.01678672 | 0.0439 | 0.0999 | 0.0717 | 0.0717 | TRUE | 0.192 |
| Tradd | 495 | 698 | 730 | 723 | 569 | 580 | 526 | 697 | 733 | 568 | 612 | 635 | 628.56 | 652 | 605 | 0.927 | -0.109 | 0.45612134 | 0.66096526 | 0.0153 | 0.0121 | 0.0138 | 0.0138 | TRUE | 0.3358 |
| Traf1 | 31 | 42 | 44 | 60 | 49 | 42 | 33 | 42 | 44 | 47 | 53 | 46 | 44.15 | 40 | 49 | 1.222 | 0.29 | 0.32234481 | 0.54362942 | 0 | 0.0815 | 0.0389 | 0.0389 | TRUE | 0.1599 |
| Traf2 | 557 | 611 | 540 | 878 | 598 | 625 | 592 | 610 | 542 | 690 | 643 | 684 | 626.97 | 581 | 672 | 1.158 | 0.211 | 0.03615384 | 0.13078403 | 9.00E-04 | 0.0122 | 0.0057 | 0.0057 | TRUE | 0.041 |
| Traf3 | 1277 | 1290 | 1350 | 2041 | 1452 | 1269 | 1356 | 1288 | 1356 | 1604 | 1562 | 1389 | 1426.02 | 1333 | 1518 | 1.14 | 0.189 | 0.0376067 | 0.13464991 | 0.0027 | 0.0092 | 0.0052 | 0.0052 | TRUE | 0.0663 |
| Traf3ip1 | 905 | 858 | 744 | 1302 | 992 | 942 | 961 | 857 | 747 | 1024 | 1067 | 1031 | 947.87 | 855 | 1041 | 1.217 | 0.284 | 0.01332282 | 0.06484511 | 0.0065 | 0.0104 | 0.0084 | 0.0084 | TRUE | 0.1454 |
| Traf3ip2 | 695 | 600 | 647 | 729 | 558 | 544 | 738 | 599 | 650 | 573 | 600 | 595 | 626.01 | 662 | 589 | 0.89 | -0.168 | 0.14064727 | 0.32593224 | 0.0044 | 0.0122 | 0.0078 | 0.0078 | TRUE | 0.1175 |
| Traf3ip3 | 2 | 1 | 0 | 2 | 2 | 0 | 2 | 1 | 0 | 2 | 2 | 0 | 1.14 | 1 | 1 | 1.213 | 0.279 | 0.8846965 | NA | 0 | 2.8937 | 1.7759 | 1.7759 | TRUE | 0.2368 |
| Traf4 | 6256 | 7170 | 6757 | 6570 | 4757 | 5244 | 6645 | 7161 | 6785 | 5165 | 5118 | 5740 | 6102.32 | 6864 | 5341 | 0.778 | -0.362 | 4.54E-06 | 0.00010167 | 0.0026 | 0.0074 | 0.0043 | 0.0043 | TRUE | 0.0519 |
| Traf5 | 61 | 56 | 47 | 82 | 81 | 60 | 65 | 56 | 47 | 64 | 87 | 66 | 64.2 | 56 | 72 | 1.292 | 0.369 | 0.16782695 | 0.36558665 | 0.0119 | 0.0582 | 0.0357 | 0.0357 | TRUE | 0.277 |
| Traf6 | 1361 | 1230 | 1230 | 2039 | 1410 | 1243 | 1446 | 1228 | 1235 | 1603 | 1517 | 1361 | 1398.29 | 1303 | 1494 | 1.147 | 0.198 | 0.07173949 | 0.20895202 | 0.0068 | 0.0093 | 0.008 | 0.008 | TRUE | 0.1097 |
| Traf7 | 8034 | 7407 | 7425 | 9213 | 6906 | 6384 | 8534 | 7397 | 7455 | 7243 | 7430 | 6988 | 7507.95 | 7795 | 7220 | 0.926 | -0.11 | 0.18995447 | 0.39508927 | 0.0036 | 0.0073 | 0.005 | 0.005 | TRUE | 0.0837 |
| Trafd1 | 790 | 733 | 695 | 1121 | 831 | 774 | 839 | 732 | 698 | 881 | 894 | 847 | 815.27 | 756 | 874 | 1.156 | 0.21 | 0.04619581 | 0.1555841 | 0.0033 | 0.0109 | 0.0067 | 0.0067 | TRUE | 0.1078 |
| Traip | 1694 | 1556 | 1511 | 1996 | 1511 | 1380 | 1799 | 1554 | 1517 | 1569 | 1626 | 1511 | 1596 | 1623 | 1569 | 0.966 | -0.049 | 0.61206071 | 0.78028154 | 0.0039 | 0.009 | 0.0062 | 0.0062 | TRUE | 0.1078 |

| | | | | | | | | | | | | | | | | | | | | | | | | | |
|---|---|---|---|---|---|---|---|---|---|---|---|---|---|---|---|---|---|---|---|---|---|---|---|---|---|
| Trak1 | 2621 | 3132 | 3378 | 3253 | 2505 | 2824 | 2784 | 3128 | 3392 | 2557 | 2695 | 3091 | 2941.26 | 3101 | 2781 | 0.896 | -0.158 | 0.16133813 | 0.35629025 | 0.0093 | 0.008 | 0.0088 | 0.0088 | TRUE | 0.1151 |
| Trak2 | 4118 | 3305 | 3228 | 4765 | 3262 | 2735 | 4374 | 3301 | 3241 | 3746 | 3510 | 2994 | 3527.58 | 3639 | 3417 | 0.939 | -0.091 | 0.52728147 | 0.71611843 | 0.0187 | 0.0078 | 0.0145 | 0.0145 | TRUE | 0.3799 |
| Tram1 | 3764 | 4061 | 3994 | 5786 | 4178 | 4158 | 3998 | 4056 | 4010 | 4548 | 4495 | 4552 | 4276.59 | 4021 | 4532 | 1.127 | 0.172 | 0.00241201 | 0.01785178 | 0 | 0.0077 | 0.0021 | 0.0021 | TRUE | 7.00E-04 |
| Tram1l1 | 52 | 90 | 85 | 189 | 126 | 105 | 55 | 90 | 85 | 149 | 136 | 115 | 104.93 | 77 | 133 | 1.733 | 0.793 | 0.00120266 | 0.01025873 | 0.0266 | 0.0383 | 0.0329 | 0.0329 | TRUE | 0.5425 |
| Tram2 | 294 | 403 | 415 | 508 | 384 | 427 | 312 | 402 | 417 | 399 | 413 | 467 | 401.9 | 377 | 426 | 1.129 | 0.176 | 0.24216335 | 0.45878684 | 0.0125 | 0.0151 | 0.0137 | 0.0137 | TRUE | 0.2572 |
| Trank1 | 380 | 469 | 473 | 585 | 444 | 425 | 404 | 468 | 475 | 460 | 478 | 465 | 458.3 | 449 | 468 | 1.041 | 0.057 | 0.61703685 | 0.7832683 | 0.0017 | 0.0141 | 0.0073 | 0.0073 | TRUE | 0.0894 |
| Trap1 | 10630 | 12241 | 12539 | 13077 | 9633 | 9444 | 11291 | 12225 | 12590 | 10280 | 10365 | 10338 | 11181.53 | 12035 | 10328 | 0.858 | -0.221 | 0.00133548 | 0.01114453 | 0.0014 | 0.0072 | 0.0033 | 0.0033 | TRUE | 0.0358 |
| Trap1a | 4667 | 5458 | 5448 | 7528 | 5285 | 5194 | 4957 | 5451 | 5470 | 5918 | 5686 | 5686 | 5528.07 | 5293 | 5763 | 1.089 | 0.123 | 0.0892685 | 0.24287437 | 0.0015 | 0.0075 | 0.0036 | 0.0036 | TRUE | 0.0374 |
| Trappc1 | 1406 | 1328 | 1262 | 2073 | 1430 | 1416 | 1493 | 1326 | 1267 | 1630 | 1539 | 1550 | 1467.52 | 1362 | 1573 | 1.155 | 0.208 | 0.02680415 | 0.10681028 | 0.0032 | 0.0091 | 0.0057 | 0.0057 | TRUE | 0.0848 |
| Trappc10 | 2577 | 2933 | 2970 | 3116 | 2285 | 2368 | 2737 | 2929 | 2982 | 2450 | 2459 | 2592 | 2691.48 | 2883 | 2500 | 0.867 | -0.206 | 0.00532042 | 0.03274268 | 0.0011 | 0.0081 | 0.0036 | 0.0036 | TRUE | 0.0236 |
| Trappc11 | 6487 | 5874 | 6296 | 8165 | 5729 | 5732 | 6890 | 5866 | 6322 | 6419 | 6164 | 6275 | 6322.65 | 6359 | 6286 | 0.989 | -0.017 | 0.84077789 | 0.91941683 | 0.0033 | 0.0074 | 0.0048 | 0.0048 | TRUE | 0.065 |
| Trappc12 | 1599 | 1723 | 1847 | 2172 | 1631 | 1624 | 1698 | 1721 | 1855 | 1707 | 1755 | 1778 | 1752.3 | 1758 | 1747 | 0.993 | -0.01 | 0.89801546 | 0.95044288 | 8.00E-04 | 0.0088 | 0.0037 | 0.0037 | TRUE | 0.028 |
| Trappc13 | 2262 | 2228 | 2196 | 3419 | 2257 | 2057 | 2403 | 2225 | 2205 | 2688 | 2428 | 2252 | 2366.77 | 2278 | 2456 | 1.079 | 0.11 | 0.24986085 | 0.46722738 | 0.0045 | 0.0083 | 0.0061 | 0.0061 | TRUE | 0.0838 |
| Trappc2 | 456 | 427 | 387 | 574 | 374 | 380 | 484 | 426 | 389 | 451 | 402 | 416 | 428.17 | 433 | 423 | 0.979 | -0.031 | 0.81078857 | 0.90283516 | 0.0056 | 0.0146 | 0.0097 | 0.0097 | TRUE | 0.1227 |
| Trappc2l | 729 | 891 | 854 | 1165 | 807 | 899 | 774 | 890 | 858 | 916 | 868 | 984 | 881.65 | 841 | 923 | 1.097 | 0.134 | 0.18955985 | 0.3947026 | 0.0033 | 0.0106 | 0.0064 | 0.0064 | TRUE | 0.056 |
| Trappc3 | 1873 | 1641 | 1470 | 2141 | 1460 | 1355 | 1989 | 1639 | 1476 | 1683 | 1571 | 1483 | 1640.27 | 1701 | 1579 | 0.929 | -0.107 | 0.40229022 | 0.61511598 | 0.0128 | 0.0089 | 0.0111 | 0.0111 | TRUE | 0.264 |
| Trappc4 | 1926 | 2106 | 2102 | 2320 | 1638 | 1732 | 2046 | 2103 | 2111 | 1824 | 1762 | 1896 | 1956.97 | 2087 | 1827 | 0.876 | -0.192 | 0.00729207 | 0.04140046 | 2.00E-04 | 0.0086 | 0.0032 | 0.0032 | TRUE | 0.0126 |
| Trappc5 | 815 | 861 | 897 | 1253 | 924 | 830 | 866 | 860 | 901 | 985 | 994 | 909 | 919 | 876 | 963 | 1.1 | 0.138 | 0.11603808 | 0.28766939 | 0 | 0.0105 | 0.0044 | 0.0044 | TRUE | 0.028 |
| Trappc6a | 354 | 699 | 722 | 631 | 450 | 562 | 376 | 698 | 725 | 496 | 484 | 615 | 565.75 | 600 | 532 | 0.886 | -0.174 | 0.45823981 | 0.66275638 | 0.0682 | 0.0127 | 0.038 | 0.038 | TRUE | 1.2461 |
| Trappc6b | 1410 | 1472 | 1442 | 2049 | 1429 | 1365 | 1498 | 1470 | 1448 | 1611 | 1538 | 1494 | 1509.7 | 1472 | 1548 | 1.052 | 0.073 | 0.3312236 | 0.55212268 | 0 | 0.0091 | 0.0034 | 0.0034 | TRUE | 0.0159 |
| Trappc8 | 6085 | 5606 | 6129 | 7299 | 5325 | 5194 | 6463 | 5599 | 6154 | 5738 | 5729 | 5686 | 5894.87 | 6072 | 5718 | 0.942 | -0.087 | 0.26764634 | 0.48739384 | 0.0024 | 0.0075 | 0.0042 | 0.0042 | TRUE | 0.0568 |
| Trappc9 | 1409 | 2151 | 2193 | 1870 | 1387 | 1712 | 1497 | 2148 | 2202 | 1470 | 1492 | 1874 | 1780.54 | 1949 | 1612 | 0.827 | -0.274 | 0.10176772 | 0.2646464 | 0.0294 | 0.0088 | 0.0197 | 0.0197 | TRUE | 0.4976 |
| Trat1 | 2 | 0 | 0 | 0 | 0 | 0 | 2 | 0 | 0 | 0 | 0 | 0 | 0.35 | 1 | 0 | 0.254 | -1.975 | 0.62446185 | NA | 10 | 9.3117 | 10 | 10 | TRUE | 0.9741 |
| Trcg1 | 0 | 0 | 0 | 1 | 0 | 0 | 0 | 0 | 0 | 1 | 0 | 0 | 0.13 | 0 | 0 | 1.859 | 0.894 | 0.82650734 | NA | 0 | 25.1515 | 10 | 10 | TRUE | 0.2563 |
| Trdmt1 | 548 | 524 | 523 | 742 | 550 | 493 | 582 | 523 | 525 | 583 | 592 | 540 | 557.55 | 543 | 572 | 1.053 | 0.074 | 0.4841737 | 0.68175944 | 0.0011 | 0.0128 | 0.0062 | 0.0062 | TRUE | 0.0446 |
| Trdn | 0 | 0 | 0 | 0 | 0 | 0 | 0 | 0 | 0 | 0 | 0 | 0 | 0 | 0 | 0 | NA | NA | NA | NA | NA | NA | NA | NA | NA | NA |
| Treh | 2 | 0 | 1 | 1 | 1 | 0 | 2 | 0 | 1 | 1 | 1 | 0 | 0.83 | 1 | 1 | 0.608 | -0.719 | 0.74058884 | NA | 0 | 3.9678 | 2.1265 | 2.1265 | TRUE | 0.1445 |
| Trem1 | 0 | 0 | 0 | 0 | 0 | 0 | 0 | 0 | 0 | 0 | 0 | 0 | 0 | 0 | 0 | NA | NA | NA | NA | NA | NA | NA | NA | NA | NA |
| Trem2 | 8 | 4 | 4 | 0 | 1 | 0 | 8 | 4 | 4 | 0 | 1 | 0 | 2.93 | 5 | 0 | 0.064 | -3.975 | 0.01128491 | NA | 0 | 1.131 | 0.6805 | 0.6805 | TRUE | 0.5204 |
| Trem3 | 0 | 0 | 0 | 0 | 0 | 0 | 0 | 0 | 0 | 0 | 0 | 0 | 0 | 0 | 0 | NA | NA | NA | NA | NA | NA | NA | NA | NA | NA |
| Treml1 | 2 | 0 | 0 | 2 | 1 | 0 | 2 | 0 | 0 | 2 | 1 | 0 | 0.8 | 1 | 1 | 1.29 | 0.367 | 0.88218031 | NA | 0 | 4.1487 | 3.1517 | 3.1517 | TRUE | 0.9743 |
| Treml2 | 304 | 276 | 249 | 296 | 233 | 182 | 323 | 276 | 250 | 233 | 251 | 199 | 255.2 | 283 | 228 | 0.805 | -0.312 | 0.05555929 | 0.17645611 | 0.0109 | 0.0198 | 0.0152 | 0.0152 | TRUE | 0.1706 |
| Treml4 | 2 | 11 | 7 | 7 | 6 | 3 | 2 | 11 | 7 | 6 | 6 | 3 | 5.9 | 7 | 5 | 0.757 | -0.403 | 0.64047319 | 0.79767036 | 0.1456 | 0.5656 | 0.3637 | 0.3637 | TRUE | 0.1895 |
| Trerf1 | 1006 | 1068 | 1209 | 1130 | 831 | 832 | 1069 | 1067 | 1214 | 888 | 894 | 911 | 1007.05 | 1117 | 898 | 0.804 | -0.315 | 0.00074326 | 0.00706608 | 0.0021 | 0.0102 | 0.0053 | 0.0053 | TRUE | 0.0702 |
| Trex1 | 202 | 213 | 218 | 259 | 222 | 225 | 215 | 213 | 219 | 204 | 239 | 246 | 222.49 | 216 | 230 | 1.062 | 0.087 | 0.55305985 | 0.73679508 | 0 | 0.0217 | 0.0111 | 0.0111 | TRUE | 0.1177 |
| Trex2 | 3 | 0 | 0 | 5 | 5 | 8 | 3 | 0 | 0 | 4 | 5 | 9 | 3.54 | 1 | 6 | 5.76 | 2.526 | 0.06182974 | NA | 0.1835 | 0.9369 | 0.7448 | 0.7448 | TRUE | 1.4181 |
| Trf | 560 | 645 | 609 | 730 | 477 | 536 | 595 | 644 | 612 | 574 | 513 | 587 | 587.39 | 617 | 558 | 0.905 | -0.144 | 0.17063286 | 0.36921683 | 0.0014 | 0.0125 | 0.0063 | 0.0063 | TRUE | 0.0558 |
| Trh | 7034 | 5350 | 6108 | 3623 | 2613 | 2442 | 7471 | 5343 | 6133 | 2848 | 2811 | 2673 | 4546.7 | 6316 | 2777 | 0.44 | -1.185 | 1.60E-20 | 4.36E-18 | 0.0147 | 0.0076 | 0.0115 | 0.0115 | TRUE | 0.2039 |
| Trhde | 0 | 0 | 0 | 0 | 2 | 0 | 0 | 0 | 0 | 0 | 2 | 0 | 0.36 | 0 | 1 | 3.775 | 1.916 | 0.63423067 | NA | 10 | 9.1927 | 10 | 10 | TRUE | 0.9743 |
| Trhr | 0 | 0 | 0 | 1 | 0 | 0 | 0 | 0 | 0 | 1 | 0 | 0 | 0.13 | 0 | 0 | 1.859 | 0.894 | 0.82650734 | NA | 0 | 25.1515 | 10 | 10 | TRUE | 0.2563 |
| Trhr2 | 0 | 1 | 0 | 1 | 0 | 0 | 0 | 1 | 0 | 1 | 0 | 0 | 0.3 | 0 | 0 | 0.954 | -0.067 | 0.98642001 | NA | 0 | 11.0817 | 9.3014 | 9.3014 | TRUE | 0.405 |
| Triap1 | 1138 | 1201 | 1156 | 1588 | 1074 | 956 | 1209 | 1199 | 1161 | 1248 | 1156 | 1046 | 1169.89 | 1190 | 1150 | 0.968 | -0.047 | 0.62714546 | 0.78999388 | 0.0033 | 0.0097 | 0.0059 | 0.0059 | TRUE | 0.074 |
| Trib1 | 328 | 325 | 310 | 470 | 332 | 274 | 348 | 325 | 311 | 369 | 357 | 300 | 335.15 | 328 | 342 | 1.045 | 0.064 | 0.63770882 | 0.79603827 | 0.0047 | 0.0167 | 0.0103 | 0.0103 | TRUE | 0.1295 |
| Trib2 | 221 | 420 | 415 | 227 | 182 | 217 | 235 | 419 | 417 | 178 | 196 | 238 | 280.45 | 357 | 204 | 0.57 | -0.811 | 0.0007282 | 0.00698182 | 0.0584 | 0.0187 | 0.0376 | 0.0376 | TRUE | 1.0192 |
| Trib3 | 376 | 382 | 366 | 596 | 432 | 356 | 399 | 382 | 368 | 469 | 465 | 390 | 411.9 | 383 | 441 | 1.154 | 0.207 | 0.09860675 | 0.25874141 | 0.0038 | 0.0149 | 0.0088 | 0.0088 | TRUE | 0.1166 |
| Tril | 17 | 16 | 6 | 35 | 14 | 9 | 18 | 16 | 6 | 28 | 15 | 10 | 15.42 | 13 | 18 | 1.327 | 0.408 | 0.51233833 | 0.70574497 | 0.2063 | 0.2206 | 0.214 | 0.214 | TRUE | 0.5912 |
| Trim10 | 7 | 8 | 5 | 6 | 6 | 5 | 7 | 8 | 5 | 5 | 6 | 5 | 6.18 | 7 | 5 | 0.809 | -0.305 | 0.68056146 | 0.82289292 | 0 | 0.5398 | 0.2332 | 0.2332 | TRUE | 0.1402 |
| Trim11 | 1028 | 1284 | 1190 | 1563 | 1243 | 1310 | 1092 | 1282 | 1195 | 1229 | 1337 | 1434 | 1261.54 | 1190 | 1333 | 1.12 | 0.163 | 0.12019172 | 0.29417501 | 0.005 | 0.0095 | 0.0071 | 0.0071 | TRUE | 0.0617 |
| Trim12a | 0 | 0 | 0 | 0 | 0 | 0 | 0 | 0 | 0 | 0 | 0 | 0 | 0 | 0 | 0 | NA | NA | NA | NA | NA | NA | NA | NA | NA | NA |
| Trim12c | 597 | 599 | 599 | 670 | 473 | 556 | 634 | 598 | 601 | 527 | 509 | 609 | 579.68 | 611 | 548 | 0.896 | -0.159 | 0.15669088 | 0.34969116 | 0.0031 | 0.0126 | 0.0073 | 0.0073 | TRUE | 0.1061 |
| Trim13 | 862 | 1062 | 1037 | 1103 | 853 | 1078 | 916 | 1061 | 1041 | 867 | 918 | 1180 | 997.07 | 1006 | 988 | 0.982 | -0.027 | 0.84824531 | 0.92405892 | 0.0159 | 0.0102 | 0.0132 | 0.0132 | TRUE | 0.3419 |
| Trim14 | 93 | 147 | 163 | 82 | 66 | 107 | 99 | 147 | 164 | 64 | 71 | 117 | 110.31 | 137 | 84 | 0.614 | -0.704 | 0.01883529 | 0.08330626 | 0.0742 | 0.0368 | 0.0551 | 0.0551 | TRUE | 1.0729 |
| Trim15 | 2 | 4 | 2 | 1 | 2 | 1 | 2 | 4 | 2 | 1 | 2 | 1 | 2.03 | 3 | 1 | 0.485 | -1.045 | 0.42847845 | NA | 0 | 1.6324 | 0.6974 | 0.6974 | TRUE | 0.208 |
| Trim16 | 245 | 198 | 192 | 273 | 190 | 168 | 260 | 198 | 193 | 215 | 204 | 184 | 208.95 | 217 | 201 | 0.929 | -0.107 | 0.54470983 | 0.72985842 | 0.0127 | 0.0227 | 0.0175 | 0.0175 | TRUE | 0.3307 |
| Trim17 | 22 | 41 | 31 | 30 | 23 | 30 | 23 | 41 | 31 | 24 | 25 | 33 | 29.44 | 32 | 27 | 0.845 | -0.243 | 0.52760686 | 0.71625197 | 0.0219 | 0.1188 | 0.0725 | 0.0725 | TRUE | 0.2218 |
| Trim2 | 2248 | 2101 | 2307 | 1417 | 1138 | 1410 | 2388 | 2098 | 2316 | 1114 | 1224 | 1543 | 1780.73 | 2267 | 1294 | 0.57 | -0.811 | 1.73E-09 | 9.95E-08 | 0.0162 | 0.0088 | 0.0125 | 0.0125 | TRUE | 0.3407 |
| Trim21 | 245 | 340 | 333 | 451 | 341 | 360 | 260 | 340 | 334 | 355 | 367 | 394 | 341.61 | 311 | 372 | 1.192 | 0.254 | 0.08217989 | 0.22887185 | 0.0085 | 0.0165 | 0.0124 | 0.0124 | TRUE | 0.2309 |
| Trim23 | 1841 | 1960 | 2020 | 2719 | 1897 | 2001 | 1955 | 1957 | 2028 | 2137 | 2041 | 2190 | 2051.69 | 1980 | 2123 | 1.072 | 0.1 | 0.15943352 | 0.35371859 | 3.00E-04 | 0.0085 | 0.0032 | 0.0032 | TRUE | 0.0135 |
| Trim24 | 8613 | 8329 | 8033 | 10195 | 7792 | 7156 | 9149 | 8318 | 8066 | 8015 | 8384 | 7833 | 8294.06 | 8511 | 8077 | 0.949 | -0.075 | 0.33441357 | 0.5548783 | 0.0026 | 0.0073 | 0.0043 | 0.0043 | TRUE | 0.0524 |
| Trim25 | 10950 | 12612 | 12382 | 14203 | 10481 | 10218 | 11631 | 12596 | 12433 | 11165 | 11277 | 11185 | 11714.47 | 12220 | 11209 | 0.917 | -0.125 | 0.04634884 | 0.15580006 | 8.00E-04 | 0.0072 | 0.0027 | 0.0027 | TRUE | 0.0217 |
| Trim26 | 2508 | 2300 | 2257 | 2443 | 1858 | 1789 | 2664 | 2297 | 2266 | 1920 | 1999 | 1958 | 2184.19 | 2409 | 1959 | 0.813 | -0.298 | 0.00120411 | 0.01026552 | 0.0037 | 0.0084 | 0.0056 | 0.0056 | TRUE | 0.1035 |
| Trim27 | 7106 | 7298 | 7556 | 9185 | 6717 | 6611 | 7548 | 7289 | 7587 | 7221 | 7227 | 7237 | 7351.3 | 7475 | 7228 | 0.967 | -0.048 | 0.38760705 | 0.60141056 | 1.00E-04 | 0.0074 | 0.0021 | 0.0021 | TRUE | 0.0058 |
| Trim28 | 84135 | 70334 | 69161 | 117317 | 81958 | 67092 | 89366 | 70243 | 69445 | 92225 | 88182 | 73443 | 80484.06 | 76351 | 84617 | 1.108 | 0.148 | 0.26031708 | 0.47980845 | 0.017 | 0.0069 | 0.0125 | 0.0125 | TRUE | 0.2723 |
| Trim29 | 215 | 278 | 279 | 102 | 97 | 139 | 228 | 278 | 280 | 80 | 104 | 152 | 187.14 | 262 | 112 | 0.426 | -1.233 | 3.29E-07 | 1.03E-05 | 0.0499 | 0.0245 | 0.0356 | 0.0356 | TRUE | 0.9539 |
| Trim3 | 810 | 891 | 881 | 1045 | 860 | 803 | 860 | 890 | 885 | 821 | 925 | 879 | 876.77 | 878 | 875 | 0.995 | -0.007 | 0.94075981 | 0.97003791 | 0.001 | 0.0107 | 0.0048 | 0.0048 | TRUE | 0.0361 |
| Trim30a | 5 | 9 | 7 | 17 | 11 | 7 | 5 | 9 | 7 | 13 | 12 | 8 | 9.03 | 7 | 11 | 1.55 | 0.632 | 0.32490044 | 0.54594394 | 0 | 0.3717 | 0.1809 | 0.1809 | TRUE | 0.234 |
| Trim30b | 0 | 1 | 0 | 0 | 0 | 0 | 0 | 1 | 0 | 0 | 0 | 0 | 0.17 | 0 | 0 | 0.49 | -1.029 | 0.80087 | NA | 0 | 19.7991 | 10 | 10 | TRUE | 0.405 |
| Trim30d | 4 | 3 | 2 | 12 | 3 | 8 | 4 | 3 | 2 | 9 | 3 | 9 | 5.11 | 3 | 7 | 2.343 | 1.228 | 0.17389568 | 0.3731541 | 0 | 0.6514 | 0.3539 | 0.3539 | TRUE | 0.5716 |
| Trim30e-ps1 | 0 | 0 | 0 | 0 | 0 | 0 | 0 | 0 | 0 | 0 | 0 | 0 | 0 | 0 | 0 | NA | NA | NA | NA | NA | NA | NA | NA | NA | NA |

| | | | | | | | | | | | | | | | | | | | | | | | | | |
|---|---|---|---|---|---|---|---|---|---|---|---|---|---|---|---|---|---|---|---|---|---|---|---|---|---|
| Trim31 | 0 | 1 | 1 | 3 | 3 | 1 | 0 | 1 | 1 | 2 | 3 | 1 | 1.45 | 1 | 2 | 3.306 | 1.725 | 0.3034607 | NA | 0 | 2.2832 | 1.0504 | 1.0504 | TRUE | 0.2212 |
| Trim32 | 630 | 712 | 673 | 1461 | 1080 | 955 | 669 | 711 | 676 | 1149 | 1162 | 1045 | 901.99 | 685 | 1119 | 1.633 | 0.707 | 1.88E-14 | 2.36E-12 | 0.0012 | 0.0106 | 0.005 | 0.005 | TRUE | 0.0384 |
| Trim33 | 6582 | 5867 | 6061 | 7404 | 5393 | 5367 | 6991 | 5859 | 6086 | 5820 | 5803 | 5875 | 6072.43 | 6312 | 5833 | 0.924 | -0.114 | 0.19706533 | 0.40371603 | 0.0043 | 0.0074 | 0.0055 | 0.0055 | TRUE | 0.1079 |
| Trim34a | 27 | 58 | 77 | 57 | 55 | 71 | 29 | 58 | 77 | 45 | 59 | 78 | 57.6 | 55 | 61 | 1.102 | 0.141 | 0.72233509 | 0.84975528 | 0.1308 | 0.0641 | 0.0954 | 0.0954 | TRUE | 0.3595 |
| Trim34b | 24 | 39 | 38 | 52 | 40 | 54 | 25 | 39 | 38 | 41 | 43 | 59 | 40.94 | 34 | 48 | 1.386 | 0.471 | 0.16025858 | 0.35474763 | 0.0204 | 0.0874 | 0.0557 | 0.0557 | TRUE | 0.3365 |
| Trim35 | 3489 | 3386 | 3369 | 4189 | 2778 | 2447 | 3706 | 3382 | 3383 | 3293 | 2989 | 2679 | 3238.51 | 3490 | 2987 | 0.856 | -0.224 | 0.02646218 | 0.10571469 | 0.0063 | 0.0079 | 0.007 | 0.007 | TRUE | 0.0988 |
| Trim36 | 243 | 289 | 266 | 345 | 211 | 267 | 258 | 289 | 267 | 271 | 227 | 292 | 267.39 | 271 | 263 | 0.972 | -0.042 | 0.77995165 | 0.88617899 | 0.0055 | 0.0192 | 0.0122 | 0.0122 | TRUE | 0.157 |
| Trim37 | 10975 | 12527 | 13132 | 13939 | 10513 | 11305 | 11657 | 12511 | 13186 | 10958 | 11311 | 12375 | 11999.72 | 12451 | 11548 | 0.927 | -0.109 | 0.1976801 | 0.40433284 | 0.0038 | 0.0072 | 0.0051 | 0.0051 | TRUE | 0.0479 |
| Trim38 | 14 | 4 | 4 | 14 | 11 | 4 | 15 | 4 | 4 | 11 | 12 | 4 | 8.35 | 8 | 9 | 1.203 | 0.266 | 0.74058829 | 0.86139812 | 0.2879 | 0.4014 | 0.3458 | 0.3458 | TRUE | 1.8778 |
| Trim39 | 691 | 582 | 592 | 989 | 808 | 571 | 734 | 581 | 594 | 777 | 869 | 625 | 696.92 | 636 | 757 | 1.19 | 0.251 | 0.10871499 | 0.27620449 | 0.0199 | 0.0116 | 0.0162 | 0.0162 | TRUE | 0.2721 |
| Trim40 | 45 | 53 | 46 | 41 | 32 | 22 | 48 | 53 | 46 | 32 | 34 | 24 | 39.61 | 49 | 30 | 0.62 | -0.691 | 0.0270496 | 0.10723237 | 0 | 0.0901 | 0.0438 | 0.0438 | TRUE | 0.1906 |
| Trim41 | 4961 | 5186 | 5239 | 7694 | 5598 | 4901 | 5269 | 5179 | 5261 | 6048 | 6023 | 5365 | 5524.29 | 5236 | 5812 | 1.11 | 0.151 | 0.0478748 | 0.15916553 | 0.002 | 0.0075 | 0.004 | 0.004 | TRUE | 0.055 |
| Trim42 | 0 | 0 | 0 | 0 | 0 | 0 | 0 | 0 | 0 | 0 | 0 | 0 | 0 | 0 | 0 | NA | NA | NA | NA | NA | NA | NA | NA | NA | NA |
| Trim43a | 12 | 6 | 3 | 141 | 98 | 33 | 13 | 6 | 3 | 111 | 105 | 36 | 45.69 | 7 | 84 | 11.708 | 3.549 | 1.28E-09 | 7.54E-08 | 0.312 | 0.079 | 0.1697 | 0.1697 | TRUE | 2.2797 |
| Trim43b | 4 | 0 | 0 | 28 | 13 | 5 | 4 | 0 | 0 | 22 | 14 | 5 | 7.62 | 1 | 14 | 10.049 | 3.329 | 0.00433197 | 0.02803607 | 1.0607 | 0.4392 | 0.5773 | 0.5773 | TRUE | 0.5293 |
| Trim43c | 1 | 2 | 0 | 7 | 3 | 1 | 1 | 2 | 0 | 6 | 3 | 1 | 2.15 | 1 | 3 | 3.266 | 1.708 | 0.24740175 | NA | 0 | 1.541 | 0.9232 | 0.9232 | TRUE | 0.1508 |
| Trim44 | 1642 | 2335 | 2548 | 3264 | 2332 | 2414 | 1744 | 2332 | 2558 | 2566 | 2509 | 2643 | 2392.01 | 2211 | 2573 | 1.163 | 0.218 | 0.12181082 | 0.29675631 | 0.0178 | 0.0083 | 0.0139 | 0.0139 | TRUE | 0.4132 |
| Trim45 | 625 | 739 | 711 | 997 | 710 | 758 | 664 | 738 | 714 | 784 | 764 | 830 | 748.88 | 705 | 793 | 1.123 | 0.167 | 0.07988933 | 0.22436154 | 8.00E-04 | 0.0113 | 0.0052 | 0.0052 | TRUE | 0.0309 |
| Trim46 | 126 | 118 | 123 | 105 | 109 | 107 | 134 | 118 | 124 | 83 | 117 | 117 | 115.36 | 125 | 106 | 0.84 | -0.252 | 0.23996185 | 0.45609524 | 0.0133 | 0.0355 | 0.0244 | 0.0244 | TRUE | 0.4047 |
| Trim47 | 163 | 326 | 353 | 176 | 161 | 200 | 173 | 326 | 354 | 138 | 173 | 219 | 230.61 | 284 | 177 | 0.62 | -0.689 | 0.01481625 | 0.07021693 | 0.0904 | 0.0212 | 0.0531 | 0.0531 | TRUE | 1.2423 |
| Trim50 | 7 | 15 | 14 | 7 | 4 | 9 | 7 | 15 | 14 | 6 | 4 | 10 | 9.36 | 12 | 7 | 0.533 | -0.907 | 0.17687831 | 0.37738204 | 0 | 0.3591 | 0.2079 | 0.2079 | TRUE | 0.438 |
| Trim52 | 201 | 287 | 277 | 312 | 331 | 353 | 213 | 287 | 278 | 245 | 356 | 386 | 294.35 | 259 | 329 | 1.265 | 0.34 | 0.10242897 | 0.26597132 | 0.0375 | 0.0181 | 0.0277 | 0.0277 | TRUE | 0.5901 |
| Trim54 | 58 | 70 | 69 | 79 | 53 | 38 | 62 | 70 | 69 | 62 | 57 | 42 | 60.25 | 67 | 54 | 0.805 | -0.313 | 0.24051855 | 0.4569331 | 0 | 0.0616 | 0.0346 | 0.0346 | TRUE | 0.2993 |
| Trim55 | 0 | 0 | 0 | 0 | 0 | 0 | 0 | 0 | 0 | 0 | 0 | 0 | 0 | 0 | 0 | NA | NA | NA | NA | NA | NA | NA | NA | NA | NA |
| Trim56 | 226 | 207 | 302 | 212 | 155 | 199 | 240 | 207 | 303 | 167 | 167 | 218 | 216.88 | 250 | 184 | 0.734 | -0.447 | 0.02597622 | 0.10433639 | 0.0265 | 0.0221 | 0.0243 | 0.0243 | TRUE | 0.359 |
| Trim58 | 0 | 0 | 0 | 2 | 0 | 1 | 0 | 0 | 0 | 2 | 0 | 1 | 0.44 | 0 | 1 | 4.782 | 2.258 | 0.50653358 | NA | 0 | 7.4188 | 6.7775 | 6.7775 | TRUE | 0.2804 |
| Trim59 | 5352 | 5869 | 5637 | 7569 | 5441 | 5566 | 5685 | 5861 | 5660 | 5950 | 5854 | 6093 | 5850.59 | 5735 | 5966 | 1.04 | 0.057 | 0.33494208 | 0.55525609 | 2.00E-04 | 0.0075 | 0.0023 | 0.0023 | TRUE | 0.0045 |
| Trim6 | 4326 | 5338 | 5456 | 8402 | 6224 | 5553 | 4595 | 5331 | 5478 | 6605 | 6697 | 6079 | 5797.46 | 5135 | 6460 | 1.258 | 0.331 | 0.00047737 | 0.00500001 | 0.0053 | 0.0075 | 0.0063 | 0.0063 | TRUE | 0.103 |
| Trim60 | 42 | 36 | 20 | 53 | 40 | 44 | 45 | 36 | 20 | 42 | 43 | 48 | 38.92 | 34 | 44 | 1.32 | 0.401 | 0.27137002 | 0.49139471 | 0.0421 | 0.0916 | 0.0695 | 0.0695 | TRUE | 0.5769 |
| Trim61 | 10 | 9 | 12 | 10 | 6 | 4 | 11 | 9 | 12 | 8 | 6 | 4 | 8.39 | 11 | 6 | 0.598 | -0.741 | 0.256595 | 0.47519694 | 0 | 0.3995 | 0.1816 | 0.1816 | TRUE | 0.1661 |
| Trim62 | 152 | 169 | 160 | 192 | 153 | 144 | 161 | 169 | 161 | 151 | 165 | 158 | 160.68 | 164 | 158 | 0.962 | -0.055 | 0.72255834 | 0.84995452 | 0 | 0.0274 | 0.0113 | 0.0113 | TRUE | 0.0167 |
| Trim63 | 360 | 436 | 399 | 257 | 218 | 248 | 382 | 435 | 401 | 202 | 235 | 271 | 321.09 | 406 | 236 | 0.579 | -0.789 | 1.61E-07 | 5.43E-06 | 0.009 | 0.0172 | 0.013 | 0.013 | TRUE | 0.1898 |
| Trim65 | 25 | 40 | 68 | 38 | 33 | 37 | 27 | 40 | 68 | 30 | 36 | 41 | 40.11 | 45 | 36 | 0.781 | -0.356 | 0.38213138 | 0.59637602 | 0.0993 | 0.089 | 0.0941 | 0.0941 | TRUE | 0.4497 |
| Trim66 | 60 | 57 | 50 | 150 | 96 | 90 | 64 | 57 | 50 | 118 | 103 | 99 | 81.76 | 57 | 107 | 1.879 | 0.91 | 6.86E-05 | 0.0010168 | 0 | 0.0472 | 0.0241 | 0.0241 | TRUE | 0.0942 |
| Trim67 | 62 | 45 | 40 | 63 | 45 | 33 | 66 | 45 | 40 | 50 | 48 | 36 | 47.5 | 50 | 45 | 0.893 | -0.164 | 0.61000557 | 0.77891178 | 0.0285 | 0.0763 | 0.0532 | 0.0532 | TRUE | 0.58 |
| Trim68 | 254 | 348 | 325 | 491 | 416 | 330 | 270 | 348 | 326 | 386 | 448 | 361 | 356.42 | 315 | 398 | 1.265 | 0.339 | 0.0249525 | 0.10156149 | 0.0113 | 0.0161 | 0.0136 | 0.0136 | TRUE | 0.1736 |
| Trim69 | 1 | 0 | 1 | 0 | 0 | 1 | 1 | 0 | 1 | 0 | 0 | 1 | 0.53 | 1 | 0 | 0.509 | -0.975 | 0.72674596 | NA | 0 | 6.2603 | 3.8683 | 3.8683 | TRUE | 0.5383 |
| Trim7 | 51 | 67 | 71 | 72 | 52 | 60 | 54 | 67 | 71 | 57 | 56 | 66 | 61.77 | 64 | 60 | 0.924 | -0.114 | 0.65246044 | 0.80502827 | 0 | 0.0602 | 0.0304 | 0.0304 | TRUE | 0.1573 |
| Trim71 | 6034 | 5196 | 5068 | 7432 | 5294 | 4013 | 6409 | 5189 | 5089 | 5842 | 5696 | 4393 | 5436.43 | 5562 | 5310 | 0.955 | -0.067 | 0.63553529 | 0.79511677 | 0.0201 | 0.0075 | 0.0141 | 0.0141 | TRUE | 0.278 |
| Trim72 | 18 | 35 | 43 | 22 | 16 | 19 | 19 | 35 | 43 | 17 | 17 | 21 | 25.43 | 32 | 18 | 0.566 | -0.821 | 0.06218558 | 0.1908525 | 0.0568 | 0.1365 | 0.0972 | 0.0972 | TRUE | 0.3011 |
| Trim75 | 0 | 0 | 0 | 1 | 1 | 2 | 0 | 0 | 0 | 1 | 1 | 2 | 0.68 | 0 | 1 | 7.126 | 2.833 | 0.25569871 | NA | 0 | 4.8859 | 3.111 | 3.111 | TRUE | 0.1691 |
| Trim8 | 798 | 1016 | 1027 | 1442 | 1203 | 869 | 848 | 1015 | 1031 | 1134 | 1294 | 951 | 1045.45 | 965 | 1126 | 1.168 | 0.224 | 0.11363938 | 0.28427491 | 0.0157 | 0.0101 | 0.0134 | 0.0134 | TRUE | 0.1855 |
| Trim9 | 0 | 1 | 2 | 2 | 1 | 1 | 0 | 1 | 2 | 2 | 1 | 1 | 1.12 | 1 | 1 | 1.247 | 0.319 | 0.8553168 | NA | 0 | 2.9354 | 1.3093 | 1.3093 | TRUE | 0.2166 |
| Triml1 | 5947 | 6351 | 6115 | 5414 | 4501 | 4457 | 6317 | 6343 | 6140 | 4256 | 4843 | 4879 | 5462.9 | 6267 | 4659 | 0.743 | -0.428 | 1.24E-07 | 4.30E-06 | 0.0027 | 0.0075 | 0.0045 | 0.0045 | TRUE | 0.0706 |
| Triml2 | 6672 | 7469 | 7284 | 9496 | 7063 | 6674 | 7087 | 7459 | 7314 | 7465 | 7599 | 7306 | 7371.7 | 7287 | 7457 | 1.023 | 0.033 | 0.57633053 | 0.75418558 | 3.00E-04 | 0.0074 | 0.0024 | 0.0024 | TRUE | 0.007 |
| Trio | 1163 | 1118 | 1245 | 1960 | 1399 | 1329 | 1235 | 1117 | 1250 | 1541 | 1505 | 1455 | 1350.47 | 1201 | 1500 | 1.25 | 0.322 | 0.00016842 | 0.0021048 | 0.0015 | 0.0093 | 0.0045 | 0.0045 | TRUE | 0.0453 |
| Triobp | 2258 | 2984 | 2986 | 2881 | 2271 | 2611 | 2398 | 2980 | 2998 | 2265 | 2443 | 2858 | 2657.2 | 2792 | 2522 | 0.903 | -0.147 | 0.25299427 | 0.47087865 | 0.0142 | 0.0081 | 0.0116 | 0.0116 | TRUE | 0.1845 |
| Trip10 | 2607 | 2522 | 2358 | 3683 | 2945 | 2709 | 2769 | 2519 | 2368 | 2895 | 3169 | 2965 | 2780.81 | 2552 | 3010 | 1.179 | 0.238 | 0.00816652 | 0.04514629 | 0.0037 | 0.0081 | 0.0055 | 0.0055 | TRUE | 0.0671 |
| Trip11 | 1180 | 1055 | 1120 | 1564 | 1127 | 1014 | 1253 | 1054 | 1125 | 1229 | 1213 | 1110 | 1163.94 | 1144 | 1184 | 1.036 | 0.051 | 0.61917114 | 0.78469067 | 0.0045 | 0.0097 | 0.0067 | 0.0067 | TRUE | 0.0836 |
| Trip12 | 32778 | 30183 | 32953 | 54837 | 37430 | 34845 | 34816 | 30144 | 33088 | 43109 | 40272 | 38143 | 36595.44 | 32683 | 40508 | 1.239 | 0.31 | 0.00037423 | 0.00411026 | 0.0045 | 0.007 | 0.0054 | 0.0054 | TRUE | 0.0565 |
| Trip13 | 2773 | 2378 | 2285 | 4542 | 3021 | 2598 | 2945 | 2375 | 2294 | 3571 | 3250 | 2844 | 2879.94 | 2538 | 3222 | 1.27 | 0.345 | 0.00851697 | 0.04667467 | 0.0156 | 0.008 | 0.012 | 0.012 | TRUE | 0.2385 |
| Trip4 | 1943 | 2320 | 2410 | 2766 | 2004 | 2014 | 2064 | 2317 | 2420 | 2174 | 2156 | 2205 | 2222.66 | 2267 | 2178 | 0.961 | -0.058 | 0.50745933 | 0.70172367 | 0.0027 | 0.0084 | 0.005 | 0.005 | TRUE | 0.0741 |
| Trip6 | 630 | 588 | 677 | 1029 | 690 | 676 | 669 | 587 | 680 | 809 | 742 | 740 | 704.58 | 645 | 764 | 1.185 | 0.245 | 0.02109182 | 0.0903477 | 0.003 | 0.0116 | 0.0067 | 0.0067 | TRUE | 0.0737 |
| Triqk | 62 | 92 | 85 | 103 | 89 | 88 | 66 | 92 | 85 | 81 | 96 | 96 | 86.02 | 81 | 91 | 1.118 | 0.161 | 0.48491004 | 0.68236871 | 0.0075 | 0.0452 | 0.0267 | 0.0267 | TRUE | 0.244 |
| Trit1 | 1819 | 2066 | 1952 | 2050 | 1531 | 1786 | 1932 | 2063 | 1960 | 1612 | 1647 | 1955 | 1861.55 | 1985 | 1738 | 0.875 | -0.193 | 0.05823203 | 0.18235677 | 0.0057 | 0.0087 | 0.0069 | 0.0069 | TRUE | 0.1432 |
| Trmt1 | 5588 | 5720 | 5717 | 6986 | 5321 | 4947 | 5935 | 5713 | 5740 | 5492 | 5725 | 5415 | 5670.12 | 5796 | 5544 | 0.956 | -0.064 | 0.29900748 | 0.51932012 | 4.00E-04 | 0.0075 | 0.0026 | 0.0026 | TRUE | 0.0099 |
| Trmt10a | 1819 | 1908 | 1775 | 2736 | 1890 | 1770 | 1932 | 1906 | 1782 | 2151 | 2034 | 1938 | 1956.97 | 1873 | 2041 | 1.09 | 0.124 | 0.12854411 | 0.3082557 | 0.0018 | 0.0086 | 0.0043 | 0.0043 | TRUE | 0.0276 |
| Trmt10b | 803 | 835 | 797 | 1248 | 856 | 770 | 853 | 834 | 800 | 981 | 921 | 843 | 872.02 | 829 | 915 | 1.105 | 0.144 | 0.1415634 | 0.32704526 | 0.0024 | 0.0107 | 0.0058 | 0.0058 | TRUE | 0.0561 |
| Trmt10c | 1854 | 1704 | 1660 | 2196 | 1611 | 1451 | 1969 | 1702 | 1667 | 1726 | 1733 | 1588 | 1730.98 | 1779 | 1682 | 0.946 | -0.08 | 0.4173043 | 0.62826684 | 0.0046 | 0.0088 | 0.0064 | 0.0064 | TRUE | 0.105 |
| Trmt11 | 640 | 447 | 453 | 864 | 603 | 539 | 680 | 446 | 455 | 679 | 649 | 590 | 583.18 | 527 | 639 | 1.214 | 0.28 | 0.11529262 | 0.28673367 | 0.0315 | 0.0126 | 0.0211 | 0.0211 | TRUE | 0.7481 |
| Trmt112 | 1369 | 1475 | 1484 | 1758 | 1285 | 1262 | 1454 | 1473 | 1490 | 1382 | 1383 | 1381 | 1427.22 | 1472 | 1382 | 0.939 | -0.091 | 0.19398807 | 0.39963165 | 0 | 0.0092 | 0.0029 | 0.0029 | TRUE | 0.0014 |
| Trmt12 | 75 | 73 | 62 | 230 | 150 | 90 | 80 | 73 | 62 | 181 | 161 | 99 | 109.26 | 72 | 147 | 2.059 | 1.042 | 0.00013605 | 0.00176644 | 0.0497 | 0.0371 | 0.0433 | 0.0433 | TRUE | 0.6382 |
| Trmt13 | 502 | 511 | 515 | 638 | 486 | 495 | 533 | 510 | 517 | 502 | 523 | 542 | 521.16 | 520 | 522 | 1.003 | 0.004 | 0.96963171 | 0.98448447 | 0 | 0.0132 | 0.0052 | 0.0052 | TRUE | 0.0148 |
| Trmt1l | 1224 | 1137 | 1255 | 1720 | 1210 | 1107 | 1300 | 1136 | 1260 | 1352 | 1302 | 1212 | 1260.26 | 1232 | 1289 | 1.047 | 0.066 | 0.48861529 | 0.68513873 | 0.0032 | 0.0095 | 0.0057 | 0.0057 | TRUE | 0.0564 |
| Trmt2a | 2104 | 1868 | 1750 | 2965 | 2058 | 1701 | 2235 | 1866 | 1757 | 2331 | 2214 | 1862 | 2044.12 | 1953 | 2136 | 1.094 | 0.13 | 0.31494277 | 0.5357288 | 0.0136 | 0.0085 | 0.0116 | 0.0116 | TRUE | 0.193 |
| Trmt2b | 575 | 579 | 632 | 746 | 615 | 667 | 611 | 578 | 635 | 586 | 662 | 730 | 633.65 | 608 | 659 | 1.083 | 0.115 | 0.33171744 | 0.55223216 | 0.0058 | 0.0121 | 0.0085 | 0.0085 | TRUE | 0.1152 |
| Trmt44 | 478 | 513 | 496 | 783 | 528 | 473 | 508 | 512 | 498 | 616 | 568 | 518 | 536.58 | 506 | 567 | 1.123 | 0.167 | 0.13036277 | 0.31086811 | 0.0024 | 0.013 | 0.0069 | 0.0069 | TRUE | 0.0684 |
| Trmt5 | 1350 | 1217 | 1210 | 1749 | 1262 | 1084 | 1434 | 1215 | 1215 | 1375 | 1358 | 1187 | 1297.28 | 1288 | 1307 | 1.015 | 0.021 | 0.84714146 | 0.9235581 | 0.0071 | 0.0094 | 0.0081 | 0.0081 | TRUE | 0.1174 |
| Trmt6 | 2571 | 2043 | 1739 | 3714 | 2353 | 1989 | 2731 | 2040 | 1746 | 2920 | 2532 | 2177 | 2357.66 | 2172 | 2543 | 1.171 | 0.228 | 0.19384658 | 0.39954891 | 0.0353 | 0.0083 | 0.0217 | 0.0217 | TRUE | 0.5364 |

| | | | | | | | | | | | | | | | | | | | | | | | | | |
|---|---|---|---|---|---|---|---|---|---|---|---|---|---|---|---|---|---|---|---|---|---|---|---|---|---|
| Trmt61a | 979 | 964 | 919 | 1437 | 1095 | 867 | 1040 | 963 | 923 | 1130 | 1178 | 949 | 1030.38 | 975 | 1086 | 1.114 | 0.156 | 0.17386359 | 0.3731541 | 0.0072 | 0.0101 | 0.0085 | 0.0085 | TRUE | 0.1425 |
| Trmt61b | 193 | 175 | 177 | 317 | 208 | 166 | 205 | 175 | 178 | 249 | 224 | 182 | 202.03 | 186 | 218 | 1.179 | 0.237 | 0.17880667 | 0.3799702 | 0.012 | 0.0232 | 0.0174 | 0.0174 | TRUE | 0.2298 |
| Trmu | 215 | 157 | 166 | 217 | 211 | 125 | 228 | 157 | 167 | 171 | 227 | 137 | 181.05 | 184 | 178 | 0.969 | -0.046 | 0.84783331 | 0.92391533 | 0.0467 | 0.0251 | 0.0356 | 0.0356 | TRUE | 0.3867 |
| Trnau1ap | 1165 | 1571 | 1553 | 1876 | 1318 | 1320 | 1237 | 1569 | 1559 | 1475 | 1418 | 1445 | 1450.6 | 1455 | 1446 | 0.994 | -0.009 | 0.93437066 | 0.96749848 | 0.0083 | 0.0092 | 0.0087 | 0.0087 | TRUE | 0.2056 |
| Trnp1 | 76 | 77 | 58 | 86 | 69 | 63 | 81 | 77 | 58 | 68 | 74 | 69 | 71.11 | 72 | 70 | 0.976 | -0.035 | 0.88564204 | 0.94387096 | 0 | 0.0532 | 0.0284 | 0.0284 | TRUE | 0.2538 |
| Trnt1 | 5625 | 5310 | 5095 | 8403 | 5970 | 5129 | 5975 | 5303 | 5116 | 6606 | 6423 | 5614 | 5839.57 | 5465 | 6214 | 1.137 | 0.186 | 0.06458633 | 0.19578237 | 0.0068 | 0.0075 | 0.0071 | 0.0071 | TRUE | 0.0868 |
| Tro | 3096 | 2949 | 2953 | 3713 | 2806 | 2621 | 3288 | 2945 | 2965 | 2919 | 3019 | 2869 | 3000.98 | 3066 | 2936 | 0.957 | -0.063 | 0.42391569 | 0.63416469 | 0.0018 | 0.008 | 0.0041 | 0.0041 | TRUE | 0.0487 |
| Troap | 1359 | 1237 | 1131 | 1301 | 1022 | 933 | 1443 | 1235 | 1136 | 1023 | 1100 | 1021 | 1159.7 | 1271 | 1048 | 0.824 | -0.279 | 0.0144854 | 0.06898058 | 0.0072 | 0.0097 | 0.0085 | 0.0085 | TRUE | 0.1675 |
| Trove2 | 1867 | 1792 | 2018 | 2522 | 1839 | 1866 | 1983 | 1790 | 2026 | 1983 | 1979 | 2043 | 1967.16 | 1933 | 2002 | 1.035 | 0.05 | 0.54086764 | 0.72674769 | 0.0018 | 0.0086 | 0.0043 | 0.0043 | TRUE | 0.0511 |
| Trp53 | 19415 | 16380 | 16794 | 24965 | 17572 | 15369 | 20622 | 16359 | 16863 | 19626 | 18906 | 16824 | 18199.95 | 17948 | 18452 | 1.028 | 0.04 | 0.72659346 | 0.85260263 | 0.0093 | 0.0071 | 0.0094 | 0.0094 | TRUE | 0.2078 |
| Trp53bp1 | 5673 | 5316 | 5808 | 6756 | 5159 | 5021 | 6026 | 5309 | 5832 | 5311 | 5551 | 5496 | 5587.46 | 5722 | 5453 | 0.953 | -0.07 | 0.36339588 | 0.58083215 | 0.0021 | 0.0075 | 0.0041 | 0.0041 | TRUE | 0.0487 |
| Trp53bp2 | 2785 | 2232 | 2214 | 3097 | 2168 | 1988 | 2958 | 2229 | 2223 | 2435 | 2333 | 2176 | 2392.3 | 2470 | 2315 | 0.937 | -0.093 | 0.47485291 | 0.67524193 | 0.0145 | 0.0083 | 0.0119 | 0.0119 | TRUE | 0.3617 |
| Trp53i11 | 4238 | 6023 | 6010 | 3204 | 2492 | 3336 | 4502 | 6015 | 6035 | 2519 | 2681 | 3652 | 4233.86 | 5517 | 2951 | 0.535 | -0.903 | 8.04E-08 | 2.99E-06 | 0.0339 | 0.0077 | 0.0202 | 0.0202 | TRUE | 0.5241 |
| Trp53i13 | 867 | 922 | 844 | 1103 | 799 | 745 | 921 | 921 | 847 | 867 | 860 | 816 | 871.91 | 896 | 848 | 0.946 | -0.08 | 0.36993222 | 0.58707239 | 0 | 0.0107 | 0.0046 | 0.0046 | TRUE | 0.0273 |
| Trp53inp1 | 15416 | 14542 | 13326 | 14589 | 10401 | 11159 | 16375 | 14523 | 13381 | 11469 | 11191 | 12215 | 13192.22 | 14760 | 11625 | 0.788 | -0.344 | 0.00032435 | 0.00363072 | 0.0062 | 0.0072 | 0.0065 | 0.0065 | TRUE | 0.112 |
| Trp53inp2 | 3184 | 3028 | 2776 | 3666 | 2572 | 2411 | 3382 | 3024 | 2787 | 2882 | 2767 | 2639 | 2913.65 | 3064 | 2763 | 0.902 | -0.149 | 0.12435252 | 0.30099106 | 0.0052 | 0.008 | 0.0064 | 0.0064 | TRUE | 0.0996 |
| Trp53rk | 223 | 158 | 140 | 264 | 175 | 130 | 237 | 158 | 141 | 208 | 188 | 142 | 178.89 | 179 | 179 | 1.008 | 0.012 | 0.96077406 | 0.98008034 | 0.0511 | 0.0253 | 0.0378 | 0.0378 | TRUE | 0.8727 |
| Trp53tg5 | 0 | 0 | 0 | 0 | 0 | 0 | 0 | 0 | 0 | 0 | 0 | 0 | 0 | 0 | 0 | NA | NA | NA | NA | NA | NA | NA | NA | NA | NA |
| Trp63 | 30 | 12 | 14 | 38 | 33 | 14 | 32 | 12 | 14 | 30 | 36 | 15 | 23.1 | 19 | 27 | 1.402 | 0.487 | 0.36398904 | 0.58146815 | 0.1773 | 0.1495 | 0.1628 | 0.1628 | TRUE | 1.1884 |
| Trp73 | 168 | 196 | 186 | 175 | 100 | 111 | 178 | 196 | 187 | 138 | 108 | 122 | 154.6 | 187 | 123 | 0.656 | -0.607 | 0.00045368 | 0.00482171 | 0 | 0.0282 | 0.0149 | 0.0149 | TRUE | 0.1358 |
| Trpa1 | 451 | 271 | 250 | 675 | 463 | 332 | 479 | 271 | 251 | 531 | 498 | 363 | 398.82 | 334 | 464 | 1.393 | 0.478 | 0.06231979 | 0.19104136 | 0.0808 | 0.0152 | 0.0448 | 0.0448 | TRUE | 1.649 |
| Trpc1 | 276 | 338 | 398 | 379 | 255 | 268 | 293 | 338 | 400 | 298 | 274 | 293 | 316.01 | 344 | 288 | 0.84 | -0.251 | 0.09900752 | 0.2595656 | 0.0101 | 0.0173 | 0.0135 | 0.0135 | TRUE | 0.2336 |
| Trpc2 | 384 | 449 | 490 | 662 | 459 | 464 | 408 | 448 | 492 | 520 | 494 | 508 | 478.42 | 449 | 507 | 1.129 | 0.175 | 0.12942077 | 0.30962138 | 0.0023 | 0.0138 | 0.0075 | 0.0075 | TRUE | 0.0796 |
| Trpc3 | 17 | 13 | 21 | 48 | 36 | 17 | 18 | 13 | 21 | 38 | 39 | 19 | 24.53 | 17 | 32 | 1.834 | 0.875 | 0.05543544 | 0.17613373 | 0.0667 | 0.1412 | 0.1055 | 0.1055 | TRUE | 0.8277 |
| Trpc4 | 1 | 1 | 0 | 0 | 1 | 3 | 1 | 1 | 0 | 0 | 1 | 3 | 1.07 | 1 | 1 | 2.065 | 1.046 | 0.61548484 | NA | 0 | 3.0854 | 2.0344 | 2.0344 | TRUE | 0.3261 |
| Trpc4ap | 5311 | 5879 | 5992 | 6690 | 5010 | 5253 | 5641 | 5871 | 6017 | 5259 | 5390 | 5750 | 5654.85 | 5843 | 5466 | 0.935 | -0.096 | 0.172286 | 0.37111375 | 0.0013 | 0.0075 | 0.0034 | 0.0034 | TRUE | 0.025 |
| Trpc5 | 0 | 1 | 0 | 0 | 0 | 0 | 0 | 1 | 0 | 0 | 0 | 0 | 0.17 | 0 | 0 | 0.49 | -1.029 | 0.80087 | NA | 0 | 19.7991 | 10 | 10 | TRUE | 0.405 |
| Trpc6 | 6 | 6 | 2 | 31 | 11 | 4 | 6 | 6 | 2 | 24 | 12 | 4 | 9.16 | 5 | 13 | 2.868 | 1.52 | 0.07089954 | 0.20739697 | 0.3698 | 0.3666 | 0.368 | 0.368 | TRUE | 0.1862 |
| Trpc7 | 8 | 12 | 6 | 12 | 12 | 11 | 8 | 12 | 6 | 9 | 13 | 12 | 10.15 | 9 | 11 | 1.288 | 0.365 | 0.54499779 | 0.73005818 | 0 | 0.3315 | 0.162 | 0.162 | TRUE | 0.3159 |
| Trpd52l3 | 1 | 0 | 1 | 10 | 7 | 1 | 1 | 0 | 1 | 8 | 8 | 1 | 3.09 | 1 | 6 | 8.115 | 3.021 | 0.04312488 | NA | 0 | 1.0723 | 0.7718 | 0.7718 | TRUE | 0.9755 |
| Trpm1 | 22 | 28 | 37 | 17 | 22 | 19 | 23 | 28 | 37 | 13 | 24 | 21 | 24.39 | 29 | 19 | 0.645 | -0.632 | 0.13826922 | 0.32279191 | 0.0268 | 0.142 | 0.0881 | 0.0881 | TRUE | 0.4797 |
| Trpm2 | 19 | 18 | 13 | 28 | 11 | 12 | 20 | 18 | 13 | 22 | 12 | 13 | 16.37 | 17 | 16 | 0.932 | -0.102 | 0.83872885 | 0.91819472 | 0 | 0.2082 | 0.1212 | 0.1212 | TRUE | 0.7176 |
| Trpm3 | 16 | 14 | 11 | 22 | 10 | 9 | 17 | 14 | 11 | 17 | 11 | 10 | 13.32 | 14 | 13 | 0.916 | -0.126 | 0.81444832 | 0.9051849 | 0 | 0.2542 | 0.1342 | 0.1342 | TRUE | 0.5264 |
| Trpm4 | 483 | 786 | 842 | 705 | 599 | 742 | 513 | 785 | 845 | 554 | 644 | 812 | 692.4 | 714 | 670 | 0.937 | -0.094 | 0.65430087 | 0.80568968 | 0.0509 | 0.0117 | 0.0301 | 0.0301 | TRUE | 0.717 |
| Trpm5 | 0 | 1 | 0 | 1 | 1 | 5 | 0 | 1 | 0 | 1 | 1 | 5 | 1.39 | 0 | 2 | 6.582 | 2.719 | 0.17135542 | NA | 0.7906 | 2.3787 | 1.6429 | 1.6429 | TRUE | 1.1742 |
| Trpm6 | 53 | 57 | 69 | 108 | 90 | 49 | 56 | 57 | 69 | 85 | 97 | 54 | 69.65 | 61 | 79 | 1.292 | 0.37 | 0.20206307 | 0.40962182 | 0.037 | 0.0542 | 0.0459 | 0.0459 | TRUE | 0.607 |
| Trpm7 | 5878 | 6417 | 6933 | 8847 | 6567 | 6741 | 6243 | 6409 | 6961 | 6955 | 7066 | 7379 | 6835.54 | 6538 | 7133 | 1.091 | 0.126 | 0.0893792 | 0.24300475 | 0.0019 | 0.0074 | 0.0038 | 0.0038 | TRUE | 0.0392 |
| Trpm8 | 0 | 0 | 0 | 1 | 0 | 0 | 0 | 0 | 0 | 1 | 0 | 0 | 0.13 | 0 | 0 | 1.859 | 0.894 | 0.82650734 | NA | 0 | 25.1515 | 10 | 10 | TRUE | 0.2563 |
| Trps1 | 1552 | 1506 | 1692 | 1982 | 1326 | 1409 | 1648 | 1504 | 1699 | 1558 | 1427 | 1542 | 1563.11 | 1617 | 1509 | 0.934 | -0.099 | 0.26292745 | 0.48235255 | 0.0025 | 0.009 | 0.005 | 0.005 | TRUE | 0.0453 |
| Trpt1 | 60 | 126 | 155 | 175 | 118 | 118 | 64 | 126 | 156 | 138 | 127 | 129 | 123.15 | 115 | 131 | 1.14 | 0.189 | 0.53334057 | 0.72124217 | 0.0866 | 0.0337 | 0.058 | 0.058 | TRUE | 0.6145 |
| Trpv1 | 2 | 1 | 0 | 7 | 3 | 4 | 2 | 1 | 0 | 6 | 3 | 4 | 2.71 | 1 | 4 | 4.277 | 2.097 | 0.11055431 | NA | 0 | 1.2246 | 0.6364 | 0.6364 | TRUE | 0.3781 |
| Trpv2 | 306 | 247 | 240 | 391 | 288 | 229 | 325 | 247 | 241 | 307 | 310 | 251 | 280.1 | 271 | 289 | 1.07 | 0.097 | 0.5737987 | 0.75221952 | 0.0174 | 0.0187 | 0.018 | 0.018 | TRUE | 0.339 |
| Trpv3 | 15 | 15 | 2 | 43 | 15 | 16 | 16 | 15 | 2 | 34 | 16 | 18 | 16.73 | 11 | 23 | 2.067 | 1.048 | 0.12956991 | 0.30972667 | 0.3759 | 0.2038 | 0.2761 | 0.2761 | TRUE | 1.9156 |
| Trpv4 | 106 | 115 | 96 | 113 | 114 | 109 | 113 | 115 | 96 | 89 | 123 | 119 | 109.11 | 108 | 110 | 1.016 | 0.023 | 0.91458823 | 0.95791138 | 0.0122 | 0.0371 | 0.0245 | 0.0245 | TRUE | 0.3208 |
| Trpv5 | 4 | 0 | 0 | 6 | 1 | 0 | 4 | 0 | 0 | 5 | 1 | 0 | 1.67 | 1 | 2 | 1.408 | 0.493 | 0.81692214 | NA | 4.2022 | 1.9754 | 2.6597 | 2.6597 | TRUE | 1.5097 |
| Trpv6 | 0 | 0 | 0 | 2 | 0 | 2 | 0 | 0 | 0 | 2 | 0 | 2 | 0.63 | 0 | 1 | 6.693 | 2.743 | 0.36333482 | NA | 0 | 5.2618 | 5.1693 | 5.1693 | TRUE | 0.2958 |
| Trrap | 9849 | 6970 | 7990 | 12190 | 8410 | 7393 | 10461 | 6961 | 8023 | 9583 | 9049 | 8093 | 8694.91 | 8482 | 8908 | 1.05 | 0.071 | 0.63806075 | 0.79613107 | 0.0207 | 0.0073 | 0.0162 | 0.0162 | TRUE | 0.509 |
| Trub1 | 2009 | 1930 | 1890 | 2326 | 1750 | 1620 | 2134 | 1928 | 1898 | 1829 | 1883 | 1773 | 1907.32 | 1987 | 1828 | 0.921 | -0.119 | 0.15077956 | 0.34172945 | 0.0018 | 0.0086 | 0.0045 | 0.0045 | TRUE | 0.0508 |
| Trub2 | 1745 | 1721 | 1533 | 2373 | 1784 | 1398 | 1853 | 1719 | 1539 | 1865 | 1919 | 1530 | 1737.81 | 1704 | 1771 | 1.04 | 0.057 | 0.63924741 | 0.79669776 | 0.0111 | 0.0088 | 0.0101 | 0.0101 | TRUE | 0.1701 |
| Try10 | 0 | 0 | 0 | 0 | 0 | 0 | 0 | 0 | 0 | 0 | 0 | 0 | 0 | 0 | 0 | NA | NA | NA | NA | NA | NA | NA | NA | NA | NA |
| Try4 | 0 | 0 | 0 | 0 | 0 | 0 | 0 | 0 | 0 | 0 | 0 | 0 | 0 | 0 | 0 | NA | NA | NA | NA | NA | NA | NA | NA | NA | NA |
| Try5 | 1 | 0 | 0 | 0 | 0 | 0 | 1 | 0 | 0 | 0 | 0 | 0 | 0.18 | 0 | 0 | 0.49 | -1.029 | 0.80087 | NA | 0 | 18.6165 | 10 | 10 | TRUE | 0.4614 |
| Tsacc | 114 | 172 | 192 | 224 | 184 | 185 | 121 | 172 | 193 | 176 | 198 | 203 | 177.04 | 162 | 192 | 1.184 | 0.244 | 0.23328265 | 0.44798903 | 0.0234 | 0.0255 | 0.0245 | 0.0245 | TRUE | 0.5063 |
| Tsc1 | 2997 | 3274 | 3454 | 3779 | 2929 | 3040 | 3183 | 3270 | 3468 | 2971 | 3151 | 3328 | 3228.54 | 3307 | 3150 | 0.952 | -0.071 | 0.37606512 | 0.59147378 | 0.0021 | 0.0079 | 0.0043 | 0.0043 | TRUE | 0.0306 |
| Tsc2 | 7101 | 6527 | 6770 | 7352 | 5443 | 5247 | 7543 | 6519 | 6798 | 5780 | 5856 | 5744 | 6373.08 | 6953 | 5793 | 0.833 | -0.263 | 0.00094788 | 0.00853793 | 0.0027 | 0.0074 | 0.0044 | 0.0044 | TRUE | 0.0671 |
| Tsc22d1 | 5188 | 5690 | 5107 | 6972 | 4705 | 4572 | 5511 | 5683 | 5128 | 5481 | 5062 | 5005 | 5311.52 | 5441 | 5183 | 0.953 | -0.07 | 0.3738245 | 0.59012712 | 0.0023 | 0.0075 | 0.0042 | 0.0042 | TRUE | 0.0312 |
| Tsc22d2 | 1326 | 1339 | 1345 | 2026 | 1425 | 1301 | 1408 | 1337 | 1351 | 1593 | 1533 | 1424 | 1441.05 | 1365 | 1517 | 1.112 | 0.153 | 0.06654184 | 0.19895694 | 0.0014 | 0.0092 | 0.0043 | 0.0043 | TRUE | 0.034 |
| Tsc22d3 | 260 | 460 | 543 | 403 | 310 | 319 | 276 | 459 | 545 | 317 | 334 | 349 | 380.06 | 427 | 333 | 0.78 | -0.359 | 0.11631649 | 0.28814159 | 0.0566 | 0.0156 | 0.035 | 0.035 | TRUE | 0.4486 |
| Tsc22d4 | 1784 | 1875 | 1903 | 2026 | 1548 | 1560 | 1895 | 1873 | 1911 | 1593 | 1666 | 1708 | 1774.04 | 1893 | 1656 | 0.874 | -0.194 | 0.00676827 | 0.0391309 | 0 | 0.0088 | 0.0031 | 0.0031 | TRUE | 0.0137 |
| Tsen15 | 848 | 1055 | 1112 | 1101 | 751 | 773 | 901 | 1054 | 1117 | 866 | 808 | 846 | 931.77 | 1024 | 840 | 0.821 | -0.285 | 0.00980898 | 0.05189777 | 0.0057 | 0.0104 | 0.0077 | 0.0077 | TRUE | 0.1311 |
| Tsen2 | 1768 | 1743 | 1752 | 1873 | 1435 | 1350 | 1878 | 1741 | 1759 | 1472 | 1544 | 1478 | 1645.34 | 1793 | 1498 | 0.836 | -0.259 | 0.00070583 | 0.00681295 | 6.00E-04 | 0.0089 | 0.0036 | 0.0036 | TRUE | 0.0208 |
| Tsen34 | 1576 | 1650 | 1680 | 2866 | 2074 | 1916 | 1674 | 1648 | 1687 | 2253 | 2231 | 2097 | 1931.77 | 1670 | 2194 | 1.315 | 0.395 | 4.40E-08 | 1.78E-06 | 4.00E-04 | 0.0086 | 0.0032 | 0.0032 | TRUE | 0.0177 |
| Tsen54 | 2457 | 2624 | 2571 | 3190 | 2508 | 2267 | 2610 | 2621 | 2582 | 2508 | 2698 | 2482 | 2583.28 | 2604 | 2563 | 0.984 | -0.023 | 0.7409798 | 0.86159073 | 7.00E-04 | 0.0082 | 0.0032 | 0.0032 | TRUE | 0.0258 |
| Tsfm | 2763 | 3219 | 3161 | 3290 | 2476 | 2489 | 2935 | 3215 | 3174 | 2586 | 2664 | 2725 | 2883.1 | 3108 | 2658 | 0.855 | -0.226 | 0.00217645 | 0.01646586 | 0.0012 | 0.008 | 0.0036 | 0.0036 | TRUE | 0.0287 |
| Tsg101 | 2686 | 3271 | 3173 | 3727 | 2675 | 2758 | 2853 | 3267 | 3186 | 2930 | 2878 | 3019 | 3022.15 | 3102 | 2942 | 0.948 | -0.076 | 0.35150187 | 0.57030012 | 0.0023 | 0.008 | 0.0045 | 0.0045 | TRUE | 0.0597 |
| Tsga10 | 105 | 108 | 96 | 100 | 74 | 94 | 112 | 108 | 96 | 79 | 80 | 103 | 96.15 | 105 | 87 | 0.824 | -0.279 | 0.19475725 | 0.40071236 | 0 | 0.0412 | 0.0229 | 0.0229 | TRUE | 0.2297 |
| Tsga13 | 0 | 0 | 0 | 0 | 0 | 0 | 0 | 0 | 0 | 0 | 0 | 0 | 0 | 0 | 0 | NA | NA | NA | NA | NA | NA | NA | NA | NA | NA |
| Tsga14 | 780 | 770 | 789 | 1322 | 989 | 808 | 828 | 769 | 792 | 1039 | 1064 | 884 | 896.26 | 796 | 996 | 1.251 | 0.323 | 0.00251719 | 0.01848285 | 0.0046 | 0.0106 | 0.0071 | 0.0071 | TRUE | 0.1123 |
| Tsga8 | 4 | 4 | 1 | 7 | 9 | 1 | 4 | 4 | 1 | 6 | 10 | 1 | 4.25 | 3 | 6 | 1.767 | 0.822 | 0.44731767 | NA | 0.3662 | 0.7812 | 0.5862 | 0.5862 | TRUE | 0.1171 |

| | | | | | | | | | | | | | | | | | | | | | | | | | |
|---|---|---|---|---|---|---|---|---|---|---|---|---|---|---|---|---|---|---|---|---|---|---|---|---|---|
| Tshb | 1 | 1 | 0 | 1 | 1 | 0 | 1 | 1 | 0 | 1 | 1 | 0 | 0.65 | 1 | 1 | 0.919 | -0.122 | 0.95812821 | NA | 0 | 5.0456 | 2.3857 | 2.3857 | TRUE | 0.2521 |
| Tshr | 5 | 4 | 3 | 16 | 16 | 7 | 5 | 4 | 3 | 13 | 17 | 8 | 8.3 | 4 | 13 | 3.052 | 1.61 | 0.02731761 | 0.10801036 | 0 | 0.404 | 0.2195 | 0.2195 | TRUE | 0.0874 |
| Tshz1 | 49 | 53 | 64 | 116 | 88 | 71 | 52 | 53 | 64 | 91 | 95 | 78 | 72.14 | 56 | 88 | 1.559 | 0.64 | 0.00734724 | 0.04158202 | 0 | 0.0526 | 0.0265 | 0.0265 | TRUE | 0.1262 |
| Tshz2 | 3 | 3 | 4 | 10 | 8 | 6 | 3 | 3 | 4 | 8 | 9 | 7 | 5.54 | 3 | 8 | 2.26 | 1.176 | 0.14424869 | 0.33139329 | 0 | 0.6016 | 0.254 | 0.254 | TRUE | 0.0404 |
| Tshz3 | 96 | 103 | 103 | 147 | 111 | 104 | 102 | 103 | 103 | 116 | 119 | 114 | 109.52 | 103 | 116 | 1.131 | 0.178 | 0.3301269 | 0.55093867 | 0 | 0.037 | 0.0149 | 0.0149 | TRUE | 0.0053 |
| Tsix | 80 | 78 | 51 | 215 | 192 | 134 | 85 | 78 | 51 | 169 | 207 | 147 | 122.73 | 71 | 174 | 2.441 | 1.288 | 2.13E-07 | 6.91E-06 | 0.0352 | 0.0337 | 0.0344 | 0.0344 | TRUE | 0.4089 |
| Tsks | 78 | 110 | 88 | 96 | 57 | 77 | 83 | 110 | 88 | 75 | 61 | 84 | 83.69 | 94 | 73 | 0.786 | -0.347 | 0.14727722 | 0.3363323 | 0.0106 | 0.0463 | 0.0291 | 0.0291 | TRUE | 0.2211 |
| Tsku | 279 | 383 | 362 | 490 | 396 | 369 | 296 | 383 | 363 | 385 | 426 | 404 | 376.26 | 347 | 405 | 1.164 | 0.219 | 0.11245617 | 0.28215727 | 0.0069 | 0.0157 | 0.0111 | 0.0111 | TRUE | 0.1866 |
| Tslp | 0 | 2 | 1 | 1 | 1 | 1 | 0 | 2 | 1 | 1 | 1 | 1 | 0.99 | 1 | 1 | 0.968 | -0.047 | 0.97991429 | NA | 0 | 3.3244 | 1.4549 | 1.4549 | TRUE | 0.1811 |
| Tsn | 4916 | 5074 | 4867 | 8181 | 5799 | 5253 | 5222 | 5067 | 4887 | 6431 | 6239 | 5750 | 5599.49 | 5059 | 6140 | 1.214 | 0.28 | 0.00020571 | 0.00250114 | 0.0019 | 0.0075 | 0.0039 | 0.0039 | TRUE | 0.0376 |
| Tsnax | 6253 | 6561 | 6268 | 8509 | 6241 | 6361 | 6642 | 6553 | 6294 | 6689 | 6715 | 6963 | 6642.54 | 6496 | 6789 | 1.045 | 0.064 | 0.2985241 | 0.51905693 | 4.00E-04 | 0.0074 | 0.0025 | 0.0025 | TRUE | 0.0091 |
| Tsnaxip1 | 33 | 22 | 20 | 17 | 14 | 19 | 35 | 22 | 20 | 13 | 15 | 21 | 21.06 | 26 | 16 | 0.635 | -0.656 | 0.15008511 | 0.34072916 | 0.0331 | 0.1634 | 0.0997 | 0.0997 | TRUE | 0.6051 |
| Tspan1 | 45 | 55 | 33 | 75 | 55 | 40 | 48 | 55 | 33 | 59 | 59 | 44 | 49.63 | 45 | 54 | 1.195 | 0.257 | 0.40882266 | 0.62124974 | 0.023 | 0.0733 | 0.0497 | 0.0497 | TRUE | 0.4384 |
| Tspan10 | 0 | 0 | 0 | 0 | 0 | 0 | 0 | 0 | 0 | 0 | 0 | 0 | 0 | 0 | 0 | NA | NA | NA | NA | NA | NA | NA | NA | NA | NA |
| Tspan11 | 44 | 61 | 57 | 33 | 27 | 27 | 47 | 61 | 57 | 26 | 29 | 30 | 41.57 | 55 | 28 | 0.51 | -0.971 | 0.00156495 | 0.01268283 | 0 | 0.0861 | 0.0412 | 0.0412 | TRUE | 0.1397 |
| Tspan12 | 113 | 130 | 142 | 164 | 99 | 103 | 120 | 130 | 143 | 129 | 107 | 113 | 123.44 | 131 | 116 | 0.89 | -0.168 | 0.37170144 | 0.58818853 | 0 | 0.0336 | 0.0174 | 0.0174 | TRUE | 0.1046 |
| Tspan13 | 2506 | 3119 | 2987 | 3265 | 2400 | 2574 | 2662 | 3115 | 2999 | 2567 | 2582 | 2818 | 2790.44 | 2925 | 2656 | 0.907 | -0.14 | 0.12952726 | 0.3096984 | 0.0041 | 0.0081 | 0.0058 | 0.0058 | TRUE | 0.0752 |
| Tspan14 | 2870 | 3543 | 3408 | 3151 | 2552 | 2747 | 3048 | 3538 | 3422 | 2477 | 2746 | 3007 | 3039.79 | 3336 | 2743 | 0.822 | -0.283 | 0.00706966 | 0.0403995 | 0.0071 | 0.008 | 0.0076 | 0.0076 | TRUE | 0.0885 |
| Tspan15 | 576 | 756 | 679 | 522 | 410 | 471 | 612 | 755 | 682 | 410 | 441 | 516 | 569.28 | 683 | 456 | 0.666 | -0.586 | 1.65E-05 | 0.00030381 | 0.0105 | 0.0127 | 0.0115 | 0.0115 | TRUE | 0.1518 |
| Tspan17 | 652 | 765 | 768 | 531 | 385 | 448 | 693 | 764 | 771 | 417 | 414 | 490 | 591.63 | 743 | 440 | 0.593 | -0.755 | 7.77E-11 | 5.81E-09 | 0.0038 | 0.0125 | 0.0079 | 0.0079 | TRUE | 0.1097 |
| Tspan18 | 88 | 104 | 109 | 129 | 88 | 115 | 93 | 104 | 109 | 101 | 95 | 126 | 104.79 | 102 | 107 | 1.047 | 0.066 | 0.75101318 | 0.86846395 | 0.004 | 0.0383 | 0.0218 | 0.0218 | TRUE | 0.2143 |
| Tspan2 | 696 | 710 | 652 | 1074 | 827 | 815 | 739 | 709 | 655 | 844 | 890 | 892 | 788.21 | 701 | 875 | 1.248 | 0.32 | 0.00072923 | 0.00698328 | 9.00E-04 | 0.0111 | 0.0052 | 0.0052 | TRUE | 0.0396 |
| Tspan3 | 3672 | 4776 | 4848 | 4449 | 3205 | 3659 | 3900 | 4770 | 4868 | 3497 | 3448 | 4005 | 4081.54 | 4513 | 3650 | 0.809 | -0.306 | 0.0077471 | 0.04335954 | 0.0102 | 0.0077 | 0.0093 | 0.0093 | TRUE | 0.1714 |
| Tspan31 | 2999 | 3100 | 3079 | 3176 | 2296 | 2381 | 3185 | 3096 | 3092 | 2497 | 2470 | 2606 | 2824.43 | 3124 | 2524 | 0.808 | -0.308 | 2.61E-06 | 6.33E-05 | 1.00E-04 | 0.0081 | 0.0027 | 0.0027 | TRUE | 0.0096 |
| Tspan32 | 4 | 5 | 9 | 12 | 6 | 5 | 4 | 5 | 9 | 9 | 6 | 5 | 6.61 | 6 | 7 | 1.18 | 0.239 | 0.74965697 | 0.8673408 | 0 | 0.5055 | 0.2532 | 0.2532 | TRUE | 0.4284 |
| Tspan33 | 277 | 331 | 310 | 305 | 255 | 254 | 294 | 331 | 311 | 240 | 274 | 278 | 288.04 | 312 | 264 | 0.844 | -0.245 | 0.06789134 | 0.2017299 | 0 | 0.0183 | 0.0095 | 0.0095 | TRUE | 0.0786 |
| Tspan4 | 1253 | 1597 | 1509 | 1554 | 1126 | 1243 | 1331 | 1595 | 1515 | 1222 | 1212 | 1361 | 1372.47 | 1480 | 1265 | 0.854 | -0.228 | 0.02950645 | 0.11361711 | 0.0053 | 0.0093 | 0.0072 | 0.0072 | TRUE | 0.0935 |
| Tspan5 | 989 | 1159 | 1199 | 1038 | 691 | 778 | 1050 | 1158 | 1204 | 816 | 743 | 852 | 970.51 | 1137 | 804 | 0.707 | -0.501 | 9.19E-07 | 2.56E-05 | 0.0036 | 0.0103 | 0.0064 | 0.0064 | TRUE | 0.0531 |
| Tspan6 | 213 | 232 | 251 | 485 | 354 | 342 | 226 | 232 | 252 | 381 | 381 | 374 | 307.75 | 237 | 379 | 1.601 | 0.679 | 2.85E-08 | 1.22E-06 | 0 | 0.0176 | 0.0073 | 0.0073 | TRUE | 0.0359 |
| Tspan7 | 976 | 1096 | 928 | 1456 | 997 | 1044 | 1037 | 1095 | 932 | 1145 | 1073 | 1143 | 1070.53 | 1021 | 1120 | 1.097 | 0.134 | 0.16764617 | 0.3654617 | 0.003 | 0.01 | 0.0059 | 0.0059 | TRUE | 0.0702 |
| Tspan8 | 226 | 181 | 136 | 233 | 147 | 132 | 240 | 181 | 137 | 183 | 158 | 144 | 173.87 | 186 | 162 | 0.874 | -0.194 | 0.40464971 | 0.61732751 | 0.0409 | 0.0259 | 0.0333 | 0.0333 | TRUE | 0.2415 |
| Tspan9 | 3109 | 3536 | 3410 | 4601 | 3270 | 3450 | 3302 | 3531 | 3424 | 3617 | 3518 | 3777 | 3528.26 | 3419 | 3637 | 1.064 | 0.089 | 0.20065303 | 0.40845374 | 8.00E-04 | 0.0078 | 0.0032 | 0.0032 | TRUE | 0.0135 |
| Tspo | 633 | 998 | 1043 | 719 | 502 | 670 | 672 | 997 | 1047 | 565 | 540 | 733 | 759.19 | 905 | 613 | 0.676 | -0.564 | 0.00331097 | 0.02286365 | 0.0379 | 0.0112 | 0.0252 | 0.0252 | TRUE | 0.6023 |
| Tspo2 | 34 | 68 | 65 | 22 | 13 | 28 | 36 | 68 | 65 | 17 | 14 | 31 | 38.54 | 56 | 21 | 0.363 | -1.462 | 0.00055959 | 0.00568676 | 0.1013 | 0.0924 | 0.0964 | 0.0964 | TRUE | 0.9097 |
| Tspy-ps | 0 | 0 | 0 | 0 | 0 | 0 | 0 | 0 | 0 | 0 | 0 | 0 | 0 | 0 | 0 | NA | NA | NA | NA | NA | NA | NA | NA | NA | NA |
| Tspyl1 | 2130 | 2092 | 2220 | 3194 | 2383 | 2229 | 2262 | 2089 | 2229 | 2511 | 2564 | 2440 | 2349.28 | 2193 | 2505 | 1.142 | 0.192 | 0.00823265 | 0.04540048 | 7.00E-04 | 0.0083 | 0.0034 | 0.0034 | TRUE | 0.021 |
| Tspyl2 | 908 | 807 | 735 | 1032 | 838 | 701 | 964 | 806 | 738 | 811 | 902 | 767 | 831.45 | 836 | 827 | 0.989 | -0.016 | 0.90256005 | 0.95205059 | 0.0115 | 0.0109 | 0.0112 | 0.0112 | TRUE | 0.2132 |
| Tspyl3 | 106 | 111 | 120 | 182 | 162 | 126 | 113 | 111 | 120 | 143 | 174 | 138 | 133.21 | 115 | 152 | 1.322 | 0.402 | 0.02942763 | 0.11340213 | 0 | 0.0316 | 0.0169 | 0.0169 | TRUE | 0.1702 |
| Tspyl4 | 279 | 207 | 199 | 412 | 269 | 187 | 296 | 207 | 200 | 324 | 289 | 205 | 253.48 | 234 | 273 | 1.167 | 0.223 | 0.32770418 | 0.54896977 | 0.0475 | 0.0199 | 0.0333 | 0.0333 | TRUE | 0.5885 |
| Tspyl5 | 79 | 75 | 91 | 159 | 100 | 103 | 84 | 75 | 91 | 125 | 108 | 113 | 99.25 | 83 | 115 | 1.384 | 0.469 | 0.02142778 | 0.09136496 | 0 | 0.0401 | 0.0196 | 0.0196 | TRUE | 0.076 |
| Tsr1 | 13358 | 12713 | 12751 | 16626 | 11840 | 11176 | 14189 | 12697 | 12803 | 13070 | 12739 | 12234 | 12955.26 | 13230 | 12681 | 0.959 | -0.061 | 0.41958037 | 0.63006965 | 0.0024 | 0.0072 | 0.004 | 0.004 | TRUE | 0.0491 |
| Tsr2 | 1078 | 1225 | 1216 | 1818 | 1285 | 1107 | 1145 | 1223 | 1221 | 1429 | 1383 | 1212 | 1268.83 | 1196 | 1341 | 1.122 | 0.166 | 0.0889048 | 0.24218996 | 0.0037 | 0.0095 | 0.006 | 0.006 | TRUE | 0.0847 |
| Tsr3 | 798 | 941 | 889 | 1373 | 1084 | 930 | 848 | 940 | 893 | 1079 | 1166 | 1018 | 990.62 | 894 | 1088 | 1.217 | 0.284 | 0.00334453 | 0.02305505 | 0.0026 | 0.0102 | 0.0057 | 0.0057 | TRUE | 0.0464 |
| Tssc1 | 1283 | 1262 | 1245 | 1885 | 1440 | 1374 | 1363 | 1260 | 1250 | 1482 | 1549 | 1504 | 1401.42 | 1291 | 1512 | 1.171 | 0.227 | 0.00432775 | 0.02802021 | 6.00E-04 | 0.0093 | 0.0039 | 0.0039 | TRUE | 0.0281 |
| Tssc4 | 694 | 694 | 670 | 1137 | 837 | 722 | 737 | 693 | 673 | 894 | 901 | 790 | 781.29 | 701 | 862 | 1.23 | 0.299 | 0.0031487 | 0.02204781 | 0.0024 | 0.0111 | 0.0061 | 0.0061 | TRUE | 0.0608 |
| Tssk1 | 0 | 0 | 0 | 1 | 1 | 0 | 0 | 0 | 0 | 1 | 1 | 0 | 0.31 | 0 | 1 | 3.32 | 1.731 | 0.65383052 | NA | 0 | 10.6224 | 8.9541 | 8.9541 | TRUE | 0.202 |
| Tssk2 | 0 | 0 | 0 | 0 | 0 | 0 | 0 | 0 | 0 | 0 | 0 | 0 | 0 | 0 | 0 | NA | NA | NA | NA | NA | NA | NA | NA | NA | NA |
| Tssk3 | 3 | 4 | 1 | 5 | 1 | 3 | 3 | 4 | 1 | 4 | 1 | 3 | 2.75 | 3 | 3 | 1.029 | 0.042 | 0.97154731 | NA | 0 | 1.2066 | 0.6154 | 0.6154 | TRUE | 0.3712 |
| Tssk4 | 0 | 0 | 0 | 0 | 0 | 1 | 0 | 0 | 0 | 0 | 0 | 1 | 0.18 | 0 | 0 | 1.859 | 0.894 | 0.82650734 | NA | 0 | 18.0643 | 10 | 10 | TRUE | 0.5383 |
| Tssk5 | 4 | 1 | 4 | 4 | 3 | 3 | 4 | 1 | 4 | 3 | 3 | 3 | 3.15 | 3 | 3 | 1.045 | 0.063 | 0.95234996 | NA | 0 | 1.0516 | 0.4908 | 0.4908 | TRUE | 0.478 |
| Tssk6 | 3 | 0 | 0 | 3 | 5 | 0 | 3 | 0 | 0 | 2 | 5 | 0 | 1.82 | 1 | 2 | 2.457 | 1.297 | 0.51788781 | NA | 3.287 | 1.8163 | 2.222 | 2.222 | TRUE | 0.3907 |
| Tst | 1523 | 1364 | 1408 | 1507 | 1158 | 1153 | 1618 | 1362 | 1414 | 1185 | 1246 | 1262 | 1347.75 | 1465 | 1231 | 0.84 | -0.251 | 0.01043332 | 0.05443413 | 0.0037 | 0.0093 | 0.0062 | 0.0062 | TRUE | 0.1003 |
| Tsta3 | 1913 | 2213 | 2252 | 2599 | 1818 | 1842 | 2032 | 2210 | 2261 | 2043 | 1956 | 2016 | 2086.48 | 2168 | 2005 | 0.925 | -0.112 | 0.15008163 | 0.34072916 | 0.0013 | 0.0085 | 0.0039 | 0.0039 | TRUE | 0.0362 |
| Tstd1 | 38 | 46 | 55 | 34 | 34 | 29 | 40 | 46 | 55 | 27 | 37 | 32 | 39.43 | 47 | 32 | 0.667 | -0.585 | 0.06829958 | 0.20245789 | 0 | 0.0905 | 0.0478 | 0.0478 | TRUE | 0.178 |
| Tstd2 | 1739 | 1843 | 1997 | 2478 | 1812 | 1824 | 1847 | 1841 | 2005 | 1948 | 1950 | 1997 | 1931.2 | 1898 | 1965 | 1.035 | 0.05 | 0.50671791 | 0.70098718 | 7.00E-04 | 0.0086 | 0.0035 | 0.0035 | TRUE | 0.0298 |
| Tstd3 | 730 | 929 | 927 | 1087 | 781 | 775 | 775 | 928 | 931 | 855 | 840 | 848 | 862.86 | 878 | 848 | 0.965 | -0.051 | 0.63076168 | 0.79261527 | 0.0041 | 0.0107 | 0.0069 | 0.0069 | TRUE | 0.1235 |
| Tsx | 72 | 41 | 42 | 87 | 80 | 86 | 76 | 41 | 42 | 68 | 86 | 94 | 68.03 | 53 | 83 | 1.556 | 0.638 | 0.04417953 | 0.15075017 | 0.0588 | 0.0553 | 0.0569 | 0.0569 | TRUE | 1.1897 |
| Ttbk1 | 78 | 73 | 92 | 46 | 47 | 52 | 83 | 73 | 92 | 36 | 51 | 57 | 65.3 | 83 | 48 | 0.574 | -0.802 | 0.0033981 | 0.02332239 | 0.0151 | 0.0574 | 0.0375 | 0.0375 | TRUE | 0.4345 |
| Ttbk2 | 1424 | 1480 | 1591 | 1858 | 1342 | 1356 | 1513 | 1478 | 1598 | 1461 | 1444 | 1484 | 1496.17 | 1530 | 1463 | 0.957 | -0.064 | 0.39631508 | 0.60970362 | 0 | 0.0091 | 0.0035 | 0.0035 | TRUE | 0.0184 |
| Ttc1 | 3824 | 4076 | 3965 | 4524 | 3322 | 3444 | 4062 | 4071 | 3981 | 3556 | 3574 | 3770 | 3835.74 | 4038 | 3633 | 0.9 | -0.153 | 0.01650136 | 0.07605478 | 3.00E-04 | 0.0078 | 0.0027 | 0.0027 | TRUE | 0.013 |
| Ttc12 | 387 | 608 | 634 | 451 | 396 | 468 | 411 | 607 | 637 | 355 | 426 | 512 | 491.3 | 552 | 431 | 0.78 | -0.359 | 0.07540768 | 0.21635982 | 0.0418 | 0.0136 | 0.0273 | 0.0273 | TRUE | 0.5627 |
| Ttc13 | 3394 | 3207 | 3143 | 3699 | 2730 | 2528 | 3605 | 3203 | 3156 | 2908 | 2937 | 2767 | 3096.04 | 3321 | 2871 | 0.865 | -0.21 | 0.01248516 | 0.06194271 | 0.0028 | 0.008 | 0.0048 | 0.0048 | TRUE | 0.0677 |
| Ttc14 | 2336 | 2360 | 2266 | 2867 | 2565 | 2415 | 2481 | 2357 | 2275 | 2254 | 2760 | 2644 | 2461.78 | 2371 | 2553 | 1.076 | 0.105 | 0.29965693 | 0.51980705 | 0.0057 | 0.0082 | 0.007 | 0.007 | TRUE | 0.1288 |
| Ttc16 | 64 | 75 | 70 | 76 | 64 | 44 | 68 | 75 | 70 | 60 | 69 | 48 | 64.99 | 71 | 59 | 0.83 | -0.269 | 0.28332325 | 0.50498071 | 0 | 0.0576 | 0.0299 | 0.0299 | TRUE | 0.1982 |
| Ttc17 | 456 | 429 | 450 | 592 | 405 | 429 | 484 | 428 | 452 | 465 | 436 | 470 | 455.9 | 455 | 457 | 1.005 | 0.008 | 0.94546366 | 0.97275189 | 0 | 0.0141 | 0.0065 | 0.0065 | TRUE | 0.0368 |
| Ttc18 | 22 | 21 | 14 | 30 | 23 | 25 | 23 | 21 | 14 | 24 | 25 | 27 | 22.35 | 19 | 25 | 1.295 | 0.373 | 0.36536203 | 0.58293715 | 0 | 0.1543 | 0.0766 | 0.0766 | TRUE | 0.3178 |
| Ttc19 | 1677 | 1707 | 1575 | 1886 | 1411 | 1409 | 1781 | 1705 | 1581 | 1483 | 1518 | 1542 | 1601.78 | 1689 | 1514 | 0.896 | -0.158 | 0.05415348 | 0.17296651 | 0.0014 | 0.009 | 0.0042 | 0.0042 | TRUE | 0.0376 |
| Ttc21a | 37 | 33 | 34 | 63 | 55 | 36 | 39 | 33 | 34 | 50 | 59 | 39 | 42.42 | 35 | 49 | 1.394 | 0.479 | 0.12344897 | 0.29946124 | 0 | 0.0846 | 0.0454 | 0.0454 | TRUE | 0.1372 |
| Ttc21b | 1604 | 1759 | 1869 | 1706 | 1154 | 1332 | 1704 | 1757 | 1877 | 1341 | 1242 | 1458 | 1563 | 1779 | 1347 | 0.757 | -0.402 | 2.22E-05 | 0.00039012 | 0.0035 | 0.009 | 0.0058 | 0.0058 | TRUE | 0.0612 |

| | | | | | | | | | | | | | | | | | | | | | | | | | |
|---|---|---|---|---|---|---|---|---|---|---|---|---|---|---|---|---|---|---|---|---|---|---|---|---|---|
| Ttc22 | 3 | 1 | 1 | 4 | 5 | 2 | 3 | 1 | 1 | 3 | 5 | 2 | 2.65 | 2 | 3 | 2.075 | 1.053 | 0.38061348 | NA | 0 | 1.2499 | 0.6048 | 0.6048 | TRUE | 0.4022 |
| Ttc23 | 422 | 460 | 500 | 527 | 394 | 396 | 448 | 459 | 502 | 414 | 424 | 433 | 446.9 | 470 | 424 | 0.901 | -0.15 | 0.16777799 | 0.36558665 | 0 | 0.0143 | 0.0063 | 0.0063 | TRUE | 0.0418 |
| Ttc23l | 0 | 0 | 1 | 0 | 0 | 0 | 0 | 0 | 1 | 0 | 0 | 0 | 0.17 | 0 | 0 | 0.49 | -1.029 | 0.80087 | NA | 0 | 19.6928 | 10 | 10 | TRUE | 0.4098 |
| Ttc24 | 1 | 3 | 1 | 5 | 1 | 4 | 1 | 3 | 1 | 4 | 1 | 4 | 2.41 | 2 | 3 | 1.86 | 0.896 | 0.47885697 | NA | 0 | 1.3751 | 0.6958 | 0.6958 | TRUE | 0.4018 |
| Ttc25 | 109 | 157 | 172 | 122 | 119 | 129 | 116 | 157 | 173 | 96 | 128 | 141 | 135.07 | 149 | 122 | 0.816 | -0.294 | 0.20986642 | 0.42015908 | 0.033 | 0.0313 | 0.0321 | 0.0321 | TRUE | 0.3815 |
| Ttc26 | 718 | 647 | 658 | 922 | 674 | 633 | 763 | 646 | 661 | 725 | 725 | 693 | 702.07 | 690 | 714 | 1.036 | 0.051 | 0.62822284 | 0.79087692 | 0.0029 | 0.0116 | 0.0066 | 0.0066 | TRUE | 0.0999 |
| Ttc27 | 3977 | 3847 | 4019 | 4897 | 3644 | 3584 | 4224 | 3842 | 4036 | 3850 | 3921 | 3923 | 3965.9 | 4034 | 3898 | 0.966 | -0.05 | 0.47146449 | 0.67293556 | 8.00E-04 | 0.0077 | 0.0032 | 0.0032 | TRUE | 0.0211 |
| Ttc28 | 522 | 479 | 503 | 664 | 458 | 430 | 554 | 478 | 505 | 522 | 493 | 471 | 503.9 | 512 | 495 | 0.967 | -0.048 | 0.67178428 | 0.81697754 | 0.0022 | 0.0134 | 0.0072 | 0.0072 | TRUE | 0.0589 |
| Ttc29 | 46 | 35 | 29 | 108 | 90 | 68 | 49 | 35 | 29 | 85 | 97 | 74 | 61.52 | 38 | 85 | 2.274 | 1.185 | 4.20E-05 | 0.00066889 | 0.0154 | 0.0605 | 0.041 | 0.041 | TRUE | 0.4631 |
| Ttc3 | 6792 | 5925 | 6249 | 7144 | 5216 | 4947 | 7214 | 5917 | 6275 | 5616 | 5612 | 5415 | 6008.29 | 6469 | 5548 | 0.858 | -0.221 | 0.01796645 | 0.08041929 | 0.0053 | 0.0075 | 0.0061 | 0.0061 | TRUE | 0.1239 |
| Ttc30a1 | 217 | 186 | 182 | 305 | 236 | 143 | 230 | 186 | 183 | 240 | 254 | 157 | 208.2 | 200 | 217 | 1.088 | 0.121 | 0.57579079 | 0.75380936 | 0.0358 | 0.0227 | 0.0291 | 0.0291 | TRUE | 0.6286 |
| Ttc30a2 | 2 | 1 | 1 | 6 | 2 | 4 | 2 | 1 | 1 | 5 | 2 | 4 | 2.56 | 1 | 4 | 2.763 | 1.466 | 0.23061062 | NA | 0 | 1.2926 | 0.5754 | 0.5754 | TRUE | 0.2016 |
| Ttc30b | 65 | 63 | 50 | 143 | 135 | 72 | 69 | 63 | 50 | 112 | 145 | 79 | 86.44 | 61 | 112 | 1.849 | 0.887 | 0.00198981 | 0.01530149 | 0.0483 | 0.045 | 0.0465 | 0.0465 | TRUE | 0.1095 |
| Ttc32 | 126 | 144 | 171 | 215 | 153 | 168 | 134 | 144 | 172 | 169 | 165 | 184 | 161.15 | 150 | 173 | 1.15 | 0.202 | 0.24252689 | 0.45909216 | 0.0032 | 0.0273 | 0.0153 | 0.0153 | TRUE | 0.1722 |
| Ttc33 | 1169 | 1120 | 1244 | 1582 | 1196 | 1195 | 1242 | 1119 | 1249 | 1244 | 1287 | 1308 | 1241.32 | 1203 | 1280 | 1.063 | 0.088 | 0.30603642 | 0.52665845 | 0.0014 | 0.0096 | 0.0046 | 0.0046 | TRUE | 0.0456 |
| Ttc34 | 3 | 3 | 6 | 9 | 9 | 8 | 3 | 3 | 6 | 7 | 10 | 9 | 6.29 | 4 | 9 | 2.075 | 1.053 | 0.1724536 | 0.37119832 | 0 | 0.5309 | 0.2462 | 0.2462 | TRUE | 0.3043 |
| Ttc36 | 4 | 1 | 0 | 2 | 0 | 0 | 4 | 1 | 0 | 2 | 0 | 0 | 1.14 | 2 | 1 | 0.326 | -1.617 | 0.48316247 | NA | 2.1979 | 2.9054 | 2.676 | 2.676 | TRUE | 0.6658 |
| Ttc37 | 4093 | 4146 | 4458 | 5418 | 3732 | 3939 | 4347 | 4141 | 4476 | 4259 | 4015 | 4312 | 4258.48 | 4321 | 4195 | 0.971 | -0.043 | 0.54868706 | 0.73344285 | 0.0011 | 0.0077 | 0.0034 | 0.0034 | TRUE | 0.017 |
| Ttc38 | 312 | 364 | 362 | 405 | 325 | 351 | 331 | 364 | 363 | 318 | 350 | 384 | 351.78 | 353 | 351 | 0.992 | -0.012 | 0.92510673 | 0.96302445 | 0.0032 | 0.0163 | 0.0092 | 0.0092 | TRUE | 0.0796 |
| Ttc39a | 165 | 249 | 291 | 234 | 174 | 163 | 175 | 249 | 292 | 184 | 187 | 178 | 210.95 | 239 | 183 | 0.767 | -0.383 | 0.06260441 | 0.19169031 | 0.0287 | 0.0225 | 0.0256 | 0.0256 | TRUE | 0.304 |
| Ttc39b | 4211 | 4447 | 4854 | 5597 | 3768 | 3813 | 4473 | 4441 | 4874 | 4400 | 4054 | 4174 | 4402.67 | 4596 | 4209 | 0.916 | -0.126 | 0.09739291 | 0.2563981 | 0.0019 | 0.0077 | 0.004 | 0.004 | TRUE | 0.0341 |
| Ttc39c | 594 | 662 | 698 | 811 | 543 | 539 | 631 | 661 | 701 | 638 | 584 | 590 | 634.12 | 664 | 604 | 0.91 | -0.136 | 0.17698698 | 0.37747649 | 0.0011 | 0.0121 | 0.0057 | 0.0057 | TRUE | 0.0293 |
| Ttc39d | 0 | 0 | 0 | 0 | 0 | 0 | 0 | 0 | 0 | 0 | 0 | 0 | 0 | 0 | 0 | NA | NA | NA | | NA | NA | NA | NA | NA | NA |
| Ttc4 | 2411 | 2428 | 2453 | 3081 | 2210 | 2184 | 2561 | 2425 | 2463 | 2422 | 2378 | 2391 | 2439.91 | 2483 | 2397 | 0.966 | -0.051 | 0.44360745 | 0.65031254 | 0 | 0.0083 | 0.0027 | 0.0027 | TRUE | 0.0091 |
| Ttc5 | 2662 | 3361 | 3269 | 3731 | 2880 | 2841 | 2828 | 3357 | 3282 | 2933 | 3099 | 3110 | 3101.37 | 3156 | 3047 | 0.965 | -0.051 | 0.58229984 | 0.7585095 | 0.0039 | 0.008 | 0.0058 | 0.0058 | TRUE | 0.1002 |
| Ttc7 | 1362 | 1639 | 1682 | 1288 | 989 | 1038 | 1447 | 1637 | 1689 | 1013 | 1064 | 1136 | 1330.89 | 1591 | 1071 | 0.673 | -0.572 | 8.94E-09 | 4.26E-07 | 0.0043 | 0.0094 | 0.0064 | 0.0064 | TRUE | 0.0754 |
| Ttc7b | 2802 | 3445 | 3589 | 4439 | 3054 | 3298 | 2976 | 3441 | 3604 | 3490 | 3286 | 3610 | 3401.03 | 3340 | 3462 | 1.036 | 0.052 | 0.59747269 | 0.77004845 | 0.0057 | 0.0079 | 0.0066 | 0.0066 | TRUE | 0.1102 |
| Ttc8 | 213 | 200 | 178 | 410 | 237 | 208 | 226 | 200 | 179 | 322 | 255 | 228 | 234.95 | 202 | 268 | 1.336 | 0.418 | 0.02346553 | 0.09736467 | 0.0195 | 0.0209 | 0.0202 | 0.0202 | TRUE | 0.3622 |
| Ttc9 | 63 | 112 | 109 | 117 | 104 | 98 | 67 | 112 | 109 | 92 | 112 | 107 | 99.9 | 96 | 104 | 1.075 | 0.105 | 0.67825621 | 0.82117578 | 0.032 | 0.0399 | 0.0361 | 0.0361 | TRUE | 0.6688 |
| Ttc9b | 41 | 45 | 40 | 73 | 66 | 60 | 44 | 45 | 40 | 57 | 71 | 66 | 53.79 | 43 | 65 | 1.503 | 0.588 | 0.02704734 | 0.10723237 | 0 | 0.0682 | 0.0314 | 0.0314 | TRUE | 0.0988 |
| Ttc9c | 3290 | 4074 | 4235 | 3798 | 2743 | 3519 | 3495 | 4069 | 4252 | 2986 | 2951 | 3852 | 3600.8 | 3939 | 3263 | 0.828 | -0.272 | 0.04057015 | 0.1423837 | 0.0169 | 0.0078 | 0.0124 | 0.0124 | TRUE | 0.3023 |
| Ttf1 | 1598 | 1481 | 1440 | 2092 | 1576 | 1501 | 1697 | 1479 | 1446 | 1645 | 1696 | 1643 | 1600.95 | 1541 | 1661 | 1.078 | 0.109 | 0.23993965 | 0.45609524 | 0.0032 | 0.009 | 0.0055 | 0.0055 | TRUE | 0.0948 |
| Ttf2 | 5777 | 6347 | 6938 | 6579 | 5154 | 5665 | 6136 | 6339 | 6966 | 5172 | 5545 | 6201 | 6060 | 6480 | 5639 | 0.87 | -0.201 | 0.0399147 | 0.14058388 | 0.0063 | 0.0074 | 0.0067 | 0.0067 | TRUE | 0.0925 |
| Tti1 | 4010 | 3812 | 3952 | 4941 | 3631 | 3719 | 4259 | 3807 | 3968 | 3884 | 3907 | 4071 | 3982.77 | 4011 | 3954 | 0.986 | -0.021 | 0.77961878 | 0.88603691 | 0.0017 | 0.0077 | 0.0038 | 0.0038 | TRUE | 0.0355 |
| Tti2 | 975 | 1019 | 1001 | 1316 | 1032 | 1037 | 1036 | 1018 | 1005 | 1035 | 1110 | 1135 | 1056.41 | 1020 | 1093 | 1.071 | 0.1 | 0.23556509 | 0.45072578 | 0 | 0.01 | 0.0041 | 0.0041 | TRUE | 0.0277 |
| Ttk | 5872 | 6026 | 6270 | 7203 | 5383 | 5579 | 6237 | 6018 | 6296 | 5662 | 5792 | 6107 | 6018.73 | 6184 | 5854 | 0.946 | -0.079 | 0.22639289 | 0.43925066 | 8.00E-04 | 0.0075 | 0.0029 | 0.0029 | TRUE | 0.0174 |
| Ttl | 790 | 869 | 798 | 1204 | 782 | 749 | 839 | 868 | 801 | 946 | 841 | 820 | 852.67 | 836 | 869 | 1.041 | 0.058 | 0.56145959 | 0.74264849 | 0.0028 | 0.0108 | 0.0061 | 0.0061 | TRUE | 0.0745 |
| Ttll1 | 1561 | 1465 | 1414 | 1561 | 1052 | 1130 | 1658 | 1463 | 1420 | 1227 | 1132 | 1237 | 1356.16 | 1514 | 1199 | 0.792 | -0.336 | 0.0005678 | 0.00575533 | 0.0036 | 0.0093 | 0.0061 | 0.0061 | TRUE | 0.0835 |
| Ttll10 | 3 | 5 | 0 | 3 | 1 | 4 | 3 | 5 | 0 | 2 | 1 | 4 | 2.67 | 3 | 2 | 0.952 | -0.071 | 0.95734325 | NA | 0.4401 | 1.2429 | 0.9063 | 0.9063 | TRUE | 0.4069 |
| Ttll11 | 15 | 14 | 17 | 17 | 6 | 11 | 16 | 14 | 17 | 13 | 6 | 12 | 13.14 | 16 | 10 | 0.685 | -0.545 | 0.31145616 | 0.53199265 | 0 | 0.2576 | 0.1307 | 0.1307 | TRUE | 0.3919 |
| Ttll12 | 4682 | 4091 | 4206 | 6060 | 4398 | 4044 | 4973 | 4086 | 4223 | 4764 | 4732 | 4427 | 4534.13 | 4427 | 4641 | 1.048 | 0.068 | 0.48895576 | 0.6853724 | 0.0056 | 0.0076 | 0.0068 | 0.0068 | TRUE | 0.1414 |
| Ttll13 | 44 | 38 | 33 | 77 | 40 | 33 | 47 | 38 | 33 | 61 | 43 | 36 | 42.92 | 39 | 47 | 1.197 | 0.259 | 0.43943133 | 0.64664868 | 0.0314 | 0.0837 | 0.0577 | 0.0577 | TRUE | 0.6167 |
| Ttll2 | 2 | 3 | 1 | 2 | 3 | 5 | 2 | 3 | 1 | 2 | 3 | 5 | 2.73 | 2 | 3 | 1.645 | 0.718 | 0.54038853 | NA | 0 | 1.2123 | 0.5954 | 0.5954 | TRUE | 0.1972 |
| Ttll3 | 165 | 228 | 203 | 268 | 232 | 183 | 175 | 228 | 204 | 211 | 250 | 200 | 211.24 | 202 | 220 | 1.087 | 0.12 | 0.47983912 | 0.67912904 | 0.0101 | 0.0225 | 0.0162 | 0.0162 | TRUE | 0.1462 |
| Ttll4 | 3019 | 2364 | 2530 | 3443 | 2458 | 2372 | 3207 | 2361 | 2540 | 2707 | 2645 | 2597 | 2675.97 | 2703 | 2650 | 0.98 | -0.029 | 0.8183062 | 0.90726388 | 0.0129 | 0.0081 | 0.0107 | 0.0107 | TRUE | 0.3225 |
| Ttll5 | 1270 | 1070 | 1166 | 1798 | 1311 | 1107 | 1349 | 1069 | 1171 | 1413 | 1411 | 1212 | 1270.69 | 1196 | 1345 | 1.125 | 0.17 | 0.15799765 | 0.35157152 | 0.0095 | 0.0095 | 0.0097 | 0.0097 | TRUE | 0.1493 |
| Ttll6 | 243 | 230 | 164 | 256 | 200 | 195 | 258 | 230 | 165 | 201 | 215 | 213 | 213.73 | 218 | 210 | 0.965 | -0.051 | 0.79085596 | 0.89242185 | 0.0217 | 0.0223 | 0.022 | 0.022 | TRUE | 0.4968 |
| Ttll7 | 255 | 157 | 177 | 299 | 218 | 186 | 271 | 157 | 178 | 235 | 235 | 204 | 213.1 | 202 | 225 | 1.114 | 0.156 | 0.4851266 | 0.68237017 | 0.0402 | 0.0224 | 0.0311 | 0.0311 | TRUE | 0.9668 |
| Ttll8 | 10 | 7 | 8 | 12 | 14 | 18 | 11 | 7 | 8 | 9 | 15 | 20 | 11.64 | 9 | 15 | 1.707 | 0.772 | 0.19282699 | 0.39842218 | 0 | 0.2899 | 0.1596 | 0.1596 | TRUE | 0.1802 |
| Ttll9 | 42 | 48 | 58 | 56 | 39 | 53 | 45 | 48 | 58 | 44 | 42 | 58 | 49.13 | 50 | 48 | 0.951 | -0.072 | 0.80471387 | 0.89931073 | 0 | 0.074 | 0.0409 | 0.0409 | TRUE | 0.2497 |
| Ttn | 168 | 71 | 102 | 373 | 238 | 123 | 178 | 71 | 102 | 293 | 256 | 135 | 172.62 | 117 | 228 | 1.949 | 0.963 | 0.00864109 | 0.04711474 | 0.1735 | 0.026 | 0.0903 | 0.0903 | TRUE | 1.0429 |
| Ttpa | 419 | 318 | 379 | 369 | 289 | 301 | 445 | 318 | 381 | 290 | 311 | 329 | 345.62 | 381 | 310 | 0.813 | -0.298 | 0.05831989 | 0.18243185 | 0.0136 | 0.0164 | 0.0149 | 0.0149 | TRUE | 0.1557 |
| Ttpal | 637 | 522 | 467 | 938 | 579 | 464 | 677 | 521 | 469 | 737 | 623 | 508 | 589.19 | 556 | 623 | 1.123 | 0.167 | 0.36621331 | 0.58366172 | 0.0335 | 0.0125 | 0.0229 | 0.0229 | TRUE | 0.23 |
| Ttr | 0 | 0 | 0 | 0 | 0 | 1 | 0 | 0 | 0 | 0 | 0 | 1 | 0.18 | 0 | 0 | 1.859 | 0.894 | 0.82650734 | NA | 0 | 18.0643 | 10 | 10 | TRUE | 0.5383 |
| Ttyh1 | 485 | 523 | 528 | 590 | 461 | 447 | 515 | 522 | 530 | 464 | 496 | 489 | 502.8 | 522 | 483 | 0.923 | -0.115 | 0.24790456 | 0.46464061 | 0 | 0.0135 | 0.0052 | 0.0052 | TRUE | 0.0152 |
| Ttyh2 | 1721 | 2283 | 2399 | 2323 | 1987 | 2533 | 1828 | 2280 | 2409 | 1826 | 2138 | 2773 | 2208.95 | 2172 | 2246 | 1.033 | 0.047 | 0.77982587 | 0.8861249 | 0.0322 | 0.0084 | 0.0201 | 0.0201 | TRUE | 0.5081 |
| Ttyh3 | 1750 | 1154 | 1068 | 1391 | 987 | 810 | 1859 | 1153 | 1072 | 1093 | 1062 | 887 | 1187.64 | 1361 | 1014 | 0.745 | -0.424 | 0.03776976 | 0.13501918 | 0.0553 | 0.0097 | 0.0292 | 0.0292 | TRUE | 1.2279 |
| Tub | 1890 | 2371 | 2306 | 2050 | 1641 | 1863 | 2008 | 2368 | 2315 | 1612 | 1766 | 2039 | 2017.91 | 2230 | 1806 | 0.809 | -0.306 | 0.00989086 | 0.05224333 | 0.0103 | 0.0085 | 0.0096 | 0.0096 | TRUE | 0.1547 |
| Tuba1a | 13499 | 12154 | 12312 | 12891 | 9078 | 8432 | 14338 | 12138 | 12363 | 10134 | 9767 | 9230 | 11328.43 | 12946 | 9710 | 0.75 | -0.415 | 6.52E-06 | 0.00013894 | 0.0053 | 0.0072 | 0.006 | 0.006 | TRUE | 0.1081 |
| Tuba1b | 15439 | 16267 | 16287 | 25039 | 17137 | 15550 | 16399 | 16246 | 16354 | 19684 | 18438 | 17022 | 17357.13 | 16333 | 18381 | 1.126 | 0.171 | 0.025485 | 0.10303752 | 0.0025 | 0.0071 | 0.0041 | 0.0041 | TRUE | 0.0512 |
| Tuba1c | 6889 | 7589 | 7230 | 10369 | 7364 | 6674 | 7317 | 7579 | 7260 | 8151 | 7923 | 7306 | 7589.41 | 7385 | 7793 | 1.055 | 0.078 | 0.27940088 | 0.5005155 | 0.0016 | 0.0073 | 0.0036 | 0.0036 | TRUE | 0.0365 |
| Tuba3a | 2242 | 2808 | 2413 | 3212 | 2637 | 2544 | 2381 | 2804 | 2423 | 2525 | 2837 | 2785 | 2625.96 | 2536 | 2716 | 1.07 | 0.098 | 0.32197713 | 0.54329981 | 0.0055 | 0.0082 | 0.0067 | 0.0067 | TRUE | 0.1039 |
| Tuba3b | 216 | 260 | 244 | 608 | 449 | 337 | 229 | 260 | 245 | 478 | 483 | 369 | 344.01 | 245 | 443 | 1.813 | 0.859 | 1.75E-08 | 7.94E-07 | 0.011 | 0.0165 | 0.0135 | 0.0135 | TRUE | 0.2441 |
| Tuba4a | 6488 | 4266 | 4175 | 4939 | 3621 | 3231 | 6891 | 4260 | 4192 | 3883 | 3896 | 3537 | 4443.25 | 5114 | 3772 | 0.738 | -0.439 | 0.01666549 | 0.07644185 | 0.0464 | 0.0076 | 0.024 | 0.024 | TRUE | 1.1252 |
| Tuba8 | 40 | 43 | 38 | 70 | 48 | 49 | 42 | 43 | 38 | 55 | 52 | 54 | 47.32 | 41 | 54 | 1.299 | 0.377 | 0.16791118 | 0.36563511 | 0 | 0.0765 | 0.0324 | 0.0324 | TRUE | 0.032 |
| Tubal3 | 0 | 0 | 1 | 0 | 0 | 0 | 0 | 0 | 1 | 0 | 0 | 0 | 0.17 | 0 | 0 | 0.49 | -1.029 | 0.80087 | NA | 0 | 19.6928 | 10 | 10 | TRUE | 0.4098 |
| Tubb1 | 26 | 22 | 18 | 20 | 9 | 14 | 28 | 22 | 18 | 16 | 10 | 15 | 18.07 | 23 | 14 | 0.607 | -0.721 | 0.12377623 | 0.30001518 | 0 | 0.1893 | 0.0997 | 0.0997 | TRUE | 0.2471 |
| Tubb2a | 1923 | 1948 | 1699 | 3037 | 2210 | 1912 | 2043 | 1945 | 1706 | 2387 | 2378 | 2093 | 2092.05 | 1898 | 2286 | 1.205 | 0.269 | 0.00976442 | 0.05171406 | 0.0061 | 0.0085 | 0.0073 | 0.0073 | TRUE | 0.0948 |
| Tubb2a-ps2 | 12 | 9 | 8 | 7 | 10 | 11 | 13 | 9 | 8 | 6 | 11 | 12 | 9.68 | 10 | 10 | 0.937 | -0.093 | 0.88338264 | 0.94254573 | 0 | 0.3473 | 0.1867 | 0.1867 | TRUE | 0.5801 |

| | | | | | | | | | | | | | | | | | | | | | | | | | |
|---|---|---|---|---|---|---|---|---|---|---|---|---|---|---|---|---|---|---|---|---|---|---|---|---|---|
| Tubb2b | 14575 | 19728 | 20459 | 17243 | 13483 | 15592 | 15481 | 19703 | 20543 | 13555 | 14507 | 17068 | 16809.44 | 18576 | 15043 | 0.81 | -0.304 | 0.02365005 | 0.09783886 | 0.0179 | 0.0071 | 0.013 | 0.013 | TRUE | 0.2597 |
| Tubb3 | 2615 | 2202 | 2163 | 2939 | 2141 | 1920 | 2778 | 2199 | 2172 | 2310 | 2304 | 2102 | 2310.73 | 2383 | 2239 | 0.94 | -0.09 | 0.44974058 | 0.65539207 | 0.0106 | 0.0083 | 0.0097 | 0.0097 | TRUE | 0.254 |
| Tubb4a | 248 | 151 | 135 | 137 | 117 | 101 | 263 | 151 | 136 | 108 | 126 | 111 | 148.99 | 183 | 115 | 0.626 | -0.676 | 0.01335947 | 0.06494505 | 0.0656 | 0.029 | 0.0467 | 0.0467 | TRUE | 1.5627 |
| Tubb4b | 24887 | 23992 | 23494 | 36957 | 24930 | 22416 | 26434 | 23961 | 23590 | 29053 | 26823 | 24538 | 25733.26 | 24662 | 26805 | 1.087 | 0.12 | 0.19100278 | 0.39611884 | 0.0055 | 0.007 | 0.0061 | 0.0061 | TRUE | 0.067 |
| Tubb5 | 74549 | 90323 | 93406 | 113018 | 83280 | 84175 | 79184 | 90206 | 93789 | 88846 | 89604 | 92143 | 88962.15 | 87726 | 90198 | 1.028 | 0.04 | 0.63440758 | 0.7944955 | 0.0033 | 0.0069 | 0.0051 | 0.0051 | TRUE | 0.0889 |
| Tubb6 | 5172 | 5508 | 5593 | 8806 | 6089 | 5476 | 5494 | 5501 | 5616 | 6923 | 6551 | 5994 | 6013.12 | 5537 | 6489 | 1.172 | 0.229 | 0.0035056 | 0.02387321 | 0.0024 | 0.0075 | 0.0043 | 0.0043 | TRUE | 0.0542 |
| Tubd1 | 505 | 526 | 525 | 718 | 469 | 427 | 536 | 525 | 527 | 564 | 505 | 467 | 520.89 | 529 | 512 | 0.969 | -0.045 | 0.69155823 | 0.83090245 | 0.0029 | 0.0132 | 0.0074 | 0.0074 | TRUE | 0.0976 |
| Tube1 | 657 | 903 | 981 | 1127 | 813 | 1114 | 698 | 902 | 985 | 886 | 875 | 1219 | 927.48 | 862 | 993 | 1.152 | 0.204 | 0.24618423 | 0.46320719 | 0.0325 | 0.0105 | 0.0213 | 0.0213 | TRUE | 0.4706 |
| Tubg1 | 4244 | 4149 | 4209 | 6148 | 4133 | 4120 | 4508 | 4144 | 4226 | 4833 | 4447 | 4510 | 4444.62 | 4293 | 4597 | 1.071 | 0.099 | 0.18329203 | 0.38575953 | 0.0017 | 0.0076 | 0.0038 | 0.0038 | TRUE | 0.025 |
| Tubg2 | 65 | 77 | 76 | 104 | 80 | 74 | 69 | 77 | 76 | 82 | 86 | 81 | 78.52 | 74 | 83 | 1.118 | 0.162 | 0.45350604 | 0.65889159 | 0 | 0.0489 | 0.0207 | 0.0207 | TRUE | 0.0315 |
| Tubgcp2 | 2025 | 2121 | 2101 | 3652 | 2350 | 2174 | 2151 | 2118 | 2110 | 2871 | 2528 | 2380 | 2359.66 | 2126 | 2593 | 1.22 | 0.287 | 0.00220506 | 0.01662318 | 0.0041 | 0.0083 | 0.0059 | 0.0059 | TRUE | 0.108 |
| Tubgcp3 | 2385 | 2262 | 2169 | 3508 | 2512 | 2417 | 2533 | 2259 | 2178 | 2758 | 2703 | 2646 | 2512.76 | 2323 | 2702 | 1.163 | 0.218 | 0.01100145 | 0.05653791 | 0.0028 | 0.0082 | 0.0049 | 0.0049 | TRUE | 0.0754 |
| Tubgcp4 | 1638 | 1930 | 1895 | 2547 | 1930 | 1885 | 1740 | 1928 | 1903 | 2002 | 2077 | 2063 | 1952.06 | 1857 | 2047 | 1.102 | 0.14 | 0.07151354 | 0.20844789 | 0.0011 | 0.0086 | 0.0039 | 0.0039 | TRUE | 0.0364 |
| Tubgcp5 | 1507 | 1426 | 1356 | 1962 | 1409 | 1456 | 1601 | 1424 | 1362 | 1542 | 1516 | 1594 | 1506.44 | 1462 | 1551 | 1.061 | 0.085 | 0.35724061 | 0.57570234 | 0.0029 | 0.0091 | 0.0055 | 0.0055 | TRUE | 0.0823 |
| Tubgcp6 | 2390 | 2247 | 2287 | 3368 | 2599 | 2253 | 2539 | 2244 | 2296 | 2648 | 2796 | 2466 | 2498.23 | 2360 | 2637 | 1.118 | 0.16 | 0.07518477 | 0.21599682 | 0.0036 | 0.0082 | 0.0055 | 0.0055 | TRUE | 0.0531 |
| Tufm | 7794 | 6073 | 6175 | 8625 | 6066 | 5669 | 8279 | 6065 | 6200 | 6780 | 6527 | 6206 | 6676.11 | 6848 | 6504 | 0.95 | -0.074 | 0.57020479 | 0.74900702 | 0.0152 | 0.0074 | 0.0121 | 0.0121 | TRUE | 0.4074 |
| Tuft1 | 373 | 536 | 495 | 607 | 513 | 497 | 396 | 535 | 497 | 477 | 552 | 544 | 500.28 | 476 | 524 | 1.099 | 0.137 | 0.34572132 | 0.56516549 | 0.0129 | 0.0135 | 0.0132 | 0.0132 | TRUE | 0.249 |
| Tug1 | 2591 | 1977 | 2019 | 3445 | 2675 | 2164 | 2752 | 1974 | 2027 | 2708 | 2878 | 2369 | 2451.5 | 2251 | 2652 | 1.178 | 0.236 | 0.10864109 | 0.27610173 | 0.0228 | 0.0082 | 0.0152 | 0.0152 | TRUE | 0.4581 |
| Tulp1 | 109 | 162 | 200 | 78 | 72 | 83 | 116 | 162 | 201 | 61 | 77 | 91 | 118.01 | 160 | 76 | 0.477 | -1.067 | 6.41E-05 | 0.00096094 | 0.0489 | 0.0348 | 0.0417 | 0.0417 | TRUE | 0.1263 |
| Tulp2 | 241 | 320 | 328 | 245 | 195 | 205 | 256 | 320 | 329 | 193 | 210 | 224 | 255.29 | 302 | 209 | 0.691 | -0.534 | 0.00072898 | 0.00698328 | 0.0086 | 0.0198 | 0.0139 | 0.0139 | TRUE | 0.1956 |
| Tulp3 | 6277 | 6871 | 6802 | 6027 | 4574 | 4865 | 6667 | 6862 | 6830 | 4738 | 4921 | 5326 | 5890.69 | 6786 | 4995 | 0.736 | -0.443 | 1.23E-09 | 7.35E-08 | 0.0017 | 0.0075 | 0.0037 | 0.0037 | TRUE | 0.0407 |
| Tulp4 | 981 | 858 | 892 | 1336 | 915 | 827 | 1042 | 857 | 896 | 1050 | 984 | 905 | 955.76 | 932 | 980 | 1.053 | 0.075 | 0.51542351 | 0.70778849 | 0.007 | 0.0104 | 0.0085 | 0.0085 | TRUE | 0.1277 |
| Tusc1 | 46 | 51 | 41 | 98 | 70 | 62 | 49 | 51 | 41 | 77 | 75 | 68 | 60.2 | 47 | 73 | 1.565 | 0.647 | 0.01048621 | 0.05463353 | 0 | 0.0616 | 0.0285 | 0.0285 | TRUE | 0.0948 |
| Tusc2 | 928 | 847 | 768 | 1184 | 927 | 746 | 986 | 846 | 771 | 931 | 997 | 817 | 891.26 | 868 | 915 | 1.055 | 0.077 | 0.55319425 | 0.73680632 | 0.0116 | 0.0106 | 0.0111 | 0.0111 | TRUE | 0.168 |
| Tusc3 | 1328 | 1688 | 1707 | 2627 | 1839 | 2014 | 1411 | 1686 | 1714 | 2065 | 1979 | 2205 | 1843.14 | 1604 | 2083 | 1.299 | 0.377 | 0.00033707 | 0.00375183 | 0.0061 | 0.0087 | 0.0074 | 0.0074 | TRUE | 0.133 |
| Tusc5 | 9 | 5 | 2 | 8 | 5 | 9 | 10 | 5 | 2 | 6 | 5 | 10 | 6.35 | 6 | 7 | 1.3 | 0.379 | 0.64610458 | 0.80125503 | 0.0966 | 0.526 | 0.3291 | 0.3291 | TRUE | 0.4102 |
| Tut1 | 2093 | 1590 | 1427 | 2614 | 1829 | 1544 | 2223 | 1588 | 1433 | 2055 | 1968 | 1690 | 1826.15 | 1748 | 1904 | 1.09 | 0.124 | 0.46122231 | 0.66530074 | 0.0315 | 0.0087 | 0.0199 | 0.0199 | TRUE | 0.6818 |
| Tvp23a | 118 | 99 | 81 | 136 | 90 | 74 | 125 | 99 | 81 | 107 | 97 | 81 | 98.38 | 102 | 95 | 0.936 | -0.095 | 0.69285262 | 0.83159723 | 0.023 | 0.0404 | 0.0318 | 0.0318 | TRUE | 0.304 |
| Tvp23b | 1801 | 2924 | 2752 | 2877 | 2149 | 2410 | 1913 | 2920 | 2763 | 2262 | 2312 | 2638 | 2468.08 | 2532 | 2404 | 0.949 | -0.075 | 0.63849359 | 0.79637583 | 0.0281 | 0.0082 | 0.0181 | 0.0181 | TRUE | 0.5542 |
| Twf1 | 4591 | 4222 | 4199 | 5983 | 4382 | 4349 | 4876 | 4217 | 4216 | 4703 | 4715 | 4761 | 4581.34 | 4436 | 4726 | 1.065 | 0.091 | 0.27996854 | 0.50109666 | 0.0034 | 0.0076 | 0.0049 | 0.0049 | TRUE | 0.0915 |
| Twf2 | 4611 | 5527 | 5837 | 4558 | 3174 | 3631 | 4898 | 5520 | 5861 | 3583 | 3415 | 3975 | 4541.9 | 5426 | 3658 | 0.674 | -0.569 | 2.10E-08 | 9.35E-07 | 0.0068 | 0.0076 | 0.0072 | 0.0072 | TRUE | 0.0884 |
| Twist1 | 56 | 152 | 151 | 166 | 129 | 137 | 59 | 152 | 152 | 130 | 139 | 150 | 130.36 | 121 | 140 | 1.153 | 0.206 | 0.5295106 | 0.71784779 | 0.115 | 0.0322 | 0.0694 | 0.0694 | TRUE | 1.9737 |
| Twist2 | 3 | 2 | 1 | 6 | 2 | 4 | 3 | 2 | 1 | 5 | 2 | 4 | 2.91 | 2 | 4 | 1.842 | 0.881 | 0.43123555 | NA | 0 | 1.1406 | 0.5253 | 0.5253 | TRUE | 0.2011 |
| Twistnb | 2888 | 2972 | 2666 | 3448 | 2459 | 2341 | 3068 | 2968 | 2677 | 2711 | 2646 | 2563 | 2771.92 | 2904 | 2640 | 0.909 | -0.137 | 0.09718836 | 0.25620253 | 0.0023 | 0.0081 | 0.0046 | 0.0046 | TRUE | 0.057 |
| Twsg1 | 4201 | 5401 | 5557 | 4959 | 3739 | 4455 | 4462 | 5394 | 5580 | 3898 | 4023 | 4877 | 4705.68 | 5145 | 4266 | 0.829 | -0.271 | 0.03176094 | 0.11986925 | 0.0142 | 0.0076 | 0.0112 | 0.0112 | TRUE | 0.1906 |
| Txk | 0 | 0 | 0 | 1 | 0 | 1 | 0 | 0 | 0 | 1 | 0 | 1 | 0.31 | 0 | 1 | 3.348 | 1.743 | 0.65064889 | NA | 0 | 10.5167 | 8.8968 | 8.8968 | TRUE | 0.2082 |
| Txlna | 7144 | 6148 | 5826 | 8428 | 6141 | 5642 | 7588 | 6140 | 5850 | 6625 | 6607 | 6176 | 6497.83 | 6526 | 6469 | 0.991 | -0.012 | 0.91286285 | 0.95691033 | 0.0087 | 0.0074 | 0.0091 | 0.0091 | TRUE | 0.2472 |
| Txlnb | 185 | 161 | 142 | 189 | 195 | 180 | 197 | 161 | 143 | 149 | 210 | 197 | 175.88 | 167 | 185 | 1.108 | 0.148 | 0.47395471 | 0.67469488 | 0.0243 | 0.0256 | 0.025 | 0.025 | TRUE | 0.345 |
| Txlng | 2709 | 2330 | 2387 | 2979 | 2140 | 2142 | 2877 | 2327 | 2397 | 2342 | 2303 | 2345 | 2431.72 | 2534 | 2330 | 0.92 | -0.121 | 0.23829404 | 0.45412925 | 0.0064 | 0.0083 | 0.0072 | 0.0072 | TRUE | 0.1704 |
| Txn1 | 14176 | 14990 | 14556 | 17067 | 12507 | 12732 | 15057 | 14971 | 14616 | 13417 | 13457 | 13937 | 14242.42 | 14881 | 13604 | 0.914 | -0.13 | 0.01917113 | 0.08408427 | 2.00E-04 | 0.0071 | 0.0021 | 0.0021 | TRUE | 0.0056 |
| Txn2 | 3856 | 3817 | 3760 | 4643 | 3183 | 3336 | 4096 | 3812 | 3775 | 3650 | 3425 | 3652 | 3734.95 | 3894 | 3576 | 0.918 | -0.123 | 0.09209433 | 0.24762974 | 0.0013 | 0.0078 | 0.0036 | 0.0036 | TRUE | 0.0248 |
| Txndc11 | 1421 | 1388 | 1229 | 1581 | 1214 | 1101 | 1509 | 1386 | 1234 | 1243 | 1306 | 1205 | 1313.98 | 1376 | 1251 | 0.909 | -0.137 | 0.18326569 | 0.38575556 | 0.0047 | 0.0094 | 0.0069 | 0.0069 | TRUE | 0.0987 |
| Txndc12 | 1321 | 1749 | 1701 | 1843 | 1322 | 1483 | 1403 | 1747 | 1708 | 1449 | 1422 | 1623 | 1558.74 | 1619 | 1498 | 0.925 | -0.113 | 0.32945549 | 0.55044196 | 0.0084 | 0.009 | 0.009 | 0.009 | TRUE | 0.1639 |
| Txndc15 | 560 | 586 | 560 | 949 | 648 | 598 | 595 | 585 | 562 | 746 | 697 | 655 | 640.03 | 581 | 699 | 1.206 | 0.27 | 0.00747629 | 0.04212792 | 0.0014 | 0.0121 | 0.0058 | 0.0058 | TRUE | 0.0422 |
| Txndc16 | 712 | 948 | 1001 | 866 | 587 | 717 | 756 | 947 | 1005 | 681 | 632 | 785 | 800.9 | 903 | 699 | 0.774 | -0.37 | 0.00979664 | 0.05184987 | 0.0148 | 0.011 | 0.0135 | 0.0135 | TRUE | 0.2388 |
| Txndc17 | 4544 | 4580 | 4619 | 6610 | 4669 | 4445 | 4827 | 4574 | 4638 | 5196 | 5024 | 4866 | 4854.03 | 4680 | 5029 | 1.075 | 0.104 | 0.11203077 | 0.28149099 | 6.00E-04 | 0.0076 | 0.0029 | 0.0029 | TRUE | 0.0105 |
| Txndc2 | 1 | 2 | 2 | 2 | 1 | 3 | 1 | 2 | 2 | 2 | 1 | 3 | 1.83 | 2 | 2 | 1.155 | 0.208 | 0.8769637 | NA | 0 | 1.8039 | 0.7505 | 0.7505 | TRUE | 0.262 |
| Txndc5 | 8820 | 9131 | 8942 | 9909 | 7370 | 7592 | 9368 | 9119 | 8979 | 7790 | 7930 | 8311 | 8582.72 | 9155 | 8010 | 0.875 | -0.193 | 0.00183706 | 0.01440068 | 6.00E-04 | 0.0073 | 0.0026 | 0.0026 | TRUE | 0.0131 |
| Txndc8 | 0 | 0 | 0 | 0 | 0 | 0 | 0 | 0 | 0 | 0 | 0 | 0 | 0 | 0 | 0 | NA | NA | NA | NA | NA | NA | NA | NA | NA | NA |
| Txndc9 | 2088 | 1952 | 1864 | 3781 | 2493 | 2181 | 2218 | 1949 | 1872 | 2972 | 2682 | 2387 | 2346.84 | 2013 | 2680 | 1.332 | 0.414 | 0.00029775 | 0.00339318 | 0.0094 | 0.0083 | 0.009 | 0.009 | TRUE | 0.1109 |
| Txnip | 19897 | 48235 | 45168 | 24788 | 19118 | 31005 | 21134 | 48173 | 45353 | 19486 | 20570 | 33940 | 31442.72 | 38220 | 24665 | 0.645 | -0.632 | 0.13511328 | 0.31796579 | 0.1288 | 0.007 | 0.0587 | 0.1288 | TRUE | 1.8722 |
| Txnl1 | 5721 | 6964 | 6791 | 8470 | 6221 | 6275 | 6077 | 6955 | 6819 | 6658 | 6693 | 6869 | 6678.57 | 6617 | 6740 | 1.019 | 0.027 | 0.73548171 | 0.85807931 | 0.0026 | 0.0074 | 0.0043 | 0.0043 | TRUE | 0.0621 |
| Txnl4a | 8507 | 7827 | 7299 | 11201 | 7735 | 7151 | 9036 | 7817 | 7329 | 8805 | 8322 | 7828 | 8189.57 | 8061 | 8318 | 1.032 | 0.046 | 0.65432913 | 0.80568968 | 0.0062 | 0.0073 | 0.0074 | 0.0074 | TRUE | 0.1367 |
| Txnl4b | 309 | 250 | 270 | 444 | 319 | 263 | 328 | 250 | 271 | 349 | 343 | 288 | 304.86 | 283 | 327 | 1.157 | 0.21 | 0.18411305 | 0.38697109 | 0.0119 | 0.0177 | 0.0147 | 0.0147 | TRUE | 0.2174 |
| Txnrd1 | 16549 | 18099 | 17835 | 19596 | 13359 | 13063 | 17578 | 18076 | 17908 | 15405 | 14373 | 14300 | 16273.27 | 17854 | 14693 | 0.823 | -0.281 | 6.99E-06 | 0.00014665 | 9.00E-04 | 0.0071 | 0.0028 | 0.0028 | TRUE | 0.0221 |
| Txnrd2 | 314 | 364 | 379 | 452 | 345 | 364 | 334 | 364 | 381 | 355 | 371 | 398 | 367.1 | 360 | 375 | 1.042 | 0.06 | 0.62414736 | 0.78828092 | 0 | 0.0159 | 0.0079 | 0.0079 | TRUE | 0.0441 |
| Txnrd3 | 805 | 1046 | 1066 | 1111 | 880 | 938 | 855 | 1045 | 1070 | 873 | 947 | 1027 | 969.51 | 990 | 949 | 0.958 | -0.063 | 0.61164282 | 0.78004614 | 0.0096 | 0.0103 | 0.0099 | 0.0099 | TRUE | 0.169 |
| Tyk2 | 1890 | 2353 | 2329 | 3036 | 2266 | 2292 | 2008 | 2350 | 2339 | 2387 | 2438 | 2509 | 2338.29 | 2232 | 2445 | 1.095 | 0.131 | 0.15044561 | 0.34117752 | 0.0035 | 0.0083 | 0.0055 | 0.0055 | TRUE | 0.0934 |
| Tymp | 17 | 23 | 27 | 10 | 23 | 20 | 18 | 23 | 27 | 8 | 25 | 22 | 20.44 | 23 | 18 | 0.786 | -0.347 | 0.50904502 | 0.70268115 | 0.1271 | 0.1681 | 0.1486 | 0.1486 | TRUE | 1.5681 |
| Tyms | 1926 | 1762 | 1698 | 3400 | 2192 | 2059 | 2046 | 1760 | 1705 | 2673 | 2358 | 2254 | 2132.6 | 1837 | 2428 | 1.323 | 0.404 | 0.00026584 | 0.00308299 | 0.0083 | 0.0084 | 0.0084 | 0.0084 | TRUE | 0.1193 |
| Tyms-ps | 0 | 1 | 2 | 0 | 1 | 2 | 0 | 1 | 2 | 0 | 1 | 2 | 1.05 | 1 | 1 | 1.042 | 0.059 | 0.97668875 | NA | 0 | 3.1584 | 1.9934 | 1.9934 | TRUE | 0.2193 |
| Tyr | 0 | 0 | 0 | 1 | 1 | 0 | 0 | 0 | 0 | 1 | 1 | 0 | 0.31 | 0 | 1 | 3.32 | 1.731 | 0.65383052 | NA | 0 | 10.6224 | 8.9541 | 8.9541 | TRUE | 0.202 |
| Tyro3 | 2748 | 3638 | 3719 | 3882 | 2829 | 3087 | 2919 | 3633 | 3734 | 3052 | 3044 | 3379 | 3293.53 | 3429 | 3158 | 0.921 | -0.119 | 0.30214251 | 0.5227503 | 0.0098 | 0.0079 | 0.0093 | 0.0093 | TRUE | 0.2055 |
| Tyrobp | 0 | 0 | 2 | 1 | 0 | 0 | 0 | 0 | 2 | 1 | 0 | 0 | 0.47 | 1 | 0 | 0.52 | -0.943 | 0.78484416 | NA | 0 | 7.0808 | 6.8441 | 6.8441 | TRUE | 0.9739 |
| Tyrp1 | 4 | 2 | 3 | 6 | 2 | 1 | 4 | 2 | 3 | 5 | 2 | 1 | 2.87 | 3 | 3 | 0.89 | -0.168 | 0.88163144 | NA | 0 | 1.1547 | 0.5699 | 0.5699 | TRUE | 0.7021 |
| Tysnd1 | 1054 | 1031 | 1007 | 1471 | 1083 | 792 | 1120 | 1030 | 1011 | 1156 | 1165 | 867 | 1058.16 | 1054 | 1063 | 1.01 | 0.014 | 0.91751813 | 0.95962175 | 0.0141 | 0.01 | 0.0122 | 0.0122 | TRUE | 0.3075 |
| Tyw1 | 2739 | 2632 | 2658 | 3152 | 2377 | 2404 | 2909 | 2629 | 2669 | 2478 | 2558 | 2632 | 2645.62 | 2736 | 2556 | 0.934 | -0.098 | 0.20243357 | 0.41001441 | 0.0014 | 0.0081 | 0.0039 | 0.0039 | TRUE | 0.0373 |
| Tyw3 | 215 | 214 | 205 | 373 | 243 | 207 | 228 | 214 | 206 | 293 | 261 | 227 | 238.2 | 216 | 260 | 1.21 | 0.275 | 0.07616075 | 0.21774648 | 0.0063 | 0.0207 | 0.0131 | 0.0131 | TRUE | 0.1436 |
| Tyw5 | 372 | 357 | 343 | 537 | 390 | 382 | 395 | 357 | 344 | 422 | 420 | 418 | 392.67 | 365 | 420 | 1.15 | 0.202 | 0.0778378 | 0.22065963 | 0 | 0.0153 | 0.0069 | 0.0069 | TRUE | 0.0574 |

| | | | | | | | | | | | | | | | | | | | | | | | | | |
|---|---|---|---|---|---|---|---|---|---|---|---|---|---|---|---|---|---|---|---|---|---|---|---|---|---|
| U05342 | 2 | 0 | 2 | 0 | 0 | 1 | 2 | 0 | 2 | 0 | 0 | 1 | 0.87 | 1 | 0 | 0.254 | -1.976 | 0.40898021 | NA | 0 | 3.7884 | 2.7571 | 2.7571 | TRUE | 0.5383 |
| U2af1 | 7537 | 8581 | 8117 | 9755 | 7247 | 7004 | 8006 | 8570 | 8150 | 7669 | 7797 | 7667 | 7976.47 | 8242 | 7711 | 0.936 | -0.096 | 0.11444127 | 0.28532528 | 5.00E-04 | 0.0073 | 0.0025 | 0.0025 | TRUE | 0.0148 |
| U2af1l4 | 654 | 702 | 625 | 897 | 697 | 654 | 695 | 701 | 628 | 705 | 750 | 716 | 699.05 | 675 | 724 | 1.073 | 0.101 | 0.29930106 | 0.51961003 | 8.00E-04 | 0.0116 | 0.0054 | 0.0054 | TRUE | 0.0439 |
| U2af2 | 14194 | 12618 | 12998 | 19483 | 13879 | 12138 | 15077 | 12602 | 13051 | 15316 | 14933 | 13287 | 14044.25 | 13577 | 14512 | 1.069 | 0.096 | 0.33967191 | 0.55980544 | 0.0071 | 0.0071 | 0.0073 | 0.0073 | TRUE | 0.1142 |
| U2surp | 9057 | 7544 | 7261 | 12696 | 8807 | 7458 | 9620 | 7534 | 7291 | 9981 | 9476 | 8164 | 8677.59 | 8148 | 9207 | 1.13 | 0.176 | 0.18093867 | 0.38254218 | 0.0174 | 0.0073 | 0.0124 | 0.0124 | TRUE | 0.3047 |
| U90926 | 0 | 0 | 0 | 0 | 0 | 0 | 0 | 0 | 0 | 0 | 0 | 0 | 0 | 0 | 0 | NA | NA | NA | NA | NA | NA | NA | NA | NA | NA |
| Uaca | 3835 | 4203 | 4345 | 4595 | 3344 | 3595 | 4073 | 4198 | 4363 | 3612 | 3598 | 3935 | 3963.22 | 4211 | 3715 | 0.882 | -0.181 | 0.01465543 | 0.06966409 | 0.0015 | 0.0077 | 0.0037 | 0.0037 | TRUE | 0.0324 |
| Uap1 | 2309 | 2486 | 2458 | 3221 | 2293 | 2425 | 2453 | 2483 | 2468 | 2532 | 2467 | 2655 | 2509.54 | 2468 | 2551 | 1.034 | 0.048 | 0.48240388 | 0.68092108 | 3.00E-04 | 0.0082 | 0.0029 | 0.0029 | TRUE | 0.0149 |
| Uap1l1 | 559 | 1226 | 1318 | 954 | 818 | 1032 | 594 | 1224 | 1323 | 750 | 880 | 1130 | 983.56 | 1047 | 920 | 0.878 | -0.188 | 0.48376968 | 0.68148223 | 0.106 | 0.0103 | 0.0508 | 0.0508 | TRUE | 1.7124 |
| Uba1 | 33896 | 33307 | 33001 | 42248 | 30647 | 30879 | 36004 | 33264 | 33136 | 33212 | 32974 | 33802 | 33732.05 | 34135 | 33329 | 0.976 | -0.034 | 0.59029704 | 0.76449681 | 0.0011 | 0.007 | 0.0029 | 0.0029 | TRUE | 0.0281 |
| Uba1y | 860 | 934 | 828 | 3099 | 2377 | 1947 | 913 | 933 | 831 | 2436 | 2558 | 2131 | 1633.78 | 892 | 2375 | 2.662 | 1.412 | 8.19E-42 | 8.62E-39 | 0.0057 | 0.0089 | 0.0071 | 0.0071 | TRUE | 0.0967 |
| Uba2 | 15058 | 17095 | 17413 | 21807 | 15820 | 16001 | 15994 | 17073 | 17484 | 17143 | 17021 | 17516 | 17038.6 | 16850 | 17227 | 1.022 | 0.032 | 0.62135636 | 0.78582572 | 0.001 | 0.0071 | 0.0029 | 0.0029 | TRUE | 0.0242 |
| Uba3 | 8730 | 8949 | 9103 | 10253 | 7427 | 8022 | 9273 | 8937 | 9140 | 8060 | 7991 | 8781 | 8697.18 | 9117 | 8277 | 0.908 | -0.139 | 0.04301329 | 0.14818819 | 0.0014 | 0.0073 | 0.0033 | 0.0033 | TRUE | 0.0345 |
| Uba5 | 1758 | 1583 | 1450 | 2554 | 1736 | 1473 | 1867 | 1581 | 1456 | 2008 | 1868 | 1612 | 1732.04 | 1635 | 1829 | 1.12 | 0.163 | 0.20634423 | 0.41526875 | 0.0135 | 0.0088 | 0.0114 | 0.0114 | TRUE | 0.1864 |
| Uba52 | 18216 | 22661 | 24536 | 25426 | 18899 | 20473 | 19349 | 22632 | 24637 | 19988 | 20334 | 22411 | 21558.35 | 22206 | 20911 | 0.942 | -0.087 | 0.4211107 | 0.63128581 | 0.0086 | 0.0071 | 0.0083 | 0.0083 | TRUE | 0.155 |
| Uba6 | 5267 | 4082 | 4610 | 6084 | 4410 | 4175 | 5594 | 4077 | 4629 | 4783 | 4745 | 4570 | 4733 | 4767 | 4699 | 0.986 | -0.02 | 0.86693653 | 0.93412739 | 0.0128 | 0.0076 | 0.0105 | 0.0105 | TRUE | 0.281 |
| Uba7 | 1 | 2 | 2 | 0 | 0 | 2 | 1 | 2 | 2 | 0 | 0 | 2 | 1.21 | 2 | 1 | 0.408 | -1.293 | 0.50120137 | NA | 0 | 2.7307 | 1.6535 | 1.6535 | TRUE | 0.9754 |
| Ubac1 | 3752 | 4034 | 3844 | 5531 | 3791 | 3827 | 3985 | 4029 | 3860 | 4348 | 4079 | 4189 | 4081.67 | 3958 | 4205 | 1.063 | 0.088 | 0.17607568 | 0.37637393 | 6.00E-04 | 0.0077 | 0.0028 | 0.0028 | TRUE | 0.0109 |
| Ubac2 | 2610 | 2688 | 2865 | 3370 | 2281 | 2388 | 2772 | 2685 | 2877 | 2649 | 2454 | 2614 | 2675.18 | 2778 | 2572 | 0.926 | -0.11 | 0.13197907 | 0.31297158 | 0.001 | 0.0081 | 0.0035 | 0.0035 | TRUE | 0.0195 |
| Ubald1 | 953 | 887 | 859 | 1178 | 875 | 832 | 1012 | 886 | 863 | 926 | 941 | 911 | 923.15 | 920 | 926 | 1.007 | 0.01 | 0.92213195 | 0.96145072 | 0.0027 | 0.0105 | 0.0059 | 0.0059 | TRUE | 0.0906 |
| Ubald2 | 339 | 688 | 682 | 438 | 367 | 441 | 360 | 687 | 685 | 344 | 395 | 483 | 492.32 | 577 | 407 | 0.705 | -0.505 | 0.03983887 | 0.14037951 | 0.0748 | 0.0136 | 0.0414 | 0.0414 | TRUE | 1.2664 |
| Ubap1 | 3152 | 2893 | 2791 | 3378 | 2474 | 2491 | 3348 | 2889 | 2802 | 2656 | 2662 | 2727 | 2847.31 | 3013 | 2682 | 0.89 | -0.168 | 0.0687938 | 0.20341504 | 0.0043 | 0.0081 | 0.0058 | 0.0058 | TRUE | 0.1145 |
| Ubap2 | 15552 | 15111 | 15793 | 18242 | 13588 | 13200 | 16519 | 15091 | 15858 | 14340 | 14620 | 14449 | 15146.34 | 15823 | 14470 | 0.914 | -0.129 | 0.04265733 | 0.14744498 | 0.001 | 0.0071 | 0.0028 | 0.0028 | TRUE | 0.02 |
| Ubap2l | 10800 | 11477 | 11975 | 15843 | 11603 | 11324 | 11472 | 11462 | 12024 | 12455 | 12484 | 12396 | 12048.73 | 11653 | 12445 | 1.068 | 0.095 | 0.09131295 | 0.2466075 | 3.00E-04 | 0.0072 | 0.0022 | 0.0022 | TRUE | 0.0095 |
| Ubash3a | 1338 | 1675 | 1759 | 1678 | 1283 | 1388 | 1421 | 1673 | 1766 | 1319 | 1380 | 1519 | 1513.2 | 1620 | 1406 | 0.867 | -0.205 | 0.07039845 | 0.20657198 | 0.0077 | 0.0091 | 0.0086 | 0.0086 | TRUE | 0.1386 |
| Ubash3b | 200 | 183 | 179 | 270 | 186 | 180 | 212 | 183 | 180 | 212 | 200 | 197 | 197.39 | 192 | 203 | 1.062 | 0.087 | 0.5661375 | 0.74614529 | 0 | 0.0236 | 0.0115 | 0.0115 | TRUE | 0.0958 |
| Ubb | 2481 | 3060 | 3035 | 3748 | 2868 | 2896 | 2635 | 3056 | 3047 | 2946 | 3086 | 3170 | 2990.18 | 2913 | 3067 | 1.053 | 0.074 | 0.40810544 | 0.62087325 | 0.0036 | 0.008 | 0.0054 | 0.0054 | TRUE | 0.0842 |
| Ubc | 459 | 515 | 536 | 535 | 380 | 396 | 488 | 514 | 538 | 421 | 409 | 433 | 467.17 | 513 | 421 | 0.82 | -0.287 | 0.00667787 | 0.03884053 | 0 | 0.014 | 0.0059 | 0.0059 | TRUE | 0.0223 |
| Ubd | 0 | 0 | 0 | 0 | 0 | 0 | 0 | 0 | 0 | 0 | 0 | 0 | 0 | 0 | 0 | NA | NA | NA | NA | NA | NA | NA | NA | NA | NA |
| Ube2a | 1470 | 1533 | 1447 | 2815 | 1980 | 1792 | 1561 | 1531 | 1453 | 2213 | 2130 | 1962 | 1808.38 | 1515 | 2102 | 1.388 | 0.473 | 2.16E-08 | 9.51E-07 | 0.0021 | 0.0087 | 0.0046 | 0.0046 | TRUE | 0.0408 |
| Ube2b | 3400 | 4546 | 4081 | 4993 | 3896 | 3891 | 3611 | 4540 | 4098 | 3925 | 4192 | 4259 | 4104.26 | 4083 | 4125 | 1.01 | 0.015 | 0.88778398 | 0.94494054 | 0.0072 | 0.0077 | 0.0074 | 0.0074 | TRUE | 0.1241 |
| Ube2c | 6930 | 9338 | 9625 | 9180 | 6667 | 7553 | 7361 | 9326 | 9665 | 7217 | 7173 | 8268 | 8168.2 | 8784 | 7553 | 0.86 | -0.218 | 0.07680317 | 0.21885087 | 0.0132 | 0.0073 | 0.0108 | 0.0108 | TRUE | 0.2451 |
| Ube2cbp | 555 | 432 | 414 | 611 | 411 | 424 | 590 | 431 | 416 | 480 | 442 | 464 | 470.55 | 479 | 462 | 0.966 | -0.05 | 0.75344476 | 0.86987269 | 0.0177 | 0.0139 | 0.0159 | 0.0159 | TRUE | 0.4723 |
| Ube2d1 | 1398 | 1312 | 1300 | 1692 | 1227 | 1178 | 1485 | 1310 | 1305 | 1330 | 1320 | 1290 | 1340.06 | 1367 | 1313 | 0.961 | -0.057 | 0.52005854 | 0.71102318 | 0.0021 | 0.0094 | 0.0049 | 0.0049 | TRUE | 0.0683 |
| Ube2d2a | 4505 | 4283 | 4389 | 6251 | 4348 | 4287 | 4785 | 4277 | 4407 | 4914 | 4678 | 4693 | 4625.77 | 4490 | 4762 | 1.061 | 0.085 | 0.25649744 | 0.47519694 | 0.0018 | 0.0076 | 0.0038 | 0.0038 | TRUE | 0.0402 |
| Ube2d2b | 1 | 3 | 0 | 1 | 3 | 2 | 1 | 3 | 0 | 1 | 3 | 2 | 1.71 | 1 | 2 | 1.496 | 0.581 | 0.709455 | NA | 0 | 1.9332 | 1.1344 | 1.1344 | TRUE | 0.5948 |
| Ube2d3 | 16180 | 18358 | 17956 | 23390 | 16457 | 15820 | 17186 | 18334 | 18030 | 18387 | 17707 | 17317 | 17826.94 | 17850 | 17804 | 0.997 | -0.004 | 0.95356801 | 0.97650213 | 8.00E-04 | 0.0071 | 0.0028 | 0.0028 | TRUE | 0.0129 |
| Ube2dnl1 | 43 | 7 | 8 | 75 | 58 | 34 | 46 | 7 | 8 | 59 | 62 | 37 | 36.55 | 20 | 53 | 2.625 | 1.392 | 0.03244587 | 0.12149483 | 0.4987 | 0.097 | 0.2708 | 0.2708 | TRUE | 6.4063 |
| Ube2dnl2 | 4 | 0 | 0 | 7 | 5 | 0 | 4 | 0 | 0 | 6 | 5 | 0 | 2.52 | 1 | 4 | 2.606 | 1.382 | 0.45894212 | NA | 4.0232 | 1.3133 | 2.0103 | 2.0103 | TRUE | 1.8997 |
| Ube2e1 | 2273 | 2740 | 2790 | 3107 | 2157 | 2292 | 2414 | 2736 | 2801 | 2442 | 2321 | 2509 | 2537.41 | 2650 | 2424 | 0.914 | -0.129 | 0.14693734 | 0.33596633 | 0.0032 | 0.0082 | 0.0053 | 0.0053 | TRUE | 0.0736 |
| Ube2e2 | 285 | 327 | 310 | 502 | 359 | 348 | 303 | 327 | 311 | 395 | 386 | 381 | 350.4 | 314 | 387 | 1.236 | 0.305 | 0.00730539 | 0.04146119 | 0 | 0.0163 | 0.0064 | 0.0064 | TRUE | 0.0151 |
| Ube2e3 | 1782 | 1757 | 1640 | 2873 | 1955 | 1768 | 1893 | 1755 | 1647 | 2259 | 2103 | 1935 | 1931.93 | 1765 | 2099 | 1.19 | 0.251 | 0.01036814 | 0.05418365 | 0.0046 | 0.0086 | 0.0064 | 0.0064 | TRUE | 0.0562 |
| Ube2f | 1416 | 1554 | 1539 | 1967 | 1280 | 1331 | 1504 | 1552 | 1545 | 1546 | 1377 | 1457 | 1496.97 | 1534 | 1460 | 0.953 | -0.07 | 0.39285666 | 0.60637095 | 0.0012 | 0.0091 | 0.0041 | 0.0041 | TRUE | 0.0331 |
| Ube2g1 | 4754 | 5275 | 5201 | 6784 | 4828 | 5179 | 5050 | 5268 | 5222 | 5333 | 5195 | 5669 | 5289.51 | 5180 | 5399 | 1.042 | 0.06 | 0.38144507 | 0.59577656 | 0.001 | 0.0075 | 0.0031 | 0.0031 | TRUE | 0.0232 |
| Ube2g2 | 1745 | 1690 | 1672 | 2194 | 1563 | 1559 | 1853 | 1688 | 1679 | 1725 | 1682 | 1707 | 1722.2 | 1740 | 1705 | 0.98 | -0.029 | 0.70857523 | 0.84126423 | 0.001 | 0.0088 | 0.0039 | 0.0039 | TRUE | 0.0391 |
| Ube2h | 3427 | 3485 | 3507 | 4292 | 3016 | 3202 | 3640 | 3481 | 3521 | 3374 | 3245 | 3505 | 3461.02 | 3547 | 3375 | 0.951 | -0.072 | 0.29042302 | 0.51139353 | 6.00E-04 | 0.0079 | 0.003 | 0.003 | TRUE | 0.0136 |
| Ube2i | 4711 | 4630 | 4613 | 6460 | 4558 | 4490 | 5004 | 4624 | 4632 | 5078 | 4904 | 4915 | 4859.56 | 4753 | 4966 | 1.045 | 0.063 | 0.35160724 | 0.5704124 | 9.00E-04 | 0.0076 | 0.0031 | 0.0031 | TRUE | 0.0259 |
| Ube2j1 | 326 | 423 | 434 | 711 | 521 | 484 | 346 | 422 | 436 | 559 | 561 | 530 | 475.64 | 401 | 550 | 1.369 | 0.453 | 0.00030591 | 0.00347104 | 0.005 | 0.0138 | 0.0092 | 0.0092 | TRUE | 0.1648 |
| Ube2j2 | 836 | 730 | 738 | 991 | 768 | 651 | 888 | 729 | 741 | 779 | 826 | 713 | 779.34 | 786 | 773 | 0.983 | -0.024 | 0.84129966 | 0.91985984 | 0.0075 | 0.0111 | 0.0091 | 0.0091 | TRUE | 0.1519 |
| Ube2k | 5294 | 6065 | 6163 | 7275 | 5399 | 5691 | 5623 | 6057 | 6188 | 5719 | 5809 | 6230 | 5937.73 | 5956 | 5919 | 0.994 | -0.009 | 0.90264857 | 0.95205059 | 0.0021 | 0.0075 | 0.004 | 0.004 | TRUE | 0.0291 |
| Ube2l3 | 5636 | 5037 | 4770 | 7636 | 5352 | 4890 | 5986 | 5030 | 4790 | 6003 | 5758 | 5353 | 5486.77 | 5269 | 5705 | 1.083 | 0.115 | 0.28222397 | 0.50364748 | 0.0085 | 0.0075 | 0.008 | 0.008 | TRUE | 0.1729 |
| Ube2l6 | 119 | 67 | 41 | 171 | 122 | 97 | 126 | 67 | 41 | 134 | 131 | 106 | 101.06 | 78 | 124 | 1.59 | 0.669 | 0.06845584 | 0.20275113 | 0.1446 | 0.0395 | 0.0866 | 0.0866 | TRUE | 1.0471 |
| Ube2m | 1991 | 2018 | 2000 | 3852 | 2747 | 2247 | 2115 | 2015 | 2008 | 3028 | 2956 | 2460 | 2430.31 | 2046 | 2815 | 1.376 | 0.461 | 7.13E-06 | 0.00014852 | 0.0058 | 0.0083 | 0.0072 | 0.0072 | TRUE | 0.1468 |
| Ube2n | 3572 | 3488 | 3387 | 4938 | 3444 | 3399 | 3794 | 3483 | 3401 | 3882 | 3706 | 3721 | 3664.44 | 3559 | 3770 | 1.059 | 0.083 | 0.27115577 | 0.49106313 | 0.0016 | 0.0078 | 0.0038 | 0.0038 | TRUE | 0.0403 |
| Ube2o | 12187 | 12357 | 12726 | 14777 | 11575 | 11511 | 12945 | 12341 | 12778 | 11617 | 12454 | 12601 | 12455.86 | 12688 | 12224 | 0.963 | -0.054 | 0.4123348 | 0.62352296 | 0.0011 | 0.0072 | 0.003 | 0.003 | TRUE | 0.0232 |
| Ube2q1 | 2720 | 3036 | 2976 | 4081 | 2875 | 2765 | 2889 | 3032 | 2988 | 3208 | 3093 | 3027 | 3039.61 | 2970 | 3109 | 1.047 | 0.067 | 0.31800949 | 0.53887305 | 4.00E-04 | 0.008 | 0.0029 | 0.0029 | TRUE | 0.0096 |
| Ube2q2 | 3088 | 3446 | 3626 | 4146 | 3214 | 3185 | 3280 | 3442 | 3641 | 3259 | 3458 | 3486 | 3427.71 | 3454 | 3401 | 0.984 | -0.023 | 0.76486303 | 0.87670202 | 0.0018 | 0.0079 | 0.0039 | 0.0039 | TRUE | 0.0272 |
| Ube2ql1 | 48 | 49 | 71 | 87 | 61 | 54 | 51 | 49 | 71 | 68 | 66 | 59 | 60.72 | 57 | 64 | 1.13 | 0.176 | 0.51279379 | 0.70604833 | 0.0075 | 0.0612 | 0.0355 | 0.0355 | TRUE | 0.4066 |
| Ube2r2 | 3344 | 4097 | 3940 | 4980 | 3589 | 3796 | 3552 | 4092 | 3956 | 3915 | 3862 | 4155 | 3921.93 | 3867 | 3977 | 1.028 | 0.04 | 0.6315264 | 0.79271265 | 0.0032 | 0.0077 | 0.0049 | 0.0049 | TRUE | 0.0615 |
| Ube2s | 8220 | 9291 | 8853 | 10743 | 7653 | 7344 | 8731 | 9279 | 8889 | 8445 | 8234 | 8039 | 8603.01 | 8966 | 8239 | 0.919 | -0.122 | 0.04934587 | 0.16288888 | 6.00E-04 | 0.0073 | 0.0027 | 0.0027 | TRUE | 0.0114 |
| Ube2t | 2622 | 3593 | 3452 | 2731 | 2090 | 2480 | 2785 | 3588 | 3466 | 2147 | 2249 | 2715 | 2824.99 | 3280 | 2370 | 0.722 | -0.469 | 0.00047143 | 0.00496025 | 0.0163 | 0.0081 | 0.0126 | 0.0126 | TRUE | 0.2114 |
| Ube2u | 13 | 11 | 11 | 25 | 15 | 19 | 14 | 11 | 11 | 20 | 16 | 21 | 15.4 | 12 | 19 | 1.583 | 0.663 | 0.16786575 | 0.36558665 | 0 | 0.2208 | 0.0978 | 0.0978 | TRUE | 0.0766 |
| Ube2v1 | 2318 | 1988 | 2103 | 2828 | 2076 | 1830 | 2462 | 1985 | 2112 | 2223 | 2234 | 2003 | 2169.87 | 2186 | 2153 | 0.985 | -0.021 | 0.84228486 | 0.92049035 | 0.0076 | 0.0084 | 0.0079 | 0.0079 | TRUE | 0.147 |
| Ube2v2 | 687 | 905 | 914 | 1250 | 873 | 996 | 730 | 904 | 918 | 983 | 939 | 1090 | 927.25 | 851 | 1004 | 1.18 | 0.239 | 0.05412365 | 0.1729472 | 0.009 | 0.0105 | 0.01 | 0.01 | TRUE | 0.1823 |
| Ube2w | 654 | 767 | 724 | 1167 | 809 | 805 | 695 | 766 | 727 | 917 | 870 | 881 | 809.45 | 729 | 889 | 1.221 | 0.287 | 0.00147873 | 0.01211262 | 0 | 0.011 | 0.0046 | 0.0046 | TRUE | 0.0231 |
| Ube2z | 12940 | 14312 | 14053 | 21251 | 15561 | 14903 | 13745 | 14294 | 14111 | 16706 | 16743 | 16314 | 15318.5 | 14050 | 16588 | 1.181 | 0.24 | 1.11E-05 | 0.00021671 | 2.00E-04 | 0.0071 | 0.0021 | 0.0021 | TRUE | 0.0044 |
| Ube3a | 4946 | 4437 | 4615 | 6359 | 4456 | 4077 | 5254 | 4431 | 4634 | 4999 | 4794 | 4463 | 4762.5 | 4773 | 4752 | 0.996 | -0.006 | 0.94978095 | 0.97481103 | 0.0054 | 0.0076 | 0.0062 | 0.0062 | TRUE | 0.0944 |
| Ube3b | 4194 | 4658 | 4926 | 6086 | 4608 | 4633 | 4455 | 4652 | 4946 | 4784 | 4958 | 5072 | 4811.13 | 4684 | 4938 | 1.054 | 0.076 | 0.29912964 | 0.51947509 | 0.0015 | 0.0076 | 0.0036 | 0.0036 | TRUE | 0.0291 |
| Ube3c | 9778 | 8295 | 8575 | 10701 | 7588 | 7161 | 10386 | 8284 | 8610 | 8412 | 8164 | 7839 | 8615.97 | 9093 | 8138 | 0.895 | -0.16 | 0.12408699 | 0.30058362 | 0.0082 | 0.0073 | 0.0077 | 0.0077 | TRUE | 0.1887 |

| | | | | | | | | | | | | | | | | | | | | | | | | | |
|---|---|---|---|---|---|---|---|---|---|---|---|---|---|---|---|---|---|---|---|---|---|---|---|---|---|
| Ube4a | 2732 | 1823 | 1679 | 3735 | 2380 | 1579 | 2902 | 1821 | 1686 | 2936 | 2561 | 1728 | 2272.29 | 2136 | 2408 | 1.128 | 0.174 | 0.45772972 | 0.66232231 | 0.0792 | 0.0084 | 0.0389 | 0.0389 | TRUE | 0.8641 |
| Ube4b | 12491 | 12210 | 13247 | 13737 | 10282 | 11310 | 13268 | 12194 | 13301 | 10799 | 11063 | 12381 | 12167.6 | 12921 | 11414 | 0.883 | -0.179 | 0.0338875 | 0.12528683 | 0.0038 | 0.0072 | 0.0051 | 0.0051 | TRUE | 0.067 |
| Ubfd1 | 4164 | 3783 | 3910 | 6268 | 4722 | 4048 | 4423 | 3778 | 3926 | 4927 | 5081 | 4431 | 4427.71 | 4042 | 4813 | 1.191 | 0.252 | 0.00900125 | 0.04865243 | 0.0056 | 0.0076 | 0.0065 | 0.0065 | TRUE | 0.0824 |
| Ubiad1 | 899 | 914 | 912 | 1207 | 850 | 766 | 955 | 913 | 916 | 949 | 915 | 839 | 914.23 | 928 | 901 | 0.972 | -0.041 | 0.65280876 | 0.80520057 | 0.0012 | 0.0105 | 0.005 | 0.005 | TRUE | 0.0427 |
| Ubl3 | 2890 | 2993 | 3054 | 3080 | 2401 | 2526 | 3070 | 2989 | 3067 | 2421 | 2583 | 2765 | 2815.84 | 3042 | 2590 | 0.851 | -0.233 | 0.00325514 | 0.02255713 | 0.0018 | 0.0081 | 0.0042 | 0.0042 | TRUE | 0.0424 |
| Ubl4 | 658 | 947 | 935 | 1148 | 746 | 861 | 699 | 946 | 939 | 902 | 803 | 943 | 871.86 | 861 | 883 | 1.025 | 0.035 | 0.80646184 | 0.90024179 | 0.0163 | 0.0107 | 0.0137 | 0.0137 | TRUE | 0.3217 |
| Ubl4b | 0 | 0 | 0 | 0 | 0 | 0 | 0 | 0 | 0 | 0 | 0 | 0 | 0 | 0 | 0 | NA | NA | NA | NA | NA | NA | NA | NA | NA | NA |
| Ubl5 | 2047 | 2084 | 2174 | 2645 | 1761 | 1928 | 2174 | 2081 | 2183 | 2079 | 1895 | 2110 | 2087.17 | 2146 | 2028 | 0.945 | -0.081 | 0.30602908 | 0.52665845 | 0.0015 | 0.0085 | 0.0041 | 0.0041 | TRUE | 0.0395 |
| Ubl7 | 1920 | 2015 | 2100 | 2546 | 1821 | 1820 | 2039 | 2012 | 2109 | 2001 | 1959 | 1992 | 2018.9 | 2053 | 1984 | 0.966 | -0.049 | 0.46549948 | 0.6680031 | 0 | 0.0085 | 0.0028 | 0.0028 | TRUE | 0.0067 |
| Ublcp1 | 1966 | 2035 | 1824 | 2597 | 1762 | 1573 | 2088 | 2032 | 1831 | 2042 | 1896 | 1722 | 1935.23 | 1984 | 1887 | 0.951 | -0.072 | 0.47501608 | 0.67537669 | 0.0049 | 0.0086 | 0.0068 | 0.0068 | TRUE | 0.0701 |
| Ubn1 | 2358 | 2812 | 2951 | 3181 | 2426 | 2407 | 2505 | 2808 | 2963 | 2501 | 2610 | 2635 | 2670.3 | 2759 | 2582 | 0.936 | -0.096 | 0.28235592 | 0.50369474 | 0.0034 | 0.0081 | 0.0054 | 0.0054 | TRUE | 0.0785 |
| Ubn2 | 2740 | 2613 | 2976 | 3610 | 2768 | 2503 | 2910 | 2610 | 2988 | 2838 | 2978 | 2740 | 2844.04 | 2836 | 2852 | 1.006 | 0.008 | 0.92442062 | 0.96286755 | 0.003 | 0.0081 | 0.0049 | 0.0049 | TRUE | 0.0594 |
| Ubox5 | 1279 | 1165 | 1163 | 1613 | 1122 | 1007 | 1359 | 1163 | 1168 | 1268 | 1207 | 1102 | 1211.22 | 1230 | 1192 | 0.97 | -0.043 | 0.6839113 | 0.82485654 | 0.0056 | 0.0096 | 0.0073 | 0.0073 | TRUE | 0.1 |
| Ubp1 | 4067 | 4299 | 4363 | 6072 | 4721 | 4374 | 4320 | 4293 | 4381 | 4773 | 5080 | 4788 | 4605.85 | 4331 | 4880 | 1.127 | 0.172 | 0.0066225 | 0.03861402 | 5.00E-04 | 0.0076 | 0.0027 | 0.0027 | TRUE | 0.0154 |
| Ubqln1 | 16860 | 16927 | 16932 | 20322 | 14593 | 14974 | 17908 | 16905 | 17002 | 15976 | 15701 | 16391 | 16647.18 | 17272 | 16023 | 0.928 | -0.108 | 0.07086034 | 0.20734866 | 7.00E-04 | 0.0071 | 0.0025 | 0.0025 | TRUE | 0.0127 |
| Ubqln2 | 2185 | 1840 | 1985 | 3699 | 2728 | 2233 | 2321 | 1838 | 1993 | 2908 | 2935 | 2444 | 2406.5 | 2051 | 2762 | 1.348 | 0.43 | 0.00039546 | 0.0042993 | 0.0116 | 0.0083 | 0.0102 | 0.0102 | TRUE | 0.1606 |
| Ubqln3 | 0 | 0 | 0 | 0 | 0 | 0 | 0 | 0 | 0 | 0 | 0 | 0 | 0 | 0 | 0 | NA | NA | NA | NA | NA | NA | NA | NA | NA | NA |
| Ubqln4 | 10194 | 11093 | 11479 | 12598 | 9499 | 10512 | 10828 | 11079 | 11526 | 9904 | 10220 | 11507 | 10843.93 | 11144 | 10544 | 0.946 | -0.08 | 0.33573551 | 0.55587029 | 0.0036 | 0.0072 | 0.0049 | 0.0049 | TRUE | 0.078 |
| Ubqlnl | 0 | 0 | 0 | 0 | 0 | 0 | 0 | 0 | 0 | 0 | 0 | 0 | 0 | 0 | 0 | NA | NA | NA | NA | NA | NA | NA | NA | NA | NA |
| Ubr1 | 2916 | 2877 | 3512 | 4110 | 2698 | 2713 | 3097 | 2873 | 3526 | 3231 | 2903 | 2970 | 3100.11 | 3165 | 3035 | 0.959 | -0.061 | 0.55355215 | 0.73700313 | 0.006 | 0.008 | 0.0072 | 0.0072 | TRUE | 0.1209 |
| Ubr2 | 3687 | 3646 | 3957 | 5204 | 3717 | 3525 | 3916 | 3641 | 3973 | 4091 | 3999 | 3859 | 3913.28 | 3843 | 3983 | 1.036 | 0.052 | 0.47012014 | 0.67198975 | 0.0011 | 0.0077 | 0.0034 | 0.0034 | TRUE | 0.0259 |
| Ubr3 | 6671 | 7534 | 8048 | 10405 | 7725 | 7783 | 7086 | 7524 | 8081 | 8180 | 8312 | 8520 | 7950.34 | 7564 | 8337 | 1.102 | 0.14 | 0.06318668 | 0.19265295 | 0.0022 | 0.0073 | 0.004 | 0.004 | TRUE | 0.0437 |
| Ubr4 | 20921 | 15672 | 20279 | 29575 | 18996 | 16479 | 22222 | 15652 | 20362 | 23250 | 20439 | 18039 | 19993.79 | 19412 | 20576 | 1.06 | 0.084 | 0.56888486 | 0.7478913 | 0.0192 | 0.0071 | 0.0156 | 0.0156 | TRUE | 0.3513 |
| Ubr5 | 38753 | 29658 | 34256 | 43809 | 31162 | 29340 | 41163 | 29620 | 34397 | 34439 | 33528 | 32117 | 34210.62 | 35060 | 33361 | 0.952 | -0.072 | 0.56087263 | 0.74256648 | 0.0127 | 0.007 | 0.0109 | 0.0109 | TRUE | 0.2838 |
| Ubr7 | 11876 | 13923 | 15090 | 12025 | 8886 | 10049 | 12614 | 13905 | 15152 | 9453 | 9561 | 11000 | 11947.58 | 13890 | 10005 | 0.72 | -0.474 | 3.87E-06 | 8.90E-05 | 0.0078 | 0.0072 | 0.0075 | 0.0075 | TRUE | 0.0925 |
| Ubtd1 | 571 | 833 | 769 | 879 | 633 | 776 | 607 | 832 | 772 | 691 | 681 | 849 | 738.68 | 737 | 740 | 1.004 | 0.006 | 0.96958236 | 0.98448447 | 0.0195 | 0.0114 | 0.0156 | 0.0156 | TRUE | 0.2821 |
| Ubtd2 | 2117 | 2116 | 2088 | 3131 | 2209 | 2195 | 2249 | 2113 | 2097 | 2461 | 2377 | 2403 | 2283.22 | 2153 | 2414 | 1.122 | 0.165 | 0.01899298 | 0.08367521 | 4.00E-04 | 0.0083 | 0.0031 | 0.0031 | TRUE | 0.0182 |
| Ubtf | 12875 | 12028 | 11928 | 16482 | 12177 | 10584 | 13676 | 12012 | 11977 | 12957 | 13102 | 11586 | 12551.56 | 12555 | 12548 | 1 | -0.001 | 0.99401496 | 0.99679577 | 0.0041 | 0.0072 | 0.0059 | 0.0059 | TRUE | 0.0745 |
| Ubtfl1 | 67 | 28 | 13 | 152 | 115 | 54 | 71 | 28 | 13 | 119 | 124 | 59 | 69.09 | 37 | 101 | 2.704 | 1.435 | 0.00694761 | 0.03986953 | 0.3626 | 0.0546 | 0.1851 | 0.1851 | TRUE | 2.8036 |
| Ubxn1 | 4764 | 5682 | 5711 | 8109 | 5970 | 5831 | 5060 | 5675 | 5734 | 6375 | 6423 | 6383 | 5941.71 | 5490 | 6394 | 1.165 | 0.22 | 0.00390976 | 0.02589866 | 0.0021 | 0.0075 | 0.004 | 0.004 | TRUE | 0.057 |
| Ubxn10 | 136 | 184 | 229 | 184 | 193 | 183 | 144 | 184 | 230 | 145 | 208 | 200 | 185.13 | 186 | 184 | 0.986 | -0.02 | 0.93120682 | 0.96618603 | 0.0402 | 0.0247 | 0.0323 | 0.0323 | TRUE | 0.2266 |
| Ubxn11 | 50 | 58 | 60 | 118 | 104 | 69 | 53 | 58 | 60 | 93 | 112 | 76 | 75.25 | 57 | 94 | 1.635 | 0.71 | 0.00447489 | 0.02874304 | 0 | 0.0507 | 0.0307 | 0.0307 | TRUE | 0.133 |
| Ubxn2a | 3567 | 3493 | 3453 | 4562 | 3027 | 2678 | 3789 | 3488 | 3467 | 3586 | 3257 | 2931 | 3419.85 | 3581 | 3258 | 0.91 | -0.136 | 0.17009943 | 0.36880651 | 0.0059 | 0.0079 | 0.0068 | 0.0068 | TRUE | 0.0952 |
| Ubxn2b | 1338 | 1185 | 1160 | 1581 | 1345 | 1280 | 1421 | 1183 | 1165 | 1243 | 1447 | 1401 | 1310.1 | 1256 | 1364 | 1.085 | 0.118 | 0.31221542 | 0.53255705 | 0.0084 | 0.0094 | 0.009 | 0.009 | TRUE | 0.1573 |
| Ubxn4 | 3558 | 3929 | 3888 | 5004 | 3735 | 3559 | 3779 | 3924 | 3904 | 3934 | 4019 | 3896 | 3909.23 | 3869 | 3950 | 1.021 | 0.03 | 0.62643103 | 0.78958445 | 0 | 0.0077 | 0.0024 | 0.0024 | TRUE | 0.005 |
| Ubxn6 | 1720 | 1850 | 1798 | 1988 | 1453 | 1441 | 1827 | 1848 | 1805 | 1563 | 1563 | 1577 | 1697.25 | 1827 | 1568 | 0.858 | -0.221 | 0.00111804 | 0.00970488 | 0 | 0.0088 | 0.0027 | 0.0027 | TRUE | 0.0013 |
| Ubxn7 | 7091 | 7430 | 8330 | 9057 | 6658 | 7110 | 7532 | 7420 | 8364 | 7120 | 7164 | 7783 | 7563.84 | 7772 | 7356 | 0.946 | -0.08 | 0.33281358 | 0.55329775 | 0.0033 | 0.0073 | 0.0047 | 0.0047 | TRUE | 0.0543 |
| Ubxn8 | 882 | 872 | 894 | 1386 | 1007 | 827 | 937 | 871 | 898 | 1090 | 1083 | 905 | 963.95 | 902 | 1026 | 1.139 | 0.188 | 0.08110187 | 0.22667707 | 0.0051 | 0.0103 | 0.0073 | 0.0073 | TRUE | 0.1248 |
| Uchl1 | 4720 | 5009 | 4967 | 5522 | 4079 | 3682 | 5013 | 5003 | 4987 | 4341 | 4389 | 4031 | 4627.27 | 5001 | 4254 | 0.851 | -0.233 | 0.00047738 | 0.00500001 | 7.00E-04 | 0.0076 | 0.003 | 0.003 | TRUE | 0.0253 |
| Uchl3 | 3598 | 3964 | 4025 | 4979 | 3544 | 3474 | 3822 | 3959 | 4042 | 3914 | 3813 | 3803 | 3892.03 | 3941 | 3843 | 0.975 | -0.036 | 0.56758145 | 0.74731774 | 3.00E-04 | 0.0077 | 0.0026 | 0.0026 | TRUE | 0.0085 |
| Uchl4 | 22 | 25 | 10 | 26 | 14 | 14 | 23 | 25 | 10 | 20 | 15 | 15 | 18.2 | 19 | 17 | 0.877 | -0.189 | 0.70981584 | 0.84193543 | 0.0672 | 0.1879 | 0.1309 | 0.1309 | TRUE | 0.9659 |
| Uchl5 | 3861 | 3890 | 3724 | 4734 | 3314 | 3261 | 4101 | 3885 | 3739 | 3721 | 3566 | 3570 | 3763.69 | 3908 | 3619 | 0.926 | -0.111 | 0.11791705 | 0.29045568 | 0.001 | 0.0078 | 0.0033 | 0.0033 | TRUE | 0.0226 |
| Uck1 | 1341 | 1336 | 1254 | 1855 | 1362 | 1320 | 1424 | 1334 | 1259 | 1458 | 1465 | 1445 | 1397.74 | 1339 | 1456 | 1.088 | 0.121 | 0.14416659 | 0.33131218 | 0.0011 | 0.0093 | 0.0042 | 0.0042 | TRUE | 0.0369 |
| Uck2 | 7010 | 9385 | 9542 | 8087 | 6432 | 7278 | 7446 | 9373 | 9581 | 6357 | 6920 | 7967 | 7940.77 | 8800 | 7081 | 0.805 | -0.314 | 0.01513101 | 0.07119521 | 0.0154 | 0.0073 | 0.0119 | 0.0119 | TRUE | 0.2212 |
| Uckl1 | 553 | 807 | 741 | 787 | 599 | 580 | 587 | 806 | 744 | 619 | 644 | 635 | 672.58 | 712 | 633 | 0.887 | -0.172 | 0.20531605 | 0.41388608 | 0.0119 | 0.0118 | 0.0118 | 0.0118 | TRUE | 0.2772 |
| Ucma | 9 | 5 | 7 | 9 | 8 | 7 | 10 | 5 | 7 | 7 | 9 | 8 | 7.49 | 7 | 8 | 1.081 | 0.112 | 0.86941695 | 0.93581138 | 0 | 0.4469 | 0.2013 | 0.2013 | TRUE | 0.2134 |
| Ucn | 0 | 2 | 4 | 0 | 0 | 0 | 0 | 2 | 4 | 0 | 0 | 0 | 1 | 2 | 0 | 0.089 | -3.486 | 0.18523826 | NA | 6.1183 | 3.2938 | 3.7366 | 3.7366 | TRUE | 0.0494 |
| Ucn2 | 3 | 2 | 3 | 5 | 2 | 3 | 3 | 2 | 3 | 4 | 2 | 3 | 2.93 | 3 | 3 | 1.157 | 0.21 | 0.84262913 | NA | 0 | 1.1324 | 0.4644 | 0.4644 | TRUE | 0.0927 |
| Ucn3 | 0 | 0 | 0 | 0 | 0 | 0 | 0 | 0 | 0 | 0 | 0 | 0 | 0 | 0 | 0 | NA | NA | NA | NA | NA | NA | NA | NA | NA | NA |
| Ucp1 | 43 | 68 | 55 | 18 | 14 | 15 | 46 | 68 | 55 | 14 | 15 | 16 | 35.74 | 56 | 15 | 0.269 | -1.895 | 2.08E-07 | 6.78E-06 | 0 | 0.0991 | 0.0548 | 0.0548 | TRUE | 0.1185 |
| Ucp2 | 9273 | 8095 | 7798 | 10168 | 7767 | 7297 | 9850 | 8085 | 7830 | 7993 | 8357 | 7988 | 8350.32 | 8588 | 8113 | 0.945 | -0.082 | 0.43211141 | 0.64176836 | 0.008 | 0.0073 | 0.0078 | 0.0078 | TRUE | 0.2016 |
| Ucp3 | 11 | 3 | 4 | 10 | 8 | 6 | 12 | 3 | 4 | 8 | 9 | 7 | 6.96 | 6 | 8 | 1.241 | 0.311 | 0.6924921 | 0.83139114 | 0.0984 | 0.4805 | 0.3009 | 0.3009 | TRUE | 1.3547 |
| Uevld | 2227 | 2738 | 2956 | 2566 | 1982 | 2229 | 2365 | 2734 | 2968 | 2017 | 2133 | 2440 | 2442.96 | 2689 | 2197 | 0.816 | -0.293 | 0.01369441 | 0.06613236 | 0.0107 | 0.0083 | 0.0097 | 0.0097 | TRUE | 0.1344 |
| Ufc1 | 1485 | 1967 | 1987 | 2339 | 1628 | 1614 | 1577 | 1964 | 1995 | 1839 | 1752 | 1767 | 1815.68 | 1845 | 1786 | 0.968 | -0.048 | 0.66887587 | 0.8150064 | 0.0081 | 0.0087 | 0.0084 | 0.0084 | TRUE | 0.1948 |
| Ufd1l | 1565 | 1463 | 1382 | 2176 | 1418 | 1453 | 1662 | 1461 | 1388 | 1711 | 1526 | 1591 | 1556.32 | 1504 | 1609 | 1.071 | 0.099 | 0.33506685 | 0.55534619 | 0.0052 | 0.009 | 0.0069 | 0.0069 | TRUE | 0.1022 |
| Ufl1 | 2390 | 2489 | 2528 | 3377 | 2356 | 2426 | 2539 | 2486 | 2538 | 2655 | 2535 | 2656 | 2568.01 | 2521 | 2615 | 1.038 | 0.053 | 0.4158976 | 0.62746671 | 0 | 0.0082 | 0.0027 | 0.0027 | TRUE | 0.0086 |
| Ufm1 | 1725 | 1541 | 1576 | 2464 | 1732 | 1568 | 1832 | 1539 | 1582 | 1937 | 1864 | 1716 | 1745.11 | 1651 | 1839 | 1.114 | 0.156 | 0.12767446 | 0.30730165 | 0.0056 | 0.0088 | 0.007 | 0.007 | TRUE | 0.1106 |
| Ufsp1 | 71 | 117 | 90 | 167 | 125 | 112 | 75 | 117 | 90 | 131 | 134 | 123 | 111.83 | 94 | 129 | 1.373 | 0.457 | 0.03697778 | 0.13297163 | 0.013 | 0.0364 | 0.0254 | 0.0254 | TRUE | 0.4282 |
| Ufsp2 | 2176 | 2448 | 2618 | 2935 | 2064 | 2266 | 2311 | 2445 | 2629 | 2307 | 2221 | 2480 | 2398.9 | 2462 | 2336 | 0.949 | -0.076 | 0.38886371 | 0.60282666 | 0.0031 | 0.0083 | 0.0052 | 0.0052 | TRUE | 0.0428 |
| Ugcg | 2929 | 3441 | 3462 | 4077 | 2931 | 3132 | 3111 | 3437 | 3476 | 3205 | 3154 | 3428 | 3301.82 | 3341 | 3262 | 0.976 | -0.035 | 0.6688063 | 0.81498979 | 0.0025 | 0.0079 | 0.0045 | 0.0045 | TRUE | 0.044 |
| Ugdh | 1347 | 1590 | 1750 | 2624 | 1898 | 1928 | 1431 | 1588 | 1757 | 2063 | 2042 | 2110 | 1831.88 | 1592 | 2072 | 1.301 | 0.38 | 0.00010179 | 0.00139866 | 0.0046 | 0.0087 | 0.0063 | 0.0063 | TRUE | 0.0994 |
| Uggt1 | 9537 | 9534 | 9944 | 11422 | 8188 | 8020 | 10130 | 9522 | 9985 | 8979 | 8810 | 8779 | 9367.42 | 9879 | 8856 | 0.897 | -0.158 | 0.00814594 | 0.04506413 | 4.00E-04 | 0.0073 | 0.0024 | 0.0024 | TRUE | 0.0122 |
| Uggt2 | 361 | 426 | 424 | 435 | 294 | 340 | 383 | 425 | 426 | 342 | 316 | 372 | 377.52 | 411 | 343 | 0.834 | -0.262 | 0.03429043 | 0.12614807 | 0.0019 | 0.0156 | 0.0083 | 0.0083 | TRUE | 0.0575 |
| Ugp2 | 1540 | 1864 | 1685 | 2960 | 2064 | 2147 | 1636 | 1862 | 1692 | 2327 | 2221 | 2350 | 2014.52 | 1730 | 2299 | 1.329 | 0.411 | 9.52E-07 | 2.63E-05 | 0.002 | 0.0085 | 0.0045 | 0.0045 | TRUE | 0.0538 |
| Ugt1a1 | 0 | 0 | 0 | 0 | 0 | 0 | 0 | 0 | 0 | 0 | 0 | 0 | 0 | 0 | 0 | NA | NA | NA | NA | NA | NA | NA | NA | NA | NA |
| Ugt1a10 | 0 | 0 | 0 | 0 | 0 | 0 | 0 | 0 | 0 | 0 | 0 | 0 | 0 | 0 | 0 | NA | NA | NA | NA | NA | NA | NA | NA | NA | NA |
| Ugt1a2 | 0 | 0 | 0 | 0 | 0 | 0 | 0 | 0 | 0 | 0 | 0 | 0 | 0 | 0 | 0 | NA | NA | NA | NA | NA | NA | NA | NA | NA | NA |
| Ugt1a5 | 0 | 0 | 0 | 0 | 0 | 0 | 0 | 0 | 0 | 0 | 0 | 0 | 0 | 0 | 0 | NA | NA | NA | NA | NA | NA | NA | NA | NA | NA |
| Ugt1a6a | 0 | 0 | 1 | 0 | 1 | 0 | 0 | 0 | 1 | 0 | 1 | 0 | 0.35 | 0 | 0 | 0.954 | -0.067 | 0.98570726 | NA | 0 | 9.5099 | 8.2019 | 8.2019 | TRUE | 0.5204 |

| | | | | | | | | | | | | | | | | | | | | | | | | | |
|---|---|---|---|---|---|---|---|---|---|---|---|---|---|---|---|---|---|---|---|---|---|---|---|---|---|
| Ugt1a6b | 0 | 0 | 0 | 0 | 0 | 0 | 0 | 0 | 0 | 0 | 0 | 0 | 0 | 0 | 0 | NA | NA | NA | NA | NA | NA | NA | NA | NA | NA |
| Ugt1a7c | 0 | 0 | 0 | 0 | 0 | 0 | 0 | 0 | 0 | 0 | 0 | 0 | 0 | 0 | 0 | NA | NA | NA | NA | NA | NA | NA | NA | NA | NA |
| Ugt1a9 | 0 | 0 | 0 | 0 | 0 | 0 | 0 | 0 | 0 | 0 | 0 | 0 | 0 | 0 | 0 | NA | NA | NA | NA | NA | NA | NA | NA | NA | NA |
| Ugt2a1 | 0 | 0 | 0 | 0 | 0 | 0 | 0 | 0 | 0 | 0 | 0 | 0 | 0 | 0 | 0 | NA | NA | NA | NA | NA | NA | NA | NA | NA | NA |
| Ugt2a2 | 0 | 0 | 0 | 0 | 0 | 0 | 0 | 0 | 0 | 0 | 0 | 0 | 0 | 0 | 0 | NA | NA | NA | NA | NA | NA | NA | NA | NA | NA |
| Ugt2a3 | 0 | 0 | 0 | 0 | 0 | 0 | 0 | 0 | 0 | 0 | 0 | 0 | 0 | 0 | 0 | NA | NA | NA | NA | NA | NA | NA | NA | NA | NA |
| Ugt2b1 | 0 | 0 | 0 | 0 | 0 | 0 | 0 | 0 | 0 | 0 | 0 | 0 | 0 | 0 | 0 | NA | NA | NA | NA | NA | NA | NA | NA | NA | NA |
| Ugt2b34 | 0 | 0 | 0 | 1 | 0 | 0 | 0 | 0 | 0 | 1 | 0 | 0 | 0.13 | 0 | 0 | 1.859 | 0.894 | 0.82650734 | NA | 0 | 25.1515 | 10 | 10 | TRUE | 0.2563 |
| Ugt2b35 | 0 | 0 | 0 | 0 | 0 | 0 | 0 | 0 | 0 | 0 | 0 | 0 | 0 | 0 | 0 | NA | NA | NA | NA | NA | NA | NA | NA | NA | NA |
| Ugt2b36 | 0 | 0 | 0 | 0 | 0 | 0 | 0 | 0 | 0 | 0 | 0 | 0 | 0 | 0 | 0 | NA | NA | NA | NA | NA | NA | NA | NA | NA | NA |
| Ugt2b37 | 0 | 0 | 0 | 0 | 0 | 0 | 0 | 0 | 0 | 0 | 0 | 0 | 0 | 0 | 0 | NA | NA | NA | NA | NA | NA | NA | NA | NA | NA |
| Ugt2b38 | 0 | 0 | 0 | 0 | 0 | 0 | 0 | 0 | 0 | 0 | 0 | 0 | 0 | 0 | 0 | NA | NA | NA | NA | NA | NA | NA | NA | NA | NA |
| Ugt2b5 | 0 | 0 | 0 | 0 | 0 | 0 | 0 | 0 | 0 | 0 | 0 | 0 | 0 | 0 | 0 | NA | NA | NA | NA | NA | NA | NA | NA | NA | NA |
| Ugt3a1 | 2 | 2 | 1 | 3 | 2 | 0 | 2 | 2 | 1 | 2 | 2 | 0 | 1.61 | 2 | 1 | 0.903 | -0.147 | 0.92235436 | NA | 0 | 2.0582 | 1.0309 | 1.0309 | TRUE | 0.4575 |
| Ugt3a2 | 1 | 0 | 0 | 1 | 0 | 2 | 1 | 0 | 0 | 1 | 0 | 2 | 0.67 | 0 | 1 | 2.693 | 1.429 | 0.58190699 | NA | 0 | 4.9025 | 3.3589 | 3.3589 | TRUE | 0.3233 |
| Ugt8a | 310 | 402 | 403 | 655 | 480 | 396 | 329 | 401 | 405 | 515 | 516 | 433 | 433.38 | 378 | 488 | 1.291 | 0.368 | 0.00794582 | 0.04412074 | 0.0088 | 0.0145 | 0.0115 | 0.0115 | TRUE | 0.1468 |
| Uhmk1 | 635 | 803 | 852 | 988 | 760 | 804 | 674 | 802 | 855 | 777 | 818 | 880 | 801.07 | 777 | 825 | 1.06 | 0.084 | 0.48781032 | 0.68458483 | 0.008 | 0.011 | 0.0093 | 0.0093 | TRUE | 0.1578 |
| Uhrf1 | 13425 | 13697 | 13867 | 19294 | 14398 | 13465 | 14260 | 13679 | 13924 | 15167 | 15491 | 14740 | 14543.55 | 13954 | 15133 | 1.084 | 0.117 | 0.04234674 | 0.14676895 | 4.00E-04 | 0.0071 | 0.0023 | 0.0023 | TRUE | 0.0063 |
| Uhrf1bp1 | 5281 | 5618 | 6029 | 6889 | 5252 | 5291 | 5609 | 5611 | 6054 | 5416 | 5651 | 5792 | 5688.68 | 5758 | 5620 | 0.976 | -0.035 | 0.61418813 | 0.781712 | 0.0013 | 0.0075 | 0.0034 | 0.0034 | TRUE | 0.0247 |
| Uhrf1bp1l | 2546 | 2578 | 2750 | 3621 | 2742 | 2776 | 2704 | 2575 | 2761 | 2847 | 2950 | 3039 | 2812.63 | 2680 | 2945 | 1.099 | 0.136 | 0.05627984 | 0.17813477 | 8.00E-04 | 0.0081 | 0.0033 | 0.0033 | TRUE | 0.0144 |
| Uhrf2 | 5061 | 5801 | 5678 | 6684 | 4871 | 4987 | 5376 | 5794 | 5701 | 5254 | 5241 | 5459 | 5470.81 | 5624 | 5318 | 0.946 | -0.081 | 0.22052949 | 0.43252954 | 8.00E-04 | 0.0075 | 0.0029 | 0.0029 | TRUE | 0.0181 |
| Uimc1 | 2543 | 2226 | 2328 | 3210 | 2256 | 2013 | 2701 | 2223 | 2338 | 2523 | 2427 | 2204 | 2402.69 | 2421 | 2385 | 0.986 | -0.021 | 0.8433366 | 0.92112915 | 0.0072 | 0.0083 | 0.0076 | 0.0076 | TRUE | 0.1243 |
| Ulbp1 | 101 | 119 | 113 | 185 | 106 | 148 | 107 | 119 | 113 | 145 | 114 | 162 | 126.85 | 113 | 140 | 1.242 | 0.312 | 0.1201176 | 0.29403924 | 0.0091 | 0.0329 | 0.0211 | 0.0211 | TRUE | 0.2674 |
| Ulk1 | 15643 | 20944 | 21566 | 20803 | 17090 | 18813 | 16616 | 20917 | 21655 | 16354 | 18388 | 20594 | 19087.07 | 19729 | 18445 | 0.935 | -0.097 | 0.45508066 | 0.66012437 | 0.0144 | 0.0071 | 0.0121 | 0.0121 | TRUE | 0.2331 |
| Ulk2 | 826 | 1374 | 1491 | 1529 | 1103 | 1281 | 877 | 1372 | 1497 | 1202 | 1187 | 1402 | 1256.28 | 1249 | 1264 | 1.011 | 0.016 | 0.93051596 | 0.9657869 | 0.0418 | 0.0095 | 0.0248 | 0.0248 | TRUE | 0.8111 |
| Ulk3 | 259 | 327 | 320 | 379 | 314 | 371 | 275 | 327 | 321 | 298 | 338 | 406 | 327.48 | 308 | 347 | 1.126 | 0.171 | 0.28504399 | 0.50627409 | 0.0139 | 0.017 | 0.0153 | 0.0153 | TRUE | 0.2486 |
| Ulk4 | 165 | 248 | 272 | 256 | 252 | 255 | 175 | 248 | 273 | 201 | 271 | 279 | 241.26 | 232 | 250 | 1.076 | 0.106 | 0.61908692 | 0.78469067 | 0.0371 | 0.0206 | 0.0287 | 0.0287 | TRUE | 0.5001 |
| Umod | 1 | 1 | 2 | 1 | 3 | 1 | 1 | 1 | 2 | 1 | 3 | 1 | 1.53 | 1 | 2 | 1.228 | 0.296 | 0.84284187 | NA | 0 | 2.1607 | 0.938 | 0.938 | TRUE | 0.4441 |
| Umodl1 | 11 | 13 | 11 | 25 | 14 | 10 | 12 | 13 | 11 | 20 | 15 | 11 | 13.56 | 12 | 15 | 1.292 | 0.37 | 0.4797869 | 0.67912904 | 0 | 0.2498 | 0.1226 | 0.1226 | TRUE | 0.2014 |
| Umps | 8341 | 7017 | 6671 | 11601 | 7953 | 6643 | 8860 | 7008 | 6698 | 9120 | 8557 | 7272 | 7919.08 | 7522 | 8316 | 1.106 | 0.145 | 0.28195131 | 0.50338874 | 0.0173 | 0.0073 | 0.013 | 0.013 | TRUE | 0.2953 |
| Unc119 | 1529 | 1845 | 1747 | 2121 | 1571 | 1698 | 1624 | 1843 | 1754 | 1667 | 1690 | 1859 | 1739.54 | 1740 | 1739 | 0.999 | -0.002 | 0.98202441 | 0.99043925 | 0.0032 | 0.0088 | 0.0054 | 0.0054 | TRUE | 0.0436 |
| Unc119b | 6255 | 6366 | 5900 | 7653 | 5551 | 5561 | 6644 | 6358 | 5924 | 6016 | 5973 | 6087 | 6167.01 | 6309 | 6025 | 0.955 | -0.066 | 0.35273904 | 0.57130829 | 0.0015 | 0.0074 | 0.0035 | 0.0035 | TRUE | 0.0347 |
| Unc13a | 306 | 602 | 683 | 511 | 488 | 545 | 325 | 601 | 686 | 402 | 525 | 597 | 522.57 | 537 | 508 | 0.944 | -0.083 | 0.7484113 | 0.86655986 | 0.0889 | 0.0132 | 0.0467 | 0.0467 | TRUE | 1.0919 |
| Unc13b | 3495 | 2449 | 2618 | 3069 | 2264 | 1913 | 3712 | 2446 | 2629 | 2413 | 2436 | 2094 | 2621.58 | 2929 | 2314 | 0.79 | -0.34 | 0.03522751 | 0.12864281 | 0.0301 | 0.0082 | 0.0184 | 0.0184 | TRUE | 0.6642 |
| Unc13c | 7 | 7 | 3 | 8 | 10 | 6 | 7 | 7 | 3 | 6 | 11 | 7 | 6.84 | 6 | 8 | 1.348 | 0.431 | 0.56507266 | 0.74551564 | 0 | 0.4884 | 0.2539 | 0.2539 | TRUE | 0.4124 |
| Unc13d | 221 | 313 | 281 | 287 | 226 | 237 | 235 | 313 | 282 | 226 | 243 | 259 | 259.62 | 277 | 243 | 0.876 | -0.191 | 0.22566412 | 0.43842997 | 0.009 | 0.0196 | 0.0141 | 0.0141 | TRUE | 0.1933 |
| Unc45a | 1923 | 1582 | 1605 | 2548 | 1787 | 1501 | 2043 | 1580 | 1612 | 2003 | 1923 | 1643 | 1800.49 | 1745 | 1856 | 1.064 | 0.09 | 0.49794905 | 0.69347924 | 0.0152 | 0.0087 | 0.0122 | 0.0122 | TRUE | 0.2687 |
| Unc45b | 39 | 65 | 47 | 85 | 60 | 51 | 41 | 65 | 47 | 67 | 65 | 56 | 56.79 | 51 | 63 | 1.22 | 0.287 | 0.30955372 | 0.52992476 | 0.0114 | 0.0649 | 0.0398 | 0.0398 | TRUE | 0.4552 |
| Unc50 | 1591 | 1782 | 1718 | 2092 | 1503 | 1575 | 1690 | 1780 | 1725 | 1645 | 1617 | 1724 | 1696.74 | 1732 | 1662 | 0.96 | -0.06 | 0.41925555 | 0.62987779 | 3.00E-04 | 0.0088 | 0.0033 | 0.0033 | TRUE | 0.0126 |
| Unc5a | 428 | 677 | 723 | 330 | 283 | 403 | 455 | 676 | 726 | 259 | 304 | 441 | 476.96 | 619 | 335 | 0.54 | -0.888 | 0.00014979 | 0.00190062 | 0.0637 | 0.0138 | 0.0373 | 0.0373 | TRUE | 0.8683 |
| Unc5b | 1516 | 1759 | 1615 | 1727 | 1310 | 1459 | 1610 | 1757 | 1622 | 1358 | 1409 | 1597 | 1558.81 | 1663 | 1455 | 0.874 | -0.194 | 0.04530491 | 0.15356738 | 0.004 | 0.009 | 0.0061 | 0.0061 | TRUE | 0.0876 |
| Unc5c | 4 | 3 | 0 | 10 | 3 | 1 | 4 | 3 | 0 | 8 | 3 | 1 | 3.24 | 2 | 4 | 1.725 | 0.786 | 0.54479563 | NA | 0.6965 | 1.0243 | 0.8839 | 0.8839 | TRUE | 0.4289 |
| Unc5cl | 60 | 43 | 42 | 72 | 54 | 54 | 64 | 43 | 42 | 57 | 58 | 59 | 53.78 | 50 | 58 | 1.169 | 0.225 | 0.42805889 | 0.63805066 | 0.0061 | 0.0682 | 0.0395 | 0.0395 | TRUE | 0.4891 |
| Unc5d | 44 | 23 | 16 | 61 | 47 | 26 | 47 | 23 | 16 | 48 | 51 | 28 | 35.46 | 29 | 42 | 1.488 | 0.573 | 0.21677934 | 0.42872021 | 0.1556 | 0.0998 | 0.1259 | 0.1259 | TRUE | 1.7422 |
| Unc79 | 14 | 21 | 22 | 15 | 20 | 12 | 15 | 21 | 22 | 12 | 22 | 13 | 17.4 | 19 | 16 | 0.794 | -0.333 | 0.49184185 | 0.6881738 | 0 | 0.1963 | 0.1112 | 0.1112 | TRUE | 0.5127 |
| Unc80 | 27 | 42 | 45 | 61 | 51 | 33 | 29 | 42 | 45 | 48 | 55 | 36 | 42.46 | 39 | 46 | 1.2 | 0.262 | 0.42799133 | 0.63802262 | 0.0212 | 0.0845 | 0.0552 | 0.0552 | TRUE | 0.3614 |
| Unc93a | 17 | 7 | 5 | 6 | 6 | 3 | 18 | 7 | 5 | 5 | 6 | 3 | 7.42 | 10 | 5 | 0.483 | -1.051 | 0.19203591 | 0.39763561 | 0.1891 | 0.4509 | 0.3153 | 0.3153 | TRUE | 1.6425 |
| Unc93b1 | 193 | 221 | 226 | 209 | 159 | 152 | 205 | 221 | 227 | 164 | 171 | 166 | 192.4 | 218 | 167 | 0.768 | -0.381 | 0.00906257 | 0.04888334 | 0 | 0.024 | 0.0101 | 0.0101 | TRUE | 0.0274 |
| Uncx | 2 | 7 | 3 | 14 | 4 | 7 | 2 | 7 | 3 | 11 | 4 | 8 | 5.85 | 4 | 8 | 1.908 | 0.932 | 0.27899961 | 0.50005946 | 0.0954 | 0.5701 | 0.3456 | 0.3456 | TRUE | 0.2687 |
| Ung | 5712 | 5671 | 5832 | 6922 | 5256 | 5102 | 6067 | 5664 | 5856 | 5442 | 5655 | 5585 | 5711.4 | 5862 | 5561 | 0.948 | -0.076 | 0.22811135 | 0.44159023 | 5.00E-04 | 0.0075 | 0.0027 | 0.0027 | TRUE | 0.0114 |
| Unk | 2550 | 3346 | 3303 | 3470 | 2743 | 2775 | 2709 | 3342 | 3317 | 2728 | 2951 | 3038 | 3013.93 | 3123 | 2906 | 0.93 | -0.104 | 0.33401359 | 0.55470805 | 0.0076 | 0.008 | 0.0081 | 0.0081 | TRUE | 0.1629 |
| Unkl | 447 | 499 | 517 | 631 | 475 | 506 | 475 | 498 | 519 | 496 | 511 | 554 | 508.88 | 497 | 520 | 1.045 | 0.063 | 0.55527181 | 0.73842723 | 0 | 0.0134 | 0.0062 | 0.0062 | TRUE | 0.0365 |
| Uox | 4 | 0 | 1 | 10 | 5 | 2 | 4 | 0 | 1 | 8 | 5 | 2 | 3.45 | 2 | 5 | 3.009 | 1.589 | 0.20723655 | NA | 0.4281 | 0.9626 | 0.7544 | 0.7544 | TRUE | 0.5569 |
| Upb1 | 46 | 69 | 50 | 74 | 42 | 52 | 49 | 69 | 50 | 58 | 45 | 57 | 54.71 | 56 | 53 | 0.956 | -0.065 | 0.81893089 | 0.90748106 | 0.0115 | 0.0671 | 0.0398 | 0.0398 | TRUE | 0.3469 |
| Upf1 | 14943 | 14775 | 14725 | 17194 | 13301 | 13074 | 15872 | 14756 | 14785 | 13517 | 14311 | 14312 | 14592.11 | 15138 | 14047 | 0.928 | -0.108 | 0.10625401 | 0.27210118 | 0.0013 | 0.0071 | 0.0032 | 0.0032 | TRUE | 0.022 |
| Upf2 | 4415 | 3832 | 4069 | 4964 | 3601 | 3815 | 4690 | 3827 | 4086 | 3902 | 3874 | 4176 | 4092.52 | 4201 | 3984 | 0.948 | -0.076 | 0.43734385 | 0.64548863 | 0.0059 | 0.0077 | 0.0067 | 0.0067 | TRUE | 0.126 |
| Upf3a | 2183 | 2633 | 2587 | 3397 | 2682 | 2363 | 2319 | 2630 | 2598 | 2670 | 2886 | 2587 | 2614.79 | 2516 | 2714 | 1.079 | 0.11 | 0.21804043 | 0.43002797 | 0.0034 | 0.0082 | 0.0053 | 0.0053 | TRUE | 0.0565 |
| Upf3b | 2357 | 2120 | 1981 | 2371 | 1762 | 1727 | 2504 | 2117 | 1989 | 1864 | 1896 | 1890 | 2043.35 | 2203 | 1883 | 0.855 | -0.226 | 0.031683 | 0.11971732 | 0.0067 | 0.0085 | 0.0075 | 0.0075 | TRUE | 0.1717 |
| Upk1a | 48 | 64 | 69 | 65 | 51 | 65 | 51 | 64 | 69 | 51 | 55 | 71 | 60.22 | 61 | 59 | 0.956 | -0.065 | 0.81338934 | 0.90450545 | 0.011 | 0.0616 | 0.0372 | 0.0372 | TRUE | 0.2646 |
| Upk1b | 6 | 6 | 4 | 1 | 6 | 3 | 6 | 6 | 4 | 1 | 6 | 3 | 4.48 | 5 | 3 | 0.623 | -0.682 | 0.48799066 | NA | 0.1179 | 0.7415 | 0.4591 | 0.4591 | TRUE | 0.4065 |
| Upk2 | 38 | 61 | 44 | 79 | 55 | 49 | 40 | 61 | 44 | 62 | 59 | 54 | 53.4 | 48 | 58 | 1.204 | 0.267 | 0.34619122 | 0.56540622 | 0 | 0.0686 | 0.0391 | 0.0391 | TRUE | 0.4084 |
| Upk3a | 1 | 8 | 6 | 4 | 4 | 6 | 1 | 8 | 6 | 3 | 4 | 7 | 4.85 | 5 | 5 | 0.914 | -0.129 | 0.89212866 | 0.94758255 | 0.1799 | 0.6863 | 0.4459 | 0.4459 | TRUE | 0.6728 |
| Upk3b | 92 | 91 | 65 | 66 | 46 | 47 | 98 | 91 | 65 | 52 | 49 | 51 | 67.78 | 85 | 51 | 0.603 | -0.73 | 0.00524711 | 0.03237773 | 0 | 0.0555 | 0.0337 | 0.0337 | TRUE | 0.3798 |
| Upk3bl | 65 | 56 | 52 | 55 | 34 | 49 | 69 | 56 | 52 | 43 | 37 | 54 | 51.77 | 59 | 45 | 0.753 | -0.41 | 0.15707706 | 0.3501158 | 0.0061 | 0.0705 | 0.0407 | 0.0407 | TRUE | 0.2304 |
| Upp1 | 14508 | 20950 | 21486 | 27485 | 20419 | 24083 | 15410 | 20923 | 21574 | 21607 | 21970 | 26363 | 21307.67 | 19302 | 23313 | 1.208 | 0.272 | 0.05968879 | 0.18520786 | 0.0218 | 0.0071 | 0.015 | 0.015 | TRUE | 0.3806 |
| Upp2 | 1 | 0 | 1 | 0 | 1 | 1 | 1 | 0 | 1 | 0 | 1 | 1 | 0.71 | 1 | 1 | 1.014 | 0.02 | 0.99311765 | NA | 0 | 4.6723 | 2.3783 | 2.3783 | TRUE | 0.3742 |
| Uprt | 74 | 81 | 104 | 98 | 68 | 76 | 79 | 81 | 104 | 77 | 73 | 83 | 82.89 | 88 | 78 | 0.883 | -0.179 | 0.43320775 | 0.64227707 | 0 | 0.0466 | 0.0254 | 0.0254 | TRUE | 0.256 |
| Uqcc | 1228 | 1324 | 1371 | 1820 | 1262 | 1285 | 1304 | 1322 | 1377 | 1431 | 1358 | 1407 | 1366.41 | 1334 | 1399 | 1.048 | 0.068 | 0.37084748 | 0.58760362 | 0 | 0.0093 | 0.0034 | 0.0034 | TRUE | 0.0092 |
| Uqcr10 | 3366 | 3546 | 3455 | 4710 | 3081 | 3307 | 3575 | 3541 | 3469 | 3703 | 3315 | 3620 | 3537.25 | 3528 | 3546 | 1.005 | 0.007 | 0.91977742 | 0.96064694 | 0.0015 | 0.0078 | 0.0037 | 0.0037 | TRUE | 0.0393 |

| | | | | | | | | | | | | | | | | | | | | | | | | | |
|---|---|---|---|---|---|---|---|---|---|---|---|---|---|---|---|---|---|---|---|---|---|---|---|---|---|
| Uqcr11 | 2223 | 2567 | 2432 | 3238 | 2118 | 2267 | 2361 | 2564 | 2442 | 2545 | 2279 | 2482 | 2445.46 | 2456 | 2435 | 0.992 | -0.012 | 0.88693895 | 0.94442566 | 0.0021 | 0.0083 | 0.0044 | 0.0044 | TRUE | 0.0381 |
| Uqcrb | 4132 | 4650 | 4586 | 5652 | 3800 | 4128 | 4389 | 4644 | 4605 | 4443 | 4089 | 4519 | 4448.04 | 4546 | 4350 | 0.957 | -0.063 | 0.38940317 | 0.6035443 | 0.0015 | 0.0076 | 0.0037 | 0.0037 | TRUE | 0.0334 |
| Uqcrc1 | 19022 | 21727 | 21855 | 24277 | 17851 | 18723 | 20205 | 21699 | 21945 | 19085 | 19207 | 20495 | 20439.15 | 21283 | 19596 | 0.921 | -0.119 | 0.08686081 | 0.23829686 | 0.0017 | 0.0071 | 0.0034 | 0.0034 | TRUE | 0.024 |
| Uqcrc2 | 8210 | 10225 | 10510 | 11947 | 8341 | 8940 | 8720 | 10212 | 10553 | 9392 | 8974 | 9786 | 9606.31 | 9828 | 9384 | 0.955 | -0.067 | 0.48353226 | 0.68146464 | 0.0054 | 0.0072 | 0.0064 | 0.0064 | TRUE | 0.1187 |
| Uqcrfs1 | 6342 | 7179 | 7054 | 9124 | 6603 | 6687 | 6736 | 7170 | 7083 | 7173 | 7104 | 7320 | 7097.66 | 6996 | 7199 | 1.029 | 0.041 | 0.50125394 | 0.69669611 | 4.00E-04 | 0.0074 | 0.0026 | 0.0026 | TRUE | 0.0129 |
| Uqcrh | 3533 | 3649 | 3548 | 4817 | 3220 | 3336 | 3753 | 3644 | 3563 | 3787 | 3465 | 3652 | 3643.76 | 3653 | 3635 | 0.995 | -0.007 | 0.92160962 | 0.96126008 | 0.0011 | 0.0078 | 0.0033 | 0.0033 | TRUE | 0.0203 |
| Uqcrq | 4472 | 4727 | 4643 | 6701 | 4651 | 4470 | 4750 | 4721 | 4662 | 5268 | 5004 | 4893 | 4883.02 | 4711 | 5055 | 1.073 | 0.102 | 0.11061501 | 0.27922853 | 6.00E-04 | 0.0076 | 0.0027 | 0.0027 | TRUE | 0.0168 |
| Urb1 | 9255 | 6668 | 7098 | 8801 | 6364 | 5781 | 9830 | 6659 | 7127 | 6919 | 6847 | 6328 | 7285.19 | 7872 | 6698 | 0.851 | -0.233 | 0.11314648 | 0.28330628 | 0.0246 | 0.0074 | 0.0154 | 0.0154 | TRUE | 0.5779 |
| Urb2 | 10932 | 11188 | 11841 | 13478 | 9720 | 9856 | 11612 | 11174 | 11890 | 10595 | 10458 | 10789 | 11086.22 | 11559 | 10614 | 0.918 | -0.123 | 0.03803986 | 0.13558524 | 5.00E-04 | 0.0072 | 0.0024 | 0.0024 | TRUE | 0.0104 |
| Urgcp | 269 | 344 | 272 | 757 | 514 | 544 | 286 | 344 | 273 | 595 | 553 | 595 | 441 | 301 | 581 | 1.932 | 0.95 | 1.61E-13 | 1.87E-11 | 0.0049 | 0.0144 | 0.0094 | 0.0094 | TRUE | 0.1761 |
| Uri1 | 12492 | 11206 | 10465 | 12392 | 9218 | 8896 | 13269 | 11192 | 10508 | 9742 | 9918 | 9738 | 10727.65 | 11656 | 9799 | 0.841 | -0.25 | 0.01342297 | 0.06516069 | 0.0075 | 0.0072 | 0.0073 | 0.0073 | TRUE | 0.179 |
| Urm1 | 1668 | 1451 | 1299 | 1916 | 1379 | 1416 | 1772 | 1449 | 1304 | 1506 | 1484 | 1550 | 1510.86 | 1508 | 1513 | 1.003 | 0.005 | 0.96836556 | 0.98419772 | 0.0118 | 0.0091 | 0.0106 | 0.0106 | TRUE | 0.2802 |
| Uroc1 | 2 | 0 | 0 | 15 | 11 | 6 | 2 | 0 | 0 | 12 | 12 | 7 | 5.39 | 1 | 10 | 14.716 | 3.879 | 0.00269933 | 0.01952032 | 0 | 0.6185 | 0.4128 | 0.4128 | TRUE | 0.9748 |
| Urod | 1741 | 1668 | 1719 | 2379 | 1751 | 1750 | 1849 | 1666 | 1726 | 1870 | 1884 | 1916 | 1818.49 | 1747 | 1890 | 1.082 | 0.113 | 0.14131844 | 0.32680725 | 9.00E-04 | 0.0087 | 0.0037 | 0.0037 | TRUE | 0.0314 |
| Uros | 221 | 160 | 162 | 430 | 314 | 151 | 235 | 160 | 163 | 338 | 338 | 165 | 233.06 | 186 | 280 | 1.512 | 0.597 | 0.03774892 | 0.13498494 | 0.0953 | 0.021 | 0.055 | 0.055 | TRUE | 1.4259 |
| Usb1 | 2545 | 2527 | 2364 | 3052 | 2109 | 2176 | 2703 | 2524 | 2374 | 2399 | 2269 | 2382 | 2441.84 | 2534 | 2350 | 0.928 | -0.108 | 0.18590871 | 0.38963646 | 0.002 | 0.0083 | 0.0044 | 0.0044 | TRUE | 0.0415 |
| Use1 | 1225 | 1584 | 1568 | 2259 | 1733 | 1502 | 1301 | 1582 | 1574 | 1776 | 1865 | 1644 | 1623.7 | 1486 | 1762 | 1.185 | 0.245 | 0.02488132 | 0.10140264 | 0.0068 | 0.0089 | 0.008 | 0.008 | TRUE | 0.1418 |
| Usf1 | 2648 | 2671 | 2731 | 3261 | 2283 | 2161 | 2813 | 2668 | 2742 | 2564 | 2456 | 2366 | 2601.31 | 2741 | 2462 | 0.899 | -0.154 | 0.03115211 | 0.1183645 | 8.00E-04 | 0.0082 | 0.0033 | 0.0033 | TRUE | 0.0162 |
| Usf2 | 1046 | 1504 | 1573 | 1581 | 1268 | 1386 | 1111 | 1502 | 1579 | 1243 | 1364 | 1517 | 1386.15 | 1397 | 1375 | 0.983 | -0.025 | 0.8702398 | 0.93630208 | 0.0215 | 0.0093 | 0.0156 | 0.0156 | TRUE | 0.3858 |
| Ush1c | 80 | 93 | 77 | 80 | 64 | 75 | 85 | 93 | 77 | 63 | 69 | 82 | 78.17 | 85 | 71 | 0.834 | -0.261 | 0.25850804 | 0.4773685 | 0 | 0.049 | 0.0257 | 0.0257 | TRUE | 0.1539 |
| Ush1g | 11 | 29 | 24 | 13 | 16 | 20 | 12 | 29 | 24 | 10 | 17 | 22 | 19.01 | 22 | 16 | 0.751 | -0.412 | 0.43717622 | 0.64537096 | 0.1162 | 0.1802 | 0.1493 | 0.1493 | TRUE | 0.5983 |
| Ush2a | 9 | 8 | 1 | 8 | 7 | 7 | 10 | 8 | 1 | 6 | 8 | 8 | 6.67 | 6 | 7 | 1.157 | 0.211 | 0.80353823 | 0.89855367 | 0.1754 | 0.5006 | 0.3664 | 0.3664 | TRUE | 1.3093 |
| Ushbp1 | 36 | 56 | 62 | 147 | 128 | 157 | 38 | 56 | 62 | 116 | 138 | 172 | 96.93 | 52 | 142 | 2.705 | 1.436 | 8.23E-08 | 3.05E-06 | 0.0356 | 0.0409 | 0.0384 | 0.0384 | TRUE | 0.2246 |
| Usmg5 | 2325 | 2449 | 2359 | 3385 | 2133 | 2153 | 2470 | 2446 | 2369 | 2661 | 2295 | 2357 | 2432.81 | 2428 | 2438 | 1.005 | 0.007 | 0.9383763 | 0.9691313 | 0.0031 | 0.0083 | 0.005 | 0.005 | TRUE | 0.0792 |
| Uso1 | 7240 | 5976 | 5900 | 9655 | 6602 | 5919 | 7690 | 5968 | 5924 | 7590 | 7103 | 6479 | 6792.55 | 6527 | 7057 | 1.081 | 0.113 | 0.36681077 | 0.5843187 | 0.0132 | 0.0074 | 0.0111 | 0.0111 | TRUE | 0.2961 |
| Usp1 | 6061 | 6112 | 6182 | 8878 | 6477 | 5736 | 6438 | 6104 | 6207 | 6979 | 6969 | 6279 | 6496.06 | 6250 | 6742 | 1.079 | 0.11 | 0.14194165 | 0.32758321 | 0.0019 | 0.0074 | 0.0039 | 0.0039 | TRUE | 0.0439 |
| Usp10 | 13997 | 14332 | 14256 | 18906 | 13998 | 13745 | 14867 | 14313 | 14315 | 14862 | 15061 | 15046 | 14744.13 | 14498 | 14990 | 1.034 | 0.048 | 0.3741862 | 0.59040264 | 2.00E-04 | 0.0071 | 0.002 | 0.002 | TRUE | 0.006 |
| Usp11 | 2799 | 3797 | 4277 | 2527 | 2048 | 2319 | 2973 | 3792 | 4295 | 1987 | 2204 | 2539 | 2964.71 | 3687 | 2243 | 0.608 | -0.717 | 2.30E-06 | 5.69E-05 | 0.0235 | 0.008 | 0.0163 | 0.0163 | TRUE | 0.2397 |
| Usp12 | 3149 | 3176 | 3263 | 4274 | 3218 | 2965 | 3345 | 3172 | 3276 | 3360 | 3462 | 3246 | 3310.17 | 3264 | 3356 | 1.028 | 0.04 | 0.55138445 | 0.73549015 | 5.00E-04 | 0.0079 | 0.003 | 0.003 | TRUE | 0.0099 |
| Usp13 | 261 | 294 | 327 | 450 | 327 | 313 | 277 | 294 | 328 | 354 | 352 | 343 | 324.57 | 300 | 350 | 1.166 | 0.221 | 0.07802781 | 0.22101584 | 0 | 0.0171 | 0.0082 | 0.0082 | TRUE | 0.0789 |
| Usp14 | 8786 | 9344 | 9710 | 11830 | 8322 | 8402 | 9332 | 9332 | 9750 | 9300 | 8954 | 9197 | 9310.86 | 9471 | 9150 | 0.966 | -0.05 | 0.39625687 | 0.60967353 | 4.00E-04 | 0.0073 | 0.0024 | 0.0024 | TRUE | 0.0081 |
| Usp15 | 9599 | 8644 | 9365 | 10401 | 7497 | 7381 | 10196 | 8633 | 9403 | 8176 | 8066 | 8080 | 8759.09 | 9411 | 8107 | 0.862 | -0.215 | 0.0091923 | 0.04938043 | 0.0033 | 0.0073 | 0.0048 | 0.0048 | TRUE | 0.065 |
| Usp16 | 2250 | 2068 | 1999 | 3253 | 2220 | 1944 | 2390 | 2065 | 2007 | 2557 | 2389 | 2128 | 2256.05 | 2154 | 2358 | 1.095 | 0.131 | 0.23383428 | 0.44850881 | 0.0081 | 0.0084 | 0.0083 | 0.0083 | TRUE | 0.1107 |
| Usp17la | 618 | 205 | 151 | 1012 | 714 | 313 | 656 | 205 | 152 | 796 | 768 | 343 | 486.53 | 338 | 636 | 1.883 | 0.913 | 0.21066334 | 0.42117366 | 0.381 | 0.0137 | 0.1525 | 0.381 | TRUE | 7.7519 |
| Usp17lb | 576 | 219 | 120 | 859 | 612 | 271 | 612 | 219 | 120 | 675 | 658 | 297 | 430.24 | 317 | 543 | 1.715 | 0.778 | 0.29564993 | 0.5165093 | 0.3964 | 0.0146 | 0.1596 | 0.3964 | TRUE | 3.0208 |
| Usp17lc | 249 | 76 | 49 | 437 | 369 | 137 | 264 | 76 | 49 | 344 | 397 | 150 | 213.35 | 130 | 297 | 2.287 | 1.193 | 0.13272979 | 0.31409066 | 0.4485 | 0.0223 | 0.195 | 0.4485 | TRUE | 3.0598 |
| Usp17ld | 102 | 53 | 45 | 171 | 126 | 63 | 108 | 53 | 45 | 134 | 136 | 69 | 90.9 | 69 | 113 | 1.647 | 0.72 | 0.06505658 | 0.19649029 | 0.1647 | 0.0431 | 0.098 | 0.098 | TRUE | 2.2251 |
| Usp17le | 969 | 438 | 299 | 1427 | 1013 | 493 | 1029 | 437 | 300 | 1122 | 1090 | 540 | 753.05 | 589 | 917 | 1.557 | 0.639 | 0.29359156 | 0.51444574 | 0.2655 | 0.0113 | 0.1086 | 0.2655 | TRUE | 3.0725 |
| Usp18 | 21 | 16 | 17 | 24 | 10 | 14 | 22 | 16 | 17 | 19 | 11 | 15 | 16.72 | 18 | 15 | 0.821 | -0.284 | 0.55376746 | 0.73717254 | 0 | 0.204 | 0.1056 | 0.1056 | TRUE | 0.2908 |
| Usp19 | 6622 | 5760 | 5647 | 8055 | 6165 | 5605 | 7034 | 5753 | 5670 | 6332 | 6633 | 6136 | 6259.57 | 6152 | 6367 | 1.035 | 0.05 | 0.63616923 | 0.79536224 | 0.0068 | 0.0074 | 0.0078 | 0.0078 | TRUE | 0.1915 |
| Usp2 | 1008 | 1044 | 1057 | 816 | 650 | 806 | 1071 | 1043 | 1061 | 641 | 699 | 882 | 899.63 | 1058 | 741 | 0.699 | -0.516 | 0.0001026 | 0.00140714 | 0.0128 | 0.0106 | 0.0116 | 0.0116 | TRUE | 0.327 |
| Usp20 | 718 | 708 | 586 | 789 | 544 | 462 | 763 | 707 | 588 | 620 | 585 | 506 | 628.24 | 686 | 570 | 0.833 | -0.264 | 0.05760361 | 0.18119747 | 0.0125 | 0.0121 | 0.0124 | 0.0124 | TRUE | 0.1834 |
| Usp21 | 1870 | 1995 | 2045 | 2562 | 1921 | 2119 | 1986 | 1992 | 2053 | 2014 | 2067 | 2320 | 2072.1 | 2010 | 2134 | 1.06 | 0.085 | 0.32259664 | 0.54376341 | 0.0024 | 0.0085 | 0.0048 | 0.0048 | TRUE | 0.07 |
| Usp22 | 19989 | 19139 | 18886 | 25018 | 17582 | 16565 | 21232 | 19114 | 18964 | 19667 | 18917 | 18133 | 19337.84 | 19770 | 18906 | 0.956 | -0.064 | 0.40589654 | 0.61859119 | 0.0028 | 0.0071 | 0.0043 | 0.0043 | TRUE | 0.0512 |
| Usp24 | 5856 | 6150 | 6902 | 9435 | 6938 | 6854 | 6220 | 6142 | 6930 | 7417 | 7465 | 7503 | 6946.2 | 6431 | 7462 | 1.16 | 0.214 | 0.0041897 | 0.027385 | 0.0021 | 0.0074 | 0.0039 | 0.0039 | TRUE | 0.0564 |
| Usp25 | 834 | 1007 | 1050 | 1291 | 953 | 1117 | 886 | 1006 | 1054 | 1015 | 1025 | 1223 | 1034.81 | 982 | 1088 | 1.107 | 0.146 | 0.21917917 | 0.43131916 | 0.0082 | 0.0101 | 0.0092 | 0.0092 | TRUE | 0.1399 |
| Usp26 | 630 | 764 | 719 | 3369 | 2737 | 2522 | 669 | 763 | 722 | 2648 | 2945 | 2761 | 1751.36 | 718 | 2785 | 3.876 | 1.954 | 4.60E-101 | 1.81E-97 | 0.0026 | 0.0088 | 0.0052 | 0.0052 | TRUE | 0.0415 |
| Usp27x | 151 | 190 | 183 | 313 | 267 | 223 | 160 | 190 | 184 | 246 | 287 | 244 | 218.56 | 178 | 259 | 1.454 | 0.54 | 0.00047926 | 0.0050125 | 0.0035 | 0.022 | 0.0125 | 0.0125 | TRUE | 0.0979 |
| Usp28 | 16456 | 24384 | 26495 | 13975 | 11361 | 15814 | 17479 | 24353 | 26604 | 10986 | 12224 | 17311 | 18159.36 | 22812 | 13507 | 0.592 | -0.756 | 0.00010564 | 0.00143882 | 0.0524 | 0.0071 | 0.0274 | 0.0274 | TRUE | 0.742 |
| Usp29 | 539 | 324 | 280 | 515 | 347 | 342 | 573 | 324 | 281 | 405 | 373 | 374 | 388.3 | 393 | 384 | 0.98 | -0.029 | 0.90542124 | 0.95303683 | 0.0716 | 0.0154 | 0.0413 | 0.0413 | TRUE | 1.8471 |
| Usp3 | 970 | 1434 | 1441 | 1374 | 1068 | 1254 | 1030 | 1432 | 1447 | 1080 | 1149 | 1373 | 1251.89 | 1303 | 1201 | 0.921 | -0.119 | 0.45162126 | 0.6571006 | 0.0248 | 0.0095 | 0.0173 | 0.0173 | TRUE | 0.4019 |
| Usp30 | 847 | 1102 | 1192 | 1118 | 885 | 1067 | 900 | 1101 | 1197 | 879 | 952 | 1168 | 1032.7 | 1066 | 1000 | 0.937 | -0.094 | 0.53479676 | 0.72222061 | 0.0204 | 0.0101 | 0.0154 | 0.0154 | TRUE | 0.258 |
| Usp31 | 2242 | 2363 | 2623 | 3236 | 2400 | 2359 | 2381 | 2360 | 2634 | 2544 | 2582 | 2582 | 2513.93 | 2458 | 2569 | 1.045 | 0.064 | 0.41017713 | 0.62167122 | 0.0014 | 0.0082 | 0.0039 | 0.0039 | TRUE | 0.0473 |
| Usp32 | 6443 | 5755 | 6219 | 7556 | 5520 | 5694 | 6844 | 5748 | 6245 | 5940 | 5939 | 6233 | 6157.97 | 6279 | 6037 | 0.962 | -0.057 | 0.51683334 | 0.70855117 | 0.0038 | 0.0074 | 0.0053 | 0.0053 | TRUE | 0.0755 |
| Usp33 | 3579 | 3766 | 3864 | 5020 | 3668 | 3614 | 3802 | 3761 | 3880 | 3946 | 3947 | 3956 | 3881.92 | 3814 | 3950 | 1.036 | 0.05 | 0.3892764 | 0.60340711 | 0 | 0.0078 | 0.0022 | 0.0022 | TRUE | 0.0028 |
| Usp34 | 10437 | 9675 | 11067 | 14527 | 10007 | 9717 | 11086 | 9663 | 11112 | 11420 | 10767 | 10637 | 10780.76 | 10620 | 10941 | 1.03 | 0.043 | 0.60976874 | 0.77891178 | 0.0031 | 0.0072 | 0.005 | 0.005 | TRUE | 0.0761 |
| Usp35 | 381 | 498 | 442 | 396 | 380 | 387 | 405 | 497 | 444 | 311 | 409 | 424 | 414.94 | 449 | 381 | 0.847 | -0.239 | 0.1333841 | 0.31535513 | 0.0167 | 0.0148 | 0.0158 | 0.0158 | TRUE | 0.3104 |
| Usp36 | 4774 | 4230 | 4222 | 6041 | 4544 | 4090 | 5071 | 4225 | 4239 | 4749 | 4889 | 4477 | 4608.31 | 4512 | 4705 | 1.043 | 0.061 | 0.53932963 | 0.72536087 | 0.0056 | 0.0076 | 0.0068 | 0.0068 | TRUE | 0.1431 |
| Usp37 | 5777 | 5430 | 5508 | 7894 | 5601 | 5252 | 6136 | 5423 | 5531 | 6206 | 6026 | 5749 | 5845.15 | 5697 | 5994 | 1.052 | 0.074 | 0.35858486 | 0.57710208 | 0.0028 | 0.0075 | 0.0045 | 0.0045 | TRUE | 0.0555 |
| Usp38 | 4544 | 4185 | 4155 | 6226 | 4311 | 3830 | 4827 | 4180 | 4172 | 4894 | 4638 | 4193 | 4483.91 | 4393 | 4575 | 1.042 | 0.059 | 0.55190746 | 0.73599865 | 0.0056 | 0.0076 | 0.0069 | 0.0069 | TRUE | 0.0908 |
| Usp39 | 5895 | 5918 | 6046 | 7292 | 5218 | 5206 | 6262 | 5910 | 6071 | 5732 | 5614 | 5699 | 5881.36 | 6081 | 5682 | 0.934 | -0.098 | 0.1025394 | 0.26612662 | 3.00E-04 | 0.0075 | 0.0024 | 0.0024 | TRUE | 0.0082 |
| Usp4 | 4565 | 4903 | 4901 | 7086 | 5062 | 5073 | 4849 | 4897 | 4921 | 5570 | 5446 | 5553 | 5206.11 | 4889 | 5523 | 1.13 | 0.176 | 0.00167829 | 0.0133822 | 0 | 0.0075 | 0.0021 | 0.0021 | TRUE | 0.0018 |
| Usp40 | 979 | 971 | 1021 | 1477 | 1053 | 955 | 1040 | 970 | 1025 | 1161 | 1133 | 1045 | 1062.38 | 1012 | 1113 | 1.101 | 0.139 | 0.11563321 | 0.28713974 | 0.0013 | 0.01 | 0.0047 | 0.0047 | TRUE | 0.0335 |
| Usp42 | 1693 | 1763 | 1848 | 2343 | 1838 | 1681 | 1798 | 1761 | 1856 | 1842 | 1978 | 1840 | 1845.69 | 1805 | 1887 | 1.045 | 0.063 | 0.39854536 | 0.61194094 | 7.00E-04 | 0.0087 | 0.0035 | 0.0035 | TRUE | 0.0213 |
| Usp43 | 893 | 1092 | 1085 | 1186 | 992 | 1165 | 949 | 1091 | 1089 | 932 | 1067 | 1275 | 1067.25 | 1043 | 1091 | 1.046 | 0.064 | 0.63797177 | 0.79611447 | 0.0147 | 0.01 | 0.0125 | 0.0125 | TRUE | 0.2586 |
| Usp44 | 2812 | 2525 | 2710 | 1908 | 1417 | 1539 | 2987 | 2522 | 2721 | 1500 | 1525 | 1685 | 2156.49 | 2743 | 1570 | 0.572 | -0.806 | 2.66E-16 | 4.24E-14 | 0.0049 | 0.0084 | 0.0065 | 0.0065 | TRUE | 0.0731 |
| Usp45 | 3553 | 3638 | 3632 | 4390 | 3173 | 3191 | 3774 | 3633 | 3647 | 3451 | 3414 | 3493 | 3568.7 | 3685 | 3453 | 0.937 | -0.094 | 0.12382643 | 0.30006463 | 0 | 0.0078 | 0.0024 | 0.0024 | TRUE | 0.0054 |
| Usp46 | 755 | 979 | 1037 | 1057 | 828 | 951 | 802 | 978 | 1041 | 831 | 891 | 1041 | 930.63 | 940 | 921 | 0.978 | -0.032 | 0.81957615 | 0.9078134 | 0.0147 | 0.0104 | 0.0127 | 0.0127 | TRUE | 0.1967 |
| Usp47 | 3759 | 4226 | 4384 | 7206 | 4874 | 4953 | 3993 | 4221 | 4402 | 5665 | 5244 | 5422 | 4824.34 | 4205 | 5444 | 1.295 | 0.373 | 4.64E-07 | 1.40E-05 | 0.0016 | 0.0076 | 0.0037 | 0.0037 | TRUE | 0.0237 |

| | | | | | | | | | | | | | | | | | | | | | | | | |
|---|---|---|---|---|---|---|---|---|---|---|---|---|---|---|---|---|---|---|---|---|---|---|---|---|
| Usp48 | 11017 | 11564 | 11184 | 15780 | 12184 | 12025 | 11702 | 11549 | 11230 | 12405 | 13109 | 13163 | 12193.08 | 11494 | 12892 | 1.122 | 0.166 | 0.00650928 | 0.03815231 | 6.00E-04 | 0.0072 | 0.0026 | 0.0026 | TRUE | 0.0135 |
| Usp49 | 1226 | 1514 | 1632 | 1943 | 1471 | 1580 | 1302 | 1512 | 1639 | 1527 | 1583 | 1730 | 1548.78 | 1484 | 1613 | 1.086 | 0.119 | 0.28878896 | 0.50977692 | 0.0077 | 0.009 | 0.0085 | 0.0085 | TRUE | 0.1381 |
| Usp5 | 12301 | 12277 | 12261 | 16202 | 12053 | 11576 | 13066 | 12261 | 12311 | 12737 | 12968 | 12672 | 12669.18 | 12546 | 12792 | 1.02 | 0.028 | 0.64053355 | 0.79767036 | 6.00E-04 | 0.0072 | 0.0025 | 0.0025 | TRUE | 0.016 |
| Usp50 | 43 | 21 | 24 | 54 | 44 | 41 | 46 | 21 | 24 | 42 | 47 | 45 | 37.57 | 30 | 45 | 1.487 | 0.572 | 0.13115792 | 0.31177432 | 0.053 | 0.0946 | 0.0757 | 0.0757 | TRUE | 1.2888 |
| Usp51 | 246 | 206 | 190 | 213 | 133 | 157 | 261 | 206 | 191 | 167 | 143 | 172 | 190.04 | 219 | 161 | 0.735 | -0.445 | 0.01517644 | 0.07132386 | 0.0137 | 0.0242 | 0.0188 | 0.0188 | TRUE | 0.3052 |
| Usp53 | 785 | 748 | 703 | 988 | 774 | 654 | 834 | 747 | 706 | 777 | 833 | 716 | 768.68 | 762 | 775 | 1.017 | 0.025 | 0.82519386 | 0.91134777 | 0.005 | 0.0112 | 0.0077 | 0.0077 | TRUE | 0.0794 |
| Usp54 | 2003 | 1825 | 1803 | 3098 | 2171 | 2211 | 2128 | 1823 | 1810 | 2435 | 2336 | 2420 | 2158.69 | 1920 | 2397 | 1.249 | 0.32 | 0.00053336 | 0.00544824 | 0.0039 | 0.0084 | 0.0057 | 0.0057 | TRUE | 0.1075 |
| Usp6nl | 1242 | 1371 | 1412 | 2140 | 1527 | 1476 | 1319 | 1369 | 1418 | 1682 | 1643 | 1616 | 1507.87 | 1369 | 1647 | 1.204 | 0.267 | 0.00037961 | 0.0041549 | 0 | 0.0091 | 0.0034 | 0.0034 | TRUE | 0.0119 |
| Usp7 | 22790 | 19881 | 19813 | 26046 | 18962 | 18413 | 24207 | 19855 | 19894 | 20475 | 20402 | 20156 | 20831.65 | 21319 | 20344 | 0.954 | -0.067 | 0.48825839 | 0.68476003 | 0.0064 | 0.0071 | 0.0068 | 0.0068 | TRUE | 0.1718 |
| Usp8 | 6699 | 6305 | 6555 | 7850 | 5843 | 5552 | 7116 | 6297 | 6582 | 6171 | 6287 | 6078 | 6421.6 | 6665 | 6179 | 0.927 | -0.109 | 0.13924512 | 0.32391948 | 0.0019 | 0.0074 | 0.0038 | 0.0038 | TRUE | 0.0427 |
| Usp9x | 21204 | 20274 | 22556 | 30756 | 20820 | 19715 | 22522 | 20248 | 22649 | 24178 | 22401 | 21581 | 22263.17 | 21806 | 22720 | 1.042 | 0.059 | 0.47195013 | 0.67314138 | 0.0034 | 0.0071 | 0.0049 | 0.0049 | TRUE | 0.0478 |
| Usp9y | 18 | 11 | 6 | 204 | 153 | 126 | 19 | 11 | 6 | 160 | 165 | 138 | 83.17 | 12 | 154 | 12.92 | 3.692 | 4.23E-27 | 2.02E-24 | 0.0049 | 0.0465 | 0.0386 | 0.0386 | TRUE | 0.9917 |
| Uspl1 | 4023 | 3749 | 3748 | 4933 | 3723 | 3421 | 4273 | 3744 | 3763 | 3878 | 4006 | 3745 | 3901.53 | 3927 | 3876 | 0.987 | -0.019 | 0.82486153 | 0.91128174 | 0.0031 | 0.0077 | 0.0048 | 0.0048 | TRUE | 0.0723 |
| Ust | 65 | 56 | 48 | 170 | 138 | 88 | 69 | 56 | 48 | 134 | 148 | 96 | 91.94 | 58 | 126 | 2.191 | 1.132 | 1.42E-05 | 0.00026634 | 0.0293 | 0.0427 | 0.0362 | 0.0362 | TRUE | 0.3987 |
| Utf1 | 13947 | 15989 | 17631 | 17387 | 13330 | 14474 | 14814 | 15968 | 17703 | 13668 | 14342 | 15844 | 15390.09 | 16162 | 14618 | 0.904 | -0.145 | 0.14228836 | 0.32804737 | 0.0069 | 0.0071 | 0.007 | 0.007 | TRUE | 0.0851 |
| Utp11l | 2089 | 2480 | 2526 | 2656 | 1872 | 1922 | 2219 | 2477 | 2536 | 2088 | 2014 | 2104 | 2239.68 | 2411 | 2069 | 0.858 | -0.221 | 0.00824533 | 0.04543864 | 0.002 | 0.0084 | 0.0046 | 0.0046 | TRUE | 0.0585 |
| Utp14a | 4361 | 4224 | 4460 | 4746 | 3459 | 3488 | 4632 | 4219 | 4478 | 3731 | 3722 | 3818 | 4099.96 | 4443 | 3757 | 0.846 | -0.242 | 0.00045193 | 0.00480966 | 8.00E-04 | 0.0077 | 0.0032 | 0.0032 | TRUE | 0.0238 |
| Utp14b | 416 | 496 | 448 | 929 | 646 | 467 | 442 | 495 | 450 | 730 | 695 | 511 | 553.94 | 462 | 645 | 1.398 | 0.483 | 0.00196208 | 0.01516213 | 0.0183 | 0.0129 | 0.0157 | 0.0157 | TRUE | 0.3855 |
| Utp15 | 3967 | 3440 | 3325 | 5120 | 3607 | 3391 | 4214 | 3436 | 3339 | 4025 | 3881 | 3712 | 3767.62 | 3663 | 3873 | 1.058 | 0.081 | 0.45982346 | 0.66404367 | 0.0078 | 0.0078 | 0.0083 | 0.0083 | TRUE | 0.2104 |
| Utp18 | 6230 | 7329 | 7244 | 8683 | 6276 | 6318 | 6617 | 7320 | 7274 | 6826 | 6753 | 6916 | 6950.86 | 7070 | 6832 | 0.966 | -0.05 | 0.48080781 | 0.68004523 | 0.0014 | 0.0074 | 0.0034 | 0.0034 | TRUE | 0.0383 |
| Utp20 | 9990 | 6971 | 7605 | 13597 | 8948 | 7658 | 10611 | 6962 | 7636 | 10689 | 9627 | 8383 | 8984.77 | 8403 | 9566 | 1.139 | 0.187 | 0.25535426 | 0.47370587 | 0.0302 | 0.0073 | 0.0194 | 0.0194 | TRUE | 0.6451 |
| Utp23 | 877 | 865 | 766 | 1210 | 812 | 738 | 932 | 864 | 769 | 951 | 874 | 808 | 866.21 | 855 | 878 | 1.028 | 0.04 | 0.73219358 | 0.85630933 | 0.0068 | 0.0107 | 0.0085 | 0.0085 | TRUE | 0.0916 |
| Utp3 | 2358 | 2358 | 2340 | 4101 | 3040 | 2556 | 2505 | 2355 | 2350 | 3224 | 3271 | 2798 | 2750.31 | 2403 | 3098 | 1.289 | 0.367 | 5.35E-05 | 0.00082104 | 0.0037 | 0.0081 | 0.0056 | 0.0056 | TRUE | 0.0863 |
| Utp6 | 7897 | 8178 | 8071 | 11332 | 7740 | 7728 | 8388 | 8167 | 8104 | 8908 | 8328 | 8460 | 8392.53 | 8220 | 8565 | 1.042 | 0.06 | 0.33644378 | 0.55663394 | 6.00E-04 | 0.0073 | 0.0027 | 0.0027 | TRUE | 0.0151 |
| Utrn | 3817 | 4242 | 5079 | 4874 | 3416 | 4094 | 4054 | 4237 | 5100 | 3832 | 3675 | 4482 | 4229.87 | 4464 | 3996 | 0.895 | -0.16 | 0.19062482 | 0.39573321 | 0.0132 | 0.0077 | 0.0105 | 0.0105 | TRUE | 0.1895 |
| Uts2 | 0 | 0 | 0 | 1 | 0 | 0 | 0 | 0 | 0 | 1 | 0 | 0 | 0.13 | 0 | 0 | 1.859 | 0.894 | 0.82650734 | NA | 0 | 25.1515 | 10 | 10 | TRUE | 0.2563 |
| Uts2d | 0 | 0 | 0 | 0 | 0 | 0 | 0 | 0 | 0 | 0 | 0 | 0 | 0 | 0 | 0 | NA | NA | NA | NA | NA | NA | NA | NA | NA | NA |
| Uts2r | 18 | 7 | 9 | 26 | 13 | 10 | 19 | 7 | 9 | 20 | 14 | 11 | 13.42 | 12 | 15 | 1.309 | 0.388 | 0.51823753 | 0.70964076 | 0.1109 | 0.2524 | 0.1842 | 0.1842 | TRUE | 1.1377 |
| Uty | 439 | 471 | 454 | 903 | 675 | 626 | 466 | 470 | 456 | 710 | 726 | 685 | 585.66 | 464 | 707 | 1.524 | 0.607 | 1.56E-10 | 1.12E-08 | 0 | 0.0125 | 0.0047 | 0.0047 | TRUE | 0.0083 |
| Uvrag | 1561 | 1699 | 1711 | 2078 | 1486 | 1434 | 1658 | 1697 | 1718 | 1634 | 1599 | 1570 | 1645.84 | 1691 | 1601 | 0.947 | -0.079 | 0.26341459 | 0.48264194 | 0 | 0.0089 | 0.003 | 0.003 | TRUE | 0.004 |
| Uvssa | 208 | 170 | 198 | 324 | 176 | 184 | 221 | 170 | 199 | 255 | 189 | 201 | 205.84 | 197 | 215 | 1.099 | 0.137 | 0.46047175 | 0.66458277 | 0.017 | 0.0229 | 0.0198 | 0.0198 | TRUE | 0.2994 |
| Uxs1 | 766 | 865 | 875 | 1109 | 747 | 763 | 814 | 864 | 879 | 872 | 804 | 835 | 844.48 | 852 | 837 | 0.983 | -0.025 | 0.78217374 | 0.88739843 | 0 | 0.0108 | 0.0046 | 0.0046 | TRUE | 0.0183 |
| Uxt | 586 | 692 | 744 | 890 | 635 | 668 | 622 | 691 | 747 | 700 | 683 | 731 | 695.78 | 687 | 705 | 1.025 | 0.036 | 0.73483132 | 0.85771731 | 0.0032 | 0.0116 | 0.0068 | 0.0068 | TRUE | 0.0788 |
| V1ra8 | 0 | 0 | 0 | 0 | 0 | 0 | 0 | 0 | 0 | 0 | 0 | 0 | 0 | 0 | 0 | NA | NA | NA | NA | NA | NA | NA | NA | NA | NA |
| V1rd18 | 0 | 0 | 0 | 0 | 0 | 0 | 0 | 0 | 0 | 0 | 0 | 0 | 0 | 0 | 0 | NA | NA | NA | NA | NA | NA | NA | NA | NA | NA |
| V1rd19 | 0 | 0 | 0 | 0 | 0 | 0 | 0 | 0 | 0 | 0 | 0 | 0 | 0 | 0 | 0 | NA | NA | NA | NA | NA | NA | NA | NA | NA | NA |
| Vac14 | 4959 | 4531 | 4416 | 6469 | 4792 | 4306 | 5267 | 4525 | 4434 | 5085 | 5156 | 4714 | 4863.59 | 4742 | 4985 | 1.051 | 0.072 | 0.44636657 | 0.65236459 | 0.005 | 0.0076 | 0.0063 | 0.0063 | TRUE | 0.1142 |
| Vamp1 | 135 | 112 | 157 | 207 | 172 | 117 | 143 | 112 | 158 | 163 | 185 | 128 | 148.13 | 138 | 159 | 1.154 | 0.206 | 0.34055073 | 0.56078159 | 0.0246 | 0.0291 | 0.0269 | 0.0269 | TRUE | 0.286 |
| Vamp2 | 878 | 1141 | 1166 | 1480 | 1261 | 1338 | 933 | 1140 | 1171 | 1163 | 1357 | 1465 | 1204.63 | 1081 | 1328 | 1.228 | 0.296 | 0.02484986 | 0.10135299 | 0.0125 | 0.0096 | 0.0117 | 0.0117 | TRUE | 0.1718 |
| Vamp3 | 3270 | 3533 | 3231 | 4982 | 3476 | 3352 | 3473 | 3528 | 3244 | 3916 | 3740 | 3669 | 3595.29 | 3415 | 3775 | 1.106 | 0.145 | 0.04515532 | 0.15319203 | 0.0011 | 0.0078 | 0.0035 | 0.0035 | TRUE | 0.0234 |
| Vamp4 | 388 | 495 | 494 | 576 | 427 | 512 | 412 | 494 | 496 | 453 | 459 | 560 | 479.2 | 467 | 491 | 1.048 | 0.068 | 0.62660384 | 0.78967605 | 0.0105 | 0.0138 | 0.012 | 0.012 | TRUE | 0.1761 |
| Vamp5 | 32 | 30 | 27 | 31 | 24 | 24 | 34 | 30 | 27 | 24 | 26 | 26 | 27.92 | 30 | 25 | 0.839 | -0.254 | 0.47396119 | 0.67469488 | 0 | 0.1249 | 0.0544 | 0.0544 | TRUE | 0.0707 |
| Vamp7 | 1205 | 1456 | 1375 | 1911 | 1387 | 1454 | 1280 | 1454 | 1381 | 1502 | 1492 | 1592 | 1450.15 | 1372 | 1529 | 1.114 | 0.156 | 0.07217171 | 0.20997827 | 0.0019 | 0.0092 | 0.0047 | 0.0047 | TRUE | 0.0408 |
| Vamp8 | 350 | 454 | 390 | 628 | 395 | 471 | 372 | 453 | 392 | 494 | 425 | 516 | 441.84 | 406 | 478 | 1.179 | 0.237 | 0.07952177 | 0.22376634 | 0.0079 | 0.0144 | 0.0109 | 0.0109 | TRUE | 0.123 |
| Vangl1 | 1720 | 1859 | 1668 | 2312 | 1831 | 1911 | 1827 | 1857 | 1675 | 1818 | 1970 | 2092 | 1872.97 | 1786 | 1960 | 1.097 | 0.133 | 0.14675712 | 0.33570026 | 0.0031 | 0.0087 | 0.0055 | 0.0055 | TRUE | 0.0498 |
| Vangl2 | 1156 | 1288 | 1319 | 1185 | 1009 | 969 | 1228 | 1286 | 1324 | 932 | 1086 | 1061 | 1152.75 | 1279 | 1026 | 0.801 | -0.321 | 0.00092396 | 0.00838471 | 0.0032 | 0.0098 | 0.0059 | 0.0059 | TRUE | 0.0804 |
| Vapa | 10741 | 13632 | 13656 | 14138 | 10649 | 11183 | 11409 | 13614 | 13712 | 11114 | 11458 | 12242 | 12258.12 | 12912 | 11605 | 0.899 | -0.154 | 0.11309711 | 0.28322762 | 0.0062 | 0.0072 | 0.0067 | 0.0067 | TRUE | 0.1267 |
| Vapb | 5025 | 5867 | 5790 | 6066 | 4521 | 4962 | 5337 | 5859 | 5814 | 4769 | 4864 | 5432 | 5345.88 | 5670 | 5022 | 0.885 | -0.176 | 0.03971287 | 0.13996678 | 0.0036 | 0.0075 | 0.0051 | 0.0051 | TRUE | 0.062 |
| Vars | 14130 | 13589 | 13965 | 19105 | 14552 | 13448 | 15009 | 13571 | 14022 | 15019 | 15657 | 14721 | 14666.54 | 14201 | 15132 | 1.066 | 0.092 | 0.19240669 | 0.39803992 | 0.0017 | 0.0071 | 0.0035 | 0.0035 | TRUE | 0.0303 |
| Vars2 | 2917 | 3185 | 2930 | 3915 | 2897 | 2879 | 3098 | 3181 | 2942 | 3078 | 3117 | 3152 | 3094.58 | 3074 | 3116 | 1.013 | 0.019 | 0.77580295 | 0.88349968 | 5.00E-04 | 0.008 | 0.003 | 0.003 | TRUE | 0.0171 |
| Vash1 | 1557 | 1745 | 1705 | 1947 | 1496 | 1370 | 1654 | 1743 | 1712 | 1531 | 1610 | 1500 | 1624.74 | 1703 | 1547 | 0.908 | -0.139 | 0.06427419 | 0.19513621 | 3.00E-04 | 0.0089 | 0.0035 | 0.0035 | TRUE | 0.0149 |
| Vash2 | 932 | 836 | 875 | 782 | 586 | 503 | 990 | 835 | 879 | 615 | 630 | 551 | 749.89 | 901 | 599 | 0.665 | -0.589 | 1.70E-07 | 5.65E-06 | 0.005 | 0.0113 | 0.0078 | 0.0078 | TRUE | 0.088 |
| Vasn | 146 | 116 | 122 | 218 | 120 | 93 | 155 | 116 | 123 | 171 | 129 | 102 | 132.62 | 131 | 134 | 1.028 | 0.04 | 0.86886903 | 0.93544206 | 0.0391 | 0.0317 | 0.0354 | 0.0354 | TRUE | 0.35 |
| Vasp | 3545 | 3282 | 3339 | 5566 | 3866 | 3576 | 3765 | 3278 | 3353 | 4376 | 4160 | 3914 | 3807.59 | 3465 | 4150 | 1.198 | 0.261 | 0.00355044 | 0.02408495 | 0.004 | 0.0078 | 0.0055 | 0.0055 | TRUE | 0.0697 |
| Vat1 | 16492 | 19715 | 19888 | 15532 | 11764 | 13092 | 17517 | 19690 | 19970 | 12210 | 12657 | 14331 | 16062.54 | 19059 | 13066 | 0.686 | -0.545 | 1.05E-08 | 4.95E-07 | 0.006 | 0.0071 | 0.0065 | 0.0065 | TRUE | 0.0876 |
| Vat1l | 96 | 46 | 49 | 158 | 95 | 57 | 102 | 46 | 49 | 124 | 102 | 62 | 80.99 | 66 | 96 | 1.472 | 0.558 | 0.14462765 | 0.33208143 | 0.1442 | 0.0476 | 0.0925 | 0.0925 | TRUE | 0.7386 |
| Vaultrc5 | 0 | 2 | 0 | 0 | 2 | 0 | 0 | 2 | 0 | 0 | 2 | 0 | 0.69 | 1 | 1 | 1.052 | 0.073 | 0.98136341 | NA | 8.094 | 4.7708 | 5.501 | 5.501 | TRUE | 0.9745 |
| Vav1 | 78 | 91 | 85 | 90 | 58 | 61 | 83 | 91 | 85 | 71 | 62 | 67 | 76.5 | 86 | 67 | 0.773 | -0.371 | 0.09155405 | 0.2469204 | 0 | 0.05 | 0.0216 | 0.0216 | TRUE | 0.0298 |
| Vav2 | 1494 | 1664 | 1624 | 1894 | 1572 | 1569 | 1587 | 1662 | 1631 | 1489 | 1691 | 1718 | 1629.54 | 1627 | 1633 | 1.003 | 0.004 | 0.96451395 | 0.98213662 | 0.0028 | 0.0089 | 0.0051 | 0.0051 | TRUE | 0.0734 |
| Vav3 | 266 | 235 | 217 | 279 | 196 | 164 | 283 | 235 | 218 | 219 | 211 | 180 | 224.14 | 245 | 203 | 0.832 | -0.266 | 0.11206454 | 0.28149099 | 0.0101 | 0.0216 | 0.0157 | 0.0157 | TRUE | 0.1966 |
| Vax1 | 5 | 5 | 4 | 11 | 10 | 4 | 5 | 5 | 4 | 9 | 11 | 4 | 6.35 | 5 | 8 | 1.669 | 0.739 | 0.33668207 | 0.55673617 | 0 | 0.5256 | 0.2576 | 0.2576 | TRUE | 0.3426 |
| Vax2 | 12 | 29 | 16 | 18 | 9 | 24 | 13 | 29 | 16 | 14 | 10 | 26 | 17.98 | 19 | 17 | 0.862 | -0.215 | 0.70338939 | 0.83766892 | 0.1559 | 0.1901 | 0.1735 | 0.1735 | TRUE | 0.8764 |
| Vax2os | 74 | 93 | 93 | 68 | 50 | 78 | 79 | 93 | 93 | 53 | 54 | 85 | 76.25 | 88 | 64 | 0.723 | -0.468 | 0.08160168 | 0.22771085 | 0.0251 | 0.0501 | 0.0386 | 0.0386 | TRUE | 0.6982 |
| Vbp1 | 6699 | 9102 | 8967 | 7820 | 5810 | 6941 | 7116 | 9090 | 9004 | 6147 | 6251 | 7598 | 7534.38 | 8403 | 6665 | 0.793 | -0.334 | 0.0102327 | 0.05367149 | 0.016 | 0.0073 | 0.0121 | 0.0121 | TRUE | 0.2193 |
| Vcam1 | 3 | 0 | 0 | 1 | 2 | 1 | 3 | 0 | 0 | 1 | 2 | 1 | 1.2 | 1 | 1 | 1.268 | 0.342 | 0.86152965 | NA | 0.7687 | 2.745 | 1.9225 | 1.9225 | TRUE | 1.443 |
| Vcan | 1249 | 1163 | 1242 | 1135 | 828 | 1171 | 1327 | 1161 | 1247 | 892 | 891 | 1282 | 1133.37 | 1245 | 1022 | 0.82 | -0.286 | 0.06846643 | 0.20275113 | 0.0252 | 0.0098 | 0.0169 | 0.0169 | TRUE | 0.5895 |
| Vcl | 7332 | 6381 | 6501 | 9753 | 6502 | 6192 | 7788 | 6373 | 6528 | 7667 | 6996 | 6778 | 7021.54 | 6896 | 7147 | 1.037 | 0.052 | 0.6205501 | 0.78523764 | 0.0069 | 0.0074 | 0.0077 | 0.0077 | TRUE | 0.1561 |
| Vcp | 23782 | 22117 | 22360 | 31558 | 21260 | 20638 | 25261 | 22088 | 22452 | 24808 | 22874 | 22592 | 23345.89 | 23267 | 23425 | 1.007 | 0.01 | 0.90724577 | 0.95354464 | 0.0033 | 0.007 | 0.0051 | 0.0051 | TRUE | 0.0687 |
| Vcpip1 | 2053 | 2083 | 2099 | 3002 | 2121 | 2017 | 2181 | 2080 | 2108 | 2360 | 2282 | 2208 | 2203.08 | 2123 | 2283 | 1.076 | 0.106 | 0.1340842 | 0.31663064 | 4.00E-04 | 0.0084 | 0.0031 | 0.0031 | TRUE | 0.0107 |

| | | | | | | | | | | | | | | | | | | | | | | | | | |
|---|---|---|---|---|---|---|---|---|---|---|---|---|---|---|---|---|---|---|---|---|---|---|---|---|---|
| Vdac1 | 31063 | 38915 | 38644 | 46606 | 35063 | 38457 | 32994 | 38865 | 38803 | 36638 | 37726 | 42097 | 37853.78 | 36887 | 38820 | 1.052 | 0.074 | 0.45589301 | 0.6607558 | 0.0063 | 0.007 | 0.007 | 0.007 | TRUE | 0.1043 |
| Vdac2 | 14162 | 15306 | 14942 | 17502 | 12503 | 12678 | 15043 | 15286 | 15003 | 13759 | 13452 | 13878 | 14403.56 | 15111 | 13696 | 0.906 | -0.142 | 0.00728781 | 0.04139117 | 1.00E-04 | 0.0071 | 0.0019 | 0.0019 | TRUE | 0.003 |
| Vdac3 | 5354 | 6103 | 6081 | 7927 | 5540 | 5776 | 5687 | 6095 | 6106 | 6232 | 5961 | 6323 | 6067.17 | 5963 | 6172 | 1.035 | 0.05 | 0.4631708 | 0.66658727 | 0.001 | 0.0074 | 0.0031 | 0.0031 | TRUE | 0.0199 |
| Vdr | 37 | 55 | 24 | 71 | 64 | 45 | 39 | 55 | 24 | 56 | 69 | 49 | 48.71 | 39 | 58 | 1.469 | 0.555 | 0.1210677 | 0.29590574 | 0.0663 | 0.0745 | 0.0708 | 0.0708 | TRUE | 0.2871 |
| Vegfa | 978 | 2160 | 2182 | 2291 | 2248 | 2514 | 1039 | 2157 | 2191 | 1801 | 2419 | 2752 | 2059.78 | 1796 | 2324 | 1.294 | 0.372 | 0.32434681 | 0.54575342 | 0.1021 | 0.0085 | 0.0475 | 0.1021 | TRUE | 1.2447 |
| Vegfb | 2161 | 2789 | 2830 | 2238 | 1854 | 1996 | 2295 | 2785 | 2842 | 1759 | 1995 | 2185 | 2310.24 | 2641 | 1980 | 0.749 | -0.417 | 0.00070777 | 0.00681501 | 0.0116 | 0.0083 | 0.0105 | 0.0105 | TRUE | 0.1586 |
| Vegfc | 409 | 365 | 392 | 615 | 395 | 340 | 434 | 365 | 394 | 483 | 425 | 372 | 412.2 | 398 | 427 | 1.077 | 0.106 | 0.45757243 | 0.66215546 | 0.0102 | 0.0149 | 0.0124 | 0.0124 | TRUE | 0.1617 |
| Veph1 | 29 | 22 | 24 | 81 | 77 | 36 | 31 | 22 | 24 | 64 | 83 | 39 | 43.8 | 26 | 62 | 2.424 | 1.278 | 0.00067351 | 0.00656934 | 0.0653 | 0.0821 | 0.0739 | 0.0739 | TRUE | 0.1243 |
| Vezf1 | 2689 | 3325 | 3519 | 3849 | 3012 | 3270 | 2856 | 3321 | 3533 | 3026 | 3241 | 3580 | 3259.4 | 3237 | 3282 | 1.014 | 0.02 | 0.86087222 | 0.93125634 | 0.0091 | 0.0079 | 0.0086 | 0.0086 | TRUE | 0.1284 |
| Vezt | 686 | 709 | 692 | 859 | 675 | 619 | 729 | 708 | 695 | 675 | 726 | 678 | 701.78 | 711 | 693 | 0.975 | -0.037 | 0.68935301 | 0.82925141 | 0 | 0.0116 | 0.0046 | 0.0046 | TRUE | 0.0203 |
| Vgf | 61 | 41 | 31 | 47 | 39 | 17 | 65 | 41 | 31 | 37 | 42 | 19 | 39.06 | 46 | 33 | 0.716 | -0.482 | 0.25895001 | 0.4778477 | 0.1205 | 0.0912 | 0.105 | 0.105 | TRUE | 0.5207 |
| Vgll1 | 2 | 2 | 2 | 11 | 10 | 0 | 2 | 2 | 2 | 9 | 11 | 0 | 4.26 | 2 | 7 | 3.187 | 1.672 | 0.1760744 | NA | 0 | 0.781 | 0.7757 | 0.7757 | TRUE | 0.83 |
| Vgll2 | 2 | 5 | 3 | 11 | 2 | 3 | 2 | 5 | 3 | 9 | 2 | 3 | 4.04 | 3 | 5 | 1.422 | 0.508 | 0.61463697 | NA | 0 | 0.8233 | 0.4813 | 0.4813 | TRUE | 1.321 |
| Vgll3 | 25 | 16 | 12 | 76 | 45 | 40 | 27 | 16 | 12 | 60 | 48 | 44 | 34.42 | 18 | 51 | 2.805 | 1.488 | 0.00016155 | 0.00202708 | 0.0339 | 0.1026 | 0.0743 | 0.0743 | TRUE | 0.7951 |
| Vgll4 | 125 | 149 | 134 | 224 | 203 | 152 | 133 | 149 | 135 | 176 | 218 | 166 | 162.84 | 139 | 187 | 1.346 | 0.429 | 0.01689165 | 0.07711 | 0.0072 | 0.0271 | 0.0169 | 0.0169 | TRUE | 0.2255 |
| Vhl | 1748 | 2135 | 2022 | 2878 | 2038 | 2026 | 1857 | 2132 | 2030 | 2262 | 2193 | 2218 | 2115.37 | 2006 | 2224 | 1.109 | 0.149 | 0.07131331 | 0.20809501 | 0.0018 | 0.0085 | 0.0044 | 0.0044 | TRUE | 0.0513 |
| Vil1 | 184 | 194 | 191 | 217 | 164 | 179 | 195 | 194 | 192 | 171 | 176 | 196 | 187.33 | 194 | 181 | 0.933 | -0.101 | 0.50060183 | 0.69649719 | 0 | 0.0245 | 0.0107 | 0.0107 | TRUE | 0.0543 |
| Vill | 1624 | 2193 | 2175 | 1877 | 1545 | 1554 | 1725 | 2190 | 2184 | 1476 | 1662 | 1701 | 1823.01 | 2033 | 1613 | 0.793 | -0.335 | 0.00620489 | 0.0369436 | 0.0106 | 0.0087 | 0.0102 | 0.0102 | TRUE | 0.212 |
| Vim | 9619 | 12498 | 13359 | 10880 | 8047 | 9086 | 10217 | 12482 | 13414 | 8553 | 8658 | 9946 | 10544.99 | 12038 | 9052 | 0.752 | -0.411 | 0.00068747 | 0.00667245 | 0.0129 | 0.0072 | 0.0105 | 0.0105 | TRUE | 0.2139 |
| Vimp | 897 | 1092 | 944 | 1471 | 940 | 942 | 953 | 1091 | 948 | 1156 | 1011 | 1031 | 1031.7 | 997 | 1066 | 1.07 | 0.098 | 0.35491679 | 0.57336348 | 0.0046 | 0.0101 | 0.0071 | 0.0071 | TRUE | 0.0807 |
| Vip | 0 | 0 | 0 | 0 | 0 | 0 | 0 | 0 | 0 | 0 | 0 | 0 | 0 | 0 | 0 | NA | NA | NA | NA | NA | NA | NA | NA | NA | NA |
| Vipas39 | 1343 | 1232 | 1223 | 2011 | 1350 | 1187 | 1427 | 1230 | 1228 | 1581 | 1453 | 1299 | 1369.62 | 1295 | 1444 | 1.116 | 0.159 | 0.16033624 | 0.3548001 | 0.0075 | 0.0093 | 0.0085 | 0.0085 | TRUE | 0.0945 |
| Vipr1 | 125 | 108 | 86 | 136 | 124 | 97 | 133 | 108 | 86 | 107 | 133 | 106 | 112.25 | 109 | 115 | 1.059 | 0.083 | 0.71871909 | 0.84765105 | 0.0218 | 0.0363 | 0.0292 | 0.0292 | TRUE | 0.2337 |
| Vipr2 | 8 | 3 | 4 | 14 | 16 | 9 | 8 | 3 | 4 | 11 | 17 | 10 | 8.93 | 5 | 13 | 2.462 | 1.3 | 0.0657501 | 0.19775182 | 0.0268 | 0.3758 | 0.2215 | 0.2215 | TRUE | 0.6257 |
| Vit | 6 | 0 | 0 | 1 | 2 | 0 | 6 | 0 | 0 | 1 | 2 | 0 | 1.55 | 2 | 1 | 0.462 | -1.114 | 0.61408358 | NA | 4.151 | 2.1298 | 2.7735 | 2.7735 | TRUE | 2.779 |
| Vkorc1 | 956 | 1155 | 1160 | 1045 | 781 | 868 | 1015 | 1154 | 1165 | 821 | 840 | 950 | 990.95 | 1111 | 870 | 0.783 | -0.353 | 0.00091471 | 0.00831042 | 0.0049 | 0.0102 | 0.0072 | 0.0072 | TRUE | 0.075 |
| Vkorc1l1 | 561 | 562 | 583 | 646 | 518 | 493 | 596 | 561 | 585 | 508 | 557 | 540 | 557.9 | 581 | 535 | 0.92 | -0.12 | 0.22820416 | 0.44159023 | 0 | 0.0128 | 0.0054 | 0.0054 | TRUE | 0.0245 |
| Vldlr | 116 | 247 | 254 | 283 | 202 | 255 | 123 | 247 | 255 | 222 | 217 | 279 | 223.98 | 208 | 239 | 1.149 | 0.2 | 0.45727899 | 0.66182576 | 0.0786 | 0.0216 | 0.0478 | 0.0478 | TRUE | 1.3806 |
| Vma21 | 859 | 1034 | 954 | 1373 | 1042 | 1025 | 912 | 1033 | 958 | 1079 | 1121 | 1122 | 1037.58 | 968 | 1107 | 1.144 | 0.194 | 0.02898565 | 0.11229714 | 0.0011 | 0.0101 | 0.0047 | 0.0047 | TRUE | 0.0413 |
| Vmac | 695 | 824 | 884 | 941 | 741 | 816 | 738 | 823 | 888 | 740 | 797 | 893 | 813.17 | 816 | 810 | 0.991 | -0.013 | 0.91431065 | 0.95780402 | 0.0076 | 0.011 | 0.0091 | 0.0091 | TRUE | 0.0953 |
| Vmn1r-ps10: | 0 | 0 | 0 | 0 | 0 | 0 | 0 | 0 | 0 | 0 | 0 | 0 | 0 | 0 | 0 | NA | NA | NA | NA | NA | NA | NA | NA | NA | NA |
| Vmn1r-ps79 | 0 | 0 | 0 | 0 | 0 | 0 | 0 | 0 | 0 | 0 | 0 | 0 | 0 | 0 | 0 | NA | NA | NA | NA | NA | NA | NA | NA | NA | NA |
| Vmn1r1 | 0 | 0 | 1 | 0 | 1 | 0 | 0 | 0 | 1 | 0 | 1 | 0 | 0.35 | 0 | 0 | 0.954 | -0.067 | 0.98570726 | NA | 0 | 9.5099 | 8.2019 | 8.2019 | TRUE | 0.5204 |
| Vmn1r10 | 0 | 0 | 0 | 0 | 0 | 0 | 0 | 0 | 0 | 0 | 0 | 0 | 0 | 0 | 0 | NA | NA | NA | NA | NA | NA | NA | NA | NA | NA |
| Vmn1r100 | 0 | 0 | 0 | 0 | 0 | 0 | 0 | 0 | 0 | 0 | 0 | 0 | 0 | 0 | 0 | NA | NA | NA | NA | NA | NA | NA | NA | NA | NA |
| Vmn1r101 | 0 | 0 | 0 | 0 | 0 | 0 | 0 | 0 | 0 | 0 | 0 | 0 | 0 | 0 | 0 | NA | NA | NA | NA | NA | NA | NA | NA | NA | NA |
| Vmn1r103 | 0 | 0 | 0 | 0 | 0 | 0 | 0 | 0 | 0 | 0 | 0 | 0 | 0 | 0 | 0 | NA | NA | NA | NA | NA | NA | NA | NA | NA | NA |
| Vmn1r104 | 0 | 0 | 0 | 0 | 0 | 0 | 0 | 0 | 0 | 0 | 0 | 0 | 0 | 0 | 0 | NA | NA | NA | NA | NA | NA | NA | NA | NA | NA |
| Vmn1r107 | 0 | 0 | 0 | 0 | 0 | 0 | 0 | 0 | 0 | 0 | 0 | 0 | 0 | 0 | 0 | NA | NA | NA | NA | NA | NA | NA | NA | NA | NA |
| Vmn1r11 | 0 | 0 | 0 | 0 | 0 | 0 | 0 | 0 | 0 | 0 | 0 | 0 | 0 | 0 | 0 | NA | NA | NA | NA | NA | NA | NA | NA | NA | NA |
| Vmn1r111 | 0 | 0 | 0 | 0 | 0 | 0 | 0 | 0 | 0 | 0 | 0 | 0 | 0 | 0 | 0 | NA | NA | NA | NA | NA | NA | NA | NA | NA | NA |
| Vmn1r112 | 0 | 0 | 0 | 0 | 0 | 0 | 0 | 0 | 0 | 0 | 0 | 0 | 0 | 0 | 0 | NA | NA | NA | NA | NA | NA | NA | NA | NA | NA |
| Vmn1r113 | 0 | 0 | 0 | 0 | 0 | 0 | 0 | 0 | 0 | 0 | 0 | 0 | 0 | 0 | 0 | NA | NA | NA | NA | NA | NA | NA | NA | NA | NA |
| Vmn1r114 | 0 | 0 | 0 | 0 | 0 | 0 | 0 | 0 | 0 | 0 | 0 | 0 | 0 | 0 | 0 | NA | NA | NA | NA | NA | NA | NA | NA | NA | NA |
| Vmn1r115 | 0 | 0 | 0 | 0 | 0 | 0 | 0 | 0 | 0 | 0 | 0 | 0 | 0 | 0 | 0 | NA | NA | NA | NA | NA | NA | NA | NA | NA | NA |
| Vmn1r116 | 0 | 0 | 0 | 0 | 0 | 0 | 0 | 0 | 0 | 0 | 0 | 0 | 0 | 0 | 0 | NA | NA | NA | NA | NA | NA | NA | NA | NA | NA |
| Vmn1r117 | 0 | 0 | 0 | 0 | 0 | 0 | 0 | 0 | 0 | 0 | 0 | 0 | 0 | 0 | 0 | NA | NA | NA | NA | NA | NA | NA | NA | NA | NA |
| Vmn1r118 | 0 | 0 | 0 | 0 | 0 | 0 | 0 | 0 | 0 | 0 | 0 | 0 | 0 | 0 | 0 | NA | NA | NA | NA | NA | NA | NA | NA | NA | NA |
| Vmn1r119 | 0 | 0 | 0 | 0 | 0 | 0 | 0 | 0 | 0 | 0 | 0 | 0 | 0 | 0 | 0 | NA | NA | NA | NA | NA | NA | NA | NA | NA | NA |
| Vmn1r12 | 0 | 0 | 0 | 0 | 0 | 0 | 0 | 0 | 0 | 0 | 0 | 0 | 0 | 0 | 0 | NA | NA | NA | NA | NA | NA | NA | NA | NA | NA |
| Vmn1r120 | 0 | 0 | 0 | 0 | 0 | 0 | 0 | 0 | 0 | 0 | 0 | 0 | 0 | 0 | 0 | NA | NA | NA | NA | NA | NA | NA | NA | NA | NA |
| Vmn1r121 | 0 | 0 | 0 | 0 | 0 | 0 | 0 | 0 | 0 | 0 | 0 | 0 | 0 | 0 | 0 | NA | NA | NA | NA | NA | NA | NA | NA | NA | NA |
| Vmn1r122 | 0 | 0 | 0 | 0 | 0 | 0 | 0 | 0 | 0 | 0 | 0 | 0 | 0 | 0 | 0 | NA | NA | NA | NA | NA | NA | NA | NA | NA | NA |
| Vmn1r123 | 0 | 0 | 0 | 0 | 0 | 0 | 0 | 0 | 0 | 0 | 0 | 0 | 0 | 0 | 0 | NA | NA | NA | NA | NA | NA | NA | NA | NA | NA |
| Vmn1r124 | 0 | 0 | 0 | 0 | 0 | 0 | 0 | 0 | 0 | 0 | 0 | 0 | 0 | 0 | 0 | NA | NA | NA | NA | NA | NA | NA | NA | NA | NA |
| Vmn1r125 | 1 | 0 | 0 | 0 | 0 | 0 | 1 | 0 | 0 | 0 | 0 | 0 | 0.18 | 0 | 0 | 0.49 | -1.029 | 0.80087 | NA | 0 | 18.6165 | 10 | 10 | TRUE | 0.4614 |
| Vmn1r126 | 0 | 0 | 0 | 0 | 0 | 0 | 0 | 0 | 0 | 0 | 0 | 0 | 0 | 0 | 0 | NA | NA | NA | NA | NA | NA | NA | NA | NA | NA |
| Vmn1r127 | 0 | 0 | 0 | 0 | 0 | 0 | 0 | 0 | 0 | 0 | 0 | 0 | 0 | 0 | 0 | NA | NA | NA | NA | NA | NA | NA | NA | NA | NA |
| Vmn1r128 | 0 | 0 | 0 | 0 | 0 | 0 | 0 | 0 | 0 | 0 | 0 | 0 | 0 | 0 | 0 | NA | NA | NA | NA | NA | NA | NA | NA | NA | NA |
| Vmn1r129 | 0 | 0 | 0 | 0 | 0 | 0 | 0 | 0 | 0 | 0 | 0 | 0 | 0 | 0 | 0 | NA | NA | NA | NA | NA | NA | NA | NA | NA | NA |
| Vmn1r13 | 0 | 0 | 0 | 0 | 0 | 0 | 0 | 0 | 0 | 0 | 0 | 0 | 0 | 0 | 0 | NA | NA | NA | NA | NA | NA | NA | NA | NA | NA |
| Vmn1r130 | 0 | 0 | 0 | 0 | 0 | 0 | 0 | 0 | 0 | 0 | 0 | 0 | 0 | 0 | 0 | NA | NA | NA | NA | NA | NA | NA | NA | NA | NA |
| Vmn1r132 | 0 | 0 | 0 | 0 | 0 | 0 | 0 | 0 | 0 | 0 | 0 | 0 | 0 | 0 | 0 | NA | NA | NA | NA | NA | NA | NA | NA | NA | NA |
| Vmn1r135 | 0 | 0 | 0 | 0 | 0 | 0 | 0 | 0 | 0 | 0 | 0 | 0 | 0 | 0 | 0 | NA | NA | NA | NA | NA | NA | NA | NA | NA | NA |
| Vmn1r137 | 0 | 0 | 0 | 0 | 0 | 0 | 0 | 0 | 0 | 0 | 0 | 0 | 0 | 0 | 0 | NA | NA | NA | NA | NA | NA | NA | NA | NA | NA |
| Vmn1r139 | 0 | 0 | 0 | 0 | 0 | 0 | 0 | 0 | 0 | 0 | 0 | 0 | 0 | 0 | 0 | NA | NA | NA | NA | NA | NA | NA | NA | NA | NA |
| Vmn1r14 | 0 | 1 | 1 | 3 | 4 | 2 | 0 | 1 | 1 | 2 | 4 | 2 | 1.81 | 1 | 3 | 4.342 | 2.118 | 0.17652896 | NA | 0 | 1.828 | 0.848 | 0.848 | TRUE | 0.2213 |
| Vmn1r142 | 0 | 0 | 0 | 0 | 0 | 0 | 0 | 0 | 0 | 0 | 0 | 0 | 0 | 0 | 0 | NA | NA | NA | NA | NA | NA | NA | NA | NA | NA |
| Vmn1r143 | 0 | 0 | 0 | 0 | 0 | 0 | 0 | 0 | 0 | 0 | 0 | 0 | 0 | 0 | 0 | NA | NA | NA | NA | NA | NA | NA | NA | NA | NA |

| | | | | | | | | | | | | | | | | | | | | | | | | | |
|---|---|---|---|---|---|---|---|---|---|---|---|---|---|---|---|---|---|---|---|---|---|---|---|---|---|
| Vmn1r148 | 0 | 0 | 0 | 0 | 0 | 0 | 0 | 0 | 0 | 0 | 0 | 0 | 0 | 0 | 0 | NA | NA | NA | NA | NA | NA | NA | NA | NA | NA |
| Vmn1r149 | 0 | 0 | 0 | 0 | 0 | 0 | 0 | 0 | 0 | 0 | 0 | 0 | 0 | 0 | 0 | NA | NA | NA | NA | NA | NA | NA | NA | NA | NA |
| Vmn1r15 | 1 | 1 | 0 | 0 | 1 | 0 | 1 | 1 | 0 | 0 | 1 | 0 | 0.52 | 1 | 0 | 0.51 | -0.972 | 0.72797553 | NA | 0 | 6.3084 | 3.8879 | 3.8879 | TRUE | 0.5204 |
| Vmn1r151 | 0 | 0 | 0 | 0 | 0 | 0 | 0 | 0 | 0 | 0 | 0 | 0 | 0 | 0 | 0 | NA | NA | NA | NA | NA | NA | NA | NA | NA | NA |
| Vmn1r152 | 0 | 0 | 0 | 0 | 0 | 0 | 0 | 0 | 0 | 0 | 0 | 0 | 0 | 0 | 0 | NA | NA | NA | NA | NA | NA | NA | NA | NA | NA |
| Vmn1r155 | 0 | 0 | 0 | 0 | 0 | 0 | 0 | 0 | 0 | 0 | 0 | 0 | 0 | 0 | 0 | NA | NA | NA | NA | NA | NA | NA | NA | NA | NA |
| Vmn1r157 | 0 | 0 | 0 | 0 | 0 | 0 | 0 | 0 | 0 | 0 | 0 | 0 | 0 | 0 | 0 | NA | NA | NA | NA | NA | NA | NA | NA | NA | NA |
| Vmn1r158 | 0 | 0 | 0 | 0 | 0 | 0 | 0 | 0 | 0 | 0 | 0 | 0 | 0 | 0 | 0 | NA | NA | NA | NA | NA | NA | NA | NA | NA | NA |
| Vmn1r159 | 0 | 0 | 0 | 0 | 0 | 0 | 0 | 0 | 0 | 0 | 0 | 0 | 0 | 0 | 0 | NA | NA | NA | NA | NA | NA | NA | NA | NA | NA |
| Vmn1r16 | 0 | 0 | 0 | 0 | 0 | 0 | 0 | 0 | 0 | 0 | 0 | 0 | 0 | 0 | 0 | NA | NA | NA | NA | NA | NA | NA | NA | NA | NA |
| Vmn1r160 | 0 | 0 | 0 | 0 | 0 | 0 | 0 | 0 | 0 | 0 | 0 | 0 | 0 | 0 | 0 | NA | NA | NA | NA | NA | NA | NA | NA | NA | NA |
| Vmn1r163 | 0 | 0 | 0 | 0 | 0 | 0 | 0 | 0 | 0 | 0 | 0 | 0 | 0 | 0 | 0 | NA | NA | NA | NA | NA | NA | NA | NA | NA | NA |
| Vmn1r165 | 0 | 0 | 0 | 0 | 0 | 0 | 0 | 0 | 0 | 0 | 0 | 0 | 0 | 0 | 0 | NA | NA | NA | NA | NA | NA | NA | NA | NA | NA |
| Vmn1r167 | 0 | 0 | 0 | 0 | 0 | 0 | 0 | 0 | 0 | 0 | 0 | 0 | 0 | 0 | 0 | NA | NA | NA | NA | NA | NA | NA | NA | NA | NA |
| Vmn1r168 | 0 | 0 | 0 | 0 | 0 | 0 | 0 | 0 | 0 | 0 | 0 | 0 | 0 | 0 | 0 | NA | NA | NA | NA | NA | NA | NA | NA | NA | NA |
| Vmn1r169 | 0 | 0 | 0 | 0 | 0 | 0 | 0 | 0 | 0 | 0 | 0 | 0 | 0 | 0 | 0 | NA | NA | NA | NA | NA | NA | NA | NA | NA | NA |
| Vmn1r17 | 0 | 0 | 0 | 0 | 0 | 0 | 0 | 0 | 0 | 0 | 0 | 0 | 0 | 0 | 0 | NA | NA | NA | NA | NA | NA | NA | NA | NA | NA |
| Vmn1r170 | 0 | 0 | 0 | 0 | 0 | 0 | 0 | 0 | 0 | 0 | 0 | 0 | 0 | 0 | 0 | NA | NA | NA | NA | NA | NA | NA | NA | NA | NA |
| Vmn1r171 | 0 | 0 | 0 | 0 | 0 | 0 | 0 | 0 | 0 | 0 | 0 | 0 | 0 | 0 | 0 | NA | NA | NA | NA | NA | NA | NA | NA | NA | NA |
| Vmn1r172 | 0 | 0 | 0 | 0 | 0 | 0 | 0 | 0 | 0 | 0 | 0 | 0 | 0 | 0 | 0 | NA | NA | NA | NA | NA | NA | NA | NA | NA | NA |
| Vmn1r173 | 0 | 0 | 0 | 0 | 0 | 0 | 0 | 0 | 0 | 0 | 0 | 0 | 0 | 0 | 0 | NA | NA | NA | NA | NA | NA | NA | NA | NA | NA |
| Vmn1r174 | 0 | 0 | 0 | 0 | 0 | 0 | 0 | 0 | 0 | 0 | 0 | 0 | 0 | 0 | 0 | NA | NA | NA | NA | NA | NA | NA | NA | NA | NA |
| Vmn1r175 | 0 | 0 | 0 | 0 | 0 | 0 | 0 | 0 | 0 | 0 | 0 | 0 | 0 | 0 | 0 | NA | NA | NA | NA | NA | NA | NA | NA | NA | NA |
| Vmn1r176 | 0 | 0 | 0 | 0 | 0 | 0 | 0 | 0 | 0 | 0 | 0 | 0 | 0 | 0 | 0 | NA | NA | NA | NA | NA | NA | NA | NA | NA | NA |
| Vmn1r177 | 0 | 0 | 0 | 0 | 0 | 0 | 0 | 0 | 0 | 0 | 0 | 0 | 0 | 0 | 0 | NA | NA | NA | NA | NA | NA | NA | NA | NA | NA |
| Vmn1r178 | 3 | 1 | 1 | 1 | 0 | 0 | 3 | 1 | 1 | 1 | 0 | 0 | 1 | 2 | 0 | 0.203 | -2.303 | 0.26642713 | NA | 0 | 3.3149 | 1.7972 | 1.7972 | TRUE | 0.4044 |
| Vmn1r179 | 0 | 0 | 0 | 0 | 0 | 0 | 0 | 0 | 0 | 0 | 0 | 0 | 0 | 0 | 0 | NA | NA | NA | NA | NA | NA | NA | NA | NA | NA |
| Vmn1r18 | 0 | 0 | 0 | 0 | 0 | 0 | 0 | 0 | 0 | 0 | 0 | 0 | 0 | 0 | 0 | NA | NA | NA | NA | NA | NA | NA | NA | NA | NA |
| Vmn1r180 | 0 | 0 | 0 | 0 | 0 | 0 | 0 | 0 | 0 | 0 | 0 | 0 | 0 | 0 | 0 | NA | NA | NA | NA | NA | NA | NA | NA | NA | NA |
| Vmn1r181 | 0 | 0 | 0 | 0 | 0 | 0 | 0 | 0 | 0 | 0 | 0 | 0 | 0 | 0 | 0 | NA | NA | NA | NA | NA | NA | NA | NA | NA | NA |
| Vmn1r183 | 0 | 0 | 0 | 0 | 0 | 0 | 0 | 0 | 0 | 0 | 0 | 0 | 0 | 0 | 0 | NA | NA | NA | NA | NA | NA | NA | NA | NA | NA |
| Vmn1r184 | 0 | 0 | 0 | 0 | 0 | 0 | 0 | 0 | 0 | 0 | 0 | 0 | 0 | 0 | 0 | NA | NA | NA | NA | NA | NA | NA | NA | NA | NA |
| Vmn1r185 | 0 | 0 | 0 | 0 | 0 | 0 | 0 | 0 | 0 | 0 | 0 | 0 | 0 | 0 | 0 | NA | NA | NA | NA | NA | NA | NA | NA | NA | NA |
| Vmn1r186 | 0 | 0 | 0 | 0 | 0 | 0 | 0 | 0 | 0 | 0 | 0 | 0 | 0 | 0 | 0 | NA | NA | NA | NA | NA | NA | NA | NA | NA | NA |
| Vmn1r187 | 0 | 0 | 0 | 0 | 0 | 0 | 0 | 0 | 0 | 0 | 0 | 0 | 0 | 0 | 0 | NA | NA | NA | NA | NA | NA | NA | NA | NA | NA |
| Vmn1r188 | 0 | 0 | 0 | 0 | 0 | 0 | 0 | 0 | 0 | 0 | 0 | 0 | 0 | 0 | 0 | NA | NA | NA | NA | NA | NA | NA | NA | NA | NA |
| Vmn1r189 | 0 | 0 | 0 | 0 | 0 | 0 | 0 | 0 | 0 | 0 | 0 | 0 | 0 | 0 | 0 | NA | NA | NA | NA | NA | NA | NA | NA | NA | NA |
| Vmn1r19 | 0 | 0 | 0 | 0 | 0 | 0 | 0 | 0 | 0 | 0 | 0 | 0 | 0 | 0 | 0 | NA | NA | NA | NA | NA | NA | NA | NA | NA | NA |
| Vmn1r191 | 0 | 0 | 0 | 0 | 0 | 0 | 0 | 0 | 0 | 0 | 0 | 0 | 0 | 0 | 0 | NA | NA | NA | NA | NA | NA | NA | NA | NA | NA |
| Vmn1r192 | 0 | 0 | 0 | 0 | 0 | 0 | 0 | 0 | 0 | 0 | 0 | 0 | 0 | 0 | 0 | NA | NA | NA | NA | NA | NA | NA | NA | NA | NA |
| Vmn1r193 | 0 | 0 | 0 | 0 | 0 | 0 | 0 | 0 | 0 | 0 | 0 | 0 | 0 | 0 | 0 | NA | NA | NA | NA | NA | NA | NA | NA | NA | NA |
| Vmn1r194 | 0 | 0 | 0 | 0 | 0 | 0 | 0 | 0 | 0 | 0 | 0 | 0 | 0 | 0 | 0 | NA | NA | NA | NA | NA | NA | NA | NA | NA | NA |
| Vmn1r195 | 0 | 0 | 0 | 0 | 0 | 0 | 0 | 0 | 0 | 0 | 0 | 0 | 0 | 0 | 0 | NA | NA | NA | NA | NA | NA | NA | NA | NA | NA |
| Vmn1r196 | 0 | 0 | 0 | 0 | 0 | 0 | 0 | 0 | 0 | 0 | 0 | 0 | 0 | 0 | 0 | NA | NA | NA | NA | NA | NA | NA | NA | NA | NA |
| Vmn1r197 | 0 | 0 | 0 | 0 | 0 | 0 | 0 | 0 | 0 | 0 | 0 | 0 | 0 | 0 | 0 | NA | NA | NA | NA | NA | NA | NA | NA | NA | NA |
| Vmn1r198 | 0 | 0 | 0 | 0 | 0 | 0 | 0 | 0 | 0 | 0 | 0 | 0 | 0 | 0 | 0 | NA | NA | NA | NA | NA | NA | NA | NA | NA | NA |
| Vmn1r199 | 0 | 0 | 1 | 0 | 0 | 0 | 0 | 0 | 1 | 0 | 0 | 0 | 0.17 | 0 | 0 | 0.49 | -1.029 | 0.80087 | NA | 0 | 19.6928 | 10 | 10 | TRUE | 0.4098 |
| Vmn1r2 | 0 | 0 | 0 | 0 | 0 | 0 | 0 | 0 | 0 | 0 | 0 | 0 | 0 | 0 | 0 | NA | NA | NA | NA | NA | NA | NA | NA | NA | NA |
| Vmn1r20 | 0 | 0 | 0 | 0 | 0 | 0 | 0 | 0 | 0 | 0 | 0 | 0 | 0 | 0 | 0 | NA | NA | NA | NA | NA | NA | NA | NA | NA | NA |
| Vmn1r200 | 0 | 0 | 0 | 0 | 0 | 0 | 0 | 0 | 0 | 0 | 0 | 0 | 0 | 0 | 0 | NA | NA | NA | NA | NA | NA | NA | NA | NA | NA |
| Vmn1r201 | 0 | 0 | 0 | 0 | 0 | 0 | 0 | 0 | 0 | 0 | 0 | 0 | 0 | 0 | 0 | NA | NA | NA | NA | NA | NA | NA | NA | NA | NA |
| Vmn1r202 | 0 | 0 | 0 | 0 | 0 | 0 | 0 | 0 | 0 | 0 | 0 | 0 | 0 | 0 | 0 | NA | NA | NA | NA | NA | NA | NA | NA | NA | NA |
| Vmn1r203 | 0 | 0 | 0 | 0 | 0 | 0 | 0 | 0 | 0 | 0 | 0 | 0 | 0 | 0 | 0 | NA | NA | NA | NA | NA | NA | NA | NA | NA | NA |
| Vmn1r204 | 0 | 0 | 0 | 0 | 0 | 0 | 0 | 0 | 0 | 0 | 0 | 0 | 0 | 0 | 0 | NA | NA | NA | NA | NA | NA | NA | NA | NA | NA |
| Vmn1r205 | 0 | 0 | 0 | 0 | 0 | 0 | 0 | 0 | 0 | 0 | 0 | 0 | 0 | 0 | 0 | NA | NA | NA | NA | NA | NA | NA | NA | NA | NA |
| Vmn1r206 | 0 | 0 | 0 | 0 | 0 | 0 | 0 | 0 | 0 | 0 | 0 | 0 | 0 | 0 | 0 | NA | NA | NA | NA | NA | NA | NA | NA | NA | NA |
| Vmn1r207-p | 0 | 0 | 0 | 0 | 0 | 0 | 0 | 0 | 0 | 0 | 0 | 0 | 0 | 0 | 0 | NA | NA | NA | NA | NA | NA | NA | NA | NA | NA |
| Vmn1r208 | 0 | 0 | 0 | 0 | 0 | 0 | 0 | 0 | 0 | 0 | 0 | 0 | 0 | 0 | 0 | NA | NA | NA | NA | NA | NA | NA | NA | NA | NA |
| Vmn1r209 | 0 | 0 | 0 | 0 | 0 | 0 | 0 | 0 | 0 | 0 | 0 | 0 | 0 | 0 | 0 | NA | NA | NA | NA | NA | NA | NA | NA | NA | NA |
| Vmn1r21 | 0 | 0 | 0 | 0 | 0 | 0 | 0 | 0 | 0 | 0 | 0 | 0 | 0 | 0 | 0 | NA | NA | NA | NA | NA | NA | NA | NA | NA | NA |
| Vmn1r210 | 1 | 0 | 0 | 0 | 1 | 0 | 1 | 0 | 0 | 0 | 1 | 0 | 0.36 | 0 | 0 | 0.954 | -0.067 | 0.98557668 | NA | 0 | 9.2518 | 8.0179 | 8.0179 | TRUE | 0.5204 |
| Vmn1r211 | 0 | 0 | 0 | 0 | 0 | 0 | 0 | 0 | 0 | 0 | 0 | 0 | 0 | 0 | 0 | NA | NA | NA | NA | NA | NA | NA | NA | NA | NA |
| Vmn1r212 | 0 | 0 | 0 | 0 | 0 | 0 | 0 | 0 | 0 | 0 | 0 | 0 | 0 | 0 | 0 | NA | NA | NA | NA | NA | NA | NA | NA | NA | NA |
| Vmn1r213 | 0 | 0 | 0 | 0 | 0 | 0 | 0 | 0 | 0 | 0 | 0 | 0 | 0 | 0 | 0 | NA | NA | NA | NA | NA | NA | NA | NA | NA | NA |
| Vmn1r214 | 0 | 0 | 0 | 0 | 0 | 0 | 0 | 0 | 0 | 0 | 0 | 0 | 0 | 0 | 0 | NA | NA | NA | NA | NA | NA | NA | NA | NA | NA |
| Vmn1r215 | 0 | 0 | 0 | 0 | 0 | 0 | 0 | 0 | 0 | 0 | 0 | 0 | 0 | 0 | 0 | NA | NA | NA | NA | NA | NA | NA | NA | NA | NA |
| Vmn1r216 | 0 | 0 | 0 | 0 | 0 | 0 | 0 | 0 | 0 | 0 | 0 | 0 | 0 | 0 | 0 | NA | NA | NA | NA | NA | NA | NA | NA | NA | NA |
| Vmn1r217 | 0 | 0 | 0 | 0 | 0 | 0 | 0 | 0 | 0 | 0 | 0 | 0 | 0 | 0 | 0 | NA | NA | NA | NA | NA | NA | NA | NA | NA | NA |
| Vmn1r218 | 0 | 0 | 0 | 0 | 0 | 0 | 0 | 0 | 0 | 0 | 0 | 0 | 0 | 0 | 0 | NA | NA | NA | NA | NA | NA | NA | NA | NA | NA |

| | | | | | | | | | | | | | | | | | | | | | | | | | | |
|---|---|---|---|---|---|---|---|---|---|---|---|---|---|---|---|---|---|---|---|---|---|---|---|---|---|---|
| Vmn1r219 | 0 | 0 | 0 | 0 | 0 | 0 | 0 | 0 | 0 | 0 | 0 | 0 | 0 | 0 | 0 | NA | NA | NA | NA | NA | NA | NA | NA | NA | NA | |
| Vmn1r22 | 0 | 0 | 0 | 0 | 0 | 0 | 0 | 0 | 0 | 0 | 0 | 0 | 0 | 0 | 0 | NA | NA | NA | NA | NA | NA | NA | NA | NA | NA | |
| Vmn1r220 | 0 | 0 | 0 | 0 | 0 | 0 | 0 | 0 | 0 | 0 | 0 | 0 | 0 | 0 | 0 | NA | NA | NA | NA | NA | NA | NA | NA | NA | NA | |
| Vmn1r221 | 0 | 0 | 0 | 0 | 0 | 0 | 0 | 0 | 0 | 0 | 0 | 0 | 0 | 0 | 0 | NA | NA | NA | NA | NA | NA | NA | NA | NA | NA | |
| Vmn1r222 | 0 | 0 | 0 | 0 | 0 | 0 | 0 | 0 | 0 | 0 | 0 | 0 | 0 | 0 | 0 | NA | NA | NA | NA | NA | NA | NA | NA | NA | NA | |
| Vmn1r223 | 0 | 0 | 0 | 0 | 0 | 0 | 0 | 0 | 0 | 0 | 0 | 0 | 0 | 0 | 0 | NA | NA | NA | NA | NA | NA | NA | NA | NA | NA | |
| Vmn1r224 | 0 | 0 | 0 | 0 | 0 | 0 | 0 | 0 | 0 | 0 | 0 | 0 | 0 | 0 | 0 | NA | NA | NA | NA | NA | NA | NA | NA | NA | NA | |
| Vmn1r225 | 0 | 0 | 0 | 0 | 0 | 0 | 0 | 0 | 0 | 0 | 0 | 0 | 0 | 0 | 0 | NA | NA | NA | NA | NA | NA | NA | NA | NA | NA | |
| Vmn1r226 | 0 | 0 | 0 | 0 | 0 | 0 | 0 | 0 | 0 | 0 | 0 | 0 | 0 | 0 | 0 | NA | NA | NA | NA | NA | NA | NA | NA | NA | NA | |
| Vmn1r227 | 2 | 1 | 1 | 1 | 1 | 0 | 2 | 1 | 1 | 1 | 1 | 0 | 1 | 1 | 1 | 0.461 | -1.117 | 0.55021106 | NA | | 0 | 3.3073 | 1.3774 | 1.3774 | TRUE | 0.1869 |
| Vmn1r228 | 0 | 1 | 0 | 0 | 0 | 0 | 0 | 1 | 0 | 0 | 0 | 0 | 0.17 | 0 | 0 | 0.49 | -1.029 | 0.80087 | NA | | 0 | 19.7991 | 10 | 10 | TRUE | 0.405 |
| Vmn1r229 | 0 | 0 | 0 | 0 | 0 | 0 | 0 | 0 | 0 | 0 | 0 | 0 | 0 | 0 | 0 | NA | NA | NA | NA | NA | NA | NA | NA | NA | NA | |
| Vmn1r23 | 0 | 0 | 0 | 0 | 0 | 0 | 0 | 0 | 0 | 0 | 0 | 0 | 0 | 0 | 0 | NA | NA | NA | NA | NA | NA | NA | NA | NA | NA | |
| Vmn1r230 | 0 | 0 | 0 | 0 | 0 | 0 | 0 | 0 | 0 | 0 | 0 | 0 | 0 | 0 | 0 | NA | NA | NA | NA | NA | NA | NA | NA | NA | NA | |
| Vmn1r231 | 0 | 0 | 0 | 0 | 0 | 0 | 0 | 0 | 0 | 0 | 0 | 0 | 0 | 0 | 0 | NA | NA | NA | NA | NA | NA | NA | NA | NA | NA | |
| Vmn1r232 | 0 | 0 | 0 | 0 | 0 | 0 | 0 | 0 | 0 | 0 | 0 | 0 | 0 | 0 | 0 | NA | NA | NA | NA | NA | NA | NA | NA | NA | NA | |
| Vmn1r233 | 0 | 0 | 0 | 0 | 0 | 0 | 0 | 0 | 0 | 0 | 0 | 0 | 0 | 0 | 0 | NA | NA | NA | NA | NA | NA | NA | NA | NA | NA | |
| Vmn1r234 | 0 | 0 | 0 | 0 | 0 | 0 | 0 | 0 | 0 | 0 | 0 | 0 | 0 | 0 | 0 | NA | NA | NA | NA | NA | NA | NA | NA | NA | NA | |
| Vmn1r235 | 0 | 0 | 0 | 0 | 0 | 0 | 0 | 0 | 0 | 0 | 0 | 0 | 0 | 0 | 0 | NA | NA | NA | NA | NA | NA | NA | NA | NA | NA | |
| Vmn1r236 | 0 | 0 | 0 | 0 | 0 | 0 | 0 | 0 | 0 | 0 | 0 | 0 | 0 | 0 | 0 | NA | NA | NA | NA | NA | NA | NA | NA | NA | NA | |
| Vmn1r237 | 0 | 0 | 0 | 0 | 0 | 0 | 0 | 0 | 0 | 0 | 0 | 0 | 0 | 0 | 0 | NA | NA | NA | NA | NA | NA | NA | NA | NA | NA | |
| Vmn1r238 | 0 | 0 | 0 | 0 | 0 | 0 | 0 | 0 | 0 | 0 | 0 | 0 | 0 | 0 | 0 | NA | NA | NA | NA | NA | NA | NA | NA | NA | NA | |
| Vmn1r24 | 0 | 0 | 0 | 0 | 0 | 0 | 0 | 0 | 0 | 0 | 0 | 0 | 0 | 0 | 0 | NA | NA | NA | NA | NA | NA | NA | NA | NA | NA | |
| Vmn1r25 | 0 | 0 | 0 | 0 | 0 | 0 | 0 | 0 | 0 | 0 | 0 | 0 | 0 | 0 | 0 | NA | NA | NA | NA | NA | NA | NA | NA | NA | NA | |
| Vmn1r26 | 0 | 0 | 0 | 0 | 0 | 0 | 0 | 0 | 0 | 0 | 0 | 0 | 0 | 0 | 0 | NA | NA | NA | NA | NA | NA | NA | NA | NA | NA | |
| Vmn1r27 | 0 | 0 | 0 | 0 | 0 | 0 | 0 | 0 | 0 | 0 | 0 | 0 | 0 | 0 | 0 | NA | NA | NA | NA | NA | NA | NA | NA | NA | NA | |
| Vmn1r28 | 0 | 0 | 0 | 0 | 0 | 0 | 0 | 0 | 0 | 0 | 0 | 0 | 0 | 0 | 0 | NA | NA | NA | NA | NA | NA | NA | NA | NA | NA | |
| Vmn1r29 | 0 | 0 | 2 | 1 | 2 | 0 | 0 | 0 | 2 | 1 | 2 | 0 | 0.82 | 1 | 1 | 1.45 | 0.536 | 0.82805985 | NA | | 0 | 4.0032 | 3.1289 | 3.1289 | TRUE | 0.6991 |
| Vmn1r3 | 0 | 0 | 0 | 0 | 0 | 0 | 0 | 0 | 0 | 0 | 0 | 0 | 0 | 0 | 0 | NA | NA | NA | NA | NA | NA | NA | NA | NA | NA | |
| Vmn1r30 | 0 | 0 | 0 | 0 | 0 | 0 | 0 | 0 | 0 | 0 | 0 | 0 | 0 | 0 | 0 | NA | NA | NA | NA | NA | NA | NA | NA | NA | NA | |
| Vmn1r31 | 0 | 0 | 0 | 0 | 0 | 0 | 0 | 0 | 0 | 0 | 0 | 0 | 0 | 0 | 0 | NA | NA | NA | NA | NA | NA | NA | NA | NA | NA | |
| Vmn1r32 | 2 | 0 | 1 | 4 | 1 | 0 | 2 | 0 | 1 | 3 | 1 | 0 | 1.22 | 1 | 1 | 1.407 | 0.492 | 0.79851906 | NA | | 0 | 2.6967 | 1.8507 | 1.8507 | TRUE | 0.4341 |
| Vmn1r33 | 0 | 0 | 0 | 0 | 0 | 0 | 0 | 0 | 0 | 0 | 0 | 0 | 0 | 0 | 0 | NA | NA | NA | NA | NA | NA | NA | NA | NA | NA | |
| Vmn1r34 | 0 | 0 | 0 | 0 | 0 | 0 | 0 | 0 | 0 | 0 | 0 | 0 | 0 | 0 | 0 | NA | NA | NA | NA | NA | NA | NA | NA | NA | NA | |
| Vmn1r35 | 0 | 0 | 0 | 0 | 0 | 0 | 0 | 0 | 0 | 0 | 0 | 0 | 0 | 0 | 0 | NA | NA | NA | NA | NA | NA | NA | NA | NA | NA | |
| Vmn1r36 | 0 | 0 | 0 | 0 | 0 | 0 | 0 | 0 | 0 | 0 | 0 | 0 | 0 | 0 | 0 | NA | NA | NA | NA | NA | NA | NA | NA | NA | NA | |
| Vmn1r37 | 0 | 0 | 0 | 0 | 0 | 0 | 0 | 0 | 0 | 0 | 0 | 0 | 0 | 0 | 0 | NA | NA | NA | NA | NA | NA | NA | NA | NA | NA | |
| Vmn1r38 | 0 | 0 | 0 | 0 | 0 | 0 | 0 | 0 | 0 | 0 | 0 | 0 | 0 | 0 | 0 | NA | NA | NA | NA | NA | NA | NA | NA | NA | NA | |
| Vmn1r39 | 0 | 0 | 0 | 0 | 0 | 0 | 0 | 0 | 0 | 0 | 0 | 0 | 0 | 0 | 0 | NA | NA | NA | NA | NA | NA | NA | NA | NA | NA | |
| Vmn1r4 | 0 | 0 | 0 | 0 | 0 | 0 | 0 | 0 | 0 | 0 | 0 | 0 | 0 | 0 | 0 | NA | NA | NA | NA | NA | NA | NA | NA | NA | NA | |
| Vmn1r40 | 0 | 0 | 0 | 0 | 0 | 0 | 0 | 0 | 0 | 0 | 0 | 0 | 0 | 0 | 0 | NA | NA | NA | NA | NA | NA | NA | NA | NA | NA | |
| Vmn1r41 | 0 | 0 | 0 | 0 | 0 | 0 | 0 | 0 | 0 | 0 | 0 | 0 | 0 | 0 | 0 | NA | NA | NA | NA | NA | NA | NA | NA | NA | NA | |
| Vmn1r42 | 0 | 0 | 0 | 0 | 0 | 0 | 0 | 0 | 0 | 0 | 0 | 0 | 0 | 0 | 0 | NA | NA | NA | NA | NA | NA | NA | NA | NA | NA | |
| Vmn1r43 | 0 | 0 | 0 | 0 | 1 | 0 | 0 | 0 | 0 | 0 | 1 | 0 | 0.18 | 0 | 0 | 1.859 | 0.894 | 0.82650734 | NA | | 0 | 18.3785 | 10 | 10 | TRUE | 0.5204 |
| Vmn1r44 | 0 | 0 | 0 | 0 | 0 | 0 | 0 | 0 | 0 | 0 | 0 | 0 | 0 | 0 | 0 | NA | NA | NA | NA | NA | NA | NA | NA | NA | NA | |
| Vmn1r45 | 0 | 0 | 0 | 0 | 0 | 0 | 0 | 0 | 0 | 0 | 0 | 0 | 0 | 0 | 0 | NA | NA | NA | NA | NA | NA | NA | NA | NA | NA | |
| Vmn1r46 | 0 | 0 | 0 | 0 | 0 | 0 | 0 | 0 | 0 | 0 | 0 | 0 | 0 | 0 | 0 | NA | NA | NA | NA | NA | NA | NA | NA | NA | NA | |
| Vmn1r47 | 0 | 0 | 0 | 0 | 0 | 0 | 0 | 0 | 0 | 0 | 0 | 0 | 0 | 0 | 0 | NA | NA | NA | NA | NA | NA | NA | NA | NA | NA | |
| Vmn1r48 | 0 | 0 | 0 | 0 | 0 | 0 | 0 | 0 | 0 | 0 | 0 | 0 | 0 | 0 | 0 | NA | NA | NA | NA | NA | NA | NA | NA | NA | NA | |
| Vmn1r49 | 0 | 0 | 0 | 0 | 0 | 0 | 0 | 0 | 0 | 0 | 0 | 0 | 0 | 0 | 0 | NA | NA | NA | NA | NA | NA | NA | NA | NA | NA | |
| Vmn1r5 | 0 | 0 | 0 | 0 | 0 | 0 | 0 | 0 | 0 | 0 | 0 | 0 | 0 | 0 | 0 | NA | NA | NA | NA | NA | NA | NA | NA | NA | NA | |
| Vmn1r50 | 0 | 0 | 0 | 0 | 0 | 0 | 0 | 0 | 0 | 0 | 0 | 0 | 0 | 0 | 0 | NA | NA | NA | NA | NA | NA | NA | NA | NA | NA | |
| Vmn1r51 | 0 | 0 | 0 | 0 | 0 | 0 | 0 | 0 | 0 | 0 | 0 | 0 | 0 | 0 | 0 | NA | NA | NA | NA | NA | NA | NA | NA | NA | NA | |
| Vmn1r52 | 0 | 0 | 0 | 0 | 0 | 0 | 0 | 0 | 0 | 0 | 0 | 0 | 0 | 0 | 0 | NA | NA | NA | NA | NA | NA | NA | NA | NA | NA | |
| Vmn1r53 | 27 | 30 | 28 | 83 | 46 | 53 | 29 | 30 | 28 | 65 | 49 | 58 | 43.25 | 29 | 57 | 1.999 | 0.999 | 0.00095186 | 0.00856405 | | 0 | 0.0831 | 0.0398 | 0.0398 | TRUE | 0.1308 |
| Vmn1r54 | 0 | 0 | 0 | 0 | 0 | 0 | 0 | 0 | 0 | 0 | 0 | 0 | 0 | 0 | 0 | NA | NA | NA | NA | NA | NA | NA | NA | NA | NA | |
| Vmn1r55 | 0 | 0 | 0 | 0 | 0 | 0 | 0 | 0 | 0 | 0 | 0 | 0 | 0 | 0 | 0 | NA | NA | NA | NA | NA | NA | NA | NA | NA | NA | |
| Vmn1r56 | 0 | 0 | 0 | 0 | 0 | 0 | 0 | 0 | 0 | 0 | 0 | 0 | 0 | 0 | 0 | NA | NA | NA | NA | NA | NA | NA | NA | NA | NA | |
| Vmn1r57 | 0 | 0 | 0 | 0 | 0 | 0 | 0 | 0 | 0 | 0 | 0 | 0 | 0 | 0 | 0 | NA | NA | NA | NA | NA | NA | NA | NA | NA | NA | |
| Vmn1r58 | 0 | 0 | 0 | 0 | 0 | 0 | 0 | 0 | 0 | 0 | 0 | 0 | 0 | 0 | 0 | NA | NA | NA | NA | NA | NA | NA | NA | NA | NA | |
| Vmn1r59 | 0 | 0 | 0 | 0 | 0 | 0 | 0 | 0 | 0 | 0 | 0 | 0 | 0 | 0 | 0 | NA | NA | NA | NA | NA | NA | NA | NA | NA | NA | |
| Vmn1r6 | 0 | 0 | 0 | 0 | 0 | 0 | 0 | 0 | 0 | 0 | 0 | 0 | 0 | 0 | 0 | NA | NA | NA | NA | NA | NA | NA | NA | NA | NA | |
| Vmn1r60 | 0 | 0 | 0 | 0 | 0 | 0 | 0 | 0 | 0 | 0 | 0 | 0 | 0 | 0 | 0 | NA | NA | NA | NA | NA | NA | NA | NA | NA | NA | |
| Vmn1r61 | 0 | 0 | 0 | 0 | 0 | 0 | 0 | 0 | 0 | 0 | 0 | 0 | 0 | 0 | 0 | NA | NA | NA | NA | NA | NA | NA | NA | NA | NA | |
| Vmn1r62 | 0 | 0 | 0 | 0 | 0 | 0 | 0 | 0 | 0 | 0 | 0 | 0 | 0 | 0 | 0 | NA | NA | NA | NA | NA | NA | NA | NA | NA | NA | |
| Vmn1r63 | 0 | 0 | 0 | 0 | 0 | 0 | 0 | 0 | 0 | 0 | 0 | 0 | 0 | 0 | 0 | NA | NA | NA | NA | NA | NA | NA | NA | NA | NA | |
| Vmn1r64 | 0 | 0 | 0 | 0 | 0 | 0 | 0 | 0 | 0 | 0 | 0 | 0 | 0 | 0 | 0 | NA | NA | NA | NA | NA | NA | NA | NA | NA | NA | |
| Vmn1r65 | 0 | 0 | 0 | 0 | 0 | 0 | 0 | 0 | 0 | 0 | 0 | 0 | 0 | 0 | 0 | NA | NA | NA | NA | NA | NA | NA | NA | NA | NA | |
| Vmn1r66 | 0 | 0 | 0 | 0 | 0 | 0 | 0 | 0 | 0 | 0 | 0 | 0 | 0 | 0 | 0 | NA | NA | NA | NA | NA | NA | NA | NA | NA | NA | |

| | | | | | | | | | | | | | | | | | | | | | | | | | |
|---|---|---|---|---|---|---|---|---|---|---|---|---|---|---|---|---|---|---|---|---|---|---|---|---|---|
| Vmn1r67 | 0 | 0 | 0 | 0 | 0 | 0 | 0 | 0 | 0 | 0 | 0 | 0 | 0 | 0 | 0 | NA | NA | NA | NA | NA | NA | NA | NA | NA | NA |
| Vmn1r68 | 0 | 0 | 0 | 0 | 0 | 0 | 0 | 0 | 0 | 0 | 0 | 0 | 0 | 0 | 0 | NA | NA | NA | NA | NA | NA | NA | NA | NA | NA |
| Vmn1r69 | 0 | 0 | 0 | 0 | 0 | 0 | 0 | 0 | 0 | 0 | 0 | 0 | 0 | 0 | 0 | NA | NA | NA | NA | NA | NA | NA | NA | NA | NA |
| Vmn1r7 | 0 | 0 | 0 | 0 | 0 | 0 | 0 | 0 | 0 | 0 | 0 | 0 | 0 | 0 | 0 | NA | NA | NA | NA | NA | NA | NA | NA | NA | NA |
| Vmn1r70 | 0 | 0 | 0 | 0 | 0 | 0 | 0 | 0 | 0 | 0 | 0 | 0 | 0 | 0 | 0 | NA | NA | NA | NA | NA | NA | NA | NA | NA | NA |
| Vmn1r71 | 0 | 0 | 0 | 0 | 0 | 0 | 0 | 0 | 0 | 0 | 0 | 0 | 0 | 0 | 0 | NA | NA | NA | NA | NA | NA | NA | NA | NA | NA |
| Vmn1r72 | 0 | 0 | 0 | 0 | 0 | 0 | 0 | 0 | 0 | 0 | 0 | 0 | 0 | 0 | 0 | NA | NA | NA | NA | NA | NA | NA | NA | NA | NA |
| Vmn1r73 | 0 | 0 | 0 | 0 | 0 | 0 | 0 | 0 | 0 | 0 | 0 | 0 | 0 | 0 | 0 | NA | NA | NA | NA | NA | NA | NA | NA | NA | NA |
| Vmn1r74 | 0 | 0 | 0 | 0 | 0 | 0 | 0 | 0 | 0 | 0 | 0 | 0 | 0 | 0 | 0 | NA | NA | NA | NA | NA | NA | NA | NA | NA | NA |
| Vmn1r75 | 0 | 0 | 0 | 0 | 0 | 0 | 0 | 0 | 0 | 0 | 0 | 0 | 0 | 0 | 0 | NA | NA | NA | NA | NA | NA | NA | NA | NA | NA |
| Vmn1r76 | 0 | 0 | 0 | 0 | 0 | 0 | 0 | 0 | 0 | 0 | 0 | 0 | 0 | 0 | 0 | NA | NA | NA | NA | NA | NA | NA | NA | NA | NA |
| Vmn1r77 | 0 | 0 | 0 | 0 | 0 | 0 | 0 | 0 | 0 | 0 | 0 | 0 | 0 | 0 | 0 | NA | NA | NA | NA | NA | NA | NA | NA | NA | NA |
| Vmn1r78 | 0 | 0 | 0 | 0 | 0 | 0 | 0 | 0 | 0 | 0 | 0 | 0 | 0 | 0 | 0 | NA | NA | NA | NA | NA | NA | NA | NA | NA | NA |
| Vmn1r79 | 0 | 0 | 0 | 0 | 0 | 0 | 0 | 0 | 0 | 0 | 0 | 0 | 0 | 0 | 0 | NA | NA | NA | NA | NA | NA | NA | NA | NA | NA |
| Vmn1r8 | 0 | 0 | 0 | 0 | 0 | 0 | 0 | 0 | 0 | 0 | 0 | 0 | 0 | 0 | 0 | NA | NA | NA | NA | NA | NA | NA | NA | NA | NA |
| Vmn1r80 | 0 | 0 | 0 | 0 | 0 | 0 | 0 | 0 | 0 | 0 | 0 | 0 | 0 | 0 | 0 | NA | NA | NA | NA | NA | NA | NA | NA | NA | NA |
| Vmn1r81 | 0 | 0 | 0 | 0 | 0 | 0 | 0 | 0 | 0 | 0 | 0 | 0 | 0 | 0 | 0 | NA | NA | NA | NA | NA | NA | NA | NA | NA | NA |
| Vmn1r82 | 0 | 1 | 0 | 0 | 0 | 0 | 0 | 1 | 0 | 0 | 0 | 0 | 0.17 | 0 | 0 | 0.49 | -1.029 | 0.80087 | NA | 0 | 19.7991 | 10 | 10 | TRUE | 0.405 |
| Vmn1r83 | 0 | 0 | 0 | 0 | 0 | 0 | 0 | 0 | 0 | 0 | 0 | 0 | 0 | 0 | 0 | NA | NA | NA | NA | NA | NA | NA | NA | NA | NA |
| Vmn1r84 | 0 | 0 | 0 | 0 | 0 | 0 | 0 | 0 | 0 | 0 | 0 | 0 | 0 | 0 | 0 | NA | NA | NA | NA | NA | NA | NA | NA | NA | NA |
| Vmn1r85 | 0 | 0 | 0 | 0 | 0 | 0 | 0 | 0 | 0 | 0 | 0 | 0 | 0 | 0 | 0 | NA | NA | NA | NA | NA | NA | NA | NA | NA | NA |
| Vmn1r86 | 0 | 0 | 0 | 0 | 0 | 0 | 0 | 0 | 0 | 0 | 0 | 0 | 0 | 0 | 0 | NA | NA | NA | NA | NA | NA | NA | NA | NA | NA |
| Vmn1r87 | 0 | 0 | 0 | 0 | 0 | 0 | 0 | 0 | 0 | 0 | 0 | 0 | 0 | 0 | 0 | NA | NA | NA | NA | NA | NA | NA | NA | NA | NA |
| Vmn1r88 | 0 | 0 | 0 | 0 | 0 | 0 | 0 | 0 | 0 | 0 | 0 | 0 | 0 | 0 | 0 | NA | NA | NA | NA | NA | NA | NA | NA | NA | NA |
| Vmn1r89 | 0 | 0 | 0 | 0 | 0 | 0 | 0 | 0 | 0 | 0 | 0 | 0 | 0 | 0 | 0 | NA | NA | NA | NA | NA | NA | NA | NA | NA | NA |
| Vmn1r9 | 0 | 0 | 0 | 0 | 0 | 0 | 0 | 0 | 0 | 0 | 0 | 0 | 0 | 0 | 0 | NA | NA | NA | NA | NA | NA | NA | NA | NA | NA |
| Vmn1r90 | 0 | 0 | 0 | 0 | 0 | 0 | 0 | 0 | 0 | 0 | 0 | 0 | 0 | 0 | 0 | NA | NA | NA | NA | NA | NA | NA | NA | NA | NA |
| Vmn1r91 | 0 | 0 | 0 | 0 | 0 | 0 | 0 | 0 | 0 | 0 | 0 | 0 | 0 | 0 | 0 | NA | NA | NA | NA | NA | NA | NA | NA | NA | NA |
| Vmn1r93 | 0 | 0 | 0 | 0 | 0 | 0 | 0 | 0 | 0 | 0 | 0 | 0 | 0 | 0 | 0 | NA | NA | NA | NA | NA | NA | NA | NA | NA | NA |
| Vmn1r94 | 0 | 0 | 0 | 0 | 0 | 0 | 0 | 0 | 0 | 0 | 0 | 0 | 0 | 0 | 0 | NA | NA | NA | NA | NA | NA | NA | NA | NA | NA |
| Vmn1r95 | 0 | 0 | 0 | 0 | 0 | 0 | 0 | 0 | 0 | 0 | 0 | 0 | 0 | 0 | 0 | NA | NA | NA | NA | NA | NA | NA | NA | NA | NA |
| Vmn2r-ps11 | 0 | 0 | 0 | 0 | 0 | 0 | 0 | 0 | 0 | 0 | 0 | 0 | 0 | 0 | 0 | NA | NA | NA | NA | NA | NA | NA | NA | NA | NA |
| Vmn2r-ps12 | 0 | 0 | 0 | 0 | 0 | 0 | 0 | 0 | 0 | 0 | 0 | 0 | 0 | 0 | 0 | NA | NA | NA | NA | NA | NA | NA | NA | NA | NA |
| Vmn2r-ps15 | 0 | 0 | 0 | 0 | 0 | 0 | 0 | 0 | 0 | 0 | 0 | 0 | 0 | 0 | 0 | NA | NA | NA | NA | NA | NA | NA | NA | NA | NA |
| Vmn2r-ps54 | 1 | 3 | 3 | 1 | 1 | 2 | 1 | 3 | 3 | 1 | 1 | 2 | 1.85 | 2 | 1 | 0.558 | -0.843 | 0.53987561 | NA | 0 | 1.7842 | 0.776 | 0.776 | TRUE | 0.2264 |
| Vmn2r-ps60 | 0 | 0 | 0 | 0 | 0 | 0 | 0 | 0 | 0 | 0 | 0 | 0 | 0 | 0 | 0 | NA | NA | NA | NA | NA | NA | NA | NA | NA | NA |
| Vmn2r1 | 8 | 3 | 15 | 5 | 14 | 6 | 8 | 3 | 15 | 4 | 15 | 7 | 8.69 | 9 | 9 | 0.951 | -0.073 | 0.92991702 | 0.96556973 | 0.3723 | 0.3862 | 0.3797 | 0.3797 | TRUE | 0.2694 |
| Vmn2r10 | 0 | 0 | 0 | 0 | 0 | 0 | 0 | 0 | 0 | 0 | 0 | 0 | 0 | 0 | 0 | NA | NA | NA | NA | NA | NA | NA | NA | NA | NA |
| Vmn2r100 | 0 | 0 | 0 | 0 | 0 | 0 | 0 | 0 | 0 | 0 | 0 | 0 | 0 | 0 | 0 | NA | NA | NA | NA | NA | NA | NA | NA | NA | NA |
| Vmn2r101 | 0 | 0 | 0 | 0 | 0 | 0 | 0 | 0 | 0 | 0 | 0 | 0 | 0 | 0 | 0 | NA | NA | NA | NA | NA | NA | NA | NA | NA | NA |
| Vmn2r102 | 0 | 0 | 0 | 0 | 0 | 0 | 0 | 0 | 0 | 0 | 0 | 0 | 0 | 0 | 0 | NA | NA | NA | NA | NA | NA | NA | NA | NA | NA |
| Vmn2r103 | 0 | 0 | 0 | 0 | 0 | 0 | 0 | 0 | 0 | 0 | 0 | 0 | 0 | 0 | 0 | NA | NA | NA | NA | NA | NA | NA | NA | NA | NA |
| Vmn2r104 | 0 | 0 | 0 | 0 | 0 | 0 | 0 | 0 | 0 | 0 | 0 | 0 | 0 | 0 | 0 | NA | NA | NA | NA | NA | NA | NA | NA | NA | NA |
| Vmn2r105 | 0 | 0 | 0 | 0 | 0 | 0 | 0 | 0 | 0 | 0 | 0 | 0 | 0 | 0 | 0 | NA | NA | NA | NA | NA | NA | NA | NA | NA | NA |
| Vmn2r106 | 0 | 0 | 0 | 0 | 0 | 0 | 0 | 0 | 0 | 0 | 0 | 0 | 0 | 0 | 0 | NA | NA | NA | NA | NA | NA | NA | NA | NA | NA |
| Vmn2r107 | 0 | 0 | 0 | 0 | 0 | 0 | 0 | 0 | 0 | 0 | 0 | 0 | 0 | 0 | 0 | NA | NA | NA | NA | NA | NA | NA | NA | NA | NA |
| Vmn2r108 | 0 | 0 | 0 | 0 | 0 | 0 | 0 | 0 | 0 | 0 | 0 | 0 | 0 | 0 | 0 | NA | NA | NA | NA | NA | NA | NA | NA | NA | NA |
| Vmn2r109 | 0 | 0 | 0 | 0 | 0 | 0 | 0 | 0 | 0 | 0 | 0 | 0 | 0 | 0 | 0 | NA | NA | NA | NA | NA | NA | NA | NA | NA | NA |
| Vmn2r11 | 0 | 0 | 0 | 0 | 0 | 0 | 0 | 0 | 0 | 0 | 0 | 0 | 0 | 0 | 0 | NA | NA | NA | NA | NA | NA | NA | NA | NA | NA |
| Vmn2r110 | 0 | 0 | 0 | 0 | 0 | 0 | 0 | 0 | 0 | 0 | 0 | 0 | 0 | 0 | 0 | NA | NA | NA | NA | NA | NA | NA | NA | NA | NA |
| Vmn2r111 | 0 | 0 | 0 | 1 | 2 | 0 | 0 | 0 | 0 | 1 | 2 | 0 | 0.49 | 0 | 1 | 5.204 | 2.38 | 0.47267317 | NA | 0 | 6.7348 | 6.4099 | 6.4099 | TRUE | 0.1222 |
| Vmn2r112 | 0 | 0 | 0 | 0 | 0 | 0 | 0 | 0 | 0 | 0 | 0 | 0 | 0 | 0 | 0 | NA | NA | NA | NA | NA | NA | NA | NA | NA | NA |
| Vmn2r113 | 2 | 0 | 1 | 0 | 1 | 0 | 2 | 0 | 1 | 0 | 1 | 0 | 0.7 | 1 | 0 | 0.337 | -1.571 | 0.54072415 | NA | 0 | 4.7083 | 3.2627 | 3.2627 | TRUE | 0.278 |
| Vmn2r114 | 0 | 0 | 0 | 0 | 0 | 0 | 0 | 0 | 0 | 0 | 0 | 0 | 0 | 0 | 0 | NA | NA | NA | NA | NA | NA | NA | NA | NA | NA |
| Vmn2r115 | 0 | 0 | 0 | 0 | 0 | 0 | 0 | 0 | 0 | 0 | 0 | 0 | 0 | 0 | 0 | NA | NA | NA | NA | NA | NA | NA | NA | NA | NA |
| Vmn2r116 | 0 | 0 | 0 | 0 | 0 | 0 | 0 | 0 | 0 | 0 | 0 | 0 | 0 | 0 | 0 | NA | NA | NA | NA | NA | NA | NA | NA | NA | NA |
| Vmn2r117 | 0 | 0 | 0 | 0 | 0 | 0 | 0 | 0 | 0 | 0 | 0 | 0 | 0 | 0 | 0 | NA | NA | NA | NA | NA | NA | NA | NA | NA | NA |
| Vmn2r118 | 0 | 0 | 0 | 0 | 0 | 0 | 0 | 0 | 0 | 0 | 0 | 0 | 0 | 0 | 0 | NA | NA | NA | NA | NA | NA | NA | NA | NA | NA |
| Vmn2r12 | 0 | 0 | 0 | 0 | 0 | 0 | 0 | 0 | 0 | 0 | 0 | 0 | 0 | 0 | 0 | NA | NA | NA | NA | NA | NA | NA | NA | NA | NA |
| Vmn2r120 | 0 | 0 | 0 | 0 | 0 | 0 | 0 | 0 | 0 | 0 | 0 | 0 | 0 | 0 | 0 | NA | NA | NA | NA | NA | NA | NA | NA | NA | NA |
| Vmn2r121 | 0 | 0 | 0 | 0 | 0 | 0 | 0 | 0 | 0 | 0 | 0 | 0 | 0 | 0 | 0 | NA | NA | NA | NA | NA | NA | NA | NA | NA | NA |
| Vmn2r122 | 0 | 0 | 0 | 0 | 0 | 0 | 0 | 0 | 0 | 0 | 0 | 0 | 0 | 0 | 0 | NA | NA | NA | NA | NA | NA | NA | NA | NA | NA |
| Vmn2r123 | 0 | 0 | 0 | 0 | 0 | 0 | 0 | 0 | 0 | 0 | 0 | 0 | 0 | 0 | 0 | NA | NA | NA | NA | NA | NA | NA | NA | NA | NA |
| Vmn2r124 | 0 | 0 | 0 | 0 | 0 | 0 | 0 | 0 | 0 | 0 | 0 | 0 | 0 | 0 | 0 | NA | NA | NA | NA | NA | NA | NA | NA | NA | NA |
| Vmn2r13 | 0 | 0 | 0 | 0 | 0 | 0 | 0 | 0 | 0 | 0 | 0 | 0 | 0 | 0 | 0 | NA | NA | NA | NA | NA | NA | NA | NA | NA | NA |
| Vmn2r14 | 0 | 0 | 0 | 0 | 0 | 0 | 0 | 0 | 0 | 0 | 0 | 0 | 0 | 0 | 0 | NA | NA | NA | NA | NA | NA | NA | NA | NA | NA |
| Vmn2r15 | 0 | 0 | 0 | 0 | 0 | 0 | 0 | 0 | 0 | 0 | 0 | 0 | 0 | 0 | 0 | NA | NA | NA | NA | NA | NA | NA | NA | NA | NA |
| Vmn2r16 | 0 | 0 | 0 | 0 | 0 | 0 | 0 | 0 | 0 | 0 | 0 | 0 | 0 | 0 | 0 | NA | NA | NA | NA | NA | NA | NA | NA | NA | NA |
| Vmn2r17 | 0 | 0 | 0 | 0 | 0 | 0 | 0 | 0 | 0 | 0 | 0 | 0 | 0 | 0 | 0 | NA | NA | NA | NA | NA | NA | NA | NA | NA | NA |

| | | | | | | | | | | | | | | | | | | | | | | | | | |
|---|---|---|---|---|---|---|---|---|---|---|---|---|---|---|---|---|---|---|---|---|---|---|---|---|---|
| Vmn2r18 | 2 | 0 | 0 | 4 | 2 | 0 | 2 | 0 | 0 | 3 | 2 | 0 | 1.24 | 1 | 2 | 2.57 | 1.362 | 0.53209554 | NA | 1.8421 | 2.6706 | 2.4182 | 2.4182 | TRUE | 0.7508 |
| Vmn2r19 | 0 | 0 | 0 | 0 | 0 | 0 | 0 | 0 | 0 | 0 | 0 | 0 | 0 | 0 | 0 | NA | NA | NA | NA | NA | NA | NA | NA | NA | NA |
| Vmn2r2 | 0 | 0 | 0 | 0 | 0 | 0 | 0 | 0 | 0 | 0 | 0 | 0 | 0 | 0 | 0 | NA | NA | NA | NA | NA | NA | NA | NA | NA | NA |
| Vmn2r20 | 0 | 0 | 0 | 0 | 0 | 0 | 0 | 0 | 0 | 0 | 0 | 0 | 0 | 0 | 0 | NA | NA | NA | NA | NA | NA | NA | NA | NA | NA |
| Vmn2r21 | 0 | 0 | 0 | 0 | 0 | 0 | 0 | 0 | 0 | 0 | 0 | 0 | 0 | 0 | 0 | NA | NA | NA | NA | NA | NA | NA | NA | NA | NA |
| Vmn2r22 | 0 | 0 | 0 | 0 | 0 | 0 | 0 | 0 | 0 | 0 | 0 | 0 | 0 | 0 | 0 | NA | NA | NA | NA | NA | NA | NA | NA | NA | NA |
| Vmn2r23 | 0 | 0 | 0 | 0 | 0 | 0 | 0 | 0 | 0 | 0 | 0 | 0 | 0 | 0 | 0 | NA | NA | NA | NA | NA | NA | NA | NA | NA | NA |
| Vmn2r24 | 0 | 0 | 0 | 0 | 0 | 0 | 0 | 0 | 0 | 0 | 0 | 0 | 0 | 0 | 0 | NA | NA | NA | NA | NA | NA | NA | NA | NA | NA |
| Vmn2r25 | 0 | 0 | 1 | 0 | 0 | 0 | 0 | 0 | 1 | 0 | 0 | 0 | 0.17 | 0 | 0 | 0.49 | -1.029 | 0.80087 | NA | 0 | 19.6928 | 10 | 10 | TRUE | 0.4098 |
| Vmn2r26 | 0 | 0 | 0 | 0 | 0 | 0 | 0 | 0 | 0 | 0 | 0 | 0 | 0 | 0 | 0 | NA | NA | NA | NA | NA | NA | NA | NA | NA | NA |
| Vmn2r27 | 4 | 4 | 4 | 4 | 2 | 2 | 4 | 4 | 4 | 3 | 2 | 2 | 3.29 | 4 | 2 | 0.619 | -0.692 | 0.48905724 | NA | 0 | 1.008 | 0.4029 | 0.4029 | TRUE | 0.0778 |
| Vmn2r28 | 0 | 0 | 0 | 0 | 1 | 0 | 0 | 0 | 0 | 0 | 1 | 0 | 0.18 | 0 | 0 | 1.859 | 0.894 | 0.82650734 | NA | 0 | 18.3785 | 10 | 10 | TRUE | 0.5204 |
| Vmn2r29 | 0 | 4 | 0 | 1 | 1 | 1 | 0 | 4 | 0 | 1 | 1 | 1 | 1.16 | 1 | 1 | 0.731 | -0.453 | 0.823294 | NA | 1.1213 | 2.8504 | 2.0817 | 2.0817 | TRUE | 1.8981 |
| Vmn2r3 | 2 | 1 | 3 | 0 | 2 | 1 | 2 | 1 | 3 | 0 | 2 | 1 | 1.56 | 2 | 1 | 0.504 | -0.988 | 0.53761549 | NA | 0 | 2.1138 | 1.1267 | 1.1267 | TRUE | 0.4308 |
| Vmn2r30 | 0 | 0 | 0 | 0 | 0 | 0 | 0 | 0 | 0 | 0 | 0 | 0 | 0 | 0 | 0 | NA | NA | NA | NA | NA | NA | NA | NA | NA | NA |
| Vmn2r31 | 0 | 0 | 0 | 0 | 0 | 0 | 0 | 0 | 0 | 0 | 0 | 0 | 0 | 0 | 0 | NA | NA | NA | NA | NA | NA | NA | NA | NA | NA |
| Vmn2r32 | 0 | 0 | 0 | 0 | 0 | 0 | 0 | 0 | 0 | 0 | 0 | 0 | 0 | 0 | 0 | NA | NA | NA | NA | NA | NA | NA | NA | NA | NA |
| Vmn2r33 | 0 | 0 | 0 | 0 | 0 | 0 | 0 | 0 | 0 | 0 | 0 | 0 | 0 | 0 | 0 | NA | NA | NA | NA | NA | NA | NA | NA | NA | NA |
| Vmn2r34 | 0 | 0 | 0 | 0 | 0 | 0 | 0 | 0 | 0 | 0 | 0 | 0 | 0 | 0 | 0 | NA | NA | NA | NA | NA | NA | NA | NA | NA | NA |
| Vmn2r35 | 0 | 0 | 0 | 0 | 0 | 0 | 0 | 0 | 0 | 0 | 0 | 0 | 0 | 0 | 0 | NA | NA | NA | NA | NA | NA | NA | NA | NA | NA |
| Vmn2r36 | 0 | 0 | 0 | 0 | 0 | 0 | 0 | 0 | 0 | 0 | 0 | 0 | 0 | 0 | 0 | NA | NA | NA | NA | NA | NA | NA | NA | NA | NA |
| Vmn2r37 | 0 | 0 | 0 | 0 | 0 | 0 | 0 | 0 | 0 | 0 | 0 | 0 | 0 | 0 | 0 | NA | NA | NA | NA | NA | NA | NA | NA | NA | NA |
| Vmn2r38 | 0 | 0 | 0 | 0 | 0 | 0 | 0 | 0 | 0 | 0 | 0 | 0 | 0 | 0 | 0 | NA | NA | NA | NA | NA | NA | NA | NA | NA | NA |
| Vmn2r39 | 0 | 0 | 0 | 0 | 0 | 0 | 0 | 0 | 0 | 0 | 0 | 0 | 0 | 0 | 0 | NA | NA | NA | NA | NA | NA | NA | NA | NA | NA |
| Vmn2r4 | 0 | 0 | 0 | 0 | 0 | 0 | 0 | 0 | 0 | 0 | 0 | 0 | 0 | 0 | 0 | NA | NA | NA | NA | NA | NA | NA | NA | NA | NA |
| Vmn2r40 | 0 | 0 | 0 | 0 | 0 | 0 | 0 | 0 | 0 | 0 | 0 | 0 | 0 | 0 | 0 | NA | NA | NA | NA | NA | NA | NA | NA | NA | NA |
| Vmn2r41 | 0 | 0 | 0 | 0 | 0 | 0 | 0 | 0 | 0 | 0 | 0 | 0 | 0 | 0 | 0 | NA | NA | NA | NA | NA | NA | NA | NA | NA | NA |
| Vmn2r42 | 0 | 0 | 0 | 0 | 0 | 0 | 0 | 0 | 0 | 0 | 0 | 0 | 0 | 0 | 0 | NA | NA | NA | NA | NA | NA | NA | NA | NA | NA |
| Vmn2r43 | 0 | 0 | 0 | 0 | 0 | 0 | 0 | 0 | 0 | 0 | 0 | 0 | 0 | 0 | 0 | NA | NA | NA | NA | NA | NA | NA | NA | NA | NA |
| Vmn2r44 | 0 | 0 | 0 | 0 | 0 | 0 | 0 | 0 | 0 | 0 | 0 | 0 | 0 | 0 | 0 | NA | NA | NA | NA | NA | NA | NA | NA | NA | NA |
| Vmn2r45 | 0 | 0 | 0 | 0 | 0 | 0 | 0 | 0 | 0 | 0 | 0 | 0 | 0 | 0 | 0 | NA | NA | NA | NA | NA | NA | NA | NA | NA | NA |
| Vmn2r46 | 0 | 0 | 0 | 0 | 0 | 0 | 0 | 0 | 0 | 0 | 0 | 0 | 0 | 0 | 0 | NA | NA | NA | NA | NA | NA | NA | NA | NA | NA |
| Vmn2r47 | 0 | 0 | 0 | 0 | 0 | 0 | 0 | 0 | 0 | 0 | 0 | 0 | 0 | 0 | 0 | NA | NA | NA | NA | NA | NA | NA | NA | NA | NA |
| Vmn2r48 | 0 | 0 | 0 | 0 | 0 | 0 | 0 | 0 | 0 | 0 | 0 | 0 | 0 | 0 | 0 | NA | NA | NA | NA | NA | NA | NA | NA | NA | NA |
| Vmn2r49 | 0 | 0 | 0 | 0 | 0 | 0 | 0 | 0 | 0 | 0 | 0 | 0 | 0 | 0 | 0 | NA | NA | NA | NA | NA | NA | NA | NA | NA | NA |
| Vmn2r5 | 0 | 0 | 0 | 0 | 0 | 0 | 0 | 0 | 0 | 0 | 0 | 0 | 0 | 0 | 0 | NA | NA | NA | NA | NA | NA | NA | NA | NA | NA |
| Vmn2r50 | 0 | 0 | 0 | 0 | 0 | 0 | 0 | 0 | 0 | 0 | 0 | 0 | 0 | 0 | 0 | NA | NA | NA | NA | NA | NA | NA | NA | NA | NA |
| Vmn2r51 | 0 | 0 | 0 | 0 | 0 | 0 | 0 | 0 | 0 | 0 | 0 | 0 | 0 | 0 | 0 | NA | NA | NA | NA | NA | NA | NA | NA | NA | NA |
| Vmn2r52 | 0 | 0 | 0 | 0 | 0 | 0 | 0 | 0 | 0 | 0 | 0 | 0 | 0 | 0 | 0 | NA | NA | NA | NA | NA | NA | NA | NA | NA | NA |
| Vmn2r53 | 0 | 0 | 0 | 0 | 0 | 0 | 0 | 0 | 0 | 0 | 0 | 0 | 0 | 0 | 0 | NA | NA | NA | NA | NA | NA | NA | NA | NA | NA |
| Vmn2r54 | 0 | 0 | 0 | 0 | 1 | 0 | 0 | 0 | 0 | 0 | 1 | 0 | 0.18 | 0 | 0 | 1.859 | 0.894 | 0.82650734 | NA | 0 | 18.3785 | 10 | 10 | TRUE | 0.5204 |
| Vmn2r55 | 0 | 0 | 0 | 0 | 0 | 0 | 0 | 0 | 0 | 0 | 0 | 0 | 0 | 0 | 0 | NA | NA | NA | NA | NA | NA | NA | NA | NA | NA |
| Vmn2r56 | 0 | 0 | 0 | 0 | 0 | 0 | 0 | 0 | 0 | 0 | 0 | 0 | 0 | 0 | 0 | NA | NA | NA | NA | NA | NA | NA | NA | NA | NA |
| Vmn2r57 | 12 | 21 | 18 | 15 | 8 | 11 | 13 | 21 | 18 | 12 | 9 | 12 | 14.04 | 17 | 11 | 0.628 | -0.672 | 0.19440332 | 0.40017348 | 0 | 0.2416 | 0.1188 | 0.1188 | TRUE | 0.2492 |
| Vmn2r58 | 0 | 1 | 0 | 0 | 0 | 0 | 0 | 1 | 0 | 0 | 0 | 0 | 0.17 | 0 | 0 | 0.49 | -1.029 | 0.80087 | NA | 0 | 19.7991 | 10 | 10 | TRUE | 0.405 |
| Vmn2r59 | 0 | 0 | 0 | 0 | 0 | 0 | 0 | 0 | 0 | 0 | 0 | 0 | 0 | 0 | 0 | NA | NA | NA | NA | NA | NA | NA | NA | NA | NA |
| Vmn2r6 | 0 | 0 | 0 | 0 | 0 | 0 | 0 | 0 | 0 | 0 | 0 | 0 | 0 | 0 | 0 | NA | NA | NA | NA | NA | NA | NA | NA | NA | NA |
| Vmn2r60 | 0 | 0 | 0 | 0 | 0 | 0 | 0 | 0 | 0 | 0 | 0 | 0 | 0 | 0 | 0 | NA | NA | NA | NA | NA | NA | NA | NA | NA | NA |
| Vmn2r61 | 0 | 0 | 0 | 0 | 0 | 1 | 0 | 0 | 0 | 0 | 0 | 1 | 0.18 | 0 | 0 | 1.859 | 0.894 | 0.82650734 | NA | 0 | 18.0643 | 10 | 10 | TRUE | 0.5383 |
| Vmn2r62 | 0 | 0 | 0 | 0 | 0 | 0 | 0 | 0 | 0 | 0 | 0 | 0 | 0 | 0 | 0 | NA | NA | NA | NA | NA | NA | NA | NA | NA | NA |
| Vmn2r63 | 0 | 0 | 0 | 0 | 0 | 0 | 0 | 0 | 0 | 0 | 0 | 0 | 0 | 0 | 0 | NA | NA | NA | NA | NA | NA | NA | NA | NA | NA |
| Vmn2r65 | 0 | 0 | 0 | 0 | 0 | 0 | 0 | 0 | 0 | 0 | 0 | 0 | 0 | 0 | 0 | NA | NA | NA | NA | NA | NA | NA | NA | NA | NA |
| Vmn2r66 | 0 | 0 | 0 | 0 | 0 | 0 | 0 | 0 | 0 | 0 | 0 | 0 | 0 | 0 | 0 | NA | NA | NA | NA | NA | NA | NA | NA | NA | NA |
| Vmn2r67 | 0 | 0 | 0 | 0 | 0 | 0 | 0 | 0 | 0 | 0 | 0 | 0 | 0 | 0 | 0 | NA | NA | NA | NA | NA | NA | NA | NA | NA | NA |
| Vmn2r68 | 0 | 0 | 0 | 0 | 0 | 0 | 0 | 0 | 0 | 0 | 0 | 0 | 0 | 0 | 0 | NA | NA | NA | NA | NA | NA | NA | NA | NA | NA |
| Vmn2r69 | 0 | 0 | 0 | 0 | 0 | 0 | 0 | 0 | 0 | 0 | 0 | 0 | 0 | 0 | 0 | NA | NA | NA | NA | NA | NA | NA | NA | NA | NA |
| Vmn2r7 | 0 | 0 | 0 | 0 | 0 | 0 | 0 | 0 | 0 | 0 | 0 | 0 | 0 | 0 | 0 | NA | NA | NA | NA | NA | NA | NA | NA | NA | NA |
| Vmn2r70 | 0 | 0 | 0 | 0 | 0 | 0 | 0 | 0 | 0 | 0 | 0 | 0 | 0 | 0 | 0 | NA | NA | NA | NA | NA | NA | NA | NA | NA | NA |
| Vmn2r71 | 0 | 0 | 0 | 0 | 0 | 0 | 0 | 0 | 0 | 0 | 0 | 0 | 0 | 0 | 0 | NA | NA | NA | NA | NA | NA | NA | NA | NA | NA |
| Vmn2r72 | 0 | 0 | 1 | 0 | 0 | 0 | 0 | 0 | 1 | 0 | 0 | 0 | 0.17 | 0 | 0 | 0.49 | -1.029 | 0.80087 | NA | 0 | 19.6928 | 10 | 10 | TRUE | 0.4098 |
| Vmn2r73 | 0 | 0 | 0 | 0 | 0 | 0 | 0 | 0 | 0 | 0 | 0 | 0 | 0 | 0 | 0 | NA | NA | NA | NA | NA | NA | NA | NA | NA | NA |
| Vmn2r74 | 0 | 0 | 0 | 0 | 0 | 0 | 0 | 0 | 0 | 0 | 0 | 0 | 0 | 0 | 0 | NA | NA | NA | NA | NA | NA | NA | NA | NA | NA |
| Vmn2r75 | 0 | 0 | 0 | 0 | 0 | 0 | 0 | 0 | 0 | 0 | 0 | 0 | 0 | 0 | 0 | NA | NA | NA | NA | NA | NA | NA | NA | NA | NA |
| Vmn2r76 | 0 | 0 | 0 | 0 | 0 | 0 | 0 | 0 | 0 | 0 | 0 | 0 | 0 | 0 | 0 | NA | NA | NA | NA | NA | NA | NA | NA | NA | NA |
| Vmn2r77 | 0 | 0 | 0 | 0 | 0 | 0 | 0 | 0 | 0 | 0 | 0 | 0 | 0 | 0 | 0 | NA | NA | NA | NA | NA | NA | NA | NA | NA | NA |
| Vmn2r78 | 1 | 0 | 0 | 0 | 0 | 0 | 1 | 0 | 0 | 0 | 0 | 0 | 0.18 | 0 | 0 | 0.49 | -1.029 | 0.80087 | NA | 0 | 18.6165 | 10 | 10 | TRUE | 0.4614 |
| Vmn2r79 | 0 | 0 | 0 | 0 | 0 | 0 | 0 | 0 | 0 | 0 | 0 | 0 | 0 | 0 | 0 | NA | NA | NA | NA | NA | NA | NA | NA | NA | NA |
| Vmn2r8 | 0 | 0 | 0 | 0 | 0 | 0 | 0 | 0 | 0 | 0 | 0 | 0 | 0 | 0 | 0 | NA | NA | NA | NA | NA | NA | NA | NA | NA | NA |
| Vmn2r80 | 0 | 0 | 0 | 0 | 0 | 0 | 0 | 0 | 0 | 0 | 0 | 0 | 0 | 0 | 0 | NA | NA | NA | NA | NA | NA | NA | NA | NA | NA |

| | | | | | | | | | | | | | | | | | | | | | | | | | |
|---|---|---|---|---|---|---|---|---|---|---|---|---|---|---|---|---|---|---|---|---|---|---|---|---|---|
| Vmn2r81 | 0 | 0 | 0 | 0 | 0 | 0 | 0 | 0 | 0 | 0 | 0 | 0 | 0 | 0 | 0 | NA | NA | NA | NA | NA | NA | NA | NA | NA | NA |
| Vmn2r82 | 0 | 0 | 0 | 0 | 0 | 0 | 0 | 0 | 0 | 0 | 0 | 0 | 0 | 0 | 0 | NA | NA | NA | NA | NA | NA | NA | NA | NA | NA |
| Vmn2r83 | 0 | 0 | 0 | 0 | 0 | 0 | 0 | 0 | 0 | 0 | 0 | 0 | 0 | 0 | 0 | NA | NA | NA | NA | NA | NA | NA | NA | NA | NA |
| Vmn2r84 | 1 | 0 | 0 | 1 | 2 | 0 | 1 | 0 | 0 | 1 | 2 | 0 | 0.67 | 0 | 1 | 2.668 | 1.416 | 0.58602256 | NA | 0 | 4.9484 | 3.3683 | 3.3683 | TRUE | 0.3203 |
| Vmn2r85 | 0 | 0 | 0 | 0 | 0 | 0 | 0 | 0 | 0 | 0 | 0 | 0 | 0 | 0 | 0 | NA | NA | NA | NA | NA | NA | NA | NA | NA | NA |
| Vmn2r86 | 0 | 0 | 0 | 0 | 0 | 0 | 0 | 0 | 0 | 0 | 0 | 0 | 0 | 0 | 0 | NA | NA | NA | NA | NA | NA | NA | NA | NA | NA |
| Vmn2r87 | 1 | 0 | 0 | 0 | 0 | 0 | 1 | 0 | 0 | 0 | 0 | 0 | 0.18 | 0 | 0 | 0.49 | -1.029 | 0.80087 | NA | 0 | 18.6165 | 10 | 10 | TRUE | 0.4614 |
| Vmn2r88 | 0 | 0 | 0 | 0 | 0 | 0 | 0 | 0 | 0 | 0 | 0 | 0 | 0 | 0 | 0 | NA | NA | NA | NA | NA | NA | NA | NA | NA | NA |
| Vmn2r89 | 0 | 1 | 0 | 1 | 0 | 1 | 0 | 1 | 0 | 1 | 0 | 1 | 0.48 | 0 | 1 | 1.723 | 0.785 | 0.78325239 | NA | 0 | 6.8716 | 4.0968 | 4.0968 | TRUE | 0.405 |
| Vmn2r9 | 0 | 0 | 0 | 0 | 0 | 0 | 0 | 0 | 0 | 0 | 0 | 0 | 0 | 0 | 0 | NA | NA | NA | NA | NA | NA | NA | NA | NA | NA |
| Vmn2r90 | 3 | 2 | 2 | 3 | 5 | 3 | 3 | 2 | 2 | 2 | 5 | 3 | 3.04 | 2 | 3 | 1.517 | 0.601 | 0.57382184 | NA | 0 | 1.0921 | 0.4753 | 0.4753 | TRUE | 0.2428 |
| Vmn2r91 | 0 | 0 | 0 | 0 | 0 | 0 | 0 | 0 | 0 | 0 | 0 | 0 | 0 | 0 | 0 | NA | NA | NA | NA | NA | NA | NA | NA | NA | NA |
| Vmn2r92 | 0 | 0 | 0 | 0 | 0 | 0 | 0 | 0 | 0 | 0 | 0 | 0 | 0 | 0 | 0 | NA | NA | NA | NA | NA | NA | NA | NA | NA | NA |
| Vmn2r93 | 0 | 0 | 0 | 0 | 0 | 0 | 0 | 0 | 0 | 0 | 0 | 0 | 0 | 0 | 0 | NA | NA | NA | NA | NA | NA | NA | NA | NA | NA |
| Vmn2r94 | 1 | 0 | 0 | 4 | 4 | 0 | 1 | 0 | 0 | 3 | 4 | 0 | 1.42 | 0 | 2 | 6.822 | 2.77 | 0.19086569 | NA | 0 | 2.3295 | 2.0355 | 2.0355 | TRUE | 0.3814 |
| Vmn2r95 | 0 | 0 | 0 | 0 | 1 | 0 | 0 | 0 | 0 | 0 | 1 | 0 | 0.18 | 0 | 0 | 1.859 | 0.894 | 0.82650734 | NA | 0 | 18.3785 | 10 | 10 | TRUE | 0.5204 |
| Vmn2r96 | 0 | 0 | 0 | 0 | 0 | 0 | 0 | 0 | 0 | 0 | 0 | 0 | 0 | 0 | 0 | NA | NA | NA | NA | NA | NA | NA | NA | NA | NA |
| Vmn2r97 | 0 | 0 | 0 | 0 | 0 | 0 | 0 | 0 | 0 | 0 | 0 | 0 | 0 | 0 | 0 | NA | NA | NA | NA | NA | NA | NA | NA | NA | NA |
| Vmn2r98 | 0 | 0 | 0 | 0 | 0 | 0 | 0 | 0 | 0 | 0 | 0 | 0 | 0 | 0 | 0 | NA | NA | NA | NA | NA | NA | NA | NA | NA | NA |
| Vmn2r99 | 0 | 0 | 0 | 0 | 0 | 0 | 0 | 0 | 0 | 0 | 0 | 0 | 0 | 0 | 0 | NA | NA | NA | NA | NA | NA | NA | NA | NA | NA |
| Vmo1 | 4 | 8 | 5 | 1 | 2 | 3 | 4 | 8 | 5 | 1 | 2 | 3 | 3.91 | 6 | 2 | 0.347 | -1.528 | 0.13754094 | NA | 0 | 0.8487 | 0.4303 | 0.4303 | TRUE | 0.3992 |
| Vmp1 | 4004 | 4539 | 4366 | 5953 | 3961 | 4068 | 4253 | 4533 | 4384 | 4680 | 4262 | 4453 | 4427.44 | 4390 | 4465 | 1.017 | 0.025 | 0.7302405 | 0.85505616 | 0.0014 | 0.0076 | 0.0035 | 0.0035 | TRUE | 0.0219 |
| Vnn1 | 6 | 3 | 1 | 83 | 46 | 31 | 6 | 3 | 1 | 65 | 49 | 34 | 26.51 | 3 | 49 | 14.564 | 3.864 | 6.35E-10 | 4.07E-08 | 0.1053 | 0.1312 | 0.1224 | 0.1224 | TRUE | 0.2715 |
| Vnn3 | 0 | 0 | 0 | 0 | 1 | 0 | 0 | 0 | 0 | 0 | 1 | 0 | 0.18 | 0 | 0 | 1.859 | 0.894 | 0.82650734 | NA | 0 | 18.3785 | 10 | 10 | TRUE | 0.5204 |
| Vopp1 | 248 | 410 | 382 | 581 | 415 | 480 | 263 | 409 | 384 | 457 | 447 | 525 | 414.19 | 352 | 476 | 1.351 | 0.434 | 0.01611167 | 0.07478319 | 0.0273 | 0.0149 | 0.021 | 0.021 | TRUE | 0.5502 |
| Vprbp | 7824 | 7402 | 7970 | 9750 | 7021 | 6985 | 8310 | 7392 | 8003 | 7665 | 7554 | 7646 | 7761.78 | 7902 | 7622 | 0.965 | -0.052 | 0.4661207 | 0.66857491 | 0.0016 | 0.0073 | 0.0035 | 0.0035 | TRUE | 0.0389 |
| Vpreb1 | 5 | 5 | 3 | 21 | 14 | 21 | 5 | 5 | 3 | 17 | 15 | 23 | 11.31 | 4 | 18 | 4.094 | 2.033 | 0.00145007 | 0.01193037 | 0 | 0.2981 | 0.1516 | 0.1516 | TRUE | 0.2521 |
| Vpreb2 | 0 | 1 | 0 | 1 | 0 | 1 | 0 | 1 | 0 | 1 | 0 | 1 | 0.48 | 0 | 1 | 1.723 | 0.785 | 0.78325239 | NA | 0 | 6.8716 | 4.0968 | 4.0968 | TRUE | 0.405 |
| Vpreb3 | 10 | 11 | 16 | 8 | 8 | 11 | 11 | 11 | 16 | 6 | 9 | 12 | 10.77 | 13 | 9 | 0.705 | -0.505 | 0.39659317 | 0.60995294 | 0 | 0.3128 | 0.1597 | 0.1597 | TRUE | 0.291 |
| Vps11 | 3420 | 2879 | 3046 | 4084 | 2938 | 2866 | 3633 | 2875 | 3059 | 3211 | 3161 | 3137 | 3179.22 | 3189 | 3170 | 0.994 | -0.008 | 0.93526618 | 0.96791458 | 0.0072 | 0.0079 | 0.0075 | 0.0075 | TRUE | 0.1799 |
| Vps13a | 3622 | 3579 | 4040 | 4369 | 3201 | 3175 | 3847 | 3574 | 4057 | 3435 | 3444 | 3476 | 3638.73 | 3826 | 3452 | 0.902 | -0.149 | 0.04997023 | 0.16412546 | 0.0016 | 0.0078 | 0.0039 | 0.0039 | TRUE | 0.0404 |
| Vps13b | 3131 | 3636 | 4156 | 4615 | 3308 | 3311 | 3326 | 3631 | 4173 | 3628 | 3559 | 3624 | 3656.94 | 3710 | 3604 | 0.971 | -0.042 | 0.67522398 | 0.8190765 | 0.0055 | 0.0078 | 0.007 | 0.007 | TRUE | 0.1451 |
| Vps13c | 4990 | 4544 | 5353 | 6152 | 4254 | 4266 | 5300 | 4538 | 5375 | 4836 | 4577 | 4670 | 4882.74 | 5071 | 4694 | 0.926 | -0.111 | 0.21921471 | 0.43131916 | 0.0043 | 0.0076 | 0.0057 | 0.0057 | TRUE | 0.1032 |
| Vps13d | 4312 | 3756 | 4358 | 4525 | 3214 | 3245 | 4580 | 3751 | 4376 | 3557 | 3458 | 3552 | 3879.1 | 4236 | 3522 | 0.832 | -0.266 | 0.00515895 | 0.03198394 | 0.0049 | 0.0078 | 0.0063 | 0.0063 | TRUE | 0.1221 |
| Vps16 | 3830 | 4503 | 4564 | 5158 | 3967 | 4051 | 4068 | 4497 | 4583 | 4055 | 4268 | 4434 | 4317.6 | 4383 | 4252 | 0.97 | -0.044 | 0.59009143 | 0.76449681 | 0.0024 | 0.0077 | 0.0046 | 0.0046 | TRUE | 0.0479 |
| Vps18 | 1250 | 1254 | 1145 | 1944 | 1325 | 1194 | 1328 | 1252 | 1150 | 1528 | 1426 | 1307 | 1331.78 | 1243 | 1420 | 1.143 | 0.193 | 0.05843915 | 0.18266785 | 0.0045 | 0.0094 | 0.0068 | 0.0068 | TRUE | 0.0583 |
| Vps25 | 3206 | 3493 | 3467 | 4983 | 3441 | 3295 | 3405 | 3488 | 3481 | 3917 | 3702 | 3607 | 3600.25 | 3458 | 3742 | 1.082 | 0.114 | 0.09192367 | 0.24742196 | 8.00E-04 | 0.0078 | 0.003 | 0.003 | TRUE | 0.0207 |
| Vps26a | 3267 | 3817 | 3926 | 4301 | 3243 | 3304 | 3470 | 3812 | 3942 | 3381 | 3489 | 3617 | 3618.57 | 3741 | 3496 | 0.934 | -0.098 | 0.22104129 | 0.43311388 | 0.0023 | 0.0078 | 0.0044 | 0.0044 | TRUE | 0.0488 |
| Vps26b | 2817 | 3197 | 3150 | 4579 | 3311 | 3456 | 2992 | 3193 | 3163 | 3600 | 3562 | 3783 | 3382.2 | 3116 | 3648 | 1.171 | 0.227 | 0.00102902 | 0.00911648 | 7.00E-04 | 0.0079 | 0.0032 | 0.0032 | TRUE | 0.0146 |
| Vps28 | 1209 | 1650 | 1744 | 1822 | 1329 | 1518 | 1284 | 1648 | 1751 | 1432 | 1430 | 1662 | 1534.52 | 1561 | 1508 | 0.966 | -0.051 | 0.71140112 | 0.84292829 | 0.0161 | 0.0091 | 0.0128 | 0.0128 | TRUE | 0.2894 |
| Vps29 | 2797 | 2830 | 2827 | 3835 | 2665 | 2595 | 2971 | 2826 | 2839 | 3015 | 2867 | 2841 | 2893.11 | 2879 | 2908 | 1.011 | 0.015 | 0.82573977 | 0.9116315 | 6.00E-04 | 0.008 | 0.003 | 0.003 | TRUE | 0.0129 |
| Vps33a | 1886 | 1766 | 1828 | 2894 | 1991 | 1898 | 2003 | 1764 | 1836 | 2275 | 2142 | 2078 | 2016.23 | 1868 | 2165 | 1.16 | 0.214 | 0.01356763 | 0.06566081 | 0.0025 | 0.0085 | 0.0049 | 0.0049 | TRUE | 0.0487 |
| Vps33b | 1180 | 1157 | 1095 | 1416 | 1124 | 1011 | 1253 | 1156 | 1099 | 1113 | 1209 | 1107 | 1156.26 | 1169 | 1143 | 0.977 | -0.033 | 0.72303936 | 0.85005881 | 0.0025 | 0.0098 | 0.0054 | 0.0054 | TRUE | 0.0469 |
| Vps35 | 18745 | 20010 | 20406 | 25394 | 18351 | 19009 | 19911 | 19984 | 20490 | 19963 | 19745 | 20808 | 20150.02 | 20128 | 20172 | 1.002 | 0.003 | 0.95663451 | 0.97766996 | 4.00E-04 | 0.0071 | 0.0023 | 0.0023 | TRUE | 0.0093 |
| Vps36 | 3050 | 3635 | 3741 | 4305 | 3184 | 3168 | 3240 | 3630 | 3756 | 3384 | 3426 | 3468 | 3484.04 | 3542 | 3426 | 0.967 | -0.048 | 0.55753882 | 0.7401315 | 0.0025 | 0.0078 | 0.0046 | 0.0046 | TRUE | 0.0677 |
| Vps37a | 1884 | 2277 | 2046 | 2776 | 2031 | 2028 | 2001 | 2274 | 2054 | 2182 | 2185 | 2220 | 2152.85 | 2110 | 2196 | 1.041 | 0.057 | 0.47972772 | 0.67912904 | 0.0017 | 0.0084 | 0.0043 | 0.0043 | TRUE | 0.0562 |
| Vps37b | 906 | 876 | 831 | 1300 | 976 | 852 | 962 | 875 | 834 | 1022 | 1050 | 933 | 946.06 | 890 | 1002 | 1.125 | 0.17 | 0.09239862 | 0.24809516 | 0.0034 | 0.0104 | 0.0063 | 0.0063 | TRUE | 0.0588 |
| Vps37c | 811 | 864 | 821 | 898 | 662 | 648 | 861 | 863 | 824 | 706 | 712 | 709 | 779.37 | 849 | 709 | 0.835 | -0.261 | 0.0022156 | 0.01666687 | 0 | 0.0111 | 0.0039 | 0.0039 | TRUE | 0.008 |
| Vps37d | 118 | 193 | 191 | 155 | 115 | 172 | 125 | 193 | 192 | 122 | 124 | 188 | 157.29 | 170 | 145 | 0.848 | -0.238 | 0.34930449 | 0.56807903 | 0.0531 | 0.0278 | 0.04 | 0.04 | TRUE | 0.7105 |
| Vps39 | 2783 | 3288 | 3356 | 3731 | 2759 | 3005 | 2956 | 3284 | 3370 | 2933 | 2969 | 3289 | 3133.43 | 3203 | 3064 | 0.956 | -0.065 | 0.47330173 | 0.67421558 | 0.0038 | 0.008 | 0.0056 | 0.0056 | TRUE | 0.0552 |
| Vps41 | 5918 | 6808 | 6874 | 7896 | 5805 | 6083 | 6286 | 6799 | 6902 | 6207 | 6246 | 6659 | 6516.54 | 6662 | 6371 | 0.956 | -0.065 | 0.37681536 | 0.59206639 | 0.0018 | 0.0074 | 0.0037 | 0.0037 | TRUE | 0.0298 |
| Vps45 | 794 | 786 | 827 | 988 | 617 | 637 | 843 | 785 | 830 | 777 | 664 | 697 | 766.1 | 819 | 713 | 0.871 | -0.199 | 0.05235152 | 0.16923389 | 0.0028 | 0.0112 | 0.0063 | 0.0063 | TRUE | 0.0763 |
| Vps4a | 2578 | 2790 | 2807 | 3399 | 2383 | 2465 | 2738 | 2786 | 2819 | 2672 | 2564 | 2698 | 2712.92 | 2781 | 2645 | 0.951 | -0.072 | 0.26662624 | 0.48632232 | 0 | 0.0081 | 0.0027 | 0.0027 | TRUE | 0.0085 |
| Vps4b | 4345 | 4246 | 3980 | 6326 | 4590 | 4479 | 4615 | 4241 | 3996 | 4973 | 4939 | 4903 | 4611.09 | 4284 | 4938 | 1.153 | 0.205 | 0.00921592 | 0.04943996 | 0.0023 | 0.0076 | 0.0043 | 0.0043 | TRUE | 0.0556 |
| Vps51 | 1100 | 1211 | 1250 | 1646 | 1190 | 1211 | 1168 | 1209 | 1255 | 1294 | 1280 | 1326 | 1255.49 | 1211 | 1300 | 1.073 | 0.102 | 0.18879775 | 0.39393017 | 0 | 0.0095 | 0.0035 | 0.0035 | TRUE | 0.0122 |
| Vps52 | 1119 | 1305 | 1296 | 2121 | 1585 | 1229 | 1189 | 1303 | 1301 | 1667 | 1705 | 1345 | 1418.55 | 1264 | 1572 | 1.244 | 0.315 | 0.0069045 | 0.03974249 | 0.0082 | 0.0092 | 0.0091 | 0.0091 | TRUE | 0.1908 |
| Vps53 | 3303 | 2590 | 2582 | 3664 | 2598 | 2141 | 3508 | 2587 | 2593 | 2880 | 2795 | 2344 | 2784.49 | 2896 | 2673 | 0.923 | -0.115 | 0.4301734 | 0.64000517 | 0.0204 | 0.0081 | 0.015 | 0.015 | TRUE | 0.4153 |
| Vps54 | 4149 | 4734 | 4691 | 7490 | 5400 | 5567 | 4407 | 4728 | 4710 | 5888 | 5810 | 6094 | 5272.87 | 4615 | 5931 | 1.285 | 0.362 | 5.20E-08 | 2.04E-06 | 8.00E-04 | 0.0075 | 0.003 | 0.003 | TRUE | 0.0189 |
| Vps72 | 2157 | 2438 | 2403 | 2808 | 2119 | 2007 | 2291 | 2435 | 2413 | 2207 | 2280 | 2197 | 2303.86 | 2380 | 2228 | 0.936 | -0.095 | 0.16864014 | 0.36646353 | 3.00E-04 | 0.0083 | 0.003 | 0.003 | TRUE | 0.0128 |
| Vps8 | 988 | 1193 | 1329 | 1564 | 1146 | 1125 | 1049 | 1191 | 1334 | 1229 | 1233 | 1231 | 1211.56 | 1191 | 1231 | 1.033 | 0.047 | 0.66737899 | 0.81394227 | 0.0063 | 0.0096 | 0.0077 | 0.0077 | TRUE | 0.1315 |
| Vps9d1 | 482 | 520 | 551 | 794 | 598 | 545 | 512 | 519 | 553 | 624 | 643 | 597 | 574.79 | 528 | 621 | 1.176 | 0.234 | 0.01776262 | 0.07970188 | 0 | 0.0126 | 0.0053 | 0.0053 | TRUE | 0.0202 |
| Vrk1 | 3223 | 3566 | 3555 | 4317 | 3064 | 3062 | 3423 | 3561 | 3570 | 3394 | 3297 | 3352 | 3432.77 | 3518 | 3348 | 0.952 | -0.072 | 0.24998165 | 0.46725653 | 0 | 0.0079 | 0.0025 | 0.0025 | TRUE | 0.0067 |
| Vrk2 | 354 | 470 | 485 | 784 | 555 | 587 | 376 | 469 | 487 | 616 | 597 | 643 | 531.4 | 444 | 619 | 1.392 | 0.477 | 0.00022039 | 0.00262741 | 0.0074 | 0.0131 | 0.0101 | 0.0101 | TRUE | 0.2063 |
| Vrk3 | 1691 | 2024 | 2103 | 2171 | 1678 | 1854 | 1796 | 2021 | 2112 | 1707 | 1805 | 2029 | 1911.79 | 1976 | 1847 | 0.934 | -0.098 | 0.3538518 | 0.57222912 | 0.0066 | 0.0086 | 0.0076 | 0.0076 | TRUE | 0.0897 |
| Vrtn | 149 | 101 | 101 | 60 | 24 | 19 | 158 | 101 | 101 | 47 | 26 | 21 | 75.72 | 120 | 31 | 0.264 | -1.919 | 8.16E-08 | 3.03E-06 | 0.1011 | 0.0504 | 0.0726 | 0.0726 | TRUE | 1.5418 |
| Vsig1 | 1 | 12 | 7 | 11 | 4 | 14 | 1 | 12 | 7 | 9 | 4 | 15 | 8.06 | 7 | 9 | 1.399 | 0.484 | 0.58471797 | 0.75998583 | 0.4649 | 0.4157 | 0.438 | 0.438 | TRUE | 0.3234 |
| Vsig10 | 987 | 1233 | 1270 | 1138 | 822 | 970 | 1048 | 1231 | 1275 | 895 | 884 | 1062 | 1065.97 | 1185 | 947 | 0.798 | -0.325 | 0.00786698 | 0.0437665 | 0.0093 | 0.01 | 0.0098 | 0.0098 | TRUE | 0.1327 |
| Vsig2 | 73 | 78 | 102 | 149 | 92 | 127 | 78 | 78 | 102 | 117 | 99 | 139 | 102.17 | 86 | 118 | 1.376 | 0.46 | 0.04605703 | 0.15528248 | 0.0169 | 0.0391 | 0.0283 | 0.0283 | TRUE | 0.2086 |
| Vsig4 | 0 | 0 | 0 | 0 | 0 | 0 | 0 | 0 | 0 | 0 | 0 | 0 | 0 | 0 | 0 | NA | NA | NA | NA | NA | NA | NA | NA | NA | NA |

| | | | | | | | | | | | | | | | | | | | | | | | | | |
|---|---|---|---|---|---|---|---|---|---|---|---|---|---|---|---|---|---|---|---|---|---|---|---|---|---|
| Vsig8 | 6 | 7 | 3 | 6 | 2 | 6 | 6 | 7 | 3 | 5 | 2 | 7 | 4.97 | 5 | 5 | 0.822 | -0.283 | 0.74737599 | 0.86571784 | 0 | 0.6699 | 0.3522 | 0.3522 | TRUE | 0.2924 |
| Vsnl1 | 24 | 10 | 10 | 15 | 8 | 7 | 25 | 10 | 10 | 12 | 9 | 8 | 12.26 | 15 | 10 | 0.624 | -0.679 | 0.2794217 | 0.5005155 | 0.125 | 0.2755 | 0.1987 | 0.1987 | TRUE | 1.5681 |
| Vstm2a | 11 | 1 | 4 | 17 | 9 | 6 | 12 | 1 | 4 | 13 | 10 | 7 | 7.72 | 6 | 10 | 1.799 | 0.847 | 0.33153645 | 0.55215482 | 0.3716 | 0.4337 | 0.4068 | 0.4068 | TRUE | 1.149 |
| Vstm2b | 33 | 27 | 18 | 45 | 35 | 22 | 35 | 27 | 18 | 35 | 38 | 24 | 29.53 | 27 | 32 | 1.218 | 0.285 | 0.47777195 | 0.67729589 | 0.0395 | 0.1184 | 0.0819 | 0.0819 | TRUE | 0.2503 |
| Vstm2l | 142 | 216 | 186 | 107 | 88 | 113 | 151 | 216 | 187 | 84 | 95 | 124 | 142.63 | 185 | 101 | 0.544 | -0.879 | 9.90E-05 | 0.00136812 | 0.028 | 0.03 | 0.0291 | 0.0291 | TRUE | 0.2174 |
| Vstm4 | 3 | 2 | 6 | 10 | 4 | 4 | 3 | 2 | 6 | 8 | 4 | 4 | 4.63 | 4 | 5 | 1.496 | 0.581 | 0.52186599 | 0.71194565 | 0 | 0.7192 | 0.3691 | 0.3691 | TRUE | 0.4606 |
| Vstm5 | 37 | 63 | 72 | 100 | 85 | 90 | 39 | 63 | 72 | 79 | 91 | 99 | 73.85 | 58 | 90 | 1.532 | 0.616 | 0.02885258 | 0.11197414 | 0.0326 | 0.0515 | 0.0429 | 0.0429 | TRUE | 0.6674 |
| Vsx1 | 12 | 15 | 18 | 10 | 5 | 7 | 13 | 15 | 18 | 8 | 5 | 8 | 11.12 | 15 | 7 | 0.459 | -1.124 | 0.0526218 | 0.16989786 | 0 | 0.3032 | 0.1394 | 0.1394 | TRUE | 0.1197 |
| Vsx2 | 22 | 14 | 6 | 16 | 9 | 9 | 23 | 14 | 6 | 13 | 10 | 10 | 12.58 | 14 | 11 | 0.747 | -0.421 | 0.5045781 | 0.69949949 | 0.1414 | 0.2688 | 0.2064 | 0.2064 | TRUE | 0.175 |
| Vta1 | 1956 | 2265 | 2083 | 2525 | 1872 | 1972 | 2078 | 2262 | 2092 | 1985 | 2014 | 2159 | 2098.17 | 2144 | 2053 | 0.957 | -0.063 | 0.42934661 | 0.63907651 | 0.0015 | 0.0085 | 0.0041 | 0.0041 | TRUE | 0.0283 |
| Vtcn1 | 0 | 0 | 2 | 7 | 5 | 3 | 0 | 0 | 2 | 6 | 5 | 3 | 2.7 | 1 | 5 | 7.004 | 2.808 | 0.06007511 | NA | 0 | 1.229 | 0.7498 | 0.7498 | TRUE | 0.9737 |
| Vti1a | 590 | 491 | 489 | 748 | 533 | 571 | 627 | 490 | 491 | 588 | 573 | 625 | 565.77 | 536 | 595 | 1.111 | 0.152 | 0.249519 | 0.466779 | 0.0091 | 0.0127 | 0.0108 | 0.0108 | TRUE | 0.2543 |
| Vti1b | 1875 | 2199 | 2064 | 2832 | 1885 | 1843 | 1992 | 2196 | 2072 | 2226 | 2028 | 2017 | 2088.69 | 2087 | 2090 | 1.002 | 0.003 | 0.96735877 | 0.98366749 | 0.0023 | 0.0085 | 0.0046 | 0.0046 | TRUE | 0.0398 |
| Vtn | 2 | 3 | 1 | 5 | 1 | 3 | 2 | 3 | 1 | 4 | 1 | 3 | 2.4 | 2 | 3 | 1.374 | 0.458 | 0.70768659 | NA | 0 | 1.3782 | 0.6501 | 0.6501 | TRUE | 0.3142 |
| Vwa1 | 58 | 72 | 78 | 115 | 82 | 59 | 62 | 72 | 78 | 90 | 88 | 65 | 75.84 | 71 | 81 | 1.151 | 0.203 | 0.41169805 | 0.62279894 | 0.0101 | 0.0503 | 0.031 | 0.031 | TRUE | 0.2799 |
| Vwa2 | 197 | 244 | 217 | 252 | 171 | 207 | 209 | 244 | 218 | 198 | 184 | 227 | 213.25 | 224 | 203 | 0.906 | -0.142 | 0.36187906 | 0.58006064 | 0.0035 | 0.0224 | 0.0128 | 0.0128 | TRUE | 0.1076 |
| Vwa3a | 3 | 1 | 4 | 3 | 1 | 3 | 3 | 1 | 4 | 2 | 1 | 3 | 2.49 | 3 | 2 | 0.821 | -0.284 | 0.81356061 | NA | 0 | 1.3318 | 0.6398 | 0.6398 | TRUE | 0.3777 |
| Vwa5a | 1177 | 2146 | 2174 | 1977 | 1550 | 1990 | 1250 | 2143 | 2183 | 1554 | 1668 | 2178 | 1829.43 | 1859 | 1800 | 0.968 | -0.047 | 0.82658096 | 0.91211327 | 0.0614 | 0.0087 | 0.0323 | 0.0323 | TRUE | 0.99 |
| Vwa5b1 | 4 | 2 | 3 | 4 | 5 | 1 | 4 | 2 | 3 | 3 | 5 | 1 | 3.15 | 3 | 3 | 1.043 | 0.06 | 0.95561084 | NA | 0 | 1.054 | 0.5299 | 0.5299 | TRUE | 0.1811 |
| Vwa5b2 | 28 | 43 | 36 | 50 | 39 | 32 | 30 | 43 | 36 | 39 | 42 | 35 | 37.52 | 36 | 39 | 1.068 | 0.095 | 0.76716425 | 0.87844595 | 0 | 0.0947 | 0.047 | 0.047 | TRUE | 0.1874 |
| Vwa7 | 12 | 14 | 24 | 9 | 15 | 11 | 13 | 14 | 24 | 7 | 16 | 12 | 14.35 | 17 | 12 | 0.683 | -0.55 | 0.33156596 | 0.55215482 | 0.0774 | 0.2365 | 0.159 | 0.159 | TRUE | 0.4297 |
| Vwa8 | 1664 | 1463 | 1533 | 1808 | 1343 | 1162 | 1767 | 1461 | 1539 | 1421 | 1445 | 1272 | 1484.36 | 1589 | 1379 | 0.868 | -0.204 | 0.05814277 | 0.18220301 | 0.0063 | 0.0091 | 0.0076 | 0.0076 | TRUE | 0.1156 |
| Vwa9 | 1361 | 1330 | 1183 | 1962 | 1319 | 1222 | 1446 | 1328 | 1188 | 1542 | 1419 | 1338 | 1376.83 | 1321 | 1433 | 1.086 | 0.119 | 0.27287812 | 0.49310652 | 0.0063 | 0.0093 | 0.0078 | 0.0078 | TRUE | 0.093 |
| Vwc2 | 22 | 13 | 12 | 24 | 15 | 7 | 23 | 13 | 12 | 19 | 16 | 8 | 15.18 | 16 | 14 | 0.893 | -0.163 | 0.77222879 | 0.88115034 | 0.0952 | 0.224 | 0.1629 | 0.1629 | TRUE | 0.7069 |
| Vwc2l | 0 | 0 | 0 | 0 | 0 | 0 | 0 | 0 | 0 | 0 | 0 | 0 | 0 | 0 | 0 | NA | NA | NA | NA | NA | NA | NA | NA | NA | NA |
| Vwce | 49 | 48 | 47 | 55 | 31 | 40 | 52 | 48 | 47 | 43 | 33 | 44 | 44.59 | 49 | 40 | 0.821 | -0.285 | 0.3244367 | 0.545815 | 0 | 0.0808 | 0.0378 | 0.0378 | TRUE | 0.1525 |
| Vwde | 1 | 3 | 0 | 9 | 2 | 2 | 1 | 3 | 0 | 7 | 2 | 2 | 2.58 | 1 | 4 | 2.861 | 1.517 | 0.27284157 | NA | 0.3829 | 1.2843 | 0.8758 | 0.8758 | TRUE | 1.1483 |
| Vwf | 1007 | 1312 | 1445 | 783 | 675 | 920 | 1070 | 1310 | 1451 | 616 | 726 | 1007 | 1029.95 | 1277 | 783 | 0.612 | -0.707 | 0.00021732 | 0.00259862 | 0.0431 | 0.0101 | 0.0253 | 0.0253 | TRUE | 0.7416 |
| Wac | 3262 | 4020 | 4076 | 4964 | 3667 | 4062 | 3465 | 4015 | 4093 | 3902 | 3945 | 4446 | 3977.77 | 3858 | 4098 | 1.062 | 0.087 | 0.3852179 | 0.59920343 | 0.0061 | 0.0077 | 0.007 | 0.007 | TRUE | 0.0963 |
| Wap | 0 | 0 | 0 | 0 | 0 | 0 | 0 | 0 | 0 | 0 | 0 | 0 | 0 | 0 | 0 | NA | NA | NA | NA | NA | NA | NA | NA | NA | NA |
| Wapal | 12389 | 11522 | 11877 | 14835 | 10597 | 10585 | 13159 | 11507 | 11926 | 11662 | 11402 | 11587 | 11873.83 | 12197 | 11550 | 0.947 | -0.079 | 0.2995311 | 0.51972817 | 0.0024 | 0.0072 | 0.004 | 0.004 | TRUE | 0.0581 |
| Wars | 5230 | 4341 | 4180 | 6871 | 4803 | 4217 | 5555 | 4335 | 4197 | 5401 | 5168 | 4616 | 4878.85 | 4696 | 5062 | 1.078 | 0.109 | 0.39646111 | 0.60986878 | 0.0136 | 0.0076 | 0.0116 | 0.0116 | TRUE | 0.3119 |
| Wars2 | 149 | 146 | 108 | 219 | 207 | 152 | 158 | 146 | 108 | 172 | 223 | 166 | 162.3 | 137 | 187 | 1.36 | 0.443 | 0.0365285 | 0.1318068 | 0.0249 | 0.0272 | 0.0261 | 0.0261 | TRUE | 0.3552 |
| Was | 90 | 71 | 69 | 112 | 93 | 69 | 96 | 71 | 69 | 88 | 100 | 76 | 83.24 | 79 | 88 | 1.12 | 0.163 | 0.50118073 | 0.69669611 | 0.0131 | 0.0465 | 0.0304 | 0.0304 | TRUE | 0.3247 |
| Wasf1 | 3174 | 2971 | 2935 | 4292 | 3026 | 2907 | 3371 | 2967 | 2947 | 3374 | 3256 | 3182 | 3182.93 | 3095 | 3271 | 1.057 | 0.08 | 0.34460124 | 0.5641524 | 0.003 | 0.0079 | 0.0048 | 0.0048 | TRUE | 0.0738 |
| Wasf2 | 6638 | 7172 | 7394 | 9379 | 6386 | 6332 | 7051 | 7163 | 7424 | 7373 | 6871 | 6931 | 7135.53 | 7213 | 7058 | 0.979 | -0.031 | 0.63760173 | 0.79603827 | 9.00E-04 | 0.0074 | 0.003 | 0.003 | TRUE | 0.0187 |
| Wasf3 | 2914 | 3641 | 3993 | 2965 | 2374 | 2800 | 3095 | 3636 | 4009 | 2331 | 2554 | 3065 | 3115.17 | 3580 | 2650 | 0.74 | -0.435 | 0.00150106 | 0.01226033 | 0.0178 | 0.008 | 0.0132 | 0.0132 | TRUE | 0.2273 |
| Wash | 1344 | 1674 | 1791 | 1810 | 1408 | 1443 | 1428 | 1672 | 1798 | 1423 | 1515 | 1580 | 1569.19 | 1633 | 1506 | 0.922 | -0.117 | 0.288517 | 0.50952666 | 0.0071 | 0.009 | 0.0082 | 0.0082 | TRUE | 0.145 |
| Wasl | 4288 | 4162 | 4227 | 5218 | 3931 | 3792 | 4555 | 4157 | 4244 | 4102 | 4230 | 4151 | 4239.67 | 4319 | 4161 | 0.963 | -0.054 | 0.43783516 | 0.64573931 | 9.00E-04 | 0.0077 | 0.0032 | 0.0032 | TRUE | 0.0278 |
| Wbp1 | 741 | 1002 | 916 | 1018 | 730 | 692 | 787 | 1001 | 920 | 800 | 785 | 758 | 841.79 | 903 | 781 | 0.865 | -0.208 | 0.07099676 | 0.20748529 | 0.0066 | 0.0108 | 0.0084 | 0.0084 | TRUE | 0.1482 |
| Wbp11 | 13121 | 13444 | 13097 | 18001 | 13222 | 13036 | 13937 | 13427 | 13151 | 14151 | 14226 | 14270 | 13860.2 | 13505 | 14216 | 1.053 | 0.074 | 0.19117184 | 0.39617243 | 3.00E-04 | 0.0071 | 0.0022 | 0.0022 | TRUE | 0.0096 |
| Wbp1l | 2007 | 1881 | 1918 | 2879 | 2196 | 1897 | 2132 | 1879 | 1926 | 2263 | 2363 | 2077 | 2106.47 | 1979 | 2234 | 1.129 | 0.176 | 0.05774672 | 0.18150285 | 0.0037 | 0.0085 | 0.0057 | 0.0057 | TRUE | 0.0552 |
| Wbp2 | 7788 | 9019 | 9268 | 9126 | 6521 | 7709 | 8272 | 9007 | 9306 | 7174 | 7016 | 8439 | 8202.45 | 8862 | 7543 | 0.851 | -0.233 | 0.01955792 | 0.08547172 | 0.007 | 0.0073 | 0.007 | 0.007 | TRUE | 0.1314 |
| Wbp2nl | 84 | 91 | 83 | 151 | 110 | 94 | 89 | 91 | 83 | 119 | 118 | 103 | 100.57 | 88 | 113 | 1.293 | 0.371 | 0.05989938 | 0.18563561 | 0 | 0.0397 | 0.0178 | 0.0178 | TRUE | 0.0614 |
| Wbp4 | 1386 | 1557 | 1463 | 2126 | 1511 | 1491 | 1472 | 1555 | 1469 | 1671 | 1626 | 1632 | 1570.89 | 1499 | 1643 | 1.097 | 0.133 | 0.06889048 | 0.20357906 | 0 | 0.009 | 0.0032 | 0.0032 | TRUE | 0.013 |
| Wbp5 | 3443 | 4381 | 4142 | 5323 | 3805 | 4128 | 3657 | 4375 | 4159 | 4185 | 4094 | 4519 | 4164.77 | 4064 | 4266 | 1.05 | 0.07 | 0.46233696 | 0.66599473 | 0.0047 | 0.0077 | 0.0062 | 0.0062 | TRUE | 0.0931 |
| Wbp7 | 6144 | 5754 | 6301 | 7598 | 5635 | 5591 | 6526 | 5747 | 6327 | 5973 | 6063 | 6120 | 6125.92 | 6200 | 6052 | 0.976 | -0.035 | 0.64388389 | 0.79975876 | 0.0018 | 0.0074 | 0.0039 | 0.0039 | TRUE | 0.0499 |
| Wbscr16 | 1525 | 1785 | 1801 | 2020 | 1547 | 1479 | 1620 | 1783 | 1808 | 1588 | 1664 | 1619 | 1680.39 | 1737 | 1624 | 0.934 | -0.098 | 0.23096755 | 0.44489489 | 0.0014 | 0.0089 | 0.0042 | 0.0042 | TRUE | 0.0418 |
| Wbscr17 | 136 | 149 | 197 | 204 | 163 | 128 | 144 | 149 | 198 | 160 | 175 | 140 | 161.16 | 164 | 158 | 0.969 | -0.045 | 0.8161539 | 0.90637793 | 0.0154 | 0.0273 | 0.0214 | 0.0214 | TRUE | 0.354 |
| Wbscr22 | 2686 | 2292 | 2108 | 3208 | 2181 | 1843 | 2853 | 2289 | 2117 | 2522 | 2347 | 2017 | 2357.44 | 2420 | 2295 | 0.949 | -0.075 | 0.58678512 | 0.76150238 | 0.0181 | 0.0083 | 0.0135 | 0.0135 | TRUE | 0.2973 |
| Wbscr25 | 3 | 1 | 1 | 3 | 4 | 1 | 3 | 1 | 1 | 2 | 4 | 1 | 2.16 | 2 | 2 | 1.503 | 0.588 | 0.65402735 | NA | 0 | 1.5337 | 0.7527 | 0.7527 | TRUE | 0.3102 |
| Wbscr27 | 324 | 494 | 478 | 515 | 413 | 426 | 344 | 493 | 480 | 405 | 444 | 466 | 438.83 | 439 | 438 | 0.997 | -0.004 | 0.97927061 | 0.98867206 | 0.0191 | 0.0144 | 0.0168 | 0.0168 | TRUE | 0.4114 |
| Wbscr28 | 1 | 0 | 0 | 0 | 0 | 0 | 1 | 0 | 0 | 0 | 0 | 0 | 0.18 | 0 | 0 | 0.49 | -1.029 | 0.80087 | NA | 0 | 18.6165 | 10 | 10 | TRUE | 0.4614 |
| Wdfy1 | 3288 | 4593 | 4894 | 3994 | 3267 | 3859 | 3492 | 4587 | 4914 | 3140 | 3515 | 4224 | 3978.79 | 4331 | 3626 | 0.837 | -0.257 | 0.09921416 | 0.25984818 | 0.0259 | 0.0077 | 0.0172 | 0.0172 | TRUE | 0.3492 |
| Wdfy2 | 476 | 523 | 574 | 612 | 476 | 429 | 506 | 522 | 576 | 481 | 512 | 470 | 511.19 | 535 | 488 | 0.911 | -0.134 | 0.2204407 | 0.43243666 | 0.0014 | 0.0133 | 0.0066 | 0.0066 | TRUE | 0.0543 |
| Wdfy3 | 2476 | 2078 | 2434 | 2859 | 2100 | 1826 | 2630 | 2075 | 2444 | 2248 | 2259 | 1999 | 2275.85 | 2383 | 2169 | 0.91 | -0.136 | 0.2308868 | 0.44485903 | 0.0087 | 0.0084 | 0.0088 | 0.0088 | TRUE | 0.1549 |
| Wdfy4 | 135 | 172 | 146 | 182 | 153 | 130 | 143 | 172 | 147 | 143 | 165 | 142 | 151.96 | 154 | 150 | 0.973 | -0.04 | 0.81851659 | 0.9072905 | 0 | 0.0286 | 0.0151 | 0.0151 | TRUE | 0.1091 |
| Wdhd1 | 4677 | 4911 | 5122 | 7024 | 5226 | 5145 | 4968 | 4905 | 5143 | 5522 | 5623 | 5632 | 5298.68 | 5005 | 5592 | 1.117 | 0.16 | 0.00687097 | 0.03957958 | 2.00E-04 | 0.0075 | 0.0023 | 0.0023 | TRUE | 0.0071 |
| Wdpcp | 70 | 47 | 44 | 95 | 59 | 45 | 74 | 47 | 44 | 75 | 63 | 49 | 58.82 | 55 | 62 | 1.14 | 0.189 | 0.54961984 | 0.73406758 | 0.0461 | 0.0629 | 0.0547 | 0.0547 | TRUE | 0.5914 |
| Wdr1 | 3897 | 3315 | 3213 | 6754 | 4845 | 4331 | 4139 | 3311 | 3226 | 5309 | 5213 | 4741 | 4323.26 | 3559 | 5088 | 1.43 | 0.516 | 9.88E-06 | 0.00019646 | 0.0115 | 0.0077 | 0.0096 | 0.0096 | TRUE | 0.2474 |
| Wdr11 | 3835 | 4350 | 4373 | 5008 | 3712 | 3770 | 4073 | 4344 | 4391 | 3937 | 3994 | 4127 | 4144.4 | 4269 | 4019 | 0.941 | -0.088 | 0.19757795 | 0.40433284 | 7.00E-04 | 0.0077 | 0.0031 | 0.0031 | TRUE | 0.0196 |
| Wdr12 | 3945 | 4516 | 4390 | 5954 | 4118 | 4063 | 4190 | 4510 | 4408 | 4681 | 4431 | 4448 | 4444.56 | 4369 | 4520 | 1.035 | 0.049 | 0.47250074 | 0.67362214 | 8.00E-04 | 0.0076 | 0.0031 | 0.0031 | TRUE | 0.0156 |
| Wdr13 | 373 | 550 | 527 | 565 | 430 | 543 | 396 | 549 | 529 | 444 | 463 | 594 | 495.98 | 491 | 500 | 1.017 | 0.024 | 0.89195173 | 0.94755432 | 0.0259 | 0.0135 | 0.0198 | 0.0198 | TRUE | 0.3332 |
| Wdr16 | 37 | 29 | 35 | 40 | 33 | 19 | 39 | 29 | 35 | 31 | 36 | 21 | 31.86 | 34 | 29 | 0.852 | -0.231 | 0.52603961 | 0.71529414 | 0 | 0.1103 | 0.0641 | 0.0641 | TRUE | 0.3803 |
| Wdr17 | 55 | 70 | 85 | 91 | 54 | 57 | 58 | 70 | 85 | 72 | 58 | 62 | 67.62 | 71 | 64 | 0.901 | -0.15 | 0.56098645 | 0.74256648 | 0.0093 | 0.0556 | 0.0332 | 0.0332 | TRUE | 0.2528 |
| Wdr18 | 5907 | 5468 | 5494 | 8084 | 6039 | 5215 | 6274 | 5461 | 5517 | 6355 | 6498 | 5709 | 5968.84 | 5751 | 6187 | 1.076 | 0.106 | 0.25660996 | 0.47519694 | 0.0052 | 0.0075 | 0.0061 | 0.0061 | TRUE | 0.0773 |
| Wdr19 | 666 | 790 | 868 | 798 | 617 | 730 | 707 | 789 | 872 | 627 | 664 | 799 | 743.04 | 789 | 697 | 0.882 | -0.182 | 0.17814033 | 0.37886047 | 0.0118 | 0.0113 | 0.0118 | 0.0118 | TRUE | 0.1934 |
| Wdr20a | 742 | 811 | 849 | 1113 | 762 | 792 | 788 | 810 | 852 | 875 | 820 | 867 | 835.39 | 817 | 854 | 1.046 | 0.065 | 0.46786837 | 0.67016801 | 0 | 0.0108 | 0.0045 | 0.0045 | TRUE | 0.0173 |
| Wdr20b | 56 | 63 | 53 | 175 | 165 | 134 | 59 | 63 | 53 | 138 | 178 | 147 | 106.23 | 58 | 154 | 2.623 | 1.391 | 1.35E-10 | 9.78E-09 | 0 | 0.0379 | 0.0219 | 0.0219 | TRUE | 0.1861 |

| | | | | | | | | | | | | | | | | | | | | | | | | | |
|---|---|---|---|---|---|---|---|---|---|---|---|---|---|---|---|---|---|---|---|---|---|---|---|---|---|
| Wdr24 | 1729 | 1749 | 1613 | 1996 | 1587 | 1457 | 1837 | 1747 | 1620 | 1569 | 1708 | 1595 | 1679.07 | 1735 | 1624 | 0.936 | -0.095 | 0.27444248 | 0.49488215 | 0.0024 | 0.0089 | 0.0049 | 0.0049 | TRUE | 0.0404 |
| Wdr25 | 278 | 351 | 368 | 417 | 298 | 301 | 295 | 351 | 370 | 328 | 321 | 329 | 332.21 | 339 | 326 | 0.963 | -0.055 | 0.68098818 | 0.82315646 | 0.0036 | 0.0168 | 0.0098 | 0.0098 | TRUE | 0.1399 |
| Wdr26 | 7549 | 8149 | 8481 | 10507 | 7707 | 7670 | 8018 | 8138 | 8516 | 8260 | 8292 | 8396 | 8270.12 | 8224 | 8316 | 1.011 | 0.016 | 0.78761669 | 0.89063883 | 3.00E-04 | 0.0073 | 0.0024 | 0.0024 | TRUE | 0.0118 |
| Wdr27 | 13 | 4 | 10 | 9 | 13 | 7 | 14 | 4 | 10 | 7 | 14 | 8 | 9.43 | 9 | 10 | 1.027 | 0.039 | 0.95595429 | 0.97741618 | 0.1246 | 0.3563 | 0.246 | 0.246 | TRUE | 0.3748 |
| Wdr3 | 5097 | 4500 | 4552 | 7045 | 4911 | 4526 | 5414 | 4494 | 4571 | 5538 | 5284 | 4954 | 5042.56 | 4826 | 5259 | 1.09 | 0.124 | 0.2164378 | 0.4284208 | 0.0068 | 0.0076 | 0.0071 | 0.0071 | TRUE | 0.1381 |
| Wdr31 | 561 | 783 | 795 | 1074 | 790 | 804 | 596 | 782 | 798 | 844 | 850 | 880 | 791.75 | 725 | 858 | 1.182 | 0.242 | 0.06669702 | 0.19928507 | 0.0114 | 0.0111 | 0.0112 | 0.0112 | TRUE | 0.2867 |
| Wdr33 | 6226 | 5432 | 5556 | 8586 | 5877 | 5104 | 6613 | 5425 | 5579 | 6750 | 6323 | 5587 | 6046.16 | 5872 | 6220 | 1.059 | 0.083 | 0.4605593 | 0.66464831 | 0.0092 | 0.0074 | 0.009 | 0.009 | TRUE | 0.1484 |
| Wdr34 | 1496 | 1881 | 2077 | 2038 | 1531 | 1640 | 1589 | 1879 | 2086 | 1602 | 1647 | 1795 | 1766.29 | 1851 | 1681 | 0.908 | -0.139 | 0.24397651 | 0.46056223 | 0.01 | 0.0088 | 0.0097 | 0.0097 | TRUE | 0.1847 |
| Wdr35 | 2515 | 2876 | 2919 | 3808 | 2757 | 2989 | 2671 | 2872 | 2931 | 2994 | 2966 | 3272 | 2951.08 | 2825 | 3077 | 1.089 | 0.123 | 0.12849716 | 0.30823683 | 0.0022 | 0.008 | 0.0044 | 0.0044 | TRUE | 0.0369 |
| Wdr36 | 9303 | 10686 | 10444 | 11772 | 8658 | 9294 | 9881 | 10672 | 10487 | 9254 | 9315 | 10174 | 9963.99 | 10347 | 9581 | 0.926 | -0.111 | 0.13318014 | 0.31506182 | 0.0021 | 0.0072 | 0.0038 | 0.0038 | TRUE | 0.0357 |
| Wdr37 | 1783 | 1878 | 1968 | 2297 | 1705 | 1742 | 1894 | 1876 | 1976 | 1806 | 1834 | 1907 | 1882.1 | 1915 | 1849 | 0.965 | -0.051 | 0.4751766 | 0.67546567 | 0 | 0.0087 | 0.0032 | 0.0032 | TRUE | 0.0094 |
| Wdr38 | 0 | 1 | 1 | 1 | 0 | 0 | 0 | 1 | 1 | 1 | 0 | 0 | 0.46 | 1 | 0 | 0.52 | -0.943 | 0.74346893 | NA | 0 | 7.0944 | 4.2167 | 4.2167 | TRUE | 0.2562 |
| Wdr4 | 2102 | 1741 | 1626 | 2379 | 1759 | 1597 | 2233 | 1739 | 1633 | 1870 | 1893 | 1748 | 1852.51 | 1868 | 1837 | 0.984 | -0.024 | 0.85522413 | 0.92800791 | 0.0143 | 0.0087 | 0.0117 | 0.0117 | TRUE | 0.3517 |
| Wdr41 | 1612 | 1782 | 1751 | 2258 | 1639 | 1644 | 1712 | 1780 | 1758 | 1775 | 1763 | 1800 | 1764.71 | 1750 | 1779 | 1.017 | 0.024 | 0.72771189 | 0.85359749 | 0 | 0.0088 | 0.0028 | 0.0028 | TRUE | 0.0043 |
| Wdr43 | 11272 | 12232 | 12703 | 13549 | 10202 | 10413 | 11973 | 12216 | 12755 | 10651 | 10977 | 11399 | 11661.8 | 12315 | 11009 | 0.894 | -0.162 | 0.01193236 | 0.05995451 | 0.001 | 0.0072 | 0.0029 | 0.0029 | TRUE | 0.012 |
| Wdr44 | 852 | 607 | 617 | 1065 | 688 | 577 | 905 | 606 | 620 | 837 | 740 | 632 | 723.3 | 710 | 736 | 1.038 | 0.054 | 0.76937163 | 0.87946546 | 0.0346 | 0.0115 | 0.0228 | 0.0228 | TRUE | 0.6782 |
| Wdr45 | 217 | 164 | 175 | 521 | 339 | 252 | 230 | 164 | 176 | 410 | 365 | 276 | 270.03 | 190 | 350 | 1.847 | 0.885 | 1.38E-05 | 0.00026131 | 0.0326 | 0.0191 | 0.0258 | 0.0258 | TRUE | 0.3002 |
| Wdr45b | 7347 | 7065 | 7081 | 9213 | 6474 | 6458 | 7804 | 7056 | 7110 | 7243 | 6966 | 7069 | 7207.87 | 7323 | 7093 | 0.969 | -0.046 | 0.5193309 | 0.71052108 | 0.0016 | 0.0074 | 0.0035 | 0.0035 | TRUE | 0.0402 |
| Wdr46 | 3493 | 2439 | 2505 | 4755 | 3394 | 2551 | 3710 | 2436 | 2515 | 3738 | 3652 | 2792 | 3140.59 | 2887 | 3394 | 1.176 | 0.234 | 0.19673654 | 0.40329202 | 0.0393 | 0.008 | 0.0233 | 0.0233 | TRUE | 0.7547 |
| Wdr47 | 1430 | 1207 | 1217 | 1787 | 1357 | 1194 | 1519 | 1205 | 1222 | 1405 | 1460 | 1307 | 1353.04 | 1315 | 1391 | 1.057 | 0.081 | 0.49531356 | 0.691335 | 0.0093 | 0.0093 | 0.0093 | 0.0093 | TRUE | 0.2193 |
| Wdr48 | 5126 | 5059 | 5158 | 6123 | 4426 | 4753 | 5445 | 5052 | 5179 | 4813 | 4762 | 5203 | 5075.8 | 5225 | 4926 | 0.943 | -0.085 | 0.24533139 | 0.4621795 | 0.0016 | 0.0076 | 0.0037 | 0.0037 | TRUE | 0.0292 |
| Wdr5 | 10766 | 11360 | 10992 | 14062 | 9885 | 10026 | 11435 | 11345 | 11037 | 11054 | 10636 | 10975 | 11080.5 | 11272 | 10888 | 0.966 | -0.05 | 0.37602885 | 0.59147378 | 2.00E-04 | 0.0072 | 0.0022 | 0.0022 | TRUE | 0.005 |
| Wdr52 | 29 | 35 | 22 | 32 | 23 | 22 | 31 | 35 | 22 | 25 | 25 | 24 | 26.97 | 29 | 25 | 0.843 | -0.246 | 0.51226512 | 0.70574497 | 0 | 0.129 | 0.0645 | 0.0645 | TRUE | 0.3077 |
| Wdr53 | 473 | 486 | 455 | 651 | 441 | 408 | 502 | 485 | 457 | 512 | 474 | 447 | 479.59 | 481 | 478 | 0.994 | -0.009 | 0.93561203 | 0.96811881 | 0.0016 | 0.0138 | 0.0069 | 0.0069 | TRUE | 0.048 |
| Wdr54 | 62 | 74 | 68 | 103 | 71 | 69 | 66 | 74 | 68 | 81 | 76 | 76 | 73.49 | 69 | 78 | 1.121 | 0.165 | 0.45918295 | 0.66378277 | 0 | 0.0517 | 0.022 | 0.022 | TRUE | 0.0299 |
| Wdr55 | 2538 | 2133 | 2086 | 3157 | 2310 | 1913 | 2696 | 2130 | 2095 | 2482 | 2485 | 2094 | 2330.32 | 2307 | 2354 | 1.021 | 0.03 | 0.81954039 | 0.9078134 | 0.0144 | 0.0083 | 0.0116 | 0.0116 | TRUE | 0.2631 |
| Wdr59 | 5174 | 5447 | 5496 | 5885 | 4310 | 4647 | 5496 | 5440 | 5519 | 4626 | 4637 | 5087 | 5134.12 | 5485 | 4783 | 0.872 | -0.198 | 0.0048518 | 0.03065811 | 0.0012 | 0.0075 | 0.0034 | 0.0034 | TRUE | 0.0373 |
| Wdr5b | 20 | 28 | 25 | 44 | 48 | 19 | 21 | 28 | 25 | 35 | 52 | 21 | 30.22 | 25 | 36 | 1.439 | 0.525 | 0.2186381 | 0.43077078 | 0.0768 | 0.1159 | 0.0969 | 0.0969 | TRUE | 0.1709 |
| Wdr6 | 6790 | 4425 | 4103 | 10167 | 6680 | 4893 | 7212 | 4419 | 4120 | 7992 | 7187 | 5356 | 6047.87 | 5250 | 6845 | 1.304 | 0.383 | 0.07831138 | 0.2214575 | 0.072 | 0.0074 | 0.0339 | 0.0339 | TRUE | 1.302 |
| Wdr60 | 455 | 426 | 419 | 698 | 528 | 470 | 483 | 425 | 421 | 549 | 568 | 514 | 493.46 | 443 | 544 | 1.228 | 0.296 | 0.00885517 | 0.048077 | 0.0019 | 0.0136 | 0.0072 | 0.0072 | TRUE | 0.0719 |
| Wdr61 | 3990 | 3982 | 3969 | 4952 | 3594 | 3519 | 4238 | 3977 | 3985 | 3893 | 3867 | 3852 | 3968.69 | 4067 | 3871 | 0.952 | -0.071 | 0.26745502 | 0.48710169 | 4.00E-04 | 0.0077 | 0.0027 | 0.0027 | TRUE | 0.0165 |
| Wdr62 | 2881 | 2579 | 2525 | 2895 | 2275 | 2092 | 3060 | 2576 | 2535 | 2276 | 2448 | 2290 | 2530.8 | 2724 | 2338 | 0.858 | -0.22 | 0.02843487 | 0.11092624 | 0.006 | 0.0082 | 0.0069 | 0.0069 | TRUE | 0.1414 |
| Wdr63 | 25 | 21 | 17 | 44 | 44 | 28 | 27 | 21 | 17 | 35 | 47 | 31 | 29.53 | 22 | 38 | 1.743 | 0.801 | 0.03718938 | 0.13361068 | 0 | 0.1185 | 0.0697 | 0.0697 | TRUE | 0.3606 |
| Wdr64 | 64 | 42 | 31 | 84 | 80 | 74 | 68 | 42 | 31 | 66 | 86 | 81 | 62.36 | 47 | 78 | 1.652 | 0.724 | 0.02805526 | 0.11003973 | 0.0629 | 0.0597 | 0.0611 | 0.0611 | TRUE | 1.0894 |
| Wdr65 | 21 | 11 | 25 | 22 | 20 | 13 | 22 | 11 | 25 | 17 | 22 | 14 | 18.57 | 19 | 18 | 0.909 | -0.138 | 0.78004203 | 0.88617899 | 0.0488 | 0.1843 | 0.1209 | 0.1209 | TRUE | 0.7873 |
| Wdr67 | 2504 | 2645 | 2747 | 2964 | 2265 | 2440 | 2660 | 2642 | 2758 | 2330 | 2437 | 2671 | 2582.93 | 2687 | 2479 | 0.922 | -0.117 | 0.15584576 | 0.34845468 | 0.0022 | 0.0082 | 0.0045 | 0.0045 | TRUE | 0.0552 |
| Wdr69 | 5 | 15 | 16 | 11 | 8 | 5 | 5 | 15 | 16 | 9 | 9 | 5 | 9.85 | 12 | 8 | 0.627 | -0.674 | 0.31608016 | 0.53684153 | 0 | 0.3414 | 0.2192 | 0.2192 | TRUE | 0.9133 |
| Wdr7 | 1711 | 1449 | 1563 | 2038 | 1423 | 1343 | 1817 | 1447 | 1569 | 1602 | 1531 | 1470 | 1572.87 | 1611 | 1534 | 0.953 | -0.07 | 0.52058201 | 0.71109881 | 0.0065 | 0.009 | 0.0079 | 0.0079 | TRUE | 0.1505 |
| Wdr70 | 1882 | 1767 | 1748 | 2165 | 1536 | 1507 | 1999 | 1765 | 1755 | 1702 | 1653 | 1650 | 1753.86 | 1840 | 1668 | 0.907 | -0.141 | 0.10317029 | 0.26735607 | 0.0021 | 0.0088 | 0.0048 | 0.0048 | TRUE | 0.0691 |
| Wdr72 | 16 | 15 | 11 | 19 | 10 | 13 | 17 | 15 | 11 | 15 | 11 | 14 | 13.82 | 14 | 13 | 0.934 | -0.099 | 0.84612109 | 0.92282842 | 0 | 0.2452 | 0.1154 | 0.1154 | TRUE | 0.1806 |
| Wdr73 | 1810 | 1711 | 1514 | 2348 | 1719 | 1663 | 1923 | 1709 | 1520 | 1846 | 1850 | 1820 | 1777.89 | 1717 | 1839 | 1.071 | 0.099 | 0.34449408 | 0.5641524 | 0.006 | 0.0088 | 0.0073 | 0.0073 | TRUE | 0.1317 |
| Wdr74 | 2857 | 2865 | 2688 | 3693 | 2738 | 2422 | 3035 | 2861 | 2699 | 2903 | 2946 | 2651 | 2849.22 | 2865 | 2833 | 0.989 | -0.016 | 0.85454181 | 0.92756327 | 0.0029 | 0.0081 | 0.0049 | 0.0049 | TRUE | 0.038 |
| Wdr75 | 6724 | 6835 | 6793 | 8802 | 6191 | 6118 | 7142 | 6826 | 6821 | 6919 | 6661 | 6697 | 6844.47 | 6930 | 6759 | 0.976 | -0.036 | 0.55341145 | 0.73694743 | 3.00E-04 | 0.0074 | 0.0025 | 0.0025 | TRUE | 0.0088 |
| Wdr76 | 3639 | 4731 | 4695 | 5124 | 3986 | 4529 | 3865 | 4725 | 4714 | 4028 | 4289 | 4958 | 4429.82 | 4435 | 4425 | 0.997 | -0.004 | 0.97571521 | 0.98697924 | 0.012 | 0.0076 | 0.01 | 0.01 | TRUE | 0.1536 |
| Wdr77 | 13815 | 12251 | 11882 | 16951 | 11859 | 11293 | 14674 | 12235 | 11931 | 13326 | 12760 | 12362 | 12881.17 | 12947 | 12816 | 0.99 | -0.015 | 0.88427237 | 0.94317602 | 0.006 | 0.0072 | 0.0071 | 0.0071 | TRUE | 0.1665 |
| Wdr78 | 42 | 34 | 32 | 43 | 31 | 22 | 45 | 34 | 32 | 34 | 33 | 24 | 33.66 | 37 | 30 | 0.829 | -0.27 | 0.4350145 | 0.64338417 | 0 | 0.1048 | 0.0566 | 0.0566 | TRUE | 0.2355 |
| Wdr8 | 830 | 818 | 781 | 1225 | 912 | 843 | 882 | 817 | 784 | 963 | 981 | 923 | 891.63 | 828 | 956 | 1.155 | 0.208 | 0.02307784 | 0.09626652 | 9.00E-04 | 0.0106 | 0.0049 | 0.0049 | TRUE | 0.0384 |
| Wdr81 | 1175 | 1246 | 1358 | 1916 | 1440 | 1196 | 1248 | 1244 | 1364 | 1506 | 1549 | 1309 | 1370.13 | 1285 | 1455 | 1.132 | 0.179 | 0.07419536 | 0.21401045 | 0.0042 | 0.0093 | 0.0065 | 0.0065 | TRUE | 0.091 |
| Wdr82 | 8413 | 8909 | 8704 | 11181 | 8219 | 7510 | 8936 | 8897 | 8740 | 8790 | 8843 | 8221 | 8737.83 | 8858 | 8618 | 0.973 | -0.039 | 0.52963386 | 0.71789091 | 7.00E-04 | 0.0073 | 0.0027 | 0.0027 | TRUE | 0.0197 |
| Wdr83 | 690 | 668 | 619 | 935 | 656 | 528 | 733 | 667 | 622 | 735 | 706 | 578 | 673.4 | 674 | 673 | 1 | 0 | 0.99965781 | 0.9997212 | 0.0099 | 0.0118 | 0.0108 | 0.0108 | TRUE | 0.1772 |
| Wdr85 | 727 | 839 | 937 | 1168 | 869 | 799 | 772 | 838 | 941 | 918 | 935 | 875 | 879.8 | 850 | 909 | 1.069 | 0.097 | 0.36334715 | 0.58083215 | 0.0043 | 0.0106 | 0.007 | 0.007 | TRUE | 0.1032 |
| Wdr86 | 289 | 341 | 379 | 323 | 206 | 206 | 307 | 341 | 381 | 254 | 222 | 225 | 288.19 | 343 | 234 | 0.683 | -0.549 | 0.00014843 | 0.00188841 | 0.0056 | 0.0183 | 0.0115 | 0.0115 | TRUE | 0.1065 |
| Wdr89 | 936 | 1105 | 1205 | 1186 | 896 | 839 | 994 | 1104 | 1210 | 932 | 964 | 918 | 1020.42 | 1103 | 938 | 0.851 | -0.233 | 0.02356142 | 0.09763248 | 0.0042 | 0.0101 | 0.0067 | 0.0067 | TRUE | 0.088 |
| Wdr90 | 910 | 1074 | 1074 | 1471 | 1249 | 1204 | 967 | 1073 | 1078 | 1156 | 1344 | 1318 | 1155.97 | 1039 | 1273 | 1.223 | 0.291 | 0.00431944 | 0.02798941 | 0.0044 | 0.0098 | 0.0066 | 0.0066 | TRUE | 0.079 |
| Wdr91 | 1816 | 1675 | 1684 | 2400 | 1707 | 1687 | 1929 | 1673 | 1691 | 1887 | 1837 | 1847 | 1810.45 | 1764 | 1857 | 1.053 | 0.074 | 0.39880454 | 0.61201859 | 0.0026 | 0.0087 | 0.005 | 0.005 | TRUE | 0.0802 |
| Wdr92 | 2336 | 2675 | 2649 | 3356 | 2451 | 2349 | 2481 | 2672 | 2660 | 2638 | 2637 | 2571 | 2609.89 | 2604 | 2615 | 1.004 | 0.006 | 0.92839107 | 0.96476961 | 5.00E-04 | 0.0082 | 0.0031 | 0.0031 | TRUE | 0.0206 |
| Wdr93 | 78 | 122 | 112 | 102 | 79 | 96 | 83 | 122 | 112 | 80 | 85 | 105 | 97.9 | 106 | 90 | 0.849 | -0.237 | 0.3180212 | 0.53887305 | 0.0197 | 0.0406 | 0.0303 | 0.0303 | TRUE | 0.3467 |
| Wdr95 | 18 | 17 | 14 | 50 | 38 | 63 | 19 | 17 | 14 | 39 | 41 | 69 | 33.22 | 17 | 50 | 2.963 | 1.567 | 0.00010512 | 0.00143422 | 0.0493 | 0.1061 | 0.0772 | 0.0772 | TRUE | 0.8936 |
| Wdr96 | 96 | 146 | 174 | 186 | 159 | 138 | 102 | 146 | 175 | 146 | 171 | 151 | 148.48 | 141 | 156 | 1.106 | 0.146 | 0.51904675 | 0.71037146 | 0.0311 | 0.0291 | 0.03 | 0.03 | TRUE | 0.362 |
| Wdsub1 | 928 | 857 | 786 | 1114 | 809 | 786 | 986 | 856 | 789 | 876 | 870 | 860 | 872.9 | 877 | 869 | 0.991 | -0.013 | 0.90776085 | 0.95354464 | 0.0052 | 0.0107 | 0.0076 | 0.0076 | TRUE | 0.1393 |
| Wdtc1 | 10648 | 9697 | 9412 | 16062 | 11214 | 10356 | 11310 | 9684 | 9451 | 12627 | 12066 | 11336 | 11078.95 | 10148 | 12010 | 1.184 | 0.243 | 0.01154563 | 0.0585335 | 0.0063 | 0.0072 | 0.0066 | 0.0066 | TRUE | 0.1225 |
| Wdyhv1 | 606 | 559 | 595 | 690 | 558 | 470 | 644 | 558 | 597 | 542 | 600 | 514 | 576.11 | 600 | 552 | 0.921 | -0.119 | 0.29882337 | 0.51926751 | 0.0037 | 0.0126 | 0.0077 | 0.0077 | TRUE | 0.0654 |
| Wee1 | 3513 | 4100 | 4236 | 4715 | 3468 | 3711 | 3731 | 4095 | 4253 | 3707 | 3731 | 4062 | 3929.95 | 4026 | 3833 | 0.952 | -0.071 | 0.4016967 | 0.61434455 | 0.0032 | 0.0077 | 0.0049 | 0.0049 | TRUE | 0.0499 |
| Wee2 | 2 | 2 | 2 | 5 | 1 | 3 | 2 | 2 | 2 | 4 | 1 | 3 | 2.4 | 2 | 3 | 1.374 | 0.458 | 0.70068582 | NA | 0 | 1.3776 | 0.6003 | 0.6003 | TRUE | 0.3135 |
| Wfdc1 | 18 | 31 | 39 | 33 | 20 | 26 | 19 | 31 | 39 | 26 | 22 | 28 | 27.53 | 30 | 25 | 0.85 | -0.235 | 0.56317472 | 0.74414927 | 0.034 | 0.1266 | 0.0828 | 0.0828 | TRUE | 0.2612 |
| Wfdc10 | 64 | 57 | 68 | 77 | 41 | 57 | 68 | 57 | 68 | 61 | 44 | 62 | 60.04 | 64 | 56 | 0.867 | -0.205 | 0.44074559 | 0.64779047 | 0 | 0.0618 | 0.0344 | 0.0344 | TRUE | 0.2612 |
| Wfdc11 | 0 | 0 | 0 | 0 | 0 | 0 | 0 | 0 | 0 | 0 | 0 | 0 | 0 | 0 | 0 | NA | NA | NA | NA | NA | NA | NA | NA | NA | NA |
| Wfdc12 | 1 | 0 | 1 | 2 | 3 | 0 | 1 | 0 | 1 | 2 | 3 | 0 | 1.14 | 1 | 2 | 2.338 | 1.225 | 0.53592605 | NA | 0 | 2.8857 | 1.7735 | 1.7735 | TRUE | 0.1009 |
| Wfdc13 | 1 | 2 | 3 | 3 | 2 | 1 | 1 | 2 | 3 | 2 | 2 | 1 | 1.95 | 2 | 2 | 0.931 | -0.103 | 0.936853 | NA | 0 | 1.6997 | 0.7066 | 0.7066 | TRUE | 0.166 |

| | | | | | | | | | | | | | | | | | | | | | | | | |
|---|---|---|---|---|---|---|---|---|---|---|---|---|---|---|---|---|---|---|---|---|---|---|---|---|
| Wfdc15a | 32 | 47 | 51 | 171 | 126 | 100 | 34 | 47 | 51 | 134 | 136 | 109 | 85.27 | 44 | 126 | 2.87 | 1.521 | 9.47E-10 | 5.83E-08 | 0.0078 | 0.0455 | 0.0291 | 0.0291 | TRUE | 0.2985 |
| Wfdc15b | 0 | 0 | 0 | 0 | 0 | 0 | 0 | 0 | 0 | 0 | 0 | 0 | 0 | 0 | 0 | NA | NA | NA | NA | NA | NA | NA | NA | NA | NA |
| Wfdc16 | 0 | 0 | 0 | 1 | 1 | 0 | 0 | 0 | 0 | 1 | 1 | 0 | 0.31 | 0 | 1 | 3.32 | 1.731 | 0.65383052 | NA | 0 | 10.6224 | 8.9541 | 8.9541 | TRUE | 0.202 |
| Wfdc17 | 0 | 0 | 0 | 0 | 0 | 0 | 0 | 0 | 0 | 0 | 0 | 0 | 0 | 0 | 0 | NA | NA | NA | NA | NA | NA | NA | NA | NA | NA |
| Wfdc18 | 0 | 0 | 0 | 0 | 0 | 0 | 0 | 0 | 0 | 0 | 0 | 0 | 0 | 0 | 0 | NA | NA | NA | NA | NA | NA | NA | NA | NA | NA |
| Wfdc2 | 1005 | 1891 | 1992 | 2154 | 1692 | 2063 | 1067 | 1889 | 2000 | 1693 | 1820 | 2258 | 1788.05 | 1652 | 1924 | 1.164 | 0.219 | 0.31594569 | 0.53679796 | 0.0655 | 0.0087 | 0.0339 | 0.0339 | TRUE | 1.1542 |
| Wfdc3 | 0 | 1 | 3 | 2 | 0 | 1 | 0 | 1 | 3 | 2 | 0 | 1 | 1.11 | 1 | 1 | 0.68 | -0.556 | 0.77748599 | NA | 0 | 2.9669 | 1.8788 | 1.8788 | TRUE | 0.3912 |
| Wfdc5 | 1 | 4 | 1 | 1 | 1 | 0 | 1 | 4 | 1 | 1 | 1 | 0 | 1.32 | 2 | 1 | 0.312 | -1.681 | 0.34405227 | NA | 0 | 2.5017 | 1.2515 | 1.2515 | TRUE | 0.6725 |
| Wfdc6a | 0 | 0 | 0 | 0 | 1 | 0 | 0 | 0 | 0 | 0 | 1 | 0 | 0.18 | 0 | 0 | 1.859 | 0.894 | 0.82650734 | NA | 0 | 18.3785 | 10 | 10 | TRUE | 0.5204 |
| Wfdc6b | 0 | 0 | 0 | 0 | 0 | 0 | 0 | 0 | 0 | 0 | 0 | 0 | 0 | 0 | 0 | NA | NA | NA | NA | NA | NA | NA | NA | NA | NA |
| Wfdc8 | 6 | 5 | 5 | 5 | 2 | 8 | 6 | 5 | 5 | 4 | 2 | 9 | 5.2 | 5 | 5 | 0.898 | -0.155 | 0.8577824 | 0.92990093 | 0 | 0.6399 | 0.3459 | 0.3459 | TRUE | 0.7924 |
| Wfdc9 | 4 | 2 | 4 | 2 | 0 | 1 | 4 | 2 | 4 | 2 | 0 | 1 | 2.15 | 3 | 1 | 0.272 | -1.877 | 0.18307362 | NA | 0 | 1.5357 | 0.7597 | 0.7597 | TRUE | 0.2789 |
| Wfikkn1 | 11 | 3 | 11 | 8 | 6 | 3 | 12 | 3 | 11 | 6 | 6 | 3 | 6.96 | 9 | 5 | 0.629 | -0.669 | 0.40603452 | 0.61874169 | 0 | 0.4803 | 0.3162 | 0.3162 | TRUE | 1.0229 |
| Wfikkn2 | 71 | 67 | 41 | 115 | 90 | 56 | 75 | 67 | 41 | 90 | 97 | 61 | 72.01 | 61 | 83 | 1.358 | 0.442 | 0.15522975 | 0.34749626 | 0.0585 | 0.0527 | 0.0554 | 0.0554 | TRUE | 0.713 |
| Wfs1 | 1330 | 1603 | 1488 | 1363 | 1158 | 1168 | 1413 | 1601 | 1494 | 1071 | 1246 | 1279 | 1350.62 | 1503 | 1199 | 0.797 | -0.328 | 0.00184466 | 0.01444641 | 0.0057 | 0.0093 | 0.0072 | 0.0072 | TRUE | 0.1062 |
| Whamm | 723 | 978 | 962 | 867 | 735 | 761 | 768 | 977 | 966 | 682 | 791 | 833 | 836.01 | 904 | 769 | 0.849 | -0.236 | 0.08241386 | 0.22928761 | 0.0131 | 0.0108 | 0.0121 | 0.0121 | TRUE | 0.2043 |
| Whrn | 764 | 937 | 813 | 706 | 667 | 765 | 812 | 936 | 816 | 555 | 718 | 837 | 778.95 | 855 | 703 | 0.822 | -0.283 | 0.07709398 | 0.21943021 | 0.0218 | 0.0111 | 0.0172 | 0.0172 | TRUE | 0.3535 |
| Whsc1 | 2571 | 3695 | 4010 | 6236 | 4356 | 4338 | 2731 | 3690 | 4026 | 4902 | 4687 | 4749 | 4130.87 | 3482 | 4779 | 1.372 | 0.457 | 0.00116008 | 0.00998171 | 0.0185 | 0.0077 | 0.014 | 0.014 | TRUE | 0.433 |
| Whsc1l1 | 2598 | 2553 | 2676 | 3263 | 2372 | 2355 | 2760 | 2550 | 2687 | 2565 | 2552 | 2578 | 2615.23 | 2666 | 2565 | 0.962 | -0.055 | 0.42081433 | 0.63116133 | 4.00E-04 | 0.0082 | 0.003 | 0.003 | TRUE | 0.0175 |
| Wibg | 1711 | 1596 | 1549 | 2506 | 1844 | 1533 | 1817 | 1594 | 1555 | 1970 | 1984 | 1678 | 1766.47 | 1655 | 1877 | 1.135 | 0.182 | 0.09381427 | 0.25057385 | 0.007 | 0.0088 | 0.0079 | 0.0079 | TRUE | 0.1033 |
| Wif1 | 121 | 96 | 99 | 123 | 112 | 95 | 129 | 96 | 99 | 97 | 121 | 104 | 107.5 | 108 | 107 | 0.99 | -0.014 | 0.94810321 | 0.97406219 | 0.0104 | 0.0375 | 0.0241 | 0.0241 | TRUE | 0.2712 |
| Wipf1 | 659 | 757 | 796 | 1048 | 730 | 761 | 700 | 756 | 799 | 824 | 785 | 833 | 782.93 | 752 | 814 | 1.083 | 0.115 | 0.23236055 | 0.44654725 | 0.0012 | 0.0111 | 0.0054 | 0.0054 | TRUE | 0.0426 |
| Wipf2 | 3064 | 3185 | 3280 | 3817 | 2853 | 2975 | 3255 | 3181 | 3293 | 3001 | 3070 | 3257 | 3175.96 | 3243 | 3109 | 0.958 | -0.061 | 0.37443494 | 0.59055878 | 7.00E-04 | 0.0079 | 0.0031 | 0.0031 | TRUE | 0.0208 |
| Wipf3 | 490 | 557 | 590 | 420 | 310 | 358 | 520 | 556 | 592 | 330 | 334 | 392 | 454.13 | 556 | 352 | 0.631 | -0.664 | 8.42E-08 | 3.10E-06 | 0.0041 | 0.0142 | 0.0087 | 0.0087 | TRUE | 0.11 |
| Wipi1 | 1185 | 1908 | 1881 | 1559 | 1309 | 1559 | 1259 | 1906 | 1889 | 1226 | 1408 | 1707 | 1565.58 | 1685 | 1447 | 0.859 | -0.22 | 0.22985242 | 0.44350775 | 0.0371 | 0.009 | 0.0236 | 0.0236 | TRUE | 0.5887 |
| Wipi2 | 3717 | 3783 | 3777 | 4866 | 3405 | 3267 | 3948 | 3778 | 3793 | 3825 | 3664 | 3576 | 3763.97 | 3840 | 3688 | 0.961 | -0.057 | 0.38794379 | 0.6017358 | 6.00E-04 | 0.0078 | 0.0029 | 0.0029 | TRUE | 0.0131 |
| Wisp1 | 48 | 41 | 33 | 45 | 30 | 17 | 51 | 41 | 33 | 35 | 32 | 19 | 35.22 | 42 | 29 | 0.696 | -0.523 | 0.16501553 | 0.36167622 | 0.0407 | 0.1004 | 0.073 | 0.073 | TRUE | 0.4075 |
| Wisp2 | 18 | 12 | 12 | 23 | 14 | 14 | 19 | 12 | 12 | 18 | 15 | 15 | 15.27 | 14 | 16 | 1.131 | 0.178 | 0.71735168 | 0.8467669 | 0 | 0.2226 | 0.1081 | 0.1081 | TRUE | 0.3526 |
| Wisp3 | 18 | 20 | 18 | 21 | 7 | 9 | 19 | 20 | 18 | 17 | 8 | 10 | 15.18 | 19 | 12 | 0.605 | -0.726 | 0.15939569 | 0.35368442 | 0 | 0.224 | 0.1227 | 0.1227 | TRUE | 0.7712 |
| Wiz | 3052 | 3163 | 3303 | 3327 | 2596 | 2836 | 3242 | 3159 | 3317 | 2615 | 2793 | 3104 | 3038.38 | 3239 | 2837 | 0.876 | -0.192 | 0.03091088 | 0.11776 | 0.0036 | 0.008 | 0.0053 | 0.0053 | TRUE | 0.0819 |
| Wls | 199 | 200 | 168 | 376 | 239 | 195 | 211 | 200 | 169 | 296 | 257 | 213 | 224.33 | 193 | 255 | 1.326 | 0.407 | 0.02339376 | 0.09714756 | 0.0159 | 0.0216 | 0.0186 | 0.0186 | TRUE | 0.1678 |
| Wnk1 | 30035 | 28873 | 33108 | 40493 | 29463 | 29090 | 31902 | 28836 | 33244 | 31832 | 31700 | 31844 | 31559.75 | 31327 | 31792 | 1.015 | 0.021 | 0.77950648 | 0.88603691 | 0.0021 | 0.007 | 0.0041 | 0.0041 | TRUE | 0.0593 |
| Wnk2 | 1107 | 1265 | 1385 | 943 | 798 | 888 | 1176 | 1263 | 1391 | 741 | 859 | 972 | 1066.98 | 1277 | 857 | 0.671 | -0.577 | 7.01E-06 | 0.00014676 | 0.0112 | 0.01 | 0.0109 | 0.0109 | TRUE | 0.1715 |
| Wnk3 | 4288 | 3293 | 3431 | 5887 | 3885 | 3427 | 4555 | 3289 | 3445 | 4628 | 4180 | 3751 | 3974.63 | 3763 | 4186 | 1.113 | 0.154 | 0.28286023 | 0.50438344 | 0.0201 | 0.0077 | 0.0146 | 0.0146 | TRUE | 0.412 |
| Wnk4 | 70 | 67 | 62 | 56 | 40 | 39 | 74 | 67 | 62 | 44 | 43 | 43 | 55.55 | 68 | 43 | 0.638 | -0.647 | 0.01207202 | 0.06036778 | 0 | 0.0662 | 0.0291 | 0.0291 | TRUE | 0.0613 |
| Wnt1 | 1 | 1 | 0 | 0 | 1 | 0 | 1 | 1 | 0 | 0 | 1 | 0 | 0.52 | 1 | 0 | 0.51 | -0.972 | 0.72797553 | NA | 0 | 6.3084 | 3.8879 | 3.8879 | TRUE | 0.5204 |
| Wnt10a | 13 | 11 | 18 | 16 | 10 | 10 | 14 | 11 | 18 | 13 | 11 | 11 | 12.86 | 14 | 12 | 0.803 | -0.316 | 0.54920407 | 0.7338626 | 0 | 0.2631 | 0.1226 | 0.1226 | TRUE | 0.2414 |
| Wnt10b | 4 | 3 | 3 | 4 | 6 | 4 | 4 | 3 | 3 | 3 | 6 | 4 | 4.04 | 3 | 4 | 1.349 | 0.432 | 0.640241 | NA | 0 | 0.8225 | 0.3596 | 0.3596 | TRUE | 0.2086 |
| Wnt11 | 77 | 76 | 85 | 115 | 67 | 81 | 82 | 76 | 85 | 90 | 72 | 89 | 82.37 | 81 | 84 | 1.036 | 0.051 | 0.81588089 | 0.90620235 | 0 | 0.0469 | 0.0229 | 0.0229 | TRUE | 0.1326 |
| Wnt16 | 24 | 21 | 27 | 14 | 16 | 29 | 25 | 21 | 27 | 11 | 17 | 32 | 22.26 | 24 | 20 | 0.804 | -0.315 | 0.52180086 | 0.71192414 | 0.0996 | 0.1549 | 0.1283 | 0.1283 | TRUE | 0.7737 |
| Wnt2 | 2 | 2 | 3 | 8 | 7 | 4 | 2 | 2 | 3 | 6 | 8 | 4 | 4.22 | 2 | 6 | 2.553 | 1.352 | 0.15095251 | NA | 0 | 0.7872 | 0.3438 | 0.3438 | TRUE | 0.1237 |
| Wnt2b | 15 | 18 | 14 | 46 | 23 | 37 | 16 | 18 | 14 | 36 | 25 | 41 | 24.9 | 16 | 34 | 2.117 | 1.082 | 0.00858731 | 0.04691342 | 0 | 0.1392 | 0.0758 | 0.0758 | TRUE | 0.3589 |
| Wnt3 | 25 | 36 | 37 | 46 | 21 | 35 | 27 | 36 | 37 | 36 | 23 | 38 | 32.79 | 33 | 32 | 0.976 | -0.035 | 0.9241923 | 0.96274037 | 0.0176 | 0.1074 | 0.0664 | 0.0664 | TRUE | 0.4463 |
| Wnt3a | 39 | 44 | 49 | 71 | 46 | 55 | 41 | 44 | 49 | 56 | 49 | 60 | 50.01 | 45 | 55 | 1.23 | 0.298 | 0.2737653 | 0.49433176 | 0 | 0.0728 | 0.0332 | 0.0332 | TRUE | 0.0627 |
| Wnt4 | 43 | 36 | 47 | 72 | 48 | 41 | 46 | 36 | 47 | 57 | 52 | 45 | 46.99 | 43 | 51 | 1.194 | 0.256 | 0.3758782 | 0.59147378 | 0 | 0.077 | 0.0387 | 0.0387 | TRUE | 0.1587 |
| Wnt5a | 62 | 73 | 87 | 53 | 47 | 70 | 66 | 73 | 87 | 42 | 51 | 77 | 65.83 | 75 | 57 | 0.741 | -0.433 | 0.1518544 | 0.34295352 | 0.0428 | 0.0569 | 0.0502 | 0.0502 | TRUE | 0.8187 |
| Wnt5b | 228 | 258 | 230 | 185 | 125 | 153 | 242 | 258 | 231 | 145 | 134 | 167 | 196.37 | 244 | 149 | 0.612 | -0.71 | 6.55E-06 | 0.00013919 | 0 | 0.0237 | 0.0125 | 0.0125 | TRUE | 0.113 |
| Wnt6 | 126 | 165 | 137 | 84 | 83 | 69 | 134 | 165 | 138 | 66 | 89 | 76 | 111.18 | 146 | 77 | 0.526 | -0.926 | 1.36E-05 | 0.00025862 | 0.0083 | 0.0365 | 0.0227 | 0.0227 | TRUE | 0.1793 |
| Wnt7a | 32 | 25 | 19 | 38 | 25 | 22 | 34 | 25 | 19 | 30 | 27 | 24 | 26.48 | 26 | 27 | 1.042 | 0.06 | 0.87910701 | 0.9402059 | 0 | 0.1313 | 0.0733 | 0.0733 | TRUE | 0.3355 |
| Wnt7b | 46 | 79 | 75 | 135 | 111 | 110 | 49 | 79 | 75 | 106 | 119 | 120 | 91.51 | 68 | 115 | 1.698 | 0.763 | 0.00168769 | 0.01344354 | 0.0153 | 0.0429 | 0.0308 | 0.0308 | TRUE | 0.5127 |
| Wnt8a | 25 | 24 | 14 | 38 | 19 | 11 | 27 | 24 | 14 | 30 | 20 | 12 | 21.16 | 22 | 21 | 0.978 | -0.032 | 0.94968 | 0.97481103 | 0.1008 | 0.1626 | 0.1329 | 0.1329 | TRUE | 0.2362 |
| Wnt8b | 11 | 7 | 12 | 21 | 10 | 8 | 12 | 7 | 12 | 17 | 11 | 9 | 11.12 | 10 | 12 | 1.189 | 0.25 | 0.67378508 | 0.81814907 | 0 | 0.303 | 0.1644 | 0.1644 | TRUE | 0.5239 |
| Wnt9a | 128 | 232 | 224 | 239 | 180 | 180 | 136 | 232 | 225 | 188 | 194 | 197 | 195.19 | 198 | 193 | 0.975 | -0.036 | 0.86840633 | 0.93526226 | 0.0357 | 0.0238 | 0.0296 | 0.0296 | TRUE | 0.7968 |
| Wnt9b | 5 | 1 | 4 | 7 | 2 | 6 | 5 | 1 | 4 | 6 | 2 | 7 | 4.09 | 3 | 5 | 1.389 | 0.474 | 0.63766152 | NA | 0 | 0.8121 | 0.4792 | 0.4792 | TRUE | 0.5779 |
| Wrap53 | 1624 | 1250 | 1132 | 2845 | 1928 | 1494 | 1725 | 1248 | 1137 | 2237 | 2074 | 1635 | 1676.06 | 1370 | 1982 | 1.447 | 0.534 | 0.00272576 | 0.01967535 | 0.0374 | 0.0089 | 0.0222 | 0.0222 | TRUE | 0.6168 |
| Wrb | 2517 | 2257 | 2168 | 3126 | 2168 | 2043 | 2673 | 2254 | 2177 | 2457 | 2333 | 2236 | 2355.15 | 2368 | 2342 | 0.989 | -0.015 | 0.88328266 | 0.94254355 | 0.0068 | 0.0083 | 0.0074 | 0.0074 | TRUE | 0.1538 |
| Wrn | 1557 | 1599 | 1644 | 2338 | 1599 | 1583 | 1654 | 1597 | 1651 | 1838 | 1720 | 1733 | 1698.79 | 1634 | 1764 | 1.08 | 0.111 | 0.13156576 | 0.31241421 | 0 | 0.0088 | 0.0033 | 0.0033 | TRUE | 0.0169 |
| Wrnip1 | 2215 | 2210 | 2202 | 3555 | 2400 | 2353 | 2353 | 2207 | 2211 | 2795 | 2582 | 2576 | 2453.93 | 2257 | 2651 | 1.175 | 0.233 | 0.00233291 | 0.01738879 | 0.0014 | 0.0082 | 0.0038 | 0.0038 | TRUE | 0.0278 |
| Wsb1 | 5200 | 6344 | 5666 | 7708 | 6470 | 6144 | 5523 | 6336 | 5689 | 6059 | 6961 | 6726 | 6215.79 | 5849 | 6582 | 1.125 | 0.17 | 0.06581669 | 0.19787232 | 0.005 | 0.0074 | 0.006 | 0.006 | TRUE | 0.0646 |
| Wsb2 | 5022 | 4658 | 4764 | 4090 | 2991 | 3029 | 5334 | 4652 | 4784 | 3215 | 3218 | 3316 | 4086.48 | 4923 | 3250 | 0.66 | -0.599 | 1.12E-13 | 1.33E-11 | 0.0025 | 0.0077 | 0.0044 | 0.0044 | TRUE | 0.0649 |
| Wscd1 | 449 | 601 | 594 | 662 | 471 | 438 | 477 | 600 | 596 | 520 | 507 | 479 | 530.04 | 558 | 502 | 0.9 | -0.151 | 0.23716813 | 0.45255895 | 0.0073 | 0.0131 | 0.0099 | 0.0099 | TRUE | 0.1871 |
| Wscd2 | 146 | 242 | 227 | 319 | 248 | 236 | 155 | 242 | 228 | 251 | 267 | 258 | 233.44 | 208 | 259 | 1.24 | 0.311 | 0.10000856 | 0.26114818 | 0.0218 | 0.021 | 0.0214 | 0.0214 | TRUE | 0.5369 |
| Wt1 | 68 | 31 | 52 | 91 | 59 | 38 | 72 | 31 | 52 | 72 | 63 | 42 | 55.34 | 52 | 59 | 1.143 | 0.192 | 0.60759386 | 0.77752295 | 0.1016 | 0.0664 | 0.0831 | 0.0831 | TRUE | 0.2273 |
| Wtap | 3961 | 3809 | 3654 | 5073 | 3625 | 3486 | 4207 | 3804 | 3669 | 3988 | 3900 | 3816 | 3897.44 | 3893 | 3901 | 1.002 | 0.003 | 0.96769627 | 0.98378919 | 0.0025 | 0.0077 | 0.0044 | 0.0044 | TRUE | 0.0604 |
| Wtip | 2833 | 3168 | 2721 | 3664 | 2704 | 2370 | 3009 | 3164 | 2732 | 2880 | 2909 | 2594 | 2881.54 | 2968 | 2794 | 0.942 | -0.087 | 0.34838653 | 0.56705391 | 0.0041 | 0.008 | 0.0058 | 0.0058 | TRUE | 0.059 |
| Wwc1 | 3770 | 3839 | 3719 | 4261 | 3380 | 3498 | 4004 | 3834 | 3734 | 3350 | 3637 | 3829 | 3731.36 | 3857 | 3605 | 0.934 | -0.098 | 0.22743338 | 0.44072728 | 0.0025 | 0.0078 | 0.0045 | 0.0045 | TRUE | 0.0474 |
| Wwc2 | 7698 | 8137 | 7847 | 11784 | 8583 | 8352 | 8177 | 8126 | 7879 | 9264 | 9235 | 9143 | 8637.23 | 8061 | 9214 | 1.143 | 0.193 | 0.00045504 | 0.0048329 | 1.00E-04 | 0.0073 | 0.0021 | 0.0021 | TRUE | 0.0047 |
| Wwox | 513 | 572 | 619 | 747 | 505 | 540 | 545 | 571 | 622 | 587 | 543 | 591 | 576.57 | 579 | 574 | 0.991 | -0.013 | 0.90199006 | 0.95205059 | 0.0015 | 0.0126 | 0.0063 | 0.0063 | TRUE | 0.048 |
| Wwp1 | 3062 | 3788 | 4297 | 4465 | 3405 | 3743 | 3252 | 3783 | 4315 | 3510 | 3664 | 4097 | 3770.17 | 3783 | 3757 | 0.993 | -0.01 | 0.93209268 | 0.96628017 | 0.0129 | 0.0078 | 0.0106 | 0.0106 | TRUE | 0.1829 |

| | | | | | | | | | | | | | | | | | | | | | | | | | |
|---|---|---|---|---|---|---|---|---|---|---|---|---|---|---|---|---|---|---|---|---|---|---|---|---|---|
| Wwp2 | 5985 | 5562 | 5729 | 7838 | 5883 | 5913 | 6357 | 5555 | 5753 | 6162 | 6330 | 6473 | 6104.76 | 5888 | 6322 | 1.074 | 0.102 | 0.19407252 | 0.39967468 | 0.0026 | 0.0074 | 0.0043 | 0.0043 | TRUE | 0.0591 |
| Wwtr1 | 2582 | 3178 | 3307 | 3263 | 2267 | 2601 | 2743 | 3174 | 3321 | 2565 | 2439 | 2847 | 2848.08 | 3079 | 2617 | 0.85 | -0.235 | 0.02757021 | 0.10873654 | 0.0074 | 0.0081 | 0.0078 | 0.0078 | TRUE | 0.1108 |
| Xab2 | 5632 | 6027 | 6239 | 8953 | 6783 | 6305 | 5982 | 6019 | 6265 | 7038 | 7298 | 6902 | 6584.01 | 6089 | 7079 | 1.163 | 0.217 | 0.00045853 | 0.00485686 | 5.00E-04 | 0.0074 | 0.0026 | 0.0026 | TRUE | 0.0088 |
| Xaf1 | 714 | 1046 | 939 | 781 | 615 | 713 | 758 | 1045 | 943 | 614 | 662 | 780 | 800.34 | 915 | 685 | 0.748 | -0.419 | 0.00585645 | 0.03526842 | 0.0185 | 0.011 | 0.0154 | 0.0154 | TRUE | 0.2669 |
| Xbp1 | 4050 | 3799 | 3310 | 4682 | 3613 | 3277 | 4302 | 3794 | 3324 | 3681 | 3887 | 3587 | 3762.45 | 3807 | 3718 | 0.977 | -0.034 | 0.75841647 | 0.87248319 | 0.0088 | 0.0078 | 0.0084 | 0.0084 | TRUE | 0.1575 |
| Xcl1 | 0 | 0 | 0 | 0 | 0 | 0 | 0 | 0 | 0 | 0 | 0 | 0 | 0 | 0 | 0 | NA | NA | NA | NA | NA | NA | NA | NA | NA | NA |
| Xcr1 | 0 | 0 | 0 | 0 | 0 | 0 | 0 | 0 | 0 | 0 | 0 | 0 | 0 | 0 | 0 | NA | NA | NA | NA | NA | NA | NA | NA | NA | NA |
| Xdh | 2 | 2 | 1 | 1 | 1 | 0 | 2 | 2 | 1 | 1 | 1 | 0 | 1.16 | 2 | 1 | 0.371 | -1.432 | 0.41986158 | NA | 0 | 2.8356 | 1.2017 | 1.2017 | TRUE | 0.1856 |
| Xiap | 2858 | 3397 | 3432 | 4451 | 3129 | 3463 | 3036 | 3393 | 3446 | 3499 | 3367 | 3791 | 3421.81 | 3292 | 3552 | 1.079 | 0.11 | 0.21709865 | 0.42919026 | 0.0037 | 0.0079 | 0.0054 | 0.0054 | TRUE | 0.056 |
| Xirp1 | 8 | 5 | 0 | 5 | 5 | 3 | 8 | 5 | 0 | 4 | 5 | 3 | 4.35 | 4 | 4 | 0.936 | -0.095 | 0.93215887 | NA | 0.5161 | 0.7647 | 0.6675 | 0.6675 | TRUE | 0.2867 |
| Xirp2 | 5 | 2 | 1 | 26 | 22 | 11 | 5 | 2 | 1 | 20 | 24 | 12 | 10.74 | 3 | 19 | 6.846 | 2.775 | 0.00034565 | 0.00383645 | 0.0651 | 0.3135 | 0.221 | 0.221 | TRUE | 0.6558 |
| Xist | 3 | 2 | 3 | 206 | 169 | 102 | 3 | 2 | 3 | 162 | 182 | 112 | 77.27 | 3 | 152 | 55.796 | 5.802 | 7.39E-24 | 2.71E-21 | 0 | 0.0495 | 0.0485 | 0.0485 | TRUE | 0.1984 |
| Xk | 608 | 727 | 863 | 817 | 617 | 678 | 646 | 726 | 867 | 642 | 664 | 742 | 714.45 | 746 | 683 | 0.914 | -0.129 | 0.34376323 | 0.56363615 | 0.0121 | 0.0115 | 0.0121 | 0.0121 | TRUE | 0.2362 |
| Xkr4 | 2 | 2 | 1 | 2 | 0 | 3 | 2 | 2 | 1 | 2 | 0 | 3 | 1.66 | 2 | 2 | 0.946 | -0.08 | 0.95764323 | NA | 0 | 1.9871 | 1.0528 | 1.0528 | TRUE | 0.1946 |
| Xkr5 | 274 | 429 | 479 | 593 | 496 | 448 | 291 | 428 | 481 | 466 | 534 | 490 | 448.45 | 400 | 497 | 1.24 | 0.311 | 0.09671311 | 0.2552922 | 0.032 | 0.0142 | 0.0229 | 0.0229 | TRUE | 0.6504 |
| Xkr6 | 13 | 10 | 8 | 31 | 19 | 17 | 14 | 10 | 8 | 24 | 20 | 19 | 15.88 | 11 | 21 | 2.008 | 1.006 | 0.04217743 | 0.14646023 | 0 | 0.2144 | 0.1053 | 0.1053 | TRUE | 0.2395 |
| Xkr7 | 12 | 21 | 28 | 31 | 21 | 21 | 13 | 21 | 28 | 24 | 23 | 23 | 21.96 | 21 | 23 | 1.13 | 0.176 | 0.68815749 | 0.82833091 | 0.0176 | 0.1569 | 0.0936 | 0.0936 | TRUE | 0.2418 |
| Xkr8 | 377 | 390 | 380 | 458 | 336 | 378 | 400 | 389 | 382 | 360 | 362 | 414 | 384.47 | 390 | 379 | 0.968 | -0.047 | 0.69059833 | 0.83034167 | 0 | 0.0155 | 0.0075 | 0.0075 | TRUE | 0.0743 |
| Xkr9 | 3 | 2 | 4 | 4 | 1 | 1 | 3 | 2 | 4 | 3 | 1 | 1 | 2.42 | 3 | 2 | 0.6 | -0.737 | 0.54219425 | NA | 0 | 1.3687 | 0.622 | 0.622 | TRUE | 0.4909 |
| Xkrx | 78 | 21 | 20 | 82 | 53 | 26 | 83 | 21 | 20 | 64 | 57 | 28 | 45.64 | 41 | 50 | 1.216 | 0.282 | 0.62545108 | 0.78901854 | 0.3799 | 0.0791 | 0.2181 | 0.2181 | TRUE | 5.7701 |
| Xlr | 39 | 30 | 30 | 133 | 116 | 72 | 41 | 30 | 30 | 105 | 125 | 79 | 68.28 | 34 | 103 | 3.043 | 1.606 | 5.21E-08 | 2.05E-06 | 0.0294 | 0.0552 | 0.0428 | 0.0428 | TRUE | 0.1115 |
| Xlr3a | 3174 | 4387 | 4272 | 3477 | 2692 | 3130 | 3371 | 4381 | 4290 | 2733 | 2896 | 3426 | 3516.38 | 4014 | 3018 | 0.752 | -0.412 | 0.00213439 | 0.01620743 | 0.0164 | 0.0078 | 0.0127 | 0.0127 | TRUE | 0.2386 |
| Xlr3b | 963 | 1252 | 1140 | 1328 | 1007 | 1002 | 1023 | 1250 | 1145 | 1044 | 1083 | 1097 | 1107.04 | 1139 | 1075 | 0.943 | -0.085 | 0.40822062 | 0.62087325 | 0.0045 | 0.0099 | 0.0067 | 0.0067 | TRUE | 0.0952 |
| Xlr3c | 821 | 1155 | 936 | 1808 | 1450 | 1378 | 872 | 1154 | 940 | 1421 | 1560 | 1508 | 1242.54 | 989 | 1496 | 1.513 | 0.598 | 1.30E-06 | 3.45E-05 | 0.0109 | 0.0096 | 0.0101 | 0.0101 | TRUE | 0.2551 |
| Xlr4a | 430 | 511 | 431 | 544 | 444 | 428 | 457 | 510 | 433 | 428 | 478 | 469 | 462.29 | 467 | 458 | 0.98 | -0.029 | 0.80609304 | 0.90019698 | 0.0033 | 0.014 | 0.0081 | 0.0081 | TRUE | 0.0787 |
| Xlr4b | 304 | 344 | 303 | 495 | 320 | 372 | 323 | 344 | 304 | 389 | 344 | 407 | 351.89 | 324 | 380 | 1.176 | 0.233 | 0.06827522 | 0.20245789 | 0.0026 | 0.0163 | 0.0089 | 0.0089 | TRUE | 0.0755 |
| Xlr4c | 148 | 204 | 171 | 263 | 223 | 221 | 157 | 204 | 172 | 207 | 240 | 242 | 203.54 | 178 | 230 | 1.289 | 0.366 | 0.02839407 | 0.11084932 | 0.0075 | 0.0231 | 0.0151 | 0.0151 | TRUE | 0.1792 |
| Xlr5a | 471 | 670 | 644 | 810 | 604 | 572 | 500 | 669 | 647 | 637 | 650 | 626 | 621.47 | 605 | 638 | 1.053 | 0.074 | 0.57879926 | 0.7558849 | 0.0104 | 0.0122 | 0.0113 | 0.0113 | TRUE | 0.2689 |
| Xlr5b | 379 | 531 | 510 | 635 | 482 | 511 | 403 | 530 | 512 | 499 | 519 | 559 | 503.69 | 482 | 526 | 1.09 | 0.124 | 0.36715255 | 0.58453273 | 0.0102 | 0.0134 | 0.0117 | 0.0117 | TRUE | 0.2379 |
| Xlr5c | 150 | 201 | 179 | 272 | 187 | 192 | 159 | 201 | 180 | 214 | 201 | 210 | 194.17 | 180 | 208 | 1.158 | 0.212 | 0.17613594 | 0.37637393 | 0 | 0.0239 | 0.0125 | 0.0125 | TRUE | 0.1101 |
| Xpa | 140 | 157 | 165 | 295 | 253 | 168 | 149 | 157 | 166 | 232 | 272 | 184 | 193.2 | 157 | 229 | 1.46 | 0.546 | 0.00352586 | 0.02397008 | 0.016 | 0.024 | 0.0199 | 0.0199 | TRUE | 0.1606 |
| Xpc | 1758 | 2101 | 2026 | 2493 | 1845 | 2077 | 1867 | 2098 | 2034 | 1960 | 1985 | 2274 | 2036.4 | 2000 | 2073 | 1.036 | 0.051 | 0.59905309 | 0.77141291 | 0.0047 | 0.0085 | 0.0063 | 0.0063 | TRUE | 0.0863 |
| Xpnpep1 | 9085 | 10803 | 11120 | 12877 | 9687 | 10333 | 9650 | 10789 | 11166 | 10123 | 10423 | 11311 | 10576.86 | 10535 | 10619 | 1.008 | 0.011 | 0.89786258 | 0.95044288 | 0.0035 | 0.0072 | 0.0055 | 0.0055 | TRUE | 0.0659 |
| Xpnpep2 | 18 | 21 | 21 | 65 | 46 | 40 | 19 | 21 | 21 | 51 | 49 | 44 | 34.26 | 20 | 48 | 2.363 | 1.241 | 0.00021046 | 0.00253352 | 0 | 0.1031 | 0.0456 | 0.0456 | TRUE | 0.0473 |
| Xpnpep3 | 771 | 734 | 813 | 1161 | 775 | 726 | 819 | 733 | 816 | 913 | 834 | 795 | 818.26 | 789 | 847 | 1.074 | 0.104 | 0.31703654 | 0.53795592 | 0.0033 | 0.0109 | 0.0065 | 0.0065 | TRUE | 0.0565 |
| Xpo1 | 23325 | 24162 | 24706 | 33854 | 24206 | 23162 | 24775 | 24131 | 24807 | 26613 | 26044 | 25354 | 25287.58 | 24571 | 26004 | 1.058 | 0.082 | 0.13893441 | 0.32357848 | 4.00E-04 | 0.007 | 0.0022 | 0.0022 | TRUE | 0.0058 |
| Xpo4 | 3707 | 3495 | 4249 | 4597 | 3309 | 3203 | 3937 | 3490 | 4266 | 3614 | 3560 | 3506 | 3729.12 | 3898 | 3560 | 0.913 | -0.131 | 0.16256173 | 0.35788551 | 0.0047 | 0.0078 | 0.006 | 0.006 | TRUE | 0.102 |
| Xpo5 | 4357 | 3290 | 3299 | 7005 | 4674 | 3770 | 4628 | 3286 | 3313 | 5507 | 5029 | 4127 | 4314.8 | 3742 | 4888 | 1.306 | 0.386 | 0.01716317 | 0.07778476 | 0.0309 | 0.0077 | 0.0186 | 0.0186 | TRUE | 0.5214 |
| Xpo6 | 8239 | 7953 | 8549 | 9078 | 6686 | 6979 | 8751 | 7943 | 8584 | 7136 | 7194 | 7640 | 7874.64 | 8426 | 7323 | 0.869 | -0.202 | 0.0054658 | 0.03340046 | 0.0018 | 0.0073 | 0.0037 | 0.0037 | TRUE | 0.0308 |
| Xpo7 | 8114 | 8401 | 8899 | 10482 | 7358 | 7863 | 8619 | 8390 | 8936 | 8240 | 7917 | 8607 | 8451.39 | 8648 | 8255 | 0.955 | -0.067 | 0.31999502 | 0.54111305 | 0.0012 | 0.0073 | 0.0032 | 0.0032 | TRUE | 0.0169 |
| Xpot | 8884 | 7823 | 7357 | 11557 | 7895 | 7562 | 9436 | 7813 | 7387 | 9085 | 8495 | 8278 | 8415.67 | 8212 | 8619 | 1.05 | 0.07 | 0.52226752 | 0.71226036 | 0.0082 | 0.0073 | 0.0085 | 0.0085 | TRUE | 0.2076 |
| Xpr1 | 2519 | 2933 | 3228 | 3811 | 2711 | 3033 | 2676 | 2929 | 3241 | 2996 | 2917 | 3320 | 3013.16 | 2949 | 3078 | 1.044 | 0.061 | 0.5471899 | 0.73237263 | 0.0058 | 0.008 | 0.0072 | 0.0072 | TRUE | 0.0915 |
| Xrcc1 | 2970 | 3157 | 3254 | 3815 | 2877 | 2888 | 3155 | 3153 | 3267 | 2999 | 3095 | 3161 | 3138.47 | 3192 | 3085 | 0.966 | -0.049 | 0.44638084 | 0.65236459 | 3.00E-04 | 0.008 | 0.0027 | 0.0027 | TRUE | 0.0074 |
| Xrcc2 | 750 | 812 | 828 | 1178 | 881 | 904 | 797 | 811 | 831 | 926 | 948 | 990 | 883.75 | 813 | 955 | 1.173 | 0.23 | 0.00650062 | 0.03815231 | 0 | 0.0106 | 0.004 | 0.004 | TRUE | 0.0122 |
| Xrcc3 | 684 | 744 | 743 | 865 | 680 | 658 | 727 | 743 | 746 | 680 | 732 | 720 | 724.59 | 739 | 711 | 0.961 | -0.057 | 0.52303154 | 0.71298212 | 0 | 0.0114 | 0.0044 | 0.0044 | TRUE | 0.0179 |
| Xrcc4 | 252 | 317 | 339 | 425 | 339 | 337 | 268 | 317 | 340 | 334 | 365 | 369 | 332.07 | 308 | 356 | 1.153 | 0.205 | 0.13989742 | 0.32462044 | 0.0054 | 0.0168 | 0.0108 | 0.0108 | TRUE | 0.1478 |
| Xrcc5 | 6040 | 7264 | 7254 | 8287 | 6036 | 6252 | 6416 | 7255 | 7284 | 6515 | 6494 | 6844 | 6801.12 | 6985 | 6618 | 0.947 | -0.078 | 0.32927334 | 0.5502542 | 0.0027 | 0.0074 | 0.0045 | 0.0045 | TRUE | 0.0619 |
| Xrcc6 | 6651 | 6613 | 6561 | 8112 | 5813 | 5713 | 7065 | 6604 | 6588 | 6377 | 6254 | 6254 | 6523.69 | 6752 | 6295 | 0.932 | -0.101 | 0.1118131 | 0.28113131 | 6.00E-04 | 0.0074 | 0.0028 | 0.0028 | TRUE | 0.0199 |
| Xrcc6bp1 | 572 | 817 | 709 | 772 | 587 | 677 | 608 | 816 | 712 | 607 | 632 | 741 | 685.83 | 712 | 660 | 0.926 | -0.111 | 0.44002418 | 0.64727303 | 0.015 | 0.0117 | 0.0134 | 0.0134 | TRUE | 0.1712 |
| Xrn1 | 2583 | 2531 | 2892 | 3213 | 2361 | 2302 | 2744 | 2528 | 2904 | 2526 | 2540 | 2520 | 2626.87 | 2725 | 2529 | 0.928 | -0.108 | 0.17942167 | 0.38055926 | 0.0018 | 0.0082 | 0.0043 | 0.0043 | TRUE | 0.0489 |
| Xrn2 | 8829 | 8992 | 9152 | 12640 | 9383 | 9890 | 9378 | 8980 | 9190 | 9937 | 10096 | 10826 | 9734.37 | 9183 | 10286 | 1.12 | 0.164 | 0.01387483 | 0.0667991 | 0.0011 | 0.0072 | 0.0031 | 0.0031 | TRUE | 0.0257 |
| Xrra1 | 46 | 33 | 41 | 62 | 42 | 42 | 49 | 33 | 41 | 49 | 45 | 46 | 43.82 | 41 | 47 | 1.141 | 0.19 | 0.52417658 | 0.71388227 | 0 | 0.0821 | 0.0414 | 0.0414 | TRUE | 0.2247 |
| Xylb | 1178 | 1003 | 954 | 1376 | 994 | 923 | 1251 | 1002 | 958 | 1082 | 1069 | 1010 | 1062.07 | 1070 | 1054 | 0.985 | -0.022 | 0.86161695 | 0.93161188 | 0.0101 | 0.01 | 0.0101 | 0.0101 | TRUE | 0.2607 |
| Xylt1 | 141 | 103 | 107 | 196 | 142 | 114 | 150 | 103 | 107 | 154 | 153 | 125 | 131.95 | 120 | 144 | 1.202 | 0.266 | 0.21949196 | 0.43164928 | 0.0201 | 0.0319 | 0.0261 | 0.0261 | TRUE | 0.4679 |
| Xylt2 | 2980 | 2792 | 2746 | 3800 | 2893 | 2497 | 3165 | 2788 | 2757 | 2987 | 3113 | 2733 | 2924.04 | 2903 | 2944 | 1.014 | 0.02 | 0.8287424 | 0.91322447 | 0.0048 | 0.008 | 0.0061 | 0.0061 | TRUE | 0.0753 |
| Yae1d1 | 1382 | 1533 | 1521 | 1646 | 1172 | 1301 | 1468 | 1531 | 1527 | 1294 | 1261 | 1424 | 1417.55 | 1509 | 1326 | 0.879 | -0.187 | 0.02821934 | 0.1103584 | 0.0013 | 0.0092 | 0.0045 | 0.0045 | TRUE | 0.0489 |
| Yaf2 | 712 | 852 | 817 | 1210 | 834 | 833 | 756 | 851 | 820 | 951 | 897 | 912 | 864.65 | 809 | 920 | 1.138 | 0.186 | 0.04406849 | 0.15054109 | 0.001 | 0.0107 | 0.005 | 0.005 | TRUE | 0.0385 |
| Yap1 | 6036 | 6427 | 6506 | 10278 | 7189 | 6928 | 6411 | 6419 | 6533 | 8080 | 7735 | 7584 | 7126.86 | 6454 | 7800 | 1.209 | 0.273 | 6.07E-06 | 0.00013039 | 5.00E-04 | 0.0074 | 0.0025 | 0.0025 | TRUE | 0.0122 |
| Yars | 5686 | 5022 | 5009 | 7891 | 5584 | 4844 | 6040 | 5016 | 5030 | 6203 | 6008 | 5303 | 5599.74 | 5362 | 5838 | 1.089 | 0.123 | 0.25741587 | 0.47607984 | 0.009 | 0.0075 | 0.0083 | 0.0083 | TRUE | 0.1491 |
| Yars2 | 1311 | 1286 | 1265 | 1835 | 1485 | 1386 | 1393 | 1284 | 1270 | 1443 | 1598 | 1517 | 1417.42 | 1316 | 1519 | 1.154 | 0.207 | 0.01711698 | 0.07766659 | 0.0019 | 0.0092 | 0.0047 | 0.0047 | TRUE | 0.0312 |
| Ybey | 238 | 302 | 258 | 264 | 215 | 207 | 253 | 302 | 259 | 208 | 231 | 227 | 246.49 | 271 | 222 | 0.816 | -0.293 | 0.0428278 | 0.14775471 | 0.0026 | 0.0203 | 0.011 | 0.011 | TRUE | 0.1087 |
| Ybx1 | 55857 | 60963 | 61304 | 70148 | 51825 | 49589 | 59330 | 60884 | 61556 | 55145 | 55761 | 54283 | 57826.39 | 60590 | 55063 | 0.909 | -0.138 | 0.00842529 | 0.04628478 | 2.00E-04 | 0.007 | 0.002 | 0.002 | TRUE | 0.0041 |
| Ybx2 | 480 | 914 | 878 | 549 | 569 | 612 | 510 | 913 | 882 | 432 | 612 | 670 | 669.66 | 768 | 571 | 0.743 | -0.429 | 0.07133926 | 0.20813222 | 0.0727 | 0.0118 | 0.0393 | 0.0393 | TRUE | 1.0226 |
| Ydjc | 1416 | 1547 | 1591 | 2028 | 1511 | 1474 | 1504 | 1545 | 1598 | 1594 | 1626 | 1614 | 1580.02 | 1549 | 1611 | 1.04 | 0.057 | 0.43242224 | 0.64190268 | 0 | 0.009 | 0.0031 | 0.0031 | TRUE | 0.0091 |
| Yeats2 | 3185 | 3381 | 3481 | 4220 | 3027 | 3008 | 3383 | 3377 | 3495 | 3317 | 3257 | 3293 | 3353.66 | 3418 | 3289 | 0.962 | -0.056 | 0.36203129 | 0.58006064 | 0 | 0.0079 | 0.0024 | 0.0024 | TRUE | 0.0047 |
| Yeats4 | 3725 | 3956 | 3824 | 5018 | 3561 | 3547 | 3957 | 3951 | 3840 | 3945 | 3831 | 3883 | 3901.02 | 3916 | 3886 | 0.993 | -0.011 | 0.85888297 | 0.9304418 | 0 | 0.0077 | 0.0023 | 0.0023 | TRUE | 0.0035 |
| Yes1 | 3890 | 3720 | 3849 | 5307 | 3835 | 4108 | 4132 | 3715 | 3865 | 4172 | 4126 | 4497 | 4084.48 | 3904 | 4265 | 1.092 | 0.128 | 0.10518117 | 0.27035349 | 0.0022 | 0.0077 | 0.0042 | 0.0042 | TRUE | 0.0317 |
| Yif1a | 760 | 1105 | 1141 | 1384 | 1042 | 1093 | 807 | 1104 | 1146 | 1088 | 1121 | 1196 | 1077.02 | 1019 | 1135 | 1.114 | 0.155 | 0.28083483 | 0.50219126 | 0.0177 | 0.01 | 0.014 | 0.014 | TRUE | 0.3928 |
| Yif1b | 2338 | 3324 | 3436 | 3181 | 2448 | 2653 | 2483 | 3320 | 3450 | 2501 | 2634 | 2904 | 2881.98 | 3084 | 2680 | 0.869 | -0.203 | 0.13955024 | 0.32415115 | 0.0169 | 0.008 | 0.0133 | 0.0133 | TRUE | 0.3525 |

| | | | | | | | | | | | | | | | | | | | | | | | | | |
|---|---|---|---|---|---|---|---|---|---|---|---|---|---|---|---|---|---|---|---|---|---|---|---|---|---|
| Yipf1 | 477 | 605 | 580 | 738 | 566 | 563 | 507 | 604 | 582 | 580 | 609 | 616 | 583.11 | 564 | 602 | 1.065 | 0.091 | 0.41158225 | 0.62279432 | 0.0028 | 0.0126 | 0.0071 | 0.0071 | TRUE | 0.093 |
| Yipf2 | 542 | 589 | 511 | 664 | 514 | 530 | 576 | 588 | 513 | 522 | 553 | 580 | 555.37 | 559 | 552 | 0.986 | -0.02 | 0.85204904 | 0.92621941 | 0.0023 | 0.0128 | 0.0069 | 0.0069 | TRUE | 0.0608 |
| Yipf3 | 1528 | 1540 | 1529 | 2058 | 1426 | 1381 | 1623 | 1538 | 1535 | 1618 | 1534 | 1512 | 1560.02 | 1565 | 1555 | 0.994 | -0.009 | 0.90744019 | 0.95354464 | 6.00E-04 | 0.009 | 0.0036 | 0.0036 | TRUE | 0.0158 |
| Yipf4 | 1706 | 2173 | 2140 | 2301 | 1744 | 1779 | 1812 | 2170 | 2149 | 1809 | 1876 | 1947 | 1960.63 | 2044 | 1877 | 0.918 | -0.123 | 0.21478979 | 0.42665801 | 0.0047 | 0.0086 | 0.0066 | 0.0066 | TRUE | 0.1185 |
| Yipf5 | 2023 | 1885 | 1778 | 2894 | 2048 | 1838 | 2149 | 1883 | 1785 | 2275 | 2204 | 2012 | 2051.2 | 1939 | 2164 | 1.116 | 0.159 | 0.12143372 | 0.29634919 | 0.006 | 0.0085 | 0.0071 | 0.0071 | TRUE | 0.1081 |
| Yipf6 | 1052 | 1174 | 1167 | 1499 | 1092 | 1114 | 1117 | 1172 | 1172 | 1178 | 1175 | 1219 | 1172.41 | 1154 | 1191 | 1.032 | 0.045 | 0.56102623 | 0.74256648 | 0 | 0.0097 | 0.0035 | 0.0035 | TRUE | 0.0091 |
| Yipf7 | 7 | 8 | 3 | 8 | 6 | 5 | 7 | 8 | 3 | 6 | 6 | 5 | 6.11 | 6 | 6 | 0.991 | -0.013 | 0.98649306 | 0.99272434 | 0 | 0.5462 | 0.2658 | 0.2658 | TRUE | 0.4839 |
| Ykt6 | 3523 | 3058 | 2897 | 4520 | 3089 | 2738 | 3742 | 3054 | 2909 | 3553 | 3324 | 2997 | 3263.17 | 3235 | 3291 | 1.018 | 0.025 | 0.83499731 | 0.91595337 | 0.0124 | 0.0079 | 0.0104 | 0.0104 | TRUE | 0.2282 |
| Ylpm1 | 4930 | 4023 | 4338 | 6577 | 4791 | 3983 | 5237 | 4018 | 4356 | 5170 | 5155 | 4360 | 4715.89 | 4537 | 4895 | 1.079 | 0.11 | 0.37701017 | 0.59213348 | 0.0125 | 0.0076 | 0.0109 | 0.0109 | TRUE | 0.2216 |
| Yme1l1 | 13578 | 15270 | 15434 | 20013 | 14157 | 14168 | 14422 | 15250 | 15497 | 15733 | 15232 | 15509 | 15273.95 | 15056 | 15491 | 1.029 | 0.041 | 0.50190159 | 0.69719851 | 7.00E-04 | 0.0071 | 0.0026 | 0.0026 | TRUE | 0.0166 |
| Yod1 | 274 | 253 | 223 | 301 | 224 | 228 | 291 | 253 | 224 | 237 | 241 | 250 | 249.14 | 256 | 243 | 0.947 | -0.078 | 0.60319136 | 0.77412127 | 0.0049 | 0.0201 | 0.0122 | 0.0122 | TRUE | 0.159 |
| Ypel1 | 296 | 354 | 383 | 427 | 305 | 334 | 314 | 354 | 385 | 336 | 328 | 366 | 347 | 351 | 343 | 0.977 | -0.033 | 0.79904688 | 0.89640568 | 0.0036 | 0.0164 | 0.0096 | 0.0096 | TRUE | 0.093 |
| Ypel2 | 1601 | 2337 | 2481 | 1955 | 1648 | 2004 | 1701 | 2334 | 2491 | 1537 | 1773 | 2194 | 2004.9 | 2175 | 1835 | 0.843 | -0.246 | 0.15868033 | 0.35263911 | 0.0342 | 0.0085 | 0.0215 | 0.0215 | TRUE | 0.4406 |
| Ypel3 | 642 | 772 | 777 | 853 | 654 | 746 | 682 | 771 | 780 | 671 | 704 | 817 | 737.33 | 744 | 731 | 0.98 | -0.03 | 0.80277199 | 0.89812791 | 0.0067 | 0.0114 | 0.0088 | 0.0088 | TRUE | 0.1253 |
| Ypel4 | 0 | 0 | 0 | 0 | 2 | 0 | 0 | 0 | 0 | 0 | 2 | 0 | 0.36 | 0 | 1 | 3.775 | 1.916 | 0.63423067 | NA | 10 | 9.1927 | 10 | 10 | TRUE | 0.9743 |
| Ypel5 | 1466 | 1361 | 1322 | 1789 | 1271 | 1314 | 1557 | 1359 | 1327 | 1406 | 1368 | 1438 | 1409.35 | 1414 | 1404 | 0.993 | -0.01 | 0.91181363 | 0.95644617 | 0.0033 | 0.0092 | 0.0057 | 0.0057 | TRUE | 0.0931 |
| Yrdc | 1996 | 2276 | 1976 | 2893 | 1972 | 1850 | 2120 | 2273 | 1984 | 2274 | 2122 | 2025 | 2133.07 | 2126 | 2140 | 1.007 | 0.011 | 0.90708316 | 0.95354464 | 0.0036 | 0.0084 | 0.0055 | 0.0055 | TRUE | 0.0446 |
| Ythdc1 | 4986 | 4937 | 4481 | 6679 | 4917 | 4338 | 5296 | 4931 | 4499 | 5251 | 5290 | 4749 | 5002.59 | 4909 | 5097 | 1.038 | 0.054 | 0.55551848 | 0.73847101 | 0.0042 | 0.0076 | 0.0059 | 0.0059 | TRUE | 0.0648 |
| Ythdc2 | 2602 | 2155 | 2107 | 3234 | 2236 | 2178 | 2764 | 2152 | 2116 | 2542 | 2406 | 2384 | 2393.99 | 2344 | 2444 | 1.043 | 0.061 | 0.6150357 | 0.78215958 | 0.0116 | 0.0083 | 0.0101 | 0.0101 | TRUE | 0.2971 |
| Ythdf1 | 8763 | 7476 | 6955 | 10246 | 7316 | 6589 | 9308 | 7466 | 6984 | 8055 | 7872 | 7213 | 7816.1 | 7919 | 7713 | 0.974 | -0.038 | 0.75439178 | 0.87032896 | 0.0112 | 0.0073 | 0.0105 | 0.0105 | TRUE | 0.2872 |
| Ythdf2 | 4523 | 4270 | 3881 | 5988 | 4395 | 3950 | 4804 | 4264 | 3897 | 4707 | 4729 | 4324 | 4454.26 | 4322 | 4587 | 1.061 | 0.086 | 0.38987582 | 0.6037429 | 0.0061 | 0.0076 | 0.007 | 0.007 | TRUE | 0.1159 |
| Ythdf3 | 5091 | 5291 | 5444 | 7708 | 5373 | 5502 | 5408 | 5284 | 5466 | 6059 | 5781 | 6023 | 5670.22 | 5386 | 5954 | 1.106 | 0.145 | 0.01590344 | 0.07405641 | 3.00E-04 | 0.0075 | 0.0024 | 0.0024 | TRUE | 0.0079 |
| Ywhab | 11722 | 10207 | 10020 | 14850 | 10879 | 9736 | 12451 | 10194 | 10061 | 11674 | 11705 | 10658 | 11123.74 | 10902 | 11346 | 1.041 | 0.058 | 0.58644225 | 0.76132745 | 0.0074 | 0.0072 | 0.008 | 0.008 | TRUE | 0.1887 |
| Ywhae | 79844 | 93188 | 93175 | 104133 | 75538 | 80326 | 84808 | 93068 | 93558 | 81861 | 81274 | 87929 | 87083.09 | 90478 | 83688 | 0.925 | -0.113 | 0.12940291 | 0.30962138 | 0.0024 | 0.0069 | 0.004 | 0.004 | TRUE | 0.0368 |
| Ywhag | 22414 | 19174 | 18037 | 33024 | 23563 | 19392 | 23808 | 19149 | 18111 | 25961 | 25352 | 21228 | 22268.12 | 20356 | 24180 | 1.188 | 0.248 | 0.05540778 | 0.1760813 | 0.0167 | 0.0071 | 0.0121 | 0.0121 | TRUE | 0.2692 |
| Ywhah | 7664 | 6997 | 6877 | 10144 | 6927 | 6715 | 8141 | 6988 | 6905 | 7974 | 7453 | 7351 | 7468.63 | 7345 | 7593 | 1.034 | 0.048 | 0.6007448 | 0.77266232 | 0.0044 | 0.0073 | 0.006 | 0.006 | TRUE | 0.1096 |
| Ywhaq | 8881 | 9700 | 9394 | 11449 | 8125 | 8127 | 9433 | 9687 | 9433 | 9000 | 8742 | 8896 | 9198.64 | 9518 | 8879 | 0.933 | -0.1 | 0.06832482 | 0.20248386 | 1.00E-04 | 0.0073 | 0.0021 | 0.0021 | TRUE | 0.003 |
| Ywhaz | 14172 | 14375 | 14115 | 20461 | 14523 | 14404 | 15053 | 14356 | 14173 | 16085 | 15626 | 15767 | 15176.78 | 14527 | 15826 | 1.089 | 0.124 | 0.03500112 | 0.12811271 | 5.00E-04 | 0.0071 | 0.0024 | 0.0024 | TRUE | 0.0122 |
| Yy1 | 4811 | 4883 | 4806 | 7260 | 5511 | 5017 | 5110 | 4877 | 4826 | 5707 | 5929 | 5492 | 5323.53 | 4938 | 5709 | 1.156 | 0.21 | 0.00190268 | 0.01479736 | 9.00E-04 | 0.0075 | 0.0031 | 0.0031 | TRUE | 0.0137 |
| Yy2 | 266 | 248 | 200 | 534 | 385 | 293 | 283 | 248 | 201 | 420 | 414 | 321 | 314.3 | 244 | 385 | 1.582 | 0.662 | 0.00020858 | 0.00252229 | 0.0218 | 0.0174 | 0.0196 | 0.0196 | TRUE | 0.2632 |
| Zadh2 | 191 | 407 | 508 | 345 | 286 | 308 | 203 | 406 | 510 | 271 | 308 | 337 | 339.25 | 373 | 305 | 0.817 | -0.291 | 0.3096567 | 0.52993766 | 0.1055 | 0.0166 | 0.056 | 0.056 | TRUE | 0.4095 |
| Zan | 121 | 131 | 98 | 79 | 62 | 60 | 129 | 131 | 98 | 62 | 67 | 66 | 92.04 | 119 | 65 | 0.543 | -0.881 | 7.66E-05 | 0.00111578 | 0 | 0.0427 | 0.0239 | 0.0239 | TRUE | 0.2381 |
| Zap70 | 208 | 264 | 274 | 448 | 382 | 478 | 221 | 264 | 275 | 352 | 411 | 523 | 341.03 | 253 | 429 | 1.689 | 0.756 | 3.10E-05 | 0.0005179 | 0.0244 | 0.0166 | 0.0206 | 0.0206 | TRUE | 0.3011 |
| Zar1 | 1 | 2 | 1 | 9 | 10 | 3 | 1 | 2 | 1 | 7 | 11 | 3 | 4.2 | 1 | 7 | 5.186 | 2.374 | 0.03215249 | NA | 0 | 0.7919 | 0.4398 | 0.4398 | TRUE | 0.068 |
| Zar1l | 3 | 6 | 1 | 4 | 2 | 2 | 3 | 6 | 1 | 3 | 2 | 2 | 2.94 | 3 | 2 | 0.743 | -0.429 | 0.70457389 | NA | 0 | 1.1257 | 0.5802 | 0.5802 | TRUE | 0.2067 |
| Zbbx | 24 | 16 | 12 | 52 | 31 | 29 | 25 | 16 | 12 | 41 | 33 | 32 | 26.58 | 18 | 35 | 1.995 | 0.996 | 0.01699276 | 0.0773408 | 0.0195 | 0.1308 | 0.083 | 0.083 | TRUE | 0.6849 |
| Zbed3 | 287 | 428 | 448 | 603 | 434 | 395 | 305 | 427 | 450 | 474 | 467 | 432 | 425.92 | 394 | 458 | 1.162 | 0.216 | 0.18803244 | 0.39300923 | 0.0194 | 0.0146 | 0.0171 | 0.0171 | TRUE | 0.4472 |
| Zbed4 | 1378 | 1331 | 1434 | 1703 | 1176 | 1216 | 1464 | 1329 | 1440 | 1339 | 1265 | 1331 | 1361.34 | 1411 | 1312 | 0.93 | -0.105 | 0.20576004 | 0.41456917 | 0.0011 | 0.0093 | 0.0042 | 0.0042 | TRUE | 0.0309 |
| Zbed6 | 3495 | 2548 | 2700 | 3437 | 2647 | 2579 | 3712 | 2545 | 2711 | 2702 | 2848 | 2823 | 2890.19 | 2989 | 2791 | 0.934 | -0.099 | 0.49407351 | 0.69039846 | 0.0205 | 0.008 | 0.0148 | 0.0148 | TRUE | 0.5431 |
| Zbp1 | 66 | 86 | 96 | 43 | 43 | 44 | 70 | 86 | 96 | 34 | 46 | 48 | 63.44 | 84 | 43 | 0.503 | -0.99 | 0.00034137 | 0.00379429 | 0.0141 | 0.0588 | 0.0375 | 0.0375 | TRUE | 0.3069 |
| Zbtb1 | 587 | 663 | 726 | 924 | 701 | 667 | 623 | 662 | 729 | 726 | 754 | 730 | 704.23 | 671 | 737 | 1.097 | 0.133 | 0.18759056 | 0.3922416 | 0.0017 | 0.0116 | 0.0059 | 0.0059 | TRUE | 0.0661 |
| Zbtb10 | 1030 | 1604 | 1549 | 3267 | 2273 | 2217 | 1094 | 1602 | 1555 | 2568 | 2446 | 2427 | 1948.68 | 1417 | 2480 | 1.75 | 0.807 | 3.44E-08 | 1.44E-06 | 0.0193 | 0.0086 | 0.0149 | 0.0149 | TRUE | 0.4772 |
| Zbtb11 | 3605 | 3220 | 3268 | 3825 | 2906 | 2843 | 3829 | 3216 | 3281 | 3007 | 3127 | 3112 | 3262.02 | 3442 | 3082 | 0.895 | -0.16 | 0.08503489 | 0.23438837 | 0.0045 | 0.0079 | 0.0059 | 0.0059 | TRUE | 0.1174 |
| Zbtb12 | 556 | 696 | 832 | 598 | 502 | 494 | 591 | 695 | 835 | 470 | 540 | 541 | 612.01 | 707 | 517 | 0.73 | -0.454 | 0.00261073 | 0.01899279 | 0.0168 | 0.0123 | 0.0147 | 0.0147 | TRUE | 0.2084 |
| Zbtb14 | 715 | 890 | 949 | 1249 | 970 | 906 | 759 | 889 | 953 | 982 | 1044 | 992 | 936.41 | 867 | 1006 | 1.159 | 0.213 | 0.05537848 | 0.17605914 | 0.0054 | 0.0104 | 0.0079 | 0.0079 | TRUE | 0.1393 |
| Zbtb16 | 20 | 11 | 16 | 31 | 29 | 22 | 21 | 11 | 16 | 24 | 31 | 24 | 21.32 | 16 | 26 | 1.65 | 0.723 | 0.10077644 | 0.26271835 | 0 | 0.1614 | 0.0894 | 0.0894 | TRUE | 0.2905 |
| Zbtb17 | 2417 | 2179 | 2200 | 2654 | 1991 | 2010 | 2567 | 2176 | 2209 | 2086 | 2142 | 2200 | 2230.22 | 2317 | 2143 | 0.925 | -0.113 | 0.22540161 | 0.43815832 | 0.0041 | 0.0084 | 0.0058 | 0.0058 | TRUE | 0.1074 |
| Zbtb18 | 1130 | 1234 | 1212 | 1627 | 1251 | 1242 | 1200 | 1232 | 1217 | 1279 | 1346 | 1360 | 1272.37 | 1216 | 1328 | 1.091 | 0.126 | 0.10062666 | 0.26242067 | 0 | 0.0095 | 0.0034 | 0.0034 | TRUE | 0.0131 |
| Zbtb2 | 244 | 271 | 267 | 494 | 328 | 279 | 259 | 271 | 268 | 388 | 353 | 305 | 307.43 | 266 | 349 | 1.315 | 0.395 | 0.00480855 | 0.03043356 | 0.0052 | 0.0176 | 0.0108 | 0.0108 | TRUE | 0.1334 |
| Zbtb20 | 53 | 81 | 68 | 98 | 64 | 66 | 56 | 81 | 68 | 77 | 69 | 72 | 70.6 | 68 | 73 | 1.062 | 0.087 | 0.7219521 | 0.84955819 | 0 | 0.0536 | 0.0293 | 0.0293 | TRUE | 0.2077 |
| Zbtb21 | 1820 | 1643 | 1638 | 2366 | 1775 | 1493 | 1933 | 1641 | 1645 | 1860 | 1910 | 1634 | 1770.47 | 1740 | 1801 | 1.036 | 0.051 | 0.63882734 | 0.79655189 | 0.0073 | 0.0088 | 0.0079 | 0.0079 | TRUE | 0.1141 |
| Zbtb22 | 806 | 820 | 711 | 1252 | 869 | 743 | 856 | 819 | 714 | 984 | 935 | 813 | 853.59 | 796 | 911 | 1.145 | 0.195 | 0.10381705 | 0.26807506 | 0.008 | 0.0108 | 0.0092 | 0.0092 | TRUE | 0.1035 |
| Zbtb24 | 818 | 777 | 757 | 1022 | 801 | 727 | 869 | 776 | 760 | 803 | 862 | 796 | 811 | 802 | 820 | 1.023 | 0.033 | 0.73852521 | 0.86001326 | 0.0023 | 0.011 | 0.0059 | 0.0059 | TRUE | 0.0634 |
| Zbtb25 | 703 | 780 | 692 | 828 | 560 | 632 | 747 | 779 | 695 | 651 | 603 | 692 | 694.3 | 740 | 649 | 0.876 | -0.191 | 0.06720839 | 0.20026653 | 0.0024 | 0.0116 | 0.0064 | 0.0064 | TRUE | 0.0435 |
| Zbtb26 | 402 | 633 | 744 | 632 | 513 | 560 | 427 | 632 | 747 | 497 | 552 | 613 | 578 | 602 | 554 | 0.919 | -0.122 | 0.54271986 | 0.7283165 | 0.0427 | 0.0126 | 0.0271 | 0.0271 | TRUE | 0.3922 |
| Zbtb3 | 222 | 301 | 303 | 357 | 274 | 251 | 236 | 301 | 304 | 281 | 295 | 275 | 281.81 | 280 | 284 | 1.01 | 0.015 | 0.91947892 | 0.960526 | 0.0066 | 0.0186 | 0.0124 | 0.0124 | TRUE | 0.2129 |
| Zbtb32 | 462 | 341 | 328 | 434 | 310 | 308 | 491 | 341 | 329 | 341 | 334 | 337 | 362.08 | 387 | 337 | 0.873 | -0.197 | 0.26338981 | 0.48264194 | 0.023 | 0.016 | 0.0195 | 0.0195 | TRUE | 0.6299 |
| Zbtb33 | 836 | 755 | 778 | 1140 | 790 | 742 | 888 | 754 | 781 | 896 | 850 | 812 | 830.27 | 808 | 853 | 1.057 | 0.08 | 0.44689943 | 0.65297364 | 0.0037 | 0.0109 | 0.0067 | 0.0067 | TRUE | 0.0891 |
| Zbtb34 | 1816 | 1869 | 1827 | 2324 | 1794 | 1748 | 1929 | 1867 | 1835 | 1827 | 1930 | 1913 | 1883.44 | 1877 | 1890 | 1.007 | 0.01 | 0.89131145 | 0.94740454 | 0 | 0.0087 | 0.0032 | 0.0032 | TRUE | 0.0107 |
| Zbtb37 | 156 | 161 | 166 | 232 | 173 | 166 | 166 | 161 | 167 | 182 | 186 | 182 | 173.9 | 165 | 183 | 1.116 | 0.158 | 0.28894717 | 0.50981931 | 0 | 0.0258 | 0.0102 | 0.0102 | TRUE | 0.0039 |
| Zbtb38 | 807 | 757 | 790 | 970 | 733 | 702 | 857 | 756 | 793 | 763 | 789 | 768 | 787.68 | 802 | 773 | 0.964 | -0.053 | 0.5724124 | 0.75108888 | 9.00E-04 | 0.0111 | 0.0051 | 0.0051 | TRUE | 0.0424 |
| Zbtb39 | 821 | 905 | 894 | 1189 | 893 | 899 | 872 | 904 | 898 | 935 | 961 | 984 | 925.53 | 891 | 960 | 1.076 | 0.106 | 0.19533649 | 0.40125647 | 0 | 0.0105 | 0.0038 | 0.0038 | TRUE | 0.0066 |
| Zbtb4 | 85 | 132 | 111 | 134 | 124 | 74 | 90 | 132 | 111 | 105 | 133 | 81 | 108.89 | 111 | 106 | 0.958 | -0.062 | 0.80720037 | 0.90068918 | 0.0376 | 0.0372 | 0.0373 | 0.0373 | TRUE | 0.2225 |
| Zbtb40 | 1426 | 1381 | 1470 | 1722 | 1366 | 1238 | 1515 | 1379 | 1476 | 1354 | 1470 | 1355 | 1424.76 | 1457 | 1393 | 0.956 | -0.065 | 0.44416896 | 0.65087811 | 0.0016 | 0.0092 | 0.0045 | 0.0045 | TRUE | 0.0277 |
| Zbtb41 | 888 | 847 | 909 | 1049 | 755 | 814 | 943 | 846 | 913 | 825 | 812 | 891 | 871.65 | 901 | 843 | 0.935 | -0.096 | 0.30905587 | 0.52957217 | 0.0016 | 0.0107 | 0.0053 | 0.0053 | TRUE | 0.0337 |
| Zbtb42 | 179 | 203 | 205 | 242 | 187 | 165 | 190 | 203 | 206 | 190 | 201 | 181 | 195.13 | 200 | 191 | 0.955 | -0.066 | 0.65058781 | 0.80417914 | 0 | 0.0238 | 0.0102 | 0.0102 | TRUE | 0.0234 |
| Zbtb43 | 2080 | 2088 | 2029 | 2789 | 2131 | 1959 | 2209 | 2085 | 2037 | 2192 | 2293 | 2144 | 2160.29 | 2110 | 2210 | 1.047 | 0.066 | 0.37747467 | 0.59245004 | 9.00E-04 | 0.0084 | 0.0036 | 0.0036 | TRUE | 0.0201 |
| Zbtb44 | 4609 | 4820 | 4530 | 6546 | 4802 | 5397 | 4896 | 4814 | 4549 | 5146 | 5167 | 5908 | 5079.74 | 4753 | 5407 | 1.137 | 0.186 | 0.03058485 | 0.11685665 | 0.0037 | 0.0076 | 0.0051 | 0.0051 | TRUE | 0.0798 |
| Zbtb45 | 929 | 869 | 798 | 1666 | 1197 | 1007 | 987 | 868 | 801 | 1310 | 1288 | 1102 | 1059.3 | 885 | 1233 | 1.394 | 0.479 | 6.47E-05 | 0.0009684 | 0.0084 | 0.01 | 0.0094 | 0.0094 | TRUE | 0.1191 |

| | | | | | | | | | | | | | | | | | | | | | | | |
|---|---|---|---|---|---|---|---|---|---|---|---|---|---|---|---|---|---|---|---|---|---|---|---|
| Zbtb46 | 372 | 386 | 396 | 442 | 359 | 320 | 395 | 386 | 398 | 347 | 386 | 350 | 377.05 | 393 | 361 | 0.919 | -0.122 | 0.28479459 | 0.50626428 | 0 | 0.0156 | 0.0067 | 0.0067 | TRUE | 0.0406 |
| Zbtb48 | 577 | 695 | 651 | 784 | 583 | 532 | 613 | 694 | 654 | 616 | 627 | 582 | 631.1 | 654 | 608 | 0.931 | -0.103 | 0.30981746 | 0.53009775 | 0.001 | 0.0121 | 0.0058 | 0.0058 | TRUE | 0.0348 |
| Zbtb49 | 218 | 228 | 219 | 264 | 230 | 177 | 232 | 228 | 220 | 208 | 247 | 194 | 221.32 | 227 | 216 | 0.954 | -0.067 | 0.66048742 | 0.80885221 | 0.0032 | 0.0218 | 0.0124 | 0.0124 | TRUE | 0.1674 |
| Zbtb5 | 688 | 706 | 727 | 773 | 646 | 648 | 731 | 705 | 730 | 608 | 695 | 709 | 696.32 | 722 | 671 | 0.927 | -0.109 | 0.29306126 | 0.51418305 | 0.0025 | 0.0116 | 0.0063 | 0.0063 | TRUE | 0.0838 |
| Zbtb6 | 572 | 592 | 615 | 770 | 547 | 462 | 608 | 591 | 618 | 605 | 589 | 506 | 585.98 | 606 | 567 | 0.937 | -0.094 | 0.39830498 | 0.6116314 | 0.0029 | 0.0125 | 0.0072 | 0.0072 | TRUE | 0.101 |
| Zbtb7a | 1078 | 1753 | 1677 | 1251 | 1186 | 1268 | 1145 | 1751 | 1684 | 983 | 1276 | 1388 | 1371.2 | 1527 | 1216 | 0.796 | -0.329 | 0.0771292 | 0.21943021 | 0.0373 | 0.0093 | 0.0243 | 0.0243 | TRUE | 0.5748 |
| Zbtb7b | 703 | 883 | 894 | 1022 | 787 | 760 | 747 | 882 | 898 | 803 | 847 | 832 | 834.73 | 842 | 827 | 0.982 | -0.027 | 0.80358507 | 0.89855367 | 0.0041 | 0.0108 | 0.007 | 0.007 | TRUE | 0.1158 |
| Zbtb7c | 473 | 354 | 315 | 401 | 274 | 229 | 502 | 354 | 316 | 315 | 295 | 251 | 338.83 | 391 | 287 | 0.736 | -0.443 | 0.02555304 | 0.10325969 | 0.0338 | 0.0166 | 0.0254 | 0.0254 | TRUE | 0.7149 |
| Zbtb8a | 627 | 724 | 736 | 817 | 636 | 626 | 666 | 723 | 739 | 642 | 684 | 685 | 689.98 | 709 | 670 | 0.944 | -0.083 | 0.39278762 | 0.6063469 | 0 | 0.0117 | 0.0053 | 0.0053 | TRUE | 0.0335 |
| Zbtb8b | 461 | 497 | 521 | 435 | 349 | 388 | 490 | 496 | 523 | 342 | 376 | 425 | 441.89 | 503 | 381 | 0.755 | -0.406 | 0.00105569 | 0.00929151 | 0.004 | 0.0144 | 0.0088 | 0.0088 | TRUE | 0.1163 |
| Zbtb8os | 738 | 822 | 785 | 1583 | 1064 | 915 | 784 | 821 | 788 | 1244 | 1145 | 1002 | 963.98 | 798 | 1130 | 1.418 | 0.504 | 3.99E-06 | 9.15E-05 | 0.0056 | 0.0103 | 0.0075 | 0.0075 | TRUE | 0.1179 |
| Zbtb9 | 1119 | 1064 | 1131 | 1450 | 1035 | 1012 | 1189 | 1063 | 1136 | 1140 | 1114 | 1108 | 1124.69 | 1129 | 1121 | 0.993 | -0.01 | 0.9041309 | 0.95248677 | 8.00E-04 | 0.0098 | 0.0043 | 0.0043 | TRUE | 0.0317 |
| Zbtbd6 | 78 | 62 | 47 | 162 | 94 | 57 | 83 | 62 | 47 | 127 | 101 | 62 | 80.47 | 64 | 97 | 1.524 | 0.607 | 0.06848713 | 0.20277005 | 0.0881 | 0.0478 | 0.0671 | 0.0671 | TRUE | 0.2184 |
| Zc2hc1a | 1106 | 1610 | 1695 | 1680 | 1173 | 1464 | 1175 | 1608 | 1702 | 1321 | 1262 | 1603 | 1445 | 1495 | 1395 | 0.933 | -0.1 | 0.53203974 | 0.7201304 | 0.0264 | 0.0092 | 0.0178 | 0.0178 | TRUE | 0.4216 |
| Zc2hc1b | 0 | 0 | 0 | 0 | 0 | 0 | 0 | 0 | 0 | 0 | 0 | 0 | 0 | 0 | 0 | NA | NA | NA | NA | NA | NA | NA | NA | NA | NA |
| Zc2hc1c | 85 | 71 | 70 | 89 | 79 | 82 | 90 | 71 | 70 | 70 | 85 | 90 | 79.37 | 77 | 82 | 1.054 | 0.075 | 0.75147216 | 0.86873993 | 0 | 0.0484 | 0.028 | 0.028 | TRUE | 0.1996 |
| Zc3h10 | 1480 | 1665 | 1742 | 1430 | 1195 | 1359 | 1572 | 1663 | 1749 | 1124 | 1286 | 1488 | 1480.26 | 1661 | 1299 | 0.781 | -0.356 | 0.00325791 | 0.02256642 | 0.0102 | 0.0091 | 0.0099 | 0.0099 | TRUE | 0.1932 |
| Zc3h11a | 9779 | 8082 | 7800 | 10504 | 7339 | 6828 | 10387 | 8072 | 7832 | 8257 | 7896 | 7474 | 8319.78 | 8764 | 7876 | 0.899 | -0.154 | 0.20814185 | 0.41756491 | 0.014 | 0.0073 | 0.0107 | 0.0107 | TRUE | 0.3207 |
| Zc3h12a | 476 | 350 | 335 | 363 | 299 | 269 | 506 | 350 | 336 | 285 | 322 | 294 | 348.84 | 397 | 300 | 0.757 | -0.402 | 0.02773774 | 0.10928796 | 0.0257 | 0.0163 | 0.0212 | 0.0212 | TRUE | 0.6528 |
| Zc3h12b | 24 | 23 | 13 | 42 | 14 | 20 | 25 | 23 | 13 | 33 | 15 | 22 | 21.91 | 20 | 23 | 1.151 | 0.203 | 0.67618035 | 0.81979608 | 0.0895 | 0.1572 | 0.1244 | 0.1244 | TRUE | 0.3593 |
| Zc3h12c | 103 | 139 | 157 | 250 | 166 | 166 | 109 | 139 | 158 | 197 | 179 | 182 | 160.45 | 135 | 186 | 1.372 | 0.457 | 0.01450772 | 0.06905059 | 0.0093 | 0.0274 | 0.0187 | 0.0187 | TRUE | 0.283 |
| Zc3h12d | 3 | 6 | 7 | 18 | 4 | 5 | 3 | 6 | 7 | 14 | 4 | 5 | 6.69 | 5 | 8 | 1.504 | 0.588 | 0.4737648 | 0.67458712 | 0.1751 | 0.4994 | 0.3318 | 0.3318 | TRUE | 1.6664 |
| Zc3h13 | 4835 | 3957 | 4109 | 5849 | 4339 | 3622 | 5136 | 3952 | 4126 | 4598 | 4668 | 3965 | 4407.46 | 4405 | 4410 | 1.002 | 0.002 | 0.98593901 | 0.99254373 | 0.0138 | 0.0077 | 0.011 | 0.011 | TRUE | 0.2566 |
| Zc3h14 | 8429 | 9130 | 9542 | 11435 | 8383 | 8875 | 8953 | 9118 | 9581 | 8989 | 9020 | 9715 | 9229.41 | 9217 | 9241 | 1.002 | 0.004 | 0.95889888 | 0.97898705 | 0.0012 | 0.0073 | 0.0033 | 0.0033 | TRUE | 0.0245 |
| Zc3h15 | 6450 | 6327 | 6296 | 9482 | 6653 | 6109 | 6851 | 6319 | 6322 | 7454 | 7158 | 6687 | 6798.53 | 6497 | 7100 | 1.093 | 0.128 | 0.09750051 | 0.2565529 | 0.0024 | 0.0074 | 0.0042 | 0.0042 | TRUE | 0.0315 |
| Zc3h18 | 10783 | 8810 | 8669 | 13274 | 9951 | 8131 | 11453 | 8799 | 8705 | 10435 | 10707 | 8901 | 9833.16 | 9652 | 10014 | 1.038 | 0.053 | 0.68736246 | 0.82764541 | 0.0143 | 0.0072 | 0.0125 | 0.0125 | TRUE | 0.3255 |
| Zc3h3 | 510 | 482 | 491 | 797 | 644 | 564 | 542 | 481 | 493 | 627 | 693 | 617 | 575.49 | 505 | 646 | 1.277 | 0.353 | 0.00114065 | 0.0098631 | 0.0021 | 0.0126 | 0.0067 | 0.0067 | TRUE | 0.0473 |
| Zc3h4 | 3485 | 3572 | 3568 | 5807 | 3937 | 3245 | 3702 | 3567 | 3583 | 4565 | 4236 | 3552 | 3867.48 | 3617 | 4118 | 1.139 | 0.187 | 0.07924744 | 0.22323462 | 0.0077 | 0.0078 | 0.008 | 0.008 | TRUE | 0.1753 |
| Zc3h6 | 1147 | 1779 | 1731 | 1234 | 921 | 1303 | 1218 | 1777 | 1738 | 970 | 991 | 1426 | 1353.41 | 1578 | 1129 | 0.715 | -0.484 | 0.01191108 | 0.05988574 | 0.0444 | 0.0093 | 0.0259 | 0.0259 | TRUE | 0.6327 |
| Zc3h7a | 1792 | 1795 | 1852 | 2122 | 1570 | 1699 | 1903 | 1793 | 1860 | 1668 | 1689 | 1860 | 1795.48 | 1852 | 1739 | 0.939 | -0.091 | 0.26583367 | 0.48549428 | 0.0014 | 0.0087 | 0.0043 | 0.0043 | TRUE | 0.0441 |
| Zc3h7b | 4452 | 4867 | 5240 | 6085 | 4545 | 4748 | 4729 | 4861 | 5262 | 4784 | 4890 | 5197 | 4953.69 | 4951 | 4957 | 1.001 | 0.002 | 0.9830742 | 0.99106917 | 0.0023 | 0.0076 | 0.0041 | 0.0041 | TRUE | 0.0369 |
| Zc3h8 | 891 | 821 | 891 | 991 | 743 | 657 | 946 | 820 | 895 | 779 | 799 | 719 | 826.44 | 887 | 766 | 0.864 | -0.211 | 0.03772793 | 0.13498494 | 0.0028 | 0.0109 | 0.0062 | 0.0062 | TRUE | 0.0522 |
| Zc3hav1 | 1601 | 2053 | 2156 | 3637 | 2779 | 2956 | 1701 | 2050 | 2165 | 2859 | 2990 | 3236 | 2500.12 | 1972 | 3028 | 1.535 | 0.618 | 4.52E-08 | 1.82E-06 | 0.0089 | 0.0082 | 0.0088 | 0.0088 | TRUE | 0.1747 |
| Zc3hav1l | 881 | 1147 | 1318 | 1851 | 1365 | 1418 | 936 | 1146 | 1323 | 1455 | 1469 | 1552 | 1313.45 | 1135 | 1492 | 1.314 | 0.394 | 0.00320243 | 0.02231751 | 0.0141 | 0.0094 | 0.0121 | 0.0121 | TRUE | 0.2811 |
| Zc3hc1 | 6172 | 6487 | 6451 | 7930 | 5620 | 5662 | 6556 | 6479 | 6477 | 6234 | 6047 | 6198 | 6331.76 | 6504 | 6160 | 0.947 | -0.078 | 0.15839974 | 0.35221777 | 0 | 0.0074 | 0.0021 | 0.0021 | TRUE | 0.0031 |
| Zc4h2 | 265 | 356 | 314 | 350 | 272 | 274 | 281 | 356 | 315 | 275 | 293 | 300 | 303.34 | 317 | 289 | 0.91 | -0.137 | 0.32787641 | 0.54902503 | 0.0046 | 0.0178 | 0.0108 | 0.0108 | TRUE | 0.1253 |
| Zcchc10 | 2809 | 2083 | 1885 | 3090 | 2211 | 1911 | 2984 | 2080 | 1893 | 2429 | 2379 | 2092 | 2309.44 | 2319 | 2300 | 0.992 | -0.011 | 0.94577158 | 0.97280132 | 0.0323 | 0.0083 | 0.0201 | 0.0201 | TRUE | 0.7612 |
| Zcchc11 | 5331 | 5162 | 5443 | 6807 | 4933 | 4778 | 5662 | 5155 | 5465 | 5351 | 5308 | 5230 | 5362.03 | 5427 | 5296 | 0.976 | -0.035 | 0.60149818 | 0.77310972 | 9.00E-04 | 0.0075 | 0.0031 | 0.0031 | TRUE | 0.0235 |
| Zcchc12 | 87 | 48 | 48 | 104 | 83 | 60 | 92 | 48 | 48 | 82 | 89 | 66 | 70.88 | 63 | 79 | 1.259 | 0.332 | 0.30455851 | 0.52512118 | 0.0697 | 0.0534 | 0.0612 | 0.0612 | TRUE | 1.4508 |
| Zcchc13 | 4 | 11 | 6 | 6 | 3 | 10 | 4 | 11 | 6 | 5 | 3 | 11 | 6.69 | 7 | 6 | 0.878 | -0.188 | 0.81777412 | 0.90711028 | 0.1545 | 0.4992 | 0.3301 | 0.3301 | TRUE | 0.922 |
| Zcchc14 | 4720 | 4944 | 4596 | 5509 | 4068 | 4016 | 5013 | 4938 | 4615 | 4331 | 4377 | 4396 | 4611.63 | 4855 | 4368 | 0.9 | -0.153 | 0.02148856 | 0.0915746 | 7.00E-04 | 0.0076 | 0.003 | 0.003 | TRUE | 0.0229 |
| Zcchc16 | 0 | 0 | 0 | 1 | 0 | 0 | 0 | 0 | 0 | 1 | 0 | 0 | 0.13 | 0 | 0 | 1.859 | 0.894 | 0.82650734 | NA | 0 | 25.1515 | 10 | 10 | TRUE | 0.2563 |
| Zcchc17 | 1484 | 1202 | 1096 | 2428 | 1607 | 1211 | 1576 | 1200 | 1100 | 1909 | 1729 | 1326 | 1473.43 | 1292 | 1655 | 1.281 | 0.357 | 0.04117071 | 0.1437557 | 0.0341 | 0.0091 | 0.0214 | 0.0214 | TRUE | 0.4427 |
| Zcchc18 | 113 | 136 | 164 | 149 | 115 | 132 | 120 | 136 | 165 | 117 | 124 | 144 | 134.31 | 140 | 128 | 0.913 | -0.131 | 0.51424394 | 0.70699577 | 0.0116 | 0.0314 | 0.0214 | 0.0214 | TRUE | 0.2428 |
| Zcchc2 | 1508 | 1839 | 1896 | 2427 | 1809 | 1927 | 1602 | 1837 | 1904 | 1908 | 1946 | 2109 | 1884.31 | 1781 | 1988 | 1.116 | 0.158 | 0.10771541 | 0.27472308 | 0.0046 | 0.0087 | 0.0064 | 0.0064 | TRUE | 0.093 |
| Zcchc24 | 1767 | 1835 | 1831 | 1792 | 1316 | 1449 | 1877 | 1833 | 1839 | 1409 | 1416 | 1586 | 1659.81 | 1850 | 1470 | 0.795 | -0.332 | 6.82E-05 | 0.00101165 | 0.0015 | 0.0089 | 0.0044 | 0.0044 | TRUE | 0.0564 |
| Zcchc3 | 740 | 781 | 748 | 1857 | 1465 | 1096 | 786 | 780 | 751 | 1460 | 1576 | 1200 | 1092.15 | 772 | 1412 | 1.829 | 0.871 | 7.50E-13 | 7.73E-11 | 0.0094 | 0.0099 | 0.0096 | 0.0096 | TRUE | 0.2053 |
| Zcchc4 | 667 | 637 | 594 | 900 | 614 | 638 | 708 | 636 | 596 | 708 | 661 | 698 | 667.94 | 647 | 689 | 1.065 | 0.092 | 0.39175153 | 0.60545861 | 0.0028 | 0.0118 | 0.0067 | 0.0067 | TRUE | 0.0806 |
| Zcchc5 | 0 | 0 | 0 | 0 | 0 | 0 | 0 | 0 | 0 | 0 | 0 | 0 | 0 | 0 | 0 | NA | NA | NA | NA | NA | NA | NA | NA | NA | NA |
| Zcchc6 | 2750 | 2618 | 2578 | 3267 | 2307 | 2325 | 2921 | 2615 | 2589 | 2568 | 2482 | 2545 | 2619.95 | 2708 | 2532 | 0.935 | -0.097 | 0.22853674 | 0.44188814 | 0.0019 | 0.0082 | 0.0043 | 0.0043 | TRUE | 0.0572 |
| Zcchc7 | 1651 | 960 | 921 | 1562 | 1082 | 829 | 1754 | 959 | 925 | 1228 | 1164 | 907 | 1156.13 | 1213 | 1100 | 0.908 | -0.14 | 0.55663389 | 0.73929959 | 0.0764 | 0.0098 | 0.0399 | 0.0399 | TRUE | 1.8276 |
| Zcchc8 | 3663 | 3660 | 3639 | 4716 | 3417 | 3411 | 3891 | 3655 | 3654 | 3707 | 3676 | 3734 | 3719.61 | 3733 | 3706 | 0.993 | -0.011 | 0.87073474 | 0.93646289 | 4.00E-04 | 0.0078 | 0.0027 | 0.0027 | TRUE | 0.0165 |
| Zcchc9 | 848 | 920 | 907 | 1071 | 781 | 892 | 901 | 919 | 911 | 842 | 840 | 976 | 898.16 | 910 | 886 | 0.973 | -0.04 | 0.68830651 | 0.82844706 | 0.0026 | 0.0106 | 0.0059 | 0.0059 | TRUE | 0.0927 |
| Zcrb1 | 819 | 924 | 838 | 1468 | 1128 | 1026 | 870 | 923 | 841 | 1154 | 1214 | 1123 | 1020.83 | 878 | 1164 | 1.325 | 0.406 | 3.50E-06 | 8.15E-05 | 8.00E-04 | 0.0101 | 0.0045 | 0.0045 | TRUE | 0.0238 |
| Zcwpw1 | 245 | 395 | 356 | 438 | 348 | 341 | 260 | 394 | 357 | 344 | 374 | 373 | 350.7 | 337 | 364 | 1.077 | 0.108 | 0.53034505 | 0.71860844 | 0.0203 | 0.0163 | 0.0183 | 0.0183 | TRUE | 0.4515 |
| Zdbf2 | 5313 | 3104 | 3182 | 6966 | 4595 | 2976 | 5643 | 3100 | 3195 | 5476 | 4944 | 3258 | 4269.36 | 3979 | 4559 | 1.146 | 0.196 | 0.42535402 | 0.63552616 | 0.0803 | 0.0077 | 0.0435 | 0.0435 | TRUE | 1.6279 |
| Zdhhc1 | 254 | 279 | 294 | 234 | 221 | 207 | 270 | 279 | 295 | 184 | 238 | 227 | 248.66 | 281 | 216 | 0.765 | -0.386 | 0.01169919 | 0.05908087 | 0.006 | 0.0202 | 0.0128 | 0.0128 | TRUE | 0.2024 |
| Zdhhc11 | 0 | 0 | 0 | 0 | 0 | 0 | 0 | 0 | 0 | 0 | 0 | 0 | 0 | 0 | 0 | NA | NA | NA | NA | NA | NA | NA | NA | NA | NA |
| Zdhhc12 | 1712 | 1892 | 1871 | 2074 | 1500 | 1616 | 1818 | 1890 | 1879 | 1630 | 1614 | 1769 | 1766.66 | 1862 | 1671 | 0.897 | -0.157 | 0.04251899 | 0.14722493 | 8.00E-04 | 0.0088 | 0.0037 | 0.0037 | TRUE | 0.0311 |
| Zdhhc13 | 2239 | 2805 | 2825 | 3079 | 2111 | 2365 | 2378 | 2801 | 2837 | 2420 | 2271 | 2589 | 2549.47 | 2672 | 2427 | 0.908 | -0.139 | 0.17459287 | 0.37396928 | 0.0061 | 0.0082 | 0.0072 | 0.0072 | TRUE | 0.112 |
| Zdhhc14 | 116 | 133 | 129 | 131 | 78 | 91 | 123 | 133 | 130 | 103 | 84 | 100 | 112.01 | 129 | 96 | 0.745 | -0.425 | 0.02610039 | 0.10464041 | 0 | 0.0363 | 0.0172 | 0.0172 | TRUE | 0.1044 |
| Zdhhc15 | 396 | 379 | 364 | 703 | 438 | 373 | 421 | 379 | 365 | 553 | 471 | 408 | 432.81 | 388 | 477 | 1.233 | 0.302 | 0.0405526 | 0.14235379 | 0.0123 | 0.0145 | 0.0133 | 0.0133 | TRUE | 0.2241 |
| Zdhhc16 | 1277 | 1362 | 1287 | 1718 | 1223 | 1211 | 1356 | 1360 | 1292 | 1351 | 1316 | 1326 | 1333.5 | 1336 | 1331 | 0.996 | -0.006 | 0.94014818 | 0.96979647 | 0 | 0.0094 | 0.0033 | 0.0033 | TRUE | 0.01 |
| Zdhhc17 | 704 | 792 | 874 | 998 | 706 | 718 | 748 | 791 | 878 | 785 | 760 | 786 | 791.08 | 806 | 777 | 0.964 | -0.053 | 0.59892584 | 0.77131202 | 0.0022 | 0.0111 | 0.0059 | 0.0059 | TRUE | 0.073 |
| Zdhhc18 | 7875 | 8128 | 7844 | 9032 | 6605 | 6450 | 8365 | 8118 | 7876 | 7100 | 7107 | 7061 | 7604.29 | 8120 | 7089 | 0.873 | -0.196 | 0.0008433 | 0.00780086 | 3.00E-04 | 0.0073 | 0.0023 | 0.0023 | TRUE | 0.0085 |
| Zdhhc19 | 1 | 2 | 1 | 4 | 1 | 2 | 1 | 2 | 1 | 3 | 1 | 2 | 1.75 | 1 | 2 | 1.601 | 0.679 | 0.62457938 | NA | 0 | 1.8942 | 0.787 | 0.787 | TRUE | 0.2166 |
| Zdhhc2 | 460 | 589 | 553 | 1214 | 888 | 776 | 489 | 588 | 555 | 954 | 955 | 849 | 731.89 | 544 | 919 | 1.691 | 0.758 | 2.38E-11 | 1.95E-09 | 0.0049 | 0.0114 | 0.0078 | 0.0078 | TRUE | 0.092 |
| Zdhhc20 | 2978 | 2819 | 3070 | 2921 | 2165 | 2381 | 3163 | 2815 | 3083 | 2296 | 2329 | 2606 | 2715.53 | 3020 | 2410 | 0.798 | -0.326 | 0.00031875 | 0.0035799 | 0.0038 | 0.0081 | 0.0055 | 0.0055 | TRUE | 0.0607 |
| Zdhhc21 | 1963 | 2490 | 2899 | 3006 | 2314 | 2464 | 2085 | 2487 | 2911 | 2363 | 2490 | 2697 | 2505.46 | 2494 | 2517 | 1.009 | 0.012 | 0.92469631 | 0.96292 | 0.0156 | 0.0082 | 0.0122 | 0.0122 | TRUE | 0.1994 |

| | | | | | | | | | | | | | | | | | | | | | | | | | |
|---|---|---|---|---|---|---|---|---|---|---|---|---|---|---|---|---|---|---|---|---|---|---|---|---|---|
| Zdhhc22 | 3 | 1 | 4 | 0 | 2 | 1 | 3 | 1 | 4 | 0 | 2 | 1 | 1.91 | 3 | 1 | 0.376 | -1.41 | 0.3552763 | NA | 0 | 1.7335 | 1.0149 | 1.0149 | TRUE | 0.5306 |
| Zdhhc23 | 105 | 93 | 99 | 160 | 124 | 114 | 112 | 93 | 99 | 126 | 133 | 125 | 114.63 | 101 | 128 | 1.264 | 0.338 | 0.06990621 | 0.20554825 | 0 | 0.0356 | 0.0162 | 0.0162 | TRUE | 0.0752 |
| Zdhhc24 | 600 | 694 | 792 | 664 | 525 | 582 | 637 | 693 | 795 | 522 | 565 | 637 | 641.6 | 708 | 575 | 0.81 | -0.304 | 0.02037306 | 0.08807968 | 0.0098 | 0.012 | 0.0108 | 0.0108 | TRUE | 0.1356 |
| Zdhhc25 | 0 | 1 | 0 | 0 | 1 | 1 | 0 | 1 | 0 | 0 | 1 | 1 | 0.53 | 0 | 1 | 1.924 | 0.944 | 0.7364225 | NA | 0 | 6.2438 | 3.9727 | 3.9727 | TRUE | 0.405 |
| Zdhhc3 | 2513 | 2168 | 2129 | 4002 | 2700 | 2411 | 2669 | 2165 | 2138 | 3146 | 2905 | 2639 | 2610.42 | 2324 | 2897 | 1.247 | 0.318 | 0.00780859 | 0.04356457 | 0.0115 | 0.0082 | 0.0099 | 0.0099 | TRUE | 0.2042 |
| Zdhhc4 | 448 | 464 | 445 | 808 | 555 | 464 | 476 | 463 | 447 | 635 | 597 | 508 | 521.06 | 462 | 580 | 1.258 | 0.331 | 0.00697384 | 0.03995331 | 0.0055 | 0.0132 | 0.0089 | 0.0089 | TRUE | 0.1368 |
| Zdhhc5 | 3580 | 2886 | 2948 | 5413 | 3695 | 3066 | 3803 | 2882 | 2960 | 4255 | 3976 | 3356 | 3538.68 | 3215 | 3862 | 1.202 | 0.265 | 0.05658096 | 0.17876766 | 0.0193 | 0.0078 | 0.0136 | 0.0136 | TRUE | 0.3104 |
| Zdhhc6 | 2320 | 1956 | 1976 | 3094 | 2166 | 1968 | 2464 | 1953 | 1984 | 2432 | 2330 | 2154 | 2219.81 | 2134 | 2305 | 1.081 | 0.112 | 0.33439412 | 0.5548783 | 0.0096 | 0.0084 | 0.0093 | 0.0093 | TRUE | 0.2215 |
| Zdhhc7 | 1755 | 1883 | 1849 | 2237 | 1561 | 1621 | 1864 | 1881 | 1857 | 1759 | 1680 | 1774 | 1802.3 | 1867 | 1738 | 0.931 | -0.104 | 0.13730093 | 0.32135635 | 0 | 0.0087 | 0.0029 | 0.0029 | TRUE | 0.0101 |
| Zdhhc8 | 2257 | 2039 | 1956 | 2215 | 1656 | 1525 | 2397 | 2036 | 1964 | 1741 | 1782 | 1669 | 1931.68 | 2132 | 1731 | 0.812 | -0.301 | 0.00314949 | 0.02204781 | 0.0057 | 0.0086 | 0.007 | 0.007 | TRUE | 0.1424 |
| Zdhhc9 | 815 | 943 | 1021 | 1398 | 1002 | 1051 | 866 | 942 | 1025 | 1099 | 1078 | 1150 | 1026.7 | 944 | 1109 | 1.174 | 0.232 | 0.0180193 | 0.08060135 | 0.0027 | 0.0101 | 0.0059 | 0.0059 | TRUE | 0.0671 |
| Zeb1 | 902 | 780 | 840 | 1177 | 799 | 726 | 958 | 779 | 843 | 925 | 860 | 795 | 860.03 | 860 | 860 | 1.001 | 0.001 | 0.99146767 | 0.99557064 | 0.0072 | 0.0107 | 0.0088 | 0.0088 | TRUE | 0.1173 |
| Zeb2 | 82 | 43 | 82 | 128 | 69 | 52 | 87 | 43 | 82 | 101 | 74 | 57 | 74.03 | 71 | 77 | 1.098 | 0.134 | 0.69634985 | 0.83373793 | 0.0941 | 0.0514 | 0.0717 | 0.0717 | TRUE | 0.5195 |
| Zer1 | 878 | 1161 | 1202 | 1106 | 823 | 969 | 933 | 1160 | 1207 | 869 | 885 | 1061 | 1019.12 | 1100 | 938 | 0.853 | -0.23 | 0.09238491 | 0.24809516 | 0.0138 | 0.0101 | 0.0124 | 0.0124 | TRUE | 0.2107 |
| Zf12 | 0 | 1 | 0 | 1 | 0 | 0 | 0 | 1 | 0 | 1 | 0 | 0 | 0.3 | 0 | 0 | 0.954 | -0.067 | 0.98642001 | NA | 0 | 11.0817 | 9.3014 | 9.3014 | TRUE | 0.405 |
| Zfa-ps | 0 | 2 | 3 | 6 | 4 | 2 | 0 | 2 | 3 | 5 | 4 | 2 | 2.7 | 2 | 4 | 2.235 | 1.161 | 0.35313685 | NA | 0 | 1.2256 | 0.6927 | 0.6927 | TRUE | 0.5391 |
| Zfand1 | 1015 | 1444 | 1503 | 1586 | 1188 | 1263 | 1078 | 1442 | 1509 | 1247 | 1278 | 1383 | 1322.83 | 1343 | 1303 | 0.969 | -0.045 | 0.74669737 | 0.86556561 | 0.0166 | 0.0094 | 0.0132 | 0.0132 | TRUE | 0.357 |
| Zfand2a | 419 | 489 | 454 | 835 | 525 | 429 | 445 | 488 | 456 | 656 | 565 | 470 | 513.36 | 463 | 564 | 1.219 | 0.286 | 0.05060358 | 0.16548201 | 0.0136 | 0.0133 | 0.0135 | 0.0135 | TRUE | 0.161 |
| Zfand2b | 605 | 666 | 635 | 739 | 583 | 554 | 643 | 665 | 638 | 581 | 627 | 606 | 626.67 | 649 | 605 | 0.932 | -0.102 | 0.27887486 | 0.49999026 | 0 | 0.0122 | 0.0048 | 0.0048 | TRUE | 0.015 |
| Zfand3 | 1808 | 2043 | 2151 | 3096 | 2333 | 2203 | 1920 | 2040 | 2160 | 2434 | 2510 | 2412 | 2246.02 | 2040 | 2452 | 1.202 | 0.265 | 0.00069103 | 0.00670286 | 0.0013 | 0.0084 | 0.0039 | 0.0039 | TRUE | 0.0318 |
| Zfand4 | 138 | 227 | 193 | 231 | 186 | 123 | 147 | 227 | 194 | 182 | 200 | 135 | 180.57 | 189 | 172 | 0.911 | -0.135 | 0.55123498 | 0.73535297 | 0.0376 | 0.0251 | 0.0312 | 0.0312 | TRUE | 0.2767 |
| Zfand5 | 3069 | 3181 | 2988 | 5051 | 3444 | 3109 | 3260 | 3177 | 3000 | 3971 | 3706 | 3403 | 3419.42 | 3146 | 3693 | 1.175 | 0.232 | 0.00794744 | 0.04412074 | 0.0034 | 0.0079 | 0.0052 | 0.0052 | TRUE | 0.0572 |
| Zfand6 | 2614 | 3103 | 2744 | 3262 | 2373 | 2514 | 2777 | 3099 | 2755 | 2564 | 2553 | 2752 | 2750.05 | 2877 | 2623 | 0.912 | -0.134 | 0.11172726 | 0.28113131 | 0.0026 | 0.0081 | 0.0047 | 0.0047 | TRUE | 0.0555 |
| Zfat | 638 | 491 | 500 | 811 | 553 | 514 | 678 | 490 | 502 | 638 | 595 | 563 | 577.55 | 557 | 599 | 1.076 | 0.106 | 0.48477406 | 0.68236113 | 0.0168 | 0.0126 | 0.0148 | 0.0148 | TRUE | 0.4211 |
| Zfc3h1 | 3812 | 3638 | 3903 | 4874 | 3895 | 3654 | 4049 | 3633 | 3919 | 3832 | 4191 | 4000 | 3937.26 | 3867 | 4008 | 1.036 | 0.051 | 0.51648501 | 0.70838447 | 0.0021 | 0.0077 | 0.0042 | 0.0042 | TRUE | 0.0341 |
| Zfhx2 | 750 | 611 | 572 | 1509 | 840 | 519 | 797 | 610 | 574 | 1186 | 904 | 568 | 773.23 | 660 | 886 | 1.343 | 0.426 | 0.08050463 | 0.22559116 | 0.0753 | 0.0112 | 0.0414 | 0.0414 | TRUE | 0.1769 |
| Zfhx2as | 6 | 3 | 3 | 6 | 6 | 4 | 6 | 3 | 3 | 5 | 6 | 4 | 4.66 | 4 | 5 | 1.258 | 0.331 | 0.70398736 | 0.83798405 | 0 | 0.7146 | 0.3283 | 0.3283 | TRUE | 0.369 |
| Zfhx3 | 2059 | 1657 | 1830 | 2334 | 1439 | 1396 | 2187 | 1655 | 1838 | 1835 | 1548 | 1528 | 1765.1 | 1893 | 1637 | 0.865 | -0.209 | 0.11417572 | 0.28497856 | 0.0148 | 0.0088 | 0.012 | 0.012 | TRUE | 0.2225 |
| Zfhx4 | 37 | 18 | 18 | 51 | 29 | 17 | 39 | 18 | 18 | 40 | 31 | 19 | 27.54 | 25 | 30 | 1.205 | 0.269 | 0.57867463 | 0.75588355 | 0.1384 | 0.1265 | 0.1323 | 0.1323 | TRUE | 0.5447 |
| Zfml | 8570 | 7115 | 7168 | 10433 | 7398 | 6763 | 9103 | 7106 | 7197 | 8202 | 7960 | 7403 | 7828.44 | 7802 | 7855 | 1.007 | 0.01 | 0.93177332 | 0.96628017 | 0.0093 | 0.0073 | 0.0095 | 0.0095 | TRUE | 0.2596 |
| Zfp1 | 1686 | 1542 | 1620 | 1713 | 1255 | 1243 | 1791 | 1540 | 1627 | 1347 | 1350 | 1361 | 1502.51 | 1653 | 1353 | 0.819 | -0.289 | 0.00107674 | 0.00942413 | 0.0024 | 0.0091 | 0.005 | 0.005 | TRUE | 0.0644 |
| Zfp101 | 348 | 319 | 310 | 500 | 323 | 312 | 370 | 319 | 311 | 393 | 348 | 342 | 346.94 | 333 | 361 | 1.085 | 0.118 | 0.37835255 | 0.5934918 | 0.0047 | 0.0164 | 0.0101 | 0.0101 | TRUE | 0.1032 |
| Zfp105 | 462 | 594 | 543 | 906 | 609 | 551 | 491 | 593 | 545 | 712 | 655 | 603 | 599.97 | 543 | 657 | 1.211 | 0.276 | 0.02343511 | 0.09729363 | 0.0062 | 0.0124 | 0.009 | 0.009 | TRUE | 0.0824 |
| Zfp106 | 40306 | 39044 | 41131 | 46888 | 33452 | 36061 | 42812 | 38994 | 41300 | 36860 | 35992 | 39474 | 39238.66 | 41035 | 37442 | 0.912 | -0.132 | 0.07011575 | 0.20597236 | 0.0022 | 0.007 | 0.0038 | 0.0038 | TRUE | 0.0276 |
| Zfp108 | 28 | 40 | 44 | 60 | 46 | 29 | 30 | 40 | 44 | 47 | 49 | 32 | 40.38 | 38 | 43 | 1.13 | 0.176 | 0.59929858 | 0.77143379 | 0.0193 | 0.0885 | 0.0563 | 0.0563 | TRUE | 0.3649 |
| Zfp109 | 212 | 194 | 229 | 255 | 166 | 165 | 225 | 194 | 230 | 200 | 179 | 181 | 201.43 | 216 | 187 | 0.865 | -0.21 | 0.17398148 | 0.37328744 | 0 | 0.0233 | 0.0121 | 0.0121 | TRUE | 0.092 |
| Zfp11 | 181 | 175 | 164 | 177 | 137 | 122 | 192 | 175 | 165 | 139 | 147 | 134 | 158.63 | 177 | 140 | 0.79 | -0.339 | 0.03760036 | 0.13464991 | 0 | 0.0277 | 0.0128 | 0.0128 | TRUE | 0.0579 |
| Zfp110 | 3429 | 4163 | 4294 | 4744 | 3398 | 3289 | 3642 | 4158 | 4312 | 3729 | 3656 | 3600 | 3849.53 | 4037 | 3662 | 0.907 | -0.141 | 0.10858043 | 0.27606395 | 0.0035 | 0.0078 | 0.0053 | 0.0053 | TRUE | 0.089 |
| Zfp111 | 609 | 610 | 605 | 1232 | 889 | 701 | 647 | 609 | 607 | 969 | 957 | 767 | 759.32 | 621 | 898 | 1.446 | 0.532 | 1.29E-05 | 0.00024697 | 0.0079 | 0.0112 | 0.0094 | 0.0094 | TRUE | 0.1888 |
| Zfp112 | 11 | 17 | 11 | 26 | 21 | 16 | 12 | 17 | 11 | 20 | 23 | 18 | 16.71 | 13 | 20 | 1.524 | 0.608 | 0.19727837 | 0.40387778 | 0 | 0.2041 | 0.0975 | 0.0975 | TRUE | 0.2622 |
| Zfp113 | 257 | 330 | 368 | 414 | 329 | 285 | 273 | 330 | 370 | 325 | 354 | 312 | 327.25 | 324 | 330 | 1.019 | 0.027 | 0.8566768 | 0.92913675 | 0.0101 | 0.017 | 0.0134 | 0.0134 | TRUE | 0.2131 |
| Zfp114 | 41 | 33 | 30 | 37 | 32 | 33 | 44 | 33 | 30 | 29 | 34 | 36 | 34.38 | 36 | 33 | 0.931 | -0.102 | 0.76129186 | 0.87423727 | 0 | 0.1027 | 0.0528 | 0.0528 | TRUE | 0.2691 |
| Zfp119a | 55 | 58 | 53 | 95 | 75 | 63 | 58 | 58 | 53 | 75 | 81 | 69 | 65.65 | 56 | 75 | 1.324 | 0.404 | 0.08859804 | 0.24146679 | 0 | 0.0571 | 0.0251 | 0.0251 | TRUE | 0.0402 |
| Zfp119b | 228 | 296 | 304 | 324 | 208 | 218 | 242 | 296 | 305 | 255 | 224 | 239 | 260.03 | 281 | 239 | 0.852 | -0.232 | 0.1234242 | 0.29946124 | 0.006 | 0.0196 | 0.0125 | 0.0125 | TRUE | 0.1619 |
| Zfp12 | 439 | 576 | 666 | 708 | 577 | 559 | 466 | 575 | 669 | 557 | 621 | 612 | 583.27 | 570 | 597 | 1.045 | 0.064 | 0.67006891 | 0.81584564 | 0.016 | 0.0126 | 0.0143 | 0.0143 | TRUE | 0.2353 |
| Zfp120 | 768 | 721 | 664 | 778 | 576 | 613 | 816 | 720 | 667 | 612 | 620 | 671 | 684.15 | 734 | 634 | 0.863 | -0.212 | 0.06409836 | 0.19484295 | 0.0051 | 0.0117 | 0.008 | 0.008 | TRUE | 0.1111 |
| Zfp128 | 81 | 113 | 115 | 90 | 74 | 93 | 86 | 113 | 115 | 71 | 80 | 102 | 94.42 | 105 | 84 | 0.798 | -0.326 | 0.17434122 | 0.37365264 | 0.0194 | 0.0418 | 0.0308 | 0.0308 | TRUE | 0.3122 |
| Zfp13 | 773 | 732 | 651 | 993 | 735 | 733 | 821 | 731 | 654 | 781 | 791 | 802 | 763.27 | 735 | 791 | 1.076 | 0.106 | 0.34467236 | 0.5641524 | 0.0051 | 0.0112 | 0.0078 | 0.0078 | TRUE | 0.1225 |
| Zfp131 | 1276 | 974 | 1020 | 1849 | 1377 | 1070 | 1355 | 973 | 1024 | 1454 | 1482 | 1171 | 1243.11 | 1117 | 1369 | 1.226 | 0.293 | 0.05981579 | 0.18552893 | 0.0239 | 0.0096 | 0.0167 | 0.0167 | TRUE | 0.4145 |
| Zfp133-ps | 22 | 16 | 31 | 22 | 12 | 18 | 23 | 16 | 31 | 17 | 13 | 20 | 20.06 | 23 | 17 | 0.709 | -0.496 | 0.28375177 | 0.50528429 | 0 | 0.1711 | 0.1032 | 0.1032 | TRUE | 0.1486 |
| Zfp14 | 200 | 167 | 154 | 185 | 133 | 145 | 212 | 167 | 155 | 145 | 143 | 159 | 163.52 | 178 | 149 | 0.838 | -0.255 | 0.16664959 | 0.36414483 | 0.0101 | 0.0271 | 0.0184 | 0.0184 | TRUE | 0.3039 |
| Zfp142 | 1460 | 1240 | 1228 | 2005 | 1468 | 1243 | 1551 | 1238 | 1233 | 1576 | 1579 | 1361 | 1423.09 | 1341 | 1505 | 1.123 | 0.168 | 0.17849387 | 0.37940773 | 0.0116 | 0.0092 | 0.0105 | 0.0105 | TRUE | 0.2251 |
| Zfp143 | 2760 | 2699 | 2756 | 3252 | 2427 | 2597 | 2932 | 2696 | 2767 | 2556 | 2611 | 2843 | 2734.17 | 2798 | 2670 | 0.954 | -0.068 | 0.39904728 | 0.61217524 | 0.002 | 0.0081 | 0.0043 | 0.0043 | TRUE | 0.0386 |
| Zfp146 | 2018 | 1692 | 1685 | 2189 | 1563 | 1456 | 2143 | 1690 | 1692 | 1721 | 1682 | 1594 | 1753.59 | 1842 | 1666 | 0.905 | -0.145 | 0.21815278 | 0.43019575 | 0.01 | 0.0088 | 0.0094 | 0.0094 | TRUE | 0.2476 |
| Zfp148 | 2018 | 2090 | 2031 | 2693 | 1897 | 1895 | 2143 | 2087 | 2039 | 2117 | 2041 | 2074 | 2083.76 | 2090 | 2077 | 0.994 | -0.008 | 0.90423437 | 0.95248677 | 0 | 0.0085 | 0.0029 | 0.0029 | TRUE | 0.006 |
| Zfp157 | 980 | 1259 | 1270 | 1319 | 997 | 1091 | 1041 | 1257 | 1275 | 1037 | 1073 | 1194 | 1146.23 | 1191 | 1101 | 0.924 | -0.114 | 0.32457512 | 0.5459314 | 0.0075 | 0.0098 | 0.0088 | 0.0088 | TRUE | 0.1453 |
| Zfp160 | 215 | 198 | 194 | 399 | 303 | 189 | 228 | 198 | 195 | 314 | 326 | 207 | 244.58 | 207 | 282 | 1.366 | 0.45 | 0.02569179 | 0.103661 | 0.0299 | 0.0204 | 0.0251 | 0.0251 | TRUE | 0.5972 |
| Zfp169 | 134 | 132 | 119 | 201 | 175 | 129 | 142 | 132 | 119 | 158 | 188 | 141 | 146.86 | 131 | 162 | 1.238 | 0.308 | 0.09970332 | 0.26073963 | 0.0075 | 0.0293 | 0.0184 | 0.0184 | TRUE | 0.1946 |
| Zfp174 | 233 | 254 | 233 | 281 | 198 | 163 | 247 | 254 | 234 | 221 | 213 | 178 | 224.58 | 245 | 204 | 0.835 | -0.26 | 0.08259287 | 0.22958314 | 0.0021 | 0.0216 | 0.0117 | 0.0117 | TRUE | 0.1271 |
| Zfp180 | 1340 | 1339 | 1221 | 1560 | 1260 | 1013 | 1423 | 1337 | 1226 | 1226 | 1356 | 1109 | 1279.59 | 1329 | 1230 | 0.926 | -0.111 | 0.31544433 | 0.53617717 | 0.0066 | 0.0095 | 0.008 | 0.008 | TRUE | 0.0936 |
| Zfp182 | 236 | 201 | 175 | 420 | 303 | 260 | 251 | 201 | 176 | 330 | 326 | 285 | 261.32 | 209 | 314 | 1.503 | 0.588 | 0.00056545 | 0.00573525 | 0.0139 | 0.0195 | 0.0169 | 0.0169 | TRUE | 0.3274 |
| Zfp184 | 712 | 700 | 744 | 820 | 650 | 670 | 756 | 699 | 747 | 645 | 699 | 733 | 713.3 | 734 | 692 | 0.942 | -0.086 | 0.388605 | 0.60260329 | 0.0017 | 0.0115 | 0.0058 | 0.0058 | TRUE | 0.0454 |
| Zfp185 | 975 | 1354 | 1365 | 1968 | 1387 | 1316 | 1036 | 1352 | 1371 | 1547 | 1492 | 1441 | 1373.08 | 1253 | 1493 | 1.192 | 0.253 | 0.04360888 | 0.14952757 | 0.0108 | 0.0093 | 0.0106 | 0.0106 | TRUE | 0.275 |
| Zfp189 | 276 | 280 | 288 | 361 | 274 | 317 | 293 | 280 | 289 | 284 | 295 | 347 | 297.93 | 287 | 309 | 1.072 | 0.1 | 0.46302823 | 0.66646126 | 0.0031 | 0.018 | 0.01 | 0.01 | TRUE | 0.1317 |
| Zfp191 | 1930 | 1913 | 1873 | 2726 | 1809 | 1736 | 2050 | 1911 | 1881 | 2143 | 1946 | 1900 | 1971.81 | 1947 | 1996 | 1.026 | 0.037 | 0.66617785 | 0.81316904 | 0.0027 | 0.0086 | 0.0049 | 0.0049 | TRUE | 0.0508 |
| Zfp2 | 888 | 1021 | 1014 | 950 | 768 | 837 | 943 | 1020 | 1018 | 747 | 826 | 916 | 911.74 | 994 | 830 | 0.834 | -0.262 | 0.01574876 | 0.07355328 | 0.0051 | 0.0105 | 0.0074 | 0.0074 | TRUE | 0.0985 |
| Zfp202 | 38 | 30 | 33 | 79 | 64 | 36 | 40 | 30 | 33 | 62 | 69 | 39 | 45.64 | 34 | 57 | 1.653 | 0.725 | 0.02939762 | 0.11339183 | 0.0322 | 0.0791 | 0.0564 | 0.0564 | TRUE | 0.5693 |
| Zfp207 | 11708 | 10414 | 10548 | 15065 | 11531 | 11148 | 12436 | 10401 | 10591 | 11843 | 12407 | 12203 | 11646.77 | 11143 | 12151 | 1.09 | 0.125 | 0.17041268 | 0.36901257 | 0.0052 | 0.0072 | 0.0059 | 0.0059 | TRUE | 0.1259 |
| Zfp212 | 1071 | 1090 | 1062 | 1435 | 1058 | 958 | 1138 | 1089 | 1066 | 1128 | 1138 | 1049 | 1101.28 | 1098 | 1105 | 1.007 | 0.011 | 0.89950428 | 0.95131641 | 5.00E-04 | 0.0099 | 0.0042 | 0.0042 | TRUE | 0.0232 |

| | | | | | | | | | | | | | | | | | | | | | | | | | |
|---|---|---|---|---|---|---|---|---|---|---|---|---|---|---|---|---|---|---|---|---|---|---|---|---|---|
| Zfp213 | 268 | 313 | 291 | 442 | 296 | 241 | 285 | 313 | 292 | 347 | 318 | 264 | 303.2 | 297 | 310 | 1.048 | 0.067 | 0.64809525 | 0.80243071 | 0.0077 | 0.0178 | 0.0125 | 0.0125 | TRUE | 0.1877 |
| Zfp217 | 5421 | 5391 | 5210 | 6907 | 5172 | 5027 | 5758 | 5384 | 5231 | 5430 | 5565 | 5503 | 5478.47 | 5458 | 5499 | 1.008 | 0.011 | 0.87351286 | 0.93766043 | 0.0011 | 0.0075 | 0.0032 | 0.0032 | TRUE | 0.0282 |
| Zfp219 | 4107 | 4489 | 4325 | 5218 | 4154 | 3887 | 4362 | 4483 | 4343 | 4102 | 4469 | 4255 | 4335.78 | 4396 | 4275 | 0.972 | -0.041 | 0.54757947 | 0.73258321 | 7.00E-04 | 0.0077 | 0.003 | 0.003 | TRUE | 0.0192 |
| Zfp229 | 2676 | 2706 | 2470 | 3938 | 2655 | 2524 | 2842 | 2703 | 2480 | 3096 | 2857 | 2763 | 2790.05 | 2675 | 2905 | 1.087 | 0.12 | 0.18051437 | 0.38190109 | 0.0035 | 0.0081 | 0.0054 | 0.0054 | TRUE | 0.0493 |
| Zfp235 | 221 | 198 | 182 | 258 | 204 | 131 | 235 | 198 | 183 | 203 | 219 | 143 | 196.82 | 205 | 188 | 0.921 | -0.118 | 0.5631082 | 0.74414265 | 0.0264 | 0.0236 | 0.025 | 0.025 | TRUE | 0.4564 |
| Zfp236 | 1586 | 1694 | 1814 | 2048 | 1527 | 1646 | 1685 | 1692 | 1821 | 1610 | 1643 | 1802 | 1708.77 | 1733 | 1685 | 0.972 | -0.041 | 0.63178081 | 0.79278407 | 0.0022 | 0.0088 | 0.0047 | 0.0047 | TRUE | 0.044 |
| Zfp239 | 493 | 528 | 488 | 660 | 515 | 402 | 524 | 527 | 490 | 519 | 554 | 440 | 509 | 514 | 504 | 0.983 | -0.025 | 0.83714796 | 0.91727183 | 0.0054 | 0.0134 | 0.009 | 0.009 | TRUE | 0.14 |
| Zfp248 | 654 | 682 | 702 | 902 | 627 | 605 | 695 | 681 | 705 | 709 | 675 | 662 | 687.77 | 694 | 682 | 0.984 | -0.023 | 0.80155993 | 0.89757347 | 0 | 0.0117 | 0.0045 | 0.0045 | TRUE | 0.0151 |
| Zfp251 | 517 | 537 | 446 | 781 | 500 | 410 | 549 | 536 | 448 | 614 | 538 | 449 | 522.34 | 511 | 534 | 1.046 | 0.065 | 0.67102507 | 0.81649493 | 0.0164 | 0.0132 | 0.0149 | 0.0149 | TRUE | 0.2172 |
| Zfp259 | 4280 | 4410 | 4472 | 5835 | 4369 | 4214 | 4546 | 4404 | 4490 | 4587 | 4701 | 4613 | 4556.91 | 4480 | 4634 | 1.034 | 0.048 | 0.4055388 | 0.61834473 | 0 | 0.0076 | 0.0022 | 0.0022 | TRUE | 0.0027 |
| Zfp26 | 799 | 772 | 801 | 1346 | 987 | 842 | 849 | 771 | 804 | 1058 | 1062 | 922 | 910.96 | 808 | 1014 | 1.256 | 0.329 | 0.00118123 | 0.01010322 | 0.0033 | 0.0105 | 0.0063 | 0.0063 | TRUE | 0.0743 |
| Zfp260 | 2120 | 2440 | 2411 | 2522 | 1903 | 1887 | 2252 | 2437 | 2421 | 1983 | 2048 | 2066 | 2200.88 | 2370 | 2032 | 0.857 | -0.223 | 0.00236715 | 0.01759411 | 8.00E-04 | 0.0084 | 0.0034 | 0.0034 | TRUE | 0.0229 |
| Zfp263 | 1195 | 1635 | 1646 | 1628 | 1286 | 1411 | 1269 | 1633 | 1653 | 1280 | 1384 | 1545 | 1460.5 | 1518 | 1403 | 0.923 | -0.115 | 0.38538674 | 0.59920343 | 0.0133 | 0.0092 | 0.012 | 0.012 | TRUE | 0.2472 |
| Zfp266 | 3208 | 3588 | 3687 | 3346 | 2552 | 2698 | 3407 | 3583 | 3702 | 2630 | 2746 | 2953 | 3170.42 | 3564 | 2776 | 0.779 | -0.361 | 6.72E-06 | 0.00014214 | 0.0021 | 0.0079 | 0.0043 | 0.0043 | TRUE | 0.0376 |
| Zfp27 | 230 | 231 | 194 | 340 | 220 | 198 | 244 | 231 | 195 | 267 | 237 | 217 | 231.75 | 223 | 240 | 1.079 | 0.11 | 0.49441022 | 0.69052608 | 0.0082 | 0.0211 | 0.0144 | 0.0144 | TRUE | 0.1374 |
| Zfp273 | 189 | 176 | 200 | 264 | 205 | 200 | 201 | 176 | 201 | 208 | 221 | 219 | 204.06 | 193 | 216 | 1.12 | 0.163 | 0.26431239 | 0.48381628 | 0 | 0.023 | 0.0105 | 0.0105 | TRUE | 0.0628 |
| Zfp275 | 752 | 953 | 970 | 1246 | 960 | 1055 | 799 | 952 | 974 | 980 | 1033 | 1155 | 981.96 | 908 | 1056 | 1.161 | 0.216 | 0.06874511 | 0.20334733 | 0.0076 | 0.0103 | 0.0091 | 0.0091 | TRUE | 0.1316 |
| Zfp276 | 319 | 286 | 300 | 531 | 413 | 352 | 339 | 286 | 301 | 417 | 444 | 385 | 362.13 | 309 | 415 | 1.348 | 0.431 | 0.00085745 | 0.00789476 | 0.0032 | 0.016 | 0.0092 | 0.0092 | TRUE | 0.0823 |
| Zfp277 | 2341 | 3072 | 3080 | 3078 | 2248 | 2499 | 2487 | 3068 | 3093 | 2420 | 2419 | 2736 | 2703.53 | 2883 | 2525 | 0.876 | -0.192 | 0.09214678 | 0.24762974 | 0.0091 | 0.0081 | 0.009 | 0.009 | TRUE | 0.1749 |
| Zfp28 | 177 | 252 | 273 | 275 | 225 | 264 | 188 | 252 | 274 | 216 | 242 | 289 | 243.51 | 238 | 249 | 1.044 | 0.062 | 0.74778433 | 0.86607612 | 0.0251 | 0.0204 | 0.0228 | 0.0228 | TRUE | 0.3685 |
| Zfp280b | 7036 | 5570 | 5169 | 6876 | 4828 | 4481 | 7473 | 5563 | 5190 | 5405 | 5195 | 4905 | 5621.94 | 6075 | 5168 | 0.851 | -0.233 | 0.09759115 | 0.25673873 | 0.0212 | 0.0075 | 0.0141 | 0.0141 | TRUE | 0.494 |
| Zfp280c | 1139 | 1220 | 1267 | 1069 | 819 | 940 | 1210 | 1218 | 1272 | 840 | 881 | 1029 | 1075.16 | 1233 | 917 | 0.742 | -0.43 | 4.61E-05 | 0.00072418 | 0.0048 | 0.01 | 0.0071 | 0.0071 | TRUE | 0.1351 |
| Zfp280d | 1547 | 1267 | 1394 | 2073 | 1498 | 1468 | 1643 | 1265 | 1400 | 1630 | 1612 | 1607 | 1526.1 | 1436 | 1616 | 1.126 | 0.171 | 0.12892815 | 0.30890254 | 0.0079 | 0.0091 | 0.0085 | 0.0085 | TRUE | 0.1909 |
| Zfp281 | 5649 | 4536 | 4228 | 8194 | 5848 | 4994 | 6000 | 4530 | 4245 | 6441 | 6292 | 5467 | 5496 | 4925 | 6067 | 1.232 | 0.301 | 0.03455927 | 0.12687822 | 0.0218 | 0.0075 | 0.0144 | 0.0144 | TRUE | 0.444 |
| Zfp282 | 367 | 462 | 476 | 730 | 528 | 528 | 390 | 461 | 478 | 574 | 568 | 578 | 508.19 | 443 | 573 | 1.293 | 0.371 | 0.00151761 | 0.01236349 | 0.003 | 0.0134 | 0.0079 | 0.0079 | TRUE | 0.1264 |
| Zfp286 | 446 | 334 | 283 | 536 | 399 | 362 | 474 | 334 | 284 | 421 | 429 | 396 | 389.73 | 364 | 415 | 1.143 | 0.193 | 0.31300264 | 0.53352184 | 0.0326 | 0.0154 | 0.0238 | 0.0238 | TRUE | 0.795 |
| Zfp287 | 197 | 264 | 202 | 282 | 199 | 181 | 209 | 264 | 203 | 222 | 214 | 198 | 218.28 | 225 | 211 | 0.939 | -0.091 | 0.57837404 | 0.75564243 | 0.0079 | 0.022 | 0.0147 | 0.0147 | TRUE | 0.2468 |
| Zfp292 | 5496 | 5710 | 6337 | 6347 | 5149 | 5428 | 5838 | 5703 | 6363 | 4990 | 5540 | 5942 | 5729.12 | 5968 | 5491 | 0.92 | -0.121 | 0.19904107 | 0.40627888 | 0.0052 | 0.0075 | 0.0062 | 0.0062 | TRUE | 0.0782 |
| Zfp296 | 1711 | 1693 | 1730 | 2280 | 1745 | 1724 | 1817 | 1691 | 1737 | 1792 | 1878 | 1887 | 1800.39 | 1748 | 1852 | 1.059 | 0.083 | 0.26603372 | 0.48569092 | 5.00E-04 | 0.0087 | 0.0034 | 0.0034 | TRUE | 0.0143 |
| Zfp3 | 121 | 166 | 155 | 237 | 189 | 155 | 129 | 166 | 156 | 186 | 203 | 170 | 168.21 | 150 | 186 | 1.242 | 0.313 | 0.0743038 | 0.21416659 | 0.0054 | 0.0265 | 0.0161 | 0.0161 | TRUE | 0.1609 |
| Zfp30 | 151 | 236 | 229 | 184 | 165 | 187 | 160 | 236 | 230 | 145 | 178 | 205 | 192.15 | 209 | 176 | 0.838 | -0.254 | 0.2354845 | 0.45062625 | 0.0317 | 0.024 | 0.0279 | 0.0279 | TRUE | 0.4421 |
| Zfp300 | 18 | 5 | 6 | 32 | 10 | 3 | 19 | 5 | 6 | 25 | 11 | 3 | 11.56 | 10 | 13 | 1.324 | 0.405 | 0.63447432 | 0.794516 | 0.5749 | 0.292 | 0.4336 | 0.4336 | TRUE | 0.3888 |
| Zfp316 | 253 | 283 | 227 | 291 | 216 | 217 | 269 | 283 | 228 | 229 | 232 | 238 | 246.33 | 260 | 233 | 0.896 | -0.158 | 0.27371032 | 0.49433176 | 0.0026 | 0.0203 | 0.011 | 0.011 | TRUE | 0.1293 |
| Zfp317 | 3344 | 3029 | 2602 | 3910 | 2882 | 2801 | 3552 | 3025 | 2613 | 3074 | 3101 | 3066 | 3071.74 | 3063 | 3080 | 1.006 | 0.008 | 0.94561706 | 0.97276996 | 0.0114 | 0.008 | 0.0099 | 0.0099 | TRUE | 0.2363 |
| Zfp318 | 3385 | 3266 | 3655 | 4884 | 3779 | 3636 | 3595 | 3262 | 3670 | 3839 | 4066 | 3980 | 3735.47 | 3509 | 3962 | 1.129 | 0.175 | 0.02490799 | 0.10143269 | 0.0019 | 0.0078 | 0.0041 | 0.0041 | TRUE | 0.0463 |
| Zfp319 | 492 | 466 | 344 | 752 | 563 | 415 | 523 | 465 | 345 | 591 | 606 | 454 | 497.43 | 444 | 550 | 1.239 | 0.31 | 0.09333572 | 0.24976093 | 0.0317 | 0.0135 | 0.0225 | 0.0225 | TRUE | 0.4412 |
| Zfp322a | 2180 | 2433 | 2738 | 3102 | 2323 | 2575 | 2316 | 2430 | 2749 | 2439 | 2499 | 2819 | 2541.89 | 2498 | 2586 | 1.035 | 0.049 | 0.63422626 | 0.7943315 | 0.0066 | 0.0082 | 0.0073 | 0.0073 | TRUE | 0.0937 |
| Zfp324 | 88 | 91 | 78 | 183 | 149 | 79 | 93 | 91 | 78 | 144 | 160 | 86 | 108.89 | 87 | 130 | 1.49 | 0.576 | 0.03066217 | 0.11709534 | 0.0461 | 0.0372 | 0.0415 | 0.0415 | TRUE | 0.8511 |
| Zfp326 | 1202 | 1258 | 1242 | 1346 | 1065 | 1033 | 1277 | 1256 | 1247 | 1058 | 1146 | 1131 | 1185.83 | 1260 | 1112 | 0.881 | -0.182 | 0.02263467 | 0.0949083 | 0 | 0.0097 | 0.0038 | 0.0038 | TRUE | 0.0222 |
| Zfp329 | 632 | 536 | 583 | 809 | 684 | 536 | 671 | 535 | 585 | 636 | 736 | 587 | 625.11 | 597 | 653 | 1.093 | 0.128 | 0.34591479 | 0.56518887 | 0.011 | 0.0122 | 0.0118 | 0.0118 | TRUE | 0.1432 |
| Zfp330 | 5290 | 6637 | 6256 | 6126 | 4560 | 5339 | 5619 | 6628 | 6282 | 4816 | 4906 | 5844 | 5682.58 | 6176 | 5189 | 0.84 | -0.252 | 0.0212107 | 0.09065994 | 0.0092 | 0.0075 | 0.0084 | 0.0084 | TRUE | 0.1486 |
| Zfp334 | 739 | 821 | 829 | 1034 | 714 | 694 | 785 | 820 | 832 | 813 | 768 | 760 | 796.34 | 812 | 780 | 0.961 | -0.057 | 0.51947856 | 0.71066145 | 0 | 0.011 | 0.0044 | 0.0044 | TRUE | 0.0167 |
| Zfp335 | 1984 | 1771 | 1709 | 2036 | 1516 | 1470 | 2107 | 1769 | 1716 | 1601 | 1631 | 1609 | 1738.82 | 1864 | 1614 | 0.866 | -0.208 | 0.04290947 | 0.14795982 | 0.0057 | 0.0088 | 0.007 | 0.007 | TRUE | 0.1571 |
| Zfp341 | 600 | 704 | 796 | 625 | 514 | 640 | 637 | 703 | 799 | 491 | 553 | 701 | 647.43 | 713 | 582 | 0.814 | -0.297 | 0.0631051 | 0.19253807 | 0.0213 | 0.012 | 0.0168 | 0.0168 | TRUE | 0.3734 |
| Zfp345 | 422 | 239 | 196 | 507 | 400 | 235 | 448 | 239 | 197 | 399 | 430 | 257 | 328.32 | 295 | 362 | 1.23 | 0.299 | 0.32848853 | 0.54950119 | 0.1259 | 0.0169 | 0.0644 | 0.0644 | TRUE | 2.3375 |
| Zfp346 | 2495 | 2692 | 2690 | 2980 | 2227 | 2178 | 2650 | 2689 | 2701 | 2343 | 2396 | 2384 | 2527.1 | 2680 | 2374 | 0.886 | -0.175 | 0.00519886 | 0.03213024 | 0 | 0.0082 | 0.0024 | 0.0024 | TRUE | 0.0017 |
| Zfp35 | 1381 | 1220 | 1297 | 1853 | 1309 | 1152 | 1467 | 1218 | 1302 | 1457 | 1408 | 1261 | 1352.29 | 1329 | 1375 | 1.035 | 0.05 | 0.64190314 | 0.79842557 | 0.0065 | 0.0093 | 0.0077 | 0.0077 | TRUE | 0.0983 |
| Zfp352 | 76 | 24 | 19 | 194 | 150 | 64 | 81 | 24 | 19 | 153 | 161 | 70 | 84.62 | 41 | 128 | 3.112 | 1.638 | 0.00166746 | 0.01331608 | 0.3623 | 0.0458 | 0.1793 | 0.1793 | TRUE | 5.2137 |
| Zfp354a | 418 | 483 | 444 | 523 | 390 | 350 | 444 | 482 | 446 | 411 | 420 | 383 | 431.01 | 457 | 405 | 0.885 | -0.176 | 0.10969728 | 0.27789037 | 0 | 0.0145 | 0.0064 | 0.0064 | TRUE | 0.0266 |
| Zfp354b | 71 | 57 | 58 | 77 | 55 | 44 | 75 | 57 | 58 | 61 | 59 | 48 | 59.74 | 63 | 56 | 0.885 | -0.177 | 0.50501356 | 0.69971255 | 0 | 0.062 | 0.0339 | 0.0339 | TRUE | 0.2291 |
| Zfp354c | 1167 | 1163 | 1281 | 1483 | 1114 | 1122 | 1240 | 1161 | 1286 | 1166 | 1199 | 1228 | 1213.32 | 1229 | 1198 | 0.974 | -0.038 | 0.64824578 | 0.80252217 | 9.00E-04 | 0.0096 | 0.0042 | 0.0042 | TRUE | 0.028 |
| Zfp358 | 187 | 231 | 212 | 269 | 195 | 175 | 199 | 231 | 213 | 211 | 210 | 192 | 209.17 | 214 | 204 | 0.955 | -0.066 | 0.65089922 | 0.80429196 | 0 | 0.0227 | 0.0107 | 0.0107 | TRUE | 0.0508 |
| Zfp36 | 370 | 575 | 724 | 700 | 513 | 766 | 393 | 574 | 727 | 550 | 552 | 839 | 605.83 | 565 | 647 | 1.145 | 0.195 | 0.41870958 | 0.62930126 | 0.074 | 0.0123 | 0.0401 | 0.0401 | TRUE | 0.2665 |
| Zfp362 | 432 | 581 | 587 | 591 | 466 | 481 | 459 | 580 | 589 | 465 | 501 | 527 | 520.17 | 543 | 498 | 0.915 | -0.128 | 0.34380163 | 0.56363615 | 0.0095 | 0.0132 | 0.0112 | 0.0112 | TRUE | 0.2125 |
| Zfp365 | 1286 | 1494 | 1468 | 1679 | 1235 | 1208 | 1366 | 1492 | 1474 | 1320 | 1329 | 1322 | 1383.85 | 1444 | 1324 | 0.916 | -0.126 | 0.1082987 | 0.27558682 | 4.00E-04 | 0.0093 | 0.0037 | 0.0037 | TRUE | 0.0267 |
| Zfp366 | 5 | 3 | 0 | 3 | 1 | 3 | 5 | 3 | 0 | 2 | 1 | 3 | 2.5 | 3 | 2 | 0.813 | -0.298 | 0.82597106 | NA | 0 | 1.3225 | 0.9191 | 0.9191 | TRUE | 0.2217 |
| Zfp367 | 647 | 1028 | 979 | 908 | 751 | 824 | 687 | 1027 | 983 | 714 | 808 | 902 | 853.46 | 899 | 808 | 0.898 | -0.155 | 0.36073045 | 0.57907793 | 0.0286 | 0.0108 | 0.0197 | 0.0197 | TRUE | 0.5039 |
| Zfp369 | 580 | 525 | 585 | 777 | 572 | 528 | 616 | 524 | 587 | 611 | 615 | 578 | 588.67 | 576 | 601 | 1.045 | 0.063 | 0.55603746 | 0.73889464 | 0.0021 | 0.0125 | 0.0067 | 0.0067 | TRUE | 0.0724 |
| Zfp36l1 | 2614 | 2301 | 2484 | 4119 | 3316 | 3123 | 2777 | 2298 | 2494 | 3238 | 3568 | 3419 | 2965.54 | 2523 | 3408 | 1.351 | 0.434 | 7.12E-06 | 0.00014852 | 0.0052 | 0.008 | 0.0064 | 0.0064 | TRUE | 0.0936 |
| Zfp36l2 | 1505 | 1835 | 2172 | 2075 | 1827 | 1841 | 1599 | 1833 | 2181 | 1631 | 1966 | 2015 | 1870.72 | 1871 | 1871 | 0.999 | -0.001 | 0.99525861 | 0.99768514 | 0.0181 | 0.0087 | 0.0136 | 0.0136 | TRUE | 0.2543 |
| Zfp36l3 | 179 | 202 | 226 | 233 | 163 | 160 | 190 | 202 | 227 | 183 | 175 | 175 | 192.08 | 206 | 178 | 0.863 | -0.213 | 0.16242021 | 0.35779288 | 0 | 0.0241 | 0.0115 | 0.0115 | TRUE | 0.0839 |
| Zfp37 | 380 | 487 | 458 | 592 | 427 | 451 | 404 | 486 | 460 | 465 | 459 | 494 | 461.4 | 450 | 473 | 1.05 | 0.07 | 0.55228497 | 0.73624731 | 0.0028 | 0.014 | 0.0079 | 0.0079 | TRUE | 0.093 |
| Zfp382 | 219 | 303 | 331 | 376 | 258 | 258 | 233 | 303 | 332 | 296 | 278 | 282 | 287.2 | 289 | 285 | 0.986 | -0.02 | 0.90194247 | 0.95205059 | 0.0132 | 0.0184 | 0.0157 | 0.0157 | TRUE | 0.3258 |
| Zfp383 | 650 | 888 | 955 | 693 | 560 | 719 | 690 | 887 | 959 | 545 | 603 | 787 | 745.09 | 845 | 645 | 0.762 | -0.393 | 0.02626065 | 0.10517595 | 0.0307 | 0.0113 | 0.0211 | 0.0211 | TRUE | 0.4367 |
| Zfp384 | 2727 | 2503 | 2531 | 3642 | 2650 | 2504 | 2897 | 2500 | 2541 | 2863 | 2851 | 2741 | 2732.17 | 2646 | 2818 | 1.065 | 0.091 | 0.29148057 | 0.5121693 | 0.0031 | 0.0081 | 0.0051 | 0.0051 | TRUE | 0.0831 |
| Zfp385a | 2406 | 2458 | 2437 | 2646 | 2019 | 2227 | 2556 | 2455 | 2447 | 2080 | 2172 | 2438 | 2357.94 | 2486 | 2230 | 0.897 | -0.157 | 0.07436663 | 0.21421286 | 0.0031 | 0.0083 | 0.0052 | 0.0052 | TRUE | 0.08 |
| Zfp385b | 43 | 25 | 20 | 44 | 56 | 37 | 46 | 25 | 20 | 35 | 60 | 41 | 37.68 | 30 | 45 | 1.489 | 0.575 | 0.17011586 | 0.36880651 | 0.1039 | 0.0943 | 0.0987 | 0.0987 | TRUE | 1.2917 |
| Zfp385c | 205 | 233 | 205 | 159 | 98 | 138 | 218 | 233 | 206 | 125 | 105 | 151 | 172.96 | 219 | 127 | 0.581 | -0.784 | 1.62E-05 | 0.0002984 | 0.008 | 0.026 | 0.0176 | 0.0176 | TRUE | 0.2569 |
| Zfp386 | 142 | 97 | 75 | 201 | 115 | 87 | 151 | 97 | 75 | 158 | 124 | 95 | 116.67 | 108 | 126 | 1.172 | 0.229 | 0.45271947 | 0.65809138 | 0.0841 | 0.0351 | 0.0584 | 0.0584 | TRUE | 0.488 |

| | | | | | | | | | | | | | | | | | | | | | | | | | |
|---|---|---|---|---|---|---|---|---|---|---|---|---|---|---|---|---|---|---|---|---|---|---|---|---|---|
| Zfp389 | 3 | 7 | 3 | 5 | 4 | 4 | 3 | 7 | 3 | 4 | 4 | 4 | 4.3 | 4 | 4 | 0.951 | -0.072 | 0.93625125 | NA | 0 | 0.773 | 0.3527 | 0.3527 | TRUE | 0.4919 |
| Zfp39 | 368 | 495 | 508 | 681 | 536 | 559 | 391 | 494 | 510 | 535 | 577 | 612 | 519.88 | 465 | 575 | 1.234 | 0.303 | 0.02692304 | 0.10714867 | 0.0101 | 0.0132 | 0.0116 | 0.0116 | TRUE | 0.2242 |
| Zfp395 | 1275 | 1894 | 2061 | 2419 | 1809 | 2146 | 1354 | 1892 | 2069 | 1902 | 1946 | 2349 | 1918.74 | 1772 | 2066 | 1.166 | 0.221 | 0.1810058 | 0.38263282 | 0.0282 | 0.0086 | 0.0191 | 0.0191 | TRUE | 0.5121 |
| Zfp397 | 1543 | 1736 | 1790 | 1876 | 1504 | 1565 | 1639 | 1734 | 1797 | 1475 | 1618 | 1713 | 1662.69 | 1723 | 1602 | 0.929 | -0.107 | 0.24736925 | 0.46413368 | 0.0034 | 0.0089 | 0.0055 | 0.0055 | TRUE | 0.0597 |
| Zfp398 | 231 | 209 | 217 | 360 | 247 | 221 | 245 | 209 | 218 | 283 | 266 | 242 | 243.78 | 224 | 264 | 1.18 | 0.238 | 0.10115595 | 0.26335947 | 0.0026 | 0.0204 | 0.0111 | 0.0111 | TRUE | 0.0755 |
| Zfp40 | 167 | 205 | 186 | 301 | 227 | 170 | 177 | 205 | 187 | 237 | 244 | 186 | 205.97 | 190 | 222 | 1.174 | 0.231 | 0.16844621 | 0.36640749 | 0.0083 | 0.0229 | 0.0154 | 0.0154 | TRUE | 0.2144 |
| Zfp407 | 1527 | 1273 | 1342 | 2169 | 1495 | 1309 | 1622 | 1271 | 1348 | 1705 | 1609 | 1433 | 1497.89 | 1414 | 1582 | 1.12 | 0.163 | 0.18615098 | 0.38985171 | 0.0113 | 0.0091 | 0.0103 | 0.0103 | TRUE | 0.1995 |
| Zfp408 | 348 | 331 | 361 | 489 | 381 | 306 | 370 | 331 | 362 | 384 | 410 | 335 | 365.33 | 354 | 376 | 1.064 | 0.089 | 0.49603446 | 0.69196835 | 0.0039 | 0.0159 | 0.0095 | 0.0095 | TRUE | 0.102 |
| Zfp41 | 1438 | 1255 | 1191 | 1582 | 1217 | 1117 | 1527 | 1253 | 1196 | 1244 | 1309 | 1223 | 1292.08 | 1325 | 1259 | 0.95 | -0.075 | 0.51962098 | 0.71079463 | 0.008 | 0.0095 | 0.0089 | 0.0089 | TRUE | 0.2125 |
| Zfp410 | 658 | 802 | 770 | 1234 | 872 | 799 | 699 | 801 | 773 | 970 | 938 | 875 | 842.66 | 758 | 928 | 1.225 | 0.293 | 0.00346876 | 0.02366321 | 0.0025 | 0.0108 | 0.006 | 0.006 | TRUE | 0.0541 |
| Zfp414 | 1359 | 1673 | 1736 | 1830 | 1435 | 1524 | 1443 | 1671 | 1743 | 1439 | 1544 | 1668 | 1584.72 | 1619 | 1550 | 0.957 | -0.064 | 0.55526022 | 0.73842723 | 0.0069 | 0.009 | 0.0078 | 0.0078 | TRUE | 0.1082 |
| Zfp418 | 262 | 267 | 213 | 339 | 251 | 224 | 278 | 267 | 214 | 266 | 270 | 245 | 256.76 | 253 | 260 | 1.031 | 0.044 | 0.77185135 | 0.88099866 | 0.0067 | 0.0197 | 0.013 | 0.013 | TRUE | 0.2041 |
| Zfp42 | 42339 | 28825 | 25211 | 50192 | 33356 | 25934 | 44972 | 28788 | 25315 | 39457 | 35889 | 28389 | 33801.44 | 33025 | 34578 | 1.047 | 0.066 | 0.74723041 | 0.86561278 | 0.0503 | 0.007 | 0.0305 | 0.0305 | TRUE | 1.2259 |
| Zfp420 | 332 | 334 | 285 | 396 | 291 | 288 | 353 | 334 | 286 | 311 | 313 | 315 | 318.67 | 324 | 313 | 0.967 | -0.049 | 0.71017331 | 0.84210612 | 0.0025 | 0.0172 | 0.0094 | 0.0094 | TRUE | 0.1196 |
| Zfp422 | 686 | 691 | 674 | 1117 | 826 | 809 | 729 | 690 | 677 | 878 | 889 | 886 | 791.32 | 699 | 884 | 1.266 | 0.34 | 8.86E-05 | 0.00125888 | 0 | 0.0111 | 0.0041 | 0.0041 | TRUE | 0.0167 |
| Zfp423 | 1335 | 1434 | 1720 | 1584 | 1093 | 1084 | 1418 | 1432 | 1727 | 1245 | 1176 | 1187 | 1364.17 | 1526 | 1203 | 0.788 | -0.343 | 0.00133471 | 0.011144 | 0.0059 | 0.0093 | 0.0075 | 0.0075 | TRUE | 0.1608 |
| Zfp426 | 565 | 452 | 402 | 813 | 494 | 381 | 600 | 451 | 404 | 639 | 532 | 417 | 507.15 | 485 | 529 | 1.093 | 0.128 | 0.52301318 | 0.71298212 | 0.0416 | 0.0134 | 0.0271 | 0.0271 | TRUE | 0.2287 |
| Zfp428 | 1529 | 1665 | 1835 | 2220 | 1592 | 1286 | 1624 | 1663 | 1843 | 1745 | 1713 | 1408 | 1665.88 | 1710 | 1622 | 0.949 | -0.075 | 0.50610121 | 0.70044124 | 0.0085 | 0.0089 | 0.0087 | 0.0087 | TRUE | 0.1596 |
| Zfp429 | 20 | 26 | 19 | 31 | 16 | 14 | 21 | 26 | 19 | 24 | 17 | 15 | 20.53 | 22 | 19 | 0.868 | -0.204 | 0.63723349 | 0.79601193 | 0 | 0.1673 | 0.0869 | 0.0869 | TRUE | 0.3953 |
| Zfp433 | 49 | 39 | 31 | 64 | 36 | 35 | 52 | 39 | 31 | 50 | 39 | 38 | 41.58 | 41 | 42 | 1.051 | 0.071 | 0.83012974 | 0.9138554 | 0.022 | 0.0861 | 0.0552 | 0.0552 | TRUE | 0.388 |
| Zfp438 | 239 | 221 | 200 | 359 | 221 | 171 | 254 | 221 | 201 | 282 | 238 | 187 | 230.43 | 225 | 236 | 1.051 | 0.072 | 0.71009486 | 0.8420764 | 0.0237 | 0.0212 | 0.0224 | 0.0224 | TRUE | 0.2067 |
| Zfp442 | 121 | 132 | 108 | 148 | 157 | 104 | 129 | 132 | 108 | 116 | 169 | 114 | 127.98 | 123 | 133 | 1.08 | 0.111 | 0.62007046 | 0.78498274 | 0.0235 | 0.0326 | 0.0281 | 0.0281 | TRUE | 0.5578 |
| Zfp444 | 1039 | 1325 | 1247 | 1288 | 1013 | 1040 | 1104 | 1323 | 1252 | 1013 | 1090 | 1138 | 1153.32 | 1226 | 1080 | 0.88 | -0.184 | 0.07958234 | 0.22381823 | 0.0052 | 0.0098 | 0.0071 | 0.0071 | TRUE | 0.0915 |
| Zfp445 | 4978 | 4209 | 4300 | 6836 | 4947 | 4480 | 5288 | 4204 | 4318 | 5374 | 5323 | 4904 | 4901.57 | 4603 | 5200 | 1.13 | 0.176 | 0.10704134 | 0.27341772 | 0.0093 | 0.0076 | 0.0084 | 0.0084 | TRUE | 0.206 |
| Zfp446 | 612 | 439 | 463 | 883 | 612 | 505 | 650 | 438 | 465 | 694 | 658 | 553 | 576.47 | 518 | 635 | 1.228 | 0.296 | 0.09079572 | 0.24567338 | 0.0289 | 0.0126 | 0.0204 | 0.0204 | TRUE | 0.5803 |
| Zfp449 | 420 | 491 | 509 | 588 | 438 | 447 | 446 | 490 | 511 | 462 | 471 | 489 | 478.4 | 482 | 474 | 0.982 | -0.026 | 0.80893358 | 0.90174222 | 0 | 0.0138 | 0.0065 | 0.0065 | TRUE | 0.0501 |
| Zfp451 | 3359 | 3746 | 3936 | 4596 | 3154 | 3271 | 3568 | 3741 | 3952 | 3613 | 3394 | 3581 | 3641.39 | 3754 | 3529 | 0.94 | -0.089 | 0.23393017 | 0.4485771 | 0.0015 | 0.0078 | 0.0037 | 0.0037 | TRUE | 0.026 |
| Zfp454 | 244 | 221 | 224 | 347 | 246 | 205 | 259 | 221 | 225 | 273 | 265 | 224 | 244.45 | 235 | 254 | 1.084 | 0.116 | 0.44449702 | 0.65105934 | 0.0051 | 0.0204 | 0.0124 | 0.0124 | TRUE | 0.1118 |
| Zfp455 | 63 | 77 | 62 | 90 | 57 | 61 | 67 | 77 | 62 | 71 | 61 | 67 | 67.49 | 69 | 66 | 0.967 | -0.048 | 0.8397163 | 0.91870182 | 0 | 0.0557 | 0.0262 | 0.0262 | TRUE | 0.0994 |
| Zfp456 | 222 | 245 | 220 | 291 | 197 | 221 | 236 | 245 | 221 | 229 | 212 | 242 | 230.67 | 234 | 228 | 0.973 | -0.039 | 0.78022876 | 0.88626345 | 0 | 0.0212 | 0.0097 | 0.0097 | TRUE | 0.0371 |
| Zfp457 | 14 | 12 | 6 | 16 | 6 | 8 | 15 | 12 | 6 | 13 | 6 | 9 | 10.11 | 11 | 9 | 0.858 | -0.221 | 0.73082844 | 0.85540824 | 0 | 0.3327 | 0.1996 | 0.1996 | TRUE | 0.4807 |
| Zfp458 | 19 | 9 | 21 | 43 | 23 | 11 | 20 | 9 | 21 | 34 | 25 | 12 | 20.14 | 17 | 24 | 1.419 | 0.505 | 0.35988769 | 0.57825474 | 0.1661 | 0.1705 | 0.1683 | 0.1683 | TRUE | 0.2131 |
| Zfp459 | 1567 | 2166 | 2256 | 2384 | 2071 | 2349 | 1664 | 2163 | 2265 | 1874 | 2228 | 2571 | 2127.77 | 2031 | 2224 | 1.095 | 0.131 | 0.40031942 | 0.61323212 | 0.0252 | 0.0085 | 0.0169 | 0.0169 | TRUE | 0.2297 |
| Zfp46 | 289 | 213 | 244 | 303 | 249 | 194 | 307 | 213 | 245 | 238 | 268 | 212 | 247.19 | 255 | 239 | 0.94 | -0.089 | 0.62629126 | 0.78958445 | 0.0197 | 0.0202 | 0.02 | 0.02 | TRUE | 0.3525 |
| Zfp462 | 8860 | 8628 | 9826 | 9975 | 7300 | 7434 | 9411 | 8617 | 9866 | 7842 | 7854 | 8138 | 8621.28 | 9298 | 7945 | 0.854 | -0.227 | 0.00310004 | 0.02177901 | 0.0024 | 0.0073 | 0.0041 | 0.0041 | TRUE | 0.0502 |
| Zfp467 | 21 | 32 | 59 | 31 | 28 | 21 | 22 | 32 | 59 | 24 | 30 | 23 | 31.83 | 38 | 26 | 0.681 | -0.555 | 0.21048372 | 0.42100851 | 0.1078 | 0.1104 | 0.109 | 0.109 | TRUE | 0.6562 |
| Zfp472 | 690 | 734 | 674 | 790 | 597 | 616 | 733 | 733 | 677 | 621 | 642 | 674 | 680.07 | 714 | 646 | 0.904 | -0.146 | 0.12769814 | 0.30730165 | 0 | 0.0117 | 0.0052 | 0.0052 | TRUE | 0.025 |
| Zfp473 | 1282 | 1612 | 1761 | 1864 | 1407 | 1518 | 1362 | 1610 | 1768 | 1465 | 1514 | 1662 | 1563.45 | 1580 | 1547 | 0.979 | -0.031 | 0.79360618 | 0.89360725 | 0.0101 | 0.009 | 0.0096 | 0.0096 | TRUE | 0.1754 |
| Zfp474 | 1 | 0 | 0 | 1 | 0 | 2 | 1 | 0 | 0 | 1 | 0 | 2 | 0.67 | 0 | 1 | 2.693 | 1.429 | 0.58190699 | NA | 0 | 4.9025 | 3.3589 | 3.3589 | TRUE | 0.3233 |
| Zfp488 | 6 | 13 | 11 | 6 | 11 | 10 | 6 | 13 | 11 | 5 | 12 | 11 | 9.65 | 10 | 9 | 0.887 | -0.173 | 0.79464464 | 0.8942658 | 0 | 0.3483 | 0.2142 | 0.2142 | TRUE | 0.77 |
| Zfp493 | 365 | 288 | 289 | 455 | 341 | 225 | 388 | 288 | 290 | 358 | 367 | 246 | 322.73 | 322 | 324 | 1.007 | 0.01 | 0.95837197 | 0.97866252 | 0.0348 | 0.0171 | 0.0258 | 0.0258 | TRUE | 0.4859 |
| Zfp503 | 8 | 22 | 15 | 41 | 24 | 29 | 8 | 22 | 15 | 32 | 26 | 32 | 22.55 | 15 | 30 | 1.969 | 0.977 | 0.03252676 | 0.12159567 | 0.0217 | 0.153 | 0.1007 | 0.1007 | TRUE | 0.3852 |
| Zfp507 | 2256 | 2413 | 2276 | 3572 | 2625 | 2492 | 2396 | 2410 | 2285 | 2808 | 2824 | 2728 | 2575.29 | 2364 | 2787 | 1.179 | 0.238 | 0.00037674 | 0.00413204 | 0 | 0.0082 | 0.0028 | 0.0028 | TRUE | 0.0103 |
| Zfp51 | 597 | 641 | 666 | 783 | 538 | 522 | 634 | 640 | 669 | 616 | 579 | 571 | 618.14 | 648 | 589 | 0.91 | -0.136 | 0.1531573 | 0.34469135 | 0 | 0.0122 | 0.0049 | 0.0049 | TRUE | 0.0199 |
| Zfp511 | 767 | 879 | 965 | 1110 | 778 | 798 | 815 | 878 | 969 | 873 | 837 | 874 | 874.12 | 887 | 861 | 0.971 | -0.043 | 0.66796122 | 0.81440046 | 0.0029 | 0.0107 | 0.0061 | 0.0061 | TRUE | 0.0775 |
| Zfp512 | 2732 | 3370 | 3564 | 3410 | 2529 | 2554 | 2902 | 3366 | 3579 | 2681 | 2721 | 2796 | 3007.27 | 3282 | 2733 | 0.832 | -0.265 | 0.00691371 | 0.0397677 | 0.0053 | 0.008 | 0.0066 | 0.0066 | TRUE | 0.1245 |
| Zfp513 | 909 | 926 | 858 | 1203 | 882 | 842 | 966 | 925 | 862 | 946 | 949 | 922 | 928.04 | 918 | 939 | 1.024 | 0.034 | 0.70136939 | 0.83650923 | 6.00E-04 | 0.0105 | 0.0046 | 0.0046 | TRUE | 0.0337 |
| Zfp516 | 4948 | 6639 | 7201 | 6474 | 5228 | 5990 | 5256 | 6630 | 7231 | 5089 | 5625 | 6557 | 6064.67 | 6372 | 5757 | 0.903 | -0.147 | 0.30430284 | 0.52512118 | 0.0201 | 0.0074 | 0.0145 | 0.0145 | TRUE | 0.2865 |
| Zfp518a | 4114 | 3922 | 3874 | 8038 | 5454 | 4617 | 4370 | 3917 | 3890 | 6319 | 5868 | 5054 | 4902.95 | 4059 | 5747 | 1.416 | 0.502 | 2.64E-06 | 6.38E-05 | 0.0081 | 0.0076 | 0.008 | 0.008 | TRUE | 0.1355 |
| Zfp518b | 3772 | 2774 | 2954 | 5935 | 4096 | 3493 | 4007 | 2770 | 2966 | 4666 | 4407 | 3824 | 3773.23 | 3248 | 4299 | 1.324 | 0.405 | 0.00748468 | 0.04216011 | 0.0256 | 0.0078 | 0.0162 | 0.0162 | TRUE | 0.5074 |
| Zfp52 | 249 | 211 | 186 | 291 | 231 | 207 | 264 | 211 | 187 | 229 | 249 | 227 | 227.65 | 221 | 235 | 1.064 | 0.089 | 0.60280246 | 0.77402918 | 0.0123 | 0.0214 | 0.0167 | 0.0167 | TRUE | 0.3277 |
| Zfp521 | 71 | 97 | 111 | 119 | 77 | 112 | 75 | 97 | 111 | 94 | 83 | 123 | 97.12 | 94 | 100 | 1.051 | 0.072 | 0.77376306 | 0.88203173 | 0.0281 | 0.0408 | 0.0346 | 0.0346 | TRUE | 0.377 |
| Zfp523 | 1172 | 1168 | 1098 | 1329 | 988 | 896 | 1245 | 1166 | 1103 | 1045 | 1063 | 981 | 1100.41 | 1171 | 1030 | 0.879 | -0.185 | 0.0409751 | 0.14332177 | 0.0018 | 0.0099 | 0.005 | 0.005 | TRUE | 0.036 |
| Zfp524 | 226 | 210 | 240 | 231 | 193 | 200 | 240 | 210 | 241 | 182 | 208 | 219 | 216.49 | 230 | 203 | 0.878 | -0.188 | 0.22118072 | 0.43313412 | 0.0034 | 0.0221 | 0.0123 | 0.0123 | TRUE | 0.0988 |
| Zfp526 | 397 | 378 | 357 | 544 | 451 | 370 | 422 | 378 | 358 | 428 | 485 | 405 | 412.6 | 386 | 439 | 1.138 | 0.187 | 0.14910979 | 0.33940823 | 0.0052 | 0.0149 | 0.0097 | 0.0097 | TRUE | 0.0941 |
| Zfp53 | 411 | 484 | 499 | 653 | 466 | 435 | 437 | 483 | 501 | 513 | 501 | 476 | 485.31 | 474 | 497 | 1.05 | 0.07 | 0.52531916 | 0.7148075 | 0.0011 | 0.0137 | 0.0067 | 0.0067 | TRUE | 0.0539 |
| Zfp532 | 3632 | 2450 | 2857 | 3703 | 2469 | 2419 | 3858 | 2447 | 2869 | 2911 | 2656 | 2648 | 2898.14 | 3058 | 2738 | 0.896 | -0.159 | 0.31988815 | 0.54101722 | 0.029 | 0.008 | 0.018 | 0.018 | TRUE | 0.5934 |
| Zfp534 | 1625 | 1045 | 1124 | 2016 | 1379 | 1137 | 1726 | 1044 | 1129 | 1585 | 1484 | 1245 | 1368.58 | 1300 | 1438 | 1.107 | 0.147 | 0.44568528 | 0.65230081 | 0.0445 | 0.0093 | 0.0259 | 0.0259 | TRUE | 0.9894 |
| Zfp536 | 36 | 35 | 28 | 74 | 54 | 47 | 38 | 35 | 28 | 58 | 58 | 51 | 44.84 | 34 | 56 | 1.66 | 0.731 | 0.01280831 | 0.06316846 | 0 | 0.0804 | 0.0383 | 0.0383 | TRUE | 0.1514 |
| Zfp54 | 162 | 206 | 173 | 269 | 180 | 146 | 172 | 206 | 174 | 211 | 194 | 160 | 186.08 | 184 | 188 | 1.027 | 0.039 | 0.82656044 | 0.91211327 | 0.01 | 0.0246 | 0.0171 | 0.0171 | TRUE | 0.1841 |
| Zfp541 | 1 | 0 | 3 | 7 | 2 | 3 | 1 | 0 | 3 | 6 | 2 | 3 | 2.5 | 1 | 4 | 2.717 | 1.442 | 0.28375683 | NA | 0 | 1.3235 | 0.7957 | 0.7957 | TRUE | 0.7234 |
| Zfp551 | 74 | 92 | 89 | 160 | 114 | 109 | 79 | 92 | 89 | 126 | 123 | 119 | 104.6 | 87 | 123 | 1.415 | 0.501 | 0.00921961 | 0.04944295 | 0 | 0.0384 | 0.0168 | 0.0168 | TRUE | 0.0604 |
| Zfp553 | 1736 | 1733 | 1571 | 2138 | 1574 | 1456 | 1844 | 1731 | 1577 | 1681 | 1694 | 1594 | 1686.7 | 1717 | 1656 | 0.965 | -0.052 | 0.56428045 | 0.74500513 | 0.0027 | 0.0089 | 0.0053 | 0.0053 | TRUE | 0.0614 |
| Zfp558 | 164 | 213 | 181 | 224 | 200 | 176 | 174 | 213 | 182 | 176 | 215 | 193 | 192.1 | 190 | 195 | 1.024 | 0.034 | 0.83768624 | 0.91756284 | 0.0058 | 0.0241 | 0.0146 | 0.0146 | TRUE | 0.1242 |
| Zfp560 | 273 | 203 | 174 | 453 | 286 | 218 | 290 | 203 | 175 | 356 | 308 | 239 | 261.65 | 223 | 301 | 1.355 | 0.439 | 0.05630575 | 0.17814528 | 0.0499 | 0.0195 | 0.0341 | 0.0341 | TRUE | 0.4863 |
| Zfp563 | 248 | 320 | 297 | 342 | 229 | 249 | 263 | 320 | 298 | 269 | 246 | 273 | 278.17 | 294 | 263 | 0.894 | -0.162 | 0.24322096 | 0.45993003 | 0.0025 | 0.0187 | 0.0102 | 0.0102 | TRUE | 0.0906 |
| Zfp566 | 419 | 343 | 342 | 577 | 411 | 368 | 445 | 343 | 343 | 454 | 442 | 403 | 404.94 | 377 | 433 | 1.15 | 0.201 | 0.16638129 | 0.36370349 | 0.0104 | 0.015 | 0.0128 | 0.0128 | TRUE | 0.2837 |
| Zfp568 | 11632 | 10559 | 10450 | 15416 | 10994 | 10001 | 12355 | 10545 | 10493 | 12119 | 11829 | 10948 | 11381.48 | 11131 | 11632 | 1.045 | 0.064 | 0.49866453 | 0.69429175 | 0.0051 | 0.0072 | 0.0063 | 0.0063 | TRUE | 0.113 |
| Zfp57 | 8183 | 6865 | 5698 | 11709 | 9189 | 7338 | 8692 | 6856 | 5721 | 9205 | 9887 | 8033 | 8065.57 | 7090 | 9042 | 1.275 | 0.351 | 0.02380379 | 0.09822957 | 0.0275 | 0.0073 | 0.0172 | 0.0172 | TRUE | 0.4738 |

| | | | | | | | | | | | | | | | | | | | | | | | | | |
|---|---|---|---|---|---|---|---|---|---|---|---|---|---|---|---|---|---|---|---|---|---|---|---|---|---|
| Zfp572 | 1 | 0 | 0 | 0 | 0 | 0 | 1 | 0 | 0 | 0 | 0 | 0 | 0.18 | 0 | 0 | 0.49 | -1.029 | 0.80087 | NA | 0 | 18.6165 | 10 | 10 | TRUE | 0.4614 |
| Zfp574 | 1333 | 1219 | 1236 | 1786 | 1341 | 1275 | 1416 | 1217 | 1241 | 1404 | 1443 | 1396 | 1352.82 | 1291 | 1414 | 1.095 | 0.131 | 0.15447114 | 0.346511 | 0.0026 | 0.0093 | 0.0054 | 0.0054 | TRUE | 0.0849 |
| Zfp575 | 96 | 104 | 119 | 92 | 72 | 75 | 102 | 104 | 119 | 72 | 77 | 82 | 92.87 | 108 | 77 | 0.711 | -0.493 | 0.01707018 | 0.07758817 | 0 | 0.0424 | 0.0197 | 0.0197 | TRUE | 0.0793 |
| Zfp579 | 105 | 107 | 75 | 134 | 120 | 89 | 112 | 107 | 75 | 105 | 129 | 97 | 104.26 | 98 | 110 | 1.13 | 0.176 | 0.45537485 | 0.6603689 | 0.0217 | 0.0385 | 0.0304 | 0.0304 | TRUE | 0.3991 |
| Zfp58 | 253 | 297 | 376 | 403 | 249 | 278 | 269 | 297 | 378 | 317 | 268 | 304 | 305.32 | 315 | 296 | 0.944 | -0.084 | 0.61666107 | 0.78321617 | 0.0161 | 0.0177 | 0.017 | 0.017 | TRUE | 0.3517 |
| Zfp580 | 324 | 371 | 439 | 385 | 334 | 346 | 344 | 371 | 441 | 303 | 359 | 379 | 366.04 | 385 | 347 | 0.898 | -0.155 | 0.31149836 | 0.53199265 | 0.0124 | 0.0159 | 0.0141 | 0.0141 | TRUE | 0.1838 |
| Zfp583 | 234 | 204 | 248 | 343 | 225 | 245 | 249 | 204 | 249 | 270 | 242 | 268 | 246.87 | 234 | 260 | 1.113 | 0.155 | 0.29562421 | 0.5165093 | 0.0037 | 0.0202 | 0.0118 | 0.0118 | TRUE | 0.1408 |
| Zfp59 | 1879 | 1559 | 1338 | 2535 | 1796 | 1330 | 1996 | 1557 | 1343 | 1993 | 1932 | 1456 | 1712.9 | 1632 | 1794 | 1.099 | 0.137 | 0.42886878 | 0.63860634 | 0.0338 | 0.0088 | 0.021 | 0.021 | TRUE | 0.4577 |
| Zfp592 | 1493 | 1412 | 1386 | 2141 | 1456 | 1369 | 1586 | 1410 | 1392 | 1683 | 1567 | 1499 | 1522.66 | 1463 | 1583 | 1.083 | 0.115 | 0.2254126 | 0.43815832 | 0.0034 | 0.0091 | 0.0058 | 0.0058 | TRUE | 0.0651 |
| Zfp593 | 1124 | 1178 | 1118 | 1857 | 1204 | 1185 | 1194 | 1176 | 1123 | 1460 | 1295 | 1297 | 1257.56 | 1164 | 1351 | 1.161 | 0.216 | 0.01696698 | 0.07731702 | 0.0023 | 0.0095 | 0.0051 | 0.0051 | TRUE | 0.0616 |
| Zfp595 | 207 | 180 | 187 | 246 | 170 | 193 | 220 | 180 | 188 | 193 | 183 | 211 | 195.83 | 196 | 196 | 1 | 0.001 | 0.99648564 | 0.99828918 | 0.0028 | 0.0237 | 0.0131 | 0.0131 | TRUE | 0.1235 |
| Zfp597 | 516 | 525 | 487 | 737 | 515 | 485 | 548 | 524 | 489 | 579 | 554 | 531 | 537.63 | 520 | 555 | 1.067 | 0.094 | 0.37040048 | 0.58736743 | 0 | 0.013 | 0.0061 | 0.0061 | TRUE | 0.0327 |
| Zfp598 | 6804 | 6274 | 6305 | 7479 | 5660 | 5517 | 7227 | 6266 | 6331 | 5879 | 6090 | 6039 | 6305.38 | 6608 | 6003 | 0.908 | -0.139 | 0.09323316 | 0.24963301 | 0.0032 | 0.0074 | 0.0048 | 0.0048 | TRUE | 0.0819 |
| Zfp599 | 900 | 800 | 715 | 1119 | 738 | 541 | 956 | 799 | 718 | 880 | 794 | 592 | 789.8 | 824 | 755 | 0.917 | -0.124 | 0.47133262 | 0.67280822 | 0.0294 | 0.0111 | 0.0202 | 0.0202 | TRUE | 0.4194 |
| Zfp60 | 1225 | 1370 | 1464 | 1753 | 1367 | 1361 | 1301 | 1368 | 1470 | 1378 | 1471 | 1490 | 1413.02 | 1380 | 1446 | 1.047 | 0.067 | 0.44629284 | 0.65236459 | 0.0021 | 0.0092 | 0.0048 | 0.0048 | TRUE | 0.0394 |
| Zfp600 | 1286 | 1052 | 1025 | 2200 | 1667 | 1345 | 1366 | 1051 | 1029 | 1729 | 1794 | 1472 | 1406.86 | 1149 | 1665 | 1.45 | 0.536 | 0.00013214 | 0.00172522 | 0.0176 | 0.0092 | 0.0134 | 0.0134 | TRUE | 0.3275 |
| Zfp605 | 367 | 341 | 347 | 513 | 353 | 322 | 390 | 341 | 348 | 403 | 380 | 352 | 369.06 | 360 | 378 | 1.055 | 0.077 | 0.53656738 | 0.72331397 | 0.0023 | 0.0158 | 0.0085 | 0.0085 | TRUE | 0.0611 |
| Zfp606 | 759 | 834 | 932 | 1031 | 764 | 817 | 806 | 833 | 936 | 810 | 822 | 894 | 850.3 | 858 | 842 | 0.981 | -0.028 | 0.78300957 | 0.88795132 | 0.0034 | 0.0108 | 0.0065 | 0.0065 | TRUE | 0.0744 |
| Zfp607 | 233 | 220 | 188 | 358 | 235 | 221 | 247 | 220 | 189 | 281 | 253 | 242 | 238.7 | 219 | 259 | 1.186 | 0.247 | 0.12110755 | 0.29595728 | 0.0077 | 0.0207 | 0.014 | 0.014 | TRUE | 0.1572 |
| Zfp608 | 124 | 105 | 92 | 60 | 38 | 48 | 132 | 105 | 92 | 47 | 41 | 53 | 78.26 | 110 | 47 | 0.428 | -1.225 | 1.45E-06 | 3.80E-05 | 0.0139 | 0.049 | 0.0315 | 0.0315 | TRUE | 0.3031 |
| Zfp609 | 7092 | 6400 | 6632 | 7789 | 5632 | 5645 | 7533 | 6392 | 6659 | 6123 | 6060 | 6179 | 6491.01 | 6861 | 6121 | 0.892 | -0.165 | 0.0516039 | 0.16750292 | 0.0036 | 0.0074 | 0.005 | 0.005 | TRUE | 0.0894 |
| Zfp61 | 298 | 464 | 416 | 495 | 486 | 416 | 317 | 463 | 418 | 389 | 523 | 455 | 427.51 | 399 | 456 | 1.14 | 0.189 | 0.29311564 | 0.51418305 | 0.027 | 0.0146 | 0.0208 | 0.0208 | TRUE | 0.3246 |
| Zfp612 | 205 | 194 | 182 | 440 | 293 | 193 | 218 | 194 | 183 | 346 | 315 | 211 | 244.44 | 198 | 291 | 1.472 | 0.558 | 0.00710452 | 0.04055462 | 0.0329 | 0.0204 | 0.0267 | 0.0267 | TRUE | 0.6328 |
| Zfp616 | 0 | 0 | 0 | 0 | 0 | 0 | 0 | 0 | 0 | 0 | 0 | 0 | 0 | 0 | 0 | NA | NA | NA | NA | NA | NA | NA | NA | NA | NA |
| Zfp617 | 406 | 340 | 314 | 588 | 378 | 312 | 431 | 340 | 315 | 462 | 407 | 342 | 382.76 | 362 | 404 | 1.117 | 0.16 | 0.35699238 | 0.57553755 | 0.0224 | 0.0155 | 0.019 | 0.019 | TRUE | 0.3154 |
| Zfp618 | 157 | 148 | 156 | 217 | 170 | 100 | 167 | 148 | 157 | 171 | 183 | 109 | 155.7 | 157 | 154 | 0.985 | -0.022 | 0.92110833 | 0.96116102 | 0.03 | 0.0281 | 0.0289 | 0.0289 | TRUE | 0.6532 |
| Zfp619 | 222 | 170 | 180 | 258 | 155 | 164 | 236 | 170 | 181 | 203 | 167 | 180 | 189.24 | 196 | 183 | 0.94 | -0.09 | 0.62983049 | 0.79195749 | 0.0155 | 0.0243 | 0.0198 | 0.0198 | TRUE | 0.3497 |
| Zfp62 | 1994 | 1529 | 1478 | 2519 | 1670 | 1184 | 2118 | 1527 | 1484 | 1980 | 1797 | 1296 | 1700.37 | 1710 | 1691 | 0.99 | -0.015 | 0.93616149 | 0.96816629 | 0.0428 | 0.0088 | 0.0249 | 0.0249 | TRUE | 0.5261 |
| Zfp622 | 3495 | 3844 | 3762 | 4930 | 3668 | 3394 | 3712 | 3839 | 3777 | 3876 | 3947 | 3715 | 3811.03 | 3776 | 3846 | 1.018 | 0.026 | 0.67919421 | 0.82180676 | 3.00E-04 | 0.0078 | 0.0027 | 0.0027 | TRUE | 0.0106 |
| Zfp623 | 288 | 367 | 394 | 507 | 386 | 365 | 306 | 367 | 396 | 399 | 415 | 400 | 380.25 | 356 | 405 | 1.135 | 0.183 | 0.17233836 | 0.37117583 | 0.0055 | 0.0156 | 0.0103 | 0.0103 | TRUE | 0.1711 |
| Zfp628 | 369 | 253 | 174 | 427 | 262 | 170 | 392 | 253 | 175 | 336 | 282 | 186 | 270.5 | 273 | 268 | 0.983 | -0.025 | 0.93443472 | 0.96749848 | 0.1181 | 0.0191 | 0.0631 | 0.0631 | TRUE | 0.4179 |
| Zfp629 | 130 | 167 | 152 | 98 | 68 | 72 | 138 | 167 | 153 | 77 | 73 | 79 | 114.42 | 153 | 76 | 0.5 | -0.999 | 2.20E-07 | 7.13E-06 | 0 | 0.0357 | 0.017 | 0.017 | TRUE | 0.0707 |
| Zfp637 | 371 | 360 | 265 | 501 | 312 | 309 | 394 | 360 | 266 | 394 | 336 | 338 | 347.91 | 340 | 356 | 1.049 | 0.069 | 0.69094751 | 0.830485 | 0.0213 | 0.0164 | 0.0189 | 0.0189 | TRUE | 0.4124 |
| Zfp639 | 1524 | 1509 | 1506 | 1971 | 1408 | 1407 | 1619 | 1507 | 1512 | 1549 | 1515 | 1540 | 1540.42 | 1546 | 1535 | 0.993 | -0.01 | 0.89395743 | 0.94875827 | 0 | 0.009 | 0.0034 | 0.0034 | TRUE | 0.0203 |
| Zfp64 | 2650 | 2912 | 3062 | 3656 | 2871 | 2607 | 2815 | 2908 | 3075 | 2874 | 3089 | 2854 | 2935.74 | 2933 | 2939 | 1.002 | 0.003 | 0.97039904 | 0.9846576 | 0.0016 | 0.008 | 0.0039 | 0.0039 | TRUE | 0.024 |
| Zfp641 | 22 | 18 | 17 | 15 | 18 | 18 | 23 | 18 | 17 | 12 | 19 | 20 | 18.21 | 19 | 17 | 0.862 | -0.215 | 0.64281513 | 0.79906055 | 0 | 0.1878 | 0.0995 | 0.0995 | TRUE | 0.4416 |
| Zfp644 | 2627 | 2342 | 2398 | 3264 | 2163 | 2041 | 2790 | 2339 | 2408 | 2566 | 2327 | 2234 | 2444.09 | 2512 | 2376 | 0.946 | -0.08 | 0.44144104 | 0.64845004 | 0.0064 | 0.0083 | 0.0073 | 0.0073 | TRUE | 0.1133 |
| Zfp646 | 3298 | 3304 | 3321 | 3960 | 2922 | 2874 | 3503 | 3300 | 3335 | 3113 | 3144 | 3146 | 3256.73 | 3379 | 3134 | 0.928 | -0.108 | 0.09099737 | 0.24600797 | 2.00E-04 | 0.0079 | 0.0027 | 0.0027 | TRUE | 0.0125 |
| Zfp647 | 323 | 344 | 365 | 381 | 255 | 243 | 343 | 344 | 366 | 300 | 274 | 266 | 315.5 | 351 | 280 | 0.799 | -0.323 | 0.00895185 | 0.04848508 | 0 | 0.0173 | 0.0078 | 0.0078 | TRUE | 0.0453 |
| Zfp648 | 0 | 0 | 2 | 1 | 0 | 0 | 0 | 0 | 2 | 1 | 0 | 0 | 0.47 | 1 | 0 | 0.52 | -0.943 | 0.78484416 | NA | 0 | 7.0808 | 6.8441 | 6.8441 | TRUE | 0.9739 |
| Zfp651 | 77 | 111 | 103 | 123 | 105 | 90 | 82 | 111 | 103 | 97 | 113 | 99 | 100.71 | 99 | 103 | 1.038 | 0.054 | 0.80058927 | 0.89705129 | 0.0049 | 0.0396 | 0.0229 | 0.0229 | TRUE | 0.2137 |
| Zfp652 | 1630 | 2183 | 2629 | 2211 | 1613 | 1847 | 1731 | 2180 | 2640 | 1738 | 1735 | 2022 | 2007.79 | 2184 | 1832 | 0.839 | -0.254 | 0.10344147 | 0.26760005 | 0.0249 | 0.0085 | 0.017 | 0.017 | TRUE | 0.1604 |
| Zfp653 | 397 | 479 | 478 | 518 | 461 | 406 | 422 | 478 | 480 | 407 | 496 | 444 | 454.61 | 460 | 449 | 0.975 | -0.037 | 0.76901264 | 0.87928573 | 0.0055 | 0.0141 | 0.0094 | 0.0094 | TRUE | 0.0961 |
| Zfp654 | 1235 | 1199 | 1220 | 1495 | 1104 | 1177 | 1312 | 1197 | 1225 | 1175 | 1188 | 1288 | 1230.96 | 1245 | 1217 | 0.977 | -0.033 | 0.70666372 | 0.8398044 | 0.0016 | 0.0096 | 0.0047 | 0.0047 | TRUE | 0.031 |
| Zfp655 | 3167 | 4755 | 5140 | 4042 | 3067 | 3665 | 3364 | 4749 | 5161 | 3178 | 3300 | 4012 | 3960.53 | 4425 | 3497 | 0.79 | -0.34 | 0.04073483 | 0.14280279 | 0.0306 | 0.0077 | 0.0196 | 0.0196 | TRUE | 0.5353 |
| Zfp658 | 332 | 243 | 197 | 324 | 203 | 155 | 353 | 243 | 198 | 255 | 218 | 170 | 239.32 | 265 | 214 | 0.813 | -0.299 | 0.2230729 | 0.43505698 | 0.0589 | 0.0207 | 0.0393 | 0.0393 | TRUE | 0.5825 |
| Zfp661 | 497 | 516 | 492 | 524 | 430 | 421 | 528 | 515 | 494 | 412 | 463 | 461 | 478.78 | 512 | 445 | 0.867 | -0.206 | 0.05917391 | 0.18419005 | 0 | 0.0138 | 0.0065 | 0.0065 | TRUE | 0.0529 |
| Zfp663 | 24 | 12 | 12 | 18 | 13 | 14 | 25 | 12 | 12 | 14 | 14 | 15 | 15.5 | 16 | 14 | 0.88 | -0.185 | 0.72507045 | 0.85138559 | 0.0453 | 0.2195 | 0.1342 | 0.1342 | TRUE | 1.056 |
| Zfp664 | 5866 | 6426 | 6506 | 7707 | 5704 | 5878 | 6231 | 6418 | 6533 | 6059 | 6137 | 6434 | 6301.89 | 6394 | 6210 | 0.971 | -0.042 | 0.50176884 | 0.69709047 | 6.00E-04 | 0.0074 | 0.0027 | 0.0027 | TRUE | 0.0121 |
| Zfp667 | 725 | 616 | 567 | 660 | 481 | 473 | 770 | 615 | 569 | 519 | 518 | 518 | 584.79 | 651 | 518 | 0.795 | -0.33 | 0.01585246 | 0.07388445 | 0.0104 | 0.0125 | 0.0118 | 0.0118 | TRUE | 0.297 |
| Zfp668 | 551 | 467 | 477 | 670 | 484 | 419 | 585 | 466 | 479 | 527 | 521 | 459 | 506.12 | 510 | 502 | 0.985 | -0.021 | 0.87251909 | 0.93737812 | 0.0087 | 0.0134 | 0.0109 | 0.0109 | TRUE | 0.1918 |
| Zfp672 | 111 | 123 | 107 | 162 | 110 | 97 | 118 | 123 | 107 | 127 | 118 | 106 | 116.68 | 116 | 117 | 1.014 | 0.02 | 0.91617363 | 0.95891503 | 0 | 0.0351 | 0.0169 | 0.0169 | TRUE | 0.0671 |
| Zfp677 | 97 | 133 | 112 | 160 | 113 | 118 | 103 | 133 | 112 | 126 | 122 | 129 | 120.81 | 116 | 126 | 1.08 | 0.111 | 0.55624337 | 0.73896547 | 0 | 0.0342 | 0.0175 | 0.0175 | TRUE | 0.1604 |
| Zfp68 | 721 | 960 | 945 | 951 | 709 | 831 | 766 | 959 | 949 | 748 | 763 | 910 | 848.93 | 891 | 807 | 0.904 | -0.145 | 0.27779577 | 0.49891315 | 0.0118 | 0.0108 | 0.0117 | 0.0117 | TRUE | 0.1793 |
| Zfp687 | 2832 | 2903 | 3105 | 3968 | 2998 | 2889 | 3008 | 2899 | 3118 | 3119 | 3226 | 3162 | 3088.76 | 3008 | 3169 | 1.053 | 0.075 | 0.26433204 | 0.48381628 | 5.00E-04 | 0.008 | 0.0029 | 0.0029 | TRUE | 0.0123 |
| Zfp688 | 338 | 474 | 435 | 541 | 418 | 474 | 359 | 473 | 437 | 425 | 450 | 519 | 443.85 | 423 | 465 | 1.096 | 0.133 | 0.36940076 | 0.58676318 | 0.0128 | 0.0143 | 0.0135 | 0.0135 | TRUE | 0.2008 |
| Zfp689 | 301 | 455 | 457 | 497 | 396 | 378 | 320 | 454 | 459 | 391 | 426 | 414 | 410.59 | 411 | 410 | 0.997 | -0.005 | 0.97766224 | 0.98797393 | 0.0183 | 0.0149 | 0.0167 | 0.0167 | TRUE | 0.4317 |
| Zfp69 | 252 | 272 | 240 | 326 | 219 | 228 | 268 | 272 | 241 | 256 | 236 | 250 | 253.63 | 260 | 247 | 0.951 | -0.072 | 0.5925822 | 0.76595693 | 0 | 0.0199 | 0.009 | 0.009 | TRUE | 0.0465 |
| Zfp691 | 178 | 167 | 216 | 216 | 165 | 207 | 189 | 167 | 217 | 170 | 178 | 227 | 191.11 | 191 | 192 | 1 | -0.001 | 0.99706887 | 0.99828918 | 0.0158 | 0.0241 | 0.0199 | 0.0199 | TRUE | 0.2751 |
| Zfp692 | 1259 | 1516 | 1533 | 1456 | 1337 | 1342 | 1337 | 1514 | 1539 | 1145 | 1439 | 1469 | 1407.13 | 1463 | 1351 | 0.922 | -0.117 | 0.34714157 | 0.56607908 | 0.0115 | 0.0092 | 0.0105 | 0.0105 | TRUE | 0.2185 |
| Zfp697 | 80 | 173 | 116 | 103 | 80 | 73 | 85 | 173 | 116 | 81 | 86 | 80 | 103.53 | 125 | 82 | 0.659 | -0.602 | 0.03339811 | 0.12391319 | 0.0565 | 0.0387 | 0.0476 | 0.0476 | TRUE | 0.2969 |
| Zfp7 | 2277 | 2426 | 2488 | 2879 | 2250 | 2207 | 2419 | 2423 | 2498 | 2263 | 2421 | 2416 | 2406.61 | 2447 | 2367 | 0.967 | -0.049 | 0.48782963 | 0.68458483 | 6.00E-04 | 0.0083 | 0.0031 | 0.0031 | TRUE | 0.0182 |
| Zfp703 | 57 | 58 | 57 | 135 | 83 | 76 | 61 | 58 | 57 | 106 | 89 | 83 | 75.72 | 59 | 93 | 1.593 | 0.672 | 0.00381678 | 0.02543232 | 0 | 0.0504 | 0.0249 | 0.0249 | TRUE | 0.1658 |
| Zfp704 | 2701 | 2396 | 2656 | 3579 | 2796 | 2619 | 2869 | 2393 | 2667 | 2814 | 3008 | 2867 | 2769.59 | 2643 | 2896 | 1.096 | 0.132 | 0.15371417 | 0.34542149 | 0.0041 | 0.0081 | 0.0058 | 0.0058 | TRUE | 0.0832 |
| Zfp706 | 8866 | 9765 | 9301 | 11757 | 8716 | 9101 | 9417 | 9752 | 9339 | 9242 | 9378 | 9962 | 9515.27 | 9503 | 9527 | 1.002 | 0.004 | 0.95586957 | 0.97741618 | 8.00E-04 | 0.0072 | 0.0029 | 0.0029 | TRUE | 0.0194 |
| Zfp707 | 316 | 340 | 322 | 486 | 317 | 273 | 336 | 340 | 323 | 382 | 341 | 299 | 336.75 | 333 | 341 | 1.026 | 0.038 | 0.78355002 | 0.88830872 | 0.0052 | 0.0167 | 0.0105 | 0.0105 | TRUE | 0.1358 |
| Zfp708 | 242 | 226 | 233 | 263 | 201 | 232 | 257 | 226 | 234 | 207 | 216 | 254 | 232.28 | 239 | 226 | 0.943 | -0.085 | 0.57284161 | 0.7512479 | 0.0039 | 0.0211 | 0.0122 | 0.0122 | TRUE | 0.1293 |
| Zfp709 | 88 | 76 | 79 | 187 | 118 | 98 | 93 | 76 | 79 | 147 | 127 | 107 | 104.99 | 83 | 127 | 1.54 | 0.623 | 0.00430101 | 0.02791204 | 0.0108 | 0.0383 | 0.0243 | 0.0243 | TRUE | 0.1403 |
| Zfp71-rs1 | 739 | 529 | 526 | 932 | 672 | 430 | 785 | 528 | 528 | 733 | 723 | 471 | 627.97 | 614 | 642 | 1.047 | 0.067 | 0.75800581 | 0.87226526 | 0.0547 | 0.0121 | 0.032 | 0.032 | TRUE | 0.6976 |

| | | | | | | | | | | | | | | | | | | | | | | | | | |
|---|---|---|---|---|---|---|---|---|---|---|---|---|---|---|---|---|---|---|---|---|---|---|---|---|---|
| Zfp710 | 15126 | 16247 | 16447 | 11508 | 9297 | 10693 | 16066 | 16226 | 16515 | 9047 | 10003 | 11705 | 13260.32 | 16269 | 10252 | 0.63 | -0.666 | 2.49E-10 | 1.75E-08 | 0.0086 | 0.0072 | 0.0079 | 0.0079 | TRUE | 0.1879 |
| Zfp711 | 88 | 107 | 92 | 196 | 127 | 90 | 93 | 107 | 92 | 154 | 137 | 99 | 113.66 | 97 | 130 | 1.335 | 0.417 | 0.06830322 | 0.20245789 | 0.0216 | 0.0359 | 0.0287 | 0.0287 | TRUE | 0.4396 |
| Zfp712 | 170 | 233 | 254 | 275 | 216 | 236 | 181 | 233 | 255 | 216 | 232 | 258 | 229.21 | 223 | 235 | 1.055 | 0.078 | 0.65923926 | 0.80795659 | 0.0146 | 0.0213 | 0.0179 | 0.0179 | TRUE | 0.2977 |
| Zfp715 | 1369 | 1359 | 1262 | 1656 | 1291 | 1230 | 1454 | 1357 | 1267 | 1302 | 1389 | 1346 | 1352.64 | 1359 | 1346 | 0.99 | -0.015 | 0.86570682 | 0.93364615 | 0.0021 | 0.0093 | 0.0049 | 0.0049 | TRUE | 0.0443 |
| Zfp719 | 469 | 457 | 384 | 757 | 550 | 352 | 498 | 456 | 386 | 595 | 592 | 385 | 485.39 | 447 | 524 | 1.175 | 0.232 | 0.22459495 | 0.43701051 | 0.0353 | 0.0137 | 0.0243 | 0.0243 | TRUE | 0.6183 |
| Zfp72 | 307 | 278 | 251 | 289 | 214 | 214 | 326 | 278 | 252 | 227 | 230 | 234 | 257.91 | 285 | 230 | 0.808 | -0.307 | 0.03932974 | 0.13899559 | 0.0053 | 0.0197 | 0.0121 | 0.0121 | TRUE | 0.1743 |
| Zfp735 | 0 | 0 | 0 | 0 | 0 | 0 | 0 | 0 | 0 | 0 | 0 | 0 | 0 | 0 | 0 | NA | NA | NA | NA | NA | NA | NA | NA | NA | NA |
| Zfp738 | 252 | 297 | 311 | 323 | 274 | 249 | 268 | 297 | 312 | 254 | 295 | 273 | 282.98 | 292 | 274 | 0.935 | -0.097 | 0.47727882 | 0.6767636 | 0.0024 | 0.0185 | 0.01 | 0.01 | TRUE | 0.0598 |
| Zfp74 | 565 | 517 | 524 | 733 | 574 | 538 | 600 | 516 | 526 | 576 | 618 | 589 | 570.89 | 547 | 594 | 1.085 | 0.118 | 0.27840933 | 0.49960996 | 0.0021 | 0.0127 | 0.0068 | 0.0068 | TRUE | 0.0819 |
| Zfp740 | 5736 | 5924 | 6245 | 6677 | 4939 | 4561 | 6093 | 5916 | 6271 | 5249 | 5314 | 4993 | 5639.23 | 6093 | 5185 | 0.851 | -0.233 | 0.00035312 | 0.00391383 | 7.00E-04 | 0.0075 | 0.0029 | 0.0029 | TRUE | 0.0127 |
| Zfp746 | 710 | 1055 | 1076 | 1246 | 957 | 992 | 754 | 1054 | 1080 | 980 | 1030 | 1086 | 997.21 | 963 | 1032 | 1.071 | 0.099 | 0.50575438 | 0.70029387 | 0.0193 | 0.0102 | 0.0149 | 0.0149 | TRUE | 0.4269 |
| Zfp747 | 351 | 361 | 285 | 401 | 270 | 261 | 373 | 361 | 286 | 315 | 291 | 286 | 318.5 | 340 | 297 | 0.876 | -0.191 | 0.19572148 | 0.40189026 | 0.0085 | 0.0172 | 0.0126 | 0.0126 | TRUE | 0.2184 |
| Zfp748 | 870 | 1008 | 1023 | 1215 | 823 | 829 | 924 | 1007 | 1027 | 955 | 885 | 907 | 951.02 | 986 | 916 | 0.93 | -0.105 | 0.24726059 | 0.46398502 | 0.0013 | 0.0104 | 0.0049 | 0.0049 | TRUE | 0.0357 |
| Zfp750 | 416 | 437 | 372 | 463 | 403 | 429 | 442 | 436 | 374 | 364 | 434 | 470 | 419.84 | 417 | 423 | 1.01 | 0.015 | 0.91938066 | 0.96048697 | 0.0105 | 0.0148 | 0.0125 | 0.0125 | TRUE | 0.1776 |
| Zfp758 | 84 | 68 | 68 | 160 | 102 | 68 | 89 | 68 | 68 | 126 | 110 | 74 | 89.23 | 75 | 103 | 1.382 | 0.467 | 0.08223291 | 0.22890531 | 0.0372 | 0.0438 | 0.0406 | 0.0406 | TRUE | 0.4405 |
| Zfp759 | 110 | 91 | 103 | 261 | 183 | 120 | 117 | 91 | 103 | 205 | 197 | 131 | 140.76 | 104 | 178 | 1.72 | 0.782 | 0.00065563 | 0.00643468 | 0.0303 | 0.0303 | 0.0303 | 0.0303 | TRUE | 0.5437 |
| Zfp760 | 752 | 647 | 706 | 1085 | 818 | 700 | 799 | 646 | 709 | 853 | 880 | 766 | 775.52 | 718 | 833 | 1.161 | 0.216 | 0.06756218 | 0.20101693 | 0.0063 | 0.0112 | 0.0087 | 0.0087 | TRUE | 0.114 |
| Zfp763 | 494 | 642 | 590 | 688 | 467 | 575 | 525 | 641 | 592 | 541 | 502 | 629 | 571.84 | 586 | 557 | 0.951 | -0.073 | 0.58381976 | 0.75929786 | 0.0098 | 0.0127 | 0.0111 | 0.0111 | TRUE | 0.1452 |
| Zfp764 | 357 | 295 | 263 | 418 | 279 | 229 | 379 | 295 | 264 | 329 | 300 | 251 | 302.89 | 313 | 293 | 0.94 | -0.089 | 0.62408321 | 0.78826303 | 0.0235 | 0.0178 | 0.0207 | 0.0207 | TRUE | 0.39 |
| Zfp768 | 77 | 99 | 96 | 122 | 74 | 73 | 82 | 99 | 96 | 96 | 80 | 80 | 88.75 | 92 | 85 | 0.925 | -0.112 | 0.60618937 | 0.77673399 | 0 | 0.044 | 0.0227 | 0.0227 | TRUE | 0.1292 |
| Zfp770 | 1889 | 1456 | 1379 | 2221 | 1610 | 1361 | 2006 | 1454 | 1385 | 1746 | 1732 | 1490 | 1635.55 | 1615 | 1656 | 1.026 | 0.037 | 0.81262968 | 0.90398098 | 0.0241 | 0.0089 | 0.0166 | 0.0166 | TRUE | 0.541 |
| Zfp771 | 612 | 545 | 525 | 708 | 568 | 412 | 650 | 544 | 527 | 557 | 611 | 451 | 556.7 | 574 | 540 | 0.941 | -0.088 | 0.56047942 | 0.74228578 | 0.0162 | 0.0128 | 0.0146 | 0.0146 | TRUE | 0.2356 |
| Zfp772 | 292 | 270 | 274 | 383 | 250 | 282 | 310 | 270 | 275 | 301 | 269 | 309 | 288.95 | 285 | 293 | 1.029 | 0.041 | 0.7596042 | 0.87323722 | 0.0017 | 0.0183 | 0.0096 | 0.0096 | TRUE | 0.0663 |
| Zfp773 | 204 | 199 | 178 | 221 | 157 | 173 | 217 | 199 | 179 | 174 | 169 | 189 | 187.7 | 198 | 177 | 0.895 | -0.16 | 0.31028903 | 0.53073186 | 0 | 0.0245 | 0.0125 | 0.0125 | TRUE | 0.0792 |
| Zfp775 | 722 | 1033 | 1055 | 896 | 751 | 814 | 767 | 1032 | 1059 | 704 | 808 | 891 | 876.89 | 953 | 801 | 0.84 | -0.251 | 0.10394242 | 0.26817827 | 0.0212 | 0.0107 | 0.0161 | 0.0161 | TRUE | 0.3457 |
| Zfp777 | 849 | 1053 | 1050 | 1528 | 1177 | 1135 | 902 | 1052 | 1054 | 1201 | 1266 | 1242 | 1119.62 | 1003 | 1236 | 1.233 | 0.302 | 0.00193602 | 0.01499011 | 0.0031 | 0.0098 | 0.0059 | 0.0059 | TRUE | 0.0917 |
| Zfp78 | 75 | 70 | 68 | 94 | 65 | 59 | 80 | 70 | 68 | 74 | 70 | 65 | 71.04 | 73 | 70 | 0.96 | -0.059 | 0.79685294 | 0.89536272 | 0 | 0.0533 | 0.024 | 0.024 | TRUE | 0.0638 |
| Zfp780b | 143 | 90 | 92 | 296 | 180 | 110 | 152 | 90 | 92 | 233 | 194 | 120 | 146.82 | 111 | 182 | 1.642 | 0.715 | 0.01771179 | 0.07951904 | 0.0913 | 0.0293 | 0.0583 | 0.0583 | TRUE | 0.3366 |
| Zfp781 | 40 | 41 | 46 | 37 | 39 | 50 | 42 | 41 | 46 | 29 | 42 | 55 | 42.57 | 43 | 42 | 0.961 | -0.057 | 0.86426357 | 0.93287105 | 0.03 | 0.0843 | 0.0576 | 0.0576 | TRUE | 0.212 |
| Zfp783 | 41 | 26 | 14 | 68 | 40 | 28 | 44 | 26 | 14 | 53 | 43 | 31 | 35.12 | 28 | 42 | 1.532 | 0.615 | 0.17907335 | 0.38024408 | 0.1452 | 0.1007 | 0.1213 | 0.1213 | TRUE | 0.4848 |
| Zfp784 | 22 | 15 | 20 | 42 | 29 | 30 | 23 | 15 | 20 | 33 | 31 | 33 | 25.91 | 19 | 32 | 1.665 | 0.736 | 0.05378628 | 0.1723792 | 0 | 0.134 | 0.0635 | 0.0635 | TRUE | 0.2204 |
| Zfp786 | 83 | 102 | 75 | 139 | 100 | 64 | 88 | 102 | 75 | 109 | 108 | 70 | 92.04 | 88 | 96 | 1.085 | 0.118 | 0.64485513 | 0.80034312 | 0.0293 | 0.0427 | 0.0362 | 0.0362 | TRUE | 0.5061 |
| Zfp787 | 294 | 292 | 270 | 290 | 185 | 198 | 312 | 292 | 271 | 228 | 199 | 217 | 253.13 | 292 | 215 | 0.737 | -0.439 | 0.00162935 | 0.01307428 | 0 | 0.0199 | 0.01 | 0.01 | TRUE | 0.0432 |
| Zfp788 | 548 | 549 | 563 | 714 | 544 | 528 | 582 | 548 | 565 | 561 | 585 | 578 | 570.04 | 565 | 575 | 1.017 | 0.024 | 0.80369621 | 0.89855367 | 0 | 0.0127 | 0.0048 | 0.0048 | TRUE | 0.0081 |
| Zfp790 | 440 | 417 | 427 | 589 | 410 | 422 | 467 | 416 | 429 | 463 | 441 | 462 | 446.45 | 437 | 455 | 1.042 | 0.059 | 0.5890819 | 0.76374843 | 0 | 0.0143 | 0.0063 | 0.0063 | TRUE | 0.0407 |
| Zfp791 | 170 | 164 | 158 | 173 | 125 | 130 | 181 | 164 | 159 | 136 | 134 | 142 | 152.63 | 168 | 137 | 0.821 | -0.285 | 0.07944775 | 0.22359919 | 0 | 0.0285 | 0.0124 | 0.0124 | TRUE | 0.0473 |
| Zfp799 | 1005 | 1091 | 1102 | 1163 | 750 | 855 | 1067 | 1090 | 1107 | 914 | 807 | 936 | 986.79 | 1088 | 886 | 0.815 | -0.296 | 0.00172022 | 0.01364755 | 0.0019 | 0.0102 | 0.0054 | 0.0054 | TRUE | 0.0704 |
| Zfp800 | 425 | 643 | 659 | 823 | 608 | 564 | 451 | 642 | 662 | 647 | 654 | 617 | 612.31 | 585 | 639 | 1.093 | 0.128 | 0.41625704 | 0.62771911 | 0.0199 | 0.0123 | 0.0162 | 0.0162 | TRUE | 0.4659 |
| Zfp804a | 0 | 0 | 0 | 0 | 0 | 0 | 0 | 0 | 0 | 0 | 0 | 0 | 0 | 0 | 0 | NA | NA | NA | NA | NA | NA | NA | NA | NA | NA |
| Zfp804b | 1 | 0 | 0 | 0 | 0 | 0 | 1 | 0 | 0 | 0 | 0 | 0 | 0.18 | 0 | 0 | 0.49 | -1.029 | 0.80087 | NA | 0 | 18.6165 | 10 | 10 | TRUE | 0.4614 |
| Zfp808 | 867 | 667 | 724 | 1229 | 835 | 722 | 921 | 666 | 727 | 966 | 898 | 790 | 828.15 | 771 | 885 | 1.148 | 0.2 | 0.17804286 | 0.37877557 | 0.0181 | 0.0109 | 0.0146 | 0.0146 | TRUE | 0.3399 |
| Zfp809 | 1400 | 1192 | 1084 | 2055 | 1449 | 1253 | 1487 | 1190 | 1088 | 1615 | 1559 | 1372 | 1385.35 | 1255 | 1515 | 1.208 | 0.272 | 0.04673544 | 0.15663537 | 0.0161 | 0.0093 | 0.0128 | 0.0128 | TRUE | 0.3122 |
| Zfp81 | 499 | 604 | 586 | 699 | 491 | 538 | 530 | 603 | 588 | 549 | 528 | 589 | 564.73 | 574 | 555 | 0.967 | -0.048 | 0.65913411 | 0.80794771 | 0.0019 | 0.0127 | 0.0067 | 0.0067 | TRUE | 0.0519 |
| Zfp810 | 260 | 298 | 306 | 390 | 258 | 274 | 276 | 298 | 307 | 307 | 278 | 300 | 294.19 | 294 | 295 | 1.004 | 0.006 | 0.96099921 | 0.98013965 | 0 | 0.0181 | 0.0082 | 0.0082 | TRUE | 0.0302 |
| Zfp811 | 467 | 475 | 487 | 541 | 407 | 383 | 496 | 474 | 489 | 425 | 438 | 419 | 456.98 | 486 | 427 | 0.879 | -0.187 | 0.06750257 | 0.20087746 | 0 | 0.0141 | 0.0053 | 0.0053 | TRUE | 0.0055 |
| Zfp819 | 427 | 380 | 384 | 425 | 310 | 267 | 454 | 380 | 386 | 334 | 334 | 292 | 363.09 | 407 | 320 | 0.789 | -0.342 | 0.01062302 | 0.05509578 | 0.0051 | 0.016 | 0.0101 | 0.0101 | TRUE | 0.1187 |
| Zfp82 | 333 | 425 | 409 | 517 | 367 | 387 | 354 | 424 | 411 | 406 | 395 | 424 | 402.29 | 396 | 408 | 1.03 | 0.042 | 0.72998468 | 0.85500295 | 0.0024 | 0.0151 | 0.0083 | 0.0083 | TRUE | 0.1004 |
| Zfp820 | 543 | 443 | 455 | 770 | 588 | 491 | 577 | 442 | 457 | 605 | 633 | 537 | 541.92 | 492 | 592 | 1.204 | 0.268 | 0.05798579 | 0.1819496 | 0.0119 | 0.013 | 0.0125 | 0.0125 | TRUE | 0.2626 |
| Zfp821 | 537 | 624 | 659 | 933 | 706 | 615 | 570 | 623 | 662 | 733 | 760 | 673 | 670.26 | 618 | 722 | 1.168 | 0.224 | 0.0375686 | 0.13461602 | 0.003 | 0.0118 | 0.0068 | 0.0068 | TRUE | 0.0538 |
| Zfp825 | 480 | 477 | 519 | 551 | 391 | 411 | 510 | 476 | 521 | 433 | 421 | 450 | 468.52 | 502 | 435 | 0.865 | -0.209 | 0.04713635 | 0.15766097 | 0 | 0.0139 | 0.0059 | 0.0059 | TRUE | 0.0242 |
| Zfp827 | 519 | 431 | 449 | 626 | 474 | 366 | 551 | 430 | 451 | 492 | 510 | 401 | 472.55 | 477 | 468 | 0.98 | -0.029 | 0.84966266 | 0.92515564 | 0.0147 | 0.0139 | 0.0143 | 0.0143 | TRUE | 0.2109 |
| Zfp828 | 5729 | 5093 | 4948 | 7006 | 5046 | 4840 | 6085 | 5086 | 4968 | 5508 | 5429 | 5298 | 5395.81 | 5380 | 5412 | 1.006 | 0.009 | 0.9299394 | 0.96556973 | 0.0063 | 0.0075 | 0.0068 | 0.0068 | TRUE | 0.1601 |
| Zfp830 | 1452 | 1165 | 1176 | 2015 | 1484 | 1326 | 1542 | 1163 | 1181 | 1584 | 1597 | 1452 | 1419.81 | 1295 | 1544 | 1.192 | 0.254 | 0.05122863 | 0.16683419 | 0.0139 | 0.0092 | 0.0115 | 0.0115 | TRUE | 0.3326 |
| Zfp831 | 36 | 23 | 26 | 49 | 30 | 35 | 38 | 23 | 26 | 39 | 32 | 38 | 32.74 | 29 | 36 | 1.255 | 0.327 | 0.35804861 | 0.57647435 | 0 | 0.1075 | 0.0604 | 0.0604 | TRUE | 0.4775 |
| Zfp839 | 644 | 912 | 918 | 1030 | 740 | 789 | 684 | 911 | 922 | 810 | 796 | 864 | 831.04 | 839 | 823 | 0.981 | -0.028 | 0.83731859 | 0.91728754 | 0.0133 | 0.0109 | 0.0122 | 0.0122 | TRUE | 0.3083 |
| Zfp84 | 517 | 504 | 504 | 665 | 434 | 391 | 549 | 503 | 506 | 523 | 467 | 428 | 496.05 | 519 | 473 | 0.912 | -0.133 | 0.26907836 | 0.4890417 | 0.0044 | 0.0135 | 0.0084 | 0.0084 | TRUE | 0.1051 |
| Zfp846 | 320 | 324 | 292 | 523 | 419 | 394 | 340 | 324 | 293 | 411 | 451 | 431 | 374.99 | 319 | 431 | 1.351 | 0.434 | 0.00032178 | 0.00360967 | 0 | 0.0157 | 0.0077 | 0.0077 | TRUE | 0.0565 |
| Zfp85-rs1 | 230 | 226 | 237 | 299 | 206 | 211 | 244 | 226 | 238 | 235 | 222 | 231 | 232.61 | 236 | 229 | 0.972 | -0.04 | 0.76273879 | 0.87490299 | 0 | 0.0211 | 0.0086 | 0.0086 | TRUE | 0.0162 |
| Zfp862 | 277 | 245 | 232 | 383 | 279 | 209 | 294 | 245 | 233 | 301 | 300 | 229 | 266.99 | 257 | 277 | 1.078 | 0.108 | 0.52824418 | 0.71693204 | 0.0156 | 0.0192 | 0.0174 | 0.0174 | TRUE | 0.2501 |
| Zfp865 | 601 | 458 | 450 | 739 | 497 | 443 | 638 | 457 | 452 | 581 | 535 | 485 | 524.71 | 516 | 534 | 1.036 | 0.051 | 0.75907363 | 0.87295942 | 0.0219 | 0.0132 | 0.0177 | 0.0177 | TRUE | 0.5012 |
| Zfp866 | 1355 | 1287 | 1396 | 1900 | 1465 | 1275 | 1439 | 1285 | 1402 | 1494 | 1576 | 1396 | 1431.98 | 1375 | 1489 | 1.082 | 0.114 | 0.21225977 | 0.4232315 | 0.0028 | 0.0092 | 0.0053 | 0.0053 | TRUE | 0.0396 |
| Zfp867 | 831 | 626 | 574 | 767 | 514 | 494 | 883 | 625 | 576 | 603 | 553 | 541 | 630.16 | 695 | 566 | 0.815 | -0.295 | 0.08418992 | 0.23262849 | 0.0273 | 0.0121 | 0.0195 | 0.0195 | TRUE | 0.6594 |
| Zfp868 | 823 | 758 | 734 | 1418 | 1022 | 839 | 874 | 757 | 737 | 1115 | 1100 | 918 | 916.83 | 789 | 1044 | 1.324 | 0.405 | 0.00088629 | 0.00810349 | 0.0082 | 0.0105 | 0.0096 | 0.0096 | TRUE | 0.1313 |
| Zfp869 | 1031 | 882 | 763 | 1344 | 1001 | 957 | 1095 | 881 | 766 | 1057 | 1077 | 1048 | 987.21 | 914 | 1061 | 1.16 | 0.215 | 0.1218933 | 0.29685061 | 0.0153 | 0.0102 | 0.0129 | 0.0129 | TRUE | 0.3566 |
| Zfp87 | 506 | 583 | 611 | 661 | 493 | 489 | 537 | 582 | 614 | 520 | 530 | 535 | 553.09 | 578 | 528 | 0.914 | -0.13 | 0.20849931 | 0.4179526 | 0 | 0.0129 | 0.0059 | 0.0059 | TRUE | 0.0431 |
| Zfp870 | 849 | 887 | 859 | 1090 | 794 | 815 | 902 | 886 | 863 | 857 | 854 | 892 | 875.58 | 884 | 868 | 0.982 | -0.026 | 0.75406656 | 0.87020836 | 0 | 0.0107 | 0.0038 | 0.0038 | TRUE | 0.007 |
| Zfp871 | 2379 | 2331 | 2681 | 3190 | 2464 | 2547 | 2527 | 2328 | 2692 | 2508 | 2651 | 2788 | 2582.31 | 2516 | 2649 | 1.053 | 0.074 | 0.40891545 | 0.62126578 | 0.0034 | 0.0082 | 0.0054 | 0.0054 | TRUE | 0.0518 |
| Zfp872 | 26 | 43 | 40 | 25 | 19 | 21 | 28 | 43 | 40 | 20 | 20 | 23 | 28.97 | 37 | 21 | 0.567 | -0.818 | 0.02864717 | 0.1114332 | 0 | 0.1206 | 0.0637 | 0.0637 | TRUE | 0.3407 |
| Zfp873 | 294 | 304 | 276 | 427 | 335 | 302 | 312 | 304 | 277 | 336 | 360 | 331 | 319.95 | 298 | 342 | 1.149 | 0.201 | 0.10426614 | 0.26870679 | 0 | 0.0172 | 0.0079 | 0.0079 | TRUE | 0.0413 |

| | | | | | | | | | | | | | | | | | | | | | | | | | |
|---|---|---|---|---|---|---|---|---|---|---|---|---|---|---|---|---|---|---|---|---|---|---|---|---|---|
| Zfp874a | 338 | 378 | 398 | 520 | 388 | 405 | 359 | 378 | 400 | 409 | 417 | 443 | 400.96 | 379 | 423 | 1.116 | 0.158 | 0.16159911 | 0.35669419 | 0 | 0.0151 | 0.0067 | 0.0067 | TRUE | 0.0268 |
| Zfp874b | 637 | 628 | 646 | 779 | 608 | 593 | 677 | 627 | 649 | 612 | 654 | 649 | 644.69 | 651 | 638 | 0.98 | -0.029 | 0.76352433 | 0.87554935 | 0 | 0.012 | 0.005 | 0.005 | TRUE | 0.0161 |
| Zfp879 | 43 | 55 | 45 | 71 | 47 | 61 | 46 | 55 | 45 | 56 | 51 | 67 | 53.16 | 49 | 58 | 1.185 | 0.245 | 0.36927062 | 0.58663742 | 0 | 0.0689 | 0.0349 | 0.0349 | TRUE | 0.1482 |
| Zfp882 | 246 | 185 | 173 | 349 | 240 | 218 | 261 | 185 | 174 | 274 | 258 | 239 | 231.83 | 207 | 257 | 1.247 | 0.318 | 0.09342486 | 0.24989155 | 0.0225 | 0.0211 | 0.0216 | 0.0216 | TRUE | 0.5793 |
| Zfp9 | 113 | 170 | 138 | 247 | 203 | 192 | 120 | 170 | 139 | 194 | 218 | 210 | 175.19 | 143 | 207 | 1.45 | 0.536 | 0.00307965 | 0.02166467 | 0.0093 | 0.0257 | 0.0177 | 0.0177 | TRUE | 0.2853 |
| Zfp90 | 531 | 547 | 518 | 655 | 446 | 474 | 564 | 546 | 520 | 515 | 480 | 519 | 524.01 | 543 | 505 | 0.929 | -0.106 | 0.29842181 | 0.51905693 | 0 | 0.0132 | 0.0056 | 0.0056 | TRUE | 0.0207 |
| Zfp91 | 1886 | 2061 | 1905 | 2567 | 1812 | 1816 | 2003 | 2058 | 1913 | 2018 | 1950 | 1988 | 1988.32 | 1991 | 1985 | 0.997 | -0.004 | 0.95257552 | 0.97590263 | 3.00E-04 | 0.0086 | 0.0032 | 0.0032 | TRUE | 0.0145 |
| Zfp92 | 21 | 29 | 19 | 50 | 18 | 23 | 22 | 29 | 19 | 39 | 19 | 25 | 25.7 | 23 | 28 | 1.205 | 0.269 | 0.53518045 | 0.72254971 | 0.0569 | 0.1351 | 0.0961 | 0.0961 | TRUE | 0.6864 |
| Zfp93 | 110 | 137 | 127 | 186 | 118 | 122 | 117 | 137 | 128 | 146 | 127 | 134 | 131.32 | 127 | 136 | 1.069 | 0.096 | 0.58818207 | 0.76283262 | 0 | 0.032 | 0.0153 | 0.0153 | TRUE | 0.0537 |
| Zfp930 | 2005 | 2239 | 2371 | 2447 | 1860 | 1797 | 2130 | 2236 | 2381 | 1924 | 2001 | 1967 | 2106.42 | 2249 | 1964 | 0.873 | -0.196 | 0.01173742 | 0.05920134 | 0.0013 | 0.0085 | 0.0039 | 0.0039 | TRUE | 0.0319 |
| Zfp931 | 77 | 68 | 70 | 101 | 66 | 71 | 82 | 68 | 70 | 79 | 71 | 78 | 74.69 | 73 | 76 | 1.04 | 0.056 | 0.80458217 | 0.89928921 | 0 | 0.051 | 0.0234 | 0.0234 | TRUE | 0.0904 |
| Zfp932 | 639 | 484 | 438 | 729 | 510 | 444 | 679 | 483 | 440 | 573 | 549 | 486 | 534.96 | 534 | 536 | 1.005 | 0.007 | 0.96912579 | 0.984481 | 0.0283 | 0.0131 | 0.0207 | 0.0207 | TRUE | 0.6548 |
| Zfp933 | 227 | 167 | 155 | 289 | 194 | 148 | 241 | 167 | 156 | 227 | 209 | 162 | 193.58 | 188 | 199 | 1.064 | 0.09 | 0.68841644 | 0.82851614 | 0.038 | 0.0239 | 0.0309 | 0.0309 | TRUE | 0.6576 |
| Zfp934 | 240 | 181 | 202 | 276 | 201 | 161 | 255 | 181 | 203 | 217 | 216 | 176 | 208 | 213 | 203 | 0.957 | -0.064 | 0.73064144 | 0.85525284 | 0.0174 | 0.0227 | 0.02 | 0.02 | TRUE | 0.3239 |
| Zfp935 | 273 | 196 | 206 | 461 | 286 | 231 | 290 | 196 | 207 | 362 | 308 | 253 | 269.26 | 231 | 308 | 1.336 | 0.418 | 0.04348639 | 0.14928338 | 0.035 | 0.0191 | 0.0271 | 0.0271 | TRUE | 0.288 |
| Zfp936 | 1262 | 649 | 512 | 1792 | 1219 | 673 | 1340 | 648 | 514 | 1409 | 1312 | 737 | 993.29 | 834 | 1153 | 1.382 | 0.466 | 0.35256275 | 0.57114007 | 0.1803 | 0.0102 | 0.0775 | 0.1803 | TRUE | 3.3451 |
| Zfp937 | 478 | 483 | 466 | 522 | 393 | 411 | 508 | 482 | 468 | 410 | 423 | 450 | 456.85 | 486 | 428 | 0.879 | -0.186 | 0.08389062 | 0.2320861 | 0 | 0.0141 | 0.0061 | 0.0061 | TRUE | 0.0234 |
| Zfp938 | 423 | 492 | 475 | 526 | 364 | 389 | 449 | 491 | 477 | 413 | 392 | 426 | 441.43 | 472 | 410 | 0.868 | -0.204 | 0.05990694 | 0.18563561 | 0 | 0.0144 | 0.0062 | 0.0062 | TRUE | 0.0212 |
| Zfp939 | 98 | 93 | 116 | 149 | 113 | 88 | 104 | 93 | 116 | 117 | 122 | 96 | 108.08 | 104 | 112 | 1.071 | 0.099 | 0.63115757 | 0.79265149 | 0 | 0.0374 | 0.0211 | 0.0211 | TRUE | 0.1359 |
| Zfp94 | 256 | 238 | 238 | 404 | 242 | 212 | 272 | 238 | 239 | 318 | 260 | 232 | 259.77 | 250 | 270 | 1.086 | 0.119 | 0.47067358 | 0.67223251 | 0.0125 | 0.0196 | 0.0159 | 0.0159 | TRUE | 0.2797 |
| Zfp940 | 551 | 583 | 583 | 738 | 519 | 556 | 585 | 582 | 585 | 580 | 558 | 609 | 583.35 | 584 | 582 | 0.997 | -0.005 | 0.95887319 | 0.97898705 | 0 | 0.0126 | 0.0049 | 0.0049 | TRUE | 0.0173 |
| Zfp941 | 862 | 781 | 709 | 935 | 690 | 656 | 916 | 780 | 712 | 735 | 742 | 718 | 767.17 | 803 | 732 | 0.912 | -0.132 | 0.2655504 | 0.48514548 | 0.0065 | 0.0112 | 0.0089 | 0.0089 | TRUE | 0.1794 |
| Zfp942 | 344 | 304 | 273 | 458 | 309 | 264 | 365 | 304 | 274 | 360 | 332 | 289 | 320.77 | 314 | 327 | 1.043 | 0.061 | 0.70361465 | 0.83769895 | 0.0136 | 0.0172 | 0.0153 | 0.0153 | TRUE | 0.2263 |
| Zfp943 | 77 | 96 | 103 | 161 | 107 | 100 | 82 | 96 | 103 | 127 | 115 | 109 | 105.37 | 94 | 117 | 1.251 | 0.324 | 0.10733604 | 0.27406574 | 0 | 0.0382 | 0.0195 | 0.0195 | TRUE | 0.1161 |
| Zfp944 | 338 | 313 | 369 | 506 | 400 | 349 | 359 | 313 | 371 | 398 | 430 | 382 | 375.39 | 348 | 403 | 1.161 | 0.216 | 0.08797221 | 0.24038421 | 0.0028 | 0.0157 | 0.0088 | 0.0088 | TRUE | 0.0883 |
| Zfp945 | 1121 | 1090 | 1174 | 1490 | 1114 | 1093 | 1191 | 1089 | 1179 | 1171 | 1199 | 1196 | 1170.75 | 1153 | 1189 | 1.031 | 0.044 | 0.58797351 | 0.76262485 | 0 | 0.0097 | 0.004 | 0.004 | TRUE | 0.0285 |
| Zfp946 | 180 | 154 | 141 | 293 | 169 | 152 | 191 | 154 | 142 | 230 | 182 | 166 | 177.52 | 162 | 193 | 1.194 | 0.256 | 0.20313177 | 0.41080946 | 0.0216 | 0.0255 | 0.0235 | 0.0235 | TRUE | 0.3307 |
| Zfp947 | 80 | 64 | 54 | 122 | 89 | 55 | 85 | 64 | 54 | 96 | 96 | 60 | 75.83 | 68 | 84 | 1.245 | 0.317 | 0.27714342 | 0.49805749 | 0.045 | 0.0503 | 0.0478 | 0.0478 | TRUE | 0.5504 |
| Zfp948 | 265 | 309 | 345 | 560 | 377 | 394 | 281 | 309 | 346 | 440 | 406 | 431 | 368.94 | 312 | 426 | 1.364 | 0.448 | 0.00046802 | 0.00493428 | 0.0028 | 0.0158 | 0.009 | 0.009 | TRUE | 0.1044 |
| Zfp949 | 386 | 402 | 408 | 551 | 402 | 399 | 410 | 401 | 410 | 433 | 433 | 437 | 420.6 | 407 | 434 | 1.067 | 0.093 | 0.36951634 | 0.58685044 | 0 | 0.0147 | 0.0053 | 0.0053 | TRUE | 0.0017 |
| Zfp951 | 1109 | 682 | 647 | 1497 | 1190 | 908 | 1178 | 681 | 650 | 1177 | 1280 | 994 | 993.31 | 836 | 1150 | 1.376 | 0.461 | 0.03732398 | 0.13397219 | 0.0634 | 0.0102 | 0.0342 | 0.0342 | TRUE | 1.516 |
| Zfp952 | 528 | 514 | 535 | 629 | 438 | 495 | 561 | 513 | 537 | 494 | 471 | 542 | 519.82 | 537 | 502 | 0.935 | -0.096 | 0.37596661 | 0.59147378 | 0.0013 | 0.0132 | 0.0066 | 0.0066 | TRUE | 0.0524 |
| Zfp953 | 289 | 243 | 210 | 451 | 319 | 257 | 307 | 243 | 211 | 355 | 343 | 281 | 289.93 | 254 | 326 | 1.29 | 0.367 | 0.04206753 | 0.1461245 | 0.0207 | 0.0183 | 0.02 | 0.02 | TRUE | 0.3754 |
| Zfp954 | 400 | 379 | 356 | 562 | 383 | 403 | 425 | 379 | 357 | 442 | 412 | 441 | 409.31 | 387 | 432 | 1.117 | 0.159 | 0.18457076 | 0.38762317 | 0.0019 | 0.015 | 0.0079 | 0.0079 | TRUE | 0.0835 |
| Zfp955a | 983 | 1052 | 1162 | 1409 | 1047 | 1029 | 1044 | 1051 | 1167 | 1108 | 1127 | 1126 | 1103.68 | 1087 | 1120 | 1.03 | 0.043 | 0.62321694 | 0.78748419 | 0.0011 | 0.0099 | 0.0045 | 0.0045 | TRUE | 0.0492 |
| Zfp955b | 904 | 706 | 639 | 1381 | 934 | 665 | 960 | 705 | 642 | 1086 | 1005 | 728 | 854.24 | 769 | 940 | 1.223 | 0.29 | 0.13521357 | 0.31810686 | 0.0422 | 0.0108 | 0.026 | 0.026 | TRUE | 0.5596 |
| Zfp956 | 278 | 271 | 290 | 439 | 348 | 266 | 295 | 271 | 291 | 345 | 374 | 291 | 311.31 | 286 | 337 | 1.18 | 0.239 | 0.09285487 | 0.2489388 | 0.0058 | 0.0175 | 0.0113 | 0.0113 | TRUE | 0.1538 |
| Zfp957 | 0 | 0 | 0 | 0 | 0 | 0 | 0 | 0 | 0 | 0 | 0 | 0 | 0 | 0 | 0 | NA | NA | NA | NA | NA | NA | NA | NA | NA | NA |
| Zfp958 | 1802 | 1710 | 1596 | 2202 | 1549 | 1485 | 1914 | 1708 | 1603 | 1731 | 1667 | 1626 | 1707.94 | 1742 | 1675 | 0.962 | -0.056 | 0.55613861 | 0.73889464 | 0.0037 | 0.0088 | 0.0059 | 0.0059 | TRUE | 0.0904 |
| Zfp959 | 274 | 248 | 229 | 369 | 238 | 167 | 291 | 248 | 230 | 290 | 256 | 183 | 249.6 | 256 | 243 | 0.951 | -0.072 | 0.72086629 | 0.8487871 | 0.0302 | 0.0201 | 0.0251 | 0.0251 | TRUE | 0.51 |
| Zfp960 | 678 | 712 | 735 | 863 | 617 | 603 | 720 | 711 | 738 | 678 | 664 | 660 | 695.27 | 723 | 667 | 0.923 | -0.115 | 0.19093884 | 0.39611884 | 0 | 0.0116 | 0.0041 | 0.0041 | TRUE | 0.0039 |
| Zfp961 | 5700 | 4869 | 4827 | 6440 | 4370 | 4130 | 6054 | 4863 | 4847 | 5063 | 4702 | 4521 | 5008.22 | 5255 | 4762 | 0.906 | -0.142 | 0.20494362 | 0.41337964 | 0.0101 | 0.0076 | 0.0088 | 0.0088 | TRUE | 0.2159 |
| Zfp963 | 168 | 120 | 120 | 208 | 114 | 92 | 178 | 120 | 120 | 164 | 123 | 101 | 134.28 | 139 | 129 | 0.929 | -0.107 | 0.67876483 | 0.82153933 | 0.0497 | 0.0314 | 0.0404 | 0.0404 | TRUE | 0.529 |
| Zfp964 | 61 | 52 | 47 | 104 | 60 | 36 | 65 | 52 | 47 | 82 | 65 | 39 | 58.27 | 55 | 62 | 1.142 | 0.191 | 0.56483925 | 0.74534144 | 0.0612 | 0.0634 | 0.0624 | 0.0624 | TRUE | 0.2721 |
| Zfp97 | 877 | 961 | 999 | 1059 | 812 | 811 | 932 | 960 | 1003 | 833 | 874 | 888 | 914.72 | 965 | 865 | 0.895 | -0.159 | 0.06467813 | 0.19591002 | 0 | 0.0105 | 0.0043 | 0.0043 | TRUE | 0.0144 |
| Zfpl1 | 679 | 820 | 795 | 1145 | 816 | 822 | 721 | 819 | 798 | 900 | 878 | 900 | 836.05 | 779 | 893 | 1.145 | 0.195 | 0.03581428 | 0.12991326 | 8.00E-04 | 0.0108 | 0.005 | 0.005 | TRUE | 0.0502 |
| Zfpm1 | 1842 | 2064 | 1782 | 2290 | 1964 | 1920 | 1957 | 2061 | 1789 | 1800 | 2113 | 2102 | 1970.38 | 1936 | 2005 | 1.035 | 0.05 | 0.62920806 | 0.79150339 | 0.0062 | 0.0086 | 0.0072 | 0.0072 | TRUE | 0.0983 |
| Zfpm2 | 5 | 7 | 1 | 23 | 9 | 3 | 5 | 7 | 1 | 18 | 10 | 3 | 7.39 | 4 | 10 | 2.367 | 1.243 | 0.17941554 | 0.38055926 | 0.4599 | 0.4526 | 0.4557 | 0.4557 | TRUE | 0.1435 |
| Zfr | 7501 | 8144 | 7931 | 9651 | 6783 | 6727 | 7967 | 8133 | 7964 | 7587 | 7298 | 7364 | 7718.86 | 8021 | 7416 | 0.925 | -0.113 | 0.04484031 | 0.15251702 | 2.00E-04 | 0.0073 | 0.0022 | 0.0022 | TRUE | 0.005 |
| Zfr2 | 30 | 112 | 86 | 52 | 45 | 71 | 32 | 112 | 86 | 41 | 48 | 78 | 66.18 | 77 | 56 | 0.723 | -0.469 | 0.29958354 | 0.51974827 | 0.2192 | 0.0567 | 0.1316 | 0.1316 | TRUE | 0.4223 |
| Zfx | 1701 | 2026 | 2036 | 2944 | 2162 | 2121 | 1807 | 2023 | 2044 | 2314 | 2326 | 2322 | 2139.47 | 1958 | 2321 | 1.185 | 0.245 | 0.00242619 | 0.01794835 | 0.0016 | 0.0084 | 0.0042 | 0.0042 | TRUE | 0.055 |
| Zfy1 | 98 | 116 | 119 | 276 | 208 | 243 | 104 | 116 | 119 | 217 | 224 | 266 | 174.37 | 113 | 236 | 2.077 | 1.054 | 8.29E-10 | 5.23E-08 | 0 | 0.0258 | 0.0147 | 0.0147 | TRUE | 0.1377 |
| Zfy2 | 17 | 12 | 6 | 66 | 83 | 62 | 18 | 12 | 6 | 52 | 89 | 68 | 40.85 | 12 | 70 | 5.803 | 2.537 | 6.84E-09 | 3.37E-07 | 0.0884 | 0.0875 | 0.0884 | 0.0884 | TRUE | 0.2529 |
| Zfyve1 | 1638 | 1825 | 1928 | 2330 | 1661 | 1708 | 1740 | 1823 | 1936 | 1832 | 1787 | 1870 | 1831.15 | 1833 | 1830 | 0.998 | -0.003 | 0.97168564 | 0.98492455 | 0.0011 | 0.0087 | 0.0039 | 0.0039 | TRUE | 0.0293 |
| Zfyve16 | 1019 | 1126 | 1223 | 1805 | 1197 | 1225 | 1082 | 1125 | 1228 | 1419 | 1288 | 1341 | 1247.12 | 1145 | 1349 | 1.179 | 0.238 | 0.00982369 | 0.05195817 | 0.0025 | 0.0095 | 0.0053 | 0.0053 | TRUE | 0.0483 |
| Zfyve19 | 312 | 421 | 423 | 634 | 457 | 367 | 331 | 420 | 425 | 498 | 492 | 402 | 428.07 | 392 | 464 | 1.184 | 0.243 | 0.10915459 | 0.2770496 | 0.0139 | 0.0146 | 0.0143 | 0.0143 | TRUE | 0.2095 |
| Zfyve20 | 4556 | 4355 | 4327 | 5935 | 4106 | 3877 | 4839 | 4349 | 4345 | 4666 | 4418 | 4244 | 4476.81 | 4511 | 4443 | 0.985 | -0.022 | 0.79073726 | 0.89242185 | 0.0029 | 0.0076 | 0.0046 | 0.0046 | TRUE | 0.0492 |
| Zfyve21 | 1642 | 2223 | 2120 | 1991 | 1580 | 1820 | 1744 | 2220 | 2129 | 1565 | 1700 | 1992 | 1891.73 | 2031 | 1752 | 0.862 | -0.214 | 0.10686134 | 0.27316322 | 0.0147 | 0.0086 | 0.0121 | 0.0121 | TRUE | 0.1843 |
| Zfyve26 | 1405 | 1513 | 1691 | 2531 | 1839 | 1848 | 1492 | 1511 | 1698 | 1990 | 1979 | 2023 | 1782.1 | 1567 | 1997 | 1.274 | 0.35 | 3.33E-05 | 0.00054957 | 0.0018 | 0.0088 | 0.0045 | 0.0045 | TRUE | 0.0644 |
| Zfyve27 | 1755 | 1913 | 1759 | 1963 | 1655 | 1603 | 1864 | 1911 | 1766 | 1543 | 1781 | 1755 | 1769.91 | 1847 | 1693 | 0.916 | -0.127 | 0.16510703 | 0.36172635 | 0.0034 | 0.0088 | 0.0055 | 0.0055 | TRUE | 0.074 |
| Zfyve28 | 847 | 1101 | 1050 | 1066 | 854 | 762 | 900 | 1100 | 1054 | 838 | 919 | 834 | 940.76 | 1018 | 864 | 0.848 | -0.238 | 0.03250549 | 0.12155199 | 0.0059 | 0.0104 | 0.0078 | 0.0078 | TRUE | 0.1226 |
| Zfyve9 | 544 | 704 | 776 | 1095 | 705 | 705 | 578 | 703 | 779 | 861 | 759 | 772 | 741.86 | 687 | 797 | 1.161 | 0.216 | 0.11021639 | 0.27862899 | 0.0114 | 0.0113 | 0.0118 | 0.0118 | TRUE | 0.2257 |
| Zg16 | 0 | 0 | 2 | 1 | 0 | 0 | 0 | 0 | 2 | 1 | 0 | 0 | 0.47 | 1 | 0 | 0.52 | -0.943 | 0.78484416 | NA | 0 | 7.0808 | 6.8441 | 6.8441 | TRUE | 0.9739 |
| Zglp1 | 447 | 502 | 443 | 378 | 281 | 289 | 475 | 501 | 445 | 297 | 302 | 316 | 389.47 | 474 | 305 | 0.644 | -0.635 | 3.87E-08 | 1.59E-06 | 0 | 0.0154 | 0.0069 | 0.0069 | TRUE | 0.0333 |
| Zgpat | 591 | 637 | 560 | 964 | 749 | 647 | 628 | 636 | 562 | 758 | 806 | 708 | 683.03 | 609 | 757 | 1.244 | 0.315 | 0.00293737 | 0.02088736 | 0.0027 | 0.0117 | 0.0066 | 0.0066 | TRUE | 0.0526 |
| Zhx1 | 494 | 486 | 450 | 608 | 431 | 364 | 525 | 485 | 452 | 478 | 464 | 398 | 467.01 | 487 | 447 | 0.918 | -0.123 | 0.32632271 | 0.54741138 | 0.0052 | 0.014 | 0.0092 | 0.0092 | TRUE | 0.1011 |
| Zhx2 | 56 | 60 | 70 | 45 | 31 | 34 | 59 | 60 | 70 | 35 | 33 | 37 | 49.27 | 63 | 35 | 0.558 | -0.841 | 0.00235374 | 0.01751096 | 0 | 0.0738 | 0.0332 | 0.0332 | TRUE | 0.084 |
| Zhx3 | 85 | 47 | 62 | 97 | 90 | 55 | 90 | 47 | 62 | 76 | 97 | 60 | 72.13 | 66 | 78 | 1.171 | 0.228 | 0.47275174 | 0.67385815 | 0.0653 | 0.0526 | 0.0586 | 0.0586 | TRUE | 0.4633 |
| Zic1 | 3 | 2 | 1 | 6 | 11 | 5 | 3 | 2 | 1 | 5 | 12 | 5 | 4.7 | 2 | 7 | 3.548 | 1.827 | 0.0664879 | 0.1989088 | 0 | 0.7075 | 0.3981 | 0.3981 | TRUE | 0.7462 |

| | | | | | | | | | | | | | | | | | | | | | | | | | |
|---|---|---|---|---|---|---|---|---|---|---|---|---|---|---|---|---|---|---|---|---|---|---|---|---|---|
| Zic2 | 449 | 255 | 279 | 1872 | 1224 | 918 | 477 | 255 | 280 | 1472 | 1317 | 1005 | 800.87 | 337 | 1265 | 3.753 | 1.908 | 7.84E-16 | 1.19E-13 | 0.0743 | 0.011 | 0.0385 | 0.0385 | TRUE | 0.8234 |
| Zic3 | 11218 | 7041 | 7337 | 10329 | 6929 | 5995 | 11915 | 7032 | 7367 | 8120 | 7455 | 6562 | 8075.34 | 8771 | 7379 | 0.841 | -0.249 | 0.19550217 | 0.40149221 | 0.0538 | 0.0073 | 0.0266 | 0.0266 | TRUE | 1.2007 |
| Zic4 | 0 | 1 | 0 | 0 | 0 | 0 | 0 | 1 | 0 | 0 | 0 | 0 | 0.17 | 0 | 0 | 0.49 | -1.029 | 0.80087 | NA | 0 | 19.7991 | 10 | 10 | TRUE | 0.405 |
| Zic5 | 437 | 356 | 385 | 773 | 540 | 440 | 464 | 356 | 387 | 608 | 581 | 482 | 479.44 | 402 | 557 | 1.387 | 0.472 | 0.00171071 | 0.01359343 | 0.0145 | 0.0138 | 0.0142 | 0.0142 | TRUE | 0.2085 |
| Zik1 | 1172 | 1121 | 1106 | 1451 | 1093 | 1083 | 1245 | 1120 | 1111 | 1141 | 1176 | 1186 | 1162.86 | 1159 | 1168 | 1.008 | 0.011 | 0.89928908 | 0.95120551 | 0.0014 | 0.0097 | 0.0046 | 0.0046 | TRUE | 0.0509 |
| Zim1 | 103 | 90 | 91 | 94 | 73 | 68 | 109 | 90 | 91 | 74 | 79 | 74 | 86.26 | 97 | 76 | 0.781 | -0.357 | 0.09453067 | 0.25193271 | 0 | 0.0451 | 0.0212 | 0.0212 | TRUE | 0.1214 |
| Zim3 | 45 | 48 | 51 | 98 | 79 | 58 | 48 | 48 | 51 | 77 | 85 | 63 | 62.08 | 49 | 75 | 1.536 | 0.619 | 0.01496643 | 0.0706643 | 0 | 0.06 | 0.0298 | 0.0298 | TRUE | 0.1567 |
| Zkscan1 | 1188 | 1090 | 1122 | 1341 | 983 | 909 | 1262 | 1089 | 1127 | 1054 | 1058 | 995 | 1097.32 | 1159 | 1036 | 0.894 | -0.162 | 0.0893851 | 0.24300475 | 0.0027 | 0.0099 | 0.0056 | 0.0056 | TRUE | 0.0718 |
| Zkscan14 | 185 | 231 | 187 | 262 | 196 | 167 | 197 | 231 | 188 | 206 | 211 | 183 | 202.44 | 205 | 200 | 0.976 | -0.035 | 0.82372514 | 0.91017184 | 0.0038 | 0.0232 | 0.0132 | 0.0132 | TRUE | 0.1324 |
| Zkscan16 | 209 | 172 | 219 | 213 | 191 | 158 | 222 | 172 | 220 | 167 | 206 | 173 | 193.26 | 205 | 182 | 0.888 | -0.171 | 0.3350617 | 0.55534619 | 0.0112 | 0.0239 | 0.0174 | 0.0174 | TRUE | 0.2157 |
| Zkscan17 | 4170 | 6090 | 6882 | 6218 | 4944 | 5906 | 4429 | 6082 | 6910 | 4888 | 5319 | 6465 | 5682.38 | 5807 | 5557 | 0.957 | -0.064 | 0.70938711 | 0.84171221 | 0.0353 | 0.0075 | 0.0208 | 0.0208 | TRUE | 0.487 |
| Zkscan2 | 234 | 296 | 310 | 402 | 295 | 278 | 249 | 296 | 311 | 316 | 317 | 304 | 298.86 | 285 | 312 | 1.096 | 0.132 | 0.33588797 | 0.55600599 | 0.003 | 0.0179 | 0.0102 | 0.0102 | TRUE | 0.1396 |
| Zkscan3 | 657 | 742 | 689 | 895 | 642 | 645 | 698 | 741 | 692 | 704 | 691 | 706 | 705.18 | 710 | 700 | 0.986 | -0.021 | 0.81786134 | 0.90711833 | 0 | 0.0116 | 0.0044 | 0.0044 | TRUE | 0.0171 |
| Zkscan4 | 23 | 43 | 54 | 37 | 35 | 28 | 24 | 43 | 54 | 29 | 38 | 31 | 36.5 | 40 | 33 | 0.797 | -0.327 | 0.39502467 | 0.60854943 | 0.059 | 0.0972 | 0.0788 | 0.0788 | TRUE | 0.3017 |
| Zkscan5 | 1560 | 1670 | 1710 | 1841 | 1412 | 1361 | 1657 | 1668 | 1717 | 1447 | 1519 | 1490 | 1583.03 | 1681 | 1485 | 0.883 | -0.179 | 0.01282678 | 0.06323977 | 0 | 0.009 | 0.0031 | 0.0031 | TRUE | 0.0063 |
| Zkscan6 | 518 | 631 | 597 | 787 | 569 | 563 | 550 | 630 | 599 | 619 | 612 | 616 | 604.5 | 593 | 616 | 1.038 | 0.053 | 0.59671959 | 0.76960188 | 0 | 0.0124 | 0.0056 | 0.0056 | TRUE | 0.0468 |
| Zkscan7 | 48 | 83 | 78 | 166 | 126 | 122 | 51 | 83 | 78 | 130 | 136 | 134 | 101.97 | 71 | 133 | 1.879 | 0.91 | 8.64E-05 | 0.00123204 | 0.0115 | 0.0392 | 0.0278 | 0.0278 | TRUE | 0.5225 |
| Zkscan8 | 897 | 1193 | 1360 | 1521 | 1142 | 1313 | 953 | 1191 | 1366 | 1196 | 1229 | 1437 | 1228.59 | 1170 | 1287 | 1.1 | 0.137 | 0.35630752 | 0.57490354 | 0.0202 | 0.0096 | 0.0151 | 0.0151 | TRUE | 0.3138 |
| Zmat1 | 27 | 51 | 71 | 67 | 52 | 46 | 29 | 51 | 71 | 53 | 56 | 50 | 51.65 | 50 | 53 | 1.052 | 0.073 | 0.83920944 | 0.91840212 | 0.075 | 0.0707 | 0.0726 | 0.0726 | TRUE | 0.2147 |
| Zmat2 | 3894 | 3754 | 3652 | 5846 | 3954 | 3635 | 4136 | 3749 | 3667 | 4596 | 4254 | 3979 | 4063.55 | 3851 | 4276 | 1.111 | 0.152 | 0.09489799 | 0.25239984 | 0.0044 | 0.0077 | 0.0057 | 0.0057 | TRUE | 0.0524 |
| Zmat3 | 6379 | 7386 | 7690 | 8461 | 6064 | 6394 | 6776 | 7376 | 7722 | 6651 | 6524 | 6999 | 7008.13 | 7291 | 6725 | 0.922 | -0.117 | 0.13926864 | 0.32392251 | 0.0026 | 0.0074 | 0.0043 | 0.0043 | TRUE | 0.0466 |
| Zmat4 | 365 | 269 | 263 | 228 | 210 | 232 | 388 | 269 | 264 | 179 | 226 | 254 | 263.26 | 307 | 220 | 0.715 | -0.485 | 0.02067389 | 0.08896825 | 0.035 | 0.0194 | 0.0277 | 0.0277 | TRUE | 0.5978 |
| Zmat5 | 588 | 547 | 567 | 675 | 457 | 449 | 625 | 546 | 569 | 531 | 492 | 492 | 542.34 | 580 | 505 | 0.871 | -0.199 | 0.06552995 | 0.19721209 | 0.0017 | 0.013 | 0.0065 | 0.0065 | TRUE | 0.0524 |
| Zmiz1 | 1317 | 868 | 1005 | 2095 | 1264 | 905 | 1399 | 867 | 1009 | 1647 | 1360 | 991 | 1212.08 | 1092 | 1333 | 1.222 | 0.289 | 0.18171788 | 0.38360117 | 0.0611 | 0.0096 | 0.0328 | 0.0328 | TRUE | 0.2593 |
| Zmiz2 | 8158 | 7067 | 6956 | 8852 | 6539 | 6427 | 8665 | 7058 | 6985 | 6959 | 7036 | 7035 | 7289.56 | 7569 | 7010 | 0.926 | -0.111 | 0.27896498 | 0.50005946 | 0.0077 | 0.0074 | 0.0074 | 0.0074 | TRUE | 0.1958 |
| Zmpste24 | 2648 | 2380 | 2316 | 3345 | 2354 | 2216 | 2813 | 2377 | 2326 | 2630 | 2533 | 2426 | 2517.2 | 2505 | 2530 | 1.01 | 0.014 | 0.88528713 | 0.94387096 | 0.0059 | 0.0082 | 0.0068 | 0.0068 | TRUE | 0.1396 |
| Zmym1 | 4725 | 3622 | 3648 | 5712 | 3993 | 3305 | 5019 | 3617 | 3663 | 4490 | 4296 | 3618 | 4117.25 | 4100 | 4135 | 1.009 | 0.013 | 0.93284347 | 0.96671764 | 0.0238 | 0.0077 | 0.0159 | 0.0159 | TRUE | 0.4679 |
| Zmym2 | 9876 | 8874 | 9133 | 11655 | 8450 | 8292 | 10490 | 8863 | 9170 | 9162 | 9092 | 9077 | 9308.99 | 9508 | 9110 | 0.958 | -0.062 | 0.47191717 | 0.67314138 | 0.0039 | 0.0073 | 0.0052 | 0.0052 | TRUE | 0.0997 |
| Zmym3 | 4601 | 3746 | 3624 | 5689 | 4074 | 3394 | 4887 | 3741 | 3639 | 4472 | 4383 | 3715 | 4139.67 | 4089 | 4190 | 1.025 | 0.036 | 0.7952634 | 0.8946193 | 0.0184 | 0.0077 | 0.0133 | 0.0133 | TRUE | 0.3546 |
| Zmym4 | 4628 | 4207 | 4521 | 6296 | 4240 | 3989 | 4916 | 4202 | 4540 | 4949 | 4562 | 4367 | 4589.15 | 4553 | 4626 | 1.017 | 0.024 | 0.79867183 | 0.89623992 | 0.0049 | 0.0076 | 0.006 | 0.006 | TRUE | 0.0593 |
| Zmym5 | 2108 | 1822 | 1712 | 2948 | 2032 | 1728 | 2239 | 1820 | 1719 | 2317 | 2186 | 1892 | 2028.85 | 1926 | 2132 | 1.107 | 0.147 | 0.26057689 | 0.47989475 | 0.0146 | 0.0085 | 0.0118 | 0.0118 | TRUE | 0.2441 |
| Zmym6 | 1114 | 1506 | 1519 | 1277 | 1031 | 1174 | 1183 | 1504 | 1525 | 1004 | 1109 | 1285 | 1268.48 | 1404 | 1133 | 0.806 | -0.311 | 0.02640403 | 0.1056142 | 0.016 | 0.0095 | 0.0134 | 0.0134 | TRUE | 0.2272 |
| Zmynd10 | 32 | 89 | 121 | 15 | 16 | 51 | 34 | 89 | 121 | 12 | 17 | 56 | 54.87 | 81 | 28 | 0.344 | -1.539 | 0.01094897 | 0.05632327 | 0.4592 | 0.0669 | 0.2399 | 0.2399 | TRUE | 0.4822 |
| Zmynd11 | 885 | 1263 | 1265 | 1967 | 1312 | 1268 | 940 | 1261 | 1270 | 1546 | 1412 | 1388 | 1302.92 | 1157 | 1449 | 1.252 | 0.324 | 0.0161798 | 0.07497251 | 0.0138 | 0.0094 | 0.0123 | 0.0123 | TRUE | 0.3218 |
| Zmynd12 | 5 | 5 | 9 | 8 | 3 | 1 | 5 | 5 | 9 | 6 | 3 | 1 | 4.99 | 6 | 3 | 0.567 | -0.819 | 0.37251819 | 0.58900822 | 0.0772 | 0.6668 | 0.3952 | 0.3952 | TRUE | 0.5942 |
| Zmynd15 | 65 | 80 | 99 | 65 | 52 | 56 | 69 | 80 | 99 | 51 | 56 | 61 | 69.45 | 83 | 56 | 0.675 | -0.567 | 0.02630969 | 0.10529217 | 0 | 0.0543 | 0.032 | 0.032 | TRUE | 0.29 |
| Zmynd19 | 3504 | 3447 | 3535 | 4417 | 3322 | 3271 | 3722 | 3443 | 3550 | 3472 | 3574 | 3581 | 3556.85 | 3572 | 3542 | 0.992 | -0.012 | 0.8597719 | 0.93089328 | 6.00E-04 | 0.0078 | 0.003 | 0.003 | TRUE | 0.0165 |
| Zmynd8 | 6515 | 7435 | 8149 | 6860 | 5423 | 5979 | 6920 | 7425 | 8182 | 5393 | 5835 | 6545 | 6716.75 | 7509 | 5924 | 0.789 | -0.342 | 0.00114369 | 0.00987062 | 0.0082 | 0.0074 | 0.0078 | 0.0078 | TRUE | 0.1024 |
| Znf41-ps | 90 | 62 | 68 | 114 | 117 | 54 | 96 | 62 | 68 | 90 | 126 | 59 | 83.4 | 75 | 92 | 1.217 | 0.284 | 0.38090161 | 0.59526263 | 0.082 | 0.0464 | 0.0634 | 0.0634 | TRUE | 0.1884 |
| Znf512b | 2345 | 2130 | 2151 | 3722 | 2645 | 2272 | 2491 | 2127 | 2160 | 2926 | 2846 | 2487 | 2506.12 | 2259 | 2753 | 1.219 | 0.286 | 0.00666129 | 0.03879683 | 0.007 | 0.0082 | 0.0076 | 0.0076 | TRUE | 0.0971 |
| Znfx1 | 1755 | 2156 | 2476 | 1943 | 1539 | 1664 | 1864 | 2153 | 2486 | 1527 | 1656 | 1822 | 1918.05 | 2168 | 1668 | 0.769 | -0.379 | 0.00323563 | 0.02247131 | 0.0134 | 0.0086 | 0.0114 | 0.0114 | TRUE | 0.1747 |
| Znhit1 | 831 | 777 | 745 | 1195 | 889 | 751 | 883 | 776 | 748 | 939 | 957 | 822 | 854.12 | 802 | 906 | 1.13 | 0.177 | 0.11790193 | 0.29045568 | 0.0054 | 0.0108 | 0.008 | 0.008 | TRUE | 0.0908 |
| Znhit2 | 352 | 366 | 372 | 613 | 478 | 359 | 374 | 366 | 374 | 482 | 514 | 393 | 417.02 | 371 | 463 | 1.249 | 0.321 | 0.0177909 | 0.07980139 | 0.0074 | 0.0148 | 0.0108 | 0.0108 | TRUE | 0.1972 |
| Znhit3 | 1745 | 1488 | 1304 | 2646 | 1601 | 1318 | 1853 | 1486 | 1309 | 2080 | 1723 | 1443 | 1649.06 | 1549 | 1749 | 1.129 | 0.175 | 0.30088414 | 0.52137383 | 0.0301 | 0.0089 | 0.02 | 0.02 | TRUE | 0.2449 |
| Znhit6 | 2059 | 2172 | 2125 | 2842 | 2186 | 1994 | 2187 | 2169 | 2134 | 2234 | 2352 | 2183 | 2209.81 | 2163 | 2256 | 1.043 | 0.061 | 0.3874997 | 0.60130316 | 3.00E-04 | 0.0084 | 0.0031 | 0.0031 | TRUE | 0.0164 |
| Znrd1 | 954 | 1028 | 1065 | 1577 | 1085 | 979 | 1013 | 1027 | 1069 | 1240 | 1167 | 1072 | 1098.02 | 1036 | 1160 | 1.12 | 0.163 | 0.07847576 | 0.22169438 | 0.0023 | 0.0099 | 0.0053 | 0.0053 | TRUE | 0.0523 |
| Znrd1as | 748 | 1018 | 924 | 1080 | 727 | 829 | 795 | 1017 | 928 | 849 | 782 | 907 | 879.61 | 913 | 846 | 0.927 | -0.11 | 0.37309434 | 0.5895055 | 0.0085 | 0.0106 | 0.0098 | 0.0098 | TRUE | 0.1528 |
| Znrf1 | 2084 | 2173 | 2089 | 2071 | 1661 | 1474 | 2214 | 2170 | 2098 | 1628 | 1787 | 1614 | 1918.34 | 2161 | 1676 | 0.776 | -0.367 | 4.25E-06 | 9.61E-05 | 0.0012 | 0.0086 | 0.004 | 0.004 | TRUE | 0.0399 |
| Znrf2 | 2382 | 2383 | 2157 | 3215 | 2300 | 2293 | 2530 | 2380 | 2166 | 2527 | 2475 | 2510 | 2431.33 | 2359 | 2504 | 1.062 | 0.087 | 0.30661594 | 0.52707865 | 0.0025 | 0.0083 | 0.0048 | 0.0048 | TRUE | 0.062 |
| Znrf3 | 663 | 704 | 732 | 1048 | 726 | 697 | 704 | 703 | 735 | 824 | 781 | 763 | 751.71 | 714 | 789 | 1.106 | 0.146 | 0.1052693 | 0.27039689 | 0 | 0.0113 | 0.0045 | 0.0045 | TRUE | 0.0182 |
| Znrf4 | 0 | 0 | 0 | 0 | 0 | 0 | 0 | 0 | 0 | 0 | 0 | 0 | 0 | 0 | 0 | NA | NA | NA | NA | NA | NA | NA | NA | NA | NA |
| Zp1 | 0 | 1 | 1 | 1 | 1 | 1 | 0 | 1 | 1 | 1 | 1 | 1 | 0.83 | 1 | 1 | 1.45 | 0.536 | 0.78654731 | NA | 0 | 3.9925 | 1.5656 | 1.5656 | TRUE | 0.2233 |
| Zp2 | 3 | 4 | 1 | 7 | 5 | 9 | 3 | 4 | 1 | 6 | 5 | 10 | 4.82 | 3 | 7 | 2.519 | 1.333 | 0.15119456 | 0.34222741 | 0 | 0.6904 | 0.363 | 0.363 | TRUE | 0.3726 |
| Zp3 | 184 | 154 | 121 | 154 | 152 | 115 | 195 | 154 | 121 | 121 | 164 | 126 | 146.87 | 157 | 137 | 0.871 | -0.199 | 0.39090562 | 0.60467415 | 0.0349 | 0.0293 | 0.0321 | 0.0321 | TRUE | 0.2279 |
| Zp3r | 2 | 1 | 2 | 7 | 2 | 2 | 2 | 1 | 2 | 6 | 2 | 2 | 2.5 | 2 | 3 | 1.968 | 0.976 | 0.42058428 | NA | 0 | 1.3269 | 0.614 | 0.614 | TRUE | 0.6396 |
| Zp4-ps | 0 | 0 | 0 | 0 | 0 | 0 | 0 | 0 | 0 | 0 | 0 | 0 | 0 | 0 | 0 | NA | NA | NA | NA | NA | NA | NA | NA | NA | NA |
| Zpbp | 19 | 31 | 26 | 41 | 26 | 26 | 20 | 31 | 26 | 32 | 28 | 28 | 27.65 | 26 | 29 | 1.15 | 0.201 | 0.58539126 | 0.76057738 | 0 | 0.126 | 0.0617 | 0.0617 | TRUE | 0.2121 |
| Zpbp2 | 12 | 9 | 15 | 14 | 15 | 6 | 13 | 9 | 15 | 11 | 16 | 7 | 11.75 | 12 | 11 | 0.917 | -0.126 | 0.8319265 | 0.91474796 | 0 | 0.2872 | 0.1668 | 0.1668 | TRUE | 0.2212 |
| Zpld1 | 0 | 0 | 0 | 0 | 0 | 0 | 0 | 0 | 0 | 0 | 0 | 0 | 0 | 0 | 0 | NA | NA | NA | NA | NA | NA | NA | NA | NA | NA |
| Zranb1 | 1328 | 1440 | 1404 | 1670 | 1222 | 1332 | 1411 | 1438 | 1410 | 1313 | 1315 | 1458 | 1390.7 | 1420 | 1362 | 0.959 | -0.061 | 0.46429228 | 0.66759211 | 0.0011 | 0.0093 | 0.0042 | 0.0042 | TRUE | 0.0452 |
| Zranb2 | 6157 | 7672 | 7274 | 8847 | 6900 | 6946 | 6540 | 7662 | 7304 | 6955 | 7424 | 7603 | 7248.01 | 7169 | 7327 | 1.022 | 0.031 | 0.71989562 | 0.8481969 | 0.0042 | 0.0074 | 0.0054 | 0.0054 | TRUE | 0.0718 |
| Zranb3 | 2282 | 2315 | 2499 | 2680 | 1974 | 2027 | 2424 | 2312 | 2509 | 2107 | 2124 | 2219 | 2282.46 | 2415 | 2150 | 0.89 | -0.168 | 0.02160323 | 0.09194449 | 8.00E-04 | 0.0083 | 0.0034 | 0.0034 | TRUE | 0.017 |
| Zrsr1 | 942 | 841 | 897 | 1490 | 1031 | 857 | 1001 | 840 | 901 | 1171 | 1109 | 938 | 993.32 | 914 | 1073 | 1.175 | 0.233 | 0.05687399 | 0.17943921 | 0.0088 | 0.0102 | 0.0098 | 0.0098 | TRUE | 0.1432 |
| Zrsr2 | 2142 | 2338 | 2399 | 3139 | 2418 | 2436 | 2275 | 2335 | 2409 | 2468 | 2602 | 2667 | 2459.14 | 2340 | 2579 | 1.102 | 0.14 | 0.0537044 | 0.17229981 | 8.00E-04 | 0.0082 | 0.0034 | 0.0034 | TRUE | 0.0176 |
| Zscan10 | 2777 | 2540 | 2670 | 3151 | 2596 | 2512 | 2950 | 2537 | 2681 | 2477 | 2793 | 2750 | 2697.89 | 2723 | 2673 | 0.982 | -0.027 | 0.77678181 | 0.88398357 | 0.0047 | 0.0081 | 0.006 | 0.006 | TRUE | 0.0645 |
| Zscan12 | 683 | 934 | 915 | 930 | 649 | 796 | 725 | 933 | 919 | 731 | 698 | 871 | 812.96 | 859 | 767 | 0.892 | -0.165 | 0.24446395 | 0.46109609 | 0.0144 | 0.011 | 0.0132 | 0.0132 | TRUE | 0.2189 |
| Zscan18 | 882 | 831 | 794 | 1009 | 883 | 774 | 937 | 830 | 797 | 793 | 950 | 847 | 859.09 | 855 | 863 | 1.01 | 0.014 | 0.90482291 | 0.95259775 | 0.0066 | 0.0107 | 0.0084 | 0.0084 | TRUE | 0.0912 |
| Zscan2 | 192 | 205 | 221 | 229 | 182 | 197 | 204 | 205 | 222 | 180 | 196 | 216 | 203.68 | 210 | 197 | 0.935 | -0.097 | 0.52134098 | 0.71160449 | 0 | 0.0231 | 0.0115 | 0.0115 | TRUE | 0.0721 |
| Zscan20 | 633 | 671 | 612 | 933 | 693 | 613 | 672 | 670 | 615 | 733 | 746 | 671 | 684.52 | 652 | 717 | 1.099 | 0.137 | 0.17139251 | 0.37034626 | 0.0013 | 0.0117 | 0.0057 | 0.0057 | TRUE | 0.0357 |

| | | | | | | | | | | | | | | | | | | | | | | | | | |
|---|---|---|---|---|---|---|---|---|---|---|---|---|---|---|---|---|---|---|---|---|---|---|---|---|---|
| Zscan21 | 1065 | 1062 | 1036 | 1493 | 1007 | 948 | 1131 | 1061 | 1040 | 1174 | 1083 | 1038 | 1087.83 | 1077 | 1098 | 1.021 | 0.029 | 0.74906674 | 0.86688385 | 0.0021 | 0.0099 | 0.0052 | 0.0052 | TRUE | 0.0446 |
| Zscan22 | 217 | 208 | 146 | 412 | 248 | 184 | 230 | 208 | 147 | 324 | 267 | 201 | 229.49 | 195 | 264 | 1.358 | 0.442 | 0.05904039 | 0.18399229 | 0.0497 | 0.0213 | 0.035 | 0.035 | TRUE | 0.1821 |
| Zscan25 | 398 | 423 | 418 | 611 | 462 | 449 | 423 | 422 | 420 | 480 | 497 | 492 | 455.64 | 422 | 490 | 1.161 | 0.215 | 0.03334951 | 0.12376199 | 0 | 0.0141 | 0.0051 | 0.0051 | TRUE | 0.0035 |
| Zscan26 | 1725 | 1653 | 1717 | 2220 | 1724 | 1780 | 1832 | 1651 | 1724 | 1745 | 1855 | 1948 | 1792.63 | 1736 | 1849 | 1.065 | 0.091 | 0.29073053 | 0.51144315 | 0.0024 | 0.0087 | 0.0048 | 0.0048 | TRUE | 0.0301 |
| Zscan29 | 410 | 412 | 375 | 630 | 448 | 357 | 435 | 411 | 377 | 495 | 482 | 391 | 431.93 | 408 | 456 | 1.12 | 0.164 | 0.23362653 | 0.44829938 | 0.0085 | 0.0145 | 0.0113 | 0.0113 | TRUE | 0.1778 |
| Zscan4a | 1334 | 376 | 207 | 1962 | 1242 | 442 | 1417 | 376 | 208 | 1542 | 1336 | 484 | 893.81 | 667 | 1121 | 1.681 | 0.749 | 0.40467595 | 0.61732751 | 0.5818 | 0.0106 | 0.2132 | 0.5818 | TRUE | 4.3592 |
| Zscan4b | 380 | 106 | 69 | 413 | 325 | 101 | 404 | 106 | 69 | 325 | 350 | 111 | 227.28 | 193 | 262 | 1.357 | 0.44 | 0.61379564 | 0.78152895 | 0.5435 | 0.0214 | 0.2255 | 0.5435 | TRUE | 8.2912 |
| Zscan4c | 1290 | 413 | 263 | 1713 | 1160 | 506 | 1370 | 412 | 264 | 1347 | 1248 | 554 | 865.89 | 682 | 1050 | 1.539 | 0.622 | 0.42555291 | 0.63570946 | 0.4373 | 0.0107 | 0.1654 | 0.4373 | TRUE | 6.3222 |
| Zscan4d | 1688 | 465 | 294 | 1907 | 1346 | 454 | 1793 | 464 | 295 | 1499 | 1448 | 497 | 999.48 | 851 | 1148 | 1.349 | 0.432 | 0.62186283 | 0.78629946 | 0.5529 | 0.0102 | 0.2025 | 0.5529 | TRUE | 7.8314 |
| Zscan4e | 775 | 208 | 162 | 891 | 594 | 251 | 823 | 208 | 163 | 700 | 639 | 275 | 467.98 | 398 | 538 | 1.353 | 0.436 | 0.59023067 | 0.76449681 | 0.4701 | 0.0139 | 0.1844 | 0.4701 | TRUE | 10.0438 |
| Zscan4f | 924 | 270 | 210 | 1061 | 831 | 314 | 981 | 270 | 211 | 834 | 894 | 344 | 588.98 | 487 | 691 | 1.417 | 0.503 | 0.52053379 | 0.71109881 | 0.4404 | 0.0125 | 0.1709 | 0.4404 | TRUE | 9.1407 |
| Zscan5b | 99 | 71 | 67 | 197 | 157 | 154 | 105 | 71 | 67 | 155 | 169 | 169 | 122.62 | 81 | 164 | 2.025 | 1.018 | 4.52E-06 | 0.00010142 | 0.0166 | 0.0338 | 0.0262 | 0.0262 | TRUE | 0.6146 |
| Zswim1 | 3482 | 5118 | 5394 | 3829 | 3097 | 3863 | 3698 | 5111 | 5416 | 3010 | 3332 | 4229 | 4132.82 | 4742 | 3524 | 0.743 | -0.429 | 0.01240665 | 0.06165017 | 0.0344 | 0.0077 | 0.0209 | 0.0209 | TRUE | 0.4512 |
| Zswim2 | 2 | 1 | 0 | 5 | 2 | 1 | 2 | 1 | 0 | 4 | 2 | 1 | 1.72 | 1 | 2 | 2.365 | 1.242 | 0.42319483 | NA | 0 | 1.926 | 1.0362 | 1.0362 | TRUE | 0.3322 |
| Zswim3 | 929 | 832 | 828 | 1080 | 811 | 735 | 987 | 831 | 831 | 849 | 873 | 805 | 862.54 | 883 | 842 | 0.954 | -0.068 | 0.52967896 | 0.71789091 | 0.0043 | 0.0107 | 0.0072 | 0.0072 | TRUE | 0.125 |
| Zswim4 | 2747 | 2288 | 2391 | 3077 | 2283 | 1960 | 2918 | 2285 | 2401 | 2419 | 2456 | 2146 | 2437.41 | 2535 | 2340 | 0.924 | -0.115 | 0.33214774 | 0.55255612 | 0.0108 | 0.0083 | 0.0097 | 0.0097 | TRUE | 0.2118 |
| Zswim5 | 745 | 729 | 843 | 1115 | 883 | 768 | 791 | 728 | 846 | 877 | 950 | 841 | 838.85 | 788 | 889 | 1.127 | 0.173 | 0.09670098 | 0.2552922 | 0.0035 | 0.0108 | 0.0066 | 0.0066 | TRUE | 0.0539 |
| Zswim6 | 1252 | 1116 | 1179 | 1494 | 1083 | 1085 | 1330 | 1115 | 1184 | 1174 | 1165 | 1188 | 1192.61 | 1210 | 1176 | 0.972 | -0.04 | 0.67547807 | 0.8192587 | 0.0032 | 0.0097 | 0.0058 | 0.0058 | TRUE | 0.0906 |
| Zswim7 | 1055 | 1383 | 1409 | 1686 | 1187 | 1336 | 1121 | 1381 | 1415 | 1325 | 1277 | 1462 | 1330.27 | 1306 | 1355 | 1.037 | 0.053 | 0.65632788 | 0.80670515 | 0.0096 | 0.0094 | 0.0095 | 0.0095 | TRUE | 0.1837 |
| Zswim8 | 773 | 1030 | 1044 | 1471 | 1184 | 1090 | 821 | 1029 | 1048 | 1156 | 1274 | 1193 | 1086.92 | 966 | 1208 | 1.25 | 0.321 | 0.00731145 | 0.04148066 | 0.0083 | 0.0099 | 0.0094 | 0.0094 | TRUE | 0.2045 |
| Zufsp | 493 | 484 | 435 | 744 | 466 | 441 | 524 | 483 | 437 | 585 | 501 | 483 | 502.14 | 481 | 523 | 1.089 | 0.123 | 0.34464368 | 0.5641524 | 0.0076 | 0.0135 | 0.0102 | 0.0102 | TRUE | 0.13 |
| Zw10 | 2430 | 2003 | 2004 | 3393 | 2385 | 2086 | 2581 | 2000 | 2012 | 2667 | 2566 | 2283 | 2351.77 | 2198 | 2505 | 1.14 | 0.19 | 0.13508615 | 0.31796579 | 0.0139 | 0.0083 | 0.0112 | 0.0112 | TRUE | 0.2812 |
| Zwilch | 5787 | 6780 | 6517 | 7737 | 5758 | 5880 | 6147 | 6771 | 6544 | 6082 | 6195 | 6437 | 6362.65 | 6487 | 6238 | 0.961 | -0.057 | 0.42016863 | 0.63064279 | 0.0014 | 0.0074 | 0.0034 | 0.0034 | TRUE | 0.0257 |
| Zwint | 4966 | 4060 | 3980 | 6373 | 4468 | 3882 | 5275 | 4055 | 3996 | 5010 | 4807 | 4249 | 4565.43 | 4442 | 4689 | 1.056 | 0.078 | 0.54802073 | 0.732964 | 0.0136 | 0.0076 | 0.012 | 0.012 | TRUE | 0.3274 |
| Zxda | 11 | 34 | 36 | 45 | 54 | 45 | 12 | 34 | 36 | 35 | 58 | 49 | 37.42 | 27 | 47 | 1.734 | 0.794 | 0.07685855 | 0.21892957 | 0.1431 | 0.0949 | 0.116 | 0.116 | TRUE | 0.9572 |
| Zxdb | 35 | 65 | 90 | 102 | 78 | 103 | 37 | 65 | 90 | 80 | 84 | 113 | 78.22 | 64 | 92 | 1.434 | 0.52 | 0.12334913 | 0.29944013 | 0.0917 | 0.049 | 0.0687 | 0.0687 | TRUE | 0.2985 |
| Zxdc | 1055 | 1384 | 1328 | 1683 | 1265 | 1308 | 1121 | 1382 | 1333 | 1323 | 1361 | 1432 | 1325.36 | 1279 | 1372 | 1.072 | 0.101 | 0.34663271 | 0.56576997 | 0.0056 | 0.0094 | 0.0075 | 0.0075 | TRUE | 0.1399 |
| Zyg11a | 2079 | 2595 | 2758 | 2764 | 2044 | 2060 | 2208 | 2592 | 2769 | 2173 | 2199 | 2255 | 2366.05 | 2523 | 2209 | 0.875 | -0.192 | 0.06173815 | 0.18977404 | 0.006 | 0.0083 | 0.0072 | 0.0072 | TRUE | 0.1441 |
| Zyg11b | 4583 | 4057 | 4274 | 4688 | 3415 | 3529 | 4868 | 4052 | 4292 | 3685 | 3674 | 3863 | 4072.33 | 4404 | 3741 | 0.849 | -0.235 | 0.0102939 | 0.05392073 | 0.0046 | 0.0077 | 0.0058 | 0.0058 | TRUE | 0.1034 |
| Zyx | 1020 | 1027 | 1071 | 1700 | 1163 | 1007 | 1083 | 1026 | 1075 | 1336 | 1251 | 1102 | 1145.76 | 1061 | 1230 | 1.16 | 0.214 | 0.03586241 | 0.13002801 | 0.0044 | 0.0098 | 0.0066 | 0.0066 | TRUE | 0.0982 |
| Zzef1 | 2380 | 1766 | 2146 | 3713 | 2413 | 2013 | 2528 | 1764 | 2155 | 2919 | 2596 | 2204 | 2360.86 | 2149 | 2573 | 1.198 | 0.26 | 0.09309878 | 0.24933842 | 0.0249 | 0.0083 | 0.0169 | 0.0169 | TRUE | 0.2362 |
| Zzz3 | 4462 | 4147 | 4215 | 5525 | 3955 | 3736 | 4739 | 4142 | 4232 | 4343 | 4255 | 4090 | 4300.28 | 4371 | 4229 | 0.968 | -0.047 | 0.56516531 | 0.74551564 | 0.0027 | 0.0077 | 0.0046 | 0.0046 | TRUE | 0.066 |

**Table S5. Genes assigned to terms related to 'Neural' identified by Gene Ontology analysis**

| GO:0021915 | GO:0001841 | GO:0014020 | GO:0001843 | GO:0090177 | GO:0061351 | GO:0061076 | GO:0001840 | GO:2000177 | GO:0090179 | GO:0090178 | GO:0003407 | GO:0001839 | GO:0014033 | GO:2000178 |
|---|---|---|---|---|---|---|---|---|---|---|---|---|---|---|
| Abl2 | Abl2 | Abl2 | Abl2 | Celsr1 | Rapgef1 | Pou4f2 | Fgf8 | Rapgef1 | Celsr1 | Celsr1 | Gnat2 | Fgf8 | Anxa6 | Rapgef1 |
| Bmp4 | Bmp4 | Bmp4 | Bmp4 | Sfrp1 | Cd24a | Casz1 | Ptch1 | Cd24a | Sfrp1 | Sfrp1 | Neurod1 | T | Bmp4 | Cd24a |
| Celsr1 | Celsr1 | Celsr1 | Celsr1 | Cthrc1 | Fgf8 | | T | Fzd9 | Ptk7 | Ptk7 | Pou4f2 | Vangl2 | Fn1 | Nf1 |
| Cobl | Cobl | Cobl | Cobl | Ptk7 | Fgfr1 | | Vangl2 | Gli1 | Vangl2 | Vangl2 | Prom1 | | Gbx2 | Spint2 |
| Enah | Enah | Enah | Enah | Vangl2 | Fzd9 | | | Nf1 | | | Megf11 | | Jag1 | Tgfb1 |
| Epha2 | Ptch1 | Ptch1 | Ptch1 | | Gbx2 | | | Ngfr | | | Thy1 | | Lama5 | |
| Fgf8 | Rara | Rara | Rara | | Kif1a | | | Notch1 | | | Rab11fip4 | | Sema3f | |
| Gbx2 | Sfrp1 | Sfrp1 | Sfrp1 | | Lef1 | | | Spint2 | | | Smarcd3 | | Sema4a | |
| Nf1 | Sox11 | Spint2 | Spint2 | | Nf1 | | | Tgfb1 | | | Casz1 | | Sema6b | |
| Notch1 | Sox4 | T | T | | Ngfr | | | Ell3 | | | | | Sema6c | |
| Ptch1 | Spint2 | Tead2 | Tead2 | | Notch1 | | | Ctsz | | | | | Sema7a | |
| Rara | T | Tgfb1 | Tgfb1 | | Ncor2 | | | Smarcd3 | | | | | Sfrp1 | |
| Sfrp1 | Tead2 | Tsc2 | Tsc2 | | Spint2 | | | Smarca1 | | | | | Sox11 | |
| Sox11 | Tgfb1 | Tulp3 | Tulp3 | | Tgfb1 | | | | | | | | Mapk3 | |
| Sox4 | Tsc2 | Kif20b | Kif20b | | Ell3 | | | | | | | | Sema4g | |
| Spint2 | Tulp3 | Luzp1 | Dlc1 | | Tacc1 | | | | | | | | | |
| T | Kif20b | Dlc1 | Med12 | | Dbn1 | | | | | | | | | |
| Tcf7 | Luzp1 | Med12 | Cthrc1 | | Ctsz | | | | | | | | | |
| Tead2 | Dlc1 | Cthrc1 | Ptk7 | | Smarcd3 | | | | | | | | | |
| Itpk1 | Med12 | Ptk7 | Vangl2 | | Smarca1 | | | | | | | | | |
| Tgfb1 | Cthrc1 | Vangl2 | | | | | | | | | | | | |
| Tsc2 | Ptk7 | | | | | | | | | | | | | |
| Tulp3 | Vangl2 | | | | | | | | | | | | | |
| Kif20b | | | | | | | | | | | | | | |
| Luzp1 | | | | | | | | | | | | | | |
| Dlc1 | | | | | | | | | | | | | | |
| Med12 | | | | | | | | | | | | | | |
| Cthrc1 | | | | | | | | | | | | | | |
| Atp6ap2 | | | | | | | | | | | | | | |
| Ptk7 | | | | | | | | | | | | | | |
| Vangl2 | | | | | | | | | | | | | | |

**Table S5. Genes assigned to terms related to 'Neuron' identified by Gene Ontology analysis**

| GO:0048699 | GO:0030182 | GO:0031175 | GO:0048666 | GO:0048812 | GO:0045664 | GO:0010975 | GO:0048667 | GO:0045665 | GO:0010976 | GO:0010977 | GO:0097485 | GO:0045666 | GO:0051402 | GO:1990138 |
|---|---|---|---|---|---|---|---|---|---|---|---|---|---|---|
| Pcsk9 | Pcsk9 | Inpp5f | Inpp5f | Unc5a | Inpp5f | Inpp5f | Unc5a | Inpp5f | Rrn3 | Inpp5f | Unc5a | Rrn3 | Pcsk9 | Apoe |
| Inpp5f | Inpp5f | Rrn3 | Rrn3 | Foxp1 | Rrn3 | Rrn3 | Foxp1 | Rap1gap | Rapgef1 | Apoe | Foxp1 | Rapgef1 | Arrb1 | Ddr1 |
| Rrn3 | Rrn3 | Unc5a | Unc5a | Adarb1 | Rapgef1 | Rapgef1 | Adarb1 | Apoe | Abl2 | Cit | Agrn | Abl2 | Adarb1 | Dpysl2 |
| Ttbk1 | Unc5a | Rapgef1 | Rapgef1 | Arhgef28 | Rap1gap | Abl2 | Arhgef28 | Cd24a | Ap2a1 | Crmp1 | Ank3 | Ap2a1 | Egln3 | Dclk1 |
| Unc5a | Rapgef1 | Foxp1 | Foxp1 | Abl2 | Abl2 | Agrn | Abl2 | Cit | Apoe | Epha4 | Boc | Apoe | Agrn | Bcl11a |
| Rapgef1 | Foxp1 | Mcf2 | Mcf2 | Agrn | Agrn | Ap2a1 | Agrn | Crmp1 | Bmp4 | Bcl11a | Apbb2 | Bmp4 | Apoe | Fn1 |
| Foxp1 | Mcf2 | Adarb1 | Adarb1 | Ank3 | Ap2a1 | Apoe | Ank3 | Dll1 | Camk2b | Ptk2 | Bmpr1b | Camk2b | Hyou1 | Impact |
| Mcf2 | Ehmt2 | Arhgef28 | Arhgef28 | Boc | Apoe | Bmp4 | Boc | Epha4 | Cd24a | Gfap | Runx3 | Cd24a | Cacna1a | Arhgap4 |
| Ehmt2 | Rap1gap | Abl2 | Abl2 | Apbb2 | Bmp4 | Cacna1a | Apbb2 | Bcl11a | Cnr1 | H2-D1 | Crmp1 | Cnr1 | Casp3 | Mag |
| Rap1gap | Adarb1 | Agrn | Agrn | Apoe | Cacna1a | Camk2b | Apoe | Ptk2 | Cobl | H2-K1 | Dpysl2 | Cobl | Cit | Map1b |
| Adarb1 | Arhgef28 | Ank3 | Ank3 | Bmpr1b | Camk2b | Cd24a | Atp2b2 | Gfap | Crabp2 | Lgals1 | Enah | Crabp2 | Coro1a | Myo5b |
| Arhgef28 | Abl2 | Boc | Boc | Cacna1a | Cd24a | Cit | Bmpr1b | H2-D1 | Dlg4 | Lrp1 | Epha4 | Dlg4 | Fgf8 | Ntn1 |
| Abl2 | Agrn | Ap2a1 | Ap2a1 | Ddr1 | Cit | Cnr1 | Cacna1a | H2-K1 | Eef2k | Arhgap4 | Epha8 | Eef2k | Fzd9 | Ntrk3 |
| Agrn | Ank3 | Apbb2 | Apbb2 | Camk2a | Cnr1 | Cobl | Camk2a | Jag1 | Enc1 | Mag | Ephb3 | Enc1 | Gclm | Pak1 |

| | | | | | | | | | | | | | | |
|---|---|---|---|---|---|---|---|---|---|---|---|---|---|---|
| Ank3 | Boc | Apoe | Apoe | Camk2b | Cobl | Crabp2 | Camk2b | Lgals1 | Epha4 | Ngfr | Ext1 | Epha4 | Glp1r | Pou4f2 |
| Boc | Ap2a1 | Atp2b2 | Atp2b2 | Runx3 | Crabp2 | Crmp1 | Runx3 | Lrp1 | Bcl11a | Ntn1 | Fgf8 | Bcl11a | Grik5 | Ptprs |
| Ap2a1 | Apbb2 | Bmp4 | Prdm1 | Cit | Crmp1 | Dpysl2 | Cit | Arhgap4 | Fgfr1 | Pmp22 | Flot1 | Fgfr1 | Lrp1 | Sema3f |
| Apbb2 | Apoe | Bmpr1b | Bmp4 | Cobl | Dpysl2 | Dlg4 | Cobl | Mag | Fn1 | Ptprs | Gap43 | Fn1 | Mag | Sema4a |
| Apoe | Atp2b2 | Cacna1a | Bmpr1b | Crabp2 | Dlg4 | Eef2k | Crabp2 | Ngfr | Ikbkb | Stmn2 | Gbx2 | Ikbkb | Mt3 | Sema6b |
| Atp2b2 | Prdm1 | Ddr1 | Cacna1a | Crmp1 | Dll1 | Enc1 | Crmp1 | Notch1 | Map1b | Sema3f | Foxd1 | Impact | Mybl2 | Sema6c |
| Prdm1 | Bmp4 | Camk2a | Ddr1 | Dpysl2 | Eef2k | Epha4 | Dpysl2 | Ntn1 | Myo5b | Sema4a | Matn2 | Map1b | Nf1 | Sema7a |
| Bmp4 | Bmpr1b | Camk2b | Camk2a | Dclk1 | Enc1 | Ephb3 | Dclk1 | Pmp22 | Neu1 | Sema6b | Ngfr | Myo5b | Ngfr | Tiam1 |
| Bmpr1b | Cacna1a | Casp3 | Camk2b | Dlg4 | Epha4 | Bcl11a | Dlg4 | Pou4f2 | Nf1 | Sema6c | Ntn1 | Neu1 | Prkcg | Nrn1l |
| Tspo | Ddr1 | Runx3 | Casp3 | Eef2k | Ephb3 | Ptk2 | Eef2k | Ptprs | Ngfr | Sema7a | Etv4 | Neurod1 | Prkci | Twf2 |
| Cacna1a | Camk2a | Cd24a | Runx3 | Enah | Bcl11a | Fgfr1 | Enah | Stmn2 | Ntn1 | Thy1 | Pou4f2 | Nf1 | Lgmn | Cpne5 |
| Ddr1 | Camk2b | Cit | Cd24a | Epha4 | Ptk2 | Fn1 | Epha4 | Sema3f | Ntrk3 | Tsc2 | Ptch1 | Ngfr | Sod1 | Sema4g |
| Camk2a | Casp3 | Cnr1 | Cit | Epha8 | Fgfr1 | Gfap | Epha8 | Sema4a | P2ry2 | Dpysl3 | Reln | Ntn1 | Sod2 | Cpne1 |
| Camk2b | Runx3 | Cobl | Cnr1 | Ephb3 | Fn1 | H2-D1 | Ephb3 | Sema6b | Pak1 | Vim | Robo1 | Ntrk3 | Srpk2 | Dbn1 |
| Casp3 | Cd24a | Crabp2 | Cobl | Bcl11a | Gfap | H2-K1 | Bcl11a | Sema6c | Palm | Klk8 | Sema3f | P2ry2 | Hdac4 | Map3k13 |
| Runx3 | Cit | Crmp1 | Crabp2 | Ext1 | H2-D1 | Ikbkb | Ext1 | Sema7a | Prkci | Sema4g | Sema4a | Pak1 | Stxbp1 | Cyfip2 |
| Cd24a | Cnr1 | Dpysl2 | Crmp1 | Ptk2 | H2-K1 | Kif13b | Ptk2 | Cntn2 | Pou4f2 | Rtn4rl2 | Sema6b | Palm | Trp73 | |
| Cdh1 | Cobl | Dclk1 | Dpysl2 | Fgf8 | Ikbkb | Lgals1 | Fgf8 | Thy1 | Reln | Lrig2 | Sema6c | Prkci | Vegfb | |
| Celsr1 | Crabp2 | Dlg4 | Dclk1 | Flot1 | Impact | Lrp1 | Flot1 | Trp73 | Robo1 | Rap1gap2 | Sema7a | Pou4f2 | Wfs1 | |
| Cit | Crmp1 | Eef2k | Dlg4 | Fn1 | Jag1 | Arhgap4 | Fn1 | Tsc2 | Stmn2 | Carm1 | Cntn2 | Rara | Axl | |
| Cnr1 | Dpysl2 | Enah | Eef2k | Gap43 | Kif13b | Mag | Gap43 | Dpysl3 | Sema7a | Ctsz | Rnf165 | Reln | Vstm2l | |
| Cobl | Dclk1 | Enc1 | Enah | Gbx2 | Lgals1 | Map1b | Gbx2 | Vim | Skil | Itm2c | Plxnb1 | Robo1 | Ctsz | |
| Crabp2 | Dlg4 | Epha4 | Enc1 | Gja1 | Lrp1 | Myo5b | Foxd1 | Klk8 | Creb3l2 | Nfatc4 | Sema4g | Stmn2 | Fam162a | |
| Crmp1 | Dll1 | Epha8 | Epha4 | Foxd1 | Rac3 | Neu1 | Kif13b | Sema4g | Plk5 | | Pla2g10 | Sema7a | Pigt | |
| Dpysl2 | Eef2k | Ephb3 | Epha8 | Impact | Arhgap4 | Nf1 | Stmn1 | Rtn4rl2 | Ppp2r5d | | Vstm2l | Skil | Trim2 | |
| Dclk1 | Enah | Bcl11a | Ephb3 | Kif13b | Mag | Ngfr | Arhgap4 | Lrig2 | Tiam1 | | Nrcam | Sox11 | | |
| Dlg4 | Enc1 | Ext1 | Bcl11a | Stmn1 | Map1b | Ntn1 | Mag | Rap1gap2 | Tsc2 | | Ccdc141 | Creb3l2 | | |
| Dll1 | Epha2 | Ptk2 | Ext1 | Lifr | Myo5b | Ntrk3 | Matn2 | Zhx2 | Dpysl3 | | Bcl11b | Plk5 | | |
| Dll3 | Epha4 | Fgf8 | Ptk2 | Arhgap4 | Neu1 | P2ry2 | Map1a | Carm1 | Tbc1d24 | | Cyfip2 | Ppp2r5d | | |
| Eef2k | Epha8 | Fgfr1 | Fgf8 | Mag | Neurod1 | Pak1 | Map1b | Ctsz | Zmynd8 | | Vangl2 | Tiam1 | | |
| Enah | Ephb3 | Flot1 | Fgfr1 | Matn2 | Nf1 | Palm | Myo5b | Itm2c | Plxnb1 | | | Tsc2 | | |
| Enc1 | Bcl11a | Fn1 | Flot1 | Map1a | Ngfr | Prkci | Ngfr | Casz1 | Pacsin1 | | | Dpysl3 | | |
| Epha2 | Ext1 | Gap43 | Fn1 | Map1b | Mycn | Pmp22 | Notch1 | Nfatc4 | Twf2 | | | Tbc1d24 | | |
| Epha4 | Ptk2 | Gbx2 | Gap43 | Myo5b | Notch1 | Pou4f2 | Ntn1 | | Cpne5 | | | Zmynd8 | | |
| Epha8 | Fgf8 | Gfap | Gbx2 | Ngfr | Ntn1 | Ptprs | Tbc1d24 | | Ankrd27 | | | Plxnb1 | | |
| Ephb3 | Fgfr1 | Gfra1 | Gfap | Notch1 | Ntrk3 | Reln | Ntrk3 | | Mapk6 | | | Pacsin1 | | |
| Bcl11a | Flot1 | Gja1 | Gfra1 | Ntn1 | P2ry2 | Robo1 | Pak1 | | Dbn1 | | | Twf2 | | |
| Ext1 | Fn1 | H2-D1 | Gja1 | Tbc1d24 | Pak1 | Stmn2 | Etv4 | | Shank3 | | | Cpne5 | | |
| Ptk2 | Gap43 | H2-K1 | Gnat2 | Ntrk3 | Palm | Sema3f | Pmp22 | | Actr2 | | | Ankrd27 | | |
| Fgf8 | Gbx2 | Foxd1 | H2-D1 | Pak1 | Prkci | Sema4a | Pou4f2 | | Dab2ip | | | Cpne1 | | |
| Fgfr1 | Gfap | Ikbkb | H2-K1 | Etv4 | Pmp22 | Sema6b | Ptch1 | | Ptk7 | | | Mapk6 | | |
| Flot1 | Gfra1 | Impact | Foxd1 | Pmp22 | Pou4f2 | Sema6c | Ptprs | | Map3k13 | | | Dbn1 | | |
| Fn1 | Gja1 | Atcay | Ikbkb | Pou4f2 | Ptprs | Sema7a | Reln | | | | | Shank3 | | |
| Fzd9 | Gnat2 | Kif13b | Impact | Ptch1 | Rara | Sfrp1 | Robo1 | | | | | Actr2 | | |
| Gap43 | Nkx6-2 | Stmn1 | Atcay | Ptprs | Reln | Skil | Sema3f | | | | | Dab2ip | | |
| Gas6 | H2-D1 | Lgals1 | Kif13b | Rac2 | Robo1 | Creb3l2 | Sema4a | | | | | Ptk7 | | |
| Gbx2 | H2-K1 | Lifr | Stmn1 | Reln | Stmn2 | Cntn2 | Sema6b | | | | | Map3k13 | | |
| Gfap | Foxd1 | Lrp1 | Lgals1 | Robo1 | Sema3f | Plk5 | Sema6c | | | | | Kdm4c | | |
| Gfra1 | Ikbkb | Rac3 | Lifr | Sema3f | Sema4a | Ppp2r5d | Sema7a | | | | | Lin28a | | |
| Gja1 | Impact | Arhgap4 | Lrp1 | Sema4a | Sema6b | Thy1 | Skil | | | | | | | |
| Gnat2 | Jag1 | Mag | Rac3 | Sema6b | Sema6c | Tiam1 | Sod1 | | | | | | | |
| Nkx6-2 | Atcay | Matn2 | Arhgap4 | Sema6c | Sema7a | Tsc2 | Stxbp1 | | | | | | | |

| | | | | | | | |
|---|---|---|---|---|---|---|---|
| H2-D1 | Kif13b | Mmp2 | Mag | Sema7a | Sfrp1 | Dpysl3 | Cntn2 |
| H2-K1 | Stmn1 | Map1a | Matn2 | Skil | Skil | Vim | Thy1 |
| Hes2 | Lgals1 | Map1b | Mmp2 | Stxbp1 | Sox11 | Tbc1d24 | Tiam1 |
| Foxd1 | Lifr | Myo5b | Map1a | Cntn2 | Creb3l2 | Zmynd8 | Tsc2 |
| Ikbkb | Lrp1 | Neu1 | Map1b | Thy1 | Cntn2 | Arhgap33 | Uchl1 |
| Il6st | Rac3 | Nf1 | Myo5b | Tiam1 | Plk5 | Plxnb1 | Vim |
| Impact | Mcoln3 | Ngfr | Neu1 | Tsc2 | Ppp2r5d | Pacsin1 | Tbc1d24 |
| Jag1 | Arhgap4 | Notch1 | Neurod1 | Uchl1 | Thy1 | Twf2 | Rnf165 |
| Atcay | Mag | Ntn1 | Nf1 | Vim | Tiam1 | Cpne5 | Arhgap33 |
| Kif13b | Matn2 | Tbc1d24 | Ngfr | Tbc1d24 | Trp73 | Ankrd27 | Plxnb1 |
| Stmn1 | Mmp2 | Ntrk3 | Notch1 | Rnf165 | Tsc2 | Klk8 | Twf2 |
| Ldlr | Map1a | P2ry2 | Ntn1 | Arhgap33 | Dpysl3 | Sema4g | Ankrd27 |
| Lef1 | Map1b | Pak1 | Tbc1d24 | Nrn1l | Vim | Rtn4rl2 | Sema4g |
| Lgals1 | Myo5b | Palm | Ntrk3 | Plxnb1 | Tbc1d24 | Lrig2 | Pla2g10 |
| Lifr | Neu1 | Etv4 | P2ry2 | Pacsin1 | Zmynd8 | Prex1 | Vstm2l |
| Lrp1 | Neurod1 | Prkci | Pak1 | Twf2 | Arhgap33 | Nrcam | Nrcam |
| Rac3 | Nf1 | Pmp22 | Palm | Cpne5 | Plxnb1 | Rap1gap2 | Adcy1 |
| Mcoln3 | Ngfr | Pou4f2 | Etv4 | Kif20b | Pacsin1 | Mapk6 | Ccdc141 |
| Arhgap4 | Mycn | Ptch1 | Prkci | Nyap1 | Twf2 | Srcin1 | Srcin1 |
| Mag | Notch1 | Ptprs | Pmp22 | Ankrd27 | Cpne5 | Dbn1 | Dbn1 |
| Matn2 | Ntn1 | Rac2 | Pou4f2 | Klk8 | Ankrd27 | Shank3 | Bcl11b |
| Mmp2 | Tbc1d24 | Reln | Ptch1 | Sema4g | Klk8 | Carm1 | Shank3 |
| Mt3 | Ntrk3 | Robo1 | Ptprs | Pla2g10 | Sema4g | Ctsz | Actr2 |
| Map1a | P2ry2 | Stmn2 | Rac2 | Cpne1 | Cpne1 | Itm2c | Map3k13 |
| Map1b | Pak1 | Sema3f | Reln | Vstm2l | Rtn4rl2 | Actr2 | Nfatc4 |
| Myo5b | Palm | Sema4a | Robo1 | Nrcam | Lrig2 | Dab2ip | Cyfip2 |
| Neu1 | Etv4 | Sema6b | Rpgr | Adcy1 | Prex1 | Ptk7 | Vangl2 |
| Neurod1 | Prkci | Sema6c | Stmn2 | Ccdc141 | Nrcam | Map3k13 | Ophn1 |
| Nf1 | Pmp22 | Sema7a | Sema3f | Srcin1 | Rap1gap2 | Nfatc4 | |
| Ngfr | Pou4f2 | Sfrp1 | Sema4a | Dbn1 | Zhx2 | | |
| Mycn | Prom1 | Skil | Sema6b | Bcl11b | Mapk6 | | |
| Notch1 | Ptch1 | Sod1 | Sema6c | Shank3 | Srcin1 | | |
| Ntn1 | Ptprs | Creb3l2 | Sema7a | Actr2 | Dbn1 | | |
| Tbc1d24 | Rac2 | Stx3 | Sfrp1 | Dab2ip | Bcl11b | | |
| Ntrk3 | Rara | Stxbp1 | Skil | Map3k13 | Shank3 | | |
| P2ry2 | Reln | Cntn2 | Sod1 | Nfatc4 | Carm1 | | |
| Pak1 | Robo1 | Plk5 | Sod2 | Cyfip2 | Ctsz | | |
| Palm | Rpgr | Ppp2r5d | Creb3l2 | Vangl2 | Itm2c | | |
| Etv4 | Stmn2 | Thy1 | Stx3 | | Actr2 | | |
| Prkci | Sema3f | Tiam1 | Stxbp1 | | Dab2ip | | |
| Pmp22 | Sema4a | Tsc2 | Cntn2 | | Casz1 | | |
| Pou4f2 | Sema6b | Tulp1 | Plk5 | | Ptk7 | | |
| Prom1 | Sema6c | Uchl1 | Ppp2r5d | | Map3k13 | | |
| Ptch1 | Sema7a | Dpysl3 | Thy1 | | Nfatc4 | | |
| Ptprs | Sfrp1 | Vim | Tiam1 | | Kdm4c | | |
| Rac2 | Skil | Tbc1d24 | Trp73 | | Lin28a | | |
| Rara | Sod1 | Rnf165 | Tsc2 | | | | |
| Reln | Sod2 | Zmynd8 | Tulp1 | | | | |
| Robo1 | Sox11 | Arhgap33 | Uchl1 | | | | |
| Rpgr | Sox4 | Nrn1l | Dpysl3 | | | | |
| Stmn2 | Stat3 | Plxnb1 | Vim | | | | |
| Sema3f | Creb3l2 | Pacsin1 | Tbc1d24 | | | | |

| | | | |
|---|---|---|---|
| Sema4a | Stx3 | Twf2 | Rnf165 |
| Sema6b | Stxbp1 | Cpne5 | Zmynd8 |
| Sema6c | Cntn2 | Kif20b | Arhgap33 |
| Sema7a | Plk5 | Nyap1 | Nrn1l |
| Sfrp1 | Ppp2r5d | Ankrd27 | Plxnb1 |
| Skil | Tgm3 | Klk8 | C1ql1 |
| Sod1 | Thy1 | Sema4g | Pacsin1 |
| Sod2 | Tiam1 | Pla2g10 | Twf2 |
| Sox11 | Trp73 | Cpne1 | Cpne5 |
| Sox4 | Tsc2 | Rtn4rl2 | Kif20b |
| Stat3 | Tulp1 | Lrig2 | Nyap1 |
| Creb3l2 | Tulp3 | Micall1 | Ankrd27 |
| Stx3 | Uchl1 | Prex1 | Klk8 |
| Stxbp1 | Dpysl3 | Vstm2l | Sema4g |
| Cntn2 | Vim | Nrcam | Pla2g10 |
| Nav1 | Wnt5b | Rap1gap2 | Cpne1 |
| Plk5 | Wnt6 | Adcy1 | Rtn4rl2 |
| Ppp2r5d | Tbc1d24 | Mapk6 | Lrig2 |
| Tgfb1 | Rnf165 | Nptx2 | Micall1 |
| Tgm3 | Zmynd8 | Ccdc141 | Naglu |
| Thy1 | Arhgap33 | Srcin1 | Prex1 |
| Tiam1 | Nrn1l | Dbn1 | Vstm2l |
| Trp73 | Plxnb1 | Slc12a5 | Nrcam |
| Tsc2 | C1ql1 | Bcl11b | Cngb1 |
| Tulp1 | Pacsin1 | Shank3 | Rap1gap2 |
| Tulp3 | Twf2 | Carm1 | Adcy1 |
| Uchl1 | Cpne5 | Trappc4 | Mapk6 |
| Dpysl3 | Kif20b | Ctsz | Nptx2 |
| Vim | Nyap1 | Itm2c | Ccdc141 |
| Wnt5b | Ankrd27 | Actr2 | Srcin1 |
| Wnt6 | Klk8 | Cthrc1 | Dbn1 |
| Tbc1d24 | Sema4g | Dab2ip | Slc12a5 |
| Rnf165 | Pla2g10 | Ptk7 | Bcl11b |
| Zmynd8 | Cpne1 | Map3k13 | Shank3 |
| Arhgap33 | Rtn4rl2 | Nfatc4 | Carm1 |
| Nrn1l | Lrig2 | Manf | Trappc4 |
| Plxnb1 | Chd5 | Cyfip2 | Ctsz |
| C1ql1 | Micall1 | Vangl2 | Itm2c |
| Pacsin1 | Naglu | Clmn | Actr2 |
| Twf2 | Prex1 | Ophn1 | Cthrc1 |
| Cpne5 | Vstm2l | Efhd1 | Dab2ip |
| Kif20b | Nrcam | | Ptk7 |
| Nyap1 | Cngb1 | | Map3k13 |
| Ankrd27 | Rap1gap2 | | Nfatc4 |
| Klk8 | Zhx2 | | Manf |
| Axl | Adcy1 | | Cyfip2 |
| Sema4g | Mapk6 | | Kcnip2 |
| Pla2g10 | Nptx2 | | Vangl2 |
| Cpne1 | Ccdc141 | | Clmn |
| Rtn4rl2 | Srcin1 | | Ophn1 |
| Ell3 | Dbn1 | | Efhd1 |
| Lrig2 | Slc12a5 | | |

| | |
|---|---|
| Chd5 | Bcl11b |
| Idh2 | Shank3 |
| Micall1 | Carm1 |
| Naglu | Trappc4 |
| Prex1 | Ctsz |
| Vstm2l | Itm2c |
| Nrcam | Actr2 |
| Chd7 | Cthrc1 |
| Cngb1 | Dab2ip |
| Rap1gap2 | Casz1 |
| Zhx2 | Ptk7 |
| Adcy1 | Map3k13 |
| Mapk6 | Gpc2 |
| Nptx2 | Nfatc4 |
| Ccdc141 | Manf |
| Srcin1 | Kdm4c |
| Dbn1 | Cyfip2 |
| Plag1 | Pigt |
| Slc12a5 | Kcnip2 |
| Bcl11b | Lin28a |
| Shank3 | Smarca1 |
| Carm1 | Vangl2 |
| Trappc4 | Clmn |
| Ctsz | Ophn1 |
| Itm2c | Efhd1 |
| Actr2 | |
| Smarcd3 | |
| Cthrc1 | |
| Dab2ip | |
| Casz1 | |
| Ptk7 | |
| Map3k13 | |
| Zcchc24 | |
| Gpc2 | |
| Nfatc4 | |
| Manf | |
| Kdm4c | |
| Cyfip2 | |
| Pigt | |
| Kcnip2 | |
| Lin28a | |
| Hes7 | |
| Smarca1 | |
| Vangl2 | |
| Clmn | |
| Ophn1 | |
| Efhd1 | |

**Table S6. Reagents used in this study**

| pCAGGS puro plasmid | Source | catalog number |
|---|---|---|
| RNF12 | MRC-PPU Reagents and Services | DU50610 |
| RNF12 S212A | MRC-PPU Reagents and Services | DU53528 |
| RNF12 S214A | MRC-PPU Reagents and Services | DU50796 |
| RNF12 S227A | MRC-PPU Reagents and Services | DU53591 |
| RNF12 S229A | MRC-PPU Reagents and Services | DU53592 |
| RNF12 S212A S214A | MRC-PPU Reagents and Services | DU53518 |
| RNF12 S227A S229A | MRC-PPU Reagents and Services | DU53514 |
| RNF12 S214A S229A | MRC-PPU Reagents and Services | DU53593 |
| RNF12 S212A S214A S227A S229A | MRC-PPU Reagents and Services | DU50797 |
| RNF12 S212E S214E S227E S229E | MRC-PPU Reagents and Services | DU50798 |
| RNF12 delta NLS | MRC-PPU Reagents and Services | DU53413 |
| HA-RNF12 | MRC-PPU Reagents and Services | DU50854 |
| HA-RNF12 S212A S214A S227A S229A | MRC-PPU Reagents and Services | DU58741 |
| RNF12 W576Y | MRC-PPU Reagents and Services | DU50800 |
| FLAG SRPK1 | MRC-PPU Reagents and Services | DU53820 |
| FLAG SRPK2 | MRC-PPU Reagents and Services | DU53821 |

| siRNA | Source | catalog number |
|---|---|---|
| ON-TARGETplus Srpk2 siRNA 06 | Horizon Discovery | J-055142-06-0010 |
| Non-targeting Pool | Horizon Discovery | D-001810-10-05 |

| Inhibitor | Target | Supplier | Catalog number | Reference |
|---|---|---|---|---|
| AZ 191 | DYRK1B | Tocris | 5232 | DOI: 10.1042/BJ20130461 |
| KH CB19 | CLK DYRK | Merck Millipore | 219511 | DOI: 10.1016/j.chembiol.2010.11.009. |
| T3 | CLK3 | Aobious | AOB8827 | DOI: 10.1038/s41467-016-0008-7 |
| SPHINX31 | SRPK1 | Axon Medchem | Axon 2714 | DOI: 10.1021/acschembio.6b01048 |
| CHIR 99021 | GSK3b | Axon Medchem | Axon 1386 | DOI: 10.2337/diabetes.52.3.588 |
| PD 0325901 | MEK | Axon Medchem | Axon 1408 | DOI: 10.1016/j.bmcl.2008.10.054 |
| VX-745 | p38 | Selleckchem | S1458 | DOI: 10.1021/ml2001455 |
| JNK-IN-8 | JNK | Selleckchem | S4901 | DOI: 10.1016/j.chembiol.2011.11.010 |
| RO-3306 | CDK1 | Sigma Aldrich | SML0569 | DOI: 10.1073/pnas.0600447103 |
| Flavopiridol | CDK7/9 | Stratech | S2679 | DOI: 10.3892/ijo.9.6.1143 |
| CCT 241533 | CHK2 | Cayman | CAY19178 | DOI: 10.1158/0008-5472.CAN-10-1252 |
| Harmine | DYRK1A | Sigma Aldrich | 286044 | DOI: 10.1016/j.abb.2010.12.024 |
| WEHI-345 | RIPK2 | Cayman | CAY23023 | DOI: 10.1038/ncomms7442 |
| IRAK-4 Ina | IRAK4 | MRC-PPU Reagents and Services | | DOI: 10.1016/j.str.2006.11.001 |

| | | | | |
|---|---|---|---|---|
| GSK 461364 | PLK1/2 | Cayman | CAY18099 | DOI: 10.1158/0008-5472.CAN-09-0945 |
| SRPIN340 | SRPK1 | Sigma Aldrich | SML1088 | DOI: 10.1073/pnas.0604616103 |

| **Immunoblot antibodies** | | | |
|---|---|---|---|
| **antigen** | **Source** | **catalog number** | **extra details** |
| RNF12 | Novus Biologicals | H00051132-M01 | |
| ERK1 | BD Biosciences | 610408 | |
| SRPK1 | BD Biosciences | 611072 | |
| SRPK2 | BD Biosciences | 611118 | |
| HA-tag | Abcam | ab9110 | |
| REX1 | Abcam | ab28141 | |
| SRPK3 | R&D Systems | MAB7230-SP | |
| FLAG | Sigma Aldrich | F1804-50UG | |
| HA-HRP | Roche | 12013819001 | |
| Synaptophysin | Cell Signaling Technologies | 5461 | D35E4 |
| beta-actin | Cell Signaling Technologies | 4970 | 13E5 |
| RNF12 (1-271) | MRC-PPU Reagents and Services | S691D | third bleed |
| RNF12 pSer212/214 QRRARpSRpSPEHRR phosphopeptide | MRC-PPU Reagents and Services | SA310 | fourth bleed |
| GST | MRC-PPU Reagents and Services | S902A | third bleed |

| **Immunofluorescence antibodies** | | | |
|---|---|---|---|
| **antigen** | **Source** | **catalog number** | **extra details** |
| RNF12 | Novus Biologicals | H00051132-M01 | |
| RNF12 (1-271) | MRC-PPU Reagents and Services | S691D | third bleed |
| Tubulin β 3 | Biolegend | 801202 | |
| MAP2 | Sigma Aldrich | M2320 | |

| **Protein** | **source** | **catalog number** |
|---|---|---|
| RNF12 | MRC-PPU Reagents and Services | DU61098 |
| RNF12 S212A S214A S227A S229A | MRC-PPU Reagents and Services | DU53249 |
| DYRK1a | MRC-PPU Reagents and Services | DU19040 |
| CLK2 | MRC-PPU Reagents and Services | DU16987 |
| GSK3beta | MRC-PPU Reagents and Services | DU899 |
| ERK1 (MAPK3) | MRC-PPU Reagents and Services | DU1509 |
| ERK2 (MAPK1) | MRC-PPU Reagents and Services | DU650 |
| JNK3 alpha 1 (SAPK1b) | MRC-PPU Reagents and Services | DU1511 |
| p38 alpha (SAPK2a) | MRC-PPU Reagents and Services | DU979 |
| CDK2 - CyclinA | MRC-PPU Reagents and Services | DU43557 |
| CDK5 - p35 | MRC-PPU Reagents and Services | DU39816 |
| CDK7 - MAT1 - Cyclin H | MRC-PPU Reagents and Services | DU49574 |
| CDK9 - Cyclin T1 | MRC-PPU Reagents and Services | DU31050 |
| SRPK1 | MRC-PPU Reagents and Services | DU967 |

| | | |
|---|---|---|
| SRPK2 | MRC-PPU Reagents and Services | DU36135 |
| SRPK3 | MRC-PPU Reagents and Services | DU967 |
| SRPK1 D497A | MRC-PPU Reagents and Services | DU66208 |
| SRPK2 D541A | MRC-PPU Reagents and Services | DU66209 |
| SRPK3 H159D | MRC-PPU Reagents and Services | DU61121 |
| SRPK3 T211M | MRC-PPU Reagents and Services | DU61140 |
| SRPK3 K270M | MRC-PPU Reagents and Services | DU61135 |
| Ube1 | MRC-PPU Reagents and Services | DU32888 |
| UBE2D1 (UbcH5a) | MRC-PPU Reagents and Services | DU4315 |
| FLAG-Ubiquitin | MRC-PPU Reagents and Services | DU46789 |
| Ubiquitin IR-800 | Walden lab (Glasgow) | |

| qPCR primers | | |
|---|---|---|
| **gene** | **primer forward (5'-3')** | **primer reverse (5'-3')** |
| Gapdh | CTCGTCCCGTAGACAAAA | TGAATTTGCCGTGAGTGG |
| Ntn1 | CGCAACTGTACCAGTGACCTCT | TTGCGGCAGTAGATGAGGACGA |
| Dll1 | ACCAAGTGCCAGTCACAGAG | TCCATCTTACACCTCAGTCGC |
| Kif1a | CACCACTATTGTCAACCCCAAA | CCCCAATGTCCCTGTAGACCT |
| Gfap | CAATGCTGGCTTCAAGGAGACACG | TCAGTTCAGCTGCCAGCGCCT |
| Unc5a | GTCTGGTGTGTGACTGTAGGCA | CCGAGCATGGAGGTTGCAGTTG |

| CRISPR Cas9 knockout | | | | |
|---|---|---|---|---|
| **gene name** | **exon targeted** | **mutation** | **sense guide RNA** | **antisense guide RNA** |
| Srpk1 | 3 | Srpk1 KO | 5’ GAG CAG GAG GAG GAG ATT CT 3’ | 5’ GCG GAG TGG GGT GCA GAG CCT 3’ |
| Srpk2 | 5 | Srpk2 KO | 5’ GAT TGA TGA CTT CAA GAT CTC 3’ | 5’ GAT GTC TTT GTT TGG GTC ACT 3’ |
| Zfp42 | 4 | Zfp42 KO | 5’ GAG GAA GAT GGC TTC CCT GA 3’ | 5’ GAA TCT CAC TTT CAT CCC GGA 3’ |

| CRISPR Cas9 knockin | | | | |
|---|---|---|---|---|
| **gene name** | **exon targeted** | **mutation** | **sense guide RNA** | **antisense guide RNA** |
| Rlim | 3 | Rlim 4xSA/y | 5’ GAG TTC GTC CTG GAG AAT AC 3’ | 5’ GTA CTT GAA GAT CAA GAA CTA 3’ |
| Rlim | 3 | Rlim deltaNLS/y | 5’ GAA GCC GGA GCC CAG AGC AT 3’ | 5’ GCT CCT CTG AGC TCT GGT GGT 3’ |
| Rlim | 3 | Rlim W576Y/y | 5’ GCA GGG CAG TCT TAT CTT CT 3’ | 5’ GTG GAA TTC TCA GAC AAC CAG 3’ |
